# Big data reveals deep associations in physical examination indicators and can help predict overall underlying health status

**DOI:** 10.1101/855809

**Authors:** Haixin Wang, Ping Shuai, Yanhui Deng, Jiyun Yang, Shanshan Zhang, Yi Yin, Lin Wang, Dongyu Li, Tao Yong, Yuping Liu, Lulin Huang

## Abstract

Because of lacking of the systematic investigation of correlations between the physical examination indicators (PEIs), currently most of them are independently used for disease warning. This results in very limited diagnostic values of general physical examination. Here, we first systematically analyzed the correlations between 221 PEIs in healthy and in 34 unhealthy states in 803,614 peoples in China. We revealed rich relevant between PEIs in healthy physical status (7,662 significant correlations, 31.5% of all). However, in disease conditions, the PEI correlations changed. We further focused on the difference of these PEIs between healthy and 35 unhealthy physical status, 1,239 significant PEI difference were discovered suggesting as candidate disease markers. Finally, we established machine learning algorithms to predict the health status by using 15%-16% PEIs by feature extraction, which reached 66%-99% precision predictions depending on the physical state. This new encyclopedia of PEI correlation provides rich information to chronic disease diagnosis. Our developed machine learning algorithms will have fundamental impact in practice of general physical examination.

---

The comprehensive primary healthcare system has had a broader impact on human health compared to clinical medical treatment^1–4^. Health examinations help those who are healthy to improve their understanding of their own physical functions and maintain their health status, and inform those as to the health benefits conferred by changing unhealthy habits and avoiding risk factors that can lead to disease^5^. Physical examinations can help minimize the distress of diseases^6^. With the population size grows and ages, people’s healthcare needs are constantly increasing, and health-care provisions are becoming more sophisticated and in parallel, more costly^7^.

Health examinations are common elements of healthcare in developed countries^8^. These checks consist of general blood examination, urine examination, blood glucose examination, blood lipid examination, renal function examination and so on. However, currently, the physical examination report is assessed mainly based on one or two independent physical examination indicators (PEIs), which can only provide very limited information for physical examiners about their healthy condition or disease diagnosis^9–11^. The correlations between PEI in different physical states (i.e. healthy, hypertension, diabetes) have not been systematically investigated, even though they are expected to provide valuable information for public health care, for example by defining a small set of easily measurable PEIs that can be used in the accurate diagnosis of a disease before the disease phenogenesis.

The recent explosion of available health data promises to transform healthcare by improving care quality and as such, improving population health while constraining escalating costs^12^. Health examination centers generate systematic big data that has the capacity to reveal otherwise undetected underlying health issues^13–14^. In clinical, there is growing investment in developing big data applications for medical care, such as those based on artificial intelligence (AI) to diagnosis diseases based on clinical images^15–17^. Although AI can save cost and improve efficiency, especially for early diagnosis and prevention of chronic diseases^18^, because of insufficient systematic analysis of PEIs in physical status, currently no prediction models were generated for physical status predictions based on PEIs.

As China’s 2009 health-care reform has made impressive progress in expansion of insurance coverage, now general physical examination industry accumulates big data^19^. By using a large dataset of general health examination of Chinese population, the present study had three main aims: to determine the correlations among PEIs in healthy and unhealthy (namely, those with underlying chronic disease) patients; to elucidate the relationship between chronic disorders and normal individuals for these PEIs to discovery candidate disease markers; to develop machine learning models that can predict individual health status using a refined set of PEIs. To address these points, we included physical examination data from 80,3614 individuals who visited one health examination center between 2013 and 2018 in China. We included data from 221 PEIs associated with 35 physical conditions, with the majority unhealthy physical states being due to chronic disease.

## Results

### Study population

The study population was mainly from the Chengdu Plain, Sichuan, P.R. China (102.54°E ∼104.53°E and 30.05°N ∼31.26°N). We included 803,614 individuals who attended the Health Management Center & Physical Examination Center of Sichuan Provincial People’s Hospital in China between 2013 and 2018. The participants represented 35 healthy states based on either a healthy status or the presence of an underlying disease condition (unhealthy status). Specifically, the study population included 711,928 healthy participants, 46,981 patients with hypertension, 11,745 patients with diabetes and 32,960 with other unhealthy status (mainly are chronic disease) (Table 1). We included 221 PEIs in our analyses, which comprised patient demographic information (age and sex) and life-style indicators (alcohol consumption, tobacco use, etc.) (Extended data Table 1).

**Table 1.** Summary of the study samples, detected correlations and the different PEIs between healthy status and unhealthy states

| Body status | Sample (N) | Age (Range) | Female % | Sig. Correlation (a) | Sig. Correlation % (b) | Different PEIs (c) |
| --- | --- | --- | --- | --- | --- | --- |
| Normal condition | 711928 | 41.4 (4~105) | 45.7 | 7662 | 31.5 | - |
| Cholecystolithiasis | 993 | 50.09 (19~97) | 47.8 | 1622 | 6.9 | 28 |
| Hypertension | 46981 | 62.0 (20~102) | 36.1 | 4413 | 18.3 | 112 |
| Hypertension+Diabetes | 8586 | 67.3 (34~99) | 67.1 | 3008 | 12.6 | 100 |
| Hypertension+Coronary | 2074 | 74.0(34~98) | 40.4 | 1920 | 7.7 | 53 |
| Hypertensive+Diabetes+Coronary | 928 | 73.5(37~98) | 35 | 2014 | 8.7 | 56 |
| Hyperlipidemia | 1722 | 65.0(29~96) | 33 | 2256 | 9.5 | 51 |
| Coronary heart disease | 1325 | 68.7(27~93) | 24.6 | 1925 | 8.3 | 36 |
| Coronary+Diabetes | 280 | 70(44~89) | 20.4 | 1335 | 6.2 | 40 |
| Rhinallergosis | 156 | 39 (18~92) | 39.7 | 1278 | 6.1 | 12 |
| Hypothyroidism | 1661 | 46.7(16~94) | 84.6 | 1703 | 7.1 | 33 |
| Hyperthyroidism | 767 | 45.2 (15~89) | 66.6 | 1234 | 5.2 | 16 |
| Cervical spondylopathy | 318 | 53.7 (27~87) | 48.7 | 1589 | 7 | 14 |
| Rheumatoid arthritis | 396 | 55.0 (24~86) | 74.7 | 1039 | 4.5 | 29 |
| Chronic rhinitis | 320 | 38.6 (21~85) | 30.6 | 969 | 4.5 | 8 |
| Nephropathy | 76 | 48.6 (23~91) | 42 | 1771 | 7.8 | 31 |
| Diabetes | 11745 | 59.3(14~96) | 24.9 | 2972 | 12.3 | 91 |
| Gout | 2138 | 51.8 (20~97) | 2 | 1813 | 7.8 | 52 |
| Parkinson's syndrome | 197 | 70.4 (40~92) | 26.4 | 1659 | 8.1 | 28 |
| Stomach trouble | 1296 | 49.5 (19~95) | 39.2 | 1523 | 6.5 | 37 |
| Chronic pharyngitis | 782 | 40.2 (18~90) | 32.7 | 1487 | 6.2 | 14 |
| Lumbar disc protrusion | 385 | 56.8 (25~91) | 35.6 | 982 | 4.3 | 12 |
| Hepatitis B | 706 | 45.3 (22~82) | 22.8 | 1084 | 4.5 | 34 |
| Hypertension+other diseases | 2409 | 64.6 (28~94) | 38.1 | 1750 | 7.1 | 67 |
| Coronary+others | 101 | 69.3(36~92) | 33.7 | 569 | 3.3 | 12 |
| Diabetes+others | 373 | 63 (29~90) | 26.8 | 1045 | 4.6 | 35 |
| Bronchial disease | 574 | 60.7 (19~95) | 33.2 | 1276 | 5.6 | 19 |
| Other disease conditions | 2777 | 51.4(17~100) | 43.8 | 2066 | 8.5 | 38 |
| Brain diseases | 257 | 69.9 (27~98) | 29.5 | 1023 | 4.7 | 25 |
| Hepatic adipose infiltration | 274 | 44.1 (21~77) | 12.4 | 1139 | 5.3 | 36 |
| Asthma | 280 | 51.0 (12~91) | 57.9 | 1388 | 5.9 | 23 |
| Other Cardiac diseases | 344 | 60.0 (26~90) | 55.9 | 749 | 3.6 | 19 |
| Heart disease | 229 | 69.7 (26~96) | 42.6 | 698 | 3.5 | 28 |
| Hepatopathy | 180 | 51.6 (25~83) | 38.7 | 702 | 3.6 | 25 |
| Pregnant | 56 | 29.7(24~36) | 100 | 1015 | 8.5 | 25 |

(a) Significant correlations, the number of correlations with $P$ values calculated by PCC adjusted by all the correlations ( $P<0.05/24,322$ PEI pairs= $2\times 10^{-6}$ ).
(b) Significant correlation %, which is the percentage of significant correlations in all the correlation pairs.
Detailed information on the correlations described in (a-b) is provided in Supplementary Table 1-35.

### PEI correlations in participants with a healthy physical status

We first aimed to explore the PEI correlations in healthy status to give a landscape. Among 221 PEIs, we found 7,662 significant correlations (*P*<0.05/ 24,322 PEI pairs=2×10^-6^) in all 24,322 PEI pairs correlations (31.5%) (Table 1, Supplementary Table 1) in those with a healthy physical status (*N*=711,928, mean age 41.4, female=45.7%). This finding suggests a wide range of correlations between PEIs (Fig.1). The top 50 correlated PEIs included sex, age, red blood cell count, prealbumin (PAB), history of alcohol intake (alcohol consumption, drinking), alkaline phosphatase level (ALP), tobacco use (smoking) and so on (Fig. 1a). Among the 221 PEIs, the number of significantly correlated PEIs also suggested rich correlations between PEIs (Fig. 1b). Of these identified correlations among PEIs in healthy status, some of them are consistent with the reported literatures, but most of them are General inspection PEIs showed rich relevance to each other or to other PEIs. For example, sex showed the richest PEI correlations (151 PEI pairs, males vs. females), including hemoglobin (Hb), creatinine, uric acid (UA), drinking, smoking, body mass index (BMI) and etc., which reflect the differences in body shape, physique and living habits between males and females (Fig. 1, Fig. 2, Supplementary Table 1). Age also showed strong PEI correlations (125 PEI pairs), such as estimated glomerular filtration rate (eGFB), systolic pressure (SBP), diastolic pressure (DBP), albumin (Alb), and low-density lipoprotein (LDL-C). These findings suggest that with increasing age, body functions systematically change (Fig. 1, Fig. 2, Supplementary Table 1). We also found 124 PEI correlations with BMI which reflects the strong influence of body shape on PEIs, including UA, high-density lipoprotein (HDL-C), SBP, and DBP (Fig. 1, Fig. 2, Supplementary Table 1). Blood pressure (BP), which has many physiological meanings, we identified a set of PEIs that correlated with blood pressure (BP), including 125 PEIs for DBP and 124 PEIs for SBP (Fig. 1, Fig. 2, Supplementary Table 1). Intraocular pressure (IOP) is an important factor for the diagnosis of glaucoma^8–9^. We found 79 PEIs that were weakly correlated with IOP of the left eye (IOP-L), including IOP of the right eye (IOP-R) SBP, DBP, Alb, BMI, TG, ApoB, drinking, and TC. Similar to IOP-L, 73 PEIs were weakly correlated with IOP-R (Fig. 1, Fig. 2, Supplementary Table 1).

**Figure 1.**
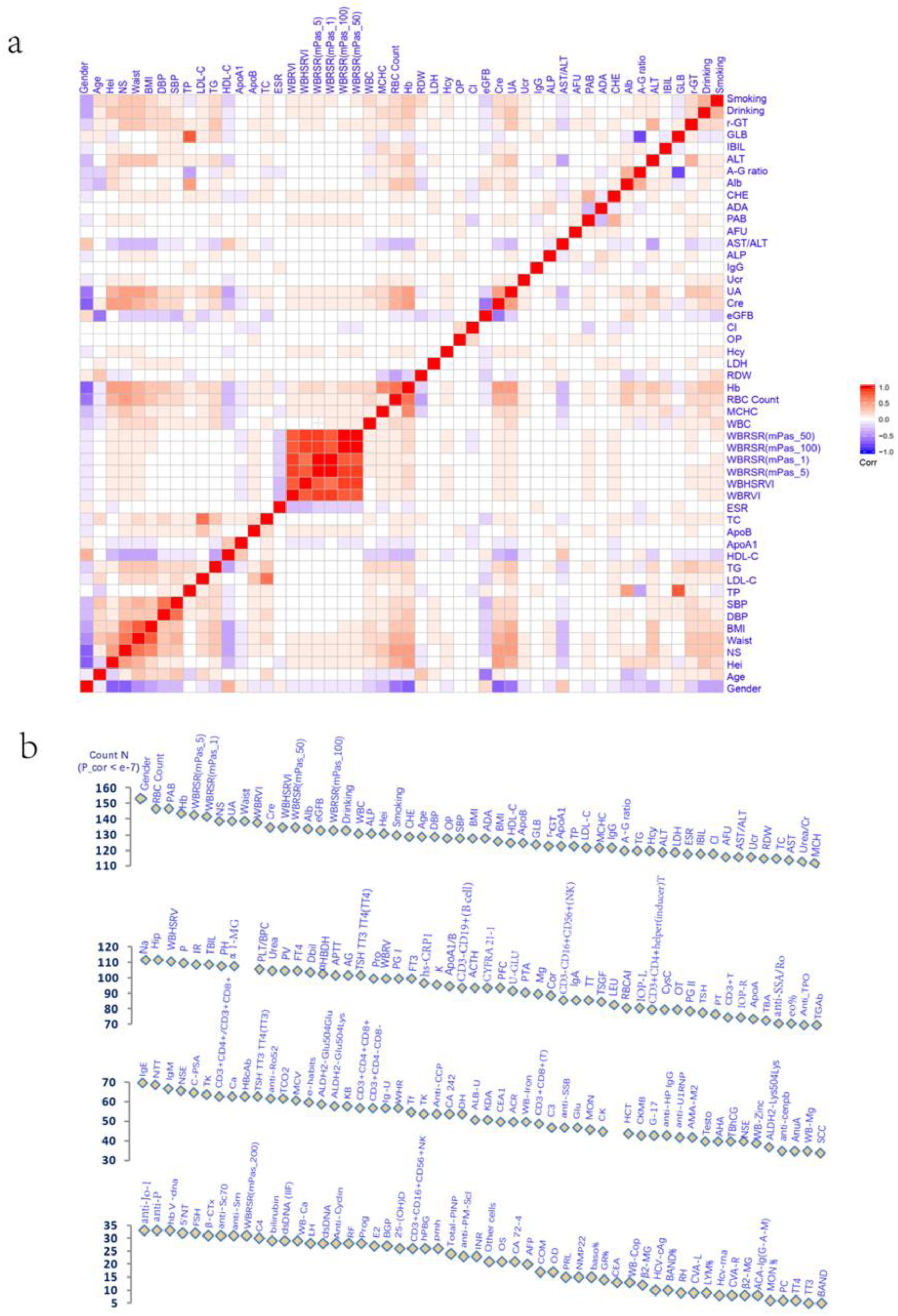
The PEI correlations detected in the healthy cohort. a, A correlation map of the top 50 correlated PEIs, each of which had >114 significant correlations with other PEIs (FDR<0.05). b, The number of statistically significant correlations detected in the healthy population of each PEI.

**Figure 2.**
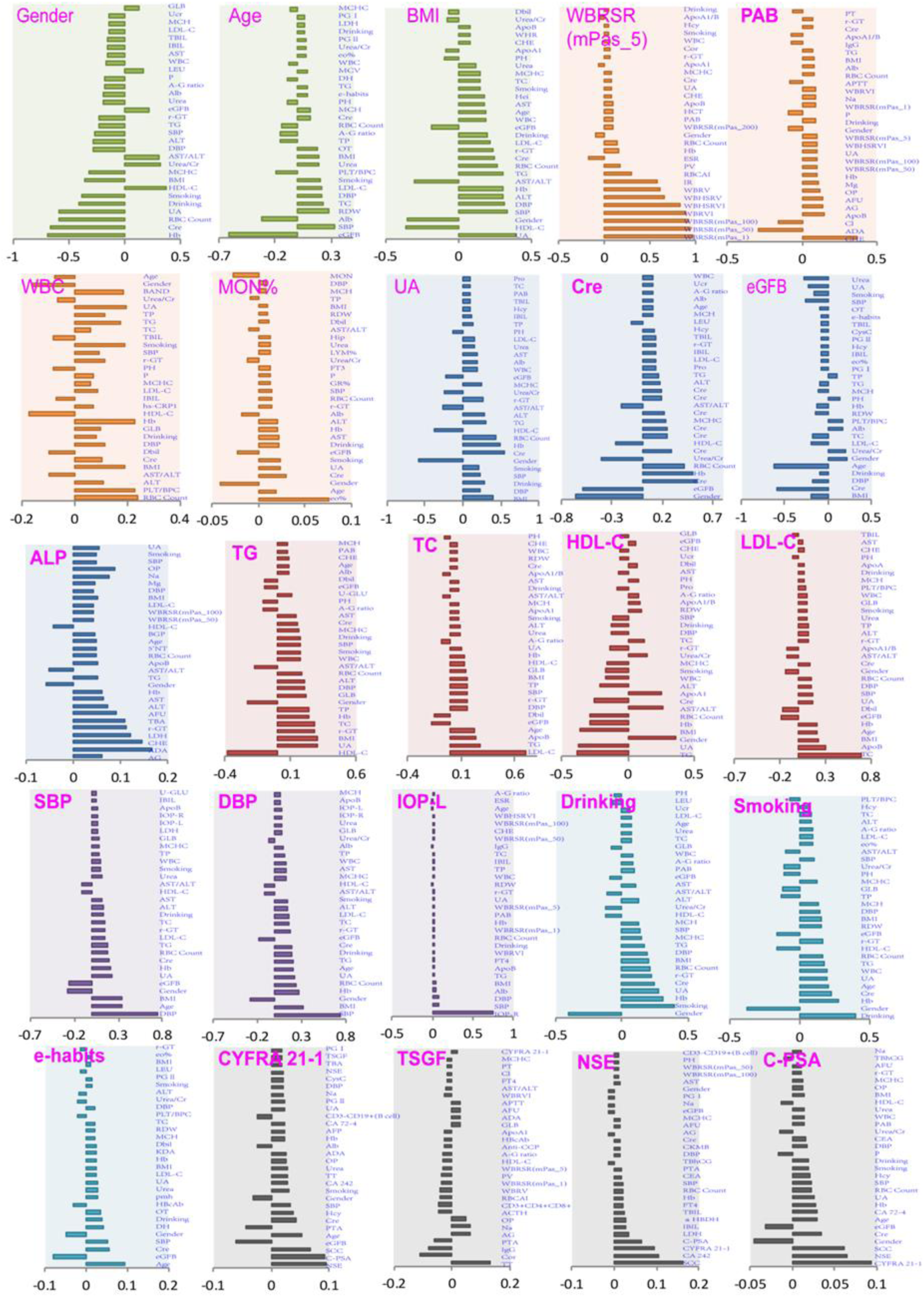
The correlation directions of typical PEIs in healthy physical conditions. The r values were calculated by the PCC method. See Extended data Table 1 for detailed PEI information.

As expected, blood lipid PEIs display many correlations. For example, 119 PEIs correlated with triglyceride (TG) (Fig. 1, Fig. 3, Supplementary Table 1). We found 122 PEIs that correlated with HDL-C, with many negative correlations, including TG, UA, and BMI (Fig. 1, Fig. 2, Supplementary Table 2). The correlation patterns between LDL and HDL showed a specific opposite trend (Fig. 1, Fig. 2, Supplementary Table 1). Out of expected, living habits have a profound impact on our body. Consistently we detected 130 PEIs that correlated with drinking, such as sex, smoking, Hb and UA (Fig. 1, Fig. 2, Supplementary Table 1). Similarly, 128 PEIs were correlated with smoking, including drinking, sex and age (Fig. 1, Fig. 2, Supplementary Table 1). We also detected 58 PEIs that weakly correlated with exercise habits (e-habits), including age, eGFB, and SBP (Fig. 1, Fig. 2, Supplementary Table 1). Tumor marker expression can indicate the occurrence and development of tumors. We detected weak correlations between several tumor markers and PEIs. For example, 88 PEIs were correlated with cytokeratin-19-fragment CYFRA21-1 (CYFRA 21-1); 83 PEIs were correlated with tumor-supplied group factors (TSGF); 64 PEIs were correlated with neuron-specific enolase (NSE); and 64 PEIs were correlated with complexed prostate special antigen (C-PSA) (Fig. 1, Fig. 2, Supplementary Table 1).

**Figure 3.**
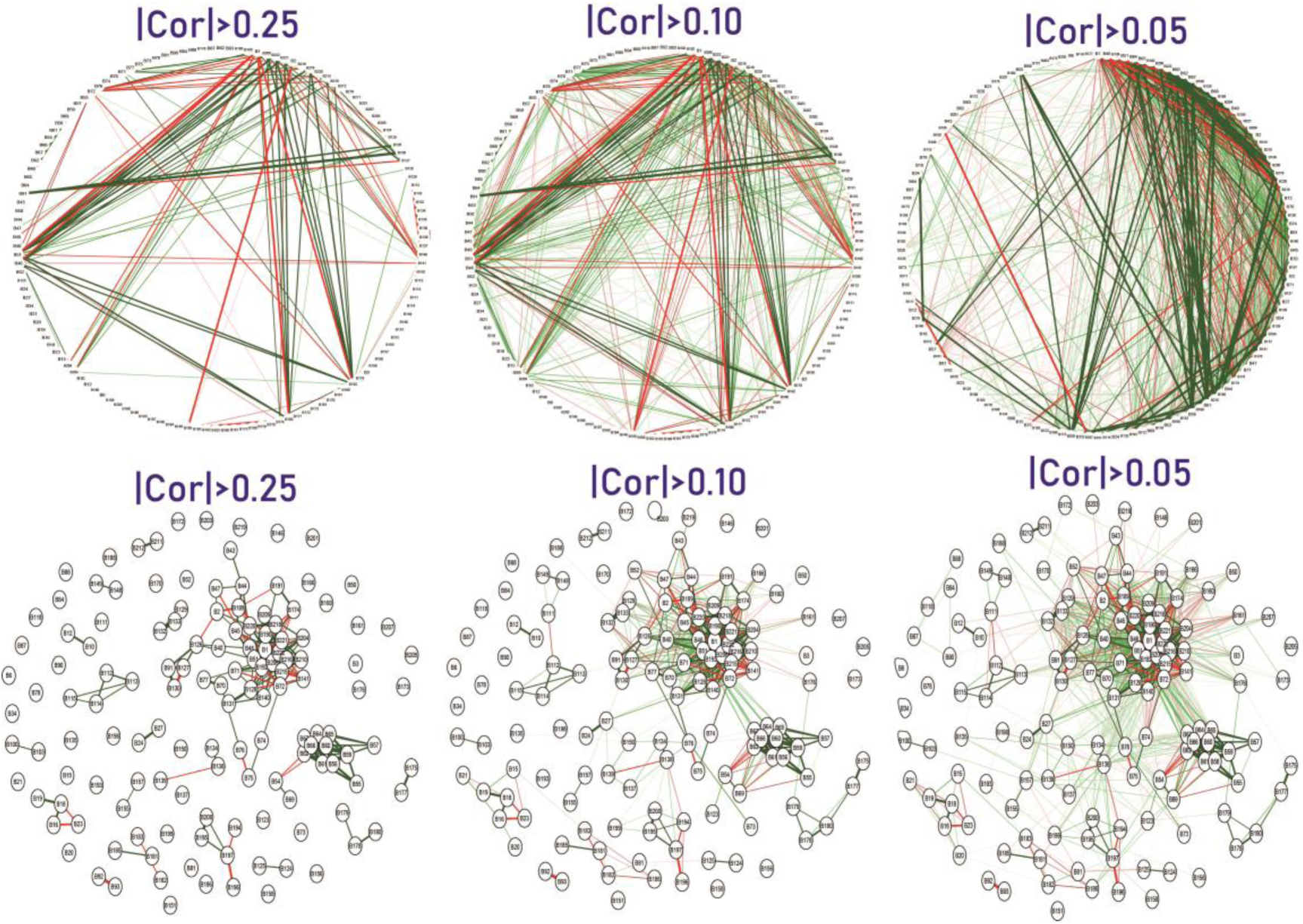
PEI networks in healthy physical status. In the weighted graphs, the green edges indicate positive weights, and the red edges indicate negative weights. The color saturation and the width of the edges correspond to the absolute weight and scale relative to the strongest weight in the graph, respectively. The circular layout shows how well the data conforms to the model while the force-oriented layout shows how each node (connected and unconnected) repulses the other, and how connected nodes attract each other. See also Supplementary Figures.

### PEI correlations in individuals with an unhealthy physical status

Next, we examined the PEI correlations in 34 unhealthy physical states. In this analysis, we also identified rich correlations in these unhealthy physical states (Table 1). Compared with the healthy physical state, we found fewer significant correlations in PEIs in those with an unhealthy physical status, which might be caused by sample size effect (Table 1, Supplementary Table 2-S35). Each unhealthy physical state has its only correlation spectrum and most of them are newly discovered in this study. For example, in the hypertension population, we found 4,413 significant correlations in the 221 PEIs of 24,322 PEI pairs (18.3%) (Supplementary Table 2). The PEI with increased correlations included monocytes (MON) (70 in hypertension vs six in healthy physical state, the same below), quantitative detection of hepatitis B virus DNA (HBV-DNA) (76 vs 33), quantitative detection of hepatitis C virus RNA (HCV-RNA) (49 vs 8), etc. (Supplementary Table 2). Those with both hypertension and coronary heart disease (hypertension+coronary) had an increased correlation of RH blood group compared with the healthy cohort (41 vs 9 in normal). Conversely, the numbers of correlations in homocysteine (Hcy) were greatly reduced in unhealthy versus healthy patients (2 vs 120). In diabetes, 10 PEI pairs increased while the remaining 195 PEI pairs decreased; the increased PEIs including MON (41 vs 6), HCV-RNA (42 vs 8), anti-Sc70 (59 vs 31) and HCV-cAg (35 vs 10) (Supplementary Table 17). These results suggest that under the unhealthy status, the PEIs have changed systematically. Each disease has its own specific PEI spectrum.

We next explored the correlation networks among the PEIs using a qgraph^8, 10^, which would show the LinkMode among PEIs. In the healthy status, we found that the PEIs showed rich interactions with both positive and negative directions (Fig. 3). In the unhealthy physical states, each of them showed its unique interaction networks with PEIs (Extended Data Figure 2 showed the network of hypertension and diabetes). These results show that there is a dependency relationship between multiple indicators in each physical state, which can be used with combination in the assessment of physical health.

### Candidate PEI markers for unhealthy physical status

To verify and discovery new candidate biomarkers or the impact of living habits for disease early diagnosis, we next calculated the difference of each of the 221 PEIs between healthy and unhealthy physical states. In total, we found 1,239 significantly different PEI pairs between healthy and 34 unhealthy physical status (*P*<0.05/34=0.0014, adjust for 34 unhealthy physical status) (Table 1, Fig. 4, Supplementary Table 36). For example, 112 PEIs were significantly different between patients with hypertension and healthy people, 100 PEIs were different between hypertension+diabetes and healthy people, and 91 PEIs were different between diabetes and the healthy people. Some of them are consistent with previous findings and the rest of them are newly discovered.

**Figure 4.**
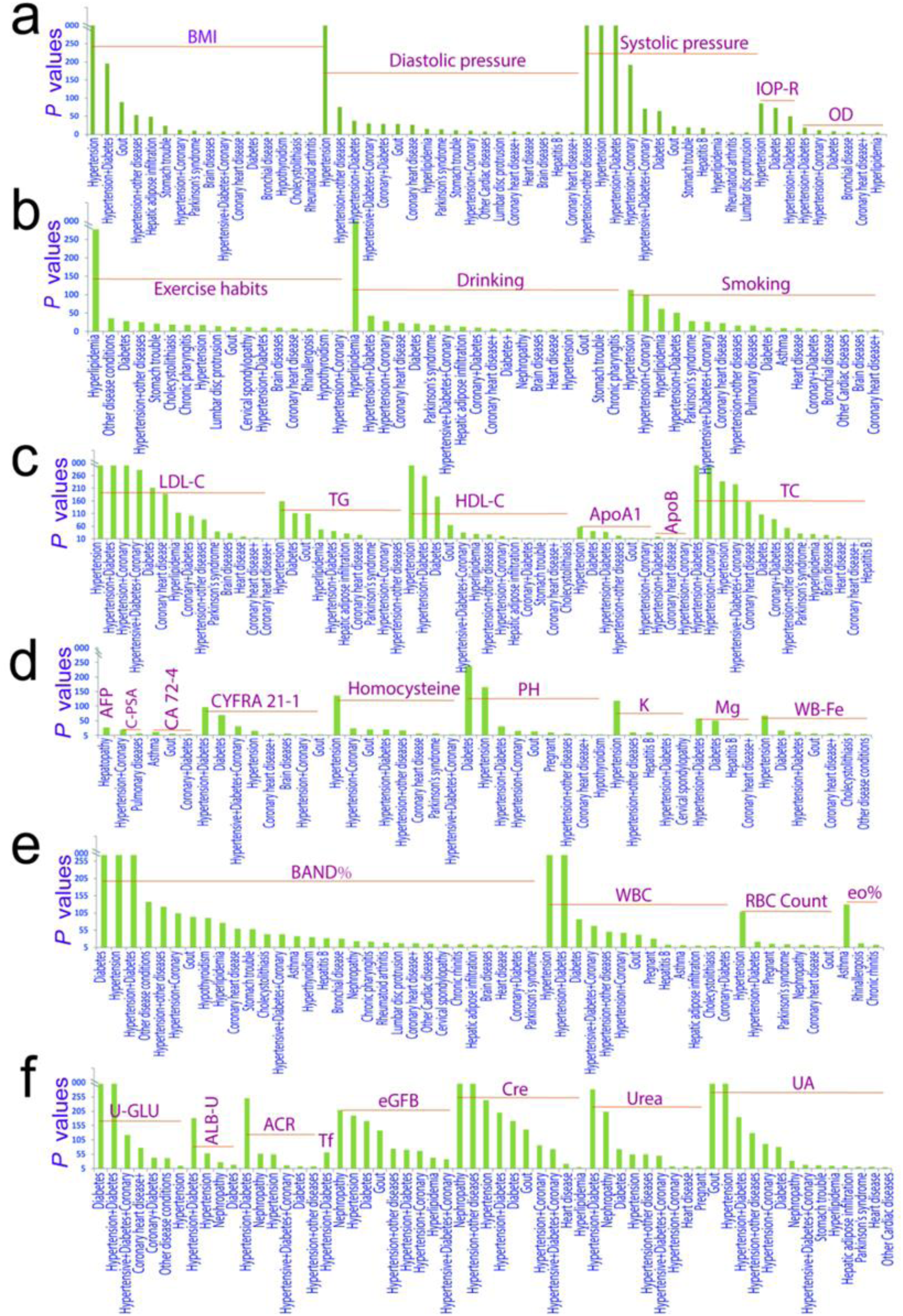
Representative candidate markers for unhealthy physical status. A linear regression model was used to compare PEIs between healthy physical states and unhealthy physical states after adjusting for sex and age. Significantly different PEIs (*P* <0.05) after Bonferroni adjustment (*P*<0.05/34 unhealthy states=1.4×10^-3^) are shown. See also Supplementary Table 36.

For many of the 221 PEI, we detected difference between healthy and unhealthy physical status, especially in PEIs involved in physique, lifestyles, blood lipids (Fig. 4, Supplementary Table 36). For example - BMI, we found differences between healthy and unhealthy physical status in 16 of the 34 unhealthy physical status, including in patients with hypertension (*P*=0) and gout (*P*=6.48×10^-90^). Exercise habits (E-habits) showed 19 differences between healthy and unhealthy status, including in hyperlipidemia (*P*=1.28×10^-277^) and diabetes (*P*=4.20×10^-29^). Dietary habits also showed differences in 10 unhealthy status, including in chronic pharyngitis (*P*=2.59×10^-19^) and cholecystolithiasis (*P*=9.43×10^-18^). We detected differences for alcohol intake habits in 20 unhealthy status, including hyperlipidemia (*P*=0), coronary heart disease (*P*=4.06×10^-24^), diabetes (*P*=1.09×10^-22^) and Parkinson’s syndrome (*P*=1.43×10^-17^). We also observed differences for smoking habits in 18 unhealthy status when compared to the unhealthy condition, including in hypertension (*P*=2.74×10^-114^), hyperlipidemia (*P*=2.69×10^-62^) and Parkinson’s syndrome (*P*=5.12×10^-29^). We found differences for IOP-R in five unhealthy status compared with healthy, including in hypertension (*P*=3.63×10^-85^) and diabetes (*P*=2.01×10^-73^); similar findings were produced for IOP-L (Fig. 4, Supplementary Table 36). For lipids PEIs, we also observed differences between 34 unhealthy and healthy status. For example, LDL-C was detected in 21 unhealthy status, including hypertension (*P*=0) and diabetes (*P*=2.95×10^-212^). HDL-C was detected in 17 unhealthy status, including in diabetes (*P*=1.92×10^-177^) (Fig. 4, Supplementary Table 36). We further conducted a detailed analysis of HDL-C and diabetes and found those with low HDL-C showed a significantly higher risk of developing diabetes than those with average values (1.26-1.75 mmol/L) in this population. Of note, those with high HDL-C levels also showed elevated risk of developing diabetes (Extended data Fig. 2).

Tumor-associated antigens also display significant differences between the healthy and unhealthy status. For example, CYFRA 21-1 was detected in 10 unhealthy status, including hypertension+diabetes (*P*=3.71×10^-97)^ and diabetes (*P*=4.52×10^-70^). CEA1 was detected in 12 unhealthy status, including hypertension+coronary (*P*=9.59×10^-29^) and diabetes (*P*=1.73×10^-18^). Alpha-fetoprotein (AFP) was detected in hepatopathy (*P*=1.08×10^-28^). C-PSA was detected in hypertension+coronary (*P*=8.38×10^-20^). Finally, the carbohydrate antigen CA724 (CA 72-4) was detected in asthma (*P*=9.92×10^-13^), gout (*P*=3.53×10^-7^) and coronary+diabetes (*P*=4.06×10^-5^) (Fig. 4, Supplementary Table 36). Among other PEIs, we also detected significant differences between the healthy and unhealthy status. For example, we found differences in urine sugar levels (U-GLU) in nine unhealthy status, including in diabetes and its associated diseases. The eosinophil rate (eo%), was found in five unhealthy status, including asthma (*P*=1.38×10^-129^) and rhinallergosis (*P*=4.05×10^-18^). Whole blood iron levels (WB-Fe) was found in 11 unhealthy status, including hypertension (*P*=2.52×10^-69^). We detected PH in 11 unhealthy status, including diabetes (*P*=1.97×10^-239^), hypertension (*P*=2.41×10^-166^), hypertension+diabetes (*P*=9.90×10^-32^) and gout (*P*=9.82×10^-15^). We found potassium (K+) in five unhealthy status, including hypertension (*P*=1.98×10^-119^) and hepatitis B (*P*=3.13×10^-10^). We also detected differences in magnesium (Mg2+) in hypertension+diabetes (*P*=3.14×10^-58^) and diabetes (*P*=5.10×10^-52^). Hcy (an indicator of cardiovascular disease) was detected in eight unhealthy status, including hypertension (*P*=1.97×10^-136^) and Parkinson’s syndrome (*P*=1.76×10^-7^) (Fig. 4, Supplementary Table 36). These results provide a set of candidate markers for chronic diseases early diagnosis.

### Machine learning to predict healthy and unhealthy physical status from PEIs

A key objective of this study was to apply PEI data and machine learning technology to develop algorithms that can predict the onset of common disease based on general physical examination. We tried three machine learning models, including kernelized support vector machine (SVM), multilayer perceptron (MLP) and random forests. Because SVM and MLP prediction models only gave very low accuracy and sensitivity in our initial training data, we excluded these models for further training. Random forests showed better performance than SVM and MLP in the initial training. However, it could not give good performance in the multi-class classification of all the physical status. Finally, we tried to use binary classification to classify each pair of healthy and unhealthy physical status (e.g. hypertension and healthy people; Parkinson’s syndrome and healthy people) and we obtained relatively better performance than the multi-class classification. Then we tried to optimize this prediction algorithm. Because the data were characterized by serious category imbalance, a random under-sampling method, was adopted that balances the data by randomly selecting the data subset of the target class. In each physical status, the top 15% or 16% representative PEIs were extracted for prediction by feature extraction. The advantage of this method is that it is usually very fast and completely independent of the model applied after feature selection.

Finally, in the random forests algorithm prediction of each pair of healthy and unhealthy physical status, the area under curve (AUC) of receiver operating characteristic curve reached 66%∼99% depending on the unhealthy physical status (average 87.6%) (Fig.5, Extended data Table 2, Extended data Table 3 and Supplementary Table 37). For classification, AUC values more than 90% indicated excellent performance, and values from 80% to 90% indicated good performance. Our algorithm provided high-precision predictions in 18 of the 34 unhealthy physical status (AUC>90%), good performance for another 9 of the unhealthy physical status (90% >AUC>80%). In our algorithm, patients with heart-related diseases showed excellent performance. For example, by extraction 30 PEI features (age, leukocyte count, monocytes, Mon%, mean corpuscular volume, red blood cell count, red cell distribution width, lymphocyte rate, platelet count, low-density lipoprotein, high-density lipoprotein, total cholesterol, carcinoembryonic antigen 1, albumin, albumin-globulin, cystatin c, glucose, urine sugar, urine creatinine, estimated glomerular filtration rate, creatinine, urea, waistline, aaist-hip Ratio, body mass index, operation history, systolic pressure, height, neck size and anamnesis, Extended data Table 2), Hypertensive+Diabetes+Coronary Heart Disease provides 99% AUC just using 909 training samples and 387 validation samples (f1-score (95%CI), 0.96(0.95-0.96); accuracy (95%CI): 0.95(0.94-0.97); specificity (95%CI): 0.95(0.94-0.95); recall (sensitivity) (95%CI): 0.95(0.94-0.97). In our algorithm, patients with Parkinson’s syndrome provides 97% AUC using 192 training samples and 83 validation samples (f1-score (95%CI), 0.91(0.90-0.91); accuracy (95%CI): 0.90(0.89-0.90); specificity (95%CI): 0.87(0.79-0.94); recall (95%CI): 0.90(0.89-0.91). For hepatic adipose infiltration, our algorithm also provided good prediction performance using 803 training samples and 115 validation samples (f1-score (95%CI), 0.82(0.78-0.87); accuracy (95%CI): 0.81(0.76-0.86); specificity (95% CI): 0.75(0.67-0.82); recall (95% CI): 0.82(0.77-0.87) and AUC (95% CI): 0.92(0.89-0.94). For chronic rhinitis, we got the lowest prediction performance in this study (AUC(95%CI):0.66(0.60-0.72)). When all unhealthy physical status were classified as one “unhealthy” status together, our algorithm also provided good predictions: f1-score (95%CI): 0.83 (0.83-0.83); accuracy (95%CI): 0.82 (0.82-0.82); specificity (95%CI): 0.81(0.81-0.81); sensitivity (95%CI): 0.84 (0.84-0.84) and AUC (95%CI): 0.9 (0.90-0.90). These results suggested that by using feature extraction of the PEIs (15-16% of all 221 PEIs) just by using small number of samples, our random forests algorithms provided good performance for majority unhealthy physical status predictions.

**Figure 5.**
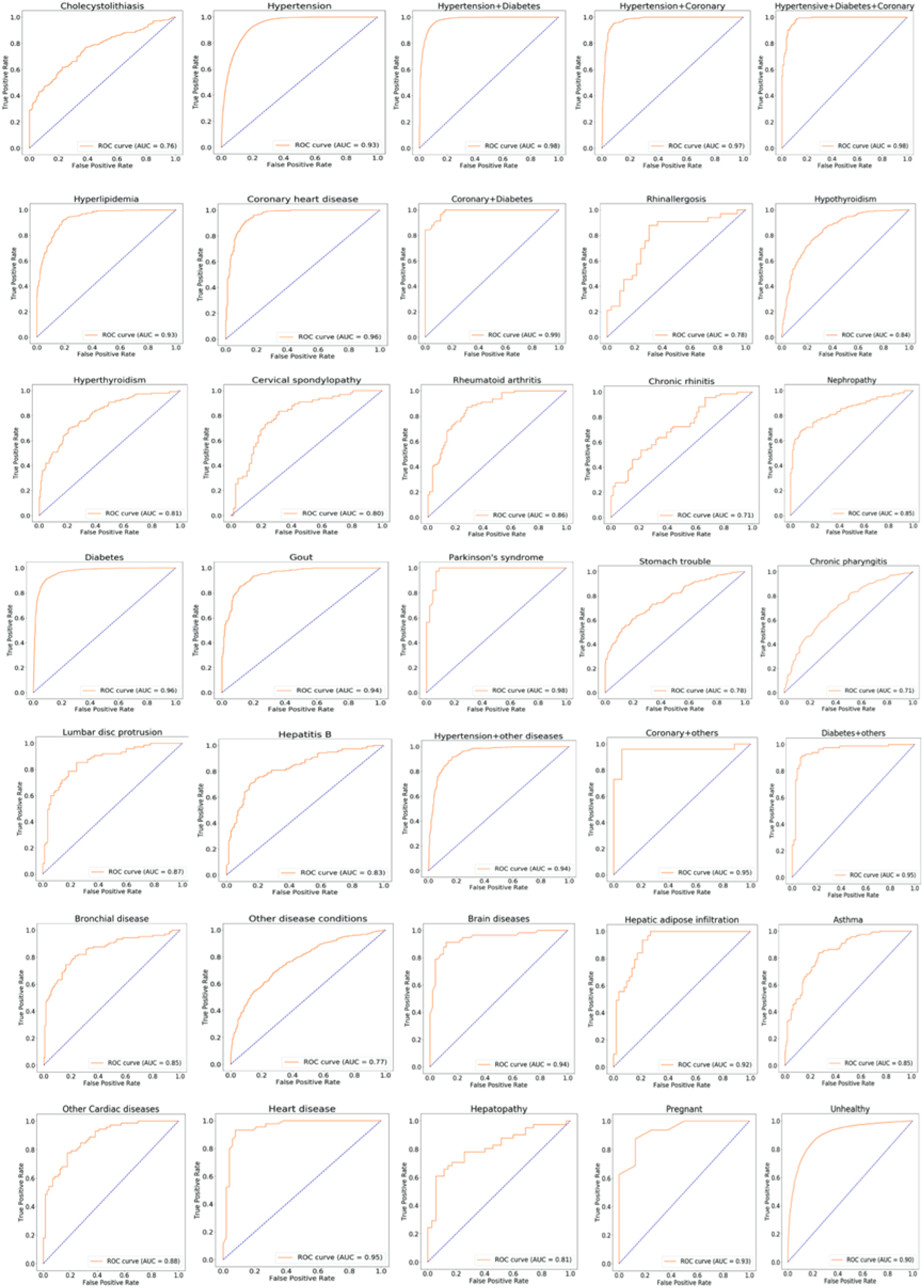
Machine learning prediction of the 35 physical status by the random forest algorism. The receiver operating characteristic curve takes the false positive rate (FPR) as the horizontal axis and the true positive rate (TPR) as the vertical axis. The horizontal axis represents the proportion of the actual negative instances in the positive class predicted by the classifier to all the negative instances, while the vertical axis represents the proportion of the actual positive instances in the positive class predicted by the classifier to all the positive instances. The AUC is the area under the ROC curve.

## Discussion

This study has produced correlation maps of 221 routine PEIs using physical examination data obtained from a Chinese population of 803,614 individuals of 35 healthy or unhealthy physical status (mainly chronic diseases). We detected a large number of correlations among PEIs in healthy or unhealthy physical states; furthermore, these correlations differed according to the 34 unhealthy physical conditions analyzed. Most of the correlations are newly observed in this study. We found that a wide range of correlations among PEIs, such as sex, age, BMI, blood lipids, blood pressures, cancer-related indicators, lifestyles including drinking, smoking, e-habits. Improving our understanding these PEI interactions will help explain disease mechanisms and pathogenesis. Our results fill the gap of systematic PEI analysis and provide rich information about how PEIs might reflect underlying health conditions. These findings provide rich information to further improve healthcare researches and clinical practice.

One of the unexpected finding from our analysis was that patients with hypertension showed more correlations between HBV-DNA and HCV-RNA to other PEIs than healthy cohort. Similarly, we found a strong correlation between hepatitis C virus and other PEIs in diabetes, suggesting that patients infected with hepatitis C may be more susceptible to diabetes. This finding implicates a phenomenon whereby viral infection can make an individual more susceptible to developing a chronic disease. For these people, antiviral therapy might be taken into consideration while treating hypertension and diabetes.

Biomarker discovery and development for clinical research, diagnostics and therapy monitoring in clinical trials are key areas of medicine and healthcare^6^. In this study, we presented many candidate markers for chronic disease. For example, we found that IOP indicators, which are considered to be a relatively independent marker for glaucoma^20^, are closely associated to hypertension, diabetes, and hypertension with diabetes. These results suggest that IOP might be affected, to some extent, by systemic diseases and might be used as one of the clinical marker of these diseases early diagnosis. Our results confirmed that low HDL-C level is a risk factor for diabetes^21^, especially in women. This result suggests that improving HDL-C level through dietary supplementation might be an effective way to prevent diabetes in patients with low HDL-C levels. However, based on our results, excessive HDL-C supplementation is also a risk factor; therefore, HDL-C supplementation should aim to bring HDL-C levels within a normal range^22^. We detected a significant increase in AFP in hepatopathy when comparing healthy cohort, which confirms AFP increase is an increased risk factor for primary liver cancer in hepatopathy^14–15^. K^+^ has significant effects on hypertension^23^ and Cl^-^, and Mg^2+^ has significant effects on diabetes, suggesting that modulation of these ions might have effects on these conditions. Living habits, such as exercise, smoking and drinking, have a more profound impact on the body than we had expected. For example, exercise, drinking or smoking history have a strong impact on hyperlipidemia^24–25^, as evidenced by comparision to healthy status. This finding suggests that by adjusting these living habits, hyperlipidemia should improve.

Because the current physical examination conclusion is generally based on a relatively independent single or several prior indicators to give advice on the results of physical examination, many of the results given are ambiguous, and the value of judging the health status of the examinees is very limited^10, 26–27^. There is an urgent need for a more accurate index system and method to judge the health status of physical examinees. In the final part of our study, we developed random forest machine learning algorithms that can predict diseases through 15%-16% of all 221 PEIs with good performance of prediction (AUC:66%∼99%; average 86%). For each disease, we defined about 30 contributed PEIs by feature extraction. In most of our prediction algorithms, only a few hundreds of samples were needed to give good prediction performance for many chronic diseases. This finding suggests machine learning on PEI data can be used to help predict the true condition of the examers, identify “at-risk” patients and indicate the most relevant follow-up physical examinations for affected individuals.

In summary, we systematically explored the correlation between various PEIs and their relationship with chronic diseases and established machine learning prediction models to predict health status. This study provides abundant information to better understand the physiological and pathological characteristics of the human body as a system. Importantly, we have identified modifiable factors and directions for disease prediction, diagnosis and treatment. Our developed machine learning algorithms can be immediately applied to clinical practice to assistant the judgment of physical examination results.

## Methods

### Study approval

The study was approved by the institutional ethics committee of Sichuan Provincial People’s Hospital and was conducted according to the Declaration of Helsinki principles. Informed consent was obtained from the participants when possible.

### Study Participants

PEI data were obtained from 803,614 Han Chinese patients visiting the Health Management Center & Physical Examination Center of Sichuan Provincial People’s Hospital in China between 2013 and 2018. The total cohort captured participants with 35 different reported health conditions, including 711,928 reported healthy participants and 91,686 unhealthy participants. The unhealthy cohort included 46,981 patients with hypertension, 11,745 with diabetes and 32,960 with other unhealthy status (Table 1).

### Detected PEIs

Only the PEIs that were recorded by the same methods were included in this study. In total, 229 PEIs were initially collected: eight PEIs that were detected in few individuals were excluded, leaving 221 PEIs for further analysis (Extended data Table 1). These PEIs included the levels of biochemical indicators and the results of blood tests. Patient lifestyles and disease conditions were also investigated during the physical examination.

### Data processing

The PEIs with string variables were converted to integer variables for data analysis. Categorized variables were digitally coded for further calculation. The mean value imputation method was used for missing data. For individuals who participated in more than one physical check, average values of each PEI were used for data analysis.

### Statistical Analyses

The Pearson correlation coefficient (PCC) method was used to calculate the correlations between two PEIs (for example, x and y) in R; this method measures the linear dependence between two variables. PCC correlation (r) (1) and *P* values (2) were calculated using the following formulae^28–30^:

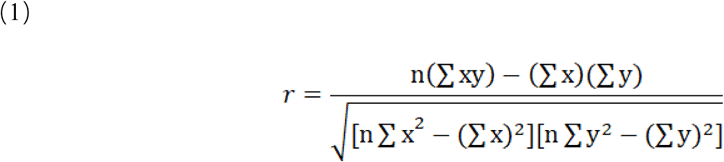

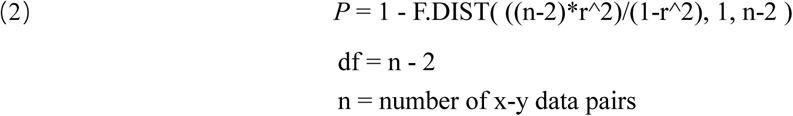

Total sample size required when using the correlation coefficient (r), when two-sieded α=0.05, β=0.20. If r=0.05, we need 3,134 samples; if r=0.10, we need 782 samples; if r=0.25, we need 123 samples; if r=0.5, we need 29 samples. The general formula for the correlation sample calculating is listed as the following (3)^31^:

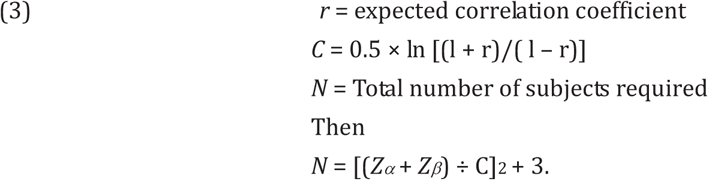

A linear regression model (lm) was used to compare PEIs between the reported healthy status and unhealthy status adjusted for sex and age in the R package^21–23^. The odds ratio of HDL-C level was calculated by using generalized linear models (glm) adjusted for age in the R package^24–25^. The correlation interaction network was conducted using qgraph^10–11^.

### Machine learning

Three machine learning models, including kernelized support vector machine (SVM)^28–29^, multilayer perceptron (MLP)^30–32^ and random forest^33^ were tested to get the prediction performance of the PEIs. By using MLP algorithm prediction in neural network to predict health and each of the 34 unhealthy status (multi classification), it could not achieve good results. We further tried prediction the healthy result is very close to zero. By using SVM algorithm prediction for making multi classification prediction, the highest F1 value of cholecystolithias is 0.70, but that of most other types of diseases is 0.00. We also tried the binary classification method, but all the results were relatively poor. When random forest algorithm is used for prediction for multi classification (health and each of the 34 unhealthy status), the F1 value of healthy status can reach 0.80-0.90, but the F1 value of unhealthy sattus is about 0.00-0.40. Then, we further chosen forest algorithm and optimized the random forest algorithm. First, due to the uneven distribution of the sample numbers of healthy and non-healthy status, and the law of large numbers^32^, we used downsampling strategy for sample randomly used. Because the data were characterized by serious category imbalance, a random under-sampling method, was adopted that balances the data by randomly selecting the data subset of the target class. Second, we used PEI feature extraction strategy to extract the most contributed PEI for each healthy and unhealthy status. Feature extraction adopts the strategy of univariate statistics in automatic feature selection. Univariate statistics select features with high confidence according to the statistical significance of the relationship between each feature and the target. This process can be achieved by using feature_selection in scikit-learn. Finally, in each healthy and non-healthy status, the top 15% or 16% representative PEIs were extracted for prediction by feature extraction. The advantage of this method is that it is usually very fast and completely independent of the model applied after feature selection. Then, the data were randomly divided such that 30% constituted the test set, and the remaining 70% were randomly divided again, with 70% as the training set for the training model and 30% as the validation set for the evaluation model. In the process of improving the generalization performance of the model by adjusting parameters, a cross-validation method with a grid search was adopted, which can be implemented by GridSearchCV provided by scikit-learn (Supplementary Table 37 and Supplementary code).

## ACKNOWLEDGMENTS

We thank all the participants in this study. This research project was supported by: the National Natural Science Foundation of China (81970839(L.H.), 81670895 (L.H.) and 81300802 (L.H.); the Department of Science and Technology of Sichuan Province, China (2015JQO057 (L.H.), 2017JQ0024 (L.H.), 2016HH0072 (L.H.) and 2013JY0195 (L.H.); the Department of Science and Technology of Sichuan Province, China (2017JZ0039 (P.S.). The author is especially indebted to Dr. Yabing Mei for taking time to make extensive comments.

## AUTHOR CONTRIBUTIONS

L.H. designed the study. P.S., Y.L., L.J., J.Y., S.Z., Y.Y., L.W., D.L. and T.Y. enrolled all the participants. L.H., H.W. and P.S. performed the data analysis. Y.D., S.Z. and Y.Y., did the machine learning prediction models. L.H. wrote the manuscript. All of authors critically revised and provided final approval for this manuscript.

## COMPETING FINANCIAL INTERESTS

The authors declare no competing interests related to this paper.

---

**Extended data Table 1.** PEIs used in this study.

| Code | PEI Name | PEI Name abbreviation | Unit |
| --- | --- | --- | --- |
| B1 | Gender | Gender | - |
| B2 | Age | Age | years |
| B3 | Homocysteine | Hcy | umol/L |
| B4 | Whole Blood Calcium | WB-Ca | mmol/L |
| B5 | Whole Blood Magnesium | WB-Mg | mmol/L |
| B6 | Iron Of Whole Blood | WB-Fe | mmol/L |
| B7 | Whole Blood Copper | WB-Cop | umol/L |
| B8 | Zinc In Whole Blood | WB-Zinc | umol/L |
| B9 | Rh Blood Group | RH | - |
| B10 | Pepsinogen I | PGI | ug/L |
| B11 | PgI/PgII | PGI/PGII | - |
| B12 | Pepsinogen II | PGII | ug/L |
| B14 | Gastrin-17 | G-17 | pmol/l |
| B15 | Cd3-Cd19+(B Cell) | CD3-CD19+(B cell) | - |
| B16 | Cd3+ Total T Lymphocyte | CD3+T | - |
| B17 | Cd3+Cd16+Cd56+Nk | CD3+CD16+CD56+NK | - |
| B18 | Cd3+Cd4+Helper(Inducer)T Cells | CD3+CD4+helper(inducer)T | - |
| B19 | Cd3+Cd4+/Cd3+Cd8+ | CD3+CD4+/CD3+CD8+ | - |
| B20 | Cd3+Cd4+Cd8+ | CD3+CD4+CD8+ | - |
| B21 | Cd3+Cd4-Cd8- | CD3+CD4-CD8- | - |
| B22 | Cd3+Cd8+(T) | CD3+CD8+(T) | - |
| B23 | Cd3-Cd16+Cd56+(Nk) | CD3-CD16+CD56+(NK) | - |
| B24 | $\alpha$ Hydroxybutyrate Dehydrogenas | αHBDH | U/L |
| B25 | Creatine Kinase | CK | U/L |
| B26 | Creatine Kinase Isoenzyme | CKMB | U/L |
| B27 | Lactic Dehydrogenase | LDH | U/L |
| B28 | Pituitary Prolactin | PRL | mIU/L |
| B29 | Estradiol E2 | E2 | pmol/L |
| B30 | Luteinizing Hormone | LH | mIU/ml |
| B31 | Follicle Stimulating Hormone | FSH | mIU/ml |
| B32 | Testosterone | Testo | nmol/L |
| B33 | Progesterone | Prog | nmol/L |
| B34 | Thymidine Kinase | TK | PM/L |
| B35 | Blood Beta-Microglobulin | β2-MG | - |
| B36 | Variant Lymphocyte Rate | V-Lym% | - |
| B37 | Band Cell | BAND | 10E-9/L |
| B38 | Band Cell% | BAND% | - |
| B39 | Other Cell Types | Other cells | - |
| B40 | Leukocyte Count | WBC | 10E-9/L |
| B41 | Monocytes | MON | - |
| B42 | Monocytes % | MON % | 10E-9/L |
| B43 | Mean Corpuscular Volume | MCV | fl |
| B44 | Mean Corpuscular Hemoglobin | MCH | pg |
| B45 | Mean Corpuscular Hemoglobin Concentration | MCHC | g/l |
| B46 | Red Blood Cell Count | RBC Count | 10E-12/L |
| B47 | Red Cell Distribution Width | RDW | fl |
| B48 | Lymphocyte Rate | LYM% | - |
| B49 | Basophilic Granulocyte Rate | baso% | - |
| B50 | Eosinophil Rate | eo% | - |
| B51 | Hemoglobin | Hb | g/L |
| B52 | Platelet Count | PLT/BPC | 10E-9/L |
| B53 | Granulocyte Rate | GR% | - |
| B54 | Erythrocyte Sedimentation Rate | ESR | mm/60min |
| B55 | Erythrocyte Deformation Index | TK | - |
| B56 | The Index Of Rigidity Of Erythrocyte | IR | - |
| B57 | Erythrocyte Aggregation Indices | RBCAI | - |
| B58 | Hematocrit | HCT | - |
| B59 | Whole Blood Reduced Viscosity | WBRV | - |
| B60 | Whole Blood Reduced Viscosity Index | WBRVI | - |
| B61 | Whole Blood High Shear Reduced Viscosity | WBHSRV | - |
| B62 | Whole Blood High Shear Reduced Viscosity Index | WBHSRVI | - |
| B63 | Shear Rate Of Whole Blood Viscosity_Mpas_1 (200) | WBRSR(mPas_200) | - |
| B64 | Shear Rate Of Whole Blood Viscosity_Mpas_3 (5) | WBRSR(mPas_5) | - |
| B65 | Shear Rate Of Whole Blood Viscosity_Mpas_4 (1) | WBRSR(mPas_1) | - |
| B66 | Shear Rate Of Whole Blood Viscosity_Mpas (100) | WBRSR(mPas_100) | - |
| B67 | Shear Rate Of Whole Blood Viscosity_Mpas (50) | WBRSR(mPas_50) | - |
| B69 | Plasma Viscosity | PV | - |
| B70 | Low-Density Lipoprotein | LDL-C | mmol/L |
| B71 | Triglyceride | TG | mmol/L |
| B72 | High-Density Lipoprotein | HDL-C | mmol/L |
| B73 | Apolipoprotein A | ApoA | mg/L |
| B74 | Apolipoprotein A1 | ApoA1 | g/L |
| B75 | Apolipoprotein A1/B | ApoA1/B | g/L |
| B76 | Apolipoprotein B | ApoB | g/L |
| B77 | Total Cholesterol | TC | mmol/L |
| B78 | Hepatitis B Virus Core Im Antibod | HBcAb | - |
| B79 | Quantitative Detection Of Hepatitis B Virus Dna | hbV-dna | IU/ml |
| B80 | Helicobacter Pylori Antibody-Igg | anti-HP IgG | - |
| B81 | Tumor Supplied Group Of Factors | TSGF | U/ML |
| B82 | Carcinoembryonic Antigen | CEA | ng/ml |
| B83 | Alpha-Fetoprotein | AFP | ng/ml |
| B84 | Complexed Prostate Special Antigen | C-PSA | ng/ml |
| B85 | Squamous Cell Carcinoma Antigen | SCC | ng/ml |
| B86 | Chorionic Gonadotropin | TBhCG | mIU/ml |
| B87 | Neuron-Specific Enolase | NSE | ng/ml |
| B88 | Carbohydrate Antigen, Ca242 | CA 242 | U/ml |
| B89 | Carbohydrate Antigen, Ca72_4 | CA 72-4 | U/ml |
| B90 | Cytokeratin-19-Fragment Cyfra21-1 | CYFRA 21-1 | ng/ml |
| B91 | Total Protein | TP | g/L |
| B92 | Aldh2-Glu504Glu | ALDH2-Glu504Glu | - |
| B93 | Aldh2-Glu504Lys | ALDH2-Glu504Lys | - |
| B94 | Aldh2-Lys504Lys | ALDH2-Lys504Lys | - |
| B95 | Anti-Cenpb | anti-cenpb | - |
| B96 | Dsdna (Iif) | dsDNA (IIF) | - |
| B97 | Anti-Dsdna Antibody | dsDNA | - |
| B98 | Jo-1 Antibody | anti-Jo-1 | - |
| B99 | Anti-Pm-Scl | anti-PM-Scl | - |
| B100 | Anti-Ro52 | anti-Ro52 | - |
| B101 | Anti-Sc70 | anti-Sc70 | - |
| B102 | Anti Sm Antibody | anti-Sm | - |
| B103 | Anti-Ssa/Ro Antibody | anti-SSA/Ro | - |
| B104 | Anti Ssb Antibody | anti-SSB | - |
| B105 | Anti-U1Rnp Antibody | anti-U1RNP | - |
| B106 | Anti-Ribosomal P Protein Antibody | anti-P | - |
| B107 | Anti-Nucleosome Antibody Systemic | AnuA | - |
| B108 | Anti-Cyclin Antibody | Anti-Cyclin | - |
| B109 | Antimitochondrial Antibody M2 | AMA-M2 | - |
| B110 | Anti-Histone Antibody | AHA | - |
| B111 | Thyroid-Stimulating Hormon | TSH | mIU/L |
| B112 | Free Thyroxine | FT4 | pmol/L |
| B113 | Free Triiodothyronine | FT3 | pmol/L |
| B114 | Total Thyroxine | TSH TT3 TT4(TT4) | nmol/L |
| B115 | Total Triiodothyronine | TSH TT3 TT4(TT3) | nmol/L |
| B116 | Total Triiodothyronine | TT3 | nmol/L |
| B117 | Total Thyroxine | TT4 | nmol/L |
| B118 | Carcinoembryonic Antigen 1 | CEA1 | ng/ml |
| B119 | Quantitative Detection Of Hepatitis C Virus Rna | Hcv-rna | - |
| B120 | Hcv Core Antigen | HCV-cAg | - |
| B121 | Complement C3 | C3 | - |
| B122 | Complement C4 | C4 | - |
| B123 | High Sensitivity C-Reactive Protein 1 | hs-CRP1 | mg/L |
| B124 | Adrenocorticotrophic Hormone | ACTH | pg/ml |
| B125 | Cortisol | Cor | ug/dl |
| B126 | Albumin | Alb | g/L |
| B127 | Albumin-Globulin | A-G ratio | - |
| B128 | Alanine Aminotransferase | ALT | U/L |
| B129 | Indirect Bilirubin | IBIL | umol/L |
| B130 | Globulin | GLB | g/L |
| B131 | Aspartate Transaminase | AST | U/L |
| B132 | Direct Bilirubin | Dbil | umol/L |
| B133 | Total Bilirubin | TBIL | umol/L |
| B134 | Cholinesterases | CHE | KU/L |
| B135 | Total Bile Acid | TBA | umol/L |
| B136 | 5-Nucleotidase | 5'NT | U/L |
| B137 | Alpha-L-Fucosidase | AFU | U/L |
| B138 | Prealbumin | PAB | mg/L |
| B139 | Adenosine Deaminase | ADA | U/L |
| B140 | R-Glutamyl Transpeptidase | r-GT | U/L |
| B141 | Ast/Alt | AST/ALT | - |
| B142 | 25-Hydroxyvitamin D | 25-(OH)D | ng/mL |
| B143 | β-Crosslaps | β-CTx | pg/mL |
| B144 | Total-Pinp | Total-PINP | ng/mL |
| B145 | Osteocalcin | BGP | ng/mL |
| B146 | Cystatin C | CysC | mg/L |
| B148 | Antithyroperoxidase Antibody | Anti_TPO | IU/ml |
| B149 | Anti-Thyroglobulin Antibodies | TGAb | IU/ml |
| B150 | Alkaline Phosphatase | ALP | U/L |
| B151 | Anti-Cyclic Peptide Containing Citrulline | Anti-CCP | Ru/ml |
| B152 | Anti Cardiolipin Antibodyig(G-A-M) | ACA-Ig(G-A-M) | u/ml |
| B153 | Glucose | Glu | mmol/L |
| B154 | Rheumatoid Factor | RF | IU/ml |
| B155 | Immunoglobulin A | IgA | g/l |
| B156 | Immunoglobulin E | IgE | IU/ml |
| B157 | Immunoglobulin G | IgG | g/l |
| B158 | Immunoglobulin M | IgM | g/l |
| B159 | Urine Beta-Microglobulin | β2-MG | - |
| B160 | Ph | PH | - |
| B161 | Leucocyte | LEU | /ul |
| B163 | Specific Gravity | SG | - |
| B164 | Calcium Oxalate Crystals | COM | HPF |
| B165 | Bilirubin | bilirubin | - |
| B166 | Protein | Pro | - |
| B167 | Phosphate Crystals | PC | - |
| B168 | Urinary Nuclear Matrix Protein 22 | NMP22 | - |
| B169 | Urate Crystal | U-CRY | - |
| B170 | Urine Sugar | U-GLU | - |
| B171 | Epithelial Cell | EC | HPF |
| B172 | Ketone Body | KB | - |
| B173 | Nitrous Acid | NTT | - |
| B174 | Urea Creatinine | Urea/Cr | - |
| B175 | Microalbuminuria | ALB-U | mg/L |
| B176 | Urine Creatinine | Ucr | umol/L |
| B177 | Ratio Of Urinary Microalbumin To Creatinin | ACR | ug/mg |
| B178 | Urinary Immunoglobulin G | Ig-U | mg/L |
| B179 | Urine Alpha-Microglobulin | α1-MG | mg/L |
| B180 | Urine Transferrin | Tf | mg/L |
| B181 | Activated Partial Thromboplastin Time | APTT | s |
| B182 | Thrombin Time | TT | s |
| B183 | Percentage Activity Of Prothrombin In Plasma | PTA | - |
| B184 | International Normalized Ratio | INR | - |
| B185 | Prothrombin Time | PT | s |
| B186 | Plasma Fibrinogen Concentration | PFC | g/l |
| B187 | Neuron-Specific Enolase | NSE | ng/ml |
| B188 | Calcium | Ca | mmol/L |
| B189 | Estimated Glomerular Filtration Rate | eGFB | ml/min |
| B190 | Creatinine | Cre | umol/L |
| B191 | Urea | Urea | mmol/L |
| B192 | Uric Acid | UA | umol/L |
| B193 | Kalium | K | mmol/L |
| B194 | Chlorine | Cl | mmol/L |
| B195 | Sodium | Na | mmol/L |
| B196 | Blood Carbon Dioxide | TCO2 | mmol/L |
| B197 | Anion Gap | AG | mmol/L |
| B198 | Phosphorus | P | mmol/L |
| B199 | Magnesium | Mg | mmol/L |
| B200 | Osmotic Pressure | OP | mOsm/L |
| B201 | Exercise Habits | e-habits | - |
| B202 | Frequency Of Exercises | e-times | - |
| B203 | Dietary Habits 1 | DH | - |
| B204 | History Of Alcohol Intake | Drinking | - |
| B205 | Known Of Drug Allergy | KDA | - |
| B206 | Waistline | Waist | cm |
| B207 | Waist-Hip Ratio | WHR | - |
| B209 | Smoking History | Smoking | - |
| B210 | Hipline | Hip | cm |
| B211 | Intra-Ocular Tension Of The Right Eye | IOP-R | mmHg |
| B212 | Intra-Ocular Tension Of The Left Eye | IOP-L | mmHg |
| B213 | Corrected Visual Acuity The Right Eye | CVA-R | - |
| B214 | Corrected Visual Acuity The Left Eye | CVA-L | - |
| B215 | Body Mass Index | BMI | - |
| B216 | Body Mass | BMI | kg |
| B218 | Diastolic Pressure | DBP | mmHg |
| B219 | Operation History | OT | - |
| B220 | Systolic Pressure | SBP | mmHg |
| B221 | Height | Hei | cm |
| B222 | Neck Size | NS | cm |
| B223 | Family History | FH | - |
| B224 | Anamnesis | pmh | - |
| B225 | Hourly Postprandial Blood Sugar | hPBG | mmol/L |
| B227 | Vision Of The Right Eye | OD | - |
| B228 | Vision Of The Left Eye | OS | - |

**Extended data Table 2.**
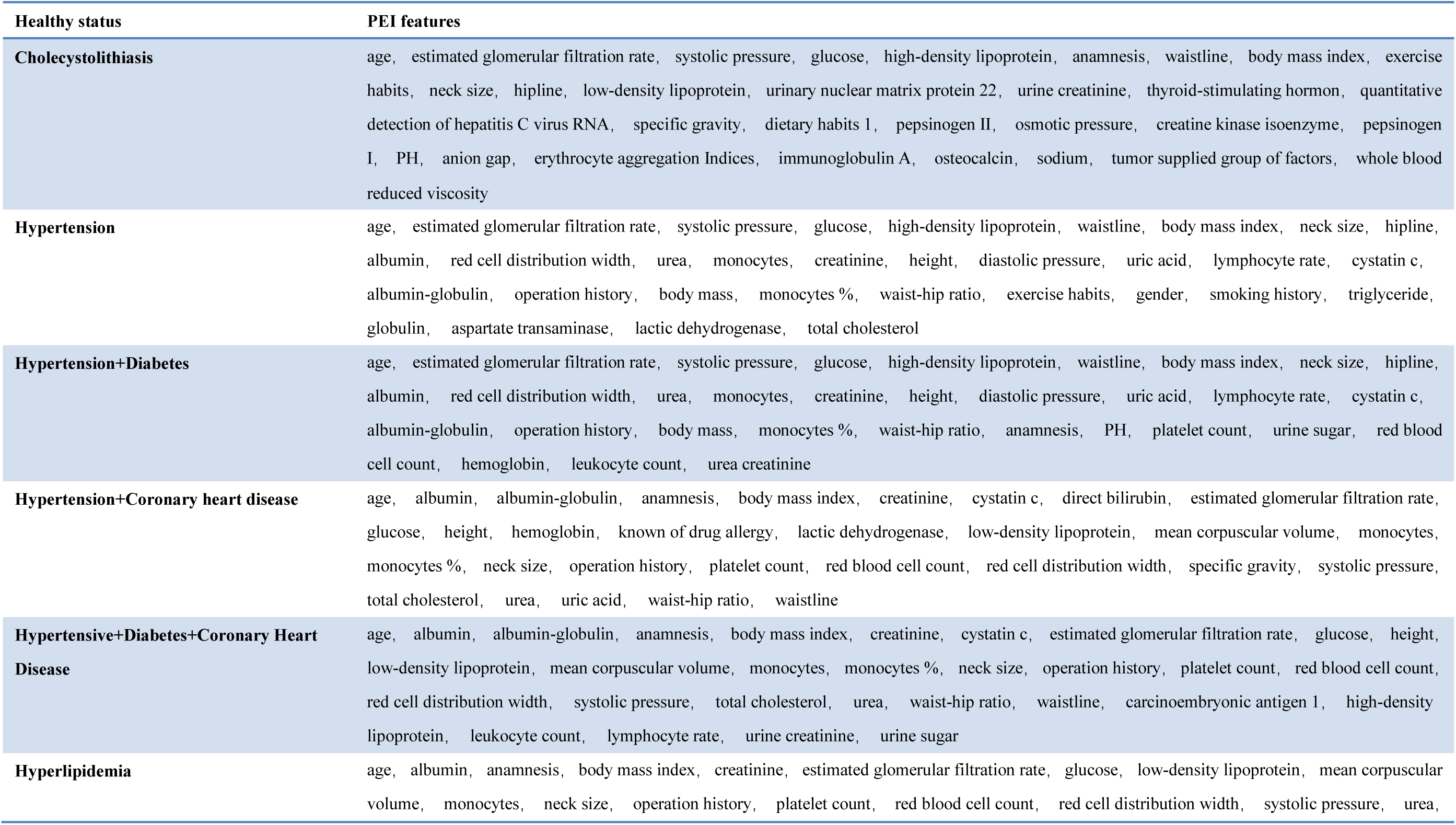
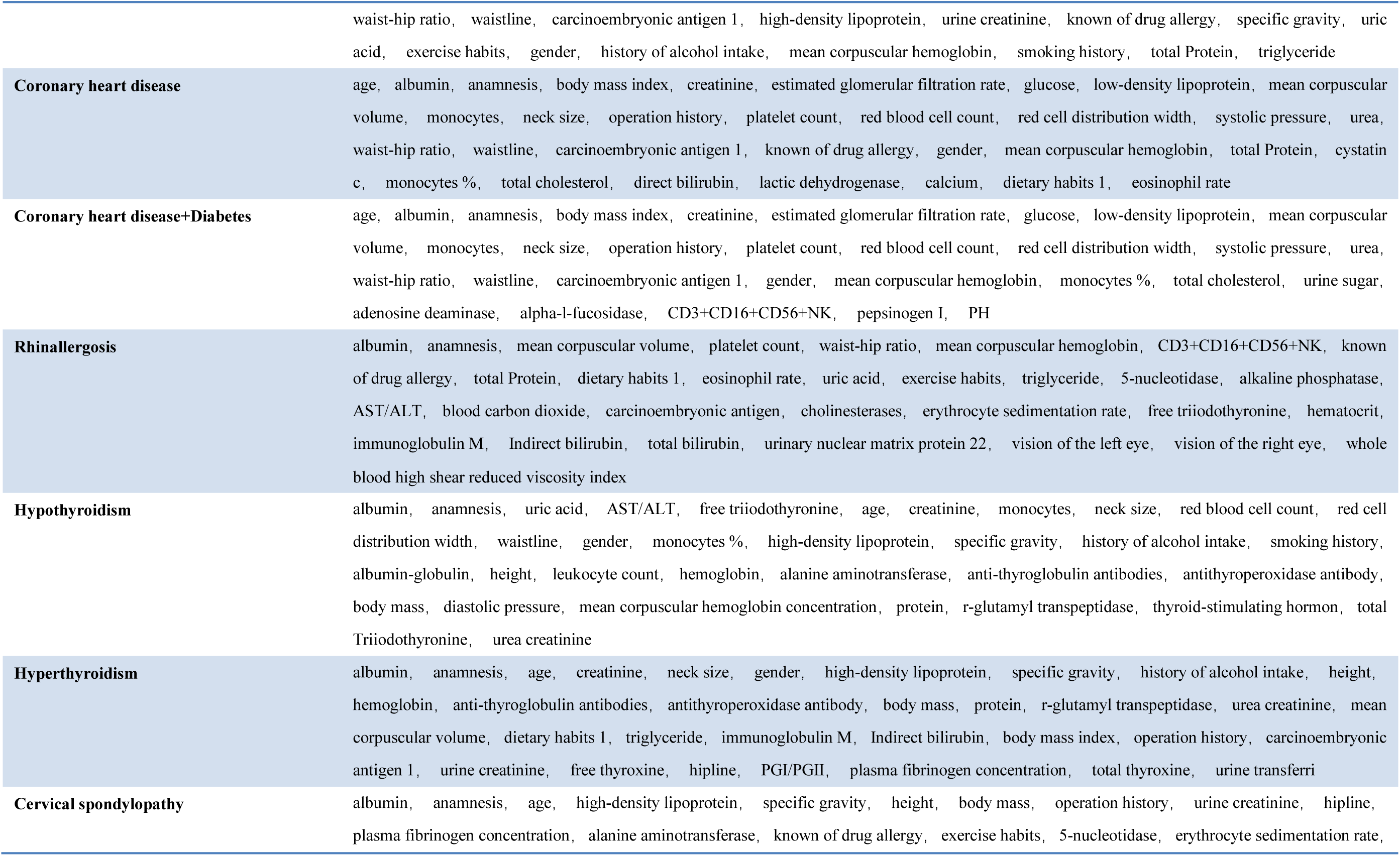
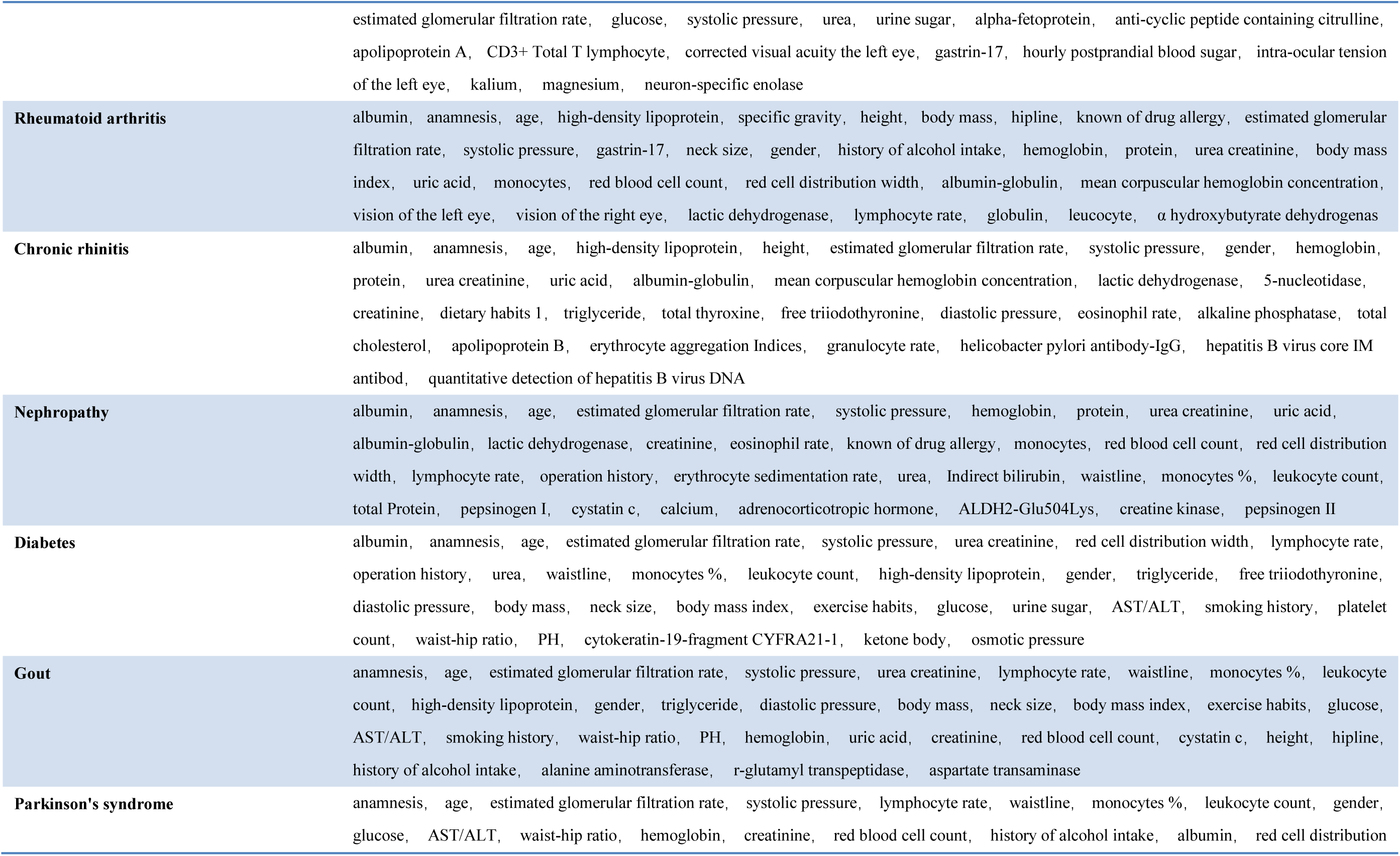
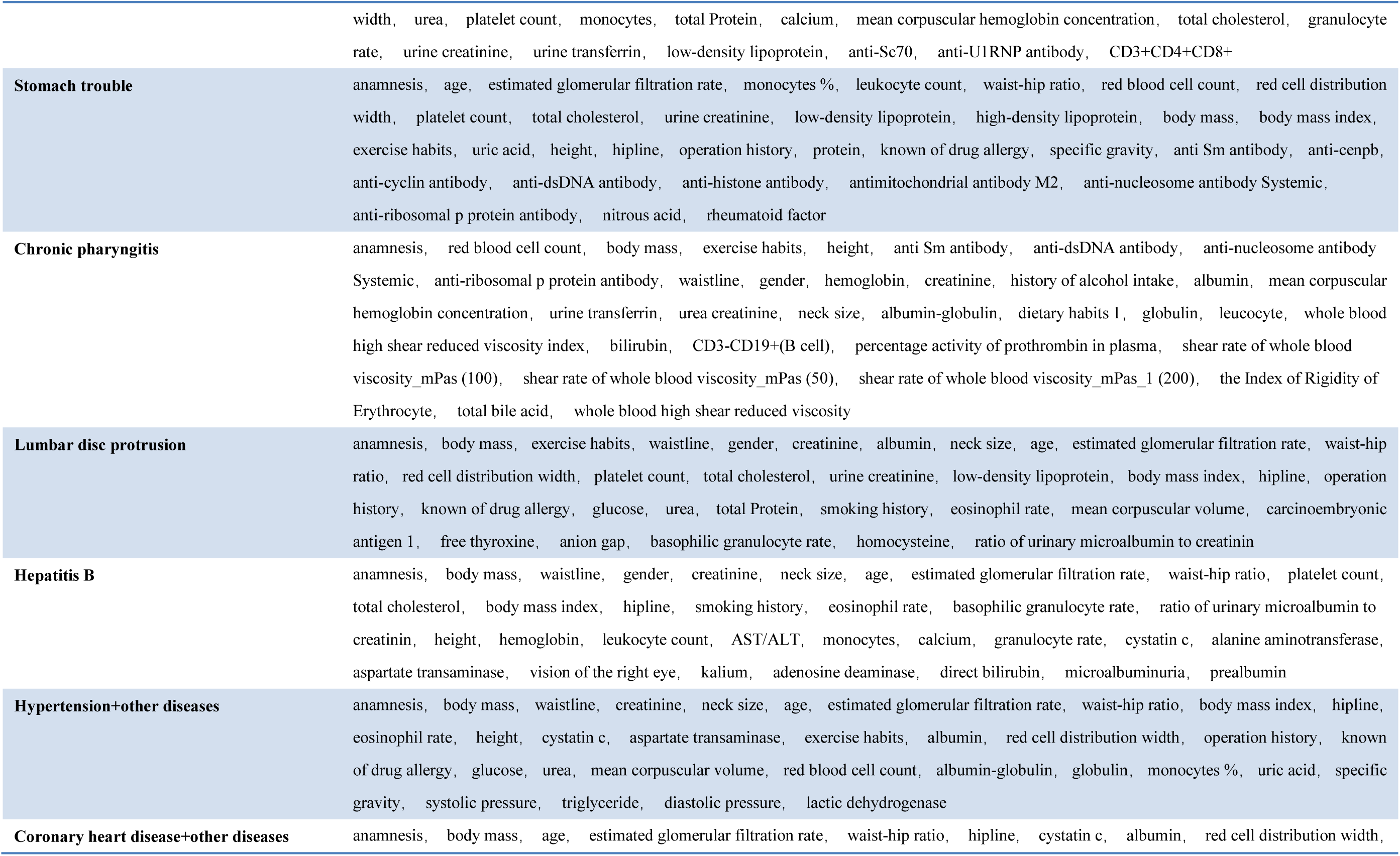
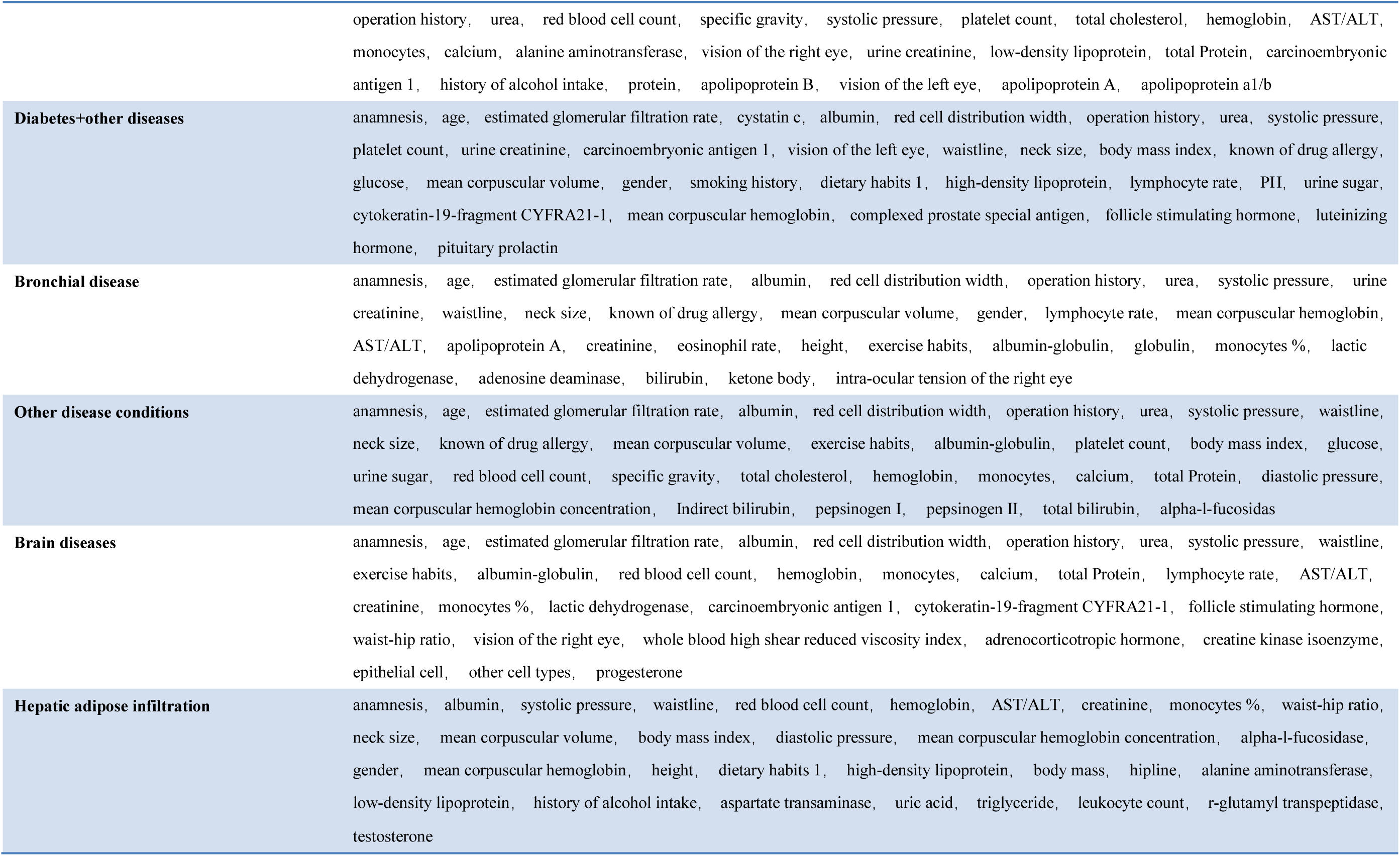
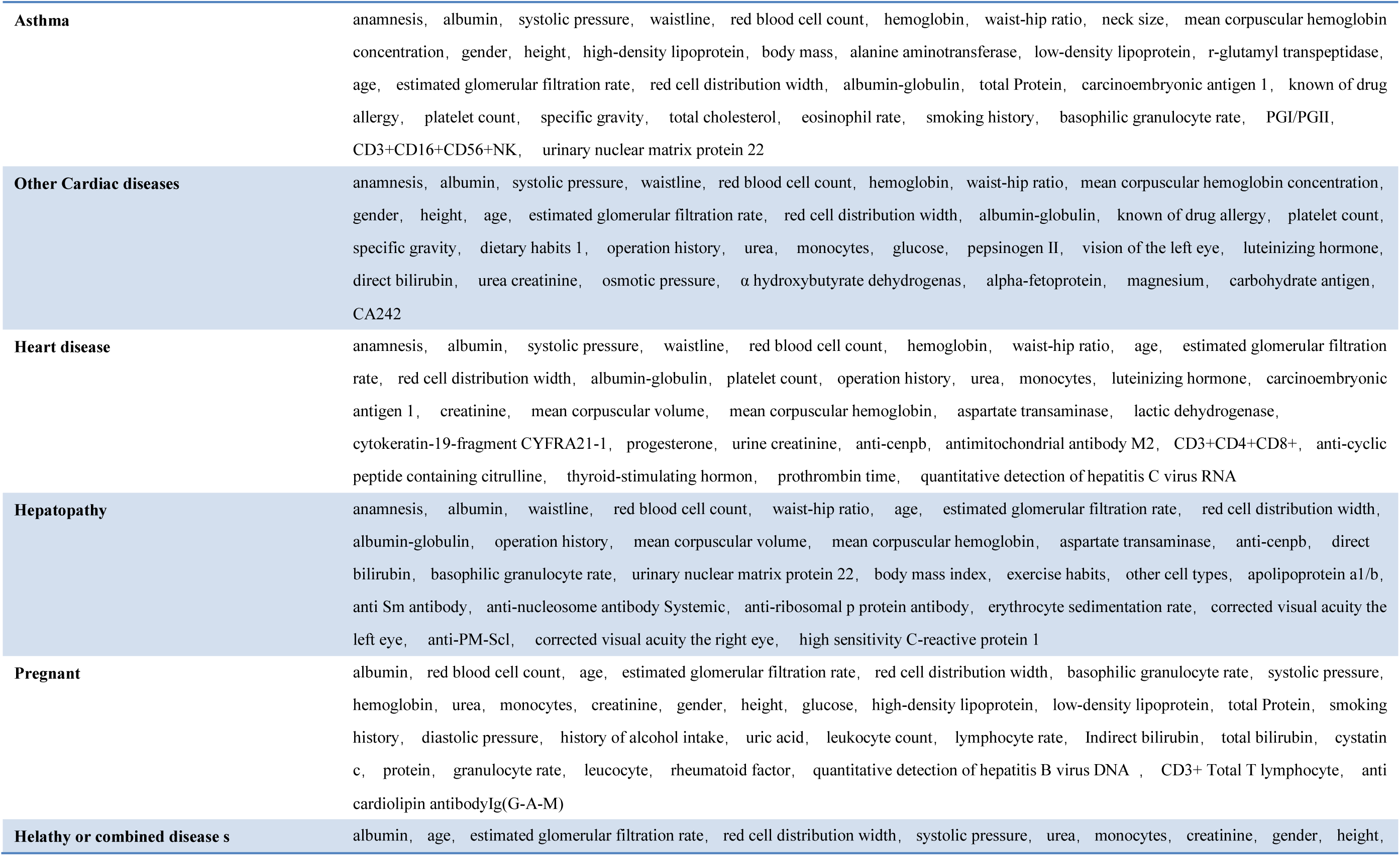
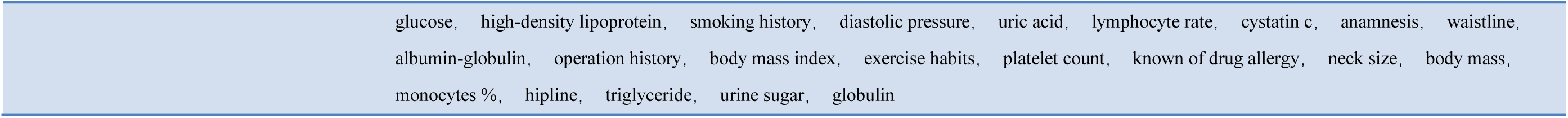
List of the 15% or 16% representative PEIs were extracted for machine learning prediction by feature extraction.

**Extended data Table 3.** Predictive Validity of Models. The number of training set and valid set samples was obtained after undersampling and data random splitting. Normal condition or disease was used to classify all kinds of diseases into disease states, undersampling with samples of healthy people, and then data division. Abbreviation: receiver operating characteristic, ROC; AUC, area under the curve.

|  | Training set (sample size) | Validation set (sample size) | F1-score (95%CI) | Accuracy (95%CI) | Specificity (95%CI) | Recall (sensitivity)(95%CI) | ROC(AUC)(95%CI) |
| --- | --- | --- | --- | --- | --- | --- | --- |
| Cholecystolithiasis | 963 | 413 | 0.69(0.66-0.71) | 0.69(0.68-0.71) | 0.73(0.70-0.76) | 0.70(0.67-0.72) | 0.77(0.75-0.79) |
| Hypertension | 46171 | 17987 | 0.86(0.86-0.86) | 0.85(0.85-0.86) | 0.82(0.82-0.82) | 0.85(0.85-0.86) | 0.92(0.92-0.93) |
| Hypertension+Diabetes | 8414 | 3065 | 0.92(0.92-0.92) | 0.92(0.92-0.92) | 0.90(0.90-0.90) | 0.92(0.92-0.92) | 0.97(0.97-0.98) |
| Hypertension+Coronary heart disease | 2032 | 871 | 0.93(0.92-0.93) | 0.93(0.92-0.93) | 0.90(0.90-0.91) | 0.93(0.92-0.93) | 0.97(0.97-0.98) |
| Hypertensive+Diabetes+Coronary Heart Disease | 909 | 389 | 0.96(0.95-0.96) | 0.95(0.94-0.97) | 0.95(0.94-0.95) | 0.95(0.94-0.97) | 0.99(0.98-0.99) |
| Hyperlipidemia | 1687 | 722 | 0.86(0.85-0.86) | 0.85(0.84-0.86) | 0.82(0.81-0.83) | 0.85(0.84-0.86) | 0.94(0.93-0.94) |
| Coronary heart disease | 1298 | 557 | 0.90(0.89-0.92) | 0.90(0.88-0.91) | 0.87(0.84-0.89) | 0.90(0.88-0.91) | 0.96(0.96-0.97) |
| Coronary heart disease+Diabetes | 274 | 118 | 0.93(0.93-0.94) | 0.94(0.92-0.96) | 0.91(0.88-0.95) | 0.94(0.92-0.95) | 0.98(0.97-1.00) |
| Rhinallergosis | 152 | 66 | 0.71(0.64-0.79) | 0.69(0.61-0.79) | 0.80(0.73-0.86) | 0.70(0.64-0.76) | 0.79(0.73-0.84) |
| Hypothyroidism | 1627 | 698 | 0.77(0.76-0.78) | 0.76(0.75-0.76) | 0.73(0.71-0.74) | 0.76(0.75-0.76) | 0.84(0.83-0.86) |
| Hyperthyroidism | 751 | 322 | 0.72(0.71-0.72) | 0.73(0.71-0.74) | 0.73(0.71-0.75) | 0.73(0.71-0.74) | 0.79(0.77-0.81) |
| Cervical spondylopathy | 311 | 134 | 0.71(0.66-0.75) | 0.71(0.69-0.73) | 0.73(0.64-0.81) | 0.71(0.69-0.73) | 0.78(0.78-0.80) |
| Rheumatoid arthritis | 387 | 167 | 0.79(0.78-0.81) | 0.78(0.77-0.78) | 0.76(0.71-0.81) | 0.78(0.77-0.79) | 0.86(0.83-0.89) |
| Chronic rhinitis | 313 | 135 | 0.61(0.58-0.64) | 0.60(0.58-0.62) | 0.57(0.56-0.57) | 0.61(0.58-0.63) | 0.66(0.60-0.72) |
| Nephropathy | 564 | 242 | 0.73(0.72-0.74) | 0.77(0.75-0.80) | 0.81(0.79-0.84) | 0.76(0.75-0.78) | 0.84(0.82-0.85) |
| Diabetes | 11545 | 4949 | 0.90(0.90-0.90) | 0.90(0.90-0.90) | 0.90(0.89-0.90) | 0.90(0.90-0.90) | 0.96(0.96-0.96) |
| Gout | 2095 | 898 | 0.88(0.88-0.88) | 0.87(0.87-0.87) | 0.85(0.84-0.87) | 0.86(0.83-0.88) | 0.94(0.94-0.94) |
| Parkinson's syndrome | 192 | 83 | 0.91(0.90-0.91) | 0.90(0.89-0.90) | 0.87(0.79-0.94) | 0.90(0.89-0.91) | 0.97(0.95-0.98) |
| Stomach trouble | 1269 | 545 | 0.68(0.68-0.69) | 0.70(0.70-0.70) | 0.71(0.70-0.73) | 0.70(0.70-0.70) | 0.77(0.75-0.78) |
| Chronic pharyngitis | 765 | 329 | 0.63(0.62-0.65) | 0.67(0.65-0.69) | 0.66(0.65-0.66) | 0.67(0.65-0.68) | 0.72(0.69-0.75) |
| Lumbar disc protrusion | 377 | 162 | 0.77(0.72-0.81) | 0.77(0.73-0.80) | 0.70(0.63-0.77) | 0.75(0.70-0.79) | 0.85(0.82-0.88) |
| Hepatitis B | 691 | 297 | 0.73(0.70-0.77) | 0.75(0.73-0.77) | 0.79(0.77-0.80) | 0.75(0.72-0.77) | 0.83(0.81-0.85) |
| Hypertension+other diseases | 2360 | 1012 | 0.86(0.85-0.88) | 0.85(0.85-0.86) | 0.82(0.81-0.83) | 0.86(0.85-0.86) | 0.93(0.93-0.94) |
| Coronary heart disease+other diseases | 98 | 43 | 0.88(0.84-0.92) | 0.87(0.84-0.89) | 0.83(0.76-0.90) | 0.86(0.83-0.88) | 0.94(0.91-0.97) |
| Diabetes+other diseases | 365 | 157 | 0.90(0.87-0.94) | 0.90(0.87-0.94) | 0.89(0.84-0.94) | 0.90(0.86-0.94) | 0.96(0.93-0.98) |
| Bronchial disease | 562 | 241 | 0.76(0.70-0.83) | 0.77(0.70-0.83) | 0.80(0.76-0.84) | 0.77(0.70-0.83) | 0.83(0.79-0.88) |
| Other disease conditions | 2720 | 1167 | 0.68(0.67-0.70) | 0.69(0.67-0.71) | 0.69(0.66-0.73) | 0.69(0.67-0.71) | 0.75(0.74-0.77) |
| Brain diseases | 251 | 108 | 0.86(0.81-0.90) | 0.86(0.82-0.91) | 0.83(0.75-0.90) | 0.87(0.82-0.91) | 0.93(0.91-0.95) |
| Hepatic adipose infiltration | 803 | 115 | 0.82(0.78-0.87) | 0.81(0.76-0.86) | 0.75(0.67-0.82) | 0.82(0.77-0.87) | 0.92(0.89-0.94) |
| Asthma | 1640 | 803 | 0.75(0.74-0.76) | 0.74(0.73-0.76) | 0.77(0.69-0.84) | 0.75(0.74-0.76) | 0.88(0.84-0.92) |
| Other Cardiac diseases | 336 | 145 | 0.79(0.78-0.81) | 0.78(0.76-0.80) | 0.80(0.75-0.86) | 0.78(0.76-0.80) | 0.88(0.84-0.92) |
| Heart disease | 224 | 96 | 0.89(0.87-0.90) | 0.89(0.85-0.92) | 0.91(0.87-0.94) | 0.89(0.86-0.92) | 0.94(0.90-0.99) |
| Hepatopathy | 176 | 76 | 0.71(0.65-0.76) | 0.72(0.66-0.78) | 0.78(0.68-0.87) | 0.73(0.68-0.78) | 0.80(0.74-0.85) |
| Pregnant | 54 | 24 | 0.83(0.76-0.90) | 0.82(0.76-0.88) | 0.85(0.79-0.92) | 0.82(0.77-0.86) | 0.91(0.90-0.93) |
| Normal or non-normal condition | 91028 | 39012 | 0.83(0.83-0.83) | 0.82(0.82-0.82) | 0.81(0.81-0.81) | 0.84(0.84-0.84) | 0.9(0.90-0.90) |

**Extended data Figure 1.**
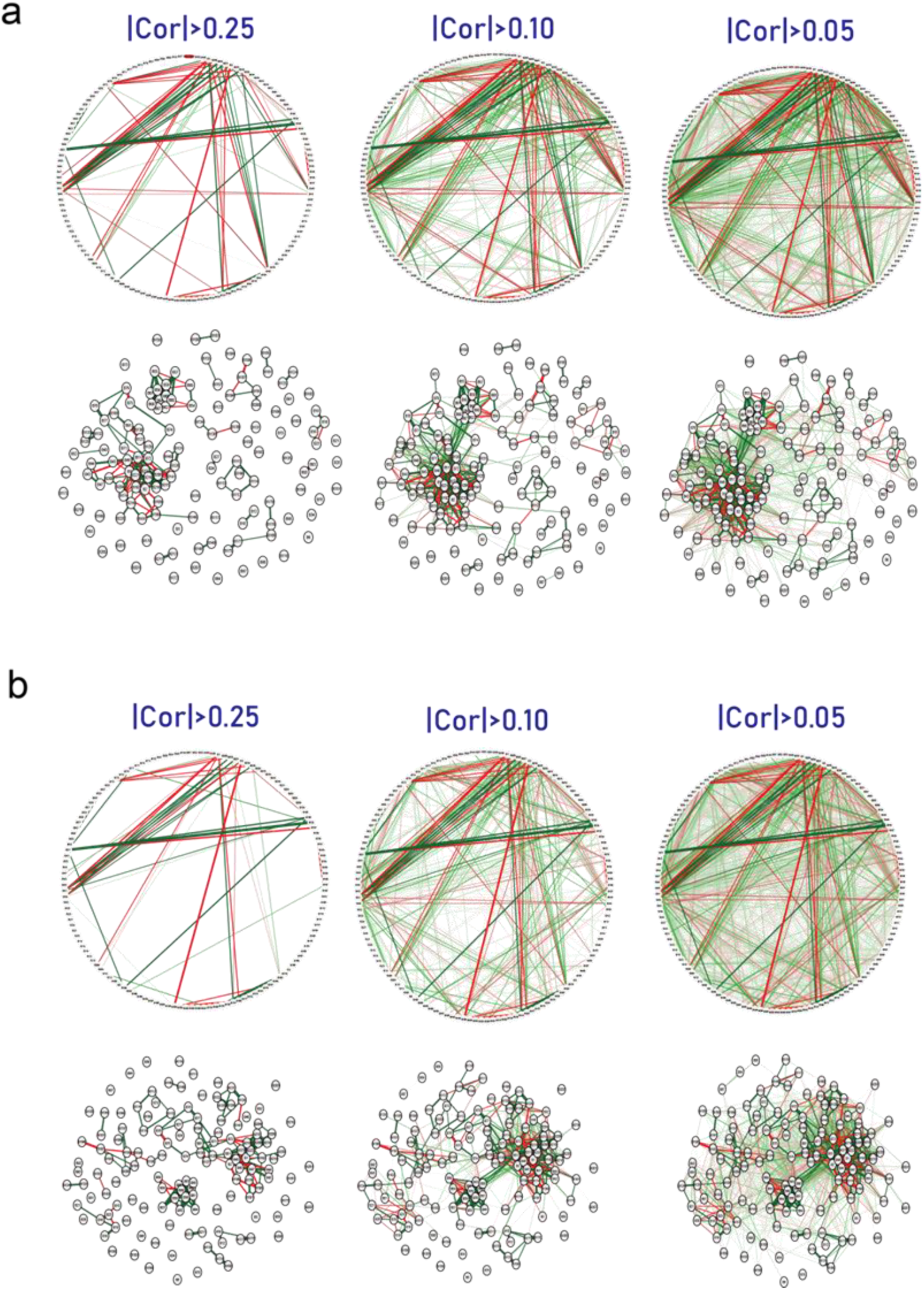
PEI networks in hypertension (a) and diabetes (b). In weighted graphs, green edges indicate positive weights, and red edges indicate negative weights. The color saturation and the width of the edges correspond to the absolute weight and scale relative to the strongest weight in the graph. At a minimum, the edge with absolute weight at this value is omitted. The circular layout is convenient to see how well the data conform to a model, but in order to show how the data clusters, another layout is more appropriate. A force-oriented layout was created by specifying layout = “spring”. In principle, what this function does is that each node (connected and unconnected) repulses each other, and connected nodes also attract each other. The full view of these figures is provided in Supplementary Figures.

**Extended data Figure 2.**
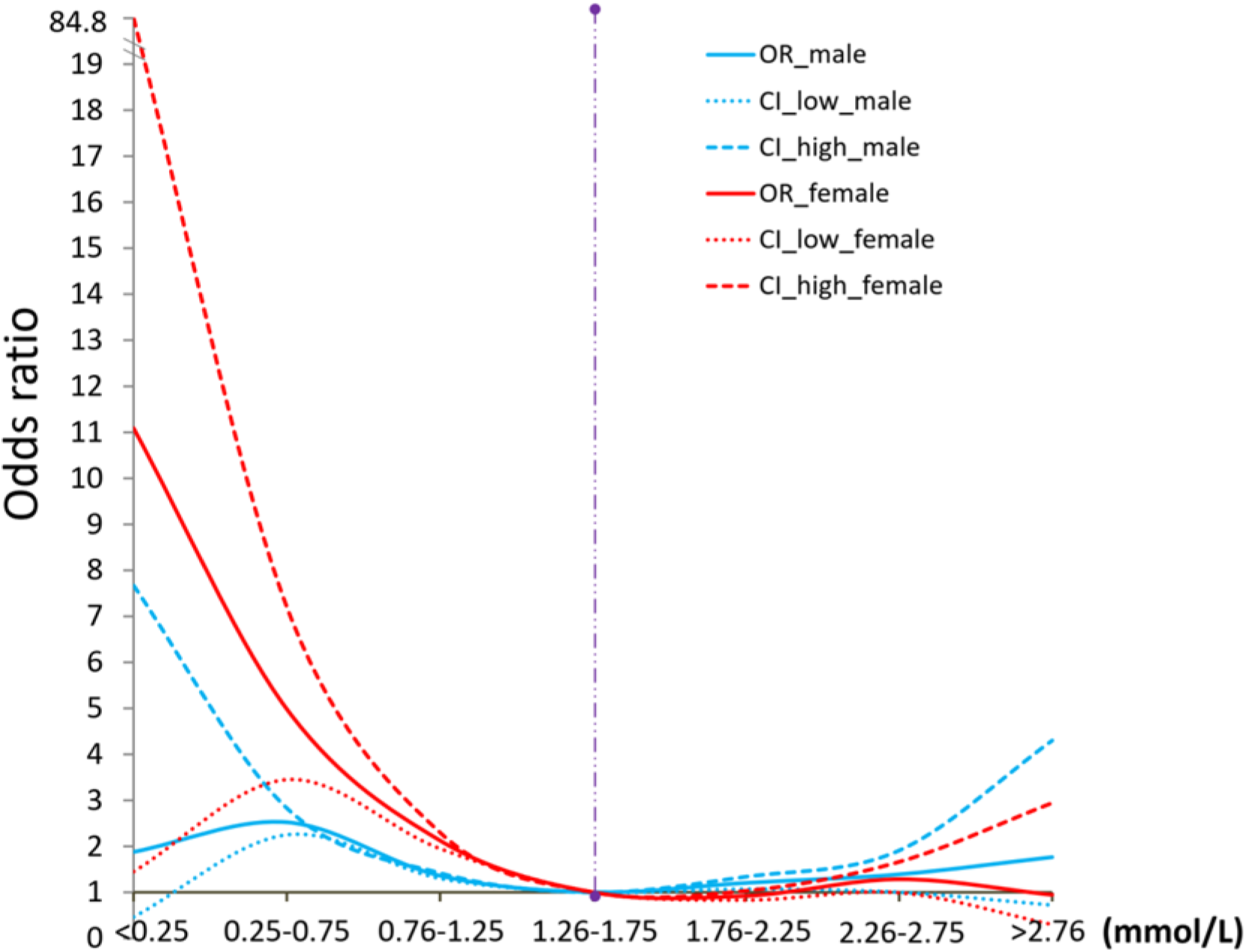
Odds ratios for HDL-C concentration in plasma from those with a normal physical status and those with diabetes. Both male and female subjects were included in this study.

## Supplementary Tables

Supplementary Table 1. PEI Correlations in Healthy status

Supplementary Table 2. PEI Correlations in Cholecystolithiasis

Supplementary Table 3. PEI Correlations in Hypertension

Supplementary Table 4. PEI Correlations in Hypertension+Diabetes

Supplementary Table 5. PEI Correlations in Hypertension+Coronary

Supplementary Table 6. PEI Correlations in Hypertensive+Diabetes+Coronary

Supplementary Table 7. PEI Correlations in Hyperlipidemia

Supplementary Table 8. PEI Correlations in Coronary heart disease

Supplementary Table 9. PEI Correlations in Coronary+Diabetes

Supplementary Table 10. PEI Correlations in Rhinallergosis

Supplementary Table 11. PEI Correlations in Hypothyroidism

Supplementary Table 12. PEI Correlations in Hyperthyroidism

Supplementary Table 13. PEI Correlations in Cervical spondylopathy

Supplementary Table 14. PEI Correlations in Rheumatoid arthritis

Supplementary Table 15. PEI Correlations in Chronic rhinitis

Supplementary Table 16. PEI Correlations in Nephropathy

Supplementary Table 17.PEI Correlations in Diabetes Supplementary Table 18. PEI Correlations in Gout

Supplementary Table 19. PEI Correlations in Parkinson’s syndrome

Supplementary Table 20. PEI Correlations in Stomach trouble

Supplementary Table 21. PEI Correlations in Chronic pharyngitis

Supplementary Table 22. PEI Correlations in Lumbar disc protrusion

Supplementary Table 23. PEI Correlations in Hepatitis B

Supplementary Table 24. PEI Correlations in Hypertension+other diseases

Supplementary Table 25. PEI Correlations in Coronary+others

Supplementary Table 26. PEI Correlations in Diabetes+others

Supplementary Table 27. PEI Correlations in Bronchial disease

Supplementary Table 28. PEI Correlations in Other disease conditions

Supplementary Table 29. PEI Correlations in Brain diseases

Supplementary Table 30. PEI Correlations in Hepatic adipose infiltration

Supplementary Table 31. PEI Correlations in Asthma

Supplementary Table 32. PEI Correlations in Other Cardiac diseases

Supplementary Table 33. PEI Correlations in Heart disease

Supplementary Table 34. PEI Correlations in Hepatopathy

Supplementary Table 35. PEI Correlations in Pregnant

Supplementary Table 36. P values of PEIs in healthy physical status vs 34 unhealthy physical status adjusted for age and sex.

Supplementary Table 37. Optimal parameter combination of machine learning

**Supplementary codes**

