## Supplementary files for "Big data reveals deep associations in physical examination indicators and can help predict overall underlying health status"

**Supplementary Tables**

Supplementary Table 1. PEI Correlations in Healthy status

Supplementary Table 2. PEI Correlations in Cholecystolithiasis

Supplementary Table 3. PEI Correlations in Hypertension

Supplementary Table 4. PEI Correlations in Hypertension+Diabetes

Supplementary Table 5. PEI Correlations in Hypertension+Coronary

Supplementary Table 6. PEI Correlations in Hypertensive+Diabetes+Coronary

Supplementary Table 7. PEI Correlations in Hyperlipidemia

Supplementary Table 8. PEI Correlations in Coronary heart disease

Supplementary Table 9. PEI Correlations in Coronary+Diabetes

Supplementary Table 10. PEI Correlations in Rhinallergosis

Supplementary Table 11. PEI Correlations in Hypothyroidism

Supplementary Table 12. PEI Correlations in Hyperthyroidism

Supplementary Table 13. PEI Correlations in Cervical spondylopathy

Supplementary Table 14. PEI Correlations in Rheumatoid arthritis

Supplementary Table 15. PEI Correlations in Chronic rhinitis

Supplementary Table 16. PEI Correlations in Nephropathy

Supplementary Table 17.PEI Correlations in Diabetes

Supplementary Table 18. PEI Correlations in Gout

Supplementary Table 19. PEI Correlations in Parkinson's syndrome

Supplementary Table 20. PEI Correlations in Stomach trouble

Supplementary Table 21. PEI Correlations in Chronic pharyngitis

Supplementary Table 22. PEI Correlations in Lumbar disc protrusion

Supplementary Table 23. PEI Correlations in Hepatitis B

Supplementary Table 24. PEI Correlations in Hypertension+other diseases

Supplementary Table 25. PEI Correlations in Coronary+others

Supplementary Table 26. PEI Correlations in Diabetes+others

Supplementary Table 27. PEI Correlations in Bronchial disease

Supplementary Table 28. PEI Correlations in Other disease conditions

Supplementary Table 29. PEI Correlations in Brain diseases

Supplementary Table 30. PEI Correlations in Hepatic adipose infiltration

Supplementary Table 31. PEI Correlations in Asthma

Supplementary Table 32. PEI Correlations in Other Cardiac diseases

Supplementary Table 33. PEI Correlations in Heart disease

Supplementary Table 34. PEI Correlations in Hepatopathy

Supplementary Table 35. PEI Correlations in Pregnant

Supplementary Table 36. P values of PEIs in healthy physical status vs 34 unhealthy physical status adjusted for age and sex.

Supplementary Table 37. Optimal parameter combination of machine learning

**Supplementary codes for machine learning**

**Supplementary Table 1. PEIs used in this study**

| Code | PEI Name | PEI Name abbreviation | Unit |
|---|---|---|---|
| B1 | Gender | Gender | - |
| B2 | Age | Age | years |
| B3 | Homocysteine | Hcy | umol/L |
| B4 | Whole Blood Calcium | WB-Ca | mmol/L |
| B5 | Whole Blood Magnesium | WB-Mg | mmol/L |
| B6 | Iron Of Whole Blood | WB-Fe | mmol/L |
| B7 | Whole Blood Copper | WB-Cop | umol/L |
| B8 | Zinc In Whole Blood | WB-Zinc | umol/L |
| B9 | Rh Blood Group | RH | - |
| B10 | Pepsinogen I | PGI | ug/L |
| B11 | PgI/PgII | PGI/PGII | - |
| B12 | Pepsinogen II | PGII | ug/L |
| B14 | Gastrin-17 | G-17 | pmol/l |
| B15 | Cd3-Cd19+(B Cell) | CD3-CD19+(B cell) | - |
| B16 | Cd3+ Total T Lymphocyte | CD3+T | - |
| B17 | Cd3+Cd16+Cd56+Nk | CD3+CD16+CD56+NK | - |
| B18 | Cd3+Cd4+Helper(Inducer)T Cells | CD3+CD4+helper(inducer)T | - |
| B19 | Cd3+Cd4+/Cd3+Cd8+ | CD3+CD4+/CD3+CD8+ | - |
| B20 | Cd3+Cd4+Cd8+ | CD3+CD4+CD8+ | - |
| B21 | Cd3+Cd4-Cd8- | CD3+CD4-CD8- | - |
| B22 | Cd3+Cd8+(T) | CD3+CD8+(T) | - |
| B23 | Cd3-Cd16+Cd56+(Nk) | CD3-CD16+CD56+(NK) | - |
| B24 | α Hydroxybutyrate Dehydrogenas | αHBDH | U/L |
| B25 | Creatine Kinase | CK | U/L |
| B26 | Creatine Kinase Isoenzyme | CKMB | U/L |
| B27 | Lactic Dehydrogenase | LDH | U/L |
| B28 | Pituitary Prolactin | PRL | mIU/L |
| B29 | Estradiol E2 | E2 | pmol/L |
| B30 | Luteinizing Hormone | LH | mIU/ml |
| B31 | Follicle Stimulating Hormone | FSH | mIU/ml |
| B32 | Testosterone | Testo | nmol/L |
| B33 | Progesterone | Prog | nmol/L |
| B34 | Thymidine Kinase | TK | PM/L |
| B35 | Blood Beta-Microglobulin | β2-MG | - |
| B36 | Variant Lymphocyte Rate | V-Lym% | - |
| B37 | Band Cell | BAND | 10E-9/L |
| B38 | Band Cell% | BAND% | - |
| B39 | Other Cell Types | Other cells | - |
| B40 | Leukocyte Count | WBC | 10E-9/L |
| B41 | Monocytes | MON | - |
| B42 | Monocytes % | MON % | 10E-9/L |

| | | | |
|---|---|---|---|
| B43 | Mean Corpuscular Volume | MCV | fl |
| B44 | Mean Corpuscular Hemoglobin | MCH | pg |
| B45 | Mean Corpuscular Hemoglobin Concentration | MCHC | g/l |
| B46 | Red Blood Cell Count | RBC Count | 10E-12/L |
| B47 | Red Cell Distribution Width | RDW | fl |
| B48 | Lymphocyte Rate | LYM% | - |
| B49 | Basophilic Granulocyte Rate | baso% | - |
| B50 | Eosinophil Rate | eo% | - |
| B51 | Hemoglobin | Hb | g/L |
| B52 | Platelet Count | PLT/BPC | 10E-9/L |
| B53 | Granulocyte Rate | GR% | - |
| B54 | Erythrocyte Sedimentation Rate | ESR | mm/60min |
| B55 | Erythrocyte Deformation Index | TK | - |
| B56 | The Index Of Rigidity Of Erythrocyte | IR | - |
| B57 | Erythrocyte Aggregation Indices | RBCAI | - |
| B58 | Hematocrit | HCT | - |
| B59 | Whole Blood Reduced Viscosity | WBRV | - |
| B60 | Whole Blood Reduced Viscosity Index | WBRVI | - |
| B61 | Whole Blood High Shear Reduced Viscosity | WBHSRV | - |
| B62 | Whole Blood High Shear Reduced Viscosity Index | WBHSRVI | - |
| B63 | Shear Rate Of Whole Blood Viscosity_Mpas_1 (200) | WBRSR(mPas_200) | - |
| B64 | Shear Rate Of Whole Blood Viscosity_Mpas_3 (5) | WBRSR(mPas_5) | - |
| B65 | Shear Rate Of Whole Blood Viscosity_Mpas_4 (1) | WBRSR(mPas_1) | - |
| B66 | Shear Rate Of Whole Blood Viscosity_Mpas (100) | WBRSR(mPas_100) | - |
| B67 | Shear Rate Of Whole Blood Viscosity_Mpas (50) | WBRSR(mPas_50) | - |
| B69 | Plasma Viscosity | PV | - |
| B70 | Low-Density Lipoprotein | LDL-C | mmol/L |
| B71 | Triglyceride | TG | mmol/L |
| B72 | High-Density Lipoprotein | HDL-C | mmol/L |
| B73 | Apolipoprotein A | ApoA | mg/L |
| B74 | Apolipoprotein A1 | ApoA1 | g/L |
| B75 | Apolipoprotein A1/B | ApoA1/B | g/L |
| B76 | Apolipoprotein B | ApoB | g/L |
| B77 | Total Cholesterol | TC | mmol/L |
| B78 | Hepatitis B Virus Core Im Antibod | HBcAb | - |
| B79 | Quantitative Detection Of Hepatitis B Virus Dna | hbV-dna | IU/ml |
| B80 | Helicobacter Pylori Antibody-Igg | anti-HP IgG | - |
| B81 | Tumor Supplied Group Of Factors | TSGF | U/ML |
| B82 | Carcinoembryonic Antigen | CEA | ng/ml |
| B83 | Alpha-Fetoprotein | AFP | ng/ml |
| B84 | Complexed Prostate Special Antigen | C-PSA | ng/ml |
| B85 | Squamous Cell Carcinoma Antigen | SCC | ng/ml |
| B86 | Chorionic Gonadotropin | TBhCG | mIU/ml |
| B87 | Neuron-Specific Enolase | NSE | ng/ml |

| | | | |
|---|---|---|---|
| B88 | Carbohydrate Antigen，Ca242 | CA 242 | U/ml |
| B89 | Carbohydrate Antigen，Ca72_4 | CA 72-4 | U/ml |
| B90 | Cytokeratin-19-Fragment Cyfra21-1 | CYFRA 21-1 | ng/ml |
| B91 | Total Protein | TP | g/L |
| B92 | Aldh2-Glu504Glu | ALDH2-Glu504Glu | - |
| B93 | Aldh2-Glu504Lys | ALDH2-Glu504Lys | - |
| B94 | Aldh2-Lys504Lys | ALDH2-Lys504Lys | - |
| B95 | Anti-Cenpb | anti-cenpb | - |
| B96 | Dsdna (Iif) | dsDNA (IIF) | - |
| B97 | Anti-Dsdna Antibody | dsDNA | - |
| B98 | Jo-1 Antibody | anti-Jo-1 | - |
| B99 | Anti-Pm-Scl | anti-PM-Scl | - |
| B100 | Anti-Ro52 | anti-Ro52 | - |
| B101 | Anti-Sc70 | anti-Sc70 | - |
| B102 | Anti Sm Antibody | anti-Sm | - |
| B103 | Anti-Ssa/Ro Antibody | anti-SSA/Ro | - |
| B104 | Anti Ssb Antibody | anti-SSB | - |
| B105 | Anti-U1Rnp Antibody | anti-U1RNP | - |
| B106 | Anti-Ribosomal P Protein Antibody | anti-P | - |
| B107 | Anti-Nucleosome Antibody Systemic | AnuA | - |
| B108 | Anti-Cyclin Antibody | Anti-Cyclin | - |
| B109 | Antimitochondrial Antibody M2 | AMA-M2 | - |
| B110 | Anti-Histone Antibody | AHA | - |
| B111 | Thyroid-Stimulating Hormon | TSH | mIU/L |
| B112 | Free Thyroxine | FT4 | pmol/L |
| B113 | Free Triiodothyronine | FT3 | pmol/L |
| B114 | Total Thyroxine | TSH TT3 TT4(TT4) | nmol/L |
| B115 | Total Triiodothyronine | TSH TT3 TT4(TT3) | nmol/L |
| B116 | Total Triiodothyronine | TT3 | nmol/L |
| B117 | Total Thyroxine | TT4 | nmol/L |
| B118 | Carcinoembryonic Antigen 1 | CEA1 | ng/ml |
| B119 | Quantitative Detection Of Hepatitis C Virus Rna | Hcv-rna | - |
| B120 | Hcv Core Antigen | HCV-cAg | - |
| B121 | Complement C3 | C3 | - |
| B122 | Complement C4 | C4 | - |
| B123 | High Sensitivity C-Reactive Protein 1 | hs-CRP1 | mg/L |
| B124 | Adrenocorticotropic Hormone | ACTH | pg/ml |
| B125 | Cortisol | Cor | ug/dl |
| B126 | Albumin | Alb | g/L |
| B127 | Albumin-Globulin | A-G ratio | - |
| B128 | Alanine Aminotransferase | ALT | U/L |
| B129 | Indirect Bilirubin | IBIL | umol/L |
| B130 | Globulin | GLB | g/L |
| B131 | Aspartate Transaminase | AST | U/L |

| | | | |
|---|---|---|---|
| B132 | Direct Bilirubin | Dbil | umol/L |
| B133 | Total Bilirubin | TBIL | umol/L |
| B134 | Cholinesterases | CHE | KU/L |
| B135 | Total Bile Acid | TBA | umol/L |
| B136 | 5-Nucleotidase | 5'NT | U/L |
| B137 | Alpha-L-Fucosidase | AFU | U/L |
| B138 | Prealbumin | PAB | mg/L |
| B139 | Adenosine Deaminase | ADA | U/L |
| B140 | R-Glutamyl Transpeptidase | r-GT | U/L |
| B141 | Ast/Alt | AST/ALT | - |
| B142 | 25-Hydroxyvitamin D | 25-(OH)D | ng/mL |
| B143 | β-Crosslaps | β-CTx | pg/mL |
| B144 | Total-Pinp | Total-PINP | ng/mL |
| B145 | Osteocalcin | BGP | ng/mL |
| B146 | Cystatin C | CysC | mg/L |
| B148 | Antithyroperoxidase Antibody | Anti_TPO | IU/ml |
| B149 | Anti-Thyroglobulin Antibodies | TGAb | IU/ml |
| B150 | Alkaline Phosphatase | ALP | U/L |
| B151 | Anti-Cyclic Peptide Containing Citrulline | Anti-CCP | Ru/ml |
| B152 | Anti Cardiolipin Antibodyig(G-A-M) | ACA-Ig(G-A-M) | u/ml |
| B153 | Glucose | Glu | mmol/L |
| B154 | Rheumatoid Factor | RF | IU/ml |
| B155 | Immunoglobulin A | IgA | g/l |
| B156 | Immunoglobulin E | IgE | IU/ml |
| B157 | Immunoglobulin G | IgG | g/l |
| B158 | Immunoglobulin M | IgM | g/l |
| B159 | Urine Beta-Microglobulin | β2-MG | - |
| B160 | Ph | PH | - |
| B161 | Leucocyte | LEU | /ul |
| B163 | Specific Gravity | SG | - |
| B164 | Calcium Oxalate Crystals | COM | HPF |
| B165 | Bilirubin | bilirubin | - |
| B166 | Protein | Pro | - |
| B167 | Phosphate Crystals | PC | - |
| B168 | Urinary Nuclear Matrix Protein 22 | NMP22 | - |
| B169 | Urate Crystal | U-CRY | - |
| B170 | Urine Sugar | U-GLU | - |
| B171 | Epithelial Cell | EC | HPF |
| B172 | Ketone Body | KB | - |
| B173 | Nitrous Acid | NTT | - |
| B174 | Urea Creatinine | Urea/Cr | - |
| B175 | Microalbuminuria | ALB-U | mg/L |
| B176 | Urine Creatinine | Ucr | umol/L |
| B177 | Ratio Of Urinary Microalbumin To Creatinin | ACR | ug/mg |

| | | | |
|---|---|---|---|
| B178 | Urinary Immunoglobulin G | Ig-U | mg/L |
| B179 | Urine Alpha-Microglobulin | α1-MG | mg/L |
| B180 | Urine Transferrin | Tf | mg/L |
| B181 | Activated Partial Thromboplastin Time | APTT | s |
| B182 | Thrombin Time | TT | s |
| B183 | Percentage Activity Of Prothrombin In Plasma | PTA | - |
| B184 | International Normalized Ratio | INR | - |
| B185 | Prothrombin Time | PT | s |
| B186 | Plasma Fibrinogen Concentration | PFC | g/l |
| B187 | Neuron-Specific Enolase | NSE | ng/ml |
| B188 | Calcium | Ca | mmol/L |
| B189 | Estimated Glomerular Filtration Rate | eGFB | ml/min |
| B190 | Creatinine | Cre | umol/L |
| B191 | Urea | Urea | mmol/L |
| B192 | Uric Acid | UA | umol/L |
| B193 | Kalium | K | mmol/L |
| B194 | Chlorine | Cl | mmol/L |
| B195 | Sodium | Na | mmol/L |
| B196 | Blood Carbon Dioxide | TCO2 | mmol/L |
| B197 | Anion Gap | AG | mmol/L |
| B198 | Phosphorus | P | mmol/L |
| B199 | Magnesium | Mg | mmol/L |
| B200 | Osmotic Pressure | OP | mOsm/L |
| B201 | Exercise Habits | e-habits | - |
| B202 | Frequency Of Exercises | e-times | - |
| B203 | Dietary Habits 1 | DH | - |
| B204 | History Of Alcohol Intake | Drinking | - |
| B205 | Known Of Drug Allergy | KDA | - |
| B206 | Waistline | Waist | cm |
| B207 | Waist-Hip Ratio | WHR | - |
| B209 | Smoking History | Smoking | - |
| B210 | Hipline | Hip | cm |
| B211 | Intra-Ocular Tension Of The Right Eye | IOP-R | mmHg |
| B212 | Intra-Ocular Tension Of The Left Eye | IOP-L | mmHg |
| B213 | Corrected Visual Acuity The Right Eye | CVA-R | - |
| B214 | Corrected Visual Acuity The Left Eye | CVA-L | - |
| B215 | Body Mass Index | BMI | - |
| B216 | Body Mass | BMI | kg |
| B218 | Diastolic Pressure | DBP | mmHg |
| B219 | Operation History | OT | - |
| B220 | Systolic Pressure | SBP | mmHg |
| B221 | Height | Hei | cm |
| B222 | Neck Size | NS | cm |
| B223 | Family History | FH | - |

| | | | |
|---|---|---|---|
| B224 | Anamnesis | pmh | - |
| B225 | Hourly Postprandial Blood Sugar | hPBG | mmol/L |
| B227 | Vision Of The Right Eye | OD | - |
| B228 | Vision Of The Left Eye | OS | - |

**Supplementary Table 1. PEI Correlations in Healthy status**

| Row | Column | Row | Column | Cor | *P* |
|---|---|---|---|---|---|
| B1 | B142 | Gender | 25-(OH)D | 0.006119 | 9.18E-08 |
| B1 | B137 | Gender | AFU | -0.03423 | 0 |
| B1 | B24 | Gender | αHBDH | 0.01309 | 0 |
| B1 | B16 | Gender | CD3+T | 0.005632 | 8.79E-07 |
| B1 | B15 | Gender | CD3-CD19+(B cell) | 0.008684 | 3.42E-14 |
| B1 | B23 | Gender | CD3-CD16+CD56+(NK) | -0.00619 | 6.67E-08 |
| B1 | B18 | Gender | CD3+CD4+helper(inducer)T | 0.007416 | 9.49E-11 |
| B1 | B54 | Gender | ESR | 0.053469 | 0 |
| B1 | B13 | Gender | PGI_PGII | -0.00659 | 8.71E-09 |
| B1 | B160 | Gender | PH | 0.069872 | 0 |
| B1 | B140 | Gender | r-GT | -0.22855 | 0 |
| B1 | B143 | Gender | β-CTx | -0.0067 | 4.93E-09 |
| B1 | B82 | Gender | CEA | -0.00569 | 6.80E-07 |
| B1 | B118 | Gender | CEA1 | -0.0265 | 0 |
| B1 | B126 | Gender | Alb | -0.1868 | 0 |
| B1 | B127 | Gender | A-G ratio | -0.17598 | 0 |
| B1 | B161 | Gender | LEU | 0.170258 | 0 |
| B1 | B40 | Gender | WBC | -0.16164 | 0 |
| B1 | B128 | Gender | ALT | -0.27333 | 0 |
| B1 | B164 | Gender | COM | 0.012681 | 0 |
| B1 | B123 | Gender | hs-CRP1 | -0.01896 | 0 |
| B1 | B111 | Gender | TSH | 0.053927 | 0 |
| B1 | B124 | Gender | ACTH | -0.01648 | 0 |
| B1 | B41 | Gender | MON％ | -0.0411 | 0 |
| B1 | B165 | Gender | bilirubin | -0.00871 | 2.78E-14 |
| B1 | B134 | Gender | CHE | -0.07687 | 0 |
| B1 | B166 | Gender | Pro | -0.17502 | 0 |
| B1 | B70 | Gender | LDL-C | -0.13992 | 0 |
| B1 | B188 | Gender | Ca | -0.0283 | 0 |
| B1 | B71 | Gender | TG | -0.23139 | 0 |
| B1 | B72 | Gender | HDL-C | 0.36582 | 0 |
| B1 | B32 | Gender | Testo | -0.01778 | 0 |
| B1 | B189 | Gender | eGFB | 0.224096 | 0 |
| B1 | B141 | Gender | AST/ALT | 0.304609 | 0 |
| B1 | B146 | Gender | CysC | -0.04504 | 0 |
| B1 | B55 | Gender | TK | 0.015659 | 0 |
| B1 | B56 | Gender | IR | -0.05577 | 0 |
| B1 | B57 | Gender | RBCAI | -0.02178 | 0 |
| B1 | B43 | Gender | MCV | -0.01009 | 0 |
| B1 | B44 | Gender | MCH | -0.13679 | 0 |
| B1 | B45 | Gender | MCHC | -0.3201 | 0 |
| B1 | B46 | Gender | RBC Count | -0.58863 | 0 |

| | | | | | |
|---|---|---|---|---|---|
| B1 | B47 | Gender | RDW | 0.067555 | 0 |
| B1 | B58 | Gender | HCT | -0.01435 | 0 |
| B1 | B181 | Gender | APTT | 0.013743 | 0 |
| B1 | B190 | Gender | Cre | -0.65885 | 0 |
| B1 | B25 | Gender | CK | -0.01605 | 0 |
| B1 | B26 | Gender | CKMB | -0.00901 | 3.77E-15 |
| B1 | B224 | Gender | pmh | -0.01316 | 0 |
| B1 | B129 | Gender | IBIL | -0.14964 | 0 |
| B1 | B150 | Gender | ALP | -0.05854 | 0 |
| B1 | B84 | Gender | C-PSA | -0.04495 | 0 |
| B1 | B222 | Gender | NS | -0.70683 | 0 |
| B1 | B100 | Gender | anti-Ro52 | 0.006769 | 3.42E-09 |
| B1 | B103 | Gender | anti-SSA/Ro | 0.008765 | 1.98E-14 |
| B1 | B148 | Gender | Anti_TPO | 0.061984 | 0 |
| B1 | B149 | Gender | TGAb | 0.055607 | 0 |
| B1 | B198 | Gender | P | 0.055717 | 0 |
| B1 | B85 | Gender | SCC | -0.02093 | 0 |
| B1 | B194 | Gender | Cl | 0.020809 | 0 |
| B1 | B199 | Gender | Mg | -0.01106 | 0 |
| B1 | B156 | Gender | IgE | -0.01045 | 0 |
| B1 | B157 | Gender | IgG | 0.032658 | 0 |
| B1 | B158 | Gender | IgM | 0.022932 | 0 |
| B1 | B195 | Gender | Na | -0.02171 | 0 |
| B1 | B2 | Gender | Age | -0.05802 | 0 |
| B1 | B176 | Gender | Ucr | -0.13356 | 0 |
| B1 | B178 | Gender | Ig-U | -0.0075 | 6.00E-11 |
| B1 | B191 | Gender | Urea | -0.18982 | 0 |
| B1 | B174 | Gender | Urea/Cr | 0.318269 | 0 |
| B1 | B192 | Gender | UA | -0.58161 | 0 |
| B1 | B170 | Gender | U-GLU | -0.05063 | 0 |
| B1 | B175 | Gender | ALB-U | -0.01348 | 0 |
| B1 | B177 | Gender | ACR | 0.006481 | 1.53E-08 |
| B1 | B179 | Gender | α1-MG | -0.04255 | 0 |
| B1 | B180 | Gender | Tf | -0.00562 | 9.21E-07 |
| B1 | B182 | Gender | TT | -0.00926 | 6.66E-16 |
| B1 | B125 | Gender | Cor | -0.00973 | 0 |
| B1 | B153 | Gender | Glu | -0.01674 | 0 |
| B1 | B138 | Gender | PAB | -0.09439 | 0 |
| B1 | B130 | Gender | GLB | 0.12852 | 0 |
| B1 | B59 | Gender | WBRV | -0.03039 | 0 |
| B1 | B60 | Gender | WBRVI | -0.09873 | 0 |
| B1 | B4 | Gender | WB-Ca | 0.008595 | 6.22E-14 |
| B1 | B61 | Gender | WBHSRV | -0.05458 | 0 |
| B1 | B62 | Gender | WBHSRVI | -0.10412 | 0 |

| | | | | | |
|---|---|---|---|---|---|
| B1 | B5 | Gender | WB-Mg | -0.01124 | 0 |
| B1 | B6 | Gender | WB-Fe | -0.01874 | 0 |
| B1 | B8 | Gender | WB-Zinc | -0.01371 | 0 |
| B1 | B63 | Gender | WBRSR(mPas_200) | -0.01144 | 0 |
| B1 | B64 | Gender | WBRSR(mPas_5) | -0.10303 | 0 |
| B1 | B65 | Gender | WBRSR(mPas_1) | -0.09742 | 0 |
| B1 | B66 | Gender | WBRSR(mPas_100) | -0.10109 | 0 |
| B1 | B67 | Gender | WBRSR(mPas_50) | -0.10189 | 0 |
| B1 | B86 | Gender | TBhCG | 0.0104 | 0 |
| B1 | B27 | Gender | LDH | -0.01558 | 0 |
| B1 | B221 | Gender | Hei | -0.66881 | 0 |
| B1 | B87 | Gender | NSE | -0.01219 | 0 |
| B1 | B187 | Gender | NSE | -0.00728 | 2.04E-10 |
| B1 | B200 | Gender | OP | -0.02806 | 0 |
| B1 | B227 | Gender | OD | -0.00874 | 2.26E-14 |
| B1 | B228 | Gender | OS | -0.00648 | 1.50E-08 |
| B1 | B50 | Gender | eo% | -0.11116 | 0 |
| B1 | B220 | Gender | SBP | -0.27128 | 0 |
| B1 | B219 | Gender | OT | 0.05683 | 0 |
| B1 | B218 | Gender | DBP | -0.27886 | 0 |
| B1 | B88 | Gender | CA 242 | 0.035479 | 0 |
| B1 | B216 | Gender | BMI | -0.61822 | 0 |
| B1 | B215 | Gender | BMI | -0.35301 | 0 |
| B1 | B131 | Gender | AST | -0.15686 | 0 |
| B1 | B3 | Gender | Hcy | -0.12242 | 0 |
| B1 | B172 | Gender | KB | 0.016762 | 0 |
| B1 | B210 | Gender | Hip | -0.3691 | 0 |
| B1 | B10 | Gender | PG I | -0.04914 | 0 |
| B1 | B12 | Gender | PG II | -0.01606 | 0 |
| B1 | B14 | Gender | G-17 | 0.008401 | 2.22E-13 |
| B1 | B209 | Gender | Smoking | -0.3766 | 0 |
| B1 | B90 | Gender | CYFRA 21-1 | -0.03144 | 0 |
| B1 | B139 | Gender | ADA | 0.02563 | 0 |
| B1 | B34 | Gender | TK | 0.010848 | 0 |
| B1 | B196 | Gender | TCO2 | -0.00938 | 2.22E-16 |
| B1 | B51 | Gender | Hb | -0.68896 | 0 |
| B1 | B185 | Gender | PT | 0.007786 | 1.06E-11 |
| B1 | B69 | Gender | PV | 0.006374 | 2.62E-08 |
| B1 | B52 | Gender | PLT/BPC | 0.095249 | 0 |
| B1 | B173 | Gender | NTT | 0.066106 | 0 |
| B1 | B207 | Gender | WHR | -0.06406 | 0 |
| B1 | B206 | Gender | Waist | -0.54132 | 0 |
| B1 | B205 | Gender | KDA | 0.052062 | 0 |
| B1 | B78 | Gender | HBcAb | 0.01236 | 0 |

| | | | | | |
|---|---|---|---|---|---|
| B1 | B197 | Gender | AG | -0.01879 | 0 |
| B1 | B204 | Gender | Drinking | -0.40366 | 0 |
| B1 | B203 | Gender | DH | -0.01991 | 0 |
| B1 | B80 | Gender | anti-HP IgG | -0.00762 | 2.96E-11 |
| B1 | B112 | Gender | FT4 | -0.02146 | 0 |
| B1 | B113 | Gender | FT3 | -0.09289 | 0 |
| B1 | B211 | Gender | IOP-R | -0.01424 | 0 |
| B1 | B201 | Gender | e-habits | -0.04823 | 0 |
| B1 | B73 | Gender | ApoA | 0.023537 | 0 |
| B1 | B74 | Gender | ApoA1 | 0.103329 | 0 |
| B1 | B75 | Gender | ApoA1/B | 0.054433 | 0 |
| B1 | B76 | Gender | ApoB | -0.05962 | 0 |
| B1 | B132 | Gender | Dbil | -0.10135 | 0 |
| B1 | B53 | Gender | GR% | 0.010359 | 0 |
| B1 | B77 | Gender | TC | -0.05594 | 0 |
| B1 | B133 | Gender | TBIL | -0.14723 | 0 |
| B1 | B135 | Gender | TBA | -0.0177 | 0 |
| B1 | B91 | Gender | TP | -0.00627 | 4.41E-08 |
| B1 | B114 | Gender | TSH TT3 TT4(TT4) | 0.025425 | 0 |
| B1 | B115 | Gender | TSH TT3 TT4(TT3) | -0.02896 | 0 |
| B1 | B212 | Gender | IOP-L | -0.01604 | 0 |
| B10 | B137 | PG I | AFU | 0.010186 | 0 |
| B10 | B24 | PG I | αHBDH | 0.017946 | 0 |
| B10 | B15 | PG I | CD3-CD19+(B cell) | -0.05491 | 0 |
| B10 | B18 | PG I | CD3+CD4+helper(inducer)T | 0.010841 | 0 |
| B10 | B19 | PG I | CD3+CD4+/CD3+CD8+ | 0.018875 | 0 |
| B10 | B20 | PG I | CD3+CD4+CD8+ | -0.00592 | 2.32E-07 |
| B10 | B21 | PG I | CD3+CD4-CD8- | 0.006365 | 2.74E-08 |
| B10 | B22 | PG I | CD3+CD8+(T) | -0.01218 | 0 |
| B10 | B92 | PG I | ALDH2-Glu504Glu | -0.0134 | 0 |
| B10 | B93 | PG I | ALDH2-Glu504Lys | 0.010022 | 0 |
| B10 | B160 | PG I | PH | -0.02943 | 0 |
| B10 | B94 | PG I | ALDH2-Lys504Lys | 0.010844 | 0 |
| B10 | B118 | PG I | CEA1 | 0.005876 | 2.90E-07 |
| B10 | B126 | PG I | Alb | -0.01971 | 0 |
| B10 | B40 | PG I | WBC | 0.019164 | 0 |
| B10 | B128 | PG I | ALT | 0.009124 | 1.78E-15 |
| B10 | B225 | PG I | hPBG | -0.00719 | 3.45E-10 |
| B10 | B123 | PG I | hs-CRP1 | 0.006447 | 1.82E-08 |
| B10 | B124 | PG I | ACTH | 0.015006 | 0 |
| B10 | B70 | PG I | LDL-C | 0.031616 | 0 |
| B10 | B71 | PG I | TG | 0.010351 | 0 |
| B10 | B189 | PG I | eGFB | -0.08535 | 0 |
| B10 | B141 | PG I | AST/ALT | -0.00774 | 1.36E-11 |

| | | | | | |
|---|---|---|---|---|---|
| B10 | B146 | PG I | CysC | 0.014413 | 0 |
| B10 | B57 | PG I | RBCAI | 0.011771 | 0 |
| B10 | B43 | PG I | MCV | 0.012397 | 0 |
| B10 | B44 | PG I | MCH | 0.018991 | 0 |
| B10 | B46 | PG I | RBC Count | 0.011124 | 0 |
| B10 | B47 | PG I | RDW | 0.035438 | 0 |
| B10 | B181 | PG I | APTT | -0.02577 | 0 |
| B10 | B190 | PG I | Cre | 0.063659 | 0 |
| B10 | B193 | PG I | K | 0.006141 | 8.26E-08 |
| B10 | B129 | PG I | IBIL | -0.0197 | 0 |
| B10 | B150 | PG I | ALP | 0.021992 | 0 |
| B10 | B84 | PG I | C-PSA | 0.008417 | 2.01E-13 |
| B10 | B222 | PG I | NS | 0.022326 | 0 |
| B10 | B100 | PG I | anti-Ro52 | -0.00932 | 4.44E-16 |
| B10 | B151 | PG I | Anti-CCP | 0.007438 | 8.39E-11 |
| B10 | B148 | PG I | Anti_TPO | -0.00902 | 3.55E-15 |
| B10 | B149 | PG I | TGAb | -0.00754 | 4.61E-11 |
| B10 | B194 | PG I | Cl | -0.00593 | 2.24E-07 |
| B10 | B199 | PG I | Mg | 0.012483 | 0 |
| B10 | B155 | PG I | IgA | -0.00851 | 1.10E-13 |
| B10 | B195 | PG I | Na | 0.020531 | 0 |
| B10 | B176 | PG I | Ucr | 0.012771 | 0 |
| B10 | B191 | PG I | Urea | 0.039168 | 0 |
| B10 | B174 | PG I | Urea/Cr | -0.01456 | 0 |
| B10 | B192 | PG I | UA | 0.026381 | 0 |
| B10 | B175 | PG I | ALB-U | 0.008551 | 8.30E-14 |
| B10 | B177 | PG I | ACR | 0.008491 | 1.23E-13 |
| B10 | B179 | PG I | α1-MG | 0.00608 | 1.11E-07 |
| B10 | B125 | PG I | Cor | 0.02443 | 0 |
| B10 | B138 | PG I | PAB | 0.02251 | 0 |
| B10 | B130 | PG I | GLB | -0.00694 | 1.36E-09 |
| B10 | B59 | PG I | WBRV | 0.010411 | 0 |
| B10 | B60 | PG I | WBRVI | 0.012409 | 0 |
| B10 | B64 | PG I | WBRSR(mPas_5) | 0.009995 | 0 |
| B10 | B65 | PG I | WBRSR(mPas_1) | 0.01276 | 0 |
| B10 | B27 | PG I | LDH | 0.012259 | 0 |
| B10 | B221 | PG I | Hei | 0.014505 | 0 |
| B10 | B87 | PG I | NSE | -0.01231 | 0 |
| B10 | B200 | PG I | OP | 0.021204 | 0 |
| B10 | B50 | PG I | eo% | 0.012345 | 0 |
| B10 | B220 | PG I | SBP | 0.022396 | 0 |
| B10 | B219 | PG I | OT | 0.005827 | 3.63E-07 |
| B10 | B218 | PG I | DBP | 0.022265 | 0 |
| B10 | B216 | PG I | BMI | 0.009571 | 0 |

| | | | | | |
|---|---|---|---|---|---|
| B10 | B131 | PG I | AST | 0.01561 | 0 |
| B10 | B12 | PG I | PG II | 0.628138 | 0 |
| B10 | B14 | PG I | G-17 | 0.085572 | 0 |
| B10 | B209 | PG I | Smoking | 0.056463 | 0 |
| B10 | B90 | PG I | CYFRA 21-1 | 0.016941 | 0 |
| B10 | B196 | PG I | TCO2 | 0.011114 | 0 |
| B10 | B51 | PG I | Hb | 0.02663 | 0 |
| B10 | B183 | PG I | PTA | -0.03943 | 0 |
| B10 | B185 | PG I | PT | -0.00743 | 8.92E-11 |
| B10 | B186 | PG I | PFC | 0.020994 | 0 |
| B10 | B52 | PG I | PLT/BPC | -0.02383 | 0 |
| B10 | B206 | PG I | Waist | 0.017401 | 0 |
| B10 | B78 | PG I | HBcAb | -0.01705 | 0 |
| B10 | B197 | PG I | AG | 0.007119 | 5.12E-10 |
| B10 | B204 | PG I | Drinking | 0.027011 | 0 |
| B10 | B80 | PG I | anti-HP IgG | 0.091035 | 0 |
| B10 | B112 | PG I | FT4 | 0.008042 | 2.20E-12 |
| B10 | B113 | PG I | FT3 | 0.014429 | 0 |
| B10 | B201 | PG I | e-habits | 0.005969 | 1.88E-07 |
| B10 | B74 | PG I | ApoA1 | 0.022909 | 0 |
| B10 | B76 | PG I | ApoB | 0.011415 | 0 |
| B10 | B132 | PG I | Dbil | -0.00683 | 2.47E-09 |
| B10 | B81 | PG I | TSGF | -0.01297 | 0 |
| B10 | B77 | PG I | TC | 0.024963 | 0 |
| B10 | B133 | PG I | TBIL | -0.01767 | 0 |
| B10 | B135 | PG I | TBA | 0.006542 | 1.12E-08 |
| B10 | B91 | PG I | TP | -0.01812 | 0 |
| B10 | B114 | PG I | TSH TT3 TT4(TT4) | 0.007412 | 9.74E-11 |
| B100 | B137 | anti-Ro52 | AFU | 0.010159 | 0 |
| B100 | B126 | anti-Ro52 | Alb | -0.00581 | 3.85E-07 |
| B100 | B121 | anti-Ro52 | C3 | -0.01934 | 0 |
| B100 | B122 | anti-Ro52 | C4 | -0.01554 | 0 |
| B100 | B124 | anti-Ro52 | ACTH | -0.00644 | 1.89E-08 |
| B100 | B181 | anti-Ro52 | APTT | 0.006038 | 1.35E-07 |
| B100 | B105 | anti-Ro52 | anti-U1RNP | 0.068691 | 0 |
| B100 | B101 | anti-Ro52 | anti-Sc70 | 0.016829 | 0 |
| B100 | B102 | anti-Ro52 | anti-Sm | 0.081385 | 0 |
| B100 | B103 | anti-Ro52 | anti-SSA/Ro | 0.304624 | 0 |
| B100 | B104 | anti-Ro52 | anti-SSB | 0.187733 | 0 |
| B100 | B106 | anti-Ro52 | anti-P | 0.051897 | 0 |
| B100 | B107 | anti-Ro52 | AnuA | 0.062463 | 0 |
| B100 | B151 | anti-Ro52 | Anti-CCP | 0.040999 | 0 |
| B100 | B149 | anti-Ro52 | TGAb | 0.006771 | 3.40E-09 |
| B100 | B108 | anti-Ro52 | Anti-Cyclin | 0.008594 | 6.26E-14 |

| | | | | | |
|---|---|---|---|---|---|
| B100 | B109 | anti-Ro52 | AMA-M2 | 0.068363 | 0 |
| B100 | B110 | anti-Ro52 | AHA | 0.013377 | 0 |
| B100 | B154 | anti-Ro52 | RF | 0.021169 | 0 |
| B100 | B194 | anti-Ro52 | Cl | 0.006234 | 5.26E-08 |
| B100 | B155 | anti-Ro52 | IgA | 0.032978 | 0 |
| B100 | B157 | anti-Ro52 | IgG | 0.063538 | 0 |
| B100 | B158 | anti-Ro52 | IgM | 0.01349 | 0 |
| B100 | B125 | anti-Ro52 | Cor | -0.00772 | 1.54E-11 |
| B100 | B138 | anti-Ro52 | PAB | -0.02044 | 0 |
| B100 | B130 | anti-Ro52 | GLB | 0.01169 | 0 |
| B100 | B139 | anti-Ro52 | ADA | 0.028859 | 0 |
| B100 | B183 | anti-Ro52 | PTA | 0.007659 | 2.29E-11 |
| B100 | B114 | anti-Ro52 | TSH TT3 TT4(TT4) | 0.011082 | 0 |
| B100 | B115 | anti-Ro52 | TSH TT3 TT4(TT3) | 0.005664 | 7.62E-07 |
| B101 | B193 | anti-Sc70 | K | 0.007466 | 7.12E-11 |
| B101 | B105 | anti-Sc70 | anti-U1RNP | 0.066968 | 0 |
| B101 | B103 | anti-Sc70 | anti-SSA/Ro | 0.023016 | 0 |
| B101 | B104 | anti-Sc70 | anti-SSB | 0.061081 | 0 |
| B101 | B108 | anti-Sc70 | Anti-Cyclin | 0.037171 | 0 |
| B101 | B109 | anti-Sc70 | AMA-M2 | 0.020519 | 0 |
| B101 | B110 | anti-Sc70 | AHA | -0.00718 | 3.67E-10 |
| B101 | B156 | anti-Sc70 | IgE | -0.00629 | 3.96E-08 |
| B101 | B157 | anti-Sc70 | IgG | 0.01654 | 0 |
| B101 | B179 | anti-Sc70 | α1-MG | -0.00657 | 9.71E-09 |
| B101 | B138 | anti-Sc70 | PAB | -0.00849 | 1.23E-13 |
| B102 | B137 | anti-Sm | AFU | -0.01489 | 0 |
| B102 | B127 | anti-Sm | A-G ratio | 0.015283 | 0 |
| B102 | B121 | anti-Sm | C3 | -0.01543 | 0 |
| B102 | B105 | anti-Sm | anti-U1RNP | 0.286285 | 0 |
| B102 | B103 | anti-Sm | anti-SSA/Ro | 0.072992 | 0 |
| B102 | B104 | anti-Sm | anti-SSB | 0.066566 | 0 |
| B102 | B106 | anti-Sm | anti-P | 0.041932 | 0 |
| B102 | B109 | anti-Sm | AMA-M2 | 0.019808 | 0 |
| B102 | B110 | anti-Sm | AHA | -0.00717 | 3.80E-10 |
| B102 | B194 | anti-Sm | Cl | -0.006 | 1.61E-07 |
| B102 | B199 | anti-Sm | Mg | 0.007209 | 3.09E-10 |
| B102 | B176 | anti-Sm | Ucr | 0.008071 | 1.83E-12 |
| B102 | B125 | anti-Sm | Cor | 0.018239 | 0 |
| B102 | B139 | anti-Sm | ADA | 0.010846 | 0 |
| B102 | B186 | anti-Sm | PFC | 0.00748 | 6.55E-11 |
| B103 | B126 | anti-SSA/Ro | Alb | -0.00704 | 8.06E-10 |
| B103 | B127 | anti-SSA/Ro | A-G ratio | -0.00704 | 8.02E-10 |
| B103 | B121 | anti-SSA/Ro | C3 | -0.01517 | 0 |
| B103 | B122 | anti-SSA/Ro | C4 | -0.01289 | 0 |

| | | | | | |
|---|---|---|---|---|---|
| B103 | B124 | anti-SSA/Ro | ACTH | -0.00751 | 5.45E-11 |
| B103 | B134 | anti-SSA/Ro | CHE | -0.00998 | 0 |
| B103 | B181 | anti-SSA/Ro | APTT | 0.012879 | 0 |
| B103 | B222 | anti-SSA/Ro | NS | -0.00681 | 2.73E-09 |
| B103 | B105 | anti-SSA/Ro | anti-U1RNP | 0.057651 | 0 |
| B103 | B104 | anti-SSA/Ro | anti-SSB | 0.22971 | 0 |
| B103 | B106 | anti-SSA/Ro | anti-P | 0.091291 | 0 |
| B103 | B107 | anti-SSA/Ro | AnuA | 0.090977 | 0 |
| B103 | B151 | anti-SSA/Ro | Anti-CCP | 0.050766 | 0 |
| B103 | B149 | anti-SSA/Ro | TGAb | 0.00767 | 2.14E-11 |
| B103 | B109 | anti-SSA/Ro | AMA-M2 | 0.040678 | 0 |
| B103 | B110 | anti-SSA/Ro | AHA | 0.007172 | 3.81E-10 |
| B103 | B154 | anti-SSA/Ro | RF | 0.011708 | 0 |
| B103 | B199 | anti-SSA/Ro | Mg | -0.00687 | 2.03E-09 |
| B103 | B155 | anti-SSA/Ro | IgA | 0.037388 | 0 |
| B103 | B157 | anti-SSA/Ro | IgG | 0.080767 | 0 |
| B103 | B158 | anti-SSA/Ro | IgM | 0.014193 | 0 |
| B103 | B195 | anti-SSA/Ro | Na | -0.01097 | 0 |
| B103 | B176 | anti-SSA/Ro | Ucr | -0.01038 | 0 |
| B103 | B192 | anti-SSA/Ro | UA | -0.00584 | 3.36E-07 |
| B103 | B138 | anti-SSA/Ro | PAB | -0.01967 | 0 |
| B103 | B130 | anti-SSA/Ro | GLB | 0.014644 | 0 |
| B103 | B221 | anti-SSA/Ro | Hei | -0.00663 | 7.15E-09 |
| B103 | B200 | anti-SSA/Ro | OP | -0.01133 | 0 |
| B103 | B139 | anti-SSA/Ro | ADA | 0.019352 | 0 |
| B103 | B185 | anti-SSA/Ro | PT | 0.00852 | 1.02E-13 |
| B103 | B197 | anti-SSA/Ro | AG | -0.01103 | 0 |
| B103 | B211 | anti-SSA/Ro | IOP-R | -0.00641 | 2.21E-08 |
| B103 | B114 | anti-SSA/Ro | TSH TT3 TT4(TT4) | 0.00721 | 3.08E-10 |
| B103 | B212 | anti-SSA/Ro | IOP-L | -0.00669 | 5.26E-09 |
| B104 | B121 | anti-SSB | C3 | -0.02096 | 0 |
| B104 | B122 | anti-SSB | C4 | -0.01066 | 0 |
| B104 | B124 | anti-SSB | ACTH | -0.00668 | 5.57E-09 |
| B104 | B181 | anti-SSB | APTT | 0.011252 | 0 |
| B104 | B105 | anti-SSB | anti-U1RNP | 0.073403 | 0 |
| B104 | B106 | anti-SSB | anti-P | 0.028919 | 0 |
| B104 | B107 | anti-SSB | AnuA | 0.045562 | 0 |
| B104 | B151 | anti-SSB | Anti-CCP | -0.00788 | 5.89E-12 |
| B104 | B108 | anti-SSB | Anti-Cyclin | -0.00587 | 2.98E-07 |
| B104 | B109 | anti-SSB | AMA-M2 | 0.056055 | 0 |
| B104 | B155 | anti-SSB | IgA | 0.018072 | 0 |
| B104 | B156 | anti-SSB | IgE | 0.009132 | 1.55E-15 |
| B104 | B157 | anti-SSB | IgG | 0.039856 | 0 |
| B104 | B179 | anti-SSB | α1-MG | -0.00642 | 2.10E-08 |

| | | | | | |
|---|---|---|---|---|---|
| B104 | B182 | anti-SSB | TT | 0.009376 | 2.22E-16 |
| B104 | B125 | anti-SSB | Cor | -0.01473 | 0 |
| B104 | B138 | anti-SSB | PAB | -0.00711 | 5.39E-10 |
| B104 | B130 | anti-SSB | GLB | 0.006943 | 1.35E-09 |
| B104 | B200 | anti-SSB | OP | -0.00738 | 1.20E-10 |
| B104 | B139 | anti-SSB | ADA | 0.00685 | 2.22E-09 |
| B104 | B196 | anti-SSB | TCO2 | -0.00599 | 1.67E-07 |
| B105 | B137 | anti-U1RNP | AFU | -0.00825 | 5.95E-13 |
| B105 | B127 | anti-U1RNP | A-G ratio | 0.006534 | 1.16E-08 |
| B105 | B121 | anti-U1RNP | C3 | -0.01244 | 0 |
| B105 | B122 | anti-U1RNP | C4 | -0.00959 | 0 |
| B105 | B111 | anti-U1RNP | TSH | 0.01013 | 0 |
| B105 | B193 | anti-U1RNP | K | 0.005837 | 3.46E-07 |
| B105 | B106 | anti-U1RNP | anti-P | 0.082975 | 0 |
| B105 | B107 | anti-U1RNP | AnuA | 0.118194 | 0 |
| B105 | B108 | anti-U1RNP | Anti-Cyclin | -0.00767 | 2.13E-11 |
| B105 | B109 | anti-U1RNP | AMA-M2 | 0.027642 | 0 |
| B105 | B110 | anti-U1RNP | AHA | 0.024614 | 0 |
| B105 | B155 | anti-U1RNP | IgA | 0.012617 | 0 |
| B105 | B156 | anti-U1RNP | IgE | -0.00645 | 1.75E-08 |
| B105 | B157 | anti-U1RNP | IgG | 0.026022 | 0 |
| B105 | B182 | anti-U1RNP | TT | 0.006256 | 4.72E-08 |
| B105 | B139 | anti-U1RNP | ADA | 0.010525 | 0 |
| B105 | B196 | anti-U1RNP | TCO2 | -0.006 | 1.64E-07 |
| B105 | B113 | anti-U1RNP | FT3 | 0.006363 | 2.77E-08 |
| B106 | B121 | anti-P | C3 | -0.0133 | 0 |
| B106 | B122 | anti-P | C4 | -0.01544 | 0 |
| B106 | B107 | anti-P | AnuA | 0.219492 | 0 |
| B106 | B108 | anti-P | Anti-Cyclin | -0.00581 | 4.00E-07 |
| B106 | B109 | anti-P | AMA-M2 | -0.00594 | 2.17E-07 |
| B106 | B110 | anti-P | AHA | 0.015638 | 0 |
| B106 | B198 | anti-P | P | -0.006 | 1.59E-07 |
| B106 | B194 | anti-P | Cl | -0.00838 | 2.61E-13 |
| B106 | B155 | anti-P | IgA | 0.006514 | 1.29E-08 |
| B106 | B156 | anti-P | IgE | 0.011679 | 0 |
| B106 | B157 | anti-P | IgG | 0.044119 | 0 |
| B106 | B195 | anti-P | Na | -0.00811 | 1.44E-12 |
| B106 | B182 | anti-P | TT | -0.01512 | 0 |
| B106 | B125 | anti-P | Cor | -0.0111 | 0 |
| B106 | B200 | anti-P | OP | -0.00578 | 4.45E-07 |
| B106 | B196 | anti-P | TCO2 | 0.008402 | 2.22E-13 |
| B106 | B186 | anti-P | PFC | 0.007512 | 5.43E-11 |
| B106 | B197 | anti-P | AG | -0.00592 | 2.37E-07 |
| B107 | B127 | AnuA | A-G ratio | -0.00624 | 5.04E-08 |

| | | | | | |
|---|---|---|---|---|---|
| B107 | B121 | AnuA | C3 | -0.02772 | 0 |
| B107 | B122 | AnuA | C4 | -0.03198 | 0 |
| B107 | B151 | AnuA | Anti-CCP | 0.013995 | 0 |
| B107 | B149 | AnuA | TGAb | 0.007849 | 7.25E-12 |
| B107 | B109 | AnuA | AMA-M2 | 0.019017 | 0 |
| B107 | B110 | AnuA | AHA | 0.078715 | 0 |
| B107 | B157 | AnuA | IgG | 0.072514 | 0 |
| B107 | B130 | AnuA | GLB | 0.009929 | 0 |
| B108 | B181 | Anti-Cyclin | APTT | 0.006197 | 6.30E-08 |
| B108 | B151 | Anti-Cyclin | Anti-CCP | 0.01202 | 0 |
| B108 | B109 | Anti-Cyclin | AMA-M2 | 0.056451 | 0 |
| B108 | B110 | Anti-Cyclin | AHA | 0.015779 | 0 |
| B108 | B198 | Anti-Cyclin | P | -0.00684 | 2.40E-09 |
| B108 | B195 | Anti-Cyclin | Na | -0.00588 | 2.88E-07 |
| B108 | B179 | Anti-Cyclin | α1-MG | -0.00695 | 1.30E-09 |
| B108 | B125 | Anti-Cyclin | Cor | 0.010466 | 0 |
| B108 | B228 | Anti-Cyclin | OS | 0.005784 | 4.42E-07 |
| B108 | B139 | Anti-Cyclin | ADA | 0.017155 | 0 |
| B109 | B225 | AMA-M2 | hPBG | 0.00827 | 5.20E-13 |
| B109 | B123 | AMA-M2 | hs-CRP1 | 0.018146 | 0 |
| B109 | B181 | AMA-M2 | APTT | -0.00573 | 5.61E-07 |
| B109 | B194 | AMA-M2 | Cl | -0.00857 | 7.11E-14 |
| B109 | B199 | AMA-M2 | Mg | -0.00817 | 9.80E-13 |
| B109 | B157 | AMA-M2 | IgG | 0.021261 | 0 |
| B109 | B158 | AMA-M2 | IgM | 0.00561 | 9.69E-07 |
| B109 | B195 | AMA-M2 | Na | -0.01041 | 0 |
| B109 | B179 | AMA-M2 | α1-MG | -0.00591 | 2.47E-07 |
| B109 | B182 | AMA-M2 | TT | -0.02179 | 0 |
| B109 | B125 | AMA-M2 | Cor | 0.023829 | 0 |
| B109 | B186 | AMA-M2 | PFC | 0.018456 | 0 |
| B11 | B12 | PGI/PGII | PG II | -0.01714 | 0 |
| B11 | B14 | PGI/PGII | G-17 | -0.0057 | 6.50E-07 |
| B110 | B121 | AHA | C3 | 0.007298 | 1.87E-10 |
| B110 | B155 | AHA | IgA | 0.019731 | 0 |
| B110 | B157 | AHA | IgG | 0.023016 | 0 |
| B110 | B158 | AHA | IgM | 0.010773 | 0 |
| B110 | B195 | AHA | Na | -0.00861 | 5.73E-14 |
| B110 | B159 | AHA | β2-MG | 0.006176 | 6.96E-08 |
| B110 | B176 | AHA | Ucr | -0.01037 | 0 |
| B110 | B179 | AHA | α1-MG | -0.0172 | 0 |
| B110 | B182 | AHA | TT | -0.00655 | 1.09E-08 |
| B110 | B138 | AHA | PAB | -0.01122 | 0 |
| B110 | B200 | AHA | OP | -0.00619 | 6.59E-08 |
| B110 | B196 | AHA | TCO2 | -0.01051 | 0 |

| | | | | | |
|---|---|---|---|---|---|
| B110 | B183 | AHA | PTA | 0.011047 | 0 |
| B110 | B112 | AHA | FT4 | 0.006071 | 1.16E-07 |
| B111 | B137 | TSH | AFU | -0.00726 | 2.38E-10 |
| B111 | B160 | TSH | PH | -0.00795 | 4.00E-12 |
| B111 | B140 | TSH | r-GT | -0.00744 | 8.24E-11 |
| B111 | B127 | TSH | A-G ratio | -0.02812 | 0 |
| B111 | B161 | TSH | LEU | 0.014382 | 0 |
| B111 | B134 | TSH | CHE | -0.01449 | 0 |
| B111 | B166 | TSH | Pro | -0.01495 | 0 |
| B111 | B189 | TSH | eGFB | -0.03173 | 0 |
| B111 | B141 | TSH | AST/ALT | 0.017824 | 0 |
| B111 | B190 | TSH | Cre | -0.00868 | 3.60E-14 |
| B111 | B193 | TSH | K | -0.00668 | 5.43E-09 |
| B111 | B129 | TSH | IBIL | -0.018 | 0 |
| B111 | B222 | TSH | NS | -0.02494 | 0 |
| B111 | B148 | TSH | Anti_TPO | 0.162953 | 0 |
| B111 | B149 | TSH | TGAb | 0.129817 | 0 |
| B111 | B198 | TSH | P | 0.014221 | 0 |
| B111 | B157 | TSH | IgG | 0.012499 | 0 |
| B111 | B176 | TSH | Ucr | -0.02336 | 0 |
| B111 | B174 | TSH | Urea/Cr | 0.009609 | 0 |
| B111 | B192 | TSH | UA | -0.02139 | 0 |
| B111 | B177 | TSH | ACR | 0.006254 | 4.75E-08 |
| B111 | B179 | TSH | α1-MG | -0.00658 | 9.10E-09 |
| B111 | B125 | TSH | Cor | 0.007793 | 1.02E-11 |
| B111 | B130 | TSH | GLB | 0.037154 | 0 |
| B111 | B221 | TSH | Hei | -0.04766 | 0 |
| B111 | B219 | TSH | OT | 0.009276 | 4.44E-16 |
| B111 | B216 | TSH | BMI | -0.01884 | 0 |
| B111 | B215 | TSH | BMI | 0.009579 | 0 |
| B111 | B131 | TSH | AST | 0.0116 | 0 |
| B111 | B172 | TSH | KB | -0.00932 | 4.44E-16 |
| B111 | B209 | TSH | Smoking | -0.03073 | 0 |
| B111 | B139 | TSH | ADA | 0.008606 | 5.77E-14 |
| B111 | B173 | TSH | NTT | 0.01005 | 0 |
| B111 | B204 | TSH | Drinking | -0.02292 | 0 |
| B111 | B112 | TSH | FT4 | -0.19912 | 0 |
| B111 | B113 | TSH | FT3 | -0.08867 | 0 |
| B111 | B132 | TSH | Dbil | -0.01877 | 0 |
| B111 | B133 | TSH | TBIL | -0.01944 | 0 |
| B111 | B135 | TSH | TBA | 0.005614 | 9.51E-07 |
| B111 | B114 | TSH | TSH TT3 TT4(TT4) | -0.10074 | 0 |
| B111 | B115 | TSH | TSH TT3 TT4(TT3) | -0.02769 | 0 |
| B112 | B140 | FT4 | r-GT | -0.01611 | 0 |

| | | | | | |
|---|---|---|---|---|---|
| B112 | B143 | FT4 | β-CTx | 0.014658 | 0 |
| B112 | B126 | FT4 | Alb | 0.05703 | 0 |
| B112 | B127 | FT4 | A-G ratio | 0.020847 | 0 |
| B112 | B128 | FT4 | ALT | 0.00599 | 1.70E-07 |
| B112 | B123 | FT4 | hs-CRP1 | 0.01995 | 0 |
| B112 | B124 | FT4 | ACTH | 0.006072 | 1.15E-07 |
| B112 | B134 | FT4 | CHE | 0.005762 | 4.89E-07 |
| B112 | B166 | FT4 | Pro | 0.024457 | 0 |
| B112 | B188 | FT4 | Ca | 0.012081 | 0 |
| B112 | B189 | FT4 | eGFB | 0.032014 | 0 |
| B112 | B141 | FT4 | AST/ALT | 0.009844 | 0 |
| B112 | B145 | FT4 | BGP | 0.013181 | 0 |
| B112 | B193 | FT4 | K | -0.00699 | 1.05E-09 |
| B112 | B129 | FT4 | IBIL | 0.05785 | 0 |
| B112 | B150 | FT4 | ALP | 0.012841 | 0 |
| B112 | B222 | FT4 | NS | -0.01936 | 0 |
| B112 | B151 | FT4 | Anti-CCP | 0.009364 | 4.44E-16 |
| B112 | B198 | FT4 | P | 0.010652 | 0 |
| B112 | B194 | FT4 | Cl | -0.01474 | 0 |
| B112 | B199 | FT4 | Mg | -0.01066 | 0 |
| B112 | B155 | FT4 | IgA | 0.006083 | 1.09E-07 |
| B112 | B157 | FT4 | IgG | 0.0109 | 0 |
| B112 | B195 | FT4 | Na | -0.01303 | 0 |
| B112 | B176 | FT4 | Ucr | 0.021399 | 0 |
| B112 | B191 | FT4 | Urea | -0.01024 | 0 |
| B112 | B170 | FT4 | U-GLU | 0.009164 | 1.33E-15 |
| B112 | B182 | FT4 | TT | -0.01947 | 0 |
| B112 | B125 | FT4 | Cor | 0.019372 | 0 |
| B112 | B138 | FT4 | PAB | -0.02082 | 0 |
| B112 | B221 | FT4 | Hei | 0.026648 | 0 |
| B112 | B200 | FT4 | OP | -0.01135 | 0 |
| B112 | B220 | FT4 | SBP | 0.013427 | 0 |
| B112 | B218 | FT4 | DBP | 0.019777 | 0 |
| B112 | B216 | FT4 | BMI | -0.03137 | 0 |
| B112 | B215 | FT4 | BMI | -0.06166 | 0 |
| B112 | B131 | FT4 | AST | 0.011017 | 0 |
| B112 | B172 | FT4 | KB | 0.027179 | 0 |
| B112 | B210 | FT4 | Hip | -0.03318 | 0 |
| B112 | B139 | FT4 | ADA | 0.012418 | 0 |
| B112 | B196 | FT4 | TCO2 | 0.007455 | 7.56E-11 |
| B112 | B183 | FT4 | PTA | 0.006364 | 2.75E-08 |
| B112 | B185 | FT4 | PT | 0.007847 | 7.34E-12 |
| B112 | B186 | FT4 | PFC | 0.019034 | 0 |
| B112 | B206 | FT4 | Waist | -0.04204 | 0 |

| | | | | | |
|---|---|---|---|---|---|
| B112 | B204 | FT4 | Drinking | -0.0097 | 0 |
| B112 | B113 | FT4 | FT3 | 0.459532 | 0 |
| B112 | B211 | FT4 | IOP-R | 0.023889 | 0 |
| B112 | B201 | FT4 | e-habits | -0.00885 | 1.09E-14 |
| B112 | B132 | FT4 | Dbil | 0.049252 | 0 |
| B112 | B117 | FT4 | TT4 | 0.013294 | 0 |
| B112 | B133 | FT4 | TBIL | 0.059506 | 0 |
| B112 | B114 | FT4 | TSH TT3 TT4(TT4) | 0.452302 | 0 |
| B112 | B115 | FT4 | TSH TT3 TT4(TT3) | 0.145866 | 0 |
| B112 | B144 | FT4 | Total-PINP | 0.01184 | 0 |
| B112 | B212 | FT4 | IOP-L | 0.023905 | 0 |
| B113 | B137 | FT3 | AFU | 0.025027 | 0 |
| B113 | B160 | FT3 | PH | -0.03034 | 0 |
| B113 | B140 | FT3 | r-GT | 0.045041 | 0 |
| B113 | B116 | FT3 | TT3 | 0.00798 | 3.23E-12 |
| B113 | B143 | FT3 | β-CTx | 0.016132 | 0 |
| B113 | B126 | FT3 | Alb | 0.060354 | 0 |
| B113 | B127 | FT3 | A-G ratio | 0.021817 | 0 |
| B113 | B161 | FT3 | LEU | -0.0113 | 0 |
| B113 | B128 | FT3 | ALT | 0.067008 | 0 |
| B113 | B123 | FT3 | hs-CRP1 | -0.00864 | 4.66E-14 |
| B113 | B124 | FT3 | ACTH | 0.009186 | 1.11E-15 |
| B113 | B134 | FT3 | CHE | 0.039874 | 0 |
| B113 | B166 | FT3 | Pro | 0.024891 | 0 |
| B113 | B188 | FT3 | Ca | 0.021535 | 0 |
| B113 | B189 | FT3 | eGFB | 0.036524 | 0 |
| B113 | B141 | FT3 | AST/ALT | -0.06346 | 0 |
| B113 | B145 | FT3 | BGP | 0.018556 | 0 |
| B113 | B146 | FT3 | CysC | 0.011272 | 0 |
| B113 | B181 | FT3 | APTT | -0.00717 | 3.80E-10 |
| B113 | B190 | FT3 | Cre | 0.024214 | 0 |
| B113 | B129 | FT3 | IBIL | -0.00789 | 5.58E-12 |
| B113 | B150 | FT3 | ALP | 0.034982 | 0 |
| B113 | B222 | FT3 | NS | 0.067587 | 0 |
| B113 | B148 | FT3 | Anti_TPO | 0.025762 | 0 |
| B113 | B149 | FT3 | TGAb | 0.016269 | 0 |
| B113 | B198 | FT3 | P | 0.00605 | 1.28E-07 |
| B113 | B194 | FT3 | Cl | -0.01603 | 0 |
| B113 | B195 | FT3 | Na | 0.015977 | 0 |
| B113 | B176 | FT3 | Ucr | 0.029288 | 0 |
| B113 | B191 | FT3 | Urea | 0.013373 | 0 |
| B113 | B192 | FT3 | UA | 0.06916 | 0 |
| B113 | B138 | FT3 | PAB | 0.015225 | 0 |
| B113 | B221 | FT3 | Hei | 0.061932 | 0 |

| | | | | | |
|---|---|---|---|---|---|
| B113 | B200 | FT3 | OP | 0.013381 | 0 |
| B113 | B220 | FT3 | SBP | 0.049233 | 0 |
| B113 | B219 | FT3 | OT | -0.00877 | 1.84E-14 |
| B113 | B218 | FT3 | DBP | 0.049017 | 0 |
| B113 | B216 | FT3 | BMI | 0.060971 | 0 |
| B113 | B215 | FT3 | BMI | 0.036898 | 0 |
| B113 | B131 | FT3 | AST | 0.039435 | 0 |
| B113 | B210 | FT3 | Hip | 0.042137 | 0 |
| B113 | B209 | FT3 | Smoking | 0.036192 | 0 |
| B113 | B139 | FT3 | ADA | 0.010447 | 0 |
| B113 | B183 | FT3 | PTA | -0.00987 | 0 |
| B113 | B186 | FT3 | PFC | 0.009477 | 0 |
| B113 | B173 | FT3 | NTT | -0.00843 | 1.81E-13 |
| B113 | B207 | FT3 | WHR | 0.006201 | 6.17E-08 |
| B113 | B206 | FT3 | Waist | 0.055377 | 0 |
| B113 | B205 | FT3 | KDA | -0.00656 | 1.03E-08 |
| B113 | B197 | FT3 | AG | 0.015931 | 0 |
| B113 | B204 | FT3 | Drinking | 0.037958 | 0 |
| B113 | B203 | FT3 | DH | 0.006819 | 2.63E-09 |
| B113 | B132 | FT3 | Dbil | 0.010858 | 0 |
| B113 | B117 | FT3 | TT4 | 0.005668 | 7.47E-07 |
| B113 | B114 | FT3 | TSH TT3 TT4(TT4) | 0.284554 | 0 |
| B113 | B115 | FT3 | TSH TT3 TT4(TT3) | 0.289797 | 0 |
| B113 | B144 | FT3 | Total-PINP | 0.016922 | 0 |
| B114 | B136 | TSH TT3 TT4(TT4) | 5'NT | 0.006893 | 1.76E-09 |
| B114 | B137 | TSH TT3 TT4(TT4) | AFU | 0.005711 | 6.16E-07 |
| B114 | B160 | TSH TT3 TT4(TT4) | PH | -0.00732 | 1.65E-10 |
| B114 | B140 | TSH TT3 TT4(TT4) | r-GT | -0.01465 | 0 |
| B114 | B143 | TSH TT3 TT4(TT4) | β-CTx | 0.005616 | 9.45E-07 |
| B114 | B127 | TSH TT3 TT4(TT4) | A-G ratio | -0.01758 | 0 |
| B114 | B161 | TSH TT3 TT4(TT4) | LEU | 0.010046 | 0 |
| B114 | B128 | TSH TT3 TT4(TT4) | ALT | 0.009947 | 0 |
| B114 | B119 | TSH TT3 TT4(TT4) | Hcv-rna | 0.007783 | 1.08E-11 |
| B114 | B123 | TSH TT3 TT4(TT4) | hs-CRP1 | 0.033665 | 0 |
| B114 | B189 | TSH TT3 TT4(TT4) | eGFB | 0.013817 | 0 |
| B114 | B141 | TSH TT3 TT4(TT4) | AST/ALT | 0.006367 | 2.72E-08 |
| B114 | B145 | TSH TT3 TT4(TT4) | BGP | 0.006877 | 1.92E-09 |
| B114 | B181 | TSH TT3 TT4(TT4) | APTT | 0.006647 | 6.52E-09 |
| B114 | B190 | TSH TT3 TT4(TT4) | Cre | -0.02782 | 0 |
| B114 | B150 | TSH TT3 TT4(TT4) | ALP | 0.027691 | 0 |
| B114 | B222 | TSH TT3 TT4(TT4) | NS | -0.03431 | 0 |
| B114 | B151 | TSH TT3 TT4(TT4) | Anti-CCP | 0.007749 | 1.33E-11 |
| B114 | B148 | TSH TT3 TT4(TT4) | Anti_TPO | 0.015698 | 0 |
| B114 | B198 | TSH TT3 TT4(TT4) | P | 0.017899 | 0 |

| | | | | | |
|---|---|---|---|---|---|
| B114 | B194 | TSH TT3 TT4(TT4) | Cl | -0.01198 | 0 |
| B114 | B199 | TSH TT3 TT4(TT4) | Mg | -0.01019 | 0 |
| B114 | B157 | TSH TT3 TT4(TT4) | IgG | 0.010137 | 0 |
| B114 | B195 | TSH TT3 TT4(TT4) | Na | 0.013731 | 0 |
| B114 | B191 | TSH TT3 TT4(TT4) | Urea | -0.01407 | 0 |
| B114 | B174 | TSH TT3 TT4(TT4) | Urea/Cr | 0.01307 | 0 |
| B114 | B192 | TSH TT3 TT4(TT4) | UA | -0.0267 | 0 |
| B114 | B170 | TSH TT3 TT4(TT4) | U-GLU | 0.006274 | 4.30E-08 |
| B114 | B182 | TSH TT3 TT4(TT4) | TT | -0.00733 | 1.55E-10 |
| B114 | B125 | TSH TT3 TT4(TT4) | Cor | 0.01055 | 0 |
| B114 | B138 | TSH TT3 TT4(TT4) | PAB | -0.06434 | 0 |
| B114 | B130 | TSH TT3 TT4(TT4) | GLB | 0.025453 | 0 |
| B114 | B221 | TSH TT3 TT4(TT4) | Hei | -0.02368 | 0 |
| B114 | B200 | TSH TT3 TT4(TT4) | OP | 0.006326 | 3.34E-08 |
| B114 | B216 | TSH TT3 TT4(TT4) | BMI | -0.03647 | 0 |
| B114 | B215 | TSH TT3 TT4(TT4) | BMI | -0.03163 | 0 |
| B114 | B131 | TSH TT3 TT4(TT4) | AST | 0.016094 | 0 |
| B114 | B172 | TSH TT3 TT4(TT4) | KB | 0.012024 | 0 |
| B114 | B210 | TSH TT3 TT4(TT4) | Hip | -0.02286 | 0 |
| B114 | B139 | TSH TT3 TT4(TT4) | ADA | 0.048708 | 0 |
| B114 | B185 | TSH TT3 TT4(TT4) | PT | 0.012444 | 0 |
| B114 | B186 | TSH TT3 TT4(TT4) | PFC | 0.041179 | 0 |
| B114 | B206 | TSH TT3 TT4(TT4) | Waist | -0.03002 | 0 |
| B114 | B197 | TSH TT3 TT4(TT4) | AG | 0.012921 | 0 |
| B114 | B204 | TSH TT3 TT4(TT4) | Drinking | -0.03478 | 0 |
| B114 | B211 | TSH TT3 TT4(TT4) | IOP-R | -0.00708 | 6.44E-10 |
| B114 | B201 | TSH TT3 TT4(TT4) | e-habits | -0.00691 | 1.60E-09 |
| B114 | B132 | TSH TT3 TT4(TT4) | Dbil | 0.014309 | 0 |
| B114 | B135 | TSH TT3 TT4(TT4) | TBA | 0.007335 | 1.52E-10 |
| B114 | B115 | TSH TT3 TT4(TT4) | TSH TT3 TT4(TT3) | 0.287072 | 0 |
| B114 | B212 | TSH TT3 TT4(TT4) | IOP-L | -0.00669 | 5.17E-09 |
| B115 | B137 | TSH TT3 TT4(TT3) | AFU | 0.023587 | 0 |
| B115 | B160 | TSH TT3 TT4(TT3) | PH | -0.00972 | 0 |
| B115 | B140 | TSH TT3 TT4(TT3) | r-GT | 0.013206 | 0 |
| B115 | B126 | TSH TT3 TT4(TT3) | Alb | 0.019396 | 0 |
| B115 | B128 | TSH TT3 TT4(TT3) | ALT | 0.028135 | 0 |
| B115 | B124 | TSH TT3 TT4(TT3) | ACTH | 0.006834 | 2.42E-09 |
| B115 | B134 | TSH TT3 TT4(TT3) | CHE | 0.036208 | 0 |
| B115 | B166 | TSH TT3 TT4(TT3) | Pro | 0.007134 | 4.70E-10 |
| B115 | B188 | TSH TT3 TT4(TT3) | Ca | 0.009319 | 4.44E-16 |
| B115 | B189 | TSH TT3 TT4(TT3) | eGFB | 0.01181 | 0 |
| B115 | B141 | TSH TT3 TT4(TT3) | AST/ALT | -0.02305 | 0 |
| B115 | B146 | TSH TT3 TT4(TT3) | CysC | 0.007701 | 1.78E-11 |
| B115 | B129 | TSH TT3 TT4(TT3) | IBIL | -0.00601 | 1.55E-07 |

| | | | | | |
|---|---|---|---|---|---|
| B115 | B150 | TSH TT3 TT4(TT3) | ALP | 0.026335 | 0 |
| B115 | B222 | TSH TT3 TT4(TT3) | NS | 0.021737 | 0 |
| B115 | B148 | TSH TT3 TT4(TT3) | Anti_TPO | 0.008556 | 8.04E-14 |
| B115 | B194 | TSH TT3 TT4(TT3) | Cl | -0.00692 | 1.50E-09 |
| B115 | B195 | TSH TT3 TT4(TT3) | Na | 0.012707 | 0 |
| B115 | B192 | TSH TT3 TT4(TT3) | UA | 0.019707 | 0 |
| B115 | B182 | TSH TT3 TT4(TT3) | TT | -0.00616 | 7.45E-08 |
| B115 | B221 | TSH TT3 TT4(TT3) | Hei | 0.015576 | 0 |
| B115 | B200 | TSH TT3 TT4(TT3) | OP | 0.011125 | 0 |
| B115 | B220 | TSH TT3 TT4(TT3) | SBP | 0.017474 | 0 |
| B115 | B218 | TSH TT3 TT4(TT3) | DBP | 0.017796 | 0 |
| B115 | B216 | TSH TT3 TT4(TT3) | BMI | 0.021331 | 0 |
| B115 | B215 | TSH TT3 TT4(TT3) | BMI | 0.017463 | 0 |
| B115 | B131 | TSH TT3 TT4(TT3) | AST | 0.018775 | 0 |
| B115 | B210 | TSH TT3 TT4(TT3) | Hip | 0.01743 | 0 |
| B115 | B209 | TSH TT3 TT4(TT3) | Smoking | 0.015248 | 0 |
| B115 | B139 | TSH TT3 TT4(TT3) | ADA | 0.01792 | 0 |
| B115 | B183 | TSH TT3 TT4(TT3) | PTA | -0.00697 | 1.14E-09 |
| B115 | B186 | TSH TT3 TT4(TT3) | PFC | 0.011657 | 0 |
| B115 | B206 | TSH TT3 TT4(TT3) | Waist | 0.024623 | 0 |
| B115 | B197 | TSH TT3 TT4(TT3) | AG | 0.010033 | 0 |
| B115 | B204 | TSH TT3 TT4(TT3) | Drinking | 0.00704 | 7.92E-10 |
| B116 | B189 | TT3 | eGFB | 0.006853 | 2.19E-09 |
| B116 | B170 | TT3 | U-GLU | 0.012643 | 0 |
| B116 | B117 | TT3 | TT4 | 0.773598 | 0 |
| B117 | B137 | TT4 | AFU | -0.00623 | 5.42E-08 |
| B117 | B190 | TT4 | Cre | -0.00566 | 7.71E-07 |
| B117 | B170 | TT4 | U-GLU | 0.009446 | 0 |
| B118 | B140 | CEA1 | r-GT | 0.018073 | 0 |
| B118 | B126 | CEA1 | Alb | -0.01669 | 0 |
| B118 | B127 | CEA1 | A-G ratio | -0.00607 | 1.18E-07 |
| B118 | B128 | CEA1 | ALT | 0.005881 | 2.83E-07 |
| B118 | B123 | CEA1 | hs-CRP1 | 0.030245 | 0 |
| B118 | B166 | CEA1 | Pro | 0.00688 | 1.90E-09 |
| B118 | B189 | CEA1 | eGFB | -0.03024 | 0 |
| B118 | B146 | CEA1 | CysC | 0.00633 | 3.26E-08 |
| B118 | B190 | CEA1 | Cre | 0.022024 | 0 |
| B118 | B150 | CEA1 | ALP | 0.009823 | 0 |
| B118 | B222 | CEA1 | NS | 0.01557 | 0 |
| B118 | B176 | CEA1 | Ucr | 0.00616 | 7.52E-08 |
| B118 | B178 | CEA1 | Ig-U | 0.005874 | 2.92E-07 |
| B118 | B191 | CEA1 | Urea | 0.011301 | 0 |
| B118 | B174 | CEA1 | Urea/Cr | -0.00668 | 5.49E-09 |
| B118 | B192 | CEA1 | UA | 0.015489 | 0 |

| | | | | | |
|---|---|---|---|---|---|
| B118 | B170 | CEA1 | U-GLU | 0.010053 | 0 |
| B118 | B179 | CEA1 | α1-MG | 0.01058 | 0 |
| B118 | B220 | CEA1 | SBP | 0.02435 | 0 |
| B118 | B218 | CEA1 | DBP | 0.014015 | 0 |
| B118 | B216 | CEA1 | BMI | 0.007085 | 6.20E-10 |
| B118 | B215 | CEA1 | BMI | 0.005894 | 2.67E-07 |
| B118 | B131 | CEA1 | AST | 0.013164 | 0 |
| B118 | B209 | CEA1 | Smoking | 0.039733 | 0 |
| B118 | B173 | CEA1 | NTT | 0.011011 | 0 |
| B118 | B206 | CEA1 | Waist | 0.01713 | 0 |
| B118 | B204 | CEA1 | Drinking | 0.017926 | 0 |
| B118 | B132 | CEA1 | Dbil | 0.007835 | 7.90E-12 |
| B119 | B120 | Hcv-rna | HCV-cAg | 0.249861 | 0 |
| B119 | B124 | Hcv-rna | ACTH | 0.011413 | 0 |
| B119 | B125 | Hcv-rna | Cor | 0.018851 | 0 |
| B119 | B138 | Hcv-rna | PAB | -0.00665 | 6.49E-09 |
| B119 | B139 | Hcv-rna | ADA | 0.0063 | 3.79E-08 |
| B119 | B196 | Hcv-rna | TCO2 | 0.00573 | 5.66E-07 |
| B119 | B135 | Hcv-rna | TBA | 0.007173 | 3.78E-10 |
| B12 | B137 | PG II | AFU | 0.012343 | 0 |
| B12 | B24 | PG II | αHBDH | 0.01839 | 0 |
| B12 | B15 | PG II | CD3-CD19+(B cell) | -0.02806 | 0 |
| B12 | B23 | PG II | CD3-CD16+CD56+(NK) | -0.00619 | 6.40E-08 |
| B12 | B18 | PG II | CD3+CD4+helper(inducer)T | 0.012764 | 0 |
| B12 | B19 | PG II | CD3+CD4+/CD3+CD8+ | 0.017442 | 0 |
| B12 | B22 | PG II | CD3+CD8+(T) | -0.01069 | 0 |
| B12 | B92 | PG II | ALDH2-Glu504Glu | -0.00967 | 0 |
| B12 | B93 | PG II | ALDH2-Glu504Lys | 0.00749 | 6.19E-11 |
| B12 | B160 | PG II | PH | -0.02213 | 0 |
| B12 | B94 | PG II | ALDH2-Lys504Lys | 0.007276 | 2.12E-10 |
| B12 | B126 | PG II | Alb | -0.03523 | 0 |
| B12 | B127 | PG II | A-G ratio | -0.0222 | 0 |
| B12 | B40 | PG II | WBC | 0.015499 | 0 |
| B12 | B124 | PG II | ACTH | 0.010602 | 0 |
| B12 | B70 | PG II | LDL-C | 0.027713 | 0 |
| B12 | B72 | PG II | HDL-C | 0.009807 | 0 |
| B12 | B189 | PG II | eGFB | -0.07519 | 0 |
| B12 | B141 | PG II | AST/ALT | 0.00632 | 3.43E-08 |
| B12 | B146 | PG II | CysC | 0.012098 | 0 |
| B12 | B43 | PG II | MCV | 0.010421 | 0 |
| B12 | B44 | PG II | MCH | 0.010982 | 0 |
| B12 | B45 | PG II | MCHC | -0.01339 | 0 |
| B12 | B46 | PG II | RBC Count | -0.0079 | 5.38E-12 |
| B12 | B47 | PG II | RDW | 0.037949 | 0 |

| | | | | | |
|---|---|---|---|---|---|
| B12 | B181 | PG II | APTT | -0.0157 | 0 |
| B12 | B190 | PG II | Cre | 0.032402 | 0 |
| B12 | B193 | PG II | K | 0.008434 | 1.79E-13 |
| B12 | B129 | PG II | IBIL | -0.02508 | 0 |
| B12 | B150 | PG II | ALP | 0.017194 | 0 |
| B12 | B84 | PG II | C-PSA | 0.007528 | 4.96E-11 |
| B12 | B109 | PG II | AMA-M2 | 0.010273 | 0 |
| B12 | B199 | PG II | Mg | 0.006445 | 1.83E-08 |
| B12 | B195 | PG II | Na | 0.013478 | 0 |
| B12 | B191 | PG II | Urea | 0.028705 | 0 |
| B12 | B125 | PG II | Cor | 0.012155 | 0 |
| B12 | B130 | PG II | GLB | 0.014081 | 0 |
| B12 | B27 | PG II | LDH | 0.013505 | 0 |
| B12 | B221 | PG II | Hei | -0.00704 | 7.98E-10 |
| B12 | B200 | PG II | OP | 0.013043 | 0 |
| B12 | B220 | PG II | SBP | 0.024386 | 0 |
| B12 | B219 | PG II | OT | 0.009089 | 2.22E-15 |
| B12 | B218 | PG II | DBP | 0.018258 | 0 |
| B12 | B131 | PG II | AST | 0.016569 | 0 |
| B12 | B210 | PG II | Hip | -0.00694 | 1.34E-09 |
| B12 | B14 | PG II | G-17 | 0.265469 | 0 |
| B12 | B209 | PG II | Smoking | 0.033967 | 0 |
| B12 | B90 | PG II | CYFRA 21-1 | 0.022581 | 0 |
| B12 | B139 | PG II | ADA | 0.010214 | 0 |
| B12 | B196 | PG II | TCO2 | 0.006708 | 4.74E-09 |
| B12 | B183 | PG II | PTA | -0.02402 | 0 |
| B12 | B185 | PG II | PT | -0.00876 | 2.07E-14 |
| B12 | B186 | PG II | PFC | 0.012034 | 0 |
| B12 | B52 | PG II | PLT/BPC | -0.01915 | 0 |
| B12 | B173 | PG II | NTT | 0.00574 | 5.41E-07 |
| B12 | B206 | PG II | Waist | 0.007856 | 6.96E-12 |
| B12 | B205 | PG II | KDA | 0.005832 | 3.55E-07 |
| B12 | B78 | PG II | HBcAb | -0.02834 | 0 |
| B12 | B204 | PG II | Drinking | 0.012233 | 0 |
| B12 | B80 | PG II | anti-HP IgG | 0.154105 | 0 |
| B12 | B112 | PG II | FT4 | 0.005975 | 1.82E-07 |
| B12 | B113 | PG II | FT3 | 0.013427 | 0 |
| B12 | B211 | PG II | IOP-R | -0.00752 | 5.16E-11 |
| B12 | B201 | PG II | e-habits | 0.013568 | 0 |
| B12 | B74 | PG II | ApoA1 | 0.020425 | 0 |
| B12 | B76 | PG II | ApoB | 0.007776 | 1.13E-11 |
| B12 | B132 | PG II | Dbil | -0.0114 | 0 |
| B12 | B77 | PG II | TC | 0.023133 | 0 |
| B12 | B133 | PG II | TBIL | -0.02321 | 0 |

| | | | | | |
|---|---|---|---|---|---|
| B12 | B135 | PG II | TBA | 0.006083 | 1.09E-07 |
| B12 | B91 | PG II | TP | -0.00982 | 0 |
| B12 | B114 | PG II | TSH TT3 TT4(TT4) | 0.008243 | 6.16E-13 |
| B12 | B212 | PG II | IOP-L | -0.00601 | 1.53E-07 |
| B120 | B140 | HCV-cAg | r-GT | 0.007076 | 6.49E-10 |
| B120 | B128 | HCV-cAg | ALT | 0.006804 | 2.84E-09 |
| B120 | B225 | HCV-cAg | hPBG | 0.008109 | 1.45E-12 |
| B120 | B194 | HCV-cAg | Cl | -0.00583 | 3.60E-07 |
| B120 | B130 | HCV-cAg | GLB | 0.006159 | 7.57E-08 |
| B120 | B131 | HCV-cAg | AST | 0.007539 | 4.65E-11 |
| B120 | B139 | HCV-cAg | ADA | 0.006177 | 6.95E-08 |
| B121 | B137 | C3 | AFU | -0.01337 | 0 |
| B121 | B122 | C3 | C4 | 0.68854 | 0 |
| B121 | B124 | C3 | ACTH | -0.00811 | 1.43E-12 |
| B121 | B134 | C3 | CHE | 0.005955 | 2.00E-07 |
| B121 | B150 | C3 | ALP | 0.006218 | 5.69E-08 |
| B121 | B194 | C3 | Cl | -0.00808 | 1.78E-12 |
| B121 | B155 | C3 | IgA | 0.014171 | 0 |
| B121 | B157 | C3 | IgG | -0.006 | 1.63E-07 |
| B121 | B195 | C3 | Na | -0.01125 | 0 |
| B121 | B125 | C3 | Cor | -0.00621 | 6.04E-08 |
| B121 | B130 | C3 | GLB | 0.007293 | 1.93E-10 |
| B121 | B200 | C3 | OP | -0.00849 | 1.23E-13 |
| B121 | B215 | C3 | BMI | 0.006331 | 3.26E-08 |
| B121 | B196 | C3 | TCO2 | -0.00869 | 3.29E-14 |
| B121 | B186 | C3 | PFC | 0.006443 | 1.85E-08 |
| B121 | B206 | C3 | Waist | 0.006079 | 1.11E-07 |
| B121 | B212 | C3 | IOP-L | 0.005631 | 8.83E-07 |
| B122 | B137 | C4 | AFU | -0.0128 | 0 |
| B122 | B134 | C4 | CHE | 0.006546 | 1.10E-08 |
| B122 | B193 | C4 | K | -0.00574 | 5.44E-07 |
| B122 | B194 | C4 | Cl | -0.00688 | 1.93E-09 |
| B122 | B155 | C4 | IgA | 0.015937 | 0 |
| B122 | B156 | C4 | IgE | -0.00675 | 3.71E-09 |
| B122 | B157 | C4 | IgG | -0.00693 | 1.43E-09 |
| B122 | B195 | C4 | Na | -0.00842 | 2.01E-13 |
| B122 | B138 | C4 | PAB | 0.006754 | 3.72E-09 |
| B122 | B200 | C4 | OP | -0.00782 | 8.86E-12 |
| B123 | B160 | hs-CRP1 | PH | -0.01379 | 0 |
| B123 | B140 | hs-CRP1 | r-GT | 0.051523 | 0 |
| B123 | B126 | hs-CRP1 | Alb | -0.04619 | 0 |
| B123 | B127 | hs-CRP1 | A-G ratio | -0.04504 | 0 |
| B123 | B128 | hs-CRP1 | ALT | 0.022421 | 0 |
| B123 | B124 | hs-CRP1 | ACTH | 0.008831 | 1.24E-14 |

| | | | | | |
|---|---|---|---|---|---|
| B123 | B165 | hs-CRP1 | bilirubin | 0.011869 | 0 |
| B123 | B166 | hs-CRP1 | Pro | 0.013669 | 0 |
| B123 | B189 | hs-CRP1 | eGFB | -0.0115 | 0 |
| B123 | B141 | hs-CRP1 | AST/ALT | -0.00895 | 5.77E-15 |
| B123 | B146 | hs-CRP1 | CysC | 0.006959 | 1.23E-09 |
| B123 | B181 | hs-CRP1 | APTT | 0.013436 | 0 |
| B123 | B190 | hs-CRP1 | Cre | 0.014177 | 0 |
| B123 | B129 | hs-CRP1 | IBIL | -0.00936 | 4.44E-16 |
| B123 | B150 | hs-CRP1 | ALP | 0.032254 | 0 |
| B123 | B222 | hs-CRP1 | NS | 0.02482 | 0 |
| B123 | B151 | hs-CRP1 | Anti-CCP | 0.03879 | 0 |
| B123 | B154 | hs-CRP1 | RF | 0.022476 | 0 |
| B123 | B194 | hs-CRP1 | Cl | -0.0153 | 0 |
| B123 | B155 | hs-CRP1 | IgA | 0.020773 | 0 |
| B123 | B156 | hs-CRP1 | IgE | 0.010865 | 0 |
| B123 | B157 | hs-CRP1 | IgG | 0.011621 | 0 |
| B123 | B176 | hs-CRP1 | Ucr | 0.024929 | 0 |
| B123 | B178 | hs-CRP1 | Ig-U | 0.053154 | 0 |
| B123 | B174 | hs-CRP1 | Urea/Cr | -0.00909 | 2.22E-15 |
| B123 | B192 | hs-CRP1 | UA | 0.023347 | 0 |
| B123 | B170 | hs-CRP1 | U-GLU | 0.019548 | 0 |
| B123 | B175 | hs-CRP1 | ALB-U | 0.033953 | 0 |
| B123 | B177 | hs-CRP1 | ACR | 0.036719 | 0 |
| B123 | B179 | hs-CRP1 | α1-MG | 0.076544 | 0 |
| B123 | B180 | hs-CRP1 | Tf | 0.018177 | 0 |
| B123 | B182 | hs-CRP1 | TT | -0.0217 | 0 |
| B123 | B125 | hs-CRP1 | Cor | 0.010643 | 0 |
| B123 | B138 | hs-CRP1 | PAB | -0.03025 | 0 |
| B123 | B130 | hs-CRP1 | GLB | 0.04401 | 0 |
| B123 | B220 | hs-CRP1 | SBP | 0.01975 | 0 |
| B123 | B218 | hs-CRP1 | DBP | 0.017932 | 0 |
| B123 | B216 | hs-CRP1 | BMI | 0.025322 | 0 |
| B123 | B215 | hs-CRP1 | BMI | 0.031161 | 0 |
| B123 | B131 | hs-CRP1 | AST | 0.02339 | 0 |
| B123 | B172 | hs-CRP1 | KB | 0.01162 | 0 |
| B123 | B210 | hs-CRP1 | Hip | 0.022239 | 0 |
| B123 | B209 | hs-CRP1 | Smoking | 0.02213 | 0 |
| B123 | B139 | hs-CRP1 | ADA | 0.034717 | 0 |
| B123 | B183 | hs-CRP1 | PTA | -0.01017 | 0 |
| B123 | B185 | hs-CRP1 | PT | 0.009813 | 0 |
| B123 | B186 | hs-CRP1 | PFC | 0.099981 | 0 |
| B123 | B173 | hs-CRP1 | NTT | 0.007088 | 6.10E-10 |
| B123 | B206 | hs-CRP1 | Waist | 0.036601 | 0 |
| B123 | B197 | hs-CRP1 | AG | 0.011258 | 0 |

| | | | | | |
|---|---|---|---|---|---|
| B123 | B204 | hs-CRP1 | Drinking | 0.013449 | 0 |
| B123 | B211 | hs-CRP1 | IOP-R | -0.00647 | 1.61E-08 |
| B123 | B132 | hs-CRP1 | Dbil | 0.007214 | 3.01E-10 |
| B123 | B133 | hs-CRP1 | TBIL | -0.00564 | 8.49E-07 |
| B123 | B212 | hs-CRP1 | IOP-L | -0.00655 | 1.05E-08 |
| B124 | B137 | ACTH | AFU | 0.017003 | 0 |
| B124 | B160 | ACTH | PH | -0.00573 | 5.64E-07 |
| B124 | B140 | ACTH | r-GT | 0.009224 | 8.88E-16 |
| B124 | B126 | ACTH | Alb | 0.006196 | 6.31E-08 |
| B124 | B128 | ACTH | ALT | 0.006318 | 3.47E-08 |
| B124 | B225 | ACTH | hPBG | -0.00585 | 3.26E-07 |
| B124 | B134 | ACTH | CHE | 0.025506 | 0 |
| B124 | B146 | ACTH | CysC | 0.007282 | 2.05E-10 |
| B124 | B181 | ACTH | APTT | -0.03804 | 0 |
| B124 | B190 | ACTH | Cre | 0.014427 | 0 |
| B124 | B129 | ACTH | IBIL | 0.007012 | 9.27E-10 |
| B124 | B150 | ACTH | ALP | 0.013387 | 0 |
| B124 | B222 | ACTH | NS | 0.013112 | 0 |
| B124 | B151 | ACTH | Anti-CCP | 0.008806 | 1.51E-14 |
| B124 | B194 | ACTH | Cl | -0.01824 | 0 |
| B124 | B155 | ACTH | IgA | 0.014279 | 0 |
| B124 | B156 | ACTH | IgE | 0.014203 | 0 |
| B124 | B195 | ACTH | Na | 0.010973 | 0 |
| B124 | B159 | ACTH | β2-MG | -0.01434 | 0 |
| B124 | B176 | ACTH | Ucr | 0.006027 | 1.43E-07 |
| B124 | B174 | ACTH | Urea/Cr | -0.0117 | 0 |
| B124 | B192 | ACTH | UA | 0.01299 | 0 |
| B124 | B179 | ACTH | α1-MG | 0.013657 | 0 |
| B124 | B182 | ACTH | TT | -0.01072 | 0 |
| B124 | B125 | ACTH | Cor | 0.374521 | 0 |
| B124 | B138 | ACTH | PAB | 0.041309 | 0 |
| B124 | B221 | ACTH | Hei | 0.010561 | 0 |
| B124 | B200 | ACTH | OP | 0.01459 | 0 |
| B124 | B220 | ACTH | SBP | 0.010108 | 0 |
| B124 | B218 | ACTH | DBP | 0.010611 | 0 |
| B124 | B216 | ACTH | BMI | 0.011081 | 0 |
| B124 | B215 | ACTH | BMI | 0.006982 | 1.09E-09 |
| B124 | B131 | ACTH | AST | 0.006621 | 7.43E-09 |
| B124 | B210 | ACTH | Hip | 0.007361 | 1.31E-10 |
| B124 | B209 | ACTH | Smoking | 0.006541 | 1.12E-08 |
| B124 | B184 | ACTH | INR | 0.013469 | 0 |
| B124 | B186 | ACTH | PFC | 0.020591 | 0 |
| B124 | B206 | ACTH | Waist | 0.00998 | 0 |
| B124 | B197 | ACTH | AG | 0.013681 | 0 |

| | | | | | |
|---|---|---|---|---|---|
| B124 | B204 | ACTH | Drinking | 0.009504 | 0 |
| B124 | B133 | ACTH | TBIL | 0.006238 | 5.15E-08 |
| B124 | B135 | ACTH | TBA | 0.017439 | 0 |
| B125 | B140 | Cor | r-GT | 0.006722 | 4.40E-09 |
| B125 | B126 | Cor | Alb | 0.009166 | 1.33E-15 |
| B125 | B127 | Cor | A-G ratio | 0.005707 | 6.28E-07 |
| B125 | B225 | Cor | hPBG | -0.01225 | 0 |
| B125 | B134 | Cor | CHE | 0.010343 | 0 |
| B125 | B181 | Cor | APTT | -0.04929 | 0 |
| B125 | B190 | Cor | Cre | 0.012807 | 0 |
| B125 | B193 | Cor | K | -0.01684 | 0 |
| B125 | B129 | Cor | IBIL | 0.006404 | 2.26E-08 |
| B125 | B150 | Cor | ALP | 0.00693 | 1.45E-09 |
| B125 | B151 | Cor | Anti-CCP | 0.01151 | 0 |
| B125 | B198 | Cor | P | 0.007489 | 6.22E-11 |
| B125 | B194 | Cor | Cl | -0.03926 | 0 |
| B125 | B156 | Cor | IgE | 0.015964 | 0 |
| B125 | B157 | Cor | IgG | 0.012758 | 0 |
| B125 | B158 | Cor | IgM | -0.00614 | 8.23E-08 |
| B125 | B195 | Cor | Na | -0.02948 | 0 |
| B125 | B176 | Cor | Ucr | 0.007507 | 5.59E-11 |
| B125 | B174 | Cor | Urea/Cr | -0.01311 | 0 |
| B125 | B179 | Cor | α1-MG | 0.019224 | 0 |
| B125 | B182 | Cor | TT | -0.05586 | 0 |
| B125 | B138 | Cor | PAB | 0.033026 | 0 |
| B125 | B221 | Cor | Hei | 0.008272 | 5.12E-13 |
| B125 | B200 | Cor | OP | -0.02224 | 0 |
| B125 | B218 | Cor | DBP | 0.006602 | 8.22E-09 |
| B125 | B215 | Cor | BMI | -0.00611 | 9.61E-08 |
| B125 | B139 | Cor | ADA | -0.0093 | 4.44E-16 |
| B125 | B196 | Cor | TCO2 | 0.011283 | 0 |
| B125 | B183 | Cor | PTA | -0.01186 | 0 |
| B125 | B185 | Cor | PT | 0.015761 | 0 |
| B125 | B186 | Cor | PFC | 0.030921 | 0 |
| B125 | B211 | Cor | IOP-R | 0.006131 | 8.67E-08 |
| B125 | B135 | Cor | TBA | 0.029553 | 0 |
| B126 | B160 | Alb | PH | -0.03543 | 0 |
| B126 | B140 | Alb | r-GT | 0.044529 | 0 |
| B126 | B127 | Alb | A-G ratio | 0.421267 | 0 |
| B126 | B161 | Alb | LEU | -0.03806 | 0 |
| B126 | B128 | Alb | ALT | 0.102731 | 0 |
| B126 | B165 | Alb | bilirubin | -0.00597 | 1.87E-07 |
| B126 | B134 | Alb | CHE | 0.0596 | 0 |
| B126 | B166 | Alb | Pro | 0.05246 | 0 |

| | | | | | |
|---|---|---|---|---|---|
| B126 | B188 | Alb | Ca | 0.109588 | 0 |
| B126 | B189 | Alb | eGFB | 0.170044 | 0 |
| B126 | B141 | Alb | AST/ALT | -0.09975 | 0 |
| B126 | B146 | Alb | CysC | -0.00731 | 1.73E-10 |
| B126 | B190 | Alb | Cre | 0.108723 | 0 |
| B126 | B224 | Alb | pmh | -0.0062 | 6.21E-08 |
| B126 | B193 | Alb | K | 0.016481 | 0 |
| B126 | B129 | Alb | IBIL | 0.142249 | 0 |
| B126 | B150 | Alb | ALP | 0.00816 | 1.05E-12 |
| B126 | B213 | Alb | CVA-R | -0.00786 | 6.77E-12 |
| B126 | B214 | Alb | CVA-L | -0.0073 | 1.86E-10 |
| B126 | B222 | Alb | NS | 0.104667 | 0 |
| B126 | B151 | Alb | Anti-CCP | -0.00588 | 2.86E-07 |
| B126 | B148 | Alb | Anti_TPO | -0.01375 | 0 |
| B126 | B149 | Alb | TGAb | -0.01091 | 0 |
| B126 | B154 | Alb | RF | -0.01162 | 0 |
| B126 | B194 | Alb | Cl | -0.05381 | 0 |
| B126 | B199 | Alb | Mg | 0.026255 | 0 |
| B126 | B155 | Alb | IgA | -0.01149 | 0 |
| B126 | B157 | Alb | IgG | -0.0135 | 0 |
| B126 | B176 | Alb | Ucr | 0.037605 | 0 |
| B126 | B191 | Alb | Urea | 0.017946 | 0 |
| B126 | B174 | Alb | Urea/Cr | -0.06306 | 0 |
| B126 | B192 | Alb | UA | 0.180228 | 0 |
| B126 | B175 | Alb | ALB-U | -0.01337 | 0 |
| B126 | B177 | Alb | ACR | -0.01653 | 0 |
| B126 | B180 | Alb | Tf | -0.00603 | 1.39E-07 |
| B126 | B138 | Alb | PAB | 0.076175 | 0 |
| B126 | B130 | Alb | GLB | -0.10615 | 0 |
| B126 | B221 | Alb | Hei | 0.203513 | 0 |
| B126 | B187 | Alb | NSE | 0.006291 | 3.97E-08 |
| B126 | B200 | Alb | OP | 0.008653 | 4.20E-14 |
| B126 | B227 | Alb | OD | -0.01228 | 0 |
| B126 | B228 | Alb | OS | -0.01074 | 0 |
| B126 | B220 | Alb | SBP | 0.031853 | 0 |
| B126 | B219 | Alb | OT | -0.05745 | 0 |
| B126 | B218 | Alb | DBP | 0.085428 | 0 |
| B126 | B216 | Alb | BMI | 0.087028 | 0 |
| B126 | B215 | Alb | BMI | -0.03157 | 0 |
| B126 | B131 | Alb | AST | 0.03819 | 0 |
| B126 | B172 | Alb | KB | 0.031786 | 0 |
| B126 | B210 | Alb | Hip | 0.027914 | 0 |
| B126 | B209 | Alb | Smoking | -0.03691 | 0 |
| B126 | B139 | Alb | ADA | -0.01979 | 0 |

| | | | | | |
|---|---|---|---|---|---|
| B126 | B196 | Alb | TCO2 | 0.006095 | 1.03E-07 |
| B126 | B186 | Alb | PFC | -0.00852 | 1.01E-13 |
| B126 | B173 | Alb | NTT | -0.03523 | 0 |
| B126 | B206 | Alb | Waist | -0.0078 | 9.66E-12 |
| B126 | B205 | Alb | KDA | -0.02335 | 0 |
| B126 | B197 | Alb | AG | 0.032192 | 0 |
| B126 | B204 | Alb | Drinking | 0.055872 | 0 |
| B126 | B203 | Alb | DH | 0.051259 | 0 |
| B126 | B211 | Alb | IOP-R | 0.0448 | 0 |
| B126 | B132 | Alb | Dbil | 0.102279 | 0 |
| B126 | B133 | Alb | TBIL | 0.14143 | 0 |
| B126 | B144 | Alb | Total-PINP | -0.00579 | 4.37E-07 |
| B126 | B212 | Alb | IOP-L | 0.038469 | 0 |
| B127 | B137 | A-G ratio | AFU | -0.00842 | 1.99E-13 |
| B127 | B160 | A-G ratio | PH | 0.013108 | 0 |
| B127 | B140 | A-G ratio | r-GT | -0.02387 | 0 |
| B127 | B161 | A-G ratio | LEU | -0.04541 | 0 |
| B127 | B128 | A-G ratio | ALT | 0.006905 | 1.66E-09 |
| B127 | B225 | A-G ratio | hPBG | -0.00843 | 1.85E-13 |
| B127 | B166 | A-G ratio | Pro | 0.037186 | 0 |
| B127 | B188 | A-G ratio | Ca | 0.041843 | 0 |
| B127 | B189 | A-G ratio | eGFB | 0.061216 | 0 |
| B127 | B146 | A-G ratio | CysC | -0.0079 | 5.43E-12 |
| B127 | B190 | A-G ratio | Cre | 0.108312 | 0 |
| B127 | B193 | A-G ratio | K | -0.01163 | 0 |
| B127 | B129 | A-G ratio | IBIL | 0.126698 | 0 |
| B127 | B150 | A-G ratio | ALP | -0.02053 | 0 |
| B127 | B222 | A-G ratio | NS | 0.079501 | 0 |
| B127 | B148 | A-G ratio | Anti_TPO | -0.03134 | 0 |
| B127 | B149 | A-G ratio | TGAb | -0.02857 | 0 |
| B127 | B154 | A-G ratio | RF | -0.01001 | 0 |
| B127 | B198 | A-G ratio | P | -0.00978 | 0 |
| B127 | B194 | A-G ratio | Cl | 0.009647 | 0 |
| B127 | B199 | A-G ratio | Mg | 0.006884 | 1.86E-09 |
| B127 | B155 | A-G ratio | IgA | -0.06381 | 0 |
| B127 | B156 | A-G ratio | IgE | -0.01008 | 0 |
| B127 | B157 | A-G ratio | IgG | -0.09351 | 0 |
| B127 | B158 | A-G ratio | IgM | -0.0148 | 0 |
| B127 | B195 | A-G ratio | Na | 0.014324 | 0 |
| B127 | B159 | A-G ratio | β2-MG | -0.01241 | 0 |
| B127 | B176 | A-G ratio | Ucr | 0.031929 | 0 |
| B127 | B178 | A-G ratio | Ig-U | -0.01067 | 0 |
| B127 | B191 | A-G ratio | Urea | 0.02214 | 0 |
| B127 | B174 | A-G ratio | Urea/Cr | -0.061 | 0 |

| | | | | | |
|---|---|---|---|---|---|
| B127 | B192 | A-G ratio | UA | 0.055539 | 0 |
| B127 | B170 | A-G ratio | U-GLU | -0.0138 | 0 |
| B127 | B175 | A-G ratio | ALB-U | -0.01229 | 0 |
| B127 | B177 | A-G ratio | ACR | -0.01589 | 0 |
| B127 | B180 | A-G ratio | Tf | -0.00683 | 2.49E-09 |
| B127 | B182 | A-G ratio | TT | -0.00634 | 3.08E-08 |
| B127 | B153 | A-G ratio | Glu | -0.00916 | 1.33E-15 |
| B127 | B138 | A-G ratio | PAB | 0.036416 | 0 |
| B127 | B130 | A-G ratio | GLB | -0.82057 | 0 |
| B127 | B221 | A-G ratio | Hei | 0.166903 | 0 |
| B127 | B200 | A-G ratio | OP | 0.011948 | 0 |
| B127 | B227 | A-G ratio | OD | -0.00638 | 2.50E-08 |
| B127 | B220 | A-G ratio | SBP | -0.0479 | 0 |
| B127 | B219 | A-G ratio | OT | -0.03425 | 0 |
| B127 | B218 | A-G ratio | DBP | -0.02345 | 0 |
| B127 | B216 | A-G ratio | BMI | 0.052086 | 0 |
| B127 | B215 | A-G ratio | BMI | -0.04869 | 0 |
| B127 | B131 | A-G ratio | AST | -0.03805 | 0 |
| B127 | B172 | A-G ratio | KB | 0.019526 | 0 |
| B127 | B209 | A-G ratio | Smoking | 0.092975 | 0 |
| B127 | B139 | A-G ratio | ADA | -0.05136 | 0 |
| B127 | B196 | A-G ratio | TCO2 | 0.005715 | 6.04E-07 |
| B127 | B185 | A-G ratio | PT | -0.00618 | 6.73E-08 |
| B127 | B186 | A-G ratio | PFC | -0.01467 | 0 |
| B127 | B173 | A-G ratio | NTT | -0.03182 | 0 |
| B127 | B206 | A-G ratio | Waist | -0.01028 | 0 |
| B127 | B205 | A-G ratio | KDA | -0.01693 | 0 |
| B127 | B204 | A-G ratio | Drinking | 0.086184 | 0 |
| B127 | B203 | A-G ratio | DH | 0.01433 | 0 |
| B127 | B211 | A-G ratio | IOP-R | 0.016402 | 0 |
| B127 | B132 | A-G ratio | Dbil | 0.124045 | 0 |
| B127 | B133 | A-G ratio | TBIL | 0.133679 | 0 |
| B127 | B212 | A-G ratio | IOP-L | 0.016316 | 0 |
| B128 | B136 | ALT | 5'NT | 0.010901 | 0 |
| B128 | B137 | ALT | AFU | 0.045256 | 0 |
| B128 | B160 | ALT | PH | -0.05802 | 0 |
| B128 | B140 | ALT | r-GT | 0.403771 | 0 |
| B128 | B161 | ALT | LEU | -0.04439 | 0 |
| B128 | B165 | ALT | bilirubin | 0.068247 | 0 |
| B128 | B134 | ALT | CHE | 0.050038 | 0 |
| B128 | B166 | ALT | Pro | 0.069432 | 0 |
| B128 | B188 | ALT | Ca | 0.021208 | 0 |
| B128 | B189 | ALT | eGFB | -0.02295 | 0 |
| B128 | B141 | ALT | AST/ALT | -0.41492 | 0 |

| | | | | | |
|---|---|---|---|---|---|
| B128 | B146 | ALT | CysC | 0.015532 | 0 |
| B128 | B181 | ALT | APTT | -0.00619 | 6.65E-08 |
| B128 | B190 | ALT | Cre | 0.1729 | 0 |
| B128 | B129 | ALT | IBIL | 0.047954 | 0 |
| B128 | B150 | ALT | ALP | 0.070974 | 0 |
| B128 | B222 | ALT | NS | 0.302839 | 0 |
| B128 | B148 | ALT | Anti_TPO | -0.017 | 0 |
| B128 | B149 | ALT | TGAb | -0.01728 | 0 |
| B128 | B198 | ALT | P | -0.01343 | 0 |
| B128 | B194 | ALT | Cl | -0.00681 | 2.78E-09 |
| B128 | B155 | ALT | IgA | 0.006064 | 1.20E-07 |
| B128 | B157 | ALT | IgG | -0.00738 | 1.14E-10 |
| B128 | B176 | ALT | Ucr | 0.047489 | 0 |
| B128 | B191 | ALT | Urea | 0.052947 | 0 |
| B128 | B174 | ALT | Urea/Cr | -0.07812 | 0 |
| B128 | B192 | ALT | UA | 0.288708 | 0 |
| B128 | B170 | ALT | U-GLU | 0.0347 | 0 |
| B128 | B175 | ALT | ALB-U | 0.008114 | 1.40E-12 |
| B128 | B179 | ALT | α1-MG | 0.019834 | 0 |
| B128 | B153 | ALT | Glu | 0.016857 | 0 |
| B128 | B138 | ALT | PAB | 0.027712 | 0 |
| B128 | B130 | ALT | GLB | 0.040043 | 0 |
| B128 | B221 | ALT | Hei | 0.189621 | 0 |
| B128 | B200 | ALT | OP | 0.008465 | 1.47E-13 |
| B128 | B220 | ALT | SBP | 0.132324 | 0 |
| B128 | B219 | ALT | OT | -0.01103 | 0 |
| B128 | B218 | ALT | DBP | 0.146832 | 0 |
| B128 | B216 | ALT | BMI | 0.338011 | 0 |
| B128 | B215 | ALT | BMI | 0.309396 | 0 |
| B128 | B131 | ALT | AST | 0.788494 | 0 |
| B128 | B172 | ALT | KB | -0.01448 | 0 |
| B128 | B210 | ALT | Hip | 0.252878 | 0 |
| B128 | B209 | ALT | Smoking | 0.086672 | 0 |
| B128 | B139 | ALT | ADA | 0.043261 | 0 |
| B128 | B173 | ALT | NTT | -0.01925 | 0 |
| B128 | B207 | ALT | WHR | 0.034004 | 0 |
| B128 | B206 | ALT | Waist | 0.322838 | 0 |
| B128 | B205 | ALT | KDA | -0.01606 | 0 |
| B128 | B197 | ALT | AG | 0.007127 | 4.91E-10 |
| B128 | B204 | ALT | Drinking | 0.120024 | 0 |
| B128 | B203 | ALT | DH | 0.019637 | 0 |
| B128 | B211 | ALT | IOP-R | 0.016359 | 0 |
| B128 | B201 | ALT | e-habits | -0.0166 | 0 |
| B128 | B132 | ALT | Dbil | 0.10045 | 0 |

| | | | | | |
|---|---|---|---|---|---|
| B128 | B133 | ALT | TBIL | 0.065192 | 0 |
| B128 | B135 | ALT | TBA | 0.034998 | 0 |
| B128 | B144 | ALT | Total-PINP | 0.00635 | 2.96E-08 |
| B128 | B212 | ALT | IOP-L | 0.015706 | 0 |
| B129 | B160 | IBIL | PH | 0.051233 | 0 |
| B129 | B140 | IBIL | r-GT | 0.030824 | 0 |
| B129 | B161 | IBIL | LEU | -0.03448 | 0 |
| B129 | B225 | IBIL | hPBG | 0.006686 | 5.31E-09 |
| B129 | B165 | IBIL | bilirubin | 0.020857 | 0 |
| B129 | B166 | IBIL | Pro | 0.062183 | 0 |
| B129 | B188 | IBIL | Ca | 0.015112 | 0 |
| B129 | B189 | IBIL | eGFB | -0.07786 | 0 |
| B129 | B141 | IBIL | AST/ALT | -0.00769 | 1.87E-11 |
| B129 | B181 | IBIL | APTT | 0.006109 | 9.65E-08 |
| B129 | B190 | IBIL | Cre | 0.133212 | 0 |
| B129 | B193 | IBIL | K | -0.02122 | 0 |
| B129 | B150 | IBIL | ALP | -0.00946 | 0 |
| B129 | B222 | IBIL | NS | 0.081814 | 0 |
| B129 | B151 | IBIL | Anti-CCP | -0.00681 | 2.83E-09 |
| B129 | B148 | IBIL | Anti_TPO | -0.00894 | 5.77E-15 |
| B129 | B149 | IBIL | TGAb | -0.00791 | 4.94E-12 |
| B129 | B198 | IBIL | P | -0.02684 | 0 |
| B129 | B194 | IBIL | Cl | -0.00836 | 2.99E-13 |
| B129 | B195 | IBIL | Na | -0.00819 | 8.70E-13 |
| B129 | B176 | IBIL | Ucr | 0.039589 | 0 |
| B129 | B191 | IBIL | Urea | -0.01121 | 0 |
| B129 | B174 | IBIL | Urea/Cr | -0.10202 | 0 |
| B129 | B192 | IBIL | UA | 0.108605 | 0 |
| B129 | B170 | IBIL | U-GLU | 0.008552 | 8.26E-14 |
| B129 | B179 | IBIL | α1-MG | 0.011436 | 0 |
| B129 | B180 | IBIL | Tf | 0.005883 | 2.81E-07 |
| B129 | B182 | IBIL | TT | -0.00579 | 4.26E-07 |
| B129 | B138 | IBIL | PAB | 0.017346 | 0 |
| B129 | B130 | IBIL | GLB | -0.09676 | 0 |
| B129 | B221 | IBIL | Hei | 0.113724 | 0 |
| B129 | B200 | IBIL | OP | -0.00813 | 1.26E-12 |
| B129 | B220 | IBIL | SBP | 0.054186 | 0 |
| B129 | B218 | IBIL | DBP | 0.052176 | 0 |
| B129 | B216 | IBIL | BMI | 0.058772 | 0 |
| B129 | B131 | IBIL | AST | 0.0704 | 0 |
| B129 | B172 | IBIL | KB | 0.029041 | 0 |
| B129 | B210 | IBIL | Hip | 0.010654 | 0 |
| B129 | B209 | IBIL | Smoking | -0.02844 | 0 |
| B129 | B196 | IBIL | TCO2 | 0.008696 | 3.15E-14 |

| | | | | | |
|---|---|---|---|---|---|
| B129 | B183 | IBIL | PTA | 0.02915 | 0 |
| B129 | B185 | IBIL | PT | 0.013682 | 0 |
| B129 | B186 | IBIL | PFC | -0.01496 | 0 |
| B129 | B173 | IBIL | NTT | -0.01453 | 0 |
| B129 | B207 | IBIL | WHR | 0.005799 | 4.13E-07 |
| B129 | B206 | IBIL | Waist | 0.028439 | 0 |
| B129 | B205 | IBIL | KDA | -0.01547 | 0 |
| B129 | B197 | IBIL | AG | -0.00615 | 7.75E-08 |
| B129 | B204 | IBIL | Drinking | 0.047248 | 0 |
| B129 | B203 | IBIL | DH | -0.02709 | 0 |
| B129 | B211 | IBIL | IOP-R | 0.020043 | 0 |
| B129 | B132 | IBIL | Dbil | 0.664802 | 0 |
| B129 | B133 | IBIL | TBIL | 0.97992 | 0 |
| B129 | B212 | IBIL | IOP-L | 0.020037 | 0 |
| B130 | B137 | GLB | AFU | 0.013476 | 0 |
| B130 | B160 | GLB | PH | -0.03737 | 0 |
| B130 | B140 | GLB | r-GT | 0.056979 | 0 |
| B130 | B143 | GLB | β-CTx | -0.00611 | 9.51E-08 |
| B130 | B161 | GLB | LEU | 0.039482 | 0 |
| B130 | B225 | GLB | hPBG | 0.011356 | 0 |
| B130 | B134 | GLB | CHE | 0.021598 | 0 |
| B130 | B166 | GLB | Pro | -0.01855 | 0 |
| B130 | B188 | GLB | Ca | 0.027329 | 0 |
| B130 | B189 | GLB | eGFB | -0.01114 | 0 |
| B130 | B141 | GLB | AST/ALT | 0.01312 | 0 |
| B130 | B146 | GLB | CysC | 0.013201 | 0 |
| B130 | B190 | GLB | Cre | -0.08038 | 0 |
| B130 | B193 | GLB | K | 0.022528 | 0 |
| B130 | B150 | GLB | ALP | 0.035468 | 0 |
| B130 | B222 | GLB | NS | -0.04728 | 0 |
| B130 | B151 | GLB | Anti-CCP | 0.005779 | 4.53E-07 |
| B130 | B148 | GLB | Anti_TPO | 0.033603 | 0 |
| B130 | B149 | GLB | TGAb | 0.031525 | 0 |
| B130 | B154 | GLB | RF | 0.008777 | 1.82E-14 |
| B130 | B198 | GLB | P | 0.01117 | 0 |
| B130 | B194 | GLB | Cl | -0.03576 | 0 |
| B130 | B155 | GLB | IgA | 0.076371 | 0 |
| B130 | B156 | GLB | IgE | 0.010498 | 0 |
| B130 | B157 | GLB | IgG | 0.114451 | 0 |
| B130 | B158 | GLB | IgM | 0.019315 | 0 |
| B130 | B195 | GLB | Na | -0.01643 | 0 |
| B130 | B176 | GLB | Ucr | -0.02393 | 0 |
| B130 | B178 | GLB | Ig-U | 0.012704 | 0 |
| B130 | B191 | GLB | Urea | -0.01586 | 0 |

| | | | | | |
|---|---|---|---|---|---|
| B130 | B174 | GLB | Urea/Cr | 0.046853 | 0 |
| B130 | B192 | GLB | UA | 0.014458 | 0 |
| B130 | B170 | GLB | U-GLU | 0.021983 | 0 |
| B130 | B175 | GLB | ALB-U | 0.012537 | 0 |
| B130 | B177 | GLB | ACR | 0.015425 | 0 |
| B130 | B180 | GLB | Tf | 0.006545 | 1.10E-08 |
| B130 | B182 | GLB | TT | 0.008607 | 5.68E-14 |
| B130 | B153 | GLB | Glu | 0.01467 | 0 |
| B130 | B138 | GLB | PAB | -0.01364 | 0 |
| B130 | B221 | GLB | Hei | -0.1159 | 0 |
| B130 | B187 | GLB | NSE | 0.007339 | 1.48E-10 |
| B130 | B200 | GLB | OP | -0.0113 | 0 |
| B130 | B220 | GLB | SBP | 0.080538 | 0 |
| B130 | B219 | GLB | OT | 0.018481 | 0 |
| B130 | B218 | GLB | DBP | 0.070546 | 0 |
| B130 | B216 | GLB | BMI | -0.02408 | 0 |
| B130 | B215 | GLB | BMI | 0.049003 | 0 |
| B130 | B131 | GLB | AST | 0.070442 | 0 |
| B130 | B172 | GLB | KB | -0.00751 | 5.41E-11 |
| B130 | B210 | GLB | Hip | 0.01748 | 0 |
| B130 | B209 | GLB | Smoking | -0.1221 | 0 |
| B130 | B139 | GLB | ADA | 0.058957 | 0 |
| B130 | B152 | GLB | ACA-Ig(G-A-M) | 0.008081 | 1.73E-12 |
| B130 | B185 | GLB | PT | 0.007247 | 2.49E-10 |
| B130 | B186 | GLB | PFC | 0.015718 | 0 |
| B130 | B173 | GLB | NTT | 0.025781 | 0 |
| B130 | B206 | GLB | Waist | 0.014334 | 0 |
| B130 | B205 | GLB | KDA | 0.011307 | 0 |
| B130 | B197 | GLB | AG | 0.01796 | 0 |
| B130 | B204 | GLB | Drinking | -0.07799 | 0 |
| B130 | B132 | GLB | Dbil | -0.09701 | 0 |
| B130 | B133 | GLB | TBIL | -0.10351 | 0 |
| B131 | B136 | AST | 5'NT | 0.009387 | 2.22E-16 |
| B131 | B137 | AST | AFU | 0.044275 | 0 |
| B131 | B160 | AST | PH | -0.0366 | 0 |
| B131 | B140 | AST | r-GT | 0.357884 | 0 |
| B131 | B161 | AST | LEU | -0.0206 | 0 |
| B131 | B165 | AST | bilirubin | 0.104042 | 0 |
| B131 | B134 | AST | CHE | 0.026954 | 0 |
| B131 | B166 | AST | Pro | 0.044375 | 0 |
| B131 | B188 | AST | Ca | 0.013735 | 0 |
| B131 | B189 | AST | eGFB | -0.05863 | 0 |
| B131 | B141 | AST | AST/ALT | -0.11653 | 0 |
| B131 | B146 | AST | CysC | 0.015066 | 0 |

| | | | | | |
|---|---|---|---|---|---|
| B131 | B190 | AST | Cre | 0.093817 | 0 |
| B131 | B224 | AST | pmh | 0.006039 | 1.35E-07 |
| B131 | B150 | AST | ALP | 0.063964 | 0 |
| B131 | B222 | AST | NS | 0.168209 | 0 |
| B131 | B148 | AST | Anti_TPO | -0.00772 | 1.57E-11 |
| B131 | B149 | AST | TGAb | -0.00838 | 2.53E-13 |
| B131 | B198 | AST | P | -0.00815 | 1.12E-12 |
| B131 | B155 | AST | IgA | 0.009449 | 0 |
| B131 | B157 | AST | IgG | 0.008003 | 2.80E-12 |
| B131 | B176 | AST | Ucr | 0.025619 | 0 |
| B131 | B191 | AST | Urea | 0.032396 | 0 |
| B131 | B174 | AST | Urea/Cr | -0.03974 | 0 |
| B131 | B192 | AST | UA | 0.179085 | 0 |
| B131 | B170 | AST | U-GLU | 0.018832 | 0 |
| B131 | B175 | AST | ALB-U | 0.007334 | 1.53E-10 |
| B131 | B179 | AST | α1-MG | 0.026263 | 0 |
| B131 | B153 | AST | Glu | 0.011646 | 0 |
| B131 | B138 | AST | PAB | 0.009596 | 0 |
| B131 | B221 | AST | Hei | 0.071748 | 0 |
| B131 | B220 | AST | SBP | 0.119283 | 0 |
| B131 | B219 | AST | OT | 0.006146 | 8.05E-08 |
| B131 | B218 | AST | DBP | 0.11835 | 0 |
| B131 | B216 | AST | BMI | 0.174074 | 0 |
| B131 | B215 | AST | BMI | 0.178589 | 0 |
| B131 | B210 | AST | Hip | 0.129439 | 0 |
| B131 | B209 | AST | Smoking | 0.06461 | 0 |
| B131 | B139 | AST | ADA | 0.060173 | 0 |
| B131 | B207 | AST | WHR | 0.022351 | 0 |
| B131 | B206 | AST | Waist | 0.190595 | 0 |
| B131 | B205 | AST | KDA | -0.00751 | 5.43E-11 |
| B131 | B204 | AST | Drinking | 0.099106 | 0 |
| B131 | B211 | AST | IOP-R | 0.008878 | 9.10E-15 |
| B131 | B132 | AST | Dbil | 0.136886 | 0 |
| B131 | B133 | AST | TBIL | 0.092784 | 0 |
| B131 | B135 | AST | TBA | 0.040404 | 0 |
| B131 | B144 | AST | Total-PINP | 0.00676 | 3.60E-09 |
| B131 | B212 | AST | IOP-L | 0.010308 | 0 |
| B132 | B142 | Dbil | 25-(OH)D | 0.005812 | 3.89E-07 |
| B132 | B160 | Dbil | PH | 0.0441 | 0 |
| B132 | B140 | Dbil | r-GT | 0.078165 | 0 |
| B132 | B161 | Dbil | LEU | -0.02185 | 0 |
| B132 | B225 | Dbil | hPBG | 0.008258 | 5.60E-13 |
| B132 | B165 | Dbil | bilirubin | 0.163582 | 0 |
| B132 | B134 | Dbil | CHE | -0.02466 | 0 |

| | | | | | |
|---|---|---|---|---|---|
| B132 | B166 | Dbil | Pro | 0.048541 | 0 |
| B132 | B188 | Dbil | Ca | 0.009162 | 1.33E-15 |
| B132 | B189 | Dbil | eGFB | -0.03356 | 0 |
| B132 | B141 | Dbil | AST/ALT | -0.00667 | 5.66E-09 |
| B132 | B181 | Dbil | APTT | 0.01019 | 0 |
| B132 | B190 | Dbil | Cre | 0.086911 | 0 |
| B132 | B193 | Dbil | K | -0.0216 | 0 |
| B132 | B150 | Dbil | ALP | 0.019648 | 0 |
| B132 | B222 | Dbil | NS | 0.017748 | 0 |
| B132 | B148 | Dbil | Anti_TPO | -0.00616 | 7.41E-08 |
| B132 | B198 | Dbil | P | -0.0249 | 0 |
| B132 | B194 | Dbil | Cl | -0.01587 | 0 |
| B132 | B155 | Dbil | IgA | 0.006187 | 6.62E-08 |
| B132 | B176 | Dbil | Ucr | 0.020863 | 0 |
| B132 | B191 | Dbil | Urea | -0.01709 | 0 |
| B132 | B174 | Dbil | Urea/Cr | -0.07553 | 0 |
| B132 | B192 | Dbil | UA | 0.041594 | 0 |
| B132 | B170 | Dbil | U-GLU | 0.009846 | 0 |
| B132 | B179 | Dbil | α1-MG | 0.013585 | 0 |
| B132 | B182 | Dbil | TT | 0.006222 | 5.57E-08 |
| B132 | B153 | Dbil | Glu | -0.00661 | 7.73E-09 |
| B132 | B221 | Dbil | Hei | 0.081533 | 0 |
| B132 | B220 | Dbil | SBP | 0.02017 | 0 |
| B132 | B219 | Dbil | OT | -0.00989 | 0 |
| B132 | B216 | Dbil | BMI | -0.01003 | 0 |
| B132 | B215 | Dbil | BMI | -0.07053 | 0 |
| B132 | B172 | Dbil | KB | 0.045898 | 0 |
| B132 | B210 | Dbil | Hip | -0.03615 | 0 |
| B132 | B209 | Dbil | Smoking | -0.03141 | 0 |
| B132 | B139 | Dbil | ADA | 0.02171 | 0 |
| B132 | B196 | Dbil | TCO2 | 0.006779 | 3.25E-09 |
| B132 | B185 | Dbil | PT | 0.016105 | 0 |
| B132 | B186 | Dbil | PFC | -0.01276 | 0 |
| B132 | B173 | Dbil | NTT | -0.00763 | 2.74E-11 |
| B132 | B206 | Dbil | Waist | -0.03988 | 0 |
| B132 | B205 | Dbil | KDA | -0.00912 | 1.78E-15 |
| B132 | B197 | Dbil | AG | 0.007783 | 1.09E-11 |
| B132 | B204 | Dbil | Drinking | 0.041136 | 0 |
| B132 | B203 | Dbil | DH | -0.00867 | 3.64E-14 |
| B132 | B201 | Dbil | e-habits | 0.02369 | 0 |
| B132 | B133 | Dbil | TBIL | 0.797354 | 0 |
| B132 | B135 | Dbil | TBA | 0.049985 | 0 |
| B132 | B212 | Dbil | IOP-L | -0.00572 | 6.04E-07 |
| B133 | B160 | TBIL | PH | 0.0529 | 0 |

| | | | | | |
|---|---|---|---|---|---|
| B133 | B140 | TBIL | r-GT | 0.045513 | 0 |
| B133 | B161 | TBIL | LEU | -0.03353 | 0 |
| B133 | B225 | TBIL | hPBG | 0.007568 | 3.92E-11 |
| B133 | B165 | TBIL | bilirubin | 0.059846 | 0 |
| B133 | B134 | TBIL | CHE | -0.0063 | 3.86E-08 |
| B133 | B166 | TBIL | Pro | 0.062817 | 0 |
| B133 | B188 | TBIL | Ca | 0.013987 | 0 |
| B133 | B189 | TBIL | eGFB | -0.07151 | 0 |
| B133 | B141 | TBIL | AST/ALT | -0.00813 | 1.27E-12 |
| B133 | B181 | TBIL | APTT | 0.007618 | 2.92E-11 |
| B133 | B190 | TBIL | Cre | 0.130135 | 0 |
| B133 | B193 | TBIL | K | -0.02267 | 0 |
| B133 | B222 | TBIL | NS | 0.070614 | 0 |
| B133 | B151 | TBIL | Anti-CCP | -0.00672 | 4.47E-09 |
| B133 | B148 | TBIL | Anti_TPO | -0.00881 | 1.40E-14 |
| B133 | B149 | TBIL | TGAb | -0.00766 | 2.33E-11 |
| B133 | B198 | TBIL | P | -0.02821 | 0 |
| B133 | B194 | TBIL | Cl | -0.01093 | 0 |
| B133 | B176 | TBIL | Ucr | 0.037389 | 0 |
| B133 | B191 | TBIL | Urea | -0.01354 | 0 |
| B133 | B174 | TBIL | Urea/Cr | -0.10204 | 0 |
| B133 | B192 | TBIL | UA | 0.098431 | 0 |
| B133 | B170 | TBIL | U-GLU | 0.009488 | 0 |
| B133 | B179 | TBIL | α1-MG | 0.012806 | 0 |
| B133 | B180 | TBIL | Tf | 0.005762 | 4.89E-07 |
| B133 | B138 | TBIL | PAB | 0.013142 | 0 |
| B133 | B221 | TBIL | Hei | 0.113077 | 0 |
| B133 | B200 | TBIL | OP | -0.00619 | 6.64E-08 |
| B133 | B220 | TBIL | SBP | 0.049086 | 0 |
| B133 | B219 | TBIL | OT | -0.00664 | 6.72E-09 |
| B133 | B218 | TBIL | DBP | 0.043096 | 0 |
| B133 | B216 | TBIL | BMI | 0.044803 | 0 |
| B133 | B215 | TBIL | BMI | -0.01987 | 0 |
| B133 | B172 | TBIL | KB | 0.035454 | 0 |
| B133 | B209 | TBIL | Smoking | -0.03117 | 0 |
| B133 | B139 | TBIL | ADA | 0.007849 | 7.23E-12 |
| B133 | B196 | TBIL | TCO2 | 0.008829 | 1.29E-14 |
| B133 | B183 | TBIL | PTA | 0.024842 | 0 |
| B133 | B185 | TBIL | PT | 0.015273 | 0 |
| B133 | B186 | TBIL | PFC | -0.0154 | 0 |
| B133 | B173 | TBIL | NTT | -0.0137 | 0 |
| B133 | B206 | TBIL | Waist | 0.012514 | 0 |
| B133 | B205 | TBIL | KDA | -0.01497 | 0 |
| B133 | B204 | TBIL | Drinking | 0.048859 | 0 |

| | | | | | |
|---|---|---|---|---|---|
| B133 | B203 | TBIL | DH | -0.02411 | 0 |
| B133 | B211 | TBIL | IOP-R | 0.014794 | 0 |
| B133 | B201 | TBIL | e-habits | 0.010167 | 0 |
| B133 | B135 | TBIL | TBA | 0.015337 | 0 |
| B133 | B212 | TBIL | IOP-L | 0.014662 | 0 |
| B134 | B136 | CHE | 5'NT | 0.02695 | 0 |
| B134 | B137 | CHE | AFU | 0.149811 | 0 |
| B134 | B160 | CHE | PH | -0.02021 | 0 |
| B134 | B140 | CHE | r-GT | 0.058452 | 0 |
| B134 | B161 | CHE | LEU | -0.00767 | 2.18E-11 |
| B134 | B166 | CHE | Pro | 0.01154 | 0 |
| B134 | B188 | CHE | Ca | 0.012583 | 0 |
| B134 | B189 | CHE | eGFB | -0.01496 | 0 |
| B134 | B141 | CHE | AST/ALT | -0.05272 | 0 |
| B134 | B146 | CHE | CysC | 0.020437 | 0 |
| B134 | B181 | CHE | APTT | -0.05503 | 0 |
| B134 | B190 | CHE | Cre | 0.046635 | 0 |
| B134 | B193 | CHE | K | 0.029672 | 0 |
| B134 | B150 | CHE | ALP | 0.144197 | 0 |
| B134 | B222 | CHE | NS | 0.0845 | 0 |
| B134 | B148 | CHE | Anti_TPO | -0.00939 | 2.22E-16 |
| B134 | B149 | CHE | TGAb | -0.01321 | 0 |
| B134 | B198 | CHE | P | -0.04383 | 0 |
| B134 | B194 | CHE | Cl | -0.09625 | 0 |
| B134 | B199 | CHE | Mg | 0.067279 | 0 |
| B134 | B155 | CHE | IgA | 0.014363 | 0 |
| B134 | B157 | CHE | IgG | -0.02899 | 0 |
| B134 | B158 | CHE | IgM | -0.01191 | 0 |
| B134 | B195 | CHE | Na | 0.072364 | 0 |
| B134 | B176 | CHE | Ucr | 0.017039 | 0 |
| B134 | B178 | CHE | Ig-U | 0.007737 | 1.43E-11 |
| B134 | B191 | CHE | Urea | 0.021847 | 0 |
| B134 | B174 | CHE | Urea/Cr | -0.01577 | 0 |
| B134 | B192 | CHE | UA | 0.081926 | 0 |
| B134 | B170 | CHE | U-GLU | 0.012457 | 0 |
| B134 | B179 | CHE | α1-MG | 0.015399 | 0 |
| B134 | B180 | CHE | Tf | 0.011231 | 0 |
| B134 | B182 | CHE | TT | 0.024748 | 0 |
| B134 | B153 | CHE | Glu | 0.006437 | 1.92E-08 |
| B134 | B138 | CHE | PAB | 0.362062 | 0 |
| B134 | B221 | CHE | Hei | 0.047744 | 0 |
| B134 | B187 | CHE | NSE | 0.006603 | 8.19E-09 |
| B134 | B200 | CHE | OP | 0.097999 | 0 |
| B134 | B220 | CHE | SBP | 0.042008 | 0 |

| | | | | | |
|---|---|---|---|---|---|
| B134 | B218 | CHE | DBP | 0.05273 | 0 |
| B134 | B216 | CHE | BMI | 0.087652 | 0 |
| B134 | B215 | CHE | BMI | 0.082421 | 0 |
| B134 | B172 | CHE | KB | -0.00706 | 7.13E-10 |
| B134 | B210 | CHE | Hip | 0.063961 | 0 |
| B134 | B209 | CHE | Smoking | 0.030482 | 0 |
| B134 | B139 | CHE | ADA | -0.0507 | 0 |
| B134 | B196 | CHE | TCO2 | 0.011728 | 0 |
| B134 | B183 | CHE | PTA | -0.00712 | 5.16E-10 |
| B134 | B184 | CHE | INR | 0.006751 | 3.77E-09 |
| B134 | B185 | CHE | PT | -0.03595 | 0 |
| B134 | B186 | CHE | PFC | 0.020694 | 0 |
| B134 | B207 | CHE | WHR | 0.009607 | 0 |
| B134 | B206 | CHE | Waist | 0.088144 | 0 |
| B134 | B197 | CHE | AG | 0.095804 | 0 |
| B134 | B204 | CHE | Drinking | 0.04913 | 0 |
| B134 | B203 | CHE | DH | 0.009168 | 1.33E-15 |
| B134 | B211 | CHE | IOP-R | 0.01937 | 0 |
| B134 | B135 | CHE | TBA | 0.014631 | 0 |
| B134 | B212 | CHE | IOP-L | 0.018614 | 0 |
| B135 | B136 | TBA | 5'NT | 0.025828 | 0 |
| B135 | B137 | TBA | AFU | 0.089927 | 0 |
| B135 | B160 | TBA | PH | -0.0072 | 3.21E-10 |
| B135 | B140 | TBA | r-GT | 0.053419 | 0 |
| B135 | B165 | TBA | bilirubin | 0.076696 | 0 |
| B135 | B141 | TBA | AST/ALT | -0.00787 | 6.33E-12 |
| B135 | B146 | TBA | CysC | 0.010812 | 0 |
| B135 | B190 | TBA | Cre | 0.0087 | 3.06E-14 |
| B135 | B193 | TBA | K | -0.02572 | 0 |
| B135 | B150 | TBA | ALP | 0.108109 | 0 |
| B135 | B222 | TBA | NS | 0.012913 | 0 |
| B135 | B198 | TBA | P | 0.007491 | 6.15E-11 |
| B135 | B199 | TBA | Mg | 0.006843 | 2.31E-09 |
| B135 | B155 | TBA | IgA | 0.009906 | 0 |
| B135 | B157 | TBA | IgG | 0.00717 | 3.86E-10 |
| B135 | B195 | TBA | Na | 0.01441 | 0 |
| B135 | B191 | TBA | Urea | 0.010688 | 0 |
| B135 | B192 | TBA | UA | 0.006981 | 1.10E-09 |
| B135 | B170 | TBA | U-GLU | 0.006035 | 1.38E-07 |
| B135 | B182 | TBA | TT | 0.008163 | 1.03E-12 |
| B135 | B138 | TBA | PAB | -0.00612 | 9.35E-08 |
| B135 | B221 | TBA | Hei | 0.008202 | 8.03E-13 |
| B135 | B200 | TBA | OP | 0.02406 | 0 |
| B135 | B220 | TBA | SBP | 0.010554 | 0 |

| | | | | | |
|---|---|---|---|---|---|
| B135 | B218 | TBA | DBP | 0.010479 | 0 |
| B135 | B216 | TBA | BMI | 0.010388 | 0 |
| B135 | B215 | TBA | BMI | 0.00833 | 3.52E-13 |
| B135 | B210 | TBA | Hip | 0.00564 | 8.50E-07 |
| B135 | B209 | TBA | Smoking | 0.014983 | 0 |
| B135 | B139 | TBA | ADA | 0.074533 | 0 |
| B135 | B206 | TBA | Waist | 0.013703 | 0 |
| B135 | B197 | TBA | AG | 0.007562 | 4.06E-11 |
| B135 | B204 | TBA | Drinking | 0.011468 | 0 |
| B136 | B137 | 5'NT | AFU | 0.010218 | 0 |
| B136 | B140 | 5'NT | r-GT | 0.031498 | 0 |
| B136 | B141 | 5'NT | AST/ALT | -0.00642 | 2.07E-08 |
| B136 | B193 | 5'NT | K | 0.012107 | 0 |
| B136 | B150 | 5'NT | ALP | 0.049173 | 0 |
| B136 | B222 | 5'NT | NS | 0.005978 | 1.80E-07 |
| B136 | B198 | 5'NT | P | 0.011103 | 0 |
| B136 | B194 | 5'NT | Cl | -0.00648 | 1.54E-08 |
| B136 | B155 | 5'NT | IgA | 0.006096 | 1.02E-07 |
| B136 | B157 | 5'NT | IgG | -0.01507 | 0 |
| B136 | B138 | 5'NT | PAB | 0.03092 | 0 |
| B136 | B216 | 5'NT | BMI | 0.007224 | 2.85E-10 |
| B136 | B215 | 5'NT | BMI | 0.006523 | 1.24E-08 |
| B136 | B210 | 5'NT | Hip | 0.006468 | 1.63E-08 |
| B136 | B139 | 5'NT | ADA | 0.017509 | 0 |
| B136 | B186 | 5'NT | PFC | 0.005963 | 1.93E-07 |
| B136 | B206 | 5'NT | Waist | 0.006954 | 1.27E-09 |
| B136 | B197 | 5'NT | AG | 0.008114 | 1.40E-12 |
| B136 | B203 | 5'NT | DH | 0.008749 | 2.20E-14 |
| B137 | B160 | AFU | PH | -0.00732 | 1.69E-10 |
| B137 | B140 | AFU | r-GT | 0.050797 | 0 |
| B137 | B165 | AFU | bilirubin | 0.008322 | 3.71E-13 |
| B137 | B166 | AFU | Pro | 0.006467 | 1.64E-08 |
| B137 | B189 | AFU | eGFB | -0.00918 | 1.11E-15 |
| B137 | B141 | AFU | AST/ALT | -0.01979 | 0 |
| B137 | B146 | AFU | CysC | 0.016412 | 0 |
| B137 | B181 | AFU | APTT | -0.02492 | 0 |
| B137 | B190 | AFU | Cre | 0.015452 | 0 |
| B137 | B193 | AFU | K | -0.01634 | 0 |
| B137 | B150 | AFU | ALP | 0.090536 | 0 |
| B137 | B222 | AFU | NS | 0.034146 | 0 |
| B137 | B154 | AFU | RF | 0.010763 | 0 |
| B137 | B198 | AFU | P | -0.05866 | 0 |
| B137 | B194 | AFU | Cl | -0.03172 | 0 |
| B137 | B155 | AFU | IgA | 0.018274 | 0 |

| | | | | | |
|---|---|---|---|---|---|
| B137 | B156 | AFU | IgE | 0.008977 | 4.44E-15 |
| B137 | B157 | AFU | IgG | -0.00788 | 5.83E-12 |
| B137 | B158 | AFU | IgM | -0.00797 | 3.35E-12 |
| B137 | B195 | AFU | Na | 0.014802 | 0 |
| B137 | B176 | AFU | Ucr | 0.008176 | 9.45E-13 |
| B137 | B178 | AFU | Ig-U | 0.014501 | 0 |
| B137 | B191 | AFU | Urea | 0.006913 | 1.58E-09 |
| B137 | B174 | AFU | Urea/Cr | -0.00734 | 1.46E-10 |
| B137 | B192 | AFU | UA | 0.028656 | 0 |
| B137 | B170 | AFU | U-GLU | 0.016133 | 0 |
| B137 | B179 | AFU | α1-MG | 0.025094 | 0 |
| B137 | B180 | AFU | Tf | 0.015975 | 0 |
| B137 | B182 | AFU | TT | 0.010393 | 0 |
| B137 | B153 | AFU | Glu | 0.006078 | 1.12E-07 |
| B137 | B138 | AFU | PAB | 0.133229 | 0 |
| B137 | B221 | AFU | Hei | 0.015779 | 0 |
| B137 | B200 | AFU | OP | 0.053294 | 0 |
| B137 | B220 | AFU | SBP | 0.029306 | 0 |
| B137 | B218 | AFU | DBP | 0.030718 | 0 |
| B137 | B216 | AFU | BMI | 0.031735 | 0 |
| B137 | B215 | AFU | BMI | 0.031007 | 0 |
| B137 | B210 | AFU | Hip | 0.022506 | 0 |
| B137 | B209 | AFU | Smoking | 0.025122 | 0 |
| B137 | B139 | AFU | ADA | 0.145506 | 0 |
| B137 | B196 | AFU | TCO2 | 0.030939 | 0 |
| B137 | B183 | AFU | PTA | 0.014871 | 0 |
| B137 | B185 | AFU | PT | -0.0133 | 0 |
| B137 | B206 | AFU | Waist | 0.037415 | 0 |
| B137 | B204 | AFU | Drinking | 0.031454 | 0 |
| B137 | B211 | AFU | IOP-R | 0.00722 | 2.91E-10 |
| B137 | B201 | AFU | e-habits | -0.00586 | 3.08E-07 |
| B137 | B212 | AFU | IOP-L | 0.008842 | 1.15E-14 |
| B138 | B160 | PAB | PH | -0.01787 | 0 |
| B138 | B140 | PAB | r-GT | 0.069815 | 0 |
| B138 | B161 | PAB | LEU | -0.0139 | 0 |
| B138 | B166 | PAB | Pro | 0.014008 | 0 |
| B138 | B188 | PAB | Ca | 0.014025 | 0 |
| B138 | B189 | PAB | eGFB | -0.01084 | 0 |
| B138 | B141 | PAB | AST/ALT | -0.04159 | 0 |
| B138 | B145 | PAB | BGP | -0.00723 | 2.75E-10 |
| B138 | B146 | PAB | CysC | 0.013909 | 0 |
| B138 | B181 | PAB | APTT | -0.08202 | 0 |
| B138 | B190 | PAB | Cre | 0.070309 | 0 |
| B138 | B224 | PAB | pmh | -0.00604 | 1.35E-07 |

| | | | | | |
|---|---|---|---|---|---|
| B138 | B193 | PAB | K | 0.019093 | 0 |
| B138 | B150 | PAB | ALP | 0.016477 | 0 |
| B138 | B222 | PAB | NS | 0.082934 | 0 |
| B138 | B151 | PAB | Anti-CCP | -0.00857 | 7.13E-14 |
| B138 | B148 | PAB | Anti_TPO | -0.01752 | 0 |
| B138 | B149 | PAB | TGAb | -0.01812 | 0 |
| B138 | B198 | PAB | P | -0.09002 | 0 |
| B138 | B194 | PAB | Cl | -0.15743 | 0 |
| B138 | B199 | PAB | Mg | 0.110038 | 0 |
| B138 | B156 | PAB | IgE | 0.009053 | 2.66E-15 |
| B138 | B157 | PAB | IgG | -0.07199 | 0 |
| B138 | B158 | PAB | IgM | -0.0257 | 0 |
| B138 | B195 | PAB | Na | 0.08734 | 0 |
| B138 | B168 | PAB | NMP22 | -0.00649 | 1.43E-08 |
| B138 | B176 | PAB | Ucr | 0.031664 | 0 |
| B138 | B191 | PAB | Urea | 0.026375 | 0 |
| B138 | B174 | PAB | Urea/Cr | -0.03228 | 0 |
| B138 | B192 | PAB | UA | 0.097115 | 0 |
| B138 | B179 | PAB | α1-MG | 0.022028 | 0 |
| B138 | B182 | PAB | TT | 0.023182 | 0 |
| B138 | B221 | PAB | Hei | 0.073079 | 0 |
| B138 | B187 | PAB | NSE | 0.009311 | 4.44E-16 |
| B138 | B200 | PAB | OP | 0.117531 | 0 |
| B138 | B220 | PAB | SBP | 0.027547 | 0 |
| B138 | B219 | PAB | OT | -0.01088 | 0 |
| B138 | B218 | PAB | DBP | 0.048837 | 0 |
| B138 | B216 | PAB | BMI | 0.075728 | 0 |
| B138 | B215 | PAB | BMI | 0.048488 | 0 |
| B138 | B172 | PAB | KB | -0.00666 | 6.12E-09 |
| B138 | B210 | PAB | Hip | 0.043556 | 0 |
| B138 | B209 | PAB | Smoking | 0.045671 | 0 |
| B138 | B139 | PAB | ADA | -0.29099 | 0 |
| B138 | B196 | PAB | TCO2 | 0.020621 | 0 |
| B138 | B183 | PAB | PTA | -0.01304 | 0 |
| B138 | B185 | PAB | PT | -0.06469 | 0 |
| B138 | B186 | PAB | PFC | -0.0241 | 0 |
| B138 | B173 | PAB | NTT | -0.01031 | 0 |
| B138 | B207 | PAB | WHR | 0.007558 | 4.15E-11 |
| B138 | B206 | PAB | Waist | 0.066945 | 0 |
| B138 | B197 | PAB | AG | 0.137579 | 0 |
| B138 | B204 | PAB | Drinking | 0.092287 | 0 |
| B138 | B203 | PAB | DH | 0.007254 | 2.41E-10 |
| B138 | B211 | PAB | IOP-R | 0.023701 | 0 |
| B138 | B144 | PAB | Total-PINP | -0.00859 | 6.31E-14 |

| | | | | | |
|---|---|---|---|---|---|
| B138 | B212 | PAB | IOP-L | 0.022233 | 0 |
| B139 | B140 | ADA | r-GT | 0.017748 | 0 |
| B139 | B161 | ADA | LEU | 0.009713 | 0 |
| B139 | B165 | ADA | bilirubin | 0.008428 | 1.87E-13 |
| B139 | B189 | ADA | eGFB | -0.0123 | 0 |
| B139 | B146 | ADA | CysC | 0.023131 | 0 |
| B139 | B181 | ADA | APTT | 0.031234 | 0 |
| B139 | B190 | ADA | Cre | -0.01636 | 0 |
| B139 | B193 | ADA | K | 0.025985 | 0 |
| B139 | B150 | ADA | ALP | 0.164595 | 0 |
| B139 | B222 | ADA | NS | -0.02197 | 0 |
| B139 | B148 | ADA | Anti_TPO | 0.012095 | 0 |
| B139 | B149 | ADA | TGAb | 0.013202 | 0 |
| B139 | B154 | ADA | RF | 0.01828 | 0 |
| B139 | B198 | ADA | P | 0.018544 | 0 |
| B139 | B194 | ADA | Cl | -0.03394 | 0 |
| B139 | B199 | ADA | Mg | -0.02925 | 0 |
| B139 | B155 | ADA | IgA | 0.054775 | 0 |
| B139 | B156 | ADA | IgE | 0.01108 | 0 |
| B139 | B157 | ADA | IgG | 0.087996 | 0 |
| B139 | B158 | ADA | IgM | 0.015894 | 0 |
| B139 | B195 | ADA | Na | 0.020693 | 0 |
| B139 | B176 | ADA | Ucr | -0.00917 | 1.33E-15 |
| B139 | B178 | ADA | Ig-U | 0.023151 | 0 |
| B139 | B174 | ADA | Urea/Cr | 0.013711 | 0 |
| B139 | B192 | ADA | UA | -0.01383 | 0 |
| B139 | B170 | ADA | U-GLU | 0.025602 | 0 |
| B139 | B175 | ADA | ALB-U | 0.007032 | 8.30E-10 |
| B139 | B177 | ADA | ACR | 0.012757 | 0 |
| B139 | B179 | ADA | α1-MG | 0.027884 | 0 |
| B139 | B180 | ADA | Tf | 0.027825 | 0 |
| B139 | B182 | ADA | TT | 0.020287 | 0 |
| B139 | B153 | ADA | Glu | 0.006449 | 1.80E-08 |
| B139 | B221 | ADA | Hei | -0.0304 | 0 |
| B139 | B200 | ADA | OP | 0.049715 | 0 |
| B139 | B220 | ADA | SBP | 0.020284 | 0 |
| B139 | B219 | ADA | OT | 0.006831 | 2.46E-09 |
| B139 | B216 | ADA | BMI | -0.01869 | 0 |
| B139 | B210 | ADA | Hip | -0.0074 | 1.03E-10 |
| B139 | B209 | ADA | Smoking | -0.01716 | 0 |
| B139 | B196 | ADA | TCO2 | -0.00854 | 9.02E-14 |
| B139 | B183 | ADA | PTA | -0.00598 | 1.82E-07 |
| B139 | B185 | ADA | PT | 0.042457 | 0 |
| B139 | B186 | ADA | PFC | 0.024961 | 0 |

| | | | | | |
|---|---|---|---|---|---|
| B139 | B173 | ADA | NTT | 0.011944 | 0 |
| B139 | B206 | ADA | Waist | -0.00711 | 5.40E-10 |
| B139 | B197 | ADA | AG | 0.040968 | 0 |
| B139 | B204 | ADA | Drinking | -0.03202 | 0 |
| B139 | B211 | ADA | IOP-R | -0.01059 | 0 |
| B139 | B201 | ADA | e-habits | 0.005907 | 2.51E-07 |
| B139 | B144 | ADA | Total-PINP | 0.006749 | 3.82E-09 |
| B139 | B212 | ADA | IOP-L | -0.00988 | 0 |
| B14 | B24 | G-17 | αHBDH | 0.007151 | 4.30E-10 |
| B14 | B19 | G-17 | CD3+CD4+/CD3+CD8+ | 0.006522 | 1.24E-08 |
| B14 | B54 | G-17 | ESR | 0.006257 | 4.70E-08 |
| B14 | B126 | G-17 | Alb | -0.01402 | 0 |
| B14 | B127 | G-17 | A-G ratio | -0.01433 | 0 |
| B14 | B40 | G-17 | WBC | 0.009536 | 0 |
| B14 | B111 | G-17 | TSH | 0.007176 | 3.73E-10 |
| B14 | B134 | G-17 | CHE | -0.00581 | 3.96E-07 |
| B14 | B189 | G-17 | eGFB | -0.01668 | 0 |
| B14 | B45 | G-17 | MCHC | -0.01208 | 0 |
| B14 | B46 | G-17 | RBC Count | -0.01296 | 0 |
| B14 | B47 | G-17 | RDW | 0.017796 | 0 |
| B14 | B129 | G-17 | IBIL | -0.00876 | 2.09E-14 |
| B14 | B150 | G-17 | ALP | 0.005781 | 4.49E-07 |
| B14 | B148 | G-17 | Anti_TPO | 0.023423 | 0 |
| B14 | B149 | G-17 | TGAb | 0.01026 | 0 |
| B14 | B176 | G-17 | Ucr | -0.00631 | 3.69E-08 |
| B14 | B191 | G-17 | Urea | 0.0081 | 1.53E-12 |
| B14 | B174 | G-17 | Urea/Cr | 0.010122 | 0 |
| B14 | B192 | G-17 | UA | -0.00913 | 1.55E-15 |
| B14 | B138 | G-17 | PAB | -0.01073 | 0 |
| B14 | B130 | G-17 | GLB | 0.012606 | 0 |
| B14 | B27 | G-17 | LDH | 0.006635 | 6.94E-09 |
| B14 | B221 | G-17 | Hei | -0.01257 | 0 |
| B14 | B220 | G-17 | SBP | 0.014123 | 0 |
| B14 | B218 | G-17 | DBP | 0.009722 | 0 |
| B14 | B88 | G-17 | CA 242 | 0.006951 | 1.29E-09 |
| B14 | B216 | G-17 | BMI | -0.00568 | 7.02E-07 |
| B14 | B131 | G-17 | AST | 0.007068 | 6.81E-10 |
| B14 | B90 | G-17 | CYFRA 21-1 | 0.009008 | 3.55E-15 |
| B14 | B139 | G-17 | ADA | 0.015703 | 0 |
| B14 | B51 | G-17 | Hb | -0.0145 | 0 |
| B14 | B78 | G-17 | HBcAb | -0.00698 | 1.13E-09 |
| B14 | B80 | G-17 | anti-HP IgG | 0.056382 | 0 |
| B14 | B133 | G-17 | TBIL | -0.00777 | 1.17E-11 |
| B140 | B160 | r-GT | PH | -0.07804 | 0 |

| | | | | | |
|---|---|---|---|---|---|
| B140 | B161 | r-GT | LEU | -0.03105 | 0 |
| B140 | B165 | r-GT | bilirubin | 0.078428 | 0 |
| B140 | B166 | r-GT | Pro | 0.067957 | 0 |
| B140 | B188 | r-GT | Ca | 0.018036 | 0 |
| B140 | B189 | r-GT | eGFB | -0.06086 | 0 |
| B140 | B141 | r-GT | AST/ALT | -0.2095 | 0 |
| B140 | B146 | r-GT | CysC | 0.017799 | 0 |
| B140 | B181 | r-GT | APTT | -0.01499 | 0 |
| B140 | B190 | r-GT | Cre | 0.13314 | 0 |
| B140 | B193 | r-GT | K | -0.00893 | 6.22E-15 |
| B140 | B150 | r-GT | ALP | 0.110765 | 0 |
| B140 | B222 | r-GT | NS | 0.251348 | 0 |
| B140 | B148 | r-GT | Anti_TPO | -0.01875 | 0 |
| B140 | B149 | r-GT | TGAb | -0.02149 | 0 |
| B140 | B198 | r-GT | P | -0.01403 | 0 |
| B140 | B194 | r-GT | Cl | -0.01384 | 0 |
| B140 | B155 | r-GT | IgA | 0.00755 | 4.35E-11 |
| B140 | B156 | r-GT | IgE | 0.006251 | 4.83E-08 |
| B140 | B157 | r-GT | IgG | -0.0148 | 0 |
| B140 | B176 | r-GT | Ucr | 0.055914 | 0 |
| B140 | B178 | r-GT | Ig-U | 0.008529 | 9.59E-14 |
| B140 | B191 | r-GT | Urea | 0.04127 | 0 |
| B140 | B174 | r-GT | Urea/Cr | -0.06255 | 0 |
| B140 | B192 | r-GT | UA | 0.262809 | 0 |
| B140 | B170 | r-GT | U-GLU | 0.068512 | 0 |
| B140 | B175 | r-GT | ALB-U | 0.013542 | 0 |
| B140 | B179 | r-GT | α1-MG | 0.033932 | 0 |
| B140 | B180 | r-GT | Tf | 0.01017 | 0 |
| B140 | B153 | r-GT | Glu | 0.027869 | 0 |
| B140 | B221 | r-GT | Hei | 0.137382 | 0 |
| B140 | B200 | r-GT | OP | 0.008215 | 7.38E-13 |
| B140 | B220 | r-GT | SBP | 0.150156 | 0 |
| B140 | B218 | r-GT | DBP | 0.177007 | 0 |
| B140 | B216 | r-GT | BMI | 0.253544 | 0 |
| B140 | B215 | r-GT | BMI | 0.23702 | 0 |
| B140 | B210 | r-GT | Hip | 0.178681 | 0 |
| B140 | B209 | r-GT | Smoking | 0.163015 | 0 |
| B140 | B185 | r-GT | PT | -0.00984 | 0 |
| B140 | B186 | r-GT | PFC | 0.005724 | 5.80E-07 |
| B140 | B173 | r-GT | NTT | -0.01211 | 0 |
| B140 | B207 | r-GT | WHR | 0.031271 | 0 |
| B140 | B206 | r-GT | Waist | 0.273163 | 0 |
| B140 | B205 | r-GT | KDA | -0.01168 | 0 |
| B140 | B197 | r-GT | AG | 0.012518 | 0 |

| | | | | | |
|---|---|---|---|---|---|
| B140 | B204 | r-GT | Drinking | 0.22511 | 0 |
| B140 | B203 | r-GT | DH | 0.01944 | 0 |
| B140 | B211 | r-GT | IOP-R | 0.02123 | 0 |
| B140 | B201 | r-GT | e-habits | -0.01188 | 0 |
| B140 | B212 | r-GT | IOP-L | 0.021965 | 0 |
| B141 | B160 | AST/ALT | PH | 0.048732 | 0 |
| B141 | B161 | AST/ALT | LEU | 0.056484 | 0 |
| B141 | B166 | AST/ALT | Pro | -0.05617 | 0 |
| B141 | B188 | AST/ALT | Ca | -0.00799 | 3.08E-12 |
| B141 | B189 | AST/ALT | eGFB | 0.041857 | 0 |
| B141 | B146 | AST/ALT | CysC | -0.01124 | 0 |
| B141 | B181 | AST/ALT | APTT | 0.009161 | 1.33E-15 |
| B141 | B190 | AST/ALT | Cre | -0.20523 | 0 |
| B141 | B150 | AST/ALT | ALP | -0.05068 | 0 |
| B141 | B222 | AST/ALT | NS | -0.31421 | 0 |
| B141 | B148 | AST/ALT | Anti_TPO | 0.01992 | 0 |
| B141 | B149 | AST/ALT | TGAb | 0.022396 | 0 |
| B141 | B198 | AST/ALT | P | 0.019157 | 0 |
| B141 | B194 | AST/ALT | Cl | 0.008969 | 4.88E-15 |
| B141 | B157 | AST/ALT | IgG | 0.023044 | 0 |
| B141 | B158 | AST/ALT | IgM | 0.007718 | 1.60E-11 |
| B141 | B195 | AST/ALT | Na | -0.01214 | 0 |
| B141 | B176 | AST/ALT | Ucr | -0.04409 | 0 |
| B141 | B191 | AST/ALT | Urea | -0.08091 | 0 |
| B141 | B174 | AST/ALT | Urea/Cr | 0.078044 | 0 |
| B141 | B192 | AST/ALT | UA | -0.2688 | 0 |
| B141 | B170 | AST/ALT | U-GLU | -0.0324 | 0 |
| B141 | B177 | AST/ALT | ACR | 0.006733 | 4.15E-09 |
| B141 | B179 | AST/ALT | α1-MG | -0.01145 | 0 |
| B141 | B182 | AST/ALT | TT | -0.00591 | 2.45E-07 |
| B141 | B153 | AST/ALT | Glu | -0.01586 | 0 |
| B141 | B221 | AST/ALT | Hei | -0.22555 | 0 |
| B141 | B200 | AST/ALT | OP | -0.01412 | 0 |
| B141 | B220 | AST/ALT | SBP | -0.1134 | 0 |
| B141 | B219 | AST/ALT | OT | 0.009352 | 4.44E-16 |
| B141 | B218 | AST/ALT | DBP | -0.12414 | 0 |
| B141 | B216 | AST/ALT | BMI | -0.34558 | 0 |
| B141 | B215 | AST/ALT | BMI | -0.30114 | 0 |
| B141 | B172 | AST/ALT | KB | 0.031638 | 0 |
| B141 | B210 | AST/ALT | Hip | -0.25943 | 0 |
| B141 | B209 | AST/ALT | Smoking | -0.09784 | 0 |
| B141 | B185 | AST/ALT | PT | 0.008626 | 5.02E-14 |
| B141 | B173 | AST/ALT | NTT | 0.026302 | 0 |
| B141 | B207 | AST/ALT | WHR | -0.03373 | 0 |

| | | | | | |
|---|---|---|---|---|---|
| B141 | B206 | AST/ALT | Waist | -0.32588 | 0 |
| B141 | B205 | AST/ALT | KDA | 0.016425 | 0 |
| B141 | B197 | AST/ALT | AG | -0.01205 | 0 |
| B141 | B204 | AST/ALT | Drinking | -0.11385 | 0 |
| B141 | B203 | AST/ALT | DH | -0.01988 | 0 |
| B141 | B211 | AST/ALT | IOP-R | -0.01289 | 0 |
| B141 | B212 | AST/ALT | IOP-L | -0.01148 | 0 |
| B142 | B143 | 25-(OH)D | β-CTx | 0.014895 | 0 |
| B142 | B189 | 25-(OH)D | eGFB | -0.01711 | 0 |
| B142 | B145 | 25-(OH)D | BGP | -0.03482 | 0 |
| B142 | B222 | 25-(OH)D | NS | -0.00739 | 1.08E-10 |
| B142 | B151 | 25-(OH)D | Anti-CCP | 0.013041 | 0 |
| B142 | B176 | 25-(OH)D | Ucr | -0.01842 | 0 |
| B142 | B182 | 25-(OH)D | TT | 0.005933 | 2.22E-07 |
| B142 | B221 | 25-(OH)D | Hei | -0.00591 | 2.48E-07 |
| B142 | B228 | 25-(OH)D | OS | 0.007436 | 8.48E-11 |
| B142 | B219 | 25-(OH)D | OT | 0.006415 | 2.13E-08 |
| B142 | B216 | 25-(OH)D | BMI | -0.00967 | 0 |
| B142 | B215 | 25-(OH)D | BMI | -0.00926 | 6.66E-16 |
| B142 | B210 | 25-(OH)D | Hip | -0.00637 | 2.64E-08 |
| B142 | B183 | 25-(OH)D | PTA | -0.01259 | 0 |
| B142 | B206 | 25-(OH)D | Waist | -0.00762 | 2.83E-11 |
| B142 | B197 | 25-(OH)D | AG | 0.005994 | 1.67E-07 |
| B142 | B201 | 25-(OH)D | e-habits | 0.008182 | 9.11E-13 |
| B143 | B145 | β-CTx | BGP | 0.579579 | 0 |
| B143 | B190 | β-CTx | Cre | 0.006473 | 1.59E-08 |
| B143 | B150 | β-CTx | ALP | 0.026408 | 0 |
| B143 | B222 | β-CTx | NS | 0.006062 | 1.21E-07 |
| B143 | B198 | β-CTx | P | 0.009819 | 0 |
| B143 | B194 | β-CTx | Cl | 0.00577 | 4.71E-07 |
| B143 | B195 | β-CTx | Na | 0.005872 | 2.95E-07 |
| B143 | B168 | β-CTx | NMP22 | 0.006052 | 1.27E-07 |
| B143 | B176 | β-CTx | Ucr | 0.027272 | 0 |
| B143 | B182 | β-CTx | TT | -0.00724 | 2.63E-10 |
| B143 | B209 | β-CTx | Smoking | 0.008219 | 7.20E-13 |
| B143 | B183 | β-CTx | PTA | 0.022752 | 0 |
| B143 | B186 | β-CTx | PFC | -0.00624 | 5.21E-08 |
| B143 | B144 | β-CTx | Total-PINP | 0.739751 | 0 |
| B144 | B145 | Total-PINP | BGP | 0.592844 | 0 |
| B144 | B150 | Total-PINP | ALP | 0.030371 | 0 |
| B144 | B148 | Total-PINP | Anti_TPO | 0.006645 | 6.56E-09 |
| B144 | B198 | Total-PINP | P | 0.008228 | 6.82E-13 |
| B144 | B194 | Total-PINP | Cl | 0.006717 | 4.51E-09 |
| B144 | B168 | Total-PINP | NMP22 | 0.010044 | 0 |

| | | | | | |
|---|---|---|---|---|---|
| B144 | B176 | Total-PINP | Ucr | 0.010411 | 0 |
| B144 | B183 | Total-PINP | PTA | 0.014922 | 0 |
| B145 | B181 | BGP | APTT | 0.005779 | 4.52E-07 |
| B145 | B150 | BGP | ALP | 0.043605 | 0 |
| B145 | B148 | BGP | Anti_TPO | 0.006281 | 4.16E-08 |
| B145 | B198 | BGP | P | 0.00786 | 6.76E-12 |
| B145 | B194 | BGP | Cl | 0.008489 | 1.25E-13 |
| B145 | B168 | BGP | NMP22 | 0.020256 | 0 |
| B145 | B182 | BGP | TT | -0.01108 | 0 |
| B145 | B183 | BGP | PTA | 0.010919 | 0 |
| B145 | B185 | BGP | PT | 0.005713 | 6.12E-07 |
| B146 | B160 | CysC | PH | -0.01925 | 0 |
| B146 | B189 | CysC | eGFB | -0.07324 | 0 |
| B146 | B190 | CysC | Cre | 0.074655 | 0 |
| B146 | B193 | CysC | K | 0.006431 | 1.97E-08 |
| B146 | B150 | CysC | ALP | 0.023331 | 0 |
| B146 | B222 | CysC | NS | 0.03703 | 0 |
| B146 | B154 | CysC | RF | 0.008555 | 8.08E-14 |
| B146 | B198 | CysC | P | -0.008 | 2.92E-12 |
| B146 | B194 | CysC | Cl | 0.007688 | 1.92E-11 |
| B146 | B199 | CysC | Mg | 0.015997 | 0 |
| B146 | B195 | CysC | Na | 0.024335 | 0 |
| B146 | B191 | CysC | Urea | 0.036488 | 0 |
| B146 | B174 | CysC | Urea/Cr | -0.01965 | 0 |
| B146 | B192 | CysC | UA | 0.046767 | 0 |
| B146 | B177 | CysC | ACR | 0.006013 | 1.53E-07 |
| B146 | B179 | CysC | α1-MG | 0.008719 | 2.71E-14 |
| B146 | B221 | CysC | Hei | 0.019733 | 0 |
| B146 | B200 | CysC | OP | 0.030452 | 0 |
| B146 | B220 | CysC | SBP | 0.031054 | 0 |
| B146 | B219 | CysC | OT | 0.006756 | 3.66E-09 |
| B146 | B218 | CysC | DBP | 0.023135 | 0 |
| B146 | B216 | CysC | BMI | 0.029109 | 0 |
| B146 | B215 | CysC | BMI | 0.025486 | 0 |
| B146 | B210 | CysC | Hip | 0.022761 | 0 |
| B146 | B209 | CysC | Smoking | 0.031166 | 0 |
| B146 | B183 | CysC | PTA | -0.00583 | 3.64E-07 |
| B146 | B206 | CysC | Waist | 0.037689 | 0 |
| B146 | B197 | CysC | AG | 0.00883 | 1.27E-14 |
| B146 | B204 | CysC | Drinking | 0.018672 | 0 |
| B146 | B201 | CysC | e-habits | 0.006621 | 7.44E-09 |
| B148 | B160 | Anti_TPO | PH | 0.007499 | 5.86E-11 |
| B148 | B161 | Anti_TPO | LEU | 0.011423 | 0 |
| B148 | B166 | Anti_TPO | Pro | -0.01072 | 0 |

| | | | | | |
|---|---|---|---|---|---|
| B148 | B189 | Anti_TPO | eGFB | 0.011652 | 0 |
| B148 | B190 | Anti_TPO | Cre | -0.04163 | 0 |
| B148 | B222 | Anti_TPO | NS | -0.0401 | 0 |
| B148 | B149 | Anti_TPO | TGAb | 0.42356 | 0 |
| B148 | B198 | Anti_TPO | P | 0.006334 | 3.21E-08 |
| B148 | B157 | Anti_TPO | IgG | 0.013488 | 0 |
| B148 | B176 | Anti_TPO | Ucr | -0.02058 | 0 |
| B148 | B191 | Anti_TPO | Urea | -0.01075 | 0 |
| B148 | B174 | Anti_TPO | Urea/Cr | 0.02317 | 0 |
| B148 | B192 | Anti_TPO | UA | -0.035 | 0 |
| B148 | B170 | Anti_TPO | U-GLU | -0.0058 | 4.12E-07 |
| B148 | B221 | Anti_TPO | Hei | -0.04204 | 0 |
| B148 | B220 | Anti_TPO | SBP | -0.01193 | 0 |
| B148 | B218 | Anti_TPO | DBP | -0.01553 | 0 |
| B148 | B216 | Anti_TPO | BMI | -0.03562 | 0 |
| B148 | B215 | Anti_TPO | BMI | -0.01727 | 0 |
| B148 | B210 | Anti_TPO | Hip | -0.01795 | 0 |
| B148 | B209 | Anti_TPO | Smoking | -0.02429 | 0 |
| B148 | B206 | Anti_TPO | Waist | -0.02874 | 0 |
| B148 | B205 | Anti_TPO | KDA | 0.005814 | 3.86E-07 |
| B148 | B197 | Anti_TPO | AG | -0.00563 | 8.71E-07 |
| B148 | B204 | Anti_TPO | Drinking | -0.03068 | 0 |
| B149 | B161 | TGAb | LEU | 0.00847 | 1.42E-13 |
| B149 | B166 | TGAb | Pro | -0.01034 | 0 |
| B149 | B189 | TGAb | eGFB | 0.009748 | 0 |
| B149 | B190 | TGAb | Cre | -0.03485 | 0 |
| B149 | B222 | TGAb | NS | -0.03983 | 0 |
| B149 | B154 | TGAb | RF | 0.006437 | 1.91E-08 |
| B149 | B198 | TGAb | P | 0.008183 | 9.03E-13 |
| B149 | B157 | TGAb | IgG | 0.015802 | 0 |
| B149 | B168 | TGAb | NMP22 | 0.015794 | 0 |
| B149 | B176 | TGAb | Ucr | -0.02114 | 0 |
| B149 | B191 | TGAb | Urea | -0.011 | 0 |
| B149 | B174 | TGAb | Urea/Cr | 0.017386 | 0 |
| B149 | B192 | TGAb | UA | -0.03396 | 0 |
| B149 | B169 | TGAb | U-CRY | 0.00561 | 9.71E-07 |
| B149 | B221 | TGAb | Hei | -0.03706 | 0 |
| B149 | B220 | TGAb | SBP | -0.01307 | 0 |
| B149 | B218 | TGAb | DBP | -0.01475 | 0 |
| B149 | B216 | TGAb | BMI | -0.03652 | 0 |
| B149 | B215 | TGAb | BMI | -0.02243 | 0 |
| B149 | B210 | TGAb | Hip | -0.02055 | 0 |
| B149 | B209 | TGAb | Smoking | -0.02615 | 0 |
| B149 | B206 | TGAb | Waist | -0.03288 | 0 |

| | | | | | |
|---|---|---|---|---|---|
| B149 | B204 | TGAb | Drinking | -0.03017 | 0 |
| B15 | B142 | CD3-CD19+(B cell) | 25-(OH)D | 0.005791 | 4.28E-07 |
| B15 | B137 | CD3-CD19+(B cell) | AFU | 0.02553 | 0 |
| B15 | B16 | CD3-CD19+(B cell) | CD3+T | -0.07039 | 0 |
| B15 | B23 | CD3-CD19+(B cell) | CD3-CD16+CD56+(NK) | -0.16096 | 0 |
| B15 | B17 | CD3-CD19+(B cell) | CD3+CD16+CD56+NK | -0.08175 | 0 |
| B15 | B18 | CD3-CD19+(B cell) | CD3+CD4+helper(inducer)T | 0.093631 | 0 |
| B15 | B19 | CD3-CD19+(B cell) | CD3+CD4+/CD3+CD8+ | 0.183395 | 0 |
| B15 | B20 | CD3-CD19+(B cell) | CD3+CD4+CD8+ | -0.03289 | 0 |
| B15 | B21 | CD3-CD19+(B cell) | CD3+CD4-CD8- | -0.05047 | 0 |
| B15 | B22 | CD3-CD19+(B cell) | CD3+CD8+(T) | -0.15853 | 0 |
| B15 | B54 | CD3-CD19+(B cell) | ESR | 0.021792 | 0 |
| B15 | B92 | CD3-CD19+(B cell) | ALDH2-Glu504Glu | 0.022409 | 0 |
| B15 | B93 | CD3-CD19+(B cell) | ALDH2-Glu504Lys | -0.01587 | 0 |
| B15 | B140 | CD3-CD19+(B cell) | r-GT | -0.00725 | 2.39E-10 |
| B15 | B94 | CD3-CD19+(B cell) | ALDH2-Lys504Lys | -0.01907 | 0 |
| B15 | B40 | CD3-CD19+(B cell) | WBC | 0.019797 | 0 |
| B15 | B121 | CD3-CD19+(B cell) | C3 | -0.01537 | 0 |
| B15 | B122 | CD3-CD19+(B cell) | C4 | -0.00935 | 4.44E-16 |
| B15 | B225 | CD3-CD19+(B cell) | hPBG | 0.009512 | 0 |
| B15 | B124 | CD3-CD19+(B cell) | ACTH | -0.00836 | 2.87E-13 |
| B15 | B70 | CD3-CD19+(B cell) | LDL-C | -0.01334 | 0 |
| B15 | B141 | CD3-CD19+(B cell) | AST/ALT | 0.005685 | 6.94E-07 |
| B15 | B145 | CD3-CD19+(B cell) | BGP | -0.01103 | 0 |
| B15 | B57 | CD3-CD19+(B cell) | RBCAI | -0.0221 | 0 |
| B15 | B44 | CD3-CD19+(B cell) | MCH | -0.00894 | 5.77E-15 |
| B15 | B47 | CD3-CD19+(B cell) | RDW | -0.00725 | 2.44E-10 |
| B15 | B181 | CD3-CD19+(B cell) | APTT | 0.023854 | 0 |
| B15 | B26 | CD3-CD19+(B cell) | CKMB | 0.005858 | 3.14E-07 |
| B15 | B193 | CD3-CD19+(B cell) | K | 0.024172 | 0 |
| B15 | B222 | CD3-CD19+(B cell) | NS | -0.01517 | 0 |
| B15 | B95 | CD3-CD19+(B cell) | anti-cenpb | -0.01857 | 0 |
| B15 | B96 | CD3-CD19+(B cell) | dsDNA (IIF) | 0.026538 | 0 |
| B15 | B98 | CD3-CD19+(B cell) | anti-Jo-1 | -0.01246 | 0 |
| B15 | B100 | CD3-CD19+(B cell) | anti-Ro52 | 0.030506 | 0 |
| B15 | B101 | CD3-CD19+(B cell) | anti-Sc70 | 0.00824 | 6.31E-13 |
| B15 | B103 | CD3-CD19+(B cell) | anti-SSA/Ro | 0.024465 | 0 |
| B15 | B104 | CD3-CD19+(B cell) | anti-SSB | 0.017121 | 0 |
| B15 | B106 | CD3-CD19+(B cell) | anti-P | 0.00906 | 2.66E-15 |
| B15 | B107 | CD3-CD19+(B cell) | AnuA | 0.018422 | 0 |
| B15 | B151 | CD3-CD19+(B cell) | Anti-CCP | -0.0139 | 0 |
| B15 | B149 | CD3-CD19+(B cell) | TGAb | 0.009658 | 0 |
| B15 | B109 | CD3-CD19+(B cell) | AMA-M2 | -0.01216 | 0 |
| B15 | B110 | CD3-CD19+(B cell) | AHA | 0.016209 | 0 |

| | | | | | |
|---|---|---|---|---|---|
| B15 | B194 | CD3-CD19+(B cell) | Cl | 0.007618 | 2.91E-11 |
| B15 | B199 | CD3-CD19+(B cell) | Mg | -0.01441 | 0 |
| B15 | B155 | CD3-CD19+(B cell) | IgA | 0.022458 | 0 |
| B15 | B156 | CD3-CD19+(B cell) | IgE | 0.013964 | 0 |
| B15 | B157 | CD3-CD19+(B cell) | IgG | 0.034429 | 0 |
| B15 | B178 | CD3-CD19+(B cell) | Ig-U | 0.013164 | 0 |
| B15 | B175 | CD3-CD19+(B cell) | ALB-U | 0.007123 | 5.02E-10 |
| B15 | B177 | CD3-CD19+(B cell) | ACR | 0.008619 | 5.28E-14 |
| B15 | B179 | CD3-CD19+(B cell) | α1-MG | 0.012505 | 0 |
| B15 | B180 | CD3-CD19+(B cell) | Tf | 0.014645 | 0 |
| B15 | B182 | CD3-CD19+(B cell) | TT | -0.01595 | 0 |
| B15 | B125 | CD3-CD19+(B cell) | Cor | -0.05394 | 0 |
| B15 | B138 | CD3-CD19+(B cell) | PAB | -0.00856 | 7.97E-14 |
| B15 | B130 | CD3-CD19+(B cell) | GLB | 0.006335 | 3.19E-08 |
| B15 | B59 | CD3-CD19+(B cell) | WBRV | -0.01324 | 0 |
| B15 | B60 | CD3-CD19+(B cell) | WBRVI | -0.01881 | 0 |
| B15 | B64 | CD3-CD19+(B cell) | WBRSR(mPas_5) | -0.01173 | 0 |
| B15 | B65 | CD3-CD19+(B cell) | WBRSR(mPas_1) | -0.01779 | 0 |
| B15 | B87 | CD3-CD19+(B cell) | NSE | 0.010713 | 0 |
| B15 | B200 | CD3-CD19+(B cell) | OP | 0.00737 | 1.24E-10 |
| B15 | B216 | CD3-CD19+(B cell) | BMI | -0.00898 | 4.44E-15 |
| B15 | B215 | CD3-CD19+(B cell) | BMI | -0.01221 | 0 |
| B15 | B210 | CD3-CD19+(B cell) | Hip | -0.00798 | 3.17E-12 |
| B15 | B90 | CD3-CD19+(B cell) | CYFRA 21-1 | -0.023 | 0 |
| B15 | B139 | CD3-CD19+(B cell) | ADA | 0.0277 | 0 |
| B15 | B34 | CD3-CD19+(B cell) | TK | 0.012498 | 0 |
| B15 | B196 | CD3-CD19+(B cell) | TCO2 | -0.00953 | 0 |
| B15 | B51 | CD3-CD19+(B cell) | Hb | -0.00823 | 6.90E-13 |
| B15 | B183 | CD3-CD19+(B cell) | PTA | 0.116881 | 0 |
| B15 | B185 | CD3-CD19+(B cell) | PT | 0.029847 | 0 |
| B15 | B186 | CD3-CD19+(B cell) | PFC | -0.02223 | 0 |
| B15 | B52 | CD3-CD19+(B cell) | PLT/BPC | 0.012683 | 0 |
| B15 | B206 | CD3-CD19+(B cell) | Waist | -0.01767 | 0 |
| B15 | B79 | CD3-CD19+(B cell) | hb V -dna | 0.021302 | 0 |
| B15 | B204 | CD3-CD19+(B cell) | Drinking | -0.01942 | 0 |
| B15 | B203 | CD3-CD19+(B cell) | DH | 0.006683 | 5.40E-09 |
| B15 | B113 | CD3-CD19+(B cell) | FT3 | -0.00796 | 3.66E-12 |
| B15 | B211 | CD3-CD19+(B cell) | IOP-R | -0.00827 | 5.27E-13 |
| B15 | B74 | CD3-CD19+(B cell) | ApoA1 | -0.01226 | 0 |
| B15 | B81 | CD3-CD19+(B cell) | TSGF | -0.01197 | 0 |
| B15 | B77 | CD3-CD19+(B cell) | TC | -0.00922 | 8.88E-16 |
| B15 | B91 | CD3-CD19+(B cell) | TP | 0.007267 | 2.23E-10 |
| B15 | B114 | CD3-CD19+(B cell) | TSH TT3 TT4(TT4) | 0.010081 | 0 |
| B15 | B144 | CD3-CD19+(B cell) | Total-PINP | -0.00697 | 1.17E-09 |

| | | | | | |
|---|---|---|---|---|---|
| B15 | B212 | CD3-CD19+(B cell) | IOP-L | -0.01333 | 0 |
| B150 | B160 | ALP | PH | -0.0353 | 0 |
| B150 | B165 | ALP | bilirubin | 0.038532 | 0 |
| B150 | B166 | ALP | Pro | 0.012606 | 0 |
| B150 | B188 | ALP | Ca | 0.006888 | 1.81E-09 |
| B150 | B189 | ALP | eGFB | -0.03756 | 0 |
| B150 | B181 | ALP | APTT | -0.01092 | 0 |
| B150 | B190 | ALP | Cre | 0.032866 | 0 |
| B150 | B224 | ALP | pmh | 0.006605 | 8.10E-09 |
| B150 | B193 | ALP | K | 0.021819 | 0 |
| B150 | B222 | ALP | NS | 0.052374 | 0 |
| B150 | B151 | ALP | Anti-CCP | 0.006612 | 7.79E-09 |
| B150 | B198 | ALP | P | -0.0366 | 0 |
| B150 | B194 | ALP | Cl | -0.02307 | 0 |
| B150 | B199 | ALP | Mg | 0.044147 | 0 |
| B150 | B155 | ALP | IgA | 0.018485 | 0 |
| B150 | B156 | ALP | IgE | 0.010177 | 0 |
| B150 | B157 | ALP | IgG | -0.00623 | 5.37E-08 |
| B150 | B195 | ALP | Na | 0.074725 | 0 |
| B150 | B168 | ALP | NMP22 | 0.006091 | 1.05E-07 |
| B150 | B176 | ALP | Ucr | 0.018873 | 0 |
| B150 | B178 | ALP | Ig-U | 0.007456 | 7.52E-11 |
| B150 | B191 | ALP | Urea | 0.028209 | 0 |
| B150 | B192 | ALP | UA | 0.053797 | 0 |
| B150 | B170 | ALP | U-GLU | 0.023972 | 0 |
| B150 | B175 | ALP | ALB-U | 0.008128 | 1.28E-12 |
| B150 | B177 | ALP | ACR | 0.009088 | 2.22E-15 |
| B150 | B179 | ALP | α1-MG | 0.033185 | 0 |
| B150 | B180 | ALP | Tf | 0.007155 | 4.20E-10 |
| B150 | B182 | ALP | TT | 0.014006 | 0 |
| B150 | B153 | ALP | Glu | 0.007377 | 1.19E-10 |
| B150 | B221 | ALP | Hei | 0.019255 | 0 |
| B150 | B200 | ALP | OP | 0.087917 | 0 |
| B150 | B220 | ALP | SBP | 0.049262 | 0 |
| B150 | B219 | ALP | OT | 0.012444 | 0 |
| B150 | B218 | ALP | DBP | 0.041892 | 0 |
| B150 | B216 | ALP | BMI | 0.047373 | 0 |
| B150 | B215 | ALP | BMI | 0.050762 | 0 |
| B150 | B172 | ALP | KB | -0.00804 | 2.30E-12 |
| B150 | B210 | ALP | Hip | 0.035404 | 0 |
| B150 | B209 | ALP | Smoking | 0.04888 | 0 |
| B150 | B183 | ALP | PTA | -0.01921 | 0 |
| B150 | B186 | ALP | PFC | 0.041455 | 0 |
| B150 | B207 | ALP | WHR | 0.008132 | 1.25E-12 |

| | | | | | |
|---|---|---|---|---|---|
| B150 | B206 | ALP | Waist | 0.06377 | 0 |
| B150 | B197 | ALP | AG | 0.060893 | 0 |
| B150 | B204 | ALP | Drinking | 0.021842 | 0 |
| B150 | B203 | ALP | DH | 0.007775 | 1.14E-11 |
| B150 | B201 | ALP | e-habits | 0.008366 | 2.79E-13 |
| B151 | B164 | Anti-CCP | COM | 0.009475 | 0 |
| B151 | B189 | Anti-CCP | eGFB | -0.00626 | 4.68E-08 |
| B151 | B181 | Anti-CCP | APTT | -0.00565 | 8.15E-07 |
| B151 | B193 | Anti-CCP | K | -0.00756 | 3.98E-11 |
| B151 | B222 | Anti-CCP | NS | 0.005799 | 4.13E-07 |
| B151 | B154 | Anti-CCP | RF | 0.125766 | 0 |
| B151 | B198 | Anti-CCP | P | 0.007564 | 4.00E-11 |
| B151 | B155 | Anti-CCP | IgA | 0.018037 | 0 |
| B151 | B157 | Anti-CCP | IgG | 0.029625 | 0 |
| B151 | B158 | Anti-CCP | IgM | 0.006149 | 7.96E-08 |
| B151 | B195 | Anti-CCP | Na | -0.00941 | 2.22E-16 |
| B151 | B176 | Anti-CCP | Ucr | -0.00715 | 4.33E-10 |
| B151 | B182 | Anti-CCP | TT | -0.01462 | 0 |
| B151 | B200 | Anti-CCP | OP | -0.00679 | 3.00E-09 |
| B151 | B183 | Anti-CCP | PTA | -0.01849 | 0 |
| B151 | B186 | Anti-CCP | PFC | 0.020491 | 0 |
| B151 | B206 | Anti-CCP | Waist | 0.006171 | 7.15E-08 |
| B151 | B197 | Anti-CCP | AG | -0.00763 | 2.70E-11 |
| B152 | B181 | ACA-Ig(G-A-M) | APTT | 0.01493 | 0 |
| B152 | B183 | ACA-Ig(G-A-M) | PTA | -0.01027 | 0 |
| B152 | B185 | ACA-Ig(G-A-M) | PT | 0.006968 | 1.18E-09 |
| B153 | B160 | Glu | PH | -0.01582 | 0 |
| B153 | B225 | Glu | hPBG | 0.008426 | 1.89E-13 |
| B153 | B166 | Glu | Pro | 0.006958 | 1.24E-09 |
| B153 | B189 | Glu | eGFB | -0.01706 | 0 |
| B153 | B222 | Glu | NS | 0.02785 | 0 |
| B153 | B191 | Glu | Urea | 0.012081 | 0 |
| B153 | B174 | Glu | Urea/Cr | 0.007863 | 6.64E-12 |
| B153 | B192 | Glu | UA | 0.009856 | 0 |
| B153 | B170 | Glu | U-GLU | 0.09198 | 0 |
| B153 | B179 | Glu | α1-MG | 0.006879 | 1.91E-09 |
| B153 | B200 | Glu | OP | 0.006002 | 1.60E-07 |
| B153 | B220 | Glu | SBP | 0.035648 | 0 |
| B153 | B219 | Glu | OT | 0.00581 | 3.92E-07 |
| B153 | B218 | Glu | DBP | 0.032365 | 0 |
| B153 | B216 | Glu | BMI | 0.02837 | 0 |
| B153 | B215 | Glu | BMI | 0.035951 | 0 |
| B153 | B172 | Glu | KB | 0.006209 | 5.93E-08 |
| B153 | B210 | Glu | Hip | 0.021159 | 0 |

| | | | | | |
|---|---|---|---|---|---|
| B153 | B209 | Glu | Smoking | 0.015658 | 0 |
| B153 | B207 | Glu | WHR | 0.005726 | 5.77E-07 |
| B153 | B206 | Glu | Waist | 0.037447 | 0 |
| B153 | B204 | Glu | Drinking | 0.01404 | 0 |
| B154 | B198 | RF | P | -0.0058 | 4.09E-07 |
| B154 | B155 | RF | IgA | 0.016506 | 0 |
| B154 | B157 | RF | IgG | 0.006442 | 1.87E-08 |
| B154 | B158 | RF | IgM | 0.012094 | 0 |
| B154 | B179 | RF | α1-MG | 0.007918 | 4.76E-12 |
| B155 | B193 | IgA | K | 0.009463 | 0 |
| B155 | B198 | IgA | P | -0.01602 | 0 |
| B155 | B194 | IgA | Cl | -0.02687 | 0 |
| B155 | B199 | IgA | Mg | -0.01319 | 0 |
| B155 | B156 | IgA | IgE | 0.063389 | 0 |
| B155 | B157 | IgA | IgG | 0.251375 | 0 |
| B155 | B158 | IgA | IgM | 0.022736 | 0 |
| B155 | B195 | IgA | Na | -0.02413 | 0 |
| B155 | B159 | IgA | β2-MG | 0.006845 | 2.29E-09 |
| B155 | B178 | IgA | Ig-U | 0.012076 | 0 |
| B155 | B174 | IgA | Urea/Cr | -0.00681 | 2.81E-09 |
| B155 | B192 | IgA | UA | 0.010352 | 0 |
| B155 | B170 | IgA | U-GLU | 0.012869 | 0 |
| B155 | B177 | IgA | ACR | 0.00711 | 5.39E-10 |
| B155 | B180 | IgA | Tf | 0.011794 | 0 |
| B155 | B182 | IgA | TT | 0.014065 | 0 |
| B155 | B200 | IgA | OP | -0.0116 | 0 |
| B155 | B220 | IgA | SBP | 0.008346 | 3.17E-13 |
| B155 | B218 | IgA | DBP | 0.008926 | 6.66E-15 |
| B155 | B215 | IgA | BMI | 0.007467 | 7.08E-11 |
| B155 | B209 | IgA | Smoking | -0.02223 | 0 |
| B155 | B185 | IgA | PT | 0.023406 | 0 |
| B155 | B206 | IgA | Waist | 0.005817 | 3.81E-07 |
| B155 | B197 | IgA | AG | 0.005918 | 2.39E-07 |
| B156 | B181 | IgE | APTT | -0.00868 | 3.60E-14 |
| B156 | B190 | IgE | Cre | 0.006802 | 2.87E-09 |
| B156 | B193 | IgE | K | 0.012508 | 0 |
| B156 | B222 | IgE | NS | 0.007884 | 5.86E-12 |
| B156 | B194 | IgE | Cl | -0.01952 | 0 |
| B156 | B157 | IgE | IgG | 0.040147 | 0 |
| B156 | B158 | IgE | IgM | -0.00831 | 4.10E-13 |
| B156 | B195 | IgE | Na | -0.01249 | 0 |
| B156 | B176 | IgE | Ucr | 0.016666 | 0 |
| B156 | B174 | IgE | Urea/Cr | -0.00615 | 7.91E-08 |
| B156 | B192 | IgE | UA | 0.010554 | 0 |

| | | | | | |
|---|---|---|---|---|---|
| B156 | B179 | IgE | α1-MG | 0.00782 | 8.65E-12 |
| B156 | B200 | IgE | OP | -0.00728 | 2.03E-10 |
| B156 | B220 | IgE | SBP | 0.007482 | 6.47E-11 |
| B156 | B218 | IgE | DBP | 0.006636 | 6.91E-09 |
| B156 | B216 | IgE | BMI | 0.007639 | 2.57E-11 |
| B156 | B215 | IgE | BMI | 0.007311 | 1.74E-10 |
| B156 | B210 | IgE | Hip | 0.005911 | 2.46E-07 |
| B156 | B209 | IgE | Smoking | 0.013895 | 0 |
| B156 | B206 | IgE | Waist | 0.007917 | 4.79E-12 |
| B156 | B204 | IgE | Drinking | 0.006697 | 5.02E-09 |
| B157 | B161 | IgG | LEU | 0.007657 | 2.31E-11 |
| B157 | B166 | IgG | Pro | -0.00936 | 4.44E-16 |
| B157 | B181 | IgG | APTT | 0.037715 | 0 |
| B157 | B190 | IgG | Cre | -0.01937 | 0 |
| B157 | B193 | IgG | K | 0.024055 | 0 |
| B157 | B222 | IgG | NS | -0.02366 | 0 |
| B157 | B198 | IgG | P | 0.010912 | 0 |
| B157 | B194 | IgG | Cl | -0.03279 | 0 |
| B157 | B199 | IgG | Mg | -0.02117 | 0 |
| B157 | B158 | IgG | IgM | 0.074957 | 0 |
| B157 | B195 | IgG | Na | -0.05446 | 0 |
| B157 | B159 | IgG | β2-MG | -0.01168 | 0 |
| B157 | B176 | IgG | Ucr | -0.01827 | 0 |
| B157 | B178 | IgG | Ig-U | 0.007715 | 1.63E-11 |
| B157 | B191 | IgG | Urea | -0.00635 | 2.98E-08 |
| B157 | B174 | IgG | Urea/Cr | 0.007228 | 2.78E-10 |
| B157 | B192 | IgG | UA | -0.0233 | 0 |
| B157 | B179 | IgG | α1-MG | -0.02382 | 0 |
| B157 | B182 | IgG | TT | 0.008872 | 9.55E-15 |
| B157 | B221 | IgG | Hei | -0.02725 | 0 |
| B157 | B187 | IgG | NSE | 0.010252 | 0 |
| B157 | B200 | IgG | OP | -0.04891 | 0 |
| B157 | B219 | IgG | OT | 0.007926 | 4.51E-12 |
| B157 | B216 | IgG | BMI | -0.01968 | 0 |
| B157 | B215 | IgG | BMI | -0.00652 | 1.25E-08 |
| B157 | B210 | IgG | Hip | -0.00892 | 7.11E-15 |
| B157 | B209 | IgG | Smoking | -0.03954 | 0 |
| B157 | B183 | IgG | PTA | 0.008681 | 3.46E-14 |
| B157 | B185 | IgG | PT | 0.046335 | 0 |
| B157 | B186 | IgG | PFC | 0.012026 | 0 |
| B157 | B206 | IgG | Waist | -0.01767 | 0 |
| B157 | B197 | IgG | AG | -0.01454 | 0 |
| B157 | B204 | IgG | Drinking | -0.03338 | 0 |
| B157 | B211 | IgG | IOP-R | -0.01581 | 0 |

| | | | | | |
|---|---|---|---|---|---|
| B157 | B212 | IgG | IOP-L | -0.0191 | 0 |
| B158 | B189 | IgM | eGFB | 0.006957 | 1.25E-09 |
| B158 | B181 | IgM | APTT | 0.024503 | 0 |
| B158 | B190 | IgM | Cre | -0.01633 | 0 |
| B158 | B222 | IgM | NS | -0.0178 | 0 |
| B158 | B198 | IgM | P | 0.008892 | 8.22E-15 |
| B158 | B195 | IgM | Na | -0.01577 | 0 |
| B158 | B174 | IgM | Urea/Cr | 0.009133 | 1.55E-15 |
| B158 | B192 | IgM | UA | -0.01955 | 0 |
| B158 | B179 | IgM | α1-MG | -0.01053 | 0 |
| B158 | B221 | IgM | Hei | -0.01809 | 0 |
| B158 | B200 | IgM | OP | -0.01387 | 0 |
| B158 | B220 | IgM | SBP | -0.00957 | 0 |
| B158 | B218 | IgM | DBP | -0.00774 | 1.41E-11 |
| B158 | B216 | IgM | BMI | -0.01809 | 0 |
| B158 | B215 | IgM | BMI | -0.01121 | 0 |
| B158 | B210 | IgM | Hip | -0.01131 | 0 |
| B158 | B209 | IgM | Smoking | -0.00775 | 1.34E-11 |
| B158 | B185 | IgM | PT | 0.006089 | 1.06E-07 |
| B158 | B206 | IgM | Waist | -0.01569 | 0 |
| B158 | B204 | IgM | Drinking | -0.00707 | 6.55E-10 |
| B158 | B211 | IgM | IOP-R | -0.00743 | 8.83E-11 |
| B158 | B212 | IgM | IOP-L | -0.00803 | 2.33E-12 |
| B159 | B193 | β2-MG | K | -0.00682 | 2.66E-09 |
| B16 | B23 | CD3+T | CD3-CD16+CD56+(NK) | -0.59249 | 0 |
| B16 | B17 | CD3+T | CD3+CD16+CD56+NK | 0.127806 | 0 |
| B16 | B18 | CD3+T | CD3+CD4+helper(inducer)T | 0.555559 | 0 |
| B16 | B19 | CD3+T | CD3+CD4+/CD3+CD8+ | -0.02047 | 0 |
| B16 | B20 | CD3+T | CD3+CD4+CD8+ | 0.16268 | 0 |
| B16 | B21 | CD3+T | CD3+CD4-CD8- | 0.186846 | 0 |
| B16 | B22 | CD3+T | CD3+CD8+(T) | 0.561987 | 0 |
| B16 | B92 | CD3+T | ALDH2-Glu504Glu | -0.00651 | 1.30E-08 |
| B16 | B94 | CD3+T | ALDH2-Lys504Lys | 0.014421 | 0 |
| B16 | B143 | CD3+T | β-CTx | 0.00563 | 8.86E-07 |
| B16 | B126 | CD3+T | Alb | -0.00633 | 3.34E-08 |
| B16 | B121 | CD3+T | C3 | 0.012266 | 0 |
| B16 | B124 | CD3+T | ACTH | -0.02896 | 0 |
| B16 | B134 | CD3+T | CHE | -0.00694 | 1.38E-09 |
| B16 | B70 | CD3+T | LDL-C | -0.00612 | 9.17E-08 |
| B16 | B145 | CD3+T | BGP | 0.006302 | 3.75E-08 |
| B16 | B55 | CD3+T | TK | 0.006621 | 7.45E-09 |
| B16 | B57 | CD3+T | RBCAI | -0.01423 | 0 |
| B16 | B46 | CD3+T | RBC Count | -0.00571 | 6.29E-07 |
| B16 | B181 | CD3+T | APTT | 0.011299 | 0 |

| | | | | | |
|---|---|---|---|---|---|
| B16 | B193 | CD3+T | K | -0.0256 | 0 |
| B16 | B150 | CD3+T | ALP | -0.01197 | 0 |
| B16 | B99 | CD3+T | anti-PM-Scl | -0.01023 | 0 |
| B16 | B105 | CD3+T | anti-U1RNP | -0.01344 | 0 |
| B16 | B96 | CD3+T | dsDNA (IIF) | -0.00581 | 3.86E-07 |
| B16 | B98 | CD3+T | anti-Jo-1 | 0.007088 | 6.10E-10 |
| B16 | B100 | CD3+T | anti-Ro52 | 0.008779 | 1.80E-14 |
| B16 | B101 | CD3+T | anti-Sc70 | -0.01083 | 0 |
| B16 | B102 | CD3+T | anti-Sm | -0.00884 | 1.20E-14 |
| B16 | B103 | CD3+T | anti-SSA/Ro | 0.015106 | 0 |
| B16 | B104 | CD3+T | anti-SSB | 0.010587 | 0 |
| B16 | B151 | CD3+T | Anti-CCP | -0.01572 | 0 |
| B16 | B198 | CD3+T | P | 0.037015 | 0 |
| B16 | B194 | CD3+T | Cl | 0.013874 | 0 |
| B16 | B199 | CD3+T | Mg | -0.00882 | 1.33E-14 |
| B16 | B155 | CD3+T | IgA | -0.02193 | 0 |
| B16 | B156 | CD3+T | IgE | -0.01746 | 0 |
| B16 | B157 | CD3+T | IgG | -0.02333 | 0 |
| B16 | B195 | CD3+T | Na | -0.01373 | 0 |
| B16 | B178 | CD3+T | Ig-U | -0.00991 | 0 |
| B16 | B177 | CD3+T | ACR | -0.00587 | 2.94E-07 |
| B16 | B179 | CD3+T | α1-MG | -0.01991 | 0 |
| B16 | B180 | CD3+T | Tf | -0.00581 | 4.00E-07 |
| B16 | B182 | CD3+T | TT | -0.02599 | 0 |
| B16 | B138 | CD3+T | PAB | -0.00868 | 3.44E-14 |
| B16 | B59 | CD3+T | WBRV | -0.01331 | 0 |
| B16 | B60 | CD3+T | WBRVI | -0.01678 | 0 |
| B16 | B61 | CD3+T | WBHSRV | -0.00599 | 1.68E-07 |
| B16 | B62 | CD3+T | WBHSRVI | -0.00868 | 3.51E-14 |
| B16 | B64 | CD3+T | WBRSR(mPas_5) | -0.01945 | 0 |
| B16 | B65 | CD3+T | WBRSR(mPas_1) | -0.0208 | 0 |
| B16 | B66 | CD3+T | WBRSR(mPas_100) | -0.01366 | 0 |
| B16 | B67 | CD3+T | WBRSR(mPas_50) | -0.01426 | 0 |
| B16 | B200 | CD3+T | OP | -0.0175 | 0 |
| B16 | B220 | CD3+T | SBP | -0.00849 | 1.22E-13 |
| B16 | B218 | CD3+T | DBP | -0.00736 | 1.32E-10 |
| B16 | B90 | CD3+T | CYFRA 21-1 | 0.007562 | 4.06E-11 |
| B16 | B139 | CD3+T | ADA | -0.00921 | 8.88E-16 |
| B16 | B35 | CD3+T | β2-MG | 0.014317 | 0 |
| B16 | B196 | CD3+T | TCO2 | -0.00753 | 4.85E-11 |
| B16 | B183 | CD3+T | PTA | 0.018023 | 0 |
| B16 | B184 | CD3+T | INR | -0.00963 | 0 |
| B16 | B186 | CD3+T | PFC | -0.01791 | 0 |
| B16 | B69 | CD3+T | PV | -0.01047 | 0 |

| | | | | | |
|---|---|---|---|---|---|
| B16 | B197 | CD3+T | AG | -0.01127 | 0 |
| B16 | B73 | CD3+T | ApoA | -0.00598 | 1.81E-07 |
| B16 | B74 | CD3+T | ApoA1 | -0.01249 | 0 |
| B16 | B75 | CD3+T | ApoA1/B | -0.00714 | 4.62E-10 |
| B16 | B76 | CD3+T | ApoB | -0.0128 | 0 |
| B16 | B91 | CD3+T | TP | -0.00685 | 2.27E-09 |
| B16 | B114 | CD3+T | TSH TT3 TT4(TT4) | -0.00606 | 1.25E-07 |
| B160 | B161 | PH | LEU | -0.01453 | 0 |
| B160 | B166 | PH | Pro | -0.00865 | 4.17E-14 |
| B160 | B188 | PH | Ca | -0.00756 | 4.12E-11 |
| B160 | B189 | PH | eGFB | 0.126966 | 0 |
| B160 | B181 | PH | APTT | 0.00575 | 5.17E-07 |
| B160 | B190 | PH | Cre | -0.09701 | 0 |
| B160 | B224 | PH | pmh | -0.01055 | 0 |
| B160 | B193 | PH | K | -0.00866 | 4.04E-14 |
| B160 | B222 | PH | NS | -0.08135 | 0 |
| B160 | B198 | PH | P | -0.0258 | 0 |
| B160 | B167 | PH | PC | 0.007371 | 1.23E-10 |
| B160 | B194 | PH | Cl | -0.01112 | 0 |
| B160 | B195 | PH | Na | -0.00805 | 2.08E-12 |
| B160 | B168 | PH | NMP22 | 0.011912 | 0 |
| B160 | B176 | PH | Ucr | -0.0486 | 0 |
| B160 | B191 | PH | Urea | -0.12436 | 0 |
| B160 | B174 | PH | Urea/Cr | -0.03313 | 0 |
| B160 | B192 | PH | UA | -0.13123 | 0 |
| B160 | B170 | PH | U-GLU | -0.04383 | 0 |
| B160 | B175 | PH | ALB-U | -0.00633 | 3.27E-08 |
| B160 | B179 | PH | α1-MG | -0.00655 | 1.08E-08 |
| B160 | B221 | PH | Hei | -0.01785 | 0 |
| B160 | B200 | PH | OP | -0.01841 | 0 |
| B160 | B220 | PH | SBP | -0.0347 | 0 |
| B160 | B219 | PH | OT | -0.01161 | 0 |
| B160 | B218 | PH | DBP | -0.05627 | 0 |
| B160 | B216 | PH | BMI | -0.07956 | 0 |
| B160 | B215 | PH | BMI | -0.09373 | 0 |
| B160 | B172 | PH | KB | -0.02695 | 0 |
| B160 | B210 | PH | Hip | -0.06148 | 0 |
| B160 | B209 | PH | Smoking | -0.11482 | 0 |
| B160 | B196 | PH | TCO2 | 0.012716 | 0 |
| B160 | B183 | PH | PTA | 0.010005 | 0 |
| B160 | B186 | PH | PFC | -0.00886 | 1.02E-14 |
| B160 | B207 | PH | WHR | -0.01404 | 0 |
| B160 | B206 | PH | Waist | -0.10629 | 0 |
| B160 | B197 | PH | AG | -0.00748 | 6.59E-11 |

| | | | | | |
|---|---|---|---|---|---|
| B160 | B204 | PH | Drinking | -0.06155 | 0 |
| B160 | B211 | PH | IOP-R | 0.012408 | 0 |
| B160 | B212 | PH | IOP-L | 0.012331 | 0 |
| B161 | B166 | LEU | Pro | 0.021578 | 0 |
| B161 | B189 | LEU | eGFB | 0.020062 | 0 |
| B161 | B190 | LEU | Cre | -0.11034 | 0 |
| B161 | B222 | LEU | NS | -0.1134 | 0 |
| B161 | B198 | LEU | P | 0.011349 | 0 |
| B161 | B168 | LEU | NMP22 | 0.011819 | 0 |
| B161 | B176 | LEU | Ucr | -0.01305 | 0 |
| B161 | B191 | LEU | Urea | -0.02035 | 0 |
| B161 | B174 | LEU | Urea/Cr | 0.062878 | 0 |
| B161 | B192 | LEU | UA | -0.09161 | 0 |
| B161 | B221 | LEU | Hei | -0.1238 | 0 |
| B161 | B220 | LEU | SBP | -0.02885 | 0 |
| B161 | B219 | LEU | OT | 0.015672 | 0 |
| B161 | B218 | LEU | DBP | -0.03639 | 0 |
| B161 | B216 | LEU | BMI | -0.09446 | 0 |
| B161 | B215 | LEU | BMI | -0.03738 | 0 |
| B161 | B172 | LEU | KB | 0.012867 | 0 |
| B161 | B210 | LEU | Hip | -0.05055 | 0 |
| B161 | B209 | LEU | Smoking | -0.05776 | 0 |
| B161 | B173 | LEU | NTT | 0.097968 | 0 |
| B161 | B207 | LEU | WHR | -0.00978 | 0 |
| B161 | B206 | LEU | Waist | -0.07318 | 0 |
| B161 | B205 | LEU | KDA | 0.008525 | 9.86E-14 |
| B161 | B204 | LEU | Drinking | -0.0646 | 0 |
| B161 | B203 | LEU | DH | -0.00594 | 2.15E-07 |
| B161 | B211 | LEU | IOP-R | -0.00771 | 1.69E-11 |
| B161 | B201 | LEU | e-habits | -0.0129 | 0 |
| B161 | B212 | LEU | IOP-L | -0.00639 | 2.42E-08 |
| B164 | B166 | COM | Pro | -0.0242 | 0 |
| B164 | B189 | COM | eGFB | 0.009408 | 2.22E-16 |
| B164 | B190 | COM | Cre | -0.01024 | 0 |
| B164 | B222 | COM | NS | -0.00878 | 1.73E-14 |
| B164 | B192 | COM | UA | -0.00845 | 1.61E-13 |
| B164 | B170 | COM | U-GLU | -0.00581 | 3.87E-07 |
| B164 | B221 | COM | Hei | -0.0076 | 3.30E-11 |
| B164 | B220 | COM | SBP | -0.00675 | 3.74E-09 |
| B164 | B216 | COM | BMI | -0.00615 | 8.06E-08 |
| B164 | B209 | COM | Smoking | -0.00988 | 0 |
| B164 | B206 | COM | Waist | -0.00657 | 9.78E-09 |
| B165 | B166 | bilirubin | Pro | 0.01089 | 0 |
| B165 | B192 | bilirubin | UA | 0.005635 | 8.65E-07 |

| | | | | | |
|---|---|---|---|---|---|
| B165 | B170 | bilirubin | U-GLU | 0.008353 | 3.05E-13 |
| B165 | B179 | bilirubin | α1-MG | 0.007469 | 6.97E-11 |
| B165 | B218 | bilirubin | DBP | 0.006481 | 1.53E-08 |
| B165 | B172 | bilirubin | KB | 0.022441 | 0 |
| B165 | B209 | bilirubin | Smoking | 0.012234 | 0 |
| B165 | B204 | bilirubin | Drinking | 0.010922 | 0 |
| B166 | B189 | Pro | eGFB | -0.02833 | 0 |
| B166 | B190 | Pro | Cre | 0.134786 | 0 |
| B166 | B193 | Pro | K | -0.01123 | 0 |
| B166 | B222 | Pro | NS | 0.125222 | 0 |
| B166 | B198 | Pro | P | -0.00563 | 8.82E-07 |
| B166 | B199 | Pro | Mg | 0.007945 | 4.02E-12 |
| B166 | B168 | Pro | NMP22 | 0.007338 | 1.49E-10 |
| B166 | B176 | Pro | Ucr | 0.091618 | 0 |
| B166 | B178 | Pro | Ig-U | 0.012436 | 0 |
| B166 | B191 | Pro | Urea | 0.075399 | 0 |
| B166 | B174 | Pro | Urea/Cr | -0.03696 | 0 |
| B166 | B192 | Pro | UA | 0.095187 | 0 |
| B166 | B170 | Pro | U-GLU | 0.052736 | 0 |
| B166 | B175 | Pro | ALB-U | 0.022481 | 0 |
| B166 | B177 | Pro | ACR | 0.013003 | 0 |
| B166 | B179 | Pro | α1-MG | 0.029629 | 0 |
| B166 | B180 | Pro | Tf | 0.009748 | 0 |
| B166 | B221 | Pro | Hei | 0.131942 | 0 |
| B166 | B200 | Pro | OP | 0.008677 | 3.55E-14 |
| B166 | B220 | Pro | SBP | 0.049179 | 0 |
| B166 | B219 | Pro | OT | -0.01407 | 0 |
| B166 | B218 | Pro | DBP | 0.056933 | 0 |
| B166 | B216 | Pro | BMI | 0.114801 | 0 |
| B166 | B215 | Pro | BMI | 0.059663 | 0 |
| B166 | B172 | Pro | KB | 0.084113 | 0 |
| B166 | B210 | Pro | Hip | 0.063535 | 0 |
| B166 | B209 | Pro | Smoking | 0.092168 | 0 |
| B166 | B173 | Pro | NTT | -0.00835 | 3.12E-13 |
| B166 | B207 | Pro | WHR | 0.010181 | 0 |
| B166 | B206 | Pro | Waist | 0.091096 | 0 |
| B166 | B205 | Pro | KDA | -0.01434 | 0 |
| B166 | B197 | Pro | AG | 0.006121 | 9.08E-08 |
| B166 | B204 | Pro | Drinking | 0.08129 | 0 |
| B166 | B203 | Pro | DH | 0.011832 | 0 |
| B167 | B178 | PC | Ig-U | -0.0084 | 2.24E-13 |
| B167 | B180 | PC | Tf | -0.01186 | 0 |
| B167 | B171 | PC | EC | -0.0494 | 0 |
| B168 | B175 | NMP22 | ALB-U | 0.010238 | 0 |

| | | | | | |
|---|---|---|---|---|---|
| B168 | B177 | NMP22 | ACR | 0.011798 | 0 |
| B168 | B219 | NMP22 | OT | 0.006474 | 1.59E-08 |
| B168 | B173 | NMP22 | NTT | 0.018297 | 0 |
| B169 | B225 | U-CRY | hPBG | -0.00619 | 6.43E-08 |
| B17 | B137 | CD3+CD16+CD56+NK | AFU | -0.01194 | 0 |
| B17 | B23 | CD3+CD16+CD56+NK | CD3-CD16+CD56+(NK) | -0.07935 | 0 |
| B17 | B18 | CD3+CD16+CD56+NK | CD3+CD4+helper(inducer)T | -0.10188 | 0 |
| B17 | B19 | CD3+CD16+CD56+NK | CD3+CD4+/CD3+CD8+ | -0.16721 | 0 |
| B17 | B20 | CD3+CD16+CD56+NK | CD3+CD4+CD8+ | 0.173318 | 0 |
| B17 | B21 | CD3+CD16+CD56+NK | CD3+CD4-CD8- | 0.239841 | 0 |
| B17 | B22 | CD3+CD16+CD56+NK | CD3+CD8+(T) | 0.198102 | 0 |
| B17 | B145 | CD3+CD16+CD56+NK | BGP | 0.005616 | 9.42E-07 |
| B17 | B55 | CD3+CD16+CD56+NK | TK | 0.006541 | 1.13E-08 |
| B17 | B96 | CD3+CD16+CD56+NK | dsDNA (IIF) | -0.0154 | 0 |
| B17 | B100 | CD3+CD16+CD56+NK | anti-Ro52 | -0.01947 | 0 |
| B17 | B103 | CD3+CD16+CD56+NK | anti-SSA/Ro | -0.00673 | 4.27E-09 |
| B17 | B110 | CD3+CD16+CD56+NK | AHA | 0.010131 | 0 |
| B17 | B199 | CD3+CD16+CD56+NK | Mg | 0.007331 | 1.55E-10 |
| B17 | B155 | CD3+CD16+CD56+NK | IgA | -0.00715 | 4.30E-10 |
| B17 | B182 | CD3+CD16+CD56+NK | TT | -0.01141 | 0 |
| B17 | B209 | CD3+CD16+CD56+NK | Smoking | -0.00575 | 5.06E-07 |
| B17 | B90 | CD3+CD16+CD56+NK | CYFRA 21-1 | -0.00854 | 9.15E-14 |
| B17 | B186 | CD3+CD16+CD56+NK | PFC | 0.005679 | 7.12E-07 |
| B17 | B69 | CD3+CD16+CD56+NK | PV | -0.01041 | 0 |
| B17 | B114 | CD3+CD16+CD56+NK | TSH TT3 TT4(TT4) | -0.00816 | 1.06E-12 |
| B170 | B225 | U-GLU | hPBG | 0.040796 | 0 |
| B170 | B189 | U-GLU | eGFB | -0.00701 | 9.55E-10 |
| B170 | B181 | U-GLU | APTT | -0.01051 | 0 |
| B170 | B222 | U-GLU | NS | 0.055219 | 0 |
| B170 | B198 | U-GLU | P | -0.00671 | 4.61E-09 |
| B170 | B194 | U-GLU | Cl | -0.02035 | 0 |
| B170 | B199 | U-GLU | Mg | -0.00804 | 2.29E-12 |
| B170 | B195 | U-GLU | Na | -0.01281 | 0 |
| B170 | B176 | U-GLU | Ucr | 0.013617 | 0 |
| B170 | B178 | U-GLU | Ig-U | 0.010876 | 0 |
| B170 | B191 | U-GLU | Urea | 0.027006 | 0 |
| B170 | B174 | U-GLU | Urea/Cr | 0.024362 | 0 |
| B170 | B192 | U-GLU | UA | -0.01435 | 0 |
| B170 | B175 | U-GLU | ALB-U | 0.018141 | 0 |
| B170 | B177 | U-GLU | ACR | 0.01435 | 0 |
| B170 | B179 | U-GLU | α1-MG | 0.035928 | 0 |
| B170 | B180 | U-GLU | Tf | 0.013807 | 0 |
| B170 | B221 | U-GLU | Hei | 0.017612 | 0 |
| B170 | B200 | U-GLU | OP | 0.018745 | 0 |

| | | | | | |
|---|---|---|---|---|---|
| B170 | B220 | U-GLU | SBP | 0.052173 | 0 |
| B170 | B218 | U-GLU | DBP | 0.051275 | 0 |
| B170 | B216 | U-GLU | BMI | 0.044254 | 0 |
| B170 | B215 | U-GLU | BMI | 0.046543 | 0 |
| B170 | B172 | U-GLU | KB | 0.076181 | 0 |
| B170 | B210 | U-GLU | Hip | 0.021164 | 0 |
| B170 | B209 | U-GLU | Smoking | 0.058272 | 0 |
| B170 | B183 | U-GLU | PTA | -0.00654 | 1.13E-08 |
| B170 | B173 | U-GLU | NTT | 0.012217 | 0 |
| B170 | B207 | U-GLU | WHR | 0.014667 | 0 |
| B170 | B206 | U-GLU | Waist | 0.062032 | 0 |
| B170 | B197 | U-GLU | AG | 0.007164 | 4.00E-10 |
| B170 | B204 | U-GLU | Drinking | 0.039986 | 0 |
| B170 | B211 | U-GLU | IOP-R | 0.012923 | 0 |
| B170 | B212 | U-GLU | IOP-L | 0.012993 | 0 |
| B172 | B225 | KB | hPBG | 0.006382 | 2.52E-08 |
| B172 | B189 | KB | eGFB | 0.041775 | 0 |
| B172 | B190 | KB | Cre | -0.02788 | 0 |
| B172 | B222 | KB | NS | -0.02974 | 0 |
| B172 | B194 | KB | Cl | -0.00647 | 1.64E-08 |
| B172 | B176 | KB | Ucr | 0.018726 | 0 |
| B172 | B191 | KB | Urea | 0.013297 | 0 |
| B172 | B174 | KB | Urea/Cr | 0.029041 | 0 |
| B172 | B192 | KB | UA | -0.01296 | 0 |
| B172 | B179 | KB | α1-MG | 0.008447 | 1.65E-13 |
| B172 | B221 | KB | Hei | -0.012 | 0 |
| B172 | B220 | KB | SBP | -0.01209 | 0 |
| B172 | B219 | KB | OT | -0.00917 | 1.11E-15 |
| B172 | B218 | KB | DBP | -0.00663 | 7.25E-09 |
| B172 | B216 | KB | BMI | -0.03734 | 0 |
| B172 | B215 | KB | BMI | -0.04178 | 0 |
| B172 | B210 | KB | Hip | -0.03227 | 0 |
| B172 | B206 | KB | Waist | -0.04336 | 0 |
| B172 | B204 | KB | Drinking | 0.005756 | 5.02E-07 |
| B172 | B203 | KB | DH | 0.006726 | 4.30E-09 |
| B173 | B189 | NTT | eGFB | -0.03015 | 0 |
| B173 | B190 | NTT | Cre | -0.03352 | 0 |
| B173 | B222 | NTT | NS | -0.0432 | 0 |
| B173 | B198 | NTT | P | 0.006416 | 2.13E-08 |
| B173 | B176 | NTT | Ucr | -0.01428 | 0 |
| B173 | B174 | NTT | Urea/Cr | 0.024008 | 0 |
| B173 | B192 | NTT | UA | -0.03545 | 0 |
| B173 | B221 | NTT | Hei | -0.05878 | 0 |
| B173 | B227 | NTT | OD | 0.007186 | 3.53E-10 |

| | | | | | |
|---|---|---|---|---|---|
| B173 | B220 | NTT | SBP | 0.007674 | 2.09E-11 |
| B173 | B219 | NTT | OT | 0.021772 | 0 |
| B173 | B218 | NTT | DBP | -0.00858 | 6.82E-14 |
| B173 | B216 | NTT | BMI | -0.0393 | 0 |
| B173 | B215 | NTT | BMI | -0.01017 | 0 |
| B173 | B210 | NTT | Hip | -0.01794 | 0 |
| B173 | B209 | NTT | Smoking | -0.02432 | 0 |
| B173 | B185 | NTT | PT | 0.010025 | 0 |
| B173 | B206 | NTT | Waist | -0.01835 | 0 |
| B173 | B205 | NTT | KDA | 0.006377 | 2.58E-08 |
| B173 | B204 | NTT | Drinking | -0.02593 | 0 |
| B173 | B203 | NTT | DH | -0.00827 | 5.25E-13 |
| B174 | B188 | Urea/Cr | Ca | -0.01131 | 0 |
| B174 | B189 | Urea/Cr | eGFB | 0.199106 | 0 |
| B174 | B190 | Urea/Cr | Cre | -0.40955 | 0 |
| B174 | B222 | Urea/Cr | NS | -0.2253 | 0 |
| B174 | B198 | Urea/Cr | P | 0.028537 | 0 |
| B174 | B194 | Urea/Cr | Cl | 0.008021 | 2.51E-12 |
| B174 | B176 | Urea/Cr | Ucr | -0.05497 | 0 |
| B174 | B191 | Urea/Cr | Urea | 0.531287 | 0 |
| B174 | B192 | Urea/Cr | UA | -0.25168 | 0 |
| B174 | B175 | Urea/Cr | ALB-U | -0.00834 | 3.37E-13 |
| B174 | B179 | Urea/Cr | α1-MG | -0.01015 | 0 |
| B174 | B221 | Urea/Cr | Hei | -0.28703 | 0 |
| B174 | B200 | Urea/Cr | OP | 0.026916 | 0 |
| B174 | B220 | Urea/Cr | SBP | -0.04791 | 0 |
| B174 | B219 | Urea/Cr | OT | 0.037463 | 0 |
| B174 | B218 | Urea/Cr | DBP | -0.072 | 0 |
| B174 | B216 | Urea/Cr | BMI | -0.20907 | 0 |
| B174 | B215 | Urea/Cr | BMI | -0.07376 | 0 |
| B174 | B210 | Urea/Cr | Hip | -0.12119 | 0 |
| B174 | B209 | Urea/Cr | Smoking | -0.10514 | 0 |
| B174 | B196 | Urea/Cr | TCO2 | -0.00864 | 4.49E-14 |
| B174 | B183 | Urea/Cr | PTA | -0.00705 | 7.64E-10 |
| B174 | B207 | Urea/Cr | WHR | -0.01304 | 0 |
| B174 | B206 | Urea/Cr | Waist | -0.12648 | 0 |
| B174 | B205 | Urea/Cr | KDA | 0.0091 | 1.78E-15 |
| B174 | B204 | Urea/Cr | Drinking | -0.12023 | 0 |
| B174 | B203 | Urea/Cr | DH | -0.0121 | 0 |
| B174 | B211 | Urea/Cr | IOP-R | -0.01156 | 0 |
| B174 | B201 | Urea/Cr | e-habits | -0.01942 | 0 |
| B174 | B212 | Urea/Cr | IOP-L | -0.00961 | 0 |
| B175 | B189 | ALB-U | eGFB | -0.02063 | 0 |
| B175 | B190 | ALB-U | Cre | 0.03468 | 0 |

| | | | | | |
|---|---|---|---|---|---|
| B175 | B222 | ALB-U | NS | 0.014193 | 0 |
| B175 | B176 | ALB-U | Ucr | 0.060978 | 0 |
| B175 | B178 | ALB-U | Ig-U | 0.050885 | 0 |
| B175 | B191 | ALB-U | Urea | 0.014406 | 0 |
| B175 | B192 | ALB-U | UA | 0.01902 | 0 |
| B175 | B177 | ALB-U | ACR | 0.68259 | 0 |
| B175 | B179 | ALB-U | α1-MG | 0.028359 | 0 |
| B175 | B180 | ALB-U | Tf | 0.068871 | 0 |
| B175 | B221 | ALB-U | Hei | 0.008149 | 1.12E-12 |
| B175 | B200 | ALB-U | OP | 0.006231 | 5.34E-08 |
| B175 | B220 | ALB-U | SBP | 0.020497 | 0 |
| B175 | B218 | ALB-U | DBP | 0.021166 | 0 |
| B175 | B216 | ALB-U | BMI | 0.011976 | 0 |
| B175 | B215 | ALB-U | BMI | 0.009864 | 0 |
| B175 | B210 | ALB-U | Hip | 0.008278 | 4.92E-13 |
| B175 | B209 | ALB-U | Smoking | 0.006233 | 5.28E-08 |
| B175 | B206 | ALB-U | Waist | 0.011881 | 0 |
| B176 | B189 | Ucr | eGFB | -0.02177 | 0 |
| B176 | B181 | Ucr | APTT | -0.00665 | 6.28E-09 |
| B176 | B190 | Ucr | Cre | 0.105648 | 0 |
| B176 | B193 | Ucr | K | -0.00622 | 5.55E-08 |
| B176 | B222 | Ucr | NS | 0.100495 | 0 |
| B176 | B194 | Ucr | Cl | -0.0101 | 0 |
| B176 | B199 | Ucr | Mg | 0.010394 | 0 |
| B176 | B195 | Ucr | Na | 0.01016 | 0 |
| B176 | B178 | Ucr | Ig-U | 0.0411 | 0 |
| B176 | B191 | Ucr | Urea | 0.031313 | 0 |
| B176 | B192 | Ucr | UA | 0.08974 | 0 |
| B176 | B177 | Ucr | ACR | -0.05089 | 0 |
| B176 | B179 | Ucr | α1-MG | 0.145167 | 0 |
| B176 | B180 | Ucr | Tf | 0.017285 | 0 |
| B176 | B221 | Ucr | Hei | 0.105982 | 0 |
| B176 | B187 | Ucr | NSE | 0.007256 | 2.37E-10 |
| B176 | B200 | Ucr | OP | 0.014097 | 0 |
| B176 | B220 | Ucr | SBP | 0.01172 | 0 |
| B176 | B219 | Ucr | OT | -0.01742 | 0 |
| B176 | B218 | Ucr | DBP | 0.033039 | 0 |
| B176 | B216 | Ucr | BMI | 0.088477 | 0 |
| B176 | B215 | Ucr | BMI | 0.041257 | 0 |
| B176 | B210 | Ucr | Hip | 0.047436 | 0 |
| B176 | B209 | Ucr | Smoking | 0.068363 | 0 |
| B176 | B183 | Ucr | PTA | 0.006981 | 1.09E-09 |
| B176 | B207 | Ucr | WHR | 0.00614 | 8.31E-08 |
| B176 | B206 | Ucr | Waist | 0.06537 | 0 |

| | | | | | |
|---|---|---|---|---|---|
| B176 | B205 | Ucr | KDA | -0.0077 | 1.81E-11 |
| B176 | B197 | Ucr | AG | 0.011435 | 0 |
| B176 | B204 | Ucr | Drinking | 0.065196 | 0 |
| B176 | B201 | Ucr | e-habits | -0.0085 | 1.15E-13 |
| B177 | B189 | ACR | eGFB | -0.0175 | 0 |
| B177 | B190 | ACR | Cre | 0.02286 | 0 |
| B177 | B199 | ACR | Mg | -0.00568 | 7.11E-07 |
| B177 | B178 | ACR | Ig-U | 0.148455 | 0 |
| B177 | B191 | ACR | Urea | 0.012404 | 0 |
| B177 | B192 | ACR | UA | 0.007429 | 8.81E-11 |
| B177 | B179 | ACR | α1-MG | 0.042284 | 0 |
| B177 | B180 | ACR | Tf | 0.242956 | 0 |
| B177 | B221 | ACR | Hei | -0.00734 | 1.49E-10 |
| B177 | B200 | ACR | OP | 0.008572 | 7.19E-14 |
| B177 | B220 | ACR | SBP | 0.021184 | 0 |
| B177 | B218 | ACR | DBP | 0.01686 | 0 |
| B177 | B186 | ACR | PFC | 0.005857 | 3.16E-07 |
| B177 | B204 | ACR | Drinking | -0.00742 | 9.09E-11 |
| B178 | B190 | Ig-U | Cre | 0.013258 | 0 |
| B178 | B222 | Ig-U | NS | 0.006285 | 4.08E-08 |
| B178 | B192 | Ig-U | UA | 0.007948 | 3.94E-12 |
| B178 | B179 | Ig-U | α1-MG | 0.330879 | 0 |
| B178 | B180 | Ig-U | Tf | 0.53879 | 0 |
| B178 | B200 | Ig-U | OP | 0.013078 | 0 |
| B178 | B220 | Ig-U | SBP | 0.010731 | 0 |
| B178 | B218 | Ig-U | DBP | 0.008388 | 2.42E-13 |
| B178 | B216 | Ig-U | BMI | 0.006632 | 7.02E-09 |
| B178 | B215 | Ig-U | BMI | 0.006287 | 4.05E-08 |
| B178 | B183 | Ig-U | PTA | 0.007051 | 7.47E-10 |
| B178 | B186 | Ig-U | PFC | 0.014591 | 0 |
| B178 | B206 | Ig-U | Waist | 0.006722 | 4.39E-09 |
| B178 | B197 | Ig-U | AG | 0.007843 | 7.52E-12 |
| B179 | B189 | α1-MG | eGFB | -0.00593 | 2.23E-07 |
| B179 | B181 | α1-MG | APTT | -0.01503 | 0 |
| B179 | B190 | α1-MG | Cre | 0.037556 | 0 |
| B179 | B193 | α1-MG | K | -0.01793 | 0 |
| B179 | B222 | α1-MG | NS | 0.028059 | 0 |
| B179 | B198 | α1-MG | P | -0.01488 | 0 |
| B179 | B194 | α1-MG | Cl | -0.00873 | 2.53E-14 |
| B179 | B199 | α1-MG | Mg | 0.007182 | 3.61E-10 |
| B179 | B195 | α1-MG | Na | 0.014559 | 0 |
| B179 | B191 | α1-MG | Urea | 0.023683 | 0 |
| B179 | B192 | α1-MG | UA | 0.023851 | 0 |
| B179 | B180 | α1-MG | Tf | 0.224128 | 0 |

| | | | | | |
|---|---|---|---|---|---|
| B179 | B221 | α1-MG | Hei | 0.025174 | 0 |
| B179 | B187 | α1-MG | NSE | 0.006498 | 1.40E-08 |
| B179 | B200 | α1-MG | OP | 0.039158 | 0 |
| B179 | B220 | α1-MG | SBP | 0.020138 | 0 |
| B179 | B218 | α1-MG | DBP | 0.019628 | 0 |
| B179 | B216 | α1-MG | BMI | 0.02086 | 0 |
| B179 | B215 | α1-MG | BMI | 0.009915 | 0 |
| B179 | B210 | α1-MG | Hip | 0.009277 | 4.44E-16 |
| B179 | B209 | α1-MG | Smoking | 0.035807 | 0 |
| B179 | B196 | α1-MG | TCO2 | -0.00596 | 1.97E-07 |
| B179 | B185 | α1-MG | PT | -0.00751 | 5.45E-11 |
| B179 | B186 | α1-MG | PFC | 0.031008 | 0 |
| B179 | B206 | α1-MG | Waist | 0.021849 | 0 |
| B179 | B197 | α1-MG | AG | 0.018984 | 0 |
| B179 | B204 | α1-MG | Drinking | 0.030057 | 0 |
| B18 | B24 | CD3+CD4+helper(inducer)T | αHBDH | -0.00737 | 1.23E-10 |
| B18 | B23 | CD3+CD4+helper(inducer)T | CD3-CD16+CD56+(NK) | -0.39092 | 0 |
| B18 | B19 | CD3+CD4+helper(inducer)T | CD3+CD4+/CD3+CD8+ | 0.716139 | 0 |
| B18 | B20 | CD3+CD4+helper(inducer)T | CD3+CD4+CD8+ | 0.099828 | 0 |
| B18 | B21 | CD3+CD4+helper(inducer)T | CD3+CD4-CD8- | -0.14759 | 0 |
| B18 | B22 | CD3+CD4+helper(inducer)T | CD3+CD8+(T) | -0.27596 | 0 |
| B18 | B92 | CD3+CD4+helper(inducer)T | ALDH2-Glu504Glu | -0.00608 | 1.10E-07 |
| B18 | B93 | CD3+CD4+helper(inducer)T | ALDH2-Glu504Lys | 0.00621 | 5.90E-08 |
| B18 | B126 | CD3+CD4+helper(inducer)T | Alb | -0.01204 | 0 |
| B18 | B121 | CD3+CD4+helper(inducer)T | C3 | 0.011777 | 0 |
| B18 | B31 | CD3+CD4+helper(inducer)T | FSH | 0.006408 | 2.21E-08 |
| B18 | B124 | CD3+CD4+helper(inducer)T | ACTH | -0.04536 | 0 |
| B18 | B134 | CD3+CD4+helper(inducer)T | CHE | -0.00799 | 3.10E-12 |
| B18 | B55 | CD3+CD4+helper(inducer)T | TK | 0.007662 | 2.25E-11 |
| B18 | B56 | CD3+CD4+helper(inducer)T | IR | -0.00736 | 1.31E-10 |
| B18 | B57 | CD3+CD4+helper(inducer)T | RBCAI | -0.00845 | 1.57E-13 |
| B18 | B46 | CD3+CD4+helper(inducer)T | RBC Count | -0.00959 | 0 |
| B18 | B181 | CD3+CD4+helper(inducer)T | APTT | -0.01699 | 0 |
| B18 | B190 | CD3+CD4+helper(inducer)T | Cre | -0.00774 | 1.37E-11 |
| B18 | B25 | CD3+CD4+helper(inducer)T | CK | -0.00649 | 1.45E-08 |
| B18 | B193 | CD3+CD4+helper(inducer)T | K | -0.02883 | 0 |
| B18 | B129 | CD3+CD4+helper(inducer)T | IBIL | -0.00953 | 0 |
| B18 | B105 | CD3+CD4+helper(inducer)T | anti-U1RNP | -0.00832 | 3.75E-13 |
| B18 | B95 | CD3+CD4+helper(inducer)T | anti-cenpb | 0.014093 | 0 |
| B18 | B98 | CD3+CD4+helper(inducer)T | anti-Jo-1 | -0.03099 | 0 |
| B18 | B100 | CD3+CD4+helper(inducer)T | anti-Ro52 | 0.009072 | 2.22E-15 |
| B18 | B101 | CD3+CD4+helper(inducer)T | anti-Sc70 | -0.00695 | 1.28E-09 |
| B18 | B102 | CD3+CD4+helper(inducer)T | anti-Sm | -0.00977 | 0 |
| B18 | B104 | CD3+CD4+helper(inducer)T | anti-SSB | 0.008914 | 7.11E-15 |

| | | | | | |
|---|---|---|---|---|---|
| B18 | B106 | CD3+CD4+helper(inducer)T | anti-P | 0.01847 | 0 |
| B18 | B107 | CD3+CD4+helper(inducer)T | AnuA | 0.005884 | 2.80E-07 |
| B18 | B109 | CD3+CD4+helper(inducer)T | AMA-M2 | 0.009827 | 0 |
| B18 | B198 | CD3+CD4+helper(inducer)T | P | 0.048553 | 0 |
| B18 | B194 | CD3+CD4+helper(inducer)T | Cl | 0.020594 | 0 |
| B18 | B199 | CD3+CD4+helper(inducer)T | Mg | -0.00868 | 3.60E-14 |
| B18 | B155 | CD3+CD4+helper(inducer)T | IgA | -0.02647 | 0 |
| B18 | B156 | CD3+CD4+helper(inducer)T | IgE | -0.00918 | 1.11E-15 |
| B18 | B157 | CD3+CD4+helper(inducer)T | IgG | -0.05305 | 0 |
| B18 | B179 | CD3+CD4+helper(inducer)T | α1-MG | -0.01711 | 0 |
| B18 | B182 | CD3+CD4+helper(inducer)T | TT | -0.01241 | 0 |
| B18 | B125 | CD3+CD4+helper(inducer)T | Cor | -0.03692 | 0 |
| B18 | B138 | CD3+CD4+helper(inducer)T | PAB | -0.01255 | 0 |
| B18 | B130 | CD3+CD4+helper(inducer)T | GLB | -0.00638 | 2.49E-08 |
| B18 | B59 | CD3+CD4+helper(inducer)T | WBRV | -0.01343 | 0 |
| B18 | B60 | CD3+CD4+helper(inducer)T | WBRVI | -0.02067 | 0 |
| B18 | B61 | CD3+CD4+helper(inducer)T | WBHSRV | -0.01432 | 0 |
| B18 | B62 | CD3+CD4+helper(inducer)T | WBHSRVI | -0.01863 | 0 |
| B18 | B64 | CD3+CD4+helper(inducer)T | WBRSR(mPas_5) | -0.02816 | 0 |
| B18 | B65 | CD3+CD4+helper(inducer)T | WBRSR(mPas_1) | -0.02673 | 0 |
| B18 | B66 | CD3+CD4+helper(inducer)T | WBRSR(mPas_100) | -0.02607 | 0 |
| B18 | B67 | CD3+CD4+helper(inducer)T | WBRSR(mPas_50) | -0.02636 | 0 |
| B18 | B27 | CD3+CD4+helper(inducer)T | LDH | -0.00586 | 3.14E-07 |
| B18 | B200 | CD3+CD4+helper(inducer)T | OP | -0.00842 | 2.02E-13 |
| B18 | B220 | CD3+CD4+helper(inducer)T | SBP | -0.00697 | 1.15E-09 |
| B18 | B209 | CD3+CD4+helper(inducer)T | Smoking | 0.016917 | 0 |
| B18 | B90 | CD3+CD4+helper(inducer)T | CYFRA 21-1 | 0.011613 | 0 |
| B18 | B139 | CD3+CD4+helper(inducer)T | ADA | -0.01047 | 0 |
| B18 | B35 | CD3+CD4+helper(inducer)T | β2-MG | 0.011826 | 0 |
| B18 | B196 | CD3+CD4+helper(inducer)T | TCO2 | -0.01246 | 0 |
| B18 | B51 | CD3+CD4+helper(inducer)T | Hb | -0.0081 | 1.51E-12 |
| B18 | B184 | CD3+CD4+helper(inducer)T | INR | -0.00825 | 5.80E-13 |
| B18 | B185 | CD3+CD4+helper(inducer)T | PT | -0.01783 | 0 |
| B18 | B69 | CD3+CD4+helper(inducer)T | PV | -0.01768 | 0 |
| B18 | B79 | CD3+CD4+helper(inducer)T | hb V -dna | -0.01 | 0 |
| B18 | B197 | CD3+CD4+helper(inducer)T | AG | -0.00575 | 5.05E-07 |
| B18 | B204 | CD3+CD4+helper(inducer)T | Drinking | 0.007366 | 1.27E-10 |
| B18 | B74 | CD3+CD4+helper(inducer)T | ApoA1 | -0.00702 | 9.02E-10 |
| B18 | B75 | CD3+CD4+helper(inducer)T | ApoA1/B | -0.00722 | 2.94E-10 |
| B18 | B132 | CD3+CD4+helper(inducer)T | Dbil | -0.00757 | 3.84E-11 |
| B18 | B81 | CD3+CD4+helper(inducer)T | TSGF | 0.016373 | 0 |
| B18 | B133 | CD3+CD4+helper(inducer)T | TBIL | -0.00969 | 0 |
| B18 | B91 | CD3+CD4+helper(inducer)T | TP | -0.01292 | 0 |
| B180 | B190 | Tf | Cre | 0.011613 | 0 |

| | | | | | |
|---|---|---|---|---|---|
| B180 | B222 | Tf | NS | 0.007631 | 2.70E-11 |
| B180 | B194 | Tf | Cl | -0.0094 | 2.22E-16 |
| B180 | B199 | Tf | Mg | -0.0093 | 4.44E-16 |
| B180 | B192 | Tf | UA | 0.01037 | 0 |
| B180 | B200 | Tf | OP | 0.012887 | 0 |
| B180 | B220 | Tf | SBP | 0.010616 | 0 |
| B180 | B218 | Tf | DBP | 0.009224 | 8.88E-16 |
| B180 | B216 | Tf | BMI | 0.007453 | 7.67E-11 |
| B180 | B215 | Tf | BMI | 0.006454 | 1.75E-08 |
| B180 | B210 | Tf | Hip | 0.006658 | 6.15E-09 |
| B180 | B183 | Tf | PTA | 0.007558 | 4.16E-11 |
| B180 | B186 | Tf | PFC | 0.011963 | 0 |
| B180 | B206 | Tf | Waist | 0.007654 | 2.36E-11 |
| B180 | B197 | Tf | AG | 0.007173 | 3.79E-10 |
| B181 | B189 | APTT | eGFB | 0.00718 | 3.65E-10 |
| B181 | B190 | APTT | Cre | -0.00816 | 1.08E-12 |
| B181 | B193 | APTT | K | 0.006432 | 1.96E-08 |
| B181 | B222 | APTT | NS | -0.01668 | 0 |
| B181 | B198 | APTT | P | 0.010186 | 0 |
| B181 | B194 | APTT | Cl | 0.0167 | 0 |
| B181 | B199 | APTT | Mg | -0.01094 | 0 |
| B181 | B191 | APTT | Urea | -0.00761 | 3.11E-11 |
| B181 | B192 | APTT | UA | -0.01176 | 0 |
| B181 | B182 | APTT | TT | 0.115298 | 0 |
| B181 | B221 | APTT | Hei | -0.00583 | 3.61E-07 |
| B181 | B200 | APTT | OP | -0.00796 | 3.72E-12 |
| B181 | B220 | APTT | SBP | -0.01241 | 0 |
| B181 | B218 | APTT | DBP | -0.01574 | 0 |
| B181 | B216 | APTT | BMI | -0.01599 | 0 |
| B181 | B215 | APTT | BMI | -0.01755 | 0 |
| B181 | B210 | APTT | Hip | -0.01203 | 0 |
| B181 | B209 | APTT | Smoking | -0.01109 | 0 |
| B181 | B183 | APTT | PTA | 0.008565 | 7.55E-14 |
| B181 | B184 | APTT | INR | -0.11264 | 0 |
| B181 | B185 | APTT | PT | 0.426257 | 0 |
| B181 | B186 | APTT | PFC | -0.1293 | 0 |
| B181 | B206 | APTT | Waist | -0.02021 | 0 |
| B181 | B197 | APTT | AG | -0.00873 | 2.44E-14 |
| B181 | B204 | APTT | Drinking | -0.01672 | 0 |
| B181 | B211 | APTT | IOP-R | -0.01183 | 0 |
| B181 | B212 | APTT | IOP-L | -0.01183 | 0 |
| B182 | B225 | TT | hPBG | 0.009232 | 8.88E-16 |
| B182 | B193 | TT | K | 0.00944 | 2.22E-16 |
| B182 | B222 | TT | NS | 0.007962 | 3.61E-12 |

| | | | | | |
|---|---|---|---|---|---|
| B182 | B194 | TT | Cl | -0.04223 | 0 |
| B182 | B199 | TT | Mg | 0.017787 | 0 |
| B182 | B195 | TT | Na | 0.051766 | 0 |
| B182 | B192 | TT | UA | 0.008455 | 1.56E-13 |
| B182 | B187 | TT | NSE | 0.006542 | 1.12E-08 |
| B182 | B200 | TT | OP | 0.044566 | 0 |
| B182 | B218 | TT | DBP | 0.006183 | 6.73E-08 |
| B182 | B216 | TT | BMI | 0.006875 | 1.95E-09 |
| B182 | B210 | TT | Hip | 0.006783 | 3.19E-09 |
| B182 | B196 | TT | TCO2 | -0.01093 | 0 |
| B182 | B183 | TT | PTA | -0.29512 | 0 |
| B182 | B185 | TT | PT | 0.012079 | 0 |
| B182 | B186 | TT | PFC | -0.20472 | 0 |
| B182 | B206 | TT | Waist | 0.008645 | 4.44E-14 |
| B182 | B197 | TT | AG | 0.066146 | 0 |
| B182 | B204 | TT | Drinking | 0.005977 | 1.81E-07 |
| B183 | B225 | PTA | hPBG | 0.007776 | 1.13E-11 |
| B183 | B190 | PTA | Cre | 0.006019 | 1.48E-07 |
| B183 | B193 | PTA | K | -0.01277 | 0 |
| B183 | B198 | PTA | P | -0.00877 | 1.95E-14 |
| B183 | B194 | PTA | Cl | 0.046794 | 0 |
| B183 | B199 | PTA | Mg | -0.02811 | 0 |
| B183 | B195 | PTA | Na | -0.01744 | 0 |
| B183 | B200 | PTA | OP | -0.00986 | 0 |
| B183 | B210 | PTA | Hip | -0.01171 | 0 |
| B183 | B185 | PTA | PT | 0.022891 | 0 |
| B183 | B186 | PTA | PFC | -0.15 | 0 |
| B183 | B206 | PTA | Waist | -0.00908 | 2.22E-15 |
| B183 | B197 | PTA | AG | -0.0444 | 0 |
| B183 | B204 | PTA | Drinking | -0.00605 | 1.29E-07 |
| B183 | B212 | PTA | IOP-L | -0.00944 | 2.22E-16 |
| B184 | B199 | INR | Mg | -0.00577 | 4.63E-07 |
| B184 | B196 | INR | TCO2 | 0.007651 | 2.39E-11 |
| B184 | B185 | INR | PT | 0.035657 | 0 |
| B184 | B186 | INR | PFC | 0.553698 | 0 |
| B184 | B197 | INR | AG | -0.00914 | 1.33E-15 |
| B185 | B222 | PT | NS | -0.00848 | 1.31E-13 |
| B185 | B199 | PT | Mg | -0.01076 | 0 |
| B185 | B195 | PT | Na | -0.00583 | 3.62E-07 |
| B185 | B219 | PT | OT | 0.009165 | 1.33E-15 |
| B185 | B218 | PT | DBP | -0.00868 | 3.42E-14 |
| B185 | B216 | PT | BMI | -0.00869 | 3.20E-14 |
| B185 | B215 | PT | BMI | -0.00841 | 2.11E-13 |
| B185 | B209 | PT | Smoking | -0.00959 | 0 |

| | | | | | |
|---|---|---|---|---|---|
| B185 | B206 | PT | Waist | -0.00888 | 8.88E-15 |
| B185 | B197 | PT | AG | -0.00907 | 2.44E-15 |
| B185 | B204 | PT | Drinking | -0.01768 | 0 |
| B185 | B211 | PT | IOP-R | -0.0064 | 2.35E-08 |
| B185 | B212 | PT | IOP-L | -0.00736 | 1.35E-10 |
| B186 | B193 | PFC | K | 0.026157 | 0 |
| B186 | B198 | PFC | P | 0.006677 | 5.56E-09 |
| B186 | B194 | PFC | Cl | -0.01295 | 0 |
| B186 | B199 | PFC | Mg | 0.011852 | 0 |
| B186 | B221 | PFC | Hei | -0.00661 | 8.05E-09 |
| B186 | B200 | PFC | OP | 0.006194 | 6.37E-08 |
| B186 | B215 | PFC | BMI | 0.006312 | 3.58E-08 |
| B186 | B209 | PFC | Smoking | 0.007221 | 2.90E-10 |
| B186 | B196 | PFC | TCO2 | 0.007351 | 1.38E-10 |
| B186 | B206 | PFC | Waist | 0.007661 | 2.26E-11 |
| B187 | B189 | NSE | eGFB | -0.00671 | 4.66E-09 |
| B187 | B190 | NSE | Cre | 0.007348 | 1.41E-10 |
| B187 | B193 | NSE | K | 0.006196 | 6.34E-08 |
| B187 | B222 | NSE | NS | 0.006182 | 6.75E-08 |
| B187 | B194 | NSE | Cl | -0.0102 | 0 |
| B187 | B192 | NSE | UA | 0.00758 | 3.64E-11 |
| B187 | B220 | NSE | SBP | 0.007197 | 3.32E-10 |
| B187 | B218 | NSE | DBP | 0.007734 | 1.45E-11 |
| B187 | B196 | NSE | TCO2 | 0.006773 | 3.35E-09 |
| B187 | B206 | NSE | Waist | 0.005979 | 1.79E-07 |
| B188 | B189 | Ca | eGFB | 0.01027 | 0 |
| B188 | B190 | Ca | Cre | 0.025429 | 0 |
| B188 | B193 | Ca | K | 0.013861 | 0 |
| B188 | B222 | Ca | NS | 0.015996 | 0 |
| B188 | B194 | Ca | Cl | -0.01254 | 0 |
| B188 | B195 | Ca | Na | 0.005674 | 7.27E-07 |
| B188 | B191 | Ca | Urea | 0.008155 | 1.08E-12 |
| B188 | B192 | Ca | UA | 0.043744 | 0 |
| B188 | B221 | Ca | Hei | 0.029559 | 0 |
| B188 | B200 | Ca | OP | 0.006021 | 1.46E-07 |
| B188 | B220 | Ca | SBP | 0.015698 | 0 |
| B188 | B218 | Ca | DBP | 0.017632 | 0 |
| B188 | B216 | Ca | BMI | 0.014775 | 0 |
| B188 | B210 | Ca | Hip | 0.007986 | 3.11E-12 |
| B188 | B197 | Ca | AG | 0.008978 | 4.44E-15 |
| B188 | B204 | Ca | Drinking | 0.010964 | 0 |
| B188 | B203 | Ca | DH | 0.011198 | 0 |
| B188 | B211 | Ca | IOP-R | 0.009369 | 4.44E-16 |
| B188 | B212 | Ca | IOP-L | 0.008669 | 3.77E-14 |

| | | | | | |
|---|---|---|---|---|---|
| B189 | B190 | eGFB | Cre | -0.58387 | 0 |
| B189 | B224 | eGFB | pmh | -0.02193 | 0 |
| B189 | B193 | eGFB | K | -0.00873 | 2.44E-14 |
| B189 | B213 | eGFB | CVA-R | -0.01819 | 0 |
| B189 | B214 | eGFB | CVA-L | -0.01487 | 0 |
| B189 | B222 | eGFB | NS | -0.20519 | 0 |
| B189 | B198 | eGFB | P | 0.019496 | 0 |
| B189 | B194 | eGFB | Cl | -0.01706 | 0 |
| B189 | B199 | eGFB | Mg | -0.02021 | 0 |
| B189 | B195 | eGFB | Na | -0.02348 | 0 |
| B189 | B191 | eGFB | Urea | -0.27207 | 0 |
| B189 | B192 | eGFB | UA | -0.23116 | 0 |
| B189 | B221 | eGFB | Hei | -0.03846 | 0 |
| B189 | B200 | eGFB | OP | -0.03049 | 0 |
| B189 | B227 | eGFB | OD | -0.02242 | 0 |
| B189 | B228 | eGFB | OS | -0.02131 | 0 |
| B189 | B220 | eGFB | SBP | -0.25659 | 0 |
| B189 | B219 | eGFB | OT | -0.10283 | 0 |
| B189 | B218 | eGFB | DBP | -0.18206 | 0 |
| B189 | B216 | eGFB | BMI | -0.15785 | 0 |
| B189 | B215 | eGFB | BMI | -0.18862 | 0 |
| B189 | B210 | eGFB | Hip | -0.11002 | 0 |
| B189 | B209 | eGFB | Smoking | -0.1587 | 0 |
| B189 | B196 | eGFB | TCO2 | -0.00923 | 8.88E-16 |
| B189 | B207 | eGFB | WHR | -0.03571 | 0 |
| B189 | B206 | eGFB | Waist | -0.24966 | 0 |
| B189 | B205 | eGFB | KDA | -0.0262 | 0 |
| B189 | B204 | eGFB | Drinking | -0.09394 | 0 |
| B189 | B203 | eGFB | DH | 0.065439 | 0 |
| B189 | B211 | eGFB | IOP-R | 0.01849 | 0 |
| B189 | B201 | eGFB | e-habits | -0.07927 | 0 |
| B189 | B212 | eGFB | IOP-L | 0.010427 | 0 |
| B19 | B23 | CD3+CD4+/CD3+CD8+ | CD3-CD16+CD56+(NK) | 0.007839 | 7.69E-12 |
| B19 | B20 | CD3+CD4+/CD3+CD8+ | CD3+CD4+CD8+ | -0.08366 | 0 |
| B19 | B21 | CD3+CD4+/CD3+CD8+ | CD3+CD4-CD8- | -0.11014 | 0 |
| B19 | B22 | CD3+CD4+/CD3+CD8+ | CD3+CD8+(T) | -0.73723 | 0 |
| B19 | B92 | CD3+CD4+/CD3+CD8+ | ALDH2-Glu504Glu | -0.01171 | 0 |
| B19 | B93 | CD3+CD4+/CD3+CD8+ | ALDH2-Glu504Lys | 0.014378 | 0 |
| B19 | B160 | CD3+CD4+/CD3+CD8+ | PH | -0.00583 | 3.57E-07 |
| B19 | B94 | CD3+CD4+/CD3+CD8+ | ALDH2-Lys504Lys | -0.00679 | 3.13E-09 |
| B19 | B126 | CD3+CD4+/CD3+CD8+ | Alb | -0.00681 | 2.77E-09 |
| B19 | B124 | CD3+CD4+/CD3+CD8+ | ACTH | -0.02373 | 0 |
| B19 | B56 | CD3+CD4+/CD3+CD8+ | IR | -0.00641 | 2.17E-08 |
| B19 | B46 | CD3+CD4+/CD3+CD8+ | RBC Count | -0.00629 | 4.01E-08 |

| | | | | | |
|---|---|---|---|---|---|
| B19 | B181 | CD3+CD4+/CD3+CD8+ | APTT | -0.02453 | 0 |
| B19 | B25 | CD3+CD4+/CD3+CD8+ | CK | -0.00587 | 3.03E-07 |
| B19 | B193 | CD3+CD4+/CD3+CD8+ | K | -0.00813 | 1.25E-12 |
| B19 | B129 | CD3+CD4+/CD3+CD8+ | IBIL | -0.00749 | 6.24E-11 |
| B19 | B150 | CD3+CD4+/CD3+CD8+ | ALP | 0.006412 | 2.16E-08 |
| B19 | B99 | CD3+CD4+/CD3+CD8+ | anti-PM-Scl | 0.00783 | 8.13E-12 |
| B19 | B105 | CD3+CD4+/CD3+CD8+ | anti-U1RNP | -0.00606 | 1.22E-07 |
| B19 | B95 | CD3+CD4+/CD3+CD8+ | anti-cenpb | 0.011873 | 0 |
| B19 | B98 | CD3+CD4+/CD3+CD8+ | anti-Jo-1 | -0.02194 | 0 |
| B19 | B100 | CD3+CD4+/CD3+CD8+ | anti-Ro52 | 0.008511 | 1.08E-13 |
| B19 | B106 | CD3+CD4+/CD3+CD8+ | anti-P | 0.026886 | 0 |
| B19 | B151 | CD3+CD4+/CD3+CD8+ | Anti-CCP | 0.025272 | 0 |
| B19 | B108 | CD3+CD4+/CD3+CD8+ | Anti-Cyclin | 0.011813 | 0 |
| B19 | B109 | CD3+CD4+/CD3+CD8+ | AMA-M2 | 0.006856 | 2.16E-09 |
| B19 | B110 | CD3+CD4+/CD3+CD8+ | AHA | -0.00769 | 1.88E-11 |
| B19 | B198 | CD3+CD4+/CD3+CD8+ | P | 0.024423 | 0 |
| B19 | B155 | CD3+CD4+/CD3+CD8+ | IgA | -0.01117 | 0 |
| B19 | B157 | CD3+CD4+/CD3+CD8+ | IgG | -0.03205 | 0 |
| B19 | B179 | CD3+CD4+/CD3+CD8+ | α1-MG | -0.00886 | 1.02E-14 |
| B19 | B180 | CD3+CD4+/CD3+CD8+ | Tf | 0.00724 | 2.59E-10 |
| B19 | B125 | CD3+CD4+/CD3+CD8+ | Cor | -0.02447 | 0 |
| B19 | B138 | CD3+CD4+/CD3+CD8+ | PAB | -0.00971 | 0 |
| B19 | B60 | CD3+CD4+/CD3+CD8+ | WBRVI | -0.00925 | 6.66E-16 |
| B19 | B61 | CD3+CD4+/CD3+CD8+ | WBHSRV | -0.00883 | 1.24E-14 |
| B19 | B62 | CD3+CD4+/CD3+CD8+ | WBHSRVI | -0.01328 | 0 |
| B19 | B64 | CD3+CD4+/CD3+CD8+ | WBRSR(mPas_5) | -0.0133 | 0 |
| B19 | B65 | CD3+CD4+/CD3+CD8+ | WBRSR(mPas_1) | -0.01116 | 0 |
| B19 | B66 | CD3+CD4+/CD3+CD8+ | WBRSR(mPas_100) | -0.01523 | 0 |
| B19 | B67 | CD3+CD4+/CD3+CD8+ | WBRSR(mPas_50) | -0.0152 | 0 |
| B19 | B209 | CD3+CD4+/CD3+CD8+ | Smoking | 0.013342 | 0 |
| B19 | B90 | CD3+CD4+/CD3+CD8+ | CYFRA 21-1 | 0.008731 | 2.49E-14 |
| B19 | B34 | CD3+CD4+/CD3+CD8+ | TK | 0.008654 | 4.17E-14 |
| B19 | B196 | CD3+CD4+/CD3+CD8+ | TCO2 | -0.00744 | 8.19E-11 |
| B19 | B183 | CD3+CD4+/CD3+CD8+ | PTA | -0.02182 | 0 |
| B19 | B184 | CD3+CD4+/CD3+CD8+ | INR | -0.01012 | 0 |
| B19 | B185 | CD3+CD4+/CD3+CD8+ | PT | -0.01025 | 0 |
| B19 | B186 | CD3+CD4+/CD3+CD8+ | PFC | 0.011992 | 0 |
| B19 | B69 | CD3+CD4+/CD3+CD8+ | PV | -0.00612 | 9.27E-08 |
| B19 | B79 | CD3+CD4+/CD3+CD8+ | hb V -dna | -0.00711 | 5.31E-10 |
| B19 | B81 | CD3+CD4+/CD3+CD8+ | TSGF | 0.011805 | 0 |
| B19 | B133 | CD3+CD4+/CD3+CD8+ | TBIL | -0.00734 | 1.47E-10 |
| B19 | B91 | CD3+CD4+/CD3+CD8+ | TP | -0.00698 | 1.08E-09 |
| B19 | B114 | CD3+CD4+/CD3+CD8+ | TSH TT3 TT4(TT4) | 0.010131 | 0 |
| B190 | B224 | Cre | pmh | 0.014467 | 0 |

| | | | | | |
|---|---|---|---|---|---|
| B190 | B222 | Cre | NS | 0.482262 | 0 |
| B190 | B198 | Cre | P | -0.03617 | 0 |
| B190 | B199 | Cre | Mg | 0.018282 | 0 |
| B190 | B195 | Cre | Na | 0.022326 | 0 |
| B190 | B191 | Cre | Urea | 0.291557 | 0 |
| B190 | B192 | Cre | UA | 0.539978 | 0 |
| B190 | B221 | Cre | Hei | 0.465767 | 0 |
| B190 | B200 | Cre | OP | 0.029514 | 0 |
| B190 | B227 | Cre | OD | 0.009489 | 0 |
| B190 | B228 | Cre | OS | 0.010538 | 0 |
| B190 | B220 | Cre | SBP | 0.194704 | 0 |
| B190 | B219 | Cre | OT | -0.01953 | 0 |
| B190 | B218 | Cre | DBP | 0.183969 | 0 |
| B190 | B216 | Cre | BMI | 0.434313 | 0 |
| B190 | B215 | Cre | BMI | 0.250718 | 0 |
| B190 | B210 | Cre | Hip | 0.268206 | 0 |
| B190 | B209 | Cre | Smoking | 0.222345 | 0 |
| B190 | B196 | Cre | TCO2 | 0.008415 | 2.03E-13 |
| B190 | B207 | Cre | WHR | 0.042384 | 0 |
| B190 | B206 | Cre | Waist | 0.362621 | 0 |
| B190 | B205 | Cre | KDA | -0.02347 | 0 |
| B190 | B197 | Cre | AG | 0.007777 | 1.12E-11 |
| B190 | B204 | Cre | Drinking | 0.241118 | 0 |
| B190 | B201 | Cre | e-habits | 0.05589 | 0 |
| B190 | B212 | Cre | IOP-L | 0.005912 | 2.45E-07 |
| B191 | B224 | Urea | pmh | 0.006897 | 1.73E-09 |
| B191 | B193 | Urea | K | 0.011089 | 0 |
| B191 | B214 | Urea | CVA-L | 0.006278 | 4.22E-08 |
| B191 | B222 | Urea | NS | 0.144281 | 0 |
| B191 | B194 | Urea | Cl | 0.008039 | 2.24E-12 |
| B191 | B199 | Urea | Mg | 0.010696 | 0 |
| B191 | B195 | Urea | Na | 0.014833 | 0 |
| B191 | B192 | Urea | UA | 0.158119 | 0 |
| B191 | B221 | Urea | Hei | 0.058377 | 0 |
| B191 | B200 | Urea | OP | 0.061755 | 0 |
| B191 | B227 | Urea | OD | 0.006875 | 1.95E-09 |
| B191 | B228 | Urea | OS | 0.011866 | 0 |
| B191 | B220 | Urea | SBP | 0.107692 | 0 |
| B191 | B219 | Urea | OT | 0.028622 | 0 |
| B191 | B218 | Urea | DBP | 0.068489 | 0 |
| B191 | B216 | Urea | BMI | 0.119375 | 0 |
| B191 | B215 | Urea | BMI | 0.121806 | 0 |
| B191 | B210 | Urea | Hip | 0.080558 | 0 |
| B191 | B209 | Urea | Smoking | 0.071088 | 0 |

| | | | | | |
|---|---|---|---|---|---|
| B191 | B207 | Urea | WHR | 0.020404 | 0 |
| B191 | B206 | Urea | Waist | 0.156584 | 0 |
| B191 | B205 | Urea | KDA | -0.00922 | 8.88E-16 |
| B191 | B197 | Urea | AG | 0.005974 | 1.83E-07 |
| B191 | B204 | Urea | Drinking | 0.069174 | 0 |
| B191 | B203 | Urea | DH | -0.01057 | 0 |
| B191 | B211 | Urea | IOP-R | -0.01017 | 0 |
| B191 | B201 | Urea | e-habits | 0.028615 | 0 |
| B192 | B222 | UA | NS | 0.514218 | 0 |
| B192 | B198 | UA | P | -0.02427 | 0 |
| B192 | B194 | UA | Cl | -0.01534 | 0 |
| B192 | B199 | UA | Mg | 0.007933 | 4.32E-12 |
| B192 | B195 | UA | Na | 0.024927 | 0 |
| B192 | B221 | UA | Hei | 0.421672 | 0 |
| B192 | B200 | UA | OP | 0.028294 | 0 |
| B192 | B220 | UA | SBP | 0.221459 | 0 |
| B192 | B219 | UA | OT | -0.02799 | 0 |
| B192 | B218 | UA | DBP | 0.227819 | 0 |
| B192 | B216 | UA | BMI | 0.519827 | 0 |
| B192 | B215 | UA | BMI | 0.391728 | 0 |
| B192 | B210 | UA | Hip | 0.358818 | 0 |
| B192 | B209 | UA | Smoking | 0.199914 | 0 |
| B192 | B207 | UA | WHR | 0.051369 | 0 |
| B192 | B206 | UA | Waist | 0.468074 | 0 |
| B192 | B205 | UA | KDA | -0.02286 | 0 |
| B192 | B197 | UA | AG | 0.02684 | 0 |
| B192 | B204 | UA | Drinking | 0.272118 | 0 |
| B192 | B203 | UA | DH | 0.036963 | 0 |
| B192 | B211 | UA | IOP-R | 0.023536 | 0 |
| B192 | B201 | UA | e-habits | 0.027226 | 0 |
| B192 | B212 | UA | IOP-L | 0.022122 | 0 |
| B193 | B222 | K | NS | -0.00656 | 1.01E-08 |
| B193 | B198 | K | P | 0.063268 | 0 |
| B193 | B194 | K | Cl | -0.01764 | 0 |
| B193 | B199 | K | Mg | 0.073608 | 0 |
| B193 | B195 | K | Na | 0.035789 | 0 |
| B193 | B200 | K | OP | 0.050483 | 0 |
| B193 | B220 | K | SBP | -0.00685 | 2.24E-09 |
| B193 | B218 | K | DBP | -0.0072 | 3.23E-10 |
| B193 | B216 | K | BMI | -0.00606 | 1.21E-07 |
| B193 | B215 | K | BMI | -0.00591 | 2.50E-07 |
| B193 | B209 | K | Smoking | 0.00773 | 1.49E-11 |
| B193 | B196 | K | TCO2 | 0.054049 | 0 |
| B193 | B206 | K | Waist | -0.00661 | 7.72E-09 |

| | | | | | |
|---|---|---|---|---|---|
| B193 | B197 | K | AG | -0.01018 | 0 |
| B193 | B203 | K | DH | 0.006021 | 1.47E-07 |
| B193 | B201 | K | e-habits | 0.006035 | 1.37E-07 |
| B194 | B222 | Cl | NS | -0.01164 | 0 |
| B194 | B198 | Cl | P | -0.03185 | 0 |
| B194 | B199 | Cl | Mg | -0.02534 | 0 |
| B194 | B195 | Cl | Na | 0.222963 | 0 |
| B194 | B221 | Cl | Hei | -0.01597 | 0 |
| B194 | B200 | Cl | OP | 0.155251 | 0 |
| B194 | B220 | Cl | SBP | -0.01109 | 0 |
| B194 | B218 | Cl | DBP | -0.01865 | 0 |
| B194 | B215 | Cl | BMI | 0.006463 | 1.68E-08 |
| B194 | B209 | Cl | Smoking | 0.007976 | 3.31E-12 |
| B194 | B196 | Cl | TCO2 | -0.15189 | 0 |
| B194 | B197 | Cl | AG | -0.3976 | 0 |
| B194 | B204 | Cl | Drinking | -0.00968 | 0 |
| B194 | B203 | Cl | DH | -0.0069 | 1.69E-09 |
| B194 | B211 | Cl | IOP-R | -0.009 | 4.00E-15 |
| B195 | B224 | Na | pmh | 0.00629 | 3.98E-08 |
| B195 | B222 | Na | NS | 0.013038 | 0 |
| B195 | B199 | Na | Mg | 0.136216 | 0 |
| B195 | B221 | Na | Hei | 0.006535 | 1.16E-08 |
| B195 | B200 | Na | OP | 0.780663 | 0 |
| B195 | B220 | Na | SBP | 0.021958 | 0 |
| B195 | B218 | Na | DBP | 0.016509 | 0 |
| B195 | B216 | Na | BMI | 0.009286 | 4.44E-16 |
| B195 | B215 | Na | BMI | 0.008044 | 2.18E-12 |
| B195 | B209 | Na | Smoking | 0.014668 | 0 |
| B195 | B196 | Na | TCO2 | 0.068238 | 0 |
| B195 | B206 | Na | Waist | 0.015191 | 0 |
| B195 | B197 | Na | AG | 0.388506 | 0 |
| B195 | B204 | Na | Drinking | 0.006236 | 5.20E-08 |
| B195 | B201 | Na | e-habits | 0.007263 | 2.29E-10 |
| B196 | B200 | TCO2 | OP | 0.057895 | 0 |
| B196 | B220 | TCO2 | SBP | 0.010271 | 0 |
| B196 | B218 | TCO2 | DBP | 0.009587 | 0 |
| B196 | B197 | TCO2 | AG | -0.65001 | 0 |
| B197 | B225 | AG | hPBG | 0.006625 | 7.31E-09 |
| B197 | B222 | AG | NS | 0.012765 | 0 |
| B197 | B198 | AG | P | 0.023818 | 0 |
| B197 | B199 | AG | Mg | 0.094062 | 0 |
| B197 | B221 | AG | Hei | 0.009948 | 0 |
| B197 | B200 | AG | OP | 0.31185 | 0 |
| B197 | B220 | AG | SBP | 0.012019 | 0 |

| | | | | | |
|---|---|---|---|---|---|
| B197 | B218 | AG | DBP | 0.014168 | 0 |
| B197 | B216 | AG | BMI | 0.008065 | 1.91E-12 |
| B197 | B206 | AG | Waist | 0.008075 | 1.79E-12 |
| B197 | B204 | AG | Drinking | 0.012224 | 0 |
| B197 | B203 | AG | DH | 0.007168 | 3.91E-10 |
| B198 | B222 | P | NS | -0.04019 | 0 |
| B198 | B199 | P | Mg | 0.010202 | 0 |
| B198 | B221 | P | Hei | -0.03138 | 0 |
| B198 | B220 | P | SBP | -0.02671 | 0 |
| B198 | B219 | P | OT | 0.007502 | 5.76E-11 |
| B198 | B218 | P | DBP | -0.02905 | 0 |
| B198 | B216 | P | BMI | -0.03492 | 0 |
| B198 | B215 | P | BMI | -0.0247 | 0 |
| B198 | B210 | P | Hip | -0.02179 | 0 |
| B198 | B209 | P | Smoking | -0.01392 | 0 |
| B198 | B206 | P | Waist | -0.03501 | 0 |
| B198 | B204 | P | Drinking | -0.02665 | 0 |
| B198 | B211 | P | IOP-R | -0.00595 | 2.03E-07 |
| B198 | B212 | P | IOP-L | -0.00673 | 4.15E-09 |
| B199 | B225 | Mg | hPBG | 0.006407 | 2.23E-08 |
| B199 | B222 | Mg | NS | 0.006601 | 8.28E-09 |
| B199 | B200 | Mg | OP | 0.123289 | 0 |
| B199 | B218 | Mg | DBP | 0.005611 | 9.64E-07 |
| B199 | B209 | Mg | Smoking | 0.008405 | 2.17E-13 |
| B199 | B206 | Mg | Waist | 0.006471 | 1.61E-08 |
| B199 | B204 | Mg | Drinking | 0.009941 | 0 |
| B2 | B142 | Age | 25-(OH)D | 0.013047 | 0 |
| B2 | B137 | Age | AFU | 0.015063 | 0 |
| B2 | B24 | Age | αHBDH | 0.05383 | 0 |
| B2 | B15 | Age | CD3-CD19+(B cell) | -0.01722 | 0 |
| B2 | B19 | Age | CD3+CD4+/CD3+CD8+ | 0.006912 | 1.60E-09 |
| B2 | B21 | Age | CD3+CD4-CD8- | -0.00789 | 5.67E-12 |
| B2 | B54 | Age | ESR | 0.030422 | 0 |
| B2 | B13 | Age | PGI_PGII | -0.00737 | 1.27E-10 |
| B2 | B160 | Age | PH | -0.0969 | 0 |
| B2 | B140 | Age | r-GT | 0.053397 | 0 |
| B2 | B82 | Age | CEA | 0.008054 | 2.04E-12 |
| B2 | B118 | Age | CEA1 | 0.040375 | 0 |
| B2 | B126 | Age | Alb | -0.3222 | 0 |
| B2 | B127 | Age | A-G ratio | -0.1507 | 0 |
| B2 | B40 | Age | WBC | -0.07639 | 0 |
| B2 | B128 | Age | ALT | -0.02705 | 0 |
| B2 | B164 | Age | COM | -0.00844 | 1.73E-13 |
| B2 | B123 | Age | hs-CRP1 | 0.017846 | 0 |

| | | | | | |
|---|---|---|---|---|---|
| B2 | B28 | Age | PRL | -0.00586 | 3.10E-07 |
| B2 | B30 | Age | LH | 0.015282 | 0 |
| B2 | B111 | Age | TSH | 0.024201 | 0 |
| B2 | B31 | Age | FSH | 0.020073 | 0 |
| B2 | B41 | Age | MON % | 0.018641 | 0 |
| B2 | B166 | Age | Pro | -0.02853 | 0 |
| B2 | B70 | Age | LDL-C | 0.211126 | 0 |
| B2 | B188 | Age | Ca | -0.03611 | 0 |
| B2 | B71 | Age | TG | 0.087963 | 0 |
| B2 | B72 | Age | HDL-C | 0.025593 | 0 |
| B2 | B189 | Age | eGFB | -0.61704 | 0 |
| B2 | B141 | Age | AST/ALT | 0.014151 | 0 |
| B2 | B146 | Age | CysC | 0.049244 | 0 |
| B2 | B56 | Age | IR | -0.00926 | 6.66E-16 |
| B2 | B43 | Age | MCV | 0.082436 | 0 |
| B2 | B44 | Age | MCH | 0.107085 | 0 |
| B2 | B45 | Age | MCHC | -0.0606 | 0 |
| B2 | B46 | Age | RBC Count | -0.14239 | 0 |
| B2 | B47 | Age | RDW | 0.275648 | 0 |
| B2 | B181 | Age | APTT | -0.00944 | 2.22E-16 |
| B2 | B190 | Age | Cre | 0.108984 | 0 |
| B2 | B25 | Age | CK | -0.0082 | 8.19E-13 |
| B2 | B26 | Age | CKMB | 0.007249 | 2.47E-10 |
| B2 | B224 | Age | pmh | 0.038327 | 0 |
| B2 | B193 | Age | K | 0.008823 | 1.33E-14 |
| B2 | B129 | Age | IBIL | 0.018501 | 0 |
| B2 | B150 | Age | ALP | 0.048589 | 0 |
| B2 | B213 | Age | CVA-R | 0.021956 | 0 |
| B2 | B214 | Age | CVA-L | 0.01748 | 0 |
| B2 | B84 | Age | C-PSA | 0.028949 | 0 |
| B2 | B222 | Age | NS | 0.093939 | 0 |
| B2 | B151 | Age | Anti-CCP | 0.008692 | 3.24E-14 |
| B2 | B154 | Age | RF | 0.006366 | 2.73E-08 |
| B2 | B48 | Age | LYM% | -0.00742 | 9.09E-11 |
| B2 | B198 | Age | P | -0.01737 | 0 |
| B2 | B194 | Age | Cl | 0.017422 | 0 |
| B2 | B199 | Age | Mg | 0.014678 | 0 |
| B2 | B157 | Age | IgG | 0.010403 | 0 |
| B2 | B195 | Age | Na | 0.025294 | 0 |
| B2 | B176 | Age | Ucr | -0.02734 | 0 |
| B2 | B191 | Age | Urea | 0.193604 | 0 |
| B2 | B174 | Age | Urea/Cr | 0.075295 | 0 |
| B2 | B192 | Age | UA | -0.01089 | 0 |
| B2 | B170 | Age | U-GLU | 0.04909 | 0 |

| | | | | | |
|---|---|---|---|---|---|
| B2 | B179 | Age | α1-MG | 0.008861 | 1.02E-14 |
| B2 | B153 | Age | Glu | 0.03657 | 0 |
| B2 | B138 | Age | PAB | -0.0189 | 0 |
| B2 | B130 | Age | GLB | 0.056443 | 0 |
| B2 | B60 | Age | WBRVI | -0.01521 | 0 |
| B2 | B61 | Age | WBHSRV | -0.00603 | 1.42E-07 |
| B2 | B62 | Age | WBHSRVI | -0.015 | 0 |
| B2 | B64 | Age | WBRSR(mPas_5) | -0.01225 | 0 |
| B2 | B65 | Age | WBRSR(mPas_1) | -0.01131 | 0 |
| B2 | B66 | Age | WBRSR(mPas_100) | -0.00985 | 0 |
| B2 | B67 | Age | WBRSR(mPas_50) | -0.00998 | 0 |
| B2 | B86 | Age | TBhCG | -0.00833 | 3.49E-13 |
| B2 | B27 | Age | LDH | 0.067283 | 0 |
| B2 | B221 | Age | Hei | -0.21056 | 0 |
| B2 | B87 | Age | NSE | 0.007843 | 7.51E-12 |
| B2 | B200 | Age | OP | 0.029237 | 0 |
| B2 | B227 | Age | OD | 0.030788 | 0 |
| B2 | B228 | Age | OS | 0.028486 | 0 |
| B2 | B50 | Age | eo% | 0.076035 | 0 |
| B2 | B220 | Age | SBP | 0.33427 | 0 |
| B2 | B219 | Age | OT | 0.178928 | 0 |
| B2 | B218 | Age | DBP | 0.212567 | 0 |
| B2 | B216 | Age | BMI | 0.019969 | 0 |
| B2 | B215 | Age | BMI | 0.185226 | 0 |
| B2 | B131 | Age | AST | 0.059654 | 0 |
| B2 | B3 | Age | Hcy | 0.022298 | 0 |
| B2 | B172 | Age | KB | -0.04448 | 0 |
| B2 | B210 | Age | Hip | 0.035208 | 0 |
| B2 | B10 | Age | PG I | 0.062429 | 0 |
| B2 | B12 | Age | PG II | 0.069004 | 0 |
| B2 | B14 | Age | G-17 | 0.0236 | 0 |
| B2 | B209 | Age | Smoking | 0.200649 | 0 |
| B2 | B90 | Age | CYFRA 21-1 | 0.051407 | 0 |
| B2 | B139 | Age | ADA | 0.024948 | 0 |
| B2 | B34 | Age | TK | -0.00717 | 3.90E-10 |
| B2 | B196 | Age | TCO2 | 0.007939 | 4.17E-12 |
| B2 | B51 | Age | Hb | -0.05999 | 0 |
| B2 | B186 | Age | PFC | 0.011036 | 0 |
| B2 | B69 | Age | PV | 0.014237 | 0 |
| B2 | B52 | Age | PLT/BPC | -0.19651 | 0 |
| B2 | B173 | Age | NTT | 0.053396 | 0 |
| B2 | B207 | Age | WHR | 0.042304 | 0 |
| B2 | B206 | Age | Waist | 0.255597 | 0 |
| B2 | B205 | Age | KDA | 0.051087 | 0 |

| | | | | | |
|---|---|---|---|---|---|
| B2 | B79 | Age | hb V -dna | -0.01185 | 0 |
| B2 | B204 | Age | Drinking | 0.067285 | 0 |
| B2 | B203 | Age | DH | -0.08575 | 0 |
| B2 | B80 | Age | anti-HP IgG | 0.027881 | 0 |
| B2 | B112 | Age | FT4 | -0.04668 | 0 |
| B2 | B113 | Age | FT3 | -0.03091 | 0 |
| B2 | B211 | Age | IOP-R | -0.03108 | 0 |
| B2 | B33 | Age | Prog | -0.00695 | 1.28E-09 |
| B2 | B201 | Age | e-habits | 0.092417 | 0 |
| B2 | B73 | Age | ApoA | 0.015942 | 0 |
| B2 | B74 | Age | ApoA1 | 0.036453 | 0 |
| B2 | B75 | Age | ApoA1/B | -0.01153 | 0 |
| B2 | B76 | Age | ApoB | 0.047252 | 0 |
| B2 | B77 | Age | TC | 0.223795 | 0 |
| B2 | B133 | Age | TBIL | 0.013788 | 0 |
| B2 | B135 | Age | TBA | 0.006842 | 2.33E-09 |
| B2 | B91 | Age | TP | -0.15174 | 0 |
| B2 | B212 | Age | IOP-L | -0.01739 | 0 |
| B20 | B142 | CD3+CD4+CD8+ | 25-(OH)D | -0.0063 | 3.79E-08 |
| B20 | B137 | CD3+CD4+CD8+ | AFU | 0.012503 | 0 |
| B20 | B23 | CD3+CD4+CD8+ | CD3-CD16+CD56+(NK) | -0.06029 | 0 |
| B20 | B21 | CD3+CD4+CD8+ | CD3+CD4-CD8- | -0.08134 | 0 |
| B20 | B22 | CD3+CD4+CD8+ | CD3+CD8+(T) | 0.205694 | 0 |
| B20 | B54 | CD3+CD4+CD8+ | ESR | 0.011212 | 0 |
| B20 | B92 | CD3+CD4+CD8+ | ALDH2-Glu504Glu | 0.018756 | 0 |
| B20 | B93 | CD3+CD4+CD8+ | ALDH2-Glu504Lys | -0.01603 | 0 |
| B20 | B94 | CD3+CD4+CD8+ | ALDH2-Lys504Lys | -0.0087 | 3.15E-14 |
| B20 | B143 | CD3+CD4+CD8+ | β-CTx | -0.00919 | 8.88E-16 |
| B20 | B124 | CD3+CD4+CD8+ | ACTH | 0.006534 | 1.17E-08 |
| B20 | B134 | CD3+CD4+CD8+ | CHE | -0.00675 | 3.83E-09 |
| B20 | B145 | CD3+CD4+CD8+ | BGP | -0.00913 | 1.78E-15 |
| B20 | B56 | CD3+CD4+CD8+ | IR | -0.0085 | 1.18E-13 |
| B20 | B181 | CD3+CD4+CD8+ | APTT | -0.0108 | 0 |
| B20 | B193 | CD3+CD4+CD8+ | K | -0.01573 | 0 |
| B20 | B150 | CD3+CD4+CD8+ | ALP | -0.00587 | 3.05E-07 |
| B20 | B105 | CD3+CD4+CD8+ | anti-U1RNP | -0.00728 | 2.07E-10 |
| B20 | B95 | CD3+CD4+CD8+ | anti-cenpb | -0.00587 | 2.94E-07 |
| B20 | B96 | CD3+CD4+CD8+ | dsDNA (IIF) | -0.00604 | 1.33E-07 |
| B20 | B97 | CD3+CD4+CD8+ | dsDNA | -0.01424 | 0 |
| B20 | B101 | CD3+CD4+CD8+ | anti-Sc70 | -0.00594 | 2.15E-07 |
| B20 | B104 | CD3+CD4+CD8+ | anti-SSB | -0.01138 | 0 |
| B20 | B107 | CD3+CD4+CD8+ | AnuA | -0.00639 | 2.38E-08 |
| B20 | B108 | CD3+CD4+CD8+ | Anti-Cyclin | -0.00715 | 4.30E-10 |
| B20 | B154 | CD3+CD4+CD8+ | RF | 0.027419 | 0 |

| | | | | | |
|---|---|---|---|---|---|
| B20 | B194 | CD3+CD4+CD8+ | Cl | 0.017549 | 0 |
| B20 | B155 | CD3+CD4+CD8+ | IgA | -0.0086 | 6.04E-14 |
| B20 | B157 | CD3+CD4+CD8+ | IgG | -0.01012 | 0 |
| B20 | B158 | CD3+CD4+CD8+ | IgM | 0.00968 | 0 |
| B20 | B195 | CD3+CD4+CD8+ | Na | -0.03264 | 0 |
| B20 | B182 | CD3+CD4+CD8+ | TT | -0.05778 | 0 |
| B20 | B125 | CD3+CD4+CD8+ | Cor | 0.031011 | 0 |
| B20 | B60 | CD3+CD4+CD8+ | WBRVI | -0.01198 | 0 |
| B20 | B62 | CD3+CD4+CD8+ | WBHSRVI | -0.01158 | 0 |
| B20 | B64 | CD3+CD4+CD8+ | WBRSR(mPas_5) | -0.00852 | 1.03E-13 |
| B20 | B65 | CD3+CD4+CD8+ | WBRSR(mPas_1) | -0.00888 | 9.10E-15 |
| B20 | B66 | CD3+CD4+CD8+ | WBRSR(mPas_100) | -0.00772 | 1.59E-11 |
| B20 | B67 | CD3+CD4+CD8+ | WBRSR(mPas_50) | -0.00825 | 6.01E-13 |
| B20 | B200 | CD3+CD4+CD8+ | OP | -0.02936 | 0 |
| B20 | B183 | CD3+CD4+CD8+ | PTA | 0.060774 | 0 |
| B20 | B184 | CD3+CD4+CD8+ | INR | -0.01143 | 0 |
| B20 | B186 | CD3+CD4+CD8+ | PFC | -0.01331 | 0 |
| B20 | B69 | CD3+CD4+CD8+ | PV | 0.009955 | 0 |
| B20 | B197 | CD3+CD4+CD8+ | AG | -0.03337 | 0 |
| B20 | B112 | CD3+CD4+CD8+ | FT4 | 0.006687 | 5.27E-09 |
| B20 | B81 | CD3+CD4+CD8+ | TSGF | -0.04496 | 0 |
| B20 | B144 | CD3+CD4+CD8+ | Total-PINP | -0.01048 | 0 |
| B200 | B225 | OP | hPBG | 0.007625 | 2.79E-11 |
| B200 | B222 | OP | NS | 0.020906 | 0 |
| B200 | B221 | OP | Hei | 0.007614 | 2.98E-11 |
| B200 | B220 | OP | SBP | 0.027335 | 0 |
| B200 | B218 | OP | DBP | 0.021643 | 0 |
| B200 | B216 | OP | BMI | 0.016159 | 0 |
| B200 | B215 | OP | BMI | 0.01661 | 0 |
| B200 | B210 | OP | Hip | 0.009388 | 2.22E-16 |
| B200 | B209 | OP | Smoking | 0.01718 | 0 |
| B200 | B206 | OP | Waist | 0.024027 | 0 |
| B200 | B204 | OP | Drinking | 0.008567 | 7.44E-14 |
| B200 | B201 | OP | e-habits | 0.00648 | 1.53E-08 |
| B201 | B224 | e-habits | pmh | 0.029061 | 0 |
| B201 | B222 | e-habits | NS | 0.038569 | 0 |
| B201 | B221 | e-habits | Hei | 0.030901 | 0 |
| B201 | B220 | e-habits | SBP | 0.051815 | 0 |
| B201 | B219 | e-habits | OT | 0.034243 | 0 |
| B201 | B218 | e-habits | DBP | 0.019519 | 0 |
| B201 | B216 | e-habits | BMI | 0.025036 | 0 |
| B201 | B215 | e-habits | BMI | 0.012022 | 0 |
| B201 | B210 | e-habits | Hip | 0.017568 | 0 |
| B201 | B209 | e-habits | Smoking | 0.015281 | 0 |

| | | | | | |
|---|---|---|---|---|---|
| B201 | B207 | e-habits | WHR | 0.00654 | 1.13E-08 |
| B201 | B206 | e-habits | Waist | 0.022283 | 0 |
| B201 | B205 | e-habits | KDA | 0.024682 | 0 |
| B201 | B204 | e-habits | Drinking | 0.038322 | 0 |
| B201 | B203 | e-habits | DH | 0.041673 | 0 |
| B201 | B212 | e-habits | IOP-L | 0.008497 | 1.18E-13 |
| B202 | B223 | e-times | FH | 0.039708 | 0 |
| B203 | B224 | DH | pmh | 0.007099 | 5.72E-10 |
| B203 | B222 | DH | NS | 0.029124 | 0 |
| B203 | B221 | DH | Hei | 0.041701 | 0 |
| B203 | B219 | DH | OT | -0.00898 | 4.44E-15 |
| B203 | B216 | DH | BMI | 0.032642 | 0 |
| B203 | B215 | DH | BMI | 0.013386 | 0 |
| B203 | B210 | DH | Hip | 0.031638 | 0 |
| B203 | B209 | DH | Smoking | 0.045301 | 0 |
| B203 | B206 | DH | Waist | 0.008631 | 4.88E-14 |
| B203 | B205 | DH | KDA | 0.007112 | 5.33E-10 |
| B203 | B204 | DH | Drinking | 0.051462 | 0 |
| B204 | B222 | Drinking | NS | 0.330087 | 0 |
| B204 | B221 | Drinking | Hei | 0.264183 | 0 |
| B204 | B220 | Drinking | SBP | 0.136096 | 0 |
| B204 | B219 | Drinking | OT | -0.02692 | 0 |
| B204 | B218 | Drinking | DBP | 0.185393 | 0 |
| B204 | B216 | Drinking | BMI | 0.290784 | 0 |
| B204 | B215 | Drinking | BMI | 0.199569 | 0 |
| B204 | B210 | Drinking | Hip | 0.175366 | 0 |
| B204 | B209 | Drinking | Smoking | 0.396509 | 0 |
| B204 | B207 | Drinking | WHR | 0.033783 | 0 |
| B204 | B206 | Drinking | Waist | 0.283569 | 0 |
| B204 | B205 | Drinking | KDA | -0.02106 | 0 |
| B204 | B211 | Drinking | IOP-R | 0.020188 | 0 |
| B204 | B212 | Drinking | IOP-L | 0.023211 | 0 |
| B205 | B224 | KDA | pmh | 0.015565 | 0 |
| B205 | B222 | KDA | NS | -0.03031 | 0 |
| B205 | B221 | KDA | Hei | -0.02989 | 0 |
| B205 | B219 | KDA | OT | 0.036333 | 0 |
| B205 | B218 | KDA | DBP | -0.00934 | 4.44E-16 |
| B205 | B216 | KDA | BMI | -0.02668 | 0 |
| B205 | B215 | KDA | BMI | -0.01448 | 0 |
| B205 | B210 | KDA | Hip | -0.01402 | 0 |
| B205 | B209 | KDA | Smoking | -0.01209 | 0 |
| B205 | B206 | KDA | Waist | -0.01379 | 0 |
| B206 | B224 | Waist | pmh | 0.013056 | 0 |
| B206 | B222 | Waist | NS | 0.740723 | 0 |

| | | | | | |
|---|---|---|---|---|---|
| B206 | B221 | Waist | Hei | 0.410046 | 0 |
| B206 | B227 | Waist | OD | 0.006603 | 8.19E-09 |
| B206 | B228 | Waist | OS | 0.006171 | 7.14E-08 |
| B206 | B220 | Waist | SBP | 0.363749 | 0 |
| B206 | B219 | Waist | OT | 0.023081 | 0 |
| B206 | B218 | Waist | DBP | 0.340036 | 0 |
| B206 | B216 | Waist | BMI | 0.852527 | 0 |
| B206 | B215 | Waist | BMI | 0.843667 | 0 |
| B206 | B210 | Waist | Hip | 0.757443 | 0 |
| B206 | B209 | Waist | Smoking | 0.256563 | 0 |
| B206 | B207 | Waist | WHR | 0.106494 | 0 |
| B206 | B211 | Waist | IOP-R | 0.032725 | 0 |
| B206 | B212 | Waist | IOP-L | 0.036407 | 0 |
| B207 | B222 | WHR | NS | 0.099593 | 0 |
| B207 | B221 | WHR | Hei | 0.032516 | 0 |
| B207 | B220 | WHR | SBP | 0.038853 | 0 |
| B207 | B219 | WHR | OT | 0.006154 | 7.74E-08 |
| B207 | B218 | WHR | DBP | 0.038459 | 0 |
| B207 | B216 | WHR | BMI | 0.076905 | 0 |
| B207 | B215 | WHR | BMI | 0.079402 | 0 |
| B207 | B210 | WHR | Hip | 0.046155 | 0 |
| B207 | B209 | WHR | Smoking | 0.03473 | 0 |
| B209 | B224 | Smoking | pmh | 0.009192 | 8.88E-16 |
| B209 | B222 | Smoking | NS | 0.286309 | 0 |
| B209 | B221 | Smoking | Hei | 0.214305 | 0 |
| B209 | B220 | Smoking | SBP | 0.09975 | 0 |
| B209 | B219 | Smoking | OT | -0.00792 | 4.63E-12 |
| B209 | B218 | Smoking | DBP | 0.142455 | 0 |
| B209 | B216 | Smoking | BMI | 0.225963 | 0 |
| B209 | B215 | Smoking | BMI | 0.150541 | 0 |
| B209 | B210 | Smoking | Hip | 0.121901 | 0 |
| B209 | B211 | Smoking | IOP-R | 0.007627 | 2.77E-11 |
| B209 | B212 | Smoking | IOP-L | 0.011704 | 0 |
| B21 | B137 | CD3+CD4-CD8- | AFU | -0.01045 | 0 |
| B21 | B23 | CD3+CD4-CD8- | CD3-CD16+CD56+(NK) | -0.09983 | 0 |
| B21 | B22 | CD3+CD4-CD8- | CD3+CD8+(T) | 0.041037 | 0 |
| B21 | B54 | CD3+CD4-CD8- | ESR | 0.005958 | 1.98E-07 |
| B21 | B92 | CD3+CD4-CD8- | ALDH2-Glu504Glu | -0.01449 | 0 |
| B21 | B93 | CD3+CD4-CD8- | ALDH2-Glu504Lys | 0.012923 | 0 |
| B21 | B126 | CD3+CD4-CD8- | Alb | 0.007522 | 5.14E-11 |
| B21 | B122 | CD3+CD4-CD8- | C4 | -0.00827 | 5.33E-13 |
| B21 | B189 | CD3+CD4-CD8- | eGFB | 0.00695 | 1.30E-09 |
| B21 | B193 | CD3+CD4-CD8- | K | 0.013068 | 0 |
| B21 | B95 | CD3+CD4-CD8- | anti-cenpb | -0.01171 | 0 |

| | | | | | |
|---|---|---|---|---|---|
| B21 | B96 | CD3+CD4-CD8- | dsDNA (IIF) | -0.00915 | 1.33E-15 |
| B21 | B97 | CD3+CD4-CD8- | dsDNA | 0.008035 | 2.30E-12 |
| B21 | B102 | CD3+CD4-CD8- | anti-Sm | 0.009223 | 8.88E-16 |
| B21 | B103 | CD3+CD4-CD8- | anti-SSA/Ro | 0.011031 | 0 |
| B21 | B107 | CD3+CD4-CD8- | AnuA | -0.00646 | 1.72E-08 |
| B21 | B108 | CD3+CD4-CD8- | Anti-Cyclin | 0.020909 | 0 |
| B21 | B109 | CD3+CD4-CD8- | AMA-M2 | -0.0088 | 1.55E-14 |
| B21 | B198 | CD3+CD4-CD8- | P | -0.00663 | 6.98E-09 |
| B21 | B194 | CD3+CD4-CD8- | Cl | -0.02203 | 0 |
| B21 | B156 | CD3+CD4-CD8- | IgE | -0.01082 | 0 |
| B21 | B157 | CD3+CD4-CD8- | IgG | 0.020038 | 0 |
| B21 | B158 | CD3+CD4-CD8- | IgM | 0.009069 | 2.44E-15 |
| B21 | B179 | CD3+CD4-CD8- | α1-MG | -0.00614 | 8.46E-08 |
| B21 | B125 | CD3+CD4-CD8- | Cor | 0.019775 | 0 |
| B21 | B62 | CD3+CD4-CD8- | WBHSRVI | -0.00658 | 9.03E-09 |
| B21 | B87 | CD3+CD4-CD8- | NSE | -0.00625 | 4.95E-08 |
| B21 | B200 | CD3+CD4-CD8- | OP | -0.00876 | 2.09E-14 |
| B21 | B215 | CD3+CD4-CD8- | BMI | -0.00622 | 5.66E-08 |
| B21 | B209 | CD3+CD4-CD8- | Smoking | -0.00711 | 5.34E-10 |
| B21 | B90 | CD3+CD4-CD8- | CYFRA 21-1 | -0.00699 | 1.02E-09 |
| B21 | B34 | CD3+CD4-CD8- | TK | 0.005942 | 2.13E-07 |
| B21 | B183 | CD3+CD4-CD8- | PTA | -0.02051 | 0 |
| B21 | B185 | CD3+CD4-CD8- | PT | 0.011668 | 0 |
| B21 | B186 | CD3+CD4-CD8- | PFC | 0.012788 | 0 |
| B21 | B69 | CD3+CD4-CD8- | PV | 0.00922 | 8.88E-16 |
| B21 | B206 | CD3+CD4-CD8- | Waist | -0.00657 | 9.69E-09 |
| B21 | B79 | CD3+CD4-CD8- | hb V -dna | 0.009423 | 2.22E-16 |
| B21 | B197 | CD3+CD4-CD8- | AG | 0.01394 | 0 |
| B21 | B204 | CD3+CD4-CD8- | Drinking | -0.0065 | 1.41E-08 |
| B21 | B33 | CD3+CD4-CD8- | Prog | 0.008449 | 1.63E-13 |
| B21 | B74 | CD3+CD4-CD8- | ApoA1 | 0.013082 | 0 |
| B21 | B75 | CD3+CD4-CD8- | ApoA1/B | 0.007037 | 8.06E-10 |
| B21 | B76 | CD3+CD4-CD8- | ApoB | -0.0089 | 7.99E-15 |
| B21 | B91 | CD3+CD4-CD8- | TP | 0.006914 | 1.57E-09 |
| B21 | B114 | CD3+CD4-CD8- | TSH TT3 TT4(TT4) | 0.007751 | 1.31E-11 |
| B210 | B222 | Hip | NS | 0.617325 | 0 |
| B210 | B221 | Hip | Hei | 0.439634 | 0 |
| B210 | B220 | Hip | SBP | 0.244922 | 0 |
| B210 | B218 | Hip | DBP | 0.229459 | 0 |
| B210 | B216 | Hip | BMI | 0.799915 | 0 |
| B210 | B215 | Hip | BMI | 0.752018 | 0 |
| B210 | B211 | Hip | IOP-R | 0.026776 | 0 |
| B210 | B212 | Hip | IOP-L | 0.026189 | 0 |
| B211 | B222 | IOP-R | NS | 0.032754 | 0 |

| | | | | | |
|---|---|---|---|---|---|
| B211 | B221 | IOP-R | Hei | 0.01198 | 0 |
| B211 | B220 | IOP-R | SBP | 0.068813 | 0 |
| B211 | B218 | IOP-R | DBP | 0.068281 | 0 |
| B211 | B216 | IOP-R | BMI | 0.030509 | 0 |
| B211 | B215 | IOP-R | BMI | 0.031521 | 0 |
| B211 | B212 | IOP-R | IOP-L | 0.742305 | 0 |
| B212 | B222 | IOP-L | NS | 0.034459 | 0 |
| B212 | B221 | IOP-L | Hei | 0.009887 | 0 |
| B212 | B228 | IOP-L | OS | 0.007073 | 6.60E-10 |
| B212 | B220 | IOP-L | SBP | 0.070111 | 0 |
| B212 | B218 | IOP-L | DBP | 0.067882 | 0 |
| B212 | B216 | IOP-L | BMI | 0.029974 | 0 |
| B212 | B215 | IOP-L | BMI | 0.032483 | 0 |
| B213 | B214 | CVA-R | CVA-L | 0.08962 | 0 |
| B213 | B227 | CVA-R | OD | 0.012842 | 0 |
| B213 | B220 | CVA-R | SBP | 0.010456 | 0 |
| B214 | B220 | CVA-L | SBP | 0.008057 | 2.01E-12 |
| B215 | B222 | BMI | NS | 0.648673 | 0 |
| B215 | B221 | BMI | Hei | 0.175979 | 0 |
| B215 | B220 | BMI | SBP | 0.333062 | 0 |
| B215 | B219 | BMI | OT | 0.020493 | 0 |
| B215 | B218 | BMI | DBP | 0.314042 | 0 |
| B215 | B216 | BMI | BMI | 0.842498 | 0 |
| B216 | B222 | BMI | NS | 0.779174 | 0 |
| B216 | B221 | BMI | Hei | 0.667687 | 0 |
| B216 | B220 | BMI | SBP | 0.309844 | 0 |
| B216 | B219 | BMI | OT | -0.02188 | 0 |
| B216 | B218 | BMI | DBP | 0.312086 | 0 |
| B218 | B222 | DBP | NS | 0.30803 | 0 |
| B218 | B221 | DBP | Hei | 0.143842 | 0 |
| B218 | B228 | DBP | OS | 0.006898 | 1.72E-09 |
| B218 | B220 | DBP | SBP | 0.737704 | 0 |
| B218 | B219 | DBP | OT | 0.021716 | 0 |
| B219 | B224 | OT | pmh | 0.040252 | 0 |
| B219 | B222 | OT | NS | -0.02176 | 0 |
| B219 | B221 | OT | Hei | -0.06766 | 0 |
| B219 | B227 | OT | OD | 0.005617 | 9.37E-07 |
| B219 | B228 | OT | OS | 0.006375 | 2.61E-08 |
| B219 | B220 | OT | SBP | 0.050604 | 0 |
| B22 | B23 | CD3+CD8+(T) | CD3-CD16+CD56+(NK) | -0.26412 | 0 |
| B22 | B92 | CD3+CD8+(T) | ALDH2-Glu504Glu | 0.012588 | 0 |
| B22 | B93 | CD3+CD8+(T) | ALDH2-Glu504Lys | -0.0151 | 0 |
| B22 | B94 | CD3+CD8+(T) | ALDH2-Lys504Lys | 0.006284 | 4.11E-08 |
| B22 | B124 | CD3+CD8+(T) | ACTH | 0.010334 | 0 |

| | | | | | |
|---|---|---|---|---|---|
| B22 | B57 | CD3+CD8+(T) | RBCAI | -0.00764 | 2.51E-11 |
| B22 | B181 | CD3+CD8+(T) | APTT | 0.027746 | 0 |
| B22 | B25 | CD3+CD8+(T) | CK | 0.009525 | 0 |
| B22 | B129 | CD3+CD8+(T) | IBIL | 0.005737 | 5.49E-07 |
| B22 | B150 | CD3+CD8+(T) | ALP | -0.00799 | 3.11E-12 |
| B22 | B99 | CD3+CD8+(T) | anti-PM-Scl | -0.00849 | 1.26E-13 |
| B22 | B95 | CD3+CD8+(T) | anti-cenpb | -0.00562 | 9.23E-07 |
| B22 | B98 | CD3+CD8+(T) | anti-Jo-1 | 0.042048 | 0 |
| B22 | B101 | CD3+CD8+(T) | anti-Sc70 | -0.00671 | 4.79E-09 |
| B22 | B102 | CD3+CD8+(T) | anti-Sm | -0.00583 | 3.55E-07 |
| B22 | B103 | CD3+CD8+(T) | anti-SSA/Ro | 0.013149 | 0 |
| B22 | B106 | CD3+CD8+(T) | anti-P | -0.01434 | 0 |
| B22 | B151 | CD3+CD8+(T) | Anti-CCP | -0.01675 | 0 |
| B22 | B108 | CD3+CD8+(T) | Anti-Cyclin | -0.01034 | 0 |
| B22 | B157 | CD3+CD8+(T) | IgG | 0.017454 | 0 |
| B22 | B195 | CD3+CD8+(T) | Na | -0.00745 | 7.79E-11 |
| B22 | B178 | CD3+CD8+(T) | Ig-U | -0.00577 | 4.62E-07 |
| B22 | B180 | CD3+CD8+(T) | Tf | -0.00568 | 6.99E-07 |
| B22 | B182 | CD3+CD8+(T) | TT | -0.01054 | 0 |
| B22 | B125 | CD3+CD8+(T) | Cor | 0.018996 | 0 |
| B22 | B62 | CD3+CD8+(T) | WBHSRVI | 0.011361 | 0 |
| B22 | B64 | CD3+CD8+(T) | WBRSR(mPas_5) | 0.007515 | 5.34E-11 |
| B22 | B66 | CD3+CD8+(T) | WBRSR(mPas_100) | 0.012206 | 0 |
| B22 | B67 | CD3+CD8+(T) | WBRSR(mPas_50) | 0.01197 | 0 |
| B22 | B209 | CD3+CD8+(T) | Smoking | -0.00662 | 7.42E-09 |
| B22 | B34 | CD3+CD8+(T) | TK | -0.00715 | 4.28E-10 |
| B22 | B35 | CD3+CD8+(T) | β2-MG | 0.006625 | 7.31E-09 |
| B22 | B183 | CD3+CD8+(T) | PTA | 0.030156 | 0 |
| B22 | B184 | CD3+CD8+(T) | INR | 0.007243 | 2.56E-10 |
| B22 | B185 | CD3+CD8+(T) | PT | 0.008826 | 1.31E-14 |
| B22 | B186 | CD3+CD8+(T) | PFC | -0.01456 | 0 |
| B22 | B197 | CD3+CD8+(T) | AG | -0.00995 | 0 |
| B22 | B74 | CD3+CD8+(T) | ApoA1 | -0.01291 | 0 |
| B22 | B81 | CD3+CD8+(T) | TSGF | -0.0075 | 5.69E-11 |
| B22 | B114 | CD3+CD8+(T) | TSH TT3 TT4(TT4) | -0.01285 | 0 |
| B220 | B225 | SBP | hPBG | 0.011394 | 0 |
| B220 | B222 | SBP | NS | 0.305004 | 0 |
| B220 | B221 | SBP | Hei | 0.116148 | 0 |
| B220 | B227 | SBP | OD | 0.014962 | 0 |
| B220 | B228 | SBP | OS | 0.017161 | 0 |
| B221 | B222 | Hei | NS | 0.546121 | 0 |
| B221 | B227 | Hei | OD | -0.00629 | 4.07E-08 |
| B222 | B224 | NS | pmh | 0.010065 | 0 |
| B227 | B228 | OD | OS | 0.12255 | 0 |

| | | | | | |
|---|---|---|---|---|---|
| B23 | B24 | CD3-CD16+CD56+(NK) | αHBDH | 0.008182 | 9.09E-13 |
| B23 | B92 | CD3-CD16+CD56+(NK) | ALDH2-Glu504Glu | 0.010913 | 0 |
| B23 | B93 | CD3-CD16+CD56+(NK) | ALDH2-Glu504Lys | -0.0069 | 1.67E-09 |
| B23 | B94 | CD3-CD16+CD56+(NK) | ALDH2-Lys504Lys | -0.0119 | 0 |
| B23 | B143 | CD3-CD16+CD56+(NK) | β-CTx | -0.0067 | 4.92E-09 |
| B23 | B126 | CD3-CD16+CD56+(NK) | Alb | 0.009537 | 0 |
| B23 | B121 | CD3-CD16+CD56+(NK) | C3 | -0.02015 | 0 |
| B23 | B122 | CD3-CD16+CD56+(NK) | C4 | 0.006095 | 1.03E-07 |
| B23 | B124 | CD3-CD16+CD56+(NK) | ACTH | 0.048086 | 0 |
| B23 | B71 | CD3-CD16+CD56+(NK) | TG | -0.00577 | 4.67E-07 |
| B23 | B55 | CD3-CD16+CD56+(NK) | TK | -0.00847 | 1.45E-13 |
| B23 | B57 | CD3-CD16+CD56+(NK) | RBCAI | 0.012698 | 0 |
| B23 | B46 | CD3-CD16+CD56+(NK) | RBC Count | 0.008581 | 6.79E-14 |
| B23 | B181 | CD3-CD16+CD56+(NK) | APTT | 0.007469 | 7.00E-11 |
| B23 | B190 | CD3-CD16+CD56+(NK) | Cre | 0.005776 | 4.59E-07 |
| B23 | B193 | CD3-CD16+CD56+(NK) | K | 0.027071 | 0 |
| B23 | B129 | CD3-CD16+CD56+(NK) | IBIL | 0.006883 | 1.86E-09 |
| B23 | B150 | CD3-CD16+CD56+(NK) | ALP | 0.007124 | 5.00E-10 |
| B23 | B105 | CD3-CD16+CD56+(NK) | anti-U1RNP | 0.024392 | 0 |
| B23 | B95 | CD3-CD16+CD56+(NK) | anti-cenpb | 0.00678 | 3.23E-09 |
| B23 | B96 | CD3-CD16+CD56+(NK) | dsDNA (IIF) | -0.00674 | 3.95E-09 |
| B23 | B98 | CD3-CD16+CD56+(NK) | anti-Jo-1 | 0.006135 | 8.51E-08 |
| B23 | B100 | CD3-CD16+CD56+(NK) | anti-Ro52 | -0.01648 | 0 |
| B23 | B101 | CD3-CD16+CD56+(NK) | anti-Sc70 | 0.020854 | 0 |
| B23 | B102 | CD3-CD16+CD56+(NK) | anti-Sm | 0.011051 | 0 |
| B23 | B103 | CD3-CD16+CD56+(NK) | anti-SSA/Ro | -0.01587 | 0 |
| B23 | B104 | CD3-CD16+CD56+(NK) | anti-SSB | -0.01724 | 0 |
| B23 | B107 | CD3-CD16+CD56+(NK) | AnuA | -0.00707 | 6.60E-10 |
| B23 | B151 | CD3-CD16+CD56+(NK) | Anti-CCP | 0.020358 | 0 |
| B23 | B108 | CD3-CD16+CD56+(NK) | Anti-Cyclin | 0.007512 | 5.45E-11 |
| B23 | B154 | CD3-CD16+CD56+(NK) | RF | 0.009043 | 2.89E-15 |
| B23 | B198 | CD3-CD16+CD56+(NK) | P | -0.05133 | 0 |
| B23 | B194 | CD3-CD16+CD56+(NK) | Cl | -0.02324 | 0 |
| B23 | B199 | CD3-CD16+CD56+(NK) | Mg | 0.010157 | 0 |
| B23 | B155 | CD3-CD16+CD56+(NK) | IgA | 0.030396 | 0 |
| B23 | B156 | CD3-CD16+CD56+(NK) | IgE | 0.012411 | 0 |
| B23 | B157 | CD3-CD16+CD56+(NK) | IgG | 0.030046 | 0 |
| B23 | B195 | CD3-CD16+CD56+(NK) | Na | 0.01918 | 0 |
| B23 | B168 | CD3-CD16+CD56+(NK) | NMP22 | 0.006306 | 3.69E-08 |
| B23 | B178 | CD3-CD16+CD56+(NK) | Ig-U | 0.008853 | 1.09E-14 |
| B23 | B179 | CD3-CD16+CD56+(NK) | α1-MG | 0.010753 | 0 |
| B23 | B180 | CD3-CD16+CD56+(NK) | Tf | 0.007669 | 2.15E-11 |
| B23 | B182 | CD3-CD16+CD56+(NK) | TT | 0.030419 | 0 |
| B23 | B125 | CD3-CD16+CD56+(NK) | Cor | 0.032065 | 0 |

| | | | | | |
|---|---|---|---|---|---|
| B23 | B130 | CD3-CD16+CD56+(NK) | GLB | 0.005762 | 4.88E-07 |
| B23 | B59 | CD3-CD16+CD56+(NK) | WBRV | 0.010434 | 0 |
| B23 | B60 | CD3-CD16+CD56+(NK) | WBRVI | 0.014656 | 0 |
| B23 | B62 | CD3-CD16+CD56+(NK) | WBHSRVI | 0.00798 | 3.24E-12 |
| B23 | B64 | CD3-CD16+CD56+(NK) | WBRSR(mPas_5) | 0.016708 | 0 |
| B23 | B65 | CD3-CD16+CD56+(NK) | WBRSR(mPas_1) | 0.018271 | 0 |
| B23 | B66 | CD3-CD16+CD56+(NK) | WBRSR(mPas_100) | 0.014156 | 0 |
| B23 | B67 | CD3-CD16+CD56+(NK) | WBRSR(mPas_50) | 0.014456 | 0 |
| B23 | B27 | CD3-CD16+CD56+(NK) | LDH | 0.006451 | 1.78E-08 |
| B23 | B200 | CD3-CD16+CD56+(NK) | OP | 0.023482 | 0 |
| B23 | B220 | CD3-CD16+CD56+(NK) | SBP | 0.009451 | 0 |
| B23 | B218 | CD3-CD16+CD56+(NK) | DBP | 0.007786 | 1.06E-11 |
| B23 | B209 | CD3-CD16+CD56+(NK) | Smoking | -0.01391 | 0 |
| B23 | B90 | CD3-CD16+CD56+(NK) | CYFRA 21-1 | -0.00665 | 6.46E-09 |
| B23 | B139 | CD3-CD16+CD56+(NK) | ADA | 0.020083 | 0 |
| B23 | B35 | CD3-CD16+CD56+(NK) | β2-MG | -0.01932 | 0 |
| B23 | B196 | CD3-CD16+CD56+(NK) | TCO2 | 0.011944 | 0 |
| B23 | B51 | CD3-CD16+CD56+(NK) | Hb | 0.005678 | 7.16E-07 |
| B23 | B183 | CD3-CD16+CD56+(NK) | PTA | -0.01266 | 0 |
| B23 | B185 | CD3-CD16+CD56+(NK) | PT | 0.008638 | 4.62E-14 |
| B23 | B186 | CD3-CD16+CD56+(NK) | PFC | 0.009359 | 4.44E-16 |
| B23 | B69 | CD3-CD16+CD56+(NK) | PV | 0.011373 | 0 |
| B23 | B197 | CD3-CD16+CD56+(NK) | AG | 0.016831 | 0 |
| B23 | B33 | CD3-CD16+CD56+(NK) | Prog | 0.006046 | 1.30E-07 |
| B23 | B73 | CD3-CD16+CD56+(NK) | ApoA | 0.006087 | 1.07E-07 |
| B23 | B74 | CD3-CD16+CD56+(NK) | ApoA1 | 0.016133 | 0 |
| B23 | B132 | CD3-CD16+CD56+(NK) | Dbil | 0.008081 | 1.73E-12 |
| B23 | B133 | CD3-CD16+CD56+(NK) | TBIL | 0.007702 | 1.76E-11 |
| B23 | B91 | CD3-CD16+CD56+(NK) | TP | 0.010834 | 0 |
| B24 | B137 | αHBDH | AFU | 0.006143 | 8.18E-08 |
| B24 | B54 | αHBDH | ESR | 0.062313 | 0 |
| B24 | B160 | αHBDH | PH | -0.013 | 0 |
| B24 | B140 | αHBDH | r-GT | 0.022478 | 0 |
| B24 | B118 | αHBDH | CEA1 | 0.011736 | 0 |
| B24 | B126 | αHBDH | Alb | 0.010663 | 0 |
| B24 | B127 | αHBDH | A-G ratio | -0.0201 | 0 |
| B24 | B161 | αHBDH | LEU | 0.01085 | 0 |
| B24 | B128 | αHBDH | ALT | 0.05181 | 0 |
| B24 | B123 | αHBDH | hs-CRP1 | 0.08152 | 0 |
| B24 | B111 | αHBDH | TSH | 0.017826 | 0 |
| B24 | B124 | αHBDH | ACTH | 0.009755 | 0 |
| B24 | B42 | αHBDH | MON | 0.007275 | 2.14E-10 |
| B24 | B165 | αHBDH | bilirubin | 0.008765 | 1.98E-14 |
| B24 | B70 | αHBDH | LDL-C | 0.02855 | 0 |

| | | | | | |
|---|---|---|---|---|---|
| B24 | B38 | αHBDH | BAND% | 0.015581 | 0 |
| B24 | B72 | αHBDH | HDL-C | 0.02477 | 0 |
| B24 | B189 | αHBDH | eGFB | -0.03468 | 0 |
| B24 | B141 | αHBDH | AST/ALT | 0.008785 | 1.73E-14 |
| B24 | B146 | αHBDH | CysC | 0.012726 | 0 |
| B24 | B43 | αHBDH | MCV | 0.011396 | 0 |
| B24 | B44 | αHBDH | MCH | 0.017836 | 0 |
| B24 | B46 | αHBDH | RBC Count | -0.01009 | 0 |
| B24 | B47 | αHBDH | RDW | 0.037417 | 0 |
| B24 | B181 | αHBDH | APTT | -0.00702 | 8.99E-10 |
| B24 | B25 | αHBDH | CK | 0.133739 | 0 |
| B24 | B26 | αHBDH | CKMB | 0.225009 | 0 |
| B24 | B193 | αHBDH | K | 0.01701 | 0 |
| B24 | B129 | αHBDH | IBIL | 0.010874 | 0 |
| B24 | B150 | αHBDH | ALP | 0.026816 | 0 |
| B24 | B222 | αHBDH | NS | -0.0061 | 1.03E-07 |
| B24 | B98 | αHBDH | anti-Jo-1 | 0.008339 | 3.32E-13 |
| B24 | B151 | αHBDH | Anti-CCP | 0.00668 | 5.46E-09 |
| B24 | B194 | αHBDH | Cl | -0.00746 | 7.40E-11 |
| B24 | B199 | αHBDH | Mg | 0.010895 | 0 |
| B24 | B157 | αHBDH | IgG | 0.007123 | 5.00E-10 |
| B24 | B195 | αHBDH | Na | 0.012793 | 0 |
| B24 | B176 | αHBDH | Ucr | -0.0124 | 0 |
| B24 | B191 | αHBDH | Urea | 0.021329 | 0 |
| B24 | B174 | αHBDH | Urea/Cr | 0.018442 | 0 |
| B24 | B175 | αHBDH | ALB-U | 0.007957 | 3.73E-12 |
| B24 | B177 | αHBDH | ACR | 0.012774 | 0 |
| B24 | B179 | αHBDH | α1-MG | 0.01516 | 0 |
| B24 | B182 | αHBDH | TT | 0.006498 | 1.40E-08 |
| B24 | B125 | αHBDH | Cor | 0.015138 | 0 |
| B24 | B138 | αHBDH | PAB | -0.0118 | 0 |
| B24 | B130 | αHBDH | GLB | 0.03306 | 0 |
| B24 | B59 | αHBDH | WBRV | 0.016313 | 0 |
| B24 | B61 | αHBDH | WBHSRV | 0.014869 | 0 |
| B24 | B64 | αHBDH | WBRSR(mPas_5) | 0.018857 | 0 |
| B24 | B65 | αHBDH | WBRSR(mPas_1) | 0.017375 | 0 |
| B24 | B66 | αHBDH | WBRSR(mPas_100) | 0.020543 | 0 |
| B24 | B67 | αHBDH | WBRSR(mPas_50) | 0.021081 | 0 |
| B24 | B27 | αHBDH | LDH | 0.723643 | 0 |
| B24 | B221 | αHBDH | Hei | -0.03526 | 0 |
| B24 | B87 | αHBDH | NSE | 0.026032 | 0 |
| B24 | B187 | αHBDH | NSE | 0.011473 | 0 |
| B24 | B200 | αHBDH | OP | 0.016147 | 0 |
| B24 | B228 | αHBDH | OS | 0.005862 | 3.10E-07 |

| | | | | | |
|---|---|---|---|---|---|
| B24 | B50 | αHBDH | eo% | 0.008694 | 3.20E-14 |
| B24 | B220 | αHBDH | SBP | 0.048447 | 0 |
| B24 | B219 | αHBDH | OT | 0.008453 | 1.59E-13 |
| B24 | B218 | αHBDH | DBP | 0.031595 | 0 |
| B24 | B215 | αHBDH | BMI | 0.020568 | 0 |
| B24 | B131 | αHBDH | AST | 0.096301 | 0 |
| B24 | B210 | αHBDH | Hip | 0.009378 | 2.22E-16 |
| B24 | B209 | αHBDH | Smoking | -0.01718 | 0 |
| B24 | B90 | αHBDH | CYFRA 21-1 | 0.010968 | 0 |
| B24 | B139 | αHBDH | ADA | 0.056551 | 0 |
| B24 | B183 | αHBDH | PTA | -0.02035 | 0 |
| B24 | B185 | αHBDH | PT | 0.016033 | 0 |
| B24 | B186 | αHBDH | PFC | 0.016409 | 0 |
| B24 | B69 | αHBDH | PV | 0.045373 | 0 |
| B24 | B52 | αHBDH | PLT/BPC | -0.02835 | 0 |
| B24 | B173 | αHBDH | NTT | 0.009923 | 0 |
| B24 | B206 | αHBDH | Waist | 0.015082 | 0 |
| B24 | B197 | αHBDH | AG | 0.009828 | 0 |
| B24 | B204 | αHBDH | Drinking | -0.01652 | 0 |
| B24 | B112 | αHBDH | FT4 | 0.010207 | 0 |
| B24 | B113 | αHBDH | FT3 | 0.007331 | 1.55E-10 |
| B24 | B211 | αHBDH | IOP-R | -0.007 | 1.00E-09 |
| B24 | B73 | αHBDH | ApoA | 0.017158 | 0 |
| B24 | B74 | αHBDH | ApoA1 | 0.064402 | 0 |
| B24 | B75 | αHBDH | ApoA1/B | 0.009231 | 8.88E-16 |
| B24 | B76 | αHBDH | ApoB | 0.021186 | 0 |
| B24 | B132 | αHBDH | Dbil | 0.013025 | 0 |
| B24 | B81 | αHBDH | TSGF | -0.01653 | 0 |
| B24 | B77 | αHBDH | TC | 0.029186 | 0 |
| B24 | B133 | αHBDH | TBIL | 0.012071 | 0 |
| B24 | B91 | αHBDH | TP | 0.034851 | 0 |
| B24 | B114 | αHBDH | TSH TT3 TT4(TT4) | 0.037439 | 0 |
| B24 | B212 | αHBDH | IOP-L | -0.00853 | 9.30E-14 |
| B25 | B54 | CK | ESR | -0.00845 | 1.66E-13 |
| B25 | B160 | CK | PH | 0.006079 | 1.11E-07 |
| B25 | B126 | CK | Alb | 0.008493 | 1.22E-13 |
| B25 | B127 | CK | A-G ratio | 0.005639 | 8.50E-07 |
| B25 | B128 | CK | ALT | 0.02272 | 0 |
| B25 | B134 | CK | CHE | 0.005941 | 2.13E-07 |
| B25 | B70 | CK | LDL-C | 0.009051 | 2.66E-15 |
| B25 | B141 | CK | AST/ALT | 0.007586 | 3.52E-11 |
| B25 | B45 | CK | MCHC | 0.007278 | 2.10E-10 |
| B25 | B46 | CK | RBC Count | 0.009968 | 0 |
| B25 | B190 | CK | Cre | 0.01507 | 0 |

| | | | | | |
|---|---|---|---|---|---|
| B25 | B26 | CK | CKMB | 0.358423 | 0 |
| B25 | B129 | CK | IBIL | 0.006719 | 4.46E-09 |
| B25 | B222 | CK | NS | 0.013682 | 0 |
| B25 | B176 | CK | Ucr | 0.006555 | 1.05E-08 |
| B25 | B174 | CK | Urea/Cr | -0.007 | 9.75E-10 |
| B25 | B192 | CK | UA | 0.014845 | 0 |
| B25 | B138 | CK | PAB | 0.005966 | 1.91E-07 |
| B25 | B60 | CK | WBRVI | 0.009676 | 0 |
| B25 | B62 | CK | WBHSRVI | 0.00799 | 3.04E-12 |
| B25 | B64 | CK | WBRSR(mPas_5) | 0.008592 | 6.31E-14 |
| B25 | B65 | CK | WBRSR(mPas_1) | 0.008944 | 5.77E-15 |
| B25 | B66 | CK | WBRSR(mPas_100) | 0.007285 | 2.01E-10 |
| B25 | B67 | CK | WBRSR(mPas_50) | 0.007587 | 3.51E-11 |
| B25 | B27 | CK | LDH | 0.232108 | 0 |
| B25 | B221 | CK | Hei | 0.016422 | 0 |
| B25 | B87 | CK | NSE | 0.010643 | 0 |
| B25 | B216 | CK | BMI | 0.01487 | 0 |
| B25 | B215 | CK | BMI | 0.007291 | 1.95E-10 |
| B25 | B131 | CK | AST | 0.086066 | 0 |
| B25 | B210 | CK | Hip | 0.008403 | 2.20E-13 |
| B25 | B51 | CK | Hb | 0.011686 | 0 |
| B25 | B206 | CK | Waist | 0.00819 | 8.67E-13 |
| B25 | B204 | CK | Drinking | 0.0079 | 5.30E-12 |
| B25 | B77 | CK | TC | 0.006485 | 1.50E-08 |
| B25 | B133 | CK | TBIL | 0.006056 | 1.24E-07 |
| B25 | B114 | CK | TSH TT3 TT4(TT4) | -0.00648 | 1.56E-08 |
| B26 | B54 | CKMB | ESR | 0.012293 | 0 |
| B26 | B140 | CKMB | r-GT | 0.006976 | 1.12E-09 |
| B26 | B118 | CKMB | CEA1 | 0.00634 | 3.10E-08 |
| B26 | B127 | CKMB | A-G ratio | -0.00898 | 4.44E-15 |
| B26 | B128 | CKMB | ALT | 0.015306 | 0 |
| B26 | B123 | CKMB | hs-CRP1 | 0.048385 | 0 |
| B26 | B189 | CKMB | eGFB | -0.00618 | 6.70E-08 |
| B26 | B55 | CKMB | TK | -0.00768 | 2.03E-11 |
| B26 | B56 | CKMB | IR | 0.009966 | 0 |
| B26 | B46 | CKMB | RBC Count | 0.012761 | 0 |
| B26 | B193 | CKMB | K | 0.010617 | 0 |
| B26 | B130 | CKMB | GLB | 0.011903 | 0 |
| B26 | B59 | CKMB | WBRV | 0.006739 | 4.03E-09 |
| B26 | B60 | CKMB | WBRVI | 0.019938 | 0 |
| B26 | B61 | CKMB | WBHSRV | 0.013575 | 0 |
| B26 | B62 | CKMB | WBHSRVI | 0.023408 | 0 |
| B26 | B64 | CKMB | WBRSR(mPas_5) | 0.025962 | 0 |
| B26 | B65 | CKMB | WBRSR(mPas_1) | 0.023548 | 0 |

| | | | | | |
|---|---|---|---|---|---|
| B26 | B66 | CKMB | WBRSR(mPas_100) | 0.027319 | 0 |
| B26 | B67 | CKMB | WBRSR(mPas_50) | 0.027183 | 0 |
| B26 | B27 | CKMB | LDH | 0.219087 | 0 |
| B26 | B87 | CKMB | NSE | 0.013491 | 0 |
| B26 | B220 | CKMB | SBP | 0.011211 | 0 |
| B26 | B218 | CKMB | DBP | 0.008327 | 3.59E-13 |
| B26 | B131 | CKMB | AST | 0.048743 | 0 |
| B26 | B90 | CKMB | CYFRA 21-1 | 0.011904 | 0 |
| B26 | B139 | CKMB | ADA | 0.01527 | 0 |
| B26 | B51 | CKMB | Hb | 0.01547 | 0 |
| B26 | B69 | CKMB | PV | 0.009481 | 0 |
| B26 | B52 | CKMB | PLT/BPC | 0.007191 | 3.42E-10 |
| B26 | B197 | CKMB | AG | 0.005844 | 3.36E-07 |
| B26 | B113 | CKMB | FT3 | 0.005731 | 5.63E-07 |
| B26 | B201 | CKMB | e-habits | 0.007185 | 3.55E-10 |
| B26 | B135 | CKMB | TBA | 0.007699 | 1.80E-11 |
| B26 | B91 | CKMB | TP | 0.007827 | 8.27E-12 |
| B26 | B114 | CKMB | TSH TT3 TT4(TT4) | 0.006286 | 4.07E-08 |
| B27 | B136 | LDH | 5'NT | 0.00686 | 2.11E-09 |
| B27 | B137 | LDH | AFU | 0.01497 | 0 |
| B27 | B54 | LDH | ESR | 0.045281 | 0 |
| B27 | B160 | LDH | PH | -0.01449 | 0 |
| B27 | B140 | LDH | r-GT | 0.061568 | 0 |
| B27 | B118 | LDH | CEA1 | 0.013925 | 0 |
| B27 | B126 | LDH | Alb | 0.012686 | 0 |
| B27 | B127 | LDH | A-G ratio | -0.02401 | 0 |
| B27 | B161 | LDH | LEU | 0.007051 | 7.48E-10 |
| B27 | B40 | LDH | WBC | 0.015275 | 0 |
| B27 | B128 | LDH | ALT | 0.124116 | 0 |
| B27 | B123 | LDH | hs-CRP1 | 0.078681 | 0 |
| B27 | B29 | LDH | E2 | -0.00567 | 7.41E-07 |
| B27 | B30 | LDH | LH | 0.007625 | 2.79E-11 |
| B27 | B111 | LDH | TSH | 0.019706 | 0 |
| B27 | B31 | LDH | FSH | 0.009738 | 0 |
| B27 | B124 | LDH | ACTH | 0.011395 | 0 |
| B27 | B165 | LDH | bilirubin | 0.020175 | 0 |
| B27 | B134 | LDH | CHE | 0.059152 | 0 |
| B27 | B166 | LDH | Pro | 0.007653 | 2.37E-11 |
| B27 | B70 | LDH | LDL-C | 0.045602 | 0 |
| B27 | B71 | LDH | TG | 0.026049 | 0 |
| B27 | B38 | LDH | BAND% | 0.009878 | 0 |
| B27 | B72 | LDH | HDL-C | 0.010474 | 0 |
| B27 | B189 | LDH | eGFB | -0.06065 | 0 |
| B27 | B141 | LDH | AST/ALT | -0.0078 | 9.68E-12 |

| | | | | | |
|---|---|---|---|---|---|
| B27 | B146 | LDH | CysC | 0.01635 | 0 |
| B27 | B55 | LDH | TK | -0.00572 | 6.03E-07 |
| B27 | B56 | LDH | IR | 0.008184 | 8.97E-13 |
| B27 | B43 | LDH | MCV | 0.012954 | 0 |
| B27 | B44 | LDH | MCH | 0.027212 | 0 |
| B27 | B45 | LDH | MCHC | 0.017473 | 0 |
| B27 | B46 | LDH | RBC Count | 0.013333 | 0 |
| B27 | B47 | LDH | RDW | 0.037852 | 0 |
| B27 | B181 | LDH | APTT | -0.01013 | 0 |
| B27 | B190 | LDH | Cre | 0.027953 | 0 |
| B27 | B193 | LDH | K | 0.022272 | 0 |
| B27 | B129 | LDH | IBIL | 0.033854 | 0 |
| B27 | B150 | LDH | ALP | 0.120096 | 0 |
| B27 | B222 | LDH | NS | 0.028986 | 0 |
| B27 | B99 | LDH | anti-PM-Scl | 0.007358 | 1.33E-10 |
| B27 | B98 | LDH | anti-Jo-1 | 0.006707 | 4.76E-09 |
| B27 | B104 | LDH | anti-SSB | -0.00566 | 7.69E-07 |
| B27 | B108 | LDH | Anti-Cyclin | 0.007766 | 1.20E-11 |
| B27 | B199 | LDH | Mg | 0.022977 | 0 |
| B27 | B157 | LDH | IgG | 0.007337 | 1.50E-10 |
| B27 | B195 | LDH | Na | 0.024814 | 0 |
| B27 | B191 | LDH | Urea | 0.028981 | 0 |
| B27 | B174 | LDH | Urea/Cr | 0.007341 | 1.46E-10 |
| B27 | B192 | LDH | UA | 0.03495 | 0 |
| B27 | B175 | LDH | ALB-U | 0.01006 | 0 |
| B27 | B177 | LDH | ACR | 0.014468 | 0 |
| B27 | B179 | LDH | α1-MG | 0.014703 | 0 |
| B27 | B125 | LDH | Cor | 0.016643 | 0 |
| B27 | B130 | LDH | GLB | 0.039367 | 0 |
| B27 | B59 | LDH | WBRV | 0.017522 | 0 |
| B27 | B60 | LDH | WBRVI | 0.01446 | 0 |
| B27 | B61 | LDH | WBHSRV | 0.02529 | 0 |
| B27 | B62 | LDH | WBHSRVI | 0.01962 | 0 |
| B27 | B64 | LDH | WBRSR(mPas_5) | 0.03544 | 0 |
| B27 | B65 | LDH | WBRSR(mPas_1) | 0.030886 | 0 |
| B27 | B66 | LDH | WBRSR(mPas_100) | 0.037791 | 0 |
| B27 | B67 | LDH | WBRSR(mPas_50) | 0.037787 | 0 |
| B27 | B221 | LDH | Hei | -0.01792 | 0 |
| B27 | B87 | LDH | NSE | 0.034266 | 0 |
| B27 | B187 | LDH | NSE | 0.013552 | 0 |
| B27 | B200 | LDH | OP | 0.028904 | 0 |
| B27 | B50 | LDH | eo% | 0.015778 | 0 |
| B27 | B220 | LDH | SBP | 0.070859 | 0 |
| B27 | B219 | LDH | OT | 0.013561 | 0 |

| | | | | | |
|---|---|---|---|---|---|
| B27 | B218 | LDH | DBP | 0.054241 | 0 |
| B27 | B216 | LDH | BMI | 0.03423 | 0 |
| B27 | B215 | LDH | BMI | 0.059159 | 0 |
| B27 | B131 | LDH | AST | 0.189361 | 0 |
| B27 | B210 | LDH | Hip | 0.035449 | 0 |
| B27 | B90 | LDH | CYFRA 21-1 | 0.010114 | 0 |
| B27 | B139 | LDH | ADA | 0.122438 | 0 |
| B27 | B152 | LDH | ACA-Ig(G-A-M) | 0.008601 | 5.95E-14 |
| B27 | B196 | LDH | TCO2 | -0.0068 | 2.98E-09 |
| B27 | B51 | LDH | Hb | 0.034384 | 0 |
| B27 | B185 | LDH | PT | 0.010929 | 0 |
| B27 | B186 | LDH | PFC | 0.015157 | 0 |
| B27 | B69 | LDH | PV | 0.047329 | 0 |
| B27 | B52 | LDH | PLT/BPC | -0.02608 | 0 |
| B27 | B173 | LDH | NTT | 0.007517 | 5.29E-11 |
| B27 | B207 | LDH | WHR | 0.006033 | 1.39E-07 |
| B27 | B206 | LDH | Waist | 0.052743 | 0 |
| B27 | B79 | LDH | hb V -dna | 0.009578 | 0 |
| B27 | B78 | LDH | HBcAb | 0.008002 | 2.82E-12 |
| B27 | B197 | LDH | AG | 0.019048 | 0 |
| B27 | B112 | LDH | FT4 | 0.010972 | 0 |
| B27 | B113 | LDH | FT3 | 0.009722 | 0 |
| B27 | B73 | LDH | ApoA | 0.014827 | 0 |
| B27 | B74 | LDH | ApoA1 | 0.037812 | 0 |
| B27 | B76 | LDH | ApoB | 0.046704 | 0 |
| B27 | B132 | LDH | Dbil | 0.022828 | 0 |
| B27 | B81 | LDH | TSGF | -0.0134 | 0 |
| B27 | B77 | LDH | TC | 0.046114 | 0 |
| B27 | B133 | LDH | TBIL | 0.032974 | 0 |
| B27 | B135 | LDH | TBA | 0.028086 | 0 |
| B27 | B91 | LDH | TP | 0.041433 | 0 |
| B27 | B114 | LDH | TSH TT3 TT4(TT4) | 0.031 | 0 |
| B27 | B115 | LDH | TSH TT3 TT4(TT3) | 0.009909 | 0 |
| B27 | B144 | LDH | Total-PINP | 0.008257 | 5.63E-13 |
| B28 | B29 | PRL | E2 | 0.203584 | 0 |
| B28 | B30 | PRL | LH | -0.12281 | 0 |
| B28 | B111 | PRL | TSH | 0.006666 | 5.89E-09 |
| B28 | B31 | PRL | FSH | -0.18198 | 0 |
| B28 | B32 | PRL | Testo | -0.03593 | 0 |
| B28 | B150 | PRL | ALP | -0.00641 | 2.23E-08 |
| B28 | B105 | PRL | anti-U1RNP | 0.012887 | 0 |
| B28 | B100 | PRL | anti-Ro52 | 0.005779 | 4.53E-07 |
| B28 | B102 | PRL | anti-Sm | 0.009847 | 0 |
| B28 | B103 | PRL | anti-SSA/Ro | 0.018653 | 0 |

| | | | | | |
|---|---|---|---|---|---|
| B28 | B195 | PRL | Na | -0.00669 | 5.29E-09 |
| B28 | B125 | PRL | Cor | 0.00562 | 9.25E-07 |
| B28 | B200 | PRL | OP | -0.00668 | 5.48E-09 |
| B28 | B33 | PRL | Prog | 0.114036 | 0 |
| B29 | B126 | E2 | Alb | -0.0078 | 9.67E-12 |
| B29 | B40 | E2 | WBC | 0.007317 | 1.68E-10 |
| B29 | B30 | E2 | LH | -0.15742 | 0 |
| B29 | B31 | E2 | FSH | -0.41511 | 0 |
| B29 | B134 | E2 | CHE | -0.0077 | 1.79E-11 |
| B29 | B70 | E2 | LDL-C | -0.00759 | 3.41E-11 |
| B29 | B32 | E2 | Testo | -0.07899 | 0 |
| B29 | B141 | E2 | AST/ALT | 0.00586 | 3.12E-07 |
| B29 | B150 | E2 | ALP | -0.01132 | 0 |
| B29 | B101 | E2 | anti-Sc70 | 0.010684 | 0 |
| B29 | B109 | E2 | AMA-M2 | 0.008358 | 2.94E-13 |
| B29 | B198 | E2 | P | -0.01532 | 0 |
| B29 | B199 | E2 | Mg | -0.00638 | 2.51E-08 |
| B29 | B195 | E2 | Na | -0.01497 | 0 |
| B29 | B191 | E2 | Urea | -0.0079 | 5.21E-12 |
| B29 | B192 | E2 | UA | -0.00573 | 5.69E-07 |
| B29 | B200 | E2 | OP | -0.01535 | 0 |
| B29 | B206 | E2 | Waist | -0.00562 | 9.36E-07 |
| B29 | B33 | E2 | Prog | 0.250751 | 0 |
| B29 | B76 | E2 | ApoB | -0.00901 | 3.77E-15 |
| B29 | B77 | E2 | TC | -0.00615 | 7.89E-08 |
| B29 | B91 | E2 | TP | -0.00605 | 1.27E-07 |
| B3 | B137 | Hcy | AFU | 0.016505 | 0 |
| B3 | B24 | Hcy | αHBDH | 0.035038 | 0 |
| B3 | B21 | Hcy | CD3+CD4-CD8- | -0.00777 | 1.16E-11 |
| B3 | B54 | Hcy | ESR | -0.01986 | 0 |
| B3 | B160 | Hcy | PH | -0.02279 | 0 |
| B3 | B140 | Hcy | r-GT | 0.04459 | 0 |
| B3 | B143 | Hcy | β-CTx | 0.00643 | 1.98E-08 |
| B3 | B118 | Hcy | CEA1 | 0.007407 | 1.00E-10 |
| B3 | B126 | Hcy | Alb | 0.027731 | 0 |
| B3 | B127 | Hcy | A-G ratio | 0.02291 | 0 |
| B3 | B161 | Hcy | LEU | -0.01247 | 0 |
| B3 | B40 | Hcy | WBC | 0.035031 | 0 |
| B3 | B128 | Hcy | ALT | 0.031564 | 0 |
| B3 | B123 | Hcy | hs-CRP1 | 0.031213 | 0 |
| B3 | B124 | Hcy | ACTH | 0.024666 | 0 |
| B3 | B41 | Hcy | MON％ | 0.005886 | 2.77E-07 |
| B3 | B134 | Hcy | CHE | 0.01948 | 0 |
| B3 | B166 | Hcy | Pro | 0.028372 | 0 |

| | | | | | |
|---|---|---|---|---|---|
| B3 | B70 | Hcy | LDL-C | 0.022399 | 0 |
| B3 | B188 | Hcy | Ca | 0.006765 | 3.50E-09 |
| B3 | B71 | Hcy | TG | 0.028893 | 0 |
| B3 | B72 | Hcy | HDL-C | -0.05307 | 0 |
| B3 | B32 | Hcy | Testo | 0.005746 | 5.26E-07 |
| B3 | B189 | Hcy | eGFB | -0.07711 | 0 |
| B3 | B141 | Hcy | AST/ALT | -0.02717 | 0 |
| B3 | B145 | Hcy | BGP | 0.005653 | 8.00E-07 |
| B3 | B146 | Hcy | CysC | 0.025888 | 0 |
| B3 | B55 | Hcy | TK | -0.00569 | 6.92E-07 |
| B3 | B56 | Hcy | IR | 0.029144 | 0 |
| B3 | B57 | Hcy | RBCAI | 0.011902 | 0 |
| B3 | B43 | Hcy | MCV | 0.010792 | 0 |
| B3 | B44 | Hcy | MCH | 0.035553 | 0 |
| B3 | B45 | Hcy | MCHC | 0.049852 | 0 |
| B3 | B46 | Hcy | RBC Count | 0.076112 | 0 |
| B3 | B47 | Hcy | RDW | 0.018916 | 0 |
| B3 | B58 | Hcy | HCT | 0.006348 | 2.99E-08 |
| B3 | B181 | Hcy | APTT | -0.00676 | 3.68E-09 |
| B3 | B190 | Hcy | Cre | 0.120071 | 0 |
| B3 | B25 | Hcy | CK | 0.015207 | 0 |
| B3 | B26 | Hcy | CKMB | 0.012352 | 0 |
| B3 | B193 | Hcy | K | -0.00964 | 0 |
| B3 | B129 | Hcy | IBIL | 0.022124 | 0 |
| B3 | B150 | Hcy | ALP | 0.041295 | 0 |
| B3 | B84 | Hcy | C-PSA | 0.019857 | 0 |
| B3 | B222 | Hcy | NS | 0.090289 | 0 |
| B3 | B148 | Hcy | Anti_TPO | -0.00845 | 1.59E-13 |
| B3 | B149 | Hcy | TGAb | -0.01164 | 0 |
| B3 | B198 | Hcy | P | -0.02322 | 0 |
| B3 | B85 | Hcy | SCC | 0.011352 | 0 |
| B3 | B194 | Hcy | Cl | -0.00859 | 6.42E-14 |
| B3 | B199 | Hcy | Mg | 0.010092 | 0 |
| B3 | B157 | Hcy | IgG | -0.01291 | 0 |
| B3 | B158 | Hcy | IgM | -0.00687 | 2.02E-09 |
| B3 | B195 | Hcy | Na | 0.02506 | 0 |
| B3 | B176 | Hcy | Ucr | 0.060897 | 0 |
| B3 | B191 | Hcy | Urea | 0.037869 | 0 |
| B3 | B174 | Hcy | Urea/Cr | -0.05401 | 0 |
| B3 | B192 | Hcy | UA | 0.099978 | 0 |
| B3 | B175 | Hcy | ALB-U | 0.013722 | 0 |
| B3 | B177 | Hcy | ACR | 0.006798 | 2.94E-09 |
| B3 | B179 | Hcy | α1-MG | 0.01148 | 0 |
| B3 | B182 | Hcy | TT | 0.018626 | 0 |

| | | | | | |
|---|---|---|---|---|---|
| B3 | B125 | Hcy | Cor | 0.022 | 0 |
| B3 | B138 | Hcy | PAB | 0.03758 | 0 |
| B3 | B130 | Hcy | GLB | -0.01379 | 0 |
| B3 | B59 | Hcy | WBRV | 0.017776 | 0 |
| B3 | B60 | Hcy | WBRVI | 0.050513 | 0 |
| B3 | B61 | Hcy | WBHSRV | 0.028567 | 0 |
| B3 | B62 | Hcy | WBHSRVI | 0.052256 | 0 |
| B3 | B64 | Hcy | WBRSR(mPas_5) | 0.051486 | 0 |
| B3 | B65 | Hcy | WBRSR(mPas_1) | 0.048881 | 0 |
| B3 | B66 | Hcy | WBRSR(mPas_100) | 0.050854 | 0 |
| B3 | B67 | Hcy | WBRSR(mPas_50) | 0.051787 | 0 |
| B3 | B86 | Hcy | TBhCG | -0.00803 | 2.38E-12 |
| B3 | B27 | Hcy | LDH | 0.045873 | 0 |
| B3 | B221 | Hcy | Hei | 0.085687 | 0 |
| B3 | B200 | Hcy | OP | 0.02482 | 0 |
| B3 | B50 | Hcy | eo% | 0.01758 | 0 |
| B3 | B220 | Hcy | SBP | 0.048104 | 0 |
| B3 | B219 | Hcy | OT | -0.00746 | 7.57E-11 |
| B3 | B218 | Hcy | DBP | 0.049028 | 0 |
| B3 | B88 | Hcy | CA 242 | -0.01 | 0 |
| B3 | B216 | Hcy | BMI | 0.079837 | 0 |
| B3 | B215 | Hcy | BMI | 0.045414 | 0 |
| B3 | B131 | Hcy | AST | 0.024822 | 0 |
| B3 | B210 | Hcy | Hip | 0.050658 | 0 |
| B3 | B10 | Hcy | PG I | 0.021548 | 0 |
| B3 | B12 | Hcy | PG II | 0.017698 | 0 |
| B3 | B14 | Hcy | G-17 | 0.011367 | 0 |
| B3 | B209 | Hcy | Smoking | 0.076504 | 0 |
| B3 | B90 | Hcy | CYFRA 21-1 | 0.036777 | 0 |
| B3 | B139 | Hcy | ADA | 0.00689 | 1.80E-09 |
| B3 | B34 | Hcy | TK | -0.00835 | 3.15E-13 |
| B3 | B51 | Hcy | Hb | 0.103389 | 0 |
| B3 | B183 | Hcy | PTA | -0.00695 | 1.29E-09 |
| B3 | B52 | Hcy | PLT/BPC | -0.01679 | 0 |
| B3 | B173 | Hcy | NTT | -0.00618 | 6.94E-08 |
| B3 | B207 | Hcy | WHR | 0.008245 | 6.08E-13 |
| B3 | B206 | Hcy | Waist | 0.073933 | 0 |
| B3 | B205 | Hcy | KDA | -0.00601 | 1.55E-07 |
| B3 | B197 | Hcy | AG | 0.021169 | 0 |
| B3 | B204 | Hcy | Drinking | 0.059746 | 0 |
| B3 | B80 | Hcy | anti-HP IgG | 0.007352 | 1.37E-10 |
| B3 | B112 | Hcy | FT4 | 0.010549 | 0 |
| B3 | B113 | Hcy | FT3 | 0.016301 | 0 |
| B3 | B73 | Hcy | ApoA | -0.00725 | 2.50E-10 |

| | | | | | |
|---|---|---|---|---|---|
| B3 | B74 | Hcy | ApoA1 | -0.03897 | 0 |
| B3 | B75 | Hcy | ApoA1/B | -0.02252 | 0 |
| B3 | B76 | Hcy | ApoB | 0.025345 | 0 |
| B3 | B132 | Hcy | Dbil | 0.023961 | 0 |
| B3 | B81 | Hcy | TSGF | 0.006291 | 3.97E-08 |
| B3 | B133 | Hcy | TBIL | 0.024054 | 0 |
| B3 | B135 | Hcy | TBA | 0.010138 | 0 |
| B3 | B115 | Hcy | TSH TT3 TT4(TT3) | 0.00749 | 6.19E-11 |
| B3 | B144 | Hcy | Total-PINP | 0.006658 | 6.15E-09 |
| B30 | B40 | LH | WBC | -0.00784 | 7.75E-12 |
| B30 | B31 | LH | FSH | 0.80422 | 0 |
| B30 | B134 | LH | CHE | 0.011506 | 0 |
| B30 | B70 | LH | LDL-C | 0.007591 | 3.42E-11 |
| B30 | B72 | LH | HDL-C | 0.008656 | 4.13E-14 |
| B30 | B32 | LH | Testo | -0.12477 | 0 |
| B30 | B189 | LH | eGFB | -0.01049 | 0 |
| B30 | B145 | LH | BGP | 0.005632 | 8.80E-07 |
| B30 | B150 | LH | ALP | 0.012699 | 0 |
| B30 | B100 | LH | anti-Ro52 | -0.00575 | 5.13E-07 |
| B30 | B101 | LH | anti-Sc70 | -0.0078 | 9.90E-12 |
| B30 | B102 | LH | anti-Sm | -0.00836 | 2.90E-13 |
| B30 | B108 | LH | Anti-Cyclin | 0.015656 | 0 |
| B30 | B198 | LH | P | 0.011728 | 0 |
| B30 | B199 | LH | Mg | 0.008257 | 5.65E-13 |
| B30 | B156 | LH | IgE | -0.00666 | 5.99E-09 |
| B30 | B195 | LH | Na | 0.017205 | 0 |
| B30 | B200 | LH | OP | 0.014623 | 0 |
| B30 | B139 | LH | ADA | 0.007147 | 4.39E-10 |
| B30 | B112 | LH | FT4 | 0.005931 | 2.24E-07 |
| B30 | B33 | LH | Prog | -0.32409 | 0 |
| B30 | B74 | LH | ApoA1 | 0.012199 | 0 |
| B30 | B77 | LH | TC | 0.010653 | 0 |
| B30 | B114 | LH | TSH TT3 TT4(TT4) | 0.006753 | 3.74E-09 |
| B31 | B40 | FSH | WBC | -0.01063 | 0 |
| B31 | B134 | FSH | CHE | 0.013394 | 0 |
| B31 | B70 | FSH | LDL-C | 0.0097 | 0 |
| B31 | B72 | FSH | HDL-C | 0.007178 | 3.68E-10 |
| B31 | B32 | FSH | Testo | -0.10966 | 0 |
| B31 | B189 | FSH | eGFB | -0.01351 | 0 |
| B31 | B145 | FSH | BGP | 0.006421 | 2.07E-08 |
| B31 | B150 | FSH | ALP | 0.017181 | 0 |
| B31 | B101 | FSH | anti-Sc70 | -0.00789 | 5.52E-12 |
| B31 | B103 | FSH | anti-SSA/Ro | -0.00626 | 4.56E-08 |
| B31 | B108 | FSH | Anti-Cyclin | 0.00692 | 1.53E-09 |

| | | | | | |
|---|---|---|---|---|---|
| B31 | B198 | FSH | P | 0.019797 | 0 |
| B31 | B194 | FSH | Cl | 0.007901 | 5.27E-12 |
| B31 | B199 | FSH | Mg | 0.009282 | 4.44E-16 |
| B31 | B156 | FSH | IgE | -0.0069 | 1.72E-09 |
| B31 | B195 | FSH | Na | 0.023315 | 0 |
| B31 | B191 | FSH | Urea | 0.007794 | 1.02E-11 |
| B31 | B200 | FSH | OP | 0.021141 | 0 |
| B31 | B219 | FSH | OT | 0.007334 | 1.53E-10 |
| B31 | B139 | FSH | ADA | 0.008362 | 2.88E-13 |
| B31 | B112 | FSH | FT4 | 0.006307 | 3.67E-08 |
| B31 | B33 | FSH | Prog | -0.29017 | 0 |
| B31 | B74 | FSH | ApoA1 | 0.007617 | 2.92E-11 |
| B31 | B76 | FSH | ApoB | 0.007694 | 1.85E-11 |
| B31 | B77 | FSH | TC | 0.012598 | 0 |
| B31 | B114 | FSH | TSH TT3 TT4(TT4) | 0.007536 | 4.72E-11 |
| B32 | B92 | Testo | ALDH2-Glu504Glu | -0.00764 | 2.59E-11 |
| B32 | B93 | Testo | ALDH2-Glu504Lys | 0.008051 | 2.08E-12 |
| B32 | B134 | Testo | CHE | 0.007504 | 5.70E-11 |
| B32 | B45 | Testo | MCHC | 0.005918 | 2.39E-07 |
| B32 | B46 | Testo | RBC Count | 0.011593 | 0 |
| B32 | B190 | Testo | Cre | 0.011358 | 0 |
| B32 | B222 | Testo | NS | 0.012226 | 0 |
| B32 | B104 | Testo | anti-SSB | 0.008184 | 8.98E-13 |
| B32 | B198 | Testo | P | -0.00828 | 4.91E-13 |
| B32 | B176 | Testo | Ucr | 0.00615 | 7.92E-08 |
| B32 | B192 | Testo | UA | 0.012265 | 0 |
| B32 | B138 | Testo | PAB | 0.015863 | 0 |
| B32 | B60 | Testo | WBRVI | 0.008318 | 3.81E-13 |
| B32 | B62 | Testo | WBHSRVI | 0.006603 | 8.17E-09 |
| B32 | B64 | Testo | WBRSR(mPas_5) | 0.009697 | 0 |
| B32 | B65 | Testo | WBRSR(mPas_1) | 0.009836 | 0 |
| B32 | B66 | Testo | WBRSR(mPas_100) | 0.00873 | 2.51E-14 |
| B32 | B67 | Testo | WBRSR(mPas_50) | 0.008997 | 4.00E-15 |
| B32 | B221 | Testo | Hei | 0.01213 | 0 |
| B32 | B200 | Testo | OP | 0.006087 | 1.07E-07 |
| B32 | B216 | Testo | BMI | 0.01088 | 0 |
| B32 | B215 | Testo | BMI | 0.00575 | 5.17E-07 |
| B32 | B209 | Testo | Smoking | 0.007902 | 5.23E-12 |
| B32 | B51 | Testo | Hb | 0.013546 | 0 |
| B32 | B186 | Testo | PFC | 0.006661 | 6.04E-09 |
| B32 | B206 | Testo | Waist | 0.009594 | 0 |
| B32 | B197 | Testo | AG | 0.00652 | 1.26E-08 |
| B32 | B204 | Testo | Drinking | 0.006203 | 6.10E-08 |
| B32 | B33 | Testo | Prog | -0.07545 | 0 |

| | | | | | |
|---|---|---|---|---|---|
| B32 | B76 | Testo | ApoB | 0.006806 | 2.82E-09 |
| B32 | B135 | Testo | TBA | 0.005869 | 3.00E-07 |
| B33 | B40 | Prog | WBC | 0.006056 | 1.24E-07 |
| B33 | B134 | Prog | CHE | -0.00677 | 3.35E-09 |
| B33 | B70 | Prog | LDL-C | -0.00801 | 2.63E-12 |
| B33 | B150 | Prog | ALP | -0.00954 | 0 |
| B33 | B101 | Prog | anti-Sc70 | 0.044794 | 0 |
| B33 | B102 | Prog | anti-Sm | 0.00676 | 3.60E-09 |
| B33 | B103 | Prog | anti-SSA/Ro | 0.012889 | 0 |
| B33 | B104 | Prog | anti-SSB | 0.006101 | 1.00E-07 |
| B33 | B107 | Prog | AnuA | 0.006444 | 1.84E-08 |
| B33 | B109 | Prog | AMA-M2 | 0.005933 | 2.22E-07 |
| B33 | B110 | Prog | AHA | 0.007451 | 7.78E-11 |
| B33 | B156 | Prog | IgE | 0.006291 | 3.97E-08 |
| B33 | B195 | Prog | Na | -0.01514 | 0 |
| B33 | B192 | Prog | UA | -0.00665 | 6.33E-09 |
| B33 | B86 | Prog | TBhCG | 0.014095 | 0 |
| B33 | B200 | Prog | OP | -0.01385 | 0 |
| B33 | B215 | Prog | BMI | -0.00564 | 8.46E-07 |
| B33 | B196 | Prog | TCO2 | -0.00826 | 5.38E-13 |
| B33 | B76 | Prog | ApoB | -0.00648 | 1.51E-08 |
| B33 | B77 | Prog | TC | -0.00718 | 3.61E-10 |
| B34 | B137 | TK | AFU | -0.01281 | 0 |
| B34 | B54 | TK | ESR | 0.007379 | 1.18E-10 |
| B34 | B93 | TK | ALDH2-Glu504Lys | 0.005921 | 2.35E-07 |
| B34 | B140 | TK | r-GT | -0.01214 | 0 |
| B34 | B143 | TK | β-CTx | -0.00597 | 1.87E-07 |
| B34 | B128 | TK | ALT | -0.00745 | 7.59E-11 |
| B34 | B123 | TK | hs-CRP1 | 0.00811 | 1.44E-12 |
| B34 | B134 | TK | CHE | -0.02192 | 0 |
| B34 | B70 | TK | LDL-C | -0.00815 | 1.12E-12 |
| B34 | B71 | TK | TG | -0.0197 | 0 |
| B34 | B72 | TK | HDL-C | 0.007167 | 3.92E-10 |
| B34 | B189 | TK | eGFB | 0.005643 | 8.38E-07 |
| B34 | B141 | TK | AST/ALT | 0.007717 | 1.61E-11 |
| B34 | B56 | TK | IR | -0.00572 | 5.86E-07 |
| B34 | B44 | TK | MCH | -0.00568 | 6.99E-07 |
| B34 | B46 | TK | RBC Count | -0.00625 | 4.92E-08 |
| B34 | B181 | TK | APTT | 0.013639 | 0 |
| B34 | B190 | TK | Cre | -0.00585 | 3.28E-07 |
| B34 | B222 | TK | NS | -0.01479 | 0 |
| B34 | B156 | TK | IgE | 0.006798 | 2.94E-09 |
| B34 | B157 | TK | IgG | 0.011607 | 0 |
| B34 | B158 | TK | IgM | 0.01064 | 0 |

| | | | | | |
|---|---|---|---|---|---|
| B34 | B195 | TK | Na | -0.00766 | 2.24E-11 |
| B34 | B168 | TK | NMP22 | 0.012893 | 0 |
| B34 | B176 | TK | Ucr | -0.00871 | 2.95E-14 |
| B34 | B192 | TK | UA | -0.01226 | 0 |
| B34 | B138 | TK | PAB | -0.02889 | 0 |
| B34 | B60 | TK | WBRVI | -0.00843 | 1.82E-13 |
| B34 | B62 | TK | WBHSRVI | -0.0098 | 0 |
| B34 | B64 | TK | WBRSR(mPas_5) | -0.0069 | 1.75E-09 |
| B34 | B65 | TK | WBRSR(mPas_1) | -0.00647 | 1.58E-08 |
| B34 | B66 | TK | WBRSR(mPas_100) | -0.00784 | 7.44E-12 |
| B34 | B67 | TK | WBRSR(mPas_50) | -0.00764 | 2.59E-11 |
| B34 | B221 | TK | Hei | -0.00591 | 2.50E-07 |
| B34 | B200 | TK | OP | -0.00717 | 3.76E-10 |
| B34 | B220 | TK | SBP | -0.00644 | 1.85E-08 |
| B34 | B218 | TK | DBP | -0.00909 | 2.22E-15 |
| B34 | B216 | TK | BMI | -0.01375 | 0 |
| B34 | B215 | TK | BMI | -0.01419 | 0 |
| B34 | B210 | TK | Hip | -0.00948 | 0 |
| B34 | B209 | TK | Smoking | -0.00872 | 2.66E-14 |
| B34 | B90 | TK | CYFRA 21-1 | -0.00682 | 2.68E-09 |
| B34 | B139 | TK | ADA | 0.01295 | 0 |
| B34 | B196 | TK | TCO2 | -0.01115 | 0 |
| B34 | B51 | TK | Hb | -0.01072 | 0 |
| B34 | B185 | TK | PT | 0.00769 | 1.90E-11 |
| B34 | B186 | TK | PFC | 0.009788 | 0 |
| B34 | B69 | TK | PV | 0.00563 | 8.86E-07 |
| B34 | B206 | TK | Waist | -0.01492 | 0 |
| B34 | B204 | TK | Drinking | -0.01436 | 0 |
| B34 | B112 | TK | FT4 | 0.005926 | 2.29E-07 |
| B34 | B75 | TK | ApoA1/B | 0.007597 | 3.30E-11 |
| B34 | B76 | TK | ApoB | -0.0164 | 0 |
| B34 | B81 | TK | TSGF | -0.01771 | 0 |
| B34 | B77 | TK | TC | -0.01251 | 0 |
| B34 | B144 | TK | Total-PINP | -0.00671 | 4.72E-09 |
| B35 | B92 | β2-MG | ALDH2-Glu504Glu | 0.011064 | 0 |
| B35 | B93 | β2-MG | ALDH2-Glu504Lys | -0.01046 | 0 |
| B35 | B124 | β2-MG | ACTH | -0.01967 | 0 |
| B35 | B199 | β2-MG | Mg | -0.00624 | 4.98E-08 |
| B35 | B159 | β2-MG | β2-MG | 0.012338 | 0 |
| B35 | B125 | β2-MG | Cor | 0.015492 | 0 |
| B35 | B138 | β2-MG | PAB | 0.00683 | 2.48E-09 |
| B35 | B139 | β2-MG | ADA | -0.00561 | 9.58E-07 |
| B36 | B44 | V-Lym% | MCH | -0.01849 | 0 |
| B36 | B45 | V-Lym% | MCHC | -0.03731 | 0 |

| | | | | | |
|---|---|---|---|---|---|
| B36 | B47 | V-Lym% | RDW | -0.00782 | 8.84E-12 |
| B37 | B40 | BAND | WBC | 0.182714 | 0 |
| B37 | B42 | BAND | MON | 0.094158 | 0 |
| B37 | B38 | BAND | BAND% | 0.460037 | 0 |
| B37 | B47 | BAND | RDW | 0.009347 | 4.44E-16 |
| B37 | B39 | BAND | Other cells | 0.042291 | 0 |
| B38 | B40 | BAND% | WBC | 0.045292 | 0 |
| B38 | B42 | BAND% | MON | 0.049296 | 0 |
| B38 | B46 | BAND% | RBC Count | -0.00677 | 3.42E-09 |
| B38 | B47 | BAND% | RDW | 0.008552 | 8.26E-14 |
| B38 | B83 | BAND% | AFP | -0.00586 | 3.12E-07 |
| B38 | B39 | BAND% | Other cells | 0.015687 | 0 |
| B38 | B86 | BAND% | TBhCG | -0.00844 | 1.68E-13 |
| B39 | B40 | Other cells | WBC | 0.019282 | 0 |
| B39 | B42 | Other cells | MON | 0.014243 | 0 |
| B39 | B70 | Other cells | LDL-C | -0.01469 | 0 |
| B39 | B72 | Other cells | HDL-C | -0.00697 | 1.18E-09 |
| B39 | B43 | Other cells | MCV | -0.05352 | 0 |
| B39 | B44 | Other cells | MCH | -0.10105 | 0 |
| B39 | B45 | Other cells | MCHC | -0.05843 | 0 |
| B39 | B46 | Other cells | RBC Count | 0.05542 | 0 |
| B39 | B47 | Other cells | RDW | -0.04076 | 0 |
| B39 | B129 | Other cells | IBIL | 0.010478 | 0 |
| B39 | B105 | Other cells | anti-U1RNP | 0.006288 | 4.03E-08 |
| B39 | B51 | Other cells | Hb | -0.04284 | 0 |
| B39 | B52 | Other cells | PLT/BPC | 0.0186 | 0 |
| B39 | B78 | Other cells | HBcAb | -0.04166 | 0 |
| B39 | B203 | Other cells | DH | 0.00628 | 4.19E-08 |
| B39 | B201 | Other cells | e-habits | 0.007322 | 1.64E-10 |
| B39 | B132 | Other cells | Dbil | 0.012957 | 0 |
| B39 | B77 | Other cells | TC | -0.01431 | 0 |
| B39 | B133 | Other cells | TBIL | 0.011857 | 0 |
| B4 | B20 | WB-Ca | CD3+CD4+CD8+ | 0.005671 | 7.39E-07 |
| B4 | B143 | WB-Ca | β-CTx | 0.007751 | 1.31E-11 |
| B4 | B124 | WB-Ca | ACTH | -0.00675 | 3.72E-09 |
| B4 | B32 | WB-Ca | Testo | -0.01267 | 0 |
| B4 | B145 | WB-Ca | BGP | 0.010033 | 0 |
| B4 | B44 | WB-Ca | MCH | -0.00607 | 1.18E-07 |
| B4 | B45 | WB-Ca | MCHC | -0.01 | 0 |
| B4 | B46 | WB-Ca | RBC Count | -0.00841 | 2.13E-13 |
| B4 | B190 | WB-Ca | Cre | -0.00783 | 8.03E-12 |
| B4 | B129 | WB-Ca | IBIL | -0.00865 | 4.44E-14 |
| B4 | B222 | WB-Ca | NS | -0.00702 | 8.92E-10 |
| B4 | B194 | WB-Ca | Cl | -0.00719 | 3.49E-10 |

| | | | | | |
|---|---|---|---|---|---|
| B4 | B192 | WB-Ca | UA | -0.0062 | 6.11E-08 |
| B4 | B125 | WB-Ca | Cor | -0.00626 | 4.72E-08 |
| B4 | B5 | WB-Ca | WB-Mg | -0.05622 | 0 |
| B4 | B6 | WB-Ca | WB-Fe | -0.15851 | 0 |
| B4 | B7 | WB-Ca | WB-Cop | 0.345601 | 0 |
| B4 | B8 | WB-Ca | WB-Zinc | 0.087553 | 0 |
| B4 | B64 | WB-Ca | WBRSR(mPas_5) | -0.00581 | 3.97E-07 |
| B4 | B66 | WB-Ca | WBRSR(mPas_100) | -0.00612 | 9.34E-08 |
| B4 | B67 | WB-Ca | WBRSR(mPas_50) | -0.00614 | 8.17E-08 |
| B4 | B221 | WB-Ca | Hei | -0.00572 | 5.82E-07 |
| B4 | B51 | WB-Ca | Hb | -0.01297 | 0 |
| B4 | B183 | WB-Ca | PTA | -0.00833 | 3.50E-13 |
| B4 | B206 | WB-Ca | Waist | -0.00564 | 8.64E-07 |
| B4 | B211 | WB-Ca | IOP-R | -0.0059 | 2.64E-07 |
| B4 | B133 | WB-Ca | TBIL | -0.0075 | 5.75E-11 |
| B4 | B144 | WB-Ca | Total-PINP | 0.007189 | 3.46E-10 |
| B40 | B137 | WBC | AFU | 0.011248 | 0 |
| B40 | B54 | WBC | ESR | 0.012157 | 0 |
| B40 | B160 | WBC | PH | -0.08048 | 0 |
| B40 | B140 | WBC | r-GT | 0.116599 | 0 |
| B40 | B118 | WBC | CEA1 | 0.014182 | 0 |
| B40 | B126 | WBC | Alb | 0.05108 | 0 |
| B40 | B127 | WBC | A-G ratio | -0.05806 | 0 |
| B40 | B161 | WBC | LEU | -0.00923 | 8.88E-16 |
| B40 | B128 | WBC | ALT | 0.107731 | 0 |
| B40 | B123 | WBC | hs-CRP1 | 0.070519 | 0 |
| B40 | B111 | WBC | TSH | -0.00692 | 1.53E-09 |
| B40 | B124 | WBC | ACTH | 0.008739 | 2.35E-14 |
| B40 | B41 | WBC | MON％ | -0.02742 | 0 |
| B40 | B42 | WBC | MON | 0.02892 | 0 |
| B40 | B165 | WBC | bilirubin | 0.006203 | 6.10E-08 |
| B40 | B134 | WBC | CHE | 0.038935 | 0 |
| B40 | B166 | WBC | Pro | 0.069601 | 0 |
| B40 | B70 | WBC | LDL-C | 0.089664 | 0 |
| B40 | B188 | WBC | Ca | 0.020938 | 0 |
| B40 | B71 | WBC | TG | 0.174823 | 0 |
| B40 | B72 | WBC | HDL-C | -0.17055 | 0 |
| B40 | B189 | WBC | eGFB | 0.022768 | 0 |
| B40 | B141 | WBC | AST/ALT | -0.09593 | 0 |
| B40 | B146 | WBC | CysC | 0.013946 | 0 |
| B40 | B56 | WBC | IR | 0.026568 | 0 |
| B40 | B57 | WBC | RBCAI | 0.007415 | 9.58E-11 |
| B40 | B43 | WBC | MCV | -0.02093 | 0 |
| B40 | B44 | WBC | MCH | -0.01164 | 0 |

| | | | | | |
|---|---|---|---|---|---|
| B40 | B45 | WBC | MCHC | 0.061404 | 0 |
| B40 | B46 | WBC | RBC Count | 0.241882 | 0 |
| B40 | B47 | WBC | RDW | -0.03174 | 0 |
| B40 | B58 | WBC | HCT | 0.008889 | 8.44E-15 |
| B40 | B181 | WBC | APTT | -0.0072 | 3.24E-10 |
| B40 | B190 | WBC | Cre | 0.103742 | 0 |
| B40 | B129 | WBC | IBIL | -0.0675 | 0 |
| B40 | B150 | WBC | ALP | 0.040977 | 0 |
| B40 | B84 | WBC | C-PSA | 0.013553 | 0 |
| B40 | B222 | WBC | NS | 0.189431 | 0 |
| B40 | B148 | WBC | Anti_TPO | -0.01727 | 0 |
| B40 | B149 | WBC | TGAb | -0.01671 | 0 |
| B40 | B48 | WBC | LYM% | -0.0282 | 0 |
| B40 | B198 | WBC | P | 0.006201 | 6.17E-08 |
| B40 | B85 | WBC | SCC | 0.014258 | 0 |
| B40 | B194 | WBC | Cl | -0.02323 | 0 |
| B40 | B155 | WBC | IgA | 0.008955 | 5.33E-15 |
| B40 | B156 | WBC | IgE | 0.011837 | 0 |
| B40 | B176 | WBC | Ucr | 0.042107 | 0 |
| B40 | B178 | WBC | Ig-U | 0.009492 | 0 |
| B40 | B191 | WBC | Urea | 0.013629 | 0 |
| B40 | B174 | WBC | Urea/Cr | -0.06166 | 0 |
| B40 | B192 | WBC | UA | 0.193542 | 0 |
| B40 | B170 | WBC | U-GLU | 0.032729 | 0 |
| B40 | B175 | WBC | ALB-U | 0.015617 | 0 |
| B40 | B177 | WBC | ACR | 0.012953 | 0 |
| B40 | B179 | WBC | α1-MG | 0.017899 | 0 |
| B40 | B125 | WBC | Cor | 0.007041 | 7.90E-10 |
| B40 | B138 | WBC | PAB | 0.034349 | 0 |
| B40 | B130 | WBC | GLB | 0.099464 | 0 |
| B40 | B59 | WBC | WBRV | 0.026457 | 0 |
| B40 | B60 | WBC | WBRVI | 0.040626 | 0 |
| B40 | B61 | WBC | WBHSRV | 0.037489 | 0 |
| B40 | B62 | WBC | WBHSRVI | 0.04414 | 0 |
| B40 | B64 | WBC | WBRSR(mPas_5) | 0.054933 | 0 |
| B40 | B65 | WBC | WBRSR(mPas_1) | 0.05045 | 0 |
| B40 | B66 | WBC | WBRSR(mPas_100) | 0.055794 | 0 |
| B40 | B67 | WBC | WBRSR(mPas_50) | 0.055989 | 0 |
| B40 | B86 | WBC | TBhCG | 0.011911 | 0 |
| B40 | B221 | WBC | Hei | 0.110518 | 0 |
| B40 | B87 | WBC | NSE | 0.008027 | 2.42E-12 |
| B40 | B49 | WBC | baso% | -0.01343 | 0 |
| B40 | B50 | WBC | eo% | 0.00839 | 2.39E-13 |
| B40 | B220 | WBC | SBP | 0.093916 | 0 |

| | | | | | |
|---|---|---|---|---|---|
| B40 | B219 | WBC | OT | -0.02403 | 0 |
| B40 | B218 | WBC | DBP | 0.116944 | 0 |
| B40 | B216 | WBC | BMI | 0.201538 | 0 |
| B40 | B215 | WBC | BMI | 0.188337 | 0 |
| B40 | B131 | WBC | AST | 0.048973 | 0 |
| B40 | B210 | WBC | Hip | 0.156867 | 0 |
| B40 | B209 | WBC | Smoking | 0.191576 | 0 |
| B40 | B90 | WBC | CYFRA 21-1 | 0.014918 | 0 |
| B40 | B139 | WBC | ADA | -0.01233 | 0 |
| B40 | B196 | WBC | TCO2 | -0.0092 | 8.88E-16 |
| B40 | B51 | WBC | Hb | 0.227264 | 0 |
| B40 | B185 | WBC | PT | -0.00731 | 1.76E-10 |
| B40 | B186 | WBC | PFC | 0.016331 | 0 |
| B40 | B69 | WBC | PV | 0.026266 | 0 |
| B40 | B52 | WBC | PLT/BPC | 0.230124 | 0 |
| B40 | B173 | WBC | NTT | -0.00846 | 1.56E-13 |
| B40 | B207 | WBC | WHR | 0.020897 | 0 |
| B40 | B206 | WBC | Waist | 0.196794 | 0 |
| B40 | B205 | WBC | KDA | -0.00612 | 9.09E-08 |
| B40 | B78 | WBC | HBcAb | -0.01808 | 0 |
| B40 | B197 | WBC | AG | 0.020173 | 0 |
| B40 | B204 | WBC | Drinking | 0.079711 | 0 |
| B40 | B203 | WBC | DH | 0.054822 | 0 |
| B40 | B80 | WBC | anti-HP IgG | 0.022348 | 0 |
| B40 | B112 | WBC | FT4 | 0.025709 | 0 |
| B40 | B113 | WBC | FT3 | 0.032676 | 0 |
| B40 | B211 | WBC | IOP-R | 0.023092 | 0 |
| B40 | B201 | WBC | e-habits | -0.01007 | 0 |
| B40 | B74 | WBC | ApoA1 | -0.0398 | 0 |
| B40 | B75 | WBC | ApoA1/B | -0.02509 | 0 |
| B40 | B76 | WBC | ApoB | 0.044966 | 0 |
| B40 | B132 | WBC | Dbil | -0.09713 | 0 |
| B40 | B53 | WBC | GR% | 0.024264 | 0 |
| B40 | B81 | WBC | TSGF | -0.00832 | 3.67E-13 |
| B40 | B77 | WBC | TC | 0.06102 | 0 |
| B40 | B133 | WBC | TBIL | -0.07995 | 0 |
| B40 | B135 | WBC | TBA | 0.006799 | 2.93E-09 |
| B40 | B91 | WBC | TP | 0.116429 | 0 |
| B40 | B114 | WBC | TSH TT3 TT4(TT4) | 0.015348 | 0 |
| B40 | B115 | WBC | TSH TT3 TT4(TT3) | 0.01437 | 0 |
| B40 | B212 | WBC | IOP-L | 0.020489 | 0 |
| B41 | B140 | MON % | r-GT | 0.014742 | 0 |
| B41 | B126 | MON % | Alb | -0.01735 | 0 |
| B41 | B127 | MON % | A-G ratio | -0.00672 | 4.47E-09 |

| | | | | | |
|---|---|---|---|---|---|
| B41 | B161 | MON％ | LEU | -0.00659 | 8.59E-09 |
| B41 | B128 | MON％ | ALT | 0.020049 | 0 |
| B41 | B123 | MON％ | hs-CRP1 | 0.006507 | 1.34E-08 |
| B41 | B166 | MON％ | Pro | 0.012656 | 0 |
| B41 | B72 | MON％ | HDL-C | -0.00755 | 4.38E-11 |
| B41 | B189 | MON％ | eGFB | -0.0223 | 0 |
| B41 | B141 | MON％ | AST/ALT | -0.01102 | 0 |
| B41 | B44 | MON％ | MCH | 0.008693 | 3.22E-14 |
| B41 | B46 | MON％ | RBC Count | 0.013774 | 0 |
| B41 | B47 | MON％ | RDW | 0.010208 | 0 |
| B41 | B190 | MON％ | Cre | 0.030156 | 0 |
| B41 | B129 | MON％ | IBIL | 0.005949 | 2.06E-07 |
| B41 | B222 | MON％ | NS | 0.026169 | 0 |
| B41 | B48 | MON％ | LYM% | 0.01232 | 0 |
| B41 | B176 | MON％ | Ucr | 0.006657 | 6.18E-09 |
| B41 | B191 | MON％ | Urea | 0.011935 | 0 |
| B41 | B174 | MON％ | Urea/Cr | -0.01235 | 0 |
| B41 | B192 | MON％ | UA | 0.023079 | 0 |
| B41 | B221 | MON％ | Hei | 0.024445 | 0 |
| B41 | B49 | MON％ | baso% | 0.007259 | 2.34E-10 |
| B41 | B50 | MON％ | eo% | 0.075646 | 0 |
| B41 | B220 | MON％ | SBP | 0.0136 | 0 |
| B41 | B218 | MON％ | DBP | 0.008313 | 3.93E-13 |
| B41 | B216 | MON％ | BMI | 0.020068 | 0 |
| B41 | B215 | MON％ | BMI | 0.009733 | 0 |
| B41 | B131 | MON％ | AST | 0.021346 | 0 |
| B41 | B210 | MON％ | Hip | 0.011905 | 0 |
| B41 | B209 | MON％ | Smoking | 0.023039 | 0 |
| B41 | B139 | MON％ | ADA | 0.006288 | 4.02E-08 |
| B41 | B51 | MON％ | Hb | 0.021044 | 0 |
| B41 | B52 | MON％ | PLT/BPC | -0.00605 | 1.29E-07 |
| B41 | B206 | MON％ | Waist | 0.020187 | 0 |
| B41 | B204 | MON％ | Drinking | 0.021816 | 0 |
| B41 | B112 | MON％ | FT4 | 0.006926 | 1.48E-09 |
| B41 | B113 | MON％ | FT3 | 0.012547 | 0 |
| B41 | B132 | MON％ | Dbil | 0.010367 | 0 |
| B41 | B53 | MON％ | GR% | 0.012906 | 0 |
| B41 | B133 | MON％ | TBIL | 0.007529 | 4.93E-11 |
| B41 | B91 | MON％ | TP | -0.00922 | 8.88E-16 |
| B42 | B228 | MON | OS | 0.006392 | 2.40E-08 |
| B43 | B54 | MCV | ESR | -0.00906 | 2.66E-15 |
| B43 | B160 | MCV | PH | -0.0118 | 0 |
| B43 | B140 | MCV | r-GT | 0.021076 | 0 |
| B43 | B126 | MCV | Alb | -0.02889 | 0 |

| | | | | | |
|---|---|---|---|---|---|
| B43 | B111 | MCV | TSH | 0.005697 | 6.57E-07 |
| B43 | B70 | MCV | LDL-C | 0.032962 | 0 |
| B43 | B72 | MCV | HDL-C | 0.034252 | 0 |
| B43 | B189 | MCV | eGFB | -0.05756 | 0 |
| B43 | B146 | MCV | CysC | 0.00567 | 7.40E-07 |
| B43 | B55 | MCV | TK | -0.01128 | 0 |
| B43 | B56 | MCV | IR | -0.00584 | 3.46E-07 |
| B43 | B57 | MCV | RBCAI | -0.00699 | 1.04E-09 |
| B43 | B44 | MCV | MCH | 0.373399 | 0 |
| B43 | B45 | MCV | MCHC | 0.127131 | 0 |
| B43 | B46 | MCV | RBC Count | -0.2282 | 0 |
| B43 | B47 | MCV | RDW | 0.227143 | 0 |
| B43 | B190 | MCV | Cre | 0.016219 | 0 |
| B43 | B129 | MCV | IBIL | -0.00588 | 2.86E-07 |
| B43 | B195 | MCV | Na | 0.00757 | 3.87E-11 |
| B43 | B191 | MCV | Urea | 0.023561 | 0 |
| B43 | B174 | MCV | Urea/Cr | 0.005755 | 5.06E-07 |
| B43 | B192 | MCV | UA | -0.01148 | 0 |
| B43 | B130 | MCV | GLB | -0.01751 | 0 |
| B43 | B59 | MCV | WBRV | -0.0143 | 0 |
| B43 | B61 | MCV | WBHSRV | -0.00723 | 2.71E-10 |
| B43 | B221 | MCV | Hei | -0.015 | 0 |
| B43 | B200 | MCV | OP | 0.006245 | 4.97E-08 |
| B43 | B49 | MCV | baso% | -0.01472 | 0 |
| B43 | B219 | MCV | OT | 0.012524 | 0 |
| B43 | B218 | MCV | DBP | 0.006055 | 1.25E-07 |
| B43 | B216 | MCV | BMI | -0.01279 | 0 |
| B43 | B215 | MCV | BMI | -0.00582 | 3.75E-07 |
| B43 | B131 | MCV | AST | 0.017726 | 0 |
| B43 | B210 | MCV | Hip | -0.01367 | 0 |
| B43 | B209 | MCV | Smoking | 0.058974 | 0 |
| B43 | B90 | MCV | CYFRA 21-1 | 0.005882 | 2.81E-07 |
| B43 | B51 | MCV | Hb | 0.09568 | 0 |
| B43 | B183 | MCV | PTA | -0.00591 | 2.43E-07 |
| B43 | B52 | MCV | PLT/BPC | -0.09612 | 0 |
| B43 | B206 | MCV | Waist | 0.006954 | 1.27E-09 |
| B43 | B205 | MCV | KDA | 0.006338 | 3.14E-08 |
| B43 | B204 | MCV | Drinking | 0.0408 | 0 |
| B43 | B112 | MCV | FT4 | -0.02414 | 0 |
| B43 | B113 | MCV | FT3 | -0.01121 | 0 |
| B43 | B201 | MCV | e-habits | 0.010841 | 0 |
| B43 | B74 | MCV | ApoA1 | 0.016164 | 0 |
| B43 | B76 | MCV | ApoB | 0.007983 | 3.17E-12 |
| B43 | B132 | MCV | Dbil | 0.008793 | 1.64E-14 |

| | | | | | |
|---|---|---|---|---|---|
| B43 | B77 | MCV | TC | 0.04107 | 0 |
| B43 | B91 | MCV | TP | -0.03279 | 0 |
| B43 | B114 | MCV | TSH TT3 TT4(TT4) | -0.00929 | 4.44E-16 |
| B44 | B137 | MCH | AFU | 0.012405 | 0 |
| B44 | B54 | MCH | ESR | -0.03269 | 0 |
| B44 | B160 | MCH | PH | -0.02149 | 0 |
| B44 | B140 | MCH | r-GT | 0.081685 | 0 |
| B44 | B118 | MCH | CEA1 | 0.011003 | 0 |
| B44 | B127 | MCH | A-G ratio | 0.052811 | 0 |
| B44 | B161 | MCH | LEU | -0.02029 | 0 |
| B44 | B128 | MCH | ALT | 0.050227 | 0 |
| B44 | B165 | MCH | bilirubin | 0.008818 | 1.38E-14 |
| B44 | B134 | MCH | CHE | 0.012462 | 0 |
| B44 | B166 | MCH | Pro | 0.024072 | 0 |
| B44 | B70 | MCH | LDL-C | 0.070308 | 0 |
| B44 | B188 | MCH | Ca | 0.007049 | 7.54E-10 |
| B44 | B71 | MCH | TG | 0.071172 | 0 |
| B44 | B72 | MCH | HDL-C | -0.00742 | 9.09E-11 |
| B44 | B189 | MCH | eGFB | -0.10424 | 0 |
| B44 | B141 | MCH | AST/ALT | -0.04654 | 0 |
| B44 | B146 | MCH | CysC | 0.012643 | 0 |
| B44 | B55 | MCH | TK | -0.01679 | 0 |
| B44 | B56 | MCH | IR | 0.010513 | 0 |
| B44 | B57 | MCH | RBCAI | -0.01136 | 0 |
| B44 | B45 | MCH | MCHC | 0.518693 | 0 |
| B44 | B46 | MCH | RBC Count | -0.28448 | 0 |
| B44 | B47 | MCH | RDW | 0.344621 | 0 |
| B44 | B58 | MCH | HCT | 0.005971 | 1.86E-07 |
| B44 | B181 | MCH | APTT | -0.00658 | 9.30E-09 |
| B44 | B190 | MCH | Cre | 0.109466 | 0 |
| B44 | B224 | MCH | pmh | 0.007293 | 1.93E-10 |
| B44 | B193 | MCH | K | -0.01153 | 0 |
| B44 | B129 | MCH | IBIL | 0.045167 | 0 |
| B44 | B150 | MCH | ALP | 0.014512 | 0 |
| B44 | B84 | MCH | C-PSA | 0.007165 | 3.96E-10 |
| B44 | B222 | MCH | NS | 0.097755 | 0 |
| B44 | B148 | MCH | Anti_TPO | -0.01537 | 0 |
| B44 | B149 | MCH | TGAb | -0.01514 | 0 |
| B44 | B198 | MCH | P | -0.01397 | 0 |
| B44 | B167 | MCH | PC | -0.01201 | 0 |
| B44 | B194 | MCH | Cl | -0.006 | 1.61E-07 |
| B44 | B157 | MCH | IgG | -0.01304 | 0 |
| B44 | B195 | MCH | Na | 0.008192 | 8.57E-13 |
| B44 | B176 | MCH | Ucr | 0.021394 | 0 |

| | | | | | |
|---|---|---|---|---|---|
| B44 | B191 | MCH | Urea | 0.045455 | 0 |
| B44 | B174 | MCH | Urea/Cr | -0.04501 | 0 |
| B44 | B192 | MCH | UA | 0.076167 | 0 |
| B44 | B170 | MCH | U-GLU | 0.013374 | 0 |
| B44 | B177 | MCH | ACR | -0.00611 | 9.39E-08 |
| B44 | B179 | MCH | α1-MG | 0.010281 | 0 |
| B44 | B182 | MCH | TT | 0.006844 | 2.30E-09 |
| B44 | B153 | MCH | Glu | 0.0065 | 1.39E-08 |
| B44 | B138 | MCH | PAB | 0.025457 | 0 |
| B44 | B130 | MCH | GLB | -0.06179 | 0 |
| B44 | B59 | MCH | WBRV | -0.01536 | 0 |
| B44 | B60 | MCH | WBRVI | 0.017757 | 0 |
| B44 | B61 | MCH | WBHSRV | 0.008423 | 1.93E-13 |
| B44 | B62 | MCH | WBHSRVI | 0.032939 | 0 |
| B44 | B64 | MCH | WBRSR(mPas_5) | 0.022069 | 0 |
| B44 | B65 | MCH | WBRSR(mPas_1) | 0.016358 | 0 |
| B44 | B66 | MCH | WBRSR(mPas_100) | 0.029397 | 0 |
| B44 | B67 | MCH | WBRSR(mPas_50) | 0.028404 | 0 |
| B44 | B221 | MCH | Hei | 0.068535 | 0 |
| B44 | B200 | MCH | OP | 0.008772 | 1.89E-14 |
| B44 | B49 | MCH | baso% | 0.329804 | 0 |
| B44 | B50 | MCH | eo% | 0.015343 | 0 |
| B44 | B220 | MCH | SBP | 0.041244 | 0 |
| B44 | B219 | MCH | OT | 0.010906 | 0 |
| B44 | B218 | MCH | DBP | 0.057045 | 0 |
| B44 | B216 | MCH | BMI | 0.070717 | 0 |
| B44 | B215 | MCH | BMI | 0.046346 | 0 |
| B44 | B131 | MCH | AST | 0.053874 | 0 |
| B44 | B172 | MCH | KB | 0.006971 | 1.16E-09 |
| B44 | B210 | MCH | Hip | 0.034118 | 0 |
| B44 | B209 | MCH | Smoking | 0.139071 | 0 |
| B44 | B90 | MCH | CYFRA 21-1 | 0.012596 | 0 |
| B44 | B51 | MCH | Hb | 0.341282 | 0 |
| B44 | B183 | MCH | PTA | -0.00672 | 4.33E-09 |
| B44 | B52 | MCH | PLT/BPC | -0.16242 | 0 |
| B44 | B173 | MCH | NTT | -0.00683 | 2.53E-09 |
| B44 | B207 | MCH | WHR | 0.011721 | 0 |
| B44 | B206 | MCH | Waist | 0.082011 | 0 |
| B44 | B197 | MCH | AG | 0.007242 | 2.57E-10 |
| B44 | B204 | MCH | Drinking | 0.122577 | 0 |
| B44 | B112 | MCH | FT4 | -0.0248 | 0 |
| B44 | B201 | MCH | e-habits | 0.02185 | 0 |
| B44 | B74 | MCH | ApoA1 | 0.006056 | 1.24E-07 |
| B44 | B75 | MCH | ApoA1/B | -0.00786 | 6.92E-12 |

| | | | | | |
|---|---|---|---|---|---|
| B44 | B76 | MCH | ApoB | 0.024311 | 0 |
| B44 | B132 | MCH | Dbil | 0.047117 | 0 |
| B44 | B77 | MCH | TC | 0.077278 | 0 |
| B44 | B133 | MCH | TBIL | 0.048796 | 0 |
| B44 | B135 | MCH | TBA | 0.006276 | 4.26E-08 |
| B44 | B91 | MCH | TP | -0.05175 | 0 |
| B44 | B114 | MCH | TSH TT3 TT4(TT4) | -0.01898 | 0 |
| B45 | B137 | MCHC | AFU | 0.016985 | 0 |
| B45 | B54 | MCHC | ESR | -0.05048 | 0 |
| B45 | B160 | MCHC | PH | -0.00706 | 7.07E-10 |
| B45 | B140 | MCHC | r-GT | 0.125507 | 0 |
| B45 | B118 | MCHC | CEA1 | 0.006781 | 3.21E-09 |
| B45 | B126 | MCHC | Alb | 0.133935 | 0 |
| B45 | B127 | MCHC | A-G ratio | 0.116677 | 0 |
| B45 | B161 | MCHC | LEU | -0.05848 | 0 |
| B45 | B128 | MCHC | ALT | 0.137613 | 0 |
| B45 | B121 | MCHC | C3 | 0.006358 | 2.84E-08 |
| B45 | B111 | MCHC | TSH | -0.02799 | 0 |
| B45 | B165 | MCHC | bilirubin | 0.005823 | 3.71E-07 |
| B45 | B134 | MCHC | CHE | 0.041957 | 0 |
| B45 | B166 | MCHC | Pro | 0.07412 | 0 |
| B45 | B70 | MCHC | LDL-C | 0.055744 | 0 |
| B45 | B188 | MCHC | Ca | 0.018315 | 0 |
| B45 | B71 | MCHC | TG | 0.166272 | 0 |
| B45 | B72 | MCHC | HDL-C | -0.15565 | 0 |
| B45 | B189 | MCHC | eGFB | -0.03656 | 0 |
| B45 | B141 | MCHC | AST/ALT | -0.15129 | 0 |
| B45 | B146 | MCHC | CysC | 0.010122 | 0 |
| B45 | B56 | MCHC | IR | 0.050953 | 0 |
| B45 | B46 | MCHC | RBC Count | 0.106713 | 0 |
| B45 | B47 | MCHC | RDW | -0.21803 | 0 |
| B45 | B58 | MCHC | HCT | 0.009149 | 1.33E-15 |
| B45 | B181 | MCHC | APTT | -0.00801 | 2.61E-12 |
| B45 | B190 | MCHC | Cre | 0.227742 | 0 |
| B45 | B193 | MCHC | K | -0.02653 | 0 |
| B45 | B129 | MCHC | IBIL | 0.155983 | 0 |
| B45 | B150 | MCHC | ALP | 0.015895 | 0 |
| B45 | B84 | MCHC | C-PSA | 0.011693 | 0 |
| B45 | B222 | MCHC | NS | 0.26076 | 0 |
| B45 | B148 | MCHC | Anti_TPO | -0.02431 | 0 |
| B45 | B149 | MCHC | TGAb | -0.02315 | 0 |
| B45 | B198 | MCHC | P | -0.01908 | 0 |
| B45 | B194 | MCHC | Cl | -0.01924 | 0 |
| B45 | B199 | MCHC | Mg | -0.00588 | 2.79E-07 |

| | | | | | |
|---|---|---|---|---|---|
| B45 | B157 | MCHC | IgG | -0.01611 | 0 |
| B45 | B158 | MCHC | IgM | -0.00791 | 5.07E-12 |
| B45 | B195 | MCHC | Na | -0.00928 | 4.44E-16 |
| B45 | B176 | MCHC | Ucr | 0.058303 | 0 |
| B45 | B191 | MCHC | Urea | 0.027176 | 0 |
| B45 | B174 | MCHC | Urea/Cr | -0.14262 | 0 |
| B45 | B192 | MCHC | UA | 0.248848 | 0 |
| B45 | B170 | MCHC | U-GLU | 0.042804 | 0 |
| B45 | B177 | MCHC | ACR | -0.00606 | 1.24E-07 |
| B45 | B179 | MCHC | α1-MG | 0.01618 | 0 |
| B45 | B153 | MCHC | Glu | 0.014506 | 0 |
| B45 | B138 | MCHC | PAB | 0.049442 | 0 |
| B45 | B130 | MCHC | GLB | -0.08154 | 0 |
| B45 | B59 | MCHC | WBRV | 0.016824 | 0 |
| B45 | B60 | MCHC | WBRVI | 0.060666 | 0 |
| B45 | B61 | MCHC | WBHSRV | 0.05142 | 0 |
| B45 | B62 | MCHC | WBHSRVI | 0.078276 | 0 |
| B45 | B63 | MCHC | WBRSR(mPas_200) | 0.008423 | 1.92E-13 |
| B45 | B64 | MCHC | WBRSR(mPas_5) | 0.069327 | 0 |
| B45 | B65 | MCHC | WBRSR(mPas_1) | 0.060529 | 0 |
| B45 | B66 | MCHC | WBRSR(mPas_100) | 0.075514 | 0 |
| B45 | B67 | MCHC | WBRSR(mPas_50) | 0.074761 | 0 |
| B45 | B221 | MCHC | Hei | 0.240264 | 0 |
| B45 | B87 | MCHC | NSE | 0.012884 | 0 |
| B45 | B49 | MCHC | baso% | -0.08102 | 0 |
| B45 | B50 | MCHC | eo% | 0.015436 | 0 |
| B45 | B220 | MCHC | SBP | 0.081719 | 0 |
| B45 | B219 | MCHC | OT | -0.02476 | 0 |
| B45 | B218 | MCHC | DBP | 0.120666 | 0 |
| B45 | B88 | MCHC | CA 242 | -0.01096 | 0 |
| B45 | B216 | MCHC | BMI | 0.239877 | 0 |
| B45 | B215 | MCHC | BMI | 0.147854 | 0 |
| B45 | B131 | MCHC | AST | 0.071171 | 0 |
| B45 | B172 | MCHC | KB | 0.009945 | 0 |
| B45 | B210 | MCHC | Hip | 0.145859 | 0 |
| B45 | B209 | MCHC | Smoking | 0.119145 | 0 |
| B45 | B90 | MCHC | CYFRA 21-1 | 0.008746 | 2.24E-14 |
| B45 | B139 | MCHC | ADA | -0.01457 | 0 |
| B45 | B51 | MCHC | Hb | 0.57014 | 0 |
| B45 | B183 | MCHC | PTA | 0.006931 | 1.44E-09 |
| B45 | B186 | MCHC | PFC | -0.00599 | 1.68E-07 |
| B45 | B52 | MCHC | PLT/BPC | -0.03727 | 0 |
| B45 | B173 | MCHC | NTT | -0.03233 | 0 |
| B45 | B207 | MCHC | WHR | 0.020458 | 0 |

| | | | | | |
|---|---|---|---|---|---|
| B45 | B206 | MCHC | Waist | 0.187266 | 0 |
| B45 | B205 | MCHC | KDA | -0.01626 | 0 |
| B45 | B197 | MCHC | AG | 0.007324 | 1.61E-10 |
| B45 | B204 | MCHC | Drinking | 0.151542 | 0 |
| B45 | B203 | MCHC | DH | 0.025009 | 0 |
| B45 | B112 | MCHC | FT4 | 0.027355 | 0 |
| B45 | B113 | MCHC | FT3 | 0.034476 | 0 |
| B45 | B211 | MCHC | IOP-R | 0.016271 | 0 |
| B45 | B201 | MCHC | e-habits | 0.009953 | 0 |
| B45 | B73 | MCHC | ApoA | -0.01328 | 0 |
| B45 | B74 | MCHC | ApoA1 | -0.05103 | 0 |
| B45 | B75 | MCHC | ApoA1/B | -0.02669 | 0 |
| B45 | B76 | MCHC | ApoB | 0.033665 | 0 |
| B45 | B132 | MCHC | Dbil | 0.093638 | 0 |
| B45 | B53 | MCHC | GR% | -0.0075 | 5.90E-11 |
| B45 | B81 | MCHC | TSGF | -0.01852 | 0 |
| B45 | B77 | MCHC | TC | 0.041651 | 0 |
| B45 | B133 | MCHC | TBIL | 0.150271 | 0 |
| B45 | B91 | MCHC | TP | 0.013391 | 0 |
| B45 | B114 | MCHC | TSH TT3 TT4(TT4) | -0.01106 | 0 |
| B45 | B115 | MCHC | TSH TT3 TT4(TT3) | 0.006286 | 4.07E-08 |
| B45 | B212 | MCHC | IOP-L | 0.013947 | 0 |
| B46 | B137 | RBC Count | AFU | 0.02665 | 0 |
| B46 | B54 | RBC Count | ESR | -0.04958 | 0 |
| B46 | B160 | RBC Count | PH | -0.05744 | 0 |
| B46 | B140 | RBC Count | r-GT | 0.163294 | 0 |
| B46 | B143 | RBC Count | β-CTx | 0.007371 | 1.23E-10 |
| B46 | B118 | RBC Count | CEA1 | 0.006055 | 1.25E-07 |
| B46 | B126 | RBC Count | Alb | 0.296364 | 0 |
| B46 | B127 | RBC Count | A-G ratio | 0.137564 | 0 |
| B46 | B161 | RBC Count | LEU | -0.10357 | 0 |
| B46 | B128 | RBC Count | ALT | 0.245399 | 0 |
| B46 | B123 | RBC Count | hs-CRP1 | 0.007345 | 1.43E-10 |
| B46 | B111 | RBC Count | TSH | -0.0519 | 0 |
| B46 | B124 | RBC Count | ACTH | 0.01248 | 0 |
| B46 | B134 | RBC Count | CHE | 0.079588 | 0 |
| B46 | B166 | RBC Count | Pro | 0.130202 | 0 |
| B46 | B70 | RBC Count | LDL-C | 0.142941 | 0 |
| B46 | B188 | RBC Count | Ca | 0.047782 | 0 |
| B46 | B71 | RBC Count | TG | 0.184238 | 0 |
| B46 | B72 | RBC Count | HDL-C | -0.28815 | 0 |
| B46 | B189 | RBC Count | eGFB | -0.05149 | 0 |
| B46 | B141 | RBC Count | AST/ALT | -0.27685 | 0 |
| B46 | B146 | RBC Count | CysC | 0.020639 | 0 |

| | | | | | |
|---|---|---|---|---|---|
| B46 | B55 | RBC Count | TK | -0.0147 | 0 |
| B46 | B56 | RBC Count | IR | 0.077533 | 0 |
| B46 | B57 | RBC Count | RBCAI | 0.035061 | 0 |
| B46 | B47 | RBC Count | RDW | -0.38034 | 0 |
| B46 | B58 | RBC Count | HCT | 0.018811 | 0 |
| B46 | B181 | RBC Count | APTT | -0.00724 | 2.66E-10 |
| B46 | B190 | RBC Count | Cre | 0.421161 | 0 |
| B46 | B193 | RBC Count | K | 0.008208 | 7.71E-13 |
| B46 | B129 | RBC Count | IBIL | 0.139534 | 0 |
| B46 | B150 | RBC Count | ALP | 0.049339 | 0 |
| B46 | B84 | RBC Count | C-PSA | 0.022902 | 0 |
| B46 | B222 | RBC Count | NS | 0.458771 | 0 |
| B46 | B100 | RBC Count | anti-Ro52 | -0.00609 | 1.08E-07 |
| B46 | B103 | RBC Count | anti-SSA/Ro | -0.00829 | 4.66E-13 |
| B46 | B148 | RBC Count | Anti_TPO | -0.04057 | 0 |
| B46 | B149 | RBC Count | TGAb | -0.03611 | 0 |
| B46 | B198 | RBC Count | P | -0.03132 | 0 |
| B46 | B85 | RBC Count | SCC | 0.015678 | 0 |
| B46 | B194 | RBC Count | Cl | -0.0326 | 0 |
| B46 | B199 | RBC Count | Mg | 0.011756 | 0 |
| B46 | B155 | RBC Count | IgA | 0.007584 | 3.56E-11 |
| B46 | B157 | RBC Count | IgG | -0.01711 | 0 |
| B46 | B158 | RBC Count | IgM | -0.01299 | 0 |
| B46 | B195 | RBC Count | Na | 0.008741 | 2.31E-14 |
| B46 | B176 | RBC Count | Ucr | 0.100774 | 0 |
| B46 | B178 | RBC Count | Ig-U | 0.005901 | 2.58E-07 |
| B46 | B191 | RBC Count | Urea | 0.078449 | 0 |
| B46 | B174 | RBC Count | Urea/Cr | -0.2371 | 0 |
| B46 | B192 | RBC Count | UA | 0.439121 | 0 |
| B46 | B170 | RBC Count | U-GLU | 0.044493 | 0 |
| B46 | B175 | RBC Count | ALB-U | 0.012359 | 0 |
| B46 | B179 | RBC Count | α1-MG | 0.026408 | 0 |
| B46 | B180 | RBC Count | Tf | 0.009733 | 0 |
| B46 | B182 | RBC Count | TT | 0.00806 | 1.97E-12 |
| B46 | B125 | RBC Count | Cor | 0.013248 | 0 |
| B46 | B153 | RBC Count | Glu | 0.009283 | 4.44E-16 |
| B46 | B138 | RBC Count | PAB | 0.080242 | 0 |
| B46 | B130 | RBC Count | GLB | -0.04154 | 0 |
| B46 | B59 | RBC Count | WBRV | 0.059226 | 0 |
| B46 | B60 | RBC Count | WBRVI | 0.134921 | 0 |
| B46 | B61 | RBC Count | WBHSRV | 0.083171 | 0 |
| B46 | B62 | RBC Count | WBHSRVI | 0.137139 | 0 |
| B46 | B63 | RBC Count | WBRSR(mPas_200) | 0.015804 | 0 |
| B46 | B64 | RBC Count | WBRSR(mPas_5) | 0.146163 | 0 |

| | | | | | |
|---|---|---|---|---|---|
| B46 | B65 | RBC Count | WBRSR(mPas_1) | 0.138743 | 0 |
| B46 | B66 | RBC Count | WBRSR(mPas_100) | 0.141169 | 0 |
| B46 | B67 | RBC Count | WBRSR(mPas_50) | 0.14281 | 0 |
| B46 | B86 | RBC Count | TBhCG | -0.01305 | 0 |
| B46 | B221 | RBC Count | Hei | 0.446902 | 0 |
| B46 | B87 | RBC Count | NSE | 0.019412 | 0 |
| B46 | B187 | RBC Count | NSE | 0.011751 | 0 |
| B46 | B200 | RBC Count | OP | 0.012996 | 0 |
| B46 | B49 | RBC Count | baso% | 0.057184 | 0 |
| B46 | B50 | RBC Count | eo% | 0.057189 | 0 |
| B46 | B220 | RBC Count | SBP | 0.1785 | 0 |
| B46 | B219 | RBC Count | OT | -0.05354 | 0 |
| B46 | B218 | RBC Count | DBP | 0.231775 | 0 |
| B46 | B88 | RBC Count | CA 242 | -0.02139 | 0 |
| B46 | B216 | RBC Count | BMI | 0.438868 | 0 |
| B46 | B215 | RBC Count | BMI | 0.268384 | 0 |
| B46 | B131 | RBC Count | AST | 0.106116 | 0 |
| B46 | B210 | RBC Count | Hip | 0.286075 | 0 |
| B46 | B209 | RBC Count | Smoking | 0.167107 | 0 |
| B46 | B90 | RBC Count | CYFRA 21-1 | 0.014758 | 0 |
| B46 | B139 | RBC Count | ADA | -0.01568 | 0 |
| B46 | B51 | RBC Count | Hb | 0.708712 | 0 |
| B46 | B185 | RBC Count | PT | -0.00735 | 1.42E-10 |
| B46 | B186 | RBC Count | PFC | -0.00577 | 4.78E-07 |
| B46 | B69 | RBC Count | PV | 0.009172 | 1.33E-15 |
| B46 | B52 | RBC Count | PLT/BPC | 0.08631 | 0 |
| B46 | B173 | RBC Count | NTT | -0.05184 | 0 |
| B46 | B207 | RBC Count | WHR | 0.039745 | 0 |
| B46 | B206 | RBC Count | Waist | 0.357086 | 0 |
| B46 | B205 | RBC Count | KDA | -0.0376 | 0 |
| B46 | B197 | RBC Count | AG | 0.022229 | 0 |
| B46 | B204 | RBC Count | Drinking | 0.208541 | 0 |
| B46 | B203 | RBC Count | DH | 0.035999 | 0 |
| B46 | B112 | RBC Count | FT4 | 0.06412 | 0 |
| B46 | B113 | RBC Count | FT3 | 0.108877 | 0 |
| B46 | B211 | RBC Count | IOP-R | 0.025597 | 0 |
| B46 | B201 | RBC Count | e-habits | 0.009379 | 2.22E-16 |
| B46 | B73 | RBC Count | ApoA | -0.01426 | 0 |
| B46 | B74 | RBC Count | ApoA1 | -0.07646 | 0 |
| B46 | B75 | RBC Count | ApoA1/B | -0.04937 | 0 |
| B46 | B76 | RBC Count | ApoB | 0.066011 | 0 |
| B46 | B132 | RBC Count | Dbil | 0.06168 | 0 |
| B46 | B53 | RBC Count | GR% | -0.00562 | 9.40E-07 |
| B46 | B81 | RBC Count | TSGF | -0.01043 | 0 |

| | | | | | |
|---|---|---|---|---|---|
| B46 | B77 | RBC Count | TC | 0.057253 | 0 |
| B46 | B133 | RBC Count | TBIL | 0.128598 | 0 |
| B46 | B135 | RBC Count | TBA | 0.007622 | 2.85E-11 |
| B46 | B91 | RBC Count | TP | 0.148389 | 0 |
| B46 | B115 | RBC Count | TSH TT3 TT4(TT3) | 0.036598 | 0 |
| B46 | B212 | RBC Count | IOP-L | 0.023185 | 0 |
| B47 | B54 | RDW | ESR | 0.016458 | 0 |
| B47 | B160 | RDW | PH | -0.04848 | 0 |
| B47 | B140 | RDW | r-GT | 0.038347 | 0 |
| B47 | B118 | RDW | CEA1 | 0.018694 | 0 |
| B47 | B126 | RDW | Alb | -0.17325 | 0 |
| B47 | B127 | RDW | A-G ratio | -0.07067 | 0 |
| B47 | B161 | RDW | LEU | 0.017629 | 0 |
| B47 | B128 | RDW | ALT | -0.02207 | 0 |
| B47 | B123 | RDW | hs-CRP1 | 0.009671 | 0 |
| B47 | B111 | RDW | TSH | 0.042407 | 0 |
| B47 | B165 | RDW | bilirubin | 0.01602 | 0 |
| B47 | B134 | RDW | CHE | -0.02627 | 0 |
| B47 | B166 | RDW | Pro | -0.01867 | 0 |
| B47 | B70 | RDW | LDL-C | 0.020896 | 0 |
| B47 | B188 | RDW | Ca | -0.02179 | 0 |
| B47 | B72 | RDW | HDL-C | 0.098941 | 0 |
| B47 | B189 | RDW | eGFB | -0.15226 | 0 |
| B47 | B141 | RDW | AST/ALT | 0.058055 | 0 |
| B47 | B146 | RDW | CysC | 0.015339 | 0 |
| B47 | B55 | RDW | TK | -0.00655 | 1.08E-08 |
| B47 | B56 | RDW | IR | -0.02324 | 0 |
| B47 | B57 | RDW | RBCAI | -0.01052 | 0 |
| B47 | B190 | RDW | Cre | -0.0384 | 0 |
| B47 | B224 | RDW | pmh | 0.011113 | 0 |
| B47 | B83 | RDW | AFP | 0.006048 | 1.29E-07 |
| B47 | B193 | RDW | K | 0.006898 | 1.72E-09 |
| B47 | B129 | RDW | IBIL | -0.06167 | 0 |
| B47 | B150 | RDW | ALP | 0.023234 | 0 |
| B47 | B213 | RDW | CVA-R | 0.007651 | 2.39E-11 |
| B47 | B214 | RDW | CVA-L | 0.008175 | 9.52E-13 |
| B47 | B222 | RDW | NS | -0.05816 | 0 |
| B47 | B154 | RDW | RF | 0.006131 | 8.69E-08 |
| B47 | B48 | RDW | LYM% | -0.00669 | 5.30E-09 |
| B47 | B198 | RDW | P | -0.0105 | 0 |
| B47 | B194 | RDW | Cl | 0.018966 | 0 |
| B47 | B199 | RDW | Mg | 0.006708 | 4.72E-09 |
| B47 | B195 | RDW | Na | 0.014894 | 0 |
| B47 | B176 | RDW | Ucr | -0.02191 | 0 |

| | | | | | |
|---|---|---|---|---|---|
| B47 | B191 | RDW | Urea | 0.053411 | 0 |
| B47 | B174 | RDW | Urea/Cr | 0.068566 | 0 |
| B47 | B192 | RDW | UA | -0.08599 | 0 |
| B47 | B170 | RDW | U-GLU | -0.01118 | 0 |
| B47 | B138 | RDW | PAB | -0.01938 | 0 |
| B47 | B130 | RDW | GLB | 0.021714 | 0 |
| B47 | B59 | RDW | WBRV | -0.02104 | 0 |
| B47 | B60 | RDW | WBRVI | -0.02872 | 0 |
| B47 | B61 | RDW | WBHSRV | -0.0261 | 0 |
| B47 | B62 | RDW | WBHSRVI | -0.02797 | 0 |
| B47 | B64 | RDW | WBRSR(mPas_5) | -0.03215 | 0 |
| B47 | B65 | RDW | WBRSR(mPas_1) | -0.03064 | 0 |
| B47 | B66 | RDW | WBRSR(mPas_100) | -0.03025 | 0 |
| B47 | B67 | RDW | WBRSR(mPas_50) | -0.03058 | 0 |
| B47 | B221 | RDW | Hei | -0.12818 | 0 |
| B47 | B87 | RDW | NSE | -0.00707 | 6.82E-10 |
| B47 | B200 | RDW | OP | 0.010244 | 0 |
| B47 | B227 | RDW | OD | 0.008208 | 7.73E-13 |
| B47 | B228 | RDW | OS | 0.010504 | 0 |
| B47 | B49 | RDW | baso% | 0.152752 | 0 |
| B47 | B50 | RDW | eo% | 0.019948 | 0 |
| B47 | B220 | RDW | SBP | 0.028335 | 0 |
| B47 | B219 | RDW | OT | 0.045741 | 0 |
| B47 | B89 | RDW | CA 72-4 | 0.006272 | 4.35E-08 |
| B47 | B216 | RDW | BMI | -0.07973 | 0 |
| B47 | B215 | RDW | BMI | -0.01221 | 0 |
| B47 | B131 | RDW | AST | 0.047312 | 0 |
| B47 | B210 | RDW | Hip | -0.05113 | 0 |
| B47 | B209 | RDW | Smoking | 0.150727 | 0 |
| B47 | B90 | RDW | CYFRA 21-1 | 0.011365 | 0 |
| B47 | B139 | RDW | ADA | 0.016705 | 0 |
| B47 | B51 | RDW | Hb | -0.15237 | 0 |
| B47 | B183 | RDW | PTA | -0.01327 | 0 |
| B47 | B186 | RDW | PFC | 0.006371 | 2.67E-08 |
| B47 | B52 | RDW | PLT/BPC | -0.20927 | 0 |
| B47 | B173 | RDW | NTT | 0.023664 | 0 |
| B47 | B205 | RDW | KDA | 0.016254 | 0 |
| B47 | B78 | RDW | HBcAb | 0.011318 | 0 |
| B47 | B204 | RDW | Drinking | 0.050244 | 0 |
| B47 | B203 | RDW | DH | -0.01473 | 0 |
| B47 | B112 | RDW | FT4 | -0.07597 | 0 |
| B47 | B113 | RDW | FT3 | -0.04452 | 0 |
| B47 | B211 | RDW | IOP-R | -0.0256 | 0 |
| B47 | B201 | RDW | e-habits | 0.021666 | 0 |

| | | | | | |
|---|---|---|---|---|---|
| B47 | B74 | RDW | ApoA1 | 0.045592 | 0 |
| B47 | B75 | RDW | ApoA1/B | 0.018419 | 0 |
| B47 | B76 | RDW | ApoB | -0.00885 | 1.11E-14 |
| B47 | B132 | RDW | Dbil | 0.012914 | 0 |
| B47 | B81 | RDW | TSGF | 0.013684 | 0 |
| B47 | B77 | RDW | TC | 0.061672 | 0 |
| B47 | B133 | RDW | TBIL | -0.04627 | 0 |
| B47 | B135 | RDW | TBA | 0.011341 | 0 |
| B47 | B91 | RDW | TP | -0.08885 | 0 |
| B47 | B114 | RDW | TSH TT3 TT4(TT4) | -0.01651 | 0 |
| B47 | B115 | RDW | TSH TT3 TT4(TT3) | -0.01184 | 0 |
| B47 | B212 | RDW | IOP-L | -0.0219 | 0 |
| B48 | B123 | LYM% | hs-CRP1 | -0.00695 | 1.29E-09 |
| B48 | B221 | LYM% | Hei | 0.006547 | 1.09E-08 |
| B48 | B50 | LYM% | eo% | 0.033887 | 0 |
| B48 | B209 | LYM% | Smoking | -0.00655 | 1.08E-08 |
| B48 | B91 | LYM% | TP | 0.005613 | 9.57E-07 |
| B49 | B70 | baso% | LDL-C | 0.007303 | 1.82E-10 |
| B49 | B189 | baso% | eGFB | -0.00966 | 0 |
| B49 | B190 | baso% | Cre | 0.006684 | 5.37E-09 |
| B49 | B167 | baso% | PC | -0.02783 | 0 |
| B49 | B50 | baso% | eo% | 0.051679 | 0 |
| B49 | B51 | baso% | Hb | -0.02445 | 0 |
| B49 | B52 | baso% | PLT/BPC | 0.006076 | 1.13E-07 |
| B49 | B77 | baso% | TC | 0.007284 | 2.02E-10 |
| B5 | B23 | WB-Mg | CD3-CD16+CD56+(NK) | -0.00707 | 6.59E-10 |
| B5 | B21 | WB-Mg | CD3+CD4-CD8- | -0.00705 | 7.56E-10 |
| B5 | B128 | WB-Mg | ALT | 0.007334 | 1.53E-10 |
| B5 | B121 | WB-Mg | C3 | -0.00699 | 1.03E-09 |
| B5 | B29 | WB-Mg | E2 | -0.00831 | 4.10E-13 |
| B5 | B134 | WB-Mg | CHE | 0.007764 | 1.22E-11 |
| B5 | B71 | WB-Mg | TG | 0.005627 | 8.99E-07 |
| B5 | B72 | WB-Mg | HDL-C | -0.00716 | 4.13E-10 |
| B5 | B32 | WB-Mg | Testo | 0.008932 | 6.22E-15 |
| B5 | B141 | WB-Mg | AST/ALT | -0.00718 | 3.59E-10 |
| B5 | B44 | WB-Mg | MCH | 0.007029 | 8.44E-10 |
| B5 | B45 | WB-Mg | MCHC | 0.008788 | 1.69E-14 |
| B5 | B46 | WB-Mg | RBC Count | 0.010355 | 0 |
| B5 | B190 | WB-Mg | Cre | 0.008 | 2.86E-12 |
| B5 | B222 | WB-Mg | NS | 0.009447 | 0 |
| B5 | B108 | WB-Mg | Anti-Cyclin | -0.00706 | 7.02E-10 |
| B5 | B157 | WB-Mg | IgG | -0.00823 | 6.51E-13 |
| B5 | B195 | WB-Mg | Na | 0.006127 | 8.84E-08 |
| B5 | B192 | WB-Mg | UA | 0.009279 | 4.44E-16 |

| | | | | | |
|---|---|---|---|---|---|
| B5 | B138 | WB-Mg | PAB | 0.008277 | 4.97E-13 |
| B5 | B6 | WB-Mg | WB-Fe | 0.723819 | 0 |
| B5 | B7 | WB-Mg | WB-Cop | 0.056887 | 0 |
| B5 | B8 | WB-Mg | WB-Zinc | 0.43236 | 0 |
| B5 | B221 | WB-Mg | Hei | 0.006693 | 5.11E-09 |
| B5 | B200 | WB-Mg | OP | 0.005814 | 3.85E-07 |
| B5 | B88 | WB-Mg | CA 242 | 0.008691 | 3.24E-14 |
| B5 | B216 | WB-Mg | BMI | 0.009643 | 0 |
| B5 | B215 | WB-Mg | BMI | 0.008224 | 6.96E-13 |
| B5 | B210 | WB-Mg | Hip | 0.006337 | 3.15E-08 |
| B5 | B51 | WB-Mg | Hb | 0.01568 | 0 |
| B5 | B206 | WB-Mg | Waist | 0.009485 | 0 |
| B5 | B204 | WB-Mg | Drinking | 0.006231 | 5.32E-08 |
| B5 | B76 | WB-Mg | ApoB | 0.008921 | 6.66E-15 |
| B50 | B137 | eo% | AFU | 0.007594 | 3.36E-11 |
| B50 | B160 | eo% | PH | -0.03242 | 0 |
| B50 | B140 | eo% | r-GT | 0.030693 | 0 |
| B50 | B118 | eo% | CEA1 | 0.009263 | 6.66E-16 |
| B50 | B126 | eo% | Alb | -0.04573 | 0 |
| B50 | B161 | eo% | LEU | -0.01743 | 0 |
| B50 | B128 | eo% | ALT | 0.029238 | 0 |
| B50 | B123 | eo% | hs-CRP1 | -0.00566 | 7.69E-07 |
| B50 | B166 | eo% | Pro | 0.014053 | 0 |
| B50 | B70 | eo% | LDL-C | 0.037368 | 0 |
| B50 | B71 | eo% | TG | 0.021696 | 0 |
| B50 | B72 | eo% | HDL-C | -0.03393 | 0 |
| B50 | B189 | eo% | eGFB | -0.08348 | 0 |
| B50 | B141 | eo% | AST/ALT | -0.03426 | 0 |
| B50 | B146 | eo% | CysC | 0.013231 | 0 |
| B50 | B190 | eo% | Cre | 0.101063 | 0 |
| B50 | B193 | eo% | K | 0.005944 | 2.11E-07 |
| B50 | B129 | eo% | IBIL | -0.01838 | 0 |
| B50 | B150 | eo% | ALP | 0.010741 | 0 |
| B50 | B222 | eo% | NS | 0.077173 | 0 |
| B50 | B85 | eo% | SCC | 0.014847 | 0 |
| B50 | B156 | eo% | IgE | 0.026791 | 0 |
| B50 | B176 | eo% | Ucr | 0.013611 | 0 |
| B50 | B191 | eo% | Urea | 0.041334 | 0 |
| B50 | B174 | eo% | Urea/Cr | -0.03721 | 0 |
| B50 | B192 | eo% | UA | 0.071782 | 0 |
| B50 | B130 | eo% | GLB | -0.01122 | 0 |
| B50 | B62 | eo% | WBHSRVI | 0.005826 | 3.64E-07 |
| B50 | B221 | eo% | Hei | 0.071713 | 0 |
| B50 | B220 | eo% | SBP | 0.030904 | 0 |

| | | | | | |
|---|---|---|---|---|---|
| B50 | B218 | eo% | DBP | 0.026646 | 0 |
| B50 | B216 | eo% | BMI | 0.066337 | 0 |
| B50 | B215 | eo% | BMI | 0.03911 | 0 |
| B50 | B131 | eo% | AST | 0.022935 | 0 |
| B50 | B172 | eo% | KB | -0.01757 | 0 |
| B50 | B210 | eo% | Hip | 0.041519 | 0 |
| B50 | B209 | eo% | Smoking | 0.095944 | 0 |
| B50 | B90 | eo% | CYFRA 21-1 | 0.014205 | 0 |
| B50 | B51 | eo% | Hb | 0.069618 | 0 |
| B50 | B173 | eo% | NTT | -0.00565 | 8.01E-07 |
| B50 | B207 | eo% | WHR | 0.008236 | 6.46E-13 |
| B50 | B206 | eo% | Waist | 0.07011 | 0 |
| B50 | B205 | eo% | KDA | 0.005884 | 2.79E-07 |
| B50 | B204 | eo% | Drinking | 0.058835 | 0 |
| B50 | B113 | eo% | FT3 | 0.008556 | 8.02E-14 |
| B50 | B211 | eo% | IOP-R | -0.009 | 4.00E-15 |
| B50 | B201 | eo% | e-habits | 0.011885 | 0 |
| B50 | B74 | eo% | ApoA1 | -0.00949 | 0 |
| B50 | B75 | eo% | ApoA1/B | -0.00914 | 1.55E-15 |
| B50 | B76 | eo% | ApoB | 0.009238 | 8.88E-16 |
| B50 | B132 | eo% | Dbil | -0.0242 | 0 |
| B50 | B53 | eo% | GR% | 0.02333 | 0 |
| B50 | B77 | eo% | TC | 0.025358 | 0 |
| B50 | B133 | eo% | TBIL | -0.02121 | 0 |
| B50 | B91 | eo% | TP | -0.038 | 0 |
| B50 | B212 | eo% | IOP-L | -0.00792 | 4.62E-12 |
| B51 | B137 | Hb | AFU | 0.036285 | 0 |
| B51 | B54 | Hb | ESR | -0.07548 | 0 |
| B51 | B160 | Hb | PH | -0.07245 | 0 |
| B51 | B140 | Hb | r-GT | 0.22511 | 0 |
| B51 | B143 | Hb | β-CTx | 0.007978 | 3.29E-12 |
| B51 | B118 | Hb | CEA1 | 0.014492 | 0 |
| B51 | B126 | Hb | Alb | 0.294475 | 0 |
| B51 | B127 | Hb | A-G ratio | 0.175962 | 0 |
| B51 | B161 | Hb | LEU | -0.11915 | 0 |
| B51 | B128 | Hb | ALT | 0.283834 | 0 |
| B51 | B164 | Hb | COM | -0.00677 | 3.35E-09 |
| B51 | B123 | Hb | hs-CRP1 | 0.006078 | 1.12E-07 |
| B51 | B111 | Hb | TSH | -0.0532 | 0 |
| B51 | B124 | Hb | ACTH | 0.01516 | 0 |
| B51 | B134 | Hb | CHE | 0.089699 | 0 |
| B51 | B166 | Hb | Pro | 0.147743 | 0 |
| B51 | B70 | Hb | LDL-C | 0.202292 | 0 |
| B51 | B188 | Hb | Ca | 0.053089 | 0 |

| | | | | | |
|---|---|---|---|---|---|
| B51 | B71 | Hb | TG | 0.239243 | 0 |
| B51 | B72 | Hb | HDL-C | -0.29033 | 0 |
| B51 | B189 | Hb | eGFB | -0.12971 | 0 |
| B51 | B141 | Hb | AST/ALT | -0.31128 | 0 |
| B51 | B146 | Hb | CysC | 0.029721 | 0 |
| B51 | B55 | Hb | TK | -0.02911 | 0 |
| B51 | B56 | Hb | IR | 0.085089 | 0 |
| B51 | B57 | Hb | RBCAI | 0.023204 | 0 |
| B51 | B58 | Hb | HCT | 0.023831 | 0 |
| B51 | B181 | Hb | APTT | -0.01224 | 0 |
| B51 | B190 | Hb | Cre | 0.502252 | 0 |
| B51 | B224 | Hb | pmh | 0.007871 | 6.32E-12 |
| B51 | B129 | Hb | IBIL | 0.169845 | 0 |
| B51 | B150 | Hb | ALP | 0.060886 | 0 |
| B51 | B84 | Hb | C-PSA | 0.027993 | 0 |
| B51 | B222 | Hb | NS | 0.532854 | 0 |
| B51 | B100 | Hb | anti-Ro52 | -0.00638 | 2.52E-08 |
| B51 | B103 | Hb | anti-SSA/Ro | -0.00931 | 4.44E-16 |
| B51 | B148 | Hb | Anti_TPO | -0.0517 | 0 |
| B51 | B149 | Hb | TGAb | -0.04743 | 0 |
| B51 | B198 | Hb | P | -0.04232 | 0 |
| B51 | B85 | Hb | SCC | 0.017093 | 0 |
| B51 | B194 | Hb | Cl | -0.03746 | 0 |
| B51 | B199 | Hb | Mg | 0.012575 | 0 |
| B51 | B156 | Hb | IgE | 0.008164 | 1.02E-12 |
| B51 | B157 | Hb | IgG | -0.02736 | 0 |
| B51 | B158 | Hb | IgM | -0.01634 | 0 |
| B51 | B195 | Hb | Na | 0.014727 | 0 |
| B51 | B176 | Hb | Ucr | 0.116492 | 0 |
| B51 | B178 | Hb | Ig-U | 0.006389 | 2.44E-08 |
| B51 | B191 | Hb | Urea | 0.111983 | 0 |
| B51 | B174 | Hb | Urea/Cr | -0.27033 | 0 |
| B51 | B192 | Hb | UA | 0.491369 | 0 |
| B51 | B170 | Hb | U-GLU | 0.055079 | 0 |
| B51 | B175 | Hb | ALB-U | 0.011217 | 0 |
| B51 | B177 | Hb | ACR | -0.00655 | 1.08E-08 |
| B51 | B179 | Hb | α1-MG | 0.034433 | 0 |
| B51 | B180 | Hb | Tf | 0.00968 | 0 |
| B51 | B182 | Hb | TT | 0.013835 | 0 |
| B51 | B125 | Hb | Cor | 0.015983 | 0 |
| B51 | B153 | Hb | Glu | 0.014654 | 0 |
| B51 | B138 | Hb | PAB | 0.100009 | 0 |
| B51 | B130 | Hb | GLB | -0.0878 | 0 |
| B51 | B59 | Hb | WBRV | 0.044083 | 0 |

| | | | | | |
|---|---|---|---|---|---|
| B51 | B60 | Hb | WBRVI | 0.147109 | 0 |
| B51 | B61 | Hb | WBHSRV | 0.08909 | 0 |
| B51 | B62 | Hb | WBHSRVI | 0.163495 | 0 |
| B51 | B63 | Hb | WBRSR(mPas_200) | 0.018274 | 0 |
| B51 | B64 | Hb | WBRSR(mPas_5) | 0.162584 | 0 |
| B51 | B65 | Hb | WBRSR(mPas_1) | 0.149934 | 0 |
| B51 | B66 | Hb | WBRSR(mPas_100) | 0.16454 | 0 |
| B51 | B67 | Hb | WBRSR(mPas_50) | 0.165173 | 0 |
| B51 | B86 | Hb | TBhCG | -0.01156 | 0 |
| B51 | B221 | Hb | Hei | 0.498251 | 0 |
| B51 | B87 | Hb | NSE | 0.020628 | 0 |
| B51 | B187 | Hb | NSE | 0.014108 | 0 |
| B51 | B200 | Hb | OP | 0.019422 | 0 |
| B51 | B220 | Hb | SBP | 0.207602 | 0 |
| B51 | B219 | Hb | OT | -0.04564 | 0 |
| B51 | B218 | Hb | DBP | 0.273879 | 0 |
| B51 | B88 | Hb | CA 242 | -0.02284 | 0 |
| B51 | B216 | Hb | BMI | 0.493654 | 0 |
| B51 | B215 | Hb | BMI | 0.305373 | 0 |
| B51 | B131 | Hb | AST | 0.147225 | 0 |
| B51 | B210 | Hb | Hip | 0.313404 | 0 |
| B51 | B209 | Hb | Smoking | 0.274759 | 0 |
| B51 | B90 | Hb | CYFRA 21-1 | 0.024156 | 0 |
| B51 | B139 | Hb | ADA | -0.01934 | 0 |
| B51 | B196 | Hb | TCO2 | 0.006322 | 3.41E-08 |
| B51 | B185 | Hb | PT | -0.01029 | 0 |
| B51 | B52 | Hb | PLT/BPC | -0.04153 | 0 |
| B51 | B173 | Hb | NTT | -0.05676 | 0 |
| B51 | B207 | Hb | WHR | 0.048574 | 0 |
| B51 | B206 | Hb | Waist | 0.420168 | 0 |
| B51 | B205 | Hb | KDA | -0.03399 | 0 |
| B51 | B197 | Hb | AG | 0.027878 | 0 |
| B51 | B204 | Hb | Drinking | 0.303026 | 0 |
| B51 | B203 | Hb | DH | 0.039617 | 0 |
| B51 | B112 | Hb | FT4 | 0.043181 | 0 |
| B51 | B113 | Hb | FT3 | 0.105316 | 0 |
| B51 | B211 | Hb | IOP-R | 0.02449 | 0 |
| B51 | B201 | Hb | e-habits | 0.024775 | 0 |
| B51 | B73 | Hb | ApoA | -0.01541 | 0 |
| B51 | B74 | Hb | ApoA1 | -0.07259 | 0 |
| B51 | B75 | Hb | ApoA1/B | -0.05641 | 0 |
| B51 | B76 | Hb | ApoB | 0.086624 | 0 |
| B51 | B132 | Hb | Dbil | 0.091654 | 0 |
| B51 | B53 | Hb | GR% | -0.00816 | 1.07E-12 |

| | | | | | |
|---|---|---|---|---|---|
| B51 | B81 | Hb | TSGF | -0.0116 | 0 |
| B51 | B77 | Hb | TC | 0.121989 | 0 |
| B51 | B133 | Hb | TBIL | 0.160915 | 0 |
| B51 | B135 | Hb | TBA | 0.012009 | 0 |
| B51 | B91 | Hb | TP | 0.107758 | 0 |
| B51 | B114 | Hb | TSH TT3 TT4(TT4) | -0.00952 | 0 |
| B51 | B115 | Hb | TSH TT3 TT4(TT3) | 0.033734 | 0 |
| B51 | B212 | Hb | IOP-L | 0.022583 | 0 |
| B52 | B54 | PLT/BPC | ESR | 0.017593 | 0 |
| B52 | B160 | PLT/BPC | PH | 0.030061 | 0 |
| B52 | B118 | PLT/BPC | CEA1 | -0.00953 | 0 |
| B52 | B126 | PLT/BPC | Alb | 0.007513 | 5.42E-11 |
| B52 | B127 | PLT/BPC | A-G ratio | -0.0868 | 0 |
| B52 | B161 | PLT/BPC | LEU | 0.018724 | 0 |
| B52 | B128 | PLT/BPC | ALT | -0.05388 | 0 |
| B52 | B123 | PLT/BPC | hs-CRP1 | 0.017242 | 0 |
| B52 | B111 | PLT/BPC | TSH | -0.02181 | 0 |
| B52 | B134 | PLT/BPC | CHE | 0.012541 | 0 |
| B52 | B166 | PLT/BPC | Pro | 0.006366 | 2.74E-08 |
| B52 | B70 | PLT/BPC | LDL-C | 0.074683 | 0 |
| B52 | B188 | PLT/BPC | Ca | 0.013505 | 0 |
| B52 | B71 | PLT/BPC | TG | -0.04154 | 0 |
| B52 | B72 | PLT/BPC | HDL-C | 0.01162 | 0 |
| B52 | B189 | PLT/BPC | eGFB | 0.159328 | 0 |
| B52 | B141 | PLT/BPC | AST/ALT | 0.053415 | 0 |
| B52 | B146 | PLT/BPC | CysC | -0.01885 | 0 |
| B52 | B190 | PLT/BPC | Cre | -0.09429 | 0 |
| B52 | B224 | PLT/BPC | pmh | -0.00984 | 0 |
| B52 | B193 | PLT/BPC | K | 0.025644 | 0 |
| B52 | B129 | PLT/BPC | IBIL | -0.08672 | 0 |
| B52 | B150 | PLT/BPC | ALP | -0.01897 | 0 |
| B52 | B214 | PLT/BPC | CVA-L | -0.00594 | 2.10E-07 |
| B52 | B222 | PLT/BPC | NS | -0.04059 | 0 |
| B52 | B148 | PLT/BPC | Anti_TPO | 0.009656 | 0 |
| B52 | B149 | PLT/BPC | TGAb | 0.007344 | 1.44E-10 |
| B52 | B198 | PLT/BPC | P | 0.031992 | 0 |
| B52 | B194 | PLT/BPC | Cl | -0.01308 | 0 |
| B52 | B155 | PLT/BPC | IgA | 0.008 | 2.86E-12 |
| B52 | B156 | PLT/BPC | IgE | 0.005981 | 1.78E-07 |
| B52 | B157 | PLT/BPC | IgG | 0.008686 | 3.38E-14 |
| B52 | B195 | PLT/BPC | Na | -0.01883 | 0 |
| B52 | B176 | PLT/BPC | Ucr | 0.006476 | 1.57E-08 |
| B52 | B191 | PLT/BPC | Urea | -0.12398 | 0 |
| B52 | B174 | PLT/BPC | Urea/Cr | -0.03316 | 0 |

| | | | | | |
|---|---|---|---|---|---|
| B52 | B192 | PLT/BPC | UA | -0.02731 | 0 |
| B52 | B170 | PLT/BPC | U-GLU | -0.02274 | 0 |
| B52 | B153 | PLT/BPC | Glu | -0.00743 | 8.95E-11 |
| B52 | B138 | PLT/BPC | PAB | 0.009739 | 0 |
| B52 | B130 | PLT/BPC | GLB | 0.1062 | 0 |
| B52 | B59 | PLT/BPC | WBRV | 0.010684 | 0 |
| B52 | B61 | PLT/BPC | WBHSRV | 0.011403 | 0 |
| B52 | B64 | PLT/BPC | WBRSR(mPas_5) | 0.009677 | 0 |
| B52 | B65 | PLT/BPC | WBRSR(mPas_1) | 0.008361 | 2.88E-13 |
| B52 | B66 | PLT/BPC | WBRSR(mPas_100) | 0.010364 | 0 |
| B52 | B67 | PLT/BPC | WBRSR(mPas_50) | 0.010379 | 0 |
| B52 | B87 | PLT/BPC | NSE | 0.009365 | 4.44E-16 |
| B52 | B187 | PLT/BPC | NSE | 0.005761 | 4.92E-07 |
| B52 | B200 | PLT/BPC | OP | -0.02153 | 0 |
| B52 | B220 | PLT/BPC | SBP | -0.03882 | 0 |
| B52 | B219 | PLT/BPC | OT | -0.02318 | 0 |
| B52 | B218 | PLT/BPC | DBP | -0.02134 | 0 |
| B52 | B88 | PLT/BPC | CA 242 | 0.006905 | 1.65E-09 |
| B52 | B89 | PLT/BPC | CA 72-4 | -0.00733 | 1.60E-10 |
| B52 | B215 | PLT/BPC | BMI | -0.01265 | 0 |
| B52 | B131 | PLT/BPC | AST | -0.06791 | 0 |
| B52 | B172 | PLT/BPC | KB | 0.006961 | 1.22E-09 |
| B52 | B210 | PLT/BPC | Hip | 0.009666 | 0 |
| B52 | B209 | PLT/BPC | Smoking | -0.07134 | 0 |
| B52 | B139 | PLT/BPC | ADA | -0.02212 | 0 |
| B52 | B183 | PLT/BPC | PTA | 0.011467 | 0 |
| B52 | B69 | PLT/BPC | PV | 0.014197 | 0 |
| B52 | B207 | PLT/BPC | WHR | -0.00771 | 1.70E-11 |
| B52 | B206 | PLT/BPC | Waist | -0.04948 | 0 |
| B52 | B205 | PLT/BPC | KDA | 0.009325 | 4.44E-16 |
| B52 | B78 | PLT/BPC | HBcAb | -0.02139 | 0 |
| B52 | B204 | PLT/BPC | Drinking | -0.05432 | 0 |
| B52 | B203 | PLT/BPC | DH | 0.025226 | 0 |
| B52 | B80 | PLT/BPC | anti-HP IgG | 0.005747 | 5.24E-07 |
| B52 | B112 | PLT/BPC | FT4 | 0.045683 | 0 |
| B52 | B211 | PLT/BPC | IOP-R | 0.019225 | 0 |
| B52 | B201 | PLT/BPC | e-habits | -0.02088 | 0 |
| B52 | B73 | PLT/BPC | ApoA | 0.00679 | 3.07E-09 |
| B52 | B74 | PLT/BPC | ApoA1 | -0.01919 | 0 |
| B52 | B75 | PLT/BPC | ApoA1/B | -0.01905 | 0 |
| B52 | B76 | PLT/BPC | ApoB | 0.03053 | 0 |
| B52 | B132 | PLT/BPC | Dbil | -0.1335 | 0 |
| B52 | B77 | PLT/BPC | TC | 0.035261 | 0 |
| B52 | B133 | PLT/BPC | TBIL | -0.10504 | 0 |

| | | | | | |
|---|---|---|---|---|---|
| B52 | B135 | PLT/BPC | TBA | -0.00786 | 6.61E-12 |
| B52 | B91 | PLT/BPC | TP | 0.095242 | 0 |
| B52 | B114 | PLT/BPC | TSH TT3 TT4(TT4) | 0.016661 | 0 |
| B52 | B212 | PLT/BPC | IOP-L | 0.015626 | 0 |
| B53 | B70 | GR% | LDL-C | -0.00676 | 3.62E-09 |
| B53 | B190 | GR% | Cre | -0.00892 | 6.66E-15 |
| B53 | B222 | GR% | NS | -0.00563 | 8.94E-07 |
| B53 | B192 | GR% | UA | -0.00612 | 9.17E-08 |
| B53 | B221 | GR% | Hei | -0.00966 | 0 |
| B53 | B216 | GR% | BMI | -0.00563 | 8.76E-07 |
| B53 | B77 | GR% | TC | -0.00576 | 4.83E-07 |
| B54 | B160 | ESR | PH | -0.00966 | 0 |
| B54 | B126 | ESR | Alb | -0.0415 | 0 |
| B54 | B127 | ESR | A-G ratio | -0.06539 | 0 |
| B54 | B161 | ESR | LEU | 0.024328 | 0 |
| B54 | B128 | ESR | ALT | -0.01122 | 0 |
| B54 | B123 | ESR | hs-CRP1 | 0.187069 | 0 |
| B54 | B111 | ESR | TSH | 0.01569 | 0 |
| B54 | B124 | ESR | ACTH | -0.00967 | 0 |
| B54 | B134 | ESR | CHE | -0.0095 | 0 |
| B54 | B166 | ESR | Pro | -0.00643 | 1.99E-08 |
| B54 | B70 | ESR | LDL-C | 0.014131 | 0 |
| B54 | B71 | ESR | TG | -0.00795 | 3.94E-12 |
| B54 | B72 | ESR | HDL-C | 0.017707 | 0 |
| B54 | B141 | ESR | AST/ALT | 0.023805 | 0 |
| B54 | B55 | ESR | TK | 0.037163 | 0 |
| B54 | B56 | ESR | IR | -0.16488 | 0 |
| B54 | B57 | ESR | RBCAI | -0.05474 | 0 |
| B54 | B58 | ESR | HCT | -0.04086 | 0 |
| B54 | B181 | ESR | APTT | 0.006208 | 5.96E-08 |
| B54 | B190 | ESR | Cre | -0.03407 | 0 |
| B54 | B193 | ESR | K | 0.011998 | 0 |
| B54 | B129 | ESR | IBIL | -0.0295 | 0 |
| B54 | B150 | ESR | ALP | 0.013957 | 0 |
| B54 | B222 | ESR | NS | -0.03538 | 0 |
| B54 | B105 | ESR | anti-U1RNP | 0.00661 | 7.90E-09 |
| B54 | B96 | ESR | dsDNA (IIF) | 0.01028 | 0 |
| B54 | B98 | ESR | anti-Jo-1 | -0.00743 | 8.83E-11 |
| B54 | B100 | ESR | anti-Ro52 | 0.013715 | 0 |
| B54 | B101 | ESR | anti-Sc70 | 0.012708 | 0 |
| B54 | B102 | ESR | anti-Sm | 0.007135 | 4.68E-10 |
| B54 | B103 | ESR | anti-SSA/Ro | 0.030426 | 0 |
| B54 | B106 | ESR | anti-P | 0.007521 | 5.17E-11 |
| B54 | B107 | ESR | AnuA | 0.018133 | 0 |

| | | | | | |
|---|---|---|---|---|---|
| B54 | B151 | ESR | Anti-CCP | 0.0299 | 0 |
| B54 | B148 | ESR | Anti_TPO | 0.014474 | 0 |
| B54 | B149 | ESR | TGAb | 0.015391 | 0 |
| B54 | B110 | ESR | AHA | 0.005961 | 1.95E-07 |
| B54 | B154 | ESR | RF | 0.013551 | 0 |
| B54 | B198 | ESR | P | 0.024557 | 0 |
| B54 | B155 | ESR | IgA | 0.054268 | 0 |
| B54 | B156 | ESR | IgE | 0.011768 | 0 |
| B54 | B157 | ESR | IgG | 0.077813 | 0 |
| B54 | B158 | ESR | IgM | 0.023271 | 0 |
| B54 | B195 | ESR | Na | -0.01939 | 0 |
| B54 | B176 | ESR | Ucr | -0.01876 | 0 |
| B54 | B178 | ESR | Ig-U | 0.006435 | 1.93E-08 |
| B54 | B174 | ESR | Urea/Cr | 0.02693 | 0 |
| B54 | B192 | ESR | UA | -0.02738 | 0 |
| B54 | B175 | ESR | ALB-U | 0.022258 | 0 |
| B54 | B177 | ESR | ACR | 0.028607 | 0 |
| B54 | B182 | ESR | TT | -0.03893 | 0 |
| B54 | B125 | ESR | Cor | -0.00622 | 5.56E-08 |
| B54 | B138 | ESR | PAB | -0.04933 | 0 |
| B54 | B130 | ESR | GLB | 0.078945 | 0 |
| B54 | B59 | ESR | WBRV | 0.027191 | 0 |
| B54 | B60 | ESR | WBRVI | -0.27842 | 0 |
| B54 | B61 | ESR | WBHSRV | -0.0464 | 0 |
| B54 | B62 | ESR | WBHSRVI | -0.29596 | 0 |
| B54 | B63 | ESR | WBRSR(mPas_200) | -0.01987 | 0 |
| B54 | B64 | ESR | WBRSR(mPas_5) | -0.16797 | 0 |
| B54 | B65 | ESR | WBRSR(mPas_1) | -0.16428 | 0 |
| B54 | B66 | ESR | WBRSR(mPas_100) | -0.15624 | 0 |
| B54 | B67 | ESR | WBRSR(mPas_50) | -0.1602 | 0 |
| B54 | B221 | ESR | Hei | -0.05114 | 0 |
| B54 | B200 | ESR | OP | -0.01117 | 0 |
| B54 | B220 | ESR | SBP | 0.011958 | 0 |
| B54 | B219 | ESR | OT | 0.007445 | 8.06E-11 |
| B54 | B89 | ESR | CA 72-4 | -0.0066 | 8.32E-09 |
| B54 | B216 | ESR | BMI | -0.03067 | 0 |
| B54 | B131 | ESR | AST | 0.005725 | 5.79E-07 |
| B54 | B210 | ESR | Hip | -0.01119 | 0 |
| B54 | B209 | ESR | Smoking | -0.02153 | 0 |
| B54 | B139 | ESR | ADA | 0.047935 | 0 |
| B54 | B196 | ESR | TCO2 | -0.00906 | 2.44E-15 |
| B54 | B183 | ESR | PTA | 0.007874 | 6.21E-12 |
| B54 | B185 | ESR | PT | 0.02255 | 0 |
| B54 | B186 | ESR | PFC | 0.105067 | 0 |

| | | | | | |
|---|---|---|---|---|---|
| B54 | B69 | ESR | PV | 0.350583 | 0 |
| B54 | B173 | ESR | NTT | 0.017116 | 0 |
| B54 | B206 | ESR | Waist | -0.01405 | 0 |
| B54 | B197 | ESR | AG | -0.00724 | 2.60E-10 |
| B54 | B204 | ESR | Drinking | -0.02964 | 0 |
| B54 | B112 | ESR | FT4 | 0.018041 | 0 |
| B54 | B113 | ESR | FT3 | -0.00738 | 1.17E-10 |
| B54 | B211 | ESR | IOP-R | -0.01557 | 0 |
| B54 | B73 | ESR | ApoA | 0.033934 | 0 |
| B54 | B74 | ESR | ApoA1 | 0.041023 | 0 |
| B54 | B76 | ESR | ApoB | 0.027026 | 0 |
| B54 | B132 | ESR | Dbil | -0.02549 | 0 |
| B54 | B77 | ESR | TC | 0.015769 | 0 |
| B54 | B133 | ESR | TBIL | -0.03023 | 0 |
| B54 | B135 | ESR | TBA | -0.00705 | 7.35E-10 |
| B54 | B91 | ESR | TP | 0.041896 | 0 |
| B54 | B114 | ESR | TSH TT3 TT4(TT4) | 0.053628 | 0 |
| B54 | B212 | ESR | IOP-L | -0.01663 | 0 |
| B55 | B137 | TK | AFU | -0.01159 | 0 |
| B55 | B126 | TK | Alb | -0.00967 | 0 |
| B55 | B124 | TK | ACTH | -0.01029 | 0 |
| B55 | B134 | TK | CHE | -0.00766 | 2.32E-11 |
| B55 | B70 | TK | LDL-C | -0.0089 | 7.55E-15 |
| B55 | B56 | TK | IR | 0.692617 | 0 |
| B55 | B57 | TK | RBCAI | 0.110263 | 0 |
| B55 | B58 | TK | HCT | -0.09445 | 0 |
| B55 | B181 | TK | APTT | -0.00712 | 4.98E-10 |
| B55 | B190 | TK | Cre | -0.01397 | 0 |
| B55 | B193 | TK | K | -0.01144 | 0 |
| B55 | B129 | TK | IBIL | -0.00915 | 1.33E-15 |
| B55 | B222 | TK | NS | -0.00735 | 1.42E-10 |
| B55 | B107 | TK | AnuA | -0.00602 | 1.46E-07 |
| B55 | B155 | TK | IgA | 0.011947 | 0 |
| B55 | B158 | TK | IgM | 0.006012 | 1.53E-07 |
| B55 | B176 | TK | Ucr | -0.00848 | 1.36E-13 |
| B55 | B174 | TK | Urea/Cr | 0.008217 | 7.30E-13 |
| B55 | B192 | TK | UA | -0.00828 | 4.98E-13 |
| B55 | B125 | TK | Cor | 0.012372 | 0 |
| B55 | B138 | TK | PAB | -0.00667 | 5.84E-09 |
| B55 | B59 | TK | WBRV | 0.524444 | 0 |
| B55 | B60 | TK | WBRVI | 0.071851 | 0 |
| B55 | B61 | TK | WBHSRV | 0.652153 | 0 |
| B55 | B62 | TK | WBHSRVI | 0.159897 | 0 |
| B55 | B64 | TK | WBRSR(mPas_5) | 0.030805 | 0 |

| | | | | | |
|---|---|---|---|---|---|
| B55 | B66 | TK | WBRSR(mPas_100) | 0.079978 | 0 |
| B55 | B67 | TK | WBRSR(mPas_50) | 0.07384 | 0 |
| B55 | B221 | TK | Hei | -0.01181 | 0 |
| B55 | B209 | TK | Smoking | -0.01075 | 0 |
| B55 | B183 | TK | PTA | -0.02015 | 0 |
| B55 | B69 | TK | PV | -0.19268 | 0 |
| B55 | B78 | TK | HBcAb | 0.007489 | 6.23E-11 |
| B55 | B204 | TK | Drinking | -0.00748 | 6.65E-11 |
| B55 | B74 | TK | ApoA1 | -0.01123 | 0 |
| B55 | B76 | TK | ApoB | -0.00924 | 8.88E-16 |
| B55 | B77 | TK | TC | -0.00571 | 6.21E-07 |
| B55 | B133 | TK | TBIL | -0.00734 | 1.49E-10 |
| B55 | B91 | TK | TP | -0.00652 | 1.23E-08 |
| B56 | B137 | IR | AFU | 0.016021 | 0 |
| B56 | B160 | IR | PH | -0.00839 | 2.33E-13 |
| B56 | B140 | IR | r-GT | 0.031289 | 0 |
| B56 | B126 | IR | Alb | 0.015421 | 0 |
| B56 | B127 | IR | A-G ratio | 0.018033 | 0 |
| B56 | B161 | IR | LEU | -0.01008 | 0 |
| B56 | B128 | IR | ALT | 0.025373 | 0 |
| B56 | B121 | IR | C3 | 0.007462 | 7.27E-11 |
| B56 | B111 | IR | TSH | -0.01306 | 0 |
| B56 | B124 | IR | ACTH | 0.014594 | 0 |
| B56 | B134 | IR | CHE | 0.038374 | 0 |
| B56 | B166 | IR | Pro | 0.012799 | 0 |
| B56 | B70 | IR | LDL-C | 0.012004 | 0 |
| B56 | B71 | IR | TG | 0.024878 | 0 |
| B56 | B72 | IR | HDL-C | -0.03063 | 0 |
| B56 | B141 | IR | AST/ALT | -0.02763 | 0 |
| B56 | B57 | IR | RBCAI | -0.07951 | 0 |
| B56 | B58 | IR | HCT | -0.01457 | 0 |
| B56 | B181 | IR | APTT | -0.02043 | 0 |
| B56 | B190 | IR | Cre | 0.03468 | 0 |
| B56 | B193 | IR | K | -0.00582 | 3.80E-07 |
| B56 | B129 | IR | IBIL | 0.013126 | 0 |
| B56 | B150 | IR | ALP | 0.018889 | 0 |
| B56 | B222 | IR | NS | 0.051298 | 0 |
| B56 | B96 | IR | dsDNA (IIF) | -0.00576 | 4.94E-07 |
| B56 | B98 | IR | anti-Jo-1 | 0.007096 | 5.82E-10 |
| B56 | B100 | IR | anti-Ro52 | -0.00729 | 1.94E-10 |
| B56 | B103 | IR | anti-SSA/Ro | -0.0172 | 0 |
| B56 | B107 | IR | AnuA | -0.01125 | 0 |
| B56 | B148 | IR | Anti_TPO | -0.01032 | 0 |
| B56 | B149 | IR | TGAb | -0.0099 | 0 |

| | | | | | |
|---|---|---|---|---|---|
| B56 | B110 | IR | AHA | -0.00881 | 1.49E-14 |
| B56 | B198 | IR | P | -0.02018 | 0 |
| B56 | B194 | IR | Cl | -0.03314 | 0 |
| B56 | B199 | IR | Mg | 0.006916 | 1.56E-09 |
| B56 | B155 | IR | IgA | 0.009784 | 0 |
| B56 | B157 | IR | IgG | -0.02632 | 0 |
| B56 | B158 | IR | IgM | -0.00994 | 0 |
| B56 | B176 | IR | Ucr | 0.023007 | 0 |
| B56 | B174 | IR | Urea/Cr | -0.02388 | 0 |
| B56 | B192 | IR | UA | 0.043865 | 0 |
| B56 | B170 | IR | U-GLU | 0.01144 | 0 |
| B56 | B177 | IR | ACR | -0.0064 | 2.25E-08 |
| B56 | B179 | IR | α1-MG | 0.019526 | 0 |
| B56 | B182 | IR | TT | 0.019082 | 0 |
| B56 | B125 | IR | Cor | 0.023453 | 0 |
| B56 | B138 | IR | PAB | 0.058108 | 0 |
| B56 | B130 | IR | GLB | -0.0134 | 0 |
| B56 | B59 | IR | WBRV | 0.616791 | 0 |
| B56 | B60 | IR | WBRVI | 0.588248 | 0 |
| B56 | B61 | IR | WBHSRV | 0.937299 | 0 |
| B56 | B62 | IR | WBHSRVI | 0.772861 | 0 |
| B56 | B63 | IR | WBRSR(mPas_200) | 0.064732 | 0 |
| B56 | B64 | IR | WBRSR(mPas_5) | 0.577484 | 0 |
| B56 | B65 | IR | WBRSR(mPas_1) | 0.496499 | 0 |
| B56 | B66 | IR | WBRSR(mPas_100) | 0.659466 | 0 |
| B56 | B67 | IR | WBRSR(mPas_50) | 0.652239 | 0 |
| B56 | B221 | IR | Hei | 0.041068 | 0 |
| B56 | B187 | IR | NSE | 0.007116 | 5.20E-10 |
| B56 | B200 | IR | OP | 0.010247 | 0 |
| B56 | B220 | IR | SBP | 0.013658 | 0 |
| B56 | B218 | IR | DBP | 0.022354 | 0 |
| B56 | B216 | IR | BMI | 0.04585 | 0 |
| B56 | B215 | IR | BMI | 0.031273 | 0 |
| B56 | B131 | IR | AST | 0.009604 | 0 |
| B56 | B210 | IR | Hip | 0.031274 | 0 |
| B56 | B209 | IR | Smoking | 0.026892 | 0 |
| B56 | B90 | IR | CYFRA 21-1 | 0.006046 | 1.31E-07 |
| B56 | B139 | IR | ADA | -0.00605 | 1.29E-07 |
| B56 | B183 | IR | PTA | -0.02766 | 0 |
| B56 | B185 | IR | PT | -0.01046 | 0 |
| B56 | B186 | IR | PFC | -0.00873 | 2.44E-14 |
| B56 | B69 | IR | PV | -0.24745 | 0 |
| B56 | B173 | IR | NTT | -0.00832 | 3.87E-13 |
| B56 | B206 | IR | Waist | 0.043409 | 0 |

| | | | | | |
|---|---|---|---|---|---|
| B56 | B78 | IR | HBcAb | 0.008689 | 3.31E-14 |
| B56 | B197 | IR | AG | 0.019582 | 0 |
| B56 | B204 | IR | Drinking | 0.028372 | 0 |
| B56 | B113 | IR | FT3 | 0.01978 | 0 |
| B56 | B211 | IR | IOP-R | 0.011658 | 0 |
| B56 | B73 | IR | ApoA | -0.01303 | 0 |
| B56 | B74 | IR | ApoA1 | -0.07043 | 0 |
| B56 | B75 | IR | ApoA1/B | -0.03871 | 0 |
| B56 | B76 | IR | ApoB | 0.048421 | 0 |
| B56 | B132 | IR | Dbil | 0.014359 | 0 |
| B56 | B77 | IR | TC | 0.006292 | 3.96E-08 |
| B56 | B133 | IR | TBIL | 0.014329 | 0 |
| B56 | B114 | IR | TSH TT3 TT4(TT4) | -0.01478 | 0 |
| B56 | B115 | IR | TSH TT3 TT4(TT3) | 0.010913 | 0 |
| B56 | B212 | IR | IOP-L | 0.010168 | 0 |
| B57 | B92 | RBCAI | ALDH2-Glu504Glu | -0.00807 | 1.82E-12 |
| B57 | B93 | RBCAI | ALDH2-Glu504Lys | 0.007726 | 1.52E-11 |
| B57 | B126 | RBCAI | Alb | 0.013638 | 0 |
| B57 | B127 | RBCAI | A-G ratio | 0.018305 | 0 |
| B57 | B161 | RBCAI | LEU | -0.00685 | 2.24E-09 |
| B57 | B123 | RBCAI | hs-CRP1 | -0.01716 | 0 |
| B57 | B111 | RBCAI | TSH | -0.00785 | 7.36E-12 |
| B57 | B124 | RBCAI | ACTH | 0.011525 | 0 |
| B57 | B134 | RBCAI | CHE | 0.006856 | 2.16E-09 |
| B57 | B71 | RBCAI | TG | -0.00735 | 1.38E-10 |
| B57 | B72 | RBCAI | HDL-C | 0.008919 | 6.88E-15 |
| B57 | B58 | RBCAI | HCT | 0.012272 | 0 |
| B57 | B181 | RBCAI | APTT | -0.01504 | 0 |
| B57 | B190 | RBCAI | Cre | 0.014577 | 0 |
| B57 | B193 | RBCAI | K | 0.00797 | 3.44E-12 |
| B57 | B129 | RBCAI | IBIL | 0.00601 | 1.55E-07 |
| B57 | B222 | RBCAI | NS | 0.013231 | 0 |
| B57 | B96 | RBCAI | dsDNA (IIF) | 0.006567 | 9.87E-09 |
| B57 | B100 | RBCAI | anti-Ro52 | -0.00727 | 2.19E-10 |
| B57 | B103 | RBCAI | anti-SSA/Ro | -0.00627 | 4.29E-08 |
| B57 | B104 | RBCAI | anti-SSB | -0.00788 | 5.88E-12 |
| B57 | B107 | RBCAI | AnuA | 0.005955 | 2.00E-07 |
| B57 | B151 | RBCAI | Anti-CCP | 0.016623 | 0 |
| B57 | B109 | RBCAI | AMA-M2 | -0.00651 | 1.31E-08 |
| B57 | B194 | RBCAI | Cl | 0.015345 | 0 |
| B57 | B158 | RBCAI | IgM | -0.00702 | 8.75E-10 |
| B57 | B176 | RBCAI | Ucr | 0.01157 | 0 |
| B57 | B174 | RBCAI | Urea/Cr | -0.01259 | 0 |
| B57 | B192 | RBCAI | UA | 0.009341 | 4.44E-16 |

| | | | | | |
|---|---|---|---|---|---|
| B57 | B125 | RBCAI | Cor | 0.025294 | 0 |
| B57 | B138 | RBCAI | PAB | 0.009412 | 2.22E-16 |
| B57 | B130 | RBCAI | GLB | -0.01903 | 0 |
| B57 | B59 | RBCAI | WBRV | 0.46308 | 0 |
| B57 | B60 | RBCAI | WBRVI | 0.453115 | 0 |
| B57 | B61 | RBCAI | WBHSRV | -0.07802 | 0 |
| B57 | B62 | RBCAI | WBHSRVI | -0.03042 | 0 |
| B57 | B64 | RBCAI | WBRSR(mPas_5) | 0.293588 | 0 |
| B57 | B65 | RBCAI | WBRSR(mPas_1) | 0.448014 | 0 |
| B57 | B67 | RBCAI | WBRSR(mPas_50) | 0.051256 | 0 |
| B57 | B86 | RBCAI | TBhCG | -0.00686 | 2.09E-09 |
| B57 | B221 | RBCAI | Hei | 0.015363 | 0 |
| B57 | B216 | RBCAI | BMI | 0.008185 | 8.96E-13 |
| B57 | B209 | RBCAI | Smoking | 0.013965 | 0 |
| B57 | B90 | RBCAI | CYFRA 21-1 | 0.006445 | 1.84E-08 |
| B57 | B139 | RBCAI | ADA | -0.02017 | 0 |
| B57 | B183 | RBCAI | PTA | -0.03768 | 0 |
| B57 | B184 | RBCAI | INR | 0.0166 | 0 |
| B57 | B186 | RBCAI | PFC | 0.00831 | 4.02E-13 |
| B57 | B69 | RBCAI | PV | 0.011574 | 0 |
| B57 | B206 | RBCAI | Waist | 0.00565 | 8.09E-07 |
| B57 | B78 | RBCAI | HBcAb | -0.01921 | 0 |
| B57 | B204 | RBCAI | Drinking | 0.013301 | 0 |
| B57 | B112 | RBCAI | FT4 | 0.01791 | 0 |
| B57 | B113 | RBCAI | FT3 | 0.006878 | 1.91E-09 |
| B57 | B211 | RBCAI | IOP-R | 0.010078 | 0 |
| B57 | B73 | RBCAI | ApoA | -0.00588 | 2.84E-07 |
| B57 | B74 | RBCAI | ApoA1 | 0.03585 | 0 |
| B57 | B75 | RBCAI | ApoA1/B | 0.016023 | 0 |
| B57 | B76 | RBCAI | ApoB | -0.01928 | 0 |
| B57 | B81 | RBCAI | TSGF | -0.04164 | 0 |
| B57 | B91 | RBCAI | TP | -0.00782 | 8.42E-12 |
| B57 | B212 | RBCAI | IOP-L | 0.012163 | 0 |
| B58 | B140 | HCT | r-GT | 0.006346 | 3.01E-08 |
| B58 | B126 | HCT | Alb | 0.005902 | 2.57E-07 |
| B58 | B128 | HCT | ALT | 0.006803 | 2.85E-09 |
| B58 | B72 | HCT | HDL-C | -0.00663 | 6.97E-09 |
| B58 | B141 | HCT | AST/ALT | -0.00603 | 1.40E-07 |
| B58 | B190 | HCT | Cre | 0.009654 | 0 |
| B58 | B222 | HCT | NS | 0.012207 | 0 |
| B58 | B194 | HCT | Cl | -0.0084 | 2.27E-13 |
| B58 | B157 | HCT | IgG | -0.00849 | 1.28E-13 |
| B58 | B174 | HCT | Urea/Cr | -0.006 | 1.61E-07 |
| B58 | B192 | HCT | UA | 0.010366 | 0 |

| | | | | | |
|---|---|---|---|---|---|
| B58 | B138 | HCT | PAB | 0.011106 | 0 |
| B58 | B59 | HCT | WBRV | -0.03671 | 0 |
| B58 | B60 | HCT | WBRVI | 0.084846 | 0 |
| B58 | B61 | HCT | WBHSRV | -0.01501 | 0 |
| B58 | B62 | HCT | WBHSRVI | 0.094695 | 0 |
| B58 | B63 | HCT | WBRSR(mPas_200) | 0.010419 | 0 |
| B58 | B64 | HCT | WBRSR(mPas_5) | 0.094997 | 0 |
| B58 | B65 | HCT | WBRSR(mPas_1) | 0.08734 | 0 |
| B58 | B66 | HCT | WBRSR(mPas_100) | 0.097165 | 0 |
| B58 | B67 | HCT | WBRSR(mPas_50) | 0.097515 | 0 |
| B58 | B221 | HCT | Hei | 0.009857 | 0 |
| B58 | B218 | HCT | DBP | 0.006593 | 8.60E-09 |
| B58 | B216 | HCT | BMI | 0.010476 | 0 |
| B58 | B215 | HCT | BMI | 0.00711 | 5.38E-10 |
| B58 | B210 | HCT | Hip | 0.007777 | 1.12E-11 |
| B58 | B209 | HCT | Smoking | 0.010598 | 0 |
| B58 | B206 | HCT | Waist | 0.010086 | 0 |
| B58 | B204 | HCT | Drinking | 0.006414 | 2.15E-08 |
| B58 | B74 | HCT | ApoA1 | -0.0104 | 0 |
| B58 | B75 | HCT | ApoA1/B | -0.00724 | 2.59E-10 |
| B58 | B76 | HCT | ApoB | 0.009337 | 4.44E-16 |
| B59 | B137 | WBRV | AFU | 0.006855 | 2.16E-09 |
| B59 | B160 | WBRV | PH | -0.00908 | 2.22E-15 |
| B59 | B140 | WBRV | r-GT | 0.020313 | 0 |
| B59 | B126 | WBRV | Alb | 0.02348 | 0 |
| B59 | B128 | WBRV | ALT | 0.013684 | 0 |
| B59 | B121 | WBRV | C3 | 0.007129 | 4.85E-10 |
| B59 | B123 | WBRV | hs-CRP1 | 0.023684 | 0 |
| B59 | B111 | WBRV | TSH | -0.00643 | 1.97E-08 |
| B59 | B124 | WBRV | ACTH | 0.014672 | 0 |
| B59 | B134 | WBRV | CHE | 0.028496 | 0 |
| B59 | B166 | WBRV | Pro | 0.008776 | 1.82E-14 |
| B59 | B70 | WBRV | LDL-C | 0.014758 | 0 |
| B59 | B71 | WBRV | TG | 0.015289 | 0 |
| B59 | B141 | WBRV | AST/ALT | -0.00744 | 8.13E-11 |
| B59 | B181 | WBRV | APTT | -0.02417 | 0 |
| B59 | B190 | WBRV | Cre | 0.020222 | 0 |
| B59 | B193 | WBRV | K | 0.009804 | 0 |
| B59 | B129 | WBRV | IBIL | 0.009836 | 0 |
| B59 | B150 | WBRV | ALP | 0.016363 | 0 |
| B59 | B222 | WBRV | NS | 0.027082 | 0 |
| B59 | B99 | WBRV | anti-PM-Scl | 0.00712 | 5.10E-10 |
| B59 | B104 | WBRV | anti-SSB | -0.00947 | 0 |
| B59 | B151 | WBRV | Anti-CCP | 0.015524 | 0 |

| | | | | | |
|---|---|---|---|---|---|
| B59 | B198 | WBRV | P | -0.00924 | 8.88E-16 |
| B59 | B194 | WBRV | Cl | -0.01838 | 0 |
| B59 | B155 | WBRV | IgA | 0.020994 | 0 |
| B59 | B156 | WBRV | IgE | 0.009207 | 8.88E-16 |
| B59 | B157 | WBRV | IgG | 0.022984 | 0 |
| B59 | B195 | WBRV | Na | -0.00922 | 8.88E-16 |
| B59 | B176 | WBRV | Ucr | 0.014442 | 0 |
| B59 | B174 | WBRV | Urea/Cr | -0.01533 | 0 |
| B59 | B192 | WBRV | UA | 0.025777 | 0 |
| B59 | B170 | WBRV | U-GLU | 0.009921 | 0 |
| B59 | B179 | WBRV | α1-MG | 0.013709 | 0 |
| B59 | B182 | WBRV | TT | -0.00645 | 1.83E-08 |
| B59 | B125 | WBRV | Cor | 0.038132 | 0 |
| B59 | B138 | WBRV | PAB | 0.031083 | 0 |
| B59 | B130 | WBRV | GLB | 0.010883 | 0 |
| B59 | B60 | WBRV | WBRVI | 0.639131 | 0 |
| B59 | B61 | WBRV | WBHSRV | 0.701324 | 0 |
| B59 | B62 | WBRV | WBHSRVI | 0.355169 | 0 |
| B59 | B63 | WBRV | WBRSR(mPas_200) | 0.044243 | 0 |
| B59 | B64 | WBRV | WBRSR(mPas_5) | 0.615759 | 0 |
| B59 | B65 | WBRV | WBRSR(mPas_1) | 0.679647 | 0 |
| B59 | B66 | WBRV | WBRSR(mPas_100) | 0.437669 | 0 |
| B59 | B67 | WBRV | WBRSR(mPas_50) | 0.469037 | 0 |
| B59 | B221 | WBRV | Hei | 0.019678 | 0 |
| B59 | B220 | WBRV | SBP | 0.008061 | 1.96E-12 |
| B59 | B218 | WBRV | DBP | 0.017004 | 0 |
| B59 | B216 | WBRV | BMI | 0.022065 | 0 |
| B59 | B215 | WBRV | BMI | 0.015087 | 0 |
| B59 | B131 | WBRV | AST | 0.011704 | 0 |
| B59 | B210 | WBRV | Hip | 0.01725 | 0 |
| B59 | B209 | WBRV | Smoking | 0.014934 | 0 |
| B59 | B90 | WBRV | CYFRA 21-1 | 0.005852 | 3.23E-07 |
| B59 | B183 | WBRV | PTA | -0.02934 | 0 |
| B59 | B184 | WBRV | INR | 0.018696 | 0 |
| B59 | B186 | WBRV | PFC | 0.030527 | 0 |
| B59 | B69 | WBRV | PV | 0.148166 | 0 |
| B59 | B206 | WBRV | Waist | 0.023277 | 0 |
| B59 | B78 | WBRV | HBcAb | -0.01004 | 0 |
| B59 | B197 | WBRV | AG | 0.00734 | 1.47E-10 |
| B59 | B204 | WBRV | Drinking | 0.015424 | 0 |
| B59 | B112 | WBRV | FT4 | 0.027051 | 0 |
| B59 | B113 | WBRV | FT3 | 0.013634 | 0 |
| B59 | B211 | WBRV | IOP-R | 0.014035 | 0 |
| B59 | B75 | WBRV | ApoA1/B | -0.00635 | 2.96E-08 |

| | | | | | |
|---|---|---|---|---|---|
| B59 | B76 | WBRV | ApoB | 0.021793 | 0 |
| B59 | B81 | WBRV | TSGF | -0.0415 | 0 |
| B59 | B77 | WBRV | TC | 0.008702 | 3.02E-14 |
| B59 | B133 | WBRV | TBIL | 0.008564 | 7.59E-14 |
| B59 | B91 | WBRV | TP | 0.023977 | 0 |
| B59 | B114 | WBRV | TSH TT3 TT4(TT4) | 0.008511 | 1.08E-13 |
| B59 | B115 | WBRV | TSH TT3 TT4(TT3) | 0.007399 | 1.05E-10 |
| B59 | B212 | WBRV | IOP-L | 0.01332 | 0 |
| B6 | B137 | WB-Fe | AFU | 0.00565 | 8.11E-07 |
| B6 | B140 | WB-Fe | r-GT | 0.005998 | 1.64E-07 |
| B6 | B126 | WB-Fe | Alb | 0.006532 | 1.18E-08 |
| B6 | B40 | WB-Fe | WBC | 0.006331 | 3.25E-08 |
| B6 | B128 | WB-Fe | ALT | 0.008575 | 7.08E-14 |
| B6 | B29 | WB-Fe | E2 | -0.00562 | 9.33E-07 |
| B6 | B134 | WB-Fe | CHE | 0.008973 | 4.88E-15 |
| B6 | B70 | WB-Fe | LDL-C | 0.006048 | 1.29E-07 |
| B6 | B72 | WB-Fe | HDL-C | -0.00726 | 2.34E-10 |
| B6 | B32 | WB-Fe | Testo | 0.014461 | 0 |
| B6 | B189 | WB-Fe | eGFB | -0.00612 | 9.28E-08 |
| B6 | B141 | WB-Fe | AST/ALT | -0.00957 | 0 |
| B6 | B145 | WB-Fe | BGP | -0.00677 | 3.40E-09 |
| B6 | B44 | WB-Fe | MCH | 0.012148 | 0 |
| B6 | B45 | WB-Fe | MCHC | 0.017503 | 0 |
| B6 | B46 | WB-Fe | RBC Count | 0.020118 | 0 |
| B6 | B190 | WB-Fe | Cre | 0.014423 | 0 |
| B6 | B129 | WB-Fe | IBIL | 0.006605 | 8.09E-09 |
| B6 | B222 | WB-Fe | NS | 0.014153 | 0 |
| B6 | B195 | WB-Fe | Na | 0.005676 | 7.23E-07 |
| B6 | B176 | WB-Fe | Ucr | 0.006342 | 3.08E-08 |
| B6 | B174 | WB-Fe | Urea/Cr | -0.00762 | 2.94E-11 |
| B6 | B192 | WB-Fe | UA | 0.014128 | 0 |
| B6 | B138 | WB-Fe | PAB | 0.012725 | 0 |
| B6 | B60 | WB-Fe | WBRVI | 0.008325 | 3.65E-13 |
| B6 | B62 | WB-Fe | WBHSRVI | 0.009671 | 0 |
| B6 | B7 | WB-Fe | WB-Cop | -0.0158 | 0 |
| B6 | B8 | WB-Fe | WB-Zinc | 0.567008 | 0 |
| B6 | B64 | WB-Fe | WBRSR(mPas_5) | 0.008891 | 8.44E-15 |
| B6 | B65 | WB-Fe | WBRSR(mPas_1) | 0.008085 | 1.68E-12 |
| B6 | B66 | WB-Fe | WBRSR(mPas_100) | 0.009065 | 2.44E-15 |
| B6 | B67 | WB-Fe | WBRSR(mPas_50) | 0.009036 | 3.11E-15 |
| B6 | B27 | WB-Fe | LDH | 0.006113 | 9.47E-08 |
| B6 | B221 | WB-Fe | Hei | 0.013266 | 0 |
| B6 | B200 | WB-Fe | OP | 0.00671 | 4.68E-09 |
| B6 | B220 | WB-Fe | SBP | 0.007201 | 3.24E-10 |

| | | | | | |
|---|---|---|---|---|---|
| B6 | B218 | WB-Fe | DBP | 0.007711 | 1.67E-11 |
| B6 | B216 | WB-Fe | BMI | 0.013835 | 0 |
| B6 | B215 | WB-Fe | BMI | 0.008937 | 6.22E-15 |
| B6 | B210 | WB-Fe | Hip | 0.008656 | 4.13E-14 |
| B6 | B209 | WB-Fe | Smoking | 0.006995 | 1.02E-09 |
| B6 | B51 | WB-Fe | Hb | 0.029163 | 0 |
| B6 | B206 | WB-Fe | Waist | 0.011707 | 0 |
| B6 | B204 | WB-Fe | Drinking | 0.007805 | 9.50E-12 |
| B6 | B113 | WB-Fe | FT3 | 0.008991 | 4.00E-15 |
| B6 | B76 | WB-Fe | ApoB | 0.007388 | 1.12E-10 |
| B6 | B133 | WB-Fe | TBIL | 0.006179 | 6.86E-08 |
| B60 | B137 | WBRVI | AFU | 0.028921 | 0 |
| B60 | B92 | WBRVI | ALDH2-Glu504Glu | -0.00861 | 5.51E-14 |
| B60 | B93 | WBRVI | ALDH2-Glu504Lys | 0.007373 | 1.22E-10 |
| B60 | B160 | WBRVI | PH | -0.00954 | 0 |
| B60 | B140 | WBRVI | r-GT | 0.042959 | 0 |
| B60 | B126 | WBRVI | Alb | 0.036866 | 0 |
| B60 | B127 | WBRVI | A-G ratio | 0.034292 | 0 |
| B60 | B161 | WBRVI | LEU | -0.02063 | 0 |
| B60 | B128 | WBRVI | ALT | 0.037697 | 0 |
| B60 | B111 | WBRVI | TSH | -0.02411 | 0 |
| B60 | B124 | WBRVI | ACTH | 0.034631 | 0 |
| B60 | B134 | WBRVI | CHE | 0.058938 | 0 |
| B60 | B166 | WBRVI | Pro | 0.021104 | 0 |
| B60 | B70 | WBRVI | LDL-C | 0.025391 | 0 |
| B60 | B188 | WBRVI | Ca | 0.006478 | 1.55E-08 |
| B60 | B71 | WBRVI | TG | 0.025983 | 0 |
| B60 | B72 | WBRVI | HDL-C | -0.03416 | 0 |
| B60 | B189 | WBRVI | eGFB | -0.00638 | 2.59E-08 |
| B60 | B141 | WBRVI | AST/ALT | -0.0389 | 0 |
| B60 | B146 | WBRVI | CysC | 0.007931 | 4.38E-12 |
| B60 | B181 | WBRVI | APTT | -0.02834 | 0 |
| B60 | B190 | WBRVI | Cre | 0.066281 | 0 |
| B60 | B193 | WBRVI | K | 0.006016 | 1.50E-07 |
| B60 | B129 | WBRVI | IBIL | 0.029247 | 0 |
| B60 | B150 | WBRVI | ALP | 0.028572 | 0 |
| B60 | B84 | WBRVI | C-PSA | 0.007706 | 1.72E-11 |
| B60 | B222 | WBRVI | NS | 0.080105 | 0 |
| B60 | B99 | WBRVI | anti-PM-Scl | 0.005801 | 4.09E-07 |
| B60 | B105 | WBRVI | anti-U1RNP | -0.01158 | 0 |
| B60 | B95 | WBRVI | anti-cenpb | -0.00661 | 7.70E-09 |
| B60 | B98 | WBRVI | anti-Jo-1 | 0.005614 | 9.51E-07 |
| B60 | B100 | WBRVI | anti-Ro52 | -0.01454 | 0 |
| B60 | B101 | WBRVI | anti-Sc70 | -0.00818 | 9.14E-13 |

| | | | | | |
|---|---|---|---|---|---|
| B60 | B103 | WBRVI | anti-SSA/Ro | -0.02398 | 0 |
| B60 | B104 | WBRVI | anti-SSB | -0.00696 | 1.23E-09 |
| B60 | B107 | WBRVI | AnuA | -0.00704 | 7.91E-10 |
| B60 | B148 | WBRVI | Anti_TPO | -0.01761 | 0 |
| B60 | B149 | WBRVI | TGAb | -0.01665 | 0 |
| B60 | B109 | WBRVI | AMA-M2 | -0.01041 | 0 |
| B60 | B110 | WBRVI | AHA | -0.01124 | 0 |
| B60 | B198 | WBRVI | P | -0.034 | 0 |
| B60 | B194 | WBRVI | Cl | -0.02887 | 0 |
| B60 | B199 | WBRVI | Mg | 0.010432 | 0 |
| B60 | B156 | WBRVI | IgE | 0.006334 | 3.21E-08 |
| B60 | B157 | WBRVI | IgG | -0.03951 | 0 |
| B60 | B158 | WBRVI | IgM | -0.02255 | 0 |
| B60 | B195 | WBRVI | Na | 0.009325 | 4.44E-16 |
| B60 | B176 | WBRVI | Ucr | 0.04342 | 0 |
| B60 | B191 | WBRVI | Urea | 0.008277 | 4.95E-13 |
| B60 | B174 | WBRVI | Urea/Cr | -0.0456 | 0 |
| B60 | B192 | WBRVI | UA | 0.069495 | 0 |
| B60 | B170 | WBRVI | U-GLU | 0.013816 | 0 |
| B60 | B179 | WBRVI | α1-MG | 0.032337 | 0 |
| B60 | B182 | WBRVI | TT | 0.025805 | 0 |
| B60 | B125 | WBRVI | Cor | 0.036431 | 0 |
| B60 | B138 | WBRVI | PAB | 0.084846 | 0 |
| B60 | B130 | WBRVI | GLB | -0.02913 | 0 |
| B60 | B61 | WBRVI | WBHSRV | 0.54836 | 0 |
| B60 | B62 | WBRVI | WBHSRVI | 0.812154 | 0 |
| B60 | B63 | WBRVI | WBRSR(mPas_200) | 0.079658 | 0 |
| B60 | B64 | WBRVI | WBRSR(mPas_5) | 0.900017 | 0 |
| B60 | B65 | WBRVI | WBRSR(mPas_1) | 0.927042 | 0 |
| B60 | B66 | WBRVI | WBRSR(mPas_100) | 0.759581 | 0 |
| B60 | B67 | WBRVI | WBRSR(mPas_50) | 0.788976 | 0 |
| B60 | B86 | WBRVI | TBhCG | -0.00729 | 1.98E-10 |
| B60 | B221 | WBRVI | Hei | 0.072385 | 0 |
| B60 | B87 | WBRVI | NSE | 0.007398 | 1.05E-10 |
| B60 | B187 | WBRVI | NSE | 0.007042 | 7.85E-10 |
| B60 | B200 | WBRVI | OP | 0.018703 | 0 |
| B60 | B228 | WBRVI | OS | 0.005861 | 3.11E-07 |
| B60 | B220 | WBRVI | SBP | 0.01503 | 0 |
| B60 | B219 | WBRVI | OT | -0.01025 | 0 |
| B60 | B218 | WBRVI | DBP | 0.034684 | 0 |
| B60 | B88 | WBRVI | CA 242 | -0.00715 | 4.32E-10 |
| B60 | B216 | WBRVI | BMI | 0.068076 | 0 |
| B60 | B215 | WBRVI | BMI | 0.038192 | 0 |
| B60 | B131 | WBRVI | AST | 0.019098 | 0 |

| | | | | | |
|---|---|---|---|---|---|
| B60 | B210 | WBRVI | Hip | 0.042245 | 0 |
| B60 | B209 | WBRVI | Smoking | 0.053253 | 0 |
| B60 | B90 | WBRVI | CYFRA 21-1 | 0.012188 | 0 |
| B60 | B139 | WBRVI | ADA | -0.02137 | 0 |
| B60 | B183 | WBRVI | PTA | -0.0443 | 0 |
| B60 | B184 | WBRVI | INR | 0.019578 | 0 |
| B60 | B185 | WBRVI | PT | -0.01533 | 0 |
| B60 | B186 | WBRVI | PFC | -0.0098 | 0 |
| B60 | B69 | WBRVI | PV | -0.15533 | 0 |
| B60 | B173 | WBRVI | NTT | -0.01236 | 0 |
| B60 | B207 | WBRVI | WHR | 0.006334 | 3.21E-08 |
| B60 | B206 | WBRVI | Waist | 0.059731 | 0 |
| B60 | B78 | WBRVI | HBcAb | -0.00926 | 6.66E-16 |
| B60 | B197 | WBRVI | AG | 0.025977 | 0 |
| B60 | B204 | WBRVI | Drinking | 0.051587 | 0 |
| B60 | B112 | WBRVI | FT4 | 0.013312 | 0 |
| B60 | B113 | WBRVI | FT3 | 0.033214 | 0 |
| B60 | B211 | WBRVI | IOP-R | 0.021535 | 0 |
| B60 | B73 | WBRVI | ApoA | -0.01808 | 0 |
| B60 | B74 | WBRVI | ApoA1 | -0.05382 | 0 |
| B60 | B75 | WBRVI | ApoA1/B | -0.03767 | 0 |
| B60 | B76 | WBRVI | ApoB | 0.05363 | 0 |
| B60 | B132 | WBRVI | Dbil | 0.01837 | 0 |
| B60 | B81 | WBRVI | TSGF | -0.02518 | 0 |
| B60 | B77 | WBRVI | TC | 0.010246 | 0 |
| B60 | B133 | WBRVI | TBIL | 0.028358 | 0 |
| B60 | B135 | WBRVI | TBA | 0.007356 | 1.34E-10 |
| B60 | B114 | WBRVI | TSH TT3 TT4(TT4) | -0.01944 | 0 |
| B60 | B115 | WBRVI | TSH TT3 TT4(TT3) | 0.017597 | 0 |
| B60 | B212 | WBRVI | IOP-L | 0.023447 | 0 |
| B61 | B137 | WBHSRV | AFU | 0.020753 | 0 |
| B61 | B160 | WBHSRV | PH | -0.01359 | 0 |
| B61 | B140 | WBHSRV | r-GT | 0.038829 | 0 |
| B61 | B126 | WBHSRV | Alb | 0.025219 | 0 |
| B61 | B161 | WBHSRV | LEU | -0.00841 | 2.14E-13 |
| B61 | B128 | WBHSRV | ALT | 0.029542 | 0 |
| B61 | B121 | WBHSRV | C3 | 0.009539 | 0 |
| B61 | B123 | WBHSRV | hs-CRP1 | 0.039522 | 0 |
| B61 | B111 | WBHSRV | TSH | -0.01098 | 0 |
| B61 | B124 | WBHSRV | ACTH | 0.022417 | 0 |
| B61 | B134 | WBHSRV | CHE | 0.051755 | 0 |
| B61 | B166 | WBHSRV | Pro | 0.013999 | 0 |
| B61 | B70 | WBHSRV | LDL-C | 0.02239 | 0 |
| B61 | B188 | WBHSRV | Ca | 0.006328 | 3.31E-08 |

| | | | | | |
|---|---|---|---|---|---|
| B61 | B71 | WBHSRV | TG | 0.035729 | 0 |
| B61 | B72 | WBHSRV | HDL-C | -0.03028 | 0 |
| B61 | B141 | WBHSRV | AST/ALT | -0.02702 | 0 |
| B61 | B181 | WBHSRV | APTT | -0.02502 | 0 |
| B61 | B190 | WBHSRV | Cre | 0.035814 | 0 |
| B61 | B129 | WBHSRV | IBIL | 0.015665 | 0 |
| B61 | B150 | WBHSRV | ALP | 0.027958 | 0 |
| B61 | B222 | WBHSRV | NS | 0.051892 | 0 |
| B61 | B105 | WBHSRV | anti-U1RNP | -0.00586 | 3.19E-07 |
| B61 | B98 | WBHSRV | anti-Jo-1 | 0.008335 | 3.40E-13 |
| B61 | B103 | WBHSRV | anti-SSA/Ro | -0.00811 | 1.45E-12 |
| B61 | B107 | WBHSRV | AnuA | -0.00999 | 0 |
| B61 | B148 | WBHSRV | Anti_TPO | -0.00781 | 9.05E-12 |
| B61 | B149 | WBHSRV | TGAb | -0.00741 | 9.89E-11 |
| B61 | B108 | WBHSRV | Anti-Cyclin | -0.00575 | 5.06E-07 |
| B61 | B198 | WBHSRV | P | -0.01801 | 0 |
| B61 | B194 | WBHSRV | Cl | -0.04476 | 0 |
| B61 | B199 | WBHSRV | Mg | 0.009468 | 0 |
| B61 | B155 | WBHSRV | IgA | 0.030972 | 0 |
| B61 | B156 | WBHSRV | IgE | 0.007946 | 4.01E-12 |
| B61 | B157 | WBHSRV | IgG | 0.008793 | 1.64E-14 |
| B61 | B195 | WBHSRV | Na | -0.00689 | 1.82E-09 |
| B61 | B176 | WBHSRV | Ucr | 0.024229 | 0 |
| B61 | B178 | WBHSRV | Ig-U | 0.006334 | 3.20E-08 |
| B61 | B174 | WBHSRV | Urea/Cr | -0.02387 | 0 |
| B61 | B192 | WBHSRV | UA | 0.048657 | 0 |
| B61 | B170 | WBHSRV | U-GLU | 0.016031 | 0 |
| B61 | B179 | WBHSRV | α1-MG | 0.024499 | 0 |
| B61 | B182 | WBHSRV | TT | 0.008588 | 6.48E-14 |
| B61 | B125 | WBHSRV | Cor | 0.03944 | 0 |
| B61 | B138 | WBHSRV | PAB | 0.062107 | 0 |
| B61 | B130 | WBHSRV | GLB | 0.018838 | 0 |
| B61 | B62 | WBHSRV | WBHSRVI | 0.721489 | 0 |
| B61 | B63 | WBHSRV | WBRSR(mPas_200) | 0.07399 | 0 |
| B61 | B64 | WBHSRV | WBRSR(mPas_5) | 0.664765 | 0 |
| B61 | B65 | WBHSRV | WBRSR(mPas_1) | 0.574121 | 0 |
| B61 | B66 | WBHSRV | WBRSR(mPas_100) | 0.742621 | 0 |
| B61 | B67 | WBHSRV | WBRSR(mPas_50) | 0.734691 | 0 |
| B61 | B221 | WBHSRV | Hei | 0.038117 | 0 |
| B61 | B87 | WBHSRV | NSE | 0.006905 | 1.65E-09 |
| B61 | B187 | WBHSRV | NSE | 0.0059 | 2.59E-07 |
| B61 | B220 | WBHSRV | SBP | 0.020445 | 0 |
| B61 | B218 | WBHSRV | DBP | 0.029571 | 0 |
| B61 | B216 | WBHSRV | BMI | 0.046847 | 0 |

| | | | | | |
|---|---|---|---|---|---|
| B61 | B215 | WBHSRV | BMI | 0.034752 | 0 |
| B61 | B131 | WBHSRV | AST | 0.016511 | 0 |
| B61 | B210 | WBHSRV | Hip | 0.033196 | 0 |
| B61 | B209 | WBHSRV | Smoking | 0.024406 | 0 |
| B61 | B183 | WBHSRV | PTA | -0.01247 | 0 |
| B61 | B184 | WBHSRV | INR | 0.006966 | 1.19E-09 |
| B61 | B185 | WBHSRV | PT | -0.00717 | 3.82E-10 |
| B61 | B186 | WBHSRV | PFC | 0.023364 | 0 |
| B61 | B69 | WBHSRV | PV | 0.051181 | 0 |
| B61 | B173 | WBHSRV | NTT | -0.00709 | 6.05E-10 |
| B61 | B206 | WBHSRV | Waist | 0.046464 | 0 |
| B61 | B78 | WBHSRV | HBcAb | 0.007553 | 4.27E-11 |
| B61 | B197 | WBHSRV | AG | 0.020864 | 0 |
| B61 | B204 | WBHSRV | Drinking | 0.025803 | 0 |
| B61 | B112 | WBHSRV | FT4 | 0.01595 | 0 |
| B61 | B113 | WBHSRV | FT3 | 0.021941 | 0 |
| B61 | B211 | WBHSRV | IOP-R | 0.013762 | 0 |
| B61 | B74 | WBHSRV | ApoA1 | -0.06033 | 0 |
| B61 | B75 | WBHSRV | ApoA1/B | -0.04257 | 0 |
| B61 | B76 | WBHSRV | ApoB | 0.071348 | 0 |
| B61 | B132 | WBHSRV | Dbil | 0.010967 | 0 |
| B61 | B81 | WBHSRV | TSGF | -0.00891 | 7.55E-15 |
| B61 | B77 | WBHSRV | TC | 0.017326 | 0 |
| B61 | B133 | WBHSRV | TBIL | 0.015523 | 0 |
| B61 | B91 | WBHSRV | TP | 0.031907 | 0 |
| B61 | B115 | WBHSRV | TSH TT3 TT4(TT3) | 0.013947 | 0 |
| B61 | B212 | WBHSRV | IOP-L | 0.010468 | 0 |
| B62 | B137 | WBHSRVI | AFU | 0.03578 | 0 |
| B62 | B160 | WBHSRVI | PH | -0.01218 | 0 |
| B62 | B140 | WBHSRVI | r-GT | 0.051505 | 0 |
| B62 | B126 | WBHSRVI | Alb | 0.033307 | 0 |
| B62 | B127 | WBHSRVI | A-G ratio | 0.025907 | 0 |
| B62 | B161 | WBHSRVI | LEU | -0.02026 | 0 |
| B62 | B128 | WBHSRVI | ALT | 0.045093 | 0 |
| B62 | B121 | WBHSRVI | C3 | 0.006815 | 2.68E-09 |
| B62 | B123 | WBHSRVI | hs-CRP1 | 0.008028 | 2.40E-12 |
| B62 | B111 | WBHSRVI | TSH | -0.02424 | 0 |
| B62 | B124 | WBHSRVI | ACTH | 0.035326 | 0 |
| B62 | B134 | WBHSRVI | CHE | 0.069042 | 0 |
| B62 | B166 | WBHSRVI | Pro | 0.022421 | 0 |
| B62 | B70 | WBHSRVI | LDL-C | 0.028079 | 0 |
| B62 | B188 | WBHSRVI | Ca | 0.007892 | 5.58E-12 |
| B62 | B71 | WBHSRVI | TG | 0.037735 | 0 |
| B62 | B72 | WBHSRVI | HDL-C | -0.04922 | 0 |

| | | | | | |
|---|---|---|---|---|---|
| B62 | B189 | WBHSRVI | eGFB | -0.00962 | 0 |
| B62 | B141 | WBHSRVI | AST/ALT | -0.04858 | 0 |
| B62 | B146 | WBHSRVI | CysC | 0.006931 | 1.44E-09 |
| B62 | B181 | WBHSRVI | APTT | -0.02509 | 0 |
| B62 | B190 | WBHSRVI | Cre | 0.069288 | 0 |
| B62 | B129 | WBHSRVI | IBIL | 0.030192 | 0 |
| B62 | B150 | WBHSRVI | ALP | 0.033746 | 0 |
| B62 | B84 | WBHSRVI | C-PSA | 0.0096 | 0 |
| B62 | B222 | WBHSRVI | NS | 0.088696 | 0 |
| B62 | B105 | WBHSRVI | anti-U1RNP | -0.0107 | 0 |
| B62 | B95 | WBHSRVI | anti-cenpb | -0.00784 | 7.66E-12 |
| B62 | B98 | WBHSRVI | anti-Jo-1 | 0.009509 | 0 |
| B62 | B100 | WBHSRVI | anti-Ro52 | -0.01204 | 0 |
| B62 | B101 | WBHSRVI | anti-Sc70 | -0.00781 | 9.33E-12 |
| B62 | B103 | WBHSRVI | anti-SSA/Ro | -0.02442 | 0 |
| B62 | B107 | WBHSRVI | AnuA | -0.01193 | 0 |
| B62 | B151 | WBHSRVI | Anti-CCP | -0.00962 | 0 |
| B62 | B148 | WBHSRVI | Anti_TPO | -0.01838 | 0 |
| B62 | B149 | WBHSRVI | TGAb | -0.01702 | 0 |
| B62 | B109 | WBHSRVI | AMA-M2 | -0.00682 | 2.59E-09 |
| B62 | B110 | WBHSRVI | AHA | -0.01277 | 0 |
| B62 | B198 | WBHSRVI | P | -0.03615 | 0 |
| B62 | B194 | WBHSRVI | Cl | -0.04633 | 0 |
| B62 | B199 | WBHSRVI | Mg | 0.014951 | 0 |
| B62 | B155 | WBHSRVI | IgA | 0.006797 | 2.95E-09 |
| B62 | B157 | WBHSRVI | IgG | -0.04388 | 0 |
| B62 | B158 | WBHSRVI | IgM | -0.02278 | 0 |
| B62 | B195 | WBHSRVI | Na | 0.009242 | 6.66E-16 |
| B62 | B176 | WBHSRVI | Ucr | 0.045665 | 0 |
| B62 | B178 | WBHSRVI | Ig-U | 0.005819 | 3.77E-07 |
| B62 | B191 | WBHSRVI | Urea | 0.010202 | 0 |
| B62 | B174 | WBHSRVI | Urea/Cr | -0.04592 | 0 |
| B62 | B192 | WBHSRVI | UA | 0.077661 | 0 |
| B62 | B170 | WBHSRVI | U-GLU | 0.016868 | 0 |
| B62 | B177 | WBHSRVI | ACR | -0.00599 | 1.72E-07 |
| B62 | B179 | WBHSRVI | α1-MG | 0.035744 | 0 |
| B62 | B180 | WBHSRVI | Tf | 0.006234 | 5.24E-08 |
| B62 | B182 | WBHSRVI | TT | 0.035255 | 0 |
| B62 | B125 | WBHSRVI | Cor | 0.033762 | 0 |
| B62 | B138 | WBHSRVI | PAB | 0.097056 | 0 |
| B62 | B130 | WBHSRVI | GLB | -0.01903 | 0 |
| B62 | B63 | WBHSRVI | WBRSR(mPas_200) | 0.093264 | 0 |
| B62 | B64 | WBHSRVI | WBRSR(mPas_5) | 0.834503 | 0 |
| B62 | B65 | WBHSRVI | WBRSR(mPas_1) | 0.739122 | 0 |

| | | | | | |
|---|---|---|---|---|---|
| B62 | B66 | WBHSRVI | WBRSR(mPas_100) | 0.907817 | 0 |
| B62 | B67 | WBHSRVI | WBRSR(mPas_50) | 0.903004 | 0 |
| B62 | B221 | WBHSRVI | Hei | 0.077001 | 0 |
| B62 | B87 | WBHSRVI | NSE | 0.010134 | 0 |
| B62 | B187 | WBHSRVI | NSE | 0.010071 | 0 |
| B62 | B200 | WBHSRVI | OP | 0.020139 | 0 |
| B62 | B220 | WBHSRVI | SBP | 0.022064 | 0 |
| B62 | B219 | WBHSRVI | OT | -0.01082 | 0 |
| B62 | B218 | WBHSRVI | DBP | 0.039489 | 0 |
| B62 | B88 | WBHSRVI | CA 242 | -0.00752 | 5.33E-11 |
| B62 | B216 | WBHSRVI | BMI | 0.078295 | 0 |
| B62 | B215 | WBHSRVI | BMI | 0.048405 | 0 |
| B62 | B131 | WBHSRVI | AST | 0.020376 | 0 |
| B62 | B210 | WBHSRVI | Hip | 0.049138 | 0 |
| B62 | B209 | WBHSRVI | Smoking | 0.053638 | 0 |
| B62 | B90 | WBHSRVI | CYFRA 21-1 | 0.009225 | 8.88E-16 |
| B62 | B139 | WBHSRVI | ADA | -0.01207 | 0 |
| B62 | B183 | WBHSRVI | PTA | -0.02575 | 0 |
| B62 | B184 | WBHSRVI | INR | 0.007307 | 1.78E-10 |
| B62 | B185 | WBHSRVI | PT | -0.01625 | 0 |
| B62 | B186 | WBHSRVI | PFC | -0.01428 | 0 |
| B62 | B69 | WBHSRVI | PV | -0.19438 | 0 |
| B62 | B173 | WBHSRVI | NTT | -0.01508 | 0 |
| B62 | B207 | WBHSRVI | WHR | 0.007381 | 1.17E-10 |
| B62 | B206 | WBHSRVI | Waist | 0.069807 | 0 |
| B62 | B78 | WBHSRVI | HBcAb | 0.005861 | 3.10E-07 |
| B62 | B197 | WBHSRVI | AG | 0.033224 | 0 |
| B62 | B204 | WBHSRVI | Drinking | 0.052742 | 0 |
| B62 | B113 | WBHSRVI | FT3 | 0.035293 | 0 |
| B62 | B211 | WBHSRVI | IOP-R | 0.017523 | 0 |
| B62 | B73 | WBHSRVI | ApoA | -0.0161 | 0 |
| B62 | B74 | WBHSRVI | ApoA1 | -0.09626 | 0 |
| B62 | B75 | WBHSRVI | ApoA1/B | -0.06082 | 0 |
| B62 | B76 | WBHSRVI | ApoB | 0.085183 | 0 |
| B62 | B132 | WBHSRVI | Dbil | 0.02261 | 0 |
| B62 | B77 | WBHSRVI | TC | 0.015431 | 0 |
| B62 | B133 | WBHSRVI | TBIL | 0.030226 | 0 |
| B62 | B135 | WBHSRVI | TBA | 0.008658 | 4.09E-14 |
| B62 | B114 | WBHSRVI | TSH TT3 TT4(TT4) | -0.02338 | 0 |
| B62 | B115 | WBHSRVI | TSH TT3 TT4(TT3) | 0.020391 | 0 |
| B62 | B212 | WBHSRVI | IOP-L | 0.017718 | 0 |
| B63 | B140 | WBRSR(mPas_200) | r-GT | 0.006179 | 6.88E-08 |
| B63 | B134 | WBRSR(mPas_200) | CHE | 0.007841 | 7.62E-12 |
| B63 | B190 | WBRSR(mPas_200) | Cre | 0.008113 | 1.41E-12 |

| | | | | | |
|---|---|---|---|---|---|
| B63 | B222 | WBRSR(mPas_200) | NS | 0.008996 | 4.00E-15 |
| B63 | B194 | WBRSR(mPas_200) | Cl | -0.00599 | 1.67E-07 |
| B63 | B192 | WBRSR(mPas_200) | UA | 0.008946 | 5.77E-15 |
| B63 | B125 | WBRSR(mPas_200) | Cor | 0.005931 | 2.25E-07 |
| B63 | B138 | WBRSR(mPas_200) | PAB | 0.01032 | 0 |
| B63 | B64 | WBRSR(mPas_200) | WBRSR(mPas_5) | 0.096483 | 0 |
| B63 | B65 | WBRSR(mPas_200) | WBRSR(mPas_1) | 0.086244 | 0 |
| B63 | B66 | WBRSR(mPas_200) | WBRSR(mPas_100) | 0.101358 | 0 |
| B63 | B67 | WBRSR(mPas_200) | WBRSR(mPas_50) | 0.100746 | 0 |
| B63 | B221 | WBRSR(mPas_200) | Hei | 0.008157 | 1.07E-12 |
| B63 | B218 | WBRSR(mPas_200) | DBP | 0.005718 | 5.98E-07 |
| B63 | B216 | WBRSR(mPas_200) | BMI | 0.008285 | 4.72E-13 |
| B63 | B69 | WBRSR(mPas_200) | PV | 0.022775 | 0 |
| B63 | B206 | WBRSR(mPas_200) | Waist | 0.007647 | 2.46E-11 |
| B63 | B74 | WBRSR(mPas_200) | ApoA1 | -0.00875 | 2.18E-14 |
| B63 | B75 | WBRSR(mPas_200) | ApoA1/B | -0.00657 | 9.89E-09 |
| B63 | B76 | WBRSR(mPas_200) | ApoB | 0.011183 | 0 |
| B64 | B136 | WBRSR(mPas_5) | 5'NT | 0.005959 | 1.97E-07 |
| B64 | B137 | WBRSR(mPas_5) | AFU | 0.037182 | 0 |
| B64 | B160 | WBRSR(mPas_5) | PH | -0.01654 | 0 |
| B64 | B140 | WBRSR(mPas_5) | r-GT | 0.055793 | 0 |
| B64 | B126 | WBRSR(mPas_5) | Alb | 0.047542 | 0 |
| B64 | B127 | WBRSR(mPas_5) | A-G ratio | 0.011442 | 0 |
| B64 | B161 | WBRSR(mPas_5) | LEU | -0.01938 | 0 |
| B64 | B128 | WBRSR(mPas_5) | ALT | 0.045941 | 0 |
| B64 | B121 | WBRSR(mPas_5) | C3 | 0.008862 | 1.02E-14 |
| B64 | B123 | WBRSR(mPas_5) | hs-CRP1 | 0.03626 | 0 |
| B64 | B111 | WBRSR(mPas_5) | TSH | -0.02258 | 0 |
| B64 | B124 | WBRSR(mPas_5) | ACTH | 0.044739 | 0 |
| B64 | B134 | WBRSR(mPas_5) | CHE | 0.080016 | 0 |
| B64 | B166 | WBRSR(mPas_5) | Pro | 0.023421 | 0 |
| B64 | B70 | WBRSR(mPas_5) | LDL-C | 0.038884 | 0 |
| B64 | B188 | WBRSR(mPas_5) | Ca | 0.010408 | 0 |
| B64 | B71 | WBRSR(mPas_5) | TG | 0.042391 | 0 |
| B64 | B72 | WBRSR(mPas_5) | HDL-C | -0.03989 | 0 |
| B64 | B189 | WBRSR(mPas_5) | eGFB | -0.00933 | 4.44E-16 |
| B64 | B141 | WBRSR(mPas_5) | AST/ALT | -0.04293 | 0 |
| B64 | B146 | WBRSR(mPas_5) | CysC | 0.007481 | 6.51E-11 |
| B64 | B181 | WBRSR(mPas_5) | APTT | -0.03365 | 0 |
| B64 | B190 | WBRSR(mPas_5) | Cre | 0.070868 | 0 |
| B64 | B193 | WBRSR(mPas_5) | K | 0.018128 | 0 |
| B64 | B129 | WBRSR(mPas_5) | IBIL | 0.033682 | 0 |
| B64 | B150 | WBRSR(mPas_5) | ALP | 0.041297 | 0 |
| B64 | B84 | WBRSR(mPas_5) | C-PSA | 0.008411 | 2.08E-13 |

| | | | | | |
|---|---|---|---|---|---|
| B64 | B222 | WBRSR(mPas_5) | NS | 0.08642 | 0 |
| B64 | B105 | WBRSR(mPas_5) | anti-U1RNP | -0.01266 | 0 |
| B64 | B95 | WBRSR(mPas_5) | anti-cenpb | -0.00702 | 8.80E-10 |
| B64 | B98 | WBRSR(mPas_5) | anti-Jo-1 | 0.008214 | 7.44E-13 |
| B64 | B100 | WBRSR(mPas_5) | anti-Ro52 | -0.0108 | 0 |
| B64 | B101 | WBRSR(mPas_5) | anti-Sc70 | -0.00828 | 4.89E-13 |
| B64 | B103 | WBRSR(mPas_5) | anti-SSA/Ro | -0.01562 | 0 |
| B64 | B104 | WBRSR(mPas_5) | anti-SSB | -0.00788 | 5.97E-12 |
| B64 | B107 | WBRSR(mPas_5) | AnuA | -0.00746 | 7.46E-11 |
| B64 | B148 | WBRSR(mPas_5) | Anti_TPO | -0.01585 | 0 |
| B64 | B149 | WBRSR(mPas_5) | TGAb | -0.01484 | 0 |
| B64 | B109 | WBRSR(mPas_5) | AMA-M2 | -0.00669 | 5.13E-09 |
| B64 | B110 | WBRSR(mPas_5) | AHA | -0.00659 | 8.79E-09 |
| B64 | B198 | WBRSR(mPas_5) | P | -0.03385 | 0 |
| B64 | B194 | WBRSR(mPas_5) | Cl | -0.04922 | 0 |
| B64 | B199 | WBRSR(mPas_5) | Mg | 0.015757 | 0 |
| B64 | B155 | WBRSR(mPas_5) | IgA | 0.023766 | 0 |
| B64 | B156 | WBRSR(mPas_5) | IgE | 0.014269 | 0 |
| B64 | B158 | WBRSR(mPas_5) | IgM | -0.01535 | 0 |
| B64 | B176 | WBRSR(mPas_5) | Ucr | 0.046864 | 0 |
| B64 | B178 | WBRSR(mPas_5) | Ig-U | 0.010458 | 0 |
| B64 | B191 | WBRSR(mPas_5) | Urea | 0.010002 | 0 |
| B64 | B174 | WBRSR(mPas_5) | Urea/Cr | -0.04719 | 0 |
| B64 | B192 | WBRSR(mPas_5) | UA | 0.079731 | 0 |
| B64 | B170 | WBRSR(mPas_5) | U-GLU | 0.020365 | 0 |
| B64 | B179 | WBRSR(mPas_5) | α1-MG | 0.040275 | 0 |
| B64 | B180 | WBRSR(mPas_5) | Tf | 0.00915 | 1.33E-15 |
| B64 | B182 | WBRSR(mPas_5) | TT | 0.019447 | 0 |
| B64 | B125 | WBRSR(mPas_5) | Cor | 0.054948 | 0 |
| B64 | B138 | WBRSR(mPas_5) | PAB | 0.096229 | 0 |
| B64 | B130 | WBRSR(mPas_5) | GLB | 0.007478 | 6.63E-11 |
| B64 | B65 | WBRSR(mPas_5) | WBRSR(mPas_1) | 0.973614 | 0 |
| B64 | B66 | WBRSR(mPas_5) | WBRSR(mPas_100) | 0.916792 | 0 |
| B64 | B67 | WBRSR(mPas_5) | WBRSR(mPas_50) | 0.934203 | 0 |
| B64 | B86 | WBRSR(mPas_5) | TBhCG | -0.00628 | 4.29E-08 |
| B64 | B221 | WBRSR(mPas_5) | Hei | 0.07358 | 0 |
| B64 | B87 | WBRSR(mPas_5) | NSE | 0.009823 | 0 |
| B64 | B187 | WBRSR(mPas_5) | NSE | 0.006979 | 1.11E-09 |
| B64 | B200 | WBRSR(mPas_5) | OP | 0.013933 | 0 |
| B64 | B228 | WBRSR(mPas_5) | OS | 0.006378 | 2.57E-08 |
| B64 | B220 | WBRSR(mPas_5) | SBP | 0.024874 | 0 |
| B64 | B219 | WBRSR(mPas_5) | OT | -0.01007 | 0 |
| B64 | B218 | WBRSR(mPas_5) | DBP | 0.045256 | 0 |
| B64 | B88 | WBRSR(mPas_5) | CA 242 | -0.00674 | 3.92E-09 |

| | | | | | |
|---|---|---|---|---|---|
| B64 | B216 | WBRSR(mPas_5) | BMI | 0.074961 | 0 |
| B64 | B215 | WBRSR(mPas_5) | BMI | 0.04648 | 0 |
| B64 | B131 | WBRSR(mPas_5) | AST | 0.027768 | 0 |
| B64 | B210 | WBRSR(mPas_5) | Hip | 0.047812 | 0 |
| B64 | B209 | WBRSR(mPas_5) | Smoking | 0.052846 | 0 |
| B64 | B90 | WBRSR(mPas_5) | CYFRA 21-1 | 0.009109 | 1.78E-15 |
| B64 | B139 | WBRSR(mPas_5) | ADA | -0.00667 | 5.77E-09 |
| B64 | B183 | WBRSR(mPas_5) | PTA | -0.02254 | 0 |
| B64 | B184 | WBRSR(mPas_5) | INR | 0.020663 | 0 |
| B64 | B185 | WBRSR(mPas_5) | PT | -0.01287 | 0 |
| B64 | B186 | WBRSR(mPas_5) | PFC | 0.023642 | 0 |
| B64 | B69 | WBRSR(mPas_5) | PV | 0.17087 | 0 |
| B64 | B173 | WBRSR(mPas_5) | NTT | -0.01267 | 0 |
| B64 | B207 | WBRSR(mPas_5) | WHR | 0.00714 | 4.57E-10 |
| B64 | B206 | WBRSR(mPas_5) | Waist | 0.068259 | 0 |
| B64 | B197 | WBRSR(mPas_5) | AG | 0.030891 | 0 |
| B64 | B204 | WBRSR(mPas_5) | Drinking | 0.050944 | 0 |
| B64 | B112 | WBRSR(mPas_5) | FT4 | 0.023595 | 0 |
| B64 | B113 | WBRSR(mPas_5) | FT3 | 0.037038 | 0 |
| B64 | B211 | WBRSR(mPas_5) | IOP-R | 0.022612 | 0 |
| B64 | B73 | WBRSR(mPas_5) | ApoA | -0.0078 | 1.00E-11 |
| B64 | B74 | WBRSR(mPas_5) | ApoA1 | -0.05925 | 0 |
| B64 | B75 | WBRSR(mPas_5) | ApoA1/B | -0.0514 | 0 |
| B64 | B76 | WBRSR(mPas_5) | ApoB | 0.09233 | 0 |
| B64 | B132 | WBRSR(mPas_5) | Dbil | 0.016638 | 0 |
| B64 | B81 | WBRSR(mPas_5) | TSGF | -0.03057 | 0 |
| B64 | B77 | WBRSR(mPas_5) | TC | 0.024593 | 0 |
| B64 | B133 | WBRSR(mPas_5) | TBIL | 0.031511 | 0 |
| B64 | B135 | WBRSR(mPas_5) | TBA | 0.00806 | 1.96E-12 |
| B64 | B91 | WBRSR(mPas_5) | TP | 0.035925 | 0 |
| B64 | B114 | WBRSR(mPas_5) | TSH TT3 TT4(TT4) | -0.00658 | 9.25E-09 |
| B64 | B115 | WBRSR(mPas_5) | TSH TT3 TT4(TT3) | 0.022762 | 0 |
| B64 | B212 | WBRSR(mPas_5) | IOP-L | 0.022229 | 0 |
| B65 | B137 | WBRSR(mPas_1) | AFU | 0.032851 | 0 |
| B65 | B92 | WBRSR(mPas_1) | ALDH2-Glu504Glu | -0.00714 | 4.44E-10 |
| B65 | B93 | WBRSR(mPas_1) | ALDH2-Glu504Lys | 0.006051 | 1.27E-07 |
| B65 | B160 | WBRSR(mPas_1) | PH | -0.015 | 0 |
| B65 | B140 | WBRSR(mPas_1) | r-GT | 0.049856 | 0 |
| B65 | B126 | WBRSR(mPas_1) | Alb | 0.045686 | 0 |
| B65 | B127 | WBRSR(mPas_1) | A-G ratio | 0.015479 | 0 |
| B65 | B161 | WBRSR(mPas_1) | LEU | -0.01872 | 0 |
| B65 | B128 | WBRSR(mPas_1) | ALT | 0.041172 | 0 |
| B65 | B121 | WBRSR(mPas_1) | C3 | 0.008024 | 2.46E-12 |
| B65 | B123 | WBRSR(mPas_1) | hs-CRP1 | 0.027025 | 0 |

| | | | | | |
|---|---|---|---|---|---|
| B65 | B111 | WBRSR(mPas_1) | TSH | -0.0215 | 0 |
| B65 | B124 | WBRSR(mPas_1) | ACTH | 0.042875 | 0 |
| B65 | B134 | WBRSR(mPas_1) | CHE | 0.07271 | 0 |
| B65 | B166 | WBRSR(mPas_1) | Pro | 0.021736 | 0 |
| B65 | B70 | WBRSR(mPas_1) | LDL-C | 0.036091 | 0 |
| B65 | B188 | WBRSR(mPas_1) | Ca | 0.009181 | 1.11E-15 |
| B65 | B71 | WBRSR(mPas_1) | TG | 0.035869 | 0 |
| B65 | B72 | WBRSR(mPas_1) | HDL-C | -0.03308 | 0 |
| B65 | B189 | WBRSR(mPas_1) | eGFB | -0.00786 | 6.61E-12 |
| B65 | B141 | WBRSR(mPas_1) | AST/ALT | -0.03799 | 0 |
| B65 | B146 | WBRSR(mPas_1) | CysC | 0.007606 | 3.13E-11 |
| B65 | B181 | WBRSR(mPas_1) | APTT | -0.03382 | 0 |
| B65 | B190 | WBRSR(mPas_1) | Cre | 0.066731 | 0 |
| B65 | B193 | WBRSR(mPas_1) | K | 0.018175 | 0 |
| B65 | B129 | WBRSR(mPas_1) | IBIL | 0.031165 | 0 |
| B65 | B150 | WBRSR(mPas_1) | ALP | 0.037442 | 0 |
| B65 | B84 | WBRSR(mPas_1) | C-PSA | 0.007593 | 3.39E-11 |
| B65 | B222 | WBRSR(mPas_1) | NS | 0.080264 | 0 |
| B65 | B99 | WBRSR(mPas_1) | anti-PM-Scl | 0.006005 | 1.58E-07 |
| B65 | B105 | WBRSR(mPas_1) | anti-U1RNP | -0.01216 | 0 |
| B65 | B95 | WBRSR(mPas_1) | anti-cenpb | -0.00661 | 7.85E-09 |
| B65 | B98 | WBRSR(mPas_1) | anti-Jo-1 | 0.006351 | 2.94E-08 |
| B65 | B100 | WBRSR(mPas_1) | anti-Ro52 | -0.01152 | 0 |
| B65 | B101 | WBRSR(mPas_1) | anti-Sc70 | -0.00812 | 1.33E-12 |
| B65 | B103 | WBRSR(mPas_1) | anti-SSA/Ro | -0.01574 | 0 |
| B65 | B104 | WBRSR(mPas_1) | anti-SSB | -0.00924 | 8.88E-16 |
| B65 | B151 | WBRSR(mPas_1) | Anti-CCP | 0.007659 | 2.28E-11 |
| B65 | B148 | WBRSR(mPas_1) | Anti_TPO | -0.0152 | 0 |
| B65 | B149 | WBRSR(mPas_1) | TGAb | -0.01432 | 0 |
| B65 | B109 | WBRSR(mPas_1) | AMA-M2 | -0.00777 | 1.16E-11 |
| B65 | B110 | WBRSR(mPas_1) | AHA | -0.0066 | 8.11E-09 |
| B65 | B198 | WBRSR(mPas_1) | P | -0.03173 | 0 |
| B65 | B194 | WBRSR(mPas_1) | Cl | -0.04093 | 0 |
| B65 | B199 | WBRSR(mPas_1) | Mg | 0.012408 | 0 |
| B65 | B155 | WBRSR(mPas_1) | IgA | 0.018506 | 0 |
| B65 | B156 | WBRSR(mPas_1) | IgE | 0.01353 | 0 |
| B65 | B158 | WBRSR(mPas_1) | IgM | -0.01543 | 0 |
| B65 | B176 | WBRSR(mPas_1) | Ucr | 0.043736 | 0 |
| B65 | B178 | WBRSR(mPas_1) | Ig-U | 0.009242 | 6.66E-16 |
| B65 | B191 | WBRSR(mPas_1) | Urea | 0.009186 | 1.11E-15 |
| B65 | B174 | WBRSR(mPas_1) | Urea/Cr | -0.04485 | 0 |
| B65 | B192 | WBRSR(mPas_1) | UA | 0.073249 | 0 |
| B65 | B170 | WBRSR(mPas_1) | U-GLU | 0.018293 | 0 |
| B65 | B179 | WBRSR(mPas_1) | α1-MG | 0.037103 | 0 |

| | | | | | |
|---|---|---|---|---|---|
| B65 | B180 | WBRSR(mPas_1) | Tf | 0.007774 | 1.14E-11 |
| B65 | B182 | WBRSR(mPas_1) | TT | 0.017173 | 0 |
| B65 | B125 | WBRSR(mPas_1) | Cor | 0.052751 | 0 |
| B65 | B138 | WBRSR(mPas_1) | PAB | 0.087897 | 0 |
| B65 | B66 | WBRSR(mPas_1) | WBRSR(mPas_100) | 0.820515 | 0 |
| B65 | B67 | WBRSR(mPas_1) | WBRSR(mPas_50) | 0.848598 | 0 |
| B65 | B86 | WBRSR(mPas_1) | TBhCG | -0.00709 | 6.18E-10 |
| B65 | B221 | WBRSR(mPas_1) | Hei | 0.069111 | 0 |
| B65 | B87 | WBRSR(mPas_1) | NSE | 0.008195 | 8.40E-13 |
| B65 | B187 | WBRSR(mPas_1) | NSE | 0.005722 | 5.87E-07 |
| B65 | B200 | WBRSR(mPas_1) | OP | 0.013399 | 0 |
| B65 | B228 | WBRSR(mPas_1) | OS | 0.006479 | 1.54E-08 |
| B65 | B220 | WBRSR(mPas_1) | SBP | 0.021115 | 0 |
| B65 | B219 | WBRSR(mPas_1) | OT | -0.00934 | 4.44E-16 |
| B65 | B218 | WBRSR(mPas_1) | DBP | 0.041347 | 0 |
| B65 | B88 | WBRSR(mPas_1) | CA 242 | -0.00625 | 4.77E-08 |
| B65 | B216 | WBRSR(mPas_1) | BMI | 0.068519 | 0 |
| B65 | B215 | WBRSR(mPas_1) | BMI | 0.041179 | 0 |
| B65 | B131 | WBRSR(mPas_1) | AST | 0.025649 | 0 |
| B65 | B210 | WBRSR(mPas_1) | Hip | 0.043492 | 0 |
| B65 | B209 | WBRSR(mPas_1) | Smoking | 0.051007 | 0 |
| B65 | B90 | WBRSR(mPas_1) | CYFRA 21-1 | 0.009764 | 0 |
| B65 | B139 | WBRSR(mPas_1) | ADA | -0.01037 | 0 |
| B65 | B183 | WBRSR(mPas_1) | PTA | -0.03007 | 0 |
| B65 | B184 | WBRSR(mPas_1) | INR | 0.024471 | 0 |
| B65 | B185 | WBRSR(mPas_1) | PT | -0.01248 | 0 |
| B65 | B186 | WBRSR(mPas_1) | PFC | 0.022678 | 0 |
| B65 | B69 | WBRSR(mPas_1) | PV | 0.153411 | 0 |
| B65 | B173 | WBRSR(mPas_1) | NTT | -0.01103 | 0 |
| B65 | B207 | WBRSR(mPas_1) | WHR | 0.006524 | 1.23E-08 |
| B65 | B206 | WBRSR(mPas_1) | Waist | 0.062136 | 0 |
| B65 | B78 | WBRSR(mPas_1) | HBcAb | -0.00964 | 0 |
| B65 | B197 | WBRSR(mPas_1) | AG | 0.027575 | 0 |
| B65 | B204 | WBRSR(mPas_1) | Drinking | 0.048958 | 0 |
| B65 | B112 | WBRSR(mPas_1) | FT4 | 0.024672 | 0 |
| B65 | B113 | WBRSR(mPas_1) | FT3 | 0.035315 | 0 |
| B65 | B211 | WBRSR(mPas_1) | IOP-R | 0.022321 | 0 |
| B65 | B73 | WBRSR(mPas_1) | ApoA | -0.00812 | 1.36E-12 |
| B65 | B74 | WBRSR(mPas_1) | ApoA1 | -0.0428 | 0 |
| B65 | B75 | WBRSR(mPas_1) | ApoA1/B | -0.04163 | 0 |
| B65 | B76 | WBRSR(mPas_1) | ApoB | 0.077727 | 0 |
| B65 | B132 | WBRSR(mPas_1) | Dbil | 0.014358 | 0 |
| B65 | B81 | WBRSR(mPas_1) | TSGF | -0.03783 | 0 |
| B65 | B77 | WBRSR(mPas_1) | TC | 0.021417 | 0 |

| | | | | | |
|---|---|---|---|---|---|
| B65 | B133 | WBRSR(mPas_1) | TBIL | 0.028883 | 0 |
| B65 | B135 | WBRSR(mPas_1) | TBA | 0.007207 | 3.13E-10 |
| B65 | B91 | WBRSR(mPas_1) | TP | 0.029112 | 0 |
| B65 | B114 | WBRSR(mPas_1) | TSH TT3 TT4(TT4) | -0.00578 | 4.47E-07 |
| B65 | B115 | WBRSR(mPas_1) | TSH TT3 TT4(TT3) | 0.02104 | 0 |
| B65 | B212 | WBRSR(mPas_1) | IOP-L | 0.022744 | 0 |
| B66 | B136 | WBRSR(mPas_100) | 5'NT | 0.006625 | 7.31E-09 |
| B66 | B137 | WBRSR(mPas_100) | AFU | 0.04083 | 0 |
| B66 | B160 | WBRSR(mPas_100) | PH | -0.01741 | 0 |
| B66 | B140 | WBRSR(mPas_100) | r-GT | 0.059224 | 0 |
| B66 | B126 | WBRSR(mPas_100) | Alb | 0.04402 | 0 |
| B66 | B161 | WBRSR(mPas_100) | LEU | -0.01836 | 0 |
| B66 | B128 | WBRSR(mPas_100) | ALT | 0.048667 | 0 |
| B66 | B121 | WBRSR(mPas_100) | C3 | 0.006945 | 1.33E-09 |
| B66 | B123 | WBRSR(mPas_100) | hs-CRP1 | 0.046471 | 0 |
| B66 | B111 | WBRSR(mPas_100) | TSH | -0.02125 | 0 |
| B66 | B124 | WBRSR(mPas_100) | ACTH | 0.041105 | 0 |
| B66 | B134 | WBRSR(mPas_100) | CHE | 0.080342 | 0 |
| B66 | B166 | WBRSR(mPas_100) | Pro | 0.023129 | 0 |
| B66 | B70 | WBRSR(mPas_100) | LDL-C | 0.039802 | 0 |
| B66 | B188 | WBRSR(mPas_100) | Ca | 0.011293 | 0 |
| B66 | B71 | WBRSR(mPas_100) | TG | 0.049005 | 0 |
| B66 | B72 | WBRSR(mPas_100) | HDL-C | -0.04752 | 0 |
| B66 | B189 | WBRSR(mPas_100) | eGFB | -0.01186 | 0 |
| B66 | B141 | WBRSR(mPas_100) | AST/ALT | -0.04645 | 0 |
| B66 | B146 | WBRSR(mPas_100) | CysC | 0.006717 | 4.50E-09 |
| B66 | B181 | WBRSR(mPas_100) | APTT | -0.02735 | 0 |
| B66 | B190 | WBRSR(mPas_100) | Cre | 0.069334 | 0 |
| B66 | B193 | WBRSR(mPas_100) | K | 0.012216 | 0 |
| B66 | B129 | WBRSR(mPas_100) | IBIL | 0.032719 | 0 |
| B66 | B150 | WBRSR(mPas_100) | ALP | 0.042493 | 0 |
| B66 | B84 | WBRSR(mPas_100) | C-PSA | 0.009589 | 0 |
| B66 | B222 | WBRSR(mPas_100) | NS | 0.088589 | 0 |
| B66 | B105 | WBRSR(mPas_100) | anti-U1RNP | -0.01138 | 0 |
| B66 | B95 | WBRSR(mPas_100) | anti-cenpb | -0.00733 | 1.57E-10 |
| B66 | B98 | WBRSR(mPas_100) | anti-Jo-1 | 0.011125 | 0 |
| B66 | B100 | WBRSR(mPas_100) | anti-Ro52 | -0.00623 | 5.38E-08 |
| B66 | B101 | WBRSR(mPas_100) | anti-Sc70 | -0.00685 | 2.23E-09 |
| B66 | B103 | WBRSR(mPas_100) | anti-SSA/Ro | -0.01275 | 0 |
| B66 | B107 | WBRSR(mPas_100) | AnuA | -0.00979 | 0 |
| B66 | B148 | WBRSR(mPas_100) | Anti_TPO | -0.01457 | 0 |
| B66 | B149 | WBRSR(mPas_100) | TGAb | -0.01364 | 0 |
| B66 | B109 | WBRSR(mPas_100) | AMA-M2 | -0.00657 | 9.88E-09 |
| B66 | B110 | WBRSR(mPas_100) | AHA | -0.00899 | 4.44E-15 |

| | | | | | |
|---|---|---|---|---|---|
| B66 | B154 | WBRSR(mPas_100) | RF | 0.006366 | 2.73E-08 |
| B66 | B198 | WBRSR(mPas_100) | P | -0.03364 | 0 |
| B66 | B194 | WBRSR(mPas_100) | Cl | -0.05712 | 0 |
| B66 | B199 | WBRSR(mPas_100) | Mg | 0.017739 | 0 |
| B66 | B155 | WBRSR(mPas_100) | IgA | 0.030033 | 0 |
| B66 | B156 | WBRSR(mPas_100) | IgE | 0.01184 | 0 |
| B66 | B157 | WBRSR(mPas_100) | IgG | -0.00731 | 1.80E-10 |
| B66 | B158 | WBRSR(mPas_100) | IgM | -0.01417 | 0 |
| B66 | B176 | WBRSR(mPas_100) | Ucr | 0.04672 | 0 |
| B66 | B178 | WBRSR(mPas_100) | Ig-U | 0.010539 | 0 |
| B66 | B191 | WBRSR(mPas_100) | Urea | 0.010656 | 0 |
| B66 | B174 | WBRSR(mPas_100) | Urea/Cr | -0.04509 | 0 |
| B66 | B192 | WBRSR(mPas_100) | UA | 0.081483 | 0 |
| B66 | B170 | WBRSR(mPas_100) | U-GLU | 0.021988 | 0 |
| B66 | B179 | WBRSR(mPas_100) | α1-MG | 0.039665 | 0 |
| B66 | B180 | WBRSR(mPas_100) | Tf | 0.010095 | 0 |
| B66 | B182 | WBRSR(mPas_100) | TT | 0.021359 | 0 |
| B66 | B125 | WBRSR(mPas_100) | Cor | 0.035863 | 0 |
| B66 | B138 | WBRSR(mPas_100) | PAB | 0.098544 | 0 |
| B66 | B130 | WBRSR(mPas_100) | GLB | 0.018211 | 0 |
| B66 | B67 | WBRSR(mPas_100) | WBRSR(mPas_50) | 0.995076 | 0 |
| B66 | B221 | WBRSR(mPas_100) | Hei | 0.072041 | 0 |
| B66 | B87 | WBRSR(mPas_100) | NSE | 0.01169 | 0 |
| B66 | B187 | WBRSR(mPas_100) | NSE | 0.008555 | 8.06E-14 |
| B66 | B200 | WBRSR(mPas_100) | OP | 0.015827 | 0 |
| B66 | B220 | WBRSR(mPas_100) | SBP | 0.029859 | 0 |
| B66 | B219 | WBRSR(mPas_100) | OT | -0.00985 | 0 |
| B66 | B218 | WBRSR(mPas_100) | DBP | 0.047247 | 0 |
| B66 | B88 | WBRSR(mPas_100) | CA 242 | -0.00688 | 1.87E-09 |
| B66 | B216 | WBRSR(mPas_100) | BMI | 0.077846 | 0 |
| B66 | B215 | WBRSR(mPas_100) | BMI | 0.051439 | 0 |
| B66 | B131 | WBRSR(mPas_100) | AST | 0.027969 | 0 |
| B66 | B210 | WBRSR(mPas_100) | Hip | 0.051031 | 0 |
| B66 | B209 | WBRSR(mPas_100) | Smoking | 0.04995 | 0 |
| B66 | B90 | WBRSR(mPas_100) | CYFRA 21-1 | 0.007157 | 4.14E-10 |
| B66 | B185 | WBRSR(mPas_100) | PT | -0.0134 | 0 |
| B66 | B186 | WBRSR(mPas_100) | PFC | 0.013827 | 0 |
| B66 | B69 | WBRSR(mPas_100) | PV | 0.146808 | 0 |
| B66 | B173 | WBRSR(mPas_100) | NTT | -0.01321 | 0 |
| B66 | B207 | WBRSR(mPas_100) | WHR | 0.007633 | 2.66E-11 |
| B66 | B206 | WBRSR(mPas_100) | Waist | 0.072342 | 0 |
| B66 | B197 | WBRSR(mPas_100) | AG | 0.034032 | 0 |
| B66 | B204 | WBRSR(mPas_100) | Drinking | 0.049401 | 0 |
| B66 | B112 | WBRSR(mPas_100) | FT4 | 0.01674 | 0 |

| | | | | | |
|---|---|---|---|---|---|
| B66 | B113 | WBRSR(mPas_100) | FT3 | 0.036482 | 0 |
| B66 | B211 | WBRSR(mPas_100) | IOP-R | 0.020073 | 0 |
| B66 | B73 | WBRSR(mPas_100) | ApoA | -0.00568 | 6.98E-07 |
| B66 | B74 | WBRSR(mPas_100) | ApoA1 | -0.0806 | 0 |
| B66 | B75 | WBRSR(mPas_100) | ApoA1/B | -0.06348 | 0 |
| B66 | B76 | WBRSR(mPas_100) | ApoB | 0.109661 | 0 |
| B66 | B132 | WBRSR(mPas_100) | Dbil | 0.01839 | 0 |
| B66 | B81 | WBRSR(mPas_100) | TSGF | -0.00995 | 0 |
| B66 | B77 | WBRSR(mPas_100) | TC | 0.02811 | 0 |
| B66 | B133 | WBRSR(mPas_100) | TBIL | 0.031198 | 0 |
| B66 | B135 | WBRSR(mPas_100) | TBA | 0.008386 | 2.46E-13 |
| B66 | B91 | WBRSR(mPas_100) | TP | 0.042931 | 0 |
| B66 | B114 | WBRSR(mPas_100) | TSH TT3 TT4(TT4) | -0.0066 | 8.31E-09 |
| B66 | B115 | WBRSR(mPas_100) | TSH TT3 TT4(TT3) | 0.023694 | 0 |
| B66 | B212 | WBRSR(mPas_100) | IOP-L | 0.018306 | 0 |
| B67 | B136 | WBRSR(mPas_50) | 5'NT | 0.006529 | 1.20E-08 |
| B67 | B137 | WBRSR(mPas_50) | AFU | 0.040761 | 0 |
| B67 | B160 | WBRSR(mPas_50) | PH | -0.0173 | 0 |
| B67 | B140 | WBRSR(mPas_50) | r-GT | 0.058994 | 0 |
| B67 | B126 | WBRSR(mPas_50) | Alb | 0.044997 | 0 |
| B67 | B161 | WBRSR(mPas_50) | LEU | -0.01876 | 0 |
| B67 | B128 | WBRSR(mPas_50) | ALT | 0.048412 | 0 |
| B67 | B121 | WBRSR(mPas_50) | C3 | 0.006855 | 2.17E-09 |
| B67 | B123 | WBRSR(mPas_50) | hs-CRP1 | 0.044888 | 0 |
| B67 | B111 | WBRSR(mPas_50) | TSH | -0.02147 | 0 |
| B67 | B124 | WBRSR(mPas_50) | ACTH | 0.041684 | 0 |
| B67 | B134 | WBRSR(mPas_50) | CHE | 0.080124 | 0 |
| B67 | B166 | WBRSR(mPas_50) | Pro | 0.023434 | 0 |
| B67 | B70 | WBRSR(mPas_50) | LDL-C | 0.040125 | 0 |
| B67 | B188 | WBRSR(mPas_50) | Ca | 0.011307 | 0 |
| B67 | B71 | WBRSR(mPas_50) | TG | 0.048274 | 0 |
| B67 | B72 | WBRSR(mPas_50) | HDL-C | -0.04675 | 0 |
| B67 | B189 | WBRSR(mPas_50) | eGFB | -0.01169 | 0 |
| B67 | B141 | WBRSR(mPas_50) | AST/ALT | -0.04613 | 0 |
| B67 | B146 | WBRSR(mPas_50) | CysC | 0.006848 | 2.25E-09 |
| B67 | B181 | WBRSR(mPas_50) | APTT | -0.02765 | 0 |
| B67 | B190 | WBRSR(mPas_50) | Cre | 0.069945 | 0 |
| B67 | B193 | WBRSR(mPas_50) | K | 0.012483 | 0 |
| B67 | B129 | WBRSR(mPas_50) | IBIL | 0.033116 | 0 |
| B67 | B150 | WBRSR(mPas_50) | ALP | 0.04256 | 0 |
| B67 | B84 | WBRSR(mPas_50) | C-PSA | 0.009475 | 0 |
| B67 | B222 | WBRSR(mPas_50) | NS | 0.088923 | 0 |
| B67 | B105 | WBRSR(mPas_50) | anti-U1RNP | -0.01162 | 0 |
| B67 | B95 | WBRSR(mPas_50) | anti-cenpb | -0.00723 | 2.76E-10 |

| | | | | | |
|---|---|---|---|---|---|
| B67 | B98 | WBRSR(mPas_50) | anti-Jo-1 | 0.010941 | 0 |
| B67 | B100 | WBRSR(mPas_50) | anti-Ro52 | -0.00649 | 1.43E-08 |
| B67 | B101 | WBRSR(mPas_50) | anti-Sc70 | -0.00696 | 1.21E-09 |
| B67 | B103 | WBRSR(mPas_50) | anti-SSA/Ro | -0.01297 | 0 |
| B67 | B107 | WBRSR(mPas_50) | AnuA | -0.00944 | 0 |
| B67 | B148 | WBRSR(mPas_50) | Anti_TPO | -0.01466 | 0 |
| B67 | B149 | WBRSR(mPas_50) | TGAb | -0.01384 | 0 |
| B67 | B109 | WBRSR(mPas_50) | AMA-M2 | -0.00704 | 7.87E-10 |
| B67 | B110 | WBRSR(mPas_50) | AHA | -0.00888 | 8.88E-15 |
| B67 | B154 | WBRSR(mPas_50) | RF | 0.006243 | 5.02E-08 |
| B67 | B198 | WBRSR(mPas_50) | P | -0.03377 | 0 |
| B67 | B194 | WBRSR(mPas_50) | Cl | -0.0557 | 0 |
| B67 | B199 | WBRSR(mPas_50) | Mg | 0.017632 | 0 |
| B67 | B155 | WBRSR(mPas_50) | IgA | 0.030083 | 0 |
| B67 | B156 | WBRSR(mPas_50) | IgE | 0.011971 | 0 |
| B67 | B157 | WBRSR(mPas_50) | IgG | -0.00791 | 5.12E-12 |
| B67 | B158 | WBRSR(mPas_50) | IgM | -0.0147 | 0 |
| B67 | B176 | WBRSR(mPas_50) | Ucr | 0.047219 | 0 |
| B67 | B178 | WBRSR(mPas_50) | Ig-U | 0.010514 | 0 |
| B67 | B191 | WBRSR(mPas_50) | Urea | 0.010449 | 0 |
| B67 | B174 | WBRSR(mPas_50) | Urea/Cr | -0.04576 | 0 |
| B67 | B192 | WBRSR(mPas_50) | UA | 0.08168 | 0 |
| B67 | B170 | WBRSR(mPas_50) | U-GLU | 0.021992 | 0 |
| B67 | B179 | WBRSR(mPas_50) | α1-MG | 0.039651 | 0 |
| B67 | B180 | WBRSR(mPas_50) | Tf | 0.009981 | 0 |
| B67 | B182 | WBRSR(mPas_50) | TT | 0.021286 | 0 |
| B67 | B125 | WBRSR(mPas_50) | Cor | 0.037367 | 0 |
| B67 | B138 | WBRSR(mPas_50) | PAB | 0.098774 | 0 |
| B67 | B130 | WBRSR(mPas_50) | GLB | 0.016803 | 0 |
| B67 | B221 | WBRSR(mPas_50) | Hei | 0.072566 | 0 |
| B67 | B87 | WBRSR(mPas_50) | NSE | 0.011523 | 0 |
| B67 | B187 | WBRSR(mPas_50) | NSE | 0.008363 | 2.86E-13 |
| B67 | B200 | WBRSR(mPas_50) | OP | 0.016273 | 0 |
| B67 | B220 | WBRSR(mPas_50) | SBP | 0.029494 | 0 |
| B67 | B219 | WBRSR(mPas_50) | OT | -0.00986 | 0 |
| B67 | B218 | WBRSR(mPas_50) | DBP | 0.047434 | 0 |
| B67 | B88 | WBRSR(mPas_50) | CA 242 | -0.00692 | 1.54E-09 |
| B67 | B216 | WBRSR(mPas_50) | BMI | 0.077873 | 0 |
| B67 | B215 | WBRSR(mPas_50) | BMI | 0.051112 | 0 |
| B67 | B131 | WBRSR(mPas_50) | AST | 0.027969 | 0 |
| B67 | B210 | WBRSR(mPas_50) | Hip | 0.051053 | 0 |
| B67 | B209 | WBRSR(mPas_50) | Smoking | 0.050672 | 0 |
| B67 | B90 | WBRSR(mPas_50) | CYFRA 21-1 | 0.007251 | 2.44E-10 |
| B67 | B183 | WBRSR(mPas_50) | PTA | -0.00722 | 2.94E-10 |

| | | | | | |
|---|---|---|---|---|---|
| B67 | B185 | WBRSR(mPas_50) | PT | -0.01391 | 0 |
| B67 | B186 | WBRSR(mPas_50) | PFC | 0.011528 | 0 |
| B67 | B69 | WBRSR(mPas_50) | PV | 0.145414 | 0 |
| B67 | B173 | WBRSR(mPas_50) | NTT | -0.01322 | 0 |
| B67 | B207 | WBRSR(mPas_50) | WHR | 0.007628 | 2.75E-11 |
| B67 | B206 | WBRSR(mPas_50) | Waist | 0.072288 | 0 |
| B67 | B197 | WBRSR(mPas_50) | AG | 0.034039 | 0 |
| B67 | B204 | WBRSR(mPas_50) | Drinking | 0.049951 | 0 |
| B67 | B112 | WBRSR(mPas_50) | FT4 | 0.017761 | 0 |
| B67 | B113 | WBRSR(mPas_50) | FT3 | 0.036806 | 0 |
| B67 | B211 | WBRSR(mPas_50) | IOP-R | 0.020957 | 0 |
| B67 | B73 | WBRSR(mPas_50) | ApoA | -0.00613 | 8.72E-08 |
| B67 | B74 | WBRSR(mPas_50) | ApoA1 | -0.07714 | 0 |
| B67 | B75 | WBRSR(mPas_50) | ApoA1/B | -0.06176 | 0 |
| B67 | B76 | WBRSR(mPas_50) | ApoB | 0.107737 | 0 |
| B67 | B132 | WBRSR(mPas_50) | Dbil | 0.018353 | 0 |
| B67 | B81 | WBRSR(mPas_50) | TSGF | -0.01239 | 0 |
| B67 | B77 | WBRSR(mPas_50) | TC | 0.02784 | 0 |
| B67 | B133 | WBRSR(mPas_50) | TBIL | 0.031508 | 0 |
| B67 | B135 | WBRSR(mPas_50) | TBA | 0.008389 | 2.40E-13 |
| B67 | B91 | WBRSR(mPas_50) | TP | 0.042331 | 0 |
| B67 | B114 | WBRSR(mPas_50) | TSH TT3 TT4(TT4) | -0.00657 | 9.46E-09 |
| B67 | B115 | WBRSR(mPas_50) | TSH TT3 TT4(TT3) | 0.023618 | 0 |
| B67 | B212 | WBRSR(mPas_50) | IOP-L | 0.01909 | 0 |
| B69 | B137 | PV | AFU | 0.009355 | 4.44E-16 |
| B69 | B160 | PV | PH | -0.01453 | 0 |
| B69 | B140 | PV | r-GT | 0.019253 | 0 |
| B69 | B126 | PV | Alb | 0.024067 | 0 |
| B69 | B127 | PV | A-G ratio | -0.05445 | 0 |
| B69 | B161 | PV | LEU | 0.005876 | 2.90E-07 |
| B69 | B128 | PV | ALT | 0.0086 | 6.00E-14 |
| B69 | B121 | PV | C3 | 0.012645 | 0 |
| B69 | B122 | PV | C4 | 0.012188 | 0 |
| B69 | B123 | PV | hs-CRP1 | 0.101694 | 0 |
| B69 | B111 | PV | TSH | 0.008035 | 2.29E-12 |
| B69 | B124 | PV | ACTH | 0.019867 | 0 |
| B69 | B134 | PV | CHE | 0.04015 | 0 |
| B69 | B70 | PV | LDL-C | 0.029419 | 0 |
| B69 | B188 | PV | Ca | 0.007664 | 2.21E-11 |
| B69 | B71 | PV | TG | 0.026669 | 0 |
| B69 | B189 | PV | eGFB | -0.00584 | 3.45E-07 |
| B69 | B181 | PV | APTT | -0.01487 | 0 |
| B69 | B193 | PV | K | 0.030719 | 0 |
| B69 | B150 | PV | ALP | 0.023857 | 0 |

| | | | | | |
|---|---|---|---|---|---|
| B69 | B102 | PV | anti-Sm | 0.009064 | 2.66E-15 |
| B69 | B103 | PV | anti-SSA/Ro | 0.026871 | 0 |
| B69 | B107 | PV | AnuA | 0.005704 | 6.36E-07 |
| B69 | B151 | PV | Anti-CCP | 0.022775 | 0 |
| B69 | B148 | PV | Anti_TPO | 0.007209 | 3.09E-10 |
| B69 | B149 | PV | TGAb | 0.00835 | 3.09E-13 |
| B69 | B109 | PV | AMA-M2 | 0.006359 | 2.83E-08 |
| B69 | B110 | PV | AHA | 0.013166 | 0 |
| B69 | B154 | PV | RF | 0.006536 | 1.15E-08 |
| B69 | B198 | PV | P | 0.006445 | 1.83E-08 |
| B69 | B194 | PV | Cl | -0.02943 | 0 |
| B69 | B199 | PV | Mg | 0.008601 | 5.95E-14 |
| B69 | B155 | PV | IgA | 0.054288 | 0 |
| B69 | B156 | PV | IgE | 0.018811 | 0 |
| B69 | B157 | PV | IgG | 0.098155 | 0 |
| B69 | B158 | PV | IgM | 0.021485 | 0 |
| B69 | B195 | PV | Na | -0.02285 | 0 |
| B69 | B178 | PV | Ig-U | 0.011914 | 0 |
| B69 | B192 | PV | UA | 0.008648 | 4.35E-14 |
| B69 | B170 | PV | U-GLU | 0.011282 | 0 |
| B69 | B175 | PV | ALB-U | 0.009763 | 0 |
| B69 | B177 | PV | ACR | 0.012657 | 0 |
| B69 | B179 | PV | α1-MG | 0.010281 | 0 |
| B69 | B180 | PV | Tf | 0.008351 | 3.08E-13 |
| B69 | B182 | PV | TT | -0.02803 | 0 |
| B69 | B125 | PV | Cor | 0.045723 | 0 |
| B69 | B138 | PV | PAB | 0.008497 | 1.18E-13 |
| B69 | B130 | PV | GLB | 0.087785 | 0 |
| B69 | B221 | PV | Hei | -0.01278 | 0 |
| B69 | B200 | PV | OP | -0.01499 | 0 |
| B69 | B220 | PV | SBP | 0.017314 | 0 |
| B69 | B218 | PV | DBP | 0.018503 | 0 |
| B69 | B215 | PV | BMI | 0.009074 | 2.22E-15 |
| B69 | B131 | PV | AST | 0.017833 | 0 |
| B69 | B209 | PV | Smoking | -0.00587 | 2.99E-07 |
| B69 | B139 | PV | ADA | 0.028199 | 0 |
| B69 | B183 | PV | PTA | 0.040337 | 0 |
| B69 | B184 | PV | INR | 0.008817 | 1.38E-14 |
| B69 | B185 | PV | PT | 0.006881 | 1.88E-09 |
| B69 | B186 | PV | PFC | 0.088952 | 0 |
| B69 | B206 | PV | Waist | 0.007343 | 1.45E-10 |
| B69 | B204 | PV | Drinking | -0.0071 | 5.65E-10 |
| B69 | B112 | PV | FT4 | 0.03458 | 0 |
| B69 | B73 | PV | ApoA | 0.027433 | 0 |

| | | | | | |
|---|---|---|---|---|---|
| B69 | B74 | PV | ApoA1 | 0.032591 | 0 |
| B69 | B75 | PV | ApoA1/B | -0.00978 | 0 |
| B69 | B76 | PV | ApoB | 0.062832 | 0 |
| B69 | B132 | PV | Dbil | -0.01076 | 0 |
| B69 | B81 | PV | TSGF | -0.03746 | 0 |
| B69 | B77 | PV | TC | 0.031198 | 0 |
| B69 | B91 | PV | TP | 0.089977 | 0 |
| B69 | B114 | PV | TSH TT3 TT4(TT4) | 0.042099 | 0 |
| B69 | B115 | PV | TSH TT3 TT4(TT3) | 0.008632 | 4.84E-14 |
| B7 | B17 | WB-Cop | CD3+CD16+CD56+NK | 0.005648 | 8.19E-07 |
| B7 | B21 | WB-Cop | CD3+CD4-CD8- | 0.00728 | 2.07E-10 |
| B7 | B92 | WB-Cop | ALDH2-Glu504Glu | 0.007169 | 3.88E-10 |
| B7 | B29 | WB-Cop | E2 | -0.00961 | 0 |
| B7 | B145 | WB-Cop | BGP | 0.005741 | 5.38E-07 |
| B7 | B100 | WB-Cop | anti-Ro52 | 0.006655 | 6.25E-09 |
| B7 | B194 | WB-Cop | Cl | -0.00624 | 5.04E-08 |
| B7 | B8 | WB-Cop | WB-Zinc | 0.12801 | 0 |
| B7 | B139 | WB-Cop | ADA | 0.006056 | 1.24E-07 |
| B7 | B183 | WB-Cop | PTA | -0.00804 | 2.29E-12 |
| B70 | B137 | LDL-C | AFU | 0.011305 | 0 |
| B70 | B160 | LDL-C | PH | -0.06247 | 0 |
| B70 | B140 | LDL-C | r-GT | 0.112318 | 0 |
| B70 | B118 | LDL-C | CEA1 | 0.009693 | 0 |
| B70 | B126 | LDL-C | Alb | 0.049628 | 0 |
| B70 | B127 | LDL-C | A-G ratio | -0.04588 | 0 |
| B70 | B161 | LDL-C | LEU | -0.01366 | 0 |
| B70 | B128 | LDL-C | ALT | 0.109537 | 0 |
| B70 | B111 | LDL-C | TSH | 0.025447 | 0 |
| B70 | B134 | LDL-C | CHE | 0.061736 | 0 |
| B70 | B166 | LDL-C | Pro | 0.018909 | 0 |
| B70 | B188 | LDL-C | Ca | 0.026635 | 0 |
| B70 | B71 | LDL-C | TG | 0.054724 | 0 |
| B70 | B72 | LDL-C | HDL-C | -0.01584 | 0 |
| B70 | B189 | LDL-C | eGFB | -0.18963 | 0 |
| B70 | B141 | LDL-C | AST/ALT | -0.12327 | 0 |
| B70 | B146 | LDL-C | CysC | 0.011112 | 0 |
| B70 | B181 | LDL-C | APTT | -0.01575 | 0 |
| B70 | B190 | LDL-C | Cre | 0.133332 | 0 |
| B70 | B224 | LDL-C | pmh | 0.008792 | 1.64E-14 |
| B70 | B193 | LDL-C | K | 0.021605 | 0 |
| B70 | B129 | LDL-C | IBIL | -0.01132 | 0 |
| B70 | B150 | LDL-C | ALP | 0.042074 | 0 |
| B70 | B84 | LDL-C | C-PSA | 0.007996 | 2.93E-12 |
| B70 | B222 | LDL-C | NS | 0.17903 | 0 |

| | | | | | |
|---|---|---|---|---|---|
| B70 | B148 | LDL-C | Anti_TPO | -0.00721 | 3.15E-10 |
| B70 | B149 | LDL-C | TGAb | -0.01223 | 0 |
| B70 | B194 | LDL-C | Cl | -0.01945 | 0 |
| B70 | B199 | LDL-C | Mg | 0.022106 | 0 |
| B70 | B155 | LDL-C | IgA | -0.00874 | 2.26E-14 |
| B70 | B157 | LDL-C | IgG | -0.01311 | 0 |
| B70 | B158 | LDL-C | IgM | -0.00662 | 7.44E-09 |
| B70 | B195 | LDL-C | Na | 0.016551 | 0 |
| B70 | B176 | LDL-C | Ucr | 0.01846 | 0 |
| B70 | B191 | LDL-C | Urea | 0.096648 | 0 |
| B70 | B174 | LDL-C | Urea/Cr | -0.01725 | 0 |
| B70 | B192 | LDL-C | UA | 0.154107 | 0 |
| B70 | B170 | LDL-C | U-GLU | 0.020398 | 0 |
| B70 | B182 | LDL-C | TT | 0.008865 | 9.99E-15 |
| B70 | B153 | LDL-C | Glu | 0.020301 | 0 |
| B70 | B138 | LDL-C | PAB | 0.0379 | 0 |
| B70 | B130 | LDL-C | GLB | 0.091565 | 0 |
| B70 | B86 | LDL-C | TBhCG | -0.00571 | 6.33E-07 |
| B70 | B221 | LDL-C | Hei | 0.032399 | 0 |
| B70 | B187 | LDL-C | NSE | 0.007472 | 6.87E-11 |
| B70 | B200 | LDL-C | OP | 0.018967 | 0 |
| B70 | B220 | LDL-C | SBP | 0.152743 | 0 |
| B70 | B219 | LDL-C | OT | 0.040882 | 0 |
| B70 | B218 | LDL-C | DBP | 0.149865 | 0 |
| B70 | B216 | LDL-C | BMI | 0.181468 | 0 |
| B70 | B215 | LDL-C | BMI | 0.220734 | 0 |
| B70 | B131 | LDL-C | AST | 0.056807 | 0 |
| B70 | B172 | LDL-C | KB | -0.01811 | 0 |
| B70 | B210 | LDL-C | Hip | 0.155064 | 0 |
| B70 | B209 | LDL-C | Smoking | 0.095043 | 0 |
| B70 | B196 | LDL-C | TCO2 | 0.007556 | 4.19E-11 |
| B70 | B183 | LDL-C | PTA | -0.0095 | 0 |
| B70 | B185 | LDL-C | PT | -0.01274 | 0 |
| B70 | B186 | LDL-C | PFC | 0.009659 | 0 |
| B70 | B207 | LDL-C | WHR | 0.029713 | 0 |
| B70 | B206 | LDL-C | Waist | 0.235114 | 0 |
| B70 | B205 | LDL-C | KDA | 0.012208 | 0 |
| B70 | B78 | LDL-C | HBcAb | 0.009177 | 1.11E-15 |
| B70 | B197 | LDL-C | AG | 0.016327 | 0 |
| B70 | B204 | LDL-C | Drinking | 0.065971 | 0 |
| B70 | B80 | LDL-C | anti-HP IgG | 0.016841 | 0 |
| B70 | B112 | LDL-C | FT4 | -0.03144 | 0 |
| B70 | B113 | LDL-C | FT3 | 0.009328 | 4.44E-16 |
| B70 | B211 | LDL-C | IOP-R | 0.010382 | 0 |

| | | | | | |
|---|---|---|---|---|---|
| B70 | B201 | LDL-C | e-habits | 0.02633 | 0 |
| B70 | B73 | LDL-C | ApoA | 0.065895 | 0 |
| B70 | B74 | LDL-C | ApoA1 | 0.031286 | 0 |
| B70 | B75 | LDL-C | ApoA1/B | -0.12256 | 0 |
| B70 | B76 | LDL-C | ApoB | 0.29875 | 0 |
| B70 | B132 | LDL-C | Dbil | -0.17898 | 0 |
| B70 | B77 | LDL-C | TC | 0.702373 | 0 |
| B70 | B133 | LDL-C | TBIL | -0.05625 | 0 |
| B70 | B91 | LDL-C | TP | 0.108741 | 0 |
| B70 | B212 | LDL-C | IOP-L | 0.012175 | 0 |
| B71 | B136 | TG | 5'NT | 0.008487 | 1.27E-13 |
| B71 | B137 | TG | AFU | 0.026811 | 0 |
| B71 | B160 | TG | PH | -0.10558 | 0 |
| B71 | B140 | TG | r-GT | 0.280684 | 0 |
| B71 | B82 | TG | CEA | 0.007695 | 1.84E-11 |
| B71 | B118 | TG | CEA1 | 0.010273 | 0 |
| B71 | B126 | TG | Alb | 0.088561 | 0 |
| B71 | B127 | TG | A-G ratio | -0.11386 | 0 |
| B71 | B161 | TG | LEU | -0.03477 | 0 |
| B71 | B128 | TG | ALT | 0.203419 | 0 |
| B71 | B120 | TG | HCV-cAg | 0.012953 | 0 |
| B71 | B121 | TG | C3 | 0.010166 | 0 |
| B71 | B122 | TG | C4 | 0.008321 | 3.73E-13 |
| B71 | B225 | TG | hPBG | 0.012083 | 0 |
| B71 | B164 | TG | COM | -0.00731 | 1.78E-10 |
| B71 | B123 | TG | hs-CRP1 | 0.010398 | 0 |
| B71 | B111 | TG | TSH | 0.012425 | 0 |
| B71 | B124 | TG | ACTH | 0.00923 | 8.88E-16 |
| B71 | B134 | TG | CHE | 0.084892 | 0 |
| B71 | B166 | TG | Pro | 0.049322 | 0 |
| B71 | B188 | TG | Ca | 0.020504 | 0 |
| B71 | B72 | TG | HDL-C | -0.38296 | 0 |
| B71 | B189 | TG | eGFB | -0.10316 | 0 |
| B71 | B141 | TG | AST/ALT | -0.17909 | 0 |
| B71 | B146 | TG | CysC | 0.037524 | 0 |
| B71 | B181 | TG | APTT | -0.01561 | 0 |
| B71 | B190 | TG | Cre | 0.155513 | 0 |
| B71 | B193 | TG | K | -0.01137 | 0 |
| B71 | B129 | TG | IBIL | -0.03346 | 0 |
| B71 | B150 | TG | ALP | 0.052395 | 0 |
| B71 | B84 | TG | C-PSA | 0.009058 | 2.66E-15 |
| B71 | B222 | TG | NS | 0.286972 | 0 |
| B71 | B148 | TG | Anti_TPO | -0.01557 | 0 |
| B71 | B149 | TG | TGAb | -0.01816 | 0 |

| | | | | | |
|---|---|---|---|---|---|
| B71 | B198 | TG | P | -0.01873 | 0 |
| B71 | B194 | TG | Cl | -0.02835 | 0 |
| B71 | B199 | TG | Mg | 0.011226 | 0 |
| B71 | B155 | TG | IgA | 0.020054 | 0 |
| B71 | B157 | TG | IgG | -0.01533 | 0 |
| B71 | B195 | TG | Na | -0.0077 | 1.74E-11 |
| B71 | B176 | TG | Ucr | 0.034027 | 0 |
| B71 | B191 | TG | Urea | 0.05374 | 0 |
| B71 | B174 | TG | Urea/Cr | -0.06696 | 0 |
| B71 | B192 | TG | UA | 0.30307 | 0 |
| B71 | B170 | TG | U-GLU | 0.104891 | 0 |
| B71 | B175 | TG | ALB-U | 0.015604 | 0 |
| B71 | B177 | TG | ACR | 0.008013 | 2.63E-12 |
| B71 | B179 | TG | α1-MG | 0.018284 | 0 |
| B71 | B153 | TG | Glu | 0.039806 | 0 |
| B71 | B138 | TG | PAB | 0.073812 | 0 |
| B71 | B130 | TG | GLB | 0.216693 | 0 |
| B71 | B221 | TG | Hei | 0.13065 | 0 |
| B71 | B87 | TG | NSE | 0.010395 | 0 |
| B71 | B187 | TG | NSE | 0.008429 | 1.86E-13 |
| B71 | B200 | TG | OP | 0.006891 | 1.79E-09 |
| B71 | B220 | TG | SBP | 0.173343 | 0 |
| B71 | B219 | TG | OT | 0.011927 | 0 |
| B71 | B218 | TG | DBP | 0.203935 | 0 |
| B71 | B88 | TG | CA 242 | -0.00853 | 9.66E-14 |
| B71 | B216 | TG | BMI | 0.29716 | 0 |
| B71 | B215 | TG | BMI | 0.300896 | 0 |
| B71 | B131 | TG | AST | 0.134622 | 0 |
| B71 | B172 | TG | KB | -0.01196 | 0 |
| B71 | B210 | TG | Hip | 0.210564 | 0 |
| B71 | B209 | TG | Smoking | 0.174543 | 0 |
| B71 | B90 | TG | CYFRA 21-1 | 0.008586 | 6.57E-14 |
| B71 | B139 | TG | ADA | -0.02604 | 0 |
| B71 | B196 | TG | TCO2 | -0.00572 | 5.92E-07 |
| B71 | B185 | TG | PT | -0.01324 | 0 |
| B71 | B173 | TG | NTT | -0.01005 | 0 |
| B71 | B207 | TG | WHR | 0.036362 | 0 |
| B71 | B206 | TG | Waist | 0.322684 | 0 |
| B71 | B197 | TG | AG | 0.018407 | 0 |
| B71 | B204 | TG | Drinking | 0.170423 | 0 |
| B71 | B203 | TG | DH | 0.025036 | 0 |
| B71 | B80 | TG | anti-HP IgG | 0.006074 | 1.14E-07 |
| B71 | B112 | TG | FT4 | -0.04249 | 0 |
| B71 | B113 | TG | FT3 | 0.02873 | 0 |

| | | | | | |
|---|---|---|---|---|---|
| B71 | B211 | TG | IOP-R | 0.024473 | 0 |
| B71 | B73 | TG | ApoA | -0.02545 | 0 |
| B71 | B74 | TG | ApoA1 | -0.071 | 0 |
| B71 | B75 | TG | ApoA1/B | -0.01134 | 0 |
| B71 | B76 | TG | ApoB | 0.052268 | 0 |
| B71 | B132 | TG | Dbil | -0.09963 | 0 |
| B71 | B77 | TG | TC | 0.277804 | 0 |
| B71 | B133 | TG | TBIL | -0.05322 | 0 |
| B71 | B135 | TG | TBA | 0.015314 | 0 |
| B71 | B91 | TG | TP | 0.239175 | 0 |
| B71 | B114 | TG | TSH TT3 TT4(TT4) | -0.0326 | 0 |
| B71 | B115 | TG | TSH TT3 TT4(TT3) | 0.006213 | 5.83E-08 |
| B71 | B212 | TG | IOP-L | 0.025153 | 0 |
| B72 | B142 | HDL-C | 25-(OH)D | 0.008602 | 5.91E-14 |
| B72 | B137 | HDL-C | AFU | -0.01326 | 0 |
| B72 | B160 | HDL-C | PH | 0.076333 | 0 |
| B72 | B140 | HDL-C | r-GT | -0.1392 | 0 |
| B72 | B118 | HDL-C | CEA1 | -0.00861 | 5.51E-14 |
| B72 | B126 | HDL-C | Alb | -0.0065 | 1.36E-08 |
| B72 | B127 | HDL-C | A-G ratio | 0.083458 | 0 |
| B72 | B161 | HDL-C | LEU | 0.05353 | 0 |
| B72 | B128 | HDL-C | ALT | -0.20615 | 0 |
| B72 | B225 | HDL-C | hPBG | -0.00976 | 0 |
| B72 | B164 | HDL-C | COM | 0.00601 | 1.55E-07 |
| B72 | B123 | HDL-C | hs-CRP1 | -0.03048 | 0 |
| B72 | B111 | HDL-C | TSH | 0.016563 | 0 |
| B72 | B124 | HDL-C | ACTH | -0.00629 | 3.89E-08 |
| B72 | B134 | HDL-C | CHE | -0.06157 | 0 |
| B72 | B166 | HDL-C | Pro | -0.08246 | 0 |
| B72 | B189 | HDL-C | eGFB | 0.057565 | 0 |
| B72 | B141 | HDL-C | AST/ALT | 0.259041 | 0 |
| B72 | B146 | HDL-C | CysC | -0.02819 | 0 |
| B72 | B190 | HDL-C | Cre | -0.25863 | 0 |
| B72 | B193 | HDL-C | K | 0.01109 | 0 |
| B72 | B129 | HDL-C | IBIL | 0.02371 | 0 |
| B72 | B150 | HDL-C | ALP | -0.04288 | 0 |
| B72 | B84 | HDL-C | C-PSA | -0.01331 | 0 |
| B72 | B222 | HDL-C | NS | -0.38355 | 0 |
| B72 | B148 | HDL-C | Anti_TPO | 0.018597 | 0 |
| B72 | B149 | HDL-C | TGAb | 0.020066 | 0 |
| B72 | B198 | HDL-C | P | 0.026812 | 0 |
| B72 | B85 | HDL-C | SCC | -0.006 | 1.62E-07 |
| B72 | B194 | HDL-C | Cl | -0.00659 | 8.57E-09 |
| B72 | B155 | HDL-C | IgA | -0.01415 | 0 |

| | | | | | |
|---|---|---|---|---|---|
| B72 | B157 | HDL-C | IgG | 0.012986 | 0 |
| B72 | B158 | HDL-C | IgM | 0.008263 | 5.43E-13 |
| B72 | B195 | HDL-C | Na | -0.01533 | 0 |
| B72 | B176 | HDL-C | Ucr | -0.06213 | 0 |
| B72 | B178 | HDL-C | Ig-U | -0.00733 | 1.57E-10 |
| B72 | B191 | HDL-C | Urea | -0.04755 | 0 |
| B72 | B174 | HDL-C | Urea/Cr | 0.142797 | 0 |
| B72 | B192 | HDL-C | UA | -0.37684 | 0 |
| B72 | B170 | HDL-C | U-GLU | -0.04628 | 0 |
| B72 | B175 | HDL-C | ALB-U | -0.01001 | 0 |
| B72 | B179 | HDL-C | α1-MG | -0.0213 | 0 |
| B72 | B180 | HDL-C | Tf | -0.00603 | 1.38E-07 |
| B72 | B182 | HDL-C | TT | -0.00903 | 3.11E-15 |
| B72 | B125 | HDL-C | Cor | 0.008015 | 2.60E-12 |
| B72 | B153 | HDL-C | Glu | -0.01956 | 0 |
| B72 | B138 | HDL-C | PAB | -0.03913 | 0 |
| B72 | B130 | HDL-C | GLB | -0.05751 | 0 |
| B72 | B86 | HDL-C | TBhCG | 0.009394 | 2.22E-16 |
| B72 | B221 | HDL-C | Hei | -0.26528 | 0 |
| B72 | B200 | HDL-C | OP | -0.01835 | 0 |
| B72 | B220 | HDL-C | SBP | -0.11651 | 0 |
| B72 | B219 | HDL-C | OT | 0.020387 | 0 |
| B72 | B218 | HDL-C | DBP | -0.12187 | 0 |
| B72 | B88 | HDL-C | CA 242 | 0.016927 | 0 |
| B72 | B216 | HDL-C | BMI | -0.40889 | 0 |
| B72 | B215 | HDL-C | BMI | -0.35792 | 0 |
| B72 | B131 | HDL-C | AST | -0.07013 | 0 |
| B72 | B172 | HDL-C | KB | 0.024672 | 0 |
| B72 | B210 | HDL-C | Hip | -0.29564 | 0 |
| B72 | B209 | HDL-C | Smoking | -0.16393 | 0 |
| B72 | B90 | HDL-C | CYFRA 21-1 | -0.00819 | 8.77E-13 |
| B72 | B196 | HDL-C | TCO2 | 0.008552 | 8.24E-14 |
| B72 | B173 | HDL-C | NTT | 0.022222 | 0 |
| B72 | B207 | HDL-C | WHR | -0.04233 | 0 |
| B72 | B206 | HDL-C | Waist | -0.39315 | 0 |
| B72 | B205 | HDL-C | KDA | 0.016929 | 0 |
| B72 | B78 | HDL-C | HBcAb | 0.008097 | 1.56E-12 |
| B72 | B197 | HDL-C | AG | -0.01151 | 0 |
| B72 | B204 | HDL-C | Drinking | -0.12023 | 0 |
| B72 | B203 | HDL-C | DH | -0.03689 | 0 |
| B72 | B80 | HDL-C | anti-HP IgG | -0.01167 | 0 |
| B72 | B112 | HDL-C | FT4 | 0.03819 | 0 |
| B72 | B113 | HDL-C | FT3 | -0.05243 | 0 |
| B72 | B73 | HDL-C | ApoA | 0.02899 | 0 |

| | | | | | |
|---|---|---|---|---|---|
| B72 | B74 | HDL-C | ApoA1 | 0.252805 | 0 |
| B72 | B75 | HDL-C | ApoA1/B | 0.086103 | 0 |
| B72 | B76 | HDL-C | ApoB | -0.03811 | 0 |
| B72 | B132 | HDL-C | Dbil | 0.068316 | 0 |
| B72 | B81 | HDL-C | TSGF | -0.02987 | 0 |
| B72 | B77 | HDL-C | TC | 0.122275 | 0 |
| B72 | B133 | HDL-C | TBIL | 0.036277 | 0 |
| B72 | B135 | HDL-C | TBA | -0.00824 | 6.13E-13 |
| B72 | B91 | HDL-C | TP | -0.05218 | 0 |
| B72 | B114 | HDL-C | TSH TT3 TT4(TT4) | 0.019044 | 0 |
| B72 | B115 | HDL-C | TSH TT3 TT4(TT3) | -0.02401 | 0 |
| B73 | B136 | ApoA | 5'NT | -0.0064 | 2.30E-08 |
| B73 | B137 | ApoA | AFU | -0.02361 | 0 |
| B73 | B140 | ApoA | r-GT | -0.02287 | 0 |
| B73 | B126 | ApoA | Alb | -0.01813 | 0 |
| B73 | B127 | ApoA | A-G ratio | -0.0166 | 0 |
| B73 | B128 | ApoA | ALT | -0.02009 | 0 |
| B73 | B123 | ApoA | hs-CRP1 | 0.014101 | 0 |
| B73 | B124 | ApoA | ACTH | -0.0064 | 2.29E-08 |
| B73 | B134 | ApoA | CHE | -0.00759 | 3.39E-11 |
| B73 | B166 | ApoA | Pro | -0.00642 | 2.05E-08 |
| B73 | B189 | ApoA | eGFB | -0.00875 | 2.20E-14 |
| B73 | B141 | ApoA | AST/ALT | 0.012548 | 0 |
| B73 | B190 | ApoA | Cre | -0.00912 | 1.78E-15 |
| B73 | B193 | ApoA | K | 0.022317 | 0 |
| B73 | B222 | ApoA | NS | -0.0229 | 0 |
| B73 | B154 | ApoA | RF | 0.008034 | 2.32E-12 |
| B73 | B198 | ApoA | P | 0.014025 | 0 |
| B73 | B199 | ApoA | Mg | 0.006078 | 1.12E-07 |
| B73 | B159 | ApoA | β2-MG | -0.00605 | 1.25E-07 |
| B73 | B176 | ApoA | Ucr | -0.0069 | 1.67E-09 |
| B73 | B174 | ApoA | Urea/Cr | 0.00622 | 5.61E-08 |
| B73 | B192 | ApoA | UA | -0.01918 | 0 |
| B73 | B170 | ApoA | U-GLU | -0.00773 | 1.46E-11 |
| B73 | B179 | ApoA | α1-MG | -0.00564 | 8.65E-07 |
| B73 | B182 | ApoA | TT | -0.00737 | 1.26E-10 |
| B73 | B138 | ApoA | PAB | -0.02642 | 0 |
| B73 | B130 | ApoA | GLB | 0.011915 | 0 |
| B73 | B221 | ApoA | Hei | -0.02417 | 0 |
| B73 | B200 | ApoA | OP | -0.00636 | 2.83E-08 |
| B73 | B218 | ApoA | DBP | -0.00699 | 1.06E-09 |
| B73 | B216 | ApoA | BMI | -0.0248 | 0 |
| B73 | B215 | ApoA | BMI | -0.01493 | 0 |
| B73 | B131 | ApoA | AST | -0.01493 | 0 |

| | | | | | |
|---|---|---|---|---|---|
| B73 | B210 | ApoA | Hip | -0.01603 | 0 |
| B73 | B209 | ApoA | Smoking | -0.01125 | 0 |
| B73 | B139 | ApoA | ADA | 0.007752 | 1.31E-11 |
| B73 | B186 | ApoA | PFC | 0.018223 | 0 |
| B73 | B206 | ApoA | Waist | -0.01855 | 0 |
| B73 | B204 | ApoA | Drinking | -0.01438 | 0 |
| B73 | B113 | ApoA | FT3 | -0.01222 | 0 |
| B73 | B74 | ApoA | ApoA1 | 0.063271 | 0 |
| B73 | B75 | ApoA | ApoA1/B | -0.08203 | 0 |
| B73 | B76 | ApoA | ApoB | 0.220151 | 0 |
| B73 | B132 | ApoA | Dbil | -0.015 | 0 |
| B73 | B77 | ApoA | TC | 0.045832 | 0 |
| B73 | B135 | ApoA | TBA | -0.02263 | 0 |
| B73 | B114 | ApoA | TSH TT3 TT4(TT4) | 0.005966 | 1.90E-07 |
| B74 | B137 | ApoA1 | AFU | -0.00935 | 4.44E-16 |
| B74 | B92 | ApoA1 | ALDH2-Glu504Glu | 0.014405 | 0 |
| B74 | B93 | ApoA1 | ALDH2-Glu504Lys | -0.01241 | 0 |
| B74 | B140 | ApoA1 | r-GT | -0.01811 | 0 |
| B74 | B94 | ApoA1 | ALDH2-Lys504Lys | -0.00643 | 1.95E-08 |
| B74 | B143 | ApoA1 | β-CTx | -0.01269 | 0 |
| B74 | B126 | ApoA1 | Alb | 0.043187 | 0 |
| B74 | B127 | ApoA1 | A-G ratio | 0.008912 | 7.11E-15 |
| B74 | B161 | ApoA1 | LEU | 0.019324 | 0 |
| B74 | B128 | ApoA1 | ALT | -0.04725 | 0 |
| B74 | B121 | ApoA1 | C3 | -0.00728 | 2.07E-10 |
| B74 | B123 | ApoA1 | hs-CRP1 | -0.05786 | 0 |
| B74 | B111 | ApoA1 | TSH | 0.019688 | 0 |
| B74 | B124 | ApoA1 | ACTH | -0.0102 | 0 |
| B74 | B165 | ApoA1 | bilirubin | -0.00809 | 1.65E-12 |
| B74 | B134 | ApoA1 | CHE | -0.06623 | 0 |
| B74 | B166 | ApoA1 | Pro | -0.02553 | 0 |
| B74 | B188 | ApoA1 | Ca | 0.007503 | 5.74E-11 |
| B74 | B141 | ApoA1 | AST/ALT | 0.053786 | 0 |
| B74 | B146 | ApoA1 | CysC | -0.00967 | 0 |
| B74 | B181 | ApoA1 | APTT | -0.0215 | 0 |
| B74 | B190 | ApoA1 | Cre | -0.07246 | 0 |
| B74 | B193 | ApoA1 | K | 0.040426 | 0 |
| B74 | B129 | ApoA1 | IBIL | -0.01964 | 0 |
| B74 | B150 | ApoA1 | ALP | -0.02016 | 0 |
| B74 | B84 | ApoA1 | C-PSA | -0.00749 | 6.01E-11 |
| B74 | B222 | ApoA1 | NS | -0.10818 | 0 |
| B74 | B95 | ApoA1 | anti-cenpb | 0.007713 | 1.65E-11 |
| B74 | B148 | ApoA1 | Anti_TPO | 0.007453 | 7.69E-11 |
| B74 | B149 | ApoA1 | TGAb | 0.007208 | 3.11E-10 |

| | | | | | |
|---|---|---|---|---|---|
| B74 | B154 | ApoA1 | RF | -0.00704 | 7.81E-10 |
| B74 | B198 | ApoA1 | P | 0.063598 | 0 |
| B74 | B194 | ApoA1 | Cl | -0.05405 | 0 |
| B74 | B199 | ApoA1 | Mg | 0.037822 | 0 |
| B74 | B155 | ApoA1 | IgA | -0.01476 | 0 |
| B74 | B157 | ApoA1 | IgG | 0.012839 | 0 |
| B74 | B176 | ApoA1 | Ucr | -0.04821 | 0 |
| B74 | B178 | ApoA1 | Ig-U | -0.00804 | 2.28E-12 |
| B74 | B174 | ApoA1 | Urea/Cr | 0.05692 | 0 |
| B74 | B192 | ApoA1 | UA | -0.09383 | 0 |
| B74 | B170 | ApoA1 | U-GLU | -0.01231 | 0 |
| B74 | B177 | ApoA1 | ACR | 0.006985 | 1.07E-09 |
| B74 | B179 | ApoA1 | α1-MG | -0.03048 | 0 |
| B74 | B180 | ApoA1 | Tf | -0.00813 | 1.28E-12 |
| B74 | B125 | ApoA1 | Cor | 0.01652 | 0 |
| B74 | B138 | ApoA1 | PAB | -0.03162 | 0 |
| B74 | B221 | ApoA1 | Hei | -0.09676 | 0 |
| B74 | B87 | ApoA1 | NSE | -0.00649 | 1.46E-08 |
| B74 | B219 | ApoA1 | OT | 0.008687 | 3.33E-14 |
| B74 | B218 | ApoA1 | DBP | -0.00764 | 2.55E-11 |
| B74 | B88 | ApoA1 | CA 242 | 0.007821 | 8.57E-12 |
| B74 | B216 | ApoA1 | BMI | -0.11399 | 0 |
| B74 | B215 | ApoA1 | BMI | -0.083 | 0 |
| B74 | B131 | ApoA1 | AST | -0.00939 | 2.22E-16 |
| B74 | B210 | ApoA1 | Hip | -0.07465 | 0 |
| B74 | B209 | ApoA1 | Smoking | -0.04632 | 0 |
| B74 | B139 | ApoA1 | ADA | 0.006078 | 1.12E-07 |
| B74 | B196 | ApoA1 | TCO2 | 0.024544 | 0 |
| B74 | B183 | ApoA1 | PTA | -0.02356 | 0 |
| B74 | B185 | ApoA1 | PT | -0.01232 | 0 |
| B74 | B173 | ApoA1 | NTT | 0.009705 | 0 |
| B74 | B207 | ApoA1 | WHR | -0.01105 | 0 |
| B74 | B206 | ApoA1 | Waist | -0.10181 | 0 |
| B74 | B78 | ApoA1 | HBcAb | 0.022863 | 0 |
| B74 | B197 | ApoA1 | AG | 0.014146 | 0 |
| B74 | B204 | ApoA1 | Drinking | -0.02259 | 0 |
| B74 | B80 | ApoA1 | anti-HP IgG | -0.01038 | 0 |
| B74 | B112 | ApoA1 | FT4 | 0.008353 | 3.04E-13 |
| B74 | B201 | ApoA1 | e-habits | -0.0062 | 6.30E-08 |
| B74 | B75 | ApoA1 | ApoA1/B | 0.295874 | 0 |
| B74 | B76 | ApoA1 | ApoB | -0.04483 | 0 |
| B74 | B81 | ApoA1 | TSGF | -0.02724 | 0 |
| B74 | B77 | ApoA1 | TC | 0.078541 | 0 |
| B74 | B133 | ApoA1 | TBIL | -0.01627 | 0 |

| | | | | | |
|---|---|---|---|---|---|
| B74 | B135 | ApoA1 | TBA | -0.0082 | 8.34E-13 |
| B74 | B91 | ApoA1 | TP | 0.030414 | 0 |
| B74 | B114 | ApoA1 | TSH TT3 TT4(TT4) | 0.01959 | 0 |
| B74 | B115 | ApoA1 | TSH TT3 TT4(TT3) | -0.00732 | 1.68E-10 |
| B74 | B212 | ApoA1 | IOP-L | -0.00776 | 1.25E-11 |
| B75 | B136 | ApoA1/B | 5'NT | -0.00585 | 3.27E-07 |
| B75 | B137 | ApoA1/B | AFU | -0.02775 | 0 |
| B75 | B160 | ApoA1/B | PH | 0.006687 | 5.29E-09 |
| B75 | B140 | ApoA1/B | r-GT | -0.02822 | 0 |
| B75 | B127 | ApoA1/B | A-G ratio | 0.041383 | 0 |
| B75 | B128 | ApoA1/B | ALT | -0.02816 | 0 |
| B75 | B123 | ApoA1/B | hs-CRP1 | -0.02291 | 0 |
| B75 | B124 | ApoA1/B | ACTH | -0.00718 | 3.71E-10 |
| B75 | B134 | ApoA1/B | CHE | -0.07473 | 0 |
| B75 | B166 | ApoA1/B | Pro | -0.01136 | 0 |
| B75 | B188 | ApoA1/B | Ca | -0.00579 | 4.26E-07 |
| B75 | B189 | ApoA1/B | eGFB | 0.018843 | 0 |
| B75 | B141 | ApoA1/B | AST/ALT | 0.062072 | 0 |
| B75 | B181 | ApoA1/B | APTT | 0.011803 | 0 |
| B75 | B190 | ApoA1/B | Cre | -0.04017 | 0 |
| B75 | B129 | ApoA1/B | IBIL | -0.00794 | 4.05E-12 |
| B75 | B150 | ApoA1/B | ALP | -0.02559 | 0 |
| B75 | B84 | ApoA1/B | C-PSA | -0.00576 | 4.84E-07 |
| B75 | B222 | ApoA1/B | NS | -0.06247 | 0 |
| B75 | B149 | ApoA1/B | TGAb | 0.006115 | 9.34E-08 |
| B75 | B198 | ApoA1/B | P | 0.023786 | 0 |
| B75 | B194 | ApoA1/B | Cl | 0.008464 | 1.47E-13 |
| B75 | B199 | ApoA1/B | Mg | -0.0179 | 0 |
| B75 | B157 | ApoA1/B | IgG | 0.013556 | 0 |
| B75 | B158 | ApoA1/B | IgM | 0.008193 | 8.51E-13 |
| B75 | B195 | ApoA1/B | Na | -0.02085 | 0 |
| B75 | B176 | ApoA1/B | Ucr | -0.02496 | 0 |
| B75 | B178 | ApoA1/B | Ig-U | -0.0059 | 2.61E-07 |
| B75 | B191 | ApoA1/B | Urea | -0.01302 | 0 |
| B75 | B174 | ApoA1/B | Urea/Cr | 0.019231 | 0 |
| B75 | B192 | ApoA1/B | UA | -0.05618 | 0 |
| B75 | B170 | ApoA1/B | U-GLU | -0.01023 | 0 |
| B75 | B179 | ApoA1/B | α1-MG | -0.01788 | 0 |
| B75 | B138 | ApoA1/B | PAB | -0.07058 | 0 |
| B75 | B221 | ApoA1/B | Hei | -0.03557 | 0 |
| B75 | B200 | ApoA1/B | OP | -0.02697 | 0 |
| B75 | B220 | ApoA1/B | SBP | -0.0266 | 0 |
| B75 | B218 | ApoA1/B | DBP | -0.02963 | 0 |
| B75 | B216 | ApoA1/B | BMI | -0.06218 | 0 |

| | | | | | |
|---|---|---|---|---|---|
| B75 | B215 | ApoA1/B | BMI | -0.0584 | 0 |
| B75 | B131 | ApoA1/B | AST | -0.0102 | 0 |
| B75 | B172 | ApoA1/B | KB | 0.00858 | 6.84E-14 |
| B75 | B210 | ApoA1/B | Hip | -0.0457 | 0 |
| B75 | B209 | ApoA1/B | Smoking | -0.02778 | 0 |
| B75 | B196 | ApoA1/B | TCO2 | -0.00754 | 4.57E-11 |
| B75 | B185 | ApoA1/B | PT | 0.007724 | 1.55E-11 |
| B75 | B186 | ApoA1/B | PFC | -0.00838 | 2.61E-13 |
| B75 | B207 | ApoA1/B | WHR | -0.00745 | 7.61E-11 |
| B75 | B206 | ApoA1/B | Waist | -0.06746 | 0 |
| B75 | B78 | ApoA1/B | HBcAb | 0.012844 | 0 |
| B75 | B197 | ApoA1/B | AG | -0.01183 | 0 |
| B75 | B204 | ApoA1/B | Drinking | -0.02216 | 0 |
| B75 | B80 | ApoA1/B | anti-HP IgG | -0.00567 | 7.30E-07 |
| B75 | B112 | ApoA1/B | FT4 | 0.018393 | 0 |
| B75 | B211 | ApoA1/B | IOP-R | -0.01258 | 0 |
| B75 | B76 | ApoA1/B | ApoB | -0.45204 | 0 |
| B75 | B132 | ApoA1/B | Dbil | 0.02571 | 0 |
| B75 | B77 | ApoA1/B | TC | -0.06904 | 0 |
| B75 | B114 | ApoA1/B | TSH TT3 TT4(TT4) | 0.014433 | 0 |
| B75 | B212 | ApoA1/B | IOP-L | -0.01466 | 0 |
| B76 | B136 | ApoB | 5'NT | 0.012015 | 0 |
| B76 | B137 | ApoB | AFU | 0.03498 | 0 |
| B76 | B160 | ApoB | PH | -0.02505 | 0 |
| B76 | B140 | ApoB | r-GT | 0.067104 | 0 |
| B76 | B118 | ApoB | CEA1 | 0.006242 | 5.04E-08 |
| B76 | B126 | ApoB | Alb | 0.031667 | 0 |
| B76 | B127 | ApoB | A-G ratio | -0.01967 | 0 |
| B76 | B128 | ApoB | ALT | 0.039652 | 0 |
| B76 | B123 | ApoB | hs-CRP1 | 0.018175 | 0 |
| B76 | B111 | ApoB | TSH | 0.020482 | 0 |
| B76 | B124 | ApoB | ACTH | 0.010691 | 0 |
| B76 | B134 | ApoB | CHE | 0.148628 | 0 |
| B76 | B166 | ApoB | Pro | 0.010341 | 0 |
| B76 | B188 | ApoB | Ca | 0.015756 | 0 |
| B76 | B189 | ApoB | eGFB | -0.04337 | 0 |
| B76 | B141 | ApoB | AST/ALT | -0.03774 | 0 |
| B76 | B146 | ApoB | CysC | 0.006304 | 3.72E-08 |
| B76 | B181 | ApoB | APTT | -0.03808 | 0 |
| B76 | B190 | ApoB | Cre | 0.046402 | 0 |
| B76 | B193 | ApoB | K | 0.043747 | 0 |
| B76 | B129 | ApoB | IBIL | 0.010727 | 0 |
| B76 | B150 | ApoB | ALP | 0.050289 | 0 |
| B76 | B84 | ApoB | C-PSA | 0.005799 | 4.14E-07 |

| | | | | | |
|---|---|---|---|---|---|
| B76 | B222 | ApoB | NS | 0.074734 | 0 |
| B76 | B97 | ApoB | dsDNA | -0.00785 | 7.33E-12 |
| B76 | B100 | ApoB | anti-Ro52 | -0.00851 | 1.08E-13 |
| B76 | B103 | ApoB | anti-SSA/Ro | -0.0101 | 0 |
| B76 | B104 | ApoB | anti-SSB | -0.0087 | 3.06E-14 |
| B76 | B198 | ApoB | P | -0.01343 | 0 |
| B76 | B194 | ApoB | Cl | -0.05611 | 0 |
| B76 | B199 | ApoB | Mg | 0.069022 | 0 |
| B76 | B156 | ApoB | IgE | 0.006927 | 1.47E-09 |
| B76 | B157 | ApoB | IgG | -0.0233 | 0 |
| B76 | B158 | ApoB | IgM | -0.00996 | 0 |
| B76 | B195 | ApoB | Na | 0.040871 | 0 |
| B76 | B176 | ApoB | Ucr | 0.025898 | 0 |
| B76 | B178 | ApoB | Ig-U | 0.008585 | 6.64E-14 |
| B76 | B191 | ApoB | Urea | 0.032004 | 0 |
| B76 | B174 | ApoB | Urea/Cr | -0.00871 | 2.82E-14 |
| B76 | B192 | ApoB | UA | 0.073097 | 0 |
| B76 | B170 | ApoB | U-GLU | 0.024138 | 0 |
| B76 | B179 | ApoB | α1-MG | 0.023902 | 0 |
| B76 | B182 | ApoB | TT | 0.014453 | 0 |
| B76 | B125 | ApoB | Cor | 0.010109 | 0 |
| B76 | B153 | ApoB | Glu | 0.009657 | 0 |
| B76 | B138 | ApoB | PAB | 0.147966 | 0 |
| B76 | B130 | ApoB | GLB | 0.036582 | 0 |
| B76 | B221 | ApoB | Hei | 0.021267 | 0 |
| B76 | B187 | ApoB | NSE | 0.008919 | 6.88E-15 |
| B76 | B200 | ApoB | OP | 0.05806 | 0 |
| B76 | B220 | ApoB | SBP | 0.058823 | 0 |
| B76 | B219 | ApoB | OT | 0.006033 | 1.38E-07 |
| B76 | B218 | ApoB | DBP | 0.063913 | 0 |
| B76 | B216 | ApoB | BMI | 0.0689 | 0 |
| B76 | B215 | ApoB | BMI | 0.078114 | 0 |
| B76 | B131 | ApoB | AST | 0.022718 | 0 |
| B76 | B172 | ApoB | KB | -0.00565 | 8.06E-07 |
| B76 | B210 | ApoB | Hip | 0.053373 | 0 |
| B76 | B209 | ApoB | Smoking | 0.041066 | 0 |
| B76 | B90 | ApoB | CYFRA 21-1 | 0.008557 | 7.95E-14 |
| B76 | B196 | ApoB | TCO2 | 0.013702 | 0 |
| B76 | B183 | ApoB | PTA | -0.01507 | 0 |
| B76 | B185 | ApoB | PT | -0.02602 | 0 |
| B76 | B186 | ApoB | PFC | 0.026238 | 0 |
| B76 | B207 | ApoB | WHR | 0.010228 | 0 |
| B76 | B206 | ApoB | Waist | 0.088554 | 0 |
| B76 | B78 | ApoB | HBcAb | -0.01376 | 0 |

| | | | | | |
|---|---|---|---|---|---|
| B76 | B197 | ApoB | AG | 0.049732 | 0 |
| B76 | B204 | ApoB | Drinking | 0.036986 | 0 |
| B76 | B112 | ApoB | FT4 | -0.03061 | 0 |
| B76 | B211 | ApoB | IOP-R | 0.024828 | 0 |
| B76 | B132 | ApoB | Dbil | -0.05236 | 0 |
| B76 | B81 | ApoB | TSGF | 0.006561 | 1.02E-08 |
| B76 | B77 | ApoB | TC | 0.231797 | 0 |
| B76 | B91 | ApoB | TP | 0.050663 | 0 |
| B76 | B114 | ApoB | TSH TT3 TT4(TT4) | -0.01662 | 0 |
| B76 | B212 | ApoB | IOP-L | 0.024532 | 0 |
| B77 | B137 | TC | AFU | 0.016095 | 0 |
| B77 | B160 | TC | PH | -0.05727 | 0 |
| B77 | B140 | TC | r-GT | 0.149644 | 0 |
| B77 | B118 | TC | CEA1 | 0.010913 | 0 |
| B77 | B126 | TC | Alb | 0.054009 | 0 |
| B77 | B127 | TC | A-G ratio | -0.08872 | 0 |
| B77 | B128 | TC | ALT | 0.084454 | 0 |
| B77 | B111 | TC | TSH | 0.032561 | 0 |
| B77 | B134 | TC | CHE | 0.057358 | 0 |
| B77 | B188 | TC | Ca | 0.028306 | 0 |
| B77 | B189 | TC | eGFB | -0.18161 | 0 |
| B77 | B141 | TC | AST/ALT | -0.07349 | 0 |
| B77 | B146 | TC | CysC | 0.013297 | 0 |
| B77 | B181 | TC | APTT | -0.01666 | 0 |
| B77 | B190 | TC | Cre | 0.068126 | 0 |
| B77 | B224 | TC | pmh | 0.006469 | 1.63E-08 |
| B77 | B193 | TC | K | 0.015774 | 0 |
| B77 | B129 | TC | IBIL | -0.01851 | 0 |
| B77 | B150 | TC | ALP | 0.033567 | 0 |
| B77 | B84 | TC | C-PSA | 0.005864 | 3.07E-07 |
| B77 | B222 | TC | NS | 0.102148 | 0 |
| B77 | B149 | TC | TGAb | -0.00756 | 4.13E-11 |
| B77 | B85 | TC | SCC | -0.00638 | 2.56E-08 |
| B77 | B194 | TC | Cl | -0.02296 | 0 |
| B77 | B199 | TC | Mg | 0.020862 | 0 |
| B77 | B157 | TC | IgG | -0.0115 | 0 |
| B77 | B191 | TC | Urea | 0.084538 | 0 |
| B77 | B174 | TC | Urea/Cr | 0.017992 | 0 |
| B77 | B192 | TC | UA | 0.095406 | 0 |
| B77 | B170 | TC | U-GLU | 0.044415 | 0 |
| B77 | B175 | TC | ALB-U | 0.00681 | 2.76E-09 |
| B77 | B153 | TC | Glu | 0.025517 | 0 |
| B77 | B138 | TC | PAB | 0.042361 | 0 |
| B77 | B130 | TC | GLB | 0.134853 | 0 |

| | | | | | |
|---|---|---|---|---|---|
| B77 | B221 | TC | Hei | -0.02776 | 0 |
| B77 | B87 | TC | NSE | 0.008264 | 5.40E-13 |
| B77 | B187 | TC | NSE | 0.007111 | 5.34E-10 |
| B77 | B200 | TC | OP | 0.009802 | 0 |
| B77 | B220 | TC | SBP | 0.149067 | 0 |
| B77 | B219 | TC | OT | 0.047469 | 0 |
| B77 | B218 | TC | DBP | 0.15024 | 0 |
| B77 | B216 | TC | BMI | 0.096645 | 0 |
| B77 | B215 | TC | BMI | 0.148328 | 0 |
| B77 | B131 | TC | AST | 0.070057 | 0 |
| B77 | B172 | TC | KB | -0.00873 | 2.53E-14 |
| B77 | B210 | TC | Hip | 0.085225 | 0 |
| B77 | B209 | TC | Smoking | 0.081294 | 0 |
| B77 | B90 | TC | CYFRA 21-1 | 0.009656 | 0 |
| B77 | B139 | TC | ADA | -0.00856 | 7.82E-14 |
| B77 | B196 | TC | TCO2 | 0.005811 | 3.91E-07 |
| B77 | B185 | TC | PT | -0.01289 | 0 |
| B77 | B186 | TC | PFC | 0.007733 | 1.47E-11 |
| B77 | B173 | TC | NTT | 0.009907 | 0 |
| B77 | B207 | TC | WHR | 0.021008 | 0 |
| B77 | B206 | TC | Waist | 0.157925 | 0 |
| B77 | B205 | TC | KDA | 0.0145 | 0 |
| B77 | B197 | TC | AG | 0.012012 | 0 |
| B77 | B204 | TC | Drinking | 0.072091 | 0 |
| B77 | B80 | TC | anti-HP IgG | 0.011782 | 0 |
| B77 | B112 | TC | FT4 | -0.03317 | 0 |
| B77 | B113 | TC | FT3 | -0.00725 | 2.42E-10 |
| B77 | B211 | TC | IOP-R | 0.016944 | 0 |
| B77 | B201 | TC | e-habits | 0.021419 | 0 |
| B77 | B132 | TC | Dbil | -0.15695 | 0 |
| B77 | B133 | TC | TBIL | -0.05626 | 0 |
| B77 | B91 | TC | TP | 0.148378 | 0 |
| B77 | B114 | TC | TSH TT3 TT4(TT4) | -0.01304 | 0 |
| B77 | B212 | TC | IOP-L | 0.019196 | 0 |
| B78 | B142 | HBcAb | 25-(OH)D | -0.0087 | 3.06E-14 |
| B78 | B160 | HBcAb | PH | -0.01911 | 0 |
| B78 | B126 | HBcAb | Alb | 0.015803 | 0 |
| B78 | B127 | HBcAb | A-G ratio | 0.007068 | 6.81E-10 |
| B78 | B161 | HBcAb | LEU | 0.006581 | 9.15E-09 |
| B78 | B128 | HBcAb | ALT | 0.01278 | 0 |
| B78 | B111 | HBcAb | TSH | 0.010068 | 0 |
| B78 | B189 | HBcAb | eGFB | 0.008232 | 6.61E-13 |
| B78 | B190 | HBcAb | Cre | -0.02051 | 0 |
| B78 | B224 | HBcAb | pmh | -0.00563 | 8.73E-07 |

| | | | | | |
|---|---|---|---|---|---|
| B78 | B151 | HBcAb | Anti-CCP | 0.008485 | 1.28E-13 |
| B78 | B110 | HBcAb | AHA | 0.008394 | 2.33E-13 |
| B78 | B157 | HBcAb | IgG | 0.020005 | 0 |
| B78 | B195 | HBcAb | Na | -0.0063 | 3.77E-08 |
| B78 | B176 | HBcAb | Ucr | -0.0088 | 1.60E-14 |
| B78 | B178 | HBcAb | Ig-U | -0.0104 | 0 |
| B78 | B191 | HBcAb | Urea | 0.015086 | 0 |
| B78 | B174 | HBcAb | Urea/Cr | 0.026904 | 0 |
| B78 | B192 | HBcAb | UA | -0.01921 | 0 |
| B78 | B175 | HBcAb | ALB-U | -0.00764 | 2.58E-11 |
| B78 | B177 | HBcAb | ACR | -0.0089 | 7.99E-15 |
| B78 | B179 | HBcAb | α1-MG | -0.01783 | 0 |
| B78 | B180 | HBcAb | Tf | -0.0096 | 0 |
| B78 | B221 | HBcAb | Hei | -0.02391 | 0 |
| B78 | B220 | HBcAb | SBP | -0.01515 | 0 |
| B78 | B216 | HBcAb | BMI | -0.00688 | 1.90E-09 |
| B78 | B215 | HBcAb | BMI | 0.008615 | 5.44E-14 |
| B78 | B131 | HBcAb | AST | 0.013822 | 0 |
| B78 | B172 | HBcAb | KB | 0.006066 | 1.18E-07 |
| B78 | B205 | HBcAb | KDA | -0.00689 | 1.84E-09 |
| B78 | B79 | HBcAb | hb V -dna | 0.010568 | 0 |
| B78 | B204 | HBcAb | Drinking | 0.007379 | 1.17E-10 |
| B78 | B203 | HBcAb | DH | -0.02271 | 0 |
| B78 | B80 | HBcAb | anti-HP IgG | -0.17537 | 0 |
| B78 | B112 | HBcAb | FT4 | -0.01793 | 0 |
| B78 | B113 | HBcAb | FT3 | 0.00607 | 1.16E-07 |
| B78 | B201 | HBcAb | e-habits | -0.03047 | 0 |
| B78 | B81 | HBcAb | TSGF | -0.02779 | 0 |
| B78 | B91 | HBcAb | TP | 0.010079 | 0 |
| B78 | B212 | HBcAb | IOP-L | -0.0066 | 8.51E-09 |
| B79 | B137 | hb V -dna | AFU | 0.012384 | 0 |
| B79 | B92 | hb V -dna | ALDH2-Glu504Glu | 0.007985 | 3.13E-12 |
| B79 | B93 | hb V -dna | ALDH2-Glu504Lys | -0.00701 | 9.39E-10 |
| B79 | B126 | hb V -dna | Alb | -0.00974 | 0 |
| B79 | B128 | hb V -dna | ALT | 0.032258 | 0 |
| B79 | B165 | hb V -dna | bilirubin | 0.00576 | 4.94E-07 |
| B79 | B134 | hb V -dna | CHE | -0.00954 | 0 |
| B79 | B189 | hb V -dna | eGFB | 0.005931 | 2.24E-07 |
| B79 | B141 | hb V -dna | AST/ALT | -0.00694 | 1.39E-09 |
| B79 | B181 | hb V -dna | APTT | 0.007989 | 3.06E-12 |
| B79 | B194 | hb V -dna | Cl | 0.011523 | 0 |
| B79 | B199 | hb V -dna | Mg | -0.00598 | 1.82E-07 |
| B79 | B157 | hb V -dna | IgG | 0.011982 | 0 |
| B79 | B195 | hb V -dna | Na | -0.00642 | 2.08E-08 |

| | | | | | |
|---|---|---|---|---|---|
| B79 | B138 | hb V -dna | PAB | -0.02168 | 0 |
| B79 | B200 | hb V -dna | OP | -0.00626 | 4.65E-08 |
| B79 | B218 | hb V -dna | DBP | -0.00567 | 7.33E-07 |
| B79 | B131 | hb V -dna | AST | 0.034175 | 0 |
| B79 | B139 | hb V -dna | ADA | 0.019321 | 0 |
| B79 | B183 | hb V -dna | PTA | 0.018016 | 0 |
| B79 | B185 | hb V -dna | PT | 0.01159 | 0 |
| B79 | B186 | hb V -dna | PFC | -0.00573 | 5.72E-07 |
| B79 | B197 | hb V -dna | AG | -0.00827 | 5.03E-13 |
| B79 | B81 | hb V -dna | TSGF | -0.01305 | 0 |
| B79 | B135 | hb V -dna | TBA | 0.012257 | 0 |
| B79 | B91 | hb V -dna | TP | -0.00588 | 2.90E-07 |
| B8 | B92 | WB-Zinc | ALDH2-Glu504Glu | -0.00588 | 2.82E-07 |
| B8 | B93 | WB-Zinc | ALDH2-Glu504Lys | 0.008628 | 4.97E-14 |
| B8 | B94 | WB-Zinc | ALDH2-Lys504Lys | -0.00732 | 1.65E-10 |
| B8 | B126 | WB-Zinc | Alb | 0.006971 | 1.15E-09 |
| B8 | B121 | WB-Zinc | C3 | -0.00606 | 1.22E-07 |
| B8 | B122 | WB-Zinc | C4 | -0.00907 | 2.22E-15 |
| B8 | B134 | WB-Zinc | CHE | 0.007705 | 1.73E-11 |
| B8 | B72 | WB-Zinc | HDL-C | -0.00571 | 6.11E-07 |
| B8 | B189 | WB-Zinc | eGFB | -0.00654 | 1.11E-08 |
| B8 | B141 | WB-Zinc | AST/ALT | -0.00706 | 6.97E-10 |
| B8 | B45 | WB-Zinc | MCHC | 0.005951 | 2.04E-07 |
| B8 | B46 | WB-Zinc | RBC Count | 0.016817 | 0 |
| B8 | B190 | WB-Zinc | Cre | 0.01131 | 0 |
| B8 | B222 | WB-Zinc | NS | 0.010093 | 0 |
| B8 | B199 | WB-Zinc | Mg | 0.006089 | 1.06E-07 |
| B8 | B195 | WB-Zinc | Na | 0.009673 | 0 |
| B8 | B192 | WB-Zinc | UA | 0.010571 | 0 |
| B8 | B138 | WB-Zinc | PAB | 0.013398 | 0 |
| B8 | B60 | WB-Zinc | WBRVI | 0.007907 | 5.07E-12 |
| B8 | B62 | WB-Zinc | WBHSRVI | 0.007446 | 7.99E-11 |
| B8 | B64 | WB-Zinc | WBRSR(mPas_5) | 0.00729 | 1.96E-10 |
| B8 | B65 | WB-Zinc | WBRSR(mPas_1) | 0.007222 | 2.88E-10 |
| B8 | B66 | WB-Zinc | WBRSR(mPas_100) | 0.006855 | 2.17E-09 |
| B8 | B67 | WB-Zinc | WBRSR(mPas_50) | 0.007001 | 9.80E-10 |
| B8 | B221 | WB-Zinc | Hei | 0.010543 | 0 |
| B8 | B200 | WB-Zinc | OP | 0.009367 | 4.44E-16 |
| B8 | B216 | WB-Zinc | BMI | 0.010381 | 0 |
| B8 | B215 | WB-Zinc | BMI | 0.006501 | 1.38E-08 |
| B8 | B210 | WB-Zinc | Hip | 0.006671 | 5.73E-09 |
| B8 | B51 | WB-Zinc | Hb | 0.017016 | 0 |
| B8 | B186 | WB-Zinc | PFC | -0.00642 | 2.10E-08 |
| B8 | B206 | WB-Zinc | Waist | 0.008994 | 4.00E-15 |

| | | | | | |
|---|---|---|---|---|---|
| B8 | B197 | WB-Zinc | AG | 0.00736 | 1.31E-10 |
| B8 | B135 | WB-Zinc | TBA | 0.008482 | 1.31E-13 |
| B80 | B160 | anti-HP IgG | PH | -0.01611 | 0 |
| B80 | B140 | anti-HP IgG | r-GT | 0.006878 | 1.92E-09 |
| B80 | B126 | anti-HP IgG | Alb | -0.03284 | 0 |
| B80 | B127 | anti-HP IgG | A-G ratio | -0.02156 | 0 |
| B80 | B161 | anti-HP IgG | LEU | -0.00897 | 4.88E-15 |
| B80 | B166 | anti-HP IgG | Pro | -0.02176 | 0 |
| B80 | B189 | anti-HP IgG | eGFB | -0.02562 | 0 |
| B80 | B141 | anti-HP IgG | AST/ALT | -0.00749 | 6.24E-11 |
| B80 | B190 | anti-HP IgG | Cre | 0.01416 | 0 |
| B80 | B129 | anti-HP IgG | IBIL | -0.01256 | 0 |
| B80 | B222 | anti-HP IgG | NS | 0.016048 | 0 |
| B80 | B174 | anti-HP IgG | Urea/Cr | -0.01101 | 0 |
| B80 | B192 | anti-HP IgG | UA | 0.006675 | 5.63E-09 |
| B80 | B130 | anti-HP IgG | GLB | 0.01159 | 0 |
| B80 | B221 | anti-HP IgG | Hei | 0.010826 | 0 |
| B80 | B220 | anti-HP IgG | SBP | 0.006017 | 1.50E-07 |
| B80 | B216 | anti-HP IgG | BMI | 0.018739 | 0 |
| B80 | B215 | anti-HP IgG | BMI | 0.016486 | 0 |
| B80 | B172 | anti-HP IgG | KB | -0.00712 | 5.06E-10 |
| B80 | B210 | anti-HP IgG | Hip | 0.016135 | 0 |
| B80 | B209 | anti-HP IgG | Smoking | 0.012143 | 0 |
| B80 | B206 | anti-HP IgG | Waist | 0.020605 | 0 |
| B80 | B203 | anti-HP IgG | DH | 0.010744 | 0 |
| B80 | B201 | anti-HP IgG | e-habits | 0.010701 | 0 |
| B80 | B132 | anti-HP IgG | Dbil | -0.01541 | 0 |
| B80 | B133 | anti-HP IgG | TBIL | -0.01419 | 0 |
| B80 | B91 | anti-HP IgG | TP | -0.01049 | 0 |
| B81 | B137 | TSGF | AFU | 0.026353 | 0 |
| B81 | B92 | TSGF | ALDH2-Glu504Glu | -0.01276 | 0 |
| B81 | B93 | TSGF | ALDH2-Glu504Lys | 0.013373 | 0 |
| B81 | B160 | TSGF | PH | 0.012419 | 0 |
| B81 | B126 | TSGF | Alb | -0.01495 | 0 |
| B81 | B127 | TSGF | A-G ratio | -0.02943 | 0 |
| B81 | B124 | TSGF | ACTH | -0.04503 | 0 |
| B81 | B134 | TSGF | CHE | 0.015693 | 0 |
| B81 | B189 | TSGF | eGFB | 0.00984 | 0 |
| B81 | B141 | TSGF | AST/ALT | -0.02069 | 0 |
| B81 | B181 | TSGF | APTT | 0.025896 | 0 |
| B81 | B129 | TSGF | IBIL | -0.01592 | 0 |
| B81 | B150 | TSGF | ALP | 0.013103 | 0 |
| B81 | B100 | TSGF | anti-Ro52 | 0.007374 | 1.21E-10 |
| B81 | B104 | TSGF | anti-SSB | 0.010724 | 0 |

| | | | | | |
|---|---|---|---|---|---|
| B81 | B151 | TSGF | Anti-CCP | -0.02794 | 0 |
| B81 | B108 | TSGF | Anti-Cyclin | -0.00727 | 2.20E-10 |
| B81 | B198 | TSGF | P | -0.0077 | 1.74E-11 |
| B81 | B194 | TSGF | Cl | -0.01902 | 0 |
| B81 | B199 | TSGF | Mg | 0.012926 | 0 |
| B81 | B155 | TSGF | IgA | 0.007658 | 2.29E-11 |
| B81 | B156 | TSGF | IgE | -0.01537 | 0 |
| B81 | B157 | TSGF | IgG | -0.07881 | 0 |
| B81 | B158 | TSGF | IgM | -0.00606 | 1.22E-07 |
| B81 | B195 | TSGF | Na | 0.059431 | 0 |
| B81 | B178 | TSGF | Ig-U | -0.00587 | 3.00E-07 |
| B81 | B192 | TSGF | UA | 0.010552 | 0 |
| B81 | B179 | TSGF | α1-MG | -0.00759 | 3.44E-11 |
| B81 | B180 | TSGF | Tf | -0.00653 | 1.18E-08 |
| B81 | B182 | TSGF | TT | 0.130388 | 0 |
| B81 | B125 | TSGF | Cor | -0.11148 | 0 |
| B81 | B138 | TSGF | PAB | 0.012995 | 0 |
| B81 | B130 | TSGF | GLB | 0.026816 | 0 |
| B81 | B187 | TSGF | NSE | 0.005641 | 8.45E-07 |
| B81 | B200 | TSGF | OP | 0.04553 | 0 |
| B81 | B220 | TSGF | SBP | 0.006292 | 3.95E-08 |
| B81 | B218 | TSGF | DBP | -0.00593 | 2.28E-07 |
| B81 | B131 | TSGF | AST | -0.00662 | 7.43E-09 |
| B81 | B90 | TSGF | CYFRA 21-1 | 0.017851 | 0 |
| B81 | B139 | TSGF | ADA | 0.026465 | 0 |
| B81 | B196 | TSGF | TCO2 | -0.01596 | 0 |
| B81 | B183 | TSGF | PTA | -0.06212 | 0 |
| B81 | B184 | TSGF | INR | -0.00743 | 8.79E-11 |
| B81 | B185 | TSGF | PT | -0.01859 | 0 |
| B81 | B186 | TSGF | PFC | -0.00648 | 1.56E-08 |
| B81 | B197 | TSGF | AG | 0.059765 | 0 |
| B81 | B112 | TSGF | FT4 | -0.01918 | 0 |
| B81 | B113 | TSGF | FT3 | 0.006417 | 2.12E-08 |
| B81 | B132 | TSGF | Dbil | 0.010915 | 0 |
| B81 | B133 | TSGF | TBIL | -0.0099 | 0 |
| B81 | B135 | TSGF | TBA | -0.00639 | 2.45E-08 |
| B81 | B91 | TSGF | TP | 0.013661 | 0 |
| B81 | B212 | TSGF | IOP-L | 0.006088 | 1.07E-07 |
| B82 | B189 | CEA | eGFB | -0.00762 | 2.94E-11 |
| B82 | B83 | CEA | AFP | 0.007914 | 4.87E-12 |
| B82 | B84 | CEA | C-PSA | 0.014847 | 0 |
| B82 | B85 | CEA | SCC | 0.03199 | 0 |
| B82 | B86 | CEA | TBhCG | 0.008162 | 1.04E-12 |
| B82 | B87 | CEA | NSE | 0.016547 | 0 |

| | | | | | |
|---|---|---|---|---|---|
| B82 | B88 | CEA | CA 242 | 0.030864 | 0 |
| B82 | B89 | CEA | CA 72-4 | 0.043844 | 0 |
| B82 | B209 | CEA | Smoking | 0.008022 | 2.50E-12 |
| B82 | B90 | CEA | CYFRA 21-1 | 0.014698 | 0 |
| B83 | B137 | AFP | AFU | 0.027567 | 0 |
| B83 | B140 | AFP | r-GT | 0.006417 | 2.12E-08 |
| B83 | B126 | AFP | Alb | -0.00826 | 5.68E-13 |
| B83 | B193 | AFP | K | 0.008847 | 1.13E-14 |
| B83 | B150 | AFP | ALP | 0.011178 | 0 |
| B83 | B84 | AFP | C-PSA | 0.006031 | 1.40E-07 |
| B83 | B85 | AFP | SCC | 0.006899 | 1.71E-09 |
| B83 | B194 | AFP | Cl | -0.00922 | 8.88E-16 |
| B83 | B155 | AFP | IgA | -0.00597 | 1.91E-07 |
| B83 | B138 | AFP | PAB | -0.00863 | 4.75E-14 |
| B83 | B86 | AFP | TBhCG | 0.184804 | 0 |
| B83 | B88 | AFP | CA 242 | 0.039676 | 0 |
| B83 | B89 | AFP | CA 72-4 | 0.017582 | 0 |
| B83 | B131 | AFP | AST | 0.007121 | 5.05E-10 |
| B83 | B90 | AFP | CYFRA 21-1 | 0.023591 | 0 |
| B83 | B139 | AFP | ADA | 0.013703 | 0 |
| B83 | B114 | AFP | TSH TT3 TT4(TT4) | 0.005627 | 8.99E-07 |
| B84 | B137 | C-PSA | AFU | 0.010664 | 0 |
| B84 | B140 | C-PSA | r-GT | 0.011086 | 0 |
| B84 | B128 | C-PSA | ALT | 0.00689 | 1.79E-09 |
| B84 | B134 | C-PSA | CHE | 0.006295 | 3.88E-08 |
| B84 | B166 | C-PSA | Pro | 0.008197 | 8.28E-13 |
| B84 | B189 | C-PSA | eGFB | -0.03205 | 0 |
| B84 | B141 | C-PSA | AST/ALT | -0.00721 | 3.03E-10 |
| B84 | B146 | C-PSA | CysC | 0.008725 | 2.58E-14 |
| B84 | B190 | C-PSA | Cre | 0.034592 | 0 |
| B84 | B129 | C-PSA | IBIL | 0.008308 | 4.08E-13 |
| B84 | B222 | C-PSA | NS | 0.029918 | 0 |
| B84 | B198 | C-PSA | P | -0.01734 | 0 |
| B84 | B85 | C-PSA | SCC | 0.060655 | 0 |
| B84 | B157 | C-PSA | IgG | -0.00566 | 7.82E-07 |
| B84 | B195 | C-PSA | Na | 0.009848 | 0 |
| B84 | B191 | C-PSA | Urea | 0.013341 | 0 |
| B84 | B174 | C-PSA | Urea/Cr | -0.01415 | 0 |
| B84 | B192 | C-PSA | UA | 0.02494 | 0 |
| B84 | B138 | C-PSA | PAB | 0.013905 | 0 |
| B84 | B130 | C-PSA | GLB | -0.00567 | 7.57E-07 |
| B84 | B86 | C-PSA | TBhCG | -0.01028 | 0 |
| B84 | B221 | C-PSA | Hei | 0.021118 | 0 |
| B84 | B87 | C-PSA | NSE | 0.065083 | 0 |

| | | | | | |
|---|---|---|---|---|---|
| B84 | B200 | C-PSA | OP | 0.012017 | 0 |
| B84 | B220 | C-PSA | SBP | 0.022236 | 0 |
| B84 | B218 | C-PSA | DBP | 0.017188 | 0 |
| B84 | B88 | C-PSA | CA 242 | 0.006693 | 5.11E-09 |
| B84 | B89 | C-PSA | CA 72-4 | 0.028595 | 0 |
| B84 | B216 | C-PSA | BMI | 0.020573 | 0 |
| B84 | B215 | C-PSA | BMI | 0.013133 | 0 |
| B84 | B131 | C-PSA | AST | 0.00696 | 1.23E-09 |
| B84 | B210 | C-PSA | Hip | 0.009997 | 0 |
| B84 | B209 | C-PSA | Smoking | 0.019676 | 0 |
| B84 | B90 | C-PSA | CYFRA 21-1 | 0.093019 | 0 |
| B84 | B206 | C-PSA | Waist | 0.023414 | 0 |
| B84 | B204 | C-PSA | Drinking | 0.018548 | 0 |
| B84 | B113 | C-PSA | FT3 | 0.006091 | 1.05E-07 |
| B84 | B133 | C-PSA | TBIL | 0.007819 | 8.69E-12 |
| B84 | B91 | C-PSA | TP | -0.00635 | 2.91E-08 |
| B85 | B166 | SCC | Pro | 0.006194 | 6.40E-08 |
| B85 | B189 | SCC | eGFB | -0.0163 | 0 |
| B85 | B146 | SCC | CysC | 0.00645 | 1.79E-08 |
| B85 | B190 | SCC | Cre | 0.028664 | 0 |
| B85 | B222 | SCC | NS | 0.010874 | 0 |
| B85 | B156 | SCC | IgE | 0.008599 | 6.00E-14 |
| B85 | B195 | SCC | Na | 0.008672 | 3.71E-14 |
| B85 | B191 | SCC | Urea | 0.007632 | 2.68E-11 |
| B85 | B174 | SCC | Urea/Cr | -0.01351 | 0 |
| B85 | B192 | SCC | UA | 0.016167 | 0 |
| B85 | B138 | SCC | PAB | 0.007245 | 2.53E-10 |
| B85 | B130 | SCC | GLB | -0.00707 | 6.68E-10 |
| B85 | B86 | SCC | TBhCG | -0.01726 | 0 |
| B85 | B221 | SCC | Hei | 0.014669 | 0 |
| B85 | B87 | SCC | NSE | 0.162894 | 0 |
| B85 | B200 | SCC | OP | 0.008372 | 2.68E-13 |
| B85 | B88 | SCC | CA 242 | 0.050806 | 0 |
| B85 | B216 | SCC | BMI | 0.008434 | 1.79E-13 |
| B85 | B209 | SCC | Smoking | 0.010487 | 0 |
| B85 | B90 | SCC | CYFRA 21-1 | 0.066678 | 0 |
| B85 | B197 | SCC | AG | 0.006871 | 1.99E-09 |
| B85 | B204 | SCC | Drinking | 0.007156 | 4.18E-10 |
| B85 | B91 | SCC | TP | -0.00808 | 1.70E-12 |
| B86 | B137 | TBhCG | AFU | 0.009367 | 4.44E-16 |
| B86 | B160 | TBhCG | PH | 0.006558 | 1.03E-08 |
| B86 | B126 | TBhCG | Alb | -0.01536 | 0 |
| B86 | B127 | TBhCG | A-G ratio | -0.00644 | 1.84E-08 |
| B86 | B134 | TBhCG | CHE | -0.00886 | 9.99E-15 |

| | | | | | |
|---|---|---|---|---|---|
| B86 | B189 | TBhCG | eGFB | 0.015692 | 0 |
| B86 | B190 | TBhCG | Cre | -0.01335 | 0 |
| B86 | B129 | TBhCG | IBIL | -0.00715 | 4.20E-10 |
| B86 | B191 | TBhCG | Urea | -0.01211 | 0 |
| B86 | B192 | TBhCG | UA | -0.0123 | 0 |
| B86 | B138 | TBhCG | PAB | -0.00669 | 5.20E-09 |
| B86 | B87 | TBhCG | NSE | -0.01372 | 0 |
| B86 | B200 | TBhCG | OP | -0.00651 | 1.33E-08 |
| B86 | B220 | TBhCG | SBP | -0.00655 | 1.09E-08 |
| B86 | B218 | TBhCG | DBP | -0.00853 | 9.81E-14 |
| B86 | B88 | TBhCG | CA 242 | 0.008172 | 9.69E-13 |
| B86 | B89 | TBhCG | CA 72-4 | 0.025218 | 0 |
| B86 | B186 | TBhCG | PFC | 0.006211 | 5.86E-08 |
| B86 | B204 | TBhCG | Drinking | -0.00567 | 7.32E-07 |
| B86 | B133 | TBhCG | TBIL | -0.00702 | 8.87E-10 |
| B86 | B91 | TBhCG | TP | -0.00892 | 6.66E-15 |
| B86 | B114 | TBhCG | TSH TT3 TT4(TT4) | 0.013783 | 0 |
| B87 | B137 | NSE | AFU | 0.013029 | 0 |
| B87 | B160 | NSE | PH | 0.011408 | 0 |
| B87 | B140 | NSE | r-GT | 0.007016 | 9.06E-10 |
| B87 | B128 | NSE | ALT | 0.005931 | 2.24E-07 |
| B87 | B134 | NSE | CHE | 0.01025 | 0 |
| B87 | B189 | NSE | eGFB | -0.01279 | 0 |
| B87 | B190 | NSE | Cre | 0.013376 | 0 |
| B87 | B129 | NSE | IBIL | 0.028079 | 0 |
| B87 | B150 | NSE | ALP | -0.00695 | 1.27E-09 |
| B87 | B222 | NSE | NS | 0.009292 | 4.44E-16 |
| B87 | B154 | NSE | RF | 0.005946 | 2.09E-07 |
| B87 | B198 | NSE | P | 0.005917 | 2.40E-07 |
| B87 | B195 | NSE | Na | -0.01278 | 0 |
| B87 | B191 | NSE | Urea | 0.006419 | 2.09E-08 |
| B87 | B192 | NSE | UA | 0.008832 | 1.24E-14 |
| B87 | B182 | NSE | TT | -0.00925 | 6.66E-16 |
| B87 | B138 | NSE | PAB | 0.00575 | 5.16E-07 |
| B87 | B200 | NSE | OP | -0.00998 | 0 |
| B87 | B220 | NSE | SBP | 0.018983 | 0 |
| B87 | B218 | NSE | DBP | 0.013515 | 0 |
| B87 | B88 | NSE | CA 242 | 0.105575 | 0 |
| B87 | B216 | NSE | BMI | 0.010012 | 0 |
| B87 | B215 | NSE | BMI | 0.010097 | 0 |
| B87 | B131 | NSE | AST | 0.01198 | 0 |
| B87 | B209 | NSE | Smoking | 0.005612 | 9.59E-07 |
| B87 | B90 | NSE | CYFRA 21-1 | 0.09353 | 0 |
| B87 | B183 | NSE | PTA | 0.015408 | 0 |

| | | | | | |
|---|---|---|---|---|---|
| B87 | B206 | NSE | Waist | 0.009991 | 0 |
| B87 | B197 | NSE | AG | -0.01326 | 0 |
| B87 | B204 | NSE | Drinking | 0.005972 | 1.85E-07 |
| B87 | B112 | NSE | FT4 | 0.021714 | 0 |
| B87 | B211 | NSE | IOP-R | 0.007611 | 3.03E-11 |
| B87 | B133 | NSE | TBIL | 0.022522 | 0 |
| B87 | B114 | NSE | TSH TT3 TT4(TT4) | -0.00575 | 5.17E-07 |
| B87 | B212 | NSE | IOP-L | 0.007074 | 6.58E-10 |
| B88 | B140 | CA 242 | r-GT | -0.01064 | 0 |
| B88 | B126 | CA 242 | Alb | -0.00712 | 4.98E-10 |
| B88 | B127 | CA 242 | A-G ratio | -0.01104 | 0 |
| B88 | B161 | CA 242 | LEU | 0.00819 | 8.63E-13 |
| B88 | B128 | CA 242 | ALT | -0.01301 | 0 |
| B88 | B189 | CA 242 | eGFB | 0.008233 | 6.59E-13 |
| B88 | B141 | CA 242 | AST/ALT | 0.017977 | 0 |
| B88 | B190 | CA 242 | Cre | -0.02436 | 0 |
| B88 | B129 | CA 242 | IBIL | -0.00592 | 2.33E-07 |
| B88 | B150 | CA 242 | ALP | -0.00745 | 7.91E-11 |
| B88 | B222 | CA 242 | NS | -0.02957 | 0 |
| B88 | B198 | CA 242 | P | 0.006346 | 3.02E-08 |
| B88 | B156 | CA 242 | IgE | -0.00601 | 1.54E-07 |
| B88 | B191 | CA 242 | Urea | -0.00792 | 4.64E-12 |
| B88 | B174 | CA 242 | Urea/Cr | 0.01112 | 0 |
| B88 | B192 | CA 242 | UA | -0.0272 | 0 |
| B88 | B170 | CA 242 | U-GLU | 0.007355 | 1.35E-10 |
| B88 | B138 | CA 242 | PAB | -0.01203 | 0 |
| B88 | B130 | CA 242 | GLB | 0.010771 | 0 |
| B88 | B221 | CA 242 | Hei | -0.02717 | 0 |
| B88 | B218 | CA 242 | DBP | -0.00692 | 1.55E-09 |
| B88 | B89 | CA 242 | CA 72-4 | -0.02408 | 0 |
| B88 | B216 | CA 242 | BMI | -0.0292 | 0 |
| B88 | B215 | CA 242 | BMI | -0.01995 | 0 |
| B88 | B210 | CA 242 | Hip | -0.02021 | 0 |
| B88 | B209 | CA 242 | Smoking | -0.01302 | 0 |
| B88 | B90 | CA 242 | CYFRA 21-1 | 0.028851 | 0 |
| B88 | B139 | CA 242 | ADA | 0.006077 | 1.12E-07 |
| B88 | B206 | CA 242 | Waist | -0.02614 | 0 |
| B88 | B204 | CA 242 | Drinking | -0.01754 | 0 |
| B88 | B113 | CA 242 | FT3 | -0.00704 | 7.80E-10 |
| B89 | B129 | CA 72-4 | IBIL | -0.01618 | 0 |
| B89 | B221 | CA 72-4 | Hei | -0.00569 | 6.89E-07 |
| B89 | B215 | CA 72-4 | BMI | 0.00846 | 1.51E-13 |
| B89 | B210 | CA 72-4 | Hip | 0.005749 | 5.18E-07 |
| B89 | B90 | CA 72-4 | CYFRA 21-1 | 0.023236 | 0 |

| | | | | | |
|---|---|---|---|---|---|
| B89 | B196 | CA 72-4 | TCO2 | -0.00562 | 9.23E-07 |
| B89 | B183 | CA 72-4 | PTA | -0.00955 | 0 |
| B89 | B206 | CA 72-4 | Waist | 0.007569 | 3.88E-11 |
| B89 | B112 | CA 72-4 | FT4 | -0.00651 | 1.30E-08 |
| B89 | B132 | CA 72-4 | Dbil | -0.00967 | 0 |
| B89 | B133 | CA 72-4 | TBIL | -0.01553 | 0 |
| B89 | B135 | CA 72-4 | TBA | 0.005729 | 5.69E-07 |
| B9 | B17 | RH | CD3+CD16+CD56+NK | -0.00578 | 4.43E-07 |
| B9 | B116 | RH | TT3 | 0.018573 | 0 |
| B9 | B46 | RH | RBC Count | -0.00625 | 4.75E-08 |
| B9 | B102 | RH | anti-Sm | 0.007495 | 6.02E-11 |
| B9 | B152 | RH | ACA-Ig(G-A-M) | 0.007909 | 5.02E-12 |
| B9 | B51 | RH | Hb | -0.00806 | 1.91E-12 |
| B9 | B69 | RH | PV | 0.009731 | 0 |
| B9 | B203 | RH | DH | -0.00761 | 3.03E-11 |
| B9 | B81 | RH | TSGF | -0.01173 | 0 |
| B90 | B137 | CYFRA 21-1 | AFU | 0.012962 | 0 |
| B90 | B92 | CYFRA 21-1 | ALDH2-Glu504Glu | -0.00725 | 2.39E-10 |
| B90 | B160 | CYFRA 21-1 | PH | -0.01224 | 0 |
| B90 | B140 | CYFRA 21-1 | r-GT | 0.006955 | 1.27E-09 |
| B90 | B94 | CYFRA 21-1 | ALDH2-Lys504Lys | 0.01121 | 0 |
| B90 | B126 | CYFRA 21-1 | Alb | -0.02435 | 0 |
| B90 | B127 | CYFRA 21-1 | A-G ratio | -0.00815 | 1.09E-12 |
| B90 | B128 | CYFRA 21-1 | ALT | 0.00655 | 1.08E-08 |
| B90 | B111 | CYFRA 21-1 | TSH | 0.007458 | 7.43E-11 |
| B90 | B166 | CYFRA 21-1 | Pro | 0.010981 | 0 |
| B90 | B189 | CYFRA 21-1 | eGFB | -0.05998 | 0 |
| B90 | B146 | CYFRA 21-1 | CysC | 0.019842 | 0 |
| B90 | B181 | CYFRA 21-1 | APTT | -0.01474 | 0 |
| B90 | B190 | CYFRA 21-1 | Cre | 0.043186 | 0 |
| B90 | B150 | CYFRA 21-1 | ALP | 0.012525 | 0 |
| B90 | B222 | CYFRA 21-1 | NS | 0.020133 | 0 |
| B90 | B198 | CYFRA 21-1 | P | 0.005995 | 1.66E-07 |
| B90 | B194 | CYFRA 21-1 | Cl | -0.00749 | 6.36E-11 |
| B90 | B199 | CYFRA 21-1 | Mg | 0.006694 | 5.09E-09 |
| B90 | B195 | CYFRA 21-1 | Na | 0.021619 | 0 |
| B90 | B191 | CYFRA 21-1 | Urea | 0.027488 | 0 |
| B90 | B174 | CYFRA 21-1 | Urea/Cr | -0.00672 | 4.43E-09 |
| B90 | B192 | CYFRA 21-1 | UA | 0.022808 | 0 |
| B90 | B170 | CYFRA 21-1 | U-GLU | 0.012828 | 0 |
| B90 | B182 | CYFRA 21-1 | TT | 0.02864 | 0 |
| B90 | B138 | CYFRA 21-1 | PAB | 0.005876 | 2.89E-07 |
| B90 | B221 | CYFRA 21-1 | Hei | 0.011906 | 0 |
| B90 | B187 | CYFRA 21-1 | NSE | 0.01929 | 0 |

| | | | | | |
|---|---|---|---|---|---|
| B90 | B200 | CYFRA 21-1 | OP | 0.024833 | 0 |
| B90 | B220 | CYFRA 21-1 | SBP | 0.032423 | 0 |
| B90 | B219 | CYFRA 21-1 | OT | 0.011971 | 0 |
| B90 | B218 | CYFRA 21-1 | DBP | 0.020761 | 0 |
| B90 | B216 | CYFRA 21-1 | BMI | 0.008105 | 1.48E-12 |
| B90 | B131 | CYFRA 21-1 | AST | 0.01325 | 0 |
| B90 | B209 | CYFRA 21-1 | Smoking | 0.029513 | 0 |
| B90 | B139 | CYFRA 21-1 | ADA | 0.024387 | 0 |
| B90 | B196 | CYFRA 21-1 | TCO2 | 0.009371 | 4.44E-16 |
| B90 | B183 | CYFRA 21-1 | PTA | -0.04368 | 0 |
| B90 | B185 | CYFRA 21-1 | PT | -0.01068 | 0 |
| B90 | B186 | CYFRA 21-1 | PFC | 0.010505 | 0 |
| B90 | B206 | CYFRA 21-1 | Waist | 0.019578 | 0 |
| B90 | B197 | CYFRA 21-1 | AG | 0.010117 | 0 |
| B90 | B204 | CYFRA 21-1 | Drinking | 0.007612 | 3.02E-11 |
| B90 | B113 | CYFRA 21-1 | FT3 | 0.005984 | 1.75E-07 |
| B90 | B201 | CYFRA 21-1 | e-habits | 0.006201 | 6.16E-08 |
| B90 | B135 | CYFRA 21-1 | TBA | 0.018988 | 0 |
| B90 | B91 | CYFRA 21-1 | TP | -0.01241 | 0 |
| B91 | B137 | TP | AFU | 0.014859 | 0 |
| B91 | B160 | TP | PH | -0.05393 | 0 |
| B91 | B140 | TP | r-GT | 0.076129 | 0 |
| B91 | B143 | TP | β-CTx | -0.00565 | 7.96E-07 |
| B91 | B118 | TP | CEA1 | -0.00818 | 9.19E-13 |
| B91 | B126 | TP | Alb | 0.529233 | 0 |
| B91 | B127 | TP | A-G ratio | -0.43892 | 0 |
| B91 | B161 | TP | LEU | 0.010063 | 0 |
| B91 | B128 | TP | ALT | 0.097752 | 0 |
| B91 | B120 | TP | HCV-cAg | 0.006235 | 5.23E-08 |
| B91 | B121 | TP | C3 | 0.008095 | 1.58E-12 |
| B91 | B122 | TP | C4 | 0.006302 | 3.76E-08 |
| B91 | B225 | TP | hPBG | 0.010314 | 0 |
| B91 | B123 | TP | hs-CRP1 | 0.008879 | 9.10E-15 |
| B91 | B111 | TP | TSH | 0.032563 | 0 |
| B91 | B134 | TP | CHE | 0.055407 | 0 |
| B91 | B166 | TP | Pro | 0.016826 | 0 |
| B91 | B188 | TP | Ca | 0.092438 | 0 |
| B91 | B189 | TP | eGFB | 0.095952 | 0 |
| B91 | B141 | TP | AST/ALT | -0.0492 | 0 |
| B91 | B146 | TP | CysC | 0.006793 | 3.01E-09 |
| B91 | B193 | TP | K | 0.029375 | 0 |
| B91 | B150 | TP | ALP | 0.035282 | 0 |
| B91 | B222 | TP | NS | 0.0246 | 0 |
| B91 | B100 | TP | anti-Ro52 | 0.006262 | 4.57E-08 |

| | | | | | |
|---|---|---|---|---|---|
| B91 | B103 | TP | anti-SSA/Ro | 0.008406 | 2.16E-13 |
| B91 | B107 | TP | AnuA | 0.00588 | 2.84E-07 |
| B91 | B148 | TP | Anti_TPO | 0.020093 | 0 |
| B91 | B149 | TP | TGAb | 0.020108 | 0 |
| B91 | B198 | TP | P | 0.008966 | 4.88E-15 |
| B91 | B194 | TP | Cl | -0.06391 | 0 |
| B91 | B199 | TP | Mg | 0.020097 | 0 |
| B91 | B155 | TP | IgA | 0.058089 | 0 |
| B91 | B156 | TP | IgE | 0.007408 | 9.97E-11 |
| B91 | B157 | TP | IgG | 0.089226 | 0 |
| B91 | B158 | TP | IgM | 0.015142 | 0 |
| B91 | B195 | TP | Na | -0.01168 | 0 |
| B91 | B178 | TP | Ig-U | 0.00743 | 8.78E-11 |
| B91 | B192 | TP | UA | 0.124138 | 0 |
| B91 | B170 | TP | U-GLU | 0.015279 | 0 |
| B91 | B182 | TP | TT | 0.00715 | 4.31E-10 |
| B91 | B153 | TP | Glu | 0.015733 | 0 |
| B91 | B138 | TP | PAB | 0.035649 | 0 |
| B91 | B130 | TP | GLB | 0.786316 | 0 |
| B91 | B221 | TP | Hei | 0.027403 | 0 |
| B91 | B187 | TP | NSE | 0.010213 | 0 |
| B91 | B220 | TP | SBP | 0.088378 | 0 |
| B91 | B219 | TP | OT | -0.01985 | 0 |
| B91 | B218 | TP | DBP | 0.11309 | 0 |
| B91 | B216 | TP | BMI | 0.033451 | 0 |
| B91 | B215 | TP | BMI | 0.022208 | 0 |
| B91 | B131 | TP | AST | 0.083612 | 0 |
| B91 | B172 | TP | KB | 0.013317 | 0 |
| B91 | B210 | TP | Hip | 0.032208 | 0 |
| B91 | B209 | TP | Smoking | -0.12697 | 0 |
| B91 | B139 | TP | ADA | 0.037977 | 0 |
| B91 | B152 | TP | ACA-Ig(G-A-M) | 0.009007 | 3.77E-15 |
| B91 | B183 | TP | PTA | -0.00755 | 4.37E-11 |
| B91 | B186 | TP | PFC | 0.008122 | 1.33E-12 |
| B91 | B206 | TP | Waist | 0.007401 | 1.03E-10 |
| B91 | B197 | TP | AG | 0.035331 | 0 |
| B91 | B204 | TP | Drinking | -0.03181 | 0 |
| B91 | B203 | TP | DH | 0.03649 | 0 |
| B91 | B112 | TP | FT4 | 0.036694 | 0 |
| B91 | B113 | TP | FT3 | 0.036929 | 0 |
| B91 | B211 | TP | IOP-R | 0.026051 | 0 |
| B91 | B132 | TP | Dbil | -0.01917 | 0 |
| B91 | B114 | TP | TSH TT3 TT4(TT4) | 0.024084 | 0 |
| B91 | B115 | TP | TSH TT3 TT4(TT3) | 0.014753 | 0 |

| | | | | | |
|---|---|---|---|---|---|
| B91 | B144 | TP | Total-PINP | -0.00772 | 1.61E-11 |
| B91 | B212 | TP | IOP-L | 0.020379 | 0 |
| B92 | B137 | ALDH2-Glu504Glu | AFU | -0.00828 | 4.83E-13 |
| B92 | B93 | ALDH2-Glu504Glu | ALDH2-Glu504Lys | -0.93834 | 0 |
| B92 | B140 | ALDH2-Glu504Glu | r-GT | 0.011946 | 0 |
| B92 | B94 | ALDH2-Glu504Glu | ALDH2-Lys504Lys | -0.2403 | 0 |
| B92 | B128 | ALDH2-Glu504Glu | ALT | 0.006264 | 4.54E-08 |
| B92 | B121 | ALDH2-Glu504Glu | C3 | 0.006753 | 3.73E-09 |
| B92 | B122 | ALDH2-Glu504Glu | C4 | -0.01556 | 0 |
| B92 | B225 | ALDH2-Glu504Glu | hPBG | 0.007462 | 7.27E-11 |
| B92 | B134 | ALDH2-Glu504Glu | CHE | -0.01334 | 0 |
| B92 | B181 | ALDH2-Glu504Glu | APTT | -0.00984 | 0 |
| B92 | B150 | ALDH2-Glu504Glu | ALP | -0.00743 | 8.78E-11 |
| B92 | B95 | ALDH2-Glu504Glu | anti-cenpb | -0.00631 | 3.52E-08 |
| B92 | B97 | ALDH2-Glu504Glu | dsDNA | -0.00819 | 8.69E-13 |
| B92 | B98 | ALDH2-Glu504Glu | anti-Jo-1 | 0.01036 | 0 |
| B92 | B103 | ALDH2-Glu504Glu | anti-SSA/Ro | 0.018127 | 0 |
| B92 | B104 | ALDH2-Glu504Glu | anti-SSB | 0.008987 | 4.44E-15 |
| B92 | B106 | ALDH2-Glu504Glu | anti-P | 0.007281 | 2.06E-10 |
| B92 | B199 | ALDH2-Glu504Glu | Mg | -0.00648 | 1.54E-08 |
| B92 | B155 | ALDH2-Glu504Glu | IgA | 0.0118 | 0 |
| B92 | B156 | ALDH2-Glu504Glu | IgE | -0.00604 | 1.34E-07 |
| B92 | B157 | ALDH2-Glu504Glu | IgG | 0.009776 | 0 |
| B92 | B158 | ALDH2-Glu504Glu | IgM | 0.010695 | 0 |
| B92 | B195 | ALDH2-Glu504Glu | Na | -0.01545 | 0 |
| B92 | B176 | ALDH2-Glu504Glu | Ucr | -0.0059 | 2.54E-07 |
| B92 | B178 | ALDH2-Glu504Glu | Ig-U | -0.00966 | 0 |
| B92 | B180 | ALDH2-Glu504Glu | Tf | -0.01261 | 0 |
| B92 | B182 | ALDH2-Glu504Glu | TT | -0.00809 | 1.67E-12 |
| B92 | B125 | ALDH2-Glu504Glu | Cor | -0.00908 | 2.22E-15 |
| B92 | B138 | ALDH2-Glu504Glu | PAB | 0.017929 | 0 |
| B92 | B200 | ALDH2-Glu504Glu | OP | -0.01385 | 0 |
| B92 | B131 | ALDH2-Glu504Glu | AST | 0.006705 | 4.80E-09 |
| B92 | B139 | ALDH2-Glu504Glu | ADA | 0.011189 | 0 |
| B92 | B183 | ALDH2-Glu504Glu | PTA | 0.020885 | 0 |
| B92 | B186 | ALDH2-Glu504Glu | PFC | -0.03249 | 0 |
| B92 | B204 | ALDH2-Glu504Glu | Drinking | 0.020353 | 0 |
| B92 | B112 | ALDH2-Glu504Glu | FT4 | -0.00756 | 4.02E-11 |
| B92 | B114 | ALDH2-Glu504Glu | TSH TT3 TT4(TT4) | -0.01562 | 0 |
| B93 | B137 | ALDH2-Glu504Lys | AFU | 0.009021 | 3.55E-15 |
| B93 | B140 | ALDH2-Glu504Lys | r-GT | -0.01092 | 0 |
| B93 | B94 | ALDH2-Glu504Lys | ALDH2-Lys504Lys | -0.10688 | 0 |
| B93 | B128 | ALDH2-Glu504Lys | ALT | -0.00583 | 3.51E-07 |
| B93 | B121 | ALDH2-Glu504Lys | C3 | -0.006 | 1.60E-07 |

| | | | | | |
|---|---|---|---|---|---|
| B93 | B122 | ALDH2-Glu504Lys | C4 | 0.013816 | 0 |
| B93 | B225 | ALDH2-Glu504Lys | hPBG | -0.00647 | 1.63E-08 |
| B93 | B124 | ALDH2-Glu504Lys | ACTH | -0.00621 | 5.98E-08 |
| B93 | B134 | ALDH2-Glu504Lys | CHE | 0.011947 | 0 |
| B93 | B181 | ALDH2-Glu504Lys | APTT | 0.010921 | 0 |
| B93 | B150 | ALDH2-Glu504Lys | ALP | 0.006742 | 3.95E-09 |
| B93 | B95 | ALDH2-Glu504Lys | anti-cenpb | 0.008522 | 1.00E-13 |
| B93 | B97 | ALDH2-Glu504Lys | dsDNA | 0.010535 | 0 |
| B93 | B98 | ALDH2-Glu504Lys | anti-Jo-1 | -0.00922 | 8.88E-16 |
| B93 | B103 | ALDH2-Glu504Lys | anti-SSA/Ro | -0.01546 | 0 |
| B93 | B104 | ALDH2-Glu504Lys | anti-SSB | -0.00807 | 1.90E-12 |
| B93 | B106 | ALDH2-Glu504Lys | anti-P | -0.00664 | 6.80E-09 |
| B93 | B199 | ALDH2-Glu504Lys | Mg | 0.006322 | 3.39E-08 |
| B93 | B155 | ALDH2-Glu504Lys | IgA | -0.01043 | 0 |
| B93 | B157 | ALDH2-Glu504Lys | IgG | -0.01046 | 0 |
| B93 | B158 | ALDH2-Glu504Lys | IgM | -0.00976 | 0 |
| B93 | B195 | ALDH2-Glu504Lys | Na | 0.016295 | 0 |
| B93 | B176 | ALDH2-Glu504Lys | Ucr | 0.006575 | 9.44E-09 |
| B93 | B178 | ALDH2-Glu504Lys | Ig-U | 0.011195 | 0 |
| B93 | B180 | ALDH2-Glu504Lys | Tf | 0.014059 | 0 |
| B93 | B182 | ALDH2-Glu504Lys | TT | 0.005969 | 1.87E-07 |
| B93 | B125 | ALDH2-Glu504Lys | Cor | 0.007738 | 1.42E-11 |
| B93 | B138 | ALDH2-Glu504Lys | PAB | -0.0157 | 0 |
| B93 | B200 | ALDH2-Glu504Lys | OP | 0.01502 | 0 |
| B93 | B131 | ALDH2-Glu504Lys | AST | -0.00597 | 1.85E-07 |
| B93 | B139 | ALDH2-Glu504Lys | ADA | -0.00956 | 0 |
| B93 | B196 | ALDH2-Glu504Lys | TCO2 | 0.006468 | 1.63E-08 |
| B93 | B183 | ALDH2-Glu504Lys | PTA | -0.0172 | 0 |
| B93 | B186 | ALDH2-Glu504Lys | PFC | 0.027449 | 0 |
| B93 | B204 | ALDH2-Glu504Lys | Drinking | -0.01737 | 0 |
| B93 | B112 | ALDH2-Glu504Lys | FT4 | 0.007532 | 4.85E-11 |
| B93 | B114 | ALDH2-Glu504Lys | TSH TT3 TT4(TT4) | 0.015977 | 0 |
| B94 | B122 | ALDH2-Lys504Lys | C4 | 0.005923 | 2.33E-07 |
| B94 | B124 | ALDH2-Lys504Lys | ACTH | 0.016307 | 0 |
| B94 | B105 | ALDH2-Lys504Lys | anti-U1RNP | -0.00739 | 1.09E-10 |
| B94 | B95 | ALDH2-Lys504Lys | anti-cenpb | -0.00577 | 4.73E-07 |
| B94 | B97 | ALDH2-Lys504Lys | dsDNA | -0.00605 | 1.26E-07 |
| B94 | B100 | ALDH2-Lys504Lys | anti-Ro52 | -0.00721 | 3.06E-10 |
| B94 | B102 | ALDH2-Lys504Lys | anti-Sm | -0.00578 | 4.44E-07 |
| B94 | B103 | ALDH2-Lys504Lys | anti-SSA/Ro | -0.00858 | 6.88E-14 |
| B94 | B108 | ALDH2-Lys504Lys | Anti-Cyclin | -0.00588 | 2.85E-07 |
| B94 | B109 | ALDH2-Lys504Lys | AMA-M2 | -0.0058 | 4.06E-07 |
| B94 | B110 | ALDH2-Lys504Lys | AHA | 0.016206 | 0 |
| B94 | B198 | ALDH2-Lys504Lys | P | 0.007055 | 7.29E-10 |

| | | | | | |
|---|---|---|---|---|---|
| B94 | B194 | ALDH2-Lys504Lys | Cl | 0.011486 | 0 |
| B94 | B156 | ALDH2-Lys504Lys | IgE | 0.018842 | 0 |
| B94 | B179 | ALDH2-Lys504Lys | α1-MG | -0.01165 | 0 |
| B94 | B182 | ALDH2-Lys504Lys | TT | 0.006328 | 3.30E-08 |
| B94 | B138 | ALDH2-Lys504Lys | PAB | -0.00628 | 4.22E-08 |
| B94 | B187 | ALDH2-Lys504Lys | NSE | 0.008087 | 1.66E-12 |
| B94 | B139 | ALDH2-Lys504Lys | ADA | -0.00587 | 3.00E-07 |
| B94 | B196 | ALDH2-Lys504Lys | TCO2 | -0.00723 | 2.79E-10 |
| B94 | B183 | ALDH2-Lys504Lys | PTA | -0.01172 | 0 |
| B94 | B185 | ALDH2-Lys504Lys | PT | 0.006959 | 1.24E-09 |
| B94 | B186 | ALDH2-Lys504Lys | PFC | 0.015994 | 0 |
| B94 | B204 | ALDH2-Lys504Lys | Drinking | -0.00974 | 0 |
| B95 | B193 | anti-cenpb | K | 0.008497 | 1.19E-13 |
| B95 | B99 | anti-cenpb | anti-PM-Scl | 0.012844 | 0 |
| B95 | B97 | anti-cenpb | dsDNA | 0.012748 | 0 |
| B95 | B98 | anti-cenpb | anti-Jo-1 | -0.00599 | 1.69E-07 |
| B95 | B100 | anti-cenpb | anti-Ro52 | 0.039815 | 0 |
| B95 | B101 | anti-cenpb | anti-Sc70 | -0.0059 | 2.57E-07 |
| B95 | B103 | anti-cenpb | anti-SSA/Ro | 0.027409 | 0 |
| B95 | B104 | anti-cenpb | anti-SSB | 0.017915 | 0 |
| B95 | B106 | anti-cenpb | anti-P | -0.00639 | 2.43E-08 |
| B95 | B151 | anti-cenpb | Anti-CCP | -0.00625 | 4.91E-08 |
| B95 | B109 | anti-cenpb | AMA-M2 | 0.079647 | 0 |
| B95 | B199 | anti-cenpb | Mg | -0.00701 | 9.37E-10 |
| B95 | B157 | anti-cenpb | IgG | 0.013932 | 0 |
| B95 | B195 | anti-cenpb | Na | 0.006494 | 1.43E-08 |
| B95 | B125 | anti-cenpb | Cor | -0.01914 | 0 |
| B95 | B138 | anti-cenpb | PAB | -0.01204 | 0 |
| B95 | B200 | anti-cenpb | OP | 0.006341 | 3.09E-08 |
| B95 | B139 | anti-cenpb | ADA | 0.010962 | 0 |
| B96 | B127 | dsDNA (IIF) | A-G ratio | -0.00618 | 6.96E-08 |
| B96 | B105 | dsDNA (IIF) | anti-U1RNP | 0.086568 | 0 |
| B96 | B97 | dsDNA (IIF) | dsDNA | 0.152215 | 0 |
| B96 | B100 | dsDNA (IIF) | anti-Ro52 | 0.103872 | 0 |
| B96 | B102 | dsDNA (IIF) | anti-Sm | -0.00693 | 1.43E-09 |
| B96 | B103 | dsDNA (IIF) | anti-SSA/Ro | 0.097177 | 0 |
| B96 | B104 | dsDNA (IIF) | anti-SSB | 0.174663 | 0 |
| B96 | B106 | dsDNA (IIF) | anti-P | 0.121867 | 0 |
| B96 | B107 | dsDNA (IIF) | AnuA | 0.602683 | 0 |
| B96 | B151 | dsDNA (IIF) | Anti-CCP | 0.009058 | 2.66E-15 |
| B96 | B109 | dsDNA (IIF) | AMA-M2 | 0.110505 | 0 |
| B96 | B110 | dsDNA (IIF) | AHA | 0.068124 | 0 |
| B96 | B155 | dsDNA (IIF) | IgA | 0.006931 | 1.44E-09 |
| B96 | B156 | dsDNA (IIF) | IgE | 0.00693 | 1.44E-09 |

| | | | | | |
|---|---|---|---|---|---|
| B96 | B157 | dsDNA (IIF) | IgG | 0.073274 | 0 |
| B96 | B179 | dsDNA (IIF) | α1-MG | 0.008467 | 1.44E-13 |
| B96 | B130 | dsDNA (IIF) | GLB | 0.009769 | 0 |
| B96 | B152 | dsDNA (IIF) | ACA-Ig(G-A-M) | -0.01188 | 0 |
| B96 | B183 | dsDNA (IIF) | PTA | 0.005672 | 7.36E-07 |
| B97 | B121 | dsDNA | C3 | -0.01542 | 0 |
| B97 | B122 | dsDNA | C4 | -0.01784 | 0 |
| B97 | B105 | dsDNA | anti-U1RNP | 0.065061 | 0 |
| B97 | B100 | dsDNA | anti-Ro52 | 0.030207 | 0 |
| B97 | B102 | dsDNA | anti-Sm | 0.048145 | 0 |
| B97 | B103 | dsDNA | anti-SSA/Ro | 0.060723 | 0 |
| B97 | B104 | dsDNA | anti-SSB | 0.04287 | 0 |
| B97 | B106 | dsDNA | anti-P | 0.050724 | 0 |
| B97 | B107 | dsDNA | AnuA | 0.177627 | 0 |
| B97 | B108 | dsDNA | Anti-Cyclin | -0.00691 | 1.57E-09 |
| B97 | B109 | dsDNA | AMA-M2 | 0.038138 | 0 |
| B97 | B110 | dsDNA | AHA | 0.0414 | 0 |
| B97 | B199 | dsDNA | Mg | 0.01165 | 0 |
| B97 | B155 | dsDNA | IgA | 0.006467 | 1.65E-08 |
| B97 | B157 | dsDNA | IgG | 0.016796 | 0 |
| B97 | B195 | dsDNA | Na | 0.010331 | 0 |
| B97 | B179 | dsDNA | α1-MG | -0.00649 | 1.48E-08 |
| B97 | B182 | dsDNA | TT | 0.007031 | 8.34E-10 |
| B97 | B125 | dsDNA | Cor | -0.00643 | 1.95E-08 |
| B97 | B197 | dsDNA | AG | 0.007758 | 1.26E-11 |
| B98 | B122 | anti-Jo-1 | C4 | 0.028752 | 0 |
| B98 | B124 | anti-Jo-1 | ACTH | 0.007464 | 7.18E-11 |
| B98 | B100 | anti-Jo-1 | anti-Ro52 | -0.00585 | 3.27E-07 |
| B98 | B103 | anti-Jo-1 | anti-SSA/Ro | 0.048558 | 0 |
| B98 | B109 | anti-Jo-1 | AMA-M2 | -0.00779 | 1.01E-11 |
| B98 | B110 | anti-Jo-1 | AHA | -0.00731 | 1.72E-10 |
| B98 | B194 | anti-Jo-1 | Cl | -0.00602 | 1.48E-07 |
| B98 | B155 | anti-Jo-1 | IgA | 0.008218 | 7.26E-13 |
| B98 | B156 | anti-Jo-1 | IgE | -0.00771 | 1.72E-11 |
| B98 | B157 | anti-Jo-1 | IgG | 0.014698 | 0 |
| B98 | B139 | anti-Jo-1 | ADA | 0.007483 | 6.44E-11 |
| B98 | B186 | anti-Jo-1 | PFC | -0.01013 | 0 |
| B99 | B137 | anti-PM-Scl | AFU | -0.00962 | 0 |
| B99 | B124 | anti-PM-Scl | ACTH | 0.015796 | 0 |
| B99 | B150 | anti-PM-Scl | ALP | 0.006979 | 1.11E-09 |
| B99 | B100 | anti-PM-Scl | anti-Ro52 | 0.045148 | 0 |
| B99 | B103 | anti-PM-Scl | anti-SSA/Ro | 0.011285 | 0 |
| B99 | B104 | anti-PM-Scl | anti-SSB | 0.042925 | 0 |
| B99 | B109 | anti-PM-Scl | AMA-M2 | -0.00584 | 3.36E-07 |

| | | | | | |
|---|---|---|---|---|---|
| B99 | B199 | anti-PM-Scl | Mg | 0.007926 | 4.51E-12 |
| B99 | B179 | anti-PM-Scl | α1-MG | -0.00703 | 8.31E-10 |
| B99 | B125 | anti-PM-Scl | Cor | 0.016071 | 0 |
| B99 | B138 | anti-PM-Scl | PAB | -0.00626 | 4.54E-08 |
| B99 | B196 | anti-PM-Scl | TCO2 | -0.00612 | 9.15E-08 |
| B99 | B183 | anti-PM-Scl | PTA | -0.00686 | 2.05E-09 |
| B99 | B186 | anti-PM-Scl | PFC | 0.007068 | 6.81E-10 |
| B99 | B212 | anti-PM-Scl | IOP-L | 0.006889 | 1.81E-09 |

| **Code** | **PEI Name** | **PEI Name abbreviation** | **Unit** |
|---|---|---|---|
| B1 | Gender | Gender | - |
| B2 | Age | Age | years |
| B3 | Homocysteine | Hcy | umol/L |
| B4 | Whole Blood Calcium | WB-Ca | mmol/L |
| B5 | Whole Blood Magnesium | WB-Mg | mmol/L |
| B6 | Iron Of Whole Blood | WB-Fe | mmol/L |
| B7 | Whole Blood Copper | WB-Cop | umol/L |
| B8 | Zinc In Whole Blood | WB-Zinc | umol/L |
| B9 | Rh Blood Group | RH | - |
| B10 | Pepsinogen I | PGI | ug/L |
| B11 | PgI/PgII | PGI/PGII | - |
| B12 | Pepsinogen II | PGII | ug/L |
| B14 | Gastrin-17 | G-17 | pmol/l |
| B15 | Cd3-Cd19+(B Cell) | CD3-CD19+(B cell) | - |
| B16 | Cd3+ Total T Lymphocyte | CD3+T | - |
| B17 | Cd3+Cd16+Cd56+Nk | CD3+CD16+CD56+NK | - |
| B18 | Cd3+Cd4+Helper(Inducer)T Cells | CD3+CD4+helper(inducer)T | - |
| B19 | Cd3+Cd4+/Cd3+Cd8+ | CD3+CD4+/CD3+CD8+ | - |
| B20 | Cd3+Cd4+Cd8+ | CD3+CD4+CD8+ | - |
| B21 | Cd3+Cd4-Cd8- | CD3+CD4-CD8- | - |
| B22 | Cd3+Cd8+(T) | CD3+CD8+(T) | - |
| B23 | Cd3-Cd16+Cd56+(Nk) | CD3-CD16+CD56+(NK) | - |
| B24 | α Hydroxybutyrate Dehydrogenas | αHBDH | U/L |
| B25 | Creatine Kinase | CK | U/L |
| B26 | Creatine Kinase Isoenzyme | CKMB | U/L |
| B27 | Lactic Dehydrogenase | LDH | U/L |
| B28 | Pituitary Prolactin | PRL | mIU/L |
| B29 | Estradiol E2 | E2 | pmol/L |
| B30 | Luteinizing Hormone | LH | mIU/ml |

| | | | |
|---|---|---|---|
| B31 | Follicle Stimulating Hormone | FSH | mIU/ml |
| B32 | Testosterone | Testo | nmol/L |
| B33 | Progesterone | Prog | nmol/L |
| B34 | Thymidine Kinase | TK | PM/L |
| B35 | Blood Beta-Microglobulin | β2-MG | - |
| B36 | Variant Lymphocyte Rate | V-Lym% | - |
| B37 | Band Cell | BAND | 10E-9/L |
| B38 | Band Cell% | BAND% | - |
| B39 | Other Cell Types | Other cells | - |
| B40 | Leukocyte Count | WBC | 10E-9/L |
| B41 | Monocytes | MON | - |
| B42 | Monocytes % | MON % | 10E-9/L |
| B43 | Mean Corpuscular Volume | MCV | fl |
| B44 | Mean Corpuscular Hemoglobin | MCH | pg |
| B45 | Mean Corpuscular Hemoglobin Concentration | MCHC | g/l |
| B46 | Red Blood Cell Count | RBC Count | 10E-12/L |
| B47 | Red Cell Distribution Width | RDW | fl |
| B48 | Lymphocyte Rate | LYM% | - |
| B49 | Basophilic Granulocyte Rate | baso% | - |
| B50 | Eosinophil Rate | eo% | - |
| B51 | Hemoglobin | Hb | g/L |
| B52 | Platelet Count | PLT/BPC | 10E-9/L |
| B53 | Granulocyte Rate | GR% | - |
| B54 | Erythrocyte Sedimentation Rate | ESR | mm/60min |
| B55 | Erythrocyte Deformation Index | TK | - |
| B56 | The Index Of Rigidity Of Erythrocyte | IR | - |
| B57 | Erythrocyte Aggregation Indices | RBCAI | - |
| B58 | Hematocrit | HCT | - |
| B59 | Whole Blood Reduced Viscosity | WBRV | - |
| B60 | Whole Blood Reduced Viscosity Index | WBRVI | - |
| B61 | Whole Blood High Shear Reduced Viscosity | WBHSRV | - |
| B62 | Whole Blood High Shear Reduced Viscosity Index | WBHSRVI | - |
| B63 | Shear Rate Of Whole Blood Viscosity_Mpas_1 (200) | WBRSR(mPas_200) | - |
| B64 | Shear Rate Of Whole Blood Viscosity_Mpas_3 (5) | WBRSR(mPas_5) | - |
| B65 | Shear Rate Of Whole Blood Viscosity_Mpas_4 (1) | WBRSR(mPas_1) | - |
| B66 | Shear Rate Of Whole Blood Viscosity_Mpas (100) | WBRSR(mPas_100) | - |
| B67 | Shear Rate Of Whole Blood Viscosity_Mpas (50) | WBRSR(mPas_50) | - |
| B69 | Plasma Viscosity | PV | - |
| B70 | Low-Density Lipoprotein | LDL-C | mmol/L |
| B71 | Triglyceride | TG | mmol/L |
| B72 | High-Density Lipoprotein | HDL-C | mmol/L |
| B73 | Apolipoprotein A | ApoA | mg/L |
| B74 | Apolipoprotein A1 | ApoA1 | g/L |
| B75 | Apolipoprotein A1/B | ApoA1/B | g/L |

| | | | |
|---|---|---|---|
| B76 | Apolipoprotein B | ApoB | g/L |
| B77 | Total Cholesterol | TC | mmol/L |
| B78 | Hepatitis B Virus Core Im Antibod | HBcAb | - |
| B79 | Quantitative Detection Of Hepatitis B Virus Dna | hbV-dna | IU/ml |
| B80 | Helicobacter Pylori Antibody-Igg | anti-HP IgG | - |
| B81 | Tumor Supplied Group Of Factors | TSGF | U/ML |
| B82 | Carcinoembryonic Antigen | CEA | ng/ml |
| B83 | Alpha-Fetoprotein | AFP | ng/ml |
| B84 | Complexed Prostate Special Antigen | C-PSA | ng/ml |
| B85 | Squamous Cell Carcinoma Antigen | SCC | ng/ml |
| B86 | Chorionic Gonadotropin | TBhCG | mIU/ml |
| B87 | Neuron-Specific Enolase | NSE | ng/ml |
| B88 | Carbohydrate Antigen，Ca242 | CA 242 | U/ml |
| B89 | Carbohydrate Antigen，Ca72_4 | CA 72-4 | U/ml |
| B90 | Cytokeratin-19-Fragment Cyfra21-1 | CYFRA 21-1 | ng/ml |
| B91 | Total Protein | TP | g/L |
| B92 | Aldh2-Glu504Glu | ALDH2-Glu504Glu | - |
| B93 | Aldh2-Glu504Lys | ALDH2-Glu504Lys | - |
| B94 | Aldh2-Lys504Lys | ALDH2-Lys504Lys | - |
| B95 | Anti-Cenpb | anti-cenpb | - |
| B96 | Dsdna (Iif) | dsDNA (IIF) | - |
| B97 | Anti-Dsdna Antibody | dsDNA | - |
| B98 | Jo-1 Antibody | anti-Jo-1 | - |
| B99 | Anti-Pm-Scl | anti-PM-Scl | - |
| B100 | Anti-Ro52 | anti-Ro52 | - |
| B101 | Anti-Sc70 | anti-Sc70 | - |
| B102 | Anti Sm Antibody | anti-Sm | - |
| B103 | Anti-Ssa/Ro Antibody | anti-SSA/Ro | - |
| B104 | Anti Ssb Antibody | anti-SSB | - |
| B105 | Anti-U1Rnp Antibody | anti-U1RNP | - |
| B106 | Anti-Ribosomal P Protein Antibody | anti-P | - |
| B107 | Anti-Nucleosome Antibody Systemic | AnuA | - |
| B108 | Anti-Cyclin Antibody | Anti-Cyclin | - |
| B109 | Antimitochondrial Antibody M2 | AMA-M2 | - |
| B110 | Anti-Histone Antibody | AHA | - |
| B111 | Thyroid-Stimulating Hormon | TSH | mIU/L |
| B112 | Free Thyroxine | FT4 | pmol/L |
| B113 | Free Triiodothyronine | FT3 | pmol/L |
| B114 | Total Thyroxine | TSH TT3 TT4(TT4) | nmol/L |
| B115 | Total Triiodothyronine | TSH TT3 TT4(TT3) | nmol/L |
| B116 | Total Triiodothyronine | TT3 | nmol/L |
| B117 | Total Thyroxine | TT4 | nmol/L |
| B118 | Carcinoembryonic Antigen 1 | CEA1 | ng/ml |
| B119 | Quantitative Detection Of Hepatitis C Virus Rna | Hcv-rna | - |

| | | | |
|---|---|---|---|
| B120 | Hcv Core Antigen | HCV-cAg | - |
| B121 | Complement C3 | C3 | - |
| B122 | Complement C4 | C4 | - |
| B123 | High Sensitivity C-Reactive Protein 1 | hs-CRP1 | mg/L |
| B124 | Adrenocorticotropic Hormone | ACTH | pg/ml |
| B125 | Cortisol | Cor | ug/dl |
| B126 | Albumin | Alb | g/L |
| B127 | Albumin-Globulin | A-G ratio | - |
| B128 | Alanine Aminotransferase | ALT | U/L |
| B129 | Indirect Bilirubin | IBIL | umol/L |
| B130 | Globulin | GLB | g/L |
| B131 | Aspartate Transaminase | AST | U/L |
| B132 | Direct Bilirubin | Dbil | umol/L |
| B133 | Total Bilirubin | TBIL | umol/L |
| B134 | Cholinesterases | CHE | KU/L |
| B135 | Total Bile Acid | TBA | umol/L |
| B136 | 5-Nucleotidase | 5'NT | U/L |
| B137 | Alpha-L-Fucosidase | AFU | U/L |
| B138 | Prealbumin | PAB | mg/L |
| B139 | Adenosine Deaminase | ADA | U/L |
| B140 | R-Glutamyl Transpeptidase | r-GT | U/L |
| B141 | Ast/Alt | AST/ALT | - |
| B142 | 25-Hydroxyvitamin D | 25-(OH)D | ng/mL |
| B143 | β-Crosslaps | β-CTx | pg/mL |
| B144 | Total-Pinp | Total-PINP | ng/mL |
| B145 | Osteocalcin | BGP | ng/mL |
| B146 | Cystatin C | CysC | mg/L |
| B148 | Antithyroperoxidase Antibody | Anti_TPO | IU/ml |
| B149 | Anti-Thyroglobulin Antibodies | TGAb | IU/ml |
| B150 | Alkaline Phosphatase | ALP | U/L |
| B151 | Anti-Cyclic Peptide Containing Citrulline | Anti-CCP | Ru/ml |
| B152 | Anti Cardiolipin Antibodyig(G-A-M) | ACA-Ig(G-A-M) | u/ml |
| B153 | Glucose | Glu | mmol/L |
| B154 | Rheumatoid Factor | RF | IU/ml |
| B155 | Immunoglobulin A | IgA | g/l |
| B156 | Immunoglobulin E | IgE | IU/ml |
| B157 | Immunoglobulin G | IgG | g/l |
| B158 | Immunoglobulin M | IgM | g/l |
| B159 | Urine Beta-Microglobulin | β2-MG | - |
| B160 | Ph | PH | - |
| B161 | Leucocyte | LEU | /ul |
| B163 | Specific Gravity | SG | - |
| B164 | Calcium Oxalate Crystals | COM | HPF |
| B165 | Bilirubin | bilirubin | - |

| | | | |
|---|---|---|---|
| B166 | Protein | Pro | - |
| B167 | Phosphate Crystals | PC | - |
| B168 | Urinary Nuclear Matrix Protein 22 | NMP22 | - |
| B169 | Urate Crystal | U-CRY | - |
| B170 | Urine Sugar | U-GLU | - |
| B171 | Epithelial Cell | EC | HPF |
| B172 | Ketone Body | KB | - |
| B173 | Nitrous Acid | NTT | - |
| B174 | Urea Creatinine | Urea/Cr | - |
| B175 | Microalbuminuria | ALB-U | mg/L |
| B176 | Urine Creatinine | Ucr | umol/L |
| B177 | Ratio Of Urinary Microalbumin To Creatinin | ACR | ug/mg |
| B178 | Urinary Immunoglobulin G | Ig-U | mg/L |
| B179 | Urine Alpha-Microglobulin | α1-MG | mg/L |
| B180 | Urine Transferrin | Tf | mg/L |
| B181 | Activated Partial Thromboplastin Time | APTT | s |
| B182 | Thrombin Time | TT | s |
| B183 | Percentage Activity Of Prothrombin In Plasma | PTA | - |
| B184 | International Normalized Ratio | INR | - |
| B185 | Prothrombin Time | PT | s |
| B186 | Plasma Fibrinogen Concentration | PFC | g/l |
| B187 | Neuron-Specific Enolase | NSE | ng/ml |
| B188 | Calcium | Ca | mmol/L |
| B189 | Estimated Glomerular Filtration Rate | eGFB | ml/min |
| B190 | Creatinine | Cre | umol/L |
| B191 | Urea | Urea | mmol/L |
| B192 | Uric Acid | UA | umol/L |
| B193 | Kalium | K | mmol/L |
| B194 | Chlorine | Cl | mmol/L |
| B195 | Sodium | Na | mmol/L |
| B196 | Blood Carbon Dioxide | TCO2 | mmol/L |
| B197 | Anion Gap | AG | mmol/L |
| B198 | Phosphorus | P | mmol/L |
| B199 | Magnesium | Mg | mmol/L |
| B200 | Osmotic Pressure | OP | mOsm/L |
| B201 | Exercise Habits | e-habits | - |
| B202 | Frequency Of Exercises | e-times | - |
| B203 | Dietary Habits 1 | DH | - |
| B204 | History Of Alcohol Intake | Drinking | - |
| B205 | Known Of Drug Allergy | KDA | - |
| B206 | Waistline | Waist | cm |
| B207 | Waist-Hip Ratio | WHR | - |
| B209 | Smoking History | Smoking | - |
| B210 | Hipline | Hip | cm |

| | | | |
|---|---|---|---|
| B211 | Intra-Ocular Tension Of The Right Eye | IOP-R | mmHg |
| B212 | Intra-Ocular Tension Of The Left Eye | IOP-L | mmHg |
| B213 | Corrected Visual Acuity The Right Eye | CVA-R | - |
| B214 | Corrected Visual Acuity The Left Eye | CVA-L | - |
| B215 | Body Mass Index | BMI | - |
| B216 | Body Mass | BMI | kg |
| B218 | Diastolic Pressure | DBP | mmHg |
| B219 | Operation History | OT | - |
| B220 | Systolic Pressure | SBP | mmHg |
| B221 | Height | Hei | cm |
| B222 | Neck Size | NS | cm |
| B223 | Family History | FH | - |
| B224 | Anamnesis | pmh | - |
| B225 | Hourly Postprandial Blood Sugar | hPBG | mmol/L |
| B227 | Vision Of The Right Eye | OD | - |
| B228 | Vision Of The Left Eye | OS | - |

**Supplementary Table 2. PEI Correlations in Cholecystolithiasis**

| Row | Column | Row | Column | Cor | *P* |
|---|---|---|---|---|---|
| B1 | B44 | Gender | MCH | -0.18202 | 9.05E-09 |
| B1 | B45 | Gender | MCHC | -0.31897 | 0 |
| B1 | B46 | Gender | RBC Count | -0.50021 | 0 |
| B1 | B51 | Gender | Hb | -0.64675 | 0 |
| B1 | B71 | Gender | TG | -0.18502 | 5.09E-09 |
| B1 | B72 | Gender | HDL-C | 0.304637 | 0 |
| B1 | B126 | Gender | Alb | -0.17113 | 6.72E-08 |
| B1 | B127 | Gender | A-G ratio | -0.17028 | 7.82E-08 |
| B1 | B128 | Gender | ALT | -0.28035 | 0 |
| B1 | B131 | Gender | AST | -0.15719 | 7.31E-07 |
| B1 | B140 | Gender | r-GT | -0.252 | 8.88E-16 |
| B1 | B141 | Gender | AST/ALT | 0.252548 | 8.88E-16 |
| B1 | B161 | Gender | LEU | 0.16185 | 3.37E-07 |
| B1 | B163 | Gender | SG | -0.37088 | 0 |
| B1 | B166 | Gender | Pro | -0.17468 | 3.55E-08 |
| B1 | B174 | Gender | Urea/Cr | 0.3788 | 0 |
| B1 | B190 | Gender | Cre | -0.51593 | 0 |
| B1 | B192 | Gender | UA | -0.562 | 0 |
| B1 | B204 | Gender | Drinking | -0.36613 | 0 |
| B1 | B206 | Gender | Waist | -0.50716 | 0 |
| B1 | B209 | Gender | Smoking | -0.39753 | 0 |
| B1 | B210 | Gender | Hip | -0.31913 | 0 |
| B1 | B215 | Gender | BMI | -0.29241 | 0 |
| B1 | B216 | Gender | BMI | -0.60175 | 0 |
| B1 | B218 | Gender | DBP | -0.19742 | 4.29E-10 |
| B1 | B221 | Gender | Hei | -0.67814 | 0 |
| B1 | B222 | Gender | NS | -0.72003 | 0 |
| B10 | B12 | PG Ⅰ | PG Ⅱ | 0.502872 | 0 |
| B10 | B189 | PG Ⅰ | eGFB | -0.23336 | 1.27E-13 |
| B10 | B190 | PG Ⅰ | Cre | 0.499029 | 0 |
| B10 | B191 | PG Ⅰ | Urea | 0.37099 | 0 |
| B100 | B101 | anti-Ro52 | anti-Sc70 | -0.34578 | 0 |
| B100 | B102 | anti-Ro52 | anti-Sm | -0.34578 | 0 |
| B100 | B103 | anti-Ro52 | anti-SSA/Ro | 0.97918 | 0 |
| B100 | B104 | anti-Ro52 | anti-SSB | 0.98189 | 0 |
| B100 | B105 | anti-Ro52 | anti-U1RNP | -0.34578 | 0 |
| B100 | B106 | anti-Ro52 | anti-P | -0.34578 | 0 |
| B100 | B107 | anti-Ro52 | AnuA | -0.34578 | 0 |
| B100 | B108 | anti-Ro52 | Anti-Cyclin | -0.34578 | 0 |
| B100 | B109 | anti-Ro52 | AMA-M2 | -0.34578 | 0 |
| B100 | B110 | anti-Ro52 | AHA | -0.34578 | 0 |
| B100 | B157 | anti-Ro52 | IgG | 0.516221 | 0 |

| | | | | | |
|---|---|---|---|---|---|
| B101 | B102 | anti-Sc70 | anti-Sm | 1 | 0 |
| B101 | B103 | anti-Sc70 | anti-SSA/Ro | -0.27591 | 0 |
| B101 | B104 | anti-Sc70 | anti-SSB | -0.32367 | 0 |
| B101 | B105 | anti-Sc70 | anti-U1RNP | 1 | 0 |
| B101 | B107 | anti-Sc70 | AnuA | 1 | 0 |
| B101 | B108 | anti-Sc70 | Anti-Cyclin | 1 | 0 |
| B101 | B109 | anti-Sc70 | AMA-M2 | 1 | 0 |
| B101 | B134 | anti-Sc70 | CHE | -0.29695 | 0 |
| B101 | B151 | anti-Sc70 | Anti-CCP | -0.23405 | 1.07E-13 |
| B101 | B155 | anti-Sc70 | IgA | -0.17398 | 4.03E-08 |
| B101 | B156 | anti-Sc70 | IgE | -0.42651 | 0 |
| B101 | B157 | anti-Sc70 | IgG | -0.21397 | 1.21E-11 |
| B101 | B175 | anti-Sc70 | ALB-U | 0.22405 | 1.19E-12 |
| B101 | B176 | anti-Sc70 | Ucr | 0.200525 | 2.24E-10 |
| B101 | B178 | anti-Sc70 | Ig-U | 0.771362 | 0 |
| B101 | B179 | anti-Sc70 | α1-MG | 0.719949 | 0 |
| B101 | B180 | anti-Sc70 | Tf | 0.779882 | 0 |
| B101 | B181 | anti-Sc70 | APTT | 0.331973 | 0 |
| B101 | B182 | anti-Sc70 | TT | 0.330901 | 0 |
| B101 | B183 | anti-Sc70 | PTA | 0.331949 | 0 |
| B101 | B184 | anti-Sc70 | INR | 0.331973 | 0 |
| B101 | B185 | anti-Sc70 | PT | 0.331264 | 0 |
| B101 | B186 | anti-Sc70 | PFC | -0.33197 | 0 |
| B101 | B194 | anti-Sc70 | Cl | -0.33683 | 0 |
| B101 | B197 | anti-Sc70 | AG | 0.254711 | 4.44E-16 |
| B102 | B103 | anti-Sm | anti-SSA/Ro | -0.27591 | 0 |
| B102 | B104 | anti-Sm | anti-SSB | -0.32367 | 0 |
| B102 | B107 | anti-Sm | AnuA | 1 | 0 |
| B102 | B134 | anti-Sm | CHE | -0.29695 | 0 |
| B102 | B151 | anti-Sm | Anti-CCP | -0.23405 | 1.07E-13 |
| B102 | B155 | anti-Sm | IgA | -0.17398 | 4.03E-08 |
| B102 | B156 | anti-Sm | IgE | -0.42651 | 0 |
| B102 | B157 | anti-Sm | IgG | -0.21397 | 1.21E-11 |
| B102 | B175 | anti-Sm | ALB-U | 0.22405 | 1.19E-12 |
| B102 | B176 | anti-Sm | Ucr | 0.200525 | 2.24E-10 |
| B102 | B178 | anti-Sm | Ig-U | 0.771362 | 0 |
| B102 | B179 | anti-Sm | α1-MG | 0.719949 | 0 |
| B102 | B180 | anti-Sm | Tf | 0.779882 | 0 |
| B102 | B181 | anti-Sm | APTT | 0.331973 | 0 |
| B102 | B182 | anti-Sm | TT | 0.330901 | 0 |
| B102 | B183 | anti-Sm | PTA | 0.331949 | 0 |
| B102 | B184 | anti-Sm | INR | 0.331973 | 0 |
| B102 | B185 | anti-Sm | PT | 0.331264 | 0 |
| B102 | B186 | anti-Sm | PFC | -0.33197 | 0 |

| | | | | | |
|---|---|---|---|---|---|
| B102 | B194 | anti-Sm | Cl | -0.33683 | 0 |
| B102 | B197 | anti-Sm | AG | 0.254711 | 4.44E-16 |
| B103 | B104 | anti-SSA/Ro | anti-SSB | 0.998747 | 0 |
| B103 | B105 | anti-SSA/Ro | anti-U1RNP | -0.27591 | 0 |
| B103 | B106 | anti-SSA/Ro | anti-P | -0.27591 | 0 |
| B103 | B107 | anti-SSA/Ro | AnuA | -0.27591 | 0 |
| B103 | B108 | anti-SSA/Ro | Anti-Cyclin | -0.27591 | 0 |
| B103 | B109 | anti-SSA/Ro | AMA-M2 | -0.27591 | 0 |
| B103 | B110 | anti-SSA/Ro | AHA | -0.27591 | 0 |
| B103 | B157 | anti-SSA/Ro | IgG | 0.524665 | 0 |
| B104 | B105 | anti-SSB | anti-U1RNP | -0.32367 | 0 |
| B104 | B106 | anti-SSB | anti-P | -0.32367 | 0 |
| B104 | B107 | anti-SSB | AnuA | -0.32367 | 0 |
| B104 | B108 | anti-SSB | Anti-Cyclin | -0.32367 | 0 |
| B104 | B109 | anti-SSB | AMA-M2 | -0.32367 | 0 |
| B104 | B110 | anti-SSB | AHA | -0.32367 | 0 |
| B104 | B157 | anti-SSB | IgG | 0.527611 | 0 |
| B105 | B106 | anti-U1RNP | anti-P | 1 | 0 |
| B105 | B107 | anti-U1RNP | AnuA | 1 | 0 |
| B105 | B109 | anti-U1RNP | AMA-M2 | 1 | 0 |
| B105 | B134 | anti-U1RNP | CHE | -0.29695 | 0 |
| B105 | B151 | anti-U1RNP | Anti-CCP | -0.23405 | 1.07E-13 |
| B105 | B155 | anti-U1RNP | IgA | -0.17398 | 4.03E-08 |
| B105 | B156 | anti-U1RNP | IgE | -0.42651 | 0 |
| B105 | B157 | anti-U1RNP | IgG | -0.21397 | 1.21E-11 |
| B105 | B175 | anti-U1RNP | ALB-U | 0.22405 | 1.19E-12 |
| B105 | B176 | anti-U1RNP | Ucr | 0.200525 | 2.24E-10 |
| B105 | B178 | anti-U1RNP | Ig-U | 0.771362 | 0 |
| B105 | B179 | anti-U1RNP | α1-MG | 0.719949 | 0 |
| B105 | B180 | anti-U1RNP | Tf | 0.779882 | 0 |
| B105 | B181 | anti-U1RNP | APTT | 0.331973 | 0 |
| B105 | B182 | anti-U1RNP | TT | 0.330901 | 0 |
| B105 | B183 | anti-U1RNP | PTA | 0.331949 | 0 |
| B105 | B184 | anti-U1RNP | INR | 0.331973 | 0 |
| B105 | B185 | anti-U1RNP | PT | 0.331264 | 0 |
| B105 | B186 | anti-U1RNP | PFC | -0.33197 | 0 |
| B105 | B194 | anti-U1RNP | Cl | -0.33683 | 0 |
| B105 | B197 | anti-U1RNP | AG | 0.254711 | 4.44E-16 |
| B106 | B107 | anti-P | AnuA | 1 | 0 |
| B106 | B108 | anti-P | Anti-Cyclin | 1 | 0 |
| B106 | B109 | anti-P | AMA-M2 | 1 | 0 |
| B106 | B110 | anti-P | AHA | 1 | 0 |
| B106 | B134 | anti-P | CHE | -0.29695 | 0 |
| B106 | B151 | anti-P | Anti-CCP | -0.23405 | 1.07E-13 |

| | | | | | |
|---|---|---|---|---|---|
| B106 | B155 | anti-P | IgA | -0.17398 | 4.03E-08 |
| B106 | B156 | anti-P | IgE | -0.42651 | 0 |
| B106 | B157 | anti-P | IgG | -0.21397 | 1.21E-11 |
| B106 | B175 | anti-P | ALB-U | 0.22405 | 1.19E-12 |
| B106 | B176 | anti-P | Ucr | 0.200525 | 2.24E-10 |
| B106 | B178 | anti-P | Ig-U | 0.771362 | 0 |
| B106 | B179 | anti-P | α1-MG | 0.719949 | 0 |
| B106 | B180 | anti-P | Tf | 0.779882 | 0 |
| B106 | B181 | anti-P | APTT | 0.331973 | 0 |
| B106 | B182 | anti-P | TT | 0.330901 | 0 |
| B106 | B183 | anti-P | PTA | 0.331949 | 0 |
| B106 | B184 | anti-P | INR | 0.331973 | 0 |
| B106 | B185 | anti-P | PT | 0.331264 | 0 |
| B106 | B186 | anti-P | PFC | -0.33197 | 0 |
| B106 | B194 | anti-P | Cl | -0.33683 | 0 |
| B106 | B197 | anti-P | AG | 0.254711 | 4.44E-16 |
| B107 | B108 | AnuA | Anti-Cyclin | 1 | 0 |
| B107 | B109 | AnuA | AMA-M2 | 1 | 0 |
| B107 | B110 | AnuA | AHA | 1 | 0 |
| B107 | B134 | AnuA | CHE | -0.29695 | 0 |
| B107 | B151 | AnuA | Anti-CCP | -0.23405 | 1.07E-13 |
| B107 | B155 | AnuA | IgA | -0.17398 | 4.03E-08 |
| B107 | B156 | AnuA | IgE | -0.42651 | 0 |
| B107 | B157 | AnuA | IgG | -0.21397 | 1.21E-11 |
| B107 | B175 | AnuA | ALB-U | 0.224049 | 1.19E-12 |
| B107 | B176 | AnuA | Ucr | 0.200525 | 2.25E-10 |
| B107 | B178 | AnuA | Ig-U | 0.771362 | 0 |
| B107 | B179 | AnuA | α1-MG | 0.719949 | 0 |
| B107 | B180 | AnuA | Tf | 0.779882 | 0 |
| B107 | B181 | AnuA | APTT | 0.331973 | 0 |
| B107 | B182 | AnuA | TT | 0.330901 | 0 |
| B107 | B183 | AnuA | PTA | 0.331949 | 0 |
| B107 | B184 | AnuA | INR | 0.331973 | 0 |
| B107 | B185 | AnuA | PT | 0.331264 | 0 |
| B107 | B186 | AnuA | PFC | -0.33197 | 0 |
| B107 | B194 | AnuA | Cl | -0.33683 | 0 |
| B107 | B197 | AnuA | AG | 0.254711 | 4.44E-16 |
| B108 | B110 | Anti-Cyclin | AHA | 1 | 0 |
| B108 | B134 | Anti-Cyclin | CHE | -0.29695 | 0 |
| B108 | B151 | Anti-Cyclin | Anti-CCP | -0.23405 | 1.07E-13 |
| B108 | B155 | Anti-Cyclin | IgA | -0.17398 | 4.03E-08 |
| B108 | B156 | Anti-Cyclin | IgE | -0.42651 | 0 |
| B108 | B157 | Anti-Cyclin | IgG | -0.21397 | 1.21E-11 |
| B108 | B175 | Anti-Cyclin | ALB-U | 0.22405 | 1.19E-12 |

| | | | | | |
|---|---|---|---|---|---|
| B108 | B176 | Anti-Cyclin | Ucr | 0.200525 | 2.24E-10 |
| B108 | B178 | Anti-Cyclin | Ig-U | 0.771362 | 0 |
| B108 | B179 | Anti-Cyclin | α1-MG | 0.719949 | 0 |
| B108 | B180 | Anti-Cyclin | Tf | 0.779882 | 0 |
| B108 | B181 | Anti-Cyclin | APTT | 0.331973 | 0 |
| B108 | B182 | Anti-Cyclin | TT | 0.330901 | 0 |
| B108 | B183 | Anti-Cyclin | PTA | 0.331949 | 0 |
| B108 | B184 | Anti-Cyclin | INR | 0.331973 | 0 |
| B108 | B185 | Anti-Cyclin | PT | 0.331264 | 0 |
| B108 | B186 | Anti-Cyclin | PFC | -0.33197 | 0 |
| B108 | B194 | Anti-Cyclin | Cl | -0.33683 | 0 |
| B108 | B197 | Anti-Cyclin | AG | 0.254711 | 4.44E-16 |
| B109 | B110 | AMA-M2 | AHA | 1 | 0 |
| B109 | B134 | AMA-M2 | CHE | -0.29695 | 0 |
| B109 | B151 | AMA-M2 | Anti-CCP | -0.23405 | 1.07E-13 |
| B109 | B155 | AMA-M2 | IgA | -0.17398 | 4.03E-08 |
| B109 | B156 | AMA-M2 | IgE | -0.42651 | 0 |
| B109 | B157 | AMA-M2 | IgG | -0.21397 | 1.21E-11 |
| B109 | B175 | AMA-M2 | ALB-U | 0.22405 | 1.19E-12 |
| B109 | B176 | AMA-M2 | Ucr | 0.200525 | 2.24E-10 |
| B109 | B178 | AMA-M2 | Ig-U | 0.771362 | 0 |
| B109 | B179 | AMA-M2 | α1-MG | 0.719949 | 0 |
| B109 | B180 | AMA-M2 | Tf | 0.779882 | 0 |
| B109 | B181 | AMA-M2 | APTT | 0.331973 | 0 |
| B109 | B182 | AMA-M2 | TT | 0.330901 | 0 |
| B109 | B183 | AMA-M2 | PTA | 0.331949 | 0 |
| B109 | B184 | AMA-M2 | INR | 0.331973 | 0 |
| B109 | B185 | AMA-M2 | PT | 0.331264 | 0 |
| B109 | B186 | AMA-M2 | PFC | -0.33197 | 0 |
| B109 | B194 | AMA-M2 | Cl | -0.33683 | 0 |
| B109 | B197 | AMA-M2 | AG | 0.254711 | 4.44E-16 |
| B11 | B12 | PGI/PGII | PG II | -0.41922 | 0 |
| B11 | B14 | PGI/PGII | G-17 | -0.1779 | 1.96E-08 |
| B11 | B95 | PGI/PGII | anti-cenpb | -0.28758 | 0 |
| B11 | B97 | PGI/PGII | dsDNA | -0.28758 | 0 |
| B11 | B98 | PGI/PGII | anti-Jo-1 | -0.28758 | 0 |
| B11 | B101 | PGI/PGII | anti-Sc70 | -0.28758 | 0 |
| B11 | B102 | PGI/PGII | anti-Sm | -0.28758 | 0 |
| B11 | B105 | PGI/PGII | anti-U1RNP | -0.28758 | 0 |
| B11 | B106 | PGI/PGII | anti-P | -0.28758 | 0 |
| B11 | B107 | PGI/PGII | AnuA | -0.28758 | 0 |
| B11 | B108 | PGI/PGII | Anti-Cyclin | -0.28758 | 0 |
| B11 | B109 | PGI/PGII | AMA-M2 | -0.28758 | 0 |
| B11 | B110 | PGI/PGII | AHA | -0.28758 | 0 |

| | | | | | |
|---|---|---|---|---|---|
| B110 | B134 | AHA | CHE | -0.29695 | 0 |
| B110 | B151 | AHA | Anti-CCP | -0.23405 | 1.07E-13 |
| B110 | B155 | AHA | IgA | -0.17398 | 4.03E-08 |
| B110 | B156 | AHA | IgE | -0.42651 | 0 |
| B110 | B157 | AHA | IgG | -0.21397 | 1.21E-11 |
| B110 | B175 | AHA | ALB-U | 0.22405 | 1.19E-12 |
| B110 | B176 | AHA | Ucr | 0.200525 | 2.24E-10 |
| B110 | B178 | AHA | Ig-U | 0.771362 | 0 |
| B110 | B179 | AHA | α1-MG | 0.719949 | 0 |
| B110 | B180 | AHA | Tf | 0.779882 | 0 |
| B110 | B181 | AHA | APTT | 0.331973 | 0 |
| B110 | B182 | AHA | TT | 0.330901 | 0 |
| B110 | B183 | AHA | PTA | 0.331949 | 0 |
| B110 | B184 | AHA | INR | 0.331973 | 0 |
| B110 | B185 | AHA | PT | 0.331264 | 0 |
| B110 | B186 | AHA | PFC | -0.33197 | 0 |
| B110 | B194 | AHA | Cl | -0.33683 | 0 |
| B110 | B197 | AHA | AG | 0.254711 | 4.44E-16 |
| B111 | B134 | TSH | CHE | 0.195932 | 5.82E-10 |
| B111 | B148 | TSH | Anti_TPO | 0.248254 | 2.89E-15 |
| B111 | B149 | TSH | TGAb | 0.182178 | 8.78E-09 |
| B112 | B114 | FT4 | TSH TT3 TT4(TT4) | 0.375623 | 0 |
| B112 | B154 | FT4 | RF | 0.160852 | 3.98E-07 |
| B113 | B115 | FT3 | TSH TT3 TT4(TT3) | 0.303551 | 0 |
| B113 | B148 | FT3 | Anti_TPO | -0.15709 | 7.43E-07 |
| B114 | B115 | TSH TT3 TT4(TT4) | TSH TT3 TT4(TT3) | 0.334011 | 0 |
| B115 | B124 | TSH TT3 TT4(TT3) | ACTH | 0.166034 | 1.65E-07 |
| B115 | B125 | TSH TT3 TT4(TT3) | Cor | 0.166034 | 1.65E-07 |
| B116 | B167 | TT3 | PC | 0.465184 | 0 |
| B116 | B202 | TT3 | e-times | 0.708765 | 0 |
| B117 | B167 | TT4 | PC | 0.576681 | 0 |
| B117 | B202 | TT4 | e-times | 0.843804 | 0 |
| B118 | B176 | CEA1 | Ucr | -0.16162 | 3.50E-07 |
| B118 | B214 | CEA1 | CVA-L | 0.168556 | 1.06E-07 |
| B118 | B220 | CEA1 | SBP | 0.163352 | 2.61E-07 |
| B118 | B228 | CEA1 | OS | 0.232578 | 1.53E-13 |
| B119 | B120 | Hcv-rna | HCV-cAg | 0.56139 | 0 |
| B119 | B121 | Hcv-rna | C3 | -0.88101 | 0 |
| B119 | B122 | Hcv-rna | C4 | -0.88101 | 0 |
| B119 | B159 | Hcv-rna | β2-MG | -0.88101 | 0 |
| B119 | B169 | Hcv-rna | U-CRY | -0.88101 | 0 |
| B119 | B195 | Hcv-rna | Na | 0.242012 | 1.44E-14 |
| B119 | B200 | Hcv-rna | OP | 0.182567 | 8.15E-09 |
| B12 | B14 | PG II | G-17 | 0.22303 | 1.52E-12 |

| | | | | | |
|---|---|---|---|---|---|
| B12 | B80 | PG II | anti-HP IgG | 0.194788 | 7.35E-10 |
| B120 | B121 | HCV-cAg | C3 | -0.49459 | 0 |
| B120 | B122 | HCV-cAg | C4 | -0.49459 | 0 |
| B120 | B154 | HCV-cAg | RF | -0.1817 | 9.62E-09 |
| B120 | B157 | HCV-cAg | IgG | -0.19536 | 6.54E-10 |
| B120 | B159 | HCV-cAg | β2-MG | -0.49459 | 0 |
| B120 | B169 | HCV-cAg | U-CRY | -0.49459 | 0 |
| B121 | B122 | C3 | C4 | 1 | 0 |
| B121 | B159 | C3 | β2-MG | 1 | 0 |
| B121 | B169 | C3 | U-CRY | 0.999999 | 0 |
| B121 | B195 | C3 | Na | -0.19632 | 5.38E-10 |
| B122 | B195 | C4 | Na | -0.19632 | 5.38E-10 |
| B123 | B138 | hs-CRP1 | PAB | -0.30676 | 0 |
| B123 | B139 | hs-CRP1 | ADA | 0.283631 | 0 |
| B123 | B146 | hs-CRP1 | CysC | -0.15627 | 8.50E-07 |
| B123 | B154 | hs-CRP1 | RF | 0.293498 | 0 |
| B123 | B168 | hs-CRP1 | NMP22 | -0.23009 | 2.82E-13 |
| B123 | B193 | hs-CRP1 | K | -0.21884 | 4.02E-12 |
| B123 | B195 | hs-CRP1 | Na | -0.27462 | 0 |
| B123 | B199 | hs-CRP1 | Mg | -0.2122 | 1.80E-11 |
| B123 | B211 | hs-CRP1 | IOP-R | 0.174237 | 3.84E-08 |
| B123 | B212 | hs-CRP1 | IOP-L | 0.159912 | 4.66E-07 |
| B124 | B125 | ACTH | Cor | 1 | 0 |
| B126 | B127 | Alb | A-G ratio | 0.519628 | 0 |
| B126 | B129 | Alb | IBIL | 0.191077 | 1.55E-09 |
| B126 | B130 | Alb | GLB | -0.20895 | 3.69E-11 |
| B126 | B133 | Alb | TBIL | 0.17225 | 5.50E-08 |
| B126 | B188 | Alb | Ca | 0.34146 | 0 |
| B126 | B189 | Alb | eGFB | 0.194454 | 7.86E-10 |
| B126 | B192 | Alb | UA | 0.157443 | 7.01E-07 |
| B126 | B221 | Alb | Hei | 0.218326 | 4.51E-12 |
| B127 | B129 | A-G ratio | IBIL | 0.21057 | 2.58E-11 |
| B127 | B130 | A-G ratio | GLB | -0.90294 | 0 |
| B127 | B133 | A-G ratio | TBIL | 0.194415 | 7.92E-10 |
| B127 | B221 | A-G ratio | Hei | 0.217709 | 5.20E-12 |
| B128 | B131 | ALT | AST | 0.777816 | 0 |
| B128 | B140 | ALT | r-GT | 0.469661 | 0 |
| B128 | B141 | ALT | AST/ALT | -0.56915 | 0 |
| B128 | B163 | ALT | SG | 0.186083 | 4.15E-09 |
| B128 | B192 | ALT | UA | 0.315074 | 0 |
| B128 | B206 | ALT | Waist | 0.34859 | 0 |
| B128 | B210 | ALT | Hip | 0.254 | 6.66E-16 |
| B128 | B215 | ALT | BMI | 0.344584 | 0 |
| B128 | B216 | ALT | BMI | 0.381513 | 0 |

| | | | | | |
|---|---|---|---|---|---|
| B128 | B221 | ALT | Hei | 0.206616 | 6.13E-11 |
| B128 | B222 | ALT | NS | 0.351156 | 0 |
| B129 | B130 | IBIL | GLB | -0.173 | 4.81E-08 |
| B129 | B132 | IBIL | Dbil | 0.677229 | 0 |
| B129 | B133 | IBIL | TBIL | 0.98049 | 0 |
| B130 | B133 | GLB | TBIL | -0.16015 | 4.48E-07 |
| B130 | B221 | GLB | Hei | -0.15737 | 7.10E-07 |
| B131 | B140 | AST | r-GT | 0.288175 | 0 |
| B131 | B141 | AST | AST/ALT | -0.16313 | 2.71E-07 |
| B131 | B192 | AST | UA | 0.219507 | 3.44E-12 |
| B131 | B206 | AST | Waist | 0.20965 | 3.17E-11 |
| B131 | B215 | AST | BMI | 0.200723 | 2.15E-10 |
| B131 | B216 | AST | BMI | 0.181482 | 1.00E-08 |
| B131 | B222 | AST | NS | 0.185155 | 4.96E-09 |
| B132 | B133 | Dbil | TBIL | 0.808538 | 0 |
| B132 | B139 | Dbil | ADA | 0.157393 | 7.07E-07 |
| B134 | B151 | CHE | Anti-CCP | 0.474144 | 0 |
| B134 | B155 | CHE | IgA | 0.525154 | 0 |
| B134 | B178 | CHE | Ig-U | -0.31521 | 0 |
| B134 | B179 | CHE | α1-MG | -0.1735 | 4.39E-08 |
| B134 | B180 | CHE | Tf | -0.34073 | 0 |
| B134 | B193 | CHE | K | -0.25664 | 4.44E-16 |
| B134 | B197 | CHE | AG | -0.26748 | 0 |
| B135 | B137 | TBA | AFU | 0.224115 | 1.17E-12 |
| B135 | B139 | TBA | ADA | 0.303365 | 0 |
| B135 | B148 | TBA | Anti_TPO | 0.164122 | 2.29E-07 |
| B135 | B153 | TBA | Glu | 0.180074 | 1.31E-08 |
| B135 | B194 | TBA | Cl | -0.23849 | 3.51E-14 |
| B135 | B197 | TBA | AG | 0.301568 | 0 |
| B135 | B198 | TBA | P | 0.160832 | 4.00E-07 |
| B135 | B199 | TBA | Mg | 0.178777 | 1.67E-08 |
| B135 | B200 | TBA | OP | 0.233061 | 1.36E-13 |
| B136 | B140 | 5'NT | r-GT | 0.267343 | 0 |
| B136 | B150 | 5'NT | ALP | 0.253871 | 6.66E-16 |
| B136 | B196 | 5'NT | TCO2 | -0.17394 | 4.06E-08 |
| B137 | B139 | AFU | ADA | 0.362117 | 0 |
| B137 | B146 | AFU | CysC | 0.216388 | 7.03E-12 |
| B137 | B150 | AFU | ALP | 0.176001 | 2.78E-08 |
| B137 | B153 | AFU | Glu | 0.273862 | 0 |
| B137 | B194 | AFU | Cl | -0.19824 | 3.61E-10 |
| B137 | B200 | AFU | OP | 0.380681 | 0 |
| B137 | B205 | AFU | KDA | 0.171724 | 6.04E-08 |
| B138 | B139 | PAB | ADA | -0.39446 | 0 |
| B138 | B150 | PAB | ALP | -0.15919 | 5.26E-07 |

| | | | | | |
|---|---|---|---|---|---|
| B138 | B156 | PAB | IgE | -0.25107 | 1.33E-15 |
| B138 | B175 | PAB | ALB-U | 0.242752 | 1.20E-14 |
| B138 | B178 | PAB | Ig-U | 0.165432 | 1.83E-07 |
| B138 | B179 | PAB | α1-MG | 0.161145 | 3.79E-07 |
| B138 | B180 | PAB | Tf | 0.165287 | 1.87E-07 |
| B138 | B194 | PAB | Cl | -0.18783 | 2.95E-09 |
| B138 | B199 | PAB | Mg | 0.23943 | 2.78E-14 |
| B139 | B150 | ADA | ALP | 0.299514 | 0 |
| B139 | B153 | ADA | Glu | 0.420627 | 0 |
| B139 | B170 | ADA | U-GLU | 0.199851 | 2.59E-10 |
| B139 | B194 | ADA | Cl | -0.29477 | 0 |
| B139 | B195 | ADA | Na | -0.16638 | 1.55E-07 |
| B139 | B200 | ADA | OP | 0.514541 | 0 |
| B139 | B205 | ADA | KDA | 0.177994 | 1.93E-08 |
| B14 | B129 | G-17 | IBIL | 0.166838 | 1.43E-07 |
| B14 | B133 | G-17 | TBIL | 0.163868 | 2.39E-07 |
| B140 | B141 | r-GT | AST/ALT | -0.34942 | 0 |
| B140 | B192 | r-GT | UA | 0.275665 | 0 |
| B140 | B204 | r-GT | Drinking | 0.172709 | 5.06E-08 |
| B140 | B206 | r-GT | Waist | 0.305629 | 0 |
| B140 | B210 | r-GT | Hip | 0.241205 | 1.78E-14 |
| B140 | B215 | r-GT | BMI | 0.283028 | 0 |
| B140 | B216 | r-GT | BMI | 0.321796 | 0 |
| B140 | B218 | r-GT | DBP | 0.161083 | 3.83E-07 |
| B140 | B221 | r-GT | Hei | 0.17885 | 1.64E-08 |
| B140 | B222 | r-GT | NS | 0.323443 | 0 |
| B141 | B163 | AST/ALT | SG | -0.18847 | 2.60E-09 |
| B141 | B192 | AST/ALT | UA | -0.23555 | 7.37E-14 |
| B141 | B206 | AST/ALT | Waist | -0.26774 | 0 |
| B141 | B210 | AST/ALT | Hip | -0.18236 | 8.48E-09 |
| B141 | B215 | AST/ALT | BMI | -0.2619 | 0 |
| B141 | B216 | AST/ALT | BMI | -0.33123 | 0 |
| B141 | B221 | AST/ALT | Hei | -0.23435 | 9.90E-14 |
| B141 | B222 | AST/ALT | NS | -0.29951 | 0 |
| B142 | B143 | 25-(OH)D | β-CTx | -1 | 0 |
| B142 | B175 | 25-(OH)D | ALB-U | -0.17475 | 3.50E-08 |
| B143 | B144 | β-CTx | Total-PINP | 0.999997 | 0 |
| B143 | B145 | β-CTx | BGP | 0.999997 | 0 |
| B143 | B175 | β-CTx | ALB-U | 0.174778 | 3.48E-08 |
| B144 | B175 | Total-PINP | ALB-U | 0.174746 | 3.50E-08 |
| B145 | B175 | BGP | ALB-U | 0.174746 | 3.50E-08 |
| B146 | B181 | CysC | APTT | -0.17173 | 6.04E-08 |
| B146 | B182 | CysC | TT | -0.17121 | 6.63E-08 |
| B146 | B183 | CysC | PTA | -0.17172 | 6.05E-08 |

| | | | | | |
|---|---|---|---|---|---|
| B146 | B184 | CysC | INR | -0.17173 | 6.04E-08 |
| B146 | B185 | CysC | PT | -0.17135 | 6.46E-08 |
| B146 | B186 | CysC | PFC | 0.171732 | 6.04E-08 |
| B146 | B189 | CysC | eGFB | -0.18361 | 6.68E-09 |
| B146 | B192 | CysC | UA | 0.174623 | 3.58E-08 |
| B146 | B195 | CysC | Na | 0.1554 | 9.79E-07 |
| B146 | B199 | CysC | Mg | 0.182408 | 8.40E-09 |
| B146 | B200 | CysC | OP | 0.226437 | 6.77E-13 |
| B146 | B206 | CysC | Waist | 0.187368 | 3.23E-09 |
| B148 | B149 | Anti_TPO | TGAb | 0.231204 | 2.14E-13 |
| B149 | B228 | TGAb | OS | 0.495633 | 0 |
| B15 | B16 | CD3-CD19+(B cell) | CD3+T | -0.68148 | 0 |
| B15 | B17 | CD3-CD19+(B cell) | CD3+CD16+CD56+NK | -0.26043 | 0 |
| B15 | B20 | CD3-CD19+(B cell) | CD3+CD4+CD8+ | 0.270452 | 0 |
| B15 | B22 | CD3-CD19+(B cell) | CD3+CD8+(T) | 0.177102 | 2.27E-08 |
| B15 | B23 | CD3-CD19+(B cell) | CD3-CD16+CD56+(NK) | 0.314987 | 0 |
| B15 | B79 | CD3-CD19+(B cell) | hb V -dna | 0.184882 | 5.23E-09 |
| B15 | B99 | CD3-CD19+(B cell) | anti-PM-Scl | 0.271389 | 0 |
| B15 | B120 | CD3-CD19+(B cell) | HCV-cAg | 0.268668 | 0 |
| B15 | B138 | CD3-CD19+(B cell) | PAB | 0.17187 | 5.89E-08 |
| B15 | B156 | CD3-CD19+(B cell) | IgE | -0.40673 | 0 |
| B15 | B157 | CD3-CD19+(B cell) | IgG | -0.31507 | 0 |
| B15 | B158 | CD3-CD19+(B cell) | IgM | -0.18491 | 5.21E-09 |
| B15 | B178 | CD3-CD19+(B cell) | Ig-U | 0.236666 | 5.57E-14 |
| B15 | B179 | CD3-CD19+(B cell) | α1-MG | 0.218243 | 4.60E-12 |
| B15 | B180 | CD3-CD19+(B cell) | Tf | 0.224059 | 1.19E-12 |
| B15 | B181 | CD3-CD19+(B cell) | APTT | -0.26043 | 0 |
| B15 | B182 | CD3-CD19+(B cell) | TT | -0.25859 | 2.22E-16 |
| B15 | B183 | CD3-CD19+(B cell) | PTA | -0.26044 | 0 |
| B15 | B184 | CD3-CD19+(B cell) | INR | -0.26043 | 0 |
| B15 | B185 | CD3-CD19+(B cell) | PT | -0.26033 | 0 |
| B15 | B186 | CD3-CD19+(B cell) | PFC | 0.260432 | 0 |
| B15 | B199 | CD3-CD19+(B cell) | Mg | 0.156325 | 8.42E-07 |
| B150 | B194 | ALP | Cl | -0.16289 | 2.83E-07 |
| B150 | B200 | ALP | OP | 0.198711 | 3.28E-10 |
| B151 | B155 | Anti-CCP | IgA | 0.204536 | 9.60E-11 |
| B151 | B156 | Anti-CCP | IgE | 0.237949 | 4.04E-14 |
| B151 | B178 | Anti-CCP | Ig-U | -0.4731 | 0 |
| B151 | B179 | Anti-CCP | α1-MG | -0.36783 | 0 |
| B151 | B180 | Anti-CCP | Tf | -0.4481 | 0 |
| B151 | B181 | Anti-CCP | APTT | 0.494682 | 0 |
| B151 | B182 | Anti-CCP | TT | 0.491607 | 0 |
| B151 | B183 | Anti-CCP | PTA | 0.49469 | 0 |
| B151 | B184 | Anti-CCP | INR | 0.494682 | 0 |

| | | | | | |
|---|---|---|---|---|---|
| B151 | B185 | Anti-CCP | PT | 0.49429 | 0 |
| B151 | B186 | Anti-CCP | PFC | -0.49468 | 0 |
| B151 | B193 | Anti-CCP | K | 0.183491 | 6.84E-09 |
| B151 | B196 | Anti-CCP | TCO2 | 0.206152 | 6.78E-11 |
| B151 | B197 | Anti-CCP | AG | -0.39127 | 0 |
| B151 | B198 | Anti-CCP | P | -0.17917 | 1.55E-08 |
| B152 | B177 | ACA-Ig(G-A-M) | ACR | 0.224473 | 1.08E-12 |
| B152 | B178 | ACA-Ig(G-A-M) | Ig-U | -0.21827 | 4.57E-12 |
| B152 | B179 | ACA-Ig(G-A-M) | α1-MG | -0.35067 | 0 |
| B152 | B180 | ACA-Ig(G-A-M) | Tf | -0.23983 | 2.51E-14 |
| B152 | B195 | ACA-Ig(G-A-M) | Na | -0.17134 | 6.47E-08 |
| B152 | B196 | ACA-Ig(G-A-M) | TCO2 | 0.189847 | 1.98E-09 |
| B152 | B197 | ACA-Ig(G-A-M) | AG | -0.17957 | 1.43E-08 |
| B152 | B200 | ACA-Ig(G-A-M) | OP | -0.17289 | 4.90E-08 |
| B152 | B211 | ACA-Ig(G-A-M) | IOP-R | -0.2388 | 3.24E-14 |
| B152 | B212 | ACA-Ig(G-A-M) | IOP-L | -0.19711 | 4.57E-10 |
| B153 | B170 | Glu | U-GLU | 0.575713 | 0 |
| B153 | B200 | Glu | OP | 0.327728 | 0 |
| B153 | B206 | Glu | Waist | 0.21512 | 9.36E-12 |
| B153 | B213 | Glu | CVA-R | 0.221537 | 2.15E-12 |
| B153 | B215 | Glu | BMI | 0.159915 | 4.66E-07 |
| B153 | B220 | Glu | SBP | 0.231718 | 1.89E-13 |
| B153 | B222 | Glu | NS | 0.183985 | 6.22E-09 |
| B154 | B155 | RF | IgA | 0.430572 | 0 |
| B154 | B157 | RF | IgG | 0.572648 | 0 |
| B154 | B158 | RF | IgM | 0.184784 | 5.33E-09 |
| B154 | B161 | RF | LEU | 0.177772 | 2.01E-08 |
| B155 | B157 | IgA | IgG | 0.360284 | 0 |
| B155 | B158 | IgA | IgM | 0.168654 | 1.04E-07 |
| B155 | B178 | IgA | Ig-U | -0.17746 | 2.13E-08 |
| B155 | B180 | IgA | Tf | -0.19472 | 7.45E-10 |
| B156 | B158 | IgE | IgM | -0.27827 | 0 |
| B156 | B175 | IgE | ALB-U | -0.15735 | 7.13E-07 |
| B156 | B176 | IgE | Ucr | -0.1792 | 1.54E-08 |
| B156 | B178 | IgE | Ig-U | -0.49302 | 0 |
| B156 | B179 | IgE | α1-MG | -0.45623 | 0 |
| B156 | B180 | IgE | Tf | -0.45119 | 0 |
| B156 | B194 | IgE | Cl | 0.316797 | 0 |
| B156 | B197 | IgE | AG | -0.27929 | 0 |
| B157 | B158 | IgG | IgM | 0.33764 | 0 |
| B157 | B181 | IgG | APTT | 0.161402 | 3.63E-07 |
| B157 | B182 | IgG | TT | 0.15879 | 5.62E-07 |
| B157 | B183 | IgG | PTA | 0.161451 | 3.60E-07 |
| B157 | B184 | IgG | INR | 0.161402 | 3.63E-07 |

| | | | | | |
|---|---|---|---|---|---|
| B157 | B185 | IgG | PT | 0.161995 | 3.28E-07 |
| B157 | B186 | IgG | PFC | -0.1614 | 3.63E-07 |
| B158 | B178 | IgM | Ig-U | 0.15856 | 5.83E-07 |
| B158 | B181 | IgM | APTT | 0.23611 | 6.39E-14 |
| B158 | B182 | IgM | TT | 0.229355 | 3.36E-13 |
| B158 | B183 | IgM | PTA | 0.236266 | 6.17E-14 |
| B158 | B184 | IgM | INR | 0.23611 | 6.39E-14 |
| B158 | B185 | IgM | PT | 0.238295 | 3.71E-14 |
| B158 | B186 | IgM | PFC | -0.23611 | 6.39E-14 |
| B159 | B169 | β2-MG | U-CRY | 1 | 0 |
| B159 | B195 | β2-MG | Na | -0.19632 | 5.38E-10 |
| B16 | B17 | CD3+T | CD3+CD16+CD56+NK | 0.868628 | 0 |
| B16 | B18 | CD3+T | CD3+CD4+helper(inducer)T | -0.46022 | 0 |
| B16 | B19 | CD3+T | CD3+CD4+/CD3+CD8+ | -0.42269 | 0 |
| B16 | B20 | CD3+T | CD3+CD4+CD8+ | 0.27631 | 0 |
| B16 | B21 | CD3+T | CD3+CD4-CD8- | 0.402159 | 0 |
| B16 | B22 | CD3+T | CD3+CD8+(T) | 0.456233 | 0 |
| B16 | B23 | CD3+T | CD3-CD16+CD56+(NK) | -0.18875 | 2.46E-09 |
| B16 | B24 | CD3+T | αHBDH | 0.177788 | 2.00E-08 |
| B16 | B56 | CD3+T | IR | -0.15844 | 5.95E-07 |
| B16 | B95 | CD3+T | anti-cenpb | 0.237656 | 4.35E-14 |
| B16 | B97 | CD3+T | dsDNA | 0.237656 | 4.35E-14 |
| B16 | B98 | CD3+T | anti-Jo-1 | 0.237656 | 4.35E-14 |
| B16 | B99 | CD3+T | anti-PM-Scl | -0.87028 | 0 |
| B16 | B101 | CD3+T | anti-Sc70 | 0.237656 | 4.35E-14 |
| B16 | B102 | CD3+T | anti-Sm | 0.237656 | 4.35E-14 |
| B16 | B105 | CD3+T | anti-U1RNP | 0.237656 | 4.35E-14 |
| B16 | B106 | CD3+T | anti-P | 0.237656 | 4.35E-14 |
| B16 | B107 | CD3+T | AnuA | 0.237656 | 4.35E-14 |
| B16 | B108 | CD3+T | Anti-Cyclin | 0.237656 | 4.35E-14 |
| B16 | B109 | CD3+T | AMA-M2 | 0.237656 | 4.35E-14 |
| B16 | B110 | CD3+T | AHA | 0.237656 | 4.35E-14 |
| B16 | B151 | CD3+T | Anti-CCP | 0.428917 | 0 |
| B16 | B157 | CD3+T | IgG | 0.275495 | 0 |
| B16 | B158 | CD3+T | IgM | 0.318008 | 0 |
| B16 | B181 | CD3+T | APTT | 0.868628 | 0 |
| B16 | B182 | CD3+T | TT | 0.863656 | 0 |
| B16 | B183 | CD3+T | PTA | 0.86863 | 0 |
| B16 | B184 | CD3+T | INR | 0.868628 | 0 |
| B16 | B185 | CD3+T | PT | 0.867747 | 0 |
| B16 | B186 | CD3+T | PFC | -0.86863 | 0 |
| B16 | B193 | CD3+T | K | 0.251283 | 1.33E-15 |
| B16 | B198 | CD3+T | P | -0.30216 | 0 |
| B16 | B199 | CD3+T | Mg | -0.25297 | 8.88E-16 |

| | | | | | |
|---|---|---|---|---|---|
| B16 | B204 | CD3+T | Drinking | -0.18172 | 9.58E-09 |
| B160 | B192 | PH | UA | -0.15825 | 6.14E-07 |
| B161 | B173 | LEU | NTT | 0.316902 | 0 |
| B163 | B166 | SG | Pro | 0.380664 | 0 |
| B163 | B176 | SG | Ucr | 0.275922 | 0 |
| B163 | B190 | SG | Cre | 0.195126 | 6.86E-10 |
| B163 | B191 | SG | Urea | 0.23807 | 3.91E-14 |
| B163 | B192 | SG | UA | 0.249288 | 2.22E-15 |
| B163 | B206 | SG | Waist | 0.229982 | 2.89E-13 |
| B163 | B209 | SG | Smoking | 0.192291 | 1.22E-09 |
| B163 | B210 | SG | Hip | 0.184912 | 5.20E-09 |
| B163 | B215 | SG | BMI | 0.161663 | 3.47E-07 |
| B163 | B216 | SG | BMI | 0.293824 | 0 |
| B163 | B221 | SG | Hei | 0.302459 | 0 |
| B163 | B222 | SG | NS | 0.321061 | 0 |
| B168 | B175 | NMP22 | ALB-U | 0.202793 | 1.39E-10 |
| B169 | B195 | U-CRY | Na | -0.19632 | 5.38E-10 |
| B17 | B18 | CD3+CD16+CD56+NK | CD3+CD4+helper(inducer)T | -0.43712 | 0 |
| B17 | B19 | CD3+CD16+CD56+NK | CD3+CD4+/CD3+CD8+ | -0.48831 | 0 |
| B17 | B20 | CD3+CD16+CD56+NK | CD3+CD4+CD8+ | 0.427657 | 0 |
| B17 | B21 | CD3+CD16+CD56+NK | CD3+CD4-CD8- | 0.568357 | 0 |
| B17 | B22 | CD3+CD16+CD56+NK | CD3+CD8+(T) | 0.582584 | 0 |
| B17 | B24 | CD3+CD16+CD56+NK | αHBDH | 0.2041 | 1.05E-10 |
| B17 | B56 | CD3+CD16+CD56+NK | IR | -0.18264 | 8.04E-09 |
| B17 | B95 | CD3+CD16+CD56+NK | anti-cenpb | 0.331973 | 0 |
| B17 | B97 | CD3+CD16+CD56+NK | dsDNA | 0.331973 | 0 |
| B17 | B98 | CD3+CD16+CD56+NK | anti-Jo-1 | 0.331973 | 0 |
| B17 | B99 | CD3+CD16+CD56+NK | anti-PM-Scl | -0.9983 | 0 |
| B17 | B101 | CD3+CD16+CD56+NK | anti-Sc70 | 0.331973 | 0 |
| B17 | B102 | CD3+CD16+CD56+NK | anti-Sm | 0.331973 | 0 |
| B17 | B105 | CD3+CD16+CD56+NK | anti-U1RNP | 0.331973 | 0 |
| B17 | B106 | CD3+CD16+CD56+NK | anti-P | 0.331973 | 0 |
| B17 | B107 | CD3+CD16+CD56+NK | AnuA | 0.331973 | 0 |
| B17 | B108 | CD3+CD16+CD56+NK | Anti-Cyclin | 0.331973 | 0 |
| B17 | B109 | CD3+CD16+CD56+NK | AMA-M2 | 0.331973 | 0 |
| B17 | B110 | CD3+CD16+CD56+NK | AHA | 0.331973 | 0 |
| B17 | B146 | CD3+CD16+CD56+NK | CysC | -0.17173 | 6.04E-08 |
| B17 | B151 | CD3+CD16+CD56+NK | Anti-CCP | 0.494682 | 0 |
| B17 | B157 | CD3+CD16+CD56+NK | IgG | 0.161402 | 3.63E-07 |
| B17 | B158 | CD3+CD16+CD56+NK | IgM | 0.23611 | 6.39E-14 |
| B17 | B181 | CD3+CD16+CD56+NK | APTT | 1 | 0 |
| B17 | B182 | CD3+CD16+CD56+NK | TT | 0.994483 | 0 |
| B17 | B183 | CD3+CD16+CD56+NK | PTA | 0.999995 | 0 |
| B17 | B185 | CD3+CD16+CD56+NK | PT | 0.998892 | 0 |

| | | | | | |
|---|---|---|---|---|---|
| B17 | B186 | CD3+CD16+CD56+NK | PFC | -1 | 0 |
| B17 | B193 | CD3+CD16+CD56+NK | K | 0.283561 | 0 |
| B17 | B194 | CD3+CD16+CD56+NK | Cl | -0.2028 | 1.39E-10 |
| B17 | B198 | CD3+CD16+CD56+NK | P | -0.36229 | 0 |
| B17 | B199 | CD3+CD16+CD56+NK | Mg | -0.26397 | 0 |
| B17 | B204 | CD3+CD16+CD56+NK | Drinking | -0.1983 | 3.57E-10 |
| B170 | B172 | U-GLU | KB | 0.186013 | 4.20E-09 |
| B170 | B213 | U-GLU | CVA-R | 0.253468 | 6.66E-16 |
| B173 | B227 | NTT | OD | 0.194413 | 7.93E-10 |
| B174 | B190 | Urea/Cr | Cre | -0.38659 | 0 |
| B174 | B191 | Urea/Cr | Urea | 0.568913 | 0 |
| B174 | B192 | Urea/Cr | UA | -0.26117 | 0 |
| B174 | B206 | Urea/Cr | Waist | -0.16993 | 8.33E-08 |
| B174 | B216 | Urea/Cr | BMI | -0.26229 | 0 |
| B174 | B221 | Urea/Cr | Hei | -0.33826 | 0 |
| B174 | B222 | Urea/Cr | NS | -0.25307 | 8.88E-16 |
| B175 | B176 | ALB-U | Ucr | 0.355126 | 0 |
| B175 | B177 | ALB-U | ACR | 0.375797 | 0 |
| B175 | B178 | ALB-U | Ig-U | 0.219825 | 3.20E-12 |
| B175 | B179 | ALB-U | α1-MG | 0.215949 | 7.76E-12 |
| B175 | B180 | ALB-U | Tf | 0.218864 | 3.99E-12 |
| B175 | B194 | ALB-U | Cl | -0.23602 | 6.55E-14 |
| B176 | B177 | Ucr | ACR | -0.18201 | 9.07E-09 |
| B176 | B178 | Ucr | Ig-U | 0.228055 | 4.60E-13 |
| B176 | B179 | Ucr | α1-MG | 0.241709 | 1.55E-14 |
| B176 | B180 | Ucr | Tf | 0.222601 | 1.68E-12 |
| B178 | B179 | Ig-U | α1-MG | 0.965659 | 0 |
| B178 | B180 | Ig-U | Tf | 0.98973 | 0 |
| B178 | B194 | Ig-U | Cl | -0.29488 | 0 |
| B178 | B197 | Ig-U | AG | 0.326446 | 0 |
| B179 | B180 | α1-MG | Tf | 0.973876 | 0 |
| B179 | B194 | α1-MG | Cl | -0.2766 | 0 |
| B179 | B197 | α1-MG | AG | 0.322681 | 0 |
| B18 | B19 | CD3+CD4+helper(inducer)T | CD3+CD4+/CD3+CD8+ | 0.98491 | 0 |
| B18 | B20 | CD3+CD4+helper(inducer)T | CD3+CD4+CD8+ | -0.70753 | 0 |
| B18 | B21 | CD3+CD4+helper(inducer)T | CD3+CD4-CD8- | -0.67214 | 0 |
| B18 | B22 | CD3+CD4+helper(inducer)T | CD3+CD8+(T) | -0.92668 | 0 |
| B18 | B23 | CD3+CD4+helper(inducer)T | CD3-CD16+CD56+(NK) | 0.242448 | 1.29E-14 |
| B18 | B79 | CD3+CD4+helper(inducer)T | hb V -dna | -0.25866 | 2.22E-16 |
| B18 | B95 | CD3+CD4+helper(inducer)T | anti-cenpb | -0.40527 | 0 |
| B18 | B97 | CD3+CD4+helper(inducer)T | dsDNA | -0.40527 | 0 |
| B18 | B98 | CD3+CD4+helper(inducer)T | anti-Jo-1 | -0.40527 | 0 |
| B18 | B99 | CD3+CD4+helper(inducer)T | anti-PM-Scl | 0.420306 | 0 |
| B18 | B101 | CD3+CD4+helper(inducer)T | anti-Sc70 | -0.40527 | 0 |

| | | | | | |
|---|---|---|---|---|---|
| B18 | B102 | CD3+CD4+helper(inducer)T | anti-Sm | -0.40527 | 0 |
| B18 | B105 | CD3+CD4+helper(inducer)T | anti-U1RNP | -0.40527 | 0 |
| B18 | B106 | CD3+CD4+helper(inducer)T | anti-P | -0.40527 | 0 |
| B18 | B107 | CD3+CD4+helper(inducer)T | AnuA | -0.40527 | 0 |
| B18 | B108 | CD3+CD4+helper(inducer)T | Anti-Cyclin | -0.40527 | 0 |
| B18 | B109 | CD3+CD4+helper(inducer)T | AMA-M2 | -0.40527 | 0 |
| B18 | B110 | CD3+CD4+helper(inducer)T | AHA | -0.40527 | 0 |
| B18 | B138 | CD3+CD4+helper(inducer)T | PAB | -0.21157 | 2.07E-11 |
| B18 | B151 | CD3+CD4+helper(inducer)T | Anti-CCP | -0.21679 | 6.41E-12 |
| B18 | B156 | CD3+CD4+helper(inducer)T | IgE | 0.746105 | 0 |
| B18 | B158 | CD3+CD4+helper(inducer)T | IgM | -0.4022 | 0 |
| B18 | B175 | CD3+CD4+helper(inducer)T | ALB-U | -0.19897 | 3.10E-10 |
| B18 | B176 | CD3+CD4+helper(inducer)T | Ucr | -0.1609 | 3.95E-07 |
| B18 | B178 | CD3+CD4+helper(inducer)T | Ig-U | -0.33027 | 0 |
| B18 | B179 | CD3+CD4+helper(inducer)T | α1-MG | -0.30473 | 0 |
| B18 | B180 | CD3+CD4+helper(inducer)T | Tf | -0.31279 | 0 |
| B18 | B181 | CD3+CD4+helper(inducer)T | APTT | -0.43712 | 0 |
| B18 | B182 | CD3+CD4+helper(inducer)T | TT | -0.43483 | 0 |
| B18 | B183 | CD3+CD4+helper(inducer)T | PTA | -0.43711 | 0 |
| B18 | B184 | CD3+CD4+helper(inducer)T | INR | -0.43712 | 0 |
| B18 | B185 | CD3+CD4+helper(inducer)T | PT | -0.43657 | 0 |
| B18 | B186 | CD3+CD4+helper(inducer)T | PFC | 0.437115 | 0 |
| B18 | B194 | CD3+CD4+helper(inducer)T | Cl | 0.241115 | 1.82E-14 |
| B18 | B198 | CD3+CD4+helper(inducer)T | P | 0.213027 | 1.50E-11 |
| B180 | B194 | Tf | Cl | -0.28563 | 0 |
| B180 | B197 | Tf | AG | 0.352806 | 0 |
| B181 | B182 | APTT | TT | 0.994483 | 0 |
| B181 | B183 | APTT | PTA | 0.999995 | 0 |
| B181 | B184 | APTT | INR | 1 | 0 |
| B181 | B185 | APTT | PT | 0.998892 | 0 |
| B181 | B186 | APTT | PFC | -1 | 0 |
| B181 | B193 | APTT | K | 0.283561 | 0 |
| B181 | B194 | APTT | Cl | -0.2028 | 1.39E-10 |
| B181 | B198 | APTT | P | -0.36229 | 0 |
| B181 | B199 | APTT | Mg | -0.26397 | 0 |
| B181 | B204 | APTT | Drinking | -0.1983 | 3.57E-10 |
| B182 | B183 | TT | PTA | 0.994163 | 0 |
| B182 | B184 | TT | INR | 0.994483 | 0 |
| B182 | B185 | TT | PT | 0.988446 | 0 |
| B182 | B186 | TT | PFC | -0.99448 | 0 |
| B182 | B193 | TT | K | 0.274866 | 0 |
| B182 | B194 | TT | Cl | -0.21348 | 1.35E-11 |
| B182 | B198 | TT | P | -0.36067 | 0 |
| B182 | B199 | TT | Mg | -0.26388 | 0 |

| | | | | | |
|---|---|---|---|---|---|
| B182 | B204 | TT | Drinking | -0.19498 | 7.06E-10 |
| B183 | B184 | PTA | INR | 0.999995 | 0 |
| B183 | B185 | PTA | PT | 0.999029 | 0 |
| B183 | B186 | PTA | PFC | -1 | 0 |
| B183 | B193 | PTA | K | 0.283764 | 0 |
| B183 | B194 | PTA | Cl | -0.20246 | 1.49E-10 |
| B183 | B198 | PTA | P | -0.36228 | 0 |
| B183 | B199 | PTA | Mg | -0.26393 | 0 |
| B183 | B204 | PTA | Drinking | -0.19836 | 3.52E-10 |
| B184 | B185 | INR | PT | 0.998892 | 0 |
| B184 | B186 | INR | PFC | -1 | 0 |
| B184 | B193 | INR | K | 0.283561 | 0 |
| B184 | B194 | INR | Cl | -0.2028 | 1.39E-10 |
| B184 | B198 | INR | P | -0.36229 | 0 |
| B184 | B199 | INR | Mg | -0.26397 | 0 |
| B184 | B204 | INR | Drinking | -0.1983 | 3.57E-10 |
| B185 | B186 | PT | PFC | -0.99889 | 0 |
| B185 | B193 | PT | K | 0.286445 | 0 |
| B185 | B194 | PT | Cl | -0.19728 | 4.41E-10 |
| B185 | B198 | PT | P | -0.36172 | 0 |
| B185 | B199 | PT | Mg | -0.26306 | 0 |
| B185 | B204 | PT | Drinking | -0.19908 | 3.04E-10 |
| B186 | B193 | PFC | K | -0.28356 | 0 |
| B186 | B194 | PFC | Cl | 0.202798 | 1.39E-10 |
| B186 | B198 | PFC | P | 0.362289 | 0 |
| B186 | B199 | PFC | Mg | 0.263966 | 0 |
| B186 | B204 | PFC | Drinking | 0.198299 | 3.57E-10 |
| B188 | B196 | Ca | TCO2 | 0.170335 | 7.74E-08 |
| B189 | B190 | eGFB | Cre | -0.51868 | 0 |
| B189 | B191 | eGFB | Urea | -0.37192 | 0 |
| B189 | B192 | eGFB | UA | -0.19602 | 5.71E-10 |
| B189 | B220 | eGFB | SBP | -0.33073 | 0 |
| B19 | B20 | CD3+CD4+/CD3+CD8+ | CD3+CD4+CD8+ | -0.75088 | 0 |
| B19 | B21 | CD3+CD4+/CD3+CD8+ | CD3+CD4-CD8- | -0.72853 | 0 |
| B19 | B22 | CD3+CD4+/CD3+CD8+ | CD3+CD8+(T) | -0.96288 | 0 |
| B19 | B23 | CD3+CD4+/CD3+CD8+ | CD3-CD16+CD56+(NK) | 0.182827 | 7.76E-09 |
| B19 | B79 | CD3+CD4+/CD3+CD8+ | hb V -dna | -0.27184 | 0 |
| B19 | B95 | CD3+CD4+/CD3+CD8+ | anti-cenpb | -0.43567 | 0 |
| B19 | B97 | CD3+CD4+/CD3+CD8+ | dsDNA | -0.43567 | 0 |
| B19 | B98 | CD3+CD4+/CD3+CD8+ | anti-Jo-1 | -0.43567 | 0 |
| B19 | B99 | CD3+CD4+/CD3+CD8+ | anti-PM-Scl | 0.470588 | 0 |
| B19 | B101 | CD3+CD4+/CD3+CD8+ | anti-Sc70 | -0.43567 | 0 |
| B19 | B102 | CD3+CD4+/CD3+CD8+ | anti-Sm | -0.43567 | 0 |
| B19 | B105 | CD3+CD4+/CD3+CD8+ | anti-U1RNP | -0.43567 | 0 |

| | | | | | |
|---|---|---|---|---|---|
| B19 | B106 | CD3+CD4+/CD3+CD8+ | anti-P | -0.43567 | 0 |
| B19 | B107 | CD3+CD4+/CD3+CD8+ | AnuA | -0.43567 | 0 |
| B19 | B108 | CD3+CD4+/CD3+CD8+ | Anti-Cyclin | -0.43567 | 0 |
| B19 | B109 | CD3+CD4+/CD3+CD8+ | AMA-M2 | -0.43567 | 0 |
| B19 | B110 | CD3+CD4+/CD3+CD8+ | AHA | -0.43567 | 0 |
| B19 | B138 | CD3+CD4+/CD3+CD8+ | PAB | -0.22136 | 2.24E-12 |
| B19 | B151 | CD3+CD4+/CD3+CD8+ | Anti-CCP | -0.24236 | 1.33E-14 |
| B19 | B156 | CD3+CD4+/CD3+CD8+ | IgE | 0.770562 | 0 |
| B19 | B158 | CD3+CD4+/CD3+CD8+ | IgM | -0.37318 | 0 |
| B19 | B175 | CD3+CD4+/CD3+CD8+ | ALB-U | -0.21233 | 1.75E-11 |
| B19 | B176 | CD3+CD4+/CD3+CD8+ | Ucr | -0.17153 | 6.26E-08 |
| B19 | B178 | CD3+CD4+/CD3+CD8+ | Ig-U | -0.34733 | 0 |
| B19 | B179 | CD3+CD4+/CD3+CD8+ | α1-MG | -0.32043 | 0 |
| B19 | B180 | CD3+CD4+/CD3+CD8+ | Tf | -0.32892 | 0 |
| B19 | B181 | CD3+CD4+/CD3+CD8+ | APTT | -0.48831 | 0 |
| B19 | B182 | CD3+CD4+/CD3+CD8+ | TT | -0.48581 | 0 |
| B19 | B183 | CD3+CD4+/CD3+CD8+ | PTA | -0.48831 | 0 |
| B19 | B184 | CD3+CD4+/CD3+CD8+ | INR | -0.48831 | 0 |
| B19 | B185 | CD3+CD4+/CD3+CD8+ | PT | -0.48769 | 0 |
| B19 | B186 | CD3+CD4+/CD3+CD8+ | PFC | 0.488313 | 0 |
| B19 | B194 | CD3+CD4+/CD3+CD8+ | Cl | 0.261796 | 0 |
| B19 | B198 | CD3+CD4+/CD3+CD8+ | P | 0.235604 | 7.26E-14 |
| B190 | B191 | Cre | Urea | 0.50424 | 0 |
| B190 | B192 | Cre | UA | 0.471329 | 0 |
| B190 | B204 | Cre | Drinking | 0.18294 | 7.59E-09 |
| B190 | B206 | Cre | Waist | 0.287713 | 0 |
| B190 | B209 | Cre | Smoking | 0.181742 | 9.54E-09 |
| B190 | B210 | Cre | Hip | 0.164916 | 2.00E-07 |
| B190 | B216 | Cre | BMI | 0.289705 | 0 |
| B190 | B221 | Cre | Hei | 0.341231 | 0 |
| B190 | B222 | Cre | NS | 0.380308 | 0 |
| B191 | B192 | Urea | UA | 0.202779 | 1.40E-10 |
| B191 | B220 | Urea | SBP | 0.163352 | 2.61E-07 |
| B192 | B204 | UA | Drinking | 0.235879 | 6.79E-14 |
| B192 | B206 | UA | Waist | 0.450732 | 0 |
| B192 | B209 | UA | Smoking | 0.16365 | 2.48E-07 |
| B192 | B210 | UA | Hip | 0.333237 | 0 |
| B192 | B215 | UA | BMI | 0.315932 | 0 |
| B192 | B216 | UA | BMI | 0.454795 | 0 |
| B192 | B218 | UA | DBP | 0.203104 | 1.30E-10 |
| B192 | B220 | UA | SBP | 0.159735 | 4.80E-07 |
| B192 | B221 | UA | Hei | 0.386722 | 0 |
| B192 | B222 | UA | NS | 0.504643 | 0 |
| B194 | B195 | Cl | Na | 0.406988 | 0 |

| | | | | | |
|---|---|---|---|---|---|
| B194 | B196 | Cl | TCO2 | -0.24427 | 7.99E-15 |
| B194 | B197 | Cl | AG | -0.26825 | 0 |
| B194 | B228 | Cl | OS | 0.162854 | 2.84E-07 |
| B195 | B196 | Na | TCO2 | 0.167158 | 1.35E-07 |
| B195 | B200 | Na | OP | 0.599878 | 0 |
| B196 | B197 | TCO2 | AG | -0.69588 | 0 |
| B196 | B198 | TCO2 | P | 0.164909 | 2.00E-07 |
| B196 | B200 | TCO2 | OP | 0.243745 | 9.33E-15 |
| B198 | B228 | P | OS | -0.15563 | 9.44E-07 |
| B2 | B43 | Age | MCV | 0.159579 | 4.93E-07 |
| B2 | B46 | Age | RBC Count | -0.18175 | 9.53E-09 |
| B2 | B47 | Age | RDW | 0.281922 | 0 |
| B2 | B52 | Age | PLT/BPC | -0.18233 | 8.52E-09 |
| B2 | B77 | Age | TC | 0.158194 | 6.20E-07 |
| B2 | B118 | Age | CEA1 | 0.192097 | 1.27E-09 |
| B2 | B126 | Age | Alb | -0.3248 | 0 |
| B2 | B127 | Age | A-G ratio | -0.19608 | 5.65E-10 |
| B2 | B141 | Age | AST/ALT | 0.164101 | 2.30E-07 |
| B2 | B153 | Age | Glu | 0.219247 | 3.65E-12 |
| B2 | B176 | Age | Ucr | -0.18007 | 1.31E-08 |
| B2 | B189 | Age | eGFB | -0.63196 | 0 |
| B2 | B190 | Age | Cre | 0.194015 | 8.59E-10 |
| B2 | B191 | Age | Urea | 0.256261 | 4.44E-16 |
| B2 | B201 | Age | e-habits | 0.195611 | 6.21E-10 |
| B2 | B206 | Age | Waist | 0.201243 | 1.93E-10 |
| B2 | B220 | Age | SBP | 0.511494 | 0 |
| B2 | B221 | Age | Hei | -0.22789 | 4.79E-13 |
| B20 | B21 | CD3+CD4+CD8+ | CD3+CD4-CD8- | 0.213121 | 1.47E-11 |
| B20 | B22 | CD3+CD4+CD8+ | CD3+CD8+(T) | 0.88645 | 0 |
| B20 | B23 | CD3+CD4+CD8+ | CD3-CD16+CD56+(NK) | 0.508941 | 0 |
| B20 | B79 | CD3+CD4+CD8+ | hb V -dna | 0.184696 | 5.42E-09 |
| B20 | B95 | CD3+CD4+CD8+ | anti-cenpb | 0.328412 | 0 |
| B20 | B97 | CD3+CD4+CD8+ | dsDNA | 0.328412 | 0 |
| B20 | B98 | CD3+CD4+CD8+ | anti-Jo-1 | 0.328412 | 0 |
| B20 | B99 | CD3+CD4+CD8+ | anti-PM-Scl | -0.41542 | 0 |
| B20 | B101 | CD3+CD4+CD8+ | anti-Sc70 | 0.328412 | 0 |
| B20 | B102 | CD3+CD4+CD8+ | anti-Sm | 0.328412 | 0 |
| B20 | B105 | CD3+CD4+CD8+ | anti-U1RNP | 0.328412 | 0 |
| B20 | B106 | CD3+CD4+CD8+ | anti-P | 0.328412 | 0 |
| B20 | B107 | CD3+CD4+CD8+ | AnuA | 0.328412 | 0 |
| B20 | B108 | CD3+CD4+CD8+ | Anti-Cyclin | 0.328412 | 0 |
| B20 | B109 | CD3+CD4+CD8+ | AMA-M2 | 0.328412 | 0 |
| B20 | B110 | CD3+CD4+CD8+ | AHA | 0.328412 | 0 |
| B20 | B120 | CD3+CD4+CD8+ | HCV-cAg | 0.23463 | 9.24E-14 |

| | | | | | |
|---|---|---|---|---|---|
| B20 | B138 | CD3+CD4+CD8+ | PAB | 0.16432 | 2.21E-07 |
| B20 | B151 | CD3+CD4+CD8+ | Anti-CCP | 0.213107 | 1.47E-11 |
| B20 | B156 | CD3+CD4+CD8+ | IgE | -0.51864 | 0 |
| B20 | B158 | CD3+CD4+CD8+ | IgM | 0.220625 | 2.66E-12 |
| B20 | B178 | CD3+CD4+CD8+ | Ig-U | 0.23702 | 5.11E-14 |
| B20 | B179 | CD3+CD4+CD8+ | α1-MG | 0.218445 | 4.39E-12 |
| B20 | B180 | CD3+CD4+CD8+ | Tf | 0.224313 | 1.12E-12 |
| B20 | B181 | CD3+CD4+CD8+ | APTT | 0.427657 | 0 |
| B20 | B182 | CD3+CD4+CD8+ | TT | 0.425658 | 0 |
| B20 | B183 | CD3+CD4+CD8+ | PTA | 0.427644 | 0 |
| B20 | B184 | CD3+CD4+CD8+ | INR | 0.427657 | 0 |
| B20 | B185 | CD3+CD4+CD8+ | PT | 0.427021 | 0 |
| B20 | B186 | CD3+CD4+CD8+ | PFC | -0.42766 | 0 |
| B204 | B206 | Drinking | Waist | 0.276212 | 0 |
| B204 | B209 | Drinking | Smoking | 0.365854 | 0 |
| B204 | B210 | Drinking | Hip | 0.189203 | 2.25E-09 |
| B204 | B215 | Drinking | BMI | 0.185739 | 4.43E-09 |
| B204 | B216 | Drinking | BMI | 0.276259 | 0 |
| B204 | B221 | Drinking | Hei | 0.241138 | 1.80E-14 |
| B204 | B222 | Drinking | NS | 0.298335 | 0 |
| B206 | B209 | Waist | Smoking | 0.230872 | 2.33E-13 |
| B206 | B210 | Waist | Hip | 0.730068 | 0 |
| B206 | B215 | Waist | BMI | 0.82773 | 0 |
| B206 | B216 | Waist | BMI | 0.840225 | 0 |
| B206 | B218 | Waist | DBP | 0.246816 | 4.00E-15 |
| B206 | B220 | Waist | SBP | 0.276708 | 0 |
| B206 | B221 | Waist | Hei | 0.392198 | 0 |
| B206 | B222 | Waist | NS | 0.778052 | 0 |
| B209 | B216 | Smoking | BMI | 0.229599 | 3.17E-13 |
| B209 | B221 | Smoking | Hei | 0.249846 | 1.78E-15 |
| B209 | B222 | Smoking | NS | 0.289161 | 0 |
| B21 | B22 | CD3+CD4-CD8- | CD3+CD8+(T) | 0.618194 | 0 |
| B21 | B23 | CD3+CD4-CD8- | CD3-CD16+CD56+(NK) | -0.59632 | 0 |
| B21 | B79 | CD3+CD4-CD8- | hb V -dna | 0.205615 | 7.61E-11 |
| B21 | B95 | CD3+CD4-CD8- | anti-cenpb | 0.396064 | 0 |
| B21 | B97 | CD3+CD4-CD8- | dsDNA | 0.396064 | 0 |
| B21 | B98 | CD3+CD4-CD8- | anti-Jo-1 | 0.396064 | 0 |
| B21 | B99 | CD3+CD4-CD8- | anti-PM-Scl | -0.55458 | 0 |
| B21 | B101 | CD3+CD4-CD8- | anti-Sc70 | 0.396064 | 0 |
| B21 | B102 | CD3+CD4-CD8- | anti-Sm | 0.396064 | 0 |
| B21 | B105 | CD3+CD4-CD8- | anti-U1RNP | 0.396064 | 0 |
| B21 | B106 | CD3+CD4-CD8- | anti-P | 0.396064 | 0 |
| B21 | B107 | CD3+CD4-CD8- | AnuA | 0.396064 | 0 |
| B21 | B108 | CD3+CD4-CD8- | Anti-Cyclin | 0.396064 | 0 |

| | | | | | |
|---|---|---|---|---|---|
| B21 | B109 | CD3+CD4-CD8- | AMA-M2 | 0.396064 | 0 |
| B21 | B110 | CD3+CD4-CD8- | AHA | 0.396064 | 0 |
| B21 | B151 | CD3+CD4-CD8- | Anti-CCP | 0.281253 | 0 |
| B21 | B156 | CD3+CD4-CD8- | IgE | -0.57335 | 0 |
| B21 | B158 | CD3+CD4-CD8- | IgM | 0.264964 | 0 |
| B21 | B175 | CD3+CD4-CD8- | ALB-U | 0.188523 | 2.57E-09 |
| B21 | B178 | CD3+CD4-CD8- | Ig-U | 0.262465 | 0 |
| B21 | B179 | CD3+CD4-CD8- | α1-MG | 0.242188 | 1.38E-14 |
| B21 | B180 | CD3+CD4-CD8- | Tf | 0.248585 | 2.66E-15 |
| B21 | B181 | CD3+CD4-CD8- | APTT | 0.568357 | 0 |
| B21 | B182 | CD3+CD4-CD8- | TT | 0.565244 | 0 |
| B21 | B183 | CD3+CD4-CD8- | PTA | 0.568354 | 0 |
| B21 | B184 | CD3+CD4-CD8- | INR | 0.568357 | 0 |
| B21 | B185 | CD3+CD4-CD8- | PT | 0.567718 | 0 |
| B21 | B186 | CD3+CD4-CD8- | PFC | -0.56836 | 0 |
| B21 | B193 | CD3+CD4-CD8- | K | 0.174499 | 3.66E-08 |
| B21 | B194 | CD3+CD4-CD8- | Cl | -0.39671 | 0 |
| B21 | B197 | CD3+CD4-CD8- | AG | 0.206597 | 6.16E-11 |
| B21 | B198 | CD3+CD4-CD8- | P | -0.32945 | 0 |
| B210 | B215 | Hip | BMI | 0.756217 | 0 |
| B210 | B216 | Hip | BMI | 0.768833 | 0 |
| B210 | B221 | Hip | Hei | 0.367737 | 0 |
| B210 | B222 | Hip | NS | 0.630005 | 0 |
| B211 | B212 | IOP-R | IOP-L | 0.826274 | 0 |
| B214 | B219 | CVA-L | OT | 0.166802 | 1.44E-07 |
| B215 | B216 | BMI | BMI | 0.82088 | 0 |
| B215 | B218 | BMI | DBP | 0.25985 | 0 |
| B215 | B220 | BMI | SBP | 0.254899 | 4.44E-16 |
| B215 | B222 | BMI | NS | 0.675684 | 0 |
| B216 | B218 | BMI | DBP | 0.24693 | 4.00E-15 |
| B216 | B221 | BMI | Hei | 0.675255 | 0 |
| B216 | B222 | BMI | NS | 0.820925 | 0 |
| B218 | B220 | DBP | SBP | 0.591999 | 0 |
| B218 | B222 | DBP | NS | 0.251425 | 1.33E-15 |
| B22 | B79 | CD3+CD8+(T) | hb V -dna | 0.242056 | 1.42E-14 |
| B22 | B95 | CD3+CD8+(T) | anti-cenpb | 0.437554 | 0 |
| B22 | B97 | CD3+CD8+(T) | dsDNA | 0.437554 | 0 |
| B22 | B98 | CD3+CD8+(T) | anti-Jo-1 | 0.437554 | 0 |
| B22 | B99 | CD3+CD8+(T) | anti-PM-Scl | -0.56652 | 0 |
| B22 | B101 | CD3+CD8+(T) | anti-Sc70 | 0.437554 | 0 |
| B22 | B102 | CD3+CD8+(T) | anti-Sm | 0.437554 | 0 |
| B22 | B105 | CD3+CD8+(T) | anti-U1RNP | 0.437554 | 0 |
| B22 | B106 | CD3+CD8+(T) | anti-P | 0.437554 | 0 |
| B22 | B107 | CD3+CD8+(T) | AnuA | 0.437553 | 0 |

| | | | | | |
|---|---|---|---|---|---|
| B22 | B108 | CD3+CD8+(T) | Anti-Cyclin | 0.437554 | 0 |
| B22 | B109 | CD3+CD8+(T) | AMA-M2 | 0.437554 | 0 |
| B22 | B110 | CD3+CD8+(T) | AHA | 0.437554 | 0 |
| B22 | B120 | CD3+CD8+(T) | HCV-cAg | 0.165354 | 1.85E-07 |
| B22 | B138 | CD3+CD8+(T) | PAB | 0.198081 | 3.74E-10 |
| B22 | B151 | CD3+CD8+(T) | Anti-CCP | 0.289299 | 0 |
| B22 | B156 | CD3+CD8+(T) | IgE | -0.68871 | 0 |
| B22 | B158 | CD3+CD8+(T) | IgM | 0.337854 | 0 |
| B22 | B175 | CD3+CD8+(T) | ALB-U | 0.207555 | 5.00E-11 |
| B22 | B176 | CD3+CD8+(T) | Ucr | 0.166895 | 1.42E-07 |
| B22 | B178 | CD3+CD8+(T) | Ig-U | 0.309797 | 0 |
| B22 | B179 | CD3+CD8+(T) | α1-MG | 0.285693 | 0 |
| B22 | B180 | CD3+CD8+(T) | Tf | 0.293302 | 0 |
| B22 | B181 | CD3+CD8+(T) | APTT | 0.582584 | 0 |
| B22 | B182 | CD3+CD8+(T) | TT | 0.579627 | 0 |
| B22 | B183 | CD3+CD8+(T) | PTA | 0.582574 | 0 |
| B22 | B184 | CD3+CD8+(T) | INR | 0.582584 | 0 |
| B22 | B185 | CD3+CD8+(T) | PT | 0.581824 | 0 |
| B22 | B186 | CD3+CD8+(T) | PFC | -0.58258 | 0 |
| B22 | B194 | CD3+CD8+(T) | Cl | -0.20885 | 3.78E-11 |
| B22 | B198 | CD3+CD8+(T) | P | -0.23615 | 6.35E-14 |
| B220 | B222 | SBP | NS | 0.200963 | 2.05E-10 |
| B221 | B222 | Hei | NS | 0.559134 | 0 |
| B23 | B75 | CD3-CD16+CD56+(NK) | ApoA1/B | 0.191126 | 1.54E-09 |
| B23 | B76 | CD3-CD16+CD56+(NK) | ApoB | -0.19713 | 4.55E-10 |
| B23 | B120 | CD3-CD16+CD56+(NK) | HCV-cAg | 0.210065 | 2.89E-11 |
| B23 | B137 | CD3-CD16+CD56+(NK) | AFU | -0.17825 | 1.84E-08 |
| B23 | B156 | CD3-CD16+CD56+(NK) | IgE | 0.207111 | 5.51E-11 |
| B23 | B158 | CD3-CD16+CD56+(NK) | IgM | -0.17704 | 2.30E-08 |
| B23 | B194 | CD3-CD16+CD56+(NK) | Cl | 0.269282 | 0 |
| B23 | B197 | CD3-CD16+CD56+(NK) | AG | -0.22991 | 2.94E-13 |
| B24 | B26 | αHBDH | CKMB | 0.499784 | 0 |
| B24 | B27 | αHBDH | LDH | 0.307643 | 0 |
| B24 | B55 | αHBDH | TK | -0.2905 | 0 |
| B24 | B58 | αHBDH | HCT | 0.291852 | 0 |
| B24 | B61 | αHBDH | WBHSRV | 0.165554 | 1.79E-07 |
| B24 | B63 | αHBDH | WBRSR(mPas_200) | 0.310512 | 0 |
| B24 | B64 | αHBDH | WBRSR(mPas_5) | 0.291219 | 0 |
| B24 | B65 | αHBDH | WBRSR(mPas_1) | 0.265227 | 0 |
| B24 | B66 | αHBDH | WBRSR(mPas_100) | 0.176597 | 2.49E-08 |
| B24 | B67 | αHBDH | WBRSR(mPas_50) | 0.182331 | 8.53E-09 |
| B24 | B69 | αHBDH | PV | 0.2073 | 5.29E-11 |
| B24 | B75 | αHBDH | ApoA1/B | 0.164022 | 2.33E-07 |
| B24 | B99 | αHBDH | anti-PM-Scl | -0.20087 | 2.09E-10 |

| | | | | | |
|---|---|---|---|---|---|
| B24 | B154 | αHBDH | RF | 0.408502 | 0 |
| B24 | B155 | αHBDH | IgA | 0.218012 | 4.85E-12 |
| B24 | B157 | αHBDH | IgG | 0.262655 | 0 |
| B24 | B181 | αHBDH | APTT | 0.2041 | 1.05E-10 |
| B24 | B182 | αHBDH | TT | 0.203193 | 1.28E-10 |
| B24 | B183 | αHBDH | PTA | 0.204092 | 1.06E-10 |
| B24 | B184 | αHBDH | INR | 0.2041 | 1.05E-10 |
| B24 | B185 | αHBDH | PT | 0.203775 | 1.13E-10 |
| B24 | B186 | αHBDH | PFC | -0.2041 | 1.05E-10 |
| B25 | B26 | CK | CKMB | 0.39795 | 0 |
| B25 | B59 | CK | WBRV | -0.17286 | 4.93E-08 |
| B25 | B123 | CK | hs-CRP1 | -0.19367 | 9.21E-10 |
| B25 | B124 | CK | ACTH | 0.325781 | 0 |
| B25 | B125 | CK | Cor | 0.325781 | 0 |
| B25 | B138 | CK | PAB | 0.165038 | 1.96E-07 |
| B25 | B154 | CK | RF | -0.17054 | 7.47E-08 |
| B25 | B168 | CK | NMP22 | 0.174902 | 3.40E-08 |
| B26 | B27 | CKMB | LDH | 0.176326 | 2.62E-08 |
| B26 | B55 | CKMB | TK | -0.25657 | 4.44E-16 |
| B26 | B58 | CKMB | HCT | 0.247271 | 3.55E-15 |
| B26 | B61 | CKMB | WBHSRV | 0.216193 | 7.34E-12 |
| B26 | B63 | CKMB | WBRSR(mPas_200) | 0.335252 | 0 |
| B26 | B64 | CKMB | WBRSR(mPas_5) | 0.247472 | 3.55E-15 |
| B26 | B65 | CKMB | WBRSR(mPas_1) | 0.189014 | 2.34E-09 |
| B26 | B69 | CKMB | PV | 0.276487 | 0 |
| B26 | B78 | CKMB | HBcAb | -0.18975 | 2.02E-09 |
| B26 | B95 | CKMB | anti-cenpb | 0.155429 | 9.74E-07 |
| B26 | B97 | CKMB | dsDNA | 0.155429 | 9.74E-07 |
| B26 | B98 | CKMB | anti-Jo-1 | 0.155429 | 9.74E-07 |
| B26 | B101 | CKMB | anti-Sc70 | 0.155429 | 9.74E-07 |
| B26 | B102 | CKMB | anti-Sm | 0.155429 | 9.74E-07 |
| B26 | B105 | CKMB | anti-U1RNP | 0.155429 | 9.74E-07 |
| B26 | B106 | CKMB | anti-P | 0.155429 | 9.74E-07 |
| B26 | B107 | CKMB | AnuA | 0.155429 | 9.74E-07 |
| B26 | B108 | CKMB | Anti-Cyclin | 0.155429 | 9.74E-07 |
| B26 | B109 | CKMB | AMA-M2 | 0.155429 | 9.74E-07 |
| B26 | B110 | CKMB | AHA | 0.155429 | 9.74E-07 |
| B26 | B134 | CKMB | CHE | -0.21222 | 1.79E-11 |
| B26 | B151 | CKMB | Anti-CCP | -0.21549 | 8.62E-12 |
| B26 | B168 | CKMB | NMP22 | 0.189556 | 2.10E-09 |
| B27 | B38 | LDH | BAND% | 0.617147 | 0 |
| B27 | B47 | LDH | RDW | 0.184635 | 5.49E-09 |
| B27 | B148 | LDH | Anti_TPO | 0.218283 | 4.56E-12 |
| B28 | B30 | PRL | LH | -0.7792 | 0 |

| | | | | | |
|---|---|---|---|---|---|
| B28 | B31 | PRL | FSH | -0.48213 | 0 |
| B28 | B32 | PRL | Testo | -0.75579 | 0 |
| B28 | B33 | PRL | Prog | -0.52949 | 0 |
| B29 | B30 | E2 | LH | -0.20972 | 3.12E-11 |
| B29 | B31 | E2 | FSH | 0.322966 | 0 |
| B29 | B33 | E2 | Prog | 0.235749 | 7.02E-14 |
| B29 | B137 | E2 | AFU | 0.188885 | 2.40E-09 |
| B29 | B150 | E2 | ALP | 0.160191 | 4.45E-07 |
| B29 | B198 | E2 | P | -0.22932 | 3.39E-13 |
| B3 | B58 | Hcy | HCT | 0.216049 | 7.59E-12 |
| B3 | B60 | Hcy | WBRVI | 0.175931 | 2.82E-08 |
| B3 | B62 | Hcy | WBHSRVI | 0.185579 | 4.57E-09 |
| B3 | B64 | Hcy | WBRSR(mPas_5) | 0.160181 | 4.46E-07 |
| B3 | B65 | Hcy | WBRSR(mPas_1) | 0.159096 | 5.34E-07 |
| B3 | B66 | Hcy | WBRSR(mPas_100) | 0.180963 | 1.11E-08 |
| B3 | B67 | Hcy | WBRSR(mPas_50) | 0.18364 | 6.64E-09 |
| B3 | B114 | Hcy | TSH TT3 TT4(TT4) | 0.155589 | 9.50E-07 |
| B30 | B31 | LH | FSH | 0.820986 | 0 |
| B30 | B32 | LH | Testo | 0.945916 | 0 |
| B30 | B33 | LH | Prog | 0.869631 | 0 |
| B30 | B95 | LH | anti-cenpb | 0.176145 | 2.71E-08 |
| B30 | B97 | LH | dsDNA | 0.176145 | 2.71E-08 |
| B30 | B98 | LH | anti-Jo-1 | 0.176145 | 2.71E-08 |
| B30 | B101 | LH | anti-Sc70 | 0.176145 | 2.71E-08 |
| B30 | B102 | LH | anti-Sm | 0.176145 | 2.71E-08 |
| B30 | B105 | LH | anti-U1RNP | 0.176145 | 2.71E-08 |
| B30 | B106 | LH | anti-P | 0.176145 | 2.71E-08 |
| B30 | B107 | LH | AnuA | 0.176145 | 2.71E-08 |
| B30 | B108 | LH | Anti-Cyclin | 0.176145 | 2.71E-08 |
| B30 | B109 | LH | AMA-M2 | 0.176145 | 2.71E-08 |
| B30 | B110 | LH | AHA | 0.176145 | 2.71E-08 |
| B30 | B178 | LH | Ig-U | 0.225766 | 7.95E-13 |
| B30 | B179 | LH | α1-MG | 0.208132 | 4.41E-11 |
| B30 | B180 | LH | Tf | 0.213701 | 1.29E-11 |
| B31 | B32 | FSH | Testo | 0.934657 | 0 |
| B31 | B33 | FSH | Prog | 0.99557 | 0 |
| B31 | B95 | FSH | anti-cenpb | 0.198669 | 3.31E-10 |
| B31 | B97 | FSH | dsDNA | 0.198669 | 3.31E-10 |
| B31 | B98 | FSH | anti-Jo-1 | 0.198669 | 3.31E-10 |
| B31 | B101 | FSH | anti-Sc70 | 0.198669 | 3.31E-10 |
| B31 | B102 | FSH | anti-Sm | 0.198669 | 3.31E-10 |
| B31 | B105 | FSH | anti-U1RNP | 0.198669 | 3.31E-10 |
| B31 | B106 | FSH | anti-P | 0.198669 | 3.31E-10 |
| B31 | B107 | FSH | AnuA | 0.198669 | 3.31E-10 |

| | | | | | |
|---|---|---|---|---|---|
| B31 | B108 | FSH | Anti-Cyclin | 0.198669 | 3.31E-10 |
| B31 | B109 | FSH | AMA-M2 | 0.198669 | 3.31E-10 |
| B31 | B110 | FSH | AHA | 0.198669 | 3.31E-10 |
| B31 | B178 | FSH | Ig-U | 0.254841 | 4.44E-16 |
| B31 | B179 | FSH | α1-MG | 0.234893 | 8.66E-14 |
| B31 | B180 | FSH | Tf | 0.241193 | 1.78E-14 |
| B32 | B33 | Testo | Prog | 0.955554 | 0 |
| B32 | B178 | Testo | Ig-U | 0.190223 | 1.84E-09 |
| B32 | B179 | Testo | α1-MG | 0.17526 | 3.19E-08 |
| B32 | B180 | Testo | Tf | 0.179988 | 1.33E-08 |
| B33 | B95 | Prog | anti-cenpb | 0.205529 | 7.75E-11 |
| B33 | B97 | Prog | dsDNA | 0.205529 | 7.75E-11 |
| B33 | B98 | Prog | anti-Jo-1 | 0.205529 | 7.75E-11 |
| B33 | B101 | Prog | anti-Sc70 | 0.205529 | 7.75E-11 |
| B33 | B102 | Prog | anti-Sm | 0.205529 | 7.75E-11 |
| B33 | B105 | Prog | anti-U1RNP | 0.205529 | 7.75E-11 |
| B33 | B106 | Prog | anti-P | 0.205529 | 7.75E-11 |
| B33 | B107 | Prog | AnuA | 0.205529 | 7.76E-11 |
| B33 | B108 | Prog | Anti-Cyclin | 0.205529 | 7.75E-11 |
| B33 | B109 | Prog | AMA-M2 | 0.205529 | 7.75E-11 |
| B33 | B110 | Prog | AHA | 0.205529 | 7.75E-11 |
| B33 | B178 | Prog | Ig-U | 0.263554 | 0 |
| B33 | B179 | Prog | α1-MG | 0.242942 | 1.13E-14 |
| B33 | B180 | Prog | Tf | 0.249452 | 2.22E-15 |
| B35 | B92 | β2-MG | ALDH2-Glu504Glu | 0.730988 | 0 |
| B35 | B94 | β2-MG | ALDH2-Lys504Lys | 0.613636 | 0 |
| B35 | B96 | β2-MG | dsDNA (IIF) | 0.940127 | 0 |
| B35 | B116 | β2-MG | TT3 | 0.837161 | 0 |
| B35 | B167 | β2-MG | PC | 0.46127 | 0 |
| B37 | B38 | BAND | BAND% | 0.63316 | 0 |
| B38 | B148 | BAND% | Anti_TPO | 0.168657 | 1.04E-07 |
| B4 | B7 | WB-Ca | WB-Cop | 1 | 0 |
| B4 | B28 | WB-Ca | PRL | 0.991516 | 0 |
| B4 | B30 | WB-Ca | LH | -0.84269 | 0 |
| B4 | B31 | WB-Ca | FSH | -0.59145 | 0 |
| B4 | B32 | WB-Ca | Testo | -0.83448 | 0 |
| B4 | B33 | WB-Ca | Prog | -0.63526 | 0 |
| B40 | B42 | WBC | MON | 0.45174 | 0 |
| B40 | B46 | WBC | RBC Count | 0.263793 | 0 |
| B40 | B48 | WBC | LYM% | -0.22809 | 4.56E-13 |
| B40 | B51 | WBC | Hb | 0.161295 | 3.70E-07 |
| B40 | B52 | WBC | PLT/BPC | 0.272471 | 0 |
| B40 | B53 | WBC | GR% | 0.215081 | 9.44E-12 |
| B40 | B72 | WBC | HDL-C | -0.19452 | 7.75E-10 |

| | | | | | |
|---|---|---|---|---|---|
| B40 | B192 | WBC | UA | 0.16159 | 3.52E-07 |
| B40 | B206 | WBC | Waist | 0.198571 | 3.38E-10 |
| B40 | B209 | WBC | Smoking | 0.198138 | 3.69E-10 |
| B40 | B215 | WBC | BMI | 0.18421 | 5.96E-09 |
| B41 | B42 | MON％ | MON | 0.808279 | 0 |
| B41 | B53 | MON％ | GR% | -0.29607 | 0 |
| B42 | B192 | MON | UA | 0.181996 | 9.09E-09 |
| B43 | B44 | MCV | MCH | 0.929391 | 0 |
| B43 | B45 | MCV | MCHC | 0.333526 | 0 |
| B43 | B46 | MCV | RBC Count | -0.53634 | 0 |
| B43 | B47 | MCV | RDW | 0.593605 | 0 |
| B43 | B51 | MCV | Hb | 0.235625 | 7.24E-14 |
| B43 | B52 | MCV | PLT/BPC | -0.19096 | 1.59E-09 |
| B44 | B45 | MCH | MCHC | 0.65614 | 0 |
| B44 | B46 | MCH | RBC Count | -0.41454 | 0 |
| B44 | B47 | MCH | RDW | 0.425203 | 0 |
| B44 | B51 | MCH | Hb | 0.40871 | 0 |
| B44 | B52 | MCH | PLT/BPC | -0.17344 | 4.44E-08 |
| B45 | B51 | MCHC | Hb | 0.568577 | 0 |
| B45 | B71 | MCHC | TG | 0.181519 | 9.95E-09 |
| B45 | B126 | MCHC | Alb | 0.161066 | 3.84E-07 |
| B45 | B153 | MCHC | Glu | 0.156185 | 8.62E-07 |
| B45 | B192 | MCHC | UA | 0.181305 | 1.04E-08 |
| B45 | B206 | MCHC | Waist | 0.185377 | 4.75E-09 |
| B45 | B210 | MCHC | Hip | 0.158053 | 6.34E-07 |
| B45 | B216 | MCHC | BMI | 0.260427 | 0 |
| B45 | B221 | MCHC | Hei | 0.257321 | 2.22E-16 |
| B45 | B222 | MCHC | NS | 0.285474 | 0 |
| B46 | B47 | RBC Count | RDW | -0.44193 | 0 |
| B46 | B51 | RBC Count | Hb | 0.652328 | 0 |
| B46 | B71 | RBC Count | TG | 0.164329 | 2.21E-07 |
| B46 | B72 | RBC Count | HDL-C | -0.25094 | 1.33E-15 |
| B46 | B113 | RBC Count | FT3 | 0.174439 | 3.70E-08 |
| B46 | B126 | RBC Count | Alb | 0.266646 | 0 |
| B46 | B127 | RBC Count | A-G ratio | 0.164363 | 2.20E-07 |
| B46 | B128 | RBC Count | ALT | 0.260605 | 0 |
| B46 | B140 | RBC Count | r-GT | 0.217416 | 5.56E-12 |
| B46 | B141 | RBC Count | AST/ALT | -0.30043 | 0 |
| B46 | B163 | RBC Count | SG | 0.244014 | 8.44E-15 |
| B46 | B174 | RBC Count | Urea/Cr | -0.25206 | 8.88E-16 |
| B46 | B190 | RBC Count | Cre | 0.218232 | 4.61E-12 |
| B46 | B192 | RBC Count | UA | 0.373959 | 0 |
| B46 | B206 | RBC Count | Waist | 0.335862 | 0 |
| B46 | B209 | RBC Count | Smoking | 0.189765 | 2.01E-09 |

| | | | | | |
|---|---|---|---|---|---|
| B46 | B210 | RBC Count | Hip | 0.231666 | 1.92E-13 |
| B46 | B215 | RBC Count | BMI | 0.248633 | 2.66E-15 |
| B46 | B216 | RBC Count | BMI | 0.39548 | 0 |
| B46 | B218 | RBC Count | DBP | 0.217319 | 5.68E-12 |
| B46 | B221 | RBC Count | Hei | 0.373874 | 0 |
| B46 | B222 | RBC Count | NS | 0.41626 | 0 |
| B47 | B52 | RDW | PLT/BPC | -0.29523 | 0 |
| B47 | B126 | RDW | Alb | -0.19528 | 6.64E-10 |
| B47 | B189 | RDW | eGFB | -0.25436 | 4.44E-16 |
| B48 | B53 | LYM% | GR% | -0.91991 | 0 |
| B49 | B50 | baso% | eo% | 0.269582 | 0 |
| B49 | B53 | baso% | GR% | -0.17335 | 4.51E-08 |
| B5 | B6 | WB-Mg | WB-Fe | -1 | 0 |
| B5 | B7 | WB-Mg | WB-Cop | -1 | 0 |
| B5 | B28 | WB-Mg | PRL | -0.99152 | 0 |
| B5 | B30 | WB-Mg | LH | 0.842692 | 0 |
| B5 | B31 | WB-Mg | FSH | 0.591453 | 0 |
| B5 | B32 | WB-Mg | Testo | 0.834483 | 0 |
| B5 | B33 | WB-Mg | Prog | 0.635264 | 0 |
| B50 | B53 | eo% | GR% | -0.28746 | 0 |
| B50 | B209 | eo% | Smoking | 0.167596 | 1.25E-07 |
| B51 | B58 | Hb | HCT | 0.173433 | 4.44E-08 |
| B51 | B64 | Hb | WBRSR(mPas_5) | 0.167636 | 1.25E-07 |
| B51 | B65 | Hb | WBRSR(mPas_1) | 0.16824 | 1.12E-07 |
| B51 | B71 | Hb | TG | 0.234713 | 9.06E-14 |
| B51 | B72 | Hb | HDL-C | -0.20101 | 2.03E-10 |
| B51 | B113 | Hb | FT3 | 0.214672 | 1.04E-11 |
| B51 | B126 | Hb | Alb | 0.281614 | 0 |
| B51 | B127 | Hb | A-G ratio | 0.210046 | 2.90E-11 |
| B51 | B128 | Hb | ALT | 0.305759 | 0 |
| B51 | B138 | Hb | PAB | 0.155302 | 9.95E-07 |
| B51 | B140 | Hb | r-GT | 0.270003 | 0 |
| B51 | B141 | Hb | AST/ALT | -0.28736 | 0 |
| B51 | B163 | Hb | SG | 0.288435 | 0 |
| B51 | B174 | Hb | Urea/Cr | -0.31558 | 0 |
| B51 | B190 | Hb | Cre | 0.327012 | 0 |
| B51 | B192 | Hb | UA | 0.42431 | 0 |
| B51 | B204 | Hb | Drinking | 0.239895 | 2.49E-14 |
| B51 | B206 | Hb | Waist | 0.388224 | 0 |
| B51 | B209 | Hb | Smoking | 0.286431 | 0 |
| B51 | B210 | Hb | Hip | 0.260215 | 0 |
| B51 | B215 | Hb | BMI | 0.266715 | 0 |
| B51 | B216 | Hb | BMI | 0.476943 | 0 |
| B51 | B218 | Hb | DBP | 0.251293 | 1.33E-15 |

| | | | | | |
|---|---|---|---|---|---|
| B51 | B221 | Hb | Hei | 0.487677 | 0 |
| B51 | B222 | Hb | NS | 0.514167 | 0 |
| B52 | B129 | PLT/BPC | IBIL | -0.20037 | 2.32E-10 |
| B52 | B132 | PLT/BPC | Dbil | -0.21673 | 6.50E-12 |
| B52 | B133 | PLT/BPC | TBIL | -0.21783 | 5.06E-12 |
| B52 | B189 | PLT/BPC | eGFB | 0.165376 | 1.84E-07 |
| B52 | B190 | PLT/BPC | Cre | -0.16758 | 1.26E-07 |
| B52 | B191 | PLT/BPC | Urea | -0.17518 | 3.24E-08 |
| B54 | B56 | ESR | IR | -0.48872 | 0 |
| B54 | B57 | ESR | RBCAI | -0.24705 | 4.00E-15 |
| B54 | B58 | ESR | HCT | -0.53734 | 0 |
| B54 | B59 | ESR | WBRV | -0.26347 | 0 |
| B54 | B60 | ESR | WBRVI | -0.53597 | 0 |
| B54 | B61 | ESR | WBHSRV | -0.45296 | 0 |
| B54 | B62 | ESR | WBHSRVI | -0.57851 | 0 |
| B54 | B63 | ESR | WBRSR(mPas_200) | -0.43781 | 0 |
| B54 | B64 | ESR | WBRSR(mPas_5) | -0.48454 | 0 |
| B54 | B65 | ESR | WBRSR(mPas_1) | -0.48085 | 0 |
| B54 | B66 | ESR | WBRSR(mPas_100) | -0.48492 | 0 |
| B54 | B67 | ESR | WBRSR(mPas_50) | -0.49435 | 0 |
| B54 | B73 | ESR | ApoA | -0.19776 | 3.99E-10 |
| B54 | B78 | ESR | HBcAb | 0.180248 | 1.26E-08 |
| B54 | B123 | ESR | hs-CRP1 | 0.257635 | 2.22E-16 |
| B54 | B138 | ESR | PAB | -0.31623 | 0 |
| B54 | B153 | ESR | Glu | 0.174341 | 3.77E-08 |
| B54 | B155 | ESR | IgA | 0.193093 | 1.04E-09 |
| B54 | B200 | ESR | OP | 0.156531 | 8.14E-07 |
| B55 | B56 | TK | IR | 0.638554 | 0 |
| B55 | B58 | TK | HCT | -0.41436 | 0 |
| B55 | B59 | TK | WBRV | -0.24051 | 2.11E-14 |
| B55 | B62 | TK | WBHSRVI | 0.191153 | 1.53E-09 |
| B55 | B63 | TK | WBRSR(mPas_200) | -0.48505 | 0 |
| B55 | B64 | TK | WBRSR(mPas_5) | -0.49097 | 0 |
| B55 | B65 | TK | WBRSR(mPas_1) | -0.46513 | 0 |
| B55 | B66 | TK | WBRSR(mPas_100) | -0.18335 | 7.02E-09 |
| B55 | B67 | TK | WBRSR(mPas_50) | -0.18682 | 3.59E-09 |
| B55 | B69 | TK | PV | -0.68421 | 0 |
| B55 | B99 | TK | anti-PM-Scl | 0.159517 | 4.98E-07 |
| B55 | B134 | TK | CHE | -0.23741 | 4.62E-14 |
| B55 | B151 | TK | Anti-CCP | -0.28809 | 0 |
| B55 | B195 | TK | Na | -0.15653 | 8.14E-07 |
| B55 | B200 | TK | OP | -0.16914 | 9.57E-08 |
| B56 | B58 | IR | HCT | 0.426213 | 0 |
| B56 | B60 | IR | WBRVI | 0.675721 | 0 |

| | | | | | |
|---|---|---|---|---|---|
| B56 | B61 | IR | WBHSRV | 0.488544 | 0 |
| B56 | B62 | IR | WBHSRVI | 0.87503 | 0 |
| B56 | B63 | IR | WBRSR(mPas_200) | 0.188419 | 2.63E-09 |
| B56 | B64 | IR | WBRSR(mPas_5) | 0.202937 | 1.35E-10 |
| B56 | B65 | IR | WBRSR(mPas_1) | 0.196917 | 4.75E-10 |
| B56 | B66 | IR | WBRSR(mPas_100) | 0.549277 | 0 |
| B56 | B67 | IR | WBRSR(mPas_50) | 0.554074 | 0 |
| B56 | B69 | IR | PV | -0.67794 | 0 |
| B56 | B99 | IR | anti-PM-Scl | 0.19261 | 1.14E-09 |
| B56 | B134 | IR | CHE | -0.21816 | 4.69E-12 |
| B56 | B139 | IR | ADA | 0.179355 | 1.50E-08 |
| B56 | B151 | IR | Anti-CCP | -0.39603 | 0 |
| B56 | B155 | IR | IgA | -0.16833 | 1.10E-07 |
| B56 | B178 | IR | Ig-U | 0.194854 | 7.25E-10 |
| B56 | B180 | IR | Tf | 0.187183 | 3.35E-09 |
| B56 | B181 | IR | APTT | -0.18264 | 8.04E-09 |
| B56 | B182 | IR | TT | -0.18154 | 9.91E-09 |
| B56 | B183 | IR | PTA | -0.18264 | 8.04E-09 |
| B56 | B184 | IR | INR | -0.18264 | 8.04E-09 |
| B56 | B185 | IR | PT | -0.18248 | 8.30E-09 |
| B56 | B186 | IR | PFC | 0.182639 | 8.04E-09 |
| B56 | B197 | IR | AG | 0.185423 | 4.71E-09 |
| B56 | B228 | IR | OS | 0.179299 | 1.51E-08 |
| B57 | B58 | RBCAI | HCT | 0.205959 | 7.07E-11 |
| B57 | B59 | RBCAI | WBRV | 0.703405 | 0 |
| B57 | B60 | RBCAI | WBRVI | 0.67358 | 0 |
| B57 | B61 | RBCAI | WBHSRV | -0.17025 | 7.86E-08 |
| B57 | B62 | RBCAI | WBHSRVI | 0.182694 | 7.96E-09 |
| B57 | B64 | RBCAI | WBRSR(mPas_5) | 0.383675 | 0 |
| B57 | B65 | RBCAI | WBRSR(mPas_1) | 0.575485 | 0 |
| B57 | B66 | RBCAI | WBRSR(mPas_100) | 0.162186 | 3.18E-07 |
| B57 | B67 | RBCAI | WBRSR(mPas_50) | 0.222449 | 1.74E-12 |
| B57 | B95 | RBCAI | anti-cenpb | -0.17305 | 4.77E-08 |
| B57 | B97 | RBCAI | dsDNA | -0.17305 | 4.77E-08 |
| B57 | B98 | RBCAI | anti-Jo-1 | -0.17305 | 4.77E-08 |
| B57 | B101 | RBCAI | anti-Sc70 | -0.17305 | 4.77E-08 |
| B57 | B102 | RBCAI | anti-Sm | -0.17305 | 4.77E-08 |
| B57 | B105 | RBCAI | anti-U1RNP | -0.17305 | 4.77E-08 |
| B57 | B106 | RBCAI | anti-P | -0.17304 | 4.77E-08 |
| B57 | B107 | RBCAI | AnuA | -0.17304 | 4.77E-08 |
| B57 | B108 | RBCAI | Anti-Cyclin | -0.17305 | 4.77E-08 |
| B57 | B109 | RBCAI | AMA-M2 | -0.17305 | 4.77E-08 |
| B57 | B110 | RBCAI | AHA | -0.17305 | 4.77E-08 |
| B57 | B134 | RBCAI | CHE | 0.162487 | 3.02E-07 |

| | | | | | |
|---|---|---|---|---|---|
| B57 | B151 | RBCAI | Anti-CCP | 0.480729 | 0 |
| B57 | B156 | RBCAI | IgE | 0.163671 | 2.47E-07 |
| B57 | B165 | RBCAI | bilirubin | 0.467975 | 0 |
| B57 | B178 | RBCAI | Ig-U | -0.25663 | 4.44E-16 |
| B57 | B179 | RBCAI | α1-MG | -0.20839 | 4.17E-11 |
| B57 | B180 | RBCAI | Tf | -0.23629 | 6.13E-14 |
| B57 | B194 | RBCAI | Cl | 0.238751 | 3.29E-14 |
| B57 | B197 | RBCAI | AG | -0.17288 | 4.92E-08 |
| B57 | B198 | RBCAI | P | -0.18639 | 3.91E-09 |
| B57 | B228 | RBCAI | OS | 0.358405 | 0 |
| B58 | B59 | HCT | WBRV | 0.162842 | 2.85E-07 |
| B58 | B60 | HCT | WBRVI | 0.718378 | 0 |
| B58 | B61 | HCT | WBHSRV | 0.510952 | 0 |
| B58 | B62 | HCT | WBHSRVI | 0.803503 | 0 |
| B58 | B63 | HCT | WBRSR(mPas_200) | 0.751168 | 0 |
| B58 | B64 | HCT | WBRSR(mPas_5) | 0.77024 | 0 |
| B58 | B65 | HCT | WBRSR(mPas_1) | 0.737146 | 0 |
| B58 | B66 | HCT | WBRSR(mPas_100) | 0.840656 | 0 |
| B58 | B67 | HCT | WBRSR(mPas_50) | 0.850548 | 0 |
| B58 | B78 | HCT | HBcAb | -0.22632 | 6.97E-13 |
| B58 | B123 | HCT | hs-CRP1 | -0.25229 | 8.88E-16 |
| B58 | B137 | HCT | AFU | 0.202362 | 1.52E-10 |
| B58 | B138 | HCT | PAB | 0.321857 | 0 |
| B58 | B146 | HCT | CysC | 0.165261 | 1.88E-07 |
| B58 | B165 | HCT | bilirubin | 0.176585 | 2.50E-08 |
| B58 | B175 | HCT | ALB-U | 0.155482 | 9.66E-07 |
| B58 | B195 | HCT | Na | 0.183866 | 6.36E-09 |
| B58 | B200 | HCT | OP | 0.274134 | 0 |
| B58 | B228 | HCT | OS | 0.282227 | 0 |
| B59 | B60 | WBRV | WBRVI | 0.447326 | 0 |
| B59 | B61 | WBRV | WBHSRV | 0.387169 | 0 |
| B59 | B63 | WBRV | WBRSR(mPas_200) | 0.443899 | 0 |
| B59 | B64 | WBRV | WBRSR(mPas_5) | 0.699172 | 0 |
| B59 | B65 | WBRV | WBRSR(mPas_1) | 0.777997 | 0 |
| B59 | B66 | WBRV | WBRSR(mPas_100) | 0.333559 | 0 |
| B59 | B67 | WBRV | WBRSR(mPas_50) | 0.3732 | 0 |
| B59 | B69 | WBRV | PV | 0.430991 | 0 |
| B59 | B124 | WBRV | ACTH | -0.15816 | 6.23E-07 |
| B59 | B125 | WBRV | Cor | -0.15816 | 6.23E-07 |
| B59 | B139 | WBRV | ADA | 0.222563 | 1.69E-12 |
| B59 | B151 | WBRV | Anti-CCP | 0.38262 | 0 |
| B59 | B165 | WBRV | bilirubin | 0.34056 | 0 |
| B59 | B178 | WBRV | Ig-U | -0.18237 | 8.46E-09 |
| B59 | B180 | WBRV | Tf | -0.16377 | 2.43E-07 |

| | | | | | |
|---|---|---|---|---|---|
| B59 | B198 | WBRV | P | -0.19208 | 1.27E-09 |
| B59 | B228 | WBRV | OS | 0.272052 | 0 |
| B6 | B7 | WB-Fe | WB-Cop | 1 | 0 |
| B6 | B28 | WB-Fe | PRL | 0.991516 | 0 |
| B6 | B30 | WB-Fe | LH | -0.84269 | 0 |
| B6 | B31 | WB-Fe | FSH | -0.59145 | 0 |
| B6 | B32 | WB-Fe | Testo | -0.83448 | 0 |
| B6 | B33 | WB-Fe | Prog | -0.63526 | 0 |
| B60 | B61 | WBRVI | WBHSRV | 0.348606 | 0 |
| B60 | B62 | WBRVI | WBHSRVI | 0.845692 | 0 |
| B60 | B63 | WBRVI | WBRSR(mPas_200) | 0.386234 | 0 |
| B60 | B64 | WBRVI | WBRSR(mPas_5) | 0.636648 | 0 |
| B60 | B65 | WBRVI | WBRSR(mPas_1) | 0.728849 | 0 |
| B60 | B66 | WBRVI | WBRSR(mPas_100) | 0.690821 | 0 |
| B60 | B67 | WBRVI | WBRSR(mPas_50) | 0.7332 | 0 |
| B60 | B69 | WBRVI | PV | -0.41041 | 0 |
| B60 | B73 | WBRVI | ApoA | 0.172278 | 5.47E-08 |
| B60 | B123 | WBRVI | hs-CRP1 | -0.16887 | 1.00E-07 |
| B60 | B138 | WBRVI | PAB | 0.180128 | 1.29E-08 |
| B60 | B139 | WBRVI | ADA | 0.160697 | 4.09E-07 |
| B60 | B146 | WBRVI | CysC | 0.171467 | 6.33E-08 |
| B60 | B165 | WBRVI | bilirubin | 0.352208 | 0 |
| B60 | B200 | WBRVI | OP | 0.158151 | 6.24E-07 |
| B60 | B228 | WBRVI | OS | 0.452061 | 0 |
| B61 | B62 | WBHSRV | WBHSRVI | 0.595976 | 0 |
| B61 | B63 | WBHSRV | WBRSR(mPas_200) | 0.857028 | 0 |
| B61 | B64 | WBHSRV | WBRSR(mPas_5) | 0.725553 | 0 |
| B61 | B65 | WBHSRV | WBRSR(mPas_1) | 0.598066 | 0 |
| B61 | B66 | WBHSRV | WBRSR(mPas_100) | 0.754845 | 0 |
| B61 | B67 | WBHSRV | WBRSR(mPas_50) | 0.739909 | 0 |
| B61 | B69 | WBHSRV | PV | 0.288437 | 0 |
| B61 | B134 | WBHSRV | CHE | -0.16228 | 3.13E-07 |
| B61 | B137 | WBHSRV | AFU | 0.23665 | 5.60E-14 |
| B61 | B139 | WBHSRV | ADA | 0.413968 | 0 |
| B61 | B151 | WBHSRV | Anti-CCP | -0.27868 | 0 |
| B61 | B153 | WBHSRV | Glu | 0.198539 | 3.40E-10 |
| B61 | B178 | WBHSRV | Ig-U | 0.164957 | 1.98E-07 |
| B61 | B180 | WBHSRV | Tf | 0.157748 | 6.67E-07 |
| B61 | B194 | WBHSRV | Cl | -0.25538 | 4.44E-16 |
| B61 | B200 | WBHSRV | OP | 0.215959 | 7.74E-12 |
| B62 | B63 | WBHSRVI | WBRSR(mPas_200) | 0.527692 | 0 |
| B62 | B64 | WBHSRVI | WBRSR(mPas_5) | 0.56514 | 0 |
| B62 | B65 | WBHSRVI | WBRSR(mPas_1) | 0.549805 | 0 |
| B62 | B66 | WBHSRVI | WBRSR(mPas_100) | 0.817151 | 0 |

| | | | | | |
|---|---|---|---|---|---|
| B62 | B67 | WBHSRVI | WBRSR(mPas_50) | 0.827887 | 0 |
| B62 | B69 | WBHSRVI | PV | -0.45789 | 0 |
| B62 | B78 | WBHSRVI | HBcAb | -0.19575 | 6.04E-10 |
| B62 | B123 | WBHSRVI | hs-CRP1 | -0.15896 | 5.46E-07 |
| B62 | B137 | WBHSRVI | AFU | 0.196296 | 5.40E-10 |
| B62 | B138 | WBHSRVI | PAB | 0.234479 | 9.59E-14 |
| B62 | B139 | WBHSRVI | ADA | 0.230163 | 2.76E-13 |
| B62 | B146 | WBHSRVI | CysC | 0.178623 | 1.71E-08 |
| B62 | B151 | WBHSRVI | Anti-CCP | -0.31497 | 0 |
| B62 | B178 | WBHSRVI | Ig-U | 0.155462 | 9.69E-07 |
| B62 | B197 | WBHSRVI | AG | 0.183952 | 6.26E-09 |
| B62 | B200 | WBHSRVI | OP | 0.235132 | 8.17E-14 |
| B62 | B228 | WBHSRVI | OS | 0.303547 | 0 |
| B63 | B64 | WBRSR(mPas_200) | WBRSR(mPas_5) | 0.911195 | 0 |
| B63 | B65 | WBRSR(mPas_200) | WBRSR(mPas_1) | 0.799695 | 0 |
| B63 | B66 | WBRSR(mPas_200) | WBRSR(mPas_100) | 0.803022 | 0 |
| B63 | B67 | WBRSR(mPas_200) | WBRSR(mPas_50) | 0.796622 | 0 |
| B63 | B69 | WBRSR(mPas_200) | PV | 0.498608 | 0 |
| B63 | B78 | WBRSR(mPas_200) | HBcAb | -0.17259 | 5.17E-08 |
| B63 | B137 | WBRSR(mPas_200) | AFU | 0.245758 | 5.33E-15 |
| B63 | B138 | WBRSR(mPas_200) | PAB | 0.18324 | 7.17E-09 |
| B63 | B139 | WBRSR(mPas_200) | ADA | 0.328778 | 0 |
| B63 | B153 | WBRSR(mPas_200) | Glu | 0.163718 | 2.45E-07 |
| B63 | B194 | WBRSR(mPas_200) | Cl | -0.21689 | 6.26E-12 |
| B63 | B200 | WBRSR(mPas_200) | OP | 0.270855 | 0 |
| B63 | B228 | WBRSR(mPas_200) | OS | 0.163511 | 2.54E-07 |
| B64 | B65 | WBRSR(mPas_5) | WBRSR(mPas_1) | 0.975674 | 0 |
| B64 | B66 | WBRSR(mPas_5) | WBRSR(mPas_100) | 0.805302 | 0 |
| B64 | B67 | WBRSR(mPas_5) | WBRSR(mPas_50) | 0.825014 | 0 |
| B64 | B69 | WBRSR(mPas_5) | PV | 0.402682 | 0 |
| B64 | B137 | WBRSR(mPas_5) | AFU | 0.204298 | 1.01E-10 |
| B64 | B138 | WBRSR(mPas_5) | PAB | 0.16638 | 1.55E-07 |
| B64 | B139 | WBRSR(mPas_5) | ADA | 0.309133 | 0 |
| B64 | B165 | WBRSR(mPas_5) | bilirubin | 0.260312 | 0 |
| B64 | B200 | WBRSR(mPas_5) | OP | 0.245156 | 6.44E-15 |
| B64 | B228 | WBRSR(mPas_5) | OS | 0.316202 | 0 |
| B65 | B66 | WBRSR(mPas_1) | WBRSR(mPas_100) | 0.75314 | 0 |
| B65 | B67 | WBRSR(mPas_1) | WBRSR(mPas_50) | 0.785242 | 0 |
| B65 | B69 | WBRSR(mPas_1) | PV | 0.315035 | 0 |
| B65 | B137 | WBRSR(mPas_1) | AFU | 0.162276 | 3.13E-07 |
| B65 | B139 | WBRSR(mPas_1) | ADA | 0.266234 | 0 |
| B65 | B151 | WBRSR(mPas_1) | Anti-CCP | 0.157704 | 6.72E-07 |
| B65 | B165 | WBRSR(mPas_1) | bilirubin | 0.344616 | 0 |
| B65 | B200 | WBRSR(mPas_1) | OP | 0.207309 | 5.28E-11 |

| | | | | | |
|---|---|---|---|---|---|
| B65 | B228 | WBRSR(mPas_1) | OS | 0.375248 | 0 |
| B66 | B67 | WBRSR(mPas_100) | WBRSR(mPas_50) | 0.997475 | 0 |
| B66 | B78 | WBRSR(mPas_100) | HBcAb | -0.18953 | 2.11E-09 |
| B66 | B137 | WBRSR(mPas_100) | AFU | 0.311448 | 0 |
| B66 | B138 | WBRSR(mPas_100) | PAB | 0.206292 | 6.58E-11 |
| B66 | B139 | WBRSR(mPas_100) | ADA | 0.431549 | 0 |
| B66 | B153 | WBRSR(mPas_100) | Glu | 0.220261 | 2.89E-12 |
| B66 | B194 | WBRSR(mPas_100) | Cl | -0.20713 | 5.48E-11 |
| B66 | B200 | WBRSR(mPas_100) | OP | 0.351182 | 0 |
| B66 | B228 | WBRSR(mPas_100) | OS | 0.238814 | 3.24E-14 |
| B67 | B78 | WBRSR(mPas_50) | HBcAb | -0.18109 | 1.08E-08 |
| B67 | B137 | WBRSR(mPas_50) | AFU | 0.301093 | 0 |
| B67 | B138 | WBRSR(mPas_50) | PAB | 0.203402 | 1.22E-10 |
| B67 | B139 | WBRSR(mPas_50) | ADA | 0.425038 | 0 |
| B67 | B153 | WBRSR(mPas_50) | Glu | 0.211123 | 2.29E-11 |
| B67 | B165 | WBRSR(mPas_50) | bilirubin | 0.164538 | 2.13E-07 |
| B67 | B194 | WBRSR(mPas_50) | Cl | -0.19282 | 1.09E-09 |
| B67 | B200 | WBRSR(mPas_50) | OP | 0.34289 | 0 |
| B67 | B228 | WBRSR(mPas_50) | OS | 0.267575 | 0 |
| B69 | B115 | PV | TSH TT3 TT4(TT3) | 0.165203 | 1.90E-07 |
| B69 | B151 | PV | Anti-CCP | 0.177477 | 2.12E-08 |
| B7 | B8 | WB-Cop | WB-Zinc | 1 | 0 |
| B7 | B28 | WB-Cop | PRL | 0.991516 | 0 |
| B7 | B30 | WB-Cop | LH | -0.84269 | 0 |
| B7 | B31 | WB-Cop | FSH | -0.59145 | 0 |
| B7 | B32 | WB-Cop | Testo | -0.83448 | 0 |
| B7 | B33 | WB-Cop | Prog | -0.63526 | 0 |
| B70 | B76 | LDL-C | ApoB | 0.185226 | 4.90E-09 |
| B70 | B77 | LDL-C | TC | 0.858077 | 0 |
| B70 | B132 | LDL-C | Dbil | -0.21757 | 5.37E-12 |
| B71 | B72 | TG | HDL-C | -0.41343 | 0 |
| B71 | B77 | TG | TC | 0.255079 | 4.44E-16 |
| B71 | B91 | TG | TP | 0.289772 | 0 |
| B71 | B128 | TG | ALT | 0.186157 | 4.09E-09 |
| B71 | B130 | TG | GLB | 0.213522 | 1.34E-11 |
| B71 | B140 | TG | r-GT | 0.20896 | 3.68E-11 |
| B71 | B141 | TG | AST/ALT | -0.17797 | 1.94E-08 |
| B71 | B153 | TG | Glu | 0.162277 | 3.13E-07 |
| B71 | B192 | TG | UA | 0.231908 | 1.81E-13 |
| B71 | B206 | TG | Waist | 0.262616 | 0 |
| B71 | B210 | TG | Hip | 0.159067 | 5.36E-07 |
| B71 | B215 | TG | BMI | 0.239496 | 2.73E-14 |
| B71 | B216 | TG | BMI | 0.255438 | 4.44E-16 |
| B71 | B218 | TG | DBP | 0.192698 | 1.12E-09 |

| | | | | | |
|---|---|---|---|---|---|
| B71 | B222 | TG | NS | 0.23114 | 2.18E-13 |
| B72 | B74 | HDL-C | ApoA1 | 0.206127 | 6.82E-11 |
| B72 | B77 | HDL-C | TC | 0.243012 | 1.11E-14 |
| B72 | B128 | HDL-C | ALT | -0.2498 | 1.78E-15 |
| B72 | B140 | HDL-C | r-GT | -0.21741 | 5.57E-12 |
| B72 | B141 | HDL-C | AST/ALT | 0.292891 | 0 |
| B72 | B163 | HDL-C | SG | -0.16849 | 1.07E-07 |
| B72 | B190 | HDL-C | Cre | -0.17455 | 3.63E-08 |
| B72 | B192 | HDL-C | UA | -0.34774 | 0 |
| B72 | B206 | HDL-C | Waist | -0.32867 | 0 |
| B72 | B209 | HDL-C | Smoking | -0.1704 | 7.65E-08 |
| B72 | B210 | HDL-C | Hip | -0.24359 | 9.55E-15 |
| B72 | B215 | HDL-C | BMI | -0.31149 | 0 |
| B72 | B216 | HDL-C | BMI | -0.35137 | 0 |
| B72 | B221 | HDL-C | Hei | -0.21381 | 1.26E-11 |
| B72 | B222 | HDL-C | NS | -0.33561 | 0 |
| B73 | B76 | ApoA | ApoB | 0.183858 | 6.37E-09 |
| B73 | B124 | ApoA | ACTH | 0.27259 | 0 |
| B73 | B125 | ApoA | Cor | 0.27259 | 0 |
| B73 | B138 | ApoA | PAB | 0.169019 | 9.77E-08 |
| B73 | B180 | ApoA | Tf | 0.155423 | 9.75E-07 |
| B73 | B196 | ApoA | TCO2 | 0.184564 | 5.56E-09 |
| B74 | B75 | ApoA1 | ApoA1/B | 0.589474 | 0 |
| B74 | B78 | ApoA1 | HBcAb | -0.16355 | 2.52E-07 |
| B74 | B123 | ApoA1 | hs-CRP1 | -0.18111 | 1.07E-08 |
| B74 | B124 | ApoA1 | ACTH | -0.16503 | 1.96E-07 |
| B74 | B125 | ApoA1 | Cor | -0.16503 | 1.96E-07 |
| B74 | B136 | ApoA1 | 5'NT | -0.15615 | 8.67E-07 |
| B74 | B139 | ApoA1 | ADA | -0.18833 | 2.67E-09 |
| B74 | B195 | ApoA1 | Na | 0.208147 | 4.40E-11 |
| B75 | B76 | ApoA1/B | ApoB | -0.74071 | 0 |
| B75 | B124 | ApoA1/B | ACTH | -0.16468 | 2.08E-07 |
| B75 | B125 | ApoA1/B | Cor | -0.16468 | 2.08E-07 |
| B75 | B138 | ApoA1/B | PAB | -0.2266 | 6.51E-13 |
| B76 | B77 | ApoB | TC | 0.213138 | 1.46E-11 |
| B76 | B138 | ApoB | PAB | 0.404497 | 0 |
| B76 | B139 | ApoB | ADA | -0.15664 | 8.00E-07 |
| B77 | B132 | TC | Dbil | -0.19753 | 4.19E-10 |
| B77 | B218 | TC | DBP | 0.159975 | 4.61E-07 |
| B78 | B123 | HBcAb | hs-CRP1 | 0.169737 | 8.61E-08 |
| B78 | B134 | HBcAb | CHE | 0.238824 | 3.24E-14 |
| B78 | B138 | HBcAb | PAB | -0.18571 | 4.46E-09 |
| B78 | B175 | HBcAb | ALB-U | -0.2974 | 0 |
| B78 | B193 | HBcAb | K | -0.19911 | 3.02E-10 |

| | | | | | |
|---|---|---|---|---|---|
| B79 | B95 | hb V -dna | anti-cenpb | 0.215035 | 9.54E-12 |
| B79 | B97 | hb V -dna | dsDNA | 0.215035 | 9.54E-12 |
| B79 | B98 | hb V -dna | anti-Jo-1 | 0.215035 | 9.54E-12 |
| B79 | B101 | hb V -dna | anti-Sc70 | 0.215035 | 9.54E-12 |
| B79 | B102 | hb V -dna | anti-Sm | 0.215035 | 9.54E-12 |
| B79 | B105 | hb V -dna | anti-U1RNP | 0.215035 | 9.54E-12 |
| B79 | B106 | hb V -dna | anti-P | 0.215035 | 9.54E-12 |
| B79 | B107 | hb V -dna | AnuA | 0.215035 | 9.54E-12 |
| B79 | B108 | hb V -dna | Anti-Cyclin | 0.215035 | 9.54E-12 |
| B79 | B109 | hb V -dna | AMA-M2 | 0.215035 | 9.54E-12 |
| B79 | B110 | hb V -dna | AHA | 0.215035 | 9.54E-12 |
| B79 | B114 | hb V -dna | TSH TT3 TT4(TT4) | 0.172125 | 5.63E-08 |
| B79 | B115 | hb V -dna | TSH TT3 TT4(TT3) | 0.155324 | 9.91E-07 |
| B79 | B119 | hb V -dna | Hcv-rna | 0.21504 | 9.53E-12 |
| B79 | B120 | hb V -dna | HCV-cAg | 0.240362 | 2.20E-14 |
| B79 | B156 | hb V -dna | IgE | -0.29079 | 0 |
| B79 | B178 | hb V -dna | Ig-U | 0.226782 | 6.24E-13 |
| B79 | B179 | hb V -dna | α1-MG | 0.263076 | 0 |
| B79 | B180 | hb V -dna | Tf | 0.258 | 2.22E-16 |
| B8 | B28 | WB-Zinc | PRL | 0.991516 | 0 |
| B8 | B30 | WB-Zinc | LH | -0.84269 | 0 |
| B8 | B31 | WB-Zinc | FSH | -0.59145 | 0 |
| B8 | B32 | WB-Zinc | Testo | -0.83448 | 0 |
| B8 | B33 | WB-Zinc | Prog | -0.63526 | 0 |
| B82 | B83 | CEA | AFP | 0.373548 | 0 |
| B82 | B87 | CEA | NSE | 0.317378 | 0 |
| B82 | B88 | CEA | CA 242 | 0.389121 | 0 |
| B82 | B89 | CEA | CA 72-4 | 0.221963 | 1.95E-12 |
| B83 | B84 | AFP | C-PSA | 0.3166 | 0 |
| B83 | B87 | AFP | NSE | 0.382715 | 0 |
| B83 | B88 | AFP | CA 242 | 0.197765 | 3.99E-10 |
| B83 | B89 | AFP | CA 72-4 | 0.423413 | 0 |
| B83 | B103 | AFP | anti-SSA/Ro | -0.1565 | 8.19E-07 |
| B83 | B104 | AFP | anti-SSB | -0.15939 | 5.09E-07 |
| B83 | B152 | AFP | ACA-Ig(G-A-M) | 0.306304 | 0 |
| B84 | B85 | C-PSA | SCC | 0.474324 | 0 |
| B84 | B89 | C-PSA | CA 72-4 | 0.166948 | 1.40E-07 |
| B84 | B90 | C-PSA | CYFRA 21-1 | 0.24454 | 7.55E-15 |
| B84 | B112 | C-PSA | FT4 | 0.156484 | 8.21E-07 |
| B85 | B87 | SCC | NSE | 0.192159 | 1.25E-09 |
| B85 | B88 | SCC | CA 242 | 0.606969 | 0 |
| B86 | B90 | TBhCG | CYFRA 21-1 | 0.459366 | 0 |
| B87 | B88 | NSE | CA 242 | 0.366829 | 0 |
| B87 | B89 | NSE | CA 72-4 | 0.308775 | 0 |

| | | | | | |
|---|---|---|---|---|---|
| B87 | B138 | NSE | PAB | -0.15756 | 6.88E-07 |
| B87 | B139 | NSE | ADA | 0.194811 | 7.31E-10 |
| B87 | B150 | NSE | ALP | 0.208125 | 4.42E-11 |
| B87 | B196 | NSE | TCO2 | 0.232135 | 1.71E-13 |
| B89 | B193 | CA 72-4 | K | 0.164057 | 2.31E-07 |
| B89 | B199 | CA 72-4 | Mg | -0.20145 | 1.85E-10 |
| B9 | B223 | RH | FH | -0.40109 | 0 |
| B91 | B126 | TP | Alb | 0.378881 | 0 |
| B91 | B127 | TP | A-G ratio | -0.55483 | 0 |
| B91 | B130 | TP | GLB | 0.82584 | 0 |
| B91 | B188 | TP | Ca | 0.225773 | 7.94E-13 |
| B92 | B94 | ALDH2-Glu504Glu | ALDH2-Lys504Lys | 0.536323 | 0 |
| B92 | B96 | ALDH2-Glu504Glu | dsDNA (IIF) | -0.27468 | 0 |
| B92 | B116 | ALDH2-Glu504Glu | TT3 | 0.633866 | 0 |
| B92 | B202 | ALDH2-Glu504Glu | e-times | 0.938612 | 0 |
| B93 | B94 | ALDH2-Glu504Lys | ALDH2-Lys504Lys | 0.938466 | 0 |
| B93 | B116 | ALDH2-Glu504Lys | TT3 | 0.525765 | 0 |
| B93 | B167 | ALDH2-Glu504Lys | PC | 0.786527 | 0 |
| B94 | B96 | ALDH2-Lys504Lys | dsDNA (IIF) | 0.745386 | 0 |
| B94 | B167 | ALDH2-Lys504Lys | PC | 0.762338 | 0 |
| B95 | B97 | anti-cenpb | dsDNA | 1 | 0 |
| B95 | B99 | anti-cenpb | anti-PM-Scl | -0.27646 | 0 |
| B95 | B100 | anti-cenpb | anti-Ro52 | -0.34578 | 0 |
| B95 | B103 | anti-cenpb | anti-SSA/Ro | -0.27591 | 0 |
| B95 | B104 | anti-cenpb | anti-SSB | -0.32367 | 0 |
| B95 | B105 | anti-cenpb | anti-U1RNP | 1 | 0 |
| B95 | B107 | anti-cenpb | AnuA | 1 | 0 |
| B95 | B108 | anti-cenpb | Anti-Cyclin | 1 | 0 |
| B95 | B109 | anti-cenpb | AMA-M2 | 1 | 0 |
| B95 | B110 | anti-cenpb | AHA | 1 | 0 |
| B95 | B134 | anti-cenpb | CHE | -0.29695 | 0 |
| B95 | B151 | anti-cenpb | Anti-CCP | -0.23405 | 1.07E-13 |
| B95 | B155 | anti-cenpb | IgA | -0.17398 | 4.03E-08 |
| B95 | B156 | anti-cenpb | IgE | -0.42651 | 0 |
| B95 | B157 | anti-cenpb | IgG | -0.21397 | 1.21E-11 |
| B95 | B175 | anti-cenpb | ALB-U | 0.22405 | 1.19E-12 |
| B95 | B176 | anti-cenpb | Ucr | 0.200525 | 2.24E-10 |
| B95 | B178 | anti-cenpb | Ig-U | 0.771362 | 0 |
| B95 | B179 | anti-cenpb | α1-MG | 0.719949 | 0 |
| B95 | B180 | anti-cenpb | Tf | 0.779882 | 0 |
| B95 | B181 | anti-cenpb | APTT | 0.331973 | 0 |
| B95 | B182 | anti-cenpb | TT | 0.330901 | 0 |
| B95 | B183 | anti-cenpb | PTA | 0.331949 | 0 |
| B95 | B184 | anti-cenpb | INR | 0.331973 | 0 |

| | | | | | |
|---|---|---|---|---|---|
| B95 | B185 | anti-cenpb | PT | 0.331264 | 0 |
| B95 | B186 | anti-cenpb | PFC | -0.33197 | 0 |
| B95 | B194 | anti-cenpb | CI | -0.33683 | 0 |
| B95 | B197 | anti-cenpb | AG | 0.254711 | 4.44E-16 |
| B96 | B116 | dsDNA (IIF) | TT3 | 0.97619 | 0 |
| B97 | B98 | dsDNA | anti-Jo-1 | 1 | 0 |
| B97 | B99 | dsDNA | anti-PM-Scl | -0.27646 | 0 |
| B97 | B100 | dsDNA | anti-Ro52 | -0.34578 | 0 |
| B97 | B102 | dsDNA | anti-Sm | 1 | 0 |
| B97 | B103 | dsDNA | anti-SSA/Ro | -0.27591 | 0 |
| B97 | B104 | dsDNA | anti-SSB | -0.32367 | 0 |
| B97 | B105 | dsDNA | anti-U1RNP | 1 | 0 |
| B97 | B106 | dsDNA | anti-P | 1 | 0 |
| B97 | B107 | dsDNA | AnuA | 1 | 0 |
| B97 | B108 | dsDNA | Anti-Cyclin | 1 | 0 |
| B97 | B109 | dsDNA | AMA-M2 | 1 | 0 |
| B97 | B110 | dsDNA | AHA | 1 | 0 |
| B97 | B134 | dsDNA | CHE | -0.29695 | 0 |
| B97 | B151 | dsDNA | Anti-CCP | -0.23405 | 1.07E-13 |
| B97 | B155 | dsDNA | IgA | -0.17398 | 4.03E-08 |
| B97 | B156 | dsDNA | IgE | -0.42651 | 0 |
| B97 | B157 | dsDNA | IgG | -0.21397 | 1.21E-11 |
| B97 | B175 | dsDNA | ALB-U | 0.22405 | 1.19E-12 |
| B97 | B176 | dsDNA | Ucr | 0.200525 | 2.24E-10 |
| B97 | B178 | dsDNA | Ig-U | 0.771362 | 0 |
| B97 | B179 | dsDNA | α1-MG | 0.719949 | 0 |
| B97 | B180 | dsDNA | Tf | 0.779882 | 0 |
| B97 | B181 | dsDNA | APTT | 0.331973 | 0 |
| B97 | B182 | dsDNA | TT | 0.330901 | 0 |
| B97 | B183 | dsDNA | PTA | 0.331949 | 0 |
| B97 | B184 | dsDNA | INR | 0.331973 | 0 |
| B97 | B185 | dsDNA | PT | 0.331264 | 0 |
| B97 | B186 | dsDNA | PFC | -0.33197 | 0 |
| B97 | B194 | dsDNA | CI | -0.33683 | 0 |
| B97 | B197 | dsDNA | AG | 0.254711 | 4.44E-16 |
| B98 | B99 | anti-Jo-1 | anti-PM-Scl | -0.27646 | 0 |
| B98 | B100 | anti-Jo-1 | anti-Ro52 | -0.34578 | 0 |
| B98 | B101 | anti-Jo-1 | anti-Sc70 | 1 | 0 |
| B98 | B102 | anti-Jo-1 | anti-Sm | 1 | 0 |
| B98 | B103 | anti-Jo-1 | anti-SSA/Ro | -0.27591 | 0 |
| B98 | B104 | anti-Jo-1 | anti-SSB | -0.32367 | 0 |
| B98 | B107 | anti-Jo-1 | AnuA | 1 | 0 |
| B98 | B108 | anti-Jo-1 | Anti-Cyclin | 1 | 0 |
| B98 | B134 | anti-Jo-1 | CHE | -0.29695 | 0 |

| | | | | | |
|---|---|---|---|---|---|
| B98 | B151 | anti-Jo-1 | Anti-CCP | -0.23405 | 1.07E-13 |
| B98 | B155 | anti-Jo-1 | IgA | -0.17398 | 4.03E-08 |
| B98 | B156 | anti-Jo-1 | IgE | -0.42651 | 0 |
| B98 | B157 | anti-Jo-1 | IgG | -0.21397 | 1.21E-11 |
| B98 | B175 | anti-Jo-1 | ALB-U | 0.22405 | 1.19E-12 |
| B98 | B176 | anti-Jo-1 | Ucr | 0.200525 | 2.24E-10 |
| B98 | B178 | anti-Jo-1 | Ig-U | 0.771362 | 0 |
| B98 | B179 | anti-Jo-1 | α1-MG | 0.719949 | 0 |
| B98 | B180 | anti-Jo-1 | Tf | 0.779882 | 0 |
| B98 | B181 | anti-Jo-1 | APTT | 0.331973 | 0 |
| B98 | B182 | anti-Jo-1 | TT | 0.330901 | 0 |
| B98 | B183 | anti-Jo-1 | PTA | 0.331949 | 0 |
| B98 | B184 | anti-Jo-1 | INR | 0.331973 | 0 |
| B98 | B185 | anti-Jo-1 | PT | 0.331264 | 0 |
| B98 | B186 | anti-Jo-1 | PFC | -0.33197 | 0 |
| B98 | B194 | anti-Jo-1 | Cl | -0.33683 | 0 |
| B98 | B197 | anti-Jo-1 | AG | 0.254711 | 4.44E-16 |
| B99 | B101 | anti-PM-Scl | anti-Sc70 | -0.27646 | 0 |
| B99 | B102 | anti-PM-Scl | anti-Sm | -0.27646 | 0 |
| B99 | B105 | anti-PM-Scl | anti-U1RNP | -0.27646 | 0 |
| B99 | B106 | anti-PM-Scl | anti-P | -0.27646 | 0 |
| B99 | B107 | anti-PM-Scl | AnuA | -0.27646 | 0 |
| B99 | B108 | anti-PM-Scl | Anti-Cyclin | -0.27646 | 0 |
| B99 | B109 | anti-PM-Scl | AMA-M2 | -0.27646 | 0 |
| B99 | B110 | anti-PM-Scl | AHA | -0.27646 | 0 |
| B99 | B146 | anti-PM-Scl | CysC | 0.174585 | 3.61E-08 |
| B99 | B151 | anti-PM-Scl | Anti-CCP | -0.51844 | 0 |
| B99 | B157 | anti-PM-Scl | IgG | -0.17765 | 2.05E-08 |
| B99 | B158 | anti-PM-Scl | IgM | -0.23263 | 1.51E-13 |
| B99 | B181 | anti-PM-Scl | APTT | -0.9983 | 0 |
| B99 | B182 | anti-PM-Scl | TT | -0.99275 | 0 |
| B99 | B183 | anti-PM-Scl | PTA | -0.9983 | 0 |
| B99 | B184 | anti-PM-Scl | INR | -0.9983 | 0 |
| B99 | B185 | anti-PM-Scl | PT | -0.99722 | 0 |
| B99 | B186 | anti-PM-Scl | PFC | 0.998302 | 0 |
| B99 | B193 | anti-PM-Scl | K | -0.2897 | 0 |
| B99 | B194 | anti-PM-Scl | Cl | 0.185809 | 4.37E-09 |
| B99 | B198 | anti-PM-Scl | P | 0.364908 | 0 |
| B99 | B199 | anti-PM-Scl | Mg | 0.264999 | 0 |
| B99 | B204 | anti-PM-Scl | Drinking | 0.200102 | 2.45E-10 |

**Supplementary Table 3. PEI Correlations in Hypertension**

| Row | Column | Row | Column | Cor | *P* |
|---|---|---|---|---|---|
| B1 | B111 | Gender | TSH | 0.071717 | 0 |
| B1 | B113 | Gender | FT3 | -0.0512 | 0 |
| B1 | B114 | Gender | TSH TT3 TT4(TT4) | 0.052299 | 0 |
| B1 | B126 | Gender | Alb | -0.07375 | 0 |
| B1 | B127 | Gender | A-G ratio | -0.1529 | 0 |
| B1 | B128 | Gender | ALT | -0.15361 | 0 |
| B1 | B129 | Gender | IBIL | -0.13299 | 0 |
| B1 | B130 | Gender | GLB | 0.138351 | 0 |
| B1 | B131 | Gender | AST | -0.04977 | 0 |
| B1 | B132 | Gender | Dbil | -0.1529 | 0 |
| B1 | B133 | Gender | TBIL | -0.14844 | 0 |
| B1 | B138 | Gender | PAB | -0.08218 | 0 |
| B1 | B139 | Gender | ADA | 0.03961 | 0 |
| B1 | B140 | Gender | r-GT | -0.14838 | 0 |
| B1 | B141 | Gender | AST/ALT | 0.192708 | 0 |
| B1 | B146 | Gender | CysC | -0.04363 | 0 |
| B1 | B148 | Gender | Anti_TPO | 0.065245 | 0 |
| B1 | B149 | Gender | TGAb | 0.056628 | 0 |
| B1 | B153 | Gender | Glu | -0.04835 | 0 |
| B1 | B160 | Gender | PH | 0.065417 | 0 |
| B1 | B161 | Gender | LEU | 0.206085 | 0 |
| B1 | B163 | Gender | SG | -0.38918 | 0 |
| B1 | B166 | Gender | Pro | -0.1573 | 0 |
| B1 | B170 | Gender | U-GLU | -0.05839 | 0 |
| B1 | B173 | Gender | NTT | 0.118155 | 0 |
| B1 | B174 | Gender | Urea/Cr | 0.411039 | 0 |
| B1 | B176 | Gender | Ucr | -0.14529 | 0 |
| B1 | B179 | Gender | α1-MG | -0.04096 | 0 |
| B1 | B188 | Gender | Ca | 0.063179 | 0 |
| B1 | B190 | Gender | Cre | -0.32238 | 0 |
| B1 | B191 | Gender | Urea | -0.06407 | 0 |
| B1 | B192 | Gender | UA | -0.40615 | 0 |
| B1 | B198 | Gender | P | 0.084699 | 0 |
| B1 | B2 | Gender | Age | 0.124492 | 0 |
| B1 | B203 | Gender | DH | -0.04512 | 0 |
| B1 | B204 | Gender | Drinking | -0.32752 | 0 |
| B1 | B205 | Gender | KDA | 0.071334 | 0 |
| B1 | B206 | Gender | Waist | -0.3719 | 0 |
| B1 | B209 | Gender | Smoking | -0.33436 | 0 |
| B1 | B210 | Gender | Hip | -0.24997 | 0 |
| B1 | B215 | Gender | BMI | -0.16036 | 0 |

| | | | | | |
|---|---|---|---|---|---|
| B1 | B216 | Gender | BMI | -0.52624 | 0 |
| B1 | B218 | Gender | DBP | -0.16885 | 0 |
| B1 | B219 | Gender | OT | 0.139931 | 0 |
| B1 | B220 | Gender | SBP | 0.058115 | 0 |
| B1 | B221 | Gender | Hei | -0.6861 | 0 |
| B1 | B222 | Gender | NS | -0.6465 | 0 |
| B1 | B24 | Gender | αHBDH | 0.051592 | 0 |
| B1 | B25 | Gender | CK | -0.04166 | 0 |
| B1 | B27 | Gender | LDH | 0.046952 | 0 |
| B1 | B3 | Gender | Hcy | -0.11243 | 0 |
| B1 | B40 | Gender | WBC | -0.14967 | 0 |
| B1 | B42 | Gender | MON | -0.24501 | 0 |
| B1 | B44 | Gender | MCH | -0.1555 | 0 |
| B1 | B45 | Gender | MCHC | -0.28992 | 0 |
| B1 | B46 | Gender | RBC Count | -0.41655 | 0 |
| B1 | B48 | Gender | LYM% | 0.123838 | 0 |
| B1 | B50 | Gender | eo% | -0.11995 | 0 |
| B1 | B51 | Gender | Hb | -0.54237 | 0 |
| B1 | B52 | Gender | PLT/BPC | 0.100778 | 0 |
| B1 | B53 | Gender | GR% | -0.04932 | 0 |
| B1 | B54 | Gender | ESR | 0.07864 | 0 |
| B1 | B56 | Gender | IR | -0.06884 | 0 |
| B1 | B58 | Gender | HCT | -0.06309 | 0 |
| B1 | B60 | Gender | WBRVI | -0.10287 | 0 |
| B1 | B61 | Gender | WBHSRV | -0.0603 | 0 |
| B1 | B62 | Gender | WBHSRVI | -0.10874 | 0 |
| B1 | B63 | Gender | WBRSR(mPas_200) | -0.09821 | 0 |
| B1 | B64 | Gender | WBRSR(mPas_5) | -0.09895 | 0 |
| B1 | B65 | Gender | WBRSR(mPas_1) | -0.09482 | 0 |
| B1 | B66 | Gender | WBRSR(mPas_100) | -0.09738 | 0 |
| B1 | B67 | Gender | WBRSR(mPas_50) | -0.09846 | 0 |
| B1 | B70 | Gender | LDL-C | 0.071795 | 0 |
| B1 | B71 | Gender | TG | -0.08957 | 0 |
| B1 | B72 | Gender | HDL-C | 0.313334 | 0 |
| B1 | B74 | Gender | ApoA1 | 0.123934 | 0 |
| B1 | B75 | Gender | ApoA1/B | 0.051883 | 0 |
| B1 | B77 | Gender | TC | 0.149732 | 0 |
| B1 | B84 | Gender | C-PSA | -0.04196 | 0 |
| B1 | B86 | Gender | TBhCG | 0.051205 | 0 |
| B1 | B91 | Gender | TP | 0.081199 | 0 |
| B1 | B157 | Gender | IgG | 0.037115 | 8.88E-16 |
| B1 | B134 | Gender | CHE | -0.03668 | 1.78E-15 |
| B1 | B49 | Gender | baso% | -0.0366 | 1.78E-15 |
| B1 | B59 | Gender | WBRV | -0.03546 | 1.38E-14 |

| | | | | | |
|---|---|---|---|---|---|
| B1 | B207 | Gender | WHR | -0.03466 | 5.24E-14 |
| B1 | B201 | Gender | e-habits | -0.03144 | 8.79E-12 |
| B1 | B137 | Gender | AFU | -0.0314 | 9.29E-12 |
| B1 | B85 | Gender | SCC | -0.0314 | 9.34E-12 |
| B1 | B144 | Gender | Total-PINP | 0.030891 | 1.99E-11 |
| B1 | B142 | Gender | 25-(OH)D | 0.030451 | 3.82E-11 |
| B1 | B73 | Gender | ApoA | 0.030437 | 3.90E-11 |
| B1 | B193 | Gender | K | 0.030064 | 6.71E-11 |
| B1 | B57 | Gender | RBCAI | -0.02981 | 9.74E-11 |
| B1 | B189 | Gender | eGFB | 0.029768 | 1.03E-10 |
| B1 | B43 | Gender | MCV | -0.0296 | 1.31E-10 |
| B1 | B31 | Gender | FSH | 0.029463 | 1.59E-10 |
| B1 | B33 | Gender | Prog | -0.02905 | 2.87E-10 |
| B1 | B143 | Gender | β-CTx | 0.027973 | 1.26E-09 |
| B1 | B69 | Gender | PV | 0.02788 | 1.43E-09 |
| B1 | B150 | Gender | ALP | 0.027667 | 1.90E-09 |
| B1 | B82 | Gender | CEA | -0.02543 | 3.40E-08 |
| B1 | B158 | Gender | IgM | 0.025113 | 4.99E-08 |
| B1 | B30 | Gender | LH | 0.024373 | 1.22E-07 |
| B1 | B29 | Gender | E2 | -0.02293 | 6.43E-07 |
| B1 | B172 | Gender | KB | -0.02257 | 9.60E-07 |
| B10 | B107 | PG I | AnuA | -0.06445 | 0 |
| B10 | B11 | PG I | PGI/PGII | 0.095873 | 0 |
| B10 | B119 | PG I | Hcv-rna | -0.05147 | 0 |
| B10 | B12 | PG I | PG II | 0.554769 | 0 |
| B10 | B126 | PG I | Alb | -0.03961 | 0 |
| B10 | B14 | PG I | G-17 | 0.05082 | 0 |
| B10 | B146 | PG I | CysC | 0.07041 | 0 |
| B10 | B15 | PG I | CD3-CD19+(B cell) | -0.04705 | 0 |
| B10 | B159 | PG I | β2-MG | 0.038497 | 0 |
| B10 | B175 | PG I | ALB-U | 0.057422 | 0 |
| B10 | B177 | PG I | ACR | 0.077714 | 0 |
| B10 | B181 | PG I | APTT | -0.04071 | 0 |
| B10 | B189 | PG I | eGFB | -0.09737 | 0 |
| B10 | B190 | PG I | Cre | 0.093365 | 0 |
| B10 | B191 | PG I | Urea | 0.085856 | 0 |
| B10 | B35 | PG I | β2-MG | 0.062654 | 0 |
| B10 | B79 | PG I | hb V -dna | -0.06371 | 0 |
| B10 | B80 | PG I | anti-HP IgG | 0.052653 | 0 |
| B10 | B183 | PG I | PTA | -0.03757 | 4.44E-16 |
| B10 | B186 | PG I | PFC | 0.037075 | 8.88E-16 |
| B10 | B184 | PG I | INR | -0.03128 | 1.12E-11 |
| B10 | B192 | PG I | UA | 0.028985 | 3.12E-10 |
| B10 | B47 | PG I | RDW | 0.028891 | 3.56E-10 |

| | | | | | |
|---|---|---|---|---|---|
| B10 | B215 | PG I | BMI | -0.02878 | 4.14E-10 |
| B10 | B200 | PG I | OP | 0.027843 | 1.50E-09 |
| B10 | B210 | PG I | Hip | -0.02655 | 8.27E-09 |
| B10 | B133 | PG I | TBIL | -0.02554 | 2.94E-08 |
| B10 | B129 | PG I | IBIL | -0.02492 | 6.32E-08 |
| B10 | B198 | PG I | P | 0.02301 | 5.88E-07 |
| B100 | B103 | anti-Ro52 | anti-SSA/Ro | 0.387877 | 0 |
| B100 | B104 | anti-Ro52 | anti-SSB | 0.329561 | 0 |
| B100 | B105 | anti-Ro52 | anti-U1RNP | 0.07212 | 0 |
| B100 | B109 | anti-Ro52 | AMA-M2 | 0.062129 | 0 |
| B100 | B157 | anti-Ro52 | IgG | 0.052587 | 0 |
| B100 | B155 | anti-Ro52 | IgA | 0.035914 | 6.22E-15 |
| B100 | B139 | anti-Ro52 | ADA | 0.032244 | 2.55E-12 |
| B100 | B138 | anti-Ro52 | PAB | -0.02619 | 1.30E-08 |
| B100 | B195 | anti-Ro52 | Na | -0.02473 | 7.94E-08 |
| B100 | B114 | anti-Ro52 | TSH TT3 TT4(TT4) | 0.024251 | 1.41E-07 |
| B100 | B182 | anti-Ro52 | TT | 0.023591 | 3.03E-07 |
| B100 | B199 | anti-Ro52 | Mg | -0.02331 | 4.19E-07 |
| B100 | B119 | anti-Ro52 | Hcv-rna | 0.0228 | 7.44E-07 |
| B101 | B103 | anti-Sc70 | anti-SSA/Ro | 0.241719 | 0 |
| B101 | B108 | anti-Sc70 | Anti-Cyclin | 0.034201 | 1.12E-13 |
| B102 | B105 | anti-Sm | anti-U1RNP | 0.340392 | 0 |
| B102 | B109 | anti-Sm | AMA-M2 | 0.238616 | 0 |
| B102 | B182 | anti-Sm | TT | -0.0466 | 0 |
| B102 | B186 | anti-Sm | PFC | 0.051299 | 0 |
| B102 | B199 | anti-Sm | Mg | -0.0379 | 2.22E-16 |
| B102 | B197 | anti-Sm | AG | -0.03514 | 2.35E-14 |
| B102 | B107 | anti-Sm | AnuA | -0.03508 | 2.60E-14 |
| B102 | B183 | anti-Sm | PTA | -0.02779 | 1.61E-09 |
| B102 | B134 | anti-Sm | CHE | -0.02434 | 1.27E-07 |
| B103 | B104 | anti-SSA/Ro | anti-SSB | 0.327209 | 0 |
| B103 | B105 | anti-SSA/Ro | anti-U1RNP | 0.08675 | 0 |
| B103 | B109 | anti-SSA/Ro | AMA-M2 | 0.074334 | 0 |
| B103 | B110 | anti-SSA/Ro | AHA | 0.12884 | 0 |
| B103 | B155 | anti-SSA/Ro | IgA | 0.041149 | 0 |
| B103 | B157 | anti-SSA/Ro | IgG | 0.075154 | 0 |
| B103 | B138 | anti-SSA/Ro | PAB | -0.03122 | 1.21E-11 |
| B103 | B139 | anti-SSA/Ro | ADA | 0.03038 | 4.24E-11 |
| B103 | B119 | anti-SSA/Ro | Hcv-rna | 0.026591 | 7.80E-09 |
| B103 | B158 | anti-SSA/Ro | IgM | 0.02415 | 1.58E-07 |
| B103 | B107 | anti-SSA/Ro | AnuA | -0.02383 | 2.30E-07 |
| B103 | B127 | anti-SSA/Ro | A-G ratio | -0.02347 | 3.49E-07 |
| B103 | B130 | anti-SSA/Ro | GLB | 0.023443 | 3.60E-07 |
| B104 | B109 | anti-SSB | AMA-M2 | 0.038708 | 0 |

| | | | | | |
|---|---|---|---|---|---|
| B104 | B155 | anti-SSB | IgA | 0.047448 | 0 |
| B104 | B157 | anti-SSB | IgG | 0.11359 | 0 |
| B104 | B158 | anti-SSB | IgM | 0.044152 | 0 |
| B104 | B107 | anti-SSB | AnuA | -0.0374 | 4.44E-16 |
| B104 | B195 | anti-SSB | Na | -0.03349 | 3.56E-13 |
| B104 | B136 | anti-SSB | 5'NT | 0.028851 | 3.77E-10 |
| B105 | B107 | anti-U1RNP | AnuA | -0.06863 | 0 |
| B105 | B109 | anti-U1RNP | AMA-M2 | 0.056055 | 0 |
| B105 | B110 | anti-U1RNP | AHA | 0.073356 | 0 |
| B105 | B151 | anti-U1RNP | Anti-CCP | 0.138415 | 0 |
| B105 | B182 | anti-U1RNP | TT | -0.03901 | 0 |
| B105 | B186 | anti-U1RNP | PFC | 0.038954 | 0 |
| B105 | B199 | anti-U1RNP | Mg | -0.03869 | 0 |
| B105 | B135 | anti-U1RNP | TBA | 0.035904 | 6.44E-15 |
| B105 | B125 | anti-U1RNP | Cor | -0.02288 | 6.78E-07 |
| B106 | B110 | anti-P | AHA | 0.034711 | 4.84E-14 |
| B106 | B212 | anti-P | IOP-L | 0.02912 | 2.59E-10 |
| B106 | B170 | anti-P | U-GLU | 0.027627 | 2.00E-09 |
| B106 | B179 | anti-P | α1-MG | 0.024371 | 1.22E-07 |
| B106 | B193 | anti-P | K | -0.02363 | 2.91E-07 |
| B106 | B178 | anti-P | Ig-U | 0.022658 | 8.72E-07 |
| B107 | B119 | AnuA | Hcv-rna | 0.139827 | 0 |
| B107 | B125 | AnuA | Cor | -0.03926 | 0 |
| B107 | B134 | AnuA | CHE | -0.05036 | 0 |
| B107 | B138 | AnuA | PAB | -0.06145 | 0 |
| B107 | B150 | AnuA | ALP | -0.0381 | 0 |
| B107 | B151 | AnuA | Anti-CCP | -0.05379 | 0 |
| B107 | B155 | AnuA | IgA | -0.05318 | 0 |
| B107 | B158 | AnuA | IgM | 0.039338 | 0 |
| B107 | B178 | AnuA | Ig-U | -0.03961 | 0 |
| B107 | B181 | AnuA | APTT | 0.106004 | 0 |
| B107 | B183 | AnuA | PTA | 0.115774 | 0 |
| B107 | B184 | AnuA | INR | 0.130397 | 0 |
| B107 | B185 | AnuA | PT | 0.044087 | 0 |
| B107 | B186 | AnuA | PFC | -0.13112 | 0 |
| B107 | B189 | AnuA | eGFB | -0.06885 | 0 |
| B107 | B193 | AnuA | K | 0.125729 | 0 |
| B107 | B194 | AnuA | Cl | 0.05476 | 0 |
| B107 | B196 | AnuA | TCO2 | -0.04576 | 0 |
| B107 | B199 | AnuA | Mg | -0.04411 | 0 |
| B107 | B152 | AnuA | ACA-Ig(G-A-M) | -0.03781 | 2.22E-16 |
| B107 | B179 | AnuA | α1-MG | -0.03691 | 1.11E-15 |
| B107 | B123 | AnuA | hs-CRP1 | -0.03361 | 2.95E-13 |
| B107 | B157 | AnuA | IgG | -0.03245 | 1.85E-12 |

| | | | | | |
|---|---|---|---|---|---|
| B107 | B218 | AnuA | DBP | -0.0303 | 4.77E-11 |
| B107 | B212 | AnuA | IOP-L | -0.02998 | 7.56E-11 |
| B107 | B195 | AnuA | Na | -0.0295 | 1.52E-10 |
| B107 | B122 | AnuA | C4 | -0.02714 | 3.81E-09 |
| B107 | B108 | AnuA | Anti-Cyclin | 0.026552 | 8.21E-09 |
| B107 | B204 | AnuA | Drinking | -0.02569 | 2.45E-08 |
| B107 | B201 | AnuA | e-habits | 0.02462 | 9.06E-08 |
| B107 | B198 | AnuA | P | 0.023786 | 2.42E-07 |
| B107 | B210 | AnuA | Hip | -0.02279 | 7.50E-07 |
| B107 | B190 | AnuA | Cre | 0.022691 | 8.40E-07 |
| B107 | B174 | AnuA | Urea/Cr | -0.02263 | 9.01E-07 |
| B108 | B123 | Anti-Cyclin | hs-CRP1 | 0.075708 | 0 |
| B108 | B176 | Anti-Cyclin | Ucr | 0.045386 | 0 |
| B108 | B181 | Anti-Cyclin | APTT | 0.039287 | 0 |
| B108 | B135 | Anti-Cyclin | TBA | 0.031622 | 6.63E-12 |
| B108 | B179 | Anti-Cyclin | α1-MG | -0.0256 | 2.72E-08 |
| B109 | B158 | AMA-M2 | IgM | 0.056779 | 0 |
| B109 | B199 | AMA-M2 | Mg | -0.04527 | 0 |
| B109 | B180 | AMA-M2 | Tf | 0.037918 | 2.22E-16 |
| B109 | B135 | AMA-M2 | TBA | 0.035508 | 1.27E-14 |
| B109 | B182 | AMA-M2 | TT | -0.03492 | 3.42E-14 |
| B109 | B155 | AMA-M2 | IgA | 0.024878 | 6.64E-08 |
| B109 | B195 | AMA-M2 | Na | -0.02437 | 1.22E-07 |
| B11 | B12 | PGI/PGII | PG II | -0.43774 | 0 |
| B11 | B14 | PGI/PGII | G-17 | -0.18533 | 0 |
| B11 | B175 | PGI/PGII | ALB-U | 0.054278 | 0 |
| B11 | B177 | PGI/PGII | ACR | 0.058894 | 0 |
| B11 | B79 | PGI/PGII | hb V -dna | 0.056905 | 0 |
| B11 | B80 | PGI/PGII | anti-HP IgG | -0.10986 | 0 |
| B11 | B139 | PGI/PGII | ADA | -0.03308 | 6.91E-13 |
| B11 | B127 | PGI/PGII | A-G ratio | 0.030782 | 2.34E-11 |
| B11 | B107 | PGI/PGII | AnuA | 0.030388 | 4.19E-11 |
| B11 | B47 | PGI/PGII | RDW | -0.02995 | 7.87E-11 |
| B11 | B146 | PGI/PGII | CysC | -0.02778 | 1.64E-09 |
| B11 | B153 | PGI/PGII | Glu | 0.026888 | 5.31E-09 |
| B11 | B183 | PGI/PGII | PTA | 0.026882 | 5.36E-09 |
| B11 | B166 | PGI/PGII | Pro | 0.026219 | 1.26E-08 |
| B11 | B78 | PGI/PGII | HBcAb | 0.025332 | 3.81E-08 |
| B11 | B137 | PGI/PGII | AFU | -0.02502 | 5.61E-08 |
| B11 | B24 | PGI/PGII | αHBDH | -0.02392 | 2.07E-07 |
| B11 | B130 | PGI/PGII | GLB | -0.02355 | 3.17E-07 |
| B11 | B201 | PGI/PGII | e-habits | -0.02342 | 3.70E-07 |
| B11 | B189 | PGI/PGII | eGFB | 0.023366 | 3.93E-07 |
| B11 | B126 | PGI/PGII | Alb | 0.023174 | 4.89E-07 |

| | | | | | |
|---|---|---|---|---|---|
| B11 | B182 | PGI/PGII | TT | -0.02271 | 8.27E-07 |
| B110 | B122 | AHA | C4 | 0.056661 | 0 |
| B110 | B138 | AHA | PAB | -0.03524 | 2.00E-14 |
| B110 | B182 | AHA | TT | -0.03475 | 4.55E-14 |
| B110 | B139 | AHA | ADA | 0.026786 | 6.06E-09 |
| B110 | B155 | AHA | IgA | 0.025445 | 3.32E-08 |
| B110 | B137 | AHA | AFU | -0.02455 | 9.81E-08 |
| B111 | B112 | TSH | FT4 | -0.21359 | 0 |
| B111 | B113 | TSH | FT3 | -0.09914 | 0 |
| B111 | B114 | TSH | TSH TT3 TT4(TT4) | -0.10954 | 0 |
| B111 | B115 | TSH | TSH TT3 TT4(TT3) | -0.04984 | 0 |
| B111 | B127 | TSH | A-G ratio | -0.04218 | 0 |
| B111 | B130 | TSH | GLB | 0.051328 | 0 |
| B111 | B148 | TSH | Anti_TPO | 0.205116 | 0 |
| B111 | B149 | TSH | TGAb | 0.186666 | 0 |
| B111 | B163 | TSH | SG | -0.05667 | 0 |
| B111 | B204 | TSH | Drinking | -0.0383 | 0 |
| B111 | B209 | TSH | Smoking | -0.04892 | 0 |
| B111 | B221 | TSH | Hei | -0.05542 | 0 |
| B111 | B176 | TSH | Ucr | -0.03703 | 8.88E-16 |
| B111 | B189 | TSH | eGFB | -0.03648 | 2.22E-15 |
| B111 | B222 | TSH | NS | -0.03588 | 6.66E-15 |
| B111 | B216 | TSH | BMI | -0.03027 | 4.95E-11 |
| B111 | B220 | TSH | SBP | 0.029313 | 1.97E-10 |
| B111 | B121 | TSH | C3 | 0.029305 | 1.99E-10 |
| B111 | B157 | TSH | IgG | 0.025477 | 3.19E-08 |
| B111 | B219 | TSH | OT | 0.023544 | 3.20E-07 |
| B112 | B113 | FT4 | FT3 | 0.273486 | 0 |
| B112 | B114 | FT4 | TSH TT3 TT4(TT4) | 0.487362 | 0 |
| B112 | B115 | FT4 | TSH TT3 TT4(TT3) | 0.173978 | 0 |
| B112 | B125 | FT4 | Cor | 0.049188 | 0 |
| B112 | B126 | FT4 | Alb | 0.044009 | 0 |
| B112 | B129 | FT4 | IBIL | 0.045532 | 0 |
| B112 | B133 | FT4 | TBIL | 0.044092 | 0 |
| B112 | B206 | FT4 | Waist | -0.04742 | 0 |
| B112 | B210 | FT4 | Hip | -0.04218 | 0 |
| B112 | B215 | FT4 | BMI | -0.05695 | 0 |
| B112 | B216 | FT4 | BMI | -0.05216 | 0 |
| B112 | B222 | FT4 | NS | -0.03918 | 0 |
| B112 | B204 | FT4 | Drinking | -0.03541 | 1.49E-14 |
| B112 | B192 | FT4 | UA | -0.0343 | 9.46E-14 |
| B112 | B189 | FT4 | eGFB | 0.034107 | 1.31E-13 |
| B112 | B187 | FT4 | NSE | 0.029288 | 2.04E-10 |
| B112 | B138 | FT4 | PAB | -0.02867 | 4.84E-10 |

| | | | | | |
|---|---|---|---|---|---|
| B112 | B132 | FT4 | Dbil | 0.027395 | 2.73E-09 |
| B112 | B139 | FT4 | ADA | 0.027216 | 3.45E-09 |
| B112 | B178 | FT4 | Ig-U | 0.026888 | 5.31E-09 |
| B112 | B149 | FT4 | TGAb | -0.02645 | 9.38E-09 |
| B112 | B153 | FT4 | Glu | 0.026426 | 9.65E-09 |
| B112 | B194 | FT4 | Cl | -0.02603 | 1.60E-08 |
| B112 | B188 | FT4 | Ca | 0.026005 | 1.65E-08 |
| B112 | B141 | FT4 | AST/ALT | 0.025541 | 2.95E-08 |
| B112 | B150 | FT4 | ALP | 0.025357 | 3.70E-08 |
| B112 | B140 | FT4 | r-GT | -0.02469 | 8.32E-08 |
| B112 | B180 | FT4 | Tf | 0.024077 | 1.73E-07 |
| B112 | B181 | FT4 | APTT | -0.0237 | 2.68E-07 |
| B112 | B182 | FT4 | TT | -0.02259 | 9.36E-07 |
| B113 | B114 | FT3 | TSH TT3 TT4(TT4) | 0.175276 | 0 |
| B113 | B115 | FT3 | TSH TT3 TT4(TT3) | 0.402342 | 0 |
| B113 | B123 | FT3 | hs-CRP1 | -0.03959 | 0 |
| B113 | B126 | FT3 | Alb | 0.072542 | 0 |
| B113 | B128 | FT3 | ALT | 0.060341 | 0 |
| B113 | B134 | FT3 | CHE | 0.069704 | 0 |
| B113 | B141 | FT3 | AST/ALT | -0.05917 | 0 |
| B113 | B163 | FT3 | SG | 0.042578 | 0 |
| B113 | B189 | FT3 | eGFB | 0.072851 | 0 |
| B113 | B216 | FT3 | BMI | 0.049145 | 0 |
| B113 | B218 | FT3 | DBP | 0.048315 | 0 |
| B113 | B221 | FT3 | Hei | 0.038869 | 0 |
| B113 | B222 | FT3 | NS | 0.048072 | 0 |
| B113 | B215 | FT3 | BMI | 0.036223 | 3.55E-15 |
| B113 | B206 | FT3 | Waist | 0.036194 | 4.00E-15 |
| B113 | B188 | FT3 | Ca | 0.035556 | 1.15E-14 |
| B113 | B131 | FT3 | AST | 0.034691 | 5.00E-14 |
| B113 | B176 | FT3 | Ucr | 0.03428 | 9.86E-14 |
| B113 | B210 | FT3 | Hip | 0.033914 | 1.79E-13 |
| B113 | B119 | FT3 | Hcv-rna | -0.0319 | 4.31E-12 |
| B113 | B137 | FT3 | AFU | 0.031692 | 5.96E-12 |
| B113 | B127 | FT3 | A-G ratio | 0.031539 | 7.53E-12 |
| B113 | B209 | FT3 | Smoking | 0.030219 | 5.36E-11 |
| B113 | B191 | FT3 | Urea | -0.02909 | 2.68E-10 |
| B113 | B190 | FT3 | Cre | -0.0288 | 4.04E-10 |
| B113 | B219 | FT3 | OT | -0.02868 | 4.76E-10 |
| B113 | B197 | FT3 | AG | 0.028251 | 8.62E-10 |
| B113 | B140 | FT3 | r-GT | 0.027917 | 1.36E-09 |
| B113 | B125 | FT3 | Cor | 0.024964 | 5.98E-08 |
| B113 | B138 | FT3 | PAB | 0.023829 | 2.30E-07 |
| B113 | B187 | FT3 | NSE | 0.02339 | 3.82E-07 |

| | | | | | |
|---|---|---|---|---|---|
| B113 | B194 | FT3 | Cl | -0.02298 | 6.08E-07 |
| B114 | B115 | TSH TT3 TT4(TT4) | TSH TT3 TT4(TT3) | 0.425803 | 0 |
| B114 | B138 | TSH TT3 TT4(TT4) | PAB | -0.07801 | 0 |
| B114 | B139 | TSH TT3 TT4(TT4) | ADA | 0.071701 | 0 |
| B114 | B150 | TSH TT3 TT4(TT4) | ALP | 0.040207 | 0 |
| B114 | B187 | TSH TT3 TT4(TT4) | NSE | 0.039982 | 0 |
| B114 | B192 | TSH TT3 TT4(TT4) | UA | -0.04623 | 0 |
| B114 | B194 | TSH TT3 TT4(TT4) | Cl | -0.04058 | 0 |
| B114 | B204 | TSH TT3 TT4(TT4) | Drinking | -0.0482 | 0 |
| B114 | B216 | TSH TT3 TT4(TT4) | BMI | -0.04677 | 0 |
| B114 | B221 | TSH TT3 TT4(TT4) | Hei | -0.04003 | 0 |
| B114 | B222 | TSH TT3 TT4(TT4) | NS | -0.04992 | 0 |
| B114 | B198 | TSH TT3 TT4(TT4) | P | 0.035952 | 5.77E-15 |
| B114 | B186 | TSH TT3 TT4(TT4) | PFC | 0.035648 | 9.99E-15 |
| B114 | B206 | TSH TT3 TT4(TT4) | Waist | -0.03558 | 1.11E-14 |
| B114 | B215 | TSH TT3 TT4(TT4) | BMI | -0.03072 | 2.58E-11 |
| B114 | B149 | TSH TT3 TT4(TT4) | TGAb | -0.02951 | 1.48E-10 |
| B114 | B178 | TSH TT3 TT4(TT4) | Ig-U | 0.026276 | 1.17E-08 |
| B114 | B130 | TSH TT3 TT4(TT4) | GLB | 0.026204 | 1.28E-08 |
| B114 | B140 | TSH TT3 TT4(TT4) | r-GT | -0.02616 | 1.36E-08 |
| B114 | B188 | TSH TT3 TT4(TT4) | Ca | 0.025904 | 1.87E-08 |
| B114 | B163 | TSH TT3 TT4(TT4) | SG | -0.02574 | 2.29E-08 |
| B114 | B197 | TSH TT3 TT4(TT4) | AG | 0.025573 | 2.83E-08 |
| B114 | B180 | TSH TT3 TT4(TT4) | Tf | 0.025213 | 4.42E-08 |
| B115 | B124 | TSH TT3 TT4(TT3) | ACTH | 0.041197 | 0 |
| B115 | B126 | TSH TT3 TT4(TT3) | Alb | 0.055101 | 0 |
| B115 | B128 | TSH TT3 TT4(TT3) | ALT | 0.054999 | 0 |
| B115 | B134 | TSH TT3 TT4(TT3) | CHE | 0.097409 | 0 |
| B115 | B137 | TSH TT3 TT4(TT3) | AFU | 0.052451 | 0 |
| B115 | B141 | TSH TT3 TT4(TT3) | AST/ALT | -0.04713 | 0 |
| B115 | B188 | TSH TT3 TT4(TT3) | Ca | 0.043938 | 0 |
| B115 | B189 | TSH TT3 TT4(TT3) | eGFB | 0.041813 | 0 |
| B115 | B215 | TSH TT3 TT4(TT3) | BMI | 0.036228 | 3.55E-15 |
| B115 | B216 | TSH TT3 TT4(TT3) | BMI | 0.035081 | 2.60E-14 |
| B115 | B125 | TSH TT3 TT4(TT3) | Cor | 0.034855 | 3.80E-14 |
| B115 | B131 | TSH TT3 TT4(TT3) | AST | 0.032543 | 1.60E-12 |
| B115 | B149 | TSH TT3 TT4(TT3) | TGAb | -0.03242 | 1.94E-12 |
| B115 | B210 | TSH TT3 TT4(TT3) | Hip | 0.031705 | 5.85E-12 |
| B115 | B206 | TSH TT3 TT4(TT3) | Waist | 0.031623 | 6.62E-12 |
| B115 | B187 | TSH TT3 TT4(TT3) | NSE | 0.031457 | 8.52E-12 |
| B115 | B218 | TSH TT3 TT4(TT3) | DBP | 0.029601 | 1.31E-10 |
| B115 | B139 | TSH TT3 TT4(TT3) | ADA | 0.02915 | 2.48E-10 |
| B115 | B222 | TSH TT3 TT4(TT3) | NS | 0.028447 | 6.59E-10 |
| B115 | B181 | TSH TT3 TT4(TT3) | APTT | -0.0265 | 8.75E-09 |

| | | | | | |
|---|---|---|---|---|---|
| B115 | B212 | TSH TT3 TT4(TT3) | IOP-L | 0.026433 | 9.57E-09 |
| B115 | B132 | TSH TT3 TT4(TT3) | Dbil | -0.02519 | 4.54E-08 |
| B115 | B211 | TSH TT3 TT4(TT3) | IOP-R | 0.02513 | 4.89E-08 |
| B115 | B191 | TSH TT3 TT4(TT3) | Urea | -0.02444 | 1.12E-07 |
| B115 | B150 | TSH TT3 TT4(TT3) | ALP | 0.024282 | 1.36E-07 |
| B115 | B133 | TSH TT3 TT4(TT3) | TBIL | -0.02421 | 1.47E-07 |
| B115 | B190 | TSH TT3 TT4(TT3) | Cre | -0.02337 | 3.91E-07 |
| B115 | B178 | TSH TT3 TT4(TT3) | Ig-U | 0.023132 | 5.12E-07 |
| B116 | B117 | TT3 | TT4 | 0.482667 | 0 |
| B119 | B121 | Hcv-rna | C3 | -0.11422 | 0 |
| B119 | B122 | Hcv-rna | C4 | -0.07205 | 0 |
| B119 | B138 | Hcv-rna | PAB | -0.0687 | 0 |
| B119 | B150 | Hcv-rna | ALP | -0.03879 | 0 |
| B119 | B181 | Hcv-rna | APTT | 0.066776 | 0 |
| B119 | B183 | Hcv-rna | PTA | 0.080739 | 0 |
| B119 | B184 | Hcv-rna | INR | 0.071933 | 0 |
| B119 | B185 | Hcv-rna | PT | 0.039204 | 0 |
| B119 | B186 | Hcv-rna | PFC | -0.06474 | 0 |
| B119 | B189 | Hcv-rna | eGFB | -0.04576 | 0 |
| B119 | B193 | Hcv-rna | K | 0.060676 | 0 |
| B119 | B194 | Hcv-rna | Cl | 0.081126 | 0 |
| B119 | B197 | Hcv-rna | AG | -0.06115 | 0 |
| B119 | B199 | Hcv-rna | Mg | -0.05295 | 0 |
| B119 | B222 | Hcv-rna | NS | -0.03932 | 0 |
| B119 | B134 | Hcv-rna | CHE | -0.03599 | 5.55E-15 |
| B119 | B125 | Hcv-rna | Cor | -0.03217 | 2.87E-12 |
| B119 | B218 | Hcv-rna | DBP | -0.03211 | 3.12E-12 |
| B119 | B158 | Hcv-rna | IgM | 0.03017 | 5.76E-11 |
| B119 | B204 | Hcv-rna | Drinking | -0.0274 | 2.71E-09 |
| B119 | B163 | Hcv-rna | SG | -0.02651 | 8.69E-09 |
| B119 | B216 | Hcv-rna | BMI | -0.02497 | 5.96E-08 |
| B119 | B221 | Hcv-rna | Hei | -0.02414 | 1.61E-07 |
| B119 | B210 | Hcv-rna | Hip | -0.02358 | 3.09E-07 |
| B119 | B146 | Hcv-rna | CysC | -0.02322 | 4.65E-07 |
| B119 | B212 | Hcv-rna | IOP-L | -0.02301 | 5.91E-07 |
| B119 | B198 | Hcv-rna | P | 0.022751 | 7.86E-07 |
| B12 | B107 | PG II | AnuA | -0.0651 | 0 |
| B12 | B126 | PG II | Alb | -0.04298 | 0 |
| B12 | B14 | PG II | G-17 | 0.172843 | 0 |
| B12 | B146 | PG II | CysC | 0.05754 | 0 |
| B12 | B15 | PG II | CD3-CD19+(B cell) | -0.05886 | 0 |
| B12 | B183 | PG II | PTA | -0.04685 | 0 |
| B12 | B186 | PG II | PFC | 0.068336 | 0 |
| B12 | B189 | PG II | eGFB | -0.0805 | 0 |

| | | | | | |
|---|---|---|---|---|---|
| B12 | B190 | PG II | Cre | 0.054095 | 0 |
| B12 | B191 | PG II | Urea | 0.055962 | 0 |
| B12 | B79 | PG II | hb V -dna | -0.07345 | 0 |
| B12 | B80 | PG II | anti-HP IgG | 0.108533 | 0 |
| B12 | B93 | PG II | ALDH2-Glu504Lys | 0.038985 | 0 |
| B12 | B99 | PG II | anti-PM-Scl | 0.040831 | 0 |
| B12 | B92 | PG II | ALDH2-Glu504Glu | -0.03682 | 1.33E-15 |
| B12 | B47 | PG II | RDW | 0.036209 | 3.77E-15 |
| B12 | B35 | PG II | β2-MG | 0.036192 | 4.00E-15 |
| B12 | B127 | PG II | A-G ratio | -0.03572 | 8.88E-15 |
| B12 | B199 | PG II | Mg | 0.033993 | 1.58E-13 |
| B12 | B184 | PG II | INR | -0.03086 | 2.09E-11 |
| B12 | B216 | PG II | BMI | -0.03046 | 3.77E-11 |
| B12 | B119 | PG II | Hcv-rna | -0.02933 | 1.91E-10 |
| B12 | B90 | PG II | CYFRA 21-1 | 0.029148 | 2.48E-10 |
| B12 | B215 | PG II | BMI | -0.02895 | 3.26E-10 |
| B12 | B197 | PG II | AG | 0.027634 | 1.98E-09 |
| B12 | B78 | PG II | HBcAb | -0.02755 | 2.22E-09 |
| B12 | B129 | PG II | IBIL | -0.02717 | 3.67E-09 |
| B12 | B195 | PG II | Na | 0.025842 | 2.02E-08 |
| B12 | B200 | PG II | OP | 0.025181 | 4.59E-08 |
| B12 | B210 | PG II | Hip | -0.02477 | 7.55E-08 |
| B12 | B133 | PG II | TBIL | -0.02469 | 8.35E-08 |
| B12 | B130 | PG II | GLB | 0.024575 | 9.56E-08 |
| B12 | B139 | PG II | ADA | 0.023653 | 2.83E-07 |
| B12 | B182 | PG II | TT | -0.02263 | 8.95E-07 |
| B120 | B158 | HCV-cAg | IgM | 0.045093 | 0 |
| B120 | B182 | HCV-cAg | TT | 0.042333 | 0 |
| B120 | B186 | HCV-cAg | PFC | 0.048963 | 0 |
| B121 | B122 | C3 | C4 | 0.509212 | 0 |
| B121 | B156 | C3 | IgE | 0.053691 | 0 |
| B121 | B195 | C3 | Na | -0.04185 | 0 |
| B121 | B155 | C3 | IgA | 0.037132 | 8.88E-16 |
| B121 | B157 | C3 | IgG | 0.03173 | 5.62E-12 |
| B121 | B196 | C3 | TCO2 | -0.03102 | 1.65E-11 |
| B121 | B194 | C3 | Cl | -0.02968 | 1.17E-10 |
| B121 | B200 | C3 | OP | -0.02826 | 8.54E-10 |
| B121 | B141 | C3 | AST/ALT | 0.027188 | 3.59E-09 |
| B121 | B197 | C3 | AG | 0.024282 | 1.36E-07 |
| B122 | B156 | C4 | IgE | 0.047871 | 0 |
| B123 | B126 | hs-CRP1 | Alb | -0.07371 | 0 |
| B123 | B127 | hs-CRP1 | A-G ratio | -0.06077 | 0 |
| B123 | B130 | hs-CRP1 | GLB | 0.04455 | 0 |
| B123 | B138 | hs-CRP1 | PAB | -0.04328 | 0 |

| | | | | | |
|---|---|---|---|---|---|
| B123 | B176 | hs-CRP1 | Ucr | 0.046302 | 0 |
| B123 | B179 | hs-CRP1 | α1-MG | 0.045318 | 0 |
| B123 | B186 | hs-CRP1 | PFC | 0.076071 | 0 |
| B123 | B190 | hs-CRP1 | Cre | 0.037079 | 8.88E-16 |
| B123 | B206 | hs-CRP1 | Waist | 0.036952 | 8.88E-16 |
| B123 | B215 | hs-CRP1 | BMI | 0.035773 | 7.99E-15 |
| B123 | B175 | hs-CRP1 | ALB-U | 0.033074 | 6.94E-13 |
| B123 | B177 | hs-CRP1 | ACR | 0.030648 | 2.86E-11 |
| B123 | B150 | hs-CRP1 | ALP | 0.029626 | 1.26E-10 |
| B123 | B192 | hs-CRP1 | UA | 0.029548 | 1.41E-10 |
| B123 | B140 | hs-CRP1 | r-GT | 0.029345 | 1.88E-10 |
| B123 | B191 | hs-CRP1 | Urea | 0.027293 | 3.12E-09 |
| B123 | B216 | hs-CRP1 | BMI | 0.025886 | 1.92E-08 |
| B123 | B210 | hs-CRP1 | Hip | 0.024025 | 1.83E-07 |
| B123 | B193 | hs-CRP1 | K | -0.02376 | 2.51E-07 |
| B123 | B188 | hs-CRP1 | Ca | -0.02343 | 3.66E-07 |
| B124 | B125 | ACTH | Cor | 0.537347 | 0 |
| B124 | B134 | ACTH | CHE | 0.044663 | 0 |
| B124 | B135 | ACTH | TBA | 0.041874 | 0 |
| B124 | B157 | ACTH | IgG | 0.0342 | 1.13E-13 |
| B124 | B183 | ACTH | PTA | -0.02977 | 1.03E-10 |
| B124 | B195 | ACTH | Na | -0.02632 | 1.11E-08 |
| B125 | B134 | Cor | CHE | 0.044108 | 0 |
| B125 | B157 | Cor | IgG | 0.038996 | 0 |
| B125 | B181 | Cor | APTT | -0.05354 | 0 |
| B125 | B183 | Cor | PTA | -0.05927 | 0 |
| B125 | B194 | Cor | Cl | -0.05055 | 0 |
| B125 | B195 | Cor | Na | -0.05959 | 0 |
| B125 | B156 | Cor | IgE | -0.03506 | 2.71E-14 |
| B125 | B184 | Cor | INR | -0.03389 | 1.86E-13 |
| B125 | B126 | Cor | Alb | 0.033369 | 4.33E-13 |
| B125 | B135 | Cor | TBA | 0.03092 | 1.91E-11 |
| B125 | B182 | Cor | TT | -0.03036 | 4.39E-11 |
| B125 | B138 | Cor | PAB | 0.03023 | 5.28E-11 |
| B125 | B200 | Cor | OP | -0.02964 | 1.24E-10 |
| B125 | B186 | Cor | PFC | 0.028299 | 8.07E-10 |
| B125 | B137 | Cor | AFU | 0.024912 | 6.37E-08 |
| B126 | B127 | Alb | A-G ratio | 0.492015 | 0 |
| B126 | B128 | Alb | ALT | 0.143325 | 0 |
| B126 | B129 | Alb | IBIL | 0.114229 | 0 |
| B126 | B130 | Alb | GLB | -0.16 | 0 |
| B126 | B131 | Alb | AST | 0.054105 | 0 |
| B126 | B132 | Alb | Dbil | 0.048942 | 0 |
| B126 | B133 | Alb | TBIL | 0.105256 | 0 |

| | | | | | |
|---|---|---|---|---|---|
| B126 | B134 | Alb | CHE | 0.091459 | 0 |
| B126 | B138 | Alb | PAB | 0.097417 | 0 |
| B126 | B141 | Alb | AST/ALT | -0.16403 | 0 |
| B126 | B146 | Alb | CysC | -0.0803 | 0 |
| B126 | B153 | Alb | Glu | 0.068232 | 0 |
| B126 | B161 | Alb | LEU | -0.0438 | 0 |
| B126 | B163 | Alb | SG | 0.049303 | 0 |
| B126 | B175 | Alb | ALB-U | -0.03942 | 0 |
| B126 | B176 | Alb | Ucr | 0.040923 | 0 |
| B126 | B177 | Alb | ACR | -0.05689 | 0 |
| B126 | B188 | Alb | Ca | 0.388354 | 0 |
| B126 | B189 | Alb | eGFB | 0.245767 | 0 |
| B126 | B190 | Alb | Cre | -0.08983 | 0 |
| B126 | B191 | Alb | Urea | -0.11203 | 0 |
| B126 | B192 | Alb | UA | 0.043903 | 0 |
| B126 | B194 | Alb | Cl | -0.05728 | 0 |
| B126 | B197 | Alb | AG | 0.038349 | 0 |
| B126 | B199 | Alb | Mg | 0.039633 | 0 |
| B126 | B203 | Alb | DH | 0.039503 | 0 |
| B126 | B204 | Alb | Drinking | 0.045174 | 0 |
| B126 | B211 | Alb | IOP-R | 0.053882 | 0 |
| B126 | B212 | Alb | IOP-L | 0.052193 | 0 |
| B126 | B216 | Alb | BMI | 0.097359 | 0 |
| B126 | B218 | Alb | DBP | 0.143532 | 0 |
| B126 | B219 | Alb | OT | -0.03991 | 0 |
| B126 | B221 | Alb | Hei | 0.117444 | 0 |
| B126 | B222 | Alb | NS | 0.089696 | 0 |
| B126 | B140 | Alb | r-GT | 0.036731 | 1.55E-15 |
| B126 | B173 | Alb | NTT | -0.03626 | 3.55E-15 |
| B126 | B215 | Alb | BMI | 0.033894 | 1.86E-13 |
| B126 | B210 | Alb | Hip | 0.032721 | 1.21E-12 |
| B126 | B172 | Alb | KB | 0.031547 | 7.44E-12 |
| B126 | B155 | Alb | IgA | -0.03118 | 1.30E-11 |
| B126 | B139 | Alb | ADA | -0.03001 | 7.28E-11 |
| B126 | B174 | Alb | Urea/Cr | -0.02897 | 3.18E-10 |
| B126 | B228 | Alb | OS | -0.02796 | 1.28E-09 |
| B126 | B205 | Alb | KDA | -0.02405 | 1.78E-07 |
| B126 | B157 | Alb | IgG | -0.02388 | 2.18E-07 |
| B127 | B129 | A-G ratio | IBIL | 0.129484 | 0 |
| B127 | B130 | A-G ratio | GLB | -0.90162 | 0 |
| B127 | B131 | A-G ratio | AST | -0.06477 | 0 |
| B127 | B132 | A-G ratio | Dbil | 0.10118 | 0 |
| B127 | B133 | A-G ratio | TBIL | 0.131652 | 0 |
| B127 | B138 | A-G ratio | PAB | 0.058497 | 0 |

| | | | | | |
|---|---|---|---|---|---|
| B127 | B139 | A-G ratio | ADA | -0.06683 | 0 |
| B127 | B141 | A-G ratio | AST/ALT | -0.11966 | 0 |
| B127 | B146 | A-G ratio | CysC | -0.06115 | 0 |
| B127 | B150 | A-G ratio | ALP | -0.04996 | 0 |
| B127 | B153 | A-G ratio | Glu | -0.06167 | 0 |
| B127 | B155 | A-G ratio | IgA | -0.08781 | 0 |
| B127 | B157 | A-G ratio | IgG | -0.11739 | 0 |
| B127 | B161 | A-G ratio | LEU | -0.06186 | 0 |
| B127 | B163 | A-G ratio | SG | 0.1003 | 0 |
| B127 | B173 | A-G ratio | NTT | -0.04711 | 0 |
| B127 | B174 | A-G ratio | Urea/Cr | -0.04501 | 0 |
| B127 | B176 | A-G ratio | Ucr | 0.043462 | 0 |
| B127 | B177 | A-G ratio | ACR | -0.03953 | 0 |
| B127 | B188 | A-G ratio | Ca | 0.041598 | 0 |
| B127 | B189 | A-G ratio | eGFB | 0.198697 | 0 |
| B127 | B190 | A-G ratio | Cre | -0.04187 | 0 |
| B127 | B191 | A-G ratio | Urea | -0.07555 | 0 |
| B127 | B192 | A-G ratio | UA | -0.0444 | 0 |
| B127 | B204 | A-G ratio | Drinking | 0.098074 | 0 |
| B127 | B209 | A-G ratio | Smoking | 0.129065 | 0 |
| B127 | B216 | A-G ratio | BMI | 0.102209 | 0 |
| B127 | B218 | A-G ratio | DBP | 0.063646 | 0 |
| B127 | B220 | A-G ratio | SBP | -0.10344 | 0 |
| B127 | B221 | A-G ratio | Hei | 0.176335 | 0 |
| B127 | B222 | A-G ratio | NS | 0.100607 | 0 |
| B127 | B128 | A-G ratio | ALT | 0.035172 | 2.22E-14 |
| B127 | B205 | A-G ratio | KDA | -0.03383 | 2.07E-13 |
| B127 | B158 | A-G ratio | IgM | -0.03363 | 2.84E-13 |
| B127 | B219 | A-G ratio | OT | -0.03349 | 3.57E-13 |
| B127 | B149 | A-G ratio | TGAb | -0.03203 | 3.56E-12 |
| B127 | B136 | A-G ratio | 5'NT | -0.03166 | 6.30E-12 |
| B127 | B175 | A-G ratio | ALB-U | -0.02874 | 4.38E-10 |
| B127 | B148 | A-G ratio | Anti_TPO | -0.02639 | 1.01E-08 |
| B127 | B212 | A-G ratio | IOP-L | 0.026111 | 1.44E-08 |
| B127 | B140 | A-G ratio | r-GT | -0.02459 | 9.40E-08 |
| B127 | B211 | A-G ratio | IOP-R | 0.023349 | 4.01E-07 |
| B127 | B186 | A-G ratio | PFC | -0.02285 | 7.07E-07 |
| B127 | B134 | A-G ratio | CHE | 0.02273 | 8.05E-07 |
| B128 | B129 | ALT | IBIL | 0.051427 | 0 |
| B128 | B131 | ALT | AST | 0.742183 | 0 |
| B128 | B132 | ALT | Dbil | 0.100488 | 0 |
| B128 | B133 | ALT | TBIL | 0.068559 | 0 |
| B128 | B134 | ALT | CHE | 0.057641 | 0 |
| B128 | B136 | ALT | 5'NT | 0.072309 | 0 |

| | | | | | |
|---|---|---|---|---|---|
| B128 | B137 | ALT | AFU | 0.05439 | 0 |
| B128 | B139 | ALT | ADA | 0.040679 | 0 |
| B128 | B140 | ALT | r-GT | 0.358489 | 0 |
| B128 | B141 | ALT | AST/ALT | -0.49844 | 0 |
| B128 | B150 | ALT | ALP | 0.04454 | 0 |
| B128 | B153 | ALT | Glu | 0.098031 | 0 |
| B128 | B160 | ALT | PH | -0.04255 | 0 |
| B128 | B163 | ALT | SG | 0.114729 | 0 |
| B128 | B166 | ALT | Pro | 0.045979 | 0 |
| B128 | B170 | ALT | U-GLU | 0.041984 | 0 |
| B128 | B174 | ALT | Urea/Cr | -0.06206 | 0 |
| B128 | B176 | ALT | Ucr | 0.067298 | 0 |
| B128 | B188 | ALT | Ca | 0.059837 | 0 |
| B128 | B189 | ALT | eGFB | 0.140244 | 0 |
| B128 | B191 | ALT | Urea | -0.05036 | 0 |
| B128 | B192 | ALT | UA | 0.175498 | 0 |
| B128 | B203 | ALT | DH | 0.041517 | 0 |
| B128 | B204 | ALT | Drinking | 0.105922 | 0 |
| B128 | B206 | ALT | Waist | 0.213473 | 0 |
| B128 | B209 | ALT | Smoking | 0.074441 | 0 |
| B128 | B210 | ALT | Hip | 0.181444 | 0 |
| B128 | B215 | ALT | BMI | 0.230384 | 0 |
| B128 | B216 | ALT | BMI | 0.266859 | 0 |
| B128 | B218 | ALT | DBP | 0.124008 | 0 |
| B128 | B219 | ALT | OT | -0.03882 | 0 |
| B128 | B220 | ALT | SBP | -0.04622 | 0 |
| B128 | B221 | ALT | Hei | 0.161585 | 0 |
| B128 | B222 | ALT | NS | 0.223495 | 0 |
| B128 | B173 | ALT | NTT | -0.03162 | 6.70E-12 |
| B128 | B212 | ALT | IOP-L | 0.03155 | 7.40E-12 |
| B128 | B161 | ALT | LEU | -0.03123 | 1.20E-11 |
| B128 | B135 | ALT | TBA | 0.03063 | 2.93E-11 |
| B128 | B211 | ALT | IOP-R | 0.029837 | 9.32E-11 |
| B128 | B201 | ALT | e-habits | -0.0297 | 1.14E-10 |
| B128 | B138 | ALT | PAB | 0.029046 | 2.87E-10 |
| B128 | B130 | ALT | GLB | 0.026745 | 6.40E-09 |
| B128 | B198 | ALT | P | -0.02488 | 6.62E-08 |
| B128 | B193 | ALT | K | -0.02461 | 9.17E-08 |
| B128 | B205 | ALT | KDA | -0.02435 | 1.25E-07 |
| B128 | B149 | ALT | TGAb | -0.02406 | 1.76E-07 |
| B128 | B148 | ALT | Anti_TPO | -0.02404 | 1.80E-07 |
| B128 | B165 | ALT | bilirubin | 0.023928 | 2.05E-07 |
| B129 | B130 | IBIL | GLB | -0.1045 | 0 |
| B129 | B131 | IBIL | AST | 0.083968 | 0 |

| | | | | | |
|---|---|---|---|---|---|
| B129 | B132 | IBIL | Dbil | 0.641139 | 0 |
| B129 | B133 | IBIL | TBIL | 0.978225 | 0 |
| B129 | B140 | IBIL | r-GT | 0.041231 | 0 |
| B129 | B160 | IBIL | PH | 0.07786 | 0 |
| B129 | B163 | IBIL | SG | 0.06853 | 0 |
| B129 | B166 | IBIL | Pro | 0.041236 | 0 |
| B129 | B174 | IBIL | Urea/Cr | -0.09968 | 0 |
| B129 | B191 | IBIL | Urea | -0.06626 | 0 |
| B129 | B192 | IBIL | UA | 0.053292 | 0 |
| B129 | B198 | IBIL | P | -0.03941 | 0 |
| B129 | B209 | IBIL | Smoking | -0.04949 | 0 |
| B129 | B216 | IBIL | BMI | 0.066142 | 0 |
| B129 | B221 | IBIL | Hei | 0.100567 | 0 |
| B129 | B222 | IBIL | NS | 0.077336 | 0 |
| B129 | B176 | IBIL | Ucr | 0.037182 | 6.66E-16 |
| B129 | B189 | IBIL | eGFB | -0.03715 | 8.88E-16 |
| B129 | B188 | IBIL | Ca | 0.036157 | 4.00E-15 |
| B129 | B183 | IBIL | PTA | 0.035687 | 9.33E-15 |
| B129 | B204 | IBIL | Drinking | 0.033949 | 1.70E-13 |
| B129 | B161 | IBIL | LEU | -0.03358 | 3.10E-13 |
| B129 | B218 | IBIL | DBP | 0.031434 | 8.82E-12 |
| B129 | B150 | IBIL | ALP | -0.03054 | 3.34E-11 |
| B129 | B146 | IBIL | CysC | -0.02854 | 5.77E-10 |
| B129 | B165 | IBIL | bilirubin | 0.025037 | 5.47E-08 |
| B129 | B193 | IBIL | K | -0.0244 | 1.17E-07 |
| B129 | B205 | IBIL | KDA | -0.02434 | 1.27E-07 |
| B129 | B206 | IBIL | Waist | 0.023996 | 1.90E-07 |
| B129 | B173 | IBIL | NTT | -0.02269 | 8.39E-07 |
| B130 | B131 | GLB | AST | 0.111932 | 0 |
| B130 | B132 | GLB | Dbil | -0.08071 | 0 |
| B130 | B133 | GLB | TBIL | -0.10599 | 0 |
| B130 | B136 | GLB | 5'NT | 0.044965 | 0 |
| B130 | B139 | GLB | ADA | 0.065882 | 0 |
| B130 | B140 | GLB | r-GT | 0.065015 | 0 |
| B130 | B141 | GLB | AST/ALT | 0.074911 | 0 |
| B130 | B146 | GLB | CysC | 0.049696 | 0 |
| B130 | B150 | GLB | ALP | 0.0536 | 0 |
| B130 | B153 | GLB | Glu | 0.104081 | 0 |
| B130 | B155 | GLB | IgA | 0.091155 | 0 |
| B130 | B157 | GLB | IgG | 0.127641 | 0 |
| B130 | B161 | GLB | LEU | 0.054145 | 0 |
| B130 | B163 | GLB | SG | -0.09084 | 0 |
| B130 | B173 | GLB | NTT | 0.042834 | 0 |
| B130 | B188 | GLB | Ca | 0.107795 | 0 |

| | | | | | |
|---|---|---|---|---|---|
| B130 | B189 | GLB | eGFB | -0.13282 | 0 |
| B130 | B191 | GLB | Urea | 0.046945 | 0 |
| B130 | B192 | GLB | UA | 0.076451 | 0 |
| B130 | B204 | GLB | Drinking | -0.08907 | 0 |
| B130 | B209 | GLB | Smoking | -0.13743 | 0 |
| B130 | B216 | GLB | BMI | -0.07934 | 0 |
| B130 | B220 | GLB | SBP | 0.110137 | 0 |
| B130 | B221 | GLB | Hei | -0.15331 | 0 |
| B130 | B222 | GLB | NS | -0.07793 | 0 |
| B130 | B174 | GLB | Urea/Cr | 0.037165 | 6.66E-16 |
| B130 | B158 | GLB | IgM | 0.036821 | 1.33E-15 |
| B130 | B194 | GLB | Cl | -0.03613 | 4.44E-15 |
| B130 | B149 | GLB | TGAb | 0.034107 | 1.31E-13 |
| B130 | B138 | GLB | PAB | -0.03135 | 1.00E-11 |
| B130 | B176 | GLB | Ucr | -0.02994 | 8.00E-11 |
| B130 | B148 | GLB | Anti_TPO | 0.029114 | 2.61E-10 |
| B130 | B205 | GLB | KDA | 0.027047 | 4.32E-09 |
| B130 | B166 | GLB | Pro | 0.02606 | 1.54E-08 |
| B130 | B193 | GLB | K | 0.025328 | 3.83E-08 |
| B130 | B170 | GLB | U-GLU | 0.024973 | 5.92E-08 |
| B130 | B219 | GLB | OT | 0.022702 | 8.30E-07 |
| B131 | B132 | AST | Dbil | 0.169738 | 0 |
| B131 | B133 | AST | TBIL | 0.113394 | 0 |
| B131 | B135 | AST | TBA | 0.048742 | 0 |
| B131 | B136 | AST | 5'NT | 0.080567 | 0 |
| B131 | B137 | AST | AFU | 0.060139 | 0 |
| B131 | B139 | AST | ADA | 0.065919 | 0 |
| B131 | B140 | AST | r-GT | 0.39731 | 0 |
| B131 | B141 | AST | AST/ALT | -0.06955 | 0 |
| B131 | B150 | AST | ALP | 0.049596 | 0 |
| B131 | B153 | AST | Glu | 0.063458 | 0 |
| B131 | B165 | AST | bilirubin | 0.066323 | 0 |
| B131 | B166 | AST | Pro | 0.038043 | 0 |
| B131 | B188 | AST | Ca | 0.051209 | 0 |
| B131 | B192 | AST | UA | 0.125298 | 0 |
| B131 | B204 | AST | Drinking | 0.089523 | 0 |
| B131 | B206 | AST | Waist | 0.088841 | 0 |
| B131 | B209 | AST | Smoking | 0.038413 | 0 |
| B131 | B210 | AST | Hip | 0.052198 | 0 |
| B131 | B215 | AST | BMI | 0.091469 | 0 |
| B131 | B216 | AST | BMI | 0.076293 | 0 |
| B131 | B218 | AST | DBP | 0.042024 | 0 |
| B131 | B222 | AST | NS | 0.07755 | 0 |
| B131 | B189 | AST | eGFB | 0.037548 | 4.44E-16 |

| | | | | | |
|---|---|---|---|---|---|
| B131 | B225 | AST | hPBG | 0.036275 | 3.33E-15 |
| B131 | B163 | AST | SG | 0.035269 | 1.91E-14 |
| B131 | B191 | AST | Urea | -0.0322 | 2.74E-12 |
| B131 | B176 | AST | Ucr | 0.028485 | 6.25E-10 |
| B131 | B179 | AST | α1-MG | 0.026553 | 8.19E-09 |
| B131 | B174 | AST | Urea/Cr | -0.02633 | 1.10E-08 |
| B132 | B133 | Dbil | TBIL | 0.786212 | 0 |
| B132 | B140 | Dbil | r-GT | 0.15977 | 0 |
| B132 | B160 | Dbil | PH | 0.047042 | 0 |
| B132 | B163 | Dbil | SG | 0.051732 | 0 |
| B132 | B165 | Dbil | bilirubin | 0.069013 | 0 |
| B132 | B166 | Dbil | Pro | 0.042883 | 0 |
| B132 | B174 | Dbil | Urea/Cr | -0.093 | 0 |
| B132 | B189 | Dbil | eGFB | -0.07582 | 0 |
| B132 | B192 | Dbil | UA | 0.04697 | 0 |
| B132 | B198 | Dbil | P | -0.04455 | 0 |
| B132 | B204 | Dbil | Drinking | 0.06815 | 0 |
| B132 | B216 | Dbil | BMI | 0.040725 | 0 |
| B132 | B221 | Dbil | Hei | 0.093605 | 0 |
| B132 | B222 | Dbil | NS | 0.069878 | 0 |
| B132 | B193 | Dbil | K | -0.03659 | 1.78E-15 |
| B132 | B161 | Dbil | LEU | -0.03411 | 1.31E-13 |
| B132 | B139 | Dbil | ADA | 0.033979 | 1.61E-13 |
| B132 | B191 | Dbil | Urea | -0.03214 | 2.97E-12 |
| B132 | B134 | Dbil | CHE | -0.03126 | 1.15E-11 |
| B132 | B190 | Dbil | Cre | 0.028026 | 1.17E-09 |
| B132 | B172 | Dbil | KB | 0.024326 | 1.29E-07 |
| B132 | B176 | Dbil | Ucr | 0.023647 | 2.85E-07 |
| B132 | B136 | Dbil | 5'NT | 0.022699 | 8.33E-07 |
| B133 | B140 | TBIL | r-GT | 0.076369 | 0 |
| B133 | B160 | TBIL | PH | 0.075491 | 0 |
| B133 | B163 | TBIL | SG | 0.069083 | 0 |
| B133 | B165 | TBIL | bilirubin | 0.038854 | 0 |
| B133 | B166 | TBIL | Pro | 0.04482 | 0 |
| B133 | B174 | TBIL | Urea/Cr | -0.10537 | 0 |
| B133 | B189 | TBIL | eGFB | -0.05037 | 0 |
| B133 | B191 | TBIL | Urea | -0.06202 | 0 |
| B133 | B192 | TBIL | UA | 0.055625 | 0 |
| B133 | B198 | TBIL | P | -0.04365 | 0 |
| B133 | B204 | TBIL | Drinking | 0.045719 | 0 |
| B133 | B209 | TBIL | Smoking | -0.0457 | 0 |
| B133 | B216 | TBIL | BMI | 0.064313 | 0 |
| B133 | B221 | TBIL | Hei | 0.10633 | 0 |
| B133 | B222 | TBIL | NS | 0.081183 | 0 |

| | | | | | |
|---|---|---|---|---|---|
| B133 | B161 | TBIL | LEU | -0.0363 | 3.11E-15 |
| B133 | B176 | TBIL | Ucr | 0.036332 | 3.11E-15 |
| B133 | B183 | TBIL | PTA | 0.032428 | 1.92E-12 |
| B133 | B188 | TBIL | Ca | 0.030583 | 3.14E-11 |
| B133 | B193 | TBIL | K | -0.02963 | 1.25E-10 |
| B133 | B206 | TBIL | Waist | 0.025291 | 4.01E-08 |
| B133 | B218 | TBIL | DBP | 0.024783 | 7.44E-08 |
| B133 | B150 | TBIL | ALP | -0.02401 | 1.88E-07 |
| B134 | B136 | CHE | 5'NT | 0.132961 | 0 |
| B134 | B137 | CHE | AFU | 0.157904 | 0 |
| B134 | B138 | CHE | PAB | 0.348863 | 0 |
| B134 | B139 | CHE | ADA | -0.10819 | 0 |
| B134 | B140 | CHE | r-GT | 0.040754 | 0 |
| B134 | B141 | CHE | AST/ALT | -0.06235 | 0 |
| B134 | B146 | CHE | CysC | -0.03974 | 0 |
| B134 | B150 | CHE | ALP | 0.044697 | 0 |
| B134 | B157 | CHE | IgG | -0.06114 | 0 |
| B134 | B181 | CHE | APTT | -0.06066 | 0 |
| B134 | B184 | CHE | INR | -0.05021 | 0 |
| B134 | B188 | CHE | Ca | 0.062607 | 0 |
| B134 | B189 | CHE | eGFB | 0.042173 | 0 |
| B134 | B192 | CHE | UA | 0.042482 | 0 |
| B134 | B193 | CHE | K | -0.04424 | 0 |
| B134 | B194 | CHE | Cl | -0.1414 | 0 |
| B134 | B195 | CHE | Na | 0.05485 | 0 |
| B134 | B197 | CHE | AG | 0.140927 | 0 |
| B134 | B199 | CHE | Mg | 0.044271 | 0 |
| B134 | B200 | CHE | OP | 0.040075 | 0 |
| B134 | B204 | CHE | Drinking | 0.045299 | 0 |
| B134 | B206 | CHE | Waist | 0.061004 | 0 |
| B134 | B210 | CHE | Hip | 0.050886 | 0 |
| B134 | B215 | CHE | BMI | 0.068069 | 0 |
| B134 | B216 | CHE | BMI | 0.076328 | 0 |
| B134 | B218 | CHE | DBP | 0.039099 | 0 |
| B134 | B221 | CHE | Hei | 0.043716 | 0 |
| B134 | B222 | CHE | NS | 0.062816 | 0 |
| B134 | B185 | CHE | PT | -0.03793 | 2.22E-16 |
| B134 | B186 | CHE | PFC | 0.036215 | 3.77E-15 |
| B134 | B163 | CHE | SG | 0.035944 | 6.00E-15 |
| B134 | B153 | CHE | Glu | 0.033335 | 4.57E-13 |
| B134 | B156 | CHE | IgE | -0.03319 | 5.77E-13 |
| B134 | B212 | CHE | IOP-L | 0.032917 | 8.89E-13 |
| B134 | B209 | CHE | Smoking | 0.027375 | 2.80E-09 |
| B134 | B211 | CHE | IOP-R | 0.023777 | 2.45E-07 |

| | | | | | |
|---|---|---|---|---|---|
| B135 | B136 | TBA | 5'NT | 0.092569 | 0 |
| B135 | B137 | TBA | AFU | 0.091105 | 0 |
| B135 | B139 | TBA | ADA | 0.101901 | 0 |
| B135 | B140 | TBA | r-GT | 0.053213 | 0 |
| B135 | B150 | TBA | ALP | 0.08296 | 0 |
| B135 | B182 | TBA | TT | 0.040025 | 0 |
| B135 | B193 | TBA | K | -0.07819 | 0 |
| B135 | B200 | TBA | OP | 0.034894 | 3.55E-14 |
| B135 | B179 | TBA | α1-MG | 0.034864 | 3.73E-14 |
| B135 | B198 | TBA | P | -0.03316 | 6.08E-13 |
| B135 | B195 | TBA | Na | 0.029378 | 1.80E-10 |
| B135 | B194 | TBA | Cl | 0.026331 | 1.09E-08 |
| B135 | B181 | TBA | APTT | 0.025157 | 4.73E-08 |
| B135 | B165 | TBA | bilirubin | 0.024574 | 9.57E-08 |
| B135 | B156 | TBA | IgE | -0.02353 | 3.25E-07 |
| B136 | B137 | 5'NT | AFU | 0.074189 | 0 |
| B136 | B138 | 5'NT | PAB | 0.073439 | 0 |
| B136 | B139 | 5'NT | ADA | 0.17969 | 0 |
| B136 | B140 | 5'NT | r-GT | 0.19426 | 0 |
| B136 | B150 | 5'NT | ALP | 0.2108 | 0 |
| B136 | B155 | 5'NT | IgA | 0.047615 | 0 |
| B136 | B157 | 5'NT | IgG | 0.055904 | 0 |
| B136 | B158 | 5'NT | IgM | 0.04148 | 0 |
| B136 | B179 | 5'NT | α1-MG | 0.090308 | 0 |
| B136 | B181 | 5'NT | APTT | 0.041144 | 0 |
| B136 | B182 | 5'NT | TT | 0.06403 | 0 |
| B136 | B194 | 5'NT | Cl | -0.04344 | 0 |
| B136 | B197 | 5'NT | AG | 0.061889 | 0 |
| B136 | B186 | 5'NT | PFC | 0.031025 | 1.63E-11 |
| B136 | B200 | 5'NT | OP | 0.028216 | 9.04E-10 |
| B136 | B193 | 5'NT | K | -0.02741 | 2.68E-09 |
| B136 | B141 | 5'NT | AST/ALT | -0.02548 | 3.18E-08 |
| B136 | B196 | 5'NT | TCO2 | -0.02535 | 3.71E-08 |
| B137 | B138 | AFU | PAB | 0.093297 | 0 |
| B137 | B139 | AFU | ADA | 0.183255 | 0 |
| B137 | B140 | AFU | r-GT | 0.068504 | 0 |
| B137 | B150 | AFU | ALP | 0.06927 | 0 |
| B137 | B156 | AFU | IgE | -0.05784 | 0 |
| B137 | B179 | AFU | α1-MG | 0.038602 | 0 |
| B137 | B193 | AFU | K | -0.04339 | 0 |
| B137 | B194 | AFU | Cl | -0.04279 | 0 |
| B137 | B197 | AFU | AG | 0.038073 | 0 |
| B137 | B198 | AFU | P | -0.05524 | 0 |
| B137 | B200 | AFU | OP | 0.067118 | 0 |

| | | | | | |
|---|---|---|---|---|---|
| B137 | B222 | AFU | NS | 0.035984 | 5.55E-15 |
| B137 | B206 | AFU | Waist | 0.035525 | 1.22E-14 |
| B137 | B216 | AFU | BMI | 0.032944 | 8.52E-13 |
| B137 | B153 | AFU | Glu | 0.031918 | 4.22E-12 |
| B137 | B204 | AFU | Drinking | 0.026258 | 1.20E-08 |
| B137 | B215 | AFU | BMI | 0.026052 | 1.55E-08 |
| B137 | B195 | AFU | Na | 0.024925 | 6.27E-08 |
| B137 | B221 | AFU | Hei | 0.02264 | 8.89E-07 |
| B138 | B139 | PAB | ADA | -0.32591 | 0 |
| B138 | B140 | PAB | r-GT | 0.064877 | 0 |
| B138 | B141 | PAB | AST/ALT | -0.04211 | 0 |
| B138 | B150 | PAB | ALP | -0.0571 | 0 |
| B138 | B157 | PAB | IgG | -0.09821 | 0 |
| B138 | B163 | PAB | SG | 0.043528 | 0 |
| B138 | B179 | PAB | α1-MG | 0.038586 | 0 |
| B138 | B181 | PAB | APTT | -0.07859 | 0 |
| B138 | B184 | PAB | INR | -0.05571 | 0 |
| B138 | B185 | PAB | PT | -0.04488 | 0 |
| B138 | B188 | PAB | Ca | 0.056489 | 0 |
| B138 | B190 | PAB | Cre | 0.039408 | 0 |
| B138 | B192 | PAB | UA | 0.087583 | 0 |
| B138 | B194 | PAB | Cl | -0.14683 | 0 |
| B138 | B197 | PAB | AG | 0.117761 | 0 |
| B138 | B198 | PAB | P | -0.04249 | 0 |
| B138 | B199 | PAB | Mg | 0.042032 | 0 |
| B138 | B204 | PAB | Drinking | 0.09791 | 0 |
| B138 | B212 | PAB | IOP-L | 0.038376 | 0 |
| B138 | B216 | PAB | BMI | 0.063435 | 0 |
| B138 | B221 | PAB | Hei | 0.08231 | 0 |
| B138 | B222 | PAB | NS | 0.075804 | 0 |
| B138 | B176 | PAB | Ucr | 0.037906 | 2.22E-16 |
| B138 | B174 | PAB | Urea/Cr | -0.03728 | 4.44E-16 |
| B138 | B218 | PAB | DBP | 0.036974 | 8.88E-16 |
| B138 | B206 | PAB | Waist | 0.036889 | 1.11E-15 |
| B138 | B211 | PAB | IOP-R | 0.034605 | 5.77E-14 |
| B138 | B209 | PAB | Smoking | 0.033975 | 1.63E-13 |
| B138 | B200 | PAB | OP | 0.032213 | 2.68E-12 |
| B138 | B186 | PAB | PFC | -0.02974 | 1.07E-10 |
| B138 | B210 | PAB | Hip | 0.026443 | 9.44E-09 |
| B138 | B161 | PAB | LEU | -0.02559 | 2.78E-08 |
| B138 | B219 | PAB | OT | -0.02417 | 1.55E-07 |
| B139 | B140 | ADA | r-GT | 0.041458 | 0 |
| B139 | B146 | ADA | CysC | 0.07773 | 0 |
| B139 | B150 | ADA | ALP | 0.182014 | 0 |

| | | | | | |
|---|---|---|---|---|---|
| B139 | B153 | ADA | Glu | 0.042726 | 0 |
| B139 | B155 | ADA | IgA | 0.084211 | 0 |
| B139 | B157 | ADA | IgG | 0.085627 | 0 |
| B139 | B179 | ADA | α1-MG | 0.03823 | 0 |
| B139 | B186 | ADA | PFC | 0.038024 | 0 |
| B139 | B193 | ADA | K | -0.05724 | 0 |
| B139 | B194 | ADA | Cl | -0.04626 | 0 |
| B139 | B195 | ADA | Na | 0.09312 | 0 |
| B139 | B197 | ADA | AG | 0.082966 | 0 |
| B139 | B200 | ADA | OP | 0.115009 | 0 |
| B139 | B221 | ADA | Hei | -0.04791 | 0 |
| B139 | B220 | ADA | SBP | 0.036531 | 2.22E-15 |
| B139 | B174 | ADA | Urea/Cr | 0.035575 | 1.13E-14 |
| B139 | B222 | ADA | NS | -0.03078 | 2.36E-11 |
| B139 | B181 | ADA | APTT | 0.029247 | 2.16E-10 |
| B139 | B182 | ADA | TT | 0.028477 | 6.32E-10 |
| B139 | B216 | ADA | BMI | -0.02787 | 1.44E-09 |
| B139 | B204 | ADA | Drinking | -0.02743 | 2.60E-09 |
| B139 | B184 | ADA | INR | 0.02641 | 9.85E-09 |
| B139 | B163 | ADA | SG | -0.02638 | 1.03E-08 |
| B139 | B185 | ADA | PT | 0.025443 | 3.33E-08 |
| B139 | B154 | ADA | RF | 0.024979 | 5.87E-08 |
| B139 | B209 | ADA | Smoking | -0.02256 | 9.67E-07 |
| B14 | B80 | G-17 | anti-HP IgG | 0.038216 | 0 |
| B14 | B139 | G-17 | ADA | 0.033056 | 7.14E-13 |
| B140 | B141 | r-GT | AST/ALT | -0.14319 | 0 |
| B140 | B150 | r-GT | ALP | 0.123542 | 0 |
| B140 | B153 | r-GT | Glu | 0.107362 | 0 |
| B140 | B160 | r-GT | PH | -0.05079 | 0 |
| B140 | B163 | r-GT | SG | 0.097937 | 0 |
| B140 | B165 | r-GT | bilirubin | 0.060442 | 0 |
| B140 | B166 | r-GT | Pro | 0.053349 | 0 |
| B140 | B170 | r-GT | U-GLU | 0.059085 | 0 |
| B140 | B174 | r-GT | Urea/Cr | -0.0802 | 0 |
| B140 | B176 | r-GT | Ucr | 0.06463 | 0 |
| B140 | B179 | r-GT | α1-MG | 0.079367 | 0 |
| B140 | B188 | r-GT | Ca | 0.041977 | 0 |
| B140 | B189 | r-GT | eGFB | 0.081265 | 0 |
| B140 | B191 | r-GT | Urea | -0.04725 | 0 |
| B140 | B192 | r-GT | UA | 0.175731 | 0 |
| B140 | B203 | r-GT | DH | 0.043248 | 0 |
| B140 | B204 | r-GT | Drinking | 0.209676 | 0 |
| B140 | B206 | r-GT | Waist | 0.134856 | 0 |
| B140 | B209 | r-GT | Smoking | 0.140665 | 0 |

| | | | | | |
|---|---|---|---|---|---|
| B140 | B210 | r-GT | Hip | 0.090138 | 0 |
| B140 | B215 | r-GT | BMI | 0.105103 | 0 |
| B140 | B216 | r-GT | BMI | 0.150006 | 0 |
| B140 | B218 | r-GT | DBP | 0.09648 | 0 |
| B140 | B221 | r-GT | Hei | 0.121214 | 0 |
| B140 | B222 | r-GT | NS | 0.160643 | 0 |
| B140 | B193 | r-GT | K | -0.03528 | 1.84E-14 |
| B140 | B201 | r-GT | e-habits | -0.02663 | 7.40E-09 |
| B140 | B219 | r-GT | OT | -0.02645 | 9.35E-09 |
| B140 | B211 | r-GT | IOP-R | 0.025775 | 2.20E-08 |
| B140 | B212 | r-GT | IOP-L | 0.025746 | 2.28E-08 |
| B140 | B182 | r-GT | TT | 0.023878 | 2.18E-07 |
| B140 | B198 | r-GT | P | -0.02349 | 3.39E-07 |
| B141 | B146 | AST/ALT | CysC | 0.041776 | 0 |
| B141 | B153 | AST/ALT | Glu | -0.0798 | 0 |
| B141 | B161 | AST/ALT | LEU | 0.053495 | 0 |
| B141 | B163 | AST/ALT | SG | -0.1362 | 0 |
| B141 | B170 | AST/ALT | U-GLU | -0.04113 | 0 |
| B141 | B173 | AST/ALT | NTT | 0.0472 | 0 |
| B141 | B174 | AST/ALT | Urea/Cr | 0.068322 | 0 |
| B141 | B176 | AST/ALT | Ucr | -0.06852 | 0 |
| B141 | B189 | AST/ALT | eGFB | -0.15748 | 0 |
| B141 | B191 | AST/ALT | Urea | 0.054503 | 0 |
| B141 | B192 | AST/ALT | UA | -0.14668 | 0 |
| B141 | B203 | AST/ALT | DH | -0.04182 | 0 |
| B141 | B204 | AST/ALT | Drinking | -0.08152 | 0 |
| B141 | B205 | AST/ALT | KDA | 0.040631 | 0 |
| B141 | B206 | AST/ALT | Waist | -0.24087 | 0 |
| B141 | B209 | AST/ALT | Smoking | -0.07697 | 0 |
| B141 | B210 | AST/ALT | Hip | -0.22459 | 0 |
| B141 | B211 | AST/ALT | IOP-R | -0.03939 | 0 |
| B141 | B215 | AST/ALT | BMI | -0.26512 | 0 |
| B141 | B216 | AST/ALT | BMI | -0.33233 | 0 |
| B141 | B218 | AST/ALT | DBP | -0.14312 | 0 |
| B141 | B219 | AST/ALT | OT | 0.048723 | 0 |
| B141 | B220 | AST/ALT | SBP | 0.089517 | 0 |
| B141 | B221 | AST/ALT | Hei | -0.23736 | 0 |
| B141 | B222 | AST/ALT | NS | -0.2639 | 0 |
| B141 | B157 | AST/ALT | IgG | 0.037204 | 6.66E-16 |
| B141 | B212 | AST/ALT | IOP-L | -0.0347 | 4.93E-14 |
| B141 | B188 | AST/ALT | Ca | -0.03463 | 5.57E-14 |
| B141 | B160 | AST/ALT | PH | 0.032853 | 9.83E-13 |
| B141 | B166 | AST/ALT | Pro | -0.02981 | 9.64E-11 |
| B141 | B224 | AST/ALT | pmh | 0.029321 | 1.95E-10 |

| | | | | | |
|---|---|---|---|---|---|
| B141 | B228 | AST/ALT | OS | 0.028921 | 3.41E-10 |
| B141 | B148 | AST/ALT | Anti_TPO | 0.024483 | 1.07E-07 |
| B141 | B149 | AST/ALT | TGAb | 0.023916 | 2.08E-07 |
| B142 | B144 | 25-(OH)D | Total-PINP | -0.04594 | 0 |
| B142 | B145 | 25-(OH)D | BGP | -0.05649 | 0 |
| B142 | B176 | 25-(OH)D | Ucr | -0.04358 | 0 |
| B142 | B189 | 25-(OH)D | eGFB | -0.03343 | 3.92E-13 |
| B142 | B216 | 25-(OH)D | BMI | -0.03208 | 3.30E-12 |
| B142 | B222 | 25-(OH)D | NS | -0.0296 | 1.30E-10 |
| B142 | B218 | 25-(OH)D | DBP | -0.028 | 1.21E-09 |
| B142 | B221 | 25-(OH)D | Hei | -0.02501 | 5.69E-08 |
| B142 | B215 | 25-(OH)D | BMI | -0.02451 | 1.03E-07 |
| B142 | B210 | 25-(OH)D | Hip | -0.02301 | 5.85E-07 |
| B143 | B144 | β-CTx | Total-PINP | 0.790642 | 0 |
| B143 | B145 | β-CTx | BGP | 0.573647 | 0 |
| B143 | B150 | β-CTx | ALP | 0.051779 | 0 |
| B143 | B176 | β-CTx | Ucr | 0.026696 | 6.81E-09 |
| B143 | B192 | β-CTx | UA | -0.02513 | 4.91E-08 |
| B143 | B183 | β-CTx | PTA | 0.023165 | 4.94E-07 |
| B144 | B145 | Total-PINP | BGP | 0.57821 | 0 |
| B144 | B150 | Total-PINP | ALP | 0.067005 | 0 |
| B144 | B177 | Total-PINP | ACR | -0.0277 | 1.82E-09 |
| B145 | B150 | BGP | ALP | 0.065247 | 0 |
| B145 | B201 | BGP | e-habits | -0.02391 | 2.10E-07 |
| B146 | B174 | CysC | Urea/Cr | -0.0557 | 0 |
| B146 | B189 | CysC | eGFB | -0.19826 | 0 |
| B146 | B190 | CysC | Cre | 0.274403 | 0 |
| B146 | B191 | CysC | Urea | 0.179065 | 0 |
| B146 | B192 | CysC | UA | 0.095235 | 0 |
| B146 | B195 | CysC | Na | 0.050335 | 0 |
| B146 | B200 | CysC | OP | 0.072231 | 0 |
| B146 | B220 | CysC | SBP | 0.039374 | 0 |
| B146 | B194 | CysC | Cl | 0.036657 | 1.78E-15 |
| B146 | B160 | CysC | PH | -0.03557 | 1.15E-14 |
| B146 | B177 | CysC | ACR | 0.032497 | 1.72E-12 |
| B146 | B218 | CysC | DBP | -0.03176 | 5.36E-12 |
| B146 | B150 | CysC | ALP | 0.027193 | 3.56E-09 |
| B146 | B228 | CysC | OS | 0.024869 | 6.71E-08 |
| B146 | B175 | CysC | ALB-U | 0.02348 | 3.45E-07 |
| B146 | B186 | CysC | PFC | 0.022735 | 8.00E-07 |
| B148 | B149 | Anti_TPO | TGAb | 0.480417 | 0 |
| B148 | B221 | Anti_TPO | Hei | -0.04687 | 0 |
| B148 | B222 | Anti_TPO | NS | -0.03963 | 0 |
| B148 | B204 | Anti_TPO | Drinking | -0.03386 | 1.95E-13 |

| | | | | | |
|---|---|---|---|---|---|
| B148 | B216 | Anti_TPO | BMI | -0.03298 | 8.07E-13 |
| B148 | B192 | Anti_TPO | UA | -0.02692 | 5.06E-09 |
| B148 | B206 | Anti_TPO | Waist | -0.02619 | 1.30E-08 |
| B148 | B163 | Anti_TPO | SG | -0.02349 | 3.41E-07 |
| B149 | B216 | TGAb | BMI | -0.04098 | 0 |
| B149 | B221 | TGAb | Hei | -0.0455 | 0 |
| B149 | B222 | TGAb | NS | -0.03972 | 0 |
| B149 | B204 | TGAb | Drinking | -0.03274 | 1.17E-12 |
| B149 | B192 | TGAb | UA | -0.02825 | 8.57E-10 |
| B149 | B206 | TGAb | Waist | -0.02753 | 2.27E-09 |
| B149 | B209 | TGAb | Smoking | -0.02705 | 4.31E-09 |
| B149 | B163 | TGAb | SG | -0.02513 | 4.88E-08 |
| B149 | B174 | TGAb | Urea/Cr | 0.022605 | 9.24E-07 |
| B15 | B100 | CD3-CD19+(B cell) | anti-Ro52 | 0.0658 | 0 |
| B15 | B103 | CD3-CD19+(B cell) | anti-SSA/Ro | 0.054423 | 0 |
| B15 | B106 | CD3-CD19+(B cell) | anti-P | 0.039246 | 0 |
| B15 | B107 | CD3-CD19+(B cell) | AnuA | 0.439816 | 0 |
| B15 | B119 | CD3-CD19+(B cell) | Hcv-rna | 0.117881 | 0 |
| B15 | B122 | CD3-CD19+(B cell) | C4 | 0.051542 | 0 |
| B15 | B125 | CD3-CD19+(B cell) | Cor | -0.0483 | 0 |
| B15 | B136 | CD3-CD19+(B cell) | 5'NT | 0.047995 | 0 |
| B15 | B138 | CD3-CD19+(B cell) | PAB | -0.09995 | 0 |
| B15 | B157 | CD3-CD19+(B cell) | IgG | 0.045303 | 0 |
| B15 | B158 | CD3-CD19+(B cell) | IgM | 0.044003 | 0 |
| B15 | B16 | CD3-CD19+(B cell) | CD3+T | -0.09467 | 0 |
| B15 | B17 | CD3-CD19+(B cell) | CD3+CD16+CD56+NK | -0.13696 | 0 |
| B15 | B18 | CD3-CD19+(B cell) | CD3+CD4+helper(inducer)T | 0.114633 | 0 |
| B15 | B181 | CD3-CD19+(B cell) | APTT | 0.099941 | 0 |
| B15 | B182 | CD3-CD19+(B cell) | TT | 0.056101 | 0 |
| B15 | B183 | CD3-CD19+(B cell) | PTA | 0.097817 | 0 |
| B15 | B184 | CD3-CD19+(B cell) | INR | 0.115354 | 0 |
| B15 | B185 | CD3-CD19+(B cell) | PT | 0.047916 | 0 |
| B15 | B186 | CD3-CD19+(B cell) | PFC | -0.12454 | 0 |
| B15 | B19 | CD3-CD19+(B cell) | CD3+CD4+/CD3+CD8+ | 0.197557 | 0 |
| B15 | B193 | CD3-CD19+(B cell) | K | 0.087969 | 0 |
| B15 | B194 | CD3-CD19+(B cell) | Cl | 0.074453 | 0 |
| B15 | B196 | CD3-CD19+(B cell) | TCO2 | -0.04079 | 0 |
| B15 | B198 | CD3-CD19+(B cell) | P | 0.045033 | 0 |
| B15 | B199 | CD3-CD19+(B cell) | Mg | -0.05346 | 0 |
| B15 | B20 | CD3-CD19+(B cell) | CD3+CD4+CD8+ | -0.09839 | 0 |
| B15 | B21 | CD3-CD19+(B cell) | CD3+CD4-CD8- | 0.070818 | 0 |
| B15 | B22 | CD3-CD19+(B cell) | CD3+CD8+(T) | -0.25419 | 0 |
| B15 | B23 | CD3-CD19+(B cell) | CD3-CD16+CD56+(NK) | -0.23244 | 0 |
| B15 | B32 | CD3-CD19+(B cell) | Testo | -0.05265 | 0 |

| | | | | | |
|---|---|---|---|---|---|
| B15 | B33 | CD3-CD19+(B cell) | Prog | 0.051788 | 0 |
| B15 | B79 | CD3-CD19+(B cell) | hb V -dna | 0.20053 | 0 |
| B15 | B92 | CD3-CD19+(B cell) | ALDH2-Glu504Glu | -0.06633 | 0 |
| B15 | B93 | CD3-CD19+(B cell) | ALDH2-Glu504Lys | 0.050768 | 0 |
| B15 | B94 | CD3-CD19+(B cell) | ALDH2-Lys504Lys | 0.091063 | 0 |
| B15 | B96 | CD3-CD19+(B cell) | dsDNA (IIF) | 0.085257 | 0 |
| B15 | B99 | CD3-CD19+(B cell) | anti-PM-Scl | -0.05178 | 0 |
| B15 | B212 | CD3-CD19+(B cell) | IOP-L | -0.03678 | 1.33E-15 |
| B15 | B29 | CD3-CD19+(B cell) | E2 | 0.036604 | 1.78E-15 |
| B15 | B189 | CD3-CD19+(B cell) | eGFB | -0.03604 | 5.11E-15 |
| B15 | B64 | CD3-CD19+(B cell) | WBRSR(mPas_5) | -0.03529 | 1.82E-14 |
| B15 | B40 | CD3-CD19+(B cell) | WBC | 0.034293 | 9.64E-14 |
| B15 | B98 | CD3-CD19+(B cell) | anti-Jo-1 | 0.033773 | 2.26E-13 |
| B15 | B134 | CD3-CD19+(B cell) | CHE | -0.03205 | 3.45E-12 |
| B15 | B69 | CD3-CD19+(B cell) | PV | -0.03188 | 4.48E-12 |
| B15 | B65 | CD3-CD19+(B cell) | WBRSR(mPas_1) | -0.03167 | 6.15E-12 |
| B15 | B42 | CD3-CD19+(B cell) | MON | 0.030648 | 2.86E-11 |
| B15 | B139 | CD3-CD19+(B cell) | ADA | 0.029525 | 1.46E-10 |
| B15 | B211 | CD3-CD19+(B cell) | IOP-R | -0.02951 | 1.48E-10 |
| B15 | B86 | CD3-CD19+(B cell) | TBhCG | 0.029504 | 1.50E-10 |
| B15 | B76 | CD3-CD19+(B cell) | ApoB | -0.02906 | 2.81E-10 |
| B15 | B222 | CD3-CD19+(B cell) | NS | -0.02854 | 5.79E-10 |
| B15 | B45 | CD3-CD19+(B cell) | MCHC | -0.02789 | 1.40E-09 |
| B15 | B104 | CD3-CD19+(B cell) | anti-SSB | 0.0262 | 1.29E-08 |
| B15 | B51 | CD3-CD19+(B cell) | Hb | -0.02615 | 1.38E-08 |
| B15 | B204 | CD3-CD19+(B cell) | Drinking | -0.02612 | 1.43E-08 |
| B15 | B197 | CD3-CD19+(B cell) | AG | -0.02496 | 6.04E-08 |
| B15 | B90 | CD3-CD19+(B cell) | CYFRA 21-1 | -0.02481 | 7.19E-08 |
| B15 | B97 | CD3-CD19+(B cell) | dsDNA | 0.024754 | 7.71E-08 |
| B15 | B81 | CD3-CD19+(B cell) | TSGF | -0.02456 | 9.78E-08 |
| B15 | B108 | CD3-CD19+(B cell) | Anti-Cyclin | -0.0238 | 2.37E-07 |
| B15 | B60 | CD3-CD19+(B cell) | WBRVI | -0.02299 | 5.99E-07 |
| B150 | B179 | ALP | α1-MG | 0.055586 | 0 |
| B150 | B186 | ALP | PFC | 0.041669 | 0 |
| B150 | B193 | ALP | K | -0.04172 | 0 |
| B150 | B195 | ALP | Na | 0.076378 | 0 |
| B150 | B197 | ALP | AG | 0.05977 | 0 |
| B150 | B199 | ALP | Mg | 0.054464 | 0 |
| B150 | B200 | ALP | OP | 0.065499 | 0 |
| B150 | B221 | ALP | Hei | -0.03081 | 2.26E-11 |
| B150 | B170 | ALP | U-GLU | 0.029376 | 1.80E-10 |
| B150 | B174 | ALP | Urea/Cr | 0.027238 | 3.36E-09 |
| B150 | B198 | ALP | P | -0.02673 | 6.51E-09 |
| B150 | B155 | ALP | IgA | 0.026715 | 6.65E-09 |

| | | | | | |
|---|---|---|---|---|---|
| B150 | B216 | ALP | BMI | -0.02626 | 1.20E-08 |
| B151 | B157 | Anti-CCP | IgG | 0.031934 | 4.12E-12 |
| B152 | B181 | ACA-Ig(G-A-M) | APTT | -0.04662 | 0 |
| B152 | B182 | ACA-Ig(G-A-M) | TT | -0.04497 | 0 |
| B152 | B183 | ACA-Ig(G-A-M) | PTA | 0.079601 | 0 |
| B152 | B156 | ACA-Ig(G-A-M) | IgE | 0.029088 | 2.70E-10 |
| B152 | B155 | ACA-Ig(G-A-M) | IgA | 0.026325 | 1.10E-08 |
| B152 | B194 | ACA-Ig(G-A-M) | Cl | 0.026038 | 1.58E-08 |
| B152 | B186 | ACA-Ig(G-A-M) | PFC | 0.022976 | 6.11E-07 |
| B153 | B160 | Glu | PH | -0.07298 | 0 |
| B153 | B163 | Glu | SG | 0.107225 | 0 |
| B153 | B166 | Glu | Pro | 0.079095 | 0 |
| B153 | B170 | Glu | U-GLU | 0.513936 | 0 |
| B153 | B172 | Glu | KB | 0.07406 | 0 |
| B153 | B174 | Glu | Urea/Cr | 0.055293 | 0 |
| B153 | B180 | Glu | Tf | 0.040778 | 0 |
| B153 | B188 | Glu | Ca | 0.069559 | 0 |
| B153 | B194 | Glu | Cl | -0.04444 | 0 |
| B153 | B206 | Glu | Waist | 0.138325 | 0 |
| B153 | B210 | Glu | Hip | 0.080769 | 0 |
| B153 | B215 | Glu | BMI | 0.118914 | 0 |
| B153 | B216 | Glu | BMI | 0.098025 | 0 |
| B153 | B220 | Glu | SBP | 0.096938 | 0 |
| B153 | B222 | Glu | NS | 0.09918 | 0 |
| B153 | B225 | Glu | hPBG | 0.061668 | 0 |
| B153 | B212 | Glu | IOP-L | 0.037197 | 6.66E-16 |
| B153 | B204 | Glu | Drinking | 0.035429 | 1.44E-14 |
| B153 | B178 | Glu | Ig-U | 0.034137 | 1.25E-13 |
| B153 | B211 | Glu | IOP-R | 0.033757 | 2.32E-13 |
| B153 | B179 | Glu | α1-MG | 0.033606 | 2.96E-13 |
| B153 | B177 | Glu | ACR | 0.033302 | 4.82E-13 |
| B153 | B191 | Glu | Urea | 0.030962 | 1.79E-11 |
| B153 | B218 | Glu | DBP | 0.030631 | 2.93E-11 |
| B153 | B195 | Glu | Na | -0.02946 | 1.60E-10 |
| B153 | B175 | Glu | ALB-U | 0.027157 | 3.74E-09 |
| B153 | B200 | Glu | OP | 0.025471 | 3.21E-08 |
| B154 | B155 | RF | IgA | 0.023975 | 1.94E-07 |
| B155 | B157 | IgA | IgG | 0.283739 | 0 |
| B155 | B158 | IgA | IgM | -0.0476 | 0 |
| B155 | B186 | IgA | PFC | 0.037988 | 0 |
| B155 | B156 | IgA | IgE | 0.033538 | 3.30E-13 |
| B155 | B178 | IgA | Ig-U | 0.023955 | 1.99E-07 |
| B155 | B209 | IgA | Smoking | -0.02386 | 2.23E-07 |
| B155 | B194 | IgA | Cl | -0.02333 | 4.08E-07 |

| | | | | | |
|---|---|---|---|---|---|
| B156 | B157 | IgE | IgG | 0.040895 | 0 |
| B156 | B193 | IgE | K | 0.046638 | 0 |
| B156 | B172 | IgE | KB | 0.025038 | 5.47E-08 |
| B157 | B193 | IgG | K | 0.046034 | 0 |
| B157 | B195 | IgG | Na | -0.06284 | 0 |
| B157 | B200 | IgG | OP | -0.03956 | 0 |
| B157 | B209 | IgG | Smoking | -0.04333 | 0 |
| B157 | B221 | IgG | Hei | -0.03735 | 4.44E-16 |
| B157 | B222 | IgG | NS | -0.0375 | 4.44E-16 |
| B157 | B198 | IgG | P | 0.035355 | 1.64E-14 |
| B157 | B184 | IgG | INR | 0.03527 | 1.89E-14 |
| B157 | B204 | IgG | Drinking | -0.03509 | 2.58E-14 |
| B157 | B197 | IgG | AG | -0.03444 | 7.53E-14 |
| B157 | B216 | IgG | BMI | -0.03124 | 1.18E-11 |
| B157 | B179 | IgG | α1-MG | -0.03123 | 1.20E-11 |
| B157 | B185 | IgG | PT | 0.02478 | 7.47E-08 |
| B157 | B194 | IgG | Cl | -0.02455 | 9.87E-08 |
| B157 | B183 | IgG | PTA | 0.024154 | 1.58E-07 |
| B159 | B186 | β2-MG | PFC | 0.034419 | 7.86E-14 |
| B159 | B182 | β2-MG | TT | -0.03112 | 1.42E-11 |
| B159 | B183 | β2-MG | PTA | -0.02425 | 1.40E-07 |
| B16 | B108 | CD3+T | Anti-Cyclin | 0.057579 | 0 |
| B16 | B138 | CD3+T | PAB | 0.044985 | 0 |
| B16 | B139 | CD3+T | ADA | -0.04762 | 0 |
| B16 | B155 | CD3+T | IgA | -0.03943 | 0 |
| B16 | B157 | CD3+T | IgG | -0.09188 | 0 |
| B16 | B17 | CD3+T | CD3+CD16+CD56+NK | 0.162667 | 0 |
| B16 | B18 | CD3+T | CD3+CD4+helper(inducer)T | 0.456512 | 0 |
| B16 | B19 | CD3+T | CD3+CD4+/CD3+CD8+ | -0.17303 | 0 |
| B16 | B20 | CD3+T | CD3+CD4+CD8+ | 0.143701 | 0 |
| B16 | B21 | CD3+T | CD3+CD4-CD8- | 0.169731 | 0 |
| B16 | B22 | CD3+T | CD3+CD8+(T) | 0.605682 | 0 |
| B16 | B23 | CD3+T | CD3-CD16+CD56+(NK) | -0.61812 | 0 |
| B16 | B30 | CD3+T | LH | 0.040437 | 0 |
| B16 | B32 | CD3+T | Testo | -0.08731 | 0 |
| B16 | B74 | CD3+T | ApoA1 | -0.04827 | 0 |
| B16 | B92 | CD3+T | ALDH2-Glu504Glu | 0.068408 | 0 |
| B16 | B93 | CD3+T | ALDH2-Glu504Lys | -0.07013 | 0 |
| B16 | B104 | CD3+T | anti-SSB | -0.03449 | 6.99E-14 |
| B16 | B54 | CD3+T | ESR | -0.0328 | 1.07E-12 |
| B16 | B105 | CD3+T | anti-U1RNP | -0.03188 | 4.49E-12 |
| B16 | B184 | CD3+T | INR | -0.03125 | 1.17E-11 |
| B16 | B81 | CD3+T | TSGF | 0.027415 | 2.66E-09 |
| B16 | B96 | CD3+T | dsDNA (IIF) | -0.02721 | 3.49E-09 |

| | | | | | |
|---|---|---|---|---|---|
| B16 | B185 | CD3+T | PT | -0.02678 | 6.11E-09 |
| B16 | B58 | CD3+T | HCT | -0.02555 | 2.92E-08 |
| B16 | B125 | CD3+T | Cor | -0.02421 | 1.47E-07 |
| B16 | B31 | CD3+T | FSH | 0.024116 | 1.65E-07 |
| B16 | B28 | CD3+T | PRL | 0.023638 | 2.88E-07 |
| B16 | B176 | CD3+T | Ucr | 0.023451 | 3.57E-07 |
| B16 | B119 | CD3+T | Hcv-rna | -0.02295 | 6.33E-07 |
| B16 | B100 | CD3+T | anti-Ro52 | -0.02276 | 7.77E-07 |
| B160 | B163 | PH | SG | -0.19168 | 0 |
| B160 | B170 | PH | U-GLU | -0.04764 | 0 |
| B160 | B174 | PH | Urea/Cr | -0.04605 | 0 |
| B160 | B176 | PH | Ucr | -0.08041 | 0 |
| B160 | B188 | PH | Ca | -0.04522 | 0 |
| B160 | B189 | PH | eGFB | 0.062311 | 0 |
| B160 | B190 | PH | Cre | -0.08959 | 0 |
| B160 | B191 | PH | Urea | -0.17333 | 0 |
| B160 | B192 | PH | UA | -0.16695 | 0 |
| B160 | B206 | PH | Waist | -0.08462 | 0 |
| B160 | B209 | PH | Smoking | -0.07087 | 0 |
| B160 | B210 | PH | Hip | -0.06277 | 0 |
| B160 | B215 | PH | BMI | -0.06142 | 0 |
| B160 | B216 | PH | BMI | -0.0676 | 0 |
| B160 | B218 | PH | DBP | 0.055755 | 0 |
| B160 | B220 | PH | SBP | 0.064866 | 0 |
| B160 | B221 | PH | Hei | -0.04052 | 0 |
| B160 | B222 | PH | NS | -0.07102 | 0 |
| B160 | B204 | PH | Drinking | -0.0373 | 4.44E-16 |
| B160 | B198 | PH | P | -0.03547 | 1.35E-14 |
| B160 | B166 | PH | Pro | -0.03405 | 1.44E-13 |
| B160 | B193 | PH | K | -0.0339 | 1.83E-13 |
| B160 | B194 | PH | Cl | -0.03133 | 1.04E-11 |
| B160 | B196 | PH | TCO2 | 0.030083 | 6.53E-11 |
| B160 | B168 | PH | NMP22 | 0.02525 | 4.22E-08 |
| B161 | B163 | LEU | SG | -0.04887 | 0 |
| B161 | B173 | LEU | NTT | 0.170839 | 0 |
| B161 | B174 | LEU | Urea/Cr | 0.099057 | 0 |
| B161 | B190 | LEU | Cre | -0.04438 | 0 |
| B161 | B192 | LEU | UA | -0.06959 | 0 |
| B161 | B204 | LEU | Drinking | -0.0628 | 0 |
| B161 | B206 | LEU | Waist | -0.04403 | 0 |
| B161 | B209 | LEU | Smoking | -0.06035 | 0 |
| B161 | B216 | LEU | BMI | -0.09582 | 0 |
| B161 | B218 | LEU | DBP | -0.0425 | 0 |
| B161 | B221 | LEU | Hei | -0.15606 | 0 |

| | | | | | |
|---|---|---|---|---|---|
| B161 | B222 | LEU | NS | -0.1216 | 0 |
| B161 | B219 | LEU | OT | 0.036818 | 1.33E-15 |
| B161 | B166 | LEU | Pro | 0.031343 | 1.01E-11 |
| B161 | B210 | LEU | Hip | -0.02821 | 9.14E-10 |
| B161 | B189 | LEU | eGFB | -0.0275 | 2.38E-09 |
| B161 | B201 | LEU | e-habits | -0.02378 | 2.44E-07 |
| B163 | B166 | SG | Pro | 0.321912 | 0 |
| B163 | B170 | SG | U-GLU | 0.119404 | 0 |
| B163 | B172 | SG | KB | 0.075096 | 0 |
| B163 | B173 | SG | NTT | -0.0539 | 0 |
| B163 | B176 | SG | Ucr | 0.285825 | 0 |
| B163 | B179 | SG | α1-MG | 0.060563 | 0 |
| B163 | B188 | SG | Ca | -0.04734 | 0 |
| B163 | B189 | SG | eGFB | 0.05877 | 0 |
| B163 | B190 | SG | Cre | 0.095175 | 0 |
| B163 | B191 | SG | Urea | 0.174321 | 0 |
| B163 | B192 | SG | UA | 0.159515 | 0 |
| B163 | B204 | SG | Drinking | 0.163182 | 0 |
| B163 | B205 | SG | KDA | -0.04552 | 0 |
| B163 | B206 | SG | Waist | 0.217039 | 0 |
| B163 | B209 | SG | Smoking | 0.18082 | 0 |
| B163 | B210 | SG | Hip | 0.159358 | 0 |
| B163 | B215 | SG | BMI | 0.151395 | 0 |
| B163 | B216 | SG | BMI | 0.2864 | 0 |
| B163 | B218 | SG | DBP | 0.095031 | 0 |
| B163 | B219 | SG | OT | -0.07261 | 0 |
| B163 | B220 | SG | SBP | -0.08025 | 0 |
| B163 | B221 | SG | Hei | 0.295616 | 0 |
| B163 | B222 | SG | NS | 0.311398 | 0 |
| B163 | B201 | SG | e-habits | -0.0319 | 4.34E-12 |
| B163 | B165 | SG | bilirubin | 0.03108 | 1.50E-11 |
| B163 | B203 | SG | DH | 0.028864 | 3.70E-10 |
| B163 | B198 | SG | P | -0.023 | 5.93E-07 |
| B163 | B178 | SG | Ig-U | 0.022821 | 7.27E-07 |
| B165 | B172 | bilirubin | KB | 0.039154 | 0 |
| B166 | B170 | Pro | U-GLU | 0.076825 | 0 |
| B166 | B172 | Pro | KB | 0.05263 | 0 |
| B166 | B174 | Pro | Urea/Cr | -0.04696 | 0 |
| B166 | B175 | Pro | ALB-U | 0.047301 | 0 |
| B166 | B176 | Pro | Ucr | 0.128767 | 0 |
| B166 | B179 | Pro | α1-MG | 0.039013 | 0 |
| B166 | B189 | Pro | eGFB | -0.0618 | 0 |
| B166 | B190 | Pro | Cre | 0.099287 | 0 |
| B166 | B191 | Pro | Urea | 0.080083 | 0 |

| | | | | | |
|---|---|---|---|---|---|
| B166 | B192 | Pro | UA | 0.08385 | 0 |
| B166 | B204 | Pro | Drinking | 0.055173 | 0 |
| B166 | B206 | Pro | Waist | 0.093623 | 0 |
| B166 | B209 | Pro | Smoking | 0.086151 | 0 |
| B166 | B210 | Pro | Hip | 0.055265 | 0 |
| B166 | B215 | Pro | BMI | 0.062949 | 0 |
| B166 | B216 | Pro | BMI | 0.104501 | 0 |
| B166 | B218 | Pro | DBP | 0.054078 | 0 |
| B166 | B221 | Pro | Hei | 0.097571 | 0 |
| B166 | B222 | Pro | NS | 0.121801 | 0 |
| B166 | B177 | Pro | ACR | 0.036813 | 1.33E-15 |
| B166 | B188 | Pro | Ca | -0.0348 | 4.20E-14 |
| B166 | B220 | Pro | SBP | 0.032763 | 1.13E-12 |
| B166 | B178 | Pro | Ig-U | 0.028269 | 8.41E-10 |
| B166 | B203 | Pro | DH | 0.024102 | 1.68E-07 |
| B168 | B183 | NMP22 | PTA | 0.037927 | 2.22E-16 |
| B168 | B173 | NMP22 | NTT | 0.031065 | 1.54E-11 |
| B168 | B193 | NMP22 | K | -0.02877 | 4.23E-10 |
| B169 | B196 | U-CRY | TCO2 | 0.023381 | 3.86E-07 |
| B17 | B136 | CD3+CD16+CD56+NK | 5'NT | -0.04081 | 0 |
| B17 | B18 | CD3+CD16+CD56+NK | CD3+CD4+helper(inducer)T | -0.16747 | 0 |
| B17 | B182 | CD3+CD16+CD56+NK | TT | -0.08322 | 0 |
| B17 | B183 | CD3+CD16+CD56+NK | PTA | 0.061179 | 0 |
| B17 | B19 | CD3+CD16+CD56+NK | CD3+CD4+/CD3+CD8+ | -0.24224 | 0 |
| B17 | B20 | CD3+CD16+CD56+NK | CD3+CD4+CD8+ | 0.072881 | 0 |
| B17 | B21 | CD3+CD16+CD56+NK | CD3+CD4-CD8- | 0.259926 | 0 |
| B17 | B22 | CD3+CD16+CD56+NK | CD3+CD8+(T) | 0.269105 | 0 |
| B17 | B23 | CD3+CD16+CD56+NK | CD3-CD16+CD56+(NK) | -0.06483 | 0 |
| B17 | B95 | CD3+CD16+CD56+NK | anti-cenpb | -0.0426 | 0 |
| B17 | B99 | CD3+CD16+CD56+NK | anti-PM-Scl | -0.06513 | 0 |
| B17 | B137 | CD3+CD16+CD56+NK | AFU | -0.03236 | 2.12E-12 |
| B17 | B186 | CD3+CD16+CD56+NK | PFC | 0.032341 | 2.19E-12 |
| B17 | B125 | CD3+CD16+CD56+NK | Cor | 0.032013 | 3.65E-12 |
| B17 | B181 | CD3+CD16+CD56+NK | APTT | 0.031655 | 6.31E-12 |
| B17 | B76 | CD3+CD16+CD56+NK | ApoB | 0.027619 | 2.02E-09 |
| B17 | B105 | CD3+CD16+CD56+NK | anti-U1RNP | 0.026864 | 5.48E-09 |
| B17 | B139 | CD3+CD16+CD56+NK | ADA | -0.02578 | 2.19E-08 |
| B17 | B135 | CD3+CD16+CD56+NK | TBA | -0.02324 | 4.55E-07 |
| B170 | B172 | U-GLU | KB | 0.091254 | 0 |
| B170 | B180 | U-GLU | Tf | 0.040283 | 0 |
| B170 | B192 | U-GLU | UA | -0.03971 | 0 |
| B170 | B204 | U-GLU | Drinking | 0.037976 | 0 |
| B170 | B206 | U-GLU | Waist | 0.055683 | 0 |
| B170 | B209 | U-GLU | Smoking | 0.046635 | 0 |

| | | | | | |
|---|---|---|---|---|---|
| B170 | B216 | U-GLU | BMI | 0.049388 | 0 |
| B170 | B221 | U-GLU | Hei | 0.03985 | 0 |
| B170 | B222 | U-GLU | NS | 0.059792 | 0 |
| B170 | B225 | U-GLU | hPBG | 0.037665 | 4.44E-16 |
| B170 | B215 | U-GLU | BMI | 0.035249 | 1.95E-14 |
| B170 | B194 | U-GLU | Cl | -0.03512 | 2.42E-14 |
| B170 | B178 | U-GLU | Ig-U | 0.03473 | 4.69E-14 |
| B170 | B218 | U-GLU | DBP | 0.034012 | 1.53E-13 |
| B170 | B220 | U-GLU | SBP | 0.031496 | 8.03E-12 |
| B170 | B179 | U-GLU | α1-MG | 0.030623 | 2.97E-11 |
| B170 | B189 | U-GLU | eGFB | 0.02934 | 1.89E-10 |
| B170 | B177 | U-GLU | ACR | 0.028679 | 4.78E-10 |
| B170 | B191 | U-GLU | Urea | 0.0263 | 1.13E-08 |
| B170 | B174 | U-GLU | Urea/Cr | 0.025653 | 2.56E-08 |
| B170 | B175 | U-GLU | ALB-U | 0.025039 | 5.46E-08 |
| B172 | B204 | KB | Drinking | 0.030439 | 3.89E-11 |
| B172 | B188 | KB | Ca | 0.028183 | 9.45E-10 |
| B172 | B176 | KB | Ucr | 0.027342 | 2.92E-09 |
| B172 | B189 | KB | eGFB | 0.026845 | 5.62E-09 |
| B173 | B174 | NTT | Urea/Cr | 0.054511 | 0 |
| B173 | B189 | NTT | eGFB | -0.05707 | 0 |
| B173 | B204 | NTT | Drinking | -0.04258 | 0 |
| B173 | B209 | NTT | Smoking | -0.04148 | 0 |
| B173 | B216 | NTT | BMI | -0.07475 | 0 |
| B173 | B221 | NTT | Hei | -0.09571 | 0 |
| B173 | B222 | NTT | NS | -0.07996 | 0 |
| B173 | B218 | NTT | DBP | -0.03622 | 3.55E-15 |
| B173 | B192 | NTT | UA | -0.03245 | 1.84E-12 |
| B173 | B206 | NTT | Waist | -0.03241 | 1.96E-12 |
| B173 | B191 | NTT | Urea | 0.030391 | 4.17E-11 |
| B173 | B176 | NTT | Ucr | -0.03036 | 4.39E-11 |
| B173 | B210 | NTT | Hip | -0.03031 | 4.67E-11 |
| B173 | B220 | NTT | SBP | 0.025759 | 2.25E-08 |
| B173 | B215 | NTT | BMI | -0.02517 | 4.67E-08 |
| B173 | B219 | NTT | OT | 0.023957 | 1.99E-07 |
| B174 | B176 | Urea/Cr | Ucr | -0.06917 | 0 |
| B174 | B189 | Urea/Cr | eGFB | 0.18244 | 0 |
| B174 | B190 | Urea/Cr | Cre | -0.32958 | 0 |
| B174 | B191 | Urea/Cr | Urea | 0.492792 | 0 |
| B174 | B192 | Urea/Cr | UA | -0.25096 | 0 |
| B174 | B198 | Urea/Cr | P | 0.042676 | 0 |
| B174 | B200 | Urea/Cr | OP | 0.046807 | 0 |
| B174 | B204 | Urea/Cr | Drinking | -0.13302 | 0 |
| B174 | B206 | Urea/Cr | Waist | -0.12498 | 0 |

| | | | | | |
|---|---|---|---|---|---|
| B174 | B209 | Urea/Cr | Smoking | -0.13893 | 0 |
| B174 | B210 | Urea/Cr | Hip | -0.10316 | 0 |
| B174 | B215 | Urea/Cr | BMI | -0.04082 | 0 |
| B174 | B216 | Urea/Cr | BMI | -0.23203 | 0 |
| B174 | B218 | Urea/Cr | DBP | -0.07828 | 0 |
| B174 | B219 | Urea/Cr | OT | 0.056075 | 0 |
| B174 | B220 | Urea/Cr | SBP | 0.041228 | 0 |
| B174 | B221 | Urea/Cr | Hei | -0.33345 | 0 |
| B174 | B222 | Urea/Cr | NS | -0.27261 | 0 |
| B174 | B188 | Urea/Cr | Ca | 0.022577 | 9.54E-07 |
| B175 | B176 | ALB-U | Ucr | 0.084365 | 0 |
| B175 | B177 | ALB-U | ACR | 0.792501 | 0 |
| B175 | B178 | ALB-U | Ig-U | 0.118585 | 0 |
| B175 | B179 | ALB-U | α1-MG | 0.057124 | 0 |
| B175 | B180 | ALB-U | Tf | 0.110939 | 0 |
| B175 | B189 | ALB-U | eGFB | -0.06568 | 0 |
| B175 | B190 | ALB-U | Cre | 0.114583 | 0 |
| B175 | B191 | ALB-U | Urea | 0.073086 | 0 |
| B175 | B192 | ALB-U | UA | 0.039827 | 0 |
| B175 | B218 | ALB-U | DBP | 0.044999 | 0 |
| B175 | B220 | ALB-U | SBP | 0.040491 | 0 |
| B175 | B187 | ALB-U | NSE | 0.033168 | 5.97E-13 |
| B175 | B198 | ALB-U | P | 0.029456 | 1.61E-10 |
| B175 | B216 | ALB-U | BMI | 0.028949 | 3.28E-10 |
| B175 | B206 | ALB-U | Waist | 0.027592 | 2.10E-09 |
| B175 | B222 | ALB-U | NS | 0.026164 | 1.35E-08 |
| B175 | B215 | ALB-U | BMI | 0.025756 | 2.25E-08 |
| B176 | B177 | Ucr | ACR | -0.059 | 0 |
| B176 | B178 | Ucr | Ig-U | 0.073756 | 0 |
| B176 | B179 | Ucr | α1-MG | 0.162156 | 0 |
| B176 | B190 | Ucr | Cre | 0.05094 | 0 |
| B176 | B192 | Ucr | UA | 0.08102 | 0 |
| B176 | B204 | Ucr | Drinking | 0.058543 | 0 |
| B176 | B206 | Ucr | Waist | 0.078324 | 0 |
| B176 | B209 | Ucr | Smoking | 0.072424 | 0 |
| B176 | B210 | Ucr | Hip | 0.06406 | 0 |
| B176 | B215 | Ucr | BMI | 0.058082 | 0 |
| B176 | B216 | Ucr | BMI | 0.119513 | 0 |
| B176 | B218 | Ucr | DBP | 0.040657 | 0 |
| B176 | B220 | Ucr | SBP | -0.04897 | 0 |
| B176 | B221 | Ucr | Hei | 0.12591 | 0 |
| B176 | B222 | Ucr | NS | 0.128483 | 0 |
| B176 | B201 | Ucr | e-habits | -0.03783 | 2.22E-16 |
| B176 | B219 | Ucr | OT | -0.0345 | 6.82E-14 |

| | | | | | |
|---|---|---|---|---|---|
| B176 | B193 | Ucr | K | -0.03398 | 1.60E-13 |
| B176 | B180 | Ucr | Tf | 0.030644 | 2.87E-11 |
| B176 | B198 | Ucr | P | -0.0236 | 3.02E-07 |
| B177 | B178 | ACR | Ig-U | 0.217913 | 0 |
| B177 | B179 | ACR | α1-MG | 0.074194 | 0 |
| B177 | B180 | ACR | Tf | 0.257501 | 0 |
| B177 | B189 | ACR | eGFB | -0.06192 | 0 |
| B177 | B190 | ACR | Cre | 0.108365 | 0 |
| B177 | B191 | ACR | Urea | 0.081464 | 0 |
| B177 | B198 | ACR | P | 0.038441 | 0 |
| B177 | B220 | ACR | SBP | 0.048348 | 0 |
| B177 | B218 | ACR | DBP | 0.031197 | 1.26E-11 |
| B177 | B192 | ACR | UA | 0.025754 | 2.26E-08 |
| B178 | B179 | Ig-U | α1-MG | 0.423824 | 0 |
| B178 | B180 | Ig-U | Tf | 0.881563 | 0 |
| B178 | B187 | Ig-U | NSE | 0.048409 | 0 |
| B178 | B193 | Ig-U | K | -0.03247 | 1.78E-12 |
| B178 | B186 | Ig-U | PFC | 0.028039 | 1.15E-09 |
| B178 | B194 | Ig-U | Cl | -0.02365 | 2.84E-07 |
| B179 | B180 | α1-MG | Tf | 0.351159 | 0 |
| B179 | B182 | α1-MG | TT | 0.058 | 0 |
| B179 | B193 | α1-MG | K | -0.0693 | 0 |
| B179 | B198 | α1-MG | P | -0.03821 | 0 |
| B179 | B225 | α1-MG | hPBG | 0.04598 | 0 |
| B179 | B181 | α1-MG | APTT | 0.03675 | 1.33E-15 |
| B179 | B183 | α1-MG | PTA | -0.03612 | 4.44E-15 |
| B179 | B209 | α1-MG | Smoking | 0.029393 | 1.76E-10 |
| B179 | B221 | α1-MG | Hei | 0.026793 | 6.01E-09 |
| B179 | B190 | α1-MG | Cre | 0.025798 | 2.14E-08 |
| B179 | B200 | α1-MG | OP | 0.025006 | 5.68E-08 |
| B179 | B191 | α1-MG | Urea | 0.022613 | 9.16E-07 |
| B18 | B104 | CD3+CD4+helper(inducer)T | anti-SSB | -0.05025 | 0 |
| B18 | B107 | CD3+CD4+helper(inducer)T | AnuA | -0.04646 | 0 |
| B18 | B108 | CD3+CD4+helper(inducer)T | Anti-Cyclin | 0.04815 | 0 |
| B18 | B110 | CD3+CD4+helper(inducer)T | AHA | 0.046148 | 0 |
| B18 | B122 | CD3+CD4+helper(inducer)T | C4 | 0.052849 | 0 |
| B18 | B125 | CD3+CD4+helper(inducer)T | Cor | -0.05464 | 0 |
| B18 | B136 | CD3+CD4+helper(inducer)T | 5'NT | 0.041816 | 0 |
| B18 | B157 | CD3+CD4+helper(inducer)T | IgG | -0.06273 | 0 |
| B18 | B183 | CD3+CD4+helper(inducer)T | PTA | -0.04519 | 0 |
| B18 | B186 | CD3+CD4+helper(inducer)T | PFC | 0.046977 | 0 |
| B18 | B19 | CD3+CD4+helper(inducer)T | CD3+CD4+/CD3+CD8+ | 0.657733 | 0 |
| B18 | B193 | CD3+CD4+helper(inducer)T | K | -0.04124 | 0 |
| B18 | B20 | CD3+CD4+helper(inducer)T | CD3+CD4+CD8+ | 0.109364 | 0 |

| | | | | | |
|---|---|---|---|---|---|
| B18 | B21 | CD3+CD4+helper(inducer)T | CD3+CD4-CD8- | -0.12315 | 0 |
| B18 | B22 | CD3+CD4+helper(inducer)T | CD3+CD8+(T) | -0.33889 | 0 |
| B18 | B23 | CD3+CD4+helper(inducer)T | CD3-CD16+CD56+(NK) | -0.34513 | 0 |
| B18 | B32 | CD3+CD4+helper(inducer)T | Testo | -0.05826 | 0 |
| B18 | B79 | CD3+CD4+helper(inducer)T | hb V -dna | -0.05031 | 0 |
| B18 | B93 | CD3+CD4+helper(inducer)T | ALDH2-Glu504Lys | -0.03976 | 0 |
| B18 | B98 | CD3+CD4+helper(inducer)T | anti-Jo-1 | -0.04621 | 0 |
| B18 | B99 | CD3+CD4+helper(inducer)T | anti-PM-Scl | 0.039911 | 0 |
| B18 | B92 | CD3+CD4+helper(inducer)T | ALDH2-Glu504Glu | 0.037955 | 2.22E-16 |
| B18 | B198 | CD3+CD4+helper(inducer)T | P | 0.036425 | 2.66E-15 |
| B18 | B156 | CD3+CD4+helper(inducer)T | IgE | 0.036155 | 4.00E-15 |
| B18 | B195 | CD3+CD4+helper(inducer)T | Na | 0.036074 | 4.88E-15 |
| B18 | B194 | CD3+CD4+helper(inducer)T | Cl | 0.03501 | 2.93E-14 |
| B18 | B97 | CD3+CD4+helper(inducer)T | dsDNA | -0.03482 | 4.02E-14 |
| B18 | B138 | CD3+CD4+helper(inducer)T | PAB | -0.03307 | 6.99E-13 |
| B18 | B196 | CD3+CD4+helper(inducer)T | TCO2 | -0.03207 | 3.35E-12 |
| B18 | B74 | CD3+CD4+helper(inducer)T | ApoA1 | -0.03191 | 4.29E-12 |
| B18 | B95 | CD3+CD4+helper(inducer)T | anti-cenpb | 0.0317 | 5.89E-12 |
| B18 | B30 | CD3+CD4+helper(inducer)T | LH | 0.026895 | 5.26E-09 |
| B18 | B197 | CD3+CD4+helper(inducer)T | AG | 0.024818 | 7.14E-08 |
| B18 | B123 | CD3+CD4+helper(inducer)T | hs-CRP1 | 0.024013 | 1.86E-07 |
| B18 | B184 | CD3+CD4+helper(inducer)T | INR | -0.02387 | 2.20E-07 |
| B18 | B28 | CD3+CD4+helper(inducer)T | PRL | 0.0229 | 6.66E-07 |
| B18 | B124 | CD3+CD4+helper(inducer)T | ACTH | -0.02279 | 7.55E-07 |
| B18 | B75 | CD3+CD4+helper(inducer)T | ApoA1/B | -0.02269 | 8.45E-07 |
| B180 | B187 | Tf | NSE | 0.053879 | 0 |
| B180 | B186 | Tf | PFC | 0.032851 | 9.87E-13 |
| B181 | B182 | APTT | TT | 0.099627 | 0 |
| B181 | B183 | APTT | PTA | 0.069529 | 0 |
| B181 | B184 | APTT | INR | 0.501398 | 0 |
| B181 | B185 | APTT | PT | 0.462767 | 0 |
| B181 | B186 | APTT | PFC | -0.04954 | 0 |
| B181 | B194 | APTT | Cl | 0.033494 | 3.54E-13 |
| B181 | B197 | APTT | AG | -0.03329 | 4.92E-13 |
| B181 | B193 | APTT | K | 0.030528 | 3.41E-11 |
| B182 | B183 | TT | PTA | -0.20101 | 0 |
| B182 | B184 | TT | INR | -0.04572 | 0 |
| B182 | B186 | TT | PFC | -0.21609 | 0 |
| B182 | B194 | TT | Cl | -0.04799 | 0 |
| B182 | B195 | TT | Na | 0.060301 | 0 |
| B182 | B197 | TT | AG | 0.071067 | 0 |
| B182 | B200 | TT | OP | 0.037293 | 4.44E-16 |
| B182 | B199 | TT | Mg | 0.027241 | 3.34E-09 |
| B182 | B198 | TT | P | -0.02371 | 2.66E-07 |

| | | | | | |
|---|---|---|---|---|---|
| B183 | B184 | PTA | INR | 0.115539 | 0 |
| B183 | B186 | PTA | PFC | -0.17544 | 0 |
| B183 | B194 | PTA | Cl | 0.045934 | 0 |
| B183 | B197 | PTA | AG | -0.07086 | 0 |
| B183 | B198 | PTA | P | -0.03137 | 9.78E-12 |
| B183 | B196 | PTA | TCO2 | 0.028905 | 3.49E-10 |
| B183 | B195 | PTA | Na | -0.02372 | 2.60E-07 |
| B183 | B200 | PTA | OP | -0.02261 | 9.19E-07 |
| B184 | B185 | INR | PT | 0.945881 | 0 |
| B184 | B186 | INR | PFC | -0.06959 | 0 |
| B184 | B193 | INR | K | 0.040335 | 0 |
| B184 | B195 | INR | Na | -0.04417 | 0 |
| B184 | B197 | INR | AG | -0.03753 | 4.44E-16 |
| B184 | B189 | INR | eGFB | -0.03644 | 2.66E-15 |
| B184 | B200 | INR | OP | -0.03575 | 8.44E-15 |
| B184 | B199 | INR | Mg | -0.02734 | 2.92E-09 |
| B184 | B194 | INR | Cl | 0.022674 | 8.56E-07 |
| B185 | B186 | PT | PFC | 0.047293 | 0 |
| B185 | B195 | PT | Na | -0.02355 | 3.19E-07 |
| B185 | B189 | PT | eGFB | -0.0233 | 4.23E-07 |
| B186 | B193 | PFC | K | -0.04717 | 0 |
| B186 | B194 | PFC | Cl | -0.06659 | 0 |
| B186 | B198 | PFC | P | 0.047584 | 0 |
| B186 | B199 | PFC | Mg | 0.03997 | 0 |
| B186 | B197 | PFC | AG | 0.036318 | 3.11E-15 |
| B187 | B225 | NSE | hPBG | -0.03264 | 1.37E-12 |
| B187 | B194 | NSE | Cl | -0.03247 | 1.80E-12 |
| B187 | B193 | NSE | K | -0.02666 | 7.15E-09 |
| B188 | B190 | Ca | Cre | -0.03868 | 0 |
| B188 | B192 | Ca | UA | 0.085267 | 0 |
| B188 | B193 | Ca | K | 0.060831 | 0 |
| B188 | B194 | Ca | Cl | -0.0447 | 0 |
| B188 | B211 | Ca | IOP-R | 0.037671 | 4.44E-16 |
| B188 | B206 | Ca | Waist | -0.03498 | 3.06E-14 |
| B188 | B212 | Ca | IOP-L | 0.033515 | 3.43E-13 |
| B188 | B222 | Ca | NS | -0.03264 | 1.37E-12 |
| B188 | B197 | Ca | AG | 0.031638 | 6.47E-12 |
| B188 | B216 | Ca | BMI | -0.03106 | 1.54E-11 |
| B188 | B218 | Ca | DBP | 0.030295 | 4.80E-11 |
| B188 | B198 | Ca | P | 0.027273 | 3.21E-09 |
| B188 | B201 | Ca | e-habits | 0.027161 | 3.71E-09 |
| B188 | B215 | Ca | BMI | -0.02569 | 2.45E-08 |
| B188 | B221 | Ca | Hei | -0.02435 | 1.24E-07 |
| B189 | B190 | eGFB | Cre | -0.50678 | 0 |

| | | | | | |
|---|---|---|---|---|---|
| B189 | B191 | eGFB | Urea | -0.42399 | 0 |
| B189 | B192 | eGFB | UA | -0.23658 | 0 |
| B189 | B193 | eGFB | K | -0.04264 | 0 |
| B189 | B203 | eGFB | DH | 0.039829 | 0 |
| B189 | B204 | eGFB | Drinking | 0.104047 | 0 |
| B189 | B209 | eGFB | Smoking | 0.068922 | 0 |
| B189 | B210 | eGFB | Hip | 0.10232 | 0 |
| B189 | B212 | eGFB | IOP-L | 0.039446 | 0 |
| B189 | B215 | eGFB | BMI | 0.098161 | 0 |
| B189 | B216 | eGFB | BMI | 0.136396 | 0 |
| B189 | B218 | eGFB | DBP | 0.187327 | 0 |
| B189 | B219 | eGFB | OT | -0.05556 | 0 |
| B189 | B220 | eGFB | SBP | -0.13501 | 0 |
| B189 | B221 | eGFB | Hei | 0.107677 | 0 |
| B189 | B222 | eGFB | NS | 0.05857 | 0 |
| B189 | B211 | eGFB | IOP-R | 0.037641 | 4.44E-16 |
| B189 | B227 | eGFB | OD | -0.03551 | 1.27E-14 |
| B189 | B228 | eGFB | OS | -0.03262 | 1.42E-12 |
| B189 | B201 | eGFB | e-habits | -0.03179 | 5.13E-12 |
| B189 | B194 | eGFB | Cl | -0.02831 | 7.89E-10 |
| B189 | B205 | eGFB | KDA | -0.0257 | 2.42E-08 |
| B189 | B224 | eGFB | pmh | 0.025693 | 2.44E-08 |
| B19 | B102 | CD3+CD4+/CD3+CD8+ | anti-Sm | 0.051509 | 0 |
| B19 | B105 | CD3+CD4+/CD3+CD8+ | anti-U1RNP | 0.039565 | 0 |
| B19 | B110 | CD3+CD4+/CD3+CD8+ | AHA | 0.077427 | 0 |
| B19 | B122 | CD3+CD4+/CD3+CD8+ | C4 | 0.058347 | 0 |
| B19 | B136 | CD3+CD4+/CD3+CD8+ | 5'NT | 0.041193 | 0 |
| B19 | B138 | CD3+CD4+/CD3+CD8+ | PAB | -0.06562 | 0 |
| B19 | B139 | CD3+CD4+/CD3+CD8+ | ADA | 0.041654 | 0 |
| B19 | B156 | CD3+CD4+/CD3+CD8+ | IgE | 0.049758 | 0 |
| B19 | B183 | CD3+CD4+/CD3+CD8+ | PTA | -0.04008 | 0 |
| B19 | B20 | CD3+CD4+/CD3+CD8+ | CD3+CD4+CD8+ | -0.08073 | 0 |
| B19 | B21 | CD3+CD4+/CD3+CD8+ | CD3+CD4-CD8- | -0.05772 | 0 |
| B19 | B22 | CD3+CD4+/CD3+CD8+ | CD3+CD8+(T) | -0.77321 | 0 |
| B19 | B23 | CD3+CD4+/CD3+CD8+ | CD3-CD16+CD56+(NK) | 0.105484 | 0 |
| B19 | B58 | CD3+CD4+/CD3+CD8+ | HCT | 0.058853 | 0 |
| B19 | B79 | CD3+CD4+/CD3+CD8+ | hb V -dna | -0.03965 | 0 |
| B19 | B98 | CD3+CD4+/CD3+CD8+ | anti-Jo-1 | -0.04966 | 0 |
| B19 | B104 | CD3+CD4+/CD3+CD8+ | anti-SSB | -0.03431 | 9.41E-14 |
| B19 | B81 | CD3+CD4+/CD3+CD8+ | TSGF | -0.03423 | 1.07E-13 |
| B19 | B54 | CD3+CD4+/CD3+CD8+ | ESR | 0.032468 | 1.80E-12 |
| B19 | B195 | CD3+CD4+/CD3+CD8+ | Na | 0.031509 | 7.87E-12 |
| B19 | B134 | CD3+CD4+/CD3+CD8+ | CHE | -0.03098 | 1.76E-11 |
| B19 | B186 | CD3+CD4+/CD3+CD8+ | PFC | 0.030434 | 3.92E-11 |

| | | | | | |
|---|---|---|---|---|---|
| B19 | B107 | CD3+CD4+/CD3+CD8+ | AnuA | -0.02983 | 9.40E-11 |
| B19 | B95 | CD3+CD4+/CD3+CD8+ | anti-cenpb | 0.028554 | 5.68E-10 |
| B19 | B56 | CD3+CD4+/CD3+CD8+ | IR | -0.02843 | 6.71E-10 |
| B19 | B155 | CD3+CD4+/CD3+CD8+ | IgA | 0.027126 | 3.89E-09 |
| B19 | B32 | CD3+CD4+/CD3+CD8+ | Testo | -0.02711 | 3.99E-09 |
| B19 | B62 | CD3+CD4+/CD3+CD8+ | WBHSRVI | -0.0265 | 8.82E-09 |
| B19 | B198 | CD3+CD4+/CD3+CD8+ | P | 0.025792 | 2.15E-08 |
| B19 | B137 | CD3+CD4+/CD3+CD8+ | AFU | -0.02503 | 5.53E-08 |
| B19 | B61 | CD3+CD4+/CD3+CD8+ | WBHSRV | -0.02429 | 1.34E-07 |
| B19 | B60 | CD3+CD4+/CD3+CD8+ | WBRVI | -0.02345 | 3.57E-07 |
| B19 | B99 | CD3+CD4+/CD3+CD8+ | anti-PM-Scl | 0.02336 | 3.96E-07 |
| B19 | B55 | CD3+CD4+/CD3+CD8+ | TK | -0.02328 | 4.33E-07 |
| B19 | B119 | CD3+CD4+/CD3+CD8+ | Hcv-rna | -0.02315 | 5.05E-07 |
| B19 | B69 | CD3+CD4+/CD3+CD8+ | PV | 0.022953 | 6.27E-07 |
| B190 | B191 | Cre | Urea | 0.524005 | 0 |
| B190 | B192 | Cre | UA | 0.34191 | 0 |
| B190 | B204 | Cre | Drinking | 0.052938 | 0 |
| B190 | B206 | Cre | Waist | 0.106204 | 0 |
| B190 | B209 | Cre | Smoking | 0.073678 | 0 |
| B190 | B210 | Cre | Hip | 0.052268 | 0 |
| B190 | B216 | Cre | BMI | 0.135614 | 0 |
| B190 | B221 | Cre | Hei | 0.196191 | 0 |
| B190 | B222 | Cre | NS | 0.191996 | 0 |
| B190 | B219 | Cre | OT | -0.03332 | 4.66E-13 |
| B190 | B224 | Cre | pmh | 0.025841 | 2.03E-08 |
| B190 | B215 | Cre | BMI | 0.024165 | 1.56E-07 |
| B191 | B192 | Urea | UA | 0.210353 | 0 |
| B191 | B200 | Urea | OP | 0.058142 | 0 |
| B191 | B204 | Urea | Drinking | -0.04011 | 0 |
| B191 | B218 | Urea | DBP | -0.07389 | 0 |
| B191 | B220 | Urea | SBP | 0.038126 | 0 |
| B191 | B221 | Urea | Hei | -0.03963 | 0 |
| B191 | B216 | Urea | BMI | -0.0314 | 9.23E-12 |
| B191 | B211 | Urea | IOP-R | -0.02825 | 8.65E-10 |
| B191 | B206 | Urea | Waist | 0.028169 | 9.64E-10 |
| B191 | B209 | Urea | Smoking | -0.02685 | 5.57E-09 |
| B191 | B212 | Urea | IOP-L | -0.02368 | 2.75E-07 |
| B192 | B203 | UA | DH | 0.046149 | 0 |
| B192 | B204 | UA | Drinking | 0.179036 | 0 |
| B192 | B206 | UA | Waist | 0.306236 | 0 |
| B192 | B209 | UA | Smoking | 0.137573 | 0 |
| B192 | B210 | UA | Hip | 0.218911 | 0 |
| B192 | B215 | UA | BMI | 0.237918 | 0 |
| B192 | B216 | UA | BMI | 0.350447 | 0 |

| | | | | | |
|---|---|---|---|---|---|
| B192 | B218 | UA | DBP | 0.072648 | 0 |
| B192 | B219 | UA | OT | -0.05602 | 0 |
| B192 | B221 | UA | Hei | 0.298414 | 0 |
| B192 | B222 | UA | NS | 0.366099 | 0 |
| B192 | B220 | UA | SBP | -0.02685 | 5.56E-09 |
| B192 | B207 | UA | WHR | 0.025659 | 2.54E-08 |
| B193 | B194 | K | Cl | 0.082736 | 0 |
| B193 | B195 | K | Na | -0.10703 | 0 |
| B193 | B196 | K | TCO2 | -0.042 | 0 |
| B193 | B197 | K | AG | -0.08886 | 0 |
| B193 | B198 | K | P | 0.14843 | 0 |
| B193 | B200 | K | OP | -0.05786 | 0 |
| B193 | B204 | K | Drinking | -0.03826 | 0 |
| B193 | B216 | K | BMI | -0.03868 | 0 |
| B193 | B222 | K | NS | -0.03788 | 2.22E-16 |
| B193 | B221 | K | Hei | -0.03075 | 2.47E-11 |
| B193 | B206 | K | Waist | -0.02847 | 6.34E-10 |
| B193 | B215 | K | BMI | -0.0273 | 3.09E-09 |
| B193 | B218 | K | DBP | -0.0271 | 4.01E-09 |
| B194 | B195 | Cl | Na | 0.311991 | 0 |
| B194 | B196 | Cl | TCO2 | -0.18314 | 0 |
| B194 | B197 | Cl | AG | -0.36172 | 0 |
| B194 | B198 | Cl | P | -0.03983 | 0 |
| B194 | B200 | Cl | OP | 0.160852 | 0 |
| B195 | B196 | Na | TCO2 | 0.075184 | 0 |
| B195 | B197 | Na | AG | 0.33613 | 0 |
| B195 | B199 | Na | Mg | 0.129475 | 0 |
| B195 | B200 | Na | OP | 0.579329 | 0 |
| B195 | B198 | Na | P | -0.03484 | 3.91E-14 |
| B195 | B207 | Na | WHR | 0.023698 | 2.68E-07 |
| B196 | B197 | TCO2 | AG | -0.64905 | 0 |
| B196 | B200 | TCO2 | OP | 0.058942 | 0 |
| B197 | B199 | AG | Mg | 0.085774 | 0 |
| B197 | B200 | AG | OP | 0.196241 | 0 |
| B198 | B216 | P | BMI | -0.03931 | 0 |
| B198 | B221 | P | Hei | -0.0539 | 0 |
| B198 | B222 | P | NS | -0.05108 | 0 |
| B198 | B204 | P | Drinking | -0.03591 | 6.22E-15 |
| B198 | B206 | P | Waist | -0.03118 | 1.30E-11 |
| B198 | B200 | P | OP | -0.02832 | 7.84E-10 |
| B199 | B200 | Mg | OP | 0.077148 | 0 |
| B2 | B107 | Age | AnuA | 0.062409 | 0 |
| B2 | B113 | Age | FT3 | -0.06627 | 0 |
| B2 | B115 | Age | TSH TT3 TT4(TT3) | -0.04231 | 0 |

| | | | | | |
|---|---|---|---|---|---|
| B2 | B119 | Age | Hcv-rna | 0.044276 | 0 |
| B2 | B12 | Age | PG II | 0.042263 | 0 |
| B2 | B126 | Age | Alb | -0.28205 | 0 |
| B2 | B127 | Age | A-G ratio | -0.20823 | 0 |
| B2 | B128 | Age | ALT | -0.26433 | 0 |
| B2 | B130 | Age | GLB | 0.128224 | 0 |
| B2 | B131 | Age | AST | -0.04948 | 0 |
| B2 | B132 | Age | Dbil | 0.082916 | 0 |
| B2 | B134 | Age | CHE | -0.08009 | 0 |
| B2 | B138 | Age | PAB | -0.07638 | 0 |
| B2 | B139 | Age | ADA | 0.049542 | 0 |
| B2 | B140 | Age | r-GT | -0.15359 | 0 |
| B2 | B141 | Age | AST/ALT | 0.337023 | 0 |
| B2 | B146 | Age | CysC | 0.121262 | 0 |
| B2 | B153 | Age | Glu | 0.044062 | 0 |
| B2 | B161 | Age | LEU | 0.06433 | 0 |
| B2 | B163 | Age | SG | -0.14265 | 0 |
| B2 | B173 | Age | NTT | 0.082897 | 0 |
| B2 | B174 | Age | Urea/Cr | 0.081079 | 0 |
| B2 | B176 | Age | Ucr | -0.0908 | 0 |
| B2 | B188 | Age | Ca | -0.07707 | 0 |
| B2 | B189 | Age | eGFB | -0.52421 | 0 |
| B2 | B190 | Age | Cre | 0.078873 | 0 |
| B2 | B191 | Age | Urea | 0.199046 | 0 |
| B2 | B192 | Age | UA | -0.07875 | 0 |
| B2 | B193 | Age | K | 0.040464 | 0 |
| B2 | B201 | Age | e-habits | 0.061812 | 0 |
| B2 | B203 | Age | DH | -0.08528 | 0 |
| B2 | B204 | Age | Drinking | -0.1954 | 0 |
| B2 | B205 | Age | KDA | 0.063246 | 0 |
| B2 | B206 | Age | Waist | -0.10544 | 0 |
| B2 | B209 | Age | Smoking | -0.15181 | 0 |
| B2 | B210 | Age | Hip | -0.20832 | 0 |
| B2 | B211 | Age | IOP-R | -0.07278 | 0 |
| B2 | B212 | Age | IOP-L | -0.06948 | 0 |
| B2 | B215 | Age | BMI | -0.22875 | 0 |
| B2 | B216 | Age | BMI | -0.37465 | 0 |
| B2 | B218 | Age | DBP | -0.3729 | 0 |
| B2 | B219 | Age | OT | 0.124826 | 0 |
| B2 | B220 | Age | SBP | 0.210202 | 0 |
| B2 | B221 | Age | Hei | -0.33981 | 0 |
| B2 | B222 | Age | NS | -0.23789 | 0 |
| B2 | B227 | Age | OD | 0.053492 | 0 |
| B2 | B228 | Age | OS | 0.068843 | 0 |

| | | | | | |
|---|---|---|---|---|---|
| B2 | B24 | Age | αHBDH | 0.056877 | 0 |
| B2 | B27 | Age | LDH | 0.06258 | 0 |
| B2 | B40 | Age | WBC | -0.05389 | 0 |
| B2 | B42 | Age | MON | 0.077687 | 0 |
| B2 | B43 | Age | MCV | 0.150143 | 0 |
| B2 | B44 | Age | MCH | 0.050639 | 0 |
| B2 | B45 | Age | MCHC | -0.16853 | 0 |
| B2 | B46 | Age | RBC Count | -0.30167 | 0 |
| B2 | B47 | Age | RDW | 0.23361 | 0 |
| B2 | B50 | Age | eo% | 0.052806 | 0 |
| B2 | B51 | Age | Hb | -0.28533 | 0 |
| B2 | B52 | Age | PLT/BPC | -0.11319 | 0 |
| B2 | B62 | Age | WBHSRVI | -0.04805 | 0 |
| B2 | B63 | Age | WBRSR(mPas_200) | -0.0402 | 0 |
| B2 | B64 | Age | WBRSR(mPas_5) | -0.03864 | 0 |
| B2 | B66 | Age | WBRSR(mPas_100) | -0.04295 | 0 |
| B2 | B67 | Age | WBRSR(mPas_50) | -0.04257 | 0 |
| B2 | B71 | Age | TG | -0.20307 | 0 |
| B2 | B72 | Age | HDL-C | 0.197179 | 0 |
| B2 | B74 | Age | ApoA1 | 0.064409 | 0 |
| B2 | B75 | Age | ApoA1/B | 0.051887 | 0 |
| B2 | B79 | Age | hb V -dna | 0.064807 | 0 |
| B2 | B86 | Age | TBhCG | 0.056099 | 0 |
| B2 | B90 | Age | CYFRA 21-1 | 0.052121 | 0 |
| B2 | B91 | Age | TP | -0.05193 | 0 |
| B2 | B15 | Age | CD3-CD19+(B cell) | 0.036713 | 1.55E-15 |
| B2 | B60 | Age | WBRVI | -0.03628 | 3.33E-15 |
| B2 | B56 | Age | IR | -0.03562 | 1.02E-14 |
| B2 | B224 | Age | pmh | 0.035453 | 1.38E-14 |
| B2 | B3 | Age | Hcy | 0.035403 | 1.51E-14 |
| B2 | B184 | Age | INR | 0.035306 | 1.78E-14 |
| B2 | B54 | Age | ESR | 0.034989 | 3.04E-14 |
| B2 | B111 | Age | TSH | 0.03405 | 1.44E-13 |
| B2 | B70 | Age | LDL-C | -0.03385 | 1.99E-13 |
| B2 | B65 | Age | WBRSR(mPas_1) | -0.03373 | 2.41E-13 |
| B2 | B142 | Age | 25-(OH)D | 0.033441 | 3.85E-13 |
| B2 | B61 | Age | WBHSRV | -0.03309 | 6.72E-13 |
| B2 | B133 | Age | TBIL | 0.032162 | 2.90E-12 |
| B2 | B77 | Age | TC | -0.03201 | 3.68E-12 |
| B2 | B149 | Age | TGAb | 0.031367 | 9.77E-12 |
| B2 | B49 | Age | baso% | -0.03115 | 1.35E-11 |
| B2 | B157 | Age | IgG | 0.030777 | 2.36E-11 |
| B2 | B118 | Age | CEA1 | 0.02965 | 1.22E-10 |
| B2 | B10 | Age | PG I | 0.028164 | 9.70E-10 |

| | | | | | |
|---|---|---|---|---|---|
| B2 | B84 | Age | C-PSA | 0.02571 | 2.39E-08 |
| B2 | B76 | Age | ApoB | -0.02562 | 2.65E-08 |
| B2 | B53 | Age | GR% | -0.02539 | 3.55E-08 |
| B2 | B172 | Age | KB | -0.02517 | 4.67E-08 |
| B2 | B48 | Age | LYM% | -0.02389 | 2.15E-07 |
| B2 | B144 | Age | Total-PINP | -0.02363 | 2.90E-07 |
| B2 | B185 | Age | PT | 0.023501 | 3.37E-07 |
| B2 | B73 | Age | ApoA | 0.023318 | 4.15E-07 |
| B20 | B107 | CD3+CD4+CD8+ | AnuA | -0.06114 | 0 |
| B20 | B21 | CD3+CD4+CD8+ | CD3+CD4-CD8- | -0.06728 | 0 |
| B20 | B22 | CD3+CD4+CD8+ | CD3+CD8+(T) | 0.219818 | 0 |
| B20 | B23 | CD3+CD4+CD8+ | CD3-CD16+CD56+(NK) | -0.082 | 0 |
| B20 | B79 | CD3+CD4+CD8+ | hb V -dna | -0.04477 | 0 |
| B20 | B138 | CD3+CD4+CD8+ | PAB | 0.033118 | 6.46E-13 |
| B20 | B199 | CD3+CD4+CD8+ | Mg | 0.031367 | 9.76E-12 |
| B20 | B187 | CD3+CD4+CD8+ | NSE | 0.028957 | 3.25E-10 |
| B20 | B136 | CD3+CD4+CD8+ | 5'NT | -0.02874 | 4.39E-10 |
| B20 | B148 | CD3+CD4+CD8+ | Anti_TPO | 0.028457 | 6.50E-10 |
| B201 | B204 | e-habits | Drinking | 0.075874 | 0 |
| B201 | B205 | e-habits | KDA | 0.045297 | 0 |
| B201 | B218 | e-habits | DBP | -0.04229 | 0 |
| B201 | B224 | e-habits | pmh | 0.062608 | 0 |
| B201 | B215 | e-habits | BMI | -0.0363 | 3.11E-15 |
| B201 | B210 | e-habits | Hip | -0.02255 | 9.81E-07 |
| B203 | B204 | DH | Drinking | 0.077104 | 0 |
| B203 | B206 | DH | Waist | 0.051822 | 0 |
| B203 | B209 | DH | Smoking | 0.096666 | 0 |
| B203 | B210 | DH | Hip | 0.057514 | 0 |
| B203 | B215 | DH | BMI | 0.06011 | 0 |
| B203 | B216 | DH | BMI | 0.073412 | 0 |
| B203 | B218 | DH | DBP | 0.049893 | 0 |
| B203 | B221 | DH | Hei | 0.049409 | 0 |
| B203 | B222 | DH | NS | 0.066148 | 0 |
| B204 | B206 | Drinking | Waist | 0.177858 | 0 |
| B204 | B209 | Drinking | Smoking | 0.396459 | 0 |
| B204 | B210 | Drinking | Hip | 0.125408 | 0 |
| B204 | B215 | Drinking | BMI | 0.115221 | 0 |
| B204 | B216 | Drinking | BMI | 0.247754 | 0 |
| B204 | B218 | Drinking | DBP | 0.126336 | 0 |
| B204 | B219 | Drinking | OT | -0.07009 | 0 |
| B204 | B220 | Drinking | SBP | -0.0445 | 0 |
| B204 | B221 | Drinking | Hei | 0.269908 | 0 |
| B204 | B222 | Drinking | NS | 0.271878 | 0 |
| B204 | B212 | Drinking | IOP-L | 0.037104 | 8.88E-16 |

| | | | | | |
|---|---|---|---|---|---|
| B204 | B211 | Drinking | IOP-R | 0.035053 | 2.73E-14 |
| B204 | B205 | Drinking | KDA | -0.02981 | 9.71E-11 |
| B205 | B216 | KDA | BMI | -0.04792 | 0 |
| B205 | B218 | KDA | DBP | -0.0665 | 0 |
| B205 | B219 | KDA | OT | 0.039138 | 0 |
| B205 | B221 | KDA | Hei | -0.04303 | 0 |
| B205 | B222 | KDA | NS | -0.0473 | 0 |
| B205 | B206 | KDA | Waist | -0.03424 | 1.05E-13 |
| B205 | B215 | KDA | BMI | -0.03161 | 6.76E-12 |
| B205 | B209 | KDA | Smoking | -0.03067 | 2.76E-11 |
| B205 | B210 | KDA | Hip | -0.02635 | 1.06E-08 |
| B205 | B220 | KDA | SBP | -0.02393 | 2.06E-07 |
| B206 | B207 | Waist | WHR | 0.073367 | 0 |
| B206 | B209 | Waist | Smoking | 0.152873 | 0 |
| B206 | B210 | Waist | Hip | 0.766366 | 0 |
| B206 | B215 | Waist | BMI | 0.797093 | 0 |
| B206 | B216 | Waist | BMI | 0.798864 | 0 |
| B206 | B218 | Waist | DBP | 0.127938 | 0 |
| B206 | B219 | Waist | OT | -0.0478 | 0 |
| B206 | B221 | Waist | Hei | 0.37801 | 0 |
| B206 | B222 | Waist | NS | 0.676928 | 0 |
| B206 | B212 | Waist | IOP-L | 0.03155 | 7.40E-12 |
| B206 | B211 | Waist | IOP-R | 0.02517 | 4.65E-08 |
| B207 | B215 | WHR | BMI | 0.041216 | 0 |
| B207 | B216 | WHR | BMI | 0.04334 | 0 |
| B207 | B222 | WHR | NS | 0.078304 | 0 |
| B207 | B209 | WHR | Smoking | 0.026258 | 1.20E-08 |
| B207 | B221 | WHR | Hei | 0.023454 | 3.55E-07 |
| B209 | B210 | Smoking | Hip | 0.091049 | 0 |
| B209 | B215 | Smoking | BMI | 0.061397 | 0 |
| B209 | B216 | Smoking | BMI | 0.209098 | 0 |
| B209 | B218 | Smoking | DBP | 0.07241 | 0 |
| B209 | B219 | Smoking | OT | -0.07171 | 0 |
| B209 | B220 | Smoking | SBP | -0.07704 | 0 |
| B209 | B221 | Smoking | Hei | 0.272708 | 0 |
| B209 | B222 | Smoking | NS | 0.244606 | 0 |
| B21 | B107 | CD3+CD4-CD8- | AnuA | 0.072979 | 0 |
| B21 | B23 | CD3+CD4-CD8- | CD3-CD16+CD56+(NK) | -0.12461 | 0 |
| B21 | B81 | CD3+CD4-CD8- | TSGF | -0.03958 | 0 |
| B21 | B98 | CD3+CD4-CD8- | anti-Jo-1 | 0.078102 | 0 |
| B21 | B125 | CD3+CD4-CD8- | Cor | 0.031368 | 9.74E-12 |
| B21 | B183 | CD3+CD4-CD8- | PTA | 0.031202 | 1.25E-11 |
| B21 | B198 | CD3+CD4-CD8- | P | -0.03037 | 4.31E-11 |
| B21 | B63 | CD3+CD4-CD8- | WBRSR(mPas_200) | 0.029096 | 2.67E-10 |

| | | | | | |
|---|---|---|---|---|---|
| B21 | B69 | CD3+CD4-CD8- | PV | 0.028026 | 1.17E-09 |
| B21 | B96 | CD3+CD4-CD8- | dsDNA (IIF) | 0.027482 | 2.43E-09 |
| B21 | B66 | CD3+CD4-CD8- | WBRSR(mPas_100) | 0.026179 | 1.32E-08 |
| B21 | B22 | CD3+CD4-CD8- | CD3+CD8+(T) | 0.025927 | 1.82E-08 |
| B21 | B95 | CD3+CD4-CD8- | anti-cenpb | -0.02585 | 2.00E-08 |
| B21 | B108 | CD3+CD4-CD8- | Anti-Cyclin | -0.02585 | 2.01E-08 |
| B21 | B61 | CD3+CD4-CD8- | WBHSRV | 0.02559 | 2.77E-08 |
| B21 | B67 | CD3+CD4-CD8- | WBRSR(mPas_50) | 0.025548 | 2.92E-08 |
| B21 | B64 | CD3+CD4-CD8- | WBRSR(mPas_5) | 0.024744 | 7.81E-08 |
| B21 | B124 | CD3+CD4-CD8- | ACTH | 0.02471 | 8.14E-08 |
| B21 | B193 | CD3+CD4-CD8- | K | 0.023407 | 3.75E-07 |
| B21 | B105 | CD3+CD4-CD8- | anti-U1RNP | 0.023282 | 4.33E-07 |
| B21 | B136 | CD3+CD4-CD8- | 5'NT | -0.02323 | 4.61E-07 |
| B21 | B181 | CD3+CD4-CD8- | APTT | -0.02321 | 4.69E-07 |
| B21 | B94 | CD3+CD4-CD8- | ALDH2-Lys504Lys | -0.02295 | 6.29E-07 |
| B210 | B215 | Hip | BMI | 0.760949 | 0 |
| B210 | B216 | Hip | BMI | 0.783482 | 0 |
| B210 | B218 | Hip | DBP | 0.142364 | 0 |
| B210 | B219 | Hip | OT | -0.03864 | 0 |
| B210 | B221 | Hip | Hei | 0.397218 | 0 |
| B210 | B222 | Hip | NS | 0.592621 | 0 |
| B210 | B220 | Hip | SBP | -0.0357 | 9.10E-15 |
| B210 | B212 | Hip | IOP-L | 0.024724 | 8.00E-08 |
| B211 | B212 | IOP-R | IOP-L | 0.73689 | 0 |
| B211 | B215 | IOP-R | BMI | 0.041315 | 0 |
| B211 | B218 | IOP-R | DBP | 0.072378 | 0 |
| B211 | B220 | IOP-R | SBP | 0.05348 | 0 |
| B211 | B222 | IOP-R | NS | 0.0314 | 9.29E-12 |
| B211 | B216 | IOP-R | BMI | 0.030142 | 6.00E-11 |
| B212 | B215 | IOP-L | BMI | 0.044938 | 0 |
| B212 | B218 | IOP-L | DBP | 0.072236 | 0 |
| B212 | B220 | IOP-L | SBP | 0.059402 | 0 |
| B212 | B222 | IOP-L | NS | 0.03076 | 2.42E-11 |
| B212 | B216 | IOP-L | BMI | 0.029562 | 1.38E-10 |
| B212 | B228 | IOP-L | OS | 0.02382 | 2.33E-07 |
| B213 | B214 | CVA-R | CVA-L | 0.046984 | 0 |
| B215 | B216 | BMI | BMI | 0.792149 | 0 |
| B215 | B218 | BMI | DBP | 0.165822 | 0 |
| B215 | B221 | BMI | Hei | 0.134591 | 0 |
| B215 | B222 | BMI | NS | 0.574264 | 0 |
| B215 | B219 | BMI | OT | -0.03688 | 1.11E-15 |
| B216 | B218 | BMI | DBP | 0.233824 | 0 |
| B216 | B219 | BMI | OT | -0.0903 | 0 |
| B216 | B220 | BMI | SBP | -0.08465 | 0 |

| | | | | | |
|---|---|---|---|---|---|
| B216 | B221 | BMI | Hei | 0.704188 | 0 |
| B216 | B222 | BMI | NS | 0.759673 | 0 |
| B216 | B228 | BMI | OS | -0.02959 | 1.33E-10 |
| B218 | B219 | DBP | OT | -0.09227 | 0 |
| B218 | B220 | DBP | SBP | 0.467863 | 0 |
| B218 | B221 | DBP | Hei | 0.187156 | 0 |
| B218 | B222 | DBP | NS | 0.194134 | 0 |
| B218 | B224 | DBP | pmh | -0.0249 | 6.46E-08 |
| B219 | B221 | OT | Hei | -0.10605 | 0 |
| B219 | B222 | OT | NS | -0.09648 | 0 |
| B219 | B224 | OT | pmh | 0.039477 | 0 |
| B22 | B110 | CD3+CD8+(T) | AHA | -0.04894 | 0 |
| B22 | B122 | CD3+CD8+(T) | C4 | -0.05449 | 0 |
| B22 | B138 | CD3+CD8+(T) | PAB | 0.071457 | 0 |
| B22 | B139 | CD3+CD8+(T) | ADA | -0.04517 | 0 |
| B22 | B155 | CD3+CD8+(T) | IgA | -0.04342 | 0 |
| B22 | B157 | CD3+CD8+(T) | IgG | -0.03846 | 0 |
| B22 | B23 | CD3+CD8+(T) | CD3-CD16+CD56+(NK) | -0.30728 | 0 |
| B22 | B32 | CD3+CD8+(T) | Testo | -0.04787 | 0 |
| B22 | B58 | CD3+CD8+(T) | HCT | -0.04081 | 0 |
| B22 | B81 | CD3+CD8+(T) | TSGF | 0.042499 | 0 |
| B22 | B102 | CD3+CD8+(T) | anti-Sm | -0.03686 | 1.33E-15 |
| B22 | B107 | CD3+CD8+(T) | AnuA | -0.03599 | 5.55E-15 |
| B22 | B105 | CD3+CD8+(T) | anti-U1RNP | -0.03585 | 7.11E-15 |
| B22 | B54 | CD3+CD8+(T) | ESR | -0.03581 | 7.55E-15 |
| B22 | B96 | CD3+CD8+(T) | dsDNA (IIF) | -0.03575 | 8.44E-15 |
| B22 | B156 | CD3+CD8+(T) | IgE | -0.03419 | 1.14E-13 |
| B22 | B195 | CD3+CD8+(T) | Na | -0.0341 | 1.32E-13 |
| B22 | B103 | CD3+CD8+(T) | anti-SSA/Ro | -0.03285 | 9.95E-13 |
| B22 | B134 | CD3+CD8+(T) | CHE | 0.031407 | 9.20E-12 |
| B22 | B95 | CD3+CD8+(T) | anti-cenpb | -0.02863 | 5.14E-10 |
| B22 | B194 | CD3+CD8+(T) | Cl | -0.02831 | 7.95E-10 |
| B22 | B93 | CD3+CD8+(T) | ALDH2-Glu504Lys | -0.02713 | 3.86E-09 |
| B22 | B92 | CD3+CD8+(T) | ALDH2-Glu504Glu | 0.02597 | 1.72E-08 |
| B22 | B97 | CD3+CD8+(T) | dsDNA | 0.02538 | 3.60E-08 |
| B22 | B98 | CD3+CD8+(T) | anti-Jo-1 | 0.025268 | 4.13E-08 |
| B22 | B185 | CD3+CD8+(T) | PT | -0.02459 | 9.36E-08 |
| B22 | B184 | CD3+CD8+(T) | INR | -0.02323 | 4.57E-07 |
| B22 | B183 | CD3+CD8+(T) | PTA | 0.023028 | 5.77E-07 |
| B22 | B99 | CD3+CD8+(T) | anti-PM-Scl | -0.02282 | 7.27E-07 |
| B220 | B221 | SBP | Hei | -0.15257 | 0 |
| B220 | B222 | SBP | NS | -0.04402 | 0 |
| B220 | B228 | SBP | OS | 0.0263 | 1.13E-08 |
| B220 | B224 | SBP | pmh | -0.02559 | 2.79E-08 |

| | | | | | |
|---|---|---|---|---|---|
| B221 | B222 | Hei | NS | 0.573975 | 0 |
| B221 | B228 | Hei | OS | -0.0337 | 2.55E-13 |
| B227 | B228 | OD | OS | 0.098378 | 0 |
| B23 | B108 | CD3-CD16+CD56+(NK) | Anti-Cyclin | -0.04184 | 0 |
| B23 | B125 | CD3-CD16+CD56+(NK) | Cor | 0.053553 | 0 |
| B23 | B136 | CD3-CD16+CD56+(NK) | 5'NT | -0.04151 | 0 |
| B23 | B157 | CD3-CD16+CD56+(NK) | IgG | 0.053737 | 0 |
| B23 | B32 | CD3-CD16+CD56+(NK) | Testo | -0.05219 | 0 |
| B23 | B74 | CD3-CD16+CD56+(NK) | ApoA1 | 0.055786 | 0 |
| B23 | B122 | CD3-CD16+CD56+(NK) | C4 | -0.03696 | 8.88E-16 |
| B23 | B33 | CD3-CD16+CD56+(NK) | Prog | 0.033704 | 2.52E-13 |
| B23 | B139 | CD3-CD16+CD56+(NK) | ADA | 0.033434 | 3.90E-13 |
| B23 | B98 | CD3-CD16+CD56+(NK) | anti-Jo-1 | -0.03253 | 1.63E-12 |
| B23 | B156 | CD3-CD16+CD56+(NK) | IgE | 0.028707 | 4.60E-10 |
| B23 | B93 | CD3-CD16+CD56+(NK) | ALDH2-Glu504Lys | 0.028576 | 5.51E-10 |
| B23 | B92 | CD3-CD16+CD56+(NK) | ALDH2-Glu504Glu | -0.02678 | 6.11E-09 |
| B23 | B184 | CD3-CD16+CD56+(NK) | INR | 0.026008 | 1.64E-08 |
| B23 | B155 | CD3-CD16+CD56+(NK) | IgA | 0.02585 | 2.00E-08 |
| B23 | B134 | CD3-CD16+CD56+(NK) | CHE | -0.02406 | 1.76E-07 |
| B23 | B99 | CD3-CD16+CD56+(NK) | anti-PM-Scl | 0.023665 | 2.79E-07 |
| B23 | B198 | CD3-CD16+CD56+(NK) | P | -0.02353 | 3.25E-07 |
| B23 | B96 | CD3-CD16+CD56+(NK) | dsDNA (IIF) | 0.023486 | 3.42E-07 |
| B23 | B76 | CD3-CD16+CD56+(NK) | ApoB | 0.022584 | 9.46E-07 |
| B24 | B114 | αHBDH | TSH TT3 TT4(TT4) | 0.053841 | 0 |
| B24 | B123 | αHBDH | hs-CRP1 | 0.058144 | 0 |
| B24 | B127 | αHBDH | A-G ratio | -0.03916 | 0 |
| B24 | B130 | αHBDH | GLB | 0.044517 | 0 |
| B24 | B131 | αHBDH | AST | 0.071963 | 0 |
| B24 | B139 | αHBDH | ADA | 0.06393 | 0 |
| B24 | B146 | αHBDH | CysC | 0.040932 | 0 |
| B24 | B150 | αHBDH | ALP | 0.038184 | 0 |
| B24 | B174 | αHBDH | Urea/Cr | 0.04713 | 0 |
| B24 | B216 | αHBDH | BMI | -0.0389 | 0 |
| B24 | B220 | αHBDH | SBP | 0.06354 | 0 |
| B24 | B221 | αHBDH | Hei | -0.07026 | 0 |
| B24 | B222 | αHBDH | NS | -0.04773 | 0 |
| B24 | B25 | αHBDH | CK | 0.133829 | 0 |
| B24 | B26 | αHBDH | CKMB | 0.273672 | 0 |
| B24 | B27 | αHBDH | LDH | 0.68828 | 0 |
| B24 | B43 | αHBDH | MCV | 0.047812 | 0 |
| B24 | B44 | αHBDH | MCH | 0.038565 | 0 |
| B24 | B46 | αHBDH | RBC Count | -0.04363 | 0 |
| B24 | B47 | αHBDH | RDW | 0.076446 | 0 |
| B24 | B52 | αHBDH | PLT/BPC | -0.04219 | 0 |

| | | | | | |
|---|---|---|---|---|---|
| B24 | B54 | αHBDH | ESR | 0.097883 | 0 |
| B24 | B69 | αHBDH | PV | 0.085441 | 0 |
| B24 | B72 | αHBDH | HDL-C | 0.059566 | 0 |
| B24 | B74 | αHBDH | ApoA1 | 0.095002 | 0 |
| B24 | B91 | αHBDH | TP | 0.037783 | 2.22E-16 |
| B24 | B177 | αHBDH | ACR | 0.037544 | 4.44E-16 |
| B24 | B209 | αHBDH | Smoking | -0.03656 | 2.00E-15 |
| B24 | B176 | αHBDH | Ucr | -0.03626 | 3.55E-15 |
| B24 | B138 | αHBDH | PAB | -0.03574 | 8.44E-15 |
| B24 | B163 | αHBDH | SG | -0.03522 | 2.09E-14 |
| B24 | B141 | αHBDH | AST/ALT | 0.03509 | 2.58E-14 |
| B24 | B186 | αHBDH | PFC | 0.033983 | 1.60E-13 |
| B24 | B204 | αHBDH | Drinking | -0.03379 | 2.21E-13 |
| B24 | B76 | αHBDH | ApoB | 0.033179 | 5.87E-13 |
| B24 | B77 | αHBDH | TC | 0.031841 | 4.75E-12 |
| B24 | B75 | αHBDH | ApoA1/B | 0.030834 | 2.17E-11 |
| B24 | B191 | αHBDH | Urea | 0.03039 | 4.17E-11 |
| B24 | B70 | αHBDH | LDL-C | 0.029993 | 7.44E-11 |
| B24 | B110 | αHBDH | AHA | 0.02784 | 1.50E-09 |
| B24 | B183 | αHBDH | PTA | -0.02727 | 3.24E-09 |
| B24 | B107 | αHBDH | AnuA | -0.02638 | 1.02E-08 |
| B24 | B128 | αHBDH | ALT | 0.025952 | 1.76E-08 |
| B24 | B192 | αHBDH | UA | -0.02579 | 2.17E-08 |
| B24 | B115 | αHBDH | TSH TT3 TT4(TT3) | 0.025409 | 3.47E-08 |
| B24 | B73 | αHBDH | ApoA | 0.025114 | 4.99E-08 |
| B24 | B175 | αHBDH | ALB-U | 0.024882 | 6.60E-08 |
| B24 | B134 | αHBDH | CHE | -0.02396 | 1.98E-07 |
| B24 | B71 | αHBDH | TG | -0.02297 | 6.15E-07 |
| B24 | B40 | αHBDH | WBC | -0.02271 | 8.23E-07 |
| B25 | B128 | CK | ALT | 0.03955 | 0 |
| B25 | B131 | CK | AST | 0.056486 | 0 |
| B25 | B175 | CK | ALB-U | 0.047233 | 0 |
| B25 | B177 | CK | ACR | 0.057008 | 0 |
| B25 | B190 | CK | Cre | 0.050177 | 0 |
| B25 | B216 | CK | BMI | 0.042996 | 0 |
| B25 | B222 | CK | NS | 0.042678 | 0 |
| B25 | B26 | CK | CKMB | 0.22824 | 0 |
| B25 | B27 | CK | LDH | 0.14616 | 0 |
| B25 | B192 | CK | UA | 0.035971 | 5.77E-15 |
| B25 | B174 | CK | Urea/Cr | -0.0353 | 1.80E-14 |
| B25 | B221 | CK | Hei | 0.033368 | 4.33E-13 |
| B25 | B215 | CK | BMI | 0.030323 | 4.61E-11 |
| B25 | B206 | CK | Waist | 0.029688 | 1.15E-10 |
| B25 | B204 | CK | Drinking | 0.025144 | 4.80E-08 |

| | | | | | |
|---|---|---|---|---|---|
| B25 | B210 | CK | Hip | 0.025134 | 4.87E-08 |
| B25 | B138 | CK | PAB | 0.023935 | 2.04E-07 |
| B26 | B131 | CKMB | AST | 0.040018 | 0 |
| B26 | B27 | CKMB | LDH | 0.218679 | 0 |
| B26 | B62 | CKMB | WBHSRVI | 0.045341 | 0 |
| B26 | B63 | CKMB | WBRSR(mPas_200) | 0.058721 | 0 |
| B26 | B64 | CKMB | WBRSR(mPas_5) | 0.051472 | 0 |
| B26 | B65 | CKMB | WBRSR(mPas_1) | 0.043826 | 0 |
| B26 | B66 | CKMB | WBRSR(mPas_100) | 0.059859 | 0 |
| B26 | B67 | CKMB | WBRSR(mPas_50) | 0.059052 | 0 |
| B26 | B61 | CKMB | WBHSRV | 0.035128 | 2.40E-14 |
| B26 | B90 | CKMB | CYFRA 21-1 | 0.033111 | 6.54E-13 |
| B26 | B60 | CKMB | WBRVI | 0.032582 | 1.51E-12 |
| B26 | B51 | CKMB | Hb | 0.031206 | 1.24E-11 |
| B26 | B69 | CKMB | PV | 0.031115 | 1.43E-11 |
| B26 | B139 | CKMB | ADA | 0.030601 | 3.06E-11 |
| B26 | B130 | CKMB | GLB | 0.030071 | 6.64E-11 |
| B26 | B177 | CKMB | ACR | 0.028668 | 4.85E-10 |
| B26 | B127 | CKMB | A-G ratio | -0.02857 | 5.54E-10 |
| B26 | B54 | CKMB | ESR | 0.027384 | 2.77E-09 |
| B26 | B220 | CKMB | SBP | 0.026253 | 1.20E-08 |
| B26 | B46 | CKMB | RBC Count | 0.024835 | 7.00E-08 |
| B26 | B58 | CKMB | HCT | 0.024703 | 8.20E-08 |
| B26 | B87 | CKMB | NSE | 0.022794 | 7.49E-07 |
| B27 | B114 | LDH | TSH TT3 TT4(TT4) | 0.050433 | 0 |
| B27 | B118 | LDH | CEA1 | 0.057966 | 0 |
| B27 | B123 | LDH | hs-CRP1 | 0.052881 | 0 |
| B27 | B127 | LDH | A-G ratio | -0.04969 | 0 |
| B27 | B128 | LDH | ALT | 0.080138 | 0 |
| B27 | B130 | LDH | GLB | 0.055008 | 0 |
| B27 | B131 | LDH | AST | 0.149293 | 0 |
| B27 | B136 | LDH | 5'NT | 0.03805 | 0 |
| B27 | B138 | LDH | PAB | -0.05853 | 0 |
| B27 | B139 | LDH | ADA | 0.140226 | 0 |
| B27 | B140 | LDH | r-GT | 0.038378 | 0 |
| B27 | B145 | LDH | BGP | -0.03975 | 0 |
| B27 | B146 | LDH | CysC | 0.041484 | 0 |
| B27 | B150 | LDH | ALP | 0.130333 | 0 |
| B27 | B163 | LDH | SG | -0.03819 | 0 |
| B27 | B176 | LDH | Ucr | -0.05749 | 0 |
| B27 | B177 | LDH | ACR | 0.055198 | 0 |
| B27 | B220 | LDH | SBP | 0.060418 | 0 |
| B27 | B221 | LDH | Hei | -0.06349 | 0 |
| B27 | B43 | LDH | MCV | 0.046474 | 0 |

| | | | | | |
|---|---|---|---|---|---|
| B27 | B44 | LDH | MCH | 0.041046 | 0 |
| B27 | B47 | LDH | RDW | 0.067166 | 0 |
| B27 | B54 | LDH | ESR | 0.065439 | 0 |
| B27 | B69 | LDH | PV | 0.074583 | 0 |
| B27 | B72 | LDH | HDL-C | 0.053515 | 0 |
| B27 | B74 | LDH | ApoA1 | 0.071928 | 0 |
| B27 | B76 | LDH | ApoB | 0.045261 | 0 |
| B27 | B91 | LDH | TP | 0.045326 | 0 |
| B27 | B186 | LDH | PFC | 0.037511 | 4.44E-16 |
| B27 | B77 | LDH | TC | 0.037235 | 6.66E-16 |
| B27 | B174 | LDH | Urea/Cr | 0.037106 | 8.88E-16 |
| B27 | B175 | LDH | ALB-U | 0.03708 | 8.88E-16 |
| B27 | B46 | LDH | RBC Count | -0.03596 | 5.77E-15 |
| B27 | B66 | LDH | WBRSR(mPas_100) | 0.03559 | 1.11E-14 |
| B27 | B67 | LDH | WBRSR(mPas_50) | 0.035018 | 2.89E-14 |
| B27 | B209 | LDH | Smoking | -0.03424 | 1.06E-13 |
| B27 | B87 | LDH | NSE | 0.034225 | 1.08E-13 |
| B27 | B70 | LDH | LDL-C | 0.033584 | 3.06E-13 |
| B27 | B52 | LDH | PLT/BPC | -0.03284 | 1.00E-12 |
| B27 | B64 | LDH | WBRSR(mPas_5) | 0.032795 | 1.08E-12 |
| B27 | B129 | LDH | IBIL | 0.032125 | 3.07E-12 |
| B27 | B63 | LDH | WBRSR(mPas_200) | 0.031866 | 4.57E-12 |
| B27 | B191 | LDH | Urea | 0.031512 | 7.84E-12 |
| B27 | B133 | LDH | TBIL | 0.03148 | 8.23E-12 |
| B27 | B204 | LDH | Drinking | -0.03086 | 2.09E-11 |
| B27 | B65 | LDH | WBRSR(mPas_1) | 0.028913 | 3.45E-10 |
| B27 | B197 | LDH | AG | 0.028819 | 3.94E-10 |
| B27 | B189 | LDH | eGFB | -0.02869 | 4.72E-10 |
| B27 | B200 | LDH | OP | 0.028562 | 5.62E-10 |
| B27 | B222 | LDH | NS | -0.02638 | 1.02E-08 |
| B27 | B135 | LDH | TBA | 0.025805 | 2.12E-08 |
| B27 | B141 | LDH | AST/ALT | 0.025356 | 3.70E-08 |
| B27 | B112 | LDH | FT4 | 0.025018 | 5.60E-08 |
| B27 | B195 | LDH | Na | 0.02494 | 6.16E-08 |
| B27 | B61 | LDH | WBHSRV | 0.024414 | 1.16E-07 |
| B27 | B215 | LDH | BMI | 0.02417 | 1.55E-07 |
| B27 | B111 | LDH | TSH | 0.023907 | 2.10E-07 |
| B27 | B109 | LDH | AMA-M2 | 0.022884 | 6.78E-07 |
| B28 | B108 | PRL | Anti-Cyclin | 0.041547 | 0 |
| B28 | B29 | PRL | E2 | 0.166889 | 0 |
| B28 | B30 | PRL | LH | -0.13841 | 0 |
| B28 | B31 | PRL | FSH | -0.24829 | 0 |
| B28 | B32 | PRL | Testo | -0.06296 | 0 |
| B28 | B33 | PRL | Prog | 0.214543 | 0 |

| | | | | | |
|---|---|---|---|---|---|
| B29 | B108 | E2 | Anti-Cyclin | -0.06542 | 0 |
| B29 | B110 | E2 | AHA | 0.041219 | 0 |
| B29 | B181 | E2 | APTT | 0.038284 | 0 |
| B29 | B198 | E2 | P | -0.03951 | 0 |
| B29 | B30 | E2 | LH | -0.40478 | 0 |
| B29 | B31 | E2 | FSH | -0.64645 | 0 |
| B29 | B32 | E2 | Testo | -0.13263 | 0 |
| B29 | B33 | E2 | Prog | 0.551986 | 0 |
| B29 | B121 | E2 | C3 | -0.03558 | 1.11E-14 |
| B29 | B139 | E2 | ADA | -0.03226 | 2.50E-12 |
| B29 | B195 | E2 | Na | -0.03047 | 3.71E-11 |
| B29 | B114 | E2 | TSH TT3 TT4(TT4) | -0.02895 | 3.29E-10 |
| B29 | B107 | E2 | AnuA | 0.027881 | 1.42E-09 |
| B29 | B199 | E2 | Mg | -0.02593 | 1.81E-08 |
| B29 | B196 | E2 | TCO2 | -0.02559 | 2.76E-08 |
| B29 | B79 | E2 | hb V -dna | 0.025005 | 5.69E-08 |
| B3 | B123 | Hcy | hs-CRP1 | 0.04207 | 0 |
| B3 | B132 | Hcy | Dbil | 0.042754 | 0 |
| B3 | B138 | Hcy | PAB | 0.038723 | 0 |
| B3 | B140 | Hcy | r-GT | 0.055992 | 0 |
| B3 | B146 | Hcy | CysC | 0.071722 | 0 |
| B3 | B163 | Hcy | SG | 0.047466 | 0 |
| B3 | B174 | Hcy | Urea/Cr | -0.06693 | 0 |
| B3 | B175 | Hcy | ALB-U | 0.063701 | 0 |
| B3 | B176 | Hcy | Ucr | 0.058348 | 0 |
| B3 | B177 | Hcy | ACR | 0.052669 | 0 |
| B3 | B189 | Hcy | eGFB | -0.12635 | 0 |
| B3 | B190 | Hcy | Cre | 0.109901 | 0 |
| B3 | B191 | Hcy | Urea | 0.076924 | 0 |
| B3 | B192 | Hcy | UA | 0.108538 | 0 |
| B3 | B204 | Hcy | Drinking | 0.052307 | 0 |
| B3 | B206 | Hcy | Waist | 0.065077 | 0 |
| B3 | B209 | Hcy | Smoking | 0.056217 | 0 |
| B3 | B216 | Hcy | BMI | 0.063781 | 0 |
| B3 | B221 | Hcy | Hei | 0.06574 | 0 |
| B3 | B222 | Hcy | NS | 0.073265 | 0 |
| B3 | B27 | Hcy | LDH | 0.040838 | 0 |
| B3 | B42 | Hcy | MON | 0.045142 | 0 |
| B3 | B44 | Hcy | MCH | 0.043172 | 0 |
| B3 | B47 | Hcy | RDW | 0.062723 | 0 |
| B3 | B48 | Hcy | LYM% | -0.0478 | 0 |
| B3 | B51 | Hcy | Hb | 0.060994 | 0 |
| B3 | B72 | Hcy | HDL-C | -0.04834 | 0 |
| B3 | B74 | Hcy | ApoA1 | -0.05874 | 0 |

| | | | | | |
|---|---|---|---|---|---|
| B3 | B79 | Hcy | hb V -dna | -0.04466 | 0 |
| B3 | B86 | Hcy | TBhCG | -0.04479 | 0 |
| B3 | B75 | Hcy | ApoA1/B | -0.03754 | 4.44E-16 |
| B3 | B43 | Hcy | MCV | 0.033264 | 5.12E-13 |
| B3 | B10 | Hcy | PG Ⅰ | 0.033214 | 5.55E-13 |
| B3 | B210 | Hcy | Hip | 0.03293 | 8.71E-13 |
| B3 | B45 | Hcy | MCHC | 0.032881 | 9.41E-13 |
| B3 | B215 | Hcy | BMI | 0.032223 | 2.64E-12 |
| B3 | B133 | Hcy | TBIL | 0.032054 | 3.42E-12 |
| B3 | B46 | Hcy | RBC Count | 0.031705 | 5.84E-12 |
| B3 | B150 | Hcy | ALP | 0.030207 | 5.45E-11 |
| B3 | B53 | Hcy | GR% | 0.029832 | 9.39E-11 |
| B3 | B193 | Hcy | K | -0.0298 | 9.87E-11 |
| B3 | B62 | Hcy | WBHSRVI | 0.029208 | 2.28E-10 |
| B3 | B24 | Hcy | αHBDH | 0.028837 | 3.84E-10 |
| B3 | B126 | Hcy | Alb | -0.02859 | 5.37E-10 |
| B3 | B66 | Hcy | WBRSR(mPas_100) | 0.027897 | 1.39E-09 |
| B3 | B52 | Hcy | PLT/BPC | -0.02765 | 1.94E-09 |
| B3 | B67 | Hcy | WBRSR(mPas_50) | 0.027478 | 2.44E-09 |
| B3 | B90 | Hcy | CYFRA 21-1 | 0.026753 | 6.33E-09 |
| B3 | B63 | Hcy | WBRSR(mPas_200) | 0.026326 | 1.10E-08 |
| B3 | B137 | Hcy | AFU | 0.026001 | 1.66E-08 |
| B3 | B50 | Hcy | eo% | 0.025885 | 1.92E-08 |
| B3 | B220 | Hcy | SBP | 0.025709 | 2.39E-08 |
| B3 | B129 | Hcy | IBIL | 0.025462 | 3.25E-08 |
| B3 | B198 | Hcy | P | -0.02541 | 3.45E-08 |
| B3 | B166 | Hcy | Pro | 0.025326 | 3.84E-08 |
| B3 | B184 | Hcy | INR | -0.02496 | 6.01E-08 |
| B3 | B119 | Hcy | Hcv-rna | -0.02384 | 2.28E-07 |
| B3 | B179 | Hcy | α1-MG | 0.0238 | 2.38E-07 |
| B3 | B12 | Hcy | PG II | 0.023772 | 2.46E-07 |
| B3 | B64 | Hcy | WBRSR(mPas_5) | 0.02331 | 4.19E-07 |
| B3 | B131 | Hcy | AST | 0.022876 | 6.84E-07 |
| B30 | B108 | LH | Anti-Cyclin | -0.04793 | 0 |
| B30 | B121 | LH | C3 | 0.055612 | 0 |
| B30 | B198 | LH | P | 0.040657 | 0 |
| B30 | B31 | LH | FSH | 0.839542 | 0 |
| B30 | B32 | LH | Testo | -0.33544 | 0 |
| B30 | B33 | LH | Prog | -0.43777 | 0 |
| B30 | B36 | LH | V-Lym% | -0.05235 | 0 |
| B30 | B115 | LH | TSH TT3 TT4(TT3) | 0.028893 | 3.55E-10 |
| B30 | B122 | LH | C4 | 0.026358 | 1.05E-08 |
| B31 | B108 | FSH | Anti-Cyclin | -0.04018 | 0 |
| B31 | B198 | FSH | P | 0.04625 | 0 |

| | | | | | |
|---|---|---|---|---|---|
| B31 | B32 | FSH | Testo | -0.23757 | 0 |
| B31 | B33 | FSH | Prog | -0.53324 | 0 |
| B31 | B36 | FSH | V-Lym% | -0.04712 | 0 |
| B31 | B115 | FSH | TSH TT3 TT4(TT3) | 0.03656 | 2.22E-15 |
| B31 | B114 | FSH | TSH TT3 TT4(TT4) | 0.031682 | 6.05E-12 |
| B31 | B121 | FSH | C3 | 0.030482 | 3.65E-11 |
| B31 | B195 | FSH | Na | 0.02868 | 4.77E-10 |
| B31 | B138 | FSH | PAB | -0.02668 | 6.92E-09 |
| B32 | B108 | Testo | Anti-Cyclin | 0.245022 | 0 |
| B32 | B123 | Testo | hs-CRP1 | 0.041894 | 0 |
| B32 | B33 | Testo | Prog | -0.16514 | 0 |
| B32 | B212 | Testo | IOP-L | 0.029421 | 1.69E-10 |
| B32 | B211 | Testo | IOP-R | 0.025055 | 5.35E-08 |
| B33 | B103 | Prog | anti-SSA/Ro | -0.04961 | 0 |
| B33 | B108 | Prog | Anti-Cyclin | -0.06411 | 0 |
| B33 | B110 | Prog | AHA | -0.05026 | 0 |
| B33 | B125 | Prog | Cor | 0.045047 | 0 |
| B33 | B181 | Prog | APTT | 0.050426 | 0 |
| B33 | B138 | Prog | PAB | 0.036088 | 4.66E-15 |
| B33 | B107 | Prog | AnuA | 0.034839 | 3.91E-14 |
| B33 | B198 | Prog | P | -0.03431 | 9.44E-14 |
| B33 | B193 | Prog | K | 0.029453 | 1.61E-10 |
| B33 | B199 | Prog | Mg | -0.02867 | 4.85E-10 |
| B33 | B93 | Prog | ALDH2-Glu504Lys | 0.024733 | 7.91E-08 |
| B33 | B92 | Prog | ALDH2-Glu504Glu | -0.02467 | 8.52E-08 |
| B33 | B137 | Prog | AFU | 0.023969 | 1.96E-07 |
| B33 | B184 | Prog | INR | 0.022756 | 7.82E-07 |
| B34 | B138 | TK | PAB | -0.03893 | 0 |
| B34 | B86 | TK | TBhCG | 0.032383 | 2.06E-12 |
| B34 | B71 | TK | TG | -0.02938 | 1.80E-10 |
| B34 | B79 | TK | hb V -dna | 0.028231 | 8.86E-10 |
| B34 | B193 | TK | K | 0.027234 | 3.37E-09 |
| B34 | B125 | TK | Cor | 0.026512 | 8.64E-09 |
| B34 | B137 | TK | AFU | -0.02614 | 1.38E-08 |
| B34 | B135 | TK | TBA | 0.025398 | 3.52E-08 |
| B34 | B134 | TK | CHE | -0.0247 | 8.23E-08 |
| B34 | B198 | TK | P | 0.024498 | 1.05E-07 |
| B34 | B81 | TK | TSGF | -0.02326 | 4.45E-07 |
| B35 | B125 | β2-MG | Cor | -0.05468 | 0 |
| B35 | B159 | β2-MG | β2-MG | 0.601897 | 0 |
| B35 | B181 | β2-MG | APTT | 0.039646 | 0 |
| B35 | B182 | β2-MG | TT | -0.06565 | 0 |
| B35 | B184 | β2-MG | INR | 0.044951 | 0 |
| B35 | B185 | β2-MG | PT | 0.063408 | 0 |

| | | | | | |
|---|---|---|---|---|---|
| B35 | B186 | β2-MG | PFC | 0.060118 | 0 |
| B35 | B124 | β2-MG | ACTH | 0.025539 | 2.95E-08 |
| B37 | B38 | BAND | BAND% | 0.892053 | 0 |
| B37 | B39 | BAND | Other cells | 0.354909 | 0 |
| B37 | B40 | BAND | WBC | 0.108625 | 0 |
| B37 | B42 | BAND | MON | 0.117027 | 0 |
| B37 | B47 | BAND | RDW | 0.085334 | 0 |
| B37 | B53 | BAND | GR% | -0.02852 | 6.00E-10 |
| B37 | B141 | BAND | AST/ALT | 0.025993 | 1.67E-08 |
| B38 | B39 | BAND% | Other cells | 0.387787 | 0 |
| B38 | B40 | BAND% | WBC | 0.082307 | 0 |
| B38 | B42 | BAND% | MON | 0.100535 | 0 |
| B38 | B47 | BAND% | RDW | 0.079057 | 0 |
| B38 | B53 | BAND% | GR% | -0.03537 | 1.60E-14 |
| B38 | B141 | BAND% | AST/ALT | 0.025358 | 3.69E-08 |
| B39 | B40 | Other cells | WBC | 0.052925 | 0 |
| B39 | B42 | Other cells | MON | 0.052823 | 0 |
| B39 | B43 | Other cells | MCV | -0.07255 | 0 |
| B39 | B44 | Other cells | MCH | -0.07916 | 0 |
| B39 | B45 | Other cells | MCHC | -0.04989 | 0 |
| B39 | B46 | Other cells | RBC Count | 0.036655 | 1.78E-15 |
| B39 | B51 | Other cells | Hb | -0.03512 | 2.44E-14 |
| B4 | B5 | WB-Ca | WB-Mg | 0.181293 | 0 |
| B4 | B6 | WB-Ca | WB-Fe | 0.070394 | 0 |
| B4 | B7 | WB-Ca | WB-Cop | 0.131195 | 0 |
| B4 | B8 | WB-Ca | WB-Zinc | -0.10782 | 0 |
| B4 | B100 | WB-Ca | anti-Ro52 | -0.03491 | 3.46E-14 |
| B40 | B123 | WBC | hs-CRP1 | 0.086941 | 0 |
| B40 | B127 | WBC | A-G ratio | -0.10491 | 0 |
| B40 | B128 | WBC | ALT | 0.061564 | 0 |
| B40 | B129 | WBC | IBIL | -0.07256 | 0 |
| B40 | B130 | WBC | GLB | 0.131906 | 0 |
| B40 | B132 | WBC | Dbil | -0.0868 | 0 |
| B40 | B133 | WBC | TBIL | -0.08189 | 0 |
| B40 | B140 | WBC | r-GT | 0.074642 | 0 |
| B40 | B141 | WBC | AST/ALT | -0.06969 | 0 |
| B40 | B153 | WBC | Glu | 0.066361 | 0 |
| B40 | B160 | WBC | PH | -0.0825 | 0 |
| B40 | B163 | WBC | SG | 0.093591 | 0 |
| B40 | B166 | WBC | Pro | 0.082814 | 0 |
| B40 | B170 | WBC | U-GLU | 0.04585 | 0 |
| B40 | B174 | WBC | Urea/Cr | -0.08639 | 0 |
| B40 | B176 | WBC | Ucr | 0.047914 | 0 |
| B40 | B188 | WBC | Ca | 0.058318 | 0 |

| | | | | | |
|---|---|---|---|---|---|
| B40 | B190 | WBC | Cre | 0.081981 | 0 |
| B40 | B192 | WBC | UA | 0.16591 | 0 |
| B40 | B203 | WBC | DH | 0.043355 | 0 |
| B40 | B204 | WBC | Drinking | 0.038285 | 0 |
| B40 | B206 | WBC | Waist | 0.163751 | 0 |
| B40 | B209 | WBC | Smoking | 0.176442 | 0 |
| B40 | B210 | WBC | Hip | 0.11635 | 0 |
| B40 | B215 | WBC | BMI | 0.1375 | 0 |
| B40 | B216 | WBC | BMI | 0.153572 | 0 |
| B40 | B221 | WBC | Hei | 0.093359 | 0 |
| B40 | B222 | WBC | NS | 0.171844 | 0 |
| B40 | B42 | WBC | MON | 0.665533 | 0 |
| B40 | B43 | WBC | MCV | -0.05869 | 0 |
| B40 | B44 | WBC | MCH | -0.04238 | 0 |
| B40 | B46 | WBC | RBC Count | 0.218918 | 0 |
| B40 | B48 | WBC | LYM% | -0.19354 | 0 |
| B40 | B49 | WBC | baso% | -0.0763 | 0 |
| B40 | B51 | WBC | Hb | 0.199471 | 0 |
| B40 | B52 | WBC | PLT/BPC | 0.252248 | 0 |
| B40 | B53 | WBC | GR% | 0.210673 | 0 |
| B40 | B60 | WBC | WBRVI | 0.039165 | 0 |
| B40 | B62 | WBC | WBHSRVI | 0.038053 | 0 |
| B40 | B63 | WBC | WBRSR(mPas_200) | 0.054885 | 0 |
| B40 | B64 | WBC | WBRSR(mPas_5) | 0.053543 | 0 |
| B40 | B65 | WBC | WBRSR(mPas_1) | 0.050719 | 0 |
| B40 | B66 | WBC | WBRSR(mPas_100) | 0.054876 | 0 |
| B40 | B67 | WBC | WBRSR(mPas_50) | 0.055465 | 0 |
| B40 | B70 | WBC | LDL-C | 0.045462 | 0 |
| B40 | B71 | WBC | TG | 0.146738 | 0 |
| B40 | B72 | WBC | HDL-C | -0.15952 | 0 |
| B40 | B77 | WBC | TC | 0.040307 | 0 |
| B40 | B91 | WBC | TP | 0.128677 | 0 |
| B40 | B61 | WBC | WBHSRV | 0.037941 | 2.22E-16 |
| B40 | B146 | WBC | CysC | 0.037369 | 4.44E-16 |
| B40 | B75 | WBC | ApoA1/B | -0.03615 | 4.22E-15 |
| B40 | B76 | WBC | ApoB | 0.036133 | 4.44E-15 |
| B40 | B74 | WBC | ApoA1 | -0.03487 | 3.71E-14 |
| B40 | B134 | WBC | CHE | 0.034822 | 4.02E-14 |
| B40 | B59 | WBC | WBRV | 0.034524 | 6.59E-14 |
| B40 | B69 | WBC | PV | 0.032804 | 1.06E-12 |
| B40 | B107 | WBC | AnuA | 0.029733 | 1.08E-10 |
| B40 | B218 | WBC | DBP | 0.02926 | 2.12E-10 |
| B40 | B194 | WBC | Cl | -0.02782 | 1.55E-09 |
| B40 | B150 | WBC | ALP | 0.027805 | 1.58E-09 |

| | | | | | |
|---|---|---|---|---|---|
| B40 | B191 | WBC | Urea | 0.026406 | 9.90E-09 |
| B40 | B56 | WBC | IR | 0.023031 | 5.74E-07 |
| B40 | B112 | WBC | FT4 | 0.022999 | 5.96E-07 |
| B40 | B83 | WBC | AFP | 0.022911 | 6.57E-07 |
| B40 | B58 | WBC | HCT | 0.022811 | 7.35E-07 |
| B41 | B42 | MON% | MON | 0.026957 | 4.85E-09 |
| B42 | B123 | MON | hs-CRP1 | 0.081788 | 0 |
| B42 | B126 | MON | Alb | -0.08972 | 0 |
| B42 | B127 | MON | A-G ratio | -0.12921 | 0 |
| B42 | B128 | MON | ALT | 0.074387 | 0 |
| B42 | B130 | MON | GLB | 0.118621 | 0 |
| B42 | B131 | MON | AST | 0.078164 | 0 |
| B42 | B140 | MON | r-GT | 0.090861 | 0 |
| B42 | B146 | MON | CysC | 0.063924 | 0 |
| B42 | B160 | MON | PH | -0.06547 | 0 |
| B42 | B163 | MON | SG | 0.126502 | 0 |
| B42 | B166 | MON | Pro | 0.084389 | 0 |
| B42 | B174 | MON | Urea/Cr | -0.11081 | 0 |
| B42 | B176 | MON | Ucr | 0.051024 | 0 |
| B42 | B189 | MON | eGFB | -0.0922 | 0 |
| B42 | B190 | MON | Cre | 0.12557 | 0 |
| B42 | B191 | MON | Urea | 0.067013 | 0 |
| B42 | B192 | MON | UA | 0.188808 | 0 |
| B42 | B204 | MON | Drinking | 0.077806 | 0 |
| B42 | B206 | MON | Waist | 0.174541 | 0 |
| B42 | B209 | MON | Smoking | 0.185912 | 0 |
| B42 | B210 | MON | Hip | 0.108602 | 0 |
| B42 | B215 | MON | BMI | 0.104645 | 0 |
| B42 | B216 | MON | BMI | 0.149288 | 0 |
| B42 | B221 | MON | Hei | 0.125438 | 0 |
| B42 | B222 | MON | NS | 0.199522 | 0 |
| B42 | B46 | MON | RBC Count | 0.152986 | 0 |
| B42 | B47 | MON | RDW | 0.098107 | 0 |
| B42 | B48 | MON | LYM% | -0.15616 | 0 |
| B42 | B50 | MON | eo% | 0.047674 | 0 |
| B42 | B51 | MON | Hb | 0.185042 | 0 |
| B42 | B52 | MON | PLT/BPC | 0.163112 | 0 |
| B42 | B62 | MON | WBHSRVI | 0.039587 | 0 |
| B42 | B63 | MON | WBRSR(mPas_200) | 0.048941 | 0 |
| B42 | B64 | MON | WBRSR(mPas_5) | 0.044422 | 0 |
| B42 | B65 | MON | WBRSR(mPas_1) | 0.041353 | 0 |
| B42 | B66 | MON | WBRSR(mPas_100) | 0.048047 | 0 |
| B42 | B67 | MON | WBRSR(mPas_50) | 0.047985 | 0 |
| B42 | B71 | MON | TG | 0.041155 | 0 |

| | | | | | |
|---|---|---|---|---|---|
| B42 | B72 | MON | HDL-C | -0.11724 | 0 |
| B42 | B91 | MON | TP | 0.053891 | 0 |
| B42 | B44 | MON | MCH | 0.037931 | 2.22E-16 |
| B42 | B107 | MON | AnuA | 0.037608 | 4.44E-16 |
| B42 | B60 | MON | WBRVI | 0.036706 | 1.55E-15 |
| B42 | B77 | MON | TC | -0.03449 | 6.99E-14 |
| B42 | B90 | MON | CYFRA 21-1 | 0.034164 | 1.19E-13 |
| B42 | B43 | MON | MCV | 0.033453 | 3.79E-13 |
| B42 | B74 | MON | ApoA1 | -0.03337 | 4.31E-13 |
| B42 | B113 | MON | FT3 | 0.032316 | 2.28E-12 |
| B42 | B129 | MON | IBIL | -0.03177 | 5.32E-12 |
| B42 | B170 | MON | U-GLU | 0.030791 | 2.31E-11 |
| B42 | B161 | MON | LEU | -0.02896 | 3.23E-10 |
| B42 | B61 | MON | WBHSRV | 0.027563 | 2.18E-09 |
| B42 | B203 | MON | DH | 0.026803 | 5.93E-09 |
| B42 | B153 | MON | Glu | 0.02628 | 1.16E-08 |
| B42 | B58 | MON | HCT | 0.025429 | 3.39E-08 |
| B42 | B118 | MON | CEA1 | 0.024677 | 8.46E-08 |
| B42 | B141 | MON | AST/ALT | -0.02466 | 8.63E-08 |
| B42 | B194 | MON | CI | -0.02455 | 9.85E-08 |
| B42 | B133 | MON | TBIL | -0.0245 | 1.05E-07 |
| B42 | B112 | MON | FT4 | 0.023316 | 4.16E-07 |
| B42 | B75 | MON | ApoA1/B | -0.02278 | 7.62E-07 |
| B43 | B112 | MCV | FT4 | -0.04219 | 0 |
| B43 | B126 | MCV | Alb | -0.05689 | 0 |
| B43 | B131 | MCV | AST | 0.068195 | 0 |
| B43 | B132 | MCV | Dbil | 0.070929 | 0 |
| B43 | B140 | MCV | r-GT | 0.063713 | 0 |
| B43 | B141 | MCV | AST/ALT | 0.074362 | 0 |
| B43 | B189 | MCV | eGFB | -0.09978 | 0 |
| B43 | B191 | MCV | Urea | 0.048248 | 0 |
| B43 | B204 | MCV | Drinking | 0.079735 | 0 |
| B43 | B209 | MCV | Smoking | 0.085334 | 0 |
| B43 | B215 | MCV | BMI | -0.05282 | 0 |
| B43 | B216 | MCV | BMI | -0.05002 | 0 |
| B43 | B218 | MCV | DBP | -0.0558 | 0 |
| B43 | B44 | MCV | MCH | 0.901898 | 0 |
| B43 | B45 | MCV | MCHC | 0.177171 | 0 |
| B43 | B46 | MCV | RBC Count | -0.53735 | 0 |
| B43 | B47 | MCV | RDW | 0.585043 | 0 |
| B43 | B51 | MCV | Hb | 0.116625 | 0 |
| B43 | B52 | MCV | PLT/BPC | -0.15828 | 0 |
| B43 | B53 | MCV | GR% | -0.05872 | 0 |
| B43 | B59 | MCV | WBRV | -0.04091 | 0 |

| | | | | | |
|---|---|---|---|---|---|
| B43 | B72 | MCV | HDL-C | 0.090648 | 0 |
| B43 | B91 | MCV | TP | -0.05757 | 0 |
| B43 | B74 | MCV | ApoA1 | 0.036531 | 2.22E-15 |
| B43 | B55 | MCV | TK | -0.0343 | 9.55E-14 |
| B43 | B160 | MCV | PH | -0.03359 | 3.03E-13 |
| B43 | B153 | MCV | Glu | -0.03187 | 4.53E-12 |
| B43 | B48 | MCV | LYM% | 0.03166 | 6.26E-12 |
| B43 | B77 | MCV | TC | 0.030979 | 1.75E-11 |
| B43 | B190 | MCV | Cre | 0.029586 | 1.34E-10 |
| B43 | B57 | MCV | RBCAI | -0.02948 | 1.56E-10 |
| B43 | B210 | MCV | Hip | -0.0291 | 2.64E-10 |
| B43 | B201 | MCV | e-habits | 0.028693 | 4.69E-10 |
| B43 | B71 | MCV | TG | -0.02708 | 4.13E-09 |
| B43 | B130 | MCV | GLB | -0.02631 | 1.12E-08 |
| B43 | B134 | MCV | CHE | -0.02545 | 3.31E-08 |
| B43 | B146 | MCV | CysC | 0.023232 | 4.58E-07 |
| B44 | B127 | MCH | A-G ratio | 0.060051 | 0 |
| B44 | B128 | MCH | ALT | 0.047466 | 0 |
| B44 | B129 | MCH | IBIL | 0.080058 | 0 |
| B44 | B130 | MCH | GLB | -0.0604 | 0 |
| B44 | B131 | MCH | AST | 0.087678 | 0 |
| B44 | B132 | MCH | Dbil | 0.114684 | 0 |
| B44 | B133 | MCH | TBIL | 0.095367 | 0 |
| B44 | B140 | MCH | r-GT | 0.103767 | 0 |
| B44 | B163 | MCH | SG | 0.063518 | 0 |
| B44 | B174 | MCH | Urea/Cr | -0.06762 | 0 |
| B44 | B190 | MCH | Cre | 0.038087 | 0 |
| B44 | B192 | MCH | UA | 0.055787 | 0 |
| B44 | B204 | MCH | Drinking | 0.1373 | 0 |
| B44 | B206 | MCH | Waist | 0.05226 | 0 |
| B44 | B209 | MCH | Smoking | 0.12945 | 0 |
| B44 | B216 | MCH | BMI | 0.059231 | 0 |
| B44 | B221 | MCH | Hei | 0.096803 | 0 |
| B44 | B222 | MCH | NS | 0.092207 | 0 |
| B44 | B45 | MCH | MCHC | 0.582185 | 0 |
| B44 | B46 | MCH | RBC Count | -0.41693 | 0 |
| B44 | B47 | MCH | RDW | 0.399959 | 0 |
| B44 | B51 | MCH | Hb | 0.312533 | 0 |
| B44 | B52 | MCH | PLT/BPC | -0.1368 | 0 |
| B44 | B53 | MCH | GR% | -0.05722 | 0 |
| B44 | B54 | MCH | ESR | -0.04097 | 0 |
| B44 | B62 | MCH | WBHSRVI | 0.040239 | 0 |
| B44 | B71 | MCH | TG | 0.053753 | 0 |
| B44 | B91 | MCH | TP | -0.04895 | 0 |

| | | | | | |
|---|---|---|---|---|---|
| B44 | B63 | MCH | WBRSR(mPas_200) | 0.037168 | 6.66E-16 |
| B44 | B66 | MCH | WBRSR(mPas_100) | 0.034801 | 4.17E-14 |
| B44 | B48 | MCH | LYM% | 0.033554 | 3.22E-13 |
| B44 | B67 | MCH | WBRSR(mPas_50) | 0.032698 | 1.26E-12 |
| B44 | B176 | MCH | Ucr | 0.032058 | 3.40E-12 |
| B44 | B112 | MCH | FT4 | -0.03116 | 1.33E-11 |
| B44 | B189 | MCH | eGFB | -0.03103 | 1.63E-11 |
| B44 | B114 | MCH | TSH TT3 TT4(TT4) | -0.02976 | 1.04E-10 |
| B44 | B58 | MCH | HCT | 0.029716 | 1.11E-10 |
| B44 | B198 | MCH | P | -0.0294 | 1.75E-10 |
| B44 | B138 | MCH | PAB | 0.029026 | 2.95E-10 |
| B44 | B219 | MCH | OT | -0.02867 | 4.85E-10 |
| B44 | B57 | MCH | RBCAI | -0.02806 | 1.11E-09 |
| B44 | B210 | MCH | Hip | 0.027905 | 1.38E-09 |
| B44 | B203 | MCH | DH | 0.02611 | 1.44E-08 |
| B44 | B166 | MCH | Pro | 0.02492 | 6.31E-08 |
| B44 | B161 | MCH | LEU | -0.02475 | 7.71E-08 |
| B44 | B172 | MCH | KB | 0.023201 | 4.74E-07 |
| B44 | B201 | MCH | e-habits | 0.022592 | 9.37E-07 |
| B45 | B126 | MCHC | Alb | 0.130591 | 0 |
| B45 | B127 | MCHC | A-G ratio | 0.125934 | 0 |
| B45 | B128 | MCHC | ALT | 0.13378 | 0 |
| B45 | B129 | MCHC | IBIL | 0.173084 | 0 |
| B45 | B130 | MCHC | GLB | -0.08613 | 0 |
| B45 | B131 | MCHC | AST | 0.064723 | 0 |
| B45 | B132 | MCHC | Dbil | 0.117655 | 0 |
| B45 | B133 | MCHC | TBIL | 0.171122 | 0 |
| B45 | B134 | MCHC | CHE | 0.04179 | 0 |
| B45 | B138 | MCHC | PAB | 0.052618 | 0 |
| B45 | B140 | MCHC | r-GT | 0.107674 | 0 |
| B45 | B141 | MCHC | AST/ALT | -0.14611 | 0 |
| B45 | B153 | MCHC | Glu | 0.108664 | 0 |
| B45 | B161 | MCHC | LEU | -0.06743 | 0 |
| B45 | B163 | MCHC | SG | 0.135622 | 0 |
| B45 | B166 | MCHC | Pro | 0.05405 | 0 |
| B45 | B170 | MCHC | U-GLU | 0.065476 | 0 |
| B45 | B173 | MCHC | NTT | -0.05233 | 0 |
| B45 | B174 | MCHC | Urea/Cr | -0.15215 | 0 |
| B45 | B176 | MCHC | Ucr | 0.070823 | 0 |
| B45 | B189 | MCHC | eGFB | 0.116434 | 0 |
| B45 | B191 | MCHC | Urea | -0.0796 | 0 |
| B45 | B192 | MCHC | UA | 0.142377 | 0 |
| B45 | B193 | MCHC | K | -0.05607 | 0 |
| B45 | B203 | MCHC | DH | 0.039856 | 0 |

| | | | | | |
|---|---|---|---|---|---|
| B45 | B204 | MCHC | Drinking | 0.153135 | 0 |
| B45 | B206 | MCHC | Waist | 0.136006 | 0 |
| B45 | B209 | MCHC | Smoking | 0.124822 | 0 |
| B45 | B210 | MCHC | Hip | 0.117365 | 0 |
| B45 | B215 | MCHC | BMI | 0.102642 | 0 |
| B45 | B216 | MCHC | BMI | 0.226577 | 0 |
| B45 | B218 | MCHC | DBP | 0.119369 | 0 |
| B45 | B219 | MCHC | OT | -0.06485 | 0 |
| B45 | B220 | MCHC | SBP | -0.04977 | 0 |
| B45 | B221 | MCHC | Hei | 0.249038 | 0 |
| B45 | B222 | MCHC | NS | 0.242206 | 0 |
| B45 | B46 | MCHC | RBC Count | 0.059778 | 0 |
| B45 | B47 | MCHC | RDW | -0.19955 | 0 |
| B45 | B51 | MCHC | Hb | 0.497198 | 0 |
| B45 | B54 | MCHC | ESR | -0.05902 | 0 |
| B45 | B56 | MCHC | IR | 0.073772 | 0 |
| B45 | B60 | MCHC | WBRVI | 0.063578 | 0 |
| B45 | B61 | MCHC | WBHSRV | 0.074713 | 0 |
| B45 | B62 | MCHC | WBHSRVI | 0.086838 | 0 |
| B45 | B63 | MCHC | WBRSR(mPas_200) | 0.084546 | 0 |
| B45 | B64 | MCHC | WBRSR(mPas_5) | 0.073205 | 0 |
| B45 | B65 | MCHC | WBRSR(mPas_1) | 0.063251 | 0 |
| B45 | B66 | MCHC | WBRSR(mPas_100) | 0.081371 | 0 |
| B45 | B67 | MCHC | WBRSR(mPas_50) | 0.080178 | 0 |
| B45 | B71 | MCHC | TG | 0.17231 | 0 |
| B45 | B72 | MCHC | HDL-C | -0.1491 | 0 |
| B45 | B74 | MCHC | ApoA1 | -0.06268 | 0 |
| B45 | B58 | MCHC | HCT | 0.037161 | 6.66E-16 |
| B45 | B146 | MCHC | CysC | -0.03349 | 3.57E-13 |
| B45 | B194 | MCHC | Cl | -0.03218 | 2.82E-12 |
| B45 | B111 | MCHC | TSH | -0.03208 | 3.31E-12 |
| B45 | B75 | MCHC | ApoA1/B | -0.03136 | 9.93E-12 |
| B45 | B160 | MCHC | PH | 0.030791 | 2.31E-11 |
| B45 | B190 | MCHC | Cre | 0.030241 | 5.19E-11 |
| B45 | B59 | MCHC | WBRV | 0.03022 | 5.36E-11 |
| B45 | B172 | MCHC | KB | 0.02822 | 8.99E-10 |
| B45 | B198 | MCHC | P | -0.02765 | 1.93E-09 |
| B45 | B114 | MCHC | TSH TT3 TT4(TT4) | -0.02683 | 5.74E-09 |
| B45 | B149 | MCHC | TGAb | -0.02551 | 3.05E-08 |
| B45 | B73 | MCHC | ApoA | -0.02481 | 7.23E-08 |
| B45 | B148 | MCHC | Anti_TPO | -0.02413 | 1.63E-07 |
| B45 | B55 | MCHC | TK | 0.023568 | 3.12E-07 |
| B45 | B195 | MCHC | Na | -0.02255 | 9.82E-07 |
| B46 | B111 | RBC Count | TSH | -0.05217 | 0 |

| | | | | | |
|---|---|---|---|---|---|
| B46 | B113 | RBC Count | FT3 | 0.093337 | 0 |
| B46 | B115 | RBC Count | TSH TT3 TT4(TT3) | 0.061776 | 0 |
| B46 | B126 | RBC Count | Alb | 0.240363 | 0 |
| B46 | B127 | RBC Count | A-G ratio | 0.128421 | 0 |
| B46 | B128 | RBC Count | ALT | 0.192564 | 0 |
| B46 | B129 | RBC Count | IBIL | 0.1317 | 0 |
| B46 | B130 | RBC Count | GLB | -0.05352 | 0 |
| B46 | B132 | RBC Count | Dbil | 0.055655 | 0 |
| B46 | B133 | RBC Count | TBIL | 0.121127 | 0 |
| B46 | B134 | RBC Count | CHE | 0.078155 | 0 |
| B46 | B138 | RBC Count | PAB | 0.062093 | 0 |
| B46 | B140 | RBC Count | r-GT | 0.085724 | 0 |
| B46 | B141 | RBC Count | AST/ALT | -0.27009 | 0 |
| B46 | B146 | RBC Count | CysC | -0.04644 | 0 |
| B46 | B153 | RBC Count | Glu | 0.059763 | 0 |
| B46 | B161 | RBC Count | LEU | -0.09926 | 0 |
| B46 | B163 | RBC Count | SG | 0.237486 | 0 |
| B46 | B166 | RBC Count | Pro | 0.087731 | 0 |
| B46 | B170 | RBC Count | U-GLU | 0.053828 | 0 |
| B46 | B173 | RBC Count | NTT | -0.06821 | 0 |
| B46 | B174 | RBC Count | Urea/Cr | -0.22316 | 0 |
| B46 | B176 | RBC Count | Ucr | 0.103055 | 0 |
| B46 | B188 | RBC Count | Ca | 0.090683 | 0 |
| B46 | B189 | RBC Count | eGFB | 0.185915 | 0 |
| B46 | B191 | RBC Count | Urea | -0.13342 | 0 |
| B46 | B192 | RBC Count | UA | 0.205474 | 0 |
| B46 | B194 | RBC Count | Cl | -0.04581 | 0 |
| B46 | B198 | RBC Count | P | -0.04127 | 0 |
| B46 | B203 | RBC Count | DH | 0.038967 | 0 |
| B46 | B204 | RBC Count | Drinking | 0.144403 | 0 |
| B46 | B205 | RBC Count | KDA | -0.04952 | 0 |
| B46 | B206 | RBC Count | Waist | 0.270529 | 0 |
| B46 | B209 | RBC Count | Smoking | 0.14454 | 0 |
| B46 | B210 | RBC Count | Hip | 0.231995 | 0 |
| B46 | B215 | RBC Count | BMI | 0.229537 | 0 |
| B46 | B216 | RBC Count | BMI | 0.377675 | 0 |
| B46 | B218 | RBC Count | DBP | 0.248801 | 0 |
| B46 | B219 | RBC Count | OT | -0.08225 | 0 |
| B46 | B221 | RBC Count | Hei | 0.351894 | 0 |
| B46 | B222 | RBC Count | NS | 0.370386 | 0 |
| B46 | B47 | RBC Count | RDW | -0.3989 | 0 |
| B46 | B48 | RBC Count | LYM% | -0.04809 | 0 |
| B46 | B49 | RBC Count | baso% | 0.066265 | 0 |
| B46 | B51 | RBC Count | Hb | 0.724535 | 0 |

| | | | | | |
|---|---|---|---|---|---|
| B46 | B52 | RBC Count | PLT/BPC | 0.092356 | 0 |
| B46 | B53 | RBC Count | GR% | 0.047564 | 0 |
| B46 | B54 | RBC Count | ESR | -0.06172 | 0 |
| B46 | B56 | RBC Count | IR | 0.104259 | 0 |
| B46 | B57 | RBC Count | RBCAI | 0.054886 | 0 |
| B46 | B58 | RBC Count | HCT | 0.092467 | 0 |
| B46 | B59 | RBC Count | WBRV | 0.085424 | 0 |
| B46 | B60 | RBC Count | WBRVI | 0.165232 | 0 |
| B46 | B61 | RBC Count | WBHSRV | 0.110139 | 0 |
| B46 | B62 | RBC Count | WBHSRVI | 0.167287 | 0 |
| B46 | B63 | RBC Count | WBRSR(mPas_200) | 0.169111 | 0 |
| B46 | B64 | RBC Count | WBRSR(mPas_5) | 0.175563 | 0 |
| B46 | B65 | RBC Count | WBRSR(mPas_1) | 0.168015 | 0 |
| B46 | B66 | RBC Count | WBRSR(mPas_100) | 0.171198 | 0 |
| B46 | B67 | RBC Count | WBRSR(mPas_50) | 0.173329 | 0 |
| B46 | B70 | RBC Count | LDL-C | 0.079357 | 0 |
| B46 | B71 | RBC Count | TG | 0.119677 | 0 |
| B46 | B72 | RBC Count | HDL-C | -0.22577 | 0 |
| B46 | B74 | RBC Count | ApoA1 | -0.08107 | 0 |
| B46 | B75 | RBC Count | ApoA1/B | -0.06913 | 0 |
| B46 | B76 | RBC Count | ApoB | 0.043581 | 0 |
| B46 | B79 | RBC Count | hb V -dna | -0.03976 | 0 |
| B46 | B91 | RBC Count | TP | 0.094642 | 0 |
| B46 | B148 | RBC Count | Anti_TPO | -0.03134 | 1.02E-11 |
| B46 | B149 | RBC Count | TGAb | -0.02866 | 4.92E-10 |
| B46 | B197 | RBC Count | AG | 0.028416 | 6.88E-10 |
| B46 | B119 | RBC Count | Hcv-rna | -0.02837 | 7.30E-10 |
| B46 | B142 | RBC Count | 25-(OH)D | -0.02731 | 3.07E-09 |
| B46 | B137 | RBC Count | AFU | 0.026927 | 5.05E-09 |
| B46 | B220 | RBC Count | SBP | -0.02616 | 1.36E-08 |
| B46 | B112 | RBC Count | FT4 | 0.026146 | 1.38E-08 |
| B46 | B55 | RBC Count | TK | -0.02556 | 2.87E-08 |
| B46 | B187 | RBC Count | NSE | 0.024664 | 8.59E-08 |
| B46 | B228 | RBC Count | OS | -0.02405 | 1.78E-07 |
| B46 | B73 | RBC Count | ApoA | -0.02389 | 2.14E-07 |
| B46 | B139 | RBC Count | ADA | -0.02375 | 2.52E-07 |
| B46 | B131 | RBC Count | AST | 0.023398 | 3.79E-07 |
| B46 | B190 | RBC Count | Cre | 0.022946 | 6.32E-07 |
| B47 | B111 | RDW | TSH | 0.038918 | 0 |
| B47 | B112 | RDW | FT4 | -0.04554 | 0 |
| B47 | B126 | RDW | Alb | -0.16833 | 0 |
| B47 | B127 | RDW | A-G ratio | -0.09883 | 0 |
| B47 | B130 | RDW | GLB | 0.054134 | 0 |
| B47 | B131 | RDW | AST | 0.092004 | 0 |

| | | | | | |
|---|---|---|---|---|---|
| B47 | B132 | RDW | Dbil | 0.08373 | 0 |
| B47 | B134 | RDW | CHE | -0.05501 | 0 |
| B47 | B140 | RDW | r-GT | 0.070781 | 0 |
| B47 | B141 | RDW | AST/ALT | 0.125786 | 0 |
| B47 | B146 | RDW | CysC | 0.058354 | 0 |
| B47 | B153 | RDW | Glu | -0.08009 | 0 |
| B47 | B160 | RDW | PH | -0.04866 | 0 |
| B47 | B188 | RDW | Ca | -0.05898 | 0 |
| B47 | B189 | RDW | eGFB | -0.17463 | 0 |
| B47 | B190 | RDW | Cre | 0.07201 | 0 |
| B47 | B191 | RDW | Urea | 0.107915 | 0 |
| B47 | B204 | RDW | Drinking | 0.038502 | 0 |
| B47 | B209 | RDW | Smoking | 0.103969 | 0 |
| B47 | B210 | RDW | Hip | -0.05137 | 0 |
| B47 | B215 | RDW | BMI | -0.06474 | 0 |
| B47 | B216 | RDW | BMI | -0.09744 | 0 |
| B47 | B218 | RDW | DBP | -0.07167 | 0 |
| B47 | B220 | RDW | SBP | 0.055261 | 0 |
| B47 | B221 | RDW | Hei | -0.07996 | 0 |
| B47 | B222 | RDW | NS | -0.05504 | 0 |
| B47 | B50 | RDW | eo% | 0.046395 | 0 |
| B47 | B51 | RDW | Hb | -0.124 | 0 |
| B47 | B52 | RDW | PLT/BPC | -0.19477 | 0 |
| B47 | B71 | RDW | TG | -0.05725 | 0 |
| B47 | B72 | RDW | HDL-C | 0.119633 | 0 |
| B47 | B74 | RDW | ApoA1 | 0.047867 | 0 |
| B47 | B75 | RDW | ApoA1/B | 0.040134 | 0 |
| B47 | B91 | RDW | TP | -0.05114 | 0 |
| B47 | B170 | RDW | U-GLU | -0.03536 | 1.62E-14 |
| B47 | B113 | RDW | FT3 | -0.0335 | 3.49E-13 |
| B47 | B211 | RDW | IOP-R | -0.03296 | 8.25E-13 |
| B47 | B129 | RDW | IBIL | -0.03278 | 1.10E-12 |
| B47 | B212 | RDW | IOP-L | -0.03277 | 1.13E-12 |
| B47 | B128 | RDW | ALT | -0.03258 | 1.51E-12 |
| B47 | B138 | RDW | PAB | -0.03161 | 6.76E-12 |
| B47 | B61 | RDW | WBHSRV | -0.02982 | 9.51E-11 |
| B47 | B194 | RDW | Cl | 0.02949 | 1.53E-10 |
| B47 | B48 | RDW | LYM% | -0.02913 | 2.55E-10 |
| B47 | B118 | RDW | CEA1 | 0.028645 | 5.02E-10 |
| B47 | B56 | RDW | IR | -0.02825 | 8.61E-10 |
| B47 | B150 | RDW | ALP | 0.026611 | 7.60E-09 |
| B47 | B115 | RDW | TSH TT3 TT4(TT3) | -0.0265 | 8.74E-09 |
| B47 | B139 | RDW | ADA | 0.025452 | 3.29E-08 |
| B47 | B228 | RDW | OS | 0.025204 | 4.46E-08 |

| | | | | | |
|---|---|---|---|---|---|
| B47 | B70 | RDW | LDL-C | -0.02515 | 4.76E-08 |
| B47 | B55 | RDW | TK | -0.02495 | 6.11E-08 |
| B47 | B174 | RDW | Urea/Cr | 0.024325 | 1.29E-07 |
| B47 | B59 | RDW | WBRV | -0.02366 | 2.79E-07 |
| B47 | B219 | RDW | OT | 0.023085 | 5.41E-07 |
| B47 | B173 | RDW | NTT | 0.022808 | 7.38E-07 |
| B48 | B123 | LYM% | hs-CRP1 | -0.07244 | 0 |
| B48 | B126 | LYM% | Alb | 0.057145 | 0 |
| B48 | B130 | LYM% | GLB | 0.060846 | 0 |
| B48 | B131 | LYM% | AST | 0.039981 | 0 |
| B48 | B132 | LYM% | Dbil | -0.05999 | 0 |
| B48 | B146 | LYM% | CysC | -0.04438 | 0 |
| B48 | B163 | LYM% | SG | -0.05837 | 0 |
| B48 | B166 | LYM% | Pro | -0.05933 | 0 |
| B48 | B174 | LYM% | Urea/Cr | 0.051211 | 0 |
| B48 | B188 | LYM% | Ca | 0.050221 | 0 |
| B48 | B189 | LYM% | eGFB | 0.087141 | 0 |
| B48 | B190 | LYM% | Cre | -0.1072 | 0 |
| B48 | B191 | LYM% | Urea | -0.07087 | 0 |
| B48 | B192 | LYM% | UA | -0.06864 | 0 |
| B48 | B206 | LYM% | Waist | -0.05255 | 0 |
| B48 | B209 | LYM% | Smoking | -0.07121 | 0 |
| B48 | B221 | LYM% | Hei | -0.04053 | 0 |
| B48 | B222 | LYM% | NS | -0.06772 | 0 |
| B48 | B49 | LYM% | baso% | 0.049793 | 0 |
| B48 | B52 | LYM% | PLT/BPC | 0.039893 | 0 |
| B48 | B53 | LYM% | GR% | -0.93767 | 0 |
| B48 | B70 | LYM% | LDL-C | 0.066832 | 0 |
| B48 | B72 | LYM% | HDL-C | 0.080074 | 0 |
| B48 | B77 | LYM% | TC | 0.09312 | 0 |
| B48 | B91 | LYM% | TP | 0.088987 | 0 |
| B48 | B127 | LYM% | A-G ratio | -0.03675 | 1.33E-15 |
| B48 | B133 | LYM% | TBIL | -0.03511 | 2.46E-14 |
| B48 | B153 | LYM% | Glu | -0.03486 | 3.73E-14 |
| B48 | B111 | LYM% | TSH | 0.033818 | 2.10E-13 |
| B48 | B74 | LYM% | ApoA1 | 0.032208 | 2.70E-12 |
| B48 | B175 | LYM% | ALB-U | -0.03081 | 2.26E-11 |
| B48 | B149 | LYM% | TGAb | 0.030121 | 6.18E-11 |
| B48 | B201 | LYM% | e-habits | 0.030119 | 6.20E-11 |
| B48 | B51 | LYM% | Hb | -0.02718 | 3.61E-09 |
| B48 | B179 | LYM% | α1-MG | -0.02709 | 4.05E-09 |
| B48 | B86 | LYM% | TBhCG | 0.026692 | 6.85E-09 |
| B48 | B198 | LYM% | P | 0.026671 | 7.04E-09 |
| B48 | B177 | LYM% | ACR | -0.02641 | 9.80E-09 |

| | | | | | |
|---|---|---|---|---|---|
| B48 | B128 | LYM% | ALT | 0.0261 | 1.46E-08 |
| B48 | B170 | LYM% | U-GLU | -0.02604 | 1.58E-08 |
| B48 | B176 | LYM% | Ucr | -0.02567 | 2.52E-08 |
| B48 | B148 | LYM% | Anti_TPO | 0.024669 | 8.54E-08 |
| B48 | B220 | LYM% | SBP | -0.02461 | 9.18E-08 |
| B48 | B204 | LYM% | Drinking | -0.02402 | 1.85E-07 |
| B48 | B140 | LYM% | r-GT | -0.02372 | 2.62E-07 |
| B48 | B129 | LYM% | IBIL | -0.02349 | 3.42E-07 |
| B49 | B140 | baso% | r-GT | 0.044011 | 0 |
| B49 | B209 | baso% | Smoking | 0.042246 | 0 |
| B49 | B221 | baso% | Hei | 0.046981 | 0 |
| B49 | B50 | baso% | eo% | 0.271707 | 0 |
| B49 | B51 | baso% | Hb | 0.06827 | 0 |
| B49 | B52 | baso% | PLT/BPC | 0.061358 | 0 |
| B49 | B53 | baso% | GR% | -0.1712 | 0 |
| B49 | B70 | baso% | LDL-C | 0.038375 | 0 |
| B49 | B77 | baso% | TC | 0.04191 | 0 |
| B49 | B204 | baso% | Drinking | 0.037725 | 2.22E-16 |
| B49 | B216 | baso% | BMI | 0.035093 | 2.55E-14 |
| B49 | B128 | baso% | ALT | 0.030514 | 3.48E-11 |
| B49 | B62 | baso% | WBHSRVI | 0.02987 | 8.88E-11 |
| B49 | B174 | baso% | Urea/Cr | -0.02968 | 1.17E-10 |
| B49 | B220 | baso% | SBP | -0.02899 | 3.10E-10 |
| B49 | B191 | baso% | Urea | -0.02898 | 3.15E-10 |
| B49 | B141 | baso% | AST/ALT | -0.02781 | 1.58E-09 |
| B49 | B123 | baso% | hs-CRP1 | -0.02746 | 2.51E-09 |
| B49 | B192 | baso% | UA | 0.026823 | 5.78E-09 |
| B49 | B66 | baso% | WBRSR(mPas_100) | 0.02578 | 2.19E-08 |
| B49 | B67 | baso% | WBRSR(mPas_50) | 0.025595 | 2.75E-08 |
| B49 | B129 | baso% | IBIL | -0.02558 | 2.81E-08 |
| B49 | B222 | baso% | NS | 0.02457 | 9.62E-08 |
| B49 | B131 | baso% | AST | 0.024335 | 1.27E-07 |
| B49 | B153 | baso% | Glu | -0.02389 | 2.15E-07 |
| B49 | B60 | baso% | WBRVI | 0.023427 | 3.66E-07 |
| B49 | B54 | baso% | ESR | -0.02292 | 6.48E-07 |
| B49 | B63 | baso% | WBRSR(mPas_200) | 0.022842 | 7.10E-07 |
| B5 | B6 | WB-Mg | WB-Fe | 0.697108 | 0 |
| B5 | B7 | WB-Mg | WB-Cop | 0.469229 | 0 |
| B5 | B8 | WB-Mg | WB-Zinc | 0.56987 | 0 |
| B5 | B107 | WB-Mg | AnuA | -0.03099 | 1.72E-11 |
| B5 | B74 | WB-Mg | ApoA1 | -0.02291 | 6.58E-07 |
| B5 | B15 | WB-Mg | CD3-CD19+(B cell) | -0.02257 | 9.57E-07 |
| B50 | B126 | eo% | Alb | -0.07138 | 0 |
| B50 | B146 | eo% | CysC | 0.04181 | 0 |

| | | | | | |
|---|---|---|---|---|---|
| B50 | B174 | eo% | Urea/Cr | -0.07158 | 0 |
| B50 | B189 | eo% | eGFB | -0.082 | 0 |
| B50 | B190 | eo% | Cre | 0.089597 | 0 |
| B50 | B191 | eo% | Urea | 0.048001 | 0 |
| B50 | B192 | eo% | UA | 0.072681 | 0 |
| B50 | B204 | eo% | Drinking | 0.04504 | 0 |
| B50 | B206 | eo% | Waist | 0.047376 | 0 |
| B50 | B209 | eo% | Smoking | 0.086929 | 0 |
| B50 | B216 | eo% | BMI | 0.051107 | 0 |
| B50 | B221 | eo% | Hei | 0.070233 | 0 |
| B50 | B222 | eo% | NS | 0.063682 | 0 |
| B50 | B53 | eo% | GR% | -0.24678 | 0 |
| B50 | B91 | eo% | TP | -0.05461 | 0 |
| B50 | B153 | eo% | Glu | -0.03586 | 6.88E-15 |
| B50 | B129 | eo% | IBIL | -0.03502 | 2.89E-14 |
| B50 | B133 | eo% | TBIL | -0.03355 | 3.24E-13 |
| B50 | B210 | eo% | Hip | 0.032014 | 3.64E-12 |
| B50 | B77 | eo% | TC | -0.03083 | 2.19E-11 |
| B50 | B156 | eo% | IgE | 0.030229 | 5.28E-11 |
| B50 | B72 | eo% | HDL-C | -0.02997 | 7.67E-11 |
| B50 | B188 | eo% | Ca | -0.02865 | 4.98E-10 |
| B50 | B52 | eo% | PLT/BPC | -0.02851 | 6.00E-10 |
| B50 | B160 | eo% | PH | -0.02777 | 1.65E-09 |
| B50 | B51 | eo% | Hb | 0.023674 | 2.76E-07 |
| B50 | B211 | eo% | IOP-R | -0.023 | 5.97E-07 |
| B50 | B220 | eo% | SBP | -0.023 | 5.98E-07 |
| B51 | B111 | Hb | TSH | -0.05688 | 0 |
| B51 | B113 | Hb | FT3 | 0.093228 | 0 |
| B51 | B115 | Hb | TSH TT3 TT4(TT3) | 0.051632 | 0 |
| B51 | B126 | Hb | Alb | 0.259903 | 0 |
| B51 | B127 | Hb | A-G ratio | 0.176466 | 0 |
| B51 | B128 | Hb | ALT | 0.2346 | 0 |
| B51 | B129 | Hb | IBIL | 0.191184 | 0 |
| B51 | B130 | Hb | GLB | -0.09754 | 0 |
| B51 | B131 | Hb | AST | 0.082942 | 0 |
| B51 | B132 | Hb | Dbil | 0.130706 | 0 |
| B51 | B133 | Hb | TBIL | 0.189252 | 0 |
| B51 | B134 | Hb | CHE | 0.081531 | 0 |
| B51 | B137 | Hb | AFU | 0.04246 | 0 |
| B51 | B138 | Hb | PAB | 0.0861 | 0 |
| B51 | B140 | Hb | r-GT | 0.159068 | 0 |
| B51 | B141 | Hb | AST/ALT | -0.2863 | 0 |
| B51 | B146 | Hb | CysC | -0.04701 | 0 |
| B51 | B148 | Hb | Anti_TPO | -0.0426 | 0 |

| | | | | | |
|---|---|---|---|---|---|
| B51 | B149 | Hb | TGAb | -0.03961 | 0 |
| B51 | B153 | Hb | Glu | 0.078938 | 0 |
| B51 | B161 | Hb | LEU | -0.1193 | 0 |
| B51 | B163 | Hb | SG | 0.293968 | 0 |
| B51 | B166 | Hb | Pro | 0.109703 | 0 |
| B51 | B170 | Hb | U-GLU | 0.070397 | 0 |
| B51 | B173 | Hb | NTT | -0.08266 | 0 |
| B51 | B174 | Hb | Urea/Cr | -0.28019 | 0 |
| B51 | B176 | Hb | Ucr | 0.131565 | 0 |
| B51 | B188 | Hb | Ca | 0.098183 | 0 |
| B51 | B189 | Hb | eGFB | 0.175889 | 0 |
| B51 | B190 | Hb | Cre | 0.049393 | 0 |
| B51 | B191 | Hb | Urea | -0.13771 | 0 |
| B51 | B192 | Hb | UA | 0.251602 | 0 |
| B51 | B194 | Hb | Cl | -0.04991 | 0 |
| B51 | B198 | Hb | P | -0.06509 | 0 |
| B51 | B203 | Hb | DH | 0.06009 | 0 |
| B51 | B204 | Hb | Drinking | 0.247168 | 0 |
| B51 | B205 | Hb | KDA | -0.04716 | 0 |
| B51 | B206 | Hb | Waist | 0.321566 | 0 |
| B51 | B209 | Hb | Smoking | 0.241912 | 0 |
| B51 | B210 | Hb | Hip | 0.265893 | 0 |
| B51 | B215 | Hb | BMI | 0.243738 | 0 |
| B51 | B216 | Hb | BMI | 0.439802 | 0 |
| B51 | B218 | Hb | DBP | 0.267471 | 0 |
| B51 | B219 | Hb | OT | -0.10647 | 0 |
| B51 | B220 | Hb | SBP | -0.04305 | 0 |
| B51 | B221 | Hb | Hei | 0.437633 | 0 |
| B51 | B222 | Hb | NS | 0.454253 | 0 |
| B51 | B54 | Hb | ESR | -0.09398 | 0 |
| B51 | B55 | Hb | TK | -0.04205 | 0 |
| B51 | B56 | Hb | IR | 0.123996 | 0 |
| B51 | B58 | Hb | HCT | 0.119468 | 0 |
| B51 | B59 | Hb | WBRV | 0.071045 | 0 |
| B51 | B60 | Hb | WBRVI | 0.182968 | 0 |
| B51 | B61 | Hb | WBHSRV | 0.128851 | 0 |
| B51 | B62 | Hb | WBHSRVI | 0.205735 | 0 |
| B51 | B63 | Hb | WBRSR(mPas_200) | 0.205492 | 0 |
| B51 | B64 | Hb | WBRSR(mPas_5) | 0.199835 | 0 |
| B51 | B65 | Hb | WBRSR(mPas_1) | 0.184509 | 0 |
| B51 | B66 | Hb | WBRSR(mPas_100) | 0.205898 | 0 |
| B51 | B67 | Hb | WBRSR(mPas_50) | 0.206341 | 0 |
| B51 | B70 | Hb | LDL-C | 0.090872 | 0 |
| B51 | B71 | Hb | TG | 0.16395 | 0 |

| | | | | | |
|---|---|---|---|---|---|
| B51 | B72 | Hb | HDL-C | -0.22883 | 0 |
| B51 | B74 | Hb | ApoA1 | -0.08345 | 0 |
| B51 | B75 | Hb | ApoA1/B | -0.07328 | 0 |
| B51 | B76 | Hb | ApoB | 0.055222 | 0 |
| B51 | B77 | Hb | TC | 0.04171 | 0 |
| B51 | B79 | Hb | hb V -dna | -0.04073 | 0 |
| B51 | B91 | Hb | TP | 0.066506 | 0 |
| B51 | B172 | Hb | KB | 0.036271 | 3.55E-15 |
| B51 | B57 | Hb | RBCAI | 0.032907 | 9.04E-13 |
| B51 | B179 | Hb | α1-MG | 0.032406 | 1.98E-12 |
| B51 | B193 | Hb | K | -0.03143 | 8.93E-12 |
| B51 | B197 | Hb | AG | 0.029954 | 7.87E-11 |
| B51 | B86 | Hb | TBhCG | -0.02957 | 1.36E-10 |
| B51 | B119 | Hb | Hcv-rna | -0.02922 | 2.26E-10 |
| B51 | B114 | Hb | TSH TT3 TT4(TT4) | -0.02759 | 2.11E-09 |
| B51 | B73 | Hb | ApoA | -0.02745 | 2.52E-09 |
| B51 | B139 | Hb | ADA | -0.0262 | 1.29E-08 |
| B51 | B228 | Hb | OS | -0.02617 | 1.33E-08 |
| B51 | B187 | Hb | NSE | 0.025624 | 2.66E-08 |
| B51 | B142 | Hb | 25-(OH)D | -0.02526 | 4.18E-08 |
| B51 | B157 | Hb | IgG | -0.02507 | 5.25E-08 |
| B51 | B160 | Hb | PH | -0.02274 | 7.92E-07 |
| B52 | B112 | PLT/BPC | FT4 | 0.038456 | 0 |
| B52 | B127 | PLT/BPC | A-G ratio | -0.0916 | 0 |
| B52 | B128 | PLT/BPC | ALT | -0.06799 | 0 |
| B52 | B129 | PLT/BPC | IBIL | -0.11232 | 0 |
| B52 | B130 | PLT/BPC | GLB | 0.100141 | 0 |
| B52 | B131 | PLT/BPC | AST | -0.08779 | 0 |
| B52 | B132 | PLT/BPC | Dbil | -0.17739 | 0 |
| B52 | B133 | PLT/BPC | TBIL | -0.1384 | 0 |
| B52 | B146 | PLT/BPC | CysC | -0.04042 | 0 |
| B52 | B160 | PLT/BPC | PH | 0.043275 | 0 |
| B52 | B188 | PLT/BPC | Ca | 0.065669 | 0 |
| B52 | B189 | PLT/BPC | eGFB | 0.097005 | 0 |
| B52 | B190 | PLT/BPC | Cre | -0.07461 | 0 |
| B52 | B191 | PLT/BPC | Urea | -0.10792 | 0 |
| B52 | B192 | PLT/BPC | UA | -0.05843 | 0 |
| B52 | B198 | PLT/BPC | P | 0.043334 | 0 |
| B52 | B220 | PLT/BPC | SBP | -0.04244 | 0 |
| B52 | B70 | PLT/BPC | LDL-C | 0.134981 | 0 |
| B52 | B77 | PLT/BPC | TC | 0.122747 | 0 |
| B52 | B91 | PLT/BPC | TP | 0.093164 | 0 |
| B52 | B134 | PLT/BPC | CHE | 0.037605 | 4.44E-16 |
| B52 | B153 | PLT/BPC | Glu | -0.03612 | 4.44E-15 |

| | | | | | |
|---|---|---|---|---|---|
| B52 | B139 | PLT/BPC | ADA | -0.03601 | 5.33E-15 |
| B52 | B76 | PLT/BPC | ApoB | 0.033663 | 2.70E-13 |
| B52 | B222 | PLT/BPC | NS | -0.0318 | 5.08E-12 |
| B52 | B141 | PLT/BPC | AST/ALT | 0.030629 | 2.94E-11 |
| B52 | B206 | PLT/BPC | Waist | -0.03046 | 3.79E-11 |
| B52 | B193 | PLT/BPC | K | 0.028477 | 6.32E-10 |
| B52 | B201 | PLT/BPC | e-habits | -0.02465 | 8.77E-08 |
| B52 | B161 | PLT/BPC | LEU | 0.023839 | 2.28E-07 |
| B52 | B195 | PLT/BPC | Na | -0.02342 | 3.71E-07 |
| B52 | B194 | PLT/BPC | Cl | -0.02277 | 7.71E-07 |
| B52 | B53 | PLT/BPC | GR% | -0.02255 | 9.81E-07 |
| B53 | B123 | GR% | hs-CRP1 | 0.066752 | 0 |
| B53 | B127 | GR% | A-G ratio | 0.05445 | 0 |
| B53 | B130 | GR% | GLB | -0.06088 | 0 |
| B53 | B131 | GR% | AST | -0.05996 | 0 |
| B53 | B132 | GR% | Dbil | 0.040137 | 0 |
| B53 | B153 | GR% | Glu | 0.047207 | 0 |
| B53 | B166 | GR% | Pro | 0.046387 | 0 |
| B53 | B190 | GR% | Cre | 0.060129 | 0 |
| B53 | B191 | GR% | Urea | 0.040244 | 0 |
| B53 | B70 | GR% | LDL-C | -0.04986 | 0 |
| B53 | B72 | GR% | HDL-C | -0.06959 | 0 |
| B53 | B77 | GR% | TC | -0.06391 | 0 |
| B53 | B91 | GR% | TP | -0.05693 | 0 |
| B53 | B163 | GR% | SG | 0.035985 | 5.55E-15 |
| B53 | B128 | GR% | ALT | -0.03404 | 1.45E-13 |
| B53 | B111 | GR% | TSH | -0.03367 | 2.68E-13 |
| B53 | B220 | GR% | SBP | 0.032696 | 1.26E-12 |
| B53 | B218 | GR% | DBP | 0.032083 | 3.27E-12 |
| B53 | B189 | GR% | eGFB | -0.03144 | 8.69E-12 |
| B53 | B133 | GR% | TBIL | 0.031112 | 1.43E-11 |
| B53 | B201 | GR% | e-habits | -0.03056 | 3.25E-11 |
| B53 | B209 | GR% | Smoking | 0.029871 | 8.87E-11 |
| B53 | B222 | GR% | NS | 0.029738 | 1.08E-10 |
| B53 | B175 | GR% | ALB-U | 0.028979 | 3.15E-10 |
| B53 | B170 | GR% | U-GLU | 0.028802 | 4.03E-10 |
| B53 | B192 | GR% | UA | 0.02636 | 1.05E-08 |
| B53 | B74 | GR% | ApoA1 | -0.02634 | 1.08E-08 |
| B53 | B188 | GR% | Ca | -0.02593 | 1.80E-08 |
| B53 | B149 | GR% | TGAb | -0.02568 | 2.49E-08 |
| B53 | B129 | GR% | IBIL | 0.025199 | 4.49E-08 |
| B53 | B179 | GR% | α1-MG | 0.024867 | 6.73E-08 |
| B53 | B206 | GR% | Waist | 0.024421 | 1.15E-07 |
| B53 | B86 | GR% | TBhCG | -0.02389 | 2.14E-07 |

| | | | | | |
|---|---|---|---|---|---|
| B53 | B177 | GR% | ACR | 0.023637 | 2.88E-07 |
| B53 | B198 | GR% | P | -0.02352 | 3.28E-07 |
| B53 | B112 | GR% | FT4 | 0.022745 | 7.92E-07 |
| B54 | B107 | ESR | AnuA | -0.05372 | 0 |
| B54 | B114 | ESR | TSH TT3 TT4(TT4) | 0.054765 | 0 |
| B54 | B121 | ESR | C3 | 0.04571 | 0 |
| B54 | B123 | ESR | hs-CRP1 | 0.149412 | 0 |
| B54 | B126 | ESR | Alb | -0.05478 | 0 |
| B54 | B127 | ESR | A-G ratio | -0.10239 | 0 |
| B54 | B129 | ESR | IBIL | -0.05183 | 0 |
| B54 | B130 | ESR | GLB | 0.102625 | 0 |
| B54 | B132 | ESR | Dbil | -0.04268 | 0 |
| B54 | B133 | ESR | TBIL | -0.05342 | 0 |
| B54 | B136 | ESR | 5'NT | 0.042129 | 0 |
| B54 | B138 | ESR | PAB | -0.05854 | 0 |
| B54 | B139 | ESR | ADA | 0.04872 | 0 |
| B54 | B141 | ESR | AST/ALT | 0.04452 | 0 |
| B54 | B155 | ESR | IgA | 0.059265 | 0 |
| B54 | B157 | ESR | IgG | 0.085224 | 0 |
| B54 | B174 | ESR | Urea/Cr | 0.048926 | 0 |
| B54 | B186 | ESR | PFC | 0.106369 | 0 |
| B54 | B198 | ESR | P | 0.051977 | 0 |
| B54 | B216 | ESR | BMI | -0.04426 | 0 |
| B54 | B221 | ESR | Hei | -0.07797 | 0 |
| B54 | B222 | ESR | NS | -0.05313 | 0 |
| B54 | B55 | ESR | TK | -0.04341 | 0 |
| B54 | B56 | ESR | IR | -0.24523 | 0 |
| B54 | B58 | ESR | HCT | -0.1416 | 0 |
| B54 | B59 | ESR | WBRV | 0.054587 | 0 |
| B54 | B60 | ESR | WBRVI | -0.27233 | 0 |
| B54 | B61 | ESR | WBHSRV | -0.08489 | 0 |
| B54 | B62 | ESR | WBHSRVI | -0.32815 | 0 |
| B54 | B63 | ESR | WBRSR(mPas_200) | -0.16293 | 0 |
| B54 | B64 | ESR | WBRSR(mPas_5) | -0.14181 | 0 |
| B54 | B65 | ESR | WBRSR(mPas_1) | -0.12822 | 0 |
| B54 | B66 | ESR | WBRSR(mPas_100) | -0.14984 | 0 |
| B54 | B67 | ESR | WBRSR(mPas_50) | -0.14885 | 0 |
| B54 | B69 | ESR | PV | 0.45622 | 0 |
| B54 | B70 | ESR | LDL-C | 0.04978 | 0 |
| B54 | B73 | ESR | ApoA | 0.081539 | 0 |
| B54 | B74 | ESR | ApoA1 | 0.087368 | 0 |
| B54 | B76 | ESR | ApoB | 0.090925 | 0 |
| B54 | B77 | ESR | TC | 0.046984 | 0 |
| B54 | B91 | ESR | TP | 0.060266 | 0 |

| | | | | | |
|---|---|---|---|---|---|
| B54 | B101 | ESR | anti-Sc70 | 0.037378 | 4.44E-16 |
| B54 | B161 | ESR | LEU | 0.035968 | 5.77E-15 |
| B54 | B111 | ESR | TSH | 0.034384 | 8.30E-14 |
| B54 | B104 | ESR | anti-SSB | 0.034269 | 1.00E-13 |
| B54 | B156 | ESR | IgE | 0.034155 | 1.21E-13 |
| B54 | B163 | ESR | SG | -0.03328 | 5.00E-13 |
| B54 | B204 | ESR | Drinking | -0.03148 | 8.27E-12 |
| B54 | B72 | ESR | HDL-C | 0.031211 | 1.23E-11 |
| B54 | B192 | ESR | UA | -0.0302 | 5.53E-11 |
| B54 | B134 | ESR | CHE | -0.03012 | 6.17E-11 |
| B54 | B122 | ESR | C4 | 0.029986 | 7.51E-11 |
| B54 | B97 | ESR | dsDNA | 0.028597 | 5.35E-10 |
| B54 | B177 | ESR | ACR | 0.027515 | 2.32E-09 |
| B54 | B75 | ESR | ApoA1/B | -0.02656 | 8.16E-09 |
| B54 | B209 | ESR | Smoking | -0.02627 | 1.17E-08 |
| B54 | B148 | ESR | Anti_TPO | 0.026241 | 1.22E-08 |
| B54 | B176 | ESR | Ucr | -0.02458 | 9.53E-08 |
| B54 | B103 | ESR | anti-SSA/Ro | 0.023952 | 2.00E-07 |
| B54 | B191 | ESR | Urea | 0.023926 | 2.06E-07 |
| B54 | B137 | ESR | AFU | -0.02375 | 2.52E-07 |
| B54 | B92 | ESR | ALDH2-Glu504Glu | -0.02373 | 2.60E-07 |
| B54 | B93 | ESR | ALDH2-Glu504Lys | 0.023647 | 2.85E-07 |
| B54 | B158 | ESR | IgM | 0.023318 | 4.15E-07 |
| B55 | B121 | TK | C3 | -0.06814 | 0 |
| B55 | B122 | TK | C4 | -0.05776 | 0 |
| B55 | B56 | TK | IR | 0.724677 | 0 |
| B55 | B57 | TK | RBCAI | -0.16718 | 0 |
| B55 | B58 | TK | HCT | -0.33158 | 0 |
| B55 | B59 | TK | WBRV | 0.384377 | 0 |
| B55 | B60 | TK | WBRVI | 0.181103 | 0 |
| B55 | B61 | TK | WBHSRV | 0.642358 | 0 |
| B55 | B62 | TK | WBHSRVI | 0.332604 | 0 |
| B55 | B63 | TK | WBRSR(mPas_200) | 0.217128 | 0 |
| B55 | B64 | TK | WBRSR(mPas_5) | 0.141433 | 0 |
| B55 | B65 | TK | WBRSR(mPas_1) | 0.091863 | 0 |
| B55 | B66 | TK | WBRSR(mPas_100) | 0.208069 | 0 |
| B55 | B67 | TK | WBRSR(mPas_50) | 0.197747 | 0 |
| B55 | B69 | TK | PV | -0.30643 | 0 |
| B55 | B74 | TK | ApoA1 | -0.04166 | 0 |
| B55 | B78 | TK | HBcAb | 0.039546 | 0 |
| B55 | B107 | TK | AnuA | 0.031246 | 1.17E-11 |
| B55 | B183 | TK | PTA | -0.03054 | 3.35E-11 |
| B55 | B199 | TK | Mg | -0.02985 | 9.17E-11 |
| B55 | B120 | TK | HCV-cAg | 0.029611 | 1.29E-10 |

| | | | | | |
|---|---|---|---|---|---|
| B55 | B198 | TK | P | 0.02934 | 1.89E-10 |
| B55 | B73 | TK | ApoA | -0.02499 | 5.79E-08 |
| B55 | B84 | TK | C-PSA | -0.02437 | 1.23E-07 |
| B55 | B100 | TK | anti-Ro52 | -0.02393 | 2.05E-07 |
| B55 | B123 | TK | hs-CRP1 | -0.02309 | 5.35E-07 |
| B55 | B81 | TK | TSGF | 0.02266 | 8.69E-07 |
| B55 | B115 | TK | TSH TT3 TT4(TT3) | -0.02259 | 9.35E-07 |
| B56 | B107 | IR | AnuA | 0.038004 | 0 |
| B56 | B121 | IR | C3 | -0.06009 | 0 |
| B56 | B122 | IR | C4 | -0.04841 | 0 |
| B56 | B129 | IR | IBIL | 0.038663 | 0 |
| B56 | B133 | IR | TBIL | 0.039896 | 0 |
| B56 | B134 | IR | CHE | 0.058681 | 0 |
| B56 | B138 | IR | PAB | 0.054626 | 0 |
| B56 | B163 | IR | SG | 0.038207 | 0 |
| B56 | B174 | IR | Urea/Cr | -0.04785 | 0 |
| B56 | B192 | IR | UA | 0.041612 | 0 |
| B56 | B194 | IR | Cl | -0.03811 | 0 |
| B56 | B206 | IR | Waist | 0.048595 | 0 |
| B56 | B210 | IR | Hip | 0.043317 | 0 |
| B56 | B215 | IR | BMI | 0.039249 | 0 |
| B56 | B216 | IR | BMI | 0.063912 | 0 |
| B56 | B221 | IR | Hei | 0.056905 | 0 |
| B56 | B222 | IR | NS | 0.062755 | 0 |
| B56 | B57 | IR | RBCAI | -0.05919 | 0 |
| B56 | B58 | IR | HCT | 0.104578 | 0 |
| B56 | B59 | IR | WBRV | 0.540399 | 0 |
| B56 | B60 | IR | WBRVI | 0.683507 | 0 |
| B56 | B61 | IR | WBHSRV | 0.9229 | 0 |
| B56 | B62 | IR | WBHSRVI | 0.882109 | 0 |
| B56 | B63 | IR | WBRSR(mPas_200) | 0.767481 | 0 |
| B56 | B64 | IR | WBRSR(mPas_5) | 0.678806 | 0 |
| B56 | B65 | IR | WBRSR(mPas_1) | 0.59463 | 0 |
| B56 | B66 | IR | WBRSR(mPas_100) | 0.755023 | 0 |
| B56 | B67 | IR | WBRSR(mPas_50) | 0.747348 | 0 |
| B56 | B69 | IR | PV | -0.29974 | 0 |
| B56 | B73 | IR | ApoA | -0.04144 | 0 |
| B56 | B74 | IR | ApoA1 | -0.0908 | 0 |
| B56 | B75 | IR | ApoA1/B | -0.05795 | 0 |
| B56 | B72 | IR | HDL-C | -0.03736 | 4.44E-16 |
| B56 | B100 | IR | anti-Ro52 | -0.03693 | 1.11E-15 |
| B56 | B141 | IR | AST/ALT | -0.0353 | 1.82E-14 |
| B56 | B157 | IR | IgG | -0.03424 | 1.06E-13 |
| B56 | B132 | IR | Dbil | 0.033201 | 5.67E-13 |

| | | | | | |
|---|---|---|---|---|---|
| B56 | B187 | IR | NSE | 0.032203 | 2.72E-12 |
| B56 | B115 | IR | TSH TT3 TT4(TT3) | 0.031836 | 4.79E-12 |
| B56 | B176 | IR | Ucr | 0.031369 | 9.73E-12 |
| B56 | B78 | IR | HBcAb | 0.029584 | 1.34E-10 |
| B56 | B120 | IR | HCV-cAg | 0.029284 | 2.05E-10 |
| B56 | B218 | IR | DBP | 0.028832 | 3.86E-10 |
| B56 | B128 | IR | ALT | 0.028802 | 4.03E-10 |
| B56 | B197 | IR | AG | 0.027877 | 1.43E-09 |
| B56 | B71 | IR | TG | 0.027832 | 1.52E-09 |
| B56 | B76 | IR | ApoB | 0.027123 | 3.90E-09 |
| B56 | B127 | IR | A-G ratio | 0.02657 | 8.02E-09 |
| B56 | B183 | IR | PTA | -0.02547 | 3.21E-08 |
| B56 | B113 | IR | FT3 | 0.025407 | 3.48E-08 |
| B56 | B140 | IR | r-GT | 0.024811 | 7.20E-08 |
| B56 | B104 | IR | anti-SSB | -0.02323 | 4.59E-07 |
| B56 | B126 | IR | Alb | 0.023188 | 4.81E-07 |
| B56 | B207 | IR | WHR | 0.022585 | 9.45E-07 |
| B57 | B121 | RBCAI | C3 | -0.06254 | 0 |
| B57 | B122 | RBCAI | C4 | -0.04609 | 0 |
| B57 | B58 | RBCAI | HCT | 0.059074 | 0 |
| B57 | B59 | RBCAI | WBRV | 0.668907 | 0 |
| B57 | B60 | RBCAI | WBRVI | 0.570792 | 0 |
| B57 | B61 | RBCAI | WBHSRV | -0.05768 | 0 |
| B57 | B64 | RBCAI | WBRSR(mPas_5) | 0.392466 | 0 |
| B57 | B65 | RBCAI | WBRSR(mPas_1) | 0.563471 | 0 |
| B57 | B66 | RBCAI | WBRSR(mPas_100) | 0.064714 | 0 |
| B57 | B67 | RBCAI | WBRSR(mPas_50) | 0.123325 | 0 |
| B57 | B76 | RBCAI | ApoB | -0.04103 | 0 |
| B57 | B81 | RBCAI | TSGF | -0.05233 | 0 |
| B57 | B75 | RBCAI | ApoA1/B | 0.035903 | 6.44E-15 |
| B57 | B135 | RBCAI | TBA | 0.032947 | 8.48E-13 |
| B57 | B92 | RBCAI | ALDH2-Glu504Glu | 0.032662 | 1.33E-12 |
| B57 | B93 | RBCAI | ALDH2-Glu504Lys | -0.03249 | 1.73E-12 |
| B57 | B102 | RBCAI | anti-Sm | 0.031577 | 7.10E-12 |
| B57 | B105 | RBCAI | anti-U1RNP | 0.03062 | 2.98E-11 |
| B57 | B189 | RBCAI | eGFB | 0.027472 | 2.46E-09 |
| B57 | B127 | RBCAI | A-G ratio | 0.027409 | 2.68E-09 |
| B57 | B112 | RBCAI | FT4 | 0.026954 | 4.87E-09 |
| B57 | B78 | RBCAI | HBcAb | -0.02614 | 1.38E-08 |
| B57 | B130 | RBCAI | GLB | -0.02601 | 1.64E-08 |
| B57 | B63 | RBCAI | WBRSR(mPas_200) | 0.025169 | 4.66E-08 |
| B57 | B74 | RBCAI | ApoA1 | 0.024914 | 6.36E-08 |
| B57 | B183 | RBCAI | PTA | -0.02479 | 7.42E-08 |
| B57 | B111 | RBCAI | TSH | -0.02477 | 7.61E-08 |

| | | | | | |
|---|---|---|---|---|---|
| B57 | B186 | RBCAI | PFC | -0.02434 | 1.26E-07 |
| B57 | B107 | RBCAI | AnuA | -0.02425 | 1.40E-07 |
| B57 | B209 | RBCAI | Smoking | 0.02407 | 1.74E-07 |
| B57 | B72 | RBCAI | HDL-C | 0.023283 | 4.32E-07 |
| B58 | B125 | HCT | Cor | 0.082765 | 0 |
| B58 | B134 | HCT | CHE | 0.04948 | 0 |
| B58 | B137 | HCT | AFU | 0.048203 | 0 |
| B58 | B138 | HCT | PAB | 0.044427 | 0 |
| B58 | B192 | HCT | UA | 0.0384 | 0 |
| B58 | B216 | HCT | BMI | 0.043763 | 0 |
| B58 | B221 | HCT | Hei | 0.045092 | 0 |
| B58 | B222 | HCT | NS | 0.050237 | 0 |
| B58 | B60 | HCT | WBRVI | 0.323094 | 0 |
| B58 | B61 | HCT | WBHSRV | 0.098564 | 0 |
| B58 | B62 | HCT | WBHSRVI | 0.361107 | 0 |
| B58 | B63 | HCT | WBRSR(mPas_200) | 0.377236 | 0 |
| B58 | B64 | HCT | WBRSR(mPas_5) | 0.359795 | 0 |
| B58 | B65 | HCT | WBRSR(mPas_1) | 0.330705 | 0 |
| B58 | B66 | HCT | WBRSR(mPas_100) | 0.368967 | 0 |
| B58 | B67 | HCT | WBRSR(mPas_50) | 0.370445 | 0 |
| B58 | B74 | HCT | ApoA1 | -0.03828 | 0 |
| B58 | B92 | HCT | ALDH2-Glu504Glu | -0.06165 | 0 |
| B58 | B93 | HCT | ALDH2-Glu504Lys | 0.066113 | 0 |
| B58 | B163 | HCT | SG | 0.037085 | 8.88E-16 |
| B58 | B206 | HCT | Waist | 0.035834 | 7.11E-15 |
| B58 | B69 | HCT | PV | 0.035241 | 2.00E-14 |
| B58 | B176 | HCT | Ucr | 0.034974 | 3.11E-14 |
| B58 | B141 | HCT | AST/ALT | -0.03471 | 4.84E-14 |
| B58 | B75 | HCT | ApoA1/B | -0.03361 | 2.95E-13 |
| B58 | B174 | HCT | Urea/Cr | -0.03197 | 3.87E-12 |
| B58 | B128 | HCT | ALT | 0.02719 | 3.57E-09 |
| B58 | B72 | HCT | HDL-C | -0.02665 | 7.21E-09 |
| B58 | B129 | HCT | IBIL | 0.025778 | 2.19E-08 |
| B58 | B133 | HCT | TBIL | 0.024961 | 6.01E-08 |
| B58 | B204 | HCT | Drinking | 0.024621 | 9.05E-08 |
| B58 | B199 | HCT | Mg | 0.024273 | 1.37E-07 |
| B58 | B209 | HCT | Smoking | 0.023267 | 4.40E-07 |
| B58 | B76 | HCT | ApoB | 0.023125 | 5.17E-07 |
| B58 | B210 | HCT | Hip | 0.023016 | 5.84E-07 |
| B58 | B113 | HCT | FT3 | 0.022847 | 7.06E-07 |
| B59 | B112 | WBRV | FT4 | 0.043356 | 0 |
| B59 | B121 | WBRV | C3 | -0.06956 | 0 |
| B59 | B122 | WBRV | C4 | -0.0546 | 0 |
| B59 | B134 | WBRV | CHE | 0.039312 | 0 |

| | | | | | |
|---|---|---|---|---|---|
| B59 | B60 | WBRV | WBRVI | 0.769684 | 0 |
| B59 | B61 | WBRV | WBHSRV | 0.64807 | 0 |
| B59 | B62 | WBRV | WBHSRVI | 0.464133 | 0 |
| B59 | B63 | WBRV | WBRSR(mPas_200) | 0.548887 | 0 |
| B59 | B64 | WBRV | WBRSR(mPas_5) | 0.760027 | 0 |
| B59 | B65 | WBRV | WBRSR(mPas_1) | 0.82402 | 0 |
| B59 | B66 | WBRV | WBRSR(mPas_100) | 0.568793 | 0 |
| B59 | B67 | WBRV | WBRSR(mPas_50) | 0.605986 | 0 |
| B59 | B69 | WBRV | PV | 0.186082 | 0 |
| B59 | B91 | WBRV | TP | 0.037321 | 4.44E-16 |
| B59 | B115 | WBRV | TSH TT3 TT4(TT3) | 0.033932 | 1.75E-13 |
| B59 | B81 | WBRV | TSGF | -0.0339 | 1.85E-13 |
| B59 | B125 | WBRV | Cor | 0.033854 | 1.98E-13 |
| B59 | B194 | WBRV | Cl | -0.0321 | 3.18E-12 |
| B59 | B189 | WBRV | eGFB | 0.031924 | 4.18E-12 |
| B59 | B126 | WBRV | Alb | 0.031858 | 4.62E-12 |
| B59 | B135 | WBRV | TBA | 0.031434 | 8.83E-12 |
| B59 | B120 | WBRV | HCV-cAg | 0.031088 | 1.49E-11 |
| B59 | B222 | WBRV | NS | 0.029152 | 2.47E-10 |
| B59 | B123 | WBRV | hs-CRP1 | 0.027931 | 1.33E-09 |
| B59 | B163 | WBRV | SG | 0.027852 | 1.48E-09 |
| B59 | B107 | WBRV | AnuA | -0.02757 | 2.16E-09 |
| B59 | B210 | WBRV | Hip | 0.027474 | 2.45E-09 |
| B59 | B216 | WBRV | BMI | 0.026799 | 5.96E-09 |
| B59 | B206 | WBRV | Waist | 0.025957 | 1.75E-08 |
| B59 | B113 | WBRV | FT3 | 0.025483 | 3.17E-08 |
| B59 | B93 | WBRV | ALDH2-Glu504Lys | -0.02544 | 3.32E-08 |
| B59 | B114 | WBRV | TSH TT3 TT4(TT4) | 0.025198 | 4.49E-08 |
| B59 | B92 | WBRV | ALDH2-Glu504Glu | 0.025102 | 5.06E-08 |
| B59 | B97 | WBRV | dsDNA | 0.025026 | 5.55E-08 |
| B59 | B129 | WBRV | IBIL | 0.024349 | 1.25E-07 |
| B59 | B138 | WBRV | PAB | 0.024052 | 1.78E-07 |
| B59 | B221 | WBRV | Hei | 0.023396 | 3.80E-07 |
| B59 | B70 | WBRV | LDL-C | 0.02328 | 4.33E-07 |
| B59 | B140 | WBRV | r-GT | 0.023001 | 5.94E-07 |
| B59 | B183 | WBRV | PTA | -0.02297 | 6.13E-07 |
| B59 | B124 | WBRV | ACTH | 0.022543 | 9.90E-07 |
| B6 | B7 | WB-Fe | WB-Cop | 0.330578 | 0 |
| B6 | B8 | WB-Fe | WB-Zinc | 0.643656 | 0 |
| B6 | B113 | WB-Fe | FT3 | -0.02996 | 7.81E-11 |
| B6 | B29 | WB-Fe | E2 | 0.023378 | 3.88E-07 |
| B6 | B33 | WB-Fe | Prog | 0.02272 | 8.14E-07 |
| B6 | B74 | WB-Fe | ApoA1 | -0.02254 | 9.90E-07 |
| B60 | B113 | WBRVI | FT3 | 0.048256 | 0 |

| | | | | | |
|---|---|---|---|---|---|
| B60 | B115 | WBRVI | TSH TT3 TT4(TT3) | 0.052176 | 0 |
| B60 | B121 | WBRVI | C3 | -0.06376 | 0 |
| B60 | B122 | WBRVI | C4 | -0.04884 | 0 |
| B60 | B127 | WBRVI | A-G ratio | 0.041825 | 0 |
| B60 | B128 | WBRVI | ALT | 0.038504 | 0 |
| B60 | B129 | WBRVI | IBIL | 0.046607 | 0 |
| B60 | B133 | WBRVI | TBIL | 0.045026 | 0 |
| B60 | B134 | WBRVI | CHE | 0.06543 | 0 |
| B60 | B138 | WBRVI | PAB | 0.064358 | 0 |
| B60 | B141 | WBRVI | AST/ALT | -0.04165 | 0 |
| B60 | B163 | WBRVI | SG | 0.064981 | 0 |
| B60 | B174 | WBRVI | Urea/Cr | -0.05655 | 0 |
| B60 | B176 | WBRVI | Ucr | 0.050442 | 0 |
| B60 | B192 | WBRVI | UA | 0.053938 | 0 |
| B60 | B204 | WBRVI | Drinking | 0.039644 | 0 |
| B60 | B206 | WBRVI | Waist | 0.06037 | 0 |
| B60 | B209 | WBRVI | Smoking | 0.040517 | 0 |
| B60 | B210 | WBRVI | Hip | 0.049517 | 0 |
| B60 | B216 | WBRVI | BMI | 0.075365 | 0 |
| B60 | B218 | WBRVI | DBP | 0.03876 | 0 |
| B60 | B221 | WBRVI | Hei | 0.077753 | 0 |
| B60 | B222 | WBRVI | NS | 0.083412 | 0 |
| B60 | B61 | WBRVI | WBHSRV | 0.642513 | 0 |
| B60 | B62 | WBRVI | WBHSRVI | 0.822324 | 0 |
| B60 | B63 | WBRVI | WBRSR(mPas_200) | 0.753455 | 0 |
| B60 | B64 | WBRVI | WBRSR(mPas_5) | 0.902817 | 0 |
| B60 | B65 | WBRVI | WBRSR(mPas_1) | 0.929779 | 0 |
| B60 | B66 | WBRVI | WBRSR(mPas_100) | 0.768266 | 0 |
| B60 | B67 | WBRVI | WBRSR(mPas_50) | 0.798028 | 0 |
| B60 | B69 | WBRVI | PV | -0.17913 | 0 |
| B60 | B73 | WBRVI | ApoA | -0.0419 | 0 |
| B60 | B74 | WBRVI | ApoA1 | -0.06563 | 0 |
| B60 | B75 | WBRVI | ApoA1/B | -0.03775 | 2.22E-16 |
| B60 | B157 | WBRVI | IgG | -0.03759 | 4.44E-16 |
| B60 | B215 | WBRVI | BMI | 0.037292 | 4.44E-16 |
| B60 | B140 | WBRVI | r-GT | 0.037051 | 8.88E-16 |
| B60 | B111 | WBRVI | TSH | -0.03626 | 3.55E-15 |
| B60 | B126 | WBRVI | Alb | 0.035748 | 8.44E-15 |
| B60 | B130 | WBRVI | GLB | -0.03344 | 3.87E-13 |
| B60 | B198 | WBRVI | P | -0.03129 | 1.10E-11 |
| B60 | B194 | WBRVI | Cl | -0.0306 | 3.09E-11 |
| B60 | B125 | WBRVI | Cor | 0.029382 | 1.79E-10 |
| B60 | B100 | WBRVI | anti-Ro52 | -0.02912 | 2.57E-10 |
| B60 | B135 | WBRVI | TBA | 0.028419 | 6.85E-10 |

| | | | | | |
|---|---|---|---|---|---|
| B60 | B132 | WBRVI | Dbil | 0.028222 | 8.97E-10 |
| B60 | B148 | WBRVI | Anti_TPO | -0.02807 | 1.11E-09 |
| B60 | B124 | WBRVI | ACTH | 0.02759 | 2.10E-09 |
| B60 | B72 | WBRVI | HDL-C | -0.02705 | 4.29E-09 |
| B60 | B149 | WBRVI | TGAb | -0.02678 | 6.09E-09 |
| B60 | B161 | WBRVI | LEU | -0.02591 | 1.85E-08 |
| B60 | B79 | WBRVI | hb V -dna | -0.02503 | 5.51E-08 |
| B60 | B104 | WBRVI | anti-SSB | -0.0246 | 9.23E-08 |
| B60 | B189 | WBRVI | eGFB | 0.024296 | 1.33E-07 |
| B60 | B139 | WBRVI | ADA | -0.02425 | 1.40E-07 |
| B60 | B183 | WBRVI | PTA | -0.0235 | 3.36E-07 |
| B60 | B92 | WBRVI | ALDH2-Glu504Glu | 0.023504 | 3.36E-07 |
| B60 | B187 | WBRVI | NSE | 0.023457 | 3.54E-07 |
| B61 | B115 | WBHSRV | TSH TT3 TT4(TT3) | 0.051519 | 0 |
| B61 | B120 | WBHSRV | HCV-cAg | 0.047139 | 0 |
| B61 | B121 | WBHSRV | C3 | -0.05693 | 0 |
| B61 | B122 | WBHSRV | C4 | -0.04158 | 0 |
| B61 | B134 | WBHSRV | CHE | 0.075461 | 0 |
| B61 | B138 | WBHSRV | PAB | 0.049545 | 0 |
| B61 | B174 | WBHSRV | Urea/Cr | -0.04272 | 0 |
| B61 | B192 | WBHSRV | UA | 0.044204 | 0 |
| B61 | B194 | WBHSRV | Cl | -0.05968 | 0 |
| B61 | B206 | WBHSRV | Waist | 0.049541 | 0 |
| B61 | B210 | WBHSRV | Hip | 0.0439 | 0 |
| B61 | B215 | WBHSRV | BMI | 0.041511 | 0 |
| B61 | B216 | WBHSRV | BMI | 0.060095 | 0 |
| B61 | B221 | WBHSRV | Hei | 0.048223 | 0 |
| B61 | B222 | WBHSRV | NS | 0.056294 | 0 |
| B61 | B62 | WBHSRV | WBHSRVI | 0.826278 | 0 |
| B61 | B63 | WBHSRV | WBRSR(mPas_200) | 0.870694 | 0 |
| B61 | B64 | WBHSRV | WBRSR(mPas_5) | 0.779342 | 0 |
| B61 | B65 | WBHSRV | WBRSR(mPas_1) | 0.684366 | 0 |
| B61 | B66 | WBHSRV | WBRSR(mPas_100) | 0.857048 | 0 |
| B61 | B67 | WBHSRV | WBRSR(mPas_50) | 0.848976 | 0 |
| B61 | B69 | WBHSRV | PV | 0.074632 | 0 |
| B61 | B71 | WBHSRV | TG | 0.0465 | 0 |
| B61 | B74 | WBHSRV | ApoA1 | -0.07042 | 0 |
| B61 | B75 | WBHSRV | ApoA1/B | -0.07439 | 0 |
| B61 | B76 | WBHSRV | ApoB | 0.072708 | 0 |
| B61 | B91 | WBHSRV | TP | 0.047119 | 0 |
| B61 | B126 | WBHSRV | Alb | 0.037248 | 4.44E-16 |
| B61 | B163 | WBHSRV | SG | 0.037581 | 4.44E-16 |
| B61 | B129 | WBHSRV | IBIL | 0.037195 | 6.66E-16 |
| B61 | B133 | WBHSRV | TBIL | 0.036478 | 2.22E-15 |

| | | | | | |
|---|---|---|---|---|---|
| B61 | B140 | WBHSRV | r-GT | 0.035153 | 2.31E-14 |
| B61 | B186 | WBHSRV | PFC | 0.034588 | 5.93E-14 |
| B61 | B125 | WBHSRV | Cor | 0.034263 | 1.01E-13 |
| B61 | B218 | WBHSRV | DBP | 0.033778 | 2.24E-13 |
| B61 | B72 | WBHSRV | HDL-C | -0.03282 | 1.04E-12 |
| B61 | B123 | WBHSRV | hs-CRP1 | 0.03224 | 2.57E-12 |
| B61 | B113 | WBHSRV | FT3 | 0.032231 | 2.60E-12 |
| B61 | B100 | WBHSRV | anti-Ro52 | -0.03206 | 3.40E-12 |
| B61 | B187 | WBHSRV | NSE | 0.030883 | 2.02E-11 |
| B61 | B176 | WBHSRV | Ucr | 0.030793 | 2.30E-11 |
| B61 | B128 | WBHSRV | ALT | 0.030396 | 4.14E-11 |
| B61 | B136 | WBHSRV | 5'NT | 0.029432 | 1.66E-10 |
| B61 | B188 | WBHSRV | Ca | 0.029218 | 2.25E-10 |
| B61 | B153 | WBHSRV | Glu | 0.028438 | 6.67E-10 |
| B61 | B141 | WBHSRV | AST/ALT | -0.02811 | 1.05E-09 |
| B61 | B197 | WBHSRV | AG | 0.027895 | 1.40E-09 |
| B61 | B130 | WBHSRV | GLB | 0.027613 | 2.04E-09 |
| B61 | B124 | WBHSRV | ACTH | 0.025077 | 5.21E-08 |
| B61 | B132 | WBHSRV | Dbil | 0.025033 | 5.50E-08 |
| B61 | B78 | WBHSRV | HBcAb | 0.024854 | 6.84E-08 |
| B61 | B73 | WBHSRV | ApoA | -0.02479 | 7.38E-08 |
| B61 | B207 | WBHSRV | WHR | 0.023616 | 2.95E-07 |
| B61 | B170 | WBHSRV | U-GLU | 0.023577 | 3.09E-07 |
| B61 | B77 | WBHSRV | TC | 0.0235 | 3.37E-07 |
| B61 | B200 | WBHSRV | OP | -0.02316 | 4.96E-07 |
| B62 | B113 | WBHSRVI | FT3 | 0.049634 | 0 |
| B62 | B115 | WBHSRVI | TSH TT3 TT4(TT3) | 0.061179 | 0 |
| B62 | B121 | WBHSRVI | C3 | -0.0498 | 0 |
| B62 | B128 | WBHSRVI | ALT | 0.045958 | 0 |
| B62 | B129 | WBHSRVI | IBIL | 0.052546 | 0 |
| B62 | B132 | WBHSRVI | Dbil | 0.040108 | 0 |
| B62 | B133 | WBHSRVI | TBIL | 0.053074 | 0 |
| B62 | B134 | WBHSRVI | CHE | 0.083916 | 0 |
| B62 | B138 | WBHSRVI | PAB | 0.074372 | 0 |
| B62 | B140 | WBHSRVI | r-GT | 0.041742 | 0 |
| B62 | B141 | WBHSRVI | AST/ALT | -0.05685 | 0 |
| B62 | B157 | WBHSRVI | IgG | -0.04256 | 0 |
| B62 | B163 | WBHSRVI | SG | 0.064706 | 0 |
| B62 | B174 | WBHSRVI | Urea/Cr | -0.06415 | 0 |
| B62 | B176 | WBHSRVI | Ucr | 0.053181 | 0 |
| B62 | B192 | WBHSRVI | UA | 0.063557 | 0 |
| B62 | B194 | WBHSRVI | Cl | -0.0488 | 0 |
| B62 | B204 | WBHSRVI | Drinking | 0.038508 | 0 |
| B62 | B206 | WBHSRVI | Waist | 0.070531 | 0 |

| | | | | | |
|---|---|---|---|---|---|
| B62 | B210 | WBHSRVI | Hip | 0.055909 | 0 |
| B62 | B215 | WBHSRVI | BMI | 0.05095 | 0 |
| B62 | B216 | WBHSRVI | BMI | 0.090127 | 0 |
| B62 | B218 | WBHSRVI | DBP | 0.043438 | 0 |
| B62 | B221 | WBHSRVI | Hei | 0.085913 | 0 |
| B62 | B222 | WBHSRVI | NS | 0.093531 | 0 |
| B62 | B63 | WBHSRVI | WBRSR(mPas_200) | 0.912713 | 0 |
| B62 | B64 | WBHSRVI | WBRSR(mPas_5) | 0.840143 | 0 |
| B62 | B65 | WBHSRVI | WBRSR(mPas_1) | 0.754901 | 0 |
| B62 | B66 | WBHSRVI | WBRSR(mPas_100) | 0.902629 | 0 |
| B62 | B67 | WBHSRVI | WBRSR(mPas_50) | 0.898357 | 0 |
| B62 | B69 | WBHSRVI | PV | -0.22649 | 0 |
| B62 | B72 | WBHSRVI | HDL-C | -0.05056 | 0 |
| B62 | B73 | WBHSRVI | ApoA | -0.0427 | 0 |
| B62 | B74 | WBHSRVI | ApoA1 | -0.10202 | 0 |
| B62 | B75 | WBHSRVI | ApoA1/B | -0.07662 | 0 |
| B62 | B76 | WBHSRVI | ApoB | 0.04825 | 0 |
| B62 | B100 | WBHSRVI | anti-Ro52 | -0.03778 | 2.22E-16 |
| B62 | B126 | WBHSRVI | Alb | 0.035816 | 7.55E-15 |
| B62 | B187 | WBHSRVI | NSE | 0.035795 | 7.77E-15 |
| B62 | B122 | WBHSRVI | C4 | -0.03502 | 2.86E-14 |
| B62 | B209 | WBHSRVI | Smoking | 0.034583 | 5.97E-14 |
| B62 | B107 | WBHSRVI | AnuA | 0.034316 | 9.28E-14 |
| B62 | B197 | WBHSRVI | AG | 0.033267 | 5.10E-13 |
| B62 | B127 | WBHSRVI | A-G ratio | 0.032687 | 1.28E-12 |
| B62 | B161 | WBHSRVI | LEU | -0.02947 | 1.57E-10 |
| B62 | B125 | WBHSRVI | Cor | 0.029424 | 1.68E-10 |
| B62 | B111 | WBHSRVI | TSH | -0.02924 | 2.20E-10 |
| B62 | B198 | WBHSRVI | P | -0.02908 | 2.73E-10 |
| B62 | B137 | WBHSRVI | AFU | 0.028718 | 4.53E-10 |
| B62 | B71 | WBHSRVI | TG | 0.02857 | 5.56E-10 |
| B62 | B148 | WBHSRVI | Anti_TPO | -0.02837 | 7.29E-10 |
| B62 | B79 | WBHSRVI | hb V -dna | -0.02827 | 8.38E-10 |
| B62 | B124 | WBHSRVI | ACTH | 0.02759 | 2.10E-09 |
| B62 | B207 | WBHSRVI | WHR | 0.02631 | 1.12E-08 |
| B62 | B149 | WBHSRVI | TGAb | -0.02417 | 1.54E-07 |
| B62 | B104 | WBHSRVI | anti-SSB | -0.02413 | 1.62E-07 |
| B62 | B101 | WBHSRVI | anti-Sc70 | -0.02412 | 1.65E-07 |
| B62 | B130 | WBHSRVI | GLB | -0.02326 | 4.43E-07 |
| B62 | B188 | WBHSRVI | Ca | 0.023256 | 4.46E-07 |
| B63 | B113 | WBRSR(mPas_200) | FT3 | 0.052978 | 0 |
| B63 | B115 | WBRSR(mPas_200) | TSH TT3 TT4(TT3) | 0.078079 | 0 |
| B63 | B121 | WBRSR(mPas_200) | C3 | -0.04495 | 0 |
| B63 | B123 | WBRSR(mPas_200) | hs-CRP1 | 0.045571 | 0 |

| | | | | | |
|---|---|---|---|---|---|
| B63 | B125 | WBRSR(mPas_200) | Cor | 0.04916 | 0 |
| B63 | B126 | WBRSR(mPas_200) | Alb | 0.04778 | 0 |
| B63 | B128 | WBRSR(mPas_200) | ALT | 0.045886 | 0 |
| B63 | B129 | WBRSR(mPas_200) | IBIL | 0.052816 | 0 |
| B63 | B133 | WBRSR(mPas_200) | TBIL | 0.050873 | 0 |
| B63 | B134 | WBRSR(mPas_200) | CHE | 0.095918 | 0 |
| B63 | B138 | WBRSR(mPas_200) | PAB | 0.06557 | 0 |
| B63 | B140 | WBRSR(mPas_200) | r-GT | 0.05149 | 0 |
| B63 | B141 | WBRSR(mPas_200) | AST/ALT | -0.04688 | 0 |
| B63 | B163 | WBRSR(mPas_200) | SG | 0.062746 | 0 |
| B63 | B174 | WBRSR(mPas_200) | Urea/Cr | -0.05981 | 0 |
| B63 | B176 | WBRSR(mPas_200) | Ucr | 0.051716 | 0 |
| B63 | B188 | WBRSR(mPas_200) | Ca | 0.045218 | 0 |
| B63 | B192 | WBRSR(mPas_200) | UA | 0.065719 | 0 |
| B63 | B194 | WBRSR(mPas_200) | Cl | -0.0678 | 0 |
| B63 | B206 | WBRSR(mPas_200) | Waist | 0.068424 | 0 |
| B63 | B210 | WBRSR(mPas_200) | Hip | 0.052874 | 0 |
| B63 | B215 | WBRSR(mPas_200) | BMI | 0.049818 | 0 |
| B63 | B216 | WBRSR(mPas_200) | BMI | 0.082537 | 0 |
| B63 | B218 | WBRSR(mPas_200) | DBP | 0.045252 | 0 |
| B63 | B221 | WBRSR(mPas_200) | Hei | 0.075091 | 0 |
| B63 | B222 | WBRSR(mPas_200) | NS | 0.083695 | 0 |
| B63 | B64 | WBRSR(mPas_200) | WBRSR(mPas_5) | 0.919882 | 0 |
| B63 | B65 | WBRSR(mPas_200) | WBRSR(mPas_1) | 0.827608 | 0 |
| B63 | B66 | WBRSR(mPas_200) | WBRSR(mPas_100) | 0.984721 | 0 |
| B63 | B67 | WBRSR(mPas_200) | WBRSR(mPas_50) | 0.979905 | 0 |
| B63 | B69 | WBRSR(mPas_200) | PV | 0.172005 | 0 |
| B63 | B71 | WBRSR(mPas_200) | TG | 0.046139 | 0 |
| B63 | B72 | WBRSR(mPas_200) | HDL-C | -0.04461 | 0 |
| B63 | B74 | WBRSR(mPas_200) | ApoA1 | -0.07945 | 0 |
| B63 | B75 | WBRSR(mPas_200) | ApoA1/B | -0.09066 | 0 |
| B63 | B76 | WBRSR(mPas_200) | ApoB | 0.092255 | 0 |
| B63 | B91 | WBRSR(mPas_200) | TP | 0.05392 | 0 |
| B63 | B137 | WBRSR(mPas_200) | AFU | 0.034305 | 9.46E-14 |
| B63 | B204 | WBRSR(mPas_200) | Drinking | 0.03272 | 1.21E-12 |
| B63 | B187 | WBRSR(mPas_200) | NSE | 0.032519 | 1.66E-12 |
| B63 | B124 | WBRSR(mPas_200) | ACTH | 0.032404 | 1.99E-12 |
| B63 | B120 | WBRSR(mPas_200) | HCV-cAg | 0.031862 | 4.60E-12 |
| B63 | B132 | WBRSR(mPas_200) | Dbil | 0.031255 | 1.16E-11 |
| B63 | B197 | WBRSR(mPas_200) | AG | 0.03115 | 1.35E-11 |
| B63 | B136 | WBRSR(mPas_200) | 5'NT | 0.03109 | 1.48E-11 |
| B63 | B70 | WBRSR(mPas_200) | LDL-C | 0.031078 | 1.51E-11 |
| B63 | B100 | WBRSR(mPas_200) | anti-Ro52 | -0.03033 | 4.54E-11 |
| B63 | B209 | WBRSR(mPas_200) | Smoking | 0.029703 | 1.13E-10 |

| | | | | | |
|---|---|---|---|---|---|
| B63 | B130 | WBRSR(mPas_200) | GLB | 0.028191 | 9.35E-10 |
| B63 | B153 | WBRSR(mPas_200) | Glu | 0.028006 | 1.20E-09 |
| B63 | B77 | WBRSR(mPas_200) | TC | 0.027264 | 3.24E-09 |
| B63 | B186 | WBRSR(mPas_200) | PFC | 0.027027 | 4.43E-09 |
| B63 | B170 | WBRSR(mPas_200) | U-GLU | 0.026994 | 4.62E-09 |
| B63 | B122 | WBRSR(mPas_200) | C4 | -0.02682 | 5.82E-09 |
| B63 | B179 | WBRSR(mPas_200) | α1-MG | 0.026387 | 1.01E-08 |
| B63 | B207 | WBRSR(mPas_200) | WHR | 0.025816 | 2.09E-08 |
| B63 | B148 | WBRSR(mPas_200) | Anti_TPO | -0.0252 | 4.49E-08 |
| B63 | B166 | WBRSR(mPas_200) | Pro | 0.024857 | 6.81E-08 |
| B63 | B200 | WBRSR(mPas_200) | OP | -0.02473 | 7.96E-08 |
| B63 | B112 | WBRSR(mPas_200) | FT4 | 0.024637 | 8.88E-08 |
| B63 | B198 | WBRSR(mPas_200) | P | -0.02448 | 1.07E-07 |
| B63 | B73 | WBRSR(mPas_200) | ApoA | -0.02332 | 4.13E-07 |
| B63 | B161 | WBRSR(mPas_200) | LEU | -0.02312 | 5.18E-07 |
| B63 | B131 | WBRSR(mPas_200) | AST | 0.023096 | 5.34E-07 |
| B64 | B113 | WBRSR(mPas_5) | FT3 | 0.056368 | 0 |
| B64 | B115 | WBRSR(mPas_5) | TSH TT3 TT4(TT3) | 0.076297 | 0 |
| B64 | B121 | WBRSR(mPas_5) | C3 | -0.05717 | 0 |
| B64 | B123 | WBRSR(mPas_5) | hs-CRP1 | 0.04101 | 0 |
| B64 | B125 | WBRSR(mPas_5) | Cor | 0.050069 | 0 |
| B64 | B126 | WBRSR(mPas_5) | Alb | 0.050453 | 0 |
| B64 | B128 | WBRSR(mPas_5) | ALT | 0.043667 | 0 |
| B64 | B129 | WBRSR(mPas_5) | IBIL | 0.048805 | 0 |
| B64 | B133 | WBRSR(mPas_5) | TBIL | 0.04599 | 0 |
| B64 | B134 | WBRSR(mPas_5) | CHE | 0.09055 | 0 |
| B64 | B138 | WBRSR(mPas_5) | PAB | 0.063768 | 0 |
| B64 | B140 | WBRSR(mPas_5) | r-GT | 0.04965 | 0 |
| B64 | B141 | WBRSR(mPas_5) | AST/ALT | -0.04113 | 0 |
| B64 | B163 | WBRSR(mPas_5) | SG | 0.065161 | 0 |
| B64 | B174 | WBRSR(mPas_5) | Urea/Cr | -0.05526 | 0 |
| B64 | B176 | WBRSR(mPas_5) | Ucr | 0.05106 | 0 |
| B64 | B188 | WBRSR(mPas_5) | Ca | 0.041777 | 0 |
| B64 | B192 | WBRSR(mPas_5) | UA | 0.060626 | 0 |
| B64 | B194 | WBRSR(mPas_5) | Cl | -0.05975 | 0 |
| B64 | B206 | WBRSR(mPas_5) | Waist | 0.066278 | 0 |
| B64 | B210 | WBRSR(mPas_5) | Hip | 0.053736 | 0 |
| B64 | B215 | WBRSR(mPas_5) | BMI | 0.045232 | 0 |
| B64 | B216 | WBRSR(mPas_5) | BMI | 0.078549 | 0 |
| B64 | B218 | WBRSR(mPas_5) | DBP | 0.045073 | 0 |
| B64 | B221 | WBRSR(mPas_5) | Hei | 0.073911 | 0 |
| B64 | B222 | WBRSR(mPas_5) | NS | 0.082499 | 0 |
| B64 | B65 | WBRSR(mPas_5) | WBRSR(mPas_1) | 0.976519 | 0 |
| B64 | B66 | WBRSR(mPas_5) | WBRSR(mPas_100) | 0.923723 | 0 |

| | | | | | |
|---|---|---|---|---|---|
| B64 | B67 | WBRSR(mPas_5) | WBRSR(mPas_50) | 0.941551 | 0 |
| B64 | B69 | WBRSR(mPas_5) | PV | 0.166934 | 0 |
| B64 | B74 | WBRSR(mPas_5) | ApoA1 | -0.05878 | 0 |
| B64 | B75 | WBRSR(mPas_5) | ApoA1/B | -0.06881 | 0 |
| B64 | B76 | WBRSR(mPas_5) | ApoB | 0.070256 | 0 |
| B64 | B91 | WBRSR(mPas_5) | TP | 0.044145 | 0 |
| B64 | B122 | WBRSR(mPas_5) | C4 | -0.03785 | 2.22E-16 |
| B64 | B204 | WBRSR(mPas_5) | Drinking | 0.035522 | 1.24E-14 |
| B64 | B71 | WBRSR(mPas_5) | TG | 0.035104 | 2.49E-14 |
| B64 | B209 | WBRSR(mPas_5) | Smoking | 0.034841 | 3.91E-14 |
| B64 | B124 | WBRSR(mPas_5) | ACTH | 0.034382 | 8.35E-14 |
| B64 | B137 | WBRSR(mPas_5) | AFU | 0.033197 | 5.70E-13 |
| B64 | B112 | WBRSR(mPas_5) | FT4 | 0.031769 | 5.30E-12 |
| B64 | B70 | WBRSR(mPas_5) | LDL-C | 0.031533 | 7.59E-12 |
| B64 | B72 | WBRSR(mPas_5) | HDL-C | -0.03127 | 1.14E-11 |
| B64 | B198 | WBRSR(mPas_5) | P | -0.0298 | 9.87E-11 |
| B64 | B135 | WBRSR(mPas_5) | TBA | 0.028384 | 7.19E-10 |
| B64 | B100 | WBRSR(mPas_5) | anti-Ro52 | -0.02753 | 2.27E-09 |
| B64 | B187 | WBRSR(mPas_5) | NSE | 0.026597 | 7.75E-09 |
| B64 | B148 | WBRSR(mPas_5) | Anti_TPO | -0.02658 | 7.96E-09 |
| B64 | B73 | WBRSR(mPas_5) | ApoA | -0.02642 | 9.72E-09 |
| B64 | B197 | WBRSR(mPas_5) | AG | 0.025996 | 1.67E-08 |
| B64 | B79 | WBRSR(mPas_5) | hb V -dna | -0.0257 | 2.42E-08 |
| B64 | B189 | WBRSR(mPas_5) | eGFB | 0.025533 | 2.97E-08 |
| B64 | B120 | WBRSR(mPas_5) | HCV-cAg | 0.02552 | 3.02E-08 |
| B64 | B179 | WBRSR(mPas_5) | α1-MG | 0.025399 | 3.51E-08 |
| B64 | B132 | WBRSR(mPas_5) | Dbil | 0.025248 | 4.23E-08 |
| B64 | B170 | WBRSR(mPas_5) | U-GLU | 0.025051 | 5.38E-08 |
| B64 | B149 | WBRSR(mPas_5) | TGAb | -0.0248 | 7.25E-08 |
| B64 | B131 | WBRSR(mPas_5) | AST | 0.024691 | 8.32E-08 |
| B64 | B153 | WBRSR(mPas_5) | Glu | 0.023709 | 2.65E-07 |
| B64 | B166 | WBRSR(mPas_5) | Pro | 0.023694 | 2.70E-07 |
| B64 | B111 | WBRSR(mPas_5) | TSH | -0.02343 | 3.65E-07 |
| B65 | B113 | WBRSR(mPas_1) | FT3 | 0.052903 | 0 |
| B65 | B115 | WBRSR(mPas_1) | TSH TT3 TT4(TT3) | 0.068335 | 0 |
| B65 | B121 | WBRSR(mPas_1) | C3 | -0.06067 | 0 |
| B65 | B122 | WBRSR(mPas_1) | C4 | -0.04203 | 0 |
| B65 | B125 | WBRSR(mPas_1) | Cor | 0.046892 | 0 |
| B65 | B126 | WBRSR(mPas_1) | Alb | 0.047416 | 0 |
| B65 | B128 | WBRSR(mPas_1) | ALT | 0.039509 | 0 |
| B65 | B129 | WBRSR(mPas_1) | IBIL | 0.045272 | 0 |
| B65 | B133 | WBRSR(mPas_1) | TBIL | 0.042214 | 0 |
| B65 | B134 | WBRSR(mPas_1) | CHE | 0.078956 | 0 |
| B65 | B138 | WBRSR(mPas_1) | PAB | 0.058028 | 0 |

| | | | | | |
|---|---|---|---|---|---|
| B65 | B140 | WBRSR(mPas_1) | r-GT | 0.046461 | 0 |
| B65 | B163 | WBRSR(mPas_1) | SG | 0.063406 | 0 |
| B65 | B174 | WBRSR(mPas_1) | Urea/Cr | -0.05092 | 0 |
| B65 | B176 | WBRSR(mPas_1) | Ucr | 0.048437 | 0 |
| B65 | B192 | WBRSR(mPas_1) | UA | 0.055621 | 0 |
| B65 | B194 | WBRSR(mPas_1) | Cl | -0.0502 | 0 |
| B65 | B206 | WBRSR(mPas_1) | Waist | 0.060982 | 0 |
| B65 | B210 | WBRSR(mPas_1) | Hip | 0.050749 | 0 |
| B65 | B215 | WBRSR(mPas_1) | BMI | 0.039817 | 0 |
| B65 | B216 | WBRSR(mPas_1) | BMI | 0.072022 | 0 |
| B65 | B218 | WBRSR(mPas_1) | DBP | 0.041465 | 0 |
| B65 | B221 | WBRSR(mPas_1) | Hei | 0.069656 | 0 |
| B65 | B222 | WBRSR(mPas_1) | NS | 0.077329 | 0 |
| B65 | B66 | WBRSR(mPas_1) | WBRSR(mPas_100) | 0.839557 | 0 |
| B65 | B67 | WBRSR(mPas_1) | WBRSR(mPas_50) | 0.867929 | 0 |
| B65 | B69 | WBRSR(mPas_1) | PV | 0.14849 | 0 |
| B65 | B74 | WBRSR(mPas_1) | ApoA1 | -0.04734 | 0 |
| B65 | B75 | WBRSR(mPas_1) | ApoA1/B | -0.05251 | 0 |
| B65 | B76 | WBRSR(mPas_1) | ApoB | 0.053361 | 0 |
| B65 | B188 | WBRSR(mPas_1) | Ca | 0.037542 | 4.44E-16 |
| B65 | B209 | WBRSR(mPas_1) | Smoking | 0.036026 | 5.33E-15 |
| B65 | B141 | WBRSR(mPas_1) | AST/ALT | -0.03544 | 1.42E-14 |
| B65 | B91 | WBRSR(mPas_1) | TP | 0.03516 | 2.26E-14 |
| B65 | B123 | WBRSR(mPas_1) | hs-CRP1 | 0.035141 | 2.35E-14 |
| B65 | B204 | WBRSR(mPas_1) | Drinking | 0.034473 | 7.19E-14 |
| B65 | B112 | WBRSR(mPas_1) | FT4 | 0.033078 | 6.89E-13 |
| B65 | B135 | WBRSR(mPas_1) | TBA | 0.032539 | 1.61E-12 |
| B65 | B124 | WBRSR(mPas_1) | ACTH | 0.032408 | 1.98E-12 |
| B65 | B70 | WBRSR(mPas_1) | LDL-C | 0.02999 | 7.47E-11 |
| B65 | B198 | WBRSR(mPas_1) | P | -0.0299 | 8.51E-11 |
| B65 | B189 | WBRSR(mPas_1) | eGFB | 0.029038 | 2.90E-10 |
| B65 | B137 | WBRSR(mPas_1) | AFU | 0.027212 | 3.47E-09 |
| B65 | B81 | WBRSR(mPas_1) | TSGF | -0.02676 | 6.27E-09 |
| B65 | B71 | WBRSR(mPas_1) | TG | 0.026303 | 1.13E-08 |
| B65 | B111 | WBRSR(mPas_1) | TSH | -0.02582 | 2.07E-08 |
| B65 | B148 | WBRSR(mPas_1) | Anti_TPO | -0.02567 | 2.50E-08 |
| B65 | B79 | WBRSR(mPas_1) | hb V -dna | -0.02538 | 3.59E-08 |
| B65 | B73 | WBRSR(mPas_1) | ApoA | -0.02518 | 4.62E-08 |
| B65 | B149 | WBRSR(mPas_1) | TGAb | -0.02466 | 8.61E-08 |
| B65 | B92 | WBRSR(mPas_1) | ALDH2-Glu504Glu | 0.024135 | 1.61E-07 |
| B65 | B131 | WBRSR(mPas_1) | AST | 0.024117 | 1.65E-07 |
| B65 | B100 | WBRSR(mPas_1) | anti-Ro52 | -0.0237 | 2.68E-07 |
| B65 | B170 | WBRSR(mPas_1) | U-GLU | 0.023418 | 3.70E-07 |
| B65 | B72 | WBRSR(mPas_1) | HDL-C | -0.02296 | 6.23E-07 |

| | | | | | |
|---|---|---|---|---|---|
| B65 | B93 | WBRSR(mPas_1) | ALDH2-Glu504Lys | -0.02273 | 8.01E-07 |
| B66 | B113 | WBRSR(mPas_100) | FT3 | 0.055542 | 0 |
| B66 | B115 | WBRSR(mPas_100) | TSH TT3 TT4(TT3) | 0.079475 | 0 |
| B66 | B121 | WBRSR(mPas_100) | C3 | -0.04616 | 0 |
| B66 | B123 | WBRSR(mPas_100) | hs-CRP1 | 0.048567 | 0 |
| B66 | B126 | WBRSR(mPas_100) | Alb | 0.048198 | 0 |
| B66 | B128 | WBRSR(mPas_100) | ALT | 0.045715 | 0 |
| B66 | B129 | WBRSR(mPas_100) | IBIL | 0.049926 | 0 |
| B66 | B133 | WBRSR(mPas_100) | TBIL | 0.048438 | 0 |
| B66 | B134 | WBRSR(mPas_100) | CHE | 0.093379 | 0 |
| B66 | B138 | WBRSR(mPas_100) | PAB | 0.063924 | 0 |
| B66 | B140 | WBRSR(mPas_100) | r-GT | 0.050725 | 0 |
| B66 | B141 | WBRSR(mPas_100) | AST/ALT | -0.04591 | 0 |
| B66 | B163 | WBRSR(mPas_100) | SG | 0.061768 | 0 |
| B66 | B174 | WBRSR(mPas_100) | Urea/Cr | -0.05755 | 0 |
| B66 | B176 | WBRSR(mPas_100) | Ucr | 0.052232 | 0 |
| B66 | B188 | WBRSR(mPas_100) | Ca | 0.045042 | 0 |
| B66 | B192 | WBRSR(mPas_100) | UA | 0.064495 | 0 |
| B66 | B194 | WBRSR(mPas_100) | Cl | -0.06489 | 0 |
| B66 | B206 | WBRSR(mPas_100) | Waist | 0.069277 | 0 |
| B66 | B210 | WBRSR(mPas_100) | Hip | 0.054859 | 0 |
| B66 | B215 | WBRSR(mPas_100) | BMI | 0.051056 | 0 |
| B66 | B216 | WBRSR(mPas_100) | BMI | 0.082359 | 0 |
| B66 | B218 | WBRSR(mPas_100) | DBP | 0.048316 | 0 |
| B66 | B221 | WBRSR(mPas_100) | Hei | 0.073435 | 0 |
| B66 | B222 | WBRSR(mPas_100) | NS | 0.085024 | 0 |
| B66 | B67 | WBRSR(mPas_100) | WBRSR(mPas_50) | 0.997581 | 0 |
| B66 | B69 | WBRSR(mPas_100) | PV | 0.167861 | 0 |
| B66 | B71 | WBRSR(mPas_100) | TG | 0.044246 | 0 |
| B66 | B72 | WBRSR(mPas_100) | HDL-C | -0.04351 | 0 |
| B66 | B74 | WBRSR(mPas_100) | ApoA1 | -0.07806 | 0 |
| B66 | B75 | WBRSR(mPas_100) | ApoA1/B | -0.09123 | 0 |
| B66 | B76 | WBRSR(mPas_100) | ApoB | 0.09386 | 0 |
| B66 | B91 | WBRSR(mPas_100) | TP | 0.054815 | 0 |
| B66 | B137 | WBRSR(mPas_100) | AFU | 0.034199 | 1.13E-13 |
| B66 | B125 | WBRSR(mPas_100) | Cor | 0.033913 | 1.80E-13 |
| B66 | B187 | WBRSR(mPas_100) | NSE | 0.033794 | 2.18E-13 |
| B66 | B204 | WBRSR(mPas_100) | Drinking | 0.032465 | 1.81E-12 |
| B66 | B120 | WBRSR(mPas_100) | HCV-cAg | 0.032331 | 2.23E-12 |
| B66 | B70 | WBRSR(mPas_100) | LDL-C | 0.03233 | 2.23E-12 |
| B66 | B136 | WBRSR(mPas_100) | 5'NT | 0.031865 | 4.58E-12 |
| B66 | B186 | WBRSR(mPas_100) | PFC | 0.031802 | 5.04E-12 |
| B66 | B79 | WBRSR(mPas_100) | hb V -dna | -0.03166 | 6.24E-12 |
| B66 | B197 | WBRSR(mPas_100) | AG | 0.031429 | 8.90E-12 |

| | | | | | |
|---|---|---|---|---|---|
| B66 | B132 | WBRSR(mPas_100) | Dbil | 0.030869 | 2.06E-11 |
| B66 | B124 | WBRSR(mPas_100) | ACTH | 0.030428 | 3.95E-11 |
| B66 | B198 | WBRSR(mPas_100) | P | -0.02981 | 9.70E-11 |
| B66 | B209 | WBRSR(mPas_100) | Smoking | 0.029513 | 1.48E-10 |
| B66 | B122 | WBRSR(mPas_100) | C4 | -0.02933 | 1.92E-10 |
| B66 | B100 | WBRSR(mPas_100) | anti-Ro52 | -0.02922 | 2.25E-10 |
| B66 | B130 | WBRSR(mPas_100) | GLB | 0.028902 | 3.51E-10 |
| B66 | B179 | WBRSR(mPas_100) | α1-MG | 0.028239 | 8.76E-10 |
| B66 | B170 | WBRSR(mPas_100) | U-GLU | 0.027646 | 1.95E-09 |
| B66 | B207 | WBRSR(mPas_100) | WHR | 0.026832 | 5.72E-09 |
| B66 | B153 | WBRSR(mPas_100) | Glu | 0.026758 | 6.29E-09 |
| B66 | B77 | WBRSR(mPas_100) | TC | 0.026739 | 6.44E-09 |
| B66 | B148 | WBRSR(mPas_100) | Anti_TPO | -0.02544 | 3.35E-08 |
| B66 | B166 | WBRSR(mPas_100) | Pro | 0.024352 | 1.25E-07 |
| B66 | B73 | WBRSR(mPas_100) | ApoA | -0.02422 | 1.46E-07 |
| B66 | B131 | WBRSR(mPas_100) | AST | 0.023475 | 3.47E-07 |
| B66 | B200 | WBRSR(mPas_100) | OP | -0.02308 | 5.42E-07 |
| B67 | B113 | WBRSR(mPas_50) | FT3 | 0.056433 | 0 |
| B67 | B115 | WBRSR(mPas_50) | TSH TT3 TT4(TT3) | 0.079669 | 0 |
| B67 | B121 | WBRSR(mPas_50) | C3 | -0.04896 | 0 |
| B67 | B123 | WBRSR(mPas_50) | hs-CRP1 | 0.048033 | 0 |
| B67 | B126 | WBRSR(mPas_50) | Alb | 0.048916 | 0 |
| B67 | B128 | WBRSR(mPas_50) | ALT | 0.045623 | 0 |
| B67 | B129 | WBRSR(mPas_50) | IBIL | 0.049766 | 0 |
| B67 | B133 | WBRSR(mPas_50) | TBIL | 0.048153 | 0 |
| B67 | B134 | WBRSR(mPas_50) | CHE | 0.092771 | 0 |
| B67 | B138 | WBRSR(mPas_50) | PAB | 0.06413 | 0 |
| B67 | B140 | WBRSR(mPas_50) | r-GT | 0.050686 | 0 |
| B67 | B141 | WBRSR(mPas_50) | AST/ALT | -0.04514 | 0 |
| B67 | B163 | WBRSR(mPas_50) | SG | 0.062689 | 0 |
| B67 | B174 | WBRSR(mPas_50) | Urea/Cr | -0.05774 | 0 |
| B67 | B176 | WBRSR(mPas_50) | Ucr | 0.052598 | 0 |
| B67 | B188 | WBRSR(mPas_50) | Ca | 0.044926 | 0 |
| B67 | B192 | WBRSR(mPas_50) | UA | 0.064208 | 0 |
| B67 | B194 | WBRSR(mPas_50) | Cl | -0.06366 | 0 |
| B67 | B206 | WBRSR(mPas_50) | Waist | 0.069321 | 0 |
| B67 | B210 | WBRSR(mPas_50) | Hip | 0.05503 | 0 |
| B67 | B215 | WBRSR(mPas_50) | BMI | 0.050357 | 0 |
| B67 | B216 | WBRSR(mPas_50) | BMI | 0.082229 | 0 |
| B67 | B218 | WBRSR(mPas_50) | DBP | 0.04837 | 0 |
| B67 | B221 | WBRSR(mPas_50) | Hei | 0.074038 | 0 |
| B67 | B222 | WBRSR(mPas_50) | NS | 0.085304 | 0 |
| B67 | B69 | WBRSR(mPas_50) | PV | 0.167044 | 0 |
| B67 | B71 | WBRSR(mPas_50) | TG | 0.042534 | 0 |

| | | | | | |
|---|---|---|---|---|---|
| B67 | B72 | WBRSR(mPas_50) | HDL-C | -0.04175 | 0 |
| B67 | B74 | WBRSR(mPas_50) | ApoA1 | -0.07585 | 0 |
| B67 | B75 | WBRSR(mPas_50) | ApoA1/B | -0.08866 | 0 |
| B67 | B76 | WBRSR(mPas_50) | ApoB | 0.090897 | 0 |
| B67 | B91 | WBRSR(mPas_50) | TP | 0.053772 | 0 |
| B67 | B125 | WBRSR(mPas_50) | Cor | 0.034612 | 5.68E-14 |
| B67 | B137 | WBRSR(mPas_50) | AFU | 0.033648 | 2.76E-13 |
| B67 | B204 | WBRSR(mPas_50) | Drinking | 0.03309 | 6.75E-13 |
| B67 | B187 | WBRSR(mPas_50) | NSE | 0.033029 | 7.45E-13 |
| B67 | B70 | WBRSR(mPas_50) | LDL-C | 0.032633 | 1.39E-12 |
| B67 | B79 | WBRSR(mPas_50) | hb V -dna | -0.03207 | 3.36E-12 |
| B67 | B122 | WBRSR(mPas_50) | C4 | -0.03158 | 7.05E-12 |
| B67 | B120 | WBRSR(mPas_50) | HCV-cAg | 0.031449 | 8.62E-12 |
| B67 | B124 | WBRSR(mPas_50) | ACTH | 0.030974 | 1.76E-11 |
| B67 | B197 | WBRSR(mPas_50) | AG | 0.030885 | 2.01E-11 |
| B67 | B136 | WBRSR(mPas_50) | 5'NT | 0.030777 | 2.36E-11 |
| B67 | B209 | WBRSR(mPas_50) | Smoking | 0.03068 | 2.72E-11 |
| B67 | B198 | WBRSR(mPas_50) | P | -0.03059 | 3.12E-11 |
| B67 | B132 | WBRSR(mPas_50) | Dbil | 0.030313 | 4.67E-11 |
| B67 | B186 | WBRSR(mPas_50) | PFC | 0.029844 | 9.23E-11 |
| B67 | B100 | WBRSR(mPas_50) | anti-Ro52 | -0.02873 | 4.43E-10 |
| B67 | B179 | WBRSR(mPas_50) | α1-MG | 0.028011 | 1.19E-09 |
| B67 | B170 | WBRSR(mPas_50) | U-GLU | 0.027737 | 1.73E-09 |
| B67 | B130 | WBRSR(mPas_50) | GLB | 0.027278 | 3.19E-09 |
| B67 | B153 | WBRSR(mPas_50) | Glu | 0.026421 | 9.71E-09 |
| B67 | B207 | WBRSR(mPas_50) | WHR | 0.026403 | 9.93E-09 |
| B67 | B77 | WBRSR(mPas_50) | TC | 0.025874 | 1.94E-08 |
| B67 | B148 | WBRSR(mPas_50) | Anti_TPO | -0.02572 | 2.35E-08 |
| B67 | B73 | WBRSR(mPas_50) | ApoA | -0.02486 | 6.80E-08 |
| B67 | B166 | WBRSR(mPas_50) | Pro | 0.024832 | 7.02E-08 |
| B67 | B131 | WBRSR(mPas_50) | AST | 0.023894 | 2.14E-07 |
| B67 | B112 | WBRSR(mPas_50) | FT4 | 0.023217 | 4.66E-07 |
| B67 | B149 | WBRSR(mPas_50) | TGAb | -0.023 | 5.92E-07 |
| B67 | B200 | WBRSR(mPas_50) | OP | -0.02267 | 8.57E-07 |
| B69 | B107 | PV | AnuA | -0.10305 | 0 |
| B69 | B109 | PV | AMA-M2 | 0.038571 | 0 |
| B69 | B112 | PV | FT4 | 0.042755 | 0 |
| B69 | B114 | PV | TSH TT3 TT4(TT4) | 0.067339 | 0 |
| B69 | B115 | PV | TSH TT3 TT4(TT3) | 0.047586 | 0 |
| B69 | B123 | PV | hs-CRP1 | 0.113282 | 0 |
| B69 | B125 | PV | Cor | 0.053793 | 0 |
| B69 | B127 | PV | A-G ratio | -0.09019 | 0 |
| B69 | B130 | PV | GLB | 0.124137 | 0 |
| B69 | B134 | PV | CHE | 0.039872 | 0 |

| | | | | | |
|---|---|---|---|---|---|
| B69 | B136 | PV | 5'NT | 0.058947 | 0 |
| B69 | B155 | PV | IgA | 0.072741 | 0 |
| B69 | B157 | PV | IgG | 0.105354 | 0 |
| B69 | B186 | PV | PFC | 0.089862 | 0 |
| B69 | B188 | PV | Ca | 0.051833 | 0 |
| B69 | B194 | PV | Cl | -0.04841 | 0 |
| B69 | B70 | PV | LDL-C | 0.06493 | 0 |
| B69 | B71 | PV | TG | 0.042896 | 0 |
| B69 | B73 | PV | ApoA | 0.049306 | 0 |
| B69 | B74 | PV | ApoA1 | 0.065902 | 0 |
| B69 | B75 | PV | ApoA1/B | -0.03842 | 0 |
| B69 | B76 | PV | ApoB | 0.118015 | 0 |
| B69 | B77 | PV | TC | 0.073771 | 0 |
| B69 | B91 | PV | TP | 0.132609 | 0 |
| B69 | B139 | PV | ADA | 0.037466 | 4.44E-16 |
| B69 | B97 | PV | dsDNA | 0.036814 | 1.33E-15 |
| B69 | B153 | PV | Glu | 0.034203 | 1.12E-13 |
| B69 | B126 | PV | Alb | 0.034169 | 1.18E-13 |
| B69 | B158 | PV | IgM | 0.033556 | 3.21E-13 |
| B69 | B111 | PV | TSH | 0.033195 | 5.72E-13 |
| B69 | B183 | PV | PTA | 0.031594 | 6.92E-12 |
| B69 | B193 | PV | K | 0.030433 | 3.92E-11 |
| B69 | B221 | PV | Hei | -0.02933 | 1.91E-10 |
| B69 | B95 | PV | anti-cenpb | 0.028662 | 4.90E-10 |
| B69 | B150 | PV | ALP | 0.027944 | 1.31E-09 |
| B69 | B120 | PV | HCV-cAg | 0.027629 | 2.00E-09 |
| B69 | B195 | PV | Na | -0.02722 | 3.46E-09 |
| B69 | B199 | PV | Mg | 0.026725 | 6.56E-09 |
| B69 | B140 | PV | r-GT | 0.025129 | 4.89E-08 |
| B69 | B81 | PV | TSGF | -0.02423 | 1.45E-07 |
| B69 | B131 | PV | AST | 0.023625 | 2.92E-07 |
| B69 | B196 | PV | TCO2 | 0.023452 | 3.56E-07 |
| B7 | B113 | WB-Cop | FT3 | 0.053273 | 0 |
| B7 | B8 | WB-Cop | WB-Zinc | 0.296204 | 0 |
| B7 | B23 | WB-Cop | CD3-CD16+CD56+(NK) | -0.03533 | 1.73E-14 |
| B7 | B100 | WB-Cop | anti-Ro52 | -0.03517 | 2.24E-14 |
| B7 | B21 | WB-Cop | CD3+CD4-CD8- | -0.02696 | 4.84E-09 |
| B7 | B107 | WB-Cop | AnuA | -0.02617 | 1.33E-08 |
| B7 | B115 | WB-Cop | TSH TT3 TT4(TT3) | 0.024463 | 1.09E-07 |
| B7 | B17 | WB-Cop | CD3+CD16+CD56+NK | 0.023483 | 3.44E-07 |
| B7 | B195 | WB-Cop | Na | -0.02283 | 7.24E-07 |
| B70 | B126 | LDL-C | Alb | 0.073653 | 0 |
| B70 | B127 | LDL-C | A-G ratio | -0.08118 | 0 |
| B70 | B129 | LDL-C | IBIL | -0.03881 | 0 |

| | | | | | |
|---|---|---|---|---|---|
| B70 | B130 | LDL-C | GLB | 0.108905 | 0 |
| B70 | B132 | LDL-C | Dbil | -0.20475 | 0 |
| B70 | B133 | LDL-C | TBIL | -0.08651 | 0 |
| B70 | B134 | LDL-C | CHE | 0.048435 | 0 |
| B70 | B188 | LDL-C | Ca | 0.105638 | 0 |
| B70 | B189 | LDL-C | eGFB | 0.043021 | 0 |
| B70 | B218 | LDL-C | DBP | 0.071081 | 0 |
| B70 | B220 | LDL-C | SBP | 0.067659 | 0 |
| B70 | B221 | LDL-C | Hei | -0.06535 | 0 |
| B70 | B71 | LDL-C | TG | -0.06821 | 0 |
| B70 | B72 | LDL-C | HDL-C | 0.123035 | 0 |
| B70 | B73 | LDL-C | ApoA | 0.068285 | 0 |
| B70 | B74 | LDL-C | ApoA1 | 0.055146 | 0 |
| B70 | B75 | LDL-C | ApoA1/B | -0.20894 | 0 |
| B70 | B76 | LDL-C | ApoB | 0.327584 | 0 |
| B70 | B77 | LDL-C | TC | 0.832036 | 0 |
| B70 | B91 | LDL-C | TP | 0.142424 | 0 |
| B70 | B111 | LDL-C | TSH | 0.03542 | 1.47E-14 |
| B70 | B194 | LDL-C | Cl | -0.0351 | 2.51E-14 |
| B70 | B174 | LDL-C | Urea/Cr | 0.03456 | 6.22E-14 |
| B70 | B222 | LDL-C | NS | -0.03233 | 2.22E-12 |
| B70 | B131 | LDL-C | AST | -0.0306 | 3.08E-11 |
| B70 | B136 | LDL-C | 5'NT | 0.029966 | 7.74E-11 |
| B70 | B153 | LDL-C | Glu | 0.029007 | 3.03E-10 |
| B70 | B161 | LDL-C | LEU | 0.028429 | 6.75E-10 |
| B70 | B190 | LDL-C | Cre | -0.02814 | 1.00E-09 |
| B70 | B215 | LDL-C | BMI | 0.027761 | 1.67E-09 |
| B70 | B119 | LDL-C | Hcv-rna | -0.0267 | 6.79E-09 |
| B70 | B150 | LDL-C | ALP | 0.025372 | 3.63E-08 |
| B70 | B146 | LDL-C | CysC | -0.02393 | 2.05E-07 |
| B70 | B86 | LDL-C | TBhCG | 0.023693 | 2.70E-07 |
| B70 | B181 | LDL-C | APTT | -0.02362 | 2.93E-07 |
| B71 | B126 | TG | Alb | 0.150119 | 0 |
| B71 | B127 | TG | A-G ratio | -0.10828 | 0 |
| B71 | B128 | TG | ALT | 0.155299 | 0 |
| B71 | B129 | TG | IBIL | -0.05472 | 0 |
| B71 | B130 | TG | GLB | 0.213029 | 0 |
| B71 | B131 | TG | AST | 0.094286 | 0 |
| B71 | B132 | TG | Dbil | -0.11575 | 0 |
| B71 | B133 | TG | TBIL | -0.07534 | 0 |
| B71 | B134 | TG | CHE | 0.079428 | 0 |
| B71 | B136 | TG | 5'NT | 0.038953 | 0 |
| B71 | B138 | TG | PAB | 0.068414 | 0 |
| B71 | B140 | TG | r-GT | 0.206844 | 0 |

| | | | | | |
|---|---|---|---|---|---|
| B71 | B141 | TG | AST/ALT | -0.14344 | 0 |
| B71 | B153 | TG | Glu | 0.189924 | 0 |
| B71 | B160 | TG | PH | -0.06649 | 0 |
| B71 | B163 | TG | SG | 0.07648 | 0 |
| B71 | B166 | TG | Pro | 0.049499 | 0 |
| B71 | B170 | TG | U-GLU | 0.1043 | 0 |
| B71 | B174 | TG | Urea/Cr | -0.05522 | 0 |
| B71 | B188 | TG | Ca | 0.089575 | 0 |
| B71 | B189 | TG | eGFB | 0.080301 | 0 |
| B71 | B192 | TG | UA | 0.213864 | 0 |
| B71 | B203 | TG | DH | 0.066315 | 0 |
| B71 | B204 | TG | Drinking | 0.112054 | 0 |
| B71 | B206 | TG | Waist | 0.156569 | 0 |
| B71 | B209 | TG | Smoking | 0.125514 | 0 |
| B71 | B210 | TG | Hip | 0.108123 | 0 |
| B71 | B211 | TG | IOP-R | 0.040032 | 0 |
| B71 | B212 | TG | IOP-L | 0.04086 | 0 |
| B71 | B215 | TG | BMI | 0.161435 | 0 |
| B71 | B216 | TG | BMI | 0.18333 | 0 |
| B71 | B218 | TG | DBP | 0.118913 | 0 |
| B71 | B221 | TG | Hei | 0.107883 | 0 |
| B71 | B222 | TG | NS | 0.162612 | 0 |
| B71 | B72 | TG | HDL-C | -0.38096 | 0 |
| B71 | B73 | TG | ApoA | -0.04001 | 0 |
| B71 | B74 | TG | ApoA1 | -0.05905 | 0 |
| B71 | B77 | TG | TC | 0.278171 | 0 |
| B71 | B91 | TG | TP | 0.282048 | 0 |
| B71 | B176 | TG | Ucr | 0.034685 | 5.04E-14 |
| B71 | B219 | TG | OT | -0.03433 | 9.08E-14 |
| B71 | B194 | TG | Cl | -0.03381 | 2.14E-13 |
| B71 | B191 | TG | Urea | -0.03153 | 7.68E-12 |
| B71 | B112 | TG | FT4 | -0.03078 | 2.33E-11 |
| B71 | B114 | TG | TSH TT3 TT4(TT4) | -0.03072 | 2.57E-11 |
| B71 | B109 | TG | AMA-M2 | 0.030415 | 4.03E-11 |
| B71 | B76 | TG | ApoB | 0.028084 | 1.08E-09 |
| B71 | B150 | TG | ALP | 0.027901 | 1.39E-09 |
| B71 | B86 | TG | TBhCG | -0.02636 | 1.05E-08 |
| B71 | B201 | TG | e-habits | -0.02625 | 1.21E-08 |
| B71 | B135 | TG | TBA | 0.025571 | 2.84E-08 |
| B71 | B157 | TG | IgG | -0.0253 | 3.97E-08 |
| B71 | B137 | TG | AFU | 0.024111 | 1.66E-07 |
| B71 | B193 | TG | K | -0.02388 | 2.17E-07 |
| B72 | B112 | HDL-C | FT4 | 0.061734 | 0 |
| B72 | B127 | HDL-C | A-G ratio | 0.05612 | 0 |

| | | | | | |
|---|---|---|---|---|---|
| B72 | B128 | HDL-C | ALT | -0.16935 | 0 |
| B72 | B130 | HDL-C | GLB | -0.06923 | 0 |
| B72 | B132 | HDL-C | Dbil | 0.061629 | 0 |
| B72 | B133 | HDL-C | TBIL | 0.040983 | 0 |
| B72 | B134 | HDL-C | CHE | -0.05291 | 0 |
| B72 | B140 | HDL-C | r-GT | -0.06139 | 0 |
| B72 | B141 | HDL-C | AST/ALT | 0.266028 | 0 |
| B72 | B153 | HDL-C | Glu | -0.11058 | 0 |
| B72 | B160 | HDL-C | PH | 0.069072 | 0 |
| B72 | B161 | HDL-C | LEU | 0.055718 | 0 |
| B72 | B163 | HDL-C | SG | -0.17009 | 0 |
| B72 | B166 | HDL-C | Pro | -0.07222 | 0 |
| B72 | B170 | HDL-C | U-GLU | -0.05759 | 0 |
| B72 | B174 | HDL-C | Urea/Cr | 0.166279 | 0 |
| B72 | B176 | HDL-C | Ucr | -0.07209 | 0 |
| B72 | B188 | HDL-C | Ca | 0.069442 | 0 |
| B72 | B190 | HDL-C | Cre | -0.12094 | 0 |
| B72 | B192 | HDL-C | UA | -0.29621 | 0 |
| B72 | B203 | HDL-C | DH | -0.04514 | 0 |
| B72 | B204 | HDL-C | Drinking | -0.06794 | 0 |
| B72 | B206 | HDL-C | Waist | -0.32944 | 0 |
| B72 | B209 | HDL-C | Smoking | -0.14148 | 0 |
| B72 | B210 | HDL-C | Hip | -0.2478 | 0 |
| B72 | B215 | HDL-C | BMI | -0.29403 | 0 |
| B72 | B216 | HDL-C | BMI | -0.38448 | 0 |
| B72 | B218 | HDL-C | DBP | -0.08424 | 0 |
| B72 | B219 | HDL-C | OT | 0.048585 | 0 |
| B72 | B220 | HDL-C | SBP | 0.071989 | 0 |
| B72 | B221 | HDL-C | Hei | -0.2918 | 0 |
| B72 | B222 | HDL-C | NS | -0.34696 | 0 |
| B72 | B73 | HDL-C | ApoA | 0.047937 | 0 |
| B72 | B74 | HDL-C | ApoA1 | 0.295675 | 0 |
| B72 | B75 | HDL-C | ApoA1/B | 0.139488 | 0 |
| B72 | B77 | HDL-C | TC | 0.253632 | 0 |
| B72 | B91 | HDL-C | TP | -0.05465 | 0 |
| B72 | B113 | HDL-C | FT3 | -0.0353 | 1.82E-14 |
| B72 | B81 | HDL-C | TSGF | -0.03486 | 3.75E-14 |
| B72 | B115 | HDL-C | TSH TT3 TT4(TT3) | -0.03454 | 6.39E-14 |
| B72 | B173 | HDL-C | NTT | 0.032614 | 1.43E-12 |
| B72 | B129 | HDL-C | IBIL | 0.030291 | 4.82E-11 |
| B72 | B205 | HDL-C | KDA | 0.029048 | 2.86E-10 |
| B72 | B193 | HDL-C | K | 0.028376 | 7.26E-10 |
| B72 | B198 | HDL-C | P | 0.028283 | 8.25E-10 |
| B72 | B114 | HDL-C | TSH TT3 TT4(TT4) | 0.027966 | 1.27E-09 |

| | | | | | |
|---|---|---|---|---|---|
| B72 | B189 | HDL-C | eGFB | 0.027176 | 3.64E-09 |
| B72 | B149 | HDL-C | TGAb | 0.026844 | 5.63E-09 |
| B72 | B123 | HDL-C | hs-CRP1 | -0.02524 | 4.30E-08 |
| B72 | B179 | HDL-C | α1-MG | -0.02305 | 5.59E-07 |
| B72 | B224 | HDL-C | pmh | 0.022967 | 6.18E-07 |
| B73 | B123 | ApoA | hs-CRP1 | 0.064069 | 0 |
| B73 | B137 | ApoA | AFU | -0.04156 | 0 |
| B73 | B193 | ApoA | K | 0.047775 | 0 |
| B73 | B74 | ApoA | ApoA1 | 0.092623 | 0 |
| B73 | B75 | ApoA | ApoA1/B | -0.10362 | 0 |
| B73 | B76 | ApoA | ApoB | 0.208747 | 0 |
| B73 | B77 | ApoA | TC | 0.054293 | 0 |
| B73 | B222 | ApoA | NS | -0.03775 | 2.22E-16 |
| B73 | B216 | ApoA | BMI | -0.03519 | 2.15E-14 |
| B73 | B206 | ApoA | Waist | -0.02793 | 1.34E-09 |
| B73 | B127 | ApoA | A-G ratio | -0.02758 | 2.13E-09 |
| B73 | B128 | ApoA | ALT | -0.02715 | 3.75E-09 |
| B73 | B140 | ApoA | r-GT | -0.02661 | 7.66E-09 |
| B73 | B126 | ApoA | Alb | -0.02624 | 1.22E-08 |
| B73 | B215 | ApoA | BMI | -0.02618 | 1.32E-08 |
| B73 | B204 | ApoA | Drinking | -0.02548 | 3.17E-08 |
| B73 | B163 | ApoA | SG | -0.02533 | 3.82E-08 |
| B73 | B135 | ApoA | TBA | -0.02522 | 4.38E-08 |
| B73 | B221 | ApoA | Hei | -0.0248 | 7.26E-08 |
| B73 | B194 | ApoA | Cl | 0.023694 | 2.70E-07 |
| B73 | B131 | ApoA | AST | -0.02355 | 3.18E-07 |
| B73 | B198 | ApoA | P | 0.023486 | 3.43E-07 |
| B74 | B123 | ApoA1 | hs-CRP1 | -0.05125 | 0 |
| B74 | B126 | ApoA1 | Alb | 0.046823 | 0 |
| B74 | B128 | ApoA1 | ALT | -0.04487 | 0 |
| B74 | B134 | ApoA1 | CHE | -0.05716 | 0 |
| B74 | B141 | ApoA1 | AST/ALT | 0.076663 | 0 |
| B74 | B163 | ApoA1 | SG | -0.0609 | 0 |
| B74 | B174 | ApoA1 | Urea/Cr | 0.075867 | 0 |
| B74 | B176 | ApoA1 | Ucr | -0.06838 | 0 |
| B74 | B188 | ApoA1 | Ca | 0.051087 | 0 |
| B74 | B190 | ApoA1 | Cre | -0.04225 | 0 |
| B74 | B192 | ApoA1 | UA | -0.07795 | 0 |
| B74 | B193 | ApoA1 | K | 0.06652 | 0 |
| B74 | B194 | ApoA1 | Cl | -0.06757 | 0 |
| B74 | B198 | ApoA1 | P | 0.096406 | 0 |
| B74 | B199 | ApoA1 | Mg | 0.046666 | 0 |
| B74 | B206 | ApoA1 | Waist | -0.10394 | 0 |
| B74 | B209 | ApoA1 | Smoking | -0.04832 | 0 |

| | | | | | |
|---|---|---|---|---|---|
| B74 | B210 | ApoA1 | Hip | -0.08062 | 0 |
| B74 | B215 | ApoA1 | BMI | -0.08402 | 0 |
| B74 | B216 | ApoA1 | BMI | -0.12861 | 0 |
| B74 | B220 | ApoA1 | SBP | 0.050341 | 0 |
| B74 | B221 | ApoA1 | Hei | -0.11156 | 0 |
| B74 | B222 | ApoA1 | NS | -0.12788 | 0 |
| B74 | B75 | ApoA1 | ApoA1/B | 0.437847 | 0 |
| B74 | B77 | ApoA1 | TC | 0.115457 | 0 |
| B74 | B86 | ApoA1 | TBhCG | 0.062068 | 0 |
| B74 | B91 | ApoA1 | TP | 0.033537 | 3.30E-13 |
| B74 | B138 | ApoA1 | PAB | 0.033471 | 3.68E-13 |
| B74 | B114 | ApoA1 | TSH TT3 TT4(TT4) | 0.033203 | 5.65E-13 |
| B74 | B181 | ApoA1 | APTT | -0.03242 | 1.94E-12 |
| B74 | B76 | ApoA1 | ApoB | 0.030574 | 3.19E-11 |
| B74 | B81 | ApoA1 | TSGF | -0.02734 | 2.94E-09 |
| B74 | B161 | ApoA1 | LEU | 0.027196 | 3.55E-09 |
| B74 | B156 | ApoA1 | IgE | 0.0266 | 7.72E-09 |
| B74 | B197 | ApoA1 | AG | 0.024398 | 1.18E-07 |
| B74 | B179 | ApoA1 | α1-MG | -0.02423 | 1.44E-07 |
| B74 | B121 | ApoA1 | C3 | 0.023857 | 2.23E-07 |
| B74 | B204 | ApoA1 | Drinking | -0.02344 | 3.59E-07 |
| B74 | B92 | ApoA1 | ALDH2-Glu504Glu | 0.023147 | 5.04E-07 |
| B75 | B123 | ApoA1/B | hs-CRP1 | -0.05699 | 0 |
| B75 | B132 | ApoA1/B | Dbil | 0.050082 | 0 |
| B75 | B134 | ApoA1/B | CHE | -0.13173 | 0 |
| B75 | B136 | ApoA1/B | 5'NT | -0.06172 | 0 |
| B75 | B137 | ApoA1/B | AFU | -0.03965 | 0 |
| B75 | B138 | ApoA1/B | PAB | -0.07594 | 0 |
| B75 | B141 | ApoA1/B | AST/ALT | 0.053494 | 0 |
| B75 | B192 | ApoA1/B | UA | -0.05347 | 0 |
| B75 | B195 | ApoA1/B | Na | -0.04498 | 0 |
| B75 | B200 | ApoA1/B | OP | -0.04001 | 0 |
| B75 | B206 | ApoA1/B | Waist | -0.06738 | 0 |
| B75 | B210 | ApoA1/B | Hip | -0.05018 | 0 |
| B75 | B215 | ApoA1/B | BMI | -0.06328 | 0 |
| B75 | B216 | ApoA1/B | BMI | -0.07063 | 0 |
| B75 | B221 | ApoA1/B | Hei | -0.04213 | 0 |
| B75 | B222 | ApoA1/B | NS | -0.06589 | 0 |
| B75 | B76 | ApoA1/B | ApoB | -0.70084 | 0 |
| B75 | B77 | ApoA1/B | TC | -0.1457 | 0 |
| B75 | B218 | ApoA1/B | DBP | -0.03583 | 7.33E-15 |
| B75 | B119 | ApoA1/B | Hcv-rna | 0.035441 | 1.42E-14 |
| B75 | B79 | ApoA1/B | hb V -dna | 0.033157 | 6.08E-13 |
| B75 | B163 | ApoA1/B | SG | -0.0328 | 1.07E-12 |

| | | | | | |
|---|---|---|---|---|---|
| B75 | B176 | ApoA1/B | Ucr | -0.03274 | 1.18E-12 |
| B75 | B153 | ApoA1/B | Glu | -0.03128 | 1.11E-11 |
| B75 | B193 | ApoA1/B | K | 0.030829 | 2.18E-11 |
| B75 | B150 | ApoA1/B | ALP | -0.03007 | 6.64E-11 |
| B75 | B128 | ApoA1/B | ALT | -0.02976 | 1.04E-10 |
| B75 | B197 | ApoA1/B | AG | -0.02949 | 1.54E-10 |
| B75 | B107 | ApoA1/B | AnuA | 0.027854 | 1.48E-09 |
| B75 | B139 | ApoA1/B | ADA | -0.02743 | 2.60E-09 |
| B75 | B86 | ApoA1/B | TBhCG | 0.026261 | 1.19E-08 |
| B75 | B224 | ApoA1/B | pmh | 0.0256 | 2.74E-08 |
| B75 | B178 | ApoA1/B | Ig-U | -0.02545 | 3.30E-08 |
| B75 | B209 | ApoA1/B | Smoking | -0.02436 | 1.24E-07 |
| B75 | B180 | ApoA1/B | Tf | -0.02335 | 3.99E-07 |
| B75 | B186 | ApoA1/B | PFC | -0.0228 | 7.44E-07 |
| B75 | B135 | ApoA1/B | TBA | 0.022654 | 8.76E-07 |
| B76 | B119 | ApoB | Hcv-rna | -0.04339 | 0 |
| B76 | B123 | ApoB | hs-CRP1 | 0.048295 | 0 |
| B76 | B130 | ApoB | GLB | 0.055502 | 0 |
| B76 | B132 | ApoB | Dbil | -0.06514 | 0 |
| B76 | B134 | ApoB | CHE | 0.169802 | 0 |
| B76 | B136 | ApoB | 5'NT | 0.105269 | 0 |
| B76 | B138 | ApoB | PAB | 0.107726 | 0 |
| B76 | B153 | ApoB | Glu | 0.044272 | 0 |
| B76 | B181 | ApoB | APTT | -0.04798 | 0 |
| B76 | B186 | ApoB | PFC | 0.044096 | 0 |
| B76 | B188 | ApoB | Ca | 0.063994 | 0 |
| B76 | B194 | ApoB | Cl | -0.09376 | 0 |
| B76 | B196 | ApoB | TCO2 | 0.039075 | 0 |
| B76 | B197 | ApoB | AG | 0.051458 | 0 |
| B76 | B199 | ApoB | Mg | 0.042456 | 0 |
| B76 | B200 | ApoB | OP | 0.040994 | 0 |
| B76 | B218 | ApoB | DBP | 0.040375 | 0 |
| B76 | B77 | ApoB | TC | 0.304537 | 0 |
| B76 | B91 | ApoB | TP | 0.072128 | 0 |
| B76 | B127 | ApoB | A-G ratio | -0.03773 | 2.22E-16 |
| B76 | B180 | ApoB | Tf | 0.037323 | 4.44E-16 |
| B76 | B126 | ApoB | Alb | 0.036771 | 1.33E-15 |
| B76 | B140 | ApoB | r-GT | 0.036426 | 2.66E-15 |
| B76 | B150 | ApoB | ALP | 0.036201 | 4.00E-15 |
| B76 | B192 | ApoB | UA | 0.033015 | 7.62E-13 |
| B76 | B184 | ApoB | INR | -0.03297 | 8.16E-13 |
| B76 | B215 | ApoB | BMI | 0.032322 | 2.26E-12 |
| B76 | B79 | ApoB | hb V -dna | -0.03098 | 1.73E-11 |
| B76 | B220 | ApoB | SBP | 0.030641 | 2.89E-11 |

| | | | | | |
|---|---|---|---|---|---|
| B76 | B178 | ApoB | Ig-U | 0.030317 | 4.65E-11 |
| B76 | B206 | ApoB | Waist | 0.029617 | 1.28E-10 |
| B76 | B170 | ApoB | U-GLU | 0.029004 | 3.04E-10 |
| B76 | B195 | ApoB | Na | 0.0278 | 1.59E-09 |
| B76 | B107 | ApoB | AnuA | -0.02747 | 2.48E-09 |
| B76 | B135 | ApoB | TBA | -0.02609 | 1.48E-08 |
| B76 | B141 | ApoB | AST/ALT | -0.02578 | 2.18E-08 |
| B76 | B137 | ApoB | AFU | 0.025776 | 2.20E-08 |
| B76 | B111 | ApoB | TSH | 0.023468 | 3.50E-07 |
| B76 | B179 | ApoB | α1-MG | 0.023151 | 5.02E-07 |
| B76 | B185 | ApoB | PT | -0.02294 | 6.35E-07 |
| B77 | B111 | TC | TSH | 0.046918 | 0 |
| B77 | B126 | TC | Alb | 0.115746 | 0 |
| B77 | B127 | TC | A-G ratio | -0.12217 | 0 |
| B77 | B129 | TC | IBIL | -0.05399 | 0 |
| B77 | B130 | TC | GLB | 0.181722 | 0 |
| B77 | B132 | TC | Dbil | -0.22625 | 0 |
| B77 | B133 | TC | TBIL | -0.10457 | 0 |
| B77 | B134 | TC | CHE | 0.060787 | 0 |
| B77 | B136 | TC | 5'NT | 0.04395 | 0 |
| B77 | B138 | TC | PAB | 0.039462 | 0 |
| B77 | B140 | TC | r-GT | 0.080469 | 0 |
| B77 | B153 | TC | Glu | 0.081489 | 0 |
| B77 | B161 | TC | LEU | 0.039898 | 0 |
| B77 | B163 | TC | SG | -0.04834 | 0 |
| B77 | B174 | TC | Urea/Cr | 0.075207 | 0 |
| B77 | B188 | TC | Ca | 0.156336 | 0 |
| B77 | B190 | TC | Cre | -0.05005 | 0 |
| B77 | B194 | TC | Cl | -0.04449 | 0 |
| B77 | B206 | TC | Waist | -0.05576 | 0 |
| B77 | B210 | TC | Hip | -0.04882 | 0 |
| B77 | B216 | TC | BMI | -0.0919 | 0 |
| B77 | B218 | TC | DBP | 0.079758 | 0 |
| B77 | B220 | TC | SBP | 0.099327 | 0 |
| B77 | B221 | TC | Hei | -0.12497 | 0 |
| B77 | B222 | TC | NS | -0.0959 | 0 |
| B77 | B91 | TC | TP | 0.233301 | 0 |
| B77 | B212 | TC | IOP-L | 0.036527 | 2.22E-15 |
| B77 | B189 | TC | eGFB | 0.034415 | 7.90E-14 |
| B77 | B211 | TC | IOP-R | 0.033107 | 6.58E-13 |
| B77 | B170 | TC | U-GLU | 0.032359 | 2.13E-12 |
| B77 | B150 | TC | ALP | 0.02919 | 2.34E-10 |
| B77 | B177 | TC | ACR | 0.026502 | 8.75E-09 |
| B77 | B181 | TC | APTT | -0.0262 | 1.29E-08 |

| | | | | | |
|---|---|---|---|---|---|
| B77 | B215 | TC | BMI | -0.02448 | 1.07E-07 |
| B77 | B184 | TC | INR | -0.02394 | 2.03E-07 |
| B77 | B185 | TC | PT | -0.02306 | 5.55E-07 |
| B78 | B80 | HBcAb | anti-HP IgG | -0.16027 | 0 |
| B78 | B180 | HBcAb | Tf | 0.024423 | 1.15E-07 |
| B79 | B100 | hb V -dna | anti-Ro52 | 0.047519 | 0 |
| B79 | B103 | hb V -dna | anti-SSA/Ro | 0.041094 | 0 |
| B79 | B107 | hb V -dna | AnuA | 0.286921 | 0 |
| B79 | B119 | hb V -dna | Hcv-rna | 0.348262 | 0 |
| B79 | B134 | hb V -dna | CHE | -0.05967 | 0 |
| B79 | B138 | hb V -dna | PAB | -0.06002 | 0 |
| B79 | B150 | hb V -dna | ALP | -0.042 | 0 |
| B79 | B157 | hb V -dna | IgG | 0.038244 | 0 |
| B79 | B176 | hb V -dna | Ucr | -0.04365 | 0 |
| B79 | B179 | hb V -dna | α1-MG | -0.0508 | 0 |
| B79 | B181 | hb V -dna | APTT | 0.076941 | 0 |
| B79 | B182 | hb V -dna | TT | -0.07047 | 0 |
| B79 | B183 | hb V -dna | PTA | 0.193192 | 0 |
| B79 | B184 | hb V -dna | INR | 0.142832 | 0 |
| B79 | B185 | hb V -dna | PT | 0.048411 | 0 |
| B79 | B186 | hb V -dna | PFC | -0.14102 | 0 |
| B79 | B187 | hb V -dna | NSE | -0.0387 | 0 |
| B79 | B189 | hb V -dna | eGFB | -0.04727 | 0 |
| B79 | B193 | hb V -dna | K | 0.106669 | 0 |
| B79 | B194 | hb V -dna | Cl | 0.073341 | 0 |
| B79 | B195 | hb V -dna | Na | -0.04419 | 0 |
| B79 | B197 | hb V -dna | AG | -0.07344 | 0 |
| B79 | B199 | hb V -dna | Mg | -0.04866 | 0 |
| B79 | B204 | hb V -dna | Drinking | -0.0383 | 0 |
| B79 | B81 | hb V -dna | TSGF | -0.12091 | 0 |
| B79 | B90 | hb V -dna | CYFRA 21-1 | -0.03818 | 0 |
| B79 | B92 | hb V -dna | ALDH2-Glu504Glu | -0.04943 | 0 |
| B79 | B94 | hb V -dna | ALDH2-Lys504Lys | 0.104349 | 0 |
| B79 | B96 | hb V -dna | dsDNA (IIF) | 0.073205 | 0 |
| B79 | B98 | hb V -dna | anti-Jo-1 | 0.05121 | 0 |
| B79 | B218 | hb V -dna | DBP | -0.03718 | 6.66E-16 |
| B79 | B200 | hb V -dna | OP | -0.03695 | 8.88E-16 |
| B79 | B222 | hb V -dna | NS | -0.03713 | 8.88E-16 |
| B79 | B137 | hb V -dna | AFU | -0.03526 | 1.91E-14 |
| B79 | B216 | hb V -dna | BMI | -0.03449 | 6.93E-14 |
| B79 | B109 | hb V -dna | AMA-M2 | 0.031588 | 6.99E-12 |
| B79 | B93 | hb V -dna | ALDH2-Glu504Lys | 0.031182 | 1.29E-11 |
| B79 | B135 | hb V -dna | TBA | 0.030896 | 1.98E-11 |
| B79 | B110 | hb V -dna | AHA | 0.030515 | 3.47E-11 |

| | | | | | |
|---|---|---|---|---|---|
| B79 | B206 | hb V -dna | Waist | -0.02991 | 8.44E-11 |
| B79 | B97 | hb V -dna | dsDNA | 0.029202 | 2.30E-10 |
| B79 | B141 | hb V -dna | AST/ALT | 0.028994 | 3.08E-10 |
| B79 | B113 | hb V -dna | FT3 | -0.02817 | 9.60E-10 |
| B79 | B105 | hb V -dna | anti-U1RNP | -0.02804 | 1.15E-09 |
| B79 | B210 | hb V -dna | Hip | -0.02791 | 1.37E-09 |
| B79 | B163 | hb V -dna | SG | -0.02721 | 3.49E-09 |
| B79 | B215 | hb V -dna | BMI | -0.027 | 4.60E-09 |
| B79 | B128 | hb V -dna | ALT | -0.02632 | 1.10E-08 |
| B79 | B95 | hb V -dna | anti-cenpb | 0.025735 | 2.31E-08 |
| B79 | B99 | hb V -dna | anti-PM-Scl | -0.02537 | 3.64E-08 |
| B79 | B198 | hb V -dna | P | 0.025237 | 4.29E-08 |
| B79 | B178 | hb V -dna | Ig-U | -0.02483 | 7.06E-08 |
| B79 | B221 | hb V -dna | Hei | -0.02473 | 7.92E-08 |
| B79 | B102 | hb V -dna | anti-Sm | -0.02322 | 4.63E-07 |
| B79 | B115 | hb V -dna | TSH TT3 TT4(TT3) | -0.02271 | 8.24E-07 |
| B8 | B113 | WB-Zinc | FT3 | -0.03535 | 1.64E-14 |
| B8 | B23 | WB-Zinc | CD3-CD16+CD56+(NK) | -0.03284 | 1.01E-12 |
| B8 | B29 | WB-Zinc | E2 | -0.03072 | 2.58E-11 |
| B8 | B33 | WB-Zinc | Prog | -0.02942 | 1.68E-10 |
| B8 | B18 | WB-Zinc | CD3+CD4+helper(inducer)T | 0.029036 | 2.91E-10 |
| B8 | B31 | WB-Zinc | FSH | 0.028594 | 5.38E-10 |
| B80 | B166 | anti-HP IgG | Pro | -0.02348 | 3.46E-07 |
| B81 | B124 | TSGF | ACTH | -0.04478 | 0 |
| B81 | B125 | TSGF | Cor | -0.09872 | 0 |
| B81 | B136 | TSGF | 5'NT | 0.047723 | 0 |
| B81 | B151 | TSGF | Anti-CCP | -0.04446 | 0 |
| B81 | B157 | TSGF | IgG | -0.05984 | 0 |
| B81 | B181 | TSGF | APTT | 0.055504 | 0 |
| B81 | B182 | TSGF | TT | 0.158345 | 0 |
| B81 | B183 | TSGF | PTA | -0.0584 | 0 |
| B81 | B184 | TSGF | INR | -0.05235 | 0 |
| B81 | B194 | TSGF | Cl | -0.03834 | 0 |
| B81 | B195 | TSGF | Na | 0.050085 | 0 |
| B81 | B197 | TSGF | AG | 0.058949 | 0 |
| B81 | B200 | TSGF | OP | 0.055709 | 0 |
| B81 | B92 | TSGF | ALDH2-Glu504Glu | 0.040004 | 0 |
| B81 | B127 | TSGF | A-G ratio | -0.03676 | 1.33E-15 |
| B81 | B93 | TSGF | ALDH2-Glu504Lys | -0.03534 | 1.69E-14 |
| B81 | B139 | TSGF | ADA | 0.035059 | 2.71E-14 |
| B81 | B137 | TSGF | AFU | 0.034232 | 1.07E-13 |
| B81 | B193 | TSGF | K | -0.03399 | 1.60E-13 |
| B81 | B179 | TSGF | α1-MG | 0.031899 | 4.34E-12 |
| B81 | B108 | TSGF | Anti-Cyclin | 0.03096 | 1.80E-11 |

| | | | | | |
|---|---|---|---|---|---|
| B81 | B189 | TSGF | eGFB | -0.03053 | 3.39E-11 |
| B81 | B130 | TSGF | GLB | 0.029515 | 1.48E-10 |
| B81 | B150 | TSGF | ALP | 0.029268 | 2.10E-10 |
| B81 | B94 | TSGF | ALDH2-Lys504Lys | -0.02901 | 3.01E-10 |
| B81 | B198 | TSGF | P | -0.02636 | 1.05E-08 |
| B81 | B112 | TSGF | FT4 | -0.02614 | 1.40E-08 |
| B81 | B224 | TSGF | pmh | -0.02591 | 1.85E-08 |
| B81 | B132 | TSGF | Dbil | 0.024364 | 1.23E-07 |
| B81 | B134 | TSGF | CHE | 0.023786 | 2.42E-07 |
| B81 | B95 | TSGF | anti-cenpb | -0.02373 | 2.59E-07 |
| B81 | B141 | TSGF | AST/ALT | -0.02353 | 3.26E-07 |
| B81 | B110 | TSGF | AHA | -0.02255 | 9.85E-07 |
| B82 | B83 | CEA | AFP | 0.119676 | 0 |
| B82 | B86 | CEA | TBhCG | -0.13499 | 0 |
| B82 | B87 | CEA | NSE | 0.092738 | 0 |
| B82 | B88 | CEA | CA 242 | 0.235054 | 0 |
| B82 | B89 | CEA | CA 72-4 | 0.115828 | 0 |
| B82 | B85 | CEA | SCC | 0.037765 | 2.22E-16 |
| B82 | B209 | CEA | Smoking | 0.033378 | 4.27E-13 |
| B82 | B84 | CEA | C-PSA | 0.030492 | 3.60E-11 |
| B83 | B85 | AFP | SCC | 0.103167 | 0 |
| B83 | B86 | AFP | TBhCG | 0.097319 | 0 |
| B83 | B87 | AFP | NSE | 0.12049 | 0 |
| B83 | B88 | AFP | CA 242 | 0.164298 | 0 |
| B83 | B90 | AFP | CYFRA 21-1 | 0.065555 | 0 |
| B83 | B138 | AFP | PAB | 0.036445 | 2.66E-15 |
| B83 | B136 | AFP | 5'NT | 0.034126 | 1.27E-13 |
| B83 | B194 | AFP | Cl | -0.02465 | 8.73E-08 |
| B83 | B140 | AFP | r-GT | 0.023599 | 3.01E-07 |
| B84 | B85 | C-PSA | SCC | 0.051642 | 0 |
| B84 | B86 | C-PSA | TBhCG | -0.12859 | 0 |
| B84 | B88 | C-PSA | CA 242 | 0.033652 | 2.74E-13 |
| B84 | B90 | C-PSA | CYFRA 21-1 | 0.024293 | 1.34E-07 |
| B84 | B189 | C-PSA | eGFB | -0.02358 | 3.08E-07 |
| B84 | B101 | C-PSA | anti-Sc70 | 0.023571 | 3.11E-07 |
| B84 | B87 | C-PSA | NSE | 0.022563 | 9.68E-07 |
| B85 | B86 | SCC | TBhCG | -0.04038 | 0 |
| B85 | B87 | SCC | NSE | 0.172992 | 0 |
| B85 | B88 | SCC | CA 242 | 0.058478 | 0 |
| B85 | B90 | SCC | CYFRA 21-1 | 0.055212 | 0 |
| B85 | B189 | SCC | eGFB | -0.03058 | 3.14E-11 |
| B85 | B190 | SCC | Cre | 0.029057 | 2.82E-10 |
| B85 | B191 | SCC | Urea | 0.023616 | 2.95E-07 |
| B86 | B119 | TBhCG | Hcv-rna | 0.044909 | 0 |

| | | | | | |
|---|---|---|---|---|---|
| B86 | B129 | TBhCG | IBIL | -0.04708 | 0 |
| B86 | B133 | TBhCG | TBIL | -0.04294 | 0 |
| B86 | B134 | TBhCG | CHE | -0.04065 | 0 |
| B86 | B137 | TBhCG | AFU | -0.03968 | 0 |
| B86 | B193 | TBhCG | K | 0.084202 | 0 |
| B86 | B198 | TBhCG | P | 0.062711 | 0 |
| B86 | B204 | TBhCG | Drinking | -0.04929 | 0 |
| B86 | B216 | TBhCG | BMI | -0.04562 | 0 |
| B86 | B221 | TBhCG | Hei | -0.04295 | 0 |
| B86 | B222 | TBhCG | NS | -0.03987 | 0 |
| B86 | B87 | TBhCG | NSE | -0.0948 | 0 |
| B86 | B90 | TBhCG | CYFRA 21-1 | 0.202399 | 0 |
| B86 | B183 | TBhCG | PTA | -0.03601 | 5.33E-15 |
| B86 | B146 | TBhCG | CysC | -0.02876 | 4.28E-10 |
| B86 | B160 | TBhCG | PH | -0.02856 | 5.63E-10 |
| B86 | B136 | TBhCG | 5'NT | 0.027922 | 1.35E-09 |
| B86 | B218 | TBhCG | DBP | -0.02731 | 3.06E-09 |
| B86 | B192 | TBhCG | UA | -0.02689 | 5.28E-09 |
| B86 | B215 | TBhCG | BMI | -0.02675 | 6.33E-09 |
| B86 | B107 | TBhCG | AnuA | 0.026372 | 1.03E-08 |
| B86 | B141 | TBhCG | AST/ALT | 0.025558 | 2.89E-08 |
| B86 | B88 | TBhCG | CA 242 | -0.02513 | 4.86E-08 |
| B86 | B128 | TBhCG | ALT | -0.02477 | 7.60E-08 |
| B86 | B163 | TBhCG | SG | -0.0244 | 1.17E-07 |
| B86 | B158 | TBhCG | IgM | 0.023634 | 2.89E-07 |
| B86 | B131 | TBhCG | AST | -0.02359 | 3.04E-07 |
| B86 | B174 | TBhCG | Urea/Cr | 0.023488 | 3.42E-07 |
| B86 | B206 | TBhCG | Waist | -0.02294 | 6.35E-07 |
| B86 | B116 | TBhCG | TT3 | 0.022733 | 8.02E-07 |
| B87 | B110 | NSE | AHA | 0.046956 | 0 |
| B87 | B88 | NSE | CA 242 | 0.147555 | 0 |
| B87 | B89 | NSE | CA 72-4 | -0.04286 | 0 |
| B87 | B90 | NSE | CYFRA 21-1 | 0.062139 | 0 |
| B87 | B129 | NSE | IBIL | 0.034702 | 4.91E-14 |
| B87 | B189 | NSE | eGFB | -0.02795 | 1.29E-09 |
| B87 | B133 | NSE | TBIL | 0.026745 | 6.39E-09 |
| B87 | B119 | NSE | Hcv-rna | 0.025621 | 2.67E-08 |
| B87 | B183 | NSE | PTA | 0.024566 | 9.67E-08 |
| B88 | B138 | CA 242 | PAB | -0.03059 | 3.10E-11 |
| B90 | B126 | CYFRA 21-1 | Alb | -0.04174 | 0 |
| B90 | B146 | CYFRA 21-1 | CysC | 0.041166 | 0 |
| B90 | B187 | CYFRA 21-1 | NSE | 0.038551 | 0 |
| B90 | B189 | CYFRA 21-1 | eGFB | -0.06843 | 0 |
| B90 | B190 | CYFRA 21-1 | Cre | 0.036128 | 4.44E-15 |

| | | | | | |
|---|---|---|---|---|---|
| B90 | B191 | CYFRA 21-1 | Urea | 0.03326 | 5.16E-13 |
| B90 | B183 | CYFRA 21-1 | PTA | -0.03189 | 4.43E-12 |
| B90 | B186 | CYFRA 21-1 | PFC | 0.030374 | 4.27E-11 |
| B90 | B108 | CYFRA 21-1 | Anti-Cyclin | 0.029725 | 1.09E-10 |
| B90 | B127 | CYFRA 21-1 | A-G ratio | -0.02693 | 5.02E-09 |
| B90 | B107 | CYFRA 21-1 | AnuA | -0.02677 | 6.17E-09 |
| B90 | B153 | CYFRA 21-1 | Glu | 0.026081 | 1.50E-08 |
| B90 | B200 | CYFRA 21-1 | OP | 0.023379 | 3.87E-07 |
| B91 | B111 | TP | TSH | 0.049474 | 0 |
| B91 | B126 | TP | Alb | 0.450521 | 0 |
| B91 | B127 | TP | A-G ratio | -0.52256 | 0 |
| B91 | B128 | TP | ALT | 0.10949 | 0 |
| B91 | B130 | TP | GLB | 0.809169 | 0 |
| B91 | B131 | TP | AST | 0.133401 | 0 |
| B91 | B132 | TP | Dbil | -0.04387 | 0 |
| B91 | B134 | TP | CHE | 0.061968 | 0 |
| B91 | B136 | TP | 5'NT | 0.048189 | 0 |
| B91 | B139 | TP | ADA | 0.041762 | 0 |
| B91 | B140 | TP | r-GT | 0.080693 | 0 |
| B91 | B150 | TP | ALP | 0.04017 | 0 |
| B91 | B153 | TP | Glu | 0.134765 | 0 |
| B91 | B155 | TP | IgA | 0.063907 | 0 |
| B91 | B157 | TP | IgG | 0.101205 | 0 |
| B91 | B163 | TP | SG | -0.05284 | 0 |
| B91 | B188 | TP | Ca | 0.32867 | 0 |
| B91 | B190 | TP | Cre | -0.03847 | 0 |
| B91 | B192 | TP | UA | 0.095304 | 0 |
| B91 | B194 | TP | Cl | -0.06678 | 0 |
| B91 | B204 | TP | Drinking | -0.05361 | 0 |
| B91 | B209 | TP | Smoking | -0.12507 | 0 |
| B91 | B218 | TP | DBP | 0.071995 | 0 |
| B91 | B220 | TP | SBP | 0.088851 | 0 |
| B91 | B221 | TP | Hei | -0.0687 | 0 |
| B91 | B197 | TP | AG | 0.037859 | 2.22E-16 |
| B91 | B215 | TP | BMI | 0.037691 | 2.22E-16 |
| B91 | B115 | TP | TSH TT3 TT4(TT3) | 0.036782 | 1.33E-15 |
| B91 | B113 | TP | FT3 | 0.035887 | 6.66E-15 |
| B91 | B133 | TP | TBIL | -0.0334 | 4.10E-13 |
| B91 | B114 | TP | TSH TT3 TT4(TT4) | 0.032605 | 1.45E-12 |
| B91 | B193 | TP | K | 0.032124 | 3.07E-12 |
| B91 | B199 | TP | Mg | 0.03114 | 1.37E-11 |
| B91 | B158 | TP | IgM | 0.030902 | 1.96E-11 |
| B91 | B141 | TP | AST/ALT | -0.02993 | 8.18E-11 |
| B91 | B138 | TP | PAB | 0.029661 | 1.20E-10 |

| | | | | | |
|---|---|---|---|---|---|
| B91 | B166 | TP | Pro | 0.029315 | 1.96E-10 |
| B91 | B149 | TP | TGAb | 0.028738 | 4.41E-10 |
| B91 | B112 | TP | FT4 | 0.026947 | 4.91E-09 |
| B91 | B129 | TP | IBIL | -0.02676 | 6.31E-09 |
| B91 | B170 | TP | U-GLU | 0.026692 | 6.85E-09 |
| B91 | B189 | TP | eGFB | 0.026213 | 1.27E-08 |
| B91 | B211 | TP | IOP-R | 0.025026 | 5.55E-08 |
| B91 | B137 | TP | AFU | 0.025021 | 5.58E-08 |
| B91 | B191 | TP | Urea | -0.02422 | 1.45E-07 |
| B91 | B160 | TP | PH | -0.02421 | 1.47E-07 |
| B91 | B148 | TP | Anti_TPO | 0.024181 | 1.53E-07 |
| B91 | B161 | TP | LEU | 0.022919 | 6.52E-07 |
| B91 | B203 | TP | DH | 0.022828 | 7.22E-07 |
| B92 | B138 | ALDH2-Glu504Glu | PAB | 0.073598 | 0 |
| B92 | B157 | ALDH2-Glu504Glu | IgG | -0.04169 | 0 |
| B92 | B193 | ALDH2-Glu504Glu | K | -0.03917 | 0 |
| B92 | B93 | ALDH2-Glu504Glu | ALDH2-Glu504Lys | -0.98319 | 0 |
| B92 | B94 | ALDH2-Glu504Glu | ALDH2-Lys504Lys | -0.17577 | 0 |
| B92 | B97 | ALDH2-Glu504Glu | dsDNA | -0.05659 | 0 |
| B92 | B108 | ALDH2-Glu504Glu | Anti-Cyclin | -0.03355 | 3.26E-13 |
| B92 | B124 | ALDH2-Glu504Glu | ACTH | -0.03311 | 6.51E-13 |
| B92 | B125 | ALDH2-Glu504Glu | Cor | -0.03224 | 2.58E-12 |
| B92 | B176 | ALDH2-Glu504Glu | Ucr | 0.031003 | 1.69E-11 |
| B92 | B99 | ALDH2-Glu504Glu | anti-PM-Scl | 0.030797 | 2.29E-11 |
| B92 | B96 | ALDH2-Glu504Glu | dsDNA (IIF) | -0.03012 | 6.17E-11 |
| B92 | B204 | ALDH2-Glu504Glu | Drinking | 0.029884 | 8.70E-11 |
| B92 | B198 | ALDH2-Glu504Glu | P | -0.02891 | 3.49E-10 |
| B92 | B135 | ALDH2-Glu504Glu | TBA | 0.026621 | 7.51E-09 |
| B92 | B107 | ALDH2-Glu504Glu | AnuA | -0.02648 | 8.95E-09 |
| B92 | B181 | ALDH2-Glu504Glu | APTT | -0.02648 | 9.04E-09 |
| B92 | B179 | ALDH2-Glu504Glu | α1-MG | 0.024268 | 1.38E-07 |
| B92 | B140 | ALDH2-Glu504Glu | r-GT | 0.024063 | 1.75E-07 |
| B93 | B138 | ALDH2-Glu504Lys | PAB | -0.06705 | 0 |
| B93 | B157 | ALDH2-Glu504Lys | IgG | 0.042141 | 0 |
| B93 | B193 | ALDH2-Glu504Lys | K | 0.038785 | 0 |
| B93 | B97 | ALDH2-Glu504Lys | dsDNA | 0.058233 | 0 |
| B93 | B108 | ALDH2-Glu504Lys | Anti-Cyclin | 0.037079 | 8.88E-16 |
| B93 | B124 | ALDH2-Glu504Lys | ACTH | 0.035234 | 2.00E-14 |
| B93 | B125 | ALDH2-Glu504Lys | Cor | 0.034073 | 1.39E-13 |
| B93 | B99 | ALDH2-Glu504Lys | anti-PM-Scl | -0.02996 | 7.83E-11 |
| B93 | B179 | ALDH2-Glu504Lys | α1-MG | -0.02955 | 1.41E-10 |
| B93 | B176 | ALDH2-Glu504Lys | Ucr | -0.02879 | 4.09E-10 |
| B93 | B198 | ALDH2-Glu504Lys | P | 0.028208 | 9.13E-10 |
| B93 | B135 | ALDH2-Glu504Lys | TBA | -0.02658 | 7.92E-09 |

| | | | | | |
|---|---|---|---|---|---|
| B93 | B204 | ALDH2-Glu504Lys | Drinking | -0.02617 | 1.34E-08 |
| B94 | B107 | ALDH2-Lys504Lys | AnuA | 0.066196 | 0 |
| B94 | B119 | ALDH2-Lys504Lys | Hcv-rna | 0.060135 | 0 |
| B94 | B138 | ALDH2-Lys504Lys | PAB | -0.04216 | 0 |
| B94 | B183 | ALDH2-Lys504Lys | PTA | 0.062508 | 0 |
| B94 | B184 | ALDH2-Lys504Lys | INR | 0.052287 | 0 |
| B94 | B96 | ALDH2-Lys504Lys | dsDNA (IIF) | 0.046009 | 0 |
| B94 | B181 | ALDH2-Lys504Lys | APTT | 0.036986 | 8.88E-16 |
| B94 | B186 | ALDH2-Lys504Lys | PFC | -0.0371 | 8.88E-16 |
| B94 | B179 | ALDH2-Lys504Lys | α1-MG | 0.025958 | 1.75E-08 |
| B94 | B180 | ALDH2-Lys504Lys | Tf | 0.02377 | 2.47E-07 |
| B94 | B204 | ALDH2-Lys504Lys | Drinking | -0.02291 | 6.59E-07 |
| B95 | B107 | anti-cenpb | AnuA | 0.043126 | 0 |
| B95 | B183 | anti-cenpb | PTA | 0.057131 | 0 |
| B95 | B134 | anti-cenpb | CHE | 0.029598 | 1.31E-10 |
| B96 | B105 | dsDNA (IIF) | anti-U1RNP | -0.05386 | 0 |
| B96 | B107 | dsDNA (IIF) | AnuA | 0.190139 | 0 |
| B96 | B119 | dsDNA (IIF) | Hcv-rna | 0.080446 | 0 |
| B96 | B122 | dsDNA (IIF) | C4 | -0.10754 | 0 |
| B96 | B124 | dsDNA (IIF) | ACTH | -0.05241 | 0 |
| B96 | B138 | dsDNA (IIF) | PAB | -0.05069 | 0 |
| B96 | B157 | dsDNA (IIF) | IgG | 0.045133 | 0 |
| B96 | B183 | dsDNA (IIF) | PTA | 0.054809 | 0 |
| B96 | B186 | dsDNA (IIF) | PFC | -0.04484 | 0 |
| B96 | B97 | dsDNA (IIF) | dsDNA | -0.37414 | 0 |
| B96 | B100 | dsDNA (IIF) | anti-Ro52 | 0.035719 | 8.88E-15 |
| B96 | B137 | dsDNA (IIF) | AFU | 0.034709 | 4.84E-14 |
| B96 | B98 | dsDNA (IIF) | anti-Jo-1 | 0.033599 | 2.99E-13 |
| B96 | B211 | dsDNA (IIF) | IOP-R | -0.03099 | 1.72E-11 |
| B96 | B114 | dsDNA (IIF) | TSH TT3 TT4(TT4) | 0.02812 | 1.03E-09 |
| B96 | B212 | dsDNA (IIF) | IOP-L | -0.02713 | 3.86E-09 |
| B96 | B109 | dsDNA (IIF) | AMA-M2 | -0.02585 | 2.01E-08 |
| B96 | B184 | dsDNA (IIF) | INR | 0.02555 | 2.91E-08 |
| B96 | B199 | dsDNA (IIF) | Mg | -0.02393 | 2.06E-07 |
| B97 | B122 | dsDNA | C4 | 0.090067 | 0 |
| B97 | B156 | dsDNA | IgE | 0.059585 | 0 |
| B97 | B109 | dsDNA | AMA-M2 | 0.034444 | 7.53E-14 |
| B97 | B158 | dsDNA | IgM | 0.030331 | 4.55E-11 |
| B97 | B157 | dsDNA | IgG | 0.02628 | 1.16E-08 |
| B97 | B211 | dsDNA | IOP-R | 0.02357 | 3.11E-07 |
| B98 | B104 | anti-Jo-1 | anti-SSB | 0.424772 | 0 |
| B98 | B107 | anti-Jo-1 | AnuA | 0.107359 | 0 |
| B98 | B136 | anti-Jo-1 | 5'NT | 0.07154 | 0 |
| B98 | B157 | anti-Jo-1 | IgG | 0.042911 | 0 |

| | | | | | |
|---|---|---|---|---|---|
| B98 | B193 | anti-Jo-1 | K | 0.048868 | 0 |
| B98 | B197 | anti-Jo-1 | AG | -0.04687 | 0 |
| B98 | B138 | anti-Jo-1 | PAB | -0.03397 | 1.65E-13 |
| B98 | B195 | anti-Jo-1 | Na | -0.03366 | 2.73E-13 |
| B98 | B196 | anti-Jo-1 | TCO2 | 0.029696 | 1.14E-10 |
| B98 | B200 | anti-Jo-1 | OP | -0.02941 | 1.72E-10 |
| B98 | B115 | anti-Jo-1 | TSH TT3 TT4(TT3) | -0.02641 | 9.84E-09 |
| B98 | B119 | anti-Jo-1 | Hcv-rna | 0.025229 | 4.33E-08 |
| B98 | B186 | anti-Jo-1 | PFC | -0.02349 | 3.41E-07 |
| B98 | B184 | anti-Jo-1 | INR | 0.023308 | 4.20E-07 |
| B98 | B135 | anti-Jo-1 | TBA | -0.02308 | 5.41E-07 |
| B99 | B181 | anti-PM-Scl | APTT | -0.05231 | 0 |
| B99 | B138 | anti-PM-Scl | PAB | -0.03019 | 5.59E-11 |
| B99 | B134 | anti-PM-Scl | CHE | -0.02956 | 1.39E-10 |
| B99 | B196 | anti-PM-Scl | TCO2 | -0.02613 | 1.41E-08 |
| B99 | B183 | anti-PM-Scl | PTA | -0.02608 | 1.49E-08 |

**Supplementary Table 4. PEI Correlations in Hypertension+Diabetes**

| Row | Column | Row | Column | Cor | *P* |
|---|---|---|---|---|---|
| B1 | B127 | Gender | A-G ratio | -0.12851 | 0 |
| B1 | B128 | Gender | ALT | -0.10755 | 0 |
| B1 | B129 | Gender | IBIL | -0.1385 | 0 |
| B1 | B130 | Gender | GLB | 0.134163 | 0 |
| B1 | B132 | Gender | Dbil | -0.14545 | 0 |
| B1 | B133 | Gender | TBIL | -0.15168 | 0 |
| B1 | B140 | Gender | r-GT | -0.0897 | 0 |
| B1 | B141 | Gender | AST/ALT | 0.153424 | 0 |
| B1 | B161 | Gender | LEU | 0.231079 | 0 |
| B1 | B163 | Gender | SG | -0.3312 | 0 |
| B1 | B166 | Gender | Pro | -0.11354 | 0 |
| B1 | B170 | Gender | U-GLU | -0.13798 | 0 |
| B1 | B173 | Gender | NTT | 0.161704 | 0 |
| B1 | B174 | Gender | Urea/Cr | 0.383518 | 0 |
| B1 | B176 | Gender | Ucr | -0.09721 | 0 |
| B1 | B190 | Gender | Cre | -0.21401 | 0 |
| B1 | B192 | Gender | UA | -0.21452 | 0 |
| B1 | B2 | Gender | Age | 0.153036 | 0 |
| B1 | B204 | Gender | Drinking | -0.26344 | 0 |
| B1 | B205 | Gender | KDA | 0.091752 | 0 |
| B1 | B206 | Gender | Waist | -0.27292 | 0 |
| B1 | B207 | Gender | WHR | -0.19793 | 0 |
| B1 | B209 | Gender | Smoking | -0.30681 | 0 |
| B1 | B210 | Gender | Hip | -0.20304 | 0 |
| B1 | B216 | Gender | BMI | -0.49449 | 0 |
| B1 | B218 | Gender | DBP | -0.16018 | 0 |
| B1 | B219 | Gender | OT | 0.117138 | 0 |
| B1 | B220 | Gender | SBP | 0.09064 | 0 |
| B1 | B221 | Gender | Hei | -0.70189 | 0 |
| B1 | B222 | Gender | NS | -0.40299 | 0 |
| B1 | B3 | Gender | Hcy | -0.09383 | 0 |
| B1 | B41 | Gender | MON % | -0.18094 | 0 |
| B1 | B42 | Gender | MON | -0.18464 | 0 |
| B1 | B44 | Gender | MCH | -0.15223 | 0 |
| B1 | B45 | Gender | MCHC | -0.26604 | 0 |
| B1 | B46 | Gender | RBC Count | -0.33072 | 0 |
| B1 | B48 | Gender | LYM% | 0.1453 | 0 |
| B1 | B50 | Gender | eo% | -0.11451 | 0 |
| B1 | B51 | Gender | Hb | -0.44565 | 0 |
| B1 | B52 | Gender | PLT/BPC | 0.131197 | 0 |
| B1 | B72 | Gender | HDL-C | 0.258304 | 0 |

| | | | | | |
|---|---|---|---|---|---|
| B1 | B74 | Gender | ApoA1 | 0.120705 | 0 |
| B1 | B77 | Gender | TC | 0.154559 | 0 |
| B1 | B91 | Gender | TP | 0.089766 | 0 |
| B1 | B70 | Gender | LDL-C | 0.087694 | 4.44E-16 |
| B1 | B215 | Gender | BMI | -0.08593 | 1.55E-15 |
| B1 | B62 | Gender | WBHSRVI | -0.07919 | 2.00E-13 |
| B1 | B60 | Gender | WBRVI | -0.07645 | 1.31E-12 |
| B1 | B149 | Gender | TGAb | 0.075437 | 2.58E-12 |
| B1 | B64 | Gender | WBRSR(mPas_5) | -0.07081 | 5.07E-11 |
| B1 | B65 | Gender | WBRSR(mPas_1) | -0.06971 | 1.00E-10 |
| B1 | B49 | Gender | baso% | -0.06812 | 2.64E-10 |
| B1 | B53 | Gender | GR% | -0.06736 | 4.16E-10 |
| B1 | B67 | Gender | WBRSR(mPas_50) | -0.06692 | 5.41E-10 |
| B1 | B66 | Gender | WBRSR(mPas_100) | -0.06601 | 9.23E-10 |
| B1 | B63 | Gender | WBRSR(mPas_200) | -0.06529 | 1.40E-09 |
| B1 | B114 | Gender | TSH TT3 TT4(TT4) | 0.064045 | 2.86E-09 |
| B1 | B148 | Gender | Anti_TPO | 0.060982 | 1.56E-08 |
| B1 | B40 | Gender | WBC | -0.0603 | 2.25E-08 |
| B1 | B201 | Gender | e-habits | -0.05701 | 1.25E-07 |
| B1 | B86 | Gender | TBhCG | 0.056362 | 1.73E-07 |
| B1 | B172 | Gender | KB | -0.05545 | 2.73E-07 |
| B1 | B56 | Gender | IR | -0.05418 | 5.09E-07 |
| B1 | B54 | Gender | ESR | 0.053542 | 6.90E-07 |
| B10 | B11 | PG I | PGI/PGII | 0.137877 | 0 |
| B10 | B12 | PG I | PG II | 0.489321 | 0 |
| B10 | B189 | PG I | eGFB | -0.1358 | 0 |
| B10 | B190 | PG I | Cre | 0.174876 | 0 |
| B10 | B191 | PG I | Urea | 0.148788 | 0 |
| B10 | B46 | PG I | RBC Count | -0.08805 | 4.44E-16 |
| B10 | B145 | PG I | BGP | -0.08574 | 1.78E-15 |
| B10 | B51 | PG I | Hb | -0.07726 | 7.55E-13 |
| B10 | B126 | PG I | Alb | -0.05962 | 3.22E-08 |
| B10 | B80 | PG I | anti-HP IgG | 0.055506 | 2.65E-07 |
| B10 | B144 | PG I | Total-PINP | -0.05335 | 7.58E-07 |
| B100 | B101 | anti-Ro52 | anti-Sc70 | 0.163075 | 0 |
| B100 | B104 | anti-Ro52 | anti-SSB | 0.163075 | 0 |
| B100 | B107 | anti-Ro52 | AnuA | 0.161154 | 0 |
| B100 | B109 | anti-Ro52 | AMA-M2 | 0.94335 | 0 |
| B100 | B136 | anti-Ro52 | 5'NT | 0.197629 | 0 |
| B100 | B138 | anti-Ro52 | PAB | -0.08191 | 2.93E-14 |
| B100 | B139 | anti-Ro52 | ADA | 0.074712 | 4.16E-12 |
| B100 | B184 | anti-Ro52 | INR | 0.072702 | 1.54E-11 |
| B100 | B198 | anti-Ro52 | P | 0.068092 | 2.69E-10 |
| B100 | B183 | anti-Ro52 | PTA | 0.065635 | 1.15E-09 |

| | | | | | |
|---|---|---|---|---|---|
| B100 | B119 | anti-Ro52 | Hcv-rna | 0.065257 | 1.43E-09 |
| B100 | B150 | anti-Ro52 | ALP | 0.053938 | 5.71E-07 |
| B100 | B140 | anti-Ro52 | r-GT | 0.053858 | 5.93E-07 |
| B100 | B161 | anti-Ro52 | LEU | 0.053185 | 8.18E-07 |
| B101 | B107 | anti-Sc70 | AnuA | 0.99526 | 0 |
| B101 | B109 | anti-Sc70 | AMA-M2 | 0.285121 | 0 |
| B101 | B115 | anti-Sc70 | TSH TT3 TT4(TT3) | 0.097149 | 0 |
| B101 | B119 | anti-Sc70 | Hcv-rna | 0.22665 | 0 |
| B101 | B124 | anti-Sc70 | ACTH | -0.1739 | 0 |
| B101 | B138 | anti-Sc70 | PAB | -0.12155 | 0 |
| B101 | B155 | anti-Sc70 | IgA | -0.11956 | 0 |
| B101 | B157 | anti-Sc70 | IgG | 0.102519 | 0 |
| B101 | B158 | anti-Sc70 | IgM | 0.152403 | 0 |
| B101 | B181 | anti-Sc70 | APTT | 0.174055 | 0 |
| B101 | B182 | anti-Sc70 | TT | 0.095817 | 0 |
| B101 | B184 | anti-Sc70 | INR | 0.270459 | 0 |
| B101 | B185 | anti-Sc70 | PT | 0.1744 | 0 |
| B101 | B186 | anti-Sc70 | PFC | -0.11045 | 0 |
| B101 | B194 | anti-Sc70 | Cl | 0.185271 | 0 |
| B101 | B195 | anti-Sc70 | Na | 0.134201 | 0 |
| B101 | B196 | anti-Sc70 | TCO2 | 0.119685 | 0 |
| B101 | B197 | anti-Sc70 | AG | -0.14837 | 0 |
| B101 | B199 | anti-Sc70 | Mg | 0.168027 | 0 |
| B101 | B200 | anti-Sc70 | OP | -0.11588 | 0 |
| B101 | B179 | anti-Sc70 | α1-MG | -0.08204 | 2.66E-14 |
| B101 | B136 | anti-Sc70 | 5'NT | -0.08078 | 6.57E-14 |
| B101 | B178 | anti-Sc70 | Ig-U | -0.07759 | 6.05E-13 |
| B101 | B139 | anti-Sc70 | ADA | -0.07705 | 8.74E-13 |
| B101 | B180 | anti-Sc70 | Tf | -0.0763 | 1.45E-12 |
| B101 | B125 | anti-Sc70 | Cor | -0.0744 | 5.11E-12 |
| B101 | B189 | anti-Sc70 | eGFB | -0.07423 | 5.72E-12 |
| B101 | B121 | anti-Sc70 | C3 | -0.07191 | 2.54E-11 |
| B101 | B212 | anti-Sc70 | IOP-L | -0.06297 | 5.22E-09 |
| B101 | B102 | anti-Sc70 | anti-Sm | -0.06065 | 1.86E-08 |
| B101 | B103 | anti-Sc70 | anti-SSA/Ro | 0.056307 | 1.78E-07 |
| B101 | B211 | anti-Sc70 | IOP-R | -0.05567 | 2.45E-07 |
| B102 | B103 | anti-Sm | anti-SSA/Ro | 0.776383 | 0 |
| B102 | B105 | anti-Sm | anti-U1RNP | 0.997977 | 0 |
| B102 | B151 | anti-Sm | Anti-CCP | 0.077406 | 6.86E-13 |
| B102 | B107 | anti-Sm | AnuA | -0.06128 | 1.33E-08 |
| B102 | B104 | anti-Sm | anti-SSB | -0.06065 | 1.86E-08 |
| B103 | B105 | anti-SSA/Ro | anti-U1RNP | 0.781397 | 0 |
| B103 | B124 | anti-SSA/Ro | ACTH | 0.207708 | 0 |
| B103 | B182 | anti-SSA/Ro | TT | -0.14512 | 0 |

| | | | | | |
|---|---|---|---|---|---|
| B103 | B151 | anti-SSA/Ro | Anti-CCP | 0.084699 | 3.77E-15 |
| B103 | B185 | anti-SSA/Ro | PT | -0.08089 | 6.06E-14 |
| B103 | B104 | anti-SSA/Ro | anti-SSB | 0.056307 | 1.78E-07 |
| B103 | B107 | anti-SSA/Ro | AnuA | 0.054593 | 4.16E-07 |
| B104 | B107 | anti-SSB | AnuA | 0.995262 | 0 |
| B104 | B109 | anti-SSB | AMA-M2 | 0.285121 | 0 |
| B104 | B115 | anti-SSB | TSH TT3 TT4(TT3) | 0.097149 | 0 |
| B104 | B119 | anti-SSB | Hcv-rna | 0.22665 | 0 |
| B104 | B124 | anti-SSB | ACTH | -0.1739 | 0 |
| B104 | B138 | anti-SSB | PAB | -0.12155 | 0 |
| B104 | B155 | anti-SSB | IgA | -0.11956 | 0 |
| B104 | B157 | anti-SSB | IgG | 0.102519 | 0 |
| B104 | B158 | anti-SSB | IgM | 0.152403 | 0 |
| B104 | B181 | anti-SSB | APTT | 0.174055 | 0 |
| B104 | B182 | anti-SSB | TT | 0.095817 | 0 |
| B104 | B184 | anti-SSB | INR | 0.270459 | 0 |
| B104 | B185 | anti-SSB | PT | 0.1744 | 0 |
| B104 | B186 | anti-SSB | PFC | -0.11045 | 0 |
| B104 | B194 | anti-SSB | Cl | 0.185271 | 0 |
| B104 | B195 | anti-SSB | Na | 0.134201 | 0 |
| B104 | B196 | anti-SSB | TCO2 | 0.119685 | 0 |
| B104 | B197 | anti-SSB | AG | -0.14837 | 0 |
| B104 | B199 | anti-SSB | Mg | 0.168027 | 0 |
| B104 | B200 | anti-SSB | OP | -0.11588 | 0 |
| B104 | B179 | anti-SSB | α1-MG | -0.08204 | 2.66E-14 |
| B104 | B136 | anti-SSB | 5'NT | -0.08078 | 6.57E-14 |
| B104 | B178 | anti-SSB | Ig-U | -0.07759 | 6.05E-13 |
| B104 | B139 | anti-SSB | ADA | -0.07705 | 8.74E-13 |
| B104 | B180 | anti-SSB | Tf | -0.0763 | 1.45E-12 |
| B104 | B125 | anti-SSB | Cor | -0.0744 | 5.11E-12 |
| B104 | B189 | anti-SSB | eGFB | -0.07423 | 5.72E-12 |
| B104 | B121 | anti-SSB | C3 | -0.07191 | 2.54E-11 |
| B104 | B212 | anti-SSB | IOP-L | -0.06297 | 5.22E-09 |
| B104 | B211 | anti-SSB | IOP-R | -0.05567 | 2.45E-07 |
| B105 | B151 | anti-U1RNP | Anti-CCP | 0.078292 | 3.74E-13 |
| B106 | B194 | anti-P | Cl | 0.085267 | 2.44E-15 |
| B106 | B197 | anti-P | AG | -0.06377 | 3.34E-09 |
| B107 | B109 | AnuA | AMA-M2 | 0.28289 | 0 |
| B107 | B115 | AnuA | TSH TT3 TT4(TT3) | 0.099071 | 0 |
| B107 | B119 | AnuA | Hcv-rna | 0.227775 | 0 |
| B107 | B124 | AnuA | ACTH | -0.17474 | 0 |
| B107 | B138 | AnuA | PAB | -0.12213 | 0 |
| B107 | B155 | AnuA | IgA | -0.12015 | 0 |
| B107 | B157 | AnuA | IgG | 0.103022 | 0 |

| | | | | | |
|---|---|---|---|---|---|
| B107 | B158 | AnuA | IgM | 0.153164 | 0 |
| B107 | B181 | AnuA | APTT | 0.174909 | 0 |
| B107 | B182 | AnuA | TT | 0.09628 | 0 |
| B107 | B184 | AnuA | INR | 0.27178 | 0 |
| B107 | B185 | AnuA | PT | 0.175248 | 0 |
| B107 | B186 | AnuA | PFC | -0.11101 | 0 |
| B107 | B194 | AnuA | Cl | 0.186205 | 0 |
| B107 | B195 | AnuA | Na | 0.13484 | 0 |
| B107 | B196 | AnuA | TCO2 | 0.120255 | 0 |
| B107 | B197 | AnuA | AG | -0.14912 | 0 |
| B107 | B199 | AnuA | Mg | 0.16892 | 0 |
| B107 | B200 | AnuA | OP | -0.11655 | 0 |
| B107 | B179 | AnuA | α1-MG | -0.08247 | 1.95E-14 |
| B107 | B136 | AnuA | 5'NT | -0.08117 | 4.97E-14 |
| B107 | B178 | AnuA | Ig-U | -0.07798 | 4.63E-13 |
| B107 | B139 | AnuA | ADA | -0.07751 | 6.40E-13 |
| B107 | B180 | AnuA | Tf | -0.07668 | 1.12E-12 |
| B107 | B125 | AnuA | Cor | -0.07477 | 4.01E-12 |
| B107 | B189 | AnuA | eGFB | -0.07365 | 8.35E-12 |
| B107 | B121 | AnuA | C3 | -0.07225 | 2.04E-11 |
| B107 | B212 | AnuA | IOP-L | -0.06206 | 8.66E-09 |
| B107 | B211 | AnuA | IOP-R | -0.0552 | 3.09E-07 |
| B108 | B151 | Anti-Cyclin | Anti-CCP | -0.10356 | 0 |
| B108 | B185 | Anti-Cyclin | PT | -0.09646 | 0 |
| B108 | B199 | Anti-Cyclin | Mg | -0.12652 | 0 |
| B108 | B145 | Anti-Cyclin | BGP | -0.0676 | 3.60E-10 |
| B108 | B195 | Anti-Cyclin | Na | -0.06676 | 5.95E-10 |
| B108 | B125 | Anti-Cyclin | Cor | -0.06471 | 1.95E-09 |
| B108 | B137 | Anti-Cyclin | AFU | 0.055259 | 3.00E-07 |
| B108 | B176 | Anti-Cyclin | Ucr | -0.05487 | 3.63E-07 |
| B108 | B144 | Anti-Cyclin | Total-PINP | -0.0548 | 3.76E-07 |
| B108 | B182 | Anti-Cyclin | TT | -0.05429 | 4.81E-07 |
| B109 | B136 | AMA-M2 | 5'NT | 0.201726 | 0 |
| B109 | B184 | AMA-M2 | INR | 0.102764 | 0 |
| B109 | B138 | AMA-M2 | PAB | -0.08797 | 4.44E-16 |
| B109 | B119 | AMA-M2 | Hcv-rna | 0.086293 | 1.11E-15 |
| B109 | B185 | AMA-M2 | PT | 0.066219 | 8.16E-10 |
| B109 | B181 | AMA-M2 | APTT | 0.06616 | 8.45E-10 |
| B109 | B124 | AMA-M2 | ACTH | -0.06593 | 9.65E-10 |
| B109 | B198 | AMA-M2 | P | 0.065495 | 1.24E-09 |
| B109 | B200 | AMA-M2 | OP | -0.06375 | 3.37E-09 |
| B109 | B150 | AMA-M2 | ALP | 0.05941 | 3.61E-08 |
| B109 | B158 | AMA-M2 | IgM | 0.05807 | 7.26E-08 |
| B109 | B139 | AMA-M2 | ADA | 0.055115 | 3.22E-07 |

| | | | | | |
|---|---|---|---|---|---|
| B109 | B161 | AMA-M2 | LEU | 0.054432 | 4.50E-07 |
| B109 | B140 | AMA-M2 | r-GT | 0.05296 | 9.11E-07 |
| B11 | B12 | PGI/PGII | PG II | -0.38597 | 0 |
| B11 | B14 | PGI/PGII | G-17 | -0.257 | 0 |
| B11 | B80 | PGI/PGII | anti-HP IgG | -0.12218 | 0 |
| B11 | B145 | PGI/PGII | BGP | -0.08747 | 4.44E-16 |
| B11 | B144 | PGI/PGII | Total-PINP | -0.06283 | 5.65E-09 |
| B11 | B149 | PGI/PGII | TGAb | -0.05882 | 4.92E-08 |
| B11 | B143 | PGI/PGII | β-CTx | -0.0581 | 7.14E-08 |
| B110 | B119 | AHA | Hcv-rna | -0.1391 | 0 |
| B110 | B150 | AHA | ALP | 0.095019 | 0 |
| B110 | B194 | AHA | Cl | -0.1925 | 0 |
| B110 | B195 | AHA | Na | -0.1366 | 0 |
| B110 | B200 | AHA | OP | -0.09559 | 0 |
| B110 | B228 | AHA | OS | 0.125856 | 0 |
| B111 | B112 | TSH | FT4 | -0.21582 | 0 |
| B111 | B113 | TSH | FT3 | -0.16747 | 0 |
| B111 | B114 | TSH | TSH TT3 TT4(TT4) | -0.10289 | 0 |
| B111 | B139 | TSH | ADA | 0.110945 | 0 |
| B111 | B148 | TSH | Anti_TPO | 0.167898 | 0 |
| B111 | B149 | TSH | TGAb | 0.181257 | 0 |
| B111 | B115 | TSH | TSH TT3 TT4(TT3) | -0.08865 | 2.22E-16 |
| B111 | B131 | TSH | AST | 0.058975 | 4.54E-08 |
| B112 | B113 | FT4 | FT3 | 0.182664 | 0 |
| B112 | B114 | FT4 | TSH TT3 TT4(TT4) | 0.522105 | 0 |
| B112 | B115 | FT4 | TSH TT3 TT4(TT3) | 0.19654 | 0 |
| B112 | B189 | FT4 | eGFB | 0.064772 | 1.89E-09 |
| B112 | B199 | FT4 | Mg | -0.05551 | 2.65E-07 |
| B113 | B114 | FT3 | TSH TT3 TT4(TT4) | 0.249349 | 0 |
| B113 | B115 | FT3 | TSH TT3 TT4(TT3) | 0.518979 | 0 |
| B113 | B126 | FT3 | Alb | 0.097472 | 0 |
| B113 | B145 | FT3 | BGP | 0.104127 | 0 |
| B113 | B189 | FT3 | eGFB | 0.093612 | 0 |
| B113 | B191 | FT3 | Urea | -0.08825 | 2.22E-16 |
| B113 | B123 | FT3 | hs-CRP1 | -0.07409 | 6.26E-12 |
| B113 | B190 | FT3 | Cre | -0.07354 | 8.92E-12 |
| B113 | B176 | FT3 | Ucr | 0.070766 | 5.22E-11 |
| B113 | B146 | FT3 | CysC | -0.06563 | 1.15E-09 |
| B113 | B134 | FT3 | CHE | 0.061747 | 1.03E-08 |
| B113 | B144 | FT3 | Total-PINP | 0.058829 | 4.90E-08 |
| B113 | B127 | FT3 | A-G ratio | 0.056024 | 2.05E-07 |
| B113 | B195 | FT3 | Na | 0.054276 | 4.85E-07 |
| B114 | B115 | TSH TT3 TT4(TT4) | TSH TT3 TT4(TT3) | 0.436263 | 0 |
| B114 | B199 | TSH TT3 TT4(TT4) | Mg | -0.06472 | 1.94E-09 |

| | | | | | |
|---|---|---|---|---|---|
| B114 | B136 | TSH TT3 TT4(TT4) | 5'NT | 0.064671 | 2.00E-09 |
| B114 | B225 | TSH TT3 TT4(TT4) | hPBG | 0.059038 | 4.39E-08 |
| B114 | B156 | TSH TT3 TT4(TT4) | IgE | 0.057742 | 8.60E-08 |
| B114 | B149 | TSH TT3 TT4(TT4) | TGAb | 0.053852 | 5.95E-07 |
| B115 | B123 | TSH TT3 TT4(TT3) | hs-CRP1 | -0.09596 | 0 |
| B115 | B200 | TSH TT3 TT4(TT3) | OP | -0.09334 | 0 |
| B115 | B191 | TSH TT3 TT4(TT3) | Urea | -0.08672 | 8.88E-16 |
| B115 | B190 | TSH TT3 TT4(TT3) | Cre | -0.0831 | 1.24E-14 |
| B115 | B146 | TSH TT3 TT4(TT3) | CysC | -0.08143 | 4.13E-14 |
| B115 | B119 | TSH TT3 TT4(TT3) | Hcv-rna | 0.079472 | 1.65E-13 |
| B115 | B196 | TSH TT3 TT4(TT3) | TCO2 | 0.078929 | 2.41E-13 |
| B115 | B136 | TSH TT3 TT4(TT3) | 5'NT | 0.070333 | 6.84E-11 |
| B115 | B199 | TSH TT3 TT4(TT3) | Mg | 0.062089 | 8.52E-09 |
| B115 | B186 | TSH TT3 TT4(TT3) | PFC | -0.06178 | 1.01E-08 |
| B115 | B189 | TSH TT3 TT4(TT3) | eGFB | 0.060903 | 1.63E-08 |
| B115 | B179 | TSH TT3 TT4(TT3) | α1-MG | -0.05746 | 9.92E-08 |
| B115 | B134 | TSH TT3 TT4(TT3) | CHE | 0.054191 | 5.05E-07 |
| B115 | B126 | TSH TT3 TT4(TT3) | Alb | 0.053753 | 6.24E-07 |
| B116 | B117 | TT3 | TT4 | -0.42507 | 0 |
| B118 | B189 | CEA1 | eGFB | -0.07469 | 4.21E-12 |
| B118 | B126 | CEA1 | Alb | -0.06618 | 8.33E-10 |
| B119 | B120 | Hcv-rna | HCV-cAg | -0.15138 | 0 |
| B119 | B124 | Hcv-rna | ACTH | -0.14992 | 0 |
| B119 | B157 | Hcv-rna | IgG | 0.096414 | 0 |
| B119 | B183 | Hcv-rna | PTA | 0.11187 | 0 |
| B119 | B184 | Hcv-rna | INR | 0.120911 | 0 |
| B119 | B194 | Hcv-rna | Cl | 0.16201 | 0 |
| B119 | B197 | Hcv-rna | AG | -0.11004 | 0 |
| B119 | B186 | Hcv-rna | PFC | -0.08131 | 4.51E-14 |
| B119 | B181 | Hcv-rna | APTT | 0.077338 | 7.18E-13 |
| B119 | B179 | Hcv-rna | α1-MG | -0.07723 | 7.74E-13 |
| B119 | B200 | Hcv-rna | OP | -0.07118 | 4.02E-11 |
| B119 | B193 | Hcv-rna | K | 0.066158 | 8.46E-10 |
| B119 | B121 | Hcv-rna | C3 | -0.06546 | 1.27E-09 |
| B119 | B182 | Hcv-rna | TT | 0.06529 | 1.40E-09 |
| B119 | B185 | Hcv-rna | PT | 0.064618 | 2.06E-09 |
| B119 | B139 | Hcv-rna | ADA | -0.06335 | 4.23E-09 |
| B119 | B211 | Hcv-rna | IOP-R | -0.06318 | 4.65E-09 |
| B119 | B212 | Hcv-rna | IOP-L | -0.06305 | 4.99E-09 |
| B119 | B151 | Hcv-rna | Anti-CCP | -0.059 | 4.47E-08 |
| B119 | B134 | Hcv-rna | CHE | 0.058918 | 4.67E-08 |
| B119 | B189 | Hcv-rna | eGFB | -0.05404 | 5.42E-07 |
| B12 | B14 | PG II | G-17 | 0.478576 | 0 |
| B12 | B189 | PG II | eGFB | -0.10346 | 0 |

| | | | | | |
|---|---|---|---|---|---|
| B12 | B190 | PG II | Cre | 0.093293 | 0 |
| B12 | B80 | PG II | anti-HP IgG | 0.126273 | 0 |
| B12 | B191 | PG II | Urea | 0.078561 | 3.10E-13 |
| B120 | B155 | HCV-cAg | IgA | 0.151734 | 0 |
| B120 | B158 | HCV-cAg | IgM | 0.354185 | 0 |
| B120 | B181 | HCV-cAg | APTT | 0.106332 | 0 |
| B120 | B184 | HCV-cAg | INR | -0.18451 | 0 |
| B120 | B185 | HCV-cAg | PT | -0.1221 | 0 |
| B120 | B186 | HCV-cAg | PFC | 0.235194 | 0 |
| B120 | B195 | HCV-cAg | Na | -0.09503 | 0 |
| B120 | B198 | HCV-cAg | P | 0.14511 | 0 |
| B120 | B199 | HCV-cAg | Mg | 0.097477 | 0 |
| B120 | B194 | HCV-cAg | Cl | -0.07945 | 1.67E-13 |
| B120 | B134 | HCV-cAg | CHE | 0.072499 | 1.75E-11 |
| B120 | B182 | HCV-cAg | TT | 0.068573 | 2.01E-10 |
| B120 | B124 | HCV-cAg | ACTH | -0.06568 | 1.12E-09 |
| B120 | B196 | HCV-cAg | TCO2 | -0.06161 | 1.11E-08 |
| B120 | B183 | HCV-cAg | PTA | -0.05564 | 2.48E-07 |
| B121 | B122 | C3 | C4 | 0.888077 | 0 |
| B121 | B134 | C3 | CHE | 0.103215 | 0 |
| B121 | B194 | C3 | Cl | -0.09202 | 0 |
| B121 | B155 | C3 | IgA | 0.056258 | 1.83E-07 |
| B122 | B134 | C4 | CHE | 0.096094 | 0 |
| B122 | B194 | C4 | Cl | -0.07536 | 2.71E-12 |
| B122 | B158 | C4 | IgM | 0.062493 | 6.81E-09 |
| B122 | B155 | C4 | IgA | 0.058841 | 4.86E-08 |
| B122 | B156 | C4 | IgE | -0.05324 | 7.98E-07 |
| B123 | B125 | hs-CRP1 | Cor | 0.142839 | 0 |
| B123 | B179 | hs-CRP1 | α1-MG | 0.197096 | 0 |
| B123 | B186 | hs-CRP1 | PFC | 0.200928 | 0 |
| B123 | B126 | hs-CRP1 | Alb | -0.08027 | 9.39E-14 |
| B123 | B127 | hs-CRP1 | A-G ratio | -0.07014 | 7.72E-11 |
| B123 | B176 | hs-CRP1 | Ucr | 0.066032 | 9.11E-10 |
| B123 | B124 | hs-CRP1 | ACTH | 0.061317 | 1.30E-08 |
| B123 | B157 | hs-CRP1 | IgG | -0.05821 | 6.74E-08 |
| B123 | B194 | hs-CRP1 | Cl | -0.05778 | 8.44E-08 |
| B123 | B150 | hs-CRP1 | ALP | 0.056859 | 1.35E-07 |
| B123 | B130 | hs-CRP1 | GLB | 0.056832 | 1.37E-07 |
| B123 | B153 | hs-CRP1 | Glu | 0.056702 | 1.46E-07 |
| B123 | B175 | hs-CRP1 | ALB-U | 0.054019 | 5.49E-07 |
| B124 | B125 | ACTH | Cor | 0.337736 | 0 |
| B124 | B182 | ACTH | TT | -0.1231 | 0 |
| B124 | B183 | ACTH | PTA | -0.12986 | 0 |
| B124 | B184 | ACTH | INR | -0.10328 | 0 |

| | | | | | |
|---|---|---|---|---|---|
| B124 | B186 | ACTH | PFC | 0.158316 | 0 |
| B124 | B181 | ACTH | APTT | -0.0658 | 1.04E-09 |
| B124 | B194 | ACTH | Cl | -0.06212 | 8.38E-09 |
| B124 | B212 | ACTH | IOP-L | 0.059668 | 3.15E-08 |
| B124 | B155 | ACTH | IgA | -0.05661 | 1.53E-07 |
| B124 | B151 | ACTH | Anti-CCP | 0.056213 | 1.87E-07 |
| B124 | B137 | ACTH | AFU | -0.05372 | 6.34E-07 |
| B125 | B159 | Cor | β2-MG | -0.21125 | 0 |
| B125 | B181 | Cor | APTT | -0.16551 | 0 |
| B125 | B184 | Cor | INR | -0.15857 | 0 |
| B125 | B186 | Cor | PFC | 0.18127 | 0 |
| B125 | B225 | Cor | hPBG | 0.164001 | 0 |
| B125 | B183 | Cor | PTA | -0.08522 | 2.66E-15 |
| B125 | B156 | Cor | IgE | -0.06822 | 2.48E-10 |
| B125 | B151 | Cor | Anti-CCP | 0.061685 | 1.06E-08 |
| B125 | B157 | Cor | IgG | -0.06168 | 1.06E-08 |
| B125 | B155 | Cor | IgA | -0.05852 | 5.76E-08 |
| B126 | B127 | Alb | A-G ratio | 0.523887 | 0 |
| B126 | B128 | Alb | ALT | 0.123123 | 0 |
| B126 | B129 | Alb | IBIL | 0.158081 | 0 |
| B126 | B130 | Alb | GLB | -0.20006 | 0 |
| B126 | B133 | Alb | TBIL | 0.150828 | 0 |
| B126 | B138 | Alb | PAB | 0.103262 | 0 |
| B126 | B141 | Alb | AST/ALT | -0.13606 | 0 |
| B126 | B146 | Alb | CysC | -0.13226 | 0 |
| B126 | B175 | Alb | ALB-U | -0.11669 | 0 |
| B126 | B177 | Alb | ACR | -0.13885 | 0 |
| B126 | B188 | Alb | Ca | 0.35527 | 0 |
| B126 | B189 | Alb | eGFB | 0.273514 | 0 |
| B126 | B190 | Alb | Cre | -0.16744 | 0 |
| B126 | B191 | Alb | Urea | -0.20664 | 0 |
| B126 | B218 | Alb | DBP | 0.143708 | 0 |
| B126 | B221 | Alb | Hei | 0.111628 | 0 |
| B126 | B132 | Alb | Dbil | 0.088526 | 2.22E-16 |
| B126 | B178 | Alb | Ig-U | -0.08629 | 1.11E-15 |
| B126 | B204 | Alb | Drinking | 0.083697 | 7.99E-15 |
| B126 | B216 | Alb | BMI | 0.081147 | 5.06E-14 |
| B126 | B180 | Alb | Tf | -0.08051 | 7.93E-14 |
| B126 | B166 | Alb | Pro | -0.07869 | 2.83E-13 |
| B126 | B134 | Alb | CHE | 0.062631 | 6.31E-09 |
| B126 | B173 | Alb | NTT | -0.0601 | 2.50E-08 |
| B126 | B131 | Alb | AST | 0.055088 | 3.26E-07 |
| B126 | B139 | Alb | ADA | -0.05504 | 3.34E-07 |
| B126 | B179 | Alb | α1-MG | -0.05308 | 8.59E-07 |

| | | | | | |
|---|---|---|---|---|---|
| B127 | B129 | A-G ratio | IBIL | 0.161975 | 0 |
| B127 | B130 | A-G ratio | GLB | -0.88609 | 0 |
| B127 | B132 | A-G ratio | Dbil | 0.13161 | 0 |
| B127 | B133 | A-G ratio | TBIL | 0.166505 | 0 |
| B127 | B141 | A-G ratio | AST/ALT | -0.12129 | 0 |
| B127 | B153 | A-G ratio | Glu | -0.11349 | 0 |
| B127 | B157 | A-G ratio | IgG | -0.09943 | 0 |
| B127 | B189 | A-G ratio | eGFB | 0.229236 | 0 |
| B127 | B190 | A-G ratio | Cre | -0.11513 | 0 |
| B127 | B191 | A-G ratio | Urea | -0.12758 | 0 |
| B127 | B204 | A-G ratio | Drinking | 0.111134 | 0 |
| B127 | B209 | A-G ratio | Smoking | 0.108884 | 0 |
| B127 | B216 | A-G ratio | BMI | 0.090908 | 0 |
| B127 | B220 | A-G ratio | SBP | -0.12114 | 0 |
| B127 | B221 | A-G ratio | Hei | 0.174514 | 0 |
| B127 | B146 | A-G ratio | CysC | -0.08795 | 4.44E-16 |
| B127 | B155 | A-G ratio | IgA | -0.08317 | 1.18E-14 |
| B127 | B192 | A-G ratio | UA | -0.07993 | 1.19E-13 |
| B127 | B163 | A-G ratio | SG | 0.077515 | 6.37E-13 |
| B127 | B150 | A-G ratio | ALP | -0.07728 | 7.49E-13 |
| B127 | B139 | A-G ratio | ADA | -0.0756 | 2.31E-12 |
| B127 | B138 | A-G ratio | PAB | 0.074009 | 6.60E-12 |
| B127 | B166 | A-G ratio | Pro | -0.06993 | 8.78E-11 |
| B127 | B140 | A-G ratio | r-GT | -0.06757 | 3.67E-10 |
| B127 | B177 | A-G ratio | ACR | -0.06107 | 1.48E-08 |
| B127 | B161 | A-G ratio | LEU | -0.05934 | 3.75E-08 |
| B127 | B188 | A-G ratio | Ca | 0.05742 | 1.01E-07 |
| B127 | B207 | A-G ratio | WHR | -0.0562 | 1.88E-07 |
| B127 | B131 | A-G ratio | AST | -0.05573 | 2.38E-07 |
| B127 | B222 | A-G ratio | NS | 0.053047 | 8.74E-07 |
| B128 | B131 | ALT | AST | 0.78266 | 0 |
| B128 | B132 | ALT | Dbil | 0.155954 | 0 |
| B128 | B133 | ALT | TBIL | 0.110425 | 0 |
| B128 | B140 | ALT | r-GT | 0.347904 | 0 |
| B128 | B141 | ALT | AST/ALT | -0.49505 | 0 |
| B128 | B163 | ALT | SG | 0.093193 | 0 |
| B128 | B189 | ALT | eGFB | 0.147465 | 0 |
| B128 | B191 | ALT | Urea | -0.10772 | 0 |
| B128 | B206 | ALT | Waist | 0.112597 | 0 |
| B128 | B215 | ALT | BMI | 0.109935 | 0 |
| B128 | B216 | ALT | BMI | 0.152465 | 0 |
| B128 | B218 | ALT | DBP | 0.105302 | 0 |
| B128 | B221 | ALT | Hei | 0.115145 | 0 |
| B128 | B210 | ALT | Hip | 0.082547 | 1.84E-14 |

| | | | | | |
|---|---|---|---|---|---|
| B128 | B129 | ALT | IBIL | 0.082491 | 1.91E-14 |
| B128 | B136 | ALT | 5'NT | 0.082324 | 2.18E-14 |
| B128 | B207 | ALT | WHR | 0.079105 | 2.13E-13 |
| B128 | B222 | ALT | NS | 0.079017 | 2.26E-13 |
| B128 | B204 | ALT | Drinking | 0.078563 | 3.10E-13 |
| B128 | B190 | ALT | Cre | -0.06806 | 2.74E-10 |
| B128 | B137 | ALT | AFU | 0.066885 | 5.52E-10 |
| B128 | B176 | ALT | Ucr | 0.063426 | 4.05E-09 |
| B128 | B139 | ALT | ADA | 0.058049 | 7.34E-08 |
| B128 | B188 | ALT | Ca | 0.057114 | 1.19E-07 |
| B128 | B192 | ALT | UA | 0.056377 | 1.72E-07 |
| B129 | B130 | IBIL | GLB | -0.12929 | 0 |
| B129 | B131 | IBIL | AST | 0.11458 | 0 |
| B129 | B132 | IBIL | Dbil | 0.62861 | 0 |
| B129 | B133 | IBIL | TBIL | 0.974239 | 0 |
| B129 | B163 | IBIL | SG | 0.091817 | 0 |
| B129 | B191 | IBIL | Urea | -0.12879 | 0 |
| B129 | B221 | IBIL | Hei | 0.128133 | 0 |
| B129 | B174 | IBIL | Urea/Cr | -0.0856 | 1.78E-15 |
| B129 | B190 | IBIL | Cre | -0.07919 | 2.01E-13 |
| B129 | B218 | IBIL | DBP | 0.071893 | 2.57E-11 |
| B129 | B216 | IBIL | BMI | 0.067074 | 4.93E-10 |
| B129 | B160 | IBIL | PH | 0.064828 | 1.83E-09 |
| B129 | B144 | IBIL | Total-PINP | -0.06166 | 1.07E-08 |
| B129 | B145 | IBIL | BGP | -0.05867 | 5.31E-08 |
| B129 | B143 | IBIL | β-CTx | -0.05317 | 8.25E-07 |
| B130 | B131 | GLB | AST | 0.095717 | 0 |
| B130 | B132 | GLB | Dbil | -0.10963 | 0 |
| B130 | B133 | GLB | TBIL | -0.13422 | 0 |
| B130 | B153 | GLB | Glu | 0.131622 | 0 |
| B130 | B155 | GLB | IgA | 0.099175 | 0 |
| B130 | B157 | GLB | IgG | 0.123394 | 0 |
| B130 | B189 | GLB | eGFB | -0.16273 | 0 |
| B130 | B192 | GLB | UA | 0.107441 | 0 |
| B130 | B204 | GLB | Drinking | -0.10191 | 0 |
| B130 | B209 | GLB | Smoking | -0.11527 | 0 |
| B130 | B220 | GLB | SBP | 0.129724 | 0 |
| B130 | B221 | GLB | Hei | -0.16547 | 0 |
| B130 | B141 | GLB | AST/ALT | 0.086842 | 8.88E-16 |
| B130 | B163 | GLB | SG | -0.08404 | 6.22E-15 |
| B130 | B216 | GLB | BMI | -0.08307 | 1.27E-14 |
| B130 | B140 | GLB | r-GT | 0.082697 | 1.64E-14 |
| B130 | B150 | GLB | ALP | 0.081325 | 4.44E-14 |
| B130 | B188 | GLB | Ca | 0.076089 | 1.67E-12 |

| | | | | | |
|---|---|---|---|---|---|
| B130 | B191 | GLB | Urea | 0.072085 | 2.28E-11 |
| B130 | B190 | GLB | Cre | 0.071282 | 3.78E-11 |
| B130 | B139 | GLB | ADA | 0.066078 | 8.86E-10 |
| B130 | B207 | GLB | WHR | 0.060884 | 1.64E-08 |
| B130 | B158 | GLB | IgM | 0.058247 | 6.63E-08 |
| B130 | B161 | GLB | LEU | 0.056633 | 1.51E-07 |
| B130 | B136 | GLB | 5'NT | 0.055338 | 2.88E-07 |
| B130 | B194 | GLB | Cl | -0.05519 | 3.11E-07 |
| B130 | B166 | GLB | Pro | 0.052955 | 9.13E-07 |
| B131 | B132 | AST | Dbil | 0.206544 | 0 |
| B131 | B133 | AST | TBIL | 0.150596 | 0 |
| B131 | B136 | AST | 5'NT | 0.11215 | 0 |
| B131 | B139 | AST | ADA | 0.090851 | 0 |
| B131 | B140 | AST | r-GT | 0.42592 | 0 |
| B131 | B141 | AST | AST/ALT | -0.10245 | 0 |
| B131 | B188 | AST | Ca | 0.080932 | 5.91E-14 |
| B131 | B191 | AST | Urea | -0.07934 | 1.81E-13 |
| B131 | B137 | AST | AFU | 0.078955 | 2.36E-13 |
| B131 | B192 | AST | UA | 0.075053 | 3.32E-12 |
| B131 | B204 | AST | Drinking | 0.070389 | 6.60E-11 |
| B131 | B215 | AST | BMI | 0.066383 | 7.41E-10 |
| B131 | B206 | AST | Waist | 0.06339 | 4.13E-09 |
| B131 | B189 | AST | eGFB | 0.060431 | 2.10E-08 |
| B131 | B190 | AST | Cre | -0.05837 | 6.22E-08 |
| B131 | B216 | AST | BMI | 0.05782 | 8.26E-08 |
| B132 | B133 | Dbil | TBIL | 0.787705 | 0 |
| B132 | B140 | Dbil | r-GT | 0.144084 | 0 |
| B132 | B191 | Dbil | Urea | -0.10764 | 0 |
| B132 | B221 | Dbil | Hei | 0.116554 | 0 |
| B132 | B174 | Dbil | Urea/Cr | -0.08106 | 5.37E-14 |
| B132 | B152 | Dbil | ACA-Ig(G-A-M) | 0.068314 | 2.35E-10 |
| B132 | B204 | Dbil | Drinking | 0.067252 | 4.44E-10 |
| B132 | B160 | Dbil | PH | 0.061508 | 1.17E-08 |
| B132 | B141 | Dbil | AST/ALT | -0.05809 | 7.19E-08 |
| B132 | B190 | Dbil | Cre | -0.05563 | 2.49E-07 |
| B132 | B135 | Dbil | TBA | 0.054602 | 4.14E-07 |
| B133 | B174 | TBIL | Urea/Cr | -0.09132 | 0 |
| B133 | B191 | TBIL | Urea | -0.1333 | 0 |
| B133 | B221 | TBIL | Hei | 0.135157 | 0 |
| B133 | B163 | TBIL | SG | 0.087178 | 4.44E-16 |
| B133 | B190 | TBIL | Cre | -0.07888 | 2.50E-13 |
| B133 | B218 | TBIL | DBP | 0.069555 | 1.10E-10 |
| B133 | B140 | TBIL | r-GT | 0.069375 | 1.23E-10 |
| B133 | B160 | TBIL | PH | 0.069055 | 1.50E-10 |

| | | | | | |
|---|---|---|---|---|---|
| B133 | B216 | TBIL | BMI | 0.066936 | 5.35E-10 |
| B133 | B152 | TBIL | ACA-Ig(G-A-M) | 0.060925 | 1.61E-08 |
| B133 | B145 | TBIL | BGP | -0.05377 | 6.20E-07 |
| B134 | B135 | CHE | TBA | -0.10557 | 0 |
| B134 | B136 | CHE | 5'NT | 0.167477 | 0 |
| B134 | B137 | CHE | AFU | 0.232748 | 0 |
| B134 | B138 | CHE | PAB | 0.250666 | 0 |
| B134 | B146 | CHE | CysC | -0.11392 | 0 |
| B134 | B184 | CHE | INR | -0.0898 | 0 |
| B134 | B194 | CHE | Cl | -0.18043 | 0 |
| B134 | B197 | CHE | AG | 0.101758 | 0 |
| B134 | B198 | CHE | P | -0.12829 | 0 |
| B134 | B151 | CHE | Anti-CCP | -0.08121 | 4.84E-14 |
| B134 | B216 | CHE | BMI | 0.0804 | 8.59E-14 |
| B134 | B185 | CHE | PT | -0.07931 | 1.85E-13 |
| B134 | B150 | CHE | ALP | 0.077365 | 7.05E-13 |
| B134 | B215 | CHE | BMI | 0.071693 | 2.92E-11 |
| B134 | B140 | CHE | r-GT | 0.071686 | 2.93E-11 |
| B134 | B186 | CHE | PFC | 0.065731 | 1.09E-09 |
| B134 | B195 | CHE | Na | -0.06561 | 1.16E-09 |
| B134 | B153 | CHE | Glu | 0.064508 | 2.19E-09 |
| B134 | B190 | CHE | Cre | -0.06344 | 4.02E-09 |
| B134 | B193 | CHE | K | -0.06181 | 9.95E-09 |
| B134 | B218 | CHE | DBP | 0.060132 | 2.46E-08 |
| B134 | B199 | CHE | Mg | -0.0587 | 5.25E-08 |
| B134 | B141 | CHE | AST/ALT | -0.05826 | 6.59E-08 |
| B134 | B206 | CHE | Waist | 0.055866 | 2.22E-07 |
| B134 | B170 | CHE | U-GLU | 0.054775 | 3.80E-07 |
| B135 | B137 | TBA | AFU | 0.122446 | 0 |
| B135 | B138 | TBA | PAB | -0.16254 | 0 |
| B135 | B139 | TBA | ADA | 0.157055 | 0 |
| B135 | B150 | TBA | ALP | 0.221218 | 0 |
| B135 | B157 | TBA | IgG | 0.134816 | 0 |
| B135 | B194 | TBA | Cl | 0.055092 | 3.26E-07 |
| B135 | B198 | TBA | P | 0.053514 | 7.00E-07 |
| B136 | B137 | 5'NT | AFU | 0.182733 | 0 |
| B136 | B139 | 5'NT | ADA | 0.184754 | 0 |
| B136 | B140 | 5'NT | r-GT | 0.310373 | 0 |
| B136 | B150 | 5'NT | ALP | 0.249265 | 0 |
| B136 | B155 | 5'NT | IgA | 0.105224 | 0 |
| B136 | B157 | 5'NT | IgG | 0.096301 | 0 |
| B136 | B182 | 5'NT | TT | 0.109363 | 0 |
| B136 | B184 | 5'NT | INR | -0.12153 | 0 |
| B136 | B193 | 5'NT | K | -0.15961 | 0 |

| | | | | | |
|---|---|---|---|---|---|
| B136 | B194 | 5'NT | Cl | -0.18477 | 0 |
| B136 | B197 | 5'NT | AG | 0.137573 | 0 |
| B136 | B198 | 5'NT | P | -0.13305 | 0 |
| B136 | B181 | 5'NT | APTT | -0.08546 | 2.22E-15 |
| B136 | B185 | 5'NT | PT | -0.06968 | 1.02E-10 |
| B136 | B199 | 5'NT | Mg | -0.06864 | 1.93E-10 |
| B136 | B195 | 5'NT | Na | -0.0677 | 3.39E-10 |
| B136 | B196 | 5'NT | TCO2 | -0.05465 | 4.04E-07 |
| B137 | B139 | AFU | ADA | 0.322652 | 0 |
| B137 | B140 | AFU | r-GT | 0.130002 | 0 |
| B137 | B150 | AFU | ALP | 0.165956 | 0 |
| B137 | B180 | AFU | Tf | 0.100361 | 0 |
| B137 | B194 | AFU | Cl | -0.17934 | 0 |
| B137 | B199 | AFU | Mg | -0.15935 | 0 |
| B137 | B200 | AFU | OP | 0.150628 | 0 |
| B137 | B155 | AFU | IgA | 0.08749 | 4.44E-16 |
| B137 | B178 | AFU | Ig-U | 0.088119 | 4.44E-16 |
| B137 | B204 | AFU | Drinking | 0.077102 | 8.43E-13 |
| B137 | B175 | AFU | ALB-U | 0.069408 | 1.21E-10 |
| B137 | B198 | AFU | P | -0.0683 | 2.38E-10 |
| B137 | B193 | AFU | K | -0.06655 | 6.72E-10 |
| B137 | B157 | AFU | IgG | 0.062453 | 6.96E-09 |
| B137 | B170 | AFU | U-GLU | 0.061713 | 1.05E-08 |
| B137 | B153 | AFU | Glu | 0.061076 | 1.48E-08 |
| B137 | B179 | AFU | α1-MG | 0.059522 | 3.40E-08 |
| B137 | B197 | AFU | AG | 0.058108 | 7.12E-08 |
| B137 | B163 | AFU | SG | 0.054735 | 3.88E-07 |
| B137 | B225 | AFU | hPBG | 0.053875 | 5.88E-07 |
| B137 | B195 | AFU | Na | -0.05377 | 6.20E-07 |
| B138 | B139 | PAB | ADA | -0.3112 | 0 |
| B138 | B150 | PAB | ALP | -0.15324 | 0 |
| B138 | B157 | PAB | IgG | -0.11377 | 0 |
| B138 | B194 | PAB | Cl | -0.14952 | 0 |
| B138 | B197 | PAB | AG | 0.129872 | 0 |
| B138 | B198 | PAB | P | -0.11826 | 0 |
| B138 | B185 | PAB | PT | -0.08273 | 1.62E-14 |
| B138 | B184 | PAB | INR | -0.07725 | 7.64E-13 |
| B138 | B155 | PAB | IgA | -0.07678 | 1.05E-12 |
| B138 | B212 | PAB | IOP-L | 0.073373 | 9.97E-12 |
| B138 | B204 | PAB | Drinking | 0.072738 | 1.50E-11 |
| B138 | B146 | PAB | CysC | -0.06913 | 1.43E-10 |
| B138 | B181 | PAB | APTT | -0.06436 | 2.39E-09 |
| B138 | B211 | PAB | IOP-R | 0.059634 | 3.21E-08 |
| B138 | B188 | PAB | Ca | 0.057983 | 7.60E-08 |

| | | | | | |
|---|---|---|---|---|---|
| B138 | B180 | PAB | Tf | 0.054446 | 4.46E-07 |
| B139 | B140 | ADA | r-GT | 0.101992 | 0 |
| B139 | B146 | ADA | CysC | 0.119859 | 0 |
| B139 | B150 | ADA | ALP | 0.259728 | 0 |
| B139 | B153 | ADA | Glu | 0.090296 | 0 |
| B139 | B155 | ADA | IgA | 0.098713 | 0 |
| B139 | B181 | ADA | APTT | -0.10043 | 0 |
| B139 | B186 | ADA | PFC | 0.091788 | 0 |
| B139 | B194 | ADA | Cl | -0.22164 | 0 |
| B139 | B197 | ADA | AG | 0.097127 | 0 |
| B139 | B199 | ADA | Mg | -0.21036 | 0 |
| B139 | B200 | ADA | OP | 0.323721 | 0 |
| B139 | B193 | ADA | K | -0.07496 | 3.54E-12 |
| B139 | B178 | ADA | Ig-U | 0.074858 | 3.78E-12 |
| B139 | B182 | ADA | TT | 0.068709 | 1.85E-10 |
| B139 | B170 | ADA | U-GLU | 0.068621 | 1.95E-10 |
| B139 | B148 | ADA | Anti_TPO | 0.068292 | 2.38E-10 |
| B139 | B180 | ADA | Tf | 0.066436 | 7.19E-10 |
| B139 | B177 | ADA | ACR | 0.065871 | 1.00E-09 |
| B139 | B195 | ADA | Na | -0.06296 | 5.25E-09 |
| B139 | B152 | ADA | ACA-Ig(G-A-M) | 0.060544 | 1.97E-08 |
| B139 | B184 | ADA | INR | -0.05885 | 4.84E-08 |
| B14 | B149 | G-17 | TGAb | 0.139228 | 0 |
| B14 | B148 | G-17 | Anti_TPO | 0.072761 | 1.48E-11 |
| B140 | B141 | r-GT | AST/ALT | -0.10962 | 0 |
| B140 | B150 | r-GT | ALP | 0.138885 | 0 |
| B140 | B194 | r-GT | Cl | -0.09249 | 0 |
| B140 | B204 | r-GT | Drinking | 0.131537 | 0 |
| B140 | B206 | r-GT | Waist | 0.090611 | 0 |
| B140 | B209 | r-GT | Smoking | 0.095859 | 0 |
| B140 | B216 | r-GT | BMI | 0.098027 | 0 |
| B140 | B153 | r-GT | Glu | 0.088684 | 2.22E-16 |
| B140 | B218 | r-GT | DBP | 0.086411 | 8.88E-16 |
| B140 | B189 | r-GT | eGFB | 0.0832 | 1.15E-14 |
| B140 | B192 | r-GT | UA | 0.082293 | 2.22E-14 |
| B140 | B163 | r-GT | SG | 0.080989 | 5.66E-14 |
| B140 | B207 | r-GT | WHR | 0.080854 | 6.22E-14 |
| B140 | B170 | r-GT | U-GLU | 0.080341 | 8.97E-14 |
| B140 | B215 | r-GT | BMI | 0.076983 | 9.14E-13 |
| B140 | B193 | r-GT | K | -0.07688 | 9.80E-13 |
| B140 | B221 | r-GT | Hei | 0.064992 | 1.66E-09 |
| B140 | B222 | r-GT | NS | 0.061405 | 1.24E-08 |
| B140 | B176 | r-GT | Ucr | 0.05717 | 1.15E-07 |
| B140 | B199 | r-GT | Mg | -0.05705 | 1.22E-07 |

| | | | | | |
|---|---|---|---|---|---|
| B140 | B197 | r-GT | AG | 0.056368 | 1.73E-07 |
| B141 | B153 | AST/ALT | Glu | -0.11327 | 0 |
| B141 | B163 | AST/ALT | SG | -0.11846 | 0 |
| B141 | B189 | AST/ALT | eGFB | -0.15906 | 0 |
| B141 | B191 | AST/ALT | Urea | 0.103493 | 0 |
| B141 | B206 | AST/ALT | Waist | -0.14821 | 0 |
| B141 | B207 | AST/ALT | WHR | -0.10625 | 0 |
| B141 | B210 | AST/ALT | Hip | -0.11328 | 0 |
| B141 | B215 | AST/ALT | BMI | -0.1385 | 0 |
| B141 | B216 | AST/ALT | BMI | -0.2135 | 0 |
| B141 | B218 | AST/ALT | DBP | -0.13248 | 0 |
| B141 | B221 | AST/ALT | Hei | -0.18679 | 0 |
| B141 | B222 | AST/ALT | NS | -0.12504 | 0 |
| B141 | B209 | AST/ALT | Smoking | -0.08424 | 5.33E-15 |
| B141 | B170 | AST/ALT | U-GLU | -0.07501 | 3.43E-12 |
| B141 | B204 | AST/ALT | Drinking | -0.06936 | 1.24E-10 |
| B141 | B176 | AST/ALT | Ucr | -0.05393 | 5.73E-07 |
| B141 | B161 | AST/ALT | LEU | 0.053676 | 6.47E-07 |
| B142 | B143 | 25-(OH)D | β-CTx | -0.15428 | 0 |
| B142 | B144 | 25-(OH)D | Total-PINP | -0.12927 | 0 |
| B142 | B151 | 25-(OH)D | Anti-CCP | -0.13191 | 0 |
| B142 | B176 | 25-(OH)D | Ucr | -0.07351 | 9.14E-12 |
| B143 | B144 | β-CTx | Total-PINP | 0.696225 | 0 |
| B143 | B145 | β-CTx | BGP | 0.62655 | 0 |
| B143 | B151 | β-CTx | Anti-CCP | 0.275707 | 0 |
| B143 | B150 | β-CTx | ALP | 0.088187 | 2.22E-16 |
| B143 | B201 | β-CTx | e-habits | -0.06803 | 2.79E-10 |
| B144 | B145 | Total-PINP | BGP | 0.63851 | 0 |
| B144 | B151 | Total-PINP | Anti-CCP | 0.246128 | 0 |
| B144 | B201 | Total-PINP | e-habits | -0.08176 | 3.26E-14 |
| B144 | B150 | Total-PINP | ALP | 0.078993 | 2.30E-13 |
| B144 | B190 | Total-PINP | Cre | 0.061521 | 1.16E-08 |
| B144 | B176 | Total-PINP | Ucr | 0.053811 | 6.07E-07 |
| B145 | B201 | BGP | e-habits | -0.10991 | 0 |
| B145 | B176 | BGP | Ucr | 0.076614 | 1.17E-12 |
| B145 | B150 | BGP | ALP | 0.075 | 3.44E-12 |
| B145 | B183 | BGP | PTA | -0.06326 | 4.44E-09 |
| B145 | B203 | BGP | DH | 0.058455 | 5.95E-08 |
| B145 | B207 | BGP | WHR | -0.05528 | 2.97E-07 |
| B145 | B175 | BGP | ALB-U | -0.05329 | 7.78E-07 |
| B146 | B189 | CysC | eGFB | -0.2364 | 0 |
| B146 | B190 | CysC | Cre | 0.360512 | 0 |
| B146 | B191 | CysC | Urea | 0.271955 | 0 |
| B146 | B192 | CysC | UA | 0.106225 | 0 |

| | | | | | |
|---|---|---|---|---|---|
| B146 | B198 | CysC | P | 0.163459 | 0 |
| B146 | B200 | CysC | OP | 0.192858 | 0 |
| B146 | B174 | CysC | Urea/Cr | -0.08355 | 8.88E-15 |
| B146 | B196 | CysC | TCO2 | -0.08133 | 4.44E-14 |
| B146 | B188 | CysC | Ca | -0.06751 | 3.81E-10 |
| B146 | B218 | CysC | DBP | -0.05997 | 2.68E-08 |
| B148 | B149 | Anti_TPO | TGAb | 0.520305 | 0 |
| B149 | B221 | TGAb | Hei | -0.07261 | 1.63E-11 |
| B15 | B100 | CD3-CD19+(B cell) | anti-Ro52 | 0.106232 | 0 |
| B15 | B101 | CD3-CD19+(B cell) | anti-Sc70 | 0.46939 | 0 |
| B15 | B103 | CD3-CD19+(B cell) | anti-SSA/Ro | 0.128249 | 0 |
| B15 | B104 | CD3-CD19+(B cell) | anti-SSB | 0.46939 | 0 |
| B15 | B107 | CD3-CD19+(B cell) | AnuA | 0.471646 | 0 |
| B15 | B109 | CD3-CD19+(B cell) | AMA-M2 | 0.177977 | 0 |
| B15 | B110 | CD3-CD19+(B cell) | AHA | -0.10072 | 0 |
| B15 | B115 | CD3-CD19+(B cell) | TSH TT3 TT4(TT3) | 0.09607 | 0 |
| B15 | B124 | CD3-CD19+(B cell) | ACTH | -0.11149 | 0 |
| B15 | B136 | CD3-CD19+(B cell) | 5'NT | -0.10737 | 0 |
| B15 | B151 | CD3-CD19+(B cell) | Anti-CCP | 0.09029 | 0 |
| B15 | B158 | CD3-CD19+(B cell) | IgM | 0.117072 | 0 |
| B15 | B16 | CD3-CD19+(B cell) | CD3+T | 0.130458 | 0 |
| B15 | B18 | CD3-CD19+(B cell) | CD3+CD4+helper(inducer)T | 0.25555 | 0 |
| B15 | B181 | CD3-CD19+(B cell) | APTT | 0.12944 | 0 |
| B15 | B184 | CD3-CD19+(B cell) | INR | 0.111378 | 0 |
| B15 | B186 | CD3-CD19+(B cell) | PFC | -0.13881 | 0 |
| B15 | B19 | CD3-CD19+(B cell) | CD3+CD4+/CD3+CD8+ | 0.123482 | 0 |
| B15 | B194 | CD3-CD19+(B cell) | Cl | 0.139788 | 0 |
| B15 | B195 | CD3-CD19+(B cell) | Na | 0.090821 | 0 |
| B15 | B199 | CD3-CD19+(B cell) | Mg | 0.168775 | 0 |
| B15 | B20 | CD3-CD19+(B cell) | CD3+CD4+CD8+ | -0.1182 | 0 |
| B15 | B200 | CD3-CD19+(B cell) | OP | -0.09499 | 0 |
| B15 | B21 | CD3-CD19+(B cell) | CD3+CD4-CD8- | 0.203413 | 0 |
| B15 | B22 | CD3-CD19+(B cell) | CD3+CD8+(T) | -0.21084 | 0 |
| B15 | B225 | CD3-CD19+(B cell) | hPBG | -0.09839 | 0 |
| B15 | B23 | CD3-CD19+(B cell) | CD3-CD16+CD56+(NK) | -0.43942 | 0 |
| B15 | B28 | CD3-CD19+(B cell) | PRL | 0.136065 | 0 |
| B15 | B29 | CD3-CD19+(B cell) | E2 | 0.106428 | 0 |
| B15 | B30 | CD3-CD19+(B cell) | LH | -0.20844 | 0 |
| B15 | B31 | CD3-CD19+(B cell) | FSH | -0.16554 | 0 |
| B15 | B33 | CD3-CD19+(B cell) | Prog | 0.15222 | 0 |
| B15 | B79 | CD3-CD19+(B cell) | hb V -dna | 0.176109 | 0 |
| B15 | B92 | CD3-CD19+(B cell) | ALDH2-Glu504Glu | -0.18591 | 0 |
| B15 | B93 | CD3-CD19+(B cell) | ALDH2-Glu504Lys | 0.178613 | 0 |
| B15 | B94 | CD3-CD19+(B cell) | ALDH2-Lys504Lys | 0.208787 | 0 |

| | | | | | |
|---|---|---|---|---|---|
| B15 | B95 | CD3-CD19+(B cell) | anti-cenpb | -0.15416 | 0 |
| B15 | B96 | CD3-CD19+(B cell) | dsDNA (IIF) | 0.146863 | 0 |
| B15 | B97 | CD3-CD19+(B cell) | dsDNA | -0.1361 | 0 |
| B15 | B98 | CD3-CD19+(B cell) | anti-Jo-1 | -0.13175 | 0 |
| B15 | B99 | CD3-CD19+(B cell) | anti-PM-Scl | 0.46939 | 0 |
| B15 | B197 | CD3-CD19+(B cell) | AG | -0.0829 | 1.42E-14 |
| B15 | B119 | CD3-CD19+(B cell) | Hcv-rna | 0.079959 | 1.17E-13 |
| B15 | B123 | CD3-CD19+(B cell) | hs-CRP1 | -0.07991 | 1.21E-13 |
| B15 | B139 | CD3-CD19+(B cell) | ADA | -0.07843 | 3.39E-13 |
| B15 | B17 | CD3-CD19+(B cell) | CD3+CD16+CD56+NK | 0.078146 | 4.13E-13 |
| B15 | B125 | CD3-CD19+(B cell) | Cor | -0.06098 | 1.56E-08 |
| B15 | B157 | CD3-CD19+(B cell) | IgG | 0.060603 | 1.91E-08 |
| B15 | B113 | CD3-CD19+(B cell) | FT3 | 0.059844 | 2.87E-08 |
| B15 | B122 | CD3-CD19+(B cell) | C4 | 0.058826 | 4.90E-08 |
| B15 | B156 | CD3-CD19+(B cell) | IgE | -0.05727 | 1.10E-07 |
| B15 | B179 | CD3-CD19+(B cell) | α1-MG | -0.05573 | 2.37E-07 |
| B15 | B177 | CD3-CD19+(B cell) | ACR | -0.05566 | 2.46E-07 |
| B15 | B178 | CD3-CD19+(B cell) | Ig-U | -0.05456 | 4.22E-07 |
| B15 | B134 | CD3-CD19+(B cell) | CHE | -0.05438 | 4.61E-07 |
| B150 | B186 | ALP | PFC | 0.105022 | 0 |
| B150 | B200 | ALP | OP | 0.110871 | 0 |
| B150 | B153 | ALP | Glu | 0.080426 | 8.44E-14 |
| B150 | B155 | ALP | IgA | 0.080022 | 1.12E-13 |
| B150 | B194 | ALP | Cl | -0.06153 | 1.16E-08 |
| B150 | B182 | ALP | TT | 0.0536 | 6.71E-07 |
| B150 | B199 | ALP | Mg | -0.05346 | 7.18E-07 |
| B151 | B152 | Anti-CCP | ACA-Ig(G-A-M) | -0.13705 | 0 |
| B151 | B183 | Anti-CCP | PTA | -0.08597 | 1.33E-15 |
| B151 | B185 | Anti-CCP | PT | 0.069497 | 1.14E-10 |
| B152 | B183 | ACA-Ig(G-A-M) | PTA | 0.060701 | 1.81E-08 |
| B152 | B182 | ACA-Ig(G-A-M) | TT | -0.05356 | 6.86E-07 |
| B153 | B160 | Glu | PH | -0.08957 | 0 |
| B153 | B163 | Glu | SG | 0.218745 | 0 |
| B153 | B166 | Glu | Pro | 0.108253 | 0 |
| B153 | B170 | Glu | U-GLU | 0.538424 | 0 |
| B153 | B172 | Glu | KB | 0.131358 | 0 |
| B153 | B174 | Glu | Urea/Cr | 0.120708 | 0 |
| B153 | B194 | Glu | Cl | -0.09555 | 0 |
| B153 | B195 | Glu | Na | -0.0936 | 0 |
| B153 | B206 | Glu | Waist | 0.104389 | 0 |
| B153 | B207 | Glu | WHR | 0.097139 | 0 |
| B153 | B218 | Glu | DBP | 0.101749 | 0 |
| B153 | B220 | Glu | SBP | 0.108417 | 0 |
| B153 | B192 | Glu | UA | -0.08673 | 8.88E-16 |

| | | | | | |
|---|---|---|---|---|---|
| B153 | B216 | Glu | BMI | 0.084777 | 3.55E-15 |
| B153 | B215 | Glu | BMI | 0.079311 | 1.85E-13 |
| B153 | B225 | Glu | hPBG | 0.073701 | 8.06E-12 |
| B153 | B188 | Glu | Ca | 0.069563 | 1.10E-10 |
| B153 | B204 | Glu | Drinking | 0.064981 | 1.67E-09 |
| B153 | B200 | Glu | OP | 0.058725 | 5.17E-08 |
| B153 | B210 | Glu | Hip | 0.058171 | 6.89E-08 |
| B154 | B183 | RF | PTA | 0.061697 | 1.06E-08 |
| B155 | B157 | IgA | IgG | 0.416146 | 0 |
| B155 | B182 | IgA | TT | 0.09156 | 0 |
| B155 | B184 | IgA | INR | -0.10921 | 0 |
| B155 | B194 | IgA | Cl | -0.08531 | 2.44E-15 |
| B155 | B195 | IgA | Na | -0.07843 | 3.39E-13 |
| B155 | B185 | IgA | PT | -0.07775 | 5.44E-13 |
| B155 | B186 | IgA | PFC | 0.06453 | 2.17E-09 |
| B155 | B197 | IgA | AG | 0.05484 | 3.68E-07 |
| B156 | B199 | IgE | Mg | -0.07509 | 3.23E-12 |
| B156 | B183 | IgE | PTA | 0.069132 | 1.43E-10 |
| B156 | B185 | IgE | PT | 0.066567 | 6.65E-10 |
| B156 | B184 | IgE | INR | 0.055061 | 3.31E-07 |
| B157 | B200 | IgG | OP | -0.12423 | 0 |
| B157 | B195 | IgG | Na | -0.08065 | 7.19E-14 |
| B157 | B199 | IgG | Mg | 0.069631 | 1.05E-10 |
| B157 | B158 | IgG | IgM | 0.068603 | 1.97E-10 |
| B157 | B180 | IgG | Tf | -0.06559 | 1.17E-09 |
| B157 | B178 | IgG | Ig-U | -0.05891 | 4.69E-08 |
| B157 | B212 | IgG | IOP-L | -0.05672 | 1.45E-07 |
| B157 | B184 | IgG | INR | 0.055664 | 2.46E-07 |
| B157 | B204 | IgG | Drinking | -0.05442 | 4.52E-07 |
| B158 | B198 | IgM | P | 0.104742 | 0 |
| B158 | B199 | IgM | Mg | 0.136894 | 0 |
| B158 | B200 | IgM | OP | -0.09107 | 0 |
| B158 | B181 | IgM | APTT | 0.075363 | 2.71E-12 |
| B158 | B186 | IgM | PFC | 0.073672 | 8.21E-12 |
| B159 | B181 | β2-MG | APTT | 0.148057 | 0 |
| B159 | B184 | β2-MG | INR | 0.153074 | 0 |
| B159 | B186 | β2-MG | PFC | -0.09161 | 0 |
| B159 | B185 | β2-MG | PT | 0.08688 | 8.88E-16 |
| B159 | B183 | β2-MG | PTA | 0.056872 | 1.34E-07 |
| B16 | B101 | CD3+T | anti-Sc70 | 0.113431 | 0 |
| B16 | B104 | CD3+T | anti-SSB | 0.113431 | 0 |
| B16 | B107 | CD3+T | AnuA | 0.113983 | 0 |
| B16 | B110 | CD3+T | AHA | -0.26064 | 0 |
| B16 | B122 | CD3+T | C4 | 0.125872 | 0 |

| | | | | | |
|---|---|---|---|---|---|
| B16 | B137 | CD3+T | AFU | -0.09521 | 0 |
| B16 | B139 | CD3+T | ADA | -0.09932 | 0 |
| B16 | B17 | CD3+T | CD3+CD16+CD56+NK | 0.229289 | 0 |
| B16 | B18 | CD3+T | CD3+CD4+helper(inducer)T | 0.463219 | 0 |
| B16 | B194 | CD3+T | Cl | 0.092997 | 0 |
| B16 | B21 | CD3+T | CD3+CD4-CD8- | 0.162433 | 0 |
| B16 | B22 | CD3+T | CD3+CD8+(T) | 0.429998 | 0 |
| B16 | B23 | CD3+T | CD3-CD16+CD56+(NK) | -0.8729 | 0 |
| B16 | B28 | CD3+T | PRL | 0.091472 | 0 |
| B16 | B96 | CD3+T | dsDNA (IIF) | 0.126874 | 0 |
| B16 | B97 | CD3+T | dsDNA | -0.2841 | 0 |
| B16 | B98 | CD3+T | anti-Jo-1 | 0.14744 | 0 |
| B16 | B99 | CD3+T | anti-PM-Scl | 0.113431 | 0 |
| B16 | B180 | CD3+T | Tf | -0.08797 | 4.44E-16 |
| B16 | B158 | CD3+T | IgM | 0.085217 | 2.66E-15 |
| B16 | B33 | CD3+T | Prog | 0.084639 | 4.00E-15 |
| B16 | B183 | CD3+T | PTA | -0.08389 | 6.88E-15 |
| B16 | B195 | CD3+T | Na | 0.076762 | 1.06E-12 |
| B16 | B54 | CD3+T | ESR | -0.0765 | 1.27E-12 |
| B16 | B199 | CD3+T | Mg | 0.074087 | 6.27E-12 |
| B16 | B200 | CD3+T | OP | -0.0697 | 1.01E-10 |
| B16 | B175 | CD3+T | ALB-U | -0.06909 | 1.47E-10 |
| B16 | B69 | CD3+T | PV | -0.06843 | 2.19E-10 |
| B16 | B30 | CD3+T | LH | -0.06748 | 3.88E-10 |
| B16 | B31 | CD3+T | FSH | -0.0668 | 5.81E-10 |
| B16 | B178 | CD3+T | Ig-U | -0.06525 | 1.44E-09 |
| B16 | B121 | CD3+T | C3 | 0.061514 | 1.17E-08 |
| B16 | B29 | CD3+T | E2 | 0.060463 | 2.06E-08 |
| B16 | B119 | CD3+T | Hcv-rna | 0.05895 | 4.59E-08 |
| B16 | B186 | CD3+T | PFC | -0.05649 | 1.63E-07 |
| B16 | B136 | CD3+T | 5'NT | 0.052897 | 9.38E-07 |
| B160 | B163 | PH | SG | -0.16413 | 0 |
| B160 | B191 | PH | Urea | -0.1198 | 0 |
| B160 | B192 | PH | UA | -0.13197 | 0 |
| B160 | B176 | PH | Ucr | -0.07849 | 3.27E-13 |
| B160 | B220 | PH | SBP | 0.068505 | 2.09E-10 |
| B160 | B174 | PH | Urea/Cr | -0.06208 | 8.57E-09 |
| B160 | B218 | PH | DBP | 0.059893 | 2.79E-08 |
| B161 | B173 | LEU | NTT | 0.213174 | 0 |
| B161 | B174 | LEU | Urea/Cr | 0.099319 | 0 |
| B161 | B216 | LEU | BMI | -0.09033 | 0 |
| B161 | B221 | LEU | Hei | -0.17237 | 0 |
| B161 | B222 | LEU | NS | -0.08426 | 5.33E-15 |
| B161 | B209 | LEU | Smoking | -0.06893 | 1.62E-10 |

| | | | | | |
|---|---|---|---|---|---|
| B161 | B204 | LEU | Drinking | -0.05952 | 3.40E-08 |
| B161 | B218 | LEU | DBP | -0.05585 | 2.23E-07 |
| B163 | B166 | SG | Pro | 0.254502 | 0 |
| B163 | B170 | SG | U-GLU | 0.282109 | 0 |
| B163 | B172 | SG | KB | 0.143978 | 0 |
| B163 | B174 | SG | Urea/Cr | 0.090974 | 0 |
| B163 | B176 | SG | Ucr | 0.288752 | 0 |
| B163 | B189 | SG | eGFB | 0.111127 | 0 |
| B163 | B191 | SG | Urea | 0.089572 | 0 |
| B163 | B204 | SG | Drinking | 0.128654 | 0 |
| B163 | B206 | SG | Waist | 0.163896 | 0 |
| B163 | B207 | SG | WHR | 0.109238 | 0 |
| B163 | B209 | SG | Smoking | 0.176425 | 0 |
| B163 | B210 | SG | Hip | 0.128076 | 0 |
| B163 | B215 | SG | BMI | 0.121051 | 0 |
| B163 | B216 | SG | BMI | 0.252181 | 0 |
| B163 | B218 | SG | DBP | 0.10627 | 0 |
| B163 | B221 | SG | Hei | 0.266263 | 0 |
| B163 | B222 | SG | NS | 0.183947 | 0 |
| B163 | B220 | SG | SBP | -0.07786 | 5.03E-13 |
| B163 | B173 | SG | NTT | -0.06804 | 2.77E-10 |
| B163 | B179 | SG | α1-MG | 0.061701 | 1.05E-08 |
| B163 | B188 | SG | Ca | -0.0577 | 8.80E-08 |
| B163 | B201 | SG | e-habits | -0.05683 | 1.37E-07 |
| B164 | B178 | COM | Ig-U | 0.078247 | 3.86E-13 |
| B166 | B170 | Pro | U-GLU | 0.149345 | 0 |
| B166 | B172 | Pro | KB | 0.103611 | 0 |
| B166 | B176 | Pro | Ucr | 0.111 | 0 |
| B166 | B189 | Pro | eGFB | -0.10901 | 0 |
| B166 | B190 | Pro | Cre | 0.123254 | 0 |
| B166 | B191 | Pro | Urea | 0.116275 | 0 |
| B166 | B207 | Pro | WHR | 0.072996 | 1.27E-11 |
| B166 | B206 | Pro | Waist | 0.06661 | 6.49E-10 |
| B166 | B175 | Pro | ALB-U | 0.059715 | 3.07E-08 |
| B166 | B177 | Pro | ACR | 0.059607 | 3.25E-08 |
| B166 | B216 | Pro | BMI | 0.057565 | 9.42E-08 |
| B166 | B188 | Pro | Ca | -0.05592 | 2.16E-07 |
| B166 | B221 | Pro | Hei | 0.054663 | 4.02E-07 |
| B166 | B222 | Pro | NS | 0.053221 | 8.04E-07 |
| B169 | B203 | U-CRY | DH | 0.065392 | 1.32E-09 |
| B17 | B101 | CD3+CD16+CD56+NK | anti-Sc70 | 0.208531 | 0 |
| B17 | B104 | CD3+CD16+CD56+NK | anti-SSB | 0.208531 | 0 |
| B17 | B107 | CD3+CD16+CD56+NK | AnuA | 0.209532 | 0 |
| B17 | B108 | CD3+CD16+CD56+NK | Anti-Cyclin | -0.15999 | 0 |

| | | | | | |
|---|---|---|---|---|---|
| B17 | B137 | CD3+CD16+CD56+NK | AFU | -0.11764 | 0 |
| B17 | B145 | CD3+CD16+CD56+NK | BGP | 0.102583 | 0 |
| B17 | B151 | CD3+CD16+CD56+NK | Anti-CCP | 0.15066 | 0 |
| B17 | B175 | CD3+CD16+CD56+NK | ALB-U | -0.11732 | 0 |
| B17 | B178 | CD3+CD16+CD56+NK | Ig-U | -0.10714 | 0 |
| B17 | B180 | CD3+CD16+CD56+NK | Tf | -0.14121 | 0 |
| B17 | B183 | CD3+CD16+CD56+NK | PTA | -0.20922 | 0 |
| B17 | B185 | CD3+CD16+CD56+NK | PT | 0.092702 | 0 |
| B17 | B19 | CD3+CD16+CD56+NK | CD3+CD4+/CD3+CD8+ | -0.16098 | 0 |
| B17 | B21 | CD3+CD16+CD56+NK | CD3+CD4-CD8- | 0.352404 | 0 |
| B17 | B22 | CD3+CD16+CD56+NK | CD3+CD8+(T) | 0.173597 | 0 |
| B17 | B23 | CD3+CD16+CD56+NK | CD3-CD16+CD56+(NK) | -0.18501 | 0 |
| B17 | B96 | CD3+CD16+CD56+NK | dsDNA (IIF) | 0.126016 | 0 |
| B17 | B99 | CD3+CD16+CD56+NK | anti-PM-Scl | 0.208531 | 0 |
| B17 | B195 | CD3+CD16+CD56+NK | Na | 0.079426 | 1.70E-13 |
| B17 | B109 | CD3+CD16+CD56+NK | AMA-M2 | 0.079056 | 2.20E-13 |
| B17 | B20 | CD3+CD16+CD56+NK | CD3+CD4+CD8+ | 0.077495 | 6.45E-13 |
| B17 | B138 | CD3+CD16+CD56+NK | PAB | -0.07598 | 1.80E-12 |
| B17 | B184 | CD3+CD16+CD56+NK | INR | 0.074858 | 3.78E-12 |
| B17 | B97 | CD3+CD16+CD56+NK | dsDNA | -0.06948 | 1.16E-10 |
| B17 | B158 | CD3+CD16+CD56+NK | IgM | 0.067803 | 3.20E-10 |
| B17 | B199 | CD3+CD16+CD56+NK | Mg | 0.06744 | 3.97E-10 |
| B17 | B194 | CD3+CD16+CD56+NK | Cl | 0.066375 | 7.45E-10 |
| B17 | B34 | CD3+CD16+CD56+NK | TK | 0.065704 | 1.10E-09 |
| B17 | B182 | CD3+CD16+CD56+NK | TT | 0.063789 | 3.30E-09 |
| B17 | B144 | CD3+CD16+CD56+NK | Total-PINP | 0.063721 | 3.43E-09 |
| B17 | B136 | CD3+CD16+CD56+NK | 5'NT | 0.058202 | 6.78E-08 |
| B17 | B69 | CD3+CD16+CD56+NK | PV | -0.05796 | 7.68E-08 |
| B17 | B57 | CD3+CD16+CD56+NK | RBCAI | 0.057363 | 1.04E-07 |
| B170 | B172 | U-GLU | KB | 0.147134 | 0 |
| B170 | B192 | U-GLU | UA | -0.11749 | 0 |
| B170 | B204 | U-GLU | Drinking | 0.109117 | 0 |
| B170 | B209 | U-GLU | Smoking | 0.097708 | 0 |
| B170 | B216 | U-GLU | BMI | 0.090597 | 0 |
| B170 | B218 | U-GLU | DBP | 0.102135 | 0 |
| B170 | B221 | U-GLU | Hei | 0.111835 | 0 |
| B170 | B194 | U-GLU | Cl | -0.08103 | 5.51E-14 |
| B170 | B207 | U-GLU | WHR | 0.080547 | 7.75E-14 |
| B170 | B206 | U-GLU | Waist | 0.076262 | 1.49E-12 |
| B170 | B222 | U-GLU | NS | 0.073803 | 7.54E-12 |
| B170 | B225 | U-GLU | hPBG | 0.068629 | 1.94E-10 |
| B170 | B200 | U-GLU | OP | 0.067096 | 4.87E-10 |
| B170 | B220 | U-GLU | SBP | 0.06175 | 1.03E-08 |
| B170 | B176 | U-GLU | Ucr | 0.059835 | 2.88E-08 |

| | | | | | |
|---|---|---|---|---|---|
| B170 | B174 | U-GLU | Urea/Cr | 0.057637 | 9.07E-08 |
| B170 | B189 | U-GLU | eGFB | 0.053327 | 7.65E-07 |
| B170 | B195 | U-GLU | Na | -0.05282 | 9.73E-07 |
| B173 | B221 | NTT | Hei | -0.12945 | 0 |
| B173 | B216 | NTT | BMI | -0.07445 | 4.93E-12 |
| B173 | B174 | NTT | Urea/Cr | 0.066804 | 5.79E-10 |
| B173 | B222 | NTT | NS | -0.05832 | 6.39E-08 |
| B173 | B204 | NTT | Drinking | -0.05462 | 4.11E-07 |
| B174 | B189 | Urea/Cr | eGFB | 0.224326 | 0 |
| B174 | B190 | Urea/Cr | Cre | -0.33012 | 0 |
| B174 | B191 | Urea/Cr | Urea | 0.323161 | 0 |
| B174 | B192 | Urea/Cr | UA | -0.18102 | 0 |
| B174 | B204 | Urea/Cr | Drinking | -0.09799 | 0 |
| B174 | B209 | Urea/Cr | Smoking | -0.11222 | 0 |
| B174 | B216 | Urea/Cr | BMI | -0.18585 | 0 |
| B174 | B221 | Urea/Cr | Hei | -0.28103 | 0 |
| B174 | B222 | Urea/Cr | NS | -0.15311 | 0 |
| B174 | B206 | Urea/Cr | Waist | -0.08563 | 1.78E-15 |
| B174 | B200 | Urea/Cr | OP | 0.076389 | 1.37E-12 |
| B174 | B218 | Urea/Cr | DBP | -0.07629 | 1.46E-12 |
| B174 | B210 | Urea/Cr | Hip | -0.07009 | 7.94E-11 |
| B174 | B207 | Urea/Cr | WHR | -0.05826 | 6.59E-08 |
| B175 | B177 | ALB-U | ACR | 0.848692 | 0 |
| B175 | B178 | ALB-U | Ig-U | 0.633189 | 0 |
| B175 | B179 | ALB-U | α1-MG | 0.298483 | 0 |
| B175 | B180 | ALB-U | Tf | 0.714224 | 0 |
| B175 | B189 | ALB-U | eGFB | -0.06798 | 2.88E-10 |
| B175 | B190 | ALB-U | Cre | 0.057694 | 8.81E-08 |
| B176 | B177 | Ucr | ACR | -0.09286 | 0 |
| B176 | B179 | Ucr | α1-MG | 0.143693 | 0 |
| B176 | B221 | Ucr | Hei | 0.092519 | 0 |
| B176 | B216 | Ucr | BMI | 0.088104 | 4.44E-16 |
| B176 | B209 | Ucr | Smoking | 0.066087 | 8.82E-10 |
| B176 | B222 | Ucr | NS | 0.062748 | 5.91E-09 |
| B176 | B201 | Ucr | e-habits | -0.05361 | 6.69E-07 |
| B177 | B178 | ACR | Ig-U | 0.534878 | 0 |
| B177 | B179 | ACR | α1-MG | 0.284178 | 0 |
| B177 | B180 | ACR | Tf | 0.558802 | 0 |
| B177 | B220 | ACR | SBP | 0.080181 | 1.00E-13 |
| B177 | B186 | ACR | PFC | 0.075358 | 2.72E-12 |
| B177 | B189 | ACR | eGFB | -0.07317 | 1.14E-11 |
| B177 | B190 | ACR | Cre | 0.069542 | 1.11E-10 |
| B177 | B191 | ACR | Urea | 0.060842 | 1.68E-08 |
| B178 | B179 | Ig-U | α1-MG | 0.53678 | 0 |

| | | | | | |
|---|---|---|---|---|---|
| B178 | B180 | Ig-U | Tf | 0.970606 | 0 |
| B178 | B200 | Ig-U | OP | 0.081967 | 2.80E-14 |
| B178 | B186 | Ig-U | PFC | 0.076338 | 1.41E-12 |
| B178 | B184 | Ig-U | INR | -0.05516 | 3.16E-07 |
| B179 | B180 | α1-MG | Tf | 0.478549 | 0 |
| B179 | B200 | α1-MG | OP | 0.072641 | 1.60E-11 |
| B179 | B186 | α1-MG | PFC | 0.062342 | 7.41E-09 |
| B18 | B110 | CD3+CD4+helper(inducer)T | AHA | -0.10595 | 0 |
| B18 | B122 | CD3+CD4+helper(inducer)T | C4 | 0.148445 | 0 |
| B18 | B124 | CD3+CD4+helper(inducer)T | ACTH | 0.140133 | 0 |
| B18 | B139 | CD3+CD4+helper(inducer)T | ADA | -0.0922 | 0 |
| B18 | B156 | CD3+CD4+helper(inducer)T | IgE | -0.13859 | 0 |
| B18 | B19 | CD3+CD4+helper(inducer)T | CD3+CD4+/CD3+CD8+ | 0.733839 | 0 |
| B18 | B22 | CD3+CD4+helper(inducer)T | CD3+CD8+(T) | -0.51924 | 0 |
| B18 | B23 | CD3+CD4+helper(inducer)T | CD3-CD16+CD56+(NK) | -0.50781 | 0 |
| B18 | B28 | CD3+CD4+helper(inducer)T | PRL | 0.253586 | 0 |
| B18 | B29 | CD3+CD4+helper(inducer)T | E2 | 0.192524 | 0 |
| B18 | B30 | CD3+CD4+helper(inducer)T | LH | -0.35772 | 0 |
| B18 | B31 | CD3+CD4+helper(inducer)T | FSH | -0.29106 | 0 |
| B18 | B33 | CD3+CD4+helper(inducer)T | Prog | 0.277885 | 0 |
| B18 | B92 | CD3+CD4+helper(inducer)T | ALDH2-Glu504Glu | 0.129659 | 0 |
| B18 | B93 | CD3+CD4+helper(inducer)T | ALDH2-Glu504Lys | -0.12926 | 0 |
| B18 | B138 | CD3+CD4+helper(inducer)T | PAB | 0.086837 | 8.88E-16 |
| B18 | B97 | CD3+CD4+helper(inducer)T | dsDNA | -0.0831 | 1.24E-14 |
| B18 | B158 | CD3+CD4+helper(inducer)T | IgM | 0.081977 | 2.80E-14 |
| B18 | B136 | CD3+CD4+helper(inducer)T | 5'NT | -0.08154 | 3.82E-14 |
| B18 | B125 | CD3+CD4+helper(inducer)T | Cor | 0.078598 | 3.03E-13 |
| B18 | B96 | CD3+CD4+helper(inducer)T | dsDNA (IIF) | 0.077981 | 4.63E-13 |
| B18 | B21 | CD3+CD4+helper(inducer)T | CD3+CD4-CD8- | -0.07692 | 9.54E-13 |
| B18 | B94 | CD3+CD4+helper(inducer)T | ALDH2-Lys504Lys | -0.07567 | 2.21E-12 |
| B18 | B200 | CD3+CD4+helper(inducer)T | OP | -0.07463 | 4.40E-12 |
| B18 | B176 | CD3+CD4+helper(inducer)T | Ucr | 0.074609 | 4.45E-12 |
| B18 | B197 | CD3+CD4+helper(inducer)T | AG | -0.07403 | 6.49E-12 |
| B18 | B157 | CD3+CD4+helper(inducer)T | IgG | -0.06592 | 9.72E-10 |
| B18 | B24 | CD3+CD4+helper(inducer)T | αHBDH | -0.06296 | 5.25E-09 |
| B18 | B186 | CD3+CD4+helper(inducer)T | PFC | -0.06036 | 2.18E-08 |
| B18 | B181 | CD3+CD4+helper(inducer)T | APTT | 0.056567 | 1.56E-07 |
| B18 | B26 | CD3+CD4+helper(inducer)T | CKMB | -0.05632 | 1.77E-07 |
| B18 | B49 | CD3+CD4+helper(inducer)T | baso% | -0.0563 | 1.79E-07 |
| B18 | B178 | CD3+CD4+helper(inducer)T | Ig-U | -0.05458 | 4.18E-07 |
| B180 | B200 | Tf | OP | 0.071903 | 2.55E-11 |
| B180 | B186 | Tf | PFC | 0.06364 | 3.59E-09 |
| B181 | B184 | APTT | INR | 0.532547 | 0 |
| B181 | B185 | APTT | PT | 0.467061 | 0 |

| | | | | | |
|---|---|---|---|---|---|
| B181 | B194 | APTT | Cl | 0.102522 | 0 |
| B181 | B199 | APTT | Mg | 0.108974 | 0 |
| B181 | B200 | APTT | OP | -0.09177 | 0 |
| B181 | B197 | APTT | AG | -0.0793 | 1.85E-13 |
| B181 | B212 | APTT | IOP-L | -0.07306 | 1.22E-11 |
| B181 | B211 | APTT | IOP-R | -0.06101 | 1.53E-08 |
| B182 | B183 | TT | PTA | -0.16684 | 0 |
| B182 | B184 | TT | INR | -0.15454 | 0 |
| B182 | B195 | TT | Na | 0.055757 | 2.34E-07 |
| B182 | B200 | TT | OP | 0.053177 | 8.22E-07 |
| B183 | B184 | PTA | INR | 0.157789 | 0 |
| B183 | B186 | PTA | PFC | -0.1174 | 0 |
| B183 | B197 | PTA | AG | -0.08912 | 0 |
| B183 | B195 | PTA | Na | -0.05808 | 7.23E-08 |
| B183 | B200 | PTA | OP | -0.05552 | 2.64E-07 |
| B184 | B185 | INR | PT | 0.81983 | 0 |
| B184 | B186 | INR | PFC | -0.25083 | 0 |
| B184 | B194 | INR | Cl | 0.118704 | 0 |
| B184 | B196 | INR | TCO2 | 0.096339 | 0 |
| B184 | B197 | INR | AG | -0.14608 | 0 |
| B184 | B199 | INR | Mg | 0.098272 | 0 |
| B184 | B200 | INR | OP | -0.09873 | 0 |
| B184 | B193 | INR | K | 0.076738 | 1.08E-12 |
| B185 | B193 | PT | K | 0.091699 | 0 |
| B185 | B197 | PT | AG | -0.09731 | 0 |
| B185 | B199 | PT | Mg | 0.077453 | 6.64E-13 |
| B185 | B196 | PT | TCO2 | 0.073127 | 1.17E-11 |
| B185 | B194 | PT | Cl | 0.070112 | 7.84E-11 |
| B185 | B212 | PT | IOP-L | -0.06101 | 1.54E-08 |
| B185 | B200 | PT | OP | -0.05685 | 1.36E-07 |
| B186 | B225 | PFC | hPBG | 0.078761 | 2.70E-13 |
| B186 | B194 | PFC | Cl | -0.07223 | 2.08E-11 |
| B186 | B197 | PFC | AG | 0.064421 | 2.31E-09 |
| B186 | B200 | PFC | OP | 0.062238 | 7.85E-09 |
| B188 | B192 | Ca | UA | 0.102715 | 0 |
| B188 | B190 | Ca | Cre | -0.08703 | 6.66E-16 |
| B188 | B193 | Ca | K | 0.055522 | 2.63E-07 |
| B188 | B191 | Ca | Urea | -0.05363 | 6.63E-07 |
| B189 | B190 | eGFB | Cre | -0.57109 | 0 |
| B189 | B191 | eGFB | Urea | -0.54279 | 0 |
| B189 | B192 | eGFB | UA | -0.28922 | 0 |
| B189 | B204 | eGFB | Drinking | 0.113189 | 0 |
| B189 | B209 | eGFB | Smoking | 0.093262 | 0 |
| B189 | B210 | eGFB | Hip | 0.091579 | 0 |

| | | | | | |
|---|---|---|---|---|---|
| B189 | B216 | eGFB | BMI | 0.138969 | 0 |
| B189 | B218 | eGFB | DBP | 0.19613 | 0 |
| B189 | B220 | eGFB | SBP | -0.14107 | 0 |
| B189 | B221 | eGFB | Hei | 0.125996 | 0 |
| B189 | B215 | eGFB | BMI | 0.081324 | 4.44E-14 |
| B189 | B194 | eGFB | Cl | -0.05901 | 4.46E-08 |
| B189 | B212 | eGFB | IOP-L | 0.056907 | 1.32E-07 |
| B189 | B207 | eGFB | WHR | -0.05672 | 1.45E-07 |
| B19 | B101 | CD3+CD4+/CD3+CD8+ | anti-Sc70 | -0.16701 | 0 |
| B19 | B104 | CD3+CD4+/CD3+CD8+ | anti-SSB | -0.16701 | 0 |
| B19 | B107 | CD3+CD4+/CD3+CD8+ | AnuA | -0.1678 | 0 |
| B19 | B122 | CD3+CD4+/CD3+CD8+ | C4 | 0.113103 | 0 |
| B19 | B124 | CD3+CD4+/CD3+CD8+ | ACTH | 0.153856 | 0 |
| B19 | B125 | CD3+CD4+/CD3+CD8+ | Cor | 0.102847 | 0 |
| B19 | B136 | CD3+CD4+/CD3+CD8+ | 5'NT | -0.10729 | 0 |
| B19 | B156 | CD3+CD4+/CD3+CD8+ | IgE | -0.12966 | 0 |
| B19 | B157 | CD3+CD4+/CD3+CD8+ | IgG | -0.11142 | 0 |
| B19 | B195 | CD3+CD4+/CD3+CD8+ | Na | -0.09127 | 0 |
| B19 | B20 | CD3+CD4+/CD3+CD8+ | CD3+CD4+CD8+ | -0.14085 | 0 |
| B19 | B22 | CD3+CD4+/CD3+CD8+ | CD3+CD8+(T) | -0.77694 | 0 |
| B19 | B23 | CD3+CD4+/CD3+CD8+ | CD3-CD16+CD56+(NK) | -0.11653 | 0 |
| B19 | B28 | CD3+CD4+/CD3+CD8+ | PRL | 0.134673 | 0 |
| B19 | B29 | CD3+CD4+/CD3+CD8+ | E2 | 0.103039 | 0 |
| B19 | B30 | CD3+CD4+/CD3+CD8+ | LH | -0.19517 | 0 |
| B19 | B31 | CD3+CD4+/CD3+CD8+ | FSH | -0.15856 | 0 |
| B19 | B33 | CD3+CD4+/CD3+CD8+ | Prog | 0.149439 | 0 |
| B19 | B92 | CD3+CD4+/CD3+CD8+ | ALDH2-Glu504Glu | 0.132475 | 0 |
| B19 | B93 | CD3+CD4+/CD3+CD8+ | ALDH2-Glu504Lys | -0.13088 | 0 |
| B19 | B94 | CD3+CD4+/CD3+CD8+ | ALDH2-Lys504Lys | -0.09499 | 0 |
| B19 | B97 | CD3+CD4+/CD3+CD8+ | dsDNA | 0.107598 | 0 |
| B19 | B99 | CD3+CD4+/CD3+CD8+ | anti-PM-Scl | -0.16701 | 0 |
| B19 | B158 | CD3+CD4+/CD3+CD8+ | IgM | 0.082 | 2.75E-14 |
| B19 | B21 | CD3+CD4+/CD3+CD8+ | CD3+CD4-CD8- | -0.0801 | 1.06E-13 |
| B19 | B197 | CD3+CD4+/CD3+CD8+ | AG | -0.07827 | 3.81E-13 |
| B19 | B183 | CD3+CD4+/CD3+CD8+ | PTA | 0.077332 | 7.21E-13 |
| B19 | B73 | CD3+CD4+/CD3+CD8+ | ApoA | 0.073079 | 1.20E-11 |
| B19 | B185 | CD3+CD4+/CD3+CD8+ | PT | -0.06846 | 2.15E-10 |
| B19 | B109 | CD3+CD4+/CD3+CD8+ | AMA-M2 | -0.06319 | 4.62E-09 |
| B19 | B138 | CD3+CD4+/CD3+CD8+ | PAB | 0.054276 | 4.85E-07 |
| B19 | B24 | CD3+CD4+/CD3+CD8+ | αHBDH | -0.05349 | 7.09E-07 |
| B190 | B191 | Cre | Urea | 0.646286 | 0 |
| B190 | B192 | Cre | UA | 0.26022 | 0 |
| B190 | B198 | Cre | P | 0.097843 | 0 |
| B190 | B221 | Cre | Hei | 0.110656 | 0 |

| | | | | | |
|---|---|---|---|---|---|
| B190 | B222 | Cre | NS | 0.077279 | 7.47E-13 |
| B190 | B200 | Cre | OP | 0.070613 | 5.74E-11 |
| B190 | B196 | Cre | TCO2 | -0.06469 | 1.97E-09 |
| B190 | B207 | Cre | WHR | 0.064554 | 2.14E-09 |
| B190 | B216 | Cre | BMI | 0.062055 | 8.68E-09 |
| B190 | B206 | Cre | Waist | 0.053887 | 5.85E-07 |
| B191 | B192 | Urea | UA | 0.270509 | 0 |
| B191 | B200 | Urea | OP | 0.10083 | 0 |
| B191 | B218 | Urea | DBP | -0.12461 | 0 |
| B191 | B198 | Urea | P | 0.075653 | 2.23E-12 |
| B191 | B204 | Urea | Drinking | -0.06806 | 2.73E-10 |
| B191 | B220 | Urea | SBP | 0.063843 | 3.20E-09 |
| B191 | B209 | Urea | Smoking | -0.053 | 8.96E-07 |
| B192 | B206 | UA | Waist | 0.181067 | 0 |
| B192 | B207 | UA | WHR | 0.147958 | 0 |
| B192 | B210 | UA | Hip | 0.11949 | 0 |
| B192 | B215 | UA | BMI | 0.136529 | 0 |
| B192 | B216 | UA | BMI | 0.195821 | 0 |
| B192 | B221 | UA | Hei | 0.156525 | 0 |
| B192 | B222 | UA | NS | 0.130143 | 0 |
| B192 | B204 | UA | Drinking | 0.065596 | 1.17E-09 |
| B193 | B195 | K | Na | -0.13299 | 0 |
| B193 | B197 | K | AG | -0.12331 | 0 |
| B193 | B199 | K | Mg | 0.1526 | 0 |
| B193 | B198 | K | P | 0.070292 | 7.01E-11 |
| B193 | B200 | K | OP | -0.06913 | 1.44E-10 |
| B194 | B195 | Cl | Na | 0.356668 | 0 |
| B194 | B196 | Cl | TCO2 | -0.12645 | 0 |
| B194 | B197 | Cl | AG | -0.40326 | 0 |
| B194 | B199 | Cl | Mg | 0.17413 | 0 |
| B195 | B196 | Na | TCO2 | 0.143094 | 0 |
| B195 | B197 | Na | AG | 0.269914 | 0 |
| B195 | B200 | Na | OP | 0.551638 | 0 |
| B195 | B199 | Na | Mg | 0.0876 | 4.44E-16 |
| B196 | B197 | TCO2 | AG | -0.63687 | 0 |
| B197 | B199 | AG | Mg | -0.09042 | 0 |
| B197 | B200 | AG | OP | 0.343288 | 0 |
| B199 | B200 | Mg | OP | -0.2002 | 0 |
| B199 | B216 | Mg | BMI | -0.06336 | 4.20E-09 |
| B199 | B204 | Mg | Drinking | -0.06252 | 6.70E-09 |
| B199 | B221 | Mg | Hei | -0.05802 | 7.45E-08 |
| B199 | B225 | Mg | hPBG | -0.05283 | 9.67E-07 |
| B2 | B126 | Age | Alb | -0.25925 | 0 |
| B2 | B127 | Age | A-G ratio | -0.22298 | 0 |

| | | | | | |
|---|---|---|---|---|---|
| B2 | B128 | Age | ALT | -0.21767 | 0 |
| B2 | B130 | Age | GLB | 0.158168 | 0 |
| B2 | B134 | Age | CHE | -0.10058 | 0 |
| B2 | B140 | Age | r-GT | -0.13694 | 0 |
| B2 | B141 | Age | AST/ALT | 0.275056 | 0 |
| B2 | B146 | Age | CysC | 0.097924 | 0 |
| B2 | B163 | Age | SG | -0.16844 | 0 |
| B2 | B170 | Age | U-GLU | -0.09517 | 0 |
| B2 | B173 | Age | NTT | 0.091594 | 0 |
| B2 | B176 | Age | Ucr | -0.09728 | 0 |
| B2 | B189 | Age | eGFB | -0.46944 | 0 |
| B2 | B190 | Age | Cre | 0.100894 | 0 |
| B2 | B191 | Age | Urea | 0.197271 | 0 |
| B2 | B199 | Age | Mg | 0.089962 | 0 |
| B2 | B203 | Age | DH | -0.09278 | 0 |
| B2 | B204 | Age | Drinking | -0.23178 | 0 |
| B2 | B206 | Age | Waist | -0.12803 | 0 |
| B2 | B209 | Age | Smoking | -0.22728 | 0 |
| B2 | B210 | Age | Hip | -0.16866 | 0 |
| B2 | B211 | Age | IOP-R | -0.09398 | 0 |
| B2 | B212 | Age | IOP-L | -0.09965 | 0 |
| B2 | B215 | Age | BMI | -0.23185 | 0 |
| B2 | B216 | Age | BMI | -0.38829 | 0 |
| B2 | B218 | Age | DBP | -0.39829 | 0 |
| B2 | B219 | Age | OT | 0.114069 | 0 |
| B2 | B220 | Age | SBP | 0.18955 | 0 |
| B2 | B221 | Age | Hei | -0.34285 | 0 |
| B2 | B222 | Age | NS | -0.17063 | 0 |
| B2 | B41 | Age | MON％ | 0.155147 | 0 |
| B2 | B42 | Age | MON | 0.101007 | 0 |
| B2 | B43 | Age | MCV | 0.13543 | 0 |
| B2 | B45 | Age | MCHC | -0.18903 | 0 |
| B2 | B46 | Age | RBC Count | -0.31174 | 0 |
| B2 | B47 | Age | RDW | 0.247659 | 0 |
| B2 | B51 | Age | Hb | -0.31442 | 0 |
| B2 | B71 | Age | TG | -0.24335 | 0 |
| B2 | B72 | Age | HDL-C | 0.165495 | 0 |
| B2 | B74 | Age | ApoA1 | 0.089655 | 0 |
| B2 | B101 | Age | anti-Sc70 | 0.084269 | 5.33E-15 |
| B2 | B104 | Age | anti-SSB | 0.084269 | 5.33E-15 |
| B2 | B99 | Age | anti-PM-Scl | 0.084269 | 5.33E-15 |
| B2 | B107 | Age | AnuA | 0.083771 | 7.55E-15 |
| B2 | B27 | Age | LDH | 0.083061 | 1.27E-14 |
| B2 | B188 | Age | Ca | -0.0827 | 1.64E-14 |

| | | | | | |
|---|---|---|---|---|---|
| B2 | B48 | Age | LYM% | -0.08084 | 6.31E-14 |
| B2 | B138 | Age | PAB | -0.07416 | 5.99E-12 |
| B2 | B113 | Age | FT3 | -0.07243 | 1.83E-11 |
| B2 | B24 | Age | αHBDH | 0.070473 | 6.26E-11 |
| B2 | B77 | Age | TC | -0.07015 | 7.63E-11 |
| B2 | B94 | Age | ALDH2-Lys504Lys | 0.069855 | 9.18E-11 |
| B2 | B62 | Age | WBHSRVI | -0.068 | 2.84E-10 |
| B2 | B227 | Age | OD | 0.067815 | 3.17E-10 |
| B2 | B118 | Age | CEA1 | 0.067565 | 3.68E-10 |
| B2 | B184 | Age | INR | 0.067126 | 4.78E-10 |
| B2 | B66 | Age | WBRSR(mPas_100) | -0.06291 | 5.40E-09 |
| B2 | B56 | Age | IR | -0.06275 | 5.92E-09 |
| B2 | B67 | Age | WBRSR(mPas_50) | -0.06227 | 7.69E-09 |
| B2 | B61 | Age | WBHSRV | -0.06038 | 2.15E-08 |
| B2 | B205 | Age | KDA | 0.058969 | 4.55E-08 |
| B2 | B63 | Age | WBRSR(mPas_200) | -0.05882 | 4.92E-08 |
| B2 | B131 | Age | AST | -0.05801 | 7.48E-08 |
| B2 | B64 | Age | WBRSR(mPas_5) | -0.05749 | 9.80E-08 |
| B2 | B79 | Age | hb V -dna | 0.057309 | 1.07E-07 |
| B2 | B153 | Age | Glu | -0.05722 | 1.12E-07 |
| B2 | B185 | Age | PT | 0.057104 | 1.19E-07 |
| B2 | B161 | Age | LEU | 0.055945 | 2.13E-07 |
| B2 | B112 | Age | FT4 | -0.05491 | 3.56E-07 |
| B2 | B65 | Age | WBRSR(mPas_1) | -0.0529 | 9.39E-07 |
| B20 | B151 | CD3+CD4+CD8+ | Anti-CCP | -0.11858 | 0 |
| B20 | B181 | CD3+CD4+CD8+ | APTT | 0.133183 | 0 |
| B20 | B21 | CD3+CD4+CD8+ | CD3+CD4-CD8- | -0.14315 | 0 |
| B20 | B22 | CD3+CD4+CD8+ | CD3+CD8+(T) | 0.160975 | 0 |
| B20 | B79 | CD3+CD4+CD8+ | hb V -dna | -0.17488 | 0 |
| B20 | B81 | CD3+CD4+CD8+ | TSGF | 0.094201 | 0 |
| B20 | B196 | CD3+CD4+CD8+ | TCO2 | -0.08851 | 2.22E-16 |
| B20 | B194 | CD3+CD4+CD8+ | Cl | 0.08322 | 1.13E-14 |
| B20 | B74 | CD3+CD4+CD8+ | ApoA1 | -0.07881 | 2.62E-13 |
| B20 | B119 | CD3+CD4+CD8+ | Hcv-rna | -0.07796 | 4.70E-13 |
| B20 | B137 | CD3+CD4+CD8+ | AFU | 0.070056 | 8.11E-11 |
| B20 | B150 | CD3+CD4+CD8+ | ALP | 0.068918 | 1.63E-10 |
| B20 | B138 | CD3+CD4+CD8+ | PAB | -0.06838 | 2.26E-10 |
| B20 | B107 | CD3+CD4+CD8+ | AnuA | -0.06729 | 4.34E-10 |
| B20 | B101 | CD3+CD4+CD8+ | anti-Sc70 | -0.06697 | 5.24E-10 |
| B20 | B104 | CD3+CD4+CD8+ | anti-SSB | -0.06697 | 5.24E-10 |
| B20 | B99 | CD3+CD4+CD8+ | anti-PM-Scl | -0.06697 | 5.24E-10 |
| B20 | B75 | CD3+CD4+CD8+ | ApoA1/B | -0.0666 | 6.52E-10 |
| B20 | B200 | CD3+CD4+CD8+ | OP | 0.065558 | 1.20E-09 |
| B20 | B182 | CD3+CD4+CD8+ | TT | -0.06463 | 2.05E-09 |

| | | | | | |
|---|---|---|---|---|---|
| B20 | B69 | CD3+CD4+CD8+ | PV | -0.05975 | 3.02E-08 |
| B20 | B55 | CD3+CD4+CD8+ | TK | 0.058464 | 5.92E-08 |
| B20 | B56 | CD3+CD4+CD8+ | IR | 0.056404 | 1.70E-07 |
| B20 | B94 | CD3+CD4+CD8+ | ALDH2-Lys504Lys | 0.055681 | 2.43E-07 |
| B201 | B204 | e-habits | Drinking | 0.075318 | 2.79E-12 |
| B202 | B223 | e-times | FH | -0.51925 | 0 |
| B203 | B209 | DH | Smoking | 0.079871 | 1.25E-13 |
| B203 | B216 | DH | BMI | 0.076689 | 1.11E-12 |
| B203 | B215 | DH | BMI | 0.06347 | 3.95E-09 |
| B203 | B204 | DH | Drinking | 0.058336 | 6.33E-08 |
| B203 | B210 | DH | Hip | 0.057032 | 1.24E-07 |
| B203 | B206 | DH | Waist | 0.055791 | 2.30E-07 |
| B204 | B206 | Drinking | Waist | 0.124616 | 0 |
| B204 | B209 | Drinking | Smoking | 0.368555 | 0 |
| B204 | B210 | Drinking | Hip | 0.097167 | 0 |
| B204 | B216 | Drinking | BMI | 0.212415 | 0 |
| B204 | B218 | Drinking | DBP | 0.118769 | 0 |
| B204 | B221 | Drinking | Hei | 0.235947 | 0 |
| B204 | B222 | Drinking | NS | 0.153695 | 0 |
| B204 | B215 | Drinking | BMI | 0.086909 | 6.66E-16 |
| B204 | B207 | Drinking | WHR | 0.085294 | 2.44E-15 |
| B204 | B220 | Drinking | SBP | -0.08456 | 4.22E-15 |
| B204 | B219 | Drinking | OT | -0.07302 | 1.25E-11 |
| B205 | B221 | KDA | Hei | -0.07142 | 3.47E-11 |
| B206 | B207 | Waist | WHR | 0.716404 | 0 |
| B206 | B209 | Waist | Smoking | 0.097959 | 0 |
| B206 | B210 | Waist | Hip | 0.740675 | 0 |
| B206 | B215 | Waist | BMI | 0.791569 | 0 |
| B206 | B216 | Waist | BMI | 0.779129 | 0 |
| B206 | B218 | Waist | DBP | 0.11334 | 0 |
| B206 | B221 | Waist | Hei | 0.33044 | 0 |
| B206 | B222 | Waist | NS | 0.442615 | 0 |
| B206 | B219 | Waist | OT | -0.06199 | 9.01E-09 |
| B207 | B215 | WHR | BMI | 0.401022 | 0 |
| B207 | B216 | WHR | BMI | 0.370225 | 0 |
| B207 | B221 | WHR | Hei | 0.126724 | 0 |
| B207 | B222 | WHR | NS | 0.250995 | 0 |
| B207 | B210 | WHR | Hip | 0.08536 | 2.22E-15 |
| B207 | B209 | WHR | Smoking | 0.083718 | 7.99E-15 |
| B209 | B216 | Smoking | BMI | 0.185295 | 0 |
| B209 | B220 | Smoking | SBP | -0.12904 | 0 |
| B209 | B221 | Smoking | Hei | 0.262865 | 0 |
| B209 | B222 | Smoking | NS | 0.145435 | 0 |
| B209 | B219 | Smoking | OT | -0.07308 | 1.20E-11 |

| | | | | | |
|---|---|---|---|---|---|
| B209 | B218 | Smoking | DBP | 0.063781 | 3.32E-09 |
| B209 | B210 | Smoking | Hip | 0.060131 | 2.46E-08 |
| B21 | B101 | CD3+CD4-CD8- | anti-Sc70 | 0.125962 | 0 |
| B21 | B104 | CD3+CD4-CD8- | anti-SSB | 0.125962 | 0 |
| B21 | B107 | CD3+CD4-CD8- | AnuA | 0.126569 | 0 |
| B21 | B138 | CD3+CD4-CD8- | PAB | -0.13664 | 0 |
| B21 | B151 | CD3+CD4-CD8- | Anti-CCP | 0.123623 | 0 |
| B21 | B183 | CD3+CD4-CD8- | PTA | -0.11587 | 0 |
| B21 | B23 | CD3+CD4-CD8- | CD3-CD16+CD56+(NK) | -0.22222 | 0 |
| B21 | B99 | CD3+CD4-CD8- | anti-PM-Scl | 0.125962 | 0 |
| B21 | B158 | CD3+CD4-CD8- | IgM | 0.082547 | 1.84E-14 |
| B21 | B180 | CD3+CD4-CD8- | Tf | -0.07897 | 2.33E-13 |
| B21 | B30 | CD3+CD4-CD8- | LH | -0.07614 | 1.61E-12 |
| B21 | B187 | CD3+CD4-CD8- | NSE | 0.068278 | 2.40E-10 |
| B21 | B22 | CD3+CD4-CD8- | CD3+CD8+(T) | -0.06512 | 1.54E-09 |
| B21 | B94 | CD3+CD4-CD8- | ALDH2-Lys504Lys | 0.06376 | 3.35E-09 |
| B21 | B81 | CD3+CD4-CD8- | TSGF | -0.06339 | 4.12E-09 |
| B21 | B31 | CD3+CD4-CD8- | FSH | -0.06209 | 8.53E-09 |
| B21 | B178 | CD3+CD4-CD8- | Ig-U | -0.06166 | 1.07E-08 |
| B21 | B175 | CD3+CD4-CD8- | ALB-U | -0.06089 | 1.64E-08 |
| B21 | B194 | CD3+CD4-CD8- | Cl | 0.060262 | 2.30E-08 |
| B21 | B156 | CD3+CD4-CD8- | IgE | -0.05747 | 9.88E-08 |
| B21 | B92 | CD3+CD4-CD8- | ALDH2-Glu504Glu | -0.05456 | 4.23E-07 |
| B21 | B139 | CD3+CD4-CD8- | ADA | -0.05419 | 5.06E-07 |
| B210 | B215 | Hip | BMI | 0.739314 | 0 |
| B210 | B216 | Hip | BMI | 0.751748 | 0 |
| B210 | B218 | Hip | DBP | 0.125106 | 0 |
| B210 | B221 | Hip | Hei | 0.350276 | 0 |
| B210 | B222 | Hip | NS | 0.400558 | 0 |
| B211 | B212 | IOP-R | IOP-L | 0.777675 | 0 |
| B211 | B218 | IOP-R | DBP | 0.074779 | 3.98E-12 |
| B211 | B215 | IOP-R | BMI | 0.053903 | 5.80E-07 |
| B212 | B218 | IOP-L | DBP | 0.077124 | 8.31E-13 |
| B212 | B215 | IOP-L | BMI | 0.058915 | 4.68E-08 |
| B215 | B216 | BMI | BMI | 0.776218 | 0 |
| B215 | B218 | BMI | DBP | 0.145497 | 0 |
| B215 | B221 | BMI | Hei | 0.095216 | 0 |
| B215 | B222 | BMI | NS | 0.381745 | 0 |
| B215 | B219 | BMI | OT | -0.05588 | 2.20E-07 |
| B216 | B218 | BMI | DBP | 0.243321 | 0 |
| B216 | B219 | BMI | OT | -0.09589 | 0 |
| B216 | B221 | BMI | Hei | 0.69376 | 0 |
| B216 | B222 | BMI | NS | 0.510631 | 0 |
| B216 | B220 | BMI | SBP | -0.07711 | 8.42E-13 |

| | | | | | |
|---|---|---|---|---|---|
| B218 | B220 | DBP | SBP | 0.411964 | 0 |
| B218 | B221 | DBP | Hei | 0.216004 | 0 |
| B218 | B222 | DBP | NS | 0.118144 | 0 |
| B218 | B219 | DBP | OT | -0.0767 | 1.11E-12 |
| B219 | B221 | OT | Hei | -0.08451 | 4.44E-15 |
| B219 | B222 | OT | NS | -0.0689 | 1.65E-10 |
| B22 | B101 | CD3+CD8+(T) | anti-Sc70 | 0.098107 | 0 |
| B22 | B104 | CD3+CD8+(T) | anti-SSB | 0.098107 | 0 |
| B22 | B107 | CD3+CD8+(T) | AnuA | 0.098568 | 0 |
| B22 | B110 | CD3+CD8+(T) | AHA | -0.11753 | 0 |
| B22 | B125 | CD3+CD8+(T) | Cor | -0.09271 | 0 |
| B22 | B136 | CD3+CD8+(T) | 5'NT | 0.146308 | 0 |
| B22 | B156 | CD3+CD8+(T) | IgE | 0.171964 | 0 |
| B22 | B183 | CD3+CD8+(T) | PTA | -0.10051 | 0 |
| B22 | B195 | CD3+CD8+(T) | Na | 0.111958 | 0 |
| B22 | B23 | CD3+CD8+(T) | CD3-CD16+CD56+(NK) | -0.22958 | 0 |
| B22 | B28 | CD3+CD8+(T) | PRL | -0.1609 | 0 |
| B22 | B29 | CD3+CD8+(T) | E2 | -0.13224 | 0 |
| B22 | B30 | CD3+CD8+(T) | LH | 0.293783 | 0 |
| B22 | B31 | CD3+CD8+(T) | FSH | 0.226987 | 0 |
| B22 | B33 | CD3+CD8+(T) | Prog | -0.19351 | 0 |
| B22 | B97 | CD3+CD8+(T) | dsDNA | -0.15601 | 0 |
| B22 | B99 | CD3+CD8+(T) | anti-PM-Scl | 0.098107 | 0 |
| B22 | B93 | CD3+CD8+(T) | ALDH2-Glu504Lys | 0.086141 | 1.33E-15 |
| B22 | B92 | CD3+CD8+(T) | ALDH2-Glu504Glu | -0.08236 | 2.13E-14 |
| B22 | B193 | CD3+CD8+(T) | K | -0.08073 | 6.79E-14 |
| B22 | B122 | CD3+CD8+(T) | C4 | -0.07901 | 2.27E-13 |
| B22 | B185 | CD3+CD8+(T) | PT | 0.075433 | 2.58E-12 |
| B22 | B73 | CD3+CD8+(T) | ApoA | -0.06667 | 6.26E-10 |
| B22 | B69 | CD3+CD8+(T) | PV | -0.06594 | 9.59E-10 |
| B22 | B124 | CD3+CD8+(T) | ACTH | -0.06342 | 4.06E-09 |
| B22 | B24 | CD3+CD8+(T) | αHBDH | 0.063367 | 4.19E-09 |
| B22 | B194 | CD3+CD8+(T) | Cl | 0.060035 | 2.59E-08 |
| B22 | B54 | CD3+CD8+(T) | ESR | -0.05953 | 3.39E-08 |
| B22 | B197 | CD3+CD8+(T) | AG | 0.053551 | 6.87E-07 |
| B22 | B138 | CD3+CD8+(T) | PAB | -0.05346 | 7.19E-07 |
| B220 | B221 | SBP | Hei | -0.13631 | 0 |
| B221 | B222 | Hei | NS | 0.376237 | 0 |
| B227 | B228 | OD | OS | 0.072631 | 1.61E-11 |
| B23 | B101 | CD3-CD16+CD56+(NK) | anti-Sc70 | -0.10311 | 0 |
| B23 | B104 | CD3-CD16+CD56+(NK) | anti-SSB | -0.10311 | 0 |
| B23 | B107 | CD3-CD16+CD56+(NK) | AnuA | -0.10362 | 0 |
| B23 | B110 | CD3-CD16+CD56+(NK) | AHA | 0.306932 | 0 |
| B23 | B122 | CD3-CD16+CD56+(NK) | C4 | -0.13445 | 0 |

| | | | | | |
|---|---|---|---|---|---|
| B23 | B139 | CD3-CD16+CD56+(NK) | ADA | 0.100519 | 0 |
| B23 | B158 | CD3-CD16+CD56+(NK) | IgM | -0.09022 | 0 |
| B23 | B183 | CD3-CD16+CD56+(NK) | PTA | 0.119259 | 0 |
| B23 | B194 | CD3-CD16+CD56+(NK) | Cl | -0.11377 | 0 |
| B23 | B199 | CD3-CD16+CD56+(NK) | Mg | -0.10709 | 0 |
| B23 | B28 | CD3-CD16+CD56+(NK) | PRL | -0.11636 | 0 |
| B23 | B30 | CD3-CD16+CD56+(NK) | LH | 0.122704 | 0 |
| B23 | B31 | CD3-CD16+CD56+(NK) | FSH | 0.107466 | 0 |
| B23 | B33 | CD3-CD16+CD56+(NK) | Prog | -0.11648 | 0 |
| B23 | B94 | CD3-CD16+CD56+(NK) | ALDH2-Lys504Lys | 0.089768 | 0 |
| B23 | B96 | CD3-CD16+CD56+(NK) | dsDNA (IIF) | -0.09927 | 0 |
| B23 | B97 | CD3-CD16+CD56+(NK) | dsDNA | 0.327534 | 0 |
| B23 | B98 | CD3-CD16+CD56+(NK) | anti-Jo-1 | -0.09138 | 0 |
| B23 | B99 | CD3-CD16+CD56+(NK) | anti-PM-Scl | -0.10311 | 0 |
| B23 | B195 | CD3-CD16+CD56+(NK) | Na | -0.08246 | 1.95E-14 |
| B23 | B29 | CD3-CD16+CD56+(NK) | E2 | -0.08198 | 2.78E-14 |
| B23 | B186 | CD3-CD16+CD56+(NK) | PFC | 0.078652 | 2.92E-13 |
| B23 | B95 | CD3-CD16+CD56+(NK) | anti-cenpb | 0.075641 | 2.25E-12 |
| B23 | B180 | CD3-CD16+CD56+(NK) | Tf | 0.074723 | 4.13E-12 |
| B23 | B54 | CD3-CD16+CD56+(NK) | ESR | 0.073136 | 1.16E-11 |
| B23 | B137 | CD3-CD16+CD56+(NK) | AFU | 0.072534 | 1.71E-11 |
| B23 | B200 | CD3-CD16+CD56+(NK) | OP | 0.072159 | 2.17E-11 |
| B23 | B178 | CD3-CD16+CD56+(NK) | Ig-U | 0.06005 | 2.57E-08 |
| B23 | B69 | CD3-CD16+CD56+(NK) | PV | 0.058126 | 7.05E-08 |
| B23 | B175 | CD3-CD16+CD56+(NK) | ALB-U | 0.057388 | 1.03E-07 |
| B23 | B121 | CD3-CD16+CD56+(NK) | C3 | -0.0545 | 4.35E-07 |
| B23 | B184 | CD3-CD16+CD56+(NK) | INR | 0.053325 | 7.66E-07 |
| B24 | B111 | αHBDH | TSH | 0.191223 | 0 |
| B24 | B146 | αHBDH | CysC | 0.114008 | 0 |
| B24 | B177 | αHBDH | ACR | 0.151699 | 0 |
| B24 | B179 | αHBDH | α1-MG | 0.093425 | 0 |
| B24 | B25 | αHBDH | CK | 0.291389 | 0 |
| B24 | B26 | αHBDH | CKMB | 0.412911 | 0 |
| B24 | B27 | αHBDH | LDH | 0.649514 | 0 |
| B24 | B37 | αHBDH | BAND | -0.13662 | 0 |
| B24 | B38 | αHBDH | BAND% | -0.19124 | 0 |
| B24 | B47 | αHBDH | RDW | 0.097283 | 0 |
| B24 | B50 | αHBDH | eo% | 0.09623 | 0 |
| B24 | B74 | αHBDH | ApoA1 | 0.105328 | 0 |
| B24 | B69 | αHBDH | PV | 0.086438 | 8.88E-16 |
| B24 | B139 | αHBDH | ADA | 0.083338 | 1.04E-14 |
| B24 | B175 | αHBDH | ALB-U | 0.077123 | 8.31E-13 |
| B24 | B131 | αHBDH | AST | 0.076123 | 1.63E-12 |
| B24 | B72 | αHBDH | HDL-C | 0.075362 | 2.71E-12 |

| | | | | | |
|---|---|---|---|---|---|
| B24 | B178 | αHBDH | Ig-U | 0.068671 | 1.89E-10 |
| B24 | B54 | αHBDH | ESR | 0.066926 | 5.38E-10 |
| B24 | B127 | αHBDH | A-G ratio | -0.0667 | 6.16E-10 |
| B24 | B199 | αHBDH | Mg | 0.065132 | 1.53E-09 |
| B24 | B123 | αHBDH | hs-CRP1 | 0.058296 | 6.46E-08 |
| B24 | B220 | αHBDH | SBP | 0.056718 | 1.45E-07 |
| B24 | B130 | αHBDH | GLB | 0.055408 | 2.79E-07 |
| B24 | B124 | αHBDH | ACTH | 0.054479 | 4.39E-07 |
| B24 | B221 | αHBDH | Hei | -0.05311 | 8.49E-07 |
| B24 | B126 | αHBDH | Alb | -0.05289 | 9.41E-07 |
| B25 | B111 | CK | TSH | 0.290853 | 0 |
| B25 | B114 | CK | TSH TT3 TT4(TT4) | -0.10133 | 0 |
| B25 | B177 | CK | ACR | 0.089616 | 0 |
| B25 | B26 | CK | CKMB | 0.427238 | 0 |
| B25 | B27 | CK | LDH | 0.242266 | 0 |
| B25 | B131 | CK | AST | 0.086275 | 1.11E-15 |
| B25 | B112 | CK | FT4 | -0.08161 | 3.64E-14 |
| B25 | B178 | CK | Ig-U | 0.071328 | 3.67E-11 |
| B25 | B87 | CK | NSE | 0.059428 | 3.57E-08 |
| B25 | B180 | CK | Tf | 0.058465 | 5.92E-08 |
| B25 | B193 | CK | K | -0.0564 | 1.70E-07 |
| B25 | B115 | CK | TSH TT3 TT4(TT3) | -0.05524 | 3.03E-07 |
| B26 | B111 | CKMB | TSH | 0.190536 | 0 |
| B26 | B27 | CKMB | LDH | 0.31559 | 0 |
| B26 | B66 | CKMB | WBRSR(mPas_100) | 0.080055 | 1.10E-13 |
| B26 | B67 | CKMB | WBRSR(mPas_50) | 0.076473 | 1.29E-12 |
| B26 | B177 | CKMB | ACR | 0.075196 | 3.02E-12 |
| B26 | B63 | CKMB | WBRSR(mPas_200) | 0.075111 | 3.20E-12 |
| B26 | B62 | CKMB | WBHSRVI | 0.074307 | 5.43E-12 |
| B26 | B64 | CKMB | WBRSR(mPas_5) | 0.072571 | 1.67E-11 |
| B26 | B131 | CKMB | AST | 0.066554 | 6.71E-10 |
| B26 | B139 | CKMB | ADA | 0.06642 | 7.26E-10 |
| B26 | B81 | CKMB | TSGF | 0.064364 | 2.38E-09 |
| B26 | B87 | CKMB | NSE | 0.06402 | 2.90E-09 |
| B26 | B65 | CKMB | WBRSR(mPas_1) | 0.061934 | 9.27E-09 |
| B26 | B60 | CKMB | WBRVI | 0.0578 | 8.35E-08 |
| B26 | B114 | CKMB | TSH TT3 TT4(TT4) | -0.05567 | 2.45E-07 |
| B27 | B111 | LDH | TSH | 0.133829 | 0 |
| B27 | B118 | LDH | CEA1 | 0.10943 | 0 |
| B27 | B131 | LDH | AST | 0.132312 | 0 |
| B27 | B139 | LDH | ADA | 0.145368 | 0 |
| B27 | B146 | LDH | CysC | 0.124898 | 0 |
| B27 | B175 | LDH | ALB-U | 0.090258 | 0 |
| B27 | B177 | LDH | ACR | 0.139935 | 0 |

| | | | | | |
|---|---|---|---|---|---|
| B27 | B37 | LDH | BAND | -0.08915 | 0 |
| B27 | B38 | LDH | BAND% | -0.12691 | 0 |
| B27 | B47 | LDH | RDW | 0.085563 | 2.00E-15 |
| B27 | B72 | LDH | HDL-C | 0.080957 | 5.77E-14 |
| B27 | B127 | LDH | A-G ratio | -0.07964 | 1.47E-13 |
| B27 | B50 | LDH | eo% | 0.079275 | 1.89E-13 |
| B27 | B74 | LDH | ApoA1 | 0.075062 | 3.30E-12 |
| B27 | B179 | LDH | α1-MG | 0.073092 | 1.19E-11 |
| B27 | B150 | LDH | ALP | 0.072969 | 1.29E-11 |
| B27 | B69 | LDH | PV | 0.067918 | 2.98E-10 |
| B27 | B126 | LDH | Alb | -0.06546 | 1.27E-09 |
| B27 | B30 | LDH | LH | -0.06461 | 2.07E-09 |
| B27 | B128 | LDH | ALT | 0.064138 | 2.71E-09 |
| B27 | B123 | LDH | hs-CRP1 | 0.063609 | 3.65E-09 |
| B27 | B33 | LDH | Prog | 0.061945 | 9.22E-09 |
| B27 | B134 | LDH | CHE | -0.06014 | 2.45E-08 |
| B27 | B130 | LDH | GLB | 0.059366 | 3.69E-08 |
| B27 | B220 | LDH | SBP | 0.05909 | 4.27E-08 |
| B27 | B178 | LDH | Ig-U | 0.056687 | 1.47E-07 |
| B27 | B190 | LDH | Cre | 0.056318 | 1.77E-07 |
| B27 | B113 | LDH | FT3 | -0.05431 | 4.77E-07 |
| B27 | B28 | LDH | PRL | 0.053804 | 6.09E-07 |
| B27 | B191 | LDH | Urea | 0.053707 | 6.38E-07 |
| B28 | B121 | PRL | C3 | 0.109718 | 0 |
| B28 | B122 | PRL | C4 | 0.127688 | 0 |
| B28 | B156 | PRL | IgE | -0.1629 | 0 |
| B28 | B29 | PRL | E2 | 0.597137 | 0 |
| B28 | B30 | PRL | LH | -0.70238 | 0 |
| B28 | B31 | PRL | FSH | -0.80939 | 0 |
| B28 | B32 | PRL | Testo | -0.26458 | 0 |
| B28 | B33 | PRL | Prog | 0.802833 | 0 |
| B28 | B49 | PRL | baso% | -0.09392 | 0 |
| B28 | B50 | PRL | eo% | -0.09579 | 0 |
| B28 | B150 | PRL | ALP | 0.075785 | 2.04E-12 |
| B28 | B113 | PRL | FT3 | 0.066672 | 6.26E-10 |
| B28 | B176 | PRL | Ucr | 0.064426 | 2.30E-09 |
| B28 | B119 | PRL | Hcv-rna | 0.063362 | 4.20E-09 |
| B28 | B79 | PRL | hb V -dna | 0.053902 | 5.81E-07 |
| B29 | B122 | E2 | C4 | 0.089603 | 0 |
| B29 | B156 | E2 | IgE | -0.12808 | 0 |
| B29 | B30 | E2 | LH | -0.40288 | 0 |
| B29 | B31 | E2 | FSH | -0.65218 | 0 |
| B29 | B33 | E2 | Prog | 0.121265 | 0 |
| B29 | B32 | E2 | Testo | -0.07931 | 1.85E-13 |

| | | | | | |
|---|---|---|---|---|---|
| B29 | B121 | E2 | C3 | 0.0725 | 1.75E-11 |
| B29 | B125 | E2 | Cor | -0.05434 | 4.71E-07 |
| B3 | B189 | Hcy | eGFB | -0.16068 | 0 |
| B3 | B190 | Hcy | Cre | 0.096202 | 0 |
| B3 | B191 | Hcy | Urea | 0.11488 | 0 |
| B3 | B192 | Hcy | UA | 0.092261 | 0 |
| B3 | B74 | Hcy | ApoA1 | -0.08943 | 0 |
| B3 | B86 | Hcy | TBhCG | -0.09846 | 0 |
| B3 | B146 | Hcy | CysC | 0.087134 | 4.44E-16 |
| B3 | B120 | Hcy | HCV-cAg | -0.0731 | 1.19E-11 |
| B3 | B115 | Hcy | TSH TT3 TT4(TT3) | -0.06292 | 5.37E-09 |
| B3 | B84 | Hcy | C-PSA | 0.061712 | 1.05E-08 |
| B3 | B199 | Hcy | Mg | -0.0557 | 2.41E-07 |
| B3 | B48 | Hcy | LYM% | -0.05443 | 4.49E-07 |
| B30 | B121 | LH | C3 | -0.12954 | 0 |
| B30 | B122 | LH | C4 | -0.16301 | 0 |
| B30 | B150 | LH | ALP | -0.09064 | 0 |
| B30 | B156 | LH | IgE | 0.243842 | 0 |
| B30 | B31 | LH | FSH | 0.885853 | 0 |
| B30 | B32 | LH | Testo | -0.26107 | 0 |
| B30 | B33 | LH | Prog | -0.60418 | 0 |
| B30 | B49 | LH | baso% | 0.114471 | 0 |
| B30 | B50 | LH | eo% | 0.115242 | 0 |
| B30 | B176 | LH | Ucr | -0.06544 | 1.28E-09 |
| B30 | B113 | LH | FT3 | -0.06425 | 2.55E-09 |
| B30 | B199 | LH | Mg | -0.05965 | 3.18E-08 |
| B30 | B203 | LH | DH | 0.057867 | 8.06E-08 |
| B30 | B158 | LH | IgM | -0.05739 | 1.03E-07 |
| B30 | B193 | LH | K | -0.05415 | 5.16E-07 |
| B30 | B155 | LH | IgA | -0.05351 | 7.00E-07 |
| B31 | B121 | FSH | C3 | -0.0968 | 0 |
| B31 | B122 | FSH | C4 | -0.1286 | 0 |
| B31 | B156 | FSH | IgE | 0.203052 | 0 |
| B31 | B32 | FSH | Testo | -0.20413 | 0 |
| B31 | B33 | FSH | Prog | -0.57595 | 0 |
| B31 | B49 | FSH | baso% | 0.091654 | 0 |
| B31 | B50 | FSH | eo% | 0.091419 | 0 |
| B31 | B150 | FSH | ALP | -0.07727 | 7.51E-13 |
| B31 | B113 | FSH | FT3 | -0.05623 | 1.86E-07 |
| B32 | B178 | Testo | Ig-U | 0.141301 | 0 |
| B32 | B33 | Testo | Prog | -0.21045 | 0 |
| B32 | B63 | Testo | WBRSR(mPas_200) | 0.121214 | 0 |
| B32 | B64 | Testo | WBRSR(mPas_5) | 0.091205 | 0 |
| B32 | B66 | Testo | WBRSR(mPas_100) | 0.122549 | 0 |

| | | | | | |
|---|---|---|---|---|---|
| B32 | B67 | Testo | WBRSR(mPas_50) | 0.118366 | 0 |
| B32 | B62 | Testo | WBHSRVI | 0.087113 | 6.66E-16 |
| B32 | B61 | Testo | WBHSRV | 0.079539 | 1.57E-13 |
| B32 | B180 | Testo | Tf | 0.076296 | 1.45E-12 |
| B32 | B69 | Testo | PV | 0.074351 | 5.28E-12 |
| B32 | B65 | Testo | WBRSR(mPas_1) | 0.069286 | 1.30E-10 |
| B32 | B200 | Testo | OP | 0.066186 | 8.32E-10 |
| B32 | B177 | Testo | ACR | 0.06038 | 2.15E-08 |
| B33 | B121 | Prog | C3 | 0.104763 | 0 |
| B33 | B122 | Prog | C4 | 0.129493 | 0 |
| B33 | B156 | Prog | IgE | -0.18581 | 0 |
| B33 | B49 | Prog | baso% | -0.09493 | 0 |
| B33 | B50 | Prog | eo% | -0.10006 | 0 |
| B33 | B176 | Prog | Ucr | 0.074584 | 4.53E-12 |
| B33 | B150 | Prog | ALP | 0.071793 | 2.74E-11 |
| B33 | B113 | Prog | FT3 | 0.061781 | 1.01E-08 |
| B34 | B73 | TK | ApoA | 0.123663 | 0 |
| B34 | B81 | TK | TSGF | -0.08875 | 2.22E-16 |
| B34 | B193 | TK | K | -0.08707 | 6.66E-16 |
| B34 | B79 | TK | hb V -dna | 0.077125 | 8.30E-13 |
| B34 | B184 | TK | INR | 0.07399 | 6.68E-12 |
| B34 | B185 | TK | PT | 0.0667 | 6.15E-10 |
| B34 | B107 | TK | AnuA | 0.059446 | 3.54E-08 |
| B34 | B101 | TK | anti-Sc70 | 0.059155 | 4.13E-08 |
| B34 | B104 | TK | anti-SSB | 0.059155 | 4.13E-08 |
| B34 | B99 | TK | anti-PM-Scl | 0.059155 | 4.13E-08 |
| B35 | B125 | β2-MG | Cor | -0.21088 | 0 |
| B35 | B159 | β2-MG | β2-MG | 0.99982 | 0 |
| B35 | B181 | β2-MG | APTT | 0.149658 | 0 |
| B35 | B184 | β2-MG | INR | 0.155095 | 0 |
| B35 | B186 | β2-MG | PFC | -0.09282 | 0 |
| B35 | B185 | β2-MG | PT | 0.088531 | 2.22E-16 |
| B35 | B74 | β2-MG | ApoA1 | -0.07883 | 2.57E-13 |
| B35 | B183 | β2-MG | PTA | 0.058419 | 6.06E-08 |
| B36 | B164 | V-Lym% | COM | -0.16995 | 0 |
| B36 | B40 | V-Lym% | WBC | -0.11107 | 0 |
| B36 | B48 | V-Lym% | LYM% | -0.07098 | 4.57E-11 |
| B36 | B53 | V-Lym% | GR% | 0.065337 | 1.36E-09 |
| B37 | B38 | BAND | BAND% | 0.955417 | 0 |
| B37 | B49 | BAND | baso% | -0.22818 | 0 |
| B37 | B50 | BAND | eo% | -0.12212 | 0 |
| B37 | B41 | BAND | MON % | 0.066526 | 6.82E-10 |
| B37 | B78 | BAND | HBcAb | -0.05477 | 3.82E-07 |
| B37 | B146 | BAND | CysC | -0.05471 | 3.93E-07 |

| | | | | | |
|---|---|---|---|---|---|
| B38 | B49 | BAND% | baso% | -0.20542 | 0 |
| B38 | B50 | BAND% | eo% | -0.14433 | 0 |
| B38 | B146 | BAND% | CysC | -0.07555 | 2.39E-12 |
| B38 | B41 | BAND% | MON % | 0.071911 | 2.54E-11 |
| B38 | B40 | BAND% | WBC | -0.06405 | 2.85E-09 |
| B39 | B44 | Other cells | MCH | -0.09796 | 0 |
| B39 | B43 | Other cells | MCV | -0.08655 | 8.88E-16 |
| B39 | B45 | Other cells | MCHC | -0.06611 | 8.69E-10 |
| B4 | B108 | WB-Ca | Anti-Cyclin | -0.17346 | 0 |
| B4 | B17 | WB-Ca | CD3+CD16+CD56+NK | 0.0987 | 0 |
| B4 | B5 | WB-Ca | WB-Mg | 0.612206 | 0 |
| B4 | B7 | WB-Ca | WB-Cop | 0.734551 | 0 |
| B4 | B8 | WB-Ca | WB-Zinc | 0.101182 | 0 |
| B4 | B185 | WB-Ca | PT | 0.083019 | 1.31E-14 |
| B4 | B54 | WB-Ca | ESR | 0.072426 | 1.83E-11 |
| B4 | B184 | WB-Ca | INR | 0.05928 | 3.86E-08 |
| B40 | B127 | WBC | A-G ratio | -0.129 | 0 |
| B40 | B129 | WBC | IBIL | -0.10871 | 0 |
| B40 | B130 | WBC | GLB | 0.144757 | 0 |
| B40 | B132 | WBC | Dbil | -0.13456 | 0 |
| B40 | B133 | WBC | TBIL | -0.12511 | 0 |
| B40 | B192 | WBC | UA | 0.091153 | 0 |
| B40 | B206 | WBC | Waist | 0.103582 | 0 |
| B40 | B207 | WBC | WHR | 0.096126 | 0 |
| B40 | B209 | WBC | Smoking | 0.132646 | 0 |
| B40 | B41 | WBC | MON % | -0.16593 | 0 |
| B40 | B42 | WBC | MON | 0.653613 | 0 |
| B40 | B46 | WBC | RBC Count | 0.145648 | 0 |
| B40 | B48 | WBC | LYM% | -0.1829 | 0 |
| B40 | B51 | WBC | Hb | 0.105616 | 0 |
| B40 | B52 | WBC | PLT/BPC | 0.278309 | 0 |
| B40 | B53 | WBC | GR% | 0.203 | 0 |
| B40 | B72 | WBC | HDL-C | -0.11698 | 0 |
| B40 | B91 | WBC | TP | 0.113365 | 0 |
| B40 | B166 | WBC | Pro | 0.084243 | 5.33E-15 |
| B40 | B153 | WBC | Glu | 0.076395 | 1.36E-12 |
| B40 | B44 | WBC | MCH | -0.07533 | 2.77E-12 |
| B40 | B43 | WBC | MCV | -0.07378 | 7.68E-12 |
| B40 | B160 | WBC | PH | -0.07083 | 5.02E-11 |
| B40 | B49 | WBC | baso% | -0.06933 | 1.27E-10 |
| B40 | B215 | WBC | BMI | 0.068189 | 2.53E-10 |
| B40 | B216 | WBC | BMI | 0.067555 | 3.71E-10 |
| B40 | B71 | WBC | TG | 0.066661 | 6.29E-10 |
| B40 | B188 | WBC | Ca | 0.062125 | 8.35E-09 |

| | | | | | |
|---|---|---|---|---|---|
| B40 | B222 | WBC | NS | 0.060479 | 2.04E-08 |
| B40 | B163 | WBC | SG | 0.05893 | 4.64E-08 |
| B40 | B210 | WBC | Hip | 0.055986 | 2.09E-07 |
| B40 | B170 | WBC | U-GLU | 0.053318 | 7.68E-07 |
| B41 | B126 | MON % | Alb | -0.12774 | 0 |
| B41 | B131 | MON % | AST | 0.091132 | 0 |
| B41 | B132 | MON % | Dbil | 0.118358 | 0 |
| B41 | B189 | MON % | eGFB | -0.1314 | 0 |
| B41 | B190 | MON % | Cre | 0.098661 | 0 |
| B41 | B42 | MON % | MON | 0.595941 | 0 |
| B41 | B43 | MON % | MCV | 0.125608 | 0 |
| B41 | B44 | MON % | MCH | 0.106274 | 0 |
| B41 | B47 | MON % | RDW | 0.129092 | 0 |
| B41 | B53 | MON % | GR% | -0.19323 | 0 |
| B41 | B71 | MON % | TG | -0.10718 | 0 |
| B41 | B221 | MON % | Hei | 0.086763 | 8.88E-16 |
| B41 | B77 | MON % | TC | -0.08664 | 8.88E-16 |
| B41 | B191 | MON % | Urea | 0.084483 | 4.44E-15 |
| B41 | B49 | MON % | baso% | 0.078031 | 4.47E-13 |
| B41 | B218 | MON % | DBP | -0.07764 | 5.84E-13 |
| B41 | B192 | MON % | UA | 0.0724 | 1.86E-11 |
| B41 | B127 | MON % | A-G ratio | -0.06936 | 1.24E-10 |
| B41 | B146 | MON % | CysC | 0.067913 | 2.99E-10 |
| B41 | B212 | MON % | IOP-L | -0.06741 | 4.05E-10 |
| B41 | B133 | MON % | TBIL | 0.06596 | 9.49E-10 |
| B41 | B174 | MON % | Urea/Cr | -0.06254 | 6.63E-09 |
| B41 | B211 | MON % | IOP-R | -0.06042 | 2.11E-08 |
| B41 | B46 | MON % | RBC Count | -0.05988 | 2.81E-08 |
| B41 | B91 | MON % | TP | -0.05336 | 7.54E-07 |
| B42 | B126 | MON | Alb | -0.11289 | 0 |
| B42 | B127 | MON | A-G ratio | -0.15774 | 0 |
| B42 | B130 | MON | GLB | 0.142905 | 0 |
| B42 | B189 | MON | eGFB | -0.10363 | 0 |
| B42 | B190 | MON | Cre | 0.10257 | 0 |
| B42 | B191 | MON | Urea | 0.089329 | 0 |
| B42 | B192 | MON | UA | 0.121954 | 0 |
| B42 | B206 | MON | Waist | 0.110888 | 0 |
| B42 | B207 | MON | WHR | 0.111857 | 0 |
| B42 | B209 | MON | Smoking | 0.132578 | 0 |
| B42 | B221 | MON | Hei | 0.090755 | 0 |
| B42 | B48 | MON | LYM% | -0.17626 | 0 |
| B42 | B51 | MON | Hb | 0.093665 | 0 |
| B42 | B52 | MON | PLT/BPC | 0.197003 | 0 |
| B42 | B163 | MON | SG | 0.087053 | 6.66E-16 |

| | | | | | |
|---|---|---|---|---|---|
| B42 | B166 | MON | Pro | 0.086494 | 8.88E-16 |
| B42 | B46 | MON | RBC Count | 0.083679 | 7.99E-15 |
| B42 | B222 | MON | NS | 0.082285 | 2.22E-14 |
| B42 | B216 | MON | BMI | 0.081001 | 5.60E-14 |
| B42 | B174 | MON | Urea/Cr | -0.07958 | 1.53E-13 |
| B42 | B72 | MON | HDL-C | -0.06856 | 2.02E-10 |
| B42 | B123 | MON | hs-CRP1 | 0.062285 | 7.64E-09 |
| B42 | B146 | MON | CysC | 0.061467 | 1.20E-08 |
| B42 | B91 | MON | TP | 0.059864 | 2.84E-08 |
| B42 | B210 | MON | Hip | 0.053353 | 7.55E-07 |
| B42 | B129 | MON | IBIL | -0.05327 | 7.85E-07 |
| B43 | B153 | MCV | Glu | -0.11351 | 0 |
| B43 | B189 | MCV | eGFB | -0.09732 | 0 |
| B43 | B44 | MCV | MCH | 0.883656 | 0 |
| B43 | B45 | MCV | MCHC | 0.114511 | 0 |
| B43 | B46 | MCV | RBC Count | -0.5334 | 0 |
| B43 | B47 | MCV | RDW | 0.581018 | 0 |
| B43 | B52 | MCV | PLT/BPC | -0.13205 | 0 |
| B43 | B53 | MCV | GR% | -0.10591 | 0 |
| B43 | B72 | MCV | HDL-C | 0.139975 | 0 |
| B43 | B141 | MCV | AST/ALT | 0.077907 | 4.87E-13 |
| B43 | B71 | MCV | TG | -0.07784 | 5.09E-13 |
| B43 | B48 | MCV | LYM% | 0.074329 | 5.35E-12 |
| B43 | B218 | MCV | DBP | -0.07137 | 3.57E-11 |
| B43 | B204 | MCV | Drinking | 0.068058 | 2.74E-10 |
| B43 | B209 | MCV | Smoking | 0.066034 | 9.09E-10 |
| B43 | B190 | MCV | Cre | 0.064276 | 2.50E-09 |
| B43 | B132 | MCV | Dbil | 0.06321 | 4.57E-09 |
| B43 | B191 | MCV | Urea | 0.063098 | 4.87E-09 |
| B43 | B112 | MCV | FT4 | -0.05783 | 8.21E-08 |
| B43 | B170 | MCV | U-GLU | -0.05608 | 2.00E-07 |
| B43 | B135 | MCV | TBA | -0.05504 | 3.34E-07 |
| B43 | B59 | MCV | WBRV | -0.05333 | 7.64E-07 |
| B44 | B132 | MCH | Dbil | 0.116382 | 0 |
| B44 | B133 | MCH | TBIL | 0.098961 | 0 |
| B44 | B204 | MCH | Drinking | 0.139269 | 0 |
| B44 | B209 | MCH | Smoking | 0.125557 | 0 |
| B44 | B45 | MCH | MCHC | 0.564308 | 0 |
| B44 | B46 | MCH | RBC Count | -0.39453 | 0 |
| B44 | B47 | MCH | RDW | 0.353494 | 0 |
| B44 | B51 | MCH | Hb | 0.268762 | 0 |
| B44 | B52 | MCH | PLT/BPC | -0.11414 | 0 |
| B44 | B53 | MCH | GR% | -0.10736 | 0 |
| B44 | B221 | MCH | Hei | 0.08494 | 3.11E-15 |

| | | | | | |
|---|---|---|---|---|---|
| B44 | B129 | MCH | IBIL | 0.082299 | 2.22E-14 |
| B44 | B48 | MCH | LYM% | 0.081524 | 3.86E-14 |
| B44 | B174 | MCH | Urea/Cr | -0.06893 | 1.62E-10 |
| B44 | B72 | MCH | HDL-C | 0.068001 | 2.84E-10 |
| B44 | B140 | MCH | r-GT | 0.061745 | 1.03E-08 |
| B44 | B131 | MCH | AST | 0.058498 | 5.82E-08 |
| B44 | B220 | MCH | SBP | -0.05745 | 9.99E-08 |
| B44 | B135 | MCH | TBA | -0.05576 | 2.35E-07 |
| B44 | B150 | MCH | ALP | -0.05378 | 6.16E-07 |
| B45 | B126 | MCHC | Alb | 0.131928 | 0 |
| B45 | B127 | MCHC | A-G ratio | 0.11621 | 0 |
| B45 | B128 | MCHC | ALT | 0.106588 | 0 |
| B45 | B129 | MCHC | IBIL | 0.189126 | 0 |
| B45 | B132 | MCHC | Dbil | 0.127479 | 0 |
| B45 | B133 | MCHC | TBIL | 0.18678 | 0 |
| B45 | B141 | MCHC | AST/ALT | -0.11864 | 0 |
| B45 | B153 | MCHC | Glu | 0.13771 | 0 |
| B45 | B163 | MCHC | SG | 0.125556 | 0 |
| B45 | B170 | MCHC | U-GLU | 0.102889 | 0 |
| B45 | B174 | MCHC | Urea/Cr | -0.10554 | 0 |
| B45 | B189 | MCHC | eGFB | 0.159608 | 0 |
| B45 | B191 | MCHC | Urea | -0.14654 | 0 |
| B45 | B204 | MCHC | Drinking | 0.16775 | 0 |
| B45 | B209 | MCHC | Smoking | 0.142795 | 0 |
| B45 | B216 | MCHC | BMI | 0.172594 | 0 |
| B45 | B218 | MCHC | DBP | 0.142266 | 0 |
| B45 | B221 | MCHC | Hei | 0.234085 | 0 |
| B45 | B222 | MCHC | NS | 0.12465 | 0 |
| B45 | B46 | MCHC | RBC Count | 0.104396 | 0 |
| B45 | B47 | MCHC | RDW | -0.28016 | 0 |
| B45 | B51 | MCHC | Hb | 0.490069 | 0 |
| B45 | B71 | MCHC | TG | 0.169519 | 0 |
| B45 | B72 | MCHC | HDL-C | -0.1025 | 0 |
| B45 | B130 | MCHC | GLB | -0.08108 | 5.33E-14 |
| B45 | B190 | MCHC | Cre | -0.07656 | 1.22E-12 |
| B45 | B161 | MCHC | LEU | -0.07556 | 2.38E-12 |
| B45 | B140 | MCHC | r-GT | 0.075504 | 2.47E-12 |
| B45 | B62 | MCHC | WBHSRVI | 0.07405 | 6.42E-12 |
| B45 | B194 | MCHC | Cl | -0.07248 | 1.77E-11 |
| B45 | B146 | MCHC | CysC | -0.07209 | 2.27E-11 |
| B45 | B66 | MCHC | WBRSR(mPas_100) | 0.071617 | 3.06E-11 |
| B45 | B63 | MCHC | WBRSR(mPas_200) | 0.070458 | 6.33E-11 |
| B45 | B67 | MCHC | WBRSR(mPas_50) | 0.06899 | 1.56E-10 |
| B45 | B206 | MCHC | Waist | 0.065881 | 9.94E-10 |

| | | | | | |
|---|---|---|---|---|---|
| B45 | B219 | MCHC | OT | -0.06495 | 1.71E-09 |
| B45 | B64 | MCHC | WBRSR(mPas_5) | 0.064298 | 2.47E-09 |
| B45 | B94 | MCHC | ALDH2-Lys504Lys | -0.06194 | 9.22E-09 |
| B45 | B120 | MCHC | HCV-cAg | 0.061839 | 9.77E-09 |
| B45 | B56 | MCHC | IR | 0.060974 | 1.56E-08 |
| B45 | B61 | MCHC | WBHSRV | 0.060143 | 2.44E-08 |
| B45 | B131 | MCHC | AST | 0.058663 | 5.34E-08 |
| B45 | B193 | MCHC | K | -0.05795 | 7.73E-08 |
| B45 | B65 | MCHC | WBRSR(mPas_1) | 0.057022 | 1.24E-07 |
| B45 | B60 | MCHC | WBRVI | 0.054571 | 4.20E-07 |
| B45 | B138 | MCHC | PAB | 0.054516 | 4.31E-07 |
| B45 | B54 | MCHC | ESR | -0.05447 | 4.41E-07 |
| B45 | B145 | MCHC | BGP | -0.05438 | 4.60E-07 |
| B45 | B220 | MCHC | SBP | -0.0536 | 6.71E-07 |
| B46 | B113 | RBC Count | FT3 | 0.096377 | 0 |
| B46 | B126 | RBC Count | Alb | 0.269637 | 0 |
| B46 | B127 | RBC Count | A-G ratio | 0.153565 | 0 |
| B46 | B128 | RBC Count | ALT | 0.166313 | 0 |
| B46 | B129 | RBC Count | IBIL | 0.186386 | 0 |
| B46 | B133 | RBC Count | TBIL | 0.169879 | 0 |
| B46 | B141 | RBC Count | AST/ALT | -0.23779 | 0 |
| B46 | B146 | RBC Count | CysC | -0.10594 | 0 |
| B46 | B153 | RBC Count | Glu | 0.144489 | 0 |
| B46 | B161 | RBC Count | LEU | -0.08949 | 0 |
| B46 | B163 | RBC Count | SG | 0.237652 | 0 |
| B46 | B170 | RBC Count | U-GLU | 0.138157 | 0 |
| B46 | B173 | RBC Count | NTT | -0.09229 | 0 |
| B46 | B174 | RBC Count | Urea/Cr | -0.11671 | 0 |
| B46 | B188 | RBC Count | Ca | 0.089184 | 0 |
| B46 | B189 | RBC Count | eGFB | 0.296256 | 0 |
| B46 | B190 | RBC Count | Cre | -0.16355 | 0 |
| B46 | B191 | RBC Count | Urea | -0.26147 | 0 |
| B46 | B204 | RBC Count | Drinking | 0.112901 | 0 |
| B46 | B206 | RBC Count | Waist | 0.1701 | 0 |
| B46 | B207 | RBC Count | WHR | 0.104117 | 0 |
| B46 | B209 | RBC Count | Smoking | 0.131196 | 0 |
| B46 | B210 | RBC Count | Hip | 0.143247 | 0 |
| B46 | B215 | RBC Count | BMI | 0.131154 | 0 |
| B46 | B216 | RBC Count | BMI | 0.300431 | 0 |
| B46 | B218 | RBC Count | DBP | 0.273798 | 0 |
| B46 | B221 | RBC Count | Hei | 0.330475 | 0 |
| B46 | B222 | RBC Count | NS | 0.195568 | 0 |
| B46 | B47 | RBC Count | RDW | -0.45902 | 0 |
| B46 | B51 | RBC Count | Hb | 0.771649 | 0 |

| | | | | | |
|---|---|---|---|---|---|
| B46 | B56 | RBC Count | IR | 0.091864 | 0 |
| B46 | B60 | RBC Count | WBRVI | 0.143157 | 0 |
| B46 | B61 | RBC Count | WBHSRV | 0.096883 | 0 |
| B46 | B62 | RBC Count | WBHSRVI | 0.147846 | 0 |
| B46 | B63 | RBC Count | WBRSR(mPas_200) | 0.149688 | 0 |
| B46 | B64 | RBC Count | WBRSR(mPas_5) | 0.15759 | 0 |
| B46 | B65 | RBC Count | WBRSR(mPas_1) | 0.151643 | 0 |
| B46 | B66 | RBC Count | WBRSR(mPas_100) | 0.151204 | 0 |
| B46 | B67 | RBC Count | WBRSR(mPas_50) | 0.153761 | 0 |
| B46 | B71 | RBC Count | TG | 0.101587 | 0 |
| B46 | B72 | RBC Count | HDL-C | -0.17157 | 0 |
| B46 | B91 | RBC Count | TP | 0.096509 | 0 |
| B46 | B176 | RBC Count | Ucr | 0.088504 | 2.22E-16 |
| B46 | B59 | RBC Count | WBRV | 0.081322 | 4.46E-14 |
| B46 | B130 | RBC Count | GLB | -0.07948 | 1.64E-13 |
| B46 | B132 | RBC Count | Dbil | 0.077029 | 8.86E-13 |
| B46 | B49 | RBC Count | baso% | 0.076532 | 1.24E-12 |
| B46 | B70 | RBC Count | LDL-C | 0.074655 | 4.32E-12 |
| B46 | B52 | RBC Count | PLT/BPC | 0.070358 | 6.73E-11 |
| B46 | B54 | RBC Count | ESR | -0.06202 | 8.83E-09 |
| B46 | B115 | RBC Count | TSH TT3 TT4(TT3) | 0.061996 | 8.96E-09 |
| B46 | B74 | RBC Count | ApoA1 | -0.0592 | 4.04E-08 |
| B46 | B219 | RBC Count | OT | -0.0591 | 4.26E-08 |
| B46 | B172 | RBC Count | KB | 0.058109 | 7.12E-08 |
| B46 | B57 | RBC Count | RBCAI | 0.055221 | 3.06E-07 |
| B47 | B126 | RDW | Alb | -0.14538 | 0 |
| B47 | B127 | RDW | A-G ratio | -0.08983 | 0 |
| B47 | B141 | RDW | AST/ALT | 0.135393 | 0 |
| B47 | B146 | RDW | CysC | 0.090543 | 0 |
| B47 | B153 | RDW | Glu | -0.16978 | 0 |
| B47 | B170 | RDW | U-GLU | -0.09271 | 0 |
| B47 | B189 | RDW | eGFB | -0.2091 | 0 |
| B47 | B190 | RDW | Cre | 0.142228 | 0 |
| B47 | B191 | RDW | Urea | 0.163624 | 0 |
| B47 | B218 | RDW | DBP | -0.11137 | 0 |
| B47 | B221 | RDW | Hei | -0.08972 | 0 |
| B47 | B51 | RDW | Hb | -0.24174 | 0 |
| B47 | B52 | RDW | PLT/BPC | -0.19784 | 0 |
| B47 | B71 | RDW | TG | -0.09953 | 0 |
| B47 | B72 | RDW | HDL-C | 0.121169 | 0 |
| B47 | B192 | RDW | UA | 0.085078 | 2.89E-15 |
| B47 | B132 | RDW | Dbil | 0.084627 | 4.00E-15 |
| B47 | B216 | RDW | BMI | -0.08005 | 1.10E-13 |
| B47 | B163 | RDW | SG | -0.06846 | 2.15E-10 |

| | | | | | |
|---|---|---|---|---|---|
| B47 | B112 | RDW | FT4 | -0.06189 | 9.52E-09 |
| B47 | B128 | RDW | ALT | -0.05737 | 1.04E-07 |
| B48 | B189 | LYM% | eGFB | 0.151675 | 0 |
| B48 | B190 | LYM% | Cre | -0.14748 | 0 |
| B48 | B191 | LYM% | Urea | -0.14239 | 0 |
| B48 | B53 | LYM% | GR% | -0.93334 | 0 |
| B48 | B72 | LYM% | HDL-C | 0.115498 | 0 |
| B48 | B77 | LYM% | TC | 0.097699 | 0 |
| B48 | B91 | LYM% | TP | 0.088635 | 2.22E-16 |
| B48 | B126 | LYM% | Alb | 0.087672 | 4.44E-16 |
| B48 | B166 | LYM% | Pro | -0.08596 | 1.33E-15 |
| B48 | B192 | LYM% | UA | -0.08485 | 3.55E-15 |
| B48 | B123 | LYM% | hs-CRP1 | -0.08364 | 8.44E-15 |
| B48 | B153 | LYM% | Glu | -0.08134 | 4.42E-14 |
| B48 | B146 | LYM% | CysC | -0.07126 | 3.83E-11 |
| B48 | B131 | LYM% | AST | 0.070372 | 6.67E-11 |
| B48 | B207 | LYM% | WHR | -0.06469 | 1.98E-09 |
| B48 | B70 | LYM% | LDL-C | 0.064664 | 2.01E-09 |
| B48 | B118 | LYM% | CEA1 | -0.05929 | 3.84E-08 |
| B48 | B163 | LYM% | SG | -0.05578 | 2.31E-07 |
| B48 | B50 | LYM% | eo% | -0.05487 | 3.63E-07 |
| B48 | B51 | LYM% | Hb | 0.053051 | 8.72E-07 |
| B49 | B50 | baso% | eo% | 0.259722 | 0 |
| B49 | B53 | baso% | GR% | -0.16402 | 0 |
| B49 | B52 | baso% | PLT/BPC | 0.083498 | 9.33E-15 |
| B49 | B51 | baso% | Hb | 0.083214 | 1.13E-14 |
| B49 | B221 | baso% | Hei | 0.067408 | 4.05E-10 |
| B49 | B156 | baso% | IgE | 0.057553 | 9.48E-08 |
| B49 | B216 | baso% | BMI | 0.057234 | 1.11E-07 |
| B5 | B108 | WB-Mg | Anti-Cyclin | -0.33692 | 0 |
| B5 | B17 | WB-Mg | CD3+CD16+CD56+NK | 0.191982 | 0 |
| B5 | B21 | WB-Mg | CD3+CD4-CD8- | 0.09406 | 0 |
| B5 | B6 | WB-Mg | WB-Fe | 0.521552 | 0 |
| B5 | B7 | WB-Mg | WB-Cop | 0.553771 | 0 |
| B5 | B8 | WB-Mg | WB-Zinc | 0.577559 | 0 |
| B5 | B145 | WB-Mg | BGP | 0.073212 | 1.11E-11 |
| B5 | B15 | WB-Mg | CD3-CD19+(B cell) | 0.072641 | 1.60E-11 |
| B5 | B54 | WB-Mg | ESR | 0.061376 | 1.26E-08 |
| B5 | B144 | WB-Mg | Total-PINP | 0.059169 | 4.10E-08 |
| B5 | B107 | WB-Mg | AnuA | 0.058458 | 5.94E-08 |
| B5 | B101 | WB-Mg | anti-Sc70 | 0.058195 | 6.81E-08 |
| B5 | B104 | WB-Mg | anti-SSB | 0.058195 | 6.81E-08 |
| B5 | B99 | WB-Mg | anti-PM-Scl | 0.058195 | 6.81E-08 |
| B5 | B195 | WB-Mg | Na | 0.056516 | 1.60E-07 |

| | | | | | |
|---|---|---|---|---|---|
| B5 | B143 | WB-Mg | β-CTx | 0.053932 | 5.72E-07 |
| B50 | B156 | eo% | IgE | 0.101561 | 0 |
| B50 | B189 | eo% | eGFB | -0.09253 | 0 |
| B50 | B190 | eo% | Cre | 0.121959 | 0 |
| B50 | B191 | eo% | Urea | 0.107642 | 0 |
| B50 | B192 | eo% | UA | 0.096102 | 0 |
| B50 | B53 | eo% | GR% | -0.23096 | 0 |
| B50 | B146 | eo% | CysC | 0.079059 | 2.20E-13 |
| B50 | B126 | eo% | Alb | -0.06685 | 5.63E-10 |
| B50 | B133 | eo% | TBIL | -0.06496 | 1.69E-09 |
| B50 | B222 | eo% | NS | 0.063061 | 4.97E-09 |
| B50 | B129 | eo% | IBIL | -0.06211 | 8.44E-09 |
| B50 | B174 | eo% | Urea/Cr | -0.06035 | 2.19E-08 |
| B50 | B209 | eo% | Smoking | 0.05863 | 5.43E-08 |
| B50 | B220 | eo% | SBP | -0.05535 | 2.87E-07 |
| B50 | B132 | eo% | Dbil | -0.05448 | 4.38E-07 |
| B51 | B113 | Hb | FT3 | 0.113284 | 0 |
| B51 | B126 | Hb | Alb | 0.296711 | 0 |
| B51 | B127 | Hb | A-G ratio | 0.187167 | 0 |
| B51 | B128 | Hb | ALT | 0.18081 | 0 |
| B51 | B129 | Hb | IBIL | 0.246651 | 0 |
| B51 | B130 | Hb | GLB | -0.11083 | 0 |
| B51 | B132 | Hb | Dbil | 0.148133 | 0 |
| B51 | B133 | Hb | TBIL | 0.238241 | 0 |
| B51 | B140 | Hb | r-GT | 0.089298 | 0 |
| B51 | B141 | Hb | AST/ALT | -0.24406 | 0 |
| B51 | B146 | Hb | CysC | -0.11172 | 0 |
| B51 | B153 | Hb | Glu | 0.133765 | 0 |
| B51 | B161 | Hb | LEU | -0.11004 | 0 |
| B51 | B163 | Hb | SG | 0.276625 | 0 |
| B51 | B170 | Hb | U-GLU | 0.146628 | 0 |
| B51 | B173 | Hb | NTT | -0.11137 | 0 |
| B51 | B174 | Hb | Urea/Cr | -0.16721 | 0 |
| B51 | B176 | Hb | Ucr | 0.095592 | 0 |
| B51 | B188 | Hb | Ca | 0.096458 | 0 |
| B51 | B189 | Hb | eGFB | 0.31089 | 0 |
| B51 | B190 | Hb | Cre | -0.16229 | 0 |
| B51 | B191 | Hb | Urea | -0.28798 | 0 |
| B51 | B204 | Hb | Drinking | 0.212042 | 0 |
| B51 | B206 | Hb | Waist | 0.190465 | 0 |
| B51 | B207 | Hb | WHR | 0.114336 | 0 |
| B51 | B209 | Hb | Smoking | 0.221467 | 0 |
| B51 | B210 | Hb | Hip | 0.163541 | 0 |
| B51 | B215 | Hb | BMI | 0.127981 | 0 |

| | | | | | |
|---|---|---|---|---|---|
| B51 | B216 | Hb | BMI | 0.343618 | 0 |
| B51 | B218 | Hb | DBP | 0.293901 | 0 |
| B51 | B221 | Hb | Hei | 0.401919 | 0 |
| B51 | B222 | Hb | NS | 0.228783 | 0 |
| B51 | B56 | Hb | IR | 0.093542 | 0 |
| B51 | B60 | Hb | WBRVI | 0.147445 | 0 |
| B51 | B61 | Hb | WBHSRV | 0.095494 | 0 |
| B51 | B62 | Hb | WBHSRVI | 0.165827 | 0 |
| B51 | B63 | Hb | WBRSR(mPas_200) | 0.163153 | 0 |
| B51 | B64 | Hb | WBRSR(mPas_5) | 0.164259 | 0 |
| B51 | B65 | Hb | WBRSR(mPas_1) | 0.15358 | 0 |
| B51 | B66 | Hb | WBRSR(mPas_100) | 0.163963 | 0 |
| B51 | B67 | Hb | WBRSR(mPas_50) | 0.164171 | 0 |
| B51 | B70 | Hb | LDL-C | 0.095641 | 0 |
| B51 | B71 | Hb | TG | 0.114917 | 0 |
| B51 | B72 | Hb | HDL-C | -0.13167 | 0 |
| B51 | B91 | Hb | TP | 0.084874 | 3.33E-15 |
| B51 | B219 | Hb | OT | -0.08324 | 1.11E-14 |
| B51 | B54 | Hb | ESR | -0.08234 | 2.13E-14 |
| B51 | B172 | Hb | KB | 0.067764 | 3.27E-10 |
| B51 | B194 | Hb | Cl | -0.06721 | 4.56E-10 |
| B51 | B77 | Hb | TC | 0.063468 | 3.96E-09 |
| B51 | B134 | Hb | CHE | 0.062304 | 7.57E-09 |
| B51 | B149 | Hb | TGAb | -0.06187 | 9.62E-09 |
| B51 | B59 | Hb | WBRV | 0.060458 | 2.07E-08 |
| B51 | B131 | Hb | AST | 0.060038 | 2.59E-08 |
| B51 | B115 | Hb | TSH TT3 TT4(TT3) | 0.059647 | 3.18E-08 |
| B51 | B220 | Hb | SBP | -0.05919 | 4.05E-08 |
| B51 | B74 | Hb | ApoA1 | -0.05671 | 1.46E-07 |
| B51 | B94 | Hb | ALDH2-Lys504Lys | -0.05465 | 4.05E-07 |
| B51 | B138 | Hb | PAB | 0.053838 | 5.99E-07 |
| B52 | B128 | PLT/BPC | ALT | -0.11509 | 0 |
| B52 | B129 | PLT/BPC | IBIL | -0.11235 | 0 |
| B52 | B130 | PLT/BPC | GLB | 0.091876 | 0 |
| B52 | B131 | PLT/BPC | AST | -0.09909 | 0 |
| B52 | B132 | PLT/BPC | Dbil | -0.17379 | 0 |
| B52 | B133 | PLT/BPC | TBIL | -0.13921 | 0 |
| B52 | B191 | PLT/BPC | Urea | -0.09191 | 0 |
| B52 | B70 | PLT/BPC | LDL-C | 0.122838 | 0 |
| B52 | B77 | PLT/BPC | TC | 0.088412 | 2.22E-16 |
| B52 | B141 | PLT/BPC | AST/ALT | 0.087308 | 4.44E-16 |
| B52 | B127 | PLT/BPC | A-G ratio | -0.08136 | 4.33E-14 |
| B52 | B188 | PLT/BPC | Ca | 0.079158 | 2.05E-13 |
| B52 | B91 | PLT/BPC | TP | 0.078394 | 3.49E-13 |

| | | | | | |
|---|---|---|---|---|---|
| B52 | B189 | PLT/BPC | eGFB | 0.077838 | 5.10E-13 |
| B52 | B192 | PLT/BPC | UA | -0.06814 | 2.61E-10 |
| B52 | B190 | PLT/BPC | Cre | -0.06706 | 4.97E-10 |
| B52 | B112 | PLT/BPC | FT4 | 0.059773 | 2.98E-08 |
| B52 | B221 | PLT/BPC | Hei | -0.05577 | 2.33E-07 |
| B53 | B189 | GR% | eGFB | -0.09097 | 0 |
| B53 | B72 | GR% | HDL-C | -0.10177 | 0 |
| B53 | B191 | GR% | Urea | 0.088457 | 2.22E-16 |
| B53 | B153 | GR% | Glu | 0.087147 | 4.44E-16 |
| B53 | B190 | GR% | Cre | 0.086134 | 1.33E-15 |
| B53 | B131 | GR% | AST | -0.08367 | 7.99E-15 |
| B53 | B123 | GR% | hs-CRP1 | 0.079446 | 1.68E-13 |
| B53 | B166 | GR% | Pro | 0.069786 | 9.58E-11 |
| B53 | B77 | GR% | TC | -0.06765 | 3.51E-10 |
| B53 | B91 | GR% | TP | -0.06538 | 1.33E-09 |
| B54 | B101 | ESR | anti-Sc70 | -0.09588 | 0 |
| B54 | B104 | ESR | anti-SSB | -0.09588 | 0 |
| B54 | B107 | ESR | AnuA | -0.09639 | 0 |
| B54 | B123 | ESR | hs-CRP1 | 0.116567 | 0 |
| B54 | B127 | ESR | A-G ratio | -0.09455 | 0 |
| B54 | B130 | ESR | GLB | 0.108206 | 0 |
| B54 | B56 | ESR | IR | -0.20063 | 0 |
| B54 | B57 | ESR | RBCAI | -0.23157 | 0 |
| B54 | B58 | ESR | HCT | -0.10093 | 0 |
| B54 | B60 | ESR | WBRVI | -0.37798 | 0 |
| B54 | B62 | ESR | WBHSRVI | -0.34325 | 0 |
| B54 | B63 | ESR | WBRSR(mPas_200) | -0.16769 | 0 |
| B54 | B64 | ESR | WBRSR(mPas_5) | -0.21453 | 0 |
| B54 | B65 | ESR | WBRSR(mPas_1) | -0.2358 | 0 |
| B54 | B66 | ESR | WBRSR(mPas_100) | -0.14693 | 0 |
| B54 | B67 | ESR | WBRSR(mPas_50) | -0.16062 | 0 |
| B54 | B69 | ESR | PV | 0.458887 | 0 |
| B54 | B76 | ESR | ApoB | 0.11205 | 0 |
| B54 | B81 | ESR | TSGF | 0.089054 | 0 |
| B54 | B99 | ESR | anti-PM-Scl | -0.09588 | 0 |
| B54 | B156 | ESR | IgE | 0.085721 | 1.78E-15 |
| B54 | B179 | ESR | α1-MG | 0.078737 | 2.75E-13 |
| B54 | B91 | ESR | TP | 0.073425 | 9.63E-12 |
| B54 | B115 | ESR | TSH TT3 TT4(TT3) | -0.07289 | 1.36E-11 |
| B54 | B155 | ESR | IgA | 0.069282 | 1.31E-10 |
| B54 | B199 | ESR | Mg | -0.06391 | 3.08E-09 |
| B54 | B77 | ESR | TC | 0.062518 | 6.72E-09 |
| B54 | B221 | ESR | Hei | -0.06062 | 1.90E-08 |
| B54 | B225 | ESR | hPBG | 0.060286 | 2.27E-08 |

| | | | | | |
|---|---|---|---|---|---|
| B54 | B157 | ESR | IgG | 0.05986 | 2.84E-08 |
| B54 | B75 | ESR | ApoA1/B | -0.05647 | 1.64E-07 |
| B54 | B195 | ESR | Na | -0.05586 | 2.23E-07 |
| B54 | B186 | ESR | PFC | 0.053457 | 7.19E-07 |
| B55 | B156 | TK | IgE | -0.09852 | 0 |
| B55 | B56 | TK | IR | 0.769071 | 0 |
| B55 | B57 | TK | RBCAI | -0.1484 | 0 |
| B55 | B58 | TK | HCT | -0.56624 | 0 |
| B55 | B59 | TK | WBRV | 0.557922 | 0 |
| B55 | B60 | TK | WBRVI | 0.226532 | 0 |
| B55 | B61 | TK | WBHSRV | 0.713446 | 0 |
| B55 | B62 | TK | WBHSRVI | 0.380229 | 0 |
| B55 | B63 | TK | WBRSR(mPas_200) | 0.284647 | 0 |
| B55 | B64 | TK | WBRSR(mPas_5) | 0.212026 | 0 |
| B55 | B65 | TK | WBRSR(mPas_1) | 0.173944 | 0 |
| B55 | B66 | TK | WBRSR(mPas_100) | 0.284059 | 0 |
| B55 | B67 | TK | WBRSR(mPas_50) | 0.276498 | 0 |
| B55 | B69 | TK | PV | -0.23762 | 0 |
| B55 | B79 | TK | hb V -dna | -0.08436 | 4.88E-15 |
| B55 | B94 | TK | ALDH2-Lys504Lys | -0.08243 | 2.00E-14 |
| B55 | B103 | TK | anti-SSA/Ro | 0.078915 | 2.43E-13 |
| B55 | B92 | TK | ALDH2-Glu504Glu | 0.071426 | 3.45E-11 |
| B55 | B107 | TK | AnuA | -0.0701 | 7.90E-11 |
| B55 | B101 | TK | anti-Sc70 | -0.06976 | 9.75E-11 |
| B55 | B104 | TK | anti-SSB | -0.06976 | 9.75E-11 |
| B55 | B99 | TK | anti-PM-Scl | -0.06976 | 9.75E-11 |
| B55 | B93 | TK | ALDH2-Glu504Lys | -0.06847 | 2.14E-10 |
| B55 | B196 | TK | TCO2 | -0.06563 | 1.15E-09 |
| B55 | B111 | TK | TSH | 0.059531 | 3.38E-08 |
| B55 | B184 | TK | INR | -0.05952 | 3.41E-08 |
| B55 | B119 | TK | Hcv-rna | -0.0583 | 6.44E-08 |
| B55 | B186 | TK | PFC | 0.056253 | 1.83E-07 |
| B55 | B75 | TK | ApoA1/B | 0.053213 | 8.07E-07 |
| B56 | B156 | IR | IgE | -0.0912 | 0 |
| B56 | B58 | IR | HCT | -0.23905 | 0 |
| B56 | B59 | IR | WBRV | 0.6836 | 0 |
| B56 | B60 | IR | WBRVI | 0.693749 | 0 |
| B56 | B61 | IR | WBHSRV | 0.932728 | 0 |
| B56 | B62 | IR | WBHSRVI | 0.8776 | 0 |
| B56 | B63 | IR | WBRSR(mPas_200) | 0.775448 | 0 |
| B56 | B64 | IR | WBRSR(mPas_5) | 0.713913 | 0 |
| B56 | B65 | IR | WBRSR(mPas_1) | 0.653555 | 0 |
| B56 | B66 | IR | WBRSR(mPas_100) | 0.765679 | 0 |
| B56 | B67 | IR | WBRSR(mPas_50) | 0.763972 | 0 |

| | | | | | |
|---|---|---|---|---|---|
| B56 | B69 | IR | PV | -0.23576 | 0 |
| B56 | B94 | IR | ALDH2-Lys504Lys | -0.0903 | 0 |
| B56 | B79 | IR | hb V -dna | -0.0855 | 2.22E-15 |
| B56 | B74 | IR | ApoA1 | -0.08205 | 2.66E-14 |
| B56 | B89 | IR | CA 72-4 | 0.08112 | 5.15E-14 |
| B56 | B92 | IR | ALDH2-Glu504Glu | 0.078982 | 2.32E-13 |
| B56 | B134 | IR | CHE | 0.078347 | 3.60E-13 |
| B56 | B184 | IR | INR | -0.07824 | 3.87E-13 |
| B56 | B93 | IR | ALDH2-Glu504Lys | -0.07577 | 2.06E-12 |
| B56 | B108 | IR | Anti-Cyclin | -0.07379 | 7.59E-12 |
| B56 | B197 | IR | AG | 0.072965 | 1.30E-11 |
| B56 | B119 | IR | Hcv-rna | -0.07174 | 2.83E-11 |
| B56 | B120 | IR | HCV-cAg | 0.071625 | 3.05E-11 |
| B56 | B103 | IR | anti-SSA/Ro | 0.069953 | 8.65E-11 |
| B56 | B125 | IR | Cor | 0.065594 | 1.17E-09 |
| B56 | B123 | IR | hs-CRP1 | 0.063685 | 3.50E-09 |
| B56 | B138 | IR | PAB | 0.056844 | 1.36E-07 |
| B57 | B58 | RBCAI | HCT | 0.13755 | 0 |
| B57 | B59 | RBCAI | WBRV | 0.577785 | 0 |
| B57 | B60 | RBCAI | WBRVI | 0.633696 | 0 |
| B57 | B62 | RBCAI | WBHSRVI | 0.126823 | 0 |
| B57 | B64 | RBCAI | WBRSR(mPas_5) | 0.40013 | 0 |
| B57 | B65 | RBCAI | WBRSR(mPas_1) | 0.549716 | 0 |
| B57 | B67 | RBCAI | WBRSR(mPas_50) | 0.129731 | 0 |
| B57 | B69 | RBCAI | PV | -0.19118 | 0 |
| B57 | B81 | RBCAI | TSGF | -0.09249 | 0 |
| B57 | B76 | RBCAI | ApoB | -0.07688 | 9.82E-13 |
| B57 | B122 | RBCAI | C4 | -0.07502 | 3.40E-12 |
| B57 | B183 | RBCAI | PTA | -0.07389 | 7.15E-12 |
| B57 | B75 | RBCAI | ApoA1/B | 0.065895 | 9.86E-10 |
| B57 | B186 | RBCAI | PFC | -0.06545 | 1.28E-09 |
| B57 | B66 | RBCAI | WBRSR(mPas_100) | 0.064675 | 2.00E-09 |
| B57 | B121 | RBCAI | C3 | -0.06156 | 1.14E-08 |
| B57 | B125 | RBCAI | Cor | 0.059704 | 3.09E-08 |
| B57 | B111 | RBCAI | TSH | -0.05952 | 3.41E-08 |
| B57 | B178 | RBCAI | Ig-U | -0.05798 | 7.61E-08 |
| B57 | B61 | RBCAI | WBHSRV | -0.05621 | 1.87E-07 |
| B57 | B156 | RBCAI | IgE | -0.05462 | 4.10E-07 |
| B57 | B136 | RBCAI | 5'NT | -0.05387 | 5.89E-07 |
| B58 | B59 | HCT | WBRV | -0.26615 | 0 |
| B58 | B61 | HCT | WBHSRV | -0.23616 | 0 |
| B58 | B64 | HCT | WBRSR(mPas_5) | 0.104823 | 0 |
| B58 | B65 | HCT | WBRSR(mPas_1) | 0.121595 | 0 |
| B58 | B69 | HCT | PV | 0.130756 | 0 |

| | | | | | |
|---|---|---|---|---|---|
| B58 | B60 | HCT | WBRVI | 0.075908 | 1.88E-12 |
| B58 | B67 | HCT | WBRSR(mPas_50) | 0.074213 | 5.78E-12 |
| B58 | B66 | HCT | WBRSR(mPas_100) | 0.067488 | 3.86E-10 |
| B58 | B63 | HCT | WBRSR(mPas_200) | 0.063436 | 4.03E-09 |
| B59 | B103 | WBRV | anti-SSA/Ro | 0.098417 | 0 |
| B59 | B123 | WBRV | hs-CRP1 | 0.08999 | 0 |
| B59 | B60 | WBRV | WBRVI | 0.764572 | 0 |
| B59 | B61 | WBRV | WBHSRV | 0.730593 | 0 |
| B59 | B62 | WBRV | WBHSRVI | 0.56759 | 0 |
| B59 | B63 | WBRV | WBRSR(mPas_200) | 0.598411 | 0 |
| B59 | B64 | WBRV | WBRSR(mPas_5) | 0.760547 | 0 |
| B59 | B65 | WBRV | WBRSR(mPas_1) | 0.80159 | 0 |
| B59 | B66 | WBRV | WBRSR(mPas_100) | 0.601513 | 0 |
| B59 | B67 | WBRV | WBRSR(mPas_50) | 0.639835 | 0 |
| B59 | B125 | WBRV | Cor | 0.087907 | 4.44E-16 |
| B59 | B156 | WBRV | IgE | -0.08283 | 1.51E-14 |
| B59 | B89 | WBRV | CA 72-4 | 0.076259 | 1.49E-12 |
| B59 | B107 | WBRV | AnuA | -0.07506 | 3.30E-12 |
| B59 | B101 | WBRV | anti-Sc70 | -0.07471 | 4.18E-12 |
| B59 | B104 | WBRV | anti-SSB | -0.07471 | 4.18E-12 |
| B59 | B99 | WBRV | anti-PM-Scl | -0.07471 | 4.18E-12 |
| B59 | B75 | WBRV | ApoA1/B | 0.069484 | 1.15E-10 |
| B59 | B194 | WBRV | Cl | -0.06765 | 3.51E-10 |
| B59 | B185 | WBRV | PT | -0.06542 | 1.30E-09 |
| B59 | B69 | WBRV | PV | 0.064509 | 2.19E-09 |
| B59 | B79 | WBRV | hb V -dna | -0.06353 | 3.82E-09 |
| B59 | B124 | WBRV | ACTH | 0.063507 | 3.87E-09 |
| B59 | B119 | WBRV | Hcv-rna | -0.0626 | 6.43E-09 |
| B59 | B184 | WBRV | INR | -0.06099 | 1.55E-08 |
| B59 | B94 | WBRV | ALDH2-Lys504Lys | -0.06022 | 2.34E-08 |
| B59 | B120 | WBRV | HCV-cAg | 0.055124 | 3.20E-07 |
| B6 | B108 | WB-Fe | Anti-Cyclin | -0.48381 | 0 |
| B6 | B145 | WB-Fe | BGP | 0.103477 | 0 |
| B6 | B15 | WB-Fe | CD3-CD19+(B cell) | 0.102909 | 0 |
| B6 | B17 | WB-Fe | CD3+CD16+CD56+NK | 0.275108 | 0 |
| B6 | B21 | WB-Fe | CD3+CD4-CD8- | 0.134671 | 0 |
| B6 | B60 | WB-Fe | WBRVI | 0.10327 | 0 |
| B6 | B62 | WB-Fe | WBHSRVI | 0.103488 | 0 |
| B6 | B63 | WB-Fe | WBRSR(mPas_200) | 0.091245 | 0 |
| B6 | B64 | WB-Fe | WBRSR(mPas_5) | 0.0979 | 0 |
| B6 | B65 | WB-Fe | WBRSR(mPas_1) | 0.098396 | 0 |
| B6 | B66 | WB-Fe | WBRSR(mPas_100) | 0.090728 | 0 |
| B6 | B67 | WB-Fe | WBRSR(mPas_50) | 0.092421 | 0 |
| B6 | B7 | WB-Fe | WB-Cop | -0.40295 | 0 |

| | | | | | |
|---|---|---|---|---|---|
| B6 | B8 | WB-Fe | WB-Zinc | 0.80767 | 0 |
| B6 | B144 | WB-Fe | Total-PINP | 0.083286 | 1.07E-14 |
| B6 | B56 | WB-Fe | IR | 0.08176 | 3.26E-14 |
| B6 | B107 | WB-Fe | AnuA | 0.081666 | 3.49E-14 |
| B6 | B101 | WB-Fe | anti-Sc70 | 0.081278 | 4.62E-14 |
| B6 | B104 | WB-Fe | anti-SSB | 0.081278 | 4.62E-14 |
| B6 | B99 | WB-Fe | anti-PM-Scl | 0.081278 | 4.62E-14 |
| B6 | B195 | WB-Fe | Na | 0.080878 | 6.13E-14 |
| B6 | B143 | WB-Fe | β-CTx | 0.075857 | 1.95E-12 |
| B6 | B151 | WB-Fe | Anti-CCP | 0.072743 | 1.50E-11 |
| B6 | B199 | WB-Fe | Mg | 0.071129 | 4.16E-11 |
| B6 | B61 | WB-Fe | WBHSRV | 0.07021 | 7.37E-11 |
| B6 | B18 | WB-Fe | CD3+CD4+helper(inducer)T | -0.06739 | 4.10E-10 |
| B6 | B57 | WB-Fe | RBCAI | 0.061066 | 1.49E-08 |
| B6 | B59 | WB-Fe | WBRV | 0.060367 | 2.17E-08 |
| B60 | B108 | WBRVI | Anti-Cyclin | -0.09926 | 0 |
| B60 | B125 | WBRVI | Cor | 0.096663 | 0 |
| B60 | B61 | WBRVI | WBHSRV | 0.607817 | 0 |
| B60 | B62 | WBRVI | WBHSRVI | 0.842493 | 0 |
| B60 | B63 | WBRVI | WBRSR(mPas_200) | 0.736307 | 0 |
| B60 | B64 | WBRVI | WBRSR(mPas_5) | 0.898168 | 0 |
| B60 | B65 | WBRVI | WBRSR(mPas_1) | 0.934267 | 0 |
| B60 | B66 | WBRVI | WBRSR(mPas_100) | 0.73146 | 0 |
| B60 | B67 | WBRVI | WBRSR(mPas_50) | 0.768998 | 0 |
| B60 | B69 | WBRVI | PV | -0.24943 | 0 |
| B60 | B156 | WBRVI | IgE | -0.07948 | 1.64E-13 |
| B60 | B185 | WBRVI | PT | -0.07145 | 3.40E-11 |
| B60 | B123 | WBRVI | hs-CRP1 | 0.070124 | 7.78E-11 |
| B60 | B183 | WBRVI | PTA | -0.0685 | 2.10E-10 |
| B60 | B134 | WBRVI | CHE | 0.066712 | 6.11E-10 |
| B60 | B221 | WBRVI | Hei | 0.064843 | 1.81E-09 |
| B60 | B103 | WBRVI | anti-SSA/Ro | 0.062054 | 8.68E-09 |
| B60 | B119 | WBRVI | Hcv-rna | -0.06167 | 1.07E-08 |
| B60 | B89 | WBRVI | CA 72-4 | 0.06115 | 1.42E-08 |
| B60 | B79 | WBRVI | hb V -dna | -0.06095 | 1.59E-08 |
| B60 | B94 | WBRVI | ALDH2-Lys504Lys | -0.05882 | 4.92E-08 |
| B60 | B129 | WBRVI | IBIL | 0.056885 | 1.33E-07 |
| B60 | B181 | WBRVI | APTT | -0.05566 | 2.46E-07 |
| B60 | B218 | WBRVI | DBP | 0.053103 | 8.51E-07 |
| B61 | B120 | WBHSRV | HCV-cAg | 0.097737 | 0 |
| B61 | B123 | WBHSRV | hs-CRP1 | 0.107367 | 0 |
| B61 | B134 | WBHSRV | CHE | 0.097792 | 0 |
| B61 | B184 | WBHSRV | INR | -0.09348 | 0 |
| B61 | B62 | WBHSRV | WBHSRVI | 0.814665 | 0 |

| | | | | | |
|---|---|---|---|---|---|
| B61 | B63 | WBHSRV | WBRSR(mPas_200) | 0.862779 | 0 |
| B61 | B64 | WBHSRV | WBRSR(mPas_5) | 0.771802 | 0 |
| B61 | B65 | WBHSRV | WBRSR(mPas_1) | 0.68934 | 0 |
| B61 | B66 | WBHSRV | WBRSR(mPas_100) | 0.853111 | 0 |
| B61 | B67 | WBHSRV | WBRSR(mPas_50) | 0.847544 | 0 |
| B61 | B69 | WBHSRV | PV | 0.117545 | 0 |
| B61 | B89 | WBHSRV | CA 72-4 | 0.085422 | 2.22E-15 |
| B61 | B194 | WBHSRV | CI | -0.08489 | 3.33E-15 |
| B61 | B74 | WBHSRV | ApoA1 | -0.08128 | 4.57E-14 |
| B61 | B197 | WBHSRV | AG | 0.080674 | 7.06E-14 |
| B61 | B94 | WBHSRV | ALDH2-Lys504Lys | -0.07887 | 2.50E-13 |
| B61 | B71 | WBHSRV | TG | 0.077863 | 5.02E-13 |
| B61 | B92 | WBHSRV | ALDH2-Glu504Glu | 0.077181 | 7.99E-13 |
| B61 | B93 | WBHSRV | ALDH2-Glu504Lys | -0.07467 | 4.27E-12 |
| B61 | B136 | WBHSRV | 5'NT | 0.074358 | 5.25E-12 |
| B61 | B186 | WBHSRV | PFC | 0.073356 | 1.01E-11 |
| B61 | B103 | WBHSRV | anti-SSA/Ro | 0.070363 | 6.71E-11 |
| B61 | B125 | WBHSRV | Cor | 0.069892 | 8.98E-11 |
| B61 | B138 | WBHSRV | PAB | 0.068275 | 2.41E-10 |
| B61 | B79 | WBHSRV | hb V -dna | -0.06661 | 6.50E-10 |
| B61 | B156 | WBHSRV | IgE | -0.06272 | 6.01E-09 |
| B61 | B91 | WBHSRV | TP | 0.059765 | 2.99E-08 |
| B61 | B185 | WBHSRV | PT | -0.05772 | 8.72E-08 |
| B61 | B121 | WBHSRV | C3 | 0.057513 | 9.67E-08 |
| B61 | B119 | WBHSRV | Hcv-rna | -0.05746 | 9.92E-08 |
| B61 | B108 | WBHSRV | Anti-Cyclin | -0.05544 | 2.75E-07 |
| B61 | B140 | WBHSRV | r-GT | 0.054347 | 4.68E-07 |
| B62 | B108 | WBHSRVI | Anti-Cyclin | -0.10395 | 0 |
| B62 | B134 | WBHSRVI | CHE | 0.111292 | 0 |
| B62 | B63 | WBHSRVI | WBRSR(mPas_200) | 0.913386 | 0 |
| B62 | B64 | WBHSRVI | WBRSR(mPas_5) | 0.875431 | 0 |
| B62 | B65 | WBHSRVI | WBRSR(mPas_1) | 0.814485 | 0 |
| B62 | B66 | WBHSRVI | WBRSR(mPas_100) | 0.899059 | 0 |
| B62 | B67 | WBHSRVI | WBRSR(mPas_50) | 0.901577 | 0 |
| B62 | B69 | WBHSRVI | PV | -0.19438 | 0 |
| B62 | B123 | WBHSRVI | hs-CRP1 | 0.084241 | 5.33E-15 |
| B62 | B74 | WBHSRVI | ApoA1 | -0.07959 | 1.51E-13 |
| B62 | B125 | WBHSRVI | Cor | 0.075447 | 2.56E-12 |
| B62 | B221 | WBHSRVI | Hei | 0.074457 | 4.92E-12 |
| B62 | B197 | WBHSRVI | AG | 0.071736 | 2.84E-11 |
| B62 | B120 | WBHSRVI | HCV-cAg | 0.070266 | 7.12E-11 |
| B62 | B184 | WBHSRVI | INR | -0.06964 | 1.05E-10 |
| B62 | B138 | WBHSRVI | PAB | 0.06812 | 2.64E-10 |
| B62 | B89 | WBHSRVI | CA 72-4 | 0.067081 | 4.91E-10 |

| | | | | | |
|---|---|---|---|---|---|
| B62 | B94 | WBHSRVI | ALDH2-Lys504Lys | -0.06681 | 5.78E-10 |
| B62 | B156 | WBHSRVI | IgE | -0.06447 | 2.24E-09 |
| B62 | B216 | WBHSRVI | BMI | 0.06101 | 1.53E-08 |
| B62 | B185 | WBHSRVI | PT | -0.06059 | 1.92E-08 |
| B62 | B218 | WBHSRVI | DBP | 0.06002 | 2.61E-08 |
| B62 | B92 | WBHSRVI | ALDH2-Glu504Glu | 0.059025 | 4.42E-08 |
| B62 | B129 | WBHSRVI | IBIL | 0.05847 | 5.90E-08 |
| B62 | B163 | WBHSRVI | SG | 0.058156 | 6.95E-08 |
| B62 | B79 | WBHSRVI | hb V -dna | -0.05716 | 1.16E-07 |
| B62 | B93 | WBHSRVI | ALDH2-Glu504Lys | -0.05667 | 1.48E-07 |
| B62 | B121 | WBHSRVI | C3 | 0.054336 | 4.71E-07 |
| B62 | B133 | WBHSRVI | TBIL | 0.053537 | 6.92E-07 |
| B62 | B204 | WBHSRVI | Drinking | 0.053141 | 8.36E-07 |
| B63 | B120 | WBRSR(mPas_200) | HCV-cAg | 0.096194 | 0 |
| B63 | B123 | WBRSR(mPas_200) | hs-CRP1 | 0.130946 | 0 |
| B63 | B134 | WBRSR(mPas_200) | CHE | 0.129242 | 0 |
| B63 | B64 | WBRSR(mPas_200) | WBRSR(mPas_5) | 0.929679 | 0 |
| B63 | B65 | WBRSR(mPas_200) | WBRSR(mPas_1) | 0.846247 | 0 |
| B63 | B66 | WBRSR(mPas_200) | WBRSR(mPas_100) | 0.982956 | 0 |
| B63 | B67 | WBRSR(mPas_200) | WBRSR(mPas_50) | 0.981738 | 0 |
| B63 | B69 | WBRSR(mPas_200) | PV | 0.206035 | 0 |
| B63 | B194 | WBRSR(mPas_200) | Cl | -0.08875 | 2.22E-16 |
| B63 | B108 | WBRSR(mPas_200) | Anti-Cyclin | -0.08382 | 7.33E-15 |
| B63 | B138 | WBRSR(mPas_200) | PAB | 0.077667 | 5.74E-13 |
| B63 | B125 | WBRSR(mPas_200) | Cor | 0.076836 | 1.01E-12 |
| B63 | B184 | WBRSR(mPas_200) | INR | -0.07625 | 1.50E-12 |
| B63 | B197 | WBRSR(mPas_200) | AG | 0.073743 | 7.84E-12 |
| B63 | B74 | WBRSR(mPas_200) | ApoA1 | -0.07333 | 1.03E-11 |
| B63 | B89 | WBRSR(mPas_200) | CA 72-4 | 0.070852 | 4.95E-11 |
| B63 | B121 | WBRSR(mPas_200) | C3 | 0.069877 | 9.06E-11 |
| B63 | B71 | WBRSR(mPas_200) | TG | 0.068413 | 2.21E-10 |
| B63 | B136 | WBRSR(mPas_200) | 5'NT | 0.066292 | 7.82E-10 |
| B63 | B129 | WBRSR(mPas_200) | IBIL | 0.063762 | 3.35E-09 |
| B63 | B91 | WBRSR(mPas_200) | TP | 0.061968 | 9.10E-09 |
| B63 | B218 | WBRSR(mPas_200) | DBP | 0.061864 | 9.64E-09 |
| B63 | B122 | WBRSR(mPas_200) | C4 | 0.059269 | 3.89E-08 |
| B63 | B221 | WBRSR(mPas_200) | Hei | 0.057851 | 8.13E-08 |
| B63 | B185 | WBRSR(mPas_200) | PT | -0.05779 | 8.39E-08 |
| B63 | B133 | WBRSR(mPas_200) | TBIL | 0.056043 | 2.03E-07 |
| B63 | B140 | WBRSR(mPas_200) | r-GT | 0.055895 | 2.19E-07 |
| B63 | B216 | WBRSR(mPas_200) | BMI | 0.054575 | 4.19E-07 |
| B64 | B123 | WBRSR(mPas_5) | hs-CRP1 | 0.12714 | 0 |
| B64 | B125 | WBRSR(mPas_5) | Cor | 0.09959 | 0 |
| B64 | B134 | WBRSR(mPas_5) | CHE | 0.098605 | 0 |

| | | | | | |
|---|---|---|---|---|---|
| B64 | B65 | WBRSR(mPas_5) | WBRSR(mPas_1) | 0.977885 | 0 |
| B64 | B66 | WBRSR(mPas_5) | WBRSR(mPas_100) | 0.924186 | 0 |
| B64 | B67 | WBRSR(mPas_5) | WBRSR(mPas_50) | 0.942781 | 0 |
| B64 | B69 | WBRSR(mPas_5) | PV | 0.132518 | 0 |
| B64 | B108 | WBRSR(mPas_5) | Anti-Cyclin | -0.088 | 4.44E-16 |
| B64 | B194 | WBRSR(mPas_5) | CI | -0.08302 | 1.31E-14 |
| B64 | B120 | WBRSR(mPas_5) | HCV-cAg | 0.078338 | 3.62E-13 |
| B64 | B184 | WBRSR(mPas_5) | INR | -0.07411 | 6.19E-12 |
| B64 | B185 | WBRSR(mPas_5) | PT | -0.07257 | 1.67E-11 |
| B64 | B89 | WBRSR(mPas_5) | CA 72-4 | 0.071265 | 3.82E-11 |
| B64 | B129 | WBRSR(mPas_5) | IBIL | 0.06548 | 1.25E-09 |
| B64 | B218 | WBRSR(mPas_5) | DBP | 0.064447 | 2.27E-09 |
| B64 | B138 | WBRSR(mPas_5) | PAB | 0.064414 | 2.32E-09 |
| B64 | B197 | WBRSR(mPas_5) | AG | 0.064342 | 2.41E-09 |
| B64 | B74 | WBRSR(mPas_5) | ApoA1 | -0.0599 | 2.78E-08 |
| B64 | B103 | WBRSR(mPas_5) | anti-SSA/Ro | 0.058724 | 5.17E-08 |
| B64 | B221 | WBRSR(mPas_5) | Hei | 0.058273 | 6.54E-08 |
| B64 | B133 | WBRSR(mPas_5) | TBIL | 0.05801 | 7.49E-08 |
| B64 | B163 | WBRSR(mPas_5) | SG | 0.052924 | 9.26E-07 |
| B65 | B123 | WBRSR(mPas_1) | hs-CRP1 | 0.117892 | 0 |
| B65 | B125 | WBRSR(mPas_1) | Cor | 0.106675 | 0 |
| B65 | B66 | WBRSR(mPas_1) | WBRSR(mPas_100) | 0.842672 | 0 |
| B65 | B67 | WBRSR(mPas_1) | WBRSR(mPas_50) | 0.8722 | 0 |
| B65 | B108 | WBRSR(mPas_1) | Anti-Cyclin | -0.0875 | 4.44E-16 |
| B65 | B134 | WBRSR(mPas_1) | CHE | 0.085842 | 1.78E-15 |
| B65 | B69 | WBRSR(mPas_1) | PV | 0.078482 | 3.28E-13 |
| B65 | B194 | WBRSR(mPas_1) | CI | -0.0773 | 7.37E-13 |
| B65 | B185 | WBRSR(mPas_1) | PT | -0.07541 | 2.63E-12 |
| B65 | B89 | WBRSR(mPas_1) | CA 72-4 | 0.068241 | 2.46E-10 |
| B65 | B103 | WBRSR(mPas_1) | anti-SSA/Ro | 0.066106 | 8.72E-10 |
| B65 | B184 | WBRSR(mPas_1) | INR | -0.06522 | 1.46E-09 |
| B65 | B120 | WBRSR(mPas_1) | HCV-cAg | 0.063634 | 3.60E-09 |
| B65 | B138 | WBRSR(mPas_1) | PAB | 0.0635 | 3.88E-09 |
| B65 | B129 | WBRSR(mPas_1) | IBIL | 0.062564 | 6.55E-09 |
| B65 | B218 | WBRSR(mPas_1) | DBP | 0.060959 | 1.58E-08 |
| B65 | B197 | WBRSR(mPas_1) | AG | 0.060829 | 1.69E-08 |
| B65 | B133 | WBRSR(mPas_1) | TBIL | 0.055626 | 2.50E-07 |
| B65 | B221 | WBRSR(mPas_1) | Hei | 0.054686 | 3.97E-07 |
| B65 | B181 | WBRSR(mPas_1) | APTT | -0.0539 | 5.81E-07 |
| B65 | B156 | WBRSR(mPas_1) | IgE | -0.05352 | 6.97E-07 |
| B66 | B120 | WBRSR(mPas_100) | HCV-cAg | 0.098271 | 0 |
| B66 | B123 | WBRSR(mPas_100) | hs-CRP1 | 0.136816 | 0 |
| B66 | B134 | WBRSR(mPas_100) | CHE | 0.125301 | 0 |
| B66 | B67 | WBRSR(mPas_100) | WBRSR(mPas_50) | 0.994633 | 0 |

| | | | | | |
|---|---|---|---|---|---|
| B66 | B69 | WBRSR(mPas_100) | PV | 0.208638 | 0 |
| B66 | B194 | WBRSR(mPas_100) | CI | -0.08653 | 8.88E-16 |
| B66 | B184 | WBRSR(mPas_100) | INR | -0.08486 | 3.33E-15 |
| B66 | B108 | WBRSR(mPas_100) | Anti-Cyclin | -0.08417 | 5.77E-15 |
| B66 | B197 | WBRSR(mPas_100) | AG | 0.079904 | 1.22E-13 |
| B66 | B138 | WBRSR(mPas_100) | PAB | 0.075709 | 2.15E-12 |
| B66 | B74 | WBRSR(mPas_100) | ApoA1 | -0.07558 | 2.35E-12 |
| B66 | B89 | WBRSR(mPas_100) | CA 72-4 | 0.071894 | 2.57E-11 |
| B66 | B185 | WBRSR(mPas_100) | PT | -0.0669 | 5.48E-10 |
| B66 | B71 | WBRSR(mPas_100) | TG | 0.066122 | 8.64E-10 |
| B66 | B218 | WBRSR(mPas_100) | DBP | 0.065769 | 1.06E-09 |
| B66 | B136 | WBRSR(mPas_100) | 5'NT | 0.065002 | 1.65E-09 |
| B66 | B91 | WBRSR(mPas_100) | TP | 0.063452 | 3.99E-09 |
| B66 | B129 | WBRSR(mPas_100) | IBIL | 0.063062 | 4.97E-09 |
| B66 | B92 | WBRSR(mPas_100) | ALDH2-Glu504Glu | 0.058545 | 5.68E-08 |
| B66 | B221 | WBRSR(mPas_100) | Hei | 0.057849 | 8.14E-08 |
| B66 | B140 | WBRSR(mPas_100) | r-GT | 0.057078 | 1.21E-07 |
| B66 | B93 | WBRSR(mPas_100) | ALDH2-Glu504Lys | -0.05688 | 1.33E-07 |
| B66 | B163 | WBRSR(mPas_100) | SG | 0.056342 | 1.75E-07 |
| B66 | B94 | WBRSR(mPas_100) | ALDH2-Lys504Lys | -0.05623 | 1.85E-07 |
| B66 | B133 | WBRSR(mPas_100) | TBIL | 0.056114 | 1.96E-07 |
| B66 | B216 | WBRSR(mPas_100) | BMI | 0.055541 | 2.61E-07 |
| B66 | B186 | WBRSR(mPas_100) | PFC | 0.05485 | 3.67E-07 |
| B67 | B120 | WBRSR(mPas_50) | HCV-cAg | 0.09574 | 0 |
| B67 | B123 | WBRSR(mPas_50) | hs-CRP1 | 0.137365 | 0 |
| B67 | B134 | WBRSR(mPas_50) | CHE | 0.123178 | 0 |
| B67 | B69 | WBRSR(mPas_50) | PV | 0.199283 | 0 |
| B67 | B108 | WBRSR(mPas_50) | Anti-Cyclin | -0.08551 | 2.22E-15 |
| B67 | B194 | WBRSR(mPas_50) | CI | -0.08526 | 2.66E-15 |
| B67 | B184 | WBRSR(mPas_50) | INR | -0.0843 | 5.11E-15 |
| B67 | B197 | WBRSR(mPas_50) | AG | 0.07886 | 2.52E-13 |
| B67 | B74 | WBRSR(mPas_50) | ApoA1 | -0.07745 | 6.64E-13 |
| B67 | B138 | WBRSR(mPas_50) | PAB | 0.073606 | 8.57E-12 |
| B67 | B89 | WBRSR(mPas_50) | CA 72-4 | 0.07261 | 1.63E-11 |
| B67 | B185 | WBRSR(mPas_50) | PT | -0.06962 | 1.06E-10 |
| B67 | B218 | WBRSR(mPas_50) | DBP | 0.06543 | 1.29E-09 |
| B67 | B129 | WBRSR(mPas_50) | IBIL | 0.064651 | 2.02E-09 |
| B67 | B71 | WBRSR(mPas_50) | TG | 0.063358 | 4.21E-09 |
| B67 | B136 | WBRSR(mPas_50) | 5'NT | 0.061505 | 1.17E-08 |
| B67 | B91 | WBRSR(mPas_50) | TP | 0.059258 | 3.91E-08 |
| B67 | B221 | WBRSR(mPas_50) | Hei | 0.058652 | 5.37E-08 |
| B67 | B92 | WBRSR(mPas_50) | ALDH2-Glu504Glu | 0.058323 | 6.37E-08 |
| B67 | B133 | WBRSR(mPas_50) | TBIL | 0.057578 | 9.35E-08 |
| B67 | B94 | WBRSR(mPas_50) | ALDH2-Lys504Lys | -0.05737 | 1.04E-07 |

| | | | | | |
|---|---|---|---|---|---|
| B67 | B93 | WBRSR(mPas_50) | ALDH2-Glu504Lys | -0.05658 | 1.55E-07 |
| B67 | B216 | WBRSR(mPas_50) | BMI | 0.056439 | 1.67E-07 |
| B67 | B163 | WBRSR(mPas_50) | SG | 0.055233 | 3.04E-07 |
| B67 | B140 | WBRSR(mPas_50) | r-GT | 0.054004 | 5.53E-07 |
| B69 | B123 | PV | hs-CRP1 | 0.109091 | 0 |
| B69 | B127 | PV | A-G ratio | -0.09493 | 0 |
| B69 | B130 | PV | GLB | 0.13039 | 0 |
| B69 | B136 | PV | 5'NT | 0.090784 | 0 |
| B69 | B155 | PV | IgA | 0.120961 | 0 |
| B69 | B156 | PV | IgE | 0.095404 | 0 |
| B69 | B157 | PV | IgG | 0.092743 | 0 |
| B69 | B194 | PV | Cl | -0.11974 | 0 |
| B69 | B91 | PV | TP | 0.127391 | 0 |
| B69 | B114 | PV | TSH TT3 TT4(TT4) | 0.087743 | 4.44E-16 |
| B69 | B195 | PV | Na | -0.08467 | 4.00E-15 |
| B69 | B71 | PV | TG | 0.08079 | 6.53E-14 |
| B69 | B77 | PV | TC | 0.078931 | 2.40E-13 |
| B69 | B180 | PV | Tf | 0.078793 | 2.65E-13 |
| B69 | B76 | PV | ApoB | 0.075008 | 3.42E-12 |
| B69 | B175 | PV | ALB-U | 0.072351 | 1.92E-11 |
| B69 | B179 | PV | α1-MG | 0.072 | 2.40E-11 |
| B69 | B137 | PV | AFU | 0.068374 | 2.27E-10 |
| B69 | B120 | PV | HCV-cAg | 0.062964 | 5.24E-09 |
| B69 | B186 | PV | PFC | 0.061503 | 1.17E-08 |
| B69 | B183 | PV | PTA | 0.05888 | 4.77E-08 |
| B69 | B178 | PV | Ig-U | 0.058423 | 6.05E-08 |
| B69 | B112 | PV | FT4 | 0.057113 | 1.19E-07 |
| B69 | B122 | PV | C4 | 0.057017 | 1.25E-07 |
| B69 | B134 | PV | CHE | 0.054747 | 3.85E-07 |
| B69 | B139 | PV | ADA | 0.054262 | 4.88E-07 |
| B69 | B108 | PV | Anti-Cyclin | 0.052944 | 9.17E-07 |
| B7 | B8 | WB-Cop | WB-Zinc | -0.22448 | 0 |
| B7 | B62 | WB-Cop | WBHSRVI | -0.08656 | 8.88E-16 |
| B7 | B56 | WB-Cop | IR | -0.08556 | 2.00E-15 |
| B7 | B54 | WB-Cop | ESR | 0.077908 | 4.87E-13 |
| B7 | B60 | WB-Cop | WBRVI | -0.07207 | 2.30E-11 |
| B7 | B61 | WB-Cop | WBHSRV | -0.06837 | 2.27E-10 |
| B7 | B63 | WB-Cop | WBRSR(mPas_200) | -0.06765 | 3.50E-10 |
| B7 | B67 | WB-Cop | WBRSR(mPas_50) | -0.06486 | 1.80E-09 |
| B7 | B66 | WB-Cop | WBRSR(mPas_100) | -0.0647 | 1.97E-09 |
| B7 | B64 | WB-Cop | WBRSR(mPas_5) | -0.06281 | 5.70E-09 |
| B7 | B55 | WB-Cop | TK | -0.06259 | 6.46E-09 |
| B7 | B108 | WB-Cop | Anti-Cyclin | 0.060669 | 1.84E-08 |
| B7 | B69 | WB-Cop | PV | 0.059004 | 4.47E-08 |

| | | | | | |
|---|---|---|---|---|---|
| B7 | B65 | WB-Cop | WBRSR(mPas_1) | -0.05882 | 4.92E-08 |
| B70 | B127 | LDL-C | A-G ratio | -0.13669 | 0 |
| B70 | B130 | LDL-C | GLB | 0.135952 | 0 |
| B70 | B132 | LDL-C | Dbil | -0.22177 | 0 |
| B70 | B133 | LDL-C | TBIL | -0.0934 | 0 |
| B70 | B220 | LDL-C | SBP | 0.099338 | 0 |
| B70 | B72 | LDL-C | HDL-C | 0.169174 | 0 |
| B70 | B75 | LDL-C | ApoA1/B | -0.19075 | 0 |
| B70 | B76 | LDL-C | ApoB | 0.285936 | 0 |
| B70 | B77 | LDL-C | TC | 0.813208 | 0 |
| B70 | B91 | LDL-C | TP | 0.120936 | 0 |
| B70 | B221 | LDL-C | Hei | -0.08556 | 2.00E-15 |
| B70 | B218 | LDL-C | DBP | 0.074103 | 6.21E-12 |
| B70 | B74 | LDL-C | ApoA1 | 0.073976 | 6.74E-12 |
| B70 | B188 | LDL-C | Ca | 0.068358 | 2.29E-10 |
| B70 | B128 | LDL-C | ALT | -0.06518 | 1.49E-09 |
| B70 | B141 | LDL-C | AST/ALT | 0.061967 | 9.11E-09 |
| B70 | B71 | LDL-C | TG | -0.05808 | 7.23E-08 |
| B70 | B153 | LDL-C | Glu | 0.053963 | 5.64E-07 |
| B70 | B131 | LDL-C | AST | -0.05383 | 6.02E-07 |
| B71 | B126 | TG | Alb | 0.127529 | 0 |
| B71 | B127 | TG | A-G ratio | -0.11896 | 0 |
| B71 | B128 | TG | ALT | 0.106461 | 0 |
| B71 | B130 | TG | GLB | 0.211958 | 0 |
| B71 | B132 | TG | Dbil | -0.10872 | 0 |
| B71 | B134 | TG | CHE | 0.126546 | 0 |
| B71 | B136 | TG | 5'NT | 0.101085 | 0 |
| B71 | B140 | TG | r-GT | 0.162746 | 0 |
| B71 | B153 | TG | Glu | 0.240642 | 0 |
| B71 | B170 | TG | U-GLU | 0.152219 | 0 |
| B71 | B189 | TG | eGFB | 0.089521 | 0 |
| B71 | B192 | TG | UA | 0.131907 | 0 |
| B71 | B194 | TG | Cl | -0.10271 | 0 |
| B71 | B204 | TG | Drinking | 0.104858 | 0 |
| B71 | B206 | TG | Waist | 0.136312 | 0 |
| B71 | B207 | TG | WHR | 0.128333 | 0 |
| B71 | B209 | TG | Smoking | 0.1193 | 0 |
| B71 | B215 | TG | BMI | 0.157026 | 0 |
| B71 | B216 | TG | BMI | 0.150521 | 0 |
| B71 | B218 | TG | DBP | 0.139098 | 0 |
| B71 | B222 | TG | NS | 0.090726 | 0 |
| B71 | B72 | TG | HDL-C | -0.34672 | 0 |
| B71 | B77 | TG | TC | 0.387895 | 0 |
| B71 | B91 | TG | TP | 0.273816 | 0 |

| | | | | | |
|---|---|---|---|---|---|
| B71 | B120 | TG | HCV-cAg | 0.088107 | 4.44E-16 |
| B71 | B188 | TG | Ca | 0.087295 | 4.44E-16 |
| B71 | B195 | TG | Na | -0.08659 | 8.88E-16 |
| B71 | B141 | TG | AST/ALT | -0.08618 | 1.33E-15 |
| B71 | B133 | TG | TBIL | -0.07921 | 1.98E-13 |
| B71 | B131 | TG | AST | 0.075802 | 2.02E-12 |
| B71 | B163 | TG | SG | 0.07553 | 2.42E-12 |
| B71 | B210 | TG | Hip | 0.072812 | 1.43E-11 |
| B71 | B74 | TG | ApoA1 | -0.07094 | 4.69E-11 |
| B71 | B172 | TG | KB | 0.069369 | 1.24E-10 |
| B71 | B155 | TG | IgA | 0.066961 | 5.27E-10 |
| B71 | B137 | TG | AFU | 0.065522 | 1.22E-09 |
| B71 | B138 | TG | PAB | 0.062954 | 5.27E-09 |
| B71 | B184 | TG | INR | -0.06161 | 1.11E-08 |
| B71 | B81 | TG | TSGF | 0.060771 | 1.75E-08 |
| B71 | B129 | TG | IBIL | -0.06024 | 2.32E-08 |
| B71 | B221 | TG | Hei | 0.05449 | 4.37E-07 |
| B72 | B128 | HDL-C | ALT | -0.09761 | 0 |
| B72 | B141 | HDL-C | AST/ALT | 0.184369 | 0 |
| B72 | B153 | HDL-C | Glu | -0.15163 | 0 |
| B72 | B163 | HDL-C | SG | -0.13851 | 0 |
| B72 | B174 | HDL-C | Urea/Cr | 0.102069 | 0 |
| B72 | B192 | HDL-C | UA | -0.20787 | 0 |
| B72 | B206 | HDL-C | Waist | -0.25577 | 0 |
| B72 | B207 | HDL-C | WHR | -0.20694 | 0 |
| B72 | B209 | HDL-C | Smoking | -0.1468 | 0 |
| B72 | B210 | HDL-C | Hip | -0.17191 | 0 |
| B72 | B215 | HDL-C | BMI | -0.22394 | 0 |
| B72 | B216 | HDL-C | BMI | -0.3107 | 0 |
| B72 | B221 | HDL-C | Hei | -0.24022 | 0 |
| B72 | B222 | HDL-C | NS | -0.20628 | 0 |
| B72 | B74 | HDL-C | ApoA1 | 0.241322 | 0 |
| B72 | B77 | HDL-C | TC | 0.234569 | 0 |
| B72 | B170 | HDL-C | U-GLU | -0.08469 | 4.00E-15 |
| B72 | B75 | HDL-C | ApoA1/B | 0.083077 | 1.24E-14 |
| B72 | B190 | HDL-C | Cre | -0.08182 | 3.11E-14 |
| B72 | B130 | HDL-C | GLB | -0.08063 | 7.31E-14 |
| B72 | B160 | HDL-C | PH | 0.079449 | 1.67E-13 |
| B72 | B218 | HDL-C | DBP | -0.07844 | 3.38E-13 |
| B72 | B220 | HDL-C | SBP | 0.065783 | 1.05E-09 |
| B72 | B91 | HDL-C | TP | -0.06518 | 1.49E-09 |
| B72 | B189 | HDL-C | eGFB | 0.060569 | 1.95E-08 |
| B72 | B127 | HDL-C | A-G ratio | 0.060502 | 2.02E-08 |
| B72 | B176 | HDL-C | Ucr | -0.05493 | 3.52E-07 |

| | | | | | |
|---|---|---|---|---|---|
| B72 | B188 | HDL-C | Ca | 0.052881 | 9.45E-07 |
| B73 | B137 | ApoA | AFU | -0.13593 | 0 |
| B73 | B74 | ApoA | ApoA1 | 0.092379 | 0 |
| B73 | B76 | ApoA | ApoB | 0.138609 | 0 |
| B73 | B75 | ApoA | ApoA1/B | -0.07176 | 2.79E-11 |
| B73 | B199 | ApoA | Mg | 0.071688 | 2.93E-11 |
| B73 | B96 | ApoA | dsDNA (IIF) | -0.06729 | 4.33E-10 |
| B73 | B122 | ApoA | C4 | 0.065783 | 1.05E-09 |
| B73 | B194 | ApoA | Cl | 0.065073 | 1.59E-09 |
| B73 | B138 | ApoA | PAB | -0.05482 | 3.72E-07 |
| B74 | B110 | ApoA1 | AHA | 0.092582 | 0 |
| B74 | B195 | ApoA1 | Na | 0.116184 | 0 |
| B74 | B196 | ApoA1 | TCO2 | 0.104174 | 0 |
| B74 | B199 | ApoA1 | Mg | 0.143379 | 0 |
| B74 | B216 | ApoA1 | BMI | -0.11385 | 0 |
| B74 | B221 | ApoA1 | Hei | -0.1083 | 0 |
| B74 | B75 | ApoA1 | ApoA1/B | 0.334482 | 0 |
| B74 | B76 | ApoA1 | ApoB | 0.139879 | 0 |
| B74 | B77 | ApoA1 | TC | 0.093033 | 0 |
| B74 | B176 | ApoA1 | Ucr | -0.08818 | 2.22E-16 |
| B74 | B86 | ApoA1 | TBhCG | 0.085716 | 1.78E-15 |
| B74 | B159 | ApoA1 | β2-MG | -0.07867 | 2.88E-13 |
| B74 | B123 | ApoA1 | hs-CRP1 | -0.07592 | 1.87E-12 |
| B74 | B95 | ApoA1 | anti-cenpb | 0.07592 | 1.87E-12 |
| B74 | B206 | ApoA1 | Waist | -0.07569 | 2.17E-12 |
| B74 | B97 | ApoA1 | dsDNA | 0.075469 | 2.52E-12 |
| B74 | B120 | ApoA1 | HCV-cAg | -0.07378 | 7.66E-12 |
| B74 | B222 | ApoA1 | NS | -0.07359 | 8.67E-12 |
| B74 | B141 | ApoA1 | AST/ALT | 0.069223 | 1.35E-10 |
| B74 | B163 | ApoA1 | SG | -0.06899 | 1.56E-10 |
| B74 | B81 | ApoA1 | TSGF | -0.0679 | 3.01E-10 |
| B74 | B146 | ApoA1 | CysC | -0.06727 | 4.40E-10 |
| B74 | B188 | ApoA1 | Ca | 0.066482 | 6.99E-10 |
| B74 | B181 | ApoA1 | APTT | -0.06315 | 4.74E-09 |
| B74 | B207 | ApoA1 | WHR | -0.06248 | 6.86E-09 |
| B74 | B215 | ApoA1 | BMI | -0.06213 | 8.34E-09 |
| B74 | B107 | ApoA1 | AnuA | 0.061968 | 9.10E-09 |
| B74 | B101 | ApoA1 | anti-Sc70 | 0.061603 | 1.11E-08 |
| B74 | B104 | ApoA1 | anti-SSB | 0.061603 | 1.11E-08 |
| B74 | B99 | ApoA1 | anti-PM-Scl | 0.061603 | 1.11E-08 |
| B74 | B209 | ApoA1 | Smoking | -0.05896 | 4.57E-08 |
| B74 | B151 | ApoA1 | Anti-CCP | 0.057065 | 1.22E-07 |
| B74 | B125 | ApoA1 | Cor | 0.056824 | 1.37E-07 |
| B74 | B180 | ApoA1 | Tf | 0.056417 | 1.69E-07 |

| | | | | | |
|---|---|---|---|---|---|
| B74 | B134 | ApoA1 | CHE | -0.05626 | 1.83E-07 |
| B74 | B115 | ApoA1 | TSH TT3 TT4(TT3) | 0.053932 | 5.72E-07 |
| B74 | B109 | ApoA1 | AMA-M2 | 0.053391 | 7.42E-07 |
| B75 | B134 | ApoA1/B | CHE | -0.09454 | 0 |
| B75 | B136 | ApoA1/B | 5'NT | 0.100135 | 0 |
| B75 | B155 | ApoA1/B | IgA | 0.10361 | 0 |
| B75 | B157 | ApoA1/B | IgG | 0.123422 | 0 |
| B75 | B76 | ApoA1/B | ApoB | -0.72783 | 0 |
| B75 | B77 | ApoA1/B | TC | -0.11978 | 0 |
| B75 | B182 | ApoA1/B | TT | 0.077833 | 5.12E-13 |
| B75 | B139 | ApoA1/B | ADA | -0.0729 | 1.35E-11 |
| B75 | B132 | ApoA1/B | Dbil | 0.071892 | 2.57E-11 |
| B75 | B113 | ApoA1/B | FT3 | 0.069754 | 9.78E-11 |
| B75 | B125 | ApoA1/B | Cor | -0.06398 | 2.96E-09 |
| B75 | B193 | ApoA1/B | K | -0.06153 | 1.16E-08 |
| B75 | B179 | ApoA1/B | α1-MG | -0.05669 | 1.47E-07 |
| B75 | B106 | ApoA1/B | anti-P | 0.055598 | 2.54E-07 |
| B76 | B134 | ApoB | CHE | 0.152328 | 0 |
| B76 | B139 | ApoB | ADA | 0.089248 | 0 |
| B76 | B77 | ApoB | TC | 0.253275 | 0 |
| B76 | B125 | ApoB | Cor | 0.083408 | 9.77E-15 |
| B76 | B132 | ApoB | Dbil | -0.08319 | 1.15E-14 |
| B76 | B193 | ApoB | K | 0.07967 | 1.43E-13 |
| B76 | B135 | ApoB | TBA | -0.07825 | 3.84E-13 |
| B76 | B122 | ApoB | C4 | 0.069023 | 1.53E-10 |
| B76 | B91 | ApoB | TP | 0.065414 | 1.30E-09 |
| B76 | B150 | ApoB | ALP | 0.06265 | 6.25E-09 |
| B76 | B121 | ApoB | C3 | 0.061943 | 9.23E-09 |
| B76 | B137 | ApoB | AFU | 0.061588 | 1.12E-08 |
| B76 | B130 | ApoB | GLB | 0.059139 | 4.16E-08 |
| B76 | B106 | ApoB | anti-P | -0.05874 | 5.13E-08 |
| B76 | B148 | ApoB | Anti_TPO | 0.058427 | 6.04E-08 |
| B76 | B157 | ApoB | IgG | -0.0579 | 7.91E-08 |
| B76 | B196 | ApoB | TCO2 | 0.057186 | 1.14E-07 |
| B76 | B181 | ApoB | APTT | -0.05584 | 2.25E-07 |
| B76 | B153 | ApoB | Glu | 0.054293 | 4.81E-07 |
| B76 | B111 | ApoB | TSH | 0.054281 | 4.84E-07 |
| B77 | B127 | TC | A-G ratio | -0.18313 | 0 |
| B77 | B130 | TC | GLB | 0.217545 | 0 |
| B77 | B132 | TC | Dbil | -0.23659 | 0 |
| B77 | B133 | TC | TBIL | -0.11404 | 0 |
| B77 | B153 | TC | Glu | 0.140766 | 0 |
| B77 | B188 | TC | Ca | 0.11624 | 0 |
| B77 | B218 | TC | DBP | 0.096675 | 0 |

| | | | | | |
|---|---|---|---|---|---|
| B77 | B220 | TC | SBP | 0.128036 | 0 |
| B77 | B221 | TC | Hei | -0.13515 | 0 |
| B77 | B91 | TC | TP | 0.223438 | 0 |
| B77 | B174 | TC | Urea/Cr | 0.082511 | 1.91E-14 |
| B77 | B134 | TC | CHE | 0.08201 | 2.71E-14 |
| B77 | B136 | TC | 5'NT | 0.081299 | 4.53E-14 |
| B77 | B170 | TC | U-GLU | 0.079813 | 1.30E-13 |
| B77 | B140 | TC | r-GT | 0.079764 | 1.35E-13 |
| B77 | B216 | TC | BMI | -0.07655 | 1.23E-12 |
| B77 | B141 | TC | AST/ALT | 0.068973 | 1.58E-10 |
| B77 | B194 | TC | Cl | -0.06234 | 7.43E-09 |
| B77 | B175 | TC | ALB-U | 0.060521 | 2.00E-08 |
| B77 | B177 | TC | ACR | 0.060118 | 2.48E-08 |
| B77 | B150 | TC | ALP | 0.05765 | 9.02E-08 |
| B77 | B129 | TC | IBIL | -0.05744 | 1.00E-07 |
| B77 | B222 | TC | NS | -0.05398 | 5.61E-07 |
| B77 | B180 | TC | Tf | 0.053855 | 5.94E-07 |
| B78 | B102 | HBcAb | anti-Sm | 0.116822 | 0 |
| B78 | B103 | HBcAb | anti-SSA/Ro | 0.098037 | 0 |
| B78 | B105 | HBcAb | anti-U1RNP | 0.118621 | 0 |
| B78 | B80 | HBcAb | anti-HP IgG | -0.10733 | 0 |
| B78 | B186 | HBcAb | PFC | -0.05974 | 3.03E-08 |
| B78 | B98 | HBcAb | anti-Jo-1 | -0.05402 | 5.50E-07 |
| B79 | B101 | hb V -dna | anti-Sc70 | 0.246321 | 0 |
| B79 | B104 | hb V -dna | anti-SSB | 0.246321 | 0 |
| B79 | B107 | hb V -dna | AnuA | 0.247549 | 0 |
| B79 | B109 | hb V -dna | AMA-M2 | 0.093831 | 0 |
| B79 | B119 | hb V -dna | Hcv-rna | 0.327036 | 0 |
| B79 | B120 | hb V -dna | HCV-cAg | -0.13367 | 0 |
| B79 | B124 | hb V -dna | ACTH | -0.09908 | 0 |
| B79 | B139 | hb V -dna | ADA | -0.12936 | 0 |
| B79 | B157 | hb V -dna | IgG | 0.129405 | 0 |
| B79 | B181 | hb V -dna | APTT | 0.111708 | 0 |
| B79 | B183 | hb V -dna | PTA | 0.158517 | 0 |
| B79 | B184 | hb V -dna | INR | 0.283293 | 0 |
| B79 | B185 | hb V -dna | PT | 0.161344 | 0 |
| B79 | B186 | hb V -dna | PFC | -0.14812 | 0 |
| B79 | B194 | hb V -dna | Cl | 0.097753 | 0 |
| B79 | B196 | hb V -dna | TCO2 | 0.130083 | 0 |
| B79 | B197 | hb V -dna | AG | -0.17039 | 0 |
| B79 | B200 | hb V -dna | OP | -0.13932 | 0 |
| B79 | B81 | hb V -dna | TSGF | -0.17262 | 0 |
| B79 | B92 | hb V -dna | ALDH2-Glu504Glu | -0.20247 | 0 |
| B79 | B93 | hb V -dna | ALDH2-Glu504Lys | 0.189629 | 0 |

| | | | | | |
|---|---|---|---|---|---|
| B79 | B94 | hb V -dna | ALDH2-Lys504Lys | 0.30032 | 0 |
| B79 | B96 | hb V -dna | dsDNA (IIF) | 0.102139 | 0 |
| B79 | B99 | hb V -dna | anti-PM-Scl | 0.246321 | 0 |
| B79 | B179 | hb V -dna | α1-MG | -0.08563 | 1.78E-15 |
| B79 | B198 | hb V -dna | P | -0.07687 | 9.85E-13 |
| B79 | B193 | hb V -dna | K | 0.074385 | 5.16E-12 |
| B79 | B115 | hb V -dna | TSH TT3 TT4(TT3) | 0.073213 | 1.11E-11 |
| B79 | B100 | hb V -dna | anti-Ro52 | 0.070984 | 4.56E-11 |
| B79 | B134 | hb V -dna | CHE | -0.06556 | 1.20E-09 |
| B79 | B125 | hb V -dna | Cor | 0.061691 | 1.06E-08 |
| B79 | B177 | hb V -dna | ACR | -0.0606 | 1.92E-08 |
| B79 | B123 | hb V -dna | hs-CRP1 | -0.05968 | 3.12E-08 |
| B79 | B178 | hb V -dna | Ig-U | -0.05762 | 9.16E-08 |
| B79 | B170 | hb V -dna | U-GLU | -0.05582 | 2.27E-07 |
| B8 | B108 | WB-Zinc | Anti-Cyclin | -0.28458 | 0 |
| B8 | B17 | WB-Zinc | CD3+CD16+CD56+NK | 0.161873 | 0 |
| B8 | B185 | WB-Zinc | PT | -0.08658 | 8.88E-16 |
| B8 | B21 | WB-Zinc | CD3+CD4-CD8- | 0.079251 | 1.92E-13 |
| B8 | B184 | WB-Zinc | INR | -0.06199 | 9.00E-09 |
| B8 | B145 | WB-Zinc | BGP | 0.061025 | 1.52E-08 |
| B8 | B15 | WB-Zinc | CD3-CD19+(B cell) | 0.060667 | 1.85E-08 |
| B81 | B125 | TSGF | Cor | -0.14959 | 0 |
| B81 | B139 | TSGF | ADA | 0.104218 | 0 |
| B81 | B151 | TSGF | Anti-CCP | -0.15147 | 0 |
| B81 | B183 | TSGF | PTA | -0.09181 | 0 |
| B81 | B184 | TSGF | INR | -0.19047 | 0 |
| B81 | B185 | TSGF | PT | -0.1652 | 0 |
| B81 | B186 | TSGF | PFC | 0.13555 | 0 |
| B81 | B197 | TSGF | AG | 0.122987 | 0 |
| B81 | B200 | TSGF | OP | 0.092473 | 0 |
| B81 | B92 | TSGF | ALDH2-Glu504Glu | 0.137371 | 0 |
| B81 | B93 | TSGF | ALDH2-Glu504Lys | -0.13212 | 0 |
| B81 | B94 | TSGF | ALDH2-Lys504Lys | -0.15214 | 0 |
| B81 | B182 | TSGF | TT | 0.087808 | 4.44E-16 |
| B81 | B134 | TSGF | CHE | 0.08556 | 2.00E-15 |
| B81 | B155 | TSGF | IgA | 0.081875 | 3.00E-14 |
| B81 | B196 | TSGF | TCO2 | -0.07777 | 5.34E-13 |
| B81 | B108 | TSGF | Anti-Cyclin | 0.076871 | 9.86E-13 |
| B81 | B98 | TSGF | anti-Jo-1 | -0.07485 | 3.81E-12 |
| B81 | B194 | TSGF | Cl | -0.07075 | 5.28E-11 |
| B81 | B199 | TSGF | Mg | -0.06621 | 8.18E-10 |
| B81 | B107 | TSGF | AnuA | -0.06472 | 1.94E-09 |
| B81 | B101 | TSGF | anti-Sc70 | -0.06439 | 2.34E-09 |
| B81 | B104 | TSGF | anti-SSB | -0.06439 | 2.34E-09 |

| | | | | | |
|---|---|---|---|---|---|
| B81 | B99 | TSGF | anti-PM-Scl | -0.06439 | 2.34E-09 |
| B81 | B178 | TSGF | Ig-U | 0.062113 | 8.40E-09 |
| B81 | B177 | TSGF | ACR | 0.056452 | 1.66E-07 |
| B82 | B83 | CEA | AFP | 0.148396 | 0 |
| B82 | B84 | CEA | C-PSA | 0.154253 | 0 |
| B82 | B85 | CEA | SCC | 0.217115 | 0 |
| B82 | B86 | CEA | TBhCG | -0.25405 | 0 |
| B82 | B87 | CEA | NSE | 0.141703 | 0 |
| B82 | B88 | CEA | CA 242 | 0.481277 | 0 |
| B82 | B90 | CEA | CYFRA 21-1 | 0.346823 | 0 |
| B82 | B89 | CEA | CA 72-4 | 0.086718 | 8.88E-16 |
| B82 | B139 | CEA | ADA | 0.053106 | 8.50E-07 |
| B83 | B106 | AFP | anti-P | 0.144115 | 0 |
| B83 | B136 | AFP | 5'NT | 0.12838 | 0 |
| B83 | B139 | AFP | ADA | 0.094992 | 0 |
| B83 | B85 | AFP | SCC | 0.317113 | 0 |
| B83 | B86 | AFP | TBhCG | 0.163733 | 0 |
| B83 | B87 | AFP | NSE | 0.250257 | 0 |
| B83 | B88 | AFP | CA 242 | 0.178464 | 0 |
| B83 | B89 | AFP | CA 72-4 | 0.079937 | 1.19E-13 |
| B83 | B150 | AFP | ALP | 0.076886 | 9.76E-13 |
| B83 | B140 | AFP | r-GT | 0.07601 | 1.76E-12 |
| B83 | B137 | AFP | AFU | 0.075179 | 3.06E-12 |
| B84 | B228 | C-PSA | OS | 0.106655 | 0 |
| B84 | B85 | C-PSA | SCC | 0.18802 | 0 |
| B84 | B86 | C-PSA | TBhCG | -0.21528 | 0 |
| B84 | B87 | C-PSA | NSE | 0.116465 | 0 |
| B84 | B88 | C-PSA | CA 242 | 0.117916 | 0 |
| B84 | B116 | C-PSA | TT3 | -0.0784 | 3.47E-13 |
| B84 | B90 | C-PSA | CYFRA 21-1 | 0.062164 | 8.17E-09 |
| B84 | B117 | C-PSA | TT4 | 0.05939 | 3.65E-08 |
| B85 | B87 | SCC | NSE | 0.28637 | 0 |
| B85 | B88 | SCC | CA 242 | 0.240651 | 0 |
| B85 | B90 | SCC | CYFRA 21-1 | 0.119799 | 0 |
| B85 | B191 | SCC | Urea | 0.05607 | 2.01E-07 |
| B85 | B135 | SCC | TBA | 0.054326 | 4.73E-07 |
| B85 | B168 | SCC | NMP22 | 0.053281 | 7.82E-07 |
| B85 | B189 | SCC | eGFB | -0.05278 | 9.89E-07 |
| B86 | B139 | TBhCG | ADA | -0.11661 | 0 |
| B86 | B199 | TBhCG | Mg | 0.118443 | 0 |
| B86 | B200 | TBhCG | OP | -0.0894 | 0 |
| B86 | B88 | TBhCG | CA 242 | -0.11668 | 0 |
| B86 | B137 | TBhCG | AFU | -0.08813 | 4.44E-16 |
| B86 | B194 | TBhCG | Cl | 0.067502 | 3.82E-10 |

| | | | | | |
|---|---|---|---|---|---|
| B86 | B115 | TBhCG | TSH TT3 TT4(TT3) | 0.056846 | 1.36E-07 |
| B86 | B89 | TBhCG | CA 72-4 | -0.05504 | 3.35E-07 |
| B86 | B129 | TBhCG | IBIL | -0.05387 | 5.89E-07 |
| B87 | B88 | NSE | CA 242 | 0.223606 | 0 |
| B87 | B90 | NSE | CYFRA 21-1 | 0.083343 | 1.02E-14 |
| B87 | B198 | NSE | P | 0.071673 | 2.95E-11 |
| B88 | B90 | CA 242 | CYFRA 21-1 | 0.191616 | 0 |
| B9 | B101 | RH | anti-Sc70 | -0.09394 | 0 |
| B9 | B104 | RH | anti-SSB | -0.09394 | 0 |
| B9 | B107 | RH | AnuA | -0.09438 | 0 |
| B9 | B125 | RH | Cor | 0.137001 | 0 |
| B9 | B131 | RH | AST | 0.093637 | 0 |
| B9 | B136 | RH | 5'NT | 0.411662 | 0 |
| B9 | B139 | RH | ADA | 0.090407 | 0 |
| B9 | B140 | RH | r-GT | 0.110302 | 0 |
| B9 | B15 | RH | CD3-CD19+(B cell) | -0.1409 | 0 |
| B9 | B18 | RH | CD3+CD4+helper(inducer)T | -0.22267 | 0 |
| B9 | B181 | RH | APTT | -0.15857 | 0 |
| B9 | B184 | RH | INR | -0.16639 | 0 |
| B9 | B185 | RH | PT | -0.10297 | 0 |
| B9 | B19 | RH | CD3+CD4+/CD3+CD8+ | -0.13906 | 0 |
| B9 | B193 | RH | K | -0.08982 | 0 |
| B9 | B22 | RH | CD3+CD8+(T) | 0.20984 | 0 |
| B9 | B99 | RH | anti-PM-Scl | -0.09394 | 0 |
| B9 | B23 | RH | CD3-CD16+CD56+(NK) | 0.065935 | 9.63E-10 |
| B9 | B150 | RH | ALP | 0.064009 | 2.91E-09 |
| B9 | B76 | RH | ApoB | 0.062375 | 7.27E-09 |
| B9 | B200 | RH | OP | 0.057444 | 1.00E-07 |
| B9 | B27 | RH | LDH | 0.057076 | 1.21E-07 |
| B9 | B183 | RH | PTA | -0.05681 | 1.38E-07 |
| B9 | B198 | RH | P | -0.05564 | 2.48E-07 |
| B90 | B190 | CYFRA 21-1 | Cre | 0.105923 | 0 |
| B90 | B189 | CYFRA 21-1 | eGFB | -0.08444 | 4.66E-15 |
| B90 | B191 | CYFRA 21-1 | Urea | 0.075234 | 2.95E-12 |
| B90 | B139 | CYFRA 21-1 | ADA | 0.072876 | 1.37E-11 |
| B90 | B199 | CYFRA 21-1 | Mg | -0.06245 | 6.98E-09 |
| B90 | B200 | CYFRA 21-1 | OP | 0.06128 | 1.33E-08 |
| B90 | B146 | CYFRA 21-1 | CysC | 0.061256 | 1.34E-08 |
| B91 | B126 | TP | Alb | 0.44473 | 0 |
| B91 | B127 | TP | A-G ratio | -0.48117 | 0 |
| B91 | B128 | TP | ALT | 0.091397 | 0 |
| B91 | B130 | TP | GLB | 0.788563 | 0 |
| B91 | B131 | TP | AST | 0.122031 | 0 |
| B91 | B153 | TP | Glu | 0.109582 | 0 |

| | | | | | |
|---|---|---|---|---|---|
| B91 | B157 | TP | IgG | 0.111989 | 0 |
| B91 | B188 | TP | Ca | 0.292489 | 0 |
| B91 | B192 | TP | UA | 0.104777 | 0 |
| B91 | B209 | TP | Smoking | -0.09538 | 0 |
| B91 | B218 | TP | DBP | 0.093728 | 0 |
| B91 | B220 | TP | SBP | 0.098682 | 0 |
| B91 | B155 | TP | IgA | 0.085224 | 2.66E-15 |
| B91 | B221 | TP | Hei | -0.08132 | 4.49E-14 |
| B91 | B194 | TP | Cl | -0.0812 | 4.88E-14 |
| B91 | B177 | TP | ACR | -0.07947 | 1.66E-13 |
| B91 | B175 | TP | ALB-U | -0.07269 | 1.55E-11 |
| B91 | B158 | TP | IgM | 0.068913 | 1.64E-10 |
| B91 | B136 | TP | 5'NT | 0.06618 | 8.35E-10 |
| B91 | B134 | TP | CHE | 0.06427 | 2.51E-09 |
| B91 | B178 | TP | Ig-U | -0.06386 | 3.17E-09 |
| B91 | B191 | TP | Urea | -0.06379 | 3.29E-09 |
| B91 | B140 | TP | r-GT | 0.062766 | 5.86E-09 |
| B91 | B180 | TP | Tf | -0.06252 | 6.73E-09 |
| B91 | B163 | TP | SG | -0.05908 | 4.29E-08 |
| B91 | B150 | TP | ALP | 0.058389 | 6.16E-08 |
| B91 | B120 | TP | HCV-cAg | 0.057837 | 8.19E-08 |
| B91 | B111 | TP | TSH | 0.052912 | 9.31E-07 |
| B92 | B101 | ALDH2-Glu504Glu | anti-Sc70 | -0.27314 | 0 |
| B92 | B104 | ALDH2-Glu504Glu | anti-SSB | -0.27314 | 0 |
| B92 | B107 | ALDH2-Glu504Glu | AnuA | -0.26218 | 0 |
| B92 | B109 | ALDH2-Glu504Glu | AMA-M2 | -0.10357 | 0 |
| B92 | B119 | ALDH2-Glu504Glu | Hcv-rna | -0.19616 | 0 |
| B92 | B124 | ALDH2-Glu504Glu | ACTH | 0.204391 | 0 |
| B92 | B138 | ALDH2-Glu504Glu | PAB | 0.101473 | 0 |
| B92 | B179 | ALDH2-Glu504Glu | α1-MG | 0.092321 | 0 |
| B92 | B181 | ALDH2-Glu504Glu | APTT | -0.12422 | 0 |
| B92 | B183 | ALDH2-Glu504Glu | PTA | -0.10756 | 0 |
| B92 | B184 | ALDH2-Glu504Glu | INR | -0.20983 | 0 |
| B92 | B185 | ALDH2-Glu504Glu | PT | -0.13279 | 0 |
| B92 | B186 | ALDH2-Glu504Glu | PFC | 0.11306 | 0 |
| B92 | B194 | ALDH2-Glu504Glu | Cl | -0.12676 | 0 |
| B92 | B197 | ALDH2-Glu504Glu | AG | 0.117697 | 0 |
| B92 | B93 | ALDH2-Glu504Glu | ALDH2-Glu504Lys | -0.99846 | 0 |
| B92 | B94 | ALDH2-Glu504Glu | ALDH2-Lys504Lys | -0.56054 | 0 |
| B92 | B96 | ALDH2-Glu504Glu | dsDNA (IIF) | 0.155462 | 0 |
| B92 | B99 | ALDH2-Glu504Glu | anti-PM-Scl | -0.27314 | 0 |
| B92 | B122 | ALDH2-Glu504Glu | C4 | -0.08021 | 9.81E-14 |
| B92 | B100 | ALDH2-Glu504Glu | anti-Ro52 | -0.07809 | 4.28E-13 |
| B92 | B134 | ALDH2-Glu504Glu | CHE | 0.077516 | 6.36E-13 |

| | | | | | |
|---|---|---|---|---|---|
| B92 | B125 | ALDH2-Glu504Glu | Cor | 0.074842 | 3.82E-12 |
| B92 | B212 | ALDH2-Glu504Glu | IOP-L | 0.067527 | 3.77E-10 |
| B92 | B121 | ALDH2-Glu504Glu | C3 | -0.05895 | 4.60E-08 |
| B92 | B155 | ALDH2-Glu504Glu | IgA | 0.054521 | 4.31E-07 |
| B92 | B200 | ALDH2-Glu504Glu | OP | 0.053737 | 6.29E-07 |
| B92 | B178 | ALDH2-Glu504Glu | Ig-U | 0.053002 | 8.93E-07 |
| B93 | B101 | ALDH2-Glu504Lys | anti-Sc70 | 0.255563 | 0 |
| B93 | B104 | ALDH2-Glu504Lys | anti-SSB | 0.255563 | 0 |
| B93 | B107 | ALDH2-Glu504Lys | AnuA | 0.245052 | 0 |
| B93 | B109 | ALDH2-Glu504Lys | AMA-M2 | 0.096901 | 0 |
| B93 | B119 | ALDH2-Glu504Lys | Hcv-rna | 0.182874 | 0 |
| B93 | B124 | ALDH2-Glu504Lys | ACTH | -0.19661 | 0 |
| B93 | B138 | ALDH2-Glu504Lys | PAB | -0.098 | 0 |
| B93 | B179 | ALDH2-Glu504Lys | α1-MG | -0.08907 | 0 |
| B93 | B181 | ALDH2-Glu504Lys | APTT | 0.116933 | 0 |
| B93 | B183 | ALDH2-Glu504Lys | PTA | 0.101137 | 0 |
| B93 | B184 | ALDH2-Glu504Lys | INR | 0.199474 | 0 |
| B93 | B185 | ALDH2-Glu504Lys | PT | 0.128631 | 0 |
| B93 | B186 | ALDH2-Glu504Lys | PFC | -0.109 | 0 |
| B93 | B194 | ALDH2-Glu504Lys | Cl | 0.122512 | 0 |
| B93 | B197 | ALDH2-Glu504Lys | AG | -0.11391 | 0 |
| B93 | B94 | ALDH2-Glu504Lys | ALDH2-Lys504Lys | 0.513678 | 0 |
| B93 | B96 | ALDH2-Glu504Lys | dsDNA (IIF) | -0.17217 | 0 |
| B93 | B99 | ALDH2-Glu504Lys | anti-PM-Scl | 0.255563 | 0 |
| B93 | B125 | ALDH2-Glu504Lys | Cor | -0.07753 | 6.28E-13 |
| B93 | B122 | ALDH2-Glu504Lys | C4 | 0.076864 | 9.91E-13 |
| B93 | B134 | ALDH2-Glu504Lys | CHE | -0.07424 | 5.67E-12 |
| B93 | B100 | ALDH2-Glu504Lys | anti-Ro52 | 0.073059 | 1.22E-11 |
| B93 | B212 | ALDH2-Glu504Lys | IOP-L | -0.06228 | 7.66E-09 |
| B93 | B121 | ALDH2-Glu504Lys | C3 | 0.056639 | 1.51E-07 |
| B93 | B155 | ALDH2-Glu504Lys | IgA | -0.05333 | 7.64E-07 |
| B94 | B100 | ALDH2-Lys504Lys | anti-Ro52 | 0.117053 | 0 |
| B94 | B101 | ALDH2-Lys504Lys | anti-Sc70 | 0.408833 | 0 |
| B94 | B104 | ALDH2-Lys504Lys | anti-SSB | 0.408833 | 0 |
| B94 | B107 | ALDH2-Lys504Lys | AnuA | 0.396363 | 0 |
| B94 | B109 | ALDH2-Lys504Lys | AMA-M2 | 0.155153 | 0 |
| B94 | B119 | ALDH2-Lys504Lys | Hcv-rna | 0.30355 | 0 |
| B94 | B122 | ALDH2-Lys504Lys | C4 | 0.093056 | 0 |
| B94 | B124 | ALDH2-Lys504Lys | ACTH | -0.22589 | 0 |
| B94 | B134 | ALDH2-Lys504Lys | CHE | -0.09048 | 0 |
| B94 | B138 | ALDH2-Lys504Lys | PAB | -0.10628 | 0 |
| B94 | B157 | ALDH2-Lys504Lys | IgG | 0.107041 | 0 |
| B94 | B179 | ALDH2-Lys504Lys | α1-MG | -0.09814 | 0 |
| B94 | B181 | ALDH2-Lys504Lys | APTT | 0.175443 | 0 |

| | | | | | |
|---|---|---|---|---|---|
| B94 | B183 | ALDH2-Lys504Lys | PTA | 0.153612 | 0 |
| B94 | B184 | ALDH2-Lys504Lys | INR | 0.267246 | 0 |
| B94 | B185 | ALDH2-Lys504Lys | PT | 0.133437 | 0 |
| B94 | B186 | ALDH2-Lys504Lys | PFC | -0.12124 | 0 |
| B94 | B194 | ALDH2-Lys504Lys | Cl | 0.13139 | 0 |
| B94 | B197 | ALDH2-Lys504Lys | AG | -0.11978 | 0 |
| B94 | B212 | ALDH2-Lys504Lys | IOP-L | -0.11453 | 0 |
| B94 | B96 | ALDH2-Lys504Lys | dsDNA (IIF) | 0.165605 | 0 |
| B94 | B99 | ALDH2-Lys504Lys | anti-PM-Scl | 0.408833 | 0 |
| B94 | B211 | ALDH2-Lys504Lys | IOP-R | -0.08718 | 4.44E-16 |
| B94 | B193 | ALDH2-Lys504Lys | K | 0.087093 | 6.66E-16 |
| B94 | B200 | ALDH2-Lys504Lys | OP | -0.06994 | 8.71E-11 |
| B94 | B121 | ALDH2-Lys504Lys | C3 | 0.066094 | 8.78E-10 |
| B94 | B196 | ALDH2-Lys504Lys | TCO2 | 0.062828 | 5.66E-09 |
| B94 | B108 | ALDH2-Lys504Lys | Anti-Cyclin | 0.059977 | 2.67E-08 |
| B94 | B178 | ALDH2-Lys504Lys | Ig-U | -0.0597 | 3.10E-08 |
| B94 | B216 | ALDH2-Lys504Lys | BMI | -0.05849 | 5.85E-08 |
| B94 | B199 | ALDH2-Lys504Lys | Mg | 0.058167 | 6.91E-08 |
| B94 | B221 | ALDH2-Lys504Lys | Hei | -0.05745 | 9.99E-08 |
| B94 | B115 | ALDH2-Lys504Lys | TSH TT3 TT4(TT3) | 0.056054 | 2.02E-07 |
| B94 | B204 | ALDH2-Lys504Lys | Drinking | -0.05566 | 2.46E-07 |
| B94 | B189 | ALDH2-Lys504Lys | eGFB | -0.05522 | 3.05E-07 |
| B94 | B95 | ALDH2-Lys504Lys | anti-cenpb | 0.054383 | 4.60E-07 |
| B95 | B195 | anti-cenpb | Na | 0.077948 | 4.73E-13 |
| B95 | B193 | anti-cenpb | K | 0.063178 | 4.65E-09 |
| B95 | B134 | anti-cenpb | CHE | -0.05634 | 1.75E-07 |
| B96 | B101 | dsDNA (IIF) | anti-Sc70 | 0.25428 | 0 |
| B96 | B104 | dsDNA (IIF) | anti-SSB | 0.254279 | 0 |
| B96 | B107 | dsDNA (IIF) | AnuA | 0.255492 | 0 |
| B96 | B109 | dsDNA (IIF) | AMA-M2 | 0.096322 | 0 |
| B96 | B110 | dsDNA (IIF) | AHA | -0.35554 | 0 |
| B96 | B119 | dsDNA (IIF) | Hcv-rna | 0.180161 | 0 |
| B96 | B182 | dsDNA (IIF) | TT | 0.121242 | 0 |
| B96 | B194 | dsDNA (IIF) | Cl | 0.162303 | 0 |
| B96 | B197 | dsDNA (IIF) | AG | -0.10253 | 0 |
| B96 | B97 | dsDNA (IIF) | dsDNA | -0.52548 | 0 |
| B96 | B99 | dsDNA (IIF) | anti-PM-Scl | 0.25428 | 0 |
| B96 | B193 | dsDNA (IIF) | K | 0.063446 | 4.01E-09 |
| B96 | B184 | dsDNA (IIF) | INR | 0.063048 | 5.00E-09 |
| B96 | B156 | dsDNA (IIF) | IgE | -0.06273 | 5.97E-09 |
| B96 | B186 | dsDNA (IIF) | PFC | -0.05522 | 3.06E-07 |
| B96 | B181 | dsDNA (IIF) | APTT | 0.054337 | 4.71E-07 |
| B96 | B195 | dsDNA (IIF) | Na | 0.053571 | 6.81E-07 |
| B96 | B103 | dsDNA (IIF) | anti-SSA/Ro | 0.05352 | 6.98E-07 |

| | | | | | |
|---|---|---|---|---|---|
| B97 | B101 | dsDNA | anti-Sc70 | -0.09071 | 0 |
| B97 | B104 | dsDNA | anti-SSB | -0.09071 | 0 |
| B97 | B107 | dsDNA | AnuA | -0.09158 | 0 |
| B97 | B110 | dsDNA | AHA | 0.939101 | 0 |
| B97 | B119 | dsDNA | Hcv-rna | -0.14071 | 0 |
| B97 | B150 | dsDNA | ALP | 0.091821 | 0 |
| B97 | B194 | dsDNA | Cl | -0.20482 | 0 |
| B97 | B195 | dsDNA | Na | -0.12632 | 0 |
| B97 | B228 | dsDNA | OS | 0.118216 | 0 |
| B97 | B99 | dsDNA | anti-PM-Scl | -0.09071 | 0 |
| B97 | B197 | dsDNA | AG | 0.06615 | 8.50E-10 |
| B97 | B200 | dsDNA | OP | -0.0649 | 1.75E-09 |
| B98 | B124 | anti-Jo-1 | ACTH | -0.10257 | 0 |
| B98 | B125 | anti-Jo-1 | Cor | 0.337758 | 0 |
| B98 | B181 | anti-Jo-1 | APTT | -0.16743 | 0 |
| B98 | B184 | anti-Jo-1 | INR | -0.16255 | 0 |
| B98 | B185 | anti-Jo-1 | PT | -0.10038 | 0 |
| B98 | B195 | anti-Jo-1 | Na | -0.10928 | 0 |
| B98 | B225 | anti-Jo-1 | hPBG | 0.175819 | 0 |
| B98 | B199 | anti-Jo-1 | Mg | -0.07898 | 2.32E-13 |
| B98 | B194 | anti-Jo-1 | Cl | -0.07863 | 2.96E-13 |
| B98 | B172 | anti-Jo-1 | KB | 0.075533 | 2.42E-12 |
| B98 | B151 | anti-Jo-1 | Anti-CCP | 0.066462 | 7.08E-10 |
| B98 | B107 | anti-Jo-1 | AnuA | -0.0653 | 1.39E-09 |
| B98 | B101 | anti-Jo-1 | anti-Sc70 | -0.0647 | 1.97E-09 |
| B98 | B104 | anti-Jo-1 | anti-SSB | -0.0647 | 1.97E-09 |
| B98 | B99 | anti-Jo-1 | anti-PM-Scl | -0.0647 | 1.97E-09 |
| B98 | B138 | anti-Jo-1 | PAB | 0.06305 | 5.00E-09 |
| B98 | B183 | anti-Jo-1 | PTA | -0.05731 | 1.07E-07 |
| B99 | B100 | anti-PM-Scl | anti-Ro52 | 0.163075 | 0 |
| B99 | B104 | anti-PM-Scl | anti-SSB | 1 | 0 |
| B99 | B107 | anti-PM-Scl | AnuA | 0.995262 | 0 |
| B99 | B109 | anti-PM-Scl | AMA-M2 | 0.285121 | 0 |
| B99 | B115 | anti-PM-Scl | TSH TT3 TT4(TT3) | 0.097149 | 0 |
| B99 | B119 | anti-PM-Scl | Hcv-rna | 0.22665 | 0 |
| B99 | B124 | anti-PM-Scl | ACTH | -0.1739 | 0 |
| B99 | B138 | anti-PM-Scl | PAB | -0.12155 | 0 |
| B99 | B155 | anti-PM-Scl | IgA | -0.11956 | 0 |
| B99 | B157 | anti-PM-Scl | IgG | 0.102519 | 0 |
| B99 | B158 | anti-PM-Scl | IgM | 0.152403 | 0 |
| B99 | B181 | anti-PM-Scl | APTT | 0.174055 | 0 |
| B99 | B182 | anti-PM-Scl | TT | 0.095817 | 0 |
| B99 | B184 | anti-PM-Scl | INR | 0.270459 | 0 |
| B99 | B185 | anti-PM-Scl | PT | 0.1744 | 0 |

| | | | | | |
|---|---|---|---|---|---|
| B99 | B186 | anti-PM-Scl | PFC | -0.11045 | 0 |
| B99 | B194 | anti-PM-Scl | Cl | 0.185271 | 0 |
| B99 | B195 | anti-PM-Scl | Na | 0.134201 | 0 |
| B99 | B196 | anti-PM-Scl | TCO2 | 0.119685 | 0 |
| B99 | B197 | anti-PM-Scl | AG | -0.14837 | 0 |
| B99 | B199 | anti-PM-Scl | Mg | 0.168027 | 0 |
| B99 | B200 | anti-PM-Scl | OP | -0.11588 | 0 |
| B99 | B179 | anti-PM-Scl | α1-MG | -0.08204 | 2.66E-14 |
| B99 | B136 | anti-PM-Scl | 5'NT | -0.08078 | 6.57E-14 |
| B99 | B178 | anti-PM-Scl | Ig-U | -0.07759 | 6.05E-13 |
| B99 | B139 | anti-PM-Scl | ADA | -0.07705 | 8.74E-13 |
| B99 | B180 | anti-PM-Scl | Tf | -0.0763 | 1.45E-12 |
| B99 | B125 | anti-PM-Scl | Cor | -0.0744 | 5.11E-12 |
| B99 | B189 | anti-PM-Scl | eGFB | -0.07423 | 5.72E-12 |
| B99 | B121 | anti-PM-Scl | C3 | -0.07191 | 2.54E-11 |
| B99 | B212 | anti-PM-Scl | IOP-L | -0.06297 | 5.22E-09 |
| B99 | B102 | anti-PM-Scl | anti-Sm | -0.06065 | 1.86E-08 |
| B99 | B103 | anti-PM-Scl | anti-SSA/Ro | 0.056307 | 1.78E-07 |
| B99 | B211 | anti-PM-Scl | IOP-R | -0.05567 | 2.45E-07 |

**Supplementary Table 5. PEI Correlations in Hypertension+Coronary**

| Row | Column | Row | Column | Cor | *P* |
|---|---|---|---|---|---|
| B1 | B132 | Gender | Dbil | -0.18785 | 0 |
| B1 | B161 | Gender | LEU | 0.205613 | 0 |
| B1 | B163 | Gender | SG | -0.26225 | 0 |
| B1 | B174 | Gender | Urea/Cr | 0.367654 | 0 |
| B1 | B190 | Gender | Cre | -0.41531 | 0 |
| B1 | B192 | Gender | UA | -0.34775 | 0 |
| B1 | B204 | Gender | Drinking | -0.21708 | 0 |
| B1 | B206 | Gender | Waist | -0.29726 | 0 |
| B1 | B207 | Gender | WHR | -0.22 | 0 |
| B1 | B209 | Gender | Smoking | -0.22765 | 0 |
| B1 | B210 | Gender | Hip | -0.18635 | 0 |
| B1 | B216 | Gender | BMI | -0.49505 | 0 |
| B1 | B221 | Gender | Hei | -0.7245 | 0 |
| B1 | B222 | Gender | NS | -0.63871 | 0 |
| B1 | B42 | Gender | MON | -0.23091 | 0 |
| B1 | B45 | Gender | MCHC | -0.2022 | 0 |
| B1 | B46 | Gender | RBC Count | -0.37808 | 0 |
| B1 | B51 | Gender | Hb | -0.46703 | 0 |
| B1 | B72 | Gender | HDL-C | 0.294963 | 0 |
| B1 | B77 | Gender | TC | 0.226815 | 0 |
| B1 | B40 | Gender | WBC | -0.17467 | 1.11E-15 |
| B1 | B173 | Gender | NTT | 0.165963 | 2.82E-14 |
| B1 | B133 | Gender | TBIL | -0.16089 | 1.69E-13 |
| B1 | B141 | Gender | AST/ALT | 0.156217 | 8.43E-13 |
| B1 | B129 | Gender | IBIL | -0.13756 | 3.16E-10 |
| B1 | B41 | Gender | MON％ | -0.1328 | 1.27E-09 |
| B1 | B219 | Gender | OT | -0.12045 | 3.75E-08 |
| B1 | B50 | Gender | eo% | -0.11904 | 5.41E-08 |
| B1 | B2 | Gender | Age | 0.118513 | 6.20E-08 |
| B1 | B130 | Gender | GLB | 0.117726 | 7.58E-08 |
| B1 | B127 | Gender | A-G ratio | -0.11674 | 9.75E-08 |
| B1 | B52 | Gender | PLT/BPC | 0.113118 | 2.40E-07 |
| B1 | B191 | Gender | Urea | -0.11259 | 2.74E-07 |
| B1 | B58 | Gender | HCT | -0.11187 | 3.26E-07 |
| B1 | B201 | Gender | e-habits | -0.11181 | 3.31E-07 |
| B1 | B3 | Gender | Hcy | -0.11145 | 3.61E-07 |
| B1 | B70 | Gender | LDL-C | 0.110486 | 4.56E-07 |
| B1 | B205 | Gender | KDA | 0.108006 | 8.21E-07 |
| B10 | B11 | PG Ⅰ | PGI/PGII | 0.20813 | 0 |
| B10 | B12 | PG Ⅰ | PG II | 0.453391 | 0 |
| B10 | B145 | PG Ⅰ | BGP | -0.13924 | 1.91E-10 |

| | | | | | |
|---|---|---|---|---|---|
| B10 | B144 | PG Ⅰ | Total-PINP | -0.1187 | 5.90E-08 |
| B10 | B189 | PG Ⅰ | eGFB | -0.10822 | 7.82E-07 |
| B10 | B143 | PG Ⅰ | β-CTx | -0.10761 | 9.02E-07 |
| B100 | B102 | anti-Ro52 | anti-Sm | -0.27973 | 0 |
| B100 | B105 | anti-Ro52 | anti-U1RNP | -0.33755 | 0 |
| B100 | B107 | anti-Ro52 | AnuA | 0.999998 | 0 |
| B100 | B108 | anti-Ro52 | Anti-Cyclin | -0.22672 | 0 |
| B100 | B109 | anti-Ro52 | AMA-M2 | -0.26181 | 0 |
| B100 | B125 | anti-Ro52 | Cor | 0.234516 | 0 |
| B100 | B154 | anti-Ro52 | RF | 0.204755 | 0 |
| B100 | B183 | anti-Ro52 | PTA | 0.273181 | 0 |
| B100 | B155 | anti-Ro52 | IgA | -0.16404 | 5.60E-14 |
| B100 | B156 | anti-Ro52 | IgE | -0.14475 | 3.52E-11 |
| B100 | B181 | anti-Ro52 | APTT | 0.13878 | 2.19E-10 |
| B100 | B195 | anti-Ro52 | Na | 0.137023 | 3.70E-10 |
| B100 | B103 | anti-Ro52 | anti-SSA/Ro | -0.13644 | 4.40E-10 |
| B100 | B137 | anti-Ro52 | AFU | -0.12881 | 3.93E-09 |
| B100 | B198 | anti-Ro52 | P | -0.12517 | 1.07E-08 |
| B100 | B139 | anti-Ro52 | ADA | 0.118712 | 5.89E-08 |
| B100 | B138 | anti-Ro52 | PAB | 0.113683 | 2.09E-07 |
| B100 | B185 | anti-Ro52 | PT | -0.10972 | 5.48E-07 |
| B101 | B102 | anti-Sc70 | anti-Sm | -0.27973 | 0 |
| B101 | B104 | anti-Sc70 | anti-SSB | 1 | 0 |
| B101 | B105 | anti-Sc70 | anti-U1RNP | -0.33755 | 0 |
| B101 | B107 | anti-Sc70 | AnuA | 0.999999 | 0 |
| B101 | B108 | anti-Sc70 | Anti-Cyclin | -0.22672 | 0 |
| B101 | B109 | anti-Sc70 | AMA-M2 | -0.26181 | 0 |
| B101 | B125 | anti-Sc70 | Cor | 0.234516 | 0 |
| B101 | B154 | anti-Sc70 | RF | 0.204755 | 0 |
| B101 | B183 | anti-Sc70 | PTA | 0.273181 | 0 |
| B101 | B155 | anti-Sc70 | IgA | -0.16404 | 5.60E-14 |
| B101 | B156 | anti-Sc70 | IgE | -0.14475 | 3.52E-11 |
| B101 | B181 | anti-Sc70 | APTT | 0.13878 | 2.19E-10 |
| B101 | B195 | anti-Sc70 | Na | 0.137023 | 3.70E-10 |
| B101 | B103 | anti-Sc70 | anti-SSA/Ro | -0.13644 | 4.40E-10 |
| B101 | B137 | anti-Sc70 | AFU | -0.12881 | 3.93E-09 |
| B101 | B198 | anti-Sc70 | P | -0.12517 | 1.07E-08 |
| B101 | B139 | anti-Sc70 | ADA | 0.118712 | 5.89E-08 |
| B101 | B138 | anti-Sc70 | PAB | 0.113683 | 2.09E-07 |
| B101 | B185 | anti-Sc70 | PT | -0.10972 | 5.48E-07 |
| B102 | B104 | anti-Sm | anti-SSB | -0.27973 | 0 |
| B102 | B105 | anti-Sm | anti-U1RNP | 0.970103 | 0 |
| B102 | B106 | anti-Sm | anti-P | -0.27973 | 0 |
| B102 | B107 | anti-Sm | AnuA | -0.27973 | 0 |

| | | | | | |
|---|---|---|---|---|---|
| B102 | B110 | anti-Sm | AHA | -0.27973 | 0 |
| B102 | B124 | anti-Sm | ACTH | -0.36402 | 0 |
| B102 | B125 | anti-Sm | Cor | -0.44287 | 0 |
| B102 | B138 | anti-Sm | PAB | -0.30825 | 0 |
| B102 | B156 | anti-Sm | IgE | 0.282049 | 0 |
| B102 | B185 | anti-Sm | PT | 0.306909 | 0 |
| B102 | B228 | anti-Sm | OS | 0.179948 | 0 |
| B102 | B120 | anti-Sm | HCV-cAg | -0.17165 | 3.55E-15 |
| B102 | B186 | anti-Sm | PFC | 0.150587 | 5.46E-12 |
| B102 | B183 | anti-Sm | PTA | -0.14799 | 1.26E-11 |
| B102 | B179 | anti-Sm | α1-MG | 0.14166 | 9.16E-11 |
| B102 | B134 | anti-Sm | CHE | -0.13424 | 8.38E-10 |
| B102 | B181 | anti-Sm | APTT | 0.125827 | 8.94E-09 |
| B102 | B182 | anti-Sm | TT | -0.11359 | 2.14E-07 |
| B103 | B158 | anti-SSA/Ro | IgM | 0.386089 | 0 |
| B103 | B178 | anti-SSA/Ro | Ig-U | 0.185639 | 0 |
| B103 | B179 | anti-SSA/Ro | α1-MG | 0.256249 | 0 |
| B103 | B180 | anti-SSA/Ro | Tf | 0.329367 | 0 |
| B103 | B104 | anti-SSA/Ro | anti-SSB | -0.13644 | 4.40E-10 |
| B103 | B106 | anti-SSA/Ro | anti-P | -0.13644 | 4.40E-10 |
| B103 | B107 | anti-SSA/Ro | AnuA | -0.13644 | 4.40E-10 |
| B103 | B110 | anti-SSA/Ro | AHA | -0.13644 | 4.40E-10 |
| B103 | B151 | anti-SSA/Ro | Anti-CCP | 0.124293 | 1.35E-08 |
| B103 | B177 | anti-SSA/Ro | ACR | 0.110605 | 4.43E-07 |
| B104 | B105 | anti-SSB | anti-U1RNP | -0.33755 | 0 |
| B104 | B106 | anti-SSB | anti-P | 1 | 0 |
| B104 | B107 | anti-SSB | AnuA | 0.999998 | 0 |
| B104 | B108 | anti-SSB | Anti-Cyclin | -0.22672 | 0 |
| B104 | B109 | anti-SSB | AMA-M2 | -0.26181 | 0 |
| B104 | B110 | anti-SSB | AHA | 1 | 0 |
| B104 | B125 | anti-SSB | Cor | 0.234516 | 0 |
| B104 | B154 | anti-SSB | RF | 0.204755 | 0 |
| B104 | B183 | anti-SSB | PTA | 0.273181 | 0 |
| B104 | B155 | anti-SSB | IgA | -0.16404 | 5.60E-14 |
| B104 | B156 | anti-SSB | IgE | -0.14475 | 3.52E-11 |
| B104 | B181 | anti-SSB | APTT | 0.13878 | 2.19E-10 |
| B104 | B195 | anti-SSB | Na | 0.137023 | 3.70E-10 |
| B104 | B137 | anti-SSB | AFU | -0.12881 | 3.93E-09 |
| B104 | B198 | anti-SSB | P | -0.12517 | 1.07E-08 |
| B104 | B139 | anti-SSB | ADA | 0.118712 | 5.89E-08 |
| B104 | B138 | anti-SSB | PAB | 0.113683 | 2.09E-07 |
| B104 | B185 | anti-SSB | PT | -0.10972 | 5.48E-07 |
| B105 | B106 | anti-U1RNP | anti-P | -0.33755 | 0 |
| B105 | B107 | anti-U1RNP | AnuA | -0.33755 | 0 |

| | | | | | |
|---|---|---|---|---|---|
| B105 | B108 | anti-U1RNP | Anti-Cyclin | 0.213195 | 0 |
| B105 | B110 | anti-U1RNP | AHA | -0.33755 | 0 |
| B105 | B124 | anti-U1RNP | ACTH | -0.27546 | 0 |
| B105 | B125 | anti-U1RNP | Cor | -0.43629 | 0 |
| B105 | B138 | anti-U1RNP | PAB | -0.27426 | 0 |
| B105 | B156 | anti-U1RNP | IgE | 0.273801 | 0 |
| B105 | B185 | anti-U1RNP | PT | 0.282689 | 0 |
| B105 | B183 | anti-U1RNP | PTA | -0.17855 | 2.22E-16 |
| B105 | B228 | anti-U1RNP | OS | 0.174314 | 1.33E-15 |
| B105 | B120 | anti-U1RNP | HCV-cAg | -0.16572 | 3.06E-14 |
| B105 | B179 | anti-U1RNP | α1-MG | 0.137064 | 3.66E-10 |
| B105 | B134 | anti-U1RNP | CHE | -0.13633 | 4.55E-10 |
| B105 | B186 | anti-U1RNP | PFC | 0.130221 | 2.65E-09 |
| B105 | B195 | anti-U1RNP | Na | 0.111044 | 3.98E-07 |
| B106 | B107 | anti-P | AnuA | 0.999999 | 0 |
| B106 | B108 | anti-P | Anti-Cyclin | -0.22672 | 0 |
| B106 | B109 | anti-P | AMA-M2 | -0.26181 | 0 |
| B106 | B125 | anti-P | Cor | 0.234516 | 0 |
| B106 | B154 | anti-P | RF | 0.204755 | 0 |
| B106 | B183 | anti-P | PTA | 0.273181 | 0 |
| B106 | B155 | anti-P | IgA | -0.16404 | 5.60E-14 |
| B106 | B156 | anti-P | IgE | -0.14475 | 3.52E-11 |
| B106 | B181 | anti-P | APTT | 0.13878 | 2.19E-10 |
| B106 | B195 | anti-P | Na | 0.137023 | 3.70E-10 |
| B106 | B137 | anti-P | AFU | -0.12881 | 3.93E-09 |
| B106 | B198 | anti-P | P | -0.12517 | 1.07E-08 |
| B106 | B139 | anti-P | ADA | 0.118712 | 5.89E-08 |
| B106 | B138 | anti-P | PAB | 0.113683 | 2.09E-07 |
| B106 | B185 | anti-P | PT | -0.10972 | 5.48E-07 |
| B107 | B108 | AnuA | Anti-Cyclin | -0.22671 | 0 |
| B107 | B109 | AnuA | AMA-M2 | -0.26181 | 0 |
| B107 | B110 | AnuA | AHA | 0.999998 | 0 |
| B107 | B125 | AnuA | Cor | 0.234516 | 0 |
| B107 | B154 | AnuA | RF | 0.204754 | 0 |
| B107 | B183 | AnuA | PTA | 0.273181 | 0 |
| B107 | B155 | AnuA | IgA | -0.16404 | 5.60E-14 |
| B107 | B156 | AnuA | IgE | -0.14475 | 3.52E-11 |
| B107 | B181 | AnuA | APTT | 0.13878 | 2.19E-10 |
| B107 | B195 | AnuA | Na | 0.137023 | 3.70E-10 |
| B107 | B137 | AnuA | AFU | -0.12881 | 3.93E-09 |
| B107 | B198 | AnuA | P | -0.12517 | 1.07E-08 |
| B107 | B139 | AnuA | ADA | 0.118712 | 5.89E-08 |
| B107 | B138 | AnuA | PAB | 0.113683 | 2.09E-07 |
| B107 | B185 | AnuA | PT | -0.10972 | 5.48E-07 |

| | | | | | |
|---|---|---|---|---|---|
| B108 | B110 | Anti-Cyclin | AHA | -0.22672 | 0 |
| B108 | B124 | Anti-Cyclin | ACTH | 0.333032 | 0 |
| B108 | B135 | Anti-Cyclin | TBA | 0.348353 | 0 |
| B108 | B181 | Anti-Cyclin | APTT | -0.31556 | 0 |
| B108 | B198 | Anti-Cyclin | P | 0.266175 | 0 |
| B108 | B184 | Anti-Cyclin | INR | -0.17385 | 1.55E-15 |
| B108 | B151 | Anti-Cyclin | Anti-CCP | -0.16694 | 2.00E-14 |
| B108 | B195 | Anti-Cyclin | Na | 0.15744 | 5.57E-13 |
| B108 | B196 | Anti-Cyclin | TCO2 | 0.141027 | 1.11E-10 |
| B108 | B183 | Anti-Cyclin | PTA | -0.13104 | 2.10E-09 |
| B108 | B137 | Anti-Cyclin | AFU | 0.12105 | 3.21E-08 |
| B108 | B182 | Anti-Cyclin | TT | 0.119197 | 5.19E-08 |
| B108 | B138 | Anti-Cyclin | PAB | 0.113653 | 2.11E-07 |
| B108 | B139 | Anti-Cyclin | ADA | -0.11059 | 4.44E-07 |
| B109 | B110 | AMA-M2 | AHA | -0.26181 | 0 |
| B109 | B151 | AMA-M2 | Anti-CCP | 0.220051 | 0 |
| B109 | B198 | AMA-M2 | P | 0.246809 | 0 |
| B109 | B120 | AMA-M2 | HCV-cAg | -0.17068 | 5.11E-15 |
| B109 | B196 | AMA-M2 | TCO2 | -0.16256 | 9.46E-14 |
| B109 | B199 | AMA-M2 | Mg | -0.15659 | 7.43E-13 |
| B109 | B197 | AMA-M2 | AG | 0.149153 | 8.69E-12 |
| B109 | B137 | AMA-M2 | AFU | 0.133206 | 1.13E-09 |
| B109 | B179 | AMA-M2 | α1-MG | -0.11733 | 8.37E-08 |
| B109 | B195 | AMA-M2 | Na | -0.11019 | 4.89E-07 |
| B11 | B12 | PGI/PGII | PG II | -0.51051 | 0 |
| B11 | B14 | PGI/PGII | G-17 | -0.22661 | 0 |
| B11 | B80 | PGI/PGII | anti-HP IgG | -0.22687 | 0 |
| B110 | B125 | AHA | Cor | 0.234516 | 0 |
| B110 | B154 | AHA | RF | 0.204755 | 0 |
| B110 | B183 | AHA | PTA | 0.273181 | 0 |
| B110 | B155 | AHA | IgA | -0.16404 | 5.60E-14 |
| B110 | B156 | AHA | IgE | -0.14475 | 3.52E-11 |
| B110 | B181 | AHA | APTT | 0.13878 | 2.19E-10 |
| B110 | B195 | AHA | Na | 0.137023 | 3.70E-10 |
| B110 | B137 | AHA | AFU | -0.12881 | 3.93E-09 |
| B110 | B198 | AHA | P | -0.12517 | 1.07E-08 |
| B110 | B139 | AHA | ADA | 0.118712 | 5.89E-08 |
| B110 | B138 | AHA | PAB | 0.113683 | 2.09E-07 |
| B110 | B185 | AHA | PT | -0.10972 | 5.48E-07 |
| B111 | B112 | TSH | FT4 | -0.18086 | 0 |
| B112 | B114 | FT4 | TSH TT3 TT4(TT4) | 0.417322 | 0 |
| B112 | B125 | FT4 | Cor | 0.115093 | 1.47E-07 |
| B113 | B114 | FT3 | TSH TT3 TT4(TT4) | 0.218855 | 0 |
| B113 | B115 | FT3 | TSH TT3 TT4(TT3) | 0.402109 | 0 |

| | | | | | |
|---|---|---|---|---|---|
| B113 | B126 | FT3 | Alb | 0.155879 | 9.45E-13 |
| B113 | B176 | FT3 | Ucr | 0.137811 | 2.93E-10 |
| B114 | B115 | TSH TT3 TT4(TT4) | TSH TT3 TT4(TT3) | 0.425703 | 0 |
| B114 | B125 | TSH TT3 TT4(TT4) | Cor | 0.127745 | 5.28E-09 |
| B115 | B134 | TSH TT3 TT4(TT3) | CHE | 0.17635 | 4.44E-16 |
| B115 | B196 | TSH TT3 TT4(TT3) | TCO2 | 0.1215 | 2.85E-08 |
| B115 | B135 | TSH TT3 TT4(TT3) | TBA | -0.11678 | 9.65E-08 |
| B116 | B117 | TT3 | TT4 | 0.542235 | 0 |
| B119 | B120 | Hcv-rna | HCV-cAg | 0.386606 | 0 |
| B119 | B125 | Hcv-rna | Cor | 0.232793 | 0 |
| B119 | B135 | Hcv-rna | TBA | -0.26329 | 0 |
| B119 | B137 | Hcv-rna | AFU | -0.22152 | 0 |
| B119 | B199 | Hcv-rna | Mg | 0.165722 | 3.06E-14 |
| B119 | B194 | Hcv-rna | Cl | -0.13295 | 1.22E-09 |
| B119 | B124 | Hcv-rna | ACTH | 0.130491 | 2.45E-09 |
| B119 | B134 | Hcv-rna | CHE | 0.109902 | 5.24E-07 |
| B12 | B84 | PG II | C-PSA | 0.18918 | 0 |
| B12 | B80 | PG II | anti-HP IgG | 0.151688 | 3.81E-12 |
| B120 | B121 | HCV-cAg | C3 | -0.38661 | 0 |
| B120 | B122 | HCV-cAg | C4 | -0.38661 | 0 |
| B120 | B152 | HCV-cAg | ACA-Ig(G-A-M) | -0.38672 | 0 |
| B120 | B169 | HCV-cAg | U-CRY | -0.38661 | 0 |
| B120 | B125 | HCV-cAg | Cor | 0.166303 | 2.49E-14 |
| B120 | B136 | HCV-cAg | 5'NT | -0.15559 | 1.04E-12 |
| B120 | B146 | HCV-cAg | CysC | -0.12751 | 5.63E-09 |
| B120 | B134 | HCV-cAg | CHE | 0.124535 | 1.27E-08 |
| B120 | B199 | HCV-cAg | Mg | 0.118731 | 5.86E-08 |
| B120 | B124 | HCV-cAg | ACTH | 0.113911 | 1.98E-07 |
| B120 | B137 | HCV-cAg | AFU | -0.10911 | 6.32E-07 |
| B121 | B125 | C3 | Cor | -0.23279 | 0 |
| B121 | B135 | C3 | TBA | 0.263295 | 0 |
| B121 | B137 | C3 | AFU | 0.221518 | 0 |
| B121 | B199 | C3 | Mg | -0.16572 | 3.06E-14 |
| B121 | B194 | C3 | Cl | 0.132954 | 1.22E-09 |
| B121 | B124 | C3 | ACTH | -0.13049 | 2.45E-09 |
| B121 | B134 | C3 | CHE | -0.1099 | 5.24E-07 |
| B122 | B125 | C4 | Cor | -0.23279 | 0 |
| B122 | B135 | C4 | TBA | 0.263296 | 0 |
| B122 | B137 | C4 | AFU | 0.221519 | 0 |
| B122 | B169 | C4 | U-CRY | 0.999992 | 0 |
| B122 | B199 | C4 | Mg | -0.16572 | 3.06E-14 |
| B122 | B194 | C4 | Cl | 0.132955 | 1.21E-09 |
| B122 | B124 | C4 | ACTH | -0.13049 | 2.45E-09 |
| B122 | B134 | C4 | CHE | -0.1099 | 5.24E-07 |

| | | | | | |
|---|---|---|---|---|---|
| B123 | B138 | hs-CRP1 | PAB | -0.19699 | 0 |
| B123 | B139 | hs-CRP1 | ADA | 0.272404 | 0 |
| B123 | B193 | hs-CRP1 | K | -0.27 | 0 |
| B123 | B194 | hs-CRP1 | Cl | -0.17772 | 4.44E-16 |
| B123 | B224 | hs-CRP1 | pmh | 0.115258 | 1.41E-07 |
| B123 | B196 | hs-CRP1 | TCO2 | 0.113578 | 2.15E-07 |
| B124 | B125 | ACTH | Cor | 0.608604 | 0 |
| B124 | B135 | ACTH | TBA | 0.215726 | 0 |
| B124 | B181 | ACTH | APTT | -0.30335 | 0 |
| B124 | B186 | ACTH | PFC | 0.178117 | 4.44E-16 |
| B124 | B199 | ACTH | Mg | 0.143514 | 5.17E-11 |
| B124 | B152 | ACTH | ACA-Ig(G-A-M) | -0.13053 | 2.43E-09 |
| B124 | B169 | ACTH | U-CRY | -0.13049 | 2.45E-09 |
| B124 | B136 | ACTH | 5'NT | -0.13028 | 2.60E-09 |
| B124 | B140 | ACTH | r-GT | -0.12026 | 3.95E-08 |
| B124 | B156 | ACTH | IgE | -0.11478 | 1.59E-07 |
| B125 | B152 | Cor | ACA-Ig(G-A-M) | -0.23286 | 0 |
| B125 | B169 | Cor | U-CRY | -0.2328 | 0 |
| B125 | B181 | Cor | APTT | -0.28438 | 0 |
| B125 | B185 | Cor | PT | -0.17457 | 1.33E-15 |
| B125 | B138 | Cor | PAB | 0.164916 | 4.09E-14 |
| B125 | B182 | Cor | TT | -0.15128 | 4.36E-12 |
| B125 | B199 | Cor | Mg | 0.147547 | 1.45E-11 |
| B125 | B194 | Cor | Cl | -0.13967 | 1.68E-10 |
| B126 | B127 | Alb | A-G ratio | 0.49192 | 0 |
| B126 | B188 | Alb | Ca | 0.278822 | 0 |
| B126 | B189 | Alb | eGFB | 0.252026 | 0 |
| B126 | B130 | Alb | GLB | -0.16719 | 1.82E-14 |
| B126 | B190 | Alb | Cre | -0.16576 | 3.02E-14 |
| B126 | B191 | Alb | Urea | -0.1619 | 1.19E-13 |
| B126 | B141 | Alb | AST/ALT | -0.15635 | 8.06E-13 |
| B126 | B216 | Alb | BMI | 0.128323 | 4.50E-09 |
| B126 | B221 | Alb | Hei | 0.123588 | 1.64E-08 |
| B126 | B134 | Alb | CHE | 0.109603 | 5.63E-07 |
| B127 | B130 | A-G ratio | GLB | -0.90608 | 0 |
| B127 | B221 | A-G ratio | Hei | 0.174007 | 1.33E-15 |
| B127 | B189 | A-G ratio | eGFB | 0.161193 | 1.53E-13 |
| B127 | B132 | A-G ratio | Dbil | 0.144548 | 3.75E-11 |
| B127 | B209 | A-G ratio | Smoking | 0.131251 | 1.98E-09 |
| B127 | B133 | A-G ratio | TBIL | 0.126218 | 8.03E-09 |
| B127 | B163 | A-G ratio | SG | 0.119485 | 4.82E-08 |
| B127 | B216 | A-G ratio | BMI | 0.11941 | 4.92E-08 |
| B127 | B129 | A-G ratio | IBIL | 0.108413 | 7.46E-07 |
| B128 | B131 | ALT | AST | 0.750504 | 0 |

| | | | | | |
|---|---|---|---|---|---|
| B128 | B140 | ALT | r-GT | 0.199166 | 0 |
| B128 | B141 | ALT | AST/ALT | -0.36467 | 0 |
| B128 | B216 | ALT | BMI | 0.112636 | 2.71E-07 |
| B129 | B132 | IBIL | Dbil | 0.666556 | 0 |
| B129 | B133 | IBIL | TBIL | 0.979071 | 0 |
| B130 | B132 | GLB | Dbil | -0.14872 | 9.99E-12 |
| B130 | B221 | GLB | Hei | -0.14855 | 1.06E-11 |
| B130 | B209 | GLB | Smoking | -0.13743 | 3.29E-10 |
| B130 | B133 | GLB | TBIL | -0.12201 | 2.49E-08 |
| B130 | B220 | GLB | SBP | 0.117971 | 7.12E-08 |
| B130 | B192 | GLB | UA | 0.11675 | 9.72E-08 |
| B130 | B163 | GLB | SG | -0.11011 | 4.99E-07 |
| B131 | B140 | AST | r-GT | 0.266777 | 0 |
| B132 | B133 | Dbil | TBIL | 0.80421 | 0 |
| B132 | B221 | Dbil | Hei | 0.127175 | 6.18E-09 |
| B133 | B221 | TBIL | Hei | 0.117291 | 8.47E-08 |
| B134 | B136 | CHE | 5'NT | -0.21374 | 0 |
| B134 | B138 | CHE | PAB | 0.386548 | 0 |
| B134 | B159 | CHE | β2-MG | -0.19401 | 0 |
| B134 | B194 | CHE | Cl | -0.15947 | 2.78E-13 |
| B134 | B139 | CHE | ADA | -0.15669 | 7.17E-13 |
| B134 | B185 | CHE | PT | -0.15028 | 6.03E-12 |
| B134 | B196 | CHE | TCO2 | 0.14751 | 1.47E-11 |
| B134 | B137 | CHE | AFU | 0.126977 | 6.53E-09 |
| B134 | B152 | CHE | ACA-Ig(G-A-M) | -0.10993 | 5.20E-07 |
| B134 | B169 | CHE | U-CRY | -0.1099 | 5.24E-07 |
| B135 | B137 | TBA | AFU | 0.19707 | 0 |
| B135 | B152 | TBA | ACA-Ig(G-A-M) | 0.263366 | 0 |
| B135 | B169 | TBA | U-CRY | 0.263298 | 0 |
| B135 | B195 | TBA | Na | 0.24905 | 0 |
| B135 | B200 | TBA | OP | 0.218785 | 0 |
| B135 | B139 | TBA | ADA | 0.172302 | 2.66E-15 |
| B135 | B159 | TBA | β2-MG | -0.15388 | 1.85E-12 |
| B135 | B196 | TBA | TCO2 | 0.145552 | 2.74E-11 |
| B135 | B138 | TBA | PAB | -0.12824 | 4.60E-09 |
| B135 | B181 | TBA | APTT | -0.10724 | 9.84E-07 |
| B136 | B198 | 5'NT | P | 0.246082 | 0 |
| B136 | B140 | 5'NT | r-GT | 0.170233 | 6.00E-15 |
| B136 | B194 | 5'NT | Cl | -0.12484 | 1.17E-08 |
| B136 | B200 | 5'NT | OP | 0.121488 | 2.86E-08 |
| B136 | B197 | 5'NT | AG | 0.114527 | 1.70E-07 |
| B136 | B199 | 5'NT | Mg | -0.10882 | 6.79E-07 |
| B137 | B152 | AFU | ACA-Ig(G-A-M) | 0.221578 | 0 |
| B137 | B169 | AFU | U-CRY | 0.221521 | 0 |

| | | | | | |
|---|---|---|---|---|---|
| B137 | B196 | AFU | TCO2 | 0.265338 | 0 |
| B137 | B185 | AFU | PT | -0.17593 | 6.66E-16 |
| B137 | B194 | AFU | Cl | -0.16059 | 1.88E-13 |
| B137 | B199 | AFU | Mg | -0.15805 | 4.53E-13 |
| B137 | B159 | AFU | β2-MG | -0.1502 | 6.19E-12 |
| B137 | B180 | AFU | Tf | 0.143721 | 4.85E-11 |
| B137 | B178 | AFU | Ig-U | 0.139119 | 1.98E-10 |
| B138 | B139 | PAB | ADA | -0.43427 | 0 |
| B138 | B178 | PAB | Ig-U | -0.24877 | 0 |
| B138 | B180 | PAB | Tf | -0.22637 | 0 |
| B138 | B185 | PAB | PT | -0.27079 | 0 |
| B138 | B184 | PAB | INR | -0.16021 | 2.15E-13 |
| B138 | B179 | PAB | α1-MG | -0.1534 | 2.17E-12 |
| B138 | B182 | PAB | TT | 0.13944 | 1.80E-10 |
| B138 | B159 | PAB | β2-MG | -0.13797 | 2.80E-10 |
| B138 | B181 | PAB | APTT | -0.12009 | 4.12E-08 |
| B138 | B193 | PAB | K | 0.119469 | 4.84E-08 |
| B138 | B186 | PAB | PFC | -0.10778 | 8.67E-07 |
| B139 | B178 | ADA | Ig-U | 0.191453 | 0 |
| B139 | B200 | ADA | OP | 0.193705 | 0 |
| B139 | B180 | ADA | Tf | 0.177265 | 4.44E-16 |
| B139 | B182 | ADA | TT | -0.17637 | 4.44E-16 |
| B139 | B193 | ADA | K | -0.17701 | 4.44E-16 |
| B139 | B197 | ADA | AG | 0.162695 | 9.02E-14 |
| B139 | B195 | ADA | Na | 0.162431 | 9.90E-14 |
| B139 | B199 | ADA | Mg | 0.142688 | 6.68E-11 |
| B139 | B186 | ADA | PFC | 0.112399 | 2.87E-07 |
| B14 | B22 | G-17 | CD3+CD8+(T) | -0.17994 | 0 |
| B14 | B19 | G-17 | CD3+CD4+/CD3+CD8+ | 0.151762 | 3.72E-12 |
| B14 | B25 | G-17 | CK | 0.116216 | 1.11E-07 |
| B140 | B150 | r-GT | ALP | 0.160021 | 2.29E-13 |
| B140 | B141 | r-GT | AST/ALT | -0.1559 | 9.38E-13 |
| B140 | B216 | r-GT | BMI | 0.141286 | 1.03E-10 |
| B140 | B184 | r-GT | INR | -0.13861 | 2.31E-10 |
| B140 | B206 | r-GT | Waist | 0.120067 | 4.15E-08 |
| B140 | B215 | r-GT | BMI | 0.113729 | 2.07E-07 |
| B140 | B192 | r-GT | UA | 0.110706 | 4.32E-07 |
| B141 | B206 | AST/ALT | Waist | -0.19087 | 0 |
| B141 | B215 | AST/ALT | BMI | -0.18206 | 0 |
| B141 | B216 | AST/ALT | BMI | -0.24878 | 0 |
| B141 | B221 | AST/ALT | Hei | -0.19006 | 0 |
| B141 | B222 | AST/ALT | NS | -0.20112 | 0 |
| B141 | B207 | AST/ALT | WHR | -0.13487 | 6.98E-10 |
| B141 | B210 | AST/ALT | Hip | -0.13485 | 7.02E-10 |

| | | | | | |
|---|---|---|---|---|---|
| B142 | B151 | 25-(OH)D | Anti-CCP | -0.18964 | 0 |
| B142 | B168 | 25-(OH)D | NMP22 | 0.40179 | 0 |
| B142 | B177 | 25-(OH)D | ACR | 0.172301 | 2.66E-15 |
| B142 | B175 | 25-(OH)D | ALB-U | 0.171105 | 4.44E-15 |
| B143 | B144 | β-CTx | Total-PINP | 0.833381 | 0 |
| B143 | B145 | β-CTx | BGP | 0.455304 | 0 |
| B143 | B151 | β-CTx | Anti-CCP | -0.15148 | 4.08E-12 |
| B144 | B145 | Total-PINP | BGP | 0.499681 | 0 |
| B144 | B177 | Total-PINP | ACR | 0.187788 | 0 |
| B144 | B175 | Total-PINP | ALB-U | 0.171765 | 3.33E-15 |
| B144 | B151 | Total-PINP | Anti-CCP | -0.11607 | 1.15E-07 |
| B146 | B190 | CysC | Cre | 0.17976 | 0 |
| B146 | B200 | CysC | OP | 0.163683 | 6.35E-14 |
| B146 | B194 | CysC | Cl | 0.146078 | 2.32E-11 |
| B146 | B189 | CysC | eGFB | -0.13956 | 1.73E-10 |
| B146 | B191 | CysC | Urea | 0.137033 | 3.69E-10 |
| B146 | B192 | CysC | UA | 0.134515 | 7.74E-10 |
| B146 | B179 | CysC | α1-MG | 0.134207 | 8.46E-10 |
| B148 | B149 | Anti_TPO | TGAb | 0.489007 | 0 |
| B15 | B100 | CD3-CD19+(B cell) | anti-Ro52 | 0.434097 | 0 |
| B15 | B101 | CD3-CD19+(B cell) | anti-Sc70 | 0.434097 | 0 |
| B15 | B102 | CD3-CD19+(B cell) | anti-Sm | -0.74637 | 0 |
| B15 | B104 | CD3-CD19+(B cell) | anti-SSB | 0.434097 | 0 |
| B15 | B105 | CD3-CD19+(B cell) | anti-U1RNP | -0.77701 | 0 |
| B15 | B106 | CD3-CD19+(B cell) | anti-P | 0.434097 | 0 |
| B15 | B107 | CD3-CD19+(B cell) | AnuA | 0.434096 | 0 |
| B15 | B108 | CD3-CD19+(B cell) | Anti-Cyclin | -0.18818 | 0 |
| B15 | B109 | CD3-CD19+(B cell) | AMA-M2 | -0.27735 | 0 |
| B15 | B110 | CD3-CD19+(B cell) | AHA | 0.434097 | 0 |
| B15 | B120 | CD3-CD19+(B cell) | HCV-cAg | 0.227238 | 0 |
| B15 | B124 | CD3-CD19+(B cell) | ACTH | 0.253964 | 0 |
| B15 | B125 | CD3-CD19+(B cell) | Cor | 0.417769 | 0 |
| B15 | B136 | CD3-CD19+(B cell) | 5'NT | -0.23975 | 0 |
| B15 | B138 | CD3-CD19+(B cell) | PAB | 0.266068 | 0 |
| B15 | B16 | CD3-CD19+(B cell) | CD3+T | 0.502518 | 0 |
| B15 | B18 | CD3-CD19+(B cell) | CD3+CD4+helper(inducer)T | 0.302353 | 0 |
| B15 | B181 | CD3-CD19+(B cell) | APTT | -0.18581 | 0 |
| B15 | B185 | CD3-CD19+(B cell) | PT | -0.31048 | 0 |
| B15 | B186 | CD3-CD19+(B cell) | PFC | -0.1991 | 0 |
| B15 | B19 | CD3-CD19+(B cell) | CD3+CD4+/CD3+CD8+ | 0.256275 | 0 |
| B15 | B198 | CD3-CD19+(B cell) | P | -0.25879 | 0 |
| B15 | B20 | CD3-CD19+(B cell) | CD3+CD4+CD8+ | 0.300325 | 0 |
| B15 | B21 | CD3-CD19+(B cell) | CD3+CD4-CD8- | 0.199171 | 0 |
| B15 | B23 | CD3-CD19+(B cell) | CD3-CD16+CD56+(NK) | -0.66868 | 0 |

| | | | | | |
|---|---|---|---|---|---|
| B15 | B55 | CD3-CD19+(B cell) | TK | 0.237744 | 0 |
| B15 | B56 | CD3-CD19+(B cell) | IR | 0.204065 | 0 |
| B15 | B60 | CD3-CD19+(B cell) | WBRVI | 0.210333 | 0 |
| B15 | B62 | CD3-CD19+(B cell) | WBHSRVI | 0.196642 | 0 |
| B15 | B69 | CD3-CD19+(B cell) | PV | -0.47332 | 0 |
| B15 | B79 | CD3-CD19+(B cell) | hb V -dna | 0.206411 | 0 |
| B15 | B92 | CD3-CD19+(B cell) | ALDH2-Glu504Glu | 0.40486 | 0 |
| B15 | B93 | CD3-CD19+(B cell) | ALDH2-Glu504Lys | -0.41027 | 0 |
| B15 | B94 | CD3-CD19+(B cell) | ALDH2-Lys504Lys | 0.335158 | 0 |
| B15 | B95 | CD3-CD19+(B cell) | anti-cenpb | 0.434097 | 0 |
| B15 | B97 | CD3-CD19+(B cell) | dsDNA | 0.434097 | 0 |
| B15 | B98 | CD3-CD19+(B cell) | anti-Jo-1 | 0.434097 | 0 |
| B15 | B99 | CD3-CD19+(B cell) | anti-PM-Scl | 0.434097 | 0 |
| B15 | B183 | CD3-CD19+(B cell) | PTA | 0.174829 | 1.11E-15 |
| B15 | B156 | CD3-CD19+(B cell) | IgE | -0.17426 | 1.33E-15 |
| B15 | B182 | CD3-CD19+(B cell) | TT | 0.174149 | 1.33E-15 |
| B15 | B54 | CD3-CD19+(B cell) | ESR | -0.15993 | 2.37E-13 |
| B15 | B197 | CD3-CD19+(B cell) | AG | -0.15328 | 2.25E-12 |
| B15 | B134 | CD3-CD19+(B cell) | CHE | 0.149955 | 6.70E-12 |
| B15 | B199 | CD3-CD19+(B cell) | Mg | 0.137353 | 3.36E-10 |
| B15 | B228 | CD3-CD19+(B cell) | OS | -0.13287 | 1.25E-09 |
| B15 | B193 | CD3-CD19+(B cell) | K | -0.12812 | 4.75E-09 |
| B15 | B151 | CD3-CD19+(B cell) | Anti-CCP | -0.12577 | 9.08E-09 |
| B15 | B158 | CD3-CD19+(B cell) | IgM | 0.122945 | 1.94E-08 |
| B15 | B57 | CD3-CD19+(B cell) | RBCAI | 0.112862 | 2.56E-07 |
| B15 | B25 | CD3-CD19+(B cell) | CK | 0.108953 | 6.57E-07 |
| B15 | B34 | CD3-CD19+(B cell) | TK | -0.10749 | 9.26E-07 |
| B150 | B176 | ALP | Ucr | -0.12326 | 1.79E-08 |
| B150 | B195 | ALP | Na | 0.117826 | 7.39E-08 |
| B150 | B197 | ALP | AG | 0.109306 | 6.04E-07 |
| B151 | B168 | Anti-CCP | NMP22 | -0.26762 | 0 |
| B151 | B181 | Anti-CCP | APTT | 0.19287 | 0 |
| B151 | B156 | Anti-CCP | IgE | 0.118772 | 5.80E-08 |
| B152 | B199 | ACA-Ig(G-A-M) | Mg | -0.16577 | 3.02E-14 |
| B152 | B194 | ACA-Ig(G-A-M) | Cl | 0.132989 | 1.20E-09 |
| B153 | B170 | Glu | U-GLU | 0.422119 | 0 |
| B153 | B206 | Glu | Waist | 0.179042 | 2.22E-16 |
| B153 | B215 | Glu | BMI | 0.170683 | 4.88E-15 |
| B153 | B210 | Glu | Hip | 0.152525 | 2.89E-12 |
| B153 | B222 | Glu | NS | 0.137939 | 2.82E-10 |
| B153 | B216 | Glu | BMI | 0.131767 | 1.71E-09 |
| B153 | B178 | Glu | Ig-U | 0.111007 | 4.02E-07 |
| B153 | B160 | Glu | PH | -0.10855 | 7.24E-07 |
| B155 | B157 | IgA | IgG | 0.237382 | 0 |

| | | | | | |
|---|---|---|---|---|---|
| B155 | B156 | IgA | IgE | 0.173605 | 1.78E-15 |
| B155 | B195 | IgA | Na | -0.12189 | 2.57E-08 |
| B156 | B157 | IgE | IgG | 0.187215 | 0 |
| B156 | B195 | IgE | Na | -0.17055 | 5.33E-15 |
| B156 | B194 | IgE | Cl | -0.15 | 6.60E-12 |
| B156 | B159 | IgE | β2-MG | 0.127301 | 5.97E-09 |
| B157 | B158 | IgG | IgM | -0.22341 | 0 |
| B157 | B182 | IgG | TT | -0.16781 | 1.44E-14 |
| B157 | B195 | IgG | Na | -0.16702 | 1.91E-14 |
| B157 | B194 | IgG | Cl | -0.13545 | 5.89E-10 |
| B157 | B159 | IgG | β2-MG | 0.11899 | 5.48E-08 |
| B158 | B159 | IgM | β2-MG | -0.48856 | 0 |
| B158 | B180 | IgM | Tf | 0.126432 | 7.58E-09 |
| B159 | B199 | β2-MG | Mg | 0.137218 | 3.50E-10 |
| B16 | B100 | CD3+T | anti-Ro52 | 0.292769 | 0 |
| B16 | B101 | CD3+T | anti-Sc70 | 0.292769 | 0 |
| B16 | B102 | CD3+T | anti-Sm | -0.82773 | 0 |
| B16 | B104 | CD3+T | anti-SSB | 0.292769 | 0 |
| B16 | B105 | CD3+T | anti-U1RNP | -0.75466 | 0 |
| B16 | B106 | CD3+T | anti-P | 0.292769 | 0 |
| B16 | B107 | CD3+T | AnuA | 0.292769 | 0 |
| B16 | B108 | CD3+T | Anti-Cyclin | 0.228855 | 0 |
| B16 | B110 | CD3+T | AHA | 0.292769 | 0 |
| B16 | B124 | CD3+T | ACTH | 0.409083 | 0 |
| B16 | B125 | CD3+T | Cor | 0.324952 | 0 |
| B16 | B136 | CD3+T | 5'NT | 0.21856 | 0 |
| B16 | B138 | CD3+T | PAB | 0.270377 | 0 |
| B16 | B156 | CD3+T | IgE | -0.22273 | 0 |
| B16 | B18 | CD3+T | CD3+CD4+helper(inducer)T | 0.80126 | 0 |
| B16 | B185 | CD3+T | PT | -0.26139 | 0 |
| B16 | B19 | CD3+T | CD3+CD4+/CD3+CD8+ | 0.308089 | 0 |
| B16 | B198 | CD3+T | P | 0.184234 | 0 |
| B16 | B20 | CD3+T | CD3+CD4+CD8+ | -0.23734 | 0 |
| B16 | B22 | CD3+T | CD3+CD8+(T) | 0.311183 | 0 |
| B16 | B23 | CD3+T | CD3-CD16+CD56+(NK) | -0.95899 | 0 |
| B16 | B55 | CD3+T | TK | 0.294705 | 0 |
| B16 | B56 | CD3+T | IR | 0.271599 | 0 |
| B16 | B62 | CD3+T | WBHSRVI | 0.262177 | 0 |
| B16 | B69 | CD3+T | PV | -0.47251 | 0 |
| B16 | B95 | CD3+T | anti-cenpb | 0.292769 | 0 |
| B16 | B97 | CD3+T | dsDNA | 0.292769 | 0 |
| B16 | B98 | CD3+T | anti-Jo-1 | 0.292769 | 0 |
| B16 | B99 | CD3+T | anti-PM-Scl | 0.292769 | 0 |
| B16 | B181 | CD3+T | APTT | -0.17941 | 2.22E-16 |

| | | | | | |
|---|---|---|---|---|---|
| B16 | B81 | CD3+T | TSGF | 0.176896 | 4.44E-16 |
| B16 | B183 | CD3+T | PTA | 0.172372 | 2.66E-15 |
| B16 | B109 | CD3+T | AMA-M2 | 0.168666 | 1.07E-14 |
| B16 | B21 | CD3+T | CD3+CD4-CD8- | -0.16279 | 8.70E-14 |
| B16 | B228 | CD3+T | OS | -0.1494 | 8.02E-12 |
| B16 | B61 | CD3+T | WBHSRV | 0.14576 | 2.56E-11 |
| B16 | B60 | CD3+T | WBRVI | 0.129529 | 3.21E-09 |
| B16 | B179 | CD3+T | α1-MG | -0.12401 | 1.46E-08 |
| B16 | B54 | CD3+T | ESR | -0.10965 | 5.56E-07 |
| B16 | B182 | CD3+T | TT | 0.107778 | 8.67E-07 |
| B160 | B163 | PH | SG | -0.22288 | 0 |
| B160 | B191 | PH | Urea | -0.22761 | 0 |
| B160 | B190 | PH | Cre | -0.17357 | 1.78E-15 |
| B160 | B192 | PH | UA | -0.17223 | 2.89E-15 |
| B161 | B173 | LEU | NTT | 0.205024 | 0 |
| B161 | B221 | LEU | Hei | -0.12644 | 7.56E-09 |
| B163 | B166 | SG | Pro | 0.320612 | 0 |
| B163 | B176 | SG | Ucr | 0.359705 | 0 |
| B163 | B191 | SG | Urea | 0.18978 | 0 |
| B163 | B216 | SG | BMI | 0.218358 | 0 |
| B163 | B221 | SG | Hei | 0.209711 | 0 |
| B163 | B222 | SG | NS | 0.199666 | 0 |
| B163 | B174 | SG | Urea/Cr | 0.130439 | 2.49E-09 |
| B163 | B210 | SG | Hip | 0.127655 | 5.41E-09 |
| B163 | B206 | SG | Waist | 0.126034 | 8.45E-09 |
| B163 | B215 | SG | BMI | 0.122943 | 1.94E-08 |
| B166 | B176 | Pro | Ucr | 0.17232 | 2.66E-15 |
| B166 | B190 | Pro | Cre | 0.130772 | 2.26E-09 |
| B166 | B191 | Pro | Urea | 0.116553 | 1.02E-07 |
| B167 | B223 | PC | FH | 0.502639 | 0 |
| B168 | B175 | NMP22 | ALB-U | 0.300219 | 0 |
| B168 | B177 | NMP22 | ACR | 0.280838 | 0 |
| B168 | B173 | NMP22 | NTT | 0.1106 | 4.43E-07 |
| B169 | B199 | U-CRY | Mg | -0.16572 | 3.06E-14 |
| B169 | B194 | U-CRY | Cl | 0.132956 | 1.21E-09 |
| B17 | B100 | CD3+CD16+CD56+NK | anti-Ro52 | 0.300783 | 0 |
| B17 | B101 | CD3+CD16+CD56+NK | anti-Sc70 | 0.300783 | 0 |
| B17 | B104 | CD3+CD16+CD56+NK | anti-SSB | 0.300783 | 0 |
| B17 | B106 | CD3+CD16+CD56+NK | anti-P | 0.300783 | 0 |
| B17 | B107 | CD3+CD16+CD56+NK | AnuA | 0.300783 | 0 |
| B17 | B110 | CD3+CD16+CD56+NK | AHA | 0.300783 | 0 |
| B17 | B137 | CD3+CD16+CD56+NK | AFU | -0.22074 | 0 |
| B17 | B151 | CD3+CD16+CD56+NK | Anti-CCP | 0.197844 | 0 |
| B17 | B181 | CD3+CD16+CD56+NK | APTT | 0.416309 | 0 |

| | | | | | |
|---|---|---|---|---|---|
| B17 | B19 | CD3+CD16+CD56+NK | CD3+CD4+/CD3+CD8+ | -0.26215 | 0 |
| B17 | B20 | CD3+CD16+CD56+NK | CD3+CD4+CD8+ | 0.390063 | 0 |
| B17 | B21 | CD3+CD16+CD56+NK | CD3+CD4-CD8- | 0.384703 | 0 |
| B17 | B22 | CD3+CD16+CD56+NK | CD3+CD8+(T) | 0.302299 | 0 |
| B17 | B25 | CD3+CD16+CD56+NK | CK | -0.18775 | 0 |
| B17 | B34 | CD3+CD16+CD56+NK | TK | 0.309032 | 0 |
| B17 | B95 | CD3+CD16+CD56+NK | anti-cenpb | 0.300783 | 0 |
| B17 | B97 | CD3+CD16+CD56+NK | dsDNA | 0.300783 | 0 |
| B17 | B98 | CD3+CD16+CD56+NK | anti-Jo-1 | 0.300783 | 0 |
| B17 | B99 | CD3+CD16+CD56+NK | anti-PM-Scl | 0.300783 | 0 |
| B17 | B185 | CD3+CD16+CD56+NK | PT | 0.176668 | 4.44E-16 |
| B17 | B18 | CD3+CD16+CD56+NK | CD3+CD4+helper(inducer)T | -0.17167 | 3.55E-15 |
| B17 | B182 | CD3+CD16+CD56+NK | TT | 0.153944 | 1.81E-12 |
| B17 | B183 | CD3+CD16+CD56+NK | PTA | 0.150694 | 5.27E-12 |
| B17 | B199 | CD3+CD16+CD56+NK | Mg | -0.13721 | 3.50E-10 |
| B17 | B134 | CD3+CD16+CD56+NK | CHE | -0.13496 | 6.80E-10 |
| B17 | B158 | CD3+CD16+CD56+NK | IgM | 0.126931 | 6.61E-09 |
| B17 | B60 | CD3+CD16+CD56+NK | WBRVI | -0.12613 | 8.22E-09 |
| B17 | B138 | CD3+CD16+CD56+NK | PAB | -0.12032 | 3.88E-08 |
| B17 | B176 | CD3+CD16+CD56+NK | Ucr | -0.11532 | 1.39E-07 |
| B170 | B228 | U-GLU | OS | 0.119773 | 4.47E-08 |
| B170 | B190 | U-GLU | Cre | 0.113328 | 2.28E-07 |
| B173 | B222 | NTT | NS | -0.11574 | 1.25E-07 |
| B174 | B189 | Urea/Cr | eGFB | 0.188401 | 0 |
| B174 | B190 | Urea/Cr | Cre | -0.37159 | 0 |
| B174 | B191 | Urea/Cr | Urea | 0.471356 | 0 |
| B174 | B216 | Urea/Cr | BMI | -0.19134 | 0 |
| B174 | B221 | Urea/Cr | Hei | -0.28253 | 0 |
| B174 | B222 | Urea/Cr | NS | -0.24473 | 0 |
| B174 | B192 | Urea/Cr | UA | -0.16102 | 1.62E-13 |
| B174 | B206 | Urea/Cr | Waist | -0.11423 | 1.83E-07 |
| B175 | B177 | ALB-U | ACR | 0.799262 | 0 |
| B176 | B214 | Ucr | CVA-L | 0.175488 | 8.88E-16 |
| B176 | B177 | Ucr | ACR | -0.16658 | 2.26E-14 |
| B178 | B179 | Ig-U | α1-MG | 0.508778 | 0 |
| B178 | B180 | Ig-U | Tf | 0.977038 | 0 |
| B178 | B197 | Ig-U | AG | -0.18247 | 0 |
| B178 | B196 | Ig-U | TCO2 | 0.132886 | 1.24E-09 |
| B178 | B200 | Ig-U | OP | 0.111103 | 3.93E-07 |
| B178 | B198 | Ig-U | P | 0.10786 | 8.50E-07 |
| B179 | B180 | α1-MG | Tf | 0.45688 | 0 |
| B179 | B194 | α1-MG | Cl | 0.187441 | 0 |
| B179 | B197 | α1-MG | AG | -0.20122 | 0 |
| B179 | B186 | α1-MG | PFC | 0.161292 | 1.47E-13 |

| | | | | | |
|---|---|---|---|---|---|
| B179 | B200 | α1-MG | OP | 0.15014 | 6.31E-12 |
| B18 | B102 | CD3+CD4+helper(inducer)T | anti-Sm | -0.73759 | 0 |
| B18 | B105 | CD3+CD4+helper(inducer)T | anti-U1RNP | -0.62695 | 0 |
| B18 | B108 | CD3+CD4+helper(inducer)T | Anti-Cyclin | 0.379543 | 0 |
| B18 | B109 | CD3+CD4+helper(inducer)T | AMA-M2 | 0.405803 | 0 |
| B18 | B124 | CD3+CD4+helper(inducer)T | ACTH | 0.392819 | 0 |
| B18 | B125 | CD3+CD4+helper(inducer)T | Cor | 0.335056 | 0 |
| B18 | B136 | CD3+CD4+helper(inducer)T | 5'NT | 0.322609 | 0 |
| B18 | B138 | CD3+CD4+helper(inducer)T | PAB | 0.261504 | 0 |
| B18 | B156 | CD3+CD4+helper(inducer)T | IgE | -0.24024 | 0 |
| B18 | B181 | CD3+CD4+helper(inducer)T | APTT | -0.30098 | 0 |
| B18 | B185 | CD3+CD4+helper(inducer)T | PT | -0.29482 | 0 |
| B18 | B19 | CD3+CD4+helper(inducer)T | CD3+CD4+/CD3+CD8+ | 0.773566 | 0 |
| B18 | B198 | CD3+CD4+helper(inducer)T | P | 0.275816 | 0 |
| B18 | B20 | CD3+CD4+helper(inducer)T | CD3+CD4+CD8+ | -0.47658 | 0 |
| B18 | B21 | CD3+CD4+helper(inducer)T | CD3+CD4-CD8- | -0.32139 | 0 |
| B18 | B22 | CD3+CD4+helper(inducer)T | CD3+CD8+(T) | -0.28261 | 0 |
| B18 | B23 | CD3+CD4+helper(inducer)T | CD3-CD16+CD56+(NK) | -0.80182 | 0 |
| B18 | B55 | CD3+CD4+helper(inducer)T | TK | 0.331785 | 0 |
| B18 | B56 | CD3+CD4+helper(inducer)T | IR | 0.273179 | 0 |
| B18 | B60 | CD3+CD4+helper(inducer)T | WBRVI | 0.245043 | 0 |
| B18 | B62 | CD3+CD4+helper(inducer)T | WBHSRVI | 0.224677 | 0 |
| B18 | B69 | CD3+CD4+helper(inducer)T | PV | -0.46907 | 0 |
| B18 | B186 | CD3+CD4+helper(inducer)T | PFC | -0.17273 | 2.22E-15 |
| B18 | B81 | CD3+CD4+helper(inducer)T | TSGF | 0.167197 | 1.82E-14 |
| B18 | B93 | CD3+CD4+helper(inducer)T | ALDH2-Glu504Lys | 0.158588 | 3.76E-13 |
| B18 | B92 | CD3+CD4+helper(inducer)T | ALDH2-Glu504Glu | -0.15649 | 7.68E-13 |
| B18 | B179 | CD3+CD4+helper(inducer)T | α1-MG | -0.15639 | 7.95E-13 |
| B18 | B54 | CD3+CD4+helper(inducer)T | ESR | -0.15537 | 1.12E-12 |
| B18 | B184 | CD3+CD4+helper(inducer)T | INR | -0.15274 | 2.70E-12 |
| B18 | B61 | CD3+CD4+helper(inducer)T | WBHSRV | 0.135248 | 6.25E-10 |
| B18 | B57 | CD3+CD4+helper(inducer)T | RBCAI | 0.135177 | 6.38E-10 |
| B18 | B228 | CD3+CD4+helper(inducer)T | OS | -0.13492 | 6.87E-10 |
| B18 | B94 | CD3+CD4+helper(inducer)T | ALDH2-Lys504Lys | -0.12963 | 3.12E-09 |
| B18 | B59 | CD3+CD4+helper(inducer)T | WBRV | 0.123764 | 1.56E-08 |
| B18 | B182 | CD3+CD4+helper(inducer)T | TT | 0.122388 | 2.25E-08 |
| B18 | B137 | CD3+CD4+helper(inducer)T | AFU | 0.12149 | 2.86E-08 |
| B18 | B139 | CD3+CD4+helper(inducer)T | ADA | -0.12038 | 3.82E-08 |
| B180 | B197 | Tf | AG | -0.17106 | 4.44E-15 |
| B180 | B196 | Tf | TCO2 | 0.131447 | 1.87E-09 |
| B180 | B200 | Tf | OP | 0.107379 | 9.52E-07 |
| B181 | B184 | APTT | INR | 0.413204 | 0 |
| B181 | B185 | APTT | PT | 0.34529 | 0 |
| B181 | B182 | APTT | TT | -0.13181 | 1.69E-09 |

| | | | | | |
|---|---|---|---|---|---|
| B182 | B183 | TT | PTA | -0.2832 | 0 |
| B182 | B184 | TT | INR | -0.21404 | 0 |
| B182 | B185 | TT | PT | -0.33012 | 0 |
| B182 | B187 | TT | NSE | -0.15901 | 3.26E-13 |
| B182 | B195 | TT | Na | 0.107256 | 9.79E-07 |
| B183 | B184 | PTA | INR | 0.201747 | 0 |
| B183 | B186 | PTA | PFC | 0.241168 | 0 |
| B184 | B185 | INR | PT | 0.857259 | 0 |
| B184 | B186 | INR | PFC | -0.15126 | 4.38E-12 |
| B186 | B193 | PFC | K | 0.110832 | 4.19E-07 |
| B189 | B190 | eGFB | Cre | -0.60751 | 0 |
| B189 | B191 | eGFB | Urea | -0.41723 | 0 |
| B189 | B192 | eGFB | UA | -0.27087 | 0 |
| B189 | B218 | eGFB | DBP | 0.169673 | 7.33E-15 |
| B189 | B220 | eGFB | SBP | -0.115 | 1.51E-07 |
| B189 | B210 | eGFB | Hip | 0.111256 | 3.79E-07 |
| B189 | B216 | eGFB | BMI | 0.109236 | 6.14E-07 |
| B19 | B100 | CD3+CD4+/CD3+CD8+ | anti-Ro52 | -0.2983 | 0 |
| B19 | B101 | CD3+CD4+/CD3+CD8+ | anti-Sc70 | -0.2983 | 0 |
| B19 | B102 | CD3+CD4+/CD3+CD8+ | anti-Sm | -0.49415 | 0 |
| B19 | B104 | CD3+CD4+/CD3+CD8+ | anti-SSB | -0.2983 | 0 |
| B19 | B105 | CD3+CD4+/CD3+CD8+ | anti-U1RNP | -0.43202 | 0 |
| B19 | B106 | CD3+CD4+/CD3+CD8+ | anti-P | -0.2983 | 0 |
| B19 | B107 | CD3+CD4+/CD3+CD8+ | AnuA | -0.2983 | 0 |
| B19 | B108 | CD3+CD4+/CD3+CD8+ | Anti-Cyclin | 0.192917 | 0 |
| B19 | B109 | CD3+CD4+/CD3+CD8+ | AMA-M2 | 0.403614 | 0 |
| B19 | B110 | CD3+CD4+/CD3+CD8+ | AHA | -0.2983 | 0 |
| B19 | B124 | CD3+CD4+/CD3+CD8+ | ACTH | 0.217394 | 0 |
| B19 | B125 | CD3+CD4+/CD3+CD8+ | Cor | 0.28324 | 0 |
| B19 | B136 | CD3+CD4+/CD3+CD8+ | 5'NT | 0.214756 | 0 |
| B19 | B139 | CD3+CD4+/CD3+CD8+ | ADA | -0.21317 | 0 |
| B19 | B156 | CD3+CD4+/CD3+CD8+ | IgE | -0.22308 | 0 |
| B19 | B181 | CD3+CD4+/CD3+CD8+ | APTT | -0.28451 | 0 |
| B19 | B185 | CD3+CD4+/CD3+CD8+ | PT | -0.23856 | 0 |
| B19 | B186 | CD3+CD4+/CD3+CD8+ | PFC | -0.26885 | 0 |
| B19 | B20 | CD3+CD4+/CD3+CD8+ | CD3+CD4+CD8+ | -0.36734 | 0 |
| B19 | B22 | CD3+CD4+/CD3+CD8+ | CD3+CD8+(T) | -0.8021 | 0 |
| B19 | B23 | CD3+CD4+/CD3+CD8+ | CD3-CD16+CD56+(NK) | -0.42518 | 0 |
| B19 | B25 | CD3+CD4+/CD3+CD8+ | CK | 0.285997 | 0 |
| B19 | B34 | CD3+CD4+/CD3+CD8+ | TK | -0.22837 | 0 |
| B19 | B54 | CD3+CD4+/CD3+CD8+ | ESR | -0.17986 | 0 |
| B19 | B55 | CD3+CD4+/CD3+CD8+ | TK | 0.312015 | 0 |
| B19 | B56 | CD3+CD4+/CD3+CD8+ | IR | 0.222585 | 0 |
| B19 | B57 | CD3+CD4+/CD3+CD8+ | RBCAI | 0.322942 | 0 |

| | | | | | |
|---|---|---|---|---|---|
| B19 | B59 | CD3+CD4+/CD3+CD8+ | WBRV | 0.248465 | 0 |
| B19 | B60 | CD3+CD4+/CD3+CD8+ | WBRVI | 0.309957 | 0 |
| B19 | B69 | CD3+CD4+/CD3+CD8+ | PV | -0.37563 | 0 |
| B19 | B95 | CD3+CD4+/CD3+CD8+ | anti-cenpb | -0.2983 | 0 |
| B19 | B97 | CD3+CD4+/CD3+CD8+ | dsDNA | -0.2983 | 0 |
| B19 | B98 | CD3+CD4+/CD3+CD8+ | anti-Jo-1 | -0.2983 | 0 |
| B19 | B99 | CD3+CD4+/CD3+CD8+ | anti-PM-Scl | -0.2983 | 0 |
| B19 | B21 | CD3+CD4+/CD3+CD8+ | CD3+CD4-CD8- | -0.17394 | 1.33E-15 |
| B19 | B65 | CD3+CD4+/CD3+CD8+ | WBRSR(mPas_1) | 0.166521 | 2.31E-14 |
| B19 | B157 | CD3+CD4+/CD3+CD8+ | IgG | -0.16115 | 1.55E-13 |
| B19 | B93 | CD3+CD4+/CD3+CD8+ | ALDH2-Glu504Lys | 0.158139 | 4.38E-13 |
| B19 | B182 | CD3+CD4+/CD3+CD8+ | TT | 0.15614 | 8.65E-13 |
| B19 | B92 | CD3+CD4+/CD3+CD8+ | ALDH2-Glu504Glu | -0.15607 | 8.86E-13 |
| B19 | B138 | CD3+CD4+/CD3+CD8+ | PAB | 0.149245 | 8.43E-12 |
| B19 | B137 | CD3+CD4+/CD3+CD8+ | AFU | 0.146634 | 1.94E-11 |
| B19 | B193 | CD3+CD4+/CD3+CD8+ | K | -0.14466 | 3.62E-11 |
| B19 | B76 | CD3+CD4+/CD3+CD8+ | ApoB | -0.14019 | 1.43E-10 |
| B19 | B62 | CD3+CD4+/CD3+CD8+ | WBHSRVI | 0.139214 | 1.93E-10 |
| B19 | B179 | CD3+CD4+/CD3+CD8+ | α1-MG | -0.13842 | 2.44E-10 |
| B19 | B194 | CD3+CD4+/CD3+CD8+ | Cl | -0.13513 | 6.46E-10 |
| B19 | B94 | CD3+CD4+/CD3+CD8+ | ALDH2-Lys504Lys | -0.12882 | 3.91E-09 |
| B19 | B71 | CD3+CD4+/CD3+CD8+ | TG | 0.115445 | 1.35E-07 |
| B19 | B184 | CD3+CD4+/CD3+CD8+ | INR | -0.11472 | 1.62E-07 |
| B19 | B198 | CD3+CD4+/CD3+CD8+ | P | 0.111437 | 3.62E-07 |
| B19 | B75 | CD3+CD4+/CD3+CD8+ | ApoA1/B | 0.109286 | 6.07E-07 |
| B19 | B197 | CD3+CD4+/CD3+CD8+ | AG | 0.108418 | 7.46E-07 |
| B19 | B58 | CD3+CD4+/CD3+CD8+ | HCT | -0.10813 | 7.98E-07 |
| B190 | B191 | Cre | Urea | 0.597297 | 0 |
| B190 | B192 | Cre | UA | 0.432332 | 0 |
| B190 | B221 | Cre | Hei | 0.26698 | 0 |
| B190 | B222 | Cre | NS | 0.274139 | 0 |
| B190 | B216 | Cre | BMI | 0.163971 | 5.73E-14 |
| B190 | B206 | Cre | Waist | 0.135017 | 6.68E-10 |
| B190 | B207 | Cre | WHR | 0.134831 | 7.05E-10 |
| B191 | B192 | Urea | UA | 0.308584 | 0 |
| B192 | B206 | UA | Waist | 0.264919 | 0 |
| B192 | B207 | UA | WHR | 0.180472 | 0 |
| B192 | B210 | UA | Hip | 0.207145 | 0 |
| B192 | B215 | UA | BMI | 0.20391 | 0 |
| B192 | B216 | UA | BMI | 0.311538 | 0 |
| B192 | B221 | UA | Hei | 0.268713 | 0 |
| B192 | B222 | UA | NS | 0.325184 | 0 |
| B193 | B194 | K | Cl | 0.36024 | 0 |
| B193 | B200 | K | OP | 0.297414 | 0 |

| | | | | | |
|---|---|---|---|---|---|
| B193 | B195 | K | Na | 0.152412 | 3.00E-12 |
| B193 | B199 | K | Mg | 0.125035 | 1.11E-08 |
| B194 | B195 | Cl | Na | 0.386468 | 0 |
| B194 | B196 | Cl | TCO2 | -0.25842 | 0 |
| B194 | B197 | Cl | AG | -0.33921 | 0 |
| B194 | B200 | Cl | OP | 0.463706 | 0 |
| B195 | B197 | Na | AG | 0.26149 | 0 |
| B195 | B198 | Na | P | -0.22239 | 0 |
| B195 | B200 | Na | OP | 0.871635 | 0 |
| B195 | B196 | Na | TCO2 | 0.138143 | 2.65E-10 |
| B196 | B197 | TCO2 | AG | -0.58307 | 0 |
| B196 | B199 | TCO2 | Mg | -0.17162 | 3.55E-15 |
| B196 | B200 | TCO2 | OP | 0.13748 | 3.23E-10 |
| B196 | B198 | TCO2 | P | -0.13341 | 1.06E-09 |
| B197 | B199 | AG | Mg | 0.139858 | 1.59E-10 |
| B198 | B200 | P | OP | -0.20584 | 0 |
| B198 | B199 | P | Mg | -0.12112 | 3.15E-08 |
| B2 | B126 | Age | Alb | -0.25659 | 0 |
| B2 | B127 | Age | A-G ratio | -0.18385 | 0 |
| B2 | B141 | Age | AST/ALT | 0.296104 | 0 |
| B2 | B163 | Age | SG | -0.19333 | 0 |
| B2 | B189 | Age | eGFB | -0.40789 | 0 |
| B2 | B191 | Age | Urea | 0.209161 | 0 |
| B2 | B210 | Age | Hip | -0.2054 | 0 |
| B2 | B215 | Age | BMI | -0.23398 | 0 |
| B2 | B216 | Age | BMI | -0.36524 | 0 |
| B2 | B218 | Age | DBP | -0.29872 | 0 |
| B2 | B220 | Age | SBP | 0.231753 | 0 |
| B2 | B221 | Age | Hei | -0.30725 | 0 |
| B2 | B222 | Age | NS | -0.21856 | 0 |
| B2 | B46 | Age | RBC Count | -0.2634 | 0 |
| B2 | B47 | Age | RDW | 0.208905 | 0 |
| B2 | B51 | Age | Hb | -0.30456 | 0 |
| B2 | B72 | Age | HDL-C | 0.229562 | 0 |
| B2 | B71 | Age | TG | -0.17959 | 2.22E-16 |
| B2 | B176 | Age | Ucr | -0.17626 | 6.66E-16 |
| B2 | B128 | Age | ALT | -0.17289 | 2.22E-15 |
| B2 | B190 | Age | Cre | 0.163174 | 7.62E-14 |
| B2 | B206 | Age | Waist | -0.14649 | 2.03E-11 |
| B2 | B74 | Age | ApoA1 | 0.141699 | 9.05E-11 |
| B2 | B41 | Age | MON％ | 0.138176 | 2.63E-10 |
| B2 | B209 | Age | Smoking | -0.13675 | 4.02E-10 |
| B2 | B94 | Age | ALDH2-Lys504Lys | 0.128759 | 3.99E-09 |
| B2 | B130 | Age | GLB | 0.110347 | 4.71E-07 |

| | | | | | |
|---|---|---|---|---|---|
| B2 | B79 | Age | hb V -dna | 0.108004 | 8.22E-07 |
| B20 | B100 | CD3+CD4+CD8+ | anti-Ro52 | 0.236801 | 0 |
| B20 | B101 | CD3+CD4+CD8+ | anti-Sc70 | 0.236801 | 0 |
| B20 | B104 | CD3+CD4+CD8+ | anti-SSB | 0.236801 | 0 |
| B20 | B106 | CD3+CD4+CD8+ | anti-P | 0.236801 | 0 |
| B20 | B107 | CD3+CD4+CD8+ | AnuA | 0.2368 | 0 |
| B20 | B108 | CD3+CD4+CD8+ | Anti-Cyclin | -0.2822 | 0 |
| B20 | B110 | CD3+CD4+CD8+ | AHA | 0.236801 | 0 |
| B20 | B136 | CD3+CD4+CD8+ | 5'NT | -0.81738 | 0 |
| B20 | B181 | CD3+CD4+CD8+ | APTT | 0.244057 | 0 |
| B20 | B198 | CD3+CD4+CD8+ | P | -0.28224 | 0 |
| B20 | B200 | CD3+CD4+CD8+ | OP | -0.18423 | 0 |
| B20 | B21 | CD3+CD4+CD8+ | CD3+CD4-CD8- | 0.31993 | 0 |
| B20 | B22 | CD3+CD4+CD8+ | CD3+CD8+(T) | 0.296521 | 0 |
| B20 | B23 | CD3+CD4+CD8+ | CD3-CD16+CD56+(NK) | 0.246043 | 0 |
| B20 | B74 | CD3+CD4+CD8+ | ApoA1 | 0.200603 | 0 |
| B20 | B92 | CD3+CD4+CD8+ | ALDH2-Glu504Glu | 0.215469 | 0 |
| B20 | B93 | CD3+CD4+CD8+ | ALDH2-Glu504Lys | -0.21833 | 0 |
| B20 | B95 | CD3+CD4+CD8+ | anti-cenpb | 0.236801 | 0 |
| B20 | B97 | CD3+CD4+CD8+ | dsDNA | 0.236801 | 0 |
| B20 | B98 | CD3+CD4+CD8+ | anti-Jo-1 | 0.236801 | 0 |
| B20 | B99 | CD3+CD4+CD8+ | anti-PM-Scl | 0.236801 | 0 |
| B20 | B94 | CD3+CD4+CD8+ | ALDH2-Lys504Lys | 0.177902 | 4.44E-16 |
| B20 | B134 | CD3+CD4+CD8+ | CHE | 0.161645 | 1.30E-13 |
| B20 | B193 | CD3+CD4+CD8+ | K | -0.15156 | 3.98E-12 |
| B20 | B120 | CD3+CD4+CD8+ | HCV-cAg | 0.147342 | 1.55E-11 |
| B20 | B151 | CD3+CD4+CD8+ | Anti-CCP | 0.143083 | 5.91E-11 |
| B20 | B183 | CD3+CD4+CD8+ | PTA | 0.118507 | 6.21E-08 |
| B20 | B34 | CD3+CD4+CD8+ | TK | 0.117674 | 7.68E-08 |
| B20 | B25 | CD3+CD4+CD8+ | CK | -0.11368 | 2.09E-07 |
| B20 | B137 | CD3+CD4+CD8+ | AFU | -0.11081 | 4.21E-07 |
| B20 | B81 | CD3+CD4+CD8+ | TSGF | -0.10981 | 5.36E-07 |
| B20 | B79 | CD3+CD4+CD8+ | hb V -dna | 0.108765 | 6.87E-07 |
| B201 | B204 | e-habits | Drinking | 0.208545 | 0 |
| B202 | B223 | e-times | FH | 0.6484 | 0 |
| B204 | B209 | Drinking | Smoking | 0.33833 | 0 |
| B204 | B221 | Drinking | Hei | 0.169848 | 6.88E-15 |
| B204 | B222 | Drinking | NS | 0.16542 | 3.42E-14 |
| B204 | B216 | Drinking | BMI | 0.14155 | 9.48E-11 |
| B205 | B222 | KDA | NS | -0.10857 | 7.18E-07 |
| B206 | B207 | Waist | WHR | 0.621222 | 0 |
| B206 | B210 | Waist | Hip | 0.754244 | 0 |
| B206 | B215 | Waist | BMI | 0.790754 | 0 |
| B206 | B216 | Waist | BMI | 0.795572 | 0 |

| | | | | | |
|---|---|---|---|---|---|
| B206 | B221 | Waist | Hei | 0.360138 | 0 |
| B206 | B222 | Waist | NS | 0.667941 | 0 |
| B207 | B215 | WHR | BMI | 0.336904 | 0 |
| B207 | B216 | WHR | BMI | 0.341698 | 0 |
| B207 | B222 | WHR | NS | 0.344144 | 0 |
| B207 | B221 | WHR | Hei | 0.157906 | 4.75E-13 |
| B207 | B210 | WHR | Hip | 0.126665 | 7.11E-09 |
| B209 | B221 | Smoking | Hei | 0.20381 | 0 |
| B209 | B222 | Smoking | NS | 0.175385 | 8.88E-16 |
| B209 | B216 | Smoking | BMI | 0.155178 | 1.20E-12 |
| B21 | B100 | CD3+CD4-CD8- | anti-Ro52 | 0.468546 | 0 |
| B21 | B101 | CD3+CD4-CD8- | anti-Sc70 | 0.468546 | 0 |
| B21 | B104 | CD3+CD4-CD8- | anti-SSB | 0.468546 | 0 |
| B21 | B105 | CD3+CD4-CD8- | anti-U1RNP | -0.19489 | 0 |
| B21 | B106 | CD3+CD4-CD8- | anti-P | 0.468546 | 0 |
| B21 | B107 | CD3+CD4-CD8- | AnuA | 0.468545 | 0 |
| B21 | B108 | CD3+CD4-CD8- | Anti-Cyclin | -0.83772 | 0 |
| B21 | B110 | CD3+CD4-CD8- | AHA | 0.468546 | 0 |
| B21 | B124 | CD3+CD4-CD8- | ACTH | -0.25657 | 0 |
| B21 | B135 | CD3+CD4-CD8- | TBA | -0.28536 | 0 |
| B21 | B151 | CD3+CD4-CD8- | Anti-CCP | 0.272545 | 0 |
| B21 | B181 | CD3+CD4-CD8- | APTT | 0.399105 | 0 |
| B21 | B183 | CD3+CD4-CD8- | PTA | 0.234255 | 0 |
| B21 | B184 | CD3+CD4-CD8- | INR | 0.179706 | 0 |
| B21 | B198 | CD3+CD4-CD8- | P | -0.24088 | 0 |
| B21 | B25 | CD3+CD4-CD8- | CK | -0.18928 | 0 |
| B21 | B79 | CD3+CD4-CD8- | hb V -dna | 0.323628 | 0 |
| B21 | B81 | CD3+CD4-CD8- | TSGF | -0.31898 | 0 |
| B21 | B92 | CD3+CD4-CD8- | ALDH2-Glu504Glu | 0.631142 | 0 |
| B21 | B93 | CD3+CD4-CD8- | ALDH2-Glu504Lys | -0.63961 | 0 |
| B21 | B94 | CD3+CD4-CD8- | ALDH2-Lys504Lys | 0.523283 | 0 |
| B21 | B95 | CD3+CD4-CD8- | anti-cenpb | 0.468546 | 0 |
| B21 | B97 | CD3+CD4-CD8- | dsDNA | 0.468546 | 0 |
| B21 | B98 | CD3+CD4-CD8- | anti-Jo-1 | 0.468546 | 0 |
| B21 | B99 | CD3+CD4-CD8- | anti-PM-Scl | 0.468546 | 0 |
| B21 | B137 | CD3+CD4-CD8- | AFU | -0.16943 | 7.99E-15 |
| B21 | B74 | CD3+CD4-CD8- | ApoA1 | 0.158204 | 4.29E-13 |
| B21 | B23 | CD3+CD4-CD8- | CD3-CD16+CD56+(NK) | 0.143438 | 5.30E-11 |
| B21 | B60 | CD3+CD4-CD8- | WBRVI | -0.12783 | 5.15E-09 |
| B21 | B155 | CD3+CD4-CD8- | IgA | -0.12504 | 1.11E-08 |
| B21 | B158 | CD3+CD4-CD8- | IgM | 0.122086 | 2.44E-08 |
| B21 | B136 | CD3+CD4-CD8- | 5'NT | 0.121436 | 2.90E-08 |
| B21 | B125 | CD3+CD4-CD8- | Cor | 0.120866 | 3.36E-08 |
| B21 | B34 | CD3+CD4-CD8- | TK | 0.11238 | 2.88E-07 |

| | | | | | |
|---|---|---|---|---|---|
| B21 | B54 | CD3+CD4-CD8- | ESR | 0.109162 | 6.25E-07 |
| B210 | B215 | Hip | BMI | 0.769689 | 0 |
| B210 | B216 | Hip | BMI | 0.779982 | 0 |
| B210 | B221 | Hip | Hei | 0.364468 | 0 |
| B210 | B222 | Hip | NS | 0.588266 | 0 |
| B211 | B212 | IOP-R | IOP-L | 0.767812 | 0 |
| B215 | B216 | BMI | BMI | 0.778045 | 0 |
| B215 | B222 | BMI | NS | 0.542868 | 0 |
| B216 | B221 | BMI | Hei | 0.695406 | 0 |
| B216 | B222 | BMI | NS | 0.765443 | 0 |
| B216 | B218 | BMI | DBP | 0.135172 | 6.39E-10 |
| B216 | B220 | BMI | SBP | -0.12272 | 2.06E-08 |
| B218 | B220 | DBP | SBP | 0.397651 | 0 |
| B218 | B221 | DBP | Hei | 0.158296 | 4.16E-13 |
| B219 | B222 | OT | NS | 0.121727 | 2.68E-08 |
| B22 | B100 | CD3+CD8+(T) | anti-Ro52 | 0.466933 | 0 |
| B22 | B101 | CD3+CD8+(T) | anti-Sc70 | 0.466933 | 0 |
| B22 | B104 | CD3+CD8+(T) | anti-SSB | 0.466933 | 0 |
| B22 | B106 | CD3+CD8+(T) | anti-P | 0.466933 | 0 |
| B22 | B107 | CD3+CD8+(T) | AnuA | 0.466932 | 0 |
| B22 | B109 | CD3+CD8+(T) | AMA-M2 | -0.29678 | 0 |
| B22 | B110 | CD3+CD8+(T) | AHA | 0.466933 | 0 |
| B22 | B139 | CD3+CD8+(T) | ADA | 0.189924 | 0 |
| B22 | B157 | CD3+CD8+(T) | IgG | 0.188819 | 0 |
| B22 | B181 | CD3+CD8+(T) | APTT | 0.182325 | 0 |
| B22 | B183 | CD3+CD8+(T) | PTA | 0.186857 | 0 |
| B22 | B186 | CD3+CD8+(T) | PFC | 0.219535 | 0 |
| B22 | B25 | CD3+CD8+(T) | CK | -0.32355 | 0 |
| B22 | B34 | CD3+CD8+(T) | TK | 0.268891 | 0 |
| B22 | B57 | CD3+CD8+(T) | RBCAI | -0.3575 | 0 |
| B22 | B59 | CD3+CD8+(T) | WBRV | -0.24882 | 0 |
| B22 | B60 | CD3+CD8+(T) | WBRVI | -0.20792 | 0 |
| B22 | B76 | CD3+CD8+(T) | ApoB | 0.188575 | 0 |
| B22 | B95 | CD3+CD8+(T) | anti-cenpb | 0.466933 | 0 |
| B22 | B97 | CD3+CD8+(T) | dsDNA | 0.466933 | 0 |
| B22 | B98 | CD3+CD8+(T) | anti-Jo-1 | 0.466933 | 0 |
| B22 | B99 | CD3+CD8+(T) | anti-PM-Scl | 0.466933 | 0 |
| B22 | B23 | CD3+CD8+(T) | CD3-CD16+CD56+(NK) | -0.17097 | 4.44E-15 |
| B22 | B136 | CD3+CD8+(T) | 5'NT | -0.16475 | 4.35E-14 |
| B22 | B75 | CD3+CD8+(T) | ApoA1/B | -0.16235 | 1.02E-13 |
| B22 | B65 | CD3+CD8+(T) | WBRSR(mPas_1) | -0.16183 | 1.22E-13 |
| B22 | B58 | CD3+CD8+(T) | HCT | 0.158728 | 3.58E-13 |
| B22 | B137 | CD3+CD8+(T) | AFU | -0.15004 | 6.52E-12 |
| B22 | B193 | CD3+CD8+(T) | K | 0.149957 | 6.70E-12 |

| | | | | | |
|---|---|---|---|---|---|
| B22 | B197 | CD3+CD8+(T) | AG | -0.14703 | 1.71E-11 |
| B22 | B93 | CD3+CD8+(T) | ALDH2-Glu504Lys | -0.14216 | 7.86E-11 |
| B22 | B92 | CD3+CD8+(T) | ALDH2-Glu504Glu | 0.140316 | 1.38E-10 |
| B22 | B194 | CD3+CD8+(T) | Cl | 0.136767 | 4.00E-10 |
| B22 | B71 | CD3+CD8+(T) | TG | -0.12231 | 2.30E-08 |
| B22 | B94 | CD3+CD8+(T) | ALDH2-Lys504Lys | 0.115412 | 1.36E-07 |
| B22 | B55 | CD3+CD8+(T) | TK | -0.10991 | 5.23E-07 |
| B220 | B221 | SBP | Hei | -0.1405 | 1.31E-10 |
| B221 | B222 | Hei | NS | 0.5974 | 0 |
| B23 | B100 | CD3-CD16+CD56+(NK) | anti-Ro52 | -0.2727 | 0 |
| B23 | B101 | CD3-CD16+CD56+(NK) | anti-Sc70 | -0.2727 | 0 |
| B23 | B102 | CD3-CD16+CD56+(NK) | anti-Sm | 0.868509 | 0 |
| B23 | B104 | CD3-CD16+CD56+(NK) | anti-SSB | -0.2727 | 0 |
| B23 | B105 | CD3-CD16+CD56+(NK) | anti-U1RNP | 0.807192 | 0 |
| B23 | B106 | CD3-CD16+CD56+(NK) | anti-P | -0.2727 | 0 |
| B23 | B107 | CD3-CD16+CD56+(NK) | AnuA | -0.2727 | 0 |
| B23 | B110 | CD3-CD16+CD56+(NK) | AHA | -0.2727 | 0 |
| B23 | B124 | CD3-CD16+CD56+(NK) | ACTH | -0.41314 | 0 |
| B23 | B125 | CD3-CD16+CD56+(NK) | Cor | -0.36293 | 0 |
| B23 | B136 | CD3-CD16+CD56+(NK) | 5'NT | -0.20635 | 0 |
| B23 | B138 | CD3-CD16+CD56+(NK) | PAB | -0.27362 | 0 |
| B23 | B156 | CD3-CD16+CD56+(NK) | IgE | 0.243486 | 0 |
| B23 | B181 | CD3-CD16+CD56+(NK) | APTT | 0.262009 | 0 |
| B23 | B185 | CD3-CD16+CD56+(NK) | PT | 0.305845 | 0 |
| B23 | B55 | CD3-CD16+CD56+(NK) | TK | -0.32081 | 0 |
| B23 | B56 | CD3-CD16+CD56+(NK) | IR | -0.29233 | 0 |
| B23 | B60 | CD3-CD16+CD56+(NK) | WBRVI | -0.18626 | 0 |
| B23 | B62 | CD3-CD16+CD56+(NK) | WBHSRVI | -0.27789 | 0 |
| B23 | B69 | CD3-CD16+CD56+(NK) | PV | 0.524321 | 0 |
| B23 | B95 | CD3-CD16+CD56+(NK) | anti-cenpb | -0.2727 | 0 |
| B23 | B97 | CD3-CD16+CD56+(NK) | dsDNA | -0.2727 | 0 |
| B23 | B98 | CD3-CD16+CD56+(NK) | anti-Jo-1 | -0.2727 | 0 |
| B23 | B99 | CD3-CD16+CD56+(NK) | anti-PM-Scl | -0.2727 | 0 |
| B23 | B108 | CD3-CD16+CD56+(NK) | Anti-Cyclin | -0.17466 | 1.11E-15 |
| B23 | B228 | CD3-CD16+CD56+(NK) | OS | 0.156942 | 6.59E-13 |
| B23 | B81 | CD3-CD16+CD56+(NK) | TSGF | -0.15577 | 9.80E-13 |
| B23 | B54 | CD3-CD16+CD56+(NK) | ESR | 0.153824 | 1.88E-12 |
| B23 | B151 | CD3-CD16+CD56+(NK) | Anti-CCP | 0.153482 | 2.11E-12 |
| B23 | B182 | CD3-CD16+CD56+(NK) | TT | -0.15255 | 2.87E-12 |
| B23 | B186 | CD3-CD16+CD56+(NK) | PFC | 0.14801 | 1.25E-11 |
| B23 | B183 | CD3-CD16+CD56+(NK) | PTA | -0.14719 | 1.63E-11 |
| B23 | B61 | CD3-CD16+CD56+(NK) | WBHSRV | -0.14426 | 4.10E-11 |
| B23 | B179 | CD3-CD16+CD56+(NK) | α1-MG | 0.124335 | 1.34E-08 |
| B23 | B93 | CD3-CD16+CD56+(NK) | ALDH2-Glu504Lys | 0.109277 | 6.08E-07 |

| | | | | | |
|---|---|---|---|---|---|
| B23 | B92 | CD3-CD16+CD56+(NK) | ALDH2-Glu504Glu | -0.10784 | 8.54E-07 |
| B24 | B25 | αHBDH | CK | 0.185369 | 0 |
| B24 | B27 | αHBDH | LDH | 0.508159 | 0 |
| B24 | B54 | αHBDH | ESR | 0.251309 | 0 |
| B24 | B58 | αHBDH | HCT | -0.26642 | 0 |
| B24 | B84 | αHBDH | C-PSA | 0.170548 | 5.33E-15 |
| B24 | B75 | αHBDH | ApoA1/B | 0.15526 | 1.16E-12 |
| B24 | B62 | αHBDH | WBHSRVI | -0.14692 | 1.77E-11 |
| B24 | B60 | αHBDH | WBRVI | -0.14216 | 7.86E-11 |
| B24 | B69 | αHBDH | PV | 0.133111 | 1.16E-09 |
| B24 | B74 | αHBDH | ApoA1 | 0.118134 | 6.83E-08 |
| B24 | B26 | αHBDH | CKMB | 0.114087 | 1.89E-07 |
| B24 | B139 | αHBDH | ADA | 0.113896 | 1.98E-07 |
| B25 | B100 | CK | anti-Ro52 | -0.20217 | 0 |
| B25 | B101 | CK | anti-Sc70 | -0.20217 | 0 |
| B25 | B104 | CK | anti-SSB | -0.20217 | 0 |
| B25 | B106 | CK | anti-P | -0.20217 | 0 |
| B25 | B107 | CK | AnuA | -0.20217 | 0 |
| B25 | B110 | CK | AHA | -0.20217 | 0 |
| B25 | B181 | CK | APTT | -0.20816 | 0 |
| B25 | B95 | CK | anti-cenpb | -0.20217 | 0 |
| B25 | B97 | CK | dsDNA | -0.20217 | 0 |
| B25 | B98 | CK | anti-Jo-1 | -0.20217 | 0 |
| B25 | B99 | CK | anti-PM-Scl | -0.20217 | 0 |
| B25 | B26 | CK | CKMB | 0.167193 | 1.82E-14 |
| B25 | B57 | CK | RBCAI | 0.16399 | 5.71E-14 |
| B25 | B139 | CK | ADA | -0.15731 | 5.82E-13 |
| B25 | B151 | CK | Anti-CCP | -0.15188 | 3.58E-12 |
| B25 | B34 | CK | TK | -0.14803 | 1.25E-11 |
| B25 | B76 | CK | ApoB | -0.14587 | 2.48E-11 |
| B25 | B27 | CK | LDH | 0.144975 | 3.28E-11 |
| B25 | B54 | CK | ESR | -0.14492 | 3.33E-11 |
| B25 | B75 | CK | ApoA1/B | 0.138462 | 2.41E-10 |
| B25 | B186 | CK | PFC | -0.13413 | 8.65E-10 |
| B25 | B108 | CK | Anti-Cyclin | 0.127529 | 5.61E-09 |
| B25 | B135 | CK | TBA | 0.108335 | 7.60E-07 |
| B26 | B123 | CKMB | hs-CRP1 | 0.251045 | 0 |
| B26 | B131 | CKMB | AST | 0.118825 | 5.72E-08 |
| B26 | B27 | CKMB | LDH | 0.111658 | 3.43E-07 |
| B27 | B150 | LDH | ALP | 0.283976 | 0 |
| B27 | B176 | LDH | Ucr | -0.21387 | 0 |
| B27 | B37 | LDH | BAND | -0.29853 | 0 |
| B27 | B38 | LDH | BAND% | -0.32609 | 0 |
| B27 | B54 | LDH | ESR | 0.147114 | 1.67E-11 |

| | | | | | |
|---|---|---|---|---|---|
| B27 | B139 | LDH | ADA | 0.128313 | 4.51E-09 |
| B27 | B84 | LDH | C-PSA | 0.12776 | 5.26E-09 |
| B27 | B58 | LDH | HCT | -0.12321 | 1.81E-08 |
| B27 | B163 | LDH | SG | -0.12298 | 1.93E-08 |
| B27 | B138 | LDH | PAB | -0.11308 | 2.43E-07 |
| B27 | B69 | LDH | PV | 0.109415 | 5.89E-07 |
| B27 | B131 | LDH | AST | 0.109219 | 6.17E-07 |
| B27 | B199 | LDH | Mg | 0.108342 | 7.59E-07 |
| B28 | B29 | PRL | E2 | 0.817691 | 0 |
| B28 | B30 | PRL | LH | 0.495176 | 0 |
| B28 | B31 | PRL | FSH | -0.29949 | 0 |
| B28 | B32 | PRL | Testo | 0.982135 | 0 |
| B28 | B33 | PRL | Prog | 0.95868 | 0 |
| B28 | B75 | PRL | ApoA1/B | -0.16648 | 2.35E-14 |
| B28 | B76 | PRL | ApoB | 0.134703 | 7.32E-10 |
| B28 | B174 | PRL | Urea/Cr | -0.12324 | 1.79E-08 |
| B28 | B156 | PRL | IgE | 0.11327 | 2.32E-07 |
| B29 | B30 | E2 | LH | 0.373514 | 0 |
| B29 | B31 | E2 | FSH | -0.6248 | 0 |
| B29 | B32 | E2 | Testo | 0.705601 | 0 |
| B29 | B33 | E2 | Prog | 0.621095 | 0 |
| B3 | B190 | Hcy | Cre | 0.14549 | 2.79E-11 |
| B3 | B189 | Hcy | eGFB | -0.13267 | 1.32E-09 |
| B30 | B31 | LH | FSH | 0.485133 | 0 |
| B30 | B32 | LH | Testo | 0.424409 | 0 |
| B30 | B33 | LH | Prog | 0.463477 | 0 |
| B30 | B200 | LH | OP | -0.13886 | 2.14E-10 |
| B30 | B195 | LH | Na | -0.13271 | 1.30E-09 |
| B31 | B32 | FSH | Testo | -0.23893 | 0 |
| B31 | B33 | FSH | Prog | -0.12165 | 2.74E-08 |
| B32 | B33 | Testo | Prog | 0.991969 | 0 |
| B32 | B75 | Testo | ApoA1/B | -0.19873 | 0 |
| B32 | B76 | Testo | ApoB | 0.157876 | 4.80E-13 |
| B32 | B156 | Testo | IgE | 0.133407 | 1.07E-09 |
| B32 | B174 | Testo | Urea/Cr | -0.11892 | 5.57E-08 |
| B32 | B155 | Testo | IgA | 0.118918 | 5.58E-08 |
| B32 | B199 | Testo | Mg | 0.114013 | 1.93E-07 |
| B32 | B137 | Testo | AFU | 0.111668 | 3.43E-07 |
| B33 | B75 | Prog | ApoA1/B | -0.21827 | 0 |
| B33 | B76 | Prog | ApoB | 0.174325 | 1.33E-15 |
| B33 | B156 | Prog | IgE | 0.147029 | 1.71E-11 |
| B33 | B155 | Prog | IgA | 0.130706 | 2.31E-09 |
| B33 | B199 | Prog | Mg | 0.126173 | 8.13E-09 |
| B33 | B137 | Prog | AFU | 0.123677 | 1.60E-08 |

| | | | | | |
|---|---|---|---|---|---|
| B33 | B174 | Prog | Urea/Cr | -0.11963 | 4.64E-08 |
| B34 | B181 | TK | APTT | 0.208739 | 0 |
| B34 | B182 | TK | TT | -0.19984 | 0 |
| B34 | B81 | TK | TSGF | -0.19421 | 0 |
| B34 | B94 | TK | ALDH2-Lys504Lys | 0.175401 | 8.88E-16 |
| B34 | B186 | TK | PFC | 0.171147 | 4.22E-15 |
| B34 | B100 | TK | anti-Ro52 | 0.170217 | 6.00E-15 |
| B34 | B101 | TK | anti-Sc70 | 0.170217 | 6.00E-15 |
| B34 | B104 | TK | anti-SSB | 0.170217 | 6.00E-15 |
| B34 | B106 | TK | anti-P | 0.170217 | 6.00E-15 |
| B34 | B107 | TK | AnuA | 0.170216 | 6.00E-15 |
| B34 | B110 | TK | AHA | 0.170217 | 6.00E-15 |
| B34 | B95 | TK | anti-cenpb | 0.170217 | 6.00E-15 |
| B34 | B97 | TK | dsDNA | 0.170217 | 6.00E-15 |
| B34 | B98 | TK | anti-Jo-1 | 0.170217 | 6.00E-15 |
| B34 | B99 | TK | anti-PM-Scl | 0.170217 | 6.00E-15 |
| B34 | B138 | TK | PAB | -0.14965 | 7.39E-12 |
| B34 | B183 | TK | PTA | 0.147684 | 1.39E-11 |
| B34 | B79 | TK | hb V -dna | 0.143124 | 5.84E-11 |
| B34 | B185 | TK | PT | 0.128982 | 3.74E-09 |
| B34 | B187 | TK | NSE | 0.120886 | 3.35E-08 |
| B34 | B194 | TK | Cl | -0.11922 | 5.16E-08 |
| B34 | B157 | TK | IgG | 0.116683 | 9.88E-08 |
| B34 | B92 | TK | ALDH2-Glu504Glu | -0.11594 | 1.19E-07 |
| B34 | B151 | TK | Anti-CCP | 0.111425 | 3.63E-07 |
| B35 | B202 | β2-MG | e-times | 0.575696 | 0 |
| B35 | B223 | β2-MG | FH | 0.99355 | 0 |
| B37 | B38 | BAND | BAND% | 0.856733 | 0 |
| B37 | B227 | BAND | OD | -0.11694 | 9.25E-08 |
| B4 | B5 | WB-Ca | WB-Mg | 0.869365 | 0 |
| B4 | B6 | WB-Ca | WB-Fe | 0.486503 | 0 |
| B4 | B7 | WB-Ca | WB-Cop | 0.49016 | 0 |
| B4 | B8 | WB-Ca | WB-Zinc | 0.290579 | 0 |
| B4 | B197 | WB-Ca | AG | -0.13429 | 8.26E-10 |
| B40 | B192 | WBC | UA | 0.220884 | 0 |
| B40 | B222 | WBC | NS | 0.214832 | 0 |
| B40 | B42 | WBC | MON | 0.674285 | 0 |
| B40 | B46 | WBC | RBC Count | 0.2594 | 0 |
| B40 | B51 | WBC | Hb | 0.223165 | 0 |
| B40 | B52 | WBC | PLT/BPC | 0.223021 | 0 |
| B40 | B72 | WBC | HDL-C | -0.1638 | 6.08E-14 |
| B40 | B216 | WBC | BMI | 0.158992 | 3.27E-13 |
| B40 | B53 | WBC | GR% | 0.156558 | 7.51E-13 |
| B40 | B206 | WBC | Waist | 0.153801 | 1.90E-12 |

| | | | | | |
|---|---|---|---|---|---|
| B40 | B41 | WBC | MON％ | -0.15136 | 4.24E-12 |
| B40 | B190 | WBC | Cre | 0.148555 | 1.05E-11 |
| B40 | B48 | WBC | LYM% | -0.13719 | 3.53E-10 |
| B40 | B210 | WBC | Hip | 0.128293 | 4.54E-09 |
| B40 | B215 | WBC | BMI | 0.122333 | 2.29E-08 |
| B40 | B221 | WBC | Hei | 0.121619 | 2.76E-08 |
| B40 | B130 | WBC | GLB | 0.119188 | 5.21E-08 |
| B40 | B43 | WBC | MCV | -0.11385 | 2.01E-07 |
| B40 | B91 | WBC | TP | 0.109876 | 5.27E-07 |
| B41 | B42 | MON％ | MON | 0.599858 | 0 |
| B41 | B53 | MON％ | GR% | -0.1587 | 3.61E-13 |
| B41 | B132 | MON％ | Dbil | 0.157136 | 6.17E-13 |
| B41 | B126 | MON％ | Alb | -0.13976 | 1.63E-10 |
| B41 | B77 | MON％ | TC | -0.12761 | 5.49E-09 |
| B41 | B71 | MON％ | TG | -0.11746 | 8.11E-08 |
| B41 | B189 | MON％ | eGFB | -0.11072 | 4.31E-07 |
| B41 | B70 | MON％ | LDL-C | -0.11045 | 4.60E-07 |
| B42 | B192 | MON | UA | 0.231015 | 0 |
| B42 | B222 | MON | NS | 0.201537 | 0 |
| B42 | B51 | MON | Hb | 0.181779 | 0 |
| B42 | B190 | MON | Cre | 0.172264 | 2.66E-15 |
| B42 | B46 | MON | RBC Count | 0.170619 | 5.11E-15 |
| B42 | B52 | MON | PLT/BPC | 0.16392 | 5.86E-14 |
| B42 | B48 | MON | LYM% | -0.15723 | 5.98E-13 |
| B42 | B77 | MON | TC | -0.14756 | 1.45E-11 |
| B42 | B221 | MON | Hei | 0.138786 | 2.19E-10 |
| B42 | B72 | MON | HDL-C | -0.13804 | 2.74E-10 |
| B42 | B216 | MON | BMI | 0.12687 | 6.72E-09 |
| B42 | B206 | MON | Waist | 0.117762 | 7.51E-08 |
| B42 | B174 | MON | Urea/Cr | -0.112 | 3.16E-07 |
| B43 | B44 | MCV | MCH | 0.88805 | 0 |
| B43 | B46 | MCV | RBC Count | -0.571 | 0 |
| B43 | B47 | MCV | RDW | 0.570257 | 0 |
| B44 | B45 | MCH | MCHC | 0.538201 | 0 |
| B44 | B46 | MCH | RBC Count | -0.5004 | 0 |
| B44 | B47 | MCH | RDW | 0.373901 | 0 |
| B44 | B51 | MCH | Hb | 0.186364 | 0 |
| B45 | B47 | MCHC | RDW | -0.22893 | 0 |
| B45 | B51 | MCHC | Hb | 0.335748 | 0 |
| B45 | B221 | MCHC | Hei | 0.169715 | 7.11E-15 |
| B45 | B129 | MCHC | IBIL | 0.161473 | 1.39E-13 |
| B45 | B222 | MCHC | NS | 0.156506 | 7.64E-13 |
| B45 | B133 | MCHC | TBIL | 0.155474 | 1.08E-12 |
| B45 | B141 | MCHC | AST/ALT | -0.14139 | 9.96E-11 |

| | | | | | |
|---|---|---|---|---|---|
| B45 | B216 | MCHC | BMI | 0.138405 | 2.45E-10 |
| B45 | B189 | MCHC | eGFB | 0.130857 | 2.21E-09 |
| B45 | B126 | MCHC | Alb | 0.123238 | 1.80E-08 |
| B45 | B204 | MCHC | Drinking | 0.113982 | 1.94E-07 |
| B46 | B126 | RBC Count | Alb | 0.225578 | 0 |
| B46 | B163 | RBC Count | SG | 0.195147 | 0 |
| B46 | B174 | RBC Count | Urea/Cr | -0.19127 | 0 |
| B46 | B189 | RBC Count | eGFB | 0.212803 | 0 |
| B46 | B206 | RBC Count | Waist | 0.214556 | 0 |
| B46 | B210 | RBC Count | Hip | 0.183985 | 0 |
| B46 | B216 | RBC Count | BMI | 0.320202 | 0 |
| B46 | B218 | RBC Count | DBP | 0.180117 | 0 |
| B46 | B221 | RBC Count | Hei | 0.332525 | 0 |
| B46 | B222 | RBC Count | NS | 0.333469 | 0 |
| B46 | B47 | RBC Count | RDW | -0.43402 | 0 |
| B46 | B51 | RBC Count | Hb | 0.747719 | 0 |
| B46 | B72 | RBC Count | HDL-C | -0.20137 | 0 |
| B46 | B141 | RBC Count | AST/ALT | -0.17868 | 2.22E-16 |
| B46 | B191 | RBC Count | Urea | -0.16303 | 8.02E-14 |
| B46 | B215 | RBC Count | BMI | 0.15867 | 3.65E-13 |
| B46 | B192 | RBC Count | UA | 0.15401 | 1.77E-12 |
| B46 | B113 | RBC Count | FT3 | 0.13859 | 2.32E-10 |
| B46 | B207 | RBC Count | WHR | 0.131557 | 1.81E-09 |
| B46 | B129 | RBC Count | IBIL | 0.11875 | 5.83E-08 |
| B46 | B209 | RBC Count | Smoking | 0.113137 | 2.39E-07 |
| B46 | B58 | RBC Count | HCT | 0.111199 | 3.84E-07 |
| B46 | B133 | RBC Count | TBIL | 0.111134 | 3.90E-07 |
| B46 | B91 | RBC Count | TP | 0.110367 | 4.69E-07 |
| B47 | B126 | RDW | Alb | -0.18679 | 0 |
| B47 | B189 | RDW | eGFB | -0.20696 | 0 |
| B47 | B51 | RDW | Hb | -0.20636 | 0 |
| B47 | B52 | RDW | PLT/BPC | -0.1741 | 1.33E-15 |
| B47 | B190 | RDW | Cre | 0.152442 | 2.97E-12 |
| B47 | B191 | RDW | Urea | 0.136881 | 3.86E-10 |
| B47 | B127 | RDW | A-G ratio | -0.11972 | 4.53E-08 |
| B47 | B216 | RDW | BMI | -0.11117 | 3.86E-07 |
| B48 | B190 | LYM% | Cre | -0.1833 | 0 |
| B48 | B53 | LYM% | GR% | -0.93078 | 0 |
| B48 | B189 | LYM% | eGFB | 0.162895 | 8.39E-14 |
| B48 | B191 | LYM% | Urea | -0.15015 | 6.29E-12 |
| B48 | B192 | LYM% | UA | -0.11768 | 7.67E-08 |
| B48 | B72 | LYM% | HDL-C | 0.110876 | 4.15E-07 |
| B49 | B50 | baso% | eo% | 0.283038 | 0 |
| B49 | B53 | baso% | GR% | -0.13412 | 8.67E-10 |

| | | | | | |
|---|---|---|---|---|---|
| B49 | B52 | baso% | PLT/BPC | 0.120076 | 4.14E-08 |
| B5 | B6 | WB-Mg | WB-Fe | 0.723597 | 0 |
| B5 | B8 | WB-Mg | WB-Zinc | 0.22807 | 0 |
| B5 | B120 | WB-Mg | HCV-cAg | 0.133361 | 1.08E-09 |
| B5 | B123 | WB-Mg | hs-CRP1 | -0.11834 | 6.47E-08 |
| B50 | B53 | eo% | GR% | -0.23932 | 0 |
| B50 | B190 | eo% | Cre | 0.134352 | 8.11E-10 |
| B51 | B126 | Hb | Alb | 0.262755 | 0 |
| B51 | B141 | Hb | AST/ALT | -0.24592 | 0 |
| B51 | B163 | Hb | SG | 0.211422 | 0 |
| B51 | B174 | Hb | Urea/Cr | -0.25847 | 0 |
| B51 | B189 | Hb | eGFB | 0.257581 | 0 |
| B51 | B191 | Hb | Urea | -0.23004 | 0 |
| B51 | B206 | Hb | Waist | 0.247425 | 0 |
| B51 | B210 | Hb | Hip | 0.224961 | 0 |
| B51 | B216 | Hb | BMI | 0.393776 | 0 |
| B51 | B218 | Hb | DBP | 0.235244 | 0 |
| B51 | B221 | Hb | Hei | 0.429786 | 0 |
| B51 | B222 | Hb | NS | 0.402375 | 0 |
| B51 | B72 | Hb | HDL-C | -0.21169 | 0 |
| B51 | B215 | Hb | BMI | 0.176317 | 4.44E-16 |
| B51 | B129 | Hb | IBIL | 0.175848 | 8.88E-16 |
| B51 | B133 | Hb | TBIL | 0.175568 | 8.88E-16 |
| B51 | B209 | Hb | Smoking | 0.164189 | 5.33E-14 |
| B51 | B192 | Hb | UA | 0.149003 | 9.12E-12 |
| B51 | B127 | Hb | A-G ratio | 0.139497 | 1.77E-10 |
| B51 | B207 | Hb | WHR | 0.137093 | 3.63E-10 |
| B51 | B58 | Hb | HCT | 0.134728 | 7.27E-10 |
| B51 | B132 | Hb | Dbil | 0.128628 | 4.13E-09 |
| B51 | B204 | Hb | Drinking | 0.127812 | 5.18E-09 |
| B51 | B113 | Hb | FT3 | 0.126148 | 8.19E-09 |
| B51 | B173 | Hb | NTT | -0.11624 | 1.10E-07 |
| B51 | B188 | Hb | Ca | 0.112424 | 2.85E-07 |
| B52 | B132 | PLT/BPC | Dbil | -0.20031 | 0 |
| B52 | B133 | PLT/BPC | TBIL | -0.14799 | 1.26E-11 |
| B52 | B70 | PLT/BPC | LDL-C | 0.135978 | 5.04E-10 |
| B52 | B129 | PLT/BPC | IBIL | -0.11737 | 8.30E-08 |
| B52 | B190 | PLT/BPC | Cre | -0.11068 | 4.35E-07 |
| B53 | B190 | GR% | Cre | 0.11915 | 5.26E-08 |
| B53 | B191 | GR% | Urea | 0.1138 | 2.03E-07 |
| B53 | B189 | GR% | eGFB | -0.11338 | 2.26E-07 |
| B53 | B153 | GR% | Glu | 0.109031 | 6.45E-07 |
| B54 | B102 | ESR | anti-Sm | 0.20821 | 0 |
| B54 | B103 | ESR | anti-SSA/Ro | 0.208423 | 0 |

| | | | | | |
|---|---|---|---|---|---|
| B54 | B105 | ESR | anti-U1RNP | 0.187345 | 0 |
| B54 | B57 | ESR | RBCAI | -0.19769 | 0 |
| B54 | B58 | ESR | HCT | -0.44649 | 0 |
| B54 | B60 | ESR | WBRVI | -0.37464 | 0 |
| B54 | B62 | ESR | WBHSRVI | -0.3423 | 0 |
| B54 | B69 | ESR | PV | 0.518071 | 0 |
| B54 | B156 | ESR | IgE | 0.167278 | 1.75E-14 |
| B54 | B56 | ESR | IR | -0.15554 | 1.06E-12 |
| B54 | B74 | ESR | ApoA1 | 0.154913 | 1.31E-12 |
| B54 | B65 | ESR | WBRSR(mPas_1) | -0.14159 | 9.37E-11 |
| B54 | B94 | ESR | ALDH2-Lys504Lys | 0.141131 | 1.08E-10 |
| B54 | B73 | ESR | ApoA | 0.138992 | 2.06E-10 |
| B54 | B93 | ESR | ALDH2-Glu504Lys | -0.11076 | 4.27E-07 |
| B55 | B102 | TK | anti-Sm | -0.36132 | 0 |
| B55 | B105 | TK | anti-U1RNP | -0.37649 | 0 |
| B55 | B109 | TK | AMA-M2 | 0.236557 | 0 |
| B55 | B156 | TK | IgE | -0.18281 | 0 |
| B55 | B56 | TK | IR | 0.874343 | 0 |
| B55 | B58 | TK | HCT | -0.32265 | 0 |
| B55 | B59 | TK | WBRV | 0.611619 | 0 |
| B55 | B60 | TK | WBRVI | 0.377932 | 0 |
| B55 | B61 | TK | WBHSRV | 0.802596 | 0 |
| B55 | B62 | TK | WBHSRVI | 0.55019 | 0 |
| B55 | B63 | TK | WBRSR(mPas_200) | 0.4281 | 0 |
| B55 | B64 | TK | WBRSR(mPas_5) | 0.369617 | 0 |
| B55 | B65 | TK | WBRSR(mPas_1) | 0.304004 | 0 |
| B55 | B66 | TK | WBRSR(mPas_100) | 0.479927 | 0 |
| B55 | B67 | TK | WBRSR(mPas_50) | 0.471573 | 0 |
| B55 | B69 | TK | PV | -0.29537 | 0 |
| B55 | B75 | TK | ApoA1/B | 0.181461 | 0 |
| B55 | B125 | TK | Cor | 0.164409 | 4.93E-14 |
| B55 | B124 | TK | ACTH | 0.145895 | 2.46E-11 |
| B55 | B76 | TK | ApoB | -0.13434 | 8.14E-10 |
| B56 | B102 | IR | anti-Sm | -0.32749 | 0 |
| B56 | B105 | IR | anti-U1RNP | -0.33102 | 0 |
| B56 | B59 | IR | WBRV | 0.657759 | 0 |
| B56 | B60 | IR | WBRVI | 0.664707 | 0 |
| B56 | B61 | IR | WBHSRV | 0.910284 | 0 |
| B56 | B62 | IR | WBHSRVI | 0.877751 | 0 |
| B56 | B63 | IR | WBRSR(mPas_200) | 0.739452 | 0 |
| B56 | B64 | IR | WBRSR(mPas_5) | 0.678443 | 0 |
| B56 | B65 | IR | WBRSR(mPas_1) | 0.58907 | 0 |
| B56 | B66 | IR | WBRSR(mPas_100) | 0.772737 | 0 |
| B56 | B67 | IR | WBRSR(mPas_50) | 0.768218 | 0 |

| | | | | | |
|---|---|---|---|---|---|
| B56 | B69 | IR | PV | -0.31188 | 0 |
| B56 | B58 | IR | HCT | 0.151207 | 4.46E-12 |
| B56 | B75 | IR | ApoA1/B | 0.142089 | 8.03E-11 |
| B56 | B76 | IR | ApoB | -0.13832 | 2.52E-10 |
| B56 | B124 | IR | ACTH | 0.129402 | 3.33E-09 |
| B56 | B73 | IR | ApoA | 0.120491 | 3.71E-08 |
| B56 | B125 | IR | Cor | 0.117044 | 9.02E-08 |
| B56 | B109 | IR | AMA-M2 | 0.11316 | 2.38E-07 |
| B57 | B100 | RBCAI | anti-Ro52 | -0.19079 | 0 |
| B57 | B101 | RBCAI | anti-Sc70 | -0.19079 | 0 |
| B57 | B104 | RBCAI | anti-SSB | -0.19079 | 0 |
| B57 | B106 | RBCAI | anti-P | -0.19079 | 0 |
| B57 | B107 | RBCAI | AnuA | -0.19079 | 0 |
| B57 | B110 | RBCAI | AHA | -0.19079 | 0 |
| B57 | B59 | RBCAI | WBRV | 0.555059 | 0 |
| B57 | B60 | RBCAI | WBRVI | 0.650941 | 0 |
| B57 | B64 | RBCAI | WBRSR(mPas_5) | 0.390919 | 0 |
| B57 | B65 | RBCAI | WBRSR(mPas_1) | 0.595561 | 0 |
| B57 | B69 | RBCAI | PV | -0.20198 | 0 |
| B57 | B95 | RBCAI | anti-cenpb | -0.19079 | 0 |
| B57 | B97 | RBCAI | dsDNA | -0.19079 | 0 |
| B57 | B98 | RBCAI | anti-Jo-1 | -0.19079 | 0 |
| B57 | B99 | RBCAI | anti-PM-Scl | -0.19079 | 0 |
| B57 | B157 | RBCAI | IgG | -0.16615 | 2.64E-14 |
| B57 | B184 | RBCAI | INR | -0.1642 | 5.28E-14 |
| B57 | B58 | RBCAI | HCT | 0.162359 | 1.01E-13 |
| B57 | B94 | RBCAI | ALDH2-Lys504Lys | -0.15736 | 5.72E-13 |
| B57 | B93 | RBCAI | ALDH2-Glu504Lys | 0.138098 | 2.69E-10 |
| B57 | B92 | RBCAI | ALDH2-Glu504Glu | -0.13435 | 8.11E-10 |
| B57 | B102 | RBCAI | anti-Sm | -0.13409 | 8.75E-10 |
| B57 | B108 | RBCAI | Anti-Cyclin | 0.129992 | 2.82E-09 |
| B57 | B156 | RBCAI | IgE | -0.12624 | 7.98E-09 |
| B57 | B109 | RBCAI | AMA-M2 | -0.11948 | 4.83E-08 |
| B57 | B183 | RBCAI | PTA | -0.11742 | 8.20E-08 |
| B57 | B185 | RBCAI | PT | -0.11273 | 2.64E-07 |
| B58 | B109 | HCT | AMA-M2 | -0.23809 | 0 |
| B58 | B60 | HCT | WBRVI | 0.553331 | 0 |
| B58 | B62 | HCT | WBHSRVI | 0.59494 | 0 |
| B58 | B63 | HCT | WBRSR(mPas_200) | 0.524321 | 0 |
| B58 | B64 | HCT | WBRSR(mPas_5) | 0.538985 | 0 |
| B58 | B65 | HCT | WBRSR(mPas_1) | 0.514321 | 0 |
| B58 | B66 | HCT | WBRSR(mPas_100) | 0.468438 | 0 |
| B58 | B67 | HCT | WBRSR(mPas_50) | 0.477662 | 0 |
| B58 | B96 | HCT | dsDNA (IIF) | 0.170963 | 4.44E-15 |

| | | | | | |
|---|---|---|---|---|---|
| B58 | B156 | HCT | IgE | 0.1666 | 2.24E-14 |
| B58 | B155 | HCT | IgA | 0.132438 | 1.41E-09 |
| B58 | B69 | HCT | PV | -0.12887 | 3.87E-09 |
| B58 | B75 | HCT | ApoA1/B | -0.12555 | 9.63E-09 |
| B58 | B197 | HCT | AG | -0.11675 | 9.72E-08 |
| B58 | B196 | HCT | TCO2 | 0.113572 | 2.15E-07 |
| B58 | B114 | HCT | TSH TT3 TT4(TT4) | -0.11023 | 4.84E-07 |
| B59 | B60 | WBRV | WBRVI | 0.717387 | 0 |
| B59 | B61 | WBRV | WBHSRV | 0.755449 | 0 |
| B59 | B62 | WBRV | WBHSRVI | 0.497084 | 0 |
| B59 | B63 | WBRV | WBRSR(mPas_200) | 0.609618 | 0 |
| B59 | B64 | WBRV | WBRSR(mPas_5) | 0.812095 | 0 |
| B59 | B65 | WBRV | WBRSR(mPas_1) | 0.84867 | 0 |
| B59 | B66 | WBRV | WBRSR(mPas_100) | 0.640775 | 0 |
| B59 | B67 | WBRV | WBRSR(mPas_50) | 0.674756 | 0 |
| B59 | B75 | WBRV | ApoA1/B | 0.180326 | 0 |
| B59 | B100 | WBRV | anti-Ro52 | -0.15982 | 2.46E-13 |
| B59 | B101 | WBRV | anti-Sc70 | -0.15982 | 2.46E-13 |
| B59 | B104 | WBRV | anti-SSB | -0.15982 | 2.46E-13 |
| B59 | B106 | WBRV | anti-P | -0.15982 | 2.46E-13 |
| B59 | B107 | WBRV | AnuA | -0.15982 | 2.46E-13 |
| B59 | B110 | WBRV | AHA | -0.15982 | 2.46E-13 |
| B59 | B95 | WBRV | anti-cenpb | -0.15982 | 2.46E-13 |
| B59 | B97 | WBRV | dsDNA | -0.15982 | 2.46E-13 |
| B59 | B98 | WBRV | anti-Jo-1 | -0.15982 | 2.46E-13 |
| B59 | B99 | WBRV | anti-PM-Scl | -0.15982 | 2.46E-13 |
| B59 | B76 | WBRV | ApoB | -0.15074 | 5.20E-12 |
| B59 | B69 | WBRV | PV | 0.115186 | 1.44E-07 |
| B59 | B73 | WBRV | ApoA | 0.112528 | 2.78E-07 |
| B6 | B7 | WB-Fe | WB-Cop | -0.23476 | 0 |
| B6 | B8 | WB-Fe | WB-Zinc | 0.590005 | 0 |
| B6 | B123 | WB-Fe | hs-CRP1 | -0.15593 | 9.27E-13 |
| B6 | B85 | WB-Fe | SCC | -0.15103 | 4.73E-12 |
| B6 | B86 | WB-Fe | TBhCG | 0.135194 | 6.35E-10 |
| B60 | B102 | WBRVI | anti-Sm | -0.30319 | 0 |
| B60 | B105 | WBRVI | anti-U1RNP | -0.27361 | 0 |
| B60 | B61 | WBRVI | WBHSRV | 0.530508 | 0 |
| B60 | B62 | WBRVI | WBHSRVI | 0.796344 | 0 |
| B60 | B63 | WBRVI | WBRSR(mPas_200) | 0.603617 | 0 |
| B60 | B64 | WBRVI | WBRSR(mPas_5) | 0.836374 | 0 |
| B60 | B65 | WBRVI | WBRSR(mPas_1) | 0.895158 | 0 |
| B60 | B66 | WBRVI | WBRSR(mPas_100) | 0.641076 | 0 |
| B60 | B67 | WBRVI | WBRSR(mPas_50) | 0.680527 | 0 |
| B60 | B69 | WBRVI | PV | -0.37162 | 0 |

| | | | | | |
|---|---|---|---|---|---|
| B60 | B93 | WBRVI | ALDH2-Glu504Lys | 0.173717 | 1.78E-15 |
| B60 | B92 | WBRVI | ALDH2-Glu504Glu | -0.1709 | 4.66E-15 |
| B60 | B94 | WBRVI | ALDH2-Lys504Lys | -0.15401 | 1.77E-12 |
| B60 | B76 | WBRVI | ApoB | -0.14009 | 1.48E-10 |
| B60 | B100 | WBRVI | anti-Ro52 | -0.12681 | 6.84E-09 |
| B60 | B101 | WBRVI | anti-Sc70 | -0.12681 | 6.84E-09 |
| B60 | B104 | WBRVI | anti-SSB | -0.12681 | 6.84E-09 |
| B60 | B106 | WBRVI | anti-P | -0.12681 | 6.84E-09 |
| B60 | B107 | WBRVI | AnuA | -0.12681 | 6.84E-09 |
| B60 | B110 | WBRVI | AHA | -0.12681 | 6.84E-09 |
| B60 | B95 | WBRVI | anti-cenpb | -0.12681 | 6.84E-09 |
| B60 | B97 | WBRVI | dsDNA | -0.12681 | 6.84E-09 |
| B60 | B98 | WBRVI | anti-Jo-1 | -0.12681 | 6.84E-09 |
| B60 | B99 | WBRVI | anti-PM-Scl | -0.12681 | 6.84E-09 |
| B60 | B125 | WBRVI | Cor | 0.126267 | 7.93E-09 |
| B61 | B62 | WBHSRV | WBHSRVI | 0.752583 | 0 |
| B61 | B63 | WBHSRV | WBRSR(mPas_200) | 0.860319 | 0 |
| B61 | B64 | WBHSRV | WBRSR(mPas_5) | 0.764671 | 0 |
| B61 | B65 | WBHSRV | WBRSR(mPas_1) | 0.646642 | 0 |
| B61 | B66 | WBHSRV | WBRSR(mPas_100) | 0.858801 | 0 |
| B61 | B67 | WBHSRV | WBRSR(mPas_50) | 0.850463 | 0 |
| B61 | B105 | WBHSRV | anti-U1RNP | -0.15511 | 1.22E-12 |
| B61 | B73 | WBHSRV | ApoA | 0.151748 | 3.73E-12 |
| B61 | B102 | WBHSRV | anti-Sm | -0.14516 | 3.10E-11 |
| B61 | B75 | WBHSRV | ApoA1/B | 0.135952 | 5.08E-10 |
| B61 | B109 | WBHSRV | AMA-M2 | 0.128237 | 4.61E-09 |
| B61 | B96 | WBHSRV | dsDNA (IIF) | 0.120998 | 3.25E-08 |
| B61 | B76 | WBHSRV | ApoB | -0.10821 | 7.82E-07 |
| B62 | B102 | WBHSRVI | anti-Sm | -0.31218 | 0 |
| B62 | B105 | WBHSRVI | anti-U1RNP | -0.30357 | 0 |
| B62 | B63 | WBHSRVI | WBRSR(mPas_200) | 0.82684 | 0 |
| B62 | B64 | WBHSRVI | WBRSR(mPas_5) | 0.780748 | 0 |
| B62 | B65 | WBHSRVI | WBRSR(mPas_1) | 0.695639 | 0 |
| B62 | B66 | WBHSRVI | WBRSR(mPas_100) | 0.830438 | 0 |
| B62 | B67 | WBHSRVI | WBRSR(mPas_50) | 0.830203 | 0 |
| B62 | B69 | WBHSRVI | PV | -0.34745 | 0 |
| B62 | B96 | WBHSRVI | dsDNA (IIF) | 0.167919 | 1.38E-14 |
| B62 | B93 | WBHSRVI | ALDH2-Glu504Lys | 0.120301 | 3.90E-08 |
| B62 | B92 | WBHSRVI | ALDH2-Glu504Glu | -0.11958 | 4.70E-08 |
| B62 | B124 | WBHSRVI | ACTH | 0.118056 | 6.97E-08 |
| B62 | B195 | WBHSRVI | Na | -0.11756 | 7.91E-08 |
| B62 | B155 | WBHSRVI | IgA | 0.116073 | 1.15E-07 |
| B63 | B64 | WBRSR(mPas_200) | WBRSR(mPas_5) | 0.896778 | 0 |
| B63 | B65 | WBRSR(mPas_200) | WBRSR(mPas_1) | 0.768705 | 0 |

| | | | | | |
|---|---|---|---|---|---|
| B63 | B66 | WBRSR(mPas_200) | WBRSR(mPas_100) | 0.953376 | 0 |
| B63 | B67 | WBRSR(mPas_200) | WBRSR(mPas_50) | 0.946992 | 0 |
| B63 | B69 | WBRSR(mPas_200) | PV | 0.224471 | 0 |
| B63 | B96 | WBRSR(mPas_200) | dsDNA (IIF) | 0.198301 | 0 |
| B63 | B182 | WBRSR(mPas_200) | TT | -0.15973 | 2.54E-13 |
| B63 | B157 | WBRSR(mPas_200) | IgG | 0.153057 | 2.43E-12 |
| B63 | B156 | WBRSR(mPas_200) | IgE | 0.14025 | 1.41E-10 |
| B63 | B155 | WBRSR(mPas_200) | IgA | 0.140135 | 1.46E-10 |
| B63 | B185 | WBRSR(mPas_200) | PT | 0.128025 | 4.89E-09 |
| B63 | B195 | WBRSR(mPas_200) | Na | -0.11785 | 7.34E-08 |
| B63 | B73 | WBRSR(mPas_200) | ApoA | 0.115192 | 1.44E-07 |
| B63 | B123 | WBRSR(mPas_200) | hs-CRP1 | 0.112521 | 2.78E-07 |
| B64 | B65 | WBRSR(mPas_5) | WBRSR(mPas_1) | 0.971511 | 0 |
| B64 | B66 | WBRSR(mPas_5) | WBRSR(mPas_100) | 0.881656 | 0 |
| B64 | B67 | WBRSR(mPas_5) | WBRSR(mPas_50) | 0.9046 | 0 |
| B64 | B96 | WBRSR(mPas_5) | dsDNA (IIF) | 0.150826 | 5.05E-12 |
| B64 | B123 | WBRSR(mPas_5) | hs-CRP1 | 0.138812 | 2.17E-10 |
| B64 | B69 | WBRSR(mPas_5) | PV | 0.13498 | 6.75E-10 |
| B64 | B93 | WBRSR(mPas_5) | ALDH2-Glu504Lys | 0.126762 | 6.92E-09 |
| B64 | B92 | WBRSR(mPas_5) | ALDH2-Glu504Glu | -0.12642 | 7.60E-09 |
| B64 | B155 | WBRSR(mPas_5) | IgA | 0.121544 | 2.82E-08 |
| B64 | B100 | WBRSR(mPas_5) | anti-Ro52 | -0.11736 | 8.32E-08 |
| B64 | B101 | WBRSR(mPas_5) | anti-Sc70 | -0.11736 | 8.32E-08 |
| B64 | B104 | WBRSR(mPas_5) | anti-SSB | -0.11736 | 8.32E-08 |
| B64 | B106 | WBRSR(mPas_5) | anti-P | -0.11736 | 8.32E-08 |
| B64 | B107 | WBRSR(mPas_5) | AnuA | -0.11736 | 8.32E-08 |
| B64 | B110 | WBRSR(mPas_5) | AHA | -0.11736 | 8.32E-08 |
| B64 | B95 | WBRSR(mPas_5) | anti-cenpb | -0.11736 | 8.32E-08 |
| B64 | B97 | WBRSR(mPas_5) | dsDNA | -0.11736 | 8.32E-08 |
| B64 | B98 | WBRSR(mPas_5) | anti-Jo-1 | -0.11736 | 8.32E-08 |
| B64 | B99 | WBRSR(mPas_5) | anti-PM-Scl | -0.11736 | 8.32E-08 |
| B64 | B182 | WBRSR(mPas_5) | TT | -0.10952 | 5.74E-07 |
| B64 | B195 | WBRSR(mPas_5) | Na | -0.10862 | 7.11E-07 |
| B65 | B66 | WBRSR(mPas_1) | WBRSR(mPas_100) | 0.768386 | 0 |
| B65 | B67 | WBRSR(mPas_1) | WBRSR(mPas_50) | 0.804901 | 0 |
| B65 | B100 | WBRSR(mPas_1) | anti-Ro52 | -0.14979 | 7.07E-12 |
| B65 | B101 | WBRSR(mPas_1) | anti-Sc70 | -0.14979 | 7.07E-12 |
| B65 | B104 | WBRSR(mPas_1) | anti-SSB | -0.14979 | 7.07E-12 |
| B65 | B106 | WBRSR(mPas_1) | anti-P | -0.14979 | 7.07E-12 |
| B65 | B107 | WBRSR(mPas_1) | AnuA | -0.14979 | 7.07E-12 |
| B65 | B110 | WBRSR(mPas_1) | AHA | -0.14979 | 7.07E-12 |
| B65 | B95 | WBRSR(mPas_1) | anti-cenpb | -0.14979 | 7.07E-12 |
| B65 | B97 | WBRSR(mPas_1) | dsDNA | -0.14979 | 7.07E-12 |
| B65 | B98 | WBRSR(mPas_1) | anti-Jo-1 | -0.14979 | 7.07E-12 |

| | | | | | |
|---|---|---|---|---|---|
| B65 | B99 | WBRSR(mPas_1) | anti-PM-Scl | -0.14979 | 7.07E-12 |
| B65 | B93 | WBRSR(mPas_1) | ALDH2-Glu504Lys | 0.143796 | 4.74E-11 |
| B65 | B92 | WBRSR(mPas_1) | ALDH2-Glu504Glu | -0.1426 | 6.87E-11 |
| B65 | B123 | WBRSR(mPas_1) | hs-CRP1 | 0.139492 | 1.77E-10 |
| B65 | B96 | WBRSR(mPas_1) | dsDNA (IIF) | 0.116376 | 1.07E-07 |
| B65 | B76 | WBRSR(mPas_1) | ApoB | -0.10936 | 5.97E-07 |
| B66 | B67 | WBRSR(mPas_100) | WBRSR(mPas_50) | 0.997582 | 0 |
| B66 | B96 | WBRSR(mPas_100) | dsDNA (IIF) | 0.195885 | 0 |
| B66 | B155 | WBRSR(mPas_100) | IgA | 0.157931 | 4.71E-13 |
| B66 | B157 | WBRSR(mPas_100) | IgG | 0.157818 | 4.89E-13 |
| B66 | B69 | WBRSR(mPas_100) | PV | 0.155539 | 1.06E-12 |
| B66 | B182 | WBRSR(mPas_100) | TT | -0.15032 | 5.95E-12 |
| B66 | B156 | WBRSR(mPas_100) | IgE | 0.129713 | 3.05E-09 |
| B66 | B73 | WBRSR(mPas_100) | ApoA | 0.126354 | 7.74E-09 |
| B66 | B195 | WBRSR(mPas_100) | Na | -0.12412 | 1.42E-08 |
| B66 | B123 | WBRSR(mPas_100) | hs-CRP1 | 0.12057 | 3.64E-08 |
| B66 | B185 | WBRSR(mPas_100) | PT | 0.120047 | 4.17E-08 |
| B67 | B96 | WBRSR(mPas_50) | dsDNA (IIF) | 0.189785 | 0 |
| B67 | B155 | WBRSR(mPas_50) | IgA | 0.15587 | 9.48E-13 |
| B67 | B69 | WBRSR(mPas_50) | PV | 0.146308 | 2.16E-11 |
| B67 | B157 | WBRSR(mPas_50) | IgG | 0.145326 | 2.94E-11 |
| B67 | B182 | WBRSR(mPas_50) | TT | -0.14039 | 1.35E-10 |
| B67 | B123 | WBRSR(mPas_50) | hs-CRP1 | 0.126393 | 7.66E-09 |
| B67 | B156 | WBRSR(mPas_50) | IgE | 0.126216 | 8.04E-09 |
| B67 | B73 | WBRSR(mPas_50) | ApoA | 0.125507 | 9.75E-09 |
| B67 | B195 | WBRSR(mPas_50) | Na | -0.12287 | 1.98E-08 |
| B67 | B185 | WBRSR(mPas_50) | PT | 0.110386 | 4.67E-07 |
| B67 | B194 | WBRSR(mPas_50) | Cl | -0.10977 | 5.41E-07 |
| B69 | B102 | PV | anti-Sm | 0.627639 | 0 |
| B69 | B105 | PV | anti-U1RNP | 0.604828 | 0 |
| B69 | B125 | PV | Cor | -0.2453 | 0 |
| B69 | B138 | PV | PAB | -0.22453 | 0 |
| B69 | B156 | PV | IgE | 0.230995 | 0 |
| B69 | B185 | PV | PT | 0.2679 | 0 |
| B69 | B124 | PV | ACTH | -0.17937 | 2.22E-16 |
| B69 | B186 | PV | PFC | 0.167264 | 1.78E-14 |
| B69 | B181 | PV | APTT | 0.155367 | 1.12E-12 |
| B69 | B184 | PV | INR | 0.137843 | 2.90E-10 |
| B69 | B139 | PV | ADA | 0.132056 | 1.57E-09 |
| B69 | B120 | PV | HCV-cAg | -0.11966 | 4.61E-08 |
| B69 | B100 | PV | anti-Ro52 | -0.11922 | 5.17E-08 |
| B69 | B101 | PV | anti-Sc70 | -0.11922 | 5.17E-08 |
| B69 | B104 | PV | anti-SSB | -0.11922 | 5.17E-08 |
| B69 | B106 | PV | anti-P | -0.11922 | 5.17E-08 |

| | | | | | |
|---|---|---|---|---|---|
| B69 | B107 | PV | AnuA | -0.11922 | 5.17E-08 |
| B69 | B110 | PV | AHA | -0.11922 | 5.17E-08 |
| B69 | B95 | PV | anti-cenpb | -0.11922 | 5.17E-08 |
| B69 | B97 | PV | dsDNA | -0.11922 | 5.17E-08 |
| B69 | B98 | PV | anti-Jo-1 | -0.11922 | 5.17E-08 |
| B69 | B99 | PV | anti-PM-Scl | -0.11922 | 5.17E-08 |
| B69 | B182 | PV | TT | -0.11496 | 1.52E-07 |
| B69 | B228 | PV | OS | 0.111954 | 3.20E-07 |
| B7 | B197 | WB-Cop | AG | -0.18106 | 0 |
| B7 | B8 | WB-Cop | WB-Zinc | 0.401874 | 0 |
| B7 | B196 | WB-Cop | TCO2 | 0.127191 | 6.15E-09 |
| B70 | B132 | LDL-C | Dbil | -0.30711 | 0 |
| B70 | B76 | LDL-C | ApoB | 0.232088 | 0 |
| B70 | B77 | LDL-C | TC | 0.894083 | 0 |
| B70 | B75 | LDL-C | ApoA1/B | -0.17666 | 4.44E-16 |
| B70 | B220 | LDL-C | SBP | 0.167237 | 1.78E-14 |
| B70 | B91 | LDL-C | TP | 0.163778 | 6.15E-14 |
| B70 | B130 | LDL-C | GLB | 0.160707 | 1.81E-13 |
| B70 | B133 | LDL-C | TBIL | -0.15675 | 7.03E-13 |
| B70 | B218 | LDL-C | DBP | 0.150072 | 6.45E-12 |
| B70 | B127 | LDL-C | A-G ratio | -0.13201 | 1.59E-09 |
| B70 | B219 | LDL-C | OT | -0.10818 | 7.88E-07 |
| B71 | B132 | TG | Dbil | -0.18427 | 0 |
| B71 | B153 | TG | Glu | 0.204701 | 0 |
| B71 | B206 | TG | Waist | 0.186522 | 0 |
| B71 | B215 | TG | BMI | 0.225808 | 0 |
| B71 | B216 | TG | BMI | 0.19205 | 0 |
| B71 | B72 | TG | HDL-C | -0.36876 | 0 |
| B71 | B77 | TG | TC | 0.281525 | 0 |
| B71 | B91 | TG | TP | 0.206669 | 0 |
| B71 | B192 | TG | UA | 0.162318 | 1.03E-13 |
| B71 | B210 | TG | Hip | 0.161448 | 1.40E-13 |
| B71 | B130 | TG | GLB | 0.142316 | 7.49E-11 |
| B71 | B126 | TG | Alb | 0.129953 | 2.85E-09 |
| B71 | B141 | TG | AST/ALT | -0.1285 | 4.28E-09 |
| B71 | B222 | TG | NS | 0.124097 | 1.43E-08 |
| B71 | B140 | TG | r-GT | 0.118755 | 5.82E-08 |
| B72 | B141 | HDL-C | AST/ALT | 0.239425 | 0 |
| B72 | B192 | HDL-C | UA | -0.33507 | 0 |
| B72 | B206 | HDL-C | Waist | -0.34279 | 0 |
| B72 | B207 | HDL-C | WHR | -0.21362 | 0 |
| B72 | B210 | HDL-C | Hip | -0.26256 | 0 |
| B72 | B215 | HDL-C | BMI | -0.32139 | 0 |
| B72 | B216 | HDL-C | BMI | -0.41338 | 0 |

| | | | | | |
|---|---|---|---|---|---|
| B72 | B221 | HDL-C | Hei | -0.29805 | 0 |
| B72 | B222 | HDL-C | NS | -0.35151 | 0 |
| B72 | B74 | HDL-C | ApoA1 | 0.248652 | 0 |
| B72 | B77 | HDL-C | TC | 0.288421 | 0 |
| B72 | B174 | HDL-C | Urea/Cr | 0.147612 | 1.42E-11 |
| B72 | B112 | HDL-C | FT4 | 0.142874 | 6.31E-11 |
| B72 | B163 | HDL-C | SG | -0.14282 | 6.41E-11 |
| B72 | B190 | HDL-C | Cre | -0.14118 | 1.06E-10 |
| B72 | B75 | HDL-C | ApoA1/B | 0.131193 | 2.01E-09 |
| B72 | B128 | HDL-C | ALT | -0.11846 | 6.29E-08 |
| B72 | B153 | HDL-C | Glu | -0.11623 | 1.11E-07 |
| B72 | B140 | HDL-C | r-GT | -0.11562 | 1.29E-07 |
| B73 | B200 | ApoA | OP | 0.12417 | 1.40E-08 |
| B73 | B196 | ApoA | TCO2 | 0.114981 | 1.52E-07 |
| B73 | B74 | ApoA | ApoA1 | 0.10857 | 7.19E-07 |
| B74 | B75 | ApoA1 | ApoA1/B | 0.462701 | 0 |
| B74 | B156 | ApoA1 | IgE | 0.165703 | 3.11E-14 |
| B74 | B195 | ApoA1 | Na | -0.16265 | 9.15E-14 |
| B74 | B216 | ApoA1 | BMI | -0.15862 | 3.72E-13 |
| B74 | B200 | ApoA1 | OP | -0.1532 | 2.31E-12 |
| B74 | B151 | ApoA1 | Anti-CCP | 0.137985 | 2.78E-10 |
| B74 | B221 | ApoA1 | Hei | -0.1347 | 7.34E-10 |
| B74 | B94 | ApoA1 | ALDH2-Lys504Lys | 0.134621 | 7.50E-10 |
| B74 | B181 | ApoA1 | APTT | 0.134583 | 7.58E-10 |
| B74 | B206 | ApoA1 | Waist | -0.12092 | 3.32E-08 |
| B74 | B136 | ApoA1 | 5'NT | -0.12007 | 4.14E-08 |
| B74 | B141 | ApoA1 | AST/ALT | 0.115233 | 1.42E-07 |
| B74 | B108 | ApoA1 | Anti-Cyclin | -0.11516 | 1.45E-07 |
| B74 | B81 | ApoA1 | TSGF | -0.11022 | 4.86E-07 |
| B74 | B222 | ApoA1 | NS | -0.10936 | 5.97E-07 |
| B75 | B76 | ApoA1/B | ApoB | -0.76249 | 0 |
| B75 | B197 | ApoA1/B | AG | -0.14704 | 1.71E-11 |
| B75 | B135 | ApoA1/B | TBA | 0.128806 | 3.93E-09 |
| B75 | B77 | ApoA1/B | TC | -0.12513 | 1.08E-08 |
| B75 | B199 | ApoA1/B | Mg | -0.12158 | 2.79E-08 |
| B75 | B139 | ApoA1/B | ADA | -0.11971 | 4.55E-08 |
| B75 | B90 | ApoA1/B | CYFRA 21-1 | 0.116256 | 1.10E-07 |
| B75 | B134 | ApoA1/B | CHE | -0.11099 | 4.04E-07 |
| B76 | B197 | ApoB | AG | 0.198419 | 0 |
| B76 | B77 | ApoB | TC | 0.215737 | 0 |
| B76 | B100 | ApoB | anti-Ro52 | 0.165585 | 3.24E-14 |
| B76 | B101 | ApoB | anti-Sc70 | 0.165585 | 3.24E-14 |
| B76 | B104 | ApoB | anti-SSB | 0.165585 | 3.24E-14 |
| B76 | B106 | ApoB | anti-P | 0.165585 | 3.24E-14 |

| | | | | | |
|---|---|---|---|---|---|
| B76 | B107 | ApoB | AnuA | 0.165585 | 3.24E-14 |
| B76 | B110 | ApoB | AHA | 0.165585 | 3.24E-14 |
| B76 | B95 | ApoB | anti-cenpb | 0.165585 | 3.24E-14 |
| B76 | B97 | ApoB | dsDNA | 0.165585 | 3.24E-14 |
| B76 | B98 | ApoB | anti-Jo-1 | 0.165585 | 3.24E-14 |
| B76 | B99 | ApoB | anti-PM-Scl | 0.165585 | 3.24E-14 |
| B76 | B134 | ApoB | CHE | 0.147647 | 1.41E-11 |
| B76 | B193 | ApoB | K | 0.135304 | 6.15E-10 |
| B76 | B194 | ApoB | Cl | -0.12598 | 8.58E-09 |
| B76 | B135 | ApoB | TBA | -0.11692 | 9.30E-08 |
| B76 | B159 | ApoB | β2-MG | 0.11375 | 2.06E-07 |
| B76 | B181 | ApoB | APTT | 0.109559 | 5.69E-07 |
| B77 | B132 | TC | Dbil | -0.31191 | 0 |
| B77 | B220 | TC | SBP | 0.192433 | 0 |
| B77 | B221 | TC | Hei | -0.19209 | 0 |
| B77 | B91 | TC | TP | 0.174294 | 1.33E-15 |
| B77 | B130 | TC | GLB | 0.159681 | 2.58E-13 |
| B77 | B133 | TC | TBIL | -0.15509 | 1.23E-12 |
| B77 | B222 | TC | NS | -0.13906 | 2.01E-10 |
| B77 | B218 | TC | DBP | 0.13478 | 7.16E-10 |
| B77 | B219 | TC | OT | -0.13141 | 1.89E-09 |
| B77 | B127 | TC | A-G ratio | -0.12371 | 1.58E-08 |
| B77 | B216 | TC | BMI | -0.12168 | 2.72E-08 |
| B77 | B174 | TC | Urea/Cr | 0.109743 | 5.44E-07 |
| B77 | B190 | TC | Cre | -0.10791 | 8.40E-07 |
| B78 | B96 | HBcAb | dsDNA (IIF) | -0.23949 | 0 |
| B78 | B80 | HBcAb | anti-HP IgG | -0.14201 | 8.22E-11 |
| B79 | B100 | hb V -dna | anti-Ro52 | 0.246533 | 0 |
| B79 | B101 | hb V -dna | anti-Sc70 | 0.246533 | 0 |
| B79 | B104 | hb V -dna | anti-SSB | 0.246533 | 0 |
| B79 | B106 | hb V -dna | anti-P | 0.246533 | 0 |
| B79 | B107 | hb V -dna | AnuA | 0.246533 | 0 |
| B79 | B108 | hb V -dna | Anti-Cyclin | -0.26259 | 0 |
| B79 | B110 | hb V -dna | AHA | 0.246533 | 0 |
| B79 | B119 | hb V -dna | Hcv-rna | 0.369822 | 0 |
| B79 | B121 | hb V -dna | C3 | -0.36982 | 0 |
| B79 | B122 | hb V -dna | C4 | -0.36982 | 0 |
| B79 | B125 | hb V -dna | Cor | 0.370172 | 0 |
| B79 | B135 | hb V -dna | TBA | -0.21863 | 0 |
| B79 | B152 | hb V -dna | ACA-Ig(G-A-M) | -0.36992 | 0 |
| B79 | B169 | hb V -dna | U-CRY | -0.36983 | 0 |
| B79 | B182 | hb V -dna | TT | -0.20476 | 0 |
| B79 | B183 | hb V -dna | PTA | 0.250114 | 0 |
| B79 | B81 | hb V -dna | TSGF | -0.33585 | 0 |

| | | | | | |
|---|---|---|---|---|---|
| B79 | B92 | hb V -dna | ALDH2-Glu504Glu | 0.306911 | 0 |
| B79 | B93 | hb V -dna | ALDH2-Glu504Lys | -0.32643 | 0 |
| B79 | B94 | hb V -dna | ALDH2-Lys504Lys | 0.618974 | 0 |
| B79 | B95 | hb V -dna | anti-cenpb | 0.246533 | 0 |
| B79 | B97 | hb V -dna | dsDNA | 0.246533 | 0 |
| B79 | B98 | hb V -dna | anti-Jo-1 | 0.246533 | 0 |
| B79 | B99 | hb V -dna | anti-PM-Scl | 0.246533 | 0 |
| B79 | B124 | hb V -dna | ACTH | 0.15953 | 2.72E-13 |
| B79 | B186 | hb V -dna | PFC | 0.142622 | 6.82E-11 |
| B79 | B120 | hb V -dna | HCV-cAg | 0.136368 | 4.50E-10 |
| B79 | B137 | hb V -dna | AFU | -0.13135 | 1.92E-09 |
| B79 | B195 | hb V -dna | Na | -0.12616 | 8.15E-09 |
| B79 | B187 | hb V -dna | NSE | 0.116356 | 1.07E-07 |
| B79 | B138 | hb V -dna | PAB | -0.11598 | 1.18E-07 |
| B81 | B108 | TSGF | Anti-Cyclin | 0.286663 | 0 |
| B81 | B182 | TSGF | TT | 0.2381 | 0 |
| B81 | B94 | TSGF | ALDH2-Lys504Lys | -0.45344 | 0 |
| B81 | B138 | TSGF | PAB | 0.176402 | 4.44E-16 |
| B81 | B157 | TSGF | IgG | -0.17373 | 1.78E-15 |
| B81 | B93 | TSGF | ALDH2-Glu504Lys | 0.163201 | 7.55E-14 |
| B81 | B125 | TSGF | Cor | -0.15 | 6.60E-12 |
| B81 | B92 | TSGF | ALDH2-Glu504Glu | -0.14722 | 1.61E-11 |
| B81 | B184 | TSGF | INR | -0.13964 | 1.69E-10 |
| B81 | B183 | TSGF | PTA | -0.13541 | 5.96E-10 |
| B81 | B181 | TSGF | APTT | -0.1228 | 2.02E-08 |
| B81 | B198 | TSGF | P | 0.117888 | 7.27E-08 |
| B81 | B102 | TSGF | anti-Sm | -0.11631 | 1.08E-07 |
| B81 | B100 | TSGF | anti-Ro52 | -0.11339 | 2.25E-07 |
| B81 | B101 | TSGF | anti-Sc70 | -0.11339 | 2.25E-07 |
| B81 | B104 | TSGF | anti-SSB | -0.11339 | 2.25E-07 |
| B81 | B106 | TSGF | anti-P | -0.11339 | 2.25E-07 |
| B81 | B107 | TSGF | AnuA | -0.11339 | 2.25E-07 |
| B81 | B110 | TSGF | AHA | -0.11339 | 2.25E-07 |
| B81 | B95 | TSGF | anti-cenpb | -0.11339 | 2.25E-07 |
| B81 | B97 | TSGF | dsDNA | -0.11339 | 2.25E-07 |
| B81 | B98 | TSGF | anti-Jo-1 | -0.11339 | 2.25E-07 |
| B81 | B99 | TSGF | anti-PM-Scl | -0.11339 | 2.25E-07 |
| B82 | B116 | CEA | TT3 | -0.20162 | 0 |
| B82 | B83 | CEA | AFP | 0.202669 | 0 |
| B82 | B86 | CEA | TBhCG | -0.41291 | 0 |
| B82 | B88 | CEA | CA 242 | 0.422788 | 0 |
| B82 | B85 | CEA | SCC | 0.151277 | 4.36E-12 |
| B82 | B87 | CEA | NSE | 0.126604 | 7.23E-09 |
| B82 | B117 | CEA | TT4 | -0.10987 | 5.28E-07 |

| | | | | | |
|---|---|---|---|---|---|
| B83 | B84 | AFP | C-PSA | 0.279306 | 0 |
| B83 | B85 | AFP | SCC | 0.251752 | 0 |
| B83 | B87 | AFP | NSE | 0.178779 | 2.22E-16 |
| B83 | B115 | AFP | TSH TT3 TT4(TT3) | -0.13693 | 3.81E-10 |
| B84 | B115 | C-PSA | TSH TT3 TT4(TT3) | -0.23336 | 0 |
| B84 | B86 | C-PSA | TBhCG | -0.202 | 0 |
| B84 | B199 | C-PSA | Mg | 0.175304 | 8.88E-16 |
| B84 | B227 | C-PSA | OD | 0.138389 | 2.47E-10 |
| B84 | B90 | C-PSA | CYFRA 21-1 | 0.135793 | 5.32E-10 |
| B84 | B200 | C-PSA | OP | 0.131914 | 1.64E-09 |
| B84 | B87 | C-PSA | NSE | 0.121875 | 2.58E-08 |
| B84 | B114 | C-PSA | TSH TT3 TT4(TT4) | -0.10763 | 8.98E-07 |
| B85 | B87 | SCC | NSE | 0.264954 | 0 |
| B85 | B90 | SCC | CYFRA 21-1 | 0.203145 | 0 |
| B85 | B117 | SCC | TT4 | -0.15783 | 4.87E-13 |
| B85 | B116 | SCC | TT3 | -0.1528 | 2.64E-12 |
| B86 | B87 | TBhCG | NSE | -0.32069 | 0 |
| B86 | B88 | TBhCG | CA 242 | -0.2504 | 0 |
| B86 | B89 | TBhCG | CA 72-4 | 0.250829 | 0 |
| B86 | B200 | TBhCG | OP | -0.16667 | 2.18E-14 |
| B86 | B195 | TBhCG | Na | -0.15102 | 4.73E-12 |
| B86 | B199 | TBhCG | Mg | -0.14759 | 1.43E-11 |
| B86 | B129 | TBhCG | IBIL | -0.13575 | 5.39E-10 |
| B86 | B133 | TBhCG | TBIL | -0.12828 | 4.56E-09 |
| B86 | B194 | TBhCG | Cl | -0.11851 | 6.20E-08 |
| B86 | B196 | TBhCG | TCO2 | 0.117592 | 7.84E-08 |
| B86 | B197 | TBhCG | AG | -0.11292 | 2.52E-07 |
| B87 | B89 | NSE | CA 72-4 | -0.17615 | 6.66E-16 |
| B87 | B88 | NSE | CA 242 | 0.175175 | 8.88E-16 |
| B87 | B129 | NSE | IBIL | 0.120344 | 3.86E-08 |
| B88 | B186 | CA 242 | PFC | 0.151794 | 3.68E-12 |
| B88 | B90 | CA 242 | CYFRA 21-1 | 0.115404 | 1.36E-07 |
| B9 | B100 | RH | anti-Ro52 | 0.300783 | 0 |
| B9 | B101 | RH | anti-Sc70 | 0.300783 | 0 |
| B9 | B104 | RH | anti-SSB | 0.300783 | 0 |
| B9 | B106 | RH | anti-P | 0.300783 | 0 |
| B9 | B107 | RH | AnuA | 0.300783 | 0 |
| B9 | B110 | RH | AHA | 0.300783 | 0 |
| B9 | B125 | RH | Cor | 0.309099 | 0 |
| B9 | B15 | RH | CD3-CD19+(B cell) | 0.250125 | 0 |
| B9 | B151 | RH | Anti-CCP | 0.197844 | 0 |
| B9 | B185 | RH | PT | -0.22196 | 0 |
| B9 | B19 | RH | CD3+CD4+/CD3+CD8+ | 0.235304 | 0 |
| B9 | B21 | RH | CD3+CD4-CD8- | 0.313829 | 0 |

| | | | | | |
|---|---|---|---|---|---|
| B9 | B22 | RH | CD3+CD8+(T) | -0.34507 | 0 |
| B9 | B57 | RH | RBCAI | 0.198628 | 0 |
| B9 | B62 | RH | WBHSRVI | -0.1838 | 0 |
| B9 | B79 | RH | hb V -dna | 0.276546 | 0 |
| B9 | B81 | RH | TSGF | -0.22435 | 0 |
| B9 | B92 | RH | ALDH2-Glu504Glu | 0.538708 | 0 |
| B9 | B93 | RH | ALDH2-Glu504Lys | -0.54594 | 0 |
| B9 | B94 | RH | ALDH2-Lys504Lys | 0.446782 | 0 |
| B9 | B95 | RH | anti-cenpb | 0.300783 | 0 |
| B9 | B97 | RH | dsDNA | 0.300783 | 0 |
| B9 | B98 | RH | anti-Jo-1 | 0.300783 | 0 |
| B9 | B99 | RH | anti-PM-Scl | 0.300783 | 0 |
| B9 | B196 | RH | TCO2 | 0.163517 | 6.75E-14 |
| B9 | B138 | RH | PAB | 0.16234 | 1.02E-13 |
| B9 | B56 | RH | IR | -0.15821 | 4.28E-13 |
| B9 | B195 | RH | Na | 0.156877 | 6.74E-13 |
| B9 | B66 | RH | WBRSR(mPas_100) | -0.15685 | 6.81E-13 |
| B9 | B183 | RH | PTA | 0.151496 | 4.06E-12 |
| B9 | B63 | RH | WBRSR(mPas_200) | -0.15148 | 4.08E-12 |
| B9 | B74 | RH | ApoA1 | 0.150469 | 5.67E-12 |
| B9 | B67 | RH | WBRSR(mPas_50) | -0.14034 | 1.37E-10 |
| B9 | B61 | RH | WBHSRV | -0.13672 | 4.05E-10 |
| B9 | B155 | RH | IgA | -0.12849 | 4.30E-09 |
| B9 | B182 | RH | TT | 0.127238 | 6.07E-09 |
| B9 | B16 | RH | CD3+T | -0.12323 | 1.80E-08 |
| B9 | B156 | RH | IgE | 0.123162 | 1.83E-08 |
| B9 | B200 | RH | OP | 0.118647 | 5.99E-08 |
| B9 | B186 | RH | PFC | -0.11427 | 1.81E-07 |
| B9 | B181 | RH | APTT | -0.11378 | 2.04E-07 |
| B90 | B157 | CYFRA 21-1 | IgG | 0.1155 | 1.33E-07 |
| B91 | B126 | TP | Alb | 0.442776 | 0 |
| B91 | B127 | TP | A-G ratio | -0.53138 | 0 |
| B91 | B130 | TP | GLB | 0.809972 | 0 |
| B91 | B188 | TP | Ca | 0.225093 | 0 |
| B91 | B209 | TP | Smoking | -0.13261 | 1.34E-09 |
| B91 | B153 | TP | Glu | 0.11681 | 9.57E-08 |
| B91 | B131 | TP | AST | 0.115072 | 1.48E-07 |
| B91 | B192 | TP | UA | 0.110114 | 4.98E-07 |
| B91 | B220 | TP | SBP | 0.108919 | 6.62E-07 |
| B92 | B100 | ALDH2-Glu504Glu | anti-Ro52 | 0.487165 | 0 |
| B92 | B101 | ALDH2-Glu504Glu | anti-Sc70 | 0.487165 | 0 |
| B92 | B104 | ALDH2-Glu504Glu | anti-SSB | 0.487165 | 0 |
| B92 | B106 | ALDH2-Glu504Glu | anti-P | 0.487165 | 0 |
| B92 | B107 | ALDH2-Glu504Glu | AnuA | 0.487164 | 0 |

| | | | | | |
|---|---|---|---|---|---|
| B92 | B108 | ALDH2-Glu504Glu | Anti-Cyclin | -0.51096 | 0 |
| B92 | B110 | ALDH2-Glu504Glu | AHA | 0.487165 | 0 |
| B92 | B135 | ALDH2-Glu504Glu | TBA | -0.19776 | 0 |
| B92 | B182 | ALDH2-Glu504Glu | TT | 0.288465 | 0 |
| B92 | B185 | ALDH2-Glu504Glu | PT | -0.19664 | 0 |
| B92 | B93 | ALDH2-Glu504Glu | ALDH2-Glu504Lys | -0.99933 | 0 |
| B92 | B94 | ALDH2-Glu504Glu | ALDH2-Lys504Lys | 0.495838 | 0 |
| B92 | B95 | ALDH2-Glu504Glu | anti-cenpb | 0.487165 | 0 |
| B92 | B97 | ALDH2-Glu504Glu | dsDNA | 0.487165 | 0 |
| B92 | B98 | ALDH2-Glu504Glu | anti-Jo-1 | 0.487165 | 0 |
| B92 | B99 | ALDH2-Glu504Glu | anti-PM-Scl | 0.487165 | 0 |
| B92 | B124 | ALDH2-Glu504Glu | ACTH | -0.17683 | 4.44E-16 |
| B92 | B183 | ALDH2-Glu504Glu | PTA | 0.167879 | 1.42E-14 |
| B92 | B198 | ALDH2-Glu504Glu | P | -0.15761 | 5.26E-13 |
| B92 | B138 | ALDH2-Glu504Glu | PAB | 0.129778 | 3.00E-09 |
| B92 | B194 | ALDH2-Glu504Glu | Cl | 0.125513 | 9.74E-09 |
| B92 | B155 | ALDH2-Glu504Glu | IgA | -0.1205 | 3.70E-08 |
| B92 | B105 | ALDH2-Glu504Glu | anti-U1RNP | -0.11636 | 1.07E-07 |
| B93 | B100 | ALDH2-Glu504Lys | anti-Ro52 | -0.49364 | 0 |
| B93 | B101 | ALDH2-Glu504Lys | anti-Sc70 | -0.49364 | 0 |
| B93 | B104 | ALDH2-Glu504Lys | anti-SSB | -0.49364 | 0 |
| B93 | B106 | ALDH2-Glu504Lys | anti-P | -0.49364 | 0 |
| B93 | B107 | ALDH2-Glu504Lys | AnuA | -0.49364 | 0 |
| B93 | B108 | ALDH2-Glu504Lys | Anti-Cyclin | 0.517823 | 0 |
| B93 | B110 | ALDH2-Glu504Lys | AHA | -0.49364 | 0 |
| B93 | B135 | ALDH2-Glu504Lys | TBA | 0.197295 | 0 |
| B93 | B182 | ALDH2-Glu504Lys | TT | -0.26748 | 0 |
| B93 | B185 | ALDH2-Glu504Lys | PT | 0.191832 | 0 |
| B93 | B94 | ALDH2-Glu504Lys | ALDH2-Lys504Lys | -0.52738 | 0 |
| B93 | B95 | ALDH2-Glu504Lys | anti-cenpb | -0.49364 | 0 |
| B93 | B97 | ALDH2-Glu504Lys | dsDNA | -0.49364 | 0 |
| B93 | B98 | ALDH2-Glu504Lys | anti-Jo-1 | -0.49364 | 0 |
| B93 | B99 | ALDH2-Glu504Lys | anti-PM-Scl | -0.49364 | 0 |
| B93 | B183 | ALDH2-Glu504Lys | PTA | -0.17845 | 2.22E-16 |
| B93 | B124 | ALDH2-Glu504Lys | ACTH | 0.167659 | 1.53E-14 |
| B93 | B198 | ALDH2-Glu504Lys | P | 0.1635 | 6.79E-14 |
| B93 | B138 | ALDH2-Glu504Lys | PAB | -0.1239 | 1.50E-08 |
| B93 | B194 | ALDH2-Glu504Lys | Cl | -0.12215 | 2.40E-08 |
| B93 | B155 | ALDH2-Glu504Lys | IgA | 0.119302 | 5.06E-08 |
| B93 | B105 | ALDH2-Glu504Lys | anti-U1RNP | 0.117893 | 7.26E-08 |
| B94 | B100 | ALDH2-Lys504Lys | anti-Ro52 | 0.402497 | 0 |
| B94 | B101 | ALDH2-Lys504Lys | anti-Sc70 | 0.402497 | 0 |
| B94 | B104 | ALDH2-Lys504Lys | anti-SSB | 0.402497 | 0 |
| B94 | B106 | ALDH2-Lys504Lys | anti-P | 0.402497 | 0 |

| | | | | | |
|---|---|---|---|---|---|
| B94 | B107 | ALDH2-Lys504Lys | AnuA | 0.402496 | 0 |
| B94 | B108 | ALDH2-Lys504Lys | Anti-Cyclin | -0.42389 | 0 |
| B94 | B110 | ALDH2-Lys504Lys | AHA | 0.402497 | 0 |
| B94 | B125 | ALDH2-Lys504Lys | Cor | 0.363801 | 0 |
| B94 | B139 | ALDH2-Lys504Lys | ADA | 0.214157 | 0 |
| B94 | B182 | ALDH2-Lys504Lys | TT | -0.34882 | 0 |
| B94 | B183 | ALDH2-Lys504Lys | PTA | 0.336162 | 0 |
| B94 | B184 | ALDH2-Lys504Lys | INR | 0.222288 | 0 |
| B94 | B198 | ALDH2-Lys504Lys | P | -0.22006 | 0 |
| B94 | B95 | ALDH2-Lys504Lys | anti-cenpb | 0.402497 | 0 |
| B94 | B97 | ALDH2-Lys504Lys | dsDNA | 0.402497 | 0 |
| B94 | B98 | ALDH2-Lys504Lys | anti-Jo-1 | 0.402497 | 0 |
| B94 | B99 | ALDH2-Lys504Lys | anti-PM-Scl | 0.402497 | 0 |
| B94 | B157 | ALDH2-Lys504Lys | IgG | 0.155732 | 9.93E-13 |
| B94 | B158 | ALDH2-Lys504Lys | IgM | 0.146643 | 1.94E-11 |
| B94 | B187 | ALDH2-Lys504Lys | NSE | 0.141212 | 1.05E-10 |
| B94 | B124 | ALDH2-Lys504Lys | ACTH | 0.126515 | 7.41E-09 |
| B94 | B186 | ALDH2-Lys504Lys | PFC | 0.125412 | 1.00E-08 |
| B94 | B156 | ALDH2-Lys504Lys | IgE | 0.125122 | 1.08E-08 |
| B94 | B154 | ALDH2-Lys504Lys | RF | 0.123335 | 1.75E-08 |
| B94 | B140 | ALDH2-Lys504Lys | r-GT | -0.1184 | 6.39E-08 |
| B95 | B100 | anti-cenpb | anti-Ro52 | 1 | 0 |
| B95 | B102 | anti-cenpb | anti-Sm | -0.27973 | 0 |
| B95 | B104 | anti-cenpb | anti-SSB | 1 | 0 |
| B95 | B105 | anti-cenpb | anti-U1RNP | -0.33755 | 0 |
| B95 | B107 | anti-cenpb | AnuA | 0.999998 | 0 |
| B95 | B108 | anti-cenpb | Anti-Cyclin | -0.22672 | 0 |
| B95 | B109 | anti-cenpb | AMA-M2 | -0.26181 | 0 |
| B95 | B110 | anti-cenpb | AHA | 1 | 0 |
| B95 | B125 | anti-cenpb | Cor | 0.234516 | 0 |
| B95 | B154 | anti-cenpb | RF | 0.204755 | 0 |
| B95 | B183 | anti-cenpb | PTA | 0.273181 | 0 |
| B95 | B96 | anti-cenpb | dsDNA (IIF) | 0.300783 | 0 |
| B95 | B97 | anti-cenpb | dsDNA | 1 | 0 |
| B95 | B155 | anti-cenpb | IgA | -0.16404 | 5.60E-14 |
| B95 | B156 | anti-cenpb | IgE | -0.14475 | 3.52E-11 |
| B95 | B181 | anti-cenpb | APTT | 0.13878 | 2.19E-10 |
| B95 | B195 | anti-cenpb | Na | 0.137023 | 3.70E-10 |
| B95 | B103 | anti-cenpb | anti-SSA/Ro | -0.13644 | 4.40E-10 |
| B95 | B137 | anti-cenpb | AFU | -0.12881 | 3.93E-09 |
| B95 | B198 | anti-cenpb | P | -0.12517 | 1.07E-08 |
| B95 | B139 | anti-cenpb | ADA | 0.118712 | 5.89E-08 |
| B95 | B138 | anti-cenpb | PAB | 0.113683 | 2.09E-07 |
| B95 | B185 | anti-cenpb | PT | -0.10972 | 5.48E-07 |

| | | | | | |
|---|---|---|---|---|---|
| B96 | B100 | dsDNA (IIF) | anti-Ro52 | 0.300783 | 0 |
| B96 | B101 | dsDNA (IIF) | anti-Sc70 | 0.300783 | 0 |
| B96 | B104 | dsDNA (IIF) | anti-SSB | 0.300783 | 0 |
| B96 | B106 | dsDNA (IIF) | anti-P | 0.300783 | 0 |
| B96 | B107 | dsDNA (IIF) | AnuA | 0.300783 | 0 |
| B96 | B110 | dsDNA (IIF) | AHA | 0.300783 | 0 |
| B96 | B97 | dsDNA (IIF) | dsDNA | 0.300783 | 0 |
| B96 | B98 | dsDNA (IIF) | anti-Jo-1 | 0.300783 | 0 |
| B96 | B99 | dsDNA (IIF) | anti-PM-Scl | 0.300783 | 0 |
| B96 | B155 | dsDNA (IIF) | IgA | 0.167849 | 1.42E-14 |
| B96 | B157 | dsDNA (IIF) | IgG | 0.153947 | 1.81E-12 |
| B96 | B151 | dsDNA (IIF) | Anti-CCP | -0.11584 | 1.22E-07 |
| B96 | B158 | dsDNA (IIF) | IgM | 0.107858 | 8.51E-07 |
| B97 | B101 | dsDNA | anti-Sc70 | 1 | 0 |
| B97 | B102 | dsDNA | anti-Sm | -0.27973 | 0 |
| B97 | B104 | dsDNA | anti-SSB | 1 | 0 |
| B97 | B105 | dsDNA | anti-U1RNP | -0.33755 | 0 |
| B97 | B106 | dsDNA | anti-P | 1 | 0 |
| B97 | B107 | dsDNA | AnuA | 0.999998 | 0 |
| B97 | B108 | dsDNA | Anti-Cyclin | -0.22672 | 0 |
| B97 | B109 | dsDNA | AMA-M2 | -0.26181 | 0 |
| B97 | B110 | dsDNA | AHA | 1 | 0 |
| B97 | B125 | dsDNA | Cor | 0.234516 | 0 |
| B97 | B154 | dsDNA | RF | 0.204755 | 0 |
| B97 | B183 | dsDNA | PTA | 0.273181 | 0 |
| B97 | B99 | dsDNA | anti-PM-Scl | 1 | 0 |
| B97 | B155 | dsDNA | IgA | -0.16404 | 5.60E-14 |
| B97 | B156 | dsDNA | IgE | -0.14475 | 3.52E-11 |
| B97 | B181 | dsDNA | APTT | 0.13878 | 2.19E-10 |
| B97 | B195 | dsDNA | Na | 0.137023 | 3.70E-10 |
| B97 | B103 | dsDNA | anti-SSA/Ro | -0.13644 | 4.40E-10 |
| B97 | B137 | dsDNA | AFU | -0.12881 | 3.93E-09 |
| B97 | B198 | dsDNA | P | -0.12517 | 1.07E-08 |
| B97 | B139 | dsDNA | ADA | 0.118712 | 5.89E-08 |
| B97 | B138 | dsDNA | PAB | 0.113683 | 2.09E-07 |
| B97 | B185 | dsDNA | PT | -0.10972 | 5.48E-07 |
| B98 | B102 | anti-Jo-1 | anti-Sm | -0.27973 | 0 |
| B98 | B105 | anti-Jo-1 | anti-U1RNP | -0.33755 | 0 |
| B98 | B107 | anti-Jo-1 | AnuA | 0.999998 | 0 |
| B98 | B108 | anti-Jo-1 | Anti-Cyclin | -0.22672 | 0 |
| B98 | B109 | anti-Jo-1 | AMA-M2 | -0.26181 | 0 |
| B98 | B125 | anti-Jo-1 | Cor | 0.234516 | 0 |
| B98 | B154 | anti-Jo-1 | RF | 0.204755 | 0 |
| B98 | B183 | anti-Jo-1 | PTA | 0.273181 | 0 |

| | | | | | |
|---|---|---|---|---|---|
| B98 | B155 | anti-Jo-1 | IgA | -0.16404 | 5.60E-14 |
| B98 | B156 | anti-Jo-1 | IgE | -0.14475 | 3.52E-11 |
| B98 | B181 | anti-Jo-1 | APTT | 0.13878 | 2.19E-10 |
| B98 | B195 | anti-Jo-1 | Na | 0.137023 | 3.70E-10 |
| B98 | B103 | anti-Jo-1 | anti-SSA/Ro | -0.13644 | 4.40E-10 |
| B98 | B137 | anti-Jo-1 | AFU | -0.12881 | 3.93E-09 |
| B98 | B198 | anti-Jo-1 | P | -0.12517 | 1.07E-08 |
| B98 | B139 | anti-Jo-1 | ADA | 0.118712 | 5.89E-08 |
| B98 | B138 | anti-Jo-1 | PAB | 0.113683 | 2.09E-07 |
| B98 | B185 | anti-Jo-1 | PT | -0.10972 | 5.48E-07 |
| B99 | B101 | anti-PM-Scl | anti-Sc70 | 1 | 0 |
| B99 | B102 | anti-PM-Scl | anti-Sm | -0.27973 | 0 |
| B99 | B104 | anti-PM-Scl | anti-SSB | 1 | 0 |
| B99 | B105 | anti-PM-Scl | anti-U1RNP | -0.33755 | 0 |
| B99 | B107 | anti-PM-Scl | AnuA | 0.999998 | 0 |
| B99 | B108 | anti-PM-Scl | Anti-Cyclin | -0.22672 | 0 |
| B99 | B109 | anti-PM-Scl | AMA-M2 | -0.26181 | 0 |
| B99 | B110 | anti-PM-Scl | AHA | 1 | 0 |
| B99 | B125 | anti-PM-Scl | Cor | 0.234516 | 0 |
| B99 | B154 | anti-PM-Scl | RF | 0.204755 | 0 |
| B99 | B183 | anti-PM-Scl | PTA | 0.273181 | 0 |
| B99 | B155 | anti-PM-Scl | IgA | -0.16404 | 5.60E-14 |
| B99 | B156 | anti-PM-Scl | IgE | -0.14475 | 3.52E-11 |
| B99 | B181 | anti-PM-Scl | APTT | 0.13878 | 2.19E-10 |
| B99 | B195 | anti-PM-Scl | Na | 0.137023 | 3.70E-10 |
| B99 | B103 | anti-PM-Scl | anti-SSA/Ro | -0.13644 | 4.40E-10 |
| B99 | B137 | anti-PM-Scl | AFU | -0.12881 | 3.93E-09 |
| B99 | B198 | anti-PM-Scl | P | -0.12517 | 1.07E-08 |
| B99 | B139 | anti-PM-Scl | ADA | 0.118712 | 5.89E-08 |
| B99 | B138 | anti-PM-Scl | PAB | 0.113683 | 2.09E-07 |
| B99 | B185 | anti-PM-Scl | PT | -0.10972 | 5.48E-07 |

**Supplementary Table 6. PEI Correlations in Hypertensive+Diabetes+Coronary**

| Row | Column | Row | Column | Cor | *P* |
|---|---|---|---|---|---|
| B1 | B161 | Gender | LEU | 0.311715 | 0 |
| B1 | B174 | Gender | Urea/Cr | 0.377869 | 0 |
| B1 | B216 | Gender | BMI | -0.46849 | 0 |
| B1 | B221 | Gender | Hei | -0.69222 | 0 |
| B1 | B222 | Gender | NS | -0.55315 | 0 |
| B1 | B46 | Gender | RBC Count | -0.32883 | 0 |
| B1 | B51 | Gender | Hb | -0.39553 | 0 |
| B1 | B173 | Gender | NTT | 0.260691 | 6.66E-16 |
| B1 | B45 | Gender | MCHC | -0.25945 | 8.88E-16 |
| B1 | B209 | Gender | Smoking | -0.25534 | 2.66E-15 |
| B1 | B72 | Gender | HDL-C | 0.22396 | 5.16E-12 |
| B1 | B205 | Gender | KDA | 0.201732 | 5.61E-10 |
| B1 | B163 | Gender | SG | -0.20051 | 7.16E-10 |
| B1 | B190 | Gender | Cre | -0.18301 | 1.97E-08 |
| B1 | B210 | Gender | Hip | -0.17261 | 1.22E-07 |
| B1 | B2 | Gender | Age | 0.172135 | 1.32E-07 |
| B1 | B204 | Gender | Drinking | -0.16505 | 4.29E-07 |
| B1 | B42 | Gender | MON | -0.1637 | 5.33E-07 |
| B1 | B50 | Gender | eo% | -0.16063 | 8.71E-07 |
| B10 | B12 | PG I | PG II | 0.621901 | 0 |
| B10 | B144 | PG I | Total-PINP | 0.289629 | 0 |
| B10 | B145 | PG I | BGP | 0.281342 | 0 |
| B10 | B175 | PG I | ALB-U | 0.271446 | 0 |
| B10 | B191 | PG I | Urea | 0.27824 | 0 |
| B10 | B11 | PG I | PGI/PGII | 0.252651 | 5.55E-15 |
| B10 | B177 | PG I | ACR | 0.235389 | 3.78E-13 |
| B10 | B190 | PG I | Cre | 0.219859 | 1.27E-11 |
| B10 | B189 | PG I | eGFB | -0.19314 | 3.00E-09 |
| B100 | B103 | anti-Ro52 | anti-SSA/Ro | 1 | 0 |
| B100 | B104 | anti-Ro52 | anti-SSB | 1 | 0 |
| B100 | B106 | anti-Ro52 | anti-P | 1 | 0 |
| B100 | B107 | anti-Ro52 | AnuA | 0.925318 | 0 |
| B100 | B109 | anti-Ro52 | AMA-M2 | 1 | 0 |
| B100 | B110 | anti-Ro52 | AHA | 0.925318 | 0 |
| B100 | B119 | anti-Ro52 | Hcv-rna | 0.433836 | 0 |
| B100 | B120 | anti-Ro52 | HCV-cAg | 0.352785 | 0 |
| B100 | B121 | anti-Ro52 | C3 | -0.43384 | 0 |
| B100 | B122 | anti-Ro52 | C4 | -0.43384 | 0 |
| B100 | B124 | anti-Ro52 | ACTH | -0.37674 | 0 |
| B100 | B125 | anti-Ro52 | Cor | -0.37674 | 0 |
| B100 | B139 | anti-Ro52 | ADA | -0.45548 | 0 |

| | | | | | |
|---|---|---|---|---|---|
| B100 | B152 | anti-Ro52 | ACA-Ig(G-A-M) | -0.43381 | 0 |
| B100 | B157 | anti-Ro52 | IgG | -0.33455 | 0 |
| B100 | B159 | anti-Ro52 | β2-MG | -0.43384 | 0 |
| B100 | B169 | anti-Ro52 | U-CRY | -0.43383 | 0 |
| B100 | B180 | anti-Ro52 | Tf | 0.440213 | 0 |
| B100 | B193 | anti-Ro52 | K | 0.3357 | 0 |
| B100 | B194 | anti-Ro52 | Cl | 0.354015 | 0 |
| B100 | B155 | anti-Ro52 | IgA | -0.24381 | 5.02E-14 |
| B100 | B137 | anti-Ro52 | AFU | -0.24337 | 5.60E-14 |
| B100 | B154 | anti-Ro52 | RF | 0.242501 | 6.93E-14 |
| B100 | B195 | anti-Ro52 | Na | 0.219271 | 1.45E-11 |
| B100 | B184 | anti-Ro52 | INR | -0.20636 | 2.21E-10 |
| B100 | B183 | anti-Ro52 | PTA | 0.202624 | 4.70E-10 |
| B100 | B178 | anti-Ro52 | Ig-U | 0.196769 | 1.49E-09 |
| B100 | B135 | anti-Ro52 | TBA | -0.18726 | 9.04E-09 |
| B100 | B196 | anti-Ro52 | TCO2 | -0.17771 | 5.04E-08 |
| B101 | B102 | anti-Sc70 | anti-Sm | 1 | 0 |
| B101 | B104 | anti-Sc70 | anti-SSB | 1 | 0 |
| B101 | B105 | anti-Sc70 | anti-U1RNP | 1 | 0 |
| B101 | B107 | anti-Sc70 | AnuA | 0.925317 | 0 |
| B101 | B108 | anti-Sc70 | Anti-Cyclin | 1 | 0 |
| B101 | B109 | anti-Sc70 | AMA-M2 | 1 | 0 |
| B101 | B110 | anti-Sc70 | AHA | 0.925318 | 0 |
| B101 | B119 | anti-Sc70 | Hcv-rna | 0.433836 | 0 |
| B101 | B120 | anti-Sc70 | HCV-cAg | 0.352785 | 0 |
| B101 | B121 | anti-Sc70 | C3 | -0.43384 | 0 |
| B101 | B122 | anti-Sc70 | C4 | -0.43384 | 0 |
| B101 | B124 | anti-Sc70 | ACTH | -0.37674 | 0 |
| B101 | B125 | anti-Sc70 | Cor | -0.37674 | 0 |
| B101 | B139 | anti-Sc70 | ADA | -0.45548 | 0 |
| B101 | B152 | anti-Sc70 | ACA-Ig(G-A-M) | -0.43381 | 0 |
| B101 | B157 | anti-Sc70 | IgG | -0.33455 | 0 |
| B101 | B159 | anti-Sc70 | β2-MG | -0.43384 | 0 |
| B101 | B169 | anti-Sc70 | U-CRY | -0.43383 | 0 |
| B101 | B180 | anti-Sc70 | Tf | 0.440213 | 0 |
| B101 | B193 | anti-Sc70 | K | 0.3357 | 0 |
| B101 | B194 | anti-Sc70 | Cl | 0.354015 | 0 |
| B101 | B155 | anti-Sc70 | IgA | -0.24381 | 5.02E-14 |
| B101 | B137 | anti-Sc70 | AFU | -0.24337 | 5.60E-14 |
| B101 | B154 | anti-Sc70 | RF | 0.242501 | 6.93E-14 |
| B101 | B195 | anti-Sc70 | Na | 0.219271 | 1.45E-11 |
| B101 | B184 | anti-Sc70 | INR | -0.20636 | 2.21E-10 |
| B101 | B183 | anti-Sc70 | PTA | 0.202624 | 4.70E-10 |
| B101 | B178 | anti-Sc70 | Ig-U | 0.196769 | 1.49E-09 |

| | | | | | |
|---|---|---|---|---|---|
| B101 | B135 | anti-Sc70 | TBA | -0.18726 | 9.04E-09 |
| B101 | B196 | anti-Sc70 | TCO2 | -0.17771 | 5.04E-08 |
| B102 | B107 | anti-Sm | AnuA | 0.925318 | 0 |
| B102 | B110 | anti-Sm | AHA | 0.925318 | 0 |
| B102 | B119 | anti-Sm | Hcv-rna | 0.433836 | 0 |
| B102 | B120 | anti-Sm | HCV-cAg | 0.352785 | 0 |
| B102 | B121 | anti-Sm | C3 | -0.43384 | 0 |
| B102 | B122 | anti-Sm | C4 | -0.43384 | 0 |
| B102 | B124 | anti-Sm | ACTH | -0.37674 | 0 |
| B102 | B125 | anti-Sm | Cor | -0.37674 | 0 |
| B102 | B139 | anti-Sm | ADA | -0.45548 | 0 |
| B102 | B152 | anti-Sm | ACA-Ig(G-A-M) | -0.43381 | 0 |
| B102 | B157 | anti-Sm | IgG | -0.33455 | 0 |
| B102 | B159 | anti-Sm | β2-MG | -0.43384 | 0 |
| B102 | B169 | anti-Sm | U-CRY | -0.43383 | 0 |
| B102 | B180 | anti-Sm | Tf | 0.440213 | 0 |
| B102 | B193 | anti-Sm | K | 0.3357 | 0 |
| B102 | B194 | anti-Sm | Cl | 0.354015 | 0 |
| B102 | B155 | anti-Sm | IgA | -0.24381 | 5.02E-14 |
| B102 | B137 | anti-Sm | AFU | -0.24337 | 5.60E-14 |
| B102 | B154 | anti-Sm | RF | 0.242501 | 6.93E-14 |
| B102 | B195 | anti-Sm | Na | 0.219271 | 1.45E-11 |
| B102 | B184 | anti-Sm | INR | -0.20636 | 2.21E-10 |
| B102 | B183 | anti-Sm | PTA | 0.202624 | 4.70E-10 |
| B102 | B178 | anti-Sm | Ig-U | 0.196769 | 1.49E-09 |
| B102 | B135 | anti-Sm | TBA | -0.18726 | 9.04E-09 |
| B102 | B196 | anti-Sm | TCO2 | -0.17771 | 5.04E-08 |
| B103 | B104 | anti-SSA/Ro | anti-SSB | 1 | 0 |
| B103 | B106 | anti-SSA/Ro | anti-P | 1 | 0 |
| B103 | B107 | anti-SSA/Ro | AnuA | 0.925318 | 0 |
| B103 | B109 | anti-SSA/Ro | AMA-M2 | 1 | 0 |
| B103 | B110 | anti-SSA/Ro | AHA | 0.925318 | 0 |
| B103 | B119 | anti-SSA/Ro | Hcv-rna | 0.433836 | 0 |
| B103 | B120 | anti-SSA/Ro | HCV-cAg | 0.352785 | 0 |
| B103 | B121 | anti-SSA/Ro | C3 | -0.43384 | 0 |
| B103 | B122 | anti-SSA/Ro | C4 | -0.43384 | 0 |
| B103 | B124 | anti-SSA/Ro | ACTH | -0.37674 | 0 |
| B103 | B125 | anti-SSA/Ro | Cor | -0.37674 | 0 |
| B103 | B139 | anti-SSA/Ro | ADA | -0.45548 | 0 |
| B103 | B152 | anti-SSA/Ro | ACA-Ig(G-A-M) | -0.43381 | 0 |
| B103 | B157 | anti-SSA/Ro | IgG | -0.33455 | 0 |
| B103 | B159 | anti-SSA/Ro | β2-MG | -0.43384 | 0 |
| B103 | B169 | anti-SSA/Ro | U-CRY | -0.43383 | 0 |
| B103 | B180 | anti-SSA/Ro | Tf | 0.440213 | 0 |

| | | | | | |
|---|---|---|---|---|---|
| B103 | B193 | anti-SSA/Ro | K | 0.3357 | 0 |
| B103 | B194 | anti-SSA/Ro | Cl | 0.354015 | 0 |
| B103 | B155 | anti-SSA/Ro | IgA | -0.24381 | 5.02E-14 |
| B103 | B137 | anti-SSA/Ro | AFU | -0.24337 | 5.60E-14 |
| B103 | B154 | anti-SSA/Ro | RF | 0.242501 | 6.93E-14 |
| B103 | B195 | anti-SSA/Ro | Na | 0.219271 | 1.45E-11 |
| B103 | B184 | anti-SSA/Ro | INR | -0.20636 | 2.21E-10 |
| B103 | B183 | anti-SSA/Ro | PTA | 0.202624 | 4.70E-10 |
| B103 | B178 | anti-SSA/Ro | Ig-U | 0.196769 | 1.49E-09 |
| B103 | B135 | anti-SSA/Ro | TBA | -0.18726 | 9.04E-09 |
| B103 | B196 | anti-SSA/Ro | TCO2 | -0.17771 | 5.04E-08 |
| B104 | B105 | anti-SSB | anti-U1RNP | 1 | 0 |
| B104 | B106 | anti-SSB | anti-P | 1 | 0 |
| B104 | B107 | anti-SSB | AnuA | 0.925318 | 0 |
| B104 | B108 | anti-SSB | Anti-Cyclin | 1 | 0 |
| B104 | B109 | anti-SSB | AMA-M2 | 1 | 0 |
| B104 | B110 | anti-SSB | AHA | 0.925318 | 0 |
| B104 | B119 | anti-SSB | Hcv-rna | 0.433836 | 0 |
| B104 | B120 | anti-SSB | HCV-cAg | 0.352785 | 0 |
| B104 | B121 | anti-SSB | C3 | -0.43384 | 0 |
| B104 | B122 | anti-SSB | C4 | -0.43384 | 0 |
| B104 | B124 | anti-SSB | ACTH | -0.37674 | 0 |
| B104 | B125 | anti-SSB | Cor | -0.37674 | 0 |
| B104 | B139 | anti-SSB | ADA | -0.45548 | 0 |
| B104 | B152 | anti-SSB | ACA-Ig(G-A-M) | -0.43381 | 0 |
| B104 | B157 | anti-SSB | IgG | -0.33455 | 0 |
| B104 | B159 | anti-SSB | β2-MG | -0.43384 | 0 |
| B104 | B169 | anti-SSB | U-CRY | -0.43383 | 0 |
| B104 | B180 | anti-SSB | Tf | 0.440213 | 0 |
| B104 | B193 | anti-SSB | K | 0.3357 | 0 |
| B104 | B194 | anti-SSB | Cl | 0.354015 | 0 |
| B104 | B155 | anti-SSB | IgA | -0.24381 | 5.02E-14 |
| B104 | B137 | anti-SSB | AFU | -0.24337 | 5.60E-14 |
| B104 | B154 | anti-SSB | RF | 0.242501 | 6.93E-14 |
| B104 | B195 | anti-SSB | Na | 0.219271 | 1.45E-11 |
| B104 | B184 | anti-SSB | INR | -0.20636 | 2.21E-10 |
| B104 | B183 | anti-SSB | PTA | 0.202624 | 4.70E-10 |
| B104 | B178 | anti-SSB | Ig-U | 0.196769 | 1.49E-09 |
| B104 | B135 | anti-SSB | TBA | -0.18726 | 9.04E-09 |
| B104 | B196 | anti-SSB | TCO2 | -0.17771 | 5.04E-08 |
| B105 | B106 | anti-U1RNP | anti-P | 1 | 0 |
| B105 | B107 | anti-U1RNP | AnuA | 0.925318 | 0 |
| B105 | B109 | anti-U1RNP | AMA-M2 | 1 | 0 |
| B105 | B110 | anti-U1RNP | AHA | 0.925318 | 0 |

| | | | | | |
|---|---|---|---|---|---|
| B105 | B119 | anti-U1RNP | Hcv-rna | 0.433836 | 0 |
| B105 | B120 | anti-U1RNP | HCV-cAg | 0.352785 | 0 |
| B105 | B121 | anti-U1RNP | C3 | -0.43384 | 0 |
| B105 | B122 | anti-U1RNP | C4 | -0.43384 | 0 |
| B105 | B124 | anti-U1RNP | ACTH | -0.37674 | 0 |
| B105 | B125 | anti-U1RNP | Cor | -0.37674 | 0 |
| B105 | B139 | anti-U1RNP | ADA | -0.45548 | 0 |
| B105 | B152 | anti-U1RNP | ACA-Ig(G-A-M) | -0.43381 | 0 |
| B105 | B157 | anti-U1RNP | IgG | -0.33455 | 0 |
| B105 | B159 | anti-U1RNP | β2-MG | -0.43384 | 0 |
| B105 | B169 | anti-U1RNP | U-CRY | -0.43383 | 0 |
| B105 | B180 | anti-U1RNP | Tf | 0.440213 | 0 |
| B105 | B193 | anti-U1RNP | K | 0.3357 | 0 |
| B105 | B194 | anti-U1RNP | Cl | 0.354015 | 0 |
| B105 | B155 | anti-U1RNP | IgA | -0.24381 | 5.02E-14 |
| B105 | B137 | anti-U1RNP | AFU | -0.24337 | 5.60E-14 |
| B105 | B154 | anti-U1RNP | RF | 0.242501 | 6.93E-14 |
| B105 | B195 | anti-U1RNP | Na | 0.219271 | 1.45E-11 |
| B105 | B184 | anti-U1RNP | INR | -0.20636 | 2.21E-10 |
| B105 | B183 | anti-U1RNP | PTA | 0.202624 | 4.70E-10 |
| B105 | B178 | anti-U1RNP | Ig-U | 0.196769 | 1.49E-09 |
| B105 | B135 | anti-U1RNP | TBA | -0.18726 | 9.04E-09 |
| B105 | B196 | anti-U1RNP | TCO2 | -0.17771 | 5.04E-08 |
| B106 | B107 | anti-P | AnuA | 0.925318 | 0 |
| B106 | B108 | anti-P | Anti-Cyclin | 1 | 0 |
| B106 | B109 | anti-P | AMA-M2 | 1 | 0 |
| B106 | B110 | anti-P | AHA | 0.925318 | 0 |
| B106 | B119 | anti-P | Hcv-rna | 0.433836 | 0 |
| B106 | B120 | anti-P | HCV-cAg | 0.352785 | 0 |
| B106 | B121 | anti-P | C3 | -0.43384 | 0 |
| B106 | B122 | anti-P | C4 | -0.43384 | 0 |
| B106 | B124 | anti-P | ACTH | -0.37674 | 0 |
| B106 | B125 | anti-P | Cor | -0.37674 | 0 |
| B106 | B139 | anti-P | ADA | -0.45548 | 0 |
| B106 | B152 | anti-P | ACA-Ig(G-A-M) | -0.43381 | 0 |
| B106 | B157 | anti-P | IgG | -0.33455 | 0 |
| B106 | B159 | anti-P | β2-MG | -0.43384 | 0 |
| B106 | B169 | anti-P | U-CRY | -0.43383 | 0 |
| B106 | B180 | anti-P | Tf | 0.440213 | 0 |
| B106 | B193 | anti-P | K | 0.3357 | 0 |
| B106 | B194 | anti-P | Cl | 0.354015 | 0 |
| B106 | B155 | anti-P | IgA | -0.24381 | 5.02E-14 |
| B106 | B137 | anti-P | AFU | -0.24337 | 5.60E-14 |
| B106 | B154 | anti-P | RF | 0.242501 | 6.93E-14 |

| | | | | | |
|---|---|---|---|---|---|
| B106 | B195 | anti-P | Na | 0.219271 | 1.45E-11 |
| B106 | B184 | anti-P | INR | -0.20636 | 2.21E-10 |
| B106 | B183 | anti-P | PTA | 0.202624 | 4.70E-10 |
| B106 | B178 | anti-P | Ig-U | 0.196769 | 1.49E-09 |
| B106 | B135 | anti-P | TBA | -0.18726 | 9.04E-09 |
| B106 | B196 | anti-P | TCO2 | -0.17771 | 5.04E-08 |
| B107 | B108 | AnuA | Anti-Cyclin | 0.925318 | 0 |
| B107 | B109 | AnuA | AMA-M2 | 0.925318 | 0 |
| B107 | B110 | AnuA | AHA | 1 | 0 |
| B107 | B119 | AnuA | Hcv-rna | 0.469108 | 0 |
| B107 | B120 | AnuA | HCV-cAg | 0.382247 | 0 |
| B107 | B121 | AnuA | C3 | -0.46911 | 0 |
| B107 | B122 | AnuA | C4 | -0.46911 | 0 |
| B107 | B124 | AnuA | ACTH | -0.40715 | 0 |
| B107 | B125 | AnuA | Cor | -0.40715 | 0 |
| B107 | B139 | AnuA | ADA | -0.41313 | 0 |
| B107 | B152 | AnuA | ACA-Ig(G-A-M) | -0.46907 | 0 |
| B107 | B155 | AnuA | IgA | -0.28136 | 0 |
| B107 | B159 | AnuA | β2-MG | -0.46911 | 0 |
| B107 | B169 | AnuA | U-CRY | -0.4691 | 0 |
| B107 | B180 | AnuA | Tf | 0.475926 | 0 |
| B107 | B194 | AnuA | Cl | 0.416382 | 0 |
| B107 | B195 | AnuA | Na | 0.287963 | 0 |
| B107 | B137 | AnuA | AFU | -0.25172 | 7.11E-15 |
| B107 | B184 | AnuA | INR | -0.22365 | 5.52E-12 |
| B107 | B183 | AnuA | PTA | 0.218414 | 1.74E-11 |
| B107 | B154 | AnuA | RF | 0.214323 | 4.20E-11 |
| B107 | B178 | AnuA | Ig-U | 0.212351 | 6.37E-11 |
| B107 | B193 | AnuA | K | 0.211512 | 7.59E-11 |
| B107 | B157 | AnuA | IgG | -0.20289 | 4.45E-10 |
| B107 | B135 | AnuA | TBA | -0.18054 | 3.06E-08 |
| B107 | B181 | AnuA | APTT | -0.16572 | 3.84E-07 |
| B108 | B110 | Anti-Cyclin | AHA | 0.925318 | 0 |
| B108 | B119 | Anti-Cyclin | Hcv-rna | 0.433836 | 0 |
| B108 | B120 | Anti-Cyclin | HCV-cAg | 0.352785 | 0 |
| B108 | B121 | Anti-Cyclin | C3 | -0.43384 | 0 |
| B108 | B122 | Anti-Cyclin | C4 | -0.43384 | 0 |
| B108 | B124 | Anti-Cyclin | ACTH | -0.37674 | 0 |
| B108 | B125 | Anti-Cyclin | Cor | -0.37674 | 0 |
| B108 | B139 | Anti-Cyclin | ADA | -0.45548 | 0 |
| B108 | B152 | Anti-Cyclin | ACA-Ig(G-A-M) | -0.43381 | 0 |
| B108 | B157 | Anti-Cyclin | IgG | -0.33455 | 0 |
| B108 | B159 | Anti-Cyclin | β2-MG | -0.43384 | 0 |
| B108 | B169 | Anti-Cyclin | U-CRY | -0.43383 | 0 |

| | | | | | |
|---|---|---|---|---|---|
| B108 | B180 | Anti-Cyclin | Tf | 0.440213 | 0 |
| B108 | B193 | Anti-Cyclin | K | 0.3357 | 0 |
| B108 | B194 | Anti-Cyclin | Cl | 0.354015 | 0 |
| B108 | B155 | Anti-Cyclin | IgA | -0.24381 | 5.02E-14 |
| B108 | B137 | Anti-Cyclin | AFU | -0.24337 | 5.60E-14 |
| B108 | B154 | Anti-Cyclin | RF | 0.242501 | 6.93E-14 |
| B108 | B195 | Anti-Cyclin | Na | 0.219271 | 1.45E-11 |
| B108 | B184 | Anti-Cyclin | INR | -0.20636 | 2.21E-10 |
| B108 | B183 | Anti-Cyclin | PTA | 0.202624 | 4.70E-10 |
| B108 | B178 | Anti-Cyclin | Ig-U | 0.196769 | 1.49E-09 |
| B108 | B135 | Anti-Cyclin | TBA | -0.18726 | 9.04E-09 |
| B108 | B196 | Anti-Cyclin | TCO2 | -0.17771 | 5.04E-08 |
| B109 | B110 | AMA-M2 | AHA | 0.925318 | 0 |
| B109 | B119 | AMA-M2 | Hcv-rna | 0.433836 | 0 |
| B109 | B120 | AMA-M2 | HCV-cAg | 0.352785 | 0 |
| B109 | B121 | AMA-M2 | C3 | -0.43384 | 0 |
| B109 | B122 | AMA-M2 | C4 | -0.43384 | 0 |
| B109 | B124 | AMA-M2 | ACTH | -0.37674 | 0 |
| B109 | B125 | AMA-M2 | Cor | -0.37674 | 0 |
| B109 | B139 | AMA-M2 | ADA | -0.45548 | 0 |
| B109 | B152 | AMA-M2 | ACA-Ig(G-A-M) | -0.43381 | 0 |
| B109 | B157 | AMA-M2 | IgG | -0.33455 | 0 |
| B109 | B159 | AMA-M2 | β2-MG | -0.43384 | 0 |
| B109 | B169 | AMA-M2 | U-CRY | -0.43383 | 0 |
| B109 | B180 | AMA-M2 | Tf | 0.440213 | 0 |
| B109 | B193 | AMA-M2 | K | 0.3357 | 0 |
| B109 | B194 | AMA-M2 | Cl | 0.354015 | 0 |
| B109 | B155 | AMA-M2 | IgA | -0.24381 | 5.02E-14 |
| B109 | B137 | AMA-M2 | AFU | -0.24337 | 5.60E-14 |
| B109 | B154 | AMA-M2 | RF | 0.242501 | 6.93E-14 |
| B109 | B195 | AMA-M2 | Na | 0.219271 | 1.45E-11 |
| B109 | B184 | AMA-M2 | INR | -0.20636 | 2.21E-10 |
| B109 | B183 | AMA-M2 | PTA | 0.202624 | 4.70E-10 |
| B109 | B178 | AMA-M2 | Ig-U | 0.196769 | 1.49E-09 |
| B109 | B135 | AMA-M2 | TBA | -0.18726 | 9.04E-09 |
| B109 | B196 | AMA-M2 | TCO2 | -0.17771 | 5.04E-08 |
| B11 | B12 | PGI/PGII | PG II | -0.40811 | 0 |
| B11 | B14 | PGI/PGII | G-17 | -0.25011 | 1.07E-14 |
| B11 | B132 | PGI/PGII | Dbil | -0.17949 | 3.68E-08 |
| B110 | B119 | AHA | Hcv-rna | 0.469108 | 0 |
| B110 | B120 | AHA | HCV-cAg | 0.382248 | 0 |
| B110 | B121 | AHA | C3 | -0.46911 | 0 |
| B110 | B122 | AHA | C4 | -0.46911 | 0 |
| B110 | B124 | AHA | ACTH | -0.40715 | 0 |

| | | | | | |
|---|---|---|---|---|---|
| B110 | B125 | AHA | Cor | -0.40715 | 0 |
| B110 | B139 | AHA | ADA | -0.41313 | 0 |
| B110 | B152 | AHA | ACA-Ig(G-A-M) | -0.46908 | 0 |
| B110 | B155 | AHA | IgA | -0.28136 | 0 |
| B110 | B159 | AHA | β2-MG | -0.46911 | 0 |
| B110 | B169 | AHA | U-CRY | -0.46911 | 0 |
| B110 | B180 | AHA | Tf | 0.475926 | 0 |
| B110 | B194 | AHA | Cl | 0.416382 | 0 |
| B110 | B195 | AHA | Na | 0.287963 | 0 |
| B110 | B137 | AHA | AFU | -0.25172 | 7.11E-15 |
| B110 | B184 | AHA | INR | -0.22365 | 5.52E-12 |
| B110 | B183 | AHA | PTA | 0.218414 | 1.74E-11 |
| B110 | B154 | AHA | RF | 0.214323 | 4.20E-11 |
| B110 | B178 | AHA | Ig-U | 0.212352 | 6.37E-11 |
| B110 | B193 | AHA | K | 0.211513 | 7.59E-11 |
| B110 | B157 | AHA | IgG | -0.20289 | 4.45E-10 |
| B110 | B135 | AHA | TBA | -0.18054 | 3.06E-08 |
| B110 | B181 | AHA | APTT | -0.16572 | 3.84E-07 |
| B111 | B228 | TSH | OS | 0.170241 | 1.82E-07 |
| B112 | B114 | FT4 | TSH TT3 TT4(TT4) | 0.49946 | 0 |
| B112 | B181 | FT4 | APTT | -0.1621 | 6.89E-07 |
| B113 | B115 | FT3 | TSH TT3 TT4(TT3) | 0.413715 | 0 |
| B113 | B185 | FT3 | PT | -0.21094 | 8.57E-11 |
| B113 | B184 | FT3 | INR | -0.2066 | 2.10E-10 |
| B113 | B181 | FT3 | APTT | -0.19625 | 1.65E-09 |
| B113 | B176 | FT3 | Ucr | 0.194952 | 2.12E-09 |
| B113 | B183 | FT3 | PTA | -0.18813 | 7.70E-09 |
| B113 | B177 | FT3 | ACR | -0.17385 | 9.84E-08 |
| B114 | B171 | TSH TT3 TT4(TT4) | EC | -0.31416 | 0 |
| B114 | B124 | TSH TT3 TT4(TT4) | ACTH | 0.244607 | 4.13E-14 |
| B114 | B125 | TSH TT3 TT4(TT4) | Cor | 0.244607 | 4.13E-14 |
| B114 | B135 | TSH TT3 TT4(TT4) | TBA | 0.219373 | 1.42E-11 |
| B114 | B151 | TSH TT3 TT4(TT4) | Anti-CCP | 0.183933 | 1.66E-08 |
| B114 | B180 | TSH TT3 TT4(TT4) | Tf | -0.1621 | 6.89E-07 |
| B115 | B134 | TSH TT3 TT4(TT3) | CHE | 0.199864 | 8.13E-10 |
| B116 | B117 | TT3 | TT4 | -0.96562 | 0 |
| B118 | B126 | CEA1 | Alb | -0.19643 | 1.59E-09 |
| B118 | B140 | CEA1 | r-GT | 0.172776 | 1.18E-07 |
| B118 | B132 | CEA1 | Dbil | 0.170568 | 1.72E-07 |
| B119 | B120 | Hcv-rna | HCV-cAg | 0.545646 | 0 |
| B119 | B122 | Hcv-rna | C4 | -1 | 0 |
| B119 | B124 | Hcv-rna | ACTH | -0.57673 | 0 |
| B119 | B125 | Hcv-rna | Cor | -0.57673 | 0 |
| B119 | B139 | Hcv-rna | ADA | -0.43647 | 0 |

| | | | | | |
|---|---|---|---|---|---|
| B119 | B151 | Hcv-rna | Anti-CCP | -0.31503 | 0 |
| B119 | B152 | Hcv-rna | ACA-Ig(G-A-M) | -0.99994 | 0 |
| B119 | B155 | Hcv-rna | IgA | -0.34006 | 0 |
| B119 | B159 | Hcv-rna | β2-MG | -1 | 0 |
| B119 | B169 | Hcv-rna | U-CRY | -1 | 0 |
| B119 | B135 | Hcv-rna | TBA | -0.25213 | 6.22E-15 |
| B119 | B184 | Hcv-rna | INR | -0.25025 | 1.02E-14 |
| B119 | B193 | Hcv-rna | K | 0.249729 | 1.15E-14 |
| B119 | B138 | Hcv-rna | PAB | 0.239223 | 1.52E-13 |
| B119 | B157 | Hcv-rna | IgG | -0.21452 | 4.03E-11 |
| B119 | B180 | Hcv-rna | Tf | 0.193367 | 2.87E-09 |
| B119 | B194 | Hcv-rna | Cl | 0.188594 | 7.06E-09 |
| B119 | B195 | Hcv-rna | Na | 0.183484 | 1.80E-08 |
| B119 | B136 | Hcv-rna | 5'NT | -0.18293 | 1.99E-08 |
| B119 | B185 | Hcv-rna | PT | -0.16585 | 3.76E-07 |
| B12 | B144 | PG II | Total-PINP | 0.352405 | 0 |
| B12 | B145 | PG II | BGP | 0.381873 | 0 |
| B12 | B175 | PG II | ALB-U | 0.36334 | 0 |
| B12 | B177 | PG II | ACR | 0.324465 | 0 |
| B12 | B190 | PG II | Cre | 0.306566 | 0 |
| B12 | B191 | PG II | Urea | 0.404415 | 0 |
| B12 | B80 | PG II | anti-HP IgG | 0.20969 | 1.11E-10 |
| B12 | B189 | PG II | eGFB | -0.20533 | 2.72E-10 |
| B12 | B17 | PG II | CD3+CD16+CD56+NK | -0.18178 | 2.45E-08 |
| B12 | B143 | PG II | β-CTx | 0.172767 | 1.19E-07 |
| B120 | B121 | HCV-cAg | C3 | -0.54565 | 0 |
| B120 | B122 | HCV-cAg | C4 | -0.54565 | 0 |
| B120 | B124 | HCV-cAg | ACTH | -0.31469 | 0 |
| B120 | B125 | HCV-cAg | Cor | -0.31469 | 0 |
| B120 | B152 | HCV-cAg | ACA-Ig(G-A-M) | -0.54562 | 0 |
| B120 | B155 | HCV-cAg | IgA | -0.33734 | 0 |
| B120 | B159 | HCV-cAg | β2-MG | -0.54565 | 0 |
| B120 | B169 | HCV-cAg | U-CRY | -0.54565 | 0 |
| B120 | B139 | HCV-cAg | ADA | -0.21722 | 2.26E-11 |
| B120 | B135 | HCV-cAg | TBA | -0.21442 | 4.12E-11 |
| B120 | B156 | HCV-cAg | IgE | 0.203616 | 3.85E-10 |
| B120 | B157 | HCV-cAg | IgG | -0.1912 | 4.33E-09 |
| B120 | B193 | HCV-cAg | K | 0.187032 | 9.43E-09 |
| B120 | B180 | HCV-cAg | Tf | 0.178888 | 4.10E-08 |
| B120 | B151 | HCV-cAg | Anti-CCP | -0.17864 | 4.29E-08 |
| B121 | B122 | C3 | C4 | 1 | 0 |
| B121 | B124 | C3 | ACTH | 0.576727 | 0 |
| B121 | B125 | C3 | Cor | 0.576727 | 0 |
| B121 | B139 | C3 | ADA | 0.436469 | 0 |

| | | | | | |
|---|---|---|---|---|---|
| B121 | B151 | C3 | Anti-CCP | 0.315028 | 0 |
| B121 | B152 | C3 | ACA-Ig(G-A-M) | 0.99995 | 0 |
| B121 | B155 | C3 | IgA | 0.340064 | 0 |
| B121 | B159 | C3 | β2-MG | 1 | 0 |
| B121 | B169 | C3 | U-CRY | 0.999998 | 0 |
| B121 | B135 | C3 | TBA | 0.252131 | 6.22E-15 |
| B121 | B184 | C3 | INR | 0.250246 | 1.02E-14 |
| B121 | B193 | C3 | K | -0.24973 | 1.15E-14 |
| B121 | B138 | C3 | PAB | -0.23922 | 1.52E-13 |
| B121 | B157 | C3 | IgG | 0.214518 | 4.03E-11 |
| B121 | B180 | C3 | Tf | -0.19337 | 2.87E-09 |
| B121 | B194 | C3 | Cl | -0.18859 | 7.06E-09 |
| B121 | B195 | C3 | Na | -0.18348 | 1.80E-08 |
| B121 | B136 | C3 | 5'NT | 0.182935 | 1.99E-08 |
| B121 | B185 | C3 | PT | 0.165846 | 3.76E-07 |
| B122 | B124 | C4 | ACTH | 0.576727 | 0 |
| B122 | B125 | C4 | Cor | 0.576727 | 0 |
| B122 | B139 | C4 | ADA | 0.436468 | 0 |
| B122 | B151 | C4 | Anti-CCP | 0.315028 | 0 |
| B122 | B152 | C4 | ACA-Ig(G-A-M) | 0.999949 | 0 |
| B122 | B155 | C4 | IgA | 0.340064 | 0 |
| B122 | B135 | C4 | TBA | 0.252131 | 6.22E-15 |
| B122 | B184 | C4 | INR | 0.250245 | 1.02E-14 |
| B122 | B193 | C4 | K | -0.24973 | 1.15E-14 |
| B122 | B138 | C4 | PAB | -0.23922 | 1.52E-13 |
| B122 | B157 | C4 | IgG | 0.214518 | 4.03E-11 |
| B122 | B180 | C4 | Tf | -0.19337 | 2.87E-09 |
| B122 | B194 | C4 | Cl | -0.18859 | 7.06E-09 |
| B122 | B195 | C4 | Na | -0.18348 | 1.80E-08 |
| B122 | B136 | C4 | 5'NT | 0.182934 | 1.99E-08 |
| B122 | B185 | C4 | PT | 0.165846 | 3.76E-07 |
| B123 | B172 | hs-CRP1 | KB | 0.234466 | 4.69E-13 |
| B124 | B135 | ACTH | TBA | 0.59701 | 0 |
| B124 | B136 | ACTH | 5'NT | 0.280012 | 0 |
| B124 | B138 | ACTH | PAB | -0.37484 | 0 |
| B124 | B139 | ACTH | ADA | 0.572505 | 0 |
| B124 | B151 | ACTH | Anti-CCP | 0.540198 | 0 |
| B124 | B152 | ACTH | ACA-Ig(G-A-M) | 0.576694 | 0 |
| B124 | B157 | ACTH | IgG | 0.346398 | 0 |
| B124 | B158 | ACTH | IgM | 0.326704 | 0 |
| B124 | B159 | ACTH | β2-MG | 0.576727 | 0 |
| B124 | B169 | ACTH | U-CRY | 0.576724 | 0 |
| B124 | B180 | ACTH | Tf | -0.33798 | 0 |
| B124 | B184 | ACTH | INR | 0.437275 | 0 |

| | | | | | |
|---|---|---|---|---|---|
| B124 | B185 | ACTH | PT | 0.305403 | 0 |
| B124 | B193 | ACTH | K | -0.32623 | 0 |
| B124 | B194 | ACTH | Cl | -0.44099 | 0 |
| B124 | B197 | ACTH | AG | 0.277445 | 0 |
| B124 | B199 | ACTH | Mg | 0.27051 | 0 |
| B124 | B183 | ACTH | PTA | -0.26493 | 2.22E-16 |
| B124 | B155 | ACTH | IgA | 0.238695 | 1.73E-13 |
| B124 | B195 | ACTH | Na | -0.18283 | 2.03E-08 |
| B124 | B198 | ACTH | P | 0.163147 | 5.83E-07 |
| B125 | B135 | Cor | TBA | 0.59701 | 0 |
| B125 | B136 | Cor | 5'NT | 0.280012 | 0 |
| B125 | B138 | Cor | PAB | -0.37484 | 0 |
| B125 | B139 | Cor | ADA | 0.572505 | 0 |
| B125 | B151 | Cor | Anti-CCP | 0.540198 | 0 |
| B125 | B152 | Cor | ACA-Ig(G-A-M) | 0.576694 | 0 |
| B125 | B157 | Cor | IgG | 0.346398 | 0 |
| B125 | B158 | Cor | IgM | 0.326704 | 0 |
| B125 | B159 | Cor | β2-MG | 0.576727 | 0 |
| B125 | B169 | Cor | U-CRY | 0.576724 | 0 |
| B125 | B180 | Cor | Tf | -0.33798 | 0 |
| B125 | B184 | Cor | INR | 0.437275 | 0 |
| B125 | B185 | Cor | PT | 0.305403 | 0 |
| B125 | B193 | Cor | K | -0.32623 | 0 |
| B125 | B194 | Cor | Cl | -0.44099 | 0 |
| B125 | B197 | Cor | AG | 0.277445 | 0 |
| B125 | B199 | Cor | Mg | 0.27051 | 0 |
| B125 | B183 | Cor | PTA | -0.26493 | 2.22E-16 |
| B125 | B155 | Cor | IgA | 0.238695 | 1.73E-13 |
| B125 | B195 | Cor | Na | -0.18283 | 2.03E-08 |
| B125 | B198 | Cor | P | 0.163147 | 5.83E-07 |
| B126 | B127 | Alb | A-G ratio | 0.537404 | 0 |
| B126 | B188 | Alb | Ca | 0.345114 | 0 |
| B126 | B189 | Alb | eGFB | 0.308957 | 0 |
| B126 | B191 | Alb | Urea | -0.26648 | 0 |
| B126 | B190 | Alb | Cre | -0.23542 | 3.75E-13 |
| B126 | B130 | Alb | GLB | -0.19366 | 2.72E-09 |
| B127 | B130 | A-G ratio | GLB | -0.901 | 0 |
| B127 | B189 | A-G ratio | eGFB | 0.231861 | 8.59E-13 |
| B127 | B129 | A-G ratio | IBIL | 0.168286 | 2.52E-07 |
| B127 | B220 | A-G ratio | SBP | -0.16383 | 5.22E-07 |
| B128 | B131 | ALT | AST | 0.740293 | 0 |
| B128 | B132 | ALT | Dbil | 0.349139 | 0 |
| B128 | B140 | ALT | r-GT | 0.434533 | 0 |
| B128 | B141 | ALT | AST/ALT | -0.54309 | 0 |

| | | | | | |
|---|---|---|---|---|---|
| B128 | B133 | ALT | TBIL | 0.255834 | 2.44E-15 |
| B128 | B216 | ALT | BMI | 0.195355 | 1.96E-09 |
| B128 | B136 | ALT | 5'NT | 0.166803 | 3.22E-07 |
| B129 | B132 | IBIL | Dbil | 0.325892 | 0 |
| B129 | B133 | IBIL | TBIL | 0.823504 | 0 |
| B129 | B130 | IBIL | GLB | -0.16241 | 6.56E-07 |
| B130 | B153 | GLB | Glu | 0.166722 | 3.26E-07 |
| B131 | B132 | AST | Dbil | 0.462077 | 0 |
| B131 | B133 | AST | TBIL | 0.349927 | 0 |
| B131 | B140 | AST | r-GT | 0.527614 | 0 |
| B132 | B133 | Dbil | TBIL | 0.80468 | 0 |
| B132 | B140 | Dbil | r-GT | 0.885761 | 0 |
| B133 | B140 | TBIL | r-GT | 0.596874 | 0 |
| B134 | B138 | CHE | PAB | 0.561042 | 0 |
| B134 | B139 | CHE | ADA | -0.23891 | 1.64E-13 |
| B134 | B157 | CHE | IgG | -0.23567 | 3.54E-13 |
| B134 | B186 | CHE | PFC | -0.23307 | 6.50E-13 |
| B134 | B171 | CHE | EC | -0.21139 | 7.79E-11 |
| B134 | B150 | CHE | ALP | -0.20191 | 5.42E-10 |
| B134 | B136 | CHE | 5'NT | -0.17349 | 1.05E-07 |
| B134 | B135 | CHE | TBA | -0.17247 | 1.25E-07 |
| B134 | B182 | CHE | TT | 0.165503 | 3.98E-07 |
| B135 | B138 | TBA | PAB | -0.29783 | 0 |
| B135 | B139 | TBA | ADA | 0.403014 | 0 |
| B135 | B151 | TBA | Anti-CCP | 0.347199 | 0 |
| B135 | B157 | TBA | IgG | 0.281903 | 0 |
| B135 | B193 | TBA | K | -0.38757 | 0 |
| B135 | B194 | TBA | Cl | -0.26644 | 0 |
| B135 | B197 | TBA | AG | 0.287397 | 0 |
| B135 | B198 | TBA | P | 0.453291 | 0 |
| B135 | B152 | TBA | ACA-Ig(G-A-M) | 0.252117 | 6.22E-15 |
| B135 | B159 | TBA | β2-MG | 0.252131 | 6.22E-15 |
| B135 | B169 | TBA | U-CRY | 0.25213 | 6.22E-15 |
| B135 | B184 | TBA | INR | 0.249488 | 1.24E-14 |
| B135 | B200 | TBA | OP | 0.224905 | 4.18E-12 |
| B135 | B180 | TBA | Tf | -0.19797 | 1.18E-09 |
| B135 | B185 | TBA | PT | 0.187019 | 9.46E-09 |
| B135 | B137 | TBA | AFU | -0.16133 | 7.79E-07 |
| B136 | B137 | 5'NT | AFU | 0.286331 | 0 |
| B136 | B150 | 5'NT | ALP | 0.291357 | 0 |
| B136 | B193 | 5'NT | K | 0.386247 | 0 |
| B136 | B196 | 5'NT | TCO2 | -0.33329 | 0 |
| B136 | B197 | 5'NT | AG | 0.289579 | 0 |
| B136 | B199 | 5'NT | Mg | 0.41058 | 0 |

| | | | | | |
|---|---|---|---|---|---|
| B136 | B159 | 5'NT | β2-MG | 0.182935 | 1.99E-08 |
| B136 | B169 | 5'NT | U-CRY | 0.182934 | 1.99E-08 |
| B136 | B152 | 5'NT | ACA-Ig(G-A-M) | 0.182924 | 2.00E-08 |
| B136 | B151 | 5'NT | Anti-CCP | 0.174557 | 8.72E-08 |
| B136 | B158 | 5'NT | IgM | 0.17024 | 1.82E-07 |
| B136 | B186 | 5'NT | PFC | -0.17017 | 1.84E-07 |
| B137 | B139 | AFU | ADA | 0.420277 | 0 |
| B137 | B193 | AFU | K | 0.484625 | 0 |
| B137 | B194 | AFU | Cl | -0.41315 | 0 |
| B137 | B195 | AFU | Na | -0.5994 | 0 |
| B137 | B200 | AFU | OP | -0.28686 | 0 |
| B137 | B150 | AFU | ALP | 0.264603 | 2.22E-16 |
| B137 | B138 | AFU | PAB | -0.20337 | 4.04E-10 |
| B137 | B180 | AFU | Tf | -0.1701 | 1.86E-07 |
| B138 | B139 | PAB | ADA | -0.55876 | 0 |
| B138 | B157 | PAB | IgG | -0.26832 | 0 |
| B138 | B151 | PAB | Anti-CCP | -0.25944 | 8.88E-16 |
| B138 | B194 | PAB | Cl | 0.257126 | 1.78E-15 |
| B138 | B159 | PAB | β2-MG | -0.23922 | 1.52E-13 |
| B138 | B169 | PAB | U-CRY | -0.23922 | 1.52E-13 |
| B138 | B152 | PAB | ACA-Ig(G-A-M) | -0.23921 | 1.53E-13 |
| B138 | B195 | PAB | Na | 0.230646 | 1.14E-12 |
| B138 | B184 | PAB | INR | -0.22658 | 2.87E-12 |
| B138 | B185 | PAB | PT | -0.20314 | 4.23E-10 |
| B138 | B182 | PAB | TT | 0.197383 | 1.32E-09 |
| B138 | B155 | PAB | IgA | -0.16648 | 3.39E-07 |
| B139 | B151 | ADA | Anti-CCP | 0.30815 | 0 |
| B139 | B152 | ADA | ACA-Ig(G-A-M) | 0.436444 | 0 |
| B139 | B155 | ADA | IgA | 0.269165 | 0 |
| B139 | B157 | ADA | IgG | 0.469032 | 0 |
| B139 | B159 | ADA | β2-MG | 0.436469 | 0 |
| B139 | B169 | ADA | U-CRY | 0.436467 | 0 |
| B139 | B180 | ADA | Tf | -0.2679 | 0 |
| B139 | B184 | ADA | INR | 0.305602 | 0 |
| B139 | B193 | ADA | K | -0.27232 | 0 |
| B139 | B194 | ADA | Cl | -0.52179 | 0 |
| B139 | B195 | ADA | Na | -0.38191 | 0 |
| B139 | B198 | ADA | P | 0.238974 | 1.62E-13 |
| B139 | B185 | ADA | PT | 0.22976 | 1.39E-12 |
| B139 | B154 | ADA | RF | -0.21187 | 7.04E-11 |
| B139 | B196 | ADA | TCO2 | 0.193911 | 2.59E-09 |
| B139 | B146 | ADA | CysC | 0.167969 | 2.65E-07 |
| B14 | B114 | G-17 | TSH TT3 TT4(TT4) | 0.177105 | 5.61E-08 |
| B141 | B216 | AST/ALT | BMI | -0.1861 | 1.12E-08 |

| | | | | | |
|---|---|---|---|---|---|
| B141 | B222 | AST/ALT | NS | -0.1616 | 7.47E-07 |
| B142 | B143 | 25-(OH)D | β-CTx | -0.33868 | 0 |
| B142 | B144 | 25-(OH)D | Total-PINP | -0.31317 | 0 |
| B142 | B151 | 25-(OH)D | Anti-CCP | -0.42731 | 0 |
| B142 | B168 | 25-(OH)D | NMP22 | -0.25766 | 1.55E-15 |
| B142 | B145 | 25-(OH)D | BGP | -0.20523 | 2.78E-10 |
| B142 | B175 | 25-(OH)D | ALB-U | -0.16265 | 6.31E-07 |
| B143 | B144 | β-CTx | Total-PINP | 0.799358 | 0 |
| B143 | B145 | β-CTx | BGP | 0.772205 | 0 |
| B143 | B151 | β-CTx | Anti-CCP | 0.278567 | 0 |
| B143 | B175 | β-CTx | ALB-U | 0.267389 | 0 |
| B143 | B191 | β-CTx | Urea | 0.212662 | 5.96E-11 |
| B143 | B177 | β-CTx | ACR | 0.19387 | 2.61E-09 |
| B143 | B190 | β-CTx | Cre | 0.177596 | 5.15E-08 |
| B144 | B145 | Total-PINP | BGP | 0.862777 | 0 |
| B144 | B175 | Total-PINP | ALB-U | 0.529697 | 0 |
| B144 | B177 | Total-PINP | ACR | 0.444951 | 0 |
| B144 | B190 | Total-PINP | Cre | 0.304735 | 0 |
| B144 | B191 | Total-PINP | Urea | 0.337366 | 0 |
| B144 | B151 | Total-PINP | Anti-CCP | 0.243696 | 5.17E-14 |
| B145 | B175 | BGP | ALB-U | 0.622157 | 0 |
| B145 | B177 | BGP | ACR | 0.545746 | 0 |
| B145 | B190 | BGP | Cre | 0.344116 | 0 |
| B145 | B191 | BGP | Urea | 0.364225 | 0 |
| B146 | B214 | CysC | CVA-L | 0.237184 | 2.47E-13 |
| B146 | B173 | CysC | NTT | 0.219377 | 1.41E-11 |
| B146 | B199 | CysC | Mg | -0.16989 | 1.93E-07 |
| B148 | B149 | Anti_TPO | TGAb | 0.204687 | 3.10E-10 |
| B15 | B100 | CD3-CD19+(B cell) | anti-Ro52 | 0.443462 | 0 |
| B15 | B101 | CD3-CD19+(B cell) | anti-Sc70 | 0.443462 | 0 |
| B15 | B102 | CD3-CD19+(B cell) | anti-Sm | 0.443462 | 0 |
| B15 | B103 | CD3-CD19+(B cell) | anti-SSA/Ro | 0.443462 | 0 |
| B15 | B104 | CD3-CD19+(B cell) | anti-SSB | 0.443462 | 0 |
| B15 | B105 | CD3-CD19+(B cell) | anti-U1RNP | 0.443462 | 0 |
| B15 | B106 | CD3-CD19+(B cell) | anti-P | 0.443462 | 0 |
| B15 | B107 | CD3-CD19+(B cell) | AnuA | 0.299599 | 0 |
| B15 | B108 | CD3-CD19+(B cell) | Anti-Cyclin | 0.443462 | 0 |
| B15 | B109 | CD3-CD19+(B cell) | AMA-M2 | 0.443462 | 0 |
| B15 | B110 | CD3-CD19+(B cell) | AHA | 0.299599 | 0 |
| B15 | B139 | CD3-CD19+(B cell) | ADA | -0.48431 | 0 |
| B15 | B157 | CD3-CD19+(B cell) | IgG | -0.41878 | 0 |
| B15 | B17 | CD3-CD19+(B cell) | CD3+CD16+CD56+NK | 0.508556 | 0 |
| B15 | B18 | CD3-CD19+(B cell) | CD3+CD4+helper(inducer) T | -0.36469 | 0 |

| | | | | | |
|---|---|---|---|---|---|
| B15 | B19 | CD3-CD19+(B cell) | CD3+CD4+/CD3+CD8+ | -0.59228 | 0 |
| B15 | B193 | CD3-CD19+(B cell) | K | 0.307483 | 0 |
| B15 | B20 | CD3-CD19+(B cell) | CD3+CD4+CD8+ | -0.61919 | 0 |
| B15 | B21 | CD3-CD19+(B cell) | CD3+CD4-CD8- | 0.602925 | 0 |
| B15 | B22 | CD3-CD19+(B cell) | CD3+CD8+(T) | 0.602957 | 0 |
| B15 | B23 | CD3-CD19+(B cell) | CD3-CD16+CD56+(NK) | -0.40702 | 0 |
| B15 | B95 | CD3-CD19+(B cell) | anti-cenpb | -0.42794 | 0 |
| B15 | B97 | CD3-CD19+(B cell) | dsDNA | 0.443462 | 0 |
| B15 | B98 | CD3-CD19+(B cell) | anti-Jo-1 | 0.443462 | 0 |
| B15 | B99 | CD3-CD19+(B cell) | anti-PM-Scl | 0.443462 | 0 |
| B15 | B135 | CD3-CD19+(B cell) | TBA | -0.264 | 4.44E-16 |
| B15 | B16 | CD3-CD19+(B cell) | CD3+T | 0.256072 | 2.22E-15 |
| B15 | B124 | CD3-CD19+(B cell) | ACTH | -0.23421 | 4.98E-13 |
| B15 | B125 | CD3-CD19+(B cell) | Cor | -0.23421 | 4.98E-13 |
| B15 | B180 | CD3-CD19+(B cell) | Tf | 0.21566 | 3.16E-11 |
| B15 | B138 | CD3-CD19+(B cell) | PAB | 0.188963 | 6.59E-09 |
| B15 | B74 | CD3-CD19+(B cell) | ApoA1 | 0.173054 | 1.13E-07 |
| B15 | B92 | CD3-CD19+(B cell) | ALDH2-Glu504Glu | -0.16403 | 5.05E-07 |
| B15 | B93 | CD3-CD19+(B cell) | ALDH2-Glu504Lys | 0.164034 | 5.05E-07 |
| B15 | B94 | CD3-CD19+(B cell) | ALDH2-Lys504Lys | 0.164034 | 5.05E-07 |
| B15 | B154 | CD3-CD19+(B cell) | RF | 0.159765 | 9.99E-07 |
| B150 | B195 | ALP | Na | -0.26468 | 2.22E-16 |
| B150 | B193 | ALP | K | 0.218778 | 1.61E-11 |
| B150 | B198 | ALP | P | -0.20699 | 1.94E-10 |
| B150 | B200 | ALP | OP | -0.20336 | 4.05E-10 |
| B150 | B181 | ALP | APTT | -0.19764 | 1.26E-09 |
| B151 | B152 | Anti-CCP | ACA-Ig(G-A-M) | 0.31501 | 0 |
| B151 | B159 | Anti-CCP | β2-MG | 0.315028 | 0 |
| B151 | B168 | Anti-CCP | NMP22 | 0.426315 | 0 |
| B151 | B169 | Anti-CCP | U-CRY | 0.315027 | 0 |
| B151 | B194 | Anti-CCP | Cl | -0.22356 | 5.63E-12 |
| B151 | B184 | Anti-CCP | INR | 0.209939 | 1.05E-10 |
| B151 | B158 | Anti-CCP | IgM | 0.195062 | 2.07E-09 |
| B151 | B180 | Anti-CCP | Tf | -0.18634 | 1.07E-08 |
| B151 | B157 | Anti-CCP | IgG | 0.173131 | 1.11E-07 |
| B151 | B154 | Anti-CCP | RF | -0.17032 | 1.79E-07 |
| B151 | B185 | Anti-CCP | PT | 0.163092 | 5.88E-07 |
| B152 | B155 | ACA-Ig(G-A-M) | IgA | 0.340045 | 0 |
| B152 | B159 | ACA-Ig(G-A-M) | β2-MG | 0.999955 | 0 |
| B152 | B169 | ACA-Ig(G-A-M) | U-CRY | 0.999985 | 0 |
| B152 | B184 | ACA-Ig(G-A-M) | INR | 0.250231 | 1.02E-14 |
| B152 | B193 | ACA-Ig(G-A-M) | K | -0.24972 | 1.15E-14 |
| B152 | B157 | ACA-Ig(G-A-M) | IgG | 0.214506 | 4.04E-11 |
| B152 | B180 | ACA-Ig(G-A-M) | Tf | -0.19336 | 2.88E-09 |

| | | | | | |
|---|---|---|---|---|---|
| B152 | B194 | ACA-Ig(G-A-M) | Cl | -0.18858 | 7.08E-09 |
| B152 | B195 | ACA-Ig(G-A-M) | Na | -0.18347 | 1.81E-08 |
| B152 | B185 | ACA-Ig(G-A-M) | PT | 0.165837 | 3.77E-07 |
| B153 | B170 | Glu | U-GLU | 0.520439 | 0 |
| B153 | B163 | Glu | SG | 0.18395 | 1.66E-08 |
| B153 | B165 | Glu | bilirubin | -0.18369 | 1.74E-08 |
| B154 | B194 | RF | Cl | 0.17494 | 8.17E-08 |
| B155 | B157 | IgA | IgG | 0.377399 | 0 |
| B155 | B158 | IgA | IgM | -0.35052 | 0 |
| B155 | B159 | IgA | β2-MG | 0.340064 | 0 |
| B155 | B169 | IgA | U-CRY | 0.340062 | 0 |
| B155 | B193 | IgA | K | -0.1968 | 1.48E-09 |
| B155 | B194 | IgA | Cl | -0.18272 | 2.07E-08 |
| B156 | B196 | IgE | TCO2 | -0.22162 | 8.65E-12 |
| B157 | B193 | IgG | K | -0.2992 | 0 |
| B157 | B159 | IgG | β2-MG | 0.214518 | 4.03E-11 |
| B157 | B169 | IgG | U-CRY | 0.214517 | 4.03E-11 |
| B157 | B184 | IgG | INR | 0.178413 | 4.46E-08 |
| B158 | B199 | IgM | Mg | 0.184385 | 1.53E-08 |
| B158 | B186 | IgM | PFC | -0.1662 | 3.55E-07 |
| B159 | B184 | β2-MG | INR | 0.250246 | 1.02E-14 |
| B159 | B193 | β2-MG | K | -0.24973 | 1.15E-14 |
| B159 | B180 | β2-MG | Tf | -0.19337 | 2.87E-09 |
| B159 | B194 | β2-MG | Cl | -0.18859 | 7.06E-09 |
| B159 | B195 | β2-MG | Na | -0.18348 | 1.80E-08 |
| B159 | B185 | β2-MG | PT | 0.165846 | 3.76E-07 |
| B16 | B100 | CD3+T | anti-Ro52 | 0.385844 | 0 |
| B16 | B101 | CD3+T | anti-Sc70 | 0.385844 | 0 |
| B16 | B102 | CD3+T | anti-Sm | 0.385844 | 0 |
| B16 | B103 | CD3+T | anti-SSA/Ro | 0.385844 | 0 |
| B16 | B104 | CD3+T | anti-SSB | 0.385844 | 0 |
| B16 | B105 | CD3+T | anti-U1RNP | 0.385844 | 0 |
| B16 | B106 | CD3+T | anti-P | 0.385844 | 0 |
| B16 | B107 | CD3+T | AnuA | 0.469506 | 0 |
| B16 | B108 | CD3+T | Anti-Cyclin | 0.385844 | 0 |
| B16 | B109 | CD3+T | AMA-M2 | 0.385844 | 0 |
| B16 | B110 | CD3+T | AHA | 0.469506 | 0 |
| B16 | B119 | CD3+T | Hcv-rna | 0.617272 | 0 |
| B16 | B120 | CD3+T | HCV-cAg | 0.336041 | 0 |
| B16 | B121 | CD3+T | C3 | -0.61727 | 0 |
| B16 | B122 | CD3+T | C4 | -0.61727 | 0 |
| B16 | B124 | CD3+T | ACTH | -0.89126 | 0 |
| B16 | B125 | CD3+T | Cor | -0.89126 | 0 |
| B16 | B135 | CD3+T | TBA | -0.53188 | 0 |

| | | | | | |
|---|---|---|---|---|---|
| B16 | B138 | CD3+T | PAB | 0.294026 | 0 |
| B16 | B139 | CD3+T | ADA | -0.66852 | 0 |
| B16 | B151 | CD3+T | Anti-CCP | -0.42758 | 0 |
| B16 | B152 | CD3+T | ACA-Ig(G-A-M) | -0.61724 | 0 |
| B16 | B155 | CD3+T | IgA | -0.30085 | 0 |
| B16 | B157 | CD3+T | IgG | -0.39327 | 0 |
| B16 | B159 | CD3+T | β2-MG | -0.61727 | 0 |
| B16 | B169 | CD3+T | U-CRY | -0.61727 | 0 |
| B16 | B18 | CD3+T | CD3+CD4+helper(inducer) T | 0.664175 | 0 |
| B16 | B183 | CD3+T | PTA | 0.293286 | 0 |
| B16 | B184 | CD3+T | INR | -0.41453 | 0 |
| B16 | B185 | CD3+T | PT | -0.27597 | 0 |
| B16 | B19 | CD3+T | CD3+CD4+/CD3+CD8+ | -0.38672 | 0 |
| B16 | B193 | CD3+T | K | 0.31108 | 0 |
| B16 | B194 | CD3+T | Cl | 0.418845 | 0 |
| B16 | B22 | CD3+T | CD3+CD8+(T) | 0.74405 | 0 |
| B16 | B23 | CD3+T | CD3-CD16+CD56+(NK) | -0.96923 | 0 |
| B16 | B24 | CD3+T | αHBDH | -0.27985 | 0 |
| B16 | B63 | CD3+T | WBRSR(mPas_200) | -0.28509 | 0 |
| B16 | B64 | CD3+T | WBRSR(mPas_5) | -0.2955 | 0 |
| B16 | B66 | CD3+T | WBRSR(mPas_100) | -0.32463 | 0 |
| B16 | B67 | CD3+T | WBRSR(mPas_50) | -0.32731 | 0 |
| B16 | B79 | CD3+T | hb V -dna | 0.444097 | 0 |
| B16 | B92 | CD3+T | ALDH2-Glu504Glu | -0.62949 | 0 |
| B16 | B93 | CD3+T | ALDH2-Glu504Lys | 0.629494 | 0 |
| B16 | B94 | CD3+T | ALDH2-Lys504Lys | 0.629494 | 0 |
| B16 | B97 | CD3+T | dsDNA | 0.385844 | 0 |
| B16 | B98 | CD3+T | anti-Jo-1 | 0.385844 | 0 |
| B16 | B99 | CD3+T | anti-PM-Scl | 0.385844 | 0 |
| B16 | B180 | CD3+T | Tf | 0.264551 | 2.22E-16 |
| B16 | B197 | CD3+T | AG | -0.26186 | 4.44E-16 |
| B16 | B61 | CD3+T | WBHSRV | -0.26192 | 4.44E-16 |
| B16 | B65 | CD3+T | WBRSR(mPas_1) | -0.25243 | 5.77E-15 |
| B16 | B199 | CD3+T | Mg | -0.24198 | 7.84E-14 |
| B16 | B195 | CD3+T | Na | 0.24074 | 1.06E-13 |
| B16 | B62 | CD3+T | WBHSRVI | -0.21168 | 7.34E-11 |
| B16 | B59 | CD3+T | WBRV | -0.20924 | 1.22E-10 |
| B16 | B198 | CD3+T | P | -0.19583 | 1.79E-09 |
| B16 | B158 | CD3+T | IgM | -0.19347 | 2.82E-09 |
| B16 | B54 | CD3+T | ESR | -0.19187 | 3.82E-09 |
| B16 | B136 | CD3+T | 5'NT | -0.17772 | 5.04E-08 |
| B16 | B60 | CD3+T | WBRVI | -0.17476 | 8.42E-08 |
| B16 | B56 | CD3+T | IR | -0.17125 | 1.53E-07 |

| | | | | | |
|---|---|---|---|---|---|
| B16 | B182 | CD3+T | TT | -0.16038 | 9.06E-07 |
| B160 | B163 | PH | SG | -0.16868 | 2.36E-07 |
| B161 | B173 | LEU | NTT | 0.312406 | 0 |
| B161 | B221 | LEU | Hei | -0.21485 | 3.75E-11 |
| B163 | B166 | SG | Pro | 0.289655 | 0 |
| B163 | B176 | SG | Ucr | 0.287433 | 0 |
| B163 | B216 | SG | BMI | 0.292263 | 0 |
| B163 | B222 | SG | NS | 0.26524 | 2.22E-16 |
| B163 | B170 | SG | U-GLU | 0.245423 | 3.38E-14 |
| B163 | B221 | SG | Hei | 0.218254 | 1.81E-11 |
| B163 | B210 | SG | Hip | 0.213886 | 4.61E-11 |
| B163 | B206 | SG | Waist | 0.196693 | 1.51E-09 |
| B163 | B215 | SG | BMI | 0.196569 | 1.55E-09 |
| B163 | B218 | SG | DBP | 0.162605 | 6.36E-07 |
| B163 | B172 | SG | KB | 0.160254 | 9.25E-07 |
| B163 | B189 | SG | eGFB | 0.160005 | 9.62E-07 |
| B166 | B170 | Pro | U-GLU | 0.19956 | 8.63E-10 |
| B169 | B184 | U-CRY | INR | 0.250244 | 1.02E-14 |
| B169 | B193 | U-CRY | K | -0.24973 | 1.15E-14 |
| B169 | B180 | U-CRY | Tf | -0.19337 | 2.87E-09 |
| B169 | B194 | U-CRY | Cl | -0.18859 | 7.06E-09 |
| B169 | B195 | U-CRY | Na | -0.18348 | 1.80E-08 |
| B169 | B185 | U-CRY | PT | 0.165845 | 3.76E-07 |
| B17 | B100 | CD3+CD16+CD56+NK | anti-Ro52 | 0.506404 | 0 |
| B17 | B101 | CD3+CD16+CD56+NK | anti-Sc70 | 0.506404 | 0 |
| B17 | B102 | CD3+CD16+CD56+NK | anti-Sm | 0.506404 | 0 |
| B17 | B103 | CD3+CD16+CD56+NK | anti-SSA/Ro | 0.506404 | 0 |
| B17 | B104 | CD3+CD16+CD56+NK | anti-SSB | 0.506404 | 0 |
| B17 | B105 | CD3+CD16+CD56+NK | anti-U1RNP | 0.506404 | 0 |
| B17 | B106 | CD3+CD16+CD56+NK | anti-P | 0.506404 | 0 |
| B17 | B107 | CD3+CD16+CD56+NK | AnuA | 0.547413 | 0 |
| B17 | B108 | CD3+CD16+CD56+NK | Anti-Cyclin | 0.506404 | 0 |
| B17 | B109 | CD3+CD16+CD56+NK | AMA-M2 | 0.506404 | 0 |
| B17 | B110 | CD3+CD16+CD56+NK | AHA | 0.547413 | 0 |
| B17 | B18 | CD3+CD16+CD56+NK | CD3+CD4+helper(inducer) T | -0.38716 | 0 |
| B17 | B181 | CD3+CD16+CD56+NK | APTT | -0.27186 | 0 |
| B17 | B20 | CD3+CD16+CD56+NK | CD3+CD4+CD8+ | -0.34185 | 0 |
| B17 | B21 | CD3+CD16+CD56+NK | CD3+CD4-CD8- | 0.977995 | 0 |
| B17 | B22 | CD3+CD16+CD56+NK | CD3+CD8+(T) | 0.269626 | 0 |
| B17 | B34 | CD3+CD16+CD56+NK | TK | 0.27812 | 0 |
| B17 | B97 | CD3+CD16+CD56+NK | dsDNA | 0.506404 | 0 |
| B17 | B98 | CD3+CD16+CD56+NK | anti-Jo-1 | 0.506404 | 0 |
| B17 | B99 | CD3+CD16+CD56+NK | anti-PM-Scl | 0.506404 | 0 |

| | | | | | |
|---|---|---|---|---|---|
| B17 | B19 | CD3+CD16+CD56+NK | CD3+CD4+/CD3+CD8+ | -0.24112 | 9.68E-14 |
| B17 | B156 | CD3+CD16+CD56+NK | IgE | -0.20062 | 7.00E-10 |
| B17 | B180 | CD3+CD16+CD56+NK | Tf | 0.182719 | 2.07E-08 |
| B17 | B183 | CD3+CD16+CD56+NK | PTA | 0.179299 | 3.81E-08 |
| B17 | B196 | CD3+CD16+CD56+NK | TCO2 | 0.173565 | 1.03E-07 |
| B17 | B73 | CD3+CD16+CD56+NK | ApoA | -0.17159 | 1.45E-07 |
| B17 | B151 | CD3+CD16+CD56+NK | Anti-CCP | 0.170202 | 1.83E-07 |
| B17 | B23 | CD3+CD16+CD56+NK | CD3-CD16+CD56+(NK) | -0.16127 | 7.87E-07 |
| B172 | B200 | KB | OP | 0.252669 | 5.55E-15 |
| B173 | B221 | NTT | Hei | -0.17758 | 5.16E-08 |
| B174 | B190 | Urea/Cr | Cre | -0.30404 | 0 |
| B174 | B191 | Urea/Cr | Urea | 0.304336 | 0 |
| B174 | B221 | Urea/Cr | Hei | -0.26461 | 2.22E-16 |
| B174 | B222 | Urea/Cr | NS | -0.18251 | 2.15E-08 |
| B174 | B189 | Urea/Cr | eGFB | 0.1653 | 4.11E-07 |
| B175 | B177 | ALB-U | ACR | 0.929329 | 0 |
| B175 | B190 | ALB-U | Cre | 0.359039 | 0 |
| B175 | B191 | ALB-U | Urea | 0.389708 | 0 |
| B176 | B204 | Ucr | Drinking | -0.21962 | 1.34E-11 |
| B176 | B177 | Ucr | ACR | -0.18957 | 5.89E-09 |
| B176 | B178 | Ucr | Ig-U | 0.182815 | 2.04E-08 |
| B176 | B179 | Ucr | α1-MG | 0.175852 | 6.97E-08 |
| B176 | B209 | Ucr | Smoking | -0.17462 | 8.63E-08 |
| B177 | B190 | ACR | Cre | 0.328991 | 0 |
| B177 | B191 | ACR | Urea | 0.349128 | 0 |
| B178 | B179 | Ig-U | α1-MG | 0.85617 | 0 |
| B178 | B180 | Ig-U | Tf | 0.786737 | 0 |
| B179 | B180 | α1-MG | Tf | 0.425214 | 0 |
| B18 | B119 | CD3+CD4+helper(inducer)T | Hcv-rna | 0.27889 | 0 |
| B18 | B121 | CD3+CD4+helper(inducer)T | C3 | -0.27889 | 0 |
| B18 | B122 | CD3+CD4+helper(inducer)T | C4 | -0.27889 | 0 |
| B18 | B124 | CD3+CD4+helper(inducer)T | ACTH | -0.46313 | 0 |
| B18 | B125 | CD3+CD4+helper(inducer)T | Cor | -0.46313 | 0 |
| B18 | B139 | CD3+CD4+helper(inducer)T | ADA | -0.30785 | 0 |
| B18 | B152 | CD3+CD4+helper(inducer)T | ACA-Ig(G-A-M) | -0.27887 | 0 |
| B18 | B159 | CD3+CD4+helper(inducer)T | β2-MG | -0.27889 | 0 |
| B18 | B169 | CD3+CD4+helper(inducer)T | U-CRY | -0.27889 | 0 |
| B18 | B19 | CD3+CD4+helper(inducer)T | CD3+CD4+/CD3+CD8+ | 0.352986 | 0 |
| B18 | B194 | CD3+CD4+helper(inducer)T | Cl | 0.271142 | 0 |
| B18 | B20 | CD3+CD4+helper(inducer)T | CD3+CD4+CD8+ | 0.356158 | 0 |
| B18 | B21 | CD3+CD4+helper(inducer)T | CD3+CD4-CD8- | -0.46846 | 0 |
| B18 | B23 | CD3+CD4+helper(inducer)T | CD3-CD16+CD56+(NK) | -0.57493 | 0 |
| B18 | B92 | CD3+CD4+helper(inducer)T | ALDH2-Glu504Glu | -0.32797 | 0 |
| B18 | B93 | CD3+CD4+helper(inducer)T | ALDH2-Glu504Lys | 0.32797 | 0 |

| | | | | | |
|---|---|---|---|---|---|
| B18 | B94 | CD3+CD4+helper(inducer)T | ALDH2-Lys504Lys | 0.32797 | 0 |
| B18 | B95 | CD3+CD4+helper(inducer)T | anti-cenpb | 0.547557 | 0 |
| B18 | B199 | CD3+CD4+helper(inducer)T | Mg | -0.26511 | 2.22E-16 |
| B18 | B135 | CD3+CD4+helper(inducer)T | TBA | -0.25813 | 1.33E-15 |
| B18 | B79 | CD3+CD4+helper(inducer)T | hb V -dna | 0.232597 | 7.25E-13 |
| B18 | B151 | CD3+CD4+helper(inducer)T | Anti-CCP | -0.23224 | 7.87E-13 |
| B18 | B184 | CD3+CD4+helper(inducer)T | INR | -0.21107 | 8.34E-11 |
| B18 | B198 | CD3+CD4+helper(inducer)T | P | -0.2061 | 2.32E-10 |
| B18 | B100 | CD3+CD4+helper(inducer)T | anti-Ro52 | -0.1955 | 1.91E-09 |
| B18 | B101 | CD3+CD4+helper(inducer)T | anti-Sc70 | -0.1955 | 1.91E-09 |
| B18 | B102 | CD3+CD4+helper(inducer)T | anti-Sm | -0.1955 | 1.91E-09 |
| B18 | B103 | CD3+CD4+helper(inducer)T | anti-SSA/Ro | -0.1955 | 1.91E-09 |
| B18 | B104 | CD3+CD4+helper(inducer)T | anti-SSB | -0.1955 | 1.91E-09 |
| B18 | B105 | CD3+CD4+helper(inducer)T | anti-U1RNP | -0.1955 | 1.91E-09 |
| B18 | B106 | CD3+CD4+helper(inducer)T | anti-P | -0.1955 | 1.91E-09 |
| B18 | B108 | CD3+CD4+helper(inducer)T | Anti-Cyclin | -0.1955 | 1.91E-09 |
| B18 | B109 | CD3+CD4+helper(inducer)T | AMA-M2 | -0.1955 | 1.91E-09 |
| B18 | B97 | CD3+CD4+helper(inducer)T | dsDNA | -0.1955 | 1.91E-09 |
| B18 | B98 | CD3+CD4+helper(inducer)T | anti-Jo-1 | -0.1955 | 1.91E-09 |
| B18 | B99 | CD3+CD4+helper(inducer)T | anti-PM-Scl | -0.1955 | 1.91E-09 |
| B18 | B67 | CD3+CD4+helper(inducer)T | WBRSR(mPas_50) | -0.17456 | 8.72E-08 |
| B18 | B64 | CD3+CD4+helper(inducer)T | WBRSR(mPas_5) | -0.17372 | 1.01E-07 |
| B18 | B195 | CD3+CD4+helper(inducer)T | Na | 0.172109 | 1.33E-07 |
| B18 | B211 | CD3+CD4+helper(inducer)T | IOP-R | -0.17061 | 1.71E-07 |
| B18 | B66 | CD3+CD4+helper(inducer)T | WBRSR(mPas_100) | -0.16933 | 2.12E-07 |
| B18 | B197 | CD3+CD4+helper(inducer)T | AG | -0.16502 | 4.31E-07 |
| B18 | B155 | CD3+CD4+helper(inducer)T | IgA | -0.16308 | 5.89E-07 |
| B18 | B65 | CD3+CD4+helper(inducer)T | WBRSR(mPas_1) | -0.1612 | 7.96E-07 |
| B180 | B194 | Tf | Cl | 0.304139 | 0 |
| B180 | B195 | Tf | Na | 0.187413 | 8.79E-09 |
| B181 | B182 | APTT | TT | 0.638126 | 0 |
| B181 | B183 | APTT | PTA | 0.524274 | 0 |
| B181 | B184 | APTT | INR | 0.822094 | 0 |
| B181 | B185 | APTT | PT | 0.78087 | 0 |
| B182 | B184 | TT | INR | 0.386807 | 0 |
| B182 | B186 | TT | PFC | -0.75055 | 0 |
| B182 | B185 | TT | PT | 0.241921 | 7.95E-14 |
| B182 | B197 | TT | AG | 0.160609 | 8.74E-07 |
| B183 | B184 | PTA | INR | 0.586316 | 0 |
| B183 | B185 | PTA | PT | 0.619629 | 0 |
| B183 | B197 | PTA | AG | -0.20533 | 2.72E-10 |
| B183 | B186 | PTA | PFC | 0.191225 | 4.31E-09 |
| B184 | B185 | INR | PT | 0.956719 | 0 |
| B184 | B194 | INR | Cl | -0.23652 | 2.90E-13 |

| | | | | | |
|---|---|---|---|---|---|
| B184 | B193 | INR | K | -0.21381 | 4.68E-11 |
| B184 | B195 | INR | Na | -0.16261 | 6.35E-07 |
| B185 | B186 | PT | PFC | 0.280722 | 0 |
| B185 | B194 | PT | Cl | -0.17552 | 7.39E-08 |
| B185 | B193 | PT | K | -0.16928 | 2.13E-07 |
| B186 | B195 | PFC | Na | -0.17553 | 7.37E-08 |
| B186 | B200 | PFC | OP | -0.1639 | 5.16E-07 |
| B189 | B190 | eGFB | Cre | -0.55214 | 0 |
| B189 | B191 | eGFB | Urea | -0.57439 | 0 |
| B189 | B192 | eGFB | UA | -0.32807 | 0 |
| B189 | B218 | eGFB | DBP | 0.196044 | 1.72E-09 |
| B189 | B203 | eGFB | DH | 0.171545 | 1.46E-07 |
| B19 | B100 | CD3+CD4+/CD3+CD8+ | anti-Ro52 | -0.58695 | 0 |
| B19 | B101 | CD3+CD4+/CD3+CD8+ | anti-Sc70 | -0.58695 | 0 |
| B19 | B102 | CD3+CD4+/CD3+CD8+ | anti-Sm | -0.58695 | 0 |
| B19 | B103 | CD3+CD4+/CD3+CD8+ | anti-SSA/Ro | -0.58695 | 0 |
| B19 | B104 | CD3+CD4+/CD3+CD8+ | anti-SSB | -0.58695 | 0 |
| B19 | B105 | CD3+CD4+/CD3+CD8+ | anti-U1RNP | -0.58695 | 0 |
| B19 | B106 | CD3+CD4+/CD3+CD8+ | anti-P | -0.58695 | 0 |
| B19 | B107 | CD3+CD4+/CD3+CD8+ | AnuA | -0.37269 | 0 |
| B19 | B108 | CD3+CD4+/CD3+CD8+ | Anti-Cyclin | -0.58695 | 0 |
| B19 | B109 | CD3+CD4+/CD3+CD8+ | AMA-M2 | -0.58695 | 0 |
| B19 | B110 | CD3+CD4+/CD3+CD8+ | AHA | -0.37269 | 0 |
| B19 | B114 | CD3+CD4+/CD3+CD8+ | TSH TT3 TT4(TT4) | 0.274132 | 0 |
| B19 | B119 | CD3+CD4+/CD3+CD8+ | Hcv-rna | -0.37435 | 0 |
| B19 | B121 | CD3+CD4+/CD3+CD8+ | C3 | 0.37435 | 0 |
| B19 | B122 | CD3+CD4+/CD3+CD8+ | C4 | 0.37435 | 0 |
| B19 | B124 | CD3+CD4+/CD3+CD8+ | ACTH | 0.63744 | 0 |
| B19 | B125 | CD3+CD4+/CD3+CD8+ | Cor | 0.63744 | 0 |
| B19 | B135 | CD3+CD4+/CD3+CD8+ | TBA | 0.410616 | 0 |
| B19 | B138 | CD3+CD4+/CD3+CD8+ | PAB | -0.35872 | 0 |
| B19 | B139 | CD3+CD4+/CD3+CD8+ | ADA | 0.410459 | 0 |
| B19 | B151 | CD3+CD4+/CD3+CD8+ | Anti-CCP | 0.355992 | 0 |
| B19 | B152 | CD3+CD4+/CD3+CD8+ | ACA-Ig(G-A-M) | 0.374329 | 0 |
| B19 | B157 | CD3+CD4+/CD3+CD8+ | IgG | 0.378775 | 0 |
| B19 | B158 | CD3+CD4+/CD3+CD8+ | IgM | 0.289804 | 0 |
| B19 | B159 | CD3+CD4+/CD3+CD8+ | β2-MG | 0.37435 | 0 |
| B19 | B169 | CD3+CD4+/CD3+CD8+ | U-CRY | 0.374348 | 0 |
| B19 | B180 | CD3+CD4+/CD3+CD8+ | Tf | -0.29745 | 0 |
| B19 | B184 | CD3+CD4+/CD3+CD8+ | INR | 0.273091 | 0 |
| B19 | B193 | CD3+CD4+/CD3+CD8+ | K | -0.45401 | 0 |
| B19 | B21 | CD3+CD4+/CD3+CD8+ | CD3+CD4-CD8- | -0.29308 | 0 |
| B19 | B22 | CD3+CD4+/CD3+CD8+ | CD3+CD8+(T) | -0.89384 | 0 |
| B19 | B23 | CD3+CD4+/CD3+CD8+ | CD3-CD16+CD56+(NK) | 0.456717 | 0 |

| | | | | | |
|---|---|---|---|---|---|
| B19 | B79 | CD3+CD4+/CD3+CD8+ | hb V -dna | -0.31675 | 0 |
| B19 | B92 | CD3+CD4+/CD3+CD8+ | ALDH2-Glu504Glu | 0.449808 | 0 |
| B19 | B93 | CD3+CD4+/CD3+CD8+ | ALDH2-Glu504Lys | -0.44981 | 0 |
| B19 | B94 | CD3+CD4+/CD3+CD8+ | ALDH2-Lys504Lys | -0.44981 | 0 |
| B19 | B95 | CD3+CD4+/CD3+CD8+ | anti-cenpb | 0.625369 | 0 |
| B19 | B97 | CD3+CD4+/CD3+CD8+ | dsDNA | -0.58695 | 0 |
| B19 | B98 | CD3+CD4+/CD3+CD8+ | anti-Jo-1 | -0.58695 | 0 |
| B19 | B99 | CD3+CD4+/CD3+CD8+ | anti-PM-Scl | -0.58695 | 0 |
| B19 | B20 | CD3+CD4+/CD3+CD8+ | CD3+CD4+CD8+ | 0.254105 | 4.00E-15 |
| B19 | B156 | CD3+CD4+/CD3+CD8+ | IgE | -0.22696 | 2.63E-12 |
| B19 | B194 | CD3+CD4+/CD3+CD8+ | Cl | -0.21328 | 5.23E-11 |
| B19 | B66 | CD3+CD4+/CD3+CD8+ | WBRSR(mPas_100) | 0.203065 | 4.30E-10 |
| B19 | B120 | CD3+CD4+/CD3+CD8+ | HCV-cAg | -0.20213 | 5.18E-10 |
| B19 | B67 | CD3+CD4+/CD3+CD8+ | WBRSR(mPas_50) | 0.198093 | 1.15E-09 |
| B19 | B24 | CD3+CD4+/CD3+CD8+ | αHBDH | 0.195601 | 1.87E-09 |
| B19 | B185 | CD3+CD4+/CD3+CD8+ | PT | 0.191902 | 3.80E-09 |
| B19 | B63 | CD3+CD4+/CD3+CD8+ | WBRSR(mPas_200) | 0.190859 | 4.62E-09 |
| B19 | B62 | CD3+CD4+/CD3+CD8+ | WBHSRVI | 0.169139 | 2.18E-07 |
| B19 | B61 | CD3+CD4+/CD3+CD8+ | WBHSRV | 0.164573 | 4.63E-07 |
| B19 | B136 | CD3+CD4+/CD3+CD8+ | 5'NT | 0.164345 | 4.80E-07 |
| B19 | B183 | CD3+CD4+/CD3+CD8+ | PTA | -0.16073 | 8.58E-07 |
| B190 | B191 | Cre | Urea | 0.619123 | 0 |
| B190 | B207 | Cre | WHR | 0.434298 | 0 |
| B190 | B192 | Cre | UA | 0.215575 | 3.21E-11 |
| B191 | B192 | Urea | UA | 0.281736 | 0 |
| B193 | B198 | K | P | -0.19924 | 9.20E-10 |
| B193 | B195 | K | Na | -0.17746 | 5.27E-08 |
| B193 | B196 | K | TCO2 | -0.16654 | 3.36E-07 |
| B193 | B197 | K | AG | 0.16062 | 8.73E-07 |
| B194 | B195 | Cl | Na | 0.497107 | 0 |
| B194 | B196 | Cl | TCO2 | -0.49512 | 0 |
| B194 | B197 | Cl | AG | -0.2978 | 0 |
| B194 | B198 | Cl | P | -0.16329 | 5.70E-07 |
| B194 | B200 | Cl | OP | 0.161571 | 7.50E-07 |
| B195 | B197 | Na | AG | 0.269477 | 0 |
| B195 | B200 | Na | OP | 0.673743 | 0 |
| B195 | B198 | Na | P | 0.198834 | 9.96E-10 |
| B196 | B197 | TCO2 | AG | -0.43419 | 0 |
| B196 | B198 | TCO2 | P | -0.18118 | 2.73E-08 |
| B197 | B198 | AG | P | 0.535825 | 0 |
| B197 | B200 | AG | OP | 0.467884 | 0 |
| B198 | B200 | P | OP | 0.285742 | 0 |
| B198 | B199 | P | Mg | -0.18502 | 1.36E-08 |
| B199 | B200 | Mg | OP | -0.34061 | 0 |

| | | | | | |
|---|---|---|---|---|---|
| B2 | B163 | Age | SG | -0.29075 | 0 |
| B2 | B189 | Age | eGFB | -0.42783 | 0 |
| B2 | B216 | Age | BMI | -0.39055 | 0 |
| B2 | B218 | Age | DBP | -0.30821 | 0 |
| B2 | B221 | Age | Hei | -0.3635 | 0 |
| B2 | B222 | Age | NS | -0.27285 | 0 |
| B2 | B46 | Age | RBC Count | -0.31889 | 0 |
| B2 | B51 | Age | Hb | -0.31657 | 0 |
| B2 | B141 | Age | AST/ALT | 0.254601 | 3.55E-15 |
| B2 | B126 | Age | Alb | -0.25378 | 4.22E-15 |
| B2 | B128 | Age | ALT | -0.23108 | 1.03E-12 |
| B2 | B47 | Age | RDW | 0.209134 | 1.25E-10 |
| B2 | B127 | Age | A-G ratio | -0.19663 | 1.53E-09 |
| B2 | B215 | Age | BMI | -0.19165 | 3.98E-09 |
| B2 | B191 | Age | Urea | 0.179213 | 3.87E-08 |
| B2 | B176 | Age | Ucr | -0.17267 | 1.21E-07 |
| B2 | B203 | Age | DH | -0.16639 | 3.44E-07 |
| B2 | B210 | Age | Hip | -0.16619 | 3.55E-07 |
| B2 | B71 | Age | TG | -0.16608 | 3.62E-07 |
| B2 | B209 | Age | Smoking | -0.16416 | 4.95E-07 |
| B2 | B220 | Age | SBP | 0.16034 | 9.12E-07 |
| B20 | B21 | CD3+CD4+CD8+ | CD3+CD4-CD8- | -0.46032 | 0 |
| B20 | B87 | CD3+CD4+CD8+ | NSE | 0.288252 | 0 |
| B20 | B22 | CD3+CD4+CD8+ | CD3+CD8+(T) | -0.24732 | 2.13E-14 |
| B20 | B199 | CD3+CD4+CD8+ | Mg | -0.19316 | 2.99E-09 |
| B20 | B74 | CD3+CD4+CD8+ | ApoA1 | -0.1605 | 8.89E-07 |
| B201 | B204 | e-habits | Drinking | 0.204584 | 3.16E-10 |
| B204 | B209 | Drinking | Smoking | 0.517493 | 0 |
| B205 | B221 | KDA | Hei | -0.17776 | 5.00E-08 |
| B206 | B207 | Waist | WHR | 0.513301 | 0 |
| B206 | B210 | Waist | Hip | 0.736295 | 0 |
| B206 | B215 | Waist | BMI | 0.794754 | 0 |
| B206 | B216 | Waist | BMI | 0.75418 | 0 |
| B206 | B222 | Waist | NS | 0.635494 | 0 |
| B206 | B221 | Waist | Hei | 0.257134 | 1.78E-15 |
| B207 | B215 | WHR | BMI | 0.30677 | 0 |
| B207 | B216 | WHR | BMI | 0.251805 | 6.88E-15 |
| B207 | B222 | WHR | NS | 0.222942 | 6.47E-12 |
| B209 | B221 | Smoking | Hei | 0.224612 | 4.46E-12 |
| B209 | B222 | Smoking | NS | 0.177199 | 5.52E-08 |
| B21 | B100 | CD3+CD4-CD8- | anti-Ro52 | 0.478849 | 0 |
| B21 | B101 | CD3+CD4-CD8- | anti-Sc70 | 0.478849 | 0 |
| B21 | B102 | CD3+CD4-CD8- | anti-Sm | 0.478849 | 0 |
| B21 | B103 | CD3+CD4-CD8- | anti-SSA/Ro | 0.478849 | 0 |

| | | | | | |
|---|---|---|---|---|---|
| B21 | B104 | CD3+CD4-CD8- | anti-SSB | 0.478849 | 0 |
| B21 | B105 | CD3+CD4-CD8- | anti-U1RNP | 0.478849 | 0 |
| B21 | B106 | CD3+CD4-CD8- | anti-P | 0.478849 | 0 |
| B21 | B107 | CD3+CD4-CD8- | AnuA | 0.517672 | 0 |
| B21 | B108 | CD3+CD4-CD8- | Anti-Cyclin | 0.478849 | 0 |
| B21 | B109 | CD3+CD4-CD8- | AMA-M2 | 0.478849 | 0 |
| B21 | B110 | CD3+CD4-CD8- | AHA | 0.517672 | 0 |
| B21 | B22 | CD3+CD4-CD8- | CD3+CD8+(T) | 0.28714 | 0 |
| B21 | B97 | CD3+CD4-CD8- | dsDNA | 0.478849 | 0 |
| B21 | B98 | CD3+CD4-CD8- | anti-Jo-1 | 0.478849 | 0 |
| B21 | B99 | CD3+CD4-CD8- | anti-PM-Scl | 0.478849 | 0 |
| B21 | B34 | CD3+CD4-CD8- | TK | 0.208689 | 1.37E-10 |
| B21 | B180 | CD3+CD4-CD8- | Tf | 0.208521 | 1.41E-10 |
| B21 | B181 | CD3+CD4-CD8- | APTT | -0.20403 | 3.54E-10 |
| B210 | B215 | Hip | BMI | 0.70934 | 0 |
| B210 | B216 | Hip | BMI | 0.731883 | 0 |
| B210 | B221 | Hip | Hei | 0.325612 | 0 |
| B210 | B222 | Hip | NS | 0.632276 | 0 |
| B211 | B212 | IOP-R | IOP-L | 0.742208 | 0 |
| B213 | B214 | CVA-R | CVA-L | -0.17209 | 1.33E-07 |
| B215 | B216 | BMI | BMI | 0.739076 | 0 |
| B215 | B222 | BMI | NS | 0.549001 | 0 |
| B216 | B221 | BMI | Hei | 0.683234 | 0 |
| B216 | B222 | BMI | NS | 0.752759 | 0 |
| B216 | B218 | BMI | DBP | 0.209781 | 1.09E-10 |
| B218 | B220 | DBP | SBP | 0.408534 | 0 |
| B218 | B221 | DBP | Hei | 0.191096 | 4.42E-09 |
| B22 | B100 | CD3+CD8+(T) | anti-Ro52 | 0.645824 | 0 |
| B22 | B101 | CD3+CD8+(T) | anti-Sc70 | 0.645824 | 0 |
| B22 | B102 | CD3+CD8+(T) | anti-Sm | 0.645824 | 0 |
| B22 | B103 | CD3+CD8+(T) | anti-SSA/Ro | 0.645824 | 0 |
| B22 | B104 | CD3+CD8+(T) | anti-SSB | 0.645824 | 0 |
| B22 | B105 | CD3+CD8+(T) | anti-U1RNP | 0.645824 | 0 |
| B22 | B106 | CD3+CD8+(T) | anti-P | 0.645824 | 0 |
| B22 | B107 | CD3+CD8+(T) | AnuA | 0.528359 | 0 |
| B22 | B108 | CD3+CD8+(T) | Anti-Cyclin | 0.645824 | 0 |
| B22 | B109 | CD3+CD8+(T) | AMA-M2 | 0.645824 | 0 |
| B22 | B110 | CD3+CD8+(T) | AHA | 0.528359 | 0 |
| B22 | B119 | CD3+CD8+(T) | Hcv-rna | 0.578055 | 0 |
| B22 | B120 | CD3+CD8+(T) | HCV-cAg | 0.313095 | 0 |
| B22 | B121 | CD3+CD8+(T) | C3 | -0.57806 | 0 |
| B22 | B122 | CD3+CD8+(T) | C4 | -0.57806 | 0 |
| B22 | B124 | CD3+CD8+(T) | ACTH | -0.83621 | 0 |
| B22 | B125 | CD3+CD8+(T) | Cor | -0.83621 | 0 |

| | | | | | |
|---|---|---|---|---|---|
| B22 | B135 | CD3+CD8+(T) | TBA | -0.52147 | 0 |
| B22 | B138 | CD3+CD8+(T) | PAB | 0.38841 | 0 |
| B22 | B139 | CD3+CD8+(T) | ADA | -0.62706 | 0 |
| B22 | B151 | CD3+CD8+(T) | Anti-CCP | -0.42439 | 0 |
| B22 | B152 | CD3+CD8+(T) | ACA-Ig(G-A-M) | -0.57802 | 0 |
| B22 | B157 | CD3+CD8+(T) | IgG | -0.46782 | 0 |
| B22 | B158 | CD3+CD8+(T) | IgM | -0.2665 | 0 |
| B22 | B159 | CD3+CD8+(T) | β2-MG | -0.57806 | 0 |
| B22 | B169 | CD3+CD8+(T) | U-CRY | -0.57805 | 0 |
| B22 | B180 | CD3+CD8+(T) | Tf | 0.330112 | 0 |
| B22 | B184 | CD3+CD8+(T) | INR | -0.37799 | 0 |
| B22 | B193 | CD3+CD8+(T) | K | 0.488858 | 0 |
| B22 | B194 | CD3+CD8+(T) | Cl | 0.330778 | 0 |
| B22 | B23 | CD3+CD8+(T) | CD3-CD16+CD56+(NK) | -0.77592 | 0 |
| B22 | B66 | CD3+CD8+(T) | WBRSR(mPas_100) | -0.29045 | 0 |
| B22 | B67 | CD3+CD8+(T) | WBRSR(mPas_50) | -0.28876 | 0 |
| B22 | B79 | CD3+CD8+(T) | hb V -dna | 0.41559 | 0 |
| B22 | B92 | CD3+CD8+(T) | ALDH2-Glu504Glu | -0.5901 | 0 |
| B22 | B93 | CD3+CD8+(T) | ALDH2-Glu504Lys | 0.590103 | 0 |
| B22 | B94 | CD3+CD8+(T) | ALDH2-Lys504Lys | 0.590103 | 0 |
| B22 | B95 | CD3+CD8+(T) | anti-cenpb | -0.39561 | 0 |
| B22 | B97 | CD3+CD8+(T) | dsDNA | 0.645824 | 0 |
| B22 | B98 | CD3+CD8+(T) | anti-Jo-1 | 0.645824 | 0 |
| B22 | B99 | CD3+CD8+(T) | anti-PM-Scl | 0.645824 | 0 |
| B22 | B63 | CD3+CD8+(T) | WBRSR(mPas_200) | -0.26266 | 4.44E-16 |
| B22 | B24 | CD3+CD8+(T) | αHBDH | -0.25883 | 1.11E-15 |
| B22 | B185 | CD3+CD8+(T) | PT | -0.25537 | 2.66E-15 |
| B22 | B183 | CD3+CD8+(T) | PTA | 0.254205 | 3.77E-15 |
| B22 | B64 | CD3+CD8+(T) | WBRSR(mPas_5) | -0.24495 | 3.80E-14 |
| B22 | B155 | CD3+CD8+(T) | IgA | -0.24277 | 6.48E-14 |
| B22 | B61 | CD3+CD8+(T) | WBHSRV | -0.23476 | 4.37E-13 |
| B22 | B114 | CD3+CD8+(T) | TSH TT3 TT4(TT4) | -0.2223 | 7.46E-12 |
| B22 | B62 | CD3+CD8+(T) | WBHSRVI | -0.2136 | 4.89E-11 |
| B22 | B197 | CD3+CD8+(T) | AG | -0.19806 | 1.16E-09 |
| B22 | B65 | CD3+CD8+(T) | WBRSR(mPas_1) | -0.19661 | 1.54E-09 |
| B22 | B136 | CD3+CD8+(T) | 5'NT | -0.17126 | 1.53E-07 |
| B22 | B156 | CD3+CD8+(T) | IgE | 0.170248 | 1.81E-07 |
| B22 | B154 | CD3+CD8+(T) | RF | 0.169713 | 1.98E-07 |
| B22 | B56 | CD3+CD8+(T) | IR | -0.16896 | 2.25E-07 |
| B220 | B221 | SBP | Hei | -0.17961 | 3.61E-08 |
| B221 | B222 | Hei | NS | 0.530881 | 0 |
| B23 | B100 | CD3-CD16+CD56+(NK) | anti-Ro52 | -0.36042 | 0 |
| B23 | B101 | CD3-CD16+CD56+(NK) | anti-Sc70 | -0.36042 | 0 |
| B23 | B102 | CD3-CD16+CD56+(NK) | anti-Sm | -0.36042 | 0 |

| | | | | | |
|---|---|---|---|---|---|
| B23 | B103 | CD3-CD16+CD56+(NK) | anti-SSA/Ro | -0.36042 | 0 |
| B23 | B104 | CD3-CD16+CD56+(NK) | anti-SSB | -0.36042 | 0 |
| B23 | B105 | CD3-CD16+CD56+(NK) | anti-U1RNP | -0.36042 | 0 |
| B23 | B106 | CD3-CD16+CD56+(NK) | anti-P | -0.36042 | 0 |
| B23 | B107 | CD3-CD16+CD56+(NK) | AnuA | -0.43092 | 0 |
| B23 | B108 | CD3-CD16+CD56+(NK) | Anti-Cyclin | -0.36042 | 0 |
| B23 | B109 | CD3-CD16+CD56+(NK) | AMA-M2 | -0.36042 | 0 |
| B23 | B110 | CD3-CD16+CD56+(NK) | AHA | -0.43092 | 0 |
| B23 | B119 | CD3-CD16+CD56+(NK) | Hcv-rna | -0.5182 | 0 |
| B23 | B120 | CD3-CD16+CD56+(NK) | HCV-cAg | -0.28136 | 0 |
| B23 | B121 | CD3-CD16+CD56+(NK) | C3 | 0.5182 | 0 |
| B23 | B122 | CD3-CD16+CD56+(NK) | C4 | 0.518199 | 0 |
| B23 | B124 | CD3-CD16+CD56+(NK) | ACTH | 0.899773 | 0 |
| B23 | B125 | CD3-CD16+CD56+(NK) | Cor | 0.899773 | 0 |
| B23 | B135 | CD3-CD16+CD56+(NK) | TBA | 0.57149 | 0 |
| B23 | B138 | CD3-CD16+CD56+(NK) | PAB | -0.29828 | 0 |
| B23 | B139 | CD3-CD16+CD56+(NK) | ADA | 0.677416 | 0 |
| B23 | B151 | CD3-CD16+CD56+(NK) | Anti-CCP | 0.441254 | 0 |
| B23 | B152 | CD3-CD16+CD56+(NK) | ACA-Ig(G-A-M) | 0.51817 | 0 |
| B23 | B157 | CD3-CD16+CD56+(NK) | IgG | 0.414368 | 0 |
| B23 | B159 | CD3-CD16+CD56+(NK) | β2-MG | 0.5182 | 0 |
| B23 | B169 | CD3-CD16+CD56+(NK) | U-CRY | 0.518197 | 0 |
| B23 | B180 | CD3-CD16+CD56+(NK) | Tf | -0.29804 | 0 |
| B23 | B183 | CD3-CD16+CD56+(NK) | PTA | -0.28655 | 0 |
| B23 | B184 | CD3-CD16+CD56+(NK) | INR | 0.414032 | 0 |
| B23 | B185 | CD3-CD16+CD56+(NK) | PT | 0.278718 | 0 |
| B23 | B193 | CD3-CD16+CD56+(NK) | K | -0.30012 | 0 |
| B23 | B194 | CD3-CD16+CD56+(NK) | Cl | -0.42043 | 0 |
| B23 | B24 | CD3-CD16+CD56+(NK) | αHBDH | 0.281761 | 0 |
| B23 | B63 | CD3-CD16+CD56+(NK) | WBRSR(mPas_200) | 0.285976 | 0 |
| B23 | B64 | CD3-CD16+CD56+(NK) | WBRSR(mPas_5) | 0.301042 | 0 |
| B23 | B66 | CD3-CD16+CD56+(NK) | WBRSR(mPas_100) | 0.326208 | 0 |
| B23 | B67 | CD3-CD16+CD56+(NK) | WBRSR(mPas_50) | 0.329607 | 0 |
| B23 | B79 | CD3-CD16+CD56+(NK) | hb V -dna | -0.44822 | 0 |
| B23 | B92 | CD3-CD16+CD56+(NK) | ALDH2-Glu504Glu | 0.635449 | 0 |
| B23 | B93 | CD3-CD16+CD56+(NK) | ALDH2-Glu504Lys | -0.63545 | 0 |
| B23 | B94 | CD3-CD16+CD56+(NK) | ALDH2-Lys504Lys | -0.63545 | 0 |
| B23 | B97 | CD3-CD16+CD56+(NK) | dsDNA | -0.36042 | 0 |
| B23 | B98 | CD3-CD16+CD56+(NK) | anti-Jo-1 | -0.36042 | 0 |
| B23 | B99 | CD3-CD16+CD56+(NK) | anti-PM-Scl | -0.36042 | 0 |
| B23 | B61 | CD3-CD16+CD56+(NK) | WBHSRV | 0.263106 | 4.44E-16 |
| B23 | B195 | CD3-CD16+CD56+(NK) | Na | -0.25958 | 8.88E-16 |
| B23 | B65 | CD3-CD16+CD56+(NK) | WBRSR(mPas_1) | 0.259751 | 8.88E-16 |
| B23 | B197 | CD3-CD16+CD56+(NK) | AG | 0.252951 | 5.33E-15 |

| | | | | | |
|---|---|---|---|---|---|
| B23 | B155 | CD3-CD16+CD56+(NK) | IgA | 0.239068 | 1.58E-13 |
| B23 | B198 | CD3-CD16+CD56+(NK) | P | 0.236039 | 3.24E-13 |
| B23 | B158 | CD3-CD16+CD56+(NK) | IgM | 0.227709 | 2.22E-12 |
| B23 | B199 | CD3-CD16+CD56+(NK) | Mg | 0.221937 | 8.07E-12 |
| B23 | B59 | CD3-CD16+CD56+(NK) | WBRV | 0.219652 | 1.33E-11 |
| B23 | B62 | CD3-CD16+CD56+(NK) | WBHSRVI | 0.20749 | 1.75E-10 |
| B23 | B54 | CD3-CD16+CD56+(NK) | ESR | 0.203202 | 4.18E-10 |
| B23 | B136 | CD3-CD16+CD56+(NK) | 5'NT | 0.182778 | 2.05E-08 |
| B23 | B60 | CD3-CD16+CD56+(NK) | WBRVI | 0.177296 | 5.42E-08 |
| B23 | B27 | CD3-CD16+CD56+(NK) | LDH | 0.169679 | 2.00E-07 |
| B23 | B56 | CD3-CD16+CD56+(NK) | IR | 0.168527 | 2.42E-07 |
| B24 | B124 | αHBDH | ACTH | 0.313291 | 0 |
| B24 | B125 | αHBDH | Cor | 0.313291 | 0 |
| B24 | B26 | αHBDH | CKMB | 0.36185 | 0 |
| B24 | B27 | αHBDH | LDH | 0.48692 | 0 |
| B24 | B135 | αHBDH | TBA | 0.241072 | 9.77E-14 |
| B24 | B138 | αHBDH | PAB | -0.23804 | 2.02E-13 |
| B24 | B139 | αHBDH | ADA | 0.191549 | 4.06E-09 |
| B24 | B151 | αHBDH | Anti-CCP | 0.177468 | 5.26E-08 |
| B24 | B119 | αHBDH | Hcv-rna | -0.17596 | 6.85E-08 |
| B24 | B121 | αHBDH | C3 | 0.175957 | 6.85E-08 |
| B24 | B122 | αHBDH | C4 | 0.175957 | 6.85E-08 |
| B24 | B159 | αHBDH | β2-MG | 0.175957 | 6.85E-08 |
| B24 | B169 | αHBDH | U-CRY | 0.175957 | 6.85E-08 |
| B24 | B152 | αHBDH | ACA-Ig(G-A-M) | 0.175947 | 6.86E-08 |
| B24 | B67 | αHBDH | WBRSR(mPas_50) | 0.168005 | 2.64E-07 |
| B24 | B92 | αHBDH | ALDH2-Glu504Glu | 0.167863 | 2.70E-07 |
| B24 | B93 | αHBDH | ALDH2-Glu504Lys | -0.16786 | 2.70E-07 |
| B24 | B94 | αHBDH | ALDH2-Lys504Lys | -0.16786 | 2.70E-07 |
| B24 | B64 | αHBDH | WBRSR(mPas_5) | 0.167796 | 2.73E-07 |
| B24 | B66 | αHBDH | WBRSR(mPas_100) | 0.164226 | 4.90E-07 |
| B24 | B157 | αHBDH | IgG | 0.163937 | 5.13E-07 |
| B25 | B87 | CK | NSE | 0.382323 | 0 |
| B25 | B82 | CK | CEA | 0.212205 | 6.57E-11 |
| B25 | B86 | CK | TBhCG | -0.20853 | 1.41E-10 |
| B25 | B26 | CK | CKMB | 0.206905 | 1.97E-10 |
| B25 | B89 | CK | CA 72-4 | 0.173526 | 1.04E-07 |
| B26 | B89 | CKMB | CA 72-4 | 0.320188 | 0 |
| B26 | B87 | CKMB | NSE | 0.252218 | 6.22E-15 |
| B26 | B155 | CKMB | IgA | 0.213051 | 5.50E-11 |
| B26 | B27 | CKMB | LDH | 0.211688 | 7.32E-11 |
| B26 | B82 | CKMB | CEA | 0.188941 | 6.62E-09 |
| B26 | B156 | CKMB | IgE | -0.18192 | 2.39E-08 |
| B26 | B124 | CKMB | ACTH | 0.179604 | 3.61E-08 |

| | | | | | |
|---|---|---|---|---|---|
| B26 | B125 | CKMB | Cor | 0.179604 | 3.61E-08 |
| B26 | B163 | CKMB | SG | 0.173039 | 1.13E-07 |
| B26 | B138 | CKMB | PAB | -0.1695 | 2.06E-07 |
| B26 | B170 | CKMB | U-GLU | 0.160381 | 9.06E-07 |
| B27 | B138 | LDH | PAB | -0.22528 | 3.84E-12 |
| B27 | B136 | LDH | 5'NT | 0.214471 | 4.07E-11 |
| B27 | B139 | LDH | ADA | 0.19657 | 1.55E-09 |
| B27 | B137 | LDH | AFU | 0.182792 | 2.04E-08 |
| B27 | B124 | LDH | ACTH | 0.178635 | 4.29E-08 |
| B27 | B125 | LDH | Cor | 0.178635 | 4.29E-08 |
| B28 | B202 | PRL | e-times | 0.823087 | 0 |
| B28 | B35 | PRL | β2-MG | 0.54386 | 0 |
| B28 | B167 | PRL | PC | 0.229018 | 1.65E-12 |
| B29 | B202 | E2 | e-times | 0.617052 | 0 |
| B29 | B31 | E2 | FSH | 0.627628 | 0 |
| B29 | B32 | E2 | Testo | 0.502928 | 0 |
| B3 | B124 | Hcy | ACTH | 0.195264 | 2.00E-09 |
| B3 | B125 | Hcy | Cor | 0.195264 | 2.00E-09 |
| B3 | B84 | Hcy | C-PSA | 0.178085 | 4.72E-08 |
| B3 | B23 | Hcy | CD3-CD16+CD56+(NK) | 0.172196 | 1.31E-07 |
| B3 | B16 | Hcy | CD3+T | -0.17097 | 1.61E-07 |
| B30 | B33 | LH | Prog | -0.23652 | 2.89E-13 |
| B31 | B167 | FSH | PC | 0.705999 | 0 |
| B31 | B35 | FSH | β2-MG | 0.652926 | 0 |
| B31 | B32 | FSH | Testo | 0.240048 | 1.25E-13 |
| B32 | B202 | Testo | e-times | 0.796437 | 0 |
| B33 | B36 | Prog | V-Lym% | -0.65133 | 0 |
| B33 | B38 | Prog | BAND% | -0.59611 | 0 |
| B33 | B223 | Prog | FH | -0.22193 | 8.08E-12 |
| B34 | B181 | TK | APTT | -0.1673 | 2.96E-07 |
| B34 | B107 | TK | AnuA | 0.166439 | 3.41E-07 |
| B34 | B110 | TK | AHA | 0.166439 | 3.41E-07 |
| B35 | B167 | β2-MG | PC | 0.464399 | 0 |
| B35 | B37 | β2-MG | BAND | 0.911291 | 0 |
| B36 | B223 | V-Lym% | FH | 0.319553 | 0 |
| B37 | B202 | BAND | e-times | 0.490137 | 0 |
| B38 | B223 | BAND% | FH | -0.75074 | 0 |
| B39 | B78 | Other cells | HBcAb | 0.522491 | 0 |
| B39 | B225 | Other cells | hPBG | -0.22102 | 9.87E-12 |
| B4 | B137 | WB-Ca | AFU | -0.31359 | 0 |
| B4 | B139 | WB-Ca | ADA | -0.44985 | 0 |
| B4 | B15 | WB-Ca | CD3-CD19+(B cell) | 0.444867 | 0 |
| B4 | B157 | WB-Ca | IgG | -0.37859 | 0 |
| B4 | B16 | WB-Ca | CD3+T | 0.298987 | 0 |

| | | | | | |
|---|---|---|---|---|---|
| B4 | B18 | WB-Ca | CD3+CD4+helper(inducer) T | 0.385709 | 0 |
| B4 | B23 | WB-Ca | CD3-CD16+CD56+(NK) | -0.33846 | 0 |
| B4 | B6 | WB-Ca | WB-Fe | -1 | 0 |
| B4 | B19 | WB-Ca | CD3+CD4+/CD3+CD8+ | 0.228677 | 1.78E-12 |
| B4 | B54 | WB-Ca | ESR | -0.2235 | 5.71E-12 |
| B4 | B59 | WB-Ca | WBRV | -0.21355 | 4.94E-11 |
| B4 | B198 | WB-Ca | P | -0.20696 | 1.95E-10 |
| B4 | B85 | WB-Ca | SCC | 0.196611 | 1.54E-09 |
| B4 | B69 | WB-Ca | PV | -0.17633 | 6.42E-08 |
| B4 | B57 | WB-Ca | RBCAI | -0.16653 | 3.36E-07 |
| B40 | B42 | WBC | MON | 0.681892 | 0 |
| B40 | B130 | WBC | GLB | 0.257887 | 1.33E-15 |
| B40 | B52 | WBC | PLT/BPC | 0.22867 | 1.79E-12 |
| B40 | B127 | WBC | A-G ratio | -0.22438 | 4.69E-12 |
| B40 | B91 | WBC | TP | 0.200133 | 7.71E-10 |
| B40 | B72 | WBC | HDL-C | -0.17722 | 5.49E-08 |
| B40 | B192 | WBC | UA | 0.176175 | 6.59E-08 |
| B40 | B71 | WBC | TG | 0.16418 | 4.93E-07 |
| B41 | B42 | MON％ | MON | 0.603712 | 0 |
| B41 | B118 | MON％ | CEA1 | 0.167424 | 2.90E-07 |
| B42 | B127 | MON | A-G ratio | -0.2144 | 4.13E-11 |
| B42 | B192 | MON | UA | 0.204855 | 3.00E-10 |
| B42 | B48 | MON | LYM% | -0.20005 | 7.83E-10 |
| B42 | B130 | MON | GLB | 0.19437 | 2.37E-09 |
| B42 | B52 | MON | PLT/BPC | 0.183399 | 1.83E-08 |
| B42 | B123 | MON | hs-CRP1 | 0.168929 | 2.26E-07 |
| B43 | B44 | MCV | MCH | 0.855515 | 0 |
| B43 | B46 | MCV | RBC Count | -0.43928 | 0 |
| B43 | B47 | MCV | RDW | 0.521707 | 0 |
| B43 | B52 | MCV | PLT/BPC | -0.16061 | 8.74E-07 |
| B44 | B45 | MCH | MCHC | 0.56285 | 0 |
| B44 | B46 | MCH | RBC Count | -0.28999 | 0 |
| B44 | B47 | MCH | RDW | 0.270455 | 0 |
| B44 | B51 | MCH | Hb | 0.274286 | 0 |
| B44 | B133 | MCH | TBIL | 0.179231 | 3.86E-08 |
| B45 | B47 | MCHC | RDW | -0.31683 | 0 |
| B45 | B51 | MCHC | Hb | 0.467885 | 0 |
| B45 | B222 | MCHC | NS | 0.206077 | 2.34E-10 |
| B45 | B221 | MCHC | Hei | 0.190497 | 4.95E-09 |
| B45 | B129 | MCHC | IBIL | 0.179708 | 3.55E-08 |
| B46 | B126 | RBC Count | Alb | 0.268896 | 0 |
| B46 | B189 | RBC Count | eGFB | 0.345415 | 0 |
| B46 | B191 | RBC Count | Urea | -0.28976 | 0 |

| | | | | | |
|---|---|---|---|---|---|
| B46 | B216 | RBC Count | BMI | 0.303708 | 0 |
| B46 | B218 | RBC Count | DBP | 0.27977 | 0 |
| B46 | B221 | RBC Count | Hei | 0.322046 | 0 |
| B46 | B47 | RBC Count | RDW | -0.44008 | 0 |
| B46 | B51 | RBC Count | Hb | 0.836771 | 0 |
| B46 | B222 | RBC Count | NS | 0.260232 | 8.88E-16 |
| B46 | B163 | RBC Count | SG | 0.257412 | 1.78E-15 |
| B46 | B141 | RBC Count | AST/ALT | -0.23897 | 1.62E-13 |
| B46 | B129 | RBC Count | IBIL | 0.218175 | 1.84E-11 |
| B46 | B190 | RBC Count | Cre | -0.20855 | 1.41E-10 |
| B46 | B170 | RBC Count | U-GLU | 0.207082 | 1.90E-10 |
| B46 | B173 | RBC Count | NTT | -0.1886 | 7.06E-09 |
| B46 | B128 | RBC Count | ALT | 0.160174 | 9.37E-07 |
| B47 | B51 | RDW | Hb | -0.28797 | 0 |
| B47 | B111 | RDW | TSH | 0.215616 | 3.19E-11 |
| B47 | B126 | RDW | Alb | -0.21213 | 6.67E-11 |
| B47 | B191 | RDW | Urea | 0.174288 | 9.13E-08 |
| B47 | B127 | RDW | A-G ratio | -0.16734 | 2.95E-07 |
| B47 | B52 | RDW | PLT/BPC | -0.16675 | 3.24E-07 |
| B47 | B170 | RDW | U-GLU | -0.16206 | 6.94E-07 |
| B48 | B53 | LYM% | GR% | -0.92051 | 0 |
| B48 | B190 | LYM% | Cre | -0.18857 | 7.09E-09 |
| B48 | B189 | LYM% | eGFB | 0.184265 | 1.57E-08 |
| B48 | B72 | LYM% | HDL-C | 0.179033 | 4.00E-08 |
| B49 | B50 | baso% | eo% | 0.253322 | 4.66E-15 |
| B5 | B137 | WB-Mg | AFU | -0.31359 | 0 |
| B5 | B139 | WB-Mg | ADA | -0.44985 | 0 |
| B5 | B15 | WB-Mg | CD3-CD19+(B cell) | 0.444867 | 0 |
| B5 | B157 | WB-Mg | IgG | -0.37859 | 0 |
| B5 | B16 | WB-Mg | CD3+T | 0.298987 | 0 |
| B5 | B18 | WB-Mg | CD3+CD4+helper(inducer) T | 0.385709 | 0 |
| B5 | B23 | WB-Mg | CD3-CD16+CD56+(NK) | -0.33846 | 0 |
| B5 | B19 | WB-Mg | CD3+CD4+/CD3+CD8+ | 0.228677 | 1.78E-12 |
| B5 | B54 | WB-Mg | ESR | -0.2235 | 5.71E-12 |
| B5 | B59 | WB-Mg | WBRV | -0.21355 | 4.94E-11 |
| B5 | B198 | WB-Mg | P | -0.20696 | 1.95E-10 |
| B5 | B85 | WB-Mg | SCC | 0.196611 | 1.54E-09 |
| B5 | B69 | WB-Mg | PV | -0.17633 | 6.42E-08 |
| B5 | B57 | WB-Mg | RBCAI | -0.16653 | 3.36E-07 |
| B50 | B132 | eo% | Dbil | 0.560793 | 0 |
| B50 | B133 | eo% | TBIL | 0.343238 | 0 |
| B50 | B140 | eo% | r-GT | 0.611773 | 0 |
| B50 | B131 | eo% | AST | 0.264732 | 2.22E-16 |

| | | | | | |
|---|---|---|---|---|---|
| B50 | B53 | eo% | GR% | -0.261 | 6.66E-16 |
| B50 | B128 | eo% | ALT | 0.221886 | 8.17E-12 |
| B50 | B190 | eo% | Cre | 0.180207 | 3.25E-08 |
| B51 | B126 | Hb | Alb | 0.27008 | 0 |
| B51 | B129 | Hb | IBIL | 0.30224 | 0 |
| B51 | B163 | Hb | SG | 0.26875 | 0 |
| B51 | B189 | Hb | eGFB | 0.355955 | 0 |
| B51 | B191 | Hb | Urea | -0.30892 | 0 |
| B51 | B216 | Hb | BMI | 0.334989 | 0 |
| B51 | B221 | Hb | Hei | 0.342946 | 0 |
| B51 | B222 | Hb | NS | 0.305232 | 0 |
| B51 | B218 | Hb | DBP | 0.265743 | 2.22E-16 |
| B51 | B133 | Hb | TBIL | 0.226019 | 3.25E-12 |
| B51 | B141 | Hb | AST/ALT | -0.22355 | 5.65E-12 |
| B51 | B209 | Hb | Smoking | 0.197105 | 1.40E-09 |
| B51 | B170 | Hb | U-GLU | 0.195129 | 2.05E-09 |
| B51 | B173 | Hb | NTT | -0.19135 | 4.21E-09 |
| B51 | B190 | Hb | Cre | -0.18418 | 1.59E-08 |
| B51 | B128 | Hb | ALT | 0.183591 | 1.77E-08 |
| B51 | B127 | Hb | A-G ratio | 0.183146 | 1.92E-08 |
| B51 | B206 | Hb | Waist | 0.161049 | 8.15E-07 |
| B51 | B210 | Hb | Hip | 0.159903 | 9.78E-07 |
| B52 | B129 | PLT/BPC | IBIL | -0.24075 | 1.05E-13 |
| B52 | B133 | PLT/BPC | TBIL | -0.23583 | 3.40E-13 |
| B53 | B140 | GR% | r-GT | -0.17998 | 3.38E-08 |
| B54 | B58 | ESR | HCT | -0.47296 | 0 |
| B54 | B59 | ESR | WBRV | 0.319805 | 0 |
| B54 | B69 | ESR | PV | 0.366541 | 0 |
| B54 | B62 | ESR | WBHSRVI | -0.24238 | 7.13E-14 |
| B54 | B139 | ESR | ADA | 0.241139 | 9.61E-14 |
| B54 | B138 | ESR | PAB | -0.18855 | 7.12E-09 |
| B54 | B55 | ESR | TK | 0.18139 | 2.63E-08 |
| B54 | B172 | ESR | KB | 0.176953 | 5.76E-08 |
| B54 | B195 | ESR | Na | -0.17465 | 8.59E-08 |
| B54 | B123 | ESR | hs-CRP1 | 0.169378 | 2.10E-07 |
| B55 | B56 | TK | IR | 0.894206 | 0 |
| B55 | B57 | TK | RBCAI | -0.26997 | 0 |
| B55 | B58 | TK | HCT | -0.46133 | 0 |
| B55 | B59 | TK | WBRV | 0.380495 | 0 |
| B55 | B61 | TK | WBHSRV | 0.804455 | 0 |
| B55 | B62 | TK | WBHSRVI | 0.570656 | 0 |
| B55 | B63 | TK | WBRSR(mPas_200) | 0.314666 | 0 |
| B55 | B66 | TK | WBRSR(mPas_100) | 0.29943 | 0 |
| B55 | B67 | TK | WBRSR(mPas_50) | 0.274019 | 0 |

| | | | | | |
|---|---|---|---|---|---|
| B55 | B69 | TK | PV | -0.57917 | 0 |
| B55 | B60 | TK | WBRVI | 0.242988 | 6.13E-14 |
| B56 | B59 | IR | WBRV | 0.43152 | 0 |
| B56 | B60 | IR | WBRVI | 0.502272 | 0 |
| B56 | B61 | IR | WBHSRV | 0.937139 | 0 |
| B56 | B62 | IR | WBHSRVI | 0.875193 | 0 |
| B56 | B63 | IR | WBRSR(mPas_200) | 0.662446 | 0 |
| B56 | B64 | IR | WBRSR(mPas_5) | 0.459886 | 0 |
| B56 | B65 | IR | WBRSR(mPas_1) | 0.334621 | 0 |
| B56 | B66 | IR | WBRSR(mPas_100) | 0.642572 | 0 |
| B56 | B67 | IR | WBRSR(mPas_50) | 0.620565 | 0 |
| B56 | B69 | IR | PV | -0.56965 | 0 |
| B56 | B57 | IR | RBCAI | -0.24282 | 6.39E-14 |
| B56 | B183 | IR | PTA | -0.21675 | 2.50E-11 |
| B56 | B182 | IR | TT | 0.199663 | 8.46E-10 |
| B56 | B124 | IR | ACTH | 0.18862 | 7.03E-09 |
| B56 | B125 | IR | Cor | 0.18862 | 7.03E-09 |
| B56 | B135 | IR | TBA | 0.180605 | 3.02E-08 |
| B56 | B186 | IR | PFC | -0.17743 | 5.30E-08 |
| B57 | B195 | RBCAI | Na | 0.329545 | 0 |
| B57 | B59 | RBCAI | WBRV | 0.67815 | 0 |
| B57 | B60 | RBCAI | WBRVI | 0.630381 | 0 |
| B57 | B61 | RBCAI | WBHSRV | -0.2691 | 0 |
| B57 | B64 | RBCAI | WBRSR(mPas_5) | 0.427094 | 0 |
| B57 | B65 | RBCAI | WBRSR(mPas_1) | 0.648145 | 0 |
| B57 | B200 | RBCAI | OP | 0.224312 | 4.77E-12 |
| B57 | B138 | RBCAI | PAB | 0.196139 | 1.69E-09 |
| B57 | B58 | RBCAI | HCT | 0.180666 | 2.99E-08 |
| B57 | B194 | RBCAI | Cl | 0.173868 | 9.82E-08 |
| B58 | B60 | HCT | WBRVI | 0.49517 | 0 |
| B58 | B62 | HCT | WBHSRVI | 0.455675 | 0 |
| B58 | B63 | HCT | WBRSR(mPas_200) | 0.585226 | 0 |
| B58 | B64 | HCT | WBRSR(mPas_5) | 0.618184 | 0 |
| B58 | B65 | HCT | WBRSR(mPas_1) | 0.544419 | 0 |
| B58 | B66 | HCT | WBRSR(mPas_100) | 0.576972 | 0 |
| B58 | B67 | HCT | WBRSR(mPas_50) | 0.595268 | 0 |
| B58 | B137 | HCT | AFU | 0.16285 | 6.11E-07 |
| B59 | B60 | WBRV | WBRVI | 0.808762 | 0 |
| B59 | B61 | WBRV | WBHSRV | 0.491196 | 0 |
| B59 | B62 | WBRV | WBHSRVI | 0.389416 | 0 |
| B59 | B63 | WBRV | WBRSR(mPas_200) | 0.416068 | 0 |
| B59 | B64 | WBRV | WBRSR(mPas_5) | 0.752807 | 0 |
| B59 | B65 | WBRV | WBRSR(mPas_1) | 0.842291 | 0 |
| B59 | B66 | WBRV | WBRSR(mPas_100) | 0.452953 | 0 |

| | | | | | |
|---|---|---|---|---|---|
| B59 | B67 | WBRV | WBRSR(mPas_50) | 0.514036 | 0 |
| B59 | B195 | WBRV | Na | 0.226093 | 3.20E-12 |
| B59 | B200 | WBRV | OP | 0.217211 | 2.26E-11 |
| B59 | B186 | WBRV | PFC | -0.20799 | 1.58E-10 |
| B59 | B183 | WBRV | PTA | -0.19786 | 1.21E-09 |
| B59 | B182 | WBRV | TT | 0.193051 | 3.05E-09 |
| B59 | B171 | WBRV | EC | 0.183523 | 1.79E-08 |
| B59 | B197 | WBRV | AG | 0.176614 | 6.11E-08 |
| B59 | B124 | WBRV | ACTH | 0.165568 | 3.94E-07 |
| B59 | B125 | WBRV | Cor | 0.165568 | 3.94E-07 |
| B6 | B137 | WB-Fe | AFU | 0.313586 | 0 |
| B6 | B139 | WB-Fe | ADA | 0.449853 | 0 |
| B6 | B15 | WB-Fe | CD3-CD19+(B cell) | -0.44487 | 0 |
| B6 | B157 | WB-Fe | IgG | 0.378593 | 0 |
| B6 | B16 | WB-Fe | CD3+T | -0.29899 | 0 |
| B6 | B18 | WB-Fe | CD3+CD4+helper(inducer) T | -0.38571 | 0 |
| B6 | B23 | WB-Fe | CD3-CD16+CD56+(NK) | 0.338457 | 0 |
| B6 | B7 | WB-Fe | WB-Cop | -1 | 0 |
| B6 | B8 | WB-Fe | WB-Zinc | 1 | 0 |
| B6 | B19 | WB-Fe | CD3+CD4+/CD3+CD8+ | -0.22868 | 1.78E-12 |
| B6 | B54 | WB-Fe | ESR | 0.223503 | 5.71E-12 |
| B6 | B59 | WB-Fe | WBRV | 0.213551 | 4.94E-11 |
| B6 | B198 | WB-Fe | P | 0.206965 | 1.95E-10 |
| B6 | B85 | WB-Fe | SCC | -0.19661 | 1.54E-09 |
| B6 | B69 | WB-Fe | PV | 0.176326 | 6.42E-08 |
| B6 | B57 | WB-Fe | RBCAI | 0.166529 | 3.36E-07 |
| B60 | B61 | WBRVI | WBHSRV | 0.465817 | 0 |
| B60 | B62 | WBRVI | WBHSRVI | 0.678396 | 0 |
| B60 | B63 | WBRVI | WBRSR(mPas_200) | 0.594514 | 0 |
| B60 | B64 | WBRVI | WBRSR(mPas_5) | 0.872737 | 0 |
| B60 | B65 | WBRVI | WBRSR(mPas_1) | 0.917133 | 0 |
| B60 | B66 | WBRVI | WBRSR(mPas_100) | 0.61004 | 0 |
| B60 | B67 | WBRVI | WBRSR(mPas_50) | 0.666928 | 0 |
| B60 | B69 | WBRVI | PV | -0.34589 | 0 |
| B60 | B183 | WBRVI | PTA | -0.23338 | 6.04E-13 |
| B60 | B171 | WBRVI | EC | 0.229996 | 1.32E-12 |
| B60 | B182 | WBRVI | TT | 0.228257 | 1.96E-12 |
| B60 | B186 | WBRVI | PFC | -0.21788 | 1.96E-11 |
| B60 | B200 | WBRVI | OP | 0.213278 | 5.24E-11 |
| B60 | B197 | WBRVI | AG | 0.208734 | 1.35E-10 |
| B60 | B195 | WBRVI | Na | 0.194516 | 2.30E-09 |
| B60 | B124 | WBRVI | ACTH | 0.163421 | 5.58E-07 |
| B60 | B125 | WBRVI | Cor | 0.163421 | 5.58E-07 |

| | | | | | |
|---|---|---|---|---|---|
| B61 | B124 | WBHSRV | ACTH | 0.268099 | 0 |
| B61 | B125 | WBHSRV | Cor | 0.268099 | 0 |
| B61 | B62 | WBHSRV | WBHSRVI | 0.851605 | 0 |
| B61 | B63 | WBHSRV | WBRSR(mPas_200) | 0.805666 | 0 |
| B61 | B64 | WBHSRV | WBRSR(mPas_5) | 0.588068 | 0 |
| B61 | B65 | WBHSRV | WBRSR(mPas_1) | 0.431316 | 0 |
| B61 | B66 | WBHSRV | WBRSR(mPas_100) | 0.799626 | 0 |
| B61 | B67 | WBHSRV | WBRSR(mPas_50) | 0.776897 | 0 |
| B61 | B69 | WBHSRV | PV | -0.25547 | 2.66E-15 |
| B61 | B183 | WBHSRV | PTA | -0.23949 | 1.43E-13 |
| B61 | B135 | WBHSRV | TBA | 0.221432 | 9.03E-12 |
| B61 | B182 | WBHSRV | TT | 0.212379 | 6.33E-11 |
| B61 | B186 | WBHSRV | PFC | -0.19619 | 1.67E-09 |
| B61 | B194 | WBHSRV | CI | -0.19584 | 1.78E-09 |
| B61 | B92 | WBHSRV | ALDH2-Glu504Glu | 0.190896 | 4.59E-09 |
| B61 | B93 | WBHSRV | ALDH2-Glu504Lys | -0.1909 | 4.59E-09 |
| B61 | B94 | WBHSRV | ALDH2-Lys504Lys | -0.1909 | 4.59E-09 |
| B62 | B63 | WBHSRVI | WBRSR(mPas_200) | 0.875575 | 0 |
| B62 | B64 | WBHSRVI | WBRSR(mPas_5) | 0.70655 | 0 |
| B62 | B65 | WBHSRVI | WBRSR(mPas_1) | 0.554739 | 0 |
| B62 | B66 | WBHSRVI | WBRSR(mPas_100) | 0.856188 | 0 |
| B62 | B67 | WBHSRVI | WBRSR(mPas_50) | 0.844693 | 0 |
| B62 | B69 | WBHSRVI | PV | -0.44835 | 0 |
| B62 | B183 | WBHSRVI | PTA | -0.25511 | 3.11E-15 |
| B62 | B124 | WBHSRVI | ACTH | 0.237892 | 2.09E-13 |
| B62 | B125 | WBHSRVI | Cor | 0.237892 | 2.09E-13 |
| B62 | B182 | WBHSRVI | TT | 0.232931 | 6.71E-13 |
| B62 | B186 | WBHSRVI | PFC | -0.20088 | 6.65E-10 |
| B62 | B194 | WBHSRVI | CI | -0.18176 | 2.46E-08 |
| B62 | B135 | WBHSRVI | TBA | 0.180606 | 3.02E-08 |
| B62 | B171 | WBHSRVI | EC | 0.176573 | 6.15E-08 |
| B62 | B92 | WBHSRVI | ALDH2-Glu504Glu | 0.170646 | 1.70E-07 |
| B62 | B93 | WBHSRVI | ALDH2-Glu504Lys | -0.17065 | 1.70E-07 |
| B62 | B94 | WBHSRVI | ALDH2-Lys504Lys | -0.17065 | 1.70E-07 |
| B62 | B197 | WBHSRVI | AG | 0.163177 | 5.80E-07 |
| B63 | B124 | WBRSR(mPas_200) | ACTH | 0.300675 | 0 |
| B63 | B125 | WBRSR(mPas_200) | Cor | 0.300675 | 0 |
| B63 | B64 | WBRSR(mPas_200) | WBRSR(mPas_5) | 0.839448 | 0 |
| B63 | B65 | WBRSR(mPas_200) | WBRSR(mPas_1) | 0.662658 | 0 |
| B63 | B66 | WBRSR(mPas_200) | WBRSR(mPas_100) | 0.994329 | 0 |
| B63 | B67 | WBRSR(mPas_200) | WBRSR(mPas_50) | 0.985617 | 0 |
| B63 | B194 | WBRSR(mPas_200) | CI | -0.26242 | 4.44E-16 |
| B63 | B183 | WBRSR(mPas_200) | PTA | -0.25655 | 2.00E-15 |
| B63 | B182 | WBRSR(mPas_200) | TT | 0.227746 | 2.20E-12 |

| | | | | | |
|---|---|---|---|---|---|
| B63 | B92 | WBRSR(mPas_200) | ALDH2-Glu504Glu | 0.21472 | 3.86E-11 |
| B63 | B93 | WBRSR(mPas_200) | ALDH2-Glu504Lys | -0.21472 | 3.86E-11 |
| B63 | B94 | WBRSR(mPas_200) | ALDH2-Lys504Lys | -0.21472 | 3.86E-11 |
| B63 | B186 | WBRSR(mPas_200) | PFC | -0.21155 | 7.53E-11 |
| B63 | B135 | WBRSR(mPas_200) | TBA | 0.202269 | 5.04E-10 |
| B63 | B137 | WBRSR(mPas_200) | AFU | 0.185975 | 1.15E-08 |
| B63 | B139 | WBRSR(mPas_200) | ADA | 0.184818 | 1.42E-08 |
| B63 | B195 | WBRSR(mPas_200) | Na | -0.1775 | 5.23E-08 |
| B64 | B124 | WBRSR(mPas_5) | ACTH | 0.279773 | 0 |
| B64 | B125 | WBRSR(mPas_5) | Cor | 0.279773 | 0 |
| B64 | B183 | WBRSR(mPas_5) | PTA | -0.26715 | 0 |
| B64 | B65 | WBRSR(mPas_5) | WBRSR(mPas_1) | 0.959934 | 0 |
| B64 | B66 | WBRSR(mPas_5) | WBRSR(mPas_100) | 0.864088 | 0 |
| B64 | B67 | WBRSR(mPas_5) | WBRSR(mPas_50) | 0.905204 | 0 |
| B64 | B182 | WBRSR(mPas_5) | TT | 0.245363 | 3.44E-14 |
| B64 | B186 | WBRSR(mPas_5) | PFC | -0.23599 | 3.28E-13 |
| B64 | B171 | WBRSR(mPas_5) | EC | 0.210717 | 8.97E-11 |
| B64 | B92 | WBRSR(mPas_5) | ALDH2-Glu504Glu | 0.199972 | 7.96E-10 |
| B64 | B93 | WBRSR(mPas_5) | ALDH2-Glu504Lys | -0.19997 | 7.96E-10 |
| B64 | B94 | WBRSR(mPas_5) | ALDH2-Lys504Lys | -0.19997 | 7.96E-10 |
| B64 | B197 | WBRSR(mPas_5) | AG | 0.198004 | 1.17E-09 |
| B64 | B139 | WBRSR(mPas_5) | ADA | 0.183668 | 1.74E-08 |
| B64 | B135 | WBRSR(mPas_5) | TBA | 0.18241 | 2.19E-08 |
| B64 | B158 | WBRSR(mPas_5) | IgM | 0.168146 | 2.58E-07 |
| B64 | B194 | WBRSR(mPas_5) | Cl | -0.16106 | 8.14E-07 |
| B65 | B66 | WBRSR(mPas_1) | WBRSR(mPas_100) | 0.696918 | 0 |
| B65 | B67 | WBRSR(mPas_1) | WBRSR(mPas_50) | 0.757138 | 0 |
| B65 | B183 | WBRSR(mPas_1) | PTA | -0.23898 | 1.61E-13 |
| B65 | B182 | WBRSR(mPas_1) | TT | 0.226011 | 3.26E-12 |
| B65 | B186 | WBRSR(mPas_1) | PFC | -0.22542 | 3.72E-12 |
| B65 | B124 | WBRSR(mPas_1) | ACTH | 0.224958 | 4.13E-12 |
| B65 | B125 | WBRSR(mPas_1) | Cor | 0.224958 | 4.13E-12 |
| B65 | B171 | WBRSR(mPas_1) | EC | 0.217537 | 2.11E-11 |
| B65 | B197 | WBRSR(mPas_1) | AG | 0.200031 | 7.87E-10 |
| B65 | B200 | WBRSR(mPas_1) | OP | 0.180409 | 3.13E-08 |
| B65 | B92 | WBRSR(mPas_1) | ALDH2-Glu504Glu | 0.161093 | 8.10E-07 |
| B65 | B93 | WBRSR(mPas_1) | ALDH2-Glu504Lys | -0.16109 | 8.10E-07 |
| B65 | B94 | WBRSR(mPas_1) | ALDH2-Lys504Lys | -0.16109 | 8.10E-07 |
| B66 | B124 | WBRSR(mPas_100) | ACTH | 0.331523 | 0 |
| B66 | B125 | WBRSR(mPas_100) | Cor | 0.331523 | 0 |
| B66 | B183 | WBRSR(mPas_100) | PTA | -0.27237 | 0 |
| B66 | B194 | WBRSR(mPas_100) | Cl | -0.28857 | 0 |
| B66 | B67 | WBRSR(mPas_100) | WBRSR(mPas_50) | 0.995929 | 0 |
| B66 | B182 | WBRSR(mPas_100) | TT | 0.238002 | 2.04E-13 |

| | | | | | |
|---|---|---|---|---|---|
| B66 | B92 | WBRSR(mPas_100) | ALDH2-Glu504Glu | 0.236138 | 3.17E-13 |
| B66 | B93 | WBRSR(mPas_100) | ALDH2-Glu504Lys | -0.23614 | 3.17E-13 |
| B66 | B94 | WBRSR(mPas_100) | ALDH2-Lys504Lys | -0.23614 | 3.17E-13 |
| B66 | B139 | WBRSR(mPas_100) | ADA | 0.219261 | 1.45E-11 |
| B66 | B186 | WBRSR(mPas_100) | PFC | -0.2151 | 3.56E-11 |
| B66 | B135 | WBRSR(mPas_100) | TBA | 0.214794 | 3.80E-11 |
| B66 | B137 | WBRSR(mPas_100) | AFU | 0.213345 | 5.16E-11 |
| B66 | B195 | WBRSR(mPas_100) | Na | -0.19773 | 1.24E-09 |
| B66 | B107 | WBRSR(mPas_100) | AnuA | -0.17391 | 9.74E-08 |
| B66 | B110 | WBRSR(mPas_100) | AHA | -0.17391 | 9.74E-08 |
| B66 | B79 | WBRSR(mPas_100) | hb V -dna | -0.16941 | 2.09E-07 |
| B66 | B100 | WBRSR(mPas_100) | anti-Ro52 | -0.16195 | 7.06E-07 |
| B66 | B101 | WBRSR(mPas_100) | anti-Sc70 | -0.16195 | 7.06E-07 |
| B66 | B102 | WBRSR(mPas_100) | anti-Sm | -0.16195 | 7.06E-07 |
| B66 | B103 | WBRSR(mPas_100) | anti-SSA/Ro | -0.16195 | 7.06E-07 |
| B66 | B104 | WBRSR(mPas_100) | anti-SSB | -0.16195 | 7.06E-07 |
| B66 | B105 | WBRSR(mPas_100) | anti-U1RNP | -0.16195 | 7.06E-07 |
| B66 | B106 | WBRSR(mPas_100) | anti-P | -0.16195 | 7.06E-07 |
| B66 | B108 | WBRSR(mPas_100) | Anti-Cyclin | -0.16195 | 7.06E-07 |
| B66 | B109 | WBRSR(mPas_100) | AMA-M2 | -0.16195 | 7.06E-07 |
| B66 | B97 | WBRSR(mPas_100) | dsDNA | -0.16195 | 7.06E-07 |
| B66 | B98 | WBRSR(mPas_100) | anti-Jo-1 | -0.16195 | 7.06E-07 |
| B66 | B99 | WBRSR(mPas_100) | anti-PM-Scl | -0.16195 | 7.06E-07 |
| B66 | B171 | WBRSR(mPas_100) | EC | 0.161569 | 7.51E-07 |
| B67 | B124 | WBRSR(mPas_50) | ACTH | 0.32942 | 0 |
| B67 | B125 | WBRSR(mPas_50) | Cor | 0.32942 | 0 |
| B67 | B183 | WBRSR(mPas_50) | PTA | -0.27743 | 0 |
| B67 | B194 | WBRSR(mPas_50) | Cl | -0.2746 | 0 |
| B67 | B182 | WBRSR(mPas_50) | TT | 0.244308 | 4.44E-14 |
| B67 | B92 | WBRSR(mPas_50) | ALDH2-Glu504Glu | 0.234699 | 4.44E-13 |
| B67 | B93 | WBRSR(mPas_50) | ALDH2-Glu504Lys | -0.2347 | 4.44E-13 |
| B67 | B94 | WBRSR(mPas_50) | ALDH2-Lys504Lys | -0.2347 | 4.44E-13 |
| B67 | B186 | WBRSR(mPas_50) | PFC | -0.22244 | 7.22E-12 |
| B67 | B139 | WBRSR(mPas_50) | ADA | 0.219542 | 1.37E-11 |
| B67 | B135 | WBRSR(mPas_50) | TBA | 0.212166 | 6.62E-11 |
| B67 | B137 | WBRSR(mPas_50) | AFU | 0.208874 | 1.32E-10 |
| B67 | B171 | WBRSR(mPas_50) | EC | 0.174182 | 9.30E-08 |
| B67 | B107 | WBRSR(mPas_50) | AnuA | -0.17231 | 1.28E-07 |
| B67 | B110 | WBRSR(mPas_50) | AHA | -0.17231 | 1.28E-07 |
| B67 | B195 | WBRSR(mPas_50) | Na | -0.16918 | 2.17E-07 |
| B67 | B79 | WBRSR(mPas_50) | hb V -dna | -0.16846 | 2.45E-07 |
| B67 | B197 | WBRSR(mPas_50) | AG | 0.167923 | 2.67E-07 |
| B67 | B100 | WBRSR(mPas_50) | anti-Ro52 | -0.1605 | 8.90E-07 |
| B67 | B101 | WBRSR(mPas_50) | anti-Sc70 | -0.1605 | 8.90E-07 |

| | | | | | |
|---|---|---|---|---|---|
| B67 | B102 | WBRSR(mPas_50) | anti-Sm | -0.1605 | 8.90E-07 |
| B67 | B103 | WBRSR(mPas_50) | anti-SSA/Ro | -0.1605 | 8.90E-07 |
| B67 | B104 | WBRSR(mPas_50) | anti-SSB | -0.1605 | 8.90E-07 |
| B67 | B105 | WBRSR(mPas_50) | anti-U1RNP | -0.1605 | 8.90E-07 |
| B67 | B106 | WBRSR(mPas_50) | anti-P | -0.1605 | 8.90E-07 |
| B67 | B108 | WBRSR(mPas_50) | Anti-Cyclin | -0.1605 | 8.90E-07 |
| B67 | B109 | WBRSR(mPas_50) | AMA-M2 | -0.1605 | 8.90E-07 |
| B67 | B97 | WBRSR(mPas_50) | dsDNA | -0.1605 | 8.90E-07 |
| B67 | B98 | WBRSR(mPas_50) | anti-Jo-1 | -0.1605 | 8.90E-07 |
| B67 | B99 | WBRSR(mPas_50) | anti-PM-Scl | -0.1605 | 8.90E-07 |
| B69 | B137 | PV | AFU | 0.243401 | 5.55E-14 |
| B69 | B195 | PV | Na | -0.20385 | 3.67E-10 |
| B69 | B139 | PV | ADA | 0.188479 | 7.21E-09 |
| B69 | B138 | PV | PAB | -0.18451 | 1.50E-08 |
| B69 | B141 | PV | AST/ALT | 0.162561 | 6.41E-07 |
| B7 | B137 | WB-Cop | AFU | -0.31359 | 0 |
| B7 | B139 | WB-Cop | ADA | -0.44985 | 0 |
| B7 | B15 | WB-Cop | CD3-CD19+(B cell) | 0.444867 | 0 |
| B7 | B157 | WB-Cop | IgG | -0.37859 | 0 |
| B7 | B16 | WB-Cop | CD3+T | 0.298987 | 0 |
| B7 | B18 | WB-Cop | CD3+CD4+helper(inducer) T | 0.385709 | 0 |
| B7 | B23 | WB-Cop | CD3-CD16+CD56+(NK) | -0.33846 | 0 |
| B7 | B8 | WB-Cop | WB-Zinc | -1 | 0 |
| B7 | B19 | WB-Cop | CD3+CD4+/CD3+CD8+ | 0.228677 | 1.78E-12 |
| B7 | B54 | WB-Cop | ESR | -0.2235 | 5.71E-12 |
| B7 | B59 | WB-Cop | WBRV | -0.21355 | 4.94E-11 |
| B7 | B198 | WB-Cop | P | -0.20696 | 1.95E-10 |
| B7 | B85 | WB-Cop | SCC | 0.196611 | 1.54E-09 |
| B7 | B69 | WB-Cop | PV | -0.17633 | 6.42E-08 |
| B7 | B57 | WB-Cop | RBCAI | -0.16653 | 3.36E-07 |
| B70 | B77 | LDL-C | TC | 0.900823 | 0 |
| B70 | B76 | LDL-C | ApoB | 0.190481 | 4.96E-09 |
| B70 | B127 | LDL-C | A-G ratio | -0.16998 | 1.90E-07 |
| B71 | B153 | TG | Glu | 0.32208 | 0 |
| B71 | B181 | TG | APTT | 0.308366 | 0 |
| B71 | B185 | TG | PT | 0.271857 | 0 |
| B71 | B72 | TG | HDL-C | -0.36076 | 0 |
| B71 | B77 | TG | TC | 0.284601 | 0 |
| B71 | B91 | TG | TP | 0.288373 | 0 |
| B71 | B184 | TG | INR | 0.265547 | 2.22E-16 |
| B71 | B130 | TG | GLB | 0.25178 | 7.11E-15 |
| B71 | B182 | TG | TT | 0.21153 | 7.57E-11 |
| B71 | B183 | TG | PTA | 0.193574 | 2.76E-09 |

| | | | | | |
|---|---|---|---|---|---|
| B71 | B127 | TG | A-G ratio | -0.17168 | 1.43E-07 |
| B71 | B207 | TG | WHR | 0.163305 | 5.68E-07 |
| B71 | B192 | TG | UA | 0.161238 | 7.91E-07 |
| B72 | B216 | HDL-C | BMI | -0.27778 | 0 |
| B72 | B222 | HDL-C | NS | -0.27838 | 0 |
| B72 | B77 | HDL-C | TC | 0.295111 | 0 |
| B72 | B206 | HDL-C | Waist | -0.2228 | 6.68E-12 |
| B72 | B192 | HDL-C | UA | -0.22272 | 6.79E-12 |
| B72 | B74 | HDL-C | ApoA1 | 0.209125 | 1.25E-10 |
| B72 | B221 | HDL-C | Hei | -0.20119 | 6.25E-10 |
| B72 | B215 | HDL-C | BMI | -0.19764 | 1.26E-09 |
| B72 | B153 | HDL-C | Glu | -0.19337 | 2.87E-09 |
| B72 | B207 | HDL-C | WHR | -0.1795 | 3.68E-08 |
| B72 | B163 | HDL-C | SG | -0.16262 | 6.35E-07 |
| B72 | B141 | HDL-C | AST/ALT | 0.1604 | 9.04E-07 |
| B73 | B178 | ApoA | Ig-U | 0.253552 | 4.44E-15 |
| B73 | B179 | ApoA | α1-MG | 0.234896 | 4.24E-13 |
| B73 | B76 | ApoA | ApoB | -0.18249 | 2.16E-08 |
| B74 | B136 | ApoA1 | 5'NT | 0.360936 | 0 |
| B74 | B139 | ApoA1 | ADA | -0.35519 | 0 |
| B74 | B75 | ApoA1 | ApoA1/B | 0.401407 | 0 |
| B74 | B76 | ApoA1 | ApoB | 0.284241 | 0 |
| B74 | B199 | ApoA1 | Mg | 0.235101 | 4.04E-13 |
| B74 | B186 | ApoA1 | PFC | -0.2128 | 5.80E-11 |
| B74 | B88 | ApoA1 | CA 242 | 0.207493 | 1.75E-10 |
| B74 | B182 | ApoA1 | TT | 0.177851 | 4.92E-08 |
| B74 | B195 | ApoA1 | Na | 0.177694 | 5.06E-08 |
| B74 | B193 | ApoA1 | K | 0.160466 | 8.94E-07 |
| B75 | B136 | ApoA1/B | 5'NT | 0.357197 | 0 |
| B75 | B76 | ApoA1/B | ApoB | -0.70347 | 0 |
| B75 | B198 | ApoA1/B | P | -0.26545 | 2.22E-16 |
| B75 | B134 | ApoA1/B | CHE | -0.25022 | 1.02E-14 |
| B75 | B138 | ApoA1/B | PAB | -0.22496 | 4.13E-12 |
| B75 | B137 | ApoA1/B | AFU | 0.201134 | 6.32E-10 |
| B75 | B135 | ApoA1/B | TBA | -0.19351 | 2.80E-09 |
| B75 | B172 | ApoA1/B | KB | -0.16914 | 2.18E-07 |
| B75 | B193 | ApoA1/B | K | 0.167982 | 2.65E-07 |
| B75 | B96 | ApoA1/B | dsDNA (IIF) | 0.162104 | 6.89E-07 |
| B76 | B138 | ApoB | PAB | 0.349614 | 0 |
| B76 | B139 | ApoB | ADA | -0.31816 | 0 |
| B76 | B96 | ApoB | dsDNA (IIF) | -0.27953 | 0 |
| B76 | B134 | ApoB | CHE | 0.229739 | 1.40E-12 |
| B76 | B77 | ApoB | TC | 0.211754 | 7.22E-11 |
| B76 | B172 | ApoB | KB | 0.181503 | 2.58E-08 |

| | | | | | |
|---|---|---|---|---|---|
| B76 | B137 | ApoB | AFU | -0.1791 | 3.95E-08 |
| B76 | B92 | ApoB | ALDH2-Glu504Glu | 0.177725 | 5.03E-08 |
| B76 | B93 | ApoB | ALDH2-Glu504Lys | -0.17773 | 5.03E-08 |
| B76 | B94 | ApoB | ALDH2-Lys504Lys | -0.17773 | 5.03E-08 |
| B76 | B86 | ApoB | TBhCG | 0.176758 | 5.96E-08 |
| B77 | B127 | TC | A-G ratio | -0.18553 | 1.24E-08 |
| B77 | B130 | TC | GLB | 0.183532 | 1.79E-08 |
| B77 | B140 | TC | r-GT | 0.182566 | 2.13E-08 |
| B79 | B100 | hb V -dna | anti-Ro52 | 0.374482 | 0 |
| B79 | B101 | hb V -dna | anti-Sc70 | 0.374482 | 0 |
| B79 | B102 | hb V -dna | anti-Sm | 0.374482 | 0 |
| B79 | B103 | hb V -dna | anti-SSA/Ro | 0.374482 | 0 |
| B79 | B104 | hb V -dna | anti-SSB | 0.374482 | 0 |
| B79 | B105 | hb V -dna | anti-U1RNP | 0.374482 | 0 |
| B79 | B106 | hb V -dna | anti-P | 0.374482 | 0 |
| B79 | B107 | hb V -dna | AnuA | 0.405152 | 0 |
| B79 | B108 | hb V -dna | Anti-Cyclin | 0.374482 | 0 |
| B79 | B109 | hb V -dna | AMA-M2 | 0.374482 | 0 |
| B79 | B110 | hb V -dna | AHA | 0.405152 | 0 |
| B79 | B119 | hb V -dna | Hcv-rna | 0.286021 | 0 |
| B79 | B121 | hb V -dna | C3 | -0.28602 | 0 |
| B79 | B122 | hb V -dna | C4 | -0.28602 | 0 |
| B79 | B124 | hb V -dna | ACTH | -0.49919 | 0 |
| B79 | B125 | hb V -dna | Cor | -0.49919 | 0 |
| B79 | B139 | hb V -dna | ADA | -0.42128 | 0 |
| B79 | B151 | hb V -dna | Anti-CCP | -0.27421 | 0 |
| B79 | B152 | hb V -dna | ACA-Ig(G-A-M) | -0.286 | 0 |
| B79 | B159 | hb V -dna | β2-MG | -0.28602 | 0 |
| B79 | B169 | hb V -dna | U-CRY | -0.28602 | 0 |
| B79 | B180 | hb V -dna | Tf | 0.32558 | 0 |
| B79 | B194 | hb V -dna | Cl | 0.397401 | 0 |
| B79 | B92 | hb V -dna | ALDH2-Glu504Glu | -0.35164 | 0 |
| B79 | B93 | hb V -dna | ALDH2-Glu504Lys | 0.351642 | 0 |
| B79 | B94 | hb V -dna | ALDH2-Lys504Lys | 0.351642 | 0 |
| B79 | B97 | hb V -dna | dsDNA | 0.374482 | 0 |
| B79 | B98 | hb V -dna | anti-Jo-1 | 0.374482 | 0 |
| B79 | B99 | hb V -dna | anti-PM-Scl | 0.374482 | 0 |
| B79 | B135 | hb V -dna | TBA | -0.25305 | 4.88E-15 |
| B79 | B195 | hb V -dna | Na | 0.249779 | 1.15E-14 |
| B79 | B184 | hb V -dna | INR | -0.21576 | 3.09E-11 |
| B79 | B199 | hb V -dna | Mg | -0.2006 | 7.03E-10 |
| B79 | B137 | hb V -dna | AFU | -0.19993 | 8.03E-10 |
| B79 | B157 | hb V -dna | IgG | -0.17002 | 1.89E-07 |
| B79 | B158 | hb V -dna | IgM | -0.16938 | 2.10E-07 |

| | | | | | |
|---|---|---|---|---|---|
| B79 | B196 | hb V -dna | TCO2 | -0.16257 | 6.40E-07 |
| B8 | B137 | WB-Zinc | AFU | 0.313586 | 0 |
| B8 | B139 | WB-Zinc | ADA | 0.449853 | 0 |
| B8 | B15 | WB-Zinc | CD3-CD19+(B cell) | -0.44487 | 0 |
| B8 | B157 | WB-Zinc | IgG | 0.378593 | 0 |
| B8 | B16 | WB-Zinc | CD3+T | -0.29899 | 0 |
| B8 | B18 | WB-Zinc | CD3+CD4+helper(inducer) T | -0.38571 | 0 |
| B8 | B23 | WB-Zinc | CD3-CD16+CD56+(NK) | 0.338457 | 0 |
| B8 | B19 | WB-Zinc | CD3+CD4+/CD3+CD8+ | -0.22868 | 1.78E-12 |
| B8 | B54 | WB-Zinc | ESR | 0.223503 | 5.71E-12 |
| B8 | B59 | WB-Zinc | WBRV | 0.213551 | 4.94E-11 |
| B8 | B198 | WB-Zinc | P | 0.206965 | 1.95E-10 |
| B8 | B85 | WB-Zinc | SCC | -0.19661 | 1.54E-09 |
| B8 | B69 | WB-Zinc | PV | 0.176326 | 6.42E-08 |
| B8 | B57 | WB-Zinc | RBCAI | 0.166529 | 3.36E-07 |
| B82 | B85 | CEA | SCC | 0.289879 | 0 |
| B82 | B86 | CEA | TBhCG | -0.47109 | 0 |
| B82 | B87 | CEA | NSE | 0.394631 | 0 |
| B82 | B88 | CEA | CA 242 | 0.243003 | 6.13E-14 |
| B82 | B89 | CEA | CA 72-4 | 0.234519 | 4.63E-13 |
| B82 | B84 | CEA | C-PSA | 0.210717 | 8.97E-11 |
| B82 | B83 | CEA | AFP | 0.191743 | 3.91E-09 |
| B83 | B85 | AFP | SCC | 0.282203 | 0 |
| B83 | B89 | AFP | CA 72-4 | 0.263046 | 4.44E-16 |
| B84 | B85 | C-PSA | SCC | 0.42895 | 0 |
| B84 | B86 | C-PSA | TBhCG | -0.23501 | 4.13E-13 |
| B84 | B117 | C-PSA | TT4 | -0.20277 | 4.56E-10 |
| B84 | B116 | C-PSA | TT3 | 0.18219 | 2.28E-08 |
| B85 | B117 | SCC | TT4 | -0.1881 | 7.74E-09 |
| B85 | B87 | SCC | NSE | 0.185814 | 1.18E-08 |
| B85 | B116 | SCC | TT3 | 0.178047 | 4.75E-08 |
| B86 | B87 | TBhCG | NSE | -0.27091 | 0 |
| B86 | B139 | TBhCG | ADA | -0.21724 | 2.25E-11 |
| B86 | B198 | TBhCG | P | -0.20754 | 1.73E-10 |
| B86 | B199 | TBhCG | Mg | 0.172344 | 1.27E-07 |
| B86 | B200 | TBhCG | OP | -0.16445 | 4.72E-07 |
| B86 | B88 | TBhCG | CA 242 | -0.16249 | 6.47E-07 |
| B87 | B88 | NSE | CA 242 | 0.202657 | 4.66E-10 |
| B87 | B89 | NSE | CA 72-4 | 0.18747 | 8.70E-09 |
| B88 | B197 | CA 242 | AG | 0.283448 | 0 |
| B88 | B198 | CA 242 | P | 0.393063 | 0 |
| B88 | B200 | CA 242 | OP | 0.283328 | 0 |
| B88 | B195 | CA 242 | Na | 0.226365 | 3.01E-12 |

| | | | | | |
|---|---|---|---|---|---|
| B88 | B135 | CA 242 | TBA | 0.205991 | 2.38E-10 |
| B90 | B154 | CYFRA 21-1 | RF | -0.1775 | 5.24E-08 |
| B90 | B197 | CYFRA 21-1 | AG | 0.16327 | 5.72E-07 |
| B91 | B126 | TP | Alb | 0.474847 | 0 |
| B91 | B127 | TP | A-G ratio | -0.45969 | 0 |
| B91 | B130 | TP | GLB | 0.771438 | 0 |
| B91 | B188 | TP | Ca | 0.319268 | 0 |
| B92 | B100 | ALDH2-Glu504Glu | anti-Ro52 | -0.53308 | 0 |
| B92 | B101 | ALDH2-Glu504Glu | anti-Sc70 | -0.53308 | 0 |
| B92 | B102 | ALDH2-Glu504Glu | anti-Sm | -0.53308 | 0 |
| B92 | B103 | ALDH2-Glu504Glu | anti-SSA/Ro | -0.53308 | 0 |
| B92 | B104 | ALDH2-Glu504Glu | anti-SSB | -0.53308 | 0 |
| B92 | B105 | ALDH2-Glu504Glu | anti-U1RNP | -0.53308 | 0 |
| B92 | B106 | ALDH2-Glu504Glu | anti-P | -0.53308 | 0 |
| B92 | B107 | ALDH2-Glu504Glu | AnuA | -0.5761 | 0 |
| B92 | B108 | ALDH2-Glu504Glu | Anti-Cyclin | -0.53308 | 0 |
| B92 | B109 | ALDH2-Glu504Glu | AMA-M2 | -0.53308 | 0 |
| B92 | B110 | ALDH2-Glu504Glu | AHA | -0.5761 | 0 |
| B92 | B119 | ALDH2-Glu504Glu | Hcv-rna | -0.4067 | 0 |
| B92 | B120 | ALDH2-Glu504Glu | HCV-cAg | -0.44528 | 0 |
| B92 | B121 | ALDH2-Glu504Glu | C3 | 0.406705 | 0 |
| B92 | B122 | ALDH2-Glu504Glu | C4 | 0.406704 | 0 |
| B92 | B124 | ALDH2-Glu504Glu | ACTH | 0.706725 | 0 |
| B92 | B125 | ALDH2-Glu504Glu | Cor | 0.706725 | 0 |
| B92 | B135 | ALDH2-Glu504Glu | TBA | 0.49474 | 0 |
| B92 | B139 | ALDH2-Glu504Glu | ADA | 0.371758 | 0 |
| B92 | B151 | ALDH2-Glu504Glu | Anti-CCP | 0.383911 | 0 |
| B92 | B152 | ALDH2-Glu504Glu | ACA-Ig(G-A-M) | 0.406681 | 0 |
| B92 | B157 | ALDH2-Glu504Glu | IgG | 0.28515 | 0 |
| B92 | B159 | ALDH2-Glu504Glu | β2-MG | 0.406704 | 0 |
| B92 | B169 | ALDH2-Glu504Glu | U-CRY | 0.406703 | 0 |
| B92 | B180 | ALDH2-Glu504Glu | Tf | -0.40795 | 0 |
| B92 | B184 | ALDH2-Glu504Glu | INR | 0.307844 | 0 |
| B92 | B193 | ALDH2-Glu504Glu | K | -0.34169 | 0 |
| B92 | B194 | ALDH2-Glu504Glu | Cl | -0.43843 | 0 |
| B92 | B198 | ALDH2-Glu504Glu | P | 0.2696 | 0 |
| B92 | B93 | ALDH2-Glu504Glu | ALDH2-Glu504Lys | -1 | 0 |
| B92 | B94 | ALDH2-Glu504Glu | ALDH2-Lys504Lys | -1 | 0 |
| B92 | B96 | ALDH2-Glu504Glu | dsDNA (IIF) | -0.70673 | 0 |
| B92 | B97 | ALDH2-Glu504Glu | dsDNA | -0.53308 | 0 |
| B92 | B98 | ALDH2-Glu504Glu | anti-Jo-1 | -0.53308 | 0 |
| B92 | B99 | ALDH2-Glu504Glu | anti-PM-Scl | -0.53308 | 0 |
| B92 | B156 | ALDH2-Glu504Glu | IgE | -0.23924 | 1.52E-13 |
| B92 | B196 | ALDH2-Glu504Glu | TCO2 | 0.229601 | 1.44E-12 |

| | | | | | |
|---|---|---|---|---|---|
| B92 | B114 | ALDH2-Glu504Glu | TSH TT3 TT4(TT4) | 0.228984 | 1.66E-12 |
| B92 | B155 | ALDH2-Glu504Glu | IgA | 0.226636 | 2.83E-12 |
| B92 | B185 | ALDH2-Glu504Glu | PT | 0.214486 | 4.05E-11 |
| B92 | B195 | ALDH2-Glu504Glu | Na | -0.20422 | 3.41E-10 |
| B92 | B183 | ALDH2-Glu504Glu | PTA | -0.18799 | 7.89E-09 |
| B92 | B154 | ALDH2-Glu504Glu | RF | -0.16625 | 3.52E-07 |
| B93 | B100 | ALDH2-Glu504Lys | anti-Ro52 | 0.533077 | 0 |
| B93 | B101 | ALDH2-Glu504Lys | anti-Sc70 | 0.533077 | 0 |
| B93 | B102 | ALDH2-Glu504Lys | anti-Sm | 0.533077 | 0 |
| B93 | B103 | ALDH2-Glu504Lys | anti-SSA/Ro | 0.533077 | 0 |
| B93 | B104 | ALDH2-Glu504Lys | anti-SSB | 0.533077 | 0 |
| B93 | B105 | ALDH2-Glu504Lys | anti-U1RNP | 0.533077 | 0 |
| B93 | B106 | ALDH2-Glu504Lys | anti-P | 0.533077 | 0 |
| B93 | B107 | ALDH2-Glu504Lys | AnuA | 0.576102 | 0 |
| B93 | B108 | ALDH2-Glu504Lys | Anti-Cyclin | 0.533077 | 0 |
| B93 | B109 | ALDH2-Glu504Lys | AMA-M2 | 0.533077 | 0 |
| B93 | B110 | ALDH2-Glu504Lys | AHA | 0.576102 | 0 |
| B93 | B119 | ALDH2-Glu504Lys | Hcv-rna | 0.406704 | 0 |
| B93 | B120 | ALDH2-Glu504Lys | HCV-cAg | 0.445278 | 0 |
| B93 | B121 | ALDH2-Glu504Lys | C3 | -0.4067 | 0 |
| B93 | B122 | ALDH2-Glu504Lys | C4 | -0.4067 | 0 |
| B93 | B124 | ALDH2-Glu504Lys | ACTH | -0.70673 | 0 |
| B93 | B125 | ALDH2-Glu504Lys | Cor | -0.70673 | 0 |
| B93 | B135 | ALDH2-Glu504Lys | TBA | -0.49474 | 0 |
| B93 | B139 | ALDH2-Glu504Lys | ADA | -0.37176 | 0 |
| B93 | B151 | ALDH2-Glu504Lys | Anti-CCP | -0.38391 | 0 |
| B93 | B152 | ALDH2-Glu504Lys | ACA-Ig(G-A-M) | -0.40668 | 0 |
| B93 | B157 | ALDH2-Glu504Lys | IgG | -0.28515 | 0 |
| B93 | B159 | ALDH2-Glu504Lys | β2-MG | -0.4067 | 0 |
| B93 | B169 | ALDH2-Glu504Lys | U-CRY | -0.4067 | 0 |
| B93 | B180 | ALDH2-Glu504Lys | Tf | 0.407948 | 0 |
| B93 | B184 | ALDH2-Glu504Lys | INR | -0.30784 | 0 |
| B93 | B193 | ALDH2-Glu504Lys | K | 0.341692 | 0 |
| B93 | B194 | ALDH2-Glu504Lys | Cl | 0.438426 | 0 |
| B93 | B198 | ALDH2-Glu504Lys | P | -0.2696 | 0 |
| B93 | B94 | ALDH2-Glu504Lys | ALDH2-Lys504Lys | 1 | 0 |
| B93 | B96 | ALDH2-Glu504Lys | dsDNA (IIF) | 0.706725 | 0 |
| B93 | B97 | ALDH2-Glu504Lys | dsDNA | 0.533077 | 0 |
| B93 | B98 | ALDH2-Glu504Lys | anti-Jo-1 | 0.533077 | 0 |
| B93 | B99 | ALDH2-Glu504Lys | anti-PM-Scl | 0.533077 | 0 |
| B93 | B156 | ALDH2-Glu504Lys | IgE | 0.23924 | 1.52E-13 |
| B93 | B196 | ALDH2-Glu504Lys | TCO2 | -0.2296 | 1.44E-12 |
| B93 | B114 | ALDH2-Glu504Lys | TSH TT3 TT4(TT4) | -0.22898 | 1.66E-12 |
| B93 | B155 | ALDH2-Glu504Lys | IgA | -0.22664 | 2.83E-12 |

| | | | | | |
|---|---|---|---|---|---|
| B93 | B185 | ALDH2-Glu504Lys | PT | -0.21449 | 4.05E-11 |
| B93 | B195 | ALDH2-Glu504Lys | Na | 0.204215 | 3.41E-10 |
| B93 | B183 | ALDH2-Glu504Lys | PTA | 0.187995 | 7.89E-09 |
| B93 | B154 | ALDH2-Glu504Lys | RF | 0.16625 | 3.52E-07 |
| B94 | B100 | ALDH2-Lys504Lys | anti-Ro52 | 0.533077 | 0 |
| B94 | B101 | ALDH2-Lys504Lys | anti-Sc70 | 0.533077 | 0 |
| B94 | B102 | ALDH2-Lys504Lys | anti-Sm | 0.533077 | 0 |
| B94 | B103 | ALDH2-Lys504Lys | anti-SSA/Ro | 0.533077 | 0 |
| B94 | B104 | ALDH2-Lys504Lys | anti-SSB | 0.533077 | 0 |
| B94 | B105 | ALDH2-Lys504Lys | anti-U1RNP | 0.533077 | 0 |
| B94 | B106 | ALDH2-Lys504Lys | anti-P | 0.533077 | 0 |
| B94 | B107 | ALDH2-Lys504Lys | AnuA | 0.576102 | 0 |
| B94 | B108 | ALDH2-Lys504Lys | Anti-Cyclin | 0.533077 | 0 |
| B94 | B109 | ALDH2-Lys504Lys | AMA-M2 | 0.533077 | 0 |
| B94 | B110 | ALDH2-Lys504Lys | AHA | 0.576102 | 0 |
| B94 | B119 | ALDH2-Lys504Lys | Hcv-rna | 0.406704 | 0 |
| B94 | B120 | ALDH2-Lys504Lys | HCV-cAg | 0.445278 | 0 |
| B94 | B121 | ALDH2-Lys504Lys | C3 | -0.4067 | 0 |
| B94 | B122 | ALDH2-Lys504Lys | C4 | -0.4067 | 0 |
| B94 | B124 | ALDH2-Lys504Lys | ACTH | -0.70673 | 0 |
| B94 | B125 | ALDH2-Lys504Lys | Cor | -0.70673 | 0 |
| B94 | B135 | ALDH2-Lys504Lys | TBA | -0.49474 | 0 |
| B94 | B139 | ALDH2-Lys504Lys | ADA | -0.37176 | 0 |
| B94 | B151 | ALDH2-Lys504Lys | Anti-CCP | -0.38391 | 0 |
| B94 | B152 | ALDH2-Lys504Lys | ACA-Ig(G-A-M) | -0.40668 | 0 |
| B94 | B157 | ALDH2-Lys504Lys | IgG | -0.28515 | 0 |
| B94 | B159 | ALDH2-Lys504Lys | β2-MG | -0.4067 | 0 |
| B94 | B169 | ALDH2-Lys504Lys | U-CRY | -0.4067 | 0 |
| B94 | B180 | ALDH2-Lys504Lys | Tf | 0.407948 | 0 |
| B94 | B184 | ALDH2-Lys504Lys | INR | -0.30784 | 0 |
| B94 | B193 | ALDH2-Lys504Lys | K | 0.341692 | 0 |
| B94 | B194 | ALDH2-Lys504Lys | Cl | 0.438426 | 0 |
| B94 | B198 | ALDH2-Lys504Lys | P | -0.2696 | 0 |
| B94 | B96 | ALDH2-Lys504Lys | dsDNA (IIF) | 0.706725 | 0 |
| B94 | B97 | ALDH2-Lys504Lys | dsDNA | 0.533077 | 0 |
| B94 | B98 | ALDH2-Lys504Lys | anti-Jo-1 | 0.533077 | 0 |
| B94 | B99 | ALDH2-Lys504Lys | anti-PM-Scl | 0.533077 | 0 |
| B94 | B156 | ALDH2-Lys504Lys | IgE | 0.23924 | 1.52E-13 |
| B94 | B196 | ALDH2-Lys504Lys | TCO2 | -0.2296 | 1.44E-12 |
| B94 | B114 | ALDH2-Lys504Lys | TSH TT3 TT4(TT4) | -0.22898 | 1.66E-12 |
| B94 | B155 | ALDH2-Lys504Lys | IgA | -0.22664 | 2.83E-12 |
| B94 | B185 | ALDH2-Lys504Lys | PT | -0.21449 | 4.05E-11 |
| B94 | B195 | ALDH2-Lys504Lys | Na | 0.204215 | 3.41E-10 |
| B94 | B183 | ALDH2-Lys504Lys | PTA | 0.187995 | 7.89E-09 |

| | | | | | |
|---|---|---|---|---|---|
| B94 | B154 | ALDH2-Lys504Lys | RF | 0.16625 | 3.52E-07 |
| B95 | B100 | anti-cenpb | anti-Ro52 | -0.34746 | 0 |
| B95 | B101 | anti-cenpb | anti-Sc70 | -0.34746 | 0 |
| B95 | B102 | anti-cenpb | anti-Sm | -0.34746 | 0 |
| B95 | B103 | anti-cenpb | anti-SSA/Ro | -0.34746 | 0 |
| B95 | B104 | anti-cenpb | anti-SSB | -0.34746 | 0 |
| B95 | B105 | anti-cenpb | anti-U1RNP | -0.34746 | 0 |
| B95 | B106 | anti-cenpb | anti-P | -0.34746 | 0 |
| B95 | B108 | anti-cenpb | Anti-Cyclin | -0.34746 | 0 |
| B95 | B109 | anti-cenpb | AMA-M2 | -0.34746 | 0 |
| B95 | B157 | anti-cenpb | IgG | 0.380048 | 0 |
| B95 | B193 | anti-cenpb | K | -0.36175 | 0 |
| B95 | B97 | anti-cenpb | dsDNA | -0.34746 | 0 |
| B95 | B98 | anti-cenpb | anti-Jo-1 | -0.34746 | 0 |
| B95 | B99 | anti-cenpb | anti-PM-Scl | -0.34746 | 0 |
| B95 | B156 | anti-cenpb | IgE | -0.2204 | 1.13E-11 |
| B95 | B114 | anti-cenpb | TSH TT3 TT4(TT4) | 0.209411 | 1.18E-10 |
| B95 | B199 | anti-cenpb | Mg | -0.19517 | 2.03E-09 |
| B95 | B139 | anti-cenpb | ADA | 0.178886 | 4.10E-08 |
| B95 | B115 | anti-cenpb | TSH TT3 TT4(TT3) | 0.16711 | 3.06E-07 |
| B96 | B100 | dsDNA (IIF) | anti-Ro52 | 0.376739 | 0 |
| B96 | B101 | dsDNA (IIF) | anti-Sc70 | 0.376739 | 0 |
| B96 | B102 | dsDNA (IIF) | anti-Sm | 0.376739 | 0 |
| B96 | B103 | dsDNA (IIF) | anti-SSA/Ro | 0.376739 | 0 |
| B96 | B104 | dsDNA (IIF) | anti-SSB | 0.376739 | 0 |
| B96 | B105 | dsDNA (IIF) | anti-U1RNP | 0.376739 | 0 |
| B96 | B106 | dsDNA (IIF) | anti-P | 0.376739 | 0 |
| B96 | B107 | dsDNA (IIF) | AnuA | 0.407146 | 0 |
| B96 | B108 | dsDNA (IIF) | Anti-Cyclin | 0.376739 | 0 |
| B96 | B109 | dsDNA (IIF) | AMA-M2 | 0.376739 | 0 |
| B96 | B110 | dsDNA (IIF) | AHA | 0.407146 | 0 |
| B96 | B120 | dsDNA (IIF) | HCV-cAg | 0.314689 | 0 |
| B96 | B138 | dsDNA (IIF) | PAB | -0.33578 | 0 |
| B96 | B97 | dsDNA (IIF) | dsDNA | 0.376739 | 0 |
| B96 | B98 | dsDNA (IIF) | anti-Jo-1 | 0.376739 | 0 |
| B96 | B99 | dsDNA (IIF) | anti-PM-Scl | 0.376739 | 0 |
| B96 | B180 | dsDNA (IIF) | Tf | 0.238636 | 1.75E-13 |
| B96 | B196 | dsDNA (IIF) | TCO2 | -0.22539 | 3.75E-12 |
| B96 | B156 | dsDNA (IIF) | IgE | 0.221822 | 8.28E-12 |
| B96 | B198 | dsDNA (IIF) | P | -0.21792 | 1.94E-11 |
| B96 | B137 | dsDNA (IIF) | AFU | -0.18171 | 2.48E-08 |
| B96 | B194 | dsDNA (IIF) | Cl | 0.178708 | 4.23E-08 |
| B97 | B100 | dsDNA | anti-Ro52 | 1 | 0 |
| B97 | B102 | dsDNA | anti-Sm | 1 | 0 |

| | | | | | |
|---|---|---|---|---|---|
| B97 | B103 | dsDNA | anti-SSA/Ro | 1 | 0 |
| B97 | B104 | dsDNA | anti-SSB | 1 | 0 |
| B97 | B105 | dsDNA | anti-U1RNP | 1 | 0 |
| B97 | B106 | dsDNA | anti-P | 1 | 0 |
| B97 | B107 | dsDNA | AnuA | 0.925318 | 0 |
| B97 | B108 | dsDNA | Anti-Cyclin | 1 | 0 |
| B97 | B109 | dsDNA | AMA-M2 | 1 | 0 |
| B97 | B110 | dsDNA | AHA | 0.925318 | 0 |
| B97 | B119 | dsDNA | Hcv-rna | 0.433836 | 0 |
| B97 | B120 | dsDNA | HCV-cAg | 0.352785 | 0 |
| B97 | B121 | dsDNA | C3 | -0.43384 | 0 |
| B97 | B122 | dsDNA | C4 | -0.43384 | 0 |
| B97 | B124 | dsDNA | ACTH | -0.37674 | 0 |
| B97 | B125 | dsDNA | Cor | -0.37674 | 0 |
| B97 | B139 | dsDNA | ADA | -0.45548 | 0 |
| B97 | B152 | dsDNA | ACA-Ig(G-A-M) | -0.43381 | 0 |
| B97 | B157 | dsDNA | IgG | -0.33455 | 0 |
| B97 | B159 | dsDNA | β2-MG | -0.43384 | 0 |
| B97 | B169 | dsDNA | U-CRY | -0.43383 | 0 |
| B97 | B180 | dsDNA | Tf | 0.440213 | 0 |
| B97 | B193 | dsDNA | K | 0.3357 | 0 |
| B97 | B194 | dsDNA | Cl | 0.354015 | 0 |
| B97 | B99 | dsDNA | anti-PM-Scl | 1 | 0 |
| B97 | B155 | dsDNA | IgA | -0.24381 | 5.02E-14 |
| B97 | B137 | dsDNA | AFU | -0.24337 | 5.60E-14 |
| B97 | B154 | dsDNA | RF | 0.242501 | 6.93E-14 |
| B97 | B195 | dsDNA | Na | 0.219271 | 1.45E-11 |
| B97 | B184 | dsDNA | INR | -0.20636 | 2.21E-10 |
| B97 | B183 | dsDNA | PTA | 0.202624 | 4.70E-10 |
| B97 | B178 | dsDNA | Ig-U | 0.196769 | 1.49E-09 |
| B97 | B135 | dsDNA | TBA | -0.18726 | 9.04E-09 |
| B97 | B196 | dsDNA | TCO2 | -0.17771 | 5.04E-08 |
| B98 | B101 | anti-Jo-1 | anti-Sc70 | 1 | 0 |
| B98 | B102 | anti-Jo-1 | anti-Sm | 1 | 0 |
| B98 | B107 | anti-Jo-1 | AnuA | 0.925317 | 0 |
| B98 | B108 | anti-Jo-1 | Anti-Cyclin | 1 | 0 |
| B98 | B110 | anti-Jo-1 | AHA | 0.925318 | 0 |
| B98 | B119 | anti-Jo-1 | Hcv-rna | 0.433836 | 0 |
| B98 | B120 | anti-Jo-1 | HCV-cAg | 0.352785 | 0 |
| B98 | B121 | anti-Jo-1 | C3 | -0.43384 | 0 |
| B98 | B122 | anti-Jo-1 | C4 | -0.43384 | 0 |
| B98 | B124 | anti-Jo-1 | ACTH | -0.37674 | 0 |
| B98 | B125 | anti-Jo-1 | Cor | -0.37674 | 0 |
| B98 | B139 | anti-Jo-1 | ADA | -0.45548 | 0 |

| | | | | | |
|---|---|---|---|---|---|
| B98 | B152 | anti-Jo-1 | ACA-Ig(G-A-M) | -0.43381 | 0 |
| B98 | B157 | anti-Jo-1 | IgG | -0.33455 | 0 |
| B98 | B159 | anti-Jo-1 | β2-MG | -0.43384 | 0 |
| B98 | B169 | anti-Jo-1 | U-CRY | -0.43383 | 0 |
| B98 | B180 | anti-Jo-1 | Tf | 0.440213 | 0 |
| B98 | B193 | anti-Jo-1 | K | 0.3357 | 0 |
| B98 | B194 | anti-Jo-1 | Cl | 0.354015 | 0 |
| B98 | B99 | anti-Jo-1 | anti-PM-Scl | 1 | 0 |
| B98 | B155 | anti-Jo-1 | IgA | -0.24381 | 5.02E-14 |
| B98 | B137 | anti-Jo-1 | AFU | -0.24337 | 5.60E-14 |
| B98 | B154 | anti-Jo-1 | RF | 0.242501 | 6.93E-14 |
| B98 | B195 | anti-Jo-1 | Na | 0.219271 | 1.45E-11 |
| B98 | B184 | anti-Jo-1 | INR | -0.20636 | 2.21E-10 |
| B98 | B183 | anti-Jo-1 | PTA | 0.202624 | 4.70E-10 |
| B98 | B178 | anti-Jo-1 | Ig-U | 0.196769 | 1.49E-09 |
| B98 | B135 | anti-Jo-1 | TBA | -0.18726 | 9.04E-09 |
| B98 | B196 | anti-Jo-1 | TCO2 | -0.17771 | 5.04E-08 |
| B99 | B101 | anti-PM-Scl | anti-Sc70 | 1 | 0 |
| B99 | B104 | anti-PM-Scl | anti-SSB | 1 | 0 |
| B99 | B106 | anti-PM-Scl | anti-P | 1 | 0 |
| B99 | B107 | anti-PM-Scl | AnuA | 0.925318 | 0 |
| B99 | B110 | anti-PM-Scl | AHA | 0.925318 | 0 |
| B99 | B119 | anti-PM-Scl | Hcv-rna | 0.433836 | 0 |
| B99 | B120 | anti-PM-Scl | HCV-cAg | 0.352785 | 0 |
| B99 | B121 | anti-PM-Scl | C3 | -0.43384 | 0 |
| B99 | B122 | anti-PM-Scl | C4 | -0.43384 | 0 |
| B99 | B124 | anti-PM-Scl | ACTH | -0.37674 | 0 |
| B99 | B125 | anti-PM-Scl | Cor | -0.37674 | 0 |
| B99 | B139 | anti-PM-Scl | ADA | -0.45548 | 0 |
| B99 | B152 | anti-PM-Scl | ACA-Ig(G-A-M) | -0.43381 | 0 |
| B99 | B157 | anti-PM-Scl | IgG | -0.33455 | 0 |
| B99 | B159 | anti-PM-Scl | β2-MG | -0.43384 | 0 |
| B99 | B169 | anti-PM-Scl | U-CRY | -0.43383 | 0 |
| B99 | B180 | anti-PM-Scl | Tf | 0.440213 | 0 |
| B99 | B193 | anti-PM-Scl | K | 0.3357 | 0 |
| B99 | B194 | anti-PM-Scl | Cl | 0.354015 | 0 |
| B99 | B155 | anti-PM-Scl | IgA | -0.24381 | 5.02E-14 |
| B99 | B137 | anti-PM-Scl | AFU | -0.24337 | 5.60E-14 |
| B99 | B154 | anti-PM-Scl | RF | 0.242501 | 6.93E-14 |
| B99 | B195 | anti-PM-Scl | Na | 0.219271 | 1.45E-11 |
| B99 | B184 | anti-PM-Scl | INR | -0.20636 | 2.21E-10 |
| B99 | B183 | anti-PM-Scl | PTA | 0.202624 | 4.70E-10 |
| B99 | B178 | anti-PM-Scl | Ig-U | 0.196769 | 1.49E-09 |
| B99 | B135 | anti-PM-Scl | TBA | -0.18726 | 9.04E-09 |

| | | | | | |
|---|---|---|---|---|---|
| B99 | B196 | anti-PM-Scl | TCO2 | -0.17771 | 5.04E-08 |

**Supplementary Table 7. PEI Correlations in Hyperlipidemia**

| Row | Column | Row | Column | Cor | *P* |
|---|---|---|---|---|---|
| B1 | B161 | Gender | LEU | 0.241648 | 0 |
| B1 | B163 | Gender | SG | -0.29208 | 0 |
| B1 | B174 | Gender | Urea/Cr | 0.405881 | 0 |
| B1 | B190 | Gender | Cre | -0.48747 | 0 |
| B1 | B192 | Gender | UA | -0.45552 | 0 |
| B1 | B204 | Gender | Drinking | -0.35298 | 0 |
| B1 | B206 | Gender | Waist | -0.44721 | 0 |
| B1 | B207 | Gender | WHR | -0.35161 | 0 |
| B1 | B209 | Gender | Smoking | -0.36934 | 0 |
| B1 | B210 | Gender | Hip | -0.28234 | 0 |
| B1 | B215 | Gender | BMI | -0.2678 | 0 |
| B1 | B216 | Gender | BMI | -0.57511 | 0 |
| B1 | B221 | Gender | Hei | -0.6864 | 0 |
| B1 | B222 | Gender | NS | -0.70613 | 0 |
| B1 | B40 | Gender | WBC | -0.19843 | 0 |
| B1 | B42 | Gender | MON | -0.28678 | 0 |
| B1 | B45 | Gender | MCHC | -0.28185 | 0 |
| B1 | B46 | Gender | RBC Count | -0.39348 | 0 |
| B1 | B51 | Gender | Hb | -0.51389 | 0 |
| B1 | B72 | Gender | HDL-C | 0.383801 | 0 |
| B1 | B41 | Gender | MON％ | -0.19516 | 4.44E-16 |
| B1 | B50 | Gender | eo% | -0.15026 | 3.69E-10 |
| B1 | B44 | Gender | MCH | -0.14784 | 7.08E-10 |
| B1 | B3 | Gender | Hcy | -0.14415 | 1.87E-09 |
| B1 | B141 | Gender | AST/ALT | 0.140733 | 4.49E-09 |
| B1 | B132 | Gender | Dbil | -0.13882 | 7.28E-09 |
| B1 | B77 | Gender | TC | 0.137485 | 1.01E-08 |
| B1 | B128 | Gender | ALT | -0.13675 | 1.22E-08 |
| B1 | B201 | Gender | e-habits | -0.13149 | 4.34E-08 |
| B1 | B133 | Gender | TBIL | -0.13009 | 6.05E-08 |
| B1 | B118 | Gender | CEA1 | -0.12936 | 7.17E-08 |
| B1 | B173 | Gender | NTT | 0.122837 | 3.17E-07 |
| B1 | B48 | Gender | LYM% | 0.119136 | 7.13E-07 |
| B10 | B12 | PG I | PG II | 0.563497 | 0 |
| B10 | B11 | PG I | PGI/PGII | 0.12008 | 5.81E-07 |
| B100 | B157 | anti-Ro52 | IgG | 0.206843 | 0 |
| B100 | B151 | anti-Ro52 | Anti-CCP | 0.193813 | 4.44E-16 |
| B100 | B178 | anti-Ro52 | Ig-U | -0.13784 | 9.29E-09 |
| B101 | B104 | anti-Sc70 | anti-SSB | 1 | 0 |
| B101 | B107 | anti-Sc70 | AnuA | 0.999999 | 0 |
| B101 | B119 | anti-Sc70 | Hcv-rna | 0.384926 | 0 |
| B101 | B120 | anti-Sc70 | HCV-cAg | 0.349051 | 0 |

| | | | | | |
|---|---|---|---|---|---|
| B101 | B121 | anti-Sc70 | C3 | -0.27501 | 0 |
| B101 | B122 | anti-Sc70 | C4 | 0.242862 | 0 |
| B101 | B125 | anti-Sc70 | Cor | 0.298639 | 0 |
| B101 | B134 | anti-Sc70 | CHE | -0.23432 | 0 |
| B101 | B151 | anti-Sc70 | Anti-CCP | 0.245648 | 0 |
| B101 | B152 | anti-Sc70 | ACA-Ig(G-A-M) | -0.38501 | 0 |
| B101 | B156 | anti-Sc70 | IgE | 0.234142 | 0 |
| B101 | B157 | anti-Sc70 | IgG | 0.328845 | 0 |
| B101 | B159 | anti-Sc70 | β2-MG | -0.38492 | 0 |
| B101 | B138 | anti-Sc70 | PAB | -0.18058 | 4.35E-14 |
| B101 | B154 | anti-Sc70 | RF | 0.158265 | 3.99E-11 |
| B101 | B178 | anti-Sc70 | Ig-U | 0.151079 | 2.96E-10 |
| B101 | B193 | anti-Sc70 | K | 0.125601 | 1.70E-07 |
| B101 | B182 | anti-Sc70 | TT | 0.124542 | 2.17E-07 |
| B101 | B183 | anti-Sc70 | PTA | -0.11862 | 7.97E-07 |
| B102 | B103 | anti-Sm | anti-SSA/Ro | 1 | 0 |
| B102 | B104 | anti-Sm | anti-SSB | 1 | 0 |
| B102 | B105 | anti-Sm | anti-U1RNP | 1 | 0 |
| B102 | B106 | anti-Sm | anti-P | 1 | 0 |
| B102 | B107 | anti-Sm | AnuA | 0.999999 | 0 |
| B102 | B108 | anti-Sm | Anti-Cyclin | 1 | 0 |
| B102 | B109 | anti-Sm | AMA-M2 | 1 | 0 |
| B102 | B110 | anti-Sm | AHA | 1 | 0 |
| B102 | B119 | anti-Sm | Hcv-rna | 0.384926 | 0 |
| B102 | B120 | anti-Sm | HCV-cAg | 0.349052 | 0 |
| B102 | B121 | anti-Sm | C3 | -0.27501 | 0 |
| B102 | B122 | anti-Sm | C4 | 0.242862 | 0 |
| B102 | B125 | anti-Sm | Cor | 0.298639 | 0 |
| B102 | B134 | anti-Sm | CHE | -0.23432 | 0 |
| B102 | B151 | anti-Sm | Anti-CCP | 0.245648 | 0 |
| B102 | B152 | anti-Sm | ACA-Ig(G-A-M) | -0.385 | 0 |
| B102 | B156 | anti-Sm | IgE | 0.234142 | 0 |
| B102 | B157 | anti-Sm | IgG | 0.328845 | 0 |
| B102 | B159 | anti-Sm | β2-MG | -0.38492 | 0 |
| B102 | B138 | anti-Sm | PAB | -0.18058 | 4.35E-14 |
| B102 | B154 | anti-Sm | RF | 0.158265 | 3.99E-11 |
| B102 | B178 | anti-Sm | Ig-U | 0.151079 | 2.96E-10 |
| B102 | B193 | anti-Sm | K | 0.125601 | 1.70E-07 |
| B102 | B182 | anti-Sm | TT | 0.124542 | 2.17E-07 |
| B102 | B183 | anti-Sm | PTA | -0.11862 | 7.97E-07 |
| B103 | B107 | anti-SSA/Ro | AnuA | 0.999999 | 0 |
| B103 | B119 | anti-SSA/Ro | Hcv-rna | 0.384926 | 0 |
| B103 | B120 | anti-SSA/Ro | HCV-cAg | 0.349051 | 0 |
| B103 | B121 | anti-SSA/Ro | C3 | -0.27501 | 0 |

| | | | | | |
|---|---|---|---|---|---|
| B103 | B122 | anti-SSA/Ro | C4 | 0.242862 | 0 |
| B103 | B125 | anti-SSA/Ro | Cor | 0.298639 | 0 |
| B103 | B134 | anti-SSA/Ro | CHE | -0.23432 | 0 |
| B103 | B151 | anti-SSA/Ro | Anti-CCP | 0.245648 | 0 |
| B103 | B152 | anti-SSA/Ro | ACA-Ig(G-A-M) | -0.385 | 0 |
| B103 | B156 | anti-SSA/Ro | IgE | 0.234142 | 0 |
| B103 | B157 | anti-SSA/Ro | IgG | 0.328845 | 0 |
| B103 | B159 | anti-SSA/Ro | β2-MG | -0.38492 | 0 |
| B103 | B138 | anti-SSA/Ro | PAB | -0.18058 | 4.35E-14 |
| B103 | B154 | anti-SSA/Ro | RF | 0.158265 | 3.99E-11 |
| B103 | B178 | anti-SSA/Ro | Ig-U | 0.151079 | 2.96E-10 |
| B103 | B193 | anti-SSA/Ro | K | 0.125601 | 1.70E-07 |
| B103 | B182 | anti-SSA/Ro | TT | 0.124542 | 2.17E-07 |
| B103 | B183 | anti-SSA/Ro | PTA | -0.11862 | 7.97E-07 |
| B104 | B106 | anti-SSB | anti-P | 1 | 0 |
| B104 | B107 | anti-SSB | AnuA | 0.999999 | 0 |
| B104 | B108 | anti-SSB | Anti-Cyclin | 1 | 0 |
| B104 | B110 | anti-SSB | AHA | 1 | 0 |
| B104 | B119 | anti-SSB | Hcv-rna | 0.384926 | 0 |
| B104 | B120 | anti-SSB | HCV-cAg | 0.349051 | 0 |
| B104 | B121 | anti-SSB | C3 | -0.27501 | 0 |
| B104 | B122 | anti-SSB | C4 | 0.242862 | 0 |
| B104 | B125 | anti-SSB | Cor | 0.298639 | 0 |
| B104 | B134 | anti-SSB | CHE | -0.23432 | 0 |
| B104 | B151 | anti-SSB | Anti-CCP | 0.245648 | 0 |
| B104 | B152 | anti-SSB | ACA-Ig(G-A-M) | -0.385 | 0 |
| B104 | B156 | anti-SSB | IgE | 0.234142 | 0 |
| B104 | B157 | anti-SSB | IgG | 0.328845 | 0 |
| B104 | B159 | anti-SSB | β2-MG | -0.38492 | 0 |
| B104 | B138 | anti-SSB | PAB | -0.18058 | 4.35E-14 |
| B104 | B154 | anti-SSB | RF | 0.158265 | 3.99E-11 |
| B104 | B178 | anti-SSB | Ig-U | 0.151079 | 2.96E-10 |
| B104 | B193 | anti-SSB | K | 0.125601 | 1.70E-07 |
| B104 | B182 | anti-SSB | TT | 0.124542 | 2.17E-07 |
| B104 | B183 | anti-SSB | PTA | -0.11862 | 7.97E-07 |
| B105 | B107 | anti-U1RNP | AnuA | 0.999999 | 0 |
| B105 | B108 | anti-U1RNP | Anti-Cyclin | 1 | 0 |
| B105 | B119 | anti-U1RNP | Hcv-rna | 0.384926 | 0 |
| B105 | B120 | anti-U1RNP | HCV-cAg | 0.349051 | 0 |
| B105 | B121 | anti-U1RNP | C3 | -0.27501 | 0 |
| B105 | B122 | anti-U1RNP | C4 | 0.242862 | 0 |
| B105 | B125 | anti-U1RNP | Cor | 0.298639 | 0 |
| B105 | B134 | anti-U1RNP | CHE | -0.23432 | 0 |
| B105 | B151 | anti-U1RNP | Anti-CCP | 0.245648 | 0 |

| | | | | | |
|---|---|---|---|---|---|
| B105 | B152 | anti-U1RNP | ACA-Ig(G-A-M) | -0.385 | 0 |
| B105 | B156 | anti-U1RNP | IgE | 0.234142 | 0 |
| B105 | B157 | anti-U1RNP | IgG | 0.328845 | 0 |
| B105 | B159 | anti-U1RNP | β2-MG | -0.38492 | 0 |
| B105 | B138 | anti-U1RNP | PAB | -0.18058 | 4.35E-14 |
| B105 | B154 | anti-U1RNP | RF | 0.158265 | 3.99E-11 |
| B105 | B178 | anti-U1RNP | Ig-U | 0.151079 | 2.96E-10 |
| B105 | B193 | anti-U1RNP | K | 0.125601 | 1.70E-07 |
| B105 | B182 | anti-U1RNP | TT | 0.124542 | 2.17E-07 |
| B105 | B183 | anti-U1RNP | PTA | -0.11862 | 7.97E-07 |
| B106 | B107 | anti-P | AnuA | 0.999999 | 0 |
| B106 | B119 | anti-P | Hcv-rna | 0.384926 | 0 |
| B106 | B120 | anti-P | HCV-cAg | 0.349051 | 0 |
| B106 | B121 | anti-P | C3 | -0.27501 | 0 |
| B106 | B122 | anti-P | C4 | 0.242862 | 0 |
| B106 | B125 | anti-P | Cor | 0.298639 | 0 |
| B106 | B134 | anti-P | CHE | -0.23432 | 0 |
| B106 | B151 | anti-P | Anti-CCP | 0.245648 | 0 |
| B106 | B152 | anti-P | ACA-Ig(G-A-M) | -0.38501 | 0 |
| B106 | B156 | anti-P | IgE | 0.234142 | 0 |
| B106 | B157 | anti-P | IgG | 0.328845 | 0 |
| B106 | B159 | anti-P | β2-MG | -0.38492 | 0 |
| B106 | B138 | anti-P | PAB | -0.18058 | 4.35E-14 |
| B106 | B154 | anti-P | RF | 0.158265 | 3.99E-11 |
| B106 | B178 | anti-P | Ig-U | 0.151079 | 2.96E-10 |
| B106 | B193 | anti-P | K | 0.125601 | 1.70E-07 |
| B106 | B182 | anti-P | TT | 0.124542 | 2.17E-07 |
| B106 | B183 | anti-P | PTA | -0.11862 | 7.97E-07 |
| B107 | B108 | AnuA | Anti-Cyclin | 0.999999 | 0 |
| B107 | B109 | AnuA | AMA-M2 | 0.999999 | 0 |
| B107 | B110 | AnuA | AHA | 0.999999 | 0 |
| B107 | B119 | AnuA | Hcv-rna | 0.384925 | 0 |
| B107 | B120 | AnuA | HCV-cAg | 0.349053 | 0 |
| B107 | B121 | AnuA | C3 | -0.27501 | 0 |
| B107 | B122 | AnuA | C4 | 0.242862 | 0 |
| B107 | B125 | AnuA | Cor | 0.298638 | 0 |
| B107 | B134 | AnuA | CHE | -0.23432 | 0 |
| B107 | B151 | AnuA | Anti-CCP | 0.245647 | 0 |
| B107 | B152 | AnuA | ACA-Ig(G-A-M) | -0.38501 | 0 |
| B107 | B156 | AnuA | IgE | 0.234142 | 0 |
| B107 | B157 | AnuA | IgG | 0.328845 | 0 |
| B107 | B159 | AnuA | β2-MG | -0.38492 | 0 |
| B107 | B138 | AnuA | PAB | -0.18058 | 4.35E-14 |
| B107 | B154 | AnuA | RF | 0.158265 | 3.99E-11 |

| | | | | | |
|---|---|---|---|---|---|
| B107 | B178 | AnuA | Ig-U | 0.151079 | 2.96E-10 |
| B107 | B193 | AnuA | K | 0.125601 | 1.70E-07 |
| B107 | B182 | AnuA | TT | 0.124542 | 2.17E-07 |
| B107 | B183 | AnuA | PTA | -0.11862 | 7.97E-07 |
| B108 | B119 | Anti-Cyclin | Hcv-rna | 0.384926 | 0 |
| B108 | B120 | Anti-Cyclin | HCV-cAg | 0.349051 | 0 |
| B108 | B121 | Anti-Cyclin | C3 | -0.27501 | 0 |
| B108 | B122 | Anti-Cyclin | C4 | 0.242862 | 0 |
| B108 | B125 | Anti-Cyclin | Cor | 0.298639 | 0 |
| B108 | B134 | Anti-Cyclin | CHE | -0.23432 | 0 |
| B108 | B151 | Anti-Cyclin | Anti-CCP | 0.245648 | 0 |
| B108 | B152 | Anti-Cyclin | ACA-Ig(G-A-M) | -0.38501 | 0 |
| B108 | B156 | Anti-Cyclin | IgE | 0.234142 | 0 |
| B108 | B157 | Anti-Cyclin | IgG | 0.328845 | 0 |
| B108 | B159 | Anti-Cyclin | β2-MG | -0.38492 | 0 |
| B108 | B138 | Anti-Cyclin | PAB | -0.18058 | 4.35E-14 |
| B108 | B154 | Anti-Cyclin | RF | 0.158265 | 3.99E-11 |
| B108 | B178 | Anti-Cyclin | Ig-U | 0.151079 | 2.96E-10 |
| B108 | B193 | Anti-Cyclin | K | 0.125601 | 1.70E-07 |
| B108 | B182 | Anti-Cyclin | TT | 0.124542 | 2.17E-07 |
| B108 | B183 | Anti-Cyclin | PTA | -0.11862 | 7.97E-07 |
| B109 | B119 | AMA-M2 | Hcv-rna | 0.384926 | 0 |
| B109 | B120 | AMA-M2 | HCV-cAg | 0.349051 | 0 |
| B109 | B121 | AMA-M2 | C3 | -0.27501 | 0 |
| B109 | B122 | AMA-M2 | C4 | 0.242862 | 0 |
| B109 | B125 | AMA-M2 | Cor | 0.298639 | 0 |
| B109 | B134 | AMA-M2 | CHE | -0.23432 | 0 |
| B109 | B151 | AMA-M2 | Anti-CCP | 0.245648 | 0 |
| B109 | B152 | AMA-M2 | ACA-Ig(G-A-M) | -0.385 | 0 |
| B109 | B156 | AMA-M2 | IgE | 0.234142 | 0 |
| B109 | B157 | AMA-M2 | IgG | 0.328845 | 0 |
| B109 | B159 | AMA-M2 | β2-MG | -0.38492 | 0 |
| B109 | B138 | AMA-M2 | PAB | -0.18058 | 4.35E-14 |
| B109 | B154 | AMA-M2 | RF | 0.158265 | 3.99E-11 |
| B109 | B178 | AMA-M2 | Ig-U | 0.151079 | 2.96E-10 |
| B109 | B193 | AMA-M2 | K | 0.125601 | 1.70E-07 |
| B109 | B182 | AMA-M2 | TT | 0.124542 | 2.17E-07 |
| B109 | B183 | AMA-M2 | PTA | -0.11862 | 7.97E-07 |
| B11 | B12 | PGI/PGII | PG II | -0.49833 | 0 |
| B11 | B14 | PGI/PGII | G-17 | -0.2062 | 0 |
| B11 | B21 | PGI/PGII | CD3+CD4-CD8- | 0.124866 | 2.01E-07 |
| B110 | B119 | AHA | Hcv-rna | 0.384926 | 0 |
| B110 | B120 | AHA | HCV-cAg | 0.349051 | 0 |
| B110 | B121 | AHA | C3 | -0.27501 | 0 |

| | | | | | |
|---|---|---|---|---|---|
| B110 | B122 | AHA | C4 | 0.242862 | 0 |
| B110 | B125 | AHA | Cor | 0.298639 | 0 |
| B110 | B134 | AHA | CHE | -0.23432 | 0 |
| B110 | B151 | AHA | Anti-CCP | 0.245648 | 0 |
| B110 | B152 | AHA | ACA-Ig(G-A-M) | -0.385 | 0 |
| B110 | B156 | AHA | IgE | 0.234142 | 0 |
| B110 | B157 | AHA | IgG | 0.328845 | 0 |
| B110 | B159 | AHA | β2-MG | -0.38492 | 0 |
| B110 | B138 | AHA | PAB | -0.18058 | 4.35E-14 |
| B110 | B154 | AHA | RF | 0.158265 | 3.99E-11 |
| B110 | B178 | AHA | Ig-U | 0.151079 | 2.96E-10 |
| B110 | B193 | AHA | K | 0.125601 | 1.70E-07 |
| B110 | B182 | AHA | TT | 0.124542 | 2.17E-07 |
| B110 | B183 | AHA | PTA | -0.11862 | 7.97E-07 |
| B111 | B112 | TSH | FT4 | -0.20619 | 0 |
| B111 | B148 | TSH | Anti_TPO | 0.289146 | 0 |
| B111 | B179 | TSH | α1-MG | -0.15679 | 6.07E-11 |
| B111 | B114 | TSH | TSH TT3 TT4(TT4) | -0.12447 | 2.20E-07 |
| B112 | B113 | FT4 | FT3 | 0.207854 | 0 |
| B112 | B114 | FT4 | TSH TT3 TT4(TT4) | 0.314788 | 0 |
| B112 | B204 | FT4 | Drinking | -0.17419 | 3.37E-13 |
| B112 | B150 | FT4 | ALP | 0.121745 | 4.04E-07 |
| B113 | B115 | FT3 | TSH TT3 TT4(TT3) | 0.29357 | 0 |
| B113 | B189 | FT3 | eGFB | 0.242024 | 0 |
| B113 | B176 | FT3 | Ucr | 0.195416 | 4.44E-16 |
| B113 | B144 | FT3 | Total-PINP | 0.172988 | 4.91E-13 |
| B113 | B191 | FT3 | Urea | -0.16866 | 1.86E-12 |
| B113 | B190 | FT3 | Cre | -0.14454 | 1.69E-09 |
| B113 | B126 | FT3 | Alb | 0.130022 | 6.14E-08 |
| B113 | B128 | FT3 | ALT | 0.124977 | 1.96E-07 |
| B114 | B115 | TSH TT3 TT4(TT4) | TSH TT3 TT4(TT3) | 0.366559 | 0 |
| B114 | B134 | TSH TT3 TT4(TT4) | CHE | -0.26418 | 0 |
| B114 | B138 | TSH TT3 TT4(TT4) | PAB | -0.24535 | 0 |
| B114 | B139 | TSH TT3 TT4(TT4) | ADA | 0.198875 | 0 |
| B114 | B124 | TSH TT3 TT4(TT4) | ACTH | 0.196371 | 2.22E-16 |
| B114 | B125 | TSH TT3 TT4(TT4) | Cor | 0.174904 | 2.69E-13 |
| B114 | B181 | TSH TT3 TT4(TT4) | APTT | 0.149107 | 5.04E-10 |
| B114 | B158 | TSH TT3 TT4(TT4) | IgM | -0.13878 | 7.34E-09 |
| B114 | B197 | TSH TT3 TT4(TT4) | AG | 0.129399 | 7.11E-08 |
| B115 | B119 | TSH TT3 TT4(TT3) | Hcv-rna | -0.26976 | 0 |
| B115 | B125 | TSH TT3 TT4(TT3) | Cor | -0.24097 | 0 |
| B115 | B152 | TSH TT3 TT4(TT3) | ACA-Ig(G-A-M) | 0.269812 | 0 |
| B115 | B159 | TSH TT3 TT4(TT3) | β2-MG | 0.269755 | 0 |
| B115 | B158 | TSH TT3 TT4(TT3) | IgM | 0.185286 | 9.10E-15 |

| | | | | | |
|---|---|---|---|---|---|
| B115 | B139 | TSH TT3 TT4(TT3) | ADA | 0.14744 | 7.87E-10 |
| B115 | B186 | TSH TT3 TT4(TT3) | PFC | 0.135541 | 1.64E-08 |
| B115 | B197 | TSH TT3 TT4(TT3) | AG | 0.127959 | 9.93E-08 |
| B115 | B183 | TSH TT3 TT4(TT3) | PTA | -0.12041 | 5.41E-07 |
| B117 | B202 | TT4 | e-times | 0.954475 | 0 |
| B117 | B223 | TT4 | FH | 0.580007 | 0 |
| B118 | B209 | CEA1 | Smoking | 0.18963 | 2.22E-15 |
| B118 | B189 | CEA1 | eGFB | -0.18487 | 1.07E-14 |
| B118 | B153 | CEA1 | Glu | 0.177394 | 1.22E-13 |
| B118 | B190 | CEA1 | Cre | 0.162684 | 1.11E-11 |
| B118 | B204 | CEA1 | Drinking | 0.14767 | 7.40E-10 |
| B118 | B220 | CEA1 | SBP | 0.128863 | 8.05E-08 |
| B118 | B206 | CEA1 | Waist | 0.126111 | 1.52E-07 |
| B118 | B191 | CEA1 | Urea | 0.124253 | 2.31E-07 |
| B118 | B222 | CEA1 | NS | 0.122242 | 3.62E-07 |
| B118 | B170 | CEA1 | U-GLU | 0.119715 | 6.29E-07 |
| B119 | B120 | Hcv-rna | HCV-cAg | 0.456823 | 0 |
| B119 | B124 | Hcv-rna | ACTH | 0.396008 | 0 |
| B119 | B125 | Hcv-rna | Cor | 0.612351 | 0 |
| B119 | B134 | Hcv-rna | CHE | -0.26306 | 0 |
| B119 | B158 | Hcv-rna | IgM | -0.49345 | 0 |
| B119 | B159 | Hcv-rna | β2-MG | -0.99999 | 0 |
| B119 | B179 | Hcv-rna | α1-MG | -0.23378 | 0 |
| B119 | B156 | Hcv-rna | IgE | 0.182861 | 2.04E-14 |
| B119 | B203 | Hcv-rna | DH | -0.1777 | 1.10E-13 |
| B119 | B155 | Hcv-rna | IgA | 0.165035 | 5.55E-12 |
| B119 | B151 | Hcv-rna | Anti-CCP | 0.155937 | 7.71E-11 |
| B119 | B138 | Hcv-rna | PAB | -0.1451 | 1.46E-09 |
| B119 | B184 | Hcv-rna | INR | 0.144909 | 1.53E-09 |
| B119 | B181 | Hcv-rna | APTT | 0.132409 | 3.49E-08 |
| B119 | B186 | Hcv-rna | PFC | -0.132 | 3.85E-08 |
| B119 | B196 | Hcv-rna | TCO2 | -0.11885 | 7.59E-07 |
| B12 | B200 | PG II | OP | -0.1234 | 2.80E-07 |
| B120 | B121 | HCV-cAg | C3 | -0.24949 | 0 |
| B120 | B122 | HCV-cAg | C4 | 0.211315 | 0 |
| B120 | B125 | HCV-cAg | Cor | 0.278836 | 0 |
| B120 | B134 | HCV-cAg | CHE | -0.21023 | 0 |
| B120 | B152 | HCV-cAg | ACA-Ig(G-A-M) | -0.45693 | 0 |
| B120 | B159 | HCV-cAg | β2-MG | -0.45682 | 0 |
| B120 | B155 | HCV-cAg | IgA | 0.18479 | 1.07E-14 |
| B120 | B124 | HCV-cAg | ACTH | 0.179863 | 5.48E-14 |
| B120 | B158 | HCV-cAg | IgM | -0.17099 | 9.12E-13 |
| B120 | B138 | HCV-cAg | PAB | -0.14604 | 1.14E-09 |
| B120 | B157 | HCV-cAg | IgG | 0.142019 | 3.24E-09 |

| | | | | | |
|---|---|---|---|---|---|
| B120 | B203 | HCV-cAg | DH | -0.13713 | 1.11E-08 |
| B120 | B199 | HCV-cAg | Mg | -0.13084 | 5.06E-08 |
| B120 | B195 | HCV-cAg | Na | -0.11851 | 8.15E-07 |
| B121 | B122 | C3 | C4 | -0.99649 | 0 |
| B121 | B134 | C3 | CHE | 0.328621 | 0 |
| B121 | B156 | C3 | IgE | 0.421915 | 0 |
| B121 | B198 | C3 | P | -0.21683 | 0 |
| B121 | B193 | C3 | K | 0.174542 | 3.02E-13 |
| B121 | B224 | C3 | pmh | 0.130934 | 4.96E-08 |
| B121 | B138 | C3 | PAB | 0.129366 | 7.16E-08 |
| B121 | B194 | C3 | Cl | -0.12125 | 4.50E-07 |
| B122 | B134 | C4 | CHE | -0.3067 | 0 |
| B122 | B156 | C4 | IgE | -0.43738 | 0 |
| B122 | B198 | C4 | P | 0.214051 | 0 |
| B122 | B193 | C4 | K | -0.17296 | 4.94E-13 |
| B122 | B224 | C4 | pmh | -0.13185 | 3.99E-08 |
| B122 | B194 | C4 | Cl | 0.127949 | 9.95E-08 |
| B123 | B186 | hs-CRP1 | PFC | 0.189738 | 2.00E-15 |
| B124 | B125 | ACTH | Cor | 0.544605 | 0 |
| B124 | B152 | ACTH | ACA-Ig(G-A-M) | -0.39609 | 0 |
| B124 | B155 | ACTH | IgA | 0.273508 | 0 |
| B124 | B158 | ACTH | IgM | -0.49415 | 0 |
| B124 | B159 | ACTH | β2-MG | -0.396 | 0 |
| B124 | B179 | ACTH | α1-MG | -0.28788 | 0 |
| B124 | B182 | ACTH | TT | 0.287552 | 0 |
| B124 | B178 | ACTH | Ig-U | -0.15307 | 1.71E-10 |
| B124 | B134 | ACTH | CHE | -0.12844 | 8.88E-08 |
| B124 | B183 | ACTH | PTA | 0.125098 | 1.91E-07 |
| B124 | B203 | ACTH | DH | -0.12088 | 4.88E-07 |
| B124 | B193 | ACTH | K | -0.11843 | 8.30E-07 |
| B125 | B134 | Cor | CHE | -0.26509 | 0 |
| B125 | B152 | Cor | ACA-Ig(G-A-M) | -0.61247 | 0 |
| B125 | B155 | Cor | IgA | 0.220856 | 0 |
| B125 | B156 | Cor | IgE | 0.217728 | 0 |
| B125 | B158 | Cor | IgM | -0.61831 | 0 |
| B125 | B159 | Cor | β2-MG | -0.61234 | 0 |
| B125 | B179 | Cor | α1-MG | -0.29925 | 0 |
| B125 | B182 | Cor | TT | 0.256708 | 0 |
| B125 | B199 | Cor | Mg | -0.19846 | 0 |
| B125 | B203 | Cor | DH | -0.1872 | 4.88E-15 |
| B125 | B184 | Cor | INR | 0.184915 | 1.02E-14 |
| B125 | B151 | Cor | Anti-CCP | 0.177814 | 1.06E-13 |
| B125 | B186 | Cor | PFC | -0.17673 | 1.51E-13 |
| B125 | B181 | Cor | APTT | 0.172666 | 5.43E-13 |

| | | | | | |
|---|---|---|---|---|---|
| B125 | B196 | Cor | TCO2 | -0.1312 | 4.65E-08 |
| B125 | B178 | Cor | Ig-U | -0.12803 | 9.76E-08 |
| B125 | B157 | Cor | IgG | 0.121293 | 4.46E-07 |
| B126 | B127 | Alb | A-G ratio | 0.472143 | 0 |
| B126 | B188 | Alb | Ca | 0.265379 | 0 |
| B126 | B189 | Alb | eGFB | 0.289856 | 0 |
| B126 | B128 | Alb | ALT | 0.188705 | 2.89E-15 |
| B126 | B141 | Alb | AST/ALT | -0.16131 | 1.66E-11 |
| B126 | B176 | Alb | Ucr | 0.158069 | 4.22E-11 |
| B126 | B163 | Alb | SG | 0.155075 | 9.82E-11 |
| B126 | B191 | Alb | Urea | -0.13778 | 9.42E-09 |
| B126 | B129 | Alb | IBIL | 0.129585 | 6.80E-08 |
| B126 | B218 | Alb | DBP | 0.121862 | 3.93E-07 |
| B126 | B190 | Alb | Cre | -0.11944 | 6.68E-07 |
| B127 | B130 | A-G ratio | GLB | -0.88154 | 0 |
| B127 | B189 | A-G ratio | eGFB | 0.19365 | 4.44E-16 |
| B127 | B221 | A-G ratio | Hei | 0.164594 | 6.33E-12 |
| B127 | B220 | A-G ratio | SBP | -0.1468 | 9.33E-10 |
| B127 | B139 | A-G ratio | ADA | -0.12677 | 1.31E-07 |
| B127 | B138 | A-G ratio | PAB | 0.126143 | 1.51E-07 |
| B128 | B131 | ALT | AST | 0.658829 | 0 |
| B128 | B140 | ALT | r-GT | 0.515257 | 0 |
| B128 | B141 | ALT | AST/ALT | -0.47902 | 0 |
| B128 | B215 | ALT | BMI | 0.22648 | 0 |
| B128 | B216 | ALT | BMI | 0.246858 | 0 |
| B128 | B222 | ALT | NS | 0.214803 | 0 |
| B128 | B171 | ALT | EC | 0.189853 | 1.78E-15 |
| B128 | B206 | ALT | Waist | 0.18928 | 2.22E-15 |
| B128 | B192 | ALT | UA | 0.174112 | 3.45E-13 |
| B128 | B163 | ALT | SG | 0.156351 | 6.87E-11 |
| B128 | B210 | ALT | Hip | 0.152881 | 1.81E-10 |
| B128 | B133 | ALT | TBIL | 0.149074 | 5.09E-10 |
| B128 | B129 | ALT | IBIL | 0.136237 | 1.38E-08 |
| B128 | B132 | ALT | Dbil | 0.134948 | 1.89E-08 |
| B128 | B221 | ALT | Hei | 0.134647 | 2.04E-08 |
| B128 | B189 | ALT | eGFB | 0.131612 | 4.22E-08 |
| B128 | B176 | ALT | Ucr | 0.129687 | 6.64E-08 |
| B129 | B132 | IBIL | Dbil | 0.556623 | 0 |
| B129 | B133 | IBIL | TBIL | 0.971653 | 0 |
| B130 | B220 | GLB | SBP | 0.129281 | 7.31E-08 |
| B130 | B221 | GLB | Hei | -0.1205 | 5.30E-07 |
| B131 | B140 | AST | r-GT | 0.472169 | 0 |
| B131 | B171 | AST | EC | 0.525899 | 0 |
| B131 | B132 | AST | Dbil | 0.145512 | 1.31E-09 |

| | | | | | |
|---|---|---|---|---|---|
| B131 | B170 | AST | U-GLU | 0.133082 | 2.97E-08 |
| B131 | B133 | AST | TBIL | 0.122571 | 3.36E-07 |
| B132 | B133 | Dbil | TBIL | 0.737123 | 0 |
| B132 | B140 | Dbil | r-GT | 0.12513 | 1.90E-07 |
| B134 | B138 | CHE | PAB | 0.464754 | 0 |
| B134 | B152 | CHE | ACA-Ig(G-A-M) | 0.263108 | 0 |
| B134 | B159 | CHE | β2-MG | 0.263053 | 0 |
| B134 | B199 | CHE | Mg | 0.20485 | 0 |
| B134 | B137 | CHE | AFU | 0.163224 | 9.48E-12 |
| B134 | B139 | CHE | ADA | -0.15955 | 2.76E-11 |
| B134 | B135 | CHE | TBA | -0.15607 | 7.43E-11 |
| B134 | B158 | CHE | IgM | 0.139672 | 5.87E-09 |
| B134 | B146 | CHE | CysC | -0.132 | 3.85E-08 |
| B134 | B179 | CHE | α1-MG | 0.128877 | 8.03E-08 |
| B134 | B180 | CHE | Tf | 0.127605 | 1.08E-07 |
| B134 | B194 | CHE | Cl | -0.12561 | 1.70E-07 |
| B134 | B196 | CHE | TCO2 | 0.119206 | 7.02E-07 |
| B135 | B137 | TBA | AFU | 0.228946 | 0 |
| B135 | B193 | TBA | K | -0.16966 | 1.38E-12 |
| B135 | B178 | TBA | Ig-U | -0.15801 | 4.29E-11 |
| B135 | B199 | TBA | Mg | -0.15718 | 5.43E-11 |
| B135 | B136 | TBA | 5'NT | -0.15428 | 1.23E-10 |
| B135 | B138 | TBA | PAB | -0.13906 | 6.85E-09 |
| B135 | B180 | TBA | Tf | -0.1385 | 7.88E-09 |
| B136 | B137 | 5'NT | AFU | -0.20896 | 0 |
| B136 | B155 | 5'NT | IgA | -0.20216 | 0 |
| B136 | B196 | 5'NT | TCO2 | -0.21339 | 0 |
| B136 | B197 | 5'NT | AG | 0.294158 | 0 |
| B136 | B157 | 5'NT | IgG | -0.19322 | 4.44E-16 |
| B136 | B139 | 5'NT | ADA | 0.187917 | 3.77E-15 |
| B136 | B183 | 5'NT | PTA | -0.17739 | 1.22E-13 |
| B136 | B181 | 5'NT | APTT | -0.17108 | 8.86E-13 |
| B136 | B194 | 5'NT | Cl | -0.14401 | 1.94E-09 |
| B136 | B199 | 5'NT | Mg | -0.14082 | 4.40E-09 |
| B136 | B158 | 5'NT | IgM | -0.13309 | 2.97E-08 |
| B136 | B198 | 5'NT | P | 0.132076 | 3.78E-08 |
| B136 | B156 | 5'NT | IgE | -0.12443 | 2.22E-07 |
| B137 | B138 | AFU | PAB | 0.298379 | 0 |
| B137 | B193 | AFU | K | -0.19264 | 8.88E-16 |
| B137 | B139 | AFU | ADA | -0.18902 | 2.66E-15 |
| B137 | B199 | AFU | Mg | 0.136712 | 1.23E-08 |
| B138 | B139 | PAB | ADA | -0.52179 | 0 |
| B138 | B155 | PAB | IgA | -0.20036 | 0 |
| B138 | B156 | PAB | IgE | -0.23202 | 0 |

| | | | | | |
|---|---|---|---|---|---|
| B138 | B199 | PAB | Mg | 0.197986 | 0 |
| B138 | B179 | PAB | α1-MG | 0.171527 | 7.73E-13 |
| B138 | B157 | PAB | IgG | -0.17115 | 8.69E-13 |
| B138 | B183 | PAB | PTA | 0.161295 | 1.67E-11 |
| B138 | B146 | PAB | CysC | -0.15942 | 2.87E-11 |
| B138 | B182 | PAB | TT | -0.15398 | 1.33E-10 |
| B138 | B193 | PAB | K | -0.14947 | 4.57E-10 |
| B138 | B152 | PAB | ACA-Ig(G-A-M) | 0.145131 | 1.45E-09 |
| B138 | B159 | PAB | β2-MG | 0.1451 | 1.46E-09 |
| B138 | B181 | PAB | APTT | -0.12974 | 6.56E-08 |
| B138 | B184 | PAB | INR | -0.11944 | 6.67E-07 |
| B139 | B155 | ADA | IgA | 0.295388 | 0 |
| B139 | B156 | ADA | IgE | 0.413322 | 0 |
| B139 | B199 | ADA | Mg | -0.21612 | 0 |
| B139 | B146 | ADA | CysC | 0.195943 | 2.22E-16 |
| B139 | B158 | ADA | IgM | 0.179299 | 6.59E-14 |
| B139 | B149 | ADA | TGAb | 0.173057 | 4.81E-13 |
| B139 | B157 | ADA | IgG | 0.153819 | 1.39E-10 |
| B139 | B148 | ADA | Anti_TPO | 0.144984 | 1.50E-09 |
| B139 | B183 | ADA | PTA | -0.12827 | 9.24E-08 |
| B14 | B47 | G-17 | RDW | 0.179684 | 5.82E-14 |
| B14 | B43 | G-17 | MCV | 0.166724 | 3.35E-12 |
| B14 | B44 | G-17 | MCH | 0.166069 | 4.07E-12 |
| B14 | B74 | G-17 | ApoA1 | -0.16416 | 7.19E-12 |
| B14 | B46 | G-17 | RBC Count | -0.13216 | 3.71E-08 |
| B140 | B171 | r-GT | EC | 0.210073 | 0 |
| B140 | B141 | r-GT | AST/ALT | -0.16922 | 1.57E-12 |
| B140 | B181 | r-GT | APTT | -0.15232 | 2.11E-10 |
| B140 | B216 | r-GT | BMI | 0.142514 | 2.85E-09 |
| B140 | B189 | r-GT | eGFB | 0.141063 | 4.13E-09 |
| B140 | B222 | r-GT | NS | 0.123564 | 2.70E-07 |
| B140 | B215 | r-GT | BMI | 0.123363 | 2.82E-07 |
| B141 | B215 | AST/ALT | BMI | -0.2117 | 0 |
| B141 | B216 | AST/ALT | BMI | -0.25324 | 0 |
| B141 | B222 | AST/ALT | NS | -0.21627 | 0 |
| B141 | B206 | AST/ALT | Waist | -0.1939 | 4.44E-16 |
| B141 | B221 | AST/ALT | Hei | -0.17506 | 2.56E-13 |
| B141 | B163 | AST/ALT | SG | -0.16773 | 2.47E-12 |
| B141 | B210 | AST/ALT | Hip | -0.15648 | 6.62E-11 |
| B141 | B192 | AST/ALT | UA | -0.14477 | 1.59E-09 |
| B141 | B176 | AST/ALT | Ucr | -0.14157 | 3.64E-09 |
| B141 | B189 | AST/ALT | eGFB | -0.12242 | 3.48E-07 |
| B142 | B151 | 25-(OH)D | Anti-CCP | -0.16875 | 1.81E-12 |
| B142 | B154 | 25-(OH)D | RF | -0.16044 | 2.13E-11 |

| | | | | | |
|---|---|---|---|---|---|
| B142 | B144 | 25-(OH)D | Total-PINP | -0.1485 | 5.94E-10 |
| B143 | B144 | β-CTx | Total-PINP | 0.824272 | 0 |
| B143 | B145 | β-CTx | BGP | 0.812161 | 0 |
| B143 | B150 | β-CTx | ALP | 0.165681 | 4.58E-12 |
| B143 | B168 | β-CTx | NMP22 | 0.149883 | 4.09E-10 |
| B143 | B151 | β-CTx | Anti-CCP | 0.125436 | 1.77E-07 |
| B144 | B145 | Total-PINP | BGP | 0.794003 | 0 |
| B144 | B151 | Total-PINP | Anti-CCP | 0.160659 | 2.01E-11 |
| B144 | B150 | Total-PINP | ALP | 0.139831 | 5.64E-09 |
| B144 | B168 | Total-PINP | NMP22 | 0.134558 | 2.08E-08 |
| B145 | B150 | BGP | ALP | 0.157256 | 5.32E-11 |
| B146 | B197 | CysC | AG | -0.13966 | 5.89E-09 |
| B146 | B189 | CysC | eGFB | -0.12072 | 5.05E-07 |
| B146 | B193 | CysC | K | 0.118104 | 8.90E-07 |
| B148 | B149 | Anti_TPO | TGAb | 0.594659 | 0 |
| B148 | B155 | Anti_TPO | IgA | 0.151166 | 2.89E-10 |
| B148 | B198 | Anti_TPO | P | 0.125236 | 1.85E-07 |
| B149 | B155 | TGAb | IgA | 0.133584 | 2.63E-08 |
| B149 | B156 | TGAb | IgE | 0.131517 | 4.32E-08 |
| B15 | B101 | CD3-CD19+(B cell) | anti-Sc70 | 0.503508 | 0 |
| B15 | B102 | CD3-CD19+(B cell) | anti-Sm | 0.503508 | 0 |
| B15 | B103 | CD3-CD19+(B cell) | anti-SSA/Ro | 0.503508 | 0 |
| B15 | B104 | CD3-CD19+(B cell) | anti-SSB | 0.503508 | 0 |
| B15 | B105 | CD3-CD19+(B cell) | anti-U1RNP | 0.503508 | 0 |
| B15 | B106 | CD3-CD19+(B cell) | anti-P | 0.503508 | 0 |
| B15 | B107 | CD3-CD19+(B cell) | AnuA | 0.503507 | 0 |
| B15 | B108 | CD3-CD19+(B cell) | Anti-Cyclin | 0.503508 | 0 |
| B15 | B109 | CD3-CD19+(B cell) | AMA-M2 | 0.503508 | 0 |
| B15 | B110 | CD3-CD19+(B cell) | AHA | 0.503508 | 0 |
| B15 | B121 | CD3-CD19+(B cell) | C3 | -0.63292 | 0 |
| B15 | B122 | CD3-CD19+(B cell) | C4 | 0.618635 | 0 |
| B15 | B134 | CD3-CD19+(B cell) | CHE | -0.22803 | 0 |
| B15 | B138 | CD3-CD19+(B cell) | PAB | -0.21923 | 0 |
| B15 | B151 | CD3-CD19+(B cell) | Anti-CCP | 0.322064 | 0 |
| B15 | B157 | CD3-CD19+(B cell) | IgG | 0.217577 | 0 |
| B15 | B17 | CD3-CD19+(B cell) | CD3+CD16+CD56+NK | -0.28992 | 0 |
| B15 | B18 | CD3-CD19+(B cell) | CD3+CD4+helper(inducer)T | 0.647539 | 0 |
| B15 | B19 | CD3-CD19+(B cell) | CD3+CD4+/CD3+CD8+ | 0.748441 | 0 |
| B15 | B21 | CD3-CD19+(B cell) | CD3+CD4-CD8- | 0.212241 | 0 |
| B15 | B22 | CD3-CD19+(B cell) | CD3+CD8+(T) | -0.6321 | 0 |
| B15 | B23 | CD3-CD19+(B cell) | CD3-CD16+CD56+(NK) | -0.35071 | 0 |
| B15 | B54 | CD3-CD19+(B cell) | ESR | 0.206402 | 0 |
| B15 | B95 | CD3-CD19+(B cell) | anti-cenpb | 0.503508 | 0 |
| B15 | B97 | CD3-CD19+(B cell) | dsDNA | 0.503508 | 0 |

| | | | | | |
|---|---|---|---|---|---|
| B15 | B98 | CD3-CD19+(B cell) | anti-Jo-1 | 0.503508 | 0 |
| B15 | B99 | CD3-CD19+(B cell) | anti-PM-Scl | 0.503508 | 0 |
| B15 | B120 | CD3-CD19+(B cell) | HCV-cAg | 0.19201 | 8.88E-16 |
| B15 | B125 | CD3-CD19+(B cell) | Cor | 0.18866 | 2.89E-15 |
| B15 | B152 | CD3-CD19+(B cell) | ACA-Ig(G-A-M) | -0.17317 | 4.63E-13 |
| B15 | B119 | CD3-CD19+(B cell) | Hcv-rna | 0.173137 | 4.69E-13 |
| B15 | B159 | CD3-CD19+(B cell) | β2-MG | -0.17314 | 4.69E-13 |
| B15 | B94 | CD3-CD19+(B cell) | ALDH2-Lys504Lys | 0.173137 | 4.69E-13 |
| B15 | B135 | CD3-CD19+(B cell) | TBA | 0.155596 | 8.49E-11 |
| B15 | B181 | CD3-CD19+(B cell) | APTT | 0.154548 | 1.14E-10 |
| B15 | B55 | CD3-CD19+(B cell) | TK | 0.145308 | 1.38E-09 |
| B15 | B79 | CD3-CD19+(B cell) | hb V -dna | 0.140311 | 5.00E-09 |
| B15 | B58 | CD3-CD19+(B cell) | HCT | -0.1392 | 6.61E-09 |
| B15 | B74 | CD3-CD19+(B cell) | ApoA1 | 0.135284 | 1.74E-08 |
| B15 | B92 | CD3-CD19+(B cell) | ALDH2-Glu504Glu | -0.12773 | 1.05E-07 |
| B15 | B183 | CD3-CD19+(B cell) | PTA | -0.1271 | 1.21E-07 |
| B15 | B67 | CD3-CD19+(B cell) | WBRSR(mPas_50) | -0.12263 | 3.32E-07 |
| B15 | B66 | CD3-CD19+(B cell) | WBRSR(mPas_100) | -0.11976 | 6.23E-07 |
| B15 | B93 | CD3-CD19+(B cell) | ALDH2-Glu504Lys | 0.118986 | 7.36E-07 |
| B151 | B156 | Anti-CCP | IgE | 0.231514 | 0 |
| B151 | B157 | Anti-CCP | IgG | 0.241795 | 0 |
| B151 | B168 | Anti-CCP | NMP22 | -0.22189 | 0 |
| B151 | B152 | Anti-CCP | ACA-Ig(G-A-M) | -0.15597 | 7.65E-11 |
| B151 | B159 | Anti-CCP | β2-MG | -0.15594 | 7.72E-11 |
| B151 | B183 | Anti-CCP | PTA | -0.12697 | 1.25E-07 |
| B151 | B193 | Anti-CCP | K | 0.123628 | 2.66E-07 |
| B151 | B197 | Anti-CCP | AG | 0.120424 | 5.39E-07 |
| B152 | B158 | ACA-Ig(G-A-M) | IgM | 0.493545 | 0 |
| B152 | B179 | ACA-Ig(G-A-M) | α1-MG | 0.233822 | 0 |
| B152 | B156 | ACA-Ig(G-A-M) | IgE | -0.1829 | 2.04E-14 |
| B152 | B203 | ACA-Ig(G-A-M) | DH | 0.177736 | 1.09E-13 |
| B152 | B155 | ACA-Ig(G-A-M) | IgA | -0.16507 | 5.49E-12 |
| B152 | B184 | ACA-Ig(G-A-M) | INR | -0.14494 | 1.52E-09 |
| B152 | B181 | ACA-Ig(G-A-M) | APTT | -0.13244 | 3.47E-08 |
| B152 | B186 | ACA-Ig(G-A-M) | PFC | 0.132025 | 3.83E-08 |
| B152 | B196 | ACA-Ig(G-A-M) | TCO2 | 0.118869 | 7.55E-07 |
| B153 | B170 | Glu | U-GLU | 0.561095 | 0 |
| B153 | B206 | Glu | Waist | 0.220321 | 0 |
| B153 | B207 | Glu | WHR | 0.20817 | 0 |
| B153 | B220 | Glu | SBP | 0.20607 | 0 |
| B153 | B215 | Glu | BMI | 0.187585 | 4.22E-15 |
| B153 | B222 | Glu | NS | 0.164376 | 6.75E-12 |
| B153 | B216 | Glu | BMI | 0.145489 | 1.32E-09 |
| B153 | B191 | Glu | Urea | 0.126274 | 1.46E-07 |

| | | | | | |
|---|---|---|---|---|---|
| B153 | B175 | Glu | ALB-U | 0.12082 | 4.94E-07 |
| B153 | B163 | Glu | SG | 0.118206 | 8.70E-07 |
| B155 | B156 | IgA | IgE | 0.43841 | 0 |
| B155 | B196 | IgA | TCO2 | 0.176142 | 1.82E-13 |
| B155 | B159 | IgA | β2-MG | -0.16503 | 5.55E-12 |
| B155 | B199 | IgA | Mg | -0.15137 | 2.73E-10 |
| B155 | B197 | IgA | AG | -0.14728 | 8.20E-10 |
| B155 | B182 | IgA | TT | 0.1399 | 5.55E-09 |
| B155 | B179 | IgA | α1-MG | -0.11872 | 7.79E-07 |
| B156 | B157 | IgE | IgG | 0.447624 | 0 |
| B156 | B159 | IgE | β2-MG | -0.18286 | 2.04E-14 |
| B156 | B182 | IgE | TT | 0.140413 | 4.87E-09 |
| B156 | B224 | IgE | pmh | 0.123276 | 2.88E-07 |
| B157 | B181 | IgG | APTT | 0.215681 | 0 |
| B157 | B158 | IgG | IgM | 0.194657 | 4.44E-16 |
| B157 | B184 | IgG | INR | 0.127724 | 1.05E-07 |
| B158 | B159 | IgM | β2-MG | 0.493441 | 0 |
| B158 | B179 | IgM | α1-MG | 0.3265 | 0 |
| B158 | B196 | IgM | TCO2 | 0.228037 | 0 |
| B158 | B178 | IgM | Ig-U | 0.18154 | 3.18E-14 |
| B158 | B183 | IgM | PTA | -0.16442 | 6.66E-12 |
| B158 | B197 | IgM | AG | -0.14663 | 9.75E-10 |
| B158 | B203 | IgM | DH | 0.143474 | 2.23E-09 |
| B158 | B198 | IgM | P | -0.13629 | 1.36E-08 |
| B158 | B182 | IgM | TT | -0.13121 | 4.64E-08 |
| B159 | B179 | β2-MG | α1-MG | 0.233773 | 0 |
| B159 | B203 | β2-MG | DH | 0.177699 | 1.10E-13 |
| B159 | B184 | β2-MG | INR | -0.14491 | 1.53E-09 |
| B159 | B181 | β2-MG | APTT | -0.13241 | 3.49E-08 |
| B159 | B186 | β2-MG | PFC | 0.131997 | 3.85E-08 |
| B159 | B196 | β2-MG | TCO2 | 0.118844 | 7.59E-07 |
| B16 | B115 | CD3+T | TSH TT3 TT4(TT3) | 0.250048 | 0 |
| B16 | B119 | CD3+T | Hcv-rna | -0.20794 | 0 |
| B16 | B124 | CD3+T | ACTH | -0.54744 | 0 |
| B16 | B125 | CD3+T | Cor | -0.30617 | 0 |
| B16 | B152 | CD3+T | ACA-Ig(G-A-M) | 0.207985 | 0 |
| B16 | B155 | CD3+T | IgA | -0.40969 | 0 |
| B16 | B158 | CD3+T | IgM | 0.321015 | 0 |
| B16 | B159 | CD3+T | β2-MG | 0.207942 | 0 |
| B16 | B17 | CD3+T | CD3+CD16+CD56+NK | 0.627785 | 0 |
| B16 | B179 | CD3+T | α1-MG | 0.286162 | 0 |
| B16 | B18 | CD3+T | CD3+CD4+helper(inducer)T | 0.634562 | 0 |
| B16 | B181 | CD3+T | APTT | 0.236034 | 0 |
| B16 | B182 | CD3+T | TT | -0.25966 | 0 |

| | | | | | |
|---|---|---|---|---|---|
| B16 | B183 | CD3+T | PTA | -0.21784 | 0 |
| B16 | B20 | CD3+T | CD3+CD4+CD8+ | -0.50689 | 0 |
| B16 | B21 | CD3+T | CD3+CD4-CD8- | 0.390532 | 0 |
| B16 | B22 | CD3+T | CD3+CD8+(T) | 0.423953 | 0 |
| B16 | B23 | CD3+T | CD3-CD16+CD56+(NK) | -0.93468 | 0 |
| B16 | B25 | CD3+T | CK | 0.201892 | 0 |
| B16 | B58 | CD3+T | HCT | 0.199656 | 0 |
| B16 | B94 | CD3+T | ALDH2-Lys504Lys | -0.20794 | 0 |
| B16 | B157 | CD3+T | IgG | 0.196417 | 2.22E-16 |
| B16 | B66 | CD3+T | WBRSR(mPas_100) | 0.187815 | 4.00E-15 |
| B16 | B63 | CD3+T | WBRSR(mPas_200) | 0.185147 | 9.55E-15 |
| B16 | B178 | CD3+T | Ig-U | 0.180736 | 4.13E-14 |
| B16 | B67 | CD3+T | WBRSR(mPas_50) | 0.176867 | 1.44E-13 |
| B16 | B62 | CD3+T | WBHSRVI | 0.174668 | 2.90E-13 |
| B16 | B79 | CD3+T | hb V -dna | -0.16971 | 1.35E-12 |
| B16 | B76 | CD3+T | ApoB | 0.161359 | 1.64E-11 |
| B16 | B92 | CD3+T | ALDH2-Glu504Glu | 0.151754 | 2.46E-10 |
| B16 | B61 | CD3+T | WBHSRV | 0.150836 | 3.16E-10 |
| B16 | B75 | CD3+T | ApoA1/B | -0.14318 | 2.40E-09 |
| B16 | B93 | CD3+T | ALDH2-Glu504Lys | -0.14124 | 3.95E-09 |
| B16 | B56 | CD3+T | IR | 0.135151 | 1.80E-08 |
| B16 | B185 | CD3+T | PT | 0.131542 | 4.29E-08 |
| B16 | B64 | CD3+T | WBRSR(mPas_5) | 0.128816 | 8.14E-08 |
| B16 | B73 | CD3+T | ApoA | -0.12621 | 1.48E-07 |
| B16 | B194 | CD3+T | CI | -0.12263 | 3.32E-07 |
| B16 | B186 | CD3+T | PFC | 0.117876 | 9.34E-07 |
| B160 | B191 | PH | Urea | -0.14621 | 1.09E-09 |
| B160 | B192 | PH | UA | -0.14607 | 1.13E-09 |
| B160 | B163 | PH | SG | -0.13911 | 6.76E-09 |
| B160 | B190 | PH | Cre | -0.13229 | 3.59E-08 |
| B161 | B221 | LEU | Hei | -0.1869 | 5.33E-15 |
| B161 | B173 | LEU | NTT | 0.16002 | 2.41E-11 |
| B161 | B222 | LEU | NS | -0.13573 | 1.56E-08 |
| B163 | B166 | SG | Pro | 0.33842 | 0 |
| B163 | B176 | SG | Ucr | 0.570032 | 0 |
| B163 | B216 | SG | BMI | 0.280659 | 0 |
| B163 | B221 | SG | Hei | 0.264325 | 0 |
| B163 | B222 | SG | NS | 0.294306 | 0 |
| B163 | B191 | SG | Urea | 0.192252 | 8.88E-16 |
| B163 | B215 | SG | BMI | 0.185957 | 7.33E-15 |
| B163 | B189 | SG | eGFB | 0.18558 | 8.44E-15 |
| B163 | B206 | SG | Waist | 0.170039 | 1.22E-12 |
| B163 | B210 | SG | Hip | 0.155204 | 9.48E-11 |
| B163 | B192 | SG | UA | 0.153848 | 1.38E-10 |

| | | | | | |
|---|---|---|---|---|---|
| B163 | B170 | SG | U-GLU | 0.150252 | 3.70E-10 |
| B163 | B220 | SG | SBP | -0.13362 | 2.61E-08 |
| B163 | B165 | SG | bilirubin | 0.132135 | 3.73E-08 |
| B164 | B213 | COM | CVA-R | -0.12949 | 6.96E-08 |
| B165 | B189 | bilirubin | eGFB | 0.169036 | 1.66E-12 |
| B166 | B176 | Pro | Ucr | 0.158293 | 3.96E-11 |
| B166 | B222 | Pro | NS | 0.13332 | 2.81E-08 |
| B166 | B175 | Pro | ALB-U | 0.12337 | 2.82E-07 |
| B166 | B216 | Pro | BMI | 0.122923 | 3.11E-07 |
| B166 | B190 | Pro | Cre | 0.122745 | 3.24E-07 |
| B167 | B173 | PC | NTT | -0.1737 | 3.93E-13 |
| B17 | B114 | CD3+CD16+CD56+NK | TSH TT3 TT4(TT4) | 0.211004 | 0 |
| B17 | B115 | CD3+CD16+CD56+NK | TSH TT3 TT4(TT3) | 0.217132 | 0 |
| B17 | B157 | CD3+CD16+CD56+NK | IgG | 0.199016 | 0 |
| B17 | B19 | CD3+CD16+CD56+NK | CD3+CD4+/CD3+CD8+ | -0.22513 | 0 |
| B17 | B198 | CD3+CD16+CD56+NK | P | 0.311371 | 0 |
| B17 | B20 | CD3+CD16+CD56+NK | CD3+CD4+CD8+ | -0.29988 | 0 |
| B17 | B21 | CD3+CD16+CD56+NK | CD3+CD4-CD8- | 0.584192 | 0 |
| B17 | B22 | CD3+CD16+CD56+NK | CD3+CD8+(T) | 0.538831 | 0 |
| B17 | B23 | CD3+CD16+CD56+NK | CD3-CD16+CD56+(NK) | -0.37322 | 0 |
| B17 | B148 | CD3+CD16+CD56+NK | Anti_TPO | 0.195192 | 4.44E-16 |
| B17 | B124 | CD3+CD16+CD56+NK | ACTH | -0.19177 | 8.88E-16 |
| B17 | B194 | CD3+CD16+CD56+NK | Cl | -0.17807 | 9.81E-14 |
| B17 | B136 | CD3+CD16+CD56+NK | 5'NT | 0.169881 | 1.28E-12 |
| B17 | B69 | CD3+CD16+CD56+NK | PV | 0.160117 | 2.35E-11 |
| B17 | B25 | CD3+CD16+CD56+NK | CK | 0.16001 | 2.42E-11 |
| B17 | B57 | CD3+CD16+CD56+NK | RBCAI | -0.14468 | 1.63E-09 |
| B17 | B156 | CD3+CD16+CD56+NK | IgE | -0.13975 | 5.76E-09 |
| B17 | B200 | CD3+CD16+CD56+NK | OP | -0.13824 | 8.40E-09 |
| B17 | B101 | CD3+CD16+CD56+NK | anti-Sc70 | 0.135702 | 1.57E-08 |
| B17 | B102 | CD3+CD16+CD56+NK | anti-Sm | 0.135702 | 1.57E-08 |
| B17 | B103 | CD3+CD16+CD56+NK | anti-SSA/Ro | 0.135702 | 1.57E-08 |
| B17 | B104 | CD3+CD16+CD56+NK | anti-SSB | 0.135702 | 1.57E-08 |
| B17 | B105 | CD3+CD16+CD56+NK | anti-U1RNP | 0.135702 | 1.57E-08 |
| B17 | B106 | CD3+CD16+CD56+NK | anti-P | 0.135702 | 1.57E-08 |
| B17 | B107 | CD3+CD16+CD56+NK | AnuA | 0.135702 | 1.57E-08 |
| B17 | B108 | CD3+CD16+CD56+NK | Anti-Cyclin | 0.135702 | 1.57E-08 |
| B17 | B109 | CD3+CD16+CD56+NK | AMA-M2 | 0.135702 | 1.57E-08 |
| B17 | B110 | CD3+CD16+CD56+NK | AHA | 0.135702 | 1.57E-08 |
| B17 | B95 | CD3+CD16+CD56+NK | anti-cenpb | 0.135702 | 1.57E-08 |
| B17 | B97 | CD3+CD16+CD56+NK | dsDNA | 0.135702 | 1.57E-08 |
| B17 | B98 | CD3+CD16+CD56+NK | anti-Jo-1 | 0.135702 | 1.57E-08 |
| B17 | B99 | CD3+CD16+CD56+NK | anti-PM-Scl | 0.135702 | 1.57E-08 |
| B17 | B113 | CD3+CD16+CD56+NK | FT3 | 0.135598 | 1.61E-08 |

| | | | | | |
|---|---|---|---|---|---|
| B17 | B183 | CD3+CD16+CD56+NK | PTA | -0.13413 | 2.31E-08 |
| B17 | B142 | CD3+CD16+CD56+NK | 25-(OH)D | 0.129874 | 6.36E-08 |
| B17 | B144 | CD3+CD16+CD56+NK | Total-PINP | 0.127245 | 1.17E-07 |
| B17 | B112 | CD3+CD16+CD56+NK | FT4 | 0.118493 | 8.19E-07 |
| B170 | B171 | U-GLU | EC | 0.13283 | 3.16E-08 |
| B174 | B189 | Urea/Cr | eGFB | 0.207076 | 0 |
| B174 | B190 | Urea/Cr | Cre | -0.40954 | 0 |
| B174 | B191 | Urea/Cr | Urea | 0.526654 | 0 |
| B174 | B192 | Urea/Cr | UA | -0.24593 | 0 |
| B174 | B216 | Urea/Cr | BMI | -0.27247 | 0 |
| B174 | B221 | Urea/Cr | Hei | -0.34514 | 0 |
| B174 | B222 | Urea/Cr | NS | -0.27831 | 0 |
| B174 | B209 | Urea/Cr | Smoking | -0.16144 | 1.60E-11 |
| B174 | B204 | Urea/Cr | Drinking | -0.15787 | 4.46E-11 |
| B174 | B206 | Urea/Cr | Waist | -0.14995 | 4.01E-10 |
| B174 | B210 | Urea/Cr | Hip | -0.14537 | 1.36E-09 |
| B174 | B200 | Urea/Cr | OP | 0.117695 | 9.71E-07 |
| B175 | B177 | ALB-U | ACR | 0.812916 | 0 |
| B175 | B190 | ALB-U | Cre | 0.246736 | 0 |
| B175 | B191 | ALB-U | Urea | 0.210051 | 0 |
| B175 | B227 | ALB-U | OD | 0.130894 | 5.00E-08 |
| B176 | B189 | Ucr | eGFB | 0.210985 | 0 |
| B176 | B220 | Ucr | SBP | -0.18682 | 5.33E-15 |
| B176 | B201 | Ucr | e-habits | -0.15891 | 3.32E-11 |
| B176 | B219 | Ucr | OT | -0.13441 | 2.16E-08 |
| B177 | B190 | ACR | Cre | 0.287134 | 0 |
| B177 | B191 | ACR | Urea | 0.221665 | 0 |
| B177 | B189 | ACR | eGFB | -0.15892 | 3.31E-11 |
| B177 | B227 | ACR | OD | 0.155517 | 8.68E-11 |
| B178 | B179 | Ig-U | α1-MG | 0.678946 | 0 |
| B178 | B180 | Ig-U | Tf | 0.826112 | 0 |
| B179 | B180 | α1-MG | Tf | 0.435963 | 0 |
| B179 | B200 | α1-MG | OP | 0.154607 | 1.12E-10 |
| B18 | B101 | CD3+CD4+helper(inducer)T | anti-Sc70 | 0.235992 | 0 |
| B18 | B102 | CD3+CD4+helper(inducer)T | anti-Sm | 0.235992 | 0 |
| B18 | B103 | CD3+CD4+helper(inducer)T | anti-SSA/Ro | 0.235992 | 0 |
| B18 | B104 | CD3+CD4+helper(inducer)T | anti-SSB | 0.235992 | 0 |
| B18 | B105 | CD3+CD4+helper(inducer)T | anti-U1RNP | 0.235992 | 0 |
| B18 | B106 | CD3+CD4+helper(inducer)T | anti-P | 0.235992 | 0 |
| B18 | B107 | CD3+CD4+helper(inducer)T | AnuA | 0.235992 | 0 |
| B18 | B108 | CD3+CD4+helper(inducer)T | Anti-Cyclin | 0.235992 | 0 |
| B18 | B109 | CD3+CD4+helper(inducer)T | AMA-M2 | 0.235992 | 0 |
| B18 | B110 | CD3+CD4+helper(inducer)T | AHA | 0.235992 | 0 |
| B18 | B121 | CD3+CD4+helper(inducer)T | C3 | -0.33585 | 0 |

| | | | | | |
|---|---|---|---|---|---|
| B18 | B122 | CD3+CD4+helper(inducer)T | C4 | 0.331524 | 0 |
| B18 | B124 | CD3+CD4+helper(inducer)T | ACTH | -0.33353 | 0 |
| B18 | B155 | CD3+CD4+helper(inducer)T | IgA | -0.27576 | 0 |
| B18 | B157 | CD3+CD4+helper(inducer)T | IgG | 0.292349 | 0 |
| B18 | B181 | CD3+CD4+helper(inducer)T | APTT | 0.446673 | 0 |
| B18 | B19 | CD3+CD4+helper(inducer)T | CD3+CD4+/CD3+CD8+ | 0.795837 | 0 |
| B18 | B20 | CD3+CD4+helper(inducer)T | CD3+CD4+CD8+ | -0.49817 | 0 |
| B18 | B21 | CD3+CD4+helper(inducer)T | CD3+CD4-CD8- | 0.313182 | 0 |
| B18 | B22 | CD3+CD4+helper(inducer)T | CD3+CD8+(T) | -0.38197 | 0 |
| B18 | B23 | CD3+CD4+helper(inducer)T | CD3-CD16+CD56+(NK) | -0.76304 | 0 |
| B18 | B95 | CD3+CD4+helper(inducer)T | anti-cenpb | 0.235992 | 0 |
| B18 | B97 | CD3+CD4+helper(inducer)T | dsDNA | 0.235992 | 0 |
| B18 | B98 | CD3+CD4+helper(inducer)T | anti-Jo-1 | 0.235992 | 0 |
| B18 | B99 | CD3+CD4+helper(inducer)T | anti-PM-Scl | 0.235992 | 0 |
| B18 | B182 | CD3+CD4+helper(inducer)T | TT | -0.185 | 1.02E-14 |
| B18 | B55 | CD3+CD4+helper(inducer)T | TK | 0.175358 | 2.33E-13 |
| B18 | B179 | CD3+CD4+helper(inducer)T | α1-MG | 0.160828 | 1.91E-11 |
| B18 | B151 | CD3+CD4+helper(inducer)T | Anti-CCP | 0.158136 | 4.14E-11 |
| B18 | B158 | CD3+CD4+helper(inducer)T | IgM | 0.156749 | 6.14E-11 |
| B18 | B56 | CD3+CD4+helper(inducer)T | IR | 0.144251 | 1.82E-09 |
| B18 | B57 | CD3+CD4+helper(inducer)T | RBCAI | -0.139 | 6.95E-09 |
| B18 | B137 | CD3+CD4+helper(inducer)T | AFU | 0.136626 | 1.25E-08 |
| B18 | B76 | CD3+CD4+helper(inducer)T | ApoB | 0.129774 | 6.51E-08 |
| B18 | B184 | CD3+CD4+helper(inducer)T | INR | 0.127746 | 1.04E-07 |
| B18 | B178 | CD3+CD4+helper(inducer)T | Ig-U | 0.127609 | 1.08E-07 |
| B18 | B73 | CD3+CD4+helper(inducer)T | ApoA | -0.123 | 3.06E-07 |
| B18 | B61 | CD3+CD4+helper(inducer)T | WBHSRV | 0.121932 | 3.87E-07 |
| B180 | B197 | Tf | AG | 0.211949 | 0 |
| B180 | B194 | Tf | Cl | -0.17615 | 1.81E-13 |
| B180 | B198 | Tf | P | -0.17111 | 8.79E-13 |
| B180 | B196 | Tf | TCO2 | -0.16875 | 1.82E-12 |
| B180 | B200 | Tf | OP | 0.129942 | 6.26E-08 |
| B180 | B199 | Tf | Mg | -0.12634 | 1.44E-07 |
| B181 | B183 | APTT | PTA | -0.26776 | 0 |
| B181 | B184 | APTT | INR | 0.494501 | 0 |
| B181 | B185 | APTT | PT | 0.377255 | 0 |
| B181 | B186 | APTT | PFC | -0.1913 | 1.33E-15 |
| B181 | B193 | APTT | K | -0.1229 | 3.12E-07 |
| B182 | B183 | TT | PTA | -0.71523 | 0 |
| B182 | B184 | TT | INR | -0.22139 | 0 |
| B182 | B185 | TT | PT | -0.32822 | 0 |
| B182 | B193 | TT | K | 0.137874 | 9.21E-09 |
| B183 | B186 | PTA | PFC | -0.28506 | 0 |
| B183 | B193 | PTA | K | -0.20269 | 0 |

| | | | | | |
|---|---|---|---|---|---|
| B184 | B185 | INR | PT | 0.889451 | 0 |
| B184 | B193 | INR | K | -0.11963 | 6.41E-07 |
| B185 | B186 | PT | PFC | 0.154056 | 1.30E-10 |
| B189 | B190 | eGFB | Cre | -0.69281 | 0 |
| B189 | B191 | eGFB | Urea | -0.42856 | 0 |
| B189 | B220 | eGFB | SBP | -0.24708 | 0 |
| B189 | B192 | eGFB | UA | -0.18601 | 7.11E-15 |
| B189 | B218 | eGFB | DBP | 0.146314 | 1.06E-09 |
| B189 | B219 | eGFB | OT | -0.13624 | 1.38E-08 |
| B189 | B227 | eGFB | OD | -0.13007 | 6.07E-08 |
| B189 | B209 | eGFB | Smoking | -0.1213 | 4.45E-07 |
| B19 | B101 | CD3+CD4+/CD3+CD8+ | anti-Sc70 | 0.414177 | 0 |
| B19 | B102 | CD3+CD4+/CD3+CD8+ | anti-Sm | 0.414177 | 0 |
| B19 | B103 | CD3+CD4+/CD3+CD8+ | anti-SSA/Ro | 0.414177 | 0 |
| B19 | B104 | CD3+CD4+/CD3+CD8+ | anti-SSB | 0.414177 | 0 |
| B19 | B105 | CD3+CD4+/CD3+CD8+ | anti-U1RNP | 0.414177 | 0 |
| B19 | B106 | CD3+CD4+/CD3+CD8+ | anti-P | 0.414177 | 0 |
| B19 | B107 | CD3+CD4+/CD3+CD8+ | AnuA | 0.414176 | 0 |
| B19 | B108 | CD3+CD4+/CD3+CD8+ | Anti-Cyclin | 0.414177 | 0 |
| B19 | B109 | CD3+CD4+/CD3+CD8+ | AMA-M2 | 0.414177 | 0 |
| B19 | B110 | CD3+CD4+/CD3+CD8+ | AHA | 0.414177 | 0 |
| B19 | B119 | CD3+CD4+/CD3+CD8+ | Hcv-rna | 0.319469 | 0 |
| B19 | B120 | CD3+CD4+/CD3+CD8+ | HCV-cAg | 0.225309 | 0 |
| B19 | B121 | CD3+CD4+/CD3+CD8+ | C3 | -0.33505 | 0 |
| B19 | B122 | CD3+CD4+/CD3+CD8+ | C4 | 0.308411 | 0 |
| B19 | B125 | CD3+CD4+/CD3+CD8+ | Cor | 0.3612 | 0 |
| B19 | B151 | CD3+CD4+/CD3+CD8+ | Anti-CCP | 0.2498 | 0 |
| B19 | B152 | CD3+CD4+/CD3+CD8+ | ACA-Ig(G-A-M) | -0.31953 | 0 |
| B19 | B157 | CD3+CD4+/CD3+CD8+ | IgG | 0.28412 | 0 |
| B19 | B159 | CD3+CD4+/CD3+CD8+ | β2-MG | -0.31947 | 0 |
| B19 | B181 | CD3+CD4+/CD3+CD8+ | APTT | 0.383382 | 0 |
| B19 | B20 | CD3+CD4+/CD3+CD8+ | CD3+CD4+CD8+ | -0.23686 | 0 |
| B19 | B21 | CD3+CD4+/CD3+CD8+ | CD3+CD4-CD8- | 0.313647 | 0 |
| B19 | B22 | CD3+CD4+/CD3+CD8+ | CD3+CD8+(T) | -0.84406 | 0 |
| B19 | B23 | CD3+CD4+/CD3+CD8+ | CD3-CD16+CD56+(NK) | -0.24149 | 0 |
| B19 | B55 | CD3+CD4+/CD3+CD8+ | TK | 0.226431 | 0 |
| B19 | B79 | CD3+CD4+/CD3+CD8+ | hb V -dna | 0.260055 | 0 |
| B19 | B92 | CD3+CD4+/CD3+CD8+ | ALDH2-Glu504Glu | -0.23408 | 0 |
| B19 | B93 | CD3+CD4+/CD3+CD8+ | ALDH2-Glu504Lys | 0.217927 | 0 |
| B19 | B94 | CD3+CD4+/CD3+CD8+ | ALDH2-Lys504Lys | 0.319469 | 0 |
| B19 | B95 | CD3+CD4+/CD3+CD8+ | anti-cenpb | 0.414177 | 0 |
| B19 | B97 | CD3+CD4+/CD3+CD8+ | dsDNA | 0.414177 | 0 |
| B19 | B98 | CD3+CD4+/CD3+CD8+ | anti-Jo-1 | 0.414177 | 0 |
| B19 | B99 | CD3+CD4+/CD3+CD8+ | anti-PM-Scl | 0.414177 | 0 |

| | | | | | |
|---|---|---|---|---|---|
| B19 | B137 | CD3+CD4+/CD3+CD8+ | AFU | 0.16206 | 1.33E-11 |
| B19 | B138 | CD3+CD4+/CD3+CD8+ | PAB | -0.154 | 1.33E-10 |
| B19 | B58 | CD3+CD4+/CD3+CD8+ | HCT | -0.15399 | 1.33E-10 |
| B19 | B199 | CD3+CD4+/CD3+CD8+ | Mg | -0.14533 | 1.37E-09 |
| B19 | B184 | CD3+CD4+/CD3+CD8+ | INR | 0.134719 | 2.00E-08 |
| B19 | B134 | CD3+CD4+/CD3+CD8+ | CHE | -0.12728 | 1.16E-07 |
| B19 | B158 | CD3+CD4+/CD3+CD8+ | IgM | -0.12247 | 3.44E-07 |
| B19 | B34 | CD3+CD4+/CD3+CD8+ | TK | 0.121468 | 4.29E-07 |
| B190 | B191 | Cre | Urea | 0.527548 | 0 |
| B190 | B192 | Cre | UA | 0.434368 | 0 |
| B190 | B204 | Cre | Drinking | 0.21324 | 0 |
| B190 | B206 | Cre | Waist | 0.257772 | 0 |
| B190 | B209 | Cre | Smoking | 0.214933 | 0 |
| B190 | B210 | Cre | Hip | 0.197378 | 0 |
| B190 | B216 | Cre | BMI | 0.309412 | 0 |
| B190 | B221 | Cre | Hei | 0.32023 | 0 |
| B190 | B222 | Cre | NS | 0.361433 | 0 |
| B190 | B215 | Cre | BMI | 0.184282 | 1.29E-14 |
| B190 | B207 | Cre | WHR | 0.177891 | 1.04E-13 |
| B190 | B227 | Cre | OD | 0.153525 | 1.51E-10 |
| B191 | B192 | Urea | UA | 0.178154 | 9.55E-14 |
| B192 | B204 | UA | Drinking | 0.232069 | 0 |
| B192 | B206 | UA | Waist | 0.38464 | 0 |
| B192 | B207 | UA | WHR | 0.261741 | 0 |
| B192 | B210 | UA | Hip | 0.270493 | 0 |
| B192 | B215 | UA | BMI | 0.31654 | 0 |
| B192 | B216 | UA | BMI | 0.424406 | 0 |
| B192 | B221 | UA | Hei | 0.350105 | 0 |
| B192 | B222 | UA | NS | 0.454056 | 0 |
| B192 | B209 | UA | Smoking | 0.182383 | 2.40E-14 |
| B193 | B200 | K | OP | -0.1371 | 1.12E-08 |
| B193 | B196 | K | TCO2 | 0.137059 | 1.13E-08 |
| B194 | B195 | Cl | Na | 0.442314 | 0 |
| B194 | B197 | Cl | AG | -0.40625 | 0 |
| B194 | B200 | Cl | OP | 0.315895 | 0 |
| B195 | B197 | Na | AG | 0.217881 | 0 |
| B195 | B200 | Na | OP | 0.839185 | 0 |
| B196 | B197 | TCO2 | AG | -0.75857 | 0 |
| B196 | B199 | TCO2 | Mg | 0.308633 | 0 |
| B197 | B199 | AG | Mg | -0.2459 | 0 |
| B197 | B200 | AG | OP | 0.201614 | 0 |
| B2 | B113 | Age | FT3 | -0.266 | 0 |
| B2 | B126 | Age | Alb | -0.36802 | 0 |
| B2 | B127 | Age | A-G ratio | -0.22343 | 0 |

| | | | | | |
|---|---|---|---|---|---|
| B2 | B128 | Age | ALT | -0.25272 | 0 |
| B2 | B141 | Age | AST/ALT | 0.257824 | 0 |
| B2 | B163 | Age | SG | -0.31725 | 0 |
| B2 | B176 | Age | Ucr | -0.39564 | 0 |
| B2 | B189 | Age | eGFB | -0.61091 | 0 |
| B2 | B191 | Age | Urea | 0.216124 | 0 |
| B2 | B216 | Age | BMI | -0.24167 | 0 |
| B2 | B218 | Age | DBP | -0.22227 | 0 |
| B2 | B219 | Age | OT | 0.204399 | 0 |
| B2 | B220 | Age | SBP | 0.402249 | 0 |
| B2 | B221 | Age | Hei | -0.28123 | 0 |
| B2 | B46 | Age | RBC Count | -0.32003 | 0 |
| B2 | B47 | Age | RDW | 0.323647 | 0 |
| B2 | B51 | Age | Hb | -0.31985 | 0 |
| B2 | B71 | Age | TG | -0.18966 | 2.22E-15 |
| B2 | B118 | Age | CEA1 | 0.183785 | 1.51E-14 |
| B2 | B70 | Age | LDL-C | -0.18161 | 3.11E-14 |
| B2 | B77 | Age | TC | -0.17607 | 1.86E-13 |
| B2 | B190 | Age | Cre | 0.175734 | 2.07E-13 |
| B2 | B140 | Age | r-GT | -0.17506 | 2.56E-13 |
| B2 | B41 | Age | MON％ | 0.169738 | 1.34E-12 |
| B2 | B45 | Age | MCHC | -0.1696 | 1.40E-12 |
| B2 | B201 | Age | e-habits | 0.16904 | 1.66E-12 |
| B2 | B222 | Age | NS | -0.16574 | 4.50E-12 |
| B2 | B52 | Age | PLT/BPC | -0.15913 | 3.12E-11 |
| B2 | B138 | Age | PAB | -0.14995 | 4.02E-10 |
| B2 | B43 | Age | MCV | 0.143808 | 2.04E-09 |
| B2 | B134 | Age | CHE | -0.14101 | 4.18E-09 |
| B2 | B72 | Age | HDL-C | 0.139257 | 6.52E-09 |
| B2 | B153 | Age | Glu | 0.13729 | 1.06E-08 |
| B2 | B91 | Age | TP | -0.13025 | 5.82E-08 |
| B2 | B210 | Age | Hip | -0.12444 | 2.22E-07 |
| B2 | B27 | Age | LDH | 0.123156 | 2.95E-07 |
| B2 | B227 | Age | OD | 0.122456 | 3.45E-07 |
| B2 | B79 | Age | hbV-dna | 0.118474 | 8.22E-07 |
| B2 | B132 | Age | Dbil | 0.118431 | 8.29E-07 |
| B2 | B146 | Age | CysC | 0.118328 | 8.48E-07 |
| B20 | B151 | CD3+CD4+CD8+ | Anti-CCP | 0.198094 | 0 |
| B20 | B181 | CD3+CD4+CD8+ | APTT | -0.45473 | 0 |
| B20 | B182 | CD3+CD4+CD8+ | TT | 0.308778 | 0 |
| B20 | B184 | CD3+CD4+CD8+ | INR | -0.21951 | 0 |
| B20 | B21 | CD3+CD4+CD8+ | CD3+CD4-CD8- | -0.29869 | 0 |
| B20 | B23 | CD3+CD4+CD8+ | CD3-CD16+CD56+(NK) | 0.402624 | 0 |
| B20 | B25 | CD3+CD4+CD8+ | CK | -0.44315 | 0 |

| | | | | | |
|---|---|---|---|---|---|
| B20 | B57 | CD3+CD4+CD8+ | RBCAI | 0.234848 | 0 |
| B20 | B155 | CD3+CD4+CD8+ | IgA | 0.196068 | 2.22E-16 |
| B20 | B125 | CD3+CD4+CD8+ | Cor | -0.18592 | 7.55E-15 |
| B20 | B101 | CD3+CD4+CD8+ | anti-Sc70 | 0.184252 | 1.29E-14 |
| B20 | B102 | CD3+CD4+CD8+ | anti-Sm | 0.184252 | 1.29E-14 |
| B20 | B103 | CD3+CD4+CD8+ | anti-SSA/Ro | 0.184252 | 1.29E-14 |
| B20 | B104 | CD3+CD4+CD8+ | anti-SSB | 0.184252 | 1.29E-14 |
| B20 | B105 | CD3+CD4+CD8+ | anti-U1RNP | 0.184252 | 1.29E-14 |
| B20 | B106 | CD3+CD4+CD8+ | anti-P | 0.184252 | 1.29E-14 |
| B20 | B107 | CD3+CD4+CD8+ | AnuA | 0.184252 | 1.29E-14 |
| B20 | B108 | CD3+CD4+CD8+ | Anti-Cyclin | 0.184252 | 1.29E-14 |
| B20 | B109 | CD3+CD4+CD8+ | AMA-M2 | 0.184252 | 1.29E-14 |
| B20 | B110 | CD3+CD4+CD8+ | AHA | 0.184252 | 1.29E-14 |
| B20 | B95 | CD3+CD4+CD8+ | anti-cenpb | 0.184252 | 1.29E-14 |
| B20 | B97 | CD3+CD4+CD8+ | dsDNA | 0.184252 | 1.29E-14 |
| B20 | B98 | CD3+CD4+CD8+ | anti-Jo-1 | 0.184252 | 1.29E-14 |
| B20 | B99 | CD3+CD4+CD8+ | anti-PM-Scl | 0.184252 | 1.29E-14 |
| B20 | B183 | CD3+CD4+CD8+ | PTA | -0.18203 | 2.71E-14 |
| B20 | B152 | CD3+CD4+CD8+ | ACA-Ig(G-A-M) | 0.169831 | 1.30E-12 |
| B20 | B119 | CD3+CD4+CD8+ | Hcv-rna | -0.1698 | 1.32E-12 |
| B20 | B159 | CD3+CD4+CD8+ | β2-MG | 0.169795 | 1.32E-12 |
| B20 | B94 | CD3+CD4+CD8+ | ALDH2-Lys504Lys | -0.1698 | 1.32E-12 |
| B20 | B193 | CD3+CD4+CD8+ | K | 0.157749 | 4.62E-11 |
| B20 | B186 | CD3+CD4+CD8+ | PFC | 0.153386 | 1.57E-10 |
| B20 | B185 | CD3+CD4+CD8+ | PT | -0.15171 | 2.49E-10 |
| B20 | B195 | CD3+CD4+CD8+ | Na | 0.150496 | 3.47E-10 |
| B20 | B156 | CD3+CD4+CD8+ | IgE | 0.142264 | 3.04E-09 |
| B20 | B55 | CD3+CD4+CD8+ | TK | -0.14081 | 4.41E-09 |
| B20 | B199 | CD3+CD4+CD8+ | Mg | 0.140415 | 4.87E-09 |
| B20 | B79 | CD3+CD4+CD8+ | hb V -dna | -0.13907 | 6.83E-09 |
| B20 | B56 | CD3+CD4+CD8+ | IR | -0.13854 | 7.80E-09 |
| B20 | B200 | CD3+CD4+CD8+ | OP | 0.131168 | 4.69E-08 |
| B20 | B61 | CD3+CD4+CD8+ | WBHSRV | -0.12741 | 1.13E-07 |
| B20 | B92 | CD3+CD4+CD8+ | ALDH2-Glu504Glu | 0.123226 | 2.91E-07 |
| B20 | B59 | CD3+CD4+CD8+ | WBRV | 0.121556 | 4.21E-07 |
| B20 | B176 | CD3+CD4+CD8+ | Ucr | -0.11831 | 8.52E-07 |
| B201 | B204 | e-habits | Drinking | 0.266674 | 0 |
| B201 | B209 | e-habits | Smoking | 0.167425 | 2.71E-12 |
| B202 | B223 | e-times | FH | 0.565128 | 0 |
| B204 | B206 | Drinking | Waist | 0.215797 | 0 |
| B204 | B209 | Drinking | Smoking | 0.473199 | 0 |
| B204 | B216 | Drinking | BMI | 0.254788 | 0 |
| B204 | B221 | Drinking | Hei | 0.278206 | 0 |
| B204 | B222 | Drinking | NS | 0.291659 | 0 |

| | | | | | |
|---|---|---|---|---|---|
| B204 | B210 | Drinking | Hip | 0.160512 | 2.09E-11 |
| B204 | B207 | Drinking | WHR | 0.15006 | 3.90E-10 |
| B204 | B215 | Drinking | BMI | 0.13695 | 1.16E-08 |
| B206 | B207 | Waist | WHR | 0.699355 | 0 |
| B206 | B209 | Waist | Smoking | 0.207289 | 0 |
| B206 | B210 | Waist | Hip | 0.739727 | 0 |
| B206 | B215 | Waist | BMI | 0.819736 | 0 |
| B206 | B216 | Waist | BMI | 0.832963 | 0 |
| B206 | B221 | Waist | Hei | 0.429444 | 0 |
| B206 | B222 | Waist | NS | 0.747318 | 0 |
| B206 | B218 | Waist | DBP | 0.150164 | 3.79E-10 |
| B206 | B220 | Waist | SBP | 0.136262 | 1.37E-08 |
| B207 | B210 | WHR | Hip | 0.257019 | 0 |
| B207 | B215 | WHR | BMI | 0.455456 | 0 |
| B207 | B216 | WHR | BMI | 0.454879 | 0 |
| B207 | B221 | WHR | Hei | 0.226686 | 0 |
| B207 | B222 | WHR | NS | 0.488376 | 0 |
| B207 | B209 | WHR | Smoking | 0.149229 | 4.88E-10 |
| B207 | B220 | WHR | SBP | 0.133858 | 2.46E-08 |
| B209 | B216 | Smoking | BMI | 0.220757 | 0 |
| B209 | B221 | Smoking | Hei | 0.260486 | 0 |
| B209 | B222 | Smoking | NS | 0.297237 | 0 |
| B209 | B210 | Smoking | Hip | 0.125563 | 1.72E-07 |
| B21 | B101 | CD3+CD4-CD8- | anti-Sc70 | 0.41225 | 0 |
| B21 | B102 | CD3+CD4-CD8- | anti-Sm | 0.41225 | 0 |
| B21 | B103 | CD3+CD4-CD8- | anti-SSA/Ro | 0.41225 | 0 |
| B21 | B104 | CD3+CD4-CD8- | anti-SSB | 0.41225 | 0 |
| B21 | B105 | CD3+CD4-CD8- | anti-U1RNP | 0.41225 | 0 |
| B21 | B106 | CD3+CD4-CD8- | anti-P | 0.41225 | 0 |
| B21 | B107 | CD3+CD4-CD8- | AnuA | 0.412249 | 0 |
| B21 | B108 | CD3+CD4-CD8- | Anti-Cyclin | 0.41225 | 0 |
| B21 | B109 | CD3+CD4-CD8- | AMA-M2 | 0.41225 | 0 |
| B21 | B110 | CD3+CD4-CD8- | AHA | 0.41225 | 0 |
| B21 | B119 | CD3+CD4-CD8- | Hcv-rna | 0.463911 | 0 |
| B21 | B120 | CD3+CD4-CD8- | HCV-cAg | 0.240167 | 0 |
| B21 | B125 | CD3+CD4-CD8- | Cor | 0.53427 | 0 |
| B21 | B134 | CD3+CD4-CD8- | CHE | -0.22053 | 0 |
| B21 | B136 | CD3+CD4-CD8- | 5'NT | 0.27851 | 0 |
| B21 | B151 | CD3+CD4-CD8- | Anti-CCP | 0.27979 | 0 |
| B21 | B152 | CD3+CD4-CD8- | ACA-Ig(G-A-M) | -0.464 | 0 |
| B21 | B157 | CD3+CD4-CD8- | IgG | 0.276216 | 0 |
| B21 | B158 | CD3+CD4-CD8- | IgM | -0.38878 | 0 |
| B21 | B159 | CD3+CD4-CD8- | β2-MG | -0.46391 | 0 |
| B21 | B184 | CD3+CD4-CD8- | INR | 0.198245 | 0 |

| | | | | | |
|---|---|---|---|---|---|
| B21 | B194 | CD3+CD4-CD8- | Cl | -0.2029 | 0 |
| B21 | B197 | CD3+CD4-CD8- | AG | 0.239999 | 0 |
| B21 | B198 | CD3+CD4-CD8- | P | 0.233654 | 0 |
| B21 | B23 | CD3+CD4-CD8- | CD3-CD16+CD56+(NK) | -0.25681 | 0 |
| B21 | B58 | CD3+CD4-CD8- | HCT | -0.28226 | 0 |
| B21 | B79 | CD3+CD4-CD8- | hb V -dna | 0.378355 | 0 |
| B21 | B92 | CD3+CD4-CD8- | ALDH2-Glu504Glu | -0.33891 | 0 |
| B21 | B93 | CD3+CD4-CD8- | ALDH2-Glu504Lys | 0.315446 | 0 |
| B21 | B94 | CD3+CD4-CD8- | ALDH2-Lys504Lys | 0.463911 | 0 |
| B21 | B95 | CD3+CD4-CD8- | anti-cenpb | 0.41225 | 0 |
| B21 | B97 | CD3+CD4-CD8- | dsDNA | 0.41225 | 0 |
| B21 | B98 | CD3+CD4-CD8- | anti-Jo-1 | 0.41225 | 0 |
| B21 | B99 | CD3+CD4-CD8- | anti-PM-Scl | 0.41225 | 0 |
| B21 | B156 | CD3+CD4-CD8- | IgE | 0.193874 | 4.44E-16 |
| B21 | B196 | CD3+CD4-CD8- | TCO2 | -0.19198 | 8.88E-16 |
| B21 | B22 | CD3+CD4-CD8- | CD3+CD8+(T) | -0.17942 | 6.35E-14 |
| B21 | B114 | CD3+CD4-CD8- | TSH TT3 TT4(TT4) | 0.178523 | 8.48E-14 |
| B21 | B183 | CD3+CD4-CD8- | PTA | -0.17256 | 5.60E-13 |
| B21 | B81 | CD3+CD4-CD8- | TSGF | 0.167413 | 2.72E-12 |
| B21 | B62 | CD3+CD4-CD8- | WBHSRVI | -0.16622 | 3.89E-12 |
| B21 | B75 | CD3+CD4-CD8- | ApoA1/B | 0.146481 | 1.01E-09 |
| B21 | B66 | CD3+CD4-CD8- | WBRSR(mPas_100) | -0.14406 | 1.91E-09 |
| B21 | B203 | CD3+CD4-CD8- | DH | -0.14174 | 3.47E-09 |
| B21 | B193 | CD3+CD4-CD8- | K | 0.141092 | 4.10E-09 |
| B21 | B67 | CD3+CD4-CD8- | WBRSR(mPas_50) | -0.14072 | 4.50E-09 |
| B21 | B55 | CD3+CD4-CD8- | TK | 0.138477 | 7.93E-09 |
| B21 | B74 | CD3+CD4-CD8- | ApoA1 | 0.137956 | 9.03E-09 |
| B21 | B63 | CD3+CD4-CD8- | WBRSR(mPas_200) | -0.12637 | 1.43E-07 |
| B21 | B181 | CD3+CD4-CD8- | APTT | 0.124483 | 2.19E-07 |
| B21 | B76 | CD3+CD4-CD8- | ApoB | -0.11945 | 6.66E-07 |
| B21 | B112 | CD3+CD4-CD8- | FT4 | 0.118207 | 8.70E-07 |
| B210 | B215 | Hip | BMI | 0.749414 | 0 |
| B210 | B216 | Hip | BMI | 0.771012 | 0 |
| B210 | B221 | Hip | Hei | 0.409209 | 0 |
| B210 | B222 | Hip | NS | 0.61652 | 0 |
| B210 | B218 | Hip | DBP | 0.158072 | 4.22E-11 |
| B211 | B212 | IOP-R | IOP-L | 0.77683 | 0 |
| B215 | B216 | BMI | BMI | 0.827869 | 0 |
| B215 | B222 | BMI | NS | 0.672988 | 0 |
| B215 | B218 | BMI | DBP | 0.193681 | 4.44E-16 |
| B215 | B221 | BMI | Hei | 0.194239 | 4.44E-16 |
| B216 | B218 | BMI | DBP | 0.24158 | 0 |
| B216 | B221 | BMI | Hei | 0.70549 | 0 |
| B216 | B222 | BMI | NS | 0.825704 | 0 |

| | | | | | |
|---|---|---|---|---|---|
| B218 | B220 | DBP | SBP | 0.461762 | 0 |
| B218 | B222 | DBP | NS | 0.196799 | 2.22E-16 |
| B218 | B221 | DBP | Hei | 0.168409 | 2.01E-12 |
| B22 | B101 | CD3+CD8+(T) | anti-Sc70 | -0.37136 | 0 |
| B22 | B102 | CD3+CD8+(T) | anti-Sm | -0.37136 | 0 |
| B22 | B103 | CD3+CD8+(T) | anti-SSA/Ro | -0.37136 | 0 |
| B22 | B104 | CD3+CD8+(T) | anti-SSB | -0.37136 | 0 |
| B22 | B105 | CD3+CD8+(T) | anti-U1RNP | -0.37136 | 0 |
| B22 | B106 | CD3+CD8+(T) | anti-P | -0.37136 | 0 |
| B22 | B107 | CD3+CD8+(T) | AnuA | -0.37136 | 0 |
| B22 | B108 | CD3+CD8+(T) | Anti-Cyclin | -0.37136 | 0 |
| B22 | B109 | CD3+CD8+(T) | AMA-M2 | -0.37136 | 0 |
| B22 | B110 | CD3+CD8+(T) | AHA | -0.37136 | 0 |
| B22 | B115 | CD3+CD8+(T) | TSH TT3 TT4(TT3) | 0.244994 | 0 |
| B22 | B119 | CD3+CD8+(T) | Hcv-rna | -0.47249 | 0 |
| B22 | B120 | CD3+CD8+(T) | HCV-cAg | -0.25295 | 0 |
| B22 | B121 | CD3+CD8+(T) | C3 | 0.211997 | 0 |
| B22 | B124 | CD3+CD8+(T) | ACTH | -0.35063 | 0 |
| B22 | B125 | CD3+CD8+(T) | Cor | -0.57676 | 0 |
| B22 | B151 | CD3+CD8+(T) | Anti-CCP | -0.22884 | 0 |
| B22 | B152 | CD3+CD8+(T) | ACA-Ig(G-A-M) | 0.47258 | 0 |
| B22 | B156 | CD3+CD8+(T) | IgE | -0.21348 | 0 |
| B22 | B158 | CD3+CD8+(T) | IgM | 0.348529 | 0 |
| B22 | B159 | CD3+CD8+(T) | β2-MG | 0.472481 | 0 |
| B22 | B179 | CD3+CD8+(T) | α1-MG | 0.208477 | 0 |
| B22 | B181 | CD3+CD8+(T) | APTT | -0.23703 | 0 |
| B22 | B23 | CD3+CD8+(T) | CD3-CD16+CD56+(NK) | -0.25854 | 0 |
| B22 | B55 | CD3+CD8+(T) | TK | -0.22602 | 0 |
| B22 | B58 | CD3+CD8+(T) | HCT | 0.284287 | 0 |
| B22 | B79 | CD3+CD8+(T) | hb V -dna | -0.38508 | 0 |
| B22 | B92 | CD3+CD8+(T) | ALDH2-Glu504Glu | 0.345549 | 0 |
| B22 | B93 | CD3+CD8+(T) | ALDH2-Glu504Lys | -0.32166 | 0 |
| B22 | B94 | CD3+CD8+(T) | ALDH2-Lys504Lys | -0.47249 | 0 |
| B22 | B95 | CD3+CD8+(T) | anti-cenpb | -0.37136 | 0 |
| B22 | B97 | CD3+CD8+(T) | dsDNA | -0.37136 | 0 |
| B22 | B98 | CD3+CD8+(T) | anti-Jo-1 | -0.37136 | 0 |
| B22 | B99 | CD3+CD8+(T) | anti-PM-Scl | -0.37136 | 0 |
| B22 | B155 | CD3+CD8+(T) | IgA | -0.18673 | 5.77E-15 |
| B22 | B138 | CD3+CD8+(T) | PAB | 0.182043 | 2.69E-14 |
| B22 | B66 | CD3+CD8+(T) | WBRSR(mPas_100) | 0.178676 | 8.06E-14 |
| B22 | B199 | CD3+CD8+(T) | Mg | 0.177844 | 1.05E-13 |
| B22 | B67 | CD3+CD8+(T) | WBRSR(mPas_50) | 0.175664 | 2.11E-13 |
| B22 | B122 | CD3+CD8+(T) | C4 | -0.1725 | 5.72E-13 |
| B22 | B81 | CD3+CD8+(T) | TSGF | -0.17079 | 9.72E-13 |

| | | | | | |
|---|---|---|---|---|---|
| B22 | B182 | CD3+CD8+(T) | TT | -0.15453 | 1.15E-10 |
| B22 | B157 | CD3+CD8+(T) | IgG | -0.15205 | 2.27E-10 |
| B22 | B75 | CD3+CD8+(T) | ApoA1/B | -0.15204 | 2.27E-10 |
| B22 | B63 | CD3+CD8+(T) | WBRSR(mPas_200) | 0.148377 | 6.13E-10 |
| B22 | B64 | CD3+CD8+(T) | WBRSR(mPas_5) | 0.14563 | 1.27E-09 |
| B22 | B203 | CD3+CD8+(T) | DH | 0.143906 | 1.99E-09 |
| B22 | B74 | CD3+CD8+(T) | ApoA1 | -0.1398 | 5.69E-09 |
| B22 | B137 | CD3+CD8+(T) | AFU | -0.13944 | 6.22E-09 |
| B22 | B62 | CD3+CD8+(T) | WBHSRVI | 0.136924 | 1.17E-08 |
| B22 | B113 | CD3+CD8+(T) | FT3 | 0.136037 | 1.45E-08 |
| B22 | B134 | CD3+CD8+(T) | CHE | 0.135434 | 1.68E-08 |
| B22 | B25 | CD3+CD8+(T) | CK | 0.124602 | 2.14E-07 |
| B220 | B221 | SBP | Hei | -0.12401 | 2.44E-07 |
| B221 | B222 | Hei | NS | 0.600798 | 0 |
| B23 | B115 | CD3-CD16+CD56+(NK) | TSH TT3 TT4(TT3) | -0.21814 | 0 |
| B23 | B119 | CD3-CD16+CD56+(NK) | Hcv-rna | 0.246572 | 0 |
| B23 | B121 | CD3-CD16+CD56+(NK) | C3 | 0.226825 | 0 |
| B23 | B122 | CD3-CD16+CD56+(NK) | C4 | -0.24755 | 0 |
| B23 | B124 | CD3-CD16+CD56+(NK) | ACTH | 0.578942 | 0 |
| B23 | B125 | CD3-CD16+CD56+(NK) | Cor | 0.353449 | 0 |
| B23 | B152 | CD3-CD16+CD56+(NK) | ACA-Ig(G-A-M) | -0.24662 | 0 |
| B23 | B155 | CD3-CD16+CD56+(NK) | IgA | 0.433744 | 0 |
| B23 | B158 | CD3-CD16+CD56+(NK) | IgM | -0.38221 | 0 |
| B23 | B159 | CD3-CD16+CD56+(NK) | β2-MG | -0.24657 | 0 |
| B23 | B179 | CD3-CD16+CD56+(NK) | α1-MG | -0.30437 | 0 |
| B23 | B181 | CD3-CD16+CD56+(NK) | APTT | -0.22842 | 0 |
| B23 | B182 | CD3-CD16+CD56+(NK) | TT | 0.249971 | 0 |
| B23 | B183 | CD3-CD16+CD56+(NK) | PTA | 0.235656 | 0 |
| B23 | B58 | CD3-CD16+CD56+(NK) | HCT | -0.20716 | 0 |
| B23 | B76 | CD3-CD16+CD56+(NK) | ApoB | -0.20225 | 0 |
| B23 | B79 | CD3-CD16+CD56+(NK) | hb V -dna | 0.201465 | 0 |
| B23 | B94 | CD3-CD16+CD56+(NK) | ALDH2-Lys504Lys | 0.246572 | 0 |
| B23 | B178 | CD3-CD16+CD56+(NK) | Ig-U | -0.18817 | 3.55E-15 |
| B23 | B157 | CD3-CD16+CD56+(NK) | IgG | -0.18045 | 4.53E-14 |
| B23 | B92 | CD3-CD16+CD56+(NK) | ALDH2-Glu504Glu | -0.17962 | 5.95E-14 |
| B23 | B62 | CD3-CD16+CD56+(NK) | WBHSRVI | -0.17669 | 1.52E-13 |
| B23 | B63 | CD3-CD16+CD56+(NK) | WBRSR(mPas_200) | -0.16723 | 2.87E-12 |
| B23 | B93 | CD3-CD16+CD56+(NK) | ALDH2-Glu504Lys | 0.167148 | 2.95E-12 |
| B23 | B66 | CD3-CD16+CD56+(NK) | WBRSR(mPas_100) | -0.16237 | 1.22E-11 |
| B23 | B73 | CD3-CD16+CD56+(NK) | ApoA | 0.159899 | 2.50E-11 |
| B23 | B186 | CD3-CD16+CD56+(NK) | PFC | -0.15544 | 8.87E-11 |
| B23 | B75 | CD3-CD16+CD56+(NK) | ApoA1/B | 0.152418 | 2.05E-10 |
| B23 | B67 | CD3-CD16+CD56+(NK) | WBRSR(mPas_50) | -0.15122 | 2.85E-10 |
| B23 | B25 | CD3-CD16+CD56+(NK) | CK | -0.1509 | 3.10E-10 |

| | | | | | |
|---|---|---|---|---|---|
| B23 | B56 | CD3-CD16+CD56+(NK) | IR | -0.13993 | 5.50E-09 |
| B23 | B61 | CD3-CD16+CD56+(NK) | WBHSRV | -0.12967 | 6.68E-08 |
| B23 | B156 | CD3-CD16+CD56+(NK) | IgE | 0.125961 | 1.57E-07 |
| B23 | B185 | CD3-CD16+CD56+(NK) | PT | -0.12525 | 1.85E-07 |
| B24 | B27 | αHBDH | LDH | 0.192947 | 6.66E-16 |
| B24 | B25 | αHBDH | CK | 0.169385 | 1.49E-12 |
| B24 | B74 | αHBDH | ApoA1 | 0.1295 | 6.94E-08 |
| B24 | B76 | αHBDH | ApoB | 0.127235 | 1.17E-07 |
| B25 | B181 | CK | APTT | 0.249118 | 0 |
| B25 | B114 | CK | TSH TT3 TT4(TT4) | 0.183203 | 1.82E-14 |
| B25 | B134 | CK | CHE | -0.16299 | 1.02E-11 |
| B25 | B196 | CK | TCO2 | 0.158775 | 3.45E-11 |
| B25 | B34 | CK | TK | 0.1442 | 1.84E-09 |
| B25 | B136 | CK | 5'NT | -0.14051 | 4.75E-09 |
| B25 | B186 | CK | PFC | -0.12898 | 7.83E-08 |
| B25 | B197 | CK | AG | -0.12639 | 1.42E-07 |
| B25 | B182 | CK | TT | -0.12426 | 2.30E-07 |
| B25 | B135 | CK | TBA | 0.12128 | 4.47E-07 |
| B27 | B38 | LDH | BAND% | 0.406565 | 0 |
| B27 | B37 | LDH | BAND | -0.17353 | 4.15E-13 |
| B27 | B150 | LDH | ALP | 0.160651 | 2.01E-11 |
| B27 | B47 | LDH | RDW | 0.147995 | 6.79E-10 |
| B27 | B176 | LDH | Ucr | -0.13186 | 3.97E-08 |
| B28 | B125 | PRL | Cor | 0.29953 | 0 |
| B28 | B29 | PRL | E2 | 0.99974 | 0 |
| B28 | B30 | PRL | LH | -0.96577 | 0 |
| B28 | B31 | PRL | FSH | -1 | 0 |
| B28 | B32 | PRL | Testo | 0.99984 | 0 |
| B28 | B33 | PRL | Prog | 0.999776 | 0 |
| B29 | B125 | E2 | Cor | 0.292876 | 0 |
| B29 | B30 | E2 | LH | -0.95961 | 0 |
| B29 | B31 | E2 | FSH | -0.99975 | 0 |
| B29 | B32 | E2 | Testo | 0.999171 | 0 |
| B29 | B33 | E2 | Prog | 0.999999 | 0 |
| B3 | B190 | Hcy | Cre | 0.185556 | 8.44E-15 |
| B3 | B189 | Hcy | eGFB | -0.14968 | 4.32E-10 |
| B3 | B222 | Hcy | NS | 0.12267 | 3.29E-07 |
| B3 | B216 | Hcy | BMI | 0.122186 | 3.66E-07 |
| B3 | B221 | Hcy | Hei | 0.120689 | 5.09E-07 |
| B30 | B125 | LH | Cor | -0.36406 | 0 |
| B30 | B31 | LH | FSH | 0.965667 | 0 |
| B30 | B32 | LH | Testo | -0.97026 | 0 |
| B30 | B33 | LH | Prog | -0.96007 | 0 |
| B30 | B143 | LH | β-CTx | 0.122239 | 3.62E-07 |

| | | | | | |
|---|---|---|---|---|---|
| B30 | B134 | LH | CHE | 0.121081 | 4.67E-07 |
| B30 | B145 | LH | BGP | 0.118718 | 7.80E-07 |
| B31 | B125 | FSH | Cor | -0.29941 | 0 |
| B31 | B32 | FSH | Testo | -0.99983 | 0 |
| B31 | B33 | FSH | Prog | -0.99978 | 0 |
| B32 | B125 | Testo | Cor | 0.304643 | 0 |
| B32 | B33 | Testo | Prog | 0.999237 | 0 |
| B33 | B125 | Prog | Cor | 0.293361 | 0 |
| B34 | B101 | TK | anti-Sc70 | 0.185289 | 9.10E-15 |
| B34 | B102 | TK | anti-Sm | 0.185289 | 9.10E-15 |
| B34 | B103 | TK | anti-SSA/Ro | 0.185289 | 9.10E-15 |
| B34 | B104 | TK | anti-SSB | 0.185289 | 9.10E-15 |
| B34 | B105 | TK | anti-U1RNP | 0.185289 | 9.10E-15 |
| B34 | B106 | TK | anti-P | 0.185289 | 9.10E-15 |
| B34 | B107 | TK | AnuA | 0.185289 | 9.10E-15 |
| B34 | B108 | TK | Anti-Cyclin | 0.185289 | 9.10E-15 |
| B34 | B109 | TK | AMA-M2 | 0.185289 | 9.10E-15 |
| B34 | B110 | TK | AHA | 0.185289 | 9.10E-15 |
| B34 | B95 | TK | anti-cenpb | 0.185289 | 9.10E-15 |
| B34 | B97 | TK | dsDNA | 0.185289 | 9.10E-15 |
| B34 | B98 | TK | anti-Jo-1 | 0.185289 | 9.10E-15 |
| B34 | B99 | TK | anti-PM-Scl | 0.185289 | 9.10E-15 |
| B34 | B193 | TK | K | 0.173809 | 3.80E-13 |
| B34 | B196 | TK | TCO2 | 0.170099 | 1.20E-12 |
| B34 | B157 | TK | IgG | 0.169298 | 1.53E-12 |
| B34 | B197 | TK | AG | -0.15786 | 4.47E-11 |
| B34 | B78 | TK | HBcAb | 0.155361 | 9.07E-11 |
| B34 | B93 | TK | ALDH2-Glu504Lys | -0.15456 | 1.13E-10 |
| B34 | B114 | TK | TSH TT3 TT4(TT4) | 0.153617 | 1.47E-10 |
| B34 | B136 | TK | 5'NT | -0.1513 | 2.78E-10 |
| B34 | B92 | TK | ALDH2-Glu504Glu | 0.14825 | 6.34E-10 |
| B34 | B54 | TK | ESR | 0.137018 | 1.14E-08 |
| B34 | B138 | TK | PAB | -0.13233 | 3.56E-08 |
| B34 | B200 | TK | OP | -0.12401 | 2.44E-07 |
| B35 | B117 | β2-MG | TT4 | 0.988317 | 0 |
| B35 | B202 | β2-MG | e-times | 0.634161 | 0 |
| B36 | B227 | V-Lym% | OD | -0.1833 | 1.78E-14 |
| B37 | B38 | BAND | BAND% | 0.355575 | 0 |
| B39 | B44 | Other cells | MCH | -0.16927 | 1.55E-12 |
| B39 | B43 | Other cells | MCV | -0.16546 | 4.89E-12 |
| B4 | B181 | WB-Ca | APTT | 0.333728 | 0 |
| B4 | B184 | WB-Ca | INR | 0.406803 | 0 |
| B4 | B185 | WB-Ca | PT | 0.230644 | 0 |
| B4 | B5 | WB-Ca | WB-Mg | 0.260266 | 0 |

| | | | | | |
|---|---|---|---|---|---|
| B4 | B6 | WB-Ca | WB-Fe | 0.849105 | 0 |
| B4 | B7 | WB-Ca | WB-Cop | 0.820412 | 0 |
| B4 | B193 | WB-Ca | K | -0.18583 | 7.55E-15 |
| B4 | B8 | WB-Ca | WB-Zinc | 0.185112 | 9.77E-15 |
| B4 | B136 | WB-Ca | 5'NT | 0.169805 | 1.31E-12 |
| B4 | B186 | WB-Ca | PFC | -0.16375 | 8.12E-12 |
| B4 | B182 | WB-Ca | TT | -0.137 | 1.14E-08 |
| B4 | B157 | WB-Ca | IgG | -0.12967 | 6.66E-08 |
| B40 | B206 | WBC | Waist | 0.200841 | 0 |
| B40 | B222 | WBC | NS | 0.215308 | 0 |
| B40 | B42 | WBC | MON | 0.678988 | 0 |
| B40 | B46 | WBC | RBC Count | 0.218887 | 0 |
| B40 | B51 | WBC | Hb | 0.218621 | 0 |
| B40 | B52 | WBC | PLT/BPC | 0.238041 | 0 |
| B40 | B72 | WBC | HDL-C | -0.19798 | 0 |
| B40 | B216 | WBC | BMI | 0.196833 | 2.22E-16 |
| B40 | B53 | WBC | GR% | 0.193612 | 4.44E-16 |
| B40 | B192 | WBC | UA | 0.189147 | 2.44E-15 |
| B40 | B48 | WBC | LYM% | -0.17902 | 7.22E-14 |
| B40 | B215 | WBC | BMI | 0.17884 | 7.64E-14 |
| B40 | B209 | WBC | Smoking | 0.172646 | 5.46E-13 |
| B40 | B207 | WBC | WHR | 0.147114 | 8.58E-10 |
| B40 | B41 | WBC | MON % | -0.13979 | 5.70E-09 |
| B40 | B128 | WBC | ALT | 0.133982 | 2.39E-08 |
| B40 | B210 | WBC | Hip | 0.131868 | 3.97E-08 |
| B40 | B221 | WBC | Hei | 0.123132 | 2.97E-07 |
| B41 | B42 | MON % | MON | 0.603857 | 0 |
| B41 | B53 | MON % | GR% | -0.21334 | 0 |
| B41 | B126 | MON % | Alb | -0.17767 | 1.11E-13 |
| B41 | B190 | MON % | Cre | 0.168484 | 1.97E-12 |
| B41 | B189 | MON % | eGFB | -0.164 | 7.54E-12 |
| B41 | B47 | MON % | RDW | 0.143145 | 2.42E-09 |
| B41 | B49 | MON % | baso% | 0.130222 | 5.86E-08 |
| B41 | B204 | MON % | Drinking | 0.121887 | 3.91E-07 |
| B41 | B209 | MON % | Smoking | 0.121759 | 4.02E-07 |
| B42 | B190 | MON | Cre | 0.207257 | 0 |
| B42 | B192 | MON | UA | 0.197821 | 0 |
| B42 | B206 | MON | Waist | 0.209904 | 0 |
| B42 | B209 | MON | Smoking | 0.222968 | 0 |
| B42 | B222 | MON | NS | 0.221241 | 0 |
| B42 | B216 | MON | BMI | 0.176438 | 1.65E-13 |
| B42 | B51 | MON | Hb | 0.173167 | 4.64E-13 |
| B42 | B207 | MON | WHR | 0.156828 | 6.00E-11 |
| B42 | B72 | MON | HDL-C | -0.15578 | 8.06E-11 |

| | | | | | |
|---|---|---|---|---|---|
| B42 | B204 | MON | Drinking | 0.148508 | 5.92E-10 |
| B42 | B215 | MON | BMI | 0.147847 | 7.06E-10 |
| B42 | B46 | MON | RBC Count | 0.146086 | 1.13E-09 |
| B42 | B48 | MON | LYM% | -0.1457 | 1.25E-09 |
| B42 | B189 | MON | eGFB | -0.13212 | 3.74E-08 |
| B42 | B221 | MON | Hei | 0.129229 | 7.40E-08 |
| B42 | B210 | MON | Hip | 0.126748 | 1.31E-07 |
| B42 | B52 | MON | PLT/BPC | 0.122605 | 3.34E-07 |
| B43 | B44 | MCV | MCH | 0.871328 | 0 |
| B43 | B46 | MCV | RBC Count | -0.54705 | 0 |
| B43 | B47 | MCV | RDW | 0.587097 | 0 |
| B43 | B52 | MCV | PLT/BPC | -0.15471 | 1.09E-10 |
| B43 | B218 | MCV | DBP | -0.12451 | 2.18E-07 |
| B43 | B189 | MCV | eGFB | -0.12381 | 2.55E-07 |
| B44 | B45 | MCH | MCHC | 0.547262 | 0 |
| B44 | B46 | MCH | RBC Count | -0.42046 | 0 |
| B44 | B47 | MCH | RDW | 0.377826 | 0 |
| B44 | B51 | MCH | Hb | 0.208038 | 0 |
| B44 | B52 | MCH | PLT/BPC | -0.13391 | 2.43E-08 |
| B45 | B216 | MCHC | BMI | 0.227695 | 0 |
| B45 | B221 | MCHC | Hei | 0.28203 | 0 |
| B45 | B222 | MCHC | NS | 0.267349 | 0 |
| B45 | B47 | MCHC | RDW | -0.24607 | 0 |
| B45 | B51 | MCHC | Hb | 0.446771 | 0 |
| B45 | B71 | MCHC | TG | 0.292903 | 0 |
| B45 | B72 | MCHC | HDL-C | -0.19357 | 4.44E-16 |
| B45 | B192 | MCHC | UA | 0.188547 | 3.11E-15 |
| B45 | B129 | MCHC | IBIL | 0.180162 | 4.97E-14 |
| B45 | B133 | MCHC | TBIL | 0.175539 | 2.20E-13 |
| B45 | B163 | MCHC | SG | 0.155057 | 9.87E-11 |
| B45 | B174 | MCHC | Urea/Cr | -0.1416 | 3.60E-09 |
| B45 | B206 | MCHC | Waist | 0.132368 | 3.52E-08 |
| B45 | B126 | MCHC | Alb | 0.126626 | 1.35E-07 |
| B45 | B141 | MCHC | AST/ALT | -0.11893 | 7.45E-07 |
| B46 | B126 | RBC Count | Alb | 0.253137 | 0 |
| B46 | B141 | RBC Count | AST/ALT | -0.21297 | 0 |
| B46 | B163 | RBC Count | SG | 0.264573 | 0 |
| B46 | B174 | RBC Count | Urea/Cr | -0.24991 | 0 |
| B46 | B189 | RBC Count | eGFB | 0.203754 | 0 |
| B46 | B192 | RBC Count | UA | 0.197544 | 0 |
| B46 | B206 | RBC Count | Waist | 0.212292 | 0 |
| B46 | B215 | RBC Count | BMI | 0.208201 | 0 |
| B46 | B216 | RBC Count | BMI | 0.351322 | 0 |
| B46 | B218 | RBC Count | DBP | 0.274319 | 0 |

| | | | | | |
|---|---|---|---|---|---|
| B46 | B221 | RBC Count | Hei | 0.352341 | 0 |
| B46 | B222 | RBC Count | NS | 0.340002 | 0 |
| B46 | B47 | RBC Count | RDW | -0.45641 | 0 |
| B46 | B51 | RBC Count | Hb | 0.788157 | 0 |
| B46 | B72 | RBC Count | HDL-C | -0.22218 | 0 |
| B46 | B113 | RBC Count | FT3 | 0.190221 | 1.78E-15 |
| B46 | B128 | RBC Count | ALT | 0.188405 | 3.11E-15 |
| B46 | B129 | RBC Count | IBIL | 0.182309 | 2.46E-14 |
| B46 | B191 | RBC Count | Urea | -0.18068 | 4.22E-14 |
| B46 | B176 | RBC Count | Ucr | 0.177226 | 1.28E-13 |
| B46 | B210 | RBC Count | Hip | 0.170653 | 1.01E-12 |
| B46 | B133 | RBC Count | TBIL | 0.164194 | 7.12E-12 |
| B46 | B71 | RBC Count | TG | 0.152478 | 2.02E-10 |
| B46 | B207 | RBC Count | WHR | 0.144876 | 1.55E-09 |
| B46 | B91 | RBC Count | TP | 0.131257 | 4.59E-08 |
| B46 | B70 | RBC Count | LDL-C | 0.128661 | 8.44E-08 |
| B46 | B209 | RBC Count | Smoking | 0.123893 | 2.50E-07 |
| B46 | B161 | RBC Count | LEU | -0.12338 | 2.81E-07 |
| B46 | B58 | RBC Count | HCT | 0.121012 | 4.74E-07 |
| B46 | B52 | RBC Count | PLT/BPC | 0.117644 | 9.81E-07 |
| B47 | B189 | RDW | eGFB | -0.26237 | 0 |
| B47 | B51 | RDW | Hb | -0.26061 | 0 |
| B47 | B52 | RDW | PLT/BPC | -0.19351 | 4.44E-16 |
| B47 | B191 | RDW | Urea | 0.175583 | 2.17E-13 |
| B47 | B126 | RDW | Alb | -0.15288 | 1.81E-10 |
| B47 | B218 | RDW | DBP | -0.14292 | 2.57E-09 |
| B47 | B190 | RDW | Cre | 0.141588 | 3.61E-09 |
| B47 | B163 | RDW | SG | -0.13306 | 2.99E-08 |
| B47 | B185 | RDW | PT | 0.132029 | 3.82E-08 |
| B47 | B221 | RDW | Hei | -0.12557 | 1.72E-07 |
| B47 | B184 | RDW | INR | 0.124922 | 1.99E-07 |
| B47 | B176 | RDW | Ucr | -0.12439 | 2.24E-07 |
| B47 | B72 | RDW | HDL-C | 0.121601 | 4.17E-07 |
| B47 | B216 | RDW | BMI | -0.12025 | 5.60E-07 |
| B48 | B53 | LYM% | GR% | -0.93483 | 0 |
| B48 | B77 | LYM% | TC | 0.158457 | 3.78E-11 |
| B48 | B192 | LYM% | UA | -0.14214 | 3.14E-09 |
| B48 | B72 | LYM% | HDL-C | 0.13705 | 1.13E-08 |
| B48 | B190 | LYM% | Cre | -0.13193 | 3.91E-08 |
| B48 | B189 | LYM% | eGFB | 0.12825 | 9.28E-08 |
| B48 | B207 | LYM% | WHR | -0.1257 | 1.67E-07 |
| B48 | B206 | LYM% | Waist | -0.12383 | 2.54E-07 |
| B48 | B153 | LYM% | Glu | -0.11912 | 7.15E-07 |
| B48 | B91 | LYM% | TP | 0.118193 | 8.73E-07 |

| | | | | | |
|---|---|---|---|---|---|
| B49 | B50 | baso% | eo% | 0.258069 | 0 |
| B49 | B53 | baso% | GR% | -0.22323 | 0 |
| B5 | B136 | WB-Mg | 5'NT | 0.328978 | 0 |
| B5 | B155 | WB-Mg | IgA | -0.20115 | 0 |
| B5 | B157 | WB-Mg | IgG | -0.2474 | 0 |
| B5 | B6 | WB-Mg | WB-Fe | 0.719574 | 0 |
| B5 | B7 | WB-Mg | WB-Cop | 0.289189 | 0 |
| B5 | B8 | WB-Mg | WB-Zinc | -0.57613 | 0 |
| B5 | B57 | WB-Mg | RBCAI | -0.1722 | 6.28E-13 |
| B5 | B138 | WB-Mg | PAB | 0.169655 | 1.38E-12 |
| B5 | B158 | WB-Mg | IgM | -0.16505 | 5.53E-12 |
| B5 | B58 | WB-Mg | HCT | 0.151616 | 2.55E-10 |
| B5 | B156 | WB-Mg | IgE | -0.12976 | 6.53E-08 |
| B5 | B195 | WB-Mg | Na | 0.125514 | 1.74E-07 |
| B5 | B137 | WB-Mg | AFU | 0.125325 | 1.81E-07 |
| B5 | B66 | WB-Mg | WBRSR(mPas_100) | 0.12474 | 2.07E-07 |
| B5 | B63 | WB-Mg | WBRSR(mPas_200) | 0.121484 | 4.27E-07 |
| B50 | B53 | eo% | GR% | -0.25287 | 0 |
| B50 | B222 | eo% | NS | 0.137345 | 1.05E-08 |
| B50 | B209 | eo% | Smoking | 0.123507 | 2.73E-07 |
| B50 | B204 | eo% | Drinking | 0.120411 | 5.41E-07 |
| B51 | B113 | Hb | FT3 | 0.199217 | 0 |
| B51 | B126 | Hb | Alb | 0.255877 | 0 |
| B51 | B128 | Hb | ALT | 0.230937 | 0 |
| B51 | B129 | Hb | IBIL | 0.210682 | 0 |
| B51 | B133 | Hb | TBIL | 0.205164 | 0 |
| B51 | B141 | Hb | AST/ALT | -0.2387 | 0 |
| B51 | B163 | Hb | SG | 0.294535 | 0 |
| B51 | B174 | Hb | Urea/Cr | -0.29336 | 0 |
| B51 | B176 | Hb | Ucr | 0.197856 | 0 |
| B51 | B192 | Hb | UA | 0.258485 | 0 |
| B51 | B206 | Hb | Waist | 0.272896 | 0 |
| B51 | B210 | Hb | Hip | 0.20239 | 0 |
| B51 | B215 | Hb | BMI | 0.22554 | 0 |
| B51 | B216 | Hb | BMI | 0.422259 | 0 |
| B51 | B218 | Hb | DBP | 0.261166 | 0 |
| B51 | B221 | Hb | Hei | 0.457378 | 0 |
| B51 | B222 | Hb | NS | 0.438025 | 0 |
| B51 | B71 | Hb | TG | 0.214931 | 0 |
| B51 | B72 | Hb | HDL-C | -0.249 | 0 |
| B51 | B207 | Hb | WHR | 0.194609 | 4.44E-16 |
| B51 | B189 | Hb | eGFB | 0.190374 | 1.78E-15 |
| B51 | B209 | Hb | Smoking | 0.188249 | 3.33E-15 |
| B51 | B204 | Hb | Drinking | 0.176515 | 1.61E-13 |

| | | | | | |
|---|---|---|---|---|---|
| B51 | B191 | Hb | Urea | -0.15622 | 7.13E-11 |
| B51 | B161 | Hb | LEU | -0.1518 | 2.43E-10 |
| B51 | B140 | Hb | r-GT | 0.143436 | 2.25E-09 |
| B51 | B127 | Hb | A-G ratio | 0.139592 | 5.99E-09 |
| B51 | B58 | Hb | HCT | 0.132908 | 3.10E-08 |
| B51 | B132 | Hb | Dbil | 0.119629 | 6.41E-07 |
| B51 | B190 | Hb | Cre | 0.118791 | 7.68E-07 |
| B52 | B70 | PLT/BPC | LDL-C | 0.165776 | 4.45E-12 |
| B52 | B132 | PLT/BPC | Dbil | -0.13971 | 5.82E-09 |
| B52 | B186 | PLT/BPC | PFC | 0.122922 | 3.11E-07 |
| B53 | B72 | GR% | HDL-C | -0.12039 | 5.44E-07 |
| B54 | B56 | ESR | IR | -0.26436 | 0 |
| B54 | B58 | ESR | HCT | -0.27016 | 0 |
| B54 | B60 | ESR | WBRVI | -0.35469 | 0 |
| B54 | B62 | ESR | WBHSRVI | -0.30281 | 0 |
| B54 | B64 | ESR | WBRSR(mPas_5) | -0.21317 | 0 |
| B54 | B65 | ESR | WBRSR(mPas_1) | -0.21145 | 0 |
| B54 | B69 | ESR | PV | 0.437617 | 0 |
| B54 | B138 | ESR | PAB | -0.19002 | 1.78E-15 |
| B54 | B78 | ESR | HBcAb | -0.18239 | 2.40E-14 |
| B54 | B157 | ESR | IgG | 0.179105 | 7.02E-14 |
| B54 | B67 | ESR | WBRSR(mPas_50) | -0.17058 | 1.04E-12 |
| B54 | B66 | ESR | WBRSR(mPas_100) | -0.16085 | 1.90E-11 |
| B54 | B121 | ESR | C3 | -0.15902 | 3.21E-11 |
| B54 | B57 | ESR | RBCAI | -0.15766 | 4.74E-11 |
| B54 | B55 | ESR | TK | -0.1545 | 1.15E-10 |
| B54 | B100 | ESR | anti-Ro52 | 0.152839 | 1.83E-10 |
| B54 | B122 | ESR | C4 | 0.151447 | 2.68E-10 |
| B54 | B134 | ESR | CHE | -0.14327 | 2.35E-09 |
| B54 | B63 | ESR | WBRSR(mPas_200) | -0.13617 | 1.40E-08 |
| B54 | B61 | ESR | WBHSRV | -0.13047 | 5.52E-08 |
| B54 | B101 | ESR | anti-Sc70 | 0.126492 | 1.39E-07 |
| B54 | B102 | ESR | anti-Sm | 0.126492 | 1.39E-07 |
| B54 | B103 | ESR | anti-SSA/Ro | 0.126492 | 1.39E-07 |
| B54 | B104 | ESR | anti-SSB | 0.126492 | 1.39E-07 |
| B54 | B105 | ESR | anti-U1RNP | 0.126492 | 1.39E-07 |
| B54 | B106 | ESR | anti-P | 0.126492 | 1.39E-07 |
| B54 | B107 | ESR | AnuA | 0.126492 | 1.39E-07 |
| B54 | B108 | ESR | Anti-Cyclin | 0.126492 | 1.39E-07 |
| B54 | B109 | ESR | AMA-M2 | 0.126492 | 1.39E-07 |
| B54 | B110 | ESR | AHA | 0.126492 | 1.39E-07 |
| B54 | B95 | ESR | anti-cenpb | 0.126492 | 1.39E-07 |
| B54 | B97 | ESR | dsDNA | 0.126492 | 1.39E-07 |
| B54 | B98 | ESR | anti-Jo-1 | 0.126492 | 1.39E-07 |

| | | | | | |
|---|---|---|---|---|---|
| B54 | B99 | ESR | anti-PM-Scl | 0.126492 | 1.39E-07 |
| B54 | B137 | ESR | AFU | -0.12109 | 4.66E-07 |
| B54 | B151 | ESR | Anti-CCP | 0.120891 | 4.87E-07 |
| B54 | B135 | ESR | TBA | 0.117881 | 9.33E-07 |
| B55 | B100 | TK | anti-Ro52 | -0.21528 | 0 |
| B55 | B156 | TK | IgE | 0.245646 | 0 |
| B55 | B157 | TK | IgG | 0.200104 | 0 |
| B55 | B181 | TK | APTT | 0.220883 | 0 |
| B55 | B56 | TK | IR | 0.794046 | 0 |
| B55 | B59 | TK | WBRV | 0.398205 | 0 |
| B55 | B60 | TK | WBRVI | 0.435229 | 0 |
| B55 | B61 | TK | WBHSRV | 0.712537 | 0 |
| B55 | B62 | TK | WBHSRVI | 0.568087 | 0 |
| B55 | B63 | TK | WBRSR(mPas_200) | 0.483142 | 0 |
| B55 | B64 | TK | WBRSR(mPas_5) | 0.39548 | 0 |
| B55 | B65 | TK | WBRSR(mPas_1) | 0.320667 | 0 |
| B55 | B66 | TK | WBRSR(mPas_100) | 0.43621 | 0 |
| B55 | B67 | TK | WBRSR(mPas_50) | 0.435939 | 0 |
| B55 | B69 | TK | PV | -0.31592 | 0 |
| B55 | B184 | TK | INR | 0.173131 | 4.69E-13 |
| B55 | B152 | TK | ACA-Ig(G-A-M) | -0.17147 | 7.88E-13 |
| B55 | B119 | TK | Hcv-rna | 0.171431 | 7.96E-13 |
| B55 | B159 | TK | β2-MG | -0.17143 | 7.97E-13 |
| B55 | B125 | TK | Cor | 0.166021 | 4.13E-12 |
| B55 | B138 | TK | PAB | -0.16195 | 1.38E-11 |
| B55 | B73 | TK | ApoA | -0.15382 | 1.39E-10 |
| B55 | B135 | TK | TBA | 0.139562 | 6.04E-09 |
| B55 | B79 | TK | hb V -dna | 0.138625 | 7.64E-09 |
| B55 | B185 | TK | PT | 0.13782 | 9.34E-09 |
| B55 | B94 | TK | ALDH2-Lys504Lys | 0.128796 | 8.18E-08 |
| B55 | B92 | TK | ALDH2-Glu504Glu | -0.12486 | 2.01E-07 |
| B55 | B58 | TK | HCT | 0.122974 | 3.07E-07 |
| B55 | B93 | TK | ALDH2-Glu504Lys | 0.118577 | 8.04E-07 |
| B56 | B100 | IR | anti-Ro52 | -0.24122 | 0 |
| B56 | B58 | IR | HCT | 0.665741 | 0 |
| B56 | B59 | IR | WBRV | 0.433104 | 0 |
| B56 | B60 | IR | WBRVI | 0.740648 | 0 |
| B56 | B61 | IR | WBHSRV | 0.924881 | 0 |
| B56 | B62 | IR | WBHSRVI | 0.949802 | 0 |
| B56 | B63 | IR | WBRSR(mPas_200) | 0.850747 | 0 |
| B56 | B64 | IR | WBRSR(mPas_5) | 0.740152 | 0 |
| B56 | B65 | IR | WBRSR(mPas_1) | 0.620014 | 0 |
| B56 | B66 | IR | WBRSR(mPas_100) | 0.828273 | 0 |
| B56 | B67 | IR | WBRSR(mPas_50) | 0.826939 | 0 |

| | | | | | |
|---|---|---|---|---|---|
| B56 | B69 | IR | PV | -0.2732 | 0 |
| B56 | B73 | IR | ApoA | -0.23342 | 0 |
| B56 | B158 | IR | IgM | 0.147033 | 8.77E-10 |
| B56 | B115 | IR | TSH TT3 TT4(TT3) | 0.143194 | 2.39E-09 |
| B56 | B137 | IR | AFU | 0.142784 | 2.66E-09 |
| B56 | B181 | IR | APTT | 0.129557 | 6.85E-08 |
| B56 | B124 | IR | ACTH | -0.11932 | 6.86E-07 |
| B57 | B100 | RBCAI | anti-Ro52 | 0.576761 | 0 |
| B57 | B121 | RBCAI | C3 | 0.272098 | 0 |
| B57 | B122 | RBCAI | C4 | -0.27038 | 0 |
| B57 | B156 | RBCAI | IgE | 0.255426 | 0 |
| B57 | B59 | RBCAI | WBRV | 0.752545 | 0 |
| B57 | B60 | RBCAI | WBRVI | 0.533065 | 0 |
| B57 | B64 | RBCAI | WBRSR(mPas_5) | 0.344074 | 0 |
| B57 | B65 | RBCAI | WBRSR(mPas_1) | 0.566048 | 0 |
| B57 | B151 | RBCAI | Anti-CCP | 0.181244 | 3.51E-14 |
| B57 | B198 | RBCAI | P | -0.15723 | 5.35E-11 |
| B57 | B93 | RBCAI | ALDH2-Glu504Lys | 0.155972 | 7.64E-11 |
| B57 | B183 | RBCAI | PTA | -0.1517 | 2.50E-10 |
| B57 | B92 | RBCAI | ALDH2-Glu504Glu | -0.15034 | 3.62E-10 |
| B57 | B182 | RBCAI | TT | 0.142384 | 2.95E-09 |
| B57 | B194 | RBCAI | Cl | -0.13052 | 5.47E-08 |
| B57 | B155 | RBCAI | IgA | 0.12532 | 1.82E-07 |
| B57 | B101 | RBCAI | anti-Sc70 | -0.12276 | 3.23E-07 |
| B57 | B102 | RBCAI | anti-Sm | -0.12276 | 3.23E-07 |
| B57 | B103 | RBCAI | anti-SSA/Ro | -0.12276 | 3.23E-07 |
| B57 | B104 | RBCAI | anti-SSB | -0.12276 | 3.23E-07 |
| B57 | B105 | RBCAI | anti-U1RNP | -0.12276 | 3.23E-07 |
| B57 | B106 | RBCAI | anti-P | -0.12276 | 3.23E-07 |
| B57 | B107 | RBCAI | AnuA | -0.12276 | 3.23E-07 |
| B57 | B108 | RBCAI | Anti-Cyclin | -0.12276 | 3.23E-07 |
| B57 | B109 | RBCAI | AMA-M2 | -0.12276 | 3.23E-07 |
| B57 | B110 | RBCAI | AHA | -0.12276 | 3.23E-07 |
| B57 | B95 | RBCAI | anti-cenpb | -0.12276 | 3.23E-07 |
| B57 | B97 | RBCAI | dsDNA | -0.12276 | 3.23E-07 |
| B57 | B98 | RBCAI | anti-Jo-1 | -0.12276 | 3.23E-07 |
| B57 | B99 | RBCAI | anti-PM-Scl | -0.12276 | 3.23E-07 |
| B57 | B138 | RBCAI | PAB | -0.11858 | 8.04E-07 |
| B58 | B119 | HCT | Hcv-rna | -0.30186 | 0 |
| B58 | B124 | HCT | ACTH | -0.26931 | 0 |
| B58 | B125 | HCT | Cor | -0.3781 | 0 |
| B58 | B138 | HCT | PAB | 0.202058 | 0 |
| B58 | B152 | HCT | ACA-Ig(G-A-M) | 0.301919 | 0 |
| B58 | B156 | HCT | IgE | -0.21054 | 0 |

| | | | | | |
|---|---|---|---|---|---|
| B58 | B157 | HCT | IgG | -0.22905 | 0 |
| B58 | B158 | HCT | IgM | 0.269345 | 0 |
| B58 | B159 | HCT | β2-MG | 0.301856 | 0 |
| B58 | B59 | HCT | WBRV | 0.262724 | 0 |
| B58 | B60 | HCT | WBRVI | 0.698997 | 0 |
| B58 | B61 | HCT | WBHSRV | 0.674185 | 0 |
| B58 | B62 | HCT | WBHSRVI | 0.854004 | 0 |
| B58 | B63 | HCT | WBRSR(mPas_200) | 0.836296 | 0 |
| B58 | B64 | HCT | WBRSR(mPas_5) | 0.759868 | 0 |
| B58 | B65 | HCT | WBRSR(mPas_1) | 0.66472 | 0 |
| B58 | B66 | HCT | WBRSR(mPas_100) | 0.841793 | 0 |
| B58 | B67 | HCT | WBRSR(mPas_50) | 0.847132 | 0 |
| B58 | B73 | HCT | ApoA | -0.22547 | 0 |
| B58 | B74 | HCT | ApoA1 | -0.20905 | 0 |
| B58 | B79 | HCT | hb V -dna | -0.24645 | 0 |
| B58 | B92 | HCT | ALDH2-Glu504Glu | 0.268601 | 0 |
| B58 | B93 | HCT | ALDH2-Glu504Lys | -0.25515 | 0 |
| B58 | B94 | HCT | ALDH2-Lys504Lys | -0.27579 | 0 |
| B58 | B100 | HCT | anti-Ro52 | -0.19354 | 4.44E-16 |
| B58 | B115 | HCT | TSH TT3 TT4(TT3) | 0.186313 | 6.44E-15 |
| B58 | B184 | HCT | INR | -0.17443 | 3.12E-13 |
| B58 | B134 | HCT | CHE | 0.171712 | 7.30E-13 |
| B58 | B137 | HCT | AFU | 0.162993 | 1.02E-11 |
| B58 | B151 | HCT | Anti-CCP | -0.16257 | 1.15E-11 |
| B58 | B198 | HCT | P | -0.15392 | 1.36E-10 |
| B58 | B179 | HCT | α1-MG | 0.153716 | 1.43E-10 |
| B58 | B155 | HCT | IgA | -0.1514 | 2.71E-10 |
| B58 | B120 | HCT | HCV-cAg | -0.14883 | 5.44E-10 |
| B58 | B222 | HCT | NS | 0.143848 | 2.02E-09 |
| B58 | B101 | HCT | anti-Sc70 | -0.14298 | 2.53E-09 |
| B58 | B102 | HCT | anti-Sm | -0.14298 | 2.53E-09 |
| B58 | B103 | HCT | anti-SSA/Ro | -0.14298 | 2.53E-09 |
| B58 | B104 | HCT | anti-SSB | -0.14298 | 2.53E-09 |
| B58 | B105 | HCT | anti-U1RNP | -0.14298 | 2.53E-09 |
| B58 | B106 | HCT | anti-P | -0.14298 | 2.53E-09 |
| B58 | B107 | HCT | AnuA | -0.14298 | 2.53E-09 |
| B58 | B108 | HCT | Anti-Cyclin | -0.14298 | 2.53E-09 |
| B58 | B109 | HCT | AMA-M2 | -0.14298 | 2.53E-09 |
| B58 | B110 | HCT | AHA | -0.14298 | 2.53E-09 |
| B58 | B95 | HCT | anti-cenpb | -0.14298 | 2.53E-09 |
| B58 | B97 | HCT | dsDNA | -0.14298 | 2.53E-09 |
| B58 | B98 | HCT | anti-Jo-1 | -0.14298 | 2.53E-09 |
| B58 | B99 | HCT | anti-PM-Scl | -0.14298 | 2.53E-09 |
| B58 | B182 | HCT | TT | -0.13956 | 6.04E-09 |

| | | | | | |
|---|---|---|---|---|---|
| B58 | B75 | HCT | ApoA1/B | -0.13409 | 2.33E-08 |
| B58 | B180 | HCT | Tf | 0.1224 | 3.49E-07 |
| B59 | B100 | WBRV | anti-Ro52 | 0.452814 | 0 |
| B59 | B121 | WBRV | C3 | 0.362512 | 0 |
| B59 | B122 | WBRV | C4 | -0.36894 | 0 |
| B59 | B151 | WBRV | Anti-CCP | 0.226704 | 0 |
| B59 | B156 | WBRV | IgE | 0.339574 | 0 |
| B59 | B194 | WBRV | Cl | -0.22172 | 0 |
| B59 | B60 | WBRV | WBRVI | 0.792257 | 0 |
| B59 | B61 | WBRV | WBHSRV | 0.560416 | 0 |
| B59 | B62 | WBRV | WBHSRVI | 0.380671 | 0 |
| B59 | B63 | WBRV | WBRSR(mPas_200) | 0.505491 | 0 |
| B59 | B64 | WBRV | WBRSR(mPas_5) | 0.787068 | 0 |
| B59 | B65 | WBRV | WBRSR(mPas_1) | 0.890295 | 0 |
| B59 | B66 | WBRV | WBRSR(mPas_100) | 0.548957 | 0 |
| B59 | B67 | WBRV | WBRSR(mPas_50) | 0.587319 | 0 |
| B59 | B69 | WBRV | PV | 0.26207 | 0 |
| B59 | B198 | WBRV | P | -0.19084 | 1.33E-15 |
| B59 | B157 | WBRV | IgG | 0.178444 | 8.70E-14 |
| B59 | B183 | WBRV | PTA | -0.13282 | 3.17E-08 |
| B59 | B138 | WBRV | PAB | -0.13092 | 4.97E-08 |
| B6 | B136 | WB-Fe | 5'NT | 0.266014 | 0 |
| B6 | B157 | WB-Fe | IgG | -0.20162 | 0 |
| B6 | B184 | WB-Fe | INR | 0.231202 | 0 |
| B6 | B7 | WB-Fe | WB-Cop | 0.674072 | 0 |
| B6 | B8 | WB-Fe | WB-Zinc | -0.25903 | 0 |
| B6 | B181 | WB-Fe | APTT | 0.189608 | 2.22E-15 |
| B6 | B155 | WB-Fe | IgA | -0.16318 | 9.61E-12 |
| B6 | B138 | WB-Fe | PAB | 0.140644 | 4.60E-09 |
| B6 | B57 | WB-Fe | RBCAI | -0.13967 | 5.88E-09 |
| B6 | B158 | WB-Fe | IgM | -0.13497 | 1.88E-08 |
| B6 | B185 | WB-Fe | PT | 0.131094 | 4.77E-08 |
| B6 | B193 | WB-Fe | K | -0.12603 | 1.55E-07 |
| B6 | B58 | WB-Fe | HCT | 0.123225 | 2.91E-07 |
| B60 | B121 | WBRVI | C3 | 0.285609 | 0 |
| B60 | B122 | WBRVI | C4 | -0.27255 | 0 |
| B60 | B158 | WBRVI | IgM | 0.25423 | 0 |
| B60 | B198 | WBRVI | P | -0.20807 | 0 |
| B60 | B61 | WBRVI | WBHSRV | 0.709266 | 0 |
| B60 | B62 | WBRVI | WBHSRVI | 0.783034 | 0 |
| B60 | B63 | WBRVI | WBRSR(mPas_200) | 0.724128 | 0 |
| B60 | B64 | WBRVI | WBRSR(mPas_5) | 0.902104 | 0 |
| B60 | B65 | WBRVI | WBRSR(mPas_1) | 0.93285 | 0 |
| B60 | B66 | WBRVI | WBRSR(mPas_100) | 0.760976 | 0 |

| | | | | | |
|---|---|---|---|---|---|
| B60 | B67 | WBRVI | WBRSR(mPas_50) | 0.79471 | 0 |
| B60 | B73 | WBRVI | ApoA | -0.20415 | 0 |
| B60 | B94 | WBRVI | ALDH2-Lys504Lys | -0.20395 | 0 |
| B60 | B125 | WBRVI | Cor | -0.19184 | 8.88E-16 |
| B60 | B194 | WBRVI | CI | -0.17581 | 2.02E-13 |
| B60 | B124 | WBRVI | ACTH | -0.16876 | 1.81E-12 |
| B60 | B152 | WBRVI | ACA-Ig(G-A-M) | 0.15707 | 5.60E-11 |
| B60 | B119 | WBRVI | Hcv-rna | -0.15704 | 5.65E-11 |
| B60 | B159 | WBRVI | β2-MG | 0.157037 | 5.66E-11 |
| B60 | B134 | WBRVI | CHE | 0.155785 | 8.05E-11 |
| B60 | B101 | WBRVI | anti-Sc70 | -0.15327 | 1.62E-10 |
| B60 | B102 | WBRVI | anti-Sm | -0.15327 | 1.62E-10 |
| B60 | B103 | WBRVI | anti-SSA/Ro | -0.15327 | 1.62E-10 |
| B60 | B104 | WBRVI | anti-SSB | -0.15327 | 1.62E-10 |
| B60 | B105 | WBRVI | anti-U1RNP | -0.15327 | 1.62E-10 |
| B60 | B106 | WBRVI | anti-P | -0.15327 | 1.62E-10 |
| B60 | B107 | WBRVI | AnuA | -0.15327 | 1.62E-10 |
| B60 | B108 | WBRVI | Anti-Cyclin | -0.15327 | 1.62E-10 |
| B60 | B109 | WBRVI | AMA-M2 | -0.15327 | 1.62E-10 |
| B60 | B110 | WBRVI | AHA | -0.15327 | 1.62E-10 |
| B60 | B95 | WBRVI | anti-cenpb | -0.15327 | 1.62E-10 |
| B60 | B97 | WBRVI | dsDNA | -0.15327 | 1.62E-10 |
| B60 | B98 | WBRVI | anti-Jo-1 | -0.15327 | 1.62E-10 |
| B60 | B99 | WBRVI | anti-PM-Scl | -0.15327 | 1.62E-10 |
| B60 | B137 | WBRVI | AFU | 0.146201 | 1.09E-09 |
| B60 | B69 | WBRVI | PV | -0.14593 | 1.17E-09 |
| B60 | B156 | WBRVI | IgE | 0.141699 | 3.51E-09 |
| B60 | B114 | WBRVI | TSH TT3 TT4(TT4) | -0.1375 | 1.01E-08 |
| B60 | B120 | WBRVI | HCV-cAg | -0.13741 | 1.03E-08 |
| B60 | B79 | WBRVI | hb V -dna | -0.1285 | 8.76E-08 |
| B60 | B100 | WBRVI | anti-Ro52 | 0.124471 | 2.20E-07 |
| B60 | B115 | WBRVI | TSH TT3 TT4(TT3) | 0.124168 | 2.36E-07 |
| B60 | B222 | WBRVI | NS | 0.118516 | 8.14E-07 |
| B61 | B62 | WBHSRV | WBHSRVI | 0.892133 | 0 |
| B61 | B63 | WBHSRV | WBRSR(mPas_200) | 0.953616 | 0 |
| B61 | B64 | WBHSRV | WBRSR(mPas_5) | 0.851012 | 0 |
| B61 | B65 | WBHSRV | WBRSR(mPas_1) | 0.728448 | 0 |
| B61 | B66 | WBHSRV | WBRSR(mPas_100) | 0.935546 | 0 |
| B61 | B67 | WBHSRV | WBRSR(mPas_50) | 0.932497 | 0 |
| B61 | B73 | WBHSRV | ApoA | -0.20364 | 0 |
| B61 | B122 | WBHSRV | C4 | -0.18217 | 2.58E-14 |
| B61 | B121 | WBHSRV | C3 | 0.180356 | 4.66E-14 |
| B61 | B93 | WBHSRV | ALDH2-Glu504Lys | -0.16561 | 4.68E-12 |
| B61 | B194 | WBHSRV | CI | -0.16299 | 1.02E-11 |

| | | | | | |
|---|---|---|---|---|---|
| B61 | B92 | WBHSRV | ALDH2-Glu504Glu | 0.162784 | 1.08E-11 |
| B61 | B181 | WBHSRV | APTT | 0.137529 | 1.00E-08 |
| B61 | B137 | WBHSRV | AFU | 0.126824 | 1.29E-07 |
| B61 | B74 | WBHSRV | ApoA1 | -0.12342 | 2.78E-07 |
| B61 | B100 | WBHSRV | anti-Ro52 | -0.11874 | 7.76E-07 |
| B62 | B100 | WBHSRVI | anti-Ro52 | -0.22557 | 0 |
| B62 | B125 | WBHSRVI | Cor | -0.22629 | 0 |
| B62 | B158 | WBHSRVI | IgM | 0.210436 | 0 |
| B62 | B63 | WBHSRVI | WBRSR(mPas_200) | 0.911803 | 0 |
| B62 | B64 | WBHSRVI | WBRSR(mPas_5) | 0.805164 | 0 |
| B62 | B65 | WBHSRVI | WBRSR(mPas_1) | 0.679506 | 0 |
| B62 | B66 | WBHSRVI | WBRSR(mPas_100) | 0.904615 | 0 |
| B62 | B67 | WBHSRVI | WBRSR(mPas_50) | 0.903074 | 0 |
| B62 | B69 | WBHSRVI | PV | -0.20917 | 0 |
| B62 | B73 | WBHSRVI | ApoA | -0.24521 | 0 |
| B62 | B124 | WBHSRVI | ACTH | -0.19197 | 8.88E-16 |
| B62 | B115 | WBHSRVI | TSH TT3 TT4(TT3) | 0.186015 | 7.11E-15 |
| B62 | B94 | WBHSRVI | ALDH2-Lys504Lys | -0.17124 | 8.46E-13 |
| B62 | B74 | WBHSRVI | ApoA1 | -0.1707 | 9.99E-13 |
| B62 | B152 | WBHSRVI | ACA-Ig(G-A-M) | 0.161981 | 1.37E-11 |
| B62 | B119 | WBHSRVI | Hcv-rna | -0.16195 | 1.38E-11 |
| B62 | B159 | WBHSRVI | β2-MG | 0.161946 | 1.38E-11 |
| B62 | B137 | WBHSRVI | AFU | 0.159364 | 2.91E-11 |
| B62 | B92 | WBHSRVI | ALDH2-Glu504Glu | 0.142957 | 2.54E-09 |
| B62 | B93 | WBHSRVI | ALDH2-Glu504Lys | -0.13443 | 2.15E-08 |
| B62 | B79 | WBHSRVI | hbⅤ-dna | -0.13255 | 3.38E-08 |
| B62 | B222 | WBHSRVI | NS | 0.128463 | 8.84E-08 |
| B62 | B179 | WBHSRVI | α1-MG | 0.12393 | 2.48E-07 |
| B62 | B134 | WBHSRVI | CHE | 0.120498 | 5.30E-07 |
| B63 | B64 | WBRSR(mPas_200) | WBRSR(mPas_5) | 0.899864 | 0 |
| B63 | B65 | WBRSR(mPas_200) | WBRSR(mPas_1) | 0.771134 | 0 |
| B63 | B66 | WBRSR(mPas_200) | WBRSR(mPas_100) | 0.992718 | 0 |
| B63 | B67 | WBRSR(mPas_200) | WBRSR(mPas_50) | 0.988302 | 0 |
| B63 | B73 | WBRSR(mPas_200) | ApoA | -0.2104 | 0 |
| B63 | B92 | WBRSR(mPas_200) | ALDH2-Glu504Glu | 0.258645 | 0 |
| B63 | B93 | WBRSR(mPas_200) | ALDH2-Glu504Lys | -0.25772 | 0 |
| B63 | B69 | WBRSR(mPas_200) | PV | 0.196525 | 2.22E-16 |
| B63 | B125 | WBRSR(mPas_200) | Cor | -0.18926 | 2.22E-15 |
| B63 | B124 | WBRSR(mPas_200) | ACTH | -0.1766 | 1.57E-13 |
| B63 | B74 | WBRSR(mPas_200) | ApoA1 | -0.16804 | 2.25E-12 |
| B63 | B121 | WBRSR(mPas_200) | C3 | 0.164956 | 5.68E-12 |
| B63 | B122 | WBRSR(mPas_200) | C4 | -0.15845 | 3.79E-11 |
| B63 | B158 | WBRSR(mPas_200) | IgM | 0.158078 | 4.21E-11 |
| B63 | B194 | WBRSR(mPas_200) | Cl | -0.15287 | 1.81E-10 |

| | | | | | |
|---|---|---|---|---|---|
| B63 | B198 | WBRSR(mPas_200) | P | -0.14133 | 3.86E-09 |
| B63 | B115 | WBRSR(mPas_200) | TSH TT3 TT4(TT3) | 0.134953 | 1.89E-08 |
| B63 | B137 | WBRSR(mPas_200) | AFU | 0.134552 | 2.08E-08 |
| B63 | B182 | WBRSR(mPas_200) | TT | -0.12244 | 3.46E-07 |
| B63 | B222 | WBRSR(mPas_200) | NS | 0.120661 | 5.12E-07 |
| B64 | B121 | WBRSR(mPas_5) | C3 | 0.304924 | 0 |
| B64 | B122 | WBRSR(mPas_5) | C4 | -0.29661 | 0 |
| B64 | B158 | WBRSR(mPas_5) | IgM | 0.206613 | 0 |
| B64 | B198 | WBRSR(mPas_5) | P | -0.22719 | 0 |
| B64 | B65 | WBRSR(mPas_5) | WBRSR(mPas_1) | 0.964521 | 0 |
| B64 | B66 | WBRSR(mPas_5) | WBRSR(mPas_100) | 0.929791 | 0 |
| B64 | B67 | WBRSR(mPas_5) | WBRSR(mPas_50) | 0.945663 | 0 |
| B64 | B69 | WBRSR(mPas_5) | PV | 0.216858 | 0 |
| B64 | B194 | WBRSR(mPas_5) | Cl | -0.19555 | 2.22E-16 |
| B64 | B73 | WBRSR(mPas_5) | ApoA | -0.19485 | 4.44E-16 |
| B64 | B92 | WBRSR(mPas_5) | ALDH2-Glu504Glu | 0.191728 | 8.88E-16 |
| B64 | B125 | WBRSR(mPas_5) | Cor | -0.19115 | 1.33E-15 |
| B64 | B93 | WBRSR(mPas_5) | ALDH2-Glu504Lys | -0.18773 | 4.00E-15 |
| B64 | B124 | WBRSR(mPas_5) | ACTH | -0.17466 | 2.91E-13 |
| B64 | B100 | WBRSR(mPas_5) | anti-Ro52 | 0.147334 | 8.10E-10 |
| B64 | B137 | WBRSR(mPas_5) | AFU | 0.141971 | 3.28E-09 |
| B64 | B101 | WBRSR(mPas_5) | anti-Sc70 | -0.13822 | 8.46E-09 |
| B64 | B102 | WBRSR(mPas_5) | anti-Sm | -0.13822 | 8.46E-09 |
| B64 | B103 | WBRSR(mPas_5) | anti-SSA/Ro | -0.13822 | 8.46E-09 |
| B64 | B104 | WBRSR(mPas_5) | anti-SSB | -0.13822 | 8.46E-09 |
| B64 | B105 | WBRSR(mPas_5) | anti-U1RNP | -0.13822 | 8.46E-09 |
| B64 | B106 | WBRSR(mPas_5) | anti-P | -0.13822 | 8.46E-09 |
| B64 | B107 | WBRSR(mPas_5) | AnuA | -0.13822 | 8.46E-09 |
| B64 | B108 | WBRSR(mPas_5) | Anti-Cyclin | -0.13822 | 8.46E-09 |
| B64 | B109 | WBRSR(mPas_5) | AMA-M2 | -0.13822 | 8.46E-09 |
| B64 | B110 | WBRSR(mPas_5) | AHA | -0.13822 | 8.46E-09 |
| B64 | B95 | WBRSR(mPas_5) | anti-cenpb | -0.13822 | 8.46E-09 |
| B64 | B97 | WBRSR(mPas_5) | dsDNA | -0.13822 | 8.46E-09 |
| B64 | B98 | WBRSR(mPas_5) | anti-Jo-1 | -0.13822 | 8.46E-09 |
| B64 | B99 | WBRSR(mPas_5) | anti-PM-Scl | -0.13822 | 8.46E-09 |
| B64 | B134 | WBRSR(mPas_5) | CHE | 0.136759 | 1.21E-08 |
| B64 | B222 | WBRSR(mPas_5) | NS | 0.120588 | 5.20E-07 |
| B64 | B156 | WBRSR(mPas_5) | IgE | 0.119644 | 6.39E-07 |
| B64 | B74 | WBRSR(mPas_5) | ApoA1 | -0.11768 | 9.75E-07 |
| B65 | B100 | WBRSR(mPas_1) | anti-Ro52 | 0.272732 | 0 |
| B65 | B121 | WBRSR(mPas_1) | C3 | 0.342131 | 0 |
| B65 | B122 | WBRSR(mPas_1) | C4 | -0.3346 | 0 |
| B65 | B158 | WBRSR(mPas_1) | IgM | 0.202338 | 0 |
| B65 | B194 | WBRSR(mPas_1) | Cl | -0.22008 | 0 |

| | | | | | |
|---|---|---|---|---|---|
| B65 | B198 | WBRSR(mPas_1) | P | -0.2364 | 0 |
| B65 | B66 | WBRSR(mPas_1) | WBRSR(mPas_100) | 0.814691 | 0 |
| B65 | B67 | WBRSR(mPas_1) | WBRSR(mPas_50) | 0.845969 | 0 |
| B65 | B69 | WBRSR(mPas_1) | PV | 0.214734 | 0 |
| B65 | B156 | WBRSR(mPas_1) | IgE | 0.167387 | 2.74E-12 |
| B65 | B73 | WBRSR(mPas_1) | ApoA | -0.16448 | 6.54E-12 |
| B65 | B125 | WBRSR(mPas_1) | Cor | -0.16161 | 1.52E-11 |
| B65 | B101 | WBRSR(mPas_1) | anti-Sc70 | -0.15047 | 3.49E-10 |
| B65 | B102 | WBRSR(mPas_1) | anti-Sm | -0.15047 | 3.49E-10 |
| B65 | B103 | WBRSR(mPas_1) | anti-SSA/Ro | -0.15047 | 3.49E-10 |
| B65 | B104 | WBRSR(mPas_1) | anti-SSB | -0.15047 | 3.49E-10 |
| B65 | B105 | WBRSR(mPas_1) | anti-U1RNP | -0.15047 | 3.49E-10 |
| B65 | B106 | WBRSR(mPas_1) | anti-P | -0.15047 | 3.49E-10 |
| B65 | B107 | WBRSR(mPas_1) | AnuA | -0.15047 | 3.49E-10 |
| B65 | B108 | WBRSR(mPas_1) | Anti-Cyclin | -0.15047 | 3.49E-10 |
| B65 | B109 | WBRSR(mPas_1) | AMA-M2 | -0.15047 | 3.49E-10 |
| B65 | B110 | WBRSR(mPas_1) | AHA | -0.15047 | 3.49E-10 |
| B65 | B95 | WBRSR(mPas_1) | anti-cenpb | -0.15047 | 3.49E-10 |
| B65 | B97 | WBRSR(mPas_1) | dsDNA | -0.15047 | 3.49E-10 |
| B65 | B98 | WBRSR(mPas_1) | anti-Jo-1 | -0.15047 | 3.49E-10 |
| B65 | B99 | WBRSR(mPas_1) | anti-PM-Scl | -0.15047 | 3.49E-10 |
| B65 | B124 | WBRSR(mPas_1) | ACTH | -0.15035 | 3.61E-10 |
| B65 | B134 | WBRSR(mPas_1) | CHE | 0.145052 | 1.48E-09 |
| B65 | B92 | WBRSR(mPas_1) | ALDH2-Glu504Glu | 0.131658 | 4.17E-08 |
| B65 | B93 | WBRSR(mPas_1) | ALDH2-Glu504Lys | -0.12698 | 1.24E-07 |
| B65 | B137 | WBRSR(mPas_1) | AFU | 0.124004 | 2.44E-07 |
| B65 | B120 | WBRSR(mPas_1) | HCV-cAg | -0.1211 | 4.65E-07 |
| B66 | B125 | WBRSR(mPas_100) | Cor | -0.21061 | 0 |
| B66 | B67 | WBRSR(mPas_100) | WBRSR(mPas_50) | 0.99705 | 0 |
| B66 | B69 | WBRSR(mPas_100) | PV | 0.214928 | 0 |
| B66 | B73 | WBRSR(mPas_100) | ApoA | -0.21067 | 0 |
| B66 | B92 | WBRSR(mPas_100) | ALDH2-Glu504Glu | 0.255946 | 0 |
| B66 | B93 | WBRSR(mPas_100) | ALDH2-Glu504Lys | -0.25343 | 0 |
| B66 | B121 | WBRSR(mPas_100) | C3 | 0.196454 | 2.22E-16 |
| B66 | B122 | WBRSR(mPas_100) | C4 | -0.18787 | 3.77E-15 |
| B66 | B124 | WBRSR(mPas_100) | ACTH | -0.18597 | 7.33E-15 |
| B66 | B74 | WBRSR(mPas_100) | ApoA1 | -0.17962 | 5.95E-14 |
| B66 | B158 | WBRSR(mPas_100) | IgM | 0.167973 | 2.30E-12 |
| B66 | B198 | WBRSR(mPas_100) | P | -0.1609 | 1.87E-11 |
| B66 | B194 | WBRSR(mPas_100) | Cl | -0.15923 | 3.03E-11 |
| B66 | B137 | WBRSR(mPas_100) | AFU | 0.143606 | 2.15E-09 |
| B66 | B115 | WBRSR(mPas_100) | TSH TT3 TT4(TT3) | 0.143202 | 2.39E-09 |
| B66 | B182 | WBRSR(mPas_100) | TT | -0.12729 | 1.16E-07 |
| B66 | B222 | WBRSR(mPas_100) | NS | 0.12359 | 2.68E-07 |

| | | | | | |
|---|---|---|---|---|---|
| B66 | B134 | WBRSR(mPas_100) | CHE | 0.120439 | 5.37E-07 |
| B66 | B101 | WBRSR(mPas_100) | anti-Sc70 | -0.11838 | 8.38E-07 |
| B66 | B102 | WBRSR(mPas_100) | anti-Sm | -0.11838 | 8.38E-07 |
| B66 | B103 | WBRSR(mPas_100) | anti-SSA/Ro | -0.11838 | 8.38E-07 |
| B66 | B104 | WBRSR(mPas_100) | anti-SSB | -0.11838 | 8.38E-07 |
| B66 | B105 | WBRSR(mPas_100) | anti-U1RNP | -0.11838 | 8.38E-07 |
| B66 | B106 | WBRSR(mPas_100) | anti-P | -0.11838 | 8.38E-07 |
| B66 | B107 | WBRSR(mPas_100) | AnuA | -0.11838 | 8.38E-07 |
| B66 | B108 | WBRSR(mPas_100) | Anti-Cyclin | -0.11838 | 8.38E-07 |
| B66 | B109 | WBRSR(mPas_100) | AMA-M2 | -0.11838 | 8.38E-07 |
| B66 | B110 | WBRSR(mPas_100) | AHA | -0.11838 | 8.38E-07 |
| B66 | B95 | WBRSR(mPas_100) | anti-cenpb | -0.11838 | 8.38E-07 |
| B66 | B97 | WBRSR(mPas_100) | dsDNA | -0.11838 | 8.38E-07 |
| B66 | B98 | WBRSR(mPas_100) | anti-Jo-1 | -0.11838 | 8.38E-07 |
| B66 | B99 | WBRSR(mPas_100) | anti-PM-Scl | -0.11838 | 8.38E-07 |
| B67 | B121 | WBRSR(mPas_50) | C3 | 0.21538 | 0 |
| B67 | B122 | WBRSR(mPas_50) | C4 | -0.20671 | 0 |
| B67 | B125 | WBRSR(mPas_50) | Cor | -0.20839 | 0 |
| B67 | B69 | WBRSR(mPas_50) | PV | 0.207615 | 0 |
| B67 | B73 | WBRSR(mPas_50) | ApoA | -0.20623 | 0 |
| B67 | B92 | WBRSR(mPas_50) | ALDH2-Glu504Glu | 0.243975 | 0 |
| B67 | B93 | WBRSR(mPas_50) | ALDH2-Glu504Lys | -0.24108 | 0 |
| B67 | B124 | WBRSR(mPas_50) | ACTH | -0.18367 | 1.58E-14 |
| B67 | B158 | WBRSR(mPas_50) | IgM | 0.174568 | 2.99E-13 |
| B67 | B74 | WBRSR(mPas_50) | ApoA1 | -0.17368 | 3.95E-13 |
| B67 | B194 | WBRSR(mPas_50) | Cl | -0.17293 | 5.00E-13 |
| B67 | B198 | WBRSR(mPas_50) | P | -0.16737 | 2.76E-12 |
| B67 | B137 | WBRSR(mPas_50) | AFU | 0.144055 | 1.91E-09 |
| B67 | B115 | WBRSR(mPas_50) | TSH TT3 TT4(TT3) | 0.140894 | 4.31E-09 |
| B67 | B134 | WBRSR(mPas_50) | CHE | 0.125187 | 1.87E-07 |
| B67 | B222 | WBRSR(mPas_50) | NS | 0.124953 | 1.97E-07 |
| B69 | B100 | PV | anti-Ro52 | 0.374087 | 0 |
| B69 | B92 | PV | ALDH2-Glu504Glu | 0.263249 | 0 |
| B69 | B93 | PV | ALDH2-Glu504Lys | -0.27866 | 0 |
| B69 | B94 | PV | ALDH2-Lys504Lys | 0.240579 | 0 |
| B69 | B151 | PV | Anti-CCP | 0.177098 | 1.34E-13 |
| B69 | B152 | PV | ACA-Ig(G-A-M) | -0.15634 | 6.89E-11 |
| B69 | B119 | PV | Hcv-rna | 0.156308 | 6.95E-11 |
| B69 | B159 | PV | β2-MG | -0.15631 | 6.95E-11 |
| B69 | B122 | PV | C4 | -0.15006 | 3.90E-10 |
| B69 | B121 | PV | C3 | 0.136921 | 1.17E-08 |
| B69 | B194 | PV | Cl | -0.13211 | 3.75E-08 |
| B69 | B79 | PV | hb V -dna | 0.127619 | 1.07E-07 |
| B69 | B158 | PV | IgM | -0.12632 | 1.45E-07 |

| | | | | | |
|---|---|---|---|---|---|
| B69 | B76 | PV | ApoB | -0.12577 | 1.64E-07 |
| B7 | B136 | WB-Cop | 5'NT | 0.317561 | 0 |
| B7 | B157 | WB-Cop | IgG | -0.23954 | 0 |
| B7 | B181 | WB-Cop | APTT | 0.254367 | 0 |
| B7 | B184 | WB-Cop | INR | 0.309115 | 0 |
| B7 | B8 | WB-Cop | WB-Zinc | 0.520338 | 0 |
| B7 | B155 | WB-Cop | IgA | -0.19441 | 4.44E-16 |
| B7 | B185 | WB-Cop | PT | 0.175143 | 2.49E-13 |
| B7 | B57 | WB-Cop | RBCAI | -0.16642 | 3.67E-12 |
| B7 | B138 | WB-Cop | PAB | 0.165365 | 5.03E-12 |
| B7 | B193 | WB-Cop | K | -0.16293 | 1.03E-11 |
| B7 | B158 | WB-Cop | IgM | -0.16002 | 2.42E-11 |
| B7 | B58 | WB-Cop | HCT | 0.146644 | 9.72E-10 |
| B7 | B156 | WB-Cop | IgE | -0.12586 | 1.61E-07 |
| B7 | B186 | WB-Cop | PFC | -0.12442 | 2.22E-07 |
| B7 | B137 | WB-Cop | AFU | 0.121204 | 4.55E-07 |
| B7 | B66 | WB-Cop | WBRSR(mPas_100) | 0.120415 | 5.40E-07 |
| B70 | B132 | LDL-C | Dbil | -0.25579 | 0 |
| B70 | B76 | LDL-C | ApoB | 0.238165 | 0 |
| B70 | B77 | LDL-C | TC | 0.801317 | 0 |
| B70 | B189 | LDL-C | eGFB | 0.15226 | 2.14E-10 |
| B70 | B75 | LDL-C | ApoA1/B | -0.15222 | 2.17E-10 |
| B70 | B72 | LDL-C | HDL-C | 0.134382 | 2.17E-08 |
| B70 | B153 | LDL-C | Glu | -0.11878 | 7.70E-07 |
| B71 | B130 | TG | GLB | 0.343881 | 0 |
| B71 | B153 | TG | Glu | 0.198564 | 0 |
| B71 | B167 | TG | PC | -0.19882 | 0 |
| B71 | B72 | TG | HDL-C | -0.40662 | 0 |
| B71 | B77 | TG | TC | 0.347883 | 0 |
| B71 | B91 | TG | TP | 0.401001 | 0 |
| B71 | B192 | TG | UA | 0.191365 | 1.11E-15 |
| B71 | B222 | TG | NS | 0.177157 | 1.31E-13 |
| B71 | B126 | TG | Alb | 0.173547 | 4.12E-13 |
| B71 | B127 | TG | A-G ratio | -0.16736 | 2.76E-12 |
| B71 | B216 | TG | BMI | 0.166489 | 3.59E-12 |
| B71 | B218 | TG | DBP | 0.16413 | 7.26E-12 |
| B71 | B140 | TG | r-GT | 0.161023 | 1.80E-11 |
| B71 | B163 | TG | SG | 0.154185 | 1.26E-10 |
| B71 | B128 | TG | ALT | 0.15071 | 3.27E-10 |
| B71 | B215 | TG | BMI | 0.141374 | 3.82E-09 |
| B71 | B141 | TG | AST/ALT | -0.14114 | 4.05E-09 |
| B71 | B170 | TG | U-GLU | 0.136568 | 1.27E-08 |
| B71 | B206 | TG | Waist | 0.136176 | 1.40E-08 |
| B71 | B132 | TG | Dbil | -0.12494 | 1.98E-07 |

| | | | | | |
|---|---|---|---|---|---|
| B71 | B166 | TG | Pro | 0.122761 | 3.22E-07 |
| B71 | B189 | TG | eGFB | 0.118108 | 8.89E-07 |
| B72 | B141 | HDL-C | AST/ALT | 0.23547 | 0 |
| B72 | B153 | HDL-C | Glu | -0.21882 | 0 |
| B72 | B163 | HDL-C | SG | -0.20164 | 0 |
| B72 | B190 | HDL-C | Cre | -0.20417 | 0 |
| B72 | B192 | HDL-C | UA | -0.35801 | 0 |
| B72 | B206 | HDL-C | Waist | -0.41942 | 0 |
| B72 | B207 | HDL-C | WHR | -0.28512 | 0 |
| B72 | B210 | HDL-C | Hip | -0.3023 | 0 |
| B72 | B215 | HDL-C | BMI | -0.3831 | 0 |
| B72 | B216 | HDL-C | BMI | -0.44293 | 0 |
| B72 | B221 | HDL-C | Hei | -0.3081 | 0 |
| B72 | B222 | HDL-C | NS | -0.46646 | 0 |
| B72 | B77 | HDL-C | TC | 0.186903 | 5.33E-15 |
| B72 | B74 | HDL-C | ApoA1 | 0.176484 | 1.63E-13 |
| B72 | B128 | HDL-C | ALT | -0.16618 | 3.94E-12 |
| B72 | B174 | HDL-C | Urea/Cr | 0.157244 | 5.33E-11 |
| B72 | B209 | HDL-C | Smoking | -0.15112 | 2.92E-10 |
| B73 | B137 | ApoA | AFU | -0.26408 | 0 |
| B73 | B193 | ApoA | K | 0.174415 | 3.14E-13 |
| B73 | B138 | ApoA | PAB | -0.14086 | 4.35E-09 |
| B73 | B134 | ApoA | CHE | -0.13723 | 1.08E-08 |
| B73 | B125 | ApoA | Cor | 0.132503 | 3.41E-08 |
| B73 | B135 | ApoA | TBA | -0.12889 | 8.00E-08 |
| B74 | B135 | ApoA1 | TBA | 0.266957 | 0 |
| B74 | B195 | ApoA1 | Na | 0.198073 | 0 |
| B74 | B197 | ApoA1 | AG | 0.324219 | 0 |
| B74 | B75 | ApoA1 | ApoA1/B | 0.508079 | 0 |
| B74 | B86 | ApoA1 | TBhCG | 0.202895 | 0 |
| B74 | B196 | ApoA1 | TCO2 | -0.17949 | 6.17E-14 |
| B74 | B76 | ApoA1 | ApoB | 0.168426 | 2.00E-12 |
| B74 | B138 | ApoA1 | PAB | -0.16585 | 4.35E-12 |
| B74 | B152 | ApoA1 | ACA-Ig(G-A-M) | -0.14057 | 4.68E-09 |
| B74 | B119 | ApoA1 | Hcv-rna | 0.140542 | 4.71E-09 |
| B74 | B159 | ApoA1 | β2-MG | -0.14054 | 4.72E-09 |
| B74 | B182 | ApoA1 | TT | 0.135859 | 1.51E-08 |
| B74 | B199 | ApoA1 | Mg | -0.12749 | 1.11E-07 |
| B74 | B183 | ApoA1 | PTA | -0.12258 | 3.36E-07 |
| B74 | B88 | ApoA1 | CA 242 | 0.120271 | 5.57E-07 |
| B74 | B200 | ApoA1 | OP | 0.119324 | 6.84E-07 |
| B75 | B119 | ApoA1/B | Hcv-rna | 0.197133 | 0 |
| B75 | B135 | ApoA1/B | TBA | 0.22158 | 0 |
| B75 | B152 | ApoA1/B | ACA-Ig(G-A-M) | -0.19717 | 0 |

| | | | | | |
|---|---|---|---|---|---|
| B75 | B159 | ApoA1/B | β2-MG | -0.19713 | 0 |
| B75 | B196 | ApoA1/B | TCO2 | -0.25946 | 0 |
| B75 | B197 | ApoA1/B | AG | 0.226304 | 0 |
| B75 | B198 | ApoA1/B | P | 0.210667 | 0 |
| B75 | B76 | ApoA1/B | ApoB | -0.70614 | 0 |
| B75 | B158 | ApoA1/B | IgM | -0.18474 | 1.11E-14 |
| B75 | B155 | ApoA1/B | IgA | 0.177159 | 1.31E-13 |
| B75 | B79 | ApoA1/B | hb V -dna | 0.17487 | 2.72E-13 |
| B75 | B199 | ApoA1/B | Mg | -0.1628 | 1.07E-11 |
| B75 | B125 | ApoA1/B | Cor | 0.159877 | 2.51E-11 |
| B75 | B179 | ApoA1/B | α1-MG | -0.14286 | 2.61E-09 |
| B75 | B156 | ApoA1/B | IgE | 0.14026 | 5.06E-09 |
| B75 | B94 | ApoA1/B | ALDH2-Lys504Lys | 0.13651 | 1.29E-08 |
| B75 | B138 | ApoA1/B | PAB | -0.13638 | 1.33E-08 |
| B75 | B180 | ApoA1/B | Tf | -0.13442 | 2.15E-08 |
| B75 | B134 | ApoA1/B | CHE | -0.1312 | 4.65E-08 |
| B75 | B124 | ApoA1/B | ACTH | 0.120616 | 5.17E-07 |
| B76 | B158 | ApoB | IgM | 0.19607 | 2.22E-16 |
| B76 | B77 | ApoB | TC | 0.188946 | 2.66E-15 |
| B76 | B152 | ApoB | ACA-Ig(G-A-M) | 0.179356 | 6.48E-14 |
| B76 | B119 | ApoB | Hcv-rna | -0.17932 | 6.55E-14 |
| B76 | B159 | ApoB | β2-MG | 0.179318 | 6.55E-14 |
| B76 | B79 | ApoB | hb V -dna | -0.16494 | 5.71E-12 |
| B76 | B125 | ApoB | Cor | -0.15986 | 2.53E-11 |
| B76 | B198 | ApoB | P | -0.15797 | 4.34E-11 |
| B76 | B155 | ApoB | IgA | -0.15678 | 6.08E-11 |
| B76 | B196 | ApoB | TCO2 | 0.151163 | 2.89E-10 |
| B76 | B124 | ApoB | ACTH | -0.14787 | 7.02E-10 |
| B76 | B115 | ApoB | TSH TT3 TT4(TT3) | 0.147397 | 7.96E-10 |
| B76 | B181 | ApoB | APTT | 0.138334 | 8.21E-09 |
| B76 | B94 | ApoB | ALDH2-Lys504Lys | -0.13617 | 1.40E-08 |
| B76 | B88 | ApoB | CA 242 | 0.123796 | 2.56E-07 |
| B77 | B130 | TC | GLB | 0.249616 | 0 |
| B77 | B132 | TC | Dbil | -0.27348 | 0 |
| B77 | B91 | TC | TP | 0.267983 | 0 |
| B77 | B127 | TC | A-G ratio | -0.16966 | 1.37E-12 |
| B77 | B206 | TC | Waist | -0.15657 | 6.46E-11 |
| B77 | B189 | TC | eGFB | 0.146918 | 9.04E-10 |
| B77 | B216 | TC | BMI | -0.13014 | 5.97E-08 |
| B77 | B222 | TC | NS | -0.1239 | 2.50E-07 |
| B78 | B79 | HBcAb | hb V -dna | 0.18978 | 2.00E-15 |
| B79 | B101 | hb V -dna | anti-Sc70 | 0.312626 | 0 |
| B79 | B102 | hb V -dna | anti-Sm | 0.312626 | 0 |
| B79 | B103 | hb V -dna | anti-SSA/Ro | 0.312626 | 0 |

| | | | | | |
|---|---|---|---|---|---|
| B79 | B104 | hb V -dna | anti-SSB | 0.312626 | 0 |
| B79 | B105 | hb V -dna | anti-U1RNP | 0.312626 | 0 |
| B79 | B106 | hb V -dna | anti-P | 0.312626 | 0 |
| B79 | B107 | hb V -dna | AnuA | 0.312626 | 0 |
| B79 | B108 | hb V -dna | Anti-Cyclin | 0.312626 | 0 |
| B79 | B109 | hb V -dna | AMA-M2 | 0.312626 | 0 |
| B79 | B110 | hb V -dna | AHA | 0.312626 | 0 |
| B79 | B119 | hb V -dna | Hcv-rna | 0.611302 | 0 |
| B79 | B120 | hb V -dna | HCV-cAg | 0.276786 | 0 |
| B79 | B124 | hb V -dna | ACTH | 0.322904 | 0 |
| B79 | B125 | hb V -dna | Cor | 0.499478 | 0 |
| B79 | B134 | hb V -dna | CHE | -0.21361 | 0 |
| B79 | B152 | hb V -dna | ACA-Ig(G-A-M) | -0.61142 | 0 |
| B79 | B158 | hb V -dna | IgM | -0.40409 | 0 |
| B79 | B159 | hb V -dna | β2-MG | -0.6113 | 0 |
| B79 | B92 | hb V -dna | ALDH2-Glu504Glu | -0.29786 | 0 |
| B79 | B93 | hb V -dna | ALDH2-Glu504Lys | 0.277306 | 0 |
| B79 | B94 | hb V -dna | ALDH2-Lys504Lys | 0.406584 | 0 |
| B79 | B95 | hb V -dna | anti-cenpb | 0.312626 | 0 |
| B79 | B97 | hb V -dna | dsDNA | 0.312626 | 0 |
| B79 | B98 | hb V -dna | anti-Jo-1 | 0.312626 | 0 |
| B79 | B99 | hb V -dna | anti-PM-Scl | 0.312626 | 0 |
| B79 | B179 | hb V -dna | α1-MG | -0.19057 | 1.55E-15 |
| B79 | B115 | hb V -dna | TSH TT3 TT4(TT3) | -0.18123 | 3.51E-14 |
| B79 | B156 | hb V -dna | IgE | 0.148676 | 5.66E-10 |
| B79 | B81 | hb V -dna | TSGF | 0.147341 | 8.08E-10 |
| B79 | B155 | hb V -dna | IgA | 0.134674 | 2.02E-08 |
| B79 | B151 | hb V -dna | Anti-CCP | 0.125644 | 1.69E-07 |
| B79 | B203 | hb V -dna | DH | -0.12321 | 2.92E-07 |
| B79 | B184 | hb V -dna | INR | 0.117779 | 9.54E-07 |
| B8 | B184 | WB-Zinc | INR | 0.210806 | 0 |
| B8 | B181 | WB-Zinc | APTT | 0.173829 | 3.77E-13 |
| B8 | B123 | WB-Zinc | hs-CRP1 | 0.130822 | 5.09E-08 |
| B8 | B185 | WB-Zinc | PT | 0.119388 | 6.75E-07 |
| B8 | B86 | WB-Zinc | TBhCG | -0.11806 | 8.98E-07 |
| B81 | B125 | TSGF | Cor | 0.22092 | 0 |
| B81 | B152 | TSGF | ACA-Ig(G-A-M) | -0.18032 | 4.75E-14 |
| B81 | B119 | TSGF | Hcv-rna | 0.180279 | 4.80E-14 |
| B81 | B159 | TSGF | β2-MG | -0.18028 | 4.80E-14 |
| B81 | B94 | TSGF | ALDH2-Lys504Lys | 0.180279 | 4.80E-14 |
| B81 | B124 | TSGF | ACTH | 0.142981 | 2.53E-09 |
| B81 | B158 | TSGF | IgM | -0.13754 | 1.00E-08 |
| B81 | B92 | TSGF | ALDH2-Glu504Glu | -0.13127 | 4.58E-08 |
| B81 | B93 | TSGF | ALDH2-Glu504Lys | 0.12215 | 3.69E-07 |

| | | | | | |
|---|---|---|---|---|---|
| B82 | B83 | CEA | AFP | 0.213956 | 0 |
| B82 | B85 | CEA | SCC | 0.338078 | 0 |
| B82 | B88 | CEA | CA 242 | 0.383507 | 0 |
| B82 | B87 | CEA | NSE | 0.180443 | 4.53E-14 |
| B82 | B138 | CEA | PAB | -0.16441 | 6.69E-12 |
| B82 | B84 | CEA | C-PSA | 0.160429 | 2.14E-11 |
| B82 | B196 | CEA | TCO2 | 0.134492 | 2.11E-08 |
| B82 | B183 | CEA | PTA | -0.13022 | 5.86E-08 |
| B82 | B181 | CEA | APTT | 0.128377 | 9.02E-08 |
| B83 | B137 | AFP | AFU | 0.210057 | 0 |
| B83 | B84 | AFP | C-PSA | 0.411867 | 0 |
| B83 | B87 | AFP | NSE | 0.404988 | 0 |
| B83 | B89 | AFP | CA 72-4 | 0.142585 | 2.80E-09 |
| B83 | B86 | AFP | TBhCG | -0.1369 | 1.17E-08 |
| B84 | B197 | C-PSA | AG | -0.1973 | 0 |
| B84 | B87 | C-PSA | NSE | 0.3458 | 0 |
| B84 | B88 | C-PSA | CA 242 | 0.273804 | 0 |
| B84 | B85 | C-PSA | SCC | 0.189116 | 2.66E-15 |
| B84 | B114 | C-PSA | TSH TT3 TT4(TT4) | -0.15354 | 1.51E-10 |
| B84 | B196 | C-PSA | TCO2 | 0.150333 | 3.62E-10 |
| B85 | B87 | SCC | NSE | 0.407628 | 0 |
| B85 | B88 | SCC | CA 242 | 0.413618 | 0 |
| B85 | B89 | SCC | CA 72-4 | 0.173786 | 3.82E-13 |
| B85 | B181 | SCC | APTT | 0.134294 | 2.22E-08 |
| B85 | B86 | SCC | TBhCG | 0.128594 | 8.57E-08 |
| B86 | B138 | TBhCG | PAB | -0.24672 | 0 |
| B86 | B197 | TBhCG | AG | 0.237617 | 0 |
| B86 | B199 | TBhCG | Mg | -0.22495 | 0 |
| B86 | B87 | TBhCG | NSE | -0.24355 | 0 |
| B86 | B88 | TBhCG | CA 242 | 0.358697 | 0 |
| B86 | B90 | TBhCG | CYFRA 21-1 | 0.236773 | 0 |
| B86 | B139 | TBhCG | ADA | 0.185733 | 7.99E-15 |
| B86 | B115 | TBhCG | TSH TT3 TT4(TT3) | 0.165253 | 5.20E-12 |
| B86 | B136 | TBhCG | 5'NT | 0.158556 | 3.67E-11 |
| B86 | B137 | TBhCG | AFU | -0.15515 | 9.63E-11 |
| B86 | B196 | TBhCG | TCO2 | -0.15265 | 1.93E-10 |
| B86 | B181 | TBhCG | APTT | 0.145662 | 1.26E-09 |
| B86 | B114 | TBhCG | TSH TT3 TT4(TT4) | 0.141454 | 3.74E-09 |
| B86 | B89 | TBhCG | CA 72-4 | 0.136049 | 1.45E-08 |
| B86 | B183 | TBhCG | PTA | -0.12326 | 2.88E-07 |
| B87 | B88 | NSE | CA 242 | 0.179368 | 6.44E-14 |
| B87 | B155 | NSE | IgA | 0.123634 | 2.65E-07 |
| B88 | B138 | CA 242 | PAB | -0.18141 | 3.31E-14 |
| B88 | B139 | CA 242 | ADA | 0.161726 | 1.47E-11 |

| | | | | | |
|---|---|---|---|---|---|
| B88 | B136 | CA 242 | 5'NT | 0.130729 | 5.20E-08 |
| B88 | B137 | CA 242 | AFU | -0.11922 | 7.00E-07 |
| B89 | B137 | CA 72-4 | AFU | 0.333274 | 0 |
| B89 | B138 | CA 72-4 | PAB | 0.153954 | 1.34E-10 |
| B89 | B139 | CA 72-4 | ADA | -0.14598 | 1.16E-09 |
| B89 | B134 | CA 72-4 | CHE | 0.122289 | 3.58E-07 |
| B9 | B35 | RH | β2-MG | 0.713974 | 0 |
| B90 | B184 | CYFRA 21-1 | INR | 0.227589 | 0 |
| B90 | B185 | CYFRA 21-1 | PT | 0.24048 | 0 |
| B90 | B123 | CYFRA 21-1 | hs-CRP1 | 0.153738 | 1.43E-10 |
| B90 | B187 | CYFRA 21-1 | NSE | 0.119755 | 6.23E-07 |
| B91 | B126 | TP | Alb | 0.499596 | 0 |
| B91 | B127 | TP | A-G ratio | -0.4909 | 0 |
| B91 | B130 | TP | GLB | 0.812667 | 0 |
| B91 | B188 | TP | Ca | 0.254618 | 0 |
| B91 | B218 | TP | DBP | 0.138361 | 8.16E-09 |
| B92 | B119 | ALDH2-Glu504Glu | Hcv-rna | -0.36478 | 0 |
| B92 | B124 | ALDH2-Glu504Glu | ACTH | -0.2894 | 0 |
| B92 | B125 | ALDH2-Glu504Glu | Cor | -0.44727 | 0 |
| B92 | B152 | ALDH2-Glu504Glu | ACA-Ig(G-A-M) | 0.364851 | 0 |
| B92 | B158 | ALDH2-Glu504Glu | IgM | 0.358725 | 0 |
| B92 | B159 | ALDH2-Glu504Glu | β2-MG | 0.364774 | 0 |
| B92 | B93 | ALDH2-Glu504Glu | ALDH2-Glu504Lys | -0.99847 | 0 |
| B92 | B101 | ALDH2-Glu504Glu | anti-Sc70 | -0.18845 | 3.11E-15 |
| B92 | B102 | ALDH2-Glu504Glu | anti-Sm | -0.18845 | 3.11E-15 |
| B92 | B103 | ALDH2-Glu504Glu | anti-SSA/Ro | -0.18845 | 3.11E-15 |
| B92 | B104 | ALDH2-Glu504Glu | anti-SSB | -0.18845 | 3.11E-15 |
| B92 | B105 | ALDH2-Glu504Glu | anti-U1RNP | -0.18845 | 3.11E-15 |
| B92 | B106 | ALDH2-Glu504Glu | anti-P | -0.18845 | 3.11E-15 |
| B92 | B107 | ALDH2-Glu504Glu | AnuA | -0.18845 | 3.11E-15 |
| B92 | B108 | ALDH2-Glu504Glu | Anti-Cyclin | -0.18845 | 3.11E-15 |
| B92 | B109 | ALDH2-Glu504Glu | AMA-M2 | -0.18845 | 3.11E-15 |
| B92 | B110 | ALDH2-Glu504Glu | AHA | -0.18845 | 3.11E-15 |
| B92 | B95 | ALDH2-Glu504Glu | anti-cenpb | -0.18845 | 3.11E-15 |
| B92 | B97 | ALDH2-Glu504Glu | dsDNA | -0.18845 | 3.11E-15 |
| B92 | B98 | ALDH2-Glu504Glu | anti-Jo-1 | -0.18845 | 3.11E-15 |
| B92 | B99 | ALDH2-Glu504Glu | anti-PM-Scl | -0.18845 | 3.11E-15 |
| B92 | B114 | ALDH2-Glu504Glu | TSH TT3 TT4(TT4) | -0.17109 | 8.84E-13 |
| B92 | B179 | ALDH2-Glu504Glu | α1-MG | 0.169721 | 1.35E-12 |
| B92 | B120 | ALDH2-Glu504Glu | HCV-cAg | -0.16747 | 2.68E-12 |
| B92 | B94 | ALDH2-Glu504Glu | ALDH2-Lys504Lys | -0.15928 | 2.98E-11 |
| B92 | B156 | ALDH2-Glu504Glu | IgE | -0.13423 | 2.25E-08 |
| B92 | B134 | ALDH2-Glu504Glu | CHE | 0.124259 | 2.31E-07 |
| B92 | B155 | ALDH2-Glu504Glu | IgA | -0.11991 | 6.03E-07 |

| | | | | | |
|---|---|---|---|---|---|
| B93 | B119 | ALDH2-Glu504Lys | Hcv-rna | 0.339553 | 0 |
| B93 | B124 | ALDH2-Glu504Lys | ACTH | 0.269373 | 0 |
| B93 | B125 | ALDH2-Glu504Lys | Cor | 0.416298 | 0 |
| B93 | B152 | ALDH2-Glu504Lys | ACA-Ig(G-A-M) | -0.33962 | 0 |
| B93 | B158 | ALDH2-Glu504Lys | IgM | -0.33373 | 0 |
| B93 | B159 | ALDH2-Glu504Lys | β2-MG | -0.33955 | 0 |
| B93 | B101 | ALDH2-Glu504Lys | anti-Sc70 | 0.175562 | 2.18E-13 |
| B93 | B102 | ALDH2-Glu504Lys | anti-Sm | 0.175562 | 2.18E-13 |
| B93 | B103 | ALDH2-Glu504Lys | anti-SSA/Ro | 0.175562 | 2.18E-13 |
| B93 | B104 | ALDH2-Glu504Lys | anti-SSB | 0.175562 | 2.18E-13 |
| B93 | B105 | ALDH2-Glu504Lys | anti-U1RNP | 0.175562 | 2.18E-13 |
| B93 | B106 | ALDH2-Glu504Lys | anti-P | 0.175562 | 2.18E-13 |
| B93 | B107 | ALDH2-Glu504Lys | AnuA | 0.175561 | 2.18E-13 |
| B93 | B108 | ALDH2-Glu504Lys | Anti-Cyclin | 0.175562 | 2.18E-13 |
| B93 | B109 | ALDH2-Glu504Lys | AMA-M2 | 0.175562 | 2.18E-13 |
| B93 | B110 | ALDH2-Glu504Lys | AHA | 0.175562 | 2.18E-13 |
| B93 | B95 | ALDH2-Glu504Lys | anti-cenpb | 0.175562 | 2.18E-13 |
| B93 | B97 | ALDH2-Glu504Lys | dsDNA | 0.175562 | 2.18E-13 |
| B93 | B98 | ALDH2-Glu504Lys | anti-Jo-1 | 0.175562 | 2.18E-13 |
| B93 | B99 | ALDH2-Glu504Lys | anti-PM-Scl | 0.175562 | 2.18E-13 |
| B93 | B114 | ALDH2-Glu504Lys | TSH TT3 TT4(TT4) | 0.166503 | 3.58E-12 |
| B93 | B179 | ALDH2-Glu504Lys | α1-MG | -0.15794 | 4.38E-11 |
| B93 | B120 | ALDH2-Glu504Lys | HCV-cAg | 0.156061 | 7.45E-11 |
| B93 | B156 | ALDH2-Glu504Lys | IgE | 0.124988 | 1.96E-07 |
| B94 | B101 | ALDH2-Lys504Lys | anti-Sc70 | 0.255109 | 0 |
| B94 | B102 | ALDH2-Lys504Lys | anti-Sm | 0.255109 | 0 |
| B94 | B103 | ALDH2-Lys504Lys | anti-SSA/Ro | 0.255109 | 0 |
| B94 | B104 | ALDH2-Lys504Lys | anti-SSB | 0.255109 | 0 |
| B94 | B105 | ALDH2-Lys504Lys | anti-U1RNP | 0.255109 | 0 |
| B94 | B106 | ALDH2-Lys504Lys | anti-P | 0.255109 | 0 |
| B94 | B107 | ALDH2-Lys504Lys | AnuA | 0.255109 | 0 |
| B94 | B108 | ALDH2-Lys504Lys | Anti-Cyclin | 0.255109 | 0 |
| B94 | B109 | ALDH2-Lys504Lys | AMA-M2 | 0.255109 | 0 |
| B94 | B110 | ALDH2-Lys504Lys | AHA | 0.255109 | 0 |
| B94 | B115 | ALDH2-Lys504Lys | TSH TT3 TT4(TT3) | -0.21858 | 0 |
| B94 | B119 | ALDH2-Lys504Lys | Hcv-rna | 0.498836 | 0 |
| B94 | B120 | ALDH2-Lys504Lys | HCV-cAg | 0.225864 | 0 |
| B94 | B124 | ALDH2-Lys504Lys | ACTH | 0.396008 | 0 |
| B94 | B125 | ALDH2-Lys504Lys | Cor | 0.612351 | 0 |
| B94 | B152 | ALDH2-Lys504Lys | ACA-Ig(G-A-M) | -0.49894 | 0 |
| B94 | B158 | ALDH2-Lys504Lys | IgM | -0.49386 | 0 |
| B94 | B159 | ALDH2-Lys504Lys | β2-MG | -0.49883 | 0 |
| B94 | B179 | ALDH2-Lys504Lys | α1-MG | -0.23286 | 0 |
| B94 | B95 | ALDH2-Lys504Lys | anti-cenpb | 0.255109 | 0 |

| | | | | | |
|---|---|---|---|---|---|
| B94 | B97 | ALDH2-Lys504Lys | dsDNA | 0.255109 | 0 |
| B94 | B98 | ALDH2-Lys504Lys | anti-Jo-1 | 0.255109 | 0 |
| B94 | B99 | ALDH2-Lys504Lys | anti-PM-Scl | 0.255109 | 0 |
| B94 | B156 | ALDH2-Lys504Lys | IgE | 0.182861 | 2.04E-14 |
| B94 | B134 | ALDH2-Lys504Lys | CHE | -0.16818 | 2.15E-12 |
| B94 | B155 | ALDH2-Lys504Lys | IgA | 0.16461 | 6.30E-12 |
| B94 | B151 | ALDH2-Lys504Lys | Anti-CCP | 0.155926 | 7.74E-11 |
| B94 | B203 | ALDH2-Lys504Lys | DH | -0.15198 | 2.31E-10 |
| B94 | B184 | ALDH2-Lys504Lys | INR | 0.144909 | 1.53E-09 |
| B94 | B181 | ALDH2-Lys504Lys | APTT | 0.132409 | 3.49E-08 |
| B94 | B186 | ALDH2-Lys504Lys | PFC | -0.132 | 3.85E-08 |
| B94 | B182 | ALDH2-Lys504Lys | TT | 0.125636 | 1.69E-07 |
| B94 | B199 | ALDH2-Lys504Lys | Mg | -0.1191 | 7.18E-07 |
| B95 | B103 | anti-cenpb | anti-SSA/Ro | 1 | 0 |
| B95 | B104 | anti-cenpb | anti-SSB | 1 | 0 |
| B95 | B105 | anti-cenpb | anti-U1RNP | 1 | 0 |
| B95 | B107 | anti-cenpb | AnuA | 0.999999 | 0 |
| B95 | B108 | anti-cenpb | Anti-Cyclin | 1 | 0 |
| B95 | B110 | anti-cenpb | AHA | 1 | 0 |
| B95 | B119 | anti-cenpb | Hcv-rna | 0.384926 | 0 |
| B95 | B120 | anti-cenpb | HCV-cAg | 0.349051 | 0 |
| B95 | B121 | anti-cenpb | C3 | -0.27501 | 0 |
| B95 | B122 | anti-cenpb | C4 | 0.242862 | 0 |
| B95 | B125 | anti-cenpb | Cor | 0.298639 | 0 |
| B95 | B134 | anti-cenpb | CHE | -0.23432 | 0 |
| B95 | B151 | anti-cenpb | Anti-CCP | 0.245648 | 0 |
| B95 | B152 | anti-cenpb | ACA-Ig(G-A-M) | -0.385 | 0 |
| B95 | B156 | anti-cenpb | IgE | 0.234142 | 0 |
| B95 | B157 | anti-cenpb | IgG | 0.328845 | 0 |
| B95 | B159 | anti-cenpb | β2-MG | -0.38492 | 0 |
| B95 | B97 | anti-cenpb | dsDNA | 1 | 0 |
| B95 | B138 | anti-cenpb | PAB | -0.18058 | 4.35E-14 |
| B95 | B154 | anti-cenpb | RF | 0.158265 | 3.99E-11 |
| B95 | B178 | anti-cenpb | Ig-U | 0.151079 | 2.96E-10 |
| B95 | B193 | anti-cenpb | K | 0.125601 | 1.70E-07 |
| B95 | B182 | anti-cenpb | TT | 0.124542 | 2.17E-07 |
| B95 | B183 | anti-cenpb | PTA | -0.11862 | 7.97E-07 |
| B96 | B116 | dsDNA (IIF) | TT3 | -0.86844 | 0 |
| B97 | B101 | dsDNA | anti-Sc70 | 1 | 0 |
| B97 | B102 | dsDNA | anti-Sm | 1 | 0 |
| B97 | B104 | dsDNA | anti-SSB | 1 | 0 |
| B97 | B106 | dsDNA | anti-P | 1 | 0 |
| B97 | B107 | dsDNA | AnuA | 0.999999 | 0 |
| B97 | B110 | dsDNA | AHA | 1 | 0 |

| | | | | | |
|---|---|---|---|---|---|
| B97 | B119 | dsDNA | Hcv-rna | 0.384926 | 0 |
| B97 | B120 | dsDNA | HCV-cAg | 0.349051 | 0 |
| B97 | B121 | dsDNA | C3 | -0.27501 | 0 |
| B97 | B122 | dsDNA | C4 | 0.242862 | 0 |
| B97 | B125 | dsDNA | Cor | 0.298639 | 0 |
| B97 | B134 | dsDNA | CHE | -0.23432 | 0 |
| B97 | B151 | dsDNA | Anti-CCP | 0.245648 | 0 |
| B97 | B152 | dsDNA | ACA-Ig(G-A-M) | -0.38501 | 0 |
| B97 | B156 | dsDNA | IgE | 0.234142 | 0 |
| B97 | B157 | dsDNA | IgG | 0.328845 | 0 |
| B97 | B159 | dsDNA | β2-MG | -0.38492 | 0 |
| B97 | B99 | dsDNA | anti-PM-Scl | 1 | 0 |
| B97 | B138 | dsDNA | PAB | -0.18058 | 4.35E-14 |
| B97 | B154 | dsDNA | RF | 0.158265 | 3.99E-11 |
| B97 | B178 | dsDNA | Ig-U | 0.151079 | 2.96E-10 |
| B97 | B193 | dsDNA | K | 0.125601 | 1.70E-07 |
| B97 | B182 | dsDNA | TT | 0.124542 | 2.17E-07 |
| B97 | B183 | dsDNA | PTA | -0.11862 | 7.97E-07 |
| B98 | B107 | anti-Jo-1 | AnuA | 0.999999 | 0 |
| B98 | B119 | anti-Jo-1 | Hcv-rna | 0.384926 | 0 |
| B98 | B120 | anti-Jo-1 | HCV-cAg | 0.349051 | 0 |
| B98 | B121 | anti-Jo-1 | C3 | -0.27501 | 0 |
| B98 | B122 | anti-Jo-1 | C4 | 0.242862 | 0 |
| B98 | B125 | anti-Jo-1 | Cor | 0.298639 | 0 |
| B98 | B134 | anti-Jo-1 | CHE | -0.23432 | 0 |
| B98 | B151 | anti-Jo-1 | Anti-CCP | 0.245648 | 0 |
| B98 | B152 | anti-Jo-1 | ACA-Ig(G-A-M) | -0.38501 | 0 |
| B98 | B156 | anti-Jo-1 | IgE | 0.234142 | 0 |
| B98 | B157 | anti-Jo-1 | IgG | 0.328845 | 0 |
| B98 | B159 | anti-Jo-1 | β2-MG | -0.38492 | 0 |
| B98 | B138 | anti-Jo-1 | PAB | -0.18058 | 4.35E-14 |
| B98 | B154 | anti-Jo-1 | RF | 0.158265 | 3.99E-11 |
| B98 | B178 | anti-Jo-1 | Ig-U | 0.151079 | 2.96E-10 |
| B98 | B193 | anti-Jo-1 | K | 0.125601 | 1.70E-07 |
| B98 | B182 | anti-Jo-1 | TT | 0.124542 | 2.17E-07 |
| B98 | B183 | anti-Jo-1 | PTA | -0.11862 | 7.97E-07 |
| B99 | B101 | anti-PM-Scl | anti-Sc70 | 1 | 0 |
| B99 | B102 | anti-PM-Scl | anti-Sm | 1 | 0 |
| B99 | B104 | anti-PM-Scl | anti-SSB | 1 | 0 |
| B99 | B107 | anti-PM-Scl | AnuA | 0.999999 | 0 |
| B99 | B108 | anti-PM-Scl | Anti-Cyclin | 1 | 0 |
| B99 | B119 | anti-PM-Scl | Hcv-rna | 0.384926 | 0 |
| B99 | B120 | anti-PM-Scl | HCV-cAg | 0.349051 | 0 |
| B99 | B121 | anti-PM-Scl | C3 | -0.27501 | 0 |

| | | | | | |
|---|---|---|---|---|---|
| B99 | B122 | anti-PM-Scl | C4 | 0.242862 | 0 |
| B99 | B125 | anti-PM-Scl | Cor | 0.298639 | 0 |
| B99 | B134 | anti-PM-Scl | CHE | -0.23432 | 0 |
| B99 | B151 | anti-PM-Scl | Anti-CCP | 0.245648 | 0 |
| B99 | B152 | anti-PM-Scl | ACA-Ig(G-A-M) | -0.38501 | 0 |
| B99 | B156 | anti-PM-Scl | IgE | 0.234142 | 0 |
| B99 | B157 | anti-PM-Scl | IgG | 0.328845 | 0 |
| B99 | B159 | anti-PM-Scl | β2-MG | -0.38492 | 0 |
| B99 | B138 | anti-PM-Scl | PAB | -0.18058 | 4.35E-14 |
| B99 | B154 | anti-PM-Scl | RF | 0.158265 | 3.99E-11 |
| B99 | B178 | anti-PM-Scl | Ig-U | 0.151079 | 2.96E-10 |
| B99 | B193 | anti-PM-Scl | K | 0.125601 | 1.70E-07 |
| B99 | B182 | anti-PM-Scl | TT | 0.124542 | 2.17E-07 |
| B99 | B183 | anti-PM-Scl | PTA | -0.11862 | 7.97E-07 |

**Supplementary Table 8. PEI Correlations in Coronary heart disease**

| Row | Column | Row | Column | Cor | *P* |
|---|---|---|---|---|---|
| B1 | B161 | Gender | LEU | 0.224483458 | 0 |
| B1 | B163 | Gender | SG | -0.30350997<br>2 | 0 |
| B1 | B174 | Gender | Urea/Cr | 0.360119848 | 0 |
| B1 | B190 | Gender | Cre | -0.44189101 | 0 |
| B1 | B192 | Gender | UA | -0.3679103 | 0 |
| B1 | B206 | Gender | Waist | -0.23172666<br>3 | 0 |
| B1 | B209 | Gender | Smoking | -0.23370356<br>2 | 0 |
| B1 | B216 | Gender | BMI | -0.43412196<br>9 | 0 |
| B1 | B221 | Gender | Hei | -0.64078416<br>5 | 0 |
| B1 | B222 | Gender | NS | -0.56397399<br>7 | 0 |
| B1 | B45 | Gender | MCHC | -0.27111803<br>1 | 0 |
| B1 | B46 | Gender | RBC Count | -0.35455233<br>8 | 0 |
| B1 | B51 | Gender | Hb | -0.47967226<br>3 | 0 |
| B1 | B72 | Gender | HDL-C | 0.353525548 | 0 |
| B1 | B77 | Gender | TC | 0.277378551 | 0 |
| B1 | B207 | Gender | WHR | -0.22094592<br>4 | 4.44E-16 |
| B1 | B42 | Gender | MON | -0.22006129<br>8 | 4.44E-16 |
| B1 | B141 | Gender | AST/ALT | 0.216882862 | 1.33E-15 |
| B1 | B205 | Gender | KDA | 0.211720674 | 6.66E-15 |
| B1 | B132 | Gender | Dbil | -0.20649327 | 3.15E-14 |
| B1 | B130 | Gender | GLB | 0.18629387 | 8.22E-12 |
| B1 | B2 | Gender | Age | 0.182663651 | 2.10E-11 |
| B1 | B219 | Gender | OT | -0.17884739<br>5 | 5.49E-11 |
| B1 | B204 | Gender | Drinking | -0.17815758<br>2 | 6.52E-11 |
| B1 | B127 | Gender | A-G ratio | -0.17775302 | 7.21E-11 |
| B1 | B128 | Gender | ALT | -0.17577808<br>3 | 1.17E-10 |
| B1 | B220 | Gender | SBP | 0.175649265 | 1.21E-10 |
| B1 | B41 | Gender | MON % | -0.17009921 | 4.63E-10 |

| | | | | | |
|---|---|---|---|---|---|
| | | | | 7 | |
| B1 | B133 | Gender | TBIL | -0.161908336 | 3.08E-09 |
| B1 | B70 | Gender | LDL-C | 0.154279223 | 1.66E-08 |
| B1 | B210 | Gender | Hip | -0.144681217 | 1.23E-07 |
| B1 | B48 | Gender | LYM% | 0.14414447 | 1.37E-07 |
| B1 | B50 | Gender | eo% | -0.143439195 | 1.58E-07 |
| B1 | B74 | Gender | ApoA1 | 0.141575684 | 2.28E-07 |
| B10 | B11 | PG Ⅰ | PGI/PGII | 0.261067925 | 0 |
| B10 | B12 | PG Ⅰ | PG Ⅱ | 0.457146812 | 0 |
| B10 | B165 | PG Ⅰ | bilirubin | 0.410825967 | 0 |
| B10 | B227 | PG Ⅰ | OD | 0.205903407 | 3.75E-14 |
| B10 | B14 | PG Ⅰ | G-17 | 0.17916346 | 5.08E-11 |
| B10 | B112 | PG Ⅰ | FT4 | 0.146588236 | 8.33E-08 |
| B100 | B119 | anti-Ro52 | Hcv-rna | 0.161376293 | 3.48E-09 |
| B100 | B197 | anti-Ro52 | AG | -0.150214952 | 3.93E-08 |
| B100 | B115 | anti-Ro52 | TSH TT3 TT4(TT3) | 0.142367131 | 1.95E-07 |
| B101 | B104 | anti-Sc70 | anti-SSB | 0.999999993 | 0 |
| B101 | B107 | anti-Sc70 | AnuA | 0.999999468 | 0 |
| B101 | B119 | anti-Sc70 | Hcv-rna | 0.470017916 | 0 |
| B101 | B120 | anti-Sc70 | HCV-cAg | 0.311176082 | 0 |
| B101 | B125 | anti-Sc70 | Cor | -0.319333509 | 0 |
| B101 | B152 | anti-Sc70 | ACA-Ig(G-A-M) | -0.263417922 | 0 |
| B101 | B154 | anti-Sc70 | RF | 0.277811861 | 0 |
| B101 | B155 | anti-Sc70 | IgA | -0.244461163 | 0 |
| B101 | B158 | anti-Sc70 | IgM | 0.292027928 | 0 |
| B101 | B159 | anti-Sc70 | β2-MG | -0.263404818 | 0 |
| B101 | B169 | anti-Sc70 | U-CRY | -0.263405908 | 0 |
| B101 | B197 | anti-Sc70 | AG | 0.240927349 | 0 |
| B101 | B199 | anti-Sc70 | Mg | 0.216390069 | 1.78E-15 |
| B101 | B137 | anti-Sc70 | AFU | 0.204688407 | 5.33E-14 |
| B101 | B134 | anti-Sc70 | CHE | 0.187404368 | 6.15E-12 |
| B101 | B200 | anti-Sc70 | OP | 0.187095702 | 6.66E-12 |
| B101 | B195 | anti-Sc70 | Na | 0.184163402 | 1.43E-11 |
| B101 | B193 | anti-Sc70 | K | 0.182545786 | 2.16E-11 |
| B101 | B108 | anti-Sc70 | Anti-Cyclin | -0.18087605 | 3.30E-11 |

| | | | | | |
|---|---|---|---|---|---|
| | | | | 3 | |
| B101 | B114 | anti-Sc70 | TSH TT3 TT4(TT4) | 0.173261898 | 2.17E-10 |
| B101 | B138 | anti-Sc70 | PAB | 0.164858125 | 1.57E-09 |
| B101 | B136 | anti-Sc70 | 5'NT | 0.151810374 | 2.81E-08 |
| B101 | B196 | anti-Sc70 | TCO2 | -0.138358662 | 4.28E-07 |
| B102 | B103 | anti-Sm | anti-SSA/Ro | 0.999999995 | 0 |
| B102 | B104 | anti-Sm | anti-SSB | 0.999999993 | 0 |
| B102 | B105 | anti-Sm | anti-U1RNP | 0.999999995 | 0 |
| B102 | B107 | anti-Sm | AnuA | 0.9999996 | 0 |
| B102 | B109 | anti-Sm | AMA-M2 | 0.999999997 | 0 |
| B102 | B110 | anti-Sm | AHA | 0.999999992 | 0 |
| B102 | B119 | anti-Sm | Hcv-rna | 0.470017904 | 0 |
| B102 | B120 | anti-Sm | HCV-cAg | 0.311176286 | 0 |
| B102 | B125 | anti-Sm | Cor | -0.319333499 | 0 |
| B102 | B152 | anti-Sm | ACA-Ig(G-A-M) | -0.26341618 | 0 |
| B102 | B154 | anti-Sm | RF | 0.277811854 | 0 |
| B102 | B155 | anti-Sm | IgA | -0.244461157 | 0 |
| B102 | B158 | anti-Sm | IgM | 0.292027921 | 0 |
| B102 | B159 | anti-Sm | β2-MG | -0.263403589 | 0 |
| B102 | B169 | anti-Sm | U-CRY | -0.263405001 | 0 |
| B102 | B197 | anti-Sm | AG | 0.240927343 | 0 |
| B102 | B199 | anti-Sm | Mg | 0.216390066 | 1.78E-15 |
| B102 | B137 | anti-Sm | AFU | 0.204688403 | 5.33E-14 |
| B102 | B134 | anti-Sm | CHE | 0.187404363 | 6.15E-12 |
| B102 | B200 | anti-Sm | OP | 0.187095702 | 6.66E-12 |
| B102 | B195 | anti-Sm | Na | 0.184163403 | 1.43E-11 |
| B102 | B193 | anti-Sm | K | 0.182545783 | 2.16E-11 |
| B102 | B108 | anti-Sm | Anti-Cyclin | -0.180876048 | 3.30E-11 |
| B102 | B114 | anti-Sm | TSH TT3 TT4(TT4) | 0.173261894 | 2.17E-10 |
| B102 | B138 | anti-Sm | PAB | 0.164858122 | 1.57E-09 |
| B102 | B136 | anti-Sm | 5'NT | 0.151810371 | 2.81E-08 |
| B102 | B196 | anti-Sm | TCO2 | -0.138358658 | 4.28E-07 |
| B103 | B104 | anti-SSA/Ro | anti-SSB | 0.999999999 | 0 |
| B103 | B107 | anti-SSA/Ro | AnuA | 0.999999435 | 0 |
| B103 | B119 | anti-SSA/Ro | Hcv-rna | 0.470017907 | 0 |
| B103 | B120 | anti-SSA/Ro | HCV-cAg | 0.311176207 | 0 |
| B103 | B125 | anti-SSA/Ro | Cor | -0.31933350 | 0 |

| | | | | | |
|---|---|---|---|---|---|
| | | | | 3 | |
| B103 | B152 | anti-SSA/Ro | ACA-Ig(G-A-M) | -0.263416228 | 0 |
| B103 | B154 | anti-SSA/Ro | RF | 0.277811855 | 0 |
| B103 | B155 | anti-SSA/Ro | IgA | -0.244461159 | 0 |
| B103 | B158 | anti-SSA/Ro | IgM | 0.292027922 | 0 |
| B103 | B159 | anti-SSA/Ro | β2-MG | -0.263404086 | 0 |
| B103 | B169 | anti-SSA/Ro | U-CRY | -0.263405449 | 0 |
| B103 | B197 | anti-SSA/Ro | AG | 0.240927345 | 0 |
| B103 | B199 | anti-SSA/Ro | Mg | 0.216390065 | 1.78E-15 |
| B103 | B137 | anti-SSA/Ro | AFU | 0.204688404 | 5.33E-14 |
| B103 | B134 | anti-SSA/Ro | CHE | 0.187404364 | 6.15E-12 |
| B103 | B200 | anti-SSA/Ro | OP | 0.187095704 | 6.66E-12 |
| B103 | B195 | anti-SSA/Ro | Na | 0.184163407 | 1.43E-11 |
| B103 | B193 | anti-SSA/Ro | K | 0.182545784 | 2.16E-11 |
| B103 | B108 | anti-SSA/Ro | Anti-Cyclin | -0.180876049 | 3.30E-11 |
| B103 | B114 | anti-SSA/Ro | TSH TT3 TT4(TT4) | 0.173261895 | 2.17E-10 |
| B103 | B138 | anti-SSA/Ro | PAB | 0.164858123 | 1.57E-09 |
| B103 | B136 | anti-SSA/Ro | 5'NT | 0.151810372 | 2.81E-08 |
| B103 | B196 | anti-SSA/Ro | TCO2 | -0.138358659 | 4.28E-07 |
| B104 | B106 | anti-SSB | anti-P | 0.99999999 | 0 |
| B104 | B107 | anti-SSB | AnuA | 0.999999493 | 0 |
| B104 | B110 | anti-SSB | AHA | 0.999999994 | 0 |
| B104 | B119 | anti-SSB | Hcv-rna | 0.470017908 | 0 |
| B104 | B120 | anti-SSB | HCV-cAg | 0.311176126 | 0 |
| B104 | B125 | anti-SSB | Cor | -0.319333503 | 0 |
| B104 | B152 | anti-SSB | ACA-Ig(G-A-M) | -0.263415582 | 0 |
| B104 | B154 | anti-SSB | RF | 0.277811855 | 0 |
| B104 | B155 | anti-SSB | IgA | -0.244461159 | 0 |
| B104 | B158 | anti-SSB | IgM | 0.292027922 | 0 |
| B104 | B159 | anti-SSB | β2-MG | -0.263404444 | 0 |
| B104 | B169 | anti-SSB | U-CRY | -0.263405697 | 0 |
| B104 | B197 | anti-SSB | AG | 0.240927345 | 0 |
| B104 | B199 | anti-SSB | Mg | 0.216390064 | 1.78E-15 |

| | | | | | |
|---|---|---|---|---|---|
| B104 | B137 | anti-SSB | AFU | 0.204688404 | 5.33E-14 |
| B104 | B134 | anti-SSB | CHE | 0.187404364 | 6.15E-12 |
| B104 | B200 | anti-SSB | OP | 0.187095705 | 6.66E-12 |
| B104 | B195 | anti-SSB | Na | 0.184163405 | 1.43E-11 |
| B104 | B193 | anti-SSB | K | 0.182545784 | 2.16E-11 |
| B104 | B108 | anti-SSB | Anti-Cyclin | -0.18087605 | 3.30E-11 |
| B104 | B114 | anti-SSB | TSH TT3 TT4(TT4) | 0.173261895 | 2.17E-10 |
| B104 | B138 | anti-SSB | PAB | 0.164858123 | 1.57E-09 |
| B104 | B136 | anti-SSB | 5'NT | 0.151810372 | 2.81E-08 |
| B104 | B196 | anti-SSB | TCO2 | -0.13835866 | 4.28E-07 |
| B105 | B107 | anti-U1RNP | AnuA | 0.999999418 | 0 |
| B105 | B119 | anti-U1RNP | Hcv-rna | 0.470017908 | 0 |
| B105 | B120 | anti-U1RNP | HCV-cAg | 0.311176192 | 0 |
| B105 | B125 | anti-U1RNP | Cor | -0.319333502 | 0 |
| B105 | B152 | anti-U1RNP | ACA-Ig(G-A-M) | -0.26341651 | 0 |
| B105 | B154 | anti-U1RNP | RF | 0.277811856 | 0 |
| B105 | B155 | anti-U1RNP | IgA | -0.244461159 | 0 |
| B105 | B158 | anti-U1RNP | IgM | 0.292027923 | 0 |
| B105 | B159 | anti-U1RNP | β2-MG | -0.263404289 | 0 |
| B105 | B169 | anti-U1RNP | U-CRY | -0.263405283 | 0 |
| B105 | B197 | anti-U1RNP | AG | 0.240927345 | 0 |
| B105 | B199 | anti-U1RNP | Mg | 0.216390065 | 1.78E-15 |
| B105 | B137 | anti-U1RNP | AFU | 0.204688404 | 5.33E-14 |
| B105 | B134 | anti-U1RNP | CHE | 0.187404365 | 6.15E-12 |
| B105 | B200 | anti-U1RNP | OP | 0.187095705 | 6.66E-12 |
| B105 | B195 | anti-U1RNP | Na | 0.184163407 | 1.43E-11 |
| B105 | B193 | anti-U1RNP | K | 0.182545784 | 2.16E-11 |
| B105 | B108 | anti-U1RNP | Anti-Cyclin | -0.18087605 | 3.30E-11 |
| B105 | B114 | anti-U1RNP | TSH TT3 TT4(TT4) | 0.173261895 | 2.17E-10 |
| B105 | B138 | anti-U1RNP | PAB | 0.164858123 | 1.57E-09 |
| B105 | B136 | anti-U1RNP | 5'NT | 0.151810372 | 2.81E-08 |
| B105 | B196 | anti-U1RNP | TCO2 | -0.138358659 | 4.28E-07 |
| B106 | B107 | anti-P | AnuA | 0.999999631 | 0 |
| B106 | B119 | anti-P | Hcv-rna | 0.470017909 | 0 |
| B106 | B120 | anti-P | HCV-cAg | 0.311176275 | 0 |
| B106 | B125 | anti-P | Cor | -0.3193335 | 0 |
| B106 | B152 | anti-P | ACA-Ig(G-A-M) | -0.263416046 | 0 |
| B106 | B154 | anti-P | RF | 0.277811857 | 0 |

| | | | | | |
|---|---|---|---|---|---|
| B106 | B155 | anti-P | IgA | -0.244461159 | 0 |
| B106 | B158 | anti-P | IgM | 0.292027924 | 0 |
| B106 | B159 | anti-P | β2-MG | -0.26340441 | 0 |
| B106 | B169 | anti-P | U-CRY | -0.263405089 | 0 |
| B106 | B197 | anti-P | AG | 0.240927346 | 0 |
| B106 | B199 | anti-P | Mg | 0.216390066 | 1.78E-15 |
| B106 | B137 | anti-P | AFU | 0.204688405 | 5.33E-14 |
| B106 | B134 | anti-P | CHE | 0.187404365 | 6.15E-12 |
| B106 | B200 | anti-P | OP | 0.187095699 | 6.66E-12 |
| B106 | B195 | anti-P | Na | 0.184163403 | 1.43E-11 |
| B106 | B193 | anti-P | K | 0.182545786 | 2.16E-11 |
| B106 | B108 | anti-P | Anti-Cyclin | -0.180876051 | 3.30E-11 |
| B106 | B114 | anti-P | TSH TT3 TT4(TT4) | 0.173261896 | 2.17E-10 |
| B106 | B138 | anti-P | PAB | 0.164858124 | 1.57E-09 |
| B106 | B136 | anti-P | 5'NT | 0.151810372 | 2.81E-08 |
| B106 | B196 | anti-P | TCO2 | -0.138358659 | 4.28E-07 |
| B107 | B109 | AnuA | AMA-M2 | 0.999999447 | 0 |
| B107 | B110 | AnuA | AHA | 0.999999467 | 0 |
| B107 | B119 | AnuA | Hcv-rna | 0.47001764 | 0 |
| B107 | B120 | AnuA | HCV-cAg | 0.311176813 | 0 |
| B107 | B125 | AnuA | Cor | -0.319333283 | 0 |
| B107 | B152 | AnuA | ACA-Ig(G-A-M) | -0.263412426 | 0 |
| B107 | B154 | AnuA | RF | 0.277811698 | 0 |
| B107 | B155 | AnuA | IgA | -0.244461016 | 0 |
| B107 | B158 | AnuA | IgM | 0.292027759 | 0 |
| B107 | B159 | AnuA | β2-MG | -0.263402253 | 0 |
| B107 | B169 | AnuA | U-CRY | -0.263401605 | 0 |
| B107 | B197 | AnuA | AG | 0.240927208 | 0 |
| B107 | B199 | AnuA | Mg | 0.21638995 | 1.78E-15 |
| B107 | B137 | AnuA | AFU | 0.204688287 | 5.33E-14 |
| B107 | B134 | AnuA | CHE | 0.187404261 | 6.15E-12 |
| B107 | B200 | AnuA | OP | 0.187095577 | 6.67E-12 |
| B107 | B195 | AnuA | Na | 0.184163278 | 1.43E-11 |
| B107 | B193 | AnuA | K | 0.182545683 | 2.16E-11 |
| B107 | B108 | AnuA | Anti-Cyclin | -0.18087594 | 3.30E-11 |

| | | | | | |
|---|---|---|---|---|---|
| | | | | 8 | |
| B107 | B114 | AnuA | TSH TT3 TT4(TT4) | 0.173261795 | 2.17E-10 |
| B107 | B138 | AnuA | PAB | 0.164858031 | 1.57E-09 |
| B107 | B136 | AnuA | 5'NT | 0.151810285 | 2.81E-08 |
| B107 | B196 | AnuA | TCO2 | -0.138358582 | 4.28E-07 |
| B108 | B151 | Anti-Cyclin | Anti-CCP | 0.277090214 | 0 |
| B108 | B156 | Anti-Cyclin | IgE | 0.452888359 | 0 |
| B108 | B158 | Anti-Cyclin | IgM | 0.228221872 | 0 |
| B108 | B179 | Anti-Cyclin | α1-MG | 0.272315859 | 0 |
| B108 | B194 | Anti-Cyclin | Cl | 0.211036689 | 8.44E-15 |
| B108 | B157 | Anti-Cyclin | IgG | 0.192922904 | 1.42E-12 |
| B108 | B109 | Anti-Cyclin | AMA-M2 | -0.180876049 | 3.30E-11 |
| B108 | B110 | Anti-Cyclin | AHA | -0.18087605 | 3.30E-11 |
| B108 | B195 | Anti-Cyclin | Na | 0.137596226 | 4.96E-07 |
| B109 | B119 | AMA-M2 | Hcv-rna | 0.470017907 | 0 |
| B109 | B120 | AMA-M2 | HCV-cAg | 0.311176226 | 0 |
| B109 | B125 | AMA-M2 | Cor | -0.319333502 | 0 |
| B109 | B152 | AMA-M2 | ACA-Ig(G-A-M) | -0.263416311 | 0 |
| B109 | B154 | AMA-M2 | RF | 0.277811855 | 0 |
| B109 | B155 | AMA-M2 | IgA | -0.244461158 | 0 |
| B109 | B158 | AMA-M2 | IgM | 0.292027922 | 0 |
| B109 | B159 | AMA-M2 | β2-MG | -0.263403875 | 0 |
| B109 | B169 | AMA-M2 | U-CRY | -0.263405201 | 0 |
| B109 | B197 | AMA-M2 | AG | 0.240927345 | 0 |
| B109 | B199 | AMA-M2 | Mg | 0.216390065 | 1.78E-15 |
| B109 | B137 | AMA-M2 | AFU | 0.204688404 | 5.33E-14 |
| B109 | B134 | AMA-M2 | CHE | 0.187404364 | 6.15E-12 |
| B109 | B200 | AMA-M2 | OP | 0.187095704 | 6.66E-12 |
| B109 | B195 | AMA-M2 | Na | 0.184163408 | 1.43E-11 |
| B109 | B193 | AMA-M2 | K | 0.182545784 | 2.16E-11 |
| B109 | B114 | AMA-M2 | TSH TT3 TT4(TT4) | 0.173261895 | 2.17E-10 |
| B109 | B138 | AMA-M2 | PAB | 0.164858123 | 1.57E-09 |
| B109 | B136 | AMA-M2 | 5'NT | 0.151810372 | 2.81E-08 |
| B109 | B196 | AMA-M2 | TCO2 | -0.138358658 | 4.28E-07 |
| B11 | B12 | PGI/PGII | PG II | -0.527260379 | 0 |

| | | | | | |
|---|---|---|---|---|---|
| B11 | B139 | PGI/PGII | ADA | 0.179276844 | 4.93E-11 |
| B11 | B153 | PGI/PGII | Glu | 0.177111101 | 8.46E-11 |
| B11 | B137 | PGI/PGII | AFU | 0.155096173 | 1.39E-08 |
| B11 | B135 | PGI/PGII | TBA | 0.147746289 | 6.57E-08 |
| B11 | B14 | PGI/PGII | G-17 | -0.14382958 | 1.46E-07 |
| B11 | B200 | PGI/PGII | OP | 0.139675073 | 3.32E-07 |
| B110 | B119 | AHA | Hcv-rna | 0.470017908 | 0 |
| B110 | B120 | AHA | HCV-cAg | 0.311176178 | 0 |
| B110 | B125 | AHA | Cor | -0.319333502 | 0 |
| B110 | B152 | AHA | ACA-Ig(G-A-M) | -0.263416304 | 0 |
| B110 | B154 | AHA | RF | 0.277811856 | 0 |
| B110 | B155 | AHA | IgA | -0.244461159 | 0 |
| B110 | B158 | AHA | IgM | 0.292027923 | 0 |
| B110 | B159 | AHA | β2-MG | -0.263404134 | 0 |
| B110 | B169 | AHA | U-CRY | -0.26340539 | 0 |
| B110 | B197 | AHA | AG | 0.240927346 | 0 |
| B110 | B199 | AHA | Mg | 0.216390065 | 1.78E-15 |
| B110 | B137 | AHA | AFU | 0.204688404 | 5.33E-14 |
| B110 | B134 | AHA | CHE | 0.187404365 | 6.15E-12 |
| B110 | B200 | AHA | OP | 0.187095703 | 6.66E-12 |
| B110 | B195 | AHA | Na | 0.184163406 | 1.43E-11 |
| B110 | B193 | AHA | K | 0.182545784 | 2.16E-11 |
| B110 | B114 | AHA | TSH TT3 TT4(TT4) | 0.173261895 | 2.17E-10 |
| B110 | B138 | AHA | PAB | 0.164858123 | 1.57E-09 |
| B110 | B136 | AHA | 5'NT | 0.151810372 | 2.81E-08 |
| B110 | B196 | AHA | TCO2 | -0.138358659 | 4.28E-07 |
| B111 | B203 | TSH | DH | 0.177199096 | 8.27E-11 |
| B111 | B112 | TSH | FT4 | -0.164129984 | 1.86E-09 |
| B111 | B115 | TSH | TSH TT3 TT4(TT3) | -0.158408039 | 6.74E-09 |
| B111 | B113 | TSH | FT3 | -0.145986901 | 9.42E-08 |
| B112 | B113 | FT4 | FT3 | 0.441320394 | 0 |
| B112 | B114 | FT4 | TSH TT3 TT4(TT4) | 0.339311682 | 0 |
| B112 | B227 | FT4 | OD | 0.18764734 | 5.77E-12 |
| B112 | B149 | FT4 | TGAb | 0.180539099 | 3.59E-11 |
| B113 | B115 | FT3 | TSH TT3 TT4(TT3) | 0.360796287 | 0 |
| B114 | B115 | TSH TT3 TT4(TT4) | TSH TT3 TT4(TT3) | 0.226830476 | 0 |

| | | | | | |
|---|---|---|---|---|---|
| B115 | B119 | TSH TT3 TT4(TT3) | Hcv-rna | 0.211830085 | 6.66E-15 |
| B115 | B194 | TSH TT3 TT4(TT3) | Cl | 0.17163294 | 3.21E-10 |
| B115 | B126 | TSH TT3 TT4(TT3) | Alb | 0.16958152 | 5.23E-10 |
| B115 | B139 | TSH TT3 TT4(TT3) | ADA | -0.168174481 | 7.28E-10 |
| B115 | B134 | TSH TT3 TT4(TT3) | CHE | 0.162051053 | 2.98E-09 |
| B115 | B120 | TSH TT3 TT4(TT3) | HCV-cAg | 0.15336224 | 2.02E-08 |
| B115 | B195 | TSH TT3 TT4(TT3) | Na | 0.145095615 | 1.13E-07 |
| B116 | B167 | TT3 | PC | 0.359380749 | 0 |
| B116 | B202 | TT3 | e-times | 0.738141684 | 0 |
| B117 | B202 | TT4 | e-times | 0.82486282 | 0 |
| B119 | B120 | Hcv-rna | HCV-cAg | 0.56199294 | 0 |
| B119 | B125 | Hcv-rna | Cor | -0.420070212 | 0 |
| B119 | B134 | Hcv-rna | CHE | 0.256589275 | 0 |
| B119 | B138 | Hcv-rna | PAB | 0.326353159 | 0 |
| B119 | B139 | Hcv-rna | ADA | -0.386084861 | 0 |
| B119 | B152 | Hcv-rna | ACA-Ig(G-A-M) | -0.701836938 | 0 |
| B119 | B153 | Hcv-rna | Glu | -0.263364678 | 0 |
| B119 | B155 | Hcv-rna | IgA | -0.297020816 | 0 |
| B119 | B159 | Hcv-rna | β2-MG | -0.701805446 | 0 |
| B119 | B169 | Hcv-rna | U-CRY | -0.701808922 | 0 |
| B119 | B195 | Hcv-rna | Na | 0.357398754 | 0 |
| B119 | B194 | Hcv-rna | Cl | 0.21770731 | 1.11E-15 |
| B119 | B158 | Hcv-rna | IgM | 0.195964093 | 6.19E-13 |
| B119 | B197 | Hcv-rna | AG | 0.163037098 | 2.39E-09 |
| B119 | B157 | Hcv-rna | IgG | -0.145898317 | 9.59E-08 |
| B12 | B84 | PG II | C-PSA | 0.224922884 | 0 |
| B12 | B111 | PG II | TSH | 0.205836484 | 3.82E-14 |
| B12 | B14 | PG II | G-17 | 0.179834582 | 4.29E-11 |
| B12 | B165 | PG II | bilirubin | 0.144643986 | 1.24E-07 |
| B12 | B83 | PG II | AFP | -0.13842571 | 4.23E-07 |
| B12 | B164 | PG II | COM | -0.136212969 | 6.47E-07 |
| B120 | B125 | HCV-cAg | Cor | -0.238811316 | 0 |
| B120 | B152 | HCV-cAg | ACA-Ig(G-A-M) | -0.39683070 | 0 |

| | | | | | |
|---|---|---|---|---|---|
| | | | | 6 | |
| B120 | B159 | HCV-cAg | β2-MG | -0.396809587 | 0 |
| B120 | B169 | HCV-cAg | U-CRY | -0.396809162 | 0 |
| B120 | B139 | HCV-cAg | ADA | -0.222618568 | 2.22E-16 |
| B120 | B195 | HCV-cAg | Na | 0.210082147 | 1.11E-14 |
| B120 | B158 | HCV-cAg | IgM | 0.209094619 | 1.47E-14 |
| B120 | B138 | HCV-cAg | PAB | 0.190474809 | 2.73E-12 |
| B120 | B134 | HCV-cAg | CHE | 0.155565561 | 1.26E-08 |
| B120 | B154 | HCV-cAg | RF | 0.154455792 | 1.60E-08 |
| B120 | B156 | HCV-cAg | IgE | -0.145518963 | 1.04E-07 |
| B120 | B194 | HCV-cAg | Cl | 0.139662235 | 3.33E-07 |
| B121 | B122 | C3 | C4 | 0.985295194 | 0 |
| B121 | B125 | C3 | Cor | -0.290116033 | 0 |
| B121 | B136 | C3 | 5'NT | 0.497710874 | 0 |
| B121 | B157 | C3 | IgG | -0.300293175 | 0 |
| B121 | B158 | C3 | IgM | -0.165119503 | 1.48E-09 |
| B121 | B124 | C3 | ACTH | -0.153965316 | 1.77E-08 |
| B121 | B155 | C3 | IgA | 0.135240936 | 7.78E-07 |
| B122 | B125 | C4 | Cor | -0.361506618 | 0 |
| B122 | B136 | C4 | 5'NT | 0.60600754 | 0 |
| B122 | B157 | C4 | IgG | -0.318776806 | 0 |
| B122 | B124 | C4 | ACTH | -0.196401587 | 5.48E-13 |
| B122 | B158 | C4 | IgM | -0.135238041 | 7.78E-07 |
| B123 | B177 | hs-CRP1 | ACR | 0.179215281 | 5.01E-11 |
| B123 | B213 | hs-CRP1 | CVA-R | 0.153029595 | 2.17E-08 |
| B124 | B125 | ACTH | Cor | 0.371832818 | 0 |
| B124 | B136 | ACTH | 5'NT | -0.246827677 | 0 |
| B124 | B152 | ACTH | ACA-Ig(G-A-M) | 0.363606783 | 0 |
| B124 | B155 | ACTH | IgA | -0.273864253 | 0 |
| B124 | B159 | ACTH | β2-MG | 0.363590621 | 0 |

| | | | | | |
|---|---|---|---|---|---|
| B124 | B169 | ACTH | U-CRY | 0.363592416 | 0 |
| B124 | B179 | ACTH | α1-MG | 0.3010837 | 0 |
| B124 | B182 | ACTH | TT | -0.250885073 | 0 |
| B124 | B199 | ACTH | Mg | -0.299417707 | 0 |
| B124 | B138 | ACTH | PAB | 0.192597437 | 1.55E-12 |
| B124 | B157 | ACTH | IgG | -0.186135025 | 8.56E-12 |
| B124 | B151 | ACTH | Anti-CCP | 0.162358138 | 2.78E-09 |
| B124 | B195 | ACTH | Na | -0.150389594 | 3.79E-08 |
| B124 | B148 | ACTH | Anti_TPO | -0.139819814 | 3.22E-07 |
| B125 | B136 | Cor | 5'NT | -0.522170462 | 0 |
| B125 | B152 | Cor | ACA-Ig(G-A-M) | 0.364010052 | 0 |
| B125 | B159 | Cor | β2-MG | 0.363993846 | 0 |
| B125 | B169 | Cor | U-CRY | 0.363995638 | 0 |
| B125 | B199 | Cor | Mg | -0.340340447 | 0 |
| B125 | B186 | Cor | PFC | 0.189790838 | 3.28E-12 |
| B125 | B181 | Cor | APTT | -0.187361305 | 6.22E-12 |
| B125 | B154 | Cor | RF | -0.174563408 | 1.58E-10 |
| B125 | B183 | Cor | PTA | -0.172140031 | 2.84E-10 |
| B125 | B151 | Cor | Anti-CCP | 0.163941182 | 1.94E-09 |
| B125 | B179 | Cor | α1-MG | 0.159304919 | 5.52E-09 |
| B125 | B196 | Cor | TCO2 | 0.153083144 | 2.14E-08 |
| B125 | B158 | Cor | IgM | -0.151570166 | 2.95E-08 |
| B125 | B138 | Cor | PAB | -0.136112463 | 6.59E-07 |
| B126 | B127 | Alb | A-G ratio | 0.514306614 | 0 |
| B126 | B188 | Alb | Ca | 0.403950605 | 0 |
| B126 | B189 | Alb | eGFB | 0.295765923 | 0 |
| B126 | B141 | Alb | AST/ALT | -0.206196546 | 3.46E-14 |
| B126 | B216 | Alb | BMI | 0.199137655 | 2.57E-13 |
| B126 | B138 | Alb | PAB | 0.189613506 | 3.43E-12 |
| B126 | B221 | Alb | Hei | 0.183003349 | 1.92E-11 |
| B126 | B130 | Alb | GLB | -0.17833783 | 6.24E-11 |

| | | | | | |
|---|---|---|---|---|---|
| | | | | 9 | |
| B126 | B222 | Alb | NS | 0.166373658 | 1.11E-09 |
| B126 | B134 | Alb | CHE | 0.164352063 | 1.77E-09 |
| B126 | B139 | Alb | ADA | -0.156857589 | 9.47E-09 |
| B126 | B128 | Alb | ALT | 0.152290919 | 2.54E-08 |
| B126 | B191 | Alb | Urea | -0.139659128 | 3.33E-07 |
| B127 | B130 | A-G ratio | GLB | -0.904469979 | 0 |
| B127 | B221 | A-G ratio | Hei | 0.225147005 | 0 |
| B127 | B189 | A-G ratio | eGFB | 0.220939194 | 4.44E-16 |
| B127 | B141 | A-G ratio | AST/ALT | -0.200629624 | 1.69E-13 |
| B127 | B220 | A-G ratio | SBP | -0.167093951 | 9.38E-10 |
| B127 | B216 | A-G ratio | BMI | 0.165588044 | 1.33E-09 |
| B127 | B222 | A-G ratio | NS | 0.158369577 | 6.80E-09 |
| B128 | B131 | ALT | AST | 0.675583817 | 0 |
| B128 | B140 | ALT | r-GT | 0.410977703 | 0 |
| B128 | B141 | ALT | AST/ALT | -0.584567589 | 0 |
| B128 | B215 | ALT | BMI | 0.235433195 | 0 |
| B128 | B216 | ALT | BMI | 0.285474834 | 0 |
| B128 | B222 | ALT | NS | 0.233340901 | 0 |
| B128 | B206 | ALT | Waist | 0.220392971 | 4.44E-16 |
| B128 | B210 | ALT | Hip | 0.18601428 | 8.84E-12 |
| B128 | B221 | ALT | Hei | 0.177055566 | 8.57E-11 |
| B128 | B189 | ALT | eGFB | 0.168733687 | 6.39E-10 |
| B128 | B207 | ALT | WHR | 0.165173271 | 1.46E-09 |
| B128 | B213 | ALT | CVA-R | 0.141679147 | 2.24E-07 |
| B129 | B132 | IBIL | Dbil | 0.67701573 | 0 |
| B129 | B133 | IBIL | TBIL | 0.978991087 | 0 |
| B129 | B221 | IBIL | Hei | 0.135874519 | 6.90E-07 |
| B130 | B221 | GLB | Hei | -0.1829539 | 1.95E-11 |
| B130 | B220 | GLB | SBP | 0.179341443 | 4.85E-11 |
| B130 | B178 | GLB | Ig-U | 0.156683974 | 9.84E-09 |
| B130 | B179 | GLB | α1-MG | 0.152370972 | 2.49E-08 |
| B130 | B180 | GLB | Tf | 0.149312848 | 4.75E-08 |
| B130 | B141 | GLB | AST/ALT | 0.143852492 | 1.45E-07 |
| B130 | B177 | GLB | ACR | 0.139366873 | 3.52E-07 |
| B130 | B218 | GLB | DBP | 0.139072717 | 3.73E-07 |
| B131 | B140 | AST | r-GT | 0.379401013 | 0 |
| B131 | B213 | AST | CVA-R | 0.319598905 | 0 |

| | | | | | |
|---|---|---|---|---|---|
| B131 | B150 | AST | ALP | 0.216519907 | 1.55E-15 |
| B132 | B133 | Dbil | TBIL | 0.812768155 | 0 |
| B133 | B221 | TBIL | Hei | 0.139478338 | 3.45E-07 |
| B134 | B138 | CHE | PAB | 0.541644109 | 0 |
| B134 | B139 | CHE | ADA | -0.36109793<br>4 | 0 |
| B134 | B195 | CHE | Na | 0.220730772 | 4.44E-16 |
| B134 | B198 | CHE | P | 0.217863877 | 1.11E-15 |
| B134 | B153 | CHE | Glu | -0.18574984<br>6 | 9.47E-12 |
| B134 | B193 | CHE | K | 0.139860482 | 3.20E-07 |
| B135 | B137 | TBA | AFU | 0.295682763 | 0 |
| B135 | B138 | TBA | PAB | -0.29049546<br>5 | 0 |
| B135 | B139 | TBA | ADA | 0.35467039 | 0 |
| B135 | B194 | TBA | Cl | 0.182016021 | 2.47E-11 |
| B135 | B197 | TBA | AG | -0.15598201<br>1 | 1.15E-08 |
| B135 | B198 | TBA | P | -0.14586436<br>8 | 9.66E-08 |
| B136 | B157 | 5'NT | IgG | -0.21489781<br>8 | 2.66E-15 |
| B136 | B150 | 5'NT | ALP | 0.206488242 | 3.18E-14 |
| B136 | B140 | 5'NT | r-GT | 0.195577999 | 6.87E-13 |
| B136 | B137 | 5'NT | AFU | 0.151863641 | 2.78E-08 |
| B137 | B139 | AFU | ADA | 0.400172241 | 0 |
| B137 | B153 | AFU | Glu | 0.273801255 | 0 |
| B137 | B200 | AFU | OP | 0.230390425 | 0 |
| B137 | B138 | AFU | PAB | -0.22231342<br>7 | 2.22E-16 |
| B137 | B150 | AFU | ALP | 0.188832341 | 4.22E-12 |
| B137 | B155 | AFU | IgA | -0.18737177<br>5 | 6.20E-12 |
| B137 | B199 | AFU | Mg | 0.169994635 | 4.74E-10 |
| B138 | B139 | PAB | ADA | -0.70927999<br>6 | 0 |
| B138 | B153 | PAB | Glu | -0.33402649<br>8 | 0 |
| B138 | B195 | PAB | Na | 0.246806345 | 0 |
| B138 | B200 | PAB | OP | -0.15339065<br>4 | 2.01E-08 |
| B138 | B170 | PAB | U-GLU | -0.14348572 | 1.56E-07 |
| B139 | B150 | ADA | ALP | 0.249067988 | 0 |
| B139 | B153 | ADA | Glu | 0.480663436 | 0 |

| | | | | | |
|---|---|---|---|---|---|
| B139 | B194 | ADA | Cl | -0.239005322 | 0 |
| B139 | B195 | ADA | Na | -0.31941111 | 0 |
| B139 | B196 | ADA | TCO2 | -0.275645108 | 0 |
| B139 | B200 | ADA | OP | 0.295684458 | 0 |
| B139 | B170 | ADA | U-GLU | 0.211702563 | 6.88E-15 |
| B139 | B146 | ADA | CysC | 0.179991052 | 4.12E-11 |
| B139 | B197 | ADA | AG | 0.148042644 | 6.18E-08 |
| B14 | B165 | G-17 | bilirubin | 0.632681548 | 0 |
| B14 | B227 | G-17 | OD | 0.248945729 | 0 |
| B14 | B123 | G-17 | hs-CRP1 | 0.155279957 | 1.34E-08 |
| B14 | B149 | G-17 | TGAb | 0.15321584 | 2.08E-08 |
| B14 | B82 | G-17 | CEA | 0.140975685 | 2.57E-07 |
| B140 | B150 | r-GT | ALP | 0.196468924 | 5.38E-13 |
| B140 | B141 | r-GT | AST/ALT | -0.192759971 | 1.48E-12 |
| B141 | B206 | AST/ALT | Waist | -0.27141692 | 0 |
| B141 | B215 | AST/ALT | BMI | -0.276153474 | 0 |
| B141 | B216 | AST/ALT | BMI | -0.360441152 | 0 |
| B141 | B221 | AST/ALT | Hei | -0.263709661 | 0 |
| B141 | B222 | AST/ALT | NS | -0.278076833 | 0 |
| B141 | B210 | AST/ALT | Hip | -0.221537982 | 4.44E-16 |
| B141 | B189 | AST/ALT | eGFB | -0.217508127 | 1.33E-15 |
| B141 | B207 | AST/ALT | WHR | -0.213152568 | 4.44E-15 |
| B142 | B145 | 25-(OH)D | BGP | 0.24374097 | 0 |
| B142 | B184 | 25-(OH)D | INR | 0.335651334 | 0 |
| B142 | B185 | 25-(OH)D | PT | 0.330334862 | 0 |
| B142 | B181 | 25-(OH)D | APTT | 0.158693999 | 6.32E-09 |
| B142 | B201 | 25-(OH)D | e-habits | 0.148795515 | 5.29E-08 |
| B143 | B144 | β-CTx | Total-PINP | 0.783055347 | 0 |
| B143 | B145 | β-CTx | BGP | 0.585470229 | 0 |
| B144 | B145 | Total-PINP | BGP | 0.670731251 | 0 |
| B144 | B150 | Total-PINP | ALP | 0.249377682 | 0 |
| B145 | B150 | BGP | ALP | 0.189591549 | 3.45E-12 |
| B145 | B184 | BGP | INR | 0.152922394 | 2.22E-08 |
| B145 | B227 | BGP | OD | 0.152383537 | 2.49E-08 |

| | | | | | |
|---|---|---|---|---|---|
| B145 | B185 | BGP | PT | 0.150577236 | 3.64E-08 |
| B146 | B190 | CysC | Cre | 0.205920689 | 3.73E-14 |
| B146 | B191 | CysC | Urea | 0.192100781 | 1.77E-12 |
| B146 | B189 | CysC | eGFB | -0.188800883 | 4.26E-12 |
| B148 | B149 | Anti_TPO | TGAb | 0.489232853 | 0 |
| B149 | B165 | TGAb | bilirubin | 0.256008926 | 0 |
| B15 | B101 | CD3-CD19+(B cell) | anti-Sc70 | 0.507137695 | 0 |
| B15 | B102 | CD3-CD19+(B cell) | anti-Sm | 0.507137683 | 0 |
| B15 | B103 | CD3-CD19+(B cell) | anti-SSA/Ro | 0.507137687 | 0 |
| B15 | B104 | CD3-CD19+(B cell) | anti-SSB | 0.507137687 | 0 |
| B15 | B105 | CD3-CD19+(B cell) | anti-U1RNP | 0.507137687 | 0 |
| B15 | B106 | CD3-CD19+(B cell) | anti-P | 0.507137689 | 0 |
| B15 | B107 | CD3-CD19+(B cell) | AnuA | 0.5071374 | 0 |
| B15 | B109 | CD3-CD19+(B cell) | AMA-M2 | 0.507137686 | 0 |
| B15 | B110 | CD3-CD19+(B cell) | AHA | 0.507137688 | 0 |
| B15 | B119 | CD3-CD19+(B cell) | Hcv-rna | 0.642361166 | 0 |
| B15 | B120 | CD3-CD19+(B cell) | HCV-cAg | 0.340519736 | 0 |
| B15 | B121 | CD3-CD19+(B cell) | C3 | -0.345639012 | 0 |
| B15 | B122 | CD3-CD19+(B cell) | C4 | -0.289895446 | 0 |
| B15 | B124 | CD3-CD19+(B cell) | ACTH | -0.268992726 | 0 |
| B15 | B125 | CD3-CD19+(B cell) | Cor | -0.531711375 | 0 |
| B15 | B152 | CD3-CD19+(B cell) | ACA-Ig(G-A-M) | -0.412326136 | 0 |
| B15 | B155 | CD3-CD19+(B cell) | IgA | -0.274865997 | 0 |
| B15 | B158 | CD3-CD19+(B cell) | IgM | 0.296134198 | 0 |
| B15 | B159 | CD3-CD19+(B cell) | β2-MG | -0.412307591 | 0 |
| B15 | B16 | CD3-CD19+(B cell) | CD3+T | 0.482134281 | 0 |
| B15 | B169 | CD3-CD19+(B cell) | U-CRY | -0.412309641 | 0 |
| B15 | B18 | CD3-CD19+(B cell) | CD3+CD4+helper(inducer) T | 0.378926472 | 0 |
| B15 | B19 | CD3-CD19+(B cell) | CD3+CD4+/CD3+CD8+ | 0.405798914 | 0 |
| B15 | B197 | CD3-CD19+(B cell) | AG | 0.241030558 | 0 |
| B15 | B22 | CD3-CD19+(B cell) | CD3+CD8+(T) | 0.42494134 | 0 |
| B15 | B28 | CD3-CD19+(B cell) | PRL | 0.237155653 | 0 |
| B15 | B29 | CD3-CD19+(B cell) | E2 | 0.237155653 | 0 |
| B15 | B30 | CD3-CD19+(B cell) | LH | -0.23715565 | 0 |

| | | | | | |
|---|---|---|---|---|---|
| | | | | 4 | |
| B15 | B31 | CD3-CD19+(B cell) | FSH | -0.237155653 | 0 |
| B15 | B32 | CD3-CD19+(B cell) | Testo | 0.237155653 | 0 |
| B15 | B33 | CD3-CD19+(B cell) | Prog | 0.237155653 | 0 |
| B15 | B79 | CD3-CD19+(B cell) | hb V -dna | 0.284326945 | 0 |
| B15 | B94 | CD3-CD19+(B cell) | ALDH2-Lys504Lys | 0.329615104 | 0 |
| B15 | B97 | CD3-CD19+(B cell) | dsDNA | 0.507137687 | 0 |
| B15 | B99 | CD3-CD19+(B cell) | anti-PM-Scl | 0.50713769 | 0 |
| B15 | B20 | CD3-CD19+(B cell) | CD3+CD4+CD8+ | 0.214252989 | 3.11E-15 |
| B15 | B92 | CD3-CD19+(B cell) | ALDH2-Glu504Glu | -0.20250422 | 9.95E-14 |
| B15 | B196 | CD3-CD19+(B cell) | TCO2 | -0.20205742 | 1.13E-13 |
| B15 | B199 | CD3-CD19+(B cell) | Mg | 0.18548082 | 1.02E-11 |
| B15 | B195 | CD3-CD19+(B cell) | Na | 0.185013722 | 1.15E-11 |
| B15 | B93 | CD3-CD19+(B cell) | ALDH2-Glu504Lys | 0.184377066 | 1.35E-11 |
| B15 | B100 | CD3-CD19+(B cell) | anti-Ro52 | 0.173732141 | 1.93E-10 |
| B15 | B179 | CD3-CD19+(B cell) | α1-MG | -0.161223942 | 3.60E-09 |
| B15 | B181 | CD3-CD19+(B cell) | APTT | 0.156465085 | 1.03E-08 |
| B15 | B183 | CD3-CD19+(B cell) | PTA | 0.143254204 | 1.64E-07 |
| B150 | B200 | ALP | OP | 0.189961631 | 3.13E-12 |
| B150 | B213 | ALP | CVA-R | 0.185020688 | 1.14E-11 |
| B150 | B153 | ALP | Glu | 0.164981702 | 1.53E-09 |
| B150 | B197 | ALP | AG | 0.136647838 | 5.95E-07 |
| B151 | B186 | Anti-CCP | PFC | 0.249789235 | 0 |
| B151 | B155 | Anti-CCP | IgA | -0.180190496 | 3.92E-11 |
| B152 | B199 | ACA-Ig(G-A-M) | Mg | -0.338684981 | 0 |
| B152 | B193 | ACA-Ig(G-A-M) | K | -0.154289782 | 1.65E-08 |
| B152 | B179 | ACA-Ig(G-A-M) | α1-MG | 0.151070363 | 3.28E-08 |
| B153 | B170 | Glu | U-GLU | 0.446395089 | 0 |
| B153 | B194 | Glu | Cl | -0.275098064 | 0 |
| B153 | B195 | Glu | Na | -0.260061503 | 0 |
| B153 | B200 | Glu | OP | 0.299517666 | 0 |
| B153 | B196 | Glu | TCO2 | -0.164818762 | 1.59E-09 |
| B154 | B158 | RF | IgM | 0.182180847 | 2.37E-11 |
| B154 | B156 | RF | IgE | -0.151376901 | 3.08E-08 |
| B155 | B157 | IgA | IgG | 0.314840025 | 0 |

| | | | | | |
|---|---|---|---|---|---|
| B155 | B200 | IgA | OP | -0.20880574<br>1 | 1.60E-14 |
| B155 | B156 | IgA | IgE | -0.20370897<br>1 | 7.06E-14 |
| B155 | B195 | IgA | Na | -0.18200276<br>1 | 2.48E-11 |
| B157 | B158 | IgG | IgM | 0.180430207 | 3.69E-11 |
| B157 | B198 | IgG | P | 0.139391526 | 3.51E-07 |
| B158 | B195 | IgM | Na | 0.204190441 | 6.15E-14 |
| B158 | B197 | IgM | AG | 0.162274287 | 2.84E-09 |
| B158 | B200 | IgM | OP | 0.157599669 | 8.05E-09 |
| B159 | B169 | β2-MG | U-CRY | 0.999993797 | 0 |
| B159 | B199 | β2-MG | Mg | -0.33866967<br>9 | 0 |
| B159 | B193 | β2-MG | K | -0.15428287<br>1 | 1.66E-08 |
| B159 | B179 | β2-MG | α1-MG | 0.151063565 | 3.29E-08 |
| B16 | B101 | CD3+T | anti-Sc70 | 0.242614894 | 0 |
| B16 | B102 | CD3+T | anti-Sm | 0.242614889 | 0 |
| B16 | B103 | CD3+T | anti-SSA/Ro | 0.242614891 | 0 |
| B16 | B104 | CD3+T | anti-SSB | 0.242614891 | 0 |
| B16 | B105 | CD3+T | anti-U1RNP | 0.242614891 | 0 |
| B16 | B106 | CD3+T | anti-P | 0.242614892 | 0 |
| B16 | B107 | CD3+T | AnuA | 0.242614751 | 0 |
| B16 | B109 | CD3+T | AMA-M2 | 0.242614891 | 0 |
| B16 | B110 | CD3+T | AHA | 0.242614891 | 0 |
| B16 | B124 | CD3+T | ACTH | -0.65613694<br>7 | 0 |
| B16 | B125 | CD3+T | Cor | -0.50033635<br>5 | 0 |
| B16 | B152 | CD3+T | ACA-Ig(G-A-M) | -0.32780374 | 0 |
| B16 | B155 | CD3+T | IgA | 0.23122311 | 0 |
| B16 | B157 | CD3+T | IgG | 0.406121345 | 0 |
| B16 | B159 | CD3+T | β2-MG | -0.32778896<br>3 | 0 |
| B16 | B169 | CD3+T | U-CRY | -0.32779059<br>2 | 0 |
| B16 | B17 | CD3+T | CD3+CD16+CD56+NK | 0.276495548 | 0 |
| B16 | B18 | CD3+T | CD3+CD4+helper(inducer)<br>T | 0.947825442 | 0 |
| B16 | B19 | CD3+T | CD3+CD4+/CD3+CD8+ | 0.410300003 | 0 |
| B16 | B199 | CD3+T | Mg | 0.365023457 | 0 |
| B16 | B20 | CD3+T | CD3+CD4+CD8+ | 0.367581861 | 0 |
| B16 | B21 | CD3+T | CD3+CD4-CD8- | 0.423992737 | 0 |

| | | | | | |
|---|---|---|---|---|---|
| B16 | B22 | CD3+T | CD3+CD8+(T) | 0.954211859 | 0 |
| B16 | B79 | CD3+T | hbⅤ-dna | 0.322801774 | 0 |
| B16 | B94 | CD3+T | ALDH2-Lys504Lys | 0.281956025 | 0 |
| B16 | B97 | CD3+T | dsDNA | 0.242614891 | 0 |
| B16 | B99 | CD3+T | anti-PM-Scl | 0.242614892 | 0 |
| B16 | B179 | CD3+T | α1-MG | -0.221710131 | 4.44E-16 |
| B16 | B182 | CD3+T | TT | 0.208505688 | 1.78E-14 |
| B16 | B23 | CD3+T | CD3-CD16+CD56+(NK) | -0.201841715 | 1.20E-13 |
| B16 | B121 | CD3+T | C3 | -0.201122037 | 1.47E-13 |
| B16 | B122 | CD3+T | C4 | -0.183889098 | 1.53E-11 |
| B16 | B158 | CD3+T | IgM | 0.180457252 | 3.67E-11 |
| B16 | B181 | CD3+T | APTT | 0.175336817 | 1.31E-10 |
| B16 | B92 | CD3+T | ALDH2-Glu504Glu | -0.17197922 | 2.95E-10 |
| B16 | B73 | CD3+T | ApoA | -0.159411806 | 5.39E-09 |
| B16 | B93 | CD3+T | ALDH2-Glu504Lys | 0.156493168 | 1.03E-08 |
| B16 | B62 | CD3+T | WBHSRVI | -0.155812207 | 1.19E-08 |
| B16 | B200 | CD3+T | OP | -0.152582386 | 2.38E-08 |
| B16 | B154 | CD3+T | RF | 0.150401011 | 3.78E-08 |
| B16 | B75 | CD3+T | ApoA1/B | 0.142991811 | 1.72E-07 |
| B16 | B58 | CD3+T | HCT | -0.136474343 | 6.15E-07 |
| B160 | B191 | PH | Urea | -0.213411298 | 4.00E-15 |
| B160 | B163 | PH | SG | -0.170275506 | 4.44E-10 |
| B160 | B190 | PH | Cre | -0.15074061 | 3.52E-08 |
| B160 | B192 | PH | UA | -0.150684659 | 3.56E-08 |
| B161 | B173 | LEU | NTT | 0.18218184 | 2.37E-11 |
| B161 | B221 | LEU | Hei | -0.168184074 | 7.27E-10 |
| B163 | B166 | SG | Pro | 0.278841086 | 0 |
| B163 | B176 | SG | Ucr | 0.300941415 | 0 |
| B163 | B191 | SG | Urea | 0.236318972 | 0 |
| B163 | B222 | SG | NS | 0.208398882 | 1.82E-14 |
| B163 | B216 | SG | BMI | 0.15652893 | 1.02E-08 |
| B163 | B190 | SG | Cre | 0.154269548 | 1.66E-08 |

| | | | | | |
|---|---|---|---|---|---|
| B163 | B221 | SG | Hei | 0.149728778 | 4.35E-08 |
| B163 | B205 | SG | KDA | -0.135474784 | 7.44E-07 |
| B163 | B188 | SG | Ca | -0.13451611 | 8.92E-07 |
| B163 | B192 | SG | UA | 0.134125508 | 9.59E-07 |
| B165 | B227 | bilirubin | OD | 0.402409468 | 0 |
| B166 | B176 | Pro | Ucr | 0.134929655 | 8.25E-07 |
| B167 | B202 | PC | e-times | 0.988683958 | 0 |
| B169 | B199 | U-CRY | Mg | -0.338671316 | 0 |
| B169 | B193 | U-CRY | K | -0.154283614 | 1.66E-08 |
| B169 | B179 | U-CRY | α1-MG | 0.151064321 | 3.29E-08 |
| B17 | B101 | CD3+CD16+CD56+NK | anti-Sc70 | 0.374917967 | 0 |
| B17 | B102 | CD3+CD16+CD56+NK | anti-Sm | 0.37491796 | 0 |
| B17 | B103 | CD3+CD16+CD56+NK | anti-SSA/Ro | 0.374917962 | 0 |
| B17 | B104 | CD3+CD16+CD56+NK | anti-SSB | 0.374917963 | 0 |
| B17 | B105 | CD3+CD16+CD56+NK | anti-U1RNP | 0.374917962 | 0 |
| B17 | B106 | CD3+CD16+CD56+NK | anti-P | 0.374917964 | 0 |
| B17 | B107 | CD3+CD16+CD56+NK | AnuA | 0.374917759 | 0 |
| B17 | B109 | CD3+CD16+CD56+NK | AMA-M2 | 0.374917962 | 0 |
| B17 | B110 | CD3+CD16+CD56+NK | AHA | 0.374917962 | 0 |
| B17 | B151 | CD3+CD16+CD56+NK | Anti-CCP | 0.304606065 | 0 |
| B17 | B186 | CD3+CD16+CD56+NK | PFC | 0.410000097 | 0 |
| B17 | B20 | CD3+CD16+CD56+NK | CD3+CD4+CD8+ | 0.350232566 | 0 |
| B17 | B21 | CD3+CD16+CD56+NK | CD3+CD4-CD8- | 0.568994217 | 0 |
| B17 | B22 | CD3+CD16+CD56+NK | CD3+CD8+(T) | 0.278575485 | 0 |
| B17 | B23 | CD3+CD16+CD56+NK | CD3-CD16+CD56+(NK) | -0.448508984 | 0 |
| B17 | B73 | CD3+CD16+CD56+NK | ApoA | -0.263927353 | 0 |
| B17 | B81 | CD3+CD16+CD56+NK | TSGF | -0.249636293 | 0 |
| B17 | B97 | CD3+CD16+CD56+NK | dsDNA | 0.374917963 | 0 |
| B17 | B99 | CD3+CD16+CD56+NK | anti-PM-Scl | 0.374917964 | 0 |
| B17 | B155 | CD3+CD16+CD56+NK | IgA | -0.217292018 | 1.33E-15 |
| B17 | B158 | CD3+CD16+CD56+NK | IgM | 0.215941608 | 1.78E-15 |
| B17 | B18 | CD3+CD16+CD56+NK | CD3+CD4+helper(inducer) T | 0.206022395 | 3.64E-14 |
| B17 | B34 | CD3+CD16+CD56+NK | TK | 0.168754566 | 6.36E-10 |
| B17 | B199 | CD3+CD16+CD56+NK | Mg | 0.160203907 | 4.52E-09 |
| B17 | B135 | CD3+CD16+CD56+NK | TBA | -0.158329219 | 6.86E-09 |

| | | | | | |
|---|---|---|---|---|---|
| B17 | B154 | CD3+CD16+CD56+NK | RF | 0.146302473 | 8.83E-08 |
| B171 | B172 | EC | KB | -0.23789271 8 | 0 |
| B173 | B178 | NTT | Ig-U | 0.264394406 | 0 |
| B173 | B180 | NTT | Tf | 0.265865618 | 0 |
| B173 | B179 | NTT | α1-MG | 0.189046291 | 3.99E-12 |
| B173 | B177 | NTT | ACR | 0.140089073 | 3.06E-07 |
| B174 | B190 | Urea/Cr | Cre | -0.42031173 | 0 |
| B174 | B191 | Urea/Cr | Urea | 0.640680359 | 0 |
| B174 | B192 | Urea/Cr | UA | -0.22462777 5 | 0 |
| B174 | B216 | Urea/Cr | BMI | -0.24781551 2 | 0 |
| B174 | B221 | Urea/Cr | Hei | -0.32784555 9 | 0 |
| B174 | B222 | Urea/Cr | NS | -0.26932264 | 0 |
| B174 | B189 | Urea/Cr | eGFB | 0.139689952 | 3.31E-07 |
| B175 | B177 | ALB-U | ACR | 0.928396406 | 0 |
| B175 | B178 | ALB-U | Ig-U | 0.492804645 | 0 |
| B175 | B179 | ALB-U | α1-MG | 0.360487579 | 0 |
| B175 | B180 | ALB-U | Tf | 0.493376972 | 0 |
| B175 | B193 | ALB-U | K | 0.216418468 | 1.78E-15 |
| B177 | B178 | ACR | Ig-U | 0.509307032 | 0 |
| B177 | B179 | ACR | α1-MG | 0.358794291 | 0 |
| B177 | B180 | ACR | Tf | 0.513929363 | 0 |
| B177 | B193 | ACR | K | 0.235759234 | 0 |
| B178 | B179 | Ig-U | α1-MG | 0.760738744 | 0 |
| B178 | B180 | Ig-U | Tf | 0.991249441 | 0 |
| B178 | B193 | Ig-U | K | 0.454901225 | 0 |
| B178 | B195 | Ig-U | Na | -0.24540237 2 | 0 |
| B178 | B194 | Ig-U | Cl | -0.2135711 | 4.00E-15 |
| B179 | B180 | α1-MG | Tf | 0.711709327 | 0 |
| B179 | B193 | α1-MG | K | 0.294970024 | 0 |
| B179 | B195 | α1-MG | Na | -0.14969423 7 | 4.38E-08 |
| B179 | B182 | α1-MG | TT | -0.14728875 2 | 7.22E-08 |
| B18 | B124 | CD3+CD4+helper(inducer) T | ACTH | -0.78648360 4 | 0 |
| B18 | B125 | CD3+CD4+helper(inducer) T | Cor | -0.61054029 2 | 0 |
| B18 | B136 | CD3+CD4+helper(inducer) T | 5'NT | 0.24812374 | 0 |

| | | | | | |
|---|---|---|---|---|---|
| B18 | B152 | CD3+CD4+helper(inducer) T | ACA-Ig(G-A-M) | -0.329073428 | 0 |
| B18 | B155 | CD3+CD4+helper(inducer) T | IgA | 0.305364428 | 0 |
| B18 | B157 | CD3+CD4+helper(inducer) T | IgG | 0.313880452 | 0 |
| B18 | B159 | CD3+CD4+helper(inducer) T | β2-MG | -0.3290587 | 0 |
| B18 | B169 | CD3+CD4+helper(inducer) T | U-CRY | -0.329060341 | 0 |
| B18 | B179 | CD3+CD4+helper(inducer) T | α1-MG | -0.246599597 | 0 |
| B18 | B19 | CD3+CD4+helper(inducer) T | CD3+CD4+/CD3+CD8+ | 0.619451408 | 0 |
| B18 | B199 | CD3+CD4+helper(inducer) T | Mg | 0.371020662 | 0 |
| B18 | B22 | CD3+CD4+helper(inducer) T | CD3+CD8+(T) | 0.826143564 | 0 |
| B18 | B79 | CD3+CD4+helper(inducer) T | hbＶ-dna | 0.283427786 | 0 |
| B18 | B94 | CD3+CD4+helper(inducer) T | ALDH2-Lys504Lys | 0.256366745 | 0 |
| B18 | B182 | CD3+CD4+helper(inducer) T | TT | 0.220417489 | 4.44E-16 |
| B18 | B28 | CD3+CD4+helper(inducer) T | PRL | -0.203632254 | 7.22E-14 |
| B18 | B29 | CD3+CD4+helper(inducer) T | E2 | -0.203632255 | 7.22E-14 |
| B18 | B30 | CD3+CD4+helper(inducer) T | LH | 0.203632255 | 7.22E-14 |
| B18 | B31 | CD3+CD4+helper(inducer) T | FSH | 0.203632255 | 7.22E-14 |
| B18 | B32 | CD3+CD4+helper(inducer) T | Testo | -0.203632255 | 7.22E-14 |
| B18 | B33 | CD3+CD4+helper(inducer) T | Prog | -0.203632255 | 7.22E-14 |
| B18 | B21 | CD3+CD4+helper(inducer) T | CD3+CD4-CD8- | 0.19356722 | 1.19E-12 |
| B18 | B20 | CD3+CD4+helper(inducer) T | CD3+CD4+CD8+ | 0.185439048 | 1.03E-11 |
| B18 | B62 | CD3+CD4+helper(inducer) T | WBHSRVI | -0.167520728 | 8.49E-10 |
| B18 | B181 | CD3+CD4+helper(inducer) T | APTT | 0.165233888 | 1.44E-09 |

| | | | | | |
|---|---|---|---|---|---|
| B18 | B101 | CD3+CD4+helper(inducer) T | anti-Sc70 | 0.157686644 | 7.90E-09 |
| B18 | B102 | CD3+CD4+helper(inducer) T | anti-Sm | 0.15768664 | 7.90E-09 |
| B18 | B103 | CD3+CD4+helper(inducer) T | anti-SSA/Ro | 0.157686641 | 7.90E-09 |
| B18 | B104 | CD3+CD4+helper(inducer) T | anti-SSB | 0.157686642 | 7.90E-09 |
| B18 | B105 | CD3+CD4+helper(inducer) T | anti-U1RNP | 0.157686641 | 7.90E-09 |
| B18 | B106 | CD3+CD4+helper(inducer) T | anti-P | 0.157686642 | 7.90E-09 |
| B18 | B107 | CD3+CD4+helper(inducer) T | AnuA | 0.157686554 | 7.90E-09 |
| B18 | B109 | CD3+CD4+helper(inducer) T | AMA-M2 | 0.157686641 | 7.90E-09 |
| B18 | B110 | CD3+CD4+helper(inducer) T | AHA | 0.157686641 | 7.90E-09 |
| B18 | B97 | CD3+CD4+helper(inducer) T | dsDNA | 0.157686642 | 7.90E-09 |
| B18 | B99 | CD3+CD4+helper(inducer) T | anti-PM-Scl | 0.157686642 | 7.90E-09 |
| B18 | B200 | CD3+CD4+helper(inducer) T | OP | -0.15655896 | 1.01E-08 |
| B18 | B92 | CD3+CD4+helper(inducer) T | ALDH2-Glu504Glu | -0.155320241 | 1.32E-08 |
| B18 | B58 | CD3+CD4+helper(inducer) T | HCT | -0.150699644 | 3.55E-08 |
| B18 | B60 | CD3+CD4+helper(inducer) T | WBRVI | -0.149121541 | 4.94E-08 |
| B18 | B67 | CD3+CD4+helper(inducer) T | WBRSR(mPas_50) | -0.143863821 | 1.45E-07 |
| B18 | B66 | CD3+CD4+helper(inducer) T | WBRSR(mPas_100) | -0.143012651 | 1.72E-07 |
| B18 | B93 | CD3+CD4+helper(inducer) T | ALDH2-Glu504Lys | 0.141256608 | 2.43E-07 |
| B18 | B89 | CD3+CD4+helper(inducer) T | CA 72-4 | 0.136273408 | 6.39E-07 |
| B18 | B63 | CD3+CD4+helper(inducer) T | WBRSR(mPas_200) | -0.134698729 | 8.61E-07 |
| B18 | B154 | CD3+CD4+helper(inducer) T | RF | 0.13415799 | 9.54E-07 |
| B18 | B64 | CD3+CD4+helper(inducer) T | WBRSR(mPas_5) | -0.134116253 | 9.61E-07 |

| | | | | | |
|---|---|---|---|---|---|
| B180 | B193 | Tf | K | 0.443930114 | 0 |
| B180 | B195 | Tf | Na | -0.252433832 | 0 |
| B180 | B194 | Tf | Cl | -0.217085801 | 1.33E-15 |
| B181 | B184 | APTT | INR | 0.900337373 | 0 |
| B181 | B185 | APTT | PT | 0.875319302 | 0 |
| B181 | B186 | APTT | PFC | 0.246760217 | 0 |
| B181 | B182 | APTT | TT | 0.221999489 | 4.44E-16 |
| B181 | B183 | APTT | PTA | 0.143778825 | 1.47E-07 |
| B182 | B183 | TT | PTA | -0.427939555 | 0 |
| B182 | B185 | TT | PT | 0.148607509 | 5.50E-08 |
| B184 | B185 | INR | PT | 0.985546294 | 0 |
| B184 | B186 | INR | PFC | 0.175931283 | 1.13E-10 |
| B185 | B186 | PT | PFC | 0.215268929 | 2.22E-15 |
| B186 | B200 | PFC | OP | 0.194446353 | 9.37E-13 |
| B189 | B190 | eGFB | Cre | -0.641453637 | 0 |
| B189 | B191 | eGFB | Urea | -0.367449171 | 0 |
| B189 | B192 | eGFB | UA | -0.23132452 | 0 |
| B189 | B220 | eGFB | SBP | -0.184079764 | 1.46E-11 |
| B189 | B216 | eGFB | BMI | 0.173584957 | 2.00E-10 |
| B189 | B221 | eGFB | Hei | 0.137424626 | 5.13E-07 |
| B19 | B101 | CD3+CD4+/CD3+CD8+ | anti-Sc70 | 0.230182118 | 0 |
| B19 | B102 | CD3+CD4+/CD3+CD8+ | anti-Sm | 0.230182114 | 0 |
| B19 | B103 | CD3+CD4+/CD3+CD8+ | anti-SSA/Ro | 0.230182115 | 0 |
| B19 | B104 | CD3+CD4+/CD3+CD8+ | anti-SSB | 0.230182115 | 0 |
| B19 | B105 | CD3+CD4+/CD3+CD8+ | anti-U1RNP | 0.230182116 | 0 |
| B19 | B106 | CD3+CD4+/CD3+CD8+ | anti-P | 0.230182116 | 0 |
| B19 | B107 | CD3+CD4+/CD3+CD8+ | AnuA | 0.230181985 | 0 |
| B19 | B109 | CD3+CD4+/CD3+CD8+ | AMA-M2 | 0.230182116 | 0 |
| B19 | B110 | CD3+CD4+/CD3+CD8+ | AHA | 0.230182116 | 0 |
| B19 | B119 | CD3+CD4+/CD3+CD8+ | Hcv-rna | 0.329072038 | 0 |
| B19 | B121 | CD3+CD4+/CD3+CD8+ | C3 | 0.496834695 | 0 |
| B19 | B122 | CD3+CD4+/CD3+CD8+ | C4 | 0.587279114 | 0 |
| B19 | B124 | CD3+CD4+/CD3+CD8+ | ACTH | -0.658268452 | 0 |
| B19 | B125 | CD3+CD4+/CD3+CD8+ | Cor | -0.856642965 | 0 |
| B19 | B136 | CD3+CD4+/CD3+CD8+ | 5'NT | 0.675576383 | 0 |
| B19 | B152 | CD3+CD4+/CD3+CD8+ | ACA-Ig(G-A-M) | -0.38314563 | 0 |

| | | | | | |
|---|---|---|---|---|---|
| | | | | 9 | |
| B19 | B159 | CD3+CD4+/CD3+CD8+ | β2-MG | -0.383128394 | 0 |
| B19 | B169 | CD3+CD4+/CD3+CD8+ | U-CRY | -0.383130275 | 0 |
| B19 | B199 | CD3+CD4+/CD3+CD8+ | Mg | 0.295285379 | 0 |
| B19 | B20 | CD3+CD4+/CD3+CD8+ | CD3+CD4+CD8+ | -0.379384457 | 0 |
| B19 | B21 | CD3+CD4+/CD3+CD8+ | CD3+CD4-CD8- | -0.418540969 | 0 |
| B19 | B23 | CD3+CD4+/CD3+CD8+ | CD3-CD16+CD56+(NK) | 0.34330761 | 0 |
| B19 | B79 | CD3+CD4+/CD3+CD8+ | hb V -dna | 0.239125914 | 0 |
| B19 | B89 | CD3+CD4+/CD3+CD8+ | CA 72-4 | 0.570267255 | 0 |
| B19 | B94 | CD3+CD4+/CD3+CD8+ | ALDH2-Lys504Lys | 0.280490472 | 0 |
| B19 | B97 | CD3+CD4+/CD3+CD8+ | dsDNA | 0.230182116 | 0 |
| B19 | B99 | CD3+CD4+/CD3+CD8+ | anti-PM-Scl | 0.230182117 | 0 |
| B19 | B120 | CD3+CD4+/CD3+CD8+ | HCV-cAg | 0.193526896 | 1.20E-12 |
| B19 | B179 | CD3+CD4+/CD3+CD8+ | α1-MG | -0.192197471 | 1.72E-12 |
| B19 | B151 | CD3+CD4+/CD3+CD8+ | Anti-CCP | -0.18353291 | 1.68E-11 |
| B19 | B92 | CD3+CD4+/CD3+CD8+ | ALDH2-Glu504Glu | -0.168576982 | 6.63E-10 |
| B19 | B196 | CD3+CD4+/CD3+CD8+ | TCO2 | -0.161404596 | 3.45E-09 |
| B19 | B186 | CD3+CD4+/CD3+CD8+ | PFC | -0.15738008 | 8.45E-09 |
| B19 | B93 | CD3+CD4+/CD3+CD8+ | ALDH2-Glu504Lys | 0.153211914 | 2.08E-08 |
| B19 | B182 | CD3+CD4+/CD3+CD8+ | TT | 0.141441176 | 2.34E-07 |
| B19 | B157 | CD3+CD4+/CD3+CD8+ | IgG | -0.140605168 | 2.76E-07 |
| B19 | B197 | CD3+CD4+/CD3+CD8+ | AG | 0.140419453 | 2.87E-07 |
| B19 | B22 | CD3+CD4+/CD3+CD8+ | CD3+CD8+(T) | 0.135714017 | 7.11E-07 |
| B190 | B191 | Cre | Urea | 0.384690868 | 0 |
| B190 | B192 | Cre | UA | 0.479894966 | 0 |
| B190 | B221 | Cre | Hei | 0.272993903 | 0 |
| B190 | B222 | Cre | NS | 0.243896826 | 0 |
| B190 | B216 | Cre | BMI | 0.170342618 | 4.37E-10 |
| B191 | B192 | Urea | UA | 0.17023 | 4.48E-10 |
| B192 | B206 | UA | Waist | 0.272153166 | 0 |
| B192 | B207 | UA | WHR | 0.231886386 | 0 |
| B192 | B216 | UA | BMI | 0.303084029 | 0 |
| B192 | B221 | UA | Hei | 0.235744855 | 0 |
| B192 | B222 | UA | NS | 0.366601292 | 0 |
| B192 | B215 | UA | BMI | 0.220440006 | 4.44E-16 |
| B192 | B210 | UA | Hip | 0.199503272 | 2.32E-13 |

| | | | | | |
|---|---|---|---|---|---|
| B193 | B200 | K | OP | 0.182490476 | 2.19E-11 |
| B193 | B194 | K | Cl | -0.144274109 | 1.33E-07 |
| B194 | B195 | Cl | Na | 0.566103596 | 0 |
| B194 | B197 | Cl | AG | -0.30582502 | 0 |
| B194 | B198 | Cl | P | -0.240789746 | 0 |
| B195 | B197 | Na | AG | 0.332558875 | 0 |
| B195 | B200 | Na | OP | 0.56360873 | 0 |
| B196 | B197 | TCO2 | AG | -0.785808366 | 0 |
| B196 | B200 | TCO2 | OP | -0.31425964 | 0 |
| B196 | B199 | TCO2 | Mg | -0.213330931 | 4.22E-15 |
| B197 | B198 | AG | P | 0.23178507 | 0 |
| B197 | B200 | AG | OP | 0.533768634 | 0 |
| B198 | B200 | P | OP | 0.162728791 | 2.56E-09 |
| B2 | B126 | Age | Alb | -0.334984772 | 0 |
| B2 | B127 | Age | A-G ratio | -0.286644723 | 0 |
| B2 | B128 | Age | ALT | -0.331923794 | 0 |
| B2 | B141 | Age | AST/ALT | 0.407976607 | 0 |
| B2 | B189 | Age | eGFB | -0.510702551 | 0 |
| B2 | B191 | Age | Urea | 0.236359411 | 0 |
| B2 | B210 | Age | Hip | -0.225075398 | 0 |
| B2 | B215 | Age | BMI | -0.295703922 | 0 |
| B2 | B216 | Age | BMI | -0.43890545 | 0 |
| B2 | B220 | Age | SBP | 0.340665275 | 0 |
| B2 | B221 | Age | Hei | -0.363615019 | 0 |
| B2 | B222 | Age | NS | -0.310235134 | 0 |
| B2 | B46 | Age | RBC Count | -0.340852168 | 0 |
| B2 | B47 | Age | RDW | 0.234245076 | 0 |
| B2 | B51 | Age | Hb | -0.363951127 | 0 |
| B2 | B72 | Age | HDL-C | 0.235957305 | 0 |
| B2 | B206 | Age | Waist | -0.19270450 | 1.50E-12 |

| | | | | | |
|---|---|---|---|---|---|
| | | | | 3 | |
| B2 | B41 | Age | MON％ | 0.18709544 | 6.67E-12 |
| B2 | B130 | Age | GLB | 0.186201654 | 8.42E-12 |
| B2 | B71 | Age | TG | -0.184210382 | 1.41E-11 |
| B2 | B146 | Age | CysC | 0.175049951 | 1.40E-10 |
| B2 | B209 | Age | Smoking | -0.157663926 | 7.94E-09 |
| B2 | B190 | Age | Cre | 0.149435749 | 4.63E-08 |
| B2 | B176 | Age | Ucr | -0.145040807 | 1.14E-07 |
| B2 | B188 | Age | Ca | -0.137746405 | 4.82E-07 |
| B2 | B45 | Age | MCHC | -0.137010904 | 5.55E-07 |
| B20 | B101 | CD3+CD4+CD8+ | anti-Sc70 | 0.249193824 | 0 |
| B20 | B102 | CD3+CD4+CD8+ | anti-Sm | 0.249193819 | 0 |
| B20 | B103 | CD3+CD4+CD8+ | anti-SSA/Ro | 0.24919382 | 0 |
| B20 | B104 | CD3+CD4+CD8+ | anti-SSB | 0.24919382 | 0 |
| B20 | B105 | CD3+CD4+CD8+ | anti-U1RNP | 0.249193821 | 0 |
| B20 | B106 | CD3+CD4+CD8+ | anti-P | 0.249193821 | 0 |
| B20 | B107 | CD3+CD4+CD8+ | AnuA | 0.249193683 | 0 |
| B20 | B109 | CD3+CD4+CD8+ | AMA-M2 | 0.24919382 | 0 |
| B20 | B110 | CD3+CD4+CD8+ | AHA | 0.249193821 | 0 |
| B20 | B121 | CD3+CD4+CD8+ | C3 | -0.252807789 | 0 |
| B20 | B122 | CD3+CD4+CD8+ | C4 | -0.307357228 | 0 |
| B20 | B136 | CD3+CD4+CD8+ | 5'NT | -0.282796289 | 0 |
| B20 | B151 | CD3+CD4+CD8+ | Anti-CCP | 0.235966863 | 0 |
| B20 | B157 | CD3+CD4+CD8+ | IgG | 0.26713563 | 0 |
| B20 | B186 | CD3+CD4+CD8+ | PFC | 0.318087921 | 0 |
| B20 | B21 | CD3+CD4+CD8+ | CD3+CD4-CD8- | 0.758675072 | 0 |
| B20 | B22 | CD3+CD4+CD8+ | CD3+CD8+(T) | 0.508350419 | 0 |
| B20 | B23 | CD3+CD4+CD8+ | CD3-CD16+CD56+(NK) | -0.452174555 | 0 |
| B20 | B89 | CD3+CD4+CD8+ | CA 72-4 | -0.331194383 | 0 |
| B20 | B97 | CD3+CD4+CD8+ | dsDNA | 0.249193821 | 0 |
| B20 | B99 | CD3+CD4+CD8+ | anti-PM-Scl | 0.249193821 | 0 |
| B20 | B125 | CD3+CD4+CD8+ | Cor | 0.214786281 | 2.66E-15 |
| B20 | B193 | CD3+CD4+CD8+ | K | 0.212907537 | 4.88E-15 |
| B20 | B79 | CD3+CD4+CD8+ | hb V -dna | 0.177150152 | 8.38E-11 |

| | | | | | |
|---|---|---|---|---|---|
| B20 | B75 | CD3+CD4+CD8+ | ApoA1/B | 0.17615186 | 1.07E-10 |
| B20 | B54 | CD3+CD4+CD8+ | ESR | -0.174007875 | 1.81E-10 |
| B20 | B73 | CD3+CD4+CD8+ | ApoA | -0.169366986 | 5.50E-10 |
| B20 | B74 | CD3+CD4+CD8+ | ApoA1 | 0.161336155 | 3.51E-09 |
| B20 | B76 | CD3+CD4+CD8+ | ApoB | -0.158444475 | 6.68E-09 |
| B201 | B204 | e-habits | Drinking | 0.200097169 | 1.96E-13 |
| B204 | B209 | Drinking | Smoking | 0.401420404 | 0 |
| B204 | B221 | Drinking | Hei | 0.147702396 | 6.63E-08 |
| B204 | B216 | Drinking | BMI | 0.14095501 | 2.58E-07 |
| B204 | B222 | Drinking | NS | 0.137894525 | 4.68E-07 |
| B205 | B222 | KDA | NS | -0.160859935 | 3.90E-09 |
| B206 | B207 | Waist | WHR | 0.798286207 | 0 |
| B206 | B210 | Waist | Hip | 0.783662237 | 0 |
| B206 | B215 | Waist | BMI | 0.819821101 | 0 |
| B206 | B216 | Waist | BMI | 0.790002746 | 0 |
| B206 | B221 | Waist | Hei | 0.295925409 | 0 |
| B206 | B222 | Waist | NS | 0.70822204 | 0 |
| B206 | B218 | Waist | DBP | 0.158191545 | 7.07E-09 |
| B207 | B210 | WHR | Hip | 0.266091147 | 0 |
| B207 | B215 | WHR | BMI | 0.535151035 | 0 |
| B207 | B216 | WHR | BMI | 0.481267621 | 0 |
| B207 | B222 | WHR | NS | 0.494331646 | 0 |
| B209 | B221 | Smoking | Hei | 0.177079016 | 8.52E-11 |
| B209 | B222 | Smoking | NS | 0.171840913 | 3.05E-10 |
| B209 | B216 | Smoking | BMI | 0.138822628 | 3.92E-07 |
| B21 | B101 | CD3+CD4-CD8- | anti-Sc70 | 0.324574798 | 0 |
| B21 | B102 | CD3+CD4-CD8- | anti-Sm | 0.32457479 | 0 |
| B21 | B103 | CD3+CD4-CD8- | anti-SSA/Ro | 0.324574792 | 0 |
| B21 | B104 | CD3+CD4-CD8- | anti-SSB | 0.324574792 | 0 |
| B21 | B105 | CD3+CD4-CD8- | anti-U1RNP | 0.324574792 | 0 |
| B21 | B106 | CD3+CD4-CD8- | anti-P | 0.324574793 | 0 |
| B21 | B107 | CD3+CD4-CD8- | AnuA | 0.32457461 | 0 |
| B21 | B109 | CD3+CD4-CD8- | AMA-M2 | 0.324574792 | 0 |
| B21 | B110 | CD3+CD4-CD8- | AHA | 0.324574793 | 0 |
| B21 | B121 | CD3+CD4-CD8- | C3 | -0.318406879 | 0 |
| B21 | B122 | CD3+CD4-CD8- | C4 | -0.387098289 | 0 |
| B21 | B125 | CD3+CD4-CD8- | Cor | 0.270447618 | 0 |
| B21 | B136 | CD3+CD4-CD8- | 5'NT | -0.34638569 | 0 |

| | | | | | |
|---|---|---|---|---|---|
| | | | | 5 | |
| B21 | B157 | CD3+CD4-CD8- | IgG | 0.368405118 | 0 |
| B21 | B22 | CD3+CD4-CD8- | CD3+CD8+(T) | 0.553800874 | 0 |
| B21 | B23 | CD3+CD4-CD8- | CD3-CD16+CD56+(NK) | -0.51747391<br>5 | 0 |
| B21 | B73 | CD3+CD4-CD8- | ApoA | -0.39452940<br>4 | 0 |
| B21 | B89 | CD3+CD4-CD8- | CA 72-4 | -0.41703482 | 0 |
| B21 | B97 | CD3+CD4-CD8- | dsDNA | 0.324574792 | 0 |
| B21 | B99 | CD3+CD4-CD8- | anti-PM-Scl | 0.324574794 | 0 |
| B21 | B186 | CD3+CD4-CD8- | PFC | 0.185171808 | 1.10E-11 |
| B21 | B81 | CD3+CD4-CD8- | TSGF | -0.18360552<br>1 | 1.65E-11 |
| B21 | B196 | CD3+CD4-CD8- | TCO2 | 0.176343476 | 1.02E-10 |
| B21 | B54 | CD3+CD4-CD8- | ESR | -0.17296043 | 2.33E-10 |
| B21 | B197 | CD3+CD4-CD8- | AG | -0.16951983 | 5.31E-10 |
| B21 | B74 | CD3+CD4-CD8- | ApoA1 | 0.164466081 | 1.72E-09 |
| B21 | B124 | CD3+CD4-CD8- | ACTH | 0.147199325 | 7.35E-08 |
| B21 | B75 | CD3+CD4-CD8- | ApoA1/B | 0.142979903 | 1.73E-07 |
| B21 | B79 | CD3+CD4-CD8- | hb V -dna | 0.140832946 | 2.64E-07 |
| B21 | B151 | CD3+CD4-CD8- | Anti-CCP | 0.136779675 | 5.80E-07 |
| B21 | B137 | CD3+CD4-CD8- | AFU | 0.134138078 | 9.57E-07 |
| B210 | B215 | Hip | BMI | 0.765432686 | 0 |
| B210 | B216 | Hip | BMI | 0.770932701 | 0 |
| B210 | B221 | Hip | Hei | 0.340482436 | 0 |
| B210 | B222 | Hip | NS | 0.636506585 | 0 |
| B210 | B218 | Hip | DBP | 0.154599328 | 1.55E-08 |
| B211 | B212 | IOP-R | IOP-L | 0.786535748 | 0 |
| B215 | B216 | BMI | BMI | 0.802099165 | 0 |
| B215 | B222 | BMI | NS | 0.636771224 | 0 |
| B215 | B218 | BMI | DBP | 0.173106707 | 2.25E-10 |
| B216 | B221 | BMI | Hei | 0.667990193 | 0 |
| B216 | B222 | BMI | NS | 0.788403369 | 0 |
| B216 | B219 | BMI | OT | 0.135723325 | 7.10E-07 |
| B218 | B220 | DBP | SBP | 0.535262534 | 0 |
| B219 | B222 | OT | NS | 0.177721691 | 7.27E-11 |
| B219 | B221 | OT | Hei | 0.141068123 | 2.52E-07 |
| B22 | B121 | CD3+CD8+(T) | C3 | -0.38242633<br>4 | 0 |
| B22 | B122 | CD3+CD8+(T) | C4 | -0.39758783<br>9 | 0 |
| B22 | B124 | CD3+CD8+(T) | ACTH | -0.49629944<br>3 | 0 |
| B22 | B125 | CD3+CD8+(T) | Cor | -0.30750141 | 0 |

| | | | | | |
|---|---|---|---|---|---|
| | | | | 5 | |
| B22 | B152 | CD3+CD8+(T) | ACA-Ig(G-A-M) | -0.24503475<br>2 | 0 |
| B22 | B157 | CD3+CD8+(T) | IgG | 0.474532321 | 0 |
| B22 | B159 | CD3+CD8+(T) | β2-MG | -0.24502373<br>7 | 0 |
| B22 | B169 | CD3+CD8+(T) | U-CRY | -0.24502495<br>1 | 0 |
| B22 | B199 | CD3+CD8+(T) | Mg | 0.310413673 | 0 |
| B22 | B23 | CD3+CD8+(T) | CD3-CD16+CD56+(NK) | -0.26240246<br>6 | 0 |
| B22 | B79 | CD3+CD8+(T) | hb V -dna | 0.296462758 | 0 |
| B22 | B89 | CD3+CD8+(T) | CA 72-4 | -0.28404371<br>5 | 0 |
| B22 | B94 | CD3+CD8+(T) | ALDH2-Lys504Lys | 0.252388323 | 0 |
| B22 | B155 | CD3+CD8+(T) | IgA | 0.215095706 | 2.44E-15 |
| B22 | B136 | CD3+CD8+(T) | 5'NT | -0.20035091<br>4 | 1.83E-13 |
| B22 | B101 | CD3+CD8+(T) | anti-Sc70 | 0.194931514 | 8.20E-13 |
| B22 | B102 | CD3+CD8+(T) | anti-Sm | 0.194931511 | 8.20E-13 |
| B22 | B103 | CD3+CD8+(T) | anti-SSA/Ro | 0.194931511 | 8.20E-13 |
| B22 | B104 | CD3+CD8+(T) | anti-SSB | 0.194931511 | 8.20E-13 |
| B22 | B105 | CD3+CD8+(T) | anti-U1RNP | 0.194931511 | 8.20E-13 |
| B22 | B106 | CD3+CD8+(T) | anti-P | 0.194931512 | 8.20E-13 |
| B22 | B109 | CD3+CD8+(T) | AMA-M2 | 0.194931511 | 8.20E-13 |
| B22 | B110 | CD3+CD8+(T) | AHA | 0.194931511 | 8.20E-13 |
| B22 | B97 | CD3+CD8+(T) | dsDNA | 0.194931511 | 8.20E-13 |
| B22 | B99 | CD3+CD8+(T) | anti-PM-Scl | 0.194931512 | 8.20E-13 |
| B22 | B107 | CD3+CD8+(T) | AnuA | 0.194931404 | 8.21E-13 |
| B22 | B179 | CD3+CD8+(T) | α1-MG | -0.19491153 | 8.25E-13 |
| B22 | B182 | CD3+CD8+(T) | TT | 0.184467613 | 1.32E-11 |
| B22 | B73 | CD3+CD8+(T) | ApoA | -0.17158804<br>4 | 3.24E-10 |
| B22 | B200 | CD3+CD8+(T) | OP | -0.16748015<br>8 | 8.57E-10 |
| B22 | B181 | CD3+CD8+(T) | APTT | 0.162745278 | 2.55E-09 |
| B22 | B75 | CD3+CD8+(T) | ApoA1/B | 0.161063512 | 3.73E-09 |
| B22 | B158 | CD3+CD8+(T) | IgM | 0.158792954 | 6.19E-09 |
| B22 | B92 | CD3+CD8+(T) | ALDH2-Glu504Glu | -0.15440384<br>2 | 1.61E-08 |
| B22 | B93 | CD3+CD8+(T) | ALDH2-Glu504Lys | 0.14053434 | 2.80E-07 |
| B22 | B156 | CD3+CD8+(T) | IgE | 0.139939941 | 3.15E-07 |
| B22 | B154 | CD3+CD8+(T) | RF | 0.137325544 | 5.23E-07 |
| B22 | B76 | CD3+CD8+(T) | ApoB | -0.13560418 | 7.26E-07 |

| | | | | | |
|---|---|---|---|---|---|
| B22 | B62 | CD3+CD8+(T) | WBHSRVI | -0.134635392 | 8.72E-07 |
| B220 | B221 | SBP | Hei | -0.224251263 | 0 |
| B221 | B222 | Hei | NS | 0.525319784 | 0 |
| B23 | B119 | CD3-CD16+CD56+(NK) | Hcv-rna | 0.358803356 | 0 |
| B23 | B121 | CD3-CD16+CD56+(NK) | C3 | 0.344805531 | 0 |
| B23 | B122 | CD3-CD16+CD56+(NK) | C4 | 0.301131931 | 0 |
| B23 | B125 | CD3-CD16+CD56+(NK) | Cor | -0.316549559 | 0 |
| B23 | B151 | CD3-CD16+CD56+(NK) | Anti-CCP | -0.229577919 | 0 |
| B23 | B152 | CD3-CD16+CD56+(NK) | ACA-Ig(G-A-M) | -0.289022693 | 0 |
| B23 | B157 | CD3-CD16+CD56+(NK) | IgG | -0.251414282 | 0 |
| B23 | B159 | CD3-CD16+CD56+(NK) | β2-MG | -0.289009752 | 0 |
| B23 | B169 | CD3-CD16+CD56+(NK) | U-CRY | -0.289011179 | 0 |
| B23 | B186 | CD3-CD16+CD56+(NK) | PFC | -0.286438094 | 0 |
| B23 | B196 | CD3-CD16+CD56+(NK) | TCO2 | -0.284498931 | 0 |
| B23 | B197 | CD3-CD16+CD56+(NK) | AG | 0.285499145 | 0 |
| B23 | B200 | CD3-CD16+CD56+(NK) | OP | 0.236855387 | 0 |
| B23 | B54 | CD3-CD16+CD56+(NK) | ESR | 0.259816148 | 0 |
| B23 | B195 | CD3-CD16+CD56+(NK) | Na | 0.203546966 | 7.39E-14 |
| B23 | B94 | CD3-CD16+CD56+(NK) | ALDH2-Lys504Lys | 0.201978503 | 1.15E-13 |
| B23 | B199 | CD3-CD16+CD56+(NK) | Mg | 0.198926974 | 2.72E-13 |
| B23 | B79 | CD3-CD16+CD56+(NK) | hbＶ-dna | 0.175422845 | 1.28E-10 |
| B23 | B69 | CD3-CD16+CD56+(NK) | PV | 0.169058269 | 5.92E-10 |
| B23 | B28 | CD3-CD16+CD56+(NK) | PRL | 0.158702426 | 6.31E-09 |
| B23 | B29 | CD3-CD16+CD56+(NK) | E2 | 0.158702426 | 6.31E-09 |
| B23 | B30 | CD3-CD16+CD56+(NK) | LH | -0.158702426 | 6.31E-09 |
| B23 | B31 | CD3-CD16+CD56+(NK) | FSH | -0.158702426 | 6.31E-09 |
| B23 | B32 | CD3-CD16+CD56+(NK) | Testo | 0.158702426 | 6.31E-09 |
| B23 | B33 | CD3-CD16+CD56+(NK) | Prog | 0.158702426 | 6.31E-09 |
| B23 | B120 | CD3-CD16+CD56+(NK) | HCV-cAg | 0.158486399 | 6.62E-09 |
| B23 | B124 | CD3-CD16+CD56+(NK) | ACTH | -0.141328748 | 2.40E-07 |
| B23 | B81 | CD3-CD16+CD56+(NK) | TSGF | 0.137786302 | 4.78E-07 |

| | | | | | |
|---|---|---|---|---|---|
| B23 | B73 | CD3-CD16+CD56+(NK) | ApoA | 0.137607704 | 4.95E-07 |
| B24 | B26 | αHBDH | CKMB | 0.390747925 | 0 |
| B24 | B27 | αHBDH | LDH | 0.623000507 | 0 |
| B24 | B28 | αHBDH | PRL | -0.25197420<br>1 | 0 |
| B24 | B29 | αHBDH | E2 | -0.25197420<br>2 | 0 |
| B24 | B30 | αHBDH | LH | 0.251974202 | 0 |
| B24 | B31 | αHBDH | FSH | 0.251974202 | 0 |
| B24 | B32 | αHBDH | Testo | -0.25197420<br>2 | 0 |
| B24 | B33 | αHBDH | Prog | -0.25197420<br>2 | 0 |
| B24 | B74 | αHBDH | ApoA1 | 0.243060392 | 0 |
| B24 | B25 | αHBDH | CK | 0.214738303 | 2.66E-15 |
| B24 | B135 | αHBDH | TBA | 0.21261425 | 5.33E-15 |
| B24 | B75 | αHBDH | ApoA1/B | 0.171687618 | 3.17E-10 |
| B24 | B125 | αHBDH | Cor | 0.141013563 | 2.55E-07 |
| B24 | B62 | αHBDH | WBHSRVI | -0.14072953<br>5 | 2.70E-07 |
| B24 | B78 | αHBDH | HBcAb | -0.13655547<br>7 | 6.06E-07 |
| B24 | B124 | αHBDH | ACTH | -0.13631737<br>8 | 6.34E-07 |
| B25 | B135 | CK | TBA | 0.250614296 | 0 |
| B25 | B26 | CK | CKMB | 0.251327875 | 0 |
| B25 | B27 | CK | LDH | 0.195073314 | 7.89E-13 |
| B25 | B76 | CK | ApoB | -0.16141268<br>2 | 3.45E-09 |
| B25 | B197 | CK | AG | -0.14675147<br>7 | 8.06E-08 |
| B25 | B69 | CK | PV | -0.13779422<br>9 | 4.78E-07 |
| B26 | B27 | CKMB | LDH | 0.266355104 | 0 |
| B26 | B73 | CKMB | ApoA | 0.27797223 | 0 |
| B26 | B199 | CKMB | Mg | -0.15323224<br>1 | 2.08E-08 |
| B26 | B79 | CKMB | hb V -dna | -0.13536716<br>5 | 7.59E-07 |
| B27 | B131 | LDH | AST | 0.198504902 | 3.06E-13 |
| B27 | B135 | LDH | TBA | 0.177443072 | 7.79E-11 |
| B27 | B74 | LDH | ApoA1 | 0.153743018 | 1.86E-08 |
| B27 | B176 | LDH | Ucr | -0.15192522<br>8 | 2.74E-08 |

| | | | | | |
|---|---|---|---|---|---|
| B27 | B28 | LDH | PRL | -0.13640785<br>7 | 6.23E-07 |
| B27 | B29 | LDH | E2 | -0.13640785<br>7 | 6.23E-07 |
| B27 | B30 | LDH | LH | 0.136407858 | 6.23E-07 |
| B27 | B31 | LDH | FSH | 0.136407857 | 6.23E-07 |
| B27 | B32 | LDH | Testo | -0.13640785<br>7 | 6.23E-07 |
| B27 | B33 | LDH | Prog | -0.13640785<br>7 | 6.23E-07 |
| B28 | B101 | PRL | anti-Sc70 | 0.265945926 | 0 |
| B28 | B102 | PRL | anti-Sm | 0.265945924 | 0 |
| B28 | B103 | PRL | anti-SSA/Ro | 0.265945922 | 0 |
| B28 | B104 | PRL | anti-SSB | 0.265945925 | 0 |
| B28 | B105 | PRL | anti-U1RNP | 0.265945923 | 0 |
| B28 | B106 | PRL | anti-P | 0.265945925 | 0 |
| B28 | B107 | PRL | AnuA | 0.265945811 | 0 |
| B28 | B109 | PRL | AMA-M2 | 0.265945924 | 0 |
| B28 | B110 | PRL | AHA | 0.265945924 | 0 |
| B28 | B119 | PRL | Hcv-rna | 0.350513941 | 0 |
| B28 | B124 | PRL | ACTH | 0.631222824 | 0 |
| B28 | B125 | PRL | Cor | -0.46651423<br>5 | 0 |
| B28 | B138 | PRL | PAB | 0.289685363 | 0 |
| B28 | B29 | PRL | E2 | 1 | 0 |
| B28 | B31 | PRL | FSH | -0.99999999<br>9 | 0 |
| B28 | B32 | PRL | Testo | 1 | 0 |
| B28 | B33 | PRL | Prog | 1 | 0 |
| B28 | B79 | PRL | hb V -dna | 0.352651028 | 0 |
| B28 | B92 | PRL | ALDH2-Glu504Glu | -0.24656697<br>1 | 0 |
| B28 | B93 | PRL | ALDH2-Glu504Lys | 0.224218726 | 0 |
| B28 | B94 | PRL | ALDH2-Lys504Lys | 0.407476705 | 0 |
| B28 | B97 | PRL | dsDNA | 0.265945925 | 0 |
| B28 | B99 | PRL | anti-PM-Scl | 0.265945924 | 0 |
| B28 | B155 | PRL | IgA | -0.21837863<br>8 | 8.88E-16 |
| B28 | B157 | PRL | IgG | -0.20448832 | 5.64E-14 |
| B28 | B148 | PRL | Anti_TPO | -0.20094067<br>6 | 1.55E-13 |
| B28 | B120 | PRL | HCV-cAg | 0.198178951 | 3.36E-13 |
| B28 | B65 | PRL | WBRSR(mPas_1) | 0.194378927 | 9.54E-13 |
| B28 | B64 | PRL | WBRSR(mPas_5) | 0.186668404 | 7.45E-12 |

| | | | | | |
|---|---|---|---|---|---|
| B28 | B134 | PRL | CHE | 0.169931466 | 4.81E-10 |
| B28 | B135 | PRL | TBA | -0.16957029<br>3 | 5.24E-10 |
| B28 | B60 | PRL | WBRVI | 0.168473569 | 6.79E-10 |
| B28 | B195 | PRL | Na | -0.16428019<br>3 | 1.80E-09 |
| B28 | B194 | PRL | Cl | -0.16336879<br>3 | 2.21E-09 |
| B28 | B186 | PRL | PFC | -0.15900762<br>4 | 5.90E-09 |
| B28 | B67 | PRL | WBRSR(mPas_50) | 0.155085019 | 1.39E-08 |
| B28 | B200 | PRL | OP | -0.15197433<br>7 | 2.71E-08 |
| B28 | B63 | PRL | WBRSR(mPas_200) | 0.151188295 | 3.20E-08 |
| B28 | B75 | PRL | ApoA1/B | -0.14855435 | 5.56E-08 |
| B28 | B66 | PRL | WBRSR(mPas_100) | 0.148437946 | 5.69E-08 |
| B28 | B183 | PRL | PTA | 0.143832465 | 1.46E-07 |
| B28 | B58 | PRL | HCT | 0.139647193 | 3.33E-07 |
| B28 | B182 | PRL | TT | -0.13856573<br>3 | 4.12E-07 |
| B29 | B101 | E2 | anti-Sc70 | 0.265945928 | 0 |
| B29 | B102 | E2 | anti-Sm | 0.265945921 | 0 |
| B29 | B103 | E2 | anti-SSA/Ro | 0.265945922 | 0 |
| B29 | B104 | E2 | anti-SSB | 0.265945923 | 0 |
| B29 | B105 | E2 | anti-U1RNP | 0.265945922 | 0 |
| B29 | B106 | E2 | anti-P | 0.265945924 | 0 |
| B29 | B107 | E2 | AnuA | 0.265945778 | 0 |
| B29 | B109 | E2 | AMA-M2 | 0.265945923 | 0 |
| B29 | B110 | E2 | AHA | 0.265945923 | 0 |
| B29 | B119 | E2 | Hcv-rna | 0.350513941 | 0 |
| B29 | B124 | E2 | ACTH | 0.631222824 | 0 |
| B29 | B125 | E2 | Cor | -0.46651423<br>6 | 0 |
| B29 | B138 | E2 | PAB | 0.289685363 | 0 |
| B29 | B31 | E2 | FSH | -1 | 0 |
| B29 | B32 | E2 | Testo | 1 | 0 |
| B29 | B79 | E2 | hb V -dna | 0.352651028 | 0 |
| B29 | B92 | E2 | ALDH2-Glu504Glu | -0.24656697<br>1 | 0 |
| B29 | B93 | E2 | ALDH2-Glu504Lys | 0.224218726 | 0 |
| B29 | B94 | E2 | ALDH2-Lys504Lys | 0.407476703 | 0 |
| B29 | B97 | E2 | dsDNA | 0.265945922 | 0 |
| B29 | B99 | E2 | anti-PM-Scl | 0.265945923 | 0 |
| B29 | B155 | E2 | IgA | -0.21837863 | 8.88E-16 |

| | | | | | |
|---|---|---|---|---|---|
| | | | | 8 | |
| B29 | B157 | E2 | IgG | -0.20448832 | 5.64E-14 |
| B29 | B148 | E2 | Anti_TPO | -0.200940676 | 1.55E-13 |
| B29 | B120 | E2 | HCV-cAg | 0.19817895 | 3.36E-13 |
| B29 | B65 | E2 | WBRSR(mPas_1) | 0.194378927 | 9.54E-13 |
| B29 | B64 | E2 | WBRSR(mPas_5) | 0.186668404 | 7.45E-12 |
| B29 | B134 | E2 | CHE | 0.169931466 | 4.81E-10 |
| B29 | B135 | E2 | TBA | -0.169570293 | 5.24E-10 |
| B29 | B60 | E2 | WBRVI | 0.168473569 | 6.79E-10 |
| B29 | B195 | E2 | Na | -0.164280193 | 1.80E-09 |
| B29 | B194 | E2 | Cl | -0.163368794 | 2.21E-09 |
| B29 | B186 | E2 | PFC | -0.159007624 | 5.90E-09 |
| B29 | B67 | E2 | WBRSR(mPas_50) | 0.155085019 | 1.39E-08 |
| B29 | B200 | E2 | OP | -0.151974338 | 2.71E-08 |
| B29 | B63 | E2 | WBRSR(mPas_200) | 0.151188295 | 3.20E-08 |
| B29 | B75 | E2 | ApoA1/B | -0.14855435 | 5.56E-08 |
| B29 | B66 | E2 | WBRSR(mPas_100) | 0.148437946 | 5.69E-08 |
| B29 | B183 | E2 | PTA | 0.143832465 | 1.46E-07 |
| B29 | B58 | E2 | HCT | 0.139647193 | 3.33E-07 |
| B29 | B182 | E2 | TT | -0.138565733 | 4.12E-07 |
| B3 | B84 | Hcy | C-PSA | 0.182945463 | 1.95E-11 |
| B3 | B14 | Hcy | G-17 | 0.145268058 | 1.09E-07 |
| B30 | B101 | LH | anti-Sc70 | -0.26594593 | 0 |
| B30 | B102 | LH | anti-Sm | -0.26594592 | 0 |
| B30 | B103 | LH | anti-SSA/Ro | -0.265945924 | 0 |
| B30 | B104 | LH | anti-SSB | -0.265945924 | 0 |
| B30 | B105 | LH | anti-U1RNP | -0.265945926 | 0 |
| B30 | B106 | LH | anti-P | -0.265945933 | 0 |
| B30 | B107 | LH | AnuA | -0.265945718 | 0 |
| B30 | B109 | LH | AMA-M2 | -0.265945925 | 0 |
| B30 | B110 | LH | AHA | -0.26594592 | 0 |

| | | | | | |
|---|---|---|---|---|---|
| | | | | 6 | |
| B30 | B119 | LH | Hcv-rna | -0.350513941 | 0 |
| B30 | B124 | LH | ACTH | -0.631222826 | 0 |
| B30 | B125 | LH | Cor | 0.466514237 | 0 |
| B30 | B138 | LH | PAB | -0.289685363 | 0 |
| B30 | B79 | LH | hb V -dna | -0.352651029 | 0 |
| B30 | B92 | LH | ALDH2-Glu504Glu | 0.246566972 | 0 |
| B30 | B93 | LH | ALDH2-Glu504Lys | -0.224218726 | 0 |
| B30 | B94 | LH | ALDH2-Lys504Lys | -0.407476706 | 0 |
| B30 | B97 | LH | dsDNA | -0.265945927 | 0 |
| B30 | B99 | LH | anti-PM-Scl | -0.265945929 | 0 |
| B30 | B155 | LH | IgA | 0.218378639 | 8.88E-16 |
| B30 | B157 | LH | IgG | 0.20448832 | 5.64E-14 |
| B30 | B148 | LH | Anti_TPO | 0.200940677 | 1.55E-13 |
| B30 | B120 | LH | HCV-cAg | -0.198178933 | 3.36E-13 |
| B30 | B65 | LH | WBRSR(mPas_1) | -0.194378928 | 9.54E-13 |
| B30 | B64 | LH | WBRSR(mPas_5) | -0.186668404 | 7.45E-12 |
| B30 | B134 | LH | CHE | -0.169931467 | 4.81E-10 |
| B30 | B135 | LH | TBA | 0.169570294 | 5.24E-10 |
| B30 | B60 | LH | WBRVI | -0.168473569 | 6.79E-10 |
| B30 | B195 | LH | Na | 0.164280192 | 1.80E-09 |
| B30 | B194 | LH | Cl | 0.163368794 | 2.21E-09 |
| B30 | B186 | LH | PFC | 0.159007625 | 5.90E-09 |
| B30 | B67 | LH | WBRSR(mPas_50) | -0.155085019 | 1.39E-08 |
| B30 | B200 | LH | OP | 0.151974335 | 2.71E-08 |
| B30 | B63 | LH | WBRSR(mPas_200) | -0.151188296 | 3.20E-08 |
| B30 | B75 | LH | ApoA1/B | 0.14855435 | 5.56E-08 |
| B30 | B66 | LH | WBRSR(mPas_100) | -0.148437946 | 5.69E-08 |

| | | | | | |
|---|---|---|---|---|---|
| B30 | B183 | LH | PTA | -0.143832465 | 1.46E-07 |
| B30 | B58 | LH | HCT | -0.139647193 | 3.33E-07 |
| B30 | B182 | LH | TT | 0.138565733 | 4.12E-07 |
| B31 | B101 | FSH | anti-Sc70 | -0.265945926 | 0 |
| B31 | B102 | FSH | anti-Sm | -0.265945917 | 0 |
| B31 | B103 | FSH | anti-SSA/Ro | -0.265945922 | 0 |
| B31 | B104 | FSH | anti-SSB | -0.265945922 | 0 |
| B31 | B105 | FSH | anti-U1RNP | -0.265945922 | 0 |
| B31 | B106 | FSH | anti-P | -0.265945921 | 0 |
| B31 | B107 | FSH | AnuA | -0.26594575 | 0 |
| B31 | B109 | FSH | AMA-M2 | -0.26594592 | 0 |
| B31 | B110 | FSH | AHA | -0.265945922 | 0 |
| B31 | B119 | FSH | Hcv-rna | -0.35051394 | 0 |
| B31 | B124 | FSH | ACTH | -0.631222824 | 0 |
| B31 | B125 | FSH | Cor | 0.466514237 | 0 |
| B31 | B138 | FSH | PAB | -0.289685363 | 0 |
| B31 | B32 | FSH | Testo | -0.999999999 | 0 |
| B31 | B33 | FSH | Prog | -1 | 0 |
| B31 | B79 | FSH | hb V -dna | -0.352651028 | 0 |
| B31 | B92 | FSH | ALDH2-Glu504Glu | 0.246566972 | 0 |
| B31 | B93 | FSH | ALDH2-Glu504Lys | -0.224218726 | 0 |
| B31 | B94 | FSH | ALDH2-Lys504Lys | -0.407476701 | 0 |
| B31 | B97 | FSH | dsDNA | -0.265945921 | 0 |
| B31 | B99 | FSH | anti-PM-Scl | -0.265945924 | 0 |
| B31 | B155 | FSH | IgA | 0.218378638 | 8.88E-16 |
| B31 | B157 | FSH | IgG | 0.20448832 | 5.64E-14 |
| B31 | B148 | FSH | Anti_TPO | 0.200940676 | 1.55E-13 |

| | | | | | |
|---|---|---|---|---|---|
| B31 | B120 | FSH | HCV-cAg | -0.198178924 | 3.36E-13 |
| B31 | B65 | FSH | WBRSR(mPas_1) | -0.194378927 | 9.54E-13 |
| B31 | B64 | FSH | WBRSR(mPas_5) | -0.186668404 | 7.45E-12 |
| B31 | B134 | FSH | CHE | -0.169931466 | 4.81E-10 |
| B31 | B135 | FSH | TBA | 0.169570293 | 5.24E-10 |
| B31 | B60 | FSH | WBRVI | -0.168473568 | 6.79E-10 |
| B31 | B195 | FSH | Na | 0.164280193 | 1.80E-09 |
| B31 | B194 | FSH | Cl | 0.163368795 | 2.21E-09 |
| B31 | B186 | FSH | PFC | 0.159007625 | 5.90E-09 |
| B31 | B67 | FSH | WBRSR(mPas_50) | -0.155085019 | 1.39E-08 |
| B31 | B200 | FSH | OP | 0.151974338 | 2.71E-08 |
| B31 | B63 | FSH | WBRSR(mPas_200) | -0.151188295 | 3.20E-08 |
| B31 | B75 | FSH | ApoA1/B | 0.14855435 | 5.56E-08 |
| B31 | B66 | FSH | WBRSR(mPas_100) | -0.148437946 | 5.69E-08 |
| B31 | B183 | FSH | PTA | -0.143832465 | 1.46E-07 |
| B31 | B58 | FSH | HCT | -0.139647193 | 3.33E-07 |
| B31 | B182 | FSH | TT | 0.138565733 | 4.12E-07 |
| B32 | B101 | Testo | anti-Sc70 | 0.265945925 | 0 |
| B32 | B102 | Testo | anti-Sm | 0.265945922 | 0 |
| B32 | B103 | Testo | anti-SSA/Ro | 0.265945922 | 0 |
| B32 | B104 | Testo | anti-SSB | 0.265945921 | 0 |
| B32 | B105 | Testo | anti-U1RNP | 0.265945922 | 0 |
| B32 | B106 | Testo | anti-P | 0.265945924 | 0 |
| B32 | B107 | Testo | AnuA | 0.265945781 | 0 |
| B32 | B109 | Testo | AMA-M2 | 0.265945922 | 0 |
| B32 | B110 | Testo | AHA | 0.265945922 | 0 |
| B32 | B119 | Testo | Hcv-rna | 0.350513941 | 0 |
| B32 | B124 | Testo | ACTH | 0.631222824 | 0 |
| B32 | B125 | Testo | Cor | -0.466514236 | 0 |
| B32 | B138 | Testo | PAB | 0.289685363 | 0 |
| B32 | B33 | Testo | Prog | 1 | 0 |
| B32 | B79 | Testo | hb V -dna | 0.352651028 | 0 |
| B32 | B92 | Testo | ALDH2-Glu504Glu | -0.24656697 | 0 |

| | | | | | |
|---|---|---|---|---|---|
| | | | | 1 | |
| B32 | B93 | Testo | ALDH2-Glu504Lys | 0.224218726 | 0 |
| B32 | B94 | Testo | ALDH2-Lys504Lys | 0.407476701 | 0 |
| B32 | B97 | Testo | dsDNA | 0.265945922 | 0 |
| B32 | B99 | Testo | anti-PM-Scl | 0.265945924 | 0 |
| B32 | B155 | Testo | IgA | -0.21837863 8 | 8.88E-16 |
| B32 | B157 | Testo | IgG | -0.20448832 | 5.64E-14 |
| B32 | B148 | Testo | Anti_TPO | -0.20094067 6 | 1.55E-13 |
| B32 | B120 | Testo | HCV-cAg | 0.198178942 | 3.36E-13 |
| B32 | B65 | Testo | WBRSR(mPas_1) | 0.194378927 | 9.54E-13 |
| B32 | B64 | Testo | WBRSR(mPas_5) | 0.186668404 | 7.45E-12 |
| B32 | B134 | Testo | CHE | 0.169931466 | 4.81E-10 |
| B32 | B135 | Testo | TBA | -0.16957029 3 | 5.24E-10 |
| B32 | B60 | Testo | WBRVI | 0.168473568 | 6.79E-10 |
| B32 | B195 | Testo | Na | -0.16428019 4 | 1.80E-09 |
| B32 | B194 | Testo | Cl | -0.16336879 4 | 2.21E-09 |
| B32 | B186 | Testo | PFC | -0.15900762 4 | 5.90E-09 |
| B32 | B67 | Testo | WBRSR(mPas_50) | 0.155085019 | 1.39E-08 |
| B32 | B200 | Testo | OP | -0.15197433 8 | 2.71E-08 |
| B32 | B63 | Testo | WBRSR(mPas_200) | 0.151188295 | 3.20E-08 |
| B32 | B75 | Testo | ApoA1/B | -0.14855435 | 5.56E-08 |
| B32 | B66 | Testo | WBRSR(mPas_100) | 0.148437946 | 5.69E-08 |
| B32 | B183 | Testo | PTA | 0.143832465 | 1.46E-07 |
| B32 | B58 | Testo | HCT | 0.139647193 | 3.33E-07 |
| B32 | B182 | Testo | TT | -0.13856573 3 | 4.12E-07 |
| B33 | B101 | Prog | anti-Sc70 | 0.265945927 | 0 |
| B33 | B102 | Prog | anti-Sm | 0.265945921 | 0 |
| B33 | B103 | Prog | anti-SSA/Ro | 0.265945922 | 0 |
| B33 | B104 | Prog | anti-SSB | 0.265945921 | 0 |
| B33 | B105 | Prog | anti-U1RNP | 0.265945922 | 0 |
| B33 | B106 | Prog | anti-P | 0.265945924 | 0 |
| B33 | B107 | Prog | AnuA | 0.265945767 | 0 |
| B33 | B109 | Prog | AMA-M2 | 0.265945922 | 0 |
| B33 | B110 | Prog | AHA | 0.265945923 | 0 |
| B33 | B119 | Prog | Hcv-rna | 0.350513941 | 0 |
| B33 | B124 | Prog | ACTH | 0.631222824 | 0 |

| | | | | | |
|---|---|---|---|---|---|
| B33 | B125 | Prog | Cor | -0.466514236 | 0 |
| B33 | B138 | Prog | PAB | 0.289685363 | 0 |
| B33 | B79 | Prog | hb V -dna | 0.352651028 | 0 |
| B33 | B92 | Prog | ALDH2-Glu504Glu | -0.246566971 | 0 |
| B33 | B93 | Prog | ALDH2-Glu504Lys | 0.224218726 | 0 |
| B33 | B94 | Prog | ALDH2-Lys504Lys | 0.407476702 | 0 |
| B33 | B97 | Prog | dsDNA | 0.265945922 | 0 |
| B33 | B99 | Prog | anti-PM-Scl | 0.265945923 | 0 |
| B33 | B155 | Prog | IgA | -0.218378638 | 8.88E-16 |
| B33 | B157 | Prog | IgG | -0.20448832 | 5.64E-14 |
| B33 | B148 | Prog | Anti_TPO | -0.200940677 | 1.55E-13 |
| B33 | B120 | Prog | HCV-cAg | 0.198178946 | 3.36E-13 |
| B33 | B65 | Prog | WBRSR(mPas_1) | 0.194378927 | 9.54E-13 |
| B33 | B64 | Prog | WBRSR(mPas_5) | 0.186668404 | 7.45E-12 |
| B33 | B134 | Prog | CHE | 0.169931466 | 4.81E-10 |
| B33 | B135 | Prog | TBA | -0.169570293 | 5.24E-10 |
| B33 | B60 | Prog | WBRVI | 0.168473568 | 6.79E-10 |
| B33 | B195 | Prog | Na | -0.164280193 | 1.80E-09 |
| B33 | B194 | Prog | Cl | -0.163368795 | 2.21E-09 |
| B33 | B186 | Prog | PFC | -0.159007624 | 5.90E-09 |
| B33 | B67 | Prog | WBRSR(mPas_50) | 0.155085019 | 1.39E-08 |
| B33 | B200 | Prog | OP | -0.151974338 | 2.71E-08 |
| B33 | B63 | Prog | WBRSR(mPas_200) | 0.151188295 | 3.20E-08 |
| B33 | B75 | Prog | ApoA1/B | -0.14855435 | 5.56E-08 |
| B33 | B66 | Prog | WBRSR(mPas_100) | 0.148437946 | 5.69E-08 |
| B33 | B183 | Prog | PTA | 0.143832465 | 1.46E-07 |
| B33 | B58 | Prog | HCT | 0.139647193 | 3.33E-07 |
| B33 | B182 | Prog | TT | -0.138565733 | 4.12E-07 |
| B35 | B116 | β2-MG | TT3 | 0.516508183 | 0 |
| B35 | B117 | β2-MG | TT4 | 0.737331664 | 0 |
| B35 | B167 | β2-MG | PC | 0.863642472 | 0 |
| B35 | B202 | β2-MG | e-times | 0.873730964 | 0 |
| B36 | B223 | V-Lym% | FH | 0.569548914 | 0 |
| B36 | B38 | V-Lym% | BAND% | -0.80450903 | 0 |

| | | | | | |
|---|---|---|---|---|---|
| | | | | 9 | |
| B37 | B202 | BAND | e-times | 0.374516878 | 0 |
| B38 | B96 | BAND% | dsDNA (IIF) | -0.491400677 | 0 |
| B4 | B121 | WB-Ca | C3 | 0.57491285 | 0 |
| B4 | B122 | WB-Ca | C4 | 0.698091611 | 0 |
| B4 | B124 | WB-Ca | ACTH | -0.264549505 | 0 |
| B4 | B125 | WB-Ca | Cor | -0.483483118 | 0 |
| B4 | B136 | WB-Ca | 5'NT | 0.805945093 | 0 |
| B4 | B157 | WB-Ca | IgG | -0.272858004 | 0 |
| B4 | B19 | WB-Ca | CD3+CD4+/CD3+CD8+ | 0.701348555 | 0 |
| B4 | B20 | WB-Ca | CD3+CD4+CD8+ | -0.410751519 | 0 |
| B4 | B21 | WB-Ca | CD3+CD4-CD8- | -0.517270745 | 0 |
| B4 | B22 | WB-Ca | CD3+CD8+(T) | -0.332756241 | 0 |
| B4 | B5 | WB-Ca | WB-Mg | 0.43691822 | 0 |
| B4 | B6 | WB-Ca | WB-Fe | 0.300260988 | 0 |
| B4 | B7 | WB-Ca | WB-Cop | -0.93813274 | 0 |
| B4 | B8 | WB-Ca | WB-Zinc | 0.324479811 | 0 |
| B4 | B89 | WB-Ca | CA 72-4 | 0.746362142 | 0 |
| B4 | B18 | WB-Ca | CD3+CD4+helper(inducer) T | 0.190045546 | 3.06E-12 |
| B4 | B73 | WB-Ca | ApoA | 0.142622581 | 1.85E-07 |
| B40 | B42 | WBC | MON | 0.634209449 | 0 |
| B40 | B46 | WBC | RBC Count | 0.262408922 | 0 |
| B40 | B51 | WBC | Hb | 0.249246219 | 0 |
| B40 | B53 | WBC | GR% | 0.243822057 | 0 |
| B40 | B72 | WBC | HDL-C | -0.216455575 | 1.78E-15 |
| B40 | B48 | WBC | LYM% | -0.201664445 | 1.26E-13 |
| B40 | B222 | WBC | NS | 0.194258527 | 9.86E-13 |
| B40 | B41 | WBC | MON % | -0.190171904 | 2.96E-12 |
| B40 | B52 | WBC | PLT/BPC | 0.179742947 | 4.39E-11 |
| B40 | B206 | WBC | Waist | 0.172756153 | 2.45E-10 |
| B40 | B216 | WBC | BMI | 0.170661309 | 4.05E-10 |
| B40 | B71 | WBC | TG | 0.165220675 | 1.45E-09 |
| B40 | B215 | WBC | BMI | 0.151523237 | 2.98E-08 |

| | | | | | |
|---|---|---|---|---|---|
| B40 | B207 | WBC | WHR | 0.147561507 | 6.82E-08 |
| B40 | B170 | WBC | U-GLU | 0.13783094 | 4.74E-07 |
| B40 | B209 | WBC | Smoking | 0.136849266 | 5.73E-07 |
| B41 | B42 | MON % | MON | 0.603700894 | 0 |
| B41 | B53 | MON % | GR% | -0.245574375 | 0 |
| B41 | B126 | MON % | Alb | -0.19133006 | 2.17E-12 |
| B41 | B49 | MON % | baso% | 0.188256349 | 4.91E-12 |
| B41 | B70 | MON % | LDL-C | -0.170680033 | 4.03E-10 |
| B41 | B77 | MON % | TC | -0.159036044 | 5.86E-09 |
| B41 | B190 | MON % | Cre | 0.15833692 | 6.84E-09 |
| B41 | B71 | MON % | TG | -0.157429315 | 8.36E-09 |
| B41 | B189 | MON % | eGFB | -0.153499782 | 1.96E-08 |
| B41 | B47 | MON % | RDW | 0.152940176 | 2.21E-08 |
| B41 | B46 | MON % | RBC Count | -0.148086955 | 6.12E-08 |
| B41 | B43 | MON % | MCV | 0.137386164 | 5.17E-07 |
| B41 | B141 | MON % | AST/ALT | 0.134252357 | 9.37E-07 |
| B42 | B190 | MON | Cre | 0.170791244 | 3.92E-10 |
| B42 | B222 | MON | NS | 0.168697384 | 6.44E-10 |
| B42 | B192 | MON | UA | 0.162230657 | 2.87E-09 |
| B42 | B48 | MON | LYM% | -0.157550357 | 8.14E-09 |
| B42 | B163 | MON | SG | 0.142397807 | 1.94E-07 |
| B42 | B72 | MON | HDL-C | -0.138360609 | 4.28E-07 |
| B43 | B44 | MCV | MCH | 0.91190825 | 0 |
| B43 | B46 | MCV | RBC Count | -0.6037356 | 0 |
| B43 | B47 | MCV | RDW | 0.598577924 | 0 |
| B43 | B52 | MCV | PLT/BPC | -0.193747024 | 1.13E-12 |
| B43 | B216 | MCV | BMI | -0.150072225 | 4.05E-08 |
| B43 | B215 | MCV | BMI | -0.142001183 | 2.10E-07 |
| B43 | B210 | MCV | Hip | -0.136861311 | 5.71E-07 |
| B44 | B45 | MCH | MCHC | 0.507600478 | 0 |
| B44 | B46 | MCH | RBC Count | -0.513487569 | 0 |

| | | | | | |
|---|---|---|---|---|---|
| B44 | B47 | MCH | RDW | 0.431554274 | 0 |
| B44 | B51 | MCH | Hb | 0.186628536 | 7.53E-12 |
| B44 | B52 | MCH | PLT/BPC | -0.179119198 | 5.13E-11 |
| B44 | B132 | MCH | Dbil | 0.165513162 | 1.35E-09 |
| B44 | B133 | MCH | TBIL | 0.141031208 | 2.54E-07 |
| B45 | B216 | MCHC | BMI | 0.258649692 | 0 |
| B45 | B221 | MCHC | Hei | 0.271294338 | 0 |
| B45 | B222 | MCHC | NS | 0.237094467 | 0 |
| B45 | B51 | MCHC | Hb | 0.427531085 | 0 |
| B45 | B47 | MCHC | RDW | -0.211573217 | 7.11E-15 |
| B45 | B129 | MCHC | IBIL | 0.170950743 | 3.78E-10 |
| B45 | B133 | MCHC | TBIL | 0.1686529 | 6.51E-10 |
| B45 | B141 | MCHC | AST/ALT | -0.16803522 | 7.53E-10 |
| B45 | B206 | MCHC | Waist | 0.163800488 | 2.01E-09 |
| B45 | B210 | MCHC | Hip | 0.160627549 | 4.11E-09 |
| B45 | B127 | MCHC | A-G ratio | 0.157398751 | 8.41E-09 |
| B45 | B174 | MCHC | Urea/Cr | -0.151117499 | 3.25E-08 |
| B45 | B192 | MCHC | UA | 0.151021164 | 3.32E-08 |
| B45 | B189 | MCHC | eGFB | 0.145495085 | 1.04E-07 |
| B45 | B215 | MCHC | BMI | 0.138447113 | 4.21E-07 |
| B46 | B126 | RBC Count | Alb | 0.28099964 | 0 |
| B46 | B141 | RBC Count | AST/ALT | -0.317598578 | 0 |
| B46 | B206 | RBC Count | Waist | 0.29708114 | 0 |
| B46 | B210 | RBC Count | Hip | 0.261828379 | 0 |
| B46 | B215 | RBC Count | BMI | 0.273541391 | 0 |
| B46 | B216 | RBC Count | BMI | 0.398383565 | 0 |
| B46 | B221 | RBC Count | Hei | 0.332222038 | 0 |
| B46 | B222 | RBC Count | NS | 0.372277133 | 0 |
| B46 | B47 | RBC Count | RDW | -0.466595773 | 0 |
| B46 | B51 | RBC Count | Hb | 0.737672818 | 0 |
| B46 | B72 | RBC Count | HDL-C | -0.259428932 | 0 |
| B46 | B207 | RBC Count | WHR | 0.211430698 | 7.55E-15 |
| B46 | B189 | RBC Count | eGFB | 0.20951006 | 1.31E-14 |
| B46 | B128 | RBC Count | ALT | 0.202473716 | 1.00E-13 |
| B46 | B174 | RBC Count | Urea/Cr | -0.188143521 | 5.06E-12 |
| B46 | B58 | RBC Count | HCT | 0.154386721 | 1.62E-08 |
| B46 | B127 | RBC Count | A-G ratio | 0.151760373 | 2.84E-08 |

| | | | | | |
|---|---|---|---|---|---|
| B46 | B218 | RBC Count | DBP | 0.149608975 | 4.46E-08 |
| B47 | B216 | RDW | BMI | -0.234024074 | 0 |
| B47 | B51 | RDW | Hb | -0.214918977 | 2.66E-15 |
| B47 | B210 | RDW | Hip | -0.212916597 | 4.88E-15 |
| B47 | B126 | RDW | Alb | -0.209195859 | 1.44E-14 |
| B47 | B141 | RDW | AST/ALT | 0.20630867 | 3.33E-14 |
| B47 | B189 | RDW | eGFB | -0.199962447 | 2.04E-13 |
| B47 | B215 | RDW | BMI | -0.196865445 | 4.83E-13 |
| B47 | B72 | RDW | HDL-C | 0.164829331 | 1.58E-09 |
| B47 | B222 | RDW | NS | -0.15584888 | 1.18E-08 |
| B47 | B52 | RDW | PLT/BPC | -0.155690732 | 1.22E-08 |
| B47 | B206 | RDW | Waist | -0.148951785 | 5.12E-08 |
| B47 | B221 | RDW | Hei | -0.14743246 | 7.01E-08 |
| B48 | B53 | LYM% | GR% | -0.929478782 | 0 |
| B49 | B50 | baso% | eo% | 0.324882463 | 0 |
| B49 | B53 | baso% | GR% | -0.1720913 | 2.87E-10 |
| B5 | B136 | WB-Mg | 5'NT | 0.254649603 | 0 |
| B5 | B6 | WB-Mg | WB-Fe | 0.716625456 | 0 |
| B5 | B7 | WB-Mg | WB-Cop | -0.68566311 | 0 |
| B5 | B89 | WB-Mg | CA 72-4 | 0.236421604 | 0 |
| B5 | B122 | WB-Mg | C4 | 0.223591072 | 2.22E-16 |
| B5 | B19 | WB-Mg | CD3+CD4+/CD3+CD8+ | 0.222940104 | 2.22E-16 |
| B5 | B8 | WB-Mg | WB-Zinc | 0.188066815 | 5.17E-12 |
| B5 | B121 | WB-Mg | C3 | 0.184459868 | 1.32E-11 |
| B5 | B21 | WB-Mg | CD3+CD4-CD8- | -0.167020631 | 9.54E-10 |
| B5 | B224 | WB-Mg | pmh | -0.16653491 | 1.07E-09 |
| B5 | B125 | WB-Mg | Cor | -0.152903259 | 2.23E-08 |
| B50 | B53 | eo% | GR% | -0.218245064 | 8.88E-16 |
| B51 | B126 | Hb | Alb | 0.321826951 | 0 |
| B51 | B128 | Hb | ALT | 0.230023707 | 0 |
| B51 | B141 | Hb | AST/ALT | -0.349557776 | 0 |

| | | | | | |
|---|---|---|---|---|---|
| B51 | B174 | Hb | Urea/Cr | -0.249715498 | 0 |
| B51 | B189 | Hb | eGFB | 0.225808539 | 0 |
| B51 | B206 | Hb | Waist | 0.314155964 | 0 |
| B51 | B207 | Hb | WHR | 0.231631883 | 0 |
| B51 | B210 | Hb | Hip | 0.271731413 | 0 |
| B51 | B215 | Hb | BMI | 0.277033953 | 0 |
| B51 | B216 | Hb | BMI | 0.448746236 | 0 |
| B51 | B221 | Hb | Hei | 0.423147517 | 0 |
| B51 | B222 | Hb | NS | 0.438021785 | 0 |
| B51 | B72 | Hb | HDL-C | -0.283097623 | 0 |
| B51 | B127 | Hb | A-G ratio | 0.219552734 | 6.66E-16 |
| B51 | B129 | Hb | IBIL | 0.216837321 | 1.33E-15 |
| B51 | B58 | Hb | HCT | 0.210507577 | 9.77E-15 |
| B51 | B133 | Hb | TBIL | 0.204100965 | 6.31E-14 |
| B51 | B163 | Hb | SG | 0.189884253 | 3.20E-12 |
| B51 | B62 | Hb | WBHSRVI | 0.182955372 | 1.94E-11 |
| B51 | B67 | Hb | WBRSR(mPas_50) | 0.163684857 | 2.06E-09 |
| B51 | B66 | Hb | WBRSR(mPas_100) | 0.163541638 | 2.13E-09 |
| B51 | B192 | Hb | UA | 0.16302205 | 2.40E-09 |
| B51 | B60 | Hb | WBRVI | 0.161235749 | 3.59E-09 |
| B51 | B63 | Hb | WBRSR(mPas_200) | 0.160288994 | 4.44E-09 |
| B51 | B205 | Hb | KDA | -0.156722573 | 9.76E-09 |
| B51 | B64 | Hb | WBRSR(mPas_5) | 0.153294916 | 2.05E-08 |
| B51 | B209 | Hb | Smoking | 0.1486724 | 5.42E-08 |
| B51 | B191 | Hb | Urea | -0.14747851 | 6.94E-08 |
| B51 | B218 | Hb | DBP | 0.139312155 | 3.56E-07 |
| B52 | B132 | PLT/BPC | Dbil | -0.178108736 | 6.60E-11 |
| B54 | B123 | ESR | hs-CRP1 | 0.247127303 | 0 |
| B54 | B58 | ESR | HCT | -0.374796276 | 0 |
| B54 | B59 | ESR | WBRV | 0.237010774 | 0 |
| B54 | B60 | ESR | WBRVI | -0.263611968 | 0 |
| B54 | B62 | ESR | WBHSRVI | -0.307051595 | 0 |
| B54 | B69 | ESR | PV | 0.638075941 | 0 |
| B54 | B74 | ESR | ApoA1 | 0.220513148 | 4.44E-16 |
| B54 | B56 | ESR | IR | -0.201497268 | 1.32E-13 |
| B54 | B73 | ESR | ApoA | 0.188117499 | 5.10E-12 |

| | | | | | |
|---|---|---|---|---|---|
| B54 | B158 | ESR | IgM | 0.166083328 | 1.19E-09 |
| B54 | B76 | ESR | ApoB | 0.160177282 | 4.55E-09 |
| B54 | B200 | ESR | OP | 0.139435769 | 3.48E-07 |
| B54 | B195 | ESR | Na | 0.135883271 | 6.88E-07 |
| B55 | B56 | TK | IR | 0.761707435 | 0 |
| B55 | B57 | TK | RBCAI | -0.25307444<br>4 | 0 |
| B55 | B58 | TK | HCT | -0.35011458 | 0 |
| B55 | B59 | TK | WBRV | 0.53418906 | 0 |
| B55 | B61 | TK | WBHSRV | 0.794237532 | 0 |
| B55 | B62 | TK | WBHSRVI | 0.383469533 | 0 |
| B55 | B63 | TK | WBRSR(mPas_200) | 0.422002715 | 0 |
| B55 | B64 | TK | WBRSR(mPas_5) | 0.317045998 | 0 |
| B55 | B66 | TK | WBRSR(mPas_100) | 0.419644574 | 0 |
| B55 | B67 | TK | WBRSR(mPas_50) | 0.413659435 | 0 |
| B55 | B60 | TK | WBRVI | 0.217166833 | 1.33E-15 |
| B55 | B65 | TK | WBRSR(mPas_1) | 0.214618741 | 2.89E-15 |
| B55 | B108 | TK | Anti-Cyclin | -0.17829343<br>7 | 6.31E-11 |
| B55 | B98 | TK | anti-Jo-1 | -0.16487048<br>5 | 1.57E-09 |
| B55 | B101 | TK | anti-Sc70 | -0.13433768<br>6 | 9.22E-07 |
| B55 | B102 | TK | anti-Sm | -0.13433768<br>1 | 9.22E-07 |
| B55 | B103 | TK | anti-SSA/Ro | -0.13433768<br>1 | 9.22E-07 |
| B55 | B104 | TK | anti-SSB | -0.13433768<br>1 | 9.22E-07 |
| B55 | B105 | TK | anti-U1RNP | -0.13433768<br>2 | 9.22E-07 |
| B55 | B106 | TK | anti-P | -0.13433768<br>3 | 9.22E-07 |
| B55 | B107 | TK | AnuA | -0.13433759<br>5 | 9.22E-07 |
| B55 | B109 | TK | AMA-M2 | -0.13433768<br>1 | 9.22E-07 |
| B55 | B110 | TK | AHA | -0.13433768<br>2 | 9.22E-07 |
| B55 | B97 | TK | dsDNA | -0.13433768<br>3 | 9.22E-07 |
| B55 | B99 | TK | anti-PM-Scl | -0.13433768<br>3 | 9.22E-07 |
| B56 | B57 | IR | RBCAI | -0.34469284 | 0 |

| | | | | | |
|---|---|---|---|---|---|
| | | | | 9 | |
| B56 | B58 | IR | HCT | 0.307733243 | 0 |
| B56 | B59 | IR | WBRV | 0.409847393 | 0 |
| B56 | B60 | IR | WBRVI | 0.653547886 | 0 |
| B56 | B61 | IR | WBHSRV | 0.937511417 | 0 |
| B56 | B62 | IR | WBHSRVI | 0.886568951 | 0 |
| B56 | B63 | IR | WBRSR(mPas_200) | 0.859402842 | 0 |
| B56 | B64 | IR | WBRSR(mPas_5) | 0.72202142 | 0 |
| B56 | B65 | IR | WBRSR(mPas_1) | 0.556585838 | 0 |
| B56 | B66 | IR | WBRSR(mPas_100) | 0.855006462 | 0 |
| B56 | B67 | IR | WBRSR(mPas_50) | 0.847969677 | 0 |
| B56 | B69 | IR | PV | -0.272716943 | 0 |
| B56 | B124 | IR | ACTH | 0.163481782 | 2.16E-09 |
| B56 | B108 | IR | Anti-Cyclin | -0.158986742 | 5.93E-09 |
| B56 | B98 | IR | anti-Jo-1 | -0.148968929 | 5.10E-08 |
| B57 | B108 | RBCAI | Anti-Cyclin | 0.414159852 | 0 |
| B57 | B59 | RBCAI | WBRV | 0.563628746 | 0 |
| B57 | B60 | RBCAI | WBRVI | 0.330543637 | 0 |
| B57 | B61 | RBCAI | WBHSRV | -0.296357109 | 0 |
| B57 | B62 | RBCAI | WBHSRVI | -0.322111796 | 0 |
| B57 | B63 | RBCAI | WBRSR(mPas_200) | -0.296003691 | 0 |
| B57 | B65 | RBCAI | WBRSR(mPas_1) | 0.428713544 | 0 |
| B57 | B66 | RBCAI | WBRSR(mPas_100) | -0.24314403 | 0 |
| B57 | B98 | RBCAI | anti-Jo-1 | 0.415567104 | 0 |
| B57 | B67 | RBCAI | WBRSR(mPas_50) | -0.194834708 | 8.42E-13 |
| B57 | B156 | RBCAI | IgE | 0.187557566 | 5.91E-12 |
| B57 | B69 | RBCAI | PV | 0.178722986 | 5.67E-11 |
| B57 | B64 | RBCAI | WBRSR(mPas_5) | 0.155777824 | 1.20E-08 |
| B57 | B58 | RBCAI | HCT | -0.145769748 | 9.84E-08 |
| B57 | B151 | RBCAI | Anti-CCP | 0.139862111 | 3.20E-07 |
| B58 | B59 | HCT | WBRV | -0.226886514 | 0 |
| B58 | B60 | HCT | WBRVI | 0.615173877 | 0 |
| B58 | B62 | HCT | WBHSRVI | 0.698521836 | 0 |
| B58 | B63 | HCT | WBRSR(mPas_200) | 0.603037355 | 0 |
| B58 | B64 | HCT | WBRSR(mPas_5) | 0.555311328 | 0 |

| | | | | | |
|---|---|---|---|---|---|
| B58 | B65 | HCT | WBRSR(mPas_1) | 0.470438242 | 0 |
| B58 | B66 | HCT | WBRSR(mPas_100) | 0.592623216 | 0 |
| B58 | B67 | HCT | WBRSR(mPas_50) | 0.591700969 | 0 |
| B58 | B69 | HCT | PV | -0.354748855 | 0 |
| B58 | B124 | HCT | ACTH | 0.184977871 | 1.16E-11 |
| B58 | B61 | HCT | WBHSRV | 0.163979844 | 1.93E-09 |
| B58 | B74 | HCT | ApoA1 | -0.146587323 | 8.33E-08 |
| B59 | B108 | WBRV | Anti-Cyclin | 0.292039538 | 0 |
| B59 | B60 | WBRV | WBRVI | 0.54531672 | 0 |
| B59 | B61 | WBRV | WBHSRV | 0.571476498 | 0 |
| B59 | B63 | WBRV | WBRSR(mPas_200) | 0.367529507 | 0 |
| B59 | B64 | WBRV | WBRSR(mPas_5) | 0.641857016 | 0 |
| B59 | B65 | WBRV | WBRSR(mPas_1) | 0.745722597 | 0 |
| B59 | B66 | WBRV | WBRSR(mPas_100) | 0.410180709 | 0 |
| B59 | B67 | WBRV | WBRSR(mPas_50) | 0.443150139 | 0 |
| B59 | B69 | WBRV | PV | 0.398961769 | 0 |
| B59 | B98 | WBRV | anti-Jo-1 | 0.301166569 | 0 |
| B59 | B62 | WBRV | WBHSRVI | 0.188248835 | 4.92E-12 |
| B59 | B151 | WBRV | Anti-CCP | 0.181822004 | 2.60E-11 |
| B59 | B101 | WBRV | anti-Sc70 | -0.181600759 | 2.75E-11 |
| B59 | B102 | WBRV | anti-Sm | -0.181600753 | 2.75E-11 |
| B59 | B103 | WBRV | anti-SSA/Ro | -0.181600755 | 2.75E-11 |
| B59 | B104 | WBRV | anti-SSB | -0.181600755 | 2.75E-11 |
| B59 | B105 | WBRV | anti-U1RNP | -0.181600756 | 2.75E-11 |
| B59 | B106 | WBRV | anti-P | -0.181600755 | 2.75E-11 |
| B59 | B107 | WBRV | AnuA | -0.181600644 | 2.75E-11 |
| B59 | B109 | WBRV | AMA-M2 | -0.181600755 | 2.75E-11 |
| B59 | B110 | WBRV | AHA | -0.181600756 | 2.75E-11 |
| B59 | B97 | WBRV | dsDNA | -0.181600756 | 2.75E-11 |
| B59 | B99 | WBRV | anti-PM-Scl | -0.181600757 | 2.75E-11 |
| B59 | B74 | WBRV | ApoA1 | 0.137628885 | 4.93E-07 |

| | | | | | |
|---|---|---|---|---|---|
| B6 | B7 | WB-Fe | WB-Cop | -0.36897988 2 | 0 |
| B6 | B8 | WB-Fe | WB-Zinc | 0.789531561 | 0 |
| B6 | B224 | WB-Fe | pmh | -0.20366862 2 | 7.15E-14 |
| B60 | B124 | WBRVI | ACTH | 0.232576514 | 0 |
| B60 | B61 | WBRVI | WBHSRV | 0.55960072 | 0 |
| B60 | B62 | WBRVI | WBHSRVI | 0.781407997 | 0 |
| B60 | B63 | WBRVI | WBRSR(mPas_200) | 0.717530896 | 0 |
| B60 | B64 | WBRVI | WBRSR(mPas_5) | 0.892227104 | 0 |
| B60 | B65 | WBRVI | WBRSR(mPas_1) | 0.909630533 | 0 |
| B60 | B66 | WBRVI | WBRSR(mPas_100) | 0.747131233 | 0 |
| B60 | B67 | WBRVI | WBRSR(mPas_50) | 0.77419617 | 0 |
| B60 | B69 | WBRVI | PV | -0.27926956 2 | 0 |
| B61 | B62 | WBHSRV | WBHSRVI | 0.763421612 | 0 |
| B61 | B63 | WBHSRV | WBRSR(mPas_200) | 0.872982607 | 0 |
| B61 | B64 | WBHSRV | WBRSR(mPas_5) | 0.764229341 | 0 |
| B61 | B65 | WBHSRV | WBRSR(mPas_1) | 0.610094759 | 0 |
| B61 | B66 | WBHSRV | WBRSR(mPas_100) | 0.873599978 | 0 |
| B61 | B67 | WBHSRV | WBRSR(mPas_50) | 0.868262785 | 0 |
| B61 | B124 | WBHSRV | ACTH | 0.137632547 | 4.93E-07 |
| B62 | B63 | WBHSRVI | WBRSR(mPas_200) | 0.920035429 | 0 |
| B62 | B64 | WBHSRVI | WBRSR(mPas_5) | 0.795649183 | 0 |
| B62 | B65 | WBHSRVI | WBRSR(mPas_1) | 0.63114906 | 0 |
| B62 | B66 | WBHSRVI | WBRSR(mPas_100) | 0.915977078 | 0 |
| B62 | B67 | WBHSRVI | WBRSR(mPas_50) | 0.910212771 | 0 |
| B62 | B69 | WBHSRVI | PV | -0.38197929 | 0 |
| B62 | B124 | WBHSRVI | ACTH | 0.211764767 | 6.66E-15 |
| B62 | B89 | WBHSRVI | CA 72-4 | 0.13866979 | 4.03E-07 |
| B62 | B216 | WBHSRVI | BMI | 0.137961155 | 4.63E-07 |
| B62 | B74 | WBHSRVI | ApoA1 | -0.13727221 1 | 5.28E-07 |
| B63 | B64 | WBRSR(mPas_200) | WBRSR(mPas_5) | 0.894228477 | 0 |
| B63 | B65 | WBRSR(mPas_200) | WBRSR(mPas_1) | 0.730646783 | 0 |
| B63 | B66 | WBRSR(mPas_200) | WBRSR(mPas_100) | 0.995694386 | 0 |
| B63 | B67 | WBRSR(mPas_200) | WBRSR(mPas_50) | 0.990528331 | 0 |
| B63 | B124 | WBRSR(mPas_200) | ACTH | 0.195205331 | 7.61E-13 |
| B64 | B65 | WBRSR(mPas_5) | WBRSR(mPas_1) | 0.95824965 | 0 |
| B64 | B66 | WBRSR(mPas_5) | WBRSR(mPas_100) | 0.918128653 | 0 |
| B64 | B67 | WBRSR(mPas_5) | WBRSR(mPas_50) | 0.93600549 | 0 |
| B64 | B124 | WBRSR(mPas_5) | ACTH | 0.215693133 | 2.22E-15 |
| B64 | B98 | WBRSR(mPas_5) | anti-Jo-1 | 0.179683023 | 4.46E-11 |
| B64 | B108 | WBRSR(mPas_5) | Anti-Cyclin | 0.175302504 | 1.32E-10 |

| | | | | | |
|---|---|---|---|---|---|
| B65 | B108 | WBRSR(mPas_1) | Anti-Cyclin | 0.283958695 | 0 |
| B65 | B66 | WBRSR(mPas_1) | WBRSR(mPas_100) | 0.767276722 | 0 |
| B65 | B67 | WBRSR(mPas_1) | WBRSR(mPas_50) | 0.797468972 | 0 |
| B65 | B98 | WBRSR(mPas_1) | anti-Jo-1 | 0.288531072 | 0 |
| B65 | B124 | WBRSR(mPas_1) | ACTH | 0.205677097 | 4.00E-14 |
| B65 | B151 | WBRSR(mPas_1) | Anti-CCP | 0.153171249 | 2.10E-08 |
| B65 | B69 | WBRSR(mPas_1) | PV | 0.139511676 | 3.42E-07 |
| B66 | B67 | WBRSR(mPas_100) | WBRSR(mPas_50) | 0.998365536 | 0 |
| B66 | B124 | WBRSR(mPas_100) | ACTH | 0.199234836 | 2.50E-13 |
| B67 | B124 | WBRSR(mPas_50) | ACTH | 0.203764568 | 6.95E-14 |
| B69 | B108 | PV | Anti-Cyclin | 0.317531981 | 0 |
| B69 | B98 | PV | anti-Jo-1 | 0.312759401 | 0 |
| B69 | B158 | PV | IgM | 0.219505486 | 6.66E-16 |
| B69 | B74 | PV | ApoA1 | 0.207803364 | 2.18E-14 |
| B69 | B76 | PV | ApoB | 0.205115559 | 4.71E-14 |
| B69 | B77 | PV | TC | 0.153431421 | 1.99E-08 |
| B69 | B156 | PV | IgE | 0.141231337 | 2.44E-07 |
| B69 | B70 | PV | LDL-C | 0.139508962 | 3.43E-07 |
| B69 | B130 | PV | GLB | 0.135626938 | 7.23E-07 |
| B7 | B121 | WB-Cop | C3 | -0.53891266 | 0 |
| B7 | B122 | WB-Cop | C4 | -0.654263108 | 0 |
| B7 | B124 | WB-Cop | ACTH | 0.24781699 | 0 |
| B7 | B125 | WB-Cop | Cor | 0.452553674 | 0 |
| B7 | B136 | WB-Cop | 5'NT | -0.754318312 | 0 |
| B7 | B157 | WB-Cop | IgG | 0.255859406 | 0 |
| B7 | B19 | WB-Cop | CD3+CD4+/CD3+CD8+ | -0.656816491 | 0 |
| B7 | B20 | WB-Cop | CD3+CD4+CD8+ | 0.385291105 | 0 |
| B7 | B21 | WB-Cop | CD3+CD4-CD8- | 0.485190855 | 0 |
| B7 | B22 | WB-Cop | CD3+CD8+(T) | 0.312174073 | 0 |
| B7 | B89 | WB-Cop | CA 72-4 | -0.698728278 | 0 |
| B7 | B18 | WB-Cop | CD3+CD4+helper(inducer) T | -0.177728809 | 7.26E-11 |
| B7 | B8 | WB-Cop | WB-Zinc | -0.16815063 | 7.33E-10 |
| B7 | B73 | WB-Cop | ApoA | -0.134558332 | 8.84E-07 |
| B70 | B132 | LDL-C | Dbil | -0.262177907 | 0 |
| B70 | B76 | LDL-C | ApoB | 0.30813815 | 0 |
| B70 | B77 | LDL-C | TC | 0.915712898 | 0 |
| B70 | B91 | LDL-C | TP | 0.219443674 | 6.66E-16 |

| | | | | | |
|---|---|---|---|---|---|
| B70 | B130 | LDL-C | GLB | 0.212089962 | 6.22E-15 |
| B70 | B75 | LDL-C | ApoA1/B | -0.206001948 | 3.64E-14 |
| B70 | B218 | LDL-C | DBP | 0.186783008 | 7.23E-12 |
| B70 | B219 | LDL-C | OT | -0.174335472 | 1.67E-10 |
| B70 | B127 | LDL-C | A-G ratio | -0.162194221 | 2.89E-09 |
| B70 | B72 | LDL-C | HDL-C | 0.160173109 | 4.55E-09 |
| B70 | B220 | LDL-C | SBP | 0.153427366 | 1.99E-08 |
| B70 | B174 | LDL-C | Urea/Cr | 0.135529844 | 7.36E-07 |
| B71 | B206 | TG | Waist | 0.234784433 | 0 |
| B71 | B215 | TG | BMI | 0.25468564 | 0 |
| B71 | B72 | TG | HDL-C | -0.393880003 | 0 |
| B71 | B216 | TG | BMI | 0.220602391 | 4.44E-16 |
| B71 | B77 | TG | TC | 0.220560996 | 4.44E-16 |
| B71 | B132 | TG | Dbil | -0.20483375 | 5.11E-14 |
| B71 | B140 | TG | r-GT | 0.199437992 | 2.36E-13 |
| B71 | B210 | TG | Hip | 0.198448452 | 3.11E-13 |
| B71 | B128 | TG | ALT | 0.197702792 | 3.83E-13 |
| B71 | B218 | TG | DBP | 0.193944721 | 1.07E-12 |
| B71 | B192 | TG | UA | 0.19330185 | 1.28E-12 |
| B71 | B222 | TG | NS | 0.189212362 | 3.82E-12 |
| B71 | B207 | TG | WHR | 0.180735669 | 3.42E-11 |
| B71 | B141 | TG | AST/ALT | -0.180237396 | 3.88E-11 |
| B71 | B126 | TG | Alb | 0.174478454 | 1.61E-10 |
| B71 | B91 | TG | TP | 0.174319095 | 1.68E-10 |
| B72 | B141 | HDL-C | AST/ALT | 0.273055824 | 0 |
| B72 | B190 | HDL-C | Cre | -0.240158005 | 0 |
| B72 | B192 | HDL-C | UA | -0.332798549 | 0 |
| B72 | B206 | HDL-C | Waist | -0.383096046 | 0 |
| B72 | B207 | HDL-C | WHR | -0.324555086 | 0 |
| B72 | B210 | HDL-C | Hip | -0.285378249 | 0 |
| B72 | B215 | HDL-C | BMI | -0.365942658 | 0 |
| B72 | B216 | HDL-C | BMI | -0.439348745 | 0 |

| | | | | | |
|---|---|---|---|---|---|
| B72 | B221 | HDL-C | Hei | -0.286991502 | 0 |
| B72 | B222 | HDL-C | NS | -0.441362524 | 0 |
| B72 | B74 | HDL-C | ApoA1 | 0.26633343 | 0 |
| B72 | B77 | HDL-C | TC | 0.377963385 | 0 |
| B72 | B128 | HDL-C | ALT | -0.212109178 | 6.22E-15 |
| B72 | B174 | HDL-C | Urea/Cr | 0.19795435 | 3.57E-13 |
| B72 | B163 | HDL-C | SG | -0.186593306 | 7.60E-12 |
| B72 | B219 | HDL-C | OT | -0.172555156 | 2.57E-10 |
| B72 | B153 | HDL-C | Glu | -0.156827332 | 9.54E-09 |
| B72 | B75 | HDL-C | ApoA1/B | 0.141943474 | 2.12E-07 |
| B72 | B209 | HDL-C | Smoking | -0.135526604 | 7.37E-07 |
| B73 | B194 | ApoA | Cl | -0.172671106 | 2.50E-10 |
| B73 | B75 | ApoA | ApoA1/B | -0.168163976 | 7.30E-10 |
| B73 | B111 | ApoA | TSH | 0.166729938 | 1.02E-09 |
| B73 | B157 | ApoA | IgG | -0.166400347 | 1.10E-09 |
| B73 | B101 | ApoA | anti-Sc70 | -0.158305153 | 6.89E-09 |
| B73 | B102 | ApoA | anti-Sm | -0.158305149 | 6.89E-09 |
| B73 | B103 | ApoA | anti-SSA/Ro | -0.15830515 | 6.89E-09 |
| B73 | B104 | ApoA | anti-SSB | -0.15830515 | 6.89E-09 |
| B73 | B105 | ApoA | anti-U1RNP | -0.158305151 | 6.89E-09 |
| B73 | B106 | ApoA | anti-P | -0.158305151 | 6.89E-09 |
| B73 | B107 | ApoA | AnuA | -0.15830506 | 6.89E-09 |
| B73 | B109 | ApoA | AMA-M2 | -0.15830515 | 6.89E-09 |
| B73 | B110 | ApoA | AHA | -0.158305151 | 6.89E-09 |
| B73 | B97 | ApoA | dsDNA | -0.158305151 | 6.89E-09 |
| B73 | B99 | ApoA | anti-PM-Scl | -0.158305151 | 6.89E-09 |
| B73 | B81 | ApoA | TSGF | 0.14816273 | 6.03E-08 |

| | | | | | |
|---|---|---|---|---|---|
| B74 | B137 | ApoA1 | AFU | -0.267435469 | 0 |
| B74 | B139 | ApoA1 | ADA | -0.293372003 | 0 |
| B74 | B75 | ApoA1 | ApoA1/B | 0.389390152 | 0 |
| B74 | B76 | ApoA1 | ApoB | 0.217387989 | 1.33E-15 |
| B74 | B134 | ApoA1 | CHE | 0.196672847 | 5.09E-13 |
| B74 | B77 | ApoA1 | TC | 0.185014843 | 1.15E-11 |
| B74 | B174 | ApoA1 | Urea/Cr | 0.181937898 | 2.52E-11 |
| B74 | B153 | ApoA1 | Glu | -0.174908648 | 1.45E-10 |
| B74 | B196 | ApoA1 | TCO2 | 0.167376948 | 8.78E-10 |
| B74 | B138 | ApoA1 | PAB | 0.164578455 | 1.68E-09 |
| B74 | B195 | ApoA1 | Na | 0.154844886 | 1.47E-08 |
| B74 | B190 | ApoA1 | Cre | -0.145576429 | 1.02E-07 |
| B74 | B115 | ApoA1 | TSH TT3 TT4(TT3) | 0.145415408 | 1.06E-07 |
| B74 | B199 | ApoA1 | Mg | -0.135489842 | 7.42E-07 |
| B75 | B76 | ApoA1/B | ApoB | -0.748127075 | 0 |
| B75 | B200 | ApoA1/B | OP | -0.183497869 | 1.69E-11 |
| B75 | B137 | ApoA1/B | AFU | -0.173175997 | 2.21E-10 |
| B75 | B197 | ApoA1/B | AG | -0.170081765 | 4.64E-10 |
| B75 | B196 | ApoA1/B | TCO2 | 0.167512986 | 8.50E-10 |
| B75 | B77 | ApoA1/B | TC | -0.158327181 | 6.86E-09 |
| B75 | B136 | ApoA1/B | 5'NT | -0.150959517 | 3.36E-08 |
| B75 | B134 | ApoA1/B | CHE | -0.139217452 | 3.63E-07 |
| B76 | B134 | ApoB | CHE | 0.275135269 | 0 |
| B76 | B77 | ApoB | TC | 0.302459998 | 0 |
| B76 | B138 | ApoB | PAB | 0.198313533 | 3.23E-13 |
| B76 | B197 | ApoB | AG | 0.168197119 | 7.25E-10 |
| B76 | B195 | ApoB | Na | 0.153325779 | 2.03E-08 |
| B76 | B136 | ApoB | 5'NT | 0.152423191 | 2.47E-08 |
| B77 | B132 | TC | Dbil | -0.264984406 | 0 |
| B77 | B91 | TC | TP | 0.227779001 | 0 |
| B77 | B130 | TC | GLB | 0.207988359 | 2.04E-14 |

| | | | | | |
|---|---|---|---|---|---|
| B77 | B222 | TC | NS | -0.200936378 | 1.55E-13 |
| B77 | B219 | TC | OT | -0.200283644 | 1.87E-13 |
| B77 | B221 | TC | Hei | -0.191115667 | 2.30E-12 |
| B77 | B218 | TC | DBP | 0.186625932 | 7.54E-12 |
| B77 | B174 | TC | Urea/Cr | 0.183180102 | 1.84E-11 |
| B77 | B190 | TC | Cre | -0.179368787 | 4.82E-11 |
| B77 | B220 | TC | SBP | 0.178102758 | 6.61E-11 |
| B77 | B216 | TC | BMI | -0.16776046 | 8.03E-10 |
| B77 | B127 | TC | A-G ratio | -0.149955459 | 4.15E-08 |
| B78 | B138 | HBcAb | PAB | 0.136705311 | 5.89E-07 |
| B79 | B101 | hb V -dna | anti-Sc70 | 0.373111398 | 0 |
| B79 | B102 | hb V -dna | anti-Sm | 0.37311139 | 0 |
| B79 | B103 | hb V -dna | anti-SSA/Ro | 0.373111392 | 0 |
| B79 | B104 | hb V -dna | anti-SSB | 0.373111392 | 0 |
| B79 | B105 | hb V -dna | anti-U1RNP | 0.373111392 | 0 |
| B79 | B106 | hb V -dna | anti-P | 0.373111394 | 0 |
| B79 | B107 | hb V -dna | AnuA | 0.37311118 | 0 |
| B79 | B109 | hb V -dna | AMA-M2 | 0.373111392 | 0 |
| B79 | B110 | hb V -dna | AHA | 0.373111393 | 0 |
| B79 | B119 | hb V -dna | Hcv-rna | 0.368999017 | 0 |
| B79 | B120 | hb V -dna | HCV-cAg | 0.275634178 | 0 |
| B79 | B125 | hb V -dna | Cor | -0.422631041 | 0 |
| B79 | B134 | hb V -dna | CHE | 0.230787727 | 0 |
| B79 | B138 | hb V -dna | PAB | 0.295011835 | 0 |
| B79 | B139 | hb V -dna | ADA | -0.366160685 | 0 |
| B79 | B153 | hb V -dna | Glu | -0.298114061 | 0 |
| B79 | B92 | hb V -dna | ALDH2-Glu504Glu | -0.2633534 | 0 |
| B79 | B93 | hb V -dna | ALDH2-Glu504Lys | 0.239714513 | 0 |
| B79 | B94 | hb V -dna | ALDH2-Lys504Lys | 0.430097024 | 0 |
| B79 | B97 | hb V -dna | dsDNA | 0.373111393 | 0 |
| B79 | B99 | hb V -dna | anti-PM-Scl | 0.373111394 | 0 |
| B79 | B154 | hb V -dna | RF | 0.192377417 | 1.64E-12 |
| B79 | B115 | hb V -dna | TSH TT3 TT4(TT3) | 0.182705061 | 2.07E-11 |
| B79 | B170 | hb V -dna | U-GLU | -0.176127598 | 1.08E-10 |
| B79 | B152 | hb V -dna | ACA-Ig(G-A-M) | -0.17331652 | 2.14E-10 |

| | | | | | |
|---|---|---|---|---|---|
| B79 | B159 | hb V -dna | β2-MG | -0.173308748 | 2.14E-10 |
| B79 | B169 | hb V -dna | U-CRY | -0.173309616 | 2.14E-10 |
| B79 | B200 | hb V -dna | OP | -0.169560974 | 5.25E-10 |
| B79 | B158 | hb V -dna | IgM | 0.160605022 | 4.13E-09 |
| B79 | B195 | hb V -dna | Na | 0.151443386 | 3.04E-08 |
| B79 | B135 | hb V -dna | TBA | -0.135570157 | 7.31E-07 |
| B8 | B224 | WB-Zinc | pmh | -0.140619096 | 2.76E-07 |
| B80 | B118 | anti-HP IgG | CEA1 | 0.147899736 | 6.36E-08 |
| B81 | B108 | TSGF | Anti-Cyclin | -0.214473127 | 3.11E-15 |
| B81 | B98 | TSGF | anti-Jo-1 | -0.210970497 | 8.44E-15 |
| B81 | B157 | TSGF | IgG | -0.171055471 | 3.68E-10 |
| B81 | B156 | TSGF | IgE | -0.163745335 | 2.03E-09 |
| B82 | B84 | CEA | C-PSA | 0.275011549 | 0 |
| B82 | B88 | CEA | CA 242 | 0.416593368 | 0 |
| B82 | B86 | CEA | TBhCG | -0.177888436 | 6.97E-11 |
| B83 | B86 | AFP | TBhCG | 0.285547501 | 0 |
| B83 | B88 | AFP | CA 242 | 0.273228492 | 0 |
| B83 | B134 | AFP | CHE | 0.136826145 | 5.75E-07 |
| B84 | B86 | C-PSA | TBhCG | -0.363799755 | 0 |
| B84 | B90 | C-PSA | CYFRA 21-1 | 0.236684117 | 0 |
| B84 | B88 | C-PSA | CA 242 | 0.203982843 | 6.53E-14 |
| B84 | B87 | C-PSA | NSE | 0.192255498 | 1.69E-12 |
| B85 | B87 | SCC | NSE | 0.214045891 | 3.55E-15 |
| B85 | B90 | SCC | CYFRA 21-1 | 0.209850278 | 1.20E-14 |
| B85 | B161 | SCC | LEU | 0.170841407 | 3.88E-10 |
| B86 | B87 | TBhCG | NSE | -0.218500154 | 8.88E-16 |
| B87 | B198 | NSE | P | 0.149873714 | 4.22E-08 |
| B87 | B88 | NSE | CA 242 | 0.149544834 | 4.52E-08 |
| B89 | B121 | CA 72-4 | C3 | 0.462065795 | 0 |
| B89 | B122 | CA 72-4 | C4 | 0.561205526 | 0 |
| B89 | B125 | CA 72-4 | Cor | -0.387140716 | 0 |

| | | | | | |
|---|---|---|---|---|---|
| B89 | B136 | CA 72-4 | 5'NT | 0.67189087 | 0 |
| B89 | B157 | CA 72-4 | IgG | -0.22597747<br>7 | 0 |
| B89 | B124 | CA 72-4 | ACTH | -0.21237059<br>9 | 5.77E-15 |
| B9 | B223 | RH | FH | 0.994134847 | 0 |
| B90 | B187 | CYFRA 21-1 | NSE | 0.143847142 | 1.45E-07 |
| B91 | B126 | TP | Alb | 0.483222986 | 0 |
| B91 | B127 | TP | A-G ratio | -0.47460826<br>7 | 0 |
| B91 | B130 | TP | GLB | 0.775273184 | 0 |
| B91 | B188 | TP | Ca | 0.351529697 | 0 |
| B91 | B218 | TP | DBP | 0.208698568 | 1.67E-14 |
| B91 | B220 | TP | SBP | 0.142673631 | 1.84E-07 |
| B92 | B125 | ALDH2-Glu504Glu | Cor | 0.294431822 | 0 |
| B92 | B93 | ALDH2-Glu504Glu | ALDH2-Glu504Lys | -0.99909976 | 0 |
| B92 | B94 | ALDH2-Glu504Glu | ALDH2-Lys504Lys | 0.338130403 | 0 |
| B92 | B101 | ALDH2-Glu504Glu | anti-Sc70 | -0.20052281<br>9 | 1.74E-13 |
| B92 | B102 | ALDH2-Glu504Glu | anti-Sm | -0.20052281<br>3 | 1.74E-13 |
| B92 | B103 | ALDH2-Glu504Glu | anti-SSA/Ro | -0.20052281<br>5 | 1.74E-13 |
| B92 | B104 | ALDH2-Glu504Glu | anti-SSB | -0.20052281<br>6 | 1.74E-13 |
| B92 | B105 | ALDH2-Glu504Glu | anti-U1RNP | -0.20052281<br>5 | 1.74E-13 |
| B92 | B106 | ALDH2-Glu504Glu | anti-P | -0.20052281<br>6 | 1.74E-13 |
| B92 | B107 | ALDH2-Glu504Glu | AnuA | -0.20052268<br>6 | 1.74E-13 |
| B92 | B109 | ALDH2-Glu504Glu | AMA-M2 | -0.20052281<br>5 | 1.74E-13 |
| B92 | B110 | ALDH2-Glu504Glu | AHA | -0.20052281<br>6 | 1.74E-13 |
| B92 | B97 | ALDH2-Glu504Glu | dsDNA | -0.20052281<br>5 | 1.74E-13 |
| B92 | B99 | ALDH2-Glu504Glu | anti-PM-Scl | -0.20052281<br>7 | 1.74E-13 |
| B92 | B199 | ALDH2-Glu504Glu | Mg | -0.19159346<br>7 | 2.02E-12 |
| B92 | B119 | ALDH2-Glu504Glu | Hcv-rna | -0.17510882<br>9 | 1.38E-10 |
| B92 | B186 | ALDH2-Glu504Glu | PFC | 0.142423506 | 1.93E-07 |

| | | | | | |
|---|---|---|---|---|---|
| B93 | B125 | ALDH2-Glu504Lys | Cor | -0.267679994 | 0 |
| B93 | B94 | ALDH2-Glu504Lys | ALDH2-Lys504Lys | -0.377749793 | 0 |
| B93 | B101 | ALDH2-Glu504Lys | anti-Sc70 | 0.182732703 | 2.06E-11 |
| B93 | B102 | ALDH2-Glu504Lys | anti-Sm | 0.182732699 | 2.06E-11 |
| B93 | B103 | ALDH2-Glu504Lys | anti-SSA/Ro | 0.182732699 | 2.06E-11 |
| B93 | B104 | ALDH2-Glu504Lys | anti-SSB | 0.182732699 | 2.06E-11 |
| B93 | B105 | ALDH2-Glu504Lys | anti-U1RNP | 0.182732699 | 2.06E-11 |
| B93 | B106 | ALDH2-Glu504Lys | anti-P | 0.182732701 | 2.06E-11 |
| B93 | B107 | ALDH2-Glu504Lys | AnuA | 0.182732597 | 2.06E-11 |
| B93 | B109 | ALDH2-Glu504Lys | AMA-M2 | 0.182732699 | 2.06E-11 |
| B93 | B110 | ALDH2-Glu504Lys | AHA | 0.182732699 | 2.06E-11 |
| B93 | B97 | ALDH2-Glu504Lys | dsDNA | 0.182732699 | 2.06E-11 |
| B93 | B99 | ALDH2-Glu504Lys | anti-PM-Scl | 0.1827327 | 2.06E-11 |
| B93 | B199 | ALDH2-Glu504Lys | Mg | 0.1822279 | 2.34E-11 |
| B93 | B119 | ALDH2-Glu504Lys | Hcv-rna | 0.159514932 | 5.27E-09 |
| B93 | B186 | ALDH2-Glu504Lys | PFC | -0.136513998 | 6.11E-07 |
| B94 | B101 | ALDH2-Lys504Lys | anti-Sc70 | 0.322847803 | 0 |
| B94 | B102 | ALDH2-Lys504Lys | anti-Sm | 0.322847808 | 0 |
| B94 | B103 | ALDH2-Lys504Lys | anti-SSA/Ro | 0.32284781 | 0 |
| B94 | B104 | ALDH2-Lys504Lys | anti-SSB | 0.322847812 | 0 |
| B94 | B105 | ALDH2-Lys504Lys | anti-U1RNP | 0.322847807 | 0 |
| B94 | B106 | ALDH2-Lys504Lys | anti-P | 0.322847814 | 0 |
| B94 | B107 | ALDH2-Lys504Lys | AnuA | 0.322847715 | 0 |
| B94 | B109 | ALDH2-Lys504Lys | AMA-M2 | 0.322847812 | 0 |
| B94 | B110 | ALDH2-Lys504Lys | AHA | 0.322847812 | 0 |
| B94 | B119 | ALDH2-Lys504Lys | Hcv-rna | 0.283227722 | 0 |
| B94 | B125 | ALDH2-Lys504Lys | Cor | -0.488025133 | 0 |
| B94 | B97 | ALDH2-Lys504Lys | dsDNA | 0.322847813 | 0 |
| B94 | B99 | ALDH2-Lys504Lys | anti-PM-Scl | 0.32284781 | 0 |
| B94 | B154 | ALDH2-Lys504Lys | RF | 0.214913133 | 2.66E-15 |
| B94 | B152 | ALDH2-Lys504Lys | ACA-Ig(G-A-M) | -0.201195032 | 1.44E-13 |
| B94 | B159 | ALDH2-Lys504Lys | β2-MG | -0.201185641 | 1.45E-13 |
| B94 | B169 | ALDH2-Lys504Lys | U-CRY | -0.201186688 | 1.45E-13 |
| B94 | B183 | ALDH2-Lys504Lys | PTA | 0.176948173 | 8.80E-11 |
| B94 | B120 | ALDH2-Lys504Lys | HCV-cAg | 0.155883595 | 1.17E-08 |
| B94 | B138 | ALDH2-Lys504Lys | PAB | 0.151653733 | 2.90E-08 |
| B94 | B196 | ALDH2-Lys504Lys | TCO2 | -0.14285837 | 1.77E-07 |

| | | | | | |
|---|---|---|---|---|---|
| | | | | 8 | |
| B94 | B199 | ALDH2-Lys504Lys | Mg | 0.139155536 | 3.67E-07 |
| B95 | B101 | anti-cenpb | anti-Sc70 | -0.236295827 | 0 |
| B95 | B102 | anti-cenpb | anti-Sm | -0.236295821 | 0 |
| B95 | B103 | anti-cenpb | anti-SSA/Ro | -0.236295824 | 0 |
| B95 | B104 | anti-cenpb | anti-SSB | -0.236295824 | 0 |
| B95 | B105 | anti-cenpb | anti-U1RNP | -0.236295824 | 0 |
| B95 | B106 | anti-cenpb | anti-P | -0.236295824 | 0 |
| B95 | B107 | anti-cenpb | AnuA | -0.236295686 | 0 |
| B95 | B109 | anti-cenpb | AMA-M2 | -0.236295823 | 0 |
| B95 | B110 | anti-cenpb | AHA | -0.236295824 | 0 |
| B95 | B97 | anti-cenpb | dsDNA | -0.236295824 | 0 |
| B95 | B99 | anti-cenpb | anti-PM-Scl | -0.236295825 | 0 |
| B95 | B146 | anti-cenpb | CysC | 0.202173583 | 1.09E-13 |
| B96 | B223 | dsDNA (IIF) | FH | 0.150618808 | 3.61E-08 |
| B97 | B101 | dsDNA | anti-Sc70 | 0.999999999 | 0 |
| B97 | B102 | dsDNA | anti-Sm | 0.999999997 | 0 |
| B97 | B104 | dsDNA | anti-SSB | 0.999999989 | 0 |
| B97 | B106 | dsDNA | anti-P | 0.999999995 | 0 |
| B97 | B107 | dsDNA | AnuA | 0.999999389 | 0 |
| B97 | B109 | dsDNA | AMA-M2 | 0.999999999 | 0 |
| B97 | B110 | dsDNA | AHA | 0.999999998 | 0 |
| B97 | B119 | dsDNA | Hcv-rna | 0.470017908 | 0 |
| B97 | B120 | dsDNA | HCV-cAg | 0.311176271 | 0 |
| B97 | B125 | dsDNA | Cor | -0.319333504 | 0 |
| B97 | B152 | dsDNA | ACA-Ig(G-A-M) | -0.263417303 | 0 |
| B97 | B154 | dsDNA | RF | 0.277811856 | 0 |
| B97 | B155 | dsDNA | IgA | -0.244461159 | 0 |
| B97 | B158 | dsDNA | IgM | 0.292027923 | 0 |
| B97 | B159 | dsDNA | β2-MG | -0.26340422 | 0 |

| | | | | | |
|---|---|---|---|---|---|
| | | | | 7 | |
| B97 | B169 | dsDNA | U-CRY | -0.263405676 | 0 |
| B97 | B197 | dsDNA | AG | 0.240927346 | 0 |
| B97 | B98 | dsDNA | anti-Jo-1 | -0.255281554 | 0 |
| B97 | B99 | dsDNA | anti-PM-Scl | 0.999999992 | 0 |
| B97 | B199 | dsDNA | Mg | 0.216390065 | 1.78E-15 |
| B97 | B137 | dsDNA | AFU | 0.204688404 | 5.33E-14 |
| B97 | B134 | dsDNA | CHE | 0.187404365 | 6.15E-12 |
| B97 | B200 | dsDNA | OP | 0.1870957 | 6.66E-12 |
| B97 | B195 | dsDNA | Na | 0.1841634 | 1.43E-11 |
| B97 | B193 | dsDNA | K | 0.182545784 | 2.16E-11 |
| B97 | B108 | dsDNA | Anti-Cyclin | -0.18087605 | 3.30E-11 |
| B97 | B114 | dsDNA | TSH TT3 TT4(TT4) | 0.173261895 | 2.17E-10 |
| B97 | B138 | dsDNA | PAB | 0.164858123 | 1.57E-09 |
| B97 | B136 | dsDNA | 5'NT | 0.151810372 | 2.81E-08 |
| B97 | B196 | dsDNA | TCO2 | -0.138358659 | 4.28E-07 |
| B98 | B101 | anti-Jo-1 | anti-Sc70 | -0.255281558 | 0 |
| B98 | B102 | anti-Jo-1 | anti-Sm | -0.255281552 | 0 |
| B98 | B103 | anti-Jo-1 | anti-SSA/Ro | -0.255281553 | 0 |
| B98 | B104 | anti-Jo-1 | anti-SSB | -0.255281554 | 0 |
| B98 | B105 | anti-Jo-1 | anti-U1RNP | -0.255281554 | 0 |
| B98 | B106 | anti-Jo-1 | anti-P | -0.255281555 | 0 |
| B98 | B107 | anti-Jo-1 | AnuA | -0.255281409 | 0 |
| B98 | B108 | anti-Jo-1 | Anti-Cyclin | 0.99709348 | 0 |
| B98 | B109 | anti-Jo-1 | AMA-M2 | -0.255281553 | 0 |
| B98 | B110 | anti-Jo-1 | AHA | -0.255281554 | 0 |
| B98 | B151 | anti-Jo-1 | Anti-CCP | 0.280418647 | 0 |
| B98 | B156 | anti-Jo-1 | IgE | 0.451351353 | 0 |
| B98 | B179 | anti-Jo-1 | α1-MG | 0.265734301 | 0 |
| B98 | B99 | anti-Jo-1 | anti-PM-Scl | -0.255281555 | 0 |
| B98 | B194 | anti-Jo-1 | Cl | 0.209835414 | 1.20E-14 |

| | | | | | |
|---|---|---|---|---|---|
| B98 | B158 | anti-Jo-1 | IgM | 0.201738653 | 1.24E-13 |
| B98 | B157 | anti-Jo-1 | IgG | 0.197601309 | 3.94E-13 |
| B98 | B154 | anti-Jo-1 | RF | -0.150311112 | 3.85E-08 |
| B99 | B101 | anti-PM-Scl | anti-Sc70 | 0.999999993 | 0 |
| B99 | B104 | anti-PM-Scl | anti-SSB | 0.999999995 | 0 |
| B99 | B107 | anti-PM-Scl | AnuA | 0.999999418 | 0 |
| B99 | B119 | anti-PM-Scl | Hcv-rna | 0.47001791 | 0 |
| B99 | B120 | anti-PM-Scl | HCV-cAg | 0.311176223 | 0 |
| B99 | B125 | anti-PM-Scl | Cor | -0.319333505 | 0 |
| B99 | B152 | anti-PM-Scl | ACA-Ig(G-A-M) | -0.263417164 | 0 |
| B99 | B154 | anti-PM-Scl | RF | 0.277811857 | 0 |
| B99 | B155 | anti-PM-Scl | IgA | -0.244461161 | 0 |
| B99 | B158 | anti-PM-Scl | IgM | 0.292027925 | 0 |
| B99 | B159 | anti-PM-Scl | β2-MG | -0.263404441 | 0 |
| B99 | B169 | anti-PM-Scl | U-CRY | -0.263405654 | 0 |
| B99 | B197 | anti-PM-Scl | AG | 0.240927347 | 0 |
| B99 | B199 | anti-PM-Scl | Mg | 0.216390066 | 1.78E-15 |
| B99 | B137 | anti-PM-Scl | AFU | 0.204688405 | 5.33E-14 |
| B99 | B134 | anti-PM-Scl | CHE | 0.187404365 | 6.15E-12 |
| B99 | B200 | anti-PM-Scl | OP | 0.187095705 | 6.66E-12 |
| B99 | B195 | anti-PM-Scl | Na | 0.184163407 | 1.43E-11 |
| B99 | B193 | anti-PM-Scl | K | 0.182545784 | 2.16E-11 |
| B99 | B108 | anti-PM-Scl | Anti-Cyclin | -0.180876051 | 3.30E-11 |
| B99 | B114 | anti-PM-Scl | TSH TT3 TT4(TT4) | 0.173261896 | 2.17E-10 |
| B99 | B138 | anti-PM-Scl | PAB | 0.164858124 | 1.57E-09 |
| B99 | B136 | anti-PM-Scl | 5'NT | 0.151810372 | 2.81E-08 |
| B99 | B196 | anti-PM-Scl | TCO2 | -0.13835866 | 4.28E-07 |

**Supplementary Table 9. PEI Correlations in Coronary+Diabetes**

| Row | Column | Row | Column | Cor | *P* |
|---|---|---|---|---|---|
| B1 | B221 | Gender | Hei | -0.57222 | 0 |
| B1 | B51 | Gender | Hb | -0.47231 | 0 |
| B1 | B222 | Gender | NS | -0.45835 | 4.44E-16 |
| B1 | B174 | Gender | Urea/Cr | 0.432889 | 3.24E-14 |
| B1 | B190 | Gender | Cre | -0.39981 | 3.59E-12 |
| B1 | B220 | Gender | SBP | 0.382085 | 3.65E-11 |
| B1 | B46 | Gender | RBC Count | -0.38091 | 4.23E-11 |
| B1 | B205 | Gender | KDA | 0.362514 | 4.03E-10 |
| B1 | B163 | Gender | SG | -0.31881 | 4.93E-08 |
| B1 | B45 | Gender | MCHC | -0.30481 | 1.97E-07 |
| B1 | B72 | Gender | HDL-C | 0.30013 | 3.09E-07 |
| B1 | B161 | Gender | LEU | 0.287662 | 9.79E-07 |
| B10 | B12 | PG Ⅰ | PG II | 0.775037 | 0 |
| B100 | B101 | anti-Ro52 | anti-Sc70 | 1 | 0 |
| B100 | B103 | anti-Ro52 | anti-SSA/Ro | 1 | 0 |
| B100 | B105 | anti-Ro52 | anti-U1RNP | 1 | 0 |
| B100 | B106 | anti-Ro52 | anti-P | 1 | 0 |
| B100 | B107 | anti-Ro52 | AnuA | 1 | 0 |
| B100 | B110 | anti-Ro52 | AHA | 1 | 0 |
| B100 | B125 | anti-Ro52 | Cor | 1 | 0 |
| B100 | B151 | anti-Ro52 | Anti-CCP | 0.508198 | 0 |
| B100 | B158 | anti-Ro52 | IgM | 0.497089 | 0 |
| B100 | B178 | anti-Ro52 | Ig-U | 0.6975 | 0 |
| B100 | B179 | anti-Ro52 | α1-MG | 0.724317 | 0 |
| B100 | B180 | anti-Ro52 | Tf | 0.705838 | 0 |
| B100 | B181 | anti-Ro52 | APTT | 1 | 0 |
| B100 | B182 | anti-Ro52 | TT | -0.99982 | 0 |
| B100 | B183 | anti-Ro52 | PTA | 0.999999 | 0 |
| B100 | B185 | anti-Ro52 | PT | 0.999261 | 0 |
| B100 | B194 | anti-Ro52 | Cl | 0.804248 | 0 |
| B100 | B197 | anti-Ro52 | AG | -0.67006 | 0 |
| B100 | B138 | anti-Ro52 | PAB | -0.41317 | 5.70E-13 |
| B100 | B137 | anti-Ro52 | AFU | -0.28849 | 9.08E-07 |
| B101 | B102 | anti-Sc70 | anti-Sm | 1 | 0 |
| B101 | B103 | anti-Sc70 | anti-SSA/Ro | 1 | 0 |
| B101 | B105 | anti-Sc70 | anti-U1RNP | 1 | 0 |
| B101 | B106 | anti-Sc70 | anti-P | 1 | 0 |
| B101 | B107 | anti-Sc70 | AnuA | 1 | 0 |
| B101 | B108 | anti-Sc70 | Anti-Cyclin | 1 | 0 |
| B101 | B109 | anti-Sc70 | AMA-M2 | 1 | 0 |
| B101 | B110 | anti-Sc70 | AHA | 1 | 0 |
| B101 | B124 | anti-Sc70 | ACTH | -1 | 0 |

| | | | | | |
|---|---|---|---|---|---|
| B101 | B125 | anti-Sc70 | Cor | 1 | 0 |
| B101 | B151 | anti-Sc70 | Anti-CCP | 0.508198 | 0 |
| B101 | B158 | anti-Sc70 | IgM | 0.497089 | 0 |
| B101 | B178 | anti-Sc70 | Ig-U | 0.6975 | 0 |
| B101 | B179 | anti-Sc70 | α1-MG | 0.724317 | 0 |
| B101 | B180 | anti-Sc70 | Tf | 0.705838 | 0 |
| B101 | B181 | anti-Sc70 | APTT | 1 | 0 |
| B101 | B182 | anti-Sc70 | TT | -0.99982 | 0 |
| B101 | B183 | anti-Sc70 | PTA | 0.999999 | 0 |
| B101 | B184 | anti-Sc70 | INR | 1 | 0 |
| B101 | B185 | anti-Sc70 | PT | 0.999261 | 0 |
| B101 | B194 | anti-Sc70 | CI | 0.804248 | 0 |
| B101 | B197 | anti-Sc70 | AG | -0.67006 | 0 |
| B101 | B138 | anti-Sc70 | PAB | -0.41317 | 5.70E-13 |
| B101 | B137 | anti-Sc70 | AFU | -0.28849 | 9.08E-07 |
| B102 | B104 | anti-Sm | anti-SSB | 1 | 0 |
| B102 | B106 | anti-Sm | anti-P | 1 | 0 |
| B102 | B107 | anti-Sm | AnuA | 1 | 0 |
| B102 | B108 | anti-Sm | Anti-Cyclin | 1 | 0 |
| B102 | B151 | anti-Sm | Anti-CCP | 0.508198 | 0 |
| B102 | B158 | anti-Sm | IgM | 0.497089 | 0 |
| B102 | B178 | anti-Sm | Ig-U | 0.6975 | 0 |
| B102 | B179 | anti-Sm | α1-MG | 0.724317 | 0 |
| B102 | B180 | anti-Sm | Tf | 0.705838 | 0 |
| B102 | B182 | anti-Sm | TT | -0.99982 | 0 |
| B102 | B183 | anti-Sm | PTA | 0.999999 | 0 |
| B102 | B185 | anti-Sm | PT | 0.999261 | 0 |
| B102 | B194 | anti-Sm | CI | 0.804248 | 0 |
| B102 | B197 | anti-Sm | AG | -0.67006 | 0 |
| B102 | B138 | anti-Sm | PAB | -0.41317 | 5.70E-13 |
| B102 | B137 | anti-Sm | AFU | -0.28849 | 9.08E-07 |
| B103 | B105 | anti-SSA/Ro | anti-U1RNP | 1 | 0 |
| B103 | B106 | anti-SSA/Ro | anti-P | 1 | 0 |
| B103 | B107 | anti-SSA/Ro | AnuA | 1 | 0 |
| B103 | B124 | anti-SSA/Ro | ACTH | -1 | 0 |
| B103 | B125 | anti-SSA/Ro | Cor | 1 | 0 |
| B103 | B151 | anti-SSA/Ro | Anti-CCP | 0.508198 | 0 |
| B103 | B158 | anti-SSA/Ro | IgM | 0.497089 | 0 |
| B103 | B178 | anti-SSA/Ro | Ig-U | 0.6975 | 0 |
| B103 | B179 | anti-SSA/Ro | α1-MG | 0.724317 | 0 |
| B103 | B180 | anti-SSA/Ro | Tf | 0.705838 | 0 |
| B103 | B181 | anti-SSA/Ro | APTT | 1 | 0 |
| B103 | B182 | anti-SSA/Ro | TT | -0.99982 | 0 |
| B103 | B183 | anti-SSA/Ro | PTA | 0.999999 | 0 |

| | | | | | |
|---|---|---|---|---|---|
| B103 | B185 | anti-SSA/Ro | PT | 0.999261 | 0 |
| B103 | B194 | anti-SSA/Ro | CI | 0.804248 | 0 |
| B103 | B197 | anti-SSA/Ro | AG | -0.67006 | 0 |
| B103 | B138 | anti-SSA/Ro | PAB | -0.41317 | 5.70E-13 |
| B103 | B137 | anti-SSA/Ro | AFU | -0.28849 | 9.08E-07 |
| B104 | B107 | anti-SSB | AnuA | 1 | 0 |
| B104 | B151 | anti-SSB | Anti-CCP | 0.508198 | 0 |
| B104 | B158 | anti-SSB | IgM | 0.497089 | 0 |
| B104 | B178 | anti-SSB | Ig-U | 0.6975 | 0 |
| B104 | B179 | anti-SSB | α1-MG | 0.724317 | 0 |
| B104 | B180 | anti-SSB | Tf | 0.705838 | 0 |
| B104 | B182 | anti-SSB | TT | -0.99982 | 0 |
| B104 | B183 | anti-SSB | PTA | 0.999999 | 0 |
| B104 | B185 | anti-SSB | PT | 0.999261 | 0 |
| B104 | B194 | anti-SSB | CI | 0.804248 | 0 |
| B104 | B197 | anti-SSB | AG | -0.67006 | 0 |
| B104 | B138 | anti-SSB | PAB | -0.41317 | 5.70E-13 |
| B104 | B137 | anti-SSB | AFU | -0.28849 | 9.08E-07 |
| B105 | B106 | anti-U1RNP | anti-P | 1 | 0 |
| B105 | B107 | anti-U1RNP | AnuA | 1 | 0 |
| B105 | B109 | anti-U1RNP | AMA-M2 | 1 | 0 |
| B105 | B110 | anti-U1RNP | AHA | 1 | 0 |
| B105 | B124 | anti-U1RNP | ACTH | -1 | 0 |
| B105 | B125 | anti-U1RNP | Cor | 1 | 0 |
| B105 | B151 | anti-U1RNP | Anti-CCP | 0.508198 | 0 |
| B105 | B158 | anti-U1RNP | IgM | 0.497089 | 0 |
| B105 | B178 | anti-U1RNP | Ig-U | 0.6975 | 0 |
| B105 | B179 | anti-U1RNP | α1-MG | 0.724317 | 0 |
| B105 | B180 | anti-U1RNP | Tf | 0.705838 | 0 |
| B105 | B181 | anti-U1RNP | APTT | 1 | 0 |
| B105 | B182 | anti-U1RNP | TT | -0.99982 | 0 |
| B105 | B183 | anti-U1RNP | PTA | 0.999999 | 0 |
| B105 | B184 | anti-U1RNP | INR | 1 | 0 |
| B105 | B185 | anti-U1RNP | PT | 0.999261 | 0 |
| B105 | B194 | anti-U1RNP | CI | 0.804248 | 0 |
| B105 | B197 | anti-U1RNP | AG | -0.67006 | 0 |
| B105 | B138 | anti-U1RNP | PAB | -0.41317 | 5.70E-13 |
| B105 | B137 | anti-U1RNP | AFU | -0.28849 | 9.08E-07 |
| B106 | B107 | anti-P | AnuA | 1 | 0 |
| B106 | B108 | anti-P | Anti-Cyclin | 1 | 0 |
| B106 | B110 | anti-P | AHA | 1 | 0 |
| B106 | B124 | anti-P | ACTH | -1 | 0 |
| B106 | B125 | anti-P | Cor | 1 | 0 |
| B106 | B151 | anti-P | Anti-CCP | 0.508198 | 0 |

| | | | | | |
|---|---|---|---|---|---|
| B106 | B158 | anti-P | IgM | 0.497089 | 0 |
| B106 | B178 | anti-P | Ig-U | 0.6975 | 0 |
| B106 | B179 | anti-P | α1-MG | 0.724317 | 0 |
| B106 | B180 | anti-P | Tf | 0.705838 | 0 |
| B106 | B181 | anti-P | APTT | 1 | 0 |
| B106 | B182 | anti-P | TT | -0.99982 | 0 |
| B106 | B183 | anti-P | PTA | 0.999999 | 0 |
| B106 | B184 | anti-P | INR | 1 | 0 |
| B106 | B185 | anti-P | PT | 0.999261 | 0 |
| B106 | B194 | anti-P | CI | 0.804248 | 0 |
| B106 | B197 | anti-P | AG | -0.67006 | 0 |
| B106 | B138 | anti-P | PAB | -0.41317 | 5.70E-13 |
| B106 | B137 | anti-P | AFU | -0.28849 | 9.08E-07 |
| B107 | B108 | AnuA | Anti-Cyclin | 1 | 0 |
| B107 | B109 | AnuA | AMA-M2 | 1 | 0 |
| B107 | B110 | AnuA | AHA | 1 | 0 |
| B107 | B124 | AnuA | ACTH | -1 | 0 |
| B107 | B125 | AnuA | Cor | 1 | 0 |
| B107 | B151 | AnuA | Anti-CCP | 0.508198 | 0 |
| B107 | B158 | AnuA | IgM | 0.497089 | 0 |
| B107 | B178 | AnuA | Ig-U | 0.6975 | 0 |
| B107 | B179 | AnuA | α1-MG | 0.724317 | 0 |
| B107 | B180 | AnuA | Tf | 0.705838 | 0 |
| B107 | B181 | AnuA | APTT | 1 | 0 |
| B107 | B182 | AnuA | TT | -0.99982 | 0 |
| B107 | B183 | AnuA | PTA | 0.999999 | 0 |
| B107 | B184 | AnuA | INR | 1 | 0 |
| B107 | B185 | AnuA | PT | 0.999261 | 0 |
| B107 | B186 | AnuA | PFC | -1 | 0 |
| B107 | B194 | AnuA | CI | 0.804248 | 0 |
| B107 | B197 | AnuA | AG | -0.67006 | 0 |
| B107 | B138 | AnuA | PAB | -0.41317 | 5.70E-13 |
| B107 | B137 | AnuA | AFU | -0.28849 | 9.08E-07 |
| B108 | B151 | Anti-Cyclin | Anti-CCP | 0.508198 | 0 |
| B108 | B158 | Anti-Cyclin | IgM | 0.497089 | 0 |
| B108 | B178 | Anti-Cyclin | Ig-U | 0.6975 | 0 |
| B108 | B179 | Anti-Cyclin | α1-MG | 0.724317 | 0 |
| B108 | B180 | Anti-Cyclin | Tf | 0.705838 | 0 |
| B108 | B182 | Anti-Cyclin | TT | -0.99982 | 0 |
| B108 | B183 | Anti-Cyclin | PTA | 0.999999 | 0 |
| B108 | B185 | Anti-Cyclin | PT | 0.999261 | 0 |
| B108 | B194 | Anti-Cyclin | CI | 0.804248 | 0 |
| B108 | B197 | Anti-Cyclin | AG | -0.67006 | 0 |
| B108 | B138 | Anti-Cyclin | PAB | -0.41317 | 5.70E-13 |

| | | | | | |
|---|---|---|---|---|---|
| B108 | B137 | Anti-Cyclin | AFU | -0.28849 | 9.08E-07 |
| B109 | B151 | AMA-M2 | Anti-CCP | 0.508198 | 0 |
| B109 | B158 | AMA-M2 | IgM | 0.497089 | 0 |
| B109 | B178 | AMA-M2 | Ig-U | 0.6975 | 0 |
| B109 | B179 | AMA-M2 | α1-MG | 0.724317 | 0 |
| B109 | B180 | AMA-M2 | Tf | 0.705838 | 0 |
| B109 | B181 | AMA-M2 | APTT | 1 | 0 |
| B109 | B182 | AMA-M2 | TT | -0.99982 | 0 |
| B109 | B183 | AMA-M2 | PTA | 0.999999 | 0 |
| B109 | B185 | AMA-M2 | PT | 0.999261 | 0 |
| B109 | B194 | AMA-M2 | Cl | 0.804248 | 0 |
| B109 | B197 | AMA-M2 | AG | -0.67006 | 0 |
| B109 | B138 | AMA-M2 | PAB | -0.41317 | 5.70E-13 |
| B109 | B137 | AMA-M2 | AFU | -0.28849 | 9.08E-07 |
| B11 | B12 | PGI/PGII | PG II | -0.57651 | 0 |
| B11 | B14 | PGI/PGII | G-17 | -0.43246 | 3.46E-14 |
| B110 | B124 | AHA | ACTH | -1 | 0 |
| B110 | B125 | AHA | Cor | 1 | 0 |
| B110 | B151 | AHA | Anti-CCP | 0.508198 | 0 |
| B110 | B158 | AHA | IgM | 0.497089 | 0 |
| B110 | B178 | AHA | Ig-U | 0.6975 | 0 |
| B110 | B179 | AHA | α1-MG | 0.724317 | 0 |
| B110 | B180 | AHA | Tf | 0.705838 | 0 |
| B110 | B181 | AHA | APTT | 1 | 0 |
| B110 | B182 | AHA | TT | -0.99982 | 0 |
| B110 | B183 | AHA | PTA | 0.999999 | 0 |
| B110 | B184 | AHA | INR | 1 | 0 |
| B110 | B185 | AHA | PT | 0.999261 | 0 |
| B110 | B194 | AHA | Cl | 0.804248 | 0 |
| B110 | B197 | AHA | AG | -0.67006 | 0 |
| B110 | B138 | AHA | PAB | -0.41317 | 5.70E-13 |
| B110 | B137 | AHA | AFU | -0.28849 | 9.08E-07 |
| B111 | B112 | TSH | FT4 | -0.33144 | 1.33E-08 |
| B111 | B194 | TSH | Cl | -0.30527 | 1.89E-07 |
| B112 | B114 | FT4 | TSH TT3 TT4(TT4) | 0.4702 | 0 |
| B112 | B113 | FT4 | FT3 | 0.386444 | 2.09E-11 |
| B112 | B148 | FT4 | Anti_TPO | 0.305724 | 1.81E-07 |
| B113 | B114 | FT3 | TSH TT3 TT4(TT4) | 0.525263 | 0 |
| B113 | B115 | FT3 | TSH TT3 TT4(TT3) | 0.631155 | 0 |
| B113 | B148 | FT3 | Anti_TPO | 0.551591 | 0 |
| B113 | B165 | FT3 | bilirubin | -0.46145 | 4.44E-16 |
| B114 | B115 | TSH TT3 TT4(TT4) | TSH TT3 TT4(TT3) | 0.79744 | 0 |
| B114 | B148 | TSH TT3 TT4(TT4) | Anti_TPO | 0.334264 | 9.81E-09 |
| B115 | B165 | TSH TT3 TT4(TT3) | bilirubin | -0.50974 | 0 |

| | | | | | |
|---|---|---|---|---|---|
| B115 | B148 | TSH TT3 TT4(TT3) | Anti_TPO | 0.428799 | 5.95E-14 |
| B119 | B152 | Hcv-rna | ACA-Ig(G-A-M) | -0.99999 | 0 |
| B119 | B169 | Hcv-rna | U-CRY | -1 | 0 |
| B119 | B120 | Hcv-rna | HCV-cAg | 0.404574 | 1.88E-12 |
| B12 | B14 | PG II | G-17 | 0.409985 | 8.92E-13 |
| B120 | B156 | HCV-cAg | IgE | -0.56004 | 0 |
| B120 | B121 | HCV-cAg | C3 | -0.40457 | 1.88E-12 |
| B120 | B122 | HCV-cAg | C4 | -0.40457 | 1.88E-12 |
| B120 | B152 | HCV-cAg | ACA-Ig(G-A-M) | -0.40457 | 1.88E-12 |
| B120 | B159 | HCV-cAg | β2-MG | -0.40457 | 1.88E-12 |
| B120 | B169 | HCV-cAg | U-CRY | -0.40457 | 1.88E-12 |
| B120 | B150 | HCV-cAg | ALP | -0.31865 | 5.01E-08 |
| B121 | B152 | C3 | ACA-Ig(G-A-M) | 0.999986 | 0 |
| B121 | B169 | C3 | U-CRY | 0.999999 | 0 |
| B122 | B152 | C4 | ACA-Ig(G-A-M) | 0.999989 | 0 |
| B122 | B169 | C4 | U-CRY | 1 | 0 |
| B124 | B125 | ACTH | Cor | -1 | 0 |
| B124 | B151 | ACTH | Anti-CCP | -0.5082 | 0 |
| B124 | B158 | ACTH | IgM | -0.49709 | 0 |
| B124 | B178 | ACTH | Ig-U | -0.6975 | 0 |
| B124 | B179 | ACTH | α1-MG | -0.72432 | 0 |
| B124 | B180 | ACTH | Tf | -0.70584 | 0 |
| B124 | B182 | ACTH | TT | 0.999822 | 0 |
| B124 | B183 | ACTH | PTA | -1 | 0 |
| B124 | B184 | ACTH | INR | -1 | 0 |
| B124 | B185 | ACTH | PT | -0.99926 | 0 |
| B124 | B194 | ACTH | Cl | -0.80425 | 0 |
| B124 | B197 | ACTH | AG | 0.670061 | 0 |
| B124 | B138 | ACTH | PAB | 0.413173 | 5.70E-13 |
| B124 | B137 | ACTH | AFU | 0.288494 | 9.08E-07 |
| B125 | B151 | Cor | Anti-CCP | 0.508198 | 0 |
| B125 | B158 | Cor | IgM | 0.497089 | 0 |
| B125 | B178 | Cor | Ig-U | 0.6975 | 0 |
| B125 | B179 | Cor | α1-MG | 0.724317 | 0 |
| B125 | B180 | Cor | Tf | 0.705838 | 0 |
| B125 | B181 | Cor | APTT | 1 | 0 |
| B125 | B182 | Cor | TT | -0.99982 | 0 |
| B125 | B183 | Cor | PTA | 0.999999 | 0 |
| B125 | B185 | Cor | PT | 0.999261 | 0 |
| B125 | B194 | Cor | Cl | 0.804248 | 0 |
| B125 | B197 | Cor | AG | -0.67006 | 0 |
| B125 | B138 | Cor | PAB | -0.41317 | 5.70E-13 |
| B125 | B137 | Cor | AFU | -0.28849 | 9.08E-07 |
| B126 | B127 | Alb | A-G ratio | 0.523231 | 0 |

| | | | | | |
|---|---|---|---|---|---|
| B126 | B188 | Alb | Ca | 0.365475 | 2.83E-10 |
| B127 | B130 | A-G ratio | GLB | -0.91855 | 0 |
| B128 | B131 | ALT | AST | 0.697656 | 0 |
| B128 | B141 | ALT | AST/ALT | -0.62781 | 0 |
| B128 | B140 | ALT | r-GT | 0.395249 | 6.61E-12 |
| B129 | B132 | IBIL | Dbil | 0.593097 | 0 |
| B129 | B133 | IBIL | TBIL | 0.970461 | 0 |
| B131 | B140 | AST | r-GT | 0.44286 | 7.11E-15 |
| B131 | B165 | AST | bilirubin | 0.291125 | 7.14E-07 |
| B132 | B133 | Dbil | TBIL | 0.769691 | 0 |
| B134 | B199 | CHE | Mg | -0.455 | 8.88E-16 |
| B134 | B200 | CHE | OP | 0.455672 | 8.88E-16 |
| B134 | B198 | CHE | P | 0.362124 | 4.22E-10 |
| B134 | B193 | CHE | K | 0.349133 | 1.90E-09 |
| B134 | B196 | CHE | TCO2 | 0.342543 | 3.98E-09 |
| B134 | B139 | CHE | ADA | 0.31029 | 1.16E-07 |
| B135 | B197 | TBA | AG | 0.382989 | 3.25E-11 |
| B135 | B139 | TBA | ADA | 0.297926 | 3.80E-07 |
| B136 | B139 | 5'NT | ADA | 0.467451 | 0 |
| B136 | B140 | 5'NT | r-GT | 0.612235 | 0 |
| B136 | B137 | 5'NT | AFU | 0.463506 | 2.22E-16 |
| B136 | B196 | 5'NT | TCO2 | 0.428994 | 5.80E-14 |
| B136 | B200 | 5'NT | OP | 0.408371 | 1.12E-12 |
| B136 | B150 | 5'NT | ALP | 0.401645 | 2.80E-12 |
| B136 | B138 | 5'NT | PAB | -0.38577 | 2.28E-11 |
| B136 | B198 | 5'NT | P | -0.3091 | 1.30E-07 |
| B137 | B139 | AFU | ADA | 0.766162 | 0 |
| B137 | B194 | AFU | Cl | -0.52484 | 0 |
| B137 | B200 | AFU | OP | 0.676973 | 0 |
| B137 | B140 | AFU | r-GT | 0.409662 | 9.33E-13 |
| B137 | B197 | AFU | AG | 0.372788 | 1.17E-10 |
| B137 | B181 | AFU | APTT | -0.28849 | 9.08E-07 |
| B137 | B184 | AFU | INR | -0.28849 | 9.08E-07 |
| B137 | B186 | AFU | PFC | 0.288494 | 9.08E-07 |
| B137 | B183 | AFU | PTA | -0.28848 | 9.09E-07 |
| B137 | B182 | AFU | TT | 0.288196 | 9.33E-07 |
| B137 | B185 | AFU | PT | -0.28778 | 9.69E-07 |
| B138 | B197 | PAB | AG | 0.419548 | 2.30E-13 |
| B138 | B181 | PAB | APTT | -0.41317 | 5.70E-13 |
| B138 | B183 | PAB | PTA | -0.41317 | 5.70E-13 |
| B138 | B184 | PAB | INR | -0.41317 | 5.70E-13 |
| B138 | B186 | PAB | PFC | 0.413173 | 5.70E-13 |
| B138 | B182 | PAB | TT | 0.413118 | 5.75E-13 |
| B138 | B185 | PAB | PT | -0.4129 | 5.92E-13 |

| | | | | | |
|---|---|---|---|---|---|
| B138 | B198 | PAB | P | 0.405393 | 1.68E-12 |
| B138 | B194 | PAB | Cl | -0.36041 | 5.17E-10 |
| B138 | B179 | PAB | α1-MG | -0.30141 | 2.73E-07 |
| B138 | B180 | PAB | Tf | -0.29384 | 5.56E-07 |
| B138 | B178 | PAB | Ig-U | -0.29042 | 7.62E-07 |
| B139 | B194 | ADA | Cl | -0.4914 | 0 |
| B139 | B200 | ADA | OP | 0.894141 | 0 |
| B139 | B140 | ADA | r-GT | 0.430385 | 4.71E-14 |
| B139 | B150 | ADA | ALP | 0.353673 | 1.13E-09 |
| B139 | B153 | ADA | Glu | 0.348209 | 2.11E-09 |
| B14 | B118 | G-17 | CEA1 | 0.398482 | 4.30E-12 |
| B14 | B80 | G-17 | anti-HP IgG | 0.343525 | 3.57E-09 |
| B140 | B200 | r-GT | OP | 0.404194 | 1.98E-12 |
| B140 | B227 | r-GT | OD | 0.337128 | 7.20E-09 |
| B140 | B153 | r-GT | Glu | 0.303585 | 2.22E-07 |
| B142 | B144 | 25-(OH)D | Total-PINP | -0.44357 | 6.22E-15 |
| B142 | B151 | 25-(OH)D | Anti-CCP | -0.37692 | 6.98E-11 |
| B142 | B168 | 25-(OH)D | NMP22 | -0.29844 | 3.62E-07 |
| B143 | B144 | β-CTx | Total-PINP | 0.912455 | 0 |
| B143 | B145 | β-CTx | BGP | 0.716386 | 0 |
| B143 | B154 | β-CTx | RF | -0.47963 | 0 |
| B143 | B175 | β-CTx | ALB-U | -0.29217 | 6.49E-07 |
| B144 | B145 | Total-PINP | BGP | 0.682211 | 0 |
| B144 | B154 | Total-PINP | RF | -0.52378 | 0 |
| B144 | B175 | Total-PINP | ALB-U | -0.37378 | 1.03E-10 |
| B144 | B151 | Total-PINP | Anti-CCP | 0.323049 | 3.19E-08 |
| B148 | B149 | Anti_TPO | TGAb | 0.448847 | 2.66E-15 |
| B15 | B156 | CD3-CD19+(B cell) | IgE | -0.75293 | 0 |
| B15 | B16 | CD3-CD19+(B cell) | CD3+T | 0.762273 | 0 |
| B15 | B22 | CD3-CD19+(B cell) | CD3+CD8+(T) | 0.934563 | 0 |
| B15 | B19 | CD3-CD19+(B cell) | CD3+CD4+/CD3+CD8+ | -0.45738 | 6.66E-16 |
| B15 | B120 | CD3-CD19+(B cell) | HCV-cAg | 0.381032 | 4.17E-11 |
| B15 | B158 | CD3-CD19+(B cell) | IgM | 0.35971 | 5.61E-10 |
| B15 | B100 | CD3-CD19+(B cell) | anti-Ro52 | 0.315889 | 6.62E-08 |
| B15 | B101 | CD3-CD19+(B cell) | anti-Sc70 | 0.315889 | 6.62E-08 |
| B15 | B102 | CD3-CD19+(B cell) | anti-Sm | 0.315889 | 6.62E-08 |
| B15 | B103 | CD3-CD19+(B cell) | anti-SSA/Ro | 0.315889 | 6.62E-08 |
| B15 | B104 | CD3-CD19+(B cell) | anti-SSB | 0.315889 | 6.62E-08 |
| B15 | B105 | CD3-CD19+(B cell) | anti-U1RNP | 0.315889 | 6.62E-08 |
| B15 | B106 | CD3-CD19+(B cell) | anti-P | 0.315889 | 6.62E-08 |
| B15 | B107 | CD3-CD19+(B cell) | AnuA | 0.315889 | 6.62E-08 |
| B15 | B108 | CD3-CD19+(B cell) | Anti-Cyclin | 0.315889 | 6.62E-08 |
| B15 | B109 | CD3-CD19+(B cell) | AMA-M2 | 0.315889 | 6.62E-08 |
| B15 | B110 | CD3-CD19+(B cell) | AHA | 0.315889 | 6.62E-08 |

| | | | | | |
|---|---|---|---|---|---|
| B15 | B124 | CD3-CD19+(B cell) | ACTH | -0.31589 | 6.62E-08 |
| B15 | B125 | CD3-CD19+(B cell) | Cor | 0.315889 | 6.62E-08 |
| B15 | B181 | CD3-CD19+(B cell) | APTT | 0.315889 | 6.62E-08 |
| B15 | B184 | CD3-CD19+(B cell) | INR | 0.315889 | 6.62E-08 |
| B15 | B186 | CD3-CD19+(B cell) | PFC | -0.31589 | 6.62E-08 |
| B15 | B20 | CD3-CD19+(B cell) | CD3+CD4+CD8+ | 0.315889 | 6.62E-08 |
| B15 | B21 | CD3-CD19+(B cell) | CD3+CD4-CD8- | -0.31589 | 6.62E-08 |
| B15 | B79 | CD3-CD19+(B cell) | hb V -dna | 0.315889 | 6.62E-08 |
| B15 | B92 | CD3-CD19+(B cell) | ALDH2-Glu504Glu | 0.315889 | 6.62E-08 |
| B15 | B93 | CD3-CD19+(B cell) | ALDH2-Glu504Lys | -0.31589 | 6.62E-08 |
| B15 | B94 | CD3-CD19+(B cell) | ALDH2-Lys504Lys | 0.315889 | 6.62E-08 |
| B15 | B95 | CD3-CD19+(B cell) | anti-cenpb | 0.315889 | 6.62E-08 |
| B15 | B97 | CD3-CD19+(B cell) | dsDNA | 0.315889 | 6.62E-08 |
| B15 | B98 | CD3-CD19+(B cell) | anti-Jo-1 | 0.315889 | 6.62E-08 |
| B15 | B99 | CD3-CD19+(B cell) | anti-PM-Scl | 0.315889 | 6.62E-08 |
| B15 | B183 | CD3-CD19+(B cell) | PTA | 0.315885 | 6.63E-08 |
| B15 | B182 | CD3-CD19+(B cell) | TT | -0.31577 | 6.70E-08 |
| B15 | B185 | CD3-CD19+(B cell) | PT | 0.315524 | 6.87E-08 |
| B150 | B193 | ALP | K | 0.344866 | 3.07E-09 |
| B150 | B200 | ALP | OP | 0.322082 | 3.53E-08 |
| B150 | B175 | ALP | ALB-U | 0.303947 | 2.14E-07 |
| B151 | B181 | Anti-CCP | APTT | 0.508198 | 0 |
| B151 | B182 | Anti-CCP | TT | -0.50801 | 0 |
| B151 | B183 | Anti-CCP | PTA | 0.508192 | 0 |
| B151 | B184 | Anti-CCP | INR | 0.508198 | 0 |
| B151 | B185 | Anti-CCP | PT | 0.507623 | 0 |
| B151 | B186 | Anti-CCP | PFC | -0.5082 | 0 |
| B151 | B158 | Anti-CCP | IgM | 0.417143 | 3.25E-13 |
| B151 | B155 | Anti-CCP | IgA | 0.401477 | 2.87E-12 |
| B151 | B194 | Anti-CCP | Cl | 0.401039 | 3.04E-12 |
| B151 | B179 | Anti-CCP | α1-MG | 0.364509 | 3.18E-10 |
| B151 | B154 | Anti-CCP | RF | -0.35782 | 7.00E-10 |
| B151 | B180 | Anti-CCP | Tf | 0.355003 | 9.71E-10 |
| B151 | B178 | Anti-CCP | Ig-U | 0.350723 | 1.59E-09 |
| B151 | B197 | Anti-CCP | AG | -0.33963 | 5.48E-09 |
| B152 | B159 | ACA-Ig(G-A-M) | β2-MG | 0.999992 | 0 |
| B152 | B169 | ACA-Ig(G-A-M) | U-CRY | 0.999992 | 0 |
| B153 | B170 | Glu | U-GLU | 0.62779 | 0 |
| B153 | B195 | Glu | Na | -0.35044 | 1.64E-09 |
| B153 | B200 | Glu | OP | 0.34853 | 2.04E-09 |
| B155 | B158 | IgA | IgM | 0.353613 | 1.14E-09 |
| B156 | B158 | IgE | IgM | -0.38655 | 2.06E-11 |
| B156 | B212 | IgE | IOP-L | 0.289585 | 8.22E-07 |
| B158 | B181 | IgM | APTT | 0.497089 | 0 |

| | | | | | |
|---|---|---|---|---|---|
| B158 | B182 | IgM | TT | -0.49688 | 0 |
| B158 | B183 | IgM | PTA | 0.497081 | 0 |
| B158 | B184 | IgM | INR | 0.497089 | 0 |
| B158 | B185 | IgM | PT | 0.496468 | 0 |
| B158 | B186 | IgM | PFC | -0.49709 | 0 |
| B158 | B194 | IgM | Cl | 0.394324 | 7.47E-12 |
| B158 | B179 | IgM | α1-MG | 0.354984 | 9.73E-10 |
| B158 | B180 | IgM | Tf | 0.345638 | 2.82E-09 |
| B158 | B178 | IgM | Ig-U | 0.341432 | 4.49E-09 |
| B158 | B197 | IgM | AG | -0.3112 | 1.06E-07 |
| B159 | B169 | β2-MG | U-CRY | 0.999997 | 0 |
| B16 | B156 | CD3+T | IgE | -0.78173 | 0 |
| B16 | B18 | CD3+T | CD3+CD4+helper(inducer)T | 0.824237 | 0 |
| B16 | B22 | CD3+T | CD3+CD8+(T) | 0.482099 | 0 |
| B16 | B23 | CD3+T | CD3-CD16+CD56+(NK) | -0.83239 | 0 |
| B16 | B120 | CD3+T | HCV-cAg | 0.378604 | 5.66E-11 |
| B16 | B100 | CD3+T | anti-Ro52 | -0.37332 | 1.09E-10 |
| B16 | B101 | CD3+T | anti-Sc70 | -0.37332 | 1.09E-10 |
| B16 | B102 | CD3+T | anti-Sm | -0.37332 | 1.09E-10 |
| B16 | B103 | CD3+T | anti-SSA/Ro | -0.37332 | 1.09E-10 |
| B16 | B104 | CD3+T | anti-SSB | -0.37332 | 1.09E-10 |
| B16 | B105 | CD3+T | anti-U1RNP | -0.37332 | 1.09E-10 |
| B16 | B106 | CD3+T | anti-P | -0.37332 | 1.09E-10 |
| B16 | B107 | CD3+T | AnuA | -0.37332 | 1.09E-10 |
| B16 | B108 | CD3+T | Anti-Cyclin | -0.37332 | 1.09E-10 |
| B16 | B109 | CD3+T | AMA-M2 | -0.37332 | 1.09E-10 |
| B16 | B110 | CD3+T | AHA | -0.37332 | 1.09E-10 |
| B16 | B124 | CD3+T | ACTH | 0.373321 | 1.09E-10 |
| B16 | B125 | CD3+T | Cor | -0.37332 | 1.09E-10 |
| B16 | B181 | CD3+T | APTT | -0.37332 | 1.09E-10 |
| B16 | B182 | CD3+T | TT | 0.373317 | 1.09E-10 |
| B16 | B183 | CD3+T | PTA | -0.37332 | 1.09E-10 |
| B16 | B184 | CD3+T | INR | -0.37332 | 1.09E-10 |
| B16 | B186 | CD3+T | PFC | 0.373321 | 1.09E-10 |
| B16 | B20 | CD3+T | CD3+CD4+CD8+ | -0.37332 | 1.09E-10 |
| B16 | B21 | CD3+T | CD3+CD4-CD8- | 0.373321 | 1.09E-10 |
| B16 | B79 | CD3+T | hbＶ-dna | -0.37332 | 1.09E-10 |
| B16 | B92 | CD3+T | ALDH2-Glu504Glu | -0.37332 | 1.09E-10 |
| B16 | B93 | CD3+T | ALDH2-Glu504Lys | 0.373321 | 1.09E-10 |
| B16 | B94 | CD3+T | ALDH2-Lys504Lys | -0.37332 | 1.09E-10 |
| B16 | B95 | CD3+T | anti-cenpb | -0.37332 | 1.09E-10 |
| B16 | B97 | CD3+T | dsDNA | -0.37332 | 1.09E-10 |
| B16 | B98 | CD3+T | anti-Jo-1 | -0.37332 | 1.09E-10 |
| B16 | B99 | CD3+T | anti-PM-Scl | -0.37332 | 1.09E-10 |

| | | | | | |
|---|---|---|---|---|---|
| B16 | B185 | CD3+T | PT | -0.37317 | 1.11E-10 |
| B16 | B81 | CD3+T | TSGF | 0.368619 | 1.94E-10 |
| B16 | B157 | CD3+T | IgG | -0.33995 | 5.29E-09 |
| B16 | B155 | CD3+T | IgA | 0.302949 | 2.36E-07 |
| B161 | B173 | LEU | NTT | 0.464776 | 2.22E-16 |
| B163 | B170 | SG | U-GLU | 0.372608 | 1.19E-10 |
| B163 | B176 | SG | Ucr | 0.3334 | 1.08E-08 |
| B163 | B166 | SG | Pro | 0.307529 | 1.52E-07 |
| B168 | B176 | NMP22 | Ucr | 0.303471 | 2.24E-07 |
| B17 | B116 | CD3+CD16+CD56+NK | TT3 | -0.98074 | 0 |
| B17 | B28 | CD3+CD16+CD56+NK | PRL | -0.68803 | 0 |
| B174 | B191 | Urea/Cr | Urea | 0.477027 | 0 |
| B174 | B190 | Urea/Cr | Cre | -0.42169 | 1.69E-13 |
| B174 | B221 | Urea/Cr | Hei | -0.34741 | 2.31E-09 |
| B175 | B177 | ALB-U | ACR | 0.743122 | 0 |
| B175 | B176 | ALB-U | Ucr | 0.334717 | 9.34E-09 |
| B178 | B179 | Ig-U | α1-MG | 0.999268 | 0 |
| B178 | B180 | Ig-U | Tf | 0.999932 | 0 |
| B178 | B181 | Ig-U | APTT | 0.6975 | 0 |
| B178 | B182 | Ig-U | TT | -0.69733 | 0 |
| B178 | B183 | Ig-U | PTA | 0.697497 | 0 |
| B178 | B184 | Ig-U | INR | 0.6975 | 0 |
| B178 | B185 | Ig-U | PT | 0.696886 | 0 |
| B178 | B186 | Ig-U | PFC | -0.6975 | 0 |
| B178 | B194 | Ig-U | Cl | 0.557176 | 0 |
| B178 | B197 | Ig-U | AG | -0.46693 | 0 |
| B179 | B180 | α1-MG | Tf | 0.999646 | 0 |
| B179 | B181 | α1-MG | APTT | 0.724317 | 0 |
| B179 | B182 | α1-MG | TT | -0.72408 | 0 |
| B179 | B183 | α1-MG | PTA | 0.72431 | 0 |
| B179 | B184 | α1-MG | INR | 0.724317 | 0 |
| B179 | B185 | α1-MG | PT | 0.723567 | 0 |
| B179 | B186 | α1-MG | PFC | -0.72432 | 0 |
| B179 | B194 | α1-MG | Cl | 0.578783 | 0 |
| B179 | B197 | α1-MG | AG | -0.48492 | 0 |
| B18 | B100 | CD3+CD4+helper(inducer)T | anti-Ro52 | -0.83301 | 0 |
| B18 | B101 | CD3+CD4+helper(inducer)T | anti-Sc70 | -0.83301 | 0 |
| B18 | B102 | CD3+CD4+helper(inducer)T | anti-Sm | -0.83301 | 0 |
| B18 | B103 | CD3+CD4+helper(inducer)T | anti-SSA/Ro | -0.83301 | 0 |
| B18 | B104 | CD3+CD4+helper(inducer)T | anti-SSB | -0.83301 | 0 |
| B18 | B105 | CD3+CD4+helper(inducer)T | anti-U1RNP | -0.83301 | 0 |
| B18 | B106 | CD3+CD4+helper(inducer)T | anti-P | -0.83301 | 0 |
| B18 | B107 | CD3+CD4+helper(inducer)T | AnuA | -0.83301 | 0 |
| B18 | B108 | CD3+CD4+helper(inducer)T | Anti-Cyclin | -0.83301 | 0 |

| | | | | | |
|---|---|---|---|---|---|
| B18 | B109 | CD3+CD4+helper(inducer)T | AMA-M2 | -0.83301 | 0 |
| B18 | B110 | CD3+CD4+helper(inducer)T | AHA | -0.83301 | 0 |
| B18 | B124 | CD3+CD4+helper(inducer)T | ACTH | 0.833011 | 0 |
| B18 | B125 | CD3+CD4+helper(inducer)T | Cor | -0.83301 | 0 |
| B18 | B156 | CD3+CD4+helper(inducer)T | IgE | -0.50694 | 0 |
| B18 | B178 | CD3+CD4+helper(inducer)T | Ig-U | -0.58245 | 0 |
| B18 | B179 | CD3+CD4+helper(inducer)T | α1-MG | -0.60473 | 0 |
| B18 | B180 | CD3+CD4+helper(inducer)T | Tf | -0.58938 | 0 |
| B18 | B181 | CD3+CD4+helper(inducer)T | APTT | -0.83301 | 0 |
| B18 | B182 | CD3+CD4+helper(inducer)T | TT | 0.832901 | 0 |
| B18 | B183 | CD3+CD4+helper(inducer)T | PTA | -0.83301 | 0 |
| B18 | B184 | CD3+CD4+helper(inducer)T | INR | -0.83301 | 0 |
| B18 | B185 | CD3+CD4+helper(inducer)T | PT | -0.83247 | 0 |
| B18 | B186 | CD3+CD4+helper(inducer)T | PFC | 0.833011 | 0 |
| B18 | B19 | CD3+CD4+helper(inducer)T | CD3+CD4+/CD3+CD8+ | 0.738519 | 0 |
| B18 | B194 | CD3+CD4+helper(inducer)T | CI | -0.65147 | 0 |
| B18 | B197 | CD3+CD4+helper(inducer)T | AG | 0.578919 | 0 |
| B18 | B20 | CD3+CD4+helper(inducer)T | CD3+CD4+CD8+ | -0.83301 | 0 |
| B18 | B21 | CD3+CD4+helper(inducer)T | CD3+CD4-CD8- | 0.833011 | 0 |
| B18 | B23 | CD3+CD4+helper(inducer)T | CD3-CD16+CD56+(NK) | -0.99989 | 0 |
| B18 | B79 | CD3+CD4+helper(inducer)T | hb V -dna | -0.83301 | 0 |
| B18 | B81 | CD3+CD4+helper(inducer)T | TSGF | 0.554403 | 0 |
| B18 | B92 | CD3+CD4+helper(inducer)T | ALDH2-Glu504Glu | -0.83301 | 0 |
| B18 | B93 | CD3+CD4+helper(inducer)T | ALDH2-Glu504Lys | 0.833011 | 0 |
| B18 | B94 | CD3+CD4+helper(inducer)T | ALDH2-Lys504Lys | -0.83301 | 0 |
| B18 | B95 | CD3+CD4+helper(inducer)T | anti-cenpb | -0.83301 | 0 |
| B18 | B97 | CD3+CD4+helper(inducer)T | dsDNA | -0.83301 | 0 |
| B18 | B98 | CD3+CD4+helper(inducer)T | anti-Jo-1 | -0.83301 | 0 |
| B18 | B99 | CD3+CD4+helper(inducer)T | anti-PM-Scl | -0.83301 | 0 |
| B18 | B151 | CD3+CD4+helper(inducer)T | Anti-CCP | -0.42623 | 8.70E-14 |
| B18 | B55 | CD3+CD4+helper(inducer)T | TK | -0.41853 | 2.66E-13 |
| B18 | B54 | CD3+CD4+helper(inducer)T | ESR | -0.41504 | 4.38E-13 |
| B18 | B138 | CD3+CD4+helper(inducer)T | PAB | 0.377217 | 6.73E-11 |
| B18 | B158 | CD3+CD4+helper(inducer)T | IgM | -0.29589 | 4.60E-07 |
| B180 | B181 | Tf | APTT | 0.705838 | 0 |
| B180 | B182 | Tf | TT | -0.70566 | 0 |
| B180 | B183 | Tf | PTA | 0.705835 | 0 |
| B180 | B184 | Tf | INR | 0.705838 | 0 |
| B180 | B185 | Tf | PT | 0.705219 | 0 |
| B180 | B186 | Tf | PFC | -0.70584 | 0 |
| B180 | B194 | Tf | CI | 0.563926 | 0 |
| B180 | B197 | Tf | AG | -0.47252 | 0 |
| B181 | B182 | APTT | TT | -0.99982 | 0 |
| B181 | B183 | APTT | PTA | 0.999999 | 0 |

| | | | | | |
|---|---|---|---|---|---|
| B181 | B185 | APTT | PT | 0.999261 | 0 |
| B181 | B194 | APTT | CI | 0.804248 | 0 |
| B181 | B197 | APTT | AG | -0.67006 | 0 |
| B182 | B183 | TT | PTA | -0.99984 | 0 |
| B182 | B184 | TT | INR | -0.99982 | 0 |
| B182 | B185 | TT | PT | -0.99981 | 0 |
| B182 | B186 | TT | PFC | 0.999822 | 0 |
| B182 | B194 | TT | CI | -0.80474 | 0 |
| B182 | B197 | TT | AG | 0.66989 | 0 |
| B183 | B184 | PTA | INR | 0.999999 | 0 |
| B183 | B185 | PTA | PT | 0.999304 | 0 |
| B183 | B186 | PTA | PFC | -1 | 0 |
| B183 | B194 | PTA | CI | 0.804286 | 0 |
| B183 | B197 | PTA | AG | -0.67006 | 0 |
| B184 | B185 | INR | PT | 0.999261 | 0 |
| B184 | B194 | INR | CI | 0.804248 | 0 |
| B184 | B197 | INR | AG | -0.67006 | 0 |
| B185 | B186 | PT | PFC | -0.99926 | 0 |
| B185 | B194 | PT | CI | 0.804938 | 0 |
| B185 | B197 | PT | AG | -0.66946 | 0 |
| B186 | B194 | PFC | CI | -0.80425 | 0 |
| B186 | B197 | PFC | AG | 0.670061 | 0 |
| B189 | B190 | eGFB | Cre | -0.72456 | 0 |
| B189 | B191 | eGFB | Urea | -0.49177 | 0 |
| B189 | B192 | eGFB | UA | -0.32414 | 2.85E-08 |
| B19 | B100 | CD3+CD4+/CD3+CD8+ | anti-Ro52 | -0.98822 | 0 |
| B19 | B101 | CD3+CD4+/CD3+CD8+ | anti-Sc70 | -0.98822 | 0 |
| B19 | B102 | CD3+CD4+/CD3+CD8+ | anti-Sm | -0.98822 | 0 |
| B19 | B103 | CD3+CD4+/CD3+CD8+ | anti-SSA/Ro | -0.98822 | 0 |
| B19 | B104 | CD3+CD4+/CD3+CD8+ | anti-SSB | -0.98822 | 0 |
| B19 | B105 | CD3+CD4+/CD3+CD8+ | anti-U1RNP | -0.98822 | 0 |
| B19 | B106 | CD3+CD4+/CD3+CD8+ | anti-P | -0.98822 | 0 |
| B19 | B107 | CD3+CD4+/CD3+CD8+ | AnuA | -0.98822 | 0 |
| B19 | B108 | CD3+CD4+/CD3+CD8+ | Anti-Cyclin | -0.98822 | 0 |
| B19 | B109 | CD3+CD4+/CD3+CD8+ | AMA-M2 | -0.98822 | 0 |
| B19 | B110 | CD3+CD4+/CD3+CD8+ | AHA | -0.98822 | 0 |
| B19 | B124 | CD3+CD4+/CD3+CD8+ | ACTH | 0.988218 | 0 |
| B19 | B125 | CD3+CD4+/CD3+CD8+ | Cor | -0.98822 | 0 |
| B19 | B151 | CD3+CD4+/CD3+CD8+ | Anti-CCP | -0.50141 | 0 |
| B19 | B158 | CD3+CD4+/CD3+CD8+ | IgM | -0.52393 | 0 |
| B19 | B178 | CD3+CD4+/CD3+CD8+ | Ig-U | -0.68889 | 0 |
| B19 | B179 | CD3+CD4+/CD3+CD8+ | α1-MG | -0.71541 | 0 |
| B19 | B180 | CD3+CD4+/CD3+CD8+ | Tf | -0.69713 | 0 |
| B19 | B181 | CD3+CD4+/CD3+CD8+ | APTT | -0.98822 | 0 |

| | | | | | |
|---|---|---|---|---|---|
| B19 | B182 | CD3+CD4+/CD3+CD8+ | TT | 0.988032 | 0 |
| B19 | B183 | CD3+CD4+/CD3+CD8+ | PTA | -0.98822 | 0 |
| B19 | B184 | CD3+CD4+/CD3+CD8+ | INR | -0.98822 | 0 |
| B19 | B185 | CD3+CD4+/CD3+CD8+ | PT | -0.98747 | 0 |
| B19 | B186 | CD3+CD4+/CD3+CD8+ | PFC | 0.988218 | 0 |
| B19 | B194 | CD3+CD4+/CD3+CD8+ | Cl | -0.79988 | 0 |
| B19 | B197 | CD3+CD4+/CD3+CD8+ | AG | 0.656425 | 0 |
| B19 | B20 | CD3+CD4+/CD3+CD8+ | CD3+CD4+CD8+ | -0.98822 | 0 |
| B19 | B21 | CD3+CD4+/CD3+CD8+ | CD3+CD4-CD8- | 0.988218 | 0 |
| B19 | B22 | CD3+CD4+/CD3+CD8+ | CD3+CD8+(T) | -0.74385 | 0 |
| B19 | B23 | CD3+CD4+/CD3+CD8+ | CD3-CD16+CD56+(NK) | -0.72863 | 0 |
| B19 | B54 | CD3+CD4+/CD3+CD8+ | ESR | -0.48943 | 0 |
| B19 | B55 | CD3+CD4+/CD3+CD8+ | TK | -0.49564 | 0 |
| B19 | B79 | CD3+CD4+/CD3+CD8+ | hb V -dna | -0.98822 | 0 |
| B19 | B81 | CD3+CD4+/CD3+CD8+ | TSGF | 0.514626 | 0 |
| B19 | B92 | CD3+CD4+/CD3+CD8+ | ALDH2-Glu504Glu | -0.98822 | 0 |
| B19 | B93 | CD3+CD4+/CD3+CD8+ | ALDH2-Glu504Lys | 0.988218 | 0 |
| B19 | B94 | CD3+CD4+/CD3+CD8+ | ALDH2-Lys504Lys | -0.98822 | 0 |
| B19 | B95 | CD3+CD4+/CD3+CD8+ | anti-cenpb | -0.98822 | 0 |
| B19 | B97 | CD3+CD4+/CD3+CD8+ | dsDNA | -0.98822 | 0 |
| B19 | B98 | CD3+CD4+/CD3+CD8+ | anti-Jo-1 | -0.98822 | 0 |
| B19 | B99 | CD3+CD4+/CD3+CD8+ | anti-PM-Scl | -0.98822 | 0 |
| B19 | B138 | CD3+CD4+/CD3+CD8+ | PAB | 0.399165 | 3.92E-12 |
| B19 | B137 | CD3+CD4+/CD3+CD8+ | AFU | 0.300417 | 3.00E-07 |
| B190 | B191 | Cre | Urea | 0.556065 | 0 |
| B190 | B192 | Cre | UA | 0.426496 | 8.39E-14 |
| B193 | B196 | K | TCO2 | 0.429013 | 5.77E-14 |
| B193 | B197 | K | AG | -0.38685 | 1.98E-11 |
| B194 | B197 | Cl | AG | -0.66428 | 0 |
| B194 | B200 | Cl | OP | -0.41438 | 4.81E-13 |
| B195 | B198 | Na | P | 0.586244 | 0 |
| B195 | B196 | Na | TCO2 | 0.458762 | 4.44E-16 |
| B196 | B197 | TCO2 | AG | -0.40744 | 1.27E-12 |
| B197 | B198 | AG | P | 0.35507 | 9.64E-10 |
| B2 | B189 | Age | eGFB | -0.57178 | 0 |
| B2 | B51 | Age | Hb | -0.43089 | 4.35E-14 |
| B2 | B220 | Age | SBP | 0.411746 | 6.97E-13 |
| B2 | B46 | Age | RBC Count | -0.37315 | 1.11E-10 |
| B2 | B221 | Age | Hei | -0.32417 | 2.85E-08 |
| B2 | B126 | Age | Alb | -0.32206 | 3.53E-08 |
| B2 | B128 | Age | ALT | -0.31066 | 1.11E-07 |
| B2 | B222 | Age | NS | -0.30976 | 1.22E-07 |
| B2 | B216 | Age | BMI | -0.30188 | 2.61E-07 |
| B20 | B101 | CD3+CD4+CD8+ | anti-Sc70 | 1 | 0 |

| | | | | | |
|---|---|---|---|---|---|
| B20 | B103 | CD3+CD4+CD8+ | anti-SSA/Ro | 1 | 0 |
| B20 | B105 | CD3+CD4+CD8+ | anti-U1RNP | 1 | 0 |
| B20 | B107 | CD3+CD4+CD8+ | AnuA | 1 | 0 |
| B20 | B110 | CD3+CD4+CD8+ | AHA | 1 | 0 |
| B20 | B124 | CD3+CD4+CD8+ | ACTH | -1 | 0 |
| B20 | B125 | CD3+CD4+CD8+ | Cor | 1 | 0 |
| B20 | B151 | CD3+CD4+CD8+ | Anti-CCP | 0.508198 | 0 |
| B20 | B158 | CD3+CD4+CD8+ | IgM | 0.497089 | 0 |
| B20 | B178 | CD3+CD4+CD8+ | Ig-U | 0.6975 | 0 |
| B20 | B179 | CD3+CD4+CD8+ | α1-MG | 0.724317 | 0 |
| B20 | B180 | CD3+CD4+CD8+ | Tf | 0.705838 | 0 |
| B20 | B182 | CD3+CD4+CD8+ | TT | -0.99982 | 0 |
| B20 | B183 | CD3+CD4+CD8+ | PTA | 0.999999 | 0 |
| B20 | B185 | CD3+CD4+CD8+ | PT | 0.999261 | 0 |
| B20 | B194 | CD3+CD4+CD8+ | Cl | 0.804248 | 0 |
| B20 | B197 | CD3+CD4+CD8+ | AG | -0.67006 | 0 |
| B20 | B22 | CD3+CD4+CD8+ | CD3+CD8+(T) | 0.632798 | 0 |
| B20 | B23 | CD3+CD4+CD8+ | CD3-CD16+CD56+(NK) | 0.824875 | 0 |
| B20 | B54 | CD3+CD4+CD8+ | ESR | 0.495827 | 0 |
| B20 | B55 | CD3+CD4+CD8+ | TK | 0.501718 | 0 |
| B20 | B79 | CD3+CD4+CD8+ | hb V -dna | 1 | 0 |
| B20 | B81 | CD3+CD4+CD8+ | TSGF | -0.54814 | 0 |
| B20 | B92 | CD3+CD4+CD8+ | ALDH2-Glu504Glu | 1 | 0 |
| B20 | B93 | CD3+CD4+CD8+ | ALDH2-Glu504Lys | -1 | 0 |
| B20 | B94 | CD3+CD4+CD8+ | ALDH2-Lys504Lys | 1 | 0 |
| B20 | B97 | CD3+CD4+CD8+ | dsDNA | 1 | 0 |
| B20 | B98 | CD3+CD4+CD8+ | anti-Jo-1 | 1 | 0 |
| B20 | B99 | CD3+CD4+CD8+ | anti-PM-Scl | 1 | 0 |
| B20 | B138 | CD3+CD4+CD8+ | PAB | -0.41317 | 5.70E-13 |
| B20 | B137 | CD3+CD4+CD8+ | AFU | -0.28849 | 9.08E-07 |
| B201 | B204 | e-habits | Drinking | 0.303398 | 2.26E-07 |
| B206 | B207 | Waist | WHR | 0.631412 | 0 |
| B206 | B210 | Waist | Hip | 0.720887 | 0 |
| B206 | B215 | Waist | BMI | 0.770044 | 0 |
| B206 | B216 | Waist | BMI | 0.756643 | 0 |
| B206 | B222 | Waist | NS | 0.669739 | 0 |
| B207 | B222 | WHR | NS | 0.353821 | 1.11E-09 |
| B21 | B105 | CD3+CD4-CD8- | anti-U1RNP | -1 | 0 |
| B21 | B107 | CD3+CD4-CD8- | AnuA | -1 | 0 |
| B21 | B110 | CD3+CD4-CD8- | AHA | -1 | 0 |
| B21 | B151 | CD3+CD4-CD8- | Anti-CCP | -0.5082 | 0 |
| B21 | B158 | CD3+CD4-CD8- | IgM | -0.49709 | 0 |
| B21 | B178 | CD3+CD4-CD8- | Ig-U | -0.6975 | 0 |
| B21 | B179 | CD3+CD4-CD8- | α1-MG | -0.72432 | 0 |

| | | | | | |
|---|---|---|---|---|---|
| B21 | B180 | CD3+CD4-CD8- | Tf | -0.70584 | 0 |
| B21 | B182 | CD3+CD4-CD8- | TT | 0.999822 | 0 |
| B21 | B183 | CD3+CD4-CD8- | PTA | -1 | 0 |
| B21 | B185 | CD3+CD4-CD8- | PT | -0.99926 | 0 |
| B21 | B194 | CD3+CD4-CD8- | Cl | -0.80425 | 0 |
| B21 | B197 | CD3+CD4-CD8- | AG | 0.670061 | 0 |
| B21 | B22 | CD3+CD4-CD8- | CD3+CD8+(T) | -0.6328 | 0 |
| B21 | B23 | CD3+CD4-CD8- | CD3-CD16+CD56+(NK) | -0.82488 | 0 |
| B21 | B54 | CD3+CD4-CD8- | ESR | -0.49583 | 0 |
| B21 | B55 | CD3+CD4-CD8- | TK | -0.50172 | 0 |
| B21 | B81 | CD3+CD4-CD8- | TSGF | 0.548139 | 0 |
| B21 | B94 | CD3+CD4-CD8- | ALDH2-Lys504Lys | -1 | 0 |
| B21 | B97 | CD3+CD4-CD8- | dsDNA | -1 | 0 |
| B21 | B98 | CD3+CD4-CD8- | anti-Jo-1 | -1 | 0 |
| B21 | B99 | CD3+CD4-CD8- | anti-PM-Scl | -1 | 0 |
| B21 | B138 | CD3+CD4-CD8- | PAB | 0.413173 | 5.70E-13 |
| B21 | B137 | CD3+CD4-CD8- | AFU | 0.288494 | 9.08E-07 |
| B210 | B215 | Hip | BMI | 0.80334 | 0 |
| B210 | B216 | Hip | BMI | 0.788474 | 0 |
| B210 | B222 | Hip | NS | 0.533358 | 0 |
| B211 | B212 | IOP-R | IOP-L | 0.672445 | 0 |
| B213 | B214 | CVA-R | CVA-L | 0.777644 | 0 |
| B215 | B216 | BMI | BMI | 0.770646 | 0 |
| B215 | B222 | BMI | NS | 0.571705 | 0 |
| B216 | B221 | BMI | Hei | 0.621133 | 0 |
| B216 | B222 | BMI | NS | 0.719286 | 0 |
| B218 | B220 | DBP | SBP | 0.472626 | 0 |
| B22 | B100 | CD3+CD8+(T) | anti-Ro52 | 0.632798 | 0 |
| B22 | B101 | CD3+CD8+(T) | anti-Sc70 | 0.632798 | 0 |
| B22 | B102 | CD3+CD8+(T) | anti-Sm | 0.632798 | 0 |
| B22 | B103 | CD3+CD8+(T) | anti-SSA/Ro | 0.632798 | 0 |
| B22 | B104 | CD3+CD8+(T) | anti-SSB | 0.632798 | 0 |
| B22 | B105 | CD3+CD8+(T) | anti-U1RNP | 0.632798 | 0 |
| B22 | B106 | CD3+CD8+(T) | anti-P | 0.632798 | 0 |
| B22 | B107 | CD3+CD8+(T) | AnuA | 0.632798 | 0 |
| B22 | B108 | CD3+CD8+(T) | Anti-Cyclin | 0.632798 | 0 |
| B22 | B109 | CD3+CD8+(T) | AMA-M2 | 0.632798 | 0 |
| B22 | B110 | CD3+CD8+(T) | AHA | 0.632798 | 0 |
| B22 | B124 | CD3+CD8+(T) | ACTH | -0.6328 | 0 |
| B22 | B125 | CD3+CD8+(T) | Cor | 0.632798 | 0 |
| B22 | B156 | CD3+CD8+(T) | IgE | -0.58943 | 0 |
| B22 | B158 | CD3+CD8+(T) | IgM | 0.479969 | 0 |
| B22 | B181 | CD3+CD8+(T) | APTT | 0.632798 | 0 |
| B22 | B182 | CD3+CD8+(T) | TT | -0.63263 | 0 |

| | | | | | |
|---|---|---|---|---|---|
| B22 | B183 | CD3+CD8+(T) | PTA | 0.632795 | 0 |
| B22 | B184 | CD3+CD8+(T) | INR | 0.632798 | 0 |
| B22 | B185 | CD3+CD8+(T) | PT | 0.632224 | 0 |
| B22 | B186 | CD3+CD8+(T) | PFC | -0.6328 | 0 |
| B22 | B194 | CD3+CD8+(T) | CI | 0.534791 | 0 |
| B22 | B79 | CD3+CD8+(T) | hb V -dna | 0.632798 | 0 |
| B22 | B92 | CD3+CD8+(T) | ALDH2-Glu504Glu | 0.632798 | 0 |
| B22 | B93 | CD3+CD8+(T) | ALDH2-Glu504Lys | -0.6328 | 0 |
| B22 | B94 | CD3+CD8+(T) | ALDH2-Lys504Lys | 0.632798 | 0 |
| B22 | B95 | CD3+CD8+(T) | anti-cenpb | 0.632798 | 0 |
| B22 | B97 | CD3+CD8+(T) | dsDNA | 0.632798 | 0 |
| B22 | B98 | CD3+CD8+(T) | anti-Jo-1 | 0.632798 | 0 |
| B22 | B99 | CD3+CD8+(T) | anti-PM-Scl | 0.632798 | 0 |
| B22 | B179 | CD3+CD8+(T) | α1-MG | 0.456435 | 8.88E-16 |
| B22 | B180 | CD3+CD8+(T) | Tf | 0.44468 | 5.33E-15 |
| B22 | B178 | CD3+CD8+(T) | Ig-U | 0.439381 | 1.20E-14 |
| B22 | B197 | CD3+CD8+(T) | AG | -0.39497 | 6.86E-12 |
| B22 | B151 | CD3+CD8+(T) | Anti-CCP | 0.31754 | 5.61E-08 |
| B22 | B55 | CD3+CD8+(T) | TK | 0.316652 | 6.13E-08 |
| B22 | B54 | CD3+CD8+(T) | ESR | 0.310941 | 1.08E-07 |
| B22 | B120 | CD3+CD8+(T) | HCV-cAg | 0.307639 | 1.50E-07 |
| B221 | B222 | Hei | NS | 0.432367 | 3.51E-14 |
| B227 | B228 | OD | OS | 0.396424 | 5.66E-12 |
| B23 | B100 | CD3-CD16+CD56+(NK) | anti-Ro52 | 0.824875 | 0 |
| B23 | B101 | CD3-CD16+CD56+(NK) | anti-Sc70 | 0.824875 | 0 |
| B23 | B102 | CD3-CD16+CD56+(NK) | anti-Sm | 0.824875 | 0 |
| B23 | B103 | CD3-CD16+CD56+(NK) | anti-SSA/Ro | 0.824875 | 0 |
| B23 | B104 | CD3-CD16+CD56+(NK) | anti-SSB | 0.824875 | 0 |
| B23 | B105 | CD3-CD16+CD56+(NK) | anti-U1RNP | 0.824875 | 0 |
| B23 | B106 | CD3-CD16+CD56+(NK) | anti-P | 0.824875 | 0 |
| B23 | B107 | CD3-CD16+CD56+(NK) | AnuA | 0.824875 | 0 |
| B23 | B108 | CD3-CD16+CD56+(NK) | Anti-Cyclin | 0.824875 | 0 |
| B23 | B109 | CD3-CD16+CD56+(NK) | AMA-M2 | 0.824875 | 0 |
| B23 | B110 | CD3-CD16+CD56+(NK) | AHA | 0.824875 | 0 |
| B23 | B124 | CD3-CD16+CD56+(NK) | ACTH | -0.82488 | 0 |
| B23 | B125 | CD3-CD16+CD56+(NK) | Cor | 0.824875 | 0 |
| B23 | B156 | CD3-CD16+CD56+(NK) | IgE | 0.516238 | 0 |
| B23 | B178 | CD3-CD16+CD56+(NK) | Ig-U | 0.576808 | 0 |
| B23 | B179 | CD3-CD16+CD56+(NK) | α1-MG | 0.598867 | 0 |
| B23 | B180 | CD3-CD16+CD56+(NK) | Tf | 0.583669 | 0 |
| B23 | B181 | CD3-CD16+CD56+(NK) | APTT | 0.824875 | 0 |
| B23 | B182 | CD3-CD16+CD56+(NK) | TT | -0.82477 | 0 |
| B23 | B183 | CD3-CD16+CD56+(NK) | PTA | 0.824877 | 0 |
| B23 | B184 | CD3-CD16+CD56+(NK) | INR | 0.824875 | 0 |

| | | | | | |
|---|---|---|---|---|---|
| B23 | B185 | CD3-CD16+CD56+(NK) | PT | 0.824344 | 0 |
| B23 | B186 | CD3-CD16+CD56+(NK) | PFC | -0.82488 | 0 |
| B23 | B194 | CD3-CD16+CD56+(NK) | CI | 0.644522 | 0 |
| B23 | B197 | CD3-CD16+CD56+(NK) | AG | -0.57392 | 0 |
| B23 | B79 | CD3-CD16+CD56+(NK) | hb V -dna | 0.824875 | 0 |
| B23 | B81 | CD3-CD16+CD56+(NK) | TSGF | -0.55207 | 0 |
| B23 | B92 | CD3-CD16+CD56+(NK) | ALDH2-Glu504Glu | 0.824875 | 0 |
| B23 | B93 | CD3-CD16+CD56+(NK) | ALDH2-Glu504Lys | -0.82488 | 0 |
| B23 | B94 | CD3-CD16+CD56+(NK) | ALDH2-Lys504Lys | 0.824875 | 0 |
| B23 | B95 | CD3-CD16+CD56+(NK) | anti-cenpb | 0.824875 | 0 |
| B23 | B97 | CD3-CD16+CD56+(NK) | dsDNA | 0.824875 | 0 |
| B23 | B98 | CD3-CD16+CD56+(NK) | anti-Jo-1 | 0.824875 | 0 |
| B23 | B99 | CD3-CD16+CD56+(NK) | anti-PM-Scl | 0.824875 | 0 |
| B23 | B151 | CD3-CD16+CD56+(NK) | Anti-CCP | 0.422155 | 1.58E-13 |
| B23 | B55 | CD3-CD16+CD56+(NK) | TK | 0.414464 | 4.75E-13 |
| B23 | B54 | CD3-CD16+CD56+(NK) | ESR | 0.411053 | 7.68E-13 |
| B23 | B138 | CD3-CD16+CD56+(NK) | PAB | -0.37458 | 9.35E-11 |
| B23 | B158 | CD3-CD16+CD56+(NK) | IgM | 0.289272 | 8.46E-07 |
| B24 | B26 | αHBDH | CKMB | 0.762881 | 0 |
| B24 | B27 | αHBDH | LDH | 0.624897 | 0 |
| B24 | B134 | αHBDH | CHE | 0.31918 | 4.75E-08 |
| B25 | B58 | CK | HCT | -0.47032 | 0 |
| B25 | B62 | CK | WBHSRVI | -0.36863 | 1.93E-10 |
| B25 | B63 | CK | WBRSR(mPas_200) | -0.3461 | 2.67E-09 |
| B25 | B66 | CK | WBRSR(mPas_100) | -0.32519 | 2.56E-08 |
| B25 | B67 | CK | WBRSR(mPas_50) | -0.3012 | 2.79E-07 |
| B26 | B27 | CKMB | LDH | 0.465097 | 2.22E-16 |
| B27 | B200 | LDH | OP | 0.451274 | 1.78E-15 |
| B27 | B139 | LDH | ADA | 0.440671 | 9.77E-15 |
| B27 | B88 | LDH | CA 242 | 0.415229 | 4.27E-13 |
| B27 | B82 | LDH | CEA | 0.390215 | 1.28E-11 |
| B27 | B134 | LDH | CHE | 0.385252 | 2.44E-11 |
| B27 | B75 | LDH | ApoA1/B | 0.368923 | 1.87E-10 |
| B27 | B87 | LDH | NSE | 0.303496 | 2.24E-07 |
| B27 | B137 | LDH | AFU | 0.300984 | 2.85E-07 |
| B28 | B202 | PRL | e-times | -0.77621 | 0 |
| B28 | B31 | PRL | FSH | -0.80403 | 0 |
| B29 | B116 | E2 | TT3 | -0.48807 | 0 |
| B29 | B223 | E2 | FH | -0.70598 | 0 |
| B29 | B202 | E2 | e-times | -0.37418 | 9.82E-11 |
| B30 | B167 | LH | PC | 0.920567 | 0 |
| B30 | B32 | LH | Testo | 0.453743 | 1.33E-15 |
| B31 | B35 | FSH | β2-MG | -0.56575 | 0 |
| B32 | B167 | Testo | PC | 0.431174 | 4.17E-14 |

| | | | | | |
|---|---|---|---|---|---|
| B33 | B116 | Prog | TT3 | -0.68362 | 0 |
| B33 | B35 | Prog | β2-MG | -0.89583 | 0 |
| B35 | B202 | β2-MG | e-times | 0.82734 | 0 |
| B36 | B167 | V-Lym% | PC | 0.777374 | 0 |
| B36 | B96 | V-Lym% | dsDNA (IIF) | 0.781452 | 0 |
| B37 | B139 | BAND | ADA | -0.49733 | 0 |
| B37 | B38 | BAND | BAND% | -1 | 0 |
| B37 | B71 | BAND | TG | -0.62386 | 0 |
| B37 | B82 | BAND | CEA | -0.521 | 0 |
| B37 | B90 | BAND | CYFRA 21-1 | -0.53967 | 0 |
| B37 | B88 | BAND | CA 242 | -0.46151 | 4.44E-16 |
| B37 | B87 | BAND | NSE | -0.44548 | 4.66E-15 |
| B37 | B200 | BAND | OP | -0.43491 | 2.40E-14 |
| B37 | B85 | BAND | SCC | 0.412518 | 6.25E-13 |
| B37 | B198 | BAND | P | 0.331499 | 1.32E-08 |
| B37 | B195 | BAND | Na | 0.32272 | 3.30E-08 |
| B37 | B194 | BAND | Cl | 0.31422 | 7.83E-08 |
| B37 | B196 | BAND | TCO2 | 0.305274 | 1.89E-07 |
| B37 | B199 | BAND | Mg | -0.30011 | 3.09E-07 |
| B38 | B139 | BAND% | ADA | 0.497328 | 0 |
| B38 | B71 | BAND% | TG | 0.623858 | 0 |
| B38 | B82 | BAND% | CEA | 0.521003 | 0 |
| B38 | B90 | BAND% | CYFRA 21-1 | 0.539673 | 0 |
| B38 | B88 | BAND% | CA 242 | 0.461513 | 4.44E-16 |
| B38 | B87 | BAND% | NSE | 0.445478 | 4.66E-15 |
| B38 | B200 | BAND% | OP | 0.434906 | 2.40E-14 |
| B38 | B85 | BAND% | SCC | -0.41252 | 6.25E-13 |
| B38 | B198 | BAND% | P | -0.3315 | 1.32E-08 |
| B38 | B195 | BAND% | Na | -0.32272 | 3.30E-08 |
| B38 | B194 | BAND% | Cl | -0.31422 | 7.83E-08 |
| B38 | B196 | BAND% | TCO2 | -0.30527 | 1.89E-07 |
| B38 | B199 | BAND% | Mg | 0.300114 | 3.09E-07 |
| B39 | B44 | Other cells | MCH | -0.31145 | 1.03E-07 |
| B39 | B43 | Other cells | MCV | -0.29903 | 3.42E-07 |
| B4 | B17 | WB-Ca | CD3+CD16+CD56+NK | -0.55141 | 0 |
| B4 | B29 | WB-Ca | E2 | -0.54621 | 0 |
| B40 | B42 | WBC | MON | 0.754101 | 0 |
| B40 | B209 | WBC | Smoking | 0.316292 | 6.36E-08 |
| B41 | B42 | MON % | MON | 0.561751 | 0 |
| B42 | B51 | MON | Hb | 0.293758 | 5.60E-07 |
| B43 | B44 | MCV | MCH | 0.89385 | 0 |
| B43 | B47 | MCV | RDW | 0.523748 | 0 |
| B43 | B46 | MCV | RBC Count | -0.46029 | 4.44E-16 |
| B44 | B45 | MCH | MCHC | 0.531605 | 0 |

| | | | | | |
|---|---|---|---|---|---|
| B44 | B51 | MCH | Hb | 0.327252 | 2.06E-08 |
| B44 | B46 | MCH | RBC Count | -0.32007 | 4.34E-08 |
| B45 | B51 | MCHC | Hb | 0.501964 | 0 |
| B45 | B47 | MCHC | RDW | -0.39545 | 6.44E-12 |
| B46 | B51 | RBC Count | Hb | 0.785559 | 0 |
| B46 | B47 | RBC Count | RDW | -0.40994 | 8.98E-13 |
| B46 | B189 | RBC Count | eGFB | 0.354091 | 1.08E-09 |
| B46 | B222 | RBC Count | NS | 0.349375 | 1.85E-09 |
| B46 | B218 | RBC Count | DBP | 0.317728 | 5.50E-08 |
| B46 | B191 | RBC Count | Urea | -0.29984 | 3.17E-07 |
| B46 | B70 | RBC Count | LDL-C | 0.290069 | 7.87E-07 |
| B46 | B126 | RBC Count | Alb | 0.289128 | 8.57E-07 |
| B48 | B53 | LYM% | GR% | -0.94198 | 0 |
| B5 | B96 | WB-Mg | dsDNA (IIF) | 0.802896 | 0 |
| B5 | B36 | WB-Mg | V-Lym% | 0.380936 | 4.22E-11 |
| B5 | B167 | WB-Mg | PC | 0.367825 | 2.13E-10 |
| B51 | B174 | Hb | Urea/Cr | -0.34296 | 3.79E-09 |
| B51 | B126 | Hb | Alb | 0.327038 | 2.11E-08 |
| B51 | B222 | Hb | NS | 0.324412 | 2.77E-08 |
| B51 | B221 | Hb | Hei | 0.314338 | 7.74E-08 |
| B51 | B128 | Hb | ALT | 0.311104 | 1.07E-07 |
| B51 | B218 | Hb | DBP | 0.311019 | 1.08E-07 |
| B51 | B141 | Hb | AST/ALT | -0.29704 | 4.13E-07 |
| B51 | B189 | Hb | eGFB | 0.295381 | 4.82E-07 |
| B52 | B132 | PLT/BPC | Dbil | -0.342 | 4.22E-09 |
| B54 | B100 | ESR | anti-Ro52 | 0.495827 | 0 |
| B54 | B101 | ESR | anti-Sc70 | 0.495827 | 0 |
| B54 | B102 | ESR | anti-Sm | 0.495827 | 0 |
| B54 | B103 | ESR | anti-SSA/Ro | 0.495827 | 0 |
| B54 | B104 | ESR | anti-SSB | 0.495827 | 0 |
| B54 | B105 | ESR | anti-U1RNP | 0.495827 | 0 |
| B54 | B106 | ESR | anti-P | 0.495827 | 0 |
| B54 | B107 | ESR | AnuA | 0.495827 | 0 |
| B54 | B108 | ESR | Anti-Cyclin | 0.495827 | 0 |
| B54 | B109 | ESR | AMA-M2 | 0.495827 | 0 |
| B54 | B110 | ESR | AHA | 0.495827 | 0 |
| B54 | B124 | ESR | ACTH | -0.49583 | 0 |
| B54 | B125 | ESR | Cor | 0.495827 | 0 |
| B54 | B181 | ESR | APTT | 0.495827 | 0 |
| B54 | B182 | ESR | TT | -0.49567 | 0 |
| B54 | B183 | ESR | PTA | 0.495823 | 0 |
| B54 | B184 | ESR | INR | 0.495827 | 0 |
| B54 | B185 | ESR | PT | 0.495321 | 0 |
| B54 | B186 | ESR | PFC | -0.49583 | 0 |

| | | | | | |
|---|---|---|---|---|---|
| B54 | B55 | ESR | TK | 0.586123 | 0 |
| B54 | B57 | ESR | RBCAI | 0.485173 | 0 |
| B54 | B59 | ESR | WBRV | 0.556399 | 0 |
| B54 | B60 | ESR | WBRVI | 0.512089 | 0 |
| B54 | B61 | ESR | WBHSRV | 0.538617 | 0 |
| B54 | B64 | ESR | WBRSR(mPas_5) | 0.527444 | 0 |
| B54 | B65 | ESR | WBRSR(mPas_1) | 0.529017 | 0 |
| B54 | B79 | ESR | hb V -dna | 0.495827 | 0 |
| B54 | B92 | ESR | ALDH2-Glu504Glu | 0.495827 | 0 |
| B54 | B93 | ESR | ALDH2-Glu504Lys | -0.49583 | 0 |
| B54 | B94 | ESR | ALDH2-Lys504Lys | 0.495827 | 0 |
| B54 | B95 | ESR | anti-cenpb | 0.495827 | 0 |
| B54 | B97 | ESR | dsDNA | 0.495827 | 0 |
| B54 | B98 | ESR | anti-Jo-1 | 0.495827 | 0 |
| B54 | B99 | ESR | anti-PM-Scl | 0.495827 | 0 |
| B54 | B56 | ESR | IR | 0.455914 | 8.88E-16 |
| B54 | B67 | ESR | WBRSR(mPas_50) | 0.439019 | 1.29E-14 |
| B54 | B69 | ESR | PV | 0.431432 | 4.04E-14 |
| B54 | B66 | ESR | WBRSR(mPas_100) | 0.401023 | 3.05E-12 |
| B54 | B194 | ESR | CI | 0.394606 | 7.20E-12 |
| B54 | B63 | ESR | WBRSR(mPas_200) | 0.389679 | 1.37E-11 |
| B54 | B134 | ESR | CHE | -0.38162 | 3.87E-11 |
| B54 | B179 | ESR | α1-MG | 0.356638 | 8.04E-10 |
| B54 | B197 | ESR | AG | -0.34931 | 1.86E-09 |
| B54 | B180 | ESR | Tf | 0.347396 | 2.31E-09 |
| B54 | B178 | ESR | Ig-U | 0.343232 | 3.68E-09 |
| B55 | B100 | TK | anti-Ro52 | 0.501718 | 0 |
| B55 | B101 | TK | anti-Sc70 | 0.501718 | 0 |
| B55 | B102 | TK | anti-Sm | 0.501718 | 0 |
| B55 | B103 | TK | anti-SSA/Ro | 0.501718 | 0 |
| B55 | B104 | TK | anti-SSB | 0.501718 | 0 |
| B55 | B105 | TK | anti-U1RNP | 0.501718 | 0 |
| B55 | B106 | TK | anti-P | 0.501718 | 0 |
| B55 | B107 | TK | AnuA | 0.501718 | 0 |
| B55 | B108 | TK | Anti-Cyclin | 0.501718 | 0 |
| B55 | B109 | TK | AMA-M2 | 0.501718 | 0 |
| B55 | B110 | TK | AHA | 0.501718 | 0 |
| B55 | B124 | TK | ACTH | -0.50172 | 0 |
| B55 | B125 | TK | Cor | 0.501718 | 0 |
| B55 | B181 | TK | APTT | 0.501718 | 0 |
| B55 | B182 | TK | TT | -0.50161 | 0 |
| B55 | B183 | TK | PTA | 0.501716 | 0 |
| B55 | B184 | TK | INR | 0.501718 | 0 |
| B55 | B185 | TK | PT | 0.501306 | 0 |

| | | | | | |
|---|---|---|---|---|---|
| B55 | B186 | TK | PFC | -0.50172 | 0 |
| B55 | B56 | TK | IR | 0.804792 | 0 |
| B55 | B57 | TK | RBCAI | 0.607399 | 0 |
| B55 | B59 | TK | WBRV | 0.701451 | 0 |
| B55 | B60 | TK | WBRVI | 0.726182 | 0 |
| B55 | B61 | TK | WBHSRV | 0.731167 | 0 |
| B55 | B62 | TK | WBHSRVI | 0.502569 | 0 |
| B55 | B64 | TK | WBRSR(mPas_5) | 0.663832 | 0 |
| B55 | B65 | TK | WBRSR(mPas_1) | 0.672237 | 0 |
| B55 | B66 | TK | WBRSR(mPas_100) | 0.494996 | 0 |
| B55 | B67 | TK | WBRSR(mPas_50) | 0.545568 | 0 |
| B55 | B79 | TK | hb V -dna | 0.501718 | 0 |
| B55 | B92 | TK | ALDH2-Glu504Glu | 0.501718 | 0 |
| B55 | B93 | TK | ALDH2-Glu504Lys | -0.50172 | 0 |
| B55 | B94 | TK | ALDH2-Lys504Lys | 0.501718 | 0 |
| B55 | B95 | TK | anti-cenpb | 0.501718 | 0 |
| B55 | B97 | TK | dsDNA | 0.501718 | 0 |
| B55 | B98 | TK | anti-Jo-1 | 0.501718 | 0 |
| B55 | B99 | TK | anti-PM-Scl | 0.501718 | 0 |
| B55 | B63 | TK | WBRSR(mPas_200) | 0.454335 | 1.11E-15 |
| B55 | B194 | TK | Cl | 0.407317 | 1.29E-12 |
| B55 | B179 | TK | α1-MG | 0.362663 | 3.96E-10 |
| B55 | B180 | TK | Tf | 0.353369 | 1.17E-09 |
| B55 | B178 | TK | Ig-U | 0.349176 | 1.89E-09 |
| B55 | B197 | TK | AG | -0.3173 | 5.74E-08 |
| B56 | B58 | IR | HCT | 0.540861 | 0 |
| B56 | B59 | IR | WBRV | 0.544279 | 0 |
| B56 | B60 | IR | WBRVI | 0.689051 | 0 |
| B56 | B61 | IR | WBHSRV | 0.947507 | 0 |
| B56 | B62 | IR | WBHSRVI | 0.917159 | 0 |
| B56 | B63 | IR | WBRSR(mPas_200) | 0.858743 | 0 |
| B56 | B64 | IR | WBRSR(mPas_5) | 0.744933 | 0 |
| B56 | B65 | IR | WBRSR(mPas_1) | 0.648998 | 0 |
| B56 | B66 | IR | WBRSR(mPas_100) | 0.855832 | 0 |
| B56 | B67 | IR | WBRSR(mPas_50) | 0.851482 | 0 |
| B56 | B57 | IR | RBCAI | 0.386609 | 2.05E-11 |
| B57 | B59 | RBCAI | WBRV | 0.979235 | 0 |
| B57 | B60 | RBCAI | WBRVI | 0.925271 | 0 |
| B57 | B61 | RBCAI | WBHSRV | 0.516753 | 0 |
| B57 | B64 | RBCAI | WBRSR(mPas_5) | 0.841729 | 0 |
| B57 | B65 | RBCAI | WBRSR(mPas_1) | 0.928554 | 0 |
| B57 | B67 | RBCAI | WBRSR(mPas_50) | 0.533653 | 0 |
| B57 | B69 | RBCAI | PV | 0.490685 | 0 |
| B57 | B66 | RBCAI | WBRSR(mPas_100) | 0.420134 | 2.12E-13 |

| | | | | | |
|---|---|---|---|---|---|
| B57 | B63 | RBCAI | WBRSR(mPas_200) | 0.290649 | 7.46E-07 |
| B58 | B61 | HCT | WBHSRV | 0.548502 | 0 |
| B58 | B62 | HCT | WBHSRVI | 0.825157 | 0 |
| B58 | B63 | HCT | WBRSR(mPas_200) | 0.791246 | 0 |
| B58 | B66 | HCT | WBRSR(mPas_100) | 0.72401 | 0 |
| B58 | B67 | HCT | WBRSR(mPas_50) | 0.645415 | 0 |
| B58 | B64 | HCT | WBRSR(mPas_5) | 0.30206 | 2.57E-07 |
| B59 | B60 | WBRV | WBRVI | 0.97241 | 0 |
| B59 | B61 | WBRV | WBHSRV | 0.671367 | 0 |
| B59 | B64 | WBRV | WBRSR(mPas_5) | 0.920091 | 0 |
| B59 | B65 | WBRV | WBRSR(mPas_1) | 0.976095 | 0 |
| B59 | B66 | WBRV | WBRSR(mPas_100) | 0.569521 | 0 |
| B59 | B67 | WBRV | WBRSR(mPas_50) | 0.669803 | 0 |
| B59 | B69 | WBRV | PV | 0.532643 | 0 |
| B59 | B63 | WBRV | WBRSR(mPas_200) | 0.453576 | 1.33E-15 |
| B59 | B62 | WBRV | WBHSRVI | 0.332709 | 1.16E-08 |
| B6 | B167 | WB-Fe | PC | 0.745356 | 0 |
| B6 | B8 | WB-Fe | WB-Zinc | -0.7746 | 0 |
| B6 | B96 | WB-Fe | dsDNA (IIF) | 0.650791 | 0 |
| B60 | B61 | WBRVI | WBHSRV | 0.787469 | 0 |
| B60 | B62 | WBRVI | WBHSRVI | 0.525773 | 0 |
| B60 | B63 | WBRVI | WBRSR(mPas_200) | 0.608836 | 0 |
| B60 | B64 | WBRVI | WBRSR(mPas_5) | 0.972042 | 0 |
| B60 | B65 | WBRVI | WBRSR(mPas_1) | 0.992859 | 0 |
| B60 | B66 | WBRVI | WBRSR(mPas_100) | 0.709654 | 0 |
| B60 | B67 | WBRVI | WBRSR(mPas_50) | 0.792962 | 0 |
| B60 | B69 | WBRVI | PV | 0.496335 | 0 |
| B61 | B62 | WBHSRV | WBHSRVI | 0.895667 | 0 |
| B61 | B63 | WBHSRV | WBRSR(mPas_200) | 0.934937 | 0 |
| B61 | B64 | WBHSRV | WBRSR(mPas_5) | 0.876001 | 0 |
| B61 | B65 | WBHSRV | WBRSR(mPas_1) | 0.783025 | 0 |
| B61 | B66 | WBHSRV | WBRSR(mPas_100) | 0.953067 | 0 |
| B61 | B67 | WBHSRV | WBRSR(mPas_50) | 0.96081 | 0 |
| B61 | B69 | WBHSRV | PV | 0.46712 | 0 |
| B62 | B63 | WBHSRVI | WBRSR(mPas_200) | 0.951452 | 0 |
| B62 | B64 | WBHSRVI | WBRSR(mPas_5) | 0.650411 | 0 |
| B62 | B65 | WBHSRVI | WBRSR(mPas_1) | 0.505226 | 0 |
| B62 | B66 | WBHSRVI | WBRSR(mPas_100) | 0.922524 | 0 |
| B62 | B67 | WBHSRVI | WBRSR(mPas_50) | 0.883256 | 0 |
| B63 | B64 | WBRSR(mPas_200) | WBRSR(mPas_5) | 0.759795 | 0 |
| B63 | B65 | WBRSR(mPas_200) | WBRSR(mPas_1) | 0.622145 | 0 |
| B63 | B66 | WBRSR(mPas_200) | WBRSR(mPas_100) | 0.988507 | 0 |
| B63 | B67 | WBRSR(mPas_200) | WBRSR(mPas_50) | 0.962291 | 0 |
| B63 | B69 | WBRSR(mPas_200) | PV | 0.524439 | 0 |

| | | | | | |
|---|---|---|---|---|---|
| B64 | B65 | WBRSR(mPas_5) | WBRSR(mPas_1) | 0.981623 | 0 |
| B64 | B66 | WBRSR(mPas_5) | WBRSR(mPas_100) | 0.842925 | 0 |
| B64 | B67 | WBRSR(mPas_5) | WBRSR(mPas_50) | 0.905252 | 0 |
| B64 | B69 | WBRSR(mPas_5) | PV | 0.624037 | 0 |
| B65 | B66 | WBRSR(mPas_1) | WBRSR(mPas_100) | 0.724895 | 0 |
| B65 | B67 | WBRSR(mPas_1) | WBRSR(mPas_50) | 0.807675 | 0 |
| B65 | B69 | WBRSR(mPas_1) | PV | 0.593781 | 0 |
| B66 | B67 | WBRSR(mPas_100) | WBRSR(mPas_50) | 0.991627 | 0 |
| B66 | B69 | WBRSR(mPas_100) | PV | 0.580898 | 0 |
| B67 | B69 | WBRSR(mPas_50) | PV | 0.610737 | 0 |
| B7 | B17 | WB-Cop | CD3+CD16+CD56+NK | 0.60404 | 0 |
| B7 | B29 | WB-Cop | E2 | 0.472377 | 0 |
| B70 | B77 | LDL-C | TC | 0.833984 | 0 |
| B71 | B82 | TG | CEA | 0.631337 | 0 |
| B71 | B87 | TG | NSE | 0.500126 | 0 |
| B71 | B88 | TG | CA 242 | 0.638937 | 0 |
| B71 | B90 | TG | CYFRA 21-1 | 0.560132 | 0 |
| B71 | B153 | TG | Glu | 0.454584 | 1.11E-15 |
| B71 | B77 | TG | TC | 0.453829 | 1.33E-15 |
| B71 | B200 | TG | OP | 0.437363 | 1.64E-14 |
| B71 | B75 | TG | ApoA1/B | 0.435793 | 2.09E-14 |
| B71 | B139 | TG | ADA | 0.433517 | 2.93E-14 |
| B71 | B195 | TG | Na | -0.40409 | 2.01E-12 |
| B71 | B85 | TG | SCC | -0.34408 | 3.35E-09 |
| B71 | B170 | TG | U-GLU | 0.317934 | 5.39E-08 |
| B71 | B140 | TG | r-GT | 0.317719 | 5.51E-08 |
| B72 | B222 | HDL-C | NS | -0.30335 | 2.27E-07 |
| B72 | B77 | HDL-C | TC | 0.300693 | 2.93E-07 |
| B73 | B76 | ApoA | ApoB | 0.432548 | 3.42E-14 |
| B73 | B74 | ApoA | ApoA1 | 0.408813 | 1.05E-12 |
| B73 | B135 | ApoA | TBA | -0.37755 | 6.46E-11 |
| B73 | B139 | ApoA | ADA | -0.35927 | 5.91E-10 |
| B73 | B200 | ApoA | OP | -0.35891 | 6.16E-10 |
| B74 | B76 | ApoA1 | ApoB | 0.526752 | 0 |
| B75 | B76 | ApoA1/B | ApoB | -0.62308 | 0 |
| B75 | B82 | ApoA1/B | CEA | 0.715016 | 0 |
| B75 | B88 | ApoA1/B | CA 242 | 0.755482 | 0 |
| B75 | B90 | ApoA1/B | CYFRA 21-1 | 0.46551 | 2.22E-16 |
| B75 | B87 | ApoA1/B | NSE | 0.429149 | 5.66E-14 |
| B75 | B195 | ApoA1/B | Na | -0.42815 | 6.57E-14 |
| B75 | B200 | ApoA1/B | OP | 0.427737 | 6.97E-14 |
| B75 | B153 | ApoA1/B | Glu | 0.296258 | 4.44E-07 |
| B76 | B200 | ApoB | OP | -0.44876 | 2.66E-15 |
| B76 | B82 | ApoB | CEA | -0.39938 | 3.81E-12 |

| | | | | | |
|---|---|---|---|---|---|
| B76 | B88 | ApoB | CA 242 | -0.36106 | 4.78E-10 |
| B76 | B139 | ApoB | ADA | -0.337 | 7.29E-09 |
| B76 | B90 | ApoB | CYFRA 21-1 | -0.33334 | 1.08E-08 |
| B76 | B137 | ApoB | AFU | -0.32044 | 4.18E-08 |
| B77 | B132 | TC | Dbil | -0.32053 | 4.14E-08 |
| B77 | B130 | TC | GLB | 0.307623 | 1.50E-07 |
| B78 | B134 | HBcAb | CHE | -0.36985 | 1.67E-10 |
| B78 | B176 | HBcAb | Ucr | 0.365831 | 2.71E-10 |
| B78 | B113 | HBcAb | FT3 | 0.339037 | 5.85E-09 |
| B79 | B101 | hb V -dna | anti-Sc70 | 1 | 0 |
| B79 | B103 | hb V -dna | anti-SSA/Ro | 1 | 0 |
| B79 | B105 | hb V -dna | anti-U1RNP | 1 | 0 |
| B79 | B106 | hb V -dna | anti-P | 1 | 0 |
| B79 | B107 | hb V -dna | AnuA | 1 | 0 |
| B79 | B110 | hb V -dna | AHA | 1 | 0 |
| B79 | B124 | hb V -dna | ACTH | -1 | 0 |
| B79 | B125 | hb V -dna | Cor | 1 | 0 |
| B79 | B151 | hb V -dna | Anti-CCP | 0.508198 | 0 |
| B79 | B158 | hb V -dna | IgM | 0.497089 | 0 |
| B79 | B178 | hb V -dna | Ig-U | 0.6975 | 0 |
| B79 | B179 | hb V -dna | α1-MG | 0.724317 | 0 |
| B79 | B180 | hb V -dna | Tf | 0.705838 | 0 |
| B79 | B182 | hb V -dna | TT | -0.99982 | 0 |
| B79 | B183 | hb V -dna | PTA | 0.999999 | 0 |
| B79 | B185 | hb V -dna | PT | 0.999261 | 0 |
| B79 | B194 | hb V -dna | Cl | 0.804248 | 0 |
| B79 | B197 | hb V -dna | AG | -0.67006 | 0 |
| B79 | B81 | hb V -dna | TSGF | -0.54814 | 0 |
| B79 | B93 | hb V -dna | ALDH2-Glu504Lys | -1 | 0 |
| B79 | B94 | hb V -dna | ALDH2-Lys504Lys | 1 | 0 |
| B79 | B97 | hb V -dna | dsDNA | 1 | 0 |
| B79 | B98 | hb V -dna | anti-Jo-1 | 1 | 0 |
| B79 | B99 | hb V -dna | anti-PM-Scl | 1 | 0 |
| B79 | B138 | hb V -dna | PAB | -0.41317 | 5.70E-13 |
| B79 | B137 | hb V -dna | AFU | -0.28849 | 9.08E-07 |
| B81 | B100 | TSGF | anti-Ro52 | -0.54814 | 0 |
| B81 | B101 | TSGF | anti-Sc70 | -0.54814 | 0 |
| B81 | B102 | TSGF | anti-Sm | -0.54814 | 0 |
| B81 | B103 | TSGF | anti-SSA/Ro | -0.54814 | 0 |
| B81 | B104 | TSGF | anti-SSB | -0.54814 | 0 |
| B81 | B105 | TSGF | anti-U1RNP | -0.54814 | 0 |
| B81 | B106 | TSGF | anti-P | -0.54814 | 0 |
| B81 | B107 | TSGF | AnuA | -0.54814 | 0 |
| B81 | B108 | TSGF | Anti-Cyclin | -0.54814 | 0 |

| | | | | | |
|---|---|---|---|---|---|
| B81 | B109 | TSGF | AMA-M2 | -0.54814 | 0 |
| B81 | B110 | TSGF | AHA | -0.54814 | 0 |
| B81 | B124 | TSGF | ACTH | 0.548139 | 0 |
| B81 | B125 | TSGF | Cor | -0.54814 | 0 |
| B81 | B181 | TSGF | APTT | -0.54814 | 0 |
| B81 | B182 | TSGF | TT | 0.548017 | 0 |
| B81 | B183 | TSGF | PTA | -0.54814 | 0 |
| B81 | B184 | TSGF | INR | -0.54814 | 0 |
| B81 | B185 | TSGF | PT | -0.54768 | 0 |
| B81 | B186 | TSGF | PFC | 0.548139 | 0 |
| B81 | B92 | TSGF | ALDH2-Glu504Glu | -0.54814 | 0 |
| B81 | B93 | TSGF | ALDH2-Glu504Lys | 0.548139 | 0 |
| B81 | B94 | TSGF | ALDH2-Lys504Lys | -0.54814 | 0 |
| B81 | B95 | TSGF | anti-cenpb | -0.54814 | 0 |
| B81 | B97 | TSGF | dsDNA | -0.54814 | 0 |
| B81 | B98 | TSGF | anti-Jo-1 | -0.54814 | 0 |
| B81 | B99 | TSGF | anti-PM-Scl | -0.54814 | 0 |
| B81 | B194 | TSGF | Cl | -0.43222 | 3.57E-14 |
| B81 | B179 | TSGF | α1-MG | -0.39616 | 5.86E-12 |
| B81 | B180 | TSGF | Tf | -0.386 | 2.21E-11 |
| B81 | B178 | TSGF | Ig-U | -0.38142 | 3.97E-11 |
| B81 | B197 | TSGF | AG | 0.373521 | 1.06E-10 |
| B81 | B156 | TSGF | IgE | -0.31286 | 8.96E-08 |
| B82 | B195 | CEA | Na | -0.51874 | 0 |
| B82 | B200 | CEA | OP | 0.484182 | 0 |
| B82 | B87 | CEA | NSE | 0.599168 | 0 |
| B82 | B88 | CEA | CA 242 | 0.858919 | 0 |
| B82 | B90 | CEA | CYFRA 21-1 | 0.542552 | 0 |
| B82 | B139 | CEA | ADA | 0.44322 | 6.66E-15 |
| B82 | B86 | CEA | TBhCG | -0.38146 | 3.95E-11 |
| B82 | B85 | CEA | SCC | -0.37542 | 8.42E-11 |
| B82 | B198 | CEA | P | -0.35544 | 9.23E-10 |
| B82 | B153 | CEA | Glu | 0.338824 | 5.98E-09 |
| B84 | B86 | C-PSA | TBhCG | -0.35365 | 1.13E-09 |
| B84 | B165 | C-PSA | bilirubin | 0.32718 | 2.08E-08 |
| B84 | B89 | C-PSA | CA 72-4 | 0.325591 | 2.45E-08 |
| B85 | B86 | SCC | TBhCG | 0.442602 | 7.33E-15 |
| B85 | B88 | SCC | CA 242 | -0.35147 | 1.46E-09 |
| B85 | B165 | SCC | bilirubin | 0.306871 | 1.62E-07 |
| B86 | B89 | TBhCG | CA 72-4 | -0.40344 | 2.20E-12 |
| B86 | B137 | TBhCG | AFU | -0.31819 | 5.25E-08 |
| B86 | B200 | TBhCG | OP | -0.29569 | 4.68E-07 |
| B87 | B88 | NSE | CA 242 | 0.621019 | 0 |
| B87 | B90 | NSE | CYFRA 21-1 | 0.591324 | 0 |

| | | | | | |
|---|---|---|---|---|---|
| B87 | B195 | NSE | Na | -0.42952 | 5.35E-14 |
| B87 | B89 | NSE | CA 72-4 | 0.42371 | 1.26E-13 |
| B87 | B139 | NSE | ADA | 0.414106 | 5.00E-13 |
| B87 | B200 | NSE | OP | 0.358038 | 6.83E-10 |
| B88 | B195 | CA 242 | Na | -0.52697 | 0 |
| B88 | B200 | CA 242 | OP | 0.496184 | 0 |
| B88 | B90 | CA 242 | CYFRA 21-1 | 0.566104 | 0 |
| B88 | B139 | CA 242 | ADA | 0.456174 | 8.88E-16 |
| B88 | B153 | CA 242 | Glu | 0.350841 | 1.57E-09 |
| B88 | B198 | CA 242 | P | -0.3175 | 5.63E-08 |
| B88 | B137 | CA 242 | AFU | 0.292215 | 6.46E-07 |
| B89 | B165 | CA 72-4 | bilirubin | 0.899226 | 0 |
| B89 | B115 | CA 72-4 | TSH TT3 TT4(TT3) | -0.43619 | 1.95E-14 |
| B89 | B113 | CA 72-4 | FT3 | -0.40306 | 2.31E-12 |
| B89 | B90 | CA 72-4 | CYFRA 21-1 | 0.326614 | 2.21E-08 |
| B9 | B29 | RH | E2 | -0.54545 | 0 |
| B9 | B31 | RH | FSH | -0.74574 | 0 |
| B9 | B33 | RH | Prog | -0.40234 | 2.55E-12 |
| B90 | B195 | CYFRA 21-1 | Na | -0.39667 | 5.47E-12 |
| B90 | B139 | CYFRA 21-1 | ADA | 0.381761 | 3.80E-11 |
| B90 | B200 | CYFRA 21-1 | OP | 0.375982 | 7.85E-11 |
| B90 | B194 | CYFRA 21-1 | Cl | -0.34712 | 2.39E-09 |
| B91 | B127 | TP | A-G ratio | -0.52419 | 0 |
| B91 | B130 | TP | GLB | 0.790798 | 0 |
| B91 | B126 | TP | Alb | 0.424231 | 1.17E-13 |
| B92 | B101 | ALDH2-Glu504Glu | anti-Sc70 | 1 | 0 |
| B92 | B103 | ALDH2-Glu504Glu | anti-SSA/Ro | 1 | 0 |
| B92 | B105 | ALDH2-Glu504Glu | anti-U1RNP | 1 | 0 |
| B92 | B106 | ALDH2-Glu504Glu | anti-P | 1 | 0 |
| B92 | B107 | ALDH2-Glu504Glu | AnuA | 1 | 0 |
| B92 | B110 | ALDH2-Glu504Glu | AHA | 1 | 0 |
| B92 | B151 | ALDH2-Glu504Glu | Anti-CCP | 0.508198 | 0 |
| B92 | B158 | ALDH2-Glu504Glu | IgM | 0.497089 | 0 |
| B92 | B178 | ALDH2-Glu504Glu | Ig-U | 0.6975 | 0 |
| B92 | B179 | ALDH2-Glu504Glu | α1-MG | 0.724317 | 0 |
| B92 | B180 | ALDH2-Glu504Glu | Tf | 0.705838 | 0 |
| B92 | B182 | ALDH2-Glu504Glu | TT | -0.99982 | 0 |
| B92 | B183 | ALDH2-Glu504Glu | PTA | 0.999999 | 0 |
| B92 | B185 | ALDH2-Glu504Glu | PT | 0.999261 | 0 |
| B92 | B194 | ALDH2-Glu504Glu | Cl | 0.804248 | 0 |
| B92 | B197 | ALDH2-Glu504Glu | AG | -0.67006 | 0 |
| B92 | B94 | ALDH2-Glu504Glu | ALDH2-Lys504Lys | 1 | 0 |
| B92 | B97 | ALDH2-Glu504Glu | dsDNA | 1 | 0 |
| B92 | B98 | ALDH2-Glu504Glu | anti-Jo-1 | 1 | 0 |

| | | | | | |
|---|---|---|---|---|---|
| B92 | B99 | ALDH2-Glu504Glu | anti-PM-Scl | 1 | 0 |
| B92 | B138 | ALDH2-Glu504Glu | PAB | -0.41317 | 5.70E-13 |
| B92 | B137 | ALDH2-Glu504Glu | AFU | -0.28849 | 9.08E-07 |
| B93 | B101 | ALDH2-Glu504Lys | anti-Sc70 | -1 | 0 |
| B93 | B103 | ALDH2-Glu504Lys | anti-SSA/Ro | -1 | 0 |
| B93 | B105 | ALDH2-Glu504Lys | anti-U1RNP | -1 | 0 |
| B93 | B106 | ALDH2-Glu504Lys | anti-P | -1 | 0 |
| B93 | B107 | ALDH2-Glu504Lys | AnuA | -1 | 0 |
| B93 | B110 | ALDH2-Glu504Lys | AHA | -1 | 0 |
| B93 | B151 | ALDH2-Glu504Lys | Anti-CCP | -0.5082 | 0 |
| B93 | B158 | ALDH2-Glu504Lys | IgM | -0.49709 | 0 |
| B93 | B178 | ALDH2-Glu504Lys | Ig-U | -0.6975 | 0 |
| B93 | B179 | ALDH2-Glu504Lys | α1-MG | -0.72432 | 0 |
| B93 | B180 | ALDH2-Glu504Lys | Tf | -0.70584 | 0 |
| B93 | B182 | ALDH2-Glu504Lys | TT | 0.999822 | 0 |
| B93 | B183 | ALDH2-Glu504Lys | PTA | -1 | 0 |
| B93 | B184 | ALDH2-Glu504Lys | INR | -1 | 0 |
| B93 | B185 | ALDH2-Glu504Lys | PT | -0.99926 | 0 |
| B93 | B194 | ALDH2-Glu504Lys | Cl | -0.80425 | 0 |
| B93 | B197 | ALDH2-Glu504Lys | AG | 0.670061 | 0 |
| B93 | B94 | ALDH2-Glu504Lys | ALDH2-Lys504Lys | -1 | 0 |
| B93 | B97 | ALDH2-Glu504Lys | dsDNA | -1 | 0 |
| B93 | B98 | ALDH2-Glu504Lys | anti-Jo-1 | -1 | 0 |
| B93 | B99 | ALDH2-Glu504Lys | anti-PM-Scl | -1 | 0 |
| B93 | B138 | ALDH2-Glu504Lys | PAB | 0.413173 | 5.70E-13 |
| B93 | B137 | ALDH2-Glu504Lys | AFU | 0.288494 | 9.08E-07 |
| B94 | B100 | ALDH2-Lys504Lys | anti-Ro52 | 1 | 0 |
| B94 | B101 | ALDH2-Lys504Lys | anti-Sc70 | 1 | 0 |
| B94 | B103 | ALDH2-Lys504Lys | anti-SSA/Ro | 1 | 0 |
| B94 | B105 | ALDH2-Lys504Lys | anti-U1RNP | 1 | 0 |
| B94 | B106 | ALDH2-Lys504Lys | anti-P | 1 | 0 |
| B94 | B107 | ALDH2-Lys504Lys | AnuA | 1 | 0 |
| B94 | B109 | ALDH2-Lys504Lys | AMA-M2 | 1 | 0 |
| B94 | B110 | ALDH2-Lys504Lys | AHA | 1 | 0 |
| B94 | B124 | ALDH2-Lys504Lys | ACTH | -1 | 0 |
| B94 | B125 | ALDH2-Lys504Lys | Cor | 1 | 0 |
| B94 | B151 | ALDH2-Lys504Lys | Anti-CCP | 0.508198 | 0 |
| B94 | B158 | ALDH2-Lys504Lys | IgM | 0.497089 | 0 |
| B94 | B178 | ALDH2-Lys504Lys | Ig-U | 0.6975 | 0 |
| B94 | B179 | ALDH2-Lys504Lys | α1-MG | 0.724317 | 0 |
| B94 | B180 | ALDH2-Lys504Lys | Tf | 0.705838 | 0 |
| B94 | B181 | ALDH2-Lys504Lys | APTT | 1 | 0 |
| B94 | B182 | ALDH2-Lys504Lys | TT | -0.99982 | 0 |
| B94 | B183 | ALDH2-Lys504Lys | PTA | 0.999999 | 0 |

| | | | | | |
|---|---|---|---|---|---|
| B94 | B184 | ALDH2-Lys504Lys | INR | 1 | 0 |
| B94 | B185 | ALDH2-Lys504Lys | PT | 0.999261 | 0 |
| B94 | B194 | ALDH2-Lys504Lys | CI | 0.804248 | 0 |
| B94 | B197 | ALDH2-Lys504Lys | AG | -0.67006 | 0 |
| B94 | B95 | ALDH2-Lys504Lys | anti-cenpb | 1 | 0 |
| B94 | B97 | ALDH2-Lys504Lys | dsDNA | 1 | 0 |
| B94 | B98 | ALDH2-Lys504Lys | anti-Jo-1 | 1 | 0 |
| B94 | B99 | ALDH2-Lys504Lys | anti-PM-Scl | 1 | 0 |
| B94 | B138 | ALDH2-Lys504Lys | PAB | -0.41317 | 5.70E-13 |
| B94 | B137 | ALDH2-Lys504Lys | AFU | -0.28849 | 9.08E-07 |
| B95 | B101 | anti-cenpb | anti-Sc70 | 1 | 0 |
| B95 | B106 | anti-cenpb | anti-P | 1 | 0 |
| B95 | B107 | anti-cenpb | AnuA | 1 | 0 |
| B95 | B109 | anti-cenpb | AMA-M2 | 1 | 0 |
| B95 | B151 | anti-cenpb | Anti-CCP | 0.508198 | 0 |
| B95 | B158 | anti-cenpb | IgM | 0.497089 | 0 |
| B95 | B178 | anti-cenpb | Ig-U | 0.6975 | 0 |
| B95 | B179 | anti-cenpb | α1-MG | 0.724317 | 0 |
| B95 | B180 | anti-cenpb | Tf | 0.705838 | 0 |
| B95 | B182 | anti-cenpb | TT | -0.99982 | 0 |
| B95 | B183 | anti-cenpb | PTA | 0.999999 | 0 |
| B95 | B185 | anti-cenpb | PT | 0.999261 | 0 |
| B95 | B194 | anti-cenpb | CI | 0.804248 | 0 |
| B95 | B197 | anti-cenpb | AG | -0.67006 | 0 |
| B95 | B98 | anti-cenpb | anti-Jo-1 | 1 | 0 |
| B95 | B138 | anti-cenpb | PAB | -0.41317 | 5.70E-13 |
| B95 | B137 | anti-cenpb | AFU | -0.28849 | 9.08E-07 |
| B96 | B167 | dsDNA (IIF) | PC | 0.781504 | 0 |
| B97 | B100 | dsDNA | anti-Ro52 | 1 | 0 |
| B97 | B103 | dsDNA | anti-SSA/Ro | 1 | 0 |
| B97 | B105 | dsDNA | anti-U1RNP | 1 | 0 |
| B97 | B106 | dsDNA | anti-P | 1 | 0 |
| B97 | B107 | dsDNA | AnuA | 1 | 0 |
| B97 | B108 | dsDNA | Anti-Cyclin | 1 | 0 |
| B97 | B109 | dsDNA | AMA-M2 | 1 | 0 |
| B97 | B110 | dsDNA | AHA | 1 | 0 |
| B97 | B124 | dsDNA | ACTH | -1 | 0 |
| B97 | B125 | dsDNA | Cor | 1 | 0 |
| B97 | B151 | dsDNA | Anti-CCP | 0.508198 | 0 |
| B97 | B158 | dsDNA | IgM | 0.497089 | 0 |
| B97 | B178 | dsDNA | Ig-U | 0.6975 | 0 |
| B97 | B179 | dsDNA | α1-MG | 0.724317 | 0 |
| B97 | B180 | dsDNA | Tf | 0.705838 | 0 |
| B97 | B181 | dsDNA | APTT | 1 | 0 |

| | | | | | |
|---|---|---|---|---|---|
| B97 | B182 | dsDNA | TT | -0.99982 | 0 |
| B97 | B183 | dsDNA | PTA | 0.999999 | 0 |
| B97 | B184 | dsDNA | INR | 1 | 0 |
| B97 | B185 | dsDNA | PT | 0.999261 | 0 |
| B97 | B186 | dsDNA | PFC | -1 | 0 |
| B97 | B194 | dsDNA | CI | 0.804248 | 0 |
| B97 | B197 | dsDNA | AG | -0.67006 | 0 |
| B97 | B138 | dsDNA | PAB | -0.41317 | 5.70E-13 |
| B97 | B137 | dsDNA | AFU | -0.28849 | 9.08E-07 |
| B98 | B100 | anti-Jo-1 | anti-Ro52 | 1 | 0 |
| B98 | B101 | anti-Jo-1 | anti-Sc70 | 1 | 0 |
| B98 | B102 | anti-Jo-1 | anti-Sm | 1 | 0 |
| B98 | B103 | anti-Jo-1 | anti-SSA/Ro | 1 | 0 |
| B98 | B104 | anti-Jo-1 | anti-SSB | 1 | 0 |
| B98 | B105 | anti-Jo-1 | anti-U1RNP | 1 | 0 |
| B98 | B107 | anti-Jo-1 | AnuA | 1 | 0 |
| B98 | B108 | anti-Jo-1 | Anti-Cyclin | 1 | 0 |
| B98 | B109 | anti-Jo-1 | AMA-M2 | 1 | 0 |
| B98 | B110 | anti-Jo-1 | AHA | 1 | 0 |
| B98 | B124 | anti-Jo-1 | ACTH | -1 | 0 |
| B98 | B125 | anti-Jo-1 | Cor | 1 | 0 |
| B98 | B151 | anti-Jo-1 | Anti-CCP | 0.508198 | 0 |
| B98 | B158 | anti-Jo-1 | IgM | 0.497089 | 0 |
| B98 | B178 | anti-Jo-1 | Ig-U | 0.6975 | 0 |
| B98 | B179 | anti-Jo-1 | α1-MG | 0.724317 | 0 |
| B98 | B180 | anti-Jo-1 | Tf | 0.705838 | 0 |
| B98 | B181 | anti-Jo-1 | APTT | 1 | 0 |
| B98 | B182 | anti-Jo-1 | TT | -0.99982 | 0 |
| B98 | B183 | anti-Jo-1 | PTA | 0.999999 | 0 |
| B98 | B184 | anti-Jo-1 | INR | 1 | 0 |
| B98 | B185 | anti-Jo-1 | PT | 0.999261 | 0 |
| B98 | B186 | anti-Jo-1 | PFC | -1 | 0 |
| B98 | B194 | anti-Jo-1 | CI | 0.804248 | 0 |
| B98 | B197 | anti-Jo-1 | AG | -0.67006 | 0 |
| B98 | B99 | anti-Jo-1 | anti-PM-Scl | 1 | 0 |
| B98 | B138 | anti-Jo-1 | PAB | -0.41317 | 5.70E-13 |
| B98 | B137 | anti-Jo-1 | AFU | -0.28849 | 9.08E-07 |
| B99 | B100 | anti-PM-Scl | anti-Ro52 | 1 | 0 |
| B99 | B103 | anti-PM-Scl | anti-SSA/Ro | 1 | 0 |
| B99 | B105 | anti-PM-Scl | anti-U1RNP | 1 | 0 |
| B99 | B107 | anti-PM-Scl | AnuA | 1 | 0 |
| B99 | B108 | anti-PM-Scl | Anti-Cyclin | 1 | 0 |
| B99 | B109 | anti-PM-Scl | AMA-M2 | 1 | 0 |
| B99 | B110 | anti-PM-Scl | AHA | 1 | 0 |

| | | | | | |
|---|---|---|---|---|---|
| B99 | B124 | anti-PM-Scl | ACTH | -1 | 0 |
| B99 | B125 | anti-PM-Scl | Cor | 1 | 0 |
| B99 | B151 | anti-PM-Scl | Anti-CCP | 0.508198 | 0 |
| B99 | B158 | anti-PM-Scl | IgM | 0.497089 | 0 |
| B99 | B178 | anti-PM-Scl | Ig-U | 0.6975 | 0 |
| B99 | B179 | anti-PM-Scl | α1-MG | 0.724317 | 0 |
| B99 | B180 | anti-PM-Scl | Tf | 0.705838 | 0 |
| B99 | B181 | anti-PM-Scl | APTT | 1 | 0 |
| B99 | B182 | anti-PM-Scl | TT | -0.99982 | 0 |
| B99 | B183 | anti-PM-Scl | PTA | 0.999999 | 0 |
| B99 | B184 | anti-PM-Scl | INR | 1 | 0 |
| B99 | B185 | anti-PM-Scl | PT | 0.999261 | 0 |
| B99 | B186 | anti-PM-Scl | PFC | -1 | 0 |
| B99 | B194 | anti-PM-Scl | Cl | 0.804248 | 0 |
| B99 | B197 | anti-PM-Scl | AG | -0.67006 | 0 |
| B99 | B138 | anti-PM-Scl | PAB | -0.41317 | 5.70E-13 |
| B99 | B137 | anti-PM-Scl | AFU | -0.28849 | 9.08E-07 |

**Supplementary Table 10. PEI Correlations in Rhinallergosis**

| Row | Column | Row | Column | Cor | *P* |
|---|---|---|---|---|---|
| B1 | B190 | Gender | Cre | -0.730474206 | 0 |
| B1 | B192 | Gender | UA | -0.672096816 | 0 |
| B1 | B206 | Gender | Waist | -0.632182121 | 0 |
| B1 | B207 | Gender | WHR | -0.642272664 | 0 |
| B1 | B216 | Gender | BMI | -0.656737407 | 0 |
| B1 | B221 | Gender | Hei | -0.645853302 | 0 |
| B1 | B222 | Gender | NS | -0.775250534 | 0 |
| B1 | B51 | Gender | Hb | -0.653142121 | 0 |
| B1 | B72 | Gender | HDL-C | 0.520393778 | 3.33E-12 |
| B1 | B46 | Gender | RBC Count | -0.486547047 | 1.20E-10 |
| B1 | B215 | Gender | BMI | -0.424098683 | 3.45E-08 |
| B1 | B210 | Gender | Hip | -0.416884126 | 6.18E-08 |
| B1 | B220 | Gender | SBP | -0.400924434 | 2.14E-07 |
| B1 | B141 | Gender | AST/ALT | 0.396517966 | 2.99E-07 |
| B1 | B128 | Gender | ALT | -0.385850465 | 6.54E-07 |
| B1 | B163 | Gender | SG | -0.384058046 | 7.44E-07 |
| B1 | B204 | Gender | Drinking | -0.381626539 | 8.86E-07 |
| B10 | B14 | PG Ⅰ | G-17 | -0.576591808 | 3.33E-15 |
| B10 | B11 | PG Ⅰ | PGI/PGII | 0.515479843 | 5.74E-12 |
| B100 | B101 | anti-Ro52 | anti-Sc70 | 0.999999998 | 0 |
| B100 | B103 | anti-Ro52 | anti-SSA/Ro | 1 | 0 |
| B100 | B105 | anti-Ro52 | anti-U1RNP | 1 | 0 |
| B100 | B106 | anti-Ro52 | anti-P | 1 | 0 |
| B100 | B109 | anti-Ro52 | AMA-M2 | 1 | 0 |
| B100 | B151 | anti-Ro52 | Anti-CCP | 0.7048221 | 0 |
| B100 | B156 | anti-Ro52 | IgE | -0.903553226 | 0 |
| B100 | B183 | anti-Ro52 | PTA | -0.935351263 | 0 |
| B100 | B155 | anti-Ro52 | IgA | 0.55642635 | 4.64E-14 |
| B100 | B175 | anti-Ro52 | ALB-U | 0.538122697 | 4.33E-13 |
| B100 | B158 | anti-Ro52 | IgM | -0.492296831 | 6.70E-11 |
| B100 | B198 | anti-Ro52 | P | -0.484721524 | 1.44E-10 |
| B100 | B185 | anti-Ro52 | PT | 0.471649437 | 5.14E-10 |
| B100 | B184 | anti-Ro52 | INR | 0.461953764 | 1.28E-09 |
| B100 | B181 | anti-Ro52 | APTT | 0.459306489 | 1.63E-09 |
| B100 | B168 | anti-Ro52 | NMP22 | 0.401609664 | 2.03E-07 |
| B100 | B157 | anti-Ro52 | IgG | 0.400110805 | 2.28E-07 |
| B100 | B176 | anti-Ro52 | Ucr | 0.387593593 | 5.77E-07 |
| B101 | B102 | anti-Sc70 | anti-Sm | 0.999999998 | 0 |
| B101 | B103 | anti-Sc70 | anti-SSA/Ro | 0.999999997 | 0 |
| B101 | B104 | anti-Sc70 | anti-SSB | 0.999999997 | 0 |
| B101 | B105 | anti-Sc70 | anti-U1RNP | 0.999999999 | 0 |
| B101 | B106 | anti-Sc70 | anti-P | 0.999999996 | 0 |

| | | | | | |
|---|---|---|---|---|---|
| B101 | B108 | anti-Sc70 | Anti-Cyclin | 0.999999998 | 0 |
| B101 | B109 | anti-Sc70 | AMA-M2 | 0.999999998 | 0 |
| B101 | B110 | anti-Sc70 | AHA | 0.999999998 | 0 |
| B101 | B151 | anti-Sc70 | Anti-CCP | 0.704822098 | 0 |
| B101 | B156 | anti-Sc70 | IgE | -0.903553223 | 0 |
| B101 | B183 | anti-Sc70 | PTA | -0.93535126 | 0 |
| B101 | B155 | anti-Sc70 | IgA | 0.556426348 | 4.64E-14 |
| B101 | B175 | anti-Sc70 | ALB-U | 0.538122696 | 4.33E-13 |
| B101 | B158 | anti-Sc70 | IgM | -0.49229683 | 6.70E-11 |
| B101 | B198 | anti-Sc70 | P | -0.484721523 | 1.44E-10 |
| B101 | B185 | anti-Sc70 | PT | 0.471649436 | 5.14E-10 |
| B101 | B184 | anti-Sc70 | INR | 0.461953763 | 1.28E-09 |
| B101 | B181 | anti-Sc70 | APTT | 0.459306487 | 1.63E-09 |
| B101 | B168 | anti-Sc70 | NMP22 | 0.401609665 | 2.03E-07 |
| B101 | B157 | anti-Sc70 | IgG | 0.400110804 | 2.28E-07 |
| B101 | B176 | anti-Sc70 | Ucr | 0.387593591 | 5.77E-07 |
| B102 | B103 | anti-Sm | anti-SSA/Ro | 0.999999999 | 0 |
| B102 | B104 | anti-Sm | anti-SSB | 0.999999998 | 0 |
| B102 | B106 | anti-Sm | anti-P | 0.999999999 | 0 |
| B102 | B108 | anti-Sm | Anti-Cyclin | 0.999999999 | 0 |
| B102 | B109 | anti-Sm | AMA-M2 | 1 | 0 |
| B102 | B151 | anti-Sm | Anti-CCP | 0.7048221 | 0 |
| B102 | B156 | anti-Sm | IgE | -0.903553226 | 0 |
| B102 | B183 | anti-Sm | PTA | -0.935351263 | 0 |
| B102 | B155 | anti-Sm | IgA | 0.55642635 | 4.64E-14 |
| B102 | B175 | anti-Sm | ALB-U | 0.538122697 | 4.33E-13 |
| B102 | B158 | anti-Sm | IgM | -0.492296831 | 6.70E-11 |
| B102 | B198 | anti-Sm | P | -0.484721524 | 1.44E-10 |
| B102 | B185 | anti-Sm | PT | 0.471649436 | 5.14E-10 |
| B102 | B184 | anti-Sm | INR | 0.461953764 | 1.28E-09 |
| B102 | B181 | anti-Sm | APTT | 0.459306489 | 1.63E-09 |
| B102 | B168 | anti-Sm | NMP22 | 0.401609664 | 2.03E-07 |
| B102 | B157 | anti-Sm | IgG | 0.400110805 | 2.28E-07 |
| B102 | B176 | anti-Sm | Ucr | 0.387593592 | 5.77E-07 |
| B103 | B105 | anti-SSA/Ro | anti-U1RNP | 1 | 0 |
| B103 | B109 | anti-SSA/Ro | AMA-M2 | 1 | 0 |
| B103 | B110 | anti-SSA/Ro | AHA | 1 | 0 |
| B103 | B151 | anti-SSA/Ro | Anti-CCP | 0.7048221 | 0 |
| B103 | B156 | anti-SSA/Ro | IgE | -0.903553226 | 0 |
| B103 | B183 | anti-SSA/Ro | PTA | -0.935351263 | 0 |
| B103 | B155 | anti-SSA/Ro | IgA | 0.55642635 | 4.64E-14 |
| B103 | B175 | anti-SSA/Ro | ALB-U | 0.538122697 | 4.33E-13 |
| B103 | B158 | anti-SSA/Ro | IgM | -0.492296831 | 6.70E-11 |
| B103 | B198 | anti-SSA/Ro | P | -0.484721524 | 1.44E-10 |

| | | | | | |
|---|---|---|---|---|---|
| B103 | B185 | anti-SSA/Ro | PT | 0.471649437 | 5.14E-10 |
| B103 | B184 | anti-SSA/Ro | INR | 0.461953764 | 1.28E-09 |
| B103 | B181 | anti-SSA/Ro | APTT | 0.459306489 | 1.63E-09 |
| B103 | B168 | anti-SSA/Ro | NMP22 | 0.401609664 | 2.03E-07 |
| B103 | B157 | anti-SSA/Ro | IgG | 0.400110805 | 2.28E-07 |
| B103 | B176 | anti-SSA/Ro | Ucr | 0.387593593 | 5.77E-07 |
| B104 | B151 | anti-SSB | Anti-CCP | 0.704822102 | 0 |
| B104 | B156 | anti-SSB | IgE | -0.903553229 | 0 |
| B104 | B183 | anti-SSB | PTA | -0.935351266 | 0 |
| B104 | B155 | anti-SSB | IgA | 0.556426351 | 4.64E-14 |
| B104 | B175 | anti-SSB | ALB-U | 0.538122699 | 4.33E-13 |
| B104 | B158 | anti-SSB | IgM | -0.492296833 | 6.69E-11 |
| B104 | B198 | anti-SSB | P | -0.484721525 | 1.44E-10 |
| B104 | B185 | anti-SSB | PT | 0.471649437 | 5.14E-10 |
| B104 | B184 | anti-SSB | INR | 0.461953765 | 1.28E-09 |
| B104 | B181 | anti-SSB | APTT | 0.45930649 | 1.63E-09 |
| B104 | B168 | anti-SSB | NMP22 | 0.401609664 | 2.03E-07 |
| B104 | B157 | anti-SSB | IgG | 0.400110806 | 2.28E-07 |
| B104 | B176 | anti-SSB | Ucr | 0.387593594 | 5.77E-07 |
| B105 | B110 | anti-U1RNP | AHA | 1 | 0 |
| B105 | B151 | anti-U1RNP | Anti-CCP | 0.7048221 | 0 |
| B105 | B156 | anti-U1RNP | IgE | -0.903553226 | 0 |
| B105 | B183 | anti-U1RNP | PTA | -0.935351263 | 0 |
| B105 | B155 | anti-U1RNP | IgA | 0.55642635 | 4.64E-14 |
| B105 | B175 | anti-U1RNP | ALB-U | 0.538122697 | 4.33E-13 |
| B105 | B158 | anti-U1RNP | IgM | -0.492296831 | 6.70E-11 |
| B105 | B198 | anti-U1RNP | P | -0.484721524 | 1.44E-10 |
| B105 | B185 | anti-U1RNP | PT | 0.471649436 | 5.14E-10 |
| B105 | B184 | anti-U1RNP | INR | 0.461953764 | 1.28E-09 |
| B105 | B181 | anti-U1RNP | APTT | 0.459306489 | 1.63E-09 |
| B105 | B168 | anti-U1RNP | NMP22 | 0.401609665 | 2.03E-07 |
| B105 | B157 | anti-U1RNP | IgG | 0.400110805 | 2.28E-07 |
| B105 | B176 | anti-U1RNP | Ucr | 0.387593592 | 5.77E-07 |
| B106 | B108 | anti-P | Anti-Cyclin | 1 | 0 |
| B106 | B110 | anti-P | AHA | 1 | 0 |
| B106 | B151 | anti-P | Anti-CCP | 0.7048221 | 0 |
| B106 | B156 | anti-P | IgE | -0.903553227 | 0 |
| B106 | B183 | anti-P | PTA | -0.935351264 | 0 |
| B106 | B155 | anti-P | IgA | 0.55642635 | 4.64E-14 |
| B106 | B175 | anti-P | ALB-U | 0.538122698 | 4.33E-13 |
| B106 | B158 | anti-P | IgM | -0.492296832 | 6.70E-11 |
| B106 | B198 | anti-P | P | -0.484721524 | 1.44E-10 |
| B106 | B185 | anti-P | PT | 0.471649437 | 5.14E-10 |
| B106 | B184 | anti-P | INR | 0.461953764 | 1.28E-09 |

| | | | | | |
|---|---|---|---|---|---|
| B106 | B181 | anti-P | APTT | 0.459306489 | 1.63E-09 |
| B106 | B168 | anti-P | NMP22 | 0.401609665 | 2.03E-07 |
| B106 | B157 | anti-P | IgG | 0.400110805 | 2.28E-07 |
| B106 | B176 | anti-P | Ucr | 0.387593593 | 5.77E-07 |
| B107 | B151 | AnuA | Anti-CCP | 0.704822115 | 0 |
| B107 | B156 | AnuA | IgE | -0.903553245 | 0 |
| B107 | B183 | AnuA | PTA | -0.935351283 | 0 |
| B107 | B155 | AnuA | IgA | 0.556426361 | 4.64E-14 |
| B107 | B175 | AnuA | ALB-U | 0.538122709 | 4.33E-13 |
| B107 | B158 | AnuA | IgM | -0.492296842 | 6.69E-11 |
| B107 | B198 | AnuA | P | -0.484721535 | 1.44E-10 |
| B107 | B185 | AnuA | PT | 0.471649445 | 5.14E-10 |
| B107 | B184 | AnuA | INR | 0.461953777 | 1.28E-09 |
| B107 | B181 | AnuA | APTT | 0.459306499 | 1.63E-09 |
| B107 | B168 | AnuA | NMP22 | 0.401609685 | 2.03E-07 |
| B107 | B157 | AnuA | IgG | 0.400110814 | 2.28E-07 |
| B107 | B176 | AnuA | Ucr | 0.387593601 | 5.77E-07 |
| B108 | B151 | Anti-Cyclin | Anti-CCP | 0.7048221 | 0 |
| B108 | B156 | Anti-Cyclin | IgE | -0.903553226 | 0 |
| B108 | B183 | Anti-Cyclin | PTA | -0.935351263 | 0 |
| B108 | B155 | Anti-Cyclin | IgA | 0.55642635 | 4.64E-14 |
| B108 | B175 | Anti-Cyclin | ALB-U | 0.538122697 | 4.33E-13 |
| B108 | B158 | Anti-Cyclin | IgM | -0.492296831 | 6.70E-11 |
| B108 | B198 | Anti-Cyclin | P | -0.484721524 | 1.44E-10 |
| B108 | B185 | Anti-Cyclin | PT | 0.471649436 | 5.14E-10 |
| B108 | B184 | Anti-Cyclin | INR | 0.461953764 | 1.28E-09 |
| B108 | B181 | Anti-Cyclin | APTT | 0.459306489 | 1.63E-09 |
| B108 | B168 | Anti-Cyclin | NMP22 | 0.401609665 | 2.03E-07 |
| B108 | B157 | Anti-Cyclin | IgG | 0.400110805 | 2.28E-07 |
| B108 | B176 | Anti-Cyclin | Ucr | 0.387593593 | 5.77E-07 |
| B109 | B110 | AMA-M2 | AHA | 1 | 0 |
| B109 | B151 | AMA-M2 | Anti-CCP | 0.7048221 | 0 |
| B109 | B156 | AMA-M2 | IgE | -0.903553226 | 0 |
| B109 | B183 | AMA-M2 | PTA | -0.935351263 | 0 |
| B109 | B155 | AMA-M2 | IgA | 0.55642635 | 4.64E-14 |
| B109 | B175 | AMA-M2 | ALB-U | 0.538122697 | 4.33E-13 |
| B109 | B158 | AMA-M2 | IgM | -0.492296831 | 6.70E-11 |
| B109 | B198 | AMA-M2 | P | -0.484721524 | 1.44E-10 |
| B109 | B185 | AMA-M2 | PT | 0.471649437 | 5.14E-10 |
| B109 | B184 | AMA-M2 | INR | 0.461953764 | 1.28E-09 |
| B109 | B181 | AMA-M2 | APTT | 0.459306489 | 1.63E-09 |
| B109 | B168 | AMA-M2 | NMP22 | 0.401609665 | 2.03E-07 |
| B109 | B157 | AMA-M2 | IgG | 0.400110805 | 2.28E-07 |
| B109 | B176 | AMA-M2 | Ucr | 0.387593592 | 5.77E-07 |

| | | | | | |
|---|---|---|---|---|---|
| B11 | B14 | PGI/PGII | G-17 | -0.674597955 | 0 |
| B11 | B12 | PGI/PGII | PG II | -0.489670856 | 8.74E-11 |
| B110 | B151 | AHA | Anti-CCP | 0.7048221 | 0 |
| B110 | B156 | AHA | IgE | -0.903553226 | 0 |
| B110 | B183 | AHA | PTA | -0.935351263 | 0 |
| B110 | B155 | AHA | IgA | 0.55642635 | 4.64E-14 |
| B110 | B175 | AHA | ALB-U | 0.538122697 | 4.33E-13 |
| B110 | B158 | AHA | IgM | -0.492296831 | 6.70E-11 |
| B110 | B198 | AHA | P | -0.484721524 | 1.44E-10 |
| B110 | B185 | AHA | PT | 0.471649437 | 5.14E-10 |
| B110 | B184 | AHA | INR | 0.461953764 | 1.28E-09 |
| B110 | B181 | AHA | APTT | 0.459306489 | 1.63E-09 |
| B110 | B168 | AHA | NMP22 | 0.401609665 | 2.03E-07 |
| B110 | B157 | AHA | IgG | 0.400110805 | 2.28E-07 |
| B110 | B176 | AHA | Ucr | 0.387593592 | 5.77E-07 |
| B111 | B148 | TSH | Anti_TPO | 0.87003874 | 0 |
| B111 | B149 | TSH | TGAb | 0.816950733 | 0 |
| B112 | B114 | FT4 | TSH TT3 TT4(TT4) | 0.437362613 | 1.14E-08 |
| B113 | B115 | FT3 | TSH TT3 TT4(TT3) | 0.466435778 | 8.41E-10 |
| B114 | B193 | TSH TT3 TT4(TT4) | K | -0.588727142 | 6.66E-16 |
| B114 | B134 | TSH TT3 TT4(TT4) | CHE | 0.536951247 | 4.98E-13 |
| B114 | B185 | TSH TT3 TT4(TT4) | PT | -0.485109177 | 1.38E-10 |
| B114 | B181 | TSH TT3 TT4(TT4) | APTT | -0.436845608 | 1.19E-08 |
| B115 | B134 | TSH TT3 TT4(TT3) | CHE | 0.456541437 | 2.10E-09 |
| B115 | B152 | TSH TT3 TT4(TT3) | ACA-Ig(G-A-M) | -0.42028932 | 4.70E-08 |
| B115 | B194 | TSH TT3 TT4(TT3) | Cl | -0.400591138 | 2.20E-07 |
| B115 | B131 | TSH TT3 TT4(TT3) | AST | 0.399243822 | 2.43E-07 |
| B115 | B197 | TSH TT3 TT4(TT3) | AG | 0.398188601 | 2.63E-07 |
| B116 | B124 | TT3 | ACTH | 0.513552591 | 7.10E-12 |
| B117 | B125 | TT4 | Cor | -0.914137926 | 0 |
| B117 | B202 | TT4 | e-times | -0.593913871 | 4.44E-16 |
| B117 | B180 | TT4 | Tf | -0.476042484 | 3.37E-10 |
| B119 | B121 | Hcv-rna | C3 | -0.999999984 | 0 |
| B119 | B169 | Hcv-rna | U-CRY | -0.99999944 | 0 |
| B119 | B179 | Hcv-rna | α1-MG | -0.999999966 | 0 |
| B121 | B169 | C3 | U-CRY | 0.999999432 | 0 |
| B122 | B159 | C4 | β2-MG | 0.999999731 | 0 |
| B122 | B169 | C4 | U-CRY | 0.999999715 | 0 |
| B123 | B156 | hs-CRP1 | IgE | -0.38988918 | 4.88E-07 |
| B125 | B223 | Cor | FH | -0.534296709 | 6.80E-13 |
| B125 | B178 | Cor | Ig-U | -0.451446949 | 3.32E-09 |
| B126 | B127 | Alb | A-G ratio | 0.545800046 | 1.73E-13 |
| B127 | B130 | A-G ratio | GLB | -0.918735946 | 0 |
| B128 | B131 | ALT | AST | 0.701116767 | 0 |

| | | | | | |
|---|---|---|---|---|---|
| B128 | B141 | ALT | AST/ALT | -0.654655193 | 0 |
| B128 | B216 | ALT | BMI | 0.609555706 | 0 |
| B128 | B215 | ALT | BMI | 0.59309447 | 4.44E-16 |
| B128 | B206 | ALT | Waist | 0.545366428 | 1.82E-13 |
| B128 | B210 | ALT | Hip | 0.525381598 | 1.90E-12 |
| B128 | B222 | ALT | NS | 0.513252177 | 7.33E-12 |
| B128 | B140 | ALT | r-GT | 0.423615491 | 3.59E-08 |
| B128 | B207 | ALT | WHR | 0.413649219 | 7.99E-08 |
| B129 | B132 | IBIL | Dbil | 0.679156728 | 0 |
| B129 | B133 | IBIL | TBIL | 0.976858959 | 0 |
| B131 | B215 | AST | BMI | 0.444540491 | 6.11E-09 |
| B131 | B206 | AST | Waist | 0.419988119 | 4.82E-08 |
| B131 | B216 | AST | BMI | 0.406567872 | 1.39E-07 |
| B131 | B207 | AST | WHR | 0.401877387 | 1.99E-07 |
| B132 | B133 | Dbil | TBIL | 0.820341578 | 0 |
| B134 | B193 | CHE | K | -0.692059016 | 0 |
| B134 | B136 | CHE | 5'NT | 0.387006046 | 6.02E-07 |
| B135 | B136 | TBA | 5'NT | -0.6808099 | 0 |
| B135 | B196 | TBA | TCO2 | 0.647416217 | 0 |
| B135 | B198 | TBA | P | -0.649487971 | 0 |
| B135 | B137 | TBA | AFU | 0.53834308 | 4.22E-13 |
| B135 | B194 | TBA | Cl | -0.533768098 | 7.23E-13 |
| B135 | B139 | TBA | ADA | 0.52284771 | 2.53E-12 |
| B136 | B138 | 5'NT | PAB | 0.519188303 | 3.81E-12 |
| B136 | B196 | 5'NT | TCO2 | -0.411191733 | 9.70E-08 |
| B137 | B139 | AFU | ADA | 0.727550678 | 0 |
| B137 | B194 | AFU | Cl | -0.799829167 | 0 |
| B137 | B198 | AFU | P | -0.586990126 | 8.88E-16 |
| B137 | B196 | AFU | TCO2 | 0.584532779 | 1.11E-15 |
| B137 | B200 | AFU | OP | 0.525971846 | 1.78E-12 |
| B137 | B197 | AFU | AG | 0.465415351 | 9.25E-10 |
| B138 | B150 | PAB | ALP | 0.412286239 | 8.90E-08 |
| B138 | B197 | PAB | AG | 0.391090098 | 4.47E-07 |
| B139 | B198 | ADA | P | -0.557346717 | 4.13E-14 |
| B139 | B194 | ADA | Cl | -0.556894487 | 4.37E-14 |
| B139 | B196 | ADA | TCO2 | 0.415506736 | 6.90E-08 |
| B14 | B44 | G-17 | MCH | -0.412427117 | 8.80E-08 |
| B141 | B216 | AST/ALT | BMI | -0.481976085 | 1.89E-10 |
| B141 | B210 | AST/ALT | Hip | -0.460257138 | 1.49E-09 |
| B141 | B222 | AST/ALT | NS | -0.451137336 | 3.41E-09 |
| B141 | B215 | AST/ALT | BMI | -0.449099606 | 4.09E-09 |
| B141 | B206 | AST/ALT | Waist | -0.419433405 | 5.04E-08 |
| B142 | B143 | 25-(OH)D | β-CTx | 0.999999291 | 0 |
| B142 | B151 | 25-(OH)D | Anti-CCP | -0.7048221 | 0 |

| | | | | | |
|---|---|---|---|---|---|
| B142 | B168 | 25-(OH)D | NMP22 | -0.401609664 | 2.03E-07 |
| B143 | B144 | β-CTx | Total-PINP | 0.999999291 | 0 |
| B143 | B151 | β-CTx | Anti-CCP | -0.704816154 | 0 |
| B143 | B168 | β-CTx | NMP22 | -0.40159333 | 2.04E-07 |
| B144 | B151 | Total-PINP | Anti-CCP | -0.7048221 | 0 |
| B144 | B168 | Total-PINP | NMP22 | -0.401609664 | 2.03E-07 |
| B145 | B171 | BGP | EC | 0.843661488 | 0 |
| B146 | B194 | CysC | Cl | -0.397608531 | 2.75E-07 |
| B148 | B149 | Anti_TPO | TGAb | 0.960804957 | 0 |
| B148 | B214 | Anti_TPO | CVA-L | -0.38989287 | 4.88E-07 |
| B15 | B100 | CD3-CD19+(B cell) | anti-Ro52 | 1 | 0 |
| B15 | B101 | CD3-CD19+(B cell) | anti-Sc70 | 0.999999997 | 0 |
| B15 | B102 | CD3-CD19+(B cell) | anti-Sm | 0.999999999 | 0 |
| B15 | B105 | CD3-CD19+(B cell) | anti-U1RNP | 1 | 0 |
| B15 | B109 | CD3-CD19+(B cell) | AMA-M2 | 1 | 0 |
| B15 | B110 | CD3-CD19+(B cell) | AHA | 1 | 0 |
| B15 | B151 | CD3-CD19+(B cell) | Anti-CCP | 0.7048221 | 0 |
| B15 | B156 | CD3-CD19+(B cell) | IgE | -0.903553226 | 0 |
| B15 | B16 | CD3-CD19+(B cell) | CD3+T | -1 | 0 |
| B15 | B18 | CD3-CD19+(B cell) | CD3+CD4+helper(inducer)T | -1 | 0 |
| B15 | B183 | CD3-CD19+(B cell) | PTA | -0.935351263 | 0 |
| B15 | B19 | CD3-CD19+(B cell) | CD3+CD4+/CD3+CD8+ | -1 | 0 |
| B15 | B20 | CD3-CD19+(B cell) | CD3+CD4+CD8+ | 1 | 0 |
| B15 | B57 | CD3-CD19+(B cell) | RBCAI | 0.673254212 | 0 |
| B15 | B90 | CD3-CD19+(B cell) | CYFRA 21-1 | -0.720217082 | 0 |
| B15 | B98 | CD3-CD19+(B cell) | anti-Jo-1 | 0.999999997 | 0 |
| B15 | B99 | CD3-CD19+(B cell) | anti-PM-Scl | 0.999999999 | 0 |
| B15 | B25 | CD3-CD19+(B cell) | CK | 0.5935606 | 4.44E-16 |
| B15 | B59 | CD3-CD19+(B cell) | WBRV | 0.571774066 | 6.44E-15 |
| B15 | B155 | CD3-CD19+(B cell) | IgA | 0.55642635 | 4.64E-14 |
| B15 | B175 | CD3-CD19+(B cell) | ALB-U | 0.538122697 | 4.33E-13 |
| B15 | B158 | CD3-CD19+(B cell) | IgM | -0.492296831 | 6.70E-11 |
| B15 | B60 | CD3-CD19+(B cell) | WBRVI | 0.490164144 | 8.32E-11 |
| B15 | B54 | CD3-CD19+(B cell) | ESR | 0.486953983 | 1.15E-10 |
| B15 | B198 | CD3-CD19+(B cell) | P | -0.484721524 | 1.44E-10 |
| B15 | B185 | CD3-CD19+(B cell) | PT | 0.471649437 | 5.14E-10 |
| B15 | B184 | CD3-CD19+(B cell) | INR | 0.461953764 | 1.28E-09 |
| B15 | B181 | CD3-CD19+(B cell) | APTT | 0.459306489 | 1.63E-09 |
| B15 | B65 | CD3-CD19+(B cell) | WBRSR(mPas_1) | 0.450002865 | 3.78E-09 |
| B15 | B58 | CD3-CD19+(B cell) | HCT | 0.423424837 | 3.64E-08 |
| B15 | B168 | CD3-CD19+(B cell) | NMP22 | 0.401609664 | 2.03E-07 |
| B15 | B157 | CD3-CD19+(B cell) | IgG | 0.400110805 | 2.28E-07 |
| B15 | B176 | CD3-CD19+(B cell) | Ucr | 0.387593593 | 5.77E-07 |
| B150 | B197 | ALP | AG | 0.444918983 | 5.91E-09 |

| | | | | | |
|---|---|---|---|---|---|
| B150 | B194 | ALP | Cl | -0.419423234 | 5.04E-08 |
| B151 | B156 | Anti-CCP | IgE | -0.636306618 | 0 |
| B151 | B183 | Anti-CCP | PTA | -0.66156937 | 0 |
| B151 | B175 | Anti-CCP | ALB-U | 0.594784271 | 2.22E-16 |
| B151 | B168 | Anti-CCP | NMP22 | 0.569802512 | 8.44E-15 |
| B151 | B155 | Anti-CCP | IgA | 0.386941597 | 6.04E-07 |
| B152 | B184 | ACA-Ig(G-A-M) | INR | 0.613320015 | 0 |
| B153 | B220 | Glu | SBP | 0.420870213 | 4.48E-08 |
| B154 | B157 | RF | IgG | 0.585357938 | 8.88E-16 |
| B155 | B156 | IgA | IgE | -0.69776581 | 0 |
| B155 | B183 | IgA | PTA | -0.5503322 | 9.90E-14 |
| B156 | B183 | IgE | PTA | 0.868039722 | 0 |
| B156 | B181 | IgE | APTT | -0.494514492 | 5.34E-11 |
| B156 | B185 | IgE | PT | -0.492402198 | 6.62E-11 |
| B156 | B175 | IgE | ALB-U | -0.484538328 | 1.46E-10 |
| B156 | B198 | IgE | P | 0.438501072 | 1.03E-08 |
| B156 | B184 | IgE | INR | -0.437642311 | 1.11E-08 |
| B157 | B158 | IgG | IgM | -0.730819563 | 0 |
| B157 | B182 | IgG | TT | -0.467473771 | 7.63E-10 |
| B157 | B186 | IgG | PFC | 0.443192413 | 6.88E-09 |
| B158 | B183 | IgM | PTA | 0.538267316 | 4.26E-13 |
| B158 | B181 | IgM | APTT | -0.509191177 | 1.14E-11 |
| B158 | B185 | IgM | PT | -0.474476901 | 3.92E-10 |
| B159 | B169 | β2-MG | U-CRY | 0.999999092 | 0 |
| B16 | B100 | CD3+T | anti-Ro52 | -1 | 0 |
| B16 | B101 | CD3+T | anti-Sc70 | -0.999999996 | 0 |
| B16 | B102 | CD3+T | anti-Sm | -0.999999999 | 0 |
| B16 | B103 | CD3+T | anti-SSA/Ro | -1 | 0 |
| B16 | B105 | CD3+T | anti-U1RNP | -1 | 0 |
| B16 | B109 | CD3+T | AMA-M2 | -1 | 0 |
| B16 | B110 | CD3+T | AHA | -1 | 0 |
| B16 | B151 | CD3+T | Anti-CCP | -0.7048221 | 0 |
| B16 | B156 | CD3+T | IgE | 0.903553226 | 0 |
| B16 | B17 | CD3+T | CD3+CD16+CD56+NK | -1 | 0 |
| B16 | B18 | CD3+T | CD3+CD4+helper(inducer)T | 1 | 0 |
| B16 | B183 | CD3+T | PTA | 0.935351263 | 0 |
| B16 | B19 | CD3+T | CD3+CD4+/CD3+CD8+ | 1 | 0 |
| B16 | B20 | CD3+T | CD3+CD4+CD8+ | -1 | 0 |
| B16 | B21 | CD3+T | CD3+CD4-CD8- | -1 | 0 |
| B16 | B23 | CD3+T | CD3-CD16+CD56+(NK) | -1 | 0 |
| B16 | B34 | CD3+T | TK | -1 | 0 |
| B16 | B57 | CD3+T | RBCAI | -0.673254212 | 0 |
| B16 | B90 | CD3+T | CYFRA 21-1 | 0.720217082 | 0 |
| B16 | B98 | CD3+T | anti-Jo-1 | -0.999999997 | 0 |

| | | | | | |
|---|---|---|---|---|---|
| B16 | B99 | CD3+T | anti-PM-Scl | -0.999999999 | 0 |
| B16 | B25 | CD3+T | CK | -0.5935606 | 4.44E-16 |
| B16 | B59 | CD3+T | WBRV | -0.571774066 | 6.44E-15 |
| B16 | B155 | CD3+T | IgA | -0.55642635 | 4.64E-14 |
| B16 | B175 | CD3+T | ALB-U | -0.538122697 | 4.33E-13 |
| B16 | B158 | CD3+T | IgM | 0.492296831 | 6.70E-11 |
| B16 | B60 | CD3+T | WBRVI | -0.490164144 | 8.32E-11 |
| B16 | B54 | CD3+T | ESR | -0.486953983 | 1.15E-10 |
| B16 | B198 | CD3+T | P | 0.484721524 | 1.44E-10 |
| B16 | B185 | CD3+T | PT | -0.471649437 | 5.14E-10 |
| B16 | B184 | CD3+T | INR | -0.461953764 | 1.28E-09 |
| B16 | B181 | CD3+T | APTT | -0.459306489 | 1.63E-09 |
| B16 | B65 | CD3+T | WBRSR(mPas_1) | -0.450002865 | 3.78E-09 |
| B16 | B58 | CD3+T | HCT | -0.423424837 | 3.64E-08 |
| B16 | B168 | CD3+T | NMP22 | -0.401609664 | 2.03E-07 |
| B16 | B157 | CD3+T | IgG | -0.400110805 | 2.28E-07 |
| B16 | B176 | CD3+T | Ucr | -0.387593593 | 5.77E-07 |
| B163 | B166 | SG | Pro | 0.442841356 | 7.09E-09 |
| B168 | B200 | NMP22 | OP | 0.535140384 | 6.16E-13 |
| B168 | B195 | NMP22 | Na | 0.480593065 | 2.16E-10 |
| B168 | B183 | NMP22 | PTA | -0.38241168 | 8.37E-07 |
| B169 | B179 | U-CRY | α1-MG | 0.999999915 | 0 |
| B169 | B187 | U-CRY | NSE | -0.999999748 | 0 |
| B17 | B100 | CD3+CD16+CD56+NK | anti-Ro52 | 1 | 0 |
| B17 | B101 | CD3+CD16+CD56+NK | anti-Sc70 | 0.999999997 | 0 |
| B17 | B102 | CD3+CD16+CD56+NK | anti-Sm | 1 | 0 |
| B17 | B105 | CD3+CD16+CD56+NK | anti-U1RNP | 1 | 0 |
| B17 | B109 | CD3+CD16+CD56+NK | AMA-M2 | 1 | 0 |
| B17 | B110 | CD3+CD16+CD56+NK | AHA | 1 | 0 |
| B17 | B151 | CD3+CD16+CD56+NK | Anti-CCP | 0.7048221 | 0 |
| B17 | B156 | CD3+CD16+CD56+NK | IgE | -0.903553226 | 0 |
| B17 | B183 | CD3+CD16+CD56+NK | PTA | -0.935351263 | 0 |
| B17 | B19 | CD3+CD16+CD56+NK | CD3+CD4+/CD3+CD8+ | -1 | 0 |
| B17 | B21 | CD3+CD16+CD56+NK | CD3+CD4-CD8- | 1 | 0 |
| B17 | B57 | CD3+CD16+CD56+NK | RBCAI | 0.673254212 | 0 |
| B17 | B90 | CD3+CD16+CD56+NK | CYFRA 21-1 | -0.720217082 | 0 |
| B17 | B98 | CD3+CD16+CD56+NK | anti-Jo-1 | 0.999999997 | 0 |
| B17 | B99 | CD3+CD16+CD56+NK | anti-PM-Scl | 0.999999999 | 0 |
| B17 | B25 | CD3+CD16+CD56+NK | CK | 0.5935606 | 4.44E-16 |
| B17 | B59 | CD3+CD16+CD56+NK | WBRV | 0.571774066 | 6.44E-15 |
| B17 | B155 | CD3+CD16+CD56+NK | IgA | 0.55642635 | 4.64E-14 |
| B17 | B175 | CD3+CD16+CD56+NK | ALB-U | 0.538122697 | 4.33E-13 |
| B17 | B158 | CD3+CD16+CD56+NK | IgM | -0.492296831 | 6.70E-11 |
| B17 | B60 | CD3+CD16+CD56+NK | WBRVI | 0.490164144 | 8.32E-11 |

| | | | | | |
|---|---|---|---|---|---|
| B17 | B54 | CD3+CD16+CD56+NK | ESR | 0.486953983 | 1.15E-10 |
| B17 | B198 | CD3+CD16+CD56+NK | P | -0.484721524 | 1.44E-10 |
| B17 | B185 | CD3+CD16+CD56+NK | PT | 0.471649437 | 5.14E-10 |
| B17 | B184 | CD3+CD16+CD56+NK | INR | 0.461953764 | 1.28E-09 |
| B17 | B181 | CD3+CD16+CD56+NK | APTT | 0.459306489 | 1.63E-09 |
| B17 | B65 | CD3+CD16+CD56+NK | WBRSR(mPas_1) | 0.450002865 | 3.78E-09 |
| B17 | B58 | CD3+CD16+CD56+NK | HCT | 0.423424837 | 3.64E-08 |
| B17 | B168 | CD3+CD16+CD56+NK | NMP22 | 0.401609664 | 2.03E-07 |
| B17 | B157 | CD3+CD16+CD56+NK | IgG | 0.400110805 | 2.28E-07 |
| B17 | B176 | CD3+CD16+CD56+NK | Ucr | 0.387593593 | 5.77E-07 |
| B170 | B173 | U-GLU | NTT | 1 | 0 |
| B174 | B191 | Urea/Cr | Urea | 0.654289272 | 0 |
| B174 | B190 | Urea/Cr | Cre | -0.398592719 | 2.56E-07 |
| B175 | B183 | ALB-U | PTA | -0.510579374 | 9.80E-12 |
| B175 | B198 | ALB-U | P | -0.382053618 | 8.59E-07 |
| B178 | B180 | Ig-U | Tf | -0.783645156 | 0 |
| B18 | B100 | CD3+CD4+helper(inducer)T | anti-Ro52 | -1 | 0 |
| B18 | B101 | CD3+CD4+helper(inducer)T | anti-Sc70 | -0.999999997 | 0 |
| B18 | B102 | CD3+CD4+helper(inducer)T | anti-Sm | -0.999999999 | 0 |
| B18 | B105 | CD3+CD4+helper(inducer)T | anti-U1RNP | -1 | 0 |
| B18 | B109 | CD3+CD4+helper(inducer)T | AMA-M2 | -1 | 0 |
| B18 | B110 | CD3+CD4+helper(inducer)T | AHA | -1 | 0 |
| B18 | B151 | CD3+CD4+helper(inducer)T | Anti-CCP | -0.7048221 | 0 |
| B18 | B156 | CD3+CD4+helper(inducer)T | IgE | 0.903553226 | 0 |
| B18 | B183 | CD3+CD4+helper(inducer)T | PTA | 0.935351263 | 0 |
| B18 | B19 | CD3+CD4+helper(inducer)T | CD3+CD4+/CD3+CD8+ | 1 | 0 |
| B18 | B20 | CD3+CD4+helper(inducer)T | CD3+CD4+CD8+ | -1 | 0 |
| B18 | B23 | CD3+CD4+helper(inducer)T | CD3-CD16+CD56+(NK) | -1 | 0 |
| B18 | B34 | CD3+CD4+helper(inducer)T | TK | -1 | 0 |
| B18 | B57 | CD3+CD4+helper(inducer)T | RBCAI | -0.673254212 | 0 |
| B18 | B90 | CD3+CD4+helper(inducer)T | CYFRA 21-1 | 0.720217082 | 0 |
| B18 | B98 | CD3+CD4+helper(inducer)T | anti-Jo-1 | -0.999999997 | 0 |
| B18 | B99 | CD3+CD4+helper(inducer)T | anti-PM-Scl | -0.999999999 | 0 |
| B18 | B25 | CD3+CD4+helper(inducer)T | CK | -0.5935606 | 4.44E-16 |
| B18 | B59 | CD3+CD4+helper(inducer)T | WBRV | -0.571774066 | 6.44E-15 |
| B18 | B155 | CD3+CD4+helper(inducer)T | IgA | -0.55642635 | 4.64E-14 |
| B18 | B175 | CD3+CD4+helper(inducer)T | ALB-U | -0.538122697 | 4.33E-13 |
| B18 | B158 | CD3+CD4+helper(inducer)T | IgM | 0.492296831 | 6.70E-11 |
| B18 | B60 | CD3+CD4+helper(inducer)T | WBRVI | -0.490164144 | 8.32E-11 |
| B18 | B54 | CD3+CD4+helper(inducer)T | ESR | -0.486953983 | 1.15E-10 |
| B18 | B198 | CD3+CD4+helper(inducer)T | P | 0.484721524 | 1.44E-10 |
| B18 | B185 | CD3+CD4+helper(inducer)T | PT | -0.471649437 | 5.14E-10 |
| B18 | B184 | CD3+CD4+helper(inducer)T | INR | -0.461953764 | 1.28E-09 |
| B18 | B181 | CD3+CD4+helper(inducer)T | APTT | -0.459306489 | 1.63E-09 |

| | | | | | |
|---|---|---|---|---|---|
| B18 | B65 | CD3+CD4+helper(inducer)T | WBRSR(mPas_1) | -0.450002865 | 3.78E-09 |
| B18 | B58 | CD3+CD4+helper(inducer)T | HCT | -0.423424837 | 3.64E-08 |
| B18 | B168 | CD3+CD4+helper(inducer)T | NMP22 | -0.401609664 | 2.03E-07 |
| B18 | B157 | CD3+CD4+helper(inducer)T | IgG | -0.400110805 | 2.28E-07 |
| B18 | B176 | CD3+CD4+helper(inducer)T | Ucr | -0.387593593 | 5.77E-07 |
| B180 | B223 | Tf | FH | -0.620684829 | 0 |
| B180 | B225 | Tf | hPBG | -0.784381436 | 0 |
| B181 | B182 | APTT | TT | 0.666986375 | 0 |
| B181 | B183 | APTT | PTA | -0.642674629 | 0 |
| B181 | B185 | APTT | PT | 0.897972248 | 0 |
| B181 | B186 | APTT | PFC | -0.615454491 | 0 |
| B181 | B184 | APTT | INR | 0.459067764 | 1.67E-09 |
| B182 | B186 | TT | PFC | -0.990845633 | 0 |
| B182 | B185 | TT | PT | 0.384800228 | 7.06E-07 |
| B183 | B185 | PTA | PT | -0.512121503 | 8.29E-12 |
| B183 | B198 | PTA | P | 0.451107418 | 3.42E-09 |
| B184 | B185 | INR | PT | 0.797196839 | 0 |
| B189 | B190 | eGFB | Cre | -0.641678937 | 0 |
| B189 | B219 | eGFB | OT | -0.458575056 | 1.74E-09 |
| B19 | B100 | CD3+CD4+/CD3+CD8+ | anti-Ro52 | -1 | 0 |
| B19 | B101 | CD3+CD4+/CD3+CD8+ | anti-Sc70 | -0.999999997 | 0 |
| B19 | B102 | CD3+CD4+/CD3+CD8+ | anti-Sm | -0.999999999 | 0 |
| B19 | B103 | CD3+CD4+/CD3+CD8+ | anti-SSA/Ro | -1 | 0 |
| B19 | B105 | CD3+CD4+/CD3+CD8+ | anti-U1RNP | -1 | 0 |
| B19 | B109 | CD3+CD4+/CD3+CD8+ | AMA-M2 | -1 | 0 |
| B19 | B110 | CD3+CD4+/CD3+CD8+ | AHA | -1 | 0 |
| B19 | B151 | CD3+CD4+/CD3+CD8+ | Anti-CCP | -0.7048221 | 0 |
| B19 | B156 | CD3+CD4+/CD3+CD8+ | IgE | 0.903553226 | 0 |
| B19 | B183 | CD3+CD4+/CD3+CD8+ | PTA | 0.935351263 | 0 |
| B19 | B20 | CD3+CD4+/CD3+CD8+ | CD3+CD4+CD8+ | -1 | 0 |
| B19 | B21 | CD3+CD4+/CD3+CD8+ | CD3+CD4-CD8- | -1 | 0 |
| B19 | B23 | CD3+CD4+/CD3+CD8+ | CD3-CD16+CD56+(NK) | -1 | 0 |
| B19 | B34 | CD3+CD4+/CD3+CD8+ | TK | -1 | 0 |
| B19 | B57 | CD3+CD4+/CD3+CD8+ | RBCAI | -0.673254212 | 0 |
| B19 | B90 | CD3+CD4+/CD3+CD8+ | CYFRA 21-1 | 0.720217082 | 0 |
| B19 | B98 | CD3+CD4+/CD3+CD8+ | anti-Jo-1 | -0.999999997 | 0 |
| B19 | B99 | CD3+CD4+/CD3+CD8+ | anti-PM-Scl | -0.999999999 | 0 |
| B19 | B25 | CD3+CD4+/CD3+CD8+ | CK | -0.5935606 | 4.44E-16 |
| B19 | B59 | CD3+CD4+/CD3+CD8+ | WBRV | -0.571774066 | 6.44E-15 |
| B19 | B155 | CD3+CD4+/CD3+CD8+ | IgA | -0.55642635 | 4.64E-14 |
| B19 | B175 | CD3+CD4+/CD3+CD8+ | ALB-U | -0.538122697 | 4.33E-13 |
| B19 | B158 | CD3+CD4+/CD3+CD8+ | IgM | 0.492296831 | 6.70E-11 |
| B19 | B60 | CD3+CD4+/CD3+CD8+ | WBRVI | -0.490164144 | 8.32E-11 |
| B19 | B54 | CD3+CD4+/CD3+CD8+ | ESR | -0.486953983 | 1.15E-10 |

| | | | | | |
|---|---|---|---|---|---|
| B19 | B198 | CD3+CD4+/CD3+CD8+ | P | 0.484721524 | 1.44E-10 |
| B19 | B185 | CD3+CD4+/CD3+CD8+ | PT | -0.471649437 | 5.14E-10 |
| B19 | B184 | CD3+CD4+/CD3+CD8+ | INR | -0.461953764 | 1.28E-09 |
| B19 | B181 | CD3+CD4+/CD3+CD8+ | APTT | -0.459306489 | 1.63E-09 |
| B19 | B65 | CD3+CD4+/CD3+CD8+ | WBRSR(mPas_1) | -0.450002865 | 3.78E-09 |
| B19 | B58 | CD3+CD4+/CD3+CD8+ | HCT | -0.423424837 | 3.64E-08 |
| B19 | B168 | CD3+CD4+/CD3+CD8+ | NMP22 | -0.401609664 | 2.03E-07 |
| B19 | B157 | CD3+CD4+/CD3+CD8+ | IgG | -0.400110805 | 2.28E-07 |
| B19 | B176 | CD3+CD4+/CD3+CD8+ | Ucr | -0.387593593 | 5.77E-07 |
| B190 | B192 | Cre | UA | 0.582511213 | 1.55E-15 |
| B190 | B222 | Cre | NS | 0.54748259 | 1.41E-13 |
| B190 | B221 | Cre | Hei | 0.449827544 | 3.84E-09 |
| B190 | B216 | Cre | BMI | 0.431898953 | 1.81E-08 |
| B190 | B207 | Cre | WHR | 0.416023127 | 6.62E-08 |
| B190 | B191 | Cre | Urea | 0.407081471 | 1.34E-07 |
| B190 | B206 | Cre | Waist | 0.399064822 | 2.47E-07 |
| B192 | B222 | UA | NS | 0.583597208 | 1.33E-15 |
| B192 | B216 | UA | BMI | 0.579396181 | 2.22E-15 |
| B192 | B206 | UA | Waist | 0.551924177 | 8.13E-14 |
| B192 | B221 | UA | Hei | 0.495853452 | 4.65E-11 |
| B192 | B207 | UA | WHR | 0.488180308 | 1.02E-10 |
| B192 | B210 | UA | Hip | 0.449990516 | 3.78E-09 |
| B192 | B215 | UA | BMI | 0.423567698 | 3.60E-08 |
| B194 | B196 | Cl | TCO2 | -0.572970007 | 5.33E-15 |
| B194 | B197 | Cl | AG | -0.517715404 | 4.49E-12 |
| B194 | B198 | Cl | P | 0.51761687 | 4.54E-12 |
| B195 | B200 | Na | OP | 0.861881501 | 0 |
| B195 | B197 | Na | AG | 0.533047188 | 7.86E-13 |
| B196 | B198 | TCO2 | P | -0.87937918 | 0 |
| B196 | B200 | TCO2 | OP | 0.418356505 | 5.49E-08 |
| B197 | B200 | AG | OP | 0.589632778 | 4.44E-16 |
| B198 | B200 | P | OP | -0.389341278 | 5.08E-07 |
| B2 | B189 | Age | eGFB | -0.660327006 | 0 |
| B2 | B219 | Age | OT | 0.48143245 | 1.99E-10 |
| B2 | B153 | Age | Glu | 0.47314488 | 4.45E-10 |
| B2 | B207 | Age | WHR | 0.446938084 | 4.95E-09 |
| B20 | B100 | CD3+CD4+CD8+ | anti-Ro52 | 1 | 0 |
| B20 | B101 | CD3+CD4+CD8+ | anti-Sc70 | 0.999999997 | 0 |
| B20 | B102 | CD3+CD4+CD8+ | anti-Sm | 1 | 0 |
| B20 | B105 | CD3+CD4+CD8+ | anti-U1RNP | 1 | 0 |
| B20 | B109 | CD3+CD4+CD8+ | AMA-M2 | 1 | 0 |
| B20 | B110 | CD3+CD4+CD8+ | AHA | 1 | 0 |
| B20 | B151 | CD3+CD4+CD8+ | Anti-CCP | 0.7048221 | 0 |
| B20 | B156 | CD3+CD4+CD8+ | IgE | -0.903553226 | 0 |

| | | | | | |
|---|---|---|---|---|---|
| B20 | B183 | CD3+CD4+CD8+ | PTA | -0.935351263 | 0 |
| B20 | B34 | CD3+CD4+CD8+ | TK | 1 | 0 |
| B20 | B57 | CD3+CD4+CD8+ | RBCAI | 0.673254212 | 0 |
| B20 | B90 | CD3+CD4+CD8+ | CYFRA 21-1 | -0.720217082 | 0 |
| B20 | B98 | CD3+CD4+CD8+ | anti-Jo-1 | 0.999999997 | 0 |
| B20 | B99 | CD3+CD4+CD8+ | anti-PM-Scl | 0.999999999 | 0 |
| B20 | B25 | CD3+CD4+CD8+ | CK | 0.5935606 | 4.44E-16 |
| B20 | B59 | CD3+CD4+CD8+ | WBRV | 0.571774066 | 6.44E-15 |
| B20 | B155 | CD3+CD4+CD8+ | IgA | 0.55642635 | 4.64E-14 |
| B20 | B175 | CD3+CD4+CD8+ | ALB-U | 0.538122697 | 4.33E-13 |
| B20 | B158 | CD3+CD4+CD8+ | IgM | -0.492296831 | 6.70E-11 |
| B20 | B60 | CD3+CD4+CD8+ | WBRVI | 0.490164144 | 8.32E-11 |
| B20 | B54 | CD3+CD4+CD8+ | ESR | 0.486953983 | 1.15E-10 |
| B20 | B198 | CD3+CD4+CD8+ | P | -0.484721524 | 1.44E-10 |
| B20 | B185 | CD3+CD4+CD8+ | PT | 0.471649437 | 5.14E-10 |
| B20 | B184 | CD3+CD4+CD8+ | INR | 0.461953764 | 1.28E-09 |
| B20 | B181 | CD3+CD4+CD8+ | APTT | 0.459306489 | 1.63E-09 |
| B20 | B65 | CD3+CD4+CD8+ | WBRSR(mPas_1) | 0.450002865 | 3.78E-09 |
| B20 | B58 | CD3+CD4+CD8+ | HCT | 0.423424837 | 3.64E-08 |
| B20 | B168 | CD3+CD4+CD8+ | NMP22 | 0.401609664 | 2.03E-07 |
| B20 | B157 | CD3+CD4+CD8+ | IgG | 0.400110805 | 2.28E-07 |
| B20 | B176 | CD3+CD4+CD8+ | Ucr | 0.387593593 | 5.77E-07 |
| B202 | B225 | e-times | hPBG | -0.873326063 | 0 |
| B202 | B223 | e-times | FH | -0.431244346 | 1.91E-08 |
| B204 | B209 | Drinking | Smoking | 0.442334442 | 7.41E-09 |
| B206 | B207 | Waist | WHR | 0.874619494 | 0 |
| B206 | B210 | Waist | Hip | 0.792914657 | 0 |
| B206 | B215 | Waist | BMI | 0.855027013 | 0 |
| B206 | B216 | Waist | BMI | 0.867953182 | 0 |
| B206 | B222 | Waist | NS | 0.770658776 | 0 |
| B206 | B221 | Waist | Hei | 0.448362523 | 4.37E-09 |
| B207 | B215 | WHR | BMI | 0.668817718 | 0 |
| B207 | B216 | WHR | BMI | 0.652972857 | 0 |
| B207 | B222 | WHR | NS | 0.68318672 | 0 |
| B207 | B210 | WHR | Hip | 0.452264295 | 3.09E-09 |
| B21 | B100 | CD3+CD4-CD8- | anti-Ro52 | 1 | 0 |
| B21 | B101 | CD3+CD4-CD8- | anti-Sc70 | 0.999999997 | 0 |
| B21 | B102 | CD3+CD4-CD8- | anti-Sm | 0.999999999 | 0 |
| B21 | B103 | CD3+CD4-CD8- | anti-SSA/Ro | 1 | 0 |
| B21 | B105 | CD3+CD4-CD8- | anti-U1RNP | 1 | 0 |
| B21 | B109 | CD3+CD4-CD8- | AMA-M2 | 1 | 0 |
| B21 | B110 | CD3+CD4-CD8- | AHA | 1 | 0 |
| B21 | B151 | CD3+CD4-CD8- | Anti-CCP | 0.7048221 | 0 |
| B21 | B156 | CD3+CD4-CD8- | IgE | -0.903553226 | 0 |

| | | | | | |
|---|---|---|---|---|---|
| B21 | B183 | CD3+CD4-CD8- | PTA | -0.935351263 | 0 |
| B21 | B23 | CD3+CD4-CD8- | CD3-CD16+CD56+(NK) | 1 | 0 |
| B21 | B57 | CD3+CD4-CD8- | RBCAI | 0.673254212 | 0 |
| B21 | B90 | CD3+CD4-CD8- | CYFRA 21-1 | -0.720217082 | 0 |
| B21 | B98 | CD3+CD4-CD8- | anti-Jo-1 | 0.999999997 | 0 |
| B21 | B99 | CD3+CD4-CD8- | anti-PM-Scl | 0.999999999 | 0 |
| B21 | B25 | CD3+CD4-CD8- | CK | 0.5935606 | 4.44E-16 |
| B21 | B59 | CD3+CD4-CD8- | WBRV | 0.571774066 | 6.44E-15 |
| B21 | B155 | CD3+CD4-CD8- | IgA | 0.55642635 | 4.64E-14 |
| B21 | B175 | CD3+CD4-CD8- | ALB-U | 0.538122697 | 4.33E-13 |
| B21 | B158 | CD3+CD4-CD8- | IgM | -0.492296831 | 6.70E-11 |
| B21 | B60 | CD3+CD4-CD8- | WBRVI | 0.490164144 | 8.32E-11 |
| B21 | B54 | CD3+CD4-CD8- | ESR | 0.486953983 | 1.15E-10 |
| B21 | B198 | CD3+CD4-CD8- | P | -0.484721524 | 1.44E-10 |
| B21 | B185 | CD3+CD4-CD8- | PT | 0.471649437 | 5.14E-10 |
| B21 | B184 | CD3+CD4-CD8- | INR | 0.461953764 | 1.28E-09 |
| B21 | B181 | CD3+CD4-CD8- | APTT | 0.459306489 | 1.63E-09 |
| B21 | B65 | CD3+CD4-CD8- | WBRSR(mPas_1) | 0.450002865 | 3.78E-09 |
| B21 | B58 | CD3+CD4-CD8- | HCT | 0.423424837 | 3.64E-08 |
| B21 | B168 | CD3+CD4-CD8- | NMP22 | 0.401609664 | 2.03E-07 |
| B21 | B157 | CD3+CD4-CD8- | IgG | 0.400110805 | 2.28E-07 |
| B21 | B176 | CD3+CD4-CD8- | Ucr | 0.387593593 | 5.77E-07 |
| B210 | B215 | Hip | BMI | 0.776312553 | 0 |
| B210 | B216 | Hip | BMI | 0.82466008 | 0 |
| B210 | B222 | Hip | NS | 0.656636781 | 0 |
| B210 | B221 | Hip | Hei | 0.473952504 | 4.12E-10 |
| B211 | B212 | IOP-R | IOP-L | 0.871437527 | 0 |
| B213 | B214 | CVA-R | CVA-L | 0.723999171 | 0 |
| B215 | B216 | BMI | BMI | 0.832590375 | 0 |
| B215 | B222 | BMI | NS | 0.659158344 | 0 |
| B216 | B221 | BMI | Hei | 0.70400546 | 0 |
| B216 | B222 | BMI | NS | 0.794399592 | 0 |
| B218 | B220 | DBP | SBP | 0.734518467 | 0 |
| B22 | B101 | CD3+CD8+(T) | anti-Sc70 | 0.999999997 | 0 |
| B22 | B102 | CD3+CD8+(T) | anti-Sm | 0.999999999 | 0 |
| B22 | B105 | CD3+CD8+(T) | anti-U1RNP | 1 | 0 |
| B22 | B106 | CD3+CD8+(T) | anti-P | 1 | 0 |
| B22 | B109 | CD3+CD8+(T) | AMA-M2 | 1 | 0 |
| B22 | B151 | CD3+CD8+(T) | Anti-CCP | 0.7048221 | 0 |
| B22 | B156 | CD3+CD8+(T) | IgE | -0.903553226 | 0 |
| B22 | B183 | CD3+CD8+(T) | PTA | -0.935351263 | 0 |
| B22 | B57 | CD3+CD8+(T) | RBCAI | 0.673254212 | 0 |
| B22 | B90 | CD3+CD8+(T) | CYFRA 21-1 | -0.720217082 | 0 |
| B22 | B98 | CD3+CD8+(T) | anti-Jo-1 | 0.999999997 | 0 |

| | | | | | |
|---|---|---|---|---|---|
| B22 | B99 | CD3+CD8+(T) | anti-PM-Scl | 0.999999999 | 0 |
| B22 | B25 | CD3+CD8+(T) | CK | 0.5935606 | 4.44E-16 |
| B22 | B59 | CD3+CD8+(T) | WBRV | 0.571774066 | 6.44E-15 |
| B22 | B155 | CD3+CD8+(T) | IgA | 0.55642635 | 4.64E-14 |
| B22 | B175 | CD3+CD8+(T) | ALB-U | 0.538122697 | 4.33E-13 |
| B22 | B158 | CD3+CD8+(T) | IgM | -0.492296831 | 6.70E-11 |
| B22 | B60 | CD3+CD8+(T) | WBRVI | 0.490164144 | 8.32E-11 |
| B22 | B54 | CD3+CD8+(T) | ESR | 0.486953983 | 1.15E-10 |
| B22 | B198 | CD3+CD8+(T) | P | -0.484721524 | 1.44E-10 |
| B22 | B185 | CD3+CD8+(T) | PT | 0.471649437 | 5.14E-10 |
| B22 | B184 | CD3+CD8+(T) | INR | 0.461953764 | 1.28E-09 |
| B22 | B181 | CD3+CD8+(T) | APTT | 0.459306489 | 1.63E-09 |
| B22 | B65 | CD3+CD8+(T) | WBRSR(mPas_1) | 0.450002865 | 3.78E-09 |
| B22 | B58 | CD3+CD8+(T) | HCT | 0.423424837 | 3.64E-08 |
| B22 | B168 | CD3+CD8+(T) | NMP22 | 0.401609665 | 2.03E-07 |
| B22 | B157 | CD3+CD8+(T) | IgG | 0.400110805 | 2.28E-07 |
| B22 | B176 | CD3+CD8+(T) | Ucr | 0.387593593 | 5.77E-07 |
| B221 | B222 | Hei | NS | 0.578653593 | 2.66E-15 |
| B227 | B228 | OD | OS | 0.719740153 | 0 |
| B23 | B100 | CD3-CD16+CD56+(NK) | anti-Ro52 | 1 | 0 |
| B23 | B101 | CD3-CD16+CD56+(NK) | anti-Sc70 | 0.999999997 | 0 |
| B23 | B102 | CD3-CD16+CD56+(NK) | anti-Sm | 0.999999999 | 0 |
| B23 | B105 | CD3-CD16+CD56+(NK) | anti-U1RNP | 1 | 0 |
| B23 | B109 | CD3-CD16+CD56+(NK) | AMA-M2 | 1 | 0 |
| B23 | B110 | CD3-CD16+CD56+(NK) | AHA | 1 | 0 |
| B23 | B151 | CD3-CD16+CD56+(NK) | Anti-CCP | 0.7048221 | 0 |
| B23 | B156 | CD3-CD16+CD56+(NK) | IgE | -0.903553226 | 0 |
| B23 | B183 | CD3-CD16+CD56+(NK) | PTA | -0.935351263 | 0 |
| B23 | B57 | CD3-CD16+CD56+(NK) | RBCAI | 0.673254212 | 0 |
| B23 | B90 | CD3-CD16+CD56+(NK) | CYFRA 21-1 | -0.720217082 | 0 |
| B23 | B98 | CD3-CD16+CD56+(NK) | anti-Jo-1 | 0.999999997 | 0 |
| B23 | B99 | CD3-CD16+CD56+(NK) | anti-PM-Scl | 0.999999999 | 0 |
| B23 | B25 | CD3-CD16+CD56+(NK) | CK | 0.5935606 | 4.44E-16 |
| B23 | B59 | CD3-CD16+CD56+(NK) | WBRV | 0.571774066 | 6.44E-15 |
| B23 | B155 | CD3-CD16+CD56+(NK) | IgA | 0.55642635 | 4.64E-14 |
| B23 | B175 | CD3-CD16+CD56+(NK) | ALB-U | 0.538122697 | 4.33E-13 |
| B23 | B158 | CD3-CD16+CD56+(NK) | IgM | -0.492296831 | 6.70E-11 |
| B23 | B60 | CD3-CD16+CD56+(NK) | WBRVI | 0.490164144 | 8.32E-11 |
| B23 | B54 | CD3-CD16+CD56+(NK) | ESR | 0.486953983 | 1.15E-10 |
| B23 | B198 | CD3-CD16+CD56+(NK) | P | -0.484721524 | 1.44E-10 |
| B23 | B185 | CD3-CD16+CD56+(NK) | PT | 0.471649437 | 5.14E-10 |
| B23 | B184 | CD3-CD16+CD56+(NK) | INR | 0.461953764 | 1.28E-09 |
| B23 | B181 | CD3-CD16+CD56+(NK) | APTT | 0.459306489 | 1.63E-09 |
| B23 | B65 | CD3-CD16+CD56+(NK) | WBRSR(mPas_1) | 0.450002865 | 3.78E-09 |

| | | | | | |
|---|---|---|---|---|---|
| B23 | B58 | CD3-CD16+CD56+(NK) | HCT | 0.423424837 | 3.64E-08 |
| B23 | B168 | CD3-CD16+CD56+(NK) | NMP22 | 0.401609664 | 2.03E-07 |
| B23 | B157 | CD3-CD16+CD56+(NK) | IgG | 0.400110805 | 2.28E-07 |
| B23 | B176 | CD3-CD16+CD56+(NK) | Ucr | 0.387593593 | 5.77E-07 |
| B24 | B26 | αHBDH | CKMB | 0.932085975 | 0 |
| B24 | B27 | αHBDH | LDH | 0.623270507 | 0 |
| B24 | B146 | αHBDH | CysC | 0.505558108 | 1.68E-11 |
| B25 | B100 | CK | anti-Ro52 | 0.5935606 | 4.44E-16 |
| B25 | B101 | CK | anti-Sc70 | 0.593560598 | 4.44E-16 |
| B25 | B102 | CK | anti-Sm | 0.5935606 | 4.44E-16 |
| B25 | B103 | CK | anti-SSA/Ro | 0.5935606 | 4.44E-16 |
| B25 | B104 | CK | anti-SSB | 0.593560601 | 4.44E-16 |
| B25 | B105 | CK | anti-U1RNP | 0.5935606 | 4.44E-16 |
| B25 | B106 | CK | anti-P | 0.5935606 | 4.44E-16 |
| B25 | B107 | CK | AnuA | 0.593560613 | 4.44E-16 |
| B25 | B108 | CK | Anti-Cyclin | 0.5935606 | 4.44E-16 |
| B25 | B109 | CK | AMA-M2 | 0.5935606 | 4.44E-16 |
| B25 | B110 | CK | AHA | 0.5935606 | 4.44E-16 |
| B25 | B34 | CK | TK | 0.5935606 | 4.44E-16 |
| B25 | B95 | CK | anti-cenpb | 0.5935606 | 4.44E-16 |
| B25 | B97 | CK | dsDNA | 0.5935606 | 4.44E-16 |
| B25 | B98 | CK | anti-Jo-1 | 0.593560598 | 4.44E-16 |
| B25 | B99 | CK | anti-PM-Scl | 0.593560599 | 4.44E-16 |
| B25 | B176 | CK | Ucr | 0.556651349 | 4.51E-14 |
| B25 | B183 | CK | PTA | -0.555299892 | 5.33E-14 |
| B25 | B199 | CK | Mg | 0.485772964 | 1.29E-10 |
| B25 | B81 | CK | TSGF | 0.465386889 | 9.28E-10 |
| B25 | B156 | CK | IgE | -0.442492479 | 7.31E-09 |
| B25 | B60 | CK | WBRVI | 0.420243874 | 4.72E-08 |
| B25 | B151 | CK | Anti-CCP | 0.418196062 | 5.56E-08 |
| B25 | B69 | CK | PV | -0.415127572 | 7.11E-08 |
| B25 | B155 | CK | IgA | 0.392225399 | 4.11E-07 |
| B25 | B146 | CK | CysC | 0.385339417 | 6.79E-07 |
| B26 | B27 | CKMB | LDH | 0.622732428 | 0 |
| B26 | B146 | CKMB | CysC | 0.4809585 | 2.08E-10 |
| B27 | B146 | LDH | CysC | 0.544013853 | 2.14E-13 |
| B28 | B117 | PRL | TT4 | -0.777777778 | 0 |
| B28 | B180 | PRL | Tf | -0.72659116 | 0 |
| B28 | B92 | PRL | ALDH2-Glu504Glu | -0.905821627 | 0 |
| B28 | B202 | PRL | e-times | -0.564218177 | 1.73E-14 |
| B28 | B178 | PRL | Ig-U | -0.486533633 | 1.20E-10 |
| B28 | B225 | PRL | hPBG | -0.45677344 | 2.05E-09 |
| B29 | B125 | E2 | Cor | -0.745601135 | 0 |
| B29 | B178 | E2 | Ig-U | -0.911407491 | 0 |

| | | | | | |
|---|---|---|---|---|---|
| B29 | B202 | E2 | e-times | -0.637193093 | 0 |
| B29 | B223 | E2 | FH | -0.603451429 | 0 |
| B29 | B225 | E2 | hPBG | -0.837707817 | 0 |
| B29 | B117 | E2 | TT4 | -0.529812943 | 1.14E-12 |
| B3 | B26 | Hcy | CKMB | 0.604655101 | 0 |
| B3 | B146 | Hcy | CysC | 0.59009587 | 4.44E-16 |
| B3 | B84 | Hcy | C-PSA | 0.581424821 | 1.78E-15 |
| B3 | B24 | Hcy | αHBDH | 0.530540732 | 1.05E-12 |
| B3 | B88 | Hcy | CA 242 | -0.514337787 | 6.51E-12 |
| B3 | B89 | Hcy | CA 72-4 | 0.512285366 | 8.15E-12 |
| B3 | B168 | Hcy | NMP22 | 0.473860973 | 4.16E-10 |
| B3 | B82 | Hcy | CEA | -0.414121816 | 7.70E-08 |
| B3 | B100 | Hcy | anti-Ro52 | 0.399542724 | 2.38E-07 |
| B3 | B101 | Hcy | anti-Sc70 | 0.399542723 | 2.38E-07 |
| B3 | B102 | Hcy | anti-Sm | 0.399542724 | 2.38E-07 |
| B3 | B103 | Hcy | anti-SSA/Ro | 0.399542724 | 2.38E-07 |
| B3 | B104 | Hcy | anti-SSB | 0.399542725 | 2.38E-07 |
| B3 | B105 | Hcy | anti-U1RNP | 0.399542724 | 2.38E-07 |
| B3 | B106 | Hcy | anti-P | 0.399542724 | 2.38E-07 |
| B3 | B107 | Hcy | AnuA | 0.399542733 | 2.38E-07 |
| B3 | B108 | Hcy | Anti-Cyclin | 0.399542724 | 2.38E-07 |
| B3 | B109 | Hcy | AMA-M2 | 0.399542724 | 2.38E-07 |
| B3 | B110 | Hcy | AHA | 0.399542724 | 2.38E-07 |
| B3 | B15 | Hcy | CD3-CD19+(B cell) | 0.399542724 | 2.38E-07 |
| B3 | B16 | Hcy | CD3+T | -0.399542724 | 2.38E-07 |
| B3 | B17 | Hcy | CD3+CD16+CD56+NK | 0.399542724 | 2.38E-07 |
| B3 | B18 | Hcy | CD3+CD4+helper(inducer)T | -0.399542724 | 2.38E-07 |
| B3 | B19 | Hcy | CD3+CD4+/CD3+CD8+ | -0.399542724 | 2.38E-07 |
| B3 | B20 | Hcy | CD3+CD4+CD8+ | 0.399542724 | 2.38E-07 |
| B3 | B21 | Hcy | CD3+CD4-CD8- | 0.399542724 | 2.38E-07 |
| B3 | B22 | Hcy | CD3+CD8+(T) | 0.399542724 | 2.38E-07 |
| B3 | B23 | Hcy | CD3-CD16+CD56+(NK) | 0.399542724 | 2.38E-07 |
| B3 | B34 | Hcy | TK | 0.399542724 | 2.38E-07 |
| B3 | B95 | Hcy | anti-cenpb | 0.399542725 | 2.38E-07 |
| B3 | B97 | Hcy | dsDNA | 0.399542724 | 2.38E-07 |
| B3 | B98 | Hcy | anti-Jo-1 | 0.399542723 | 2.38E-07 |
| B3 | B99 | Hcy | anti-PM-Scl | 0.399542724 | 2.38E-07 |
| B3 | B27 | Hcy | LDH | 0.39547999 | 3.23E-07 |
| B30 | B145 | LH | BGP | 0.89597867 | 0 |
| B30 | B171 | LH | EC | 0.666972969 | 0 |
| B30 | B35 | LH | β2-MG | 0.688247202 | 0 |
| B30 | B38 | LH | BAND% | 0.79381612 | 0 |
| B30 | B94 | LH | ALDH2-Lys504Lys | 0.535303379 | 6.04E-13 |
| B30 | B33 | LH | Prog | 0.480761974 | 2.12E-10 |

| | | | | | |
|---|---|---|---|---|---|
| B31 | B117 | FSH | TT4 | -0.635052896 | 0 |
| B31 | B225 | FSH | hPBG | -0.817629818 | 0 |
| B31 | B180 | FSH | Tf | -0.583030591 | 1.33E-15 |
| B31 | B125 | FSH | Cor | -0.471404521 | 5.26E-10 |
| B31 | B92 | FSH | ALDH2-Glu504Glu | -0.416025147 | 6.62E-08 |
| B32 | B145 | Testo | BGP | 0.597614305 | 2.22E-16 |
| B32 | B38 | Testo | BAND% | 0.5143445 | 6.51E-12 |
| B32 | B94 | Testo | ALDH2-Lys504Lys | 0.433554985 | 1.57E-08 |
| B33 | B94 | Prog | ALDH2-Lys504Lys | 0.724791515 | 0 |
| B33 | B37 | Prog | BAND | 0.549442256 | 1.11E-13 |
| B33 | B93 | Prog | ALDH2-Glu504Lys | 0.545132838 | 1.87E-13 |
| B33 | B36 | Prog | V-Lym% | 0.507237499 | 1.40E-11 |
| B34 | B100 | TK | anti-Ro52 | 1 | 0 |
| B34 | B101 | TK | anti-Sc70 | 0.999999997 | 0 |
| B34 | B102 | TK | anti-Sm | 1 | 0 |
| B34 | B105 | TK | anti-U1RNP | 1 | 0 |
| B34 | B109 | TK | AMA-M2 | 1 | 0 |
| B34 | B110 | TK | AHA | 1 | 0 |
| B34 | B151 | TK | Anti-CCP | 0.7048221 | 0 |
| B34 | B156 | TK | IgE | -0.903553226 | 0 |
| B34 | B183 | TK | PTA | -0.935351263 | 0 |
| B34 | B57 | TK | RBCAI | 0.673254212 | 0 |
| B34 | B90 | TK | CYFRA 21-1 | -0.720217082 | 0 |
| B34 | B98 | TK | anti-Jo-1 | 0.999999997 | 0 |
| B34 | B99 | TK | anti-PM-Scl | 0.999999999 | 0 |
| B34 | B59 | TK | WBRV | 0.571774066 | 6.44E-15 |
| B34 | B155 | TK | IgA | 0.55642635 | 4.64E-14 |
| B34 | B175 | TK | ALB-U | 0.538122697 | 4.33E-13 |
| B34 | B158 | TK | IgM | -0.492296831 | 6.70E-11 |
| B34 | B60 | TK | WBRVI | 0.490164144 | 8.32E-11 |
| B34 | B54 | TK | ESR | 0.486953983 | 1.15E-10 |
| B34 | B198 | TK | P | -0.484721524 | 1.44E-10 |
| B34 | B185 | TK | PT | 0.471649437 | 5.14E-10 |
| B34 | B184 | TK | INR | 0.461953764 | 1.28E-09 |
| B34 | B181 | TK | APTT | 0.459306489 | 1.63E-09 |
| B34 | B65 | TK | WBRSR(mPas_1) | 0.450002865 | 3.78E-09 |
| B34 | B58 | TK | HCT | 0.423424837 | 3.64E-08 |
| B34 | B168 | TK | NMP22 | 0.401609664 | 2.03E-07 |
| B34 | B157 | TK | IgG | 0.400110805 | 2.28E-07 |
| B34 | B176 | TK | Ucr | 0.387593593 | 5.77E-07 |
| B35 | B124 | β2-MG | ACTH | 0.951189731 | 0 |
| B35 | B171 | β2-MG | EC | 0.945905303 | 0 |
| B35 | B36 | β2-MG | V-Lym% | 0.818934617 | 0 |
| B35 | B38 | β2-MG | BAND% | 0.967805339 | 0 |

| | | | | | |
|---|---|---|---|---|---|
| B35 | B94 | β2-MG | ALDH2-Lys504Lys | -0.684210526 | 0 |
| B35 | B116 | β2-MG | TT3 | 0.449921271 | 3.80E-09 |
| B35 | B93 | β2-MG | ALDH2-Glu504Lys | 0.390199486 | 4.77E-07 |
| B36 | B124 | V-Lym% | ACTH | 0.778962199 | 0 |
| B36 | B37 | V-Lym% | BAND | -0.738548946 | 0 |
| B36 | B145 | V-Lym% | BGP | -0.583874208 | 1.33E-15 |
| B36 | B38 | V-Lym% | BAND% | 0.502518908 | 2.32E-11 |
| B36 | B171 | V-Lym% | EC | 0.447811076 | 4.59E-09 |
| B37 | B38 | BAND | BAND% | -0.748455199 | 0 |
| B38 | B94 | BAND% | ALDH2-Lys504Lys | 0.718049122 | 0 |
| B38 | B124 | BAND% | ACTH | 0.534522484 | 6.62E-13 |
| B39 | B78 | Other cells | HBcAb | 0.888189409 | 0 |
| B39 | B184 | Other cells | INR | 0.410087608 | 1.06E-07 |
| B4 | B33 | WB-Ca | Prog | 0.638273567 | 0 |
| B4 | B36 | WB-Ca | V-Lym% | 0.994729463 | 0 |
| B4 | B37 | WB-Ca | BAND | 0.606091527 | 0 |
| B4 | B9 | WB-Ca | RH | 0.785714286 | 0 |
| B4 | B94 | WB-Ca | ALDH2-Lys504Lys | 0.55618773 | 4.77E-14 |
| B4 | B171 | WB-Ca | EC | 0.416496564 | 6.37E-08 |
| B40 | B42 | WBC | MON | 0.662710799 | 0 |
| B40 | B52 | WBC | PLT/BPC | 0.396185433 | 3.06E-07 |
| B41 | B42 | MON % | MON | 0.681625394 | 0 |
| B43 | B44 | MCV | MCH | 0.927882323 | 0 |
| B43 | B47 | MCV | RDW | 0.553916452 | 6.35E-14 |
| B43 | B46 | MCV | RBC Count | -0.427523178 | 2.60E-08 |
| B43 | B51 | MCV | Hb | 0.383605289 | 7.69E-07 |
| B44 | B45 | MCH | MCHC | 0.665965631 | 0 |
| B44 | B51 | MCH | Hb | 0.534867766 | 6.36E-13 |
| B45 | B51 | MCHC | Hb | 0.593785636 | 4.44E-16 |
| B45 | B221 | MCHC | Hei | 0.386854209 | 6.08E-07 |
| B46 | B51 | RBC Count | Hb | 0.617771182 | 0 |
| B46 | B216 | RBC Count | BMI | 0.426842781 | 2.75E-08 |
| B46 | B222 | RBC Count | NS | 0.397583531 | 2.76E-07 |
| B46 | B72 | RBC Count | HDL-C | -0.391832687 | 4.23E-07 |
| B47 | B52 | RDW | PLT/BPC | -0.409858048 | 1.08E-07 |
| B48 | B53 | LYM% | GR% | -0.901082556 | 0 |
| B5 | B116 | WB-Mg | TT3 | 0.980580676 | 0 |
| B5 | B30 | WB-Mg | LH | 0.6 | 0 |
| B5 | B7 | WB-Mg | WB-Cop | 0.896717661 | 0 |
| B5 | B171 | WB-Mg | EC | 0.533578375 | 7.39E-13 |
| B5 | B124 | WB-Mg | ACTH | 0.480079359 | 2.27E-10 |
| B5 | B9 | WB-Mg | RH | 0.459619408 | 1.58E-09 |
| B50 | B53 | eo% | GR% | -0.42464886 | 3.30E-08 |
| B51 | B216 | Hb | BMI | 0.60870164 | 0 |

| | | | | | |
|---|---|---|---|---|---|
| B51 | B222 | Hb | NS | 0.596050737 | 2.22E-16 |
| B51 | B221 | Hb | Hei | 0.555502135 | 5.20E-14 |
| B51 | B206 | Hb | Waist | 0.505548499 | 1.68E-11 |
| B51 | B128 | Hb | ALT | 0.47684554 | 3.11E-10 |
| B51 | B192 | Hb | UA | 0.436363781 | 1.24E-08 |
| B51 | B207 | Hb | WHR | 0.43259909 | 1.70E-08 |
| B51 | B215 | Hb | BMI | 0.422192602 | 4.03E-08 |
| B51 | B210 | Hb | Hip | 0.420201984 | 4.73E-08 |
| B51 | B190 | Hb | Cre | 0.413015653 | 8.40E-08 |
| B51 | B163 | Hb | SG | 0.409572191 | 1.10E-07 |
| B51 | B141 | Hb | AST/ALT | -0.388311016 | 5.47E-07 |
| B54 | B59 | ESR | WBRV | 0.653536948 | 0 |
| B54 | B69 | ESR | PV | 0.775779336 | 0 |
| B54 | B182 | ESR | TT | -0.563927758 | 1.78E-14 |
| B54 | B186 | ESR | PFC | 0.532587861 | 8.30E-13 |
| B54 | B100 | ESR | anti-Ro52 | 0.486953983 | 1.15E-10 |
| B54 | B101 | ESR | anti-Sc70 | 0.486953981 | 1.15E-10 |
| B54 | B102 | ESR | anti-Sm | 0.486953982 | 1.15E-10 |
| B54 | B103 | ESR | anti-SSA/Ro | 0.486953983 | 1.15E-10 |
| B54 | B104 | ESR | anti-SSB | 0.486953984 | 1.15E-10 |
| B54 | B105 | ESR | anti-U1RNP | 0.486953982 | 1.15E-10 |
| B54 | B106 | ESR | anti-P | 0.486953983 | 1.15E-10 |
| B54 | B107 | ESR | AnuA | 0.486953993 | 1.15E-10 |
| B54 | B108 | ESR | Anti-Cyclin | 0.486953983 | 1.15E-10 |
| B54 | B109 | ESR | AMA-M2 | 0.486953983 | 1.15E-10 |
| B54 | B110 | ESR | AHA | 0.486953982 | 1.15E-10 |
| B54 | B95 | ESR | anti-cenpb | 0.486953983 | 1.15E-10 |
| B54 | B97 | ESR | dsDNA | 0.486953983 | 1.15E-10 |
| B54 | B98 | ESR | anti-Jo-1 | 0.486953981 | 1.15E-10 |
| B54 | B99 | ESR | anti-PM-Scl | 0.486953982 | 1.15E-10 |
| B54 | B198 | ESR | P | -0.413539252 | 8.06E-08 |
| B54 | B175 | ESR | ALB-U | 0.40546693 | 1.51E-07 |
| B54 | B157 | ESR | IgG | 0.396311532 | 3.03E-07 |
| B54 | B156 | ESR | IgE | -0.395455498 | 3.23E-07 |
| B54 | B74 | ESR | ApoA1 | 0.391156952 | 4.44E-07 |
| B55 | B56 | TK | IR | 0.754875937 | 0 |
| B55 | B59 | TK | WBRV | 0.642649442 | 0 |
| B55 | B61 | TK | WBHSRV | 0.725202895 | 0 |
| B55 | B62 | TK | WBHSRVI | 0.648399763 | 0 |
| B55 | B63 | TK | WBRSR(mPas_200) | 0.635797941 | 0 |
| B55 | B64 | TK | WBRSR(mPas_5) | 0.615046872 | 0 |
| B55 | B66 | TK | WBRSR(mPas_100) | 0.631759993 | 0 |
| B55 | B67 | TK | WBRSR(mPas_50) | 0.631564019 | 0 |
| B55 | B60 | TK | WBRVI | 0.589817092 | 4.44E-16 |

| | | | | | |
|---|---|---|---|---|---|
| B55 | B65 | TK | WBRSR(mPas_1) | 0.593692492 | 4.44E-16 |
| B55 | B57 | TK | RBCAI | -0.478585737 | 2.63E-10 |
| B55 | B58 | TK | HCT | 0.469338227 | 6.40E-10 |
| B55 | B82 | TK | CEA | -0.448671862 | 4.25E-09 |
| B55 | B85 | TK | SCC | 0.409745881 | 1.09E-07 |
| B56 | B181 | IR | APTT | 0.631205858 | 0 |
| B56 | B58 | IR | HCT | 0.928594544 | 0 |
| B56 | B59 | IR | WBRV | 0.791785242 | 0 |
| B56 | B60 | IR | WBRVI | 0.955359382 | 0 |
| B56 | B61 | IR | WBHSRV | 0.950615995 | 0 |
| B56 | B62 | IR | WBHSRVI | 0.988681151 | 0 |
| B56 | B63 | IR | WBRSR(mPas_200) | 0.944744621 | 0 |
| B56 | B64 | IR | WBRSR(mPas_5) | 0.94920596 | 0 |
| B56 | B65 | IR | WBRSR(mPas_1) | 0.938865572 | 0 |
| B56 | B66 | IR | WBRSR(mPas_100) | 0.951796663 | 0 |
| B56 | B67 | IR | WBRSR(mPas_50) | 0.952262054 | 0 |
| B56 | B185 | IR | PT | 0.556329141 | 4.71E-14 |
| B56 | B57 | IR | RBCAI | -0.486331958 | 1.22E-10 |
| B56 | B182 | IR | TT | 0.439752697 | 9.26E-09 |
| B56 | B186 | IR | PFC | -0.409324358 | 1.12E-07 |
| B56 | B183 | IR | PTA | -0.399697336 | 2.35E-07 |
| B57 | B100 | RBCAI | anti-Ro52 | 0.673254212 | 0 |
| B57 | B101 | RBCAI | anti-Sc70 | 0.67325421 | 0 |
| B57 | B102 | RBCAI | anti-Sm | 0.673254212 | 0 |
| B57 | B103 | RBCAI | anti-SSA/Ro | 0.673254212 | 0 |
| B57 | B104 | RBCAI | anti-SSB | 0.673254213 | 0 |
| B57 | B105 | RBCAI | anti-U1RNP | 0.673254212 | 0 |
| B57 | B106 | RBCAI | anti-P | 0.673254212 | 0 |
| B57 | B107 | RBCAI | AnuA | 0.673254228 | 0 |
| B57 | B108 | RBCAI | Anti-Cyclin | 0.673254212 | 0 |
| B57 | B109 | RBCAI | AMA-M2 | 0.673254212 | 0 |
| B57 | B110 | RBCAI | AHA | 0.673254212 | 0 |
| B57 | B95 | RBCAI | anti-cenpb | 0.673254212 | 0 |
| B57 | B97 | RBCAI | dsDNA | 0.673254212 | 0 |
| B57 | B98 | RBCAI | anti-Jo-1 | 0.67325421 | 0 |
| B57 | B99 | RBCAI | anti-PM-Scl | 0.673254212 | 0 |
| B57 | B156 | RBCAI | IgE | -0.580427933 | 2.00E-15 |
| B57 | B90 | RBCAI | CYFRA 21-1 | -0.579403025 | 2.22E-15 |
| B57 | B61 | RBCAI | WBHSRV | -0.561143131 | 2.55E-14 |
| B57 | B183 | RBCAI | PTA | -0.554065171 | 6.24E-14 |
| B57 | B66 | RBCAI | WBRSR(mPas_100) | -0.533783385 | 7.22E-13 |
| B57 | B63 | RBCAI | WBRSR(mPas_200) | -0.522143781 | 2.74E-12 |
| B57 | B67 | RBCAI | WBRSR(mPas_50) | -0.520820913 | 3.18E-12 |
| B57 | B151 | RBCAI | Anti-CCP | 0.470864386 | 5.53E-10 |

| | | | | | |
|---|---|---|---|---|---|
| B57 | B62 | RBCAI | WBHSRVI | -0.467385839 | 7.69E-10 |
| B57 | B182 | RBCAI | TT | -0.44356452 | 6.66E-09 |
| B57 | B64 | RBCAI | WBRSR(mPas_5) | -0.43388162 | 1.53E-08 |
| B57 | B186 | RBCAI | PFC | 0.421300532 | 4.33E-08 |
| B57 | B157 | RBCAI | IgG | 0.396282707 | 3.04E-07 |
| B58 | B181 | HCT | APTT | 0.801570407 | 0 |
| B58 | B185 | HCT | PT | 0.712066125 | 0 |
| B58 | B59 | HCT | WBRV | 0.761164617 | 0 |
| B58 | B60 | HCT | WBRVI | 0.979121342 | 0 |
| B58 | B61 | HCT | WBHSRV | 0.879164402 | 0 |
| B58 | B62 | HCT | WBHSRVI | 0.969210789 | 0 |
| B58 | B63 | HCT | WBRSR(mPas_200) | 0.921718972 | 0 |
| B58 | B64 | HCT | WBRSR(mPas_5) | 0.947871464 | 0 |
| B58 | B65 | HCT | WBRSR(mPas_1) | 0.955395958 | 0 |
| B58 | B66 | HCT | WBRSR(mPas_100) | 0.929961591 | 0 |
| B58 | B67 | HCT | WBRSR(mPas_50) | 0.932370716 | 0 |
| B58 | B183 | HCT | PTA | -0.567117229 | 1.18E-14 |
| B58 | B182 | HCT | TT | 0.506162736 | 1.58E-11 |
| B58 | B186 | HCT | PFC | -0.470353637 | 5.81E-10 |
| B58 | B156 | HCT | IgE | -0.444802333 | 5.98E-09 |
| B58 | B100 | HCT | anti-Ro52 | 0.423424837 | 3.64E-08 |
| B58 | B101 | HCT | anti-Sc70 | 0.423424835 | 3.64E-08 |
| B58 | B102 | HCT | anti-Sm | 0.423424836 | 3.64E-08 |
| B58 | B103 | HCT | anti-SSA/Ro | 0.423424837 | 3.64E-08 |
| B58 | B104 | HCT | anti-SSB | 0.423424838 | 3.64E-08 |
| B58 | B105 | HCT | anti-U1RNP | 0.423424837 | 3.64E-08 |
| B58 | B106 | HCT | anti-P | 0.423424837 | 3.64E-08 |
| B58 | B107 | HCT | AnuA | 0.423424846 | 3.64E-08 |
| B58 | B108 | HCT | Anti-Cyclin | 0.423424837 | 3.64E-08 |
| B58 | B109 | HCT | AMA-M2 | 0.423424837 | 3.64E-08 |
| B58 | B110 | HCT | AHA | 0.423424837 | 3.64E-08 |
| B58 | B95 | HCT | anti-cenpb | 0.423424837 | 3.64E-08 |
| B58 | B97 | HCT | dsDNA | 0.423424837 | 3.64E-08 |
| B58 | B98 | HCT | anti-Jo-1 | 0.423424836 | 3.64E-08 |
| B58 | B99 | HCT | anti-PM-Scl | 0.423424836 | 3.64E-08 |
| B58 | B158 | HCT | IgM | -0.4200254 | 4.80E-08 |
| B59 | B60 | WBRV | WBRVI | 0.785802654 | 0 |
| B59 | B61 | WBRV | WBHSRV | 0.883529281 | 0 |
| B59 | B62 | WBRV | WBHSRVI | 0.766621119 | 0 |
| B59 | B63 | WBRV | WBRSR(mPas_200) | 0.875785951 | 0 |
| B59 | B64 | WBRV | WBRSR(mPas_5) | 0.892262875 | 0 |
| B59 | B65 | WBRV | WBRSR(mPas_1) | 0.90995803 | 0 |
| B59 | B66 | WBRV | WBRSR(mPas_100) | 0.856566108 | 0 |
| B59 | B67 | WBRV | WBRSR(mPas_50) | 0.862799664 | 0 |

| | | | | | |
|---|---|---|---|---|---|
| B59 | B183 | WBRV | PTA | -0.581174313 | 1.78E-15 |
| B59 | B100 | WBRV | anti-Ro52 | 0.571774066 | 6.44E-15 |
| B59 | B101 | WBRV | anti-Sc70 | 0.571774064 | 6.44E-15 |
| B59 | B102 | WBRV | anti-Sm | 0.571774066 | 6.44E-15 |
| B59 | B103 | WBRV | anti-SSA/Ro | 0.571774066 | 6.44E-15 |
| B59 | B104 | WBRV | anti-SSB | 0.571774068 | 6.44E-15 |
| B59 | B105 | WBRV | anti-U1RNP | 0.571774066 | 6.44E-15 |
| B59 | B106 | WBRV | anti-P | 0.571774066 | 6.44E-15 |
| B59 | B107 | WBRV | AnuA | 0.571774077 | 6.44E-15 |
| B59 | B108 | WBRV | Anti-Cyclin | 0.571774066 | 6.44E-15 |
| B59 | B109 | WBRV | AMA-M2 | 0.571774066 | 6.44E-15 |
| B59 | B110 | WBRV | AHA | 0.571774066 | 6.44E-15 |
| B59 | B95 | WBRV | anti-cenpb | 0.571774067 | 6.44E-15 |
| B59 | B97 | WBRV | dsDNA | 0.571774066 | 6.44E-15 |
| B59 | B98 | WBRV | anti-Jo-1 | 0.571774065 | 6.44E-15 |
| B59 | B99 | WBRV | anti-PM-Scl | 0.571774066 | 6.44E-15 |
| B59 | B156 | WBRV | IgE | -0.533428991 | 7.52E-13 |
| B59 | B69 | WBRV | PV | 0.508980562 | 1.17E-11 |
| B59 | B181 | WBRV | APTT | 0.426967726 | 2.72E-08 |
| B59 | B185 | WBRV | PT | 0.40828593 | 1.22E-07 |
| B59 | B151 | WBRV | Anti-CCP | 0.404533446 | 1.63E-07 |
| B6 | B180 | WB-Fe | Tf | -0.637792804 | 0 |
| B6 | B29 | WB-Fe | E2 | -0.561951487 | 2.31E-14 |
| B6 | B28 | WB-Fe | PRL | -0.510688231 | 9.69E-12 |
| B60 | B181 | WBRVI | APTT | 0.78301838 | 0 |
| B60 | B183 | WBRVI | PTA | -0.615571695 | 0 |
| B60 | B185 | WBRVI | PT | 0.70219741 | 0 |
| B60 | B61 | WBRVI | WBHSRV | 0.875192264 | 0 |
| B60 | B62 | WBRVI | WBHSRVI | 0.971532803 | 0 |
| B60 | B63 | WBRVI | WBRSR(mPas_200) | 0.899303498 | 0 |
| B60 | B64 | WBRVI | WBRSR(mPas_5) | 0.932383701 | 0 |
| B60 | B65 | WBRVI | WBRSR(mPas_1) | 0.946566336 | 0 |
| B60 | B66 | WBRVI | WBRSR(mPas_100) | 0.906716902 | 0 |
| B60 | B67 | WBRVI | WBRSR(mPas_50) | 0.910525265 | 0 |
| B60 | B156 | WBRVI | IgE | -0.500143067 | 2.98E-11 |
| B60 | B100 | WBRVI | anti-Ro52 | 0.490164144 | 8.32E-11 |
| B60 | B101 | WBRVI | anti-Sc70 | 0.490164143 | 8.32E-11 |
| B60 | B102 | WBRVI | anti-Sm | 0.490164144 | 8.32E-11 |
| B60 | B103 | WBRVI | anti-SSA/Ro | 0.490164144 | 8.32E-11 |
| B60 | B104 | WBRVI | anti-SSB | 0.490164146 | 8.32E-11 |
| B60 | B105 | WBRVI | anti-U1RNP | 0.490164144 | 8.32E-11 |
| B60 | B106 | WBRVI | anti-P | 0.490164145 | 8.32E-11 |
| B60 | B107 | WBRVI | AnuA | 0.490164156 | 8.32E-11 |
| B60 | B108 | WBRVI | Anti-Cyclin | 0.490164144 | 8.32E-11 |

| | | | | | |
|---|---|---|---|---|---|
| B60 | B109 | WBRVI | AMA-M2 | 0.490164144 | 8.32E-11 |
| B60 | B110 | WBRVI | AHA | 0.490164144 | 8.32E-11 |
| B60 | B95 | WBRVI | anti-cenpb | 0.490164145 | 8.32E-11 |
| B60 | B97 | WBRVI | dsDNA | 0.490164145 | 8.32E-11 |
| B60 | B98 | WBRVI | anti-Jo-1 | 0.490164143 | 8.32E-11 |
| B60 | B99 | WBRVI | anti-PM-Scl | 0.490164144 | 8.32E-11 |
| B60 | B182 | WBRVI | TT | 0.439252619 | 9.67E-09 |
| B60 | B158 | WBRVI | IgM | -0.435136474 | 1.37E-08 |
| B60 | B186 | WBRVI | PFC | -0.407031957 | 1.34E-07 |
| B60 | B184 | WBRVI | INR | 0.380086499 | 9.88E-07 |
| B61 | B62 | WBHSRV | WBHSRVI | 0.938423877 | 0 |
| B61 | B63 | WBHSRV | WBRSR(mPas_200) | 0.991329285 | 0 |
| B61 | B64 | WBHSRV | WBRSR(mPas_5) | 0.97941114 | 0 |
| B61 | B65 | WBHSRV | WBRSR(mPas_1) | 0.960754975 | 0 |
| B61 | B66 | WBHSRV | WBRSR(mPas_100) | 0.989051386 | 0 |
| B61 | B67 | WBHSRV | WBRSR(mPas_50) | 0.988877746 | 0 |
| B61 | B181 | WBHSRV | APTT | 0.484147635 | 1.52E-10 |
| B61 | B185 | WBHSRV | PT | 0.428670374 | 2.36E-08 |
| B61 | B69 | WBHSRV | PV | 0.414931192 | 7.22E-08 |
| B62 | B181 | WBHSRVI | APTT | 0.697129785 | 0 |
| B62 | B185 | WBHSRVI | PT | 0.612879625 | 0 |
| B62 | B63 | WBHSRVI | WBRSR(mPas_200) | 0.951716347 | 0 |
| B62 | B64 | WBHSRVI | WBRSR(mPas_5) | 0.960325259 | 0 |
| B62 | B65 | WBHSRVI | WBRSR(mPas_1) | 0.95186254 | 0 |
| B62 | B66 | WBHSRVI | WBRSR(mPas_100) | 0.961165043 | 0 |
| B62 | B67 | WBHSRVI | WBRSR(mPas_50) | 0.961520804 | 0 |
| B62 | B182 | WBHSRVI | TT | 0.498003339 | 3.72E-11 |
| B62 | B186 | WBHSRVI | PFC | -0.463805643 | 1.08E-09 |
| B62 | B183 | WBHSRVI | PTA | -0.427831783 | 2.54E-08 |
| B63 | B64 | WBRSR(mPas_200) | WBRSR(mPas_5) | 0.994351325 | 0 |
| B63 | B65 | WBRSR(mPas_200) | WBRSR(mPas_1) | 0.980095382 | 0 |
| B63 | B66 | WBRSR(mPas_200) | WBRSR(mPas_100) | 0.999001218 | 0 |
| B63 | B67 | WBRSR(mPas_200) | WBRSR(mPas_50) | 0.999186937 | 0 |
| B63 | B181 | WBRSR(mPas_200) | APTT | 0.558198963 | 3.71E-14 |
| B63 | B185 | WBRSR(mPas_200) | PT | 0.49416294 | 5.53E-11 |
| B63 | B69 | WBRSR(mPas_200) | PV | 0.404772855 | 1.60E-07 |
| B64 | B181 | WBRSR(mPas_5) | APTT | 0.611871437 | 0 |
| B64 | B65 | WBRSR(mPas_5) | WBRSR(mPas_1) | 0.995399827 | 0 |
| B64 | B66 | WBRSR(mPas_5) | WBRSR(mPas_100) | 0.993341621 | 0 |
| B64 | B67 | WBRSR(mPas_5) | WBRSR(mPas_50) | 0.99499354 | 0 |
| B64 | B185 | WBRSR(mPas_5) | PT | 0.547257711 | 1.44E-13 |
| B64 | B183 | WBRSR(mPas_5) | PTA | -0.465573535 | 9.12E-10 |
| B65 | B181 | WBRSR(mPas_1) | APTT | 0.645768069 | 0 |
| B65 | B66 | WBRSR(mPas_1) | WBRSR(mPas_100) | 0.977776782 | 0 |

| | | | | | |
|---|---|---|---|---|---|
| B65 | B67 | WBRSR(mPas_1) | WBRSR(mPas_50) | 0.980873646 | 0 |
| B65 | B185 | WBRSR(mPas_1) | PT | 0.583001602 | 1.33E-15 |
| B65 | B183 | WBRSR(mPas_1) | PTA | -0.544523134 | 2.01E-13 |
| B65 | B156 | WBRSR(mPas_1) | IgE | -0.451685312 | 3.25E-09 |
| B65 | B100 | WBRSR(mPas_1) | anti-Ro52 | 0.450002865 | 3.78E-09 |
| B65 | B101 | WBRSR(mPas_1) | anti-Sc70 | 0.450002864 | 3.78E-09 |
| B65 | B102 | WBRSR(mPas_1) | anti-Sm | 0.450002865 | 3.78E-09 |
| B65 | B103 | WBRSR(mPas_1) | anti-SSA/Ro | 0.450002865 | 3.78E-09 |
| B65 | B104 | WBRSR(mPas_1) | anti-SSB | 0.450002866 | 3.78E-09 |
| B65 | B105 | WBRSR(mPas_1) | anti-U1RNP | 0.450002865 | 3.78E-09 |
| B65 | B106 | WBRSR(mPas_1) | anti-P | 0.450002865 | 3.78E-09 |
| B65 | B107 | WBRSR(mPas_1) | AnuA | 0.450002874 | 3.78E-09 |
| B65 | B108 | WBRSR(mPas_1) | Anti-Cyclin | 0.450002865 | 3.78E-09 |
| B65 | B109 | WBRSR(mPas_1) | AMA-M2 | 0.450002865 | 3.78E-09 |
| B65 | B110 | WBRSR(mPas_1) | AHA | 0.450002865 | 3.78E-09 |
| B65 | B95 | WBRSR(mPas_1) | anti-cenpb | 0.450002866 | 3.78E-09 |
| B65 | B97 | WBRSR(mPas_1) | dsDNA | 0.450002865 | 3.78E-09 |
| B65 | B98 | WBRSR(mPas_1) | anti-Jo-1 | 0.450002864 | 3.78E-09 |
| B65 | B99 | WBRSR(mPas_1) | anti-PM-Scl | 0.450002865 | 3.78E-09 |
| B66 | B67 | WBRSR(mPas_100) | WBRSR(mPas_50) | 0.999876245 | 0 |
| B66 | B181 | WBRSR(mPas_100) | APTT | 0.567743662 | 1.09E-14 |
| B66 | B185 | WBRSR(mPas_100) | PT | 0.501063211 | 2.70E-11 |
| B66 | B182 | WBRSR(mPas_100) | TT | 0.38978406 | 4.91E-07 |
| B67 | B181 | WBRSR(mPas_50) | APTT | 0.572046227 | 6.22E-15 |
| B67 | B185 | WBRSR(mPas_50) | PT | 0.505954957 | 1.61E-11 |
| B67 | B182 | WBRSR(mPas_50) | TT | 0.383312356 | 7.85E-07 |
| B69 | B176 | PV | Ucr | -0.437773342 | 1.10E-08 |
| B69 | B74 | PV | ApoA1 | 0.400587806 | 2.20E-07 |
| B7 | B124 | WB-Cop | ACTH | 0.978399181 | 0 |
| B7 | B33 | WB-Cop | Prog | 0.703849392 | 0 |
| B7 | B94 | WB-Cop | ALDH2-Lys504Lys | 0.844926112 | 0 |
| B7 | B36 | WB-Cop | V-Lym% | 0.571601983 | 6.66E-15 |
| B7 | B30 | WB-Cop | LH | 0.480384461 | 2.21E-10 |
| B7 | B37 | WB-Cop | BAND | 0.400320385 | 2.24E-07 |
| B70 | B77 | LDL-C | TC | 0.787478139 | 0 |
| B71 | B140 | TG | r-GT | 0.474847367 | 3.78E-10 |
| B71 | B72 | TG | HDL-C | -0.462187317 | 1.25E-09 |
| B71 | B207 | TG | WHR | 0.413425066 | 8.14E-08 |
| B71 | B77 | TG | TC | 0.381330446 | 9.04E-07 |
| B72 | B216 | HDL-C | BMI | -0.517135185 | 4.79E-12 |
| B72 | B206 | HDL-C | Waist | -0.476238126 | 3.30E-10 |
| B72 | B222 | HDL-C | NS | -0.453377628 | 2.79E-09 |
| B72 | B215 | HDL-C | BMI | -0.443219935 | 6.86E-09 |
| B72 | B192 | HDL-C | UA | -0.431655671 | 1.84E-08 |

| | | | | | |
|---|---|---|---|---|---|
| B72 | B141 | HDL-C | AST/ALT | 0.423385755 | 3.65E-08 |
| B72 | B207 | HDL-C | WHR | -0.409716864 | 1.09E-07 |
| B72 | B210 | HDL-C | Hip | -0.38937128 | 5.06E-07 |
| B72 | B190 | HDL-C | Cre | -0.386151352 | 6.40E-07 |
| B72 | B128 | HDL-C | ALT | -0.382198207 | 8.50E-07 |
| B73 | B193 | ApoA | K | -0.544186079 | 2.10E-13 |
| B73 | B134 | ApoA | CHE | 0.497565324 | 3.89E-11 |
| B73 | B75 | ApoA | ApoA1/B | 0.449204682 | 4.05E-09 |
| B73 | B114 | ApoA | TSH TT3 TT4(TT4) | 0.446470887 | 5.16E-09 |
| B74 | B198 | ApoA1 | P | -0.622342131 | 0 |
| B74 | B135 | ApoA1 | TBA | 0.595853834 | 2.22E-16 |
| B74 | B79 | ApoA1 | hb V -dna | -0.566667787 | 1.24E-14 |
| B74 | B75 | ApoA1 | ApoA1/B | 0.534624838 | 6.54E-13 |
| B74 | B196 | ApoA1 | TCO2 | 0.502442416 | 2.34E-11 |
| B74 | B194 | ApoA1 | Cl | -0.449298461 | 4.02E-09 |
| B74 | B150 | ApoA1 | ALP | 0.388595461 | 5.36E-07 |
| B75 | B76 | ApoA1/B | ApoB | -0.800440872 | 0 |
| B75 | B198 | ApoA1/B | P | -0.551897781 | 8.17E-14 |
| B75 | B135 | ApoA1/B | TBA | 0.509493311 | 1.10E-11 |
| B75 | B196 | ApoA1/B | TCO2 | 0.45010983 | 3.74E-09 |
| B75 | B199 | ApoA1/B | Mg | 0.437005778 | 1.17E-08 |
| B75 | B197 | ApoA1/B | AG | -0.415681505 | 6.80E-08 |
| B76 | B199 | ApoB | Mg | -0.62222023 | 0 |
| B76 | B197 | ApoB | AG | 0.565743498 | 1.42E-14 |
| B76 | B138 | ApoB | PAB | 0.480328932 | 2.22E-10 |
| B76 | B87 | ApoB | NSE | 0.447438573 | 4.74E-09 |
| B76 | B150 | ApoB | ALP | 0.393469514 | 3.75E-07 |
| B76 | B85 | ApoB | SCC | -0.380725394 | 9.44E-07 |
| B79 | B135 | hb V -dna | TBA | -0.454374906 | 2.55E-09 |
| B79 | B112 | hb V -dna | FT4 | -0.387441038 | 5.83E-07 |
| B8 | B125 | WB-Zinc | Cor | 0.603022689 | 0 |
| B8 | B180 | WB-Zinc | Tf | -0.824322106 | 0 |
| B8 | B29 | WB-Zinc | E2 | -0.691714464 | 0 |
| B8 | B96 | WB-Zinc | dsDNA (IIF) | 0.639602149 | 0 |
| B82 | B86 | CEA | TBhCG | -0.598505808 | 0 |
| B82 | B87 | CEA | NSE | -0.639696363 | 0 |
| B82 | B89 | CEA | CA 72-4 | -0.790443854 | 0 |
| B82 | B197 | CEA | AG | -0.525462331 | 1.88E-12 |
| B82 | B84 | CEA | C-PSA | -0.506954954 | 1.45E-11 |
| B82 | B115 | CEA | TSH TT3 TT4(TT3) | -0.439790272 | 9.23E-09 |
| B82 | B88 | CEA | CA 242 | 0.412229427 | 8.94E-08 |
| B83 | B90 | AFP | CYFRA 21-1 | 0.552268908 | 7.79E-14 |
| B83 | B200 | AFP | OP | 0.530158413 | 1.10E-12 |
| B83 | B195 | AFP | Na | 0.511071315 | 9.30E-12 |

| | | | | | |
|---|---|---|---|---|---|
| B84 | B85 | C-PSA | SCC | -0.637762019 | 0 |
| B84 | B86 | C-PSA | TBhCG | 0.60622951 | 0 |
| B84 | B88 | C-PSA | CA 242 | -0.922835511 | 0 |
| B84 | B89 | C-PSA | CA 72-4 | 0.790532091 | 0 |
| B84 | B197 | C-PSA | AG | 0.582805651 | 1.33E-15 |
| B84 | B168 | C-PSA | NMP22 | 0.568536022 | 9.77E-15 |
| B84 | B200 | C-PSA | OP | 0.444693619 | 6.03E-09 |
| B85 | B197 | SCC | AG | -0.646464351 | 0 |
| B85 | B88 | SCC | CA 242 | 0.816293757 | 0 |
| B85 | B115 | SCC | TSH TT3 TT4(TT3) | -0.537041643 | 4.92E-13 |
| B85 | B194 | SCC | Cl | 0.440621886 | 8.59E-09 |
| B86 | B168 | TBhCG | NMP22 | 0.600219258 | 0 |
| B86 | B89 | TBhCG | CA 72-4 | 0.925788656 | 0 |
| B86 | B90 | TBhCG | CYFRA 21-1 | 0.599785707 | 0 |
| B86 | B200 | TBhCG | OP | 0.522105903 | 2.75E-12 |
| B86 | B195 | TBhCG | Na | 0.517652052 | 4.52E-12 |
| B86 | B88 | TBhCG | CA 242 | -0.435820159 | 1.30E-08 |
| B87 | B115 | NSE | TSH TT3 TT4(TT3) | 0.625799513 | 0 |
| B87 | B197 | NSE | AG | 0.555096708 | 5.48E-14 |
| B87 | B194 | NSE | Cl | -0.437011322 | 1.17E-08 |
| B88 | B197 | CA 242 | AG | -0.748593379 | 0 |
| B88 | B89 | CA 242 | CA 72-4 | -0.619992658 | 0 |
| B88 | B200 | CA 242 | OP | -0.525664669 | 1.84E-12 |
| B88 | B168 | CA 242 | NMP22 | -0.482809704 | 1.74E-10 |
| B88 | B194 | CA 242 | Cl | 0.413401505 | 8.15E-08 |
| B88 | B195 | CA 242 | Na | -0.395165241 | 3.30E-07 |
| B89 | B168 | CA 72-4 | NMP22 | 0.569013526 | 9.33E-15 |
| B89 | B90 | CA 72-4 | CYFRA 21-1 | 0.484806446 | 1.42E-10 |
| B89 | B197 | CA 72-4 | AG | 0.462778319 | 1.18E-09 |
| B89 | B200 | CA 72-4 | OP | 0.448130234 | 4.46E-09 |
| B89 | B195 | CA 72-4 | Na | 0.38574515 | 6.59E-07 |
| B9 | B124 | RH | ACTH | 0.6172134 | 0 |
| B9 | B36 | RH | V-Lym% | 0.957427108 | 0 |
| B9 | B94 | RH | ALDH2-Lys504Lys | 0.405553553 | 1.50E-07 |
| B90 | B100 | CYFRA 21-1 | anti-Ro52 | -0.720217082 | 0 |
| B90 | B101 | CYFRA 21-1 | anti-Sc70 | -0.72021708 | 0 |
| B90 | B102 | CYFRA 21-1 | anti-Sm | -0.720217082 | 0 |
| B90 | B103 | CYFRA 21-1 | anti-SSA/Ro | -0.720217082 | 0 |
| B90 | B104 | CYFRA 21-1 | anti-SSB | -0.720217084 | 0 |
| B90 | B105 | CYFRA 21-1 | anti-U1RNP | -0.720217082 | 0 |
| B90 | B106 | CYFRA 21-1 | anti-P | -0.720217082 | 0 |
| B90 | B107 | CYFRA 21-1 | AnuA | -0.720217097 | 0 |
| B90 | B108 | CYFRA 21-1 | Anti-Cyclin | -0.720217082 | 0 |
| B90 | B109 | CYFRA 21-1 | AMA-M2 | -0.720217082 | 0 |

| | | | | | |
|---|---|---|---|---|---|
| B90 | B110 | CYFRA 21-1 | AHA | -0.720217082 | 0 |
| B90 | B156 | CYFRA 21-1 | IgE | 0.651658316 | 0 |
| B90 | B183 | CYFRA 21-1 | PTA | 0.66976743 | 0 |
| B90 | B95 | CYFRA 21-1 | anti-cenpb | -0.720217083 | 0 |
| B90 | B97 | CYFRA 21-1 | dsDNA | -0.720217082 | 0 |
| B90 | B98 | CYFRA 21-1 | anti-Jo-1 | -0.72021708 | 0 |
| B90 | B99 | CYFRA 21-1 | anti-PM-Scl | -0.720217081 | 0 |
| B90 | B151 | CYFRA 21-1 | Anti-CCP | -0.513118653 | 7.44E-12 |
| B90 | B155 | CYFRA 21-1 | IgA | -0.409556575 | 1.10E-07 |
| B90 | B175 | CYFRA 21-1 | ALB-U | -0.404773635 | 1.60E-07 |
| B91 | B130 | TP | GLB | 0.769458428 | 0 |
| B91 | B127 | TP | A-G ratio | -0.493930493 | 5.66E-11 |
| B91 | B126 | TP | Alb | 0.433703152 | 1.55E-08 |
| B92 | B117 | ALDH2-Glu504Glu | TT4 | -0.603881085 | 0 |
| B92 | B180 | ALDH2-Glu504Glu | Tf | -0.944689404 | 0 |
| B92 | B202 | ALDH2-Glu504Glu | e-times | -0.877579922 | 0 |
| B93 | B145 | ALDH2-Glu504Lys | BGP | 0.657375735 | 0 |
| B93 | B171 | ALDH2-Glu504Lys | EC | 0.870863572 | 0 |
| B94 | B171 | ALDH2-Lys504Lys | EC | 0.612056372 | 0 |
| B95 | B101 | anti-cenpb | anti-Sc70 | 0.999999996 | 0 |
| B95 | B106 | anti-cenpb | anti-P | 0.999999998 | 0 |
| B95 | B151 | anti-cenpb | Anti-CCP | 0.704822101 | 0 |
| B95 | B156 | anti-cenpb | IgE | -0.903553227 | 0 |
| B95 | B183 | anti-cenpb | PTA | -0.935351264 | 0 |
| B95 | B98 | anti-cenpb | anti-Jo-1 | 0.999999998 | 0 |
| B95 | B155 | anti-cenpb | IgA | 0.556426351 | 4.64E-14 |
| B95 | B175 | anti-cenpb | ALB-U | 0.538122698 | 4.33E-13 |
| B95 | B158 | anti-cenpb | IgM | -0.492296832 | 6.70E-11 |
| B95 | B198 | anti-cenpb | P | -0.484721525 | 1.44E-10 |
| B95 | B185 | anti-cenpb | PT | 0.471649437 | 5.14E-10 |
| B95 | B184 | anti-cenpb | INR | 0.461953765 | 1.28E-09 |
| B95 | B181 | anti-cenpb | APTT | 0.459306489 | 1.63E-09 |
| B95 | B168 | anti-cenpb | NMP22 | 0.401609665 | 2.03E-07 |
| B95 | B157 | anti-cenpb | IgG | 0.400110805 | 2.28E-07 |
| B95 | B176 | anti-cenpb | Ucr | 0.387593593 | 5.77E-07 |
| B97 | B101 | dsDNA | anti-Sc70 | 0.999999998 | 0 |
| B97 | B109 | dsDNA | AMA-M2 | 1 | 0 |
| B97 | B151 | dsDNA | Anti-CCP | 0.7048221 | 0 |
| B97 | B156 | dsDNA | IgE | -0.903553226 | 0 |
| B97 | B183 | dsDNA | PTA | -0.935351263 | 0 |
| B97 | B98 | dsDNA | anti-Jo-1 | 0.999999998 | 0 |
| B97 | B155 | dsDNA | IgA | 0.55642635 | 4.64E-14 |
| B97 | B175 | dsDNA | ALB-U | 0.538122697 | 4.33E-13 |
| B97 | B158 | dsDNA | IgM | -0.492296831 | 6.70E-11 |

| | | | | | |
|---|---|---|---|---|---|
| B97 | B198 | dsDNA | P | -0.484721524 | 1.44E-10 |
| B97 | B185 | dsDNA | PT | 0.471649437 | 5.14E-10 |
| B97 | B184 | dsDNA | INR | 0.461953764 | 1.28E-09 |
| B97 | B181 | dsDNA | APTT | 0.459306489 | 1.63E-09 |
| B97 | B168 | dsDNA | NMP22 | 0.401609664 | 2.03E-07 |
| B97 | B157 | dsDNA | IgG | 0.400110805 | 2.28E-07 |
| B97 | B176 | dsDNA | Ucr | 0.387593593 | 5.77E-07 |
| B98 | B100 | anti-Jo-1 | anti-Ro52 | 0.999999997 | 0 |
| B98 | B101 | anti-Jo-1 | anti-Sc70 | 1 | 0 |
| B98 | B102 | anti-Jo-1 | anti-Sm | 0.999999999 | 0 |
| B98 | B103 | anti-Jo-1 | anti-SSA/Ro | 0.999999997 | 0 |
| B98 | B104 | anti-Jo-1 | anti-SSB | 0.999999998 | 0 |
| B98 | B105 | anti-Jo-1 | anti-U1RNP | 0.999999998 | 0 |
| B98 | B106 | anti-Jo-1 | anti-P | 0.999999998 | 0 |
| B98 | B108 | anti-Jo-1 | Anti-Cyclin | 0.999999998 | 0 |
| B98 | B109 | anti-Jo-1 | AMA-M2 | 0.999999997 | 0 |
| B98 | B110 | anti-Jo-1 | AHA | 0.999999997 | 0 |
| B98 | B151 | anti-Jo-1 | Anti-CCP | 0.704822098 | 0 |
| B98 | B156 | anti-Jo-1 | IgE | -0.903553224 | 0 |
| B98 | B183 | anti-Jo-1 | PTA | -0.935351261 | 0 |
| B98 | B99 | anti-Jo-1 | anti-PM-Scl | 0.999999997 | 0 |
| B98 | B155 | anti-Jo-1 | IgA | 0.556426348 | 4.64E-14 |
| B98 | B175 | anti-Jo-1 | ALB-U | 0.538122696 | 4.33E-13 |
| B98 | B158 | anti-Jo-1 | IgM | -0.49229683 | 6.70E-11 |
| B98 | B198 | anti-Jo-1 | P | -0.484721523 | 1.44E-10 |
| B98 | B185 | anti-Jo-1 | PT | 0.471649435 | 5.14E-10 |
| B98 | B184 | anti-Jo-1 | INR | 0.461953763 | 1.28E-09 |
| B98 | B181 | anti-Jo-1 | APTT | 0.459306488 | 1.63E-09 |
| B98 | B168 | anti-Jo-1 | NMP22 | 0.401609666 | 2.03E-07 |
| B98 | B157 | anti-Jo-1 | IgG | 0.400110804 | 2.28E-07 |
| B98 | B176 | anti-Jo-1 | Ucr | 0.387593591 | 5.77E-07 |
| B99 | B100 | anti-PM-Scl | anti-Ro52 | 0.999999999 | 0 |
| B99 | B103 | anti-PM-Scl | anti-SSA/Ro | 0.999999999 | 0 |
| B99 | B108 | anti-PM-Scl | Anti-Cyclin | 1 | 0 |
| B99 | B110 | anti-PM-Scl | AHA | 1 | 0 |
| B99 | B151 | anti-PM-Scl | Anti-CCP | 0.7048221 | 0 |
| B99 | B156 | anti-PM-Scl | IgE | -0.903553225 | 0 |
| B99 | B183 | anti-PM-Scl | PTA | -0.935351262 | 0 |
| B99 | B155 | anti-PM-Scl | IgA | 0.55642635 | 4.64E-14 |
| B99 | B175 | anti-PM-Scl | ALB-U | 0.538122697 | 4.33E-13 |
| B99 | B158 | anti-PM-Scl | IgM | -0.492296831 | 6.70E-11 |
| B99 | B198 | anti-PM-Scl | P | -0.484721524 | 1.44E-10 |
| B99 | B185 | anti-PM-Scl | PT | 0.471649437 | 5.14E-10 |
| B99 | B184 | anti-PM-Scl | INR | 0.461953764 | 1.28E-09 |

| | | | | | |
|---|---|---|---|---|---|
| B99 | B181 | anti-PM-Scl | APTT | 0.459306489 | 1.63E-09 |
| B99 | B168 | anti-PM-Scl | NMP22 | 0.401609666 | 2.03E-07 |
| B99 | B157 | anti-PM-Scl | IgG | 0.400110805 | 2.28E-07 |
| B99 | B176 | anti-PM-Scl | Ucr | 0.387593592 | 5.77E-07 |

**Supplementary Table 11. PEI Correlations in Hypothyroidism**

| Row | Column | Row | Column | Cor | *P* |
|---|---|---|---|---|---|
| B1 | B163 | Gender | SG | -0.2932978 | 0 |
| B1 | B174 | Gender | Urea/Cr | 0.26229326 | 0 |
| B1 | B189 | Gender | eGFB | 0.2216233 | 0 |
| B1 | B190 | Gender | Cre | -0.3224045 | 0 |
| B1 | B192 | Gender | UA | -0.4492026 | 0 |
| B1 | B204 | Gender | Drinking | -0.4458063 | 0 |
| B1 | B206 | Gender | Waist | -0.4121516 | 0 |
| B1 | B207 | Gender | WHR | -0.4001778 | 0 |
| B1 | B209 | Gender | Smoking | -0.4627832 | 0 |
| B1 | B210 | Gender | Hip | -0.2378583 | 0 |
| B1 | B216 | Gender | BMI | -0.5070998 | 0 |
| B1 | B221 | Gender | Hei | -0.5772929 | 0 |
| B1 | B222 | Gender | NS | -0.6517958 | 0 |
| B1 | B42 | Gender | MON | -0.2016853 | 0 |
| B1 | B45 | Gender | MCHC | -0.2103364 | 0 |
| B1 | B46 | Gender | RBC Count | -0.392497 | 0 |
| B1 | B51 | Gender | Hb | -0.4968018 | 0 |
| B1 | B72 | Gender | HDL-C | 0.2333749 | 0 |
| B1 | B215 | Gender | BMI | -0.1986516 | 4.44E-16 |
| B1 | B128 | Gender | ALT | -0.1914466 | 3.55E-15 |
| B1 | B218 | Gender | DBP | -0.1818718 | 8.08E-14 |
| B1 | B166 | Gender | Pro | -0.1762402 | 4.69E-13 |
| B1 | B84 | Gender | C-PSA | -0.1620117 | 3.10E-11 |
| B1 | B141 | Gender | AST/ALT | 0.15541088 | 1.92E-10 |
| B1 | B94 | Gender | ALDH2-Lys504Lys | 0.15240508 | 4.29E-10 |
| B1 | B140 | Gender | r-GT | -0.1519831 | 4.80E-10 |
| B1 | B133 | Gender | TBIL | -0.1502997 | 7.46E-10 |
| B1 | B71 | Gender | TG | -0.1458477 | 2.35E-09 |
| B1 | B129 | Gender | IBIL | -0.1446851 | 3.15E-09 |
| B1 | B41 | Gender | MON％ | -0.1399206 | 1.03E-08 |
| B1 | B2 | Gender | Age | -0.1381718 | 1.57E-08 |
| B1 | B220 | Gender | SBP | -0.135408 | 3.03E-08 |
| B1 | B191 | Gender | Urea | -0.1341079 | 4.10E-08 |
| B1 | B44 | Gender | MCH | -0.1321422 | 6.47E-08 |
| B1 | B132 | Gender | Dbil | -0.1304053 | 9.63E-08 |
| B1 | B118 | Gender | CEA1 | -0.1234945 | 4.45E-07 |
| B10 | B11 | PGⅠ | PGI/PGII | 0.30500541 | 0 |
| B10 | B12 | PGⅠ | PGⅡ | 0.49137292 | 0 |
| B10 | B14 | PGⅠ | G-17 | -0.1742848 | 8.52E-13 |
| B100 | B101 | anti-Ro52 | anti-Sc70 | -0.3185303 | 0 |
| B100 | B102 | anti-Ro52 | anti-Sm | -0.3185303 | 0 |
| B100 | B103 | anti-Ro52 | anti-SSA/Ro | 0.70410455 | 0 |

| | | | | | |
|---|---|---|---|---|---|
| B100 | B104 | anti-Ro52 | anti-SSB | 0.56083724 | 0 |
| B100 | B105 | anti-Ro52 | anti-U1RNP | -0.3185303 | 0 |
| B100 | B107 | anti-Ro52 | AnuA | -0.3185301 | 0 |
| B100 | B108 | anti-Ro52 | Anti-Cyclin | -0.3185303 | 0 |
| B100 | B157 | anti-Ro52 | IgG | 0.20041427 | 0 |
| B100 | B180 | anti-Ro52 | Tf | -0.2434742 | 0 |
| B100 | B196 | anti-Ro52 | TCO2 | 0.23141832 | 0 |
| B100 | B197 | anti-Ro52 | AG | -0.255022 | 0 |
| B100 | B194 | anti-Ro52 | Cl | 0.17604091 | 4.98E-13 |
| B100 | B178 | anti-Ro52 | Ig-U | -0.1593497 | 6.52E-11 |
| B100 | B179 | anti-Ro52 | α1-MG | -0.1550516 | 2.11E-10 |
| B100 | B193 | anti-Ro52 | K | -0.1504767 | 7.13E-10 |
| B100 | B138 | anti-Ro52 | PAB | -0.136584 | 2.29E-08 |
| B100 | B134 | anti-Ro52 | CHE | -0.1343484 | 3.88E-08 |
| B100 | B155 | anti-Ro52 | IgA | 0.13268723 | 5.71E-08 |
| B100 | B136 | anti-Ro52 | 5'NT | -0.128511 | 1.48E-07 |
| B101 | B103 | anti-Sc70 | anti-SSA/Ro | -0.3344827 | 0 |
| B101 | B107 | anti-Sc70 | AnuA | 0.99999939 | 0 |
| B101 | B125 | anti-Sc70 | Cor | 0.26774054 | 0 |
| B101 | B135 | anti-Sc70 | TBA | 0.24449504 | 0 |
| B101 | B156 | anti-Sc70 | IgE | 0.29730773 | 0 |
| B101 | B178 | anti-Sc70 | Ig-U | 0.34819819 | 0 |
| B101 | B179 | anti-Sc70 | α1-MG | 0.34866525 | 0 |
| B101 | B180 | anti-Sc70 | Tf | 0.42439565 | 0 |
| B101 | B121 | anti-Sc70 | C3 | -0.1954528 | 8.88E-16 |
| B101 | B122 | anti-Sc70 | C4 | -0.195453 | 8.88E-16 |
| B101 | B159 | anti-Sc70 | β2-MG | -0.1954519 | 8.88E-16 |
| B101 | B169 | anti-Sc70 | U-CRY | -0.1954546 | 8.88E-16 |
| B101 | B184 | anti-Sc70 | INR | 0.1931741 | 2.00E-15 |
| B101 | B106 | anti-Sc70 | anti-P | -0.19276 | 2.22E-15 |
| B101 | B154 | anti-Sc70 | RF | -0.1785002 | 2.33E-13 |
| B101 | B139 | anti-Sc70 | ADA | 0.16693577 | 7.57E-12 |
| B101 | B119 | anti-Sc70 | Hcv-rna | 0.16323416 | 2.19E-11 |
| B101 | B151 | anti-Sc70 | Anti-CCP | 0.15767897 | 1.03E-10 |
| B101 | B104 | anti-Sc70 | anti-SSB | -0.1552047 | 2.03E-10 |
| B101 | B158 | anti-Sc70 | IgM | 0.14873463 | 1.12E-09 |
| B101 | B181 | anti-Sc70 | APTT | -0.1409302 | 8.01E-09 |
| B101 | B137 | anti-Sc70 | AFU | 0.13753181 | 1.83E-08 |
| B101 | B138 | anti-Sc70 | PAB | 0.12878441 | 1.39E-07 |
| B102 | B103 | anti-Sm | anti-SSA/Ro | -0.3344827 | 0 |
| B102 | B105 | anti-Sm | anti-U1RNP | 0.99999999 | 0 |
| B102 | B107 | anti-Sm | AnuA | 0.99999949 | 0 |
| B102 | B108 | anti-Sm | Anti-Cyclin | 0.99999999 | 0 |
| B102 | B125 | anti-Sm | Cor | 0.26774053 | 0 |

| | | | | | |
|---|---|---|---|---|---|
| B102 | B135 | anti-Sm | TBA | 0.24449504 | 0 |
| B102 | B156 | anti-Sm | IgE | 0.29730772 | 0 |
| B102 | B178 | anti-Sm | Ig-U | 0.34819818 | 0 |
| B102 | B179 | anti-Sm | α1-MG | 0.34866524 | 0 |
| B102 | B180 | anti-Sm | Tf | 0.42439564 | 0 |
| B102 | B121 | anti-Sm | C3 | -0.1954523 | 8.88E-16 |
| B102 | B122 | anti-Sm | C4 | -0.1954531 | 8.88E-16 |
| B102 | B159 | anti-Sm | β2-MG | -0.1954505 | 8.88E-16 |
| B102 | B169 | anti-Sm | U-CRY | -0.1954535 | 8.88E-16 |
| B102 | B184 | anti-Sm | INR | 0.1931741 | 2.00E-15 |
| B102 | B106 | anti-Sm | anti-P | -0.19276 | 2.22E-15 |
| B102 | B154 | anti-Sm | RF | -0.1785001 | 2.33E-13 |
| B102 | B139 | anti-Sm | ADA | 0.16693576 | 7.57E-12 |
| B102 | B119 | anti-Sm | Hcv-rna | 0.16323415 | 2.19E-11 |
| B102 | B151 | anti-Sm | Anti-CCP | 0.15767896 | 1.03E-10 |
| B102 | B104 | anti-Sm | anti-SSB | -0.1552047 | 2.03E-10 |
| B102 | B158 | anti-Sm | IgM | 0.14873462 | 1.12E-09 |
| B102 | B181 | anti-Sm | APTT | -0.1409302 | 8.01E-09 |
| B102 | B137 | anti-Sm | AFU | 0.1375318 | 1.83E-08 |
| B102 | B138 | anti-Sm | PAB | 0.12878441 | 1.39E-07 |
| B103 | B104 | anti-SSA/Ro | anti-SSB | 0.61143268 | 0 |
| B103 | B105 | anti-SSA/Ro | anti-U1RNP | -0.3344827 | 0 |
| B103 | B107 | anti-SSA/Ro | AnuA | -0.3344825 | 0 |
| B103 | B108 | anti-SSA/Ro | Anti-Cyclin | -0.3344827 | 0 |
| B103 | B180 | anti-SSA/Ro | Tf | -0.209224 | 0 |
| B103 | B199 | anti-SSA/Ro | Mg | 0.19617216 | 6.66E-16 |
| B103 | B157 | anti-SSA/Ro | IgG | 0.18682514 | 1.64E-14 |
| B103 | B138 | anti-SSA/Ro | PAB | -0.1508511 | 6.46E-10 |
| B103 | B194 | anti-SSA/Ro | Cl | 0.15074574 | 6.64E-10 |
| B103 | B176 | anti-SSA/Ro | Ucr | 0.1425359 | 5.39E-09 |
| B103 | B186 | anti-SSA/Ro | PFC | -0.1362468 | 2.48E-08 |
| B103 | B197 | anti-SSA/Ro | AG | -0.1337711 | 4.44E-08 |
| B103 | B125 | anti-SSA/Ro | Cor | -0.1329353 | 5.39E-08 |
| B103 | B156 | anti-SSA/Ro | IgE | -0.1328801 | 5.46E-08 |
| B103 | B198 | anti-SSA/Ro | P | 0.12514342 | 3.11E-07 |
| B104 | B155 | anti-SSB | IgA | 0.21776694 | 0 |
| B104 | B157 | anti-SSB | IgG | 0.33826943 | 0 |
| B104 | B194 | anti-SSB | Cl | 0.20569998 | 0 |
| B104 | B198 | anti-SSB | P | 0.27117125 | 0 |
| B104 | B197 | anti-SSB | AG | -0.1847552 | 3.20E-14 |
| B104 | B105 | anti-SSB | anti-U1RNP | -0.1552047 | 2.03E-10 |
| B104 | B107 | anti-SSB | AnuA | -0.1552046 | 2.03E-10 |
| B104 | B108 | anti-SSB | Anti-Cyclin | -0.1552047 | 2.03E-10 |
| B104 | B158 | anti-SSB | IgM | 0.12687822 | 2.13E-07 |

| | | | | | |
|---|---|---|---|---|---|
| B105 | B107 | anti-U1RNP | AnuA | 0.99999928 | 0 |
| B105 | B108 | anti-U1RNP | Anti-Cyclin | 1 | 0 |
| B105 | B125 | anti-U1RNP | Cor | 0.26774053 | 0 |
| B105 | B135 | anti-U1RNP | TBA | 0.24449504 | 0 |
| B105 | B156 | anti-U1RNP | IgE | 0.29730773 | 0 |
| B105 | B178 | anti-U1RNP | Ig-U | 0.34819819 | 0 |
| B105 | B179 | anti-U1RNP | α1-MG | 0.34866524 | 0 |
| B105 | B180 | anti-U1RNP | Tf | 0.42439565 | 0 |
| B105 | B121 | anti-U1RNP | C3 | -0.1954525 | 8.88E-16 |
| B105 | B122 | anti-U1RNP | C4 | -0.195453 | 8.88E-16 |
| B105 | B159 | anti-U1RNP | β2-MG | -0.1954514 | 8.88E-16 |
| B105 | B169 | anti-U1RNP | U-CRY | -0.195454 | 8.88E-16 |
| B105 | B184 | anti-U1RNP | INR | 0.1931741 | 2.00E-15 |
| B105 | B106 | anti-U1RNP | anti-P | -0.19276 | 2.22E-15 |
| B105 | B154 | anti-U1RNP | RF | -0.1785002 | 2.33E-13 |
| B105 | B139 | anti-U1RNP | ADA | 0.16693577 | 7.57E-12 |
| B105 | B119 | anti-U1RNP | Hcv-rna | 0.16323416 | 2.19E-11 |
| B105 | B151 | anti-U1RNP | Anti-CCP | 0.15767896 | 1.03E-10 |
| B105 | B158 | anti-U1RNP | IgM | 0.14873462 | 1.12E-09 |
| B105 | B181 | anti-U1RNP | APTT | -0.1409302 | 8.01E-09 |
| B105 | B137 | anti-U1RNP | AFU | 0.1375318 | 1.83E-08 |
| B105 | B138 | anti-U1RNP | PAB | 0.12878441 | 1.39E-07 |
| B106 | B157 | anti-P | IgG | 0.3091613 | 0 |
| B106 | B178 | anti-P | Ig-U | -0.2486824 | 0 |
| B106 | B179 | anti-P | α1-MG | -0.2580017 | 0 |
| B106 | B180 | anti-P | Tf | -0.2392454 | 0 |
| B106 | B107 | anti-P | AnuA | -0.1927599 | 2.22E-15 |
| B106 | B108 | anti-P | Anti-Cyclin | -0.19276 | 2.22E-15 |
| B107 | B108 | AnuA | Anti-Cyclin | 0.99999926 | 0 |
| B107 | B125 | AnuA | Cor | 0.26774036 | 0 |
| B107 | B135 | AnuA | TBA | 0.24449486 | 0 |
| B107 | B156 | AnuA | IgE | 0.29730752 | 0 |
| B107 | B178 | AnuA | Ig-U | 0.34819794 | 0 |
| B107 | B179 | AnuA | α1-MG | 0.348665 | 0 |
| B107 | B180 | AnuA | Tf | 0.42439535 | 0 |
| B107 | B121 | AnuA | C3 | -0.1954527 | 8.88E-16 |
| B107 | B122 | AnuA | C4 | -0.1954528 | 8.88E-16 |
| B107 | B159 | AnuA | β2-MG | -0.1954492 | 8.88E-16 |
| B107 | B169 | AnuA | U-CRY | -0.1954536 | 8.88E-16 |
| B107 | B184 | AnuA | INR | 0.19317398 | 2.00E-15 |
| B107 | B154 | AnuA | RF | -0.1785 | 2.33E-13 |
| B107 | B139 | AnuA | ADA | 0.16693565 | 7.57E-12 |
| B107 | B119 | AnuA | Hcv-rna | 0.16323404 | 2.19E-11 |
| B107 | B151 | AnuA | Anti-CCP | 0.15767885 | 1.03E-10 |

| | | | | | |
|---|---|---|---|---|---|
| B107 | B158 | AnuA | IgM | 0.14873452 | 1.12E-09 |
| B107 | B181 | AnuA | APTT | -0.1409301 | 8.01E-09 |
| B107 | B137 | AnuA | AFU | 0.1375317 | 1.83E-08 |
| B107 | B138 | AnuA | PAB | 0.12878432 | 1.39E-07 |
| B108 | B125 | Anti-Cyclin | Cor | 0.26774053 | 0 |
| B108 | B135 | Anti-Cyclin | TBA | 0.24449504 | 0 |
| B108 | B156 | Anti-Cyclin | IgE | 0.29730773 | 0 |
| B108 | B178 | Anti-Cyclin | Ig-U | 0.34819819 | 0 |
| B108 | B179 | Anti-Cyclin | α1-MG | 0.34866524 | 0 |
| B108 | B180 | Anti-Cyclin | Tf | 0.42439565 | 0 |
| B108 | B121 | Anti-Cyclin | C3 | -0.1954526 | 8.88E-16 |
| B108 | B122 | Anti-Cyclin | C4 | -0.1954532 | 8.88E-16 |
| B108 | B159 | Anti-Cyclin | β2-MG | -0.1954512 | 8.88E-16 |
| B108 | B169 | Anti-Cyclin | U-CRY | -0.1954547 | 8.88E-16 |
| B108 | B184 | Anti-Cyclin | INR | 0.19317409 | 2.00E-15 |
| B108 | B154 | Anti-Cyclin | RF | -0.1785002 | 2.33E-13 |
| B108 | B139 | Anti-Cyclin | ADA | 0.16693576 | 7.57E-12 |
| B108 | B119 | Anti-Cyclin | Hcv-rna | 0.16323416 | 2.19E-11 |
| B108 | B151 | Anti-Cyclin | Anti-CCP | 0.15767896 | 1.03E-10 |
| B108 | B158 | Anti-Cyclin | IgM | 0.14873462 | 1.12E-09 |
| B108 | B181 | Anti-Cyclin | APTT | -0.1409302 | 8.01E-09 |
| B108 | B137 | Anti-Cyclin | AFU | 0.1375318 | 1.83E-08 |
| B108 | B138 | Anti-Cyclin | PAB | 0.12878441 | 1.39E-07 |
| B109 | B194 | AMA-M2 | Cl | -0.2564007 | 0 |
| B109 | B200 | AMA-M2 | OP | -0.1646234 | 1.47E-11 |
| B109 | B137 | AMA-M2 | AFU | 0.15891578 | 7.36E-11 |
| B109 | B135 | AMA-M2 | TBA | 0.13196973 | 6.74E-08 |
| B109 | B195 | AMA-M2 | Na | -0.1265361 | 2.29E-07 |
| B11 | B12 | PGI/PGII | PG II | -0.4359501 | 0 |
| B11 | B14 | PGI/PGII | G-17 | -0.1861793 | 2.02E-14 |
| B110 | B145 | AHA | BGP | -0.2385151 | 0 |
| B110 | B182 | AHA | TT | -0.3000954 | 0 |
| B110 | B183 | AHA | PTA | 0.36807865 | 0 |
| B110 | B143 | AHA | β-CTx | 0.14045352 | 9.01E-09 |
| B110 | B151 | AHA | Anti-CCP | -0.1362611 | 2.47E-08 |
| B110 | B194 | AHA | Cl | 0.13034051 | 9.78E-08 |
| B110 | B155 | AHA | IgA | -0.1261841 | 2.48E-07 |
| B110 | B199 | AHA | Mg | 0.12473497 | 3.40E-07 |
| B111 | B112 | TSH | FT4 | -0.4603166 | 0 |
| B111 | B113 | TSH | FT3 | -0.3467826 | 0 |
| B111 | B114 | TSH | TSH TT3 TT4(TT4) | -0.3598568 | 0 |
| B111 | B115 | TSH | TSH TT3 TT4(TT3) | -0.3332098 | 0 |
| B111 | B148 | TSH | Anti_TPO | 0.20599664 | 0 |
| B111 | B149 | TSH | TGAb | 0.17080419 | 2.43E-12 |

| | | | | | |
|---|---|---|---|---|---|
| B111 | B185 | TSH | PT | -0.1606495 | 4.54E-11 |
| B111 | B183 | TSH | PTA | 0.1563651 | 1.48E-10 |
| B111 | B184 | TSH | INR | -0.1355076 | 2.95E-08 |
| B111 | B204 | TSH | Drinking | 0.12919253 | 1.27E-07 |
| B112 | B113 | FT4 | FT3 | 0.21784508 | 0 |
| B112 | B114 | FT4 | TSH TT3 TT4(TT4) | 0.60802314 | 0 |
| B112 | B115 | FT4 | TSH TT3 TT4(TT3) | 0.15174551 | 5.11E-10 |
| B112 | B185 | FT4 | PT | 0.13167384 | 7.21E-08 |
| B112 | B184 | FT4 | INR | 0.12662952 | 2.25E-07 |
| B113 | B115 | FT3 | TSH TT3 TT4(TT3) | 0.47695754 | 0 |
| B113 | B148 | FT3 | Anti_TPO | -0.1686624 | 4.57E-12 |
| B113 | B149 | FT3 | TGAb | -0.1518481 | 4.97E-10 |
| B113 | B114 | FT3 | TSH TT3 TT4(TT4) | 0.13572553 | 2.81E-08 |
| B113 | B165 | FT3 | bilirubin | 0.125624 | 2.80E-07 |
| B114 | B115 | TSH TT3 TT4(TT4) | TSH TT3 TT4(TT3) | 0.21064944 | 0 |
| B114 | B185 | TSH TT3 TT4(TT4) | PT | 0.16052999 | 4.70E-11 |
| B114 | B184 | TSH TT3 TT4(TT4) | INR | 0.15017374 | 7.71E-10 |
| B114 | B183 | TSH TT3 TT4(TT4) | PTA | -0.148593 | 1.16E-09 |
| B115 | B149 | TSH TT3 TT4(TT3) | TGAb | -0.1579154 | 9.69E-11 |
| B115 | B148 | TSH TT3 TT4(TT3) | Anti_TPO | -0.1490629 | 1.03E-09 |
| B115 | B136 | TSH TT3 TT4(TT3) | 5'NT | 0.12655294 | 2.28E-07 |
| B116 | B117 | TT3 | TT4 | 1 | 0 |
| B116 | B149 | TT3 | TGAb | -0.1660342 | 9.83E-12 |
| B117 | B149 | TT4 | TGAb | -0.1660342 | 9.83E-12 |
| B118 | B189 | CEA1 | eGFB | -0.1956132 | 8.88E-16 |
| B118 | B209 | CEA1 | Smoking | 0.15684453 | 1.30E-10 |
| B118 | B190 | CEA1 | Cre | 0.12426915 | 3.76E-07 |
| B119 | B120 | Hcv-rna | HCV-cAg | 0.46415872 | 0 |
| B119 | B121 | Hcv-rna | C3 | -0.8446383 | 0 |
| B119 | B122 | Hcv-rna | C4 | -0.8446403 | 0 |
| B119 | B124 | Hcv-rna | ACTH | 0.27490152 | 0 |
| B119 | B137 | Hcv-rna | AFU | 0.28984077 | 0 |
| B119 | B159 | Hcv-rna | β2-MG | -0.8446325 | 0 |
| B119 | B169 | Hcv-rna | U-CRY | -0.8446447 | 0 |
| B119 | B138 | Hcv-rna | PAB | 0.17784475 | 2.86E-13 |
| B119 | B125 | Hcv-rna | Cor | 0.17164912 | 1.89E-12 |
| B119 | B197 | Hcv-rna | AG | -0.1705621 | 2.61E-12 |
| B119 | B195 | Hcv-rna | Na | -0.170224 | 2.88E-12 |
| B119 | B198 | Hcv-rna | P | -0.1635085 | 2.03E-11 |
| B119 | B199 | Hcv-rna | Mg | 0.15267172 | 4.00E-10 |
| B12 | B26 | PG II | CKMB | 0.25070277 | 0 |
| B12 | B80 | PG II | anti-HP IgG | 0.12484326 | 3.32E-07 |
| B120 | B121 | HCV-cAg | C3 | -0.392049 | 0 |
| B120 | B122 | HCV-cAg | C4 | -0.3920502 | 0 |

| | | | | | |
|---|---|---|---|---|---|
| B120 | B159 | HCV-cAg | β2-MG | -0.3920467 | 0 |
| B120 | B169 | HCV-cAg | U-CRY | -0.3920506 | 0 |
| B120 | B137 | HCV-cAg | AFU | 0.176001 | 5.04E-13 |
| B121 | B159 | C3 | β2-MG | 0.99999606 | 0 |
| B121 | B198 | C3 | P | 0.20598207 | 0 |
| B121 | B137 | C3 | AFU | -0.1935644 | 1.78E-15 |
| B121 | B197 | C3 | AG | 0.15863487 | 7.95E-11 |
| B121 | B138 | C3 | PAB | -0.1496715 | 8.79E-10 |
| B121 | B194 | C3 | Cl | -0.1468888 | 1.80E-09 |
| B121 | B195 | C3 | Na | 0.145285 | 2.71E-09 |
| B121 | B199 | C3 | Mg | -0.12976 | 1.12E-07 |
| B121 | B124 | C3 | ACTH | -0.1247748 | 3.37E-07 |
| B121 | B158 | C3 | IgM | 0.12327742 | 4.66E-07 |
| B122 | B159 | C4 | β2-MG | 0.99999854 | 0 |
| B122 | B169 | C4 | U-CRY | 0.99999759 | 0 |
| B122 | B198 | C4 | P | 0.20598257 | 0 |
| B122 | B137 | C4 | AFU | -0.1935648 | 1.78E-15 |
| B122 | B197 | C4 | AG | 0.15863524 | 7.95E-11 |
| B122 | B138 | C4 | PAB | -0.1496719 | 8.79E-10 |
| B122 | B194 | C4 | Cl | -0.1468892 | 1.80E-09 |
| B122 | B195 | C4 | Na | 0.1452853 | 2.71E-09 |
| B122 | B199 | C4 | Mg | -0.1297603 | 1.12E-07 |
| B122 | B124 | C4 | ACTH | -0.1247752 | 3.37E-07 |
| B122 | B158 | C4 | IgM | 0.12327771 | 4.66E-07 |
| B123 | B182 | hs-CRP1 | TT | -0.1544025 | 2.52E-10 |
| B123 | B186 | hs-CRP1 | PFC | 0.14537558 | 2.65E-09 |
| B124 | B125 | ACTH | Cor | 0.7162258 | 0 |
| B124 | B138 | ACTH | PAB | 0.23392131 | 0 |
| B124 | B156 | ACTH | IgE | 0.34882002 | 0 |
| B124 | B137 | ACTH | AFU | 0.1800311 | 1.44E-13 |
| B124 | B134 | ACTH | CHE | 0.17422837 | 8.67E-13 |
| B124 | B186 | ACTH | PFC | 0.17157802 | 1.93E-12 |
| B124 | B182 | ACTH | TT | -0.1305272 | 9.37E-08 |
| B124 | B178 | ACTH | Ig-U | 0.12911021 | 1.29E-07 |
| B124 | B196 | ACTH | TCO2 | -0.1273804 | 1.90E-07 |
| B124 | B159 | ACTH | β2-MG | -0.124774 | 3.37E-07 |
| B124 | B169 | ACTH | U-CRY | -0.1247758 | 3.37E-07 |
| B125 | B178 | Cor | Ig-U | 0.20612019 | 0 |
| B125 | B184 | Cor | INR | 0.23769401 | 0 |
| B125 | B179 | Cor | α1-MG | 0.19706423 | 4.44E-16 |
| B125 | B138 | Cor | PAB | 0.17968426 | 1.61E-13 |
| B125 | B135 | Cor | TBA | 0.17483294 | 7.21E-13 |
| B125 | B199 | Cor | Mg | -0.1652123 | 1.25E-11 |
| B125 | B198 | Cor | P | -0.1364226 | 2.38E-08 |

| | | | | | |
|---|---|---|---|---|---|
| B125 | B180 | Cor | Tf | 0.13548544 | 2.97E-08 |
| B125 | B137 | Cor | AFU | 0.12187326 | 6.30E-07 |
| B126 | B127 | Alb | A-G ratio | 0.44329327 | 0 |
| B126 | B188 | Alb | Ca | 0.40099163 | 0 |
| B126 | B189 | Alb | eGFB | 0.17998443 | 1.47E-13 |
| B127 | B130 | A-G ratio | GLB | -0.8983219 | 0 |
| B128 | B131 | ALT | AST | 0.75778207 | 0 |
| B128 | B140 | ALT | r-GT | 0.42162058 | 0 |
| B128 | B141 | ALT | AST/ALT | -0.5828154 | 0 |
| B128 | B192 | ALT | UA | 0.25670266 | 0 |
| B128 | B206 | ALT | Waist | 0.31240565 | 0 |
| B128 | B207 | ALT | WHR | 0.28200355 | 0 |
| B128 | B210 | ALT | Hip | 0.2053167 | 0 |
| B128 | B215 | ALT | BMI | 0.29178949 | 0 |
| B128 | B216 | ALT | BMI | 0.28344158 | 0 |
| B128 | B222 | ALT | NS | 0.28410545 | 0 |
| B128 | B204 | ALT | Drinking | 0.15001302 | 8.05E-10 |
| B128 | B220 | ALT | SBP | 0.14170685 | 6.62E-09 |
| B128 | B218 | ALT | DBP | 0.1222007 | 5.87E-07 |
| B129 | B132 | IBIL | Dbil | 0.68400614 | 0 |
| B129 | B133 | IBIL | TBIL | 0.98330863 | 0 |
| B129 | B210 | IBIL | Hip | -0.1315539 | 7.41E-08 |
| B130 | B188 | GLB | Ca | 0.12483094 | 3.33E-07 |
| B131 | B140 | AST | r-GT | 0.37009095 | 0 |
| B131 | B189 | AST | eGFB | -0.1844174 | 3.57E-14 |
| B131 | B207 | AST | WHR | 0.179269 | 1.83E-13 |
| B131 | B206 | AST | Waist | 0.17864496 | 2.23E-13 |
| B131 | B222 | AST | NS | 0.15559956 | 1.82E-10 |
| B131 | B220 | AST | SBP | 0.15519191 | 2.03E-10 |
| B131 | B215 | AST | BMI | 0.15487939 | 2.21E-10 |
| B131 | B192 | AST | UA | 0.14221622 | 5.84E-09 |
| B131 | B141 | AST | AST/ALT | -0.1422078 | 5.85E-09 |
| B131 | B216 | AST | BMI | 0.12281652 | 5.15E-07 |
| B132 | B133 | Dbil | TBIL | 0.80515208 | 0 |
| B132 | B210 | Dbil | Hip | -0.157025 | 1.24E-10 |
| B132 | B215 | Dbil | BMI | -0.1565175 | 1.42E-10 |
| B133 | B210 | TBIL | Hip | -0.1458042 | 2.38E-09 |
| B134 | B136 | CHE | 5'NT | 0.38546351 | 0 |
| B134 | B138 | CHE | PAB | 0.34423965 | 0 |
| B134 | B150 | CHE | ALP | 0.27536691 | 0 |
| B134 | B193 | CHE | K | 0.1663667 | 8.93E-12 |
| B134 | B152 | CHE | ACA-Ig(G-A-M) | 0.15267483 | 3.99E-10 |
| B134 | B196 | CHE | TCO2 | -0.1360788 | 2.58E-08 |
| B134 | B137 | CHE | AFU | 0.12392262 | 4.06E-07 |

| | | | | | |
|---|---|---|---|---|---|
| B135 | B179 | TBA | α1-MG | 0.2005377 | 0 |
| B135 | B138 | TBA | PAB | 0.18955785 | 6.66E-15 |
| B135 | B178 | TBA | Ig-U | 0.17747746 | 3.20E-13 |
| B135 | B180 | TBA | Tf | 0.17299896 | 1.26E-12 |
| B135 | B184 | TBA | INR | 0.17266041 | 1.39E-12 |
| B135 | B152 | TBA | ACA-Ig(G-A-M) | 0.15787505 | 9.80E-11 |
| B135 | B182 | TBA | TT | 0.14796166 | 1.37E-09 |
| B135 | B146 | TBA | CysC | 0.13438664 | 3.85E-08 |
| B136 | B138 | 5'NT | PAB | 0.45183805 | 0 |
| B136 | B150 | 5'NT | ALP | 0.21157264 | 0 |
| B136 | B158 | 5'NT | IgM | 0.23370719 | 0 |
| B136 | B194 | 5'NT | Cl | -0.2328053 | 0 |
| B136 | B197 | 5'NT | AG | 0.26326577 | 0 |
| B136 | B199 | 5'NT | Mg | 0.18253344 | 6.55E-14 |
| B136 | B181 | 5'NT | APTT | -0.1770606 | 3.64E-13 |
| B136 | B200 | 5'NT | OP | 0.16775007 | 5.97E-12 |
| B136 | B196 | 5'NT | TCO2 | -0.1538344 | 2.93E-10 |
| B136 | B178 | 5'NT | Ig-U | -0.1301533 | 1.02E-07 |
| B137 | B199 | AFU | Mg | 0.20873923 | 0 |
| B137 | B159 | AFU | β2-MG | -0.1935631 | 1.78E-15 |
| B137 | B169 | AFU | U-CRY | -0.1935659 | 1.78E-15 |
| B137 | B156 | AFU | IgE | 0.12932412 | 1.23E-07 |
| B138 | B196 | PAB | TCO2 | -0.1844382 | 3.55E-14 |
| B138 | B197 | PAB | AG | 0.17592922 | 5.16E-13 |
| B138 | B169 | PAB | U-CRY | -0.1496727 | 8.79E-10 |
| B138 | B159 | PAB | β2-MG | -0.1496705 | 8.80E-10 |
| B138 | B200 | PAB | OP | 0.14220554 | 5.85E-09 |
| B138 | B150 | PAB | ALP | 0.12548806 | 2.89E-07 |
| B139 | B194 | ADA | Cl | -0.2738199 | 0 |
| B139 | B157 | ADA | IgG | 0.18196727 | 7.84E-14 |
| B139 | B155 | ADA | IgA | 0.17581362 | 5.34E-13 |
| B139 | B156 | ADA | IgE | 0.1473967 | 1.58E-09 |
| B139 | B197 | ADA | AG | 0.1417439 | 6.56E-09 |
| B139 | B146 | ADA | CysC | 0.13391247 | 4.30E-08 |
| B140 | B141 | r-GT | AST/ALT | -0.2263782 | 0 |
| B140 | B206 | r-GT | Waist | 0.22706987 | 0 |
| B140 | B207 | r-GT | WHR | 0.20669139 | 0 |
| B140 | B215 | r-GT | BMI | 0.20147701 | 0 |
| B140 | B216 | r-GT | BMI | 0.20757204 | 0 |
| B140 | B222 | r-GT | NS | 0.22381615 | 0 |
| B140 | B192 | r-GT | UA | 0.17330042 | 1.15E-12 |
| B140 | B220 | r-GT | SBP | 0.15157381 | 5.34E-10 |
| B140 | B210 | r-GT | Hip | 0.15146866 | 5.49E-10 |
| B140 | B218 | r-GT | DBP | 0.13277535 | 5.59E-08 |

| | | | | | |
|---|---|---|---|---|---|
| B140 | B166 | r-GT | Pro | 0.1302435 | 9.99E-08 |
| B140 | B153 | r-GT | Glu | 0.12025963 | 8.86E-07 |
| B141 | B192 | AST/ALT | UA | -0.2167204 | 0 |
| B141 | B206 | AST/ALT | Waist | -0.2860861 | 0 |
| B141 | B207 | AST/ALT | WHR | -0.2552961 | 0 |
| B141 | B215 | AST/ALT | BMI | -0.2737384 | 0 |
| B141 | B216 | AST/ALT | BMI | -0.2650887 | 0 |
| B141 | B222 | AST/ALT | NS | -0.2362241 | 0 |
| B141 | B210 | AST/ALT | Hip | -0.1987911 | 4.44E-16 |
| B141 | B153 | AST/ALT | Glu | -0.1270299 | 2.05E-07 |
| B142 | B143 | 25-(OH)D | β-CTx | -0.2344676 | 0 |
| B142 | B144 | 25-(OH)D | Total-PINP | -0.2117557 | 0 |
| B142 | B151 | 25-(OH)D | Anti-CCP | -0.2711152 | 0 |
| B143 | B144 | β-CTx | Total-PINP | 0.75244221 | 0 |
| B143 | B145 | β-CTx | BGP | 0.3875566 | 0 |
| B144 | B145 | Total-PINP | BGP | 0.47539046 | 0 |
| B146 | B189 | CysC | eGFB | -0.193011 | 2.22E-15 |
| B148 | B149 | Anti_TPO | TGAb | 0.51298912 | 0 |
| B15 | B101 | CD3-CD19+(B cell) | anti-Sc70 | 0.68607375 | 0 |
| B15 | B102 | CD3-CD19+(B cell) | anti-Sm | 0.68607372 | 0 |
| B15 | B105 | CD3-CD19+(B cell) | anti-U1RNP | 0.68607373 | 0 |
| B15 | B107 | CD3-CD19+(B cell) | AnuA | 0.68607325 | 0 |
| B15 | B108 | CD3-CD19+(B cell) | Anti-Cyclin | 0.68607373 | 0 |
| B15 | B125 | CD3-CD19+(B cell) | Cor | 0.27874228 | 0 |
| B15 | B151 | CD3-CD19+(B cell) | Anti-CCP | 0.22904968 | 0 |
| B15 | B154 | CD3-CD19+(B cell) | RF | -0.3821013 | 0 |
| B15 | B156 | CD3-CD19+(B cell) | IgE | 0.23808324 | 0 |
| B15 | B157 | CD3-CD19+(B cell) | IgG | 0.25580479 | 0 |
| B15 | B16 | CD3-CD19+(B cell) | CD3+T | 0.22107974 | 0 |
| B15 | B17 | CD3-CD19+(B cell) | CD3+CD16+CD56+NK | 0.54898906 | 0 |
| B15 | B181 | CD3-CD19+(B cell) | APTT | -0.3211687 | 0 |
| B15 | B19 | CD3-CD19+(B cell) | CD3+CD4+/CD3+CD8+ | -0.2489407 | 0 |
| B15 | B20 | CD3-CD19+(B cell) | CD3+CD4+CD8+ | -0.3255677 | 0 |
| B15 | B22 | CD3-CD19+(B cell) | CD3+CD8+(T) | 0.49604469 | 0 |
| B15 | B23 | CD3-CD19+(B cell) | CD3-CD16+CD56+(NK) | -0.4664635 | 0 |
| B15 | B79 | CD3-CD19+(B cell) | hb V -dna | 0.2744815 | 0 |
| B15 | B92 | CD3-CD19+(B cell) | ALDH2-Glu504Glu | -0.2088476 | 0 |
| B15 | B94 | CD3-CD19+(B cell) | ALDH2-Lys504Lys | 0.31321827 | 0 |
| B15 | B97 | CD3-CD19+(B cell) | dsDNA | 0.68607373 | 0 |
| B15 | B98 | CD3-CD19+(B cell) | anti-Jo-1 | 0.68607375 | 0 |
| B15 | B99 | CD3-CD19+(B cell) | anti-PM-Scl | 0.68607374 | 0 |
| B15 | B93 | CD3-CD19+(B cell) | ALDH2-Glu504Lys | 0.19453401 | 1.33E-15 |
| B15 | B139 | CD3-CD19+(B cell) | ADA | 0.19232211 | 2.66E-15 |
| B15 | B81 | CD3-CD19+(B cell) | TSGF | -0.174058 | 9.13E-13 |

| | | | | | |
|---|---|---|---|---|---|
| B15 | B155 | CD3-CD19+(B cell) | IgA | 0.16077006 | 4.39E-11 |
| B15 | B135 | CD3-CD19+(B cell) | TBA | 0.15984746 | 5.68E-11 |
| B15 | B121 | CD3-CD19+(B cell) | C3 | -0.1555827 | 1.83E-10 |
| B15 | B122 | CD3-CD19+(B cell) | C4 | -0.1555831 | 1.83E-10 |
| B15 | B159 | CD3-CD19+(B cell) | β2-MG | -0.1555816 | 1.83E-10 |
| B15 | B169 | CD3-CD19+(B cell) | U-CRY | -0.1555839 | 1.83E-10 |
| B15 | B158 | CD3-CD19+(B cell) | IgM | 0.15520381 | 2.03E-10 |
| B15 | B180 | CD3-CD19+(B cell) | Tf | 0.1547707 | 2.28E-10 |
| B15 | B124 | CD3-CD19+(B cell) | ACTH | 0.13291003 | 5.42E-08 |
| B15 | B21 | CD3-CD19+(B cell) | CD3+CD4-CD8- | 0.13224869 | 6.32E-08 |
| B15 | B119 | CD3-CD19+(B cell) | Hcv-rna | 0.13006042 | 1.04E-07 |
| B15 | B109 | CD3-CD19+(B cell) | AMA-M2 | 0.12123064 | 7.22E-07 |
| B151 | B155 | Anti-CCP | IgA | 0.13274621 | 5.63E-08 |
| B153 | B170 | Glu | U-GLU | 0.25446642 | 0 |
| B153 | B220 | Glu | SBP | 0.21926049 | 0 |
| B153 | B206 | Glu | Waist | 0.18354816 | 4.73E-14 |
| B153 | B215 | Glu | BMI | 0.1772303 | 3.46E-13 |
| B153 | B207 | Glu | WHR | 0.1641641 | 1.68E-11 |
| B153 | B189 | Glu | eGFB | -0.1378502 | 1.69E-08 |
| B153 | B222 | Glu | NS | 0.1286131 | 1.44E-07 |
| B153 | B218 | Glu | DBP | 0.12777927 | 1.74E-07 |
| B153 | B216 | Glu | BMI | 0.12770034 | 1.77E-07 |
| B153 | B210 | Glu | Hip | 0.12643148 | 2.35E-07 |
| B154 | B157 | RF | IgG | -0.1869768 | 1.55E-14 |
| B155 | B157 | IgA | IgG | 0.22446037 | 0 |
| B155 | B158 | IgA | IgM | 0.14418595 | 3.57E-09 |
| B155 | B185 | IgA | PT | 0.12377331 | 4.19E-07 |
| B156 | B186 | IgE | PFC | 0.22805413 | 0 |
| B156 | B157 | IgE | IgG | 0.19026357 | 5.33E-15 |
| B156 | B182 | IgE | TT | -0.1783868 | 2.42E-13 |
| B157 | B158 | IgG | IgM | 0.17608389 | 4.92E-13 |
| B157 | B198 | IgG | P | 0.12021953 | 8.94E-07 |
| B158 | B186 | IgM | PFC | 0.15027247 | 7.52E-10 |
| B158 | B181 | IgM | APTT | -0.126161 | 2.49E-07 |
| B158 | B159 | IgM | β2-MG | 0.12327657 | 4.66E-07 |
| B158 | B169 | IgM | U-CRY | 0.12327834 | 4.66E-07 |
| B159 | B169 | β2-MG | U-CRY | 0.99998655 | 0 |
| B159 | B198 | β2-MG | P | 0.20598064 | 0 |
| B159 | B197 | β2-MG | AG | 0.15863377 | 7.95E-11 |
| B159 | B194 | β2-MG | Cl | -0.1468877 | 1.80E-09 |
| B159 | B195 | β2-MG | Na | 0.14528391 | 2.71E-09 |
| B159 | B199 | β2-MG | Mg | -0.1297591 | 1.12E-07 |
| B16 | B125 | CD3+T | Cor | 0.22845464 | 0 |
| B16 | B18 | CD3+T | CD3+CD4+helper(inducer)T | 0.83173009 | 0 |

| | | | | | |
|---|---|---|---|---|---|
| B16 | B181 | CD3+T | APTT | -0.2887294 | 0 |
| B16 | B182 | CD3+T | TT | -0.2364736 | 0 |
| B16 | B19 | CD3+T | CD3+CD4+/CD3+CD8+ | 0.27711508 | 0 |
| B16 | B193 | CD3+T | K | -0.2073335 | 0 |
| B16 | B20 | CD3+T | CD3+CD4+CD8+ | 0.38236656 | 0 |
| B16 | B21 | CD3+T | CD3+CD4-CD8- | -0.5489844 | 0 |
| B16 | B22 | CD3+T | CD3+CD8+(T) | 0.27411402 | 0 |
| B16 | B23 | CD3+T | CD3-CD16+CD56+(NK) | -0.9431253 | 0 |
| B16 | B92 | CD3+T | ALDH2-Glu504Glu | 0.26987876 | 0 |
| B16 | B93 | CD3+T | ALDH2-Glu504Lys | -0.275253 | 0 |
| B16 | B199 | CD3+T | Mg | -0.1853228 | 2.66E-14 |
| B16 | B124 | CD3+T | ACTH | 0.1688048 | 4.38E-12 |
| B16 | B137 | CD3+T | AFU | -0.1529407 | 3.72E-10 |
| B16 | B155 | CD3+T | IgA | 0.14782886 | 1.42E-09 |
| B16 | B81 | CD3+T | TSGF | -0.1409645 | 7.95E-09 |
| B16 | B103 | CD3+T | anti-SSA/Ro | -0.1328397 | 5.51E-08 |
| B16 | B198 | CD3+T | P | -0.1297596 | 1.12E-07 |
| B16 | B121 | CD3+T | C3 | 0.12429716 | 3.74E-07 |
| B16 | B122 | CD3+T | C4 | 0.12429742 | 3.74E-07 |
| B16 | B159 | CD3+T | β2-MG | 0.12429628 | 3.74E-07 |
| B16 | B169 | CD3+T | U-CRY | 0.12429797 | 3.74E-07 |
| B16 | B123 | CD3+T | hs-CRP1 | 0.12358195 | 4.37E-07 |
| B16 | B151 | CD3+T | Anti-CCP | 0.12212929 | 5.96E-07 |
| B16 | B179 | CD3+T | α1-MG | 0.12004249 | 9.27E-07 |
| B160 | B191 | PH | Urea | -0.1491062 | 1.02E-09 |
| B160 | B163 | PH | SG | -0.1391751 | 1.23E-08 |
| B160 | B192 | PH | UA | -0.122748 | 5.22E-07 |
| B161 | B173 | LEU | NTT | 0.14966118 | 8.82E-10 |
| B163 | B165 | SG | bilirubin | 0.24317059 | 0 |
| B163 | B166 | SG | Pro | 0.38615875 | 0 |
| B163 | B176 | SG | Ucr | 0.25809519 | 0 |
| B163 | B191 | SG | Urea | 0.23945331 | 0 |
| B163 | B222 | SG | NS | 0.23067029 | 0 |
| B163 | B192 | SG | UA | 0.20016109 | 2.22E-16 |
| B163 | B216 | SG | BMI | 0.19605707 | 8.88E-16 |
| B163 | B221 | SG | Hei | 0.17495834 | 6.94E-13 |
| B163 | B190 | SG | Cre | 0.15593696 | 1.66E-10 |
| B163 | B170 | SG | U-GLU | 0.14640215 | 2.04E-09 |
| B163 | B206 | SG | Waist | 0.14140143 | 7.14E-09 |
| B163 | B207 | SG | WHR | 0.13660341 | 2.28E-08 |
| B163 | B209 | SG | Smoking | 0.13362513 | 4.59E-08 |
| B165 | B192 | bilirubin | UA | 0.12041241 | 8.58E-07 |
| B166 | B222 | Pro | NS | 0.12913172 | 1.28E-07 |
| B166 | B192 | Pro | UA | 0.12377949 | 4.18E-07 |

| | | | | | |
|---|---|---|---|---|---|
| B167 | B223 | PC | FH | 0.7144725 | 0 |
| B169 | B198 | U-CRY | P | 0.20598362 | 0 |
| B169 | B197 | U-CRY | AG | 0.15863605 | 7.95E-11 |
| B169 | B194 | U-CRY | Cl | -0.14689 | 1.80E-09 |
| B169 | B195 | U-CRY | Na | 0.14528582 | 2.71E-09 |
| B169 | B199 | U-CRY | Mg | -0.1297609 | 1.11E-07 |
| B17 | B101 | CD3+CD16+CD56+NK | anti-Sc70 | 0.39230095 | 0 |
| B17 | B102 | CD3+CD16+CD56+NK | anti-Sm | 0.39230094 | 0 |
| B17 | B105 | CD3+CD16+CD56+NK | anti-U1RNP | 0.39230095 | 0 |
| B17 | B107 | CD3+CD16+CD56+NK | AnuA | 0.39230068 | 0 |
| B17 | B108 | CD3+CD16+CD56+NK | Anti-Cyclin | 0.39230095 | 0 |
| B17 | B110 | CD3+CD16+CD56+NK | AHA | -0.2129242 | 0 |
| B17 | B154 | CD3+CD16+CD56+NK | RF | -0.3823145 | 0 |
| B17 | B157 | CD3+CD16+CD56+NK | IgG | 0.24330987 | 0 |
| B17 | B158 | CD3+CD16+CD56+NK | IgM | 0.25456847 | 0 |
| B17 | B18 | CD3+CD16+CD56+NK | CD3+CD4+helper(inducer)T | -0.2164953 | 0 |
| B17 | B181 | CD3+CD16+CD56+NK | APTT | -0.2714965 | 0 |
| B17 | B183 | CD3+CD16+CD56+NK | PTA | -0.2653238 | 0 |
| B17 | B19 | CD3+CD16+CD56+NK | CD3+CD4+/CD3+CD8+ | -0.207906 | 0 |
| B17 | B21 | CD3+CD16+CD56+NK | CD3+CD4-CD8- | 0.5053338 | 0 |
| B17 | B22 | CD3+CD16+CD56+NK | CD3+CD8+(T) | 0.27752058 | 0 |
| B17 | B79 | CD3+CD16+CD56+NK | hb V -dna | 0.24014783 | 0 |
| B17 | B94 | CD3+CD16+CD56+NK | ALDH2-Lys504Lys | 0.32890827 | 0 |
| B17 | B97 | CD3+CD16+CD56+NK | dsDNA | 0.39230095 | 0 |
| B17 | B98 | CD3+CD16+CD56+NK | anti-Jo-1 | 0.39230096 | 0 |
| B17 | B99 | CD3+CD16+CD56+NK | anti-PM-Scl | 0.39230095 | 0 |
| B17 | B23 | CD3+CD16+CD56+NK | CD3-CD16+CD56+(NK) | -0.1965728 | 6.66E-16 |
| B17 | B180 | CD3+CD16+CD56+NK | Tf | 0.1843055 | 3.71E-14 |
| B17 | B185 | CD3+CD16+CD56+NK | PT | -0.1784952 | 2.34E-13 |
| B17 | B198 | CD3+CD16+CD56+NK | P | 0.17148247 | 1.98E-12 |
| B17 | B176 | CD3+CD16+CD56+NK | Ucr | -0.1607652 | 4.40E-11 |
| B17 | B151 | CD3+CD16+CD56+NK | Anti-CCP | 0.13811787 | 1.59E-08 |
| B17 | B184 | CD3+CD16+CD56+NK | INR | -0.1329725 | 5.34E-08 |
| B17 | B195 | CD3+CD16+CD56+NK | Na | -0.1295715 | 1.16E-07 |
| B172 | B184 | KB | INR | 0.24815098 | 0 |
| B172 | B185 | KB | PT | 0.23891924 | 0 |
| B172 | B182 | KB | TT | 0.17339109 | 1.12E-12 |
| B172 | B186 | KB | PFC | -0.1552569 | 2.00E-10 |
| B174 | B189 | Urea/Cr | eGFB | 0.20199201 | 0 |
| B174 | B190 | Urea/Cr | Cre | -0.297067 | 0 |
| B174 | B191 | Urea/Cr | Urea | 0.67017031 | 0 |
| B174 | B192 | Urea/Cr | UA | -0.2006114 | 0 |
| B174 | B221 | Urea/Cr | Hei | -0.209151 | 0 |
| B174 | B222 | Urea/Cr | NS | -0.1796417 | 1.63E-13 |

| | | | | | |
|---|---|---|---|---|---|
| B174 | B216 | Urea/Cr | BMI | -0.1481332 | 1.31E-09 |
| B175 | B176 | ALB-U | Ucr | 0.24162513 | 0 |
| B175 | B177 | ALB-U | ACR | 0.95123285 | 0 |
| B178 | B179 | Ig-U | α1-MG | 0.91023718 | 0 |
| B178 | B180 | Ig-U | Tf | 0.81852579 | 0 |
| B179 | B180 | α1-MG | Tf | 0.83172071 | 0 |
| B179 | B194 | α1-MG | Cl | 0.14185911 | 6.38E-09 |
| B179 | B184 | α1-MG | INR | 0.13444585 | 3.79E-08 |
| B18 | B101 | CD3+CD4+helper(inducer)T | anti-Sc70 | -0.2490083 | 0 |
| B18 | B102 | CD3+CD4+helper(inducer)T | anti-Sm | -0.2490083 | 0 |
| B18 | B105 | CD3+CD4+helper(inducer)T | anti-U1RNP | -0.2490083 | 0 |
| B18 | B107 | CD3+CD4+helper(inducer)T | AnuA | -0.2490081 | 0 |
| B18 | B108 | CD3+CD4+helper(inducer)T | Anti-Cyclin | -0.2490083 | 0 |
| B18 | B154 | CD3+CD4+helper(inducer)T | RF | 0.20885673 | 0 |
| B18 | B182 | CD3+CD4+helper(inducer)T | TT | -0.2521065 | 0 |
| B18 | B19 | CD3+CD4+helper(inducer)T | CD3+CD4+/CD3+CD8+ | 0.70927038 | 0 |
| B18 | B20 | CD3+CD4+helper(inducer)T | CD3+CD4+CD8+ | 0.3555002 | 0 |
| B18 | B21 | CD3+CD4+helper(inducer)T | CD3+CD4-CD8- | -0.5792123 | 0 |
| B18 | B22 | CD3+CD4+helper(inducer)T | CD3+CD8+(T) | -0.2339493 | 0 |
| B18 | B23 | CD3+CD4+helper(inducer)T | CD3-CD16+CD56+(NK) | -0.7459191 | 0 |
| B18 | B92 | CD3+CD4+helper(inducer)T | ALDH2-Glu504Glu | 0.26552008 | 0 |
| B18 | B93 | CD3+CD4+helper(inducer)T | ALDH2-Glu504Lys | -0.2587733 | 0 |
| B18 | B94 | CD3+CD4+helper(inducer)T | ALDH2-Lys504Lys | -0.218704 | 0 |
| B18 | B97 | CD3+CD4+helper(inducer)T | dsDNA | -0.2490083 | 0 |
| B18 | B98 | CD3+CD4+helper(inducer)T | anti-Jo-1 | -0.2490083 | 0 |
| B18 | B99 | CD3+CD4+helper(inducer)T | anti-PM-Scl | -0.2490083 | 0 |
| B18 | B110 | CD3+CD4+helper(inducer)T | AHA | 0.19792037 | 4.44E-16 |
| B18 | B79 | CD3+CD4+helper(inducer)T | hb V -dna | -0.1938597 | 1.55E-15 |
| B18 | B181 | CD3+CD4+helper(inducer)T | APTT | -0.1828706 | 5.86E-14 |
| B18 | B137 | CD3+CD4+helper(inducer)T | AFU | -0.1731982 | 1.18E-12 |
| B18 | B193 | CD3+CD4+helper(inducer)T | K | -0.1726687 | 1.39E-12 |
| B18 | B121 | CD3+CD4+helper(inducer)T | C3 | 0.1571251 | 1.20E-10 |
| B18 | B122 | CD3+CD4+helper(inducer)T | C4 | 0.15712546 | 1.20E-10 |
| B18 | B159 | CD3+CD4+helper(inducer)T | β2-MG | 0.15712402 | 1.20E-10 |
| B18 | B169 | CD3+CD4+helper(inducer)T | U-CRY | 0.15712627 | 1.20E-10 |
| B18 | B180 | CD3+CD4+helper(inducer)T | Tf | -0.1505544 | 6.98E-10 |
| B18 | B194 | CD3+CD4+helper(inducer)T | Cl | 0.14208777 | 6.03E-09 |
| B18 | B58 | CD3+CD4+helper(inducer)T | HCT | -0.1418406 | 6.41E-09 |
| B18 | B67 | CD3+CD4+helper(inducer)T | WBRSR(mPas_50) | -0.1359838 | 2.64E-08 |
| B18 | B66 | CD3+CD4+helper(inducer)T | WBRSR(mPas_100) | -0.1347897 | 3.50E-08 |
| B18 | B158 | CD3+CD4+helper(inducer)T | IgM | -0.1335538 | 4.67E-08 |
| B18 | B64 | CD3+CD4+helper(inducer)T | WBRSR(mPas_5) | -0.1333288 | 4.92E-08 |
| B18 | B119 | CD3+CD4+helper(inducer)T | Hcv-rna | -0.1324646 | 6.01E-08 |
| B18 | B134 | CD3+CD4+helper(inducer)T | CHE | -0.1316451 | 7.26E-08 |

| | | | | | |
|---|---|---|---|---|---|
| B18 | B62 | CD3+CD4+helper(inducer)T | WBHSRVI | -0.1315191 | 7.47E-08 |
| B18 | B60 | CD3+CD4+helper(inducer)T | WBRVI | -0.1290275 | 1.32E-07 |
| B18 | B63 | CD3+CD4+helper(inducer)T | WBRSR(mPas_200) | -0.1277382 | 1.76E-07 |
| B18 | B65 | CD3+CD4+helper(inducer)T | WBRSR(mPas_1) | -0.127401 | 1.89E-07 |
| B18 | B123 | CD3+CD4+helper(inducer)T | hs-CRP1 | 0.12537802 | 2.96E-07 |
| B18 | B139 | CD3+CD4+helper(inducer)T | ADA | -0.1228832 | 5.08E-07 |
| B180 | B193 | Tf | K | 0.13640703 | 2.39E-08 |
| B181 | B183 | APTT | PTA | 0.2596008 | 0 |
| B181 | B184 | APTT | INR | 0.2258926 | 0 |
| B181 | B185 | APTT | PT | 0.3180133 | 0 |
| B181 | B186 | APTT | PFC | -0.3325534 | 0 |
| B181 | B193 | APTT | K | 0.15080935 | 6.53E-10 |
| B181 | B182 | APTT | TT | 0.13098257 | 8.45E-08 |
| B182 | B184 | TT | INR | 0.60046692 | 0 |
| B182 | B185 | TT | PT | 0.55537741 | 0 |
| B182 | B186 | TT | PFC | -0.5722458 | 0 |
| B183 | B193 | PTA | K | 0.15886036 | 7.47E-11 |
| B184 | B185 | INR | PT | 0.91166924 | 0 |
| B184 | B186 | INR | PFC | -0.529424 | 0 |
| B184 | B199 | INR | Mg | -0.145119 | 2.83E-09 |
| B185 | B186 | PT | PFC | -0.4293931 | 0 |
| B185 | B194 | PT | Cl | -0.1269233 | 2.10E-07 |
| B188 | B192 | Ca | UA | 0.1255809 | 2.83E-07 |
| B189 | B190 | eGFB | Cre | -0.5104604 | 0 |
| B189 | B191 | eGFB | Urea | -0.3514174 | 0 |
| B189 | B192 | eGFB | UA | -0.2812525 | 0 |
| B189 | B206 | eGFB | Waist | -0.249505 | 0 |
| B189 | B207 | eGFB | WHR | -0.2844988 | 0 |
| B189 | B220 | eGFB | SBP | -0.318098 | 0 |
| B189 | B222 | eGFB | NS | -0.1742918 | 8.50E-13 |
| B189 | B219 | eGFB | OT | -0.1540347 | 2.78E-10 |
| B189 | B215 | eGFB | BMI | -0.1534254 | 3.27E-10 |
| B189 | B227 | eGFB | OD | 0.14160343 | 6.79E-09 |
| B189 | B201 | eGFB | e-habits | -0.1364556 | 2.36E-08 |
| B189 | B216 | eGFB | BMI | -0.1351661 | 3.20E-08 |
| B189 | B209 | eGFB | Smoking | -0.1348556 | 3.44E-08 |
| B189 | B218 | eGFB | DBP | -0.1337971 | 4.41E-08 |
| B19 | B101 | CD3+CD4+/CD3+CD8+ | anti-Sc70 | -0.3766767 | 0 |
| B19 | B102 | CD3+CD4+/CD3+CD8+ | anti-Sm | -0.3766767 | 0 |
| B19 | B103 | CD3+CD4+/CD3+CD8+ | anti-SSA/Ro | 0.20841984 | 0 |
| B19 | B105 | CD3+CD4+/CD3+CD8+ | anti-U1RNP | -0.3766767 | 0 |
| B19 | B107 | CD3+CD4+/CD3+CD8+ | AnuA | -0.3766764 | 0 |
| B19 | B108 | CD3+CD4+/CD3+CD8+ | Anti-Cyclin | -0.3766767 | 0 |
| B19 | B124 | CD3+CD4+/CD3+CD8+ | ACTH | -0.2809772 | 0 |

| | | | | | |
|---|---|---|---|---|---|
| B19 | B125 | CD3+CD4+/CD3+CD8+ | Cor | -0.4470642 | 0 |
| B19 | B182 | CD3+CD4+/CD3+CD8+ | TT | -0.2088825 | 0 |
| B19 | B184 | CD3+CD4+/CD3+CD8+ | INR | -0.2466338 | 0 |
| B19 | B22 | CD3+CD4+/CD3+CD8+ | CD3+CD8+(T) | -0.8014754 | 0 |
| B19 | B23 | CD3+CD4+/CD3+CD8+ | CD3-CD16+CD56+(NK) | -0.2720074 | 0 |
| B19 | B60 | CD3+CD4+/CD3+CD8+ | WBRVI | -0.2018404 | 0 |
| B19 | B61 | CD3+CD4+/CD3+CD8+ | WBHSRV | -0.2170218 | 0 |
| B19 | B62 | CD3+CD4+/CD3+CD8+ | WBHSRVI | -0.2549625 | 0 |
| B19 | B63 | CD3+CD4+/CD3+CD8+ | WBRSR(mPas_200) | -0.2539727 | 0 |
| B19 | B64 | CD3+CD4+/CD3+CD8+ | WBRSR(mPas_5) | -0.2283135 | 0 |
| B19 | B65 | CD3+CD4+/CD3+CD8+ | WBRSR(mPas_1) | -0.2037966 | 0 |
| B19 | B66 | CD3+CD4+/CD3+CD8+ | WBRSR(mPas_100) | -0.252677 | 0 |
| B19 | B67 | CD3+CD4+/CD3+CD8+ | WBRSR(mPas_50) | -0.250167 | 0 |
| B19 | B79 | CD3+CD4+/CD3+CD8+ | hb V -dna | -0.3398078 | 0 |
| B19 | B81 | CD3+CD4+/CD3+CD8+ | TSGF | 0.26057891 | 0 |
| B19 | B92 | CD3+CD4+/CD3+CD8+ | ALDH2-Glu504Glu | 0.26478486 | 0 |
| B19 | B93 | CD3+CD4+/CD3+CD8+ | ALDH2-Glu504Lys | -0.2558123 | 0 |
| B19 | B94 | CD3+CD4+/CD3+CD8+ | ALDH2-Lys504Lys | -0.2532826 | 0 |
| B19 | B97 | CD3+CD4+/CD3+CD8+ | dsDNA | -0.3766767 | 0 |
| B19 | B98 | CD3+CD4+/CD3+CD8+ | anti-Jo-1 | -0.3766767 | 0 |
| B19 | B99 | CD3+CD4+/CD3+CD8+ | anti-PM-Scl | -0.3766767 | 0 |
| B19 | B56 | CD3+CD4+/CD3+CD8+ | IR | -0.1987692 | 4.44E-16 |
| B19 | B110 | CD3+CD4+/CD3+CD8+ | AHA | 0.19561198 | 8.88E-16 |
| B19 | B139 | CD3+CD4+/CD3+CD8+ | ADA | -0.1848643 | 3.11E-14 |
| B19 | B156 | CD3+CD4+/CD3+CD8+ | IgE | -0.1820605 | 7.59E-14 |
| B19 | B135 | CD3+CD4+/CD3+CD8+ | TBA | -0.1769752 | 3.74E-13 |
| B19 | B180 | CD3+CD4+/CD3+CD8+ | Tf | -0.1722681 | 1.57E-12 |
| B19 | B58 | CD3+CD4+/CD3+CD8+ | HCT | -0.1638544 | 1.84E-11 |
| B19 | B154 | CD3+CD4+/CD3+CD8+ | RF | 0.161248 | 3.84E-11 |
| B19 | B54 | CD3+CD4+/CD3+CD8+ | ESR | 0.15166946 | 5.21E-10 |
| B19 | B186 | CD3+CD4+/CD3+CD8+ | PFC | -0.1465907 | 1.95E-09 |
| B19 | B185 | CD3+CD4+/CD3+CD8+ | PT | -0.1411652 | 7.57E-09 |
| B19 | B158 | CD3+CD4+/CD3+CD8+ | IgM | -0.1398954 | 1.03E-08 |
| B19 | B194 | CD3+CD4+/CD3+CD8+ | Cl | 0.13742147 | 1.88E-08 |
| B19 | B183 | CD3+CD4+/CD3+CD8+ | PTA | -0.1354881 | 2.97E-08 |
| B19 | B176 | CD3+CD4+/CD3+CD8+ | Ucr | 0.13237712 | 6.13E-08 |
| B19 | B138 | CD3+CD4+/CD3+CD8+ | PAB | -0.1296882 | 1.13E-07 |
| B19 | B178 | CD3+CD4+/CD3+CD8+ | Ig-U | -0.1257828 | 2.71E-07 |
| B190 | B191 | Cre | Urea | 0.36251539 | 0 |
| B190 | B192 | Cre | UA | 0.29951046 | 0 |
| B190 | B221 | Cre | Hei | 0.20424686 | 0 |
| B190 | B222 | Cre | NS | 0.19710431 | 4.44E-16 |
| B190 | B207 | Cre | WHR | 0.19272671 | 2.22E-15 |
| B190 | B206 | Cre | Waist | 0.18019546 | 1.37E-13 |

| | | | | | |
|---|---|---|---|---|---|
| B190 | B216 | Cre | BMI | 0.17528842 | 6.27E-13 |
| B190 | B209 | Cre | Smoking | 0.14881289 | 1.10E-09 |
| B190 | B204 | Cre | Drinking | 0.13190647 | 6.84E-08 |
| B191 | B207 | Urea | WHR | 0.17178095 | 1.81E-12 |
| B191 | B192 | Urea | UA | 0.16536293 | 1.19E-11 |
| B191 | B206 | Urea | Waist | 0.13968118 | 1.09E-08 |
| B191 | B220 | Urea | SBP | 0.12830593 | 1.55E-07 |
| B192 | B204 | UA | Drinking | 0.25408153 | 0 |
| B192 | B206 | UA | Waist | 0.40844848 | 0 |
| B192 | B207 | UA | WHR | 0.37303285 | 0 |
| B192 | B209 | UA | Smoking | 0.21173004 | 0 |
| B192 | B210 | UA | Hip | 0.26899673 | 0 |
| B192 | B215 | UA | BMI | 0.32737937 | 0 |
| B192 | B216 | UA | BMI | 0.41380461 | 0 |
| B192 | B221 | UA | Hei | 0.24620425 | 0 |
| B192 | B222 | UA | NS | 0.43409989 | 0 |
| B192 | B220 | UA | SBP | 0.15287483 | 3.79E-10 |
| B192 | B218 | UA | DBP | 0.15282335 | 3.84E-10 |
| B193 | B198 | K | P | 0.234722 | 0 |
| B194 | B197 | Cl | AG | -0.4469758 | 0 |
| B194 | B198 | Cl | P | 0.19987365 | 2.22E-16 |
| B194 | B200 | Cl | OP | 0.14093205 | 8.01E-09 |
| B194 | B195 | Cl | Na | 0.11970862 | 9.95E-07 |
| B195 | B197 | Na | AG | 0.5228123 | 0 |
| B195 | B199 | Na | Mg | 0.24124103 | 0 |
| B195 | B200 | Na | OP | 0.93268589 | 0 |
| B196 | B197 | TCO2 | AG | -0.7124255 | 0 |
| B196 | B198 | TCO2 | P | 0.13291871 | 5.41E-08 |
| B196 | B200 | TCO2 | OP | -0.1248134 | 3.34E-07 |
| B197 | B200 | AG | OP | 0.50093376 | 0 |
| B197 | B198 | AG | P | -0.1507546 | 6.63E-10 |
| B198 | B199 | P | Mg | 0.16646766 | 8.67E-12 |
| B198 | B200 | P | OP | 0.12580041 | 2.69E-07 |
| B199 | B200 | Mg | OP | 0.22541906 | 0 |
| B2 | B118 | Age | CEA1 | 0.23330884 | 0 |
| B2 | B126 | Age | Alb | -0.2347371 | 0 |
| B2 | B131 | Age | AST | 0.21198826 | 0 |
| B2 | B153 | Age | Glu | 0.21592789 | 0 |
| B2 | B189 | Age | eGFB | -0.7026477 | 0 |
| B2 | B191 | Age | Urea | 0.25907329 | 0 |
| B2 | B206 | Age | Waist | 0.31243783 | 0 |
| B2 | B207 | Age | WHR | 0.36671545 | 0 |
| B2 | B215 | Age | BMI | 0.20533556 | 0 |
| B2 | B219 | Age | OT | 0.25046376 | 0 |

| | | | | | |
|---|---|---|---|---|---|
| B2 | B220 | Age | SBP | 0.44748115 | 0 |
| B2 | B227 | Age | OD | -0.2168498 | 0 |
| B2 | B47 | Age | RDW | 0.21934585 | 0 |
| B2 | B77 | Age | TC | 0.26450423 | 0 |
| B2 | B70 | Age | LDL-C | 0.18746997 | 1.33E-14 |
| B2 | B43 | Age | MCV | 0.18394795 | 4.17E-14 |
| B2 | B52 | Age | PLT/BPC | -0.1771751 | 3.52E-13 |
| B2 | B222 | Age | NS | 0.1679627 | 5.61E-12 |
| B2 | B146 | Age | CysC | 0.16708907 | 7.24E-12 |
| B2 | B190 | Age | Cre | 0.16612201 | 9.58E-12 |
| B2 | B44 | Age | MCH | 0.16390638 | 1.81E-11 |
| B2 | B218 | Age | DBP | 0.15990331 | 5.59E-11 |
| B2 | B201 | Age | e-habits | 0.15882785 | 7.54E-11 |
| B2 | B91 | Age | TP | -0.1537764 | 2.98E-10 |
| B2 | B221 | Age | Hei | -0.1483478 | 1.24E-09 |
| B2 | B27 | Age | LDH | 0.1366668 | 2.25E-08 |
| B2 | B71 | Age | TG | 0.13506117 | 3.28E-08 |
| B2 | B192 | Age | UA | 0.13317816 | 5.10E-08 |
| B2 | B53 | Age | GR% | -0.1292266 | 1.26E-07 |
| B2 | B224 | Age | pmh | 0.12840583 | 1.51E-07 |
| B2 | B205 | Age | KDA | 0.12405589 | 3.94E-07 |
| B20 | B119 | CD3+CD4+CD8+ | Hcv-rna | -0.35532 | 0 |
| B20 | B121 | CD3+CD4+CD8+ | C3 | 0.42068475 | 0 |
| B20 | B122 | CD3+CD4+CD8+ | C4 | 0.42068574 | 0 |
| B20 | B159 | CD3+CD4+CD8+ | β2-MG | 0.42068185 | 0 |
| B20 | B169 | CD3+CD4+CD8+ | U-CRY | 0.42068791 | 0 |
| B20 | B21 | CD3+CD4+CD8+ | CD3+CD4-CD8- | -0.3885244 | 0 |
| B20 | B22 | CD3+CD4+CD8+ | CD3+CD8+(T) | 0.28043463 | 0 |
| B20 | B23 | CD3+CD4+CD8+ | CD3-CD16+CD56+(NK) | -0.2176143 | 0 |
| B20 | B101 | CD3+CD4+CD8+ | anti-Sc70 | -0.1993207 | 2.22E-16 |
| B20 | B102 | CD3+CD4+CD8+ | anti-Sm | -0.1993207 | 2.22E-16 |
| B20 | B105 | CD3+CD4+CD8+ | anti-U1RNP | -0.1993207 | 2.22E-16 |
| B20 | B107 | CD3+CD4+CD8+ | AnuA | -0.1993206 | 2.22E-16 |
| B20 | B108 | CD3+CD4+CD8+ | Anti-Cyclin | -0.1993207 | 2.22E-16 |
| B20 | B97 | CD3+CD4+CD8+ | dsDNA | -0.1993207 | 2.22E-16 |
| B20 | B98 | CD3+CD4+CD8+ | anti-Jo-1 | -0.1993207 | 2.22E-16 |
| B20 | B99 | CD3+CD4+CD8+ | anti-PM-Scl | -0.1993207 | 2.22E-16 |
| B20 | B120 | CD3+CD4+CD8+ | HCV-cAg | -0.1648898 | 1.37E-11 |
| B20 | B195 | CD3+CD4+CD8+ | Na | 0.16161289 | 3.47E-11 |
| B20 | B200 | CD3+CD4+CD8+ | OP | 0.15040277 | 7.27E-10 |
| B20 | B82 | CD3+CD4+CD8+ | CEA | -0.1381712 | 1.57E-08 |
| B20 | B125 | CD3+CD4+CD8+ | Cor | 0.13707246 | 2.04E-08 |
| B20 | B211 | CD3+CD4+CD8+ | IOP-R | -0.1310827 | 8.26E-08 |
| B20 | B193 | CD3+CD4+CD8+ | K | -0.1304016 | 9.64E-08 |

| | | | | | |
|---|---|---|---|---|---|
| B20 | B150 | CD3+CD4+CD8+ | ALP | -0.1271455 | 2.00E-07 |
| B202 | B223 | e-times | FH | 0.55316819 | 0 |
| B204 | B206 | Drinking | Waist | 0.23636881 | 0 |
| B204 | B207 | Drinking | WHR | 0.21517746 | 0 |
| B204 | B209 | Drinking | Smoking | 0.39260708 | 0 |
| B204 | B216 | Drinking | BMI | 0.29967354 | 0 |
| B204 | B221 | Drinking | Hei | 0.28766813 | 0 |
| B204 | B222 | Drinking | NS | 0.36235843 | 0 |
| B204 | B210 | Drinking | Hip | 0.14969493 | 8.74E-10 |
| B204 | B215 | Drinking | BMI | 0.14885704 | 1.09E-09 |
| B206 | B207 | Waist | WHR | 0.84733896 | 0 |
| B206 | B210 | Waist | Hip | 0.75456699 | 0 |
| B206 | B215 | Waist | BMI | 0.80663512 | 0 |
| B206 | B216 | Waist | BMI | 0.80867433 | 0 |
| B206 | B218 | Waist | DBP | 0.24041706 | 0 |
| B206 | B220 | Waist | SBP | 0.30103419 | 0 |
| B206 | B221 | Waist | Hei | 0.26945363 | 0 |
| B206 | B222 | Waist | NS | 0.73499351 | 0 |
| B206 | B209 | Waist | Smoking | 0.17946521 | 1.73E-13 |
| B207 | B210 | WHR | Hip | 0.29529312 | 0 |
| B207 | B215 | WHR | BMI | 0.55197834 | 0 |
| B207 | B216 | WHR | BMI | 0.53007933 | 0 |
| B207 | B220 | WHR | SBP | 0.27374197 | 0 |
| B207 | B222 | WHR | NS | 0.58570468 | 0 |
| B207 | B218 | WHR | DBP | 0.19239514 | 2.66E-15 |
| B207 | B209 | WHR | Smoking | 0.17729491 | 3.39E-13 |
| B207 | B221 | WHR | Hei | 0.13708164 | 2.03E-08 |
| B209 | B216 | Smoking | BMI | 0.2129169 | 0 |
| B209 | B221 | Smoking | Hei | 0.2570443 | 0 |
| B209 | B222 | Smoking | NS | 0.30425923 | 0 |
| B21 | B125 | CD3+CD4-CD8- | Cor | -0.2154162 | 0 |
| B21 | B154 | CD3+CD4-CD8- | RF | -0.4944637 | 0 |
| B21 | B22 | CD3+CD4-CD8- | CD3+CD8+(T) | -0.2237589 | 0 |
| B21 | B23 | CD3+CD4-CD8- | CD3-CD16+CD56+(NK) | 0.41984298 | 0 |
| B21 | B94 | CD3+CD4-CD8- | ALDH2-Lys504Lys | 0.1854506 | 2.58E-14 |
| B21 | B121 | CD3+CD4-CD8- | C3 | -0.1824873 | 6.64E-14 |
| B21 | B122 | CD3+CD4-CD8- | C4 | -0.1824877 | 6.64E-14 |
| B21 | B169 | CD3+CD4-CD8- | U-CRY | -0.1824887 | 6.64E-14 |
| B21 | B159 | CD3+CD4-CD8- | β2-MG | -0.1824861 | 6.66E-14 |
| B21 | B103 | CD3+CD4-CD8- | anti-SSA/Ro | 0.17367892 | 1.02E-12 |
| B21 | B93 | CD3+CD4-CD8- | ALDH2-Glu504Lys | -0.1549293 | 2.18E-10 |
| B21 | B158 | CD3+CD4-CD8- | IgM | 0.15427737 | 2.60E-10 |
| B21 | B119 | CD3+CD4-CD8- | Hcv-rna | 0.15415878 | 2.69E-10 |
| B21 | B185 | CD3+CD4-CD8- | PT | -0.14756 | 1.52E-09 |

| | | | | | |
|---|---|---|---|---|---|
| B21 | B199 | CD3+CD4-CD8- | Mg | 0.14749427 | 1.54E-09 |
| B21 | B76 | CD3+CD4-CD8- | ApoB | 0.14379419 | 3.94E-09 |
| B21 | B137 | CD3+CD4-CD8- | AFU | 0.14015812 | 9.68E-09 |
| B21 | B92 | CD3+CD4-CD8- | ALDH2-Glu504Glu | 0.13931925 | 1.19E-08 |
| B21 | B184 | CD3+CD4-CD8- | INR | -0.1375689 | 1.81E-08 |
| B21 | B101 | CD3+CD4-CD8- | anti-Sc70 | 0.13497247 | 3.35E-08 |
| B21 | B102 | CD3+CD4-CD8- | anti-Sm | 0.13497246 | 3.35E-08 |
| B21 | B105 | CD3+CD4-CD8- | anti-U1RNP | 0.13497247 | 3.35E-08 |
| B21 | B107 | CD3+CD4-CD8- | AnuA | 0.13497237 | 3.35E-08 |
| B21 | B108 | CD3+CD4-CD8- | Anti-Cyclin | 0.13497247 | 3.35E-08 |
| B21 | B97 | CD3+CD4-CD8- | dsDNA | 0.13497247 | 3.35E-08 |
| B21 | B98 | CD3+CD4-CD8- | anti-Jo-1 | 0.13497247 | 3.35E-08 |
| B21 | B99 | CD3+CD4-CD8- | anti-PM-Scl | 0.13497247 | 3.35E-08 |
| B21 | B124 | CD3+CD4-CD8- | ACTH | -0.1344394 | 3.80E-08 |
| B21 | B193 | CD3+CD4-CD8- | K | 0.13076749 | 8.87E-08 |
| B21 | B110 | CD3+CD4-CD8- | AHA | -0.127386 | 1.90E-07 |
| B210 | B215 | Hip | BMI | 0.77540194 | 0 |
| B210 | B216 | Hip | BMI | 0.80062573 | 0 |
| B210 | B220 | Hip | SBP | 0.20414046 | 0 |
| B210 | B221 | Hip | Hei | 0.31099488 | 0 |
| B210 | B222 | Hip | NS | 0.59711062 | 0 |
| B210 | B218 | Hip | DBP | 0.19192024 | 3.11E-15 |
| B211 | B212 | IOP-R | IOP-L | 0.80360747 | 0 |
| B215 | B216 | BMI | BMI | 0.81248972 | 0 |
| B215 | B218 | BMI | DBP | 0.23199854 | 0 |
| B215 | B220 | BMI | SBP | 0.28464517 | 0 |
| B215 | B222 | BMI | NS | 0.62409917 | 0 |
| B216 | B218 | BMI | DBP | 0.2354885 | 0 |
| B216 | B220 | BMI | SBP | 0.22370123 | 0 |
| B216 | B221 | BMI | Hei | 0.5869468 | 0 |
| B216 | B222 | BMI | NS | 0.77106421 | 0 |
| B218 | B220 | DBP | SBP | 0.67987439 | 0 |
| B218 | B222 | DBP | NS | 0.22244743 | 0 |
| B22 | B101 | CD3+CD8+(T) | anti-Sc70 | 0.54445175 | 0 |
| B22 | B102 | CD3+CD8+(T) | anti-Sm | 0.54445174 | 0 |
| B22 | B103 | CD3+CD8+(T) | anti-SSA/Ro | -0.2708374 | 0 |
| B22 | B105 | CD3+CD8+(T) | anti-U1RNP | 0.54445174 | 0 |
| B22 | B107 | CD3+CD8+(T) | AnuA | 0.54445136 | 0 |
| B22 | B108 | CD3+CD8+(T) | Anti-Cyclin | 0.54445175 | 0 |
| B22 | B124 | CD3+CD8+(T) | ACTH | 0.31369181 | 0 |
| B22 | B125 | CD3+CD8+(T) | Cor | 0.56655073 | 0 |
| B22 | B135 | CD3+CD8+(T) | TBA | 0.21874534 | 0 |
| B22 | B139 | CD3+CD8+(T) | ADA | 0.20612261 | 0 |
| B22 | B156 | CD3+CD8+(T) | IgE | 0.23751085 | 0 |

| | | | | | |
|---|---|---|---|---|---|
| B22 | B178 | CD3+CD8+(T) | Ig-U | 0.20636177 | 0 |
| B22 | B184 | CD3+CD8+(T) | INR | 0.3136025 | 0 |
| B22 | B199 | CD3+CD8+(T) | Mg | -0.2189041 | 0 |
| B22 | B23 | CD3+CD8+(T) | CD3-CD16+CD56+(NK) | -0.2552459 | 0 |
| B22 | B56 | CD3+CD8+(T) | IR | 0.22036796 | 0 |
| B22 | B61 | CD3+CD8+(T) | WBHSRV | 0.23672109 | 0 |
| B22 | B62 | CD3+CD8+(T) | WBHSRVI | 0.22626464 | 0 |
| B22 | B63 | CD3+CD8+(T) | WBRSR(mPas_200) | 0.2348534 | 0 |
| B22 | B66 | CD3+CD8+(T) | WBRSR(mPas_100) | 0.22150684 | 0 |
| B22 | B67 | CD3+CD8+(T) | WBRSR(mPas_50) | 0.21766041 | 0 |
| B22 | B79 | CD3+CD8+(T) | hb V -dna | 0.38791829 | 0 |
| B22 | B81 | CD3+CD8+(T) | TSGF | -0.2973793 | 0 |
| B22 | B94 | CD3+CD8+(T) | ALDH2-Lys504Lys | 0.36184291 | 0 |
| B22 | B97 | CD3+CD8+(T) | dsDNA | 0.54445175 | 0 |
| B22 | B98 | CD3+CD8+(T) | anti-Jo-1 | 0.54445176 | 0 |
| B22 | B99 | CD3+CD8+(T) | anti-PM-Scl | 0.54445175 | 0 |
| B22 | B64 | CD3+CD8+(T) | WBRSR(mPas_5) | 0.19535705 | 8.88E-16 |
| B22 | B151 | CD3+CD8+(T) | Anti-CCP | 0.18298007 | 5.68E-14 |
| B22 | B92 | CD3+CD8+(T) | ALDH2-Glu504Glu | -0.180599 | 1.21E-13 |
| B22 | B180 | CD3+CD8+(T) | Tf | 0.1793568 | 1.79E-13 |
| B22 | B110 | CD3+CD8+(T) | AHA | -0.1751273 | 6.59E-13 |
| B22 | B65 | CD3+CD8+(T) | WBRSR(mPas_1) | 0.17016129 | 2.94E-12 |
| B22 | B185 | CD3+CD8+(T) | PT | 0.16869386 | 4.53E-12 |
| B22 | B60 | CD3+CD8+(T) | WBRVI | 0.16677528 | 7.93E-12 |
| B22 | B93 | CD3+CD8+(T) | ALDH2-Glu504Lys | 0.16241715 | 2.76E-11 |
| B22 | B54 | CD3+CD8+(T) | ESR | -0.1533458 | 3.34E-10 |
| B22 | B154 | CD3+CD8+(T) | RF | -0.1450794 | 2.85E-09 |
| B22 | B59 | CD3+CD8+(T) | WBRV | 0.13614773 | 2.54E-08 |
| B22 | B179 | CD3+CD8+(T) | α1-MG | 0.12909632 | 1.30E-07 |
| B22 | B176 | CD3+CD8+(T) | Ucr | -0.128163 | 1.60E-07 |
| B22 | B157 | CD3+CD8+(T) | IgG | 0.12535191 | 2.97E-07 |
| B22 | B194 | CD3+CD8+(T) | Cl | -0.1210922 | 7.43E-07 |
| B22 | B186 | CD3+CD8+(T) | PFC | 0.11974744 | 9.87E-07 |
| B220 | B222 | SBP | NS | 0.23071909 | 0 |
| B221 | B222 | Hei | NS | 0.44767716 | 0 |
| B23 | B125 | CD3-CD16+CD56+(NK) | Cor | -0.2244492 | 0 |
| B23 | B181 | CD3-CD16+CD56+(NK) | APTT | 0.40902807 | 0 |
| B23 | B182 | CD3-CD16+CD56+(NK) | TT | 0.28792473 | 0 |
| B23 | B193 | CD3-CD16+CD56+(NK) | K | 0.20930706 | 0 |
| B23 | B93 | CD3-CD16+CD56+(NK) | ALDH2-Glu504Lys | 0.19171153 | 3.11E-15 |
| B23 | B101 | CD3-CD16+CD56+(NK) | anti-Sc70 | -0.1913519 | 3.55E-15 |
| B23 | B102 | CD3-CD16+CD56+(NK) | anti-Sm | -0.1913519 | 3.55E-15 |
| B23 | B105 | CD3-CD16+CD56+(NK) | anti-U1RNP | -0.1913519 | 3.55E-15 |
| B23 | B107 | CD3-CD16+CD56+(NK) | AnuA | -0.1913517 | 3.55E-15 |

| | | | | | |
|---|---|---|---|---|---|
| B23 | B108 | CD3-CD16+CD56+(NK) | Anti-Cyclin | -0.1913519 | 3.55E-15 |
| B23 | B97 | CD3-CD16+CD56+(NK) | dsDNA | -0.1913519 | 3.55E-15 |
| B23 | B98 | CD3-CD16+CD56+(NK) | anti-Jo-1 | -0.1913519 | 3.55E-15 |
| B23 | B99 | CD3-CD16+CD56+(NK) | anti-PM-Scl | -0.1913519 | 3.55E-15 |
| B23 | B92 | CD3-CD16+CD56+(NK) | ALDH2-Glu504Glu | -0.1883342 | 9.99E-15 |
| B23 | B124 | CD3-CD16+CD56+(NK) | ACTH | -0.1762374 | 4.69E-13 |
| B23 | B151 | CD3-CD16+CD56+(NK) | Anti-CCP | -0.1604408 | 4.82E-11 |
| B23 | B81 | CD3-CD16+CD56+(NK) | TSGF | 0.15744556 | 1.10E-10 |
| B23 | B155 | CD3-CD16+CD56+(NK) | IgA | -0.1410096 | 7.86E-09 |
| B23 | B183 | CD3-CD16+CD56+(NK) | PTA | 0.13279114 | 5.57E-08 |
| B23 | B137 | CD3-CD16+CD56+(NK) | AFU | 0.12814204 | 1.60E-07 |
| B23 | B199 | CD3-CD16+CD56+(NK) | Mg | 0.12556406 | 2.84E-07 |
| B24 | B25 | αHBDH | CK | 0.37894246 | 0 |
| B24 | B26 | αHBDH | CKMB | 0.20819467 | 0 |
| B24 | B27 | αHBDH | LDH | 0.71620766 | 0 |
| B24 | B74 | αHBDH | ApoA1 | 0.24628549 | 0 |
| B24 | B139 | αHBDH | ADA | 0.17842462 | 2.39E-13 |
| B24 | B58 | αHBDH | HCT | -0.1451626 | 2.79E-09 |
| B24 | B62 | αHBDH | WBHSRVI | -0.1266801 | 2.22E-07 |
| B24 | B75 | αHBDH | ApoA1/B | 0.12584779 | 2.67E-07 |
| B24 | B135 | αHBDH | TBA | 0.12259315 | 5.40E-07 |
| B24 | B73 | αHBDH | ApoA | 0.12188524 | 6.28E-07 |
| B24 | B146 | αHBDH | CysC | 0.12121236 | 7.25E-07 |
| B25 | B27 | CK | LDH | 0.33677399 | 0 |
| B25 | B26 | CK | CKMB | 0.18468397 | 3.29E-14 |
| B25 | B73 | CK | ApoA | 0.15178039 | 5.06E-10 |
| B25 | B110 | CK | AHA | 0.14879016 | 1.11E-09 |
| B25 | B76 | CK | ApoB | 0.14859236 | 1.16E-09 |
| B25 | B54 | CK | ESR | 0.13187335 | 6.89E-08 |
| B25 | B75 | CK | ApoA1/B | -0.1251203 | 3.13E-07 |
| B25 | B69 | CK | PV | 0.125099 | 3.14E-07 |
| B25 | B174 | CK | Urea/Cr | -0.1241716 | 3.84E-07 |
| B25 | B136 | CK | 5'NT | 0.12378375 | 4.18E-07 |
| B26 | B27 | CKMB | LDH | 0.16573268 | 1.07E-11 |
| B27 | B74 | LDH | ApoA1 | 0.19968895 | 2.22E-16 |
| B27 | B136 | LDH | 5'NT | 0.17038957 | 2.74E-12 |
| B27 | B197 | LDH | AG | 0.16620122 | 9.36E-12 |
| B27 | B196 | LDH | TCO2 | -0.1649933 | 1.33E-11 |
| B27 | B139 | LDH | ADA | 0.15770432 | 1.03E-10 |
| B27 | B220 | LDH | SBP | 0.15756077 | 1.07E-10 |
| B27 | B131 | LDH | AST | 0.15403198 | 2.78E-10 |
| B27 | B150 | LDH | ALP | 0.14442936 | 3.36E-09 |
| B27 | B189 | LDH | eGFB | -0.1369454 | 2.10E-08 |
| B27 | B134 | LDH | CHE | 0.13030329 | 9.86E-08 |

| | | | | | |
|---|---|---|---|---|---|
| B28 | B30 | PRL | LH | 0.24065489 | 0 |
| B28 | B75 | PRL | ApoA1/B | -0.1433276 | 4.43E-09 |
| B28 | B33 | PRL | Prog | 0.13727069 | 1.94E-08 |
| B29 | B30 | E2 | LH | -0.5142006 | 0 |
| B29 | B31 | E2 | FSH | -0.6130249 | 0 |
| B29 | B32 | E2 | Testo | 0.35861151 | 0 |
| B29 | B33 | E2 | Prog | 0.31193541 | 0 |
| B29 | B73 | E2 | ApoA | -0.1706354 | 2.55E-12 |
| B29 | B194 | E2 | Cl | -0.1605977 | 4.61E-11 |
| B29 | B198 | E2 | P | -0.1426122 | 5.29E-09 |
| B29 | B74 | E2 | ApoA1 | -0.1351961 | 3.18E-08 |
| B29 | B136 | E2 | 5'NT | 0.12149497 | 6.83E-07 |
| B3 | B146 | Hcy | CysC | 0.15122392 | 5.86E-10 |
| B3 | B25 | Hcy | CK | 0.14644839 | 2.02E-09 |
| B3 | B24 | Hcy | αHBDH | 0.14599729 | 2.26E-09 |
| B3 | B125 | Hcy | Cor | 0.12567722 | 2.77E-07 |
| B3 | B199 | Hcy | Mg | -0.1253773 | 2.96E-07 |
| B3 | B50 | Hcy | eo% | 0.12403057 | 3.96E-07 |
| B30 | B31 | LH | FSH | 0.91366083 | 0 |
| B30 | B32 | LH | Testo | -0.3882613 | 0 |
| B30 | B33 | LH | Prog | -0.3496809 | 0 |
| B30 | B73 | LH | ApoA | 0.15979677 | 5.76E-11 |
| B30 | B161 | LH | LEU | 0.12596742 | 2.60E-07 |
| B31 | B32 | FSH | Testo | -0.5250149 | 0 |
| B31 | B33 | FSH | Prog | -0.4395265 | 0 |
| B31 | B73 | FSH | ApoA | 0.19195932 | 3.11E-15 |
| B31 | B194 | FSH | Cl | 0.161053 | 4.06E-11 |
| B31 | B193 | FSH | K | 0.13629079 | 2.45E-08 |
| B31 | B198 | FSH | P | 0.1350396 | 3.30E-08 |
| B32 | B33 | Testo | Prog | 0.81738285 | 0 |
| B32 | B194 | Testo | Cl | -0.1898532 | 6.22E-15 |
| B32 | B198 | Testo | P | -0.1700053 | 3.08E-12 |
| B32 | B73 | Testo | ApoA | -0.1603355 | 4.96E-11 |
| B32 | B136 | Testo | 5'NT | 0.14600284 | 2.26E-09 |
| B32 | B74 | Testo | ApoA1 | -0.1383853 | 1.49E-08 |
| B32 | B134 | Testo | CHE | 0.12860182 | 1.45E-07 |
| B32 | B193 | Testo | K | -0.1247751 | 3.37E-07 |
| B32 | B138 | Testo | PAB | 0.12041603 | 8.57E-07 |
| B33 | B73 | Prog | ApoA | -0.1782381 | 2.53E-13 |
| B33 | B194 | Prog | Cl | -0.16369 | 1.93E-11 |
| B33 | B198 | Prog | P | -0.1431525 | 4.63E-09 |
| B33 | B74 | Prog | ApoA1 | -0.1371911 | 1.98E-08 |
| B33 | B136 | Prog | 5'NT | 0.12109845 | 7.42E-07 |
| B34 | B145 | TK | BGP | 0.25358268 | 0 |

| | | | | | |
|---|---|---|---|---|---|
| B34 | B144 | TK | Total-PINP | 0.15182729 | 5.00E-10 |
| B34 | B120 | TK | HCV-cAg | -0.1443877 | 3.40E-09 |
| B34 | B87 | TK | NSE | 0.13813469 | 1.58E-08 |
| B34 | B88 | TK | CA 242 | 0.12077938 | 7.94E-07 |
| B35 | B202 | β2-MG | e-times | 0.64908514 | 0 |
| B37 | B38 | BAND | BAND% | 0.86093306 | 0 |
| B39 | B103 | Other cells | anti-SSA/Ro | 0.2056132 | 0 |
| B39 | B104 | Other cells | anti-SSB | 0.32782863 | 0 |
| B39 | B40 | Other cells | WBC | 0.22612436 | 0 |
| B39 | B100 | Other cells | anti-Ro52 | 0.18875952 | 8.66E-15 |
| B4 | B5 | WB-Ca | WB-Mg | -0.4113526 | 0 |
| B4 | B7 | WB-Ca | WB-Cop | -0.693474 | 0 |
| B4 | B73 | WB-Ca | ApoA | 0.14510162 | 2.84E-09 |
| B40 | B41 | WBC | MON % | -0.2214992 | 0 |
| B40 | B42 | WBC | MON | 0.59888889 | 0 |
| B40 | B52 | WBC | PLT/BPC | 0.20458388 | 0 |
| B40 | B53 | WBC | GR% | 0.20478241 | 0 |
| B40 | B46 | WBC | RBC Count | 0.1752712 | 6.31E-13 |
| B40 | B48 | WBC | LYM% | -0.1666593 | 8.20E-12 |
| B40 | B51 | WBC | Hb | 0.15871524 | 7.77E-11 |
| B40 | B71 | WBC | TG | 0.158238 | 8.87E-11 |
| B40 | B206 | WBC | Waist | 0.13139554 | 7.69E-08 |
| B40 | B130 | WBC | GLB | 0.13030496 | 9.86E-08 |
| B40 | B207 | WBC | WHR | 0.12711118 | 2.02E-07 |
| B40 | B49 | WBC | baso% | -0.126765 | 2.18E-07 |
| B40 | B72 | WBC | HDL-C | -0.1228655 | 5.10E-07 |
| B40 | B222 | WBC | NS | 0.12114672 | 7.35E-07 |
| B41 | B42 | MON % | MON | 0.57259544 | 0 |
| B41 | B53 | MON % | GR% | -0.2556312 | 0 |
| B42 | B51 | MON | Hb | 0.22902169 | 0 |
| B42 | B192 | MON | UA | 0.19246873 | 2.66E-15 |
| B42 | B46 | MON | RBC Count | 0.19238613 | 2.66E-15 |
| B42 | B222 | MON | NS | 0.17337443 | 1.12E-12 |
| B42 | B128 | MON | ALT | 0.16174711 | 3.34E-11 |
| B42 | B206 | MON | Waist | 0.15158477 | 5.33E-10 |
| B42 | B165 | MON | bilirubin | 0.14980292 | 8.50E-10 |
| B42 | B131 | MON | AST | 0.14406329 | 3.69E-09 |
| B42 | B207 | MON | WHR | 0.14222487 | 5.83E-09 |
| B42 | B127 | MON | A-G ratio | -0.1406375 | 8.61E-09 |
| B42 | B48 | MON | LYM% | -0.1388265 | 1.34E-08 |
| B42 | B130 | MON | GLB | 0.13593893 | 2.67E-08 |
| B42 | B209 | MON | Smoking | 0.12804176 | 1.64E-07 |
| B42 | B71 | MON | TG | 0.12654257 | 2.29E-07 |
| B42 | B216 | MON | BMI | 0.12399961 | 3.99E-07 |

| | | | | | |
|---|---|---|---|---|---|
| B42 | B52 | MON | PLT/BPC | 0.1200956 | 9.17E-07 |
| B43 | B44 | MCV | MCH | 0.93623723 | 0 |
| B43 | B45 | MCV | MCHC | 0.36960758 | 0 |
| B43 | B46 | MCV | RBC Count | -0.4941775 | 0 |
| B43 | B47 | MCV | RDW | 0.44056572 | 0 |
| B43 | B51 | MCV | Hb | 0.3384339 | 0 |
| B43 | B52 | MCV | PLT/BPC | -0.2855881 | 0 |
| B43 | B189 | MCV | eGFB | -0.159232 | 6.74E-11 |
| B44 | B45 | MCH | MCHC | 0.67023677 | 0 |
| B44 | B46 | MCH | RBC Count | -0.3852863 | 0 |
| B44 | B47 | MCH | RDW | 0.26655108 | 0 |
| B44 | B51 | MCH | Hb | 0.49013207 | 0 |
| B44 | B52 | MCH | PLT/BPC | -0.2509054 | 0 |
| B44 | B189 | MCH | eGFB | -0.1587505 | 7.70E-11 |
| B44 | B54 | MCH | ESR | -0.12352 | 4.43E-07 |
| B45 | B47 | MCHC | RDW | -0.2369142 | 0 |
| B45 | B51 | MCHC | Hb | 0.59078055 | 0 |
| B45 | B192 | MCHC | UA | 0.18212777 | 7.46E-14 |
| B45 | B133 | MCHC | TBIL | 0.15866193 | 7.89E-11 |
| B45 | B129 | MCHC | IBIL | 0.15654765 | 1.41E-10 |
| B45 | B54 | MCHC | ESR | -0.1375583 | 1.81E-08 |
| B45 | B132 | MCHC | Dbil | 0.12573918 | 2.73E-07 |
| B45 | B58 | MCHC | HCT | 0.12246231 | 5.55E-07 |
| B45 | B222 | MCHC | NS | 0.12220407 | 5.87E-07 |
| B46 | B126 | RBC Count | Alb | 0.21662443 | 0 |
| B46 | B128 | RBC Count | ALT | 0.21981334 | 0 |
| B46 | B163 | RBC Count | SG | 0.21549941 | 0 |
| B46 | B192 | RBC Count | UA | 0.30753199 | 0 |
| B46 | B204 | RBC Count | Drinking | 0.20483257 | 0 |
| B46 | B216 | RBC Count | BMI | 0.2405709 | 0 |
| B46 | B218 | RBC Count | DBP | 0.22324214 | 0 |
| B46 | B221 | RBC Count | Hei | 0.26869054 | 0 |
| B46 | B222 | RBC Count | NS | 0.27278347 | 0 |
| B46 | B47 | RBC Count | RDW | -0.4225922 | 0 |
| B46 | B51 | RBC Count | Hb | 0.60307934 | 0 |
| B46 | B72 | RBC Count | HDL-C | -0.1994591 | 2.22E-16 |
| B46 | B141 | RBC Count | AST/ALT | -0.1907583 | 4.44E-15 |
| B46 | B206 | RBC Count | Waist | 0.17426625 | 8.57E-13 |
| B46 | B188 | RBC Count | Ca | 0.16354405 | 2.01E-11 |
| B46 | B209 | RBC Count | Smoking | 0.15511162 | 2.08E-10 |
| B46 | B207 | RBC Count | WHR | 0.15509774 | 2.09E-10 |
| B46 | B91 | RBC Count | TP | 0.15333906 | 3.35E-10 |
| B46 | B165 | RBC Count | bilirubin | 0.15257404 | 4.10E-10 |
| B46 | B71 | RBC Count | TG | 0.14454777 | 3.26E-09 |

| | | | | | |
|---|---|---|---|---|---|
| B46 | B174 | RBC Count | Urea/Cr | -0.1437866 | 3.95E-09 |
| B46 | B52 | RBC Count | PLT/BPC | 0.12983587 | 1.10E-07 |
| B46 | B140 | RBC Count | r-GT | 0.12565857 | 2.78E-07 |
| B46 | B166 | RBC Count | Pro | 0.12250693 | 5.50E-07 |
| B46 | B176 | RBC Count | Ucr | 0.12220677 | 5.87E-07 |
| B47 | B52 | RDW | PLT/BPC | -0.2181827 | 0 |
| B47 | B189 | RDW | eGFB | -0.1690943 | 4.02E-12 |
| B47 | B112 | RDW | FT4 | -0.1679779 | 5.58E-12 |
| B47 | B51 | RDW | Hb | -0.1593123 | 6.59E-11 |
| B47 | B72 | RDW | HDL-C | 0.15837428 | 8.54E-11 |
| B47 | B111 | RDW | TSH | 0.15436545 | 2.54E-10 |
| B47 | B118 | RDW | CEA1 | 0.14702622 | 1.74E-09 |
| B47 | B126 | RDW | Alb | -0.1290886 | 1.30E-07 |
| B47 | B77 | RDW | TC | 0.12655077 | 2.28E-07 |
| B48 | B53 | LYM% | GR% | -0.9507621 | 0 |
| B49 | B50 | baso% | eo% | 0.28341193 | 0 |
| B49 | B53 | baso% | GR% | -0.1802886 | 1.33E-13 |
| B49 | B177 | baso% | ACR | 0.124816 | 3.34E-07 |
| B5 | B6 | WB-Mg | WB-Fe | 0.66484304 | 0 |
| B5 | B7 | WB-Mg | WB-Cop | -0.31341 | 0 |
| B5 | B8 | WB-Mg | WB-Zinc | -0.8114666 | 0 |
| B5 | B58 | WB-Mg | HCT | 0.19481305 | 1.11E-15 |
| B5 | B73 | WB-Mg | ApoA | -0.170532 | 2.63E-12 |
| B5 | B62 | WB-Mg | WBHSRVI | 0.15725577 | 1.16E-10 |
| B5 | B57 | WB-Mg | RBCAI | -0.1488251 | 1.10E-09 |
| B5 | B63 | WB-Mg | WBRSR(mPas_200) | 0.14249442 | 5.45E-09 |
| B5 | B66 | WB-Mg | WBRSR(mPas_100) | 0.13047711 | 9.48E-08 |
| B5 | B43 | WB-Mg | MCV | 0.12412284 | 3.89E-07 |
| B5 | B67 | WB-Mg | WBRSR(mPas_50) | 0.12228193 | 5.77E-07 |
| B50 | B53 | eo% | GR% | -0.2863998 | 0 |
| B50 | B190 | eo% | Cre | 0.14876266 | 1.11E-09 |
| B50 | B189 | eo% | eGFB | -0.1310091 | 8.40E-08 |
| B50 | B156 | eo% | IgE | 0.12739729 | 1.89E-07 |
| B51 | B128 | Hb | ALT | 0.22238779 | 0 |
| B51 | B141 | Hb | AST/ALT | -0.2438774 | 0 |
| B51 | B163 | Hb | SG | 0.2223385 | 0 |
| B51 | B192 | Hb | UA | 0.37124237 | 0 |
| B51 | B204 | Hb | Drinking | 0.23985776 | 0 |
| B51 | B206 | Hb | Waist | 0.22830562 | 0 |
| B51 | B207 | Hb | WHR | 0.22753752 | 0 |
| B51 | B209 | Hb | Smoking | 0.21747925 | 0 |
| B51 | B216 | Hb | BMI | 0.27422621 | 0 |
| B51 | B218 | Hb | DBP | 0.23300799 | 0 |
| B51 | B221 | Hb | Hei | 0.29564228 | 0 |

| | | | | | |
|---|---|---|---|---|---|
| B51 | B222 | Hb | NS | 0.326488 | 0 |
| B51 | B126 | Hb | Alb | 0.19138501 | 3.55E-15 |
| B51 | B174 | Hb | Urea/Cr | -0.1867712 | 1.67E-14 |
| B51 | B71 | Hb | TG | 0.18378248 | 4.40E-14 |
| B51 | B190 | Hb | Cre | 0.1827569 | 6.08E-14 |
| B51 | B165 | Hb | bilirubin | 0.17767473 | 3.01E-13 |
| B51 | B58 | Hb | HCT | 0.1735626 | 1.06E-12 |
| B51 | B188 | Hb | Ca | 0.17130596 | 2.09E-12 |
| B51 | B133 | Hb | TBIL | 0.15303946 | 3.62E-10 |
| B51 | B129 | Hb | IBIL | 0.15256254 | 4.11E-10 |
| B51 | B84 | Hb | C-PSA | 0.15087125 | 6.43E-10 |
| B51 | B72 | Hb | HDL-C | -0.1446793 | 3.16E-09 |
| B51 | B189 | Hb | eGFB | -0.1377287 | 1.74E-08 |
| B51 | B166 | Hb | Pro | 0.13339419 | 4.85E-08 |
| B51 | B220 | Hb | SBP | 0.13154106 | 7.43E-08 |
| B51 | B55 | Hb | TK | -0.1255524 | 2.85E-07 |
| B51 | B62 | Hb | WBHSRVI | 0.11987586 | 9.60E-07 |
| B51 | B210 | Hb | Hip | 0.11975353 | 9.85E-07 |
| B52 | B189 | PLT/BPC | eGFB | 0.17042872 | 2.71E-12 |
| B52 | B132 | PLT/BPC | Dbil | -0.1667448 | 8.00E-12 |
| B52 | B133 | PLT/BPC | TBIL | -0.1295119 | 1.18E-07 |
| B54 | B106 | ESR | anti-P | 0.30162618 | 0 |
| B54 | B55 | ESR | TK | 0.28770607 | 0 |
| B54 | B58 | ESR | HCT | -0.4381305 | 0 |
| B54 | B59 | ESR | WBRV | 0.34791283 | 0 |
| B54 | B60 | ESR | WBRVI | -0.2507557 | 0 |
| B54 | B62 | ESR | WBHSRVI | -0.3395407 | 0 |
| B54 | B69 | ESR | PV | 0.45805377 | 0 |
| B54 | B61 | ESR | WBHSRV | 0.16868532 | 4.54E-12 |
| B54 | B195 | ESR | Na | -0.1574253 | 1.11E-10 |
| B54 | B157 | ESR | IgG | 0.13825467 | 1.54E-08 |
| B54 | B179 | ESR | α1-MG | -0.1298473 | 1.09E-07 |
| B54 | B101 | ESR | anti-Sc70 | -0.1295783 | 1.16E-07 |
| B54 | B102 | ESR | anti-Sm | -0.1295783 | 1.16E-07 |
| B54 | B105 | ESR | anti-U1RNP | -0.1295783 | 1.16E-07 |
| B54 | B107 | ESR | AnuA | -0.1295782 | 1.16E-07 |
| B54 | B108 | ESR | Anti-Cyclin | -0.1295783 | 1.16E-07 |
| B54 | B97 | ESR | dsDNA | -0.1295783 | 1.16E-07 |
| B54 | B98 | ESR | anti-Jo-1 | -0.1295783 | 1.16E-07 |
| B54 | B99 | ESR | anti-PM-Scl | -0.1295783 | 1.16E-07 |
| B54 | B178 | ESR | Ig-U | -0.1272658 | 1.95E-07 |
| B54 | B182 | ESR | TT | -0.1264486 | 2.34E-07 |
| B54 | B94 | ESR | ALDH2-Lys504Lys | -0.1262799 | 2.43E-07 |
| B54 | B200 | ESR | OP | -0.1262065 | 2.46E-07 |

| | | | | | |
|---|---|---|---|---|---|
| B54 | B180 | ESR | Tf | -0.1250915 | 3.15E-07 |
| B55 | B56 | TK | IR | 0.76740052 | 0 |
| B55 | B58 | TK | HCT | -0.601382 | 0 |
| B55 | B59 | TK | WBRV | 0.57771287 | 0 |
| B55 | B61 | TK | WBHSRV | 0.6670247 | 0 |
| B55 | B83 | TK | AFP | 0.32591476 | 0 |
| B55 | B86 | TK | TBhCG | 0.32517757 | 0 |
| B55 | B69 | TK | PV | -0.158778 | 7.64E-11 |
| B56 | B110 | IR | AHA | -0.225677 | 0 |
| B56 | B59 | IR | WBRV | 0.63603961 | 0 |
| B56 | B60 | IR | WBRVI | 0.55503376 | 0 |
| B56 | B61 | IR | WBHSRV | 0.91354725 | 0 |
| B56 | B62 | IR | WBHSRVI | 0.69412865 | 0 |
| B56 | B63 | IR | WBRSR(mPas_200) | 0.55159827 | 0 |
| B56 | B64 | IR | WBRSR(mPas_5) | 0.49750478 | 0 |
| B56 | B65 | IR | WBRSR(mPas_1) | 0.44968532 | 0 |
| B56 | B66 | IR | WBRSR(mPas_100) | 0.55608854 | 0 |
| B56 | B67 | IR | WBRSR(mPas_50) | 0.55146005 | 0 |
| B56 | B106 | IR | anti-P | -0.1411722 | 7.55E-09 |
| B56 | B86 | IR | TBhCG | 0.1370312 | 2.06E-08 |
| B56 | B83 | IR | AFP | 0.13675002 | 2.20E-08 |
| B56 | B69 | IR | PV | -0.1363823 | 2.40E-08 |
| B56 | B138 | IR | PAB | 0.13044292 | 9.55E-08 |
| B56 | B156 | IR | IgE | 0.12682716 | 2.15E-07 |
| B56 | B125 | IR | Cor | 0.1204944 | 8.43E-07 |
| B57 | B156 | RBCAI | IgE | -0.2574271 | 0 |
| B57 | B59 | RBCAI | WBRV | 0.59050293 | 0 |
| B57 | B60 | RBCAI | WBRVI | 0.60112625 | 0 |
| B57 | B64 | RBCAI | WBRSR(mPas_5) | 0.47379047 | 0 |
| B57 | B65 | RBCAI | WBRSR(mPas_1) | 0.60800108 | 0 |
| B57 | B66 | RBCAI | WBRSR(mPas_100) | 0.21228259 | 0 |
| B57 | B67 | RBCAI | WBRSR(mPas_50) | 0.25977591 | 0 |
| B57 | B94 | RBCAI | ALDH2-Lys504Lys | 0.18262461 | 6.35E-14 |
| B57 | B58 | RBCAI | HCT | 0.17995291 | 1.48E-13 |
| B57 | B63 | RBCAI | WBRSR(mPas_200) | 0.17518772 | 6.47E-13 |
| B57 | B69 | RBCAI | PV | 0.1664757 | 8.65E-12 |
| B57 | B81 | RBCAI | TSGF | -0.15115 | 5.97E-10 |
| B57 | B92 | RBCAI | ALDH2-Glu504Glu | -0.1493841 | 9.48E-10 |
| B57 | B83 | RBCAI | AFP | -0.143516 | 4.23E-09 |
| B57 | B86 | RBCAI | TBhCG | -0.1430331 | 4.77E-09 |
| B57 | B93 | RBCAI | ALDH2-Glu504Lys | 0.14178767 | 6.49E-09 |
| B57 | B73 | RBCAI | ApoA | 0.13275584 | 5.62E-08 |
| B57 | B180 | RBCAI | Tf | 0.12185428 | 6.32E-07 |
| B58 | B60 | HCT | WBRVI | 0.68272539 | 0 |

| | | | | | |
|---|---|---|---|---|---|
| B58 | B62 | HCT | WBHSRVI | 0.74137645 | 0 |
| B58 | B63 | HCT | WBRSR(mPas_200) | 0.74075723 | 0 |
| B58 | B64 | HCT | WBRSR(mPas_5) | 0.70454054 | 0 |
| B58 | B65 | HCT | WBRSR(mPas_1) | 0.66263434 | 0 |
| B58 | B66 | HCT | WBRSR(mPas_100) | 0.72854253 | 0 |
| B58 | B67 | HCT | WBRSR(mPas_50) | 0.72685471 | 0 |
| B58 | B83 | HCT | AFP | -0.3044795 | 0 |
| B58 | B86 | HCT | TBhCG | -0.30212 | 0 |
| B58 | B110 | HCT | AHA | -0.1690959 | 4.02E-12 |
| B58 | B135 | HCT | TBA | -0.1525845 | 4.09E-10 |
| B58 | B106 | HCT | anti-P | -0.1506943 | 6.73E-10 |
| B58 | B137 | HCT | AFU | 0.13482864 | 3.47E-08 |
| B58 | B134 | HCT | CHE | 0.12228113 | 5.77E-07 |
| B59 | B60 | WBRV | WBRVI | 0.59185858 | 0 |
| B59 | B61 | WBRV | WBHSRV | 0.78707406 | 0 |
| B59 | B62 | WBRV | WBHSRVI | 0.33717856 | 0 |
| B59 | B63 | WBRV | WBRSR(mPas_200) | 0.47442326 | 0 |
| B59 | B64 | WBRV | WBRSR(mPas_5) | 0.62582685 | 0 |
| B59 | B65 | WBRV | WBRSR(mPas_1) | 0.67581168 | 0 |
| B59 | B66 | WBRV | WBRSR(mPas_100) | 0.50582408 | 0 |
| B59 | B67 | WBRV | WBRSR(mPas_50) | 0.53154798 | 0 |
| B59 | B69 | WBRV | PV | 0.39518276 | 0 |
| B59 | B81 | WBRV | TSGF | -0.1254075 | 2.94E-07 |
| B6 | B7 | WB-Fe | WB-Cop | -0.5035603 | 0 |
| B6 | B8 | WB-Fe | WB-Zinc | -0.426845 | 0 |
| B6 | B43 | WB-Fe | MCV | 0.1512009 | 5.89E-10 |
| B6 | B73 | WB-Fe | ApoA | -0.1353497 | 3.07E-08 |
| B6 | B44 | WB-Fe | MCH | 0.13217182 | 6.43E-08 |
| B60 | B110 | WBRVI | AHA | -0.2235675 | 0 |
| B60 | B61 | WBRVI | WBHSRV | 0.58782718 | 0 |
| B60 | B62 | WBRVI | WBHSRVI | 0.85054902 | 0 |
| B60 | B63 | WBRVI | WBRSR(mPas_200) | 0.81309301 | 0 |
| B60 | B64 | WBRVI | WBRSR(mPas_5) | 0.91619329 | 0 |
| B60 | B65 | WBRVI | WBRSR(mPas_1) | 0.93500705 | 0 |
| B60 | B66 | WBRVI | WBRSR(mPas_100) | 0.82839077 | 0 |
| B60 | B67 | WBRVI | WBRSR(mPas_50) | 0.84926039 | 0 |
| B60 | B106 | WBRVI | anti-P | -0.1783984 | 2.41E-13 |
| B60 | B83 | WBRVI | AFP | -0.1707495 | 2.47E-12 |
| B60 | B86 | WBRVI | TBhCG | -0.1690013 | 4.14E-12 |
| B60 | B176 | WBRVI | Ucr | 0.12913794 | 1.28E-07 |
| B60 | B81 | WBRVI | TSGF | -0.1226502 | 5.34E-07 |
| B60 | B182 | WBRVI | TT | 0.12079139 | 7.92E-07 |
| B61 | B110 | WBHSRV | AHA | -0.210857 | 0 |
| B61 | B62 | WBHSRV | WBHSRVI | 0.67569473 | 0 |

| | | | | | |
|---|---|---|---|---|---|
| B61 | B63 | WBHSRV | WBRSR(mPas_200) | 0.71753223 | 0 |
| B61 | B64 | WBHSRV | WBRSR(mPas_5) | 0.67612387 | 0 |
| B61 | B65 | WBHSRV | WBRSR(mPas_1) | 0.62606014 | 0 |
| B61 | B66 | WBHSRV | WBRSR(mPas_100) | 0.72655937 | 0 |
| B61 | B67 | WBHSRV | WBRSR(mPas_50) | 0.72369145 | 0 |
| B61 | B69 | WBHSRV | PV | 0.2683029 | 0 |
| B61 | B124 | WBHSRV | ACTH | 0.1653959 | 1.18E-11 |
| B61 | B156 | WBHSRV | IgE | 0.15147843 | 5.48E-10 |
| B61 | B136 | WBHSRV | 5'NT | 0.14400214 | 3.74E-09 |
| B61 | B138 | WBHSRV | PAB | 0.13516117 | 3.21E-08 |
| B61 | B125 | WBHSRV | Cor | 0.13060085 | 9.21E-08 |
| B62 | B110 | WBHSRVI | AHA | -0.2699924 | 0 |
| B62 | B63 | WBHSRVI | WBRSR(mPas_200) | 0.89690615 | 0 |
| B62 | B64 | WBHSRVI | WBRSR(mPas_5) | 0.82822294 | 0 |
| B62 | B65 | WBHSRVI | WBRSR(mPas_1) | 0.76267785 | 0 |
| B62 | B66 | WBHSRVI | WBRSR(mPas_100) | 0.89131569 | 0 |
| B62 | B67 | WBHSRVI | WBRSR(mPas_50) | 0.88595002 | 0 |
| B62 | B106 | WBHSRVI | anti-P | -0.1995472 | 2.22E-16 |
| B62 | B124 | WBHSRVI | ACTH | 0.16135319 | 3.73E-11 |
| B62 | B134 | WBHSRVI | CHE | 0.15683409 | 1.30E-10 |
| B62 | B138 | WBHSRVI | PAB | 0.15145405 | 5.51E-10 |
| B62 | B156 | WBHSRVI | IgE | 0.13959064 | 1.11E-08 |
| B62 | B176 | WBHSRVI | Ucr | 0.13086889 | 8.67E-08 |
| B62 | B83 | WBHSRVI | AFP | -0.1254669 | 2.90E-07 |
| B62 | B86 | WBHSRVI | TBhCG | -0.1238598 | 4.11E-07 |
| B62 | B94 | WBHSRVI | ALDH2-Lys504Lys | -0.1215686 | 6.72E-07 |
| B63 | B110 | WBRSR(mPas_200) | AHA | -0.2287775 | 0 |
| B63 | B64 | WBRSR(mPas_200) | WBRSR(mPas_5) | 0.9481393 | 0 |
| B63 | B65 | WBRSR(mPas_200) | WBRSR(mPas_1) | 0.8852284 | 0 |
| B63 | B66 | WBRSR(mPas_200) | WBRSR(mPas_100) | 0.99750844 | 0 |
| B63 | B67 | WBRSR(mPas_200) | WBRSR(mPas_50) | 0.99424194 | 0 |
| B63 | B69 | WBRSR(mPas_200) | PV | 0.40563884 | 0 |
| B63 | B124 | WBRSR(mPas_200) | ACTH | 0.1953651 | 8.88E-16 |
| B63 | B83 | WBRSR(mPas_200) | AFP | -0.1806054 | 1.21E-13 |
| B63 | B86 | WBRSR(mPas_200) | TBhCG | -0.1783281 | 2.46E-13 |
| B63 | B134 | WBRSR(mPas_200) | CHE | 0.15606211 | 1.61E-10 |
| B63 | B156 | WBRSR(mPas_200) | IgE | 0.15169416 | 5.18E-10 |
| B63 | B138 | WBRSR(mPas_200) | PAB | 0.14778129 | 1.43E-09 |
| B63 | B136 | WBRSR(mPas_200) | 5'NT | 0.14613328 | 2.19E-09 |
| B63 | B176 | WBRSR(mPas_200) | Ucr | 0.12562372 | 2.80E-07 |
| B64 | B110 | WBRSR(mPas_5) | AHA | -0.2113925 | 0 |
| B64 | B65 | WBRSR(mPas_5) | WBRSR(mPas_1) | 0.98673087 | 0 |
| B64 | B66 | WBRSR(mPas_5) | WBRSR(mPas_100) | 0.96050739 | 0 |
| B64 | B67 | WBRSR(mPas_5) | WBRSR(mPas_50) | 0.97294383 | 0 |

| | | | | | |
|---|---|---|---|---|---|
| B64 | B69 | WBRSR(mPas_5) | PV | 0.42600654 | 0 |
| B64 | B83 | WBRSR(mPas_5) | AFP | -0.2004771 | 0 |
| B64 | B86 | WBRSR(mPas_5) | TBhCG | -0.1983891 | 4.44E-16 |
| B64 | B124 | WBRSR(mPas_5) | ACTH | 0.14551352 | 2.56E-09 |
| B64 | B136 | WBRSR(mPas_5) | 5'NT | 0.14095179 | 7.97E-09 |
| B64 | B134 | WBRSR(mPas_5) | CHE | 0.13456285 | 3.69E-08 |
| B64 | B176 | WBRSR(mPas_5) | Ucr | 0.12949469 | 1.18E-07 |
| B64 | B81 | WBRSR(mPas_5) | TSGF | -0.1233974 | 4.54E-07 |
| B65 | B66 | WBRSR(mPas_1) | WBRSR(mPas_100) | 0.90354225 | 0 |
| B65 | B67 | WBRSR(mPas_1) | WBRSR(mPas_50) | 0.92348346 | 0 |
| B65 | B69 | WBRSR(mPas_1) | PV | 0.41515823 | 0 |
| B65 | B83 | WBRSR(mPas_1) | AFP | -0.2031649 | 0 |
| B65 | B86 | WBRSR(mPas_1) | TBhCG | -0.2010401 | 0 |
| B65 | B110 | WBRSR(mPas_1) | AHA | -0.1913515 | 3.55E-15 |
| B65 | B81 | WBRSR(mPas_1) | TSGF | -0.1426656 | 5.22E-09 |
| B65 | B136 | WBRSR(mPas_1) | 5'NT | 0.13042773 | 9.58E-08 |
| B65 | B176 | WBRSR(mPas_1) | Ucr | 0.12414468 | 3.87E-07 |
| B66 | B110 | WBRSR(mPas_100) | AHA | -0.2313186 | 0 |
| B66 | B67 | WBRSR(mPas_100) | WBRSR(mPas_50) | 0.99876546 | 0 |
| B66 | B69 | WBRSR(mPas_100) | PV | 0.41801796 | 0 |
| B66 | B124 | WBRSR(mPas_100) | ACTH | 0.19496486 | 1.11E-15 |
| B66 | B83 | WBRSR(mPas_100) | AFP | -0.1820668 | 7.59E-14 |
| B66 | B86 | WBRSR(mPas_100) | TBhCG | -0.179821 | 1.54E-13 |
| B66 | B134 | WBRSR(mPas_100) | CHE | 0.16561322 | 1.11E-11 |
| B66 | B136 | WBRSR(mPas_100) | 5'NT | 0.14838872 | 1.23E-09 |
| B66 | B156 | WBRSR(mPas_100) | IgE | 0.14227701 | 5.75E-09 |
| B66 | B138 | WBRSR(mPas_100) | PAB | 0.1392241 | 1.21E-08 |
| B66 | B94 | WBRSR(mPas_100) | ALDH2-Lys504Lys | -0.1294645 | 1.19E-07 |
| B66 | B176 | WBRSR(mPas_100) | Ucr | 0.12913357 | 1.28E-07 |
| B66 | B135 | WBRSR(mPas_100) | TBA | -0.1200933 | 9.18E-07 |
| B67 | B110 | WBRSR(mPas_50) | AHA | -0.2302073 | 0 |
| B67 | B69 | WBRSR(mPas_50) | PV | 0.42056381 | 0 |
| B67 | B124 | WBRSR(mPas_50) | ACTH | 0.1873715 | 1.38E-14 |
| B67 | B83 | WBRSR(mPas_50) | AFP | -0.1867696 | 1.67E-14 |
| B67 | B86 | WBRSR(mPas_50) | TBhCG | -0.1845326 | 3.46E-14 |
| B67 | B134 | WBRSR(mPas_50) | CHE | 0.16191715 | 3.18E-11 |
| B67 | B136 | WBRSR(mPas_50) | 5'NT | 0.14803636 | 1.34E-09 |
| B67 | B138 | WBRSR(mPas_50) | PAB | 0.13265649 | 5.75E-08 |
| B67 | B176 | WBRSR(mPas_50) | Ucr | 0.13006388 | 1.04E-07 |
| B67 | B156 | WBRSR(mPas_50) | IgE | 0.12605989 | 2.55E-07 |
| B67 | B135 | WBRSR(mPas_50) | TBA | -0.1200514 | 9.26E-07 |
| B69 | B106 | PV | anti-P | 0.44939311 | 0 |
| B69 | B83 | PV | AFP | -0.1726844 | 1.38E-12 |
| B69 | B86 | PV | TBhCG | -0.17027 | 2.84E-12 |

| | | | | | |
|---|---|---|---|---|---|
| B69 | B91 | PV | TP | 0.15399597 | 2.81E-10 |
| B69 | B157 | PV | IgG | 0.14419912 | 3.56E-09 |
| B69 | B158 | PV | IgM | 0.13009175 | 1.03E-07 |
| B7 | B8 | WB-Cop | WB-Zinc | 0.59563632 | 0 |
| B7 | B199 | WB-Cop | Mg | 0.15130497 | 5.73E-10 |
| B7 | B193 | WB-Cop | K | 0.12835022 | 1.53E-07 |
| B70 | B132 | LDL-C | Dbil | -0.2105849 | 0 |
| B70 | B189 | LDL-C | eGFB | -0.2012124 | 0 |
| B70 | B75 | LDL-C | ApoA1/B | -0.2012752 | 0 |
| B70 | B76 | LDL-C | ApoB | 0.26972037 | 0 |
| B70 | B77 | LDL-C | TC | 0.8956847 | 0 |
| B70 | B206 | LDL-C | Waist | 0.19450766 | 1.33E-15 |
| B70 | B215 | LDL-C | BMI | 0.17318074 | 1.19E-12 |
| B70 | B207 | LDL-C | WHR | 0.17292974 | 1.28E-12 |
| B70 | B71 | LDL-C | TG | 0.1714046 | 2.03E-12 |
| B70 | B218 | LDL-C | DBP | 0.16494534 | 1.34E-11 |
| B70 | B192 | LDL-C | UA | 0.16383743 | 1.85E-11 |
| B70 | B220 | LDL-C | SBP | 0.14773866 | 1.45E-09 |
| B70 | B188 | LDL-C | Ca | 0.14573799 | 2.42E-09 |
| B70 | B111 | LDL-C | TSH | 0.14220449 | 5.86E-09 |
| B70 | B210 | LDL-C | Hip | 0.13932095 | 1.19E-08 |
| B70 | B153 | LDL-C | Glu | 0.13377521 | 4.44E-08 |
| B70 | B216 | LDL-C | BMI | 0.12913532 | 1.28E-07 |
| B70 | B222 | LDL-C | NS | 0.124241 | 3.79E-07 |
| B71 | B128 | TG | ALT | 0.28167523 | 0 |
| B71 | B131 | TG | AST | 0.20975923 | 0 |
| B71 | B132 | TG | Dbil | -0.2087297 | 0 |
| B71 | B140 | TG | r-GT | 0.24722868 | 0 |
| B71 | B192 | TG | UA | 0.33577712 | 0 |
| B71 | B206 | TG | Waist | 0.32678238 | 0 |
| B71 | B207 | TG | WHR | 0.31759929 | 0 |
| B71 | B215 | TG | BMI | 0.32901905 | 0 |
| B71 | B216 | TG | BMI | 0.28291637 | 0 |
| B71 | B218 | TG | DBP | 0.21392909 | 0 |
| B71 | B222 | TG | NS | 0.26516514 | 0 |
| B71 | B72 | TG | HDL-C | -0.4327736 | 0 |
| B71 | B77 | TG | TC | 0.29751064 | 0 |
| B71 | B91 | TG | TP | 0.21158108 | 0 |
| B71 | B130 | TG | GLB | 0.19833445 | 4.44E-16 |
| B71 | B210 | TG | Hip | 0.19710925 | 4.44E-16 |
| B71 | B141 | TG | AST/ALT | -0.1849412 | 3.02E-14 |
| B71 | B220 | TG | SBP | 0.16191552 | 3.18E-11 |
| B71 | B153 | TG | Glu | 0.15559304 | 1.83E-10 |
| B71 | B134 | TG | CHE | 0.14327275 | 4.49E-09 |

| | | | | | |
|---|---|---|---|---|---|
| B71 | B189 | TG | eGFB | -0.1405343 | 8.83E-09 |
| B71 | B127 | TG | A-G ratio | -0.1404083 | 9.11E-09 |
| B71 | B150 | TG | ALP | 0.12355328 | 4.39E-07 |
| B71 | B74 | TG | ApoA1 | -0.1212573 | 7.18E-07 |
| B72 | B192 | HDL-C | UA | -0.3283451 | 0 |
| B72 | B206 | HDL-C | Waist | -0.3080484 | 0 |
| B72 | B207 | HDL-C | WHR | -0.2657916 | 0 |
| B72 | B210 | HDL-C | Hip | -0.2313509 | 0 |
| B72 | B215 | HDL-C | BMI | -0.3106156 | 0 |
| B72 | B216 | HDL-C | BMI | -0.3559139 | 0 |
| B72 | B222 | HDL-C | NS | -0.3279948 | 0 |
| B72 | B74 | HDL-C | ApoA1 | 0.27341745 | 0 |
| B72 | B77 | HDL-C | TC | 0.25295145 | 0 |
| B72 | B132 | HDL-C | Dbil | 0.18189763 | 8.02E-14 |
| B72 | B221 | HDL-C | Hei | -0.1796147 | 1.65E-13 |
| B72 | B141 | HDL-C | AST/ALT | 0.16387398 | 1.83E-11 |
| B72 | B75 | HDL-C | ApoA1/B | 0.16252453 | 2.68E-11 |
| B72 | B128 | HDL-C | ALT | -0.1612631 | 3.83E-11 |
| B72 | B133 | HDL-C | TBIL | 0.13856448 | 1.42E-08 |
| B72 | B140 | HDL-C | r-GT | -0.1237507 | 4.21E-07 |
| B72 | B134 | HDL-C | CHE | -0.1207147 | 8.05E-07 |
| B73 | B193 | ApoA | K | 0.14004429 | 9.95E-09 |
| B73 | B90 | ApoA | CYFRA 21-1 | 0.13390042 | 4.31E-08 |
| B73 | B137 | ApoA | AFU | 0.12650831 | 2.31E-07 |
| B73 | B76 | ApoA | ApoB | 0.12155575 | 6.74E-07 |
| B74 | B75 | ApoA1 | ApoA1/B | 0.5877379 | 0 |
| B74 | B138 | ApoA1 | PAB | -0.1527352 | 3.93E-10 |
| B74 | B198 | ApoA1 | P | 0.13739703 | 1.89E-08 |
| B74 | B200 | ApoA1 | OP | 0.12849125 | 1.48E-07 |
| B75 | B134 | ApoA1/B | CHE | -0.2295618 | 0 |
| B75 | B76 | ApoA1/B | ApoB | -0.7877488 | 0 |
| B75 | B150 | ApoA1/B | ALP | -0.1497107 | 8.71E-10 |
| B75 | B136 | ApoA1/B | 5'NT | -0.147721 | 1.46E-09 |
| B75 | B77 | ApoA1/B | TC | -0.1389592 | 1.30E-08 |
| B76 | B134 | ApoB | CHE | 0.25921783 | 0 |
| B76 | B136 | ApoB | 5'NT | 0.22690049 | 0 |
| B76 | B77 | ApoB | TC | 0.23831296 | 0 |
| B76 | B197 | ApoB | AG | 0.18747623 | 1.33E-14 |
| B76 | B198 | ApoB | P | 0.16337052 | 2.11E-11 |
| B76 | B200 | ApoB | OP | 0.15583669 | 1.71E-10 |
| B76 | B193 | ApoB | K | 0.15140962 | 5.58E-10 |
| B76 | B196 | ApoB | TCO2 | -0.136988 | 2.08E-08 |
| B76 | B155 | ApoB | IgA | -0.1311275 | 8.17E-08 |
| B76 | B150 | ApoB | ALP | 0.1244281 | 3.64E-07 |

| | | | | | |
|---|---|---|---|---|---|
| B76 | B211 | ApoB | IOP-R | 0.12347154 | 4.47E-07 |
| B76 | B156 | ApoB | IgE | 0.12064584 | 8.17E-07 |
| B77 | B189 | TC | eGFB | -0.2661557 | 0 |
| B77 | B132 | TC | Dbil | -0.1883932 | 9.77E-15 |
| B77 | B188 | TC | Ca | 0.17920973 | 1.87E-13 |
| B77 | B111 | TC | TSH | 0.17458105 | 7.79E-13 |
| B77 | B220 | TC | SBP | 0.17456856 | 7.82E-13 |
| B77 | B218 | TC | DBP | 0.16083988 | 4.31E-11 |
| B77 | B207 | TC | WHR | 0.14853057 | 1.18E-09 |
| B77 | B206 | TC | Waist | 0.14482368 | 3.04E-09 |
| B77 | B140 | TC | r-GT | 0.14460224 | 3.22E-09 |
| B77 | B91 | TC | TP | 0.13515235 | 3.21E-08 |
| B77 | B153 | TC | Glu | 0.13152716 | 7.46E-08 |
| B77 | B131 | TC | AST | 0.12958146 | 1.16E-07 |
| B77 | B215 | TC | BMI | 0.12923958 | 1.25E-07 |
| B77 | B191 | TC | Urea | 0.12490735 | 3.28E-07 |
| B78 | B154 | HBcAb | RF | 0.2299056 | 0 |
| B78 | B80 | HBcAb | anti-HP IgG | -0.1464926 | 1.99E-09 |
| B79 | B101 | hb V -dna | anti-Sc70 | 0.36623607 | 0 |
| B79 | B102 | hb V -dna | anti-Sm | 0.36623606 | 0 |
| B79 | B105 | hb V -dna | anti-U1RNP | 0.36623606 | 0 |
| B79 | B107 | hb V -dna | AnuA | 0.3662358 | 0 |
| B79 | B108 | hb V -dna | Anti-Cyclin | 0.36623606 | 0 |
| B79 | B119 | hb V -dna | Hcv-rna | 0.41630606 | 0 |
| B79 | B121 | hb V -dna | C3 | -0.2446759 | 0 |
| B79 | B122 | hb V -dna | C4 | -0.2446765 | 0 |
| B79 | B124 | hb V -dna | ACTH | 0.25750111 | 0 |
| B79 | B125 | hb V -dna | Cor | 0.41391598 | 0 |
| B79 | B137 | hb V -dna | AFU | 0.21246597 | 0 |
| B79 | B159 | hb V -dna | β2-MG | -0.2446742 | 0 |
| B79 | B169 | hb V -dna | U-CRY | -0.2446778 | 0 |
| B79 | B178 | hb V -dna | Ig-U | 0.28435161 | 0 |
| B79 | B179 | hb V -dna | α1-MG | 0.31682626 | 0 |
| B79 | B180 | hb V -dna | Tf | 0.43471341 | 0 |
| B79 | B81 | hb V -dna | TSGF | -0.2984419 | 0 |
| B79 | B94 | hb V -dna | ALDH2-Lys504Lys | 0.5218244 | 0 |
| B79 | B97 | hb V -dna | dsDNA | 0.36623606 | 0 |
| B79 | B98 | hb V -dna | anti-Jo-1 | 0.36623607 | 0 |
| B79 | B99 | hb V -dna | anti-PM-Scl | 0.36623607 | 0 |
| B79 | B197 | hb V -dna | AG | -0.1977522 | 4.44E-16 |
| B79 | B120 | hb V -dna | HCV-cAg | 0.18647142 | 1.84E-14 |
| B79 | B135 | hb V -dna | TBA | 0.18402383 | 4.06E-14 |
| B79 | B196 | hb V -dna | TCO2 | 0.17510968 | 6.63E-13 |
| B79 | B181 | hb V -dna | APTT | -0.1717766 | 1.81E-12 |

| | | | | | |
|---|---|---|---|---|---|
| B79 | B184 | hb V -dna | INR | 0.15953293 | 6.20E-11 |
| B79 | B92 | hb V -dna | ALDH2-Glu504Glu | -0.1584854 | 8.28E-11 |
| B79 | B176 | hb V -dna | Ucr | -0.1406633 | 8.56E-09 |
| B79 | B93 | hb V -dna | ALDH2-Glu504Lys | 0.12949829 | 1.18E-07 |
| B79 | B183 | hb V -dna | PTA | 0.12645038 | 2.34E-07 |
| B8 | B58 | WB-Zinc | HCT | -0.1854144 | 2.60E-14 |
| B8 | B62 | WB-Zinc | WBHSRVI | -0.1331586 | 5.12E-08 |
| B81 | B125 | TSGF | Cor | -0.3300932 | 0 |
| B81 | B179 | TSGF | α1-MG | -0.228452 | 0 |
| B81 | B180 | TSGF | Tf | -0.2456794 | 0 |
| B81 | B92 | TSGF | ALDH2-Glu504Glu | 0.20304701 | 0 |
| B81 | B94 | TSGF | ALDH2-Lys504Lys | -0.2487227 | 0 |
| B81 | B120 | TSGF | HCV-cAg | 0.19769959 | 4.44E-16 |
| B81 | B181 | TSGF | APTT | 0.1972075 | 4.44E-16 |
| B81 | B93 | TSGF | ALDH2-Glu504Lys | -0.1926902 | 2.22E-15 |
| B81 | B156 | TSGF | IgE | 0.18574095 | 2.33E-14 |
| B81 | B199 | TSGF | Mg | 0.15309279 | 3.57E-10 |
| B81 | B135 | TSGF | TBA | -0.1529992 | 3.66E-10 |
| B81 | B184 | TSGF | INR | -0.1417861 | 6.49E-09 |
| B81 | B101 | TSGF | anti-Sc70 | -0.1382216 | 1.55E-08 |
| B81 | B102 | TSGF | anti-Sm | -0.1382216 | 1.55E-08 |
| B81 | B105 | TSGF | anti-U1RNP | -0.1382216 | 1.55E-08 |
| B81 | B107 | TSGF | AnuA | -0.1382215 | 1.55E-08 |
| B81 | B108 | TSGF | Anti-Cyclin | -0.1382216 | 1.55E-08 |
| B81 | B97 | TSGF | dsDNA | -0.1382216 | 1.55E-08 |
| B81 | B98 | TSGF | anti-Jo-1 | -0.1382216 | 1.55E-08 |
| B81 | B99 | TSGF | anti-PM-Scl | -0.1382216 | 1.55E-08 |
| B81 | B178 | TSGF | Ig-U | -0.13477 | 3.51E-08 |
| B81 | B124 | TSGF | ACTH | -0.1279605 | 1.67E-07 |
| B82 | B83 | CEA | AFP | 0.24636568 | 0 |
| B82 | B84 | CEA | C-PSA | 0.53629725 | 0 |
| B82 | B85 | CEA | SCC | 0.35187384 | 0 |
| B82 | B86 | CEA | TBhCG | 0.22014833 | 0 |
| B82 | B89 | CEA | CA 72-4 | 0.2157008 | 0 |
| B82 | B90 | CEA | CYFRA 21-1 | 0.23288259 | 0 |
| B82 | B114 | CEA | TSH TT3 TT4(TT4) | -0.1236645 | 4.29E-07 |
| B83 | B86 | AFP | TBhCG | 0.99709962 | 0 |
| B84 | B85 | C-PSA | SCC | 0.28709107 | 0 |
| B84 | B90 | C-PSA | CYFRA 21-1 | 0.25868345 | 0 |
| B84 | B114 | C-PSA | TSH TT3 TT4(TT4) | -0.1384693 | 1.46E-08 |
| B84 | B192 | C-PSA | UA | 0.12574519 | 2.73E-07 |
| B84 | B209 | C-PSA | Smoking | 0.12508138 | 3.15E-07 |
| B85 | B87 | SCC | NSE | 0.24316567 | 0 |
| B85 | B90 | SCC | CYFRA 21-1 | 0.13721847 | 1.97E-08 |

| | | | | | |
|---|---|---|---|---|---|
| B9 | B137 | RH | AFU | -0.193383 | 1.78E-15 |
| B9 | B150 | RH | ALP | 0.16205818 | 3.06E-11 |
| B9 | B138 | RH | PAB | 0.15292227 | 3.74E-10 |
| B9 | B134 | RH | CHE | 0.13876508 | 1.36E-08 |
| B90 | B109 | CYFRA 21-1 | AMA-M2 | 0.15610618 | 1.59E-10 |
| B90 | B137 | CYFRA 21-1 | AFU | 0.12300729 | 4.94E-07 |
| B91 | B126 | TP | Alb | 0.54395769 | 0 |
| B91 | B127 | TP | A-G ratio | -0.4913383 | 0 |
| B91 | B130 | TP | GLB | 0.80438174 | 0 |
| B91 | B188 | TP | Ca | 0.34357935 | 0 |
| B91 | B192 | TP | UA | 0.1231953 | 4.75E-07 |
| B92 | B125 | ALDH2-Glu504Glu | Cor | -0.2093103 | 0 |
| B92 | B184 | ALDH2-Glu504Glu | INR | -0.2139305 | 0 |
| B92 | B225 | ALDH2-Glu504Glu | hPBG | -0.216276 | 0 |
| B92 | B93 | ALDH2-Glu504Glu | ALDH2-Glu504Lys | -0.9983775 | 0 |
| B92 | B94 | ALDH2-Glu504Glu | ALDH2-Lys504Lys | -0.4507836 | 0 |
| B92 | B180 | ALDH2-Glu504Glu | Tf | -0.1656057 | 1.11E-11 |
| B92 | B176 | ALDH2-Glu504Glu | Ucr | 0.16159977 | 3.48E-11 |
| B92 | B135 | ALDH2-Glu504Glu | TBA | -0.1554307 | 1.91E-10 |
| B92 | B196 | ALDH2-Glu504Glu | TCO2 | -0.1489041 | 1.07E-09 |
| B92 | B179 | ALDH2-Glu504Glu | α1-MG | -0.1473162 | 1.62E-09 |
| B92 | B186 | ALDH2-Glu504Glu | PFC | 0.1430673 | 4.73E-09 |
| B92 | B183 | ALDH2-Glu504Glu | PTA | -0.1382339 | 1.54E-08 |
| B92 | B197 | ALDH2-Glu504Glu | AG | 0.13043988 | 9.56E-08 |
| B92 | B101 | ALDH2-Glu504Glu | anti-Sc70 | -0.126844 | 2.14E-07 |
| B92 | B102 | ALDH2-Glu504Glu | anti-Sm | -0.126844 | 2.14E-07 |
| B92 | B105 | ALDH2-Glu504Glu | anti-U1RNP | -0.126844 | 2.14E-07 |
| B92 | B107 | ALDH2-Glu504Glu | AnuA | -0.1268439 | 2.14E-07 |
| B92 | B108 | ALDH2-Glu504Glu | Anti-Cyclin | -0.126844 | 2.14E-07 |
| B92 | B97 | ALDH2-Glu504Glu | dsDNA | -0.126844 | 2.14E-07 |
| B92 | B98 | ALDH2-Glu504Glu | anti-Jo-1 | -0.126844 | 2.14E-07 |
| B92 | B99 | ALDH2-Glu504Glu | anti-PM-Scl | -0.126844 | 2.14E-07 |
| B93 | B184 | ALDH2-Glu504Lys | INR | 0.20301985 | 0 |
| B93 | B225 | ALDH2-Glu504Lys | hPBG | 0.20530349 | 0 |
| B93 | B94 | ALDH2-Glu504Lys | ALDH2-Lys504Lys | 0.39922455 | 0 |
| B93 | B125 | ALDH2-Glu504Lys | Cor | 0.19870874 | 4.44E-16 |
| B93 | B176 | ALDH2-Glu504Lys | Ucr | -0.1565437 | 1.41E-10 |
| B93 | B135 | ALDH2-Glu504Lys | TBA | 0.14768882 | 1.47E-09 |
| B93 | B180 | ALDH2-Glu504Lys | Tf | 0.14559318 | 2.51E-09 |
| B93 | B196 | ALDH2-Glu504Lys | TCO2 | 0.14120421 | 7.49E-09 |
| B93 | B186 | ALDH2-Glu504Lys | PFC | -0.1357127 | 2.82E-08 |
| B93 | B179 | ALDH2-Glu504Lys | α1-MG | 0.13423319 | 3.99E-08 |
| B93 | B183 | ALDH2-Glu504Lys | PTA | 0.13110273 | 8.22E-08 |
| B93 | B197 | ALDH2-Glu504Lys | AG | -0.1236942 | 4.26E-07 |

| | | | | | |
|---|---|---|---|---|---|
| B94 | B101 | ALDH2-Lys504Lys | anti-Sc70 | 0.41886119 | 0 |
| B94 | B102 | ALDH2-Lys504Lys | anti-Sm | 0.4188612 | 0 |
| B94 | B105 | ALDH2-Lys504Lys | anti-U1RNP | 0.41886119 | 0 |
| B94 | B107 | ALDH2-Lys504Lys | AnuA | 0.41886096 | 0 |
| B94 | B108 | ALDH2-Lys504Lys | Anti-Cyclin | 0.41886118 | 0 |
| B94 | B125 | ALDH2-Lys504Lys | Cor | 0.25522481 | 0 |
| B94 | B178 | ALDH2-Lys504Lys | Ig-U | 0.22866047 | 0 |
| B94 | B179 | ALDH2-Lys504Lys | α1-MG | 0.26775524 | 0 |
| B94 | B180 | ALDH2-Lys504Lys | Tf | 0.38416472 | 0 |
| B94 | B184 | ALDH2-Lys504Lys | INR | 0.26203444 | 0 |
| B94 | B225 | ALDH2-Lys504Lys | hPBG | 0.26400212 | 0 |
| B94 | B97 | ALDH2-Lys504Lys | dsDNA | 0.4188612 | 0 |
| B94 | B98 | ALDH2-Lys504Lys | anti-Jo-1 | 0.4188612 | 0 |
| B94 | B99 | ALDH2-Lys504Lys | anti-PM-Scl | 0.4188612 | 0 |
| B94 | B135 | ALDH2-Lys504Lys | TBA | 0.18747681 | 1.33E-14 |
| B94 | B196 | ALDH2-Lys504Lys | TCO2 | 0.18404199 | 4.04E-14 |
| B94 | B156 | ALDH2-Lys504Lys | IgE | -0.1822061 | 7.26E-14 |
| B94 | B186 | ALDH2-Lys504Lys | PFC | -0.1761472 | 4.83E-13 |
| B94 | B183 | ALDH2-Lys504Lys | PTA | 0.1705894 | 2.59E-12 |
| B94 | B197 | ALDH2-Lys504Lys | AG | -0.1612302 | 3.86E-11 |
| B94 | B158 | ALDH2-Lys504Lys | IgM | 0.15737891 | 1.12E-10 |
| B94 | B222 | ALDH2-Lys504Lys | NS | -0.1511486 | 5.98E-10 |
| B94 | B176 | ALDH2-Lys504Lys | Ucr | -0.1479972 | 1.36E-09 |
| B94 | B134 | ALDH2-Lys504Lys | CHE | -0.1296921 | 1.13E-07 |
| B94 | B199 | ALDH2-Lys504Lys | Mg | -0.1290765 | 1.30E-07 |
| B94 | B193 | ALDH2-Lys504Lys | K | 0.12704368 | 2.05E-07 |
| B94 | B209 | ALDH2-Lys504Lys | Smoking | -0.1228171 | 5.15E-07 |
| B94 | B216 | ALDH2-Lys504Lys | BMI | -0.1205447 | 8.34E-07 |
| B95 | B100 | anti-cenpb | anti-Ro52 | 0.56791038 | 0 |
| B95 | B178 | anti-cenpb | Ig-U | -0.2626861 | 0 |
| B95 | B179 | anti-cenpb | α1-MG | -0.2551582 | 0 |
| B95 | B180 | anti-cenpb | Tf | -0.2197689 | 0 |
| B95 | B196 | anti-cenpb | TCO2 | 0.31117951 | 0 |
| B95 | B197 | anti-cenpb | AG | -0.2057588 | 0 |
| B95 | B101 | anti-cenpb | anti-Sc70 | -0.1861109 | 2.07E-14 |
| B95 | B102 | anti-cenpb | anti-Sm | -0.1861109 | 2.07E-14 |
| B95 | B105 | anti-cenpb | anti-U1RNP | -0.1861109 | 2.07E-14 |
| B95 | B107 | anti-cenpb | AnuA | -0.1861107 | 2.07E-14 |
| B95 | B108 | anti-cenpb | Anti-Cyclin | -0.1861109 | 2.07E-14 |
| B95 | B97 | anti-cenpb | dsDNA | -0.1861109 | 2.07E-14 |
| B95 | B98 | anti-cenpb | anti-Jo-1 | -0.1861109 | 2.07E-14 |
| B95 | B99 | anti-cenpb | anti-PM-Scl | -0.1861109 | 2.07E-14 |
| B95 | B224 | anti-cenpb | pmh | 0.15893548 | 7.32E-11 |
| B95 | B137 | anti-cenpb | AFU | -0.1250606 | 3.17E-07 |

| | | | | | |
|---|---|---|---|---|---|
| B97 | B100 | dsDNA | anti-Ro52 | -0.3185303 | 0 |
| B97 | B101 | dsDNA | anti-Sc70 | 1 | 0 |
| B97 | B102 | dsDNA | anti-Sm | 1 | 0 |
| B97 | B103 | dsDNA | anti-SSA/Ro | -0.3344827 | 0 |
| B97 | B107 | dsDNA | AnuA | 0.99999919 | 0 |
| B97 | B125 | dsDNA | Cor | 0.26774053 | 0 |
| B97 | B135 | dsDNA | TBA | 0.24449504 | 0 |
| B97 | B156 | dsDNA | IgE | 0.29730773 | 0 |
| B97 | B178 | dsDNA | Ig-U | 0.34819819 | 0 |
| B97 | B179 | dsDNA | α1-MG | 0.34866524 | 0 |
| B97 | B180 | dsDNA | Tf | 0.42439565 | 0 |
| B97 | B99 | dsDNA | anti-PM-Scl | 0.99999999 | 0 |
| B97 | B121 | dsDNA | C3 | -0.1954527 | 8.88E-16 |
| B97 | B122 | dsDNA | C4 | -0.1954533 | 8.88E-16 |
| B97 | B159 | dsDNA | β2-MG | -0.1954515 | 8.88E-16 |
| B97 | B169 | dsDNA | U-CRY | -0.1954547 | 8.88E-16 |
| B97 | B184 | dsDNA | INR | 0.1931741 | 2.00E-15 |
| B97 | B106 | dsDNA | anti-P | -0.19276 | 2.22E-15 |
| B97 | B154 | dsDNA | RF | -0.1785002 | 2.33E-13 |
| B97 | B139 | dsDNA | ADA | 0.16693577 | 7.57E-12 |
| B97 | B119 | dsDNA | Hcv-rna | 0.16323416 | 2.19E-11 |
| B97 | B151 | dsDNA | Anti-CCP | 0.15767896 | 1.03E-10 |
| B97 | B104 | dsDNA | anti-SSB | -0.1552047 | 2.03E-10 |
| B97 | B158 | dsDNA | IgM | 0.14873462 | 1.12E-09 |
| B97 | B181 | dsDNA | APTT | -0.1409302 | 8.01E-09 |
| B97 | B137 | dsDNA | AFU | 0.1375318 | 1.83E-08 |
| B97 | B138 | dsDNA | PAB | 0.12878441 | 1.39E-07 |
| B98 | B100 | anti-Jo-1 | anti-Ro52 | -0.3185303 | 0 |
| B98 | B103 | anti-Jo-1 | anti-SSA/Ro | -0.3344827 | 0 |
| B98 | B107 | anti-Jo-1 | AnuA | 0.99999916 | 0 |
| B98 | B125 | anti-Jo-1 | Cor | 0.26774054 | 0 |
| B98 | B135 | anti-Jo-1 | TBA | 0.24449505 | 0 |
| B98 | B156 | anti-Jo-1 | IgE | 0.29730774 | 0 |
| B98 | B178 | anti-Jo-1 | Ig-U | 0.3481982 | 0 |
| B98 | B179 | anti-Jo-1 | α1-MG | 0.34866525 | 0 |
| B98 | B180 | anti-Jo-1 | Tf | 0.42439566 | 0 |
| B98 | B121 | anti-Jo-1 | C3 | -0.1954526 | 8.88E-16 |
| B98 | B122 | anti-Jo-1 | C4 | -0.1954531 | 8.88E-16 |
| B98 | B159 | anti-Jo-1 | β2-MG | -0.1954518 | 8.88E-16 |
| B98 | B169 | anti-Jo-1 | U-CRY | -0.1954546 | 8.88E-16 |
| B98 | B184 | anti-Jo-1 | INR | 0.1931741 | 2.00E-15 |
| B98 | B106 | anti-Jo-1 | anti-P | -0.19276 | 2.22E-15 |
| B98 | B154 | anti-Jo-1 | RF | -0.1785002 | 2.33E-13 |
| B98 | B139 | anti-Jo-1 | ADA | 0.16693577 | 7.57E-12 |

| | | | | | |
|---|---|---|---|---|---|
| B98 | B119 | anti-Jo-1 | Hcv-rna | 0.16323416 | 2.19E-11 |
| B98 | B151 | anti-Jo-1 | Anti-CCP | 0.15767897 | 1.03E-10 |
| B98 | B104 | anti-Jo-1 | anti-SSB | -0.1552047 | 2.03E-10 |
| B98 | B158 | anti-Jo-1 | IgM | 0.14873463 | 1.12E-09 |
| B98 | B181 | anti-Jo-1 | APTT | -0.1409302 | 8.01E-09 |
| B98 | B137 | anti-Jo-1 | AFU | 0.13753181 | 1.83E-08 |
| B98 | B138 | anti-Jo-1 | PAB | 0.12878442 | 1.39E-07 |
| B99 | B100 | anti-PM-Scl | anti-Ro52 | -0.3185303 | 0 |
| B99 | B101 | anti-PM-Scl | anti-Sc70 | 1 | 0 |
| B99 | B103 | anti-PM-Scl | anti-SSA/Ro | -0.3344827 | 0 |
| B99 | B107 | anti-PM-Scl | AnuA | 0.99999928 | 0 |
| B99 | B108 | anti-PM-Scl | Anti-Cyclin | 0.99999999 | 0 |
| B99 | B125 | anti-PM-Scl | Cor | 0.26774054 | 0 |
| B99 | B135 | anti-PM-Scl | TBA | 0.24449504 | 0 |
| B99 | B156 | anti-PM-Scl | IgE | 0.29730773 | 0 |
| B99 | B178 | anti-PM-Scl | Ig-U | 0.34819819 | 0 |
| B99 | B179 | anti-PM-Scl | α1-MG | 0.34866524 | 0 |
| B99 | B180 | anti-PM-Scl | Tf | 0.42439565 | 0 |
| B99 | B121 | anti-PM-Scl | C3 | -0.1954526 | 8.88E-16 |
| B99 | B122 | anti-PM-Scl | C4 | -0.1954533 | 8.88E-16 |
| B99 | B159 | anti-PM-Scl | β2-MG | -0.1954515 | 8.88E-16 |
| B99 | B169 | anti-PM-Scl | U-CRY | -0.1954547 | 8.88E-16 |
| B99 | B184 | anti-PM-Scl | INR | 0.1931741 | 2.00E-15 |
| B99 | B106 | anti-PM-Scl | anti-P | -0.19276 | 2.22E-15 |
| B99 | B154 | anti-PM-Scl | RF | -0.1785002 | 2.33E-13 |
| B99 | B139 | anti-PM-Scl | ADA | 0.16693577 | 7.57E-12 |
| B99 | B119 | anti-PM-Scl | Hcv-rna | 0.16323416 | 2.19E-11 |
| B99 | B151 | anti-PM-Scl | Anti-CCP | 0.15767896 | 1.03E-10 |
| B99 | B104 | anti-PM-Scl | anti-SSB | -0.1552047 | 2.03E-10 |
| B99 | B158 | anti-PM-Scl | IgM | 0.14873463 | 1.12E-09 |
| B99 | B181 | anti-PM-Scl | APTT | -0.1409302 | 8.01E-09 |
| B99 | B137 | anti-PM-Scl | AFU | 0.1375318 | 1.83E-08 |
| B99 | B138 | anti-PM-Scl | PAB | 0.12878441 | 1.39E-07 |

**Supplementary Table 12. PEI Correlations in Hyperthyroidism**

| Row | Column | Row | Column | Cor | *P* |
|---|---|---|---|---|---|
| B1 | B141 | Gender | AST/ALT | 0.325837418 | 0 |
| B1 | B163 | Gender | SG | -0.47370468 | 0 |
| B1 | B174 | Gender | Urea/Cr | 0.321845394 | 0 |
| B1 | B190 | Gender | Cre | -0.66322191 | 0 |
| B1 | B192 | Gender | UA | -0.53069188 | 0 |
| B1 | B204 | Gender | Drinking | -0.39666751 | 0 |
| B1 | B206 | Gender | Waist | -0.52645017 | 0 |
| B1 | B209 | Gender | Smoking | -0.49301441 | 0 |
| B1 | B210 | Gender | Hip | -0.44400475 | 0 |
| B1 | B215 | Gender | BMI | -0.32879682 | 0 |
| B1 | B216 | Gender | BMI | -0.62804215 | 0 |
| B1 | B221 | Gender | Hei | -0.67020916 | 0 |
| B1 | B222 | Gender | NS | -0.74772688 | 0 |
| B1 | B42 | Gender | MON | -0.2940062 | 0 |
| B1 | B46 | Gender | RBC Count | -0.57046018 | 0 |
| B1 | B51 | Gender | Hb | -0.67753058 | 0 |
| B1 | B45 | Gender | MCHC | -0.29144508 | 2.22E-16 |
| B1 | B72 | Gender | HDL-C | 0.285079822 | 8.88E-16 |
| B1 | B128 | Gender | ALT | -0.27368421 | 1.20E-14 |
| B1 | B127 | Gender | A-G ratio | -0.24488045 | 6.16E-12 |
| B1 | B218 | Gender | DBP | -0.24054071 | 1.48E-11 |
| B1 | B166 | Gender | Pro | -0.22575594 | 2.55E-10 |
| B1 | B71 | Gender | TG | -0.20873412 | 5.35E-09 |
| B1 | B130 | Gender | GLB | 0.207500561 | 6.61E-09 |
| B1 | B41 | Gender | MON％ | -0.20374665 | 1.25E-08 |
| B1 | B191 | Gender | Urea | -0.19603457 | 4.41E-08 |
| B10 | B12 | PG Ⅰ | PG II | 0.614289527 | 0 |
| B100 | B103 | anti-Ro52 | anti-SSA/Ro | -0.66109405 | 0 |
| B100 | B104 | anti-Ro52 | anti-SSB | 0.999999994 | 0 |
| B100 | B106 | anti-Ro52 | anti-P | 0.999999994 | 0 |
| B100 | B107 | anti-Ro52 | AnuA | 0.99999973 | 0 |
| B100 | B109 | anti-Ro52 | AMA-M2 | 1 | 0 |
| B100 | B171 | anti-Ro52 | EC | -0.30105168 | 0 |
| B100 | B173 | anti-Ro52 | NTT | -0.29660015 | 0 |
| B100 | B179 | anti-Ro52 | α1-MG | 0.285253143 | 8.88E-16 |
| B100 | B158 | anti-Ro52 | IgM | 0.281658571 | 1.78E-15 |
| B100 | B139 | anti-Ro52 | ADA | -0.22290662 | 4.32E-10 |
| B100 | B199 | anti-Ro52 | Mg | 0.207016524 | 7.18E-09 |
| B100 | B135 | anti-Ro52 | TBA | 0.178387519 | 6.62E-07 |
| B100 | B193 | anti-Ro52 | K | -0.17780783 | 7.20E-07 |
| B100 | B157 | anti-Ro52 | IgG | -0.17568129 | 9.80E-07 |
| B101 | B102 | anti-Sc70 | anti-Sm | 0.999999988 | 0 |

| | | | | | |
|---|---|---|---|---|---|
| B101 | B103 | anti-Sc70 | anti-SSA/Ro | -0.66109406 | 0 |
| B101 | B105 | anti-Sc70 | anti-U1RNP | 0.999999999 | 0 |
| B101 | B107 | anti-Sc70 | AnuA | 0.999999522 | 0 |
| B101 | B108 | anti-Sc70 | Anti-Cyclin | 0.999999989 | 0 |
| B101 | B171 | anti-Sc70 | EC | -0.30105168 | 0 |
| B101 | B173 | anti-Sc70 | NTT | -0.29660015 | 0 |
| B101 | B179 | anti-Sc70 | α1-MG | 0.285253148 | 8.88E-16 |
| B101 | B158 | anti-Sc70 | IgM | 0.281658576 | 1.78E-15 |
| B101 | B139 | anti-Sc70 | ADA | -0.22290662 | 4.32E-10 |
| B101 | B199 | anti-Sc70 | Mg | 0.207016527 | 7.18E-09 |
| B101 | B135 | anti-Sc70 | TBA | 0.178387522 | 6.62E-07 |
| B101 | B193 | anti-Sc70 | K | -0.17780784 | 7.20E-07 |
| B101 | B157 | anti-Sc70 | IgG | -0.17568129 | 9.80E-07 |
| B102 | B103 | anti-Sm | anti-SSA/Ro | -0.66109406 | 0 |
| B102 | B107 | anti-Sm | AnuA | 0.999999833 | 0 |
| B102 | B171 | anti-Sm | EC | -0.30105168 | 0 |
| B102 | B173 | anti-Sm | NTT | -0.29660015 | 0 |
| B102 | B179 | anti-Sm | α1-MG | 0.285253146 | 8.88E-16 |
| B102 | B158 | anti-Sm | IgM | 0.281658574 | 1.78E-15 |
| B102 | B139 | anti-Sm | ADA | -0.22290662 | 4.32E-10 |
| B102 | B199 | anti-Sm | Mg | 0.207016527 | 7.18E-09 |
| B102 | B135 | anti-Sm | TBA | 0.178387521 | 6.62E-07 |
| B102 | B193 | anti-Sm | K | -0.17780784 | 7.20E-07 |
| B102 | B157 | anti-Sm | IgG | -0.17568129 | 9.80E-07 |
| B103 | B104 | anti-SSA/Ro | anti-SSB | -0.66109405 | 0 |
| B103 | B105 | anti-SSA/Ro | anti-U1RNP | -0.66109405 | 0 |
| B103 | B106 | anti-SSA/Ro | anti-P | -0.66109404 | 0 |
| B103 | B107 | anti-SSA/Ro | AnuA | -0.66109387 | 0 |
| B103 | B108 | anti-SSA/Ro | Anti-Cyclin | -0.66109405 | 0 |
| B103 | B109 | anti-SSA/Ro | AMA-M2 | -0.66109405 | 0 |
| B103 | B110 | anti-SSA/Ro | AHA | -0.66109405 | 0 |
| B103 | B139 | anti-SSA/Ro | ADA | 0.414422377 | 0 |
| B103 | B173 | anti-SSA/Ro | NTT | 0.327126499 | 0 |
| B103 | B179 | anti-SSA/Ro | α1-MG | -0.39454072 | 0 |
| B103 | B199 | anti-SSA/Ro | Mg | -0.27510782 | 8.66E-15 |
| B103 | B134 | anti-SSA/Ro | CHE | -0.25678747 | 5.14E-13 |
| B103 | B149 | anti-SSA/Ro | TGAb | 0.247924341 | 3.31E-12 |
| B103 | B150 | anti-SSA/Ro | ALP | 0.233329181 | 6.07E-11 |
| B103 | B198 | anti-SSA/Ro | P | 0.221774954 | 5.32E-10 |
| B103 | B197 | anti-SSA/Ro | AG | 0.212922277 | 2.59E-09 |
| B103 | B196 | anti-SSA/Ro | TCO2 | -0.20859179 | 5.49E-09 |
| B103 | B157 | anti-SSA/Ro | IgG | 0.204832735 | 1.04E-08 |
| B103 | B193 | anti-SSA/Ro | K | 0.199046014 | 2.71E-08 |
| B104 | B105 | anti-SSB | anti-U1RNP | 0.999999992 | 0 |

| | | | | | |
|---|---|---|---|---|---|
| B104 | B106 | anti-SSB | anti-P | 0.999999998 | 0 |
| B104 | B107 | anti-SSB | AnuA | 0.999999775 | 0 |
| B104 | B108 | anti-SSB | Anti-Cyclin | 0.99999999 | 0 |
| B104 | B109 | anti-SSB | AMA-M2 | 0.999999996 | 0 |
| B104 | B110 | anti-SSB | AHA | 0.999999993 | 0 |
| B104 | B171 | anti-SSB | EC | -0.30105168 | 0 |
| B104 | B173 | anti-SSB | NTT | -0.29660015 | 0 |
| B104 | B179 | anti-SSB | α1-MG | 0.285253141 | 8.88E-16 |
| B104 | B158 | anti-SSB | IgM | 0.281658569 | 1.78E-15 |
| B104 | B139 | anti-SSB | ADA | -0.22290661 | 4.32E-10 |
| B104 | B199 | anti-SSB | Mg | 0.207016523 | 7.18E-09 |
| B104 | B135 | anti-SSB | TBA | 0.178387518 | 6.62E-07 |
| B104 | B193 | anti-SSB | K | -0.17780783 | 7.20E-07 |
| B104 | B157 | anti-SSB | IgG | -0.17568129 | 9.80E-07 |
| B105 | B106 | anti-U1RNP | anti-P | 0.999999987 | 0 |
| B105 | B107 | anti-U1RNP | AnuA | 0.999999711 | 0 |
| B105 | B109 | anti-U1RNP | AMA-M2 | 0.999999996 | 0 |
| B105 | B171 | anti-U1RNP | EC | -0.30105168 | 0 |
| B105 | B173 | anti-U1RNP | NTT | -0.29660015 | 0 |
| B105 | B179 | anti-U1RNP | α1-MG | 0.285253143 | 8.88E-16 |
| B105 | B158 | anti-U1RNP | IgM | 0.281658571 | 1.78E-15 |
| B105 | B139 | anti-U1RNP | ADA | -0.22290662 | 4.32E-10 |
| B105 | B199 | anti-U1RNP | Mg | 0.207016524 | 7.18E-09 |
| B105 | B135 | anti-U1RNP | TBA | 0.178387519 | 6.62E-07 |
| B105 | B193 | anti-U1RNP | K | -0.17780783 | 7.20E-07 |
| B105 | B157 | anti-U1RNP | IgG | -0.17568129 | 9.80E-07 |
| B106 | B108 | anti-P | Anti-Cyclin | 0.999999993 | 0 |
| B106 | B109 | anti-P | AMA-M2 | 0.999999993 | 0 |
| B106 | B110 | anti-P | AHA | 0.999999992 | 0 |
| B106 | B171 | anti-P | EC | -0.30105167 | 0 |
| B106 | B173 | anti-P | NTT | -0.29660014 | 0 |
| B106 | B179 | anti-P | α1-MG | 0.285253141 | 8.88E-16 |
| B106 | B158 | anti-P | IgM | 0.281658568 | 1.78E-15 |
| B106 | B139 | anti-P | ADA | -0.22290661 | 4.32E-10 |
| B106 | B199 | anti-P | Mg | 0.207016524 | 7.18E-09 |
| B106 | B135 | anti-P | TBA | 0.178387517 | 6.62E-07 |
| B106 | B193 | anti-P | K | -0.17780783 | 7.20E-07 |
| B106 | B157 | anti-P | IgG | -0.17568129 | 9.80E-07 |
| B107 | B108 | AnuA | Anti-Cyclin | 0.999999687 | 0 |
| B107 | B109 | AnuA | AMA-M2 | 0.999999735 | 0 |
| B107 | B110 | AnuA | AHA | 0.99999977 | 0 |
| B107 | B171 | AnuA | EC | -0.3010516 | 0 |
| B107 | B173 | AnuA | NTT | -0.29660007 | 0 |
| B107 | B179 | AnuA | α1-MG | 0.285253067 | 8.88E-16 |

| | | | | | |
|---|---|---|---|---|---|
| B107 | B158 | AnuA | IgM | 0.281658499 | 1.78E-15 |
| B107 | B139 | AnuA | ADA | -0.22290656 | 4.32E-10 |
| B107 | B199 | AnuA | Mg | 0.207016466 | 7.18E-09 |
| B107 | B135 | AnuA | TBA | 0.178387472 | 6.62E-07 |
| B107 | B193 | AnuA | K | -0.17780779 | 7.20E-07 |
| B107 | B157 | AnuA | IgG | -0.17568124 | 9.80E-07 |
| B108 | B110 | Anti-Cyclin | AHA | 0.999999997 | 0 |
| B108 | B171 | Anti-Cyclin | EC | -0.30105168 | 0 |
| B108 | B173 | Anti-Cyclin | NTT | -0.29660015 | 0 |
| B108 | B179 | Anti-Cyclin | α1-MG | 0.285253144 | 8.88E-16 |
| B108 | B158 | Anti-Cyclin | IgM | 0.281658572 | 1.78E-15 |
| B108 | B139 | Anti-Cyclin | ADA | -0.22290662 | 4.32E-10 |
| B108 | B199 | Anti-Cyclin | Mg | 0.207016524 | 7.18E-09 |
| B108 | B135 | Anti-Cyclin | TBA | 0.178387519 | 6.62E-07 |
| B108 | B193 | Anti-Cyclin | K | -0.17780784 | 7.20E-07 |
| B108 | B157 | Anti-Cyclin | IgG | -0.17568129 | 9.80E-07 |
| B109 | B171 | AMA-M2 | EC | -0.30105168 | 0 |
| B109 | B173 | AMA-M2 | NTT | -0.29660015 | 0 |
| B109 | B179 | AMA-M2 | α1-MG | 0.285253143 | 8.88E-16 |
| B109 | B158 | AMA-M2 | IgM | 0.281658571 | 1.78E-15 |
| B109 | B139 | AMA-M2 | ADA | -0.22290662 | 4.32E-10 |
| B109 | B199 | AMA-M2 | Mg | 0.207016524 | 7.18E-09 |
| B109 | B135 | AMA-M2 | TBA | 0.178387519 | 6.62E-07 |
| B109 | B193 | AMA-M2 | K | -0.17780783 | 7.20E-07 |
| B109 | B157 | AMA-M2 | IgG | -0.17568129 | 9.80E-07 |
| B11 | B12 | PGI/PGII | PG II | -0.4519117 | 0 |
| B11 | B14 | PGI/PGII | G-17 | -0.26219205 | 1.59E-13 |
| B11 | B196 | PGI/PGII | TCO2 | 0.254617295 | 8.16E-13 |
| B11 | B197 | PGI/PGII | AG | -0.19682067 | 3.89E-08 |
| B110 | B171 | AHA | EC | -0.30105168 | 0 |
| B110 | B173 | AHA | NTT | -0.29660015 | 0 |
| B110 | B179 | AHA | α1-MG | 0.285253144 | 8.88E-16 |
| B110 | B158 | AHA | IgM | 0.281658572 | 1.78E-15 |
| B110 | B139 | AHA | ADA | -0.22290662 | 4.32E-10 |
| B110 | B199 | AHA | Mg | 0.207016524 | 7.18E-09 |
| B110 | B135 | AHA | TBA | 0.178387519 | 6.62E-07 |
| B110 | B193 | AHA | K | -0.17780783 | 7.20E-07 |
| B110 | B157 | AHA | IgG | -0.17568129 | 9.80E-07 |
| B111 | B123 | TSH | hs-CRP1 | 0.355789066 | 0 |
| B111 | B132 | TSH | Dbil | 0.345918066 | 0 |
| B111 | B165 | TSH | bilirubin | 0.297198777 | 0 |
| B111 | B112 | TSH | FT4 | -0.27704928 | 5.55E-15 |
| B111 | B133 | TSH | TBIL | 0.194883831 | 5.31E-08 |
| B111 | B131 | TSH | AST | 0.179420476 | 5.69E-07 |

| | | | | | |
|---|---|---|---|---|---|
| B111 | B130 | TSH | GLB | 0.178108961 | 6.89E-07 |
| B112 | B113 | FT4 | FT3 | 0.855980391 | 0 |
| B112 | B114 | FT4 | TSH TT3 TT4(TT4) | 0.591225565 | 0 |
| B112 | B115 | FT4 | TSH TT3 TT4(TT3) | 0.543938823 | 0 |
| B112 | B174 | FT4 | Urea/Cr | 0.293490531 | 0 |
| B112 | B145 | FT4 | BGP | 0.280214981 | 2.66E-15 |
| B112 | B143 | FT4 | β-CTx | 0.253618464 | 1.01E-12 |
| B112 | B144 | FT4 | Total-PINP | 0.252549567 | 1.26E-12 |
| B112 | B149 | FT4 | TGAb | 0.232679893 | 6.88E-11 |
| B112 | B148 | FT4 | Anti_TPO | 0.196468051 | 4.12E-08 |
| B112 | B190 | FT4 | Cre | -0.17598587 | 9.38E-07 |
| B113 | B114 | FT3 | TSH TT3 TT4(TT4) | 0.469429751 | 0 |
| B113 | B115 | FT3 | TSH TT3 TT4(TT3) | 0.571696482 | 0 |
| B113 | B174 | FT3 | Urea/Cr | 0.300519373 | 0 |
| B113 | B145 | FT3 | BGP | 0.274975697 | 8.88E-15 |
| B113 | B144 | FT3 | Total-PINP | 0.251369205 | 1.62E-12 |
| B113 | B143 | FT3 | β-CTx | 0.246912795 | 4.07E-12 |
| B113 | B149 | FT3 | TGAb | 0.225933301 | 2.47E-10 |
| B113 | B137 | FT3 | AFU | -0.20818622 | 5.88E-09 |
| B113 | B199 | FT3 | Mg | -0.17567927 | 9.80E-07 |
| B114 | B115 | TSH TT3 TT4(TT4) | TSH TT3 TT4(TT3) | 0.798713449 | 0 |
| B114 | B137 | TSH TT3 TT4(TT4) | AFU | -0.26521343 | 8.19E-14 |
| B114 | B199 | TSH TT3 TT4(TT4) | Mg | -0.23710561 | 2.91E-11 |
| B114 | B138 | TSH TT3 TT4(TT4) | PAB | -0.23682215 | 3.08E-11 |
| B114 | B149 | TSH TT3 TT4(TT4) | TGAb | 0.210291181 | 4.09E-09 |
| B114 | B148 | TSH TT3 TT4(TT4) | Anti_TPO | 0.193822857 | 6.28E-08 |
| B114 | B174 | TSH TT3 TT4(TT4) | Urea/Cr | 0.190761985 | 1.02E-07 |
| B115 | B137 | TSH TT3 TT4(TT3) | AFU | -0.29855747 | 0 |
| B115 | B199 | TSH TT3 TT4(TT3) | Mg | -0.27913543 | 3.55E-15 |
| B115 | B138 | TSH TT3 TT4(TT3) | PAB | -0.26681252 | 5.73E-14 |
| B115 | B149 | TSH TT3 TT4(TT3) | TGAb | 0.202115724 | 1.63E-08 |
| B115 | B174 | TSH TT3 TT4(TT3) | Urea/Cr | 0.193125853 | 7.02E-08 |
| B117 | B167 | TT4 | PC | 0.295954092 | 0 |
| B117 | B223 | TT4 | FH | 0.848365617 | 0 |
| B118 | B123 | CEA1 | hs-CRP1 | 0.199335416 | 2.58E-08 |
| B118 | B132 | CEA1 | Dbil | 0.182257836 | 3.74E-07 |
| B119 | B120 | Hcv-rna | HCV-cAg | 0.374988321 | 0 |
| B119 | B122 | Hcv-rna | C4 | -0.99999947 | 0 |
| B119 | B152 | Hcv-rna | ACA-Ig(G-A-M) | -0.99996911 | 0 |
| B119 | B169 | Hcv-rna | U-CRY | -0.99999274 | 0 |
| B12 | B80 | PG II | anti-HP IgG | 0.289830134 | 2.22E-16 |
| B12 | B14 | PG II | G-17 | 0.288217638 | 4.44E-16 |
| B120 | B121 | HCV-cAg | C3 | -0.37498915 | 0 |
| B120 | B122 | HCV-cAg | C4 | -0.37498851 | 0 |

| | | | | | |
|---|---|---|---|---|---|
| B120 | B152 | HCV-cAg | ACA-Ig(G-A-M) | -0.37497813 | 0 |
| B120 | B159 | HCV-cAg | β2-MG | -0.37499177 | 0 |
| B120 | B169 | HCV-cAg | U-CRY | -0.37498701 | 0 |
| B121 | B122 | C3 | C4 | 0.999999539 | 0 |
| B121 | B152 | C3 | ACA-Ig(G-A-M) | 0.999974627 | 0 |
| B121 | B169 | C3 | U-CRY | 0.999995258 | 0 |
| B122 | B152 | C4 | ACA-Ig(G-A-M) | 0.999973235 | 0 |
| B122 | B159 | C4 | β2-MG | 0.999998584 | 0 |
| B123 | B131 | hs-CRP1 | AST | 0.458078796 | 0 |
| B123 | B132 | hs-CRP1 | Dbil | 0.730082968 | 0 |
| B123 | B133 | hs-CRP1 | TBIL | 0.358254947 | 0 |
| B123 | B140 | hs-CRP1 | r-GT | 0.47808406 | 0 |
| B123 | B165 | hs-CRP1 | bilirubin | 0.630486503 | 0 |
| B123 | B126 | hs-CRP1 | Alb | -0.2865132 | 4.44E-16 |
| B123 | B188 | hs-CRP1 | Ca | -0.20872702 | 5.36E-09 |
| B123 | B130 | hs-CRP1 | GLB | 0.190756397 | 1.02E-07 |
| B124 | B125 | ACTH | Cor | -0.65419532 | 0 |
| B125 | B156 | Cor | IgE | 0.192336905 | 7.95E-08 |
| B126 | B127 | Alb | A-G ratio | 0.455174888 | 0 |
| B126 | B188 | Alb | Ca | 0.316346988 | 0 |
| B126 | B132 | Alb | Dbil | -0.22890682 | 1.41E-10 |
| B126 | B165 | Alb | bilirubin | -0.22361441 | 3.79E-10 |
| B126 | B131 | Alb | AST | -0.18627118 | 2.04E-07 |
| B127 | B130 | A-G ratio | GLB | -0.90397717 | 0 |
| B127 | B221 | A-G ratio | Hei | 0.235292855 | 4.15E-11 |
| B127 | B222 | A-G ratio | NS | 0.200839332 | 2.02E-08 |
| B127 | B131 | A-G ratio | AST | -0.18723712 | 1.76E-07 |
| B128 | B131 | ALT | AST | 0.670298342 | 0 |
| B128 | B140 | ALT | r-GT | 0.347307831 | 0 |
| B128 | B141 | ALT | AST/ALT | -0.51382551 | 0 |
| B128 | B192 | ALT | UA | 0.278868919 | 3.55E-15 |
| B128 | B216 | ALT | BMI | 0.240341182 | 1.54E-11 |
| B128 | B206 | ALT | Waist | 0.23418686 | 5.14E-11 |
| B128 | B222 | ALT | NS | 0.219701999 | 7.75E-10 |
| B128 | B221 | ALT | Hei | 0.185089978 | 2.44E-07 |
| B128 | B215 | ALT | BMI | 0.184140007 | 2.82E-07 |
| B128 | B163 | ALT | SG | 0.18282922 | 3.43E-07 |
| B129 | B132 | IBIL | Dbil | 0.477419469 | 0 |
| B129 | B133 | IBIL | TBIL | 0.937383712 | 0 |
| B130 | B131 | GLB | AST | 0.238106936 | 2.39E-11 |
| B130 | B221 | GLB | Hei | -0.20738672 | 6.74E-09 |
| B130 | B132 | GLB | Dbil | 0.197243876 | 3.63E-08 |
| B131 | B132 | AST | Dbil | 0.495840231 | 0 |
| B131 | B133 | AST | TBIL | 0.302475673 | 0 |

| | | | | | |
|---|---|---|---|---|---|
| B131 | B140 | AST | r-GT | 0.489017331 | 0 |
| B131 | B165 | AST | bilirubin | 0.36657622 | 0 |
| B131 | B177 | AST | ACR | 0.192064909 | 8.30E-08 |
| B131 | B173 | AST | NTT | 0.190951132 | 9.88E-08 |
| B131 | B175 | AST | ALB-U | 0.177045324 | 8.05E-07 |
| B132 | B133 | Dbil | TBIL | 0.753504186 | 0 |
| B132 | B140 | Dbil | r-GT | 0.446441165 | 0 |
| B132 | B165 | Dbil | bilirubin | 0.596053471 | 0 |
| B133 | B165 | TBIL | bilirubin | 0.306463593 | 0 |
| B133 | B140 | TBIL | r-GT | 0.280763062 | 2.22E-15 |
| B133 | B228 | TBIL | OS | -0.18106055 | 4.46E-07 |
| B134 | B196 | CHE | TCO2 | 0.282874961 | 1.33E-15 |
| B134 | B139 | CHE | ADA | -0.27347846 | 1.27E-14 |
| B134 | B138 | CHE | PAB | 0.222921388 | 4.31E-10 |
| B134 | B200 | CHE | OP | 0.192582802 | 7.65E-08 |
| B134 | B195 | CHE | Na | 0.186369659 | 2.01E-07 |
| B135 | B136 | TBA | 5'NT | 0.401503455 | 0 |
| B135 | B198 | TBA | P | -0.63309753 | 0 |
| B135 | B200 | TBA | OP | 0.278195447 | 4.22E-15 |
| B135 | B194 | TBA | Cl | -0.2432697 | 8.54E-12 |
| B135 | B199 | TBA | Mg | 0.19495745 | 5.24E-08 |
| B136 | B198 | 5'NT | P | -0.43803806 | 0 |
| B136 | B140 | 5'NT | r-GT | 0.248439558 | 2.97E-12 |
| B137 | B199 | AFU | Mg | 0.385124044 | 0 |
| B137 | B138 | AFU | PAB | 0.198378621 | 3.02E-08 |
| B138 | B139 | PAB | ADA | -0.34770626 | 0 |
| B138 | B199 | PAB | Mg | 0.406632552 | 0 |
| B138 | B193 | PAB | K | 0.276765411 | 5.77E-15 |
| B138 | B171 | PAB | EC | 0.224112653 | 3.46E-10 |
| B138 | B196 | PAB | TCO2 | 0.217145528 | 1.23E-09 |
| B138 | B158 | PAB | IgM | -0.21660017 | 1.35E-09 |
| B138 | B195 | PAB | Na | 0.204054261 | 1.18E-08 |
| B138 | B198 | PAB | P | -0.18646453 | 1.98E-07 |
| B138 | B200 | PAB | OP | 0.179681458 | 5.47E-07 |
| B139 | B193 | ADA | K | 0.333193234 | 0 |
| B139 | B198 | ADA | P | 0.306729588 | 0 |
| B139 | B196 | ADA | TCO2 | -0.28133897 | 2.00E-15 |
| B139 | B197 | ADA | AG | 0.271529477 | 1.98E-14 |
| B139 | B150 | ADA | ALP | 0.228197706 | 1.61E-10 |
| B139 | B149 | ADA | TGAb | 0.183985416 | 2.88E-07 |
| B14 | B34 | G-17 | TK | -0.28662686 | 4.44E-16 |
| B14 | B85 | G-17 | SCC | -0.18765533 | 1.65E-07 |
| B140 | B165 | r-GT | bilirubin | 0.344221954 | 0 |
| B140 | B192 | r-GT | UA | 0.209507568 | 4.69E-09 |

| | | | | | |
|---|---|---|---|---|---|
| B140 | B141 | r-GT | AST/ALT | -0.19107973 | 9.69E-08 |
| B140 | B206 | r-GT | Waist | 0.182439968 | 3.63E-07 |
| B140 | B145 | r-GT | BGP | 0.177427656 | 7.61E-07 |
| B141 | B216 | AST/ALT | BMI | -0.32112622 | 0 |
| B141 | B206 | AST/ALT | Waist | -0.2870654 | 4.44E-16 |
| B141 | B222 | AST/ALT | NS | -0.28167751 | 1.78E-15 |
| B141 | B221 | AST/ALT | Hei | -0.27761484 | 4.88E-15 |
| B141 | B192 | AST/ALT | UA | -0.24857344 | 2.89E-12 |
| B141 | B210 | AST/ALT | Hip | -0.22565886 | 2.60E-10 |
| B141 | B215 | AST/ALT | BMI | -0.21826877 | 1.00E-09 |
| B141 | B218 | AST/ALT | DBP | -0.17814574 | 6.85E-07 |
| B142 | B151 | 25-(OH)D | Anti-CCP | -0.37661338 | 0 |
| B142 | B143 | 25-(OH)D | β-CTx | 0.21126171 | 3.46E-09 |
| B142 | B173 | 25-(OH)D | NTT | 0.208036193 | 6.03E-09 |
| B142 | B165 | 25-(OH)D | bilirubin | -0.20672209 | 7.55E-09 |
| B142 | B168 | 25-(OH)D | NMP22 | -0.18575873 | 2.20E-07 |
| B143 | B144 | β-CTx | Total-PINP | 0.895902823 | 0 |
| B143 | B145 | β-CTx | BGP | 0.883209723 | 0 |
| B143 | B149 | β-CTx | TGAb | 0.184058537 | 2.85E-07 |
| B144 | B145 | Total-PINP | BGP | 0.939814318 | 0 |
| B144 | B168 | Total-PINP | NMP22 | -0.2406817 | 1.44E-11 |
| B144 | B149 | Total-PINP | TGAb | 0.1857808 | 2.20E-07 |
| B145 | B149 | BGP | TGAb | 0.205113059 | 9.90E-09 |
| B148 | B149 | Anti_TPO | TGAb | 0.636893318 | 0 |
| B149 | B185 | TGAb | PT | 0.203950226 | 1.20E-08 |
| B149 | B184 | TGAb | INR | 0.200193229 | 2.25E-08 |
| B149 | B150 | TGAb | ALP | 0.178468955 | 6.54E-07 |
| B15 | B16 | CD3-CD19+(B cell) | CD3+T | -0.45069499 | 0 |
| B15 | B17 | CD3-CD19+(B cell) | CD3+CD16+CD56+NK | -0.63205007 | 0 |
| B15 | B19 | CD3-CD19+(B cell) | CD3+CD4+/CD3+CD8+ | 0.353797565 | 0 |
| B15 | B197 | CD3-CD19+(B cell) | AG | 0.340023329 | 0 |
| B15 | B20 | CD3-CD19+(B cell) | CD3+CD4+CD8+ | -0.5312214 | 0 |
| B15 | B22 | CD3-CD19+(B cell) | CD3+CD8+(T) | -0.34427819 | 0 |
| B15 | B100 | CD3-CD19+(B cell) | anti-Ro52 | 0.261346919 | 1.92E-13 |
| B15 | B101 | CD3-CD19+(B cell) | anti-Sc70 | 0.261346923 | 1.92E-13 |
| B15 | B102 | CD3-CD19+(B cell) | anti-Sm | 0.261346922 | 1.92E-13 |
| B15 | B104 | CD3-CD19+(B cell) | anti-SSB | 0.261346918 | 1.92E-13 |
| B15 | B105 | CD3-CD19+(B cell) | anti-U1RNP | 0.261346919 | 1.92E-13 |
| B15 | B106 | CD3-CD19+(B cell) | anti-P | 0.261346917 | 1.92E-13 |
| B15 | B107 | CD3-CD19+(B cell) | AnuA | 0.26134685 | 1.92E-13 |
| B15 | B108 | CD3-CD19+(B cell) | Anti-Cyclin | 0.261346919 | 1.92E-13 |
| B15 | B109 | CD3-CD19+(B cell) | AMA-M2 | 0.261346918 | 1.92E-13 |
| B15 | B110 | CD3-CD19+(B cell) | AHA | 0.261346919 | 1.92E-13 |
| B15 | B95 | CD3-CD19+(B cell) | anti-cenpb | 0.261346917 | 1.92E-13 |

| | | | | | |
|---|---|---|---|---|---|
| B15 | B97 | CD3-CD19+(B cell) | dsDNA | 0.261346916 | 1.92E-13 |
| B15 | B98 | CD3-CD19+(B cell) | anti-Jo-1 | 0.261346924 | 1.92E-13 |
| B15 | B99 | CD3-CD19+(B cell) | anti-PM-Scl | 0.26134692 | 1.92E-13 |
| B15 | B138 | CD3-CD19+(B cell) | PAB | -0.24367094 | 7.88E-12 |
| B15 | B200 | CD3-CD19+(B cell) | OP | 0.227728421 | 1.76E-10 |
| B15 | B158 | CD3-CD19+(B cell) | IgM | 0.215666118 | 1.60E-09 |
| B15 | B135 | CD3-CD19+(B cell) | TBA | 0.208351037 | 5.72E-09 |
| B15 | B196 | CD3-CD19+(B cell) | TCO2 | -0.20504964 | 1.00E-08 |
| B15 | B195 | CD3-CD19+(B cell) | Na | 0.20434864 | 1.13E-08 |
| B15 | B164 | CD3-CD19+(B cell) | COM | -0.19221959 | 8.10E-08 |
| B15 | B57 | CD3-CD19+(B cell) | RBCAI | -0.17640898 | 8.82E-07 |
| B150 | B195 | ALP | Na | -0.23252218 | 7.09E-11 |
| B150 | B200 | ALP | OP | -0.18560995 | 2.25E-07 |
| B151 | B154 | Anti-CCP | RF | 0.316113776 | 0 |
| B151 | B168 | Anti-CCP | NMP22 | 0.28186116 | 1.78E-15 |
| B151 | B175 | Anti-CCP | ALB-U | 0.259268795 | 3.02E-13 |
| B151 | B165 | Anti-CCP | bilirubin | 0.224129164 | 3.45E-10 |
| B151 | B177 | Anti-CCP | ACR | 0.20742436 | 6.70E-09 |
| B151 | B173 | Anti-CCP | NTT | -0.18297974 | 3.35E-07 |
| B152 | B159 | ACA-Ig(G-A-M) | β2-MG | 0.999978385 | 0 |
| B153 | B170 | Glu | U-GLU | 0.309788707 | 0 |
| B153 | B220 | Glu | SBP | 0.246192983 | 4.72E-12 |
| B153 | B206 | Glu | Waist | 0.195266557 | 4.99E-08 |
| B154 | B173 | RF | NTT | -0.26646964 | 6.20E-14 |
| B154 | B156 | RF | IgE | 0.255610792 | 6.61E-13 |
| B154 | B157 | RF | IgG | -0.24618281 | 4.73E-12 |
| B154 | B183 | RF | PTA | 0.195567188 | 4.76E-08 |
| B154 | B158 | RF | IgM | 0.189602015 | 1.22E-07 |
| B154 | B182 | RF | TT | -0.18158411 | 4.13E-07 |
| B155 | B157 | IgA | IgG | 0.490069448 | 0 |
| B155 | B158 | IgA | IgM | 0.509141775 | 0 |
| B156 | B193 | IgE | K | -0.33072155 | 0 |
| B156 | B173 | IgE | NTT | -0.2543743 | 8.60E-13 |
| B157 | B173 | IgG | NTT | 0.302574725 | 0 |
| B158 | B183 | IgM | PTA | 0.243232037 | 8.61E-12 |
| B158 | B182 | IgM | TT | -0.22234724 | 4.79E-10 |
| B158 | B186 | IgM | PFC | 0.213477182 | 2.35E-09 |
| B16 | B19 | CD3+T | CD3+CD4+/CD3+CD8+ | -0.43358127 | 0 |
| B16 | B197 | CD3+T | AG | -0.32256386 | 0 |
| B16 | B22 | CD3+T | CD3+CD8+(T) | 0.486163841 | 0 |
| B16 | B23 | CD3+T | CD3-CD16+CD56+(NK) | -0.7960172 | 0 |
| B16 | B200 | CD3+T | OP | -0.28591855 | 6.66E-16 |
| B16 | B138 | CD3+T | PAB | 0.28464763 | 8.88E-16 |
| B16 | B21 | CD3+T | CD3+CD4-CD8- | 0.280958834 | 2.22E-15 |

| | | | | | |
|---|---|---|---|---|---|
| B16 | B135 | CD3+T | TBA | -0.25857081 | 3.50E-13 |
| B16 | B18 | CD3+T | CD3+CD4+helper(inducer)T | 0.254300173 | 8.73E-13 |
| B16 | B156 | CD3+T | IgE | -0.25085664 | 1.80E-12 |
| B16 | B195 | CD3+T | Na | -0.25013814 | 2.09E-12 |
| B16 | B20 | CD3+T | CD3+CD4+CD8+ | -0.22609141 | 2.40E-10 |
| B16 | B171 | CD3+T | EC | 0.215295352 | 1.71E-09 |
| B16 | B196 | CD3+T | TCO2 | 0.188559511 | 1.43E-07 |
| B16 | B73 | CD3+T | ApoA | 0.180488832 | 4.86E-07 |
| B163 | B166 | SG | Pro | 0.421919185 | 0 |
| B163 | B190 | SG | Cre | 0.342284255 | 0 |
| B163 | B191 | SG | Urea | 0.315939404 | 0 |
| B163 | B216 | SG | BMI | 0.307904842 | 0 |
| B163 | B221 | SG | Hei | 0.344306291 | 0 |
| B163 | B222 | SG | NS | 0.357734843 | 0 |
| B163 | B206 | SG | Waist | 0.260057163 | 2.54E-13 |
| B163 | B192 | SG | UA | 0.226465053 | 2.23E-10 |
| B163 | B176 | SG | Ucr | 0.219086702 | 8.66E-10 |
| B163 | B209 | SG | Smoking | 0.205932493 | 8.63E-09 |
| B163 | B210 | SG | Hip | 0.196416373 | 4.15E-08 |
| B166 | B190 | Pro | Cre | 0.182432046 | 3.64E-07 |
| B166 | B221 | Pro | Hei | 0.176044221 | 9.30E-07 |
| B166 | B222 | Pro | NS | 0.175549928 | 9.99E-07 |
| B167 | B223 | PC | FH | 0.293940161 | 0 |
| B17 | B18 | CD3+CD16+CD56+NK | CD3+CD4+helper(inducer)T | -0.33616565 | 0 |
| B17 | B19 | CD3+CD16+CD56+NK | CD3+CD4+/CD3+CD8+ | -0.58678119 | 0 |
| B17 | B20 | CD3+CD16+CD56+NK | CD3+CD4+CD8+ | 0.709306717 | 0 |
| B17 | B21 | CD3+CD16+CD56+NK | CD3+CD4-CD8- | -0.3723861 | 0 |
| B17 | B22 | CD3+CD16+CD56+NK | CD3+CD8+(T) | 0.631627392 | 0 |
| B17 | B23 | CD3+CD16+CD56+NK | CD3-CD16+CD56+(NK) | 0.33844168 | 0 |
| B17 | B34 | CD3+CD16+CD56+NK | TK | 0.341748786 | 0 |
| B17 | B193 | CD3+CD16+CD56+NK | K | 0.256591294 | 5.36E-13 |
| B17 | B164 | CD3+CD16+CD56+NK | COM | 0.253880236 | 9.54E-13 |
| B17 | B158 | CD3+CD16+CD56+NK | IgM | -0.2065253 | 7.80E-09 |
| B17 | B138 | CD3+CD16+CD56+NK | PAB | 0.179565929 | 5.57E-07 |
| B173 | B193 | NTT | K | 0.202207165 | 1.61E-08 |
| B174 | B190 | Urea/Cr | Cre | -0.54785908 | 0 |
| B174 | B191 | Urea/Cr | Urea | 0.619619055 | 0 |
| B174 | B221 | Urea/Cr | Hei | -0.27470626 | 9.55E-15 |
| B174 | B222 | Urea/Cr | NS | -0.26617815 | 6.62E-14 |
| B174 | B189 | Urea/Cr | eGFB | 0.25026561 | 2.04E-12 |
| B174 | B192 | Urea/Cr | UA | -0.23433287 | 5.00E-11 |
| B174 | B216 | Urea/Cr | BMI | -0.229973 | 1.15E-10 |
| B174 | B209 | Urea/Cr | Smoking | -0.20506916 | 9.98E-09 |
| B174 | B210 | Urea/Cr | Hip | -0.19853367 | 2.95E-08 |

| | | | | | |
|---|---|---|---|---|---|
| B175 | B176 | ALB-U | Ucr | 0.553118943 | 0 |
| B175 | B177 | ALB-U | ACR | 0.684695063 | 0 |
| B176 | B207 | Ucr | WHR | -0.18387939 | 2.93E-07 |
| B177 | B178 | ACR | Ig-U | 0.399738107 | 0 |
| B177 | B179 | ACR | α1-MG | 0.311138845 | 0 |
| B177 | B180 | ACR | Tf | 0.388771067 | 0 |
| B177 | B207 | ACR | WHR | 0.198561867 | 2.93E-08 |
| B178 | B179 | Ig-U | α1-MG | 0.935848691 | 0 |
| B178 | B180 | Ig-U | Tf | 0.998283348 | 0 |
| B178 | B195 | Ig-U | Na | -0.3476602 | 0 |
| B178 | B198 | Ig-U | P | -0.33257487 | 0 |
| B178 | B194 | Ig-U | Cl | -0.25785697 | 4.09E-13 |
| B178 | B200 | Ig-U | OP | -0.22327163 | 4.04E-10 |
| B179 | B180 | α1-MG | Tf | 0.950139408 | 0 |
| B179 | B195 | α1-MG | Na | -0.29981706 | 0 |
| B179 | B198 | α1-MG | P | -0.37886954 | 0 |
| B179 | B207 | α1-MG | WHR | -0.26676426 | 5.82E-14 |
| B179 | B194 | α1-MG | Cl | -0.24895877 | 2.67E-12 |
| B18 | B100 | CD3+CD4+helper(inducer)T | anti-Ro52 | -0.3406733 | 0 |
| B18 | B101 | CD3+CD4+helper(inducer)T | anti-Sc70 | -0.34067331 | 0 |
| B18 | B102 | CD3+CD4+helper(inducer)T | anti-Sm | -0.34067331 | 0 |
| B18 | B103 | CD3+CD4+helper(inducer)T | anti-SSA/Ro | 0.656567531 | 0 |
| B18 | B104 | CD3+CD4+helper(inducer)T | anti-SSB | -0.3406733 | 0 |
| B18 | B105 | CD3+CD4+helper(inducer)T | anti-U1RNP | -0.3406733 | 0 |
| B18 | B106 | CD3+CD4+helper(inducer)T | anti-P | -0.3406733 | 0 |
| B18 | B107 | CD3+CD4+helper(inducer)T | AnuA | -0.34067322 | 0 |
| B18 | B108 | CD3+CD4+helper(inducer)T | Anti-Cyclin | -0.34067331 | 0 |
| B18 | B109 | CD3+CD4+helper(inducer)T | AMA-M2 | -0.3406733 | 0 |
| B18 | B110 | CD3+CD4+helper(inducer)T | AHA | -0.34067331 | 0 |
| B18 | B139 | CD3+CD4+helper(inducer)T | ADA | 0.332512628 | 0 |
| B18 | B19 | CD3+CD4+helper(inducer)T | CD3+CD4+/CD3+CD8+ | 0.655565755 | 0 |
| B18 | B22 | CD3+CD4+helper(inducer)T | CD3+CD8+(T) | -0.51131103 | 0 |
| B18 | B23 | CD3+CD4+helper(inducer)T | CD3-CD16+CD56+(NK) | -0.57814357 | 0 |
| B18 | B95 | CD3+CD4+helper(inducer)T | anti-cenpb | -0.3406733 | 0 |
| B18 | B97 | CD3+CD4+helper(inducer)T | dsDNA | -0.3406733 | 0 |
| B18 | B98 | CD3+CD4+helper(inducer)T | anti-Jo-1 | -0.34067331 | 0 |
| B18 | B99 | CD3+CD4+helper(inducer)T | anti-PM-Scl | -0.3406733 | 0 |
| B18 | B179 | CD3+CD4+helper(inducer)T | α1-MG | -0.22299576 | 4.25E-10 |
| B18 | B194 | CD3+CD4+helper(inducer)T | Cl | -0.19652713 | 4.08E-08 |
| B180 | B195 | Tf | Na | -0.34103068 | 0 |
| B180 | B198 | Tf | P | -0.33805831 | 0 |
| B180 | B194 | Tf | Cl | -0.26918076 | 3.38E-14 |
| B180 | B200 | Tf | OP | -0.21576324 | 1.57E-09 |
| B181 | B182 | APTT | TT | -0.32584637 | 0 |

| | | | | | |
|---|---|---|---|---|---|
| B181 | B184 | APTT | INR | 0.578421572 | 0 |
| B181 | B185 | APTT | PT | 0.688605045 | 0 |
| B181 | B186 | APTT | PFC | -0.79579335 | 0 |
| B181 | B183 | APTT | PTA | 0.240714021 | 1.43E-11 |
| B182 | B183 | TT | PTA | -0.30485813 | 0 |
| B182 | B184 | TT | INR | -0.64040194 | 0 |
| B182 | B185 | TT | PT | -0.53603432 | 0 |
| B183 | B184 | PTA | INR | 0.432610688 | 0 |
| B183 | B185 | PTA | PT | 0.380206094 | 0 |
| B183 | B186 | PTA | PFC | -0.240197 | 1.58E-11 |
| B184 | B185 | INR | PT | 0.977961883 | 0 |
| B184 | B186 | INR | PFC | -0.54523131 | 0 |
| B185 | B186 | PT | PFC | -0.64453424 | 0 |
| B189 | B190 | eGFB | Cre | -0.60814849 | 0 |
| B189 | B191 | eGFB | Urea | -0.26906692 | 3.46E-14 |
| B189 | B206 | eGFB | Waist | -0.23357619 | 5.79E-11 |
| B189 | B215 | eGFB | BMI | -0.22373509 | 3.71E-10 |
| B189 | B192 | eGFB | UA | -0.20398801 | 1.20E-08 |
| B189 | B220 | eGFB | SBP | -0.2038483 | 1.22E-08 |
| B189 | B222 | eGFB | NS | -0.19974384 | 2.42E-08 |
| B19 | B100 | CD3+CD4+/CD3+CD8+ | anti-Ro52 | -0.29869143 | 0 |
| B19 | B101 | CD3+CD4+/CD3+CD8+ | anti-Sc70 | -0.29869143 | 0 |
| B19 | B102 | CD3+CD4+/CD3+CD8+ | anti-Sm | -0.29869143 | 0 |
| B19 | B103 | CD3+CD4+/CD3+CD8+ | anti-SSA/Ro | 0.406731378 | 0 |
| B19 | B104 | CD3+CD4+/CD3+CD8+ | anti-SSB | -0.29869142 | 0 |
| B19 | B105 | CD3+CD4+/CD3+CD8+ | anti-U1RNP | -0.29869143 | 0 |
| B19 | B106 | CD3+CD4+/CD3+CD8+ | anti-P | -0.29869142 | 0 |
| B19 | B107 | CD3+CD4+/CD3+CD8+ | AnuA | -0.29869135 | 0 |
| B19 | B108 | CD3+CD4+/CD3+CD8+ | Anti-Cyclin | -0.29869143 | 0 |
| B19 | B109 | CD3+CD4+/CD3+CD8+ | AMA-M2 | -0.29869143 | 0 |
| B19 | B110 | CD3+CD4+/CD3+CD8+ | AHA | -0.29869143 | 0 |
| B19 | B138 | CD3+CD4+/CD3+CD8+ | PAB | -0.31885961 | 0 |
| B19 | B139 | CD3+CD4+/CD3+CD8+ | ADA | 0.299503674 | 0 |
| B19 | B156 | CD3+CD4+/CD3+CD8+ | IgE | 0.316035596 | 0 |
| B19 | B20 | CD3+CD4+/CD3+CD8+ | CD3+CD4+CD8+ | -0.30558848 | 0 |
| B19 | B22 | CD3+CD4+/CD3+CD8+ | CD3+CD8+(T) | -0.96286334 | 0 |
| B19 | B95 | CD3+CD4+/CD3+CD8+ | anti-cenpb | -0.29869142 | 0 |
| B19 | B97 | CD3+CD4+/CD3+CD8+ | dsDNA | -0.29869142 | 0 |
| B19 | B98 | CD3+CD4+/CD3+CD8+ | anti-Jo-1 | -0.29869143 | 0 |
| B19 | B99 | CD3+CD4+/CD3+CD8+ | anti-PM-Scl | -0.29869143 | 0 |
| B19 | B34 | CD3+CD4+/CD3+CD8+ | TK | -0.25553254 | 6.72E-13 |
| B19 | B171 | CD3+CD4+/CD3+CD8+ | EC | -0.23138245 | 8.82E-11 |
| B19 | B197 | CD3+CD4+/CD3+CD8+ | AG | 0.22668673 | 2.14E-10 |
| B19 | B55 | CD3+CD4+/CD3+CD8+ | TK | 0.206082632 | 8.41E-09 |

| | | | | | |
|---|---|---|---|---|---|
| B19 | B158 | CD3+CD4+/CD3+CD8+ | IgM | 0.191339518 | 9.30E-08 |
| B19 | B193 | CD3+CD4+/CD3+CD8+ | K | -0.18294699 | 3.37E-07 |
| B190 | B192 | Cre | UA | 0.515739845 | 0 |
| B190 | B204 | Cre | Drinking | 0.341853085 | 0 |
| B190 | B206 | Cre | Waist | 0.414481993 | 0 |
| B190 | B209 | Cre | Smoking | 0.361782862 | 0 |
| B190 | B210 | Cre | Hip | 0.379291746 | 0 |
| B190 | B215 | Cre | BMI | 0.321941187 | 0 |
| B190 | B216 | Cre | BMI | 0.496673548 | 0 |
| B190 | B221 | Cre | Hei | 0.436858453 | 0 |
| B190 | B222 | Cre | NS | 0.549195986 | 0 |
| B190 | B191 | Cre | Urea | 0.257192711 | 4.71E-13 |
| B191 | B206 | Urea | Waist | 0.200793639 | 2.03E-08 |
| B192 | B206 | UA | Waist | 0.499450653 | 0 |
| B192 | B210 | UA | Hip | 0.404256929 | 0 |
| B192 | B215 | UA | BMI | 0.402815787 | 0 |
| B192 | B216 | UA | BMI | 0.525555577 | 0 |
| B192 | B221 | UA | Hei | 0.392422595 | 0 |
| B192 | B222 | UA | NS | 0.523938062 | 0 |
| B192 | B204 | UA | Drinking | 0.278164105 | 4.44E-15 |
| B192 | B209 | UA | Smoking | 0.203185537 | 1.37E-08 |
| B194 | B195 | Cl | Na | 0.309969928 | 0 |
| B194 | B197 | Cl | AG | -0.24054021 | 1.48E-11 |
| B194 | B198 | Cl | P | 0.187256757 | 1.75E-07 |
| B194 | B200 | Cl | OP | 0.181089696 | 4.44E-07 |
| B195 | B197 | Na | AG | 0.377406652 | 0 |
| B195 | B200 | Na | OP | 0.896157741 | 0 |
| B196 | B197 | TCO2 | AG | -0.82834408 | 0 |
| B196 | B199 | TCO2 | Mg | -0.19397322 | 6.14E-08 |
| B197 | B200 | AG | OP | 0.294735475 | 0 |
| B197 | B199 | AG | Mg | 0.187959654 | 1.57E-07 |
| B198 | B199 | P | Mg | -0.31088665 | 0 |
| B2 | B153 | Age | Glu | 0.311101739 | 0 |
| B2 | B189 | Age | eGFB | -0.64933678 | 0 |
| B2 | B220 | Age | SBP | 0.346908745 | 0 |
| B2 | B43 | Age | MCV | 0.298284629 | 0 |
| B2 | B47 | Age | RDW | 0.348998023 | 0 |
| B2 | B52 | Age | PLT/BPC | -0.26919932 | 3.38E-14 |
| B2 | B126 | Age | Alb | -0.26525038 | 8.13E-14 |
| B2 | B219 | Age | OT | 0.257108465 | 4.80E-13 |
| B2 | B46 | Age | RBC Count | -0.2362646 | 3.43E-11 |
| B2 | B77 | Age | TC | 0.233953394 | 5.38E-11 |
| B2 | B227 | Age | OD | -0.23127994 | 9.00E-11 |
| B2 | B44 | Age | MCH | 0.224507026 | 3.22E-10 |

| | | | | | |
|---|---|---|---|---|---|
| B2 | B221 | Age | Hei | -0.22333977 | 3.99E-10 |
| B2 | B228 | Age | OS | -0.21859666 | 9.46E-10 |
| B2 | B206 | Age | Waist | 0.215382494 | 1.68E-09 |
| B2 | B191 | Age | Urea | 0.21042167 | 4.00E-09 |
| B2 | B70 | Age | LDL-C | 0.177942803 | 7.06E-07 |
| B20 | B164 | CD3+CD4+CD8+ | COM | 0.348935773 | 0 |
| B20 | B21 | CD3+CD4+CD8+ | CD3+CD4-CD8- | -0.63374153 | 0 |
| B20 | B22 | CD3+CD4+CD8+ | CD3+CD8+(T) | 0.308015767 | 0 |
| B20 | B23 | CD3+CD4+CD8+ | CD3-CD16+CD56+(NK) | 0.574050822 | 0 |
| B20 | B161 | CD3+CD4+CD8+ | LEU | 0.229360173 | 1.30E-10 |
| B20 | B87 | CD3+CD4+CD8+ | NSE | -0.2123356 | 2.87E-09 |
| B204 | B209 | Drinking | Smoking | 0.404511543 | 0 |
| B204 | B216 | Drinking | BMI | 0.318800804 | 0 |
| B204 | B221 | Drinking | Hei | 0.302570354 | 0 |
| B204 | B222 | Drinking | NS | 0.33296055 | 0 |
| B204 | B206 | Drinking | Waist | 0.262150261 | 1.61E-13 |
| B204 | B210 | Drinking | Hip | 0.247691965 | 3.47E-12 |
| B204 | B215 | Drinking | BMI | 0.191955168 | 8.44E-08 |
| B206 | B210 | Waist | Hip | 0.776324217 | 0 |
| B206 | B215 | Waist | BMI | 0.796460944 | 0 |
| B206 | B216 | Waist | BMI | 0.831090816 | 0 |
| B206 | B220 | Waist | SBP | 0.30696212 | 0 |
| B206 | B221 | Waist | Hei | 0.425655252 | 0 |
| B206 | B222 | Waist | NS | 0.743938554 | 0 |
| B206 | B218 | Waist | DBP | 0.270565002 | 2.46E-14 |
| B206 | B209 | Waist | Smoking | 0.24369015 | 7.85E-12 |
| B206 | B207 | Waist | WHR | 0.22007991 | 7.24E-10 |
| B209 | B222 | Smoking | NS | 0.360957639 | 0 |
| B209 | B221 | Smoking | Hei | 0.283611433 | 1.33E-15 |
| B209 | B216 | Smoking | BMI | 0.268769807 | 3.69E-14 |
| B209 | B210 | Smoking | Hip | 0.182583255 | 3.56E-07 |
| B21 | B198 | CD3+CD4-CD8- | P | 0.308452975 | 0 |
| B21 | B23 | CD3+CD4-CD8- | CD3-CD16+CD56+(NK) | -0.44158433 | 0 |
| B21 | B22 | CD3+CD4-CD8- | CD3+CD8+(T) | -0.25172786 | 1.50E-12 |
| B21 | B164 | CD3+CD4-CD8- | COM | -0.20863464 | 5.44E-09 |
| B21 | B197 | CD3+CD4-CD8- | AG | -0.20191072 | 1.69E-08 |
| B21 | B194 | CD3+CD4-CD8- | CI | 0.188569851 | 1.43E-07 |
| B210 | B215 | Hip | BMI | 0.751987374 | 0 |
| B210 | B216 | Hip | BMI | 0.837949596 | 0 |
| B210 | B221 | Hip | Hei | 0.494110207 | 0 |
| B210 | B222 | Hip | NS | 0.668783266 | 0 |
| B210 | B218 | Hip | DBP | 0.263154711 | 1.29E-13 |
| B210 | B220 | Hip | SBP | 0.210200031 | 4.16E-09 |
| B211 | B212 | IOP-R | IOP-L | 0.797642859 | 0 |

| | | | | | |
|---|---|---|---|---|---|
| B213 | B214 | CVA-R | CVA-L | 0.742748637 | 0 |
| B213 | B228 | CVA-R | OS | -0.17916958 | 5.90E-07 |
| B214 | B228 | CVA-L | OS | -0.18665127 | 1.92E-07 |
| B215 | B216 | BMI | BMI | 0.82273833 | 0 |
| B215 | B222 | BMI | NS | 0.639314006 | 0 |
| B215 | B218 | BMI | DBP | 0.282006954 | 1.78E-15 |
| B215 | B220 | BMI | SBP | 0.28184145 | 1.78E-15 |
| B216 | B218 | BMI | DBP | 0.326969218 | 0 |
| B216 | B221 | BMI | Hei | 0.681320176 | 0 |
| B216 | B222 | BMI | NS | 0.799814215 | 0 |
| B216 | B220 | BMI | SBP | 0.267816065 | 4.57E-14 |
| B218 | B220 | DBP | SBP | 0.668038492 | 0 |
| B218 | B222 | DBP | NS | 0.28462791 | 8.88E-16 |
| B218 | B221 | DBP | Hei | 0.208134483 | 5.93E-09 |
| B22 | B138 | CD3+CD8+(T) | PAB | 0.342578812 | 0 |
| B22 | B156 | CD3+CD8+(T) | IgE | -0.34681166 | 0 |
| B22 | B34 | CD3+CD8+(T) | TK | 0.288694411 | 4.44E-16 |
| B22 | B171 | CD3+CD8+(T) | EC | 0.258805011 | 3.33E-13 |
| B22 | B139 | CD3+CD8+(T) | ADA | -0.25006074 | 2.13E-12 |
| B22 | B103 | CD3+CD8+(T) | anti-SSA/Ro | -0.23316021 | 6.27E-11 |
| B22 | B55 | CD3+CD8+(T) | TK | -0.22739041 | 1.88E-10 |
| B22 | B158 | CD3+CD8+(T) | IgM | -0.22063491 | 6.54E-10 |
| B22 | B193 | CD3+CD8+(T) | K | 0.189641328 | 1.21E-07 |
| B22 | B100 | CD3+CD8+(T) | anti-Ro52 | 0.18416581 | 2.80E-07 |
| B22 | B101 | CD3+CD8+(T) | anti-Sc70 | 0.184165812 | 2.80E-07 |
| B22 | B102 | CD3+CD8+(T) | anti-Sm | 0.184165812 | 2.80E-07 |
| B22 | B104 | CD3+CD8+(T) | anti-SSB | 0.184165809 | 2.80E-07 |
| B22 | B105 | CD3+CD8+(T) | anti-U1RNP | 0.18416581 | 2.80E-07 |
| B22 | B106 | CD3+CD8+(T) | anti-P | 0.184165809 | 2.80E-07 |
| B22 | B107 | CD3+CD8+(T) | AnuA | 0.184165762 | 2.80E-07 |
| B22 | B108 | CD3+CD8+(T) | Anti-Cyclin | 0.18416581 | 2.80E-07 |
| B22 | B109 | CD3+CD8+(T) | AMA-M2 | 0.18416581 | 2.80E-07 |
| B22 | B110 | CD3+CD8+(T) | AHA | 0.184165811 | 2.80E-07 |
| B22 | B95 | CD3+CD8+(T) | anti-cenpb | 0.184165809 | 2.80E-07 |
| B22 | B97 | CD3+CD8+(T) | dsDNA | 0.184165809 | 2.80E-07 |
| B22 | B98 | CD3+CD8+(T) | anti-Jo-1 | 0.184165812 | 2.80E-07 |
| B22 | B99 | CD3+CD8+(T) | anti-PM-Scl | 0.18416581 | 2.80E-07 |
| B220 | B222 | SBP | NS | 0.2501207 | 2.10E-12 |
| B221 | B222 | Hei | NS | 0.576417327 | 0 |
| B227 | B228 | OD | OS | 0.823836625 | 0 |
| B23 | B103 | CD3-CD16+CD56+(NK) | anti-SSA/Ro | -0.39446951 | 0 |
| B23 | B135 | CD3-CD16+CD56+(NK) | TBA | 0.219896019 | 7.48E-10 |
| B23 | B200 | CD3-CD16+CD56+(NK) | OP | 0.200343204 | 2.19E-08 |
| B23 | B164 | CD3-CD16+CD56+(NK) | COM | 0.197564396 | 3.45E-08 |

| | | | | | |
|---|---|---|---|---|---|
| B24 | B26 | αHBDH | CKMB | 0.528244613 | 0 |
| B24 | B27 | αHBDH | LDH | 0.816539224 | 0 |
| B24 | B132 | αHBDH | Dbil | 0.257256845 | 4.65E-13 |
| B24 | B47 | αHBDH | RDW | 0.243139576 | 8.77E-12 |
| B24 | B25 | αHBDH | CK | 0.225481449 | 2.68E-10 |
| B24 | B131 | αHBDH | AST | 0.210214701 | 4.15E-09 |
| B25 | B26 | CK | CKMB | 0.326845338 | 0 |
| B25 | B27 | CK | LDH | 0.236478099 | 3.29E-11 |
| B26 | B27 | CKMB | LDH | 0.46005364 | 0 |
| B26 | B149 | CKMB | TGAb | 0.21118749 | 3.51E-09 |
| B26 | B132 | CKMB | Dbil | 0.185090845 | 2.44E-07 |
| B26 | B131 | CKMB | AST | 0.175960727 | 9.41E-07 |
| B27 | B131 | LDH | AST | 0.228606685 | 1.49E-10 |
| B27 | B47 | LDH | RDW | 0.186705104 | 1.91E-07 |
| B28 | B139 | PRL | ADA | 0.346967777 | 0 |
| B28 | B29 | PRL | E2 | 0.793082169 | 0 |
| B28 | B31 | PRL | FSH | -0.57555326 | 0 |
| B28 | B92 | PRL | ALDH2-Glu504Glu | 0.538568014 | 0 |
| B28 | B93 | PRL | ALDH2-Glu504Lys | -0.53856801 | 0 |
| B28 | B94 | PRL | ALDH2-Lys504Lys | -0.53856801 | 0 |
| B28 | B138 | PRL | PAB | -0.28030079 | 2.66E-15 |
| B28 | B33 | PRL | Prog | 0.253116099 | 1.12E-12 |
| B28 | B39 | PRL | Other cells | -0.2203986 | 6.83E-10 |
| B28 | B156 | PRL | IgE | -0.21567959 | 1.59E-09 |
| B28 | B30 | PRL | LH | -0.17884605 | 6.19E-07 |
| B29 | B138 | E2 | PAB | -0.35912027 | 0 |
| B29 | B139 | E2 | ADA | 0.444585851 | 0 |
| B29 | B31 | E2 | FSH | -0.47535163 | 0 |
| B29 | B92 | E2 | ALDH2-Glu504Glu | 0.689359117 | 0 |
| B29 | B93 | E2 | ALDH2-Glu504Lys | -0.68935912 | 0 |
| B29 | B94 | E2 | ALDH2-Lys504Lys | -0.68935911 | 0 |
| B29 | B156 | E2 | IgE | -0.27588767 | 7.11E-15 |
| B29 | B33 | E2 | Prog | 0.241390664 | 1.25E-11 |
| B29 | B198 | E2 | P | 0.204213031 | 1.15E-08 |
| B29 | B194 | E2 | Cl | 0.185923017 | 2.15E-07 |
| B29 | B32 | E2 | Testo | -0.17901784 | 6.03E-07 |
| B29 | B39 | E2 | Other cells | -0.17623682 | 9.04E-07 |
| B3 | B123 | Hcy | hs-CRP1 | 0.337662309 | 0 |
| B3 | B132 | Hcy | Dbil | 0.320430719 | 0 |
| B3 | B24 | Hcy | αHBDH | 0.252263156 | 1.34E-12 |
| B3 | B165 | Hcy | bilirubin | 0.227965571 | 1.69E-10 |
| B3 | B27 | Hcy | LDH | 0.227044226 | 2.01E-10 |
| B3 | B47 | Hcy | RDW | 0.209005801 | 5.11E-09 |
| B3 | B131 | Hcy | AST | 0.177071292 | 8.02E-07 |

| | | | | | |
|---|---|---|---|---|---|
| B30 | B31 | LH | FSH | 0.872222492 | 0 |
| B30 | B33 | LH | Prog | -0.29809925 | 0 |
| B30 | B173 | LH | NTT | 0.210520952 | 3.93E-09 |
| B31 | B33 | FSH | Prog | -0.36839696 | 0 |
| B31 | B92 | FSH | ALDH2-Glu504Glu | -0.29669961 | 0 |
| B31 | B93 | FSH | ALDH2-Glu504Lys | 0.296699612 | 0 |
| B31 | B94 | FSH | ALDH2-Lys504Lys | 0.29669961 | 0 |
| B31 | B173 | FSH | NTT | 0.214448826 | 1.98E-09 |
| B31 | B139 | FSH | ADA | -0.1916835 | 8.81E-08 |
| B31 | B39 | FSH | Other cells | 0.175835174 | 9.58E-07 |
| B32 | B33 | Testo | Prog | 0.779485975 | 0 |
| B32 | B92 | Testo | ALDH2-Glu504Glu | -0.25930635 | 2.99E-13 |
| B32 | B93 | Testo | ALDH2-Glu504Lys | 0.25930635 | 2.99E-13 |
| B32 | B94 | Testo | ALDH2-Lys504Lys | 0.259306348 | 2.99E-13 |
| B33 | B173 | Prog | NTT | -0.25504185 | 7.46E-13 |
| B34 | B193 | TK | K | 0.377638141 | 0 |
| B34 | B85 | TK | SCC | 0.227962206 | 1.69E-10 |
| B34 | B156 | TK | IgE | -0.22710487 | 1.98E-10 |
| B34 | B87 | TK | NSE | 0.205898221 | 8.68E-09 |
| B34 | B145 | TK | BGP | -0.19871512 | 2.86E-08 |
| B34 | B197 | TK | AG | -0.18494693 | 2.49E-07 |
| B34 | B144 | TK | Total-PINP | -0.18299442 | 3.35E-07 |
| B34 | B143 | TK | β-CTx | -0.17755774 | 7.47E-07 |
| B35 | B167 | β2-MG | PC | 0.480572312 | 0 |
| B35 | B37 | β2-MG | BAND | 0.846787565 | 0 |
| B35 | B96 | β2-MG | dsDNA (IIF) | 0.771415561 | 0 |
| B36 | B116 | V-Lym% | TT3 | 0.317363237 | 0 |
| B39 | B79 | Other cells | hb V -dna | 0.356399017 | 0 |
| B39 | B75 | Other cells | ApoA1/B | -0.26985239 | 2.89E-14 |
| B39 | B76 | Other cells | ApoB | 0.209304084 | 4.85E-09 |
| B39 | B154 | Other cells | RF | 0.205463888 | 9.34E-09 |
| B39 | B145 | Other cells | BGP | -0.20135933 | 1.85E-08 |
| B39 | B175 | Other cells | ALB-U | 0.194924582 | 5.27E-08 |
| B39 | B144 | Other cells | Total-PINP | -0.18331082 | 3.19E-07 |
| B39 | B143 | Other cells | β-CTx | -0.18045603 | 4.88E-07 |
| B4 | B5 | WB-Ca | WB-Mg | 0.945885434 | 0 |
| B4 | B6 | WB-Ca | WB-Fe | 0.973106048 | 0 |
| B4 | B112 | WB-Ca | FT4 | 0.207471537 | 6.64E-09 |
| B4 | B7 | WB-Ca | WB-Cop | 0.187889218 | 1.59E-07 |
| B40 | B42 | WBC | MON | 0.647252762 | 0 |
| B40 | B48 | WBC | LYM% | -0.32131988 | 0 |
| B40 | B53 | WBC | GR% | 0.331446569 | 0 |
| B40 | B206 | WBC | Waist | 0.241098441 | 1.32E-11 |
| B40 | B216 | WBC | BMI | 0.226747536 | 2.12E-10 |

| | | | | | |
|---|---|---|---|---|---|
| B40 | B51 | WBC | Hb | 0.222124606 | 4.99E-10 |
| B40 | B52 | WBC | PLT/BPC | 0.211818268 | 3.14E-09 |
| B40 | B215 | WBC | BMI | 0.208182217 | 5.88E-09 |
| B40 | B209 | WBC | Smoking | 0.204589552 | 1.08E-08 |
| B40 | B222 | WBC | NS | 0.203134206 | 1.38E-08 |
| B40 | B210 | WBC | Hip | 0.202738425 | 1.47E-08 |
| B40 | B46 | WBC | RBC Count | 0.186061119 | 2.10E-07 |
| B40 | B72 | WBC | HDL-C | -0.17884575 | 6.19E-07 |
| B41 | B42 | MON% | MON | 0.649491808 | 0 |
| B41 | B53 | MON% | GR% | -0.2668274 | 5.73E-14 |
| B42 | B209 | MON | Smoking | 0.287527218 | 4.44E-16 |
| B42 | B51 | MON | Hb | 0.263168132 | 1.29E-13 |
| B42 | B123 | MON | hs-CRP1 | 0.245543474 | 5.39E-12 |
| B42 | B72 | MON | HDL-C | -0.23858274 | 2.18E-11 |
| B42 | B206 | MON | Waist | 0.226147804 | 2.37E-10 |
| B42 | B216 | MON | BMI | 0.218936942 | 8.90E-10 |
| B42 | B48 | MON | LYM% | -0.21852349 | 9.59E-10 |
| B42 | B222 | MON | NS | 0.213869195 | 2.19E-09 |
| B42 | B140 | MON | r-GT | 0.210472505 | 3.97E-09 |
| B42 | B221 | MON | Hei | 0.197582296 | 3.44E-08 |
| B42 | B46 | MON | RBC Count | 0.193686931 | 6.42E-08 |
| B42 | B192 | MON | UA | 0.191917234 | 8.49E-08 |
| B42 | B58 | MON | HCT | 0.185809038 | 2.19E-07 |
| B42 | B131 | MON | AST | 0.181649002 | 4.09E-07 |
| B42 | B69 | MON | PV | -0.17832111 | 6.68E-07 |
| B42 | B190 | MON | Cre | 0.178243601 | 6.76E-07 |
| B42 | B210 | MON | Hip | 0.176675268 | 8.49E-07 |
| B43 | B189 | MCV | eGFB | -0.30806825 | 0 |
| B43 | B44 | MCV | MCH | 0.907616211 | 0 |
| B43 | B46 | MCV | RBC Count | -0.51234081 | 0 |
| B43 | B47 | MCV | RDW | 0.495853842 | 0 |
| B43 | B45 | MCV | MCHC | 0.26266172 | 1.44E-13 |
| B43 | B77 | MCV | TC | 0.225659139 | 2.60E-10 |
| B43 | B52 | MCV | PLT/BPC | -0.19805459 | 3.18E-08 |
| B43 | B70 | MCV | LDL-C | 0.196797276 | 3.90E-08 |
| B43 | B51 | MCV | Hb | 0.192251837 | 8.06E-08 |
| B44 | B45 | MCH | MCHC | 0.641376328 | 0 |
| B44 | B46 | MCH | RBC Count | -0.35310216 | 0 |
| B44 | B47 | MCH | RDW | 0.305218724 | 0 |
| B44 | B51 | MCH | Hb | 0.408788772 | 0 |
| B44 | B189 | MCH | eGFB | -0.2722022 | 1.69E-14 |
| B44 | B190 | MCH | Cre | 0.226866291 | 2.07E-10 |
| B44 | B52 | MCH | PLT/BPC | -0.20531425 | 9.57E-09 |
| B44 | B209 | MCH | Smoking | 0.191724636 | 8.76E-08 |

| | | | | | |
|---|---|---|---|---|---|
| B44 | B77 | MCH | TC | 0.179652839 | 5.50E-07 |
| B45 | B51 | MCHC | Hb | 0.59082528 | 0 |
| B45 | B190 | MCHC | Cre | 0.23081079 | 9.84E-11 |
| B45 | B141 | MCHC | AST/ALT | -0.22459243 | 3.17E-10 |
| B45 | B47 | MCHC | RDW | -0.21950682 | 8.03E-10 |
| B45 | B209 | MCHC | Smoking | 0.194897274 | 5.30E-08 |
| B45 | B222 | MCHC | NS | 0.194455548 | 5.68E-08 |
| B45 | B221 | MCHC | Hei | 0.181863172 | 3.96E-07 |
| B45 | B216 | MCHC | BMI | 0.179303976 | 5.79E-07 |
| B46 | B141 | RBC Count | AST/ALT | -0.33055272 | 0 |
| B46 | B163 | RBC Count | SG | 0.31583356 | 0 |
| B46 | B190 | RBC Count | Cre | 0.313722025 | 0 |
| B46 | B192 | RBC Count | UA | 0.420686272 | 0 |
| B46 | B206 | RBC Count | Waist | 0.337568754 | 0 |
| B46 | B210 | RBC Count | Hip | 0.294039978 | 0 |
| B46 | B216 | RBC Count | BMI | 0.429131945 | 0 |
| B46 | B221 | RBC Count | Hei | 0.462019395 | 0 |
| B46 | B222 | RBC Count | NS | 0.444174895 | 0 |
| B46 | B47 | RBC Count | RDW | -0.41399313 | 0 |
| B46 | B51 | RBC Count | Hb | 0.699524485 | 0 |
| B46 | B218 | RBC Count | DBP | 0.282896869 | 1.33E-15 |
| B46 | B126 | RBC Count | Alb | 0.274651449 | 9.77E-15 |
| B46 | B127 | RBC Count | A-G ratio | 0.239004654 | 2.00E-11 |
| B46 | B128 | RBC Count | ALT | 0.228760499 | 1.45E-10 |
| B46 | B209 | RBC Count | Smoking | 0.225547888 | 2.65E-10 |
| B46 | B215 | RBC Count | BMI | 0.222769936 | 4.43E-10 |
| B46 | B72 | RBC Count | HDL-C | -0.21922088 | 8.45E-10 |
| B46 | B204 | RBC Count | Drinking | 0.213161635 | 2.48E-09 |
| B47 | B189 | RDW | eGFB | -0.30716828 | 0 |
| B47 | B111 | RDW | TSH | 0.263590151 | 1.17E-13 |
| B47 | B112 | RDW | FT4 | -0.24843181 | 2.98E-12 |
| B47 | B132 | RDW | Dbil | 0.244548562 | 6.59E-12 |
| B47 | B77 | RDW | TC | 0.238709201 | 2.12E-11 |
| B47 | B131 | RDW | AST | 0.23310689 | 6.34E-11 |
| B47 | B123 | RDW | hs-CRP1 | 0.212386531 | 2.84E-09 |
| B47 | B130 | RDW | GLB | 0.20564873 | 9.05E-09 |
| B47 | B127 | RDW | A-G ratio | -0.1983568 | 3.03E-08 |
| B47 | B113 | RDW | FT3 | -0.19658721 | 4.04E-08 |
| B47 | B221 | RDW | Hei | -0.19543667 | 4.86E-08 |
| B47 | B70 | RDW | LDL-C | 0.187865683 | 1.60E-07 |
| B47 | B52 | RDW | PLT/BPC | -0.18620796 | 2.06E-07 |
| B47 | B141 | RDW | AST/ALT | 0.179583091 | 5.55E-07 |
| B48 | B53 | LYM% | GR% | -0.94590657 | 0 |
| B49 | B50 | baso% | eo% | 0.29776153 | 0 |

| | | | | | |
|---|---|---|---|---|---|
| B49 | B53 | baso% | GR% | -0.17979503 | 5.38E-07 |
| B49 | B190 | baso% | Cre | 0.175637588 | 9.86E-07 |
| B5 | B6 | WB-Mg | WB-Fe | 0.995198051 | 0 |
| B5 | B7 | WB-Mg | WB-Cop | 0.496443154 | 0 |
| B5 | B8 | WB-Mg | WB-Zinc | 0.274549432 | 9.77E-15 |
| B5 | B78 | WB-Mg | HBcAb | -0.20806435 | 6.00E-09 |
| B5 | B112 | WB-Mg | FT4 | 0.18829976 | 1.49E-07 |
| B50 | B165 | eo% | bilirubin | -0.27234808 | 1.64E-14 |
| B50 | B53 | eo% | GR% | -0.25028149 | 2.03E-12 |
| B51 | B141 | Hb | AST/ALT | -0.35465245 | 0 |
| B51 | B163 | Hb | SG | 0.338470429 | 0 |
| B51 | B190 | Hb | Cre | 0.485907 | 0 |
| B51 | B192 | Hb | UA | 0.41466518 | 0 |
| B51 | B204 | Hb | Drinking | 0.324514643 | 0 |
| B51 | B206 | Hb | Waist | 0.369265867 | 0 |
| B51 | B209 | Hb | Smoking | 0.375460985 | 0 |
| B51 | B210 | Hb | Hip | 0.317918819 | 0 |
| B51 | B216 | Hb | BMI | 0.468117623 | 0 |
| B51 | B221 | Hb | Hei | 0.483059479 | 0 |
| B51 | B222 | Hb | NS | 0.511494621 | 0 |
| B51 | B126 | Hb | Alb | 0.27936192 | 3.11E-15 |
| B51 | B174 | Hb | Urea/Cr | -0.27960223 | 3.11E-15 |
| B51 | B128 | Hb | ALT | 0.278145203 | 4.44E-15 |
| B51 | B218 | Hb | DBP | 0.274620616 | 9.77E-15 |
| B51 | B215 | Hb | BMI | 0.253499979 | 1.03E-12 |
| B51 | B127 | Hb | A-G ratio | 0.251201082 | 1.68E-12 |
| B51 | B71 | Hb | TG | 0.209891715 | 4.39E-09 |
| B51 | B166 | Hb | Pro | 0.179036881 | 6.02E-07 |
| B51 | B62 | Hb | WBHSRVI | 0.176479625 | 8.73E-07 |
| B52 | B189 | PLT/BPC | eGFB | 0.176329148 | 8.92E-07 |
| B54 | B56 | ESR | IR | -0.51592531 | 0 |
| B54 | B58 | ESR | HCT | -0.35865271 | 0 |
| B54 | B60 | ESR | WBRVI | -0.5224656 | 0 |
| B54 | B61 | ESR | WBHSRV | -0.33423326 | 0 |
| B54 | B62 | ESR | WBHSRVI | -0.51142634 | 0 |
| B54 | B63 | ESR | WBRSR(mPas_200) | -0.32152764 | 0 |
| B54 | B64 | ESR | WBRSR(mPas_5) | -0.40125561 | 0 |
| B54 | B65 | ESR | WBRSR(mPas_1) | -0.40252247 | 0 |
| B54 | B66 | ESR | WBRSR(mPas_100) | -0.3475621 | 0 |
| B54 | B67 | ESR | WBRSR(mPas_50) | -0.36395032 | 0 |
| B54 | B69 | ESR | PV | 0.585339698 | 0 |
| B54 | B79 | ESR | hb V -dna | -0.28361006 | 1.33E-15 |
| B54 | B182 | ESR | TT | -0.27019359 | 2.69E-14 |
| B54 | B156 | ESR | IgE | 0.258110147 | 3.87E-13 |

| | | | | | |
|---|---|---|---|---|---|
| B54 | B111 | ESR | TSH | 0.23665683 | 3.18E-11 |
| B54 | B59 | ESR | WBRV | -0.228548 | 1.51E-10 |
| B54 | B125 | ESR | Cor | 0.224203067 | 3.40E-10 |
| B54 | B55 | ESR | TK | -0.22241864 | 4.73E-10 |
| B54 | B81 | ESR | TSGF | 0.221713142 | 5.38E-10 |
| B54 | B75 | ESR | ApoA1/B | 0.196319313 | 4.22E-08 |
| B54 | B57 | ESR | RBCAI | -0.19616996 | 4.32E-08 |
| B55 | B56 | TK | IR | 0.558048855 | 0 |
| B55 | B58 | TK | HCT | -0.41201262 | 0 |
| B55 | B59 | TK | WBRV | 0.537112439 | 0 |
| B55 | B61 | TK | WBHSRV | 0.500025604 | 0 |
| B55 | B69 | TK | PV | -0.26369065 | 1.15E-13 |
| B55 | B60 | TK | WBRVI | 0.210334699 | 4.06E-09 |
| B55 | B136 | TK | 5'NT | -0.19632295 | 4.21E-08 |
| B55 | B182 | TK | TT | 0.178224324 | 6.78E-07 |
| B56 | B182 | IR | TT | 0.296731276 | 0 |
| B56 | B58 | IR | HCT | 0.506579134 | 0 |
| B56 | B59 | IR | WBRV | 0.46812105 | 0 |
| B56 | B60 | IR | WBRVI | 0.766175403 | 0 |
| B56 | B61 | IR | WBHSRV | 0.911990931 | 0 |
| B56 | B62 | IR | WBHSRVI | 0.893615015 | 0 |
| B56 | B63 | IR | WBRSR(mPas_200) | 0.766215074 | 0 |
| B56 | B64 | IR | WBRSR(mPas_5) | 0.773183109 | 0 |
| B56 | B65 | IR | WBRSR(mPas_1) | 0.678534872 | 0 |
| B56 | B66 | IR | WBRSR(mPas_100) | 0.796728953 | 0 |
| B56 | B67 | IR | WBRSR(mPas_50) | 0.803873014 | 0 |
| B56 | B69 | IR | PV | -0.43057919 | 0 |
| B57 | B100 | RBCAI | anti-Ro52 | -0.30683775 | 0 |
| B57 | B101 | RBCAI | anti-Sc70 | -0.30683776 | 0 |
| B57 | B102 | RBCAI | anti-Sm | -0.30683775 | 0 |
| B57 | B104 | RBCAI | anti-SSB | -0.30683775 | 0 |
| B57 | B105 | RBCAI | anti-U1RNP | -0.30683775 | 0 |
| B57 | B106 | RBCAI | anti-P | -0.30683775 | 0 |
| B57 | B107 | RBCAI | AnuA | -0.30683767 | 0 |
| B57 | B108 | RBCAI | Anti-Cyclin | -0.30683775 | 0 |
| B57 | B109 | RBCAI | AMA-M2 | -0.30683775 | 0 |
| B57 | B110 | RBCAI | AHA | -0.30683775 | 0 |
| B57 | B136 | RBCAI | 5'NT | 0.447927009 | 0 |
| B57 | B59 | RBCAI | WBRV | 0.789469281 | 0 |
| B57 | B60 | RBCAI | WBRVI | 0.570550029 | 0 |
| B57 | B65 | RBCAI | WBRSR(mPas_1) | 0.518240627 | 0 |
| B57 | B95 | RBCAI | anti-cenpb | -0.30683775 | 0 |
| B57 | B97 | RBCAI | dsDNA | -0.30683775 | 0 |
| B57 | B98 | RBCAI | anti-Jo-1 | -0.30683776 | 0 |

| | | | | | |
|---|---|---|---|---|---|
| B57 | B99 | RBCAI | anti-PM-Scl | -0.30683775 | 0 |
| B57 | B64 | RBCAI | WBRSR(mPas_5) | 0.263118609 | 1.30E-13 |
| B57 | B69 | RBCAI | PV | -0.2290386 | 1.38E-10 |
| B57 | B171 | RBCAI | EC | 0.194946944 | 5.25E-08 |
| B58 | B134 | HCT | CHE | 0.308802914 | 0 |
| B58 | B60 | HCT | WBRVI | 0.592404372 | 0 |
| B58 | B61 | HCT | WBHSRV | 0.4948534 | 0 |
| B58 | B62 | HCT | WBHSRVI | 0.830221429 | 0 |
| B58 | B63 | HCT | WBRSR(mPas_200) | 0.836169256 | 0 |
| B58 | B64 | HCT | WBRSR(mPas_5) | 0.763537211 | 0 |
| B58 | B65 | HCT | WBRSR(mPas_1) | 0.632635154 | 0 |
| B58 | B66 | HCT | WBRSR(mPas_100) | 0.834728621 | 0 |
| B58 | B67 | HCT | WBRSR(mPas_50) | 0.828352232 | 0 |
| B58 | B194 | HCT | CI | -0.2399335 | 1.67E-11 |
| B59 | B60 | WBRV | WBRVI | 0.695987962 | 0 |
| B59 | B61 | WBRV | WBHSRV | 0.504052707 | 0 |
| B59 | B64 | WBRV | WBRSR(mPas_5) | 0.557786328 | 0 |
| B59 | B65 | WBRV | WBRSR(mPas_1) | 0.722047272 | 0 |
| B59 | B66 | WBRV | WBRSR(mPas_100) | 0.299545247 | 0 |
| B59 | B67 | WBRV | WBRSR(mPas_50) | 0.357311238 | 0 |
| B59 | B136 | WBRV | 5'NT | 0.287708939 | 4.44E-16 |
| B59 | B62 | WBRV | WBHSRVI | 0.251927711 | 1.44E-12 |
| B59 | B63 | WBRV | WBRSR(mPas_200) | 0.243832076 | 7.62E-12 |
| B59 | B100 | WBRV | anti-Ro52 | -0.19831357 | 3.05E-08 |
| B59 | B101 | WBRV | anti-Sc70 | -0.19831358 | 3.05E-08 |
| B59 | B102 | WBRV | anti-Sm | -0.19831358 | 3.05E-08 |
| B59 | B104 | WBRV | anti-SSB | -0.19831357 | 3.05E-08 |
| B59 | B105 | WBRV | anti-U1RNP | -0.19831357 | 3.05E-08 |
| B59 | B106 | WBRV | anti-P | -0.19831357 | 3.05E-08 |
| B59 | B107 | WBRV | AnuA | -0.19831352 | 3.05E-08 |
| B59 | B108 | WBRV | Anti-Cyclin | -0.19831357 | 3.05E-08 |
| B59 | B109 | WBRV | AMA-M2 | -0.19831357 | 3.05E-08 |
| B59 | B110 | WBRV | AHA | -0.19831357 | 3.05E-08 |
| B59 | B95 | WBRV | anti-cenpb | -0.19831357 | 3.05E-08 |
| B59 | B97 | WBRV | dsDNA | -0.19831357 | 3.05E-08 |
| B59 | B98 | WBRV | anti-Jo-1 | -0.19831358 | 3.05E-08 |
| B59 | B99 | WBRV | anti-PM-Scl | -0.19831357 | 3.05E-08 |
| B6 | B7 | WB-Fe | WB-Cop | 0.409091112 | 0 |
| B6 | B112 | WB-Fe | FT4 | 0.196252109 | 4.26E-08 |
| B6 | B8 | WB-Fe | WB-Zinc | 0.1791106 | 5.95E-07 |
| B60 | B136 | WBRVI | 5'NT | 0.291883456 | 0 |
| B60 | B61 | WBRVI | WBHSRV | 0.64707978 | 0 |
| B60 | B62 | WBRVI | WBHSRVI | 0.794327568 | 0 |
| B60 | B63 | WBRVI | WBRSR(mPas_200) | 0.64631546 | 0 |

| | | | | | |
|---|---|---|---|---|---|
| B60 | B64 | WBRVI | WBRSR(mPas_5) | 0.875924544 | 0 |
| B60 | B65 | WBRVI | WBRSR(mPas_1) | 0.923559518 | 0 |
| B60 | B66 | WBRVI | WBRSR(mPas_100) | 0.70802383 | 0 |
| B60 | B67 | WBRVI | WBRSR(mPas_50) | 0.751748454 | 0 |
| B60 | B69 | WBRVI | PV | -0.44620318 | 0 |
| B60 | B140 | WBRVI | r-GT | 0.183459676 | 3.12E-07 |
| B61 | B62 | WBHSRV | WBHSRVI | 0.814460802 | 0 |
| B61 | B63 | WBHSRV | WBRSR(mPas_200) | 0.868088572 | 0 |
| B61 | B64 | WBHSRV | WBRSR(mPas_5) | 0.819182142 | 0 |
| B61 | B65 | WBHSRV | WBRSR(mPas_1) | 0.707044118 | 0 |
| B61 | B66 | WBHSRV | WBRSR(mPas_100) | 0.876040536 | 0 |
| B61 | B67 | WBHSRV | WBRSR(mPas_50) | 0.876449711 | 0 |
| B61 | B182 | WBHSRV | TT | 0.245742045 | 5.17E-12 |
| B61 | B194 | WBHSRV | CI | -0.199195 | 2.64E-08 |
| B61 | B135 | WBHSRV | TBA | 0.178682635 | 6.34E-07 |
| B62 | B63 | WBHSRVI | WBRSR(mPas_200) | 0.903372348 | 0 |
| B62 | B64 | WBHSRVI | WBRSR(mPas_5) | 0.872379257 | 0 |
| B62 | B65 | WBHSRVI | WBRSR(mPas_1) | 0.742711315 | 0 |
| B62 | B66 | WBHSRVI | WBRSR(mPas_100) | 0.926674077 | 0 |
| B62 | B67 | WBHSRVI | WBRSR(mPas_50) | 0.927557645 | 0 |
| B62 | B69 | WBHSRVI | PV | -0.38662854 | 0 |
| B62 | B134 | WBHSRVI | CHE | 0.262956287 | 1.35E-13 |
| B62 | B182 | WBHSRVI | TT | 0.261038308 | 2.05E-13 |
| B63 | B64 | WBRSR(mPas_200) | WBRSR(mPas_5) | 0.900815154 | 0 |
| B63 | B65 | WBRSR(mPas_200) | WBRSR(mPas_1) | 0.749793961 | 0 |
| B63 | B66 | WBRSR(mPas_200) | WBRSR(mPas_100) | 0.993599967 | 0 |
| B63 | B67 | WBRSR(mPas_200) | WBRSR(mPas_50) | 0.985915058 | 0 |
| B63 | B194 | WBRSR(mPas_200) | CI | -0.29140005 | 2.22E-16 |
| B63 | B134 | WBRSR(mPas_200) | CHE | 0.235777911 | 3.78E-11 |
| B63 | B182 | WBRSR(mPas_200) | TT | 0.196087308 | 4.38E-08 |
| B63 | B75 | WBRSR(mPas_200) | ApoA1/B | -0.18513784 | 2.42E-07 |
| B63 | B135 | WBRSR(mPas_200) | TBA | 0.180969263 | 4.52E-07 |
| B64 | B65 | WBRSR(mPas_5) | WBRSR(mPas_1) | 0.9581316 | 0 |
| B64 | B66 | WBRSR(mPas_5) | WBRSR(mPas_100) | 0.928233561 | 0 |
| B64 | B67 | WBRSR(mPas_5) | WBRSR(mPas_50) | 0.950267496 | 0 |
| B64 | B136 | WBRSR(mPas_5) | 5'NT | 0.239347537 | 1.87E-11 |
| B64 | B194 | WBRSR(mPas_5) | CI | -0.22785677 | 1.72E-10 |
| B64 | B134 | WBRSR(mPas_5) | CHE | 0.183007916 | 3.34E-07 |
| B65 | B136 | WBRSR(mPas_1) | 5'NT | 0.349219406 | 0 |
| B65 | B66 | WBRSR(mPas_1) | WBRSR(mPas_100) | 0.792158501 | 0 |
| B65 | B67 | WBRSR(mPas_1) | WBRSR(mPas_50) | 0.831694194 | 0 |
| B65 | B194 | WBRSR(mPas_1) | CI | -0.1756314 | 9.87E-07 |
| B66 | B67 | WBRSR(mPas_100) | WBRSR(mPas_50) | 0.997466292 | 0 |
| B66 | B194 | WBRSR(mPas_100) | CI | -0.2767149 | 6.00E-15 |

| | | | | | |
|---|---|---|---|---|---|
| B66 | B134 | WBRSR(mPas_100) | CHE | 0.23729014 | 2.81E-11 |
| B66 | B182 | WBRSR(mPas_100) | TT | 0.206312012 | 8.09E-09 |
| B67 | B194 | WBRSR(mPas_50) | Cl | -0.27041492 | 2.55E-14 |
| B67 | B134 | WBRSR(mPas_50) | CHE | 0.230598136 | 1.02E-10 |
| B67 | B182 | WBRSR(mPas_50) | TT | 0.203578505 | 1.28E-08 |
| B69 | B194 | PV | Cl | -0.22186866 | 5.23E-10 |
| B69 | B193 | PV | K | 0.197544637 | 3.46E-08 |
| B69 | B111 | PV | TSH | 0.192719907 | 7.48E-08 |
| B69 | B135 | PV | TBA | 0.186472086 | 1.98E-07 |
| B69 | B199 | PV | Mg | 0.184837771 | 2.53E-07 |
| B69 | B154 | PV | RF | 0.179113353 | 5.95E-07 |
| B7 | B78 | WB-Cop | HBcAb | -0.38833911 | 0 |
| B7 | B8 | WB-Cop | WB-Zinc | 0.971010112 | 0 |
| B7 | B156 | WB-Cop | IgE | 0.280583073 | 2.44E-15 |
| B7 | B81 | WB-Cop | TSGF | 0.246173808 | 4.74E-12 |
| B7 | B39 | WB-Cop | Other cells | 0.205068515 | 9.98E-09 |
| B7 | B154 | WB-Cop | RF | 0.186777445 | 1.89E-07 |
| B7 | B157 | WB-Cop | IgG | -0.17984912 | 5.34E-07 |
| B70 | B77 | LDL-C | TC | 0.856266163 | 0 |
| B70 | B76 | LDL-C | ApoB | 0.286174136 | 6.66E-16 |
| B70 | B189 | LDL-C | eGFB | -0.26471085 | 9.15E-14 |
| B70 | B91 | LDL-C | TP | 0.240744983 | 1.42E-11 |
| B70 | B112 | LDL-C | FT4 | -0.22529031 | 2.78E-10 |
| B70 | B190 | LDL-C | Cre | 0.208000529 | 6.07E-09 |
| B70 | B215 | LDL-C | BMI | 0.203266405 | 1.35E-08 |
| B70 | B206 | LDL-C | Waist | 0.188324445 | 1.49E-07 |
| B70 | B126 | LDL-C | Alb | 0.180665376 | 4.73E-07 |
| B70 | B132 | LDL-C | Dbil | -0.17980337 | 5.38E-07 |
| B70 | B75 | LDL-C | ApoA1/B | -0.17940029 | 5.70E-07 |
| B71 | B206 | TG | Waist | 0.349446424 | 0 |
| B71 | B215 | TG | BMI | 0.326707897 | 0 |
| B71 | B216 | TG | BMI | 0.302178237 | 0 |
| B71 | B77 | TG | TC | 0.336817947 | 0 |
| B71 | B222 | TG | NS | 0.289636968 | 2.22E-16 |
| B71 | B192 | TG | UA | 0.281079757 | 2.22E-15 |
| B71 | B190 | TG | Cre | 0.235700919 | 3.83E-11 |
| B71 | B72 | TG | HDL-C | -0.23231163 | 7.38E-11 |
| B71 | B210 | TG | Hip | 0.219454251 | 8.11E-10 |
| B71 | B189 | TG | eGFB | -0.21414075 | 2.09E-09 |
| B71 | B218 | TG | DBP | 0.194575543 | 5.57E-08 |
| B71 | B141 | TG | AST/ALT | -0.19148117 | 9.10E-08 |
| B72 | B206 | HDL-C | Waist | -0.32780986 | 0 |
| B72 | B216 | HDL-C | BMI | -0.34192246 | 0 |
| B72 | B222 | HDL-C | NS | -0.33862857 | 0 |

| | | | | | |
|---|---|---|---|---|---|
| B72 | B74 | HDL-C | ApoA1 | 0.370311791 | 0 |
| B72 | B77 | HDL-C | TC | 0.37901466 | 0 |
| B72 | B192 | HDL-C | UA | -0.27106903 | 2.20E-14 |
| B72 | B215 | HDL-C | BMI | -0.26851474 | 3.91E-14 |
| B72 | B221 | HDL-C | Hei | -0.2580383 | 3.93E-13 |
| B72 | B92 | HDL-C | ALDH2-Glu504Glu | 0.253697794 | 9.92E-13 |
| B72 | B93 | HDL-C | ALDH2-Glu504Lys | -0.25369779 | 9.92E-13 |
| B72 | B94 | HDL-C | ALDH2-Lys504Lys | -0.25369779 | 9.92E-13 |
| B72 | B210 | HDL-C | Hip | -0.24167173 | 1.18E-11 |
| B72 | B209 | HDL-C | Smoking | -0.18955615 | 1.23E-07 |
| B72 | B163 | HDL-C | SG | -0.1885427 | 1.44E-07 |
| B72 | B128 | HDL-C | ALT | -0.18135433 | 4.27E-07 |
| B72 | B141 | HDL-C | AST/ALT | 0.180641879 | 4.75E-07 |
| B73 | B199 | ApoA | Mg | 0.259326793 | 2.98E-13 |
| B73 | B76 | ApoA | ApoB | 0.247317466 | 3.75E-12 |
| B73 | B74 | ApoA | ApoA1 | 0.229985178 | 1.15E-10 |
| B73 | B90 | ApoA | CYFRA 21-1 | 0.206090199 | 8.40E-09 |
| B73 | B88 | ApoA | CA 242 | 0.181000756 | 4.50E-07 |
| B74 | B123 | ApoA1 | hs-CRP1 | -0.43072709 | 0 |
| B74 | B132 | ApoA1 | Dbil | -0.36553098 | 0 |
| B74 | B165 | ApoA1 | bilirubin | -0.30147568 | 0 |
| B74 | B75 | ApoA1 | ApoA1/B | 0.459245211 | 0 |
| B74 | B92 | ApoA1 | ALDH2-Glu504Glu | 0.370511268 | 0 |
| B74 | B93 | ApoA1 | ALDH2-Glu504Lys | -0.37051127 | 0 |
| B74 | B94 | ApoA1 | ALDH2-Lys504Lys | -0.37051127 | 0 |
| B74 | B140 | ApoA1 | r-GT | -0.25445298 | 8.45E-13 |
| B74 | B131 | ApoA1 | AST | -0.24842291 | 2.98E-12 |
| B74 | B77 | ApoA1 | TC | 0.230158945 | 1.11E-10 |
| B74 | B133 | ApoA1 | TBIL | -0.21086364 | 3.71E-09 |
| B74 | B126 | ApoA1 | Alb | 0.191339507 | 9.30E-08 |
| B74 | B188 | ApoA1 | Ca | 0.187653052 | 1.65E-07 |
| B75 | B115 | ApoA1/B | TSH TT3 TT4(TT3) | 0.337589066 | 0 |
| B75 | B76 | ApoA1/B | ApoB | -0.67368507 | 0 |
| B75 | B79 | ApoA1/B | hb V -dna | -0.28232143 | 1.55E-15 |
| B75 | B114 | ApoA1/B | TSH TT3 TT4(TT4) | 0.269998612 | 2.80E-14 |
| B75 | B194 | ApoA1/B | Cl | 0.263410166 | 1.22E-13 |
| B75 | B112 | ApoA1/B | FT4 | 0.227636917 | 1.79E-10 |
| B75 | B125 | ApoA1/B | Cor | 0.222613812 | 4.56E-10 |
| B75 | B123 | ApoA1/B | hs-CRP1 | -0.22246438 | 4.69E-10 |
| B75 | B81 | ApoA1/B | TSGF | 0.221442027 | 5.65E-10 |
| B75 | B193 | ApoA1/B | K | -0.22016569 | 7.13E-10 |
| B75 | B113 | ApoA1/B | FT3 | 0.21791379 | 1.07E-09 |
| B75 | B138 | ApoA1/B | PAB | -0.21286129 | 2.62E-09 |
| B75 | B179 | ApoA1/B | α1-MG | -0.21175179 | 3.18E-09 |

| | | | | | |
|---|---|---|---|---|---|
| B75 | B211 | ApoA1/B | IOP-R | -0.20337682 | 1.33E-08 |
| B75 | B92 | ApoA1/B | ALDH2-Glu504Glu | 0.197189135 | 3.66E-08 |
| B75 | B93 | ApoA1/B | ALDH2-Glu504Lys | -0.19718914 | 3.66E-08 |
| B75 | B94 | ApoA1/B | ALDH2-Lys504Lys | -0.19718913 | 3.66E-08 |
| B75 | B199 | ApoA1/B | Mg | -0.18009334 | 5.15E-07 |
| B76 | B77 | ApoB | TC | 0.300966954 | 0 |
| B76 | B194 | ApoB | Cl | -0.25268159 | 1.23E-12 |
| B76 | B138 | ApoB | PAB | 0.242222012 | 1.05E-11 |
| B76 | B115 | ApoB | TSH TT3 TT4(TT3) | -0.21571632 | 1.58E-09 |
| B76 | B196 | ApoB | TCO2 | 0.214237139 | 2.06E-09 |
| B76 | B212 | ApoB | IOP-L | 0.195312401 | 4.96E-08 |
| B76 | B114 | ApoB | TSH TT3 TT4(TT4) | -0.19457567 | 5.57E-08 |
| B76 | B112 | ApoB | FT4 | -0.18448263 | 2.67E-07 |
| B77 | B189 | TC | eGFB | -0.32715349 | 0 |
| B77 | B91 | TC | TP | 0.266581427 | 6.04E-14 |
| B77 | B112 | TC | FT4 | -0.22598048 | 2.45E-10 |
| B77 | B219 | TC | OT | 0.201647151 | 1.77E-08 |
| B77 | B126 | TC | Alb | 0.199562231 | 2.49E-08 |
| B77 | B132 | TC | Dbil | -0.19398516 | 6.12E-08 |
| B77 | B190 | TC | Cre | 0.186497552 | 1.97E-07 |
| B77 | B113 | TC | FT3 | -0.18636171 | 2.01E-07 |
| B78 | B143 | HBcAb | β-CTx | 0.341589039 | 0 |
| B78 | B144 | HBcAb | Total-PINP | 0.353602701 | 0 |
| B78 | B145 | HBcAb | BGP | 0.382640924 | 0 |
| B78 | B83 | HBcAb | AFP | 0.226027971 | 2.42E-10 |
| B78 | B79 | HBcAb | hb V -dna | 0.217102793 | 1.24E-09 |
| B79 | B81 | hb V -dna | TSGF | -0.26701442 | 5.48E-14 |
| B79 | B125 | hb V -dna | Cor | -0.26055636 | 2.28E-13 |
| B79 | B156 | hb V -dna | IgE | -0.24314368 | 8.76E-12 |
| B8 | B78 | WB-Zinc | HBcAb | -0.37287466 | 0 |
| B8 | B156 | WB-Zinc | IgE | 0.269452115 | 3.18E-14 |
| B8 | B81 | WB-Zinc | TSGF | 0.2363008 | 3.41E-11 |
| B8 | B39 | WB-Zinc | Other cells | 0.195631161 | 4.71E-08 |
| B8 | B154 | WB-Zinc | RF | 0.177590414 | 7.43E-07 |
| B81 | B156 | TSGF | IgE | 0.314083323 | 0 |
| B81 | B125 | TSGF | Cor | 0.211265663 | 3.46E-09 |
| B82 | B84 | CEA | C-PSA | 0.417378722 | 0 |
| B82 | B85 | CEA | SCC | 0.285853664 | 6.66E-16 |
| B82 | B87 | CEA | NSE | 0.237482575 | 2.70E-11 |
| B82 | B89 | CEA | CA 72-4 | 0.225910439 | 2.48E-10 |
| B82 | B88 | CEA | CA 242 | 0.221611937 | 5.48E-10 |
| B82 | B86 | CEA | TBhCG | 0.213257356 | 2.44E-09 |
| B83 | B85 | AFP | SCC | 0.400347479 | 0 |
| B83 | B86 | AFP | TBhCG | 0.282371158 | 1.55E-15 |

| | | | | | |
|---|---|---|---|---|---|
| B83 | B89 | AFP | CA 72-4 | 0.28111785 | 2.22E-15 |
| B84 | B85 | C-PSA | SCC | 0.467840608 | 0 |
| B84 | B86 | C-PSA | TBhCG | 0.67051875 | 0 |
| B84 | B87 | C-PSA | NSE | 0.444282084 | 0 |
| B84 | B89 | C-PSA | CA 72-4 | 0.288266636 | 4.44E-16 |
| B85 | B86 | SCC | TBhCG | 0.42387002 | 0 |
| B85 | B87 | SCC | NSE | 0.299251518 | 0 |
| B85 | B89 | SCC | CA 72-4 | 0.359136884 | 0 |
| B85 | B183 | SCC | PTA | -0.18768255 | 1.64E-07 |
| B86 | B87 | TBhCG | NSE | 0.481942871 | 0 |
| B86 | B89 | TBhCG | CA 72-4 | 0.430080469 | 0 |
| B86 | B196 | TBhCG | TCO2 | 0.176797185 | 8.34E-07 |
| B90 | B120 | CYFRA 21-1 | HCV-cAg | -0.23305973 | 6.40E-11 |
| B91 | B126 | TP | Alb | 0.523184698 | 0 |
| B91 | B127 | TP | A-G ratio | -0.49132356 | 0 |
| B91 | B130 | TP | GLB | 0.784033798 | 0 |
| B91 | B188 | TP | Ca | 0.239317977 | 1.88E-11 |
| B91 | B209 | TP | Smoking | -0.17672909 | 8.42E-07 |
| B92 | B93 | ALDH2-Glu504Glu | ALDH2-Glu504Lys | -1 | 0 |
| B92 | B94 | ALDH2-Glu504Glu | ALDH2-Lys504Lys | -0.99999999 | 0 |
| B92 | B156 | ALDH2-Glu504Glu | IgE | -0.19910174 | 2.68E-08 |
| B93 | B94 | ALDH2-Glu504Lys | ALDH2-Lys504Lys | 0.999999994 | 0 |
| B93 | B156 | ALDH2-Glu504Lys | IgE | 0.199101741 | 2.68E-08 |
| B94 | B156 | ALDH2-Lys504Lys | IgE | 0.19910174 | 2.68E-08 |
| B95 | B100 | anti-cenpb | anti-Ro52 | 0.999999993 | 0 |
| B95 | B103 | anti-cenpb | anti-SSA/Ro | -0.66109405 | 0 |
| B95 | B104 | anti-cenpb | anti-SSB | 0.999999993 | 0 |
| B95 | B105 | anti-cenpb | anti-U1RNP | 0.999999995 | 0 |
| B95 | B107 | anti-cenpb | AnuA | 0.999999767 | 0 |
| B95 | B108 | anti-cenpb | Anti-Cyclin | 0.999999997 | 0 |
| B95 | B109 | anti-cenpb | AMA-M2 | 0.999999993 | 0 |
| B95 | B110 | anti-cenpb | AHA | 0.999999995 | 0 |
| B95 | B171 | anti-cenpb | EC | -0.30105168 | 0 |
| B95 | B173 | anti-cenpb | NTT | -0.29660015 | 0 |
| B95 | B97 | anti-cenpb | dsDNA | 0.999999997 | 0 |
| B95 | B179 | anti-cenpb | α1-MG | 0.285253141 | 8.88E-16 |
| B95 | B158 | anti-cenpb | IgM | 0.281658569 | 1.78E-15 |
| B95 | B139 | anti-cenpb | ADA | -0.22290661 | 4.32E-10 |
| B95 | B199 | anti-cenpb | Mg | 0.207016523 | 7.18E-09 |
| B95 | B135 | anti-cenpb | TBA | 0.178387518 | 6.62E-07 |
| B95 | B193 | anti-cenpb | K | -0.17780783 | 7.20E-07 |
| B95 | B157 | anti-cenpb | IgG | -0.17568129 | 9.80E-07 |
| B97 | B100 | dsDNA | anti-Ro52 | 0.999999993 | 0 |
| B97 | B102 | dsDNA | anti-Sm | 0.999999996 | 0 |

| | | | | | |
|---|---|---|---|---|---|
| B97 | B103 | dsDNA | anti-SSA/Ro | -0.66109405 | 0 |
| B97 | B104 | dsDNA | anti-SSB | 0.999999997 | 0 |
| B97 | B105 | dsDNA | anti-U1RNP | 0.999999992 | 0 |
| B97 | B106 | dsDNA | anti-P | 0.999999995 | 0 |
| B97 | B107 | dsDNA | AnuA | 0.999999837 | 0 |
| B97 | B108 | dsDNA | Anti-Cyclin | 0.999999997 | 0 |
| B97 | B109 | dsDNA | AMA-M2 | 0.99999999 | 0 |
| B97 | B110 | dsDNA | AHA | 0.999999995 | 0 |
| B97 | B171 | dsDNA | EC | -0.30105167 | 0 |
| B97 | B173 | dsDNA | NTT | -0.29660014 | 0 |
| B97 | B99 | dsDNA | anti-PM-Scl | 0.999999997 | 0 |
| B97 | B179 | dsDNA | α1-MG | 0.285253141 | 8.88E-16 |
| B97 | B158 | dsDNA | IgM | 0.281658569 | 1.78E-15 |
| B97 | B139 | dsDNA | ADA | -0.22290661 | 4.32E-10 |
| B97 | B199 | dsDNA | Mg | 0.207016523 | 7.18E-09 |
| B97 | B135 | dsDNA | TBA | 0.178387517 | 6.62E-07 |
| B97 | B193 | dsDNA | K | -0.17780783 | 7.20E-07 |
| B97 | B157 | dsDNA | IgG | -0.17568129 | 9.80E-07 |
| B98 | B101 | anti-Jo-1 | anti-Sc70 | 0.999999965 | 0 |
| B98 | B102 | anti-Jo-1 | anti-Sm | 0.999999992 | 0 |
| B98 | B103 | anti-Jo-1 | anti-SSA/Ro | -0.66109407 | 0 |
| B98 | B107 | anti-Jo-1 | AnuA | 0.999999555 | 0 |
| B98 | B108 | anti-Jo-1 | Anti-Cyclin | 0.999999998 | 0 |
| B98 | B171 | anti-Jo-1 | EC | -0.30105168 | 0 |
| B98 | B173 | anti-Jo-1 | NTT | -0.29660015 | 0 |
| B98 | B99 | anti-Jo-1 | anti-PM-Scl | 0.999999999 | 0 |
| B98 | B179 | anti-Jo-1 | α1-MG | 0.285253149 | 8.88E-16 |
| B98 | B158 | anti-Jo-1 | IgM | 0.281658576 | 1.78E-15 |
| B98 | B139 | anti-Jo-1 | ADA | -0.22290662 | 4.32E-10 |
| B98 | B199 | anti-Jo-1 | Mg | 0.207016526 | 7.18E-09 |
| B98 | B135 | anti-Jo-1 | TBA | 0.178387522 | 6.62E-07 |
| B98 | B193 | anti-Jo-1 | K | -0.17780784 | 7.20E-07 |
| B98 | B157 | anti-Jo-1 | IgG | -0.17568129 | 9.80E-07 |
| B99 | B101 | anti-PM-Scl | anti-Sc70 | 0.999999987 | 0 |
| B99 | B103 | anti-PM-Scl | anti-SSA/Ro | -0.66109405 | 0 |
| B99 | B104 | anti-PM-Scl | anti-SSB | 0.999999997 | 0 |
| B99 | B106 | anti-PM-Scl | anti-P | 0.999999988 | 0 |
| B99 | B107 | anti-PM-Scl | AnuA | 0.999999707 | 0 |
| B99 | B109 | anti-PM-Scl | AMA-M2 | 0.999999999 | 0 |
| B99 | B171 | anti-PM-Scl | EC | -0.30105168 | 0 |
| B99 | B173 | anti-PM-Scl | NTT | -0.29660015 | 0 |
| B99 | B179 | anti-PM-Scl | α1-MG | 0.285253144 | 8.88E-16 |
| B99 | B158 | anti-PM-Scl | IgM | 0.281658572 | 1.78E-15 |
| B99 | B139 | anti-PM-Scl | ADA | -0.22290662 | 4.32E-10 |

| | | | | | |
|---|---|---|---|---|---|
| B99 | B199 | anti-PM-Scl | Mg | 0.207016524 | 7.18E-09 |
| B99 | B135 | anti-PM-Scl | TBA | 0.17838752 | 6.62E-07 |
| B99 | B193 | anti-PM-Scl | K | -0.17780784 | 7.20E-07 |
| B99 | B157 | anti-PM-Scl | IgG | -0.17568129 | 9.80E-07 |

**Supplementary Table 13. PEI Correlations in Cervical spondylopathy**

| Row | Column | Row | Column | Cor | *P* |
|---|---|---|---|---|---|
| B1 | B190 | Gender | Cre | -0.66340451 | 0 |
| B1 | B192 | Gender | UA | -0.60573664 | 0 |
| B1 | B216 | Gender | BMI | -0.52700024 | 0 |
| B1 | B221 | Gender | Hei | -0.5609765 | 0 |
| B1 | B222 | Gender | NS | -0.6893586 | 0 |
| B1 | B46 | Gender | RBC Count | -0.45117102 | 0 |
| B1 | B51 | Gender | Hb | -0.58557155 | 0 |
| B1 | B206 | Gender | Waist | -0.43154684 | 8.88E-16 |
| B1 | B163 | Gender | SG | -0.40707091 | 4.02E-14 |
| B1 | B209 | Gender | Smoking | -0.38209899 | 1.71E-12 |
| B1 | B174 | Gender | Urea/Cr | 0.358302493 | 4.57E-11 |
| B1 | B72 | Gender | HDL-C | 0.335272777 | 8.59E-10 |
| B1 | B204 | Gender | Drinking | -0.3245909 | 3.09E-09 |
| B1 | B128 | Gender | ALT | -0.30172694 | 4.07E-08 |
| B1 | B215 | Gender | BMI | -0.28635324 | 2.04E-07 |
| B1 | B45 | Gender | MCHC | -0.28570787 | 2.18E-07 |
| B1 | B218 | Gender | DBP | -0.28486881 | 2.37E-07 |
| B1 | B210 | Gender | Hip | -0.27962808 | 4.01E-07 |
| B1 | B207 | Gender | WHR | -0.27884504 | 4.34E-07 |
| B10 | B12 | PG I | PG II | 0.619206966 | 0 |
| B10 | B14 | PG I | G-17 | 0.463184577 | 0 |
| B10 | B55 | PG I | TK | 0.289058078 | 1.55E-07 |
| B10 | B61 | PG I | WBHSRV | 0.275319006 | 6.13E-07 |
| B100 | B103 | anti-Ro52 | anti-SSA/Ro | 1 | 0 |
| B100 | B105 | anti-Ro52 | anti-U1RNP | 1 | 0 |
| B100 | B106 | anti-Ro52 | anti-P | 0.999999999 | 0 |
| B100 | B107 | anti-Ro52 | AnuA | 0.999999986 | 0 |
| B100 | B151 | anti-Ro52 | Anti-CCP | 0.665704468 | 0 |
| B100 | B156 | anti-Ro52 | IgE | 0.614367186 | 0 |
| B100 | B178 | anti-Ro52 | Ig-U | 0.524633851 | 0 |
| B100 | B180 | anti-Ro52 | Tf | 0.531153079 | 0 |
| B100 | B184 | anti-Ro52 | INR | 0.452826162 | 0 |
| B100 | B193 | anti-Ro52 | K | 0.459514228 | 0 |
| B100 | B154 | anti-Ro52 | RF | 0.419728548 | 5.33E-15 |
| B100 | B199 | anti-Ro52 | Mg | 0.402787859 | 7.82E-14 |
| B100 | B179 | anti-Ro52 | α1-MG | 0.397881945 | 1.66E-13 |
| B100 | B183 | anti-Ro52 | PTA | 0.375742668 | 4.22E-12 |
| B100 | B157 | anti-Ro52 | IgG | 0.340998994 | 4.24E-10 |
| B100 | B158 | anti-Ro52 | IgM | -0.28134467 | 3.38E-07 |
| B100 | B119 | anti-Ro52 | Hcv-rna | 0.281326061 | 3.39E-07 |
| B100 | B121 | anti-Ro52 | C3 | -0.28132607 | 3.39E-07 |
| B100 | B122 | anti-Ro52 | C4 | -0.28132616 | 3.39E-07 |

| | | | | | |
|---|---|---|---|---|---|
| B100 | B152 | anti-Ro52 | ACA-Ig(G-A-M) | -0.2813268 | 3.39E-07 |
| B100 | B159 | anti-Ro52 | β2-MG | -0.28132641 | 3.39E-07 |
| B100 | B169 | anti-Ro52 | U-CRY | -0.28132577 | 3.39E-07 |
| B101 | B102 | anti-Sc70 | anti-Sm | 0.999999998 | 0 |
| B101 | B106 | anti-Sc70 | anti-P | 0.999999999 | 0 |
| B101 | B107 | anti-Sc70 | AnuA | 0.999999968 | 0 |
| B101 | B108 | anti-Sc70 | Anti-Cyclin | 1 | 0 |
| B101 | B151 | anti-Sc70 | Anti-CCP | 0.665704468 | 0 |
| B101 | B156 | anti-Sc70 | IgE | 0.614367187 | 0 |
| B101 | B178 | anti-Sc70 | Ig-U | 0.524633852 | 0 |
| B101 | B180 | anti-Sc70 | Tf | 0.53115308 | 0 |
| B101 | B184 | anti-Sc70 | INR | 0.452826163 | 0 |
| B101 | B193 | anti-Sc70 | K | 0.459514229 | 0 |
| B101 | B154 | anti-Sc70 | RF | 0.419728548 | 5.33E-15 |
| B101 | B199 | anti-Sc70 | Mg | 0.40278786 | 7.82E-14 |
| B101 | B179 | anti-Sc70 | α1-MG | 0.397881946 | 1.66E-13 |
| B101 | B183 | anti-Sc70 | PTA | 0.375742668 | 4.22E-12 |
| B101 | B157 | anti-Sc70 | IgG | 0.340998994 | 4.24E-10 |
| B101 | B158 | anti-Sc70 | IgM | -0.28134467 | 3.38E-07 |
| B101 | B119 | anti-Sc70 | Hcv-rna | 0.281326061 | 3.39E-07 |
| B101 | B121 | anti-Sc70 | C3 | -0.28132608 | 3.39E-07 |
| B101 | B122 | anti-Sc70 | C4 | -0.28132624 | 3.39E-07 |
| B101 | B152 | anti-Sc70 | ACA-Ig(G-A-M) | -0.28132699 | 3.39E-07 |
| B101 | B159 | anti-Sc70 | β2-MG | -0.28132642 | 3.39E-07 |
| B101 | B169 | anti-Sc70 | U-CRY | -0.28132573 | 3.39E-07 |
| B102 | B104 | anti-Sm | anti-SSB | 0.999999999 | 0 |
| B102 | B106 | anti-Sm | anti-P | 1 | 0 |
| B102 | B107 | anti-Sm | AnuA | 0.999999998 | 0 |
| B102 | B108 | anti-Sm | Anti-Cyclin | 0.999999999 | 0 |
| B102 | B151 | anti-Sm | Anti-CCP | 0.665704468 | 0 |
| B102 | B156 | anti-Sm | IgE | 0.614367187 | 0 |
| B102 | B178 | anti-Sm | Ig-U | 0.524633852 | 0 |
| B102 | B180 | anti-Sm | Tf | 0.531153079 | 0 |
| B102 | B184 | anti-Sm | INR | 0.452826163 | 0 |
| B102 | B193 | anti-Sm | K | 0.459514229 | 0 |
| B102 | B154 | anti-Sm | RF | 0.419728548 | 5.33E-15 |
| B102 | B199 | anti-Sm | Mg | 0.40278786 | 7.82E-14 |
| B102 | B179 | anti-Sm | α1-MG | 0.397881945 | 1.66E-13 |
| B102 | B183 | anti-Sm | PTA | 0.375742668 | 4.22E-12 |
| B102 | B157 | anti-Sm | IgG | 0.340998994 | 4.24E-10 |
| B102 | B158 | anti-Sm | IgM | -0.28134467 | 3.38E-07 |
| B102 | B119 | anti-Sm | Hcv-rna | 0.281326061 | 3.39E-07 |
| B102 | B121 | anti-Sm | C3 | -0.28132608 | 3.39E-07 |
| B102 | B122 | anti-Sm | C4 | -0.28132619 | 3.39E-07 |

| | | | | | |
|---|---|---|---|---|---|
| B102 | B152 | anti-Sm | ACA-Ig(G-A-M) | -0.28132698 | 3.39E-07 |
| B102 | B159 | anti-Sm | β2-MG | -0.28132649 | 3.39E-07 |
| B102 | B169 | anti-Sm | U-CRY | -0.28132578 | 3.39E-07 |
| B103 | B105 | anti-SSA/Ro | anti-U1RNP | 1 | 0 |
| B103 | B106 | anti-SSA/Ro | anti-P | 1 | 0 |
| B103 | B107 | anti-SSA/Ro | AnuA | 0.999999991 | 0 |
| B103 | B151 | anti-SSA/Ro | Anti-CCP | 0.665704467 | 0 |
| B103 | B156 | anti-SSA/Ro | IgE | 0.614367186 | 0 |
| B103 | B178 | anti-SSA/Ro | Ig-U | 0.524633851 | 0 |
| B103 | B180 | anti-SSA/Ro | Tf | 0.531153079 | 0 |
| B103 | B184 | anti-SSA/Ro | INR | 0.452826162 | 0 |
| B103 | B193 | anti-SSA/Ro | K | 0.459514228 | 0 |
| B103 | B154 | anti-SSA/Ro | RF | 0.419728548 | 5.33E-15 |
| B103 | B199 | anti-SSA/Ro | Mg | 0.402787859 | 7.82E-14 |
| B103 | B179 | anti-SSA/Ro | α1-MG | 0.397881945 | 1.66E-13 |
| B103 | B183 | anti-SSA/Ro | PTA | 0.375742668 | 4.22E-12 |
| B103 | B157 | anti-SSA/Ro | IgG | 0.340998994 | 4.24E-10 |
| B103 | B158 | anti-SSA/Ro | IgM | -0.28134467 | 3.38E-07 |
| B103 | B119 | anti-SSA/Ro | Hcv-rna | 0.281326061 | 3.39E-07 |
| B103 | B121 | anti-SSA/Ro | C3 | -0.28132607 | 3.39E-07 |
| B103 | B122 | anti-SSA/Ro | C4 | -0.28132616 | 3.39E-07 |
| B103 | B152 | anti-SSA/Ro | ACA-Ig(G-A-M) | -0.28132677 | 3.39E-07 |
| B103 | B159 | anti-SSA/Ro | β2-MG | -0.28132639 | 3.39E-07 |
| B103 | B169 | anti-SSA/Ro | U-CRY | -0.28132575 | 3.39E-07 |
| B104 | B107 | anti-SSB | AnuA | 0.999999989 | 0 |
| B104 | B151 | anti-SSB | Anti-CCP | 0.665704468 | 0 |
| B104 | B156 | anti-SSB | IgE | 0.614367186 | 0 |
| B104 | B178 | anti-SSB | Ig-U | 0.524633852 | 0 |
| B104 | B180 | anti-SSB | Tf | 0.531153079 | 0 |
| B104 | B184 | anti-SSB | INR | 0.452826162 | 0 |
| B104 | B193 | anti-SSB | K | 0.459514229 | 0 |
| B104 | B154 | anti-SSB | RF | 0.419728548 | 5.33E-15 |
| B104 | B199 | anti-SSB | Mg | 0.40278786 | 7.82E-14 |
| B104 | B179 | anti-SSB | α1-MG | 0.397881945 | 1.66E-13 |
| B104 | B183 | anti-SSB | PTA | 0.375742668 | 4.22E-12 |
| B104 | B157 | anti-SSB | IgG | 0.340998994 | 4.24E-10 |
| B104 | B158 | anti-SSB | IgM | -0.28134467 | 3.38E-07 |
| B104 | B119 | anti-SSB | Hcv-rna | 0.281326061 | 3.39E-07 |
| B104 | B121 | anti-SSB | C3 | -0.28132606 | 3.39E-07 |
| B104 | B122 | anti-SSB | C4 | -0.28132617 | 3.39E-07 |
| B104 | B152 | anti-SSB | ACA-Ig(G-A-M) | -0.28132684 | 3.39E-07 |
| B104 | B159 | anti-SSB | β2-MG | -0.28132639 | 3.39E-07 |
| B104 | B169 | anti-SSB | U-CRY | -0.28132574 | 3.39E-07 |
| B105 | B106 | anti-U1RNP | anti-P | 0.999999999 | 0 |

| | | | | | |
|---|---|---|---|---|---|
| B105 | B107 | anti-U1RNP | AnuA | 0.99999999 | 0 |
| B105 | B109 | anti-U1RNP | AMA-M2 | 1 | 0 |
| B105 | B110 | anti-U1RNP | AHA | 1 | 0 |
| B105 | B151 | anti-U1RNP | Anti-CCP | 0.665704467 | 0 |
| B105 | B156 | anti-U1RNP | IgE | 0.614367186 | 0 |
| B105 | B178 | anti-U1RNP | Ig-U | 0.524633851 | 0 |
| B105 | B180 | anti-U1RNP | Tf | 0.531153079 | 0 |
| B105 | B184 | anti-U1RNP | INR | 0.452826163 | 0 |
| B105 | B193 | anti-U1RNP | K | 0.459514228 | 0 |
| B105 | B154 | anti-U1RNP | RF | 0.419728548 | 5.33E-15 |
| B105 | B199 | anti-U1RNP | Mg | 0.402787859 | 7.82E-14 |
| B105 | B179 | anti-U1RNP | α1-MG | 0.397881945 | 1.66E-13 |
| B105 | B183 | anti-U1RNP | PTA | 0.375742668 | 4.22E-12 |
| B105 | B157 | anti-U1RNP | IgG | 0.340998994 | 4.24E-10 |
| B105 | B158 | anti-U1RNP | IgM | -0.28134467 | 3.38E-07 |
| B105 | B119 | anti-U1RNP | Hcv-rna | 0.281326061 | 3.39E-07 |
| B105 | B121 | anti-U1RNP | C3 | -0.28132607 | 3.39E-07 |
| B105 | B122 | anti-U1RNP | C4 | -0.28132617 | 3.39E-07 |
| B105 | B152 | anti-U1RNP | ACA-Ig(G-A-M) | -0.28132675 | 3.39E-07 |
| B105 | B159 | anti-U1RNP | β2-MG | -0.2813264 | 3.39E-07 |
| B105 | B169 | anti-U1RNP | U-CRY | -0.28132577 | 3.39E-07 |
| B106 | B107 | anti-P | AnuA | 0.99999998 | 0 |
| B106 | B108 | anti-P | Anti-Cyclin | 1 | 0 |
| B106 | B110 | anti-P | AHA | 0.999999999 | 0 |
| B106 | B151 | anti-P | Anti-CCP | 0.665704467 | 0 |
| B106 | B156 | anti-P | IgE | 0.614367186 | 0 |
| B106 | B178 | anti-P | Ig-U | 0.524633851 | 0 |
| B106 | B180 | anti-P | Tf | 0.531153079 | 0 |
| B106 | B184 | anti-P | INR | 0.452826162 | 0 |
| B106 | B193 | anti-P | K | 0.459514229 | 0 |
| B106 | B154 | anti-P | RF | 0.419728548 | 5.33E-15 |
| B106 | B199 | anti-P | Mg | 0.402787859 | 7.82E-14 |
| B106 | B179 | anti-P | α1-MG | 0.397881945 | 1.66E-13 |
| B106 | B183 | anti-P | PTA | 0.375742668 | 4.22E-12 |
| B106 | B157 | anti-P | IgG | 0.340998994 | 4.24E-10 |
| B106 | B158 | anti-P | IgM | -0.28134467 | 3.38E-07 |
| B106 | B119 | anti-P | Hcv-rna | 0.281326061 | 3.39E-07 |
| B106 | B121 | anti-P | C3 | -0.28132609 | 3.39E-07 |
| B106 | B122 | anti-P | C4 | -0.28132622 | 3.39E-07 |
| B106 | B152 | anti-P | ACA-Ig(G-A-M) | -0.28132684 | 3.39E-07 |
| B106 | B159 | anti-P | β2-MG | -0.2813264 | 3.39E-07 |
| B106 | B169 | anti-P | U-CRY | -0.28132571 | 3.39E-07 |
| B107 | B108 | AnuA | Anti-Cyclin | 0.999999987 | 0 |
| B107 | B109 | AnuA | AMA-M2 | 0.999999986 | 0 |

| | | | | | |
|---|---|---|---|---|---|
| B107 | B110 | AnuA | AHA | 0.999999991 | 0 |
| B107 | B151 | AnuA | Anti-CCP | 0.66570446 | 0 |
| B107 | B156 | AnuA | IgE | 0.614367179 | 0 |
| B107 | B178 | AnuA | Ig-U | 0.524633845 | 0 |
| B107 | B180 | AnuA | Tf | 0.531153072 | 0 |
| B107 | B184 | AnuA | INR | 0.452826157 | 0 |
| B107 | B193 | AnuA | K | 0.459514223 | 0 |
| B107 | B154 | AnuA | RF | 0.419728543 | 5.33E-15 |
| B107 | B199 | AnuA | Mg | 0.402787855 | 7.82E-14 |
| B107 | B179 | AnuA | α1-MG | 0.39788194 | 1.66E-13 |
| B107 | B183 | AnuA | PTA | 0.375742664 | 4.22E-12 |
| B107 | B157 | AnuA | IgG | 0.34099899 | 4.24E-10 |
| B107 | B158 | AnuA | IgM | -0.28134466 | 3.38E-07 |
| B107 | B119 | AnuA | Hcv-rna | 0.281326058 | 3.39E-07 |
| B107 | B121 | AnuA | C3 | -0.28132601 | 3.39E-07 |
| B107 | B122 | AnuA | C4 | -0.28132645 | 3.39E-07 |
| B107 | B152 | AnuA | ACA-Ig(G-A-M) | -0.28132692 | 3.39E-07 |
| B107 | B159 | AnuA | β2-MG | -0.28132645 | 3.39E-07 |
| B107 | B169 | AnuA | U-CRY | -0.28132579 | 3.39E-07 |
| B108 | B110 | Anti-Cyclin | AHA | 1 | 0 |
| B108 | B151 | Anti-Cyclin | Anti-CCP | 0.665704468 | 0 |
| B108 | B156 | Anti-Cyclin | IgE | 0.614367186 | 0 |
| B108 | B178 | Anti-Cyclin | Ig-U | 0.524633851 | 0 |
| B108 | B180 | Anti-Cyclin | Tf | 0.531153079 | 0 |
| B108 | B184 | Anti-Cyclin | INR | 0.452826163 | 0 |
| B108 | B193 | Anti-Cyclin | K | 0.459514229 | 0 |
| B108 | B154 | Anti-Cyclin | RF | 0.419728548 | 5.33E-15 |
| B108 | B199 | Anti-Cyclin | Mg | 0.402787859 | 7.82E-14 |
| B108 | B179 | Anti-Cyclin | α1-MG | 0.397881945 | 1.66E-13 |
| B108 | B183 | Anti-Cyclin | PTA | 0.375742668 | 4.22E-12 |
| B108 | B157 | Anti-Cyclin | IgG | 0.340998994 | 4.24E-10 |
| B108 | B158 | Anti-Cyclin | IgM | -0.28134467 | 3.38E-07 |
| B108 | B119 | Anti-Cyclin | Hcv-rna | 0.281326061 | 3.39E-07 |
| B108 | B121 | Anti-Cyclin | C3 | -0.28132607 | 3.39E-07 |
| B108 | B122 | Anti-Cyclin | C4 | -0.28132616 | 3.39E-07 |
| B108 | B152 | Anti-Cyclin | ACA-Ig(G-A-M) | -0.28132677 | 3.39E-07 |
| B108 | B159 | Anti-Cyclin | β2-MG | -0.28132642 | 3.39E-07 |
| B108 | B169 | Anti-Cyclin | U-CRY | -0.28132576 | 3.39E-07 |
| B109 | B151 | AMA-M2 | Anti-CCP | 0.665704468 | 0 |
| B109 | B156 | AMA-M2 | IgE | 0.614367186 | 0 |
| B109 | B178 | AMA-M2 | Ig-U | 0.524633851 | 0 |
| B109 | B180 | AMA-M2 | Tf | 0.531153079 | 0 |
| B109 | B184 | AMA-M2 | INR | 0.452826162 | 0 |
| B109 | B193 | AMA-M2 | K | 0.459514229 | 0 |

| | | | | | |
|---|---|---|---|---|---|
| B109 | B154 | AMA-M2 | RF | 0.419728548 | 5.33E-15 |
| B109 | B199 | AMA-M2 | Mg | 0.402787859 | 7.82E-14 |
| B109 | B179 | AMA-M2 | α1-MG | 0.397881945 | 1.66E-13 |
| B109 | B183 | AMA-M2 | PTA | 0.375742668 | 4.22E-12 |
| B109 | B157 | AMA-M2 | IgG | 0.340998994 | 4.24E-10 |
| B109 | B158 | AMA-M2 | IgM | -0.28134467 | 3.38E-07 |
| B109 | B119 | AMA-M2 | Hcv-rna | 0.281326061 | 3.39E-07 |
| B109 | B121 | AMA-M2 | C3 | -0.28132607 | 3.39E-07 |
| B109 | B122 | AMA-M2 | C4 | -0.28132617 | 3.39E-07 |
| B109 | B152 | AMA-M2 | ACA-Ig(G-A-M) | -0.28132684 | 3.39E-07 |
| B109 | B159 | AMA-M2 | β2-MG | -0.28132642 | 3.39E-07 |
| B109 | B169 | AMA-M2 | U-CRY | -0.28132578 | 3.39E-07 |
| B11 | B12 | PGI/PGII | PG II | -0.49176828 | 0 |
| B110 | B151 | AHA | Anti-CCP | 0.665704467 | 0 |
| B110 | B156 | AHA | IgE | 0.614367186 | 0 |
| B110 | B178 | AHA | Ig-U | 0.524633851 | 0 |
| B110 | B180 | AHA | Tf | 0.531153079 | 0 |
| B110 | B184 | AHA | INR | 0.452826162 | 0 |
| B110 | B193 | AHA | K | 0.459514228 | 0 |
| B110 | B154 | AHA | RF | 0.419728548 | 5.33E-15 |
| B110 | B199 | AHA | Mg | 0.402787859 | 7.82E-14 |
| B110 | B179 | AHA | α1-MG | 0.397881945 | 1.66E-13 |
| B110 | B183 | AHA | PTA | 0.375742668 | 4.22E-12 |
| B110 | B157 | AHA | IgG | 0.340998994 | 4.24E-10 |
| B110 | B158 | AHA | IgM | -0.28134467 | 3.38E-07 |
| B110 | B119 | AHA | Hcv-rna | 0.281326061 | 3.39E-07 |
| B110 | B121 | AHA | C3 | -0.28132607 | 3.39E-07 |
| B110 | B122 | AHA | C4 | -0.28132617 | 3.39E-07 |
| B110 | B152 | AHA | ACA-Ig(G-A-M) | -0.28132685 | 3.39E-07 |
| B110 | B159 | AHA | β2-MG | -0.28132641 | 3.39E-07 |
| B110 | B169 | AHA | U-CRY | -0.28132575 | 3.39E-07 |
| B113 | B115 | FT3 | TSH TT3 TT4(TT3) | 0.449146784 | 0 |
| B114 | B115 | TSH TT3 TT4(TT4) | TSH TT3 TT4(TT3) | 0.578239605 | 0 |
| B114 | B195 | TSH TT3 TT4(TT4) | Na | 0.545521849 | 0 |
| B114 | B135 | TSH TT3 TT4(TT4) | TBA | 0.373579392 | 5.72E-12 |
| B114 | B197 | TSH TT3 TT4(TT4) | AG | 0.370917378 | 8.28E-12 |
| B114 | B200 | TSH TT3 TT4(TT4) | OP | 0.354853258 | 7.21E-11 |
| B114 | B187 | TSH TT3 TT4(TT4) | NSE | 0.310615769 | 1.53E-08 |
| B115 | B195 | TSH TT3 TT4(TT3) | Na | 0.404865188 | 5.66E-14 |
| B115 | B187 | TSH TT3 TT4(TT3) | NSE | 0.300315282 | 4.74E-08 |
| B115 | B119 | TSH TT3 TT4(TT3) | Hcv-rna | 0.274628703 | 6.56E-07 |
| B115 | B121 | TSH TT3 TT4(TT3) | C3 | -0.27462871 | 6.56E-07 |
| B115 | B122 | TSH TT3 TT4(TT3) | C4 | -0.2746288 | 6.56E-07 |
| B115 | B152 | TSH TT3 TT4(TT3) | ACA-Ig(G-A-M) | -0.27462943 | 6.56E-07 |

| | | | | | |
|---|---|---|---|---|---|
| B115 | B159 | TSH TT3 TT4(TT3) | β2-MG | -0.27462903 | 6.56E-07 |
| B115 | B169 | TSH TT3 TT4(TT3) | U-CRY | -0.27462841 | 6.56E-07 |
| B117 | B167 | TT4 | PC | 0.955636965 | 0 |
| B119 | B148 | Hcv-rna | Anti_TPO | -0.45472746 | 0 |
| B119 | B169 | Hcv-rna | U-CRY | -0.99999893 | 0 |
| B119 | B120 | Hcv-rna | HCV-cAg | 0.357558491 | 5.05E-11 |
| B12 | B14 | PG II | G-17 | 0.411104796 | 2.13E-14 |
| B120 | B121 | HCV-cAg | C3 | -0.35755848 | 5.05E-11 |
| B120 | B122 | HCV-cAg | C4 | -0.35755871 | 5.05E-11 |
| B120 | B152 | HCV-cAg | ACA-Ig(G-A-M) | -0.35755786 | 5.05E-11 |
| B120 | B159 | HCV-cAg | β2-MG | -0.3575586 | 5.05E-11 |
| B120 | B169 | HCV-cAg | U-CRY | -0.35755781 | 5.05E-11 |
| B120 | B135 | HCV-cAg | TBA | -0.31644216 | 7.94E-09 |
| B121 | B148 | C3 | Anti_TPO | 0.454727474 | 0 |
| B121 | B169 | C3 | U-CRY | 0.999998943 | 0 |
| B122 | B148 | C4 | Anti_TPO | 0.454727616 | 0 |
| B122 | B169 | C4 | U-CRY | 0.999999453 | 0 |
| B123 | B186 | hs-CRP1 | PFC | 0.448626831 | 0 |
| B123 | B177 | hs-CRP1 | ACR | 0.405253439 | 5.33E-14 |
| B123 | B194 | hs-CRP1 | Cl | -0.36515016 | 1.83E-11 |
| B123 | B181 | hs-CRP1 | APTT | -0.34261594 | 3.46E-10 |
| B123 | B199 | hs-CRP1 | Mg | -0.31923125 | 5.77E-09 |
| B124 | B181 | ACTH | APTT | -0.27280423 | 7.83E-07 |
| B125 | B187 | Cor | NSE | 0.698307366 | 0 |
| B125 | B198 | Cor | P | 0.474811123 | 0 |
| B125 | B199 | Cor | Mg | -0.34906423 | 1.53E-10 |
| B125 | B193 | Cor | K | -0.29978458 | 5.02E-08 |
| B125 | B205 | Cor | KDA | 0.295118666 | 8.24E-08 |
| B126 | B127 | Alb | A-G ratio | 0.442527588 | 0 |
| B126 | B188 | Alb | Ca | 0.397133176 | 1.86E-13 |
| B127 | B130 | A-G ratio | GLB | -0.89766643 | 0 |
| B128 | B131 | ALT | AST | 0.664598327 | 0 |
| B128 | B140 | ALT | r-GT | 0.462344814 | 0 |
| B128 | B141 | ALT | AST/ALT | -0.56501874 | 0 |
| B128 | B192 | ALT | UA | 0.384959383 | 1.13E-12 |
| B128 | B222 | ALT | NS | 0.35357227 | 8.52E-11 |
| B128 | B216 | ALT | BMI | 0.341640439 | 3.91E-10 |
| B128 | B163 | ALT | SG | 0.275409353 | 6.08E-07 |
| B129 | B132 | IBIL | Dbil | 0.651186167 | 0 |
| B129 | B133 | IBIL | TBIL | 0.978419962 | 0 |
| B131 | B140 | AST | r-GT | 0.447765374 | 0 |
| B131 | B192 | AST | UA | 0.3086149 | 1.92E-08 |
| B132 | B133 | Dbil | TBIL | 0.79386761 | 0 |
| B132 | B135 | Dbil | TBA | 0.282973 | 2.87E-07 |

| | | | | | |
|---|---|---|---|---|---|
| B134 | B135 | CHE | TBA | -0.41380577 | 1.38E-14 |
| B134 | B137 | CHE | AFU | 0.383388064 | 1.42E-12 |
| B135 | B197 | TBA | AG | 0.559320366 | 0 |
| B135 | B200 | TBA | OP | 0.515839739 | 0 |
| B135 | B198 | TBA | P | -0.43932369 | 2.22E-16 |
| B135 | B193 | TBA | K | -0.38830528 | 6.92E-13 |
| B135 | B195 | TBA | Na | 0.355944846 | 6.24E-11 |
| B135 | B196 | TBA | TCO2 | -0.32056282 | 4.94E-09 |
| B135 | B146 | TBA | CysC | 0.314629861 | 9.76E-09 |
| B135 | B194 | TBA | Cl | -0.31140224 | 1.40E-08 |
| B136 | B150 | 5'NT | ALP | 0.601534733 | 0 |
| B136 | B153 | 5'NT | Glu | 0.47586359 | 0 |
| B136 | B140 | 5'NT | r-GT | 0.339940577 | 4.83E-10 |
| B137 | B138 | AFU | PAB | 0.365113135 | 1.83E-11 |
| B137 | B197 | AFU | AG | 0.327625183 | 2.16E-09 |
| B137 | B199 | AFU | Mg | -0.29147372 | 1.21E-07 |
| B137 | B195 | AFU | Na | 0.287105454 | 1.89E-07 |
| B138 | B200 | PAB | OP | 0.385728684 | 1.01E-12 |
| B138 | B198 | PAB | P | -0.32840378 | 1.97E-09 |
| B138 | B176 | PAB | Ucr | 0.282787253 | 2.93E-07 |
| B139 | B200 | ADA | OP | 0.393560818 | 3.18E-13 |
| B139 | B150 | ADA | ALP | 0.286990572 | 1.91E-07 |
| B139 | B197 | ADA | AG | 0.281263565 | 3.41E-07 |
| B14 | B61 | G-17 | WBHSRV | 0.437839677 | 2.22E-16 |
| B14 | B55 | G-17 | TK | 0.424371674 | 2.44E-15 |
| B14 | B56 | G-17 | IR | 0.421106508 | 4.22E-15 |
| B14 | B59 | G-17 | WBRV | 0.35101503 | 1.19E-10 |
| B14 | B93 | G-17 | ALDH2-Glu504Lys | 0.349638306 | 1.42E-10 |
| B14 | B92 | G-17 | ALDH2-Glu504Glu | -0.3328717 | 1.15E-09 |
| B14 | B62 | G-17 | WBHSRVI | 0.307975416 | 2.06E-08 |
| B14 | B66 | G-17 | WBRSR(mPas_100) | 0.30545146 | 2.71E-08 |
| B14 | B63 | G-17 | WBRSR(mPas_200) | 0.304947563 | 2.87E-08 |
| B14 | B185 | G-17 | PT | -0.30015817 | 4.82E-08 |
| B14 | B67 | G-17 | WBRSR(mPas_50) | 0.296447487 | 7.16E-08 |
| B14 | B22 | G-17 | CD3+CD8+(T) | -0.27937151 | 4.12E-07 |
| B14 | B19 | G-17 | CD3+CD4+/CD3+CD8+ | 0.272123793 | 8.36E-07 |
| B14 | B64 | G-17 | WBRSR(mPas_5) | 0.270725485 | 9.56E-07 |
| B140 | B192 | r-GT | UA | 0.303854592 | 3.23E-08 |
| B140 | B141 | r-GT | AST/ALT | -0.29477788 | 8.54E-08 |
| B140 | B150 | r-GT | ALP | 0.294217624 | 9.05E-08 |
| B140 | B153 | r-GT | Glu | 0.292676552 | 1.06E-07 |
| B140 | B222 | r-GT | NS | 0.286448991 | 2.02E-07 |
| B141 | B216 | AST/ALT | BMI | -0.38337395 | 1.42E-12 |
| B141 | B221 | AST/ALT | Hei | -0.34669278 | 2.07E-10 |

| | | | | | |
|---|---|---|---|---|---|
| B141 | B192 | AST/ALT | UA | -0.31191796 | 1.33E-08 |
| B141 | B222 | AST/ALT | NS | -0.31147964 | 1.39E-08 |
| B141 | B210 | AST/ALT | Hip | -0.31108149 | 1.46E-08 |
| B141 | B206 | AST/ALT | Waist | -0.2737038 | 7.17E-07 |
| B142 | B144 | 25-(OH)D | Total-PINP | -0.74438861 | 0 |
| B142 | B151 | 25-(OH)D | Anti-CCP | -0.5231091 | 0 |
| B142 | B168 | 25-(OH)D | NMP22 | 0.913089867 | 0 |
| B142 | B154 | 25-(OH)D | RF | -0.27263202 | 7.96E-07 |
| B143 | B145 | β-CTx | BGP | 0.724602378 | 0 |
| B143 | B144 | β-CTx | Total-PINP | 0.4290153 | 1.11E-15 |
| B144 | B145 | Total-PINP | BGP | 0.607179761 | 0 |
| B144 | B168 | Total-PINP | NMP22 | -0.75829061 | 0 |
| B144 | B151 | Total-PINP | Anti-CCP | 0.389128721 | 6.13E-13 |
| B148 | B152 | Anti_TPO | ACA-Ig(G-A-M) | 0.454728667 | 0 |
| B148 | B159 | Anti_TPO | β2-MG | 0.454728001 | 0 |
| B148 | B169 | Anti_TPO | U-CRY | 0.454726971 | 0 |
| B15 | B100 | CD3-CD19+(B cell) | anti-Ro52 | 0.487717811 | 0 |
| B15 | B101 | CD3-CD19+(B cell) | anti-Sc70 | 0.487717811 | 0 |
| B15 | B102 | CD3-CD19+(B cell) | anti-Sm | 0.487717811 | 0 |
| B15 | B103 | CD3-CD19+(B cell) | anti-SSA/Ro | 0.487717811 | 0 |
| B15 | B104 | CD3-CD19+(B cell) | anti-SSB | 0.487717811 | 0 |
| B15 | B105 | CD3-CD19+(B cell) | anti-U1RNP | 0.48771781 | 0 |
| B15 | B106 | CD3-CD19+(B cell) | anti-P | 0.487717811 | 0 |
| B15 | B107 | CD3-CD19+(B cell) | AnuA | 0.487717805 | 0 |
| B15 | B108 | CD3-CD19+(B cell) | Anti-Cyclin | 0.487717811 | 0 |
| B15 | B109 | CD3-CD19+(B cell) | AMA-M2 | 0.487717811 | 0 |
| B15 | B110 | CD3-CD19+(B cell) | AHA | 0.487717811 | 0 |
| B15 | B135 | CD3-CD19+(B cell) | TBA | -0.69234486 | 0 |
| B15 | B151 | CD3-CD19+(B cell) | Anti-CCP | 0.49437704 | 0 |
| B15 | B156 | CD3-CD19+(B cell) | IgE | 0.452397262 | 0 |
| B15 | B16 | CD3-CD19+(B cell) | CD3+T | -0.6910061 | 0 |
| B15 | B178 | CD3-CD19+(B cell) | Ig-U | 0.485411482 | 0 |
| B15 | B180 | CD3-CD19+(B cell) | Tf | 0.441073948 | 0 |
| B15 | B193 | CD3-CD19+(B cell) | K | 0.507089668 | 0 |
| B15 | B195 | CD3-CD19+(B cell) | Na | -0.46655867 | 0 |
| B15 | B197 | CD3-CD19+(B cell) | AG | -0.55984251 | 0 |
| B15 | B20 | CD3-CD19+(B cell) | CD3+CD4+CD8+ | 0.455674377 | 0 |
| B15 | B200 | CD3-CD19+(B cell) | OP | -0.52288446 | 0 |
| B15 | B22 | CD3-CD19+(B cell) | CD3+CD8+(T) | -0.58622157 | 0 |
| B15 | B57 | CD3-CD19+(B cell) | RBCAI | 0.524412223 | 0 |
| B15 | B94 | CD3-CD19+(B cell) | ALDH2-Lys504Lys | 0.466351143 | 0 |
| B15 | B95 | CD3-CD19+(B cell) | anti-cenpb | 0.487717811 | 0 |
| B15 | B97 | CD3-CD19+(B cell) | dsDNA | 0.48771781 | 0 |
| B15 | B98 | CD3-CD19+(B cell) | anti-Jo-1 | 0.48771781 | 0 |

| | | | | | |
|---|---|---|---|---|---|
| B15 | B99 | CD3-CD19+(B cell) | anti-PM-Scl | 0.48771781 | 0 |
| B15 | B199 | CD3-CD19+(B cell) | Mg | 0.422765115 | 3.11E-15 |
| B15 | B179 | CD3-CD19+(B cell) | α1-MG | 0.41387299 | 1.38E-14 |
| B15 | B23 | CD3-CD19+(B cell) | CD3-CD16+CD56+(NK) | 0.409203162 | 2.89E-14 |
| B15 | B79 | CD3-CD19+(B cell) | hb V -dna | 0.394760849 | 2.65E-13 |
| B15 | B194 | CD3-CD19+(B cell) | Cl | 0.386462279 | 9.07E-13 |
| B15 | B114 | CD3-CD19+(B cell) | TSH TT3 TT4(TT4) | -0.38624108 | 9.37E-13 |
| B15 | B55 | CD3-CD19+(B cell) | TK | -0.34066035 | 4.42E-10 |
| B15 | B90 | CD3-CD19+(B cell) | CYFRA 21-1 | -0.33801211 | 6.13E-10 |
| B15 | B154 | CD3-CD19+(B cell) | RF | 0.324653455 | 3.07E-09 |
| B15 | B184 | CD3-CD19+(B cell) | INR | 0.316267256 | 8.10E-09 |
| B15 | B56 | CD3-CD19+(B cell) | IR | -0.31003622 | 1.64E-08 |
| B15 | B198 | CD3-CD19+(B cell) | P | 0.30964857 | 1.71E-08 |
| B15 | B17 | CD3-CD19+(B cell) | CD3+CD16+CD56+NK | 0.308075605 | 2.03E-08 |
| B15 | B19 | CD3-CD19+(B cell) | CD3+CD4+/CD3+CD8+ | 0.300363731 | 4.71E-08 |
| B15 | B83 | CD3-CD19+(B cell) | AFP | -0.28677498 | 1.96E-07 |
| B15 | B183 | CD3-CD19+(B cell) | PTA | 0.279638139 | 4.01E-07 |
| B150 | B153 | ALP | Glu | 0.359739508 | 3.78E-11 |
| B150 | B200 | ALP | OP | 0.298955366 | 5.48E-08 |
| B151 | B154 | Anti-CCP | RF | 0.50928063 | 0 |
| B151 | B156 | Anti-CCP | IgE | 0.622239933 | 0 |
| B151 | B178 | Anti-CCP | Ig-U | 0.460682227 | 0 |
| B151 | B184 | Anti-CCP | INR | 0.46257367 | 0 |
| B151 | B180 | Anti-CCP | Tf | 0.420935839 | 4.44E-15 |
| B151 | B168 | Anti-CCP | NMP22 | -0.3972012 | 1.84E-13 |
| B151 | B193 | Anti-CCP | K | 0.380584508 | 2.12E-12 |
| B151 | B183 | Anti-CCP | PTA | 0.378651096 | 2.80E-12 |
| B151 | B157 | Anti-CCP | IgG | 0.358453581 | 4.48E-11 |
| B151 | B179 | Anti-CCP | α1-MG | 0.336905115 | 7.03E-10 |
| B151 | B199 | Anti-CCP | Mg | 0.325633257 | 2.73E-09 |
| B151 | B158 | Anti-CCP | IgM | -0.29343417 | 9.83E-08 |
| B153 | B170 | Glu | U-GLU | 0.444498444 | 0 |
| B153 | B220 | Glu | SBP | 0.314211484 | 1.02E-08 |
| B153 | B215 | Glu | BMI | 0.298798194 | 5.57E-08 |
| B153 | B206 | Glu | Waist | 0.289727228 | 1.44E-07 |
| B154 | B156 | RF | IgE | 0.608316769 | 0 |
| B154 | B182 | RF | TT | -0.37603067 | 4.05E-12 |
| B154 | B184 | RF | INR | 0.37338812 | 5.87E-12 |
| B154 | B178 | RF | Ig-U | 0.339237894 | 5.27E-10 |
| B154 | B180 | RF | Tf | 0.333644662 | 1.05E-09 |
| B154 | B199 | RF | Mg | 0.285067747 | 2.33E-07 |
| B154 | B183 | RF | PTA | 0.279655876 | 4.00E-07 |
| B154 | B179 | RF | α1-MG | 0.275954542 | 5.76E-07 |
| B155 | B182 | IgA | TT | 0.584830956 | 0 |

| | | | | | |
|---|---|---|---|---|---|
| B155 | B157 | IgA | IgG | 0.415779691 | 1.02E-14 |
| B155 | B158 | IgA | IgM | -0.35239011 | 9.94E-11 |
| B155 | B178 | IgA | Ig-U | 0.306070311 | 2.54E-08 |
| B155 | B180 | IgA | Tf | 0.28150289 | 3.33E-07 |
| B156 | B178 | IgE | Ig-U | 0.453416948 | 0 |
| B156 | B182 | IgE | TT | -0.50835567 | 0 |
| B156 | B183 | IgE | PTA | 0.451718941 | 0 |
| B156 | B184 | IgE | INR | 0.664103269 | 0 |
| B156 | B180 | IgE | Tf | 0.414679398 | 1.20E-14 |
| B156 | B185 | IgE | PT | 0.373203868 | 6.02E-12 |
| B156 | B179 | IgE | α1-MG | 0.318785879 | 6.07E-09 |
| B156 | B193 | IgE | K | 0.318606915 | 6.20E-09 |
| B157 | B158 | IgG | IgM | -0.5827547 | 0 |
| B157 | B178 | IgG | Ig-U | 0.381510969 | 1.86E-12 |
| B157 | B184 | IgG | INR | 0.375728659 | 4.23E-12 |
| B157 | B180 | IgG | Tf | 0.349876956 | 1.38E-10 |
| B157 | B179 | IgG | α1-MG | 0.34099007 | 4.24E-10 |
| B157 | B181 | IgG | APTT | -0.28118696 | 3.44E-07 |
| B158 | B182 | IgM | TT | -0.34987854 | 1.37E-10 |
| B158 | B179 | IgM | α1-MG | -0.31302641 | 1.17E-08 |
| B158 | B178 | IgM | Ig-U | -0.27745067 | 4.98E-07 |
| B159 | B169 | β2-MG | U-CRY | 0.999997962 | 0 |
| B16 | B125 | CD3+T | Cor | 0.529680402 | 0 |
| B16 | B193 | CD3+T | K | -0.44076975 | 0 |
| B16 | B194 | CD3+T | Cl | -0.45596137 | 0 |
| B16 | B199 | CD3+T | Mg | -0.59099587 | 0 |
| B16 | B23 | CD3+T | CD3-CD16+CD56+(NK) | -0.94028915 | 0 |
| B16 | B34 | CD3+T | TK | -0.4869878 | 0 |
| B16 | B57 | CD3+T | RBCAI | -0.43841022 | 2.22E-16 |
| B16 | B22 | CD3+T | CD3+CD8+(T) | 0.433309079 | 4.44E-16 |
| B16 | B90 | CD3+T | CYFRA 21-1 | 0.424036526 | 2.66E-15 |
| B16 | B17 | CD3+T | CD3+CD16+CD56+NK | -0.42247717 | 3.33E-15 |
| B16 | B197 | CD3+T | AG | 0.41368817 | 1.42E-14 |
| B16 | B179 | CD3+T | α1-MG | -0.40382079 | 6.66E-14 |
| B16 | B83 | CD3+T | AFP | 0.393170664 | 3.37E-13 |
| B16 | B137 | CD3+T | AFU | 0.375717207 | 4.23E-12 |
| B16 | B181 | CD3+T | APTT | -0.34946051 | 1.45E-10 |
| B16 | B195 | CD3+T | Na | 0.345751944 | 2.33E-10 |
| B16 | B178 | CD3+T | Ig-U | -0.3411854 | 4.14E-10 |
| B16 | B19 | CD3+T | CD3+CD4+/CD3+CD8+ | -0.33639561 | 7.48E-10 |
| B16 | B187 | CD3+T | NSE | 0.318768392 | 6.08E-09 |
| B16 | B200 | CD3+T | OP | 0.317753517 | 6.84E-09 |
| B16 | B114 | CD3+T | TSH TT3 TT4(TT4) | 0.315217087 | 9.13E-09 |
| B16 | B186 | CD3+T | PFC | 0.310625606 | 1.53E-08 |

| | | | | | |
|---|---|---|---|---|---|
| B16 | B180 | CD3+T | Tf | -0.30805621 | 2.04E-08 |
| B16 | B151 | CD3+T | Anti-CCP | -0.28802629 | 1.72E-07 |
| B16 | B135 | CD3+T | TBA | 0.279041904 | 4.25E-07 |
| B16 | B60 | CD3+T | WBRVI | -0.2790053 | 4.27E-07 |
| B16 | B154 | CD3+T | RF | -0.27239318 | 8.14E-07 |
| B161 | B173 | LEU | NTT | 0.370335845 | 8.97E-12 |
| B163 | B222 | SG | NS | 0.351398056 | 1.13E-10 |
| B163 | B166 | SG | Pro | 0.348904827 | 1.56E-10 |
| B163 | B216 | SG | BMI | 0.321288519 | 4.55E-09 |
| B163 | B176 | SG | Ucr | 0.308315868 | 1.98E-08 |
| B163 | B190 | SG | Cre | 0.299142756 | 5.37E-08 |
| B163 | B191 | SG | Urea | 0.289274626 | 1.51E-07 |
| B163 | B221 | SG | Hei | 0.275924621 | 5.78E-07 |
| B163 | B192 | SG | UA | 0.270463551 | 9.80E-07 |
| B17 | B123 | CD3+CD16+CD56+NK | hs-CRP1 | -0.50744763 | 0 |
| B17 | B181 | CD3+CD16+CD56+NK | APTT | 0.574618957 | 0 |
| B17 | B186 | CD3+CD16+CD56+NK | PFC | -0.83312517 | 0 |
| B17 | B193 | CD3+CD16+CD56+NK | K | 0.608847387 | 0 |
| B17 | B194 | CD3+CD16+CD56+NK | Cl | 0.73597467 | 0 |
| B17 | B197 | CD3+CD16+CD56+NK | AG | -0.460943 | 0 |
| B17 | B199 | CD3+CD16+CD56+NK | Mg | 0.69970468 | 0 |
| B17 | B20 | CD3+CD16+CD56+NK | CD3+CD4+CD8+ | -0.67631864 | 0 |
| B17 | B83 | CD3+CD16+CD56+NK | AFP | -0.93046442 | 0 |
| B17 | B84 | CD3+CD16+CD56+NK | C-PSA | 0.465903548 | 0 |
| B17 | B89 | CD3+CD16+CD56+NK | CA 72-4 | 0.517486102 | 0 |
| B17 | B23 | CD3+CD16+CD56+NK | CD3-CD16+CD56+(NK) | 0.436946596 | 4.44E-16 |
| B17 | B69 | CD3+CD16+CD56+NK | PV | -0.41143938 | 2.02E-14 |
| B17 | B100 | CD3+CD16+CD56+NK | anti-Ro52 | 0.40501587 | 5.53E-14 |
| B17 | B101 | CD3+CD16+CD56+NK | anti-Sc70 | 0.405015869 | 5.53E-14 |
| B17 | B102 | CD3+CD16+CD56+NK | anti-Sm | 0.40501587 | 5.53E-14 |
| B17 | B103 | CD3+CD16+CD56+NK | anti-SSA/Ro | 0.40501587 | 5.53E-14 |
| B17 | B104 | CD3+CD16+CD56+NK | anti-SSB | 0.40501587 | 5.53E-14 |
| B17 | B105 | CD3+CD16+CD56+NK | anti-U1RNP | 0.40501587 | 5.53E-14 |
| B17 | B106 | CD3+CD16+CD56+NK | anti-P | 0.40501587 | 5.53E-14 |
| B17 | B107 | CD3+CD16+CD56+NK | AnuA | 0.405015865 | 5.53E-14 |
| B17 | B108 | CD3+CD16+CD56+NK | Anti-Cyclin | 0.40501587 | 5.53E-14 |
| B17 | B109 | CD3+CD16+CD56+NK | AMA-M2 | 0.40501587 | 5.53E-14 |
| B17 | B110 | CD3+CD16+CD56+NK | AHA | 0.40501587 | 5.53E-14 |
| B17 | B95 | CD3+CD16+CD56+NK | anti-cenpb | 0.40501587 | 5.53E-14 |
| B17 | B97 | CD3+CD16+CD56+NK | dsDNA | 0.40501587 | 5.53E-14 |
| B17 | B98 | CD3+CD16+CD56+NK | anti-Jo-1 | 0.405015869 | 5.53E-14 |
| B17 | B99 | CD3+CD16+CD56+NK | anti-PM-Scl | 0.40501587 | 5.53E-14 |
| B17 | B18 | CD3+CD16+CD56+NK | CD3+CD4+helper(inducer)T | -0.39882617 | 1.43E-13 |
| B17 | B151 | CD3+CD16+CD56+NK | Anti-CCP | 0.397040687 | 1.88E-13 |

| | | | | | |
|---|---|---|---|---|---|
| B17 | B86 | CD3+CD16+CD56+NK | TBhCG | 0.374513269 | 5.02E-12 |
| B17 | B85 | CD3+CD16+CD56+NK | SCC | 0.358793438 | 4.29E-11 |
| B17 | B156 | CD3+CD16+CD56+NK | IgE | 0.288804758 | 1.59E-07 |
| B174 | B190 | Urea/Cr | Cre | -0.5055585 | 0 |
| B174 | B191 | Urea/Cr | Urea | 0.627824022 | 0 |
| B174 | B221 | Urea/Cr | Hei | -0.30774658 | 2.11E-08 |
| B174 | B192 | Urea/Cr | UA | -0.30044165 | 4.67E-08 |
| B175 | B177 | ALB-U | ACR | 0.87197617 | 0 |
| B175 | B196 | ALB-U | TCO2 | -0.27382525 | 7.09E-07 |
| B178 | B179 | Ig-U | α1-MG | 0.830582277 | 0 |
| B178 | B180 | Ig-U | Tf | 0.956429859 | 0 |
| B178 | B184 | Ig-U | INR | 0.480135305 | 0 |
| B179 | B180 | α1-MG | Tf | 0.8899402 | 0 |
| B18 | B134 | CD3+CD4+helper(inducer)T | CHE | 0.473265022 | 0 |
| B18 | B135 | CD3+CD4+helper(inducer)T | TBA | -0.79886481 | 0 |
| B18 | B19 | CD3+CD4+helper(inducer)T | CD3+CD4+/CD3+CD8+ | 0.878428861 | 0 |
| B18 | B20 | CD3+CD4+helper(inducer)T | CD3+CD4+CD8+ | 0.677781207 | 0 |
| B18 | B22 | CD3+CD4+helper(inducer)T | CD3+CD8+(T) | -0.86073875 | 0 |
| B18 | B151 | CD3+CD4+helper(inducer)T | Anti-CCP | -0.4049726 | 5.57E-14 |
| B18 | B100 | CD3+CD4+helper(inducer)T | anti-Ro52 | -0.40395404 | 6.53E-14 |
| B18 | B101 | CD3+CD4+helper(inducer)T | anti-Sc70 | -0.40395405 | 6.53E-14 |
| B18 | B102 | CD3+CD4+helper(inducer)T | anti-Sm | -0.40395405 | 6.53E-14 |
| B18 | B103 | CD3+CD4+helper(inducer)T | anti-SSA/Ro | -0.40395404 | 6.53E-14 |
| B18 | B104 | CD3+CD4+helper(inducer)T | anti-SSB | -0.40395404 | 6.53E-14 |
| B18 | B105 | CD3+CD4+helper(inducer)T | anti-U1RNP | -0.40395404 | 6.53E-14 |
| B18 | B106 | CD3+CD4+helper(inducer)T | anti-P | -0.40395404 | 6.53E-14 |
| B18 | B107 | CD3+CD4+helper(inducer)T | AnuA | -0.40395404 | 6.53E-14 |
| B18 | B108 | CD3+CD4+helper(inducer)T | Anti-Cyclin | -0.40395404 | 6.53E-14 |
| B18 | B109 | CD3+CD4+helper(inducer)T | AMA-M2 | -0.40395404 | 6.53E-14 |
| B18 | B110 | CD3+CD4+helper(inducer)T | AHA | -0.40395404 | 6.53E-14 |
| B18 | B95 | CD3+CD4+helper(inducer)T | anti-cenpb | -0.40395404 | 6.53E-14 |
| B18 | B97 | CD3+CD4+helper(inducer)T | dsDNA | -0.40395404 | 6.53E-14 |
| B18 | B98 | CD3+CD4+helper(inducer)T | anti-Jo-1 | -0.40395404 | 6.53E-14 |
| B18 | B99 | CD3+CD4+helper(inducer)T | anti-PM-Scl | -0.40395404 | 6.53E-14 |
| B18 | B21 | CD3+CD4+helper(inducer)T | CD3+CD4-CD8- | -0.39952873 | 1.29E-13 |
| B18 | B198 | CD3+CD4+helper(inducer)T | P | 0.37543057 | 4.41E-12 |
| B18 | B83 | CD3+CD4+helper(inducer)T | AFP | 0.371069618 | 8.10E-12 |
| B18 | B57 | CD3+CD4+helper(inducer)T | RBCAI | -0.35720577 | 5.29E-11 |
| B18 | B156 | CD3+CD4+helper(inducer)T | IgE | -0.35467844 | 7.37E-11 |
| B18 | B43 | CD3+CD4+helper(inducer)T | MCV | 0.313130611 | 1.16E-08 |
| B18 | B120 | CD3+CD4+helper(inducer)T | HCV-cAg | 0.311789821 | 1.34E-08 |
| B18 | B44 | CD3+CD4+helper(inducer)T | MCH | 0.30363349 | 3.31E-08 |
| B18 | B75 | CD3+CD4+helper(inducer)T | ApoA1/B | -0.29582146 | 7.65E-08 |
| B18 | B178 | CD3+CD4+helper(inducer)T | Ig-U | -0.28254846 | 3.00E-07 |

| | | | | | |
|---|---|---|---|---|---|
| B18 | B196 | CD3+CD4+helper(inducer)T | TCO2 | 0.271722228 | 8.69E-07 |
| B180 | B184 | Tf | INR | 0.443717828 | 0 |
| B181 | B183 | APTT | PTA | -0.57113504 | 0 |
| B181 | B186 | APTT | PFC | -0.86875573 | 0 |
| B181 | B194 | APTT | Cl | 0.534408118 | 0 |
| B181 | B199 | APTT | Mg | 0.4512556 | 0 |
| B181 | B193 | APTT | K | 0.37700598 | 3.53E-12 |
| B181 | B185 | APTT | PT | 0.290252331 | 1.37E-07 |
| B182 | B183 | TT | PTA | -0.52237114 | 0 |
| B182 | B184 | TT | INR | -0.42501378 | 2.22E-15 |
| B183 | B185 | PTA | PT | -0.31950466 | 5.59E-09 |
| B183 | B186 | PTA | PFC | 0.302397815 | 3.78E-08 |
| B184 | B185 | INR | PT | 0.828685871 | 0 |
| B186 | B193 | PFC | K | -0.48508307 | 0 |
| B186 | B194 | PFC | Cl | -0.63646627 | 0 |
| B186 | B199 | PFC | Mg | -0.5547354 | 0 |
| B186 | B197 | PFC | AG | 0.381366391 | 1.90E-12 |
| B187 | B198 | NSE | P | 0.334342977 | 9.62E-10 |
| B189 | B190 | eGFB | Cre | -0.57422485 | 0 |
| B189 | B220 | eGFB | SBP | -0.35133577 | 1.14E-10 |
| B189 | B191 | eGFB | Urea | -0.29900574 | 5.45E-08 |
| B19 | B135 | CD3+CD4+/CD3+CD8+ | TBA | -0.7153298 | 0 |
| B19 | B20 | CD3+CD4+/CD3+CD8+ | CD3+CD4+CD8+ | 0.54932376 | 0 |
| B19 | B21 | CD3+CD4+/CD3+CD8+ | CD3+CD4-CD8- | -0.53069429 | 0 |
| B19 | B22 | CD3+CD4+/CD3+CD8+ | CD3+CD8+(T) | -0.93000006 | 0 |
| B19 | B34 | CD3+CD4+/CD3+CD8+ | TK | 0.488457537 | 0 |
| B19 | B134 | CD3+CD4+/CD3+CD8+ | CHE | 0.374747732 | 4.85E-12 |
| B19 | B100 | CD3+CD4+/CD3+CD8+ | anti-Ro52 | -0.355961 | 6.23E-11 |
| B19 | B101 | CD3+CD4+/CD3+CD8+ | anti-Sc70 | -0.355961 | 6.23E-11 |
| B19 | B102 | CD3+CD4+/CD3+CD8+ | anti-Sm | -0.355961 | 6.23E-11 |
| B19 | B103 | CD3+CD4+/CD3+CD8+ | anti-SSA/Ro | -0.355961 | 6.23E-11 |
| B19 | B104 | CD3+CD4+/CD3+CD8+ | anti-SSB | -0.355961 | 6.23E-11 |
| B19 | B105 | CD3+CD4+/CD3+CD8+ | anti-U1RNP | -0.355961 | 6.23E-11 |
| B19 | B106 | CD3+CD4+/CD3+CD8+ | anti-P | -0.355961 | 6.23E-11 |
| B19 | B107 | CD3+CD4+/CD3+CD8+ | AnuA | -0.355961 | 6.23E-11 |
| B19 | B108 | CD3+CD4+/CD3+CD8+ | Anti-Cyclin | -0.355961 | 6.23E-11 |
| B19 | B109 | CD3+CD4+/CD3+CD8+ | AMA-M2 | -0.355961 | 6.23E-11 |
| B19 | B110 | CD3+CD4+/CD3+CD8+ | AHA | -0.355961 | 6.23E-11 |
| B19 | B95 | CD3+CD4+/CD3+CD8+ | anti-cenpb | -0.355961 | 6.23E-11 |
| B19 | B97 | CD3+CD4+/CD3+CD8+ | dsDNA | -0.355961 | 6.23E-11 |
| B19 | B98 | CD3+CD4+/CD3+CD8+ | anti-Jo-1 | -0.355961 | 6.23E-11 |
| B19 | B99 | CD3+CD4+/CD3+CD8+ | anti-PM-Scl | -0.355961 | 6.23E-11 |
| B19 | B125 | CD3+CD4+/CD3+CD8+ | Cor | -0.34450879 | 2.73E-10 |
| B19 | B151 | CD3+CD4+/CD3+CD8+ | Anti-CCP | -0.3422218 | 3.63E-10 |

| | | | | | |
|---|---|---|---|---|---|
| B19 | B114 | CD3+CD4+/CD3+CD8+ | TSH TT3 TT4(TT4) | -0.32596209 | 2.63E-09 |
| B19 | B156 | CD3+CD4+/CD3+CD8+ | IgE | -0.30011663 | 4.84E-08 |
| B19 | B94 | CD3+CD4+/CD3+CD8+ | ALDH2-Lys504Lys | -0.29887977 | 5.53E-08 |
| B19 | B124 | CD3+CD4+/CD3+CD8+ | ACTH | -0.29652432 | 7.10E-08 |
| B19 | B120 | CD3+CD4+/CD3+CD8+ | HCV-cAg | 0.283067379 | 2.85E-07 |
| B190 | B192 | Cre | UA | 0.582171275 | 0 |
| B190 | B222 | Cre | NS | 0.437410458 | 2.22E-16 |
| B190 | B221 | Cre | Hei | 0.309129824 | 1.81E-08 |
| B190 | B191 | Cre | Urea | 0.308474079 | 1.95E-08 |
| B190 | B216 | Cre | BMI | 0.299070398 | 5.41E-08 |
| B190 | B206 | Cre | Waist | 0.276771921 | 5.32E-07 |
| B192 | B222 | UA | NS | 0.529607775 | 0 |
| B192 | B216 | UA | BMI | 0.438804968 | 2.22E-16 |
| B192 | B221 | UA | Hei | 0.407618392 | 3.69E-14 |
| B192 | B206 | UA | Waist | 0.376271794 | 3.92E-12 |
| B192 | B215 | UA | BMI | 0.284548048 | 2.45E-07 |
| B192 | B207 | UA | WHR | 0.28229958 | 3.08E-07 |
| B193 | B194 | K | Cl | 0.63801903 | 0 |
| B193 | B199 | K | Mg | 0.658969849 | 0 |
| B193 | B197 | K | AG | -0.35518322 | 6.90E-11 |
| B194 | B197 | Cl | AG | -0.44933149 | 0 |
| B194 | B199 | Cl | Mg | 0.610651521 | 0 |
| B194 | B196 | Cl | TCO2 | -0.31642884 | 7.95E-09 |
| B195 | B197 | Na | AG | 0.743430912 | 0 |
| B195 | B200 | Na | OP | 0.673850214 | 0 |
| B195 | B196 | Na | TCO2 | -0.3719814 | 7.14E-12 |
| B195 | B198 | Na | P | -0.28710205 | 1.89E-07 |
| B196 | B197 | TCO2 | AG | -0.60457206 | 0 |
| B197 | B199 | AG | Mg | -0.58307264 | 0 |
| B197 | B200 | AG | OP | 0.631187863 | 0 |
| B197 | B198 | AG | P | -0.2834388 | 2.74E-07 |
| B198 | B200 | P | OP | -0.42463839 | 2.22E-15 |
| B198 | B199 | P | Mg | -0.29993671 | 4.93E-08 |
| B2 | B189 | Age | eGFB | -0.57056915 | 0 |
| B2 | B220 | Age | SBP | 0.429708939 | 8.88E-16 |
| B2 | B47 | Age | RDW | 0.382583998 | 1.59E-12 |
| B2 | B221 | Age | Hei | -0.34415099 | 2.85E-10 |
| B2 | B191 | Age | Urea | 0.287772538 | 1.77E-07 |
| B2 | B126 | Age | Alb | -0.2798959 | 3.91E-07 |
| B20 | B135 | CD3+CD4+CD8+ | TBA | -0.61526221 | 0 |
| B20 | B181 | CD3+CD4+CD8+ | APTT | -0.53217528 | 0 |
| B20 | B186 | CD3+CD4+CD8+ | PFC | 0.608919991 | 0 |
| B20 | B22 | CD3+CD4+CD8+ | CD3+CD8+(T) | -0.61335159 | 0 |
| B20 | B83 | CD3+CD4+CD8+ | AFP | 0.629204477 | 0 |

| | | | | | |
|---|---|---|---|---|---|
| B20 | B198 | CD3+CD4+CD8+ | P | 0.426301522 | 1.78E-15 |
| B20 | B69 | CD3+CD4+CD8+ | PV | 0.42644748 | 1.78E-15 |
| B20 | B123 | CD3+CD4+CD8+ | hs-CRP1 | 0.368698624 | 1.12E-11 |
| B20 | B89 | CD3+CD4+CD8+ | CA 72-4 | -0.35944365 | 3.93E-11 |
| B20 | B134 | CD3+CD4+CD8+ | CHE | 0.355083574 | 6.99E-11 |
| B20 | B199 | CD3+CD4+CD8+ | Mg | -0.3494611 | 1.45E-10 |
| B20 | B194 | CD3+CD4+CD8+ | Cl | -0.3372181 | 6.77E-10 |
| B20 | B184 | CD3+CD4+CD8+ | INR | 0.327657303 | 2.15E-09 |
| B20 | B84 | CD3+CD4+CD8+ | C-PSA | -0.32202775 | 4.17E-09 |
| B20 | B196 | CD3+CD4+CD8+ | TCO2 | 0.312444725 | 1.25E-08 |
| B20 | B155 | CD3+CD4+CD8+ | IgA | 0.304328384 | 3.07E-08 |
| B20 | B94 | CD3+CD4+CD8+ | ALDH2-Lys504Lys | 0.287993637 | 1.73E-07 |
| B20 | B62 | CD3+CD4+CD8+ | WBHSRVI | -0.28247101 | 3.02E-07 |
| B20 | B34 | CD3+CD4+CD8+ | TK | -0.27670661 | 5.35E-07 |
| B204 | B209 | Drinking | Smoking | 0.482504471 | 0 |
| B204 | B221 | Drinking | Hei | 0.291909612 | 1.15E-07 |
| B206 | B207 | Waist | WHR | 0.560327769 | 0 |
| B206 | B210 | Waist | Hip | 0.686047819 | 0 |
| B206 | B215 | Waist | BMI | 0.886917023 | 0 |
| B206 | B216 | Waist | BMI | 0.832474216 | 0 |
| B206 | B222 | Waist | NS | 0.758881315 | 0 |
| B206 | B218 | Waist | DBP | 0.353625314 | 8.46E-11 |
| B206 | B220 | Waist | SBP | 0.317914676 | 6.71E-09 |
| B206 | B221 | Waist | Hei | 0.287568639 | 1.80E-07 |
| B207 | B215 | WHR | BMI | 0.437764615 | 2.22E-16 |
| B207 | B222 | WHR | NS | 0.433147548 | 4.44E-16 |
| B207 | B216 | WHR | BMI | 0.375596485 | 4.31E-12 |
| B209 | B222 | Smoking | NS | 0.30124367 | 4.29E-08 |
| B21 | B100 | CD3+CD4-CD8- | anti-Ro52 | 0.638459265 | 0 |
| B21 | B101 | CD3+CD4-CD8- | anti-Sc70 | 0.638459266 | 0 |
| B21 | B102 | CD3+CD4-CD8- | anti-Sm | 0.638459266 | 0 |
| B21 | B103 | CD3+CD4-CD8- | anti-SSA/Ro | 0.638459265 | 0 |
| B21 | B104 | CD3+CD4-CD8- | anti-SSB | 0.638459265 | 0 |
| B21 | B105 | CD3+CD4-CD8- | anti-U1RNP | 0.638459265 | 0 |
| B21 | B106 | CD3+CD4-CD8- | anti-P | 0.638459265 | 0 |
| B21 | B107 | CD3+CD4-CD8- | AnuA | 0.638459257 | 0 |
| B21 | B108 | CD3+CD4-CD8- | Anti-Cyclin | 0.638459265 | 0 |
| B21 | B109 | CD3+CD4-CD8- | AMA-M2 | 0.638459265 | 0 |
| B21 | B110 | CD3+CD4-CD8- | AHA | 0.638459265 | 0 |
| B21 | B151 | CD3+CD4-CD8- | Anti-CCP | 0.646429098 | 0 |
| B21 | B156 | CD3+CD4-CD8- | IgE | 0.595568957 | 0 |
| B21 | B34 | CD3+CD4-CD8- | TK | -0.57098546 | 0 |
| B21 | B79 | CD3+CD4-CD8- | hb V -dna | 0.577643304 | 0 |
| B21 | B95 | CD3+CD4-CD8- | anti-cenpb | 0.638459265 | 0 |

| | | | | | |
|---|---|---|---|---|---|
| B21 | B97 | CD3+CD4-CD8- | dsDNA | 0.638459264 | 0 |
| B21 | B98 | CD3+CD4-CD8- | anti-Jo-1 | 0.638459264 | 0 |
| B21 | B99 | CD3+CD4-CD8- | anti-PM-Scl | 0.638459264 | 0 |
| B21 | B138 | CD3+CD4-CD8- | PAB | 0.420899121 | 4.44E-15 |
| B21 | B157 | CD3+CD4-CD8- | IgG | 0.41633003 | 9.33E-15 |
| B21 | B184 | CD3+CD4-CD8- | INR | 0.408907601 | 3.02E-14 |
| B21 | B158 | CD3+CD4-CD8- | IgM | -0.40258619 | 8.06E-14 |
| B21 | B57 | CD3+CD4-CD8- | RBCAI | 0.366276945 | 1.57E-11 |
| B21 | B183 | CD3+CD4-CD8- | PTA | 0.359717767 | 3.79E-11 |
| B21 | B124 | CD3+CD4-CD8- | ACTH | 0.358018145 | 4.75E-11 |
| B21 | B39 | CD3+CD4-CD8- | Other cells | 0.330375603 | 1.55E-09 |
| B21 | B81 | CD3+CD4-CD8- | TSGF | -0.32897941 | 1.84E-09 |
| B21 | B176 | CD3+CD4-CD8- | Ucr | 0.323618957 | 3.46E-09 |
| B21 | B22 | CD3+CD4-CD8- | CD3+CD8+(T) | 0.314857621 | 9.51E-09 |
| B21 | B55 | CD3+CD4-CD8- | TK | -0.31154544 | 1.38E-08 |
| B21 | B76 | CD3+CD4-CD8- | ApoB | 0.30567245 | 2.65E-08 |
| B21 | B23 | CD3+CD4-CD8- | CD3-CD16+CD56+(NK) | -0.30163881 | 4.11E-08 |
| B21 | B178 | CD3+CD4-CD8- | Ig-U | 0.294274351 | 9.00E-08 |
| B21 | B56 | CD3+CD4-CD8- | IR | -0.27800056 | 4.71E-07 |
| B21 | B193 | CD3+CD4-CD8- | K | 0.275202711 | 6.20E-07 |
| B210 | B215 | Hip | BMI | 0.705810086 | 0 |
| B210 | B216 | Hip | BMI | 0.746899498 | 0 |
| B210 | B222 | Hip | NS | 0.570596761 | 0 |
| B210 | B221 | Hip | Hei | 0.377718456 | 3.19E-12 |
| B210 | B218 | Hip | DBP | 0.273096994 | 7.61E-07 |
| B211 | B212 | IOP-R | IOP-L | 0.757291722 | 0 |
| B215 | B216 | BMI | BMI | 0.836550562 | 0 |
| B215 | B222 | BMI | NS | 0.716129486 | 0 |
| B215 | B218 | BMI | DBP | 0.380879002 | 2.03E-12 |
| B215 | B220 | BMI | SBP | 0.331588326 | 1.34E-09 |
| B216 | B221 | BMI | Hei | 0.64852655 | 0 |
| B216 | B222 | BMI | NS | 0.811615675 | 0 |
| B216 | B218 | BMI | DBP | 0.359576853 | 3.86E-11 |
| B218 | B220 | DBP | SBP | 0.651986042 | 0 |
| B218 | B222 | DBP | NS | 0.350128823 | 1.33E-10 |
| B22 | B135 | CD3+CD8+(T) | TBA | 0.901971578 | 0 |
| B22 | B197 | CD3+CD8+(T) | AG | 0.448072081 | 0 |
| B22 | B200 | CD3+CD8+(T) | OP | 0.396108173 | 2.17E-13 |
| B22 | B114 | CD3+CD8+(T) | TSH TT3 TT4(TT4) | 0.385310802 | 1.07E-12 |
| B22 | B134 | CD3+CD8+(T) | CHE | -0.38524714 | 1.08E-12 |
| B22 | B193 | CD3+CD8+(T) | K | -0.34691664 | 2.01E-10 |
| B22 | B43 | CD3+CD8+(T) | MCV | -0.33956108 | 5.06E-10 |
| B22 | B120 | CD3+CD8+(T) | HCV-cAg | -0.33684183 | 7.09E-10 |
| B22 | B44 | CD3+CD8+(T) | MCH | -0.32746101 | 2.20E-09 |

| | | | | | |
|---|---|---|---|---|---|
| B22 | B198 | CD3+CD8+(T) | P | -0.31283238 | 1.20E-08 |
| B22 | B194 | CD3+CD8+(T) | Cl | -0.28980735 | 1.43E-07 |
| B22 | B34 | CD3+CD8+(T) | TK | -0.27858946 | 4.45E-07 |
| B22 | B195 | CD3+CD8+(T) | Na | 0.27395709 | 7.00E-07 |
| B221 | B222 | Hei | NS | 0.483491593 | 0 |
| B23 | B125 | CD3-CD16+CD56+(NK) | Cor | -0.6030885 | 0 |
| B23 | B181 | CD3-CD16+CD56+(NK) | APTT | 0.47414844 | 0 |
| B23 | B199 | CD3-CD16+CD56+(NK) | Mg | 0.574732147 | 0 |
| B23 | B34 | CD3-CD16+CD56+(NK) | TK | 0.667636574 | 0 |
| B23 | B194 | CD3-CD16+CD56+(NK) | Cl | 0.422330795 | 3.55E-15 |
| B23 | B83 | CD3-CD16+CD56+(NK) | AFP | -0.40659546 | 4.33E-14 |
| B23 | B90 | CD3-CD16+CD56+(NK) | CYFRA 21-1 | -0.38279534 | 1.54E-12 |
| B23 | B187 | CD3-CD16+CD56+(NK) | NSE | -0.37413165 | 5.29E-12 |
| B23 | B186 | CD3-CD16+CD56+(NK) | PFC | -0.35562442 | 6.51E-11 |
| B23 | B137 | CD3-CD16+CD56+(NK) | AFU | -0.35548526 | 6.63E-11 |
| B23 | B193 | CD3-CD16+CD56+(NK) | K | 0.351958081 | 1.05E-10 |
| B23 | B57 | CD3-CD16+CD56+(NK) | RBCAI | 0.315029594 | 9.33E-09 |
| B23 | B76 | CD3-CD16+CD56+(NK) | ApoB | -0.31095916 | 1.48E-08 |
| B23 | B179 | CD3-CD16+CD56+(NK) | α1-MG | 0.308065852 | 2.04E-08 |
| B24 | B26 | αHBDH | CKMB | 0.549364557 | 0 |
| B24 | B27 | αHBDH | LDH | 0.662601313 | 0 |
| B24 | B90 | αHBDH | CYFRA 21-1 | 0.275351663 | 6.11E-07 |
| B25 | B39 | CK | Other cells | 0.447485202 | 0 |
| B25 | B57 | CK | RBCAI | 0.364212542 | 2.07E-11 |
| B25 | B26 | CK | CKMB | 0.312810015 | 1.20E-08 |
| B25 | B69 | CK | PV | 0.292006289 | 1.14E-07 |
| B26 | B27 | CKMB | LDH | 0.450162754 | 0 |
| B27 | B213 | LDH | CVA-R | 0.494212905 | 0 |
| B27 | B176 | LDH | Ucr | -0.28583905 | 2.15E-07 |
| B28 | B164 | PRL | COM | -0.58343433 | 0 |
| B28 | B30 | PRL | LH | 1 | 0 |
| B28 | B32 | PRL | Testo | -1 | 0 |
| B29 | B164 | E2 | COM | 0.58343433 | 0 |
| B29 | B30 | E2 | LH | -1 | 0 |
| B29 | B32 | E2 | Testo | 1 | 0 |
| B3 | B85 | Hcy | SCC | 0.328987096 | 1.84E-09 |
| B3 | B84 | Hcy | C-PSA | 0.272195382 | 8.30E-07 |
| B30 | B164 | LH | COM | -0.58343433 | 0 |
| B30 | B31 | LH | FSH | 1 | 0 |
| B30 | B32 | LH | Testo | -1 | 0 |
| B30 | B33 | LH | Prog | -1 | 0 |
| B31 | B164 | FSH | COM | -0.58343433 | 0 |
| B31 | B32 | FSH | Testo | -1 | 0 |
| B32 | B164 | Testo | COM | 0.58343433 | 0 |

| | | | | | |
|---|---|---|---|---|---|
| B32 | B33 | Testo | Prog | 1 | 0 |
| B33 | B164 | Prog | COM | 0.58343433 | 0 |
| B34 | B124 | TK | ACTH | -0.5210076 | 0 |
| B34 | B125 | TK | Cor | -0.52169085 | 0 |
| B34 | B181 | TK | APTT | 0.557683407 | 0 |
| B34 | B79 | TK | hb V -dna | -0.35037167 | 1.29E-10 |
| B34 | B187 | TK | NSE | -0.34684582 | 2.03E-10 |
| B34 | B94 | TK | ALDH2-Lys504Lys | -0.34608381 | 2.23E-10 |
| B34 | B199 | TK | Mg | 0.333453356 | 1.07E-09 |
| B34 | B158 | TK | IgM | 0.318130245 | 6.55E-09 |
| B34 | B69 | TK | PV | -0.31217438 | 1.29E-08 |
| B34 | B92 | TK | ALDH2-Glu504Glu | 0.293175219 | 1.01E-07 |
| B34 | B93 | TK | ALDH2-Glu504Lys | -0.27604734 | 5.71E-07 |
| B34 | B198 | TK | P | -0.27539219 | 6.09E-07 |
| B35 | B116 | β2-MG | TT3 | -0.90093746 | 0 |
| B35 | B202 | β2-MG | e-times | -0.54272042 | 0 |
| B35 | B37 | β2-MG | BAND | -0.38376129 | 1.34E-12 |
| B36 | B167 | V-Lym% | PC | 0.707174992 | 0 |
| B36 | B96 | V-Lym% | dsDNA (IIF) | 0.679873454 | 0 |
| B39 | B57 | Other cells | RBCAI | 0.425113928 | 2.22E-15 |
| B39 | B180 | Other cells | Tf | 0.378815166 | 2.73E-12 |
| B39 | B60 | Other cells | WBRVI | 0.373841177 | 5.51E-12 |
| B39 | B78 | Other cells | HBcAb | 0.366008462 | 1.62E-11 |
| B39 | B65 | Other cells | WBRSR(mPas_1) | 0.357706376 | 4.95E-11 |
| B39 | B179 | Other cells | α1-MG | 0.354896324 | 7.17E-11 |
| B39 | B178 | Other cells | Ig-U | 0.349114533 | 1.52E-10 |
| B39 | B64 | Other cells | WBRSR(mPas_5) | 0.326679533 | 2.42E-09 |
| B39 | B59 | Other cells | WBRV | 0.302040361 | 3.93E-08 |
| B39 | B151 | Other cells | Anti-CCP | 0.280584068 | 3.65E-07 |
| B39 | B100 | Other cells | anti-Ro52 | 0.275282728 | 6.15E-07 |
| B39 | B101 | Other cells | anti-Sc70 | 0.275282729 | 6.15E-07 |
| B39 | B102 | Other cells | anti-Sm | 0.275282729 | 6.15E-07 |
| B39 | B103 | Other cells | anti-SSA/Ro | 0.275282728 | 6.15E-07 |
| B39 | B104 | Other cells | anti-SSB | 0.275282729 | 6.15E-07 |
| B39 | B105 | Other cells | anti-U1RNP | 0.275282728 | 6.15E-07 |
| B39 | B106 | Other cells | anti-P | 0.275282728 | 6.15E-07 |
| B39 | B107 | Other cells | AnuA | 0.275282725 | 6.15E-07 |
| B39 | B108 | Other cells | Anti-Cyclin | 0.275282729 | 6.15E-07 |
| B39 | B109 | Other cells | AMA-M2 | 0.275282728 | 6.15E-07 |
| B39 | B110 | Other cells | AHA | 0.275282728 | 6.15E-07 |
| B39 | B95 | Other cells | anti-cenpb | 0.275282729 | 6.15E-07 |
| B39 | B97 | Other cells | dsDNA | 0.275282728 | 6.15E-07 |
| B39 | B98 | Other cells | anti-Jo-1 | 0.275282728 | 6.15E-07 |
| B39 | B99 | Other cells | anti-PM-Scl | 0.275282728 | 6.15E-07 |

| | | | | | |
|---|---|---|---|---|---|
| B4 | B137 | WB-Ca | AFU | 0.452850717 | 0 |
| B4 | B138 | WB-Ca | PAB | 0.52045028 | 0 |
| B4 | B16 | WB-Ca | CD3+T | 0.659409978 | 0 |
| B4 | B21 | WB-Ca | CD3+CD4-CD8- | 0.514334406 | 0 |
| B4 | B23 | WB-Ca | CD3-CD16+CD56+(NK) | -0.62509392 | 0 |
| B4 | B5 | WB-Ca | WB-Mg | 1 | 0 |
| B4 | B6 | WB-Ca | WB-Fe | 1 | 0 |
| B4 | B7 | WB-Ca | WB-Cop | 1 | 0 |
| B4 | B76 | WB-Ca | ApoB | 0.443255337 | 0 |
| B4 | B8 | WB-Ca | WB-Zinc | 1 | 0 |
| B4 | B94 | WB-Ca | ALDH2-Lys504Lys | -0.49762845 | 0 |
| B4 | B15 | WB-Ca | CD3-CD19+(B cell) | -0.39394978 | 3.00E-13 |
| B4 | B179 | WB-Ca | α1-MG | -0.38747121 | 7.82E-13 |
| B4 | B178 | WB-Ca | Ig-U | -0.38072155 | 2.08E-12 |
| B4 | B57 | WB-Ca | RBCAI | -0.3714923 | 7.64E-12 |
| B4 | B85 | WB-Ca | SCC | -0.36823304 | 1.20E-11 |
| B4 | B92 | WB-Ca | ALDH2-Glu504Glu | 0.364342135 | 2.04E-11 |
| B4 | B75 | WB-Ca | ApoA1/B | -0.35273649 | 9.50E-11 |
| B4 | B180 | WB-Ca | Tf | -0.34443905 | 2.75E-10 |
| B4 | B93 | WB-Ca | ALDH2-Glu504Lys | -0.33929588 | 5.23E-10 |
| B4 | B90 | WB-Ca | CYFRA 21-1 | 0.312457974 | 1.25E-08 |
| B4 | B176 | WB-Ca | Ucr | 0.295282965 | 8.09E-08 |
| B4 | B200 | WB-Ca | OP | 0.291810316 | 1.16E-07 |
| B4 | B134 | WB-Ca | CHE | 0.29147904 | 1.21E-07 |
| B4 | B195 | WB-Ca | Na | 0.290527589 | 1.33E-07 |
| B4 | B198 | WB-Ca | P | -0.27659081 | 5.42E-07 |
| B40 | B42 | WBC | MON | 0.622408679 | 0 |
| B41 | B42 | MON % | MON | 0.702632156 | 0 |
| B41 | B190 | MON % | Cre | 0.271142705 | 9.18E-07 |
| B42 | B190 | MON | Cre | 0.330006996 | 1.62E-09 |
| B43 | B44 | MCV | MCH | 0.902288532 | 0 |
| B43 | B46 | MCV | RBC Count | -0.61867737 | 0 |
| B43 | B47 | MCV | RDW | 0.508239105 | 0 |
| B43 | B135 | MCV | TBA | -0.38774998 | 7.51E-13 |
| B44 | B45 | MCH | MCHC | 0.533881408 | 0 |
| B44 | B46 | MCH | RBC Count | -0.48676848 | 0 |
| B44 | B135 | MCH | TBA | -0.35367891 | 8.40E-11 |
| B44 | B47 | MCH | RDW | 0.314817957 | 9.55E-09 |
| B45 | B51 | MCHC | Hb | 0.514496546 | 0 |
| B45 | B222 | MCHC | NS | 0.327193704 | 2.27E-09 |
| B45 | B221 | MCHC | Hei | 0.32026516 | 5.12E-09 |
| B45 | B216 | MCHC | BMI | 0.317173426 | 7.31E-09 |
| B45 | B71 | MCHC | TG | 0.292574968 | 1.08E-07 |
| B45 | B47 | MCHC | RDW | -0.27814951 | 4.65E-07 |

| | | | | | |
|---|---|---|---|---|---|
| B46 | B51 | RBC Count | Hb | 0.722785008 | 0 |
| B46 | B47 | RBC Count | RDW | -0.43915076 | 2.22E-16 |
| B46 | B222 | RBC Count | NS | 0.386499263 | 9.02E-13 |
| B46 | B190 | RBC Count | Cre | 0.338915235 | 5.49E-10 |
| B46 | B216 | RBC Count | BMI | 0.334120218 | 9.88E-10 |
| B46 | B72 | RBC Count | HDL-C | -0.32221178 | 4.08E-09 |
| B46 | B218 | RBC Count | DBP | 0.305989548 | 2.56E-08 |
| B46 | B192 | RBC Count | UA | 0.303840612 | 3.24E-08 |
| B46 | B221 | RBC Count | Hei | 0.303109262 | 3.50E-08 |
| B48 | B53 | LYM% | GR% | -0.93400012 | 0 |
| B49 | B50 | baso% | eo% | 0.296701494 | 6.97E-08 |
| B5 | B137 | WB-Mg | AFU | 0.452850717 | 0 |
| B5 | B138 | WB-Mg | PAB | 0.52045028 | 0 |
| B5 | B16 | WB-Mg | CD3+T | 0.659409978 | 0 |
| B5 | B21 | WB-Mg | CD3+CD4-CD8- | 0.514334406 | 0 |
| B5 | B23 | WB-Mg | CD3-CD16+CD56+(NK) | -0.62509392 | 0 |
| B5 | B7 | WB-Mg | WB-Cop | 1 | 0 |
| B5 | B76 | WB-Mg | ApoB | 0.443255337 | 0 |
| B5 | B8 | WB-Mg | WB-Zinc | 1 | 0 |
| B5 | B94 | WB-Mg | ALDH2-Lys504Lys | -0.49762844 | 0 |
| B5 | B15 | WB-Mg | CD3-CD19+(B cell) | -0.39394978 | 3.00E-13 |
| B5 | B179 | WB-Mg | α1-MG | -0.38747121 | 7.82E-13 |
| B5 | B178 | WB-Mg | Ig-U | -0.38072155 | 2.08E-12 |
| B5 | B57 | WB-Mg | RBCAI | -0.37149229 | 7.64E-12 |
| B5 | B85 | WB-Mg | SCC | -0.36823304 | 1.20E-11 |
| B5 | B92 | WB-Mg | ALDH2-Glu504Glu | 0.364342135 | 2.04E-11 |
| B5 | B75 | WB-Mg | ApoA1/B | -0.35273649 | 9.50E-11 |
| B5 | B180 | WB-Mg | Tf | -0.34443905 | 2.75E-10 |
| B5 | B93 | WB-Mg | ALDH2-Glu504Lys | -0.33929588 | 5.23E-10 |
| B5 | B90 | WB-Mg | CYFRA 21-1 | 0.312457974 | 1.25E-08 |
| B5 | B176 | WB-Mg | Ucr | 0.295282965 | 8.09E-08 |
| B5 | B200 | WB-Mg | OP | 0.291810317 | 1.16E-07 |
| B5 | B134 | WB-Mg | CHE | 0.29147904 | 1.21E-07 |
| B5 | B195 | WB-Mg | Na | 0.29052759 | 1.33E-07 |
| B5 | B198 | WB-Mg | P | -0.27659081 | 5.42E-07 |
| B50 | B53 | eo% | GR% | -0.28346989 | 2.73E-07 |
| B51 | B216 | Hb | BMI | 0.454530105 | 0 |
| B51 | B221 | Hb | Hei | 0.457770411 | 0 |
| B51 | B222 | Hb | NS | 0.513800953 | 0 |
| B51 | B190 | Hb | Cre | 0.402104869 | 8.68E-14 |
| B51 | B192 | Hb | UA | 0.395251083 | 2.46E-13 |
| B51 | B163 | Hb | SG | 0.391164038 | 4.54E-13 |
| B51 | B72 | Hb | HDL-C | -0.34565108 | 2.36E-10 |
| B51 | B174 | Hb | Urea/Cr | -0.33483236 | 9.06E-10 |

| | | | | | |
|---|---|---|---|---|---|
| B51 | B206 | Hb | Waist | 0.328333112 | 1.98E-09 |
| B51 | B218 | Hb | DBP | 0.306676962 | 2.37E-08 |
| B51 | B141 | Hb | AST/ALT | -0.29085825 | 1.29E-07 |
| B51 | B58 | Hb | HCT | 0.274760562 | 6.47E-07 |
| B52 | B70 | PLT/BPC | LDL-C | 0.276891526 | 5.26E-07 |
| B54 | B175 | ESR | ALB-U | 0.606123921 | 0 |
| B54 | B177 | ESR | ACR | 0.680428239 | 0 |
| B54 | B58 | ESR | HCT | -0.56901985 | 0 |
| B54 | B62 | ESR | WBHSRVI | -0.5342105 | 0 |
| B54 | B69 | ESR | PV | 0.464598039 | 0 |
| B54 | B60 | ESR | WBRVI | -0.41043548 | 2.35E-14 |
| B54 | B155 | ESR | IgA | 0.392378335 | 3.79E-13 |
| B54 | B123 | ESR | hs-CRP1 | 0.377115972 | 3.48E-12 |
| B54 | B63 | ESR | WBRSR(mPas_200) | -0.338929 | 5.48E-10 |
| B54 | B56 | ESR | IR | -0.32708725 | 2.30E-09 |
| B54 | B66 | ESR | WBRSR(mPas_100) | -0.32209425 | 4.14E-09 |
| B54 | B67 | ESR | WBRSR(mPas_50) | -0.31642585 | 7.96E-09 |
| B55 | B100 | TK | anti-Ro52 | -0.48238707 | 0 |
| B55 | B101 | TK | anti-Sc70 | -0.48238707 | 0 |
| B55 | B102 | TK | anti-Sm | -0.48238707 | 0 |
| B55 | B103 | TK | anti-SSA/Ro | -0.48238707 | 0 |
| B55 | B104 | TK | anti-SSB | -0.48238707 | 0 |
| B55 | B105 | TK | anti-U1RNP | -0.48238707 | 0 |
| B55 | B106 | TK | anti-P | -0.48238707 | 0 |
| B55 | B107 | TK | AnuA | -0.48238706 | 0 |
| B55 | B108 | TK | Anti-Cyclin | -0.48238707 | 0 |
| B55 | B109 | TK | AMA-M2 | -0.48238707 | 0 |
| B55 | B110 | TK | AHA | -0.48238707 | 0 |
| B55 | B151 | TK | Anti-CCP | -0.47494673 | 0 |
| B55 | B156 | TK | IgE | -0.58443017 | 0 |
| B55 | B184 | TK | INR | -0.75603691 | 0 |
| B55 | B185 | TK | PT | -0.66802379 | 0 |
| B55 | B56 | TK | IR | 0.913873027 | 0 |
| B55 | B59 | TK | WBRV | 0.53584953 | 0 |
| B55 | B61 | TK | WBHSRV | 0.866322572 | 0 |
| B55 | B62 | TK | WBHSRVI | 0.620980942 | 0 |
| B55 | B63 | TK | WBRSR(mPas_200) | 0.489437186 | 0 |
| B55 | B66 | TK | WBRSR(mPas_100) | 0.487134729 | 0 |
| B55 | B67 | TK | WBRSR(mPas_50) | 0.455711952 | 0 |
| B55 | B79 | TK | hb V -dna | -0.50011517 | 0 |
| B55 | B92 | TK | ALDH2-Glu504Glu | -0.55736182 | 0 |
| B55 | B93 | TK | ALDH2-Glu504Lys | 0.592658712 | 0 |
| B55 | B94 | TK | ALDH2-Lys504Lys | -0.56067198 | 0 |
| B55 | B95 | TK | anti-cenpb | -0.48238707 | 0 |

| | | | | | |
|---|---|---|---|---|---|
| B55 | B97 | TK | dsDNA | -0.48238707 | 0 |
| B55 | B98 | TK | anti-Jo-1 | -0.48238707 | 0 |
| B55 | B99 | TK | anti-PM-Scl | -0.48238706 | 0 |
| B55 | B178 | TK | Ig-U | -0.43948821 | 2.22E-16 |
| B55 | B180 | TK | Tf | -0.40928355 | 2.84E-14 |
| B55 | B57 | TK | RBCAI | -0.34866845 | 1.61E-10 |
| B55 | B81 | TK | TSGF | 0.331992741 | 1.28E-09 |
| B55 | B154 | TK | RF | -0.32796641 | 2.07E-09 |
| B55 | B64 | TK | WBRSR(mPas_5) | 0.317538557 | 7.01E-09 |
| B55 | B182 | TK | TT | 0.298260569 | 5.90E-08 |
| B55 | B183 | TK | PTA | -0.29021103 | 1.37E-07 |
| B56 | B100 | IR | anti-Ro52 | -0.45070213 | 0 |
| B56 | B101 | IR | anti-Sc70 | -0.45070213 | 0 |
| B56 | B102 | IR | anti-Sm | -0.45070213 | 0 |
| B56 | B103 | IR | anti-SSA/Ro | -0.45070213 | 0 |
| B56 | B104 | IR | anti-SSB | -0.45070213 | 0 |
| B56 | B105 | IR | anti-U1RNP | -0.45070213 | 0 |
| B56 | B106 | IR | anti-P | -0.45070213 | 0 |
| B56 | B107 | IR | AnuA | -0.45070213 | 0 |
| B56 | B108 | IR | Anti-Cyclin | -0.45070213 | 0 |
| B56 | B109 | IR | AMA-M2 | -0.45070213 | 0 |
| B56 | B110 | IR | AHA | -0.45070213 | 0 |
| B56 | B151 | IR | Anti-CCP | -0.4439355 | 0 |
| B56 | B156 | IR | IgE | -0.50552841 | 0 |
| B56 | B178 | IR | Ig-U | -0.4474319 | 0 |
| B56 | B184 | IR | INR | -0.73734556 | 0 |
| B56 | B185 | IR | PT | -0.68086459 | 0 |
| B56 | B58 | IR | HCT | 0.520497407 | 0 |
| B56 | B59 | IR | WBRV | 0.702513463 | 0 |
| B56 | B60 | IR | WBRVI | 0.555886674 | 0 |
| B56 | B61 | IR | WBHSRV | 0.966021408 | 0 |
| B56 | B62 | IR | WBHSRVI | 0.880283392 | 0 |
| B56 | B63 | IR | WBRSR(mPas_200) | 0.761781409 | 0 |
| B56 | B64 | IR | WBRSR(mPas_5) | 0.599241935 | 0 |
| B56 | B65 | IR | WBRSR(mPas_1) | 0.530191926 | 0 |
| B56 | B66 | IR | WBRSR(mPas_100) | 0.756306412 | 0 |
| B56 | B67 | IR | WBRSR(mPas_50) | 0.731033291 | 0 |
| B56 | B79 | IR | hb V -dna | -0.51325512 | 0 |
| B56 | B92 | IR | ALDH2-Glu504Glu | -0.61637245 | 0 |
| B56 | B93 | IR | ALDH2-Glu504Lys | 0.65265477 | 0 |
| B56 | B94 | IR | ALDH2-Lys504Lys | -0.57061899 | 0 |
| B56 | B95 | IR | anti-cenpb | -0.45070213 | 0 |
| B56 | B97 | IR | dsDNA | -0.45070213 | 0 |
| B56 | B98 | IR | anti-Jo-1 | -0.45070213 | 0 |

| | | | | | |
|---|---|---|---|---|---|
| B56 | B99 | IR | anti-PM-Scl | -0.45070213 | 0 |
| B56 | B180 | IR | Tf | -0.41817552 | 6.88E-15 |
| B56 | B81 | IR | TSGF | 0.340114654 | 4.73E-10 |
| B57 | B100 | RBCAI | anti-Ro52 | 0.484457407 | 0 |
| B57 | B101 | RBCAI | anti-Sc70 | 0.484457408 | 0 |
| B57 | B102 | RBCAI | anti-Sm | 0.484457407 | 0 |
| B57 | B103 | RBCAI | anti-SSA/Ro | 0.484457407 | 0 |
| B57 | B104 | RBCAI | anti-SSB | 0.484457407 | 0 |
| B57 | B105 | RBCAI | anti-U1RNP | 0.484457407 | 0 |
| B57 | B106 | RBCAI | anti-P | 0.484457407 | 0 |
| B57 | B107 | RBCAI | AnuA | 0.484457401 | 0 |
| B57 | B108 | RBCAI | Anti-Cyclin | 0.484457407 | 0 |
| B57 | B109 | RBCAI | AMA-M2 | 0.484457407 | 0 |
| B57 | B110 | RBCAI | AHA | 0.484457407 | 0 |
| B57 | B151 | RBCAI | Anti-CCP | 0.495287936 | 0 |
| B57 | B178 | RBCAI | Ig-U | 0.517143354 | 0 |
| B57 | B179 | RBCAI | α1-MG | 0.48953876 | 0 |
| B57 | B180 | RBCAI | Tf | 0.468563991 | 0 |
| B57 | B58 | RBCAI | HCT | 0.458750924 | 0 |
| B57 | B59 | RBCAI | WBRV | 0.553994626 | 0 |
| B57 | B60 | RBCAI | WBRVI | 0.703072189 | 0 |
| B57 | B64 | RBCAI | WBRSR(mPas_5) | 0.607991695 | 0 |
| B57 | B65 | RBCAI | WBRSR(mPas_1) | 0.677527692 | 0 |
| B57 | B94 | RBCAI | ALDH2-Lys504Lys | 0.455319477 | 0 |
| B57 | B95 | RBCAI | anti-cenpb | 0.484457407 | 0 |
| B57 | B97 | RBCAI | dsDNA | 0.484457406 | 0 |
| B57 | B98 | RBCAI | anti-Jo-1 | 0.484457407 | 0 |
| B57 | B99 | RBCAI | anti-PM-Scl | 0.484457406 | 0 |
| B57 | B79 | RBCAI | hb V -dna | 0.438467924 | 2.22E-16 |
| B57 | B69 | RBCAI | PV | 0.434666678 | 4.44E-16 |
| B57 | B67 | RBCAI | WBRSR(mPas_50) | 0.425345731 | 2.22E-15 |
| B57 | B158 | RBCAI | IgM | -0.41064455 | 2.29E-14 |
| B57 | B66 | RBCAI | WBRSR(mPas_100) | 0.376110437 | 4.01E-12 |
| B57 | B156 | RBCAI | IgE | 0.371925905 | 7.20E-12 |
| B57 | B63 | RBCAI | WBRSR(mPas_200) | 0.36780602 | 1.27E-11 |
| B57 | B157 | RBCAI | IgG | 0.362721666 | 2.53E-11 |
| B57 | B90 | RBCAI | CYFRA 21-1 | -0.35833631 | 4.55E-11 |
| B57 | B155 | RBCAI | IgA | 0.354419713 | 7.63E-11 |
| B57 | B154 | RBCAI | RF | 0.314934631 | 9.43E-09 |
| B57 | B184 | RBCAI | INR | 0.270775909 | 9.51E-07 |
| B58 | B59 | HCT | WBRV | 0.611133444 | 0 |
| B58 | B60 | HCT | WBRVI | 0.853327718 | 0 |
| B58 | B61 | HCT | WBHSRV | 0.542280492 | 0 |
| B58 | B62 | HCT | WBHSRVI | 0.815656052 | 0 |

| | | | | | |
|---|---|---|---|---|---|
| B58 | B63 | HCT | WBRSR(mPas_200) | 0.811170945 | 0 |
| B58 | B64 | HCT | WBRSR(mPas_5) | 0.78572606 | 0 |
| B58 | B65 | HCT | WBRSR(mPas_1) | 0.787191 | 0 |
| B58 | B66 | HCT | WBRSR(mPas_100) | 0.802778804 | 0 |
| B58 | B67 | HCT | WBRSR(mPas_50) | 0.808796848 | 0 |
| B58 | B92 | HCT | ALDH2-Glu504Glu | -0.44063063 | 0 |
| B58 | B93 | HCT | ALDH2-Glu504Lys | 0.453504359 | 0 |
| B58 | B155 | HCT | IgA | -0.42360857 | 2.66E-15 |
| B58 | B185 | HCT | PT | -0.29544211 | 7.96E-08 |
| B59 | B185 | WBRV | PT | -0.55635177 | 0 |
| B59 | B60 | WBRV | WBRVI | 0.856565903 | 0 |
| B59 | B61 | WBRV | WBHSRV | 0.82222459 | 0 |
| B59 | B62 | WBRV | WBHSRVI | 0.738334888 | 0 |
| B59 | B63 | WBRV | WBRSR(mPas_200) | 0.871901034 | 0 |
| B59 | B64 | WBRV | WBRSR(mPas_5) | 0.911434055 | 0 |
| B59 | B65 | WBRV | WBRSR(mPas_1) | 0.879366271 | 0 |
| B59 | B66 | WBRV | WBRSR(mPas_100) | 0.879377723 | 0 |
| B59 | B67 | WBRV | WBRSR(mPas_50) | 0.891242559 | 0 |
| B59 | B92 | WBRV | ALDH2-Glu504Glu | -0.66502903 | 0 |
| B59 | B93 | WBRV | ALDH2-Glu504Lys | 0.67686024 | 0 |
| B59 | B69 | WBRV | PV | 0.438705305 | 2.22E-16 |
| B59 | B184 | WBRV | INR | -0.38577432 | 1.00E-12 |
| B59 | B158 | WBRV | IgM | -0.35187844 | 1.06E-10 |
| B59 | B124 | WBRV | ACTH | 0.285872389 | 2.14E-07 |
| B59 | B179 | WBRV | α1-MG | 0.28438483 | 2.49E-07 |
| B6 | B137 | WB-Fe | AFU | 0.452850717 | 0 |
| B6 | B138 | WB-Fe | PAB | 0.52045028 | 0 |
| B6 | B16 | WB-Fe | CD3+T | 0.659409978 | 0 |
| B6 | B21 | WB-Fe | CD3+CD4-CD8- | 0.514334406 | 0 |
| B6 | B23 | WB-Fe | CD3-CD16+CD56+(NK) | -0.62509392 | 0 |
| B6 | B7 | WB-Fe | WB-Cop | 1 | 0 |
| B6 | B76 | WB-Fe | ApoB | 0.443255337 | 0 |
| B6 | B8 | WB-Fe | WB-Zinc | 1 | 0 |
| B6 | B94 | WB-Fe | ALDH2-Lys504Lys | -0.49762844 | 0 |
| B6 | B15 | WB-Fe | CD3-CD19+(B cell) | -0.39394978 | 3.00E-13 |
| B6 | B179 | WB-Fe | α1-MG | -0.38747121 | 7.82E-13 |
| B6 | B178 | WB-Fe | Ig-U | -0.38072155 | 2.08E-12 |
| B6 | B57 | WB-Fe | RBCAI | -0.37149229 | 7.64E-12 |
| B6 | B85 | WB-Fe | SCC | -0.36823304 | 1.20E-11 |
| B6 | B92 | WB-Fe | ALDH2-Glu504Glu | 0.364342135 | 2.04E-11 |
| B6 | B75 | WB-Fe | ApoA1/B | -0.35273649 | 9.50E-11 |
| B6 | B180 | WB-Fe | Tf | -0.34443905 | 2.75E-10 |
| B6 | B93 | WB-Fe | ALDH2-Glu504Lys | -0.33929588 | 5.23E-10 |
| B6 | B90 | WB-Fe | CYFRA 21-1 | 0.312457974 | 1.25E-08 |

| | | | | | |
|---|---|---|---|---|---|
| B6 | B176 | WB-Fe | Ucr | 0.295282965 | 8.09E-08 |
| B6 | B200 | WB-Fe | OP | 0.291810317 | 1.16E-07 |
| B6 | B134 | WB-Fe | CHE | 0.29147904 | 1.21E-07 |
| B6 | B195 | WB-Fe | Na | 0.29052759 | 1.33E-07 |
| B6 | B198 | WB-Fe | P | -0.27659081 | 5.42E-07 |
| B60 | B61 | WBRVI | WBHSRV | 0.625919965 | 0 |
| B60 | B62 | WBRVI | WBHSRVI | 0.813815846 | 0 |
| B60 | B63 | WBRVI | WBRSR(mPas_200) | 0.882598303 | 0 |
| B60 | B64 | WBRVI | WBRSR(mPas_5) | 0.946030663 | 0 |
| B60 | B65 | WBRVI | WBRSR(mPas_1) | 0.965621928 | 0 |
| B60 | B66 | WBRVI | WBRSR(mPas_100) | 0.88344806 | 0 |
| B60 | B67 | WBRVI | WBRSR(mPas_50) | 0.903585544 | 0 |
| B60 | B92 | WBRVI | ALDH2-Glu504Glu | -0.46509955 | 0 |
| B60 | B93 | WBRVI | ALDH2-Glu504Lys | 0.471235996 | 0 |
| B60 | B185 | WBRVI | PT | -0.34038153 | 4.57E-10 |
| B60 | B199 | WBRVI | Mg | 0.273326099 | 7.44E-07 |
| B61 | B156 | WBHSRV | IgE | -0.44240159 | 0 |
| B61 | B184 | WBHSRV | INR | -0.65004671 | 0 |
| B61 | B185 | WBHSRV | PT | -0.67389552 | 0 |
| B61 | B62 | WBHSRV | WBHSRVI | 0.86833035 | 0 |
| B61 | B63 | WBHSRV | WBRSR(mPas_200) | 0.851806091 | 0 |
| B61 | B64 | WBHSRV | WBRSR(mPas_5) | 0.723031509 | 0 |
| B61 | B65 | WBHSRV | WBRSR(mPas_1) | 0.646731945 | 0 |
| B61 | B66 | WBHSRV | WBRSR(mPas_100) | 0.851208047 | 0 |
| B61 | B67 | WBHSRV | WBRSR(mPas_50) | 0.831425963 | 0 |
| B61 | B92 | WBHSRV | ALDH2-Glu504Glu | -0.65447069 | 0 |
| B61 | B93 | WBHSRV | ALDH2-Glu504Lys | 0.687104331 | 0 |
| B61 | B94 | WBHSRV | ALDH2-Lys504Lys | -0.50009272 | 0 |
| B61 | B79 | WBHSRV | hb V -dna | -0.41338243 | 1.47E-14 |
| B61 | B100 | WBHSRV | anti-Ro52 | -0.40094254 | 1.04E-13 |
| B61 | B101 | WBHSRV | anti-Sc70 | -0.40094254 | 1.04E-13 |
| B61 | B102 | WBHSRV | anti-Sm | -0.40094254 | 1.04E-13 |
| B61 | B103 | WBHSRV | anti-SSA/Ro | -0.40094254 | 1.04E-13 |
| B61 | B104 | WBHSRV | anti-SSB | -0.40094254 | 1.04E-13 |
| B61 | B105 | WBHSRV | anti-U1RNP | -0.40094254 | 1.04E-13 |
| B61 | B106 | WBHSRV | anti-P | -0.40094254 | 1.04E-13 |
| B61 | B107 | WBHSRV | AnuA | -0.40094254 | 1.04E-13 |
| B61 | B108 | WBHSRV | Anti-Cyclin | -0.40094254 | 1.04E-13 |
| B61 | B109 | WBHSRV | AMA-M2 | -0.40094254 | 1.04E-13 |
| B61 | B110 | WBHSRV | AHA | -0.40094254 | 1.04E-13 |
| B61 | B95 | WBHSRV | anti-cenpb | -0.40094254 | 1.04E-13 |
| B61 | B97 | WBHSRV | dsDNA | -0.40094254 | 1.04E-13 |
| B61 | B98 | WBHSRV | anti-Jo-1 | -0.40094254 | 1.04E-13 |
| B61 | B99 | WBHSRV | anti-PM-Scl | -0.40094254 | 1.04E-13 |

| | | | | | |
|---|---|---|---|---|---|
| B61 | B151 | WBHSRV | Anti-CCP | -0.39008491 | 5.32E-13 |
| B61 | B178 | WBHSRV | Ig-U | -0.35003561 | 1.35E-10 |
| B61 | B180 | WBHSRV | Tf | -0.32960749 | 1.70E-09 |
| B61 | B81 | WBHSRV | TSGF | 0.286518233 | 2.01E-07 |
| B62 | B184 | WBHSRVI | INR | -0.54090551 | 0 |
| B62 | B185 | WBHSRVI | PT | -0.50594891 | 0 |
| B62 | B63 | WBHSRVI | WBRSR(mPas_200) | 0.92009837 | 0 |
| B62 | B64 | WBHSRVI | WBRSR(mPas_5) | 0.807806445 | 0 |
| B62 | B65 | WBHSRVI | WBRSR(mPas_1) | 0.77760547 | 0 |
| B62 | B66 | WBHSRVI | WBRSR(mPas_100) | 0.912968149 | 0 |
| B62 | B67 | WBHSRVI | WBRSR(mPas_50) | 0.900415643 | 0 |
| B62 | B92 | WBHSRVI | ALDH2-Glu504Glu | -0.48695632 | 0 |
| B62 | B93 | WBHSRVI | ALDH2-Glu504Lys | 0.515923919 | 0 |
| B62 | B94 | WBHSRVI | ALDH2-Lys504Lys | -0.45625559 | 0 |
| B62 | B79 | WBHSRVI | hb V -dna | -0.41150801 | 2.00E-14 |
| B62 | B178 | WBHSRVI | Ig-U | -0.35914237 | 4.09E-11 |
| B62 | B155 | WBHSRVI | IgA | -0.34348134 | 3.10E-10 |
| B62 | B180 | WBHSRVI | Tf | -0.33648776 | 7.40E-10 |
| B62 | B100 | WBHSRVI | anti-Ro52 | -0.30946045 | 1.74E-08 |
| B62 | B101 | WBHSRVI | anti-Sc70 | -0.30946045 | 1.74E-08 |
| B62 | B102 | WBHSRVI | anti-Sm | -0.30946045 | 1.74E-08 |
| B62 | B103 | WBHSRVI | anti-SSA/Ro | -0.30946045 | 1.74E-08 |
| B62 | B104 | WBHSRVI | anti-SSB | -0.30946045 | 1.74E-08 |
| B62 | B105 | WBHSRVI | anti-U1RNP | -0.30946045 | 1.74E-08 |
| B62 | B106 | WBHSRVI | anti-P | -0.30946045 | 1.74E-08 |
| B62 | B107 | WBHSRVI | AnuA | -0.30946045 | 1.74E-08 |
| B62 | B108 | WBHSRVI | Anti-Cyclin | -0.30946045 | 1.74E-08 |
| B62 | B109 | WBHSRVI | AMA-M2 | -0.30946045 | 1.74E-08 |
| B62 | B110 | WBHSRVI | AHA | -0.30946045 | 1.74E-08 |
| B62 | B95 | WBHSRVI | anti-cenpb | -0.30946045 | 1.74E-08 |
| B62 | B97 | WBHSRVI | dsDNA | -0.30946045 | 1.74E-08 |
| B62 | B98 | WBHSRVI | anti-Jo-1 | -0.30946045 | 1.74E-08 |
| B62 | B99 | WBHSRVI | anti-PM-Scl | -0.30946045 | 1.74E-08 |
| B62 | B156 | WBHSRVI | IgE | -0.30840065 | 1.96E-08 |
| B62 | B151 | WBHSRVI | Anti-CCP | -0.30488052 | 2.89E-08 |
| B62 | B81 | WBHSRVI | TSGF | 0.270867887 | 9.43E-07 |
| B63 | B185 | WBRSR(mPas_200) | PT | -0.45548261 | 0 |
| B63 | B64 | WBRSR(mPas_200) | WBRSR(mPas_5) | 0.949475069 | 0 |
| B63 | B65 | WBRSR(mPas_200) | WBRSR(mPas_1) | 0.913976765 | 0 |
| B63 | B66 | WBRSR(mPas_200) | WBRSR(mPas_100) | 0.99861291 | 0 |
| B63 | B67 | WBRSR(mPas_200) | WBRSR(mPas_50) | 0.996425316 | 0 |
| B63 | B92 | WBRSR(mPas_200) | ALDH2-Glu504Glu | -0.50859719 | 0 |
| B63 | B93 | WBRSR(mPas_200) | ALDH2-Glu504Lys | 0.529174643 | 0 |
| B63 | B184 | WBRSR(mPas_200) | INR | -0.35989678 | 3.70E-11 |

| | | | | | |
|---|---|---|---|---|---|
| B63 | B69 | WBRSR(mPas_200) | PV | 0.338866835 | 5.52E-10 |
| B63 | B94 | WBRSR(mPas_200) | ALDH2-Lys504Lys | -0.30274265 | 3.65E-08 |
| B64 | B65 | WBRSR(mPas_5) | WBRSR(mPas_1) | 0.984730042 | 0 |
| B64 | B66 | WBRSR(mPas_5) | WBRSR(mPas_100) | 0.954939113 | 0 |
| B64 | B67 | WBRSR(mPas_5) | WBRSR(mPas_50) | 0.968542421 | 0 |
| B64 | B69 | WBRSR(mPas_5) | PV | 0.452306682 | 0 |
| B64 | B92 | WBRSR(mPas_5) | ALDH2-Glu504Glu | -0.49238576 | 0 |
| B64 | B93 | WBRSR(mPas_5) | ALDH2-Glu504Lys | 0.501674797 | 0 |
| B64 | B185 | WBRSR(mPas_5) | PT | -0.36976331 | 9.71E-12 |
| B65 | B66 | WBRSR(mPas_1) | WBRSR(mPas_100) | 0.919015368 | 0 |
| B65 | B67 | WBRSR(mPas_1) | WBRSR(mPas_50) | 0.937672755 | 0 |
| B65 | B93 | WBRSR(mPas_1) | ALDH2-Glu504Lys | 0.420343261 | 4.88E-15 |
| B65 | B92 | WBRSR(mPas_1) | ALDH2-Glu504Glu | -0.41298964 | 1.58E-14 |
| B65 | B69 | WBRSR(mPas_1) | PV | 0.410797816 | 2.22E-14 |
| B65 | B185 | WBRSR(mPas_1) | PT | -0.3078096 | 2.09E-08 |
| B66 | B185 | WBRSR(mPas_100) | PT | -0.45463781 | 0 |
| B66 | B67 | WBRSR(mPas_100) | WBRSR(mPas_50) | 0.99850823 | 0 |
| B66 | B92 | WBRSR(mPas_100) | ALDH2-Glu504Glu | -0.51032169 | 0 |
| B66 | B93 | WBRSR(mPas_100) | ALDH2-Glu504Lys | 0.530620321 | 0 |
| B66 | B184 | WBRSR(mPas_100) | INR | -0.36097927 | 3.20E-11 |
| B66 | B69 | WBRSR(mPas_100) | PV | 0.357887479 | 4.83E-11 |
| B66 | B94 | WBRSR(mPas_100) | ALDH2-Lys504Lys | -0.29750912 | 6.40E-08 |
| B67 | B185 | WBRSR(mPas_50) | PT | -0.44165738 | 0 |
| B67 | B92 | WBRSR(mPas_50) | ALDH2-Glu504Glu | -0.50979087 | 0 |
| B67 | B93 | WBRSR(mPas_50) | ALDH2-Glu504Lys | 0.528155573 | 0 |
| B67 | B69 | WBRSR(mPas_50) | PV | 0.375601019 | 4.30E-12 |
| B67 | B184 | WBRSR(mPas_50) | INR | -0.3355482 | 8.30E-10 |
| B69 | B123 | PV | hs-CRP1 | 0.744715887 | 0 |
| B69 | B177 | PV | ACR | 0.474885813 | 0 |
| B69 | B181 | PV | APTT | -0.44739694 | 0 |
| B69 | B186 | PV | PFC | 0.44573522 | 0 |
| B69 | B83 | PV | AFP | 0.382809593 | 1.54E-12 |
| B69 | B194 | PV | Cl | -0.38129091 | 1.92E-12 |
| B69 | B157 | PV | IgG | 0.340155443 | 4.70E-10 |
| B69 | B175 | PV | ALB-U | 0.33261087 | 1.19E-09 |
| B69 | B178 | PV | Ig-U | 0.331914848 | 1.29E-09 |
| B69 | B179 | PV | α1-MG | 0.329419199 | 1.74E-09 |
| B69 | B79 | PV | hb V -dna | 0.327917634 | 2.08E-09 |
| B69 | B180 | PV | Tf | 0.30190306 | 3.99E-08 |
| B69 | B184 | PV | INR | 0.292347037 | 1.10E-07 |
| B69 | B155 | PV | IgA | 0.273204023 | 7.53E-07 |
| B7 | B137 | WB-Cop | AFU | 0.452850717 | 0 |
| B7 | B138 | WB-Cop | PAB | 0.52045028 | 0 |
| B7 | B16 | WB-Cop | CD3+T | 0.659409978 | 0 |

| | | | | | |
|---|---|---|---|---|---|
| B7 | B21 | WB-Cop | CD3+CD4-CD8- | 0.514334406 | 0 |
| B7 | B23 | WB-Cop | CD3-CD16+CD56+(NK) | -0.62509392 | 0 |
| B7 | B76 | WB-Cop | ApoB | 0.443255337 | 0 |
| B7 | B8 | WB-Cop | WB-Zinc | 1 | 0 |
| B7 | B94 | WB-Cop | ALDH2-Lys504Lys | -0.49762844 | 0 |
| B7 | B15 | WB-Cop | CD3-CD19+(B cell) | -0.39394978 | 3.00E-13 |
| B7 | B179 | WB-Cop | α1-MG | -0.38747121 | 7.82E-13 |
| B7 | B178 | WB-Cop | Ig-U | -0.38072155 | 2.08E-12 |
| B7 | B57 | WB-Cop | RBCAI | -0.3714923 | 7.64E-12 |
| B7 | B85 | WB-Cop | SCC | -0.36823304 | 1.20E-11 |
| B7 | B92 | WB-Cop | ALDH2-Glu504Glu | 0.364342135 | 2.04E-11 |
| B7 | B75 | WB-Cop | ApoA1/B | -0.35273649 | 9.50E-11 |
| B7 | B180 | WB-Cop | Tf | -0.34443905 | 2.75E-10 |
| B7 | B93 | WB-Cop | ALDH2-Glu504Lys | -0.33929588 | 5.23E-10 |
| B7 | B90 | WB-Cop | CYFRA 21-1 | 0.312457974 | 1.25E-08 |
| B7 | B176 | WB-Cop | Ucr | 0.295282965 | 8.09E-08 |
| B7 | B200 | WB-Cop | OP | 0.291810316 | 1.16E-07 |
| B7 | B134 | WB-Cop | CHE | 0.29147904 | 1.21E-07 |
| B7 | B195 | WB-Cop | Na | 0.29052759 | 1.33E-07 |
| B7 | B198 | WB-Cop | P | -0.27659081 | 5.42E-07 |
| B70 | B77 | LDL-C | TC | 0.755437877 | 0 |
| B71 | B136 | TG | 5'NT | 0.443158291 | 0 |
| B71 | B72 | TG | HDL-C | -0.47491012 | 0 |
| B71 | B150 | TG | ALP | 0.378885339 | 2.70E-12 |
| B71 | B77 | TG | TC | 0.367596049 | 1.31E-11 |
| B71 | B153 | TG | Glu | 0.359626941 | 3.84E-11 |
| B71 | B91 | TG | TP | 0.352810433 | 9.41E-11 |
| B71 | B140 | TG | r-GT | 0.343501692 | 3.10E-10 |
| B71 | B206 | TG | Waist | 0.302559475 | 3.72E-08 |
| B71 | B222 | TG | NS | 0.299361347 | 5.25E-08 |
| B71 | B215 | TG | BMI | 0.28916963 | 1.53E-07 |
| B71 | B216 | TG | BMI | 0.28368688 | 2.67E-07 |
| B72 | B206 | HDL-C | Waist | -0.47757782 | 0 |
| B72 | B216 | HDL-C | BMI | -0.51852346 | 0 |
| B72 | B222 | HDL-C | NS | -0.50554224 | 0 |
| B72 | B215 | HDL-C | BMI | -0.42961007 | 8.88E-16 |
| B72 | B192 | HDL-C | UA | -0.40379493 | 6.68E-14 |
| B72 | B221 | HDL-C | Hei | -0.35641631 | 5.87E-11 |
| B72 | B210 | HDL-C | Hip | -0.3411968 | 4.13E-10 |
| B72 | B227 | HDL-C | OD | 0.281476334 | 3.34E-07 |
| B72 | B141 | HDL-C | AST/ALT | 0.278233133 | 4.61E-07 |
| B73 | B124 | ApoA | ACTH | 0.562557594 | 0 |
| B73 | B76 | ApoA | ApoB | 0.282507549 | 3.01E-07 |
| B74 | B75 | ApoA1 | ApoA1/B | 0.547071489 | 0 |

| | | | | | |
|---|---|---|---|---|---|
| B74 | B198 | ApoA1 | P | 0.29589142 | 7.59E-08 |
| B75 | B76 | ApoA1/B | ApoB | -0.71059925 | 0 |
| B75 | B134 | ApoA1/B | CHE | -0.42764818 | 1.33E-15 |
| B75 | B135 | ApoA1/B | TBA | 0.341571462 | 3.94E-10 |
| B75 | B124 | ApoA1/B | ACTH | -0.3282914 | 1.99E-09 |
| B76 | B124 | ApoB | ACTH | 0.425023673 | 2.22E-15 |
| B76 | B134 | ApoB | CHE | 0.39705872 | 1.88E-13 |
| B76 | B85 | ApoB | SCC | -0.33870682 | 5.63E-10 |
| B76 | B138 | ApoB | PAB | 0.330205465 | 1.59E-09 |
| B77 | B150 | TC | ALP | 0.309442637 | 1.75E-08 |
| B77 | B136 | TC | 5'NT | 0.298928159 | 5.50E-08 |
| B77 | B91 | TC | TP | 0.276405283 | 5.51E-07 |
| B77 | B153 | TC | Glu | 0.272670836 | 7.93E-07 |
| B79 | B100 | hb V -dna | anti-Ro52 | 0.725623807 | 0 |
| B79 | B101 | hb V -dna | anti-Sc70 | 0.725623808 | 0 |
| B79 | B102 | hb V -dna | anti-Sm | 0.725623808 | 0 |
| B79 | B103 | hb V -dna | anti-SSA/Ro | 0.725623807 | 0 |
| B79 | B104 | hb V -dna | anti-SSB | 0.725623808 | 0 |
| B79 | B105 | hb V -dna | anti-U1RNP | 0.725623807 | 0 |
| B79 | B106 | hb V -dna | anti-P | 0.725623807 | 0 |
| B79 | B107 | hb V -dna | AnuA | 0.725623799 | 0 |
| B79 | B108 | hb V -dna | Anti-Cyclin | 0.725623807 | 0 |
| B79 | B109 | hb V -dna | AMA-M2 | 0.725623807 | 0 |
| B79 | B110 | hb V -dna | AHA | 0.725623807 | 0 |
| B79 | B119 | hb V -dna | Hcv-rna | 0.629446092 | 0 |
| B79 | B121 | hb V -dna | C3 | -0.62944611 | 0 |
| B79 | B122 | hb V -dna | C4 | -0.62944631 | 0 |
| B79 | B151 | hb V -dna | Anti-CCP | 0.549392706 | 0 |
| B79 | B152 | hb V -dna | ACA-Ig(G-A-M) | -0.62944776 | 0 |
| B79 | B156 | hb V -dna | IgE | 0.541949567 | 0 |
| B79 | B157 | hb V -dna | IgG | 0.440503925 | 0 |
| B79 | B159 | hb V -dna | β2-MG | -0.62944684 | 0 |
| B79 | B169 | hb V -dna | U-CRY | -0.62944542 | 0 |
| B79 | B178 | hb V -dna | Ig-U | 0.498776722 | 0 |
| B79 | B180 | hb V -dna | Tf | 0.456474597 | 0 |
| B79 | B184 | hb V -dna | INR | 0.573640307 | 0 |
| B79 | B95 | hb V -dna | anti-cenpb | 0.725623808 | 0 |
| B79 | B97 | hb V -dna | dsDNA | 0.725623806 | 0 |
| B79 | B98 | hb V -dna | anti-Jo-1 | 0.725623806 | 0 |
| B79 | B99 | hb V -dna | anti-PM-Scl | 0.725623806 | 0 |
| B79 | B94 | hb V -dna | ALDH2-Lys504Lys | 0.439368322 | 2.22E-16 |
| B79 | B155 | hb V -dna | IgA | 0.36157392 | 2.96E-11 |
| B79 | B179 | hb V -dna | α1-MG | 0.352195286 | 1.02E-10 |
| B79 | B81 | hb V -dna | TSGF | -0.35064557 | 1.25E-10 |

| | | | | | |
|---|---|---|---|---|---|
| B79 | B158 | hb V -dna | IgM | -0.31427123 | 1.02E-08 |
| B79 | B185 | hb V -dna | PT | 0.313267756 | 1.14E-08 |
| B79 | B183 | hb V -dna | PTA | 0.306927622 | 2.31E-08 |
| B79 | B181 | hb V -dna | APTT | -0.30643365 | 2.44E-08 |
| B8 | B137 | WB-Zinc | AFU | 0.452850717 | 0 |
| B8 | B138 | WB-Zinc | PAB | 0.52045028 | 0 |
| B8 | B16 | WB-Zinc | CD3+T | 0.659409978 | 0 |
| B8 | B21 | WB-Zinc | CD3+CD4-CD8- | 0.514334406 | 0 |
| B8 | B23 | WB-Zinc | CD3-CD16+CD56+(NK) | -0.62509392 | 0 |
| B8 | B76 | WB-Zinc | ApoB | 0.443255337 | 0 |
| B8 | B94 | WB-Zinc | ALDH2-Lys504Lys | -0.49762845 | 0 |
| B8 | B15 | WB-Zinc | CD3-CD19+(B cell) | -0.39394978 | 3.00E-13 |
| B8 | B179 | WB-Zinc | α1-MG | -0.38747121 | 7.82E-13 |
| B8 | B178 | WB-Zinc | Ig-U | -0.38072155 | 2.08E-12 |
| B8 | B57 | WB-Zinc | RBCAI | -0.3714923 | 7.64E-12 |
| B8 | B85 | WB-Zinc | SCC | -0.36823304 | 1.20E-11 |
| B8 | B92 | WB-Zinc | ALDH2-Glu504Glu | 0.364342135 | 2.04E-11 |
| B8 | B75 | WB-Zinc | ApoA1/B | -0.35273649 | 9.50E-11 |
| B8 | B180 | WB-Zinc | Tf | -0.34443905 | 2.75E-10 |
| B8 | B93 | WB-Zinc | ALDH2-Glu504Lys | -0.33929588 | 5.23E-10 |
| B8 | B90 | WB-Zinc | CYFRA 21-1 | 0.312457974 | 1.25E-08 |
| B8 | B176 | WB-Zinc | Ucr | 0.295282965 | 8.09E-08 |
| B8 | B200 | WB-Zinc | OP | 0.291810316 | 1.16E-07 |
| B8 | B134 | WB-Zinc | CHE | 0.29147904 | 1.21E-07 |
| B8 | B195 | WB-Zinc | Na | 0.29052759 | 1.33E-07 |
| B8 | B198 | WB-Zinc | P | -0.27659081 | 5.42E-07 |
| B80 | B227 | anti-HP IgG | OD | 0.322921151 | 3.76E-09 |
| B81 | B100 | TSGF | anti-Ro52 | -0.43125203 | 8.88E-16 |
| B81 | B101 | TSGF | anti-Sc70 | -0.43125203 | 8.88E-16 |
| B81 | B102 | TSGF | anti-Sm | -0.43125203 | 8.88E-16 |
| B81 | B103 | TSGF | anti-SSA/Ro | -0.43125203 | 8.88E-16 |
| B81 | B104 | TSGF | anti-SSB | -0.43125203 | 8.88E-16 |
| B81 | B105 | TSGF | anti-U1RNP | -0.43125203 | 8.88E-16 |
| B81 | B106 | TSGF | anti-P | -0.43125203 | 8.88E-16 |
| B81 | B107 | TSGF | AnuA | -0.43125203 | 8.88E-16 |
| B81 | B108 | TSGF | Anti-Cyclin | -0.43125203 | 8.88E-16 |
| B81 | B109 | TSGF | AMA-M2 | -0.43125203 | 8.88E-16 |
| B81 | B110 | TSGF | AHA | -0.43125203 | 8.88E-16 |
| B81 | B95 | TSGF | anti-cenpb | -0.43125203 | 8.88E-16 |
| B81 | B97 | TSGF | dsDNA | -0.43125203 | 8.88E-16 |
| B81 | B98 | TSGF | anti-Jo-1 | -0.43125203 | 8.88E-16 |
| B81 | B99 | TSGF | anti-PM-Scl | -0.43125203 | 8.88E-16 |
| B81 | B180 | TSGF | Tf | -0.34985053 | 1.38E-10 |
| B81 | B184 | TSGF | INR | -0.34630417 | 2.17E-10 |

| | | | | | |
|---|---|---|---|---|---|
| B81 | B178 | TSGF | Ig-U | -0.34367174 | 3.03E-10 |
| B81 | B151 | TSGF | Anti-CCP | -0.32207176 | 4.15E-09 |
| B81 | B158 | TSGF | IgM | 0.315768681 | 8.58E-09 |
| B81 | B94 | TSGF | ALDH2-Lys504Lys | -0.29245471 | 1.09E-07 |
| B81 | B157 | TSGF | IgG | -0.2706861 | 9.59E-07 |
| B82 | B88 | CEA | CA 242 | 0.460104663 | 0 |
| B82 | B200 | CEA | OP | 0.433875678 | 4.44E-16 |
| B82 | B87 | CEA | NSE | -0.32878394 | 1.88E-09 |
| B82 | B195 | CEA | Na | 0.29463942 | 8.66E-08 |
| B83 | B123 | AFP | hs-CRP1 | 0.452510401 | 0 |
| B83 | B181 | AFP | APTT | -0.53464925 | 0 |
| B83 | B186 | AFP | PFC | 0.775203324 | 0 |
| B83 | B193 | AFP | K | -0.59387799 | 0 |
| B83 | B194 | AFP | Cl | -0.68821901 | 0 |
| B83 | B199 | AFP | Mg | -0.65113069 | 0 |
| B83 | B197 | AFP | AG | 0.412532719 | 1.69E-14 |
| B83 | B89 | AFP | CA 72-4 | -0.40780974 | 3.57E-14 |
| B83 | B100 | AFP | anti-Ro52 | -0.37696214 | 3.55E-12 |
| B83 | B101 | AFP | anti-Sc70 | -0.37696214 | 3.55E-12 |
| B83 | B102 | AFP | anti-Sm | -0.37696214 | 3.55E-12 |
| B83 | B103 | AFP | anti-SSA/Ro | -0.37696214 | 3.55E-12 |
| B83 | B104 | AFP | anti-SSB | -0.37696214 | 3.55E-12 |
| B83 | B105 | AFP | anti-U1RNP | -0.37696214 | 3.55E-12 |
| B83 | B106 | AFP | anti-P | -0.37696214 | 3.55E-12 |
| B83 | B107 | AFP | AnuA | -0.37696214 | 3.55E-12 |
| B83 | B108 | AFP | Anti-Cyclin | -0.37696214 | 3.55E-12 |
| B83 | B109 | AFP | AMA-M2 | -0.37696214 | 3.55E-12 |
| B83 | B110 | AFP | AHA | -0.37696214 | 3.55E-12 |
| B83 | B95 | AFP | anti-cenpb | -0.37696214 | 3.55E-12 |
| B83 | B97 | AFP | dsDNA | -0.37696214 | 3.55E-12 |
| B83 | B98 | AFP | anti-Jo-1 | -0.37696214 | 3.55E-12 |
| B83 | B99 | AFP | anti-PM-Scl | -0.37696214 | 3.55E-12 |
| B83 | B151 | AFP | Anti-CCP | -0.3695418 | 1.00E-11 |
| B83 | B86 | AFP | TBhCG | -0.34216834 | 3.66E-10 |
| B84 | B85 | C-PSA | SCC | 0.476247364 | 0 |
| B84 | B86 | C-PSA | TBhCG | 0.682517908 | 0 |
| B84 | B88 | C-PSA | CA 242 | -0.46121772 | 0 |
| B84 | B89 | C-PSA | CA 72-4 | 0.811662665 | 0 |
| B84 | B186 | C-PSA | PFC | -0.38735061 | 7.96E-13 |
| B84 | B194 | C-PSA | Cl | 0.340332318 | 4.60E-10 |
| B84 | B199 | C-PSA | Mg | 0.319516186 | 5.58E-09 |
| B84 | B193 | C-PSA | K | 0.277510201 | 4.95E-07 |
| B85 | B186 | SCC | PFC | -0.34642373 | 2.14E-10 |
| B85 | B87 | SCC | NSE | 0.328513462 | 1.94E-09 |

| | | | | | |
|---|---|---|---|---|---|
| B85 | B138 | SCC | PAB | -0.29993395 | 4.94E-08 |
| B85 | B200 | SCC | OP | -0.29597409 | 7.52E-08 |
| B86 | B89 | TBhCG | CA 72-4 | 0.887323378 | 0 |
| B86 | B88 | TBhCG | CA 242 | -0.34742837 | 1.88E-10 |
| B86 | B186 | TBhCG | PFC | -0.31065351 | 1.53E-08 |
| B86 | B193 | TBhCG | K | 0.301843309 | 4.02E-08 |
| B86 | B87 | TBhCG | NSE | -0.2920283 | 1.14E-07 |
| B86 | B194 | TBhCG | Cl | 0.281758987 | 3.25E-07 |
| B87 | B89 | NSE | CA 72-4 | -0.49080507 | 0 |
| B87 | B161 | NSE | LEU | 0.362926624 | 2.47E-11 |
| B88 | B111 | CA 242 | TSH | 0.382893533 | 1.52E-12 |
| B88 | B89 | CA 242 | CA 72-4 | -0.38272012 | 1.56E-12 |
| B89 | B186 | CA 72-4 | PFC | -0.43003128 | 8.88E-16 |
| B89 | B194 | CA 72-4 | Cl | 0.377360764 | 3.36E-12 |
| B89 | B199 | CA 72-4 | Mg | 0.3532433 | 8.89E-11 |
| B89 | B193 | CA 72-4 | K | 0.306669234 | 2.37E-08 |
| B89 | B181 | CA 72-4 | APTT | 0.298815402 | 5.56E-08 |
| B9 | B38 | RH | BAND% | -0.50251891 | 0 |
| B90 | B100 | CYFRA 21-1 | anti-Ro52 | -0.38294936 | 1.51E-12 |
| B90 | B101 | CYFRA 21-1 | anti-Sc70 | -0.38294936 | 1.51E-12 |
| B90 | B102 | CYFRA 21-1 | anti-Sm | -0.38294936 | 1.51E-12 |
| B90 | B103 | CYFRA 21-1 | anti-SSA/Ro | -0.38294936 | 1.51E-12 |
| B90 | B104 | CYFRA 21-1 | anti-SSB | -0.38294936 | 1.51E-12 |
| B90 | B105 | CYFRA 21-1 | anti-U1RNP | -0.38294936 | 1.51E-12 |
| B90 | B106 | CYFRA 21-1 | anti-P | -0.38294936 | 1.51E-12 |
| B90 | B107 | CYFRA 21-1 | AnuA | -0.38294936 | 1.51E-12 |
| B90 | B108 | CYFRA 21-1 | Anti-Cyclin | -0.38294936 | 1.51E-12 |
| B90 | B109 | CYFRA 21-1 | AMA-M2 | -0.38294936 | 1.51E-12 |
| B90 | B110 | CYFRA 21-1 | AHA | -0.38294936 | 1.51E-12 |
| B90 | B95 | CYFRA 21-1 | anti-cenpb | -0.38294936 | 1.51E-12 |
| B90 | B97 | CYFRA 21-1 | dsDNA | -0.38294936 | 1.51E-12 |
| B90 | B98 | CYFRA 21-1 | anti-Jo-1 | -0.38294936 | 1.51E-12 |
| B90 | B99 | CYFRA 21-1 | anti-PM-Scl | -0.38294936 | 1.51E-12 |
| B90 | B178 | CYFRA 21-1 | Ig-U | -0.36222547 | 2.71E-11 |
| B90 | B180 | CYFRA 21-1 | Tf | -0.36213819 | 2.74E-11 |
| B90 | B179 | CYFRA 21-1 | α1-MG | -0.35593868 | 6.25E-11 |
| B90 | B134 | CYFRA 21-1 | CHE | 0.354298641 | 7.75E-11 |
| B90 | B199 | CYFRA 21-1 | Mg | -0.30469114 | 2.95E-08 |
| B90 | B154 | CYFRA 21-1 | RF | -0.28547013 | 2.23E-07 |
| B90 | B151 | CYFRA 21-1 | Anti-CCP | -0.28382542 | 2.64E-07 |
| B91 | B126 | TP | Alb | 0.536914334 | 0 |
| B91 | B127 | TP | A-G ratio | -0.49207434 | 0 |
| B91 | B130 | TP | GLB | 0.797848216 | 0 |
| B91 | B188 | TP | Ca | 0.331280484 | 1.39E-09 |

| | | | | | |
|---|---|---|---|---|---|
| B92 | B184 | ALDH2-Glu504Glu | INR | 0.443743864 | 0 |
| B92 | B185 | ALDH2-Glu504Glu | PT | 0.635909382 | 0 |
| B92 | B93 | ALDH2-Glu504Glu | ALDH2-Glu504Lys | -0.99848738 | 0 |
| B92 | B124 | ALDH2-Glu504Glu | ACTH | -0.37261755 | 6.54E-12 |
| B92 | B196 | ALDH2-Glu504Glu | TCO2 | -0.28557816 | 2.21E-07 |
| B92 | B137 | ALDH2-Glu504Glu | AFU | 0.285309647 | 2.27E-07 |
| B93 | B184 | ALDH2-Glu504Lys | INR | -0.47172352 | 0 |
| B93 | B185 | ALDH2-Glu504Lys | PT | -0.65556721 | 0 |
| B93 | B124 | ALDH2-Glu504Lys | ACTH | 0.365312443 | 1.79E-11 |
| B93 | B196 | ALDH2-Glu504Lys | TCO2 | 0.284593763 | 2.44E-07 |
| B93 | B137 | ALDH2-Glu504Lys | AFU | -0.27710825 | 5.15E-07 |
| B94 | B178 | ALDH2-Lys504Lys | Ig-U | 0.561320416 | 0 |
| B94 | B180 | ALDH2-Lys504Lys | Tf | 0.514330776 | 0 |
| B94 | B184 | ALDH2-Lys504Lys | INR | 0.444188575 | 0 |
| B94 | B100 | ALDH2-Lys504Lys | anti-Ro52 | 0.399120174 | 1.37E-13 |
| B94 | B101 | ALDH2-Lys504Lys | anti-Sc70 | 0.399120174 | 1.37E-13 |
| B94 | B102 | ALDH2-Lys504Lys | anti-Sm | 0.399120174 | 1.37E-13 |
| B94 | B103 | ALDH2-Lys504Lys | anti-SSA/Ro | 0.399120174 | 1.37E-13 |
| B94 | B104 | ALDH2-Lys504Lys | anti-SSB | 0.399120176 | 1.37E-13 |
| B94 | B105 | ALDH2-Lys504Lys | anti-U1RNP | 0.399120174 | 1.37E-13 |
| B94 | B106 | ALDH2-Lys504Lys | anti-P | 0.399120174 | 1.37E-13 |
| B94 | B107 | ALDH2-Lys504Lys | AnuA | 0.399120178 | 1.37E-13 |
| B94 | B108 | ALDH2-Lys504Lys | Anti-Cyclin | 0.399120175 | 1.37E-13 |
| B94 | B109 | ALDH2-Lys504Lys | AMA-M2 | 0.399120173 | 1.37E-13 |
| B94 | B110 | ALDH2-Lys504Lys | AHA | 0.399120173 | 1.37E-13 |
| B94 | B95 | ALDH2-Lys504Lys | anti-cenpb | 0.399120174 | 1.37E-13 |
| B94 | B97 | ALDH2-Lys504Lys | dsDNA | 0.399120174 | 1.37E-13 |
| B94 | B98 | ALDH2-Lys504Lys | anti-Jo-1 | 0.399120174 | 1.37E-13 |
| B94 | B99 | ALDH2-Lys504Lys | anti-PM-Scl | 0.399120175 | 1.37E-13 |
| B94 | B156 | ALDH2-Lys504Lys | IgE | 0.388677474 | 6.55E-13 |
| B94 | B151 | ALDH2-Lys504Lys | Anti-CCP | 0.388283322 | 6.95E-13 |
| B94 | B179 | ALDH2-Lys504Lys | α1-MG | 0.351375372 | 1.13E-10 |
| B94 | B204 | ALDH2-Lys504Lys | Drinking | -0.31798911 | 6.65E-09 |
| B94 | B181 | ALDH2-Lys504Lys | APTT | -0.31639598 | 7.98E-09 |
| B94 | B154 | ALDH2-Lys504Lys | RF | 0.304899056 | 2.88E-08 |
| B94 | B125 | ALDH2-Lys504Lys | Cor | 0.296747287 | 6.93E-08 |
| B94 | B198 | ALDH2-Lys504Lys | P | 0.296433808 | 7.17E-08 |
| B95 | B101 | anti-cenpb | anti-Sc70 | 0.999999999 | 0 |
| B95 | B106 | anti-cenpb | anti-P | 0.999999999 | 0 |
| B95 | B107 | anti-cenpb | AnuA | 0.99999999 | 0 |
| B95 | B108 | anti-cenpb | Anti-Cyclin | 1 | 0 |
| B95 | B109 | anti-cenpb | AMA-M2 | 1 | 0 |
| B95 | B151 | anti-cenpb | Anti-CCP | 0.665704468 | 0 |
| B95 | B156 | anti-cenpb | IgE | 0.614367186 | 0 |

| | | | | | |
|---|---|---|---|---|---|
| B95 | B178 | anti-cenpb | Ig-U | 0.524633852 | 0 |
| B95 | B180 | anti-cenpb | Tf | 0.531153079 | 0 |
| B95 | B184 | anti-cenpb | INR | 0.452826163 | 0 |
| B95 | B193 | anti-cenpb | K | 0.459514229 | 0 |
| B95 | B98 | anti-cenpb | anti-Jo-1 | 0.999999999 | 0 |
| B95 | B154 | anti-cenpb | RF | 0.419728548 | 5.33E-15 |
| B95 | B199 | anti-cenpb | Mg | 0.402787859 | 7.82E-14 |
| B95 | B179 | anti-cenpb | α1-MG | 0.397881945 | 1.66E-13 |
| B95 | B183 | anti-cenpb | PTA | 0.375742668 | 4.22E-12 |
| B95 | B157 | anti-cenpb | IgG | 0.340998994 | 4.24E-10 |
| B95 | B158 | anti-cenpb | IgM | -0.28134467 | 3.38E-07 |
| B95 | B119 | anti-cenpb | Hcv-rna | 0.281326061 | 3.39E-07 |
| B95 | B121 | anti-cenpb | C3 | -0.28132607 | 3.39E-07 |
| B95 | B122 | anti-cenpb | C4 | -0.2813262 | 3.39E-07 |
| B95 | B152 | anti-cenpb | ACA-Ig(G-A-M) | -0.28132685 | 3.39E-07 |
| B95 | B159 | anti-cenpb | β2-MG | -0.28132641 | 3.39E-07 |
| B95 | B169 | anti-cenpb | U-CRY | -0.28132578 | 3.39E-07 |
| B96 | B167 | dsDNA (IIF) | PC | 0.948223744 | 0 |
| B97 | B100 | dsDNA | anti-Ro52 | 0.999999999 | 0 |
| B97 | B103 | dsDNA | anti-SSA/Ro | 0.999999999 | 0 |
| B97 | B105 | dsDNA | anti-U1RNP | 0.999999999 | 0 |
| B97 | B108 | dsDNA | Anti-Cyclin | 0.999999999 | 0 |
| B97 | B109 | dsDNA | AMA-M2 | 0.999999999 | 0 |
| B97 | B110 | dsDNA | AHA | 0.999999999 | 0 |
| B97 | B151 | dsDNA | Anti-CCP | 0.665704467 | 0 |
| B97 | B156 | dsDNA | IgE | 0.614367185 | 0 |
| B97 | B178 | dsDNA | Ig-U | 0.524633851 | 0 |
| B97 | B180 | dsDNA | Tf | 0.531153078 | 0 |
| B97 | B184 | dsDNA | INR | 0.452826162 | 0 |
| B97 | B193 | dsDNA | K | 0.459514228 | 0 |
| B97 | B99 | dsDNA | anti-PM-Scl | 1 | 0 |
| B97 | B154 | dsDNA | RF | 0.419728547 | 5.33E-15 |
| B97 | B199 | dsDNA | Mg | 0.402787859 | 7.82E-14 |
| B97 | B179 | dsDNA | α1-MG | 0.397881945 | 1.66E-13 |
| B97 | B183 | dsDNA | PTA | 0.375742667 | 4.22E-12 |
| B97 | B157 | dsDNA | IgG | 0.340998994 | 4.24E-10 |
| B97 | B158 | dsDNA | IgM | -0.28134467 | 3.38E-07 |
| B97 | B119 | dsDNA | Hcv-rna | 0.281326061 | 3.39E-07 |
| B97 | B121 | dsDNA | C3 | -0.28132606 | 3.39E-07 |
| B97 | B122 | dsDNA | C4 | -0.28132615 | 3.39E-07 |
| B97 | B152 | dsDNA | ACA-Ig(G-A-M) | -0.28132666 | 3.39E-07 |
| B97 | B159 | dsDNA | β2-MG | -0.28132637 | 3.39E-07 |
| B97 | B169 | dsDNA | U-CRY | -0.2813257 | 3.39E-07 |
| B98 | B100 | anti-Jo-1 | anti-Ro52 | 0.999999999 | 0 |

| | | | | | |
|---|---|---|---|---|---|
| B98 | B101 | anti-Jo-1 | anti-Sc70 | 0.999999998 | 0 |
| B98 | B102 | anti-Jo-1 | anti-Sm | 0.999999999 | 0 |
| B98 | B103 | anti-Jo-1 | anti-SSA/Ro | 0.999999999 | 0 |
| B98 | B104 | anti-Jo-1 | anti-SSB | 0.999999999 | 0 |
| B98 | B105 | anti-Jo-1 | anti-U1RNP | 0.999999999 | 0 |
| B98 | B107 | anti-Jo-1 | AnuA | 0.999999995 | 0 |
| B98 | B109 | anti-Jo-1 | AMA-M2 | 0.999999997 | 0 |
| B98 | B110 | anti-Jo-1 | AHA | 0.999999999 | 0 |
| B98 | B151 | anti-Jo-1 | Anti-CCP | 0.665704467 | 0 |
| B98 | B156 | anti-Jo-1 | IgE | 0.614367185 | 0 |
| B98 | B178 | anti-Jo-1 | Ig-U | 0.524633851 | 0 |
| B98 | B180 | anti-Jo-1 | Tf | 0.531153078 | 0 |
| B98 | B184 | anti-Jo-1 | INR | 0.452826162 | 0 |
| B98 | B193 | anti-Jo-1 | K | 0.459514228 | 0 |
| B98 | B99 | anti-Jo-1 | anti-PM-Scl | 1 | 0 |
| B98 | B154 | anti-Jo-1 | RF | 0.419728547 | 5.33E-15 |
| B98 | B199 | anti-Jo-1 | Mg | 0.402787859 | 7.82E-14 |
| B98 | B179 | anti-Jo-1 | α1-MG | 0.397881945 | 1.66E-13 |
| B98 | B183 | anti-Jo-1 | PTA | 0.375742667 | 4.22E-12 |
| B98 | B157 | anti-Jo-1 | IgG | 0.340998993 | 4.24E-10 |
| B98 | B158 | anti-Jo-1 | IgM | -0.28134467 | 3.38E-07 |
| B98 | B119 | anti-Jo-1 | Hcv-rna | 0.28132606 | 3.39E-07 |
| B98 | B121 | anti-Jo-1 | C3 | -0.28132608 | 3.39E-07 |
| B98 | B122 | anti-Jo-1 | C4 | -0.28132621 | 3.39E-07 |
| B98 | B152 | anti-Jo-1 | ACA-Ig(G-A-M) | -0.28132689 | 3.39E-07 |
| B98 | B159 | anti-Jo-1 | β2-MG | -0.28132646 | 3.39E-07 |
| B98 | B169 | anti-Jo-1 | U-CRY | -0.28132575 | 3.39E-07 |
| B99 | B100 | anti-PM-Scl | anti-Ro52 | 0.999999999 | 0 |
| B99 | B103 | anti-PM-Scl | anti-SSA/Ro | 0.999999999 | 0 |
| B99 | B105 | anti-PM-Scl | anti-U1RNP | 0.999999999 | 0 |
| B99 | B107 | anti-PM-Scl | AnuA | 0.99999999 | 0 |
| B99 | B109 | anti-PM-Scl | AMA-M2 | 0.999999999 | 0 |
| B99 | B110 | anti-PM-Scl | AHA | 1 | 0 |
| B99 | B151 | anti-PM-Scl | Anti-CCP | 0.665704467 | 0 |
| B99 | B156 | anti-PM-Scl | IgE | 0.614367185 | 0 |
| B99 | B178 | anti-PM-Scl | Ig-U | 0.524633851 | 0 |
| B99 | B180 | anti-PM-Scl | Tf | 0.531153078 | 0 |
| B99 | B184 | anti-PM-Scl | INR | 0.452826162 | 0 |
| B99 | B193 | anti-PM-Scl | K | 0.459514228 | 0 |
| B99 | B154 | anti-PM-Scl | RF | 0.419728547 | 5.33E-15 |
| B99 | B199 | anti-PM-Scl | Mg | 0.402787859 | 7.82E-14 |
| B99 | B179 | anti-PM-Scl | α1-MG | 0.397881945 | 1.66E-13 |
| B99 | B183 | anti-PM-Scl | PTA | 0.375742667 | 4.22E-12 |
| B99 | B157 | anti-PM-Scl | IgG | 0.340998993 | 4.24E-10 |

| | | | | | |
|---|---|---|---|---|---|
| B99 | B158 | anti-PM-Scl | IgM | -0.28134467 | 3.38E-07 |
| B99 | B119 | anti-PM-Scl | Hcv-rna | 0.28132606 | 3.39E-07 |
| B99 | B121 | anti-PM-Scl | C3 | -0.28132607 | 3.39E-07 |
| B99 | B122 | anti-PM-Scl | C4 | -0.28132615 | 3.39E-07 |
| B99 | B152 | anti-PM-Scl | ACA-Ig(G-A-M) | -0.28132668 | 3.39E-07 |
| B99 | B159 | anti-PM-Scl | β2-MG | -0.28132635 | 3.39E-07 |
| B99 | B169 | anti-PM-Scl | U-CRY | -0.28132572 | 3.39E-07 |

**Supplementary Table 14. PEI Correlations in Rheumatoid arthritis**

| Row | Column | Row | Column | Cor | *P* |
|---|---|---|---|---|---|
| B1 | B190 | Gender | Cre | -0.55999211 | 0 |
| B1 | B192 | Gender | UA | -0.50031825 | 0 |
| B1 | B206 | Gender | Waist | -0.42215999 | 0 |
| B1 | B207 | Gender | WHR | -0.44331733 | 0 |
| B1 | B209 | Gender | Smoking | -0.51914003 | 0 |
| B1 | B216 | Gender | BMI | -0.44426835 | 0 |
| B1 | B221 | Gender | Hei | -0.46529749 | 0 |
| B1 | B222 | Gender | NS | -0.63605267 | 0 |
| B1 | B51 | Gender | Hb | -0.45575253 | 0 |
| B1 | B72 | Gender | HDL-C | 0.409170049 | 0 |
| B1 | B46 | Gender | RBC Count | -0.39632222 | 2.22E-16 |
| B1 | B163 | Gender | SG | -0.36107627 | 1.22E-13 |
| B1 | B174 | Gender | Urea/Cr | 0.320705999 | 6.35E-11 |
| B1 | B204 | Gender | Drinking | -0.30083701 | 9.98E-10 |
| B1 | B40 | Gender | WBC | -0.28584033 | 6.98E-09 |
| B1 | B2 | Gender | Age | -0.26007114 | 1.52E-07 |
| B1 | B42 | Gender | MON | -0.24437571 | 8.55E-07 |
| B10 | B14 | PG Ⅰ | G-17 | 0.472619969 | 0 |
| B10 | B12 | PG Ⅰ | PG II | 0.352387385 | 5.08E-13 |
| B10 | B191 | PG Ⅰ | Urea | 0.251331733 | 4.04E-07 |
| B100 | B120 | anti-Ro52 | HCV-cAg | -0.40546819 | 0 |
| B100 | B204 | anti-Ro52 | Drinking | 0.359976665 | 1.47E-13 |
| B100 | B224 | anti-Ro52 | pmh | 0.308194415 | 3.69E-10 |
| B100 | B103 | anti-Ro52 | anti-SSA/Ro | 0.294488569 | 2.30E-09 |
| B100 | B101 | anti-Ro52 | anti-Sc70 | -0.24927397 | 5.05E-07 |
| B100 | B102 | anti-Ro52 | anti-Sm | -0.24927397 | 5.05E-07 |
| B100 | B105 | anti-Ro52 | anti-U1RNP | -0.24927397 | 5.05E-07 |
| B100 | B106 | anti-Ro52 | anti-P | -0.24927397 | 5.05E-07 |
| B100 | B107 | anti-Ro52 | AnuA | -0.24927397 | 5.05E-07 |
| B100 | B108 | anti-Ro52 | Anti-Cyclin | -0.24927397 | 5.05E-07 |
| B100 | B109 | anti-Ro52 | AMA-M2 | -0.24927397 | 5.05E-07 |
| B100 | B110 | anti-Ro52 | AHA | -0.24927397 | 5.05E-07 |
| B101 | B102 | anti-Sc70 | anti-Sm | 0.999999999 | 0 |
| B101 | B103 | anti-Sc70 | anti-SSA/Ro | -0.67410508 | 0 |
| B101 | B106 | anti-Sc70 | anti-P | 1 | 0 |
| B101 | B107 | anti-Sc70 | AnuA | 0.999999967 | 0 |
| B101 | B121 | anti-Sc70 | C3 | 0.329995431 | 1.63E-11 |
| B101 | B122 | anti-Sc70 | C4 | 0.329969907 | 1.64E-11 |
| B101 | B104 | anti-Sc70 | anti-SSB | -0.31611978 | 1.22E-10 |
| B101 | B155 | anti-Sc70 | IgA | 0.2516837 | 3.88E-07 |
| B101 | B157 | anti-Sc70 | IgG | -0.25117744 | 4.11E-07 |
| B102 | B103 | anti-Sm | anti-SSA/Ro | -0.67410508 | 0 |

| | | | | | |
|---|---|---|---|---|---|
| B102 | B108 | anti-Sm | Anti-Cyclin | 1 | 0 |
| B102 | B121 | anti-Sm | C3 | 0.329995431 | 1.63E-11 |
| B102 | B122 | anti-Sm | C4 | 0.329969907 | 1.64E-11 |
| B102 | B104 | anti-Sm | anti-SSB | -0.31611978 | 1.22E-10 |
| B102 | B155 | anti-Sm | IgA | 0.251683699 | 3.88E-07 |
| B102 | B157 | anti-Sm | IgG | -0.25117744 | 4.11E-07 |
| B103 | B104 | anti-SSA/Ro | anti-SSB | 0.52279203 | 0 |
| B103 | B105 | anti-SSA/Ro | anti-U1RNP | -0.67410508 | 0 |
| B103 | B106 | anti-SSA/Ro | anti-P | -0.67410508 | 0 |
| B103 | B107 | anti-SSA/Ro | AnuA | -0.67410508 | 0 |
| B103 | B108 | anti-SSA/Ro | Anti-Cyclin | -0.67410508 | 0 |
| B103 | B109 | anti-SSA/Ro | AMA-M2 | -0.67410508 | 0 |
| B103 | B110 | anti-SSA/Ro | AHA | -0.67410508 | 0 |
| B103 | B121 | anti-SSA/Ro | C3 | -0.53057894 | 0 |
| B103 | B122 | anti-SSA/Ro | C4 | -0.53058105 | 0 |
| B103 | B157 | anti-SSA/Ro | IgG | 0.320995151 | 6.09E-11 |
| B104 | B105 | anti-SSB | anti-U1RNP | -0.31611978 | 1.22E-10 |
| B104 | B106 | anti-SSB | anti-P | -0.31611978 | 1.22E-10 |
| B104 | B107 | anti-SSB | AnuA | -0.31611979 | 1.22E-10 |
| B104 | B108 | anti-SSB | Anti-Cyclin | -0.31611978 | 1.22E-10 |
| B104 | B109 | anti-SSB | AMA-M2 | -0.31611978 | 1.22E-10 |
| B104 | B110 | anti-SSB | AHA | -0.31611978 | 1.22E-10 |
| B104 | B173 | anti-SSB | NTT | 0.313147017 | 1.86E-10 |
| B105 | B106 | anti-U1RNP | anti-P | 0.999999999 | 0 |
| B105 | B108 | anti-U1RNP | Anti-Cyclin | 1 | 0 |
| B105 | B109 | anti-U1RNP | AMA-M2 | 1 | 0 |
| B105 | B110 | anti-U1RNP | AHA | 1 | 0 |
| B105 | B121 | anti-U1RNP | C3 | 0.32999543 | 1.63E-11 |
| B105 | B122 | anti-U1RNP | C4 | 0.329969907 | 1.64E-11 |
| B105 | B155 | anti-U1RNP | IgA | 0.251683699 | 3.88E-07 |
| B105 | B157 | anti-U1RNP | IgG | -0.25117744 | 4.11E-07 |
| B106 | B107 | anti-P | AnuA | 0.999999993 | 0 |
| B106 | B108 | anti-P | Anti-Cyclin | 1 | 0 |
| B106 | B109 | anti-P | AMA-M2 | 1 | 0 |
| B106 | B110 | anti-P | AHA | 0.999999999 | 0 |
| B106 | B121 | anti-P | C3 | 0.32999543 | 1.63E-11 |
| B106 | B122 | anti-P | C4 | 0.329969907 | 1.64E-11 |
| B106 | B155 | anti-P | IgA | 0.251683699 | 3.88E-07 |
| B106 | B157 | anti-P | IgG | -0.25117744 | 4.11E-07 |
| B107 | B121 | AnuA | C3 | 0.329995432 | 1.63E-11 |
| B107 | B122 | AnuA | C4 | 0.329969906 | 1.64E-11 |
| B107 | B155 | AnuA | IgA | 0.251683699 | 3.88E-07 |
| B107 | B157 | AnuA | IgG | -0.25117744 | 4.11E-07 |
| B108 | B110 | Anti-Cyclin | AHA | 1 | 0 |

| | | | | | |
|---|---|---|---|---|---|
| B108 | B121 | Anti-Cyclin | C3 | 0.32999543 | 1.63E-11 |
| B108 | B122 | Anti-Cyclin | C4 | 0.329969907 | 1.64E-11 |
| B108 | B155 | Anti-Cyclin | IgA | 0.251683699 | 3.88E-07 |
| B108 | B157 | Anti-Cyclin | IgG | -0.25117744 | 4.11E-07 |
| B109 | B121 | AMA-M2 | C3 | 0.32999543 | 1.63E-11 |
| B109 | B122 | AMA-M2 | C4 | 0.329969907 | 1.64E-11 |
| B109 | B155 | AMA-M2 | IgA | 0.251683699 | 3.88E-07 |
| B109 | B157 | AMA-M2 | IgG | -0.25117744 | 4.11E-07 |
| B11 | B12 | PGI/PGII | PG II | -0.56076253 | 0 |
| B11 | B14 | PGI/PGII | G-17 | -0.31267385 | 1.98E-10 |
| B11 | B149 | PGI/PGII | TGAb | -0.255659 | 2.50E-07 |
| B110 | B121 | AHA | C3 | 0.32999543 | 1.63E-11 |
| B110 | B122 | AHA | C4 | 0.329969907 | 1.64E-11 |
| B110 | B155 | AHA | IgA | 0.251683699 | 3.88E-07 |
| B110 | B157 | AHA | IgG | -0.25117744 | 4.11E-07 |
| B112 | B114 | FT4 | TSH TT3 TT4(TT4) | 0.373660874 | 1.44E-14 |
| B113 | B115 | FT3 | TSH TT3 TT4(TT3) | 0.527283159 | 0 |
| B113 | B134 | FT3 | CHE | -0.28482608 | 7.93E-09 |
| B113 | B196 | FT3 | TCO2 | -0.26216811 | 1.20E-07 |
| B114 | B198 | TSH TT3 TT4(TT4) | P | -0.50617096 | 0 |
| B114 | B115 | TSH TT3 TT4(TT4) | TSH TT3 TT4(TT3) | 0.302610103 | 7.87E-10 |
| B114 | B194 | TSH TT3 TT4(TT4) | Cl | 0.246790828 | 6.60E-07 |
| B115 | B134 | TSH TT3 TT4(TT3) | CHE | -0.38636903 | 1.55E-15 |
| B115 | B196 | TSH TT3 TT4(TT3) | TCO2 | -0.37464821 | 1.20E-14 |
| B115 | B200 | TSH TT3 TT4(TT3) | OP | 0.324210269 | 3.82E-11 |
| B115 | B138 | TSH TT3 TT4(TT3) | PAB | -0.269816 | 4.93E-08 |
| B116 | B117 | TT3 | TT4 | -0.52774674 | 0 |
| B119 | B120 | Hcv-rna | HCV-cAg | 0.405656033 | 0 |
| B119 | B124 | Hcv-rna | ACTH | 0.635038757 | 0 |
| B119 | B169 | Hcv-rna | U-CRY | -0.99999452 | 0 |
| B119 | B179 | Hcv-rna | α1-MG | 0.413808817 | 0 |
| B119 | B198 | Hcv-rna | P | -0.28987342 | 4.18E-09 |
| B119 | B150 | Hcv-rna | ALP | -0.2452647 | 7.77E-07 |
| B12 | B14 | PG II | G-17 | 0.412678199 | 0 |
| B12 | B149 | PG II | TGAb | 0.352155232 | 5.27E-13 |
| B120 | B152 | HCV-cAg | ACA-Ig(G-A-M) | -0.40566747 | 0 |
| B120 | B159 | HCV-cAg | β2-MG | -0.40565655 | 0 |
| B120 | B169 | HCV-cAg | U-CRY | -0.40565295 | 0 |
| B120 | B224 | HCV-cAg | pmh | -0.28144004 | 1.21E-08 |
| B120 | B139 | HCV-cAg | ADA | -0.27077664 | 4.40E-08 |
| B120 | B124 | HCV-cAg | ACTH | 0.261410263 | 1.31E-07 |
| B121 | B122 | C3 | C4 | 0.99998187 | 0 |
| B121 | B158 | C3 | IgM | 0.346244154 | 1.35E-12 |
| B121 | B205 | C3 | KDA | -0.29766287 | 1.52E-09 |

| | | | | | |
|---|---|---|---|---|---|
| B122 | B158 | C4 | IgM | 0.34629298 | 1.34E-12 |
| B122 | B205 | C4 | KDA | -0.29757726 | 1.54E-09 |
| B123 | B184 | hs-CRP1 | INR | -0.39849014 | 0 |
| B123 | B186 | hs-CRP1 | PFC | 0.608178075 | 0 |
| B123 | B150 | hs-CRP1 | ALP | 0.348181955 | 9.95E-13 |
| B123 | B156 | hs-CRP1 | IgE | 0.336060521 | 6.55E-12 |
| B123 | B125 | hs-CRP1 | Cor | -0.32273278 | 4.74E-11 |
| B123 | B179 | hs-CRP1 | α1-MG | -0.30907698 | 3.26E-10 |
| B123 | B157 | hs-CRP1 | IgG | 0.284074071 | 8.71E-09 |
| B123 | B181 | hs-CRP1 | APTT | 0.275773746 | 2.42E-08 |
| B124 | B125 | ACTH | Cor | 0.720676179 | 0 |
| B124 | B152 | ACTH | ACA-Ig(G-A-M) | -0.63506467 | 0 |
| B124 | B159 | ACTH | β2-MG | -0.63504099 | 0 |
| B124 | B169 | ACTH | U-CRY | -0.63503528 | 0 |
| B124 | B181 | ACTH | APTT | 0.459914164 | 0 |
| B124 | B198 | ACTH | P | -0.40091199 | 0 |
| B124 | B194 | ACTH | Cl | -0.38626289 | 1.55E-15 |
| B124 | B184 | ACTH | INR | -0.36417276 | 7.28E-14 |
| B124 | B179 | ACTH | α1-MG | 0.317382298 | 1.02E-10 |
| B124 | B193 | ACTH | K | -0.30289233 | 7.58E-10 |
| B124 | B186 | ACTH | PFC | 0.25856407 | 1.81E-07 |
| B124 | B200 | ACTH | OP | -0.2544029 | 2.88E-07 |
| B124 | B197 | ACTH | AG | 0.248827415 | 5.30E-07 |
| B124 | B158 | ACTH | IgM | 0.244928772 | 8.06E-07 |
| B125 | B135 | Cor | TBA | 0.421042482 | 0 |
| B125 | B194 | Cor | Cl | -0.4046803 | 0 |
| B125 | B181 | Cor | APTT | 0.347934134 | 1.03E-12 |
| B125 | B197 | Cor | AG | 0.340093677 | 3.53E-12 |
| B125 | B182 | Cor | TT | 0.332019545 | 1.21E-11 |
| B125 | B150 | Cor | ALP | -0.30968441 | 3.00E-10 |
| B125 | B179 | Cor | α1-MG | 0.284717588 | 8.04E-09 |
| B125 | B157 | Cor | IgG | -0.28079399 | 1.31E-08 |
| B125 | B199 | Cor | Mg | 0.27373739 | 3.09E-08 |
| B125 | B156 | Cor | IgE | -0.26598509 | 7.73E-08 |
| B125 | B198 | Cor | P | -0.25927097 | 1.67E-07 |
| B126 | B127 | Alb | A-G ratio | 0.682301807 | 0 |
| B126 | B130 | Alb | GLB | -0.45441572 | 0 |
| B126 | B188 | Alb | Ca | 0.509760393 | 0 |
| B127 | B130 | A-G ratio | GLB | -0.92384873 | 0 |
| B128 | B131 | ALT | AST | 0.760359693 | 0 |
| B128 | B140 | ALT | r-GT | 0.498060348 | 0 |
| B128 | B141 | ALT | AST/ALT | -0.60651243 | 0 |
| B129 | B132 | IBIL | Dbil | 0.656486907 | 0 |
| B129 | B133 | IBIL | TBIL | 0.976739304 | 0 |

| | | | | | |
|---|---|---|---|---|---|
| B131 | B140 | AST | r-GT | 0.382482716 | 3.11E-15 |
| B132 | B133 | Dbil | TBIL | 0.802818964 | 0 |
| B134 | B137 | CHE | AFU | -0.45322181 | 0 |
| B134 | B138 | CHE | PAB | 0.56125778 | 0 |
| B134 | B199 | CHE | Mg | 0.394573486 | 4.44E-16 |
| B134 | B225 | CHE | hPBG | 0.33387409 | 9.12E-12 |
| B134 | B193 | CHE | K | 0.332965914 | 1.05E-11 |
| B134 | B136 | CHE | 5'NT | 0.304357936 | 6.22E-10 |
| B135 | B139 | TBA | ADA | 0.40827134 | 0 |
| B135 | B194 | TBA | Cl | -0.5196916 | 0 |
| B135 | B225 | TBA | hPBG | -0.71502761 | 0 |
| B135 | B199 | TBA | Mg | 0.387811206 | 1.11E-15 |
| B135 | B138 | TBA | PAB | -0.30689384 | 4.40E-10 |
| B135 | B193 | TBA | K | -0.29575183 | 1.95E-09 |
| B136 | B197 | 5'NT | AG | 0.460095273 | 0 |
| B136 | B193 | 5'NT | K | 0.347352851 | 1.14E-12 |
| B136 | B195 | 5'NT | Na | 0.337856724 | 4.98E-12 |
| B136 | B196 | 5'NT | TCO2 | -0.2949023 | 2.18E-09 |
| B136 | B200 | 5'NT | OP | 0.246873773 | 6.55E-07 |
| B137 | B200 | AFU | OP | 0.294808692 | 2.21E-09 |
| B137 | B225 | AFU | hPBG | -0.28381012 | 9.00E-09 |
| B138 | B139 | PAB | ADA | -0.52537711 | 0 |
| B138 | B225 | PAB | hPBG | 0.324885593 | 3.47E-11 |
| B138 | B155 | PAB | IgA | 0.244593766 | 8.35E-07 |
| B139 | B225 | ADA | hPBG | -0.53197929 | 0 |
| B139 | B196 | ADA | TCO2 | 0.285957264 | 6.88E-09 |
| B140 | B141 | r-GT | AST/ALT | -0.26439342 | 9.29E-08 |
| B141 | B216 | AST/ALT | BMI | -0.26432881 | 9.36E-08 |
| B141 | B215 | AST/ALT | BMI | -0.24831923 | 5.60E-07 |
| B142 | B143 | 25-(OH)D | β-CTx | 0.303805838 | 6.70E-10 |
| B143 | B144 | β-CTx | Total-PINP | 0.792209477 | 0 |
| B143 | B145 | β-CTx | BGP | 0.549546646 | 0 |
| B143 | B151 | β-CTx | Anti-CCP | 0.299942955 | 1.12E-09 |
| B144 | B145 | Total-PINP | BGP | 0.682277615 | 0 |
| B144 | B151 | Total-PINP | Anti-CCP | 0.248161398 | 5.70E-07 |
| B145 | B199 | BGP | Mg | 0.313957594 | 1.66E-10 |
| B145 | B195 | BGP | Na | 0.264662274 | 9.01E-08 |
| B146 | B154 | CysC | RF | 0.471073182 | 0 |
| B146 | B190 | CysC | Cre | 0.250445881 | 4.45E-07 |
| B148 | B149 | Anti_TPO | TGAb | 0.511349684 | 0 |
| B15 | B138 | CD3-CD19+(B cell) | PAB | 0.541531759 | 0 |
| B15 | B139 | CD3-CD19+(B cell) | ADA | -0.40195918 | 0 |
| B15 | B16 | CD3-CD19+(B cell) | CD3+T | 0.838584399 | 0 |
| B15 | B17 | CD3-CD19+(B cell) | CD3+CD16+CD56+NK | 0.789619196 | 0 |

| | | | | | |
|---|---|---|---|---|---|
| B15 | B18 | CD3-CD19+(B cell) | CD3+CD4+helper(inducer)T | -0.94804226 | 0 |
| B15 | B19 | CD3-CD19+(B cell) | CD3+CD4+/CD3+CD8+ | -0.99206548 | 0 |
| B15 | B21 | CD3-CD19+(B cell) | CD3+CD4-CD8- | 0.982074448 | 0 |
| B15 | B22 | CD3-CD19+(B cell) | CD3+CD8+(T) | 0.949664592 | 0 |
| B15 | B23 | CD3-CD19+(B cell) | CD3-CD16+CD56+(NK) | -0.99578801 | 0 |
| B15 | B194 | CD3-CD19+(B cell) | Cl | -0.36052421 | 1.34E-13 |
| B15 | B200 | CD3-CD19+(B cell) | OP | -0.35562353 | 3.00E-13 |
| B15 | B58 | CD3-CD19+(B cell) | HCT | 0.346334463 | 1.33E-12 |
| B15 | B195 | CD3-CD19+(B cell) | Na | -0.34623246 | 1.35E-12 |
| B15 | B193 | CD3-CD19+(B cell) | K | -0.33387629 | 9.12E-12 |
| B15 | B196 | CD3-CD19+(B cell) | TCO2 | 0.295524667 | 2.01E-09 |
| B15 | B69 | CD3-CD19+(B cell) | PV | 0.288098295 | 5.24E-09 |
| B15 | B63 | CD3-CD19+(B cell) | WBRSR(mPas_200) | 0.272329153 | 3.66E-08 |
| B15 | B54 | CD3-CD19+(B cell) | ESR | -0.26915028 | 5.33E-08 |
| B15 | B66 | CD3-CD19+(B cell) | WBRSR(mPas_100) | 0.263523657 | 1.03E-07 |
| B15 | B67 | CD3-CD19+(B cell) | WBRSR(mPas_50) | 0.262726839 | 1.13E-07 |
| B15 | B101 | CD3-CD19+(B cell) | anti-Sc70 | 0.256331751 | 2.32E-07 |
| B15 | B102 | CD3-CD19+(B cell) | anti-Sm | 0.25633175 | 2.32E-07 |
| B15 | B105 | CD3-CD19+(B cell) | anti-U1RNP | 0.25633175 | 2.32E-07 |
| B15 | B106 | CD3-CD19+(B cell) | anti-P | 0.25633175 | 2.32E-07 |
| B15 | B107 | CD3-CD19+(B cell) | AnuA | 0.256331751 | 2.32E-07 |
| B15 | B108 | CD3-CD19+(B cell) | Anti-Cyclin | 0.25633175 | 2.32E-07 |
| B15 | B109 | CD3-CD19+(B cell) | AMA-M2 | 0.25633175 | 2.32E-07 |
| B15 | B110 | CD3-CD19+(B cell) | AHA | 0.25633175 | 2.32E-07 |
| B15 | B95 | CD3-CD19+(B cell) | anti-cenpb | 0.25633175 | 2.32E-07 |
| B15 | B97 | CD3-CD19+(B cell) | dsDNA | 0.25633175 | 2.32E-07 |
| B15 | B98 | CD3-CD19+(B cell) | anti-Jo-1 | 0.25633175 | 2.32E-07 |
| B15 | B99 | CD3-CD19+(B cell) | anti-PM-Scl | 0.25633175 | 2.32E-07 |
| B15 | B120 | CD3-CD19+(B cell) | HCV-cAg | 0.243972609 | 8.92E-07 |
| B150 | B186 | ALP | PFC | 0.437443385 | 0 |
| B150 | B179 | ALP | α1-MG | -0.35266648 | 4.85E-13 |
| B150 | B193 | ALP | K | 0.263881379 | 9.86E-08 |
| B150 | B184 | ALP | INR | -0.24543065 | 7.64E-07 |
| B150 | B152 | ALP | ACA-Ig(G-A-M) | 0.245274696 | 7.77E-07 |
| B150 | B159 | ALP | β2-MG | 0.245265556 | 7.77E-07 |
| B150 | B169 | ALP | U-CRY | 0.24526336 | 7.78E-07 |
| B152 | B179 | ACA-Ig(G-A-M) | α1-MG | -0.41382569 | 0 |
| B152 | B198 | ACA-Ig(G-A-M) | P | 0.289885257 | 4.18E-09 |
| B155 | B158 | IgA | IgM | 0.45923468 | 0 |
| B155 | B156 | IgA | IgE | 0.388569918 | 8.88E-16 |
| B155 | B157 | IgA | IgG | 0.339180494 | 4.06E-12 |
| B155 | B196 | IgA | TCO2 | 0.25963957 | 1.60E-07 |
| B156 | B157 | IgE | IgG | 0.451869186 | 0 |
| B156 | B158 | IgE | IgM | 0.463155439 | 0 |

| | | | | | |
|---|---|---|---|---|---|
| B156 | B186 | IgE | PFC | 0.415572851 | 0 |
| B156 | B184 | IgE | INR | -0.25881671 | 1.76E-07 |
| B157 | B158 | IgG | IgM | 0.579199818 | 0 |
| B157 | B186 | IgG | PFC | 0.384795593 | 2.00E-15 |
| B158 | B186 | IgM | PFC | 0.374448466 | 1.24E-14 |
| B158 | B184 | IgM | INR | -0.32803584 | 2.18E-11 |
| B158 | B181 | IgM | APTT | 0.314247899 | 1.59E-10 |
| B159 | B169 | β2-MG | U-CRY | 0.999993991 | 0 |
| B159 | B179 | β2-MG | α1-MG | -0.41381027 | 0 |
| B159 | B198 | β2-MG | P | 0.289874433 | 4.18E-09 |
| B16 | B139 | CD3+T | ADA | -0.59452548 | 0 |
| B16 | B18 | CD3+T | CD3+CD4+helper(inducer)T | -0.62169737 | 0 |
| B16 | B19 | CD3+T | CD3+CD4+/CD3+CD8+ | -0.90042055 | 0 |
| B16 | B195 | CD3+T | Na | -0.45702251 | 0 |
| B16 | B20 | CD3+T | CD3+CD4+CD8+ | 0.675954982 | 0 |
| B16 | B200 | CD3+T | OP | -0.4149932 | 0 |
| B16 | B21 | CD3+T | CD3+CD4-CD8- | 0.720866417 | 0 |
| B16 | B22 | CD3+T | CD3+CD8+(T) | 0.967033733 | 0 |
| B16 | B23 | CD3+T | CD3-CD16+CD56+(NK) | -0.7851046 | 0 |
| B16 | B120 | CD3+T | HCV-cAg | 0.381169441 | 4.00E-15 |
| B16 | B138 | CD3+T | PAB | 0.379237697 | 5.33E-15 |
| B16 | B17 | CD3+T | CD3+CD16+CD56+NK | 0.327891948 | 2.23E-11 |
| B16 | B194 | CD3+T | Cl | -0.32721974 | 2.46E-11 |
| B16 | B224 | CD3+T | pmh | -0.2840429 | 8.74E-09 |
| B16 | B150 | CD3+T | ALP | -0.25323696 | 3.27E-07 |
| B16 | B50 | CD3+T | eo% | -0.24338643 | 9.49E-07 |
| B161 | B173 | LEU | NTT | 0.354971264 | 3.34E-13 |
| B163 | B166 | SG | Pro | 0.347718911 | 1.07E-12 |
| B163 | B176 | SG | Ucr | 0.303747965 | 6.75E-10 |
| B163 | B222 | SG | NS | 0.302508433 | 7.98E-10 |
| B163 | B190 | SG | Cre | 0.290334571 | 3.94E-09 |
| B163 | B216 | SG | BMI | 0.255713074 | 2.49E-07 |
| B163 | B165 | SG | bilirubin | 0.245244822 | 7.79E-07 |
| B167 | B223 | PC | FH | 0.286125156 | 6.73E-09 |
| B169 | B179 | U-CRY | α1-MG | -0.41380655 | 0 |
| B169 | B198 | U-CRY | P | 0.289871835 | 4.18E-09 |
| B17 | B138 | CD3+CD16+CD56+NK | PAB | 0.511946769 | 0 |
| B17 | B18 | CD3+CD16+CD56+NK | CD3+CD4+helper(inducer)T | -0.94380489 | 0 |
| B17 | B19 | CD3+CD16+CD56+NK | CD3+CD4+/CD3+CD8+ | -0.70621114 | 0 |
| B17 | B20 | CD3+CD16+CD56+NK | CD3+CD4+CD8+ | -0.474561 | 0 |
| B17 | B21 | CD3+CD16+CD56+NK | CD3+CD4-CD8- | 0.891123886 | 0 |
| B17 | B22 | CD3+CD16+CD56+NK | CD3+CD8+(T) | 0.557652708 | 0 |
| B17 | B23 | CD3+CD16+CD56+NK | CD3-CD16+CD56+(NK) | -0.84255132 | 0 |
| B17 | B58 | CD3+CD16+CD56+NK | HCT | 0.442692608 | 0 |

| | | | | | |
|---|---|---|---|---|---|
| B17 | B69 | CD3+CD16+CD56+NK | PV | 0.362902569 | 9.02E-14 |
| B17 | B54 | CD3+CD16+CD56+NK | ESR | -0.34493555 | 1.66E-12 |
| B17 | B63 | CD3+CD16+CD56+NK | WBRSR(mPas_200) | 0.344786351 | 1.70E-12 |
| B17 | B196 | CD3+CD16+CD56+NK | TCO2 | 0.343004147 | 2.25E-12 |
| B17 | B66 | CD3+CD16+CD56+NK | WBRSR(mPas_100) | 0.333404254 | 9.79E-12 |
| B17 | B67 | CD3+CD16+CD56+NK | WBRSR(mPas_50) | 0.332738217 | 1.08E-11 |
| B17 | B101 | CD3+CD16+CD56+NK | anti-Sc70 | 0.329940539 | 1.65E-11 |
| B17 | B102 | CD3+CD16+CD56+NK | anti-Sm | 0.32994054 | 1.65E-11 |
| B17 | B105 | CD3+CD16+CD56+NK | anti-U1RNP | 0.329940539 | 1.65E-11 |
| B17 | B106 | CD3+CD16+CD56+NK | anti-P | 0.32994054 | 1.65E-11 |
| B17 | B107 | CD3+CD16+CD56+NK | AnuA | 0.329940543 | 1.65E-11 |
| B17 | B108 | CD3+CD16+CD56+NK | Anti-Cyclin | 0.329940539 | 1.65E-11 |
| B17 | B109 | CD3+CD16+CD56+NK | AMA-M2 | 0.329940539 | 1.65E-11 |
| B17 | B110 | CD3+CD16+CD56+NK | AHA | 0.329940539 | 1.65E-11 |
| B17 | B95 | CD3+CD16+CD56+NK | anti-cenpb | 0.329940539 | 1.65E-11 |
| B17 | B97 | CD3+CD16+CD56+NK | dsDNA | 0.32994054 | 1.65E-11 |
| B17 | B98 | CD3+CD16+CD56+NK | anti-Jo-1 | 0.329940539 | 1.65E-11 |
| B17 | B99 | CD3+CD16+CD56+NK | anti-PM-Scl | 0.32994054 | 1.65E-11 |
| B17 | B193 | CD3+CD16+CD56+NK | K | -0.31452771 | 1.53E-10 |
| B17 | B55 | CD3+CD16+CD56+NK | TK | -0.30842143 | 3.57E-10 |
| B17 | B76 | CD3+CD16+CD56+NK | ApoB | -0.26519772 | 8.47E-08 |
| B17 | B199 | CD3+CD16+CD56+NK | Mg | -0.26297417 | 1.09E-07 |
| B17 | B62 | CD3+CD16+CD56+NK | WBHSRVI | 0.259444526 | 1.64E-07 |
| B17 | B194 | CD3+CD16+CD56+NK | Cl | -0.25664255 | 2.24E-07 |
| B17 | B134 | CD3+CD16+CD56+NK | CHE | 0.249506934 | 4.93E-07 |
| B174 | B190 | Urea/Cr | Cre | -0.42236983 | 0 |
| B174 | B191 | Urea/Cr | Urea | 0.67114201 | 0 |
| B174 | B221 | Urea/Cr | Hei | -0.27859334 | 1.72E-08 |
| B174 | B192 | Urea/Cr | UA | -0.25431785 | 2.91E-07 |
| B175 | B177 | ALB-U | ACR | 0.920396331 | 0 |
| B175 | B178 | ALB-U | Ig-U | 0.982665144 | 0 |
| B175 | B179 | ALB-U | α1-MG | 0.586600138 | 0 |
| B175 | B180 | ALB-U | Tf | 0.983615954 | 0 |
| B176 | B179 | Ucr | α1-MG | 0.29384277 | 2.51E-09 |
| B177 | B178 | ACR | Ig-U | 0.885455395 | 0 |
| B177 | B179 | ACR | α1-MG | 0.489960637 | 0 |
| B177 | B180 | ACR | Tf | 0.886170264 | 0 |
| B178 | B179 | Ig-U | α1-MG | 0.642894384 | 0 |
| B178 | B180 | Ig-U | Tf | 0.999202816 | 0 |
| B179 | B180 | α1-MG | Tf | 0.638036378 | 0 |
| B179 | B186 | α1-MG | PFC | -0.42182008 | 0 |
| B179 | B184 | α1-MG | INR | 0.262389914 | 1.17E-07 |
| B18 | B138 | CD3+CD4+helper(inducer)T | PAB | -0.557126 | 0 |
| B18 | B19 | CD3+CD4+helper(inducer)T | CD3+CD4+/CD3+CD8+ | 0.900522177 | 0 |

| | | | | | |
|---|---|---|---|---|---|
| B18 | B21 | CD3+CD4+helper(inducer)T | CD3+CD4-CD8- | -0.99101622 | 0 |
| B18 | B22 | CD3+CD4+helper(inducer)T | CD3+CD8+(T) | -0.80065754 | 0 |
| B18 | B23 | CD3+CD4+helper(inducer)T | CD3-CD16+CD56+(NK) | 0.973218359 | 0 |
| B18 | B58 | CD3+CD4+helper(inducer)T | HCT | -0.41607884 | 0 |
| B18 | B69 | CD3+CD4+helper(inducer)T | PV | -0.34334058 | 2.13E-12 |
| B18 | B193 | CD3+CD4+helper(inducer)T | K | 0.34291616 | 2.28E-12 |
| B18 | B196 | CD3+CD4+helper(inducer)T | TCO2 | -0.33702351 | 5.66E-12 |
| B18 | B194 | CD3+CD4+helper(inducer)T | Cl | 0.32725667 | 2.45E-11 |
| B18 | B63 | CD3+CD4+helper(inducer)T | WBRSR(mPas_200) | -0.32545372 | 3.19E-11 |
| B18 | B54 | CD3+CD4+helper(inducer)T | ESR | 0.323818833 | 4.05E-11 |
| B18 | B66 | CD3+CD4+helper(inducer)T | WBRSR(mPas_100) | -0.31480929 | 1.47E-10 |
| B18 | B67 | CD3+CD4+helper(inducer)T | WBRSR(mPas_50) | -0.31403476 | 1.64E-10 |
| B18 | B101 | CD3+CD4+helper(inducer)T | anti-Sc70 | -0.30913959 | 3.24E-10 |
| B18 | B102 | CD3+CD4+helper(inducer)T | anti-Sm | -0.30913959 | 3.24E-10 |
| B18 | B105 | CD3+CD4+helper(inducer)T | anti-U1RNP | -0.30913959 | 3.24E-10 |
| B18 | B106 | CD3+CD4+helper(inducer)T | anti-P | -0.30913959 | 3.24E-10 |
| B18 | B107 | CD3+CD4+helper(inducer)T | AnuA | -0.30913959 | 3.24E-10 |
| B18 | B108 | CD3+CD4+helper(inducer)T | Anti-Cyclin | -0.30913959 | 3.24E-10 |
| B18 | B109 | CD3+CD4+helper(inducer)T | AMA-M2 | -0.30913959 | 3.24E-10 |
| B18 | B110 | CD3+CD4+helper(inducer)T | AHA | -0.30913959 | 3.24E-10 |
| B18 | B95 | CD3+CD4+helper(inducer)T | anti-cenpb | -0.30913959 | 3.24E-10 |
| B18 | B97 | CD3+CD4+helper(inducer)T | dsDNA | -0.30913959 | 3.24E-10 |
| B18 | B98 | CD3+CD4+helper(inducer)T | anti-Jo-1 | -0.30913959 | 3.24E-10 |
| B18 | B99 | CD3+CD4+helper(inducer)T | anti-PM-Scl | -0.30913959 | 3.24E-10 |
| B18 | B55 | CD3+CD4+helper(inducer)T | TK | 0.289457111 | 4.41E-09 |
| B18 | B200 | CD3+CD4+helper(inducer)T | OP | 0.268951268 | 5.46E-08 |
| B18 | B62 | CD3+CD4+helper(inducer)T | WBHSRVI | -0.24468841 | 8.27E-07 |
| B181 | B186 | APTT | PFC | 0.651464386 | 0 |
| B181 | B183 | APTT | PTA | -0.30408668 | 6.45E-10 |
| B181 | B194 | APTT | Cl | -0.28521455 | 7.55E-09 |
| B181 | B184 | APTT | INR | -0.28506102 | 7.70E-09 |
| B182 | B183 | TT | PTA | -0.82693263 | 0 |
| B182 | B185 | TT | PT | -0.70056512 | 0 |
| B182 | B184 | TT | INR | -0.39703352 | 2.22E-16 |
| B183 | B184 | PTA | INR | 0.508207886 | 0 |
| B183 | B185 | PTA | PT | 0.640585688 | 0 |
| B183 | B186 | PTA | PFC | -0.39160653 | 4.44E-16 |
| B184 | B185 | INR | PT | 0.765610785 | 0 |
| B184 | B186 | INR | PFC | -0.77319186 | 0 |
| B185 | B186 | PT | PFC | -0.32909928 | 1.86E-11 |
| B189 | B190 | eGFB | Cre | -0.67626556 | 0 |
| B189 | B207 | eGFB | WHR | -0.33813669 | 4.77E-12 |
| B189 | B191 | eGFB | Urea | -0.33230102 | 1.16E-11 |
| B189 | B192 | eGFB | UA | -0.32157249 | 5.61E-11 |

| | | | | | |
|---|---|---|---|---|---|
| B189 | B206 | eGFB | Waist | -0.29074816 | 3.74E-09 |
| B189 | B220 | eGFB | SBP | -0.25472313 | 2.78E-07 |
| B19 | B138 | CD3+CD4+/CD3+CD8+ | PAB | -0.51995363 | 0 |
| B19 | B139 | CD3+CD4+/CD3+CD8+ | ADA | 0.458183768 | 0 |
| B19 | B21 | CD3+CD4+/CD3+CD8+ | CD3+CD4-CD8- | -0.95058435 | 0 |
| B19 | B22 | CD3+CD4+/CD3+CD8+ | CD3+CD8+(T) | -0.98151428 | 0 |
| B19 | B23 | CD3+CD4+/CD3+CD8+ | CD3-CD16+CD56+(NK) | 0.976360002 | 0 |
| B19 | B195 | CD3+CD4+/CD3+CD8+ | Na | 0.38195102 | 3.33E-15 |
| B19 | B200 | CD3+CD4+/CD3+CD8+ | OP | 0.379750627 | 4.88E-15 |
| B19 | B194 | CD3+CD4+/CD3+CD8+ | Cl | 0.363407674 | 8.30E-14 |
| B19 | B193 | CD3+CD4+/CD3+CD8+ | K | 0.320799571 | 6.27E-11 |
| B19 | B58 | CD3+CD4+/CD3+CD8+ | HCT | -0.30891429 | 3.34E-10 |
| B19 | B20 | CD3+CD4+/CD3+CD8+ | CD3+CD4+CD8+ | -0.28805848 | 5.27E-09 |
| B19 | B120 | CD3+CD4+/CD3+CD8+ | HCV-cAg | -0.28278736 | 1.02E-08 |
| B19 | B196 | CD3+CD4+/CD3+CD8+ | TCO2 | -0.27071277 | 4.43E-08 |
| B19 | B69 | CD3+CD4+/CD3+CD8+ | PV | -0.25806676 | 1.91E-07 |
| B19 | B150 | CD3+CD4+/CD3+CD8+ | ALP | 0.249046708 | 5.18E-07 |
| B19 | B63 | CD3+CD4+/CD3+CD8+ | WBRSR(mPas_200) | -0.24358338 | 9.29E-07 |
| B190 | B192 | Cre | UA | 0.537050478 | 0 |
| B190 | B222 | Cre | NS | 0.453394612 | 0 |
| B190 | B206 | Cre | Waist | 0.345205753 | 1.59E-12 |
| B190 | B191 | Cre | Urea | 0.34427762 | 1.84E-12 |
| B190 | B216 | Cre | BMI | 0.340215146 | 3.47E-12 |
| B190 | B207 | Cre | WHR | 0.335337932 | 7.31E-12 |
| B190 | B209 | Cre | Smoking | 0.302514972 | 7.97E-10 |
| B190 | B221 | Cre | Hei | 0.278041825 | 1.84E-08 |
| B192 | B206 | UA | Waist | 0.460315903 | 0 |
| B192 | B207 | UA | WHR | 0.45202991 | 0 |
| B192 | B216 | UA | BMI | 0.416081734 | 0 |
| B192 | B222 | UA | NS | 0.466348841 | 0 |
| B192 | B215 | UA | BMI | 0.315669814 | 1.30E-10 |
| B192 | B209 | UA | Smoking | 0.291109405 | 3.57E-09 |
| B192 | B221 | UA | Hei | 0.275813577 | 2.41E-08 |
| B193 | B199 | K | Mg | 0.3332757 | 9.99E-12 |
| B193 | B196 | K | TCO2 | -0.30302026 | 7.45E-10 |
| B193 | B194 | K | Cl | 0.256142315 | 2.37E-07 |
| B194 | B197 | Cl | AG | -0.45405849 | 0 |
| B194 | B225 | Cl | hPBG | 0.338073995 | 4.82E-12 |
| B194 | B200 | Cl | OP | 0.251846916 | 3.82E-07 |
| B195 | B196 | Na | TCO2 | -0.41410286 | 0 |
| B195 | B197 | Na | AG | 0.628311413 | 0 |
| B195 | B200 | Na | OP | 0.925017526 | 0 |
| B196 | B197 | TCO2 | AG | -0.71900391 | 0 |
| B196 | B200 | TCO2 | OP | -0.51404569 | 0 |

| | | | | | |
|---|---|---|---|---|---|
| B196 | B225 | TCO2 | hPBG | -0.26429093 | 9.40E-08 |
| B197 | B200 | AG | OP | 0.609953103 | 0 |
| B197 | B199 | AG | Mg | 0.266981699 | 6.88E-08 |
| B198 | B200 | P | OP | 0.278056523 | 1.83E-08 |
| B2 | B189 | Age | eGFB | -0.60382522 | 0 |
| B2 | B207 | Age | WHR | 0.441091561 | 0 |
| B2 | B220 | Age | SBP | 0.481872534 | 0 |
| B2 | B206 | Age | Waist | 0.351577234 | 5.78E-13 |
| B2 | B190 | Age | Cre | 0.323069148 | 4.52E-11 |
| B2 | B191 | Age | Urea | 0.293251984 | 2.71E-09 |
| B2 | B77 | Age | TC | 0.288099109 | 5.24E-09 |
| B2 | B70 | Age | LDL-C | 0.287293842 | 5.81E-09 |
| B2 | B192 | Age | UA | 0.254008025 | 3.01E-07 |
| B20 | B139 | CD3+CD4+CD8+ | ADA | -0.53254946 | 0 |
| B20 | B22 | CD3+CD4+CD8+ | CD3+CD8+(T) | 0.466010006 | 0 |
| B20 | B120 | CD3+CD4+CD8+ | HCV-cAg | 0.360016524 | 1.46E-13 |
| B20 | B195 | CD3+CD4+CD8+ | Na | -0.35900494 | 1.72E-13 |
| B20 | B76 | CD3+CD4+CD8+ | ApoB | 0.277363315 | 1.99E-08 |
| B20 | B224 | CD3+CD4+CD8+ | pmh | -0.27629216 | 2.27E-08 |
| B20 | B200 | CD3+CD4+CD8+ | OP | -0.27021318 | 4.71E-08 |
| B204 | B209 | Drinking | Smoking | 0.468109181 | 0 |
| B204 | B222 | Drinking | NS | 0.329524412 | 1.75E-11 |
| B206 | B207 | Waist | WHR | 0.855610835 | 0 |
| B206 | B210 | Waist | Hip | 0.708112018 | 0 |
| B206 | B215 | Waist | BMI | 0.795076864 | 0 |
| B206 | B216 | Waist | BMI | 0.81079271 | 0 |
| B206 | B222 | Waist | NS | 0.734711855 | 0 |
| B206 | B209 | Waist | Smoking | 0.323086486 | 4.50E-11 |
| B206 | B221 | Waist | Hei | 0.312395222 | 2.06E-10 |
| B207 | B215 | WHR | BMI | 0.570095537 | 0 |
| B207 | B216 | WHR | BMI | 0.531534844 | 0 |
| B207 | B222 | WHR | NS | 0.594068499 | 0 |
| B207 | B209 | WHR | Smoking | 0.31257051 | 2.01E-10 |
| B207 | B220 | WHR | SBP | 0.287447681 | 5.70E-09 |
| B207 | B210 | WHR | Hip | 0.245233412 | 7.80E-07 |
| B209 | B222 | Smoking | NS | 0.401900359 | 0 |
| B209 | B216 | Smoking | BMI | 0.312062684 | 2.16E-10 |
| B209 | B221 | Smoking | Hei | 0.258429821 | 1.84E-07 |
| B21 | B138 | CD3+CD4-CD8- | PAB | 0.557734144 | 0 |
| B21 | B22 | CD3+CD4-CD8- | CD3+CD8+(T) | 0.873592269 | 0 |
| B21 | B23 | CD3+CD4-CD8- | CD3-CD16+CD56+(NK) | -0.99522008 | 0 |
| B21 | B58 | CD3+CD4-CD8- | HCT | 0.392109696 | 4.44E-16 |
| B21 | B194 | CD3+CD4-CD8- | Cl | -0.34544965 | 1.53E-12 |
| B21 | B193 | CD3+CD4-CD8- | K | -0.34352527 | 2.07E-12 |

| | | | | | |
|---|---|---|---|---|---|
| B21 | B69 | CD3+CD4-CD8- | PV | 0.324532548 | 3.65E-11 |
| B21 | B196 | CD3+CD4-CD8- | TCO2 | 0.323911733 | 3.99E-11 |
| B21 | B200 | CD3+CD4-CD8- | OP | -0.30884485 | 3.37E-10 |
| B21 | B63 | CD3+CD4-CD8- | WBRSR(mPas_200) | 0.307305945 | 4.16E-10 |
| B21 | B139 | CD3+CD4-CD8- | ADA | -0.30567541 | 5.20E-10 |
| B21 | B54 | CD3+CD4-CD8- | ESR | -0.30500097 | 5.70E-10 |
| B21 | B66 | CD3+CD4-CD8- | WBRSR(mPas_100) | 0.297297696 | 1.60E-09 |
| B21 | B67 | CD3+CD4-CD8- | WBRSR(mPas_50) | 0.296503841 | 1.77E-09 |
| B21 | B101 | CD3+CD4-CD8- | anti-Sc70 | 0.290915191 | 3.66E-09 |
| B21 | B102 | CD3+CD4-CD8- | anti-Sm | 0.290915191 | 3.66E-09 |
| B21 | B105 | CD3+CD4-CD8- | anti-U1RNP | 0.29091519 | 3.66E-09 |
| B21 | B106 | CD3+CD4-CD8- | anti-P | 0.29091519 | 3.66E-09 |
| B21 | B107 | CD3+CD4-CD8- | AnuA | 0.290915192 | 3.66E-09 |
| B21 | B108 | CD3+CD4-CD8- | Anti-Cyclin | 0.29091519 | 3.66E-09 |
| B21 | B109 | CD3+CD4-CD8- | AMA-M2 | 0.29091519 | 3.66E-09 |
| B21 | B110 | CD3+CD4-CD8- | AHA | 0.290915191 | 3.66E-09 |
| B21 | B95 | CD3+CD4-CD8- | anti-cenpb | 0.290915191 | 3.66E-09 |
| B21 | B97 | CD3+CD4-CD8- | dsDNA | 0.29091519 | 3.66E-09 |
| B21 | B98 | CD3+CD4-CD8- | anti-Jo-1 | 0.29091519 | 3.66E-09 |
| B21 | B99 | CD3+CD4-CD8- | anti-PM-Scl | 0.29091519 | 3.66E-09 |
| B21 | B195 | CD3+CD4-CD8- | Na | -0.28235494 | 1.08E-08 |
| B21 | B55 | CD3+CD4-CD8- | TK | -0.27260032 | 3.54E-08 |
| B210 | B215 | Hip | BMI | 0.720045334 | 0 |
| B210 | B216 | Hip | BMI | 0.794202744 | 0 |
| B210 | B222 | Hip | NS | 0.560520694 | 0 |
| B210 | B221 | Hip | Hei | 0.398160285 | 2.22E-16 |
| B211 | B212 | IOP-R | IOP-L | 0.834584827 | 0 |
| B213 | B214 | CVA-R | CVA-L | 0.787837042 | 0 |
| B215 | B216 | BMI | BMI | 0.790990592 | 0 |
| B215 | B222 | BMI | NS | 0.611567091 | 0 |
| B215 | B220 | BMI | SBP | 0.247324141 | 6.24E-07 |
| B216 | B221 | BMI | Hei | 0.628805017 | 0 |
| B216 | B222 | BMI | NS | 0.766049413 | 0 |
| B218 | B220 | DBP | SBP | 0.65328488 | 0 |
| B22 | B138 | CD3+CD8+(T) | PAB | 0.471212757 | 0 |
| B22 | B139 | CD3+CD8+(T) | ADA | -0.52977116 | 0 |
| B22 | B195 | CD3+CD8+(T) | Na | -0.42465178 | 0 |
| B22 | B200 | CD3+CD8+(T) | OP | -0.40487278 | 0 |
| B22 | B23 | CD3+CD8+(T) | CD3-CD16+CD56+(NK) | -0.91694242 | 0 |
| B22 | B194 | CD3+CD8+(T) | Cl | -0.35668982 | 2.52E-13 |
| B22 | B120 | CD3+CD8+(T) | HCV-cAg | 0.33323241 | 1.01E-11 |
| B22 | B193 | CD3+CD8+(T) | K | -0.29108758 | 3.58E-09 |
| B22 | B150 | CD3+CD8+(T) | ALP | -0.25719724 | 2.11E-07 |
| B22 | B224 | CD3+CD8+(T) | pmh | -0.24555225 | 7.54E-07 |

| | | | | | |
|---|---|---|---|---|---|
| B221 | B222 | Hei | NS | 0.466854263 | 0 |
| B23 | B138 | CD3-CD16+CD56+(NK) | PAB | -0.5518536 | 0 |
| B23 | B58 | CD3-CD16+CD56+(NK) | HCT | -0.37016111 | 2.64E-14 |
| B23 | B139 | CD3-CD16+CD56+(NK) | ADA | 0.356937282 | 2.42E-13 |
| B23 | B194 | CD3-CD16+CD56+(NK) | Cl | 0.354816717 | 3.42E-13 |
| B23 | B193 | CD3-CD16+CD56+(NK) | K | 0.340074529 | 3.54E-12 |
| B23 | B200 | CD3-CD16+CD56+(NK) | OP | 0.334472674 | 8.34E-12 |
| B23 | B195 | CD3-CD16+CD56+(NK) | Na | 0.316722193 | 1.12E-10 |
| B23 | B196 | CD3-CD16+CD56+(NK) | TCO2 | -0.31066448 | 2.62E-10 |
| B23 | B69 | CD3-CD16+CD56+(NK) | PV | -0.30711889 | 4.27E-10 |
| B23 | B63 | CD3-CD16+CD56+(NK) | WBRSR(mPas_200) | -0.29056975 | 3.83E-09 |
| B23 | B54 | CD3-CD16+CD56+(NK) | ESR | 0.287801618 | 5.45E-09 |
| B23 | B66 | CD3-CD16+CD56+(NK) | WBRSR(mPas_100) | -0.28113953 | 1.25E-08 |
| B23 | B67 | CD3-CD16+CD56+(NK) | WBRSR(mPas_50) | -0.28034056 | 1.38E-08 |
| B23 | B101 | CD3-CD16+CD56+(NK) | anti-Sc70 | -0.27430891 | 2.89E-08 |
| B23 | B102 | CD3-CD16+CD56+(NK) | anti-Sm | -0.27430891 | 2.89E-08 |
| B23 | B105 | CD3-CD16+CD56+(NK) | anti-U1RNP | -0.27430891 | 2.89E-08 |
| B23 | B106 | CD3-CD16+CD56+(NK) | anti-P | -0.27430891 | 2.89E-08 |
| B23 | B107 | CD3-CD16+CD56+(NK) | AnuA | -0.27430891 | 2.89E-08 |
| B23 | B108 | CD3-CD16+CD56+(NK) | Anti-Cyclin | -0.27430891 | 2.89E-08 |
| B23 | B109 | CD3-CD16+CD56+(NK) | AMA-M2 | -0.27430891 | 2.89E-08 |
| B23 | B110 | CD3-CD16+CD56+(NK) | AHA | -0.27430891 | 2.89E-08 |
| B23 | B95 | CD3-CD16+CD56+(NK) | anti-cenpb | -0.27430891 | 2.89E-08 |
| B23 | B97 | CD3-CD16+CD56+(NK) | dsDNA | -0.27430891 | 2.89E-08 |
| B23 | B98 | CD3-CD16+CD56+(NK) | anti-Jo-1 | -0.27430891 | 2.89E-08 |
| B23 | B99 | CD3-CD16+CD56+(NK) | anti-PM-Scl | -0.27430891 | 2.89E-08 |
| B23 | B55 | CD3-CD16+CD56+(NK) | TK | 0.25720035 | 2.11E-07 |
| B24 | B26 | αHBDH | CKMB | 0.479133327 | 0 |
| B24 | B27 | αHBDH | LDH | 0.659238805 | 0 |
| B24 | B83 | αHBDH | AFP | 0.370990967 | 2.29E-14 |
| B24 | B197 | αHBDH | AG | -0.30969944 | 3.00E-10 |
| B24 | B186 | αHBDH | PFC | 0.299148049 | 1.25E-09 |
| B24 | B82 | αHBDH | CEA | 0.281316787 | 1.23E-08 |
| B24 | B137 | αHBDH | AFU | -0.26887839 | 5.51E-08 |
| B24 | B194 | αHBDH | Cl | 0.258831474 | 1.75E-07 |
| B25 | B121 | CK | C3 | -0.48440609 | 0 |
| B25 | B122 | CK | C4 | -0.48449447 | 0 |
| B25 | B26 | CK | CKMB | 0.324867569 | 3.48E-11 |
| B25 | B103 | CK | anti-SSA/Ro | 0.273474939 | 3.19E-08 |
| B25 | B123 | CK | hs-CRP1 | -0.26481006 | 8.85E-08 |
| B26 | B83 | CKMB | AFP | 0.558016035 | 0 |
| B26 | B89 | CKMB | CA 72-4 | 0.348999923 | 8.74E-13 |
| B26 | B137 | CKMB | AFU | -0.34207839 | 2.60E-12 |
| B26 | B198 | CKMB | P | -0.34120883 | 2.97E-12 |

| | | | | | |
|---|---|---|---|---|---|
| B26 | B27 | CKMB | LDH | 0.313361575 | 1.80E-10 |
| B27 | B150 | LDH | ALP | 0.31363401 | 1.73E-10 |
| B27 | B131 | LDH | AST | 0.289146308 | 4.59E-09 |
| B27 | B137 | LDH | AFU | -0.28297433 | 9.99E-09 |
| B27 | B83 | LDH | AFP | 0.271085381 | 4.24E-08 |
| B28 | B119 | PRL | Hcv-rna | 0.518975263 | 0 |
| B28 | B120 | PRL | HCV-cAg | 0.493886247 | 0 |
| B28 | B152 | PRL | ACA-Ig(G-A-M) | -0.51899644 | 0 |
| B28 | B159 | PRL | β2-MG | -0.51897708 | 0 |
| B28 | B169 | PRL | U-CRY | -0.51897242 | 0 |
| B28 | B29 | PRL | E2 | 0.987541777 | 0 |
| B28 | B30 | PRL | LH | -0.60218313 | 0 |
| B28 | B31 | PRL | FSH | -0.95902679 | 0 |
| B28 | B32 | PRL | Testo | 0.989482462 | 0 |
| B28 | B33 | PRL | Prog | 0.987111234 | 0 |
| B28 | B79 | PRL | hb V -dna | 0.364365765 | 7.06E-14 |
| B28 | B124 | PRL | ACTH | 0.33177432 | 1.25E-11 |
| B28 | B74 | PRL | ApoA1 | -0.29252425 | 2.97E-09 |
| B28 | B82 | PRL | CEA | -0.27788024 | 1.87E-08 |
| B28 | B198 | PRL | P | -0.27243987 | 3.61E-08 |
| B29 | B119 | E2 | Hcv-rna | 0.559040365 | 0 |
| B29 | B120 | E2 | HCV-cAg | 0.462455791 | 0 |
| B29 | B152 | E2 | ACA-Ig(G-A-M) | -0.55906316 | 0 |
| B29 | B159 | E2 | β2-MG | -0.55904232 | 0 |
| B29 | B169 | E2 | U-CRY | -0.55903731 | 0 |
| B29 | B30 | E2 | LH | -0.71864792 | 0 |
| B29 | B31 | E2 | FSH | -0.98147259 | 0 |
| B29 | B32 | E2 | Testo | 0.983565549 | 0 |
| B29 | B33 | E2 | Prog | 0.99995357 | 0 |
| B29 | B79 | E2 | hb V -dna | 0.392690684 | 4.44E-16 |
| B29 | B124 | E2 | ACTH | 0.357185788 | 2.33E-13 |
| B29 | B74 | E2 | ApoA1 | -0.33132461 | 1.34E-11 |
| B29 | B198 | E2 | P | -0.30754605 | 4.03E-10 |
| B29 | B89 | E2 | CA 72-4 | -0.2907363 | 3.74E-09 |
| B29 | B199 | E2 | Mg | 0.258339383 | 1.85E-07 |
| B29 | B82 | E2 | CEA | -0.25093223 | 4.22E-07 |
| B30 | B119 | LH | Hcv-rna | -0.44171801 | 0 |
| B30 | B152 | LH | ACA-Ig(G-A-M) | 0.441736005 | 0 |
| B30 | B159 | LH | β2-MG | 0.441719543 | 0 |
| B30 | B169 | LH | U-CRY | 0.441715588 | 0 |
| B30 | B199 | LH | Mg | -0.41704917 | 0 |
| B30 | B31 | LH | FSH | 0.726380565 | 0 |
| B30 | B32 | LH | Testo | -0.60998655 | 0 |
| B30 | B33 | LH | Prog | -0.71907902 | 0 |

| | | | | | |
|---|---|---|---|---|---|
| B30 | B89 | LH | CA 72-4 | 0.598506088 | 0 |
| B30 | B134 | LH | CHE | -0.39526767 | 4.44E-16 |
| B30 | B74 | LH | ApoA1 | 0.372418957 | 1.78E-14 |
| B30 | B198 | LH | P | 0.344529151 | 1.77E-12 |
| B30 | B79 | LH | hb V -dna | -0.3104985 | 2.68E-10 |
| B30 | B136 | LH | 5'NT | -0.28838035 | 5.06E-09 |
| B30 | B124 | LH | ACTH | -0.28199939 | 1.13E-08 |
| B30 | B75 | LH | ApoA1/B | 0.28192704 | 1.14E-08 |
| B31 | B119 | FSH | Hcv-rna | -0.70691575 | 0 |
| B31 | B120 | FSH | HCV-cAg | -0.48278222 | 0 |
| B31 | B124 | FSH | ACTH | -0.45077245 | 0 |
| B31 | B152 | FSH | ACA-Ig(G-A-M) | 0.706944568 | 0 |
| B31 | B159 | FSH | β2-MG | 0.706918217 | 0 |
| B31 | B169 | FSH | U-CRY | 0.706911881 | 0 |
| B31 | B32 | FSH | Testo | -0.98286426 | 0 |
| B31 | B33 | FSH | Prog | -0.983266 | 0 |
| B31 | B79 | FSH | hb V -dna | -0.49743246 | 0 |
| B31 | B74 | FSH | ApoA1 | 0.362114053 | 1.03E-13 |
| B31 | B198 | FSH | P | 0.331866734 | 1.23E-11 |
| B31 | B179 | FSH | α1-MG | -0.29169203 | 3.31E-09 |
| B31 | B89 | FSH | CA 72-4 | 0.258507562 | 1.82E-07 |
| B32 | B119 | Testo | Hcv-rna | 0.636880971 | 0 |
| B32 | B120 | Testo | HCV-cAg | 0.51621376 | 0 |
| B32 | B124 | Testo | ACTH | 0.406428052 | 0 |
| B32 | B152 | Testo | ACA-Ig(G-A-M) | -0.63690695 | 0 |
| B32 | B159 | Testo | β2-MG | -0.63688319 | 0 |
| B32 | B169 | Testo | U-CRY | -0.63687748 | 0 |
| B32 | B33 | Testo | Prog | 0.984531319 | 0 |
| B32 | B79 | Testo | hb V -dna | 0.447846554 | 0 |
| B32 | B74 | Testo | ApoA1 | -0.31785229 | 9.55E-11 |
| B32 | B198 | Testo | P | -0.29271769 | 2.90E-09 |
| B32 | B179 | Testo | α1-MG | 0.262652216 | 1.14E-07 |
| B32 | B82 | Testo | CEA | -0.25024475 | 4.55E-07 |
| B33 | B119 | Prog | Hcv-rna | 0.567004201 | 0 |
| B33 | B120 | Prog | HCV-cAg | 0.464144194 | 0 |
| B33 | B152 | Prog | ACA-Ig(G-A-M) | -0.56702733 | 0 |
| B33 | B159 | Prog | β2-MG | -0.56700618 | 0 |
| B33 | B169 | Prog | U-CRY | -0.5670011 | 0 |
| B33 | B79 | Prog | hb V -dna | 0.398328638 | 0 |
| B33 | B124 | Prog | ACTH | 0.36222891 | 1.01E-13 |
| B33 | B74 | Prog | ApoA1 | -0.3330175 | 1.04E-11 |
| B33 | B198 | Prog | P | -0.3089025 | 3.34E-10 |
| B33 | B89 | Prog | CA 72-4 | -0.28887158 | 4.75E-09 |
| B33 | B199 | Prog | Mg | 0.256019282 | 2.41E-07 |

| | | | | | |
|---|---|---|---|---|---|
| B33 | B82 | Prog | CEA | -0.24920207 | 5.09E-07 |
| B34 | B92 | TK | ALDH2-Glu504Glu | -0.32122147 | 5.90E-11 |
| B34 | B93 | TK | ALDH2-Glu504Lys | 0.321221468 | 5.90E-11 |
| B34 | B94 | TK | ALDH2-Lys504Lys | -0.32122147 | 5.90E-11 |
| B34 | B178 | TK | Ig-U | -0.29566325 | 1.98E-09 |
| B34 | B180 | TK | Tf | -0.29263795 | 2.93E-09 |
| B34 | B175 | TK | ALB-U | -0.28595836 | 6.88E-09 |
| B34 | B125 | TK | Cor | 0.281889163 | 1.14E-08 |
| B34 | B177 | TK | ACR | -0.26981638 | 4.93E-08 |
| B34 | B124 | TK | ACTH | 0.247741848 | 5.96E-07 |
| B34 | B74 | TK | ApoA1 | -0.24752028 | 6.11E-07 |
| B35 | B116 | β2-MG | TT3 | -0.82869432 | 0 |
| B36 | B167 | V-Lym% | PC | 0.555103175 | 0 |
| B36 | B223 | V-Lym% | FH | 0.694876798 | 0 |
| B36 | B37 | V-Lym% | BAND | 0.457495711 | 0 |
| B36 | B96 | V-Lym% | dsDNA (IIF) | 0.646996639 | 0 |
| B37 | B167 | BAND | PC | -0.36629484 | 5.11E-14 |
| B39 | B78 | Other cells | HBcAb | 0.85253165 | 0 |
| B39 | B101 | Other cells | anti-Sc70 | 0.362171569 | 1.02E-13 |
| B39 | B102 | Other cells | anti-Sm | 0.362171569 | 1.02E-13 |
| B39 | B105 | Other cells | anti-U1RNP | 0.362171569 | 1.02E-13 |
| B39 | B106 | Other cells | anti-P | 0.362171568 | 1.02E-13 |
| B39 | B107 | Other cells | AnuA | 0.36217157 | 1.02E-13 |
| B39 | B108 | Other cells | Anti-Cyclin | 0.362171569 | 1.02E-13 |
| B39 | B109 | Other cells | AMA-M2 | 0.362171569 | 1.02E-13 |
| B39 | B110 | Other cells | AHA | 0.362171568 | 1.02E-13 |
| B39 | B95 | Other cells | anti-cenpb | 0.362171569 | 1.02E-13 |
| B39 | B97 | Other cells | dsDNA | 0.362171568 | 1.02E-13 |
| B39 | B98 | Other cells | anti-Jo-1 | 0.362171568 | 1.02E-13 |
| B39 | B99 | Other cells | anti-PM-Scl | 0.362171568 | 1.02E-13 |
| B39 | B103 | Other cells | anti-SSA/Ro | -0.34545047 | 1.53E-12 |
| B4 | B167 | WB-Ca | PC | 0.493215425 | 0 |
| B4 | B223 | WB-Ca | FH | 0.664291572 | 0 |
| B40 | B42 | WBC | MON | 0.68466748 | 0 |
| B40 | B53 | WBC | GR% | 0.372367616 | 1.80E-14 |
| B40 | B52 | WBC | PLT/BPC | 0.355249222 | 3.19E-13 |
| B40 | B46 | WBC | RBC Count | 0.329626635 | 1.72E-11 |
| B40 | B48 | WBC | LYM% | -0.30429244 | 6.27E-10 |
| B40 | B207 | WBC | WHR | 0.300971259 | 9.80E-10 |
| B40 | B127 | WBC | A-G ratio | -0.26915614 | 5.33E-08 |
| B40 | B192 | WBC | UA | 0.260679515 | 1.42E-07 |
| B40 | B206 | WBC | Waist | 0.259165419 | 1.69E-07 |
| B41 | B42 | MON % | MON | 0.530601006 | 0 |
| B41 | B53 | MON % | GR% | -0.24846944 | 5.51E-07 |

| | | | | | |
|---|---|---|---|---|---|
| B43 | B44 | MCV | MCH | 0.934518255 | 0 |
| B43 | B46 | MCV | RBC Count | -0.49702277 | 0 |
| B43 | B47 | MCV | RDW | 0.40859818 | 0 |
| B43 | B45 | MCV | MCHC | 0.344825273 | 1.69E-12 |
| B43 | B52 | MCV | PLT/BPC | -0.33223689 | 1.17E-11 |
| B43 | B51 | MCV | Hb | 0.289795631 | 4.22E-09 |
| B44 | B45 | MCH | MCHC | 0.654671834 | 0 |
| B44 | B51 | MCH | Hb | 0.44491138 | 0 |
| B44 | B46 | MCH | RBC Count | -0.38240145 | 3.11E-15 |
| B44 | B52 | MCH | PLT/BPC | -0.3459287 | 1.42E-12 |
| B45 | B51 | MCHC | Hb | 0.565733803 | 0 |
| B45 | B126 | MCHC | Alb | 0.30093388 | 9.85E-10 |
| B45 | B127 | MCHC | A-G ratio | 0.276715287 | 2.16E-08 |
| B45 | B216 | MCHC | BMI | 0.251628801 | 3.91E-07 |
| B46 | B47 | RBC Count | RDW | -0.40633826 | 0 |
| B46 | B51 | RBC Count | Hb | 0.646167053 | 0 |
| B46 | B222 | RBC Count | NS | 0.321774554 | 5.45E-11 |
| B46 | B192 | RBC Count | UA | 0.314311344 | 1.58E-10 |
| B46 | B216 | RBC Count | BMI | 0.311100188 | 2.47E-10 |
| B46 | B206 | RBC Count | Waist | 0.301873891 | 8.69E-10 |
| B46 | B207 | RBC Count | WHR | 0.279122672 | 1.61E-08 |
| B46 | B209 | RBC Count | Smoking | 0.253730806 | 3.10E-07 |
| B46 | B221 | RBC Count | Hei | 0.250655007 | 4.35E-07 |
| B46 | B174 | RBC Count | Urea/Cr | -0.24306124 | 9.82E-07 |
| B47 | B126 | RDW | Alb | -0.25766426 | 2.00E-07 |
| B48 | B53 | LYM% | GR% | -0.91990002 | 0 |
| B48 | B126 | LYM% | Alb | 0.29223409 | 3.09E-09 |
| B48 | B127 | LYM% | A-G ratio | 0.252148646 | 3.69E-07 |
| B49 | B50 | baso% | eo% | 0.279490587 | 1.54E-08 |
| B49 | B53 | baso% | GR% | -0.24883922 | 5.30E-07 |
| B5 | B167 | WB-Mg | PC | -0.76302703 | 0 |
| B5 | B37 | WB-Mg | BAND | -0.7200823 | 0 |
| B5 | B6 | WB-Mg | WB-Fe | 0.470013944 | 0 |
| B5 | B96 | WB-Mg | dsDNA (IIF) | 0.272772363 | 3.47E-08 |
| B50 | B53 | eo% | GR% | -0.35766757 | 2.15E-13 |
| B51 | B216 | Hb | BMI | 0.438923747 | 0 |
| B51 | B222 | Hb | NS | 0.431144862 | 0 |
| B51 | B221 | Hb | Hei | 0.390115374 | 8.88E-16 |
| B51 | B206 | Hb | Waist | 0.366627036 | 4.82E-14 |
| B51 | B192 | Hb | UA | 0.354544678 | 3.58E-13 |
| B51 | B190 | Hb | Cre | 0.327858086 | 2.24E-11 |
| B51 | B209 | Hb | Smoking | 0.315703007 | 1.30E-10 |
| B51 | B126 | Hb | Alb | 0.310592825 | 2.65E-10 |
| B51 | B163 | Hb | SG | 0.309535078 | 3.06E-10 |

| | | | | | |
|---|---|---|---|---|---|
| B51 | B210 | Hb | Hip | 0.299720425 | 1.16E-09 |
| B51 | B207 | Hb | WHR | 0.283782393 | 9.03E-09 |
| B51 | B141 | Hb | AST/ALT | -0.2743576 | 2.87E-08 |
| B51 | B215 | Hb | BMI | 0.26238389 | 1.17E-07 |
| B51 | B129 | Hb | IBIL | 0.259200107 | 1.68E-07 |
| B53 | B126 | GR% | Alb | -0.26113841 | 1.35E-07 |
| B53 | B127 | GR% | A-G ratio | -0.25138198 | 4.02E-07 |
| B54 | B55 | ESR | TK | 0.451299091 | 0 |
| B54 | B58 | ESR | HCT | -0.60175735 | 0 |
| B54 | B60 | ESR | WBRVI | -0.32791475 | 2.22E-11 |
| B54 | B69 | ESR | PV | 0.319171 | 7.92E-11 |
| B55 | B114 | TK | TSH TT3 TT4(TT4) | 0.419044692 | 0 |
| B55 | B56 | TK | IR | 0.806816132 | 0 |
| B55 | B57 | TK | RBCAI | -0.69504111 | 0 |
| B55 | B58 | TK | HCT | -0.70229403 | 0 |
| B55 | B60 | TK | WBRVI | -0.68152992 | 0 |
| B55 | B61 | TK | WBHSRV | 0.683682579 | 0 |
| B55 | B64 | TK | WBRSR(mPas_5) | -0.45071559 | 0 |
| B55 | B65 | TK | WBRSR(mPas_1) | -0.61250009 | 0 |
| B55 | B59 | TK | WBRV | -0.3576802 | 2.14E-13 |
| B55 | B196 | TK | TCO2 | -0.26519127 | 8.47E-08 |
| B55 | B156 | TK | IgE | 0.262350619 | 1.18E-07 |
| B55 | B62 | TK | WBHSRVI | 0.245437659 | 7.63E-07 |
| B56 | B114 | IR | TSH TT3 TT4(TT4) | 0.442262537 | 0 |
| B56 | B156 | IR | IgE | 0.40650249 | 0 |
| B56 | B57 | IR | RBCAI | -0.80071054 | 0 |
| B56 | B59 | IR | WBRV | -0.41684703 | 0 |
| B56 | B60 | IR | WBRVI | -0.51013545 | 0 |
| B56 | B61 | IR | WBHSRV | 0.93849919 | 0 |
| B56 | B62 | IR | WBHSRVI | 0.770195153 | 0 |
| B56 | B63 | IR | WBRSR(mPas_200) | 0.659357974 | 0 |
| B56 | B66 | IR | WBRSR(mPas_100) | 0.630403687 | 0 |
| B56 | B67 | IR | WBRSR(mPas_50) | 0.543434419 | 0 |
| B56 | B65 | IR | WBRSR(mPas_1) | -0.3896099 | 8.88E-16 |
| B57 | B114 | RBCAI | TSH TT3 TT4(TT4) | -0.41166058 | 0 |
| B57 | B156 | RBCAI | IgE | -0.46874507 | 0 |
| B57 | B59 | RBCAI | WBRV | 0.834843898 | 0 |
| B57 | B60 | RBCAI | WBRVI | 0.876672785 | 0 |
| B57 | B61 | RBCAI | WBHSRV | -0.70751906 | 0 |
| B57 | B62 | RBCAI | WBHSRVI | -0.54463124 | 0 |
| B57 | B63 | RBCAI | WBRSR(mPas_200) | -0.40915147 | 0 |
| B57 | B64 | RBCAI | WBRSR(mPas_5) | 0.494550505 | 0 |
| B57 | B65 | RBCAI | WBRSR(mPas_1) | 0.775376649 | 0 |
| B57 | B66 | RBCAI | WBRSR(mPas_100) | -0.35165239 | 5.71E-13 |

| | | | | | |
|---|---|---|---|---|---|
| B57 | B58 | RBCAI | HCT | 0.259632505 | 1.60E-07 |
| B58 | B60 | HCT | WBRVI | 0.596779414 | 0 |
| B58 | B62 | HCT | WBHSRVI | 0.506520018 | 0 |
| B58 | B63 | HCT | WBRSR(mPas_200) | 0.544878434 | 0 |
| B58 | B64 | HCT | WBRSR(mPas_5) | 0.702149714 | 0 |
| B58 | B65 | HCT | WBRSR(mPas_1) | 0.611138697 | 0 |
| B58 | B66 | HCT | WBRSR(mPas_100) | 0.53474365 | 0 |
| B58 | B67 | HCT | WBRSR(mPas_50) | 0.586139119 | 0 |
| B58 | B69 | HCT | PV | 0.271401283 | 4.09E-08 |
| B58 | B76 | HCT | ApoB | -0.24701832 | 6.44E-07 |
| B59 | B156 | WBRV | IgE | -0.40218827 | 0 |
| B59 | B60 | WBRV | WBRVI | 0.847648301 | 0 |
| B59 | B64 | WBRV | WBRSR(mPas_5) | 0.726194067 | 0 |
| B59 | B65 | WBRV | WBRSR(mPas_1) | 0.879048689 | 0 |
| B59 | B69 | WBRV | PV | 0.358782974 | 1.79E-13 |
| B59 | B62 | WBRV | WBHSRVI | -0.27282226 | 3.45E-08 |
| B59 | B114 | WBRV | TSH TT3 TT4(TT4) | -0.26532188 | 8.35E-08 |
| B6 | B167 | WB-Fe | PC | 0.418405566 | 0 |
| B6 | B8 | WB-Fe | WB-Zinc | 0.487565206 | 0 |
| B60 | B64 | WBRVI | WBRSR(mPas_5) | 0.814614693 | 0 |
| B60 | B65 | WBRVI | WBRSR(mPas_1) | 0.954959449 | 0 |
| B60 | B61 | WBRVI | WBHSRV | -0.37522913 | 1.11E-14 |
| B60 | B156 | WBRVI | IgE | -0.35946041 | 1.60E-13 |
| B60 | B114 | WBRVI | TSH TT3 TT4(TT4) | -0.33465557 | 8.11E-12 |
| B61 | B62 | WBHSRV | WBHSRVI | 0.803279548 | 0 |
| B61 | B63 | WBHSRV | WBRSR(mPas_200) | 0.813287253 | 0 |
| B61 | B66 | WBHSRV | WBRSR(mPas_100) | 0.806137567 | 0 |
| B61 | B67 | WBHSRV | WBRSR(mPas_50) | 0.742007867 | 0 |
| B61 | B69 | WBHSRV | PV | 0.551329051 | 0 |
| B61 | B114 | WBHSRV | TSH TT3 TT4(TT4) | 0.395300314 | 4.44E-16 |
| B61 | B156 | WBHSRV | IgE | 0.37416385 | 1.33E-14 |
| B61 | B92 | WBHSRV | ALDH2-Glu504Glu | -0.25647477 | 2.29E-07 |
| B61 | B93 | WBHSRV | ALDH2-Glu504Lys | 0.256474772 | 2.29E-07 |
| B61 | B94 | WBHSRV | ALDH2-Lys504Lys | -0.25647477 | 2.29E-07 |
| B62 | B63 | WBHSRVI | WBRSR(mPas_200) | 0.933823504 | 0 |
| B62 | B66 | WBHSRVI | WBRSR(mPas_100) | 0.90306135 | 0 |
| B62 | B67 | WBHSRVI | WBRSR(mPas_50) | 0.858426388 | 0 |
| B62 | B69 | WBHSRVI | PV | 0.39388528 | 4.44E-16 |
| B62 | B64 | WBHSRVI | WBRSR(mPas_5) | 0.378789292 | 5.77E-15 |
| B62 | B156 | WBHSRVI | IgE | 0.378506969 | 6.22E-15 |
| B62 | B114 | WBHSRVI | TSH TT3 TT4(TT4) | 0.273966345 | 3.01E-08 |
| B63 | B64 | WBRSR(mPas_200) | WBRSR(mPas_5) | 0.583769506 | 0 |
| B63 | B66 | WBRSR(mPas_200) | WBRSR(mPas_100) | 0.995022667 | 0 |
| B63 | B67 | WBRSR(mPas_200) | WBRSR(mPas_50) | 0.976946727 | 0 |

| | | | | | |
|---|---|---|---|---|---|
| B63 | B69 | WBRSR(mPas_200) | PV | 0.69196705 | 0 |
| B63 | B156 | WBRSR(mPas_200) | IgE | 0.305017718 | 5.69E-10 |
| B63 | B65 | WBRSR(mPas_200) | WBRSR(mPas_1) | 0.250687028 | 4.33E-07 |
| B64 | B65 | WBRSR(mPas_5) | WBRSR(mPas_1) | 0.931877813 | 0 |
| B64 | B66 | WBRSR(mPas_5) | WBRSR(mPas_100) | 0.636368702 | 0 |
| B64 | B67 | WBRSR(mPas_5) | WBRSR(mPas_50) | 0.73330688 | 0 |
| B64 | B69 | WBRSR(mPas_5) | PV | 0.653689838 | 0 |
| B65 | B67 | WBRSR(mPas_1) | WBRSR(mPas_50) | 0.437092901 | 0 |
| B65 | B69 | WBRSR(mPas_1) | PV | 0.454704271 | 0 |
| B65 | B66 | WBRSR(mPas_1) | WBRSR(mPas_100) | 0.31356686 | 1.75E-10 |
| B65 | B156 | WBRSR(mPas_1) | IgE | -0.30898422 | 3.31E-10 |
| B65 | B114 | WBRSR(mPas_1) | TSH TT3 TT4(TT4) | -0.28713521 | 5.93E-09 |
| B66 | B67 | WBRSR(mPas_100) | WBRSR(mPas_50) | 0.991053055 | 0 |
| B66 | B69 | WBRSR(mPas_100) | PV | 0.741290403 | 0 |
| B66 | B156 | WBRSR(mPas_100) | IgE | 0.27547767 | 2.51E-08 |
| B67 | B69 | WBRSR(mPas_50) | PV | 0.770733747 | 0 |
| B69 | B92 | PV | ALDH2-Glu504Glu | -0.31454597 | 1.53E-10 |
| B69 | B93 | PV | ALDH2-Glu504Lys | 0.314545972 | 1.53E-10 |
| B69 | B94 | PV | ALDH2-Lys504Lys | -0.31454597 | 1.53E-10 |
| B69 | B125 | PV | Cor | 0.276845225 | 2.12E-08 |
| B69 | B171 | PV | EC | -0.25647596 | 2.29E-07 |
| B69 | B124 | PV | ACTH | 0.243668891 | 9.21E-07 |
| B7 | B116 | WB-Cop | TT3 | -0.78920046 | 0 |
| B7 | B9 | WB-Cop | RH | -0.82105805 | 0 |
| B70 | B77 | LDL-C | TC | 0.861614842 | 0 |
| B70 | B76 | LDL-C | ApoB | 0.272023742 | 3.79E-08 |
| B70 | B206 | LDL-C | Waist | 0.249142193 | 5.12E-07 |
| B71 | B72 | TG | HDL-C | -0.36193992 | 1.06E-13 |
| B71 | B206 | TG | Waist | 0.343114007 | 2.21E-12 |
| B71 | B192 | TG | UA | 0.335175835 | 7.49E-12 |
| B71 | B207 | TG | WHR | 0.327018726 | 2.54E-11 |
| B71 | B77 | TG | TC | 0.320338174 | 6.70E-11 |
| B71 | B215 | TG | BMI | 0.275122326 | 2.62E-08 |
| B71 | B222 | TG | NS | 0.272877008 | 3.43E-08 |
| B71 | B216 | TG | BMI | 0.256053129 | 2.40E-07 |
| B72 | B192 | HDL-C | UA | -0.41787824 | 0 |
| B72 | B77 | HDL-C | TC | 0.413839498 | 0 |
| B72 | B222 | HDL-C | NS | -0.34242985 | 2.46E-12 |
| B72 | B207 | HDL-C | WHR | -0.32312084 | 4.48E-11 |
| B72 | B206 | HDL-C | Waist | -0.31304036 | 1.88E-10 |
| B72 | B216 | HDL-C | BMI | -0.30247797 | 8.01E-10 |
| B72 | B209 | HDL-C | Smoking | -0.26423414 | 9.46E-08 |
| B72 | B190 | HDL-C | Cre | -0.25796966 | 1.93E-07 |
| B72 | B174 | HDL-C | Urea/Cr | 0.251248857 | 4.07E-07 |

| | | | | | |
|---|---|---|---|---|---|
| B72 | B221 | HDL-C | Hei | -0.24767399 | 6.01E-07 |
| B73 | B123 | ApoA | hs-CRP1 | 0.462947668 | 0 |
| B73 | B186 | ApoA | PFC | 0.471328571 | 0 |
| B73 | B184 | ApoA | INR | -0.35081722 | 6.53E-13 |
| B73 | B156 | ApoA | IgE | 0.27386286 | 3.05E-08 |
| B73 | B125 | ApoA | Cor | -0.26637777 | 7.38E-08 |
| B73 | B76 | ApoA | ApoB | 0.257039762 | 2.15E-07 |
| B74 | B124 | ApoA1 | ACTH | -0.42943326 | 0 |
| B74 | B179 | ApoA1 | α1-MG | -0.42334969 | 0 |
| B74 | B75 | ApoA1 | ApoA1/B | 0.534878825 | 0 |
| B74 | B125 | ApoA1 | Cor | -0.37398506 | 1.38E-14 |
| B74 | B119 | ApoA1 | Hcv-rna | -0.33221058 | 1.17E-11 |
| B74 | B152 | ApoA1 | ACA-Ig(G-A-M) | 0.332224108 | 1.17E-11 |
| B74 | B159 | ApoA1 | β2-MG | 0.332211733 | 1.17E-11 |
| B74 | B169 | ApoA1 | U-CRY | 0.332208757 | 1.17E-11 |
| B74 | B198 | ApoA1 | P | 0.308605739 | 3.48E-10 |
| B74 | B176 | ApoA1 | Ucr | -0.27894199 | 1.64E-08 |
| B74 | B92 | ApoA1 | ALDH2-Glu504Glu | 0.269738274 | 4.98E-08 |
| B74 | B93 | ApoA1 | ALDH2-Glu504Lys | -0.26973827 | 4.98E-08 |
| B74 | B94 | ApoA1 | ALDH2-Lys504Lys | 0.269738272 | 4.98E-08 |
| B74 | B123 | ApoA1 | hs-CRP1 | 0.255078303 | 2.67E-07 |
| B74 | B150 | ApoA1 | ALP | 0.248687551 | 5.38E-07 |
| B74 | B199 | ApoA1 | Mg | -0.24673255 | 6.64E-07 |
| B75 | B76 | ApoA1/B | ApoB | -0.8840219 | 0 |
| B75 | B148 | ApoA1/B | Anti_TPO | 0.268566099 | 5.71E-08 |
| B76 | B148 | ApoB | Anti_TPO | -0.26189131 | 1.24E-07 |
| B76 | B77 | ApoB | TC | 0.243668058 | 9.21E-07 |
| B77 | B188 | TC | Ca | 0.273228542 | 3.29E-08 |
| B78 | B103 | HBcAb | anti-SSA/Ro | -0.26463462 | 9.04E-08 |
| B78 | B104 | HBcAb | anti-SSB | -0.25947482 | 1.63E-07 |
| B79 | B119 | hb V -dna | Hcv-rna | 0.706210978 | 0 |
| B79 | B125 | hb V -dna | Cor | -0.59665199 | 0 |
| B79 | B135 | hb V -dna | TBA | -0.47547986 | 0 |
| B79 | B152 | hb V -dna | ACA-Ig(G-A-M) | -0.70623977 | 0 |
| B79 | B159 | hb V -dna | β2-MG | -0.70621344 | 0 |
| B79 | B169 | hb V -dna | U-CRY | -0.70620711 | 0 |
| B79 | B181 | hb V -dna | APTT | -0.43488164 | 0 |
| B79 | B92 | hb V -dna | ALDH2-Glu504Glu | 0.706211306 | 0 |
| B79 | B93 | hb V -dna | ALDH2-Glu504Lys | -0.70621131 | 0 |
| B79 | B94 | hb V -dna | ALDH2-Lys504Lys | 0.7062113 | 0 |
| B79 | B184 | hb V -dna | INR | 0.355785889 | 2.92E-13 |
| B79 | B197 | hb V -dna | AG | -0.33404829 | 8.89E-12 |
| B79 | B81 | hb V -dna | TSGF | -0.31632257 | 1.19E-10 |
| B79 | B194 | hb V -dna | Cl | 0.291576852 | 3.36E-09 |

| | | | | | |
|---|---|---|---|---|---|
| B79 | B120 | hb V -dna | HCV-cAg | 0.282787295 | 1.02E-08 |
| B79 | B196 | hb V -dna | TCO2 | 0.270712789 | 4.43E-08 |
| B79 | B186 | hb V -dna | PFC | -0.26904517 | 5.40E-08 |
| B79 | B199 | hb V -dna | Mg | -0.26150129 | 1.30E-07 |
| B79 | B179 | hb V -dna | α1-MG | 0.260742568 | 1.41E-07 |
| B8 | B96 | WB-Zinc | dsDNA (IIF) | 0.545705156 | 0 |
| B81 | B92 | TSGF | ALDH2-Glu504Glu | -0.44998593 | 0 |
| B81 | B93 | TSGF | ALDH2-Glu504Lys | 0.449985933 | 0 |
| B81 | B94 | TSGF | ALDH2-Lys504Lys | -0.44998593 | 0 |
| B81 | B125 | TSGF | Cor | 0.396522209 | 2.22E-16 |
| B81 | B124 | TSGF | ACTH | 0.349211559 | 8.45E-13 |
| B81 | B181 | TSGF | APTT | 0.27684532 | 2.12E-08 |
| B82 | B85 | CEA | SCC | 0.458328619 | 0 |
| B82 | B86 | CEA | TBhCG | -0.53661017 | 0 |
| B82 | B87 | CEA | NSE | 0.510663492 | 0 |
| B82 | B88 | CEA | CA 242 | 0.628465589 | 0 |
| B82 | B83 | CEA | AFP | 0.331068091 | 1.39E-11 |
| B82 | B194 | CEA | Cl | 0.294499471 | 2.30E-09 |
| B83 | B89 | AFP | CA 72-4 | 0.488725542 | 0 |
| B83 | B137 | AFP | AFU | -0.33369399 | 9.38E-12 |
| B83 | B84 | AFP | C-PSA | 0.263477615 | 1.03E-07 |
| B83 | B198 | AFP | P | -0.25243085 | 3.58E-07 |
| B85 | B194 | SCC | Cl | 0.282207732 | 1.10E-08 |
| B85 | B197 | SCC | AG | -0.26085688 | 1.39E-07 |
| B85 | B224 | SCC | pmh | 0.252097865 | 3.71E-07 |
| B87 | B88 | NSE | CA 242 | 0.643778248 | 0 |
| B87 | B90 | NSE | CYFRA 21-1 | 0.525918948 | 0 |
| B88 | B90 | CA 242 | CYFRA 21-1 | 0.429367372 | 0 |
| B89 | B134 | CA 72-4 | CHE | -0.30686154 | 4.42E-10 |
| B89 | B199 | CA 72-4 | Mg | -0.28069651 | 1.33E-08 |
| B90 | B151 | CYFRA 21-1 | Anti-CCP | 0.249408012 | 4.98E-07 |
| B91 | B127 | TP | A-G ratio | -0.51909936 | 0 |
| B91 | B130 | TP | GLB | 0.766534869 | 0 |
| B91 | B188 | TP | Ca | 0.283822478 | 8.99E-09 |
| B92 | B124 | ALDH2-Glu504Glu | ACTH | -0.77295096 | 0 |
| B92 | B125 | ALDH2-Glu504Glu | Cor | -0.87884206 | 0 |
| B92 | B135 | ALDH2-Glu504Glu | TBA | -0.43789181 | 0 |
| B92 | B181 | ALDH2-Glu504Glu | APTT | -0.61649705 | 0 |
| B92 | B184 | ALDH2-Glu504Glu | INR | 0.503368653 | 0 |
| B92 | B194 | ALDH2-Glu504Glu | Cl | 0.460793281 | 0 |
| B92 | B94 | ALDH2-Glu504Glu | ALDH2-Lys504Lys | 0.999999992 | 0 |
| B92 | B197 | ALDH2-Glu504Glu | AG | -0.38983807 | 8.88E-16 |
| B92 | B186 | ALDH2-Glu504Glu | PFC | -0.38244907 | 3.11E-15 |
| B92 | B158 | ALDH2-Glu504Glu | IgM | -0.32866191 | 1.99E-11 |

| | | | | | |
|---|---|---|---|---|---|
| B92 | B199 | ALDH2-Glu504Glu | Mg | -0.31517355 | 1.40E-10 |
| B92 | B198 | ALDH2-Glu504Glu | P | 0.281615372 | 1.18E-08 |
| B93 | B124 | ALDH2-Glu504Lys | ACTH | 0.772950962 | 0 |
| B93 | B125 | ALDH2-Glu504Lys | Cor | 0.878842064 | 0 |
| B93 | B135 | ALDH2-Glu504Lys | TBA | 0.437891811 | 0 |
| B93 | B181 | ALDH2-Glu504Lys | APTT | 0.616497047 | 0 |
| B93 | B184 | ALDH2-Glu504Lys | INR | -0.50336865 | 0 |
| B93 | B194 | ALDH2-Glu504Lys | Cl | -0.46079328 | 0 |
| B93 | B94 | ALDH2-Glu504Lys | ALDH2-Lys504Lys | -0.99999999 | 0 |
| B93 | B197 | ALDH2-Glu504Lys | AG | 0.389838067 | 8.88E-16 |
| B93 | B186 | ALDH2-Glu504Lys | PFC | 0.382449069 | 3.11E-15 |
| B93 | B158 | ALDH2-Glu504Lys | IgM | 0.328661908 | 1.99E-11 |
| B93 | B199 | ALDH2-Glu504Lys | Mg | 0.315173549 | 1.40E-10 |
| B93 | B198 | ALDH2-Glu504Lys | P | -0.28161537 | 1.18E-08 |
| B94 | B124 | ALDH2-Lys504Lys | ACTH | -0.77295096 | 0 |
| B94 | B125 | ALDH2-Lys504Lys | Cor | -0.87884206 | 0 |
| B94 | B135 | ALDH2-Lys504Lys | TBA | -0.43789181 | 0 |
| B94 | B181 | ALDH2-Lys504Lys | APTT | -0.61649704 | 0 |
| B94 | B184 | ALDH2-Lys504Lys | INR | 0.503368649 | 0 |
| B94 | B194 | ALDH2-Lys504Lys | Cl | 0.460793278 | 0 |
| B94 | B197 | ALDH2-Lys504Lys | AG | -0.38983806 | 8.88E-16 |
| B94 | B186 | ALDH2-Lys504Lys | PFC | -0.38244907 | 3.11E-15 |
| B94 | B158 | ALDH2-Lys504Lys | IgM | -0.32866191 | 1.99E-11 |
| B94 | B199 | ALDH2-Lys504Lys | Mg | -0.31517355 | 1.40E-10 |
| B94 | B198 | ALDH2-Lys504Lys | P | 0.28161537 | 1.18E-08 |
| B95 | B103 | anti-cenpb | anti-SSA/Ro | -0.67410508 | 0 |
| B95 | B106 | anti-cenpb | anti-P | 0.999999999 | 0 |
| B95 | B108 | anti-cenpb | Anti-Cyclin | 1 | 0 |
| B95 | B109 | anti-cenpb | AMA-M2 | 1 | 0 |
| B95 | B97 | anti-cenpb | dsDNA | 1 | 0 |
| B95 | B98 | anti-cenpb | anti-Jo-1 | 0.999999998 | 0 |
| B95 | B99 | anti-cenpb | anti-PM-Scl | 1 | 0 |
| B95 | B121 | anti-cenpb | C3 | 0.32999543 | 1.63E-11 |
| B95 | B122 | anti-cenpb | C4 | 0.329969907 | 1.64E-11 |
| B95 | B104 | anti-cenpb | anti-SSB | -0.31611978 | 1.22E-10 |
| B95 | B155 | anti-cenpb | IgA | 0.251683699 | 3.88E-07 |
| B95 | B157 | anti-cenpb | IgG | -0.25117744 | 4.11E-07 |
| B95 | B100 | anti-cenpb | anti-Ro52 | -0.24927397 | 5.05E-07 |
| B97 | B103 | dsDNA | anti-SSA/Ro | -0.67410508 | 0 |
| B97 | B105 | dsDNA | anti-U1RNP | 0.999999998 | 0 |
| B97 | B108 | dsDNA | Anti-Cyclin | 0.999999999 | 0 |
| B97 | B109 | dsDNA | AMA-M2 | 0.999999999 | 0 |
| B97 | B110 | dsDNA | AHA | 0.999999998 | 0 |
| B97 | B99 | dsDNA | anti-PM-Scl | 0.999999999 | 0 |

| | | | | | |
|---|---|---|---|---|---|
| B97 | B121 | dsDNA | C3 | 0.32999543 | 1.63E-11 |
| B97 | B122 | dsDNA | C4 | 0.329969907 | 1.64E-11 |
| B97 | B104 | dsDNA | anti-SSB | -0.31611978 | 1.22E-10 |
| B97 | B155 | dsDNA | IgA | 0.251683699 | 3.88E-07 |
| B97 | B157 | dsDNA | IgG | -0.25117744 | 4.11E-07 |
| B97 | B100 | dsDNA | anti-Ro52 | -0.24927397 | 5.05E-07 |
| B98 | B101 | anti-Jo-1 | anti-Sc70 | 0.999999997 | 0 |
| B98 | B102 | anti-Jo-1 | anti-Sm | 0.999999999 | 0 |
| B98 | B103 | anti-Jo-1 | anti-SSA/Ro | -0.67410508 | 0 |
| B98 | B105 | anti-Jo-1 | anti-U1RNP | 0.999999998 | 0 |
| B98 | B109 | anti-Jo-1 | AMA-M2 | 0.999999997 | 0 |
| B98 | B110 | anti-Jo-1 | AHA | 0.999999999 | 0 |
| B98 | B121 | anti-Jo-1 | C3 | 0.32999543 | 1.63E-11 |
| B98 | B122 | anti-Jo-1 | C4 | 0.329969906 | 1.64E-11 |
| B98 | B104 | anti-Jo-1 | anti-SSB | -0.31611978 | 1.22E-10 |
| B98 | B155 | anti-Jo-1 | IgA | 0.251683699 | 3.88E-07 |
| B98 | B157 | anti-Jo-1 | IgG | -0.25117744 | 4.11E-07 |
| B98 | B100 | anti-Jo-1 | anti-Ro52 | -0.24927397 | 5.05E-07 |
| B99 | B103 | anti-PM-Scl | anti-SSA/Ro | -0.67410508 | 0 |
| B99 | B105 | anti-PM-Scl | anti-U1RNP | 0.999999999 | 0 |
| B99 | B106 | anti-PM-Scl | anti-P | 1 | 0 |
| B99 | B109 | anti-PM-Scl | AMA-M2 | 0.999999998 | 0 |
| B99 | B110 | anti-PM-Scl | AHA | 0.999999999 | 0 |
| B99 | B121 | anti-PM-Scl | C3 | 0.32999543 | 1.63E-11 |
| B99 | B122 | anti-PM-Scl | C4 | 0.329969907 | 1.64E-11 |
| B99 | B104 | anti-PM-Scl | anti-SSB | -0.31611978 | 1.22E-10 |
| B99 | B155 | anti-PM-Scl | IgA | 0.251683699 | 3.88E-07 |
| B99 | B157 | anti-PM-Scl | IgG | -0.25117744 | 4.11E-07 |
| B99 | B100 | anti-PM-Scl | anti-Ro52 | -0.24927397 | 5.05E-07 |

**Supplementary Table 15. PEI Correlations in Chronic rhinitis**

| Row | Column | Row | Column | Cor | *P* |
|---|---|---|---|---|---|
| B1 | B163 | Gender | SG | -0.54351 | 0 |
| B1 | B190 | Gender | Cre | -0.68471 | 0 |
| B1 | B192 | Gender | UA | -0.51787 | 0 |
| B1 | B206 | Gender | Waist | -0.58224 | 0 |
| B1 | B207 | Gender | WHR | -0.52517 | 0 |
| B1 | B210 | Gender | Hip | -0.46255 | 0 |
| B1 | B216 | Gender | BMI | -0.6616 | 0 |
| B1 | B221 | Gender | Hei | -0.66505 | 0 |
| B1 | B222 | Gender | NS | -0.70413 | 0 |
| B1 | B46 | Gender | RBC Count | -0.61346 | 0 |
| B1 | B51 | Gender | Hb | -0.69961 | 0 |
| B1 | B72 | Gender | HDL-C | 0.442135 | 0 |
| B1 | B215 | Gender | BMI | -0.40381 | 5.55E-14 |
| B1 | B141 | Gender | AST/ALT | 0.341137 | 3.67E-10 |
| B1 | B204 | Gender | Drinking | -0.32342 | 3.16E-09 |
| B1 | B174 | Gender | Urea/Cr | 0.305185 | 2.53E-08 |
| B1 | B128 | Gender | ALT | -0.30009 | 4.40E-08 |
| B1 | B191 | Gender | Urea | -0.28565 | 2.01E-07 |
| B1 | B45 | Gender | MCHC | -0.28059 | 3.35E-07 |
| B1 | B218 | Gender | DBP | -0.27417 | 6.32E-07 |
| B10 | B12 | PG Ⅰ | PG II | 0.579791 | 0 |
| B10 | B11 | PG Ⅰ | PGI/PGII | 0.270641 | 8.90E-07 |
| B100 | B103 | anti-Ro52 | anti-SSA/Ro | 1 | 0 |
| B100 | B105 | anti-Ro52 | anti-U1RNP | 1 | 0 |
| B100 | B106 | anti-Ro52 | anti-P | 1 | 0 |
| B100 | B107 | anti-Ro52 | AnuA | 1 | 0 |
| B100 | B151 | anti-Ro52 | Anti-CCP | -0.62933 | 0 |
| B100 | B154 | anti-Ro52 | RF | 0.392478 | 3.15E-13 |
| B100 | B178 | anti-Ro52 | Ig-U | 0.288523 | 1.49E-07 |
| B101 | B102 | anti-Sc70 | anti-Sm | 1 | 0 |
| B101 | B106 | anti-Sc70 | anti-P | 1 | 0 |
| B101 | B107 | anti-Sc70 | AnuA | 1 | 0 |
| B101 | B108 | anti-Sc70 | Anti-Cyclin | 1 | 0 |
| B101 | B151 | anti-Sc70 | Anti-CCP | -0.62933 | 0 |
| B101 | B154 | anti-Sc70 | RF | 0.392478 | 3.15E-13 |
| B101 | B178 | anti-Sc70 | Ig-U | 0.288523 | 1.49E-07 |
| B102 | B104 | anti-Sm | anti-SSB | 1 | 0 |
| B102 | B106 | anti-Sm | anti-P | 1 | 0 |
| B102 | B107 | anti-Sm | AnuA | 1 | 0 |
| B102 | B108 | anti-Sm | Anti-Cyclin | 1 | 0 |
| B102 | B151 | anti-Sm | Anti-CCP | -0.62933 | 0 |
| B102 | B154 | anti-Sm | RF | 0.392478 | 3.15E-13 |

| | | | | | |
|---|---|---|---|---|---|
| B102 | B178 | anti-Sm | Ig-U | 0.288523 | 1.49E-07 |
| B103 | B105 | anti-SSA/Ro | anti-U1RNP | 1 | 0 |
| B103 | B106 | anti-SSA/Ro | anti-P | 1 | 0 |
| B103 | B107 | anti-SSA/Ro | AnuA | 1 | 0 |
| B103 | B151 | anti-SSA/Ro | Anti-CCP | -0.62933 | 0 |
| B103 | B154 | anti-SSA/Ro | RF | 0.392478 | 3.15E-13 |
| B103 | B178 | anti-SSA/Ro | Ig-U | 0.288523 | 1.49E-07 |
| B104 | B107 | anti-SSB | AnuA | 1 | 0 |
| B104 | B151 | anti-SSB | Anti-CCP | -0.62933 | 0 |
| B104 | B154 | anti-SSB | RF | 0.392478 | 3.15E-13 |
| B104 | B178 | anti-SSB | Ig-U | 0.288523 | 1.49E-07 |
| B105 | B106 | anti-U1RNP | anti-P | 1 | 0 |
| B105 | B107 | anti-U1RNP | AnuA | 1 | 0 |
| B105 | B109 | anti-U1RNP | AMA-M2 | 1 | 0 |
| B105 | B110 | anti-U1RNP | AHA | 1 | 0 |
| B105 | B151 | anti-U1RNP | Anti-CCP | -0.62933 | 0 |
| B105 | B154 | anti-U1RNP | RF | 0.392478 | 3.15E-13 |
| B105 | B178 | anti-U1RNP | Ig-U | 0.288523 | 1.49E-07 |
| B106 | B107 | anti-P | AnuA | 1 | 0 |
| B106 | B108 | anti-P | Anti-Cyclin | 1 | 0 |
| B106 | B110 | anti-P | AHA | 1 | 0 |
| B106 | B151 | anti-P | Anti-CCP | -0.62933 | 0 |
| B106 | B154 | anti-P | RF | 0.392478 | 3.15E-13 |
| B106 | B178 | anti-P | Ig-U | 0.288523 | 1.49E-07 |
| B107 | B108 | AnuA | Anti-Cyclin | 1 | 0 |
| B107 | B109 | AnuA | AMA-M2 | 1 | 0 |
| B107 | B110 | AnuA | AHA | 1 | 0 |
| B107 | B151 | AnuA | Anti-CCP | -0.62933 | 0 |
| B107 | B154 | AnuA | RF | 0.392478 | 3.15E-13 |
| B107 | B178 | AnuA | Ig-U | 0.288523 | 1.49E-07 |
| B108 | B110 | Anti-Cyclin | AHA | 1 | 0 |
| B108 | B151 | Anti-Cyclin | Anti-CCP | -0.62933 | 0 |
| B108 | B154 | Anti-Cyclin | RF | 0.392478 | 3.15E-13 |
| B108 | B178 | Anti-Cyclin | Ig-U | 0.288523 | 1.49E-07 |
| B109 | B151 | AMA-M2 | Anti-CCP | -0.62933 | 0 |
| B109 | B154 | AMA-M2 | RF | 0.392478 | 3.15E-13 |
| B109 | B178 | AMA-M2 | Ig-U | 0.288523 | 1.49E-07 |
| B11 | B12 | PGI/PGII | PG II | -0.50773 | 0 |
| B11 | B85 | PGI/PGII | SCC | 0.371621 | 6.45E-12 |
| B11 | B200 | PGI/PGII | OP | -0.29386 | 8.57E-08 |
| B11 | B195 | PGI/PGII | Na | -0.28343 | 2.52E-07 |
| B11 | B87 | PGI/PGII | NSE | -0.27192 | 7.87E-07 |
| B11 | B90 | PGI/PGII | CYFRA 21-1 | -0.27026 | 9.23E-07 |
| B110 | B151 | AHA | Anti-CCP | -0.62933 | 0 |

| | | | | | |
|---|---|---|---|---|---|
| B110 | B154 | AHA | RF | 0.392478 | 3.15E-13 |
| B110 | B178 | AHA | Ig-U | 0.288523 | 1.49E-07 |
| B112 | B113 | FT4 | FT3 | 0.870398 | 0 |
| B112 | B114 | FT4 | TSH TT3 TT4(TT4) | 0.553524 | 0 |
| B112 | B115 | FT4 | TSH TT3 TT4(TT3) | 0.492649 | 0 |
| B113 | B115 | FT3 | TSH TT3 TT4(TT3) | 0.394364 | 2.37E-13 |
| B113 | B114 | FT3 | TSH TT3 TT4(TT4) | 0.362545 | 2.25E-11 |
| B114 | B115 | TSH TT3 TT4(TT4) | TSH TT3 TT4(TT3) | 0.878867 | 0 |
| B114 | B148 | TSH TT3 TT4(TT4) | Anti_TPO | 0.380804 | 1.75E-12 |
| B115 | B148 | TSH TT3 TT4(TT3) | Anti_TPO | 0.466372 | 0 |
| B116 | B142 | TT3 | 25-(OH)D | -0.73366 | 0 |
| B117 | B145 | TT4 | BGP | -0.82624 | 0 |
| B117 | B171 | TT4 | EC | 0.59222 | 0 |
| B118 | B170 | CEA1 | U-GLU | 0.283509 | 2.50E-07 |
| B119 | B120 | Hcv-rna | HCV-cAg | 0.492851 | 0 |
| B119 | B121 | Hcv-rna | C3 | -0.81521 | 0 |
| B119 | B122 | Hcv-rna | C4 | -0.81521 | 0 |
| B119 | B143 | Hcv-rna | β-CTx | -0.81522 | 0 |
| B119 | B159 | Hcv-rna | β2-MG | -0.81521 | 0 |
| B119 | B169 | Hcv-rna | U-CRY | -0.8152 | 0 |
| B120 | B121 | HCV-cAg | C3 | -0.40178 | 7.64E-14 |
| B120 | B122 | HCV-cAg | C4 | -0.40178 | 7.64E-14 |
| B120 | B143 | HCV-cAg | β-CTx | -0.40178 | 7.64E-14 |
| B120 | B159 | HCV-cAg | β2-MG | -0.40178 | 7.64E-14 |
| B120 | B169 | HCV-cAg | U-CRY | -0.40178 | 7.64E-14 |
| B121 | B169 | C3 | U-CRY | 0.999999 | 0 |
| B122 | B169 | C4 | U-CRY | 0.999999 | 0 |
| B123 | B154 | hs-CRP1 | RF | 0.406961 | 3.40E-14 |
| B124 | B180 | ACTH | Tf | -0.49456 | 0 |
| B124 | B187 | ACTH | NSE | -0.6133 | 0 |
| B124 | B185 | ACTH | PT | -0.38174 | 1.53E-12 |
| B124 | B179 | ACTH | α1-MG | 0.351614 | 9.60E-11 |
| B124 | B186 | ACTH | PFC | -0.31672 | 6.90E-09 |
| B124 | B183 | ACTH | PTA | -0.30721 | 2.02E-08 |
| B124 | B156 | ACTH | IgE | -0.2954 | 7.27E-08 |
| B125 | B180 | Cor | Tf | 0.494563 | 0 |
| B125 | B187 | Cor | NSE | 0.613304 | 0 |
| B125 | B185 | Cor | PT | 0.381739 | 1.53E-12 |
| B125 | B179 | Cor | α1-MG | -0.35161 | 9.60E-11 |
| B125 | B186 | Cor | PFC | 0.316722 | 6.90E-09 |
| B125 | B183 | Cor | PTA | 0.307213 | 2.02E-08 |
| B125 | B156 | Cor | IgE | 0.295403 | 7.27E-08 |
| B126 | B127 | Alb | A-G ratio | 0.431677 | 4.44E-16 |
| B126 | B188 | Alb | Ca | 0.434188 | 4.44E-16 |

| | | | | | |
|---|---|---|---|---|---|
| B127 | B130 | A-G ratio | GLB | -0.82835 | 0 |
| B128 | B131 | ALT | AST | 0.56791 | 0 |
| B128 | B140 | ALT | r-GT | 0.545343 | 0 |
| B128 | B141 | ALT | AST/ALT | -0.51182 | 0 |
| B128 | B216 | ALT | BMI | 0.383826 | 1.13E-12 |
| B128 | B222 | ALT | NS | 0.358311 | 3.97E-11 |
| B128 | B206 | ALT | Waist | 0.357793 | 4.25E-11 |
| B128 | B215 | ALT | BMI | 0.348559 | 1.43E-10 |
| B128 | B192 | ALT | UA | 0.348024 | 1.53E-10 |
| B128 | B210 | ALT | Hip | 0.318888 | 5.37E-09 |
| B128 | B207 | ALT | WHR | 0.281273 | 3.13E-07 |
| B129 | B132 | IBIL | Dbil | 0.718275 | 0 |
| B129 | B133 | IBIL | TBIL | 0.98139 | 0 |
| B131 | B184 | AST | INR | -0.39726 | 1.53E-13 |
| B132 | B133 | Dbil | TBIL | 0.838407 | 0 |
| B134 | B138 | CHE | PAB | 0.3848 | 9.81E-13 |
| B134 | B198 | CHE | P | -0.35725 | 4.57E-11 |
| B134 | B137 | CHE | AFU | 0.337531 | 5.75E-10 |
| B134 | B196 | CHE | TCO2 | 0.319225 | 5.17E-09 |
| B134 | B135 | CHE | TBA | -0.28957 | 1.34E-07 |
| B135 | B187 | TBA | NSE | -0.48054 | 0 |
| B136 | B195 | 5'NT | Na | -0.46029 | 0 |
| B136 | B197 | 5'NT | AG | -0.37993 | 1.99E-12 |
| B136 | B200 | 5'NT | OP | -0.31923 | 5.17E-09 |
| B136 | B196 | 5'NT | TCO2 | 0.288084 | 1.56E-07 |
| B137 | B197 | AFU | AG | 0.394698 | 2.25E-13 |
| B137 | B139 | AFU | ADA | 0.365337 | 1.54E-11 |
| B137 | B194 | AFU | Cl | -0.32216 | 3.67E-09 |
| B137 | B193 | AFU | K | -0.27409 | 6.38E-07 |
| B138 | B196 | PAB | TCO2 | 0.41056 | 1.91E-14 |
| B138 | B139 | PAB | ADA | -0.39385 | 2.56E-13 |
| B139 | B197 | ADA | AG | 0.413938 | 1.11E-14 |
| B139 | B196 | ADA | TCO2 | -0.36808 | 1.06E-11 |
| B14 | B193 | G-17 | K | 0.580689 | 0 |
| B14 | B200 | G-17 | OP | 0.388098 | 6.04E-13 |
| B14 | B195 | G-17 | Na | 0.282204 | 2.85E-07 |
| B140 | B206 | r-GT | Waist | 0.338973 | 4.81E-10 |
| B140 | B141 | r-GT | AST/ALT | -0.33526 | 7.61E-10 |
| B140 | B207 | r-GT | WHR | 0.324825 | 2.68E-09 |
| B140 | B216 | r-GT | BMI | 0.32162 | 3.91E-09 |
| B140 | B192 | r-GT | UA | 0.310201 | 1.45E-08 |
| B140 | B222 | r-GT | NS | 0.293377 | 9.01E-08 |
| B140 | B215 | r-GT | BMI | 0.2907 | 1.19E-07 |
| B141 | B192 | AST/ALT | UA | -0.36141 | 2.62E-11 |

| | | | | | |
|---|---|---|---|---|---|
| B141 | B216 | AST/ALT | BMI | -0.35223 | 8.86E-11 |
| B141 | B206 | AST/ALT | Waist | -0.34176 | 3.39E-10 |
| B141 | B173 | AST/ALT | NTT | 0.320652 | 4.38E-09 |
| B141 | B222 | AST/ALT | NS | -0.31748 | 6.32E-09 |
| B141 | B215 | AST/ALT | BMI | -0.30309 | 3.18E-08 |
| B141 | B207 | AST/ALT | WHR | -0.3026 | 3.35E-08 |
| B141 | B210 | AST/ALT | Hip | -0.27142 | 8.26E-07 |
| B142 | B223 | 25-(OH)D | FH | -0.66667 | 0 |
| B144 | B171 | Total-PINP | EC | 0.601929 | 0 |
| B145 | B167 | BGP | PC | 0.830105 | 0 |
| B145 | B171 | BGP | EC | 0.43093 | 6.66E-16 |
| B146 | B194 | CysC | Cl | 0.427149 | 1.33E-15 |
| B146 | B196 | CysC | TCO2 | -0.27633 | 5.12E-07 |
| B149 | B161 | TGAb | LEU | 0.310471 | 1.40E-08 |
| B149 | B168 | TGAb | NMP22 | -0.3058 | 2.36E-08 |
| B15 | B100 | CD3-CD19+(B cell) | anti-Ro52 | 0.486695 | 0 |
| B15 | B101 | CD3-CD19+(B cell) | anti-Sc70 | 0.486695 | 0 |
| B15 | B102 | CD3-CD19+(B cell) | anti-Sm | 0.486695 | 0 |
| B15 | B103 | CD3-CD19+(B cell) | anti-SSA/Ro | 0.486695 | 0 |
| B15 | B104 | CD3-CD19+(B cell) | anti-SSB | 0.486695 | 0 |
| B15 | B105 | CD3-CD19+(B cell) | anti-U1RNP | 0.486695 | 0 |
| B15 | B106 | CD3-CD19+(B cell) | anti-P | 0.486695 | 0 |
| B15 | B107 | CD3-CD19+(B cell) | AnuA | 0.486695 | 0 |
| B15 | B108 | CD3-CD19+(B cell) | Anti-Cyclin | 0.486695 | 0 |
| B15 | B109 | CD3-CD19+(B cell) | AMA-M2 | 0.486695 | 0 |
| B15 | B110 | CD3-CD19+(B cell) | AHA | 0.486695 | 0 |
| B15 | B139 | CD3-CD19+(B cell) | ADA | -0.48595 | 0 |
| B15 | B16 | CD3-CD19+(B cell) | CD3+T | -0.99728 | 0 |
| B15 | B17 | CD3-CD19+(B cell) | CD3+CD16+CD56+NK | 0.995002 | 0 |
| B15 | B18 | CD3-CD19+(B cell) | CD3+CD4+helper(inducer)T | 0.606256 | 0 |
| B15 | B19 | CD3-CD19+(B cell) | CD3+CD4+/CD3+CD8+ | 0.727366 | 0 |
| B15 | B194 | CD3-CD19+(B cell) | Cl | -0.44496 | 0 |
| B15 | B20 | CD3-CD19+(B cell) | CD3+CD4+CD8+ | 0.995002 | 0 |
| B15 | B21 | CD3-CD19+(B cell) | CD3+CD4-CD8- | 0.995002 | 0 |
| B15 | B22 | CD3-CD19+(B cell) | CD3+CD8+(T) | -0.7891 | 0 |
| B15 | B23 | CD3-CD19+(B cell) | CD3-CD16+CD56+(NK) | 0.777566 | 0 |
| B15 | B95 | CD3-CD19+(B cell) | anti-cenpb | 0.486695 | 0 |
| B15 | B97 | CD3-CD19+(B cell) | dsDNA | 0.486695 | 0 |
| B15 | B98 | CD3-CD19+(B cell) | anti-Jo-1 | 0.486695 | 0 |
| B15 | B99 | CD3-CD19+(B cell) | anti-PM-Scl | 0.486695 | 0 |
| B15 | B196 | CD3-CD19+(B cell) | TCO2 | 0.436496 | 2.22E-16 |
| B15 | B198 | CD3-CD19+(B cell) | P | -0.42113 | 3.55E-15 |
| B15 | B200 | CD3-CD19+(B cell) | OP | -0.38913 | 5.18E-13 |
| B15 | B168 | CD3-CD19+(B cell) | NMP22 | 0.347119 | 1.72E-10 |

| | | | | | |
|---|---|---|---|---|---|
| B15 | B138 | CD3-CD19+(B cell) | PAB | 0.326999 | 2.07E-09 |
| B15 | B146 | CD3-CD19+(B cell) | CysC | -0.32395 | 2.97E-09 |
| B15 | B156 | CD3-CD19+(B cell) | IgE | 0.28827 | 1.53E-07 |
| B15 | B193 | CD3-CD19+(B cell) | K | -0.28606 | 1.93E-07 |
| B151 | B154 | Anti-CCP | RF | -0.34423 | 2.48E-10 |
| B154 | B156 | RF | IgE | 0.400827 | 8.84E-14 |
| B154 | B184 | RF | INR | 0.351678 | 9.52E-11 |
| B154 | B182 | RF | TT | -0.29816 | 5.42E-08 |
| B155 | B198 | IgA | P | -0.29516 | 7.47E-08 |
| B156 | B184 | IgE | INR | 0.371871 | 6.23E-12 |
| B156 | B186 | IgE | PFC | 0.290467 | 1.22E-07 |
| B156 | B182 | IgE | TT | -0.27192 | 7.87E-07 |
| B159 | B169 | β2-MG | U-CRY | 0.999998 | 0 |
| B16 | B100 | CD3+T | anti-Ro52 | -0.51477 | 0 |
| B16 | B101 | CD3+T | anti-Sc70 | -0.51477 | 0 |
| B16 | B102 | CD3+T | anti-Sm | -0.51477 | 0 |
| B16 | B103 | CD3+T | anti-SSA/Ro | -0.51477 | 0 |
| B16 | B104 | CD3+T | anti-SSB | -0.51477 | 0 |
| B16 | B105 | CD3+T | anti-U1RNP | -0.51477 | 0 |
| B16 | B106 | CD3+T | anti-P | -0.51477 | 0 |
| B16 | B107 | CD3+T | AnuA | -0.51477 | 0 |
| B16 | B108 | CD3+T | Anti-Cyclin | -0.51477 | 0 |
| B16 | B109 | CD3+T | AMA-M2 | -0.51477 | 0 |
| B16 | B110 | CD3+T | AHA | -0.51477 | 0 |
| B16 | B139 | CD3+T | ADA | 0.481512 | 0 |
| B16 | B17 | CD3+T | CD3+CD16+CD56+NK | -0.98495 | 0 |
| B16 | B18 | CD3+T | CD3+CD4+helper(inducer)T | -0.66317 | 0 |
| B16 | B19 | CD3+T | CD3+CD4+/CD3+CD8+ | -0.77593 | 0 |
| B16 | B194 | CD3+T | Cl | 0.440277 | 0 |
| B16 | B20 | CD3+T | CD3+CD4+CD8+ | -0.98495 | 0 |
| B16 | B21 | CD3+T | CD3+CD4-CD8- | -0.98495 | 0 |
| B16 | B22 | CD3+T | CD3+CD8+(T) | 0.832191 | 0 |
| B16 | B23 | CD3+T | CD3-CD16+CD56+(NK) | -0.82176 | 0 |
| B16 | B95 | CD3+T | anti-cenpb | -0.51477 | 0 |
| B16 | B97 | CD3+T | dsDNA | -0.51477 | 0 |
| B16 | B98 | CD3+T | anti-Jo-1 | -0.51477 | 0 |
| B16 | B99 | CD3+T | anti-PM-Scl | -0.51477 | 0 |
| B16 | B196 | CD3+T | TCO2 | -0.43209 | 4.44E-16 |
| B16 | B198 | CD3+T | P | 0.416855 | 7.11E-15 |
| B16 | B200 | CD3+T | OP | 0.38504 | 9.47E-13 |
| B16 | B168 | CD3+T | NMP22 | -0.34303 | 2.89E-10 |
| B16 | B138 | CD3+T | PAB | -0.32382 | 3.02E-09 |
| B16 | B146 | CD3+T | CysC | 0.320622 | 4.39E-09 |
| B16 | B193 | CD3+T | K | 0.28301 | 2.63E-07 |

| | | | | | |
|---|---|---|---|---|---|
| B16 | B156 | CD3+T | IgE | -0.27834 | 4.19E-07 |
| B161 | B173 | LEU | NTT | 0.413424 | 1.20E-14 |
| B163 | B166 | SG | Pro | 0.467708 | 0 |
| B163 | B190 | SG | Cre | 0.497867 | 0 |
| B163 | B216 | SG | BMI | 0.380977 | 1.71E-12 |
| B163 | B222 | SG | NS | 0.376617 | 3.19E-12 |
| B163 | B192 | SG | UA | 0.3593 | 3.48E-11 |
| B163 | B221 | SG | Hei | 0.359242 | 3.51E-11 |
| B163 | B191 | SG | Urea | 0.34924 | 1.31E-10 |
| B163 | B206 | SG | Waist | 0.298967 | 4.97E-08 |
| B163 | B207 | SG | WHR | 0.278147 | 4.27E-07 |
| B168 | B198 | NMP22 | P | -0.31963 | 4.93E-09 |
| B17 | B100 | CD3+CD16+CD56+NK | anti-Ro52 | 0.444401 | 0 |
| B17 | B101 | CD3+CD16+CD56+NK | anti-Sc70 | 0.444401 | 0 |
| B17 | B102 | CD3+CD16+CD56+NK | anti-Sm | 0.444401 | 0 |
| B17 | B103 | CD3+CD16+CD56+NK | anti-SSA/Ro | 0.444401 | 0 |
| B17 | B104 | CD3+CD16+CD56+NK | anti-SSB | 0.444401 | 0 |
| B17 | B105 | CD3+CD16+CD56+NK | anti-U1RNP | 0.444401 | 0 |
| B17 | B106 | CD3+CD16+CD56+NK | anti-P | 0.444401 | 0 |
| B17 | B107 | CD3+CD16+CD56+NK | AnuA | 0.444401 | 0 |
| B17 | B108 | CD3+CD16+CD56+NK | Anti-Cyclin | 0.444401 | 0 |
| B17 | B109 | CD3+CD16+CD56+NK | AMA-M2 | 0.444401 | 0 |
| B17 | B110 | CD3+CD16+CD56+NK | AHA | 0.444401 | 0 |
| B17 | B139 | CD3+CD16+CD56+NK | ADA | -0.48774 | 0 |
| B17 | B18 | CD3+CD16+CD56+NK | CD3+CD4+helper(inducer)T | 0.523814 | 0 |
| B17 | B19 | CD3+CD16+CD56+NK | CD3+CD4+/CD3+CD8+ | 0.655205 | 0 |
| B17 | B194 | CD3+CD16+CD56+NK | Cl | -0.44745 | 0 |
| B17 | B22 | CD3+CD16+CD56+NK | CD3+CD8+(T) | -0.72382 | 0 |
| B17 | B23 | CD3+CD16+CD56+NK | CD3-CD16+CD56+(NK) | 0.710891 | 0 |
| B17 | B95 | CD3+CD16+CD56+NK | anti-cenpb | 0.444401 | 0 |
| B17 | B97 | CD3+CD16+CD56+NK | dsDNA | 0.444401 | 0 |
| B17 | B98 | CD3+CD16+CD56+NK | anti-Jo-1 | 0.444401 | 0 |
| B17 | B99 | CD3+CD16+CD56+NK | anti-PM-Scl | 0.444401 | 0 |
| B17 | B196 | CD3+CD16+CD56+NK | TCO2 | 0.438683 | 2.22E-16 |
| B17 | B198 | CD3+CD16+CD56+NK | P | -0.42328 | 2.44E-15 |
| B17 | B200 | CD3+CD16+CD56+NK | OP | -0.39131 | 3.75E-13 |
| B17 | B168 | CD3+CD16+CD56+NK | NMP22 | 0.349653 | 1.24E-10 |
| B17 | B138 | CD3+CD16+CD56+NK | PAB | 0.328466 | 1.74E-09 |
| B17 | B146 | CD3+CD16+CD56+NK | CysC | -0.32566 | 2.43E-09 |
| B17 | B156 | CD3+CD16+CD56+NK | IgE | 0.299231 | 4.83E-08 |
| B17 | B193 | CD3+CD16+CD56+NK | K | -0.28772 | 1.62E-07 |
| B17 | B195 | CD3+CD16+CD56+NK | Na | -0.2707 | 8.85E-07 |
| B170 | B173 | U-GLU | NTT | 0.311905 | 1.19E-08 |
| B174 | B191 | Urea/Cr | Urea | 0.628182 | 0 |

| | | | | | |
|---|---|---|---|---|---|
| B174 | B190 | Urea/Cr | Cre | -0.41186 | 1.55E-14 |
| B174 | B221 | Urea/Cr | Hei | -0.30146 | 3.80E-08 |
| B175 | B177 | ALB-U | ACR | 0.425819 | 1.55E-15 |
| B175 | B176 | ALB-U | Ucr | 0.372752 | 5.51E-12 |
| B175 | B186 | ALB-U | PFC | -0.30854 | 1.74E-08 |
| B175 | B185 | ALB-U | PT | 0.28369 | 2.45E-07 |
| B176 | B177 | Ucr | ACR | -0.44914 | 0 |
| B176 | B179 | Ucr | α1-MG | 0.320637 | 4.39E-09 |
| B177 | B179 | ACR | α1-MG | -0.38681 | 7.31E-13 |
| B178 | B180 | Ig-U | Tf | 0.713474 | 0 |
| B179 | B180 | α1-MG | Tf | -0.28444 | 2.27E-07 |
| B18 | B100 | CD3+CD4+helper(inducer)T | anti-Ro52 | 0.612527 | 0 |
| B18 | B101 | CD3+CD4+helper(inducer)T | anti-Sc70 | 0.612527 | 0 |
| B18 | B102 | CD3+CD4+helper(inducer)T | anti-Sm | 0.612527 | 0 |
| B18 | B103 | CD3+CD4+helper(inducer)T | anti-SSA/Ro | 0.612527 | 0 |
| B18 | B104 | CD3+CD4+helper(inducer)T | anti-SSB | 0.612527 | 0 |
| B18 | B105 | CD3+CD4+helper(inducer)T | anti-U1RNP | 0.612527 | 0 |
| B18 | B106 | CD3+CD4+helper(inducer)T | anti-P | 0.612527 | 0 |
| B18 | B107 | CD3+CD4+helper(inducer)T | AnuA | 0.612527 | 0 |
| B18 | B108 | CD3+CD4+helper(inducer)T | Anti-Cyclin | 0.612527 | 0 |
| B18 | B109 | CD3+CD4+helper(inducer)T | AMA-M2 | 0.612527 | 0 |
| B18 | B110 | CD3+CD4+helper(inducer)T | AHA | 0.612527 | 0 |
| B18 | B178 | CD3+CD4+helper(inducer)T | Ig-U | 0.55863 | 0 |
| B18 | B180 | CD3+CD4+helper(inducer)T | Tf | 0.447747 | 0 |
| B18 | B19 | CD3+CD4+helper(inducer)T | CD3+CD4+/CD3+CD8+ | 0.986724 | 0 |
| B18 | B20 | CD3+CD4+helper(inducer)T | CD3+CD4+CD8+ | 0.523814 | 0 |
| B18 | B21 | CD3+CD4+helper(inducer)T | CD3+CD4-CD8- | 0.523814 | 0 |
| B18 | B22 | CD3+CD4+helper(inducer)T | CD3+CD8+(T) | -0.9669 | 0 |
| B18 | B23 | CD3+CD4+helper(inducer)T | CD3-CD16+CD56+(NK) | 0.97147 | 0 |
| B18 | B92 | CD3+CD4+helper(inducer)T | ALDH2-Glu504Glu | -0.59906 | 0 |
| B18 | B93 | CD3+CD4+helper(inducer)T | ALDH2-Glu504Lys | 0.599057 | 0 |
| B18 | B94 | CD3+CD4+helper(inducer)T | ALDH2-Lys504Lys | 0.599057 | 0 |
| B18 | B95 | CD3+CD4+helper(inducer)T | anti-cenpb | 0.612527 | 0 |
| B18 | B97 | CD3+CD4+helper(inducer)T | dsDNA | 0.612527 | 0 |
| B18 | B98 | CD3+CD4+helper(inducer)T | anti-Jo-1 | 0.612527 | 0 |
| B18 | B99 | CD3+CD4+helper(inducer)T | anti-PM-Scl | 0.612527 | 0 |
| B18 | B201 | CD3+CD4+helper(inducer)T | e-habits | -0.30709 | 2.05E-08 |
| B18 | B154 | CD3+CD4+helper(inducer)T | RF | 0.289714 | 1.32E-07 |
| B18 | B24 | CD3+CD4+helper(inducer)T | αHBDH | 0.285775 | 1.98E-07 |
| B180 | B187 | Tf | NSE | 0.301135 | 3.93E-08 |
| B181 | B183 | APTT | PTA | 0.738304 | 0 |
| B181 | B184 | APTT | INR | 0.448265 | 0 |
| B181 | B185 | APTT | PT | 0.705202 | 0 |
| B181 | B186 | APTT | PFC | -0.50427 | 0 |

| | | | | | |
|---|---|---|---|---|---|
| B181 | B182 | APTT | TT | 0.288457 | 1.51E-07 |
| B182 | B186 | TT | PFC | -0.81694 | 0 |
| B182 | B184 | TT | INR | -0.36012 | 3.12E-11 |
| B183 | B184 | PTA | INR | 0.785348 | 0 |
| B183 | B185 | PTA | PT | 0.817879 | 0 |
| B184 | B185 | INR | PT | 0.778069 | 0 |
| B185 | B186 | PT | PFC | -0.36982 | 8.30E-12 |
| B187 | B227 | NSE | OD | 0.338432 | 5.14E-10 |
| B189 | B190 | eGFB | Cre | -0.52861 | 0 |
| B19 | B100 | CD3+CD4+/CD3+CD8+ | anti-Ro52 | 0.627951 | 0 |
| B19 | B101 | CD3+CD4+/CD3+CD8+ | anti-Sc70 | 0.627951 | 0 |
| B19 | B102 | CD3+CD4+/CD3+CD8+ | anti-Sm | 0.627951 | 0 |
| B19 | B103 | CD3+CD4+/CD3+CD8+ | anti-SSA/Ro | 0.627951 | 0 |
| B19 | B104 | CD3+CD4+/CD3+CD8+ | anti-SSB | 0.627951 | 0 |
| B19 | B105 | CD3+CD4+/CD3+CD8+ | anti-U1RNP | 0.627951 | 0 |
| B19 | B106 | CD3+CD4+/CD3+CD8+ | anti-P | 0.627951 | 0 |
| B19 | B107 | CD3+CD4+/CD3+CD8+ | AnuA | 0.627951 | 0 |
| B19 | B108 | CD3+CD4+/CD3+CD8+ | Anti-Cyclin | 0.627951 | 0 |
| B19 | B109 | CD3+CD4+/CD3+CD8+ | AMA-M2 | 0.627951 | 0 |
| B19 | B110 | CD3+CD4+/CD3+CD8+ | AHA | 0.627951 | 0 |
| B19 | B178 | CD3+CD4+/CD3+CD8+ | Ig-U | 0.494595 | 0 |
| B19 | B20 | CD3+CD4+/CD3+CD8+ | CD3+CD4+CD8+ | 0.655205 | 0 |
| B19 | B21 | CD3+CD4+/CD3+CD8+ | CD3+CD4-CD8- | 0.655205 | 0 |
| B19 | B22 | CD3+CD4+/CD3+CD8+ | CD3+CD8+(T) | -0.9955 | 0 |
| B19 | B23 | CD3+CD4+/CD3+CD8+ | CD3-CD16+CD56+(NK) | 0.99709 | 0 |
| B19 | B92 | CD3+CD4+/CD3+CD8+ | ALDH2-Glu504Glu | -0.53043 | 0 |
| B19 | B93 | CD3+CD4+/CD3+CD8+ | ALDH2-Glu504Lys | 0.53043 | 0 |
| B19 | B94 | CD3+CD4+/CD3+CD8+ | ALDH2-Lys504Lys | 0.53043 | 0 |
| B19 | B95 | CD3+CD4+/CD3+CD8+ | anti-cenpb | 0.627951 | 0 |
| B19 | B97 | CD3+CD4+/CD3+CD8+ | dsDNA | 0.627951 | 0 |
| B19 | B98 | CD3+CD4+/CD3+CD8+ | anti-Jo-1 | 0.627951 | 0 |
| B19 | B99 | CD3+CD4+/CD3+CD8+ | anti-PM-Scl | 0.627951 | 0 |
| B19 | B180 | CD3+CD4+/CD3+CD8+ | Tf | 0.395889 | 1.88E-13 |
| B19 | B139 | CD3+CD4+/CD3+CD8+ | ADA | -0.32443 | 2.81E-09 |
| B19 | B201 | CD3+CD4+/CD3+CD8+ | e-habits | -0.31482 | 8.58E-09 |
| B19 | B194 | CD3+CD4+/CD3+CD8+ | Cl | -0.29124 | 1.13E-07 |
| B19 | B196 | CD3+CD4+/CD3+CD8+ | TCO2 | 0.287466 | 1.67E-07 |
| B19 | B154 | CD3+CD4+/CD3+CD8+ | RF | 0.286866 | 1.77E-07 |
| B19 | B198 | CD3+CD4+/CD3+CD8+ | P | -0.27709 | 4.75E-07 |
| B190 | B192 | Cre | UA | 0.498771 | 0 |
| B190 | B216 | Cre | BMI | 0.487162 | 0 |
| B190 | B221 | Cre | Hei | 0.467129 | 0 |
| B190 | B222 | Cre | NS | 0.508297 | 0 |
| B190 | B206 | Cre | Waist | 0.405061 | 4.57E-14 |

| | | | | | |
|---|---|---|---|---|---|
| B190 | B191 | Cre | Urea | 0.401101 | 8.48E-14 |
| B190 | B207 | Cre | WHR | 0.361009 | 2.77E-11 |
| B190 | B210 | Cre | Hip | 0.329611 | 1.51E-09 |
| B190 | B215 | Cre | BMI | 0.313742 | 9.70E-09 |
| B192 | B206 | UA | Waist | 0.466739 | 0 |
| B192 | B216 | UA | BMI | 0.524312 | 0 |
| B192 | B222 | UA | NS | 0.531205 | 0 |
| B192 | B210 | UA | Hip | 0.423064 | 2.66E-15 |
| B192 | B221 | UA | Hei | 0.420098 | 4.00E-15 |
| B192 | B215 | UA | BMI | 0.395487 | 2.00E-13 |
| B192 | B207 | UA | WHR | 0.366589 | 1.30E-11 |
| B192 | B204 | UA | Drinking | 0.3147 | 8.70E-09 |
| B192 | B218 | UA | DBP | 0.272358 | 7.54E-07 |
| B193 | B195 | K | Na | 0.563333 | 0 |
| B193 | B200 | K | OP | 0.659558 | 0 |
| B193 | B197 | K | AG | 0.274893 | 5.89E-07 |
| B194 | B196 | Cl | TCO2 | -0.51003 | 0 |
| B194 | B199 | Cl | Mg | 0.375295 | 3.85E-12 |
| B195 | B197 | Na | AG | 0.490043 | 0 |
| B195 | B200 | Na | OP | 0.937903 | 0 |
| B195 | B198 | Na | P | -0.33241 | 1.08E-09 |
| B196 | B197 | TCO2 | AG | -0.6518 | 0 |
| B196 | B199 | TCO2 | Mg | -0.29047 | 1.22E-07 |
| B197 | B200 | AG | OP | 0.440121 | 0 |
| B198 | B200 | P | OP | -0.28446 | 2.27E-07 |
| B2 | B189 | Age | eGFB | -0.50502 | 0 |
| B2 | B47 | Age | RDW | 0.339152 | 4.70E-10 |
| B2 | B126 | Age | Alb | -0.3306 | 1.34E-09 |
| B2 | B207 | Age | WHR | 0.31187 | 1.20E-08 |
| B2 | B153 | Age | Glu | 0.27308 | 7.03E-07 |
| B20 | B100 | CD3+CD4+CD8+ | anti-Ro52 | 0.444401 | 0 |
| B20 | B101 | CD3+CD4+CD8+ | anti-Sc70 | 0.444401 | 0 |
| B20 | B102 | CD3+CD4+CD8+ | anti-Sm | 0.444401 | 0 |
| B20 | B103 | CD3+CD4+CD8+ | anti-SSA/Ro | 0.444401 | 0 |
| B20 | B104 | CD3+CD4+CD8+ | anti-SSB | 0.444401 | 0 |
| B20 | B105 | CD3+CD4+CD8+ | anti-U1RNP | 0.444401 | 0 |
| B20 | B106 | CD3+CD4+CD8+ | anti-P | 0.444401 | 0 |
| B20 | B107 | CD3+CD4+CD8+ | AnuA | 0.444401 | 0 |
| B20 | B108 | CD3+CD4+CD8+ | Anti-Cyclin | 0.444401 | 0 |
| B20 | B109 | CD3+CD4+CD8+ | AMA-M2 | 0.444401 | 0 |
| B20 | B110 | CD3+CD4+CD8+ | AHA | 0.444401 | 0 |
| B20 | B139 | CD3+CD4+CD8+ | ADA | -0.48774 | 0 |
| B20 | B194 | CD3+CD4+CD8+ | Cl | -0.44745 | 0 |
| B20 | B22 | CD3+CD4+CD8+ | CD3+CD8+(T) | -0.72382 | 0 |

| | | | | | |
|---|---|---|---|---|---|
| B20 | B23 | CD3+CD4+CD8+ | CD3-CD16+CD56+(NK) | 0.710891 | 0 |
| B20 | B95 | CD3+CD4+CD8+ | anti-cenpb | 0.444401 | 0 |
| B20 | B97 | CD3+CD4+CD8+ | dsDNA | 0.444401 | 0 |
| B20 | B98 | CD3+CD4+CD8+ | anti-Jo-1 | 0.444401 | 0 |
| B20 | B99 | CD3+CD4+CD8+ | anti-PM-Scl | 0.444401 | 0 |
| B20 | B196 | CD3+CD4+CD8+ | TCO2 | 0.438683 | 2.22E-16 |
| B20 | B198 | CD3+CD4+CD8+ | P | -0.42328 | 2.44E-15 |
| B20 | B200 | CD3+CD4+CD8+ | OP | -0.39131 | 3.75E-13 |
| B20 | B168 | CD3+CD4+CD8+ | NMP22 | 0.349653 | 1.24E-10 |
| B20 | B138 | CD3+CD4+CD8+ | PAB | 0.328466 | 1.74E-09 |
| B20 | B146 | CD3+CD4+CD8+ | CysC | -0.32566 | 2.43E-09 |
| B20 | B156 | CD3+CD4+CD8+ | IgE | 0.299231 | 4.83E-08 |
| B20 | B193 | CD3+CD4+CD8+ | K | -0.28772 | 1.62E-07 |
| B20 | B195 | CD3+CD4+CD8+ | Na | -0.2707 | 8.85E-07 |
| B204 | B222 | Drinking | NS | 0.308284 | 1.79E-08 |
| B204 | B206 | Drinking | Waist | 0.282212 | 2.85E-07 |
| B204 | B207 | Drinking | WHR | 0.278885 | 3.97E-07 |
| B204 | B209 | Drinking | Smoking | 0.27652 | 5.02E-07 |
| B204 | B216 | Drinking | BMI | 0.270157 | 9.33E-07 |
| B206 | B207 | Waist | WHR | 0.871097 | 0 |
| B206 | B210 | Waist | Hip | 0.804774 | 0 |
| B206 | B215 | Waist | BMI | 0.82188 | 0 |
| B206 | B216 | Waist | BMI | 0.846927 | 0 |
| B206 | B221 | Waist | Hei | 0.442832 | 0 |
| B206 | B222 | Waist | NS | 0.779891 | 0 |
| B206 | B220 | Waist | SBP | 0.334219 | 8.65E-10 |
| B206 | B218 | Waist | DBP | 0.329201 | 1.59E-09 |
| B207 | B215 | WHR | BMI | 0.613408 | 0 |
| B207 | B216 | WHR | BMI | 0.610514 | 0 |
| B207 | B222 | WHR | NS | 0.623749 | 0 |
| B207 | B210 | WHR | Hip | 0.413063 | 1.29E-14 |
| B207 | B218 | WHR | DBP | 0.302349 | 3.45E-08 |
| B207 | B221 | WHR | Hei | 0.298915 | 5.00E-08 |
| B21 | B100 | CD3+CD4-CD8- | anti-Ro52 | 0.444401 | 0 |
| B21 | B101 | CD3+CD4-CD8- | anti-Sc70 | 0.444401 | 0 |
| B21 | B102 | CD3+CD4-CD8- | anti-Sm | 0.444401 | 0 |
| B21 | B103 | CD3+CD4-CD8- | anti-SSA/Ro | 0.444401 | 0 |
| B21 | B104 | CD3+CD4-CD8- | anti-SSB | 0.444401 | 0 |
| B21 | B105 | CD3+CD4-CD8- | anti-U1RNP | 0.444401 | 0 |
| B21 | B106 | CD3+CD4-CD8- | anti-P | 0.444401 | 0 |
| B21 | B107 | CD3+CD4-CD8- | AnuA | 0.444401 | 0 |
| B21 | B108 | CD3+CD4-CD8- | Anti-Cyclin | 0.444401 | 0 |
| B21 | B109 | CD3+CD4-CD8- | AMA-M2 | 0.444401 | 0 |
| B21 | B110 | CD3+CD4-CD8- | AHA | 0.444401 | 0 |

| | | | | | |
|---|---|---|---|---|---|
| B21 | B139 | CD3+CD4-CD8- | ADA | -0.48774 | 0 |
| B21 | B194 | CD3+CD4-CD8- | Cl | -0.44745 | 0 |
| B21 | B22 | CD3+CD4-CD8- | CD3+CD8+(T) | -0.72382 | 0 |
| B21 | B23 | CD3+CD4-CD8- | CD3-CD16+CD56+(NK) | 0.710891 | 0 |
| B21 | B95 | CD3+CD4-CD8- | anti-cenpb | 0.444401 | 0 |
| B21 | B97 | CD3+CD4-CD8- | dsDNA | 0.444401 | 0 |
| B21 | B98 | CD3+CD4-CD8- | anti-Jo-1 | 0.444401 | 0 |
| B21 | B99 | CD3+CD4-CD8- | anti-PM-Scl | 0.444401 | 0 |
| B21 | B196 | CD3+CD4-CD8- | TCO2 | 0.438683 | 2.22E-16 |
| B21 | B198 | CD3+CD4-CD8- | P | -0.42328 | 2.44E-15 |
| B21 | B200 | CD3+CD4-CD8- | OP | -0.39131 | 3.75E-13 |
| B21 | B168 | CD3+CD4-CD8- | NMP22 | 0.349653 | 1.24E-10 |
| B21 | B138 | CD3+CD4-CD8- | PAB | 0.328466 | 1.74E-09 |
| B21 | B146 | CD3+CD4-CD8- | CysC | -0.32566 | 2.43E-09 |
| B21 | B156 | CD3+CD4-CD8- | IgE | 0.299231 | 4.83E-08 |
| B21 | B193 | CD3+CD4-CD8- | K | -0.28772 | 1.62E-07 |
| B21 | B195 | CD3+CD4-CD8- | Na | -0.2707 | 8.85E-07 |
| B210 | B215 | Hip | BMI | 0.784583 | 0 |
| B210 | B216 | Hip | BMI | 0.837874 | 0 |
| B210 | B221 | Hip | Hei | 0.470654 | 0 |
| B210 | B222 | Hip | NS | 0.697292 | 0 |
| B210 | B220 | Hip | SBP | 0.339506 | 4.50E-10 |
| B211 | B212 | IOP-R | IOP-L | 0.542019 | 0 |
| B211 | B227 | IOP-R | OD | 0.558799 | 0 |
| B212 | B225 | IOP-L | hPBG | 0.278502 | 4.13E-07 |
| B213 | B214 | CVA-R | CVA-L | 0.766352 | 0 |
| B215 | B216 | BMI | BMI | 0.824366 | 0 |
| B215 | B222 | BMI | NS | 0.724876 | 0 |
| B215 | B220 | BMI | SBP | 0.358288 | 3.98E-11 |
| B215 | B218 | BMI | DBP | 0.297426 | 5.86E-08 |
| B216 | B221 | BMI | Hei | 0.69468 | 0 |
| B216 | B222 | BMI | NS | 0.823286 | 0 |
| B216 | B220 | BMI | SBP | 0.36009 | 3.13E-11 |
| B216 | B218 | BMI | DBP | 0.333962 | 8.93E-10 |
| B218 | B220 | DBP | SBP | 0.679355 | 0 |
| B218 | B222 | DBP | NS | 0.29657 | 6.42E-08 |
| B22 | B100 | CD3+CD8+(T) | anti-Ro52 | -0.62926 | 0 |
| B22 | B101 | CD3+CD8+(T) | anti-Sc70 | -0.62926 | 0 |
| B22 | B102 | CD3+CD8+(T) | anti-Sm | -0.62926 | 0 |
| B22 | B103 | CD3+CD8+(T) | anti-SSA/Ro | -0.62926 | 0 |
| B22 | B104 | CD3+CD8+(T) | anti-SSB | -0.62926 | 0 |
| B22 | B105 | CD3+CD8+(T) | anti-U1RNP | -0.62926 | 0 |
| B22 | B106 | CD3+CD8+(T) | anti-P | -0.62926 | 0 |
| B22 | B107 | CD3+CD8+(T) | AnuA | -0.62926 | 0 |

| | | | | | |
|---|---|---|---|---|---|
| B22 | B108 | CD3+CD8+(T) | Anti-Cyclin | -0.62926 | 0 |
| B22 | B109 | CD3+CD8+(T) | AMA-M2 | -0.62926 | 0 |
| B22 | B110 | CD3+CD8+(T) | AHA | -0.62926 | 0 |
| B22 | B178 | CD3+CD8+(T) | Ig-U | -0.45119 | 0 |
| B22 | B23 | CD3+CD8+(T) | CD3-CD16+CD56+(NK) | -0.99983 | 0 |
| B22 | B92 | CD3+CD8+(T) | ALDH2-Glu504Glu | 0.483911 | 0 |
| B22 | B93 | CD3+CD8+(T) | ALDH2-Glu504Lys | -0.48391 | 0 |
| B22 | B94 | CD3+CD8+(T) | ALDH2-Lys504Lys | -0.48391 | 0 |
| B22 | B95 | CD3+CD8+(T) | anti-cenpb | -0.62926 | 0 |
| B22 | B97 | CD3+CD8+(T) | dsDNA | -0.62926 | 0 |
| B22 | B98 | CD3+CD8+(T) | anti-Jo-1 | -0.62926 | 0 |
| B22 | B99 | CD3+CD8+(T) | anti-PM-Scl | -0.62926 | 0 |
| B22 | B180 | CD3+CD8+(T) | Tf | -0.3608 | 2.85E-11 |
| B22 | B139 | CD3+CD8+(T) | ADA | 0.357475 | 4.44E-11 |
| B22 | B194 | CD3+CD8+(T) | Cl | 0.322112 | 3.69E-09 |
| B22 | B196 | CD3+CD8+(T) | TCO2 | -0.31756 | 6.26E-09 |
| B22 | B201 | CD3+CD8+(T) | e-habits | 0.315475 | 7.96E-09 |
| B22 | B198 | CD3+CD8+(T) | P | 0.306156 | 2.27E-08 |
| B22 | B200 | CD3+CD8+(T) | OP | 0.28173 | 2.99E-07 |
| B22 | B154 | CD3+CD8+(T) | RF | -0.28169 | 3.00E-07 |
| B220 | B222 | SBP | NS | 0.313356 | 1.01E-08 |
| B221 | B222 | Hei | NS | 0.524854 | 0 |
| B23 | B100 | CD3-CD16+CD56+(NK) | anti-Ro52 | 0.62945 | 0 |
| B23 | B101 | CD3-CD16+CD56+(NK) | anti-Sc70 | 0.62945 | 0 |
| B23 | B102 | CD3-CD16+CD56+(NK) | anti-Sm | 0.62945 | 0 |
| B23 | B103 | CD3-CD16+CD56+(NK) | anti-SSA/Ro | 0.62945 | 0 |
| B23 | B104 | CD3-CD16+CD56+(NK) | anti-SSB | 0.62945 | 0 |
| B23 | B105 | CD3-CD16+CD56+(NK) | anti-U1RNP | 0.62945 | 0 |
| B23 | B106 | CD3-CD16+CD56+(NK) | anti-P | 0.62945 | 0 |
| B23 | B107 | CD3-CD16+CD56+(NK) | AnuA | 0.62945 | 0 |
| B23 | B108 | CD3-CD16+CD56+(NK) | Anti-Cyclin | 0.62945 | 0 |
| B23 | B109 | CD3-CD16+CD56+(NK) | AMA-M2 | 0.62945 | 0 |
| B23 | B110 | CD3-CD16+CD56+(NK) | AHA | 0.62945 | 0 |
| B23 | B178 | CD3-CD16+CD56+(NK) | Ig-U | 0.460014 | 0 |
| B23 | B92 | CD3-CD16+CD56+(NK) | ALDH2-Glu504Glu | -0.49337 | 0 |
| B23 | B93 | CD3-CD16+CD56+(NK) | ALDH2-Glu504Lys | 0.493366 | 0 |
| B23 | B94 | CD3-CD16+CD56+(NK) | ALDH2-Lys504Lys | 0.493366 | 0 |
| B23 | B95 | CD3-CD16+CD56+(NK) | anti-cenpb | 0.62945 | 0 |
| B23 | B97 | CD3-CD16+CD56+(NK) | dsDNA | 0.62945 | 0 |
| B23 | B98 | CD3-CD16+CD56+(NK) | anti-Jo-1 | 0.62945 | 0 |
| B23 | B99 | CD3-CD16+CD56+(NK) | anti-PM-Scl | 0.62945 | 0 |
| B23 | B180 | CD3-CD16+CD56+(NK) | Tf | 0.367927 | 1.08E-11 |
| B23 | B139 | CD3-CD16+CD56+(NK) | ADA | -0.35126 | 1.01E-10 |
| B23 | B194 | CD3-CD16+CD56+(NK) | Cl | -0.31629 | 7.25E-09 |

| | | | | | |
|---|---|---|---|---|---|
| B23 | B201 | CD3-CD16+CD56+(NK) | e-habits | -0.31557 | 7.87E-09 |
| B23 | B196 | CD3-CD16+CD56+(NK) | TCO2 | 0.311892 | 1.20E-08 |
| B23 | B198 | CD3-CD16+CD56+(NK) | P | -0.30068 | 4.13E-08 |
| B23 | B154 | CD3-CD16+CD56+(NK) | RF | 0.282906 | 2.65E-07 |
| B23 | B200 | CD3-CD16+CD56+(NK) | OP | -0.27664 | 4.96E-07 |
| B24 | B27 | αHBDH | LDH | 0.595163 | 0 |
| B24 | B124 | αHBDH | ACTH | 0.424802 | 1.78E-15 |
| B24 | B125 | αHBDH | Cor | -0.4248 | 1.78E-15 |
| B24 | B178 | αHBDH | Ig-U | 0.310727 | 1.36E-08 |
| B24 | B198 | αHBDH | P | -0.30587 | 2.34E-08 |
| B24 | B79 | αHBDH | hb V -dna | -0.2867 | 1.80E-07 |
| B25 | B119 | CK | Hcv-rna | -0.56336 | 0 |
| B25 | B26 | CK | CKMB | 0.636663 | 0 |
| B25 | B34 | CK | TK | -0.48711 | 0 |
| B25 | B79 | CK | hb V -dna | -0.50801 | 0 |
| B25 | B138 | CK | PAB | -0.40851 | 2.66E-14 |
| B26 | B119 | CKMB | Hcv-rna | -0.38715 | 6.95E-13 |
| B26 | B138 | CKMB | PAB | -0.38599 | 8.24E-13 |
| B26 | B79 | CKMB | hb V -dna | -0.30901 | 1.65E-08 |
| B27 | B139 | LDH | ADA | 0.335072 | 7.79E-10 |
| B28 | B202 | PRL | e-times | -0.86658 | 0 |
| B28 | B29 | PRL | E2 | -0.95229 | 0 |
| B28 | B33 | PRL | Prog | -0.38656 | 7.59E-13 |
| B29 | B116 | E2 | TT3 | -0.51232 | 0 |
| B29 | B202 | E2 | e-times | -0.53182 | 0 |
| B29 | B35 | E2 | β2-MG | -0.88589 | 0 |
| B29 | B37 | E2 | BAND | 0.456633 | 0 |
| B29 | B38 | E2 | BAND% | 0.611577 | 0 |
| B30 | B145 | LH | BGP | 0.531162 | 0 |
| B30 | B167 | LH | PC | 0.919239 | 0 |
| B30 | B171 | LH | EC | 0.686199 | 0 |
| B30 | B32 | LH | Testo | 0.436564 | 2.22E-16 |
| B30 | B144 | LH | Total-PINP | 0.4 | 1.00E-13 |
| B31 | B35 | FSH | β2-MG | -0.8761 | 0 |
| B32 | B144 | Testo | Total-PINP | -0.60634 | 0 |
| B33 | B116 | Prog | TT3 | -0.67488 | 0 |
| B33 | B35 | Prog | β2-MG | -0.92984 | 0 |
| B34 | B124 | TK | ACTH | -0.52035 | 0 |
| B34 | B125 | TK | Cor | 0.520349 | 0 |
| B34 | B79 | TK | hb V -dna | 0.578798 | 0 |
| B34 | B89 | TK | CA 72-4 | -0.42459 | 2.00E-15 |
| B34 | B136 | TK | 5'NT | 0.38399 | 1.10E-12 |
| B34 | B92 | TK | ALDH2-Glu504Glu | -0.36369 | 1.93E-11 |
| B34 | B93 | TK | ALDH2-Glu504Lys | 0.363687 | 1.93E-11 |

| | | | | | |
|---|---|---|---|---|---|
| B34 | B94 | TK | ALDH2-Lys504Lys | 0.363687 | 1.93E-11 |
| B34 | B74 | TK | ApoA1 | 0.336984 | 6.15E-10 |
| B34 | B82 | TK | CEA | -0.3261 | 2.30E-09 |
| B34 | B187 | TK | NSE | 0.316734 | 6.89E-09 |
| B34 | B75 | TK | ApoA1/B | 0.303227 | 3.13E-08 |
| B34 | B119 | TK | Hcv-rna | 0.280834 | 3.27E-07 |
| B35 | B116 | β2-MG | TT3 | -0.89474 | 0 |
| B35 | B202 | β2-MG | e-times | -0.62439 | 0 |
| B35 | B142 | β2-MG | 25-(OH)D | -0.42146 | 3.11E-15 |
| B35 | B37 | β2-MG | BAND | -0.39736 | 1.51E-13 |
| B36 | B118 | V-Lym% | CEA1 | -0.53073 | 0 |
| B36 | B170 | V-Lym% | U-GLU | -0.50361 | 0 |
| B36 | B42 | V-Lym% | MON | -0.2793 | 3.81E-07 |
| B36 | B153 | V-Lym% | Glu | -0.27699 | 4.79E-07 |
| B39 | B151 | Other cells | Anti-CCP | -0.32477 | 2.70E-09 |
| B4 | B29 | WB-Ca | E2 | -0.49751 | 0 |
| B4 | B35 | WB-Ca | β2-MG | -0.36345 | 1.99E-11 |
| B40 | B42 | WBC | MON | 0.562033 | 0 |
| B41 | B42 | MON % | MON | 0.712927 | 0 |
| B41 | B190 | MON % | Cre | 0.284264 | 2.31E-07 |
| B42 | B192 | MON | UA | 0.314032 | 9.38E-09 |
| B42 | B190 | MON | Cre | 0.27533 | 5.64E-07 |
| B43 | B44 | MCV | MCH | 0.86322 | 0 |
| B43 | B46 | MCV | RBC Count | -0.60398 | 0 |
| B43 | B47 | MCV | RDW | 0.634344 | 0 |
| B44 | B45 | MCH | MCHC | 0.495116 | 0 |
| B44 | B46 | MCH | RBC Count | -0.47162 | 0 |
| B44 | B47 | MCH | RDW | 0.39738 | 1.50E-13 |
| B45 | B51 | MCHC | Hb | 0.480258 | 0 |
| B45 | B71 | MCHC | TG | 0.34837 | 1.46E-10 |
| B45 | B47 | MCHC | RDW | -0.30649 | 2.19E-08 |
| B46 | B221 | RBC Count | Hei | 0.479235 | 0 |
| B46 | B47 | RBC Count | RDW | -0.49734 | 0 |
| B46 | B51 | RBC Count | Hb | 0.756488 | 0 |
| B46 | B190 | RBC Count | Cre | 0.437282 | 2.22E-16 |
| B46 | B216 | RBC Count | BMI | 0.437987 | 2.22E-16 |
| B46 | B72 | RBC Count | HDL-C | -0.40877 | 2.55E-14 |
| B46 | B222 | RBC Count | NS | 0.39825 | 1.31E-13 |
| B46 | B163 | RBC Count | SG | 0.389887 | 4.64E-13 |
| B46 | B192 | RBC Count | UA | 0.387733 | 6.38E-13 |
| B46 | B206 | RBC Count | Waist | 0.364965 | 1.62E-11 |
| B46 | B207 | RBC Count | WHR | 0.326339 | 2.24E-09 |
| B46 | B141 | RBC Count | AST/ALT | -0.32109 | 4.16E-09 |
| B46 | B126 | RBC Count | Alb | 0.299126 | 4.89E-08 |

| | | | | | |
|---|---|---|---|---|---|
| B46 | B210 | RBC Count | Hip | 0.296716 | 6.33E-08 |
| B46 | B128 | RBC Count | ALT | 0.284418 | 2.28E-07 |
| B47 | B72 | RDW | HDL-C | 0.274493 | 6.13E-07 |
| B48 | B53 | LYM% | GR% | -0.88873 | 0 |
| B5 | B144 | WB-Mg | Total-PINP | -0.97991 | 0 |
| B5 | B96 | WB-Mg | dsDNA (IIF) | 0.835672 | 0 |
| B5 | B171 | WB-Mg | EC | 0.335754 | 7.16E-10 |
| B50 | B53 | eo% | GR% | -0.39532 | 2.05E-13 |
| B51 | B190 | Hb | Cre | 0.564461 | 0 |
| B51 | B192 | Hb | UA | 0.441822 | 0 |
| B51 | B216 | Hb | BMI | 0.494474 | 0 |
| B51 | B221 | Hb | Hei | 0.541014 | 0 |
| B51 | B222 | Hb | NS | 0.501976 | 0 |
| B51 | B163 | Hb | SG | 0.435476 | 4.44E-16 |
| B51 | B206 | Hb | Waist | 0.405196 | 4.49E-14 |
| B51 | B126 | Hb | Alb | 0.395553 | 1.98E-13 |
| B51 | B72 | Hb | HDL-C | -0.39046 | 4.26E-13 |
| B51 | B207 | Hb | WHR | 0.364503 | 1.72E-11 |
| B51 | B210 | Hb | Hip | 0.320818 | 4.29E-09 |
| B51 | B141 | Hb | AST/ALT | -0.32034 | 4.54E-09 |
| B51 | B174 | Hb | Urea/Cr | -0.30505 | 2.56E-08 |
| B51 | B204 | Hb | Drinking | 0.289919 | 1.29E-07 |
| B51 | B70 | Hb | LDL-C | 0.277341 | 4.63E-07 |
| B54 | B56 | ESR | IR | -0.68814 | 0 |
| B54 | B57 | ESR | RBCAI | -0.46707 | 0 |
| B54 | B58 | ESR | HCT | -0.60691 | 0 |
| B54 | B59 | ESR | WBRV | -0.53409 | 0 |
| B54 | B60 | ESR | WBRVI | -0.74095 | 0 |
| B54 | B61 | ESR | WBHSRV | -0.4915 | 0 |
| B54 | B62 | ESR | WBHSRVI | -0.67502 | 0 |
| B54 | B63 | ESR | WBRSR(mPas_200) | -0.47071 | 0 |
| B54 | B64 | ESR | WBRSR(mPas_5) | -0.57069 | 0 |
| B54 | B65 | ESR | WBRSR(mPas_1) | -0.60121 | 0 |
| B54 | B66 | ESR | WBRSR(mPas_100) | -0.4871 | 0 |
| B54 | B67 | ESR | WBRSR(mPas_50) | -0.50751 | 0 |
| B54 | B69 | ESR | PV | 0.425817 | 1.55E-15 |
| B54 | B55 | ESR | TK | -0.41953 | 4.44E-15 |
| B54 | B81 | ESR | TSGF | 0.368934 | 9.37E-12 |
| B54 | B157 | ESR | IgG | 0.308752 | 1.70E-08 |
| B54 | B184 | ESR | INR | 0.283097 | 2.60E-07 |
| B55 | B56 | TK | IR | 0.667938 | 0 |
| B55 | B61 | TK | WBHSRV | 0.490637 | 0 |
| B55 | B62 | TK | WBHSRVI | 0.450018 | 0 |
| B55 | B187 | TK | NSE | -0.43143 | 6.66E-16 |

| | | | | | |
|---|---|---|---|---|---|
| B55 | B135 | TK | TBA | 0.390374 | 4.31E-13 |
| B55 | B69 | TK | PV | -0.38609 | 8.12E-13 |
| B55 | B124 | TK | ACTH | 0.310149 | 1.46E-08 |
| B55 | B125 | TK | Cor | -0.31015 | 1.46E-08 |
| B56 | B58 | IR | HCT | 0.779295 | 0 |
| B56 | B59 | IR | WBRV | 0.525351 | 0 |
| B56 | B60 | IR | WBRVI | 0.812865 | 0 |
| B56 | B61 | IR | WBHSRV | 0.876785 | 0 |
| B56 | B62 | IR | WBHSRVI | 0.964233 | 0 |
| B56 | B63 | IR | WBRSR(mPas_200) | 0.806988 | 0 |
| B56 | B64 | IR | WBRSR(mPas_5) | 0.781872 | 0 |
| B56 | B65 | IR | WBRSR(mPas_1) | 0.725993 | 0 |
| B56 | B66 | IR | WBRSR(mPas_100) | 0.830883 | 0 |
| B56 | B67 | IR | WBRSR(mPas_50) | 0.82884 | 0 |
| B56 | B184 | IR | INR | -0.32694 | 2.08E-09 |
| B56 | B69 | IR | PV | -0.30563 | 2.41E-08 |
| B56 | B182 | IR | TT | 0.298984 | 4.96E-08 |
| B56 | B154 | IR | RF | -0.29661 | 6.40E-08 |
| B57 | B135 | RBCAI | TBA | -0.54596 | 0 |
| B57 | B187 | RBCAI | NSE | 0.442304 | 0 |
| B57 | B59 | RBCAI | WBRV | 0.725881 | 0 |
| B57 | B60 | RBCAI | WBRVI | 0.626906 | 0 |
| B57 | B64 | RBCAI | WBRSR(mPas_5) | 0.451388 | 0 |
| B57 | B65 | RBCAI | WBRSR(mPas_1) | 0.594047 | 0 |
| B57 | B58 | RBCAI | HCT | 0.373437 | 5.00E-12 |
| B57 | B90 | RBCAI | CYFRA 21-1 | -0.31114 | 1.30E-08 |
| B58 | B59 | HCT | WBRV | 0.662019 | 0 |
| B58 | B60 | HCT | WBRVI | 0.893316 | 0 |
| B58 | B61 | HCT | WBHSRV | 0.78676 | 0 |
| B58 | B62 | HCT | WBHSRVI | 0.913695 | 0 |
| B58 | B63 | HCT | WBRSR(mPas_200) | 0.886588 | 0 |
| B58 | B64 | HCT | WBRSR(mPas_5) | 0.925693 | 0 |
| B58 | B65 | HCT | WBRSR(mPas_1) | 0.904395 | 0 |
| B58 | B66 | HCT | WBRSR(mPas_100) | 0.906289 | 0 |
| B58 | B67 | HCT | WBRSR(mPas_50) | 0.916304 | 0 |
| B59 | B60 | WBRV | WBRVI | 0.799996 | 0 |
| B59 | B61 | WBRV | WBHSRV | 0.67875 | 0 |
| B59 | B62 | WBRV | WBHSRVI | 0.591439 | 0 |
| B59 | B63 | WBRV | WBRSR(mPas_200) | 0.726293 | 0 |
| B59 | B64 | WBRV | WBRSR(mPas_5) | 0.862532 | 0 |
| B59 | B65 | WBRV | WBRSR(mPas_1) | 0.91201 | 0 |
| B59 | B66 | WBRV | WBRSR(mPas_100) | 0.724987 | 0 |
| B59 | B67 | WBRV | WBRSR(mPas_50) | 0.750626 | 0 |
| B59 | B135 | WBRV | TBA | -0.42729 | 1.33E-15 |

| | | | | | |
|---|---|---|---|---|---|
| B59 | B187 | WBRV | NSE | 0.306149 | 2.27E-08 |
| B59 | B69 | WBRV | PV | 0.279913 | 3.59E-07 |
| B59 | B151 | WBRV | Anti-CCP | -0.27375 | 6.59E-07 |
| B6 | B167 | WB-Fe | PC | 0.532901 | 0 |
| B60 | B61 | WBRVI | WBHSRV | 0.719481 | 0 |
| B60 | B62 | WBRVI | WBHSRVI | 0.890608 | 0 |
| B60 | B63 | WBRVI | WBRSR(mPas_200) | 0.766732 | 0 |
| B60 | B64 | WBRVI | WBRSR(mPas_5) | 0.90263 | 0 |
| B60 | B65 | WBRVI | WBRSR(mPas_1) | 0.933549 | 0 |
| B60 | B66 | WBRVI | WBRSR(mPas_100) | 0.801374 | 0 |
| B60 | B67 | WBRVI | WBRSR(mPas_50) | 0.824893 | 0 |
| B60 | B157 | WBRVI | IgG | -0.28724 | 1.71E-07 |
| B60 | B154 | WBRVI | RF | -0.27245 | 7.47E-07 |
| B61 | B62 | WBHSRV | WBHSRVI | 0.879602 | 0 |
| B61 | B63 | WBHSRV | WBRSR(mPas_200) | 0.966437 | 0 |
| B61 | B64 | WBHSRV | WBRSR(mPas_5) | 0.890929 | 0 |
| B61 | B65 | WBHSRV | WBRSR(mPas_1) | 0.813323 | 0 |
| B61 | B66 | WBHSRV | WBRSR(mPas_100) | 0.9625 | 0 |
| B61 | B67 | WBHSRV | WBRSR(mPas_50) | 0.954863 | 0 |
| B62 | B63 | WBHSRVI | WBRSR(mPas_200) | 0.877479 | 0 |
| B62 | B64 | WBHSRVI | WBRSR(mPas_5) | 0.878532 | 0 |
| B62 | B65 | WBHSRVI | WBRSR(mPas_1) | 0.833025 | 0 |
| B62 | B66 | WBHSRVI | WBRSR(mPas_100) | 0.90436 | 0 |
| B62 | B67 | WBHSRVI | WBRSR(mPas_50) | 0.906996 | 0 |
| B62 | B154 | WBHSRVI | RF | -0.30279 | 3.28E-08 |
| B62 | B184 | WBHSRVI | INR | -0.28636 | 1.87E-07 |
| B62 | B182 | WBHSRVI | TT | 0.281505 | 3.06E-07 |
| B63 | B64 | WBRSR(mPas_200) | WBRSR(mPas_5) | 0.95202 | 0 |
| B63 | B65 | WBRSR(mPas_200) | WBRSR(mPas_1) | 0.888221 | 0 |
| B63 | B66 | WBRSR(mPas_200) | WBRSR(mPas_100) | 0.996585 | 0 |
| B63 | B67 | WBRSR(mPas_200) | WBRSR(mPas_50) | 0.993896 | 0 |
| B64 | B65 | WBRSR(mPas_5) | WBRSR(mPas_1) | 0.985809 | 0 |
| B64 | B66 | WBRSR(mPas_5) | WBRSR(mPas_100) | 0.961735 | 0 |
| B64 | B67 | WBRSR(mPas_5) | WBRSR(mPas_50) | 0.973528 | 0 |
| B65 | B66 | WBRSR(mPas_1) | WBRSR(mPas_100) | 0.902179 | 0 |
| B65 | B67 | WBRSR(mPas_1) | WBRSR(mPas_50) | 0.921486 | 0 |
| B66 | B67 | WBRSR(mPas_100) | WBRSR(mPas_50) | 0.998857 | 0 |
| B69 | B139 | PV | ADA | 0.338534 | 5.08E-10 |
| B69 | B135 | PV | TBA | -0.30747 | 1.96E-08 |
| B69 | B154 | PV | RF | 0.293379 | 9.01E-08 |
| B69 | B187 | PV | NSE | 0.275264 | 5.68E-07 |
| B7 | B202 | WB-Cop | e-times | 0.7698 | 0 |
| B7 | B28 | WB-Cop | PRL | -0.92916 | 0 |
| B70 | B77 | LDL-C | TC | 0.830768 | 0 |

| | | | | | |
|---|---|---|---|---|---|
| B70 | B216 | LDL-C | BMI | 0.311031 | 1.32E-08 |
| B70 | B222 | LDL-C | NS | 0.307548 | 1.95E-08 |
| B70 | B206 | LDL-C | Waist | 0.305998 | 2.31E-08 |
| B70 | B215 | LDL-C | BMI | 0.277502 | 4.56E-07 |
| B71 | B130 | TG | GLB | 0.603397 | 0 |
| B71 | B91 | TG | TP | 0.633478 | 0 |
| B71 | B72 | TG | HDL-C | -0.36347 | 1.98E-11 |
| B71 | B77 | TG | TC | 0.272361 | 7.54E-07 |
| B72 | B192 | HDL-C | UA | -0.46586 | 0 |
| B72 | B206 | HDL-C | Waist | -0.4574 | 0 |
| B72 | B216 | HDL-C | BMI | -0.47092 | 0 |
| B72 | B222 | HDL-C | NS | -0.45727 | 0 |
| B72 | B207 | HDL-C | WHR | -0.40508 | 4.57E-14 |
| B72 | B215 | HDL-C | BMI | -0.39834 | 1.30E-13 |
| B72 | B210 | HDL-C | Hip | -0.36331 | 2.03E-11 |
| B72 | B141 | HDL-C | AST/ALT | 0.357732 | 4.29E-11 |
| B72 | B163 | HDL-C | SG | -0.33369 | 9.23E-10 |
| B72 | B221 | HDL-C | Hei | -0.32672 | 2.14E-09 |
| B72 | B190 | HDL-C | Cre | -0.32029 | 4.56E-09 |
| B72 | B128 | HDL-C | ALT | -0.30503 | 2.57E-08 |
| B72 | B74 | HDL-C | ApoA1 | 0.285498 | 2.04E-07 |
| B73 | B197 | ApoA | AG | -0.4638 | 0 |
| B73 | B196 | ApoA | TCO2 | 0.433761 | 4.44E-16 |
| B73 | B80 | ApoA | anti-HP IgG | 0.358822 | 3.71E-11 |
| B73 | B199 | ApoA | Mg | -0.33982 | 4.33E-10 |
| B73 | B76 | ApoA | ApoB | 0.318984 | 5.31E-09 |
| B73 | B79 | ApoA | hb V -dna | -0.31396 | 9.46E-09 |
| B73 | B124 | ApoA | ACTH | 0.312959 | 1.06E-08 |
| B73 | B125 | ApoA | Cor | -0.31296 | 1.06E-08 |
| B73 | B195 | ApoA | Na | -0.29358 | 8.82E-08 |
| B73 | B120 | ApoA | HCV-cAg | -0.29049 | 1.22E-07 |
| B74 | B111 | ApoA1 | TSH | 0.568371 | 0 |
| B74 | B75 | ApoA1 | ApoA1/B | 0.743359 | 0 |
| B74 | B179 | ApoA1 | α1-MG | -0.3193 | 5.13E-09 |
| B74 | B86 | ApoA1 | TBhCG | 0.29397 | 8.46E-08 |
| B75 | B111 | ApoA1/B | TSH | 0.531163 | 0 |
| B75 | B76 | ApoA1/B | ApoB | -0.71769 | 0 |
| B76 | B136 | ApoB | 5'NT | 0.305357 | 2.48E-08 |
| B76 | B89 | ApoB | CA 72-4 | 0.277113 | 4.74E-07 |
| B78 | B179 | HBcAb | α1-MG | 0.448025 | 0 |
| B78 | B151 | HBcAb | Anti-CCP | -0.29176 | 1.07E-07 |
| B79 | B124 | hb V -dna | ACTH | -0.58967 | 0 |
| B79 | B125 | hb V -dna | Cor | 0.589668 | 0 |
| B79 | B92 | hb V -dna | ALDH2-Glu504Glu | -0.41376 | 1.15E-14 |

| | | | | | |
|---|---|---|---|---|---|
| B79 | B93 | hb V -dna | ALDH2-Glu504Lys | 0.413758 | 1.15E-14 |
| B79 | B94 | hb V -dna | ALDH2-Lys504Lys | 0.413758 | 1.15E-14 |
| B79 | B187 | hb V -dna | NSE | 0.359986 | 3.17E-11 |
| B79 | B119 | hb V -dna | Hcv-rna | 0.310037 | 1.47E-08 |
| B79 | B180 | hb V -dna | Tf | 0.286223 | 1.89E-07 |
| B79 | B136 | hb V -dna | 5'NT | 0.277688 | 4.47E-07 |
| B8 | B145 | WB-Zinc | BGP | -0.36148 | 2.60E-11 |
| B81 | B124 | TSGF | ACTH | 0.328442 | 1.74E-09 |
| B81 | B125 | TSGF | Cor | -0.32844 | 1.74E-09 |
| B82 | B84 | CEA | C-PSA | 0.568462 | 0 |
| B82 | B89 | CEA | CA 72-4 | 0.451178 | 0 |
| B82 | B194 | CEA | Cl | 0.391901 | 3.43E-13 |
| B82 | B196 | CEA | TCO2 | -0.27632 | 5.12E-07 |
| B84 | B199 | C-PSA | Mg | 0.369544 | 8.61E-12 |
| B84 | B196 | C-PSA | TCO2 | -0.27 | 9.47E-07 |
| B85 | B88 | SCC | CA 242 | 0.325437 | 2.49E-09 |
| B86 | B90 | TBhCG | CYFRA 21-1 | 0.64706 | 0 |
| B86 | B88 | TBhCG | CA 242 | 0.424259 | 2.22E-15 |
| B86 | B199 | TBhCG | Mg | -0.28822 | 1.54E-07 |
| B87 | B137 | NSE | AFU | 0.48171 | 0 |
| B87 | B120 | NSE | HCV-cAg | -0.35053 | 1.11E-10 |
| B87 | B194 | NSE | Cl | -0.28456 | 2.24E-07 |
| B88 | B157 | CA 242 | IgG | 0.329408 | 1.55E-09 |
| B90 | B135 | CYFRA 21-1 | TBA | 0.337916 | 5.48E-10 |
| B91 | B126 | TP | Alb | 0.452619 | 0 |
| B91 | B127 | TP | A-G ratio | -0.53508 | 0 |
| B91 | B130 | TP | GLB | 0.881668 | 0 |
| B92 | B124 | ALDH2-Glu504Glu | ACTH | 0.705998 | 0 |
| B92 | B125 | ALDH2-Glu504Glu | Cor | -0.706 | 0 |
| B92 | B180 | ALDH2-Glu504Glu | Tf | -0.72526 | 0 |
| B92 | B94 | ALDH2-Glu504Glu | ALDH2-Lys504Lys | -1 | 0 |
| B92 | B187 | ALDH2-Glu504Glu | NSE | -0.43196 | 4.44E-16 |
| B92 | B100 | ALDH2-Glu504Glu | anti-Ro52 | -0.30974 | 1.52E-08 |
| B92 | B101 | ALDH2-Glu504Glu | anti-Sc70 | -0.30974 | 1.52E-08 |
| B92 | B102 | ALDH2-Glu504Glu | anti-Sm | -0.30974 | 1.52E-08 |
| B92 | B103 | ALDH2-Glu504Glu | anti-SSA/Ro | -0.30974 | 1.52E-08 |
| B92 | B104 | ALDH2-Glu504Glu | anti-SSB | -0.30974 | 1.52E-08 |
| B92 | B105 | ALDH2-Glu504Glu | anti-U1RNP | -0.30974 | 1.52E-08 |
| B92 | B106 | ALDH2-Glu504Glu | anti-P | -0.30974 | 1.52E-08 |
| B92 | B107 | ALDH2-Glu504Glu | AnuA | -0.30974 | 1.52E-08 |
| B92 | B108 | ALDH2-Glu504Glu | Anti-Cyclin | -0.30974 | 1.52E-08 |
| B92 | B109 | ALDH2-Glu504Glu | AMA-M2 | -0.30974 | 1.52E-08 |
| B92 | B110 | ALDH2-Glu504Glu | AHA | -0.30974 | 1.52E-08 |
| B92 | B95 | ALDH2-Glu504Glu | anti-cenpb | -0.30974 | 1.52E-08 |

| | | | | | |
|---|---|---|---|---|---|
| B92 | B97 | ALDH2-Glu504Glu | dsDNA | -0.30974 | 1.52E-08 |
| B92 | B98 | ALDH2-Glu504Glu | anti-Jo-1 | -0.30974 | 1.52E-08 |
| B92 | B99 | ALDH2-Glu504Glu | anti-PM-Scl | -0.30974 | 1.52E-08 |
| B92 | B178 | ALDH2-Glu504Glu | Ig-U | -0.3068 | 2.12E-08 |
| B92 | B154 | ALDH2-Glu504Glu | RF | -0.28724 | 1.71E-07 |
| B93 | B124 | ALDH2-Glu504Lys | ACTH | -0.706 | 0 |
| B93 | B125 | ALDH2-Glu504Lys | Cor | 0.705998 | 0 |
| B93 | B180 | ALDH2-Glu504Lys | Tf | 0.725264 | 0 |
| B93 | B94 | ALDH2-Glu504Lys | ALDH2-Lys504Lys | 1 | 0 |
| B93 | B187 | ALDH2-Glu504Lys | NSE | 0.431963 | 4.44E-16 |
| B93 | B100 | ALDH2-Glu504Lys | anti-Ro52 | 0.309737 | 1.52E-08 |
| B93 | B101 | ALDH2-Glu504Lys | anti-Sc70 | 0.309737 | 1.52E-08 |
| B93 | B102 | ALDH2-Glu504Lys | anti-Sm | 0.309737 | 1.52E-08 |
| B93 | B103 | ALDH2-Glu504Lys | anti-SSA/Ro | 0.309737 | 1.52E-08 |
| B93 | B104 | ALDH2-Glu504Lys | anti-SSB | 0.309737 | 1.52E-08 |
| B93 | B105 | ALDH2-Glu504Lys | anti-U1RNP | 0.309737 | 1.52E-08 |
| B93 | B106 | ALDH2-Glu504Lys | anti-P | 0.309737 | 1.52E-08 |
| B93 | B107 | ALDH2-Glu504Lys | AnuA | 0.309737 | 1.52E-08 |
| B93 | B108 | ALDH2-Glu504Lys | Anti-Cyclin | 0.309737 | 1.52E-08 |
| B93 | B109 | ALDH2-Glu504Lys | AMA-M2 | 0.309737 | 1.52E-08 |
| B93 | B110 | ALDH2-Glu504Lys | AHA | 0.309737 | 1.52E-08 |
| B93 | B95 | ALDH2-Glu504Lys | anti-cenpb | 0.309737 | 1.52E-08 |
| B93 | B97 | ALDH2-Glu504Lys | dsDNA | 0.309737 | 1.52E-08 |
| B93 | B98 | ALDH2-Glu504Lys | anti-Jo-1 | 0.309737 | 1.52E-08 |
| B93 | B99 | ALDH2-Glu504Lys | anti-PM-Scl | 0.309737 | 1.52E-08 |
| B93 | B178 | ALDH2-Glu504Lys | Ig-U | 0.306795 | 2.12E-08 |
| B93 | B154 | ALDH2-Glu504Lys | RF | 0.287243 | 1.71E-07 |
| B94 | B124 | ALDH2-Lys504Lys | ACTH | -0.706 | 0 |
| B94 | B125 | ALDH2-Lys504Lys | Cor | 0.705998 | 0 |
| B94 | B180 | ALDH2-Lys504Lys | Tf | 0.725264 | 0 |
| B94 | B187 | ALDH2-Lys504Lys | NSE | 0.431963 | 4.44E-16 |
| B94 | B100 | ALDH2-Lys504Lys | anti-Ro52 | 0.309737 | 1.52E-08 |
| B94 | B101 | ALDH2-Lys504Lys | anti-Sc70 | 0.309737 | 1.52E-08 |
| B94 | B102 | ALDH2-Lys504Lys | anti-Sm | 0.309737 | 1.52E-08 |
| B94 | B103 | ALDH2-Lys504Lys | anti-SSA/Ro | 0.309737 | 1.52E-08 |
| B94 | B104 | ALDH2-Lys504Lys | anti-SSB | 0.309737 | 1.52E-08 |
| B94 | B105 | ALDH2-Lys504Lys | anti-U1RNP | 0.309737 | 1.52E-08 |
| B94 | B106 | ALDH2-Lys504Lys | anti-P | 0.309737 | 1.52E-08 |
| B94 | B107 | ALDH2-Lys504Lys | AnuA | 0.309737 | 1.52E-08 |
| B94 | B108 | ALDH2-Lys504Lys | Anti-Cyclin | 0.309737 | 1.52E-08 |
| B94 | B109 | ALDH2-Lys504Lys | AMA-M2 | 0.309737 | 1.52E-08 |
| B94 | B110 | ALDH2-Lys504Lys | AHA | 0.309737 | 1.52E-08 |
| B94 | B95 | ALDH2-Lys504Lys | anti-cenpb | 0.309737 | 1.52E-08 |
| B94 | B97 | ALDH2-Lys504Lys | dsDNA | 0.309737 | 1.52E-08 |

| | | | | | |
|---|---|---|---|---|---|
| B94 | B98 | ALDH2-Lys504Lys | anti-Jo-1 | 0.309737 | 1.52E-08 |
| B94 | B99 | ALDH2-Lys504Lys | anti-PM-Scl | 0.309737 | 1.52E-08 |
| B94 | B178 | ALDH2-Lys504Lys | Ig-U | 0.306795 | 2.12E-08 |
| B94 | B154 | ALDH2-Lys504Lys | RF | 0.287243 | 1.71E-07 |
| B95 | B101 | anti-cenpb | anti-Sc70 | 1 | 0 |
| B95 | B106 | anti-cenpb | anti-P | 1 | 0 |
| B95 | B107 | anti-cenpb | AnuA | 1 | 0 |
| B95 | B108 | anti-cenpb | Anti-Cyclin | 1 | 0 |
| B95 | B109 | anti-cenpb | AMA-M2 | 1 | 0 |
| B95 | B151 | anti-cenpb | Anti-CCP | -0.62933 | 0 |
| B95 | B98 | anti-cenpb | anti-Jo-1 | 1 | 0 |
| B95 | B154 | anti-cenpb | RF | 0.392478 | 3.15E-13 |
| B95 | B178 | anti-cenpb | Ig-U | 0.288523 | 1.49E-07 |
| B96 | B167 | dsDNA (IIF) | PC | 0.954786 | 0 |
| B97 | B100 | dsDNA | anti-Ro52 | 1 | 0 |
| B97 | B103 | dsDNA | anti-SSA/Ro | 1 | 0 |
| B97 | B105 | dsDNA | anti-U1RNP | 1 | 0 |
| B97 | B108 | dsDNA | Anti-Cyclin | 1 | 0 |
| B97 | B109 | dsDNA | AMA-M2 | 1 | 0 |
| B97 | B110 | dsDNA | AHA | 1 | 0 |
| B97 | B151 | dsDNA | Anti-CCP | -0.62933 | 0 |
| B97 | B99 | dsDNA | anti-PM-Scl | 1 | 0 |
| B97 | B154 | dsDNA | RF | 0.392478 | 3.15E-13 |
| B97 | B178 | dsDNA | Ig-U | 0.288523 | 1.49E-07 |
| B98 | B100 | anti-Jo-1 | anti-Ro52 | 1 | 0 |
| B98 | B101 | anti-Jo-1 | anti-Sc70 | 1 | 0 |
| B98 | B102 | anti-Jo-1 | anti-Sm | 1 | 0 |
| B98 | B103 | anti-Jo-1 | anti-SSA/Ro | 1 | 0 |
| B98 | B104 | anti-Jo-1 | anti-SSB | 1 | 0 |
| B98 | B105 | anti-Jo-1 | anti-U1RNP | 1 | 0 |
| B98 | B107 | anti-Jo-1 | AnuA | 1 | 0 |
| B98 | B109 | anti-Jo-1 | AMA-M2 | 1 | 0 |
| B98 | B110 | anti-Jo-1 | AHA | 1 | 0 |
| B98 | B151 | anti-Jo-1 | Anti-CCP | -0.62933 | 0 |
| B98 | B99 | anti-Jo-1 | anti-PM-Scl | 1 | 0 |
| B98 | B154 | anti-Jo-1 | RF | 0.392478 | 3.15E-13 |
| B98 | B178 | anti-Jo-1 | Ig-U | 0.288523 | 1.49E-07 |
| B99 | B100 | anti-PM-Scl | anti-Ro52 | 1 | 0 |
| B99 | B103 | anti-PM-Scl | anti-SSA/Ro | 1 | 0 |
| B99 | B105 | anti-PM-Scl | anti-U1RNP | 1 | 0 |
| B99 | B106 | anti-PM-Scl | anti-P | 1 | 0 |
| B99 | B107 | anti-PM-Scl | AnuA | 1 | 0 |
| B99 | B109 | anti-PM-Scl | AMA-M2 | 1 | 0 |
| B99 | B110 | anti-PM-Scl | AHA | 1 | 0 |

| | | | | | |
|---|---|---|---|---|---|
| B99 | B151 | anti-PM-Scl | Anti-CCP | -0.62933 | 0 |
| B99 | B154 | anti-PM-Scl | RF | 0.392478 | 3.15E-13 |
| B99 | B178 | anti-PM-Scl | Ig-U | 0.288523 | 1.49E-07 |

**Supplementary Table 16. PEI Correlations in Nephropathy**

| Row | Column | Row | Column | Cor | *P* |
|---|---|---|---|---|---|
| B1 | B192 | Gender | UA | -0.43349803 | 0 |
| B1 | B206 | Gender | Waist | -0.50917524 | 0 |
| B1 | B207 | Gender | WHR | -0.44638652 | 0 |
| B1 | B209 | Gender | Smoking | -0.33960109 | 0 |
| B1 | B210 | Gender | Hip | -0.3802898 | 0 |
| B1 | B216 | Gender | BMI | -0.58729438 | 0 |
| B1 | B221 | Gender | Hei | -0.65373775 | 0 |
| B1 | B222 | Gender | NS | -0.60058387 | 0 |
| B1 | B46 | Gender | RBC Count | -0.43865444 | 0 |
| B1 | B51 | Gender | Hb | -0.54324201 | 0 |
| B1 | B174 | Gender | Urea/Cr | 0.331987769 | 2.22E-16 |
| B1 | B42 | Gender | MON | -0.32035903 | 3.11E-15 |
| B1 | B72 | Gender | HDL-C | 0.315452713 | 8.88E-15 |
| B1 | B215 | Gender | BMI | -0.30264881 | 1.15E-13 |
| B1 | B163 | Gender | SG | -0.29787673 | 2.88E-13 |
| B1 | B45 | Gender | MCHC | -0.29130371 | 9.94E-13 |
| B1 | B141 | Gender | AST/ALT | 0.265651016 | 9.26E-11 |
| B1 | B204 | Gender | Drinking | -0.25697298 | 3.87E-10 |
| B1 | B40 | Gender | WBC | -0.23056764 | 2.18E-08 |
| B1 | B140 | Gender | r-GT | -0.22805989 | 3.12E-08 |
| B1 | B128 | Gender | ALT | -0.21822647 | 1.22E-07 |
| B10 | B12 | PG I | PG II | 0.62879399 | 0 |
| B10 | B145 | PG I | BGP | -0.31589593 | 8.22E-15 |
| B10 | B11 | PG I | PGI/PGII | 0.28877045 | 1.59E-12 |
| B10 | B144 | PG I | Total-PINP | -0.27012187 | 4.35E-11 |
| B10 | B189 | PG I | eGFB | -0.23350567 | 1.42E-08 |
| B10 | B191 | PG I | Urea | 0.213518371 | 2.30E-07 |
| B10 | B143 | PG I | β-CTx | -0.21027781 | 3.53E-07 |
| B10 | B228 | PG I | OS | 0.209178274 | 4.07E-07 |
| B100 | B103 | anti-Ro52 | anti-SSA/Ro | 1 | 0 |
| B100 | B104 | anti-Ro52 | anti-SSB | 0.999999998 | 0 |
| B100 | B105 | anti-Ro52 | anti-U1RNP | 1 | 0 |
| B100 | B106 | anti-Ro52 | anti-P | 0.999999998 | 0 |
| B100 | B107 | anti-Ro52 | AnuA | 0.999999975 | 0 |
| B100 | B124 | anti-Ro52 | ACTH | -0.40646943 | 0 |
| B100 | B125 | anti-Ro52 | Cor | 0.406469422 | 0 |
| B100 | B156 | anti-Ro52 | IgE | 0.379583052 | 0 |
| B100 | B178 | anti-Ro52 | Ig-U | -0.39099799 | 0 |
| B100 | B180 | anti-Ro52 | Tf | -0.37436038 | 0 |
| B100 | B184 | anti-Ro52 | INR | 0.310198726 | 2.58E-14 |
| B100 | B185 | anti-Ro52 | PT | 0.293544549 | 6.54E-13 |
| B100 | B214 | anti-Ro52 | CVA-L | -0.28581501 | 2.73E-12 |

| | | | | | |
|---|---|---|---|---|---|
| B100 | B154 | anti-Ro52 | RF | 0.285144149 | 3.08E-12 |
| B100 | B175 | anti-Ro52 | ALB-U | -0.25006036 | 1.16E-09 |
| B100 | B183 | anti-Ro52 | PTA | -0.24874269 | 1.43E-09 |
| B100 | B177 | anti-Ro52 | ACR | -0.24423985 | 2.86E-09 |
| B100 | B196 | anti-Ro52 | TCO2 | 0.244131093 | 2.91E-09 |
| B100 | B199 | anti-Ro52 | Mg | 0.234108713 | 1.30E-08 |
| B100 | B195 | anti-Ro52 | Na | 0.228135163 | 3.09E-08 |
| B100 | B182 | anti-Ro52 | TT | 0.219111153 | 1.08E-07 |
| B100 | B151 | anti-Ro52 | Anti-CCP | 0.209182419 | 4.07E-07 |
| B101 | B102 | anti-Sc70 | anti-Sm | 0.999999998 | 0 |
| B101 | B107 | anti-Sc70 | AnuA | 0.999999862 | 0 |
| B101 | B124 | anti-Sc70 | ACTH | -0.40646945 | 0 |
| B101 | B125 | anti-Sc70 | Cor | 0.406469428 | 0 |
| B101 | B156 | anti-Sc70 | IgE | 0.379583056 | 0 |
| B101 | B178 | anti-Sc70 | Ig-U | -0.390998 | 0 |
| B101 | B180 | anti-Sc70 | Tf | -0.37436039 | 0 |
| B101 | B184 | anti-Sc70 | INR | 0.310198725 | 2.58E-14 |
| B101 | B185 | anti-Sc70 | PT | 0.293544547 | 6.54E-13 |
| B101 | B214 | anti-Sc70 | CVA-L | -0.28581502 | 2.73E-12 |
| B101 | B154 | anti-Sc70 | RF | 0.285144152 | 3.08E-12 |
| B101 | B175 | anti-Sc70 | ALB-U | -0.25006036 | 1.16E-09 |
| B101 | B183 | anti-Sc70 | PTA | -0.24874269 | 1.43E-09 |
| B101 | B177 | anti-Sc70 | ACR | -0.24423985 | 2.86E-09 |
| B101 | B196 | anti-Sc70 | TCO2 | 0.244131094 | 2.91E-09 |
| B101 | B199 | anti-Sc70 | Mg | 0.234108715 | 1.30E-08 |
| B101 | B195 | anti-Sc70 | Na | 0.228135159 | 3.09E-08 |
| B101 | B182 | anti-Sc70 | TT | 0.219111154 | 1.08E-07 |
| B101 | B151 | anti-Sc70 | Anti-CCP | 0.209182422 | 4.07E-07 |
| B102 | B104 | anti-Sm | anti-SSB | 1 | 0 |
| B102 | B107 | anti-Sm | AnuA | 0.999999997 | 0 |
| B102 | B124 | anti-Sm | ACTH | -0.40646944 | 0 |
| B102 | B125 | anti-Sm | Cor | 0.406469426 | 0 |
| B102 | B156 | anti-Sm | IgE | 0.379583054 | 0 |
| B102 | B178 | anti-Sm | Ig-U | -0.39099799 | 0 |
| B102 | B180 | anti-Sm | Tf | -0.37436039 | 0 |
| B102 | B184 | anti-Sm | INR | 0.310198727 | 2.58E-14 |
| B102 | B185 | anti-Sm | PT | 0.293544548 | 6.54E-13 |
| B102 | B214 | anti-Sm | CVA-L | -0.28581501 | 2.73E-12 |
| B102 | B154 | anti-Sm | RF | 0.28514415 | 3.08E-12 |
| B102 | B175 | anti-Sm | ALB-U | -0.25006036 | 1.16E-09 |
| B102 | B183 | anti-Sm | PTA | -0.24874269 | 1.43E-09 |
| B102 | B177 | anti-Sm | ACR | -0.24423985 | 2.86E-09 |
| B102 | B196 | anti-Sm | TCO2 | 0.244131093 | 2.91E-09 |
| B102 | B199 | anti-Sm | Mg | 0.234108714 | 1.30E-08 |

| | | | | | |
|---|---|---|---|---|---|
| B102 | B195 | anti-Sm | Na | 0.228135159 | 3.09E-08 |
| B102 | B182 | anti-Sm | TT | 0.219111153 | 1.08E-07 |
| B102 | B151 | anti-Sm | Anti-CCP | 0.20918242 | 4.07E-07 |
| B103 | B104 | anti-SSA/Ro | anti-SSB | 0.999999998 | 0 |
| B103 | B105 | anti-SSA/Ro | anti-U1RNP | 1 | 0 |
| B103 | B106 | anti-SSA/Ro | anti-P | 0.999999998 | 0 |
| B103 | B107 | anti-SSA/Ro | AnuA | 0.999999985 | 0 |
| B103 | B108 | anti-SSA/Ro | Anti-Cyclin | 1 | 0 |
| B103 | B124 | anti-SSA/Ro | ACTH | -0.40646943 | 0 |
| B103 | B125 | anti-SSA/Ro | Cor | 0.406469422 | 0 |
| B103 | B156 | anti-SSA/Ro | IgE | 0.379583052 | 0 |
| B103 | B178 | anti-SSA/Ro | Ig-U | -0.39099799 | 0 |
| B103 | B180 | anti-SSA/Ro | Tf | -0.37436038 | 0 |
| B103 | B184 | anti-SSA/Ro | INR | 0.310198726 | 2.58E-14 |
| B103 | B185 | anti-SSA/Ro | PT | 0.293544549 | 6.54E-13 |
| B103 | B214 | anti-SSA/Ro | CVA-L | -0.28581501 | 2.73E-12 |
| B103 | B154 | anti-SSA/Ro | RF | 0.285144149 | 3.08E-12 |
| B103 | B175 | anti-SSA/Ro | ALB-U | -0.25006036 | 1.16E-09 |
| B103 | B183 | anti-SSA/Ro | PTA | -0.24874269 | 1.43E-09 |
| B103 | B177 | anti-SSA/Ro | ACR | -0.24423985 | 2.86E-09 |
| B103 | B196 | anti-SSA/Ro | TCO2 | 0.244131093 | 2.91E-09 |
| B103 | B199 | anti-SSA/Ro | Mg | 0.234108713 | 1.30E-08 |
| B103 | B195 | anti-SSA/Ro | Na | 0.228135162 | 3.09E-08 |
| B103 | B182 | anti-SSA/Ro | TT | 0.219111153 | 1.08E-07 |
| B103 | B151 | anti-SSA/Ro | Anti-CCP | 0.209182419 | 4.07E-07 |
| B104 | B105 | anti-SSB | anti-U1RNP | 0.999999997 | 0 |
| B104 | B107 | anti-SSB | AnuA | 0.999999974 | 0 |
| B104 | B108 | anti-SSB | Anti-Cyclin | 0.999999998 | 0 |
| B104 | B109 | anti-SSB | AMA-M2 | 0.999999998 | 0 |
| B104 | B110 | anti-SSB | AHA | 0.999999999 | 0 |
| B104 | B124 | anti-SSB | ACTH | -0.40646942 | 0 |
| B104 | B125 | anti-SSB | Cor | 0.406469423 | 0 |
| B104 | B156 | anti-SSB | IgE | 0.379583052 | 0 |
| B104 | B178 | anti-SSB | Ig-U | -0.39099799 | 0 |
| B104 | B180 | anti-SSB | Tf | -0.37436038 | 0 |
| B104 | B184 | anti-SSB | INR | 0.310198725 | 2.58E-14 |
| B104 | B185 | anti-SSB | PT | 0.293544549 | 6.54E-13 |
| B104 | B214 | anti-SSB | CVA-L | -0.28581501 | 2.73E-12 |
| B104 | B154 | anti-SSB | RF | 0.285144148 | 3.08E-12 |
| B104 | B175 | anti-SSB | ALB-U | -0.25006036 | 1.16E-09 |
| B104 | B183 | anti-SSB | PTA | -0.24874269 | 1.43E-09 |
| B104 | B177 | anti-SSB | ACR | -0.24423985 | 2.86E-09 |
| B104 | B196 | anti-SSB | TCO2 | 0.244131092 | 2.91E-09 |
| B104 | B199 | anti-SSB | Mg | 0.234108713 | 1.30E-08 |

| | | | | | |
|---|---|---|---|---|---|
| B104 | B195 | anti-SSB | Na | 0.228135162 | 3.09E-08 |
| B104 | B182 | anti-SSB | TT | 0.219111153 | 1.08E-07 |
| B104 | B151 | anti-SSB | Anti-CCP | 0.209182419 | 4.07E-07 |
| B105 | B106 | anti-U1RNP | anti-P | 0.999999995 | 0 |
| B105 | B107 | anti-U1RNP | AnuA | 0.999999973 | 0 |
| B105 | B108 | anti-U1RNP | Anti-Cyclin | 1 | 0 |
| B105 | B109 | anti-U1RNP | AMA-M2 | 0.999999998 | 0 |
| B105 | B124 | anti-U1RNP | ACTH | -0.40646943 | 0 |
| B105 | B125 | anti-U1RNP | Cor | 0.406469422 | 0 |
| B105 | B156 | anti-U1RNP | IgE | 0.379583052 | 0 |
| B105 | B178 | anti-U1RNP | Ig-U | -0.39099799 | 0 |
| B105 | B180 | anti-U1RNP | Tf | -0.37436038 | 0 |
| B105 | B184 | anti-U1RNP | INR | 0.310198726 | 2.58E-14 |
| B105 | B185 | anti-U1RNP | PT | 0.293544549 | 6.54E-13 |
| B105 | B214 | anti-U1RNP | CVA-L | -0.28581501 | 2.73E-12 |
| B105 | B154 | anti-U1RNP | RF | 0.285144149 | 3.08E-12 |
| B105 | B175 | anti-U1RNP | ALB-U | -0.25006036 | 1.16E-09 |
| B105 | B183 | anti-U1RNP | PTA | -0.24874269 | 1.43E-09 |
| B105 | B177 | anti-U1RNP | ACR | -0.24423985 | 2.86E-09 |
| B105 | B196 | anti-U1RNP | TCO2 | 0.244131093 | 2.91E-09 |
| B105 | B199 | anti-U1RNP | Mg | 0.234108713 | 1.30E-08 |
| B105 | B195 | anti-U1RNP | Na | 0.228135162 | 3.09E-08 |
| B105 | B182 | anti-U1RNP | TT | 0.219111153 | 1.08E-07 |
| B105 | B151 | anti-U1RNP | Anti-CCP | 0.209182419 | 4.07E-07 |
| B106 | B108 | anti-P | Anti-Cyclin | 0.999999999 | 0 |
| B106 | B109 | anti-P | AMA-M2 | 0.999999998 | 0 |
| B106 | B110 | anti-P | AHA | 0.999999997 | 0 |
| B106 | B124 | anti-P | ACTH | -0.40646943 | 0 |
| B106 | B125 | anti-P | Cor | 0.406469426 | 0 |
| B106 | B156 | anti-P | IgE | 0.379583052 | 0 |
| B106 | B178 | anti-P | Ig-U | -0.39099799 | 0 |
| B106 | B180 | anti-P | Tf | -0.37436038 | 0 |
| B106 | B184 | anti-P | INR | 0.310198726 | 2.58E-14 |
| B106 | B185 | anti-P | PT | 0.293544547 | 6.54E-13 |
| B106 | B214 | anti-P | CVA-L | -0.28581501 | 2.73E-12 |
| B106 | B154 | anti-P | RF | 0.285144149 | 3.08E-12 |
| B106 | B175 | anti-P | ALB-U | -0.25006036 | 1.16E-09 |
| B106 | B183 | anti-P | PTA | -0.24874269 | 1.43E-09 |
| B106 | B177 | anti-P | ACR | -0.24423985 | 2.86E-09 |
| B106 | B196 | anti-P | TCO2 | 0.244131092 | 2.91E-09 |
| B106 | B199 | anti-P | Mg | 0.234108713 | 1.30E-08 |
| B106 | B195 | anti-P | Na | 0.228135162 | 3.09E-08 |
| B106 | B182 | anti-P | TT | 0.219111154 | 1.08E-07 |
| B106 | B151 | anti-P | Anti-CCP | 0.209182419 | 4.07E-07 |

| | | | | | |
|---|---|---|---|---|---|
| B107 | B108 | AnuA | Anti-Cyclin | 0.999999962 | 0 |
| B107 | B109 | AnuA | AMA-M2 | 0.999999972 | 0 |
| B107 | B110 | AnuA | AHA | 0.999999987 | 0 |
| B107 | B124 | AnuA | ACTH | -0.40646949 | 0 |
| B107 | B125 | AnuA | Cor | 0.406469424 | 0 |
| B107 | B156 | AnuA | IgE | 0.379583045 | 0 |
| B107 | B178 | AnuA | Ig-U | -0.39099798 | 0 |
| B107 | B180 | AnuA | Tf | -0.37436038 | 0 |
| B107 | B184 | AnuA | INR | 0.31019871 | 2.58E-14 |
| B107 | B185 | AnuA | PT | 0.29354453 | 6.54E-13 |
| B107 | B214 | AnuA | CVA-L | -0.28581501 | 2.73E-12 |
| B107 | B154 | AnuA | RF | 0.285144144 | 3.08E-12 |
| B107 | B175 | AnuA | ALB-U | -0.25006035 | 1.16E-09 |
| B107 | B183 | AnuA | PTA | -0.24874269 | 1.43E-09 |
| B107 | B177 | AnuA | ACR | -0.24423984 | 2.86E-09 |
| B107 | B196 | AnuA | TCO2 | 0.244131087 | 2.91E-09 |
| B107 | B199 | AnuA | Mg | 0.234108706 | 1.30E-08 |
| B107 | B195 | AnuA | Na | 0.228135131 | 3.09E-08 |
| B107 | B182 | AnuA | TT | 0.219111147 | 1.08E-07 |
| B107 | B151 | AnuA | Anti-CCP | 0.209182416 | 4.07E-07 |
| B108 | B110 | Anti-Cyclin | AHA | 0.999999999 | 0 |
| B108 | B124 | Anti-Cyclin | ACTH | -0.40646943 | 0 |
| B108 | B125 | Anti-Cyclin | Cor | 0.406469423 | 0 |
| B108 | B156 | Anti-Cyclin | IgE | 0.379583053 | 0 |
| B108 | B178 | Anti-Cyclin | Ig-U | -0.39099799 | 0 |
| B108 | B180 | Anti-Cyclin | Tf | -0.37436038 | 0 |
| B108 | B184 | Anti-Cyclin | INR | 0.310198725 | 2.58E-14 |
| B108 | B185 | Anti-Cyclin | PT | 0.293544549 | 6.54E-13 |
| B108 | B214 | Anti-Cyclin | CVA-L | -0.28581501 | 2.73E-12 |
| B108 | B154 | Anti-Cyclin | RF | 0.285144149 | 3.08E-12 |
| B108 | B175 | Anti-Cyclin | ALB-U | -0.25006036 | 1.16E-09 |
| B108 | B183 | Anti-Cyclin | PTA | -0.24874269 | 1.43E-09 |
| B108 | B177 | Anti-Cyclin | ACR | -0.24423985 | 2.86E-09 |
| B108 | B196 | Anti-Cyclin | TCO2 | 0.244131093 | 2.91E-09 |
| B108 | B199 | Anti-Cyclin | Mg | 0.234108713 | 1.30E-08 |
| B108 | B195 | Anti-Cyclin | Na | 0.22813516 | 3.09E-08 |
| B108 | B182 | Anti-Cyclin | TT | 0.219111153 | 1.08E-07 |
| B108 | B151 | Anti-Cyclin | Anti-CCP | 0.20918242 | 4.07E-07 |
| B109 | B124 | AMA-M2 | ACTH | -0.40646943 | 0 |
| B109 | B125 | AMA-M2 | Cor | 0.406469423 | 0 |
| B109 | B156 | AMA-M2 | IgE | 0.379583052 | 0 |
| B109 | B178 | AMA-M2 | Ig-U | -0.39099799 | 0 |
| B109 | B180 | AMA-M2 | Tf | -0.37436038 | 0 |
| B109 | B184 | AMA-M2 | INR | 0.310198725 | 2.58E-14 |

| | | | | | |
|---|---|---|---|---|---|
| B109 | B185 | AMA-M2 | PT | 0.293544549 | 6.54E-13 |
| B109 | B214 | AMA-M2 | CVA-L | -0.28581501 | 2.73E-12 |
| B109 | B154 | AMA-M2 | RF | 0.285144149 | 3.08E-12 |
| B109 | B175 | AMA-M2 | ALB-U | -0.25006036 | 1.16E-09 |
| B109 | B183 | AMA-M2 | PTA | -0.24874269 | 1.43E-09 |
| B109 | B177 | AMA-M2 | ACR | -0.24423985 | 2.86E-09 |
| B109 | B196 | AMA-M2 | TCO2 | 0.244131093 | 2.91E-09 |
| B109 | B199 | AMA-M2 | Mg | 0.234108713 | 1.30E-08 |
| B109 | B195 | AMA-M2 | Na | 0.228135162 | 3.09E-08 |
| B109 | B182 | AMA-M2 | TT | 0.219111153 | 1.08E-07 |
| B109 | B151 | AMA-M2 | Anti-CCP | 0.209182419 | 4.07E-07 |
| B11 | B12 | PGI/PGII | PG II | -0.40370454 | 0 |
| B11 | B80 | PGI/PGII | anti-HP IgG | -0.28218686 | 5.26E-12 |
| B11 | B14 | PGI/PGII | G-17 | -0.25087086 | 1.02E-09 |
| B11 | B175 | PGI/PGII | ALB-U | 0.219229463 | 1.07E-07 |
| B110 | B124 | AHA | ACTH | -0.40646943 | 0 |
| B110 | B125 | AHA | Cor | 0.406469423 | 0 |
| B110 | B156 | AHA | IgE | 0.379583053 | 0 |
| B110 | B178 | AHA | Ig-U | -0.39099799 | 0 |
| B110 | B180 | AHA | Tf | -0.37436038 | 0 |
| B110 | B184 | AHA | INR | 0.310198725 | 2.58E-14 |
| B110 | B185 | AHA | PT | 0.293544548 | 6.54E-13 |
| B110 | B214 | AHA | CVA-L | -0.28581501 | 2.73E-12 |
| B110 | B154 | AHA | RF | 0.285144149 | 3.08E-12 |
| B110 | B175 | AHA | ALB-U | -0.25006036 | 1.16E-09 |
| B110 | B183 | AHA | PTA | -0.24874269 | 1.43E-09 |
| B110 | B177 | AHA | ACR | -0.24423985 | 2.86E-09 |
| B110 | B196 | AHA | TCO2 | 0.244131093 | 2.91E-09 |
| B110 | B199 | AHA | Mg | 0.234108713 | 1.30E-08 |
| B110 | B195 | AHA | Na | 0.228135162 | 3.09E-08 |
| B110 | B182 | AHA | TT | 0.219111152 | 1.08E-07 |
| B110 | B151 | AHA | Anti-CCP | 0.20918242 | 4.07E-07 |
| B112 | B114 | FT4 | TSH TT3 TT4(TT4) | 0.381034742 | 0 |
| B112 | B113 | FT4 | FT3 | 0.221525213 | 7.79E-08 |
| B112 | B115 | FT4 | TSH TT3 TT4(TT3) | 0.216804003 | 1.48E-07 |
| B112 | B227 | FT4 | OD | 0.21434938 | 2.06E-07 |
| B113 | B115 | FT3 | TSH TT3 TT4(TT3) | 0.532646094 | 0 |
| B113 | B114 | FT3 | TSH TT3 TT4(TT4) | 0.299745193 | 2.01E-13 |
| B114 | B115 | TSH TT3 TT4(TT4) | TSH TT3 TT4(TT3) | 0.539614912 | 0 |
| B117 | B167 | TT4 | PC | -0.34434328 | 0 |
| B118 | B189 | CEA1 | eGFB | -0.34663862 | 0 |
| B118 | B191 | CEA1 | Urea | 0.371629149 | 0 |
| B118 | B190 | CEA1 | Cre | 0.330611796 | 4.44E-16 |
| B118 | B220 | CEA1 | SBP | 0.215207727 | 1.84E-07 |

| | | | | | |
|---|---|---|---|---|---|
| B119 | B120 | Hcv-rna | HCV-cAg | 0.34921508 | 0 |
| B119 | B134 | Hcv-rna | CHE | 0.384008425 | 0 |
| B119 | B139 | Hcv-rna | ADA | -0.34453101 | 0 |
| B119 | B169 | Hcv-rna | U-CRY | -0.99999344 | 0 |
| B12 | B14 | PG II | G-17 | 0.35424304 | 0 |
| B12 | B80 | PG II | anti-HP IgG | 0.302441466 | 1.19E-13 |
| B120 | B121 | HCV-cAg | C3 | -0.34921535 | 0 |
| B120 | B122 | HCV-cAg | C4 | -0.34921542 | 0 |
| B120 | B152 | HCV-cAg | ACA-Ig(G-A-M) | -0.34922522 | 0 |
| B120 | B159 | HCV-cAg | β2-MG | -0.34921688 | 0 |
| B120 | B169 | HCV-cAg | U-CRY | -0.34921278 | 0 |
| B120 | B157 | HCV-cAg | IgG | -0.30379152 | 9.19E-14 |
| B121 | B134 | C3 | CHE | -0.38400851 | 0 |
| B121 | B139 | C3 | ADA | 0.344531076 | 0 |
| B121 | B169 | C3 | U-CRY | 0.999994911 | 0 |
| B122 | B134 | C4 | CHE | -0.3840085 | 0 |
| B122 | B139 | C4 | ADA | 0.344531069 | 0 |
| B122 | B169 | C4 | U-CRY | 0.999996977 | 0 |
| B124 | B151 | ACTH | Anti-CCP | -0.37877757 | 0 |
| B124 | B182 | ACTH | TT | 0.335197768 | 0 |
| B124 | B184 | ACTH | INR | -0.70503348 | 0 |
| B124 | B185 | ACTH | PT | -0.7059417 | 0 |
| B124 | B186 | ACTH | PFC | -0.38380101 | 0 |
| B124 | B222 | ACTH | NS | 0.597437357 | 0 |
| B124 | B156 | ACTH | IgE | -0.30619534 | 5.73E-14 |
| B124 | B183 | ACTH | PTA | -0.29856274 | 2.53E-13 |
| B124 | B155 | ACTH | IgA | -0.29356881 | 6.51E-13 |
| B124 | B181 | ACTH | APTT | 0.275574994 | 1.69E-11 |
| B124 | B135 | ACTH | TBA | -0.23820546 | 7.12E-09 |
| B124 | B196 | ACTH | TCO2 | -0.2154783 | 1.77E-07 |
| B124 | B198 | ACTH | P | 0.205188628 | 6.81E-07 |
| B125 | B151 | Cor | Anti-CCP | 0.378777569 | 0 |
| B125 | B182 | Cor | TT | -0.33519777 | 0 |
| B125 | B184 | Cor | INR | 0.705033474 | 0 |
| B125 | B185 | Cor | PT | 0.705941694 | 0 |
| B125 | B186 | Cor | PFC | 0.383801003 | 0 |
| B125 | B222 | Cor | NS | -0.59743735 | 0 |
| B125 | B156 | Cor | IgE | 0.306195333 | 5.73E-14 |
| B125 | B183 | Cor | PTA | 0.298562739 | 2.53E-13 |
| B125 | B155 | Cor | IgA | 0.293568811 | 6.51E-13 |
| B125 | B181 | Cor | APTT | -0.27557499 | 1.69E-11 |
| B125 | B135 | Cor | TBA | 0.238205454 | 7.12E-09 |
| B125 | B196 | Cor | TCO2 | 0.2154783 | 1.77E-07 |
| B125 | B198 | Cor | P | -0.20518863 | 6.81E-07 |

| | | | | | |
|---|---|---|---|---|---|
| B126 | B127 | Alb | A-G ratio | 0.533434841 | 0 |
| B126 | B188 | Alb | Ca | 0.390742561 | 0 |
| B126 | B189 | Alb | eGFB | 0.260646454 | 2.12E-10 |
| B126 | B191 | Alb | Urea | -0.23468199 | 1.20E-08 |
| B126 | B133 | Alb | TBIL | 0.209921394 | 3.70E-07 |
| B127 | B130 | A-G ratio | GLB | -0.85894528 | 0 |
| B127 | B132 | A-G ratio | Dbil | 0.238337418 | 6.98E-09 |
| B127 | B221 | A-G ratio | Hei | 0.220895566 | 8.50E-08 |
| B127 | B133 | A-G ratio | TBIL | 0.216240676 | 1.60E-07 |
| B128 | B131 | ALT | AST | 0.745136276 | 0 |
| B128 | B140 | ALT | r-GT | 0.487747558 | 0 |
| B128 | B141 | ALT | AST/ALT | -0.61226664 | 0 |
| B128 | B215 | ALT | BMI | 0.344283981 | 0 |
| B128 | B216 | ALT | BMI | 0.369281074 | 0 |
| B128 | B206 | ALT | Waist | 0.327033137 | 8.88E-16 |
| B128 | B222 | ALT | NS | 0.295046584 | 4.93E-13 |
| B128 | B210 | ALT | Hip | 0.292615173 | 7.78E-13 |
| B128 | B171 | ALT | EC | 0.286224673 | 2.53E-12 |
| B128 | B207 | ALT | WHR | 0.248227074 | 1.55E-09 |
| B128 | B150 | ALT | ALP | 0.216526036 | 1.54E-07 |
| B129 | B132 | IBIL | Dbil | 0.658446552 | 0 |
| B129 | B133 | IBIL | TBIL | 0.980188567 | 0 |
| B129 | B191 | IBIL | Urea | -0.26424896 | 1.17E-10 |
| B129 | B190 | IBIL | Cre | -0.22712479 | 3.56E-08 |
| B130 | B209 | GLB | Smoking | -0.21147888 | 3.01E-07 |
| B131 | B140 | AST | r-GT | 0.351456762 | 0 |
| B131 | B215 | AST | BMI | 0.249747954 | 1.22E-09 |
| B131 | B206 | AST | Waist | 0.242588886 | 3.68E-09 |
| B131 | B150 | AST | ALP | 0.232474552 | 1.65E-08 |
| B131 | B171 | AST | EC | 0.230055338 | 2.34E-08 |
| B131 | B216 | AST | BMI | 0.225417775 | 4.53E-08 |
| B131 | B207 | AST | WHR | 0.222043321 | 7.25E-08 |
| B131 | B141 | AST | AST/ALT | -0.20918703 | 4.07E-07 |
| B132 | B133 | Dbil | TBIL | 0.794349882 | 0 |
| B133 | B191 | TBIL | Urea | -0.24931043 | 1.31E-09 |
| B133 | B190 | TBIL | Cre | -0.21488195 | 1.92E-07 |
| B134 | B152 | CHE | ACA-Ig(G-A-M) | -0.38402169 | 0 |
| B134 | B159 | CHE | β2-MG | -0.38400957 | 0 |
| B134 | B169 | CHE | U-CRY | -0.38400591 | 0 |
| B134 | B168 | CHE | NMP22 | 0.245481544 | 2.37E-09 |
| B134 | B199 | CHE | Mg | 0.238839525 | 6.48E-09 |
| B134 | B137 | CHE | AFU | 0.226556351 | 3.86E-08 |
| B134 | B136 | CHE | 5'NT | 0.210973385 | 3.22E-07 |
| B135 | B198 | TBA | P | -0.25761973 | 3.48E-10 |

| | | | | | |
|---|---|---|---|---|---|
| B135 | B193 | TBA | K | -0.21673915 | 1.50E-07 |
| B135 | B138 | TBA | PAB | -0.20556661 | 6.49E-07 |
| B135 | B179 | TBA | α1-MG | -0.20399075 | 7.93E-07 |
| B136 | B157 | 5'NT | IgG | 0.368989751 | 0 |
| B136 | B194 | 5'NT | Cl | -0.4308252 | 0 |
| B136 | B150 | 5'NT | ALP | 0.211963561 | 2.83E-07 |
| B136 | B196 | 5'NT | TCO2 | 0.204928896 | 7.04E-07 |
| B137 | B139 | AFU | ADA | 0.340306645 | 0 |
| B137 | B197 | AFU | AG | -0.29191801 | 8.86E-13 |
| B137 | B196 | AFU | TCO2 | 0.289516375 | 1.38E-12 |
| B138 | B196 | PAB | TCO2 | -0.33857247 | 0 |
| B138 | B197 | PAB | AG | 0.406007493 | 0 |
| B138 | B200 | PAB | OP | 0.433357268 | 0 |
| B138 | B199 | PAB | Mg | 0.319981334 | 3.55E-15 |
| B138 | B195 | PAB | Na | 0.23133961 | 1.95E-08 |
| B138 | B193 | PAB | K | 0.213070952 | 2.44E-07 |
| B138 | B198 | PAB | P | 0.209503675 | 3.90E-07 |
| B139 | B152 | ADA | ACA-Ig(G-A-M) | 0.344542885 | 0 |
| B139 | B159 | ADA | β2-MG | 0.344532027 | 0 |
| B139 | B169 | ADA | U-CRY | 0.344528752 | 0 |
| B139 | B200 | ADA | OP | 0.410790525 | 0 |
| B139 | B195 | ADA | Na | 0.286401434 | 2.45E-12 |
| B139 | B197 | ADA | AG | 0.239979399 | 5.46E-09 |
| B140 | B141 | r-GT | AST/ALT | -0.34086476 | 0 |
| B140 | B216 | r-GT | BMI | 0.254972784 | 5.33E-10 |
| B140 | B206 | r-GT | Waist | 0.22964843 | 2.49E-08 |
| B140 | B215 | r-GT | BMI | 0.215775129 | 1.70E-07 |
| B141 | B216 | AST/ALT | BMI | -0.34302606 | 0 |
| B141 | B215 | AST/ALT | BMI | -0.29103407 | 1.04E-12 |
| B141 | B206 | AST/ALT | Waist | -0.27733172 | 1.25E-11 |
| B141 | B210 | AST/ALT | Hip | -0.26359225 | 1.31E-10 |
| B141 | B222 | AST/ALT | NS | -0.26334455 | 1.36E-10 |
| B141 | B221 | AST/ALT | Hei | -0.21875998 | 1.14E-07 |
| B142 | B143 | 25-(OH)D | β-CTx | -0.69450994 | 0 |
| B142 | B144 | 25-(OH)D | Total-PINP | -0.62122397 | 0 |
| B142 | B145 | 25-(OH)D | BGP | -0.51416787 | 0 |
| B143 | B144 | β-CTx | Total-PINP | 0.631584686 | 0 |
| B143 | B145 | β-CTx | BGP | 0.861494257 | 0 |
| B143 | B183 | β-CTx | PTA | -0.33777889 | 0 |
| B143 | B182 | β-CTx | TT | 0.324075635 | 1.33E-15 |
| B143 | B186 | β-CTx | PFC | -0.30837782 | 3.73E-14 |
| B144 | B145 | Total-PINP | BGP | 0.752932347 | 0 |
| B144 | B151 | Total-PINP | Anti-CCP | -0.21415919 | 2.11E-07 |
| B145 | B183 | BGP | PTA | -0.20717357 | 5.28E-07 |

| | | | | | |
|---|---|---|---|---|---|
| B146 | B189 | CysC | eGFB | -0.26951401 | 4.82E-11 |
| B146 | B191 | CysC | Urea | 0.235881935 | 1.00E-08 |
| B146 | B192 | CysC | UA | 0.230651556 | 2.15E-08 |
| B146 | B200 | CysC | OP | 0.215661798 | 1.73E-07 |
| B146 | B213 | CysC | CVA-R | 0.209657489 | 3.83E-07 |
| B148 | B149 | Anti_TPO | TGAb | 0.465801936 | 0 |
| B15 | B100 | CD3-CD19+(B cell) | anti-Ro52 | 0.652229026 | 0 |
| B15 | B101 | CD3-CD19+(B cell) | anti-Sc70 | 0.652229033 | 0 |
| B15 | B102 | CD3-CD19+(B cell) | anti-Sm | 0.652229029 | 0 |
| B15 | B103 | CD3-CD19+(B cell) | anti-SSA/Ro | 0.652229026 | 0 |
| B15 | B104 | CD3-CD19+(B cell) | anti-SSB | 0.652229025 | 0 |
| B15 | B105 | CD3-CD19+(B cell) | anti-U1RNP | 0.652229026 | 0 |
| B15 | B106 | CD3-CD19+(B cell) | anti-P | 0.652229025 | 0 |
| B15 | B107 | CD3-CD19+(B cell) | AnuA | 0.652229014 | 0 |
| B15 | B108 | CD3-CD19+(B cell) | Anti-Cyclin | 0.652229027 | 0 |
| B15 | B109 | CD3-CD19+(B cell) | AMA-M2 | 0.652229026 | 0 |
| B15 | B110 | CD3-CD19+(B cell) | AHA | 0.652229027 | 0 |
| B15 | B124 | CD3-CD19+(B cell) | ACTH | -0.83524692 | 0 |
| B15 | B125 | CD3-CD19+(B cell) | Cor | 0.835246914 | 0 |
| B15 | B151 | CD3-CD19+(B cell) | Anti-CCP | 0.409716483 | 0 |
| B15 | B156 | CD3-CD19+(B cell) | IgE | 0.379800516 | 0 |
| B15 | B16 | CD3-CD19+(B cell) | CD3+T | 0.454281176 | 0 |
| B15 | B17 | CD3-CD19+(B cell) | CD3+CD16+CD56+NK | -0.44494478 | 0 |
| B15 | B18 | CD3-CD19+(B cell) | CD3+CD4+helper(inducer)T | 0.539497712 | 0 |
| B15 | B184 | CD3-CD19+(B cell) | INR | 0.616597635 | 0 |
| B15 | B185 | CD3-CD19+(B cell) | PT | 0.598813108 | 0 |
| B15 | B20 | CD3-CD19+(B cell) | CD3+CD4+CD8+ | -0.82864423 | 0 |
| B15 | B222 | CD3-CD19+(B cell) | NS | -0.47002389 | 0 |
| B15 | B23 | CD3-CD19+(B cell) | CD3-CD16+CD56+(NK) | -0.81883527 | 0 |
| B15 | B54 | CD3-CD19+(B cell) | ESR | -0.34659567 | 0 |
| B15 | B79 | CD3-CD19+(B cell) | hb V -dna | 0.414678647 | 0 |
| B15 | B81 | CD3-CD19+(B cell) | TSGF | -0.35217837 | 0 |
| B15 | B92 | CD3-CD19+(B cell) | ALDH2-Glu504Glu | -0.83524691 | 0 |
| B15 | B93 | CD3-CD19+(B cell) | ALDH2-Glu504Lys | 0.835246914 | 0 |
| B15 | B94 | CD3-CD19+(B cell) | ALDH2-Lys504Lys | 0.835246903 | 0 |
| B15 | B95 | CD3-CD19+(B cell) | anti-cenpb | 0.652229026 | 0 |
| B15 | B97 | CD3-CD19+(B cell) | dsDNA | 0.652229024 | 0 |
| B15 | B98 | CD3-CD19+(B cell) | anti-Jo-1 | 0.652229028 | 0 |
| B15 | B99 | CD3-CD19+(B cell) | anti-PM-Scl | -0.83130518 | 0 |
| B15 | B61 | CD3-CD19+(B cell) | WBHSRV | 0.330241289 | 4.44E-16 |
| B15 | B181 | CD3-CD19+(B cell) | APTT | -0.31150533 | 2.00E-14 |
| B15 | B59 | CD3-CD19+(B cell) | WBRV | 0.300466174 | 1.75E-13 |
| B15 | B26 | CD3-CD19+(B cell) | CKMB | 0.2857997 | 2.74E-12 |
| B15 | B63 | CD3-CD19+(B cell) | WBRSR(mPas_200) | 0.271664623 | 3.34E-11 |

| | | | | | |
|---|---|---|---|---|---|
| B15 | B56 | CD3-CD19+(B cell) | IR | 0.270368299 | 4.17E-11 |
| B15 | B64 | CD3-CD19+(B cell) | WBRSR(mPas_5) | 0.267958684 | 6.28E-11 |
| B15 | B199 | CD3-CD19+(B cell) | Mg | 0.257813304 | 3.37E-10 |
| B15 | B66 | CD3-CD19+(B cell) | WBRSR(mPas_100) | 0.251794324 | 8.85E-10 |
| B15 | B67 | CD3-CD19+(B cell) | WBRSR(mPas_50) | 0.247365883 | 1.77E-09 |
| B15 | B24 | CD3-CD19+(B cell) | αHBDH | 0.245164308 | 2.49E-09 |
| B15 | B34 | CD3-CD19+(B cell) | TK | 0.241561047 | 4.30E-09 |
| B15 | B65 | CD3-CD19+(B cell) | WBRSR(mPas_1) | 0.235824074 | 1.01E-08 |
| B15 | B19 | CD3-CD19+(B cell) | CD3+CD4+/CD3+CD8+ | 0.233339949 | 1.46E-08 |
| B15 | B55 | CD3-CD19+(B cell) | TK | 0.228292721 | 3.02E-08 |
| B15 | B75 | CD3-CD19+(B cell) | ApoA1/B | 0.220264005 | 9.26E-08 |
| B15 | B25 | CD3-CD19+(B cell) | CK | 0.219376316 | 1.05E-07 |
| B15 | B62 | CD3-CD19+(B cell) | WBHSRVI | 0.218843553 | 1.13E-07 |
| B15 | B74 | CD3-CD19+(B cell) | ApoA1 | 0.210073482 | 3.62E-07 |
| B15 | B115 | CD3-CD19+(B cell) | TSH TT3 TT4(TT3) | 0.203401932 | 8.54E-07 |
| B15 | B154 | CD3-CD19+(B cell) | RF | 0.202360097 | 9.74E-07 |
| B150 | B171 | ALP | EC | 0.55447731 | 0 |
| B151 | B154 | Anti-CCP | RF | 0.383871842 | 0 |
| B151 | B184 | Anti-CCP | INR | 0.289700823 | 1.34E-12 |
| B151 | B185 | Anti-CCP | PT | 0.274223957 | 2.14E-11 |
| B151 | B179 | Anti-CCP | α1-MG | 0.254707981 | 5.56E-10 |
| B151 | B183 | Anti-CCP | PTA | -0.23152826 | 1.90E-08 |
| B151 | B182 | Anti-CCP | TT | 0.203834308 | 8.09E-07 |
| B153 | B206 | Glu | Waist | 0.246662232 | 1.97E-09 |
| B153 | B170 | Glu | U-GLU | 0.223875976 | 5.62E-08 |
| B153 | B207 | Glu | WHR | 0.213606064 | 2.28E-07 |
| B153 | B215 | Glu | BMI | 0.208121713 | 4.67E-07 |
| B154 | B156 | RF | IgE | 0.335003609 | 0 |
| B154 | B179 | RF | α1-MG | 0.576643114 | 0 |
| B154 | B181 | RF | APTT | -0.23364593 | 1.39E-08 |
| B154 | B184 | RF | INR | 0.219717427 | 9.99E-08 |
| B154 | B185 | RF | PT | 0.214250122 | 2.09E-07 |
| B154 | B180 | RF | Tf | 0.20305682 | 8.92E-07 |
| B155 | B156 | IgA | IgE | -0.34407235 | 0 |
| B155 | B157 | IgA | IgG | 0.387559626 | 0 |
| B155 | B185 | IgA | PT | 0.274364903 | 2.09E-11 |
| B155 | B186 | IgA | PFC | 0.273605913 | 2.39E-11 |
| B155 | B184 | IgA | INR | 0.267849691 | 6.40E-11 |
| B155 | B182 | IgA | TT | -0.26078147 | 2.08E-10 |
| B155 | B183 | IgA | PTA | 0.249001839 | 1.37E-09 |
| B155 | B158 | IgA | IgM | -0.21052242 | 3.42E-07 |
| B156 | B158 | IgE | IgM | 0.425746685 | 0 |
| B156 | B181 | IgE | APTT | -0.42726479 | 0 |
| B156 | B184 | IgE | INR | 0.466622582 | 0 |

| | | | | | |
|---|---|---|---|---|---|
| B156 | B185 | IgE | PT | 0.463727144 | 0 |
| B156 | B157 | IgE | IgG | -0.29749835 | 3.10E-13 |
| B156 | B228 | IgE | OS | 0.273104352 | 2.60E-11 |
| B157 | B228 | IgG | OS | 0.204309826 | 7.61E-07 |
| B157 | B158 | IgG | IgM | -0.20288561 | 9.12E-07 |
| B158 | B182 | IgM | TT | -0.46541891 | 0 |
| B158 | B183 | IgM | PTA | 0.470475157 | 0 |
| B158 | B186 | IgM | PFC | 0.459501084 | 0 |
| B159 | B169 | β2-MG | U-CRY | 0.999998858 | 0 |
| B16 | B124 | CD3+T | ACTH | -0.75337271 | 0 |
| B16 | B125 | CD3+T | Cor | 0.753372702 | 0 |
| B16 | B155 | CD3+T | IgA | 0.462805037 | 0 |
| B16 | B18 | CD3+T | CD3+CD4+helper(inducer)T | 0.378820343 | 0 |
| B16 | B182 | CD3+T | TT | -0.48271116 | 0 |
| B16 | B183 | CD3+T | PTA | 0.464678714 | 0 |
| B16 | B184 | CD3+T | INR | 0.515717161 | 0 |
| B16 | B185 | CD3+T | PT | 0.527423099 | 0 |
| B16 | B186 | CD3+T | PFC | 0.508227595 | 0 |
| B16 | B19 | CD3+T | CD3+CD4+/CD3+CD8+ | -0.52588974 | 0 |
| B16 | B20 | CD3+T | CD3+CD4+CD8+ | -0.78557986 | 0 |
| B16 | B21 | CD3+T | CD3+CD4-CD8- | 0.863099247 | 0 |
| B16 | B22 | CD3+T | CD3+CD8+(T) | 0.723172611 | 0 |
| B16 | B222 | CD3+T | NS | -0.47156642 | 0 |
| B16 | B23 | CD3+T | CD3-CD16+CD56+(NK) | -0.76223585 | 0 |
| B16 | B69 | CD3+T | PV | 0.390184986 | 0 |
| B16 | B74 | CD3+T | ApoA1 | 0.390530891 | 0 |
| B16 | B79 | CD3+T | hb V -dna | 0.379051246 | 0 |
| B16 | B92 | CD3+T | ALDH2-Glu504Glu | -0.7533727 | 0 |
| B16 | B93 | CD3+T | ALDH2-Glu504Lys | 0.753372702 | 0 |
| B16 | B94 | CD3+T | ALDH2-Lys504Lys | 0.753372693 | 0 |
| B16 | B99 | CD3+T | anti-PM-Scl | -0.75951936 | 0 |
| B16 | B75 | CD3+T | ApoA1/B | 0.326082305 | 8.88E-16 |
| B16 | B81 | CD3+T | TSGF | -0.31507139 | 9.77E-15 |
| B16 | B54 | CD3+T | ESR | 0.289292399 | 1.44E-12 |
| B16 | B178 | CD3+T | Ig-U | 0.272972563 | 2.66E-11 |
| B16 | B198 | CD3+T | P | -0.27142261 | 3.48E-11 |
| B16 | B17 | CD3+T | CD3+CD16+CD56+NK | 0.265138075 | 1.01E-10 |
| B16 | B180 | CD3+T | Tf | 0.257158098 | 3.75E-10 |
| B16 | B76 | CD3+T | ApoB | -0.238906 | 6.42E-09 |
| B16 | B151 | CD3+T | Anti-CCP | 0.229140954 | 2.67E-08 |
| B16 | B34 | CD3+T | TK | 0.224723113 | 4.99E-08 |
| B16 | B139 | CD3+T | ADA | 0.204714508 | 7.23E-07 |
| B16 | B135 | CD3+T | TBA | 0.203067699 | 8.91E-07 |
| B160 | B163 | PH | SG | -0.20369665 | 8.23E-07 |

| | | | | | |
|---|---|---|---|---|---|
| B163 | B216 | SG | BMI | 0.272146839 | 3.07E-11 |
| B163 | B176 | SG | Ucr | 0.270882013 | 3.82E-11 |
| B163 | B221 | SG | Hei | 0.256265842 | 4.33E-10 |
| B163 | B210 | SG | Hip | 0.217041574 | 1.44E-07 |
| B163 | B206 | SG | Waist | 0.203244506 | 8.71E-07 |
| B164 | B172 | COM | KB | -0.21502012 | 1.88E-07 |
| B167 | B223 | PC | FH | -0.22048193 | 8.99E-08 |
| B168 | B175 | NMP22 | ALB-U | 0.524405949 | 0 |
| B168 | B177 | NMP22 | ACR | 0.367054024 | 0 |
| B168 | B194 | NMP22 | Cl | 0.373754818 | 0 |
| B168 | B196 | NMP22 | TCO2 | -0.47114471 | 0 |
| B168 | B199 | NMP22 | Mg | -0.25232796 | 8.13E-10 |
| B168 | B200 | NMP22 | OP | 0.23812539 | 7.21E-09 |
| B17 | B100 | CD3+CD16+CD56+NK | anti-Ro52 | -0.40646942 | 0 |
| B17 | B101 | CD3+CD16+CD56+NK | anti-Sc70 | -0.40646943 | 0 |
| B17 | B102 | CD3+CD16+CD56+NK | anti-Sm | -0.40646942 | 0 |
| B17 | B103 | CD3+CD16+CD56+NK | anti-SSA/Ro | -0.40646942 | 0 |
| B17 | B104 | CD3+CD16+CD56+NK | anti-SSB | -0.40646942 | 0 |
| B17 | B105 | CD3+CD16+CD56+NK | anti-U1RNP | -0.40646942 | 0 |
| B17 | B106 | CD3+CD16+CD56+NK | anti-P | -0.40646942 | 0 |
| B17 | B107 | CD3+CD16+CD56+NK | AnuA | -0.40646941 | 0 |
| B17 | B108 | CD3+CD16+CD56+NK | Anti-Cyclin | -0.40646942 | 0 |
| B17 | B109 | CD3+CD16+CD56+NK | AMA-M2 | -0.40646942 | 0 |
| B17 | B110 | CD3+CD16+CD56+NK | AHA | -0.40646942 | 0 |
| B17 | B143 | CD3+CD16+CD56+NK | β-CTx | -0.37235035 | 0 |
| B17 | B151 | CD3+CD16+CD56+NK | Anti-CCP | -0.37877757 | 0 |
| B17 | B158 | CD3+CD16+CD56+NK | IgM | 0.43544915 | 0 |
| B17 | B18 | CD3+CD16+CD56+NK | CD3+CD4+helper(inducer)T | -0.54777916 | 0 |
| B17 | B182 | CD3+CD16+CD56+NK | TT | -0.8705931 | 0 |
| B17 | B183 | CD3+CD16+CD56+NK | PTA | 0.907361612 | 0 |
| B17 | B186 | CD3+CD16+CD56+NK | PFC | 0.828469925 | 0 |
| B17 | B19 | CD3+CD16+CD56+NK | CD3+CD4+/CD3+CD8+ | -0.64029469 | 0 |
| B17 | B21 | CD3+CD16+CD56+NK | CD3+CD4-CD8- | 0.62943436 | 0 |
| B17 | B22 | CD3+CD16+CD56+NK | CD3+CD8+(T) | 0.509389507 | 0 |
| B17 | B54 | CD3+CD16+CD56+NK | ESR | 0.803575963 | 0 |
| B17 | B55 | CD3+CD16+CD56+NK | TK | -0.5857551 | 0 |
| B17 | B56 | CD3+CD16+CD56+NK | IR | -0.68004681 | 0 |
| B17 | B59 | CD3+CD16+CD56+NK | WBRV | -0.41330162 | 0 |
| B17 | B60 | CD3+CD16+CD56+NK | WBRVI | -0.42620366 | 0 |
| B17 | B61 | CD3+CD16+CD56+NK | WBHSRV | -0.57716015 | 0 |
| B17 | B62 | CD3+CD16+CD56+NK | WBHSRVI | -0.53629716 | 0 |
| B17 | B63 | CD3+CD16+CD56+NK | WBRSR(mPas_200) | -0.37946254 | 0 |
| B17 | B64 | CD3+CD16+CD56+NK | WBRSR(mPas_5) | -0.38585494 | 0 |
| B17 | B65 | CD3+CD16+CD56+NK | WBRSR(mPas_1) | -0.33674866 | 0 |

| | | | | | |
|---|---|---|---|---|---|
| B17 | B66 | CD3+CD16+CD56+NK | WBRSR(mPas_100) | -0.37015563 | 0 |
| B17 | B67 | CD3+CD16+CD56+NK | WBRSR(mPas_50) | -0.36438948 | 0 |
| B17 | B69 | CD3+CD16+CD56+NK | PV | 0.517730344 | 0 |
| B17 | B95 | CD3+CD16+CD56+NK | anti-cenpb | -0.40646942 | 0 |
| B17 | B97 | CD3+CD16+CD56+NK | dsDNA | -0.40646942 | 0 |
| B17 | B98 | CD3+CD16+CD56+NK | anti-Jo-1 | -0.40646942 | 0 |
| B17 | B20 | CD3+CD16+CD56+NK | CD3+CD4+CD8+ | 0.296585411 | 3.68E-13 |
| B17 | B26 | CD3+CD16+CD56+NK | CKMB | -0.2956089 | 4.43E-13 |
| B17 | B74 | CD3+CD16+CD56+NK | ApoA1 | 0.258922692 | 2.82E-10 |
| B17 | B199 | CD3+CD16+CD56+NK | Mg | -0.23214646 | 1.73E-08 |
| B17 | B145 | CD3+CD16+CD56+NK | BGP | -0.22795763 | 3.16E-08 |
| B17 | B195 | CD3+CD16+CD56+NK | Na | -0.22586999 | 4.25E-08 |
| B17 | B24 | CD3+CD16+CD56+NK | αHBDH | -0.20645747 | 5.79E-07 |
| B170 | B178 | U-GLU | Ig-U | 0.379812166 | 0 |
| B170 | B180 | U-GLU | Tf | 0.358300232 | 0 |
| B170 | B214 | U-GLU | CVA-L | 0.537378487 | 0 |
| B170 | B171 | U-GLU | EC | 0.315904388 | 8.22E-15 |
| B170 | B191 | U-GLU | Urea | 0.270426178 | 4.13E-11 |
| B170 | B190 | U-GLU | Cre | 0.251391471 | 9.43E-10 |
| B170 | B189 | U-GLU | eGFB | -0.22030846 | 9.21E-08 |
| B174 | B189 | Urea/Cr | eGFB | 0.444390195 | 0 |
| B174 | B190 | Urea/Cr | Cre | -0.50823022 | 0 |
| B174 | B218 | Urea/Cr | DBP | -0.23673986 | 8.85E-09 |
| B174 | B220 | Urea/Cr | SBP | -0.2217208 | 7.58E-08 |
| B174 | B221 | Urea/Cr | Hei | -0.20973392 | 3.79E-07 |
| B175 | B177 | ALB-U | ACR | 0.78821998 | 0 |
| B175 | B178 | ALB-U | Ig-U | 0.531428512 | 0 |
| B175 | B180 | ALB-U | Tf | 0.503037511 | 0 |
| B175 | B196 | ALB-U | TCO2 | -0.37329977 | 0 |
| B175 | B214 | ALB-U | CVA-L | 0.375198032 | 0 |
| B175 | B179 | ALB-U | α1-MG | 0.245759422 | 2.27E-09 |
| B175 | B199 | ALB-U | Mg | -0.22703287 | 3.61E-08 |
| B175 | B200 | ALB-U | OP | 0.220908555 | 8.48E-08 |
| B176 | B177 | Ucr | ACR | -0.2361647 | 9.64E-09 |
| B176 | B183 | Ucr | PTA | 0.204552702 | 7.38E-07 |
| B177 | B178 | ACR | Ig-U | 0.438258891 | 0 |
| B177 | B180 | ACR | Tf | 0.436152295 | 0 |
| B177 | B227 | ACR | OD | 0.551575426 | 0 |
| B177 | B214 | ACR | CVA-L | 0.307753073 | 4.22E-14 |
| B177 | B196 | ACR | TCO2 | -0.29803566 | 2.79E-13 |
| B177 | B179 | ACR | α1-MG | 0.263292543 | 1.37E-10 |
| B178 | B179 | Ig-U | α1-MG | 0.480327637 | 0 |
| B178 | B180 | Ig-U | Tf | 0.95290513 | 0 |
| B178 | B214 | Ig-U | CVA-L | 0.709359068 | 0 |

| | | | | | |
|---|---|---|---|---|---|
| B178 | B193 | Ig-U | K | 0.326065457 | 8.88E-16 |
| B178 | B194 | Ig-U | Cl | -0.26272476 | 1.51E-10 |
| B178 | B195 | Ig-U | Na | -0.21846639 | 1.18E-07 |
| B179 | B180 | α1-MG | Tf | 0.718842707 | 0 |
| B179 | B214 | α1-MG | CVA-L | 0.329486073 | 4.44E-16 |
| B179 | B211 | α1-MG | IOP-R | 0.227892216 | 3.19E-08 |
| B18 | B124 | CD3+CD4+helper(inducer)T | ACTH | -0.42491543 | 0 |
| B18 | B125 | CD3+CD4+helper(inducer)T | Cor | 0.424915423 | 0 |
| B18 | B151 | CD3+CD4+helper(inducer)T | Anti-CCP | 0.413833135 | 0 |
| B18 | B178 | CD3+CD4+helper(inducer)T | Ig-U | -0.41914635 | 0 |
| B18 | B180 | CD3+CD4+helper(inducer)T | Tf | -0.39705923 | 0 |
| B18 | B182 | CD3+CD4+helper(inducer)T | TT | 0.334565536 | 0 |
| B18 | B183 | CD3+CD4+helper(inducer)T | PTA | -0.37038676 | 0 |
| B18 | B184 | CD3+CD4+helper(inducer)T | INR | 0.334531609 | 0 |
| B18 | B19 | CD3+CD4+helper(inducer)T | CD3+CD4+/CD3+CD8+ | 0.536969528 | 0 |
| B18 | B195 | CD3+CD4+helper(inducer)T | Na | 0.379093156 | 0 |
| B18 | B20 | CD3+CD4+helper(inducer)T | CD3+CD4+CD8+ | -0.7770318 | 0 |
| B18 | B23 | CD3+CD4+helper(inducer)T | CD3-CD16+CD56+(NK) | -0.41854658 | 0 |
| B18 | B54 | CD3+CD4+helper(inducer)T | ESR | -0.38513283 | 0 |
| B18 | B55 | CD3+CD4+helper(inducer)T | TK | 0.38654304 | 0 |
| B18 | B56 | CD3+CD4+helper(inducer)T | IR | 0.392670534 | 0 |
| B18 | B61 | CD3+CD4+helper(inducer)T | WBHSRV | 0.385916118 | 0 |
| B18 | B92 | CD3+CD4+helper(inducer)T | ALDH2-Glu504Glu | -0.42491542 | 0 |
| B18 | B93 | CD3+CD4+helper(inducer)T | ALDH2-Glu504Lys | 0.424915423 | 0 |
| B18 | B94 | CD3+CD4+helper(inducer)T | ALDH2-Lys504Lys | 0.424915418 | 0 |
| B18 | B99 | CD3+CD4+helper(inducer)T | anti-PM-Scl | -0.42377117 | 0 |
| B18 | B22 | CD3+CD4+helper(inducer)T | CD3+CD8+(T) | -0.33038903 | 4.44E-16 |
| B18 | B185 | CD3+CD4+helper(inducer)T | PT | 0.312191009 | 1.73E-14 |
| B18 | B59 | CD3+CD4+helper(inducer)T | WBRV | 0.300263475 | 1.82E-13 |
| B18 | B214 | CD3+CD4+helper(inducer)T | CVA-L | -0.29510209 | 4.88E-13 |
| B18 | B186 | CD3+CD4+helper(inducer)T | PFC | -0.29074932 | 1.10E-12 |
| B18 | B198 | CD3+CD4+helper(inducer)T | P | -0.28418725 | 3.67E-12 |
| B18 | B62 | CD3+CD4+helper(inducer)T | WBHSRVI | 0.273006733 | 2.65E-11 |
| B18 | B155 | CD3+CD4+helper(inducer)T | IgA | 0.268411022 | 5.82E-11 |
| B18 | B137 | CD3+CD4+helper(inducer)T | AFU | -0.26296997 | 1.45E-10 |
| B18 | B100 | CD3+CD4+helper(inducer)T | anti-Ro52 | 0.262550685 | 1.55E-10 |
| B18 | B101 | CD3+CD4+helper(inducer)T | anti-Sc70 | 0.262550687 | 1.55E-10 |
| B18 | B102 | CD3+CD4+helper(inducer)T | anti-Sm | 0.262550686 | 1.55E-10 |
| B18 | B103 | CD3+CD4+helper(inducer)T | anti-SSA/Ro | 0.262550685 | 1.55E-10 |
| B18 | B104 | CD3+CD4+helper(inducer)T | anti-SSB | 0.262550684 | 1.55E-10 |
| B18 | B105 | CD3+CD4+helper(inducer)T | anti-U1RNP | 0.262550684 | 1.55E-10 |
| B18 | B106 | CD3+CD4+helper(inducer)T | anti-P | 0.262550685 | 1.55E-10 |
| B18 | B107 | CD3+CD4+helper(inducer)T | AnuA | 0.262550677 | 1.55E-10 |
| B18 | B108 | CD3+CD4+helper(inducer)T | Anti-Cyclin | 0.262550684 | 1.55E-10 |

| | | | | | |
|---|---|---|---|---|---|
| B18 | B109 | CD3+CD4+helper(inducer)T | AMA-M2 | 0.262550685 | 1.55E-10 |
| B18 | B110 | CD3+CD4+helper(inducer)T | AHA | 0.262550684 | 1.55E-10 |
| B18 | B95 | CD3+CD4+helper(inducer)T | anti-cenpb | 0.262550684 | 1.55E-10 |
| B18 | B97 | CD3+CD4+helper(inducer)T | dsDNA | 0.262550684 | 1.55E-10 |
| B18 | B98 | CD3+CD4+helper(inducer)T | anti-Jo-1 | 0.262550684 | 1.55E-10 |
| B18 | B222 | CD3+CD4+helper(inducer)T | NS | -0.25566035 | 4.78E-10 |
| B18 | B63 | CD3+CD4+helper(inducer)T | WBRSR(mPas_200) | 0.240817608 | 4.82E-09 |
| B18 | B26 | CD3+CD4+helper(inducer)T | CKMB | 0.240114699 | 5.35E-09 |
| B18 | B64 | CD3+CD4+helper(inducer)T | WBRSR(mPas_5) | 0.236774775 | 8.81E-09 |
| B18 | B193 | CD3+CD4+helper(inducer)T | K | -0.23655411 | 9.10E-09 |
| B18 | B175 | CD3+CD4+helper(inducer)T | ALB-U | -0.23556364 | 1.05E-08 |
| B18 | B66 | CD3+CD4+helper(inducer)T | WBRSR(mPas_100) | 0.231726743 | 1.84E-08 |
| B18 | B158 | CD3+CD4+helper(inducer)T | IgM | -0.22952431 | 2.53E-08 |
| B18 | B179 | CD3+CD4+helper(inducer)T | α1-MG | -0.22843223 | 2.96E-08 |
| B18 | B67 | CD3+CD4+helper(inducer)T | WBRSR(mPas_50) | 0.227767443 | 3.25E-08 |
| B18 | B181 | CD3+CD4+helper(inducer)T | APTT | -0.21822306 | 1.22E-07 |
| B18 | B60 | CD3+CD4+helper(inducer)T | WBRVI | 0.21721023 | 1.40E-07 |
| B18 | B79 | CD3+CD4+helper(inducer)T | hb V -dna | 0.211405199 | 3.04E-07 |
| B18 | B65 | CD3+CD4+helper(inducer)T | WBRSR(mPas_1) | 0.20432608 | 7.60E-07 |
| B18 | B143 | CD3+CD4+helper(inducer)T | β-CTx | 0.203978528 | 7.94E-07 |
| B180 | B214 | Tf | CVA-L | 0.672841433 | 0 |
| B180 | B193 | Tf | K | 0.307148502 | 4.75E-14 |
| B180 | B194 | Tf | Cl | -0.2496517 | 1.24E-09 |
| B180 | B195 | Tf | Na | -0.20650891 | 5.75E-07 |
| B181 | B184 | APTT | INR | -0.87239029 | 0 |
| B181 | B185 | APTT | PT | -0.86582407 | 0 |
| B181 | B186 | APTT | PFC | -0.34008634 | 0 |
| B181 | B182 | APTT | TT | 0.273535486 | 2.42E-11 |
| B182 | B183 | TT | PTA | -0.99645978 | 0 |
| B182 | B184 | TT | INR | -0.43657144 | 0 |
| B182 | B185 | TT | PT | -0.47387643 | 0 |
| B182 | B186 | TT | PFC | -0.9965913 | 0 |
| B182 | B222 | TT | NS | 0.237818819 | 7.54E-09 |
| B183 | B184 | PTA | INR | 0.363510654 | 0 |
| B183 | B185 | PTA | PT | 0.401663282 | 0 |
| B183 | B186 | PTA | PFC | 0.987003714 | 0 |
| B183 | B222 | PTA | NS | -0.21685055 | 1.47E-07 |
| B184 | B185 | INR | PT | 0.998619253 | 0 |
| B184 | B186 | INR | PFC | 0.507494613 | 0 |
| B184 | B222 | INR | NS | -0.42896966 | 0 |
| B185 | B186 | PT | PFC | 0.542524014 | 0 |
| B185 | B222 | PT | NS | -0.43031764 | 0 |
| B186 | B222 | PFC | NS | -0.2664064 | 8.16E-11 |
| B189 | B190 | eGFB | Cre | -0.67346969 | 0 |

| | | | | | |
|---|---|---|---|---|---|
| B189 | B191 | eGFB | Urea | -0.72840104 | 0 |
| B189 | B220 | eGFB | SBP | -0.3646374 | 0 |
| B189 | B192 | eGFB | UA | -0.23525281 | 1.10E-08 |
| B19 | B100 | CD3+CD4+/CD3+CD8+ | anti-Ro52 | 0.605358308 | 0 |
| B19 | B101 | CD3+CD4+/CD3+CD8+ | anti-Sc70 | 0.605358314 | 0 |
| B19 | B102 | CD3+CD4+/CD3+CD8+ | anti-Sm | 0.605358311 | 0 |
| B19 | B103 | CD3+CD4+/CD3+CD8+ | anti-SSA/Ro | 0.605358308 | 0 |
| B19 | B104 | CD3+CD4+/CD3+CD8+ | anti-SSB | 0.605358307 | 0 |
| B19 | B105 | CD3+CD4+/CD3+CD8+ | anti-U1RNP | 0.605358308 | 0 |
| B19 | B106 | CD3+CD4+/CD3+CD8+ | anti-P | 0.605358308 | 0 |
| B19 | B107 | CD3+CD4+/CD3+CD8+ | AnuA | 0.605358296 | 0 |
| B19 | B108 | CD3+CD4+/CD3+CD8+ | Anti-Cyclin | 0.605358308 | 0 |
| B19 | B109 | CD3+CD4+/CD3+CD8+ | AMA-M2 | 0.605358308 | 0 |
| B19 | B110 | CD3+CD4+/CD3+CD8+ | AHA | 0.605358309 | 0 |
| B19 | B175 | CD3+CD4+/CD3+CD8+ | ALB-U | -0.40542736 | 0 |
| B19 | B178 | CD3+CD4+/CD3+CD8+ | Ig-U | -0.73568162 | 0 |
| B19 | B180 | CD3+CD4+/CD3+CD8+ | Tf | -0.69710211 | 0 |
| B19 | B182 | CD3+CD4+/CD3+CD8+ | TT | 0.599522562 | 0 |
| B19 | B183 | CD3+CD4+/CD3+CD8+ | PTA | -0.6183569 | 0 |
| B19 | B186 | CD3+CD4+/CD3+CD8+ | PFC | -0.57863106 | 0 |
| B19 | B21 | CD3+CD4+/CD3+CD8+ | CD3+CD4-CD8- | -0.50894277 | 0 |
| B19 | B214 | CD3+CD4+/CD3+CD8+ | CVA-L | -0.52163543 | 0 |
| B19 | B22 | CD3+CD4+/CD3+CD8+ | CD3+CD8+(T) | -0.93899878 | 0 |
| B19 | B54 | CD3+CD4+/CD3+CD8+ | ESR | -0.54646247 | 0 |
| B19 | B56 | CD3+CD4+/CD3+CD8+ | IR | 0.417478694 | 0 |
| B19 | B61 | CD3+CD4+/CD3+CD8+ | WBHSRV | 0.338003249 | 0 |
| B19 | B62 | CD3+CD4+/CD3+CD8+ | WBHSRVI | 0.351149389 | 0 |
| B19 | B69 | CD3+CD4+/CD3+CD8+ | PV | -0.3698906 | 0 |
| B19 | B95 | CD3+CD4+/CD3+CD8+ | anti-cenpb | 0.605358308 | 0 |
| B19 | B97 | CD3+CD4+/CD3+CD8+ | dsDNA | 0.605358306 | 0 |
| B19 | B98 | CD3+CD4+/CD3+CD8+ | anti-Jo-1 | 0.60535831 | 0 |
| B19 | B179 | CD3+CD4+/CD3+CD8+ | α1-MG | -0.32957816 | 4.44E-16 |
| B19 | B55 | CD3+CD4+/CD3+CD8+ | TK | 0.331308818 | 4.44E-16 |
| B19 | B74 | CD3+CD4+/CD3+CD8+ | ApoA1 | -0.32922047 | 4.44E-16 |
| B19 | B177 | CD3+CD4+/CD3+CD8+ | ACR | -0.32371297 | 1.78E-15 |
| B19 | B158 | CD3+CD4+/CD3+CD8+ | IgM | -0.29211704 | 8.54E-13 |
| B19 | B60 | CD3+CD4+/CD3+CD8+ | WBRVI | 0.282001969 | 5.43E-12 |
| B19 | B195 | CD3+CD4+/CD3+CD8+ | Na | 0.277529717 | 1.20E-11 |
| B19 | B170 | CD3+CD4+/CD3+CD8+ | U-GLU | -0.27449904 | 2.04E-11 |
| B19 | B194 | CD3+CD4+/CD3+CD8+ | Cl | 0.262741193 | 1.50E-10 |
| B19 | B64 | CD3+CD4+/CD3+CD8+ | WBRSR(mPas_5) | 0.245021913 | 2.54E-09 |
| B19 | B143 | CD3+CD4+/CD3+CD8+ | β-CTx | 0.238524395 | 6.79E-09 |
| B19 | B63 | CD3+CD4+/CD3+CD8+ | WBRSR(mPas_200) | 0.237303952 | 8.14E-09 |
| B19 | B59 | CD3+CD4+/CD3+CD8+ | WBRV | 0.236504903 | 9.17E-09 |

| | | | | | |
|---|---|---|---|---|---|
| B19 | B66 | CD3+CD4+/CD3+CD8+ | WBRSR(mPas_100) | 0.2314018 | 1.93E-08 |
| B19 | B67 | CD3+CD4+/CD3+CD8+ | WBRSR(mPas_50) | 0.22784213 | 3.22E-08 |
| B19 | B199 | CD3+CD4+/CD3+CD8+ | Mg | 0.223154118 | 6.22E-08 |
| B19 | B65 | CD3+CD4+/CD3+CD8+ | WBRSR(mPas_1) | 0.215037007 | 1.88E-07 |
| B190 | B191 | Cre | Urea | 0.815616248 | 0 |
| B190 | B220 | Cre | SBP | 0.248092077 | 1.58E-09 |
| B191 | B220 | Urea | SBP | 0.23326252 | 1.47E-08 |
| B191 | B192 | Urea | UA | 0.231235139 | 1.98E-08 |
| B192 | B206 | UA | Waist | 0.393839569 | 0 |
| B192 | B207 | UA | WHR | 0.336076201 | 0 |
| B192 | B216 | UA | BMI | 0.372091011 | 0 |
| B192 | B222 | UA | NS | 0.355803644 | 0 |
| B192 | B210 | UA | Hip | 0.306335748 | 5.60E-14 |
| B192 | B215 | UA | BMI | 0.293314865 | 6.83E-13 |
| B192 | B221 | UA | Hei | 0.276260362 | 1.50E-11 |
| B193 | B198 | K | P | 0.58890567 | 0 |
| B193 | B199 | K | Mg | 0.386893118 | 0 |
| B193 | B214 | K | CVA-L | 0.226592124 | 3.84E-08 |
| B193 | B197 | K | AG | -0.21657109 | 1.53E-07 |
| B194 | B196 | Cl | TCO2 | -0.54914653 | 0 |
| B194 | B200 | Cl | OP | 0.328745048 | 4.44E-16 |
| B194 | B195 | Cl | Na | 0.298206348 | 2.70E-13 |
| B194 | B197 | Cl | AG | -0.29356449 | 6.51E-13 |
| B194 | B199 | Cl | Mg | -0.2152783 | 1.82E-07 |
| B195 | B197 | Na | AG | 0.550662561 | 0 |
| B195 | B200 | Na | OP | 0.833230506 | 0 |
| B195 | B196 | Na | TCO2 | -0.31168659 | 1.91E-14 |
| B196 | B197 | TCO2 | AG | -0.50344039 | 0 |
| B196 | B200 | TCO2 | OP | -0.60328111 | 0 |
| B196 | B199 | TCO2 | Mg | 0.21551309 | 1.76E-07 |
| B196 | B222 | TCO2 | NS | -0.2147912 | 1.94E-07 |
| B197 | B200 | AG | OP | 0.670758646 | 0 |
| B197 | B198 | AG | P | -0.20223696 | 9.89E-07 |
| B2 | B189 | Age | eGFB | -0.47906498 | 0 |
| B2 | B191 | Age | Urea | 0.394728268 | 0 |
| B2 | B207 | Age | WHR | 0.36556453 | 0 |
| B2 | B220 | Age | SBP | 0.372568358 | 0 |
| B2 | B118 | Age | CEA1 | 0.307884034 | 4.11E-14 |
| B2 | B47 | Age | RDW | 0.297654469 | 3.01E-13 |
| B2 | B153 | Age | Glu | 0.285334594 | 2.98E-12 |
| B2 | B206 | Age | Waist | 0.277603813 | 1.19E-11 |
| B2 | B126 | Age | Alb | -0.27213687 | 3.08E-11 |
| B2 | B46 | Age | RBC Count | -0.2680526 | 6.18E-11 |
| B2 | B52 | Age | PLT/BPC | -0.26649189 | 8.04E-11 |

| | | | | | |
|---|---|---|---|---|---|
| B2 | B50 | Age | eo% | 0.253110227 | 7.18E-10 |
| B2 | B43 | Age | MCV | 0.239641258 | 5.75E-09 |
| B2 | B219 | Age | OT | 0.219315005 | 1.05E-07 |
| B2 | B221 | Age | Hei | -0.21780607 | 1.30E-07 |
| B20 | B124 | CD3+CD4+CD8+ | ACTH | 0.832497393 | 0 |
| B20 | B125 | CD3+CD4+CD8+ | Cor | -0.83249739 | 0 |
| B20 | B151 | CD3+CD4+CD8+ | Anti-CCP | -0.42924206 | 0 |
| B20 | B155 | CD3+CD4+CD8+ | IgA | -0.37711648 | 0 |
| B20 | B184 | CD3+CD4+CD8+ | INR | -0.60628594 | 0 |
| B20 | B185 | CD3+CD4+CD8+ | PT | -0.59471724 | 0 |
| B20 | B21 | CD3+CD4+CD8+ | CD3+CD4-CD8- | -0.53323707 | 0 |
| B20 | B222 | CD3+CD4+CD8+ | NS | 0.494031374 | 0 |
| B20 | B23 | CD3+CD4+CD8+ | CD3-CD16+CD56+(NK) | 0.826375671 | 0 |
| B20 | B79 | CD3+CD4+CD8+ | hb V -dna | -0.4157639 | 0 |
| B20 | B81 | CD3+CD4+CD8+ | TSGF | 0.349757509 | 0 |
| B20 | B92 | CD3+CD4+CD8+ | ALDH2-Glu504Glu | 0.832497387 | 0 |
| B20 | B93 | CD3+CD4+CD8+ | ALDH2-Glu504Lys | -0.83249739 | 0 |
| B20 | B94 | CD3+CD4+CD8+ | ALDH2-Lys504Lys | -0.83249738 | 0 |
| B20 | B99 | CD3+CD4+CD8+ | anti-PM-Scl | 0.83330331 | 0 |
| B20 | B198 | CD3+CD4+CD8+ | P | 0.31771807 | 5.77E-15 |
| B20 | B181 | CD3+CD4+CD8+ | APTT | 0.284838305 | 3.26E-12 |
| B20 | B61 | CD3+CD4+CD8+ | WBHSRV | -0.27433594 | 2.10E-11 |
| B20 | B100 | CD3+CD4+CD8+ | anti-Ro52 | -0.26923869 | 5.05E-11 |
| B20 | B101 | CD3+CD4+CD8+ | anti-Sc70 | -0.26923869 | 5.05E-11 |
| B20 | B102 | CD3+CD4+CD8+ | anti-Sm | -0.26923869 | 5.05E-11 |
| B20 | B103 | CD3+CD4+CD8+ | anti-SSA/Ro | -0.26923869 | 5.05E-11 |
| B20 | B104 | CD3+CD4+CD8+ | anti-SSB | -0.26923868 | 5.05E-11 |
| B20 | B105 | CD3+CD4+CD8+ | anti-U1RNP | -0.26923869 | 5.05E-11 |
| B20 | B106 | CD3+CD4+CD8+ | anti-P | -0.26923868 | 5.05E-11 |
| B20 | B107 | CD3+CD4+CD8+ | AnuA | -0.26923868 | 5.05E-11 |
| B20 | B108 | CD3+CD4+CD8+ | Anti-Cyclin | -0.26923869 | 5.05E-11 |
| B20 | B109 | CD3+CD4+CD8+ | AMA-M2 | -0.26923869 | 5.05E-11 |
| B20 | B110 | CD3+CD4+CD8+ | AHA | -0.26923869 | 5.05E-11 |
| B20 | B95 | CD3+CD4+CD8+ | anti-cenpb | -0.26923869 | 5.05E-11 |
| B20 | B97 | CD3+CD4+CD8+ | dsDNA | -0.26923868 | 5.05E-11 |
| B20 | B98 | CD3+CD4+CD8+ | anti-Jo-1 | -0.26923869 | 5.05E-11 |
| B20 | B75 | CD3+CD4+CD8+ | ApoA1/B | -0.25841831 | 3.06E-10 |
| B20 | B22 | CD3+CD4+CD8+ | CD3+CD8+(T) | -0.25722029 | 3.71E-10 |
| B20 | B59 | CD3+CD4+CD8+ | WBRV | -0.25289791 | 7.43E-10 |
| B20 | B195 | CD3+CD4+CD8+ | Na | -0.2506836 | 1.05E-09 |
| B20 | B34 | CD3+CD4+CD8+ | TK | -0.24410426 | 2.92E-09 |
| B20 | B26 | CD3+CD4+CD8+ | CKMB | -0.24074918 | 4.87E-09 |
| B20 | B74 | CD3+CD4+CD8+ | ApoA1 | -0.23434061 | 1.26E-08 |
| B20 | B76 | CD3+CD4+CD8+ | ApoB | 0.223229251 | 6.15E-08 |

| | | | | | |
|---|---|---|---|---|---|
| B20 | B55 | CD3+CD4+CD8+ | TK | -0.21582733 | 1.69E-07 |
| B20 | B24 | CD3+CD4+CD8+ | αHBDH | -0.21160912 | 2.96E-07 |
| B20 | B56 | CD3+CD4+CD8+ | IR | -0.20320397 | 8.76E-07 |
| B20 | B25 | CD3+CD4+CD8+ | CK | -0.20304093 | 8.94E-07 |
| B204 | B209 | Drinking | Smoking | 0.426604408 | 0 |
| B204 | B221 | Drinking | Hei | 0.229257234 | 2.63E-08 |
| B206 | B207 | Waist | WHR | 0.859220488 | 0 |
| B206 | B210 | Waist | Hip | 0.772854052 | 0 |
| B206 | B215 | Waist | BMI | 0.833088268 | 0 |
| B206 | B216 | Waist | BMI | 0.82407878 | 0 |
| B206 | B221 | Waist | Hei | 0.353370253 | 0 |
| B206 | B222 | Waist | NS | 0.64698412 | 0 |
| B206 | B220 | Waist | SBP | 0.21336123 | 2.35E-07 |
| B207 | B210 | WHR | Hip | 0.348849707 | 0 |
| B207 | B215 | WHR | BMI | 0.597711834 | 0 |
| B207 | B216 | WHR | BMI | 0.55069757 | 0 |
| B207 | B222 | WHR | NS | 0.495892349 | 0 |
| B207 | B220 | WHR | SBP | 0.245745841 | 2.27E-09 |
| B209 | B221 | Smoking | Hei | 0.225333213 | 4.58E-08 |
| B21 | B124 | CD3+CD4-CD8- | ACTH | -0.56643612 | 0 |
| B21 | B125 | CD3+CD4-CD8- | Cor | 0.56643611 | 0 |
| B21 | B155 | CD3+CD4-CD8- | IgA | 0.458627742 | 0 |
| B21 | B182 | CD3+CD4-CD8- | TT | -0.73698786 | 0 |
| B21 | B183 | CD3+CD4-CD8- | PTA | 0.739300257 | 0 |
| B21 | B184 | CD3+CD4-CD8- | INR | 0.360430627 | 0 |
| B21 | B185 | CD3+CD4-CD8- | PT | 0.387057116 | 0 |
| B21 | B186 | CD3+CD4-CD8- | PFC | 0.738012851 | 0 |
| B21 | B22 | CD3+CD4-CD8- | CD3+CD8+(T) | 0.622364293 | 0 |
| B21 | B222 | CD3+CD4-CD8- | NS | -0.3711591 | 0 |
| B21 | B23 | CD3+CD4-CD8- | CD3-CD16+CD56+(NK) | -0.57822826 | 0 |
| B21 | B54 | CD3+CD4-CD8- | ESR | 0.544072057 | 0 |
| B21 | B55 | CD3+CD4-CD8- | TK | -0.34040641 | 0 |
| B21 | B56 | CD3+CD4-CD8- | IR | -0.42680809 | 0 |
| B21 | B62 | CD3+CD4-CD8- | WBHSRVI | -0.35654054 | 0 |
| B21 | B69 | CD3+CD4-CD8- | PV | 0.520122765 | 0 |
| B21 | B74 | CD3+CD4-CD8- | ApoA1 | 0.367844536 | 0 |
| B21 | B92 | CD3+CD4-CD8- | ALDH2-Glu504Glu | -0.56643611 | 0 |
| B21 | B93 | CD3+CD4-CD8- | ALDH2-Glu504Lys | 0.56643611 | 0 |
| B21 | B94 | CD3+CD4-CD8- | ALDH2-Lys504Lys | 0.566436103 | 0 |
| B21 | B99 | CD3+CD4-CD8- | anti-PM-Scl | -0.57228414 | 0 |
| B21 | B158 | CD3+CD4-CD8- | IgM | 0.321049934 | 2.66E-15 |
| B21 | B61 | CD3+CD4-CD8- | WBHSRV | -0.29292944 | 7.34E-13 |
| B21 | B79 | CD3+CD4-CD8- | hb V -dna | 0.285630917 | 2.82E-12 |
| B21 | B75 | CD3+CD4-CD8- | ApoA1/B | 0.281225309 | 6.24E-12 |

| | | | | | |
|---|---|---|---|---|---|
| B21 | B60 | CD3+CD4-CD8- | WBRVI | -0.27399405 | 2.23E-11 |
| B21 | B100 | CD3+CD4-CD8- | anti-Ro52 | -0.24309786 | 3.41E-09 |
| B21 | B101 | CD3+CD4-CD8- | anti-Sc70 | -0.24309786 | 3.41E-09 |
| B21 | B102 | CD3+CD4-CD8- | anti-Sm | -0.24309786 | 3.41E-09 |
| B21 | B103 | CD3+CD4-CD8- | anti-SSA/Ro | -0.24309786 | 3.41E-09 |
| B21 | B104 | CD3+CD4-CD8- | anti-SSB | -0.24309786 | 3.41E-09 |
| B21 | B105 | CD3+CD4-CD8- | anti-U1RNP | -0.24309786 | 3.41E-09 |
| B21 | B106 | CD3+CD4-CD8- | anti-P | -0.24309786 | 3.41E-09 |
| B21 | B107 | CD3+CD4-CD8- | AnuA | -0.24309785 | 3.41E-09 |
| B21 | B108 | CD3+CD4-CD8- | Anti-Cyclin | -0.24309786 | 3.41E-09 |
| B21 | B109 | CD3+CD4-CD8- | AMA-M2 | -0.24309786 | 3.41E-09 |
| B21 | B110 | CD3+CD4-CD8- | AHA | -0.24309786 | 3.41E-09 |
| B21 | B95 | CD3+CD4-CD8- | anti-cenpb | -0.24309786 | 3.41E-09 |
| B21 | B97 | CD3+CD4-CD8- | dsDNA | -0.24309785 | 3.41E-09 |
| B21 | B98 | CD3+CD4-CD8- | anti-Jo-1 | -0.24309786 | 3.41E-09 |
| B21 | B81 | CD3+CD4-CD8- | TSGF | -0.23656498 | 9.08E-09 |
| B21 | B143 | CD3+CD4-CD8- | β-CTx | -0.23436602 | 1.26E-08 |
| B21 | B198 | CD3+CD4-CD8- | P | -0.22461718 | 5.07E-08 |
| B21 | B135 | CD3+CD4-CD8- | TBA | 0.214165414 | 2.11E-07 |
| B210 | B215 | Hip | BMI | 0.791524842 | 0 |
| B210 | B216 | Hip | BMI | 0.825726288 | 0 |
| B210 | B221 | Hip | Hei | 0.414697493 | 0 |
| B210 | B222 | Hip | NS | 0.573116707 | 0 |
| B211 | B212 | IOP-R | IOP-L | 0.850240995 | 0 |
| B215 | B216 | BMI | BMI | 0.82722191 | 0 |
| B215 | B222 | BMI | NS | 0.559510828 | 0 |
| B216 | B221 | BMI | Hei | 0.665239 | 0 |
| B216 | B222 | BMI | NS | 0.699313278 | 0 |
| B218 | B220 | DBP | SBP | 0.6478366 | 0 |
| B22 | B100 | CD3+CD8+(T) | anti-Ro52 | -0.40441919 | 0 |
| B22 | B101 | CD3+CD8+(T) | anti-Sc70 | -0.40441919 | 0 |
| B22 | B102 | CD3+CD8+(T) | anti-Sm | -0.40441919 | 0 |
| B22 | B103 | CD3+CD8+(T) | anti-SSA/Ro | -0.40441919 | 0 |
| B22 | B104 | CD3+CD8+(T) | anti-SSB | -0.40441919 | 0 |
| B22 | B105 | CD3+CD8+(T) | anti-U1RNP | -0.40441919 | 0 |
| B22 | B106 | CD3+CD8+(T) | anti-P | -0.40441919 | 0 |
| B22 | B107 | CD3+CD8+(T) | AnuA | -0.40441918 | 0 |
| B22 | B108 | CD3+CD8+(T) | Anti-Cyclin | -0.40441919 | 0 |
| B22 | B109 | CD3+CD8+(T) | AMA-M2 | -0.40441919 | 0 |
| B22 | B110 | CD3+CD8+(T) | AHA | -0.40441919 | 0 |
| B22 | B124 | CD3+CD8+(T) | ACTH | -0.42712165 | 0 |
| B22 | B125 | CD3+CD8+(T) | Cor | 0.427121644 | 0 |
| B22 | B175 | CD3+CD8+(T) | ALB-U | 0.405092814 | 0 |
| B22 | B178 | CD3+CD8+(T) | Ig-U | 0.740596897 | 0 |

| | | | | | |
|---|---|---|---|---|---|
| B22 | B179 | CD3+CD8+(T) | α1-MG | 0.361502717 | 0 |
| B22 | B180 | CD3+CD8+(T) | Tf | 0.700614733 | 0 |
| B22 | B182 | CD3+CD8+(T) | TT | -0.58631489 | 0 |
| B22 | B183 | CD3+CD8+(T) | PTA | 0.589259569 | 0 |
| B22 | B186 | CD3+CD8+(T) | PFC | 0.585639363 | 0 |
| B22 | B214 | CD3+CD8+(T) | CVA-L | 0.522225531 | 0 |
| B22 | B23 | CD3+CD8+(T) | CD3-CD16+CD56+(NK) | -0.43799411 | 0 |
| B22 | B54 | CD3+CD8+(T) | ESR | 0.446551758 | 0 |
| B22 | B56 | CD3+CD8+(T) | IR | -0.33909859 | 0 |
| B22 | B69 | CD3+CD8+(T) | PV | 0.409025446 | 0 |
| B22 | B74 | CD3+CD8+(T) | ApoA1 | 0.411419791 | 0 |
| B22 | B92 | CD3+CD8+(T) | ALDH2-Glu504Glu | -0.42712164 | 0 |
| B22 | B93 | CD3+CD8+(T) | ALDH2-Glu504Lys | 0.427121644 | 0 |
| B22 | B94 | CD3+CD8+(T) | ALDH2-Lys504Lys | 0.427121638 | 0 |
| B22 | B95 | CD3+CD8+(T) | anti-cenpb | -0.40441919 | 0 |
| B22 | B97 | CD3+CD8+(T) | dsDNA | -0.40441919 | 0 |
| B22 | B98 | CD3+CD8+(T) | anti-Jo-1 | -0.40441919 | 0 |
| B22 | B99 | CD3+CD8+(T) | anti-PM-Scl | -0.43428023 | 0 |
| B22 | B177 | CD3+CD8+(T) | ACR | 0.325491868 | 1.11E-15 |
| B22 | B185 | CD3+CD8+(T) | PT | 0.291779891 | 9.09E-13 |
| B22 | B158 | CD3+CD8+(T) | IgM | 0.290792956 | 1.09E-12 |
| B22 | B62 | CD3+CD8+(T) | WBHSRVI | -0.28827656 | 1.74E-12 |
| B22 | B170 | CD3+CD8+(T) | U-GLU | 0.27789519 | 1.13E-11 |
| B22 | B75 | CD3+CD8+(T) | ApoA1/B | 0.275943246 | 1.59E-11 |
| B22 | B194 | CD3+CD8+(T) | Cl | -0.27247256 | 2.90E-11 |
| B22 | B184 | CD3+CD8+(T) | INR | 0.270218899 | 4.27E-11 |
| B22 | B222 | CD3+CD8+(T) | NS | -0.26792658 | 6.31E-11 |
| B22 | B55 | CD3+CD8+(T) | TK | -0.26421988 | 1.18E-10 |
| B22 | B155 | CD3+CD8+(T) | IgA | 0.245026172 | 2.54E-09 |
| B22 | B61 | CD3+CD8+(T) | WBHSRV | -0.23500065 | 1.14E-08 |
| B22 | B60 | CD3+CD8+(T) | WBRVI | -0.22357283 | 5.86E-08 |
| B22 | B79 | CD3+CD8+(T) | hb V -dna | 0.216802857 | 1.48E-07 |
| B22 | B195 | CD3+CD8+(T) | Na | -0.20867311 | 4.35E-07 |
| B22 | B139 | CD3+CD8+(T) | ADA | 0.204702256 | 7.24E-07 |
| B221 | B222 | Hei | NS | 0.48476734 | 0 |
| B23 | B100 | CD3-CD16+CD56+(NK) | anti-Ro52 | -0.38898959 | 0 |
| B23 | B101 | CD3-CD16+CD56+(NK) | anti-Sc70 | -0.38898959 | 0 |
| B23 | B102 | CD3-CD16+CD56+(NK) | anti-Sm | -0.38898959 | 0 |
| B23 | B103 | CD3-CD16+CD56+(NK) | anti-SSA/Ro | -0.38898959 | 0 |
| B23 | B104 | CD3-CD16+CD56+(NK) | anti-SSB | -0.38898959 | 0 |
| B23 | B105 | CD3-CD16+CD56+(NK) | anti-U1RNP | -0.38898959 | 0 |
| B23 | B106 | CD3-CD16+CD56+(NK) | anti-P | -0.38898959 | 0 |
| B23 | B107 | CD3-CD16+CD56+(NK) | AnuA | -0.38898958 | 0 |
| B23 | B108 | CD3-CD16+CD56+(NK) | Anti-Cyclin | -0.38898959 | 0 |

| | | | | | |
|---|---|---|---|---|---|
| B23 | B109 | CD3-CD16+CD56+(NK) | AMA-M2 | -0.38898959 | 0 |
| B23 | B110 | CD3-CD16+CD56+(NK) | AHA | -0.38898959 | 0 |
| B23 | B124 | CD3-CD16+CD56+(NK) | ACTH | 0.999545676 | 0 |
| B23 | B125 | CD3-CD16+CD56+(NK) | Cor | -0.99954567 | 0 |
| B23 | B151 | CD3-CD16+CD56+(NK) | Anti-CCP | -0.37681798 | 0 |
| B23 | B182 | CD3-CD16+CD56+(NK) | TT | 0.351772154 | 0 |
| B23 | B184 | CD3-CD16+CD56+(NK) | INR | -0.70353199 | 0 |
| B23 | B185 | CD3-CD16+CD56+(NK) | PT | -0.70523844 | 0 |
| B23 | B186 | CD3-CD16+CD56+(NK) | PFC | -0.39954346 | 0 |
| B23 | B222 | CD3-CD16+CD56+(NK) | NS | 0.598768851 | 0 |
| B23 | B69 | CD3-CD16+CD56+(NK) | PV | -0.3519771 | 0 |
| B23 | B74 | CD3-CD16+CD56+(NK) | ApoA1 | -0.3950888 | 0 |
| B23 | B79 | CD3-CD16+CD56+(NK) | hb V -dna | -0.49876789 | 0 |
| B23 | B81 | CD3-CD16+CD56+(NK) | TSGF | 0.420157296 | 0 |
| B23 | B92 | CD3-CD16+CD56+(NK) | ALDH2-Glu504Glu | 0.999545669 | 0 |
| B23 | B93 | CD3-CD16+CD56+(NK) | ALDH2-Glu504Lys | -0.99954567 | 0 |
| B23 | B94 | CD3-CD16+CD56+(NK) | ALDH2-Lys504Lys | -0.99954566 | 0 |
| B23 | B95 | CD3-CD16+CD56+(NK) | anti-cenpb | -0.38898959 | 0 |
| B23 | B97 | CD3-CD16+CD56+(NK) | dsDNA | -0.38898959 | 0 |
| B23 | B98 | CD3-CD16+CD56+(NK) | anti-Jo-1 | -0.38898959 | 0 |
| B23 | B99 | CD3-CD16+CD56+(NK) | anti-PM-Scl | 0.9996962 | 0 |
| B23 | B75 | CD3-CD16+CD56+(NK) | ApoA1/B | -0.32738135 | 8.88E-16 |
| B23 | B183 | CD3-CD16+CD56+(NK) | PTA | -0.3158576 | 8.22E-15 |
| B23 | B156 | CD3-CD16+CD56+(NK) | IgE | -0.29812646 | 2.75E-13 |
| B23 | B155 | CD3-CD16+CD56+(NK) | IgA | -0.29782065 | 2.91E-13 |
| B23 | B34 | CD3-CD16+CD56+(NK) | TK | -0.29250989 | 7.94E-13 |
| B23 | B181 | CD3-CD16+CD56+(NK) | APTT | 0.271965706 | 3.17E-11 |
| B23 | B135 | CD3-CD16+CD56+(NK) | TBA | -0.23991083 | 5.52E-09 |
| B23 | B196 | CD3-CD16+CD56+(NK) | TCO2 | -0.21559493 | 1.75E-07 |
| B23 | B76 | CD3-CD16+CD56+(NK) | ApoB | 0.214256913 | 2.09E-07 |
| B23 | B198 | CD3-CD16+CD56+(NK) | P | 0.203892484 | 8.03E-07 |
| B23 | B25 | CD3-CD16+CD56+(NK) | CK | -0.20220769 | 9.93E-07 |
| B24 | B27 | αHBDH | LDH | 0.458458122 | 0 |
| B24 | B25 | αHBDH | CK | 0.322018208 | 2.22E-15 |
| B24 | B55 | αHBDH | TK | 0.239735348 | 5.67E-09 |
| B24 | B26 | αHBDH | CKMB | 0.235619288 | 1.04E-08 |
| B24 | B138 | αHBDH | PAB | 0.227140758 | 3.55E-08 |
| B24 | B200 | αHBDH | OP | 0.215873872 | 1.68E-07 |
| B25 | B59 | CK | WBRV | 0.263097217 | 1.42E-10 |
| B25 | B124 | CK | ACTH | -0.20438884 | 7.54E-07 |
| B25 | B125 | CK | Cor | 0.204388838 | 7.54E-07 |
| B25 | B92 | CK | ALDH2-Glu504Glu | -0.20438884 | 7.54E-07 |
| B25 | B93 | CK | ALDH2-Glu504Lys | 0.204388838 | 7.54E-07 |
| B25 | B94 | CK | ALDH2-Lys504Lys | 0.204388836 | 7.54E-07 |

| | | | | | |
|---|---|---|---|---|---|
| B25 | B99 | CK | anti-PM-Scl | -0.20383367 | 8.09E-07 |
| B25 | B131 | CK | AST | 0.202595682 | 9.45E-07 |
| B26 | B59 | CKMB | WBRV | 0.317894879 | 5.33E-15 |
| B26 | B60 | CKMB | WBRVI | 0.291272723 | 1.00E-12 |
| B26 | B64 | CKMB | WBRSR(mPas_5) | 0.285864822 | 2.70E-12 |
| B26 | B65 | CKMB | WBRSR(mPas_1) | 0.275550784 | 1.70E-11 |
| B26 | B62 | CKMB | WBHSRVI | 0.274385079 | 2.08E-11 |
| B26 | B56 | CKMB | IR | 0.272549085 | 2.87E-11 |
| B26 | B61 | CKMB | WBHSRV | 0.265540922 | 9.43E-11 |
| B26 | B57 | CKMB | RBCAI | 0.253122368 | 7.17E-10 |
| B26 | B67 | CKMB | WBRSR(mPas_50) | 0.24997216 | 1.18E-09 |
| B26 | B63 | CKMB | WBRSR(mPas_200) | 0.246931097 | 1.89E-09 |
| B26 | B66 | CKMB | WBRSR(mPas_100) | 0.246347908 | 2.07E-09 |
| B26 | B183 | CKMB | PTA | -0.21327234 | 2.38E-07 |
| B27 | B171 | LDH | EC | 0.27279742 | 2.75E-11 |
| B27 | B150 | LDH | ALP | 0.223598845 | 5.84E-08 |
| B27 | B195 | LDH | Na | 0.223211962 | 6.17E-08 |
| B27 | B200 | LDH | OP | 0.208015456 | 4.74E-07 |
| B28 | B29 | PRL | E2 | -1 | 0 |
| B28 | B33 | PRL | Prog | 1 | 0 |
| B29 | B30 | E2 | LH | 1 | 0 |
| B29 | B31 | E2 | FSH | 1 | 0 |
| B29 | B32 | E2 | Testo | -1 | 0 |
| B29 | B33 | E2 | Prog | -1 | 0 |
| B3 | B34 | Hcy | TK | -0.3685843 | 0 |
| B3 | B25 | Hcy | CK | 0.275430547 | 1.74E-11 |
| B3 | B24 | Hcy | αHBDH | 0.245422863 | 2.39E-09 |
| B3 | B187 | Hcy | NSE | 0.223196847 | 6.18E-08 |
| B3 | B26 | Hcy | CKMB | -0.22135475 | 7.97E-08 |
| B30 | B31 | LH | FSH | 1 | 0 |
| B30 | B32 | LH | Testo | -1 | 0 |
| B31 | B32 | FSH | Testo | -1 | 0 |
| B32 | B33 | Testo | Prog | 1 | 0 |
| B34 | B168 | TK | NMP22 | -0.61157281 | 0 |
| B34 | B187 | TK | NSE | -0.46404793 | 0 |
| B34 | B175 | TK | ALB-U | -0.31284161 | 1.51E-14 |
| B34 | B194 | TK | Cl | -0.30956245 | 2.93E-14 |
| B34 | B99 | TK | anti-PM-Scl | -0.29256328 | 7.86E-13 |
| B34 | B124 | TK | ACTH | -0.29249118 | 7.97E-13 |
| B34 | B125 | TK | Cor | 0.292491173 | 7.97E-13 |
| B34 | B92 | TK | ALDH2-Glu504Glu | -0.29249117 | 7.97E-13 |
| B34 | B93 | TK | ALDH2-Glu504Lys | 0.292491173 | 7.97E-13 |
| B34 | B94 | TK | ALDH2-Lys504Lys | 0.29249117 | 7.97E-13 |
| B34 | B222 | TK | NS | -0.25069532 | 1.05E-09 |

| | | | | | |
|---|---|---|---|---|---|
| B34 | B75 | TK | ApoA1/B | 0.246941664 | 1.89E-09 |
| B34 | B196 | TK | TCO2 | 0.234275141 | 1.27E-08 |
| B34 | B76 | TK | ApoB | -0.23240432 | 1.67E-08 |
| B34 | B199 | TK | Mg | 0.231537355 | 1.89E-08 |
| B34 | B177 | TK | ACR | -0.21522983 | 1.83E-07 |
| B34 | B185 | TK | PT | 0.203712926 | 8.21E-07 |
| B34 | B184 | TK | INR | 0.203465657 | 8.47E-07 |
| B36 | B116 | V-Lym% | TT3 | -0.86772183 | 0 |
| B37 | B38 | BAND | BAND% | 1 | 0 |
| B37 | B40 | BAND | WBC | 0.281639001 | 5.80E-12 |
| B37 | B120 | BAND | HCV-cAg | -0.24671753 | 1.96E-09 |
| B38 | B40 | BAND% | WBC | 0.281639001 | 5.80E-12 |
| B38 | B120 | BAND% | HCV-cAg | -0.24671753 | 1.96E-09 |
| B39 | B43 | Other cells | MCV | -0.24445933 | 2.77E-09 |
| B39 | B44 | Other cells | MCH | -0.23359143 | 1.41E-08 |
| B4 | B167 | WB-Ca | PC | -0.2754307 | 1.74E-11 |
| B40 | B42 | WBC | MON | 0.696732425 | 0 |
| B40 | B52 | WBC | PLT/BPC | 0.258148603 | 3.19E-10 |
| B40 | B46 | WBC | RBC Count | 0.236079721 | 9.76E-09 |
| B40 | B53 | WBC | GR% | 0.219781448 | 9.90E-08 |
| B40 | B192 | WBC | UA | 0.21830067 | 1.21E-07 |
| B40 | B206 | WBC | Waist | 0.210519238 | 3.42E-07 |
| B40 | B48 | WBC | LYM% | -0.20685096 | 5.50E-07 |
| B41 | B42 | MON % | MON | 0.587866152 | 0 |
| B41 | B53 | MON % | GR% | -0.23999598 | 5.45E-09 |
| B42 | B206 | MON | Waist | 0.27128583 | 3.56E-11 |
| B42 | B192 | MON | UA | 0.269185335 | 5.10E-11 |
| B42 | B216 | MON | BMI | 0.246710793 | 1.96E-09 |
| B42 | B207 | MON | WHR | 0.228121414 | 3.09E-08 |
| B42 | B210 | MON | Hip | 0.21223217 | 2.73E-07 |
| B42 | B215 | MON | BMI | 0.211968848 | 2.83E-07 |
| B42 | B222 | MON | NS | 0.211844155 | 2.87E-07 |
| B42 | B174 | MON | Urea/Cr | -0.20370888 | 8.22E-07 |
| B43 | B44 | MCV | MCH | 0.920034243 | 0 |
| B43 | B46 | MCV | RBC Count | -0.49321115 | 0 |
| B43 | B47 | MCV | RDW | 0.520019028 | 0 |
| B43 | B52 | MCV | PLT/BPC | -0.28699424 | 2.20E-12 |
| B43 | B45 | MCV | MCHC | 0.279455112 | 8.56E-12 |
| B44 | B45 | MCH | MCHC | 0.631003592 | 0 |
| B44 | B51 | MCH | Hb | 0.350876571 | 0 |
| B44 | B46 | MCH | RBC Count | -0.33174042 | 4.44E-16 |
| B44 | B47 | MCH | RDW | 0.311043455 | 2.18E-14 |
| B44 | B52 | MCH | PLT/BPC | -0.27412894 | 2.18E-11 |
| B44 | B209 | MCH | Smoking | 0.210458423 | 3.45E-07 |

| | | | | | |
|---|---|---|---|---|---|
| B45 | B51 | MCHC | Hb | 0.588066406 | 0 |
| B45 | B47 | MCHC | RDW | -0.28288721 | 4.64E-12 |
| B45 | B216 | MCHC | BMI | 0.265399475 | 9.66E-11 |
| B45 | B221 | MCHC | Hei | 0.264590033 | 1.11E-10 |
| B45 | B71 | MCHC | TG | 0.261938875 | 1.72E-10 |
| B45 | B163 | MCHC | SG | 0.241391791 | 4.42E-09 |
| B45 | B222 | MCHC | NS | 0.209321937 | 4.00E-07 |
| B46 | B189 | RBC Count | eGFB | 0.391448458 | 0 |
| B46 | B190 | RBC Count | Cre | -0.35280409 | 0 |
| B46 | B191 | RBC Count | Urea | -0.47116127 | 0 |
| B46 | B216 | RBC Count | BMI | 0.383190143 | 0 |
| B46 | B221 | RBC Count | Hei | 0.42939358 | 0 |
| B46 | B47 | RBC Count | RDW | -0.51256639 | 0 |
| B46 | B51 | RBC Count | Hb | 0.760141745 | 0 |
| B46 | B129 | RBC Count | IBIL | 0.306084828 | 5.86E-14 |
| B46 | B222 | RBC Count | NS | 0.290559643 | 1.14E-12 |
| B46 | B133 | RBC Count | TBIL | 0.288423696 | 1.69E-12 |
| B46 | B126 | RBC Count | Alb | 0.285396418 | 2.94E-12 |
| B46 | B210 | RBC Count | Hip | 0.278737559 | 9.72E-12 |
| B46 | B72 | RBC Count | HDL-C | -0.26482068 | 1.06E-10 |
| B46 | B128 | RBC Count | ALT | 0.262405821 | 1.59E-10 |
| B46 | B206 | RBC Count | Waist | 0.258332451 | 3.10E-10 |
| B46 | B141 | RBC Count | AST/ALT | -0.24951905 | 1.27E-09 |
| B46 | B163 | RBC Count | SG | 0.242010822 | 4.02E-09 |
| B46 | B140 | RBC Count | r-GT | 0.213069522 | 2.44E-07 |
| B47 | B191 | RDW | Urea | 0.301904911 | 1.33E-13 |
| B47 | B51 | RDW | Hb | -0.28954655 | 1.38E-12 |
| B47 | B190 | RDW | Cre | 0.289462301 | 1.40E-12 |
| B47 | B189 | RDW | eGFB | -0.28520949 | 3.05E-12 |
| B47 | B170 | RDW | U-GLU | 0.267145552 | 7.20E-11 |
| B47 | B118 | RDW | CEA1 | 0.257795435 | 3.38E-10 |
| B47 | B52 | RDW | PLT/BPC | -0.23437405 | 1.25E-08 |
| B47 | B214 | RDW | CVA-L | 0.206588736 | 5.69E-07 |
| B48 | B53 | LYM% | GR% | -0.92322274 | 0 |
| B48 | B189 | LYM% | eGFB | 0.323514843 | 1.78E-15 |
| B48 | B191 | LYM% | Urea | -0.29843564 | 2.59E-13 |
| B48 | B190 | LYM% | Cre | -0.25012339 | 1.15E-09 |
| B49 | B50 | baso% | eo% | 0.27492726 | 1.90E-11 |
| B5 | B116 | WB-Mg | TT3 | -0.92518727 | 0 |
| B5 | B7 | WB-Mg | WB-Cop | -0.49039971 | 0 |
| B5 | B9 | WB-Mg | RH | 0.990867389 | 0 |
| B50 | B53 | eo% | GR% | -0.22897236 | 2.74E-08 |
| B50 | B189 | eo% | eGFB | -0.21470485 | 1.97E-07 |
| B50 | B191 | eo% | Urea | 0.204394788 | 7.53E-07 |

| | | | | | |
|---|---|---|---|---|---|
| B51 | B141 | Hb | AST/ALT | -0.3357761 | 0 |
| B51 | B189 | Hb | eGFB | 0.353179472 | 0 |
| B51 | B191 | Hb | Urea | -0.42291926 | 0 |
| B51 | B210 | Hb | Hip | 0.348640533 | 0 |
| B51 | B216 | Hb | BMI | 0.461791423 | 0 |
| B51 | B221 | Hb | Hei | 0.480219036 | 0 |
| B51 | B222 | Hb | NS | 0.363019592 | 0 |
| B51 | B206 | Hb | Waist | 0.333377848 | 2.22E-16 |
| B51 | B190 | Hb | Cre | -0.32674297 | 8.88E-16 |
| B51 | B163 | Hb | SG | 0.316789546 | 6.66E-15 |
| B51 | B128 | Hb | ALT | 0.29195266 | 8.81E-13 |
| B51 | B126 | Hb | Alb | 0.276074693 | 1.55E-11 |
| B51 | B209 | Hb | Smoking | 0.275162846 | 1.82E-11 |
| B51 | B129 | Hb | IBIL | 0.272377092 | 2.95E-11 |
| B51 | B133 | Hb | TBIL | 0.26679118 | 7.65E-11 |
| B51 | B204 | Hb | Drinking | 0.262740639 | 1.50E-10 |
| B51 | B72 | Hb | HDL-C | -0.25617492 | 4.40E-10 |
| B51 | B215 | Hb | BMI | 0.253247653 | 7.02E-10 |
| B51 | B140 | Hb | r-GT | 0.211889611 | 2.86E-07 |
| B51 | B207 | Hb | WHR | 0.211816579 | 2.88E-07 |
| B51 | B71 | Hb | TG | 0.211457273 | 3.02E-07 |
| B52 | B70 | PLT/BPC | LDL-C | 0.266237083 | 8.39E-11 |
| B53 | B189 | GR% | eGFB | -0.21752314 | 1.35E-07 |
| B54 | B100 | ESR | anti-Ro52 | -0.35777954 | 0 |
| B54 | B101 | ESR | anti-Sc70 | -0.35777955 | 0 |
| B54 | B102 | ESR | anti-Sm | -0.35777954 | 0 |
| B54 | B103 | ESR | anti-SSA/Ro | -0.35777954 | 0 |
| B54 | B104 | ESR | anti-SSB | -0.35777954 | 0 |
| B54 | B105 | ESR | anti-U1RNP | -0.35777954 | 0 |
| B54 | B106 | ESR | anti-P | -0.35777954 | 0 |
| B54 | B107 | ESR | AnuA | -0.35777954 | 0 |
| B54 | B108 | ESR | Anti-Cyclin | -0.35777954 | 0 |
| B54 | B109 | ESR | AMA-M2 | -0.35777954 | 0 |
| B54 | B110 | ESR | AHA | -0.35777954 | 0 |
| B54 | B158 | ESR | IgM | 0.368120618 | 0 |
| B54 | B182 | ESR | TT | -0.79117819 | 0 |
| B54 | B183 | ESR | PTA | 0.806082601 | 0 |
| B54 | B186 | ESR | PFC | 0.769830444 | 0 |
| B54 | B55 | ESR | TK | -0.34927235 | 0 |
| B54 | B56 | ESR | IR | -0.46472201 | 0 |
| B54 | B62 | ESR | WBHSRVI | -0.41634492 | 0 |
| B54 | B69 | ESR | PV | 0.619740956 | 0 |
| B54 | B95 | ESR | anti-cenpb | -0.35777954 | 0 |
| B54 | B97 | ESR | dsDNA | -0.35777954 | 0 |

| | | | | | |
|---|---|---|---|---|---|
| B54 | B98 | ESR | anti-Jo-1 | -0.35777954 | 0 |
| B54 | B60 | ESR | WBRVI | -0.32618767 | 8.88E-16 |
| B54 | B61 | ESR | WBHSRV | -0.31880603 | 4.44E-15 |
| B54 | B74 | ESR | ApoA1 | 0.301790878 | 1.36E-13 |
| B54 | B143 | ESR | β-CTx | -0.29915105 | 2.26E-13 |
| B54 | B151 | ESR | Anti-CCP | -0.246847 | 1.92E-09 |
| B54 | B58 | ESR | HCT | -0.23295808 | 1.54E-08 |
| B54 | B63 | ESR | WBRSR(mPas_200) | -0.22081241 | 8.59E-08 |
| B54 | B66 | ESR | WBRSR(mPas_100) | -0.21787073 | 1.28E-07 |
| B54 | B67 | ESR | WBRSR(mPas_50) | -0.21057443 | 3.39E-07 |
| B55 | B182 | TK | TT | 0.487150171 | 0 |
| B55 | B183 | TK | PTA | -0.51160066 | 0 |
| B55 | B186 | TK | PFC | -0.45926206 | 0 |
| B55 | B56 | TK | IR | 0.714137659 | 0 |
| B55 | B58 | TK | HCT | -0.36073573 | 0 |
| B55 | B61 | TK | WBHSRV | 0.574572044 | 0 |
| B55 | B69 | TK | PV | -0.50307192 | 0 |
| B55 | B151 | TK | Anti-CCP | 0.297930415 | 2.85E-13 |
| B55 | B74 | TK | ApoA1 | -0.27045061 | 4.11E-11 |
| B55 | B158 | TK | IgM | -0.2505835 | 1.07E-09 |
| B55 | B62 | TK | WBHSRVI | 0.246396039 | 2.06E-09 |
| B55 | B57 | TK | RBCAI | -0.24490815 | 2.59E-09 |
| B55 | B59 | TK | WBRV | 0.234964473 | 1.15E-08 |
| B55 | B143 | TK | β-CTx | 0.218014285 | 1.26E-07 |
| B55 | B176 | TK | Ucr | -0.21225045 | 2.72E-07 |
| B56 | B182 | IR | TT | 0.59259083 | 0 |
| B56 | B183 | IR | PTA | -0.61744996 | 0 |
| B56 | B186 | IR | PFC | -0.56398298 | 0 |
| B56 | B58 | IR | HCT | 0.387009813 | 0 |
| B56 | B59 | IR | WBRV | 0.757864902 | 0 |
| B56 | B60 | IR | WBRVI | 0.702598253 | 0 |
| B56 | B61 | IR | WBHSRV | 0.944259308 | 0 |
| B56 | B62 | IR | WBHSRVI | 0.853468086 | 0 |
| B56 | B63 | IR | WBRSR(mPas_200) | 0.726666053 | 0 |
| B56 | B64 | IR | WBRSR(mPas_5) | 0.655446283 | 0 |
| B56 | B65 | IR | WBRSR(mPas_1) | 0.626336903 | 0 |
| B56 | B66 | IR | WBRSR(mPas_100) | 0.72880854 | 0 |
| B56 | B67 | IR | WBRSR(mPas_50) | 0.72301139 | 0 |
| B56 | B69 | IR | PV | -0.34793021 | 0 |
| B56 | B57 | IR | RBCAI | 0.320873383 | 2.89E-15 |
| B56 | B158 | IR | IgM | -0.30316947 | 1.04E-13 |
| B56 | B151 | IR | Anti-CCP | 0.281169474 | 6.31E-12 |
| B56 | B198 | IR | P | -0.26617891 | 8.48E-11 |
| B56 | B74 | IR | ApoA1 | -0.25483061 | 5.46E-10 |

| | | | | | |
|---|---|---|---|---|---|
| B56 | B143 | IR | β-CTx | 0.25328871 | 6.98E-10 |
| B56 | B100 | IR | anti-Ro52 | 0.233241509 | 1.48E-08 |
| B56 | B101 | IR | anti-Sc70 | 0.233241511 | 1.48E-08 |
| B56 | B102 | IR | anti-Sm | 0.23324151 | 1.48E-08 |
| B56 | B103 | IR | anti-SSA/Ro | 0.233241509 | 1.48E-08 |
| B56 | B104 | IR | anti-SSB | 0.233241509 | 1.48E-08 |
| B56 | B105 | IR | anti-U1RNP | 0.233241509 | 1.48E-08 |
| B56 | B106 | IR | anti-P | 0.233241509 | 1.48E-08 |
| B56 | B107 | IR | AnuA | 0.233241502 | 1.48E-08 |
| B56 | B108 | IR | Anti-Cyclin | 0.233241509 | 1.48E-08 |
| B56 | B109 | IR | AMA-M2 | 0.233241509 | 1.48E-08 |
| B56 | B110 | IR | AHA | 0.233241509 | 1.48E-08 |
| B56 | B95 | IR | anti-cenpb | 0.233241509 | 1.48E-08 |
| B56 | B97 | IR | dsDNA | 0.233241508 | 1.48E-08 |
| B56 | B98 | IR | anti-Jo-1 | 0.233241509 | 1.48E-08 |
| B56 | B199 | IR | Mg | 0.210690696 | 3.34E-07 |
| B57 | B58 | RBCAI | HCT | 0.776911499 | 0 |
| B57 | B59 | RBCAI | WBRV | 0.78698974 | 0 |
| B57 | B60 | RBCAI | WBRVI | 0.851811229 | 0 |
| B57 | B62 | RBCAI | WBHSRVI | 0.61872553 | 0 |
| B57 | B63 | RBCAI | WBRSR(mPas_200) | 0.581037781 | 0 |
| B57 | B64 | RBCAI | WBRSR(mPas_5) | 0.770537663 | 0 |
| B57 | B65 | RBCAI | WBRSR(mPas_1) | 0.817218372 | 0 |
| B57 | B66 | RBCAI | WBRSR(mPas_100) | 0.593696259 | 0 |
| B57 | B67 | RBCAI | WBRSR(mPas_50) | 0.62169477 | 0 |
| B57 | B61 | RBCAI | WBHSRV | 0.318120976 | 5.33E-15 |
| B58 | B59 | HCT | WBRV | 0.693900477 | 0 |
| B58 | B60 | HCT | WBRVI | 0.880199687 | 0 |
| B58 | B61 | HCT | WBHSRV | 0.477872163 | 0 |
| B58 | B62 | HCT | WBHSRVI | 0.803910992 | 0 |
| B58 | B63 | HCT | WBRSR(mPas_200) | 0.83180133 | 0 |
| B58 | B64 | HCT | WBRSR(mPas_5) | 0.850019694 | 0 |
| B58 | B65 | HCT | WBRSR(mPas_1) | 0.895921359 | 0 |
| B58 | B66 | HCT | WBRSR(mPas_100) | 0.836337631 | 0 |
| B58 | B67 | HCT | WBRSR(mPas_50) | 0.84730172 | 0 |
| B59 | B60 | WBRV | WBRVI | 0.923377477 | 0 |
| B59 | B61 | WBRV | WBHSRV | 0.812608034 | 0 |
| B59 | B62 | WBRV | WBHSRVI | 0.867018297 | 0 |
| B59 | B63 | WBRV | WBRSR(mPas_200) | 0.862669752 | 0 |
| B59 | B64 | WBRV | WBRSR(mPas_5) | 0.925952013 | 0 |
| B59 | B65 | WBRV | WBRSR(mPas_1) | 0.927849972 | 0 |
| B59 | B66 | WBRV | WBRSR(mPas_100) | 0.866392596 | 0 |
| B59 | B67 | WBRV | WBRSR(mPas_50) | 0.878546793 | 0 |
| B59 | B183 | WBRV | PTA | -0.32583742 | 8.88E-16 |

| | | | | | |
|---|---|---|---|---|---|
| B59 | B182 | WBRV | TT | 0.304350344 | 8.24E-14 |
| B59 | B186 | WBRV | PFC | -0.27881009 | 9.59E-12 |
| B59 | B151 | WBRV | Anti-CCP | 0.221080688 | 8.28E-08 |
| B59 | B199 | WBRV | Mg | 0.21523034 | 1.83E-07 |
| B59 | B100 | WBRV | anti-Ro52 | 0.211045998 | 3.19E-07 |
| B59 | B101 | WBRV | anti-Sc70 | 0.211046 | 3.19E-07 |
| B59 | B102 | WBRV | anti-Sm | 0.211045999 | 3.19E-07 |
| B59 | B103 | WBRV | anti-SSA/Ro | 0.211045998 | 3.19E-07 |
| B59 | B104 | WBRV | anti-SSB | 0.211045998 | 3.19E-07 |
| B59 | B105 | WBRV | anti-U1RNP | 0.211045998 | 3.19E-07 |
| B59 | B106 | WBRV | anti-P | 0.211045998 | 3.19E-07 |
| B59 | B107 | WBRV | AnuA | 0.211045995 | 3.19E-07 |
| B59 | B108 | WBRV | Anti-Cyclin | 0.211045998 | 3.19E-07 |
| B59 | B109 | WBRV | AMA-M2 | 0.211045998 | 3.19E-07 |
| B59 | B110 | WBRV | AHA | 0.211045998 | 3.19E-07 |
| B59 | B95 | WBRV | anti-cenpb | 0.211045998 | 3.19E-07 |
| B59 | B97 | WBRV | dsDNA | 0.211045998 | 3.19E-07 |
| B59 | B98 | WBRV | anti-Jo-1 | 0.211045999 | 3.19E-07 |
| B6 | B167 | WB-Fe | PC | 0.477921261 | 0 |
| B6 | B8 | WB-Fe | WB-Zinc | 0.795810417 | 0 |
| B60 | B182 | WBRVI | TT | 0.383477833 | 0 |
| B60 | B183 | WBRVI | PTA | -0.39718131 | 0 |
| B60 | B186 | WBRVI | PFC | -0.3671487 | 0 |
| B60 | B61 | WBRVI | WBHSRV | 0.722472145 | 0 |
| B60 | B62 | WBRVI | WBHSRVI | 0.937353682 | 0 |
| B60 | B63 | WBRVI | WBRSR(mPas_200) | 0.885705212 | 0 |
| B60 | B64 | WBRVI | WBRSR(mPas_5) | 0.949092527 | 0 |
| B60 | B65 | WBRVI | WBRSR(mPas_1) | 0.968480531 | 0 |
| B60 | B66 | WBRVI | WBRSR(mPas_100) | 0.895167163 | 0 |
| B60 | B67 | WBRVI | WBRSR(mPas_50) | 0.908560847 | 0 |
| B61 | B182 | WBHSRV | TT | 0.452614787 | 0 |
| B61 | B183 | WBHSRV | PTA | -0.47987026 | 0 |
| B61 | B186 | WBHSRV | PFC | -0.42109411 | 0 |
| B61 | B62 | WBHSRV | WBHSRVI | 0.879129407 | 0 |
| B61 | B63 | WBHSRV | WBRSR(mPas_200) | 0.86155592 | 0 |
| B61 | B64 | WBHSRV | WBRSR(mPas_5) | 0.758212846 | 0 |
| B61 | B65 | WBHSRV | WBRSR(mPas_1) | 0.726168206 | 0 |
| B61 | B66 | WBHSRV | WBRSR(mPas_100) | 0.858340074 | 0 |
| B61 | B67 | WBHSRV | WBRSR(mPas_50) | 0.84908748 | 0 |
| B61 | B151 | WBHSRV | Anti-CCP | 0.284724488 | 3.33E-12 |
| B61 | B100 | WBHSRV | anti-Ro52 | 0.245220545 | 2.46E-09 |
| B61 | B101 | WBHSRV | anti-Sc70 | 0.245220547 | 2.46E-09 |
| B61 | B102 | WBHSRV | anti-Sm | 0.245220546 | 2.46E-09 |
| B61 | B103 | WBHSRV | anti-SSA/Ro | 0.245220545 | 2.46E-09 |

| | | | | | |
|---|---|---|---|---|---|
| B61 | B104 | WBHSRV | anti-SSB | 0.245220544 | 2.46E-09 |
| B61 | B105 | WBHSRV | anti-U1RNP | 0.245220544 | 2.46E-09 |
| B61 | B106 | WBHSRV | anti-P | 0.245220544 | 2.46E-09 |
| B61 | B107 | WBHSRV | AnuA | 0.24522054 | 2.46E-09 |
| B61 | B108 | WBHSRV | Anti-Cyclin | 0.245220545 | 2.46E-09 |
| B61 | B109 | WBHSRV | AMA-M2 | 0.245220544 | 2.46E-09 |
| B61 | B110 | WBHSRV | AHA | 0.245220545 | 2.46E-09 |
| B61 | B95 | WBHSRV | anti-cenpb | 0.245220544 | 2.46E-09 |
| B61 | B97 | WBHSRV | dsDNA | 0.245220543 | 2.46E-09 |
| B61 | B98 | WBHSRV | anti-Jo-1 | 0.245220545 | 2.46E-09 |
| B61 | B199 | WBHSRV | Mg | 0.243969786 | 2.98E-09 |
| B61 | B158 | WBHSRV | IgM | -0.23473425 | 1.19E-08 |
| B61 | B198 | WBHSRV | P | -0.22783765 | 3.22E-08 |
| B61 | B143 | WBHSRV | β-CTx | 0.215040939 | 1.88E-07 |
| B61 | B136 | WBHSRV | 5'NT | 0.209920227 | 3.70E-07 |
| B62 | B182 | WBHSRVI | TT | 0.482965524 | 0 |
| B62 | B183 | WBHSRVI | PTA | -0.50059357 | 0 |
| B62 | B186 | WBHSRVI | PFC | -0.46259818 | 0 |
| B62 | B63 | WBHSRVI | WBRSR(mPas_200) | 0.941711911 | 0 |
| B62 | B64 | WBHSRVI | WBRSR(mPas_5) | 0.908791364 | 0 |
| B62 | B65 | WBHSRVI | WBRSR(mPas_1) | 0.907202534 | 0 |
| B62 | B66 | WBHSRVI | WBRSR(mPas_100) | 0.947568545 | 0 |
| B62 | B67 | WBHSRVI | WBRSR(mPas_50) | 0.948994576 | 0 |
| B62 | B158 | WBHSRVI | IgM | -0.24594181 | 2.21E-09 |
| B62 | B100 | WBHSRVI | anti-Ro52 | 0.21353833 | 2.30E-07 |
| B62 | B101 | WBHSRVI | anti-Sc70 | 0.213538332 | 2.30E-07 |
| B62 | B102 | WBHSRVI | anti-Sm | 0.213538331 | 2.30E-07 |
| B62 | B103 | WBHSRVI | anti-SSA/Ro | 0.21353833 | 2.30E-07 |
| B62 | B104 | WBHSRVI | anti-SSB | 0.21353833 | 2.30E-07 |
| B62 | B105 | WBHSRVI | anti-U1RNP | 0.21353833 | 2.30E-07 |
| B62 | B106 | WBHSRVI | anti-P | 0.21353833 | 2.30E-07 |
| B62 | B107 | WBHSRVI | AnuA | 0.213538325 | 2.30E-07 |
| B62 | B108 | WBHSRVI | Anti-Cyclin | 0.21353833 | 2.30E-07 |
| B62 | B109 | WBHSRVI | AMA-M2 | 0.21353833 | 2.30E-07 |
| B62 | B110 | WBHSRVI | AHA | 0.21353833 | 2.30E-07 |
| B62 | B95 | WBHSRVI | anti-cenpb | 0.21353833 | 2.30E-07 |
| B62 | B97 | WBHSRVI | dsDNA | 0.213538329 | 2.30E-07 |
| B62 | B98 | WBHSRVI | anti-Jo-1 | 0.213538331 | 2.30E-07 |
| B62 | B199 | WBHSRVI | Mg | 0.210385671 | 3.48E-07 |
| B62 | B198 | WBHSRVI | P | -0.20959437 | 3.86E-07 |
| B63 | B64 | WBRSR(mPas_200) | WBRSR(mPas_5) | 0.939822626 | 0 |
| B63 | B65 | WBRSR(mPas_200) | WBRSR(mPas_1) | 0.937762619 | 0 |
| B63 | B66 | WBRSR(mPas_200) | WBRSR(mPas_100) | 0.998580515 | 0 |
| B63 | B67 | WBRSR(mPas_200) | WBRSR(mPas_50) | 0.997281508 | 0 |

| | | | | | |
|---|---|---|---|---|---|
| B63 | B183 | WBRSR(mPas_200) | PTA | -0.30475621 | 7.62E-14 |
| B63 | B182 | WBRSR(mPas_200) | TT | 0.285137132 | 3.09E-12 |
| B63 | B186 | WBRSR(mPas_200) | PFC | -0.26261349 | 1.54E-10 |
| B63 | B199 | WBRSR(mPas_200) | Mg | 0.23328489 | 1.47E-08 |
| B63 | B69 | WBRSR(mPas_200) | PV | 0.217443282 | 1.36E-07 |
| B63 | B100 | WBRSR(mPas_200) | anti-Ro52 | 0.215055274 | 1.88E-07 |
| B63 | B101 | WBRSR(mPas_200) | anti-Sc70 | 0.215055276 | 1.88E-07 |
| B63 | B102 | WBRSR(mPas_200) | anti-Sm | 0.215055275 | 1.88E-07 |
| B63 | B103 | WBRSR(mPas_200) | anti-SSA/Ro | 0.215055274 | 1.88E-07 |
| B63 | B104 | WBRSR(mPas_200) | anti-SSB | 0.215055274 | 1.88E-07 |
| B63 | B105 | WBRSR(mPas_200) | anti-U1RNP | 0.215055274 | 1.88E-07 |
| B63 | B106 | WBRSR(mPas_200) | anti-P | 0.215055274 | 1.88E-07 |
| B63 | B107 | WBRSR(mPas_200) | AnuA | 0.215055268 | 1.88E-07 |
| B63 | B108 | WBRSR(mPas_200) | Anti-Cyclin | 0.215055274 | 1.88E-07 |
| B63 | B109 | WBRSR(mPas_200) | AMA-M2 | 0.215055274 | 1.88E-07 |
| B63 | B110 | WBRSR(mPas_200) | AHA | 0.215055274 | 1.88E-07 |
| B63 | B95 | WBRSR(mPas_200) | anti-cenpb | 0.215055274 | 1.88E-07 |
| B63 | B97 | WBRSR(mPas_200) | dsDNA | 0.215055273 | 1.88E-07 |
| B63 | B98 | WBRSR(mPas_200) | anti-Jo-1 | 0.215055274 | 1.88E-07 |
| B63 | B197 | WBRSR(mPas_200) | AG | 0.210459004 | 3.45E-07 |
| B64 | B65 | WBRSR(mPas_5) | WBRSR(mPas_1) | 0.986180234 | 0 |
| B64 | B66 | WBRSR(mPas_5) | WBRSR(mPas_100) | 0.946195383 | 0 |
| B64 | B67 | WBRSR(mPas_5) | WBRSR(mPas_50) | 0.955187913 | 0 |
| B64 | B183 | WBRSR(mPas_5) | PTA | -0.32242606 | 2.22E-15 |
| B64 | B182 | WBRSR(mPas_5) | TT | 0.30477369 | 7.59E-14 |
| B64 | B186 | WBRSR(mPas_5) | PFC | -0.28391203 | 3.85E-12 |
| B64 | B100 | WBRSR(mPas_5) | anti-Ro52 | 0.217682444 | 1.32E-07 |
| B64 | B101 | WBRSR(mPas_5) | anti-Sc70 | 0.217682446 | 1.32E-07 |
| B64 | B102 | WBRSR(mPas_5) | anti-Sm | 0.217682444 | 1.32E-07 |
| B64 | B103 | WBRSR(mPas_5) | anti-SSA/Ro | 0.217682444 | 1.32E-07 |
| B64 | B104 | WBRSR(mPas_5) | anti-SSB | 0.217682443 | 1.32E-07 |
| B64 | B105 | WBRSR(mPas_5) | anti-U1RNP | 0.217682444 | 1.32E-07 |
| B64 | B106 | WBRSR(mPas_5) | anti-P | 0.217682443 | 1.32E-07 |
| B64 | B107 | WBRSR(mPas_5) | AnuA | 0.217682439 | 1.32E-07 |
| B64 | B108 | WBRSR(mPas_5) | Anti-Cyclin | 0.217682444 | 1.32E-07 |
| B64 | B109 | WBRSR(mPas_5) | AMA-M2 | 0.217682444 | 1.32E-07 |
| B64 | B110 | WBRSR(mPas_5) | AHA | 0.217682444 | 1.32E-07 |
| B64 | B95 | WBRSR(mPas_5) | anti-cenpb | 0.217682444 | 1.32E-07 |
| B64 | B97 | WBRSR(mPas_5) | dsDNA | 0.217682443 | 1.32E-07 |
| B64 | B98 | WBRSR(mPas_5) | anti-Jo-1 | 0.217682444 | 1.32E-07 |
| B64 | B199 | WBRSR(mPas_5) | Mg | 0.203869795 | 8.05E-07 |
| B65 | B66 | WBRSR(mPas_1) | WBRSR(mPas_100) | 0.943457258 | 0 |
| B65 | B67 | WBRSR(mPas_1) | WBRSR(mPas_50) | 0.954484131 | 0 |
| B65 | B183 | WBRSR(mPas_1) | PTA | -0.28356081 | 4.11E-12 |

| | | | | | |
|---|---|---|---|---|---|
| B65 | B182 | WBRSR(mPas_1) | TT | 0.268600127 | 5.63E-11 |
| B65 | B186 | WBRSR(mPas_1) | PFC | -0.25072599 | 1.05E-09 |
| B66 | B67 | WBRSR(mPas_100) | WBRSR(mPas_50) | 0.999257543 | 0 |
| B66 | B183 | WBRSR(mPas_100) | PTA | -0.3058221 | 6.17E-14 |
| B66 | B182 | WBRSR(mPas_100) | TT | 0.288188238 | 1.77E-12 |
| B66 | B186 | WBRSR(mPas_100) | PFC | -0.2676089 | 6.66E-11 |
| B66 | B199 | WBRSR(mPas_100) | Mg | 0.220083691 | 9.50E-08 |
| B66 | B69 | WBRSR(mPas_100) | PV | 0.203804006 | 8.12E-07 |
| B66 | B197 | WBRSR(mPas_100) | AG | 0.203238147 | 8.72E-07 |
| B67 | B183 | WBRSR(mPas_50) | PTA | -0.30223605 | 1.24E-13 |
| B67 | B182 | WBRSR(mPas_50) | TT | 0.285112482 | 3.10E-12 |
| B67 | B186 | WBRSR(mPas_50) | PFC | -0.26504143 | 1.03E-10 |
| B67 | B199 | WBRSR(mPas_50) | Mg | 0.21758675 | 1.33E-07 |
| B67 | B197 | WBRSR(mPas_50) | AG | 0.203696023 | 8.23E-07 |
| B69 | B124 | PV | ACTH | -0.34296199 | 0 |
| B69 | B125 | PV | Cor | 0.342961985 | 0 |
| B69 | B182 | PV | TT | -0.5891094 | 0 |
| B69 | B183 | PV | PTA | 0.591356415 | 0 |
| B69 | B186 | PV | PFC | 0.587227626 | 0 |
| B69 | B74 | PV | ApoA1 | 0.492944696 | 0 |
| B69 | B92 | PV | ALDH2-Glu504Glu | -0.34296199 | 0 |
| B69 | B93 | PV | ALDH2-Glu504Lys | 0.342961985 | 0 |
| B69 | B94 | PV | ALDH2-Lys504Lys | 0.34296198 | 0 |
| B69 | B99 | PV | anti-PM-Scl | -0.34541515 | 0 |
| B69 | B158 | PV | IgM | 0.291501681 | 9.58E-13 |
| B69 | B138 | PV | PAB | 0.280031596 | 7.73E-12 |
| B69 | B185 | PV | PT | 0.278472137 | 1.02E-11 |
| B69 | B184 | PV | INR | 0.256721627 | 4.03E-10 |
| B69 | B176 | PV | Ucr | 0.223778407 | 5.70E-08 |
| B7 | B116 | WB-Cop | TT3 | -0.52991409 | 0 |
| B7 | B202 | WB-Cop | e-times | 0.361157559 | 0 |
| B7 | B9 | WB-Cop | RH | 0.571832802 | 0 |
| B70 | B77 | LDL-C | TC | 0.819840031 | 0 |
| B70 | B132 | LDL-C | Dbil | -0.27984858 | 7.98E-12 |
| B71 | B72 | TG | HDL-C | -0.38508766 | 0 |
| B71 | B140 | TG | r-GT | 0.320220727 | 3.33E-15 |
| B71 | B77 | TG | TC | 0.299871532 | 1.97E-13 |
| B71 | B130 | TG | GLB | 0.257649531 | 3.46E-10 |
| B71 | B91 | TG | TP | 0.245286358 | 2.44E-09 |
| B72 | B206 | HDL-C | Waist | -0.42956682 | 0 |
| B72 | B207 | HDL-C | WHR | -0.37349804 | 0 |
| B72 | B215 | HDL-C | BMI | -0.38945447 | 0 |
| B72 | B216 | HDL-C | BMI | -0.42539111 | 0 |
| B72 | B210 | HDL-C | Hip | -0.3296776 | 4.44E-16 |

| | | | | | |
|---|---|---|---|---|---|
| B72 | B222 | HDL-C | NS | -0.32517469 | 1.33E-15 |
| B72 | B192 | HDL-C | UA | -0.28821983 | 1.76E-12 |
| B72 | B141 | HDL-C | AST/ALT | 0.259922741 | 2.39E-10 |
| B72 | B221 | HDL-C | Hei | -0.23962559 | 5.76E-09 |
| B72 | B153 | HDL-C | Glu | -0.2064031 | 5.83E-07 |
| B73 | B119 | ApoA | Hcv-rna | -0.30853729 | 3.60E-14 |
| B73 | B121 | ApoA | C3 | 0.308537355 | 3.60E-14 |
| B73 | B122 | ApoA | C4 | 0.308537349 | 3.60E-14 |
| B73 | B152 | ApoA | ACA-Ig(G-A-M) | 0.308547929 | 3.60E-14 |
| B73 | B159 | ApoA | β2-MG | 0.308538208 | 3.60E-14 |
| B73 | B169 | ApoA | U-CRY | 0.308535268 | 3.60E-14 |
| B73 | B139 | ApoA | ADA | 0.277854849 | 1.14E-11 |
| B73 | B198 | ApoA | P | 0.221554865 | 7.76E-08 |
| B74 | B124 | ApoA1 | ACTH | -0.39026158 | 0 |
| B74 | B125 | ApoA1 | Cor | 0.39026158 | 0 |
| B74 | B182 | ApoA1 | TT | -0.35606129 | 0 |
| B74 | B183 | ApoA1 | PTA | 0.350170401 | 0 |
| B74 | B186 | ApoA1 | PFC | 0.362779097 | 0 |
| B74 | B75 | ApoA1 | ApoA1/B | 0.334445524 | 0 |
| B74 | B84 | ApoA1 | C-PSA | -0.39253316 | 0 |
| B74 | B89 | ApoA1 | CA 72-4 | -0.42396847 | 0 |
| B74 | B92 | ApoA1 | ALDH2-Glu504Glu | -0.39026158 | 0 |
| B74 | B93 | ApoA1 | ALDH2-Glu504Lys | 0.39026158 | 0 |
| B74 | B94 | ApoA1 | ALDH2-Lys504Lys | 0.390261575 | 0 |
| B74 | B99 | ApoA1 | anti-PM-Scl | -0.392132 | 0 |
| B74 | B194 | ApoA1 | Cl | -0.32511755 | 1.33E-15 |
| B74 | B196 | ApoA1 | TCO2 | 0.301954345 | 1.31E-13 |
| B74 | B185 | ApoA1 | PT | 0.270748746 | 3.90E-11 |
| B74 | B222 | ApoA1 | NS | -0.26749385 | 6.79E-11 |
| B74 | B184 | ApoA1 | INR | 0.25753661 | 3.53E-10 |
| B74 | B137 | ApoA1 | AFU | -0.2494217 | 1.28E-09 |
| B74 | B200 | ApoA1 | OP | -0.20675597 | 5.57E-07 |
| B75 | B76 | ApoA1/B | ApoB | -0.81851747 | 0 |
| B75 | B99 | ApoA1/B | anti-PM-Scl | -0.32721789 | 8.88E-16 |
| B75 | B124 | ApoA1/B | ACTH | -0.32530829 | 1.11E-15 |
| B75 | B125 | ApoA1/B | Cor | 0.325308291 | 1.11E-15 |
| B75 | B92 | ApoA1/B | ALDH2-Glu504Glu | -0.32530829 | 1.11E-15 |
| B75 | B93 | ApoA1/B | ALDH2-Glu504Lys | 0.325308291 | 1.11E-15 |
| B75 | B94 | ApoA1/B | ALDH2-Lys504Lys | 0.325308287 | 1.11E-15 |
| B75 | B196 | ApoA1/B | TCO2 | 0.31401784 | 1.20E-14 |
| B75 | B138 | ApoA1/B | PAB | -0.31094313 | 2.22E-14 |
| B75 | B137 | ApoA1/B | AFU | -0.27089387 | 3.81E-11 |
| B75 | B222 | ApoA1/B | NS | -0.26223916 | 1.63E-10 |
| B75 | B168 | ApoA1/B | NMP22 | -0.2482871 | 1.53E-09 |

| | | | | | |
|---|---|---|---|---|---|
| B75 | B194 | ApoA1/B | CI | -0.24043974 | 5.10E-09 |
| B75 | B185 | ApoA1/B | PT | 0.23115874 | 2.00E-08 |
| B75 | B89 | ApoA1/B | CA 72-4 | -0.22233223 | 6.97E-08 |
| B75 | B184 | ApoA1/B | INR | 0.221594822 | 7.71E-08 |
| B75 | B200 | ApoA1/B | OP | -0.22100286 | 8.37E-08 |
| B75 | B186 | ApoA1/B | PFC | 0.219501953 | 1.03E-07 |
| B75 | B182 | ApoA1/B | TT | -0.21121311 | 3.12E-07 |
| B75 | B135 | ApoA1/B | TBA | 0.204270315 | 7.65E-07 |
| B76 | B138 | ApoB | PAB | 0.349324961 | 0 |
| B76 | B198 | ApoB | P | 0.293203788 | 6.97E-13 |
| B76 | B168 | ApoB | NMP22 | 0.272071541 | 3.11E-11 |
| B76 | B77 | ApoB | TC | 0.215795464 | 1.70E-07 |
| B76 | B99 | ApoB | anti-PM-Scl | 0.215743281 | 1.71E-07 |
| B76 | B124 | ApoB | ACTH | 0.213961145 | 2.17E-07 |
| B76 | B125 | ApoB | Cor | -0.21396114 | 2.17E-07 |
| B76 | B92 | ApoB | ALDH2-Glu504Glu | 0.213961144 | 2.17E-07 |
| B76 | B93 | ApoB | ALDH2-Glu504Lys | -0.21396114 | 2.17E-07 |
| B76 | B94 | ApoB | ALDH2-Lys504Lys | -0.21396114 | 2.17E-07 |
| B77 | B132 | TC | Dbil | -0.30290975 | 1.09E-13 |
| B79 | B124 | hb V -dna | ACTH | -0.49888327 | 0 |
| B79 | B125 | hb V -dna | Cor | 0.498883269 | 0 |
| B79 | B179 | hb V -dna | α1-MG | -0.39051216 | 0 |
| B79 | B184 | hb V -dna | INR | 0.349710655 | 0 |
| B79 | B185 | hb V -dna | PT | 0.350150262 | 0 |
| B79 | B92 | hb V -dna | ALDH2-Glu504Glu | -0.49888327 | 0 |
| B79 | B93 | hb V -dna | ALDH2-Glu504Lys | 0.498883269 | 0 |
| B79 | B94 | hb V -dna | ALDH2-Lys504Lys | 0.498883262 | 0 |
| B79 | B99 | hb V -dna | anti-PM-Scl | -0.49891786 | 0 |
| B79 | B222 | hb V -dna | NS | -0.28310868 | 4.45E-12 |
| B79 | B154 | hb V -dna | RF | -0.25720213 | 3.72E-10 |
| B79 | B138 | hb V -dna | PAB | 0.253355248 | 6.91E-10 |
| B79 | B81 | hb V -dna | TSGF | -0.21176151 | 2.90E-07 |
| B8 | B117 | WB-Zinc | TT4 | 0.463586325 | 0 |
| B80 | B88 | anti-HP IgG | CA 242 | 0.239949172 | 5.49E-09 |
| B81 | B124 | TSGF | ACTH | 0.420405592 | 0 |
| B81 | B125 | TSGF | Cor | -0.42040559 | 0 |
| B81 | B92 | TSGF | ALDH2-Glu504Glu | 0.420405589 | 0 |
| B81 | B93 | TSGF | ALDH2-Glu504Lys | -0.42040559 | 0 |
| B81 | B94 | TSGF | ALDH2-Lys504Lys | -0.42040558 | 0 |
| B81 | B99 | TSGF | anti-PM-Scl | 0.42034409 | 0 |
| B81 | B185 | TSGF | PT | -0.29782812 | 2.91E-13 |
| B81 | B184 | TSGF | INR | -0.29743936 | 3.13E-13 |
| B81 | B222 | TSGF | NS | 0.237970755 | 7.37E-09 |
| B81 | B136 | TSGF | 5'NT | -0.22769712 | 3.28E-08 |

| | | | | | |
|---|---|---|---|---|---|
| B82 | B85 | CEA | SCC | 0.459651936 | 0 |
| B82 | B89 | CEA | CA 72-4 | 0.321021387 | 2.89E-15 |
| B82 | B87 | CEA | NSE | 0.314272057 | 1.15E-14 |
| B82 | B137 | CEA | AFU | 0.29890651 | 2.37E-13 |
| B82 | B195 | CEA | Na | -0.28629432 | 2.50E-12 |
| B82 | B200 | CEA | OP | -0.23327554 | 1.47E-08 |
| B82 | B198 | CEA | P | 0.212704732 | 2.56E-07 |
| B82 | B228 | CEA | OS | -0.20642959 | 5.81E-07 |
| B82 | B193 | CEA | K | 0.205160771 | 6.83E-07 |
| B83 | B86 | AFP | TBhCG | 0.450014916 | 0 |
| B83 | B85 | AFP | SCC | 0.227197828 | 3.52E-08 |
| B83 | B84 | AFP | C-PSA | 0.212653522 | 2.58E-07 |
| B83 | B197 | AFP | AG | 0.209137343 | 4.09E-07 |
| B84 | B135 | C-PSA | TBA | 0.387659969 | 0 |
| B84 | B89 | C-PSA | CA 72-4 | 0.740224407 | 0 |
| B84 | B193 | C-PSA | K | -0.29379477 | 6.24E-13 |
| B84 | B137 | C-PSA | AFU | 0.271075249 | 3.69E-11 |
| B84 | B204 | C-PSA | Drinking | 0.249547862 | 1.26E-09 |
| B84 | B85 | C-PSA | SCC | 0.231823274 | 1.82E-08 |
| B84 | B198 | C-PSA | P | -0.21965615 | 1.01E-07 |
| B85 | B195 | SCC | Na | -0.25292917 | 7.39E-10 |
| B85 | B155 | SCC | IgA | 0.241751522 | 4.18E-09 |
| B86 | B88 | TBhCG | CA 242 | 0.393453325 | 0 |
| B86 | B137 | TBhCG | AFU | -0.25282543 | 7.51E-10 |
| B86 | B89 | TBhCG | CA 72-4 | -0.21379696 | 2.22E-07 |
| B86 | B90 | TBhCG | CYFRA 21-1 | 0.209173999 | 4.08E-07 |
| B86 | B228 | TBhCG | OS | -0.20755757 | 5.02E-07 |
| B87 | B228 | NSE | OS | -0.24528477 | 2.44E-09 |
| B88 | B138 | CA 242 | PAB | -0.33836412 | 0 |
| B88 | B197 | CA 242 | AG | -0.35036834 | 0 |
| B88 | B196 | CA 242 | TCO2 | 0.295791543 | 4.28E-13 |
| B88 | B200 | CA 242 | OP | -0.25322983 | 7.04E-10 |
| B88 | B199 | CA 242 | Mg | -0.23134013 | 1.95E-08 |
| B89 | B135 | CA 72-4 | TBA | 0.338403278 | 0 |
| B89 | B137 | CA 72-4 | AFU | 0.486872212 | 0 |
| B89 | B204 | CA 72-4 | Drinking | 0.224046981 | 5.49E-08 |
| B9 | B202 | RH | e-times | -0.83333333 | 0 |
| B90 | B189 | CYFRA 21-1 | eGFB | -0.22127561 | 8.06E-08 |
| B91 | B126 | TP | Alb | 0.619420802 | 0 |
| B91 | B130 | TP | GLB | 0.722411878 | 0 |
| B91 | B188 | TP | Ca | 0.318762105 | 4.44E-15 |
| B91 | B127 | TP | A-G ratio | -0.30689329 | 5.00E-14 |
| B92 | B100 | ALDH2-Glu504Glu | anti-Ro52 | -0.40646942 | 0 |
| B92 | B101 | ALDH2-Glu504Glu | anti-Sc70 | -0.40646943 | 0 |

| | | | | | |
|---|---|---|---|---|---|
| B92 | B102 | ALDH2-Glu504Glu | anti-Sm | -0.40646943 | 0 |
| B92 | B103 | ALDH2-Glu504Glu | anti-SSA/Ro | -0.40646942 | 0 |
| B92 | B104 | ALDH2-Glu504Glu | anti-SSB | -0.40646942 | 0 |
| B92 | B105 | ALDH2-Glu504Glu | anti-U1RNP | -0.40646942 | 0 |
| B92 | B106 | ALDH2-Glu504Glu | anti-P | -0.40646942 | 0 |
| B92 | B107 | ALDH2-Glu504Glu | AnuA | -0.40646943 | 0 |
| B92 | B108 | ALDH2-Glu504Glu | Anti-Cyclin | -0.40646942 | 0 |
| B92 | B109 | ALDH2-Glu504Glu | AMA-M2 | -0.40646942 | 0 |
| B92 | B110 | ALDH2-Glu504Glu | AHA | -0.40646942 | 0 |
| B92 | B125 | ALDH2-Glu504Glu | Cor | -1 | 0 |
| B92 | B151 | ALDH2-Glu504Glu | Anti-CCP | -0.37877757 | 0 |
| B92 | B182 | ALDH2-Glu504Glu | TT | 0.335197767 | 0 |
| B92 | B184 | ALDH2-Glu504Glu | INR | -0.70503348 | 0 |
| B92 | B185 | ALDH2-Glu504Glu | PT | -0.7059417 | 0 |
| B92 | B186 | ALDH2-Glu504Glu | PFC | -0.383801 | 0 |
| B92 | B222 | ALDH2-Glu504Glu | NS | 0.597437353 | 0 |
| B92 | B94 | ALDH2-Glu504Glu | ALDH2-Lys504Lys | -0.99999999 | 0 |
| B92 | B95 | ALDH2-Glu504Glu | anti-cenpb | -0.40646942 | 0 |
| B92 | B97 | ALDH2-Glu504Glu | dsDNA | -0.40646942 | 0 |
| B92 | B98 | ALDH2-Glu504Glu | anti-Jo-1 | -0.40646942 | 0 |
| B92 | B99 | ALDH2-Glu504Glu | anti-PM-Scl | 0.999935487 | 0 |
| B92 | B156 | ALDH2-Glu504Glu | IgE | -0.30619533 | 5.73E-14 |
| B92 | B183 | ALDH2-Glu504Glu | PTA | -0.29856274 | 2.53E-13 |
| B92 | B155 | ALDH2-Glu504Glu | IgA | -0.29356881 | 6.51E-13 |
| B92 | B181 | ALDH2-Glu504Glu | APTT | 0.275574992 | 1.69E-11 |
| B92 | B135 | ALDH2-Glu504Glu | TBA | -0.23820545 | 7.12E-09 |
| B92 | B196 | ALDH2-Glu504Glu | TCO2 | -0.2154783 | 1.77E-07 |
| B92 | B198 | ALDH2-Glu504Glu | P | 0.205188626 | 6.81E-07 |
| B93 | B100 | ALDH2-Glu504Lys | anti-Ro52 | 0.406469422 | 0 |
| B93 | B101 | ALDH2-Glu504Lys | anti-Sc70 | 0.406469425 | 0 |
| B93 | B102 | ALDH2-Glu504Lys | anti-Sm | 0.406469424 | 0 |
| B93 | B103 | ALDH2-Glu504Lys | anti-SSA/Ro | 0.406469422 | 0 |
| B93 | B104 | ALDH2-Glu504Lys | anti-SSB | 0.406469423 | 0 |
| B93 | B105 | ALDH2-Glu504Lys | anti-U1RNP | 0.406469423 | 0 |
| B93 | B106 | ALDH2-Glu504Lys | anti-P | 0.406469423 | 0 |
| B93 | B107 | ALDH2-Glu504Lys | AnuA | 0.406469411 | 0 |
| B93 | B108 | ALDH2-Glu504Lys | Anti-Cyclin | 0.406469423 | 0 |
| B93 | B109 | ALDH2-Glu504Lys | AMA-M2 | 0.406469423 | 0 |
| B93 | B110 | ALDH2-Glu504Lys | AHA | 0.406469423 | 0 |
| B93 | B151 | ALDH2-Glu504Lys | Anti-CCP | 0.378777569 | 0 |
| B93 | B182 | ALDH2-Glu504Lys | TT | -0.33519777 | 0 |
| B93 | B184 | ALDH2-Glu504Lys | INR | 0.705033474 | 0 |
| B93 | B185 | ALDH2-Glu504Lys | PT | 0.705941693 | 0 |
| B93 | B186 | ALDH2-Glu504Lys | PFC | 0.383801003 | 0 |

| | | | | | |
|---|---|---|---|---|---|
| B93 | B222 | ALDH2-Glu504Lys | NS | -0.59743735 | 0 |
| B93 | B94 | ALDH2-Glu504Lys | ALDH2-Lys504Lys | 0.999999988 | 0 |
| B93 | B95 | ALDH2-Glu504Lys | anti-cenpb | 0.406469422 | 0 |
| B93 | B97 | ALDH2-Glu504Lys | dsDNA | 0.406469422 | 0 |
| B93 | B98 | ALDH2-Glu504Lys | anti-Jo-1 | 0.406469423 | 0 |
| B93 | B99 | ALDH2-Glu504Lys | anti-PM-Scl | -0.99993549 | 0 |
| B93 | B156 | ALDH2-Glu504Lys | IgE | 0.306195333 | 5.73E-14 |
| B93 | B183 | ALDH2-Glu504Lys | PTA | 0.298562739 | 2.53E-13 |
| B93 | B155 | ALDH2-Glu504Lys | IgA | 0.293568811 | 6.51E-13 |
| B93 | B181 | ALDH2-Glu504Lys | APTT | -0.27557499 | 1.69E-11 |
| B93 | B135 | ALDH2-Glu504Lys | TBA | 0.238205454 | 7.12E-09 |
| B93 | B196 | ALDH2-Glu504Lys | TCO2 | 0.2154783 | 1.77E-07 |
| B93 | B198 | ALDH2-Glu504Lys | P | -0.20518863 | 6.81E-07 |
| B94 | B100 | ALDH2-Lys504Lys | anti-Ro52 | 0.406469418 | 0 |
| B94 | B101 | ALDH2-Lys504Lys | anti-Sc70 | 0.406469422 | 0 |
| B94 | B102 | ALDH2-Lys504Lys | anti-Sm | 0.406469417 | 0 |
| B94 | B103 | ALDH2-Lys504Lys | anti-SSA/Ro | 0.406469417 | 0 |
| B94 | B104 | ALDH2-Lys504Lys | anti-SSB | 0.406469437 | 0 |
| B94 | B105 | ALDH2-Lys504Lys | anti-U1RNP | 0.406469421 | 0 |
| B94 | B106 | ALDH2-Lys504Lys | anti-P | 0.406469431 | 0 |
| B94 | B107 | ALDH2-Lys504Lys | AnuA | 0.406469516 | 0 |
| B94 | B108 | ALDH2-Lys504Lys | Anti-Cyclin | 0.406469423 | 0 |
| B94 | B109 | ALDH2-Lys504Lys | AMA-M2 | 0.406469418 | 0 |
| B94 | B110 | ALDH2-Lys504Lys | AHA | 0.406469416 | 0 |
| B94 | B124 | ALDH2-Lys504Lys | ACTH | -0.99999999 | 0 |
| B94 | B125 | ALDH2-Lys504Lys | Cor | 0.999999992 | 0 |
| B94 | B151 | ALDH2-Lys504Lys | Anti-CCP | 0.378777564 | 0 |
| B94 | B182 | ALDH2-Lys504Lys | TT | -0.33519776 | 0 |
| B94 | B184 | ALDH2-Lys504Lys | INR | 0.705033464 | 0 |
| B94 | B185 | ALDH2-Lys504Lys | PT | 0.705941685 | 0 |
| B94 | B186 | ALDH2-Lys504Lys | PFC | 0.383800999 | 0 |
| B94 | B222 | ALDH2-Lys504Lys | NS | -0.59743734 | 0 |
| B94 | B95 | ALDH2-Lys504Lys | anti-cenpb | 0.406469425 | 0 |
| B94 | B97 | ALDH2-Lys504Lys | dsDNA | 0.406469427 | 0 |
| B94 | B98 | ALDH2-Lys504Lys | anti-Jo-1 | 0.406469424 | 0 |
| B94 | B99 | ALDH2-Lys504Lys | anti-PM-Scl | -0.99993547 | 0 |
| B94 | B156 | ALDH2-Lys504Lys | IgE | 0.306195329 | 5.73E-14 |
| B94 | B183 | ALDH2-Lys504Lys | PTA | 0.298562735 | 2.53E-13 |
| B94 | B155 | ALDH2-Lys504Lys | IgA | 0.293568807 | 6.51E-13 |
| B94 | B181 | ALDH2-Lys504Lys | APTT | -0.27557499 | 1.69E-11 |
| B94 | B135 | ALDH2-Lys504Lys | TBA | 0.238205451 | 7.12E-09 |
| B94 | B196 | ALDH2-Lys504Lys | TCO2 | 0.215478297 | 1.77E-07 |
| B94 | B198 | ALDH2-Lys504Lys | P | -0.20518862 | 6.81E-07 |
| B95 | B100 | anti-cenpb | anti-Ro52 | 0.999999999 | 0 |

| | | | | | |
|---|---|---|---|---|---|
| B95 | B103 | anti-cenpb | anti-SSA/Ro | 0.999999999 | 0 |
| B95 | B104 | anti-cenpb | anti-SSB | 0.999999997 | 0 |
| B95 | B105 | anti-cenpb | anti-U1RNP | 0.999999999 | 0 |
| B95 | B107 | anti-cenpb | AnuA | 0.999999983 | 0 |
| B95 | B108 | anti-cenpb | Anti-Cyclin | 0.999999999 | 0 |
| B95 | B109 | anti-cenpb | AMA-M2 | 0.999999998 | 0 |
| B95 | B110 | anti-cenpb | AHA | 1 | 0 |
| B95 | B124 | anti-cenpb | ACTH | -0.40646943 | 0 |
| B95 | B125 | anti-cenpb | Cor | 0.406469423 | 0 |
| B95 | B156 | anti-cenpb | IgE | 0.379583052 | 0 |
| B95 | B178 | anti-cenpb | Ig-U | -0.39099799 | 0 |
| B95 | B180 | anti-cenpb | Tf | -0.37436038 | 0 |
| B95 | B97 | anti-cenpb | dsDNA | 0.999999999 | 0 |
| B95 | B99 | anti-cenpb | anti-PM-Scl | -0.39606511 | 0 |
| B95 | B184 | anti-cenpb | INR | 0.310198725 | 2.58E-14 |
| B95 | B185 | anti-cenpb | PT | 0.293544549 | 6.54E-13 |
| B95 | B214 | anti-cenpb | CVA-L | -0.28581501 | 2.73E-12 |
| B95 | B154 | anti-cenpb | RF | 0.285144149 | 3.08E-12 |
| B95 | B175 | anti-cenpb | ALB-U | -0.25006036 | 1.16E-09 |
| B95 | B183 | anti-cenpb | PTA | -0.24874269 | 1.43E-09 |
| B95 | B177 | anti-cenpb | ACR | -0.24423985 | 2.86E-09 |
| B95 | B196 | anti-cenpb | TCO2 | 0.244131093 | 2.91E-09 |
| B95 | B199 | anti-cenpb | Mg | 0.234108713 | 1.30E-08 |
| B95 | B195 | anti-cenpb | Na | 0.228135163 | 3.09E-08 |
| B95 | B182 | anti-cenpb | TT | 0.219111153 | 1.08E-07 |
| B95 | B151 | anti-cenpb | Anti-CCP | 0.209182419 | 4.07E-07 |
| B96 | B225 | dsDNA (IIF) | hPBG | -0.36115756 | 0 |
| B97 | B100 | dsDNA | anti-Ro52 | 0.999999997 | 0 |
| B97 | B103 | dsDNA | anti-SSA/Ro | 0.999999996 | 0 |
| B97 | B105 | dsDNA | anti-U1RNP | 0.999999996 | 0 |
| B97 | B108 | dsDNA | Anti-Cyclin | 0.999999998 | 0 |
| B97 | B109 | dsDNA | AMA-M2 | 0.999999996 | 0 |
| B97 | B110 | dsDNA | AHA | 0.999999996 | 0 |
| B97 | B124 | dsDNA | ACTH | -0.40646943 | 0 |
| B97 | B125 | dsDNA | Cor | 0.406469422 | 0 |
| B97 | B156 | dsDNA | IgE | 0.379583051 | 0 |
| B97 | B178 | dsDNA | Ig-U | -0.39099799 | 0 |
| B97 | B180 | dsDNA | Tf | -0.37436038 | 0 |
| B97 | B99 | dsDNA | anti-PM-Scl | -0.3960651 | 0 |
| B97 | B184 | dsDNA | INR | 0.310198724 | 2.58E-14 |
| B97 | B185 | dsDNA | PT | 0.293544547 | 6.54E-13 |
| B97 | B214 | dsDNA | CVA-L | -0.28581501 | 2.73E-12 |
| B97 | B154 | dsDNA | RF | 0.285144148 | 3.08E-12 |
| B97 | B175 | dsDNA | ALB-U | -0.25006036 | 1.16E-09 |

| | | | | | |
|---|---|---|---|---|---|
| B97 | B183 | dsDNA | PTA | -0.24874269 | 1.43E-09 |
| B97 | B177 | dsDNA | ACR | -0.24423985 | 2.86E-09 |
| B97 | B196 | dsDNA | TCO2 | 0.244131092 | 2.91E-09 |
| B97 | B199 | dsDNA | Mg | 0.234108712 | 1.30E-08 |
| B97 | B195 | dsDNA | Na | 0.228135161 | 3.09E-08 |
| B97 | B182 | dsDNA | TT | 0.219111152 | 1.08E-07 |
| B97 | B151 | dsDNA | Anti-CCP | 0.209182419 | 4.07E-07 |
| B98 | B101 | anti-Jo-1 | anti-Sc70 | 0.999999989 | 0 |
| B98 | B102 | anti-Jo-1 | anti-Sm | 0.999999996 | 0 |
| B98 | B105 | anti-Jo-1 | anti-U1RNP | 0.999999999 | 0 |
| B98 | B107 | anti-Jo-1 | AnuA | 0.99999994 | 0 |
| B98 | B109 | anti-Jo-1 | AMA-M2 | 0.999999999 | 0 |
| B98 | B110 | anti-Jo-1 | AHA | 0.999999999 | 0 |
| B98 | B124 | anti-Jo-1 | ACTH | -0.40646943 | 0 |
| B98 | B125 | anti-Jo-1 | Cor | 0.406469423 | 0 |
| B98 | B156 | anti-Jo-1 | IgE | 0.379583053 | 0 |
| B98 | B178 | anti-Jo-1 | Ig-U | -0.39099799 | 0 |
| B98 | B180 | anti-Jo-1 | Tf | -0.37436038 | 0 |
| B98 | B99 | anti-Jo-1 | anti-PM-Scl | -0.39606511 | 0 |
| B98 | B184 | anti-Jo-1 | INR | 0.310198724 | 2.58E-14 |
| B98 | B185 | anti-Jo-1 | PT | 0.293544545 | 6.54E-13 |
| B98 | B214 | anti-Jo-1 | CVA-L | -0.28581501 | 2.73E-12 |
| B98 | B154 | anti-Jo-1 | RF | 0.28514415 | 3.08E-12 |
| B98 | B175 | anti-Jo-1 | ALB-U | -0.25006036 | 1.16E-09 |
| B98 | B183 | anti-Jo-1 | PTA | -0.24874269 | 1.43E-09 |
| B98 | B177 | anti-Jo-1 | ACR | -0.24423985 | 2.86E-09 |
| B98 | B196 | anti-Jo-1 | TCO2 | 0.244131093 | 2.91E-09 |
| B98 | B199 | anti-Jo-1 | Mg | 0.234108713 | 1.30E-08 |
| B98 | B195 | anti-Jo-1 | Na | 0.228135157 | 3.09E-08 |
| B98 | B182 | anti-Jo-1 | TT | 0.219111153 | 1.08E-07 |
| B98 | B151 | anti-Jo-1 | Anti-CCP | 0.20918242 | 4.07E-07 |
| B99 | B100 | anti-PM-Scl | anti-Ro52 | -0.39606511 | 0 |
| B99 | B101 | anti-PM-Scl | anti-Sc70 | -0.39606511 | 0 |
| B99 | B102 | anti-PM-Scl | anti-Sm | -0.39606511 | 0 |
| B99 | B103 | anti-PM-Scl | anti-SSA/Ro | -0.39606511 | 0 |
| B99 | B104 | anti-PM-Scl | anti-SSB | -0.39606511 | 0 |
| B99 | B105 | anti-PM-Scl | anti-U1RNP | -0.39606511 | 0 |
| B99 | B106 | anti-PM-Scl | anti-P | -0.39606511 | 0 |
| B99 | B107 | anti-PM-Scl | AnuA | -0.3960651 | 0 |
| B99 | B108 | anti-PM-Scl | Anti-Cyclin | -0.39606511 | 0 |
| B99 | B109 | anti-PM-Scl | AMA-M2 | -0.39606511 | 0 |
| B99 | B110 | anti-PM-Scl | AHA | -0.39606511 | 0 |
| B99 | B124 | anti-PM-Scl | ACTH | 0.999935495 | 0 |
| B99 | B125 | anti-PM-Scl | Cor | -0.99993549 | 0 |

| | | | | | |
|---|---|---|---|---|---|
| B99 | B151 | anti-PM-Scl | Anti-CCP | -0.37806662 | 0 |
| B99 | B182 | anti-PM-Scl | TT | 0.339593995 | 0 |
| B99 | B184 | anti-PM-Scl | INR | -0.70469429 | 0 |
| B99 | B185 | anti-PM-Scl | PT | -0.70581409 | 0 |
| B99 | B186 | anti-PM-Scl | PFC | -0.3879812 | 0 |
| B99 | B222 | anti-PM-Scl | NS | 0.59809483 | 0 |
| B99 | B183 | anti-PM-Scl | PTA | -0.30314459 | 1.04E-13 |
| B99 | B156 | anti-PM-Scl | IgE | -0.30300386 | 1.07E-13 |
| B99 | B155 | anti-PM-Scl | IgA | -0.29630482 | 3.89E-13 |
| B99 | B181 | anti-PM-Scl | APTT | 0.274662507 | 1.99E-11 |
| B99 | B135 | anti-PM-Scl | TBA | -0.23732104 | 8.12E-09 |
| B99 | B196 | anti-PM-Scl | TCO2 | -0.21351821 | 2.30E-07 |
| B99 | B198 | anti-PM-Scl | P | 0.206251861 | 5.94E-07 |

**Supplementary Table 17.PEI Correlations in Diabetes**

| Row | Column | Row | Column | Cor | *P* |
|---|---|---|---|---|---|
| B1 | B126 | Gender | Alb | -0.094115831 | 0 |
| B1 | B127 | Gender | A-G ratio | -0.145731403 | 0 |
| B1 | B128 | Gender | ALT | -0.081382156 | 0 |
| B1 | B129 | Gender | IBIL | -0.099340521 | 0 |
| B1 | B130 | Gender | GLB | 0.124953029 | 0 |
| B1 | B132 | Gender | Dbil | -0.11101206 | 0 |
| B1 | B133 | Gender | TBIL | -0.11099021 | 0 |
| B1 | B140 | Gender | r-GT | -0.09344472 | 0 |
| B1 | B141 | Gender | AST/ALT | 0.121756985 | 0 |
| B1 | B148 | Gender | Anti_TPO | 0.081956341 | 0 |
| B1 | B161 | Gender | LEU | 0.225163153 | 0 |
| B1 | B163 | Gender | SG | -0.333109931 | 0 |
| B1 | B166 | Gender | Pro | -0.154635363 | 0 |
| B1 | B170 | Gender | U-GLU | -0.144332905 | 0 |
| B1 | B173 | Gender | NTT | 0.157765641 | 0 |
| B1 | B174 | Gender | Urea/Cr | 0.364584609 | 0 |
| B1 | B176 | Gender | Ucr | -0.166167979 | 0 |
| B1 | B190 | Gender | Cre | -0.368049155 | 0 |
| B1 | B191 | Gender | Urea | -0.081734077 | 0 |
| B1 | B192 | Gender | UA | -0.262956124 | 0 |
| B1 | B204 | Gender | Drinking | -0.260139157 | 0 |
| B1 | B206 | Gender | Waist | -0.288783125 | 0 |
| B1 | B209 | Gender | Smoking | -0.308687138 | 0 |
| B1 | B210 | Gender | Hip | -0.198150138 | 0 |
| B1 | B215 | Gender | BMI | -0.100119361 | 0 |
| B1 | B216 | Gender | BMI | -0.458499075 | 0 |
| B1 | B218 | Gender | DBP | -0.113132465 | 0 |
| B1 | B219 | Gender | OT | 0.125745637 | 0 |
| B1 | B220 | Gender | SBP | 0.076914388 | 0 |
| B1 | B221 | Gender | Hei | -0.642868778 | 0 |
| B1 | B222 | Gender | NS | -0.608769157 | 0 |
| B1 | B3 | Gender | Hcy | -0.105404684 | 0 |
| B1 | B40 | Gender | WBC | -0.115820861 | 0 |
| B1 | B41 | Gender | MON % | -0.175471487 | 0 |
| B1 | B42 | Gender | MON | -0.218767418 | 0 |
| B1 | B44 | Gender | MCH | -0.140983187 | 0 |
| B1 | B45 | Gender | MCHC | -0.26266774 | 0 |
| B1 | B46 | Gender | RBC Count | -0.376448818 | 0 |
| B1 | B48 | Gender | LYM% | 0.08463569 | 0 |
| B1 | B50 | Gender | eo% | -0.096821769 | 0 |
| B1 | B51 | Gender | Hb | -0.49464612 | 0 |
| B1 | B52 | Gender | PLT/BPC | 0.093423037 | 0 |

| | | | | | |
|---|---|---|---|---|---|
| B1 | B58 | Gender | HCT | -0.14135428 | 0 |
| B1 | B60 | Gender | WBRVI | -0.110166995 | 0 |
| B1 | B62 | Gender | WBHSRVI | -0.108805252 | 0 |
| B1 | B63 | Gender | WBRSR(mPas_200) | -0.093735344 | 0 |
| B1 | B65 | Gender | WBRSR(mPas_1) | -0.097584756 | 0 |
| B1 | B66 | Gender | WBRSR(mPas_100) | -0.092345803 | 0 |
| B1 | B67 | Gender | WBRSR(mPas_50) | -0.095110163 | 0 |
| B1 | B72 | Gender | HDL-C | 0.274746234 | 0 |
| B1 | B74 | Gender | ApoA1 | 0.138504758 | 0 |
| B1 | B77 | Gender | TC | 0.097068786 | 0 |
| B1 | B54 | Gender | ESR | 0.073699162 | 1.11E-15 |
| B1 | B2 | Gender | Age | 0.070981707 | 1.22E-14 |
| B1 | B113 | Gender | FT3 | -0.067836736 | 1.70E-13 |
| B1 | B149 | Gender | TGAb | 0.063304689 | 6.10E-12 |
| B1 | B56 | Gender | IR | -0.062668328 | 9.90E-12 |
| B1 | B172 | Gender | KB | -0.060667653 | 4.39E-11 |
| B1 | B75 | Gender | ApoA1/B | 0.060581649 | 4.68E-11 |
| B1 | B71 | Gender | TG | -0.060249282 | 5.96E-11 |
| B1 | B138 | Gender | PAB | -0.059426716 | 1.08E-10 |
| B1 | B198 | Gender | P | 0.057966866 | 3.05E-10 |
| B1 | B153 | Gender | Glu | -0.05697617 | 6.07E-10 |
| B1 | B27 | Gender | LDH | 0.056583363 | 7.95E-10 |
| B1 | B24 | Gender | αHBDH | 0.056579087 | 7.98E-10 |
| B1 | B91 | Gender | TP | 0.05577717 | 1.38E-09 |
| B1 | B25 | Gender | CK | -0.054689005 | 2.85E-09 |
| B1 | B114 | Gender | TSH TT3 TT4(TT4) | 0.054494968 | 3.25E-09 |
| B1 | B205 | Gender | KDA | 0.053043632 | 8.37E-09 |
| B1 | B61 | Gender | WBHSRV | -0.05200241 | 1.63E-08 |
| B1 | B111 | Gender | TSH | 0.05193736 | 1.70E-08 |
| B1 | B84 | Gender | C-PSA | -0.047667019 | 2.26E-07 |
| B1 | B49 | Gender | baso% | -0.046567763 | 4.27E-07 |
| B1 | B189 | Gender | eGFB | 0.046243131 | 5.13E-07 |
| B1 | B179 | Gender | α1-MG | -0.045323533 | 8.59E-07 |
| B10 | B11 | PG Ⅰ | PGI/PGII | 0.165564845 | 0 |
| B10 | B119 | PG Ⅰ | Hcv-rna | -0.092059861 | 0 |
| B10 | B12 | PG Ⅰ | PG II | 0.543757539 | 0 |
| B10 | B143 | PG Ⅰ | β-CTx | -0.096524665 | 0 |
| B10 | B145 | PG Ⅰ | BGP | -0.106710914 | 0 |
| B10 | B15 | PG Ⅰ | CD3-CD19+(B cell) | -0.077186201 | 0 |
| B10 | B175 | PG Ⅰ | ALB-U | 0.289761478 | 0 |
| B10 | B177 | PG Ⅰ | ACR | 0.235078333 | 0 |
| B10 | B189 | PG Ⅰ | eGFB | -0.09912661 | 0 |
| B10 | B190 | PG Ⅰ | Cre | 0.102115992 | 0 |
| B10 | B191 | PG Ⅰ | Urea | 0.085957831 | 0 |

| | | | | | |
|---|---|---|---|---|---|
| B10 | B80 | PG I | anti-HP IgG | 0.083290261 | 0 |
| B10 | B144 | PG I | Total-PINP | -0.07363458 | 1.33E-15 |
| B10 | B123 | PG I | hs-CRP1 | 0.068479424 | 1.00E-13 |
| B10 | B186 | PG I | PFC | 0.068204143 | 1.26E-13 |
| B10 | B107 | PG I | AnuA | -0.067446002 | 2.33E-13 |
| B10 | B101 | PG I | anti-Sc70 | -0.066956741 | 3.47E-13 |
| B10 | B104 | PG I | anti-SSB | -0.066956729 | 3.47E-13 |
| B10 | B98 | PG I | anti-Jo-1 | -0.066956723 | 3.47E-13 |
| B10 | B151 | PG I | Anti-CCP | 0.065144244 | 1.47E-12 |
| B10 | B18 | PG I | CD3+CD4+helper(inducer)T | 0.062996367 | 7.72E-12 |
| B10 | B125 | PG I | Cor | 0.056888008 | 6.45E-10 |
| B10 | B47 | PG I | RDW | 0.054618425 | 2.99E-09 |
| B10 | B16 | PG I | CD3+T | 0.052641652 | 1.08E-08 |
| B10 | B24 | PG I | αHBDH | 0.050742543 | 3.58E-08 |
| B10 | B25 | PG I | CK | 0.050692469 | 3.69E-08 |
| B10 | B79 | PG I | hb V -dna | -0.047537137 | 2.44E-07 |
| B10 | B126 | PG I | Alb | -0.046722878 | 3.90E-07 |
| B10 | B192 | PG I | UA | 0.046519656 | 4.38E-07 |
| B10 | B120 | PG I | HCV-cAg | -0.045399169 | 8.24E-07 |
| B100 | B101 | anti-Ro52 | anti-Sc70 | 0.09482229 | 0 |
| B100 | B104 | anti-Ro52 | anti-SSB | 0.094822278 | 0 |
| B100 | B107 | anti-Ro52 | AnuA | 0.092893593 | 0 |
| B100 | B157 | anti-Ro52 | IgG | 0.079460027 | 0 |
| B100 | B182 | anti-Ro52 | TT | -0.06027973 | 5.83E-11 |
| B100 | B193 | anti-Ro52 | K | 0.052154086 | 1.48E-08 |
| B100 | B124 | anti-Ro52 | ACTH | -0.050831261 | 3.39E-08 |
| B100 | B137 | anti-Ro52 | AFU | -0.046961108 | 3.41E-07 |
| B100 | B119 | anti-Ro52 | Hcv-rna | -0.045990699 | 5.92E-07 |
| B101 | B102 | anti-Sc70 | anti-Sm | 0.110763142 | 0 |
| B101 | B103 | anti-Sc70 | anti-SSA/Ro | 0.094458136 | 0 |
| B101 | B106 | anti-Sc70 | anti-P | 0.110785287 | 0 |
| B101 | B107 | anti-Sc70 | AnuA | 0.994375158 | 0 |
| B101 | B119 | anti-Sc70 | Hcv-rna | 0.150439903 | 0 |
| B101 | B120 | anti-Sc70 | HCV-cAg | 0.126183922 | 0 |
| B101 | B121 | anti-Sc70 | C3 | -0.094750552 | 0 |
| B101 | B122 | anti-Sc70 | C4 | -0.086964903 | 0 |
| B101 | B134 | anti-Sc70 | CHE | -0.099159467 | 0 |
| B101 | B137 | anti-Sc70 | AFU | -0.077017833 | 0 |
| B101 | B138 | anti-Sc70 | PAB | -0.07599806 | 0 |
| B101 | B139 | anti-Sc70 | ADA | -0.086100026 | 0 |
| B101 | B151 | anti-Sc70 | Anti-CCP | -0.123677072 | 0 |
| B101 | B155 | anti-Sc70 | IgA | -0.07859844 | 0 |
| B101 | B157 | anti-Sc70 | IgG | 0.143501857 | 0 |
| B101 | B181 | anti-Sc70 | APTT | 0.126973236 | 0 |

| | | | | | |
|---|---|---|---|---|---|
| B101 | B184 | anti-Sc70 | INR | 0.140026311 | 0 |
| B101 | B185 | anti-Sc70 | PT | 0.076376763 | 0 |
| B101 | B186 | anti-Sc70 | PFC | -0.08709384 | 0 |
| B101 | B194 | anti-Sc70 | Cl | 0.12305288 | 0 |
| B101 | B195 | anti-Sc70 | Na | 0.122660759 | 0 |
| B101 | B198 | anti-Sc70 | P | -0.094561464 | 0 |
| B101 | B199 | anti-Sc70 | Mg | 0.114914253 | 0 |
| B101 | B200 | anti-Sc70 | OP | -0.165117545 | 0 |
| B101 | B183 | anti-Sc70 | PTA | 0.072236214 | 4.00E-15 |
| B101 | B152 | anti-Sc70 | ACA-Ig(G-A-M) | 0.066754191 | 4.08E-13 |
| B101 | B179 | anti-Sc70 | α1-MG | -0.065045271 | 1.59E-12 |
| B101 | B212 | anti-Sc70 | IOP-L | -0.061301285 | 2.75E-11 |
| B101 | B124 | anti-Sc70 | ACTH | -0.054617221 | 2.99E-09 |
| B101 | B193 | anti-Sc70 | K | 0.05201209 | 1.62E-08 |
| B101 | B112 | anti-Sc70 | FT4 | -0.048806322 | 1.16E-07 |
| B101 | B220 | anti-Sc70 | SBP | 0.04795154 | 1.92E-07 |
| B101 | B150 | anti-Sc70 | ALP | -0.047571286 | 2.39E-07 |
| B101 | B189 | anti-Sc70 | eGFB | -0.047256567 | 2.87E-07 |
| B101 | B211 | anti-Sc70 | IOP-R | -0.046863291 | 3.60E-07 |
| B102 | B104 | anti-Sm | anti-SSB | 0.110763147 | 0 |
| B102 | B107 | anti-Sm | AnuA | 0.109310812 | 0 |
| B102 | B140 | anti-Sm | r-GT | 0.118774779 | 0 |
| B102 | B155 | anti-Sm | IgA | 0.127839175 | 0 |
| B102 | B179 | anti-Sm | α1-MG | 0.098510446 | 0 |
| B102 | B157 | anti-Sm | IgG | 0.06964853 | 3.77E-14 |
| B102 | B158 | anti-Sm | IgM | 0.059000807 | 1.47E-10 |
| B102 | B115 | anti-Sm | TSH TT3 TT4(TT3) | -0.054272634 | 3.76E-09 |
| B102 | B178 | anti-Sm | Ig-U | 0.050999672 | 3.05E-08 |
| B103 | B104 | anti-SSA/Ro | anti-SSB | 0.094458122 | 0 |
| B103 | B107 | anti-SSA/Ro | AnuA | 0.092371516 | 0 |
| B103 | B108 | anti-SSA/Ro | Anti-Cyclin | 0.626575473 | 0 |
| B103 | B141 | anti-SSA/Ro | AST/ALT | 0.068280597 | 1.18E-13 |
| B103 | B157 | anti-SSA/Ro | IgG | 0.066327519 | 5.75E-13 |
| B104 | B106 | anti-SSB | anti-P | 0.110785259 | 0 |
| B104 | B107 | anti-SSB | AnuA | 0.994376258 | 0 |
| B104 | B119 | anti-SSB | Hcv-rna | 0.150439883 | 0 |
| B104 | B120 | anti-SSB | HCV-cAg | 0.12618479 | 0 |
| B104 | B121 | anti-SSB | C3 | -0.094750482 | 0 |
| B104 | B122 | anti-SSB | C4 | -0.086964863 | 0 |
| B104 | B134 | anti-SSB | CHE | -0.099159449 | 0 |
| B104 | B137 | anti-SSB | AFU | -0.07701782 | 0 |
| B104 | B138 | anti-SSB | PAB | -0.075998044 | 0 |
| B104 | B139 | anti-SSB | ADA | -0.08610001 | 0 |
| B104 | B151 | anti-SSB | Anti-CCP | -0.123677049 | 0 |

| | | | | | |
|---|---|---|---|---|---|
| B104 | B155 | anti-SSB | IgA | -0.078598424 | 0 |
| B104 | B157 | anti-SSB | IgG | 0.143501842 | 0 |
| B104 | B181 | anti-SSB | APTT | 0.126973217 | 0 |
| B104 | B184 | anti-SSB | INR | 0.140026295 | 0 |
| B104 | B185 | anti-SSB | PT | 0.076376755 | 0 |
| B104 | B186 | anti-SSB | PFC | -0.087093817 | 0 |
| B104 | B194 | anti-SSB | Cl | 0.123052885 | 0 |
| B104 | B195 | anti-SSB | Na | 0.1226608 | 0 |
| B104 | B198 | anti-SSB | P | -0.094561437 | 0 |
| B104 | B199 | anti-SSB | Mg | 0.114914249 | 0 |
| B104 | B200 | anti-SSB | OP | -0.165117511 | 0 |
| B104 | B183 | anti-SSB | PTA | 0.072236204 | 4.00E-15 |
| B104 | B152 | anti-SSB | ACA-Ig(G-A-M) | 0.06675424 | 4.08E-13 |
| B104 | B179 | anti-SSB | α1-MG | -0.06504526 | 1.59E-12 |
| B104 | B212 | anti-SSB | IOP-L | -0.061301273 | 2.75E-11 |
| B104 | B124 | anti-SSB | ACTH | -0.054617211 | 2.99E-09 |
| B104 | B193 | anti-SSB | K | 0.052012096 | 1.62E-08 |
| B104 | B112 | anti-SSB | FT4 | -0.048806311 | 1.16E-07 |
| B104 | B220 | anti-SSB | SBP | 0.047951534 | 1.92E-07 |
| B104 | B150 | anti-SSB | ALP | -0.047571276 | 2.39E-07 |
| B104 | B189 | anti-SSB | eGFB | -0.047256558 | 2.87E-07 |
| B104 | B211 | anti-SSB | IOP-R | -0.046863283 | 3.60E-07 |
| B105 | B119 | anti-U1RNP | Hcv-rna | -0.084963313 | 0 |
| B105 | B125 | anti-U1RNP | Cor | 0.109565661 | 0 |
| B105 | B196 | anti-U1RNP | TCO2 | 0.056127644 | 1.09E-09 |
| B105 | B179 | anti-U1RNP | α1-MG | 0.051636505 | 2.05E-08 |
| B105 | B158 | anti-U1RNP | IgM | 0.050233638 | 4.89E-08 |
| B106 | B107 | anti-P | AnuA | 0.109332722 | 0 |
| B106 | B157 | anti-P | IgG | 0.08368434 | 0 |
| B106 | B123 | anti-P | hs-CRP1 | 0.073384458 | 1.55E-15 |
| B106 | B155 | anti-P | IgA | 0.069168591 | 5.66E-14 |
| B107 | B119 | AnuA | Hcv-rna | 0.151385073 | 0 |
| B107 | B120 | AnuA | HCV-cAg | 0.122875342 | 0 |
| B107 | B121 | AnuA | C3 | -0.095282408 | 0 |
| B107 | B122 | AnuA | C4 | -0.087444475 | 0 |
| B107 | B134 | AnuA | CHE | -0.10458179 | 0 |
| B107 | B138 | AnuA | PAB | -0.084602301 | 0 |
| B107 | B139 | AnuA | ADA | -0.086218483 | 0 |
| B107 | B151 | AnuA | Anti-CCP | -0.124321867 | 0 |
| B107 | B157 | AnuA | IgG | 0.145517111 | 0 |
| B107 | B181 | AnuA | APTT | 0.127680792 | 0 |
| B107 | B184 | AnuA | INR | 0.140811983 | 0 |
| B107 | B185 | AnuA | PT | 0.076786554 | 0 |
| B107 | B186 | AnuA | PFC | -0.08761144 | 0 |

| | | | | | |
|---|---|---|---|---|---|
| B107 | B194 | AnuA | Cl | 0.124650008 | 0 |
| B107 | B195 | AnuA | Na | 0.124763441 | 0 |
| B107 | B198 | AnuA | P | -0.093804267 | 0 |
| B107 | B199 | AnuA | Mg | 0.108597029 | 0 |
| B107 | B200 | AnuA | OP | -0.16232695 | 0 |
| B107 | B137 | AnuA | AFU | -0.073949171 | 8.88E-16 |
| B107 | B183 | AnuA | PTA | 0.072651555 | 2.89E-15 |
| B107 | B155 | AnuA | IgA | -0.072107587 | 4.66E-15 |
| B107 | B152 | AnuA | ACA-Ig(G-A-M) | 0.067136596 | 3.00E-13 |
| B107 | B179 | AnuA | α1-MG | -0.063648249 | 4.69E-12 |
| B107 | B212 | AnuA | IOP-L | -0.06337118 | 5.80E-12 |
| B107 | B124 | AnuA | ACTH | -0.054937091 | 2.42E-09 |
| B107 | B150 | AnuA | ALP | -0.050769582 | 3.52E-08 |
| B107 | B220 | AnuA | SBP | 0.048932252 | 1.07E-07 |
| B107 | B211 | AnuA | IOP-R | -0.04795949 | 1.91E-07 |
| B107 | B112 | AnuA | FT4 | -0.047045475 | 3.25E-07 |
| B107 | B189 | AnuA | eGFB | -0.04678867 | 3.76E-07 |
| B107 | B193 | AnuA | K | 0.046693334 | 3.97E-07 |
| B108 | B109 | Anti-Cyclin | AMA-M2 | 0.146688438 | 0 |
| B109 | B182 | AMA-M2 | TT | 0.102773861 | 0 |
| B109 | B195 | AMA-M2 | Na | -0.063625262 | 4.77E-12 |
| B109 | B119 | AMA-M2 | Hcv-rna | -0.054104454 | 4.20E-09 |
| B109 | B204 | AMA-M2 | Drinking | 0.050062658 | 5.43E-08 |
| B109 | B155 | AMA-M2 | IgA | 0.047715273 | 2.20E-07 |
| B109 | B157 | AMA-M2 | IgG | -0.047386031 | 2.67E-07 |
| B109 | B151 | AMA-M2 | Anti-CCP | -0.045329198 | 8.56E-07 |
| B11 | B12 | PGI/PGII | PG II | -0.498291336 | 0 |
| B11 | B14 | PGI/PGII | G-17 | -0.234989476 | 0 |
| B11 | B175 | PGI/PGII | ALB-U | 0.11913415 | 0 |
| B11 | B177 | PGI/PGII | ACR | 0.097007614 | 0 |
| B11 | B80 | PGI/PGII | anti-HP IgG | -0.119502957 | 0 |
| B11 | B34 | PGI/PGII | TK | 0.070837419 | 1.38E-14 |
| B11 | B145 | PGI/PGII | BGP | -0.051394021 | 2.39E-08 |
| B110 | B119 | AHA | Hcv-rna | -0.138791252 | 0 |
| B110 | B134 | AHA | CHE | 0.070314478 | 2.18E-14 |
| B110 | B138 | AHA | PAB | 0.069955001 | 2.93E-14 |
| B110 | B120 | AHA | HCV-cAg | -0.06702877 | 3.27E-13 |
| B110 | B197 | AHA | AG | 0.054032248 | 4.40E-09 |
| B110 | B196 | AHA | TCO2 | -0.045970141 | 5.98E-07 |
| B111 | B112 | TSH | FT4 | -0.219643397 | 0 |
| B111 | B113 | TSH | FT3 | -0.130398835 | 0 |
| B111 | B148 | TSH | Anti_TPO | 0.200964715 | 0 |
| B111 | B149 | TSH | TGAb | 0.132053942 | 0 |
| B111 | B189 | TSH | eGFB | -0.050036872 | 5.52E-08 |

| | | | | | |
|---|---|---|---|---|---|
| B111 | B198 | TSH | P | 0.049236109 | 8.95E-08 |
| B111 | B114 | TSH | TSH TT3 TT4(TT4) | -0.045289237 | 8.76E-07 |
| B112 | B113 | FT4 | FT3 | 0.250976763 | 0 |
| B112 | B114 | FT4 | TSH TT3 TT4(TT4) | 0.452266153 | 0 |
| B112 | B115 | FT4 | TSH TT3 TT4(TT3) | 0.087293655 | 0 |
| B112 | B153 | FT4 | Glu | 0.073258724 | 1.78E-15 |
| B112 | B188 | FT4 | Ca | 0.06195375 | 1.69E-11 |
| B112 | B120 | FT4 | HCV-cAg | -0.059124283 | 1.34E-10 |
| B112 | B189 | FT4 | eGFB | 0.057253095 | 5.01E-10 |
| B112 | B150 | FT4 | ALP | 0.054767309 | 2.71E-09 |
| B112 | B194 | FT4 | Cl | -0.050226563 | 4.91E-08 |
| B113 | B114 | FT3 | TSH TT3 TT4(TT4) | 0.104717097 | 0 |
| B113 | B115 | FT3 | TSH TT3 TT4(TT3) | 0.358200366 | 0 |
| B113 | B126 | FT3 | Alb | 0.092343097 | 0 |
| B113 | B145 | FT3 | BGP | 0.080801966 | 0 |
| B113 | B149 | FT3 | TGAb | 0.064735221 | 2.02E-12 |
| B113 | B189 | FT3 | eGFB | 0.062780944 | 9.09E-12 |
| B113 | B222 | FT3 | NS | 0.059984732 | 7.22E-11 |
| B113 | B123 | FT3 | hs-CRP1 | -0.052543674 | 1.15E-08 |
| B113 | B209 | FT3 | Smoking | 0.051274653 | 2.57E-08 |
| B113 | B216 | FT3 | BMI | 0.050526912 | 4.09E-08 |
| B113 | B188 | FT3 | Ca | 0.049440475 | 7.92E-08 |
| B113 | B134 | FT3 | CHE | 0.049283165 | 8.70E-08 |
| B113 | B141 | FT3 | AST/ALT | -0.049236072 | 8.95E-08 |
| B113 | B206 | FT3 | Waist | 0.046644196 | 4.08E-07 |
| B113 | B144 | FT3 | Total-PINP | 0.046410547 | 4.67E-07 |
| B113 | B128 | FT3 | ALT | 0.046112207 | 5.52E-07 |
| B113 | B119 | FT3 | Hcv-rna | -0.045600254 | 7.36E-07 |
| B113 | B143 | FT3 | β-CTx | 0.04538352 | 8.31E-07 |
| B114 | B115 | TSH TT3 TT4(TT4) | TSH TT3 TT4(TT3) | 0.325671044 | 0 |
| B114 | B150 | TSH TT3 TT4(TT4) | ALP | 0.059692141 | 8.93E-11 |
| B114 | B139 | TSH TT3 TT4(TT4) | ADA | 0.057049229 | 5.77E-10 |
| B114 | B186 | TSH TT3 TT4(TT4) | PFC | 0.054746578 | 2.75E-09 |
| B114 | B123 | TSH TT3 TT4(TT4) | hs-CRP1 | 0.05320768 | 7.53E-09 |
| B114 | B197 | TSH TT3 TT4(TT4) | AG | 0.052924042 | 9.04E-09 |
| B114 | B204 | TSH TT3 TT4(TT4) | Drinking | -0.048426736 | 1.45E-07 |
| B114 | B188 | TSH TT3 TT4(TT4) | Ca | 0.048390098 | 1.48E-07 |
| B114 | B194 | TSH TT3 TT4(TT4) | Cl | -0.047844555 | 2.04E-07 |
| B114 | B120 | TSH TT3 TT4(TT4) | HCV-cAg | -0.047232993 | 2.91E-07 |
| B115 | B134 | TSH TT3 TT4(TT3) | CHE | 0.076412018 | 0 |
| B115 | B126 | TSH TT3 TT4(TT3) | Alb | 0.057481141 | 4.28E-10 |
| B115 | B125 | TSH TT3 TT4(TT3) | Cor | -0.045195609 | 9.22E-07 |
| B116 | B117 | TT3 | TT4 | 0.594770853 | 0 |
| B118 | B150 | CEA1 | ALP | 0.229329471 | 0 |

| | | | | | |
|---|---|---|---|---|---|
| B118 | B219 | CEA1 | OT | 0.067693087 | 1.91E-13 |
| B119 | B120 | Hcv-rna | HCV-cAg | 0.505246111 | 0 |
| B119 | B151 | Hcv-rna | Anti-CCP | -0.096563033 | 0 |
| B119 | B183 | Hcv-rna | PTA | 0.129101314 | 0 |
| B119 | B186 | Hcv-rna | PFC | -0.08750318 | 0 |
| B119 | B194 | Hcv-rna | CI | 0.129000852 | 0 |
| B119 | B200 | Hcv-rna | OP | -0.114737538 | 0 |
| B119 | B184 | Hcv-rna | INR | 0.069489405 | 4.33E-14 |
| B119 | B199 | Hcv-rna | Mg | 0.068321963 | 1.14E-13 |
| B119 | B169 | Hcv-rna | U-CRY | -0.065438676 | 1.16E-12 |
| B119 | B137 | Hcv-rna | AFU | -0.065284508 | 1.31E-12 |
| B119 | B182 | Hcv-rna | TT | -0.05705892 | 5.73E-10 |
| B119 | B135 | Hcv-rna | TBA | -0.055938192 | 1.23E-09 |
| B119 | B121 | Hcv-rna | C3 | -0.054494779 | 3.25E-09 |
| B119 | B156 | Hcv-rna | IgE | 0.05295574 | 8.86E-09 |
| B119 | B181 | Hcv-rna | APTT | 0.049894481 | 6.01E-08 |
| B119 | B146 | Hcv-rna | CysC | -0.047113163 | 3.12E-07 |
| B119 | B196 | Hcv-rna | TCO2 | -0.045127055 | 9.58E-07 |
| B12 | B14 | PG II | G-17 | 0.208731128 | 0 |
| B12 | B80 | PG II | anti-HP IgG | 0.166607769 | 0 |
| B12 | B119 | PG II | Hcv-rna | -0.069196564 | 5.53E-14 |
| B12 | B189 | PG II | eGFB | -0.068373592 | 1.09E-13 |
| B12 | B186 | PG II | PFC | 0.064367821 | 2.69E-12 |
| B12 | B25 | PG II | CK | 0.062167831 | 1.44E-11 |
| B12 | B190 | PG II | Cre | 0.052747596 | 1.01E-08 |
| B12 | B47 | PG II | RDW | 0.052281871 | 1.36E-08 |
| B12 | B15 | PG II | CD3-CD19+(B cell) | -0.049355211 | 8.33E-08 |
| B12 | B183 | PG II | PTA | -0.046603752 | 4.18E-07 |
| B12 | B151 | PG II | Anti-CCP | 0.045095972 | 9.75E-07 |
| B120 | B121 | HCV-cAg | C3 | -0.091990789 | 0 |
| B120 | B134 | HCV-cAg | CHE | -0.082257943 | 0 |
| B120 | B183 | HCV-cAg | PTA | 0.080303431 | 0 |
| B120 | B200 | HCV-cAg | OP | -0.085758236 | 0 |
| B120 | B224 | HCV-cAg | pmh | -0.134184958 | 0 |
| B120 | B194 | HCV-cAg | CI | 0.074118626 | 8.88E-16 |
| B120 | B122 | HCV-cAg | C4 | -0.057383392 | 4.58E-10 |
| B120 | B163 | HCV-cAg | SG | -0.054082243 | 4.26E-09 |
| B120 | B189 | HCV-cAg | eGFB | -0.053829236 | 5.03E-09 |
| B120 | B151 | HCV-cAg | Anti-CCP | -0.053189104 | 7.62E-09 |
| B120 | B146 | HCV-cAg | CysC | -0.050164775 | 5.10E-08 |
| B120 | B155 | HCV-cAg | IgA | -0.046519238 | 4.39E-07 |
| B120 | B222 | HCV-cAg | NS | -0.046093749 | 5.58E-07 |
| B120 | B126 | HCV-cAg | Alb | -0.04592541 | 6.14E-07 |
| B121 | B122 | C3 | C4 | 0.915742313 | 0 |

| | | | | | |
|---|---|---|---|---|---|
| B121 | B228 | C3 | OS | -0.07228222 | 4.00E-15 |
| B121 | B227 | C3 | OD | -0.071143069 | 1.07E-14 |
| B121 | B157 | C3 | IgG | -0.05553465 | 1.62E-09 |
| B121 | B134 | C3 | CHE | 0.053480788 | 6.31E-09 |
| B122 | B228 | C4 | OS | -0.056848148 | 6.63E-10 |
| B122 | B227 | C4 | OD | -0.055364842 | 1.82E-09 |
| B122 | B134 | C4 | CHE | 0.053231354 | 7.42E-09 |
| B123 | B136 | hs-CRP1 | 5'NT | 0.07927594 | 0 |
| B123 | B138 | hs-CRP1 | PAB | -0.099772647 | 0 |
| B123 | B139 | hs-CRP1 | ADA | 0.088487414 | 0 |
| B123 | B169 | hs-CRP1 | U-CRY | 0.206797891 | 0 |
| B123 | B175 | hs-CRP1 | ALB-U | 0.11435067 | 0 |
| B123 | B177 | hs-CRP1 | ACR | 0.106257616 | 0 |
| B123 | B127 | hs-CRP1 | A-G ratio | -0.074321807 | 6.66E-16 |
| B123 | B130 | hs-CRP1 | GLB | 0.069655581 | 3.77E-14 |
| B123 | B155 | hs-CRP1 | IgA | 0.066164588 | 6.54E-13 |
| B123 | B126 | hs-CRP1 | Alb | -0.06456283 | 2.31E-12 |
| B123 | B194 | hs-CRP1 | Cl | -0.057760662 | 3.52E-10 |
| B123 | B186 | hs-CRP1 | PFC | 0.053372158 | 6.77E-09 |
| B123 | B153 | hs-CRP1 | Glu | 0.050554763 | 4.02E-08 |
| B123 | B195 | hs-CRP1 | Na | -0.0495951 | 7.21E-08 |
| B123 | B150 | hs-CRP1 | ALP | 0.049145903 | 9.45E-08 |
| B123 | B199 | hs-CRP1 | Mg | -0.048795907 | 1.16E-07 |
| B124 | B125 | ACTH | Cor | 0.476475727 | 0 |
| B124 | B196 | ACTH | TCO2 | 0.056503972 | 8.40E-10 |
| B124 | B183 | ACTH | PTA | -0.056326563 | 9.48E-10 |
| B124 | B181 | ACTH | APTT | -0.052687478 | 1.05E-08 |
| B124 | B200 | ACTH | OP | 0.047216808 | 2.94E-07 |
| B125 | B151 | Cor | Anti-CCP | 0.144695375 | 0 |
| B125 | B181 | Cor | APTT | -0.0793659 | 0 |
| B125 | B182 | Cor | TT | -0.100135988 | 0 |
| B125 | B186 | Cor | PFC | 0.20583999 | 0 |
| B125 | B138 | Cor | PAB | 0.073112152 | 1.78E-15 |
| B125 | B139 | Cor | ADA | -0.063431229 | 5.54E-12 |
| B125 | B183 | Cor | PTA | -0.059372734 | 1.12E-10 |
| B125 | B196 | Cor | TCO2 | 0.05502892 | 2.28E-09 |
| B125 | B197 | Cor | AG | -0.054352161 | 3.57E-09 |
| B125 | B150 | Cor | ALP | -0.048918453 | 1.08E-07 |
| B125 | B136 | Cor | 5'NT | -0.048551751 | 1.35E-07 |
| B126 | B127 | Alb | A-G ratio | 0.496530408 | 0 |
| B126 | B128 | Alb | ALT | 0.08210408 | 0 |
| B126 | B129 | Alb | IBIL | 0.124681794 | 0 |
| B126 | B130 | Alb | GLB | -0.16367333 | 0 |
| B126 | B133 | Alb | TBIL | 0.110072174 | 0 |

| | | | | | |
|---|---|---|---|---|---|
| B126 | B134 | Alb | CHE | 0.089039799 | 0 |
| B126 | B138 | Alb | PAB | 0.107937531 | 0 |
| B126 | B141 | Alb | AST/ALT | -0.138423483 | 0 |
| B126 | B188 | Alb | Ca | 0.416879599 | 0 |
| B126 | B189 | Alb | eGFB | 0.23961717 | 0 |
| B126 | B192 | Alb | UA | 0.089274033 | 0 |
| B126 | B216 | Alb | BMI | 0.112134079 | 0 |
| B126 | B218 | Alb | DBP | 0.149477192 | 0 |
| B126 | B221 | Alb | Hei | 0.115950089 | 0 |
| B126 | B222 | Alb | NS | 0.108031258 | 0 |
| B126 | B177 | Alb | ACR | -0.075055765 | 4.44E-16 |
| B126 | B175 | Alb | ALB-U | -0.073506476 | 1.33E-15 |
| B126 | B191 | Alb | Urea | -0.069758486 | 3.46E-14 |
| B126 | B163 | Alb | SG | 0.069270557 | 5.20E-14 |
| B126 | B190 | Alb | Cre | -0.066156656 | 6.59E-13 |
| B126 | B160 | Alb | PH | -0.061022312 | 3.38E-11 |
| B126 | B184 | Alb | INR | -0.058615524 | 1.93E-10 |
| B126 | B161 | Alb | LEU | -0.055797572 | 1.36E-09 |
| B126 | B224 | Alb | pmh | 0.05557047 | 1.58E-09 |
| B126 | B139 | Alb | ADA | -0.055221416 | 2.00E-09 |
| B126 | B215 | Alb | BMI | 0.054934495 | 2.42E-09 |
| B126 | B185 | Alb | PT | -0.054869158 | 2.53E-09 |
| B126 | B181 | Alb | APTT | -0.054810476 | 2.63E-09 |
| B126 | B229 | Alb | slit-lamp | -0.054292416 | 3.71E-09 |
| B126 | B194 | Alb | Cl | -0.049733852 | 6.63E-08 |
| B126 | B212 | Alb | IOP-L | 0.047480987 | 2.52E-07 |
| B126 | B204 | Alb | Drinking | 0.046158788 | 5.38E-07 |
| B127 | B129 | A-G ratio | IBIL | 0.133510943 | 0 |
| B127 | B130 | A-G ratio | GLB | -0.896251792 | 0 |
| B127 | B131 | A-G ratio | AST | -0.081930425 | 0 |
| B127 | B132 | A-G ratio | Dbil | 0.077190354 | 0 |
| B127 | B133 | A-G ratio | TBIL | 0.128491623 | 0 |
| B127 | B139 | A-G ratio | ADA | -0.082066925 | 0 |
| B127 | B141 | A-G ratio | AST/ALT | -0.080931356 | 0 |
| B127 | B146 | A-G ratio | CysC | -0.079347961 | 0 |
| B127 | B153 | A-G ratio | Glu | -0.111847188 | 0 |
| B127 | B155 | A-G ratio | IgA | -0.091089436 | 0 |
| B127 | B157 | A-G ratio | IgG | -0.095370226 | 0 |
| B127 | B189 | A-G ratio | eGFB | 0.165972164 | 0 |
| B127 | B204 | A-G ratio | Drinking | 0.085767361 | 0 |
| B127 | B209 | A-G ratio | Smoking | 0.112339717 | 0 |
| B127 | B216 | A-G ratio | BMI | 0.080188298 | 0 |
| B127 | B220 | A-G ratio | SBP | -0.133506395 | 0 |
| B127 | B221 | A-G ratio | Hei | 0.168866173 | 0 |

| | | | | | |
|---|---|---|---|---|---|
| B127 | B222 | A-G ratio | NS | 0.079935365 | 0 |
| B127 | B191 | A-G ratio | Urea | -0.068733355 | 8.13E-14 |
| B127 | B138 | A-G ratio | PAB | 0.067750802 | 1.82E-13 |
| B127 | B161 | A-G ratio | LEU | -0.0662527 | 6.10E-13 |
| B127 | B140 | A-G ratio | r-GT | -0.065700294 | 9.46E-13 |
| B127 | B177 | A-G ratio | ACR | -0.063837669 | 4.05E-12 |
| B127 | B170 | A-G ratio | U-GLU | -0.061425674 | 2.51E-11 |
| B127 | B150 | A-G ratio | ALP | -0.060091012 | 6.69E-11 |
| B127 | B163 | A-G ratio | SG | 0.052559799 | 1.14E-08 |
| B127 | B178 | A-G ratio | Ig-U | -0.049222459 | 9.03E-08 |
| B127 | B188 | A-G ratio | Ca | 0.047726818 | 2.19E-07 |
| B127 | B158 | A-G ratio | IgM | -0.045165572 | 9.38E-07 |
| B128 | B131 | ALT | AST | 0.839704026 | 0 |
| B128 | B132 | ALT | Dbil | 0.247248388 | 0 |
| B128 | B133 | ALT | TBIL | 0.12031803 | 0 |
| B128 | B140 | ALT | r-GT | 0.325376302 | 0 |
| B128 | B141 | ALT | AST/ALT | -0.384997413 | 0 |
| B128 | B165 | ALT | bilirubin | 0.188321606 | 0 |
| B128 | B189 | ALT | eGFB | 0.094108084 | 0 |
| B128 | B192 | ALT | UA | 0.09701503 | 0 |
| B128 | B206 | ALT | Waist | 0.122039507 | 0 |
| B128 | B210 | ALT | Hip | 0.096240229 | 0 |
| B128 | B215 | ALT | BMI | 0.136293864 | 0 |
| B128 | B216 | ALT | BMI | 0.146439356 | 0 |
| B128 | B222 | ALT | NS | 0.118980841 | 0 |
| B128 | B221 | ALT | Hei | 0.073578064 | 1.33E-15 |
| B128 | B129 | ALT | IBIL | 0.062747158 | 9.32E-12 |
| B128 | B136 | ALT | 5'NT | 0.062321566 | 1.29E-11 |
| B128 | B139 | ALT | ADA | 0.056767454 | 7.01E-10 |
| B128 | B218 | ALT | DBP | 0.054642375 | 2.94E-09 |
| B128 | B137 | ALT | AFU | 0.053685543 | 5.52E-09 |
| B128 | B130 | ALT | GLB | 0.051499334 | 2.23E-08 |
| B128 | B163 | ALT | SG | 0.051309503 | 2.52E-08 |
| B128 | B150 | ALT | ALP | 0.045893614 | 6.25E-07 |
| B129 | B130 | IBIL | GLB | -0.111101093 | 0 |
| B129 | B131 | IBIL | AST | 0.083140716 | 0 |
| B129 | B132 | IBIL | Dbil | 0.621324441 | 0 |
| B129 | B133 | IBIL | TBIL | 0.974978086 | 0 |
| B129 | B163 | IBIL | SG | 0.078786703 | 0 |
| B129 | B209 | IBIL | Smoking | -0.082572874 | 0 |
| B129 | B221 | IBIL | Hei | 0.067318384 | 2.59E-13 |
| B129 | B174 | IBIL | Urea/Cr | -0.063507679 | 5.22E-12 |
| B129 | B176 | IBIL | Ucr | 0.057214812 | 5.15E-10 |
| B129 | B224 | IBIL | pmh | -0.053874612 | 4.88E-09 |

| | | | | | |
|---|---|---|---|---|---|
| B129 | B150 | IBIL | ALP | -0.053483127 | 6.30E-09 |
| B130 | B131 | GLB | AST | 0.102089095 | 0 |
| B130 | B133 | GLB | TBIL | -0.102203578 | 0 |
| B130 | B139 | GLB | ADA | 0.077513083 | 0 |
| B130 | B140 | GLB | r-GT | 0.106473556 | 0 |
| B130 | B146 | GLB | CysC | 0.093925074 | 0 |
| B130 | B153 | GLB | Glu | 0.140254837 | 0 |
| B130 | B155 | GLB | IgA | 0.10697984 | 0 |
| B130 | B157 | GLB | IgG | 0.092856584 | 0 |
| B130 | B188 | GLB | Ca | 0.114431464 | 0 |
| B130 | B189 | GLB | eGFB | -0.086292637 | 0 |
| B130 | B209 | GLB | Smoking | -0.119421375 | 0 |
| B130 | B220 | GLB | SBP | 0.137007578 | 0 |
| B130 | B221 | GLB | Hei | -0.142748976 | 0 |
| B130 | B150 | GLB | ALP | 0.072110054 | 4.66E-15 |
| B130 | B194 | GLB | Cl | -0.071977091 | 5.33E-15 |
| B130 | B204 | GLB | Drinking | -0.069548831 | 4.13E-14 |
| B130 | B170 | GLB | U-GLU | 0.067434873 | 2.35E-13 |
| B130 | B158 | GLB | IgM | 0.065111123 | 1.51E-12 |
| B130 | B192 | GLB | UA | 0.064563227 | 2.31E-12 |
| B130 | B161 | GLB | LEU | 0.057030845 | 5.85E-10 |
| B130 | B136 | GLB | 5'NT | 0.054550053 | 3.13E-09 |
| B130 | B218 | GLB | DBP | 0.052587325 | 1.12E-08 |
| B130 | B191 | GLB | Urea | 0.052380858 | 1.28E-08 |
| B130 | B177 | GLB | ACR | 0.049941042 | 5.85E-08 |
| B130 | B216 | GLB | BMI | -0.049299447 | 8.62E-08 |
| B130 | B222 | GLB | NS | -0.048735008 | 1.21E-07 |
| B130 | B174 | GLB | Urea/Cr | 0.048609457 | 1.30E-07 |
| B130 | B160 | GLB | PH | -0.048555592 | 1.34E-07 |
| B130 | B215 | GLB | BMI | 0.048364492 | 1.50E-07 |
| B130 | B178 | GLB | Ig-U | 0.048167638 | 1.69E-07 |
| B130 | B182 | GLB | TT | 0.047805577 | 2.09E-07 |
| B130 | B132 | GLB | Dbil | -0.047741825 | 2.17E-07 |
| B131 | B132 | AST | Dbil | 0.2919091 | 0 |
| B131 | B133 | AST | TBIL | 0.149206675 | 0 |
| B131 | B140 | AST | r-GT | 0.348564231 | 0 |
| B131 | B165 | AST | bilirubin | 0.223779562 | 0 |
| B131 | B192 | AST | UA | 0.072350186 | 3.77E-15 |
| B131 | B139 | AST | ADA | 0.065993605 | 7.50E-13 |
| B131 | B135 | AST | TBA | 0.064288285 | 2.86E-12 |
| B131 | B137 | AST | AFU | 0.06187508 | 1.80E-11 |
| B131 | B136 | AST | 5'NT | 0.058423798 | 2.21E-10 |
| B131 | B141 | AST | AST/ALT | -0.056980201 | 6.05E-10 |
| B131 | B215 | AST | BMI | 0.054348067 | 3.58E-09 |

| | | | | | |
|---|---|---|---|---|---|
| B132 | B133 | Dbil | TBIL | 0.779759999 | 0 |
| B132 | B140 | Dbil | r-GT | 0.12597885 | 0 |
| B132 | B165 | Dbil | bilirubin | 0.148546435 | 0 |
| B132 | B184 | Dbil | INR | 0.096828849 | 0 |
| B132 | B185 | Dbil | PT | 0.092662611 | 0 |
| B132 | B209 | Dbil | Smoking | -0.06818649 | 1.27E-13 |
| B132 | B221 | Dbil | Hei | 0.06088012 | 3.76E-11 |
| B132 | B189 | Dbil | eGFB | -0.059501141 | 1.02E-10 |
| B132 | B174 | Dbil | Urea/Cr | -0.056286775 | 9.74E-10 |
| B132 | B181 | Dbil | APTT | 0.054627412 | 2.97E-09 |
| B132 | B224 | Dbil | pmh | -0.053638956 | 5.69E-09 |
| B132 | B204 | Dbil | Drinking | 0.051280468 | 2.56E-08 |
| B133 | B209 | TBIL | Smoking | -0.08531537 | 0 |
| B133 | B163 | TBIL | SG | 0.07384683 | 8.88E-16 |
| B133 | B221 | TBIL | Hei | 0.071172547 | 1.04E-14 |
| B133 | B174 | TBIL | Urea/Cr | -0.066592236 | 4.65E-13 |
| B133 | B165 | TBIL | bilirubin | 0.063742577 | 4.36E-12 |
| B133 | B140 | TBIL | r-GT | 0.059590572 | 9.60E-11 |
| B133 | B224 | TBIL | pmh | -0.058312892 | 2.39E-10 |
| B133 | B176 | TBIL | Ucr | 0.055235976 | 1.98E-09 |
| B133 | B150 | TBIL | ALP | -0.048028292 | 1.83E-07 |
| B134 | B136 | CHE | 5'NT | 0.125452723 | 0 |
| B134 | B137 | CHE | AFU | 0.097672439 | 0 |
| B134 | B138 | CHE | PAB | 0.313210998 | 0 |
| B134 | B188 | CHE | Ca | 0.077861455 | 0 |
| B134 | B194 | CHE | Cl | -0.214421049 | 0 |
| B134 | B195 | CHE | Na | -0.118117354 | 0 |
| B134 | B197 | CHE | AG | 0.110449858 | 0 |
| B134 | B200 | CHE | OP | 0.096078236 | 0 |
| B134 | B150 | CHE | ALP | 0.072914716 | 2.22E-15 |
| B134 | B157 | CHE | IgG | -0.070322681 | 2.13E-14 |
| B134 | B215 | CHE | BMI | 0.069432181 | 4.53E-14 |
| B134 | B199 | CHE | Mg | -0.068350396 | 1.11E-13 |
| B134 | B139 | CHE | ADA | -0.065697086 | 9.49E-13 |
| B134 | B216 | CHE | BMI | 0.060116591 | 6.56E-11 |
| B134 | B141 | CHE | AST/ALT | -0.05777616 | 3.48E-10 |
| B134 | B155 | CHE | IgA | 0.056598163 | 7.87E-10 |
| B134 | B206 | CHE | Waist | 0.05619327 | 1.04E-09 |
| B134 | B193 | CHE | K | 0.055374471 | 1.81E-09 |
| B134 | B222 | CHE | NS | 0.053809738 | 5.09E-09 |
| B134 | B152 | CHE | ACA-Ig(G-A-M) | 0.052158333 | 1.47E-08 |
| B134 | B212 | CHE | IOP-L | 0.050404366 | 4.40E-08 |
| B134 | B198 | CHE | P | 0.048882316 | 1.11E-07 |
| B134 | B184 | CHE | INR | -0.04685353 | 3.62E-07 |

| | | | | | |
|---|---|---|---|---|---|
| B134 | B153 | CHE | Glu | 0.045594984 | 7.39E-07 |
| B135 | B137 | TBA | AFU | 0.133413143 | 0 |
| B135 | B139 | TBA | ADA | 0.075523914 | 2.22E-16 |
| B135 | B196 | TBA | TCO2 | 0.072782408 | 2.66E-15 |
| B135 | B198 | TBA | P | -0.068972742 | 6.66E-14 |
| B135 | B138 | TBA | PAB | -0.0604717 | 5.07E-11 |
| B135 | B150 | TBA | ALP | 0.059859099 | 7.91E-11 |
| B135 | B181 | TBA | APTT | 0.05681133 | 6.80E-10 |
| B135 | B200 | TBA | OP | 0.047021979 | 3.29E-07 |
| B135 | B197 | TBA | AG | -0.046537838 | 4.34E-07 |
| B135 | B199 | TBA | Mg | -0.046147107 | 5.42E-07 |
| B136 | B137 | 5'NT | AFU | 0.120122202 | 0 |
| B136 | B139 | 5'NT | ADA | 0.183702751 | 0 |
| B136 | B140 | 5'NT | r-GT | 0.189492724 | 0 |
| B136 | B150 | 5'NT | ALP | 0.295049958 | 0 |
| B136 | B186 | 5'NT | PFC | 0.112363583 | 0 |
| B136 | B194 | 5'NT | Cl | -0.121414941 | 0 |
| B136 | B195 | 5'NT | Na | -0.093533445 | 0 |
| B136 | B197 | 5'NT | AG | 0.07528829 | 4.44E-16 |
| B136 | B151 | 5'NT | Anti-CCP | 0.0568385 | 6.68E-10 |
| B136 | B193 | 5'NT | K | -0.05677711 | 6.96E-10 |
| B136 | B196 | 5'NT | TCO2 | -0.055731544 | 1.42E-09 |
| B136 | B153 | 5'NT | Glu | 0.055388062 | 1.79E-09 |
| B136 | B146 | 5'NT | CysC | 0.055007977 | 2.31E-09 |
| B136 | B157 | 5'NT | IgG | -0.049884745 | 6.05E-08 |
| B136 | B182 | 5'NT | TT | 0.049800079 | 6.37E-08 |
| B136 | B138 | 5'NT | PAB | -0.046453392 | 4.55E-07 |
| B136 | B176 | 5'NT | Ucr | 0.045660257 | 7.12E-07 |
| B136 | B170 | 5'NT | U-GLU | 0.045121179 | 9.61E-07 |
| B137 | B139 | AFU | ADA | 0.259208 | 0 |
| B137 | B150 | AFU | ALP | 0.118204812 | 0 |
| B137 | B194 | AFU | Cl | -0.109059676 | 0 |
| B137 | B196 | AFU | TCO2 | 0.079779426 | 0 |
| B137 | B199 | AFU | Mg | -0.107625741 | 0 |
| B137 | B200 | AFU | OP | 0.274833152 | 0 |
| B137 | B170 | AFU | U-GLU | 0.06062916 | 4.52E-11 |
| B137 | B204 | AFU | Drinking | 0.058225686 | 2.54E-10 |
| B137 | B140 | AFU | r-GT | 0.057116368 | 5.51E-10 |
| B137 | B153 | AFU | Glu | 0.055658507 | 1.49E-09 |
| B137 | B152 | AFU | ACA-Ig(G-A-M) | -0.048898976 | 1.10E-07 |
| B138 | B139 | PAB | ADA | -0.330954142 | 0 |
| B138 | B150 | PAB | ALP | -0.129373925 | 0 |
| B138 | B157 | PAB | IgG | -0.086745142 | 0 |
| B138 | B188 | PAB | Ca | 0.076837548 | 0 |

| | | | | | |
|---|---|---|---|---|---|
| B138 | B192 | PAB | UA | 0.090189344 | 0 |
| B138 | B194 | PAB | Cl | -0.087842202 | 0 |
| B138 | B197 | PAB | AG | 0.155706778 | 0 |
| B138 | B204 | PAB | Drinking | 0.068601289 | 9.06E-14 |
| B138 | B200 | PAB | OP | 0.06756558 | 2.12E-13 |
| B138 | B158 | PAB | IgM | -0.065143227 | 1.47E-12 |
| B138 | B181 | PAB | APTT | -0.064004162 | 3.57E-12 |
| B138 | B184 | PAB | INR | -0.063642543 | 4.71E-12 |
| B138 | B199 | PAB | Mg | 0.061965747 | 1.68E-11 |
| B138 | B195 | PAB | Na | 0.060735557 | 4.18E-11 |
| B138 | B196 | PAB | TCO2 | -0.060624179 | 4.53E-11 |
| B138 | B216 | PAB | BMI | 0.058962694 | 1.51E-10 |
| B138 | B222 | PAB | NS | 0.056616658 | 7.77E-10 |
| B138 | B185 | PAB | PT | -0.055984128 | 1.20E-09 |
| B138 | B221 | PAB | Hei | 0.053967319 | 4.59E-09 |
| B138 | B148 | PAB | Anti_TPO | -0.047222166 | 2.93E-07 |
| B139 | B150 | ADA | ALP | 0.219149164 | 0 |
| B139 | B155 | ADA | IgA | 0.087067794 | 0 |
| B139 | B182 | ADA | TT | 0.09025426 | 0 |
| B139 | B186 | ADA | PFC | 0.233212149 | 0 |
| B139 | B194 | ADA | Cl | -0.236119898 | 0 |
| B139 | B195 | ADA | Na | -0.161971685 | 0 |
| B139 | B199 | ADA | Mg | -0.152224563 | 0 |
| B139 | B200 | ADA | OP | 0.166402871 | 0 |
| B139 | B153 | ADA | Glu | 0.064719289 | 2.05E-12 |
| B139 | B197 | ADA | AG | 0.062796135 | 8.98E-12 |
| B139 | B140 | ADA | r-GT | 0.058974423 | 1.49E-10 |
| B139 | B212 | ADA | IOP-L | 0.05858885 | 1.97E-10 |
| B139 | B170 | ADA | U-GLU | 0.05512893 | 2.13E-09 |
| B139 | B157 | ADA | IgG | 0.046540767 | 4.33E-07 |
| B14 | B93 | G-17 | ALDH2-Glu504Lys | 0.078152312 | 0 |
| B14 | B92 | G-17 | ALDH2-Glu504Glu | -0.073164982 | 1.78E-15 |
| B14 | B81 | G-17 | TSGF | 0.069349715 | 4.86E-14 |
| B14 | B20 | G-17 | CD3+CD4+CD8+ | 0.065915704 | 7.98E-13 |
| B14 | B80 | G-17 | anti-HP IgG | 0.055219277 | 2.00E-09 |
| B14 | B138 | G-17 | PAB | -0.054891173 | 2.50E-09 |
| B14 | B150 | G-17 | ALP | 0.05217293 | 1.46E-08 |
| B14 | B23 | G-17 | CD3-CD16+CD56+(NK) | -0.049572104 | 7.31E-08 |
| B140 | B141 | r-GT | AST/ALT | -0.13284524 | 0 |
| B140 | B150 | r-GT | ALP | 0.083221662 | 0 |
| B140 | B153 | r-GT | Glu | 0.118641463 | 0 |
| B140 | B163 | r-GT | SG | 0.091717671 | 0 |
| B140 | B165 | r-GT | bilirubin | 0.158598418 | 0 |
| B140 | B170 | r-GT | U-GLU | 0.114717807 | 0 |

| | | | | | |
|---|---|---|---|---|---|
| B140 | B176 | r-GT | Ucr | 0.095602896 | 0 |
| B140 | B189 | r-GT | eGFB | 0.098150686 | 0 |
| B140 | B192 | r-GT | UA | 0.079262781 | 0 |
| B140 | B204 | r-GT | Drinking | 0.133214396 | 0 |
| B140 | B206 | r-GT | Waist | 0.123096261 | 0 |
| B140 | B210 | r-GT | Hip | 0.083332826 | 0 |
| B140 | B215 | r-GT | BMI | 0.106164571 | 0 |
| B140 | B216 | r-GT | BMI | 0.129096772 | 0 |
| B140 | B218 | r-GT | DBP | 0.094087423 | 0 |
| B140 | B221 | r-GT | Hei | 0.082797095 | 0 |
| B140 | B222 | r-GT | NS | 0.136083199 | 0 |
| B140 | B209 | r-GT | Smoking | 0.072512824 | 3.11E-15 |
| B140 | B172 | r-GT | KB | 0.065102988 | 1.52E-12 |
| B140 | B179 | r-GT | α1-MG | 0.062152566 | 1.46E-11 |
| B140 | B191 | r-GT | Urea | -0.053177863 | 7.68E-09 |
| B140 | B186 | r-GT | PFC | 0.046365428 | 4.79E-07 |
| B140 | B166 | r-GT | Pro | 0.046008361 | 5.86E-07 |
| B140 | B174 | r-GT | Urea/Cr | -0.045116866 | 9.63E-07 |
| B141 | B153 | AST/ALT | Glu | -0.108662443 | 0 |
| B141 | B163 | AST/ALT | SG | -0.108888174 | 0 |
| B141 | B170 | AST/ALT | U-GLU | -0.086535331 | 0 |
| B141 | B189 | AST/ALT | eGFB | -0.136937964 | 0 |
| B141 | B192 | AST/ALT | UA | -0.089405788 | 0 |
| B141 | B206 | AST/ALT | Waist | -0.154261034 | 0 |
| B141 | B210 | AST/ALT | Hip | -0.11163458 | 0 |
| B141 | B215 | AST/ALT | BMI | -0.167306221 | 0 |
| B141 | B216 | AST/ALT | BMI | -0.200008335 | 0 |
| B141 | B221 | AST/ALT | Hei | -0.128382674 | 0 |
| B141 | B222 | AST/ALT | NS | -0.157542093 | 0 |
| B141 | B218 | AST/ALT | DBP | -0.072881924 | 2.22E-15 |
| B141 | B220 | AST/ALT | SBP | 0.065201611 | 1.40E-12 |
| B141 | B176 | AST/ALT | Ucr | -0.052622017 | 1.10E-08 |
| B141 | B188 | AST/ALT | Ca | -0.050867977 | 3.31E-08 |
| B141 | B160 | AST/ALT | PH | 0.048446202 | 1.43E-07 |
| B141 | B224 | AST/ALT | pmh | 0.045315626 | 8.63E-07 |
| B142 | B143 | 25-(OH)D | β-CTx | -0.162459542 | 0 |
| B142 | B144 | 25-(OH)D | Total-PINP | -0.1066258 | 0 |
| B142 | B151 | 25-(OH)D | Anti-CCP | -0.108794785 | 0 |
| B142 | B173 | 25-(OH)D | NTT | 0.053591593 | 5.87E-09 |
| B142 | B176 | 25-(OH)D | Ucr | -0.052830888 | 9.60E-09 |
| B142 | B145 | 25-(OH)D | BGP | -0.052121802 | 1.51E-08 |
| B143 | B144 | β-CTx | Total-PINP | 0.830006212 | 0 |
| B143 | B145 | β-CTx | BGP | 0.606028461 | 0 |
| B143 | B150 | β-CTx | ALP | 0.077129894 | 0 |

| | | | | | |
|---|---|---|---|---|---|
| B143 | B151 | β-CTx | Anti-CCP | 0.083865835 | 0 |
| B143 | B167 | β-CTx | PC | 0.066050433 | 7.17E-13 |
| B143 | B168 | β-CTx | NMP22 | -0.064405684 | 2.61E-12 |
| B143 | B183 | β-CTx | PTA | -0.053781522 | 5.19E-09 |
| B143 | B176 | β-CTx | Ucr | 0.048652493 | 1.27E-07 |
| B144 | B145 | Total-PINP | BGP | 0.670825076 | 0 |
| B144 | B150 | Total-PINP | ALP | 0.0981478 | 0 |
| B144 | B151 | Total-PINP | Anti-CCP | 0.140267004 | 0 |
| B144 | B183 | Total-PINP | PTA | -0.076652745 | 0 |
| B144 | B167 | Total-PINP | PC | 0.069520285 | 4.22E-14 |
| B144 | B168 | Total-PINP | NMP22 | -0.062763537 | 9.21E-12 |
| B145 | B150 | BGP | ALP | 0.10807254 | 0 |
| B145 | B167 | BGP | PC | 0.083742913 | 0 |
| B145 | B183 | BGP | PTA | -0.117629279 | 0 |
| B145 | B182 | BGP | TT | 0.056722196 | 7.23E-10 |
| B145 | B203 | BGP | DH | 0.047105472 | 3.14E-07 |
| B146 | B182 | CysC | TT | 0.105068777 | 0 |
| B146 | B189 | CysC | eGFB | -0.160940935 | 0 |
| B146 | B190 | CysC | Cre | 0.123408848 | 0 |
| B146 | B191 | CysC | Urea | 0.070706355 | 1.55E-14 |
| B146 | B155 | CysC | IgA | 0.070225714 | 2.33E-14 |
| B146 | B174 | CysC | Urea/Cr | -0.063664422 | 4.63E-12 |
| B146 | B192 | CysC | UA | 0.062961692 | 7.92E-12 |
| B146 | B163 | CysC | SG | -0.050207711 | 4.97E-08 |
| B146 | B152 | CysC | ACA-Ig(G-A-M) | -0.04955728 | 7.38E-08 |
| B146 | B200 | CysC | OP | 0.047764802 | 2.14E-07 |
| B146 | B198 | CysC | P | -0.046794083 | 3.75E-07 |
| B148 | B149 | Anti_TPO | TGAb | 0.392114459 | 0 |
| B148 | B222 | Anti_TPO | NS | -0.061901786 | 1.76E-11 |
| B148 | B181 | Anti_TPO | APTT | 0.056091393 | 1.11E-09 |
| B148 | B221 | Anti_TPO | Hei | -0.053556313 | 6.01E-09 |
| B148 | B216 | Anti_TPO | BMI | -0.052076628 | 1.55E-08 |
| B148 | B176 | Anti_TPO | Ucr | -0.048352617 | 1.52E-07 |
| B149 | B222 | TGAb | NS | -0.053118678 | 7.98E-09 |
| B149 | B221 | TGAb | Hei | -0.051338743 | 2.47E-08 |
| B15 | B101 | CD3-CD19+(B cell) | anti-Sc70 | 0.364725366 | 0 |
| B15 | B103 | CD3-CD19+(B cell) | anti-SSA/Ro | 0.078369868 | 0 |
| B15 | B104 | CD3-CD19+(B cell) | anti-SSB | 0.364725304 | 0 |
| B15 | B105 | CD3-CD19+(B cell) | anti-U1RNP | -0.102732383 | 0 |
| B15 | B107 | CD3-CD19+(B cell) | AnuA | 0.366829111 | 0 |
| B15 | B119 | CD3-CD19+(B cell) | Hcv-rna | 0.119864426 | 0 |
| B15 | B120 | CD3-CD19+(B cell) | HCV-cAg | 0.128562754 | 0 |
| B15 | B138 | CD3-CD19+(B cell) | PAB | -0.08742482 | 0 |
| B15 | B155 | CD3-CD19+(B cell) | IgA | 0.11782687 | 0 |

| | | | | | |
|---|---|---|---|---|---|
| B15 | B157 | CD3-CD19+(B cell) | IgG | 0.181660918 | 0 |
| B15 | B158 | CD3-CD19+(B cell) | IgM | 0.113373509 | 0 |
| B15 | B16 | CD3-CD19+(B cell) | CD3+T | -0.349822511 | 0 |
| B15 | B17 | CD3-CD19+(B cell) | CD3+CD16+CD56+NK | -0.162919801 | 0 |
| B15 | B181 | CD3-CD19+(B cell) | APTT | 0.141318463 | 0 |
| B15 | B183 | CD3-CD19+(B cell) | PTA | 0.132189333 | 0 |
| B15 | B184 | CD3-CD19+(B cell) | INR | 0.115478801 | 0 |
| B15 | B19 | CD3-CD19+(B cell) | CD3+CD4+/CD3+CD8+ | 0.143914863 | 0 |
| B15 | B20 | CD3-CD19+(B cell) | CD3+CD4+CD8+ | -0.16236123 | 0 |
| B15 | B200 | CD3-CD19+(B cell) | OP | -0.107138405 | 0 |
| B15 | B21 | CD3-CD19+(B cell) | CD3+CD4-CD8- | -0.172248326 | 0 |
| B15 | B22 | CD3-CD19+(B cell) | CD3+CD8+(T) | -0.29538826 | 0 |
| B15 | B79 | CD3-CD19+(B cell) | hb V -dna | 0.140413696 | 0 |
| B15 | B96 | CD3-CD19+(B cell) | dsDNA (IIF) | 0.125350753 | 0 |
| B15 | B98 | CD3-CD19+(B cell) | anti-Jo-1 | 0.364725276 | 0 |
| B15 | B194 | CD3-CD19+(B cell) | Cl | 0.070840085 | 1.38E-14 |
| B15 | B151 | CD3-CD19+(B cell) | Anti-CCP | -0.070351524 | 2.09E-14 |
| B15 | B90 | CD3-CD19+(B cell) | CYFRA 21-1 | -0.069675854 | 3.69E-14 |
| B15 | B57 | CD3-CD19+(B cell) | RBCAI | -0.06839042 | 1.08E-13 |
| B15 | B106 | CD3-CD19+(B cell) | anti-P | 0.067893497 | 1.62E-13 |
| B15 | B102 | CD3-CD19+(B cell) | anti-Sm | 0.067885497 | 1.63E-13 |
| B15 | B136 | CD3-CD19+(B cell) | 5'NT | -0.0670597 | 3.19E-13 |
| B15 | B94 | CD3-CD19+(B cell) | ALDH2-Lys504Lys | 0.066075702 | 7.03E-13 |
| B15 | B60 | CD3-CD19+(B cell) | WBRVI | -0.064078324 | 3.37E-12 |
| B15 | B31 | CD3-CD19+(B cell) | FSH | -0.063543004 | 5.08E-12 |
| B15 | B95 | CD3-CD19+(B cell) | anti-cenpb | 0.061040919 | 3.34E-11 |
| B15 | B59 | CD3-CD19+(B cell) | WBRV | -0.059444159 | 1.07E-10 |
| B15 | B65 | CD3-CD19+(B cell) | WBRSR(mPas_1) | -0.059387507 | 1.11E-10 |
| B15 | B212 | CD3-CD19+(B cell) | IOP-L | -0.057154511 | 5.37E-10 |
| B15 | B122 | CD3-CD19+(B cell) | C4 | -0.056472209 | 8.58E-10 |
| B15 | B185 | CD3-CD19+(B cell) | PT | 0.0551429 | 2.11E-09 |
| B15 | B93 | CD3-CD19+(B cell) | ALDH2-Glu504Lys | -0.053655378 | 5.63E-09 |
| B15 | B115 | CD3-CD19+(B cell) | TSH TT3 TT4(TT3) | 0.051206779 | 2.68E-08 |
| B15 | B139 | CD3-CD19+(B cell) | ADA | -0.048830774 | 1.14E-07 |
| B15 | B29 | CD3-CD19+(B cell) | E2 | 0.048602482 | 1.31E-07 |
| B15 | B18 | CD3-CD19+(B cell) | CD3+CD4+helper(inducer)T | -0.048359918 | 1.51E-07 |
| B15 | B186 | CD3-CD19+(B cell) | PFC | -0.048299975 | 1.56E-07 |
| B15 | B124 | CD3-CD19+(B cell) | ACTH | -0.04772244 | 2.19E-07 |
| B15 | B54 | CD3-CD19+(B cell) | ESR | 0.046293491 | 4.99E-07 |
| B150 | B153 | ALP | Glu | 0.086222518 | 0 |
| B150 | B170 | ALP | U-GLU | 0.088043359 | 0 |
| B150 | B194 | ALP | Cl | -0.107517107 | 0 |
| B150 | B200 | ALP | OP | 0.104668445 | 0 |
| B150 | B186 | ALP | PFC | 0.067007756 | 3.33E-13 |

| | | | | | |
|---|---|---|---|---|---|
| B150 | B195 | ALP | Na | -0.06334839 | 5.90E-12 |
| B150 | B189 | ALP | eGFB | 0.055251277 | 1.96E-09 |
| B150 | B199 | ALP | Mg | -0.050215676 | 4.94E-08 |
| B150 | B174 | ALP | Urea/Cr | 0.047516478 | 2.47E-07 |
| B150 | B190 | ALP | Cre | -0.04693737 | 3.45E-07 |
| B151 | B157 | Anti-CCP | IgG | 0.081069552 | 0 |
| B151 | B183 | Anti-CCP | PTA | -0.168138053 | 0 |
| B151 | B186 | Anti-CCP | PFC | 0.082836286 | 0 |
| B151 | B152 | Anti-CCP | ACA-Ig(G-A-M) | 0.073986997 | 8.88E-16 |
| B151 | B182 | Anti-CCP | TT | -0.060587963 | 4.65E-11 |
| B151 | B155 | Anti-CCP | IgA | 0.059781288 | 8.37E-11 |
| B151 | B194 | Anti-CCP | Cl | -0.05417684 | 4.00E-09 |
| B151 | B156 | Anti-CCP | IgE | 0.053201996 | 7.56E-09 |
| B151 | B158 | Anti-CCP | IgM | 0.048588034 | 1.32E-07 |
| B151 | B168 | Anti-CCP | NMP22 | -0.047072045 | 3.20E-07 |
| B151 | B179 | Anti-CCP | α1-MG | 0.045126717 | 9.58E-07 |
| B152 | B200 | ACA-Ig(G-A-M) | OP | -0.048322506 | 1.54E-07 |
| B153 | B160 | Glu | PH | -0.140239199 | 0 |
| B153 | B163 | Glu | SG | 0.333646115 | 0 |
| B153 | B166 | Glu | Pro | 0.104487774 | 0 |
| B153 | B170 | Glu | U-GLU | 0.640971164 | 0 |
| B153 | B172 | Glu | KB | 0.201293372 | 0 |
| B153 | B174 | Glu | Urea/Cr | 0.172239274 | 0 |
| B153 | B188 | Glu | Ca | 0.084574579 | 0 |
| B153 | B189 | Glu | eGFB | 0.129067827 | 0 |
| B153 | B190 | Glu | Cre | -0.093903693 | 0 |
| B153 | B192 | Glu | UA | -0.130506112 | 0 |
| B153 | B194 | Glu | Cl | -0.076018155 | 0 |
| B153 | B195 | Glu | Na | -0.092657492 | 0 |
| B153 | B206 | Glu | Waist | 0.084258837 | 0 |
| B153 | B218 | Glu | DBP | 0.081509543 | 0 |
| B153 | B222 | Glu | NS | 0.077359746 | 0 |
| B153 | B216 | Glu | BMI | 0.073923182 | 8.88E-16 |
| B153 | B209 | Glu | Smoking | 0.066005309 | 7.43E-13 |
| B153 | B204 | Glu | Drinking | 0.064682439 | 2.11E-12 |
| B153 | B191 | Glu | Urea | 0.061801296 | 1.90E-11 |
| B153 | B215 | Glu | BMI | 0.058528481 | 2.05E-10 |
| B153 | B179 | Glu | α1-MG | 0.052869654 | 9.36E-09 |
| B153 | B200 | Glu | OP | 0.049741185 | 6.60E-08 |
| B153 | B219 | Glu | OT | -0.049455532 | 7.85E-08 |
| B153 | B221 | Glu | Hei | 0.049205663 | 9.12E-08 |
| B153 | B225 | Glu | hPBG | 0.048174817 | 1.68E-07 |
| B154 | B157 | RF | IgG | 0.082065732 | 0 |
| B155 | B157 | IgA | IgG | 0.188006599 | 0 |

| | | | | | |
|---|---|---|---|---|---|
| B155 | B182 | IgA | TT | 0.084108475 | 0 |
| B155 | B194 | IgA | Cl | -0.084358584 | 0 |
| B155 | B156 | IgA | IgE | 0.071273225 | 9.55E-15 |
| B155 | B195 | IgA | Na | -0.068368276 | 1.10E-13 |
| B155 | B178 | IgA | Ig-U | 0.056733773 | 7.17E-10 |
| B155 | B186 | IgA | PFC | 0.04868215 | 1.25E-07 |
| B155 | B180 | IgA | Tf | 0.047242642 | 2.90E-07 |
| B155 | B170 | IgA | U-GLU | 0.046623004 | 4.13E-07 |
| B156 | B158 | IgE | IgM | -0.059891125 | 7.73E-11 |
| B157 | B158 | IgG | IgM | 0.098018984 | 0 |
| B157 | B200 | IgG | OP | -0.128973452 | 0 |
| B157 | B184 | IgG | INR | 0.049714243 | 6.71E-08 |
| B157 | B199 | IgG | Mg | 0.047843578 | 2.04E-07 |
| B159 | B186 | β2-MG | PFC | 0.052221081 | 1.42E-08 |
| B159 | B181 | β2-MG | APTT | -0.045325652 | 8.58E-07 |
| B16 | B101 | CD3+T | anti-Sc70 | -0.121056924 | 0 |
| B16 | B104 | CD3+T | anti-SSB | -0.121056889 | 0 |
| B16 | B107 | CD3+T | AnuA | -0.130646618 | 0 |
| B16 | B108 | CD3+T | Anti-Cyclin | 0.086971045 | 0 |
| B16 | B121 | CD3+T | C3 | 0.232797363 | 0 |
| B16 | B122 | CD3+T | C4 | 0.223536338 | 0 |
| B16 | B152 | CD3+T | ACA-Ig(G-A-M) | -0.086388218 | 0 |
| B16 | B155 | CD3+T | IgA | -0.08634152 | 0 |
| B16 | B157 | CD3+T | IgG | -0.107651359 | 0 |
| B16 | B17 | CD3+T | CD3+CD16+CD56+NK | 0.202889622 | 0 |
| B16 | B18 | CD3+T | CD3+CD4+helper(inducer)T | 0.392382531 | 0 |
| B16 | B19 | CD3+T | CD3+CD4+/CD3+CD8+ | -0.21752396 | 0 |
| B16 | B193 | CD3+T | K | -0.077229238 | 0 |
| B16 | B20 | CD3+T | CD3+CD4+CD8+ | 0.200174466 | 0 |
| B16 | B21 | CD3+T | CD3+CD4-CD8- | 0.320225356 | 0 |
| B16 | B22 | CD3+T | CD3+CD8+(T) | 0.573832381 | 0 |
| B16 | B23 | CD3+T | CD3-CD16+CD56+(NK) | -0.844743191 | 0 |
| B16 | B95 | CD3+T | anti-cenpb | -0.092187338 | 0 |
| B16 | B98 | CD3+T | anti-Jo-1 | -0.121056872 | 0 |
| B16 | B99 | CD3+T | anti-PM-Scl | 0.076350756 | 0 |
| B16 | B105 | CD3+T | anti-U1RNP | 0.074761967 | 4.44E-16 |
| B16 | B181 | CD3+T | APTT | -0.072450739 | 3.55E-15 |
| B16 | B79 | CD3+T | hb V -dna | -0.070922453 | 1.29E-14 |
| B16 | B74 | CD3+T | ApoA1 | -0.06854504 | 9.48E-14 |
| B16 | B124 | CD3+T | ACTH | 0.066998221 | 3.35E-13 |
| B16 | B31 | CD3+T | FSH | 0.061877937 | 1.79E-11 |
| B16 | B200 | CD3+T | OP | 0.059190576 | 1.28E-10 |
| B16 | B212 | CD3+T | IOP-L | 0.051394329 | 2.39E-08 |
| B16 | B184 | CD3+T | INR | -0.050790733 | 3.47E-08 |

| | | | | | |
|---|---|---|---|---|---|
| B16 | B29 | CD3+T | E2 | -0.048486981 | 1.40E-07 |
| B160 | B163 | PH | SG | -0.132155739 | 0 |
| B160 | B170 | PH | U-GLU | -0.124360712 | 0 |
| B160 | B191 | PH | Urea | -0.119409062 | 0 |
| B160 | B192 | PH | UA | -0.090003147 | 0 |
| B160 | B188 | PH | Ca | -0.067424428 | 2.38E-13 |
| B160 | B174 | PH | Urea/Cr | -0.062897824 | 8.32E-12 |
| B160 | B172 | PH | KB | -0.058652864 | 1.88E-10 |
| B160 | B206 | PH | Waist | -0.058403517 | 2.24E-10 |
| B161 | B173 | LEU | NTT | 0.199866777 | 0 |
| B161 | B216 | LEU | BMI | -0.089633497 | 0 |
| B161 | B221 | LEU | Hei | -0.154354806 | 0 |
| B161 | B222 | LEU | NS | -0.115783754 | 0 |
| B161 | B174 | LEU | Urea/Cr | 0.074197503 | 8.88E-16 |
| B161 | B209 | LEU | Smoking | -0.067723103 | 1.86E-13 |
| B161 | B204 | LEU | Drinking | -0.06408317 | 3.35E-12 |
| B161 | B163 | LEU | SG | -0.061504113 | 2.37E-11 |
| B161 | B189 | LEU | eGFB | -0.046003161 | 5.87E-07 |
| B163 | B166 | SG | Pro | 0.303321381 | 0 |
| B163 | B170 | SG | U-GLU | 0.389458195 | 0 |
| B163 | B172 | SG | KB | 0.201680095 | 0 |
| B163 | B174 | SG | Urea/Cr | 0.127218059 | 0 |
| B163 | B176 | SG | Ucr | 0.265943159 | 0 |
| B163 | B189 | SG | eGFB | 0.135366891 | 0 |
| B163 | B191 | SG | Urea | 0.182995106 | 0 |
| B163 | B204 | SG | Drinking | 0.125087882 | 0 |
| B163 | B206 | SG | Waist | 0.150292224 | 0 |
| B163 | B209 | SG | Smoking | 0.158015157 | 0 |
| B163 | B210 | SG | Hip | 0.100734538 | 0 |
| B163 | B215 | SG | BMI | 0.097317456 | 0 |
| B163 | B216 | SG | BMI | 0.220839347 | 0 |
| B163 | B220 | SG | SBP | -0.106016069 | 0 |
| B163 | B221 | SG | Hei | 0.246427884 | 0 |
| B163 | B222 | SG | NS | 0.254602945 | 0 |
| B163 | B179 | SG | α1-MG | 0.075084982 | 4.44E-16 |
| B163 | B219 | SG | OT | -0.068119746 | 1.35E-13 |
| B163 | B173 | SG | NTT | -0.063540094 | 5.10E-12 |
| B163 | B218 | SG | DBP | 0.057447711 | 4.38E-10 |
| B163 | B201 | SG | e-habits | -0.05215532 | 1.48E-08 |
| B163 | B200 | SG | OP | 0.046728348 | 3.89E-07 |
| B165 | B176 | bilirubin | Ucr | 0.060276704 | 5.84E-11 |
| B165 | B179 | bilirubin | α1-MG | 0.05987408 | 7.83E-11 |
| B166 | B170 | Pro | U-GLU | 0.179955831 | 0 |
| B166 | B172 | Pro | KB | 0.135624847 | 0 |

| | | | | | |
|---|---|---|---|---|---|
| B166 | B176 | Pro | Ucr | 0.135112922 | 0 |
| B166 | B190 | Pro | Cre | 0.097176899 | 0 |
| B166 | B191 | Pro | Urea | 0.103857261 | 0 |
| B166 | B209 | Pro | Smoking | 0.078659842 | 0 |
| B166 | B216 | Pro | BMI | 0.078991785 | 0 |
| B166 | B221 | Pro | Hei | 0.088779781 | 0 |
| B166 | B222 | Pro | NS | 0.113190445 | 0 |
| B166 | B206 | Pro | Waist | 0.072724461 | 2.66E-15 |
| B166 | B218 | Pro | DBP | 0.057365523 | 4.64E-10 |
| B169 | B171 | U-CRY | EC | -0.120950806 | 0 |
| B169 | B204 | U-CRY | Drinking | 0.058364318 | 2.30E-10 |
| B17 | B134 | CD3+CD16+CD56+NK | CHE | -0.081946334 | 0 |
| B17 | B164 | CD3+CD16+CD56+NK | COM | -0.075856114 | 0 |
| B17 | B18 | CD3+CD16+CD56+NK | CD3+CD4+helper(inducer)T | -0.129103371 | 0 |
| B17 | B183 | CD3+CD16+CD56+NK | PTA | 0.140396682 | 0 |
| B17 | B19 | CD3+CD16+CD56+NK | CD3+CD4+/CD3+CD8+ | -0.160473332 | 0 |
| B17 | B20 | CD3+CD16+CD56+NK | CD3+CD4+CD8+ | 0.267605298 | 0 |
| B17 | B21 | CD3+CD16+CD56+NK | CD3+CD4-CD8- | 0.470964736 | 0 |
| B17 | B22 | CD3+CD16+CD56+NK | CD3+CD8+(T) | 0.163610009 | 0 |
| B17 | B23 | CD3+CD16+CD56+NK | CD3-CD16+CD56+(NK) | -0.096667457 | 0 |
| B17 | B99 | CD3+CD16+CD56+NK | anti-PM-Scl | -0.065419678 | 1.18E-12 |
| B17 | B95 | CD3+CD16+CD56+NK | anti-cenpb | -0.063250528 | 6.36E-12 |
| B17 | B107 | CD3+CD16+CD56+NK | AnuA | 0.058628164 | 1.91E-10 |
| B17 | B101 | CD3+CD16+CD56+NK | anti-Sc70 | 0.058304011 | 2.40E-10 |
| B17 | B104 | CD3+CD16+CD56+NK | anti-SSB | 0.058304007 | 2.40E-10 |
| B17 | B98 | CD3+CD16+CD56+NK | anti-Jo-1 | 0.058304011 | 2.40E-10 |
| B17 | B115 | CD3+CD16+CD56+NK | TSH TT3 TT4(TT3) | -0.046853523 | 3.62E-07 |
| B17 | B155 | CD3+CD16+CD56+NK | IgA | -0.045452664 | 8.00E-07 |
| B170 | B172 | U-GLU | KB | 0.210319242 | 0 |
| B170 | B174 | U-GLU | Urea/Cr | 0.116547319 | 0 |
| B170 | B189 | U-GLU | eGFB | 0.13451806 | 0 |
| B170 | B192 | U-GLU | UA | -0.138146692 | 0 |
| B170 | B204 | U-GLU | Drinking | 0.08905184 | 0 |
| B170 | B206 | U-GLU | Waist | 0.083020258 | 0 |
| B170 | B209 | U-GLU | Smoking | 0.11807666 | 0 |
| B170 | B216 | U-GLU | BMI | 0.094470728 | 0 |
| B170 | B221 | U-GLU | Hei | 0.114378847 | 0 |
| B170 | B222 | U-GLU | NS | 0.122354753 | 0 |
| B170 | B194 | U-GLU | Cl | -0.071757384 | 6.22E-15 |
| B170 | B218 | U-GLU | DBP | 0.066176283 | 6.48E-13 |
| B170 | B195 | U-GLU | Na | -0.064524204 | 2.38E-12 |
| B170 | B191 | U-GLU | Urea | 0.063293842 | 6.15E-12 |
| B170 | B176 | U-GLU | Ucr | 0.061080558 | 3.24E-11 |
| B170 | B219 | U-GLU | OT | -0.06104322 | 3.33E-11 |

| | | | | | |
|---|---|---|---|---|---|
| B170 | B179 | U-GLU | α1-MG | 0.060395714 | 5.36E-11 |
| B170 | B200 | U-GLU | OP | 0.060131358 | 6.49E-11 |
| B170 | B201 | U-GLU | e-habits | -0.05239895 | 1.27E-08 |
| B170 | B190 | U-GLU | Cre | -0.049058728 | 9.96E-08 |
| B172 | B189 | KB | eGFB | 0.093614076 | 0 |
| B172 | B176 | KB | Ucr | 0.074149032 | 8.88E-16 |
| B172 | B174 | KB | Urea/Cr | 0.067991362 | 1.50E-13 |
| B172 | B221 | KB | Hei | 0.04621509 | 5.21E-07 |
| B173 | B216 | NTT | BMI | -0.076610987 | 0 |
| B173 | B221 | NTT | Hei | -0.119898592 | 0 |
| B173 | B222 | NTT | NS | -0.085497027 | 0 |
| B173 | B174 | NTT | Urea/Cr | 0.07110658 | 1.11E-14 |
| B173 | B209 | NTT | Smoking | -0.045541253 | 7.61E-07 |
| B174 | B189 | Urea/Cr | eGFB | 0.248752006 | 0 |
| B174 | B190 | Urea/Cr | Cre | -0.385194229 | 0 |
| B174 | B191 | Urea/Cr | Urea | 0.55512888 | 0 |
| B174 | B192 | Urea/Cr | UA | -0.22358618 | 0 |
| B174 | B200 | Urea/Cr | OP | 0.101306641 | 0 |
| B174 | B204 | Urea/Cr | Drinking | -0.098143515 | 0 |
| B174 | B206 | Urea/Cr | Waist | -0.112561934 | 0 |
| B174 | B209 | Urea/Cr | Smoking | -0.122874024 | 0 |
| B174 | B210 | Urea/Cr | Hip | -0.107627107 | 0 |
| B174 | B216 | Urea/Cr | BMI | -0.191321997 | 0 |
| B174 | B221 | Urea/Cr | Hei | -0.271923902 | 0 |
| B174 | B222 | Urea/Cr | NS | -0.247527922 | 0 |
| B174 | B176 | Urea/Cr | Ucr | -0.069218551 | 5.42E-14 |
| B174 | B218 | Urea/Cr | DBP | -0.057036721 | 5.82E-10 |
| B175 | B177 | ALB-U | ACR | 0.908810884 | 0 |
| B175 | B178 | ALB-U | Ig-U | 0.103266206 | 0 |
| B175 | B180 | ALB-U | Tf | 0.078293447 | 0 |
| B175 | B191 | ALB-U | Urea | 0.09415116 | 0 |
| B175 | B190 | ALB-U | Cre | 0.063517338 | 5.18E-12 |
| B175 | B176 | ALB-U | Ucr | 0.057979175 | 3.02E-10 |
| B175 | B189 | ALB-U | eGFB | -0.053580747 | 5.91E-09 |
| B176 | B179 | Ucr | α1-MG | 0.171220983 | 0 |
| B176 | B216 | Ucr | BMI | 0.120432541 | 0 |
| B176 | B221 | Ucr | Hei | 0.143257973 | 0 |
| B176 | B222 | Ucr | NS | 0.131589345 | 0 |
| B176 | B206 | Ucr | Waist | 0.071764604 | 6.22E-15 |
| B176 | B209 | Ucr | Smoking | 0.064310872 | 2.81E-12 |
| B176 | B183 | Ucr | PTA | 0.053252037 | 7.32E-09 |
| B176 | B220 | Ucr | SBP | -0.052606567 | 1.11E-08 |
| B176 | B210 | Ucr | Hip | 0.052232885 | 1.41E-08 |
| B176 | B190 | Ucr | Cre | 0.050311354 | 4.66E-08 |

| | | | | | |
|---|---|---|---|---|---|
| B177 | B178 | ACR | Ig-U | 0.234648217 | 0 |
| B177 | B180 | ACR | Tf | 0.192755825 | 0 |
| B177 | B191 | ACR | Urea | 0.111725163 | 0 |
| B177 | B190 | ACR | Cre | 0.072210587 | 4.22E-15 |
| B177 | B179 | ACR | α1-MG | 0.071085783 | 1.11E-14 |
| B177 | B189 | ACR | eGFB | -0.061182539 | 3.01E-11 |
| B178 | B179 | Ig-U | α1-MG | 0.555096349 | 0 |
| B178 | B180 | Ig-U | Tf | 0.903851033 | 0 |
| B179 | B180 | α1-MG | Tf | 0.414259968 | 0 |
| B179 | B194 | α1-MG | Cl | -0.048845335 | 1.13E-07 |
| B179 | B195 | α1-MG | Na | -0.048602002 | 1.31E-07 |
| B18 | B101 | CD3+CD4+helper(inducer)T | anti-Sc70 | -0.197470932 | 0 |
| B18 | B104 | CD3+CD4+helper(inducer)T | anti-SSB | -0.197470885 | 0 |
| B18 | B107 | CD3+CD4+helper(inducer)T | AnuA | -0.201640354 | 0 |
| B18 | B121 | CD3+CD4+helper(inducer)T | C3 | 0.147428078 | 0 |
| B18 | B122 | CD3+CD4+helper(inducer)T | C4 | 0.122256862 | 0 |
| B18 | B136 | CD3+CD4+helper(inducer)T | 5'NT | 0.151891212 | 0 |
| B18 | B151 | CD3+CD4+helper(inducer)T | Anti-CCP | 0.102288522 | 0 |
| B18 | B152 | CD3+CD4+helper(inducer)T | ACA-Ig(G-A-M) | -0.127254575 | 0 |
| B18 | B183 | CD3+CD4+helper(inducer)T | PTA | -0.088345938 | 0 |
| B18 | B186 | CD3+CD4+helper(inducer)T | PFC | 0.111643493 | 0 |
| B18 | B19 | CD3+CD4+helper(inducer)T | CD3+CD4+/CD3+CD8+ | 0.694187689 | 0 |
| B18 | B20 | CD3+CD4+helper(inducer)T | CD3+CD4+CD8+ | 0.0848274 | 0 |
| B18 | B22 | CD3+CD4+helper(inducer)T | CD3+CD8+(T) | -0.386989251 | 0 |
| B18 | B23 | CD3+CD4+helper(inducer)T | CD3-CD16+CD56+(NK) | -0.425732317 | 0 |
| B18 | B98 | CD3+CD4+helper(inducer)T | anti-Jo-1 | -0.197470859 | 0 |
| B18 | B157 | CD3+CD4+helper(inducer)T | IgG | -0.071749602 | 6.22E-15 |
| B18 | B181 | CD3+CD4+helper(inducer)T | APTT | -0.071780357 | 6.22E-15 |
| B18 | B184 | CD3+CD4+helper(inducer)T | INR | -0.06602756 | 7.30E-13 |
| B18 | B139 | CD3+CD4+helper(inducer)T | ADA | 0.06484659 | 1.85E-12 |
| B18 | B90 | CD3+CD4+helper(inducer)T | CYFRA 21-1 | 0.063332835 | 5.97E-12 |
| B18 | B193 | CD3+CD4+helper(inducer)T | K | -0.063320408 | 6.03E-12 |
| B18 | B195 | CD3+CD4+helper(inducer)T | Na | -0.062598832 | 1.04E-11 |
| B18 | B150 | CD3+CD4+helper(inducer)T | ALP | 0.059568421 | 9.76E-11 |
| B18 | B97 | CD3+CD4+helper(inducer)T | dsDNA | -0.058544256 | 2.03E-10 |
| B18 | B138 | CD3+CD4+helper(inducer)T | PAB | 0.055197958 | 2.03E-09 |
| B18 | B79 | CD3+CD4+helper(inducer)T | hb V -dna | -0.05506369 | 2.22E-09 |
| B18 | B105 | CD3+CD4+helper(inducer)T | anti-U1RNP | -0.052166115 | 1.47E-08 |
| B18 | B212 | CD3+CD4+helper(inducer)T | IOP-L | 0.051449045 | 2.30E-08 |
| B18 | B198 | CD3+CD4+helper(inducer)T | P | 0.051251653 | 2.61E-08 |
| B18 | B99 | CD3+CD4+helper(inducer)T | anti-PM-Scl | 0.049236401 | 8.95E-08 |
| B18 | B211 | CD3+CD4+helper(inducer)T | IOP-R | 0.047825521 | 2.06E-07 |
| B18 | B81 | CD3+CD4+helper(inducer)T | TSGF | -0.046445918 | 4.57E-07 |
| B181 | B184 | APTT | INR | 0.436917944 | 0 |

| | | | | | |
|---|---|---|---|---|---|
| B181 | B185 | APTT | PT | 0.428607543 | 0 |
| B181 | B182 | APTT | TT | 0.074023973 | 8.88E-16 |
| B181 | B200 | APTT | OP | -0.07346535 | 1.33E-15 |
| B181 | B194 | APTT | Cl | 0.06875106 | 7.99E-14 |
| B182 | B183 | TT | PTA | -0.276339651 | 0 |
| B182 | B185 | TT | PT | 0.100800231 | 0 |
| B182 | B194 | TT | Cl | -0.097638303 | 0 |
| B182 | B197 | TT | AG | 0.101209687 | 0 |
| B182 | B186 | TT | PFC | -0.073903993 | 8.88E-16 |
| B182 | B196 | TT | TCO2 | -0.047225884 | 2.92E-07 |
| B183 | B186 | PTA | PFC | -0.085042357 | 0 |
| B183 | B194 | PTA | Cl | 0.092475005 | 0 |
| B183 | B197 | PTA | AG | -0.081439247 | 0 |
| B183 | B184 | PTA | INR | 0.055354143 | 1.83E-09 |
| B183 | B200 | PTA | OP | -0.054160204 | 4.05E-09 |
| B183 | B193 | PTA | K | -0.049972583 | 5.74E-08 |
| B184 | B185 | INR | PT | 0.973513942 | 0 |
| B184 | B186 | INR | PFC | -0.09220699 | 0 |
| B184 | B200 | INR | OP | -0.070742593 | 1.51E-14 |
| B184 | B194 | INR | Cl | 0.061223508 | 2.92E-11 |
| B186 | B194 | PFC | Cl | -0.154511136 | 0 |
| B186 | B195 | PFC | Na | -0.092159925 | 0 |
| B188 | B192 | Ca | UA | 0.114766602 | 0 |
| B188 | B218 | Ca | DBP | 0.092044258 | 0 |
| B188 | B194 | Ca | Cl | -0.070090217 | 2.62E-14 |
| B188 | B193 | Ca | K | 0.064848587 | 1.85E-12 |
| B188 | B198 | Ca | P | 0.050726133 | 3.61E-08 |
| B188 | B197 | Ca | AG | 0.046422952 | 4.63E-07 |
| B189 | B190 | eGFB | Cre | -0.600987795 | 0 |
| B189 | B191 | eGFB | Urea | -0.359454859 | 0 |
| B189 | B192 | eGFB | UA | -0.179633416 | 0 |
| B189 | B215 | eGFB | BMI | 0.081141672 | 0 |
| B189 | B216 | eGFB | BMI | 0.111549753 | 0 |
| B189 | B218 | eGFB | DBP | 0.113710447 | 0 |
| B189 | B220 | eGFB | SBP | -0.204870922 | 0 |
| B189 | B221 | eGFB | Hei | 0.081997218 | 0 |
| B189 | B224 | eGFB | pmh | 0.086356055 | 0 |
| B189 | B229 | eGFB | slit-lamp | -0.107611735 | 0 |
| B189 | B211 | eGFB | IOP-R | 0.071277508 | 9.55E-15 |
| B189 | B219 | eGFB | OT | -0.069751418 | 3.46E-14 |
| B189 | B201 | eGFB | e-habits | -0.064518827 | 2.39E-12 |
| B189 | B212 | eGFB | IOP-L | 0.060898353 | 3.71E-11 |
| B189 | B210 | eGFB | Hip | 0.060789971 | 4.01E-11 |
| B189 | B204 | eGFB | Drinking | 0.055105482 | 2.16E-09 |

| | | | | | |
|---|---|---|---|---|---|
| B189 | B194 | eGFB | Cl | -0.054325464 | 3.63E-09 |
| B189 | B209 | eGFB | Smoking | 0.048259435 | 1.60E-07 |
| B189 | B203 | eGFB | DH | 0.047583291 | 2.38E-07 |
| B189 | B228 | eGFB | OS | -0.046106431 | 5.54E-07 |
| B19 | B101 | CD3+CD4+/CD3+CD8+ | anti-Sc70 | -0.118157459 | 0 |
| B19 | B104 | CD3+CD4+/CD3+CD8+ | anti-SSB | -0.11815744 | 0 |
| B19 | B105 | CD3+CD4+/CD3+CD8+ | anti-U1RNP | -0.087571282 | 0 |
| B19 | B107 | CD3+CD4+/CD3+CD8+ | AnuA | -0.116900042 | 0 |
| B19 | B136 | CD3+CD4+/CD3+CD8+ | 5'NT | 0.098884887 | 0 |
| B19 | B151 | CD3+CD4+/CD3+CD8+ | Anti-CCP | 0.110896153 | 0 |
| B19 | B155 | CD3+CD4+/CD3+CD8+ | IgA | 0.087237159 | 0 |
| B19 | B183 | CD3+CD4+/CD3+CD8+ | PTA | -0.08769574 | 0 |
| B19 | B20 | CD3+CD4+/CD3+CD8+ | CD3+CD4+CD8+ | -0.147350557 | 0 |
| B19 | B22 | CD3+CD4+/CD3+CD8+ | CD3+CD8+(T) | -0.814433793 | 0 |
| B19 | B23 | CD3+CD4+/CD3+CD8+ | CD3-CD16+CD56+(NK) | 0.107331521 | 0 |
| B19 | B98 | CD3+CD4+/CD3+CD8+ | anti-Jo-1 | -0.11815743 | 0 |
| B19 | B81 | CD3+CD4+/CD3+CD8+ | TSGF | -0.072414697 | 3.55E-15 |
| B19 | B186 | CD3+CD4+/CD3+CD8+ | PFC | 0.069122463 | 5.86E-14 |
| B19 | B125 | CD3+CD4+/CD3+CD8+ | Cor | -0.066606152 | 4.60E-13 |
| B19 | B100 | CD3+CD4+/CD3+CD8+ | anti-Ro52 | -0.064984891 | 1.66E-12 |
| B19 | B139 | CD3+CD4+/CD3+CD8+ | ADA | 0.062921508 | 8.17E-12 |
| B19 | B95 | CD3+CD4+/CD3+CD8+ | anti-cenpb | 0.058890016 | 1.59E-10 |
| B19 | B138 | CD3+CD4+/CD3+CD8+ | PAB | 0.056418903 | 8.90E-10 |
| B19 | B97 | CD3+CD4+/CD3+CD8+ | dsDNA | -0.05525604 | 1.96E-09 |
| B19 | B90 | CD3+CD4+/CD3+CD8+ | CYFRA 21-1 | 0.053858349 | 4.93E-09 |
| B19 | B195 | CD3+CD4+/CD3+CD8+ | Na | -0.050141173 | 5.18E-08 |
| B19 | B108 | CD3+CD4+/CD3+CD8+ | Anti-Cyclin | -0.046654459 | 4.06E-07 |
| B19 | B31 | CD3+CD4+/CD3+CD8+ | FSH | -0.045374983 | 8.35E-07 |
| B190 | B191 | Cre | Urea | 0.423101839 | 0 |
| B190 | B192 | Cre | UA | 0.291791173 | 0 |
| B190 | B206 | Cre | Waist | 0.115795574 | 0 |
| B190 | B209 | Cre | Smoking | 0.086883773 | 0 |
| B190 | B216 | Cre | BMI | 0.142250794 | 0 |
| B190 | B221 | Cre | Hei | 0.214858621 | 0 |
| B190 | B222 | Cre | NS | 0.222205557 | 0 |
| B190 | B210 | Cre | Hip | 0.075329681 | 2.22E-16 |
| B190 | B204 | Cre | Drinking | 0.056802964 | 6.84E-10 |
| B191 | B192 | Urea | UA | 0.114912694 | 0 |
| B191 | B200 | Urea | OP | 0.084707916 | 0 |
| B191 | B218 | Urea | DBP | -0.069561218 | 4.09E-14 |
| B192 | B204 | UA | Drinking | 0.09021393 | 0 |
| B192 | B206 | UA | Waist | 0.231345842 | 0 |
| B192 | B210 | UA | Hip | 0.168825735 | 0 |
| B192 | B215 | UA | BMI | 0.191150847 | 0 |

| | | | | | |
|---|---|---|---|---|---|
| B192 | B216 | UA | BMI | 0.272067628 | 0 |
| B192 | B218 | UA | DBP | 0.087018164 | 0 |
| B192 | B221 | UA | Hei | 0.216265314 | 0 |
| B192 | B222 | UA | NS | 0.259196604 | 0 |
| B193 | B198 | K | P | 0.09989593 | 0 |
| B193 | B200 | K | OP | 0.075349943 | 2.22E-16 |
| B193 | B196 | K | TCO2 | 0.067878171 | 1.64E-13 |
| B193 | B197 | K | AG | -0.047136766 | 3.08E-07 |
| B194 | B195 | Cl | Na | 0.405944577 | 0 |
| B194 | B196 | Cl | TCO2 | -0.140723475 | 0 |
| B194 | B197 | Cl | AG | -0.350415461 | 0 |
| B194 | B198 | Cl | P | -0.143142857 | 0 |
| B194 | B199 | Cl | Mg | 0.135354924 | 0 |
| B194 | B212 | Cl | IOP-L | -0.069322102 | 4.97E-14 |
| B195 | B196 | Na | TCO2 | 0.090227828 | 0 |
| B195 | B197 | Na | AG | 0.27330461 | 0 |
| B195 | B199 | Na | Mg | 0.153128498 | 0 |
| B195 | B200 | Na | OP | 0.503896135 | 0 |
| B196 | B197 | TCO2 | AG | -0.676805732 | 0 |
| B196 | B200 | TCO2 | OP | 0.07754742 | 0 |
| B197 | B198 | AG | P | 0.081443521 | 0 |
| B197 | B200 | AG | OP | 0.272443199 | 0 |
| B198 | B199 | P | Mg | -0.076367456 | 0 |
| B199 | B200 | Mg | OP | -0.138126876 | 0 |
| B199 | B216 | Mg | BMI | -0.046565876 | 4.27E-07 |
| B2 | B126 | Age | Alb | -0.280277706 | 0 |
| B2 | B127 | Age | A-G ratio | -0.199873842 | 0 |
| B2 | B128 | Age | ALT | -0.145854177 | 0 |
| B2 | B130 | Age | GLB | 0.110150619 | 0 |
| B2 | B132 | Age | Dbil | 0.085484865 | 0 |
| B2 | B134 | Age | CHE | -0.079644869 | 0 |
| B2 | B140 | Age | r-GT | -0.148916375 | 0 |
| B2 | B141 | Age | AST/ALT | 0.229420774 | 0 |
| B2 | B146 | Age | CysC | 0.113967838 | 0 |
| B2 | B153 | Age | Glu | -0.115690049 | 0 |
| B2 | B163 | Age | SG | -0.180787447 | 0 |
| B2 | B170 | Age | U-GLU | -0.12598849 | 0 |
| B2 | B172 | Age | KB | -0.100113 | 0 |
| B2 | B176 | Age | Ucr | -0.086718773 | 0 |
| B2 | B188 | Age | Ca | -0.109955382 | 0 |
| B2 | B189 | Age | eGFB | -0.612158214 | 0 |
| B2 | B190 | Age | Cre | 0.178043621 | 0 |
| B2 | B191 | Age | Urea | 0.175274314 | 0 |
| B2 | B201 | Age | e-habits | 0.08462389 | 0 |

| | | | | | |
|---|---|---|---|---|---|
| B2 | B203 | Age | DH | -0.082187271 | 0 |
| B2 | B204 | Age | Drinking | -0.12243786 | 0 |
| B2 | B209 | Age | Smoking | -0.112135824 | 0 |
| B2 | B210 | Age | Hip | -0.120107044 | 0 |
| B2 | B211 | Age | IOP-R | -0.104765576 | 0 |
| B2 | B212 | Age | IOP-L | -0.09160839 | 0 |
| B2 | B215 | Age | BMI | -0.1659732 | 0 |
| B2 | B216 | Age | BMI | -0.285598708 | 0 |
| B2 | B218 | Age | DBP | -0.171378565 | 0 |
| B2 | B219 | Age | OT | 0.122648313 | 0 |
| B2 | B220 | Age | SBP | 0.304827433 | 0 |
| B2 | B221 | Age | Hei | -0.261514504 | 0 |
| B2 | B222 | Age | NS | -0.179151668 | 0 |
| B2 | B228 | Age | OS | 0.081009733 | 0 |
| B2 | B229 | Age | slit-lamp | 0.160003398 | 0 |
| B2 | B24 | Age | αHBDH | 0.080775895 | 0 |
| B2 | B27 | Age | LDH | 0.089671163 | 0 |
| B2 | B41 | Age | MON％ | 0.192804193 | 0 |
| B2 | B42 | Age | MON | 0.082570733 | 0 |
| B2 | B43 | Age | MCV | 0.169653754 | 0 |
| B2 | B45 | Age | MCHC | -0.186510455 | 0 |
| B2 | B46 | Age | RBC Count | -0.314116779 | 0 |
| B2 | B47 | Age | RDW | 0.279217127 | 0 |
| B2 | B51 | Age | Hb | -0.29423118 | 0 |
| B2 | B52 | Age | PLT/BPC | -0.111379054 | 0 |
| B2 | B71 | Age | TG | -0.204349791 | 0 |
| B2 | B72 | Age | HDL-C | 0.170604297 | 0 |
| B2 | B74 | Age | ApoA1 | 0.088782319 | 0 |
| B2 | B107 | Age | AnuA | 0.074304398 | 6.66E-16 |
| B2 | B101 | Age | anti-Sc70 | 0.073704559 | 1.11E-15 |
| B2 | B104 | Age | anti-SSB | 0.073704548 | 1.11E-15 |
| B2 | B98 | Age | anti-Jo-1 | 0.073704542 | 1.11E-15 |
| B2 | B77 | Age | TC | -0.071973686 | 5.33E-15 |
| B2 | B62 | Age | WBHSRVI | -0.070959006 | 1.24E-14 |
| B2 | B227 | Age | OD | 0.070306356 | 2.18E-14 |
| B2 | B91 | Age | TP | -0.070206482 | 2.35E-14 |
| B2 | B40 | Age | WBC | -0.070005689 | 2.80E-14 |
| B2 | B67 | Age | WBRSR(mPas_50) | -0.069652443 | 3.77E-14 |
| B2 | B173 | Age | NTT | 0.068655774 | 8.66E-14 |
| B2 | B58 | Age | HCT | -0.067993272 | 1.49E-13 |
| B2 | B66 | Age | WBRSR(mPas_100) | -0.067371825 | 2.48E-13 |
| B2 | B138 | Age | PAB | -0.067344004 | 2.54E-13 |
| B2 | B161 | Age | LEU | 0.066058255 | 7.12E-13 |
| B2 | B63 | Age | WBRSR(mPas_200) | -0.064216068 | 3.03E-12 |

| | | | | | |
|---|---|---|---|---|---|
| B2 | B50 | Age | eo% | 0.06396994 | 3.66E-12 |
| B2 | B60 | Age | WBRVI | -0.063914522 | 3.82E-12 |
| B2 | B65 | Age | WBRSR(mPas_1) | -0.063320932 | 6.03E-12 |
| B2 | B44 | Age | MCH | 0.060260138 | 5.91E-11 |
| B2 | B194 | Age | Cl | 0.056525333 | 8.28E-10 |
| B2 | B75 | Age | ApoA1/B | 0.054339355 | 3.60E-09 |
| B2 | B195 | Age | Na | 0.053532077 | 6.10E-09 |
| B2 | B54 | Age | ESR | 0.053138867 | 7.87E-09 |
| B2 | B120 | Age | HCV-cAg | 0.053026929 | 8.46E-09 |
| B2 | B192 | Age | UA | -0.05145116 | 2.30E-08 |
| B2 | B61 | Age | WBHSRV | -0.05139253 | 2.39E-08 |
| B2 | B149 | Age | TGAb | 0.051089577 | 2.88E-08 |
| B2 | B3 | Age | Hcy | 0.050868853 | 3.31E-08 |
| B2 | B56 | Age | IR | -0.050529834 | 4.08E-08 |
| B2 | B157 | Age | IgG | 0.047481921 | 2.52E-07 |
| B2 | B12 | Age | PG II | 0.046050872 | 5.72E-07 |
| B2 | B73 | Age | ApoA | 0.045961625 | 6.01E-07 |
| B20 | B101 | CD3+CD4+CD8+ | anti-Sc70 | -0.076216462 | 0 |
| B20 | B104 | CD3+CD4+CD8+ | anti-SSB | -0.076216446 | 0 |
| B20 | B107 | CD3+CD4+CD8+ | AnuA | -0.076651767 | 0 |
| B20 | B151 | CD3+CD4+CD8+ | Anti-CCP | -0.090690405 | 0 |
| B20 | B152 | CD3+CD4+CD8+ | ACA-Ig(G-A-M) | -0.619483501 | 0 |
| B20 | B157 | CD3+CD4+CD8+ | IgG | -0.10821899 | 0 |
| B20 | B183 | CD3+CD4+CD8+ | PTA | 0.120973521 | 0 |
| B20 | B21 | CD3+CD4+CD8+ | CD3+CD4-CD8- | -0.126969142 | 0 |
| B20 | B22 | CD3+CD4+CD8+ | CD3+CD8+(T) | 0.271773292 | 0 |
| B20 | B23 | CD3+CD4+CD8+ | CD3-CD16+CD56+(NK) | -0.138265877 | 0 |
| B20 | B92 | CD3+CD4+CD8+ | ALDH2-Glu504Glu | -0.106746622 | 0 |
| B20 | B93 | CD3+CD4+CD8+ | ALDH2-Glu504Lys | 0.110915296 | 0 |
| B20 | B98 | CD3+CD4+CD8+ | anti-Jo-1 | -0.076216438 | 0 |
| B20 | B197 | CD3+CD4+CD8+ | AG | -0.074303061 | 6.66E-16 |
| B20 | B158 | CD3+CD4+CD8+ | IgM | -0.055405176 | 1.77E-09 |
| B20 | B99 | CD3+CD4+CD8+ | anti-PM-Scl | 0.055207931 | 2.02E-09 |
| B20 | B155 | CD3+CD4+CD8+ | IgA | -0.050827877 | 3.39E-08 |
| B200 | B211 | OP | IOP-R | 0.071957658 | 5.33E-15 |
| B200 | B212 | OP | IOP-L | 0.061805779 | 1.89E-11 |
| B201 | B204 | e-habits | Drinking | 0.084321341 | 0 |
| B201 | B224 | e-habits | pmh | 0.103459958 | 0 |
| B201 | B205 | e-habits | KDA | 0.045230456 | 9.05E-07 |
| B202 | B223 | e-times | FH | -0.32389084 | 0 |
| B203 | B209 | DH | Smoking | 0.065362113 | 1.24E-12 |
| B203 | B204 | DH | Drinking | 0.054800607 | 2.65E-09 |
| B204 | B206 | Drinking | Waist | 0.128894686 | 0 |
| B204 | B209 | Drinking | Smoking | 0.308817863 | 0 |

| | | | | | |
|---|---|---|---|---|---|
| B204 | B210 | Drinking | Hip | 0.085604909 | 0 |
| B204 | B216 | Drinking | BMI | 0.179387249 | 0 |
| B204 | B218 | Drinking | DBP | 0.099073617 | 0 |
| B204 | B221 | Drinking | Hei | 0.20834939 | 0 |
| B204 | B222 | Drinking | NS | 0.220359991 | 0 |
| B204 | B215 | Drinking | BMI | 0.07371782 | 1.11E-15 |
| B204 | B219 | Drinking | OT | -0.046096375 | 5.57E-07 |
| B205 | B222 | KDA | NS | -0.045964827 | 6.00E-07 |
| B206 | B207 | Waist | WHR | 0.100657118 | 0 |
| B206 | B209 | Waist | Smoking | 0.105557222 | 0 |
| B206 | B210 | Waist | Hip | 0.758147537 | 0 |
| B206 | B215 | Waist | BMI | 0.802340738 | 0 |
| B206 | B216 | Waist | BMI | 0.794663956 | 0 |
| B206 | B218 | Waist | DBP | 0.181866181 | 0 |
| B206 | B220 | Waist | SBP | 0.125140313 | 0 |
| B206 | B221 | Waist | Hei | 0.334200237 | 0 |
| B206 | B222 | Waist | NS | 0.668027611 | 0 |
| B207 | B215 | WHR | BMI | 0.057116629 | 5.51E-10 |
| B207 | B222 | WHR | NS | 0.051021284 | 3.01E-08 |
| B207 | B216 | WHR | BMI | 0.049948128 | 5.82E-08 |
| B209 | B216 | Smoking | BMI | 0.157130246 | 0 |
| B209 | B220 | Smoking | SBP | -0.099276573 | 0 |
| B209 | B221 | Smoking | Hei | 0.257746335 | 0 |
| B209 | B222 | Smoking | NS | 0.218265917 | 0 |
| B209 | B219 | Smoking | OT | -0.065935942 | 7.85E-13 |
| B209 | B210 | Smoking | Hip | 0.045488774 | 7.84E-07 |
| B21 | B124 | CD3+CD4-CD8- | ACTH | 0.243600281 | 0 |
| B21 | B152 | CD3+CD4-CD8- | ACA-Ig(G-A-M) | 0.080320743 | 0 |
| B21 | B195 | CD3+CD4-CD8- | Na | 0.081637995 | 0 |
| B21 | B200 | CD3+CD4-CD8- | OP | 0.098837166 | 0 |
| B21 | B23 | CD3+CD4-CD8- | CD3-CD16+CD56+(NK) | -0.216209984 | 0 |
| B21 | B92 | CD3+CD4-CD8- | ALDH2-Glu504Glu | -0.094508941 | 0 |
| B21 | B93 | CD3+CD4-CD8- | ALDH2-Glu504Lys | 0.100000257 | 0 |
| B21 | B138 | CD3+CD4-CD8- | PAB | 0.07289794 | 2.22E-15 |
| B21 | B81 | CD3+CD4-CD8- | TSGF | 0.056979074 | 6.06E-10 |
| B21 | B181 | CD3+CD4-CD8- | APTT | -0.054587795 | 3.05E-09 |
| B21 | B183 | CD3+CD4-CD8- | PTA | 0.050949292 | 3.15E-08 |
| B210 | B215 | Hip | BMI | 0.733358186 | 0 |
| B210 | B216 | Hip | BMI | 0.766699379 | 0 |
| B210 | B218 | Hip | DBP | 0.154114398 | 0 |
| B210 | B221 | Hip | Hei | 0.37331522 | 0 |
| B210 | B222 | Hip | NS | 0.585630193 | 0 |
| B210 | B220 | Hip | SBP | 0.059045035 | 1.42E-10 |
| B211 | B212 | IOP-R | IOP-L | 0.778622209 | 0 |

| | | | | | |
|---|---|---|---|---|---|
| B211 | B218 | IOP-R | DBP | 0.090513666 | 0 |
| B211 | B220 | IOP-R | SBP | 0.046055986 | 5.70E-07 |
| B211 | B213 | IOP-R | CVA-R | 0.045450561 | 8.01E-07 |
| B212 | B218 | IOP-L | DBP | 0.079192678 | 0 |
| B215 | B216 | BMI | BMI | 0.789955773 | 0 |
| B215 | B218 | BMI | DBP | 0.209655418 | 0 |
| B215 | B220 | BMI | SBP | 0.116393268 | 0 |
| B215 | B221 | BMI | Hei | 0.090992087 | 0 |
| B215 | B222 | BMI | NS | 0.572796595 | 0 |
| B216 | B218 | BMI | DBP | 0.222228047 | 0 |
| B216 | B221 | BMI | Hei | 0.676936 | 0 |
| B216 | B222 | BMI | NS | 0.751353162 | 0 |
| B216 | B219 | BMI | OT | -0.068432705 | 1.04E-13 |
| B218 | B220 | DBP | SBP | 0.565182993 | 0 |
| B218 | B221 | DBP | Hei | 0.113168952 | 0 |
| B218 | B222 | DBP | NS | 0.198624464 | 0 |
| B218 | B219 | DBP | OT | -0.051912388 | 1.72E-08 |
| B219 | B221 | OT | Hei | -0.08668641 | 0 |
| B219 | B222 | OT | NS | -0.073854247 | 8.88E-16 |
| B22 | B105 | CD3+CD8+(T) | anti-U1RNP | 0.119124995 | 0 |
| B22 | B121 | CD3+CD8+(T) | C3 | 0.078085501 | 0 |
| B22 | B122 | CD3+CD8+(T) | C4 | 0.084715223 | 0 |
| B22 | B155 | CD3+CD8+(T) | IgA | -0.10307405 | 0 |
| B22 | B157 | CD3+CD8+(T) | IgG | -0.077784129 | 0 |
| B22 | B23 | CD3+CD8+(T) | CD3-CD16+CD56+(NK) | -0.403679279 | 0 |
| B22 | B29 | CD3+CD8+(T) | E2 | -0.076020754 | 0 |
| B22 | B31 | CD3+CD8+(T) | FSH | 0.096638542 | 0 |
| B22 | B95 | CD3+CD8+(T) | anti-cenpb | -0.097588698 | 0 |
| B22 | B97 | CD3+CD8+(T) | dsDNA | 0.082936797 | 0 |
| B22 | B108 | CD3+CD8+(T) | Anti-Cyclin | 0.074398434 | 6.66E-16 |
| B22 | B152 | CD3+CD8+(T) | ACA-Ig(G-A-M) | -0.071091482 | 1.11E-14 |
| B22 | B33 | CD3+CD8+(T) | Prog | -0.062278175 | 1.33E-11 |
| B22 | B100 | CD3+CD8+(T) | anti-Ro52 | 0.060180791 | 6.27E-11 |
| B22 | B138 | CD3+CD8+(T) | PAB | -0.049556428 | 7.38E-08 |
| B22 | B81 | CD3+CD8+(T) | TSGF | 0.048231735 | 1.63E-07 |
| B22 | B136 | CD3+CD8+(T) | 5'NT | -0.046765216 | 3.81E-07 |
| B22 | B99 | CD3+CD8+(T) | anti-PM-Scl | 0.046625354 | 4.13E-07 |
| B220 | B221 | SBP | Hei | -0.132365631 | 0 |
| B221 | B222 | Hei | NS | 0.540482297 | 0 |
| B227 | B228 | OD | OS | 0.129484288 | 0 |
| B227 | B229 | OD | slit-lamp | 0.057928754 | 3.13E-10 |
| B23 | B101 | CD3-CD16+CD56+(NK) | anti-Sc70 | 0.15682674 | 0 |
| B23 | B104 | CD3-CD16+CD56+(NK) | anti-SSB | 0.15682672 | 0 |
| B23 | B107 | CD3-CD16+CD56+(NK) | AnuA | 0.15770509 | 0 |

| | | | | | |
|---|---|---|---|---|---|
| B23 | B121 | CD3-CD16+CD56+(NK) | C3 | -0.280423681 | 0 |
| B23 | B122 | CD3-CD16+CD56+(NK) | C4 | -0.229017703 | 0 |
| B23 | B152 | CD3-CD16+CD56+(NK) | ACA-Ig(G-A-M) | 0.093059967 | 0 |
| B23 | B193 | CD3-CD16+CD56+(NK) | K | 0.116517702 | 0 |
| B23 | B98 | CD3-CD16+CD56+(NK) | anti-Jo-1 | 0.156826712 | 0 |
| B23 | B99 | CD3-CD16+CD56+(NK) | anti-PM-Scl | -0.07978312 | 0 |
| B23 | B157 | CD3-CD16+CD56+(NK) | IgG | 0.074199427 | 8.88E-16 |
| B23 | B96 | CD3-CD16+CD56+(NK) | dsDNA (IIF) | 0.068083546 | 1.39E-13 |
| B23 | B74 | CD3-CD16+CD56+(NK) | ApoA1 | 0.067697953 | 1.90E-13 |
| B23 | B79 | CD3-CD16+CD56+(NK) | hb V -dna | 0.066744169 | 4.11E-13 |
| B23 | B124 | CD3-CD16+CD56+(NK) | ACTH | -0.06468781 | 2.10E-12 |
| B23 | B200 | CD3-CD16+CD56+(NK) | OP | -0.064655652 | 2.15E-12 |
| B23 | B109 | CD3-CD16+CD56+(NK) | AMA-M2 | -0.06437644 | 2.67E-12 |
| B23 | B212 | CD3-CD16+CD56+(NK) | IOP-L | -0.057255267 | 5.01E-10 |
| B23 | B211 | CD3-CD16+CD56+(NK) | IOP-R | -0.050981744 | 3.08E-08 |
| B23 | B181 | CD3-CD16+CD56+(NK) | APTT | 0.048434541 | 1.44E-07 |
| B23 | B120 | CD3-CD16+CD56+(NK) | HCV-cAg | 0.048244172 | 1.62E-07 |
| B23 | B184 | CD3-CD16+CD56+(NK) | INR | 0.048133873 | 1.72E-07 |
| B23 | B72 | CD3-CD16+CD56+(NK) | HDL-C | 0.046666751 | 4.03E-07 |
| B23 | B199 | CD3-CD16+CD56+(NK) | Mg | 0.04628988 | 5.00E-07 |
| B24 | B127 | αHBDH | A-G ratio | -0.076129079 | 0 |
| B24 | B130 | αHBDH | GLB | 0.076669917 | 0 |
| B24 | B131 | αHBDH | AST | 0.092969362 | 0 |
| B24 | B175 | αHBDH | ALB-U | 0.07595509 | 0 |
| B24 | B25 | αHBDH | CK | 0.210571384 | 0 |
| B24 | B26 | αHBDH | CKMB | 0.147753188 | 0 |
| B24 | B27 | αHBDH | LDH | 0.728811674 | 0 |
| B24 | B69 | αHBDH | PV | 0.08128962 | 0 |
| B24 | B74 | αHBDH | ApoA1 | 0.134522921 | 0 |
| B24 | B75 | αHBDH | ApoA1/B | 0.090421668 | 0 |
| B24 | B47 | αHBDH | RDW | 0.070799167 | 1.42E-14 |
| B24 | B177 | αHBDH | ACR | 0.068391164 | 1.08E-13 |
| B24 | B221 | αHBDH | Hei | -0.065254851 | 1.35E-12 |
| B24 | B72 | αHBDH | HDL-C | 0.064932343 | 1.73E-12 |
| B24 | B220 | αHBDH | SBP | 0.063032389 | 7.51E-12 |
| B24 | B141 | αHBDH | AST/ALT | 0.06057441 | 4.70E-11 |
| B24 | B58 | αHBDH | HCT | -0.058775737 | 1.72E-10 |
| B24 | B91 | αHBDH | TP | 0.056987147 | 6.02E-10 |
| B24 | B148 | αHBDH | Anti_TPO | 0.05349364 | 6.26E-09 |
| B24 | B176 | αHBDH | Ucr | -0.051195714 | 2.70E-08 |
| B24 | B79 | αHBDH | hb V -dna | 0.050120695 | 5.24E-08 |
| B24 | B46 | αHBDH | RBC Count | -0.049655117 | 6.95E-08 |
| B24 | B216 | αHBDH | BMI | -0.045276851 | 8.82E-07 |
| B25 | B123 | CK | hs-CRP1 | -0.079284703 | 0 |

| | | | | | |
|---|---|---|---|---|---|
| B25 | B26 | CK | CKMB | 0.212429696 | 0 |
| B25 | B27 | CK | LDH | 0.202650293 | 0 |
| B25 | B89 | CK | CA 72-4 | 0.084298682 | 0 |
| B25 | B189 | CK | eGFB | -0.068685058 | 8.44E-14 |
| B25 | B190 | CK | Cre | 0.065132473 | 1.48E-12 |
| B25 | B192 | CK | UA | 0.061866514 | 1.81E-11 |
| B25 | B114 | CK | TSH TT3 TT4(TT4) | -0.060276063 | 5.85E-11 |
| B25 | B57 | CK | RBCAI | 0.059830928 | 8.07E-11 |
| B25 | B153 | CK | Glu | -0.055983642 | 1.20E-09 |
| B25 | B131 | CK | AST | 0.052128438 | 1.50E-08 |
| B25 | B78 | CK | HBcAb | -0.047006696 | 3.32E-07 |
| B26 | B27 | CKMB | LDH | 0.120903843 | 0 |
| B26 | B145 | CKMB | BGP | -0.066410145 | 5.38E-13 |
| B26 | B183 | CKMB | PTA | 0.056880813 | 6.48E-10 |
| B26 | B34 | CKMB | TK | 0.049610911 | 7.14E-08 |
| B26 | B115 | CKMB | TSH TT3 TT4(TT3) | 0.047255881 | 2.87E-07 |
| B27 | B118 | LDH | CEA1 | 0.092661653 | 0 |
| B27 | B127 | LDH | A-G ratio | -0.098499235 | 0 |
| B27 | B130 | LDH | GLB | 0.09907058 | 0 |
| B27 | B131 | LDH | AST | 0.146680648 | 0 |
| B27 | B138 | LDH | PAB | -0.076617135 | 0 |
| B27 | B139 | LDH | ADA | 0.124426123 | 0 |
| B27 | B150 | LDH | ALP | 0.087000161 | 0 |
| B27 | B221 | LDH | Hei | -0.075959801 | 0 |
| B27 | B69 | LDH | PV | 0.078289937 | 0 |
| B27 | B74 | LDH | ApoA1 | 0.105386965 | 0 |
| B27 | B47 | LDH | RDW | 0.072314603 | 4.00E-15 |
| B27 | B91 | LDH | TP | 0.071572186 | 7.55E-15 |
| B27 | B220 | LDH | SBP | 0.071471671 | 7.99E-15 |
| B27 | B177 | LDH | ACR | 0.066245888 | 6.13E-13 |
| B27 | B72 | LDH | HDL-C | 0.065725413 | 9.28E-13 |
| B27 | B185 | LDH | PT | 0.065616449 | 1.01E-12 |
| B27 | B75 | LDH | ApoA1/B | 0.065397427 | 1.20E-12 |
| B27 | B184 | LDH | INR | 0.06427292 | 2.90E-12 |
| B27 | B186 | LDH | PFC | 0.063551745 | 5.05E-12 |
| B27 | B128 | LDH | ALT | 0.060366427 | 5.47E-11 |
| B27 | B176 | LDH | Ucr | -0.060271734 | 5.86E-11 |
| B27 | B175 | LDH | ALB-U | 0.057887602 | 3.22E-10 |
| B27 | B189 | LDH | eGFB | -0.056129562 | 1.08E-09 |
| B27 | B141 | LDH | AST/ALT | 0.055527494 | 1.63E-09 |
| B27 | B140 | LDH | r-GT | 0.053382282 | 6.73E-09 |
| B27 | B182 | LDH | TT | 0.047298375 | 2.80E-07 |
| B27 | B46 | LDH | RBC Count | -0.045788862 | 6.63E-07 |
| B27 | B41 | LDH | MON % | 0.045485835 | 7.85E-07 |

| | | | | | |
|---|---|---|---|---|---|
| B28 | B29 | PRL | E2 | -0.149561775 | 0 |
| B28 | B30 | PRL | LH | -0.274985206 | 0 |
| B28 | B31 | PRL | FSH | -0.160989545 | 0 |
| B28 | B33 | PRL | Prog | -0.187837538 | 0 |
| B28 | B32 | PRL | Testo | -0.072268188 | 4.00E-15 |
| B28 | B34 | PRL | TK | -0.049478224 | 7.74E-08 |
| B28 | B74 | PRL | ApoA1 | -0.047815822 | 2.08E-07 |
| B29 | B125 | E2 | Cor | 0.079989177 | 0 |
| B29 | B30 | E2 | LH | -0.30263891 | 0 |
| B29 | B31 | E2 | FSH | -0.538598758 | 0 |
| B29 | B32 | E2 | Testo | -0.316313048 | 0 |
| B29 | B33 | E2 | Prog | 0.484462645 | 0 |
| B29 | B173 | E2 | NTT | -0.059514368 | 1.01E-10 |
| B29 | B205 | E2 | KDA | -0.056261805 | 9.91E-10 |
| B29 | B75 | E2 | ApoA1/B | 0.046001842 | 5.88E-07 |
| B3 | B174 | Hcy | Urea/Cr | -0.091592138 | 0 |
| B3 | B189 | Hcy | eGFB | -0.13845564 | 0 |
| B3 | B190 | Hcy | Cre | 0.13098271 | 0 |
| B3 | B192 | Hcy | UA | 0.105928358 | 0 |
| B3 | B222 | Hcy | NS | 0.077971958 | 0 |
| B3 | B176 | Hcy | Ucr | 0.061766407 | 1.95E-11 |
| B3 | B47 | Hcy | RDW | 0.061394701 | 2.57E-11 |
| B3 | B221 | Hcy | Hei | 0.058812131 | 1.68E-10 |
| B3 | B216 | Hcy | BMI | 0.057876111 | 3.25E-10 |
| B3 | B206 | Hcy | Waist | 0.057519036 | 4.17E-10 |
| B3 | B191 | Hcy | Urea | 0.054173917 | 4.01E-09 |
| B3 | B25 | Hcy | CK | 0.052271406 | 1.37E-08 |
| B3 | B209 | Hcy | Smoking | 0.051347345 | 2.46E-08 |
| B3 | B111 | Hcy | TSH | 0.049773674 | 6.47E-08 |
| B3 | B175 | Hcy | ALB-U | 0.047437637 | 2.59E-07 |
| B3 | B42 | Hcy | MON | 0.046492692 | 4.45E-07 |
| B3 | B44 | Hcy | MCH | 0.046244644 | 5.13E-07 |
| B3 | B146 | Hcy | CysC | 0.045570546 | 7.49E-07 |
| B30 | B31 | LH | FSH | 0.79088348 | 0 |
| B30 | B32 | LH | Testo | -0.280747392 | 0 |
| B30 | B33 | LH | Prog | -0.066973591 | 3.42E-13 |
| B31 | B32 | FSH | Testo | -0.340823656 | 0 |
| B31 | B33 | FSH | Prog | -0.164004706 | 0 |
| B31 | B125 | FSH | Cor | -0.073056 | 2.00E-15 |
| B31 | B76 | FSH | ApoB | 0.056184083 | 1.04E-09 |
| B31 | B75 | FSH | ApoA1/B | -0.054132056 | 4.12E-09 |
| B31 | B173 | FSH | NTT | 0.050347441 | 4.56E-08 |
| B31 | B77 | FSH | TC | 0.045898194 | 6.23E-07 |
| B32 | B121 | Testo | C3 | -0.139848725 | 0 |

| | | | | | |
|---|---|---|---|---|---|
| B32 | B33 | Testo | Prog | -0.263255959 | 0 |
| B33 | B205 | Prog | KDA | -0.067687919 | 1.92E-13 |
| B34 | B173 | TK | NTT | 0.069329408 | 4.95E-14 |
| B34 | B81 | TK | TSGF | -0.066630976 | 4.51E-13 |
| B34 | B107 | TK | AnuA | 0.063873882 | 3.94E-12 |
| B34 | B101 | TK | anti-Sc70 | 0.063484371 | 5.32E-12 |
| B34 | B104 | TK | anti-SSB | 0.06348436 | 5.32E-12 |
| B34 | B98 | TK | anti-Jo-1 | 0.063484355 | 5.32E-12 |
| B34 | B151 | TK | Anti-CCP | 0.061352403 | 2.65E-11 |
| B34 | B200 | TK | OP | -0.060107859 | 6.61E-11 |
| B34 | B74 | TK | ApoA1 | 0.059219447 | 1.25E-10 |
| B34 | B139 | TK | ADA | -0.057178566 | 5.28E-10 |
| B34 | B134 | TK | CHE | -0.052784828 | 9.89E-09 |
| B35 | B159 | β2-MG | β2-MG | 0.978773458 | 0 |
| B35 | B186 | β2-MG | PFC | 0.052893004 | 9.22E-09 |
| B35 | B181 | β2-MG | APTT | -0.046001934 | 5.88E-07 |
| B36 | B184 | V-Lym% | INR | -0.15327275 | 0 |
| B36 | B185 | V-Lym% | PT | -0.15135838 | 0 |
| B36 | B37 | V-Lym% | BAND | 0.100138926 | 0 |
| B36 | B181 | V-Lym% | APTT | -0.075707442 | 2.22E-16 |
| B36 | B38 | V-Lym% | BAND% | 0.05933494 | 1.15E-10 |
| B36 | B66 | V-Lym% | WBRSR(mPas_100) | -0.050862045 | 3.32E-08 |
| B36 | B67 | V-Lym% | WBRSR(mPas_50) | -0.050370818 | 4.50E-08 |
| B36 | B63 | V-Lym% | WBRSR(mPas_200) | -0.049772319 | 6.48E-08 |
| B36 | B62 | V-Lym% | WBHSRVI | -0.049243912 | 8.91E-08 |
| B36 | B58 | V-Lym% | HCT | -0.047979356 | 1.89E-07 |
| B36 | B132 | V-Lym% | Dbil | -0.047698413 | 2.22E-07 |
| B36 | B179 | V-Lym% | α1-MG | -0.04678942 | 3.76E-07 |
| B37 | B38 | BAND | BAND% | 0.636249318 | 0 |
| B37 | B90 | BAND | CYFRA 21-1 | -0.060907317 | 3.68E-11 |
| B38 | B85 | BAND% | SCC | 0.234301572 | 0 |
| B38 | B199 | BAND% | Mg | -0.064146063 | 3.19E-12 |
| B38 | B193 | BAND% | K | -0.060427835 | 5.23E-11 |
| B38 | B139 | BAND% | ADA | 0.048312543 | 1.55E-07 |
| B38 | B137 | BAND% | AFU | 0.047866937 | 2.02E-07 |
| B38 | B138 | BAND% | PAB | -0.045339952 | 8.51E-07 |
| B38 | B41 | BAND% | MON % | 0.045169874 | 9.36E-07 |
| B39 | B43 | Other cells | MCV | -0.102353391 | 0 |
| B39 | B44 | Other cells | MCH | -0.110183498 | 0 |
| B39 | B45 | Other cells | MCHC | -0.064005031 | 3.56E-12 |
| B39 | B51 | Other cells | Hb | -0.051214403 | 2.67E-08 |
| B39 | B46 | Other cells | RBC Count | 0.049799956 | 6.37E-08 |
| B4 | B121 | WB-Ca | C3 | 0.10960052 | 0 |
| B4 | B122 | WB-Ca | C4 | -0.118566899 | 0 |

| | | | | | |
|---|---|---|---|---|---|
| B4 | B29 | WB-Ca | E2 | 0.21663564 | 0 |
| B4 | B32 | WB-Ca | Testo | -0.298579259 | 0 |
| B4 | B33 | WB-Ca | Prog | 0.169363392 | 0 |
| B4 | B5 | WB-Ca | WB-Mg | 0.394819998 | 0 |
| B4 | B6 | WB-Ca | WB-Fe | 0.264356179 | 0 |
| B4 | B7 | WB-Ca | WB-Cop | 0.671242091 | 0 |
| B4 | B8 | WB-Ca | WB-Zinc | 0.541498414 | 0 |
| B4 | B158 | WB-Ca | IgM | 0.071468044 | 7.99E-15 |
| B40 | B127 | WBC | A-G ratio | -0.101486507 | 0 |
| B40 | B129 | WBC | IBIL | -0.109268794 | 0 |
| B40 | B130 | WBC | GLB | 0.124703251 | 0 |
| B40 | B132 | WBC | Dbil | -0.126962692 | 0 |
| B40 | B133 | WBC | TBIL | -0.12314777 | 0 |
| B40 | B166 | WBC | Pro | 0.077854342 | 0 |
| B40 | B188 | WBC | Ca | 0.078175987 | 0 |
| B40 | B192 | WBC | UA | 0.092047831 | 0 |
| B40 | B206 | WBC | Waist | 0.143707779 | 0 |
| B40 | B209 | WBC | Smoking | 0.216585951 | 0 |
| B40 | B210 | WBC | Hip | 0.098087442 | 0 |
| B40 | B215 | WBC | BMI | 0.10463833 | 0 |
| B40 | B216 | WBC | BMI | 0.13179181 | 0 |
| B40 | B221 | WBC | Hei | 0.089958957 | 0 |
| B40 | B222 | WBC | NS | 0.13703924 | 0 |
| B40 | B41 | WBC | MON % | -0.141830021 | 0 |
| B40 | B42 | WBC | MON | 0.663961179 | 0 |
| B40 | B46 | WBC | RBC Count | 0.177861575 | 0 |
| B40 | B48 | WBC | LYM% | -0.189527578 | 0 |
| B40 | B51 | WBC | Hb | 0.160154653 | 0 |
| B40 | B52 | WBC | PLT/BPC | 0.245250551 | 0 |
| B40 | B53 | WBC | GR% | 0.198158985 | 0 |
| B40 | B71 | WBC | TG | 0.104308302 | 0 |
| B40 | B72 | WBC | HDL-C | -0.167792479 | 0 |
| B40 | B91 | WBC | TP | 0.116256906 | 0 |
| B40 | B174 | WBC | Urea/Cr | -0.068921795 | 6.95E-14 |
| B40 | B218 | WBC | DBP | 0.068745407 | 8.04E-14 |
| B40 | B163 | WBC | SG | 0.066495476 | 5.02E-13 |
| B40 | B170 | WBC | U-GLU | 0.066100914 | 6.88E-13 |
| B40 | B153 | WBC | Glu | 0.06569149 | 9.53E-13 |
| B40 | B112 | WBC | FT4 | 0.06110839 | 3.17E-11 |
| B40 | B49 | WBC | baso% | -0.060875785 | 3.77E-11 |
| B40 | B123 | WBC | hs-CRP1 | 0.060684544 | 4.34E-11 |
| B40 | B160 | WBC | PH | -0.055681985 | 1.47E-09 |
| B40 | B43 | WBC | MCV | -0.054753461 | 2.73E-09 |
| B40 | B63 | WBC | WBRSR(mPas_200) | 0.053524124 | 6.13E-09 |

| | | | | | |
|---|---|---|---|---|---|
| B40 | B190 | WBC | Cre | 0.052740616 | 1.02E-08 |
| B40 | B175 | WBC | ALB-U | 0.051208785 | 2.68E-08 |
| B40 | B67 | WBC | WBRSR(mPas_50) | 0.050422333 | 4.36E-08 |
| B40 | B66 | WBC | WBRSR(mPas_100) | 0.049828261 | 6.26E-08 |
| B40 | B47 | WBC | RDW | -0.047130375 | 3.09E-07 |
| B40 | B70 | WBC | LDL-C | 0.045320456 | 8.61E-07 |
| B40 | B58 | WBC | HCT | 0.045250533 | 8.95E-07 |
| B41 | B126 | MON％ | Alb | -0.127435929 | 0 |
| B41 | B131 | MON％ | AST | 0.10239394 | 0 |
| B41 | B174 | MON％ | Urea/Cr | -0.084372858 | 0 |
| B41 | B189 | MON％ | eGFB | -0.15937412 | 0 |
| B41 | B190 | MON％ | Cre | 0.104549872 | 0 |
| B41 | B42 | MON％ | MON | 0.602822676 | 0 |
| B41 | B43 | MON％ | MCV | 0.088418762 | 0 |
| B41 | B47 | MON％ | RDW | 0.090152574 | 0 |
| B41 | B49 | MON％ | baso% | 0.140395366 | 0 |
| B41 | B53 | MON％ | GR% | -0.23774016 | 0 |
| B41 | B71 | MON％ | TG | -0.097556242 | 0 |
| B41 | B132 | MON％ | Dbil | 0.07431835 | 6.66E-16 |
| B41 | B222 | MON％ | NS | 0.074004478 | 8.88E-16 |
| B41 | B50 | MON％ | eo% | 0.072128338 | 4.44E-15 |
| B41 | B204 | MON％ | Drinking | 0.071651361 | 6.88E-15 |
| B41 | B44 | MON％ | MCH | 0.071372686 | 8.88E-15 |
| B41 | B221 | MON％ | Hei | 0.069479735 | 4.35E-14 |
| B41 | B127 | MON％ | A-G ratio | -0.069316774 | 5.00E-14 |
| B41 | B77 | MON％ | TC | -0.06568153 | 9.61E-13 |
| B41 | B188 | MON％ | Ca | -0.062432344 | 1.18E-11 |
| B41 | B128 | MON％ | ALT | 0.059478616 | 1.04E-10 |
| B41 | B192 | MON％ | UA | 0.053503328 | 6.22E-09 |
| B41 | B91 | MON％ | TP | -0.052242078 | 1.40E-08 |
| B41 | B191 | MON％ | Urea | 0.050484286 | 4.19E-08 |
| B41 | B211 | MON％ | IOP-R | -0.04601865 | 5.82E-07 |
| B41 | B153 | MON％ | Glu | -0.045873451 | 6.32E-07 |
| B41 | B209 | MON％ | Smoking | 0.045498446 | 7.79E-07 |
| B42 | B126 | MON | Alb | -0.08133427 | 0 |
| B42 | B127 | MON | A-G ratio | -0.132128633 | 0 |
| B42 | B130 | MON | GLB | 0.121286734 | 0 |
| B42 | B163 | MON | SG | 0.084812057 | 0 |
| B42 | B166 | MON | Pro | 0.09140822 | 0 |
| B42 | B174 | MON | Urea/Cr | -0.119221791 | 0 |
| B42 | B189 | MON | eGFB | -0.090819524 | 0 |
| B42 | B190 | MON | Cre | 0.117624577 | 0 |
| B42 | B192 | MON | UA | 0.106859808 | 0 |
| B42 | B206 | MON | Waist | 0.144033603 | 0 |

| | | | | | |
|---|---|---|---|---|---|
| B42 | B209 | MON | Smoking | 0.206885525 | 0 |
| B42 | B210 | MON | Hip | 0.088160566 | 0 |
| B42 | B216 | MON | BMI | 0.120948741 | 0 |
| B42 | B221 | MON | Hei | 0.12275573 | 0 |
| B42 | B222 | MON | NS | 0.163983355 | 0 |
| B42 | B46 | MON | RBC Count | 0.134115177 | 0 |
| B42 | B48 | MON | LYM% | -0.151214074 | 0 |
| B42 | B50 | MON | eo% | 0.083840711 | 0 |
| B42 | B51 | MON | Hb | 0.151598896 | 0 |
| B42 | B52 | MON | PLT/BPC | 0.179647665 | 0 |
| B42 | B72 | MON | HDL-C | -0.111700728 | 0 |
| B42 | B133 | MON | TBIL | -0.067873241 | 1.65E-13 |
| B42 | B129 | MON | IBIL | -0.067246094 | 2.74E-13 |
| B42 | B123 | MON | hs-CRP1 | 0.067238439 | 2.76E-13 |
| B42 | B215 | MON | BMI | 0.062538137 | 1.09E-11 |
| B42 | B49 | MON | baso% | 0.060361128 | 5.49E-11 |
| B42 | B91 | MON | TP | 0.060208879 | 6.14E-11 |
| B42 | B176 | MON | Ucr | 0.060101441 | 6.64E-11 |
| B42 | B58 | MON | HCT | 0.05845065 | 2.17E-10 |
| B42 | B204 | MON | Drinking | 0.058003213 | 2.97E-10 |
| B42 | B175 | MON | ALB-U | 0.055884526 | 1.28E-09 |
| B42 | B132 | MON | Dbil | -0.050620569 | 3.86E-08 |
| B42 | B131 | MON | AST | 0.050496852 | 4.16E-08 |
| B42 | B63 | MON | WBRSR(mPas_200) | 0.047903505 | 1.97E-07 |
| B42 | B128 | MON | ALT | 0.046676403 | 4.01E-07 |
| B42 | B113 | MON | FT3 | 0.045650049 | 7.16E-07 |
| B43 | B153 | MCV | Glu | -0.098815555 | 0 |
| B43 | B170 | MCV | U-GLU | -0.086500696 | 0 |
| B43 | B189 | MCV | eGFB | -0.14480876 | 0 |
| B43 | B190 | MCV | Cre | 0.083709184 | 0 |
| B43 | B44 | MCV | MCH | 0.897794217 | 0 |
| B43 | B45 | MCV | MCHC | 0.152351087 | 0 |
| B43 | B46 | MCV | RBC Count | -0.570217837 | 0 |
| B43 | B47 | MCV | RDW | 0.591627875 | 0 |
| B43 | B51 | MCV | Hb | 0.099558711 | 0 |
| B43 | B52 | MCV | PLT/BPC | -0.152501837 | 0 |
| B43 | B72 | MCV | HDL-C | 0.108876808 | 0 |
| B43 | B141 | MCV | AST/ALT | 0.066922039 | 3.57E-13 |
| B43 | B218 | MCV | DBP | -0.064816164 | 1.90E-12 |
| B43 | B53 | MCV | GR% | -0.06130889 | 2.74E-11 |
| B43 | B209 | MCV | Smoking | 0.057505264 | 4.21E-10 |
| B43 | B204 | MCV | Drinking | 0.054195769 | 3.95E-09 |
| B43 | B191 | MCV | Urea | 0.052329755 | 1.32E-08 |
| B43 | B71 | MCV | TG | -0.051319815 | 2.50E-08 |

| | | | | | |
|---|---|---|---|---|---|
| B43 | B91 | MCV | TP | -0.04778489 | 2.11E-07 |
| B43 | B74 | MCV | ApoA1 | 0.046040026 | 5.75E-07 |
| B44 | B190 | MCH | Cre | 0.076675183 | 0 |
| B44 | B204 | MCH | Drinking | 0.104348881 | 0 |
| B44 | B209 | MCH | Smoking | 0.105713195 | 0 |
| B44 | B221 | MCH | Hei | 0.090851106 | 0 |
| B44 | B222 | MCH | NS | 0.080465509 | 0 |
| B44 | B45 | MCH | MCHC | 0.569629807 | 0 |
| B44 | B46 | MCH | RBC Count | -0.435420966 | 0 |
| B44 | B47 | MCH | RDW | 0.376964073 | 0 |
| B44 | B51 | MCH | Hb | 0.318036093 | 0 |
| B44 | B52 | MCH | PLT/BPC | -0.136725816 | 0 |
| B44 | B174 | MCH | Urea/Cr | -0.075175465 | 4.44E-16 |
| B44 | B127 | MCH | A-G ratio | 0.0742044 | 8.88E-16 |
| B44 | B140 | MCH | r-GT | 0.06646858 | 5.13E-13 |
| B44 | B130 | MCH | GLB | -0.062663687 | 9.93E-12 |
| B44 | B53 | MCH | GR% | -0.062423195 | 1.19E-11 |
| B44 | B163 | MCH | SG | 0.060935431 | 3.61E-11 |
| B44 | B189 | MCH | eGFB | -0.060327101 | 5.63E-11 |
| B44 | B216 | MCH | BMI | 0.059852516 | 7.95E-11 |
| B44 | B71 | MCH | TG | 0.057506358 | 4.20E-10 |
| B44 | B48 | MCH | LYM% | 0.05062827 | 3.84E-08 |
| B44 | B206 | MCH | Waist | 0.048209327 | 1.65E-07 |
| B44 | B58 | MCH | HCT | 0.047220119 | 2.93E-07 |
| B44 | B133 | MCH | TBIL | 0.046547555 | 4.32E-07 |
| B45 | B126 | MCHC | Alb | 0.142988772 | 0 |
| B45 | B127 | MCHC | A-G ratio | 0.123346222 | 0 |
| B45 | B129 | MCHC | IBIL | 0.134700637 | 0 |
| B45 | B133 | MCHC | TBIL | 0.12234495 | 0 |
| B45 | B140 | MCHC | r-GT | 0.105072106 | 0 |
| B45 | B141 | MCHC | AST/ALT | -0.113487444 | 0 |
| B45 | B153 | MCHC | Glu | 0.194164666 | 0 |
| B45 | B163 | MCHC | SG | 0.188015215 | 0 |
| B45 | B170 | MCHC | U-GLU | 0.17499088 | 0 |
| B45 | B174 | MCHC | Urea/Cr | -0.095105755 | 0 |
| B45 | B176 | MCHC | Ucr | 0.086769777 | 0 |
| B45 | B189 | MCHC | eGFB | 0.135339201 | 0 |
| B45 | B204 | MCHC | Drinking | 0.120784207 | 0 |
| B45 | B206 | MCHC | Waist | 0.119906216 | 0 |
| B45 | B209 | MCHC | Smoking | 0.12046989 | 0 |
| B45 | B210 | MCHC | Hip | 0.095928119 | 0 |
| B45 | B215 | MCHC | BMI | 0.098669666 | 0 |
| B45 | B216 | MCHC | BMI | 0.214611498 | 0 |
| B45 | B218 | MCHC | DBP | 0.130829347 | 0 |

| | | | | | |
|---|---|---|---|---|---|
| B45 | B221 | MCHC | Hei | 0.230614221 | 0 |
| B45 | B222 | MCHC | NS | 0.229288133 | 0 |
| B45 | B46 | MCHC | RBC Count | 0.086124142 | 0 |
| B45 | B47 | MCHC | RDW | -0.268816332 | 0 |
| B45 | B51 | MCHC | Hb | 0.533707614 | 0 |
| B45 | B56 | MCHC | IR | 0.078536959 | 0 |
| B45 | B58 | MCHC | HCT | 0.079855331 | 0 |
| B45 | B61 | MCHC | WBHSRV | 0.082584522 | 0 |
| B45 | B62 | MCHC | WBHSRVI | 0.096971964 | 0 |
| B45 | B63 | MCHC | WBRSR(mPas_200) | 0.097210755 | 0 |
| B45 | B66 | MCHC | WBRSR(mPas_100) | 0.094116339 | 0 |
| B45 | B67 | MCHC | WBRSR(mPas_50) | 0.094062944 | 0 |
| B45 | B71 | MCHC | TG | 0.22345131 | 0 |
| B45 | B72 | MCHC | HDL-C | -0.157699289 | 0 |
| B45 | B74 | MCHC | ApoA1 | -0.080357247 | 0 |
| B45 | B172 | MCHC | KB | 0.074705853 | 4.44E-16 |
| B45 | B60 | MCHC | WBRVI | 0.074570645 | 4.44E-16 |
| B45 | B65 | MCHC | WBRSR(mPas_1) | 0.07480544 | 4.44E-16 |
| B45 | B130 | MCHC | GLB | -0.071403454 | 8.44E-15 |
| B45 | B128 | MCHC | ALT | 0.068507481 | 9.77E-14 |
| B45 | B219 | MCHC | OT | -0.066128233 | 6.74E-13 |
| B45 | B192 | MCHC | UA | 0.061328429 | 2.70E-11 |
| B45 | B77 | MCHC | TC | 0.06049398 | 4.99E-11 |
| B45 | B166 | MCHC | Pro | 0.059811598 | 8.19E-11 |
| B45 | B191 | MCHC | Urea | -0.055545876 | 1.61E-09 |
| B45 | B54 | MCHC | ESR | -0.053101174 | 8.07E-09 |
| B45 | B161 | MCHC | LEU | -0.052875891 | 9.32E-09 |
| B45 | B193 | MCHC | K | -0.052239464 | 1.40E-08 |
| B45 | B195 | MCHC | Na | -0.051929011 | 1.70E-08 |
| B45 | B132 | MCHC | Dbil | 0.051831169 | 1.81E-08 |
| B45 | B194 | MCHC | Cl | -0.050748315 | 3.56E-08 |
| B45 | B173 | MCHC | NTT | -0.05020249 | 4.98E-08 |
| B45 | B138 | MCHC | PAB | 0.049120376 | 9.60E-08 |
| B46 | B113 | RBC Count | FT3 | 0.112664261 | 0 |
| B46 | B126 | RBC Count | Alb | 0.259401385 | 0 |
| B46 | B127 | RBC Count | A-G ratio | 0.125704856 | 0 |
| B46 | B128 | RBC Count | ALT | 0.110708627 | 0 |
| B46 | B129 | RBC Count | IBIL | 0.127155621 | 0 |
| B46 | B133 | RBC Count | TBIL | 0.113562609 | 0 |
| B46 | B134 | RBC Count | CHE | 0.080069597 | 0 |
| B46 | B141 | RBC Count | AST/ALT | -0.219488539 | 0 |
| B46 | B153 | RBC Count | Glu | 0.181772643 | 0 |
| B46 | B161 | RBC Count | LEU | -0.114222203 | 0 |
| B46 | B163 | RBC Count | SG | 0.263860701 | 0 |

| | | | | | |
|---|---|---|---|---|---|
| B46 | B166 | RBC Count | Pro | 0.085708078 | 0 |
| B46 | B170 | RBC Count | U-GLU | 0.214888807 | 0 |
| B46 | B172 | RBC Count | KB | 0.078636194 | 0 |
| B46 | B173 | RBC Count | NTT | -0.081473889 | 0 |
| B46 | B174 | RBC Count | Urea/Cr | -0.121679266 | 0 |
| B46 | B176 | RBC Count | Ucr | 0.106512371 | 0 |
| B46 | B188 | RBC Count | Ca | 0.137765394 | 0 |
| B46 | B189 | RBC Count | eGFB | 0.255249351 | 0 |
| B46 | B191 | RBC Count | Urea | -0.118320979 | 0 |
| B46 | B204 | RBC Count | Drinking | 0.114404004 | 0 |
| B46 | B206 | RBC Count | Waist | 0.185161952 | 0 |
| B46 | B209 | RBC Count | Smoking | 0.150952968 | 0 |
| B46 | B210 | RBC Count | Hip | 0.160655826 | 0 |
| B46 | B215 | RBC Count | BMI | 0.147919282 | 0 |
| B46 | B216 | RBC Count | BMI | 0.303312551 | 0 |
| B46 | B218 | RBC Count | DBP | 0.210504141 | 0 |
| B46 | B219 | RBC Count | OT | -0.089569333 | 0 |
| B46 | B221 | RBC Count | Hei | 0.320245019 | 0 |
| B46 | B222 | RBC Count | NS | 0.328511036 | 0 |
| B46 | B47 | RBC Count | RDW | -0.492306441 | 0 |
| B46 | B51 | RBC Count | Hb | 0.706125505 | 0 |
| B46 | B52 | RBC Count | PLT/BPC | 0.079324574 | 0 |
| B46 | B56 | RBC Count | IR | 0.095371956 | 0 |
| B46 | B58 | RBC Count | HCT | 0.189896129 | 0 |
| B46 | B59 | RBC Count | WBRV | 0.077253968 | 0 |
| B46 | B60 | RBC Count | WBRVI | 0.155714266 | 0 |
| B46 | B61 | RBC Count | WBHSRV | 0.101297768 | 0 |
| B46 | B62 | RBC Count | WBHSRVI | 0.158071783 | 0 |
| B46 | B63 | RBC Count | WBRSR(mPas_200) | 0.158602515 | 0 |
| B46 | B65 | RBC Count | WBRSR(mPas_1) | 0.157218462 | 0 |
| B46 | B66 | RBC Count | WBRSR(mPas_100) | 0.159722255 | 0 |
| B46 | B67 | RBC Count | WBRSR(mPas_50) | 0.163021804 | 0 |
| B46 | B70 | RBC Count | LDL-C | 0.11141218 | 0 |
| B46 | B71 | RBC Count | TG | 0.124388999 | 0 |
| B46 | B72 | RBC Count | HDL-C | -0.189425615 | 0 |
| B46 | B74 | RBC Count | ApoA1 | -0.087549647 | 0 |
| B46 | B75 | RBC Count | ApoA1/B | -0.083397811 | 0 |
| B46 | B77 | RBC Count | TC | 0.078782338 | 0 |
| B46 | B91 | RBC Count | TP | 0.115776261 | 0 |
| B46 | B54 | RBC Count | ESR | -0.0730571 | 2.00E-15 |
| B46 | B115 | RBC Count | TSH TT3 TT4(TT3) | 0.071420428 | 8.44E-15 |
| B46 | B140 | RBC Count | r-GT | 0.071338924 | 8.88E-15 |
| B46 | B192 | RBC Count | UA | 0.067996337 | 1.49E-13 |
| B46 | B64 | RBC Count | WBRSR(mPas_5) | 0.067310591 | 2.60E-13 |

| | | | | | |
|---|---|---|---|---|---|
| B46 | B111 | RBC Count | TSH | -0.06339006 | 5.72E-12 |
| B46 | B120 | RBC Count | HCV-cAg | -0.059702572 | 8.86E-11 |
| B46 | B194 | RBC Count | Cl | -0.057381567 | 4.58E-10 |
| B46 | B149 | RBC Count | TGAb | -0.055732947 | 1.42E-09 |
| B46 | B76 | RBC Count | ApoB | 0.049984274 | 5.69E-08 |
| B46 | B220 | RBC Count | SBP | -0.049657145 | 6.95E-08 |
| B46 | B138 | RBC Count | PAB | 0.049487208 | 7.70E-08 |
| B46 | B49 | RBC Count | baso% | 0.048780382 | 1.18E-07 |
| B46 | B160 | RBC Count | PH | -0.048468951 | 1.41E-07 |
| B46 | B146 | RBC Count | CysC | -0.046037039 | 5.76E-07 |
| B46 | B130 | RBC Count | GLB | -0.045619322 | 7.29E-07 |
| B47 | B126 | RDW | Alb | -0.112121616 | 0 |
| B47 | B132 | RDW | Dbil | 0.086906834 | 0 |
| B47 | B134 | RDW | CHE | -0.082631165 | 0 |
| B47 | B141 | RDW | AST/ALT | 0.10766492 | 0 |
| B47 | B153 | RDW | Glu | -0.169621273 | 0 |
| B47 | B163 | RDW | SG | -0.105215667 | 0 |
| B47 | B170 | RDW | U-GLU | -0.142170104 | 0 |
| B47 | B189 | RDW | eGFB | -0.223101323 | 0 |
| B47 | B190 | RDW | Cre | 0.117341416 | 0 |
| B47 | B191 | RDW | Urea | 0.104349989 | 0 |
| B47 | B216 | RDW | BMI | -0.117881337 | 0 |
| B47 | B218 | RDW | DBP | -0.104486299 | 0 |
| B47 | B221 | RDW | Hei | -0.105031873 | 0 |
| B47 | B222 | RDW | NS | -0.081704669 | 0 |
| B47 | B51 | RDW | Hb | -0.2323588 | 0 |
| B47 | B52 | RDW | PLT/BPC | -0.197411689 | 0 |
| B47 | B71 | RDW | TG | -0.076688601 | 0 |
| B47 | B72 | RDW | HDL-C | 0.123079108 | 0 |
| B47 | B131 | RDW | AST | 0.073683409 | 1.33E-15 |
| B47 | B74 | RDW | ApoA1 | 0.07308625 | 2.00E-15 |
| B47 | B146 | RDW | CysC | 0.072011869 | 5.11E-15 |
| B47 | B181 | RDW | APTT | 0.069824473 | 3.26E-14 |
| B47 | B215 | RDW | BMI | -0.069085877 | 6.06E-14 |
| B47 | B188 | RDW | Ca | -0.063992672 | 3.60E-12 |
| B47 | B127 | RDW | A-G ratio | -0.062653012 | 1.00E-11 |
| B47 | B75 | RDW | ApoA1/B | 0.059242622 | 1.23E-10 |
| B47 | B70 | RDW | LDL-C | -0.05894451 | 1.53E-10 |
| B47 | B195 | RDW | Na | 0.055233984 | 1.98E-09 |
| B47 | B63 | RDW | WBRSR(mPas_200) | -0.054151344 | 4.07E-09 |
| B47 | B111 | RDW | TSH | 0.053131109 | 7.91E-09 |
| B47 | B219 | RDW | OT | 0.052366028 | 1.29E-08 |
| B47 | B112 | RDW | FT4 | -0.051417514 | 2.35E-08 |
| B47 | B77 | RDW | TC | -0.051052733 | 2.95E-08 |

| | | | | | |
|---|---|---|---|---|---|
| B47 | B66 | RDW | WBRSR(mPas_100) | -0.050063877 | 5.43E-08 |
| B47 | B67 | RDW | WBRSR(mPas_50) | -0.049888568 | 6.04E-08 |
| B47 | B61 | RDW | WBHSRV | -0.049852997 | 6.17E-08 |
| B47 | B194 | RDW | CI | 0.049238529 | 8.94E-08 |
| B47 | B62 | RDW | WBHSRVI | -0.048702522 | 1.23E-07 |
| B47 | B209 | RDW | Smoking | 0.047540906 | 2.44E-07 |
| B47 | B211 | RDW | IOP-R | -0.046586234 | 4.22E-07 |
| B47 | B210 | RDW | Hip | -0.046339526 | 4.86E-07 |
| B48 | B132 | LYM% | Dbil | -0.078065014 | 0 |
| B48 | B153 | LYM% | Glu | -0.076931307 | 0 |
| B48 | B53 | LYM% | GR% | -0.936046118 | 0 |
| B48 | B70 | LYM% | LDL-C | 0.086981462 | 0 |
| B48 | B72 | LYM% | HDL-C | 0.093117081 | 0 |
| B48 | B77 | LYM% | TC | 0.095053731 | 0 |
| B48 | B189 | LYM% | eGFB | 0.073837522 | 8.88E-16 |
| B48 | B91 | LYM% | TP | 0.070963105 | 1.24E-14 |
| B48 | B190 | LYM% | Cre | -0.070041914 | 2.71E-14 |
| B48 | B170 | LYM% | U-GLU | -0.065605497 | 1.02E-12 |
| B48 | B166 | LYM% | Pro | -0.065068366 | 1.56E-12 |
| B48 | B123 | LYM% | hs-CRP1 | -0.061906109 | 1.76E-11 |
| B48 | B209 | LYM% | Smoking | -0.058535218 | 2.04E-10 |
| B48 | B133 | LYM% | TBIL | -0.058492225 | 2.10E-10 |
| B48 | B163 | LYM% | SG | -0.050395877 | 4.43E-08 |
| B48 | B126 | LYM% | Alb | 0.047490697 | 2.51E-07 |
| B48 | B130 | LYM% | GLB | 0.046758052 | 3.83E-07 |
| B48 | B129 | LYM% | IBIL | -0.045525317 | 7.68E-07 |
| B49 | B50 | baso% | eo% | 0.269393443 | 0 |
| B49 | B52 | baso% | PLT/BPC | 0.130099514 | 0 |
| B49 | B53 | baso% | GR% | -0.176206167 | 0 |
| B49 | B129 | baso% | IBIL | -0.053622052 | 5.75E-09 |
| B49 | B133 | baso% | TBIL | -0.050984789 | 3.08E-08 |
| B49 | B140 | baso% | r-GT | 0.049124954 | 9.57E-08 |
| B49 | B221 | baso% | Hei | 0.047672072 | 2.26E-07 |
| B5 | B121 | WB-Mg | C3 | 0.08115709 | 0 |
| B5 | B122 | WB-Mg | C4 | -0.12528822 | 0 |
| B5 | B29 | WB-Mg | E2 | 0.129770899 | 0 |
| B5 | B32 | WB-Mg | Testo | -0.107503107 | 0 |
| B5 | B33 | WB-Mg | Prog | 0.104072151 | 0 |
| B5 | B6 | WB-Mg | WB-Fe | 0.79870952 | 0 |
| B5 | B7 | WB-Mg | WB-Cop | 0.173416642 | 0 |
| B5 | B8 | WB-Mg | WB-Zinc | 0.247679333 | 0 |
| B5 | B120 | WB-Mg | HCV-cAg | 0.07522903 | 4.44E-16 |
| B5 | B158 | WB-Mg | IgM | 0.069078232 | 6.08E-14 |
| B5 | B31 | WB-Mg | FSH | -0.047174083 | 3.01E-07 |

| | | | | | |
|---|---|---|---|---|---|
| B50 | B189 | eo% | eGFB | -0.097600767 | 0 |
| B50 | B190 | eo% | Cre | 0.108493101 | 0 |
| B50 | B209 | eo% | Smoking | 0.077767835 | 0 |
| B50 | B53 | eo% | GR% | -0.274165774 | 0 |
| B50 | B174 | eo% | Urea/Cr | -0.069160686 | 5.68E-14 |
| B50 | B221 | eo% | Hei | 0.056772474 | 6.99E-10 |
| B50 | B222 | eo% | NS | 0.055797769 | 1.36E-09 |
| B50 | B126 | eo% | Alb | -0.055787289 | 1.37E-09 |
| B50 | B192 | eo% | UA | 0.050572856 | 3.97E-08 |
| B51 | B113 | Hb | FT3 | 0.122136469 | 0 |
| B51 | B126 | Hb | Alb | 0.301834817 | 0 |
| B51 | B127 | Hb | A-G ratio | 0.185868129 | 0 |
| B51 | B128 | Hb | ALT | 0.115005147 | 0 |
| B51 | B129 | Hb | IBIL | 0.161785317 | 0 |
| B51 | B130 | Hb | GLB | -0.093637461 | 0 |
| B51 | B133 | Hb | TBIL | 0.148437205 | 0 |
| B51 | B134 | Hb | CHE | 0.08213438 | 0 |
| B51 | B138 | Hb | PAB | 0.082075221 | 0 |
| B51 | B140 | Hb | r-GT | 0.120091935 | 0 |
| B51 | B141 | Hb | AST/ALT | -0.228096437 | 0 |
| B51 | B153 | Hb | Glu | 0.194668395 | 0 |
| B51 | B161 | Hb | LEU | -0.133341374 | 0 |
| B51 | B163 | Hb | SG | 0.326585828 | 0 |
| B51 | B166 | Hb | Pro | 0.105692674 | 0 |
| B51 | B170 | Hb | U-GLU | 0.229908083 | 0 |
| B51 | B172 | Hb | KB | 0.105994055 | 0 |
| B51 | B173 | Hb | NTT | -0.10111626 | 0 |
| B51 | B174 | Hb | Urea/Cr | -0.183102719 | 0 |
| B51 | B176 | Hb | Ucr | 0.141931297 | 0 |
| B51 | B188 | Hb | Ca | 0.146929079 | 0 |
| B51 | B189 | Hb | eGFB | 0.228573598 | 0 |
| B51 | B191 | Hb | Urea | -0.114093403 | 0 |
| B51 | B192 | Hb | UA | 0.099470693 | 0 |
| B51 | B204 | Hb | Drinking | 0.190379845 | 0 |
| B51 | B206 | Hb | Waist | 0.234186005 | 0 |
| B51 | B209 | Hb | Smoking | 0.23400538 | 0 |
| B51 | B210 | Hb | Hip | 0.194389784 | 0 |
| B51 | B215 | Hb | BMI | 0.167169044 | 0 |
| B51 | B216 | Hb | BMI | 0.368640463 | 0 |
| B51 | B218 | Hb | DBP | 0.226877114 | 0 |
| B51 | B219 | Hb | OT | -0.099221969 | 0 |
| B51 | B220 | Hb | SBP | -0.075887805 | 0 |
| B51 | B221 | Hb | Hei | 0.404961516 | 0 |
| B51 | B222 | Hb | NS | 0.405348081 | 0 |

| | | | | | |
|---|---|---|---|---|---|
| B51 | B54 | Hb | ESR | -0.099207097 | 0 |
| B51 | B56 | Hb | IR | 0.114996174 | 0 |
| B51 | B58 | Hb | HCT | 0.239239442 | 0 |
| B51 | B60 | Hb | WBRVI | 0.180629736 | 0 |
| B51 | B61 | Hb | WBHSRV | 0.118775471 | 0 |
| B51 | B62 | Hb | WBHSRVI | 0.197758628 | 0 |
| B51 | B63 | Hb | WBRSR(mPas_200) | 0.193914093 | 0 |
| B51 | B64 | Hb | WBRSR(mPas_5) | 0.081197629 | 0 |
| B51 | B65 | Hb | WBRSR(mPas_1) | 0.178672381 | 0 |
| B51 | B66 | Hb | WBRSR(mPas_100) | 0.193770894 | 0 |
| B51 | B67 | Hb | WBRSR(mPas_50) | 0.19626502 | 0 |
| B51 | B70 | Hb | LDL-C | 0.119802536 | 0 |
| B51 | B71 | Hb | TG | 0.172292337 | 0 |
| B51 | B72 | Hb | HDL-C | -0.184904026 | 0 |
| B51 | B74 | Hb | ApoA1 | -0.091694927 | 0 |
| B51 | B75 | Hb | ApoA1/B | -0.084940664 | 0 |
| B51 | B77 | Hb | TC | 0.110936707 | 0 |
| B51 | B91 | Hb | TP | 0.098133227 | 0 |
| B51 | B59 | Hb | WBRV | 0.070064012 | 2.66E-14 |
| B51 | B132 | Hb | Dbil | 0.067347681 | 2.53E-13 |
| B51 | B115 | Hb | TSH TT3 TT4(TT3) | 0.066393207 | 5.45E-13 |
| B51 | B120 | Hb | HCV-cAg | -0.064756174 | 1.99E-12 |
| B51 | B111 | Hb | TSH | -0.063820249 | 4.11E-12 |
| B51 | B194 | Hb | Cl | -0.062943852 | 8.03E-12 |
| B51 | B160 | Hb | PH | -0.062861438 | 8.55E-12 |
| B51 | B149 | Hb | TGAb | -0.060510307 | 4.93E-11 |
| B51 | B146 | Hb | CysC | -0.055351494 | 1.83E-09 |
| B51 | B76 | Hb | ApoB | 0.05474641 | 2.75E-09 |
| B51 | B148 | Hb | Anti_TPO | -0.052187312 | 1.45E-08 |
| B51 | B181 | Hb | APTT | -0.049735756 | 6.62E-08 |
| B52 | B128 | PLT/BPC | ALT | -0.084912246 | 0 |
| B52 | B129 | PLT/BPC | IBIL | -0.136158986 | 0 |
| B52 | B131 | PLT/BPC | AST | -0.09726427 | 0 |
| B52 | B132 | PLT/BPC | Dbil | -0.197921746 | 0 |
| B52 | B133 | PLT/BPC | TBIL | -0.164783279 | 0 |
| B52 | B189 | PLT/BPC | eGFB | 0.085478191 | 0 |
| B52 | B70 | PLT/BPC | LDL-C | 0.137317376 | 0 |
| B52 | B77 | PLT/BPC | TC | 0.096381882 | 0 |
| B52 | B191 | PLT/BPC | Urea | -0.07362695 | 1.33E-15 |
| B52 | B130 | PLT/BPC | GLB | 0.065085741 | 1.54E-12 |
| B52 | B127 | PLT/BPC | A-G ratio | -0.064145855 | 3.20E-12 |
| B52 | B188 | PLT/BPC | Ca | 0.059679099 | 9.01E-11 |
| B52 | B91 | PLT/BPC | TP | 0.058069376 | 2.84E-10 |
| B52 | B140 | PLT/BPC | r-GT | -0.054710173 | 2.81E-09 |

| | | | | | |
|---|---|---|---|---|---|
| B52 | B190 | PLT/BPC | Cre | -0.054168358 | 4.03E-09 |
| B52 | B146 | PLT/BPC | CysC | -0.051922656 | 1.71E-08 |
| B52 | B160 | PLT/BPC | PH | 0.051336104 | 2.47E-08 |
| B52 | B198 | PLT/BPC | P | 0.050502334 | 4.15E-08 |
| B52 | B76 | PLT/BPC | ApoB | 0.047896387 | 1.98E-07 |
| B53 | B153 | GR% | Glu | 0.089163779 | 0 |
| B53 | B70 | GR% | LDL-C | -0.080694557 | 0 |
| B53 | B72 | GR% | HDL-C | -0.085374836 | 0 |
| B53 | B77 | GR% | TC | -0.074293358 | 6.66E-16 |
| B53 | B170 | GR% | U-GLU | 0.068371016 | 1.09E-13 |
| B53 | B132 | GR% | Dbil | 0.068159078 | 1.30E-13 |
| B53 | B133 | GR% | TBIL | 0.058145546 | 2.69E-10 |
| B53 | B123 | GR% | hs-CRP1 | 0.057715202 | 3.63E-10 |
| B53 | B130 | GR% | GLB | -0.052472822 | 1.21E-08 |
| B53 | B127 | GR% | A-G ratio | 0.051401265 | 2.37E-08 |
| B53 | B91 | GR% | TP | -0.049195527 | 9.17E-08 |
| B53 | B129 | GR% | IBIL | 0.048640001 | 1.28E-07 |
| B53 | B166 | GR% | Pro | 0.048413031 | 1.46E-07 |
| B53 | B131 | GR% | AST | -0.047107094 | 3.13E-07 |
| B54 | B123 | ESR | hs-CRP1 | 0.244807951 | 0 |
| B54 | B126 | ESR | Alb | -0.081643366 | 0 |
| B54 | B127 | ESR | A-G ratio | -0.114987873 | 0 |
| B54 | B130 | ESR | GLB | 0.11054229 | 0 |
| B54 | B157 | ESR | IgG | 0.096084749 | 0 |
| B54 | B169 | ESR | U-CRY | 0.246208596 | 0 |
| B54 | B56 | ESR | IR | -0.199916917 | 0 |
| B54 | B58 | ESR | HCT | -0.350231687 | 0 |
| B54 | B60 | ESR | WBRVI | -0.286059723 | 0 |
| B54 | B62 | ESR | WBHSRVI | -0.292029735 | 0 |
| B54 | B63 | ESR | WBRSR(mPas_200) | -0.116392973 | 0 |
| B54 | B64 | ESR | WBRSR(mPas_5) | -0.076014261 | 0 |
| B54 | B65 | ESR | WBRSR(mPas_1) | -0.135158048 | 0 |
| B54 | B66 | ESR | WBRSR(mPas_100) | -0.112081862 | 0 |
| B54 | B67 | ESR | WBRSR(mPas_50) | -0.121095955 | 0 |
| B54 | B69 | ESR | PV | 0.448322343 | 0 |
| B54 | B73 | ESR | ApoA | 0.092168364 | 0 |
| B54 | B158 | ESR | IgM | 0.072164678 | 4.44E-15 |
| B54 | B114 | ESR | TSH TT3 TT4(TT4) | 0.071535348 | 7.55E-15 |
| B54 | B155 | ESR | IgA | 0.06927927 | 5.15E-14 |
| B54 | B57 | ESR | RBCAI | -0.069192949 | 5.55E-14 |
| B54 | B221 | ESR | Hei | -0.066748431 | 4.10E-13 |
| B54 | B76 | ESR | ApoB | 0.066231117 | 6.21E-13 |
| B54 | B178 | ESR | Ig-U | 0.064653744 | 2.15E-12 |
| B54 | B177 | ESR | ACR | 0.063544149 | 5.08E-12 |

| | | | | | |
|---|---|---|---|---|---|
| B54 | B198 | ESR | P | 0.062959186 | 7.94E-12 |
| B54 | B225 | ESR | hPBG | 0.062284994 | 1.32E-11 |
| B54 | B81 | ESR | TSGF | 0.061050998 | 3.31E-11 |
| B54 | B186 | ESR | PFC | 0.060866511 | 3.79E-11 |
| B54 | B179 | ESR | α1-MG | 0.054018879 | 4.44E-09 |
| B54 | B149 | ESR | TGAb | 0.052552358 | 1.15E-08 |
| B54 | B61 | ESR | WBHSRV | -0.050489828 | 4.18E-08 |
| B54 | B91 | ESR | TP | 0.050348253 | 4.56E-08 |
| B54 | B138 | ESR | PAB | -0.050104502 | 5.29E-08 |
| B54 | B161 | ESR | LEU | 0.048937517 | 1.07E-07 |
| B54 | B113 | ESR | FT3 | -0.048672961 | 1.25E-07 |
| B54 | B133 | ESR | TBIL | -0.048536007 | 1.36E-07 |
| B54 | B129 | ESR | IBIL | -0.046878323 | 3.57E-07 |
| B54 | B189 | ESR | eGFB | -0.045867834 | 6.34E-07 |
| B54 | B228 | ESR | OS | 0.045504221 | 7.77E-07 |
| B54 | B222 | ESR | NS | -0.045166428 | 9.37E-07 |
| B55 | B225 | TK | hPBG | -0.082671369 | 0 |
| B55 | B56 | TK | IR | 0.775074768 | 0 |
| B55 | B57 | TK | RBCAI | -0.23792157 | 0 |
| B55 | B58 | TK | HCT | -0.312551121 | 0 |
| B55 | B59 | TK | WBRV | 0.351651082 | 0 |
| B55 | B60 | TK | WBRVI | 0.208431345 | 0 |
| B55 | B61 | TK | WBHSRV | 0.688541884 | 0 |
| B55 | B62 | TK | WBHSRVI | 0.433244248 | 0 |
| B55 | B63 | TK | WBRSR(mPas_200) | 0.30976653 | 0 |
| B55 | B64 | TK | WBRSR(mPas_5) | 0.12253939 | 0 |
| B55 | B65 | TK | WBRSR(mPas_1) | 0.122661275 | 0 |
| B55 | B66 | TK | WBRSR(mPas_100) | 0.30068824 | 0 |
| B55 | B67 | TK | WBRSR(mPas_50) | 0.286252195 | 0 |
| B55 | B69 | TK | PV | -0.313416745 | 0 |
| B55 | B151 | TK | Anti-CCP | 0.064907049 | 1.77E-12 |
| B55 | B79 | TK | hb V -dna | -0.057809473 | 3.40E-10 |
| B55 | B193 | TK | K | -0.05672078 | 7.24E-10 |
| B55 | B89 | TK | CA 72-4 | 0.048711479 | 1.22E-07 |
| B55 | B103 | TK | anti-SSA/Ro | 0.04870457 | 1.23E-07 |
| B55 | B109 | TK | AMA-M2 | 0.047772848 | 2.13E-07 |
| B55 | B74 | TK | ApoA1 | -0.045695663 | 6.98E-07 |
| B56 | B57 | IR | RBCAI | -0.141194676 | 0 |
| B56 | B58 | IR | HCT | 0.300382332 | 0 |
| B56 | B59 | IR | WBRV | 0.509755968 | 0 |
| B56 | B60 | IR | WBRVI | 0.64522822 | 0 |
| B56 | B61 | IR | WBHSRV | 0.933391859 | 0 |
| B56 | B62 | IR | WBHSRVI | 0.899064993 | 0 |
| B56 | B63 | IR | WBRSR(mPas_200) | 0.786600881 | 0 |

| | | | | | |
|---|---|---|---|---|---|
| B56 | B64 | IR | WBRSR(mPas_5) | 0.312846549 | 0 |
| B56 | B65 | IR | WBRSR(mPas_1) | 0.566920959 | 0 |
| B56 | B66 | IR | WBRSR(mPas_100) | 0.779997135 | 0 |
| B56 | B67 | IR | WBRSR(mPas_50) | 0.772053309 | 0 |
| B56 | B69 | IR | PV | -0.262163244 | 0 |
| B56 | B74 | IR | ApoA1 | -0.117575555 | 0 |
| B56 | B75 | IR | ApoA1/B | -0.075896927 | 0 |
| B56 | B222 | IR | NS | 0.067911693 | 1.60E-13 |
| B56 | B115 | IR | TSH TT3 TT4(TT3) | 0.06756149 | 2.12E-13 |
| B56 | B225 | IR | hPBG | -0.063533328 | 5.12E-12 |
| B56 | B72 | IR | HDL-C | -0.063005482 | 7.66E-12 |
| B56 | B113 | IR | FT3 | 0.061720267 | 2.02E-11 |
| B56 | B216 | IR | BMI | 0.060411553 | 5.30E-11 |
| B56 | B79 | IR | hb V -dna | -0.057941754 | 3.10E-10 |
| B56 | B221 | IR | Hei | 0.05624792 | 1.00E-09 |
| B56 | B134 | IR | CHE | 0.055727695 | 1.42E-09 |
| B56 | B138 | IR | PAB | 0.053987243 | 4.53E-09 |
| B56 | B193 | IR | K | -0.051597388 | 2.10E-08 |
| B56 | B71 | IR | TG | 0.051511708 | 2.22E-08 |
| B56 | B93 | IR | ALDH2-Glu504Lys | 0.04916909 | 9.32E-08 |
| B56 | B183 | IR | PTA | -0.046506423 | 4.42E-07 |
| B56 | B92 | IR | ALDH2-Glu504Glu | -0.04638091 | 4.74E-07 |
| B56 | B163 | IR | SG | 0.045458492 | 7.97E-07 |
| B57 | B101 | RBCAI | anti-Sc70 | -0.089345873 | 0 |
| B57 | B104 | RBCAI | anti-SSB | -0.089345853 | 0 |
| B57 | B107 | RBCAI | AnuA | -0.089834094 | 0 |
| B57 | B151 | RBCAI | Anti-CCP | 0.078976618 | 0 |
| B57 | B58 | RBCAI | HCT | 0.127011153 | 0 |
| B57 | B59 | RBCAI | WBRV | 0.662912778 | 0 |
| B57 | B60 | RBCAI | WBRVI | 0.575528984 | 0 |
| B57 | B61 | RBCAI | WBHSRV | -0.134435496 | 0 |
| B57 | B64 | RBCAI | WBRSR(mPas_5) | 0.156620023 | 0 |
| B57 | B65 | RBCAI | WBRSR(mPas_1) | 0.563182789 | 0 |
| B57 | B98 | RBCAI | anti-Jo-1 | -0.089345835 | 0 |
| B57 | B67 | RBCAI | WBRSR(mPas_50) | 0.066097832 | 6.90E-13 |
| B57 | B81 | RBCAI | TSGF | -0.063863969 | 3.97E-12 |
| B57 | B96 | RBCAI | dsDNA (IIF) | -0.057843974 | 3.32E-10 |
| B57 | B139 | RBCAI | ADA | -0.055668188 | 1.48E-09 |
| B57 | B182 | RBCAI | TT | -0.054229418 | 3.87E-09 |
| B57 | B122 | RBCAI | C4 | 0.053521543 | 6.14E-09 |
| B57 | B74 | RBCAI | ApoA1 | 0.053480014 | 6.31E-09 |
| B57 | B169 | RBCAI | U-CRY | -0.052713952 | 1.03E-08 |
| B57 | B121 | RBCAI | C3 | 0.051566721 | 2.14E-08 |
| B57 | B103 | RBCAI | anti-SSA/Ro | 0.049026346 | 1.02E-07 |

| | | | | | |
|---|---|---|---|---|---|
| B57 | B124 | RBCAI | ACTH | 0.048431806 | 1.45E-07 |
| B57 | B125 | RBCAI | Cor | 0.047226438 | 2.92E-07 |
| B57 | B149 | RBCAI | TGAb | -0.046883797 | 3.56E-07 |
| B57 | B62 | RBCAI | WBHSRVI | -0.046518717 | 4.39E-07 |
| B58 | B113 | HCT | FT3 | 0.080612134 | 0 |
| B58 | B115 | HCT | TSH TT3 TT4(TT3) | 0.114591975 | 0 |
| B58 | B163 | HCT | SG | 0.087783891 | 0 |
| B58 | B176 | HCT | Ucr | 0.087129652 | 0 |
| B58 | B216 | HCT | BMI | 0.090537613 | 0 |
| B58 | B221 | HCT | Hei | 0.115263443 | 0 |
| B58 | B222 | HCT | NS | 0.108541603 | 0 |
| B58 | B59 | HCT | WBRV | 0.139562565 | 0 |
| B58 | B60 | HCT | WBRVI | 0.628987801 | 0 |
| B58 | B61 | HCT | WBHSRV | 0.295032259 | 0 |
| B58 | B62 | HCT | WBHSRVI | 0.674090963 | 0 |
| B58 | B63 | HCT | WBRSR(mPas_200) | 0.653473838 | 0 |
| B58 | B64 | HCT | WBRSR(mPas_5) | 0.255802165 | 0 |
| B58 | B65 | HCT | WBRSR(mPas_1) | 0.601724336 | 0 |
| B58 | B66 | HCT | WBRSR(mPas_100) | 0.642327938 | 0 |
| B58 | B67 | HCT | WBRSR(mPas_50) | 0.651953783 | 0 |
| B58 | B74 | HCT | ApoA1 | -0.119367381 | 0 |
| B58 | B75 | HCT | ApoA1/B | -0.105407226 | 0 |
| B58 | B151 | HCT | Anti-CCP | -0.074881307 | 4.44E-16 |
| B58 | B141 | HCT | AST/ALT | -0.072424788 | 3.55E-15 |
| B58 | B134 | HCT | CHE | 0.072310382 | 4.00E-15 |
| B58 | B174 | HCT | Urea/Cr | -0.065821637 | 8.60E-13 |
| B58 | B93 | HCT | ALDH2-Glu504Lys | 0.06570175 | 9.45E-13 |
| B58 | B73 | HCT | ApoA | -0.064832566 | 1.87E-12 |
| B58 | B149 | HCT | TGAb | -0.062189163 | 1.42E-11 |
| B58 | B138 | HCT | PAB | 0.061771351 | 1.94E-11 |
| B58 | B129 | HCT | IBIL | 0.061399628 | 2.56E-11 |
| B58 | B76 | HCT | ApoB | 0.06118701 | 3.00E-11 |
| B58 | B92 | HCT | ALDH2-Glu504Glu | -0.061126078 | 3.13E-11 |
| B58 | B69 | HCT | PV | -0.061071021 | 3.26E-11 |
| B58 | B133 | HCT | TBIL | 0.060574353 | 4.70E-11 |
| B58 | B126 | HCT | Alb | 0.060195948 | 6.20E-11 |
| B58 | B109 | HCT | AMA-M2 | -0.059955292 | 7.38E-11 |
| B58 | B72 | HCT | HDL-C | -0.05593056 | 1.24E-09 |
| B58 | B140 | HCT | r-GT | 0.054289745 | 3.72E-09 |
| B58 | B170 | HCT | U-GLU | 0.053332329 | 6.95E-09 |
| B58 | B206 | HCT | Waist | 0.052063061 | 1.57E-08 |
| B58 | B194 | HCT | Cl | -0.051251159 | 2.61E-08 |
| B58 | B121 | HCT | C3 | 0.05080651 | 3.44E-08 |
| B58 | B122 | HCT | C4 | 0.050738681 | 3.59E-08 |

| | | | | | |
|---|---|---|---|---|---|
| B58 | B158 | HCT | IgM | -0.048702096 | 1.23E-07 |
| B58 | B157 | HCT | IgG | -0.046713445 | 3.93E-07 |
| B58 | B209 | HCT | Smoking | 0.046399077 | 4.70E-07 |
| B58 | B148 | HCT | Anti_TPO | -0.046369962 | 4.77E-07 |
| B58 | B135 | HCT | TBA | -0.046129522 | 5.47E-07 |
| B58 | B188 | HCT | Ca | 0.045999283 | 5.89E-07 |
| B59 | B101 | WBRV | anti-Sc70 | -0.11411111 | 0 |
| B59 | B104 | WBRV | anti-SSB | -0.114111086 | 0 |
| B59 | B107 | WBRV | AnuA | -0.114759396 | 0 |
| B59 | B151 | WBRV | Anti-CCP | 0.112397994 | 0 |
| B59 | B60 | WBRV | WBRVI | 0.785799809 | 0 |
| B59 | B61 | WBRV | WBHSRV | 0.611515191 | 0 |
| B59 | B62 | WBRV | WBHSRVI | 0.463789829 | 0 |
| B59 | B63 | WBRV | WBRSR(mPas_200) | 0.555752914 | 0 |
| B59 | B64 | WBRV | WBRSR(mPas_5) | 0.355789932 | 0 |
| B59 | B65 | WBRV | WBRSR(mPas_1) | 0.858960685 | 0 |
| B59 | B66 | WBRV | WBRSR(mPas_100) | 0.585542809 | 0 |
| B59 | B67 | WBRV | WBRSR(mPas_50) | 0.624003462 | 0 |
| B59 | B69 | WBRV | PV | 0.218073663 | 0 |
| B59 | B98 | WBRV | anti-Jo-1 | -0.114111077 | 0 |
| B59 | B79 | WBRV | hb V -dna | -0.063884817 | 3.91E-12 |
| B59 | B126 | WBRV | Alb | 0.06232146 | 1.29E-11 |
| B59 | B103 | WBRV | anti-SSA/Ro | 0.060291572 | 5.78E-11 |
| B59 | B91 | WBRV | TP | 0.059412547 | 1.09E-10 |
| B59 | B122 | WBRV | C4 | 0.057249238 | 5.03E-10 |
| B59 | B121 | WBRV | C3 | 0.056456775 | 8.67E-10 |
| B59 | B138 | WBRV | PAB | 0.052621272 | 1.10E-08 |
| B59 | B71 | WBRV | TG | 0.050591072 | 3.93E-08 |
| B59 | B155 | WBRV | IgA | 0.05056007 | 4.00E-08 |
| B59 | B81 | WBRV | TSGF | -0.049477178 | 7.74E-08 |
| B59 | B119 | WBRV | Hcv-rna | -0.047811335 | 2.08E-07 |
| B59 | B96 | WBRV | dsDNA (IIF) | -0.04629968 | 4.97E-07 |
| B6 | B121 | WB-Fe | C3 | 0.076920775 | 0 |
| B6 | B122 | WB-Fe | C4 | -0.120202929 | 0 |
| B6 | B29 | WB-Fe | E2 | 0.101721114 | 0 |
| B6 | B33 | WB-Fe | Prog | 0.079296262 | 0 |
| B6 | B7 | WB-Fe | WB-Cop | 0.204859048 | 0 |
| B6 | B8 | WB-Fe | WB-Zinc | 0.452188515 | 0 |
| B6 | B158 | WB-Fe | IgM | 0.069983885 | 2.84E-14 |
| B6 | B31 | WB-Fe | FSH | -0.066460489 | 5.16E-13 |
| B60 | B113 | WBRVI | FT3 | 0.080698983 | 0 |
| B60 | B115 | WBRVI | TSH TT3 TT4(TT3) | 0.081858543 | 0 |
| B60 | B121 | WBRVI | C3 | 0.079277373 | 0 |
| B60 | B122 | WBRVI | C4 | 0.07987547 | 0 |

| | | | | | |
|---|---|---|---|---|---|
| B60 | B138 | WBRVI | PAB | 0.077708007 | 0 |
| B60 | B216 | WBRVI | BMI | 0.076916315 | 0 |
| B60 | B221 | WBRVI | Hei | 0.088764886 | 0 |
| B60 | B222 | WBRVI | NS | 0.091341397 | 0 |
| B60 | B61 | WBRVI | WBHSRV | 0.609444255 | 0 |
| B60 | B62 | WBRVI | WBHSRVI | 0.785896305 | 0 |
| B60 | B63 | WBRVI | WBRSR(mPas_200) | 0.713149422 | 0 |
| B60 | B64 | WBRVI | WBRSR(mPas_5) | 0.393546231 | 0 |
| B60 | B65 | WBRVI | WBRSR(mPas_1) | 0.936734209 | 0 |
| B60 | B66 | WBRVI | WBRSR(mPas_100) | 0.736405294 | 0 |
| B60 | B67 | WBRVI | WBRSR(mPas_50) | 0.772515329 | 0 |
| B60 | B69 | WBRVI | PV | -0.165123437 | 0 |
| B60 | B74 | WBRVI | ApoA1 | -0.082089169 | 0 |
| B60 | B75 | WBRVI | ApoA1/B | -0.079619208 | 0 |
| B60 | B107 | WBRVI | AnuA | -0.074500957 | 4.44E-16 |
| B60 | B101 | WBRVI | anti-Sc70 | -0.074076783 | 8.88E-16 |
| B60 | B104 | WBRVI | anti-SSB | -0.07407677 | 8.88E-16 |
| B60 | B98 | WBRVI | anti-Jo-1 | -0.07407676 | 8.88E-16 |
| B60 | B149 | WBRVI | TGAb | -0.07189099 | 5.77E-15 |
| B60 | B126 | WBRVI | Alb | 0.070940443 | 1.27E-14 |
| B60 | B163 | WBRVI | SG | 0.06855059 | 9.46E-14 |
| B60 | B79 | WBRVI | hb V -dna | -0.06107626 | 3.25E-11 |
| B60 | B93 | WBRVI | ALDH2-Glu504Lys | 0.05580541 | 1.35E-09 |
| B60 | B119 | WBRVI | Hcv-rna | -0.053447919 | 6.44E-09 |
| B60 | B174 | WBRVI | Urea/Cr | -0.052970313 | 8.78E-09 |
| B60 | B183 | WBRVI | PTA | -0.052513768 | 1.18E-08 |
| B60 | B127 | WBRVI | A-G ratio | 0.052490549 | 1.19E-08 |
| B60 | B148 | WBRVI | Anti_TPO | -0.052365942 | 1.29E-08 |
| B60 | B92 | WBRVI | ALDH2-Glu504Glu | -0.050876542 | 3.29E-08 |
| B60 | B140 | WBRVI | r-GT | 0.050471716 | 4.23E-08 |
| B60 | B189 | WBRVI | eGFB | 0.050190516 | 5.02E-08 |
| B60 | B72 | WBRVI | HDL-C | -0.045315173 | 8.63E-07 |
| B61 | B134 | WBHSRV | CHE | 0.096456711 | 0 |
| B61 | B194 | WBHSRV | Cl | -0.076093983 | 0 |
| B61 | B62 | WBHSRV | WBHSRVI | 0.852723349 | 0 |
| B61 | B63 | WBHSRV | WBRSR(mPas_200) | 0.890740982 | 0 |
| B61 | B64 | WBHSRV | WBRSR(mPas_5) | 0.35105402 | 0 |
| B61 | B65 | WBHSRV | WBRSR(mPas_1) | 0.653336027 | 0 |
| B61 | B66 | WBHSRV | WBRSR(mPas_100) | 0.881938447 | 0 |
| B61 | B67 | WBHSRV | WBRSR(mPas_50) | 0.873282996 | 0 |
| B61 | B69 | WBHSRV | PV | 0.082171251 | 0 |
| B61 | B74 | WBHSRV | ApoA1 | -0.088885863 | 0 |
| B61 | B75 | WBHSRV | ApoA1/B | -0.08158683 | 0 |
| B61 | B76 | WBHSRV | ApoB | 0.082420283 | 0 |

| | | | | | |
|---|---|---|---|---|---|
| B61 | B71 | WBHSRV | TG | 0.074249136 | 6.66E-16 |
| B61 | B115 | WBHSRV | TSH TT3 TT4(TT3) | 0.070658843 | 1.60E-14 |
| B61 | B91 | WBHSRV | TP | 0.064788235 | 1.94E-12 |
| B61 | B222 | WBHSRV | NS | 0.062015606 | 1.62E-11 |
| B61 | B72 | WBHSRV | HDL-C | -0.059274041 | 1.21E-10 |
| B61 | B155 | WBHSRV | IgA | 0.058269483 | 2.46E-10 |
| B61 | B216 | WBHSRV | BMI | 0.057621864 | 3.88E-10 |
| B61 | B113 | WBHSRV | FT3 | 0.057031646 | 5.84E-10 |
| B61 | B153 | WBHSRV | Glu | 0.056088825 | 1.11E-09 |
| B61 | B163 | WBHSRV | SG | 0.055588968 | 1.56E-09 |
| B61 | B138 | WBHSRV | PAB | 0.053520739 | 6.15E-09 |
| B61 | B79 | WBHSRV | hb V -dna | -0.052448889 | 1.23E-08 |
| B61 | B169 | WBHSRV | U-CRY | 0.052328214 | 1.32E-08 |
| B61 | B126 | WBHSRV | Alb | 0.052045374 | 1.58E-08 |
| B61 | B170 | WBHSRV | U-GLU | 0.049665625 | 6.91E-08 |
| B61 | B136 | WBHSRV | 5'NT | 0.047630259 | 2.31E-07 |
| B61 | B225 | WBHSRV | hPBG | -0.046278054 | 5.03E-07 |
| B61 | B119 | WBHSRV | Hcv-rna | -0.045946822 | 6.06E-07 |
| B62 | B113 | WBHSRVI | FT3 | 0.081091733 | 0 |
| B62 | B115 | WBHSRVI | TSH TT3 TT4(TT3) | 0.100258595 | 0 |
| B62 | B134 | WBHSRVI | CHE | 0.076367889 | 0 |
| B62 | B216 | WBHSRVI | BMI | 0.088017092 | 0 |
| B62 | B221 | WBHSRVI | Hei | 0.092581311 | 0 |
| B62 | B222 | WBHSRVI | NS | 0.099876142 | 0 |
| B62 | B63 | WBHSRVI | WBRSR(mPas_200) | 0.909473602 | 0 |
| B62 | B64 | WBHSRVI | WBRSR(mPas_5) | 0.362673667 | 0 |
| B62 | B65 | WBHSRVI | WBRSR(mPas_1) | 0.720667082 | 0 |
| B62 | B66 | WBHSRVI | WBRSR(mPas_100) | 0.904679297 | 0 |
| B62 | B67 | WBHSRVI | WBRSR(mPas_50) | 0.903029051 | 0 |
| B62 | B69 | WBHSRVI | PV | -0.204910842 | 0 |
| B62 | B74 | WBHSRVI | ApoA1 | -0.14422454 | 0 |
| B62 | B75 | WBHSRVI | ApoA1/B | -0.107114919 | 0 |
| B62 | B163 | WBHSRVI | SG | 0.07487523 | 4.44E-16 |
| B62 | B72 | WBHSRVI | HDL-C | -0.072646707 | 2.89E-15 |
| B62 | B93 | WBHSRVI | ALDH2-Glu504Lys | 0.071172039 | 1.04E-14 |
| B62 | B138 | WBHSRVI | PAB | 0.068163209 | 1.30E-13 |
| B62 | B92 | WBHSRVI | ALDH2-Glu504Glu | -0.066463352 | 5.15E-13 |
| B62 | B194 | WBHSRVI | Cl | -0.062125225 | 1.49E-11 |
| B62 | B176 | WBHSRVI | Ucr | 0.058075347 | 2.82E-10 |
| B62 | B141 | WBHSRVI | AST/ALT | -0.057039726 | 5.81E-10 |
| B62 | B126 | WBHSRVI | Alb | 0.056820998 | 6.76E-10 |
| B62 | B76 | WBHSRVI | ApoB | 0.056724688 | 7.22E-10 |
| B62 | B149 | WBHSRVI | TGAb | -0.056627307 | 7.72E-10 |
| B62 | B71 | WBHSRVI | TG | 0.053375855 | 6.75E-09 |

| | | | | | |
|---|---|---|---|---|---|
| B62 | B170 | WBHSRVI | U-GLU | 0.052962475 | 8.82E-09 |
| B62 | B206 | WBHSRVI | Waist | 0.052833404 | 9.58E-09 |
| B62 | B189 | WBHSRVI | eGFB | 0.05205632 | 1.57E-08 |
| B62 | B121 | WBHSRVI | C3 | 0.05150632 | 2.22E-08 |
| B62 | B225 | WBHSRVI | hPBG | -0.050178476 | 5.06E-08 |
| B62 | B122 | WBHSRVI | C4 | 0.050142212 | 5.17E-08 |
| B62 | B79 | WBHSRVI | hb V -dna | -0.049336727 | 8.43E-08 |
| B62 | B153 | WBHSRVI | Glu | 0.048190866 | 1.67E-07 |
| B62 | B140 | WBHSRVI | r-GT | 0.048071624 | 1.79E-07 |
| B62 | B119 | WBHSRVI | Hcv-rna | -0.047009962 | 3.31E-07 |
| B62 | B158 | WBHSRVI | IgM | -0.045349833 | 8.47E-07 |
| B63 | B115 | WBRSR(mPas_200) | TSH TT3 TT4(TT3) | 0.101099603 | 0 |
| B63 | B134 | WBRSR(mPas_200) | CHE | 0.119188758 | 0 |
| B63 | B163 | WBRSR(mPas_200) | SG | 0.084123802 | 0 |
| B63 | B194 | WBRSR(mPas_200) | Cl | -0.091291067 | 0 |
| B63 | B216 | WBRSR(mPas_200) | BMI | 0.079617979 | 0 |
| B63 | B221 | WBRSR(mPas_200) | Hei | 0.076337826 | 0 |
| B63 | B222 | WBRSR(mPas_200) | NS | 0.088144747 | 0 |
| B63 | B64 | WBRSR(mPas_200) | WBRSR(mPas_5) | 0.390968174 | 0 |
| B63 | B65 | WBRSR(mPas_200) | WBRSR(mPas_1) | 0.789908572 | 0 |
| B63 | B66 | WBRSR(mPas_200) | WBRSR(mPas_100) | 0.988423574 | 0 |
| B63 | B67 | WBRSR(mPas_200) | WBRSR(mPas_50) | 0.98180803 | 0 |
| B63 | B69 | WBRSR(mPas_200) | PV | 0.197184265 | 0 |
| B63 | B71 | WBRSR(mPas_200) | TG | 0.075964603 | 0 |
| B63 | B74 | WBRSR(mPas_200) | ApoA1 | -0.102318738 | 0 |
| B63 | B75 | WBRSR(mPas_200) | ApoA1/B | -0.107086927 | 0 |
| B63 | B76 | WBRSR(mPas_200) | ApoB | 0.101458342 | 0 |
| B63 | B91 | WBRSR(mPas_200) | TP | 0.078363945 | 0 |
| B63 | B113 | WBRSR(mPas_200) | FT3 | 0.071414942 | 8.44E-15 |
| B63 | B153 | WBRSR(mPas_200) | Glu | 0.069880742 | 3.11E-14 |
| B63 | B170 | WBRSR(mPas_200) | U-GLU | 0.067911233 | 1.60E-13 |
| B63 | B126 | WBRSR(mPas_200) | Alb | 0.066730471 | 4.16E-13 |
| B63 | B72 | WBRSR(mPas_200) | HDL-C | -0.065022232 | 1.61E-12 |
| B63 | B138 | WBRSR(mPas_200) | PAB | 0.062092525 | 1.53E-11 |
| B63 | B140 | WBRSR(mPas_200) | r-GT | 0.060763479 | 4.09E-11 |
| B63 | B176 | WBRSR(mPas_200) | Ucr | 0.057278491 | 4.93E-10 |
| B63 | B155 | WBRSR(mPas_200) | IgA | 0.055210975 | 2.02E-09 |
| B63 | B206 | WBRSR(mPas_200) | Waist | 0.054654735 | 2.92E-09 |
| B63 | B188 | WBRSR(mPas_200) | Ca | 0.051948027 | 1.68E-08 |
| B63 | B169 | WBRSR(mPas_200) | U-CRY | 0.051901977 | 1.73E-08 |
| B63 | B189 | WBRSR(mPas_200) | eGFB | 0.04870529 | 1.23E-07 |
| B63 | B136 | WBRSR(mPas_200) | 5'NT | 0.047948542 | 1.92E-07 |
| B63 | B121 | WBRSR(mPas_200) | C3 | 0.047762356 | 2.14E-07 |
| B63 | B141 | WBRSR(mPas_200) | AST/ALT | -0.047062785 | 3.21E-07 |

| | | | | | |
|---|---|---|---|---|---|
| B63 | B137 | WBRSR(mPas_200) | AFU | 0.046919648 | 3.49E-07 |
| B63 | B122 | WBRSR(mPas_200) | C4 | 0.046915729 | 3.50E-07 |
| B63 | B200 | WBRSR(mPas_200) | OP | 0.046099663 | 5.56E-07 |
| B63 | B77 | WBRSR(mPas_200) | TC | 0.045725469 | 6.87E-07 |
| B64 | B65 | WBRSR(mPas_5) | WBRSR(mPas_1) | 0.419994928 | 0 |
| B64 | B66 | WBRSR(mPas_5) | WBRSR(mPas_100) | 0.397213832 | 0 |
| B64 | B67 | WBRSR(mPas_5) | WBRSR(mPas_50) | 0.404102466 | 0 |
| B64 | B69 | WBRSR(mPas_5) | PV | 0.070994638 | 1.20E-14 |
| B64 | B212 | WBRSR(mPas_5) | IOP-L | 0.057986342 | 3.01E-10 |
| B64 | B74 | WBRSR(mPas_5) | ApoA1 | -0.054839599 | 2.58E-09 |
| B64 | B71 | WBRSR(mPas_5) | TG | 0.049970051 | 5.74E-08 |
| B65 | B101 | WBRSR(mPas_1) | anti-Sc70 | -0.098362646 | 0 |
| B65 | B104 | WBRSR(mPas_1) | anti-SSB | -0.098362621 | 0 |
| B65 | B107 | WBRSR(mPas_1) | AnuA | -0.098940169 | 0 |
| B65 | B115 | WBRSR(mPas_1) | TSH TT3 TT4(TT3) | 0.083426361 | 0 |
| B65 | B121 | WBRSR(mPas_1) | C3 | 0.078612353 | 0 |
| B65 | B122 | WBRSR(mPas_1) | C4 | 0.079433888 | 0 |
| B65 | B126 | WBRSR(mPas_1) | Alb | 0.081666121 | 0 |
| B65 | B163 | WBRSR(mPas_1) | SG | 0.077036711 | 0 |
| B65 | B221 | WBRSR(mPas_1) | Hei | 0.075868383 | 0 |
| B65 | B222 | WBRSR(mPas_1) | NS | 0.084217439 | 0 |
| B65 | B66 | WBRSR(mPas_1) | WBRSR(mPas_100) | 0.81116858 | 0 |
| B65 | B67 | WBRSR(mPas_1) | WBRSR(mPas_50) | 0.843204812 | 0 |
| B65 | B69 | WBRSR(mPas_1) | PV | 0.168921668 | 0 |
| B65 | B75 | WBRSR(mPas_1) | ApoA1/B | -0.08840585 | 0 |
| B65 | B98 | WBRSR(mPas_1) | anti-Jo-1 | -0.098362615 | 0 |
| B65 | B138 | WBRSR(mPas_1) | PAB | 0.075772533 | 2.22E-16 |
| B65 | B134 | WBRSR(mPas_1) | CHE | 0.074672685 | 4.44E-16 |
| B65 | B113 | WBRSR(mPas_1) | FT3 | 0.074144837 | 8.88E-16 |
| B65 | B216 | WBRSR(mPas_1) | BMI | 0.072875326 | 2.22E-15 |
| B65 | B76 | WBRSR(mPas_1) | ApoB | 0.068028003 | 1.45E-13 |
| B65 | B194 | WBRSR(mPas_1) | Cl | -0.064779012 | 1.95E-12 |
| B65 | B91 | WBRSR(mPas_1) | TP | 0.064134774 | 3.22E-12 |
| B65 | B140 | WBRSR(mPas_1) | r-GT | 0.062848057 | 8.64E-12 |
| B65 | B74 | WBRSR(mPas_1) | ApoA1 | -0.059214483 | 1.26E-10 |
| B65 | B71 | WBRSR(mPas_1) | TG | 0.058538898 | 2.04E-10 |
| B65 | B119 | WBRSR(mPas_1) | Hcv-rna | -0.058472494 | 2.13E-10 |
| B65 | B149 | WBRSR(mPas_1) | TGAb | -0.054295867 | 3.70E-09 |
| B65 | B79 | WBRSR(mPas_1) | hb V -dna | -0.053660156 | 5.61E-09 |
| B65 | B151 | WBRSR(mPas_1) | Anti-CCP | 0.051909055 | 1.73E-08 |
| B65 | B189 | WBRSR(mPas_1) | eGFB | 0.051183913 | 2.72E-08 |
| B65 | B206 | WBRSR(mPas_1) | Waist | 0.046382138 | 4.74E-07 |
| B65 | B170 | WBRSR(mPas_1) | U-GLU | 0.046208899 | 5.23E-07 |
| B66 | B115 | WBRSR(mPas_100) | TSH TT3 TT4(TT3) | 0.092859308 | 0 |

| | | | | | |
|---|---|---|---|---|---|
| B66 | B134 | WBRSR(mPas_100) | CHE | 0.115471939 | 0 |
| B66 | B163 | WBRSR(mPas_100) | SG | 0.08473142 | 0 |
| B66 | B194 | WBRSR(mPas_100) | Cl | -0.101968754 | 0 |
| B66 | B216 | WBRSR(mPas_100) | BMI | 0.080150602 | 0 |
| B66 | B222 | WBRSR(mPas_100) | NS | 0.089336006 | 0 |
| B66 | B67 | WBRSR(mPas_100) | WBRSR(mPas_50) | 0.995685571 | 0 |
| B66 | B69 | WBRSR(mPas_100) | PV | 0.196696789 | 0 |
| B66 | B71 | WBRSR(mPas_100) | TG | 0.076478425 | 0 |
| B66 | B74 | WBRSR(mPas_100) | ApoA1 | -0.103903428 | 0 |
| B66 | B75 | WBRSR(mPas_100) | ApoA1/B | -0.110236131 | 0 |
| B66 | B76 | WBRSR(mPas_100) | ApoB | 0.103017599 | 0 |
| B66 | B91 | WBRSR(mPas_100) | TP | 0.081683453 | 0 |
| B66 | B221 | WBRSR(mPas_100) | Hei | 0.075055063 | 4.44E-16 |
| B66 | B113 | WBRSR(mPas_100) | FT3 | 0.071075659 | 1.13E-14 |
| B66 | B153 | WBRSR(mPas_100) | Glu | 0.070869222 | 1.35E-14 |
| B66 | B126 | WBRSR(mPas_100) | Alb | 0.07053051 | 1.80E-14 |
| B66 | B170 | WBRSR(mPas_100) | U-GLU | 0.068040332 | 1.44E-13 |
| B66 | B72 | WBRSR(mPas_100) | HDL-C | -0.064895193 | 1.78E-12 |
| B66 | B140 | WBRSR(mPas_100) | r-GT | 0.063990728 | 3.60E-12 |
| B66 | B138 | WBRSR(mPas_100) | PAB | 0.062830917 | 8.75E-12 |
| B66 | B176 | WBRSR(mPas_100) | Ucr | 0.059768242 | 8.45E-11 |
| B66 | B107 | WBRSR(mPas_100) | AnuA | -0.058762501 | 1.74E-10 |
| B66 | B101 | WBRSR(mPas_100) | anti-Sc70 | -0.058397304 | 2.25E-10 |
| B66 | B104 | WBRSR(mPas_100) | anti-SSB | -0.05839729 | 2.25E-10 |
| B66 | B98 | WBRSR(mPas_100) | anti-Jo-1 | -0.058397281 | 2.25E-10 |
| B66 | B155 | WBRSR(mPas_100) | IgA | 0.057647391 | 3.81E-10 |
| B66 | B137 | WBRSR(mPas_100) | AFU | 0.054696442 | 2.84E-09 |
| B66 | B121 | WBRSR(mPas_100) | C3 | 0.054571272 | 3.09E-09 |
| B66 | B119 | WBRSR(mPas_100) | Hcv-rna | -0.054516964 | 3.20E-09 |
| B66 | B206 | WBRSR(mPas_100) | Waist | 0.054414958 | 3.42E-09 |
| B66 | B195 | WBRSR(mPas_100) | Na | -0.053747121 | 5.30E-09 |
| B66 | B122 | WBRSR(mPas_100) | C4 | 0.053510408 | 6.19E-09 |
| B66 | B200 | WBRSR(mPas_100) | OP | 0.052784406 | 9.89E-09 |
| B66 | B189 | WBRSR(mPas_100) | eGFB | 0.05264557 | 1.08E-08 |
| B66 | B188 | WBRSR(mPas_100) | Ca | 0.052620942 | 1.10E-08 |
| B66 | B169 | WBRSR(mPas_100) | U-CRY | 0.051448593 | 2.31E-08 |
| B66 | B136 | WBRSR(mPas_100) | 5'NT | 0.048272551 | 1.59E-07 |
| B66 | B139 | WBRSR(mPas_100) | ADA | 0.046431227 | 4.61E-07 |
| B66 | B141 | WBRSR(mPas_100) | AST/ALT | -0.045666885 | 7.09E-07 |
| B66 | B77 | WBRSR(mPas_100) | TC | 0.045476233 | 7.89E-07 |
| B66 | B215 | WBRSR(mPas_100) | BMI | 0.045102686 | 9.71E-07 |
| B67 | B115 | WBRSR(mPas_50) | TSH TT3 TT4(TT3) | 0.095229392 | 0 |
| B67 | B134 | WBRSR(mPas_50) | CHE | 0.110965534 | 0 |
| B67 | B163 | WBRSR(mPas_50) | SG | 0.085349223 | 0 |

| | | | | | |
|---|---|---|---|---|---|
| B67 | B194 | WBRSR(mPas_50) | Cl | -0.099740244 | 0 |
| B67 | B216 | WBRSR(mPas_50) | BMI | 0.081931614 | 0 |
| B67 | B221 | WBRSR(mPas_50) | Hei | 0.077023522 | 0 |
| B67 | B222 | WBRSR(mPas_50) | NS | 0.091549969 | 0 |
| B67 | B69 | WBRSR(mPas_50) | PV | 0.193785873 | 0 |
| B67 | B74 | WBRSR(mPas_50) | ApoA1 | -0.098386281 | 0 |
| B67 | B75 | WBRSR(mPas_50) | ApoA1/B | -0.109147567 | 0 |
| B67 | B76 | WBRSR(mPas_50) | ApoB | 0.101298724 | 0 |
| B67 | B91 | WBRSR(mPas_50) | TP | 0.080674939 | 0 |
| B67 | B71 | WBRSR(mPas_50) | TG | 0.075459524 | 2.22E-16 |
| B67 | B113 | WBRSR(mPas_50) | FT3 | 0.074294026 | 6.66E-16 |
| B67 | B126 | WBRSR(mPas_50) | Alb | 0.073030332 | 2.22E-15 |
| B67 | B153 | WBRSR(mPas_50) | Glu | 0.068763717 | 7.90E-14 |
| B67 | B107 | WBRSR(mPas_50) | AnuA | -0.068128322 | 1.34E-13 |
| B67 | B101 | WBRSR(mPas_50) | anti-Sc70 | -0.06771171 | 1.88E-13 |
| B67 | B104 | WBRSR(mPas_50) | anti-SSB | -0.067711705 | 1.88E-13 |
| B67 | B98 | WBRSR(mPas_50) | anti-Jo-1 | -0.067711696 | 1.88E-13 |
| B67 | B170 | WBRSR(mPas_50) | U-GLU | 0.066792384 | 3.96E-13 |
| B67 | B140 | WBRSR(mPas_50) | r-GT | 0.065096093 | 1.52E-12 |
| B67 | B138 | WBRSR(mPas_50) | PAB | 0.063981678 | 3.63E-12 |
| B67 | B119 | WBRSR(mPas_50) | Hcv-rna | -0.063583573 | 4.93E-12 |
| B67 | B72 | WBRSR(mPas_50) | HDL-C | -0.062816789 | 8.84E-12 |
| B67 | B176 | WBRSR(mPas_50) | Ucr | 0.061152109 | 3.07E-11 |
| B67 | B121 | WBRSR(mPas_50) | C3 | 0.058554502 | 2.01E-10 |
| B67 | B122 | WBRSR(mPas_50) | C4 | 0.057241018 | 5.05E-10 |
| B67 | B155 | WBRSR(mPas_50) | IgA | 0.055760831 | 1.39E-09 |
| B67 | B206 | WBRSR(mPas_50) | Waist | 0.055495994 | 1.66E-09 |
| B67 | B189 | WBRSR(mPas_50) | eGFB | 0.054733927 | 2.77E-09 |
| B67 | B93 | WBRSR(mPas_50) | ALDH2-Glu504Lys | 0.053183687 | 7.65E-09 |
| B67 | B195 | WBRSR(mPas_50) | Na | -0.05230896 | 1.34E-08 |
| B67 | B200 | WBRSR(mPas_50) | OP | 0.052185264 | 1.45E-08 |
| B67 | B188 | WBRSR(mPas_50) | Ca | 0.052131711 | 1.50E-08 |
| B67 | B137 | WBRSR(mPas_50) | AFU | 0.049914771 | 5.94E-08 |
| B67 | B92 | WBRSR(mPas_50) | ALDH2-Glu504Glu | -0.048233212 | 1.63E-07 |
| B67 | B169 | WBRSR(mPas_50) | U-CRY | 0.048011982 | 1.85E-07 |
| B67 | B79 | WBRSR(mPas_50) | hb V -dna | -0.047519418 | 2.47E-07 |
| B67 | B215 | WBRSR(mPas_50) | BMI | 0.045801364 | 6.58E-07 |
| B67 | B77 | WBRSR(mPas_50) | TC | 0.045482751 | 7.86E-07 |
| B67 | B141 | WBRSR(mPas_50) | AST/ALT | -0.045272588 | 8.84E-07 |
| B69 | B112 | PV | FT4 | 0.099558059 | 0 |
| B69 | B114 | PV | TSH TT3 TT4(TT4) | 0.125110158 | 0 |
| B69 | B123 | PV | hs-CRP1 | 0.122879592 | 0 |
| B69 | B127 | PV | A-G ratio | -0.095747942 | 0 |
| B69 | B130 | PV | GLB | 0.128995252 | 0 |

| | | | | | |
|---|---|---|---|---|---|
| B69 | B134 | PV | CHE | 0.096685686 | 0 |
| B69 | B137 | PV | AFU | 0.081384011 | 0 |
| B69 | B155 | PV | IgA | 0.119749939 | 0 |
| B69 | B169 | PV | U-CRY | 0.118836842 | 0 |
| B69 | B194 | PV | Cl | -0.086756772 | 0 |
| B69 | B225 | PV | hPBG | 0.119323817 | 0 |
| B69 | B76 | PV | ApoB | 0.117379527 | 0 |
| B69 | B91 | PV | TP | 0.132600678 | 0 |
| B69 | B107 | PV | AnuA | -0.074515974 | 4.44E-16 |
| B69 | B74 | PV | ApoA1 | 0.074690883 | 4.44E-16 |
| B69 | B101 | PV | anti-Sc70 | -0.074053718 | 8.88E-16 |
| B69 | B104 | PV | anti-SSB | -0.074053681 | 8.88E-16 |
| B69 | B98 | PV | anti-Jo-1 | -0.07405368 | 8.88E-16 |
| B69 | B186 | PV | PFC | 0.070480477 | 1.87E-14 |
| B69 | B77 | PV | TC | 0.066115786 | 6.80E-13 |
| B69 | B193 | PV | K | 0.066102642 | 6.87E-13 |
| B69 | B139 | PV | ADA | 0.065393881 | 1.21E-12 |
| B69 | B195 | PV | Na | -0.060163897 | 6.34E-11 |
| B69 | B70 | PV | LDL-C | 0.05730199 | 4.85E-10 |
| B69 | B157 | PV | IgG | 0.057150561 | 5.38E-10 |
| B69 | B183 | PV | PTA | 0.057063291 | 5.72E-10 |
| B69 | B153 | PV | Glu | 0.056463767 | 8.63E-10 |
| B69 | B158 | PV | IgM | 0.05583549 | 1.32E-09 |
| B69 | B149 | PV | TGAb | 0.051421585 | 2.34E-08 |
| B69 | B178 | PV | Ig-U | 0.05025957 | 4.81E-08 |
| B69 | B71 | PV | TG | 0.050097348 | 5.32E-08 |
| B69 | B188 | PV | Ca | 0.046945028 | 3.44E-07 |
| B69 | B200 | PV | OP | 0.046896027 | 3.54E-07 |
| B69 | B120 | PV | HCV-cAg | -0.045717485 | 6.90E-07 |
| B7 | B121 | WB-Cop | C3 | 0.097168104 | 0 |
| B7 | B225 | WB-Cop | hPBG | -0.097725112 | 0 |
| B7 | B30 | WB-Cop | LH | 0.098269296 | 0 |
| B7 | B31 | WB-Cop | FSH | 0.141782054 | 0 |
| B7 | B32 | WB-Cop | Testo | -0.321364297 | 0 |
| B7 | B8 | WB-Cop | WB-Zinc | 0.516049696 | 0 |
| B7 | B199 | WB-Cop | Mg | 0.074715044 | 4.44E-16 |
| B7 | B73 | WB-Cop | ApoA | -0.050366802 | 4.51E-08 |
| B7 | B145 | WB-Cop | BGP | 0.049241053 | 8.93E-08 |
| B70 | B127 | LDL-C | A-G ratio | -0.080155635 | 0 |
| B70 | B130 | LDL-C | GLB | 0.079114444 | 0 |
| B70 | B132 | LDL-C | Dbil | -0.193891721 | 0 |
| B70 | B133 | LDL-C | TBIL | -0.083532725 | 0 |
| B70 | B153 | LDL-C | Glu | 0.103459652 | 0 |
| B70 | B188 | LDL-C | Ca | 0.100903243 | 0 |

| | | | | | |
|---|---|---|---|---|---|
| B70 | B71 | LDL-C | TG | -0.109448753 | 0 |
| B70 | B72 | LDL-C | HDL-C | 0.162053691 | 0 |
| B70 | B75 | LDL-C | ApoA1/B | -0.216267769 | 0 |
| B70 | B76 | LDL-C | ApoB | 0.3435673 | 0 |
| B70 | B77 | LDL-C | TC | 0.768825024 | 0 |
| B70 | B91 | LDL-C | TP | 0.095355898 | 0 |
| B70 | B170 | LDL-C | U-GLU | 0.073428275 | 1.33E-15 |
| B70 | B73 | LDL-C | ApoA | 0.072914027 | 2.22E-15 |
| B70 | B74 | LDL-C | ApoA1 | 0.067765514 | 1.80E-13 |
| B70 | B131 | LDL-C | AST | -0.067520235 | 2.20E-13 |
| B70 | B128 | LDL-C | ALT | -0.050220943 | 4.93E-08 |
| B70 | B224 | LDL-C | pmh | 0.049607339 | 7.16E-08 |
| B70 | B194 | LDL-C | Cl | -0.047442826 | 2.58E-07 |
| B70 | B174 | LDL-C | Urea/Cr | 0.047060781 | 3.22E-07 |
| B71 | B126 | TG | Alb | 0.144334351 | 0 |
| B71 | B127 | TG | A-G ratio | -0.170447692 | 0 |
| B71 | B130 | TG | GLB | 0.295010615 | 0 |
| B71 | B132 | TG | Dbil | -0.105377729 | 0 |
| B71 | B133 | TG | TBIL | -0.089686784 | 0 |
| B71 | B134 | TG | CHE | 0.103611104 | 0 |
| B71 | B140 | TG | r-GT | 0.210337057 | 0 |
| B71 | B141 | TG | AST/ALT | -0.108565264 | 0 |
| B71 | B153 | TG | Glu | 0.260256697 | 0 |
| B71 | B155 | TG | IgA | 0.099970045 | 0 |
| B71 | B160 | TG | PH | -0.078679054 | 0 |
| B71 | B163 | TG | SG | 0.142244591 | 0 |
| B71 | B170 | TG | U-GLU | 0.190837358 | 0 |
| B71 | B172 | TG | KB | 0.097222481 | 0 |
| B71 | B188 | TG | Ca | 0.105098757 | 0 |
| B71 | B189 | TG | eGFB | 0.131240762 | 0 |
| B71 | B192 | TG | UA | 0.128357746 | 0 |
| B71 | B206 | TG | Waist | 0.169956995 | 0 |
| B71 | B209 | TG | Smoking | 0.09312183 | 0 |
| B71 | B210 | TG | Hip | 0.099393872 | 0 |
| B71 | B215 | TG | BMI | 0.164092939 | 0 |
| B71 | B216 | TG | BMI | 0.178035122 | 0 |
| B71 | B218 | TG | DBP | 0.125197492 | 0 |
| B71 | B221 | TG | Hei | 0.0898335 | 0 |
| B71 | B222 | TG | NS | 0.157153064 | 0 |
| B71 | B72 | TG | HDL-C | -0.390988068 | 0 |
| B71 | B74 | TG | ApoA1 | -0.076337755 | 0 |
| B71 | B77 | TG | TC | 0.368895533 | 0 |
| B71 | B91 | TG | TP | 0.353277114 | 0 |
| B71 | B129 | TG | IBIL | -0.075082563 | 4.44E-16 |

| | | | | | |
|---|---|---|---|---|---|
| B71 | B204 | TG | Drinking | 0.072087068 | 4.88E-15 |
| B71 | B195 | TG | Na | -0.069659144 | 3.75E-14 |
| B71 | B128 | TG | ALT | 0.067850883 | 1.68E-13 |
| B71 | B146 | TG | CysC | 0.065126188 | 1.49E-12 |
| B71 | B194 | TG | Cl | -0.062857322 | 8.58E-12 |
| B71 | B136 | TG | 5'NT | 0.057297471 | 4.86E-10 |
| B71 | B102 | TG | anti-Sm | 0.056752863 | 7.08E-10 |
| B71 | B75 | TG | ApoA1/B | 0.055899481 | 1.27E-09 |
| B71 | B158 | TG | IgM | 0.055466611 | 1.70E-09 |
| B71 | B166 | TG | Pro | 0.053805879 | 5.10E-09 |
| B71 | B203 | TG | DH | 0.052155316 | 1.48E-08 |
| B71 | B157 | TG | IgG | -0.050846023 | 3.35E-08 |
| B71 | B138 | TG | PAB | 0.046897102 | 3.53E-07 |
| B71 | B73 | TG | ApoA | -0.046081602 | 5.62E-07 |
| B72 | B128 | HDL-C | ALT | -0.097281317 | 0 |
| B72 | B130 | HDL-C | GLB | -0.088078455 | 0 |
| B72 | B140 | HDL-C | r-GT | -0.085479312 | 0 |
| B72 | B141 | HDL-C | AST/ALT | 0.205687758 | 0 |
| B72 | B153 | HDL-C | Glu | -0.142058007 | 0 |
| B72 | B160 | HDL-C | PH | 0.080080238 | 0 |
| B72 | B163 | HDL-C | SG | -0.158800161 | 0 |
| B72 | B166 | HDL-C | Pro | -0.07789154 | 0 |
| B72 | B170 | HDL-C | U-GLU | -0.124272573 | 0 |
| B72 | B174 | HDL-C | Urea/Cr | 0.09691832 | 0 |
| B72 | B176 | HDL-C | Ucr | -0.077434229 | 0 |
| B72 | B190 | HDL-C | Cre | -0.091882831 | 0 |
| B72 | B192 | HDL-C | UA | -0.246736566 | 0 |
| B72 | B206 | HDL-C | Waist | -0.327524924 | 0 |
| B72 | B209 | HDL-C | Smoking | -0.156353541 | 0 |
| B72 | B210 | HDL-C | Hip | -0.216681615 | 0 |
| B72 | B215 | HDL-C | BMI | -0.284922067 | 0 |
| B72 | B216 | HDL-C | BMI | -0.358554539 | 0 |
| B72 | B221 | HDL-C | Hei | -0.249455634 | 0 |
| B72 | B222 | HDL-C | NS | -0.343134082 | 0 |
| B72 | B74 | HDL-C | ApoA1 | 0.291568243 | 0 |
| B72 | B75 | HDL-C | ApoA1/B | 0.098247729 | 0 |
| B72 | B77 | HDL-C | TC | 0.194759352 | 0 |
| B72 | B91 | HDL-C | TP | -0.085798698 | 0 |
| B72 | B132 | HDL-C | Dbil | 0.072190856 | 4.44E-15 |
| B72 | B134 | HDL-C | CHE | -0.071645082 | 7.11E-15 |
| B72 | B133 | HDL-C | TBIL | 0.069152433 | 5.73E-14 |
| B72 | B218 | HDL-C | DBP | -0.067117514 | 3.04E-13 |
| B72 | B204 | HDL-C | Drinking | -0.062707382 | 9.61E-12 |
| B72 | B129 | HDL-C | IBIL | 0.061069012 | 3.27E-11 |

| | | | | | |
|---|---|---|---|---|---|
| B72 | B127 | HDL-C | A-G ratio | 0.060934892 | 3.61E-11 |
| B72 | B224 | HDL-C | pmh | 0.056894516 | 6.42E-10 |
| B72 | B173 | HDL-C | NTT | 0.046467151 | 4.52E-07 |
| B73 | B74 | ApoA | ApoA1 | 0.086018594 | 0 |
| B73 | B75 | ApoA | ApoA1/B | -0.097656444 | 0 |
| B73 | B76 | ApoA | ApoB | 0.187797781 | 0 |
| B73 | B101 | ApoA | anti-Sc70 | 0.065274017 | 1.32E-12 |
| B73 | B104 | ApoA | anti-SSB | 0.065274007 | 1.33E-12 |
| B73 | B98 | ApoA | anti-Jo-1 | 0.065274001 | 1.33E-12 |
| B73 | B107 | ApoA | AnuA | 0.064284177 | 2.87E-12 |
| B73 | B216 | ApoA | BMI | -0.055411355 | 1.76E-09 |
| B73 | B176 | ApoA | Ucr | -0.05195353 | 1.68E-08 |
| B73 | B157 | ApoA | IgG | 0.050615216 | 3.87E-08 |
| B73 | B82 | ApoA | CEA | 0.047151567 | 3.05E-07 |
| B73 | B195 | ApoA | Na | 0.047045685 | 3.24E-07 |
| B73 | B77 | ApoA | TC | 0.046639708 | 4.09E-07 |
| B73 | B128 | ApoA | ALT | -0.045666716 | 7.10E-07 |
| B74 | B123 | ApoA1 | hs-CRP1 | -0.079441446 | 0 |
| B74 | B125 | ApoA1 | Cor | 0.079449645 | 0 |
| B74 | B134 | ApoA1 | CHE | -0.096663616 | 0 |
| B74 | B139 | ApoA1 | ADA | -0.099207914 | 0 |
| B74 | B141 | ApoA1 | AST/ALT | 0.077366827 | 0 |
| B74 | B176 | ApoA1 | Ucr | -0.096818876 | 0 |
| B74 | B192 | ApoA1 | UA | -0.076749588 | 0 |
| B74 | B194 | ApoA1 | Cl | 0.07753901 | 0 |
| B74 | B195 | ApoA1 | Na | 0.162083565 | 0 |
| B74 | B199 | ApoA1 | Mg | 0.149927374 | 0 |
| B74 | B206 | ApoA1 | Waist | -0.130229686 | 0 |
| B74 | B210 | ApoA1 | Hip | -0.087407815 | 0 |
| B74 | B215 | ApoA1 | BMI | -0.099616583 | 0 |
| B74 | B216 | ApoA1 | BMI | -0.15836851 | 0 |
| B74 | B221 | ApoA1 | Hei | -0.139123444 | 0 |
| B74 | B222 | ApoA1 | NS | -0.151911821 | 0 |
| B74 | B75 | ApoA1 | ApoA1/B | 0.393379872 | 0 |
| B74 | B77 | ApoA1 | TC | 0.094235032 | 0 |
| B74 | B102 | ApoA1 | anti-Sm | -0.073823914 | 8.88E-16 |
| B74 | B163 | ApoA1 | SG | -0.069673626 | 3.71E-14 |
| B74 | B86 | ApoA1 | TBhCG | 0.069560979 | 4.09E-14 |
| B74 | B220 | ApoA1 | SBP | 0.062345354 | 1.26E-11 |
| B74 | B82 | ApoA1 | CEA | -0.058237444 | 2.52E-10 |
| B74 | B183 | ApoA1 | PTA | -0.05670156 | 7.33E-10 |
| B74 | B170 | ApoA1 | U-GLU | -0.055138699 | 2.12E-09 |
| B74 | B114 | ApoA1 | TSH TT3 TT4(TT4) | 0.054138379 | 4.11E-09 |
| B74 | B140 | ApoA1 | r-GT | -0.053608524 | 5.81E-09 |

| | | | | | |
|---|---|---|---|---|---|
| B74 | B128 | ApoA1 | ALT | -0.052266716 | 1.38E-08 |
| B74 | B209 | ApoA1 | Smoking | -0.051323032 | 2.49E-08 |
| B74 | B198 | ApoA1 | P | 0.050208663 | 4.97E-08 |
| B74 | B200 | ApoA1 | OP | -0.047164478 | 3.03E-07 |
| B75 | B134 | ApoA1/B | CHE | -0.091490506 | 0 |
| B75 | B138 | ApoA1/B | PAB | -0.101417206 | 0 |
| B75 | B146 | ApoA1/B | CysC | 0.11226343 | 0 |
| B75 | B171 | ApoA1/B | EC | 0.07954179 | 0 |
| B75 | B194 | ApoA1/B | Cl | 0.082527504 | 0 |
| B75 | B216 | ApoA1/B | BMI | -0.078718469 | 0 |
| B75 | B76 | ApoA1/B | ApoB | -0.711004615 | 0 |
| B75 | B77 | ApoA1/B | TC | -0.135837292 | 0 |
| B75 | B109 | ApoA1/B | AMA-M2 | 0.075299169 | 2.22E-16 |
| B75 | B200 | ApoA1/B | OP | -0.074482189 | 4.44E-16 |
| B75 | B137 | ApoA1/B | AFU | -0.071895138 | 5.55E-15 |
| B75 | B158 | ApoA1/B | IgM | 0.069402702 | 4.66E-14 |
| B75 | B206 | ApoA1/B | Waist | -0.068667704 | 8.57E-14 |
| B75 | B139 | ApoA1/B | ADA | -0.067309358 | 2.61E-13 |
| B75 | B222 | ApoA1/B | NS | -0.065384471 | 1.21E-12 |
| B75 | B163 | ApoA1/B | SG | -0.06253318 | 1.10E-11 |
| B75 | B215 | ApoA1/B | BMI | -0.059181193 | 1.29E-10 |
| B75 | B136 | ApoA1/B | 5'NT | 0.058321915 | 2.37E-10 |
| B75 | B221 | ApoA1/B | Hei | -0.057436619 | 4.41E-10 |
| B75 | B153 | ApoA1/B | Glu | -0.056479103 | 8.54E-10 |
| B75 | B170 | ApoA1/B | U-GLU | -0.055141351 | 2.11E-09 |
| B75 | B92 | ApoA1/B | ALDH2-Glu504Glu | 0.054863977 | 2.54E-09 |
| B75 | B197 | ApoA1/B | AG | -0.054292396 | 3.71E-09 |
| B75 | B107 | ApoA1/B | AnuA | 0.054242091 | 3.83E-09 |
| B75 | B93 | ApoA1/B | ALDH2-Glu504Lys | -0.053650079 | 5.65E-09 |
| B75 | B192 | ApoA1/B | UA | -0.05143709 | 2.32E-08 |
| B75 | B101 | ApoA1/B | anti-Sc70 | 0.050898859 | 3.25E-08 |
| B75 | B104 | ApoA1/B | anti-SSB | 0.050898852 | 3.25E-08 |
| B75 | B98 | ApoA1/B | anti-Jo-1 | 0.050898849 | 3.25E-08 |
| B75 | B141 | ApoA1/B | AST/ALT | 0.046850855 | 3.63E-07 |
| B75 | B210 | ApoA1/B | Hip | -0.046716418 | 3.92E-07 |
| B76 | B134 | ApoB | CHE | 0.108901416 | 0 |
| B76 | B138 | ApoB | PAB | 0.103439021 | 0 |
| B76 | B153 | ApoB | Glu | 0.092747163 | 0 |
| B76 | B186 | ApoB | PFC | 0.112691772 | 0 |
| B76 | B188 | ApoB | Ca | 0.07887161 | 0 |
| B76 | B193 | ApoB | K | 0.080104357 | 0 |
| B76 | B194 | ApoB | Cl | -0.131747111 | 0 |
| B76 | B77 | ApoB | TC | 0.299567428 | 0 |
| B76 | B197 | ApoB | AG | 0.074956056 | 4.44E-16 |

| | | | | | |
|---|---|---|---|---|---|
| B76 | B198 | ApoB | P | 0.07098824 | 1.22E-14 |
| B76 | B200 | ApoB | OP | 0.059767376 | 8.45E-11 |
| B76 | B125 | ApoB | Cor | 0.05738315 | 4.58E-10 |
| B76 | B170 | ApoB | U-GLU | 0.056855316 | 6.60E-10 |
| B76 | B137 | ApoB | AFU | 0.056539967 | 8.19E-10 |
| B76 | B102 | ApoB | anti-Sm | -0.055757772 | 1.40E-09 |
| B76 | B91 | ApoB | TP | 0.055178774 | 2.06E-09 |
| B76 | B158 | ApoB | IgM | -0.052352474 | 1.30E-08 |
| B76 | B124 | ApoB | ACTH | 0.052003306 | 1.63E-08 |
| B76 | B82 | ApoB | CEA | -0.048062365 | 1.80E-07 |
| B76 | B146 | ApoB | CysC | -0.047888218 | 1.99E-07 |
| B76 | B114 | ApoB | TSH TT3 TT4(TT4) | 0.047030433 | 3.27E-07 |
| B76 | B130 | ApoB | GLB | 0.046926908 | 3.47E-07 |
| B76 | B152 | ApoB | ACA-Ig(G-A-M) | -0.045805831 | 6.56E-07 |
| B76 | B139 | ApoB | ADA | 0.045223082 | 9.08E-07 |
| B77 | B126 | TC | Alb | 0.081175462 | 0 |
| B77 | B127 | TC | A-G ratio | -0.171319895 | 0 |
| B77 | B130 | TC | GLB | 0.224383411 | 0 |
| B77 | B132 | TC | Dbil | -0.212450572 | 0 |
| B77 | B133 | TC | TBIL | -0.106954061 | 0 |
| B77 | B140 | TC | r-GT | 0.10339738 | 0 |
| B77 | B153 | TC | Glu | 0.209926558 | 0 |
| B77 | B170 | TC | U-GLU | 0.144531364 | 0 |
| B77 | B174 | TC | Urea/Cr | 0.101091116 | 0 |
| B77 | B188 | TC | Ca | 0.147893911 | 0 |
| B77 | B218 | TC | DBP | 0.079736366 | 0 |
| B77 | B221 | TC | Hei | -0.07660691 | 0 |
| B77 | B91 | TC | TP | 0.251456689 | 0 |
| B77 | B172 | TC | KB | 0.075234873 | 4.44E-16 |
| B77 | B134 | TC | CHE | 0.073256538 | 1.78E-15 |
| B77 | B163 | TC | SG | 0.071787702 | 6.22E-15 |
| B77 | B189 | TC | eGFB | 0.066619262 | 4.55E-13 |
| B77 | B194 | TC | Cl | -0.065504099 | 1.11E-12 |
| B77 | B129 | TC | IBIL | -0.05866248 | 1.87E-10 |
| B77 | B190 | TC | Cre | -0.057180163 | 5.27E-10 |
| B77 | B155 | TC | IgA | 0.049882631 | 6.06E-08 |
| B77 | B186 | TC | PFC | 0.045272617 | 8.84E-07 |
| B78 | B80 | HBcAb | anti-HP IgG | -0.153278898 | 0 |
| B79 | B101 | hb V -dna | anti-Sc70 | 0.151868235 | 0 |
| B79 | B104 | hb V -dna | anti-SSB | 0.151868212 | 0 |
| B79 | B107 | hb V -dna | AnuA | 0.152763725 | 0 |
| B79 | B119 | hb V -dna | Hcv-rna | 0.175708939 | 0 |
| B79 | B120 | hb V -dna | HCV-cAg | 0.094452661 | 0 |
| B79 | B183 | hb V -dna | PTA | 0.114761011 | 0 |

| | | | | | |
|---|---|---|---|---|---|
| B79 | B194 | hb V -dna | Cl | 0.092498578 | 0 |
| B79 | B96 | hb V -dna | dsDNA (IIF) | 0.096173693 | 0 |
| B79 | B98 | hb V -dna | anti-Jo-1 | 0.151868199 | 0 |
| B79 | B184 | hb V -dna | INR | 0.074237677 | 6.66E-16 |
| B79 | B200 | hb V -dna | OP | -0.071797333 | 6.22E-15 |
| B79 | B151 | hb V -dna | Anti-CCP | -0.070611287 | 1.69E-14 |
| B79 | B81 | hb V -dna | TSGF | -0.066664796 | 4.39E-13 |
| B79 | B139 | hb V -dna | ADA | -0.065985981 | 7.55E-13 |
| B79 | B186 | hb V -dna | PFC | -0.060679446 | 4.35E-11 |
| B79 | B181 | hb V -dna | APTT | 0.055792245 | 1.36E-09 |
| B79 | B134 | hb V -dna | CHE | -0.046517858 | 4.39E-07 |
| B79 | B199 | hb V -dna | Mg | 0.045650017 | 7.16E-07 |
| B8 | B121 | WB-Zinc | C3 | 0.089388548 | 0 |
| B8 | B29 | WB-Zinc | E2 | 0.082192714 | 0 |
| B8 | B32 | WB-Zinc | Testo | -0.134610195 | 0 |
| B8 | B33 | WB-Zinc | Prog | 0.066928423 | 3.55E-13 |
| B8 | B225 | WB-Zinc | hPBG | -0.057703248 | 3.66E-10 |
| B8 | B158 | WB-Zinc | IgM | 0.049426961 | 7.98E-08 |
| B8 | B199 | WB-Zinc | Mg | 0.045465751 | 7.94E-07 |
| B81 | B125 | TSGF | Cor | -0.174954138 | 0 |
| B81 | B151 | TSGF | Anti-CCP | -0.196814723 | 0 |
| B81 | B182 | TSGF | TT | 0.140326643 | 0 |
| B81 | B183 | TSGF | PTA | -0.076580236 | 0 |
| B81 | B200 | TSGF | OP | 0.088790423 | 0 |
| B81 | B224 | TSGF | pmh | -0.090850265 | 0 |
| B81 | B92 | TSGF | ALDH2-Glu504Glu | -0.092459827 | 0 |
| B81 | B93 | TSGF | ALDH2-Glu504Lys | 0.099021157 | 0 |
| B81 | B120 | TSGF | HCV-cAg | 0.074072739 | 8.88E-16 |
| B81 | B199 | TSGF | Mg | -0.069519591 | 4.22E-14 |
| B81 | B139 | TSGF | ADA | 0.065181758 | 1.42E-12 |
| B81 | B197 | TSGF | AG | 0.064033698 | 3.48E-12 |
| B81 | B225 | TSGF | hPBG | 0.056284062 | 9.76E-10 |
| B81 | B137 | TSGF | AFU | 0.055441613 | 1.73E-09 |
| B81 | B97 | TSGF | dsDNA | 0.053919468 | 4.74E-09 |
| B81 | B194 | TSGF | Cl | -0.053539139 | 6.07E-09 |
| B81 | B150 | TSGF | ALP | 0.052953486 | 8.87E-09 |
| B81 | B184 | TSGF | INR | -0.051195913 | 2.70E-08 |
| B81 | B90 | TSGF | CYFRA 21-1 | 0.048875626 | 1.11E-07 |
| B82 | B83 | CEA | AFP | 0.096281849 | 0 |
| B82 | B86 | CEA | TBhCG | -0.210502962 | 0 |
| B82 | B87 | CEA | NSE | 0.090440249 | 0 |
| B82 | B88 | CEA | CA 242 | 0.073866838 | 8.88E-16 |
| B82 | B84 | CEA | C-PSA | 0.059125828 | 1.34E-10 |
| B82 | B158 | CEA | IgM | 0.057796231 | 3.43E-10 |

| | | | | | |
|---|---|---|---|---|---|
| B82 | B200 | CEA | OP | 0.048065342 | 1.79E-07 |
| B83 | B86 | AFP | TBhCG | 0.173201303 | 0 |
| B83 | B88 | AFP | CA 242 | 0.217450849 | 0 |
| B83 | B87 | AFP | NSE | 0.075698757 | 2.22E-16 |
| B83 | B89 | AFP | CA 72-4 | 0.075094521 | 4.44E-16 |
| B83 | B84 | AFP | C-PSA | 0.068926846 | 6.93E-14 |
| B83 | B94 | AFP | ALDH2-Lys504Lys | 0.061073997 | 3.26E-11 |
| B84 | B85 | C-PSA | SCC | 0.118102432 | 0 |
| B84 | B86 | C-PSA | TBhCG | -0.139311842 | 0 |
| B84 | B87 | C-PSA | NSE | 0.100954594 | 0 |
| B84 | B94 | C-PSA | ALDH2-Lys504Lys | 0.071283119 | 9.33E-15 |
| B84 | B89 | C-PSA | CA 72-4 | 0.046930328 | 3.47E-07 |
| B85 | B87 | SCC | NSE | 0.290527243 | 0 |
| B85 | B88 | SCC | CA 242 | 0.061349732 | 2.66E-11 |
| B85 | B199 | SCC | Mg | -0.051367007 | 2.43E-08 |
| B85 | B193 | SCC | K | -0.046069583 | 5.66E-07 |
| B86 | B200 | TBhCG | OP | -0.115188777 | 0 |
| B86 | B88 | TBhCG | CA 242 | -0.107925564 | 0 |
| B86 | B90 | TBhCG | CYFRA 21-1 | 0.128352641 | 0 |
| B86 | B137 | TBhCG | AFU | -0.07458666 | 4.44E-16 |
| B86 | B120 | TBhCG | HCV-cAg | 0.057240876 | 5.06E-10 |
| B86 | B194 | TBhCG | Cl | 0.053952146 | 4.64E-09 |
| B86 | B157 | TBhCG | IgG | 0.05333158 | 6.95E-09 |
| B86 | B199 | TBhCG | Mg | 0.053328787 | 6.96E-09 |
| B86 | B196 | TBhCG | TCO2 | -0.049842749 | 6.21E-08 |
| B86 | B139 | TBhCG | ADA | -0.045423342 | 8.13E-07 |
| B87 | B88 | NSE | CA 242 | 0.094446888 | 0 |
| B87 | B90 | NSE | CYFRA 21-1 | 0.069503017 | 4.26E-14 |
| B87 | B195 | NSE | Na | -0.066095455 | 6.91E-13 |
| B87 | B89 | NSE | CA 72-4 | -0.056399657 | 9.02E-10 |
| B87 | B200 | NSE | OP | -0.047714745 | 2.20E-07 |
| B88 | B141 | CA 242 | AST/ALT | 0.060721523 | 4.22E-11 |
| B89 | B194 | CA 72-4 | Cl | -0.075710975 | 2.22E-16 |
| B89 | B200 | CA 72-4 | OP | 0.060352427 | 5.53E-11 |
| B9 | B114 | RH | TSH TT3 TT4(TT4) | 0.081332765 | 0 |
| B9 | B225 | RH | hPBG | 0.243515901 | 0 |
| B9 | B54 | RH | ESR | 0.143382696 | 0 |
| B9 | B55 | RH | TK | -0.176463496 | 0 |
| B9 | B56 | RH | IR | -0.126896331 | 0 |
| B9 | B60 | RH | WBRVI | -0.086911842 | 0 |
| B9 | B62 | RH | WBHSRVI | -0.107836829 | 0 |
| B9 | B69 | RH | PV | 0.379407812 | 0 |
| B9 | B138 | RH | PAB | -0.074160599 | 8.88E-16 |
| B9 | B61 | RH | WBHSRV | -0.070483414 | 1.87E-14 |

| | | | | | |
|---|---|---|---|---|---|
| B9 | B195 | RH | Na | -0.059638298 | 9.28E-11 |
| B9 | B112 | RH | FT4 | 0.057006722 | 5.94E-10 |
| B9 | B58 | RH | HCT | -0.055383562 | 1.80E-09 |
| B9 | B120 | RH | HCV-cAg | -0.05035805 | 4.53E-08 |
| B9 | B76 | RH | ApoB | 0.046020968 | 5.82E-07 |
| B90 | B101 | CYFRA 21-1 | anti-Sc70 | -0.076437177 | 0 |
| B90 | B104 | CYFRA 21-1 | anti-SSB | -0.076437165 | 0 |
| B90 | B107 | CYFRA 21-1 | AnuA | -0.077046669 | 0 |
| B90 | B98 | CYFRA 21-1 | anti-Jo-1 | -0.076437158 | 0 |
| B90 | B180 | CYFRA 21-1 | Tf | 0.066898625 | 3.63E-13 |
| B90 | B178 | CYFRA 21-1 | Ig-U | 0.05387905 | 4.87E-09 |
| B90 | B139 | CYFRA 21-1 | ADA | 0.05377233 | 5.22E-09 |
| B91 | B126 | TP | Alb | 0.457050592 | 0 |
| B91 | B127 | TP | A-G ratio | -0.50783895 | 0 |
| B91 | B128 | TP | ALT | 0.096102783 | 0 |
| B91 | B130 | TP | GLB | 0.802627208 | 0 |
| B91 | B131 | TP | AST | 0.095758937 | 0 |
| B91 | B134 | TP | CHE | 0.089473854 | 0 |
| B91 | B140 | TP | r-GT | 0.112589958 | 0 |
| B91 | B153 | TP | Glu | 0.131613789 | 0 |
| B91 | B155 | TP | IgA | 0.098592673 | 0 |
| B91 | B160 | TP | PH | -0.08073028 | 0 |
| B91 | B188 | TP | Ca | 0.355299162 | 0 |
| B91 | B192 | TP | UA | 0.112184252 | 0 |
| B91 | B194 | TP | Cl | -0.094912786 | 0 |
| B91 | B209 | TP | Smoking | -0.110643648 | 0 |
| B91 | B215 | TP | BMI | 0.076861204 | 0 |
| B91 | B218 | TP | DBP | 0.137834606 | 0 |
| B91 | B220 | TP | SBP | 0.099451205 | 0 |
| B91 | B157 | TP | IgG | 0.06994301 | 2.95E-14 |
| B91 | B189 | TP | eGFB | 0.067118159 | 3.04E-13 |
| B91 | B158 | TP | IgM | 0.064035354 | 3.48E-12 |
| B91 | B150 | TP | ALP | 0.058804677 | 1.69E-10 |
| B91 | B221 | TP | Hei | -0.058570019 | 1.99E-10 |
| B91 | B146 | TP | CysC | 0.058469234 | 2.14E-10 |
| B91 | B193 | TP | K | 0.057858693 | 3.29E-10 |
| B91 | B206 | TP | Waist | 0.057810155 | 3.40E-10 |
| B91 | B141 | TP | AST/ALT | -0.050245866 | 4.85E-08 |
| B91 | B170 | TP | U-GLU | 0.049892136 | 6.02E-08 |
| B91 | B113 | TP | FT3 | 0.04922465 | 9.02E-08 |
| B91 | B136 | TP | 5'NT | 0.045175871 | 9.32E-07 |
| B92 | B105 | ALDH2-Glu504Glu | anti-U1RNP | -0.145717171 | 0 |
| B92 | B110 | ALDH2-Glu504Glu | AHA | 0.102279197 | 0 |
| B92 | B124 | ALDH2-Glu504Glu | ACTH | -0.110018095 | 0 |

| | | | | | |
|---|---|---|---|---|---|
| B92 | B136 | ALDH2-Glu504Glu | 5'NT | 0.083463433 | 0 |
| B92 | B151 | ALDH2-Glu504Glu | Anti-CCP | 0.117496035 | 0 |
| B92 | B152 | ALDH2-Glu504Glu | ACA-Ig(G-A-M) | 0.139511273 | 0 |
| B92 | B182 | ALDH2-Glu504Glu | TT | -0.091365818 | 0 |
| B92 | B93 | ALDH2-Glu504Glu | ALDH2-Glu504Lys | -0.97474339 | 0 |
| B92 | B94 | ALDH2-Glu504Glu | ALDH2-Lys504Lys | -0.144917887 | 0 |
| B92 | B100 | ALDH2-Glu504Glu | anti-Ro52 | 0.057224876 | 5.11E-10 |
| B92 | B137 | ALDH2-Glu504Glu | AFU | 0.055903674 | 1.26E-09 |
| B92 | B103 | ALDH2-Glu504Glu | anti-SSA/Ro | 0.052785333 | 9.88E-09 |
| B92 | B109 | ALDH2-Glu504Glu | AMA-M2 | 0.050483177 | 4.20E-08 |
| B92 | B155 | ALDH2-Glu504Glu | IgA | 0.047577304 | 2.39E-07 |
| B92 | B119 | ALDH2-Glu504Glu | Hcv-rna | 0.04695853 | 3.41E-07 |
| B93 | B105 | ALDH2-Glu504Lys | anti-U1RNP | 0.149285432 | 0 |
| B93 | B110 | ALDH2-Glu504Lys | AHA | -0.097208169 | 0 |
| B93 | B124 | ALDH2-Glu504Lys | ACTH | 0.11583898 | 0 |
| B93 | B136 | ALDH2-Glu504Lys | 5'NT | -0.076959737 | 0 |
| B93 | B151 | ALDH2-Glu504Lys | Anti-CCP | -0.112410094 | 0 |
| B93 | B152 | ALDH2-Glu504Lys | ACA-Ig(G-A-M) | -0.145529993 | 0 |
| B93 | B182 | ALDH2-Glu504Lys | TT | 0.093363001 | 0 |
| B93 | B94 | ALDH2-Glu504Lys | ALDH2-Lys504Lys | -0.079710436 | 0 |
| B93 | B119 | ALDH2-Glu504Lys | Hcv-rna | -0.060103777 | 6.63E-11 |
| B93 | B100 | ALDH2-Glu504Lys | anti-Ro52 | -0.058565935 | 2.00E-10 |
| B93 | B103 | ALDH2-Glu504Lys | anti-SSA/Ro | -0.054771942 | 2.70E-09 |
| B93 | B109 | ALDH2-Glu504Lys | AMA-M2 | -0.049797055 | 6.38E-08 |
| B93 | B199 | ALDH2-Glu504Lys | Mg | -0.046951906 | 3.42E-07 |
| B94 | B101 | ALDH2-Lys504Lys | anti-Sc70 | 0.102576415 | 0 |
| B94 | B104 | ALDH2-Lys504Lys | anti-SSB | 0.102576387 | 0 |
| B94 | B107 | ALDH2-Lys504Lys | AnuA | 0.101287667 | 0 |
| B94 | B98 | ALDH2-Lys504Lys | anti-Jo-1 | 0.102576385 | 0 |
| B94 | B137 | ALDH2-Lys504Lys | AFU | -0.067181458 | 2.89E-13 |
| B94 | B155 | ALDH2-Lys504Lys | IgA | -0.05784485 | 3.32E-10 |
| B94 | B119 | ALDH2-Lys504Lys | Hcv-rna | 0.057711704 | 3.64E-10 |
| B94 | B96 | ALDH2-Lys504Lys | dsDNA (IIF) | 0.056920392 | 6.31E-10 |
| B94 | B129 | ALDH2-Lys504Lys | IBIL | 0.056276987 | 9.81E-10 |
| B94 | B133 | ALDH2-Lys504Lys | TBIL | 0.053958967 | 4.62E-09 |
| B94 | B183 | ALDH2-Lys504Lys | PTA | 0.052131299 | 1.50E-08 |
| B94 | B138 | ALDH2-Lys504Lys | PAB | -0.04808649 | 1.77E-07 |
| B95 | B156 | anti-cenpb | IgE | 0.117210394 | 0 |
| B95 | B151 | anti-cenpb | Anti-CCP | -0.056659501 | 7.55E-10 |
| B96 | B100 | dsDNA (IIF) | anti-Ro52 | -0.158886648 | 0 |
| B96 | B101 | dsDNA (IIF) | anti-Sc70 | 0.255629686 | 0 |
| B96 | B104 | dsDNA (IIF) | anti-SSB | 0.255629382 | 0 |
| B96 | B107 | dsDNA (IIF) | AnuA | 0.257072523 | 0 |
| B96 | B110 | dsDNA (IIF) | AHA | -0.388032589 | 0 |

| | | | | | |
|---|---|---|---|---|---|
| B96 | B119 | dsDNA (IIF) | Hcv-rna | 0.245887127 | 0 |
| B96 | B120 | dsDNA (IIF) | HCV-cAg | 0.124392454 | 0 |
| B96 | B136 | dsDNA (IIF) | 5'NT | 0.077369236 | 0 |
| B96 | B138 | dsDNA (IIF) | PAB | -0.08524016 | 0 |
| B96 | B151 | dsDNA (IIF) | Anti-CCP | -0.143996124 | 0 |
| B96 | B182 | dsDNA (IIF) | TT | 0.082065864 | 0 |
| B96 | B183 | dsDNA (IIF) | PTA | 0.099899779 | 0 |
| B96 | B97 | dsDNA (IIF) | dsDNA | -0.374252168 | 0 |
| B96 | B98 | dsDNA (IIF) | anti-Jo-1 | 0.255629647 | 0 |
| B96 | B194 | dsDNA (IIF) | CI | 0.070448405 | 1.93E-14 |
| B96 | B157 | dsDNA (IIF) | IgG | 0.064585887 | 2.27E-12 |
| B96 | B103 | dsDNA (IIF) | anti-SSA/Ro | 0.059933996 | 7.49E-11 |
| B96 | B108 | dsDNA (IIF) | Anti-Cyclin | -0.048755251 | 1.19E-07 |
| B96 | B106 | dsDNA (IIF) | anti-P | 0.047403322 | 2.64E-07 |
| B96 | B102 | dsDNA (IIF) | anti-Sm | 0.047397693 | 2.65E-07 |
| B96 | B156 | dsDNA (IIF) | IgE | 0.04567699 | 7.05E-07 |
| B96 | B181 | dsDNA (IIF) | APTT | 0.045325472 | 8.58E-07 |
| B97 | B108 | dsDNA | Anti-Cyclin | 0.097070545 | 0 |
| B97 | B110 | dsDNA | AHA | 0.62863137 | 0 |
| B97 | B119 | dsDNA | Hcv-rna | -0.087078606 | 0 |
| B97 | B194 | dsDNA | CI | -0.067327526 | 2.57E-13 |
| B97 | B197 | dsDNA | AG | 0.064312235 | 2.81E-12 |
| B97 | B182 | dsDNA | TT | 0.049089537 | 9.78E-08 |
| B98 | B100 | anti-Jo-1 | anti-Ro52 | 0.094822272 | 0 |
| B98 | B102 | anti-Jo-1 | anti-Sm | 0.110763166 | 0 |
| B98 | B103 | anti-Jo-1 | anti-SSA/Ro | 0.094458113 | 0 |
| B98 | B106 | anti-Jo-1 | anti-P | 0.110785278 | 0 |
| B98 | B107 | anti-Jo-1 | AnuA | 0.994376517 | 0 |
| B98 | B119 | anti-Jo-1 | Hcv-rna | 0.150439869 | 0 |
| B98 | B120 | anti-Jo-1 | HCV-cAg | 0.126184869 | 0 |
| B98 | B121 | anti-Jo-1 | C3 | -0.094750439 | 0 |
| B98 | B122 | anti-Jo-1 | C4 | -0.086964846 | 0 |
| B98 | B134 | anti-Jo-1 | CHE | -0.099159435 | 0 |
| B98 | B137 | anti-Jo-1 | AFU | -0.077017813 | 0 |
| B98 | B138 | anti-Jo-1 | PAB | -0.075998036 | 0 |
| B98 | B139 | anti-Jo-1 | ADA | -0.086100003 | 0 |
| B98 | B151 | anti-Jo-1 | Anti-CCP | -0.123677037 | 0 |
| B98 | B155 | anti-Jo-1 | IgA | -0.078598417 | 0 |
| B98 | B157 | anti-Jo-1 | IgG | 0.143501834 | 0 |
| B98 | B181 | anti-Jo-1 | APTT | 0.126973213 | 0 |
| B98 | B184 | anti-Jo-1 | INR | 0.14002629 | 0 |
| B98 | B185 | anti-Jo-1 | PT | 0.076376745 | 0 |
| B98 | B186 | anti-Jo-1 | PFC | -0.087093807 | 0 |
| B98 | B194 | anti-Jo-1 | CI | 0.123052886 | 0 |

| | | | | | |
|---|---|---|---|---|---|
| B98 | B195 | anti-Jo-1 | Na | 0.122660778 | 0 |
| B98 | B198 | anti-Jo-1 | P | -0.094561432 | 0 |
| B98 | B199 | anti-Jo-1 | Mg | 0.114914237 | 0 |
| B98 | B200 | anti-Jo-1 | OP | -0.165117493 | 0 |
| B98 | B183 | anti-Jo-1 | PTA | 0.072236199 | 4.00E-15 |
| B98 | B152 | anti-Jo-1 | ACA-Ig(G-A-M) | 0.066754252 | 4.08E-13 |
| B98 | B179 | anti-Jo-1 | α1-MG | -0.065045254 | 1.59E-12 |
| B98 | B212 | anti-Jo-1 | IOP-L | -0.061301266 | 2.75E-11 |
| B98 | B124 | anti-Jo-1 | ACTH | -0.054617204 | 2.99E-09 |
| B98 | B193 | anti-Jo-1 | K | 0.052012094 | 1.62E-08 |
| B98 | B112 | anti-Jo-1 | FT4 | -0.048806306 | 1.16E-07 |
| B98 | B220 | anti-Jo-1 | SBP | 0.04795153 | 1.92E-07 |
| B98 | B150 | anti-Jo-1 | ALP | -0.047571272 | 2.39E-07 |
| B98 | B189 | anti-Jo-1 | eGFB | -0.047256553 | 2.87E-07 |
| B98 | B211 | anti-Jo-1 | IOP-R | -0.046863277 | 3.60E-07 |
| B99 | B144 | anti-PM-Scl | Total-PINP | -0.082035674 | 0 |
| B99 | B151 | anti-PM-Scl | Anti-CCP | -0.091708305 | 0 |
| B99 | B156 | anti-PM-Scl | IgE | 0.114668502 | 0 |
| B99 | B183 | anti-PM-Scl | PTA | 0.151475652 | 0 |
| B99 | B186 | anti-PM-Scl | PFC | 0.179491963 | 0 |
| B99 | B142 | anti-PM-Scl | 25-(OH)D | 0.075406501 | 2.22E-16 |
| B99 | B145 | anti-PM-Scl | BGP | -0.074472354 | 4.44E-16 |
| B99 | B195 | anti-PM-Scl | Na | -0.070154419 | 2.49E-14 |
| B99 | B143 | anti-PM-Scl | β-CTx | -0.058635233 | 1.90E-10 |
| B99 | B200 | anti-PM-Scl | OP | -0.048446452 | 1.43E-07 |
| B99 | B176 | anti-PM-Scl | Ucr | 0.045864181 | 6.35E-07 |

**Supplementary Table 18. PEI Correlations in Gout**

| Row | Column | Row | Column | Cor | *P* |
|---|---|---|---|---|---|
| B1 | B173 | Gender | NTT | 0.197349 | 0 |
| B1 | B2 | Gender | Age | 0.199052 | 0 |
| B1 | B216 | Gender | BMI | -0.21324 | 0 |
| B1 | B221 | Gender | Hei | -0.31705 | 0 |
| B1 | B222 | Gender | NS | -0.24648 | 0 |
| B1 | B46 | Gender | RBC Count | -0.19193 | 0 |
| B1 | B51 | Gender | Hb | -0.27093 | 0 |
| B1 | B174 | Gender | Urea/Cr | 0.155788 | 4.37E-13 |
| B1 | B72 | Gender | HDL-C | 0.155435 | 4.94E-13 |
| B1 | B205 | Gender | KDA | 0.145782 | 1.26E-11 |
| B1 | B206 | Gender | Waist | -0.1345 | 4.27E-10 |
| B1 | B141 | Gender | AST/ALT | 0.129381 | 1.92E-09 |
| B1 | B161 | Gender | LEU | 0.125387 | 5.98E-09 |
| B1 | B90 | Gender | CYFRA 21-1 | 0.115531 | 8.47E-08 |
| B1 | B189 | Gender | eGFB | -0.11494 | 9.85E-08 |
| B1 | B171 | Gender | EC | 0.11188 | 2.14E-07 |
| B1 | B220 | Gender | SBP | 0.107442 | 6.38E-07 |
| B1 | B163 | Gender | SG | -0.10641 | 8.16E-07 |
| B10 | B12 | PG I | PG II | 0.52179 | 0 |
| B10 | B142 | PG I | 25-(OH)D | 0.189238 | 0 |
| B10 | B145 | PG I | BGP | 0.313082 | 0 |
| B10 | B190 | PG I | Cre | 0.366439 | 0 |
| B10 | B191 | PG I | Urea | 0.213685 | 0 |
| B10 | B189 | PG I | eGFB | -0.17564 | 4.44E-16 |
| B10 | B11 | PG I | PGI/PGII | 0.142092 | 4.12E-11 |
| B10 | B184 | PG I | INR | -0.1229 | 1.19E-08 |
| B10 | B113 | PG I | FT3 | -0.11385 | 1.30E-07 |
| B10 | B228 | PG I | OS | 0.113161 | 1.55E-07 |
| B10 | B14 | PG I | G-17 | -0.11024 | 3.22E-07 |
| B10 | B79 | PG I | hb V -dna | -0.10939 | 3.98E-07 |
| B10 | B80 | PG I | anti-HP IgG | 0.106035 | 8.93E-07 |
| B10 | B170 | PG I | U-GLU | 0.105871 | 9.28E-07 |
| B100 | B102 | anti-Ro52 | anti-Sm | 1 | 0 |
| B100 | B103 | anti-Ro52 | anti-SSA/Ro | 1 | 0 |
| B100 | B107 | anti-Ro52 | AnuA | 0.999999 | 0 |
| B100 | B151 | anti-Ro52 | Anti-CCP | 0.515326 | 0 |
| B100 | B154 | anti-Ro52 | RF | 0.202149 | 0 |
| B100 | B155 | anti-Ro52 | IgA | -0.18934 | 0 |
| B100 | B179 | anti-Ro52 | α1-MG | -0.20209 | 0 |
| B100 | B184 | anti-Ro52 | INR | 0.183231 | 0 |
| B100 | B138 | anti-Ro52 | PAB | -0.16705 | 7.55E-15 |
| B100 | B180 | anti-Ro52 | Tf | -0.16223 | 4.44E-14 |

| | | | | | |
|---|---|---|---|---|---|
| B100 | B178 | anti-Ro52 | Ig-U | -0.16208 | 4.71E-14 |
| B100 | B136 | anti-Ro52 | 5'NT | -0.16186 | 5.11E-14 |
| B100 | B139 | anti-Ro52 | ADA | -0.16171 | 5.37E-14 |
| B100 | B134 | anti-Ro52 | CHE | -0.14467 | 1.81E-11 |
| B100 | B177 | anti-Ro52 | ACR | -0.13048 | 1.40E-09 |
| B100 | B137 | anti-Ro52 | AFU | -0.12035 | 2.38E-08 |
| B100 | B182 | anti-Ro52 | TT | -0.11888 | 3.53E-08 |
| B100 | B168 | anti-Ro52 | NMP22 | 0.115313 | 8.96E-08 |
| B101 | B104 | anti-Sc70 | anti-SSB | 1 | 0 |
| B101 | B107 | anti-Sc70 | AnuA | 0.999999 | 0 |
| B101 | B151 | anti-Sc70 | Anti-CCP | 0.515326 | 0 |
| B101 | B154 | anti-Sc70 | RF | 0.202149 | 0 |
| B101 | B155 | anti-Sc70 | IgA | -0.18934 | 0 |
| B101 | B179 | anti-Sc70 | α1-MG | -0.20209 | 0 |
| B101 | B184 | anti-Sc70 | INR | 0.183231 | 0 |
| B101 | B138 | anti-Sc70 | PAB | -0.16705 | 7.55E-15 |
| B101 | B180 | anti-Sc70 | Tf | -0.16223 | 4.44E-14 |
| B101 | B178 | anti-Sc70 | Ig-U | -0.16208 | 4.71E-14 |
| B101 | B136 | anti-Sc70 | 5'NT | -0.16186 | 5.11E-14 |
| B101 | B139 | anti-Sc70 | ADA | -0.16171 | 5.37E-14 |
| B101 | B134 | anti-Sc70 | CHE | -0.14467 | 1.81E-11 |
| B101 | B177 | anti-Sc70 | ACR | -0.13048 | 1.40E-09 |
| B101 | B137 | anti-Sc70 | AFU | -0.12035 | 2.38E-08 |
| B101 | B182 | anti-Sc70 | TT | -0.11888 | 3.53E-08 |
| B101 | B168 | anti-Sc70 | NMP22 | 0.115313 | 8.96E-08 |
| B102 | B103 | anti-Sm | anti-SSA/Ro | 1 | 0 |
| B102 | B104 | anti-Sm | anti-SSB | 1 | 0 |
| B102 | B105 | anti-Sm | anti-U1RNP | 1 | 0 |
| B102 | B106 | anti-Sm | anti-P | 1 | 0 |
| B102 | B107 | anti-Sm | AnuA | 0.999999 | 0 |
| B102 | B109 | anti-Sm | AMA-M2 | 1 | 0 |
| B102 | B110 | anti-Sm | AHA | 1 | 0 |
| B102 | B151 | anti-Sm | Anti-CCP | 0.515326 | 0 |
| B102 | B154 | anti-Sm | RF | 0.202149 | 0 |
| B102 | B155 | anti-Sm | IgA | -0.18934 | 0 |
| B102 | B179 | anti-Sm | α1-MG | -0.20209 | 0 |
| B102 | B184 | anti-Sm | INR | 0.183231 | 0 |
| B102 | B138 | anti-Sm | PAB | -0.16705 | 7.55E-15 |
| B102 | B180 | anti-Sm | Tf | -0.16223 | 4.44E-14 |
| B102 | B178 | anti-Sm | Ig-U | -0.16208 | 4.71E-14 |
| B102 | B136 | anti-Sm | 5'NT | -0.16186 | 5.11E-14 |
| B102 | B139 | anti-Sm | ADA | -0.16171 | 5.37E-14 |
| B102 | B134 | anti-Sm | CHE | -0.14467 | 1.81E-11 |
| B102 | B177 | anti-Sm | ACR | -0.13048 | 1.40E-09 |

| | | | | | |
|---|---|---|---|---|---|
| B102 | B137 | anti-Sm | AFU | -0.12035 | 2.38E-08 |
| B102 | B182 | anti-Sm | TT | -0.11888 | 3.53E-08 |
| B102 | B168 | anti-Sm | NMP22 | 0.115313 | 8.96E-08 |
| B103 | B104 | anti-SSA/Ro | anti-SSB | 1 | 0 |
| B103 | B107 | anti-SSA/Ro | AnuA | 0.999999 | 0 |
| B103 | B151 | anti-SSA/Ro | Anti-CCP | 0.515326 | 0 |
| B103 | B154 | anti-SSA/Ro | RF | 0.202149 | 0 |
| B103 | B155 | anti-SSA/Ro | IgA | -0.18934 | 0 |
| B103 | B179 | anti-SSA/Ro | α1-MG | -0.20209 | 0 |
| B103 | B184 | anti-SSA/Ro | INR | 0.183231 | 0 |
| B103 | B138 | anti-SSA/Ro | PAB | -0.16705 | 7.55E-15 |
| B103 | B180 | anti-SSA/Ro | Tf | -0.16223 | 4.44E-14 |
| B103 | B178 | anti-SSA/Ro | Ig-U | -0.16208 | 4.71E-14 |
| B103 | B136 | anti-SSA/Ro | 5'NT | -0.16186 | 5.11E-14 |
| B103 | B139 | anti-SSA/Ro | ADA | -0.16171 | 5.37E-14 |
| B103 | B134 | anti-SSA/Ro | CHE | -0.14467 | 1.81E-11 |
| B103 | B177 | anti-SSA/Ro | ACR | -0.13048 | 1.40E-09 |
| B103 | B137 | anti-SSA/Ro | AFU | -0.12035 | 2.38E-08 |
| B103 | B182 | anti-SSA/Ro | TT | -0.11888 | 3.53E-08 |
| B103 | B168 | anti-SSA/Ro | NMP22 | 0.115313 | 8.96E-08 |
| B104 | B106 | anti-SSB | anti-P | 1 | 0 |
| B104 | B107 | anti-SSB | AnuA | 0.999999 | 0 |
| B104 | B109 | anti-SSB | AMA-M2 | 1 | 0 |
| B104 | B110 | anti-SSB | AHA | 1 | 0 |
| B104 | B151 | anti-SSB | Anti-CCP | 0.515326 | 0 |
| B104 | B154 | anti-SSB | RF | 0.202149 | 0 |
| B104 | B155 | anti-SSB | IgA | -0.18934 | 0 |
| B104 | B179 | anti-SSB | α1-MG | -0.20209 | 0 |
| B104 | B184 | anti-SSB | INR | 0.183231 | 0 |
| B104 | B138 | anti-SSB | PAB | -0.16705 | 7.55E-15 |
| B104 | B180 | anti-SSB | Tf | -0.16223 | 4.44E-14 |
| B104 | B178 | anti-SSB | Ig-U | -0.16208 | 4.71E-14 |
| B104 | B136 | anti-SSB | 5'NT | -0.16186 | 5.11E-14 |
| B104 | B139 | anti-SSB | ADA | -0.16171 | 5.37E-14 |
| B104 | B134 | anti-SSB | CHE | -0.14467 | 1.81E-11 |
| B104 | B177 | anti-SSB | ACR | -0.13048 | 1.40E-09 |
| B104 | B137 | anti-SSB | AFU | -0.12035 | 2.38E-08 |
| B104 | B182 | anti-SSB | TT | -0.11888 | 3.53E-08 |
| B104 | B168 | anti-SSB | NMP22 | 0.115313 | 8.96E-08 |
| B105 | B107 | anti-U1RNP | AnuA | 0.999999 | 0 |
| B105 | B151 | anti-U1RNP | Anti-CCP | 0.515326 | 0 |
| B105 | B154 | anti-U1RNP | RF | 0.202149 | 0 |
| B105 | B155 | anti-U1RNP | IgA | -0.18934 | 0 |
| B105 | B179 | anti-U1RNP | α1-MG | -0.20209 | 0 |

| | | | | | |
|---|---|---|---|---|---|
| B105 | B184 | anti-U1RNP | INR | 0.183231 | 0 |
| B105 | B138 | anti-U1RNP | PAB | -0.16705 | 7.55E-15 |
| B105 | B180 | anti-U1RNP | Tf | -0.16223 | 4.44E-14 |
| B105 | B178 | anti-U1RNP | Ig-U | -0.16208 | 4.71E-14 |
| B105 | B136 | anti-U1RNP | 5'NT | -0.16186 | 5.11E-14 |
| B105 | B139 | anti-U1RNP | ADA | -0.16171 | 5.37E-14 |
| B105 | B134 | anti-U1RNP | CHE | -0.14467 | 1.81E-11 |
| B105 | B177 | anti-U1RNP | ACR | -0.13048 | 1.40E-09 |
| B105 | B137 | anti-U1RNP | AFU | -0.12035 | 2.38E-08 |
| B105 | B182 | anti-U1RNP | TT | -0.11888 | 3.53E-08 |
| B105 | B168 | anti-U1RNP | NMP22 | 0.115313 | 8.96E-08 |
| B106 | B107 | anti-P | AnuA | 0.999999 | 0 |
| B106 | B151 | anti-P | Anti-CCP | 0.515326 | 0 |
| B106 | B154 | anti-P | RF | 0.202149 | 0 |
| B106 | B155 | anti-P | IgA | -0.18934 | 0 |
| B106 | B179 | anti-P | α1-MG | -0.20209 | 0 |
| B106 | B184 | anti-P | INR | 0.183231 | 0 |
| B106 | B138 | anti-P | PAB | -0.16705 | 7.55E-15 |
| B106 | B180 | anti-P | Tf | -0.16223 | 4.44E-14 |
| B106 | B178 | anti-P | Ig-U | -0.16208 | 4.71E-14 |
| B106 | B136 | anti-P | 5'NT | -0.16186 | 5.11E-14 |
| B106 | B139 | anti-P | ADA | -0.16171 | 5.37E-14 |
| B106 | B134 | anti-P | CHE | -0.14467 | 1.81E-11 |
| B106 | B177 | anti-P | ACR | -0.13048 | 1.40E-09 |
| B106 | B137 | anti-P | AFU | -0.12035 | 2.38E-08 |
| B106 | B182 | anti-P | TT | -0.11888 | 3.53E-08 |
| B106 | B168 | anti-P | NMP22 | 0.115313 | 8.96E-08 |
| B107 | B109 | AnuA | AMA-M2 | 0.999999 | 0 |
| B107 | B110 | AnuA | AHA | 0.999999 | 0 |
| B107 | B151 | AnuA | Anti-CCP | 0.515325 | 0 |
| B107 | B154 | AnuA | RF | 0.202149 | 0 |
| B107 | B155 | AnuA | IgA | -0.18934 | 0 |
| B107 | B179 | AnuA | α1-MG | -0.20209 | 0 |
| B107 | B184 | AnuA | INR | 0.18323 | 0 |
| B107 | B138 | AnuA | PAB | -0.16705 | 7.55E-15 |
| B107 | B180 | AnuA | Tf | -0.16223 | 4.44E-14 |
| B107 | B178 | AnuA | Ig-U | -0.16208 | 4.71E-14 |
| B107 | B136 | AnuA | 5'NT | -0.16186 | 5.11E-14 |
| B107 | B139 | AnuA | ADA | -0.16171 | 5.37E-14 |
| B107 | B134 | AnuA | CHE | -0.14467 | 1.81E-11 |
| B107 | B177 | AnuA | ACR | -0.13048 | 1.40E-09 |
| B107 | B137 | AnuA | AFU | -0.12035 | 2.38E-08 |
| B107 | B182 | AnuA | TT | -0.11888 | 3.53E-08 |
| B107 | B168 | AnuA | NMP22 | 0.115313 | 8.96E-08 |

| | | | | | |
|---|---|---|---|---|---|
| B108 | B151 | Anti-Cyclin | Anti-CCP | -0.18535 | 0 |
| B109 | B151 | AMA-M2 | Anti-CCP | 0.515326 | 0 |
| B109 | B154 | AMA-M2 | RF | 0.202149 | 0 |
| B109 | B155 | AMA-M2 | IgA | -0.18934 | 0 |
| B109 | B179 | AMA-M2 | α1-MG | -0.20209 | 0 |
| B109 | B184 | AMA-M2 | INR | 0.183231 | 0 |
| B109 | B138 | AMA-M2 | PAB | -0.16705 | 7.55E-15 |
| B109 | B180 | AMA-M2 | Tf | -0.16223 | 4.44E-14 |
| B109 | B178 | AMA-M2 | Ig-U | -0.16208 | 4.71E-14 |
| B109 | B136 | AMA-M2 | 5'NT | -0.16186 | 5.11E-14 |
| B109 | B139 | AMA-M2 | ADA | -0.16171 | 5.37E-14 |
| B109 | B134 | AMA-M2 | CHE | -0.14467 | 1.81E-11 |
| B109 | B177 | AMA-M2 | ACR | -0.13048 | 1.40E-09 |
| B109 | B137 | AMA-M2 | AFU | -0.12035 | 2.38E-08 |
| B109 | B182 | AMA-M2 | TT | -0.11888 | 3.53E-08 |
| B109 | B168 | AMA-M2 | NMP22 | 0.115313 | 8.96E-08 |
| B11 | B12 | PGI/PGII | PG II | -0.58079 | 0 |
| B11 | B14 | PGI/PGII | G-17 | -0.25438 | 0 |
| B11 | B95 | PGI/PGII | anti-cenpb | 0.160113 | 9.55E-14 |
| B11 | B78 | PGI/PGII | HBcAb | 0.110943 | 2.71E-07 |
| B110 | B151 | AHA | Anti-CCP | 0.515326 | 0 |
| B110 | B154 | AHA | RF | 0.202149 | 0 |
| B110 | B155 | AHA | IgA | -0.18934 | 0 |
| B110 | B179 | AHA | α1-MG | -0.20209 | 0 |
| B110 | B184 | AHA | INR | 0.183231 | 0 |
| B110 | B138 | AHA | PAB | -0.16705 | 7.55E-15 |
| B110 | B180 | AHA | Tf | -0.16223 | 4.44E-14 |
| B110 | B178 | AHA | Ig-U | -0.16208 | 4.71E-14 |
| B110 | B136 | AHA | 5'NT | -0.16186 | 5.11E-14 |
| B110 | B139 | AHA | ADA | -0.16171 | 5.37E-14 |
| B110 | B134 | AHA | CHE | -0.14467 | 1.81E-11 |
| B110 | B177 | AHA | ACR | -0.13048 | 1.40E-09 |
| B110 | B137 | AHA | AFU | -0.12035 | 2.38E-08 |
| B110 | B182 | AHA | TT | -0.11888 | 3.53E-08 |
| B110 | B168 | AHA | NMP22 | 0.115313 | 8.96E-08 |
| B111 | B148 | TSH | Anti_TPO | 0.265086 | 0 |
| B111 | B149 | TSH | TGAb | 0.126116 | 4.87E-09 |
| B112 | B114 | FT4 | TSH TT3 TT4(TT4) | 0.435524 | 0 |
| B112 | B113 | FT4 | FT3 | 0.134126 | 4.78E-10 |
| B113 | B115 | FT3 | TSH TT3 TT4(TT3) | 0.431164 | 0 |
| B113 | B126 | FT3 | Alb | 0.144021 | 2.23E-11 |
| B113 | B189 | FT3 | eGFB | 0.135898 | 2.80E-10 |
| B113 | B191 | FT3 | Urea | -0.12767 | 3.14E-09 |
| B113 | B143 | FT3 | β-CTx | 0.127583 | 3.22E-09 |

| | | | | | |
|---|---|---|---|---|---|
| B113 | B190 | FT3 | Cre | -0.1257 | 5.48E-09 |
| B113 | B128 | FT3 | ALT | 0.115534 | 8.46E-08 |
| B114 | B115 | TSH TT3 TT4(TT4) | TSH TT3 TT4(TT3) | 0.349557 | 0 |
| B115 | B184 | TSH TT3 TT4(TT3) | INR | 0.142754 | 3.34E-11 |
| B115 | B136 | TSH TT3 TT4(TT3) | 5'NT | -0.11862 | 3.77E-08 |
| B115 | B124 | TSH TT3 TT4(TT3) | ACTH | 0.112888 | 1.66E-07 |
| B116 | B117 | TT3 | TT4 | -0.54528 | 0 |
| B118 | B153 | CEA1 | Glu | 0.14473 | 1.77E-11 |
| B118 | B209 | CEA1 | Smoking | 0.137579 | 1.68E-10 |
| B118 | B189 | CEA1 | eGFB | -0.11693 | 5.89E-08 |
| B119 | B120 | Hcv-rna | HCV-cAg | 0.441708 | 0 |
| B119 | B121 | Hcv-rna | C3 | -0.96054 | 0 |
| B119 | B122 | Hcv-rna | C4 | -0.96054 | 0 |
| B119 | B136 | Hcv-rna | 5'NT | -0.4056 | 0 |
| B119 | B139 | Hcv-rna | ADA | -0.20019 | 0 |
| B119 | B152 | Hcv-rna | ACA-Ig(G-A-M) | -0.96066 | 0 |
| B119 | B159 | Hcv-rna | β2-MG | -0.96053 | 0 |
| B119 | B169 | Hcv-rna | U-CRY | -0.96055 | 0 |
| B119 | B150 | Hcv-rna | ALP | -0.15995 | 1.01E-13 |
| B119 | B124 | Hcv-rna | ACTH | -0.14518 | 1.53E-11 |
| B119 | B125 | Hcv-rna | Cor | -0.10566 | 9.76E-07 |
| B12 | B14 | PG II | G-17 | 0.290906 | 0 |
| B12 | B145 | PG II | BGP | 0.266819 | 0 |
| B12 | B190 | PG II | Cre | 0.261205 | 0 |
| B12 | B80 | PG II | anti-HP IgG | 0.144971 | 1.64E-11 |
| B12 | B144 | PG II | Total-PINP | 0.138577 | 1.24E-10 |
| B12 | B142 | PG II | 25-(OH)D | 0.118326 | 4.08E-08 |
| B12 | B191 | PG II | Urea | 0.118168 | 4.26E-08 |
| B12 | B189 | PG II | eGFB | -0.11336 | 1.48E-07 |
| B12 | B78 | PG II | HBcAb | -0.1081 | 5.44E-07 |
| B120 | B121 | HCV-cAg | C3 | -0.42428 | 0 |
| B120 | B122 | HCV-cAg | C4 | -0.42428 | 0 |
| B120 | B136 | HCV-cAg | 5'NT | -0.20379 | 0 |
| B120 | B152 | HCV-cAg | ACA-Ig(G-A-M) | -0.42434 | 0 |
| B120 | B159 | HCV-cAg | β2-MG | -0.42427 | 0 |
| B120 | B169 | HCV-cAg | U-CRY | -0.42428 | 0 |
| B120 | B139 | HCV-cAg | ADA | -0.16596 | 1.13E-14 |
| B120 | B199 | HCV-cAg | Mg | 0.159547 | 1.17E-13 |
| B120 | B137 | HCV-cAg | AFU | -0.13281 | 7.06E-10 |
| B120 | B155 | HCV-cAg | IgA | -0.11743 | 5.17E-08 |
| B120 | B150 | HCV-cAg | ALP | -0.11175 | 2.21E-07 |
| B120 | B154 | HCV-cAg | RF | 0.109959 | 3.45E-07 |
| B121 | B136 | C3 | 5'NT | 0.3568 | 0 |
| B121 | B159 | C3 | β2-MG | 0.999993 | 0 |

| | | | | | |
|---|---|---|---|---|---|
| B121 | B139 | C3 | ADA | 0.172255 | 1.11E-15 |
| B121 | B199 | C3 | Mg | -0.13768 | 1.63E-10 |
| B121 | B150 | C3 | ALP | 0.120131 | 2.52E-08 |
| B122 | B136 | C4 | 5'NT | 0.356801 | 0 |
| B122 | B159 | C4 | β2-MG | 0.999996 | 0 |
| B122 | B169 | C4 | U-CRY | 0.999996 | 0 |
| B122 | B139 | C4 | ADA | 0.172256 | 1.11E-15 |
| B122 | B199 | C4 | Mg | -0.13768 | 1.63E-10 |
| B122 | B150 | C4 | ALP | 0.120132 | 2.52E-08 |
| B123 | B150 | hs-CRP1 | ALP | 0.163517 | 2.80E-14 |
| B123 | B127 | hs-CRP1 | A-G ratio | -0.14253 | 3.58E-11 |
| B123 | B126 | hs-CRP1 | Alb | -0.13522 | 3.44E-10 |
| B123 | B130 | hs-CRP1 | GLB | 0.120957 | 2.02E-08 |
| B123 | B200 | hs-CRP1 | OP | 0.10973 | 3.65E-07 |
| B124 | B125 | ACTH | Cor | 0.746288 | 0 |
| B124 | B182 | ACTH | TT | -0.18866 | 0 |
| B124 | B136 | ACTH | 5'NT | 0.15864 | 1.61E-13 |
| B124 | B184 | ACTH | INR | 0.15807 | 1.97E-13 |
| B124 | B137 | ACTH | AFU | 0.1423 | 3.86E-11 |
| B124 | B186 | ACTH | PFC | -0.13275 | 7.18E-10 |
| B124 | B156 | ACTH | IgE | 0.130961 | 1.22E-09 |
| B124 | B134 | ACTH | CHE | -0.11854 | 3.85E-08 |
| B124 | B138 | ACTH | PAB | -0.1158 | 7.90E-08 |
| B124 | B200 | ACTH | OP | -0.11496 | 9.82E-08 |
| B124 | B139 | ACTH | ADA | 0.1101 | 3.34E-07 |
| B125 | B156 | Cor | IgE | 0.184667 | 0 |
| B125 | B182 | Cor | TT | -0.17987 | 0 |
| B125 | B158 | Cor | IgM | -0.17453 | 4.44E-16 |
| B125 | B184 | Cor | INR | 0.162412 | 4.17E-14 |
| B125 | B181 | Cor | APTT | -0.14647 | 1.01E-11 |
| B125 | B136 | Cor | 5'NT | 0.139389 | 9.61E-11 |
| B125 | B170 | Cor | U-GLU | -0.13011 | 1.56E-09 |
| B125 | B200 | Cor | OP | -0.12373 | 9.49E-09 |
| B125 | B134 | Cor | CHE | -0.11772 | 4.78E-08 |
| B125 | B186 | Cor | PFC | -0.11252 | 1.82E-07 |
| B126 | B127 | Alb | A-G ratio | 0.465257 | 0 |
| B126 | B188 | Alb | Ca | 0.400211 | 0 |
| B126 | B189 | Alb | eGFB | 0.253012 | 0 |
| B126 | B128 | Alb | ALT | 0.175921 | 2.22E-16 |
| B126 | B221 | Alb | Hei | 0.14781 | 6.50E-12 |
| B126 | B130 | Alb | GLB | -0.13133 | 1.09E-09 |
| B126 | B131 | Alb | AST | 0.129134 | 2.07E-09 |
| B126 | B129 | Alb | IBIL | 0.120864 | 2.07E-08 |
| B126 | B191 | Alb | Urea | -0.11654 | 6.52E-08 |

| | | | | | |
|---|---|---|---|---|---|
| B126 | B141 | Alb | AST/ALT | -0.11155 | 2.33E-07 |
| B126 | B133 | Alb | TBIL | 0.110813 | 2.80E-07 |
| B126 | B138 | Alb | PAB | 0.109512 | 3.86E-07 |
| B126 | B134 | Alb | CHE | 0.105877 | 9.27E-07 |
| B127 | B130 | A-G ratio | GLB | -0.90385 | 0 |
| B127 | B189 | A-G ratio | eGFB | 0.180015 | 0 |
| B127 | B220 | A-G ratio | SBP | -0.14559 | 1.34E-11 |
| B127 | B129 | A-G ratio | IBIL | 0.144632 | 1.83E-11 |
| B127 | B133 | A-G ratio | TBIL | 0.144248 | 2.07E-11 |
| B127 | B157 | A-G ratio | IgG | -0.11744 | 5.15E-08 |
| B127 | B221 | A-G ratio | Hei | 0.115779 | 7.94E-08 |
| B127 | B191 | A-G ratio | Urea | -0.11025 | 3.21E-07 |
| B127 | B207 | A-G ratio | WHR | -0.10651 | 7.96E-07 |
| B128 | B131 | ALT | AST | 0.760334 | 0 |
| B128 | B140 | ALT | r-GT | 0.412251 | 0 |
| B128 | B141 | ALT | AST/ALT | -0.4532 | 0 |
| B128 | B189 | ALT | eGFB | 0.219143 | 0 |
| B128 | B206 | ALT | Waist | 0.230216 | 0 |
| B128 | B210 | ALT | Hip | 0.25656 | 0 |
| B128 | B215 | ALT | BMI | 0.281255 | 0 |
| B128 | B216 | ALT | BMI | 0.301971 | 0 |
| B128 | B222 | ALT | NS | 0.236792 | 0 |
| B128 | B165 | ALT | bilirubin | 0.172994 | 8.88E-16 |
| B128 | B188 | ALT | Ca | 0.161693 | 5.42E-14 |
| B128 | B192 | ALT | UA | 0.141124 | 5.59E-11 |
| B128 | B221 | ALT | Hei | 0.137736 | 1.60E-10 |
| B128 | B201 | ALT | e-habits | -0.12433 | 8.04E-09 |
| B128 | B191 | ALT | Urea | -0.11562 | 8.29E-08 |
| B129 | B132 | IBIL | Dbil | 0.642909 | 0 |
| B129 | B133 | IBIL | TBIL | 0.980634 | 0 |
| B129 | B209 | IBIL | Smoking | -0.1301 | 1.56E-09 |
| B129 | B130 | IBIL | GLB | -0.11836 | 4.05E-08 |
| B130 | B188 | GLB | Ca | 0.17468 | 4.44E-16 |
| B130 | B192 | GLB | UA | 0.1374 | 1.77E-10 |
| B130 | B220 | GLB | SBP | 0.136169 | 2.58E-10 |
| B130 | B157 | GLB | IgG | 0.125951 | 5.11E-09 |
| B130 | B133 | GLB | TBIL | -0.1207 | 2.17E-08 |
| B130 | B194 | GLB | Cl | -0.11621 | 7.11E-08 |
| B130 | B139 | GLB | ADA | 0.114167 | 1.20E-07 |
| B131 | B140 | AST | r-GT | 0.44901 | 0 |
| B131 | B165 | AST | bilirubin | 0.470627 | 0 |
| B131 | B171 | AST | EC | -0.23179 | 0 |
| B131 | B132 | AST | Dbil | 0.16828 | 4.88E-15 |
| B131 | B215 | AST | BMI | 0.163463 | 2.84E-14 |

| | | | | | |
|---|---|---|---|---|---|
| B131 | B216 | AST | BMI | 0.155156 | 5.44E-13 |
| B131 | B189 | AST | eGFB | 0.149556 | 3.64E-12 |
| B131 | B210 | AST | Hip | 0.145613 | 1.33E-11 |
| B131 | B206 | AST | Waist | 0.14272 | 3.37E-11 |
| B131 | B222 | AST | NS | 0.140645 | 6.49E-11 |
| B131 | B188 | AST | Ca | 0.128356 | 2.58E-09 |
| B131 | B192 | AST | UA | 0.117567 | 4.98E-08 |
| B132 | B133 | Dbil | TBIL | 0.780379 | 0 |
| B132 | B165 | Dbil | bilirubin | 0.26692 | 0 |
| B132 | B171 | Dbil | EC | -0.12826 | 2.66E-09 |
| B132 | B140 | Dbil | r-GT | 0.109176 | 4.19E-07 |
| B133 | B209 | TBIL | Smoking | -0.12085 | 2.08E-08 |
| B134 | B137 | CHE | AFU | 0.266932 | 0 |
| B134 | B138 | CHE | PAB | 0.458846 | 0 |
| B134 | B194 | CHE | Cl | -0.26199 | 0 |
| B134 | B151 | CHE | Anti-CCP | -0.17429 | 4.44E-16 |
| B134 | B197 | CHE | AG | 0.172822 | 8.88E-16 |
| B134 | B180 | CHE | Tf | 0.130445 | 1.41E-09 |
| B134 | B178 | CHE | Ig-U | 0.127649 | 3.16E-09 |
| B134 | B136 | CHE | 5'NT | 0.124771 | 7.10E-09 |
| B134 | B188 | CHE | Ca | 0.110124 | 3.32E-07 |
| B134 | B177 | CHE | ACR | 0.108286 | 5.20E-07 |
| B135 | B181 | TBA | APTT | -0.21345 | 0 |
| B135 | B184 | TBA | INR | -0.23031 | 0 |
| B135 | B185 | TBA | PT | -0.18867 | 0 |
| B135 | B186 | TBA | PFC | -0.19192 | 0 |
| B135 | B194 | TBA | Cl | 0.132819 | 7.04E-10 |
| B136 | B137 | 5'NT | AFU | 0.196401 | 0 |
| B136 | B139 | 5'NT | ADA | 0.467818 | 0 |
| B136 | B140 | 5'NT | r-GT | 0.193235 | 0 |
| B136 | B150 | 5'NT | ALP | 0.310564 | 0 |
| B136 | B152 | 5'NT | ACA-Ig(G-A-M) | 0.356844 | 0 |
| B136 | B159 | 5'NT | β2-MG | 0.356797 | 0 |
| B136 | B169 | 5'NT | U-CRY | 0.356803 | 0 |
| B136 | B194 | 5'NT | Cl | -0.20529 | 0 |
| B136 | B197 | 5'NT | AG | 0.147599 | 6.97E-12 |
| B136 | B155 | 5'NT | IgA | 0.13919 | 1.02E-10 |
| B136 | B153 | 5'NT | Glu | 0.121977 | 1.53E-08 |
| B137 | B139 | AFU | ADA | 0.172696 | 8.88E-16 |
| B137 | B193 | AFU | K | 0.153066 | 1.12E-12 |
| B137 | B200 | AFU | OP | 0.144061 | 2.20E-11 |
| B137 | B138 | AFU | PAB | 0.141162 | 5.52E-11 |
| B137 | B150 | AFU | ALP | 0.139329 | 9.79E-11 |
| B138 | B139 | PAB | ADA | -0.34197 | 0 |

| | | | | | |
|---|---|---|---|---|---|
| B138 | B194 | PAB | Cl | -0.184 | 0 |
| B138 | B151 | PAB | Anti-CCP | -0.16967 | 2.89E-15 |
| B138 | B154 | PAB | RF | -0.16791 | 5.55E-15 |
| B138 | B184 | PAB | INR | -0.14871 | 4.83E-12 |
| B138 | B204 | PAB | Drinking | 0.138707 | 1.19E-10 |
| B138 | B186 | PAB | PFC | 0.125605 | 5.63E-09 |
| B138 | B168 | PAB | NMP22 | -0.1229 | 1.19E-08 |
| B138 | B185 | PAB | PT | -0.12256 | 1.31E-08 |
| B138 | B199 | PAB | Mg | -0.11578 | 7.94E-08 |
| B139 | B150 | ADA | ALP | 0.210676 | 0 |
| B139 | B153 | ADA | Glu | 0.196521 | 0 |
| B139 | B155 | ADA | IgA | 0.348334 | 0 |
| B139 | B194 | ADA | Cl | -0.20383 | 0 |
| B139 | B200 | ADA | OP | 0.200351 | 0 |
| B139 | B152 | ADA | ACA-Ig(G-A-M) | 0.172276 | 8.88E-16 |
| B139 | B159 | ADA | β2-MG | 0.172253 | 1.11E-15 |
| B139 | B169 | ADA | U-CRY | 0.172256 | 1.11E-15 |
| B139 | B193 | ADA | K | 0.171471 | 1.33E-15 |
| B139 | B146 | ADA | CysC | 0.154034 | 8.02E-13 |
| B139 | B178 | ADA | Ig-U | 0.151848 | 1.69E-12 |
| B139 | B158 | ADA | IgM | 0.143684 | 2.48E-11 |
| B139 | B180 | ADA | Tf | 0.138775 | 1.16E-10 |
| B139 | B157 | ADA | IgG | 0.136464 | 2.36E-10 |
| B139 | B177 | ADA | ACR | 0.128348 | 2.59E-09 |
| B139 | B211 | ADA | IOP-R | -0.12676 | 4.07E-09 |
| B139 | B156 | ADA | IgE | 0.126456 | 4.43E-09 |
| B139 | B195 | ADA | Na | -0.12537 | 6.00E-09 |
| B139 | B212 | ADA | IOP-L | -0.11607 | 7.36E-08 |
| B139 | B140 | ADA | r-GT | 0.107715 | 5.97E-07 |
| B14 | B137 | G-17 | AFU | -0.12045 | 2.32E-08 |
| B140 | B165 | r-GT | bilirubin | 0.219125 | 0 |
| B140 | B171 | r-GT | EC | -0.23243 | 0 |
| B140 | B141 | r-GT | AST/ALT | -0.16891 | 3.77E-15 |
| B140 | B153 | r-GT | Glu | 0.164056 | 2.31E-14 |
| B140 | B204 | r-GT | Drinking | 0.148915 | 4.51E-12 |
| B140 | B206 | r-GT | Waist | 0.142633 | 3.47E-11 |
| B140 | B207 | r-GT | WHR | 0.140223 | 7.41E-11 |
| B140 | B218 | r-GT | DBP | 0.136132 | 2.61E-10 |
| B140 | B222 | r-GT | NS | 0.134212 | 4.65E-10 |
| B140 | B215 | r-GT | BMI | 0.134042 | 4.90E-10 |
| B140 | B189 | r-GT | eGFB | 0.127893 | 2.95E-09 |
| B140 | B216 | r-GT | BMI | 0.115681 | 8.15E-08 |
| B140 | B192 | r-GT | UA | 0.11549 | 8.56E-08 |
| B141 | B215 | AST/ALT | BMI | -0.18492 | 0 |

| | | | | | |
|---|---|---|---|---|---|
| B141 | B216 | AST/ALT | BMI | -0.19482 | 0 |
| B141 | B222 | AST/ALT | NS | -0.16749 | 6.44E-15 |
| B141 | B206 | AST/ALT | Waist | -0.15449 | 6.85E-13 |
| B141 | B210 | AST/ALT | Hip | -0.14479 | 1.74E-11 |
| B141 | B189 | AST/ALT | eGFB | -0.13605 | 2.67E-10 |
| B142 | B145 | 25-(OH)D | BGP | 0.446098 | 0 |
| B142 | B190 | 25-(OH)D | Cre | 0.361902 | 0 |
| B142 | B191 | 25-(OH)D | Urea | 0.164629 | 1.87E-14 |
| B142 | B144 | 25-(OH)D | Total-PINP | 0.139228 | 1.01E-10 |
| B143 | B144 | β-CTx | Total-PINP | 0.717468 | 0 |
| B143 | B145 | β-CTx | BGP | 0.375246 | 0 |
| B143 | B150 | β-CTx | ALP | 0.140821 | 6.15E-11 |
| B143 | B183 | β-CTx | PTA | -0.12919 | 2.03E-09 |
| B143 | B190 | β-CTx | Cre | 0.11988 | 2.70E-08 |
| B143 | B181 | β-CTx | APTT | 0.11209 | 2.03E-07 |
| B144 | B145 | Total-PINP | BGP | 0.665015 | 0 |
| B144 | B190 | Total-PINP | Cre | 0.357676 | 0 |
| B144 | B150 | Total-PINP | ALP | 0.144274 | 2.05E-11 |
| B144 | B191 | Total-PINP | Urea | 0.127327 | 3.46E-09 |
| B144 | B168 | Total-PINP | NMP22 | 0.123382 | 1.04E-08 |
| B145 | B190 | BGP | Cre | 0.674411 | 0 |
| B145 | B191 | BGP | Urea | 0.267285 | 0 |
| B145 | B150 | BGP | ALP | 0.157241 | 2.63E-13 |
| B145 | B177 | BGP | ACR | 0.130896 | 1.24E-09 |
| B145 | B153 | BGP | Glu | 0.106159 | 8.67E-07 |
| B146 | B189 | CysC | eGFB | -0.23059 | 0 |
| B146 | B191 | CysC | Urea | 0.185908 | 0 |
| B146 | B193 | CysC | K | 0.173411 | 6.66E-16 |
| B146 | B200 | CysC | OP | 0.170872 | 1.78E-15 |
| B146 | B190 | CysC | Cre | 0.153313 | 1.03E-12 |
| B148 | B149 | Anti_TPO | TGAb | 0.460503 | 0 |
| B15 | B100 | CD3-CD19+(B cell) | anti-Ro52 | 0.543077 | 0 |
| B15 | B101 | CD3-CD19+(B cell) | anti-Sc70 | 0.543077 | 0 |
| B15 | B102 | CD3-CD19+(B cell) | anti-Sm | 0.543077 | 0 |
| B15 | B103 | CD3-CD19+(B cell) | anti-SSA/Ro | 0.543077 | 0 |
| B15 | B104 | CD3-CD19+(B cell) | anti-SSB | 0.543077 | 0 |
| B15 | B105 | CD3-CD19+(B cell) | anti-U1RNP | 0.543077 | 0 |
| B15 | B106 | CD3-CD19+(B cell) | anti-P | 0.543077 | 0 |
| B15 | B107 | CD3-CD19+(B cell) | AnuA | 0.543077 | 0 |
| B15 | B109 | CD3-CD19+(B cell) | AMA-M2 | 0.543077 | 0 |
| B15 | B110 | CD3-CD19+(B cell) | AHA | 0.543077 | 0 |
| B15 | B151 | CD3-CD19+(B cell) | Anti-CCP | 0.383362 | 0 |
| B15 | B155 | CD3-CD19+(B cell) | IgA | -0.22513 | 0 |
| B15 | B184 | CD3-CD19+(B cell) | INR | 0.217601 | 0 |

| | | | | | |
|---|---|---|---|---|---|
| B15 | B19 | CD3-CD19+(B cell) | CD3+CD4+/CD3+CD8+ | 0.256073 | 0 |
| B15 | B21 | CD3-CD19+(B cell) | CD3+CD4-CD8- | 0.289396 | 0 |
| B15 | B22 | CD3-CD19+(B cell) | CD3+CD8+(T) | -0.31069 | 0 |
| B15 | B79 | CD3-CD19+(B cell) | hb V -dna | 0.398288 | 0 |
| B15 | B98 | CD3-CD19+(B cell) | anti-Jo-1 | 0.543077 | 0 |
| B15 | B23 | CD3-CD19+(B cell) | CD3-CD16+CD56+(NK) | -0.17194 | 1.33E-15 |
| B15 | B81 | CD3-CD19+(B cell) | TSGF | -0.16037 | 8.70E-14 |
| B15 | B134 | CD3-CD19+(B cell) | CHE | -0.15924 | 1.30E-13 |
| B15 | B95 | CD3-CD19+(B cell) | anti-cenpb | -0.15618 | 3.81E-13 |
| B15 | B186 | CD3-CD19+(B cell) | PFC | -0.15361 | 9.27E-13 |
| B15 | B185 | CD3-CD19+(B cell) | PT | 0.149086 | 4.26E-12 |
| B15 | B55 | CD3-CD19+(B cell) | TK | 0.148132 | 5.84E-12 |
| B15 | B154 | CD3-CD19+(B cell) | RF | 0.142829 | 3.26E-11 |
| B15 | B16 | CD3-CD19+(B cell) | CD3+T | -0.14128 | 5.32E-11 |
| B15 | B18 | CD3-CD19+(B cell) | CD3+CD4+helper(inducer)T | 0.13981 | 8.43E-11 |
| B15 | B138 | CD3-CD19+(B cell) | PAB | -0.13461 | 4.13E-10 |
| B15 | B158 | CD3-CD19+(B cell) | IgM | 0.133261 | 6.18E-10 |
| B15 | B59 | CD3-CD19+(B cell) | WBRV | 0.131688 | 9.83E-10 |
| B15 | B96 | CD3-CD19+(B cell) | dsDNA (IIF) | 0.128691 | 2.35E-09 |
| B15 | B34 | CD3-CD19+(B cell) | TK | 0.115741 | 8.02E-08 |
| B15 | B183 | CD3-CD19+(B cell) | PTA | 0.111816 | 2.18E-07 |
| B15 | B74 | CD3-CD19+(B cell) | ApoA1 | 0.110752 | 2.84E-07 |
| B15 | B139 | CD3-CD19+(B cell) | ADA | -0.11013 | 3.31E-07 |
| B15 | B76 | CD3-CD19+(B cell) | ApoB | -0.10917 | 4.19E-07 |
| B15 | B199 | CD3-CD19+(B cell) | Mg | 0.109035 | 4.33E-07 |
| B15 | B179 | CD3-CD19+(B cell) | α1-MG | 0.109013 | 4.36E-07 |
| B15 | B39 | CD3-CD19+(B cell) | Other cells | 0.108847 | 4.54E-07 |
| B15 | B198 | CD3-CD19+(B cell) | P | 0.105802 | 9.44E-07 |
| B150 | B197 | ALP | AG | 0.183022 | 0 |
| B150 | B177 | ALP | ACR | 0.14637 | 1.04E-11 |
| B150 | B152 | ALP | ACA-Ig(G-A-M) | 0.120146 | 2.51E-08 |
| B150 | B159 | ALP | β2-MG | 0.12013 | 2.52E-08 |
| B150 | B169 | ALP | U-CRY | 0.120132 | 2.52E-08 |
| B150 | B179 | ALP | α1-MG | 0.114021 | 1.25E-07 |
| B150 | B195 | ALP | Na | 0.113629 | 1.38E-07 |
| B150 | B200 | ALP | OP | 0.110724 | 2.86E-07 |
| B151 | B156 | Anti-CCP | IgE | 0.229055 | 0 |
| B151 | B157 | Anti-CCP | IgG | 0.199735 | 0 |
| B151 | B178 | Anti-CCP | Ig-U | -0.20025 | 0 |
| B151 | B180 | Anti-CCP | Tf | -0.21751 | 0 |
| B151 | B184 | Anti-CCP | INR | 0.213522 | 0 |
| B151 | B185 | Anti-CCP | PT | 0.230987 | 0 |
| B151 | B177 | Anti-CCP | ACR | -0.17088 | 1.78E-15 |
| B151 | B182 | Anti-CCP | TT | -0.16922 | 3.33E-15 |

| | | | | | |
|---|---|---|---|---|---|
| B151 | B168 | Anti-CCP | NMP22 | 0.159464 | 1.20E-13 |
| B151 | B158 | Anti-CCP | IgM | 0.13844 | 1.29E-10 |
| B151 | B183 | Anti-CCP | PTA | -0.13602 | 2.70E-10 |
| B151 | B186 | Anti-CCP | PFC | 0.128376 | 2.57E-09 |
| B151 | B175 | Anti-CCP | ALB-U | -0.11519 | 9.25E-08 |
| B151 | B154 | Anti-CCP | RF | 0.113304 | 1.50E-07 |
| B152 | B199 | ACA-Ig(G-A-M) | Mg | -0.1377 | 1.62E-10 |
| B153 | B170 | Glu | U-GLU | 0.51785 | 0 |
| B153 | B174 | Glu | Urea/Cr | 0.175379 | 4.44E-16 |
| B153 | B191 | Glu | Urea | 0.174841 | 4.44E-16 |
| B153 | B207 | Glu | WHR | 0.162389 | 4.22E-14 |
| B153 | B225 | Glu | hPBG | 0.146547 | 9.83E-12 |
| B153 | B155 | Glu | IgA | 0.140941 | 5.92E-11 |
| B153 | B194 | Glu | Cl | -0.11619 | 7.15E-08 |
| B153 | B163 | Glu | SG | 0.112376 | 1.89E-07 |
| B153 | B195 | Glu | Na | -0.10747 | 6.34E-07 |
| B155 | B157 | IgA | IgG | 0.298076 | 0 |
| B155 | B195 | IgA | Na | -0.21014 | 0 |
| B155 | B193 | IgA | K | 0.142415 | 3.72E-11 |
| B155 | B183 | IgA | PTA | -0.1284 | 2.55E-09 |
| B155 | B194 | IgA | Cl | -0.123 | 1.16E-08 |
| B156 | B200 | IgE | OP | 0.172424 | 8.88E-16 |
| B156 | B186 | IgE | PFC | 0.138584 | 1.23E-10 |
| B156 | B199 | IgE | Mg | 0.134034 | 4.91E-10 |
| B156 | B198 | IgE | P | 0.132989 | 6.70E-10 |
| B156 | B194 | IgE | Cl | -0.11966 | 2.86E-08 |
| B157 | B158 | IgG | IgM | 0.252545 | 0 |
| B157 | B185 | IgG | PT | 0.166174 | 1.07E-14 |
| B157 | B198 | IgG | P | 0.139143 | 1.04E-10 |
| B158 | B175 | IgM | ALB-U | 0.185747 | 0 |
| B158 | B177 | IgM | ACR | 0.255403 | 0 |
| B158 | B178 | IgM | Ig-U | 0.29157 | 0 |
| B158 | B180 | IgM | Tf | 0.289897 | 0 |
| B158 | B196 | IgM | TCO2 | 0.129165 | 2.05E-09 |
| B158 | B194 | IgM | Cl | -0.12044 | 2.32E-08 |
| B159 | B169 | β2-MG | U-CRY | 0.99999 | 0 |
| B159 | B199 | β2-MG | Mg | -0.13768 | 1.63E-10 |
| B16 | B139 | CD3+T | ADA | -0.1944 | 0 |
| B16 | B155 | CD3+T | IgA | -0.296 | 0 |
| B16 | B17 | CD3+T | CD3+CD16+CD56+NK | 0.431422 | 0 |
| B16 | B18 | CD3+T | CD3+CD4+helper(inducer)T | 0.366978 | 0 |
| B16 | B181 | CD3+T | APTT | -0.1775 | 0 |
| B16 | B182 | CD3+T | TT | -0.17739 | 0 |
| B16 | B184 | CD3+T | INR | -0.18554 | 0 |

| | | | | | |
|---|---|---|---|---|---|
| B16 | B19 | CD3+T | CD3+CD4+/CD3+CD8+ | -0.17958 | 0 |
| B16 | B20 | CD3+T | CD3+CD4+CD8+ | 0.28689 | 0 |
| B16 | B21 | CD3+T | CD3+CD4-CD8- | 0.231653 | 0 |
| B16 | B22 | CD3+T | CD3+CD8+(T) | 0.549452 | 0 |
| B16 | B23 | CD3+T | CD3-CD16+CD56+(NK) | -0.92488 | 0 |
| B16 | B79 | CD3+T | hb V -dna | -0.20385 | 0 |
| B16 | B95 | CD3+T | anti-cenpb | 0.230853 | 0 |
| B16 | B96 | CD3+T | dsDNA (IIF) | -0.25684 | 0 |
| B16 | B97 | CD3+T | dsDNA | 0.254635 | 0 |
| B16 | B186 | CD3+T | PFC | 0.169588 | 2.89E-15 |
| B16 | B93 | CD3+T | ALDH2-Glu504Lys | 0.144202 | 2.10E-11 |
| B16 | B92 | CD3+T | ALDH2-Glu504Glu | -0.13723 | 1.87E-10 |
| B16 | B81 | CD3+T | TSGF | 0.129754 | 1.73E-09 |
| B16 | B183 | CD3+T | PTA | 0.109258 | 4.10E-07 |
| B160 | B192 | PH | UA | -0.18671 | 0 |
| B160 | B191 | PH | Urea | -0.11132 | 2.47E-07 |
| B161 | B165 | LEU | bilirubin | 0.166126 | 1.07E-14 |
| B161 | B173 | LEU | NTT | 0.152705 | 1.26E-12 |
| B163 | B166 | SG | Pro | 0.2803 | 0 |
| B163 | B174 | SG | Urea/Cr | 0.296326 | 0 |
| B163 | B176 | SG | Ucr | 0.312757 | 0 |
| B163 | B191 | SG | Urea | 0.168564 | 4.44E-15 |
| B163 | B215 | SG | BMI | 0.114606 | 1.07E-07 |
| B163 | B222 | SG | NS | 0.113234 | 1.52E-07 |
| B163 | B189 | SG | eGFB | 0.109465 | 3.90E-07 |
| B163 | B206 | SG | Waist | 0.105688 | 9.70E-07 |
| B165 | B171 | bilirubin | EC | -0.46317 | 0 |
| B166 | B191 | Pro | Urea | 0.111776 | 2.20E-07 |
| B167 | B223 | PC | FH | 0.486655 | 0 |
| B168 | B183 | NMP22 | PTA | -0.16319 | 3.15E-14 |
| B168 | B176 | NMP22 | Ucr | -0.1339 | 5.11E-10 |
| B169 | B199 | U-CRY | Mg | -0.13768 | 1.63E-10 |
| B17 | B108 | CD3+CD16+CD56+NK | Anti-Cyclin | -0.268 | 0 |
| B17 | B155 | CD3+CD16+CD56+NK | IgA | -0.31566 | 0 |
| B17 | B19 | CD3+CD16+CD56+NK | CD3+CD4+/CD3+CD8+ | -0.19552 | 0 |
| B17 | B21 | CD3+CD16+CD56+NK | CD3+CD4-CD8- | 0.222373 | 0 |
| B17 | B22 | CD3+CD16+CD56+NK | CD3+CD8+(T) | 0.332682 | 0 |
| B17 | B23 | CD3+CD16+CD56+NK | CD3-CD16+CD56+(NK) | -0.39435 | 0 |
| B17 | B34 | CD3+CD16+CD56+NK | TK | -0.199 | 0 |
| B17 | B96 | CD3+CD16+CD56+NK | dsDNA (IIF) | -0.52137 | 0 |
| B17 | B97 | CD3+CD16+CD56+NK | dsDNA | 0.536786 | 0 |
| B17 | B183 | CD3+CD16+CD56+NK | PTA | 0.173597 | 6.66E-16 |
| B17 | B139 | CD3+CD16+CD56+NK | ADA | -0.16721 | 7.11E-15 |
| B17 | B157 | CD3+CD16+CD56+NK | IgG | -0.1556 | 4.66E-13 |

| | | | | | |
|---|---|---|---|---|---|
| B17 | B20 | CD3+CD16+CD56+NK | CD3+CD4+CD8+ | 0.155131 | 5.49E-13 |
| B17 | B181 | CD3+CD16+CD56+NK | APTT | -0.13465 | 4.09E-10 |
| B170 | B174 | U-GLU | Urea/Cr | 0.147783 | 6.56E-12 |
| B173 | B175 | NTT | ALB-U | 0.284926 | 0 |
| B173 | B177 | NTT | ACR | 0.133471 | 5.81E-10 |
| B174 | B190 | Urea/Cr | Cre | -0.25801 | 0 |
| B174 | B191 | Urea/Cr | Urea | 0.598731 | 0 |
| B174 | B192 | Urea/Cr | UA | -0.16422 | 2.15E-14 |
| B174 | B189 | Urea/Cr | eGFB | 0.132158 | 8.56E-10 |
| B174 | B221 | Urea/Cr | Hei | -0.1109 | 2.74E-07 |
| B175 | B177 | ALB-U | ACR | 0.854515 | 0 |
| B175 | B178 | ALB-U | Ig-U | 0.617139 | 0 |
| B175 | B179 | ALB-U | α1-MG | 0.196688 | 0 |
| B175 | B180 | ALB-U | Tf | 0.622532 | 0 |
| B175 | B184 | ALB-U | INR | -0.20123 | 0 |
| B175 | B185 | ALB-U | PT | -0.1819 | 0 |
| B176 | B179 | Ucr | α1-MG | 0.126182 | 4.79E-09 |
| B176 | B177 | Ucr | ACR | -0.12324 | 1.08E-08 |
| B177 | B178 | ACR | Ig-U | 0.847147 | 0 |
| B177 | B179 | ACR | α1-MG | 0.256054 | 0 |
| B177 | B180 | ACR | Tf | 0.85633 | 0 |
| B177 | B184 | ACR | INR | -0.27858 | 0 |
| B177 | B185 | ACR | PT | -0.25144 | 0 |
| B177 | B182 | ACR | TT | 0.138508 | 1.26E-10 |
| B177 | B194 | ACR | Cl | -0.12846 | 2.51E-09 |
| B177 | B190 | ACR | Cre | 0.126493 | 4.38E-09 |
| B177 | B181 | ACR | APTT | -0.11447 | 1.11E-07 |
| B178 | B179 | Ig-U | α1-MG | 0.363659 | 0 |
| B178 | B180 | Ig-U | Tf | 0.985744 | 0 |
| B178 | B184 | Ig-U | INR | -0.30245 | 0 |
| B178 | B185 | Ig-U | PT | -0.27617 | 0 |
| B178 | B182 | Ig-U | TT | 0.16505 | 1.60E-14 |
| B178 | B181 | Ig-U | APTT | -0.13753 | 1.70E-10 |
| B178 | B194 | Ig-U | Cl | -0.13544 | 3.22E-10 |
| B179 | B180 | α1-MG | Tf | 0.280985 | 0 |
| B18 | B124 | CD3+CD4+helper(inducer)T | ACTH | -0.18217 | 0 |
| B18 | B125 | CD3+CD4+helper(inducer)T | Cor | -0.19398 | 0 |
| B18 | B155 | CD3+CD4+helper(inducer)T | IgA | -0.25057 | 0 |
| B18 | B177 | CD3+CD4+helper(inducer)T | ACR | 0.183474 | 0 |
| B18 | B178 | CD3+CD4+helper(inducer)T | Ig-U | 0.223638 | 0 |
| B18 | B180 | CD3+CD4+helper(inducer)T | Tf | 0.218407 | 0 |
| B18 | B182 | CD3+CD4+helper(inducer)T | TT | 0.24047 | 0 |
| B18 | B186 | CD3+CD4+helper(inducer)T | PFC | -0.17767 | 0 |
| B18 | B19 | CD3+CD4+helper(inducer)T | CD3+CD4+/CD3+CD8+ | 0.778255 | 0 |

| | | | | | |
|---|---|---|---|---|---|
| B18 | B194 | CD3+CD4+helper(inducer)T | Cl | 0.196813 | 0 |
| B18 | B20 | CD3+CD4+helper(inducer)T | CD3+CD4+CD8+ | 0.254991 | 0 |
| B18 | B22 | CD3+CD4+helper(inducer)T | CD3+CD8+(T) | -0.51834 | 0 |
| B18 | B23 | CD3+CD4+helper(inducer)T | CD3-CD16+CD56+(NK) | -0.44286 | 0 |
| B18 | B95 | CD3+CD4+helper(inducer)T | anti-cenpb | 0.376973 | 0 |
| B18 | B179 | CD3+CD4+helper(inducer)T | α1-MG | 0.16184 | 5.13E-14 |
| B18 | B58 | CD3+CD4+helper(inducer)T | HCT | -0.14908 | 4.26E-12 |
| B18 | B69 | CD3+CD4+helper(inducer)T | PV | -0.14644 | 1.02E-11 |
| B18 | B175 | CD3+CD4+helper(inducer)T | ALB-U | 0.137925 | 1.51E-10 |
| B18 | B97 | CD3+CD4+helper(inducer)T | dsDNA | 0.130825 | 1.27E-09 |
| B18 | B79 | CD3+CD4+helper(inducer)T | hb V -dna | 0.126711 | 4.12E-09 |
| B18 | B157 | CD3+CD4+helper(inducer)T | IgG | -0.12379 | 9.32E-09 |
| B18 | B96 | CD3+CD4+helper(inducer)T | dsDNA (IIF) | -0.12095 | 2.02E-08 |
| B18 | B136 | CD3+CD4+helper(inducer)T | 5'NT | -0.10957 | 3.80E-07 |
| B18 | B21 | CD3+CD4+helper(inducer)T | CD3+CD4-CD8- | -0.1079 | 5.71E-07 |
| B18 | B137 | CD3+CD4+helper(inducer)T | AFU | -0.10729 | 6.61E-07 |
| B18 | B65 | CD3+CD4+helper(inducer)T | WBRSR(mPas_1) | -0.10594 | 9.13E-07 |
| B180 | B184 | Tf | INR | -0.31935 | 0 |
| B180 | B185 | Tf | PT | -0.29245 | 0 |
| B180 | B182 | Tf | TT | 0.164907 | 1.69E-14 |
| B180 | B194 | Tf | Cl | -0.14136 | 5.18E-11 |
| B180 | B181 | Tf | APTT | -0.13852 | 1.26E-10 |
| B181 | B183 | APTT | PTA | -0.34116 | 0 |
| B181 | B184 | APTT | INR | 0.276555 | 0 |
| B181 | B185 | APTT | PT | 0.290331 | 0 |
| B181 | B197 | APTT | AG | 0.179292 | 0 |
| B181 | B186 | APTT | PFC | 0.159345 | 1.25E-13 |
| B181 | B196 | APTT | TCO2 | -0.10862 | 4.79E-07 |
| B182 | B183 | TT | PTA | -0.19872 | 0 |
| B182 | B186 | TT | PFC | -0.53003 | 0 |
| B182 | B200 | TT | OP | 0.1507 | 2.49E-12 |
| B182 | B185 | TT | PT | 0.138951 | 1.10E-10 |
| B182 | B194 | TT | Cl | 0.127884 | 2.95E-09 |
| B182 | B196 | TT | TCO2 | -0.12346 | 1.02E-08 |
| B182 | B198 | TT | P | -0.11492 | 9.92E-08 |
| B182 | B195 | TT | Na | 0.108637 | 4.77E-07 |
| B183 | B200 | PTA | OP | -0.1742 | 4.44E-16 |
| B183 | B185 | PTA | PT | -0.17193 | 1.33E-15 |
| B183 | B195 | PTA | Na | -0.1441 | 2.17E-11 |
| B183 | B197 | PTA | AG | -0.13453 | 4.24E-10 |
| B184 | B185 | INR | PT | 0.897575 | 0 |
| B184 | B186 | INR | PFC | -0.26294 | 0 |
| B184 | B213 | INR | CVA-R | -0.13903 | 1.07E-10 |
| B185 | B213 | PT | CVA-R | -0.19654 | 0 |

| | | | | | |
|---|---|---|---|---|---|
| B185 | B186 | PT | PFC | -0.15728 | 2.60E-13 |
| B186 | B194 | PFC | Cl | -0.28201 | 0 |
| B186 | B197 | PFC | AG | 0.132614 | 7.48E-10 |
| B186 | B198 | PFC | P | 0.12731 | 3.48E-09 |
| B186 | B199 | PFC | Mg | 0.121343 | 1.82E-08 |
| B188 | B192 | Ca | UA | 0.131552 | 1.02E-09 |
| B189 | B190 | eGFB | Cre | -0.60994 | 0 |
| B189 | B191 | eGFB | Urea | -0.49041 | 0 |
| B189 | B210 | eGFB | Hip | 0.185834 | 0 |
| B189 | B216 | eGFB | BMI | 0.216253 | 0 |
| B189 | B220 | eGFB | SBP | -0.22437 | 0 |
| B189 | B221 | eGFB | Hei | 0.230743 | 0 |
| B189 | B222 | eGFB | NS | 0.148828 | 4.64E-12 |
| B189 | B224 | eGFB | pmh | 0.146668 | 9.45E-12 |
| B189 | B215 | eGFB | BMI | 0.114425 | 1.13E-07 |
| B189 | B201 | eGFB | e-habits | -0.1137 | 1.36E-07 |
| B19 | B182 | CD3+CD4+/CD3+CD8+ | TT | 0.480585 | 0 |
| B19 | B186 | CD3+CD4+/CD3+CD8+ | PFC | -0.29975 | 0 |
| B19 | B22 | CD3+CD4+/CD3+CD8+ | CD3+CD8+(T) | -0.87802 | 0 |
| B19 | B79 | CD3+CD4+/CD3+CD8+ | hb V -dna | 0.205462 | 0 |
| B19 | B95 | CD3+CD4+/CD3+CD8+ | anti-cenpb | 0.293217 | 0 |
| B19 | B125 | CD3+CD4+/CD3+CD8+ | Cor | -0.17251 | 8.88E-16 |
| B19 | B194 | CD3+CD4+/CD3+CD8+ | Cl | 0.171408 | 1.33E-15 |
| B19 | B58 | CD3+CD4+/CD3+CD8+ | HCT | -0.14835 | 5.44E-12 |
| B19 | B178 | CD3+CD4+/CD3+CD8+ | Ig-U | 0.14784 | 6.43E-12 |
| B19 | B124 | CD3+CD4+/CD3+CD8+ | ACTH | -0.14655 | 9.83E-12 |
| B19 | B100 | CD3+CD4+/CD3+CD8+ | anti-Ro52 | 0.145819 | 1.25E-11 |
| B19 | B101 | CD3+CD4+/CD3+CD8+ | anti-Sc70 | 0.145819 | 1.25E-11 |
| B19 | B102 | CD3+CD4+/CD3+CD8+ | anti-Sm | 0.145819 | 1.25E-11 |
| B19 | B103 | CD3+CD4+/CD3+CD8+ | anti-SSA/Ro | 0.145819 | 1.25E-11 |
| B19 | B104 | CD3+CD4+/CD3+CD8+ | anti-SSB | 0.145819 | 1.25E-11 |
| B19 | B105 | CD3+CD4+/CD3+CD8+ | anti-U1RNP | 0.145819 | 1.25E-11 |
| B19 | B106 | CD3+CD4+/CD3+CD8+ | anti-P | 0.145819 | 1.25E-11 |
| B19 | B107 | CD3+CD4+/CD3+CD8+ | AnuA | 0.145819 | 1.25E-11 |
| B19 | B109 | CD3+CD4+/CD3+CD8+ | AMA-M2 | 0.145819 | 1.25E-11 |
| B19 | B110 | CD3+CD4+/CD3+CD8+ | AHA | 0.145819 | 1.25E-11 |
| B19 | B98 | CD3+CD4+/CD3+CD8+ | anti-Jo-1 | 0.145819 | 1.25E-11 |
| B19 | B69 | CD3+CD4+/CD3+CD8+ | PV | -0.14424 | 2.07E-11 |
| B19 | B180 | CD3+CD4+/CD3+CD8+ | Tf | 0.142458 | 3.67E-11 |
| B19 | B179 | CD3+CD4+/CD3+CD8+ | α1-MG | 0.131803 | 9.50E-10 |
| B19 | B134 | CD3+CD4+/CD3+CD8+ | CHE | -0.12675 | 4.08E-09 |
| B19 | B136 | CD3+CD4+/CD3+CD8+ | 5'NT | -0.12402 | 8.76E-09 |
| B19 | B196 | CD3+CD4+/CD3+CD8+ | TCO2 | -0.11929 | 3.16E-08 |
| B19 | B177 | CD3+CD4+/CD3+CD8+ | ACR | 0.117986 | 4.46E-08 |

| | | | | | |
|---|---|---|---|---|---|
| B19 | B34 | CD3+CD4+/CD3+CD8+ | TK | 0.117803 | 4.68E-08 |
| B19 | B199 | CD3+CD4+/CD3+CD8+ | Mg | -0.11174 | 2.22E-07 |
| B19 | B183 | CD3+CD4+/CD3+CD8+ | PTA | -0.10705 | 7.01E-07 |
| B190 | B191 | Cre | Urea | 0.538521 | 0 |
| B190 | B192 | Cre | UA | 0.160632 | 7.90E-14 |
| B191 | B221 | Urea | Hei | -0.15715 | 2.72E-13 |
| B191 | B220 | Urea | SBP | 0.141028 | 5.76E-11 |
| B191 | B216 | Urea | BMI | -0.106 | 8.99E-07 |
| B192 | B216 | UA | BMI | 0.120963 | 2.02E-08 |
| B192 | B210 | UA | Hip | 0.118368 | 4.04E-08 |
| B192 | B215 | UA | BMI | 0.117919 | 4.54E-08 |
| B193 | B225 | K | hPBG | -0.12358 | 9.88E-09 |
| B193 | B197 | K | AG | -0.11605 | 7.40E-08 |
| B194 | B195 | Cl | Na | 0.259279 | 0 |
| B194 | B197 | Cl | AG | -0.35658 | 0 |
| B194 | B198 | Cl | P | -0.13494 | 3.74E-10 |
| B194 | B200 | Cl | OP | 0.131204 | 1.13E-09 |
| B195 | B197 | Na | AG | 0.328625 | 0 |
| B195 | B200 | Na | OP | 0.799789 | 0 |
| B196 | B197 | TCO2 | AG | -0.7558 | 0 |
| B197 | B200 | AG | OP | 0.349954 | 0 |
| B198 | B199 | P | Mg | 0.137335 | 1.81E-10 |
| B198 | B211 | P | IOP-R | 0.107849 | 5.78E-07 |
| B2 | B126 | Age | Alb | -0.31866 | 0 |
| B2 | B128 | Age | ALT | -0.30383 | 0 |
| B2 | B141 | Age | AST/ALT | 0.249818 | 0 |
| B2 | B153 | Age | Glu | 0.188041 | 0 |
| B2 | B188 | Age | Ca | -0.22551 | 0 |
| B2 | B189 | Age | eGFB | -0.54367 | 0 |
| B2 | B190 | Age | Cre | 0.188143 | 0 |
| B2 | B191 | Age | Urea | 0.329761 | 0 |
| B2 | B210 | Age | Hip | -0.23714 | 0 |
| B2 | B215 | Age | BMI | -0.19184 | 0 |
| B2 | B216 | Age | BMI | -0.36546 | 0 |
| B2 | B219 | Age | OT | 0.200148 | 0 |
| B2 | B220 | Age | SBP | 0.339373 | 0 |
| B2 | B221 | Age | Hei | -0.39785 | 0 |
| B2 | B222 | Age | NS | -0.23449 | 0 |
| B2 | B43 | Age | MCV | 0.191156 | 0 |
| B2 | B46 | Age | RBC Count | -0.37899 | 0 |
| B2 | B47 | Age | RDW | 0.32684 | 0 |
| B2 | B51 | Age | Hb | -0.34638 | 0 |
| B2 | B52 | Age | PLT/BPC | -0.21291 | 0 |
| B2 | B118 | Age | CEA1 | 0.174673 | 4.44E-16 |

| | | | | | |
|---|---|---|---|---|---|
| B2 | B201 | Age | e-habits | 0.175471 | 4.44E-16 |
| B2 | B192 | Age | UA | -0.17155 | 1.33E-15 |
| B2 | B41 | Age | MON％ | 0.168479 | 4.44E-15 |
| B2 | B127 | Age | A-G ratio | -0.15692 | 2.95E-13 |
| B2 | B174 | Age | Urea/Cr | 0.150566 | 2.60E-12 |
| B2 | B72 | Age | HDL-C | 0.147433 | 7.36E-12 |
| B2 | B176 | Age | Ucr | -0.14661 | 9.63E-12 |
| B2 | B71 | Age | TG | -0.14589 | 1.22E-11 |
| B2 | B91 | Age | TP | -0.1435 | 2.63E-11 |
| B2 | B45 | Age | MCHC | -0.14294 | 3.15E-11 |
| B2 | B146 | Age | CysC | 0.138818 | 1.15E-10 |
| B2 | B140 | Age | r-GT | -0.13253 | 7.66E-10 |
| B2 | B207 | Age | WHR | 0.129597 | 1.81E-09 |
| B2 | B113 | Age | FT3 | -0.12883 | 2.26E-09 |
| B2 | B227 | Age | OD | 0.122895 | 1.19E-08 |
| B2 | B50 | Age | eo% | 0.120319 | 2.40E-08 |
| B2 | B132 | Age | Dbil | 0.119485 | 3.00E-08 |
| B2 | B131 | Age | AST | -0.1188 | 3.60E-08 |
| B2 | B173 | Age | NTT | 0.115965 | 7.57E-08 |
| B2 | B203 | Age | DH | -0.11481 | 1.02E-07 |
| B20 | B151 | CD3+CD4+CD8+ | Anti-CCP | -0.22041 | 0 |
| B20 | B175 | CD3+CD4+CD8+ | ALB-U | 0.209198 | 0 |
| B20 | B177 | CD3+CD4+CD8+ | ACR | 0.289813 | 0 |
| B20 | B178 | CD3+CD4+CD8+ | Ig-U | 0.345782 | 0 |
| B20 | B180 | CD3+CD4+CD8+ | Tf | 0.343288 | 0 |
| B20 | B187 | CD3+CD4+CD8+ | NSE | 0.215175 | 0 |
| B20 | B21 | CD3+CD4+CD8+ | CD3+CD4-CD8- | -0.29285 | 0 |
| B20 | B23 | CD3+CD4+CD8+ | CD3-CD16+CD56+(NK) | -0.24682 | 0 |
| B20 | B22 | CD3+CD4+CD8+ | CD3+CD8+(T) | 0.176904 | 2.22E-16 |
| B20 | B182 | CD3+CD4+CD8+ | TT | -0.16873 | 4.00E-15 |
| B20 | B57 | CD3+CD4+CD8+ | RBCAI | -0.15653 | 3.38E-13 |
| B20 | B93 | CD3+CD4+CD8+ | ALDH2-Glu504Lys | 0.147618 | 6.92E-12 |
| B20 | B92 | CD3+CD4+CD8+ | ALDH2-Glu504Glu | -0.14113 | 5.58E-11 |
| B20 | B100 | CD3+CD4+CD8+ | anti-Ro52 | -0.13084 | 1.26E-09 |
| B20 | B101 | CD3+CD4+CD8+ | anti-Sc70 | -0.13084 | 1.26E-09 |
| B20 | B102 | CD3+CD4+CD8+ | anti-Sm | -0.13084 | 1.26E-09 |
| B20 | B103 | CD3+CD4+CD8+ | anti-SSA/Ro | -0.13084 | 1.26E-09 |
| B20 | B104 | CD3+CD4+CD8+ | anti-SSB | -0.13084 | 1.26E-09 |
| B20 | B105 | CD3+CD4+CD8+ | anti-U1RNP | -0.13084 | 1.26E-09 |
| B20 | B106 | CD3+CD4+CD8+ | anti-P | -0.13084 | 1.26E-09 |
| B20 | B107 | CD3+CD4+CD8+ | AnuA | -0.13084 | 1.26E-09 |
| B20 | B109 | CD3+CD4+CD8+ | AMA-M2 | -0.13084 | 1.26E-09 |
| B20 | B110 | CD3+CD4+CD8+ | AHA | -0.13084 | 1.26E-09 |
| B20 | B98 | CD3+CD4+CD8+ | anti-Jo-1 | -0.13084 | 1.26E-09 |

| | | | | | |
|---|---|---|---|---|---|
| B20 | B95 | CD3+CD4+CD8+ | anti-cenpb | 0.130806 | 1.27E-09 |
| B20 | B155 | CD3+CD4+CD8+ | IgA | -0.12988 | 1.67E-09 |
| B20 | B194 | CD3+CD4+CD8+ | CI | 0.126949 | 3.85E-09 |
| B20 | B179 | CD3+CD4+CD8+ | α1-MG | 0.126533 | 4.34E-09 |
| B20 | B59 | CD3+CD4+CD8+ | WBRV | -0.12145 | 1.77E-08 |
| B20 | B65 | CD3+CD4+CD8+ | WBRSR(mPas_1) | -0.10707 | 6.97E-07 |
| B20 | B60 | CD3+CD4+CD8+ | WBRVI | -0.10591 | 9.21E-07 |
| B200 | B225 | OP | hPBG | 0.111251 | 2.51E-07 |
| B202 | B223 | e-times | FH | 0.663315 | 0 |
| B204 | B209 | Drinking | Smoking | 0.264054 | 0 |
| B204 | B218 | Drinking | DBP | 0.128448 | 2.52E-09 |
| B206 | B207 | Waist | WHR | 0.692646 | 0 |
| B206 | B210 | Waist | Hip | 0.763584 | 0 |
| B206 | B215 | Waist | BMI | 0.821678 | 0 |
| B206 | B216 | Waist | BMI | 0.785472 | 0 |
| B206 | B218 | Waist | DBP | 0.211682 | 0 |
| B206 | B221 | Waist | Hei | 0.244496 | 0 |
| B206 | B222 | Waist | NS | 0.704433 | 0 |
| B206 | B220 | Waist | SBP | 0.153438 | 9.83E-13 |
| B207 | B215 | WHR | BMI | 0.456985 | 0 |
| B207 | B216 | WHR | BMI | 0.33809 | 0 |
| B207 | B222 | WHR | NS | 0.374975 | 0 |
| B207 | B218 | WHR | DBP | 0.145193 | 1.53E-11 |
| B207 | B220 | WHR | SBP | 0.133579 | 5.62E-10 |
| B207 | B210 | WHR | Hip | 0.123942 | 8.94E-09 |
| B207 | B227 | WHR | OD | -0.10854 | 4.89E-07 |
| B21 | B100 | CD3+CD4-CD8- | anti-Ro52 | 0.430525 | 0 |
| B21 | B101 | CD3+CD4-CD8- | anti-Sc70 | 0.430525 | 0 |
| B21 | B102 | CD3+CD4-CD8- | anti-Sm | 0.430525 | 0 |
| B21 | B103 | CD3+CD4-CD8- | anti-SSA/Ro | 0.430525 | 0 |
| B21 | B104 | CD3+CD4-CD8- | anti-SSB | 0.430525 | 0 |
| B21 | B105 | CD3+CD4-CD8- | anti-U1RNP | 0.430525 | 0 |
| B21 | B106 | CD3+CD4-CD8- | anti-P | 0.430525 | 0 |
| B21 | B107 | CD3+CD4-CD8- | AnuA | 0.430525 | 0 |
| B21 | B109 | CD3+CD4-CD8- | AMA-M2 | 0.430525 | 0 |
| B21 | B110 | CD3+CD4-CD8- | AHA | 0.430525 | 0 |
| B21 | B134 | CD3+CD4-CD8- | CHE | -0.19064 | 0 |
| B21 | B136 | CD3+CD4-CD8- | 5'NT | -0.18748 | 0 |
| B21 | B151 | CD3+CD4-CD8- | Anti-CCP | 0.470513 | 0 |
| B21 | B155 | CD3+CD4-CD8- | IgA | -0.21552 | 0 |
| B21 | B177 | CD3+CD4-CD8- | ACR | -0.24372 | 0 |
| B21 | B178 | CD3+CD4-CD8- | Ig-U | -0.30118 | 0 |
| B21 | B180 | CD3+CD4-CD8- | Tf | -0.29628 | 0 |
| B21 | B23 | CD3+CD4-CD8- | CD3-CD16+CD56+(NK) | -0.22839 | 0 |

| | | | | | |
|---|---|---|---|---|---|
| B21 | B57 | CD3+CD4-CD8- | RBCAI | 0.290548 | 0 |
| B21 | B59 | CD3+CD4-CD8- | WBRV | 0.181062 | 0 |
| B21 | B92 | CD3+CD4-CD8- | ALDH2-Glu504Glu | 0.193088 | 0 |
| B21 | B93 | CD3+CD4-CD8- | ALDH2-Glu504Lys | -0.20823 | 0 |
| B21 | B96 | CD3+CD4-CD8- | dsDNA (IIF) | 0.210376 | 0 |
| B21 | B98 | CD3+CD4-CD8- | anti-Jo-1 | 0.430525 | 0 |
| B21 | B74 | CD3+CD4-CD8- | ApoA1 | 0.174004 | 4.44E-16 |
| B21 | B175 | CD3+CD4-CD8- | ALB-U | -0.17184 | 1.33E-15 |
| B21 | B97 | CD3+CD4-CD8- | dsDNA | -0.1717 | 1.33E-15 |
| B21 | B139 | CD3+CD4-CD8- | ADA | -0.16823 | 4.88E-15 |
| B21 | B39 | CD3+CD4-CD8- | Other cells | 0.149238 | 4.05E-12 |
| B21 | B195 | CD3+CD4-CD8- | Na | 0.134637 | 4.10E-10 |
| B21 | B60 | CD3+CD4-CD8- | WBRVI | 0.123103 | 1.13E-08 |
| B21 | B184 | CD3+CD4-CD8- | INR | 0.113619 | 1.38E-07 |
| B21 | B185 | CD3+CD4-CD8- | PT | 0.11186 | 2.15E-07 |
| B21 | B154 | CD3+CD4-CD8- | RF | 0.107111 | 6.90E-07 |
| B210 | B215 | Hip | BMI | 0.752439 | 0 |
| B210 | B216 | Hip | BMI | 0.818078 | 0 |
| B210 | B221 | Hip | Hei | 0.400676 | 0 |
| B210 | B222 | Hip | NS | 0.658059 | 0 |
| B210 | B218 | Hip | DBP | 0.167866 | 5.55E-15 |
| B211 | B212 | IOP-R | IOP-L | 0.782797 | 0 |
| B215 | B216 | BMI | BMI | 0.839713 | 0 |
| B215 | B218 | BMI | DBP | 0.22617 | 0 |
| B215 | B222 | BMI | NS | 0.70265 | 0 |
| B215 | B220 | BMI | SBP | 0.130964 | 1.22E-09 |
| B216 | B218 | BMI | DBP | 0.236781 | 0 |
| B216 | B221 | BMI | Hei | 0.6066 | 0 |
| B216 | B222 | BMI | NS | 0.714356 | 0 |
| B218 | B220 | DBP | SBP | 0.61174 | 0 |
| B218 | B222 | DBP | NS | 0.196469 | 0 |
| B218 | B221 | DBP | Hei | 0.106213 | 8.56E-07 |
| B22 | B100 | CD3+CD8+(T) | anti-Ro52 | -0.23463 | 0 |
| B22 | B101 | CD3+CD8+(T) | anti-Sc70 | -0.23463 | 0 |
| B22 | B102 | CD3+CD8+(T) | anti-Sm | -0.23463 | 0 |
| B22 | B103 | CD3+CD8+(T) | anti-SSA/Ro | -0.23463 | 0 |
| B22 | B104 | CD3+CD8+(T) | anti-SSB | -0.23463 | 0 |
| B22 | B105 | CD3+CD8+(T) | anti-U1RNP | -0.23463 | 0 |
| B22 | B106 | CD3+CD8+(T) | anti-P | -0.23463 | 0 |
| B22 | B107 | CD3+CD8+(T) | AnuA | -0.23463 | 0 |
| B22 | B109 | CD3+CD8+(T) | AMA-M2 | -0.23463 | 0 |
| B22 | B110 | CD3+CD8+(T) | AHA | -0.23463 | 0 |
| B22 | B125 | CD3+CD8+(T) | Cor | 0.204521 | 0 |
| B22 | B182 | CD3+CD8+(T) | TT | -0.40882 | 0 |

| | | | | | |
|---|---|---|---|---|---|
| B22 | B186 | CD3+CD8+(T) | PFC | 0.335278 | 0 |
| B22 | B23 | CD3+CD8+(T) | CD3-CD16+CD56+(NK) | -0.41275 | 0 |
| B22 | B79 | CD3+CD8+(T) | hb V -dna | -0.30208 | 0 |
| B22 | B96 | CD3+CD8+(T) | dsDNA (IIF) | -0.18471 | 0 |
| B22 | B98 | CD3+CD8+(T) | anti-Jo-1 | -0.23463 | 0 |
| B22 | B124 | CD3+CD8+(T) | ACTH | 0.168677 | 4.00E-15 |
| B22 | B97 | CD3+CD8+(T) | dsDNA | 0.164637 | 1.87E-14 |
| B22 | B134 | CD3+CD8+(T) | CHE | 0.163464 | 2.84E-14 |
| B22 | B69 | CD3+CD8+(T) | PV | 0.161548 | 5.71E-14 |
| B22 | B95 | CD3+CD8+(T) | anti-cenpb | -0.15298 | 1.15E-12 |
| B22 | B34 | CD3+CD8+(T) | TK | -0.14805 | 5.99E-12 |
| B22 | B93 | CD3+CD8+(T) | ALDH2-Glu504Lys | 0.147206 | 7.93E-12 |
| B22 | B181 | CD3+CD8+(T) | APTT | -0.14152 | 4.93E-11 |
| B22 | B199 | CD3+CD8+(T) | Mg | 0.133554 | 5.66E-10 |
| B22 | B92 | CD3+CD8+(T) | ALDH2-Glu504Glu | -0.13031 | 1.47E-09 |
| B22 | B136 | CD3+CD8+(T) | 5'NT | 0.115202 | 9.22E-08 |
| B22 | B179 | CD3+CD8+(T) | α1-MG | -0.11095 | 2.71E-07 |
| B22 | B196 | CD3+CD8+(T) | TCO2 | 0.109885 | 3.52E-07 |
| B22 | B198 | CD3+CD8+(T) | P | 0.108773 | 4.62E-07 |
| B22 | B184 | CD3+CD8+(T) | INR | -0.10841 | 5.04E-07 |
| B221 | B222 | Hei | NS | 0.300426 | 0 |
| B227 | B228 | OD | OS | 0.178031 | 0 |
| B23 | B139 | CD3-CD16+CD56+(NK) | ADA | 0.220338 | 0 |
| B23 | B155 | CD3-CD16+CD56+(NK) | IgA | 0.338873 | 0 |
| B23 | B181 | CD3-CD16+CD56+(NK) | APTT | 0.192138 | 0 |
| B23 | B95 | CD3-CD16+CD56+(NK) | anti-cenpb | -0.18496 | 0 |
| B23 | B96 | CD3-CD16+CD56+(NK) | dsDNA (IIF) | 0.236755 | 0 |
| B23 | B97 | CD3-CD16+CD56+(NK) | dsDNA | -0.24028 | 0 |
| B23 | B183 | CD3-CD16+CD56+(NK) | PTA | -0.16857 | 4.44E-15 |
| B23 | B184 | CD3-CD16+CD56+(NK) | INR | 0.155288 | 5.20E-13 |
| B23 | B93 | CD3-CD16+CD56+(NK) | ALDH2-Glu504Lys | -0.15493 | 5.89E-13 |
| B23 | B180 | CD3-CD16+CD56+(NK) | Tf | -0.15466 | 6.46E-13 |
| B23 | B178 | CD3-CD16+CD56+(NK) | Ig-U | -0.15351 | 9.58E-13 |
| B23 | B92 | CD3-CD16+CD56+(NK) | ALDH2-Glu504Glu | 0.149132 | 4.20E-12 |
| B23 | B177 | CD3-CD16+CD56+(NK) | ACR | -0.13467 | 4.05E-10 |
| B23 | B194 | CD3-CD16+CD56+(NK) | Cl | -0.11871 | 3.69E-08 |
| B23 | B175 | CD3-CD16+CD56+(NK) | ALB-U | -0.10827 | 5.22E-07 |
| B24 | B26 | αHBDH | CKMB | 0.745974 | 0 |
| B24 | B27 | αHBDH | LDH | 0.749091 | 0 |
| B24 | B187 | αHBDH | NSE | 0.162835 | 3.57E-14 |
| B25 | B134 | CK | CHE | 0.153832 | 8.59E-13 |
| B25 | B26 | CK | CKMB | 0.125215 | 6.28E-09 |
| B25 | B27 | CK | LDH | 0.109676 | 3.70E-07 |
| B26 | B187 | CKMB | NSE | 0.208135 | 0 |

| | | | | | |
|---|---|---|---|---|---|
| B26 | B27 | CKMB | LDH | 0.58036 | 0 |
| B27 | B131 | LDH | AST | 0.194758 | 0 |
| B27 | B187 | LDH | NSE | 0.154736 | 6.29E-13 |
| B27 | B74 | LDH | ApoA1 | 0.121677 | 1.66E-08 |
| B28 | B223 | PRL | FH | 0.8001 | 0 |
| B28 | B31 | PRL | FSH | 0.465857 | 0 |
| B28 | B33 | PRL | Prog | 0.592945 | 0 |
| B28 | B37 | PRL | BAND | 0.537895 | 0 |
| B29 | B167 | E2 | PC | 0.391686 | 0 |
| B29 | B223 | E2 | FH | 0.934898 | 0 |
| B29 | B31 | E2 | FSH | 0.957969 | 0 |
| B29 | B33 | E2 | Prog | 0.601912 | 0 |
| B29 | B35 | E2 | β2-MG | 0.875836 | 0 |
| B29 | B37 | E2 | BAND | 0.201596 | 0 |
| B3 | B189 | Hcy | eGFB | -0.17359 | 6.66E-16 |
| B3 | B181 | Hcy | APTT | -0.16143 | 5.95E-14 |
| B3 | B183 | Hcy | PTA | 0.153556 | 9.44E-13 |
| B3 | B168 | Hcy | NMP22 | -0.1265 | 4.38E-09 |
| B3 | B39 | Hcy | Other cells | -0.12185 | 1.59E-08 |
| B30 | B38 | LH | BAND% | -0.22943 | 0 |
| B31 | B223 | FSH | FH | 0.968552 | 0 |
| B31 | B33 | FSH | Prog | 0.571819 | 0 |
| B31 | B35 | FSH | β2-MG | 0.466086 | 0 |
| B32 | B116 | Testo | TT3 | -0.5652 | 0 |
| B33 | B167 | Prog | PC | 0.951019 | 0 |
| B33 | B202 | Prog | e-times | 0.738119 | 0 |
| B33 | B37 | Prog | BAND | 0.481417 | 0 |
| B34 | B151 | TK | Anti-CCP | 0.207686 | 0 |
| B34 | B182 | TK | TT | -0.18944 | 0 |
| B34 | B187 | TK | NSE | 0.251039 | 0 |
| B34 | B79 | TK | hb V -dna | 0.260002 | 0 |
| B34 | B81 | TK | TSGF | -0.24345 | 0 |
| B34 | B138 | TK | PAB | -0.16138 | 6.06E-14 |
| B34 | B139 | TK | ADA | 0.159873 | 1.04E-13 |
| B34 | B124 | TK | ACTH | 0.129572 | 1.82E-09 |
| B34 | B57 | TK | RBCAI | 0.126079 | 4.93E-09 |
| B34 | B134 | TK | CHE | -0.12498 | 6.70E-09 |
| B34 | B184 | TK | INR | 0.124926 | 6.80E-09 |
| B34 | B180 | TK | Tf | -0.11271 | 1.74E-07 |
| B34 | B100 | TK | anti-Ro52 | 0.11125 | 2.51E-07 |
| B34 | B101 | TK | anti-Sc70 | 0.11125 | 2.51E-07 |
| B34 | B102 | TK | anti-Sm | 0.11125 | 2.51E-07 |
| B34 | B103 | TK | anti-SSA/Ro | 0.11125 | 2.51E-07 |
| B34 | B104 | TK | anti-SSB | 0.11125 | 2.51E-07 |

| | | | | | |
|---|---|---|---|---|---|
| B34 | B105 | TK | anti-U1RNP | 0.11125 | 2.51E-07 |
| B34 | B106 | TK | anti-P | 0.11125 | 2.51E-07 |
| B34 | B107 | TK | AnuA | 0.11125 | 2.51E-07 |
| B34 | B109 | TK | AMA-M2 | 0.11125 | 2.51E-07 |
| B34 | B110 | TK | AHA | 0.11125 | 2.51E-07 |
| B34 | B98 | TK | anti-Jo-1 | 0.11125 | 2.51E-07 |
| B34 | B178 | TK | Ig-U | -0.10718 | 6.80E-07 |
| B35 | B202 | β2-MG | e-times | 0.56841 | 0 |
| B35 | B223 | β2-MG | FH | 0.993533 | 0 |
| B36 | B210 | V-Lym% | Hip | -0.15085 | 2.36E-12 |
| B36 | B216 | V-Lym% | BMI | -0.13598 | 2.73E-10 |
| B36 | B215 | V-Lym% | BMI | -0.12518 | 6.33E-09 |
| B37 | B202 | BAND | e-times | 0.505238 | 0 |
| B38 | B116 | BAND% | TT3 | -0.5945 | 0 |
| B38 | B117 | BAND% | TT4 | -0.50246 | 0 |
| B39 | B151 | Other cells | Anti-CCP | 0.177151 | 0 |
| B39 | B168 | Other cells | NMP22 | 0.23584 | 0 |
| B39 | B74 | Other cells | ApoA1 | 0.231465 | 0 |
| B39 | B134 | Other cells | CHE | -0.17154 | 1.33E-15 |
| B39 | B78 | Other cells | HBcAb | 0.162422 | 4.15E-14 |
| B39 | B69 | Other cells | PV | -0.15713 | 2.74E-13 |
| B39 | B100 | Other cells | anti-Ro52 | 0.143633 | 2.52E-11 |
| B39 | B101 | Other cells | anti-Sc70 | 0.143633 | 2.52E-11 |
| B39 | B102 | Other cells | anti-Sm | 0.143633 | 2.52E-11 |
| B39 | B103 | Other cells | anti-SSA/Ro | 0.143633 | 2.52E-11 |
| B39 | B104 | Other cells | anti-SSB | 0.143633 | 2.52E-11 |
| B39 | B105 | Other cells | anti-U1RNP | 0.143633 | 2.52E-11 |
| B39 | B106 | Other cells | anti-P | 0.143633 | 2.52E-11 |
| B39 | B107 | Other cells | AnuA | 0.143633 | 2.52E-11 |
| B39 | B109 | Other cells | AMA-M2 | 0.143633 | 2.52E-11 |
| B39 | B110 | Other cells | AHA | 0.143633 | 2.52E-11 |
| B39 | B98 | Other cells | anti-Jo-1 | 0.143633 | 2.52E-11 |
| B39 | B155 | Other cells | IgA | -0.12631 | 4.61E-09 |
| B39 | B93 | Other cells | ALDH2-Glu504Lys | -0.1261 | 4.90E-09 |
| B39 | B154 | Other cells | RF | 0.123622 | 9.77E-09 |
| B39 | B182 | Other cells | TT | -0.1225 | 1.33E-08 |
| B39 | B79 | Other cells | hb V -dna | 0.115625 | 8.27E-08 |
| B39 | B92 | Other cells | ALDH2-Glu504Glu | 0.115602 | 8.32E-08 |
| B39 | B76 | Other cells | ApoB | -0.11536 | 8.84E-08 |
| B39 | B180 | Other cells | Tf | -0.10926 | 4.10E-07 |
| B4 | B6 | WB-Ca | WB-Fe | 0.353538 | 0 |
| B4 | B7 | WB-Ca | WB-Cop | -0.54963 | 0 |
| B4 | B8 | WB-Ca | WB-Zinc | -0.53111 | 0 |
| B4 | B5 | WB-Ca | WB-Mg | 0.156836 | 3.04E-13 |

| | | | | | |
|---|---|---|---|---|---|
| B4 | B69 | WB-Ca | PV | -0.15122 | 2.08E-12 |
| B4 | B138 | WB-Ca | PAB | -0.15063 | 2.54E-12 |
| B4 | B57 | WB-Ca | RBCAI | 0.123981 | 8.84E-09 |
| B4 | B55 | WB-Ca | TK | 0.116115 | 7.28E-08 |
| B4 | B54 | WB-Ca | ESR | -0.11578 | 7.94E-08 |
| B4 | B60 | WB-Ca | WBRVI | 0.107332 | 6.55E-07 |
| B40 | B130 | WBC | GLB | 0.192721 | 0 |
| B40 | B206 | WBC | Waist | 0.190365 | 0 |
| B40 | B215 | WBC | BMI | 0.187506 | 0 |
| B40 | B222 | WBC | NS | 0.179452 | 0 |
| B40 | B42 | WBC | MON | 0.685871 | 0 |
| B40 | B48 | WBC | LYM% | -0.26685 | 0 |
| B40 | B52 | WBC | PLT/BPC | 0.318541 | 0 |
| B40 | B53 | WBC | GR% | 0.289333 | 0 |
| B40 | B216 | WBC | BMI | 0.176113 | 2.22E-16 |
| B40 | B91 | WBC | TP | 0.175345 | 4.44E-16 |
| B40 | B192 | WBC | UA | 0.169852 | 2.66E-15 |
| B40 | B127 | WBC | A-G ratio | -0.16291 | 3.46E-14 |
| B40 | B210 | WBC | Hip | 0.158335 | 1.79E-13 |
| B40 | B209 | WBC | Smoking | 0.149895 | 3.26E-12 |
| B40 | B133 | WBC | TBIL | -0.13509 | 3.57E-10 |
| B40 | B129 | WBC | IBIL | -0.13011 | 1.56E-09 |
| B40 | B41 | WBC | MON % | -0.12681 | 4.00E-09 |
| B40 | B72 | WBC | HDL-C | -0.12261 | 1.29E-08 |
| B40 | B46 | WBC | RBC Count | 0.122492 | 1.33E-08 |
| B40 | B207 | WBC | WHR | 0.119158 | 3.27E-08 |
| B40 | B128 | WBC | ALT | 0.119066 | 3.35E-08 |
| B40 | B70 | WBC | LDL-C | 0.116829 | 6.05E-08 |
| B40 | B131 | WBC | AST | 0.1125 | 1.83E-07 |
| B40 | B132 | WBC | Dbil | -0.11211 | 2.02E-07 |
| B40 | B218 | WBC | DBP | 0.111505 | 2.35E-07 |
| B41 | B42 | MON % | MON | 0.606691 | 0 |
| B41 | B53 | MON % | GR% | -0.22074 | 0 |
| B41 | B43 | MON % | MCV | 0.147615 | 6.93E-12 |
| B41 | B189 | MON % | eGFB | -0.12323 | 1.09E-08 |
| B41 | B49 | MON % | baso% | 0.12162 | 1.69E-08 |
| B41 | B126 | MON % | Alb | -0.11126 | 2.50E-07 |
| B41 | B44 | MON % | MCH | 0.11106 | 2.63E-07 |
| B41 | B71 | MON % | TG | -0.10742 | 6.41E-07 |
| B42 | B48 | MON | LYM% | -0.20559 | 0 |
| B42 | B52 | MON | PLT/BPC | 0.21798 | 0 |
| B42 | B130 | MON | GLB | 0.149678 | 3.50E-12 |
| B42 | B209 | MON | Smoking | 0.147621 | 6.91E-12 |
| B42 | B127 | MON | A-G ratio | -0.14508 | 1.58E-11 |

| | | | | | |
|---|---|---|---|---|---|
| B42 | B206 | MON | Waist | 0.13319 | 6.31E-10 |
| B42 | B192 | MON | UA | 0.122866 | 1.20E-08 |
| B42 | B123 | MON | hs-CRP1 | 0.122112 | 1.48E-08 |
| B42 | B131 | MON | AST | 0.117006 | 5.77E-08 |
| B42 | B222 | MON | NS | 0.115975 | 7.55E-08 |
| B42 | B215 | MON | BMI | 0.114505 | 1.10E-07 |
| B42 | B207 | MON | WHR | 0.109687 | 3.69E-07 |
| B43 | B44 | MCV | MCH | 0.905641 | 0 |
| B43 | B46 | MCV | RBC Count | -0.5788 | 0 |
| B43 | B47 | MCV | RDW | 0.592229 | 0 |
| B43 | B45 | MCV | MCHC | 0.163271 | 3.06E-14 |
| B43 | B52 | MCV | PLT/BPC | -0.1532 | 1.07E-12 |
| B43 | B209 | MCV | Smoking | 0.148336 | 5.46E-12 |
| B43 | B204 | MCV | Drinking | 0.125274 | 6.17E-09 |
| B43 | B91 | MCV | TP | -0.11405 | 1.24E-07 |
| B43 | B126 | MCV | Alb | -0.11201 | 2.08E-07 |
| B43 | B189 | MCV | eGFB | -0.10793 | 5.66E-07 |
| B44 | B45 | MCH | MCHC | 0.563821 | 0 |
| B44 | B46 | MCH | RBC Count | -0.48749 | 0 |
| B44 | B47 | MCH | RDW | 0.395266 | 0 |
| B44 | B51 | MCH | Hb | 0.259642 | 0 |
| B44 | B209 | MCH | Smoking | 0.133029 | 6.62E-10 |
| B44 | B204 | MCH | Drinking | 0.130415 | 1.43E-09 |
| B45 | B47 | MCHC | RDW | -0.24151 | 0 |
| B45 | B51 | MCHC | Hb | 0.452649 | 0 |
| B45 | B71 | MCHC | TG | 0.17585 | 2.22E-16 |
| B45 | B129 | MCHC | IBIL | 0.135317 | 3.34E-10 |
| B45 | B126 | MCHC | Alb | 0.124104 | 8.55E-09 |
| B45 | B133 | MCHC | TBIL | 0.122101 | 1.48E-08 |
| B45 | B191 | MCHC | Urea | -0.12137 | 1.81E-08 |
| B45 | B146 | MCHC | CysC | -0.12049 | 2.29E-08 |
| B45 | B189 | MCHC | eGFB | 0.118453 | 3.95E-08 |
| B45 | B221 | MCHC | Hei | 0.110938 | 2.71E-07 |
| B45 | B216 | MCHC | BMI | 0.107639 | 6.08E-07 |
| B46 | B126 | RBC Count | Alb | 0.276184 | 0 |
| B46 | B189 | RBC Count | eGFB | 0.286395 | 0 |
| B46 | B191 | RBC Count | Urea | -0.20863 | 0 |
| B46 | B216 | RBC Count | BMI | 0.206324 | 0 |
| B46 | B221 | RBC Count | Hei | 0.194624 | 0 |
| B46 | B47 | RBC Count | RDW | -0.46517 | 0 |
| B46 | B51 | RBC Count | Hb | 0.709072 | 0 |
| B46 | B52 | RBC Count | PLT/BPC | 0.187099 | 0 |
| B46 | B58 | RBC Count | HCT | 0.173088 | 8.88E-16 |
| B46 | B128 | RBC Count | ALT | 0.170292 | 2.22E-15 |

| | | | | | |
|---|---|---|---|---|---|
| B46 | B91 | RBC Count | TP | 0.170499 | 2.22E-15 |
| B46 | B70 | RBC Count | LDL-C | 0.157834 | 2.14E-13 |
| B46 | B210 | RBC Count | Hip | 0.157699 | 2.24E-13 |
| B46 | B222 | RBC Count | NS | 0.157162 | 2.71E-13 |
| B46 | B188 | RBC Count | Ca | 0.154371 | 7.14E-13 |
| B46 | B163 | RBC Count | SG | 0.149283 | 3.99E-12 |
| B46 | B64 | RBC Count | WBRSR(mPas_5) | 0.139166 | 1.03E-10 |
| B46 | B60 | RBC Count | WBRVI | 0.137814 | 1.56E-10 |
| B46 | B190 | RBC Count | Cre | -0.13779 | 1.57E-10 |
| B46 | B67 | RBC Count | WBRSR(mPas_50) | 0.137753 | 1.59E-10 |
| B46 | B63 | RBC Count | WBRSR(mPas_200) | 0.136 | 2.72E-10 |
| B46 | B65 | RBC Count | WBRSR(mPas_1) | 0.135827 | 2.86E-10 |
| B46 | B66 | RBC Count | WBRSR(mPas_100) | 0.135789 | 2.90E-10 |
| B46 | B62 | RBC Count | WBHSRVI | 0.134457 | 4.32E-10 |
| B46 | B77 | RBC Count | TC | 0.132318 | 8.17E-10 |
| B46 | B215 | RBC Count | BMI | 0.131751 | 9.65E-10 |
| B46 | B141 | RBC Count | AST/ALT | -0.12463 | 7.39E-09 |
| B46 | B113 | RBC Count | FT3 | 0.11832 | 4.09E-08 |
| B46 | B218 | RBC Count | DBP | 0.114927 | 9.90E-08 |
| B46 | B220 | RBC Count | SBP | -0.11158 | 2.31E-07 |
| B46 | B129 | RBC Count | IBIL | 0.111089 | 2.61E-07 |
| B47 | B126 | RDW | Alb | -0.18514 | 0 |
| B47 | B189 | RDW | eGFB | -0.2309 | 0 |
| B47 | B209 | RDW | Smoking | 0.196354 | 0 |
| B47 | B51 | RDW | Hb | -0.20934 | 0 |
| B47 | B52 | RDW | PLT/BPC | -0.25592 | 0 |
| B47 | B221 | RDW | Hei | -0.17194 | 1.33E-15 |
| B47 | B216 | RDW | BMI | -0.16121 | 6.44E-14 |
| B47 | B191 | RDW | Urea | 0.158292 | 1.82E-13 |
| B47 | B146 | RDW | CysC | 0.120731 | 2.15E-08 |
| B47 | B190 | RDW | Cre | 0.120224 | 2.46E-08 |
| B47 | B128 | RDW | ALT | -0.1139 | 1.29E-07 |
| B47 | B222 | RDW | NS | -0.11177 | 2.21E-07 |
| B47 | B48 | RDW | LYM% | -0.10787 | 5.75E-07 |
| B48 | B53 | LYM% | GR% | -0.93792 | 0 |
| B48 | B189 | LYM% | eGFB | 0.148116 | 5.88E-12 |
| B48 | B123 | LYM% | hs-CRP1 | -0.12406 | 8.65E-09 |
| B48 | B190 | LYM% | Cre | -0.11302 | 1.61E-07 |
| B48 | B191 | LYM% | Urea | -0.1124 | 1.88E-07 |
| B49 | B50 | baso% | eo% | 0.353469 | 0 |
| B49 | B53 | baso% | GR% | -0.22723 | 0 |
| B49 | B220 | baso% | SBP | -0.10979 | 3.60E-07 |
| B5 | B55 | WB-Mg | TK | -0.18723 | 0 |
| B5 | B6 | WB-Mg | WB-Fe | 0.979292 | 0 |

| | | | | | |
|---|---|---|---|---|---|
| B5 | B7 | WB-Mg | WB-Cop | 0.738866 | 0 |
| B5 | B8 | WB-Mg | WB-Zinc | 0.75352 | 0 |
| B5 | B56 | WB-Mg | IR | -0.14009 | 7.71E-11 |
| B5 | B57 | WB-Mg | RBCAI | 0.122308 | 1.40E-08 |
| B5 | B61 | WB-Mg | WBHSRV | -0.12075 | 2.14E-08 |
| B50 | B53 | eo% | GR% | -0.29545 | 0 |
| B51 | B126 | Hb | Alb | 0.269323 | 0 |
| B51 | B189 | Hb | eGFB | 0.283821 | 0 |
| B51 | B191 | Hb | Urea | -0.23686 | 0 |
| B51 | B216 | Hb | BMI | 0.20485 | 0 |
| B51 | B221 | Hb | Hei | 0.211425 | 0 |
| B51 | B58 | Hb | HCT | 0.207483 | 0 |
| B51 | B60 | Hb | WBRVI | 0.178362 | 0 |
| B51 | B62 | Hb | WBHSRVI | 0.181131 | 0 |
| B51 | B63 | Hb | WBRSR(mPas_200) | 0.179446 | 0 |
| B51 | B64 | Hb | WBRSR(mPas_5) | 0.18056 | 0 |
| B51 | B66 | Hb | WBRSR(mPas_100) | 0.180484 | 0 |
| B51 | B67 | Hb | WBRSR(mPas_50) | 0.18224 | 0 |
| B51 | B70 | Hb | LDL-C | 0.192289 | 0 |
| B51 | B77 | Hb | TC | 0.178525 | 0 |
| B51 | B128 | Hb | ALT | 0.174666 | 4.44E-16 |
| B51 | B65 | Hb | WBRSR(mPas_1) | 0.174404 | 4.44E-16 |
| B51 | B222 | Hb | NS | 0.166236 | 1.02E-14 |
| B51 | B188 | Hb | Ca | 0.155038 | 5.67E-13 |
| B51 | B210 | Hb | Hip | 0.153416 | 9.90E-13 |
| B51 | B190 | Hb | Cre | -0.14751 | 7.18E-12 |
| B51 | B163 | Hb | SG | 0.1461 | 1.14E-11 |
| B51 | B220 | Hb | SBP | -0.1376 | 1.67E-10 |
| B51 | B52 | Hb | PLT/BPC | 0.1362 | 2.56E-10 |
| B51 | B141 | Hb | AST/ALT | -0.13533 | 3.33E-10 |
| B51 | B129 | Hb | IBIL | 0.133356 | 6.01E-10 |
| B51 | B174 | Hb | Urea/Cr | -0.13141 | 1.07E-09 |
| B51 | B61 | Hb | WBHSRV | 0.129583 | 1.81E-09 |
| B51 | B127 | Hb | A-G ratio | 0.129469 | 1.88E-09 |
| B51 | B218 | Hb | DBP | 0.127316 | 3.47E-09 |
| B51 | B91 | Hb | TP | 0.125732 | 5.43E-09 |
| B51 | B215 | Hb | BMI | 0.124531 | 7.59E-09 |
| B51 | B56 | Hb | IR | 0.121948 | 1.54E-08 |
| B51 | B146 | Hb | CysC | -0.12101 | 1.99E-08 |
| B51 | B204 | Hb | Drinking | 0.118766 | 3.63E-08 |
| B51 | B131 | Hb | AST | 0.115211 | 9.20E-08 |
| B51 | B140 | Hb | r-GT | 0.114726 | 1.04E-07 |
| B51 | B113 | Hb | FT3 | 0.114386 | 1.14E-07 |
| B51 | B206 | Hb | Waist | 0.111658 | 2.27E-07 |

| | | | | | |
|---|---|---|---|---|---|
| B51 | B133 | Hb | TBIL | 0.110865 | 2.76E-07 |
| B51 | B71 | Hb | TG | 0.107409 | 6.43E-07 |
| B52 | B132 | PLT/BPC | Dbil | -0.21423 | 0 |
| B52 | B133 | PLT/BPC | TBIL | -0.19255 | 0 |
| B52 | B70 | PLT/BPC | LDL-C | 0.211723 | 0 |
| B52 | B129 | PLT/BPC | IBIL | -0.16869 | 4.00E-15 |
| B52 | B91 | PLT/BPC | TP | 0.167166 | 7.11E-15 |
| B52 | B130 | PLT/BPC | GLB | 0.164453 | 2.00E-14 |
| B52 | B77 | PLT/BPC | TC | 0.164444 | 2.00E-14 |
| B52 | B188 | PLT/BPC | Ca | 0.151489 | 1.91E-12 |
| B52 | B189 | PLT/BPC | eGFB | 0.141367 | 5.18E-11 |
| B52 | B127 | PLT/BPC | A-G ratio | -0.13512 | 3.54E-10 |
| B52 | B191 | PLT/BPC | Urea | -0.11437 | 1.14E-07 |
| B52 | B216 | PLT/BPC | BMI | 0.112651 | 1.77E-07 |
| B53 | B123 | GR% | hs-CRP1 | 0.108245 | 5.25E-07 |
| B54 | B123 | ESR | hs-CRP1 | 0.270262 | 0 |
| B54 | B56 | ESR | IR | -0.21747 | 0 |
| B54 | B57 | ESR | RBCAI | -0.21879 | 0 |
| B54 | B58 | ESR | HCT | -0.43636 | 0 |
| B54 | B60 | ESR | WBRVI | -0.41107 | 0 |
| B54 | B62 | ESR | WBHSRVI | -0.35339 | 0 |
| B54 | B63 | ESR | WBRSR(mPas_200) | -0.25608 | 0 |
| B54 | B64 | ESR | WBRSR(mPas_5) | -0.30861 | 0 |
| B54 | B65 | ESR | WBRSR(mPas_1) | -0.33322 | 0 |
| B54 | B66 | ESR | WBRSR(mPas_100) | -0.26718 | 0 |
| B54 | B67 | ESR | WBRSR(mPas_50) | -0.27815 | 0 |
| B54 | B69 | ESR | PV | 0.227737 | 0 |
| B54 | B199 | ESR | Mg | 0.159758 | 1.08E-13 |
| B54 | B119 | ESR | Hcv-rna | -0.15942 | 1.22E-13 |
| B54 | B138 | ESR | PAB | -0.1367 | 2.19E-10 |
| B54 | B59 | ESR | WBRV | -0.1347 | 4.02E-10 |
| B54 | B61 | ESR | WBHSRV | -0.1327 | 7.30E-10 |
| B54 | B136 | ESR | 5'NT | 0.12248 | 1.34E-08 |
| B54 | B154 | ESR | RF | 0.107328 | 6.55E-07 |
| B55 | B177 | TK | ACR | 0.210282 | 0 |
| B55 | B178 | TK | Ig-U | 0.239813 | 0 |
| B55 | B180 | TK | Tf | 0.241012 | 0 |
| B55 | B56 | TK | IR | 0.746151 | 0 |
| B55 | B57 | TK | RBCAI | -0.24988 | 0 |
| B55 | B58 | TK | HCT | -0.21691 | 0 |
| B55 | B59 | TK | WBRV | 0.30944 | 0 |
| B55 | B60 | TK | WBRVI | 0.180952 | 0 |
| B55 | B61 | TK | WBHSRV | 0.603365 | 0 |
| B55 | B62 | TK | WBHSRVI | 0.364342 | 0 |

| | | | | | |
|---|---|---|---|---|---|
| B55 | B63 | TK | WBRSR(mPas_200) | 0.186586 | 0 |
| B55 | B69 | TK | PV | -0.43858 | 0 |
| B55 | B94 | TK | ALDH2-Lys504Lys | -0.28531 | 0 |
| B55 | B95 | TK | anti-cenpb | -0.31287 | 0 |
| B55 | B186 | TK | PFC | -0.16373 | 2.58E-14 |
| B55 | B66 | TK | WBRSR(mPas_100) | 0.157591 | 2.33E-13 |
| B55 | B175 | TK | ALB-U | 0.153068 | 1.12E-12 |
| B55 | B100 | TK | anti-Ro52 | 0.146439 | 1.02E-11 |
| B55 | B101 | TK | anti-Sc70 | 0.146439 | 1.02E-11 |
| B55 | B102 | TK | anti-Sm | 0.146439 | 1.02E-11 |
| B55 | B103 | TK | anti-SSA/Ro | 0.146439 | 1.02E-11 |
| B55 | B104 | TK | anti-SSB | 0.146439 | 1.02E-11 |
| B55 | B105 | TK | anti-U1RNP | 0.146439 | 1.02E-11 |
| B55 | B106 | TK | anti-P | 0.146439 | 1.02E-11 |
| B55 | B107 | TK | AnuA | 0.146439 | 1.02E-11 |
| B55 | B109 | TK | AMA-M2 | 0.146439 | 1.02E-11 |
| B55 | B110 | TK | AHA | 0.146439 | 1.02E-11 |
| B55 | B98 | TK | anti-Jo-1 | 0.146439 | 1.02E-11 |
| B55 | B67 | TK | WBRSR(mPas_50) | 0.142974 | 3.11E-11 |
| B55 | B138 | TK | PAB | -0.12432 | 8.05E-09 |
| B55 | B139 | TK | ADA | 0.119287 | 3.16E-08 |
| B55 | B193 | TK | K | -0.1187 | 3.69E-08 |
| B55 | B154 | TK | RF | 0.111235 | 2.52E-07 |
| B56 | B186 | IR | PFC | -0.19508 | 0 |
| B56 | B58 | IR | HCT | 0.445908 | 0 |
| B56 | B59 | IR | WBRV | 0.492516 | 0 |
| B56 | B60 | IR | WBRVI | 0.677232 | 0 |
| B56 | B61 | IR | WBHSRV | 0.915901 | 0 |
| B56 | B62 | IR | WBHSRVI | 0.889844 | 0 |
| B56 | B63 | IR | WBRSR(mPas_200) | 0.737512 | 0 |
| B56 | B64 | IR | WBRSR(mPas_5) | 0.631525 | 0 |
| B56 | B65 | IR | WBRSR(mPas_1) | 0.561728 | 0 |
| B56 | B66 | IR | WBRSR(mPas_100) | 0.725865 | 0 |
| B56 | B67 | IR | WBRSR(mPas_50) | 0.716321 | 0 |
| B56 | B69 | IR | PV | -0.33887 | 0 |
| B56 | B94 | IR | ALDH2-Lys504Lys | -0.28942 | 0 |
| B56 | B95 | IR | anti-cenpb | -0.25956 | 0 |
| B56 | B178 | IR | Ig-U | 0.176512 | 2.22E-16 |
| B56 | B180 | IR | Tf | 0.171554 | 1.33E-15 |
| B56 | B57 | IR | RBCAI | -0.16049 | 8.33E-14 |
| B56 | B177 | IR | ACR | 0.143452 | 2.67E-11 |
| B56 | B182 | IR | TT | 0.125805 | 5.32E-09 |
| B57 | B151 | RBCAI | Anti-CCP | 0.199153 | 0 |
| B57 | B177 | RBCAI | ACR | -0.23667 | 0 |

| | | | | | |
|---|---|---|---|---|---|
| B57 | B178 | RBCAI | Ig-U | -0.26907 | 0 |
| B57 | B180 | RBCAI | Tf | -0.27577 | 0 |
| B57 | B59 | RBCAI | WBRV | 0.616942 | 0 |
| B57 | B60 | RBCAI | WBRVI | 0.498016 | 0 |
| B57 | B64 | RBCAI | WBRSR(mPas_5) | 0.305186 | 0 |
| B57 | B65 | RBCAI | WBRSR(mPas_1) | 0.458436 | 0 |
| B57 | B79 | RBCAI | hb V -dna | 0.192693 | 0 |
| B57 | B92 | RBCAI | ALDH2-Glu504Glu | 0.246637 | 0 |
| B57 | B93 | RBCAI | ALDH2-Glu504Lys | -0.24416 | 0 |
| B57 | B134 | RBCAI | CHE | -0.17109 | 1.78E-15 |
| B57 | B175 | RBCAI | ALB-U | -0.17103 | 1.78E-15 |
| B57 | B61 | RBCAI | WBHSRV | -0.15911 | 1.36E-13 |
| B57 | B74 | RBCAI | ApoA1 | 0.158926 | 1.45E-13 |
| B57 | B58 | RBCAI | HCT | 0.147854 | 6.40E-12 |
| B57 | B136 | RBCAI | 5'NT | -0.13896 | 1.10E-10 |
| B57 | B95 | RBCAI | anti-cenpb | 0.137265 | 1.85E-10 |
| B57 | B81 | RBCAI | TSGF | -0.13438 | 4.43E-10 |
| B57 | B100 | RBCAI | anti-Ro52 | 0.12682 | 4.00E-09 |
| B57 | B101 | RBCAI | anti-Sc70 | 0.12682 | 4.00E-09 |
| B57 | B102 | RBCAI | anti-Sm | 0.12682 | 4.00E-09 |
| B57 | B103 | RBCAI | anti-SSA/Ro | 0.12682 | 4.00E-09 |
| B57 | B104 | RBCAI | anti-SSB | 0.12682 | 4.00E-09 |
| B57 | B105 | RBCAI | anti-U1RNP | 0.12682 | 4.00E-09 |
| B57 | B106 | RBCAI | anti-P | 0.12682 | 4.00E-09 |
| B57 | B107 | RBCAI | AnuA | 0.12682 | 4.00E-09 |
| B57 | B109 | RBCAI | AMA-M2 | 0.12682 | 4.00E-09 |
| B57 | B110 | RBCAI | AHA | 0.12682 | 4.00E-09 |
| B57 | B98 | RBCAI | anti-Jo-1 | 0.12682 | 4.00E-09 |
| B57 | B156 | RBCAI | IgE | 0.123727 | 9.49E-09 |
| B57 | B184 | RBCAI | INR | 0.109634 | 3.74E-07 |
| B57 | B76 | RBCAI | ApoB | -0.10874 | 4.66E-07 |
| B58 | B59 | HCT | WBRV | 0.336157 | 0 |
| B58 | B60 | HCT | WBRVI | 0.754827 | 0 |
| B58 | B61 | HCT | WBHSRV | 0.514014 | 0 |
| B58 | B62 | HCT | WBHSRVI | 0.785546 | 0 |
| B58 | B63 | HCT | WBRSR(mPas_200) | 0.809686 | 0 |
| B58 | B64 | HCT | WBRSR(mPas_5) | 0.81681 | 0 |
| B58 | B65 | HCT | WBRSR(mPas_1) | 0.781154 | 0 |
| B58 | B66 | HCT | WBRSR(mPas_100) | 0.824853 | 0 |
| B58 | B67 | HCT | WBRSR(mPas_50) | 0.832537 | 0 |
| B58 | B95 | HCT | anti-cenpb | -0.17675 | 2.22E-16 |
| B58 | B94 | HCT | ALDH2-Lys504Lys | -0.17301 | 8.88E-16 |
| B58 | B200 | HCT | OP | -0.1507 | 2.49E-12 |
| B58 | B76 | HCT | ApoB | 0.135028 | 3.64E-10 |

| | | | | | |
|---|---|---|---|---|---|
| B58 | B138 | HCT | PAB | 0.125525 | 5.75E-09 |
| B58 | B134 | HCT | CHE | 0.118067 | 4.37E-08 |
| B58 | B154 | HCT | RF | -0.11699 | 5.80E-08 |
| B58 | B96 | HCT | dsDNA (IIF) | -0.11646 | 6.66E-08 |
| B58 | B123 | HCT | hs-CRP1 | -0.11554 | 8.45E-08 |
| B58 | B97 | HCT | dsDNA | 0.108424 | 5.03E-07 |
| B58 | B112 | HCT | FT4 | -0.10608 | 8.84E-07 |
| B59 | B60 | WBRV | WBRVI | 0.785497 | 0 |
| B59 | B61 | WBRV | WBHSRV | 0.613473 | 0 |
| B59 | B62 | WBRV | WBHSRVI | 0.486963 | 0 |
| B59 | B63 | WBRV | WBRSR(mPas_200) | 0.573869 | 0 |
| B59 | B64 | WBRV | WBRSR(mPas_5) | 0.753314 | 0 |
| B59 | B65 | WBRV | WBRSR(mPas_1) | 0.80912 | 0 |
| B59 | B66 | WBRV | WBRSR(mPas_100) | 0.539254 | 0 |
| B59 | B67 | WBRV | WBRSR(mPas_50) | 0.566957 | 0 |
| B59 | B69 | WBRV | PV | 0.184179 | 0 |
| B59 | B79 | WBRV | hb V -dna | 0.246479 | 0 |
| B59 | B81 | WBRV | TSGF | -0.18755 | 0 |
| B59 | B95 | WBRV | anti-cenpb | -0.18405 | 0 |
| B59 | B92 | WBRV | ALDH2-Glu504Glu | 0.171252 | 1.55E-15 |
| B59 | B151 | WBRV | Anti-CCP | 0.170825 | 1.78E-15 |
| B59 | B93 | WBRV | ALDH2-Glu504Lys | -0.13271 | 7.28E-10 |
| B59 | B100 | WBRV | anti-Ro52 | 0.132041 | 8.86E-10 |
| B59 | B101 | WBRV | anti-Sc70 | 0.132041 | 8.86E-10 |
| B59 | B102 | WBRV | anti-Sm | 0.132041 | 8.86E-10 |
| B59 | B103 | WBRV | anti-SSA/Ro | 0.132041 | 8.86E-10 |
| B59 | B104 | WBRV | anti-SSB | 0.132041 | 8.86E-10 |
| B59 | B105 | WBRV | anti-U1RNP | 0.132041 | 8.86E-10 |
| B59 | B106 | WBRV | anti-P | 0.132041 | 8.86E-10 |
| B59 | B107 | WBRV | AnuA | 0.132041 | 8.86E-10 |
| B59 | B109 | WBRV | AMA-M2 | 0.132041 | 8.86E-10 |
| B59 | B110 | WBRV | AHA | 0.132041 | 8.86E-10 |
| B59 | B98 | WBRV | anti-Jo-1 | 0.132041 | 8.86E-10 |
| B59 | B157 | WBRV | IgG | 0.131594 | 1.01E-09 |
| B59 | B94 | WBRV | ALDH2-Lys504Lys | -0.1072 | 6.76E-07 |
| B6 | B7 | WB-Fe | WB-Cop | 0.58714 | 0 |
| B6 | B8 | WB-Fe | WB-Zinc | 0.604817 | 0 |
| B6 | B55 | WB-Fe | TK | -0.15353 | 9.53E-13 |
| B6 | B57 | WB-Fe | RBCAI | 0.141258 | 5.36E-11 |
| B6 | B56 | WB-Fe | IR | -0.1152 | 9.22E-08 |
| B6 | B61 | WB-Fe | WBHSRV | -0.10754 | 6.23E-07 |
| B60 | B61 | WBRVI | WBHSRV | 0.673647 | 0 |
| B60 | B62 | WBRVI | WBHSRVI | 0.828752 | 0 |
| B60 | B63 | WBRVI | WBRSR(mPas_200) | 0.772941 | 0 |

| | | | | | |
|---|---|---|---|---|---|
| B60 | B64 | WBRVI | WBRSR(mPas_5) | 0.895554 | 0 |
| B60 | B65 | WBRVI | WBRSR(mPas_1) | 0.919423 | 0 |
| B60 | B66 | WBRVI | WBRSR(mPas_100) | 0.775566 | 0 |
| B60 | B67 | WBRVI | WBRSR(mPas_50) | 0.799645 | 0 |
| B60 | B79 | WBRVI | hb V -dna | 0.188594 | 0 |
| B60 | B92 | WBRVI | ALDH2-Glu504Glu | 0.191785 | 0 |
| B60 | B94 | WBRVI | ALDH2-Lys504Lys | -0.21076 | 0 |
| B60 | B186 | WBRVI | PFC | -0.15217 | 1.51E-12 |
| B60 | B69 | WBRVI | PV | -0.13174 | 9.67E-10 |
| B60 | B123 | WBRVI | hs-CRP1 | -0.11166 | 2.27E-07 |
| B60 | B93 | WBRVI | ALDH2-Glu504Lys | -0.11135 | 2.45E-07 |
| B60 | B180 | WBRVI | Tf | -0.10791 | 5.70E-07 |
| B61 | B178 | WBHSRV | Ig-U | 0.187723 | 0 |
| B61 | B180 | WBHSRV | Tf | 0.180812 | 0 |
| B61 | B62 | WBHSRV | WBHSRVI | 0.877202 | 0 |
| B61 | B63 | WBHSRV | WBRSR(mPas_200) | 0.888884 | 0 |
| B61 | B64 | WBHSRV | WBRSR(mPas_5) | 0.786869 | 0 |
| B61 | B65 | WBHSRV | WBRSR(mPas_1) | 0.704353 | 0 |
| B61 | B66 | WBHSRV | WBRSR(mPas_100) | 0.863648 | 0 |
| B61 | B67 | WBHSRV | WBRSR(mPas_50) | 0.85361 | 0 |
| B61 | B94 | WBHSRV | ALDH2-Lys504Lys | -0.21531 | 0 |
| B61 | B95 | WBHSRV | anti-cenpb | -0.29066 | 0 |
| B61 | B177 | WBHSRV | ACR | 0.150198 | 2.94E-12 |
| B61 | B179 | WBHSRV | α1-MG | 0.113746 | 1.34E-07 |
| B61 | B185 | WBHSRV | PT | -0.11196 | 2.10E-07 |
| B61 | B186 | WBHSRV | PFC | -0.11168 | 2.25E-07 |
| B61 | B175 | WBHSRV | ALB-U | 0.107165 | 6.81E-07 |
| B62 | B63 | WBHSRVI | WBRSR(mPas_200) | 0.915553 | 0 |
| B62 | B64 | WBHSRVI | WBRSR(mPas_5) | 0.843102 | 0 |
| B62 | B65 | WBHSRVI | WBRSR(mPas_1) | 0.774109 | 0 |
| B62 | B66 | WBHSRVI | WBRSR(mPas_100) | 0.918245 | 0 |
| B62 | B67 | WBHSRVI | WBRSR(mPas_50) | 0.914977 | 0 |
| B62 | B94 | WBHSRVI | ALDH2-Lys504Lys | -0.23703 | 0 |
| B62 | B95 | WBHSRVI | anti-cenpb | -0.17651 | 2.22E-16 |
| B62 | B69 | WBHSRVI | PV | -0.16371 | 2.60E-14 |
| B62 | B186 | WBHSRVI | PFC | -0.15666 | 3.23E-13 |
| B62 | B76 | WBHSRVI | ApoB | 0.11614 | 7.24E-08 |
| B62 | B96 | WBHSRVI | dsDNA (IIF) | -0.11602 | 7.46E-08 |
| B62 | B97 | WBHSRVI | dsDNA | 0.111749 | 2.22E-07 |
| B62 | B200 | WBHSRVI | OP | -0.11115 | 2.57E-07 |
| B62 | B179 | WBHSRVI | α1-MG | 0.110527 | 3.00E-07 |
| B62 | B182 | WBHSRVI | TT | 0.109214 | 4.15E-07 |
| B63 | B64 | WBRSR(mPas_200) | WBRSR(mPas_5) | 0.93914 | 0 |
| B63 | B65 | WBRSR(mPas_200) | WBRSR(mPas_1) | 0.864196 | 0 |

| | | | | | |
|---|---|---|---|---|---|
| B63 | B66 | WBRSR(mPas_200) | WBRSR(mPas_100) | 0.987862 | 0 |
| B63 | B67 | WBRSR(mPas_200) | WBRSR(mPas_50) | 0.984948 | 0 |
| B63 | B69 | WBRSR(mPas_200) | PV | 0.236214 | 0 |
| B63 | B95 | WBRSR(mPas_200) | anti-cenpb | -0.20037 | 0 |
| B63 | B94 | WBRSR(mPas_200) | ALDH2-Lys504Lys | -0.13555 | 3.11E-10 |
| B63 | B134 | WBRSR(mPas_200) | CHE | 0.128182 | 2.71E-09 |
| B63 | B179 | WBRSR(mPas_200) | α1-MG | 0.126436 | 4.46E-09 |
| B63 | B200 | WBRSR(mPas_200) | OP | -0.1247 | 7.24E-09 |
| B63 | B194 | WBRSR(mPas_200) | Cl | -0.10649 | 8.00E-07 |
| B64 | B65 | WBRSR(mPas_5) | WBRSR(mPas_1) | 0.980815 | 0 |
| B64 | B66 | WBRSR(mPas_5) | WBRSR(mPas_100) | 0.929001 | 0 |
| B64 | B67 | WBRSR(mPas_5) | WBRSR(mPas_50) | 0.942854 | 0 |
| B64 | B69 | WBRSR(mPas_5) | PV | 0.249702 | 0 |
| B64 | B95 | WBRSR(mPas_5) | anti-cenpb | -0.15077 | 2.43E-12 |
| B64 | B94 | WBRSR(mPas_5) | ALDH2-Lys504Lys | -0.13057 | 1.36E-09 |
| B64 | B179 | WBRSR(mPas_5) | α1-MG | 0.117353 | 5.27E-08 |
| B64 | B92 | WBRSR(mPas_5) | ALDH2-Glu504Glu | 0.112646 | 1.77E-07 |
| B64 | B200 | WBRSR(mPas_5) | OP | -0.10566 | 9.75E-07 |
| B65 | B66 | WBRSR(mPas_1) | WBRSR(mPas_100) | 0.853898 | 0 |
| B65 | B67 | WBRSR(mPas_1) | WBRSR(mPas_50) | 0.87526 | 0 |
| B65 | B69 | WBRSR(mPas_1) | PV | 0.225325 | 0 |
| B65 | B92 | WBRSR(mPas_1) | ALDH2-Glu504Glu | 0.1373 | 1.83E-10 |
| B65 | B94 | WBRSR(mPas_1) | ALDH2-Lys504Lys | -0.127 | 3.80E-09 |
| B65 | B95 | WBRSR(mPas_1) | anti-cenpb | -0.11917 | 3.26E-08 |
| B65 | B79 | WBRSR(mPas_1) | hb V -dna | 0.116857 | 6.00E-08 |
| B65 | B180 | WBRSR(mPas_1) | Tf | -0.10582 | 9.40E-07 |
| B66 | B67 | WBRSR(mPas_100) | WBRSR(mPas_50) | 0.998672 | 0 |
| B66 | B69 | WBRSR(mPas_100) | PV | 0.209454 | 0 |
| B66 | B95 | WBRSR(mPas_100) | anti-cenpb | -0.18968 | 0 |
| B66 | B134 | WBRSR(mPas_100) | CHE | 0.136984 | 2.01E-10 |
| B66 | B94 | WBRSR(mPas_100) | ALDH2-Lys504Lys | -0.13237 | 8.04E-10 |
| B66 | B179 | WBRSR(mPas_100) | α1-MG | 0.128558 | 2.44E-09 |
| B66 | B200 | WBRSR(mPas_100) | OP | -0.12482 | 7.01E-09 |
| B66 | B194 | WBRSR(mPas_100) | Cl | -0.10809 | 5.46E-07 |
| B67 | B69 | WBRSR(mPas_50) | PV | 0.207898 | 0 |
| B67 | B95 | WBRSR(mPas_50) | anti-cenpb | -0.18328 | 0 |
| B67 | B94 | WBRSR(mPas_50) | ALDH2-Lys504Lys | -0.13193 | 9.16E-10 |
| B67 | B134 | WBRSR(mPas_50) | CHE | 0.127993 | 2.86E-09 |
| B67 | B179 | WBRSR(mPas_50) | α1-MG | 0.127195 | 3.59E-09 |
| B67 | B200 | WBRSR(mPas_50) | OP | -0.12305 | 1.14E-08 |
| B69 | B186 | PV | PFC | 0.210589 | 0 |
| B69 | B94 | PV | ALDH2-Lys504Lys | 0.320399 | 0 |
| B69 | B123 | PV | hs-CRP1 | 0.164287 | 2.11E-14 |
| B69 | B92 | PV | ALDH2-Glu504Glu | -0.1509 | 2.32E-12 |

| | | | | | |
|---|---|---|---|---|---|
| B69 | B194 | PV | Cl | -0.13509 | 3.58E-10 |
| B69 | B134 | PV | CHE | 0.131572 | 1.02E-09 |
| B69 | B157 | PV | IgG | 0.121863 | 1.58E-08 |
| B69 | B154 | PV | RF | -0.12147 | 1.76E-08 |
| B7 | B138 | WB-Cop | PAB | 0.19077 | 0 |
| B7 | B55 | WB-Cop | TK | -0.2376 | 0 |
| B7 | B8 | WB-Cop | WB-Zinc | 0.999758 | 0 |
| B7 | B56 | WB-Cop | IR | -0.17671 | 2.22E-16 |
| B7 | B69 | WB-Cop | PV | 0.165764 | 1.22E-14 |
| B7 | B198 | WB-Cop | P | 0.128969 | 2.17E-09 |
| B7 | B61 | WB-Cop | WBHSRV | -0.12486 | 6.93E-09 |
| B7 | B39 | WB-Cop | Other cells | -0.10996 | 3.45E-07 |
| B70 | B132 | LDL-C | Dbil | -0.20177 | 0 |
| B70 | B72 | LDL-C | HDL-C | 0.222557 | 0 |
| B70 | B76 | LDL-C | ApoB | 0.298246 | 0 |
| B70 | B77 | LDL-C | TC | 0.750692 | 0 |
| B70 | B71 | LDL-C | TG | -0.15886 | 1.49E-13 |
| B70 | B188 | LDL-C | Ca | 0.135521 | 3.14E-10 |
| B70 | B75 | LDL-C | ApoA1/B | -0.13308 | 6.53E-10 |
| B70 | B133 | LDL-C | TBIL | -0.12414 | 8.47E-09 |
| B70 | B219 | LDL-C | OT | -0.11895 | 3.46E-08 |
| B70 | B189 | LDL-C | eGFB | 0.112969 | 1.63E-07 |
| B71 | B130 | TG | GLB | 0.288629 | 0 |
| B71 | B140 | TG | r-GT | 0.248799 | 0 |
| B71 | B153 | TG | Glu | 0.222141 | 0 |
| B71 | B72 | TG | HDL-C | -0.39138 | 0 |
| B71 | B77 | TG | TC | 0.38665 | 0 |
| B71 | B91 | TG | TP | 0.350315 | 0 |
| B71 | B128 | TG | ALT | 0.170269 | 2.22E-15 |
| B71 | B127 | TG | A-G ratio | -0.16256 | 3.95E-14 |
| B71 | B126 | TG | Alb | 0.158779 | 1.53E-13 |
| B71 | B170 | TG | U-GLU | 0.158045 | 1.99E-13 |
| B71 | B132 | TG | Dbil | -0.15199 | 1.61E-12 |
| B71 | B141 | TG | AST/ALT | -0.15083 | 2.38E-12 |
| B71 | B215 | TG | BMI | 0.140346 | 7.13E-11 |
| B71 | B87 | TG | NSE | 0.139199 | 1.02E-10 |
| B71 | B216 | TG | BMI | 0.133276 | 6.15E-10 |
| B71 | B222 | TG | NS | 0.129557 | 1.83E-09 |
| B71 | B192 | TG | UA | 0.129167 | 2.05E-09 |
| B71 | B206 | TG | Waist | 0.122759 | 1.24E-08 |
| B71 | B134 | TG | CHE | 0.122505 | 1.33E-08 |
| B71 | B188 | TG | Ca | 0.117507 | 5.06E-08 |
| B71 | B133 | TG | TBIL | -0.11682 | 6.06E-08 |
| B71 | B218 | TG | DBP | 0.106075 | 8.84E-07 |

| | | | | | |
|---|---|---|---|---|---|
| B71 | B209 | TG | Smoking | 0.105586 | 9.93E-07 |
| B72 | B141 | HDL-C | AST/ALT | 0.213444 | 0 |
| B72 | B192 | HDL-C | UA | -0.17706 | 0 |
| B72 | B206 | HDL-C | Waist | -0.18391 | 0 |
| B72 | B215 | HDL-C | BMI | -0.20353 | 0 |
| B72 | B216 | HDL-C | BMI | -0.23106 | 0 |
| B72 | B222 | HDL-C | NS | -0.19198 | 0 |
| B72 | B74 | HDL-C | ApoA1 | 0.207013 | 0 |
| B72 | B77 | HDL-C | TC | 0.181467 | 0 |
| B72 | B128 | HDL-C | ALT | -0.17486 | 4.44E-16 |
| B72 | B130 | HDL-C | GLB | -0.16671 | 8.44E-15 |
| B72 | B210 | HDL-C | Hip | -0.15765 | 2.29E-13 |
| B72 | B91 | HDL-C | TP | -0.14932 | 3.94E-12 |
| B72 | B127 | HDL-C | A-G ratio | 0.139746 | 8.60E-11 |
| B72 | B221 | HDL-C | Hei | -0.13612 | 2.62E-10 |
| B72 | B207 | HDL-C | WHR | -0.12212 | 1.48E-08 |
| B72 | B132 | HDL-C | Dbil | 0.118982 | 3.43E-08 |
| B72 | B174 | HDL-C | Urea/Cr | 0.115973 | 7.56E-08 |
| B73 | B108 | ApoA | Anti-Cyclin | 0.183351 | 0 |
| B73 | B76 | ApoA | ApoB | 0.191304 | 0 |
| B73 | B198 | ApoA | P | -0.13222 | 8.42E-10 |
| B73 | B199 | ApoA | Mg | 0.124261 | 8.18E-09 |
| B73 | B74 | ApoA | ApoA1 | 0.11194 | 2.11E-07 |
| B74 | B168 | ApoA1 | NMP22 | 0.226674 | 0 |
| B74 | B75 | ApoA1 | ApoA1/B | 0.218778 | 0 |
| B74 | B79 | ApoA1 | hb V -dna | 0.202274 | 0 |
| B74 | B151 | ApoA1 | Anti-CCP | 0.17551 | 4.44E-16 |
| B74 | B123 | ApoA1 | hs-CRP1 | -0.1571 | 2.77E-13 |
| B74 | B194 | ApoA1 | Cl | -0.13891 | 1.11E-10 |
| B74 | B100 | ApoA1 | anti-Ro52 | 0.138487 | 1.27E-10 |
| B74 | B101 | ApoA1 | anti-Sc70 | 0.138487 | 1.27E-10 |
| B74 | B102 | ApoA1 | anti-Sm | 0.138487 | 1.27E-10 |
| B74 | B103 | ApoA1 | anti-SSA/Ro | 0.138487 | 1.27E-10 |
| B74 | B104 | ApoA1 | anti-SSB | 0.138487 | 1.27E-10 |
| B74 | B105 | ApoA1 | anti-U1RNP | 0.138487 | 1.27E-10 |
| B74 | B106 | ApoA1 | anti-P | 0.138487 | 1.27E-10 |
| B74 | B107 | ApoA1 | AnuA | 0.138487 | 1.27E-10 |
| B74 | B109 | ApoA1 | AMA-M2 | 0.138487 | 1.27E-10 |
| B74 | B110 | ApoA1 | AHA | 0.138487 | 1.27E-10 |
| B74 | B98 | ApoA1 | anti-Jo-1 | 0.138487 | 1.27E-10 |
| B74 | B146 | ApoA1 | CysC | -0.13415 | 4.74E-10 |
| B74 | B199 | ApoA1 | Mg | 0.126364 | 4.55E-09 |
| B74 | B81 | ApoA1 | TSGF | -0.12183 | 1.60E-08 |
| B74 | B176 | ApoA1 | Ucr | -0.12091 | 2.04E-08 |

| | | | | | |
|---|---|---|---|---|---|
| B74 | B135 | ApoA1 | TBA | 0.116758 | 6.16E-08 |
| B74 | B134 | ApoA1 | CHE | -0.11459 | 1.08E-07 |
| B74 | B182 | ApoA1 | TT | -0.1109 | 2.74E-07 |
| B74 | B119 | ApoA1 | Hcv-rna | 0.110587 | 2.96E-07 |
| B74 | B139 | ApoA1 | ADA | -0.11013 | 3.31E-07 |
| B74 | B200 | ApoA1 | OP | -0.10863 | 4.78E-07 |
| B75 | B76 | ApoA1/B | ApoB | -0.56477 | 0 |
| B75 | B197 | ApoA1/B | AG | -0.16327 | 3.06E-14 |
| B75 | B196 | ApoA1/B | TCO2 | 0.149064 | 4.29E-12 |
| B75 | B92 | ApoA1/B | ALDH2-Glu504Glu | 0.113677 | 1.36E-07 |
| B75 | B93 | ApoA1/B | ALDH2-Glu504Lys | -0.11295 | 1.64E-07 |
| B76 | B134 | ApoB | CHE | 0.226718 | 0 |
| B76 | B77 | ApoB | TC | 0.245747 | 0 |
| B76 | B100 | ApoB | anti-Ro52 | -0.16316 | 3.20E-14 |
| B76 | B101 | ApoB | anti-Sc70 | -0.16316 | 3.20E-14 |
| B76 | B102 | ApoB | anti-Sm | -0.16316 | 3.20E-14 |
| B76 | B103 | ApoB | anti-SSA/Ro | -0.16316 | 3.20E-14 |
| B76 | B104 | ApoB | anti-SSB | -0.16316 | 3.20E-14 |
| B76 | B105 | ApoB | anti-U1RNP | -0.16316 | 3.20E-14 |
| B76 | B106 | ApoB | anti-P | -0.16316 | 3.20E-14 |
| B76 | B107 | ApoB | AnuA | -0.16316 | 3.20E-14 |
| B76 | B109 | ApoB | AMA-M2 | -0.16316 | 3.20E-14 |
| B76 | B110 | ApoB | AHA | -0.16316 | 3.20E-14 |
| B76 | B98 | ApoB | anti-Jo-1 | -0.16316 | 3.20E-14 |
| B76 | B136 | ApoB | 5'NT | 0.139906 | 8.18E-11 |
| B76 | B199 | ApoB | Mg | 0.139684 | 8.77E-11 |
| B76 | B180 | ApoB | Tf | 0.128489 | 2.49E-09 |
| B76 | B79 | ApoB | hb V -dna | -0.12634 | 4.58E-09 |
| B76 | B151 | ApoB | Anti-CCP | -0.12329 | 1.07E-08 |
| B76 | B225 | ApoB | hPBG | -0.1201 | 2.54E-08 |
| B76 | B138 | ApoB | PAB | 0.119512 | 2.98E-08 |
| B76 | B178 | ApoB | Ig-U | 0.119257 | 3.19E-08 |
| B76 | B198 | ApoB | P | 0.109976 | 3.44E-07 |
| B77 | B130 | TC | GLB | 0.177243 | 0 |
| B77 | B132 | TC | Dbil | -0.24498 | 0 |
| B77 | B140 | TC | r-GT | 0.184579 | 0 |
| B77 | B188 | TC | Ca | 0.189241 | 0 |
| B77 | B91 | TC | TP | 0.244698 | 0 |
| B77 | B133 | TC | TBIL | -0.15348 | 9.68E-13 |
| B77 | B126 | TC | Alb | 0.148318 | 5.50E-12 |
| B77 | B218 | TC | DBP | 0.129182 | 2.04E-09 |
| B77 | B153 | TC | Glu | 0.12381 | 9.27E-09 |
| B77 | B134 | TC | CHE | 0.117791 | 4.70E-08 |
| B77 | B129 | TC | IBIL | -0.11142 | 2.40E-07 |

| | | | | | |
|---|---|---|---|---|---|
| B78 | B80 | HBcAb | anti-HP IgG | -0.13441 | 4.39E-10 |
| B78 | B148 | HBcAb | Anti_TPO | 0.1094 | 3.96E-07 |
| B78 | B168 | HBcAb | NMP22 | -0.10896 | 4.41E-07 |
| B79 | B100 | hb V -dna | anti-Ro52 | 0.432914 | 0 |
| B79 | B101 | hb V -dna | anti-Sc70 | 0.432914 | 0 |
| B79 | B102 | hb V -dna | anti-Sm | 0.432914 | 0 |
| B79 | B103 | hb V -dna | anti-SSA/Ro | 0.432914 | 0 |
| B79 | B104 | hb V -dna | anti-SSB | 0.432914 | 0 |
| B79 | B105 | hb V -dna | anti-U1RNP | 0.432914 | 0 |
| B79 | B106 | hb V -dna | anti-P | 0.432914 | 0 |
| B79 | B107 | hb V -dna | AnuA | 0.432914 | 0 |
| B79 | B109 | hb V -dna | AMA-M2 | 0.432914 | 0 |
| B79 | B110 | hb V -dna | AHA | 0.432914 | 0 |
| B79 | B119 | hb V -dna | Hcv-rna | 0.180068 | 0 |
| B79 | B139 | hb V -dna | ADA | -0.2014 | 0 |
| B79 | B183 | hb V -dna | PTA | 0.236315 | 0 |
| B79 | B184 | hb V -dna | INR | 0.314077 | 0 |
| B79 | B186 | hb V -dna | PFC | -0.25821 | 0 |
| B79 | B200 | hb V -dna | OP | -0.19086 | 0 |
| B79 | B81 | hb V -dna | TSGF | -0.46465 | 0 |
| B79 | B95 | hb V -dna | anti-cenpb | -0.21009 | 0 |
| B79 | B98 | hb V -dna | anti-Jo-1 | 0.432914 | 0 |
| B79 | B180 | hb V -dna | Tf | -0.17392 | 4.44E-16 |
| B79 | B136 | hb V -dna | 5'NT | -0.16469 | 1.82E-14 |
| B79 | B178 | hb V -dna | Ig-U | -0.1646 | 1.87E-14 |
| B79 | B155 | hb V -dna | IgA | -0.16212 | 4.62E-14 |
| B79 | B177 | hb V -dna | ACR | -0.16166 | 5.46E-14 |
| B79 | B151 | hb V -dna | Anti-CCP | 0.159468 | 1.20E-13 |
| B79 | B154 | hb V -dna | RF | 0.142126 | 4.07E-11 |
| B79 | B134 | hb V -dna | CHE | -0.13925 | 1.00E-10 |
| B79 | B93 | hb V -dna | ALDH2-Glu504Lys | -0.1369 | 2.07E-10 |
| B79 | B120 | hb V -dna | HCV-cAg | 0.124416 | 7.84E-09 |
| B79 | B121 | hb V -dna | C3 | -0.12289 | 1.19E-08 |
| B79 | B122 | hb V -dna | C4 | -0.12289 | 1.19E-08 |
| B79 | B152 | hb V -dna | ACA-Ig(G-A-M) | -0.1229 | 1.19E-08 |
| B79 | B159 | hb V -dna | β2-MG | -0.12289 | 1.19E-08 |
| B79 | B169 | hb V -dna | U-CRY | -0.12289 | 1.19E-08 |
| B79 | B182 | hb V -dna | TT | -0.12033 | 2.39E-08 |
| B79 | B194 | hb V -dna | CI | 0.117039 | 5.72E-08 |
| B79 | B175 | hb V -dna | ALB-U | -0.11039 | 3.10E-07 |
| B79 | B185 | hb V -dna | PT | 0.108564 | 4.86E-07 |
| B79 | B124 | hb V -dna | ACTH | 0.107977 | 5.60E-07 |
| B79 | B114 | hb V -dna | TSH TT3 TT4(TT4) | 0.10737 | 6.49E-07 |
| B8 | B138 | WB-Zinc | PAB | 0.189517 | 0 |

| | | | | | |
|---|---|---|---|---|---|
| B8 | B55 | WB-Zinc | TK | -0.23792 | 0 |
| B8 | B56 | WB-Zinc | IR | -0.17698 | 2.22E-16 |
| B8 | B69 | WB-Zinc | PV | 0.16414 | 2.22E-14 |
| B8 | B198 | WB-Zinc | P | 0.128177 | 2.72E-09 |
| B8 | B61 | WB-Zinc | WBHSRV | -0.12576 | 5.39E-09 |
| B8 | B39 | WB-Zinc | Other cells | -0.10918 | 4.18E-07 |
| B81 | B124 | TSGF | ACTH | -0.33798 | 0 |
| B81 | B125 | TSGF | Cor | -0.29212 | 0 |
| B81 | B151 | TSGF | Anti-CCP | -0.25093 | 0 |
| B81 | B180 | TSGF | Tf | 0.186537 | 0 |
| B81 | B184 | TSGF | INR | -0.32668 | 0 |
| B81 | B185 | TSGF | PT | -0.19702 | 0 |
| B81 | B186 | TSGF | PFC | 0.280145 | 0 |
| B81 | B95 | TSGF | anti-cenpb | 0.181144 | 0 |
| B81 | B100 | TSGF | anti-Ro52 | -0.169 | 3.55E-15 |
| B81 | B101 | TSGF | anti-Sc70 | -0.169 | 3.55E-15 |
| B81 | B102 | TSGF | anti-Sm | -0.169 | 3.55E-15 |
| B81 | B103 | TSGF | anti-SSA/Ro | -0.169 | 3.55E-15 |
| B81 | B104 | TSGF | anti-SSB | -0.169 | 3.55E-15 |
| B81 | B105 | TSGF | anti-U1RNP | -0.169 | 3.55E-15 |
| B81 | B106 | TSGF | anti-P | -0.169 | 3.55E-15 |
| B81 | B107 | TSGF | AnuA | -0.169 | 3.55E-15 |
| B81 | B109 | TSGF | AMA-M2 | -0.169 | 3.55E-15 |
| B81 | B110 | TSGF | AHA | -0.169 | 3.55E-15 |
| B81 | B98 | TSGF | anti-Jo-1 | -0.169 | 3.55E-15 |
| B81 | B177 | TSGF | ACR | 0.166724 | 8.44E-15 |
| B81 | B183 | TSGF | PTA | -0.16441 | 2.02E-14 |
| B81 | B178 | TSGF | Ig-U | 0.162227 | 4.46E-14 |
| B81 | B134 | TSGF | CHE | 0.143061 | 3.03E-11 |
| B81 | B200 | TSGF | OP | 0.13829 | 1.35E-10 |
| B81 | B195 | TSGF | Na | 0.124547 | 7.56E-09 |
| B81 | B158 | TSGF | IgM | 0.114768 | 1.03E-07 |
| B81 | B175 | TSGF | ALB-U | 0.114224 | 1.19E-07 |
| B81 | B182 | TSGF | TT | 0.113898 | 1.29E-07 |
| B81 | B157 | TSGF | IgG | -0.11353 | 1.41E-07 |
| B82 | B171 | CEA | EC | 0.214837 | 0 |
| B82 | B88 | CEA | CA 242 | 0.247137 | 0 |
| B82 | B89 | CEA | CA 72-4 | 0.208496 | 0 |
| B82 | B90 | CEA | CYFRA 21-1 | 0.20831 | 0 |
| B82 | B85 | CEA | SCC | 0.157344 | 2.54E-13 |
| B82 | B139 | CEA | ADA | 0.150439 | 2.71E-12 |
| B82 | B86 | CEA | TBhCG | -0.1383 | 1.35E-10 |
| B82 | B196 | CEA | TCO2 | -0.12354 | 1.00E-08 |
| B83 | B85 | AFP | SCC | 0.213946 | 0 |

| | | | | | |
|---|---|---|---|---|---|
| B83 | B88 | AFP | CA 242 | 0.210351 | 0 |
| B83 | B86 | AFP | TBhCG | 0.119853 | 2.72E-08 |
| B84 | B86 | C-PSA | TBhCG | -0.18623 | 0 |
| B84 | B87 | C-PSA | NSE | 0.187159 | 0 |
| B84 | B85 | C-PSA | SCC | 0.120947 | 2.03E-08 |
| B84 | B214 | C-PSA | CVA-L | -0.10639 | 8.20E-07 |
| B85 | B86 | SCC | TBhCG | -0.18031 | 0 |
| B85 | B87 | SCC | NSE | 0.268744 | 0 |
| B85 | B135 | SCC | TBA | 0.111537 | 2.34E-07 |
| B86 | B88 | TBhCG | CA 242 | 0.221088 | 0 |
| B86 | B89 | TBhCG | CA 72-4 | -0.15858 | 1.64E-13 |
| B86 | B90 | TBhCG | CYFRA 21-1 | 0.135285 | 3.37E-10 |
| B86 | B134 | TBhCG | CHE | -0.1118 | 2.19E-07 |
| B9 | B203 | RH | DH | -0.1135 | 1.43E-07 |
| B90 | B171 | CYFRA 21-1 | EC | 0.421429 | 0 |
| B90 | B173 | CYFRA 21-1 | NTT | 0.157556 | 2.36E-13 |
| B90 | B180 | CYFRA 21-1 | Tf | 0.11875 | 3.65E-08 |
| B90 | B177 | CYFRA 21-1 | ACR | 0.116657 | 6.32E-08 |
| B90 | B178 | CYFRA 21-1 | Ig-U | 0.115124 | 9.41E-08 |
| B91 | B126 | TP | Alb | 0.461429 | 0 |
| B91 | B127 | TP | A-G ratio | -0.53948 | 0 |
| B91 | B128 | TP | ALT | 0.192433 | 0 |
| B91 | B130 | TP | GLB | 0.818879 | 0 |
| B91 | B188 | TP | Ca | 0.387985 | 0 |
| B91 | B192 | TP | UA | 0.17607 | 2.22E-16 |
| B91 | B131 | TP | AST | 0.166223 | 1.02E-14 |
| B91 | B194 | TP | Cl | -0.1534 | 9.96E-13 |
| B91 | B209 | TP | Smoking | -0.12178 | 1.62E-08 |
| B91 | B140 | TP | r-GT | 0.120184 | 2.49E-08 |
| B92 | B134 | ALDH2-Glu504Glu | CHE | -0.18605 | 0 |
| B92 | B156 | ALDH2-Glu504Glu | IgE | 0.219798 | 0 |
| B92 | B158 | ALDH2-Glu504Glu | IgM | -0.27413 | 0 |
| B92 | B177 | ALDH2-Glu504Glu | ACR | -0.23579 | 0 |
| B92 | B178 | ALDH2-Glu504Glu | Ig-U | -0.27693 | 0 |
| B92 | B180 | ALDH2-Glu504Glu | Tf | -0.27511 | 0 |
| B92 | B93 | ALDH2-Glu504Glu | ALDH2-Glu504Lys | -0.91865 | 0 |
| B92 | B94 | ALDH2-Glu504Glu | ALDH2-Lys504Lys | -0.27933 | 0 |
| B92 | B175 | ALDH2-Glu504Glu | ALB-U | -0.17234 | 8.88E-16 |
| B92 | B151 | ALDH2-Glu504Glu | Anti-CCP | 0.146032 | 1.16E-11 |
| B93 | B134 | ALDH2-Glu504Lys | CHE | 0.204007 | 0 |
| B93 | B156 | ALDH2-Glu504Lys | IgE | -0.22429 | 0 |
| B93 | B158 | ALDH2-Glu504Lys | IgM | 0.283291 | 0 |
| B93 | B175 | ALDH2-Glu504Lys | ALB-U | 0.183381 | 0 |
| B93 | B177 | ALDH2-Glu504Lys | ACR | 0.251184 | 0 |

| | | | | | |
|---|---|---|---|---|---|
| B93 | B178 | ALDH2-Glu504Lys | Ig-U | 0.294139 | 0 |
| B93 | B180 | ALDH2-Glu504Lys | Tf | 0.292446 | 0 |
| B93 | B151 | ALDH2-Glu504Lys | Anti-CCP | -0.15821 | 1.87E-13 |
| B93 | B94 | ALDH2-Glu504Lys | ALDH2-Lys504Lys | -0.12274 | 1.25E-08 |
| B94 | B123 | ALDH2-Lys504Lys | hs-CRP1 | 0.122551 | 1.31E-08 |
| B95 | B151 | anti-cenpb | Anti-CCP | -0.22926 | 0 |
| B95 | B100 | anti-cenpb | anti-Ro52 | -0.16372 | 2.58E-14 |
| B95 | B101 | anti-cenpb | anti-Sc70 | -0.16372 | 2.58E-14 |
| B95 | B102 | anti-cenpb | anti-Sm | -0.16372 | 2.58E-14 |
| B95 | B103 | anti-cenpb | anti-SSA/Ro | -0.16372 | 2.58E-14 |
| B95 | B104 | anti-cenpb | anti-SSB | -0.16372 | 2.58E-14 |
| B95 | B105 | anti-cenpb | anti-U1RNP | -0.16372 | 2.58E-14 |
| B95 | B106 | anti-cenpb | anti-P | -0.16372 | 2.58E-14 |
| B95 | B107 | anti-cenpb | AnuA | -0.16372 | 2.58E-14 |
| B95 | B109 | anti-cenpb | AMA-M2 | -0.16372 | 2.58E-14 |
| B95 | B110 | anti-cenpb | AHA | -0.16372 | 2.58E-14 |
| B95 | B98 | anti-cenpb | anti-Jo-1 | -0.16372 | 2.58E-14 |
| B95 | B199 | anti-cenpb | Mg | -0.14657 | 9.77E-12 |
| B95 | B198 | anti-cenpb | P | -0.14451 | 1.90E-11 |
| B96 | B100 | dsDNA (IIF) | anti-Ro52 | 0.194966 | 0 |
| B96 | B101 | dsDNA (IIF) | anti-Sc70 | 0.194965 | 0 |
| B96 | B102 | dsDNA (IIF) | anti-Sm | 0.194966 | 0 |
| B96 | B103 | dsDNA (IIF) | anti-SSA/Ro | 0.194966 | 0 |
| B96 | B104 | dsDNA (IIF) | anti-SSB | 0.194966 | 0 |
| B96 | B105 | dsDNA (IIF) | anti-U1RNP | 0.194966 | 0 |
| B96 | B106 | dsDNA (IIF) | anti-P | 0.194965 | 0 |
| B96 | B107 | dsDNA (IIF) | AnuA | 0.194966 | 0 |
| B96 | B109 | dsDNA (IIF) | AMA-M2 | 0.194966 | 0 |
| B96 | B110 | dsDNA (IIF) | AHA | 0.194966 | 0 |
| B96 | B151 | dsDNA (IIF) | Anti-CCP | 0.244519 | 0 |
| B96 | B183 | dsDNA (IIF) | PTA | -0.30177 | 0 |
| B96 | B97 | dsDNA (IIF) | dsDNA | -0.99548 | 0 |
| B96 | B98 | dsDNA (IIF) | anti-Jo-1 | 0.194965 | 0 |
| B96 | B196 | dsDNA (IIF) | TCO2 | 0.15097 | 2.27E-12 |
| B96 | B182 | dsDNA (IIF) | TT | -0.11846 | 3.94E-08 |
| B97 | B151 | dsDNA | Anti-CCP | -0.19813 | 0 |
| B97 | B183 | dsDNA | PTA | 0.303052 | 0 |
| B97 | B196 | dsDNA | TCO2 | -0.15258 | 1.32E-12 |
| B97 | B182 | dsDNA | TT | 0.108651 | 4.76E-07 |
| B98 | B107 | anti-Jo-1 | AnuA | 0.999999 | 0 |
| B98 | B151 | anti-Jo-1 | Anti-CCP | 0.515326 | 0 |
| B98 | B154 | anti-Jo-1 | RF | 0.202149 | 0 |
| B98 | B155 | anti-Jo-1 | IgA | -0.18934 | 0 |
| B98 | B179 | anti-Jo-1 | α1-MG | -0.20209 | 0 |

| | | | | | |
|---|---|---|---|---|---|
| B98 | B184 | anti-Jo-1 | INR | 0.183231 | 0 |
| B98 | B138 | anti-Jo-1 | PAB | -0.16705 | 7.55E-15 |
| B98 | B180 | anti-Jo-1 | Tf | -0.16223 | 4.44E-14 |
| B98 | B178 | anti-Jo-1 | Ig-U | -0.16208 | 4.71E-14 |
| B98 | B136 | anti-Jo-1 | 5'NT | -0.16186 | 5.11E-14 |
| B98 | B139 | anti-Jo-1 | ADA | -0.16171 | 5.37E-14 |
| B98 | B134 | anti-Jo-1 | CHE | -0.14467 | 1.81E-11 |
| B98 | B177 | anti-Jo-1 | ACR | -0.13048 | 1.40E-09 |
| B98 | B137 | anti-Jo-1 | AFU | -0.12035 | 2.38E-08 |
| B98 | B182 | anti-Jo-1 | TT | -0.11888 | 3.53E-08 |
| B98 | B168 | anti-Jo-1 | NMP22 | 0.115313 | 8.96E-08 |

**Supplementary Table 19. PEI Correlations in Parkinson's syndrome**

| Row | Column | Row | Column | Cor | *P* |
|---|---|---|---|---|---|
| B1 | B221 | Gender | Hei | -0.5884125 | 0 |
| B1 | B190 | Gender | Cre | -0.4526746 | 2.41E-11 |
| B1 | B216 | Gender | BMI | -0.3901144 | 1.45E-08 |
| B10 | B12 | PG I | PG II | 0.67638998 | 0 |
| B10 | B142 | PG I | 25-(OH)D | 0.57963807 | 0 |
| B100 | B101 | anti-Ro52 | anti-Sc70 | 1 | 0 |
| B100 | B105 | anti-Ro52 | anti-U1RNP | 1 | 0 |
| B100 | B106 | anti-Ro52 | anti-P | 1 | 0 |
| B100 | B107 | anti-Ro52 | AnuA | 0.99999994 | 0 |
| B100 | B109 | anti-Ro52 | AMA-M2 | 1 | 0 |
| B100 | B110 | anti-Ro52 | AHA | 1 | 0 |
| B100 | B134 | anti-Ro52 | CHE | 0.99587065 | 0 |
| B100 | B135 | anti-Ro52 | TBA | 0.72594937 | 0 |
| B100 | B136 | anti-Ro52 | 5'NT | 0.77267967 | 0 |
| B100 | B137 | anti-Ro52 | AFU | -0.8999613 | 0 |
| B100 | B151 | anti-Ro52 | Anti-CCP | 1 | 0 |
| B100 | B156 | anti-Ro52 | IgE | 0.94663186 | 0 |
| B100 | B175 | anti-Ro52 | ALB-U | -0.8425779 | 0 |
| B100 | B177 | anti-Ro52 | ACR | -0.5594521 | 0 |
| B100 | B182 | anti-Ro52 | TT | -0.9646716 | 0 |
| B100 | B183 | anti-Ro52 | PTA | 0.70776873 | 0 |
| B100 | B185 | anti-Ro52 | PT | 0.83292135 | 0 |
| B100 | B186 | anti-Ro52 | PFC | -0.7053006 | 0 |
| B100 | B194 | anti-Ro52 | Cl | 0.65638677 | 0 |
| B100 | B195 | anti-Ro52 | Na | 0.60796773 | 0 |
| B100 | B157 | anti-Ro52 | IgG | 0.54290776 | 2.22E-16 |
| B100 | B138 | anti-Ro52 | PAB | -0.5256581 | 2.22E-15 |
| B100 | B155 | anti-Ro52 | IgA | 0.4155488 | 1.26E-09 |
| B100 | B154 | anti-Ro52 | RF | 0.35447914 | 3.21E-07 |
| B101 | B102 | anti-Sc70 | anti-Sm | 0.99999999 | 0 |
| B101 | B103 | anti-Sc70 | anti-SSA/Ro | 1 | 0 |
| B101 | B104 | anti-Sc70 | anti-SSB | 1 | 0 |
| B101 | B105 | anti-Sc70 | anti-U1RNP | 1 | 0 |
| B101 | B106 | anti-Sc70 | anti-P | 0.99999999 | 0 |
| B101 | B107 | anti-Sc70 | AnuA | 0.99999998 | 0 |
| B101 | B108 | anti-Sc70 | Anti-Cyclin | 0.99999999 | 0 |
| B101 | B109 | anti-Sc70 | AMA-M2 | 1 | 0 |
| B101 | B110 | anti-Sc70 | AHA | 1 | 0 |
| B101 | B134 | anti-Sc70 | CHE | 0.99587064 | 0 |
| B101 | B135 | anti-Sc70 | TBA | 0.72594937 | 0 |
| B101 | B136 | anti-Sc70 | 5'NT | 0.77267967 | 0 |
| B101 | B137 | anti-Sc70 | AFU | -0.8999613 | 0 |

| | | | | | |
|---|---|---|---|---|---|
| B101 | B151 | anti-Sc70 | Anti-CCP | 0.99999999 | 0 |
| B101 | B156 | anti-Sc70 | IgE | 0.94663186 | 0 |
| B101 | B175 | anti-Sc70 | ALB-U | -0.8425779 | 0 |
| B101 | B177 | anti-Sc70 | ACR | -0.5594521 | 0 |
| B101 | B182 | anti-Sc70 | TT | -0.9646716 | 0 |
| B101 | B183 | anti-Sc70 | PTA | 0.70776872 | 0 |
| B101 | B185 | anti-Sc70 | PT | 0.83292135 | 0 |
| B101 | B186 | anti-Sc70 | PFC | -0.7053006 | 0 |
| B101 | B194 | anti-Sc70 | Cl | 0.65638676 | 0 |
| B101 | B195 | anti-Sc70 | Na | 0.60796772 | 0 |
| B101 | B157 | anti-Sc70 | IgG | 0.54290776 | 2.22E-16 |
| B101 | B138 | anti-Sc70 | PAB | -0.5256581 | 2.22E-15 |
| B101 | B155 | anti-Sc70 | IgA | 0.4155488 | 1.26E-09 |
| B101 | B154 | anti-Sc70 | RF | 0.35447914 | 3.21E-07 |
| B102 | B103 | anti-Sm | anti-SSA/Ro | 1 | 0 |
| B102 | B104 | anti-Sm | anti-SSB | 1 | 0 |
| B102 | B106 | anti-Sm | anti-P | 1 | 0 |
| B102 | B107 | anti-Sm | AnuA | 0.99999999 | 0 |
| B102 | B108 | anti-Sm | Anti-Cyclin | 1 | 0 |
| B102 | B134 | anti-Sm | CHE | 0.99587065 | 0 |
| B102 | B135 | anti-Sm | TBA | 0.72594937 | 0 |
| B102 | B136 | anti-Sm | 5'NT | 0.77267967 | 0 |
| B102 | B137 | anti-Sm | AFU | -0.8999613 | 0 |
| B102 | B156 | anti-Sm | IgE | 0.94663186 | 0 |
| B102 | B175 | anti-Sm | ALB-U | -0.8425779 | 0 |
| B102 | B177 | anti-Sm | ACR | -0.5594521 | 0 |
| B102 | B182 | anti-Sm | TT | -0.9646716 | 0 |
| B102 | B183 | anti-Sm | PTA | 0.70776873 | 0 |
| B102 | B185 | anti-Sm | PT | 0.83292135 | 0 |
| B102 | B186 | anti-Sm | PFC | -0.7053006 | 0 |
| B102 | B194 | anti-Sm | Cl | 0.65638677 | 0 |
| B102 | B195 | anti-Sm | Na | 0.60796773 | 0 |
| B102 | B157 | anti-Sm | IgG | 0.54290776 | 2.22E-16 |
| B102 | B138 | anti-Sm | PAB | -0.5256581 | 2.22E-15 |
| B102 | B155 | anti-Sm | IgA | 0.4155488 | 1.26E-09 |
| B102 | B154 | anti-Sm | RF | 0.35447914 | 3.21E-07 |
| B103 | B105 | anti-SSA/Ro | anti-U1RNP | 1 | 0 |
| B103 | B107 | anti-SSA/Ro | AnuA | 0.99999995 | 0 |
| B103 | B110 | anti-SSA/Ro | AHA | 1 | 0 |
| B103 | B134 | anti-SSA/Ro | CHE | 0.99587065 | 0 |
| B103 | B135 | anti-SSA/Ro | TBA | 0.72594937 | 0 |
| B103 | B136 | anti-SSA/Ro | 5'NT | 0.77267967 | 0 |
| B103 | B137 | anti-SSA/Ro | AFU | -0.8999613 | 0 |
| B103 | B151 | anti-SSA/Ro | Anti-CCP | 1 | 0 |

| | | | | | |
|---|---|---|---|---|---|
| B103 | B156 | anti-SSA/Ro | IgE | 0.94663186 | 0 |
| B103 | B175 | anti-SSA/Ro | ALB-U | -0.8425779 | 0 |
| B103 | B177 | anti-SSA/Ro | ACR | -0.5594521 | 0 |
| B103 | B182 | anti-SSA/Ro | TT | -0.9646716 | 0 |
| B103 | B183 | anti-SSA/Ro | PTA | 0.70776873 | 0 |
| B103 | B185 | anti-SSA/Ro | PT | 0.83292135 | 0 |
| B103 | B186 | anti-SSA/Ro | PFC | -0.7053006 | 0 |
| B103 | B194 | anti-SSA/Ro | Cl | 0.65638677 | 0 |
| B103 | B195 | anti-SSA/Ro | Na | 0.60796773 | 0 |
| B103 | B157 | anti-SSA/Ro | IgG | 0.54290776 | 2.22E-16 |
| B103 | B138 | anti-SSA/Ro | PAB | -0.5256581 | 2.22E-15 |
| B103 | B155 | anti-SSA/Ro | IgA | 0.4155488 | 1.26E-09 |
| B103 | B154 | anti-SSA/Ro | RF | 0.35447914 | 3.21E-07 |
| B104 | B107 | anti-SSB | AnuA | 0.99999997 | 0 |
| B104 | B134 | anti-SSB | CHE | 0.99587065 | 0 |
| B104 | B135 | anti-SSB | TBA | 0.72594938 | 0 |
| B104 | B136 | anti-SSB | 5'NT | 0.77267967 | 0 |
| B104 | B137 | anti-SSB | AFU | -0.8999613 | 0 |
| B104 | B156 | anti-SSB | IgE | 0.94663187 | 0 |
| B104 | B175 | anti-SSB | ALB-U | -0.8425779 | 0 |
| B104 | B177 | anti-SSB | ACR | -0.5594521 | 0 |
| B104 | B182 | anti-SSB | TT | -0.9646716 | 0 |
| B104 | B183 | anti-SSB | PTA | 0.70776873 | 0 |
| B104 | B185 | anti-SSB | PT | 0.83292134 | 0 |
| B104 | B186 | anti-SSB | PFC | -0.7053006 | 0 |
| B104 | B194 | anti-SSB | Cl | 0.65638677 | 0 |
| B104 | B195 | anti-SSB | Na | 0.60796773 | 0 |
| B104 | B157 | anti-SSB | IgG | 0.54290776 | 2.22E-16 |
| B104 | B138 | anti-SSB | PAB | -0.5256581 | 2.22E-15 |
| B104 | B155 | anti-SSB | IgA | 0.4155488 | 1.26E-09 |
| B104 | B154 | anti-SSB | RF | 0.35447914 | 3.21E-07 |
| B105 | B107 | anti-U1RNP | AnuA | 0.99999996 | 0 |
| B105 | B110 | anti-U1RNP | AHA | 1 | 0 |
| B105 | B134 | anti-U1RNP | CHE | 0.99587065 | 0 |
| B105 | B135 | anti-U1RNP | TBA | 0.72594937 | 0 |
| B105 | B136 | anti-U1RNP | 5'NT | 0.77267967 | 0 |
| B105 | B137 | anti-U1RNP | AFU | -0.8999613 | 0 |
| B105 | B151 | anti-U1RNP | Anti-CCP | 1 | 0 |
| B105 | B156 | anti-U1RNP | IgE | 0.94663186 | 0 |
| B105 | B175 | anti-U1RNP | ALB-U | -0.8425779 | 0 |
| B105 | B177 | anti-U1RNP | ACR | -0.5594521 | 0 |
| B105 | B182 | anti-U1RNP | TT | -0.9646716 | 0 |
| B105 | B183 | anti-U1RNP | PTA | 0.70776872 | 0 |
| B105 | B185 | anti-U1RNP | PT | 0.83292135 | 0 |

| | | | | | |
|---|---|---|---|---|---|
| B105 | B186 | anti-U1RNP | PFC | -0.7053006 | 0 |
| B105 | B194 | anti-U1RNP | Cl | 0.65638677 | 0 |
| B105 | B195 | anti-U1RNP | Na | 0.60796773 | 0 |
| B105 | B157 | anti-U1RNP | IgG | 0.54290776 | 2.22E-16 |
| B105 | B138 | anti-U1RNP | PAB | -0.5256581 | 2.22E-15 |
| B105 | B155 | anti-U1RNP | IgA | 0.4155488 | 1.26E-09 |
| B105 | B154 | anti-U1RNP | RF | 0.35447914 | 3.21E-07 |
| B106 | B107 | anti-P | AnuA | 0.99999994 | 0 |
| B106 | B108 | anti-P | Anti-Cyclin | 1 | 0 |
| B106 | B110 | anti-P | AHA | 1 | 0 |
| B106 | B134 | anti-P | CHE | 0.99587065 | 0 |
| B106 | B135 | anti-P | TBA | 0.72594937 | 0 |
| B106 | B136 | anti-P | 5'NT | 0.77267967 | 0 |
| B106 | B137 | anti-P | AFU | -0.8999613 | 0 |
| B106 | B156 | anti-P | IgE | 0.94663186 | 0 |
| B106 | B175 | anti-P | ALB-U | -0.8425779 | 0 |
| B106 | B177 | anti-P | ACR | -0.5594521 | 0 |
| B106 | B182 | anti-P | TT | -0.9646716 | 0 |
| B106 | B183 | anti-P | PTA | 0.70776873 | 0 |
| B106 | B185 | anti-P | PT | 0.83292135 | 0 |
| B106 | B186 | anti-P | PFC | -0.7053006 | 0 |
| B106 | B194 | anti-P | Cl | 0.65638677 | 0 |
| B106 | B195 | anti-P | Na | 0.60796773 | 0 |
| B106 | B157 | anti-P | IgG | 0.54290776 | 2.22E-16 |
| B106 | B138 | anti-P | PAB | -0.5256581 | 2.22E-15 |
| B106 | B155 | anti-P | IgA | 0.4155488 | 1.26E-09 |
| B106 | B154 | anti-P | RF | 0.35447914 | 3.21E-07 |
| B107 | B108 | AnuA | Anti-Cyclin | 0.99999995 | 0 |
| B107 | B109 | AnuA | AMA-M2 | 0.99999995 | 0 |
| B107 | B110 | AnuA | AHA | 0.99999995 | 0 |
| B107 | B134 | AnuA | CHE | 0.9958706 | 0 |
| B107 | B135 | AnuA | TBA | 0.72594933 | 0 |
| B107 | B136 | AnuA | 5'NT | 0.77267963 | 0 |
| B107 | B137 | AnuA | AFU | -0.8999613 | 0 |
| B107 | B151 | AnuA | Anti-CCP | 0.99999995 | 0 |
| B107 | B156 | AnuA | IgE | 0.94663181 | 0 |
| B107 | B175 | AnuA | ALB-U | -0.8425778 | 0 |
| B107 | B177 | AnuA | ACR | -0.5594521 | 0 |
| B107 | B182 | AnuA | TT | -0.9646715 | 0 |
| B107 | B183 | AnuA | PTA | 0.70776869 | 0 |
| B107 | B185 | AnuA | PT | 0.83292129 | 0 |
| B107 | B186 | AnuA | PFC | -0.7053006 | 0 |
| B107 | B194 | AnuA | Cl | 0.65638674 | 0 |
| B107 | B195 | AnuA | Na | 0.60796771 | 0 |

| | | | | | |
|---|---|---|---|---|---|
| B107 | B157 | AnuA | IgG | 0.54290773 | 2.22E-16 |
| B107 | B138 | AnuA | PAB | -0.5256581 | 2.22E-15 |
| B107 | B155 | AnuA | IgA | 0.41554878 | 1.26E-09 |
| B107 | B154 | AnuA | RF | 0.35447912 | 3.21E-07 |
| B108 | B134 | Anti-Cyclin | CHE | 0.99587065 | 0 |
| B108 | B135 | Anti-Cyclin | TBA | 0.72594937 | 0 |
| B108 | B136 | Anti-Cyclin | 5'NT | 0.77267967 | 0 |
| B108 | B137 | Anti-Cyclin | AFU | -0.8999613 | 0 |
| B108 | B156 | Anti-Cyclin | IgE | 0.94663186 | 0 |
| B108 | B175 | Anti-Cyclin | ALB-U | -0.8425779 | 0 |
| B108 | B177 | Anti-Cyclin | ACR | -0.5594521 | 0 |
| B108 | B182 | Anti-Cyclin | TT | -0.9646716 | 0 |
| B108 | B183 | Anti-Cyclin | PTA | 0.70776873 | 0 |
| B108 | B185 | Anti-Cyclin | PT | 0.83292135 | 0 |
| B108 | B186 | Anti-Cyclin | PFC | -0.7053006 | 0 |
| B108 | B194 | Anti-Cyclin | Cl | 0.65638677 | 0 |
| B108 | B195 | Anti-Cyclin | Na | 0.60796773 | 0 |
| B108 | B157 | Anti-Cyclin | IgG | 0.54290776 | 2.22E-16 |
| B108 | B138 | Anti-Cyclin | PAB | -0.5256581 | 2.22E-15 |
| B108 | B155 | Anti-Cyclin | IgA | 0.4155488 | 1.26E-09 |
| B108 | B154 | Anti-Cyclin | RF | 0.35447914 | 3.21E-07 |
| B109 | B110 | AMA-M2 | AHA | 1 | 0 |
| B109 | B134 | AMA-M2 | CHE | 0.99587065 | 0 |
| B109 | B135 | AMA-M2 | TBA | 0.72594937 | 0 |
| B109 | B136 | AMA-M2 | 5'NT | 0.77267967 | 0 |
| B109 | B137 | AMA-M2 | AFU | -0.8999613 | 0 |
| B109 | B151 | AMA-M2 | Anti-CCP | 1 | 0 |
| B109 | B156 | AMA-M2 | IgE | 0.94663186 | 0 |
| B109 | B175 | AMA-M2 | ALB-U | -0.8425779 | 0 |
| B109 | B177 | AMA-M2 | ACR | -0.5594521 | 0 |
| B109 | B182 | AMA-M2 | TT | -0.9646716 | 0 |
| B109 | B183 | AMA-M2 | PTA | 0.70776873 | 0 |
| B109 | B185 | AMA-M2 | PT | 0.83292135 | 0 |
| B109 | B186 | AMA-M2 | PFC | -0.7053006 | 0 |
| B109 | B194 | AMA-M2 | Cl | 0.65638677 | 0 |
| B109 | B195 | AMA-M2 | Na | 0.60796773 | 0 |
| B109 | B157 | AMA-M2 | IgG | 0.54290776 | 2.22E-16 |
| B109 | B138 | AMA-M2 | PAB | -0.5256581 | 2.22E-15 |
| B109 | B155 | AMA-M2 | IgA | 0.4155488 | 1.26E-09 |
| B109 | B154 | AMA-M2 | RF | 0.35447914 | 3.21E-07 |
| B11 | B12 | PGI/PGII | PG II | -0.6945325 | 0 |
| B11 | B145 | PGI/PGII | BGP | 0.40460736 | 3.70E-09 |
| B11 | B142 | PGI/PGII | 25-(OH)D | -0.3660731 | 1.22E-07 |
| B110 | B134 | AHA | CHE | 0.99587065 | 0 |

| | | | | | |
|---|---|---|---|---|---|
| B110 | B135 | AHA | TBA | 0.72594937 | 0 |
| B110 | B136 | AHA | 5'NT | 0.77267967 | 0 |
| B110 | B137 | AHA | AFU | -0.8999613 | 0 |
| B110 | B151 | AHA | Anti-CCP | 1 | 0 |
| B110 | B156 | AHA | IgE | 0.94663186 | 0 |
| B110 | B175 | AHA | ALB-U | -0.8425779 | 0 |
| B110 | B177 | AHA | ACR | -0.5594521 | 0 |
| B110 | B182 | AHA | TT | -0.9646716 | 0 |
| B110 | B183 | AHA | PTA | 0.70776872 | 0 |
| B110 | B185 | AHA | PT | 0.83292135 | 0 |
| B110 | B186 | AHA | PFC | -0.7053006 | 0 |
| B110 | B194 | AHA | Cl | 0.65638677 | 0 |
| B110 | B195 | AHA | Na | 0.60796773 | 0 |
| B110 | B157 | AHA | IgG | 0.54290776 | 2.22E-16 |
| B110 | B138 | AHA | PAB | -0.5256581 | 2.22E-15 |
| B110 | B155 | AHA | IgA | 0.4155488 | 1.26E-09 |
| B110 | B154 | AHA | RF | 0.35447914 | 3.21E-07 |
| B112 | B114 | FT4 | TSH TT3 TT4(TT4) | 0.51297311 | 1.29E-14 |
| B112 | B149 | FT4 | TGAb | 0.3602718 | 1.99E-07 |
| B113 | B115 | FT3 | TSH TT3 TT4(TT3) | 0.47294376 | 2.26E-12 |
| B113 | B211 | FT3 | IOP-R | 0.34964909 | 4.75E-07 |
| B114 | B115 | TSH TT3 TT4(TT4) | TSH TT3 TT4(TT3) | 0.56985335 | 0 |
| B115 | B211 | TSH TT3 TT4(TT3) | IOP-R | 0.4409951 | 8.83E-11 |
| B115 | B149 | TSH TT3 TT4(TT3) | TGAb | 0.41178631 | 1.84E-09 |
| B116 | B179 | TT3 | α1-MG | -0.622543 | 0 |
| B116 | B223 | TT3 | FH | -0.7727273 | 0 |
| B116 | B117 | TT3 | TT4 | -0.3856946 | 2.18E-08 |
| B117 | B122 | TT4 | C4 | -0.8414523 | 0 |
| B117 | B159 | TT4 | β2-MG | -0.5660381 | 0 |
| B117 | B202 | TT4 | e-times | -0.3411615 | 9.30E-07 |
| B119 | B167 | Hcv-rna | PC | 0.8040844 | 0 |
| B12 | B142 | PG II | 25-(OH)D | 0.75332958 | 0 |
| B12 | B145 | PG II | BGP | -0.4411932 | 8.64E-11 |
| B12 | B144 | PG II | Total-PINP | 0.36381913 | 1.48E-07 |
| B120 | B131 | HCV-cAg | AST | -0.4461124 | 5.03E-11 |
| B120 | B209 | HCV-cAg | Smoking | -0.412515 | 1.71E-09 |
| B120 | B204 | HCV-cAg | Drinking | -0.4093236 | 2.34E-09 |
| B121 | B152 | C3 | ACA-Ig(G-A-M) | 0.49131376 | 2.29E-13 |
| B122 | B178 | C4 | Ig-U | -0.8861469 | 0 |
| B122 | B180 | C4 | Tf | -0.6522994 | 0 |
| B122 | B202 | C4 | e-times | -0.6477503 | 0 |
| B122 | B223 | C4 | FH | -0.4759973 | 1.56E-12 |
| B123 | B228 | hs-CRP1 | OS | 0.40985095 | 2.22E-09 |
| B123 | B146 | hs-CRP1 | CysC | 0.34682003 | 5.95E-07 |

| | | | | | |
|---|---|---|---|---|---|
| B125 | B178 | Cor | Ig-U | -0.5809475 | 0 |
| B126 | B127 | Alb | A-G ratio | 0.61525173 | 0 |
| B126 | B188 | Alb | Ca | 0.3830731 | 2.76E-08 |
| B127 | B130 | A-G ratio | GLB | -0.9139504 | 0 |
| B128 | B141 | ALT | AST/ALT | -0.5888712 | 0 |
| B128 | B131 | ALT | AST | 0.5182149 | 6.22E-15 |
| B129 | B132 | IBIL | Dbil | 0.68837685 | 0 |
| B129 | B133 | IBIL | TBIL | 0.97550016 | 0 |
| B131 | B209 | AST | Smoking | 0.35996303 | 2.04E-07 |
| B132 | B133 | Dbil | TBIL | 0.8309871 | 0 |
| B134 | B135 | CHE | TBA | 0.7601172 | 0 |
| B134 | B136 | CHE | 5'NT | 0.74510134 | 0 |
| B134 | B137 | CHE | AFU | -0.9350155 | 0 |
| B134 | B138 | CHE | PAB | -0.5582544 | 0 |
| B134 | B151 | CHE | Anti-CCP | 0.99587065 | 0 |
| B134 | B156 | CHE | IgE | 0.94286898 | 0 |
| B134 | B175 | CHE | ALB-U | -0.8381456 | 0 |
| B134 | B177 | CHE | ACR | -0.5561599 | 0 |
| B134 | B182 | CHE | TT | -0.9608075 | 0 |
| B134 | B183 | CHE | PTA | 0.70516629 | 0 |
| B134 | B185 | CHE | PT | 0.8292311 | 0 |
| B134 | B186 | CHE | PFC | -0.7027095 | 0 |
| B134 | B194 | CHE | Cl | 0.64959669 | 0 |
| B134 | B195 | CHE | Na | 0.592523 | 0 |
| B134 | B157 | CHE | IgG | 0.54100487 | 2.22E-16 |
| B134 | B155 | CHE | IgA | 0.41368243 | 1.52E-09 |
| B134 | B154 | CHE | RF | 0.3537353 | 3.41E-07 |
| B134 | B197 | CHE | AG | -0.3457015 | 6.51E-07 |
| B135 | B137 | TBA | AFU | -0.7797145 | 0 |
| B135 | B151 | TBA | Anti-CCP | 0.72594937 | 0 |
| B135 | B156 | TBA | IgE | 0.68692404 | 0 |
| B135 | B175 | TBA | ALB-U | -0.6135136 | 0 |
| B135 | B182 | TBA | TT | -0.7000716 | 0 |
| B135 | B185 | TBA | PT | 0.60514422 | 0 |
| B135 | B194 | TBA | Cl | 0.87428431 | 0 |
| B135 | B195 | TBA | Na | 0.81467257 | 0 |
| B135 | B193 | TBA | K | 0.53717818 | 4.44E-16 |
| B135 | B183 | TBA | PTA | 0.5131845 | 1.24E-14 |
| B135 | B186 | TBA | PFC | -0.5113906 | 1.60E-14 |
| B135 | B177 | TBA | ACR | -0.4080348 | 2.65E-09 |
| B135 | B157 | TBA | IgG | 0.39346745 | 1.07E-08 |
| B136 | B137 | 5'NT | AFU | -0.631078 | 0 |
| B136 | B138 | 5'NT | PAB | -0.7401846 | 0 |
| B136 | B151 | 5'NT | Anti-CCP | 0.77267967 | 0 |

| | | | | | |
|---|---|---|---|---|---|
| B136 | B156 | 5'NT | IgE | 0.73199083 | 0 |
| B136 | B175 | 5'NT | ALB-U | -0.6474702 | 0 |
| B136 | B182 | 5'NT | TT | -0.7458297 | 0 |
| B136 | B183 | 5'NT | PTA | 0.5480788 | 0 |
| B136 | B185 | 5'NT | PT | 0.64264114 | 0 |
| B136 | B186 | 5'NT | PFC | -0.5461759 | 0 |
| B136 | B193 | 5'NT | K | -0.651189 | 0 |
| B136 | B199 | 5'NT | Mg | 0.73501634 | 0 |
| B136 | B200 | 5'NT | OP | -0.7465021 | 0 |
| B136 | B177 | 5'NT | ACR | -0.4285959 | 3.32E-10 |
| B136 | B157 | 5'NT | IgG | 0.42076446 | 7.45E-10 |
| B136 | B139 | 5'NT | ADA | 0.4183331 | 9.53E-10 |
| B136 | B168 | 5'NT | NMP22 | -0.3433451 | 7.84E-07 |
| B137 | B138 | AFU | PAB | 0.7032151 | 0 |
| B137 | B151 | AFU | Anti-CCP | -0.8999613 | 0 |
| B137 | B156 | AFU | IgE | -0.8527687 | 0 |
| B137 | B175 | AFU | ALB-U | 0.75282988 | 0 |
| B137 | B182 | AFU | TT | 0.86885093 | 0 |
| B137 | B183 | AFU | PTA | -0.6387981 | 0 |
| B137 | B185 | AFU | PT | -0.7481606 | 0 |
| B137 | B186 | AFU | PFC | 0.63658331 | 0 |
| B137 | B197 | AFU | AG | 0.64495658 | 0 |
| B137 | B194 | AFU | Cl | -0.505512 | 3.55E-14 |
| B137 | B177 | AFU | ACR | 0.49786147 | 9.81E-14 |
| B137 | B157 | AFU | IgG | -0.4905371 | 2.53E-13 |
| B137 | B200 | AFU | OP | 0.43821038 | 1.19E-10 |
| B137 | B195 | AFU | Na | -0.4178292 | 1.00E-09 |
| B137 | B199 | AFU | Mg | -0.3788377 | 4.03E-08 |
| B137 | B155 | AFU | IgA | -0.3731164 | 6.67E-08 |
| B137 | B139 | AFU | ADA | -0.3468208 | 5.95E-07 |
| B138 | B139 | PAB | ADA | -0.8829535 | 0 |
| B138 | B193 | PAB | K | 0.75419338 | 0 |
| B138 | B196 | PAB | TCO2 | -0.7552739 | 0 |
| B138 | B197 | PAB | AG | 0.8132278 | 0 |
| B138 | B198 | PAB | P | -0.7751862 | 0 |
| B138 | B199 | PAB | Mg | -0.9118865 | 0 |
| B138 | B200 | PAB | OP | 0.93778012 | 0 |
| B138 | B151 | PAB | Anti-CCP | -0.5256581 | 2.22E-15 |
| B138 | B182 | PAB | TT | 0.50864903 | 2.33E-14 |
| B138 | B156 | PAB | IgE | -0.4995152 | 7.90E-14 |
| B138 | B185 | PAB | PT | -0.4345517 | 1.77E-10 |
| B138 | B168 | PAB | NMP22 | 0.43148148 | 2.45E-10 |
| B138 | B175 | PAB | ALB-U | 0.43044462 | 2.73E-10 |
| B138 | B183 | PAB | PTA | -0.3762317 | 5.08E-08 |

| | | | | | |
|---|---|---|---|---|---|
| B138 | B186 | PAB | PFC | 0.37494895 | 5.68E-08 |
| B138 | B150 | PAB | ALP | -0.3584529 | 2.32E-07 |
| B139 | B193 | ADA | K | -0.818546 | 0 |
| B139 | B194 | ADA | Cl | -0.6315007 | 0 |
| B139 | B195 | ADA | Na | -0.7070958 | 0 |
| B139 | B196 | ADA | TCO2 | 0.95664785 | 0 |
| B139 | B197 | ADA | AG | -0.8318764 | 0 |
| B139 | B198 | ADA | P | 0.97970375 | 0 |
| B139 | B199 | ADA | Mg | 0.89933733 | 0 |
| B139 | B200 | ADA | OP | -0.9067134 | 0 |
| B139 | B168 | ADA | NMP22 | -0.4867197 | 4.11E-13 |
| B139 | B150 | ADA | ALP | 0.3571463 | 2.58E-07 |
| B142 | B143 | 25-(OH)D | β-CTx | -0.6027455 | 0 |
| B142 | B145 | 25-(OH)D | BGP | -0.6752535 | 0 |
| B143 | B144 | β-CTx | Total-PINP | 0.57865926 | 0 |
| B143 | B145 | β-CTx | BGP | 0.90389414 | 0 |
| B143 | B150 | β-CTx | ALP | 0.51264301 | 1.33E-14 |
| B143 | B227 | β-CTx | OD | 0.44879934 | 3.73E-11 |
| B143 | B228 | β-CTx | OS | 0.39593796 | 8.46E-09 |
| B144 | B150 | Total-PINP | ALP | 0.42169042 | 6.77E-10 |
| B144 | B145 | Total-PINP | BGP | 0.36636611 | 1.19E-07 |
| B145 | B227 | BGP | OD | 0.51035541 | 1.84E-14 |
| B145 | B150 | BGP | ALP | 0.49244215 | 1.98E-13 |
| B145 | B228 | BGP | OS | 0.4518248 | 2.66E-11 |
| B148 | B149 | Anti_TPO | TGAb | 0.44226072 | 7.69E-11 |
| B15 | B100 | CD3-CD19+(B cell) | anti-Ro52 | 1 | 0 |
| B15 | B101 | CD3-CD19+(B cell) | anti-Sc70 | 0.99999999 | 0 |
| B15 | B102 | CD3-CD19+(B cell) | anti-Sm | 1 | 0 |
| B15 | B105 | CD3-CD19+(B cell) | anti-U1RNP | 1 | 0 |
| B15 | B107 | CD3-CD19+(B cell) | AnuA | 0.99999995 | 0 |
| B15 | B110 | CD3-CD19+(B cell) | AHA | 1 | 0 |
| B15 | B134 | CD3-CD19+(B cell) | CHE | 0.99587065 | 0 |
| B15 | B135 | CD3-CD19+(B cell) | TBA | 0.72594937 | 0 |
| B15 | B136 | CD3-CD19+(B cell) | 5'NT | 0.77267967 | 0 |
| B15 | B137 | CD3-CD19+(B cell) | AFU | -0.8999613 | 0 |
| B15 | B151 | CD3-CD19+(B cell) | Anti-CCP | 1 | 0 |
| B15 | B156 | CD3-CD19+(B cell) | IgE | 0.94663186 | 0 |
| B15 | B16 | CD3-CD19+(B cell) | CD3+T | 1 | 0 |
| B15 | B175 | CD3-CD19+(B cell) | ALB-U | -0.8425779 | 0 |
| B15 | B177 | CD3-CD19+(B cell) | ACR | -0.5594521 | 0 |
| B15 | B18 | CD3-CD19+(B cell) | CD3+CD4+helper(inducer)T | -1 | 0 |
| B15 | B182 | CD3-CD19+(B cell) | TT | -0.9646716 | 0 |
| B15 | B183 | CD3-CD19+(B cell) | PTA | 0.70776873 | 0 |
| B15 | B185 | CD3-CD19+(B cell) | PT | 0.83292135 | 0 |

| | | | | | |
|---|---|---|---|---|---|
| B15 | B186 | CD3-CD19+(B cell) | PFC | -0.7053006 | 0 |
| B15 | B19 | CD3-CD19+(B cell) | CD3+CD4+/CD3+CD8+ | -1 | 0 |
| B15 | B194 | CD3-CD19+(B cell) | Cl | 0.65638677 | 0 |
| B15 | B195 | CD3-CD19+(B cell) | Na | 0.60796773 | 0 |
| B15 | B20 | CD3-CD19+(B cell) | CD3+CD4+CD8+ | -1 | 0 |
| B15 | B21 | CD3-CD19+(B cell) | CD3+CD4-CD8- | 1 | 0 |
| B15 | B22 | CD3-CD19+(B cell) | CD3+CD8+(T) | 1 | 0 |
| B15 | B23 | CD3-CD19+(B cell) | CD3-CD16+CD56+(NK) | -1 | 0 |
| B15 | B24 | CD3-CD19+(B cell) | αHBDH | -0.5434727 | 0 |
| B15 | B73 | CD3-CD19+(B cell) | ApoA | -0.8899831 | 0 |
| B15 | B97 | CD3-CD19+(B cell) | dsDNA | 1 | 0 |
| B15 | B98 | CD3-CD19+(B cell) | anti-Jo-1 | 1 | 0 |
| B15 | B99 | CD3-CD19+(B cell) | anti-PM-Scl | 1 | 0 |
| B15 | B157 | CD3-CD19+(B cell) | IgG | 0.54290776 | 2.22E-16 |
| B15 | B138 | CD3-CD19+(B cell) | PAB | -0.5256581 | 2.22E-15 |
| B15 | B25 | CD3-CD19+(B cell) | CK | -0.5058335 | 3.42E-14 |
| B15 | B54 | CD3-CD19+(B cell) | ESR | -0.4962338 | 1.21E-13 |
| B15 | B87 | CD3-CD19+(B cell) | NSE | -0.4925773 | 1.95E-13 |
| B15 | B59 | CD3-CD19+(B cell) | WBRV | 0.46137883 | 8.89E-12 |
| B15 | B27 | CD3-CD19+(B cell) | LDH | -0.4611981 | 9.08E-12 |
| B15 | B55 | CD3-CD19+(B cell) | TK | 0.43493806 | 1.70E-10 |
| B15 | B155 | CD3-CD19+(B cell) | IgA | 0.4155488 | 1.26E-09 |
| B15 | B69 | CD3-CD19+(B cell) | PV | 0.39065855 | 1.38E-08 |
| B15 | B154 | CD3-CD19+(B cell) | RF | 0.35447914 | 3.21E-07 |
| B15 | B76 | CD3-CD19+(B cell) | ApoB | -0.3492309 | 4.91E-07 |
| B150 | B200 | ALP | OP | -0.383354 | 2.69E-08 |
| B150 | B199 | ALP | Mg | 0.38273217 | 2.85E-08 |
| B150 | B193 | ALP | K | -0.3598245 | 2.07E-07 |
| B151 | B156 | Anti-CCP | IgE | 0.94663186 | 0 |
| B151 | B175 | Anti-CCP | ALB-U | -0.8425779 | 0 |
| B151 | B177 | Anti-CCP | ACR | -0.5594521 | 0 |
| B151 | B182 | Anti-CCP | TT | -0.9646716 | 0 |
| B151 | B183 | Anti-CCP | PTA | 0.70776873 | 0 |
| B151 | B185 | Anti-CCP | PT | 0.83292135 | 0 |
| B151 | B186 | Anti-CCP | PFC | -0.7053006 | 0 |
| B151 | B194 | Anti-CCP | Cl | 0.65638677 | 0 |
| B151 | B195 | Anti-CCP | Na | 0.60796773 | 0 |
| B151 | B157 | Anti-CCP | IgG | 0.54290776 | 2.22E-16 |
| B151 | B155 | Anti-CCP | IgA | 0.4155488 | 1.26E-09 |
| B151 | B154 | Anti-CCP | RF | 0.35447914 | 3.21E-07 |
| B152 | B169 | ACA-Ig(G-A-M) | U-CRY | 0.83075222 | 0 |
| B154 | B156 | RF | IgE | 0.45040025 | 3.12E-11 |
| B154 | B157 | RF | IgG | 0.38796148 | 1.77E-08 |
| B155 | B157 | IgA | IgG | 0.83320521 | 0 |

| | | | | | |
|---|---|---|---|---|---|
| B155 | B156 | IgA | IgE | 0.48128688 | 8.12E-13 |
| B155 | B182 | IgA | TT | -0.4013165 | 5.08E-09 |
| B155 | B175 | IgA | ALB-U | -0.3465536 | 6.08E-07 |
| B155 | B185 | IgA | PT | 0.34517764 | 6.78E-07 |
| B156 | B157 | IgE | IgG | 0.69191476 | 0 |
| B156 | B175 | IgE | ALB-U | -0.8010866 | 0 |
| B156 | B182 | IgE | TT | -0.9127534 | 0 |
| B156 | B183 | IgE | PTA | 0.66882874 | 0 |
| B156 | B185 | IgE | PT | 0.78938459 | 0 |
| B156 | B186 | IgE | PFC | -0.6664883 | 0 |
| B156 | B194 | IgE | Cl | 0.62000123 | 0 |
| B156 | B195 | IgE | Na | 0.57396159 | 0 |
| B156 | B177 | IgE | ACR | -0.5331766 | 8.88E-16 |
| B157 | B182 | IgG | TT | -0.5227173 | 3.33E-15 |
| B157 | B175 | IgG | ALB-U | -0.4655063 | 5.48E-12 |
| B157 | B185 | IgG | PT | 0.45432183 | 2.00E-11 |
| B157 | B183 | IgG | PTA | 0.38154377 | 3.17E-08 |
| B157 | B186 | IgG | PFC | -0.3801944 | 3.57E-08 |
| B157 | B203 | IgG | DH | -0.3733204 | 6.55E-08 |
| B157 | B194 | IgG | Cl | 0.35321257 | 3.56E-07 |
| B158 | B203 | IgM | DH | -0.4316781 | 2.40E-10 |
| B159 | B179 | β2-MG | α1-MG | -0.8598892 | 0 |
| B159 | B180 | β2-MG | Tf | -0.8711401 | 0 |
| B16 | B101 | CD3+T | anti-Sc70 | 1 | 0 |
| B16 | B105 | CD3+T | anti-U1RNP | 1 | 0 |
| B16 | B107 | CD3+T | AnuA | 0.99999995 | 0 |
| B16 | B109 | CD3+T | AMA-M2 | 1 | 0 |
| B16 | B110 | CD3+T | AHA | 1 | 0 |
| B16 | B134 | CD3+T | CHE | 0.99587065 | 0 |
| B16 | B135 | CD3+T | TBA | 0.72594937 | 0 |
| B16 | B136 | CD3+T | 5'NT | 0.77267967 | 0 |
| B16 | B137 | CD3+T | AFU | -0.8999613 | 0 |
| B16 | B151 | CD3+T | Anti-CCP | 1 | 0 |
| B16 | B156 | CD3+T | IgE | 0.94663186 | 0 |
| B16 | B17 | CD3+T | CD3+CD16+CD56+NK | 1 | 0 |
| B16 | B175 | CD3+T | ALB-U | -0.8425779 | 0 |
| B16 | B177 | CD3+T | ACR | -0.5594521 | 0 |
| B16 | B18 | CD3+T | CD3+CD4+helper(inducer)T | -1 | 0 |
| B16 | B182 | CD3+T | TT | -0.9646716 | 0 |
| B16 | B183 | CD3+T | PTA | 0.70776873 | 0 |
| B16 | B185 | CD3+T | PT | 0.83292135 | 0 |
| B16 | B186 | CD3+T | PFC | -0.7053006 | 0 |
| B16 | B19 | CD3+T | CD3+CD4+/CD3+CD8+ | -1 | 0 |
| B16 | B194 | CD3+T | Cl | 0.65638677 | 0 |

| | | | | | |
|---|---|---|---|---|---|
| B16 | B195 | CD3+T | Na | 0.60796773 | 0 |
| B16 | B20 | CD3+T | CD3+CD4+CD8+ | -1 | 0 |
| B16 | B21 | CD3+T | CD3+CD4-CD8- | 1 | 0 |
| B16 | B22 | CD3+T | CD3+CD8+(T) | 1 | 0 |
| B16 | B23 | CD3+T | CD3-CD16+CD56+(NK) | -1 | 0 |
| B16 | B24 | CD3+T | αHBDH | -0.5434727 | 0 |
| B16 | B34 | CD3+T | TK | 1 | 0 |
| B16 | B73 | CD3+T | ApoA | -0.8899831 | 0 |
| B16 | B97 | CD3+T | dsDNA | 1 | 0 |
| B16 | B98 | CD3+T | anti-Jo-1 | 1 | 0 |
| B16 | B99 | CD3+T | anti-PM-Scl | 1 | 0 |
| B16 | B157 | CD3+T | IgG | 0.54290776 | 2.22E-16 |
| B16 | B138 | CD3+T | PAB | -0.5256581 | 2.22E-15 |
| B16 | B25 | CD3+T | CK | -0.5058335 | 3.42E-14 |
| B16 | B54 | CD3+T | ESR | -0.4962338 | 1.21E-13 |
| B16 | B87 | CD3+T | NSE | -0.4925773 | 1.95E-13 |
| B16 | B59 | CD3+T | WBRV | 0.46137883 | 8.89E-12 |
| B16 | B27 | CD3+T | LDH | -0.4611981 | 9.08E-12 |
| B16 | B55 | CD3+T | TK | 0.43493806 | 1.70E-10 |
| B16 | B155 | CD3+T | IgA | 0.4155488 | 1.26E-09 |
| B16 | B69 | CD3+T | PV | 0.39065855 | 1.38E-08 |
| B16 | B154 | CD3+T | RF | 0.35447914 | 3.21E-07 |
| B16 | B76 | CD3+T | ApoB | -0.3492309 | 4.91E-07 |
| B163 | B166 | SG | Pro | 0.45774864 | 1.35E-11 |
| B163 | B176 | SG | Ucr | 0.42382255 | 5.44E-10 |
| B167 | B225 | PC | hPBG | 0.72577474 | 0 |
| B168 | B193 | NMP22 | K | 0.57272213 | 0 |
| B168 | B199 | NMP22 | Mg | -0.5479018 | 0 |
| B168 | B200 | NMP22 | OP | 0.5351726 | 4.44E-16 |
| B168 | B198 | NMP22 | P | -0.4579483 | 1.32E-11 |
| B168 | B195 | NMP22 | Na | 0.43945817 | 1.04E-10 |
| B168 | B194 | NMP22 | Cl | 0.42529046 | 4.68E-10 |
| B168 | B196 | NMP22 | TCO2 | -0.3976822 | 7.18E-09 |
| B17 | B100 | CD3+CD16+CD56+NK | anti-Ro52 | 1 | 0 |
| B17 | B101 | CD3+CD16+CD56+NK | anti-Sc70 | 0.99999999 | 0 |
| B17 | B105 | CD3+CD16+CD56+NK | anti-U1RNP | 1 | 0 |
| B17 | B107 | CD3+CD16+CD56+NK | AnuA | 0.99999994 | 0 |
| B17 | B109 | CD3+CD16+CD56+NK | AMA-M2 | 1 | 0 |
| B17 | B110 | CD3+CD16+CD56+NK | AHA | 1 | 0 |
| B17 | B134 | CD3+CD16+CD56+NK | CHE | 0.99587065 | 0 |
| B17 | B135 | CD3+CD16+CD56+NK | TBA | 0.72594937 | 0 |
| B17 | B136 | CD3+CD16+CD56+NK | 5'NT | 0.77267967 | 0 |
| B17 | B137 | CD3+CD16+CD56+NK | AFU | -0.8999613 | 0 |
| B17 | B156 | CD3+CD16+CD56+NK | IgE | 0.94663186 | 0 |

| | | | | | |
|---|---|---|---|---|---|
| B17 | B175 | CD3+CD16+CD56+NK | ALB-U | -0.8425779 | 0 |
| B17 | B177 | CD3+CD16+CD56+NK | ACR | -0.5594521 | 0 |
| B17 | B18 | CD3+CD16+CD56+NK | CD3+CD4+helper(inducer)T | -1 | 0 |
| B17 | B182 | CD3+CD16+CD56+NK | TT | -0.9646716 | 0 |
| B17 | B183 | CD3+CD16+CD56+NK | PTA | 0.70776873 | 0 |
| B17 | B185 | CD3+CD16+CD56+NK | PT | 0.83292135 | 0 |
| B17 | B186 | CD3+CD16+CD56+NK | PFC | -0.7053006 | 0 |
| B17 | B194 | CD3+CD16+CD56+NK | Cl | 0.65638677 | 0 |
| B17 | B195 | CD3+CD16+CD56+NK | Na | 0.60796773 | 0 |
| B17 | B20 | CD3+CD16+CD56+NK | CD3+CD4+CD8+ | -1 | 0 |
| B17 | B24 | CD3+CD16+CD56+NK | αHBDH | -0.5434727 | 0 |
| B17 | B73 | CD3+CD16+CD56+NK | ApoA | -0.8899831 | 0 |
| B17 | B97 | CD3+CD16+CD56+NK | dsDNA | 1 | 0 |
| B17 | B98 | CD3+CD16+CD56+NK | anti-Jo-1 | 1 | 0 |
| B17 | B99 | CD3+CD16+CD56+NK | anti-PM-Scl | 1 | 0 |
| B17 | B157 | CD3+CD16+CD56+NK | IgG | 0.54290776 | 2.22E-16 |
| B17 | B138 | CD3+CD16+CD56+NK | PAB | -0.5256581 | 2.22E-15 |
| B17 | B25 | CD3+CD16+CD56+NK | CK | -0.5058335 | 3.42E-14 |
| B17 | B54 | CD3+CD16+CD56+NK | ESR | -0.4962338 | 1.21E-13 |
| B17 | B87 | CD3+CD16+CD56+NK | NSE | -0.4925773 | 1.95E-13 |
| B17 | B59 | CD3+CD16+CD56+NK | WBRV | 0.46137883 | 8.89E-12 |
| B17 | B27 | CD3+CD16+CD56+NK | LDH | -0.4611981 | 9.08E-12 |
| B17 | B55 | CD3+CD16+CD56+NK | TK | 0.43493806 | 1.70E-10 |
| B17 | B155 | CD3+CD16+CD56+NK | IgA | 0.4155488 | 1.26E-09 |
| B17 | B69 | CD3+CD16+CD56+NK | PV | 0.39065855 | 1.38E-08 |
| B17 | B154 | CD3+CD16+CD56+NK | RF | 0.35447914 | 3.21E-07 |
| B17 | B76 | CD3+CD16+CD56+NK | ApoB | -0.3492309 | 4.91E-07 |
| B171 | B205 | EC | KDA | -0.5068589 | 2.98E-14 |
| B171 | B172 | EC | KB | -0.3677829 | 1.06E-07 |
| B174 | B191 | Urea/Cr | Urea | 0.64295741 | 0 |
| B175 | B177 | ALB-U | ACR | 0.74204904 | 0 |
| B175 | B182 | ALB-U | TT | 0.81565176 | 0 |
| B175 | B183 | ALB-U | PTA | -0.6039678 | 0 |
| B175 | B185 | ALB-U | PT | -0.6958339 | 0 |
| B175 | B186 | ALB-U | PFC | 0.60191483 | 0 |
| B175 | B194 | ALB-U | Cl | -0.5618991 | 0 |
| B175 | B195 | ALB-U | Na | -0.5224373 | 3.55E-15 |
| B175 | B176 | ALB-U | Ucr | 0.51449291 | 1.04E-14 |
| B176 | B227 | Ucr | OD | -0.3631175 | 1.57E-07 |
| B177 | B182 | ACR | TT | 0.5426149 | 2.22E-16 |
| B177 | B185 | ACR | PT | -0.4598307 | 1.06E-11 |
| B177 | B183 | ACR | PTA | -0.4038122 | 4.00E-09 |
| B177 | B186 | ACR | PFC | 0.40245882 | 4.56E-09 |
| B177 | B194 | ACR | Cl | -0.3763283 | 5.03E-08 |

| | | | | | |
|---|---|---|---|---|---|
| B177 | B195 | ACR | Na | -0.3506158 | 4.39E-07 |
| B178 | B179 | Ig-U | α1-MG | -0.860946 | 0 |
| B178 | B180 | Ig-U | Tf | -0.6831301 | 0 |
| B178 | B202 | Ig-U | e-times | -0.9238698 | 0 |
| B178 | B223 | Ig-U | FH | -0.9574271 | 0 |
| B179 | B223 | α1-MG | FH | -0.9684002 | 0 |
| B18 | B100 | CD3+CD4+helper(inducer)T | anti-Ro52 | -1 | 0 |
| B18 | B101 | CD3+CD4+helper(inducer)T | anti-Sc70 | -1 | 0 |
| B18 | B102 | CD3+CD4+helper(inducer)T | anti-Sm | -1 | 0 |
| B18 | B103 | CD3+CD4+helper(inducer)T | anti-SSA/Ro | -1 | 0 |
| B18 | B105 | CD3+CD4+helper(inducer)T | anti-U1RNP | -1 | 0 |
| B18 | B106 | CD3+CD4+helper(inducer)T | anti-P | -1 | 0 |
| B18 | B107 | CD3+CD4+helper(inducer)T | AnuA | -0.9999999 | 0 |
| B18 | B108 | CD3+CD4+helper(inducer)T | Anti-Cyclin | -1 | 0 |
| B18 | B109 | CD3+CD4+helper(inducer)T | AMA-M2 | -1 | 0 |
| B18 | B110 | CD3+CD4+helper(inducer)T | AHA | -1 | 0 |
| B18 | B134 | CD3+CD4+helper(inducer)T | CHE | -0.9958706 | 0 |
| B18 | B135 | CD3+CD4+helper(inducer)T | TBA | -0.7259494 | 0 |
| B18 | B136 | CD3+CD4+helper(inducer)T | 5'NT | -0.7726797 | 0 |
| B18 | B137 | CD3+CD4+helper(inducer)T | AFU | 0.89996133 | 0 |
| B18 | B151 | CD3+CD4+helper(inducer)T | Anti-CCP | -1 | 0 |
| B18 | B156 | CD3+CD4+helper(inducer)T | IgE | -0.9466319 | 0 |
| B18 | B175 | CD3+CD4+helper(inducer)T | ALB-U | 0.84257789 | 0 |
| B18 | B177 | CD3+CD4+helper(inducer)T | ACR | 0.55945214 | 0 |
| B18 | B182 | CD3+CD4+helper(inducer)T | TT | 0.96467157 | 0 |
| B18 | B183 | CD3+CD4+helper(inducer)T | PTA | -0.7077687 | 0 |
| B18 | B185 | CD3+CD4+helper(inducer)T | PT | -0.8329214 | 0 |
| B18 | B186 | CD3+CD4+helper(inducer)T | PFC | 0.70530063 | 0 |
| B18 | B19 | CD3+CD4+helper(inducer)T | CD3+CD4+/CD3+CD8+ | 1 | 0 |
| B18 | B194 | CD3+CD4+helper(inducer)T | Cl | -0.6563868 | 0 |
| B18 | B195 | CD3+CD4+helper(inducer)T | Na | -0.6079677 | 0 |
| B18 | B20 | CD3+CD4+helper(inducer)T | CD3+CD4+CD8+ | 1 | 0 |
| B18 | B21 | CD3+CD4+helper(inducer)T | CD3+CD4-CD8- | -1 | 0 |
| B18 | B22 | CD3+CD4+helper(inducer)T | CD3+CD8+(T) | -1 | 0 |
| B18 | B23 | CD3+CD4+helper(inducer)T | CD3-CD16+CD56+(NK) | 1 | 0 |
| B18 | B24 | CD3+CD4+helper(inducer)T | αHBDH | 0.54347267 | 0 |
| B18 | B34 | CD3+CD4+helper(inducer)T | TK | -1 | 0 |
| B18 | B73 | CD3+CD4+helper(inducer)T | ApoA | 0.88998306 | 0 |
| B18 | B97 | CD3+CD4+helper(inducer)T | dsDNA | -1 | 0 |
| B18 | B98 | CD3+CD4+helper(inducer)T | anti-Jo-1 | -1 | 0 |
| B18 | B99 | CD3+CD4+helper(inducer)T | anti-PM-Scl | -1 | 0 |
| B18 | B157 | CD3+CD4+helper(inducer)T | IgG | -0.5429078 | 2.22E-16 |
| B18 | B138 | CD3+CD4+helper(inducer)T | PAB | 0.52565811 | 2.22E-15 |
| B18 | B25 | CD3+CD4+helper(inducer)T | CK | 0.50583348 | 3.42E-14 |

| | | | | | |
|---|---|---|---|---|---|
| B18 | B54 | CD3+CD4+helper(inducer)T | ESR | 0.4962338 | 1.21E-13 |
| B18 | B87 | CD3+CD4+helper(inducer)T | NSE | 0.49257731 | 1.95E-13 |
| B18 | B59 | CD3+CD4+helper(inducer)T | WBRV | -0.4613788 | 8.89E-12 |
| B18 | B27 | CD3+CD4+helper(inducer)T | LDH | 0.46119814 | 9.08E-12 |
| B18 | B55 | CD3+CD4+helper(inducer)T | TK | -0.4349381 | 1.70E-10 |
| B18 | B155 | CD3+CD4+helper(inducer)T | IgA | -0.4155488 | 1.26E-09 |
| B18 | B69 | CD3+CD4+helper(inducer)T | PV | -0.3906585 | 1.38E-08 |
| B18 | B154 | CD3+CD4+helper(inducer)T | RF | -0.3544791 | 3.21E-07 |
| B18 | B76 | CD3+CD4+helper(inducer)T | ApoB | 0.34923093 | 4.91E-07 |
| B180 | B223 | Tf | FH | -0.4077178 | 2.74E-09 |
| B181 | B183 | APTT | PTA | -0.702824 | 0 |
| B181 | B184 | APTT | INR | -0.9806196 | 0 |
| B181 | B185 | APTT | PT | 0.55763379 | 0 |
| B181 | B186 | APTT | PFC | 0.70530063 | 0 |
| B182 | B183 | TT | PTA | -0.8688809 | 0 |
| B182 | B185 | TT | PT | -0.6577016 | 0 |
| B182 | B186 | TT | PFC | 0.86714921 | 0 |
| B182 | B194 | TT | Cl | -0.6320898 | 0 |
| B182 | B195 | TT | Na | -0.585214 | 0 |
| B183 | B184 | PTA | INR | 0.54983152 | 0 |
| B183 | B186 | PTA | PFC | -0.9999939 | 0 |
| B183 | B194 | PTA | Cl | 0.46159931 | 8.67E-12 |
| B183 | B195 | PTA | Na | 0.4268813 | 3.97E-10 |
| B184 | B185 | INR | PT | -0.7094585 | 0 |
| B184 | B186 | INR | PFC | -0.5527413 | 0 |
| B185 | B194 | PT | Cl | 0.54904566 | 0 |
| B185 | B195 | PT | Na | 0.50906775 | 2.20E-14 |
| B186 | B194 | PFC | Cl | -0.4599689 | 1.05E-11 |
| B186 | B195 | PFC | Na | -0.4253689 | 4.64E-10 |
| B189 | B190 | eGFB | Cre | -0.5659166 | 0 |
| B19 | B100 | CD3+CD4+/CD3+CD8+ | anti-Ro52 | -1 | 0 |
| B19 | B101 | CD3+CD4+/CD3+CD8+ | anti-Sc70 | -1 | 0 |
| B19 | B103 | CD3+CD4+/CD3+CD8+ | anti-SSA/Ro | -1 | 0 |
| B19 | B105 | CD3+CD4+/CD3+CD8+ | anti-U1RNP | -1 | 0 |
| B19 | B107 | CD3+CD4+/CD3+CD8+ | AnuA | -0.9999999 | 0 |
| B19 | B109 | CD3+CD4+/CD3+CD8+ | AMA-M2 | -1 | 0 |
| B19 | B110 | CD3+CD4+/CD3+CD8+ | AHA | -1 | 0 |
| B19 | B134 | CD3+CD4+/CD3+CD8+ | CHE | -0.9958706 | 0 |
| B19 | B135 | CD3+CD4+/CD3+CD8+ | TBA | -0.7259494 | 0 |
| B19 | B136 | CD3+CD4+/CD3+CD8+ | 5'NT | -0.7726797 | 0 |
| B19 | B137 | CD3+CD4+/CD3+CD8+ | AFU | 0.89996133 | 0 |
| B19 | B156 | CD3+CD4+/CD3+CD8+ | IgE | -0.9466319 | 0 |
| B19 | B175 | CD3+CD4+/CD3+CD8+ | ALB-U | 0.84257789 | 0 |
| B19 | B177 | CD3+CD4+/CD3+CD8+ | ACR | 0.55945214 | 0 |

| | | | | | |
|---|---|---|---|---|---|
| B19 | B182 | CD3+CD4+/CD3+CD8+ | TT | 0.96467157 | 0 |
| B19 | B183 | CD3+CD4+/CD3+CD8+ | PTA | -0.7077687 | 0 |
| B19 | B185 | CD3+CD4+/CD3+CD8+ | PT | -0.8329214 | 0 |
| B19 | B186 | CD3+CD4+/CD3+CD8+ | PFC | 0.70530063 | 0 |
| B19 | B194 | CD3+CD4+/CD3+CD8+ | Cl | -0.6563868 | 0 |
| B19 | B195 | CD3+CD4+/CD3+CD8+ | Na | -0.6079677 | 0 |
| B19 | B20 | CD3+CD4+/CD3+CD8+ | CD3+CD4+CD8+ | 1 | 0 |
| B19 | B23 | CD3+CD4+/CD3+CD8+ | CD3-CD16+CD56+(NK) | 1 | 0 |
| B19 | B24 | CD3+CD4+/CD3+CD8+ | αHBDH | 0.54347267 | 0 |
| B19 | B34 | CD3+CD4+/CD3+CD8+ | TK | -1 | 0 |
| B19 | B73 | CD3+CD4+/CD3+CD8+ | ApoA | 0.88998306 | 0 |
| B19 | B97 | CD3+CD4+/CD3+CD8+ | dsDNA | -1 | 0 |
| B19 | B98 | CD3+CD4+/CD3+CD8+ | anti-Jo-1 | -1 | 0 |
| B19 | B99 | CD3+CD4+/CD3+CD8+ | anti-PM-Scl | -1 | 0 |
| B19 | B157 | CD3+CD4+/CD3+CD8+ | IgG | -0.5429078 | 2.22E-16 |
| B19 | B138 | CD3+CD4+/CD3+CD8+ | PAB | 0.52565811 | 2.22E-15 |
| B19 | B25 | CD3+CD4+/CD3+CD8+ | CK | 0.50583348 | 3.42E-14 |
| B19 | B54 | CD3+CD4+/CD3+CD8+ | ESR | 0.4962338 | 1.21E-13 |
| B19 | B87 | CD3+CD4+/CD3+CD8+ | NSE | 0.49257732 | 1.95E-13 |
| B19 | B59 | CD3+CD4+/CD3+CD8+ | WBRV | -0.4613788 | 8.89E-12 |
| B19 | B27 | CD3+CD4+/CD3+CD8+ | LDH | 0.46119814 | 9.08E-12 |
| B19 | B55 | CD3+CD4+/CD3+CD8+ | TK | -0.4349381 | 1.70E-10 |
| B19 | B155 | CD3+CD4+/CD3+CD8+ | IgA | -0.4155488 | 1.26E-09 |
| B19 | B69 | CD3+CD4+/CD3+CD8+ | PV | -0.3906585 | 1.38E-08 |
| B19 | B154 | CD3+CD4+/CD3+CD8+ | RF | -0.3544791 | 3.21E-07 |
| B19 | B76 | CD3+CD4+/CD3+CD8+ | ApoB | 0.34923093 | 4.91E-07 |
| B190 | B191 | Cre | Urea | 0.54747899 | 0 |
| B190 | B192 | Cre | UA | 0.39863496 | 6.56E-09 |
| B193 | B194 | K | Cl | 0.69729759 | 0 |
| B193 | B195 | K | Na | 0.71908567 | 0 |
| B193 | B196 | K | TCO2 | -0.6445795 | 0 |
| B193 | B198 | K | P | -0.7551014 | 0 |
| B193 | B199 | K | Mg | -0.9572497 | 0 |
| B193 | B200 | K | OP | 0.93525396 | 0 |
| B193 | B197 | K | AG | 0.39034638 | 1.42E-08 |
| B194 | B195 | Cl | Na | 0.99367955 | 0 |
| B194 | B196 | Cl | TCO2 | -0.644287 | 0 |
| B194 | B198 | Cl | P | -0.7229074 | 0 |
| B194 | B199 | Cl | Mg | -0.5417082 | 2.22E-16 |
| B194 | B200 | Cl | OP | 0.49750996 | 1.03E-13 |
| B195 | B196 | Na | TCO2 | -0.7260189 | 0 |
| B195 | B198 | Na | P | -0.7943549 | 0 |
| B195 | B199 | Na | Mg | -0.5922389 | 0 |
| B195 | B200 | Na | OP | 0.55414857 | 0 |

| | | | | | |
|---|---|---|---|---|---|
| B196 | B197 | TCO2 | AG | -0.880457 | 0 |
| B196 | B198 | TCO2 | P | 0.98681293 | 0 |
| B196 | B199 | TCO2 | Mg | 0.73365355 | 0 |
| B196 | B200 | TCO2 | OP | -0.7447443 | 0 |
| B197 | B198 | AG | P | -0.8144355 | 0 |
| B197 | B199 | AG | Mg | -0.600791 | 0 |
| B197 | B200 | AG | OP | 0.64240867 | 0 |
| B198 | B199 | P | Mg | 0.81171164 | 0 |
| B198 | B200 | P | OP | -0.8142551 | 0 |
| B199 | B200 | Mg | OP | -0.9976635 | 0 |
| B2 | B189 | Age | eGFB | -0.4886758 | 3.21E-13 |
| B20 | B100 | CD3+CD4+CD8+ | anti-Ro52 | -1 | 0 |
| B20 | B101 | CD3+CD4+CD8+ | anti-Sc70 | -1 | 0 |
| B20 | B102 | CD3+CD4+CD8+ | anti-Sm | -1 | 0 |
| B20 | B103 | CD3+CD4+CD8+ | anti-SSA/Ro | -1 | 0 |
| B20 | B105 | CD3+CD4+CD8+ | anti-U1RNP | -1 | 0 |
| B20 | B107 | CD3+CD4+CD8+ | AnuA | -0.9999999 | 0 |
| B20 | B109 | CD3+CD4+CD8+ | AMA-M2 | -1 | 0 |
| B20 | B110 | CD3+CD4+CD8+ | AHA | -1 | 0 |
| B20 | B134 | CD3+CD4+CD8+ | CHE | -0.9958706 | 0 |
| B20 | B135 | CD3+CD4+CD8+ | TBA | -0.7259494 | 0 |
| B20 | B136 | CD3+CD4+CD8+ | 5'NT | -0.7726797 | 0 |
| B20 | B137 | CD3+CD4+CD8+ | AFU | 0.89996133 | 0 |
| B20 | B151 | CD3+CD4+CD8+ | Anti-CCP | -1 | 0 |
| B20 | B156 | CD3+CD4+CD8+ | IgE | -0.9466319 | 0 |
| B20 | B175 | CD3+CD4+CD8+ | ALB-U | 0.84257789 | 0 |
| B20 | B177 | CD3+CD4+CD8+ | ACR | 0.55945214 | 0 |
| B20 | B182 | CD3+CD4+CD8+ | TT | 0.96467157 | 0 |
| B20 | B183 | CD3+CD4+CD8+ | PTA | -0.7077687 | 0 |
| B20 | B185 | CD3+CD4+CD8+ | PT | -0.8329214 | 0 |
| B20 | B186 | CD3+CD4+CD8+ | PFC | 0.70530063 | 0 |
| B20 | B194 | CD3+CD4+CD8+ | Cl | -0.6563868 | 0 |
| B20 | B195 | CD3+CD4+CD8+ | Na | -0.6079677 | 0 |
| B20 | B21 | CD3+CD4+CD8+ | CD3+CD4-CD8- | -1 | 0 |
| B20 | B22 | CD3+CD4+CD8+ | CD3+CD8+(T) | -1 | 0 |
| B20 | B23 | CD3+CD4+CD8+ | CD3-CD16+CD56+(NK) | 1 | 0 |
| B20 | B24 | CD3+CD4+CD8+ | αHBDH | 0.54347267 | 0 |
| B20 | B34 | CD3+CD4+CD8+ | TK | -1 | 0 |
| B20 | B73 | CD3+CD4+CD8+ | ApoA | 0.88998306 | 0 |
| B20 | B97 | CD3+CD4+CD8+ | dsDNA | -1 | 0 |
| B20 | B98 | CD3+CD4+CD8+ | anti-Jo-1 | -1 | 0 |
| B20 | B99 | CD3+CD4+CD8+ | anti-PM-Scl | -1 | 0 |
| B20 | B157 | CD3+CD4+CD8+ | IgG | -0.5429078 | 2.22E-16 |
| B20 | B138 | CD3+CD4+CD8+ | PAB | 0.52565811 | 2.22E-15 |

| | | | | | |
|---|---|---|---|---|---|
| B20 | B25 | CD3+CD4+CD8+ | CK | 0.50583348 | 3.42E-14 |
| B20 | B54 | CD3+CD4+CD8+ | ESR | 0.4962338 | 1.21E-13 |
| B20 | B87 | CD3+CD4+CD8+ | NSE | 0.49257732 | 1.95E-13 |
| B20 | B59 | CD3+CD4+CD8+ | WBRV | -0.4613788 | 8.89E-12 |
| B20 | B27 | CD3+CD4+CD8+ | LDH | 0.46119814 | 9.08E-12 |
| B20 | B55 | CD3+CD4+CD8+ | TK | -0.4349381 | 1.70E-10 |
| B20 | B155 | CD3+CD4+CD8+ | IgA | -0.4155488 | 1.26E-09 |
| B20 | B69 | CD3+CD4+CD8+ | PV | -0.3906585 | 1.38E-08 |
| B20 | B154 | CD3+CD4+CD8+ | RF | -0.3544791 | 3.21E-07 |
| B20 | B76 | CD3+CD4+CD8+ | ApoB | 0.34923093 | 4.91E-07 |
| B202 | B223 | e-times | FH | -0.4082483 | 2.60E-09 |
| B206 | B207 | Waist | WHR | 0.78940176 | 0 |
| B206 | B210 | Waist | Hip | 0.65414645 | 0 |
| B206 | B215 | Waist | BMI | 0.76917143 | 0 |
| B206 | B216 | Waist | BMI | 0.70845241 | 0 |
| B207 | B215 | WHR | BMI | 0.44271885 | 7.31E-11 |
| B207 | B216 | WHR | BMI | 0.34324468 | 7.90E-07 |
| B21 | B100 | CD3+CD4-CD8- | anti-Ro52 | 1 | 0 |
| B21 | B101 | CD3+CD4-CD8- | anti-Sc70 | 0.99999999 | 0 |
| B21 | B103 | CD3+CD4-CD8- | anti-SSA/Ro | 1 | 0 |
| B21 | B105 | CD3+CD4-CD8- | anti-U1RNP | 1 | 0 |
| B21 | B107 | CD3+CD4-CD8- | AnuA | 0.99999994 | 0 |
| B21 | B109 | CD3+CD4-CD8- | AMA-M2 | 1 | 0 |
| B21 | B110 | CD3+CD4-CD8- | AHA | 1 | 0 |
| B21 | B134 | CD3+CD4-CD8- | CHE | 0.99587065 | 0 |
| B21 | B135 | CD3+CD4-CD8- | TBA | 0.72594937 | 0 |
| B21 | B136 | CD3+CD4-CD8- | 5'NT | 0.77267967 | 0 |
| B21 | B137 | CD3+CD4-CD8- | AFU | -0.8999613 | 0 |
| B21 | B156 | CD3+CD4-CD8- | IgE | 0.94663186 | 0 |
| B21 | B175 | CD3+CD4-CD8- | ALB-U | -0.8425779 | 0 |
| B21 | B177 | CD3+CD4-CD8- | ACR | -0.5594521 | 0 |
| B21 | B182 | CD3+CD4-CD8- | TT | -0.9646716 | 0 |
| B21 | B183 | CD3+CD4-CD8- | PTA | 0.70776873 | 0 |
| B21 | B185 | CD3+CD4-CD8- | PT | 0.83292135 | 0 |
| B21 | B186 | CD3+CD4-CD8- | PFC | -0.7053006 | 0 |
| B21 | B194 | CD3+CD4-CD8- | Cl | 0.65638677 | 0 |
| B21 | B195 | CD3+CD4-CD8- | Na | 0.60796773 | 0 |
| B21 | B22 | CD3+CD4-CD8- | CD3+CD8+(T) | 1 | 0 |
| B21 | B24 | CD3+CD4-CD8- | αHBDH | -0.5434727 | 0 |
| B21 | B73 | CD3+CD4-CD8- | ApoA | -0.8899831 | 0 |
| B21 | B97 | CD3+CD4-CD8- | dsDNA | 1 | 0 |
| B21 | B98 | CD3+CD4-CD8- | anti-Jo-1 | 1 | 0 |
| B21 | B99 | CD3+CD4-CD8- | anti-PM-Scl | 1 | 0 |
| B21 | B157 | CD3+CD4-CD8- | IgG | 0.54290776 | 2.22E-16 |

| | | | | | |
|---|---|---|---|---|---|
| B21 | B138 | CD3+CD4-CD8- | PAB | -0.5256581 | 2.22E-15 |
| B21 | B25 | CD3+CD4-CD8- | CK | -0.5058335 | 3.42E-14 |
| B21 | B54 | CD3+CD4-CD8- | ESR | -0.4962338 | 1.21E-13 |
| B21 | B87 | CD3+CD4-CD8- | NSE | -0.4925773 | 1.95E-13 |
| B21 | B59 | CD3+CD4-CD8- | WBRV | 0.46137883 | 8.89E-12 |
| B21 | B27 | CD3+CD4-CD8- | LDH | -0.4611981 | 9.08E-12 |
| B21 | B55 | CD3+CD4-CD8- | TK | 0.43493806 | 1.70E-10 |
| B21 | B155 | CD3+CD4-CD8- | IgA | 0.4155488 | 1.26E-09 |
| B21 | B69 | CD3+CD4-CD8- | PV | 0.39065855 | 1.38E-08 |
| B21 | B154 | CD3+CD4-CD8- | RF | 0.35447914 | 3.21E-07 |
| B21 | B76 | CD3+CD4-CD8- | ApoB | -0.3492309 | 4.91E-07 |
| B210 | B215 | Hip | BMI | 0.70752414 | 0 |
| B210 | B216 | Hip | BMI | 0.73313141 | 0 |
| B211 | B212 | IOP-R | IOP-L | 0.9067832 | 0 |
| B213 | B214 | CVA-R | CVA-L | 0.90359812 | 0 |
| B215 | B216 | BMI | BMI | 0.77156757 | 0 |
| B216 | B221 | BMI | Hei | 0.62157938 | 0 |
| B218 | B220 | DBP | SBP | 0.53267146 | 8.88E-16 |
| B22 | B100 | CD3+CD8+(T) | anti-Ro52 | 1 | 0 |
| B22 | B101 | CD3+CD8+(T) | anti-Sc70 | 0.99999999 | 0 |
| B22 | B103 | CD3+CD8+(T) | anti-SSA/Ro | 1 | 0 |
| B22 | B105 | CD3+CD8+(T) | anti-U1RNP | 1 | 0 |
| B22 | B107 | CD3+CD8+(T) | AnuA | 0.99999995 | 0 |
| B22 | B109 | CD3+CD8+(T) | AMA-M2 | 1 | 0 |
| B22 | B110 | CD3+CD8+(T) | AHA | 1 | 0 |
| B22 | B134 | CD3+CD8+(T) | CHE | 0.99587065 | 0 |
| B22 | B135 | CD3+CD8+(T) | TBA | 0.72594937 | 0 |
| B22 | B136 | CD3+CD8+(T) | 5'NT | 0.77267967 | 0 |
| B22 | B137 | CD3+CD8+(T) | AFU | -0.8999613 | 0 |
| B22 | B156 | CD3+CD8+(T) | IgE | 0.94663186 | 0 |
| B22 | B175 | CD3+CD8+(T) | ALB-U | -0.8425779 | 0 |
| B22 | B177 | CD3+CD8+(T) | ACR | -0.5594521 | 0 |
| B22 | B182 | CD3+CD8+(T) | TT | -0.9646716 | 0 |
| B22 | B183 | CD3+CD8+(T) | PTA | 0.70776873 | 0 |
| B22 | B185 | CD3+CD8+(T) | PT | 0.83292135 | 0 |
| B22 | B186 | CD3+CD8+(T) | PFC | -0.7053006 | 0 |
| B22 | B194 | CD3+CD8+(T) | Cl | 0.65638677 | 0 |
| B22 | B195 | CD3+CD8+(T) | Na | 0.60796773 | 0 |
| B22 | B24 | CD3+CD8+(T) | αHBDH | -0.5434727 | 0 |
| B22 | B73 | CD3+CD8+(T) | ApoA | -0.8899831 | 0 |
| B22 | B97 | CD3+CD8+(T) | dsDNA | 1 | 0 |
| B22 | B98 | CD3+CD8+(T) | anti-Jo-1 | 1 | 0 |
| B22 | B99 | CD3+CD8+(T) | anti-PM-Scl | 1 | 0 |
| B22 | B157 | CD3+CD8+(T) | IgG | 0.54290776 | 2.22E-16 |

| | | | | | |
|---|---|---|---|---|---|
| B22 | B138 | CD3+CD8+(T) | PAB | -0.5256581 | 2.22E-15 |
| B22 | B25 | CD3+CD8+(T) | CK | -0.5058335 | 3.42E-14 |
| B22 | B54 | CD3+CD8+(T) | ESR | -0.4962338 | 1.21E-13 |
| B22 | B87 | CD3+CD8+(T) | NSE | -0.4925773 | 1.95E-13 |
| B22 | B59 | CD3+CD8+(T) | WBRV | 0.46137883 | 8.89E-12 |
| B22 | B27 | CD3+CD8+(T) | LDH | -0.4611981 | 9.08E-12 |
| B22 | B55 | CD3+CD8+(T) | TK | 0.43493806 | 1.70E-10 |
| B22 | B155 | CD3+CD8+(T) | IgA | 0.4155488 | 1.26E-09 |
| B22 | B69 | CD3+CD8+(T) | PV | 0.39065855 | 1.38E-08 |
| B22 | B154 | CD3+CD8+(T) | RF | 0.35447914 | 3.21E-07 |
| B22 | B76 | CD3+CD8+(T) | ApoB | -0.3492309 | 4.91E-07 |
| B227 | B228 | OD | OS | 0.56241612 | 0 |
| B23 | B100 | CD3-CD16+CD56+(NK) | anti-Ro52 | -1 | 0 |
| B23 | B101 | CD3-CD16+CD56+(NK) | anti-Sc70 | -1 | 0 |
| B23 | B103 | CD3-CD16+CD56+(NK) | anti-SSA/Ro | -1 | 0 |
| B23 | B105 | CD3-CD16+CD56+(NK) | anti-U1RNP | -1 | 0 |
| B23 | B107 | CD3-CD16+CD56+(NK) | AnuA | -0.9999999 | 0 |
| B23 | B109 | CD3-CD16+CD56+(NK) | AMA-M2 | -1 | 0 |
| B23 | B110 | CD3-CD16+CD56+(NK) | AHA | -1 | 0 |
| B23 | B134 | CD3-CD16+CD56+(NK) | CHE | -0.9958706 | 0 |
| B23 | B135 | CD3-CD16+CD56+(NK) | TBA | -0.7259494 | 0 |
| B23 | B136 | CD3-CD16+CD56+(NK) | 5'NT | -0.7726797 | 0 |
| B23 | B137 | CD3-CD16+CD56+(NK) | AFU | 0.89996133 | 0 |
| B23 | B156 | CD3-CD16+CD56+(NK) | IgE | -0.9466319 | 0 |
| B23 | B175 | CD3-CD16+CD56+(NK) | ALB-U | 0.84257789 | 0 |
| B23 | B177 | CD3-CD16+CD56+(NK) | ACR | 0.55945214 | 0 |
| B23 | B182 | CD3-CD16+CD56+(NK) | TT | 0.96467157 | 0 |
| B23 | B183 | CD3-CD16+CD56+(NK) | PTA | -0.7077687 | 0 |
| B23 | B185 | CD3-CD16+CD56+(NK) | PT | -0.8329214 | 0 |
| B23 | B186 | CD3-CD16+CD56+(NK) | PFC | 0.70530063 | 0 |
| B23 | B194 | CD3-CD16+CD56+(NK) | Cl | -0.6563868 | 0 |
| B23 | B195 | CD3-CD16+CD56+(NK) | Na | -0.6079677 | 0 |
| B23 | B24 | CD3-CD16+CD56+(NK) | αHBDH | 0.54347267 | 0 |
| B23 | B34 | CD3-CD16+CD56+(NK) | TK | -1 | 0 |
| B23 | B73 | CD3-CD16+CD56+(NK) | ApoA | 0.88998306 | 0 |
| B23 | B97 | CD3-CD16+CD56+(NK) | dsDNA | -1 | 0 |
| B23 | B98 | CD3-CD16+CD56+(NK) | anti-Jo-1 | -1 | 0 |
| B23 | B99 | CD3-CD16+CD56+(NK) | anti-PM-Scl | -1 | 0 |
| B23 | B157 | CD3-CD16+CD56+(NK) | IgG | -0.5429078 | 2.22E-16 |
| B23 | B138 | CD3-CD16+CD56+(NK) | PAB | 0.52565811 | 2.22E-15 |
| B23 | B25 | CD3-CD16+CD56+(NK) | CK | 0.50583348 | 3.42E-14 |
| B23 | B54 | CD3-CD16+CD56+(NK) | ESR | 0.4962338 | 1.21E-13 |
| B23 | B87 | CD3-CD16+CD56+(NK) | NSE | 0.49257732 | 1.95E-13 |
| B23 | B59 | CD3-CD16+CD56+(NK) | WBRV | -0.4613788 | 8.89E-12 |

| | | | | | |
|---|---|---|---|---|---|
| B23 | B27 | CD3-CD16+CD56+(NK) | LDH | 0.46119814 | 9.08E-12 |
| B23 | B55 | CD3-CD16+CD56+(NK) | TK | -0.4349381 | 1.70E-10 |
| B23 | B155 | CD3-CD16+CD56+(NK) | IgA | -0.4155488 | 1.26E-09 |
| B23 | B69 | CD3-CD16+CD56+(NK) | PV | -0.3906585 | 1.38E-08 |
| B23 | B154 | CD3-CD16+CD56+(NK) | RF | -0.3544791 | 3.21E-07 |
| B23 | B76 | CD3-CD16+CD56+(NK) | ApoB | 0.34923093 | 4.91E-07 |
| B24 | B100 | αHBDH | anti-Ro52 | -0.5434727 | 0 |
| B24 | B101 | αHBDH | anti-Sc70 | -0.5434727 | 0 |
| B24 | B102 | αHBDH | anti-Sm | -0.5434727 | 0 |
| B24 | B103 | αHBDH | anti-SSA/Ro | -0.5434727 | 0 |
| B24 | B104 | αHBDH | anti-SSB | -0.5434727 | 0 |
| B24 | B105 | αHBDH | anti-U1RNP | -0.5434727 | 0 |
| B24 | B106 | αHBDH | anti-P | -0.5434727 | 0 |
| B24 | B107 | αHBDH | AnuA | -0.5434726 | 0 |
| B24 | B108 | αHBDH | Anti-Cyclin | -0.5434727 | 0 |
| B24 | B109 | αHBDH | AMA-M2 | -0.5434727 | 0 |
| B24 | B110 | αHBDH | AHA | -0.5434727 | 0 |
| B24 | B151 | αHBDH | Anti-CCP | -0.5434727 | 0 |
| B24 | B156 | αHBDH | IgE | -0.6618361 | 0 |
| B24 | B157 | αHBDH | IgG | -0.5468235 | 0 |
| B24 | B26 | αHBDH | CKMB | 0.55595691 | 0 |
| B24 | B27 | αHBDH | LDH | 0.71453442 | 0 |
| B24 | B34 | αHBDH | TK | -0.5434727 | 0 |
| B24 | B95 | αHBDH | anti-cenpb | -0.5434727 | 0 |
| B24 | B97 | αHBDH | dsDNA | -0.5434727 | 0 |
| B24 | B98 | αHBDH | anti-Jo-1 | -0.5434727 | 0 |
| B24 | B99 | αHBDH | anti-PM-Scl | -0.5434727 | 0 |
| B24 | B134 | αHBDH | CHE | -0.5415168 | 2.22E-16 |
| B24 | B185 | αHBDH | PT | -0.5062435 | 3.22E-14 |
| B24 | B137 | αHBDH | AFU | 0.499828 | 7.57E-14 |
| B24 | B182 | αHBDH | TT | 0.49876767 | 8.70E-14 |
| B24 | B136 | αHBDH | 5'NT | -0.480534 | 8.92E-13 |
| B24 | B73 | αHBDH | ApoA | 0.44514687 | 5.60E-11 |
| B24 | B175 | αHBDH | ALB-U | 0.41339969 | 1.56E-09 |
| B24 | B146 | αHBDH | CysC | 0.3866617 | 2.00E-08 |
| B24 | B138 | αHBDH | PAB | 0.36806473 | 1.03E-07 |
| B24 | B155 | αHBDH | IgA | -0.3527175 | 3.71E-07 |
| B24 | B154 | αHBDH | RF | -0.3511323 | 4.21E-07 |
| B24 | B81 | αHBDH | TSGF | -0.3493362 | 4.87E-07 |
| B24 | B158 | αHBDH | IgM | -0.3473875 | 5.69E-07 |
| B25 | B187 | CK | NSE | -0.5615505 | 0 |
| B25 | B73 | CK | ApoA | 0.55678127 | 0 |
| B25 | B100 | CK | anti-Ro52 | -0.5058335 | 3.42E-14 |
| B25 | B101 | CK | anti-Sc70 | -0.5058335 | 3.42E-14 |

| | | | | | |
|---|---|---|---|---|---|
| B25 | B102 | CK | anti-Sm | -0.5058335 | 3.42E-14 |
| B25 | B103 | CK | anti-SSA/Ro | -0.5058335 | 3.42E-14 |
| B25 | B104 | CK | anti-SSB | -0.5058335 | 3.42E-14 |
| B25 | B105 | CK | anti-U1RNP | -0.5058335 | 3.42E-14 |
| B25 | B106 | CK | anti-P | -0.5058335 | 3.42E-14 |
| B25 | B107 | CK | AnuA | -0.5058335 | 3.42E-14 |
| B25 | B108 | CK | Anti-Cyclin | -0.5058335 | 3.42E-14 |
| B25 | B109 | CK | AMA-M2 | -0.5058335 | 3.42E-14 |
| B25 | B110 | CK | AHA | -0.5058335 | 3.42E-14 |
| B25 | B151 | CK | Anti-CCP | -0.5058335 | 3.42E-14 |
| B25 | B34 | CK | TK | -0.5058335 | 3.42E-14 |
| B25 | B95 | CK | anti-cenpb | -0.5058335 | 3.42E-14 |
| B25 | B97 | CK | dsDNA | -0.5058335 | 3.42E-14 |
| B25 | B98 | CK | anti-Jo-1 | -0.5058335 | 3.42E-14 |
| B25 | B99 | CK | anti-PM-Scl | -0.5058335 | 3.42E-14 |
| B25 | B134 | CK | CHE | -0.5037433 | 4.51E-14 |
| B25 | B185 | CK | PT | -0.4884962 | 3.28E-13 |
| B25 | B156 | CK | IgE | -0.4643675 | 6.27E-12 |
| B25 | B137 | CK | AFU | 0.4605033 | 9.84E-12 |
| B25 | B182 | CK | TT | 0.45598211 | 1.66E-11 |
| B25 | B175 | CK | ALB-U | 0.45123571 | 2.84E-11 |
| B25 | B136 | CK | 5'NT | -0.425489 | 4.58E-10 |
| B25 | B76 | CK | ApoB | 0.39654702 | 7.99E-09 |
| B25 | B90 | CK | CYFRA 21-1 | 0.39272739 | 1.14E-08 |
| B26 | B81 | CKMB | TSGF | -0.6292755 | 0 |
| B26 | B27 | CKMB | LDH | 0.47082084 | 2.91E-12 |
| B27 | B156 | LDH | IgE | -0.5539065 | 0 |
| B27 | B185 | LDH | PT | -0.4938459 | 1.65E-13 |
| B27 | B137 | LDH | AFU | 0.47795935 | 1.22E-12 |
| B27 | B134 | LDH | CHE | -0.4724875 | 2.38E-12 |
| B27 | B100 | LDH | anti-Ro52 | -0.4611981 | 9.08E-12 |
| B27 | B101 | LDH | anti-Sc70 | -0.4611981 | 9.08E-12 |
| B27 | B102 | LDH | anti-Sm | -0.4611981 | 9.08E-12 |
| B27 | B103 | LDH | anti-SSA/Ro | -0.4611981 | 9.08E-12 |
| B27 | B104 | LDH | anti-SSB | -0.4611981 | 9.08E-12 |
| B27 | B105 | LDH | anti-U1RNP | -0.4611981 | 9.08E-12 |
| B27 | B106 | LDH | anti-P | -0.4611981 | 9.08E-12 |
| B27 | B107 | LDH | AnuA | -0.4611981 | 9.08E-12 |
| B27 | B108 | LDH | Anti-Cyclin | -0.4611981 | 9.08E-12 |
| B27 | B109 | LDH | AMA-M2 | -0.4611981 | 9.08E-12 |
| B27 | B110 | LDH | AHA | -0.4611981 | 9.08E-12 |
| B27 | B151 | LDH | Anti-CCP | -0.4611981 | 9.08E-12 |
| B27 | B34 | LDH | TK | -0.4611981 | 9.08E-12 |
| B27 | B95 | LDH | anti-cenpb | -0.4611981 | 9.08E-12 |

| | | | | | |
|---|---|---|---|---|---|
| B27 | B97 | LDH | dsDNA | -0.4611981 | 9.08E-12 |
| B27 | B98 | LDH | anti-Jo-1 | -0.4611981 | 9.08E-12 |
| B27 | B99 | LDH | anti-PM-Scl | -0.4611981 | 9.08E-12 |
| B27 | B157 | LDH | IgG | -0.4511152 | 2.88E-11 |
| B27 | B73 | LDH | ApoA | 0.42330771 | 5.74E-10 |
| B27 | B182 | LDH | TT | 0.39267743 | 1.15E-08 |
| B27 | B135 | LDH | TBA | -0.3890563 | 1.60E-08 |
| B27 | B175 | LDH | ALB-U | 0.36241671 | 1.66E-07 |
| B28 | B122 | PRL | C4 | -0.9944903 | 0 |
| B28 | B202 | PRL | e-times | -0.4885042 | 3.28E-13 |
| B28 | B31 | PRL | FSH | -0.4349588 | 1.69E-10 |
| B28 | B159 | PRL | β2-MG | -0.3836829 | 2.61E-08 |
| B29 | B117 | E2 | TT4 | -0.5617468 | 0 |
| B29 | B178 | E2 | Ig-U | -0.8406728 | 0 |
| B29 | B223 | E2 | FH | -0.6062755 | 0 |
| B29 | B125 | E2 | Cor | -0.4961389 | 1.23E-13 |
| B29 | B122 | E2 | C4 | -0.462179 | 8.10E-12 |
| B29 | B202 | E2 | e-times | -0.3413944 | 9.13E-07 |
| B30 | B124 | LH | ACTH | 0.72112526 | 0 |
| B30 | B167 | LH | PC | 0.78194744 | 0 |
| B30 | B169 | LH | U-CRY | 0.58797473 | 0 |
| B30 | B225 | LH | hPBG | 0.57906602 | 0 |
| B30 | B96 | LH | dsDNA (IIF) | 0.80582296 | 0 |
| B30 | B33 | LH | Prog | 0.50434162 | 4.17E-14 |
| B30 | B121 | LH | C3 | 0.44474959 | 5.85E-11 |
| B30 | B152 | LH | ACA-Ig(G-A-M) | 0.42446505 | 5.09E-10 |
| B30 | B35 | LH | β2-MG | 0.34875597 | 5.10E-07 |
| B31 | B178 | FSH | Ig-U | -0.7830553 | 0 |
| B31 | B179 | FSH | α1-MG | -0.3771571 | 4.68E-08 |
| B32 | B152 | Testo | ACA-Ig(G-A-M) | 0.64270505 | 0 |
| B32 | B167 | Testo | PC | 0.53692484 | 4.44E-16 |
| B32 | B119 | Testo | Hcv-rna | 0.43173289 | 2.38E-10 |
| B32 | B36 | Testo | V-Lym% | 0.34517263 | 6.78E-07 |
| B33 | B152 | Prog | ACA-Ig(G-A-M) | 0.56647641 | 0 |
| B33 | B167 | Prog | PC | 0.90979927 | 0 |
| B33 | B169 | Prog | U-CRY | 0.9057953 | 0 |
| B33 | B36 | Prog | V-Lym% | 0.78749923 | 0 |
| B34 | B101 | TK | anti-Sc70 | 0.99999999 | 0 |
| B34 | B103 | TK | anti-SSA/Ro | 1 | 0 |
| B34 | B105 | TK | anti-U1RNP | 1 | 0 |
| B34 | B107 | TK | AnuA | 0.99999995 | 0 |
| B34 | B109 | TK | AMA-M2 | 1 | 0 |
| B34 | B110 | TK | AHA | 1 | 0 |
| B34 | B134 | TK | CHE | 0.99587065 | 0 |

| | | | | | |
|---|---|---|---|---|---|
| B34 | B135 | TK | TBA | 0.72594937 | 0 |
| B34 | B136 | TK | 5'NT | 0.77267967 | 0 |
| B34 | B137 | TK | AFU | -0.8999613 | 0 |
| B34 | B156 | TK | IgE | 0.94663186 | 0 |
| B34 | B175 | TK | ALB-U | -0.8425779 | 0 |
| B34 | B177 | TK | ACR | -0.5594521 | 0 |
| B34 | B182 | TK | TT | -0.9646716 | 0 |
| B34 | B183 | TK | PTA | 0.70776873 | 0 |
| B34 | B185 | TK | PT | 0.83292135 | 0 |
| B34 | B186 | TK | PFC | -0.7053006 | 0 |
| B34 | B194 | TK | Cl | 0.65638677 | 0 |
| B34 | B195 | TK | Na | 0.60796773 | 0 |
| B34 | B73 | TK | ApoA | -0.8899831 | 0 |
| B34 | B97 | TK | dsDNA | 1 | 0 |
| B34 | B98 | TK | anti-Jo-1 | 1 | 0 |
| B34 | B99 | TK | anti-PM-Scl | 1 | 0 |
| B34 | B157 | TK | IgG | 0.54290776 | 2.22E-16 |
| B34 | B138 | TK | PAB | -0.5256581 | 2.22E-15 |
| B34 | B54 | TK | ESR | -0.4962338 | 1.21E-13 |
| B34 | B87 | TK | NSE | -0.4925773 | 1.95E-13 |
| B34 | B59 | TK | WBRV | 0.46137883 | 8.89E-12 |
| B34 | B55 | TK | TK | 0.43493806 | 1.70E-10 |
| B34 | B155 | TK | IgA | 0.4155488 | 1.26E-09 |
| B34 | B69 | TK | PV | 0.39065855 | 1.38E-08 |
| B34 | B154 | TK | RF | 0.35447914 | 3.21E-07 |
| B34 | B76 | TK | ApoB | -0.3492309 | 4.91E-07 |
| B35 | B121 | β2-MG | C3 | 0.83813325 | 0 |
| B35 | B124 | β2-MG | ACTH | 0.98036761 | 0 |
| B35 | B152 | β2-MG | ACA-Ig(G-A-M) | 0.77193813 | 0 |
| B35 | B36 | β2-MG | V-Lym% | 0.90282575 | 0 |
| B35 | B96 | β2-MG | dsDNA (IIF) | 0.62123653 | 0 |
| B35 | B225 | β2-MG | hPBG | 0.41208169 | 1.78E-09 |
| B36 | B121 | V-Lym% | C3 | 0.91146543 | 0 |
| B36 | B124 | V-Lym% | ACTH | 0.79455216 | 0 |
| B36 | B167 | V-Lym% | PC | 0.81989159 | 0 |
| B36 | B225 | V-Lym% | hPBG | 0.82158384 | 0 |
| B36 | B79 | V-Lym% | hb V -dna | 0.68353656 | 0 |
| B37 | B116 | BAND | TT3 | -0.9293204 | 0 |
| B37 | B159 | BAND | β2-MG | -0.5573641 | 0 |
| B38 | B121 | BAND% | C3 | 0.92450033 | 0 |
| B38 | B124 | BAND% | ACTH | 0.81236239 | 0 |
| B38 | B152 | BAND% | ACA-Ig(G-A-M) | 0.75337824 | 0 |
| B38 | B167 | BAND% | PC | 0.81649658 | 0 |
| B39 | B63 | Other cells | WBRSR(mPas_200) | 0.49025076 | 2.62E-13 |

| | | | | | |
|---|---|---|---|---|---|
| B39 | B66 | Other cells | WBRSR(mPas_100) | 0.48693479 | 4.00E-13 |
| B39 | B54 | Other cells | ESR | -0.4843094 | 5.57E-13 |
| B39 | B67 | Other cells | WBRSR(mPas_50) | 0.48406257 | 5.75E-13 |
| B39 | B61 | Other cells | WBHSRV | 0.47276047 | 2.31E-12 |
| B39 | B64 | Other cells | WBRSR(mPas_5) | 0.46834527 | 3.92E-12 |
| B39 | B65 | Other cells | WBRSR(mPas_1) | 0.45306892 | 2.31E-11 |
| B39 | B62 | Other cells | WBHSRVI | 0.45100747 | 2.91E-11 |
| B39 | B58 | Other cells | HCT | 0.4424835 | 7.50E-11 |
| B39 | B56 | Other cells | IR | 0.4297528 | 2.94E-10 |
| B39 | B78 | Other cells | HBcAb | 0.4284827 | 3.36E-10 |
| B39 | B74 | Other cells | ApoA1 | -0.4262477 | 4.24E-10 |
| B39 | B183 | Other cells | PTA | 0.41644263 | 1.15E-09 |
| B39 | B186 | Other cells | PFC | -0.4164452 | 1.15E-09 |
| B39 | B60 | Other cells | WBRVI | 0.41381654 | 1.50E-09 |
| B39 | B75 | Other cells | ApoA1/B | -0.4105087 | 2.08E-09 |
| B39 | B111 | Other cells | TSH | -0.3673593 | 1.10E-07 |
| B39 | B182 | Other cells | TT | -0.3611201 | 1.86E-07 |
| B39 | B59 | Other cells | WBRV | 0.359333 | 2.15E-07 |
| B4 | B152 | WB-Ca | ACA-Ig(G-A-M) | 0.81446296 | 0 |
| B4 | B167 | WB-Ca | PC | 0.58969198 | 0 |
| B4 | B32 | WB-Ca | Testo | -0.6575959 | 0 |
| B4 | B5 | WB-Ca | WB-Mg | -0.5726371 | 0 |
| B4 | B7 | WB-Ca | WB-Cop | -0.7878386 | 0 |
| B4 | B169 | WB-Ca | U-CRY | -0.5333333 | 6.66E-16 |
| B4 | B33 | WB-Ca | Prog | 0.50321961 | 4.84E-14 |
| B40 | B42 | WBC | MON | 0.75898123 | 0 |
| B41 | B42 | MON % | MON | 0.69378836 | 0 |
| B42 | B206 | MON | Waist | 0.37407334 | 6.13E-08 |
| B43 | B44 | MCV | MCH | 0.94114514 | 0 |
| B43 | B45 | MCV | MCHC | 0.57510223 | 0 |
| B43 | B51 | MCV | Hb | 0.43913003 | 1.08E-10 |
| B43 | B47 | MCV | RDW | 0.43439941 | 1.80E-10 |
| B43 | B52 | MCV | PLT/BPC | -0.3672965 | 1.10E-07 |
| B44 | B45 | MCH | MCHC | 0.81380347 | 0 |
| B44 | B51 | MCH | Hb | 0.55689489 | 0 |
| B44 | B52 | MCH | PLT/BPC | -0.3627382 | 1.62E-07 |
| B45 | B51 | MCHC | Hb | 0.62432614 | 0 |
| B46 | B51 | RBC Count | Hb | 0.70694135 | 0 |
| B46 | B47 | RBC Count | RDW | -0.4141126 | 1.46E-09 |
| B46 | B221 | RBC Count | Hei | 0.38817567 | 1.74E-08 |
| B46 | B216 | RBC Count | BMI | 0.34607777 | 6.32E-07 |
| B48 | B53 | LYM% | GR% | -0.9526454 | 0 |
| B48 | B188 | LYM% | Ca | 0.34612004 | 6.29E-07 |
| B5 | B121 | WB-Mg | C3 | 0.86629616 | 0 |

| | | | | | |
|---|---|---|---|---|---|
| B5 | B152 | WB-Mg | ACA-Ig(G-A-M) | 0.70594748 | 0 |
| B5 | B30 | WB-Mg | LH | 0.57391391 | 0 |
| B5 | B7 | WB-Mg | WB-Cop | 0.55367873 | 0 |
| B5 | B79 | WB-Mg | hbＶ-dna | 0.98793535 | 0 |
| B5 | B96 | WB-Mg | dsDNA (IIF) | 0.75341058 | 0 |
| B5 | B35 | WB-Mg | β2-MG | 0.51485078 | 9.99E-15 |
| B5 | B124 | WB-Mg | ACTH | 0.38060907 | 3.44E-08 |
| B5 | B36 | WB-Mg | V-Lym% | 0.37067287 | 8.24E-08 |
| B51 | B216 | Hb | BMI | 0.4397555 | 1.01E-10 |
| B51 | B206 | Hb | Waist | 0.38318501 | 2.73E-08 |
| B54 | B181 | ESR | APTT | 0.70451315 | 0 |
| B54 | B182 | ESR | TT | 0.66497989 | 0 |
| B54 | B183 | ESR | PTA | -0.8507131 | 0 |
| B54 | B184 | ESR | INR | -0.5929311 | 0 |
| B54 | B186 | ESR | PFC | 0.8512306 | 0 |
| B54 | B55 | ESR | TK | -0.7604313 | 0 |
| B54 | B56 | ESR | IR | -0.814675 | 0 |
| B54 | B57 | ESR | RBCAI | -0.6510524 | 0 |
| B54 | B58 | ESR | HCT | -0.6530979 | 0 |
| B54 | B59 | ESR | WBRV | -0.7425358 | 0 |
| B54 | B60 | ESR | WBRVI | -0.7898201 | 0 |
| B54 | B61 | ESR | WBHSRV | -0.7909694 | 0 |
| B54 | B62 | ESR | WBHSRVI | -0.7606478 | 0 |
| B54 | B63 | ESR | WBRSR(mPas_200) | -0.7295881 | 0 |
| B54 | B64 | ESR | WBRSR(mPas_5) | -0.7652412 | 0 |
| B54 | B65 | ESR | WBRSR(mPas_1) | -0.7776913 | 0 |
| B54 | B66 | ESR | WBRSR(mPas_100) | -0.737745 | 0 |
| B54 | B67 | ESR | WBRSR(mPas_50) | -0.7420049 | 0 |
| B54 | B74 | ESR | ApoA1 | 0.56598279 | 0 |
| B54 | B100 | ESR | anti-Ro52 | -0.4962338 | 1.21E-13 |
| B54 | B101 | ESR | anti-Sc70 | -0.4962338 | 1.21E-13 |
| B54 | B102 | ESR | anti-Sm | -0.4962338 | 1.21E-13 |
| B54 | B103 | ESR | anti-SSA/Ro | -0.4962338 | 1.21E-13 |
| B54 | B104 | ESR | anti-SSB | -0.4962338 | 1.21E-13 |
| B54 | B105 | ESR | anti-U1RNP | -0.4962338 | 1.21E-13 |
| B54 | B106 | ESR | anti-P | -0.4962338 | 1.21E-13 |
| B54 | B107 | ESR | AnuA | -0.4962338 | 1.21E-13 |
| B54 | B108 | ESR | Anti-Cyclin | -0.4962338 | 1.21E-13 |
| B54 | B109 | ESR | AMA-M2 | -0.4962338 | 1.21E-13 |
| B54 | B110 | ESR | AHA | -0.4962338 | 1.21E-13 |
| B54 | B151 | ESR | Anti-CCP | -0.4962338 | 1.21E-13 |
| B54 | B95 | ESR | anti-cenpb | -0.4962338 | 1.21E-13 |
| B54 | B97 | ESR | dsDNA | -0.4962338 | 1.21E-13 |
| B54 | B98 | ESR | anti-Jo-1 | -0.4962338 | 1.21E-13 |

| | | | | | |
|---|---|---|---|---|---|
| B54 | B99 | ESR | anti-PM-Scl | -0.4962338 | 1.21E-13 |
| B54 | B134 | ESR | CHE | -0.494664 | 1.49E-13 |
| B54 | B156 | ESR | IgE | -0.4680026 | 4.08E-12 |
| B54 | B137 | ESR | AFU | 0.44933625 | 3.51E-11 |
| B54 | B175 | ESR | ALB-U | 0.42951945 | 3.01E-10 |
| B54 | B136 | ESR | 5'NT | -0.3852267 | 2.27E-08 |
| B54 | B69 | ESR | PV | 0.36275748 | 1.62E-07 |
| B54 | B135 | ESR | TBA | -0.3593128 | 2.16E-07 |
| B55 | B183 | TK | PTA | 0.6166667 | 0 |
| B55 | B186 | TK | PFC | -0.6166705 | 0 |
| B55 | B56 | TK | IR | 0.80112158 | 0 |
| B55 | B57 | TK | RBCAI | 0.96330497 | 0 |
| B55 | B59 | TK | WBRV | 0.91922695 | 0 |
| B55 | B60 | TK | WBRVI | 0.78114819 | 0 |
| B55 | B61 | TK | WBHSRV | 0.74515971 | 0 |
| B55 | B62 | TK | WBHSRVI | 0.65319386 | 0 |
| B55 | B63 | TK | WBRSR(mPas_200) | 0.59563891 | 0 |
| B55 | B64 | TK | WBRSR(mPas_5) | 0.69229608 | 0 |
| B55 | B65 | TK | WBRSR(mPas_1) | 0.74189858 | 0 |
| B55 | B66 | TK | WBRSR(mPas_100) | 0.60693324 | 0 |
| B55 | B67 | TK | WBRSR(mPas_50) | 0.61968262 | 0 |
| B55 | B182 | TK | TT | -0.5347453 | 4.44E-16 |
| B55 | B69 | TK | PV | -0.4957344 | 1.29E-13 |
| B55 | B100 | TK | anti-Ro52 | 0.43493806 | 1.70E-10 |
| B55 | B101 | TK | anti-Sc70 | 0.43493806 | 1.70E-10 |
| B55 | B102 | TK | anti-Sm | 0.43493806 | 1.70E-10 |
| B55 | B103 | TK | anti-SSA/Ro | 0.43493806 | 1.70E-10 |
| B55 | B104 | TK | anti-SSB | 0.43493806 | 1.70E-10 |
| B55 | B105 | TK | anti-U1RNP | 0.43493806 | 1.70E-10 |
| B55 | B106 | TK | anti-P | 0.43493806 | 1.70E-10 |
| B55 | B107 | TK | AnuA | 0.43493803 | 1.70E-10 |
| B55 | B108 | TK | Anti-Cyclin | 0.43493806 | 1.70E-10 |
| B55 | B109 | TK | AMA-M2 | 0.43493806 | 1.70E-10 |
| B55 | B110 | TK | AHA | 0.43493806 | 1.70E-10 |
| B55 | B151 | TK | Anti-CCP | 0.43493806 | 1.70E-10 |
| B55 | B181 | TK | APTT | -0.4349381 | 1.70E-10 |
| B55 | B95 | TK | anti-cenpb | 0.43493806 | 1.70E-10 |
| B55 | B97 | TK | dsDNA | 0.43493806 | 1.70E-10 |
| B55 | B98 | TK | anti-Jo-1 | 0.43493806 | 1.70E-10 |
| B55 | B99 | TK | anti-PM-Scl | 0.43493806 | 1.70E-10 |
| B55 | B134 | TK | CHE | 0.43359558 | 1.96E-10 |
| B55 | B58 | TK | HCT | 0.42020106 | 7.89E-10 |
| B55 | B156 | TK | IgE | 0.41007228 | 2.17E-09 |
| B55 | B137 | TK | AFU | -0.3940246 | 1.01E-08 |

| | | | | | |
|---|---|---|---|---|---|
| B55 | B73 | TK | ApoA | -0.3874641 | 1.86E-08 |
| B55 | B74 | TK | ApoA1 | -0.3845328 | 2.42E-08 |
| B55 | B175 | TK | ALB-U | -0.3772589 | 4.64E-08 |
| B55 | B76 | TK | ApoB | -0.3449933 | 6.88E-07 |
| B55 | B184 | TK | INR | 0.34085921 | 9.52E-07 |
| B56 | B181 | IR | APTT | -0.6570427 | 0 |
| B56 | B183 | IR | PTA | 0.62949899 | 0 |
| B56 | B184 | IR | INR | 0.59811784 | 0 |
| B56 | B186 | IR | PFC | -0.6305474 | 0 |
| B56 | B57 | IR | RBCAI | 0.7694476 | 0 |
| B56 | B58 | IR | HCT | 0.87323074 | 0 |
| B56 | B59 | IR | WBRV | 0.90792924 | 0 |
| B56 | B60 | IR | WBRVI | 0.9961008 | 0 |
| B56 | B61 | IR | WBHSRV | 0.98268847 | 0 |
| B56 | B62 | IR | WBHSRVI | 0.975749 | 0 |
| B56 | B63 | IR | WBRSR(mPas_200) | 0.9495959 | 0 |
| B56 | B64 | IR | WBRSR(mPas_5) | 0.9815716 | 0 |
| B56 | B65 | IR | WBRSR(mPas_1) | 0.99092567 | 0 |
| B56 | B66 | IR | WBRSR(mPas_100) | 0.95582613 | 0 |
| B56 | B67 | IR | WBRSR(mPas_50) | 0.96058792 | 0 |
| B56 | B74 | IR | ApoA1 | -0.5155095 | 9.10E-15 |
| B56 | B69 | IR | PV | -0.4072757 | 2.86E-09 |
| B56 | B182 | IR | TT | -0.3976137 | 7.23E-09 |
| B57 | B59 | RBCAI | WBRV | 0.8831367 | 0 |
| B57 | B60 | RBCAI | WBRVI | 0.77143701 | 0 |
| B57 | B61 | RBCAI | WBHSRV | 0.69105429 | 0 |
| B57 | B62 | RBCAI | WBHSRVI | 0.63129197 | 0 |
| B57 | B63 | RBCAI | WBRSR(mPas_200) | 0.55009294 | 0 |
| B57 | B64 | RBCAI | WBRSR(mPas_5) | 0.6587629 | 0 |
| B57 | B65 | RBCAI | WBRSR(mPas_1) | 0.7137132 | 0 |
| B57 | B66 | RBCAI | WBRSR(mPas_100) | 0.56548383 | 0 |
| B57 | B67 | RBCAI | WBRSR(mPas_50) | 0.57981889 | 0 |
| B57 | B69 | RBCAI | PV | -0.605056 | 0 |
| B57 | B186 | RBCAI | PFC | -0.4435997 | 6.64E-11 |
| B57 | B183 | RBCAI | PTA | 0.4433398 | 6.83E-11 |
| B57 | B58 | RBCAI | HCT | 0.41042073 | 2.10E-09 |
| B57 | B181 | RBCAI | APTT | -0.3651438 | 1.32E-07 |
| B57 | B182 | RBCAI | TT | -0.3479426 | 5.44E-07 |
| B58 | B181 | HCT | APTT | -0.7147756 | 0 |
| B58 | B184 | HCT | INR | 0.71429448 | 0 |
| B58 | B59 | HCT | WBRV | 0.60873034 | 0 |
| B58 | B60 | HCT | WBRVI | 0.88895257 | 0 |
| B58 | B61 | HCT | WBHSRV | 0.86997308 | 0 |
| B58 | B62 | HCT | WBHSRVI | 0.95798433 | 0 |

| | | | | | |
|---|---|---|---|---|---|
| B58 | B63 | HCT | WBRSR(mPas_200) | 0.94079034 | 0 |
| B58 | B64 | HCT | WBRSR(mPas_5) | 0.91808312 | 0 |
| B58 | B65 | HCT | WBRSR(mPas_1) | 0.89455223 | 0 |
| B58 | B66 | HCT | WBRSR(mPas_100) | 0.94468267 | 0 |
| B58 | B67 | HCT | WBRSR(mPas_50) | 0.94223823 | 0 |
| B58 | B75 | HCT | ApoA1/B | -0.5380056 | 4.44E-16 |
| B58 | B74 | HCT | ApoA1 | -0.4968059 | 1.13E-13 |
| B58 | B186 | HCT | PFC | -0.4557492 | 1.70E-11 |
| B58 | B185 | HCT | PT | -0.4552359 | 1.80E-11 |
| B58 | B183 | HCT | PTA | 0.4538113 | 2.12E-11 |
| B58 | B76 | HCT | ApoB | 0.44332002 | 6.85E-11 |
| B59 | B183 | WBRV | PTA | 0.58680325 | 0 |
| B59 | B186 | WBRV | PFC | -0.5865723 | 0 |
| B59 | B60 | WBRV | WBRVI | 0.88976184 | 0 |
| B59 | B61 | WBRV | WBHSRV | 0.9091563 | 0 |
| B59 | B62 | WBRV | WBHSRVI | 0.80830879 | 0 |
| B59 | B63 | WBRV | WBRSR(mPas_200) | 0.80768499 | 0 |
| B59 | B64 | WBRV | WBRSR(mPas_5) | 0.8686294 | 0 |
| B59 | B65 | WBRV | WBRSR(mPas_1) | 0.89860009 | 0 |
| B59 | B66 | WBRV | WBRSR(mPas_100) | 0.81149494 | 0 |
| B59 | B67 | WBRV | WBRSR(mPas_50) | 0.82004773 | 0 |
| B59 | B182 | WBRV | TT | -0.5421359 | 2.22E-16 |
| B59 | B100 | WBRV | anti-Ro52 | 0.46137883 | 8.89E-12 |
| B59 | B101 | WBRV | anti-Sc70 | 0.46137883 | 8.89E-12 |
| B59 | B102 | WBRV | anti-Sm | 0.46137883 | 8.89E-12 |
| B59 | B103 | WBRV | anti-SSA/Ro | 0.46137883 | 8.89E-12 |
| B59 | B104 | WBRV | anti-SSB | 0.46137883 | 8.89E-12 |
| B59 | B105 | WBRV | anti-U1RNP | 0.46137883 | 8.89E-12 |
| B59 | B106 | WBRV | anti-P | 0.46137883 | 8.89E-12 |
| B59 | B107 | WBRV | AnuA | 0.4613788 | 8.89E-12 |
| B59 | B108 | WBRV | Anti-Cyclin | 0.46137883 | 8.89E-12 |
| B59 | B109 | WBRV | AMA-M2 | 0.46137883 | 8.89E-12 |
| B59 | B110 | WBRV | AHA | 0.46137883 | 8.89E-12 |
| B59 | B151 | WBRV | Anti-CCP | 0.46137883 | 8.89E-12 |
| B59 | B95 | WBRV | anti-cenpb | 0.46137883 | 8.89E-12 |
| B59 | B97 | WBRV | dsDNA | 0.46137883 | 8.89E-12 |
| B59 | B98 | WBRV | anti-Jo-1 | 0.46137883 | 8.89E-12 |
| B59 | B99 | WBRV | anti-PM-Scl | 0.46137883 | 8.89E-12 |
| B59 | B134 | WBRV | CHE | 0.46019875 | 1.02E-11 |
| B59 | B156 | WBRV | IgE | 0.43411153 | 1.85E-10 |
| B59 | B137 | WBRV | AFU | -0.4193756 | 8.58E-10 |
| B59 | B175 | WBRV | ALB-U | -0.4059984 | 3.24E-09 |
| B59 | B74 | WBRV | ApoA1 | -0.4000496 | 5.74E-09 |
| B59 | B181 | WBRV | APTT | -0.3660408 | 1.23E-07 |

| | | | | | |
|---|---|---|---|---|---|
| B59 | B136 | WBRV | 5'NT | 0.35921626 | 2.17E-07 |
| B6 | B202 | WB-Fe | e-times | -0.6579517 | 0 |
| B6 | B223 | WB-Fe | FH | -0.6446584 | 0 |
| B60 | B181 | WBRVI | APTT | -0.6528694 | 0 |
| B60 | B183 | WBRVI | PTA | 0.57609407 | 0 |
| B60 | B184 | WBRVI | INR | 0.60792614 | 0 |
| B60 | B186 | WBRVI | PFC | -0.5773063 | 0 |
| B60 | B61 | WBRVI | WBHSRV | 0.97100709 | 0 |
| B60 | B62 | WBRVI | WBHSRVI | 0.98044547 | 0 |
| B60 | B63 | WBRVI | WBRSR(mPas_200) | 0.94557104 | 0 |
| B60 | B64 | WBRVI | WBRSR(mPas_5) | 0.97905199 | 0 |
| B60 | B65 | WBRVI | WBRSR(mPas_1) | 0.98845295 | 0 |
| B60 | B66 | WBRVI | WBRSR(mPas_100) | 0.95331708 | 0 |
| B60 | B67 | WBRVI | WBRSR(mPas_50) | 0.95822534 | 0 |
| B60 | B74 | WBRVI | ApoA1 | -0.5064491 | 3.13E-14 |
| B60 | B69 | WBRVI | PV | -0.4485225 | 3.85E-11 |
| B61 | B181 | WBHSRV | APTT | -0.5753182 | 0 |
| B61 | B183 | WBHSRV | PTA | 0.63276579 | 0 |
| B61 | B186 | WBHSRV | PFC | -0.6334022 | 0 |
| B61 | B62 | WBHSRV | WBHSRVI | 0.96781151 | 0 |
| B61 | B63 | WBHSRV | WBRSR(mPas_200) | 0.97791193 | 0 |
| B61 | B64 | WBHSRV | WBRSR(mPas_5) | 0.9927148 | 0 |
| B61 | B65 | WBHSRV | WBRSR(mPas_1) | 0.99405316 | 0 |
| B61 | B66 | WBHSRV | WBRSR(mPas_100) | 0.97911985 | 0 |
| B61 | B67 | WBHSRV | WBRSR(mPas_50) | 0.98176382 | 0 |
| B61 | B184 | WBHSRV | INR | 0.50125812 | 6.26E-14 |
| B61 | B74 | WBHSRV | ApoA1 | -0.499535 | 7.86E-14 |
| B61 | B182 | WBHSRV | TT | -0.45892 | 1.18E-11 |
| B62 | B181 | WBHSRVI | APTT | -0.687653 | 0 |
| B62 | B183 | WBHSRVI | PTA | 0.56319712 | 0 |
| B62 | B184 | WBHSRVI | INR | 0.65232076 | 0 |
| B62 | B186 | WBHSRVI | PFC | -0.5646244 | 0 |
| B62 | B63 | WBHSRVI | WBRSR(mPas_200) | 0.98087955 | 0 |
| B62 | B64 | WBHSRVI | WBRSR(mPas_5) | 0.98838129 | 0 |
| B62 | B65 | WBHSRVI | WBRSR(mPas_1) | 0.98291548 | 0 |
| B62 | B66 | WBHSRVI | WBRSR(mPas_100) | 0.98578069 | 0 |
| B62 | B67 | WBHSRVI | WBRSR(mPas_50) | 0.987364 | 0 |
| B62 | B74 | WBHSRVI | ApoA1 | -0.5183462 | 6.22E-15 |
| B62 | B75 | WBHSRVI | ApoA1/B | -0.4057026 | 3.33E-09 |
| B62 | B69 | WBHSRVI | PV | -0.3675874 | 1.07E-07 |
| B63 | B181 | WBRSR(mPas_200) | APTT | -0.5927634 | 0 |
| B63 | B183 | WBRSR(mPas_200) | PTA | 0.56485083 | 0 |
| B63 | B186 | WBRSR(mPas_200) | PFC | -0.5658072 | 0 |
| B63 | B64 | WBRSR(mPas_200) | WBRSR(mPas_5) | 0.9905868 | 0 |

| | | | | | |
|---|---|---|---|---|---|
| B63 | B65 | WBRSR(mPas_200) | WBRSR(mPas_1) | 0.97751843 | 0 |
| B63 | B66 | WBRSR(mPas_200) | WBRSR(mPas_100) | 0.9995702 | 0 |
| B63 | B67 | WBRSR(mPas_200) | WBRSR(mPas_50) | 0.99908551 | 0 |
| B63 | B184 | WBRSR(mPas_200) | INR | 0.54044693 | 2.22E-16 |
| B63 | B74 | WBRSR(mPas_200) | ApoA1 | -0.4953809 | 1.35E-13 |
| B63 | B75 | WBRSR(mPas_200) | ApoA1/B | -0.4128093 | 1.66E-09 |
| B63 | B182 | WBRSR(mPas_200) | TT | -0.3545545 | 3.19E-07 |
| B64 | B181 | WBRSR(mPas_5) | APTT | -0.599035 | 0 |
| B64 | B183 | WBRSR(mPas_5) | PTA | 0.57919979 | 0 |
| B64 | B184 | WBRSR(mPas_5) | INR | 0.54385909 | 0 |
| B64 | B186 | WBRSR(mPas_5) | PFC | -0.5801374 | 0 |
| B64 | B65 | WBRSR(mPas_5) | WBRSR(mPas_1) | 0.99712989 | 0 |
| B64 | B66 | WBRSR(mPas_5) | WBRSR(mPas_100) | 0.99299018 | 0 |
| B64 | B67 | WBRSR(mPas_5) | WBRSR(mPas_50) | 0.99489446 | 0 |
| B64 | B74 | WBRSR(mPas_5) | ApoA1 | -0.5007834 | 6.66E-14 |
| B64 | B182 | WBRSR(mPas_5) | TT | -0.3696756 | 8.98E-08 |
| B64 | B75 | WBRSR(mPas_5) | ApoA1/B | -0.3528035 | 3.68E-07 |
| B65 | B181 | WBRSR(mPas_1) | APTT | -0.5925496 | 0 |
| B65 | B183 | WBRSR(mPas_1) | PTA | 0.58527702 | 0 |
| B65 | B186 | WBRSR(mPas_1) | PFC | -0.5861618 | 0 |
| B65 | B66 | WBRSR(mPas_1) | WBRSR(mPas_100) | 0.98119267 | 0 |
| B65 | B67 | WBRSR(mPas_1) | WBRSR(mPas_50) | 0.9844047 | 0 |
| B65 | B184 | WBRSR(mPas_1) | INR | 0.53457019 | 4.44E-16 |
| B65 | B74 | WBRSR(mPas_1) | ApoA1 | -0.4979341 | 9.73E-14 |
| B65 | B182 | WBRSR(mPas_1) | TT | -0.3824413 | 2.92E-08 |
| B66 | B181 | WBRSR(mPas_100) | APTT | -0.6034778 | 0 |
| B66 | B183 | WBRSR(mPas_100) | PTA | 0.5635467 | 0 |
| B66 | B184 | WBRSR(mPas_100) | INR | 0.5533869 | 0 |
| B66 | B186 | WBRSR(mPas_100) | PFC | -0.5645601 | 0 |
| B66 | B67 | WBRSR(mPas_100) | WBRSR(mPas_50) | 0.99984321 | 0 |
| B66 | B74 | WBRSR(mPas_100) | ApoA1 | -0.4999715 | 7.42E-14 |
| B66 | B75 | WBRSR(mPas_100) | ApoA1/B | -0.4103097 | 2.12E-09 |
| B66 | B182 | WBRSR(mPas_100) | TT | -0.3453277 | 6.70E-07 |
| B67 | B181 | WBRSR(mPas_50) | APTT | -0.604755 | 0 |
| B67 | B183 | WBRSR(mPas_50) | PTA | 0.56625552 | 0 |
| B67 | B184 | WBRSR(mPas_50) | INR | 0.55414051 | 0 |
| B67 | B186 | WBRSR(mPas_50) | PFC | -0.5672658 | 0 |
| B67 | B74 | WBRSR(mPas_50) | ApoA1 | -0.5004926 | 6.93E-14 |
| B67 | B75 | WBRSR(mPas_50) | ApoA1/B | -0.4030873 | 4.29E-09 |
| B67 | B182 | WBRSR(mPas_50) | TT | -0.3481174 | 5.37E-07 |
| B69 | B181 | PV | APTT | 0.62712023 | 0 |
| B69 | B184 | PV | INR | -0.6908789 | 0 |
| B69 | B185 | PV | PT | 0.67133229 | 0 |
| B69 | B100 | PV | anti-Ro52 | 0.39065855 | 1.38E-08 |

| | | | | | |
|---|---|---|---|---|---|
| B69 | B101 | PV | anti-Sc70 | 0.39065855 | 1.38E-08 |
| B69 | B102 | PV | anti-Sm | 0.39065855 | 1.38E-08 |
| B69 | B103 | PV | anti-SSA/Ro | 0.39065855 | 1.38E-08 |
| B69 | B104 | PV | anti-SSB | 0.39065855 | 1.38E-08 |
| B69 | B105 | PV | anti-U1RNP | 0.39065855 | 1.38E-08 |
| B69 | B106 | PV | anti-P | 0.39065855 | 1.38E-08 |
| B69 | B107 | PV | AnuA | 0.39065852 | 1.38E-08 |
| B69 | B108 | PV | Anti-Cyclin | 0.39065855 | 1.38E-08 |
| B69 | B109 | PV | AMA-M2 | 0.39065855 | 1.38E-08 |
| B69 | B110 | PV | AHA | 0.39065855 | 1.38E-08 |
| B69 | B151 | PV | Anti-CCP | 0.39065855 | 1.38E-08 |
| B69 | B95 | PV | anti-cenpb | 0.39065855 | 1.38E-08 |
| B69 | B97 | PV | dsDNA | 0.39065855 | 1.38E-08 |
| B69 | B98 | PV | anti-Jo-1 | 0.39065855 | 1.38E-08 |
| B69 | B99 | PV | anti-PM-Scl | 0.39065855 | 1.38E-08 |
| B69 | B134 | PV | CHE | 0.38893947 | 1.62E-08 |
| B69 | B156 | PV | IgE | 0.37019608 | 8.59E-08 |
| B69 | B137 | PV | AFU | -0.3509711 | 4.27E-07 |
| B69 | B73 | PV | ApoA | -0.3470427 | 5.85E-07 |
| B7 | B121 | WB-Cop | C3 | 0.80119275 | 0 |
| B7 | B124 | WB-Cop | ACTH | 0.98287319 | 0 |
| B7 | B152 | WB-Cop | ACA-Ig(G-A-M) | 0.96249804 | 0 |
| B7 | B225 | WB-Cop | hPBG | 0.7878386 | 0 |
| B7 | B36 | WB-Cop | V-Lym% | 0.71919495 | 0 |
| B7 | B79 | WB-Cop | hb V -dna | 0.83062779 | 0 |
| B7 | B33 | WB-Cop | Prog | 0.50972893 | 2.00E-14 |
| B7 | B30 | WB-Cop | LH | 0.47375712 | 2.05E-12 |
| B70 | B77 | LDL-C | TC | 0.86594405 | 0 |
| B71 | B77 | TG | TC | 0.38284501 | 2.82E-08 |
| B71 | B72 | TG | HDL-C | -0.3488228 | 5.07E-07 |
| B73 | B100 | ApoA | anti-Ro52 | -0.8899831 | 0 |
| B73 | B101 | ApoA | anti-Sc70 | -0.8899831 | 0 |
| B73 | B102 | ApoA | anti-Sm | -0.8899831 | 0 |
| B73 | B103 | ApoA | anti-SSA/Ro | -0.8899831 | 0 |
| B73 | B104 | ApoA | anti-SSB | -0.8899831 | 0 |
| B73 | B105 | ApoA | anti-U1RNP | -0.8899831 | 0 |
| B73 | B106 | ApoA | anti-P | -0.8899831 | 0 |
| B73 | B107 | ApoA | AnuA | -0.889983 | 0 |
| B73 | B108 | ApoA | Anti-Cyclin | -0.8899831 | 0 |
| B73 | B109 | ApoA | AMA-M2 | -0.8899831 | 0 |
| B73 | B110 | ApoA | AHA | -0.8899831 | 0 |
| B73 | B134 | ApoA | CHE | -0.891605 | 0 |
| B73 | B135 | ApoA | TBA | -0.6260967 | 0 |
| B73 | B136 | ApoA | 5'NT | -0.7176583 | 0 |

| | | | | | |
|---|---|---|---|---|---|
| B73 | B137 | ApoA | AFU | 0.8328283 | 0 |
| B73 | B138 | ApoA | PAB | 0.55047737 | 0 |
| B73 | B151 | ApoA | Anti-CCP | -0.8899831 | 0 |
| B73 | B156 | ApoA | IgE | -0.8436641 | 0 |
| B73 | B175 | ApoA | ALB-U | 0.75073721 | 0 |
| B73 | B182 | ApoA | TT | 0.80718772 | 0 |
| B73 | B185 | ApoA | PT | -0.8491549 | 0 |
| B73 | B76 | ApoA | ApoB | 0.6159375 | 0 |
| B73 | B95 | ApoA | anti-cenpb | -0.8899831 | 0 |
| B73 | B97 | ApoA | dsDNA | -0.8899831 | 0 |
| B73 | B98 | ApoA | anti-Jo-1 | -0.8899831 | 0 |
| B73 | B99 | ApoA | anti-PM-Scl | -0.8899831 | 0 |
| B73 | B194 | ApoA | Cl | -0.5217645 | 3.77E-15 |
| B73 | B75 | ApoA | ApoA1/B | -0.4923165 | 2.02E-13 |
| B73 | B183 | ApoA | PTA | -0.4921996 | 2.05E-13 |
| B73 | B186 | ApoA | PFC | 0.48952275 | 2.88E-13 |
| B73 | B157 | ApoA | IgG | -0.4859115 | 4.55E-13 |
| B73 | B177 | ApoA | ACR | 0.47925085 | 1.04E-12 |
| B73 | B195 | ApoA | Na | -0.4707593 | 2.94E-12 |
| B73 | B87 | ApoA | NSE | 0.43291055 | 2.11E-10 |
| B73 | B200 | ApoA | OP | 0.37122074 | 7.86E-08 |
| B73 | B184 | ApoA | INR | 0.36976565 | 8.91E-08 |
| B73 | B155 | ApoA | IgA | -0.3686187 | 9.83E-08 |
| B74 | B75 | ApoA1 | ApoA1/B | 0.74490634 | 0 |
| B74 | B181 | ApoA1 | APTT | 0.50357844 | 4.62E-14 |
| B74 | B186 | ApoA1 | PFC | 0.46733768 | 4.41E-12 |
| B74 | B183 | ApoA1 | PTA | -0.466479 | 4.89E-12 |
| B74 | B184 | ApoA1 | INR | -0.4628202 | 7.52E-12 |
| B74 | B78 | ApoA1 | HBcAb | 0.41408976 | 1.46E-09 |
| B75 | B76 | ApoA1/B | ApoB | -0.8260416 | 0 |
| B75 | B184 | ApoA1/B | INR | -0.4585034 | 1.24E-11 |
| B75 | B181 | ApoA1/B | APTT | 0.43762465 | 1.27E-10 |
| B75 | B185 | ApoA1/B | PT | 0.36842814 | 1.00E-07 |
| B76 | B185 | ApoB | PT | -0.4660818 | 5.12E-12 |
| B76 | B137 | ApoB | AFU | 0.40143306 | 5.03E-09 |
| B76 | B175 | ApoB | ALB-U | 0.38411957 | 2.51E-08 |
| B76 | B184 | ApoB | INR | 0.3803051 | 3.54E-08 |
| B76 | B138 | ApoB | PAB | 0.3647266 | 1.37E-07 |
| B76 | B134 | ApoB | CHE | -0.3634936 | 1.52E-07 |
| B76 | B100 | ApoB | anti-Ro52 | -0.3492309 | 4.91E-07 |
| B76 | B101 | ApoB | anti-Sc70 | -0.3492309 | 4.91E-07 |
| B76 | B102 | ApoB | anti-Sm | -0.3492309 | 4.91E-07 |
| B76 | B103 | ApoB | anti-SSA/Ro | -0.3492309 | 4.91E-07 |
| B76 | B104 | ApoB | anti-SSB | -0.3492309 | 4.91E-07 |

| | | | | | |
|---|---|---|---|---|---|
| B76 | B105 | ApoB | anti-U1RNP | -0.3492309 | 4.91E-07 |
| B76 | B106 | ApoB | anti-P | -0.3492309 | 4.91E-07 |
| B76 | B107 | ApoB | AnuA | -0.3492309 | 4.91E-07 |
| B76 | B108 | ApoB | Anti-Cyclin | -0.3492309 | 4.91E-07 |
| B76 | B109 | ApoB | AMA-M2 | -0.3492309 | 4.91E-07 |
| B76 | B110 | ApoB | AHA | -0.3492309 | 4.91E-07 |
| B76 | B151 | ApoB | Anti-CCP | -0.3492309 | 4.91E-07 |
| B76 | B95 | ApoB | anti-cenpb | -0.3492309 | 4.91E-07 |
| B76 | B97 | ApoB | dsDNA | -0.3492309 | 4.91E-07 |
| B76 | B98 | ApoB | anti-Jo-1 | -0.3492309 | 4.91E-07 |
| B76 | B99 | ApoB | anti-PM-Scl | -0.3492309 | 4.91E-07 |
| B79 | B119 | hb V -dna | Hcv-rna | 0.88148256 | 0 |
| B79 | B121 | hb V -dna | C3 | 0.50128041 | 6.26E-14 |
| B79 | B167 | hb V -dna | PC | 0.47434165 | 1.91E-12 |
| B79 | B152 | hb V -dna | ACA-Ig(G-A-M) | 0.36275617 | 1.62E-07 |
| B8 | B9 | WB-Zinc | RH | -0.5773503 | 0 |
| B8 | B180 | WB-Zinc | Tf | -0.48795 | 3.52E-13 |
| B82 | B115 | CEA | TSH TT3 TT4(TT3) | -0.7338155 | 0 |
| B82 | B89 | CEA | CA 72-4 | -0.4287289 | 3.27E-10 |
| B82 | B86 | CEA | TBhCG | -0.4220244 | 6.55E-10 |
| B82 | B84 | CEA | C-PSA | 0.41647654 | 1.15E-09 |
| B82 | B85 | CEA | SCC | -0.4064426 | 3.10E-09 |
| B82 | B114 | CEA | TSH TT3 TT4(TT4) | -0.3953005 | 8.98E-09 |
| B82 | B113 | CEA | FT3 | -0.3596843 | 2.09E-07 |
| B83 | B138 | AFP | PAB | 0.55583554 | 0 |
| B83 | B196 | AFP | TCO2 | -0.5526982 | 0 |
| B83 | B197 | AFP | AG | 0.64049096 | 0 |
| B83 | B86 | AFP | TBhCG | 0.6010308 | 0 |
| B83 | B139 | AFP | ADA | -0.5402317 | 2.22E-16 |
| B83 | B198 | AFP | P | -0.5154863 | 9.10E-15 |
| B83 | B137 | AFP | AFU | 0.45432937 | 2.00E-11 |
| B83 | B200 | AFP | OP | 0.44525968 | 5.53E-11 |
| B83 | B199 | AFP | Mg | -0.4177161 | 1.01E-09 |
| B83 | B89 | AFP | CA 72-4 | 0.34240149 | 8.44E-07 |
| B84 | B85 | C-PSA | SCC | -0.4956825 | 1.30E-13 |
| B84 | B86 | C-PSA | TBhCG | -0.4285581 | 3.33E-10 |
| B84 | B88 | C-PSA | CA 242 | 0.3409648 | 9.44E-07 |
| B85 | B136 | SCC | 5'NT | 0.55439053 | 0 |
| B85 | B199 | SCC | Mg | 0.52864486 | 1.33E-15 |
| B85 | B200 | SCC | OP | -0.5243132 | 2.66E-15 |
| B85 | B89 | SCC | CA 72-4 | 0.52252943 | 3.55E-15 |
| B85 | B193 | SCC | K | -0.5205941 | 4.44E-15 |
| B85 | B115 | SCC | TSH TT3 TT4(TT3) | 0.47075121 | 2.94E-12 |
| B85 | B138 | SCC | PAB | -0.4596135 | 1.09E-11 |

| | | | | | |
|---|---|---|---|---|---|
| B85 | B86 | SCC | TBhCG | 0.43468873 | 1.74E-10 |
| B85 | B139 | SCC | ADA | 0.3577526 | 2.45E-07 |
| B86 | B89 | TBhCG | CA 72-4 | 0.88280151 | 0 |
| B86 | B115 | TBhCG | TSH TT3 TT4(TT3) | 0.48680264 | 4.07E-13 |
| B86 | B195 | TBhCG | Na | 0.4183629 | 9.50E-10 |
| B86 | B196 | TBhCG | TCO2 | -0.4091453 | 2.38E-09 |
| B86 | B198 | TBhCG | P | -0.4072232 | 2.87E-09 |
| B86 | B194 | TBhCG | Cl | 0.39794082 | 7.01E-09 |
| B86 | B139 | TBhCG | ADA | -0.3531067 | 3.59E-07 |
| B86 | B156 | TBhCG | IgE | 0.34154869 | 9.02E-07 |
| B87 | B135 | NSE | TBA | -0.5260749 | 2.00E-15 |
| B87 | B134 | NSE | CHE | -0.5059075 | 3.38E-14 |
| B87 | B137 | NSE | AFU | 0.49983998 | 7.55E-14 |
| B87 | B100 | NSE | anti-Ro52 | -0.4925773 | 1.95E-13 |
| B87 | B101 | NSE | anti-Sc70 | -0.4925773 | 1.95E-13 |
| B87 | B102 | NSE | anti-Sm | -0.4925773 | 1.95E-13 |
| B87 | B103 | NSE | anti-SSA/Ro | -0.4925773 | 1.95E-13 |
| B87 | B104 | NSE | anti-SSB | -0.4925773 | 1.95E-13 |
| B87 | B105 | NSE | anti-U1RNP | -0.4925773 | 1.95E-13 |
| B87 | B106 | NSE | anti-P | -0.4925773 | 1.95E-13 |
| B87 | B107 | NSE | AnuA | -0.4925773 | 1.95E-13 |
| B87 | B108 | NSE | Anti-Cyclin | -0.4925773 | 1.95E-13 |
| B87 | B109 | NSE | AMA-M2 | -0.4925773 | 1.95E-13 |
| B87 | B110 | NSE | AHA | -0.4925773 | 1.95E-13 |
| B87 | B151 | NSE | Anti-CCP | -0.4925773 | 1.95E-13 |
| B87 | B95 | NSE | anti-cenpb | -0.4925773 | 1.95E-13 |
| B87 | B97 | NSE | dsDNA | -0.4925773 | 1.95E-13 |
| B87 | B98 | NSE | anti-Jo-1 | -0.4925773 | 1.95E-13 |
| B87 | B99 | NSE | anti-PM-Scl | -0.4925773 | 1.95E-13 |
| B87 | B182 | NSE | TT | 0.47565736 | 1.62E-12 |
| B87 | B194 | NSE | Cl | -0.4509043 | 2.95E-11 |
| B87 | B156 | NSE | IgE | -0.4183922 | 9.47E-10 |
| B87 | B195 | NSE | Na | -0.4155717 | 1.26E-09 |
| B87 | B175 | NSE | ALB-U | 0.41118766 | 1.95E-09 |
| B87 | B185 | NSE | PT | -0.4092657 | 2.35E-09 |
| B87 | B183 | NSE | PTA | -0.3499233 | 4.65E-07 |
| B87 | B186 | NSE | PFC | 0.34871213 | 5.12E-07 |
| B87 | B88 | NSE | CA 242 | 0.34247015 | 8.39E-07 |
| B88 | B138 | CA 242 | PAB | 0.36376613 | 1.49E-07 |
| B88 | B197 | CA 242 | AG | 0.35612414 | 2.81E-07 |
| B89 | B115 | CA 72-4 | TSH TT3 TT4(TT3) | 0.50779814 | 2.62E-14 |
| B89 | B194 | CA 72-4 | Cl | 0.3553629 | 2.99E-07 |
| B89 | B195 | CA 72-4 | Na | 0.3549576 | 3.09E-07 |
| B9 | B117 | RH | TT4 | -0.5685352 | 0 |

| | | | | | |
|---|---|---|---|---|---|
| B9 | B159 | RH | β2-MG | -0.7288109 | 0 |
| B9 | B179 | RH | α1-MG | -0.6117753 | 0 |
| B9 | B31 | RH | FSH | -0.5753965 | 0 |
| B9 | B178 | RH | Ig-U | -0.4811252 | 8.29E-13 |
| B9 | B29 | RH | E2 | -0.3698001 | 8.88E-08 |
| B90 | B187 | CYFRA 21-1 | NSE | -0.3683835 | 1.00E-07 |
| B91 | B130 | TP | GLB | 0.75083538 | 0 |
| B91 | B127 | TP | A-G ratio | -0.4677821 | 4.19E-12 |
| B91 | B126 | TP | Alb | 0.37398769 | 6.18E-08 |
| B91 | B188 | TP | Ca | 0.35068708 | 4.37E-07 |
| B92 | B93 | ALDH2-Glu504Glu | ALDH2-Glu504Lys | -1 | 0 |
| B92 | B94 | ALDH2-Glu504Glu | ALDH2-Lys504Lys | -1 | 0 |
| B92 | B209 | ALDH2-Glu504Glu | Smoking | 0.46333027 | 7.08E-12 |
| B92 | B161 | ALDH2-Glu504Glu | LEU | 0.35775145 | 2.45E-07 |
| B93 | B94 | ALDH2-Glu504Lys | ALDH2-Lys504Lys | 1 | 0 |
| B93 | B209 | ALDH2-Glu504Lys | Smoking | -0.4633303 | 7.08E-12 |
| B93 | B161 | ALDH2-Glu504Lys | LEU | -0.3577514 | 2.45E-07 |
| B94 | B209 | ALDH2-Lys504Lys | Smoking | -0.4633303 | 7.08E-12 |
| B94 | B161 | ALDH2-Lys504Lys | LEU | -0.3577514 | 2.45E-07 |
| B95 | B101 | anti-cenpb | anti-Sc70 | 0.99999999 | 0 |
| B95 | B106 | anti-cenpb | anti-P | 1 | 0 |
| B95 | B107 | anti-cenpb | AnuA | 0.99999995 | 0 |
| B95 | B134 | anti-cenpb | CHE | 0.99587065 | 0 |
| B95 | B135 | anti-cenpb | TBA | 0.72594937 | 0 |
| B95 | B136 | anti-cenpb | 5'NT | 0.77267967 | 0 |
| B95 | B137 | anti-cenpb | AFU | -0.8999613 | 0 |
| B95 | B156 | anti-cenpb | IgE | 0.94663187 | 0 |
| B95 | B175 | anti-cenpb | ALB-U | -0.8425779 | 0 |
| B95 | B177 | anti-cenpb | ACR | -0.5594521 | 0 |
| B95 | B182 | anti-cenpb | TT | -0.9646716 | 0 |
| B95 | B183 | anti-cenpb | PTA | 0.70776873 | 0 |
| B95 | B185 | anti-cenpb | PT | 0.83292135 | 0 |
| B95 | B186 | anti-cenpb | PFC | -0.7053006 | 0 |
| B95 | B194 | anti-cenpb | Cl | 0.65638677 | 0 |
| B95 | B195 | anti-cenpb | Na | 0.60796773 | 0 |
| B95 | B98 | anti-cenpb | anti-Jo-1 | 1 | 0 |
| B95 | B157 | anti-cenpb | IgG | 0.54290776 | 2.22E-16 |
| B95 | B138 | anti-cenpb | PAB | -0.5256581 | 2.22E-15 |
| B95 | B155 | anti-cenpb | IgA | 0.4155488 | 1.26E-09 |
| B95 | B154 | anti-cenpb | RF | 0.35447914 | 3.21E-07 |
| B96 | B152 | dsDNA (IIF) | ACA-Ig(G-A-M) | 0.77354494 | 0 |
| B96 | B167 | dsDNA (IIF) | PC | 0.90267093 | 0 |
| B97 | B100 | dsDNA | anti-Ro52 | 1 | 0 |
| B97 | B101 | dsDNA | anti-Sc70 | 1 | 0 |

| | | | | | |
|---|---|---|---|---|---|
| B97 | B103 | dsDNA | anti-SSA/Ro | 1 | 0 |
| B97 | B106 | dsDNA | anti-P | 1 | 0 |
| B97 | B107 | dsDNA | AnuA | 0.99999999 | 0 |
| B97 | B108 | dsDNA | Anti-Cyclin | 1 | 0 |
| B97 | B109 | dsDNA | AMA-M2 | 1 | 0 |
| B97 | B134 | dsDNA | CHE | 0.99587065 | 0 |
| B97 | B135 | dsDNA | TBA | 0.72594937 | 0 |
| B97 | B136 | dsDNA | 5'NT | 0.77267967 | 0 |
| B97 | B137 | dsDNA | AFU | -0.8999613 | 0 |
| B97 | B151 | dsDNA | Anti-CCP | 1 | 0 |
| B97 | B156 | dsDNA | IgE | 0.94663186 | 0 |
| B97 | B175 | dsDNA | ALB-U | -0.8425779 | 0 |
| B97 | B177 | dsDNA | ACR | -0.5594521 | 0 |
| B97 | B182 | dsDNA | TT | -0.9646716 | 0 |
| B97 | B183 | dsDNA | PTA | 0.70776872 | 0 |
| B97 | B185 | dsDNA | PT | 0.83292135 | 0 |
| B97 | B186 | dsDNA | PFC | -0.7053006 | 0 |
| B97 | B194 | dsDNA | Cl | 0.65638677 | 0 |
| B97 | B195 | dsDNA | Na | 0.60796773 | 0 |
| B97 | B98 | dsDNA | anti-Jo-1 | 1 | 0 |
| B97 | B157 | dsDNA | IgG | 0.54290776 | 2.22E-16 |
| B97 | B138 | dsDNA | PAB | -0.5256581 | 2.22E-15 |
| B97 | B155 | dsDNA | IgA | 0.4155488 | 1.26E-09 |
| B97 | B154 | dsDNA | RF | 0.35447914 | 3.21E-07 |
| B98 | B100 | anti-Jo-1 | anti-Ro52 | 1 | 0 |
| B98 | B101 | anti-Jo-1 | anti-Sc70 | 1 | 0 |
| B98 | B102 | anti-Jo-1 | anti-Sm | 1 | 0 |
| B98 | B103 | anti-Jo-1 | anti-SSA/Ro | 1 | 0 |
| B98 | B104 | anti-Jo-1 | anti-SSB | 1 | 0 |
| B98 | B105 | anti-Jo-1 | anti-U1RNP | 1 | 0 |
| B98 | B106 | anti-Jo-1 | anti-P | 1 | 0 |
| B98 | B107 | anti-Jo-1 | AnuA | 0.99999999 | 0 |
| B98 | B108 | anti-Jo-1 | Anti-Cyclin | 1 | 0 |
| B98 | B109 | anti-Jo-1 | AMA-M2 | 1 | 0 |
| B98 | B110 | anti-Jo-1 | AHA | 1 | 0 |
| B98 | B134 | anti-Jo-1 | CHE | 0.99587065 | 0 |
| B98 | B135 | anti-Jo-1 | TBA | 0.72594937 | 0 |
| B98 | B136 | anti-Jo-1 | 5'NT | 0.77267967 | 0 |
| B98 | B137 | anti-Jo-1 | AFU | -0.8999613 | 0 |
| B98 | B151 | anti-Jo-1 | Anti-CCP | 1 | 0 |
| B98 | B156 | anti-Jo-1 | IgE | 0.94663186 | 0 |
| B98 | B175 | anti-Jo-1 | ALB-U | -0.8425779 | 0 |
| B98 | B177 | anti-Jo-1 | ACR | -0.5594521 | 0 |
| B98 | B182 | anti-Jo-1 | TT | -0.9646716 | 0 |

| | | | | | |
|---|---|---|---|---|---|
| B98 | B183 | anti-Jo-1 | PTA | 0.70776872 | 0 |
| B98 | B185 | anti-Jo-1 | PT | 0.83292135 | 0 |
| B98 | B186 | anti-Jo-1 | PFC | -0.7053006 | 0 |
| B98 | B194 | anti-Jo-1 | Cl | 0.65638676 | 0 |
| B98 | B195 | anti-Jo-1 | Na | 0.60796772 | 0 |
| B98 | B99 | anti-Jo-1 | anti-PM-Scl | 1 | 0 |
| B98 | B157 | anti-Jo-1 | IgG | 0.54290776 | 2.22E-16 |
| B98 | B138 | anti-Jo-1 | PAB | -0.5256581 | 2.22E-15 |
| B98 | B155 | anti-Jo-1 | IgA | 0.4155488 | 1.26E-09 |
| B98 | B154 | anti-Jo-1 | RF | 0.35447914 | 3.21E-07 |
| B99 | B100 | anti-PM-Scl | anti-Ro52 | 1 | 0 |
| B99 | B103 | anti-PM-Scl | anti-SSA/Ro | 1 | 0 |
| B99 | B107 | anti-PM-Scl | AnuA | 0.99999996 | 0 |
| B99 | B108 | anti-PM-Scl | Anti-Cyclin | 1 | 0 |
| B99 | B110 | anti-PM-Scl | AHA | 1 | 0 |
| B99 | B134 | anti-PM-Scl | CHE | 0.99587065 | 0 |
| B99 | B135 | anti-PM-Scl | TBA | 0.72594937 | 0 |
| B99 | B136 | anti-PM-Scl | 5'NT | 0.77267967 | 0 |
| B99 | B137 | anti-PM-Scl | AFU | -0.8999613 | 0 |
| B99 | B151 | anti-PM-Scl | Anti-CCP | 1 | 0 |
| B99 | B156 | anti-PM-Scl | IgE | 0.94663186 | 0 |
| B99 | B175 | anti-PM-Scl | ALB-U | -0.8425779 | 0 |
| B99 | B177 | anti-PM-Scl | ACR | -0.5594521 | 0 |
| B99 | B182 | anti-PM-Scl | TT | -0.9646716 | 0 |
| B99 | B183 | anti-PM-Scl | PTA | 0.70776872 | 0 |
| B99 | B185 | anti-PM-Scl | PT | 0.83292135 | 0 |
| B99 | B186 | anti-PM-Scl | PFC | -0.7053006 | 0 |
| B99 | B194 | anti-PM-Scl | Cl | 0.65638677 | 0 |
| B99 | B195 | anti-PM-Scl | Na | 0.60796773 | 0 |
| B99 | B157 | anti-PM-Scl | IgG | 0.54290776 | 2.22E-16 |
| B99 | B138 | anti-PM-Scl | PAB | -0.5256581 | 2.22E-15 |
| B99 | B155 | anti-PM-Scl | IgA | 0.4155488 | 1.26E-09 |
| B99 | B154 | anti-PM-Scl | RF | 0.35447914 | 3.21E-07 |

**Supplementary Table 20. PEI Correlations in Stomach trouble**

| Row | Column | Row | Column | Cor | *P* |
|---|---|---|---|---|---|
| B1 | B141 | Gender | AST/ALT | 0.288281095 | 0 |
| B1 | B163 | Gender | SG | -0.450465071 | 0 |
| B1 | B174 | Gender | Urea/Cr | 0.405477266 | 0 |
| B1 | B190 | Gender | Cre | -0.668176029 | 0 |
| B1 | B192 | Gender | UA | -0.51432267 | 0 |
| B1 | B204 | Gender | Drinking | -0.264402556 | 0 |
| B1 | B206 | Gender | Waist | -0.420316169 | 0 |
| B1 | B207 | Gender | WHR | -0.386479818 | 0 |
| B1 | B209 | Gender | Smoking | -0.340681016 | 0 |
| B1 | B210 | Gender | Hip | -0.283929011 | 0 |
| B1 | B216 | Gender | BMI | -0.547444353 | 0 |
| B1 | B221 | Gender | Hei | -0.693211922 | 0 |
| B1 | B222 | Gender | NS | -0.714435561 | 0 |
| B1 | B42 | Gender | MON | -0.255943168 | 0 |
| B1 | B45 | Gender | MCHC | -0.282533449 | 0 |
| B1 | B46 | Gender | RBC Count | -0.575045329 | 0 |
| B1 | B51 | Gender | Hb | -0.64780062 | 0 |
| B1 | B72 | Gender | HDL-C | 0.380275552 | 0 |
| B1 | B74 | Gender | ApoA1 | 0.226720748 | 0 |
| B1 | B161 | Gender | LEU | 0.212026257 | 1.22E-14 |
| B1 | B58 | Gender | HCT | -0.208947095 | 2.98E-14 |
| B1 | B215 | Gender | BMI | -0.203484251 | 1.40E-13 |
| B1 | B128 | Gender | ALT | -0.19565865 | 1.20E-12 |
| B1 | B71 | Gender | TG | -0.192110862 | 3.07E-12 |
| B1 | B138 | Gender | PAB | -0.183889906 | 2.55E-11 |
| B1 | B62 | Gender | WBHSRVI | -0.179912356 | 6.85E-11 |
| B1 | B127 | Gender | A-G ratio | -0.178403723 | 9.91E-11 |
| B1 | B176 | Gender | Ucr | -0.17518921 | 2.15E-10 |
| B1 | B64 | Gender | WBRSR(mPas_5) | -0.173692082 | 3.08E-10 |
| B1 | B63 | Gender | WBRSR(mPas_200) | -0.173190146 | 3.47E-10 |
| B1 | B67 | Gender | WBRSR(mPas_50) | -0.17285503 | 3.75E-10 |
| B1 | B66 | Gender | WBRSR(mPas_100) | -0.170093079 | 7.17E-10 |
| B1 | B3 | Gender | Hcy | -0.168737241 | 9.80E-10 |
| B1 | B40 | Gender | WBC | -0.167281943 | 1.37E-09 |
| B1 | B218 | Gender | DBP | -0.163327195 | 3.34E-09 |
| B1 | B41 | Gender | MON％ | -0.161447156 | 5.07E-09 |
| B1 | B60 | Gender | WBRVI | -0.161023584 | 5.56E-09 |
| B1 | B140 | Gender | r-GT | -0.160628306 | 6.07E-09 |
| B1 | B166 | Gender | Pro | -0.158841054 | 8.96E-09 |
| B1 | B130 | Gender | GLB | 0.156483384 | 1.49E-08 |
| B1 | B65 | Gender | WBRSR(mPas_1) | -0.155038588 | 2.02E-08 |
| B10 | B12 | PG Ⅰ | PG II | 0.713996542 | 0 |

| | | | | | |
|---|---|---|---|---|---|
| B10 | B14 | PG Ⅰ | G-17 | 0.405899693 | 0 |
| B10 | B111 | PG Ⅰ | TSH | 0.181441456 | 4.70E-11 |
| B10 | B124 | PG Ⅰ | ACTH | -0.165130772 | 2.23E-09 |
| B10 | B106 | PG Ⅰ | anti-P | 0.149366692 | 6.58E-08 |
| B10 | B158 | PG Ⅰ | IgM | 0.146370901 | 1.20E-07 |
| B10 | B135 | PG Ⅰ | TBA | 0.144615475 | 1.71E-07 |
| B100 | B151 | anti-Ro52 | Anti-CCP | -0.181694243 | 4.41E-11 |
| B100 | B185 | anti-Ro52 | PT | -0.176872259 | 1.44E-10 |
| B100 | B186 | anti-Ro52 | PFC | 0.144570335 | 1.72E-07 |
| B101 | B104 | anti-Sc70 | anti-SSB | 0.999999997 | 0 |
| B101 | B107 | anti-Sc70 | AnuA | 0.999999794 | 0 |
| B101 | B151 | anti-Sc70 | Anti-CCP | 0.245131194 | 0 |
| B101 | B154 | anti-Sc70 | RF | 0.408504393 | 0 |
| B101 | B183 | anti-Sc70 | PTA | 0.262201702 | 0 |
| B101 | B185 | anti-Sc70 | PT | 0.238563374 | 0 |
| B101 | B186 | anti-Sc70 | PFC | 0.238316949 | 0 |
| B101 | B184 | anti-Sc70 | INR | 0.211610709 | 1.38E-14 |
| B101 | B106 | anti-Sc70 | anti-P | -0.180934714 | 5.32E-11 |
| B101 | B180 | anti-Sc70 | Tf | 0.175916569 | 1.81E-10 |
| B101 | B178 | anti-Sc70 | Ig-U | 0.167849824 | 1.20E-09 |
| B101 | B156 | anti-Sc70 | IgE | 0.166233311 | 1.74E-09 |
| B101 | B120 | anti-Sc70 | HCV-cAg | 0.162135295 | 4.35E-09 |
| B101 | B175 | anti-Sc70 | ALB-U | 0.155247957 | 1.94E-08 |
| B102 | B103 | anti-Sm | anti-SSA/Ro | 0.999999998 | 0 |
| B102 | B104 | anti-Sm | anti-SSB | 0.999999997 | 0 |
| B102 | B105 | anti-Sm | anti-U1RNP | 0.999999998 | 0 |
| B102 | B107 | anti-Sm | AnuA | 0.999999848 | 0 |
| B102 | B108 | anti-Sm | Anti-Cyclin | 0.999999997 | 0 |
| B102 | B109 | anti-Sm | AMA-M2 | 0.999999999 | 0 |
| B102 | B110 | anti-Sm | AHA | 0.999999997 | 0 |
| B102 | B151 | anti-Sm | Anti-CCP | 0.245131192 | 0 |
| B102 | B154 | anti-Sm | RF | 0.408504389 | 0 |
| B102 | B183 | anti-Sm | PTA | 0.2622017 | 0 |
| B102 | B185 | anti-Sm | PT | 0.238563374 | 0 |
| B102 | B186 | anti-Sm | PFC | 0.238316947 | 0 |
| B102 | B184 | anti-Sm | INR | 0.211610709 | 1.38E-14 |
| B102 | B106 | anti-Sm | anti-P | -0.180934712 | 5.32E-11 |
| B102 | B180 | anti-Sm | Tf | 0.175916567 | 1.81E-10 |
| B102 | B178 | anti-Sm | Ig-U | 0.167849823 | 1.20E-09 |
| B102 | B156 | anti-Sm | IgE | 0.16623331 | 1.74E-09 |
| B102 | B120 | anti-Sm | HCV-cAg | 0.16213538 | 4.35E-09 |
| B102 | B175 | anti-Sm | ALB-U | 0.155247955 | 1.94E-08 |
| B103 | B104 | anti-SSA/Ro | anti-SSB | 1 | 0 |
| B103 | B107 | anti-SSA/Ro | AnuA | 0.999999785 | 0 |

| | | | | | |
|---|---|---|---|---|---|
| B103 | B151 | anti-SSA/Ro | Anti-CCP | 0.245131193 | 0 |
| B103 | B154 | anti-SSA/Ro | RF | 0.40850439 | 0 |
| B103 | B183 | anti-SSA/Ro | PTA | 0.262201701 | 0 |
| B103 | B185 | anti-SSA/Ro | PT | 0.238563373 | 0 |
| B103 | B186 | anti-SSA/Ro | PFC | 0.238316947 | 0 |
| B103 | B184 | anti-SSA/Ro | INR | 0.211610708 | 1.38E-14 |
| B103 | B106 | anti-SSA/Ro | anti-P | -0.180934713 | 5.32E-11 |
| B103 | B180 | anti-SSA/Ro | Tf | 0.175916568 | 1.81E-10 |
| B103 | B178 | anti-SSA/Ro | Ig-U | 0.167849823 | 1.20E-09 |
| B103 | B156 | anti-SSA/Ro | IgE | 0.16623331 | 1.74E-09 |
| B103 | B120 | anti-SSA/Ro | HCV-cAg | 0.162135349 | 4.35E-09 |
| B103 | B175 | anti-SSA/Ro | ALB-U | 0.155247956 | 1.94E-08 |
| B104 | B107 | anti-SSB | AnuA | 0.999999807 | 0 |
| B104 | B108 | anti-SSB | Anti-Cyclin | 0.999999999 | 0 |
| B104 | B110 | anti-SSB | AHA | 0.999999998 | 0 |
| B104 | B151 | anti-SSB | Anti-CCP | 0.245131193 | 0 |
| B104 | B154 | anti-SSB | RF | 0.40850439 | 0 |
| B104 | B183 | anti-SSB | PTA | 0.262201701 | 0 |
| B104 | B185 | anti-SSB | PT | 0.238563375 | 0 |
| B104 | B186 | anti-SSB | PFC | 0.238316948 | 0 |
| B104 | B184 | anti-SSB | INR | 0.211610708 | 1.38E-14 |
| B104 | B106 | anti-SSB | anti-P | -0.180934713 | 5.32E-11 |
| B104 | B180 | anti-SSB | Tf | 0.175916568 | 1.81E-10 |
| B104 | B178 | anti-SSB | Ig-U | 0.167849823 | 1.20E-09 |
| B104 | B156 | anti-SSB | IgE | 0.16623331 | 1.74E-09 |
| B104 | B120 | anti-SSB | HCV-cAg | 0.162135317 | 4.35E-09 |
| B104 | B175 | anti-SSB | ALB-U | 0.155247956 | 1.94E-08 |
| B105 | B107 | anti-U1RNP | AnuA | 0.999999778 | 0 |
| B105 | B108 | anti-U1RNP | Anti-Cyclin | 1 | 0 |
| B105 | B151 | anti-U1RNP | Anti-CCP | 0.245131193 | 0 |
| B105 | B154 | anti-U1RNP | RF | 0.40850439 | 0 |
| B105 | B183 | anti-U1RNP | PTA | 0.262201701 | 0 |
| B105 | B185 | anti-U1RNP | PT | 0.238563374 | 0 |
| B105 | B186 | anti-U1RNP | PFC | 0.238316948 | 0 |
| B105 | B184 | anti-U1RNP | INR | 0.211610709 | 1.38E-14 |
| B105 | B106 | anti-U1RNP | anti-P | -0.180934713 | 5.32E-11 |
| B105 | B180 | anti-U1RNP | Tf | 0.175916568 | 1.81E-10 |
| B105 | B178 | anti-U1RNP | Ig-U | 0.167849823 | 1.20E-09 |
| B105 | B156 | anti-U1RNP | IgE | 0.16623331 | 1.74E-09 |
| B105 | B120 | anti-U1RNP | HCV-cAg | 0.162135343 | 4.35E-09 |
| B105 | B175 | anti-U1RNP | ALB-U | 0.155247956 | 1.94E-08 |
| B106 | B121 | anti-P | C3 | 0.230409022 | 0 |
| B106 | B122 | anti-P | C4 | 0.230409091 | 0 |
| B106 | B159 | anti-P | β2-MG | 0.230408802 | 0 |

| | | | | | |
|---|---|---|---|---|---|
| B106 | B169 | anti-P | U-CRY | 0.230409274 | 0 |
| B106 | B119 | anti-P | Hcv-rna | -0.208066902 | 3.84E-14 |
| B106 | B107 | anti-P | AnuA | -0.180934669 | 5.32E-11 |
| B106 | B108 | anti-P | Anti-Cyclin | -0.180934713 | 5.32E-11 |
| B106 | B109 | anti-P | AMA-M2 | -0.180934713 | 5.32E-11 |
| B106 | B110 | anti-P | AHA | -0.180934713 | 5.32E-11 |
| B106 | B197 | anti-P | AG | -0.14758207 | 9.45E-08 |
| B107 | B108 | AnuA | Anti-Cyclin | 0.999999772 | 0 |
| B107 | B109 | AnuA | AMA-M2 | 0.999999789 | 0 |
| B107 | B110 | AnuA | AHA | 0.999999797 | 0 |
| B107 | B151 | AnuA | Anti-CCP | 0.245131139 | 0 |
| B107 | B154 | AnuA | RF | 0.408504302 | 0 |
| B107 | B183 | AnuA | PTA | 0.262201644 | 0 |
| B107 | B185 | AnuA | PT | 0.238563341 | 0 |
| B107 | B186 | AnuA | PFC | 0.238316901 | 0 |
| B107 | B184 | AnuA | INR | 0.211610669 | 1.38E-14 |
| B107 | B180 | AnuA | Tf | 0.175916531 | 1.81E-10 |
| B107 | B178 | AnuA | Ig-U | 0.167849787 | 1.20E-09 |
| B107 | B156 | AnuA | IgE | 0.166233274 | 1.74E-09 |
| B107 | B120 | AnuA | HCV-cAg | 0.162135608 | 4.35E-09 |
| B107 | B175 | AnuA | ALB-U | 0.155247922 | 1.94E-08 |
| B108 | B151 | Anti-Cyclin | Anti-CCP | 0.245131193 | 0 |
| B108 | B154 | Anti-Cyclin | RF | 0.408504391 | 0 |
| B108 | B183 | Anti-Cyclin | PTA | 0.262201701 | 0 |
| B108 | B185 | Anti-Cyclin | PT | 0.238563373 | 0 |
| B108 | B186 | Anti-Cyclin | PFC | 0.238316948 | 0 |
| B108 | B184 | Anti-Cyclin | INR | 0.211610707 | 1.38E-14 |
| B108 | B180 | Anti-Cyclin | Tf | 0.175916568 | 1.81E-10 |
| B108 | B178 | Anti-Cyclin | Ig-U | 0.167849823 | 1.20E-09 |
| B108 | B156 | Anti-Cyclin | IgE | 0.16623331 | 1.74E-09 |
| B108 | B120 | Anti-Cyclin | HCV-cAg | 0.162135334 | 4.35E-09 |
| B108 | B175 | Anti-Cyclin | ALB-U | 0.155247956 | 1.94E-08 |
| B109 | B151 | AMA-M2 | Anti-CCP | 0.245131193 | 0 |
| B109 | B154 | AMA-M2 | RF | 0.40850439 | 0 |
| B109 | B183 | AMA-M2 | PTA | 0.262201701 | 0 |
| B109 | B185 | AMA-M2 | PT | 0.238563374 | 0 |
| B109 | B186 | AMA-M2 | PFC | 0.238316948 | 0 |
| B109 | B184 | AMA-M2 | INR | 0.211610708 | 1.38E-14 |
| B109 | B180 | AMA-M2 | Tf | 0.175916568 | 1.81E-10 |
| B109 | B178 | AMA-M2 | Ig-U | 0.167849823 | 1.20E-09 |
| B109 | B156 | AMA-M2 | IgE | 0.16623331 | 1.74E-09 |
| B109 | B120 | AMA-M2 | HCV-cAg | 0.162135357 | 4.35E-09 |
| B109 | B175 | AMA-M2 | ALB-U | 0.155247956 | 1.94E-08 |
| B11 | B12 | PGI/PGII | PG II | -0.462402177 | 0 |

| | | | | | |
|---|---|---|---|---|---|
| B11 | B14 | PGI/PGII | G-17 | -0.230440213 | 0 |
| B110 | B151 | AHA | Anti-CCP | 0.245131193 | 0 |
| B110 | B154 | AHA | RF | 0.40850439 | 0 |
| B110 | B183 | AHA | PTA | 0.262201701 | 0 |
| B110 | B185 | AHA | PT | 0.238563374 | 0 |
| B110 | B186 | AHA | PFC | 0.238316948 | 0 |
| B110 | B184 | AHA | INR | 0.211610709 | 1.38E-14 |
| B110 | B180 | AHA | Tf | 0.175916568 | 1.81E-10 |
| B110 | B178 | AHA | Ig-U | 0.167849823 | 1.20E-09 |
| B110 | B156 | AHA | IgE | 0.16623331 | 1.74E-09 |
| B110 | B120 | AHA | HCV-cAg | 0.162135336 | 4.35E-09 |
| B110 | B175 | AHA | ALB-U | 0.155247956 | 1.94E-08 |
| B111 | B124 | TSH | ACTH | -0.338994548 | 0 |
| B111 | B149 | TSH | TGAb | 0.275920931 | 0 |
| B111 | B182 | TSH | TT | 0.244233031 | 0 |
| B111 | B184 | TSH | INR | -0.221572913 | 6.66E-16 |
| B111 | B112 | TSH | FT4 | -0.207951086 | 3.97E-14 |
| B111 | B185 | TSH | PT | -0.191377893 | 3.72E-12 |
| B111 | B186 | TSH | PFC | 0.175457112 | 2.02E-10 |
| B111 | B115 | TSH | TSH TT3 TT4(TT3) | -0.168868554 | 9.51E-10 |
| B111 | B158 | TSH | IgM | 0.161573223 | 4.93E-09 |
| B111 | B197 | TSH | AG | 0.140398708 | 3.88E-07 |
| B111 | B181 | TSH | APTT | -0.137152489 | 7.19E-07 |
| B111 | B194 | TSH | Cl | -0.135909937 | 9.07E-07 |
| B112 | B113 | FT4 | FT3 | 0.228394037 | 0 |
| B112 | B114 | FT4 | TSH TT3 TT4(TT4) | 0.424266068 | 0 |
| B112 | B115 | FT4 | TSH TT3 TT4(TT3) | 0.197004455 | 8.33E-13 |
| B113 | B115 | FT3 | TSH TT3 TT4(TT3) | 0.434349407 | 0 |
| B113 | B126 | FT3 | Alb | 0.172937139 | 3.68E-10 |
| B113 | B114 | FT3 | TSH TT3 TT4(TT4) | 0.14271622 | 2.48E-07 |
| B114 | B115 | TSH TT3 TT4(TT4) | TSH TT3 TT4(TT3) | 0.41521003 | 0 |
| B114 | B197 | TSH TT3 TT4(TT4) | AG | 0.173448597 | 3.26E-10 |
| B114 | B196 | TSH TT3 TT4(TT4) | TCO2 | -0.136021987 | 8.88E-07 |
| B115 | B182 | TSH TT3 TT4(TT3) | TT | -0.240896422 | 0 |
| B115 | B124 | TSH TT3 TT4(TT3) | ACTH | 0.182457292 | 3.65E-11 |
| B115 | B197 | TSH TT3 TT4(TT3) | AG | 0.1716552 | 4.98E-10 |
| B115 | B187 | TSH TT3 TT4(TT3) | NSE | 0.143309748 | 2.21E-07 |
| B115 | B134 | TSH TT3 TT4(TT3) | CHE | 0.141292054 | 3.27E-07 |
| B116 | B167 | TT3 | PC | 0.388432439 | 0 |
| B116 | B202 | TT3 | e-times | 0.727058904 | 0 |
| B117 | B202 | TT4 | e-times | 0.823378176 | 0 |
| B118 | B145 | CEA1 | BGP | 0.202275952 | 1.97E-13 |
| B118 | B143 | CEA1 | β-CTx | 0.174951708 | 2.28E-10 |
| B118 | B142 | CEA1 | 25-(OH)D | -0.158511814 | 9.62E-09 |

| | | | | | |
|---|---|---|---|---|---|
| B119 | B120 | Hcv-rna | HCV-cAg | 0.457134822 | 0 |
| B119 | B121 | Hcv-rna | C3 | -0.903827141 | 0 |
| B119 | B122 | Hcv-rna | C4 | -0.903827389 | 0 |
| B119 | B159 | Hcv-rna | β2-MG | -0.903826308 | 0 |
| B119 | B169 | Hcv-rna | U-CRY | -0.903828091 | 0 |
| B12 | B14 | PG II | G-17 | 0.590228123 | 0 |
| B12 | B106 | PG II | anti-P | 0.158087179 | 1.05E-08 |
| B12 | B179 | PG II | α1-MG | 0.135577153 | 9.65E-07 |
| B120 | B121 | HCV-cAg | C3 | -0.413172874 | 0 |
| B120 | B122 | HCV-cAg | C4 | -0.413172914 | 0 |
| B120 | B159 | HCV-cAg | β2-MG | -0.413173515 | 0 |
| B120 | B169 | HCV-cAg | U-CRY | -0.41317328 | 0 |
| B120 | B154 | HCV-cAg | RF | 0.182885646 | 3.28E-11 |
| B120 | B157 | HCV-cAg | IgG | -0.179845414 | 6.96E-11 |
| B121 | B159 | C3 | β2-MG | 0.999999588 | 0 |
| B121 | B136 | C3 | 5'NT | 0.149009817 | 7.07E-08 |
| B122 | B136 | C4 | 5'NT | 0.149009857 | 7.07E-08 |
| B124 | B125 | ACTH | Cor | 0.499102959 | 0 |
| B124 | B182 | ACTH | TT | -0.382877227 | 0 |
| B124 | B187 | ACTH | NSE | 0.474656038 | 0 |
| B124 | B158 | ACTH | IgM | -0.194200416 | 1.77E-12 |
| B124 | B194 | ACTH | Cl | 0.142027482 | 2.83E-07 |
| B125 | B181 | Cor | APTT | -0.250463799 | 0 |
| B125 | B184 | Cor | INR | -0.277959219 | 0 |
| B125 | B185 | Cor | PT | -0.229989722 | 0 |
| B125 | B187 | Cor | NSE | 0.219026543 | 1.55E-15 |
| B125 | B182 | Cor | TT | -0.183324604 | 2.94E-11 |
| B126 | B127 | Alb | A-G ratio | 0.493533119 | 0 |
| B126 | B188 | Alb | Ca | 0.435028811 | 0 |
| B126 | B189 | Alb | eGFB | 0.236991149 | 0 |
| B126 | B194 | Alb | Cl | -0.15840501 | 9.85E-09 |
| B126 | B221 | Alb | Hei | 0.156317207 | 1.54E-08 |
| B126 | B219 | Alb | OT | -0.137258153 | 7.05E-07 |
| B126 | B130 | Alb | GLB | -0.135619908 | 9.57E-07 |
| B127 | B130 | A-G ratio | GLB | -0.902708025 | 0 |
| B127 | B221 | A-G ratio | Hei | 0.171047922 | 5.74E-10 |
| B127 | B157 | A-G ratio | IgG | -0.14766863 | 9.28E-08 |
| B127 | B155 | A-G ratio | IgA | -0.14273869 | 2.47E-07 |
| B128 | B131 | ALT | AST | 0.826263925 | 0 |
| B128 | B140 | ALT | r-GT | 0.699101052 | 0 |
| B128 | B141 | ALT | AST/ALT | -0.452187711 | 0 |
| B128 | B192 | ALT | UA | 0.231941563 | 0 |
| B128 | B206 | ALT | Waist | 0.22833966 | 0 |
| B128 | B215 | ALT | BMI | 0.229248355 | 0 |

| | | | | | |
|---|---|---|---|---|---|
| B128 | B216 | ALT | BMI | 0.272635403 | 0 |
| B128 | B222 | ALT | NS | 0.257391731 | 0 |
| B128 | B136 | ALT | 5'NT | 0.21862218 | 1.78E-15 |
| B128 | B210 | ALT | Hip | 0.197404441 | 7.48E-13 |
| B128 | B207 | ALT | WHR | 0.173507891 | 3.22E-10 |
| B128 | B132 | ALT | Dbil | 0.169601728 | 8.03E-10 |
| B128 | B221 | ALT | Hei | 0.162565115 | 3.96E-09 |
| B128 | B163 | ALT | SG | 0.141951516 | 2.88E-07 |
| B129 | B132 | IBIL | Dbil | 0.739252044 | 0 |
| B129 | B133 | IBIL | TBIL | 0.984775832 | 0 |
| B130 | B155 | GLB | IgA | 0.153924068 | 2.56E-08 |
| B130 | B157 | GLB | IgG | 0.149647878 | 6.21E-08 |
| B130 | B221 | GLB | Hei | -0.137424551 | 6.83E-07 |
| B131 | B140 | AST | r-GT | 0.62403962 | 0 |
| B131 | B132 | AST | Dbil | 0.15387167 | 2.59E-08 |
| B131 | B192 | AST | UA | 0.139070446 | 5.00E-07 |
| B131 | B222 | AST | NS | 0.137732227 | 6.45E-07 |
| B131 | B136 | AST | 5'NT | 0.137282918 | 7.02E-07 |
| B132 | B133 | Dbil | TBIL | 0.844976861 | 0 |
| B132 | B152 | Dbil | ACA-Ig(G-A-M) | 0.142126881 | 2.78E-07 |
| B132 | B215 | Dbil | BMI | -0.141092585 | 3.40E-07 |
| B132 | B140 | Dbil | r-GT | 0.138443492 | 5.64E-07 |
| B134 | B138 | CHE | PAB | 0.428592745 | 0 |
| B134 | B194 | CHE | Cl | -0.256981747 | 0 |
| B134 | B137 | CHE | AFU | 0.21377627 | 7.33E-15 |
| B134 | B197 | CHE | AG | 0.182741347 | 3.40E-11 |
| B134 | B181 | CHE | APTT | -0.169563814 | 8.10E-10 |
| B134 | B185 | CHE | PT | -0.161453697 | 5.06E-09 |
| B134 | B199 | CHE | Mg | 0.155111553 | 1.99E-08 |
| B134 | B152 | CHE | ACA-Ig(G-A-M) | -0.135824652 | 9.21E-07 |
| B135 | B137 | TBA | AFU | 0.191177328 | 3.93E-12 |
| B135 | B158 | TBA | IgM | 0.186121189 | 1.45E-11 |
| B135 | B196 | TBA | TCO2 | -0.163329071 | 3.34E-09 |
| B135 | B197 | TBA | AG | 0.151700317 | 4.07E-08 |
| B135 | B138 | TBA | PAB | 0.14095629 | 3.49E-07 |
| B136 | B140 | 5'NT | r-GT | 0.233257108 | 0 |
| B136 | B150 | 5'NT | ALP | 0.294844881 | 0 |
| B136 | B173 | 5'NT | NTT | 0.238022784 | 0 |
| B136 | B137 | 5'NT | AFU | 0.205643481 | 7.66E-14 |
| B136 | B146 | 5'NT | CysC | 0.165558723 | 2.03E-09 |
| B136 | B159 | 5'NT | β2-MG | 0.149009679 | 7.07E-08 |
| B136 | B169 | 5'NT | U-CRY | 0.149009973 | 7.07E-08 |
| B136 | B186 | 5'NT | PFC | 0.143588203 | 2.09E-07 |
| B136 | B197 | 5'NT | AG | 0.13701965 | 7.37E-07 |

| | | | | | |
|---|---|---|---|---|---|
| B137 | B138 | AFU | PAB | 0.205721538 | 7.51E-14 |
| B137 | B181 | AFU | APTT | -0.182578491 | 3.54E-11 |
| B137 | B199 | AFU | Mg | 0.141699292 | 3.02E-07 |
| B138 | B139 | PAB | ADA | -0.343400283 | 0 |
| B138 | B194 | PAB | Cl | -0.272546899 | 0 |
| B138 | B197 | PAB | AG | 0.208955115 | 2.98E-14 |
| B138 | B199 | PAB | Mg | 0.20833191 | 3.55E-14 |
| B138 | B222 | PAB | NS | 0.185288214 | 1.79E-11 |
| B138 | B192 | PAB | UA | 0.16411141 | 2.80E-09 |
| B138 | B187 | PAB | NSE | 0.163484677 | 3.23E-09 |
| B138 | B221 | PAB | Hei | 0.155563828 | 1.81E-08 |
| B138 | B216 | PAB | BMI | 0.15488113 | 2.09E-08 |
| B138 | B181 | PAB | APTT | -0.147987722 | 8.70E-08 |
| B138 | B200 | PAB | OP | 0.137148023 | 7.20E-07 |
| B139 | B150 | ADA | ALP | 0.248175318 | 0 |
| B139 | B157 | ADA | IgG | 0.186868368 | 1.20E-11 |
| B139 | B198 | ADA | P | 0.170580089 | 6.40E-10 |
| B14 | B179 | G-17 | α1-MG | 0.229103494 | 0 |
| B14 | B168 | G-17 | NMP22 | 0.149292046 | 6.68E-08 |
| B140 | B141 | r-GT | AST/ALT | -0.244699614 | 0 |
| B140 | B192 | r-GT | UA | 0.222026411 | 6.66E-16 |
| B140 | B216 | r-GT | BMI | 0.207039058 | 5.15E-14 |
| B140 | B222 | r-GT | NS | 0.19537748 | 1.29E-12 |
| B140 | B206 | r-GT | Waist | 0.189475626 | 6.12E-12 |
| B140 | B207 | r-GT | WHR | 0.170527273 | 6.48E-10 |
| B140 | B215 | r-GT | BMI | 0.160213106 | 6.65E-09 |
| B140 | B190 | r-GT | Cre | 0.155571044 | 1.81E-08 |
| B140 | B221 | r-GT | Hei | 0.140891539 | 3.53E-07 |
| B141 | B192 | AST/ALT | UA | -0.230507133 | 0 |
| B141 | B206 | AST/ALT | Waist | -0.315142936 | 0 |
| B141 | B207 | AST/ALT | WHR | -0.245762512 | 0 |
| B141 | B210 | AST/ALT | Hip | -0.267847529 | 0 |
| B141 | B215 | AST/ALT | BMI | -0.280419413 | 0 |
| B141 | B216 | AST/ALT | BMI | -0.363508067 | 0 |
| B141 | B221 | AST/ALT | Hei | -0.26032595 | 0 |
| B141 | B222 | AST/ALT | NS | -0.350169359 | 0 |
| B141 | B163 | AST/ALT | SG | -0.192803686 | 2.56E-12 |
| B141 | B158 | AST/ALT | IgM | 0.158294638 | 1.01E-08 |
| B141 | B190 | AST/ALT | Cre | -0.15751133 | 1.19E-08 |
| B141 | B209 | AST/ALT | Smoking | -0.137033924 | 7.35E-07 |
| B142 | B143 | 25-(OH)D | β-CTx | -0.400278458 | 0 |
| B142 | B144 | 25-(OH)D | Total-PINP | -0.261468837 | 0 |
| B142 | B145 | 25-(OH)D | BGP | -0.415404856 | 0 |
| B142 | B183 | 25-(OH)D | PTA | -0.226396514 | 0 |

| | | | | | |
|---|---|---|---|---|---|
| B143 | B144 | β-CTx | Total-PINP | 0.696094382 | 0 |
| B143 | B145 | β-CTx | BGP | 0.537881946 | 0 |
| B144 | B145 | Total-PINP | BGP | 0.532953464 | 0 |
| B146 | B189 | CysC | eGFB | -0.195030558 | 1.42E-12 |
| B146 | B190 | CysC | Cre | 0.154364145 | 2.33E-08 |
| B148 | B149 | Anti_TPO | TGAb | 0.35700594 | 0 |
| B15 | B101 | CD3-CD19+(B cell) | anti-Sc70 | 0.674901875 | 0 |
| B15 | B102 | CD3-CD19+(B cell) | anti-Sm | 0.674901869 | 0 |
| B15 | B103 | CD3-CD19+(B cell) | anti-SSA/Ro | 0.674901871 | 0 |
| B15 | B104 | CD3-CD19+(B cell) | anti-SSB | 0.674901871 | 0 |
| B15 | B105 | CD3-CD19+(B cell) | anti-U1RNP | 0.674901871 | 0 |
| B15 | B107 | CD3-CD19+(B cell) | AnuA | 0.674901725 | 0 |
| B15 | B108 | CD3-CD19+(B cell) | Anti-Cyclin | 0.674901871 | 0 |
| B15 | B109 | CD3-CD19+(B cell) | AMA-M2 | 0.674901871 | 0 |
| B15 | B110 | CD3-CD19+(B cell) | AHA | 0.674901871 | 0 |
| B15 | B151 | CD3-CD19+(B cell) | Anti-CCP | 0.345989276 | 0 |
| B15 | B154 | CD3-CD19+(B cell) | RF | 0.367553232 | 0 |
| B15 | B16 | CD3-CD19+(B cell) | CD3+T | -0.259868512 | 0 |
| B15 | B17 | CD3-CD19+(B cell) | CD3+CD16+CD56+NK | 0.263869071 | 0 |
| B15 | B183 | CD3-CD19+(B cell) | PTA | 0.249052868 | 0 |
| B15 | B184 | CD3-CD19+(B cell) | INR | 0.248350885 | 0 |
| B15 | B185 | CD3-CD19+(B cell) | PT | 0.259251001 | 0 |
| B15 | B21 | CD3-CD19+(B cell) | CD3+CD4-CD8- | 0.491879365 | 0 |
| B15 | B22 | CD3-CD19+(B cell) | CD3+CD8+(T) | -0.316218689 | 0 |
| B15 | B94 | CD3-CD19+(B cell) | ALDH2-Lys504Lys | 0.311160983 | 0 |
| B15 | B95 | CD3-CD19+(B cell) | anti-cenpb | 0.674901869 | 0 |
| B15 | B97 | CD3-CD19+(B cell) | dsDNA | 0.674901871 | 0 |
| B15 | B98 | CD3-CD19+(B cell) | anti-Jo-1 | 0.674901877 | 0 |
| B15 | B99 | CD3-CD19+(B cell) | anti-PM-Scl | 0.674901872 | 0 |
| B15 | B20 | CD3-CD19+(B cell) | CD3+CD4+CD8+ | 0.223319045 | 4.44E-16 |
| B15 | B79 | CD3-CD19+(B cell) | hb V -dna | 0.222402755 | 4.44E-16 |
| B15 | B100 | CD3-CD19+(B cell) | anti-Ro52 | -0.202278973 | 1.97E-13 |
| B15 | B156 | CD3-CD19+(B cell) | IgE | 0.165715334 | 1.95E-09 |
| B15 | B120 | CD3-CD19+(B cell) | HCV-cAg | 0.165296779 | 2.15E-09 |
| B15 | B181 | CD3-CD19+(B cell) | APTT | 0.154309277 | 2.36E-08 |
| B15 | B39 | CD3-CD19+(B cell) | Other cells | 0.149427373 | 6.50E-08 |
| B15 | B134 | CD3-CD19+(B cell) | CHE | -0.141028671 | 3.44E-07 |
| B15 | B125 | CD3-CD19+(B cell) | Cor | -0.137919057 | 6.22E-07 |
| B15 | B175 | CD3-CD19+(B cell) | ALB-U | 0.136999367 | 7.40E-07 |
| B15 | B186 | CD3-CD19+(B cell) | PFC | 0.136120803 | 8.72E-07 |
| B150 | B197 | ALP | AG | 0.205235085 | 8.59E-14 |
| B150 | B200 | ALP | OP | 0.193976571 | 1.88E-12 |
| B150 | B195 | ALP | Na | 0.165402165 | 2.10E-09 |
| B151 | B186 | Anti-CCP | PFC | 0.201862241 | 2.21E-13 |

| | | | | | |
|---|---|---|---|---|---|
| B151 | B183 | Anti-CCP | PTA | 0.180455625 | 5.99E-11 |
| B151 | B185 | Anti-CCP | PT | 0.170160307 | 7.05E-10 |
| B151 | B157 | Anti-CCP | IgG | 0.15373802 | 2.66E-08 |
| B151 | B213 | Anti-CCP | CVA-R | -0.151386051 | 4.34E-08 |
| B151 | B214 | Anti-CCP | CVA-L | -0.14932916 | 6.63E-08 |
| B152 | B181 | ACA-Ig(G-A-M) | APTT | 0.56647583 | 0 |
| B152 | B182 | ACA-Ig(G-A-M) | TT | 0.415817369 | 0 |
| B152 | B185 | ACA-Ig(G-A-M) | PT | 0.316134073 | 0 |
| B152 | B186 | ACA-Ig(G-A-M) | PFC | -0.260540702 | 0 |
| B152 | B197 | ACA-Ig(G-A-M) | AG | 0.23009694 | 0 |
| B152 | B213 | ACA-Ig(G-A-M) | CVA-R | -0.192685943 | 2.64E-12 |
| B152 | B196 | ACA-Ig(G-A-M) | TCO2 | -0.179566925 | 7.46E-11 |
| B153 | B170 | Glu | U-GLU | 0.441877098 | 0 |
| B153 | B220 | Glu | SBP | 0.201567696 | 2.39E-13 |
| B153 | B215 | Glu | BMI | 0.188784398 | 7.31E-12 |
| B153 | B206 | Glu | Waist | 0.16671821 | 1.56E-09 |
| B153 | B207 | Glu | WHR | 0.140352239 | 3.92E-07 |
| B154 | B184 | RF | INR | 0.227208499 | 0 |
| B154 | B185 | RF | PT | 0.188756088 | 7.37E-12 |
| B154 | B175 | RF | ALB-U | 0.182990095 | 3.19E-11 |
| B154 | B183 | RF | PTA | 0.142143629 | 2.77E-07 |
| B155 | B157 | IgA | IgG | 0.35056202 | 0 |
| B155 | B184 | IgA | INR | 0.14205761 | 2.82E-07 |
| B157 | B185 | IgG | PT | 0.241751501 | 0 |
| B157 | B187 | IgG | NSE | -0.215678245 | 4.22E-15 |
| B157 | B184 | IgG | INR | 0.192929637 | 2.48E-12 |
| B158 | B182 | IgM | TT | 0.275426001 | 0 |
| B158 | B184 | IgM | INR | -0.168217367 | 1.11E-09 |
| B159 | B169 | β2-MG | U-CRY | 0.999997651 | 0 |
| B16 | B18 | CD3+T | CD3+CD4+helper(inducer)T | 0.551722064 | 0 |
| B16 | B184 | CD3+T | INR | -0.295101544 | 0 |
| B16 | B185 | CD3+T | PT | -0.318490222 | 0 |
| B16 | B186 | CD3+T | PFC | 0.2969627 | 0 |
| B16 | B22 | CD3+T | CD3+CD8+(T) | 0.435812992 | 0 |
| B16 | B23 | CD3+T | CD3-CD16+CD56+(NK) | -0.937099421 | 0 |
| B16 | B32 | CD3+T | Testo | -0.294425872 | 0 |
| B16 | B20 | CD3+T | CD3+CD4+CD8+ | 0.222257748 | 4.44E-16 |
| B16 | B199 | CD3+T | Mg | -0.210936141 | 1.69E-14 |
| B16 | B30 | CD3+T | LH | 0.189737805 | 5.71E-12 |
| B16 | B60 | CD3+T | WBRVI | -0.145535361 | 1.42E-07 |
| B16 | B28 | CD3+T | PRL | 0.143943973 | 1.95E-07 |
| B16 | B194 | CD3+T | Cl | -0.13986133 | 4.30E-07 |
| B16 | B31 | CD3+T | FSH | 0.136649513 | 7.90E-07 |
| B160 | B163 | PH | SG | -0.20390124 | 1.25E-13 |

| | | | | | |
|---|---|---|---|---|---|
| B161 | B221 | LEU | Hei | -0.165573129 | 2.02E-09 |
| B161 | B222 | LEU | NS | -0.13610936 | 8.74E-07 |
| B163 | B166 | SG | Pro | 0.362033642 | 0 |
| B163 | B176 | SG | Ucr | 0.421389947 | 0 |
| B163 | B190 | SG | Cre | 0.32733717 | 0 |
| B163 | B191 | SG | Urea | 0.243912776 | 0 |
| B163 | B192 | SG | UA | 0.283248798 | 0 |
| B163 | B216 | SG | BMI | 0.315183833 | 0 |
| B163 | B221 | SG | Hei | 0.384439869 | 0 |
| B163 | B222 | SG | NS | 0.351611691 | 0 |
| B163 | B206 | SG | Waist | 0.219698643 | 1.33E-15 |
| B163 | B210 | SG | Hip | 0.191940854 | 3.21E-12 |
| B163 | B209 | SG | Smoking | 0.172448768 | 4.13E-10 |
| B163 | B207 | SG | WHR | 0.170771598 | 6.12E-10 |
| B163 | B175 | SG | ALB-U | 0.144633158 | 1.70E-07 |
| B166 | B190 | Pro | Cre | 0.169010363 | 9.21E-10 |
| B166 | B176 | Pro | Ucr | 0.138508132 | 5.57E-07 |
| B167 | B202 | PC | e-times | 0.992145034 | 0 |
| B17 | B100 | CD3+CD16+CD56+NK | anti-Ro52 | 0.251170494 | 0 |
| B17 | B101 | CD3+CD16+CD56+NK | anti-Sc70 | 0.245729931 | 0 |
| B17 | B102 | CD3+CD16+CD56+NK | anti-Sm | 0.245729929 | 0 |
| B17 | B103 | CD3+CD16+CD56+NK | anti-SSA/Ro | 0.24572993 | 0 |
| B17 | B104 | CD3+CD16+CD56+NK | anti-SSB | 0.24572993 | 0 |
| B17 | B105 | CD3+CD16+CD56+NK | anti-U1RNP | 0.24572993 | 0 |
| B17 | B107 | CD3+CD16+CD56+NK | AnuA | 0.245729879 | 0 |
| B17 | B108 | CD3+CD16+CD56+NK | Anti-Cyclin | 0.24572993 | 0 |
| B17 | B109 | CD3+CD16+CD56+NK | AMA-M2 | 0.24572993 | 0 |
| B17 | B110 | CD3+CD16+CD56+NK | AHA | 0.24572993 | 0 |
| B17 | B18 | CD3+CD16+CD56+NK | CD3+CD4+helper(inducer)T | -0.358313314 | 0 |
| B17 | B186 | CD3+CD16+CD56+NK | PFC | 0.244668066 | 0 |
| B17 | B19 | CD3+CD16+CD56+NK | CD3+CD4+/CD3+CD8+ | -0.319093451 | 0 |
| B17 | B20 | CD3+CD16+CD56+NK | CD3+CD4+CD8+ | 0.391795163 | 0 |
| B17 | B21 | CD3+CD16+CD56+NK | CD3+CD4-CD8- | 0.362309822 | 0 |
| B17 | B22 | CD3+CD16+CD56+NK | CD3+CD8+(T) | 0.34390122 | 0 |
| B17 | B95 | CD3+CD16+CD56+NK | anti-cenpb | 0.245729929 | 0 |
| B17 | B97 | CD3+CD16+CD56+NK | dsDNA | 0.24572993 | 0 |
| B17 | B98 | CD3+CD16+CD56+NK | anti-Jo-1 | 0.245729933 | 0 |
| B17 | B99 | CD3+CD16+CD56+NK | anti-PM-Scl | 0.24572993 | 0 |
| B17 | B197 | CD3+CD16+CD56+NK | AG | 0.179345078 | 7.87E-11 |
| B17 | B181 | CD3+CD16+CD56+NK | APTT | 0.158713366 | 9.21E-09 |
| B17 | B185 | CD3+CD16+CD56+NK | PT | -0.145591884 | 1.41E-07 |
| B17 | B184 | CD3+CD16+CD56+NK | INR | -0.141896265 | 2.91E-07 |
| B17 | B151 | CD3+CD16+CD56+NK | Anti-CCP | 0.140441492 | 3.85E-07 |
| B171 | B173 | EC | NTT | 0.171452162 | 5.22E-10 |

| | | | | | |
|---|---|---|---|---|---|
| B174 | B190 | Urea/Cr | Cre | -0.532472142 | 0 |
| B174 | B191 | Urea/Cr | Urea | 0.65589605 | 0 |
| B174 | B192 | Urea/Cr | UA | -0.304796245 | 0 |
| B174 | B216 | Urea/Cr | BMI | -0.238016282 | 0 |
| B174 | B221 | Urea/Cr | Hei | -0.36219778 | 0 |
| B174 | B222 | Urea/Cr | NS | -0.302253173 | 0 |
| B174 | B189 | Urea/Cr | eGFB | 0.18867252 | 7.53E-12 |
| B174 | B209 | Urea/Cr | Smoking | -0.169148816 | 8.92E-10 |
| B175 | B176 | ALB-U | Ucr | 0.275299763 | 0 |
| B175 | B177 | ALB-U | ACR | 0.700588462 | 0 |
| B175 | B228 | ALB-U | OS | 0.347096065 | 0 |
| B176 | B179 | Ucr | α1-MG | 0.194611741 | 1.58E-12 |
| B176 | B221 | Ucr | Hei | 0.185809973 | 1.57E-11 |
| B176 | B190 | Ucr | Cre | 0.164971169 | 2.31E-09 |
| B176 | B192 | Ucr | UA | 0.13734985 | 6.93E-07 |
| B176 | B220 | Ucr | SBP | -0.136384657 | 8.30E-07 |
| B177 | B178 | ACR | Ig-U | 0.361221563 | 0 |
| B177 | B180 | ACR | Tf | 0.386863179 | 0 |
| B177 | B228 | ACR | OS | 0.302040237 | 0 |
| B178 | B179 | Ig-U | α1-MG | 0.553489869 | 0 |
| B178 | B180 | Ig-U | Tf | 0.958001036 | 0 |
| B179 | B180 | α1-MG | Tf | 0.448734916 | 0 |
| B18 | B19 | CD3+CD4+helper(inducer)T | CD3+CD4+/CD3+CD8+ | 0.751791706 | 0 |
| B18 | B22 | CD3+CD4+helper(inducer)T | CD3+CD8+(T) | -0.4790627 | 0 |
| B18 | B23 | CD3+CD4+helper(inducer)T | CD3-CD16+CD56+(NK) | -0.555843487 | 0 |
| B18 | B183 | CD3+CD4+helper(inducer)T | PTA | 0.167078336 | 1.43E-09 |
| B18 | B137 | CD3+CD4+helper(inducer)T | AFU | -0.160921767 | 5.69E-09 |
| B18 | B186 | CD3+CD4+helper(inducer)T | PFC | 0.157221281 | 1.27E-08 |
| B18 | B199 | CD3+CD4+helper(inducer)T | Mg | -0.143893303 | 1.97E-07 |
| B181 | B182 | APTT | TT | 0.343617051 | 0 |
| B181 | B185 | APTT | PT | 0.443007578 | 0 |
| B181 | B186 | APTT | PFC | -0.21193434 | 1.24E-14 |
| B181 | B184 | APTT | INR | 0.195500513 | 1.25E-12 |
| B181 | B183 | APTT | PTA | 0.143873776 | 1.98E-07 |
| B182 | B184 | TT | INR | -0.318744621 | 0 |
| B182 | B186 | TT | PFC | -0.332348695 | 0 |
| B182 | B194 | TT | Cl | -0.151591465 | 4.16E-08 |
| B182 | B197 | TT | AG | 0.139794638 | 4.36E-07 |
| B183 | B197 | PTA | AG | -0.220845287 | 8.88E-16 |
| B183 | B184 | PTA | INR | 0.203224461 | 1.51E-13 |
| B183 | B196 | PTA | TCO2 | 0.164197588 | 2.75E-09 |
| B183 | B185 | PTA | PT | 0.155425338 | 1.86E-08 |
| B183 | B198 | PTA | P | -0.149923448 | 5.87E-08 |
| B184 | B185 | INR | PT | 0.871635958 | 0 |

| | | | | | |
|---|---|---|---|---|---|
| B184 | B187 | INR | NSE | -0.318836057 | 0 |
| B184 | B194 | INR | Cl | 0.281033615 | 0 |
| B185 | B186 | PT | PFC | -0.237334604 | 0 |
| B185 | B187 | PT | NSE | -0.26160798 | 0 |
| B185 | B194 | PT | Cl | 0.175531602 | 1.98E-10 |
| B187 | B198 | NSE | P | 0.165762339 | 1.93E-09 |
| B188 | B194 | Ca | Cl | -0.137422524 | 6.83E-07 |
| B189 | B190 | eGFB | Cre | -0.510883261 | 0 |
| B189 | B207 | eGFB | WHR | -0.238324101 | 0 |
| B189 | B220 | eGFB | SBP | -0.301611705 | 0 |
| B189 | B191 | eGFB | Urea | -0.19587604 | 1.13E-12 |
| B189 | B206 | eGFB | Waist | -0.166668673 | 1.57E-09 |
| B189 | B209 | eGFB | Smoking | -0.140898329 | 3.53E-07 |
| B19 | B20 | CD3+CD4+/CD3+CD8+ | CD3+CD4+CD8+ | -0.25578808 | 0 |
| B19 | B22 | CD3+CD4+/CD3+CD8+ | CD3+CD8+(T) | -0.756859768 | 0 |
| B19 | B92 | CD3+CD4+/CD3+CD8+ | ALDH2-Glu504Glu | -0.241522119 | 0 |
| B19 | B93 | CD3+CD4+/CD3+CD8+ | ALDH2-Glu504Lys | 0.241712526 | 0 |
| B19 | B21 | CD3+CD4+/CD3+CD8+ | CD3+CD4-CD8- | -0.215820735 | 4.00E-15 |
| B19 | B61 | CD3+CD4+/CD3+CD8+ | WBHSRV | -0.179779223 | 7.08E-11 |
| B19 | B55 | CD3+CD4+/CD3+CD8+ | TK | -0.177869654 | 1.13E-10 |
| B19 | B56 | CD3+CD4+/CD3+CD8+ | IR | -0.157793285 | 1.12E-08 |
| B19 | B197 | CD3+CD4+/CD3+CD8+ | AG | -0.152247277 | 3.63E-08 |
| B190 | B191 | Cre | Urea | 0.227289316 | 0 |
| B190 | B192 | Cre | UA | 0.50349945 | 0 |
| B190 | B206 | Cre | Waist | 0.327046221 | 0 |
| B190 | B207 | Cre | WHR | 0.288148066 | 0 |
| B190 | B209 | Cre | Smoking | 0.233120626 | 0 |
| B190 | B210 | Cre | Hip | 0.235620596 | 0 |
| B190 | B216 | Cre | BMI | 0.408300506 | 0 |
| B190 | B221 | Cre | Hei | 0.474958449 | 0 |
| B190 | B222 | Cre | NS | 0.507992252 | 0 |
| B190 | B215 | Cre | BMI | 0.183755464 | 2.63E-11 |
| B190 | B204 | Cre | Drinking | 0.164171808 | 2.77E-09 |
| B192 | B206 | UA | Waist | 0.460447015 | 0 |
| B192 | B207 | UA | WHR | 0.386238439 | 0 |
| B192 | B210 | UA | Hip | 0.355967959 | 0 |
| B192 | B215 | UA | BMI | 0.359020477 | 0 |
| B192 | B216 | UA | BMI | 0.516931532 | 0 |
| B192 | B221 | UA | Hei | 0.423190857 | 0 |
| B192 | B222 | UA | NS | 0.529408581 | 0 |
| B192 | B209 | UA | Smoking | 0.201425517 | 2.49E-13 |
| B192 | B204 | UA | Drinking | 0.175733575 | 1.89E-10 |
| B192 | B218 | UA | DBP | 0.141548048 | 3.11E-07 |
| B193 | B198 | K | P | 0.167615556 | 1.27E-09 |

| | | | | | |
|---|---|---|---|---|---|
| B193 | B199 | K | Mg | 0.14440759 | 1.78E-07 |
| B194 | B195 | Cl | Na | 0.402223538 | 0 |
| B194 | B197 | Cl | AG | -0.305310966 | 0 |
| B194 | B200 | Cl | OP | 0.362460397 | 0 |
| B194 | B196 | Cl | TCO2 | -0.205598445 | 7.77E-14 |
| B195 | B197 | Na | AG | 0.262584482 | 0 |
| B195 | B200 | Na | OP | 0.940919452 | 0 |
| B195 | B199 | Na | Mg | 0.159194767 | 8.30E-09 |
| B196 | B197 | TCO2 | AG | -0.702913417 | 0 |
| B197 | B200 | AG | OP | 0.252013073 | 0 |
| B197 | B199 | AG | Mg | 0.145233022 | 1.51E-07 |
| B198 | B222 | P | NS | -0.136665768 | 7.88E-07 |
| B2 | B126 | Age | Alb | -0.346418275 | 0 |
| B2 | B189 | Age | eGFB | -0.599263316 | 0 |
| B2 | B207 | Age | WHR | 0.272648042 | 0 |
| B2 | B220 | Age | SBP | 0.425539156 | 0 |
| B2 | B221 | Age | Hei | -0.288198603 | 0 |
| B2 | B46 | Age | RBC Count | -0.278672758 | 0 |
| B2 | B47 | Age | RDW | 0.284977256 | 0 |
| B2 | B153 | Age | Glu | 0.223828772 | 4.44E-16 |
| B2 | B43 | Age | MCV | 0.218309389 | 1.78E-15 |
| B2 | B52 | Age | PLT/BPC | -0.211854907 | 1.29E-14 |
| B2 | B219 | Age | OT | 0.209158856 | 2.80E-14 |
| B2 | B77 | Age | TC | 0.200657019 | 3.08E-13 |
| B2 | B51 | Age | Hb | -0.200468379 | 3.24E-13 |
| B2 | B163 | Age | SG | -0.195298758 | 1.32E-12 |
| B2 | B188 | Age | Ca | -0.191243196 | 3.86E-12 |
| B2 | B201 | Age | e-habits | 0.190947769 | 4.17E-12 |
| B2 | B191 | Age | Urea | 0.180973888 | 5.27E-11 |
| B2 | B91 | Age | TP | -0.17933365 | 7.90E-11 |
| B2 | B127 | Age | A-G ratio | -0.174946843 | 2.28E-10 |
| B2 | B118 | Age | CEA1 | 0.166945367 | 1.48E-09 |
| B2 | B176 | Age | Ucr | -0.16126662 | 5.28E-09 |
| B2 | B41 | Age | MON％ | 0.161121182 | 5.45E-09 |
| B2 | B206 | Age | Waist | 0.1515724 | 4.18E-08 |
| B2 | B40 | Age | WBC | -0.150456956 | 5.26E-08 |
| B2 | B205 | Age | KDA | 0.149432884 | 6.49E-08 |
| B2 | B128 | Age | ALT | -0.149177223 | 6.84E-08 |
| B2 | B44 | Age | MCH | 0.144931578 | 1.60E-07 |
| B2 | B203 | Age | DH | -0.137994397 | 6.14E-07 |
| B20 | B106 | CD3+CD4+CD8+ | anti-P | 0.354764068 | 0 |
| B20 | B186 | CD3+CD4+CD8+ | PFC | 0.260081503 | 0 |
| B20 | B21 | CD3+CD4+CD8+ | CD3+CD4-CD8- | 0.233318759 | 0 |
| B20 | B22 | CD3+CD4+CD8+ | CD3+CD8+(T) | 0.263262393 | 0 |

| | | | | | |
|---|---|---|---|---|---|
| B20 | B23 | CD3+CD4+CD8+ | CD3-CD16+CD56+(NK) | -0.269592999 | 0 |
| B20 | B94 | CD3+CD4+CD8+ | ALDH2-Lys504Lys | 0.211435752 | 1.47E-14 |
| B20 | B196 | CD3+CD4+CD8+ | TCO2 | -0.18574348 | 1.59E-11 |
| B20 | B114 | CD3+CD4+CD8+ | TSH TT3 TT4(TT4) | 0.163914048 | 2.93E-09 |
| B20 | B185 | CD3+CD4+CD8+ | PT | -0.153364605 | 2.88E-08 |
| B20 | B59 | CD3+CD4+CD8+ | WBRV | 0.147885371 | 8.88E-08 |
| B204 | B209 | Drinking | Smoking | 0.4619582 | 0 |
| B204 | B222 | Drinking | NS | 0.260480217 | 0 |
| B204 | B216 | Drinking | BMI | 0.204981524 | 9.24E-14 |
| B204 | B221 | Drinking | Hei | 0.202246894 | 1.98E-13 |
| B204 | B206 | Drinking | Waist | 0.19226929 | 2.95E-12 |
| B204 | B207 | Drinking | WHR | 0.183407007 | 2.88E-11 |
| B206 | B207 | Waist | WHR | 0.837345332 | 0 |
| B206 | B210 | Waist | Hip | 0.760672339 | 0 |
| B206 | B215 | Waist | BMI | 0.840520955 | 0 |
| B206 | B216 | Waist | BMI | 0.832598971 | 0 |
| B206 | B218 | Waist | DBP | 0.27866219 | 0 |
| B206 | B220 | Waist | SBP | 0.262232432 | 0 |
| B206 | B221 | Waist | Hei | 0.332167966 | 0 |
| B206 | B222 | Waist | NS | 0.731626226 | 0 |
| B206 | B209 | Waist | Smoking | 0.202362773 | 1.92E-13 |
| B207 | B210 | WHR | Hip | 0.290379149 | 0 |
| B207 | B215 | WHR | BMI | 0.598750669 | 0 |
| B207 | B216 | WHR | BMI | 0.55287281 | 0 |
| B207 | B218 | WHR | DBP | 0.24890712 | 0 |
| B207 | B220 | WHR | SBP | 0.285432236 | 0 |
| B207 | B222 | WHR | NS | 0.540004086 | 0 |
| B207 | B209 | WHR | Smoking | 0.216691007 | 3.11E-15 |
| B207 | B221 | WHR | Hei | 0.170254337 | 6.90E-10 |
| B209 | B221 | Smoking | Hei | 0.244031726 | 0 |
| B209 | B222 | Smoking | NS | 0.289999832 | 0 |
| B209 | B216 | Smoking | BMI | 0.217704387 | 2.22E-15 |
| B21 | B101 | CD3+CD4-CD8- | anti-Sc70 | 0.464474834 | 0 |
| B21 | B102 | CD3+CD4-CD8- | anti-Sm | 0.46447483 | 0 |
| B21 | B103 | CD3+CD4-CD8- | anti-SSA/Ro | 0.464474831 | 0 |
| B21 | B104 | CD3+CD4-CD8- | anti-SSB | 0.464474831 | 0 |
| B21 | B105 | CD3+CD4-CD8- | anti-U1RNP | 0.464474831 | 0 |
| B21 | B107 | CD3+CD4-CD8- | AnuA | 0.464474732 | 0 |
| B21 | B108 | CD3+CD4-CD8- | Anti-Cyclin | 0.464474832 | 0 |
| B21 | B109 | CD3+CD4-CD8- | AMA-M2 | 0.464474831 | 0 |
| B21 | B110 | CD3+CD4-CD8- | AHA | 0.464474832 | 0 |
| B21 | B151 | CD3+CD4-CD8- | Anti-CCP | 0.383786356 | 0 |
| B21 | B154 | CD3+CD4-CD8- | RF | 0.23122473 | 0 |
| B21 | B186 | CD3+CD4-CD8- | PFC | 0.448507205 | 0 |

| | | | | | |
|---|---|---|---|---|---|
| B21 | B23 | CD3+CD4-CD8- | CD3-CD16+CD56+(NK) | -0.256825773 | 0 |
| B21 | B94 | CD3+CD4-CD8- | ALDH2-Lys504Lys | 0.260254796 | 0 |
| B21 | B95 | CD3+CD4-CD8- | anti-cenpb | 0.46447483 | 0 |
| B21 | B97 | CD3+CD4-CD8- | dsDNA | 0.464474831 | 0 |
| B21 | B98 | CD3+CD4-CD8- | anti-Jo-1 | 0.464474836 | 0 |
| B21 | B99 | CD3+CD4-CD8- | anti-PM-Scl | 0.464474832 | 0 |
| B21 | B32 | CD3+CD4-CD8- | Testo | -0.218668991 | 1.78E-15 |
| B21 | B183 | CD3+CD4-CD8- | PTA | 0.211978393 | 1.24E-14 |
| B21 | B69 | CD3+CD4-CD8- | PV | 0.210355773 | 2.00E-14 |
| B21 | B157 | CD3+CD4-CD8- | IgG | 0.203284981 | 1.49E-13 |
| B21 | B64 | CD3+CD4-CD8- | WBRSR(mPas_5) | 0.19149194 | 3.61E-12 |
| B21 | B65 | CD3+CD4-CD8- | WBRSR(mPas_1) | 0.183847971 | 2.57E-11 |
| B21 | B63 | CD3+CD4-CD8- | WBRSR(mPas_200) | 0.180971034 | 5.28E-11 |
| B21 | B81 | CD3+CD4-CD8- | TSGF | -0.180142806 | 6.47E-11 |
| B21 | B59 | CD3+CD4-CD8- | WBRV | 0.174025471 | 2.84E-10 |
| B21 | B66 | CD3+CD4-CD8- | WBRSR(mPas_100) | 0.17316407 | 3.49E-10 |
| B21 | B67 | CD3+CD4-CD8- | WBRSR(mPas_50) | 0.172629788 | 3.96E-10 |
| B21 | B79 | CD3+CD4-CD8- | hb V -dna | 0.164592253 | 2.52E-09 |
| B21 | B185 | CD3+CD4-CD8- | PT | 0.159057507 | 8.55E-09 |
| B21 | B61 | CD3+CD4-CD8- | WBHSRV | 0.138993534 | 5.08E-07 |
| B21 | B30 | CD3+CD4-CD8- | LH | 0.137088526 | 7.28E-07 |
| B21 | B184 | CD3+CD4-CD8- | INR | 0.135674018 | 9.48E-07 |
| B210 | B215 | Hip | BMI | 0.771726297 | 0 |
| B210 | B216 | Hip | BMI | 0.813353746 | 0 |
| B210 | B221 | Hip | Hei | 0.391187851 | 0 |
| B210 | B222 | Hip | NS | 0.648188405 | 0 |
| B210 | B218 | Hip | DBP | 0.188843468 | 7.20E-12 |
| B211 | B212 | IOP-R | IOP-L | 0.793501632 | 0 |
| B213 | B214 | CVA-R | CVA-L | 0.708256306 | 0 |
| B215 | B216 | BMI | BMI | 0.81597087 | 0 |
| B215 | B218 | BMI | DBP | 0.242289413 | 0 |
| B215 | B222 | BMI | NS | 0.635265297 | 0 |
| B215 | B220 | BMI | SBP | 0.208089197 | 3.82E-14 |
| B216 | B218 | BMI | DBP | 0.230211871 | 0 |
| B216 | B221 | BMI | Hei | 0.647030189 | 0 |
| B216 | B222 | BMI | NS | 0.823500667 | 0 |
| B218 | B220 | DBP | SBP | 0.635287821 | 0 |
| B218 | B222 | DBP | NS | 0.227795347 | 0 |
| B22 | B184 | CD3+CD8+(T) | INR | -0.263896403 | 0 |
| B22 | B185 | CD3+CD8+(T) | PT | -0.23771706 | 0 |
| B22 | B23 | CD3+CD8+(T) | CD3-CD16+CD56+(NK) | -0.314896833 | 0 |
| B22 | B94 | CD3+CD8+(T) | ALDH2-Lys504Lys | -0.175563777 | 1.97E-10 |
| B22 | B32 | CD3+CD8+(T) | Testo | -0.175084157 | 2.21E-10 |
| B22 | B55 | CD3+CD8+(T) | TK | 0.173493221 | 3.23E-10 |

| | | | | | |
|---|---|---|---|---|---|
| B22 | B182 | CD3+CD8+(T) | TT | 0.161963254 | 4.52E-09 |
| B22 | B194 | CD3+CD8+(T) | CI | -0.152295401 | 3.60E-08 |
| B22 | B183 | CD3+CD8+(T) | PTA | -0.137876016 | 6.27E-07 |
| B22 | B61 | CD3+CD8+(T) | WBHSRV | 0.135406355 | 9.96E-07 |
| B220 | B222 | SBP | NS | 0.152821396 | 3.22E-08 |
| B221 | B222 | Hei | NS | 0.588341242 | 0 |
| B23 | B184 | CD3-CD16+CD56+(NK) | INR | 0.261529346 | 0 |
| B23 | B185 | CD3-CD16+CD56+(NK) | PT | 0.283195788 | 0 |
| B23 | B186 | CD3-CD16+CD56+(NK) | PFC | -0.32256466 | 0 |
| B23 | B32 | CD3-CD16+CD56+(NK) | Testo | 0.303691376 | 0 |
| B23 | B199 | CD3-CD16+CD56+(NK) | Mg | 0.215313794 | 4.66E-15 |
| B23 | B30 | CD3-CD16+CD56+(NK) | LH | -0.195188257 | 1.36E-12 |
| B23 | B100 | CD3-CD16+CD56+(NK) | anti-Ro52 | 0.156490367 | 1.49E-08 |
| B23 | B28 | CD3-CD16+CD56+(NK) | PRL | -0.148273845 | 8.21E-08 |
| B23 | B31 | CD3-CD16+CD56+(NK) | FSH | -0.140889153 | 3.53E-07 |
| B23 | B60 | CD3-CD16+CD56+(NK) | WBRVI | 0.136973013 | 7.44E-07 |
| B24 | B26 | αHBDH | CKMB | 0.271337345 | 0 |
| B24 | B27 | αHBDH | LDH | 0.680903985 | 0 |
| B24 | B139 | αHBDH | ADA | 0.224000042 | 4.44E-16 |
| B24 | B60 | αHBDH | WBRVI | -0.204720961 | 9.95E-14 |
| B24 | B25 | αHBDH | CK | 0.203038105 | 1.59E-13 |
| B24 | B62 | αHBDH | WBHSRVI | -0.201860292 | 2.21E-13 |
| B24 | B54 | αHBDH | ESR | 0.19177501 | 3.36E-12 |
| B24 | B64 | αHBDH | WBRSR(mPas_5) | -0.190647025 | 4.51E-12 |
| B24 | B66 | αHBDH | WBRSR(mPas_100) | -0.189116592 | 6.71E-12 |
| B24 | B67 | αHBDH | WBRSR(mPas_50) | -0.185534942 | 1.68E-11 |
| B24 | B56 | αHBDH | IR | -0.169317933 | 8.57E-10 |
| B24 | B58 | αHBDH | HCT | -0.165251898 | 2.17E-09 |
| B24 | B63 | αHBDH | WBRSR(mPas_200) | -0.164366144 | 2.65E-09 |
| B24 | B74 | αHBDH | ApoA1 | 0.15584343 | 1.71E-08 |
| B24 | B61 | αHBDH | WBHSRV | -0.151444155 | 4.29E-08 |
| B24 | B65 | αHBDH | WBRSR(mPas_1) | -0.145841124 | 1.34E-07 |
| B24 | B46 | αHBDH | RBC Count | -0.14345603 | 2.15E-07 |
| B24 | B51 | αHBDH | Hb | -0.140112405 | 4.10E-07 |
| B24 | B221 | αHBDH | Hei | -0.138757383 | 5.31E-07 |
| B25 | B26 | CK | CKMB | 0.233535823 | 0 |
| B25 | B27 | CK | LDH | 0.171533202 | 5.12E-10 |
| B25 | B175 | CK | ALB-U | 0.15372555 | 2.67E-08 |
| B25 | B176 | CK | Ucr | 0.152961433 | 3.13E-08 |
| B26 | B27 | CKMB | LDH | 0.284549963 | 0 |
| B26 | B82 | CKMB | CEA | 0.202518037 | 1.84E-13 |
| B27 | B139 | LDH | ADA | 0.232586445 | 0 |
| B27 | B54 | LDH | ESR | 0.182293868 | 3.80E-11 |
| B27 | B150 | LDH | ALP | 0.166082383 | 1.80E-09 |

| | | | | | |
|---|---|---|---|---|---|
| B27 | B60 | LDH | WBRVI | -0.165640129 | 1.99E-09 |
| B27 | B62 | LDH | WBHSRVI | -0.153989811 | 2.52E-08 |
| B27 | B47 | LDH | RDW | 0.152760739 | 3.27E-08 |
| B27 | B64 | LDH | WBRSR(mPas_5) | -0.143383242 | 2.18E-07 |
| B27 | B177 | LDH | ACR | 0.13935118 | 4.74E-07 |
| B28 | B30 | PRL | LH | 0.378860843 | 0 |
| B28 | B32 | PRL | Testo | -0.460808302 | 0 |
| B28 | B33 | PRL | Prog | -0.20580213 | 7.33E-14 |
| B28 | B186 | PRL | PFC | 0.14456181 | 1.73E-07 |
| B28 | B114 | PRL | TSH TT3 TT4(TT4) | -0.143987333 | 1.93E-07 |
| B29 | B30 | E2 | LH | 0.510538264 | 0 |
| B29 | B33 | E2 | Prog | 0.709845936 | 0 |
| B29 | B157 | E2 | IgG | 0.175998661 | 1.77E-10 |
| B29 | B31 | E2 | FSH | -0.171989235 | 4.60E-10 |
| B29 | B32 | E2 | Testo | -0.159571705 | 7.65E-09 |
| B3 | B135 | Hcy | TBA | 0.18426613 | 2.32E-11 |
| B3 | B190 | Hcy | Cre | 0.17870498 | 9.21E-11 |
| B3 | B60 | Hcy | WBRVI | 0.177206072 | 1.33E-10 |
| B3 | B74 | Hcy | ApoA1 | -0.17307059 | 3.57E-10 |
| B3 | B62 | Hcy | WBHSRVI | 0.150851475 | 4.85E-08 |
| B3 | B65 | Hcy | WBRSR(mPas_1) | 0.136944156 | 7.48E-07 |
| B30 | B31 | LH | FSH | 0.701125252 | 0 |
| B30 | B32 | LH | Testo | -0.507088105 | 0 |
| B30 | B186 | LH | PFC | 0.189066102 | 6.80E-12 |
| B30 | B94 | LH | ALDH2-Lys504Lys | 0.16954094 | 8.14E-10 |
| B30 | B182 | LH | TT | -0.168942592 | 9.35E-10 |
| B31 | B32 | FSH | Testo | -0.484932022 | 0 |
| B31 | B33 | FSH | Prog | -0.410659509 | 0 |
| B31 | B186 | FSH | PFC | 0.137983321 | 6.15E-07 |
| B32 | B182 | Testo | TT | 0.264886715 | 0 |
| B32 | B186 | Testo | PFC | -0.298056013 | 0 |
| B32 | B33 | Testo | Prog | -0.250365146 | 0 |
| B32 | B94 | Testo | ALDH2-Lys504Lys | -0.270219534 | 0 |
| B32 | B92 | Testo | ALDH2-Glu504Glu | 0.176377679 | 1.62E-10 |
| B32 | B79 | Testo | hb V -dna | -0.170160729 | 7.05E-10 |
| B32 | B101 | Testo | anti-Sc70 | -0.164661188 | 2.48E-09 |
| B32 | B102 | Testo | anti-Sm | -0.164661187 | 2.48E-09 |
| B32 | B103 | Testo | anti-SSA/Ro | -0.164661187 | 2.48E-09 |
| B32 | B104 | Testo | anti-SSB | -0.164661187 | 2.48E-09 |
| B32 | B105 | Testo | anti-U1RNP | -0.164661187 | 2.48E-09 |
| B32 | B107 | Testo | AnuA | -0.164661151 | 2.48E-09 |
| B32 | B108 | Testo | Anti-Cyclin | -0.164661187 | 2.48E-09 |
| B32 | B109 | Testo | AMA-M2 | -0.164661187 | 2.48E-09 |
| B32 | B110 | Testo | AHA | -0.164661187 | 2.48E-09 |

| | | | | | |
|---|---|---|---|---|---|
| B32 | B95 | Testo | anti-cenpb | -0.164661187 | 2.48E-09 |
| B32 | B97 | Testo | dsDNA | -0.164661187 | 2.48E-09 |
| B32 | B98 | Testo | anti-Jo-1 | -0.164661189 | 2.48E-09 |
| B32 | B99 | Testo | anti-PM-Scl | -0.164661187 | 2.48E-09 |
| B32 | B93 | Testo | ALDH2-Glu504Lys | -0.161768628 | 4.72E-09 |
| B32 | B184 | Testo | INR | -0.157990765 | 1.08E-08 |
| B32 | B81 | Testo | TSGF | 0.151669686 | 4.10E-08 |
| B32 | B120 | Testo | HCV-cAg | -0.136399987 | 8.28E-07 |
| B35 | B116 | β2-MG | TT3 | 0.5873967 | 0 |
| B35 | B117 | β2-MG | TT4 | 0.750104188 | 0 |
| B35 | B167 | β2-MG | PC | 0.831387125 | 0 |
| B35 | B202 | β2-MG | e-times | 0.91001895 | 0 |
| B36 | B79 | V-Lym% | hb V -dna | 0.172510684 | 4.07E-10 |
| B37 | B202 | BAND | e-times | 0.378438803 | 0 |
| B38 | B96 | BAND% | dsDNA (IIF) | -0.47457646 | 0 |
| B39 | B175 | Other cells | ALB-U | 0.200496912 | 3.22E-13 |
| B39 | B154 | Other cells | RF | 0.186706125 | 1.25E-11 |
| B39 | B101 | Other cells | anti-Sc70 | 0.177313825 | 1.29E-10 |
| B39 | B102 | Other cells | anti-Sm | 0.177313823 | 1.29E-10 |
| B39 | B103 | Other cells | anti-SSA/Ro | 0.177313824 | 1.29E-10 |
| B39 | B104 | Other cells | anti-SSB | 0.177313824 | 1.29E-10 |
| B39 | B105 | Other cells | anti-U1RNP | 0.177313824 | 1.29E-10 |
| B39 | B107 | Other cells | AnuA | 0.177313785 | 1.29E-10 |
| B39 | B108 | Other cells | Anti-Cyclin | 0.177313824 | 1.29E-10 |
| B39 | B109 | Other cells | AMA-M2 | 0.177313824 | 1.29E-10 |
| B39 | B110 | Other cells | AHA | 0.177313824 | 1.29E-10 |
| B39 | B95 | Other cells | anti-cenpb | 0.177313824 | 1.29E-10 |
| B39 | B97 | Other cells | dsDNA | 0.177313824 | 1.29E-10 |
| B39 | B98 | Other cells | anti-Jo-1 | 0.177313825 | 1.29E-10 |
| B39 | B99 | Other cells | anti-PM-Scl | 0.177313824 | 1.29E-10 |
| B39 | B176 | Other cells | Ucr | 0.175830311 | 1.85E-10 |
| B39 | B79 | Other cells | hb V -dna | 0.149876976 | 5.93E-08 |
| B4 | B6 | WB-Ca | WB-Fe | -0.275545162 | 0 |
| B4 | B7 | WB-Ca | WB-Cop | 0.234796878 | 0 |
| B4 | B8 | WB-Ca | WB-Zinc | -0.48913571 | 0 |
| B40 | B136 | WBC | 5'NT | 0.245432164 | 0 |
| B40 | B206 | WBC | Waist | 0.244794807 | 0 |
| B40 | B216 | WBC | BMI | 0.241422003 | 0 |
| B40 | B222 | WBC | NS | 0.235686853 | 0 |
| B40 | B42 | WBC | MON | 0.65125604 | 0 |
| B40 | B46 | WBC | RBC Count | 0.302274994 | 0 |
| B40 | B48 | WBC | LYM% | -0.313286886 | 0 |
| B40 | B51 | WBC | Hb | 0.261269669 | 0 |
| B40 | B52 | WBC | PLT/BPC | 0.231094107 | 0 |

| | | | | | |
|---|---|---|---|---|---|
| B40 | B53 | WBC | GR% | 0.328225332 | 0 |
| B40 | B72 | WBC | HDL-C | -0.233398869 | 0 |
| B40 | B192 | WBC | UA | 0.224054773 | 4.44E-16 |
| B40 | B209 | WBC | Smoking | 0.218009096 | 2.22E-15 |
| B40 | B210 | WBC | Hip | 0.206120667 | 6.71E-14 |
| B40 | B215 | WBC | BMI | 0.198060733 | 6.26E-13 |
| B40 | B207 | WBC | WHR | 0.193024831 | 2.41E-12 |
| B40 | B71 | WBC | TG | 0.190775151 | 4.36E-12 |
| B40 | B128 | WBC | ALT | 0.18977775 | 5.65E-12 |
| B40 | B141 | WBC | AST/ALT | -0.173600249 | 3.15E-10 |
| B40 | B150 | WBC | ALP | 0.165780958 | 1.93E-09 |
| B40 | B190 | WBC | Cre | 0.151752682 | 4.03E-08 |
| B40 | B221 | WBC | Hei | 0.150979542 | 4.73E-08 |
| B40 | B140 | WBC | r-GT | 0.150315829 | 5.42E-08 |
| B40 | B41 | WBC | MON％ | -0.143067922 | 2.31E-07 |
| B40 | B163 | WBC | SG | 0.140173603 | 4.05E-07 |
| B41 | B42 | MON％ | MON | 0.616546868 | 0 |
| B41 | B53 | MON％ | GR% | -0.24051102 | 0 |
| B41 | B123 | MON％ | hs-CRP1 | 0.162569075 | 3.96E-09 |
| B41 | B190 | MON％ | Cre | 0.13672157 | 7.79E-07 |
| B42 | B192 | MON | UA | 0.244211143 | 0 |
| B42 | B222 | MON | NS | 0.247774621 | 0 |
| B42 | B46 | MON | RBC Count | 0.262694882 | 0 |
| B42 | B51 | MON | Hb | 0.253374741 | 0 |
| B42 | B48 | MON | LYM% | -0.217562689 | 2.22E-15 |
| B42 | B206 | MON | Waist | 0.216722012 | 3.11E-15 |
| B42 | B190 | MON | Cre | 0.209129264 | 2.84E-14 |
| B42 | B207 | MON | WHR | 0.200234496 | 3.46E-13 |
| B42 | B216 | MON | BMI | 0.199941344 | 3.75E-13 |
| B42 | B209 | MON | Smoking | 0.197223 | 7.86E-13 |
| B42 | B72 | MON | HDL-C | -0.192339229 | 2.89E-12 |
| B42 | B123 | MON | hs-CRP1 | 0.183574954 | 2.76E-11 |
| B42 | B52 | MON | PLT/BPC | 0.177518836 | 1.23E-10 |
| B42 | B163 | MON | SG | 0.173153319 | 3.50E-10 |
| B42 | B221 | MON | Hei | 0.169093219 | 9.03E-10 |
| B42 | B174 | MON | Urea/Cr | -0.153892147 | 2.58E-08 |
| B42 | B140 | MON | r-GT | 0.150621478 | 5.09E-08 |
| B42 | B210 | MON | Hip | 0.142788593 | 2.44E-07 |
| B42 | B128 | MON | ALT | 0.138542169 | 5.53E-07 |
| B42 | B141 | MON | AST/ALT | -0.135787628 | 9.28E-07 |
| B42 | B166 | MON | Pro | 0.135775601 | 9.30E-07 |
| B43 | B44 | MCV | MCH | 0.910797697 | 0 |
| B43 | B45 | MCV | MCHC | 0.228381056 | 0 |
| B43 | B46 | MCV | RBC Count | -0.513447224 | 0 |

| | | | | | |
|---|---|---|---|---|---|
| B43 | B47 | MCV | RDW | 0.51212674 | 0 |
| B43 | B52 | MCV | PLT/BPC | -0.201604644 | 2.37E-13 |
| B43 | B189 | MCV | eGFB | -0.158064548 | 1.06E-08 |
| B44 | B45 | MCH | MCHC | 0.607647972 | 0 |
| B44 | B46 | MCH | RBC Count | -0.376068164 | 0 |
| B44 | B47 | MCH | RDW | 0.310372623 | 0 |
| B44 | B51 | MCH | Hb | 0.318473774 | 0 |
| B44 | B52 | MCH | PLT/BPC | -0.160591928 | 6.12E-09 |
| B45 | B222 | MCHC | NS | 0.243578795 | 0 |
| B45 | B47 | MCHC | RDW | -0.271997659 | 0 |
| B45 | B51 | MCHC | Hb | 0.537040765 | 0 |
| B45 | B221 | MCHC | Hei | 0.219595708 | 1.33E-15 |
| B45 | B190 | MCHC | Cre | 0.201017184 | 2.79E-13 |
| B45 | B216 | MCHC | BMI | 0.1951275 | 1.38E-12 |
| B45 | B174 | MCHC | Urea/Cr | -0.192546937 | 2.74E-12 |
| B45 | B192 | MCHC | UA | 0.179393462 | 7.78E-11 |
| B45 | B133 | MCHC | TBIL | 0.167800607 | 1.22E-09 |
| B45 | B129 | MCHC | IBIL | 0.164395026 | 2.63E-09 |
| B45 | B141 | MCHC | AST/ALT | -0.152710685 | 3.30E-08 |
| B45 | B58 | MCHC | HCT | 0.145350213 | 1.48E-07 |
| B45 | B132 | MCHC | Dbil | 0.144205959 | 1.85E-07 |
| B46 | B126 | RBC Count | Alb | 0.30848828 | 0 |
| B46 | B141 | RBC Count | AST/ALT | -0.296496334 | 0 |
| B46 | B163 | RBC Count | SG | 0.374244858 | 0 |
| B46 | B174 | RBC Count | Urea/Cr | -0.291150756 | 0 |
| B46 | B190 | RBC Count | Cre | 0.388319065 | 0 |
| B46 | B192 | RBC Count | UA | 0.407305556 | 0 |
| B46 | B206 | RBC Count | Waist | 0.351681191 | 0 |
| B46 | B207 | RBC Count | WHR | 0.252195261 | 0 |
| B46 | B210 | RBC Count | Hip | 0.326411216 | 0 |
| B46 | B215 | RBC Count | BMI | 0.255964835 | 0 |
| B46 | B216 | RBC Count | BMI | 0.483451673 | 0 |
| B46 | B221 | RBC Count | Hei | 0.506036088 | 0 |
| B46 | B222 | RBC Count | NS | 0.511737517 | 0 |
| B46 | B47 | RBC Count | RDW | -0.449249703 | 0 |
| B46 | B51 | RBC Count | Hb | 0.7508312 | 0 |
| B46 | B72 | RBC Count | HDL-C | -0.345497015 | 0 |
| B46 | B128 | RBC Count | ALT | 0.208840809 | 3.06E-14 |
| B46 | B58 | RBC Count | HCT | 0.206876417 | 5.40E-14 |
| B46 | B64 | RBC Count | WBRSR(mPas_5) | 0.201195089 | 2.65E-13 |
| B46 | B67 | RBC Count | WBRSR(mPas_50) | 0.198122392 | 6.16E-13 |
| B46 | B63 | RBC Count | WBRSR(mPas_200) | 0.194871716 | 1.48E-12 |
| B46 | B66 | RBC Count | WBRSR(mPas_100) | 0.194386132 | 1.68E-12 |
| B46 | B52 | RBC Count | PLT/BPC | 0.193994844 | 1.87E-12 |

| | | | | | |
|---|---|---|---|---|---|
| B46 | B218 | RBC Count | DBP | 0.191885706 | 3.26E-12 |
| B46 | B62 | RBC Count | WBHSRVI | 0.179832348 | 6.99E-11 |
| B46 | B127 | RBC Count | A-G ratio | 0.179374919 | 7.82E-11 |
| B46 | B65 | RBC Count | WBRSR(mPas_1) | 0.177873232 | 1.13E-10 |
| B46 | B60 | RBC Count | WBRVI | 0.177777914 | 1.15E-10 |
| B46 | B138 | RBC Count | PAB | 0.172530842 | 4.05E-10 |
| B46 | B113 | RBC Count | FT3 | 0.171202282 | 5.53E-10 |
| B46 | B209 | RBC Count | Smoking | 0.170253036 | 6.90E-10 |
| B46 | B71 | RBC Count | TG | 0.167946927 | 1.18E-09 |
| B46 | B74 | RBC Count | ApoA1 | -0.160318647 | 6.50E-09 |
| B46 | B176 | RBC Count | Ucr | 0.159188237 | 8.31E-09 |
| B46 | B188 | RBC Count | Ca | 0.152569501 | 3.40E-08 |
| B46 | B166 | RBC Count | Pro | 0.151591302 | 4.16E-08 |
| B46 | B205 | RBC Count | KDA | -0.144660535 | 1.69E-07 |
| B46 | B204 | RBC Count | Drinking | 0.14378009 | 2.01E-07 |
| B46 | B61 | RBC Count | WBHSRV | 0.14367217 | 2.06E-07 |
| B46 | B91 | RBC Count | TP | 0.14137053 | 3.22E-07 |
| B47 | B51 | RDW | Hb | -0.25385996 | 0 |
| B47 | B52 | RDW | PLT/BPC | -0.269658784 | 0 |
| B47 | B126 | RDW | Alb | -0.214508521 | 5.77E-15 |
| B47 | B189 | RDW | eGFB | -0.18192983 | 4.16E-11 |
| B47 | B221 | RDW | Hei | -0.154896192 | 2.09E-08 |
| B47 | B118 | RDW | CEA1 | 0.150217854 | 5.53E-08 |
| B47 | B72 | RDW | HDL-C | 0.137357616 | 6.92E-07 |
| B48 | B53 | LYM% | GR% | -0.936689098 | 0 |
| B48 | B72 | LYM% | HDL-C | 0.13566412 | 9.49E-07 |
| B49 | B50 | baso% | eo% | 0.274896448 | 0 |
| B49 | B53 | baso% | GR% | -0.185478883 | 1.71E-11 |
| B5 | B6 | WB-Mg | WB-Fe | 0.892903209 | 0 |
| B5 | B7 | WB-Mg | WB-Cop | 0.385240284 | 0 |
| B5 | B8 | WB-Mg | WB-Zinc | 0.74990012 | 0 |
| B50 | B53 | eo% | GR% | -0.291037197 | 0 |
| B50 | B180 | eo% | Tf | 0.202473375 | 1.86E-13 |
| B50 | B178 | eo% | Ig-U | 0.195379032 | 1.29E-12 |
| B51 | B126 | Hb | Alb | 0.275546914 | 0 |
| B51 | B128 | Hb | ALT | 0.23516589 | 0 |
| B51 | B138 | Hb | PAB | 0.22853928 | 0 |
| B51 | B141 | Hb | AST/ALT | -0.33384143 | 0 |
| B51 | B163 | Hb | SG | 0.365116271 | 0 |
| B51 | B174 | Hb | Urea/Cr | -0.356106339 | 0 |
| B51 | B190 | Hb | Cre | 0.476655916 | 0 |
| B51 | B192 | Hb | UA | 0.451570634 | 0 |
| B51 | B206 | Hb | Waist | 0.378154999 | 0 |
| B51 | B207 | Hb | WHR | 0.29001424 | 0 |

| | | | | | |
|---|---|---|---|---|---|
| B51 | B209 | Hb | Smoking | 0.240721228 | 0 |
| B51 | B210 | Hb | Hip | 0.322588553 | 0 |
| B51 | B215 | Hb | BMI | 0.262683156 | 0 |
| B51 | B216 | Hb | BMI | 0.502810325 | 0 |
| B51 | B221 | Hb | Hei | 0.529709474 | 0 |
| B51 | B222 | Hb | NS | 0.562070825 | 0 |
| B51 | B58 | Hb | HCT | 0.265339086 | 0 |
| B51 | B72 | Hb | HDL-C | -0.321660845 | 0 |
| B51 | B63 | Hb | WBRSR(mPas_200) | 0.219833761 | 1.33E-15 |
| B51 | B67 | Hb | WBRSR(mPas_50) | 0.217991926 | 2.22E-15 |
| B51 | B66 | Hb | WBRSR(mPas_100) | 0.215867393 | 4.00E-15 |
| B51 | B127 | Hb | A-G ratio | 0.210841306 | 1.73E-14 |
| B51 | B218 | Hb | DBP | 0.206589392 | 5.86E-14 |
| B51 | B62 | Hb | WBHSRVI | 0.201025022 | 2.78E-13 |
| B51 | B71 | Hb | TG | 0.20024142 | 3.45E-13 |
| B51 | B64 | Hb | WBRSR(mPas_5) | 0.199488952 | 4.24E-13 |
| B51 | B204 | Hb | Drinking | 0.19561824 | 1.21E-12 |
| B51 | B65 | Hb | WBRSR(mPas_1) | 0.17625944 | 1.67E-10 |
| B51 | B140 | Hb | r-GT | 0.167234359 | 1.38E-09 |
| B51 | B176 | Hb | Ucr | 0.165594161 | 2.01E-09 |
| B51 | B113 | Hb | FT3 | 0.159677266 | 7.47E-09 |
| B51 | B60 | Hb | WBRVI | 0.158070759 | 1.06E-08 |
| B51 | B188 | Hb | Ca | 0.155461137 | 1.85E-08 |
| B51 | B161 | Hb | LEU | -0.153574659 | 2.75E-08 |
| B51 | B74 | Hb | ApoA1 | -0.147993546 | 8.69E-08 |
| B51 | B166 | Hb | Pro | 0.144472646 | 1.76E-07 |
| B51 | B54 | Hb | ESR | -0.136165271 | 8.65E-07 |
| B52 | B189 | PLT/BPC | eGFB | 0.162925535 | 3.65E-09 |
| B52 | B91 | PLT/BPC | TP | 0.152892209 | 3.18E-08 |
| B52 | B130 | PLT/BPC | GLB | 0.138603152 | 5.47E-07 |
| B52 | B188 | PLT/BPC | Ca | 0.135761613 | 9.32E-07 |
| B54 | B111 | ESR | TSH | 0.38524413 | 0 |
| B54 | B123 | ESR | hs-CRP1 | 0.250264794 | 0 |
| B54 | B186 | ESR | PFC | 0.231639101 | 0 |
| B54 | B56 | ESR | IR | -0.372277237 | 0 |
| B54 | B58 | ESR | HCT | -0.462224327 | 0 |
| B54 | B60 | ESR | WBRVI | -0.340536786 | 0 |
| B54 | B61 | ESR | WBHSRV | -0.232031141 | 0 |
| B54 | B62 | ESR | WBHSRVI | -0.476214458 | 0 |
| B54 | B63 | ESR | WBRSR(mPas_200) | -0.299640651 | 0 |
| B54 | B66 | ESR | WBRSR(mPas_100) | -0.287804452 | 0 |
| B54 | B67 | ESR | WBRSR(mPas_50) | -0.267983736 | 0 |
| B54 | B69 | ESR | PV | 0.443158176 | 0 |
| B54 | B64 | ESR | WBRSR(mPas_5) | -0.225882835 | 2.22E-16 |

| | | | | | |
|---|---|---|---|---|---|
| B54 | B139 | ESR | ADA | 0.201929383 | 2.16E-13 |
| B54 | B138 | ESR | PAB | -0.197538997 | 7.21E-13 |
| B54 | B65 | ESR | WBRSR(mPas_1) | -0.188782092 | 7.32E-12 |
| B54 | B124 | ESR | ACTH | -0.163142384 | 3.48E-09 |
| B54 | B177 | ESR | ACR | 0.137625736 | 6.58E-07 |
| B55 | B184 | TK | INR | -0.237599127 | 0 |
| B55 | B56 | TK | IR | 0.77413984 | 0 |
| B55 | B58 | TK | HCT | -0.25109206 | 0 |
| B55 | B60 | TK | WBRVI | 0.245259816 | 0 |
| B55 | B61 | TK | WBHSRV | 0.626817113 | 0 |
| B55 | B62 | TK | WBHSRVI | 0.441573115 | 0 |
| B55 | B69 | TK | PV | -0.600259537 | 0 |
| B55 | B59 | TK | WBRV | 0.21948415 | 1.33E-15 |
| B55 | B183 | TK | PTA | -0.197419248 | 7.45E-13 |
| B55 | B63 | TK | WBRSR(mPas_200) | 0.190879647 | 4.24E-12 |
| B55 | B57 | TK | RBCAI | -0.188174015 | 8.56E-12 |
| B55 | B182 | TK | TT | 0.187137904 | 1.12E-11 |
| B55 | B185 | TK | PT | -0.18235854 | 3.74E-11 |
| B55 | B66 | TK | WBRSR(mPas_100) | 0.168648055 | 1.00E-09 |
| B55 | B194 | TK | CI | -0.153938691 | 2.55E-08 |
| B55 | B139 | TK | ADA | -0.140942623 | 3.50E-07 |
| B55 | B67 | TK | WBRSR(mPas_50) | 0.13584164 | 9.18E-07 |
| B56 | B138 | IR | PAB | 0.302252027 | 0 |
| B56 | B139 | IR | ADA | -0.252731706 | 0 |
| B56 | B58 | IR | HCT | 0.392617744 | 0 |
| B56 | B59 | IR | WBRV | 0.274002331 | 0 |
| B56 | B60 | IR | WBRVI | 0.620872013 | 0 |
| B56 | B61 | IR | WBHSRV | 0.921239624 | 0 |
| B56 | B62 | IR | WBHSRVI | 0.907280116 | 0 |
| B56 | B63 | IR | WBRSR(mPas_200) | 0.717820478 | 0 |
| B56 | B64 | IR | WBRSR(mPas_5) | 0.54248513 | 0 |
| B56 | B65 | IR | WBRSR(mPas_1) | 0.41680794 | 0 |
| B56 | B66 | IR | WBRSR(mPas_100) | 0.690440233 | 0 |
| B56 | B67 | IR | WBRSR(mPas_50) | 0.665115123 | 0 |
| B56 | B69 | IR | PV | -0.527823476 | 0 |
| B56 | B184 | IR | INR | -0.223365186 | 4.44E-16 |
| B56 | B194 | IR | CI | -0.19652294 | 9.49E-13 |
| B56 | B57 | IR | RBCAI | -0.193232829 | 2.28E-12 |
| B56 | B182 | IR | TT | 0.183455295 | 2.84E-11 |
| B56 | B185 | IR | PT | -0.182941686 | 3.23E-11 |
| B57 | B182 | RBCAI | TT | -0.363792676 | 0 |
| B57 | B184 | RBCAI | INR | 0.387494986 | 0 |
| B57 | B185 | RBCAI | PT | 0.283912784 | 0 |
| B57 | B59 | RBCAI | WBRV | 0.817270796 | 0 |

| | | | | | |
|---|---|---|---|---|---|
| B57 | B60 | RBCAI | WBRVI | 0.562441675 | 0 |
| B57 | B64 | RBCAI | WBRSR(mPas_5) | 0.401606404 | 0 |
| B57 | B65 | RBCAI | WBRSR(mPas_1) | 0.592864193 | 0 |
| B57 | B81 | RBCAI | TSGF | -0.251902412 | 0 |
| B57 | B69 | RBCAI | PV | 0.220098155 | 1.11E-15 |
| B57 | B79 | RBCAI | hb V -dna | 0.194935763 | 1.45E-12 |
| B57 | B187 | RBCAI | NSE | -0.192350326 | 2.88E-12 |
| B57 | B158 | RBCAI | IgM | -0.184579669 | 2.14E-11 |
| B57 | B62 | RBCAI | WBHSRVI | -0.147947908 | 8.77E-08 |
| B57 | B89 | RBCAI | CA 72-4 | -0.14045483 | 3.84E-07 |
| B57 | B61 | RBCAI | WBHSRV | -0.138647127 | 5.42E-07 |
| B58 | B138 | HCT | PAB | 0.407953671 | 0 |
| B58 | B60 | HCT | WBRVI | 0.613200793 | 0 |
| B58 | B61 | HCT | WBHSRV | 0.474983577 | 0 |
| B58 | B62 | HCT | WBHSRVI | 0.734695877 | 0 |
| B58 | B63 | HCT | WBRSR(mPas_200) | 0.807096209 | 0 |
| B58 | B64 | HCT | WBRSR(mPas_5) | 0.745743871 | 0 |
| B58 | B65 | HCT | WBRSR(mPas_1) | 0.658352723 | 0 |
| B58 | B66 | HCT | WBRSR(mPas_100) | 0.780654482 | 0 |
| B58 | B67 | HCT | WBRSR(mPas_50) | 0.792760975 | 0 |
| B58 | B154 | HCT | RF | 0.22435181 | 4.44E-16 |
| B58 | B134 | HCT | CHE | 0.218788508 | 1.78E-15 |
| B58 | B141 | HCT | AST/ALT | -0.191708402 | 3.42E-12 |
| B58 | B137 | HCT | AFU | 0.191015244 | 4.10E-12 |
| B58 | B111 | HCT | TSH | -0.182718711 | 3.42E-11 |
| B58 | B222 | HCT | NS | 0.17133949 | 5.36E-10 |
| B58 | B183 | HCT | PTA | 0.169739828 | 7.78E-10 |
| B58 | B192 | HCT | UA | 0.16254911 | 3.97E-09 |
| B58 | B124 | HCT | ACTH | 0.153544745 | 2.77E-08 |
| B58 | B221 | HCT | Hei | 0.150810751 | 4.89E-08 |
| B58 | B139 | HCT | ADA | -0.139247698 | 4.84E-07 |
| B59 | B60 | WBRV | WBRVI | 0.74679259 | 0 |
| B59 | B61 | WBRV | WBHSRV | 0.406470797 | 0 |
| B59 | B62 | WBRV | WBHSRVI | 0.237063866 | 0 |
| B59 | B63 | WBRV | WBRSR(mPas_200) | 0.350090995 | 0 |
| B59 | B64 | WBRV | WBRSR(mPas_5) | 0.687475249 | 0 |
| B59 | B65 | WBRV | WBRSR(mPas_1) | 0.78089975 | 0 |
| B59 | B66 | WBRV | WBRSR(mPas_100) | 0.42230951 | 0 |
| B59 | B67 | WBRV | WBRSR(mPas_50) | 0.471813848 | 0 |
| B59 | B69 | WBRV | PV | 0.218350732 | 1.78E-15 |
| B59 | B182 | WBRV | TT | -0.203769011 | 1.30E-13 |
| B59 | B81 | WBRV | TSGF | -0.202836758 | 1.68E-13 |
| B59 | B184 | WBRV | INR | 0.192894182 | 2.50E-12 |
| B59 | B187 | WBRV | NSE | -0.175731566 | 1.89E-10 |

| | | | | | |
|---|---|---|---|---|---|
| B59 | B94 | WBRV | ALDH2-Lys504Lys | 0.171496045 | 5.17E-10 |
| B59 | B79 | WBRV | hb V -dna | 0.16091389 | 5.70E-09 |
| B59 | B89 | WBRV | CA 72-4 | -0.160541259 | 6.19E-09 |
| B59 | B185 | WBRV | PT | 0.147025078 | 1.06E-07 |
| B6 | B8 | WB-Fe | WB-Zinc | 0.753428247 | 0 |
| B60 | B138 | WBRVI | PAB | 0.340067549 | 0 |
| B60 | B61 | WBRVI | WBHSRV | 0.626515945 | 0 |
| B60 | B62 | WBRVI | WBHSRVI | 0.726908683 | 0 |
| B60 | B63 | WBRVI | WBRSR(mPas_200) | 0.676612949 | 0 |
| B60 | B64 | WBRVI | WBRSR(mPas_5) | 0.87297347 | 0 |
| B60 | B65 | WBRVI | WBRSR(mPas_1) | 0.895411234 | 0 |
| B60 | B66 | WBRVI | WBRSR(mPas_100) | 0.710874973 | 0 |
| B60 | B67 | WBRVI | WBRSR(mPas_50) | 0.74956117 | 0 |
| B60 | B139 | WBRVI | ADA | -0.200471337 | 3.24E-13 |
| B60 | B69 | WBRVI | PV | -0.192712104 | 2.62E-12 |
| B60 | B184 | WBRVI | INR | 0.177879287 | 1.13E-10 |
| B60 | B111 | WBRVI | TSH | -0.172169706 | 4.41E-10 |
| B60 | B182 | WBRVI | TT | -0.163178056 | 3.45E-09 |
| B60 | B81 | WBRVI | TSGF | -0.153029191 | 3.09E-08 |
| B61 | B138 | WBHSRV | PAB | 0.334388075 | 0 |
| B61 | B184 | WBHSRV | INR | -0.233420774 | 0 |
| B61 | B194 | WBHSRV | CI | -0.258943677 | 0 |
| B61 | B62 | WBHSRV | WBHSRVI | 0.886228194 | 0 |
| B61 | B63 | WBHSRV | WBRSR(mPas_200) | 0.876790379 | 0 |
| B61 | B64 | WBHSRV | WBRSR(mPas_5) | 0.715172246 | 0 |
| B61 | B65 | WBHSRV | WBRSR(mPas_1) | 0.57194659 | 0 |
| B61 | B66 | WBHSRV | WBRSR(mPas_100) | 0.853089802 | 0 |
| B61 | B67 | WBHSRV | WBRSR(mPas_50) | 0.832289118 | 0 |
| B61 | B139 | WBHSRV | ADA | -0.220834083 | 8.88E-16 |
| B61 | B182 | WBHSRV | TT | 0.181838422 | 4.25E-11 |
| B61 | B185 | WBHSRV | PT | -0.178608957 | 9.43E-11 |
| B61 | B69 | WBHSRV | PV | -0.165251214 | 2.17E-09 |
| B61 | B121 | WBHSRV | C3 | 0.1637908 | 3.01E-09 |
| B61 | B122 | WBHSRV | C4 | 0.16379084 | 3.01E-09 |
| B61 | B159 | WBHSRV | β2-MG | 0.163790645 | 3.01E-09 |
| B61 | B169 | WBHSRV | U-CRY | 0.16379097 | 3.01E-09 |
| B61 | B125 | WBHSRV | Cor | 0.163602955 | 3.14E-09 |
| B61 | B71 | WBHSRV | TG | 0.153750352 | 2.66E-08 |
| B61 | B119 | WBHSRV | Hcv-rna | -0.149519016 | 6.38E-08 |
| B62 | B138 | WBHSRVI | PAB | 0.41005784 | 0 |
| B62 | B139 | WBHSRVI | ADA | -0.257550943 | 0 |
| B62 | B63 | WBHSRVI | WBRSR(mPas_200) | 0.890126658 | 0 |
| B62 | B64 | WBHSRVI | WBRSR(mPas_5) | 0.731363897 | 0 |
| B62 | B65 | WBHSRVI | WBRSR(mPas_1) | 0.599832241 | 0 |

| | | | | | |
|---|---|---|---|---|---|
| B62 | B66 | WBHSRVI | WBRSR(mPas_100) | 0.864445185 | 0 |
| B62 | B67 | WBHSRVI | WBRSR(mPas_50) | 0.849951589 | 0 |
| B62 | B69 | WBHSRVI | PV | -0.371702065 | 0 |
| B62 | B194 | WBHSRVI | CI | -0.173238373 | 3.43E-10 |
| B62 | B134 | WBHSRVI | CHE | 0.163167622 | 3.46E-09 |
| B62 | B141 | WBHSRVI | AST/ALT | -0.160782834 | 5.87E-09 |
| B62 | B184 | WBHSRVI | INR | -0.147623545 | 9.37E-08 |
| B62 | B221 | WBHSRVI | Hei | 0.136448672 | 8.20E-07 |
| B63 | B138 | WBRSR(mPas_200) | PAB | 0.415569963 | 0 |
| B63 | B64 | WBRSR(mPas_200) | WBRSR(mPas_5) | 0.873119014 | 0 |
| B63 | B65 | WBRSR(mPas_200) | WBRSR(mPas_1) | 0.734496704 | 0 |
| B63 | B66 | WBRSR(mPas_200) | WBRSR(mPas_100) | 0.976595493 | 0 |
| B63 | B67 | WBRSR(mPas_200) | WBRSR(mPas_50) | 0.970024988 | 0 |
| B63 | B194 | WBRSR(mPas_200) | CI | -0.217049075 | 2.66E-15 |
| B63 | B134 | WBRSR(mPas_200) | CHE | 0.212149687 | 1.18E-14 |
| B63 | B154 | WBRSR(mPas_200) | RF | 0.195020983 | 1.42E-12 |
| B63 | B139 | WBRSR(mPas_200) | ADA | -0.193822458 | 1.95E-12 |
| B63 | B125 | WBRSR(mPas_200) | Cor | 0.181638839 | 4.47E-11 |
| B63 | B141 | WBRSR(mPas_200) | AST/ALT | -0.158640612 | 9.36E-09 |
| B63 | B71 | WBRSR(mPas_200) | TG | 0.149455454 | 6.46E-08 |
| B63 | B179 | WBRSR(mPas_200) | α1-MG | 0.141185294 | 3.34E-07 |
| B63 | B121 | WBRSR(mPas_200) | C3 | 0.139714024 | 4.43E-07 |
| B63 | B122 | WBRSR(mPas_200) | C4 | 0.139714058 | 4.43E-07 |
| B63 | B159 | WBRSR(mPas_200) | β2-MG | 0.139713886 | 4.43E-07 |
| B63 | B169 | WBRSR(mPas_200) | U-CRY | 0.139714162 | 4.43E-07 |
| B63 | B192 | WBRSR(mPas_200) | UA | 0.135745894 | 9.35E-07 |
| B64 | B138 | WBRSR(mPas_5) | PAB | 0.385133326 | 0 |
| B64 | B65 | WBRSR(mPas_5) | WBRSR(mPas_1) | 0.947280406 | 0 |
| B64 | B66 | WBRSR(mPas_5) | WBRSR(mPas_100) | 0.899228055 | 0 |
| B64 | B67 | WBRSR(mPas_5) | WBRSR(mPas_50) | 0.927957988 | 0 |
| B64 | B154 | WBRSR(mPas_5) | RF | 0.192419327 | 2.83E-12 |
| B64 | B69 | WBRSR(mPas_5) | PV | 0.187217873 | 1.09E-11 |
| B64 | B134 | WBRSR(mPas_5) | CHE | 0.175925669 | 1.81E-10 |
| B64 | B139 | WBRSR(mPas_5) | ADA | -0.170226772 | 6.95E-10 |
| B64 | B125 | WBRSR(mPas_5) | Cor | 0.151026236 | 4.68E-08 |
| B64 | B111 | WBRSR(mPas_5) | TSH | -0.140524284 | 3.79E-07 |
| B64 | B194 | WBRSR(mPas_5) | CI | -0.140514774 | 3.80E-07 |
| B65 | B138 | WBRSR(mPas_1) | PAB | 0.346219868 | 0 |
| B65 | B66 | WBRSR(mPas_1) | WBRSR(mPas_100) | 0.777252028 | 0 |
| B65 | B67 | WBRSR(mPas_1) | WBRSR(mPas_50) | 0.816816007 | 0 |
| B65 | B69 | WBRSR(mPas_1) | PV | 0.206991515 | 5.24E-14 |
| B65 | B182 | WBRSR(mPas_1) | TT | -0.189350548 | 6.32E-12 |
| B65 | B81 | WBRSR(mPas_1) | TSGF | -0.17772906 | 1.17E-10 |
| B65 | B184 | WBRSR(mPas_1) | INR | 0.173109759 | 3.53E-10 |

| | | | | | |
|---|---|---|---|---|---|
| B65 | B154 | WBRSR(mPas_1) | RF | 0.16885467 | 9.54E-10 |
| B65 | B111 | WBRSR(mPas_1) | TSH | -0.167143081 | 1.41E-09 |
| B65 | B134 | WBRSR(mPas_1) | CHE | 0.154295923 | 2.37E-08 |
| B65 | B124 | WBRSR(mPas_1) | ACTH | 0.146920497 | 1.08E-07 |
| B65 | B139 | WBRSR(mPas_1) | ADA | -0.145404973 | 1.46E-07 |
| B65 | B125 | WBRSR(mPas_1) | Cor | 0.144707672 | 1.68E-07 |
| B65 | B79 | WBRSR(mPas_1) | hb V -dna | 0.137012422 | 7.38E-07 |
| B65 | B187 | WBRSR(mPas_1) | NSE | -0.136264832 | 8.49E-07 |
| B66 | B138 | WBRSR(mPas_100) | PAB | 0.419631865 | 0 |
| B66 | B67 | WBRSR(mPas_100) | WBRSR(mPas_50) | 0.991595412 | 0 |
| B66 | B134 | WBRSR(mPas_100) | CHE | 0.211591919 | 1.38E-14 |
| B66 | B194 | WBRSR(mPas_100) | Cl | -0.205853422 | 7.22E-14 |
| B66 | B139 | WBRSR(mPas_100) | ADA | -0.192217157 | 2.99E-12 |
| B66 | B154 | WBRSR(mPas_100) | RF | 0.160499025 | 6.24E-09 |
| B66 | B71 | WBRSR(mPas_100) | TG | 0.155269928 | 1.93E-08 |
| B66 | B125 | WBRSR(mPas_100) | Cor | 0.150601076 | 5.11E-08 |
| B66 | B121 | WBRSR(mPas_100) | C3 | 0.141097939 | 3.39E-07 |
| B66 | B122 | WBRSR(mPas_100) | C4 | 0.141097977 | 3.39E-07 |
| B66 | B159 | WBRSR(mPas_100) | β2-MG | 0.141097814 | 3.39E-07 |
| B66 | B169 | WBRSR(mPas_100) | U-CRY | 0.141098104 | 3.39E-07 |
| B67 | B138 | WBRSR(mPas_50) | PAB | 0.418106231 | 0 |
| B67 | B134 | WBRSR(mPas_50) | CHE | 0.211401173 | 1.47E-14 |
| B67 | B194 | WBRSR(mPas_50) | Cl | -0.195404633 | 1.28E-12 |
| B67 | B139 | WBRSR(mPas_50) | ADA | -0.186804558 | 1.22E-11 |
| B67 | B154 | WBRSR(mPas_50) | RF | 0.171090364 | 5.68E-10 |
| B67 | B71 | WBRSR(mPas_50) | TG | 0.144380984 | 1.79E-07 |
| B67 | B125 | WBRSR(mPas_50) | Cor | 0.143573141 | 2.10E-07 |
| B69 | B186 | PV | PFC | 0.229574886 | 0 |
| B69 | B193 | PV | K | 0.203130687 | 1.55E-13 |
| B69 | B123 | PV | hs-CRP1 | 0.200787293 | 2.97E-13 |
| B69 | B91 | PV | TP | 0.174172199 | 2.75E-10 |
| B69 | B157 | PV | IgG | 0.173274063 | 3.40E-10 |
| B69 | B183 | PV | PTA | 0.170266708 | 6.88E-10 |
| B69 | B130 | PV | GLB | 0.167983484 | 1.17E-09 |
| B69 | B139 | PV | ADA | 0.166932334 | 1.48E-09 |
| B69 | B154 | PV | RF | 0.147921208 | 8.82E-08 |
| B69 | B151 | PV | Anti-CCP | 0.136703356 | 7.82E-07 |
| B7 | B119 | WB-Cop | Hcv-rna | -0.278536748 | 0 |
| B7 | B8 | WB-Cop | WB-Zinc | 0.321995531 | 0 |
| B7 | B79 | WB-Cop | hb V -dna | -0.160505078 | 6.24E-09 |
| B70 | B132 | LDL-C | Dbil | -0.247648694 | 0 |
| B70 | B75 | LDL-C | ApoA1/B | -0.35883879 | 0 |
| B70 | B76 | LDL-C | ApoB | 0.449632255 | 0 |
| B70 | B77 | LDL-C | TC | 0.876193493 | 0 |

| | | | | | |
|---|---|---|---|---|---|
| B70 | B215 | LDL-C | BMI | 0.202679437 | 1.76E-13 |
| B70 | B206 | LDL-C | Waist | 0.197073292 | 8.18E-13 |
| B70 | B207 | LDL-C | WHR | 0.178062033 | 1.08E-10 |
| B70 | B133 | LDL-C | TBIL | -0.147307781 | 9.98E-08 |
| B70 | B210 | LDL-C | Hip | 0.145240518 | 1.51E-07 |
| B70 | B220 | LDL-C | SBP | 0.141976596 | 2.86E-07 |
| B70 | B218 | LDL-C | DBP | 0.136742123 | 7.76E-07 |
| B71 | B192 | TG | UA | 0.303084391 | 0 |
| B71 | B206 | TG | Waist | 0.364543524 | 0 |
| B71 | B207 | TG | WHR | 0.334693774 | 0 |
| B71 | B210 | TG | Hip | 0.239689673 | 0 |
| B71 | B215 | TG | BMI | 0.311406899 | 0 |
| B71 | B216 | TG | BMI | 0.32325163 | 0 |
| B71 | B222 | TG | NS | 0.292164294 | 0 |
| B71 | B72 | TG | HDL-C | -0.452117759 | 0 |
| B71 | B77 | TG | TC | 0.235603399 | 0 |
| B71 | B218 | TG | DBP | 0.21619909 | 3.55E-15 |
| B71 | B140 | TG | r-GT | 0.202436799 | 1.88E-13 |
| B71 | B130 | TG | GLB | 0.187702547 | 9.67E-12 |
| B71 | B138 | TG | PAB | 0.185466977 | 1.71E-11 |
| B71 | B91 | TG | TP | 0.183893064 | 2.55E-11 |
| B71 | B141 | TG | AST/ALT | -0.182640226 | 3.48E-11 |
| B71 | B134 | TG | CHE | 0.165028243 | 2.28E-09 |
| B71 | B132 | TG | Dbil | -0.155864718 | 1.70E-08 |
| B71 | B128 | TG | ALT | 0.140953205 | 3.49E-07 |
| B71 | B127 | TG | A-G ratio | -0.139795822 | 4.36E-07 |
| B71 | B158 | TG | IgM | 0.139698583 | 4.44E-07 |
| B71 | B221 | TG | Hei | 0.13804899 | 6.07E-07 |
| B72 | B141 | HDL-C | AST/ALT | 0.321239329 | 0 |
| B72 | B163 | HDL-C | SG | -0.254790081 | 0 |
| B72 | B190 | HDL-C | Cre | -0.282046815 | 0 |
| B72 | B192 | HDL-C | UA | -0.424188336 | 0 |
| B72 | B206 | HDL-C | Waist | -0.470948753 | 0 |
| B72 | B207 | HDL-C | WHR | -0.384575988 | 0 |
| B72 | B210 | HDL-C | Hip | -0.370456555 | 0 |
| B72 | B215 | HDL-C | BMI | -0.427116815 | 0 |
| B72 | B216 | HDL-C | BMI | -0.51160132 | 0 |
| B72 | B221 | HDL-C | Hei | -0.326568099 | 0 |
| B72 | B222 | HDL-C | NS | -0.487605372 | 0 |
| B72 | B74 | HDL-C | ApoA1 | 0.375364799 | 0 |
| B72 | B77 | HDL-C | TC | 0.23375088 | 0 |
| B72 | B75 | HDL-C | ApoA1/B | 0.209088956 | 2.86E-14 |
| B72 | B209 | HDL-C | Smoking | -0.20505881 | 9.02E-14 |
| B72 | B128 | HDL-C | ALT | -0.172315774 | 4.26E-10 |

| | | | | | |
|---|---|---|---|---|---|
| B72 | B140 | HDL-C | r-GT | -0.153362542 | 2.88E-08 |
| B72 | B174 | HDL-C | Urea/Cr | 0.14335728 | 2.19E-07 |
| B73 | B76 | ApoA | ApoB | 0.238086792 | 0 |
| B73 | B187 | ApoA | NSE | 0.20654978 | 5.93E-14 |
| B73 | B75 | ApoA | ApoA1/B | -0.197991216 | 6.38E-13 |
| B73 | B197 | ApoA | AG | -0.145737077 | 1.37E-07 |
| B73 | B193 | ApoA | K | 0.139111996 | 4.96E-07 |
| B73 | B171 | ApoA | EC | 0.137970262 | 6.16E-07 |
| B74 | B198 | ApoA1 | P | 0.234699653 | 0 |
| B74 | B222 | ApoA1 | NS | -0.260359389 | 0 |
| B74 | B75 | ApoA1 | ApoA1/B | 0.526213607 | 0 |
| B74 | B206 | ApoA1 | Waist | -0.220085751 | 1.11E-15 |
| B74 | B216 | ApoA1 | BMI | -0.214996447 | 5.11E-15 |
| B74 | B207 | ApoA1 | WHR | -0.211190939 | 1.55E-14 |
| B74 | B221 | ApoA1 | Hei | -0.181210625 | 4.97E-11 |
| B74 | B192 | ApoA1 | UA | -0.177088315 | 1.36E-10 |
| B74 | B190 | ApoA1 | Cre | -0.15807726 | 1.06E-08 |
| B74 | B176 | ApoA1 | Ucr | -0.157229171 | 1.27E-08 |
| B74 | B215 | ApoA1 | BMI | -0.147944605 | 8.78E-08 |
| B74 | B77 | ApoA1 | TC | 0.13850072 | 5.58E-07 |
| B75 | B134 | ApoA1/B | CHE | -0.274896121 | 0 |
| B75 | B76 | ApoA1/B | ApoB | -0.8009069 | 0 |
| B75 | B77 | ApoA1/B | TC | -0.269924705 | 0 |
| B75 | B206 | ApoA1/B | Waist | -0.197403282 | 7.48E-13 |
| B75 | B207 | ApoA1/B | WHR | -0.187285337 | 1.08E-11 |
| B75 | B85 | ApoA1/B | SCC | 0.183197234 | 3.03E-11 |
| B75 | B138 | ApoA1/B | PAB | -0.177958333 | 1.10E-10 |
| B75 | B222 | ApoA1/B | NS | -0.176930481 | 1.42E-10 |
| B75 | B185 | ApoA1/B | PT | 0.169795556 | 7.68E-10 |
| B75 | B142 | ApoA1/B | 25-(OH)D | 0.168013333 | 1.16E-09 |
| B75 | B215 | ApoA1/B | BMI | -0.160412728 | 6.36E-09 |
| B75 | B216 | ApoA1/B | BMI | -0.154662107 | 2.19E-08 |
| B75 | B136 | ApoA1/B | 5'NT | -0.15022629 | 5.52E-08 |
| B75 | B184 | ApoA1/B | INR | 0.138028162 | 6.10E-07 |
| B76 | B134 | ApoB | CHE | 0.266670348 | 0 |
| B76 | B77 | ApoB | TC | 0.423770989 | 0 |
| B76 | B185 | ApoB | PT | -0.19420346 | 1.77E-12 |
| B76 | B184 | ApoB | INR | -0.168965076 | 9.30E-10 |
| B76 | B187 | ApoB | NSE | 0.16711576 | 1.42E-09 |
| B76 | B170 | ApoB | U-GLU | 0.164015796 | 2.87E-09 |
| B76 | B142 | ApoB | 25-(OH)D | -0.151201646 | 4.51E-08 |
| B76 | B138 | ApoB | PAB | 0.148303156 | 8.16E-08 |
| B76 | B136 | ApoB | 5'NT | 0.145892386 | 1.33E-07 |
| B76 | B153 | ApoB | Glu | 0.141963873 | 2.87E-07 |

| | | | | | |
|---|---|---|---|---|---|
| B77 | B132 | TC | Dbil | -0.245179015 | 0 |
| B77 | B220 | TC | SBP | 0.178842242 | 8.91E-11 |
| B77 | B218 | TC | DBP | 0.155808055 | 1.72E-08 |
| B77 | B133 | TC | TBIL | -0.153054104 | 3.07E-08 |
| B77 | B153 | TC | Glu | 0.147761034 | 9.11E-08 |
| B77 | B221 | TC | Hei | -0.137369979 | 6.90E-07 |
| B78 | B80 | HBcAb | anti-HP IgG | -0.208222588 | 3.69E-14 |
| B79 | B101 | hb V -dna | anti-Sc70 | 0.275819602 | 0 |
| B79 | B102 | hb V -dna | anti-Sm | 0.2758196 | 0 |
| B79 | B103 | hb V -dna | anti-SSA/Ro | 0.2758196 | 0 |
| B79 | B104 | hb V -dna | anti-SSB | 0.2758196 | 0 |
| B79 | B105 | hb V -dna | anti-U1RNP | 0.275819601 | 0 |
| B79 | B106 | hb V -dna | anti-P | -0.23104087 | 0 |
| B79 | B107 | hb V -dna | AnuA | 0.275819541 | 0 |
| B79 | B108 | hb V -dna | Anti-Cyclin | 0.275819601 | 0 |
| B79 | B109 | hb V -dna | AMA-M2 | 0.2758196 | 0 |
| B79 | B110 | hb V -dna | AHA | 0.275819601 | 0 |
| B79 | B119 | hb V -dna | Hcv-rna | 0.413649889 | 0 |
| B79 | B121 | hb V -dna | C3 | -0.341586912 | 0 |
| B79 | B122 | hb V -dna | C4 | -0.341587007 | 0 |
| B79 | B124 | hb V -dna | ACTH | 0.244881075 | 0 |
| B79 | B154 | hb V -dna | RF | 0.282277846 | 0 |
| B79 | B159 | hb V -dna | β2-MG | -0.341586596 | 0 |
| B79 | B169 | hb V -dna | U-CRY | -0.341587275 | 0 |
| B79 | B182 | hb V -dna | TT | -0.33913423 | 0 |
| B79 | B183 | hb V -dna | PTA | 0.306230761 | 0 |
| B79 | B184 | hb V -dna | INR | 0.375059659 | 0 |
| B79 | B185 | hb V -dna | PT | 0.252290611 | 0 |
| B79 | B81 | hb V -dna | TSGF | -0.475644893 | 0 |
| B79 | B94 | hb V -dna | ALDH2-Lys504Lys | 0.328245185 | 0 |
| B79 | B95 | hb V -dna | anti-cenpb | 0.2758196 | 0 |
| B79 | B97 | hb V -dna | dsDNA | 0.275819601 | 0 |
| B79 | B98 | hb V -dna | anti-Jo-1 | 0.275819603 | 0 |
| B79 | B99 | hb V -dna | anti-PM-Scl | 0.275819601 | 0 |
| B79 | B120 | hb V -dna | HCV-cAg | 0.220068665 | 1.11E-15 |
| B79 | B111 | hb V -dna | TSH | -0.21460228 | 5.77E-15 |
| B79 | B180 | hb V -dna | Tf | 0.190018864 | 5.31E-12 |
| B79 | B186 | hb V -dna | PFC | 0.17604641 | 1.75E-10 |
| B79 | B187 | hb V -dna | NSE | -0.171964843 | 4.63E-10 |
| B79 | B178 | hb V -dna | Ig-U | 0.158082696 | 1.06E-08 |
| B79 | B175 | hb V -dna | ALB-U | 0.139320407 | 4.77E-07 |
| B80 | B228 | anti-HP IgG | OS | 0.136629338 | 7.93E-07 |
| B81 | B124 | TSGF | ACTH | -0.288973453 | 0 |
| B81 | B182 | TSGF | TT | 0.350817422 | 0 |

| | | | | | |
|---|---|---|---|---|---|
| B81 | B183 | TSGF | PTA | -0.309852904 | 0 |
| B81 | B184 | TSGF | INR | -0.444328148 | 0 |
| B81 | B185 | TSGF | PT | -0.320852425 | 0 |
| B81 | B94 | TSGF | ALDH2-Lys504Lys | -0.365796729 | 0 |
| B81 | B187 | TSGF | NSE | 0.217893595 | 2.22E-15 |
| B81 | B154 | TSGF | RF | -0.211581001 | 1.40E-14 |
| B81 | B157 | TSGF | IgG | -0.206033753 | 6.86E-14 |
| B81 | B111 | TSGF | TSH | 0.205667248 | 7.59E-14 |
| B81 | B194 | TSGF | Cl | -0.194815418 | 1.50E-12 |
| B81 | B101 | TSGF | anti-Sc70 | -0.1507631 | 4.94E-08 |
| B81 | B102 | TSGF | anti-Sm | -0.150763099 | 4.94E-08 |
| B81 | B103 | TSGF | anti-SSA/Ro | -0.150763099 | 4.94E-08 |
| B81 | B104 | TSGF | anti-SSB | -0.150763099 | 4.94E-08 |
| B81 | B105 | TSGF | anti-U1RNP | -0.150763099 | 4.94E-08 |
| B81 | B107 | TSGF | AnuA | -0.150763066 | 4.94E-08 |
| B81 | B108 | TSGF | Anti-Cyclin | -0.150763099 | 4.94E-08 |
| B81 | B109 | TSGF | AMA-M2 | -0.150763099 | 4.94E-08 |
| B81 | B110 | TSGF | AHA | -0.150763099 | 4.94E-08 |
| B81 | B95 | TSGF | anti-cenpb | -0.150763099 | 4.94E-08 |
| B81 | B97 | TSGF | dsDNA | -0.150763099 | 4.94E-08 |
| B81 | B98 | TSGF | anti-Jo-1 | -0.150763101 | 4.94E-08 |
| B81 | B99 | TSGF | anti-PM-Scl | -0.1507631 | 4.94E-08 |
| B81 | B180 | TSGF | Tf | -0.148936769 | 7.18E-08 |
| B81 | B186 | TSGF | PFC | -0.143100988 | 2.30E-07 |
| B82 | B83 | CEA | AFP | 0.430750682 | 0 |
| B82 | B84 | CEA | C-PSA | 0.267686111 | 0 |
| B82 | B88 | CEA | CA 242 | 0.286458888 | 0 |
| B82 | B89 | CEA | CA 72-4 | 0.18324508 | 3.00E-11 |
| B82 | B136 | CEA | 5'NT | 0.183083506 | 3.12E-11 |
| B82 | B85 | CEA | SCC | 0.157780472 | 1.13E-08 |
| B82 | B90 | CEA | CYFRA 21-1 | 0.152279048 | 3.61E-08 |
| B83 | B84 | AFP | C-PSA | 0.316945666 | 0 |
| B83 | B87 | AFP | NSE | 0.316223881 | 0 |
| B83 | B88 | AFP | CA 242 | 0.185820439 | 1.56E-11 |
| B83 | B90 | AFP | CYFRA 21-1 | 0.15662503 | 1.44E-08 |
| B84 | B87 | C-PSA | NSE | 0.280281732 | 0 |
| B84 | B90 | C-PSA | CYFRA 21-1 | 0.303571082 | 0 |
| B84 | B85 | C-PSA | SCC | 0.154825574 | 2.12E-08 |
| B85 | B142 | SCC | 25-(OH)D | 0.297124931 | 0 |
| B85 | B87 | SCC | NSE | 0.237423069 | 0 |
| B85 | B193 | SCC | K | 0.150993953 | 4.71E-08 |
| B86 | B90 | TBhCG | CYFRA 21-1 | 0.33395619 | 0 |
| B87 | B89 | NSE | CA 72-4 | 0.228019879 | 0 |
| B87 | B90 | NSE | CYFRA 21-1 | 0.243898209 | 0 |

| | | | | | |
|---|---|---|---|---|---|
| B87 | B183 | NSE | PTA | 0.179824734 | 7.00E-11 |
| B87 | B186 | NSE | PFC | -0.176363968 | 1.63E-10 |
| B87 | B114 | NSE | TSH TT3 TT4(TT4) | -0.166315403 | 1.71E-09 |
| B89 | B139 | CA 72-4 | ADA | -0.167582453 | 1.28E-09 |
| B89 | B200 | CA 72-4 | OP | -0.137455161 | 6.79E-07 |
| B9 | B38 | RH | BAND% | -0.889566007 | 0 |
| B91 | B126 | TP | Alb | 0.521996113 | 0 |
| B91 | B127 | TP | A-G ratio | -0.461938452 | 0 |
| B91 | B130 | TP | GLB | 0.774226888 | 0 |
| B91 | B188 | TP | Ca | 0.369561897 | 0 |
| B91 | B194 | TP | Cl | -0.192585745 | 2.71E-12 |
| B91 | B209 | TP | Smoking | -0.157811554 | 1.12E-08 |
| B91 | B134 | TP | CHE | 0.146789789 | 1.11E-07 |
| B91 | B218 | TP | DBP | 0.143393037 | 2.17E-07 |
| B92 | B124 | ALDH2-Glu504Glu | ACTH | -0.402157471 | 0 |
| B92 | B187 | ALDH2-Glu504Glu | NSE | -0.395942131 | 0 |
| B92 | B93 | ALDH2-Glu504Glu | ALDH2-Glu504Lys | -0.998815144 | 0 |
| B92 | B156 | ALDH2-Glu504Glu | IgE | 0.135663845 | 9.49E-07 |
| B93 | B124 | ALDH2-Glu504Lys | ACTH | 0.398333283 | 0 |
| B93 | B187 | ALDH2-Glu504Lys | NSE | 0.406475462 | 0 |
| B93 | B94 | ALDH2-Glu504Lys | ALDH2-Lys504Lys | -0.179207557 | 8.14E-11 |
| B93 | B185 | ALDH2-Glu504Lys | PT | -0.138770861 | 5.30E-07 |
| B93 | B184 | ALDH2-Glu504Lys | INR | -0.136205687 | 8.58E-07 |
| B94 | B101 | ALDH2-Lys504Lys | anti-Sc70 | 0.317599287 | 0 |
| B94 | B102 | ALDH2-Lys504Lys | anti-Sm | 0.31759929 | 0 |
| B94 | B103 | ALDH2-Lys504Lys | anti-SSA/Ro | 0.31759929 | 0 |
| B94 | B104 | ALDH2-Lys504Lys | anti-SSB | 0.317599292 | 0 |
| B94 | B105 | ALDH2-Lys504Lys | anti-U1RNP | 0.317599289 | 0 |
| B94 | B107 | ALDH2-Lys504Lys | AnuA | 0.31759926 | 0 |
| B94 | B108 | ALDH2-Lys504Lys | Anti-Cyclin | 0.317599286 | 0 |
| B94 | B109 | ALDH2-Lys504Lys | AMA-M2 | 0.317599291 | 0 |
| B94 | B110 | ALDH2-Lys504Lys | AHA | 0.317599291 | 0 |
| B94 | B184 | ALDH2-Lys504Lys | INR | 0.311095331 | 0 |
| B94 | B185 | ALDH2-Lys504Lys | PT | 0.242473524 | 0 |
| B94 | B186 | ALDH2-Lys504Lys | PFC | 0.272851393 | 0 |
| B94 | B187 | ALDH2-Lys504Lys | NSE | -0.276047807 | 0 |
| B94 | B95 | ALDH2-Lys504Lys | anti-cenpb | 0.317599291 | 0 |
| B94 | B97 | ALDH2-Lys504Lys | dsDNA | 0.317599292 | 0 |
| B94 | B98 | ALDH2-Lys504Lys | anti-Jo-1 | 0.317599291 | 0 |
| B94 | B99 | ALDH2-Lys504Lys | anti-PM-Scl | 0.31759929 | 0 |
| B94 | B196 | ALDH2-Lys504Lys | TCO2 | -0.214604094 | 5.77E-15 |
| B94 | B151 | ALDH2-Lys504Lys | Anti-CCP | 0.162508253 | 4.01E-09 |
| B94 | B138 | ALDH2-Lys504Lys | PAB | -0.15808044 | 1.06E-08 |
| B94 | B182 | ALDH2-Lys504Lys | TT | -0.156549692 | 1.47E-08 |

| | | | | | |
|---|---|---|---|---|---|
| B94 | B154 | ALDH2-Lys504Lys | RF | 0.153855054 | 2.60E-08 |
| B94 | B120 | ALDH2-Lys504Lys | HCV-cAg | 0.138924809 | 5.14E-07 |
| B94 | B131 | ALDH2-Lys504Lys | AST | -0.13878668 | 5.28E-07 |
| B95 | B103 | anti-cenpb | anti-SSA/Ro | 0.999999998 | 0 |
| B95 | B104 | anti-cenpb | anti-SSB | 0.999999998 | 0 |
| B95 | B105 | anti-cenpb | anti-U1RNP | 0.999999998 | 0 |
| B95 | B107 | anti-cenpb | AnuA | 0.999999816 | 0 |
| B95 | B108 | anti-cenpb | Anti-Cyclin | 0.999999998 | 0 |
| B95 | B109 | anti-cenpb | AMA-M2 | 1 | 0 |
| B95 | B110 | anti-cenpb | AHA | 0.999999998 | 0 |
| B95 | B151 | anti-cenpb | Anti-CCP | 0.245131192 | 0 |
| B95 | B154 | anti-cenpb | RF | 0.408504389 | 0 |
| B95 | B183 | anti-cenpb | PTA | 0.2622017 | 0 |
| B95 | B185 | anti-cenpb | PT | 0.238563374 | 0 |
| B95 | B186 | anti-cenpb | PFC | 0.238316947 | 0 |
| B95 | B97 | anti-cenpb | dsDNA | 0.999999998 | 0 |
| B95 | B184 | anti-cenpb | INR | 0.211610708 | 1.38E-14 |
| B95 | B106 | anti-cenpb | anti-P | -0.180934712 | 5.32E-11 |
| B95 | B180 | anti-cenpb | Tf | 0.175916567 | 1.81E-10 |
| B95 | B178 | anti-cenpb | Ig-U | 0.167849823 | 1.20E-09 |
| B95 | B156 | anti-cenpb | IgE | 0.16623331 | 1.74E-09 |
| B95 | B120 | anti-cenpb | HCV-cAg | 0.162135381 | 4.35E-09 |
| B95 | B175 | anti-cenpb | ALB-U | 0.155247955 | 1.94E-08 |
| B96 | B223 | dsDNA (IIF) | FH | 0.247720179 | 0 |
| B97 | B101 | dsDNA | anti-Sc70 | 1 | 0 |
| B97 | B102 | dsDNA | anti-Sm | 0.999999999 | 0 |
| B97 | B103 | dsDNA | anti-SSA/Ro | 1 | 0 |
| B97 | B104 | dsDNA | anti-SSB | 0.999999996 | 0 |
| B97 | B107 | dsDNA | AnuA | 0.999999769 | 0 |
| B97 | B109 | dsDNA | AMA-M2 | 0.999999999 | 0 |
| B97 | B110 | dsDNA | AHA | 0.999999999 | 0 |
| B97 | B151 | dsDNA | Anti-CCP | 0.245131193 | 0 |
| B97 | B154 | dsDNA | RF | 0.40850439 | 0 |
| B97 | B183 | dsDNA | PTA | 0.262201701 | 0 |
| B97 | B185 | dsDNA | PT | 0.238563374 | 0 |
| B97 | B186 | dsDNA | PFC | 0.238316948 | 0 |
| B97 | B99 | dsDNA | anti-PM-Scl | 0.999999997 | 0 |
| B97 | B184 | dsDNA | INR | 0.211610708 | 1.38E-14 |
| B97 | B106 | dsDNA | anti-P | -0.180934713 | 5.32E-11 |
| B97 | B180 | dsDNA | Tf | 0.175916568 | 1.81E-10 |
| B97 | B178 | dsDNA | Ig-U | 0.167849823 | 1.20E-09 |
| B97 | B156 | dsDNA | IgE | 0.16623331 | 1.74E-09 |
| B97 | B120 | dsDNA | HCV-cAg | 0.162135376 | 4.35E-09 |
| B97 | B175 | dsDNA | ALB-U | 0.155247956 | 1.94E-08 |

| | | | | | |
|---|---|---|---|---|---|
| B98 | B101 | anti-Jo-1 | anti-Sc70 | 0.999999997 | 0 |
| B98 | B107 | anti-Jo-1 | AnuA | 0.999999729 | 0 |
| B98 | B151 | anti-Jo-1 | Anti-CCP | 0.245131195 | 0 |
| B98 | B154 | anti-Jo-1 | RF | 0.408504395 | 0 |
| B98 | B183 | anti-Jo-1 | PTA | 0.262201703 | 0 |
| B98 | B185 | anti-Jo-1 | PT | 0.238563374 | 0 |
| B98 | B186 | anti-Jo-1 | PFC | 0.23831695 | 0 |
| B98 | B184 | anti-Jo-1 | INR | 0.211610708 | 1.38E-14 |
| B98 | B106 | anti-Jo-1 | anti-P | -0.180934716 | 5.32E-11 |
| B98 | B180 | anti-Jo-1 | Tf | 0.17591657 | 1.81E-10 |
| B98 | B178 | anti-Jo-1 | Ig-U | 0.167849825 | 1.20E-09 |
| B98 | B156 | anti-Jo-1 | IgE | 0.166233312 | 1.74E-09 |
| B98 | B120 | anti-Jo-1 | HCV-cAg | 0.162135356 | 4.35E-09 |
| B98 | B175 | anti-Jo-1 | ALB-U | 0.155247957 | 1.94E-08 |
| B99 | B101 | anti-PM-Scl | anti-Sc70 | 0.999999997 | 0 |
| B99 | B104 | anti-PM-Scl | anti-SSB | 0.999999998 | 0 |
| B99 | B107 | anti-PM-Scl | AnuA | 0.999999778 | 0 |
| B99 | B108 | anti-PM-Scl | Anti-Cyclin | 0.999999999 | 0 |
| B99 | B151 | anti-PM-Scl | Anti-CCP | 0.245131193 | 0 |
| B99 | B154 | anti-PM-Scl | RF | 0.408504391 | 0 |
| B99 | B183 | anti-PM-Scl | PTA | 0.262201701 | 0 |
| B99 | B185 | anti-PM-Scl | PT | 0.238563375 | 0 |
| B99 | B186 | anti-PM-Scl | PFC | 0.238316948 | 0 |
| B99 | B184 | anti-PM-Scl | INR | 0.211610708 | 1.38E-14 |
| B99 | B106 | anti-PM-Scl | anti-P | -0.180934714 | 5.32E-11 |
| B99 | B180 | anti-PM-Scl | Tf | 0.175916568 | 1.81E-10 |
| B99 | B178 | anti-PM-Scl | Ig-U | 0.167849823 | 1.20E-09 |
| B99 | B156 | anti-PM-Scl | IgE | 0.16623331 | 1.74E-09 |
| B99 | B120 | anti-PM-Scl | HCV-cAg | 0.162135354 | 4.35E-09 |
| B99 | B175 | anti-PM-Scl | ALB-U | 0.155247956 | 1.94E-08 |

**Supplementary Table 21. PEI Correlations in Chronic pharyngitis**

| Row | Column | Row | Column | Cor | *P* |
|---|---|---|---|---|---|
| B1 | B128 | Gender | ALT | -0.299152219 | 0 |
| B1 | B141 | Gender | AST/ALT | 0.386867417 | 0 |
| B1 | B163 | Gender | SG | -0.441283164 | 0 |
| B1 | B174 | Gender | Urea/Cr | 0.428065906 | 0 |
| B1 | B190 | Gender | Cre | -0.692641759 | 0 |
| B1 | B192 | Gender | UA | -0.561546526 | 0 |
| B1 | B204 | Gender | Drinking | -0.31947499 | 0 |
| B1 | B206 | Gender | Waist | -0.517122049 | 0 |
| B1 | B207 | Gender | WHR | -0.433249874 | 0 |
| B1 | B210 | Gender | Hip | -0.435629588 | 0 |
| B1 | B215 | Gender | BMI | -0.367858787 | 0 |
| B1 | B216 | Gender | BMI | -0.642432674 | 0 |
| B1 | B220 | Gender | SBP | -0.29410124 | 0 |
| B1 | B221 | Gender | Hei | -0.672566166 | 0 |
| B1 | B222 | Gender | NS | -0.736477497 | 0 |
| B1 | B45 | Gender | MCHC | -0.354952446 | 0 |
| B1 | B46 | Gender | RBC Count | -0.623295932 | 0 |
| B1 | B51 | Gender | Hb | -0.718548228 | 0 |
| B1 | B72 | Gender | HDL-C | 0.375654965 | 0 |
| B1 | B209 | Gender | Smoking | -0.288861505 | 2.22E-16 |
| B1 | B161 | Gender | LEU | 0.243139806 | 5.50E-12 |
| B1 | B218 | Gender | DBP | -0.233601702 | 3.75E-11 |
| B1 | B126 | Gender | Alb | -0.23010353 | 7.42E-11 |
| B1 | B71 | Gender | TG | -0.210696778 | 2.69E-09 |
| B1 | B140 | Gender | r-GT | -0.200927675 | 1.45E-08 |
| B1 | B42 | Gender | MON | -0.195034111 | 3.84E-08 |
| B1 | B127 | Gender | A-G ratio | -0.194011088 | 4.53E-08 |
| B1 | B176 | Gender | Ucr | -0.191121419 | 7.21E-08 |
| B1 | B50 | Gender | eo% | -0.179359949 | 4.45E-07 |
| B10 | B12 | PG I | PG II | 0.726027849 | 0 |
| B100 | B103 | anti-Ro52 | anti-SSA/Ro | 1 | 0 |
| B100 | B104 | anti-Ro52 | anti-SSB | 0.999999999 | 0 |
| B100 | B106 | anti-Ro52 | anti-P | 0.999999999 | 0 |
| B100 | B107 | anti-Ro52 | AnuA | 0.999999947 | 0 |
| B100 | B109 | anti-Ro52 | AMA-M2 | 1 | 0 |
| B100 | B183 | anti-Ro52 | PTA | 0.303357647 | 0 |
| B100 | B154 | anti-Ro52 | RF | 0.265694332 | 4.22E-14 |
| B100 | B180 | anti-Ro52 | Tf | 0.249915727 | 1.34E-12 |
| B100 | B186 | anti-Ro52 | PFC | -0.231641262 | 5.50E-11 |
| B100 | B113 | anti-Ro52 | FT3 | -0.200835622 | 1.47E-08 |
| B100 | B114 | anti-Ro52 | TSH TT3 TT4(TT4) | 0.189036316 | 1.00E-07 |
| B101 | B102 | anti-Sc70 | anti-Sm | 0.999999998 | 0 |

| | | | | | |
|---|---|---|---|---|---|
| B101 | B107 | anti-Sc70 | AnuA | 0.999999911 | 0 |
| B101 | B108 | anti-Sc70 | Anti-Cyclin | 0.999999998 | 0 |
| B101 | B183 | anti-Sc70 | PTA | 0.303357647 | 0 |
| B101 | B154 | anti-Sc70 | RF | 0.265694333 | 4.22E-14 |
| B101 | B180 | anti-Sc70 | Tf | 0.249915728 | 1.34E-12 |
| B101 | B186 | anti-Sc70 | PFC | -0.231641263 | 5.50E-11 |
| B101 | B113 | anti-Sc70 | FT3 | -0.200835622 | 1.47E-08 |
| B101 | B114 | anti-Sc70 | TSH TT3 TT4(TT4) | 0.189036317 | 1.00E-07 |
| B102 | B107 | anti-Sm | AnuA | 0.999999966 | 0 |
| B102 | B183 | anti-Sm | PTA | 0.303357647 | 0 |
| B102 | B154 | anti-Sm | RF | 0.265694333 | 4.22E-14 |
| B102 | B180 | anti-Sm | Tf | 0.249915728 | 1.34E-12 |
| B102 | B186 | anti-Sm | PFC | -0.231641262 | 5.50E-11 |
| B102 | B113 | anti-Sm | FT3 | -0.200835622 | 1.47E-08 |
| B102 | B114 | anti-Sm | TSH TT3 TT4(TT4) | 0.189036317 | 1.00E-07 |
| B103 | B104 | anti-SSA/Ro | anti-SSB | 0.999999999 | 0 |
| B103 | B106 | anti-SSA/Ro | anti-P | 0.999999998 | 0 |
| B103 | B107 | anti-SSA/Ro | AnuA | 0.999999948 | 0 |
| B103 | B109 | anti-SSA/Ro | AMA-M2 | 1 | 0 |
| B103 | B183 | anti-SSA/Ro | PTA | 0.303357647 | 0 |
| B103 | B154 | anti-SSA/Ro | RF | 0.265694332 | 4.22E-14 |
| B103 | B180 | anti-SSA/Ro | Tf | 0.249915727 | 1.34E-12 |
| B103 | B186 | anti-SSA/Ro | PFC | -0.231641262 | 5.50E-11 |
| B103 | B113 | anti-SSA/Ro | FT3 | -0.200835622 | 1.47E-08 |
| B103 | B114 | anti-SSA/Ro | TSH TT3 TT4(TT4) | 0.189036316 | 1.00E-07 |
| B104 | B106 | anti-SSB | anti-P | 1 | 0 |
| B104 | B107 | anti-SSB | AnuA | 0.999999956 | 0 |
| B104 | B108 | anti-SSB | Anti-Cyclin | 0.999999998 | 0 |
| B104 | B109 | anti-SSB | AMA-M2 | 0.999999999 | 0 |
| B104 | B183 | anti-SSB | PTA | 0.303357646 | 0 |
| B104 | B154 | anti-SSB | RF | 0.265694332 | 4.22E-14 |
| B104 | B180 | anti-SSB | Tf | 0.249915727 | 1.34E-12 |
| B104 | B186 | anti-SSB | PFC | -0.231641261 | 5.50E-11 |
| B104 | B113 | anti-SSB | FT3 | -0.200835621 | 1.47E-08 |
| B104 | B114 | anti-SSB | TSH TT3 TT4(TT4) | 0.189036316 | 1.00E-07 |
| B105 | B187 | anti-U1RNP | NSE | 0.482869646 | 0 |
| B105 | B151 | anti-U1RNP | Anti-CCP | -0.235832021 | 2.41E-11 |
| B105 | B185 | anti-U1RNP | PT | -0.218960132 | 6.07E-10 |
| B106 | B107 | anti-P | AnuA | 0.999999999 | 0 |
| B106 | B108 | anti-P | Anti-Cyclin | 0.999999999 | 0 |
| B106 | B109 | anti-P | AMA-M2 | 0.999999999 | 0 |
| B106 | B183 | anti-P | PTA | 0.303357646 | 0 |
| B106 | B154 | anti-P | RF | 0.265694332 | 4.22E-14 |
| B106 | B180 | anti-P | Tf | 0.249915727 | 1.34E-12 |

| | | | | | |
|---|---|---|---|---|---|
| B106 | B186 | anti-P | PFC | -0.231641261 | 5.50E-11 |
| B106 | B113 | anti-P | FT3 | -0.200835621 | 1.47E-08 |
| B106 | B114 | anti-P | TSH TT3 TT4(TT4) | 0.189036316 | 1.00E-07 |
| B107 | B108 | AnuA | Anti-Cyclin | 0.999999939 | 0 |
| B107 | B109 | AnuA | AMA-M2 | 0.999999948 | 0 |
| B107 | B183 | AnuA | PTA | 0.303357631 | 0 |
| B107 | B154 | AnuA | RF | 0.265694319 | 4.22E-14 |
| B107 | B180 | AnuA | Tf | 0.249915714 | 1.34E-12 |
| B107 | B186 | AnuA | PFC | -0.231641248 | 5.50E-11 |
| B107 | B113 | AnuA | FT3 | -0.200835611 | 1.47E-08 |
| B107 | B114 | AnuA | TSH TT3 TT4(TT4) | 0.189036307 | 1.00E-07 |
| B108 | B183 | Anti-Cyclin | PTA | 0.303357647 | 0 |
| B108 | B154 | Anti-Cyclin | RF | 0.265694332 | 4.22E-14 |
| B108 | B180 | Anti-Cyclin | Tf | 0.249915727 | 1.34E-12 |
| B108 | B186 | Anti-Cyclin | PFC | -0.231641262 | 5.50E-11 |
| B108 | B113 | Anti-Cyclin | FT3 | -0.200835622 | 1.47E-08 |
| B108 | B114 | Anti-Cyclin | TSH TT3 TT4(TT4) | 0.189036316 | 1.00E-07 |
| B109 | B183 | AMA-M2 | PTA | 0.303357647 | 0 |
| B109 | B154 | AMA-M2 | RF | 0.265694332 | 4.22E-14 |
| B109 | B180 | AMA-M2 | Tf | 0.249915727 | 1.34E-12 |
| B109 | B186 | AMA-M2 | PFC | -0.231641262 | 5.50E-11 |
| B109 | B113 | AMA-M2 | FT3 | -0.200835622 | 1.47E-08 |
| B109 | B114 | AMA-M2 | TSH TT3 TT4(TT4) | 0.189036316 | 1.00E-07 |
| B11 | B12 | PGI/PGII | PG II | -0.349642644 | 0 |
| B110 | B112 | AHA | FT4 | 0.381633105 | 0 |
| B110 | B113 | AHA | FT3 | 0.884648676 | 0 |
| B110 | B157 | AHA | IgG | 0.19087427 | 7.50E-08 |
| B110 | B219 | AHA | OT | 0.183190295 | 2.49E-07 |
| B111 | B148 | TSH | Anti_TPO | 0.349192519 | 0 |
| B111 | B149 | TSH | TGAb | 0.56751198 | 0 |
| B112 | B113 | FT4 | FT3 | 0.399897943 | 0 |
| B112 | B114 | FT4 | TSH TT3 TT4(TT4) | 0.34330476 | 0 |
| B114 | B115 | TSH TT3 TT4(TT4) | TSH TT3 TT4(TT3) | 0.491973428 | 0 |
| B114 | B157 | TSH TT3 TT4(TT4) | IgG | -0.184693081 | 1.98E-07 |
| B114 | B151 | TSH TT3 TT4(TT4) | Anti-CCP | 0.17791932 | 5.52E-07 |
| B115 | B134 | TSH TT3 TT4(TT3) | CHE | 0.242786786 | 5.91E-12 |
| B116 | B202 | TT3 | e-times | 0.565685425 | 0 |
| B117 | B223 | TT4 | FH | 0.849962166 | 0 |
| B119 | B120 | Hcv-rna | HCV-cAg | 0.456301003 | 0 |
| B119 | B121 | Hcv-rna | C3 | -0.912279795 | 0 |
| B119 | B122 | Hcv-rna | C4 | -0.912279454 | 0 |
| B119 | B152 | Hcv-rna | ACA-Ig(G-A-M) | -0.912266489 | 0 |
| B119 | B159 | Hcv-rna | β2-MG | -0.912280104 | 0 |
| B119 | B169 | Hcv-rna | U-CRY | -0.912277027 | 0 |

| | | | | | |
|---|---|---|---|---|---|
| B12 | B14 | PG II | G-17 | 0.259540045 | 1.67E-13 |
| B12 | B80 | PG II | anti-HP IgG | 0.18135 | 3.30E-07 |
| B120 | B121 | HCV-cAg | C3 | -0.416276184 | 0 |
| B120 | B122 | HCV-cAg | C4 | -0.416275876 | 0 |
| B120 | B152 | HCV-cAg | ACA-Ig(G-A-M) | -0.416270575 | 0 |
| B120 | B159 | HCV-cAg | β2-MG | -0.416277401 | 0 |
| B120 | B169 | HCV-cAg | U-CRY | -0.416275283 | 0 |
| B120 | B142 | HCV-cAg | 25-(OH)D | 0.225748525 | 1.71E-10 |
| B120 | B193 | HCV-cAg | K | 0.201187623 | 1.39E-08 |
| B121 | B122 | C3 | C4 | 0.99999979 | 0 |
| B121 | B152 | C3 | ACA-Ig(G-A-M) | 0.999987965 | 0 |
| B121 | B169 | C3 | U-CRY | 0.999998161 | 0 |
| B121 | B193 | C3 | K | -0.191898068 | 6.37E-08 |
| B122 | B152 | C4 | ACA-Ig(G-A-M) | 0.999987377 | 0 |
| B122 | B159 | C4 | β2-MG | 0.999999365 | 0 |
| B122 | B193 | C4 | K | -0.191897988 | 6.37E-08 |
| B124 | B125 | ACTH | Cor | 0.617120173 | 0 |
| B124 | B181 | ACTH | APTT | -0.302719173 | 0 |
| B124 | B185 | ACTH | PT | -0.373547171 | 0 |
| B124 | B138 | ACTH | PAB | 0.235073268 | 2.80E-11 |
| B124 | B157 | ACTH | IgG | -0.22643384 | 1.50E-10 |
| B124 | B198 | ACTH | P | -0.22503458 | 1.96E-10 |
| B124 | B139 | ACTH | ADA | -0.203942291 | 8.69E-09 |
| B124 | B184 | ACTH | INR | -0.203538486 | 9.31E-09 |
| B124 | B151 | ACTH | Anti-CCP | 0.199878778 | 1.73E-08 |
| B124 | B158 | ACTH | IgM | -0.197531706 | 2.55E-08 |
| B124 | B134 | ACTH | CHE | 0.18839342 | 1.11E-07 |
| B124 | B197 | ACTH | AG | -0.18055587 | 3.72E-07 |
| B124 | B182 | ACTH | TT | 0.179885384 | 4.11E-07 |
| B125 | B151 | Cor | Anti-CCP | 0.276939551 | 3.11E-15 |
| B125 | B138 | Cor | PAB | 0.274663214 | 5.33E-15 |
| B125 | B198 | Cor | P | -0.270307851 | 1.47E-14 |
| B125 | B157 | Cor | IgG | -0.225426761 | 1.82E-10 |
| B125 | B186 | Cor | PFC | 0.206003848 | 6.10E-09 |
| B125 | B136 | Cor | 5'NT | 0.182529953 | 2.76E-07 |
| B125 | B134 | Cor | CHE | 0.18075934 | 3.61E-07 |
| B126 | B127 | Alb | A-G ratio | 0.536668913 | 0 |
| B126 | B188 | Alb | Ca | 0.44867175 | 0 |
| B126 | B221 | Alb | Hei | 0.266180301 | 3.77E-14 |
| B126 | B189 | Alb | eGFB | 0.201965325 | 1.22E-08 |
| B126 | B141 | Alb | AST/ALT | -0.195275348 | 3.69E-08 |
| B126 | B192 | Alb | UA | 0.188437205 | 1.10E-07 |
| B127 | B130 | A-G ratio | GLB | -0.894754046 | 0 |
| B127 | B221 | A-G ratio | Hei | 0.209191949 | 3.51E-09 |

| | | | | | |
|---|---|---|---|---|---|
| B128 | B131 | ALT | AST | 0.802032879 | 0 |
| B128 | B140 | ALT | r-GT | 0.537385578 | 0 |
| B128 | B141 | ALT | AST/ALT | -0.540137365 | 0 |
| B128 | B192 | ALT | UA | 0.360063875 | 0 |
| B128 | B206 | ALT | Waist | 0.374726357 | 0 |
| B128 | B210 | ALT | Hip | 0.349516654 | 0 |
| B128 | B215 | ALT | BMI | 0.357040675 | 0 |
| B128 | B216 | ALT | BMI | 0.381538963 | 0 |
| B128 | B222 | ALT | NS | 0.381685851 | 0 |
| B128 | B207 | ALT | WHR | 0.277075049 | 3.11E-15 |
| B128 | B136 | ALT | 5'NT | 0.25718538 | 2.80E-13 |
| B128 | B190 | ALT | Cre | 0.209893063 | 3.10E-09 |
| B128 | B221 | ALT | Hei | 0.19640389 | 3.07E-08 |
| B128 | B218 | ALT | DBP | 0.189918251 | 8.74E-08 |
| B128 | B163 | ALT | SG | 0.17400216 | 9.81E-07 |
| B129 | B132 | IBIL | Dbil | 0.735466394 | 0 |
| B129 | B133 | IBIL | TBIL | 0.980468757 | 0 |
| B130 | B209 | GLB | Smoking | -0.178273105 | 5.23E-07 |
| B131 | B140 | AST | r-GT | 0.463991689 | 0 |
| B131 | B215 | AST | BMI | 0.265458502 | 4.44E-14 |
| B131 | B206 | AST | Waist | 0.254918406 | 4.58E-13 |
| B131 | B222 | AST | NS | 0.244500266 | 4.15E-12 |
| B131 | B192 | AST | UA | 0.240997208 | 8.52E-12 |
| B131 | B216 | AST | BMI | 0.222378213 | 3.22E-10 |
| B131 | B210 | AST | Hip | 0.219938746 | 5.07E-10 |
| B131 | B136 | AST | 5'NT | 0.217667004 | 7.70E-10 |
| B131 | B207 | AST | WHR | 0.204292165 | 8.18E-09 |
| B131 | B141 | AST | AST/ALT | -0.194012769 | 4.53E-08 |
| B131 | B218 | AST | DBP | 0.186597794 | 1.47E-07 |
| B132 | B133 | Dbil | TBIL | 0.847315999 | 0 |
| B134 | B138 | CHE | PAB | 0.509099709 | 0 |
| B134 | B139 | CHE | ADA | -0.226724218 | 1.42E-10 |
| B134 | B136 | CHE | 5'NT | 0.212396294 | 1.99E-09 |
| B135 | B196 | TBA | TCO2 | 0.191353529 | 6.95E-08 |
| B136 | B138 | 5'NT | PAB | 0.312320481 | 0 |
| B136 | B140 | 5'NT | r-GT | 0.319023549 | 0 |
| B136 | B185 | 5'NT | PT | 0.286484499 | 4.44E-16 |
| B136 | B184 | 5'NT | INR | 0.236558336 | 2.09E-11 |
| B136 | B196 | 5'NT | TCO2 | -0.214742229 | 1.31E-09 |
| B136 | B197 | 5'NT | AG | 0.19067633 | 7.75E-08 |
| B137 | B183 | AFU | PTA | -0.231379393 | 5.79E-11 |
| B137 | B138 | AFU | PAB | 0.216462197 | 9.58E-10 |
| B137 | B151 | AFU | Anti-CCP | -0.21592445 | 1.06E-09 |
| B137 | B177 | AFU | ACR | -0.181781811 | 3.09E-07 |

| | | | | | |
|---|---|---|---|---|---|
| B137 | B175 | AFU | ALB-U | -0.178199624 | 5.29E-07 |
| B138 | B139 | PAB | ADA | -0.417199295 | 0 |
| B138 | B196 | PAB | TCO2 | -0.317224784 | 0 |
| B138 | B197 | PAB | AG | 0.280862298 | 1.33E-15 |
| B138 | B157 | PAB | IgG | -0.233008458 | 4.21E-11 |
| B138 | B199 | PAB | Mg | 0.211690295 | 2.26E-09 |
| B138 | B140 | PAB | r-GT | 0.194104311 | 4.46E-08 |
| B138 | B200 | PAB | OP | 0.183233519 | 2.48E-07 |
| B139 | B193 | ADA | K | -0.221113855 | 4.08E-10 |
| B139 | B150 | ADA | ALP | 0.207091129 | 5.05E-09 |
| B14 | B155 | G-17 | IgA | 0.292795134 | 0 |
| B14 | B186 | G-17 | PFC | 0.431903311 | 0 |
| B14 | B20 | G-17 | CD3+CD4+CD8+ | 0.573634766 | 0 |
| B14 | B54 | G-17 | ESR | 0.5421719 | 0 |
| B14 | B15 | G-17 | CD3-CD19+(B cell) | -0.237966754 | 1.57E-11 |
| B14 | B23 | G-17 | CD3-CD16+CD56+(NK) | 0.231535193 | 5.62E-11 |
| B14 | B61 | G-17 | WBHSRV | 0.230305237 | 7.13E-11 |
| B14 | B184 | G-17 | INR | 0.221221624 | 4.00E-10 |
| B14 | B151 | G-17 | Anti-CCP | -0.220973996 | 4.19E-10 |
| B14 | B55 | G-17 | TK | 0.216321049 | 9.83E-10 |
| B14 | B17 | G-17 | CD3+CD16+CD56+NK | -0.216282856 | 9.90E-10 |
| B14 | B69 | G-17 | PV | 0.209829238 | 3.13E-09 |
| B14 | B95 | G-17 | anti-cenpb | -0.204768909 | 7.54E-09 |
| B14 | B182 | G-17 | TT | 0.203991645 | 8.61E-09 |
| B14 | B100 | G-17 | anti-Ro52 | -0.200493915 | 1.56E-08 |
| B14 | B101 | G-17 | anti-Sc70 | -0.200493916 | 1.56E-08 |
| B14 | B102 | G-17 | anti-Sm | -0.200493916 | 1.56E-08 |
| B14 | B103 | G-17 | anti-SSA/Ro | -0.200493915 | 1.56E-08 |
| B14 | B104 | G-17 | anti-SSB | -0.200493915 | 1.56E-08 |
| B14 | B106 | G-17 | anti-P | -0.200493915 | 1.56E-08 |
| B14 | B107 | G-17 | AnuA | -0.200493904 | 1.56E-08 |
| B14 | B108 | G-17 | Anti-Cyclin | -0.200493915 | 1.56E-08 |
| B14 | B109 | G-17 | AMA-M2 | -0.200493915 | 1.56E-08 |
| B14 | B97 | G-17 | dsDNA | -0.200493915 | 1.56E-08 |
| B14 | B98 | G-17 | anti-Jo-1 | -0.200493916 | 1.56E-08 |
| B14 | B63 | G-17 | WBRSR(mPas_200) | 0.183568891 | 2.35E-07 |
| B14 | B183 | G-17 | PTA | -0.181836375 | 3.06E-07 |
| B14 | B56 | G-17 | IR | 0.174150901 | 9.60E-07 |
| B140 | B192 | r-GT | UA | 0.29220989 | 0 |
| B140 | B204 | r-GT | Drinking | 0.293305393 | 0 |
| B140 | B206 | r-GT | Waist | 0.317972494 | 0 |
| B140 | B216 | r-GT | BMI | 0.296002292 | 0 |
| B140 | B222 | r-GT | NS | 0.304424012 | 0 |
| B140 | B215 | r-GT | BMI | 0.288001857 | 2.22E-16 |

| | | | | | |
|---|---|---|---|---|---|
| B140 | B141 | r-GT | AST/ALT | -0.279484163 | 1.78E-15 |
| B140 | B210 | r-GT | Hip | 0.263738736 | 6.55E-14 |
| B140 | B207 | r-GT | WHR | 0.254760761 | 4.74E-13 |
| B140 | B179 | r-GT | α1-MG | 0.222755701 | 3.00E-10 |
| B140 | B176 | r-GT | Ucr | 0.184544865 | 2.02E-07 |
| B140 | B163 | r-GT | SG | 0.17421942 | 9.50E-07 |
| B141 | B192 | AST/ALT | UA | -0.326972241 | 0 |
| B141 | B206 | AST/ALT | Waist | -0.314118636 | 0 |
| B141 | B210 | AST/ALT | Hip | -0.313480935 | 0 |
| B141 | B216 | AST/ALT | BMI | -0.38180981 | 0 |
| B141 | B221 | AST/ALT | Hei | -0.307792592 | 0 |
| B141 | B222 | AST/ALT | NS | -0.369653044 | 0 |
| B141 | B215 | AST/ALT | BMI | -0.280511552 | 1.33E-15 |
| B141 | B190 | AST/ALT | Cre | -0.275349586 | 4.44E-15 |
| B141 | B207 | AST/ALT | WHR | -0.222251392 | 3.30E-10 |
| B141 | B163 | AST/ALT | SG | -0.213457236 | 1.65E-09 |
| B142 | B143 | 25-(OH)D | β-CTx | -0.700073542 | 0 |
| B142 | B144 | 25-(OH)D | Total-PINP | -0.595897486 | 0 |
| B142 | B145 | 25-(OH)D | BGP | -0.247441696 | 2.25E-12 |
| B143 | B144 | β-CTx | Total-PINP | 0.667602128 | 0 |
| B143 | B145 | β-CTx | BGP | 0.716911585 | 0 |
| B143 | B182 | β-CTx | TT | -0.301644576 | 0 |
| B143 | B183 | β-CTx | PTA | 0.366886468 | 0 |
| B143 | B227 | β-CTx | OD | -0.207855915 | 4.43E-09 |
| B144 | B145 | Total-PINP | BGP | 0.564559252 | 0 |
| B144 | B182 | Total-PINP | TT | -0.344601053 | 0 |
| B144 | B183 | Total-PINP | PTA | 0.419564221 | 0 |
| B144 | B227 | Total-PINP | OD | 0.30914475 | 0 |
| B144 | B195 | Total-PINP | Na | 0.177284009 | 6.06E-07 |
| B145 | B182 | BGP | TT | -0.422368527 | 0 |
| B145 | B183 | BGP | PTA | 0.510896128 | 0 |
| B145 | B195 | BGP | Na | 0.218704756 | 6.37E-10 |
| B145 | B151 | BGP | Anti-CCP | -0.210285955 | 2.89E-09 |
| B145 | B227 | BGP | OD | -0.199981217 | 1.70E-08 |
| B145 | B200 | BGP | OP | 0.186141521 | 1.58E-07 |
| B145 | B168 | BGP | NMP22 | -0.177008554 | 6.31E-07 |
| B146 | B194 | CysC | Cl | 0.187531705 | 1.27E-07 |
| B148 | B149 | Anti_TPO | TGAb | 0.453264965 | 0 |
| B148 | B156 | Anti_TPO | IgE | 0.256599064 | 3.18E-13 |
| B148 | B213 | Anti_TPO | CVA-R | 0.198204039 | 2.28E-08 |
| B15 | B100 | CD3-CD19+(B cell) | anti-Ro52 | 0.557179139 | 0 |
| B15 | B101 | CD3-CD19+(B cell) | anti-Sc70 | 0.55717914 | 0 |
| B15 | B102 | CD3-CD19+(B cell) | anti-Sm | 0.55717914 | 0 |
| B15 | B103 | CD3-CD19+(B cell) | anti-SSA/Ro | 0.557179139 | 0 |

| | | | | | |
|---|---|---|---|---|---|
| B15 | B104 | CD3-CD19+(B cell) | anti-SSB | 0.557179138 | 0 |
| B15 | B106 | CD3-CD19+(B cell) | anti-P | 0.557179138 | 0 |
| B15 | B107 | CD3-CD19+(B cell) | AnuA | 0.557179109 | 0 |
| B15 | B108 | CD3-CD19+(B cell) | Anti-Cyclin | 0.557179139 | 0 |
| B15 | B109 | CD3-CD19+(B cell) | AMA-M2 | 0.557179138 | 0 |
| B15 | B151 | CD3-CD19+(B cell) | Anti-CCP | 0.38396744 | 0 |
| B15 | B16 | CD3-CD19+(B cell) | CD3+T | -0.571055694 | 0 |
| B15 | B183 | CD3-CD19+(B cell) | PTA | 0.451035983 | 0 |
| B15 | B22 | CD3-CD19+(B cell) | CD3+CD8+(T) | -0.424318568 | 0 |
| B15 | B79 | CD3-CD19+(B cell) | hb V -dna | 0.379295293 | 0 |
| B15 | B94 | CD3-CD19+(B cell) | ALDH2-Lys504Lys | 0.396852137 | 0 |
| B15 | B95 | CD3-CD19+(B cell) | anti-cenpb | 0.570368504 | 0 |
| B15 | B97 | CD3-CD19+(B cell) | dsDNA | 0.557179138 | 0 |
| B15 | B98 | CD3-CD19+(B cell) | anti-Jo-1 | 0.557179141 | 0 |
| B15 | B99 | CD3-CD19+(B cell) | anti-PM-Scl | -0.321014353 | 0 |
| B15 | B154 | CD3-CD19+(B cell) | RF | 0.268837501 | 2.04E-14 |
| B15 | B21 | CD3-CD19+(B cell) | CD3+CD4-CD8- | 0.264531661 | 5.48E-14 |
| B15 | B181 | CD3-CD19+(B cell) | APTT | 0.262104581 | 9.46E-14 |
| B15 | B19 | CD3-CD19+(B cell) | CD3+CD4+/CD3+CD8+ | 0.249079624 | 1.60E-12 |
| B15 | B54 | CD3-CD19+(B cell) | ESR | -0.248117395 | 1.95E-12 |
| B15 | B18 | CD3-CD19+(B cell) | CD3+CD4+helper(inducer)T | -0.244831417 | 3.88E-12 |
| B15 | B186 | CD3-CD19+(B cell) | PFC | -0.227800546 | 1.16E-10 |
| B15 | B157 | CD3-CD19+(B cell) | IgG | 0.206506459 | 5.59E-09 |
| B15 | B81 | CD3-CD19+(B cell) | TSGF | -0.205176337 | 7.03E-09 |
| B15 | B105 | CD3-CD19+(B cell) | anti-U1RNP | -0.197761603 | 2.45E-08 |
| B15 | B155 | CD3-CD19+(B cell) | IgA | -0.193474043 | 4.94E-08 |
| B15 | B17 | CD3-CD19+(B cell) | CD3+CD16+CD56+NK | 0.193140408 | 5.22E-08 |
| B15 | B180 | CD3-CD19+(B cell) | Tf | 0.189390499 | 9.50E-08 |
| B150 | B151 | ALP | Anti-CCP | 0.233134839 | 4.11E-11 |
| B151 | B183 | Anti-CCP | PTA | 0.232454245 | 4.69E-11 |
| B151 | B155 | Anti-CCP | IgA | -0.217451569 | 8.01E-10 |
| B151 | B198 | Anti-CCP | P | -0.203447773 | 9.45E-09 |
| B151 | B175 | Anti-CCP | ALB-U | 0.199239267 | 1.92E-08 |
| B151 | B184 | Anti-CCP | INR | 0.194373274 | 4.27E-08 |
| B151 | B185 | Anti-CCP | PT | 0.175776162 | 7.57E-07 |
| B152 | B159 | ACA-Ig(G-A-M) | β2-MG | 0.999989574 | 0 |
| B152 | B193 | ACA-Ig(G-A-M) | K | -0.19189528 | 6.37E-08 |
| B153 | B170 | Glu | U-GLU | 0.427975514 | 0 |
| B153 | B220 | Glu | SBP | 0.271165398 | 1.20E-14 |
| B153 | B215 | Glu | BMI | 0.21416732 | 1.45E-09 |
| B153 | B206 | Glu | Waist | 0.199593646 | 1.81E-08 |
| B153 | B207 | Glu | WHR | 0.193424659 | 4.98E-08 |
| B154 | B183 | RF | PTA | 0.193786023 | 4.70E-08 |
| B154 | B180 | RF | Tf | 0.179333401 | 4.47E-07 |

| | | | | | |
|---|---|---|---|---|---|
| B155 | B157 | IgA | IgG | 0.350316281 | 0 |
| B155 | B158 | IgA | IgM | 0.296848723 | 0 |
| B155 | B186 | IgA | PFC | 0.308536614 | 0 |
| B156 | B157 | IgE | IgG | 0.339104788 | 0 |
| B156 | B182 | IgE | TT | -0.180798438 | 3.58E-07 |
| B157 | B158 | IgG | IgM | 0.471991441 | 0 |
| B157 | B184 | IgG | INR | 0.265047422 | 4.88E-14 |
| B157 | B181 | IgG | APTT | 0.250002141 | 1.31E-12 |
| B157 | B185 | IgG | PT | 0.224304645 | 2.25E-10 |
| B158 | B185 | IgM | PT | 0.266954593 | 3.15E-14 |
| B158 | B184 | IgM | INR | 0.203601868 | 9.21E-09 |
| B158 | B173 | IgM | NTT | 0.202569889 | 1.10E-08 |
| B159 | B193 | β2-MG | K | -0.191898144 | 6.37E-08 |
| B16 | B100 | CD3+T | anti-Ro52 | -0.33269279 | 0 |
| B16 | B101 | CD3+T | anti-Sc70 | -0.332692792 | 0 |
| B16 | B102 | CD3+T | anti-Sm | -0.332692791 | 0 |
| B16 | B103 | CD3+T | anti-SSA/Ro | -0.33269279 | 0 |
| B16 | B104 | CD3+T | anti-SSB | -0.33269279 | 0 |
| B16 | B105 | CD3+T | anti-U1RNP | 0.323599696 | 0 |
| B16 | B106 | CD3+T | anti-P | -0.332692789 | 0 |
| B16 | B107 | CD3+T | AnuA | -0.332692776 | 0 |
| B16 | B108 | CD3+T | Anti-Cyclin | -0.332692791 | 0 |
| B16 | B109 | CD3+T | AMA-M2 | -0.33269279 | 0 |
| B16 | B137 | CD3+T | AFU | 0.300773829 | 0 |
| B16 | B151 | CD3+T | Anti-CCP | -0.38475453 | 0 |
| B16 | B18 | CD3+T | CD3+CD4+helper(inducer)T | 0.370874414 | 0 |
| B16 | B183 | CD3+T | PTA | -0.419894463 | 0 |
| B16 | B184 | CD3+T | INR | -0.426024407 | 0 |
| B16 | B185 | CD3+T | PT | -0.322401586 | 0 |
| B16 | B19 | CD3+T | CD3+CD4+/CD3+CD8+ | -0.367794589 | 0 |
| B16 | B22 | CD3+T | CD3+CD8+(T) | 0.710139839 | 0 |
| B16 | B23 | CD3+T | CD3-CD16+CD56+(NK) | -0.88076915 | 0 |
| B16 | B69 | CD3+T | PV | -0.32170309 | 0 |
| B16 | B95 | CD3+T | anti-cenpb | -0.340606322 | 0 |
| B16 | B97 | CD3+T | dsDNA | -0.332692789 | 0 |
| B16 | B98 | CD3+T | anti-Jo-1 | -0.332692792 | 0 |
| B16 | B181 | CD3+T | APTT | -0.28308849 | 6.66E-16 |
| B16 | B21 | CD3+T | CD3+CD4-CD8- | -0.234627635 | 3.06E-11 |
| B16 | B154 | CD3+T | RF | -0.207738054 | 4.52E-09 |
| B16 | B90 | CD3+T | CYFRA 21-1 | 0.205104584 | 7.12E-09 |
| B16 | B157 | CD3+T | IgG | -0.196065845 | 3.24E-08 |
| B16 | B17 | CD3+T | CD3+CD16+CD56+NK | -0.193907451 | 4.61E-08 |
| B16 | B93 | CD3+T | ALDH2-Glu504Lys | 0.18079323 | 3.59E-07 |
| B16 | B92 | CD3+T | ALDH2-Glu504Glu | -0.178662426 | 4.94E-07 |

| | | | | | |
|---|---|---|---|---|---|
| B160 | B192 | PH | UA | -0.18565297 | 1.71E-07 |
| B161 | B190 | LEU | Cre | -0.175349689 | 8.06E-07 |
| B163 | B166 | SG | Pro | 0.353721947 | 0 |
| B163 | B176 | SG | Ucr | 0.347490161 | 0 |
| B163 | B190 | SG | Cre | 0.319131503 | 0 |
| B163 | B192 | SG | UA | 0.322477621 | 0 |
| B163 | B216 | SG | BMI | 0.379237529 | 0 |
| B163 | B221 | SG | Hei | 0.37670135 | 0 |
| B163 | B222 | SG | NS | 0.380834332 | 0 |
| B163 | B210 | SG | Hip | 0.28464355 | 4.44E-16 |
| B163 | B191 | SG | Urea | 0.278361493 | 2.22E-15 |
| B163 | B215 | SG | BMI | 0.235467353 | 2.59E-11 |
| B163 | B206 | SG | Waist | 0.228015741 | 1.11E-10 |
| B163 | B204 | SG | Drinking | 0.17704138 | 6.28E-07 |
| B166 | B222 | Pro | NS | 0.17654802 | 6.76E-07 |
| B167 | B172 | PC | KB | -0.357393683 | 0 |
| B169 | B193 | U-CRY | K | -0.191897471 | 6.37E-08 |
| B17 | B105 | CD3+CD16+CD56+NK | anti-U1RNP | -0.427546056 | 0 |
| B17 | B151 | CD3+CD16+CD56+NK | Anti-CCP | 0.33097868 | 0 |
| B17 | B194 | CD3+CD16+CD56+NK | Cl | -0.299910382 | 0 |
| B17 | B21 | CD3+CD16+CD56+NK | CD3+CD4-CD8- | 0.414246893 | 0 |
| B17 | B187 | CD3+CD16+CD56+NK | NSE | -0.279034 | 1.78E-15 |
| B17 | B95 | CD3+CD16+CD56+NK | anti-cenpb | 0.247653181 | 2.16E-12 |
| B17 | B100 | CD3+CD16+CD56+NK | anti-Ro52 | 0.241917819 | 7.07E-12 |
| B17 | B101 | CD3+CD16+CD56+NK | anti-Sc70 | 0.241917819 | 7.07E-12 |
| B17 | B102 | CD3+CD16+CD56+NK | anti-Sm | 0.241917819 | 7.07E-12 |
| B17 | B103 | CD3+CD16+CD56+NK | anti-SSA/Ro | 0.241917819 | 7.07E-12 |
| B17 | B104 | CD3+CD16+CD56+NK | anti-SSB | 0.241917818 | 7.07E-12 |
| B17 | B106 | CD3+CD16+CD56+NK | anti-P | 0.241917818 | 7.07E-12 |
| B17 | B107 | CD3+CD16+CD56+NK | AnuA | 0.241917807 | 7.07E-12 |
| B17 | B108 | CD3+CD16+CD56+NK | Anti-Cyclin | 0.241917819 | 7.07E-12 |
| B17 | B109 | CD3+CD16+CD56+NK | AMA-M2 | 0.241917819 | 7.07E-12 |
| B17 | B97 | CD3+CD16+CD56+NK | dsDNA | 0.241917818 | 7.07E-12 |
| B17 | B98 | CD3+CD16+CD56+NK | anti-Jo-1 | 0.241917819 | 7.07E-12 |
| B17 | B23 | CD3+CD16+CD56+NK | CD3-CD16+CD56+(NK) | 0.239214369 | 1.22E-11 |
| B17 | B182 | CD3+CD16+CD56+NK | TT | -0.215417089 | 1.16E-09 |
| B17 | B60 | CD3+CD16+CD56+NK | WBRVI | -0.210335528 | 2.87E-09 |
| B17 | B20 | CD3+CD16+CD56+NK | CD3+CD4+CD8+ | -0.204261143 | 8.23E-09 |
| B17 | B183 | CD3+CD16+CD56+NK | PTA | 0.203445217 | 9.45E-09 |
| B17 | B57 | CD3+CD16+CD56+NK | RBCAI | -0.197791609 | 2.44E-08 |
| B17 | B114 | CD3+CD16+CD56+NK | TSH TT3 TT4(TT4) | 0.194550309 | 4.15E-08 |
| B17 | B65 | CD3+CD16+CD56+NK | WBRSR(mPas_1) | -0.191202962 | 7.12E-08 |
| B17 | B90 | CD3+CD16+CD56+NK | CYFRA 21-1 | -0.185282325 | 1.81E-07 |
| B170 | B173 | U-GLU | NTT | 0.279412438 | 1.78E-15 |

| | | | | | |
|---|---|---|---|---|---|
| B171 | B227 | EC | OD | 0.601890639 | 0 |
| B173 | B181 | NTT | APTT | 0.409561683 | 0 |
| B173 | B186 | NTT | PFC | -0.247394568 | 2.28E-12 |
| B174 | B190 | Urea/Cr | Cre | -0.521386572 | 0 |
| B174 | B191 | Urea/Cr | Urea | 0.635582933 | 0 |
| B174 | B221 | Urea/Cr | Hei | -0.375996222 | 0 |
| B174 | B222 | Urea/Cr | NS | -0.291302581 | 0 |
| B174 | B216 | Urea/Cr | BMI | -0.274252938 | 5.77E-15 |
| B174 | B192 | Urea/Cr | UA | -0.26941679 | 1.80E-14 |
| B174 | B189 | Urea/Cr | eGFB | 0.209719715 | 3.20E-09 |
| B174 | B210 | Urea/Cr | Hip | -0.179962215 | 4.07E-07 |
| B175 | B177 | ALB-U | ACR | 0.798504647 | 0 |
| B175 | B176 | ALB-U | Ucr | 0.223885098 | 2.43E-10 |
| B176 | B179 | Ucr | α1-MG | 0.238966877 | 1.29E-11 |
| B176 | B178 | Ucr | Ig-U | 0.201506205 | 1.31E-08 |
| B176 | B221 | Ucr | Hei | 0.199959256 | 1.70E-08 |
| B176 | B192 | Ucr | UA | 0.188023774 | 1.18E-07 |
| B176 | B216 | Ucr | BMI | 0.185515788 | 1.74E-07 |
| B176 | B177 | Ucr | ACR | -0.176911196 | 6.40E-07 |
| B177 | B180 | ACR | Tf | 0.205459678 | 6.70E-09 |
| B178 | B179 | Ig-U | α1-MG | 0.769767517 | 0 |
| B178 | B180 | Ig-U | Tf | 0.652338799 | 0 |
| B178 | B204 | Ig-U | Drinking | 0.187300184 | 1.32E-07 |
| B179 | B204 | α1-MG | Drinking | 0.201415361 | 1.33E-08 |
| B179 | B180 | α1-MG | Tf | 0.186226223 | 1.56E-07 |
| B18 | B105 | CD3+CD4+helper(inducer)T | anti-U1RNP | -0.388855086 | 0 |
| B18 | B182 | CD3+CD4+helper(inducer)T | TT | 0.455251155 | 0 |
| B18 | B19 | CD3+CD4+helper(inducer)T | CD3+CD4+/CD3+CD8+ | 0.639249923 | 0 |
| B18 | B21 | CD3+CD4+helper(inducer)T | CD3+CD4-CD8- | -0.380106743 | 0 |
| B18 | B22 | CD3+CD4+helper(inducer)T | CD3+CD8+(T) | -0.336042871 | 0 |
| B18 | B23 | CD3+CD4+helper(inducer)T | CD3-CD16+CD56+(NK) | -0.319409168 | 0 |
| B18 | B57 | CD3+CD4+helper(inducer)T | RBCAI | -0.277742763 | 2.66E-15 |
| B18 | B157 | CD3+CD4+helper(inducer)T | IgG | -0.251966774 | 8.65E-13 |
| B18 | B65 | CD3+CD4+helper(inducer)T | WBRSR(mPas_1) | -0.249291906 | 1.53E-12 |
| B18 | B155 | CD3+CD4+helper(inducer)T | IgA | -0.245055865 | 3.70E-12 |
| B18 | B69 | CD3+CD4+helper(inducer)T | PV | -0.238129692 | 1.52E-11 |
| B18 | B183 | CD3+CD4+helper(inducer)T | PTA | -0.233563697 | 3.77E-11 |
| B18 | B64 | CD3+CD4+helper(inducer)T | WBRSR(mPas_5) | -0.231660146 | 5.48E-11 |
| B18 | B58 | CD3+CD4+helper(inducer)T | HCT | -0.230950658 | 6.29E-11 |
| B18 | B59 | CD3+CD4+helper(inducer)T | WBRV | -0.224536684 | 2.15E-10 |
| B18 | B60 | CD3+CD4+helper(inducer)T | WBRVI | -0.222759513 | 3.00E-10 |
| B18 | B20 | CD3+CD4+helper(inducer)T | CD3+CD4+CD8+ | 0.200141188 | 1.65E-08 |
| B18 | B135 | CD3+CD4+helper(inducer)T | TBA | -0.19782611 | 2.43E-08 |
| B18 | B187 | CD3+CD4+helper(inducer)T | NSE | -0.174806766 | 8.72E-07 |

| | | | | | |
|---|---|---|---|---|---|
| B18 | B151 | CD3+CD4+helper(inducer)T | Anti-CCP | -0.174268561 | 9.44E-07 |
| B181 | B184 | APTT | INR | 0.394581264 | 0 |
| B181 | B185 | APTT | PT | 0.538861032 | 0 |
| B181 | B186 | APTT | PFC | -0.612005737 | 0 |
| B181 | B183 | APTT | PTA | 0.245428766 | 3.43E-12 |
| B181 | B195 | APTT | Na | 0.179325635 | 4.47E-07 |
| B181 | B200 | APTT | OP | 0.175021824 | 8.45E-07 |
| B182 | B183 | TT | PTA | -0.373495973 | 0 |
| B182 | B184 | TT | INR | -0.181411027 | 3.27E-07 |
| B183 | B184 | PTA | INR | 0.287993467 | 2.22E-16 |
| B183 | B186 | PTA | PFC | -0.281025589 | 1.11E-15 |
| B183 | B195 | PTA | Na | 0.209143585 | 3.53E-09 |
| B183 | B200 | PTA | OP | 0.190137057 | 8.44E-08 |
| B183 | B185 | PTA | PT | 0.186880247 | 1.41E-07 |
| B184 | B185 | INR | PT | 0.864744874 | 0 |
| B185 | B186 | PT | PFC | -0.254019191 | 5.57E-13 |
| B185 | B187 | PT | NSE | -0.197356614 | 2.62E-08 |
| B186 | B187 | PFC | NSE | -0.191072512 | 7.27E-08 |
| B188 | B192 | Ca | UA | 0.20445841 | 7.95E-09 |
| B189 | B190 | eGFB | Cre | -0.588973033 | 0 |
| B189 | B191 | eGFB | Urea | -0.262561353 | 8.53E-14 |
| B189 | B207 | eGFB | WHR | -0.255929155 | 3.68E-13 |
| B189 | B206 | eGFB | Waist | -0.216866913 | 8.90E-10 |
| B189 | B220 | eGFB | SBP | -0.202559677 | 1.10E-08 |
| B189 | B215 | eGFB | BMI | -0.197489262 | 2.57E-08 |
| B19 | B105 | CD3+CD4+/CD3+CD8+ | anti-U1RNP | -0.441387899 | 0 |
| B19 | B125 | CD3+CD4+/CD3+CD8+ | Cor | -0.361212192 | 0 |
| B19 | B182 | CD3+CD4+/CD3+CD8+ | TT | 0.505622461 | 0 |
| B19 | B22 | CD3+CD4+/CD3+CD8+ | CD3+CD8+(T) | -0.857945412 | 0 |
| B19 | B23 | CD3+CD4+/CD3+CD8+ | CD3-CD16+CD56+(NK) | 0.292669875 | 0 |
| B19 | B58 | CD3+CD4+/CD3+CD8+ | HCT | -0.331078777 | 0 |
| B19 | B124 | CD3+CD4+/CD3+CD8+ | ACTH | -0.286861504 | 4.44E-16 |
| B19 | B65 | CD3+CD4+/CD3+CD8+ | WBRSR(mPas_1) | -0.260710396 | 1.29E-13 |
| B19 | B64 | CD3+CD4+/CD3+CD8+ | WBRSR(mPas_5) | -0.247517372 | 2.22E-12 |
| B19 | B185 | CD3+CD4+/CD3+CD8+ | PT | 0.245354735 | 3.48E-12 |
| B19 | B60 | CD3+CD4+/CD3+CD8+ | WBRVI | -0.244183195 | 4.44E-12 |
| B19 | B187 | CD3+CD4+/CD3+CD8+ | NSE | -0.23635864 | 2.17E-11 |
| B19 | B34 | CD3+CD4+/CD3+CD8+ | TK | 0.233861299 | 3.56E-11 |
| B19 | B184 | CD3+CD4+/CD3+CD8+ | INR | 0.232697052 | 4.48E-11 |
| B19 | B67 | CD3+CD4+/CD3+CD8+ | WBRSR(mPas_50) | -0.219522862 | 5.48E-10 |
| B19 | B66 | CD3+CD4+/CD3+CD8+ | WBRSR(mPas_100) | -0.206233504 | 5.86E-09 |
| B19 | B135 | CD3+CD4+/CD3+CD8+ | TBA | -0.202003996 | 1.21E-08 |
| B19 | B63 | CD3+CD4+/CD3+CD8+ | WBRSR(mPas_200) | -0.197603282 | 2.52E-08 |
| B19 | B62 | CD3+CD4+/CD3+CD8+ | WBHSRVI | -0.190723121 | 7.69E-08 |

| | | | | | |
|---|---|---|---|---|---|
| B19 | B156 | CD3+CD4+/CD3+CD8+ | IgE | -0.190560569 | 7.89E-08 |
| B19 | B57 | CD3+CD4+/CD3+CD8+ | RBCAI | -0.182263723 | 2.87E-07 |
| B19 | B145 | CD3+CD4+/CD3+CD8+ | BGP | -0.181966431 | 3.00E-07 |
| B190 | B191 | Cre | Urea | 0.289199776 | 0 |
| B190 | B192 | Cre | UA | 0.49398729 | 0 |
| B190 | B206 | Cre | Waist | 0.344796041 | 0 |
| B190 | B210 | Cre | Hip | 0.325857418 | 0 |
| B190 | B216 | Cre | BMI | 0.478556232 | 0 |
| B190 | B221 | Cre | Hei | 0.515974739 | 0 |
| B190 | B222 | Cre | NS | 0.522143591 | 0 |
| B190 | B215 | Cre | BMI | 0.264225817 | 5.86E-14 |
| B190 | B207 | Cre | WHR | 0.261856151 | 9.99E-14 |
| B190 | B220 | Cre | SBP | 0.230317757 | 7.12E-11 |
| B190 | B209 | Cre | Smoking | 0.188657205 | 1.07E-07 |
| B190 | B204 | Cre | Drinking | 0.186627256 | 1.47E-07 |
| B192 | B206 | UA | Waist | 0.474454295 | 0 |
| B192 | B207 | UA | WHR | 0.350414625 | 0 |
| B192 | B210 | UA | Hip | 0.446272172 | 0 |
| B192 | B215 | UA | BMI | 0.420663793 | 0 |
| B192 | B216 | UA | BMI | 0.56156704 | 0 |
| B192 | B221 | UA | Hei | 0.433889653 | 0 |
| B192 | B222 | UA | NS | 0.560119524 | 0 |
| B192 | B204 | UA | Drinking | 0.234168771 | 3.35E-11 |
| B192 | B220 | UA | SBP | 0.227876639 | 1.14E-10 |
| B192 | B218 | UA | DBP | 0.207273338 | 4.90E-09 |
| B192 | B209 | UA | Smoking | 0.190147255 | 8.43E-08 |
| B193 | B198 | K | P | 0.19365931 | 4.80E-08 |
| B194 | B196 | Cl | TCO2 | -0.409287728 | 0 |
| B194 | B197 | Cl | AG | -0.239631347 | 1.12E-11 |
| B194 | B198 | Cl | P | -0.238104451 | 1.53E-11 |
| B194 | B200 | Cl | OP | 0.227558124 | 1.21E-10 |
| B194 | B195 | Cl | Na | 0.223089432 | 2.82E-10 |
| B195 | B197 | Na | AG | 0.58451561 | 0 |
| B195 | B200 | Na | OP | 0.910352656 | 0 |
| B195 | B199 | Na | Mg | 0.207024703 | 5.11E-09 |
| B195 | B196 | Na | TCO2 | -0.204096426 | 8.46E-09 |
| B196 | B197 | TCO2 | AG | -0.626612152 | 0 |
| B196 | B200 | TCO2 | OP | -0.227164832 | 1.31E-10 |
| B197 | B200 | AG | OP | 0.548080151 | 0 |
| B199 | B200 | Mg | OP | 0.245547176 | 3.34E-12 |
| B2 | B126 | Age | Alb | -0.377697704 | 0 |
| B2 | B189 | Age | eGFB | -0.524627562 | 0 |
| B2 | B207 | Age | WHR | 0.348633601 | 0 |
| B2 | B220 | Age | SBP | 0.291253845 | 0 |

| | | | | | |
|---|---|---|---|---|---|
| B2 | B221 | Age | Hei | -0.301782548 | 0 |
| B2 | B47 | Age | RDW | 0.319632049 | 0 |
| B2 | B77 | Age | TC | 0.282230055 | 8.88E-16 |
| B2 | B153 | Age | Glu | 0.279261389 | 1.78E-15 |
| B2 | B127 | Age | A-G ratio | -0.265863523 | 4.04E-14 |
| B2 | B46 | Age | RBC Count | -0.238947265 | 1.29E-11 |
| B2 | B206 | Age | Waist | 0.223288159 | 2.72E-10 |
| B2 | B41 | Age | MON % | 0.208777774 | 3.77E-09 |
| B2 | B215 | Age | BMI | 0.197976315 | 2.37E-08 |
| B2 | B43 | Age | MCV | 0.196048179 | 3.25E-08 |
| B2 | B218 | Age | DBP | 0.188542175 | 1.09E-07 |
| B2 | B227 | Age | OD | 0.186140979 | 1.58E-07 |
| B2 | B203 | Age | DH | -0.176959989 | 6.36E-07 |
| B2 | B70 | Age | LDL-C | 0.176483577 | 6.82E-07 |
| B20 | B143 | CD3+CD4+CD8+ | β-CTx | 0.322484287 | 0 |
| B20 | B144 | CD3+CD4+CD8+ | Total-PINP | 0.368326337 | 0 |
| B20 | B145 | CD3+CD4+CD8+ | BGP | 0.451956223 | 0 |
| B20 | B21 | CD3+CD4+CD8+ | CD3+CD4-CD8- | -0.313370076 | 0 |
| B20 | B54 | CD3+CD4+CD8+ | ESR | 0.443094666 | 0 |
| B20 | B69 | CD3+CD4+CD8+ | PV | 0.397611061 | 0 |
| B20 | B94 | CD3+CD4+CD8+ | ALDH2-Lys504Lys | 0.281979786 | 8.88E-16 |
| B20 | B99 | CD3+CD4+CD8+ | anti-PM-Scl | -0.271617379 | 1.07E-14 |
| B20 | B157 | CD3+CD4+CD8+ | IgG | 0.254166674 | 5.39E-13 |
| B20 | B73 | CD3+CD4+CD8+ | ApoA | 0.24484916 | 3.87E-12 |
| B20 | B184 | CD3+CD4+CD8+ | INR | 0.244438259 | 4.21E-12 |
| B20 | B151 | CD3+CD4+CD8+ | Anti-CCP | -0.237596722 | 1.69E-11 |
| B20 | B81 | CD3+CD4+CD8+ | TSGF | -0.233276573 | 3.99E-11 |
| B20 | B183 | CD3+CD4+CD8+ | PTA | 0.221546533 | 3.77E-10 |
| B20 | B194 | CD3+CD4+CD8+ | CI | 0.212944491 | 1.81E-09 |
| B20 | B57 | CD3+CD4+CD8+ | RBCAI | -0.195801789 | 3.39E-08 |
| B20 | B79 | CD3+CD4+CD8+ | hb V -dna | 0.192373672 | 5.90E-08 |
| B20 | B55 | CD3+CD4+CD8+ | TK | 0.179982014 | 4.05E-07 |
| B20 | B185 | CD3+CD4+CD8+ | PT | 0.175038574 | 8.43E-07 |
| B204 | B209 | Drinking | Smoking | 0.411475327 | 0 |
| B204 | B216 | Drinking | BMI | 0.298114986 | 0 |
| B204 | B222 | Drinking | NS | 0.325221837 | 0 |
| B204 | B206 | Drinking | Waist | 0.287198748 | 2.22E-16 |
| B204 | B207 | Drinking | WHR | 0.259384344 | 1.73E-13 |
| B204 | B215 | Drinking | BMI | 0.248920744 | 1.65E-12 |
| B204 | B210 | Drinking | Hip | 0.210995531 | 2.55E-09 |
| B204 | B221 | Drinking | Hei | 0.202377515 | 1.13E-08 |
| B206 | B207 | Waist | WHR | 0.868317335 | 0 |
| B206 | B210 | Waist | Hip | 0.780031382 | 0 |
| B206 | B215 | Waist | BMI | 0.849163676 | 0 |

| | | | | | |
|---|---|---|---|---|---|
| B206 | B216 | Waist | BMI | 0.845246634 | 0 |
| B206 | B218 | Waist | DBP | 0.309642193 | 0 |
| B206 | B220 | Waist | SBP | 0.344969563 | 0 |
| B206 | B221 | Waist | Hei | 0.36878975 | 0 |
| B206 | B222 | Waist | NS | 0.774386702 | 0 |
| B206 | B209 | Waist | Smoking | 0.262261022 | 9.13E-14 |
| B207 | B210 | WHR | Hip | 0.370827318 | 0 |
| B207 | B215 | WHR | BMI | 0.642866082 | 0 |
| B207 | B216 | WHR | BMI | 0.58098797 | 0 |
| B207 | B220 | WHR | SBP | 0.301220765 | 0 |
| B207 | B222 | WHR | NS | 0.593055278 | 0 |
| B207 | B218 | WHR | DBP | 0.282501563 | 8.88E-16 |
| B207 | B209 | WHR | Smoking | 0.265025489 | 4.88E-14 |
| B207 | B221 | WHR | Hei | 0.177104868 | 6.22E-07 |
| B209 | B222 | Smoking | NS | 0.286296055 | 4.44E-16 |
| B209 | B216 | Smoking | BMI | 0.234666658 | 3.04E-11 |
| B209 | B221 | Smoking | Hei | 0.189123625 | 9.91E-08 |
| B21 | B124 | CD3+CD4-CD8- | ACTH | -0.475898498 | 0 |
| B21 | B187 | CD3+CD4-CD8- | NSE | -0.280672046 | 1.33E-15 |
| B21 | B186 | CD3+CD4-CD8- | PFC | 0.231490083 | 5.67E-11 |
| B21 | B139 | CD3+CD4-CD8- | ADA | 0.227943455 | 1.12E-10 |
| B21 | B182 | CD3+CD4-CD8- | TT | -0.221808146 | 3.59E-10 |
| B21 | B79 | CD3+CD4-CD8- | hb V -dna | -0.220312972 | 4.73E-10 |
| B21 | B81 | CD3+CD4-CD8- | TSGF | 0.220299213 | 4.75E-10 |
| B21 | B181 | CD3+CD4-CD8- | APTT | 0.217867078 | 7.42E-10 |
| B21 | B58 | CD3+CD4-CD8- | HCT | -0.216415815 | 9.67E-10 |
| B21 | B22 | CD3+CD4-CD8- | CD3+CD8+(T) | -0.195310033 | 3.67E-08 |
| B21 | B195 | CD3+CD4-CD8- | Na | 0.186094363 | 1.59E-07 |
| B21 | B65 | CD3+CD4-CD8- | WBRSR(mPas_1) | -0.18598095 | 1.62E-07 |
| B21 | B138 | CD3+CD4-CD8- | PAB | -0.185211699 | 1.83E-07 |
| B21 | B63 | CD3+CD4-CD8- | WBRSR(mPas_200) | -0.175807391 | 7.54E-07 |
| B21 | B66 | CD3+CD4-CD8- | WBRSR(mPas_100) | -0.174380898 | 9.28E-07 |
| B210 | B215 | Hip | BMI | 0.778057733 | 0 |
| B210 | B216 | Hip | BMI | 0.852741271 | 0 |
| B210 | B221 | Hip | Hei | 0.47777625 | 0 |
| B210 | B222 | Hip | NS | 0.707001769 | 0 |
| B210 | B220 | Hip | SBP | 0.269042413 | 1.95E-14 |
| B210 | B218 | Hip | DBP | 0.221098714 | 4.09E-10 |
| B211 | B212 | IOP-R | IOP-L | 0.785831133 | 0 |
| B215 | B216 | BMI | BMI | 0.828664958 | 0 |
| B215 | B218 | BMI | DBP | 0.305139919 | 0 |
| B215 | B220 | BMI | SBP | 0.322245705 | 0 |
| B215 | B222 | BMI | NS | 0.710141156 | 0 |
| B216 | B218 | BMI | DBP | 0.299832948 | 0 |

| | | | | | |
|---|---|---|---|---|---|
| B216 | B220 | BMI | SBP | 0.32811071 | 0 |
| B216 | B221 | BMI | Hei | 0.662583091 | 0 |
| B216 | B222 | BMI | NS | 0.845263857 | 0 |
| B218 | B220 | DBP | SBP | 0.694839566 | 0 |
| B218 | B222 | DBP | NS | 0.312499805 | 0 |
| B22 | B105 | CD3+CD8+(T) | anti-U1RNP | 0.620879295 | 0 |
| B22 | B182 | CD3+CD8+(T) | TT | -0.422620173 | 0 |
| B22 | B184 | CD3+CD8+(T) | INR | -0.299222857 | 0 |
| B22 | B185 | CD3+CD8+(T) | PT | -0.30760225 | 0 |
| B22 | B187 | CD3+CD8+(T) | NSE | 0.339651048 | 0 |
| B22 | B23 | CD3+CD8+(T) | CD3-CD16+CD56+(NK) | -0.608088344 | 0 |
| B22 | B58 | CD3+CD8+(T) | HCT | 0.33465583 | 0 |
| B22 | B60 | CD3+CD8+(T) | WBRVI | 0.26635191 | 3.64E-14 |
| B22 | B181 | CD3+CD8+(T) | APTT | -0.257760716 | 2.47E-13 |
| B22 | B125 | CD3+CD8+(T) | Cor | 0.247355746 | 2.29E-12 |
| B22 | B65 | CD3+CD8+(T) | WBRSR(mPas_1) | 0.2426069 | 6.14E-12 |
| B22 | B151 | CD3+CD8+(T) | Anti-CCP | -0.237490546 | 1.73E-11 |
| B22 | B95 | CD3+CD8+(T) | anti-cenpb | -0.23394099 | 3.50E-11 |
| B22 | B156 | CD3+CD8+(T) | IgE | 0.233306819 | 3.97E-11 |
| B22 | B100 | CD3+CD8+(T) | anti-Ro52 | -0.228533397 | 1.00E-10 |
| B22 | B101 | CD3+CD8+(T) | anti-Sc70 | -0.228533398 | 1.00E-10 |
| B22 | B102 | CD3+CD8+(T) | anti-Sm | -0.228533397 | 1.00E-10 |
| B22 | B103 | CD3+CD8+(T) | anti-SSA/Ro | -0.228533397 | 1.00E-10 |
| B22 | B104 | CD3+CD8+(T) | anti-SSB | -0.228533396 | 1.00E-10 |
| B22 | B106 | CD3+CD8+(T) | anti-P | -0.228533396 | 1.00E-10 |
| B22 | B107 | CD3+CD8+(T) | AnuA | -0.228533384 | 1.00E-10 |
| B22 | B108 | CD3+CD8+(T) | Anti-Cyclin | -0.228533397 | 1.00E-10 |
| B22 | B109 | CD3+CD8+(T) | AMA-M2 | -0.228533397 | 1.00E-10 |
| B22 | B97 | CD3+CD8+(T) | dsDNA | -0.228533396 | 1.00E-10 |
| B22 | B98 | CD3+CD8+(T) | anti-Jo-1 | -0.228533398 | 1.00E-10 |
| B22 | B64 | CD3+CD8+(T) | WBRSR(mPas_5) | 0.227809042 | 1.15E-10 |
| B22 | B62 | CD3+CD8+(T) | WBHSRVI | 0.222017934 | 3.45E-10 |
| B22 | B124 | CD3+CD8+(T) | ACTH | 0.221316712 | 3.93E-10 |
| B22 | B67 | CD3+CD8+(T) | WBRSR(mPas_50) | 0.202386605 | 1.13E-08 |
| B22 | B90 | CD3+CD8+(T) | CYFRA 21-1 | 0.199729138 | 1.77E-08 |
| B22 | B137 | CD3+CD8+(T) | AFU | 0.187626076 | 1.25E-07 |
| B22 | B148 | CD3+CD8+(T) | Anti_TPO | 0.18568736 | 1.70E-07 |
| B22 | B66 | CD3+CD8+(T) | WBRSR(mPas_100) | 0.184317923 | 2.10E-07 |
| B220 | B222 | SBP | NS | 0.361485003 | 0 |
| B221 | B222 | Hei | NS | 0.558332467 | 0 |
| B23 | B137 | CD3-CD16+CD56+(NK) | AFU | -0.314529233 | 0 |
| B23 | B151 | CD3-CD16+CD56+(NK) | Anti-CCP | 0.325999171 | 0 |
| B23 | B184 | CD3-CD16+CD56+(NK) | INR | 0.402730437 | 0 |
| B23 | B185 | CD3-CD16+CD56+(NK) | PT | 0.362763151 | 0 |

| | | | | | |
|---|---|---|---|---|---|
| B23 | B99 | CD3-CD16+CD56+(NK) | anti-PM-Scl | 0.437176143 | 0 |
| B23 | B69 | CD3-CD16+CD56+(NK) | PV | 0.286194259 | 4.44E-16 |
| B23 | B183 | CD3-CD16+CD56+(NK) | PTA | 0.263487194 | 6.93E-14 |
| B23 | B105 | CD3-CD16+CD56+(NK) | anti-U1RNP | -0.255529546 | 4.01E-13 |
| B23 | B92 | CD3-CD16+CD56+(NK) | ALDH2-Glu504Glu | 0.240539946 | 9.36E-12 |
| B23 | B93 | CD3-CD16+CD56+(NK) | ALDH2-Glu504Lys | -0.2319945 | 5.13E-11 |
| B23 | B34 | CD3-CD16+CD56+(NK) | TK | 0.224937916 | 1.99E-10 |
| B23 | B90 | CD3-CD16+CD56+(NK) | CYFRA 21-1 | -0.187022747 | 1.38E-07 |
| B24 | B177 | αHBDH | ACR | 0.331694741 | 0 |
| B24 | B25 | αHBDH | CK | 0.313009545 | 0 |
| B24 | B26 | αHBDH | CKMB | 0.344737528 | 0 |
| B24 | B27 | αHBDH | LDH | 0.718646622 | 0 |
| B24 | B175 | αHBDH | ALB-U | 0.260793351 | 1.27E-13 |
| B24 | B139 | αHBDH | ADA | 0.178843727 | 4.81E-07 |
| B25 | B37 | CK | BAND | 0.484318998 | 0 |
| B25 | B38 | CK | BAND% | 0.484318998 | 0 |
| B25 | B26 | CK | CKMB | 0.227189848 | 1.30E-10 |
| B25 | B27 | CK | LDH | 0.226881551 | 1.38E-10 |
| B25 | B123 | CK | hs-CRP1 | 0.187742355 | 1.23E-07 |
| B25 | B175 | CK | ALB-U | 0.175302674 | 8.11E-07 |
| B26 | B27 | CKMB | LDH | 0.234659131 | 3.04E-11 |
| B26 | B37 | CKMB | BAND | 0.218731025 | 6.33E-10 |
| B26 | B38 | CKMB | BAND% | 0.218731025 | 6.33E-10 |
| B26 | B175 | CKMB | ALB-U | 0.218401358 | 6.73E-10 |
| B26 | B177 | CKMB | ACR | 0.203495373 | 9.37E-09 |
| B26 | B95 | CKMB | anti-cenpb | 0.180023431 | 4.03E-07 |
| B26 | B100 | CKMB | anti-Ro52 | 0.175977521 | 7.35E-07 |
| B26 | B101 | CKMB | anti-Sc70 | 0.175977521 | 7.35E-07 |
| B26 | B102 | CKMB | anti-Sm | 0.175977521 | 7.35E-07 |
| B26 | B103 | CKMB | anti-SSA/Ro | 0.175977521 | 7.35E-07 |
| B26 | B104 | CKMB | anti-SSB | 0.17597752 | 7.35E-07 |
| B26 | B106 | CKMB | anti-P | 0.17597752 | 7.35E-07 |
| B26 | B107 | CKMB | AnuA | 0.175977512 | 7.35E-07 |
| B26 | B108 | CKMB | Anti-Cyclin | 0.175977521 | 7.35E-07 |
| B26 | B109 | CKMB | AMA-M2 | 0.17597752 | 7.35E-07 |
| B26 | B97 | CKMB | dsDNA | 0.17597752 | 7.35E-07 |
| B26 | B98 | CKMB | anti-Jo-1 | 0.175977521 | 7.35E-07 |
| B27 | B139 | LDH | ADA | 0.286750271 | 4.44E-16 |
| B27 | B177 | LDH | ACR | 0.244276693 | 4.35E-12 |
| B27 | B175 | LDH | ALB-U | 0.185098052 | 1.86E-07 |
| B27 | B135 | LDH | TBA | 0.174368741 | 9.30E-07 |
| B28 | B29 | PRL | E2 | 0.95409213 | 0 |
| B28 | B30 | PRL | LH | 0.952018403 | 0 |
| B28 | B31 | PRL | FSH | 0.304234591 | 0 |

| | | | | | |
|---|---|---|---|---|---|
| B28 | B32 | PRL | Testo | 0.355839615 | 0 |
| B28 | B33 | PRL | Prog | -0.392575683 | 0 |
| B29 | B30 | E2 | LH | 0.871911635 | 0 |
| B29 | B142 | E2 | 25-(OH)D | 0.189964639 | 8.67E-08 |
| B3 | B78 | Hcy | HBcAb | 0.179102046 | 4.62E-07 |
| B30 | B32 | LH | Testo | 0.476205574 | 0 |
| B30 | B33 | LH | Prog | -0.571425584 | 0 |
| B30 | B31 | LH | FSH | 0.282768103 | 8.88E-16 |
| B31 | B32 | FSH | Testo | 0.565904576 | 0 |
| B31 | B33 | FSH | Prog | -0.433340972 | 0 |
| B32 | B33 | Testo | Prog | -0.964288889 | 0 |
| B34 | B124 | TK | ACTH | -0.643108258 | 0 |
| B34 | B125 | TK | Cor | -0.541321054 | 0 |
| B34 | B185 | TK | PT | 0.487647182 | 0 |
| B34 | B57 | TK | RBCAI | -0.437963891 | 0 |
| B34 | B58 | TK | HCT | -0.433964412 | 0 |
| B34 | B59 | TK | WBRV | -0.297956386 | 0 |
| B34 | B60 | TK | WBRVI | -0.376098618 | 0 |
| B34 | B64 | TK | WBRSR(mPas_5) | -0.311052254 | 0 |
| B34 | B65 | TK | WBRSR(mPas_1) | -0.355455809 | 0 |
| B34 | B138 | TK | PAB | -0.284778181 | 4.44E-16 |
| B34 | B79 | TK | hb V -dna | -0.26871628 | 2.11E-14 |
| B34 | B158 | TK | IgM | 0.267081024 | 3.06E-14 |
| B34 | B62 | TK | WBHSRVI | -0.251503731 | 9.55E-13 |
| B34 | B67 | TK | WBRSR(mPas_50) | -0.242078194 | 6.84E-12 |
| B34 | B184 | TK | INR | 0.240812332 | 8.85E-12 |
| B34 | B137 | TK | AFU | -0.237169712 | 1.85E-11 |
| B34 | B134 | TK | CHE | -0.235331787 | 2.66E-11 |
| B34 | B66 | TK | WBRSR(mPas_100) | -0.232506742 | 4.65E-11 |
| B34 | B63 | TK | WBRSR(mPas_200) | -0.226519946 | 1.48E-10 |
| B34 | B198 | TK | P | 0.215426026 | 1.16E-09 |
| B34 | B139 | TK | ADA | 0.203143793 | 9.95E-09 |
| B34 | B81 | TK | TSGF | 0.197205408 | 2.69E-08 |
| B34 | B161 | TK | LEU | 0.196889751 | 2.83E-08 |
| B34 | B115 | TK | TSH TT3 TT4(TT3) | -0.175911114 | 7.42E-07 |
| B35 | B96 | β2-MG | dsDNA (IIF) | 0.789542034 | 0 |
| B36 | B141 | V-Lym% | AST/ALT | -0.18112199 | 3.41E-07 |
| B37 | B38 | BAND | BAND% | 1 | 0 |
| B37 | B58 | BAND | HCT | 0.344235736 | 0 |
| B37 | B67 | BAND | WBRSR(mPas_50) | 0.279440945 | 1.78E-15 |
| B37 | B66 | BAND | WBRSR(mPas_100) | 0.276008 | 4.00E-15 |
| B37 | B63 | BAND | WBRSR(mPas_200) | 0.269200079 | 1.89E-14 |
| B37 | B64 | BAND | WBRSR(mPas_5) | 0.26577794 | 4.13E-14 |
| B37 | B62 | BAND | WBHSRVI | 0.244847499 | 3.87E-12 |

| | | | | | |
|---|---|---|---|---|---|
| B37 | B65 | BAND | WBRSR(mPas_1) | 0.243151823 | 5.49E-12 |
| B37 | B60 | BAND | WBRVI | 0.213065003 | 1.77E-09 |
| B37 | B194 | BAND | CI | -0.194797425 | 3.99E-08 |
| B38 | B58 | BAND% | HCT | 0.344235736 | 0 |
| B38 | B67 | BAND% | WBRSR(mPas_50) | 0.279440945 | 1.78E-15 |
| B38 | B66 | BAND% | WBRSR(mPas_100) | 0.276008 | 4.00E-15 |
| B38 | B63 | BAND% | WBRSR(mPas_200) | 0.269200079 | 1.89E-14 |
| B38 | B64 | BAND% | WBRSR(mPas_5) | 0.26577794 | 4.13E-14 |
| B38 | B62 | BAND% | WBHSRVI | 0.244847499 | 3.87E-12 |
| B38 | B65 | BAND% | WBRSR(mPas_1) | 0.243151823 | 5.49E-12 |
| B38 | B60 | BAND% | WBRVI | 0.213065003 | 1.77E-09 |
| B38 | B194 | BAND% | CI | -0.194797425 | 3.99E-08 |
| B39 | B44 | Other cells | MCH | -0.17437613 | 9.29E-07 |
| B39 | B43 | Other cells | MCV | -0.174042068 | 9.75E-07 |
| B4 | B6 | WB-Ca | WB-Fe | -0.999999998 | 0 |
| B4 | B7 | WB-Ca | WB-Cop | -1 | 0 |
| B4 | B205 | WB-Ca | KDA | -0.192890215 | 5.43E-08 |
| B40 | B42 | WBC | MON | 0.677195259 | 0 |
| B40 | B53 | WBC | GR% | 0.286547516 | 4.44E-16 |
| B40 | B46 | WBC | RBC Count | 0.281882721 | 8.88E-16 |
| B40 | B48 | WBC | LYM% | -0.282757343 | 8.88E-16 |
| B40 | B51 | WBC | Hb | 0.223816942 | 2.46E-10 |
| B40 | B52 | WBC | PLT/BPC | 0.217725238 | 7.62E-10 |
| B40 | B192 | WBC | UA | 0.212970447 | 1.80E-09 |
| B40 | B209 | WBC | Smoking | 0.203388166 | 9.55E-09 |
| B40 | B72 | WBC | HDL-C | -0.197382869 | 2.61E-08 |
| B40 | B216 | WBC | BMI | 0.180263228 | 3.89E-07 |
| B40 | B210 | WBC | Hip | 0.174919547 | 8.58E-07 |
| B41 | B42 | MON % | MON | 0.653829943 | 0 |
| B41 | B53 | MON % | GR% | -0.229543803 | 8.27E-11 |
| B41 | B126 | MON % | Alb | -0.190060043 | 8.54E-08 |
| B41 | B49 | MON % | baso% | 0.186862815 | 1.41E-07 |
| B41 | B189 | MON % | eGFB | -0.186492054 | 1.50E-07 |
| B42 | B209 | MON | Smoking | 0.21804926 | 7.18E-10 |
| B42 | B48 | MON | LYM% | -0.211026607 | 2.54E-09 |
| B42 | B46 | MON | RBC Count | 0.20798999 | 4.32E-09 |
| B42 | B192 | MON | UA | 0.20581804 | 6.30E-09 |
| B42 | B51 | MON | Hb | 0.196192235 | 3.18E-08 |
| B42 | B210 | MON | Hip | 0.195149276 | 3.77E-08 |
| B42 | B206 | MON | Waist | 0.188918275 | 1.02E-07 |
| B42 | B216 | MON | BMI | 0.185821474 | 1.66E-07 |
| B42 | B222 | MON | NS | 0.184196869 | 2.14E-07 |
| B42 | B190 | MON | Cre | 0.177641405 | 5.75E-07 |
| B43 | B44 | MCV | MCH | 0.912715389 | 0 |

| | | | | | |
|---|---|---|---|---|---|
| B43 | B46 | MCV | RBC Count | -0.584656695 | 0 |
| B43 | B47 | MCV | RDW | 0.600365707 | 0 |
| B43 | B45 | MCV | MCHC | 0.177252647 | 6.09E-07 |
| B43 | B52 | MCV | PLT/BPC | -0.175641351 | 7.72E-07 |
| B44 | B45 | MCH | MCHC | 0.562175863 | 0 |
| B44 | B46 | MCH | RBC Count | -0.438299978 | 0 |
| B44 | B47 | MCH | RDW | 0.427841701 | 0 |
| B44 | B51 | MCH | Hb | 0.341578227 | 0 |
| B45 | B51 | MCHC | Hb | 0.579582529 | 0 |
| B45 | B192 | MCHC | UA | 0.284820606 | 4.44E-16 |
| B45 | B221 | MCHC | Hei | 0.285379412 | 4.44E-16 |
| B45 | B222 | MCHC | NS | 0.271679074 | 1.07E-14 |
| B45 | B216 | MCHC | BMI | 0.257570069 | 2.57E-13 |
| B45 | B141 | MCHC | AST/ALT | -0.241620916 | 7.51E-12 |
| B45 | B190 | MCHC | Cre | 0.240294481 | 9.83E-12 |
| B45 | B126 | MCHC | Alb | 0.195459292 | 3.58E-08 |
| B45 | B47 | MCHC | RDW | -0.191240317 | 7.08E-08 |
| B45 | B128 | MCHC | ALT | 0.177944027 | 5.50E-07 |
| B45 | B174 | MCHC | Urea/Cr | -0.175766554 | 7.58E-07 |
| B45 | B127 | MCHC | A-G ratio | 0.174088508 | 9.69E-07 |
| B46 | B126 | RBC Count | Alb | 0.322566551 | 0 |
| B46 | B128 | RBC Count | ALT | 0.335467764 | 0 |
| B46 | B141 | RBC Count | AST/ALT | -0.346983707 | 0 |
| B46 | B163 | RBC Count | SG | 0.323812834 | 0 |
| B46 | B174 | RBC Count | Urea/Cr | -0.337659535 | 0 |
| B46 | B190 | RBC Count | Cre | 0.443736162 | 0 |
| B46 | B192 | RBC Count | UA | 0.432529277 | 0 |
| B46 | B206 | RBC Count | Waist | 0.355749545 | 0 |
| B46 | B210 | RBC Count | Hip | 0.350209865 | 0 |
| B46 | B216 | RBC Count | BMI | 0.475650886 | 0 |
| B46 | B221 | RBC Count | Hei | 0.47694067 | 0 |
| B46 | B222 | RBC Count | NS | 0.522741129 | 0 |
| B46 | B47 | RBC Count | RDW | -0.515795955 | 0 |
| B46 | B51 | RBC Count | Hb | 0.687959087 | 0 |
| B46 | B72 | RBC Count | HDL-C | -0.29372975 | 0 |
| B46 | B215 | RBC Count | BMI | 0.284514697 | 4.44E-16 |
| B46 | B207 | RBC Count | WHR | 0.255032494 | 4.47E-13 |
| B46 | B218 | RBC Count | DBP | 0.245329207 | 3.50E-12 |
| B46 | B220 | RBC Count | SBP | 0.226265405 | 1.55E-10 |
| B46 | B188 | RBC Count | Ca | 0.216999822 | 8.69E-10 |
| B46 | B131 | RBC Count | AST | 0.209058389 | 3.59E-09 |
| B46 | B140 | RBC Count | r-GT | 0.203236643 | 9.80E-09 |
| B46 | B91 | RBC Count | TP | 0.18543373 | 1.76E-07 |
| B46 | B204 | RBC Count | Drinking | 0.182009452 | 2.98E-07 |

| | | | | | |
|---|---|---|---|---|---|
| B47 | B52 | RDW | PLT/BPC | -0.260180089 | 1.45E-13 |
| B47 | B126 | RDW | Alb | -0.223088782 | 2.82E-10 |
| B47 | B51 | RDW | Hb | -0.198752835 | 2.08E-08 |
| B47 | B221 | RDW | Hei | -0.176641344 | 6.66E-07 |
| B48 | B53 | LYM% | GR% | -0.936162853 | 0 |
| B49 | B50 | baso% | eo% | 0.333382653 | 0 |
| B49 | B53 | baso% | GR% | -0.181832687 | 3.06E-07 |
| B5 | B205 | WB-Mg | KDA | 0.192890215 | 5.43E-08 |
| B50 | B53 | eo% | GR% | -0.259096142 | 1.84E-13 |
| B51 | B126 | Hb | Alb | 0.32724754 | 0 |
| B51 | B128 | Hb | ALT | 0.329695618 | 0 |
| B51 | B141 | Hb | AST/ALT | -0.369168009 | 0 |
| B51 | B163 | Hb | SG | 0.34821928 | 0 |
| B51 | B174 | Hb | Urea/Cr | -0.373252816 | 0 |
| B51 | B190 | Hb | Cre | 0.541438032 | 0 |
| B51 | B192 | Hb | UA | 0.482555172 | 0 |
| B51 | B206 | Hb | Waist | 0.424542056 | 0 |
| B51 | B207 | Hb | WHR | 0.329908476 | 0 |
| B51 | B210 | Hb | Hip | 0.388025373 | 0 |
| B51 | B215 | Hb | BMI | 0.32329867 | 0 |
| B51 | B216 | Hb | BMI | 0.547036702 | 0 |
| B51 | B221 | Hb | Hei | 0.552109393 | 0 |
| B51 | B222 | Hb | NS | 0.588832698 | 0 |
| B51 | B220 | Hb | SBP | 0.275963924 | 4.00E-15 |
| B51 | B218 | Hb | DBP | 0.265732421 | 4.17E-14 |
| B51 | B72 | Hb | HDL-C | -0.24674843 | 2.60E-12 |
| B51 | B204 | Hb | Drinking | 0.239872895 | 1.07E-11 |
| B51 | B131 | Hb | AST | 0.23390377 | 3.53E-11 |
| B51 | B58 | Hb | HCT | 0.233231784 | 4.03E-11 |
| B51 | B140 | Hb | r-GT | 0.216161257 | 1.01E-09 |
| B51 | B71 | Hb | TG | 0.214912662 | 1.27E-09 |
| B51 | B209 | Hb | Smoking | 0.21383064 | 1.54E-09 |
| B51 | B188 | Hb | Ca | 0.213243418 | 1.71E-09 |
| B51 | B176 | Hb | Ucr | 0.206294026 | 5.80E-09 |
| B51 | B70 | Hb | LDL-C | 0.205413933 | 6.75E-09 |
| B51 | B127 | Hb | A-G ratio | 0.204011411 | 8.58E-09 |
| B51 | B161 | Hb | LEU | -0.192763159 | 5.54E-08 |
| B54 | B155 | ESR | IgA | 0.318712821 | 0 |
| B54 | B184 | ESR | INR | 0.406360992 | 0 |
| B54 | B185 | ESR | PT | 0.371545897 | 0 |
| B54 | B186 | ESR | PFC | 0.353214691 | 0 |
| B54 | B69 | ESR | PV | 0.347259397 | 0 |
| B54 | B157 | ESR | IgG | 0.264999655 | 4.93E-14 |
| B54 | B151 | ESR | Anti-CCP | -0.238850133 | 1.32E-11 |

| | | | | | |
|---|---|---|---|---|---|
| B54 | B60 | ESR | WBRVI | -0.225428378 | 1.82E-10 |
| B54 | B58 | ESR | HCT | -0.213531805 | 1.63E-09 |
| B54 | B156 | ESR | IgE | 0.21225049 | 2.04E-09 |
| B54 | B62 | ESR | WBHSRVI | -0.204487738 | 7.91E-09 |
| B54 | B158 | ESR | IgM | 0.198088834 | 2.32E-08 |
| B54 | B136 | ESR | 5'NT | 0.195714641 | 3.43E-08 |
| B54 | B92 | ESR | ALDH2-Glu504Glu | -0.173957802 | 9.87E-07 |
| B55 | B124 | TK | ACTH | -0.305372382 | 0 |
| B55 | B56 | TK | IR | 0.771723097 | 0 |
| B55 | B57 | TK | RBCAI | -0.373479788 | 0 |
| B55 | B58 | TK | HCT | -0.475624687 | 0 |
| B55 | B59 | TK | WBRV | 0.394313063 | 0 |
| B55 | B61 | TK | WBHSRV | 0.70932351 | 0 |
| B55 | B62 | TK | WBHSRVI | 0.439160804 | 0 |
| B55 | B63 | TK | WBRSR(mPas_200) | 0.352622224 | 0 |
| B55 | B66 | TK | WBRSR(mPas_100) | 0.342066769 | 0 |
| B55 | B67 | TK | WBRSR(mPas_50) | 0.307741015 | 0 |
| B55 | B95 | TK | anti-cenpb | -0.274565173 | 5.33E-15 |
| B55 | B100 | TK | anti-Ro52 | -0.266165031 | 3.77E-14 |
| B55 | B101 | TK | anti-Sc70 | -0.266165032 | 3.77E-14 |
| B55 | B102 | TK | anti-Sm | -0.266165032 | 3.77E-14 |
| B55 | B103 | TK | anti-SSA/Ro | -0.266165031 | 3.77E-14 |
| B55 | B104 | TK | anti-SSB | -0.266165031 | 3.77E-14 |
| B55 | B106 | TK | anti-P | -0.26616503 | 3.77E-14 |
| B55 | B107 | TK | AnuA | -0.266165016 | 3.77E-14 |
| B55 | B108 | TK | Anti-Cyclin | -0.266165032 | 3.77E-14 |
| B55 | B109 | TK | AMA-M2 | -0.266165031 | 3.77E-14 |
| B55 | B97 | TK | dsDNA | -0.266165031 | 3.77E-14 |
| B55 | B98 | TK | anti-Jo-1 | -0.266165032 | 3.77E-14 |
| B55 | B125 | TK | Cor | -0.234043037 | 3.43E-11 |
| B55 | B176 | TK | Ucr | -0.188804769 | 1.04E-07 |
| B55 | B69 | TK | PV | -0.18281205 | 2.64E-07 |
| B55 | B64 | TK | WBRSR(mPas_5) | 0.175337971 | 8.07E-07 |
| B56 | B59 | IR | WBRV | 0.689867427 | 0 |
| B56 | B60 | IR | WBRVI | 0.653636514 | 0 |
| B56 | B61 | IR | WBHSRV | 0.966441759 | 0 |
| B56 | B62 | IR | WBHSRVI | 0.905543796 | 0 |
| B56 | B63 | IR | WBRSR(mPas_200) | 0.830932307 | 0 |
| B56 | B64 | IR | WBRSR(mPas_5) | 0.708624252 | 0 |
| B56 | B65 | IR | WBRSR(mPas_1) | 0.628797998 | 0 |
| B56 | B66 | IR | WBRSR(mPas_100) | 0.832835184 | 0 |
| B56 | B67 | IR | WBRSR(mPas_50) | 0.813381192 | 0 |
| B56 | B95 | IR | anti-cenpb | -0.296485243 | 0 |
| B56 | B100 | IR | anti-Ro52 | -0.283989218 | 4.44E-16 |

| | | | | | |
|---|---|---|---|---|---|
| B56 | B101 | IR | anti-Sc70 | -0.283989219 | 4.44E-16 |
| B56 | B102 | IR | anti-Sm | -0.283989219 | 4.44E-16 |
| B56 | B103 | IR | anti-SSA/Ro | -0.283989218 | 4.44E-16 |
| B56 | B104 | IR | anti-SSB | -0.283989218 | 4.44E-16 |
| B56 | B106 | IR | anti-P | -0.283989218 | 4.44E-16 |
| B56 | B107 | IR | AnuA | -0.283989204 | 4.44E-16 |
| B56 | B108 | IR | Anti-Cyclin | -0.283989219 | 4.44E-16 |
| B56 | B109 | IR | AMA-M2 | -0.283989218 | 4.44E-16 |
| B56 | B97 | IR | dsDNA | -0.283989218 | 4.44E-16 |
| B56 | B98 | IR | anti-Jo-1 | -0.28398922 | 4.44E-16 |
| B56 | B158 | IR | IgM | -0.26080318 | 1.26E-13 |
| B56 | B185 | IR | PT | -0.206153358 | 5.94E-09 |
| B56 | B150 | IR | ALP | 0.204204746 | 8.31E-09 |
| B56 | B184 | IR | INR | -0.191410888 | 6.89E-08 |
| B56 | B181 | IR | APTT | -0.187597072 | 1.26E-07 |
| B56 | B204 | IR | Drinking | 0.183869453 | 2.25E-07 |
| B57 | B124 | RBCAI | ACTH | 0.335560965 | 0 |
| B57 | B154 | RBCAI | RF | 0.307687921 | 0 |
| B57 | B58 | RBCAI | HCT | 0.455692068 | 0 |
| B57 | B59 | RBCAI | WBRV | 0.559120969 | 0 |
| B57 | B60 | RBCAI | WBRVI | 0.607175886 | 0 |
| B57 | B64 | RBCAI | WBRSR(mPas_5) | 0.444909035 | 0 |
| B57 | B65 | RBCAI | WBRSR(mPas_1) | 0.577905259 | 0 |
| B57 | B79 | RBCAI | hb V -dna | 0.303349198 | 0 |
| B57 | B185 | RBCAI | PT | -0.287271858 | 2.22E-16 |
| B57 | B105 | RBCAI | anti-U1RNP | 0.27681532 | 3.11E-15 |
| B57 | B187 | RBCAI | NSE | 0.223179236 | 2.78E-10 |
| B57 | B125 | RBCAI | Cor | 0.215637662 | 1.11E-09 |
| B57 | B67 | RBCAI | WBRSR(mPas_50) | 0.202378155 | 1.13E-08 |
| B57 | B81 | RBCAI | TSGF | -0.200019595 | 1.69E-08 |
| B57 | B139 | RBCAI | ADA | -0.188405587 | 1.11E-07 |
| B57 | B180 | RBCAI | Tf | 0.185026304 | 1.88E-07 |
| B58 | B105 | HCT | anti-U1RNP | 0.317279384 | 0 |
| B58 | B124 | HCT | ACTH | 0.509002346 | 0 |
| B58 | B125 | HCT | Cor | 0.311176091 | 0 |
| B58 | B185 | HCT | PT | -0.4291117 | 0 |
| B58 | B59 | HCT | WBRV | 0.324169457 | 0 |
| B58 | B60 | HCT | WBRVI | 0.703485505 | 0 |
| B58 | B62 | HCT | WBHSRVI | 0.568228652 | 0 |
| B58 | B63 | HCT | WBRSR(mPas_200) | 0.579294029 | 0 |
| B58 | B64 | HCT | WBRSR(mPas_5) | 0.693997248 | 0 |
| B58 | B65 | HCT | WBRSR(mPas_1) | 0.714047964 | 0 |
| B58 | B66 | HCT | WBRSR(mPas_100) | 0.600464608 | 0 |
| B58 | B67 | HCT | WBRSR(mPas_50) | 0.624886355 | 0 |

| | | | | | |
|---|---|---|---|---|---|
| B58 | B181 | HCT | APTT | -0.283099146 | 6.66E-16 |
| B58 | B198 | HCT | P | -0.268292248 | 2.33E-14 |
| B58 | B187 | HCT | NSE | 0.253116971 | 6.76E-13 |
| B58 | B79 | HCT | hb V -dna | 0.20898354 | 3.64E-09 |
| B58 | B176 | HCT | Ucr | 0.205235009 | 6.96E-09 |
| B58 | B158 | HCT | IgM | -0.205055471 | 7.18E-09 |
| B58 | B61 | HCT | WBHSRV | 0.202089524 | 1.19E-08 |
| B58 | B184 | HCT | INR | -0.199871017 | 1.73E-08 |
| B59 | B154 | WBRV | RF | 0.299990083 | 0 |
| B59 | B60 | WBRV | WBRVI | 0.840180174 | 0 |
| B59 | B61 | WBRV | WBHSRV | 0.754850752 | 0 |
| B59 | B62 | WBRV | WBHSRVI | 0.706535625 | 0 |
| B59 | B63 | WBRV | WBRSR(mPas_200) | 0.760030786 | 0 |
| B59 | B64 | WBRV | WBRSR(mPas_5) | 0.862578936 | 0 |
| B59 | B65 | WBRV | WBRSR(mPas_1) | 0.881461941 | 0 |
| B59 | B66 | WBRV | WBRSR(mPas_100) | 0.77214225 | 0 |
| B59 | B67 | WBRV | WBRSR(mPas_50) | 0.790191109 | 0 |
| B59 | B79 | WBRV | hb V -dna | 0.248408248 | 1.84E-12 |
| B59 | B69 | WBRV | PV | 0.241895977 | 7.10E-12 |
| B59 | B151 | WBRV | Anti-CCP | 0.213419046 | 1.66E-09 |
| B59 | B124 | WBRV | ACTH | 0.209223224 | 3.49E-09 |
| B59 | B95 | WBRV | anti-cenpb | -0.206415535 | 5.68E-09 |
| B59 | B100 | WBRV | anti-Ro52 | -0.204106921 | 8.45E-09 |
| B59 | B101 | WBRV | anti-Sc70 | -0.204106922 | 8.45E-09 |
| B59 | B102 | WBRV | anti-Sm | -0.204106922 | 8.45E-09 |
| B59 | B103 | WBRV | anti-SSA/Ro | -0.204106921 | 8.45E-09 |
| B59 | B104 | WBRV | anti-SSB | -0.204106921 | 8.45E-09 |
| B59 | B106 | WBRV | anti-P | -0.20410692 | 8.45E-09 |
| B59 | B107 | WBRV | AnuA | -0.204106909 | 8.45E-09 |
| B59 | B108 | WBRV | Anti-Cyclin | -0.204106921 | 8.45E-09 |
| B59 | B109 | WBRV | AMA-M2 | -0.204106921 | 8.45E-09 |
| B59 | B97 | WBRV | dsDNA | -0.204106921 | 8.45E-09 |
| B59 | B98 | WBRV | anti-Jo-1 | -0.204106922 | 8.45E-09 |
| B59 | B185 | WBRV | PT | -0.191079978 | 7.26E-08 |
| B59 | B81 | WBRV | TSGF | -0.186253303 | 1.55E-07 |
| B6 | B7 | WB-Fe | WB-Cop | 0.999999999 | 0 |
| B6 | B8 | WB-Fe | WB-Zinc | -0.999999995 | 0 |
| B6 | B205 | WB-Fe | KDA | 0.192890215 | 5.43E-08 |
| B60 | B105 | WBRVI | anti-U1RNP | 0.336905642 | 0 |
| B60 | B124 | WBRVI | ACTH | 0.308689344 | 0 |
| B60 | B185 | WBRVI | PT | -0.407710966 | 0 |
| B60 | B61 | WBRVI | WBHSRV | 0.648935909 | 0 |
| B60 | B62 | WBRVI | WBHSRVI | 0.848288612 | 0 |
| B60 | B63 | WBRVI | WBRSR(mPas_200) | 0.794777447 | 0 |

| | | | | | |
|---|---|---|---|---|---|
| B60 | B64 | WBRVI | WBRSR(mPas_5) | 0.937126641 | 0 |
| B60 | B65 | WBRVI | WBRSR(mPas_1) | 0.964417394 | 0 |
| B60 | B66 | WBRVI | WBRSR(mPas_100) | 0.826416523 | 0 |
| B60 | B67 | WBRVI | WBRSR(mPas_50) | 0.852583245 | 0 |
| B60 | B187 | WBRVI | NSE | 0.249173084 | 1.57E-12 |
| B60 | B95 | WBRVI | anti-cenpb | -0.234050591 | 3.43E-11 |
| B60 | B100 | WBRVI | anti-Ro52 | -0.229155835 | 8.91E-11 |
| B60 | B101 | WBRVI | anti-Sc70 | -0.229155835 | 8.91E-11 |
| B60 | B102 | WBRVI | anti-Sm | -0.229155835 | 8.91E-11 |
| B60 | B103 | WBRVI | anti-SSA/Ro | -0.229155835 | 8.91E-11 |
| B60 | B104 | WBRVI | anti-SSB | -0.229155834 | 8.91E-11 |
| B60 | B106 | WBRVI | anti-P | -0.229155834 | 8.91E-11 |
| B60 | B107 | WBRVI | AnuA | -0.229155822 | 8.91E-11 |
| B60 | B108 | WBRVI | Anti-Cyclin | -0.229155835 | 8.91E-11 |
| B60 | B109 | WBRVI | AMA-M2 | -0.229155835 | 8.91E-11 |
| B60 | B97 | WBRVI | dsDNA | -0.229155834 | 8.91E-11 |
| B60 | B98 | WBRVI | anti-Jo-1 | -0.229155836 | 8.91E-11 |
| B60 | B79 | WBRVI | hb V -dna | 0.214199533 | 1.44E-09 |
| B60 | B158 | WBRVI | IgM | -0.211896164 | 2.18E-09 |
| B60 | B181 | WBRVI | APTT | -0.209397849 | 3.38E-09 |
| B60 | B198 | WBRVI | P | -0.19287408 | 5.45E-08 |
| B60 | B184 | WBRVI | INR | -0.17804322 | 5.42E-07 |
| B61 | B62 | WBHSRV | WBHSRVI | 0.888963931 | 0 |
| B61 | B63 | WBHSRV | WBRSR(mPas_200) | 0.899840779 | 0 |
| B61 | B64 | WBHSRV | WBRSR(mPas_5) | 0.768595727 | 0 |
| B61 | B65 | WBHSRV | WBRSR(mPas_1) | 0.68574917 | 0 |
| B61 | B66 | WBHSRV | WBRSR(mPas_100) | 0.892244204 | 0 |
| B61 | B67 | WBHSRV | WBRSR(mPas_50) | 0.871204407 | 0 |
| B61 | B95 | WBHSRV | anti-cenpb | -0.245481705 | 3.39E-12 |
| B61 | B158 | WBHSRV | IgM | -0.243747366 | 4.85E-12 |
| B61 | B150 | WBHSRV | ALP | 0.235724056 | 2.46E-11 |
| B61 | B100 | WBHSRV | anti-Ro52 | -0.233207454 | 4.05E-11 |
| B61 | B101 | WBHSRV | anti-Sc70 | -0.233207455 | 4.05E-11 |
| B61 | B102 | WBHSRV | anti-Sm | -0.233207454 | 4.05E-11 |
| B61 | B103 | WBHSRV | anti-SSA/Ro | -0.233207454 | 4.05E-11 |
| B61 | B104 | WBHSRV | anti-SSB | -0.233207454 | 4.05E-11 |
| B61 | B106 | WBHSRV | anti-P | -0.233207453 | 4.05E-11 |
| B61 | B107 | WBHSRV | AnuA | -0.23320744 | 4.05E-11 |
| B61 | B108 | WBHSRV | Anti-Cyclin | -0.233207454 | 4.05E-11 |
| B61 | B109 | WBHSRV | AMA-M2 | -0.233207454 | 4.05E-11 |
| B61 | B97 | WBHSRV | dsDNA | -0.233207454 | 4.05E-11 |
| B61 | B98 | WBHSRV | anti-Jo-1 | -0.233207455 | 4.05E-11 |
| B61 | B76 | WBHSRV | ApoB | 0.182261002 | 2.87E-07 |
| B61 | B204 | WBHSRV | Drinking | 0.179642979 | 4.26E-07 |

| | | | | | |
|---|---|---|---|---|---|
| B62 | B185 | WBHSRVI | PT | -0.342207572 | 0 |
| B62 | B63 | WBHSRVI | WBRSR(mPas_200) | 0.936975634 | 0 |
| B62 | B64 | WBHSRVI | WBRSR(mPas_5) | 0.885835493 | 0 |
| B62 | B65 | WBHSRVI | WBRSR(mPas_1) | 0.828799379 | 0 |
| B62 | B66 | WBHSRVI | WBRSR(mPas_100) | 0.94999456 | 0 |
| B62 | B67 | WBHSRVI | WBRSR(mPas_50) | 0.943933428 | 0 |
| B62 | B158 | WBHSRVI | IgM | -0.284181219 | 4.44E-16 |
| B62 | B181 | WBHSRVI | APTT | -0.271663449 | 1.07E-14 |
| B62 | B95 | WBHSRVI | anti-cenpb | -0.267429446 | 2.84E-14 |
| B62 | B100 | WBHSRVI | anti-Ro52 | -0.253859813 | 5.76E-13 |
| B62 | B101 | WBHSRVI | anti-Sc70 | -0.253859814 | 5.76E-13 |
| B62 | B102 | WBHSRVI | anti-Sm | -0.253859814 | 5.76E-13 |
| B62 | B103 | WBHSRVI | anti-SSA/Ro | -0.253859813 | 5.76E-13 |
| B62 | B104 | WBHSRVI | anti-SSB | -0.253859813 | 5.76E-13 |
| B62 | B106 | WBHSRVI | anti-P | -0.253859812 | 5.76E-13 |
| B62 | B107 | WBHSRVI | AnuA | -0.253859799 | 5.76E-13 |
| B62 | B108 | WBHSRVI | Anti-Cyclin | -0.253859813 | 5.76E-13 |
| B62 | B109 | WBHSRVI | AMA-M2 | -0.253859813 | 5.76E-13 |
| B62 | B97 | WBHSRVI | dsDNA | -0.253859813 | 5.76E-13 |
| B62 | B98 | WBHSRVI | anti-Jo-1 | -0.253859814 | 5.76E-13 |
| B62 | B184 | WBHSRVI | INR | -0.242049351 | 6.88E-12 |
| B62 | B124 | WBHSRVI | ACTH | 0.223997284 | 2.38E-10 |
| B62 | B105 | WBHSRVI | anti-U1RNP | 0.192960096 | 5.37E-08 |
| B62 | B150 | WBHSRVI | ALP | 0.176886357 | 6.43E-07 |
| B63 | B64 | WBRSR(mPas_200) | WBRSR(mPas_5) | 0.931616404 | 0 |
| B63 | B65 | WBRSR(mPas_200) | WBRSR(mPas_1) | 0.865296258 | 0 |
| B63 | B66 | WBRSR(mPas_200) | WBRSR(mPas_100) | 0.996559234 | 0 |
| B63 | B67 | WBRSR(mPas_200) | WBRSR(mPas_50) | 0.989942684 | 0 |
| B63 | B69 | WBRSR(mPas_200) | PV | 0.31296031 | 0 |
| B63 | B124 | WBRSR(mPas_200) | ACTH | 0.255550767 | 4.00E-13 |
| B63 | B158 | WBRSR(mPas_200) | IgM | -0.254232831 | 5.32E-13 |
| B63 | B181 | WBRSR(mPas_200) | APTT | -0.205384071 | 6.79E-09 |
| B63 | B185 | WBRSR(mPas_200) | PT | -0.204716193 | 7.61E-09 |
| B63 | B95 | WBRSR(mPas_200) | anti-cenpb | -0.198161644 | 2.30E-08 |
| B63 | B135 | WBRSR(mPas_200) | TBA | 0.19744672 | 2.58E-08 |
| B63 | B150 | WBRSR(mPas_200) | ALP | 0.194980459 | 3.87E-08 |
| B63 | B100 | WBRSR(mPas_200) | anti-Ro52 | -0.184037297 | 2.19E-07 |
| B63 | B101 | WBRSR(mPas_200) | anti-Sc70 | -0.184037298 | 2.19E-07 |
| B63 | B102 | WBRSR(mPas_200) | anti-Sm | -0.184037298 | 2.19E-07 |
| B63 | B103 | WBRSR(mPas_200) | anti-SSA/Ro | -0.184037297 | 2.19E-07 |
| B63 | B104 | WBRSR(mPas_200) | anti-SSB | -0.184037297 | 2.19E-07 |
| B63 | B106 | WBRSR(mPas_200) | anti-P | -0.184037297 | 2.19E-07 |
| B63 | B107 | WBRSR(mPas_200) | AnuA | -0.184037287 | 2.19E-07 |
| B63 | B108 | WBRSR(mPas_200) | Anti-Cyclin | -0.184037298 | 2.19E-07 |

| | | | | | |
|---|---|---|---|---|---|
| B63 | B109 | WBRSR(mPas_200) | AMA-M2 | -0.184037297 | 2.19E-07 |
| B63 | B97 | WBRSR(mPas_200) | dsDNA | -0.184037297 | 2.19E-07 |
| B63 | B98 | WBRSR(mPas_200) | anti-Jo-1 | -0.184037298 | 2.19E-07 |
| B64 | B124 | WBRSR(mPas_5) | ACTH | 0.316135739 | 0 |
| B64 | B65 | WBRSR(mPas_5) | WBRSR(mPas_1) | 0.982181376 | 0 |
| B64 | B66 | WBRSR(mPas_5) | WBRSR(mPas_100) | 0.946279255 | 0 |
| B64 | B67 | WBRSR(mPas_5) | WBRSR(mPas_50) | 0.966023379 | 0 |
| B64 | B185 | WBRSR(mPas_5) | PT | -0.285883262 | 4.44E-16 |
| B64 | B69 | WBRSR(mPas_5) | PV | 0.252732778 | 7.34E-13 |
| B64 | B105 | WBRSR(mPas_5) | anti-U1RNP | 0.25114612 | 1.03E-12 |
| B64 | B158 | WBRSR(mPas_5) | IgM | -0.224345447 | 2.23E-10 |
| B64 | B79 | WBRSR(mPas_5) | hb V -dna | 0.223610399 | 2.56E-10 |
| B64 | B187 | WBRSR(mPas_5) | NSE | 0.202994659 | 1.02E-08 |
| B64 | B95 | WBRSR(mPas_5) | anti-cenpb | -0.198196252 | 2.28E-08 |
| B64 | B135 | WBRSR(mPas_5) | TBA | 0.191027255 | 7.32E-08 |
| B64 | B100 | WBRSR(mPas_5) | anti-Ro52 | -0.1902175 | 8.33E-08 |
| B64 | B101 | WBRSR(mPas_5) | anti-Sc70 | -0.1902175 | 8.33E-08 |
| B64 | B102 | WBRSR(mPas_5) | anti-Sm | -0.1902175 | 8.33E-08 |
| B64 | B103 | WBRSR(mPas_5) | anti-SSA/Ro | -0.1902175 | 8.33E-08 |
| B64 | B104 | WBRSR(mPas_5) | anti-SSB | -0.190217499 | 8.33E-08 |
| B64 | B106 | WBRSR(mPas_5) | anti-P | -0.190217499 | 8.33E-08 |
| B64 | B107 | WBRSR(mPas_5) | AnuA | -0.190217488 | 8.33E-08 |
| B64 | B108 | WBRSR(mPas_5) | Anti-Cyclin | -0.1902175 | 8.33E-08 |
| B64 | B109 | WBRSR(mPas_5) | AMA-M2 | -0.190217499 | 8.33E-08 |
| B64 | B97 | WBRSR(mPas_5) | dsDNA | -0.190217499 | 8.33E-08 |
| B64 | B98 | WBRSR(mPas_5) | anti-Jo-1 | -0.190217501 | 8.33E-08 |
| B65 | B105 | WBRSR(mPas_1) | anti-U1RNP | 0.290119821 | 0 |
| B65 | B124 | WBRSR(mPas_1) | ACTH | 0.348670898 | 0 |
| B65 | B185 | WBRSR(mPas_1) | PT | -0.314415293 | 0 |
| B65 | B66 | WBRSR(mPas_1) | WBRSR(mPas_100) | 0.88527639 | 0 |
| B65 | B67 | WBRSR(mPas_1) | WBRSR(mPas_50) | 0.908535568 | 0 |
| B65 | B79 | WBRSR(mPas_1) | hb V -dna | 0.239020497 | 1.27E-11 |
| B65 | B187 | WBRSR(mPas_1) | NSE | 0.228508067 | 1.01E-10 |
| B65 | B69 | WBRSR(mPas_1) | PV | 0.222044831 | 3.43E-10 |
| B65 | B95 | WBRSR(mPas_1) | anti-cenpb | -0.2110399 | 2.53E-09 |
| B65 | B100 | WBRSR(mPas_1) | anti-Ro52 | -0.205559188 | 6.59E-09 |
| B65 | B101 | WBRSR(mPas_1) | anti-Sc70 | -0.205559189 | 6.59E-09 |
| B65 | B102 | WBRSR(mPas_1) | anti-Sm | -0.205559188 | 6.59E-09 |
| B65 | B103 | WBRSR(mPas_1) | anti-SSA/Ro | -0.205559188 | 6.59E-09 |
| B65 | B104 | WBRSR(mPas_1) | anti-SSB | -0.205559187 | 6.59E-09 |
| B65 | B106 | WBRSR(mPas_1) | anti-P | -0.205559187 | 6.59E-09 |
| B65 | B107 | WBRSR(mPas_1) | AnuA | -0.205559176 | 6.59E-09 |
| B65 | B108 | WBRSR(mPas_1) | Anti-Cyclin | -0.205559188 | 6.59E-09 |
| B65 | B109 | WBRSR(mPas_1) | AMA-M2 | -0.205559188 | 6.59E-09 |

| | | | | | |
|---|---|---|---|---|---|
| B65 | B97 | WBRSR(mPas_1) | dsDNA | -0.205559187 | 6.59E-09 |
| B65 | B98 | WBRSR(mPas_1) | anti-Jo-1 | -0.205559189 | 6.59E-09 |
| B65 | B158 | WBRSR(mPas_1) | IgM | -0.198267764 | 2.26E-08 |
| B65 | B181 | WBRSR(mPas_1) | APTT | -0.181351991 | 3.30E-07 |
| B65 | B125 | WBRSR(mPas_1) | Cor | 0.175314498 | 8.10E-07 |
| B66 | B67 | WBRSR(mPas_100) | WBRSR(mPas_50) | 0.996105112 | 0 |
| B66 | B69 | WBRSR(mPas_100) | PV | 0.278380347 | 2.22E-15 |
| B66 | B124 | WBRSR(mPas_100) | ACTH | 0.255681634 | 3.89E-13 |
| B66 | B158 | WBRSR(mPas_100) | IgM | -0.245931476 | 3.09E-12 |
| B66 | B185 | WBRSR(mPas_100) | PT | -0.233450865 | 3.86E-11 |
| B66 | B95 | WBRSR(mPas_100) | anti-cenpb | -0.220011667 | 5.00E-10 |
| B66 | B100 | WBRSR(mPas_100) | anti-Ro52 | -0.207932201 | 4.37E-09 |
| B66 | B101 | WBRSR(mPas_100) | anti-Sc70 | -0.207932202 | 4.37E-09 |
| B66 | B102 | WBRSR(mPas_100) | anti-Sm | -0.207932202 | 4.37E-09 |
| B66 | B103 | WBRSR(mPas_100) | anti-SSA/Ro | -0.207932201 | 4.37E-09 |
| B66 | B104 | WBRSR(mPas_100) | anti-SSB | -0.207932201 | 4.37E-09 |
| B66 | B106 | WBRSR(mPas_100) | anti-P | -0.2079322 | 4.37E-09 |
| B66 | B107 | WBRSR(mPas_100) | AnuA | -0.207932191 | 4.37E-09 |
| B66 | B108 | WBRSR(mPas_100) | Anti-Cyclin | -0.207932202 | 4.37E-09 |
| B66 | B109 | WBRSR(mPas_100) | AMA-M2 | -0.207932201 | 4.37E-09 |
| B66 | B97 | WBRSR(mPas_100) | dsDNA | -0.207932201 | 4.37E-09 |
| B66 | B98 | WBRSR(mPas_100) | anti-Jo-1 | -0.207932202 | 4.37E-09 |
| B66 | B181 | WBRSR(mPas_100) | APTT | -0.207883867 | 4.40E-09 |
| B66 | B150 | WBRSR(mPas_100) | ALP | 0.199401805 | 1.87E-08 |
| B66 | B135 | WBRSR(mPas_100) | TBA | 0.185634163 | 1.71E-07 |
| B67 | B124 | WBRSR(mPas_50) | ACTH | 0.269661283 | 1.69E-14 |
| B67 | B69 | WBRSR(mPas_50) | PV | 0.269448532 | 1.78E-14 |
| B67 | B158 | WBRSR(mPas_50) | IgM | -0.248520204 | 1.80E-12 |
| B67 | B185 | WBRSR(mPas_50) | PT | -0.246672075 | 2.65E-12 |
| B67 | B95 | WBRSR(mPas_50) | anti-cenpb | -0.211975796 | 2.15E-09 |
| B67 | B100 | WBRSR(mPas_50) | anti-Ro52 | -0.200446376 | 1.57E-08 |
| B67 | B101 | WBRSR(mPas_50) | anti-Sc70 | -0.200446377 | 1.57E-08 |
| B67 | B102 | WBRSR(mPas_50) | anti-Sm | -0.200446376 | 1.57E-08 |
| B67 | B103 | WBRSR(mPas_50) | anti-SSA/Ro | -0.200446376 | 1.57E-08 |
| B67 | B104 | WBRSR(mPas_50) | anti-SSB | -0.200446375 | 1.57E-08 |
| B67 | B106 | WBRSR(mPas_50) | anti-P | -0.200446375 | 1.57E-08 |
| B67 | B107 | WBRSR(mPas_50) | AnuA | -0.200446366 | 1.57E-08 |
| B67 | B108 | WBRSR(mPas_50) | Anti-Cyclin | -0.200446376 | 1.57E-08 |
| B67 | B109 | WBRSR(mPas_50) | AMA-M2 | -0.200446376 | 1.57E-08 |
| B67 | B97 | WBRSR(mPas_50) | dsDNA | -0.200446376 | 1.57E-08 |
| B67 | B98 | WBRSR(mPas_50) | anti-Jo-1 | -0.200446377 | 1.57E-08 |
| B67 | B150 | WBRSR(mPas_50) | ALP | 0.196400596 | 3.07E-08 |
| B67 | B181 | WBRSR(mPas_50) | APTT | -0.196014035 | 3.27E-08 |
| B67 | B135 | WBRSR(mPas_50) | TBA | 0.192641652 | 5.65E-08 |

| | | | | | |
|---|---|---|---|---|---|
| B67 | B105 | WBRSR(mPas_50) | anti-U1RNP | 0.17918521 | 4.57E-07 |
| B69 | B145 | PV | BGP | 0.313313686 | 0 |
| B69 | B183 | PV | PTA | 0.466849468 | 0 |
| B69 | B184 | PV | INR | 0.400227969 | 0 |
| B69 | B185 | PV | PT | 0.303097019 | 0 |
| B69 | B195 | PV | Na | 0.288183574 | 2.22E-16 |
| B69 | B144 | PV | Total-PINP | 0.257535265 | 2.59E-13 |
| B69 | B193 | PV | K | 0.25140834 | 9.74E-13 |
| B69 | B154 | PV | RF | 0.245301061 | 3.52E-12 |
| B69 | B182 | PV | TT | -0.233743771 | 3.64E-11 |
| B69 | B81 | PV | TSGF | -0.227559863 | 1.21E-10 |
| B69 | B143 | PV | β-CTx | 0.225115274 | 1.93E-10 |
| B69 | B157 | PV | IgG | 0.211511882 | 2.33E-09 |
| B69 | B199 | PV | Mg | 0.208858274 | 3.72E-09 |
| B69 | B135 | PV | TBA | 0.200272204 | 1.62E-08 |
| B69 | B137 | PV | AFU | -0.194373385 | 4.27E-08 |
| B69 | B197 | PV | AG | 0.192250873 | 6.02E-08 |
| B69 | B200 | PV | OP | 0.183573286 | 2.35E-07 |
| B69 | B151 | PV | Anti-CCP | 0.180836425 | 3.56E-07 |
| B7 | B8 | WB-Cop | WB-Zinc | -0.999999998 | 0 |
| B7 | B205 | WB-Cop | KDA | 0.192890215 | 5.43E-08 |
| B70 | B76 | LDL-C | ApoB | 0.367887918 | 0 |
| B70 | B77 | LDL-C | TC | 0.840429168 | 0 |
| B70 | B75 | LDL-C | ApoA1/B | -0.265245318 | 4.66E-14 |
| B70 | B206 | LDL-C | Waist | 0.211141812 | 2.49E-09 |
| B70 | B207 | LDL-C | WHR | 0.210062898 | 3.01E-09 |
| B70 | B132 | LDL-C | Dbil | -0.209892632 | 3.10E-09 |
| B70 | B215 | LDL-C | BMI | 0.186391197 | 1.52E-07 |
| B71 | B140 | TG | r-GT | 0.293270739 | 0 |
| B71 | B192 | TG | UA | 0.35093929 | 0 |
| B71 | B206 | TG | Waist | 0.340460704 | 0 |
| B71 | B207 | TG | WHR | 0.305047265 | 0 |
| B71 | B215 | TG | BMI | 0.304815785 | 0 |
| B71 | B216 | TG | BMI | 0.296084175 | 0 |
| B71 | B222 | TG | NS | 0.308350522 | 0 |
| B71 | B72 | TG | HDL-C | -0.423714625 | 0 |
| B71 | B77 | TG | TC | 0.299894169 | 0 |
| B71 | B128 | TG | ALT | 0.273720645 | 6.66E-15 |
| B71 | B210 | TG | Hip | 0.253351111 | 6.43E-13 |
| B71 | B91 | TG | TP | 0.247974658 | 2.01E-12 |
| B71 | B141 | TG | AST/ALT | -0.227705385 | 1.18E-10 |
| B71 | B130 | TG | GLB | 0.212227856 | 2.05E-09 |
| B71 | B131 | TG | AST | 0.192120713 | 6.15E-08 |
| B72 | B141 | HDL-C | AST/ALT | 0.321838288 | 0 |

| | | | | | |
|---|---|---|---|---|---|
| B72 | B192 | HDL-C | UA | -0.43686957 | 0 |
| B72 | B206 | HDL-C | Waist | -0.325949446 | 0 |
| B72 | B210 | HDL-C | Hip | -0.295259401 | 0 |
| B72 | B216 | HDL-C | BMI | -0.387061611 | 0 |
| B72 | B221 | HDL-C | Hei | -0.306929789 | 0 |
| B72 | B222 | HDL-C | NS | -0.396443639 | 0 |
| B72 | B74 | HDL-C | ApoA1 | 0.295094627 | 0 |
| B72 | B215 | HDL-C | BMI | -0.287630145 | 2.22E-16 |
| B72 | B190 | HDL-C | Cre | -0.251129128 | 1.03E-12 |
| B72 | B207 | HDL-C | WHR | -0.248614234 | 1.76E-12 |
| B72 | B128 | HDL-C | ALT | -0.232012999 | 5.12E-11 |
| B72 | B163 | HDL-C | SG | -0.204667309 | 7.67E-09 |
| B72 | B77 | HDL-C | TC | 0.194209799 | 4.39E-08 |
| B72 | B75 | HDL-C | ApoA1/B | 0.193896076 | 4.62E-08 |
| B72 | B140 | HDL-C | r-GT | -0.174131222 | 9.63E-07 |
| B73 | B183 | ApoA | PTA | 0.255005492 | 4.50E-13 |
| B73 | B182 | ApoA | TT | -0.233730937 | 3.65E-11 |
| B73 | B145 | ApoA | BGP | 0.217576325 | 7.83E-10 |
| B73 | B144 | ApoA | Total-PINP | 0.191327793 | 6.98E-08 |
| B74 | B75 | ApoA1 | ApoA1/B | 0.572581556 | 0 |
| B74 | B183 | ApoA1 | PTA | 0.193897966 | 4.62E-08 |
| B74 | B157 | ApoA1 | IgG | 0.178168534 | 5.32E-07 |
| B75 | B76 | ApoA1/B | ApoB | -0.776013984 | 0 |
| B75 | B77 | ApoA1/B | TC | -0.200507209 | 1.55E-08 |
| B75 | B94 | ApoA1/B | ALDH2-Lys504Lys | 0.197675503 | 2.49E-08 |
| B75 | B138 | ApoA1/B | PAB | -0.181504163 | 3.22E-07 |
| B76 | B77 | ApoB | TC | 0.323903823 | 0 |
| B76 | B138 | ApoB | PAB | 0.220765431 | 4.35E-10 |
| B76 | B186 | ApoB | PFC | 0.215703258 | 1.10E-09 |
| B76 | B105 | ApoB | anti-U1RNP | -0.183069477 | 2.54E-07 |
| B76 | B94 | ApoB | ALDH2-Lys504Lys | -0.18122839 | 3.36E-07 |
| B76 | B95 | ApoB | anti-cenpb | -0.179346709 | 4.46E-07 |
| B76 | B197 | ApoB | AG | 0.176046888 | 7.27E-07 |
| B76 | B100 | ApoB | anti-Ro52 | -0.174991237 | 8.49E-07 |
| B76 | B101 | ApoB | anti-Sc70 | -0.174991238 | 8.49E-07 |
| B76 | B102 | ApoB | anti-Sm | -0.174991238 | 8.49E-07 |
| B76 | B103 | ApoB | anti-SSA/Ro | -0.174991237 | 8.49E-07 |
| B76 | B104 | ApoB | anti-SSB | -0.174991237 | 8.49E-07 |
| B76 | B106 | ApoB | anti-P | -0.174991237 | 8.49E-07 |
| B76 | B107 | ApoB | AnuA | -0.174991228 | 8.49E-07 |
| B76 | B108 | ApoB | Anti-Cyclin | -0.174991237 | 8.49E-07 |
| B76 | B109 | ApoB | AMA-M2 | -0.174991237 | 8.49E-07 |
| B76 | B97 | ApoB | dsDNA | -0.174991237 | 8.49E-07 |
| B76 | B98 | ApoB | anti-Jo-1 | -0.174991238 | 8.49E-07 |

| | | | | | |
|---|---|---|---|---|---|
| B77 | B207 | TC | WHR | 0.246565504 | 2.71E-12 |
| B77 | B132 | TC | Dbil | -0.22518733 | 1.90E-10 |
| B77 | B206 | TC | Waist | 0.217655363 | 7.71E-10 |
| B77 | B189 | TC | eGFB | -0.205604121 | 6.53E-09 |
| B77 | B218 | TC | DBP | 0.200427058 | 1.57E-08 |
| B77 | B91 | TC | TP | 0.200303531 | 1.61E-08 |
| B77 | B215 | TC | BMI | 0.192791729 | 5.52E-08 |
| B77 | B130 | TC | GLB | 0.176535014 | 6.77E-07 |
| B78 | B80 | HBcAb | anti-HP IgG | -0.203013034 | 1.02E-08 |
| B79 | B100 | hb V -dna | anti-Ro52 | 0.336014371 | 0 |
| B79 | B101 | hb V -dna | anti-Sc70 | 0.336014372 | 0 |
| B79 | B102 | hb V -dna | anti-Sm | 0.336014372 | 0 |
| B79 | B103 | hb V -dna | anti-SSA/Ro | 0.336014371 | 0 |
| B79 | B104 | hb V -dna | anti-SSB | 0.336014371 | 0 |
| B79 | B106 | hb V -dna | anti-P | 0.336014371 | 0 |
| B79 | B107 | hb V -dna | AnuA | 0.336014354 | 0 |
| B79 | B108 | hb V -dna | Anti-Cyclin | 0.336014372 | 0 |
| B79 | B109 | hb V -dna | AMA-M2 | 0.336014371 | 0 |
| B79 | B119 | hb V -dna | Hcv-rna | 0.402716043 | 0 |
| B79 | B121 | hb V -dna | C3 | -0.29218057 | 0 |
| B79 | B122 | hb V -dna | C4 | -0.292180462 | 0 |
| B79 | B152 | hb V -dna | ACA-Ig(G-A-M) | -0.292176298 | 0 |
| B79 | B159 | hb V -dna | β2-MG | -0.292180671 | 0 |
| B79 | B169 | hb V -dna | U-CRY | -0.292179685 | 0 |
| B79 | B94 | hb V -dna | ALDH2-Lys504Lys | 0.346468664 | 0 |
| B79 | B95 | hb V -dna | anti-cenpb | 0.344174569 | 0 |
| B79 | B97 | hb V -dna | dsDNA | 0.336014371 | 0 |
| B79 | B98 | hb V -dna | anti-Jo-1 | 0.336014373 | 0 |
| B79 | B99 | hb V -dna | anti-PM-Scl | -0.304021285 | 0 |
| B79 | B81 | hb V -dna | TSGF | -0.281522886 | 8.88E-16 |
| B79 | B183 | hb V -dna | PTA | 0.274665357 | 5.33E-15 |
| B79 | B180 | hb V -dna | Tf | 0.263794956 | 6.48E-14 |
| B79 | B178 | hb V -dna | Ig-U | 0.220016447 | 5.00E-10 |
| B79 | B154 | hb V -dna | RF | 0.219619201 | 5.38E-10 |
| B79 | B124 | hb V -dna | ACTH | 0.209049403 | 3.59E-09 |
| B79 | B157 | hb V -dna | IgG | 0.19220795 | 6.06E-08 |
| B79 | B186 | hb V -dna | PFC | -0.184477399 | 2.05E-07 |
| B79 | B193 | hb V -dna | K | 0.180062021 | 4.00E-07 |
| B8 | B205 | WB-Zinc | KDA | -0.192890215 | 5.43E-08 |
| B81 | B156 | TSGF | IgE | 0.293443566 | 0 |
| B81 | B184 | TSGF | INR | -0.313953748 | 0 |
| B81 | B92 | TSGF | ALDH2-Glu504Glu | 0.286400039 | 4.44E-16 |
| B81 | B151 | TSGF | Anti-CCP | -0.283532381 | 6.66E-16 |
| B81 | B93 | TSGF | ALDH2-Glu504Lys | -0.274073576 | 6.22E-15 |

| | | | | | |
|---|---|---|---|---|---|
| B81 | B95 | TSGF | anti-cenpb | -0.218709781 | 6.36E-10 |
| B81 | B100 | TSGF | anti-Ro52 | -0.213808513 | 1.55E-09 |
| B81 | B101 | TSGF | anti-Sc70 | -0.213808514 | 1.55E-09 |
| B81 | B102 | TSGF | anti-Sm | -0.213808513 | 1.55E-09 |
| B81 | B103 | TSGF | anti-SSA/Ro | -0.213808513 | 1.55E-09 |
| B81 | B104 | TSGF | anti-SSB | -0.213808513 | 1.55E-09 |
| B81 | B106 | TSGF | anti-P | -0.213808513 | 1.55E-09 |
| B81 | B107 | TSGF | AnuA | -0.213808501 | 1.55E-09 |
| B81 | B108 | TSGF | Anti-Cyclin | -0.213808513 | 1.55E-09 |
| B81 | B109 | TSGF | AMA-M2 | -0.213808513 | 1.55E-09 |
| B81 | B97 | TSGF | dsDNA | -0.213808513 | 1.55E-09 |
| B81 | B98 | TSGF | anti-Jo-1 | -0.213808514 | 1.55E-09 |
| B81 | B94 | TSGF | ALDH2-Lys504Lys | -0.209912476 | 3.09E-09 |
| B81 | B124 | TSGF | ACTH | -0.187166643 | 1.35E-07 |
| B81 | B154 | TSGF | RF | -0.181329632 | 3.31E-07 |
| B81 | B183 | TSGF | PTA | -0.175706245 | 7.65E-07 |
| B81 | B125 | TSGF | Cor | -0.174460211 | 9.18E-07 |
| B82 | B84 | CEA | C-PSA | 0.330933801 | 0 |
| B82 | B89 | CEA | CA 72-4 | 0.536228532 | 0 |
| B82 | B87 | CEA | NSE | -0.261629474 | 1.05E-13 |
| B83 | B84 | AFP | C-PSA | 0.335808578 | 0 |
| B83 | B90 | AFP | CYFRA 21-1 | 0.347505194 | 0 |
| B83 | B88 | AFP | CA 242 | 0.259918435 | 1.54E-13 |
| B83 | B87 | AFP | NSE | 0.180518868 | 3.74E-07 |
| B84 | B89 | C-PSA | CA 72-4 | 0.196471909 | 3.03E-08 |
| B84 | B86 | C-PSA | TBhCG | -0.180105737 | 3.98E-07 |
| B85 | B89 | SCC | CA 72-4 | -0.300609802 | 0 |
| B85 | B87 | SCC | NSE | 0.17631228 | 7.00E-07 |
| B86 | B90 | TBhCG | CYFRA 21-1 | 0.33992062 | 0 |
| B86 | B87 | TBhCG | NSE | 0.228246045 | 1.06E-10 |
| B87 | B88 | NSE | CA 242 | 0.203310535 | 9.67E-09 |
| B89 | B194 | CA 72-4 | Cl | -0.17665135 | 6.66E-07 |
| B90 | B105 | CYFRA 21-1 | anti-U1RNP | 0.244161853 | 4.46E-12 |
| B90 | B137 | CYFRA 21-1 | AFU | 0.205252162 | 6.94E-09 |
| B91 | B126 | TP | Alb | 0.528669874 | 0 |
| B91 | B127 | TP | A-G ratio | -0.41022617 | 0 |
| B91 | B130 | TP | GLB | 0.750113882 | 0 |
| B91 | B188 | TP | Ca | 0.367140474 | 0 |
| B92 | B134 | ALDH2-Glu504Glu | CHE | -0.357668372 | 0 |
| B92 | B93 | ALDH2-Glu504Glu | ALDH2-Glu504Lys | -0.998790507 | 0 |
| B92 | B184 | ALDH2-Glu504Glu | INR | -0.282688142 | 8.88E-16 |
| B92 | B125 | ALDH2-Glu504Glu | Cor | -0.249661031 | 1.41E-12 |
| B92 | B185 | ALDH2-Glu504Glu | PT | -0.245732482 | 3.22E-12 |
| B92 | B99 | ALDH2-Glu504Glu | anti-PM-Scl | 0.227165462 | 1.31E-10 |

| | | | | | |
|---|---|---|---|---|---|
| B92 | B182 | ALDH2-Glu504Glu | TT | 0.211984537 | 2.14E-09 |
| B92 | B136 | ALDH2-Glu504Glu | 5'NT | -0.189750448 | 8.97E-08 |
| B92 | B199 | ALDH2-Glu504Glu | Mg | -0.187833951 | 1.21E-07 |
| B92 | B155 | ALDH2-Glu504Glu | IgA | 0.178746436 | 4.88E-07 |
| B93 | B134 | ALDH2-Glu504Lys | CHE | 0.366707612 | 0 |
| B93 | B184 | ALDH2-Glu504Lys | INR | 0.278967328 | 1.78E-15 |
| B93 | B125 | ALDH2-Glu504Lys | Cor | 0.26350544 | 6.91E-14 |
| B93 | B185 | ALDH2-Glu504Lys | PT | 0.24643839 | 2.78E-12 |
| B93 | B182 | ALDH2-Glu504Lys | TT | -0.213433133 | 1.65E-09 |
| B93 | B99 | ALDH2-Glu504Lys | anti-PM-Scl | -0.209669035 | 3.22E-09 |
| B93 | B136 | ALDH2-Glu504Lys | 5'NT | 0.206811916 | 5.30E-09 |
| B93 | B199 | ALDH2-Glu504Lys | Mg | 0.191056334 | 7.29E-08 |
| B93 | B155 | ALDH2-Glu504Lys | IgA | -0.178748517 | 4.88E-07 |
| B94 | B100 | ALDH2-Lys504Lys | anti-Ro52 | 0.358379014 | 0 |
| B94 | B101 | ALDH2-Lys504Lys | anti-Sc70 | 0.358379015 | 0 |
| B94 | B102 | ALDH2-Lys504Lys | anti-Sm | 0.358379014 | 0 |
| B94 | B103 | ALDH2-Lys504Lys | anti-SSA/Ro | 0.358379013 | 0 |
| B94 | B104 | ALDH2-Lys504Lys | anti-SSB | 0.358379018 | 0 |
| B94 | B106 | ALDH2-Lys504Lys | anti-P | 0.35837902 | 0 |
| B94 | B107 | ALDH2-Lys504Lys | AnuA | 0.358379039 | 0 |
| B94 | B108 | ALDH2-Lys504Lys | Anti-Cyclin | 0.358379014 | 0 |
| B94 | B109 | ALDH2-Lys504Lys | AMA-M2 | 0.358379015 | 0 |
| B94 | B125 | ALDH2-Lys504Lys | Cor | -0.313956197 | 0 |
| B94 | B136 | ALDH2-Lys504Lys | 5'NT | -0.37077141 | 0 |
| B94 | B138 | ALDH2-Lys504Lys | PAB | -0.343112676 | 0 |
| B94 | B140 | ALDH2-Lys504Lys | r-GT | -0.307718322 | 0 |
| B94 | B157 | ALDH2-Lys504Lys | IgG | 0.331147348 | 0 |
| B94 | B95 | ALDH2-Lys504Lys | anti-cenpb | 0.366978571 | 0 |
| B94 | B97 | ALDH2-Lys504Lys | dsDNA | 0.358379018 | 0 |
| B94 | B98 | ALDH2-Lys504Lys | anti-Jo-1 | 0.358379017 | 0 |
| B94 | B99 | ALDH2-Lys504Lys | anti-PM-Scl | -0.322501245 | 0 |
| B94 | B134 | ALDH2-Lys504Lys | CHE | -0.231628987 | 5.51E-11 |
| B94 | B124 | ALDH2-Lys504Lys | ACTH | -0.207114017 | 5.03E-09 |
| B94 | B197 | ALDH2-Lys504Lys | AG | -0.197760616 | 2.45E-08 |
| B94 | B115 | ALDH2-Lys504Lys | TSH TT3 TT4(TT3) | -0.197218136 | 2.68E-08 |
| B94 | B196 | ALDH2-Lys504Lys | TCO2 | 0.196683235 | 2.93E-08 |
| B94 | B128 | ALDH2-Lys504Lys | ALT | -0.188135599 | 1.16E-07 |
| B94 | B154 | ALDH2-Lys504Lys | RF | 0.178253435 | 5.25E-07 |
| B95 | B100 | anti-cenpb | anti-Ro52 | 0.977375647 | 0 |
| B95 | B101 | anti-cenpb | anti-Sc70 | 0.977375651 | 0 |
| B95 | B102 | anti-cenpb | anti-Sm | 0.977375651 | 0 |
| B95 | B103 | anti-cenpb | anti-SSA/Ro | 0.977375647 | 0 |
| B95 | B104 | anti-cenpb | anti-SSB | 0.977375647 | 0 |
| B95 | B106 | anti-cenpb | anti-P | 0.97737565 | 0 |

| | | | | | |
|---|---|---|---|---|---|
| B95 | B107 | anti-cenpb | AnuA | 0.977375603 | 0 |
| B95 | B108 | anti-cenpb | Anti-Cyclin | 0.977375648 | 0 |
| B95 | B109 | anti-cenpb | AMA-M2 | 0.977375647 | 0 |
| B95 | B183 | anti-cenpb | PTA | 0.310620149 | 0 |
| B95 | B97 | anti-cenpb | dsDNA | 0.977375648 | 0 |
| B95 | B98 | anti-cenpb | anti-Jo-1 | 0.977375651 | 0 |
| B95 | B154 | anti-cenpb | RF | 0.246503164 | 2.74E-12 |
| B95 | B186 | anti-cenpb | PFC | -0.236951973 | 1.93E-11 |
| B95 | B180 | anti-cenpb | Tf | 0.233809747 | 3.60E-11 |
| B95 | B113 | anti-cenpb | FT3 | -0.197019877 | 2.77E-08 |
| B95 | B114 | anti-cenpb | TSH TT3 TT4(TT4) | 0.193644947 | 4.81E-08 |
| B95 | B145 | anti-cenpb | BGP | -0.176984845 | 6.34E-07 |
| B97 | B100 | dsDNA | anti-Ro52 | 0.999999999 | 0 |
| B97 | B102 | dsDNA | anti-Sm | 0.999999999 | 0 |
| B97 | B103 | dsDNA | anti-SSA/Ro | 0.999999998 | 0 |
| B97 | B104 | dsDNA | anti-SSB | 0.999999999 | 0 |
| B97 | B106 | dsDNA | anti-P | 0.999999999 | 0 |
| B97 | B107 | dsDNA | AnuA | 0.999999966 | 0 |
| B97 | B108 | dsDNA | Anti-Cyclin | 1 | 0 |
| B97 | B109 | dsDNA | AMA-M2 | 0.999999998 | 0 |
| B97 | B183 | dsDNA | PTA | 0.303357646 | 0 |
| B97 | B154 | dsDNA | RF | 0.265694332 | 4.22E-14 |
| B97 | B180 | dsDNA | Tf | 0.249915727 | 1.34E-12 |
| B97 | B186 | dsDNA | PFC | -0.231641261 | 5.50E-11 |
| B97 | B113 | dsDNA | FT3 | -0.200835621 | 1.47E-08 |
| B97 | B114 | dsDNA | TSH TT3 TT4(TT4) | 0.189036316 | 1.00E-07 |
| B98 | B101 | anti-Jo-1 | anti-Sc70 | 0.999999994 | 0 |
| B98 | B102 | anti-Jo-1 | anti-Sm | 0.999999999 | 0 |
| B98 | B107 | anti-Jo-1 | AnuA | 0.999999915 | 0 |
| B98 | B108 | anti-Jo-1 | Anti-Cyclin | 1 | 0 |
| B98 | B183 | anti-Jo-1 | PTA | 0.303357648 | 0 |
| B98 | B154 | anti-Jo-1 | RF | 0.265694333 | 4.22E-14 |
| B98 | B180 | anti-Jo-1 | Tf | 0.249915728 | 1.34E-12 |
| B98 | B186 | anti-Jo-1 | PFC | -0.231641263 | 5.50E-11 |
| B98 | B113 | anti-Jo-1 | FT3 | -0.200835622 | 1.47E-08 |
| B98 | B114 | anti-Jo-1 | TSH TT3 TT4(TT4) | 0.189036317 | 1.00E-07 |
| B99 | B179 | anti-PM-Scl | α1-MG | 0.295188708 | 0 |

**Supplementary Table 22. PEI Correlations in Lumbar disc protrusion**

| Row | Column | Row | Column | Cor | *P* |
|---|---|---|---|---|---|
| B1 | B163 | Gender | SG | -0.40657 | 0 |
| B1 | B190 | Gender | Cre | -0.61112 | 0 |
| B1 | B192 | Gender | UA | -0.48687 | 0 |
| B1 | B206 | Gender | Waist | -0.40553 | 0 |
| B1 | B216 | Gender | BMI | -0.56496 | 0 |
| B1 | B221 | Gender | Hei | -0.71851 | 0 |
| B1 | B222 | Gender | NS | -0.69925 | 0 |
| B1 | B46 | Gender | RBC Count | -0.49097 | 0 |
| B1 | B51 | Gender | Hb | -0.64268 | 0 |
| B1 | B174 | Gender | Urea/Cr | 0.398284 | 4.44E-16 |
| B1 | B209 | Gender | Smoking | -0.37896 | 1.35E-14 |
| B1 | B207 | Gender | WHR | -0.36588 | 1.22E-13 |
| B1 | B45 | Gender | MCHC | -0.3168 | 2.01E-10 |
| B1 | B204 | Gender | Drinking | -0.31395 | 2.98E-10 |
| B1 | B210 | Gender | Hip | -0.29217 | 5.16E-09 |
| B1 | B72 | Gender | HDL-C | 0.290799 | 6.12E-09 |
| B10 | B12 | PG I | PG II | 0.642372 | 0 |
| B10 | B80 | PG I | anti-HP IgG | 0.290278 | 6.53E-09 |
| B10 | B14 | PG I | G-17 | 0.250453 | 6.43E-07 |
| B100 | B103 | anti-Ro52 | anti-SSA/Ro | 1 | 0 |
| B100 | B104 | anti-Ro52 | anti-SSB | 1 | 0 |
| B100 | B105 | anti-Ro52 | anti-U1RNP | 1 | 0 |
| B100 | B106 | anti-Ro52 | anti-P | 1 | 0 |
| B100 | B151 | anti-Ro52 | Anti-CCP | -0.70526 | 0 |
| B100 | B178 | anti-Ro52 | Ig-U | 0.453154 | 0 |
| B100 | B180 | anti-Ro52 | Tf | 0.496456 | 0 |
| B101 | B102 | anti-Sc70 | anti-Sm | 1 | 0 |
| B101 | B106 | anti-Sc70 | anti-P | 1 | 0 |
| B101 | B107 | anti-Sc70 | AnuA | 1 | 0 |
| B101 | B151 | anti-Sc70 | Anti-CCP | -0.70526 | 0 |
| B101 | B178 | anti-Sc70 | Ig-U | 0.453154 | 0 |
| B101 | B180 | anti-Sc70 | Tf | 0.496456 | 0 |
| B102 | B104 | anti-Sm | anti-SSB | 1 | 0 |
| B102 | B108 | anti-Sm | Anti-Cyclin | 1 | 0 |
| B102 | B151 | anti-Sm | Anti-CCP | -0.70526 | 0 |
| B102 | B178 | anti-Sm | Ig-U | 0.453154 | 0 |
| B102 | B180 | anti-Sm | Tf | 0.496456 | 0 |
| B103 | B104 | anti-SSA/Ro | anti-SSB | 1 | 0 |
| B103 | B105 | anti-SSA/Ro | anti-U1RNP | 1 | 0 |
| B103 | B106 | anti-SSA/Ro | anti-P | 1 | 0 |
| B103 | B151 | anti-SSA/Ro | Anti-CCP | -0.70526 | 0 |
| B103 | B178 | anti-SSA/Ro | Ig-U | 0.453154 | 0 |

| | | | | | |
|---|---|---|---|---|---|
| B103 | B180 | anti-SSA/Ro | Tf | 0.496456 | 0 |
| B104 | B105 | anti-SSB | anti-U1RNP | 1 | 0 |
| B104 | B107 | anti-SSB | AnuA | 1 | 0 |
| B104 | B109 | anti-SSB | AMA-M2 | 1 | 0 |
| B104 | B151 | anti-SSB | Anti-CCP | -0.70526 | 0 |
| B104 | B178 | anti-SSB | Ig-U | 0.453154 | 0 |
| B104 | B180 | anti-SSB | Tf | 0.496456 | 0 |
| B105 | B106 | anti-U1RNP | anti-P | 1 | 0 |
| B105 | B108 | anti-U1RNP | Anti-Cyclin | 1 | 0 |
| B105 | B109 | anti-U1RNP | AMA-M2 | 1 | 0 |
| B105 | B110 | anti-U1RNP | AHA | 1 | 0 |
| B105 | B151 | anti-U1RNP | Anti-CCP | -0.70526 | 0 |
| B105 | B178 | anti-U1RNP | Ig-U | 0.453154 | 0 |
| B105 | B180 | anti-U1RNP | Tf | 0.496456 | 0 |
| B106 | B107 | anti-P | AnuA | 1 | 0 |
| B106 | B108 | anti-P | Anti-Cyclin | 1 | 0 |
| B106 | B109 | anti-P | AMA-M2 | 1 | 0 |
| B106 | B110 | anti-P | AHA | 1 | 0 |
| B106 | B151 | anti-P | Anti-CCP | -0.70526 | 0 |
| B106 | B178 | anti-P | Ig-U | 0.453154 | 0 |
| B106 | B180 | anti-P | Tf | 0.496456 | 0 |
| B107 | B109 | AnuA | AMA-M2 | 1 | 0 |
| B107 | B151 | AnuA | Anti-CCP | -0.70526 | 0 |
| B107 | B178 | AnuA | Ig-U | 0.453154 | 0 |
| B107 | B180 | AnuA | Tf | 0.496456 | 0 |
| B108 | B110 | Anti-Cyclin | AHA | 1 | 0 |
| B108 | B151 | Anti-Cyclin | Anti-CCP | -0.70526 | 0 |
| B108 | B178 | Anti-Cyclin | Ig-U | 0.453154 | 0 |
| B108 | B180 | Anti-Cyclin | Tf | 0.496456 | 0 |
| B109 | B151 | AMA-M2 | Anti-CCP | -0.70526 | 0 |
| B109 | B178 | AMA-M2 | Ig-U | 0.453154 | 0 |
| B109 | B180 | AMA-M2 | Tf | 0.496456 | 0 |
| B11 | B12 | PGI/PGII | PG II | -0.46109 | 0 |
| B110 | B151 | AHA | Anti-CCP | -0.70526 | 0 |
| B110 | B178 | AHA | Ig-U | 0.453154 | 0 |
| B110 | B180 | AHA | Tf | 0.496456 | 0 |
| B112 | B114 | FT4 | TSH TT3 TT4(TT4) | 0.378698 | 1.42E-14 |
| B113 | B115 | FT3 | TSH TT3 TT4(TT3) | 0.333009 | 2.01E-11 |
| B114 | B115 | TSH TT3 TT4(TT4) | TSH TT3 TT4(TT3) | 0.492134 | 0 |
| B116 | B117 | TT3 | TT4 | -0.59259 | 0 |
| B119 | B120 | Hcv-rna | HCV-cAg | 0.426588 | 0 |
| B119 | B121 | Hcv-rna | C3 | -0.70525 | 0 |
| B119 | B122 | Hcv-rna | C4 | -0.70525 | 0 |
| B119 | B138 | Hcv-rna | PAB | 0.462648 | 0 |

| | | | | | |
|---|---|---|---|---|---|
| B119 | B152 | Hcv-rna | ACA-Ig(G-A-M) | -0.70526 | 0 |
| B119 | B159 | Hcv-rna | β2-MG | -0.70525 | 0 |
| B119 | B169 | Hcv-rna | U-CRY | -0.70525 | 0 |
| B119 | B123 | Hcv-rna | hs-CRP1 | -0.39754 | 4.44E-16 |
| B119 | B193 | Hcv-rna | K | -0.25579 | 3.63E-07 |
| B12 | B14 | PG II | G-17 | 0.41275 | 0 |
| B12 | B145 | PG II | BGP | 0.27629 | 3.56E-08 |
| B12 | B144 | PG II | Total-PINP | 0.272514 | 5.54E-08 |
| B120 | B121 | HCV-cAg | C3 | -0.30085 | 1.70E-09 |
| B120 | B122 | HCV-cAg | C4 | -0.30085 | 1.70E-09 |
| B120 | B152 | HCV-cAg | ACA-Ig(G-A-M) | -0.30086 | 1.70E-09 |
| B120 | B159 | HCV-cAg | β2-MG | -0.30085 | 1.70E-09 |
| B120 | B169 | HCV-cAg | U-CRY | -0.30085 | 1.70E-09 |
| B121 | B169 | C3 | U-CRY | 0.999998 | 0 |
| B121 | B138 | C3 | PAB | -0.27702 | 3.27E-08 |
| B122 | B169 | C4 | U-CRY | 0.999999 | 0 |
| B122 | B138 | C4 | PAB | -0.27703 | 3.27E-08 |
| B123 | B193 | hs-CRP1 | K | 0.462036 | 0 |
| B123 | B138 | hs-CRP1 | PAB | -0.33853 | 8.90E-12 |
| B123 | B125 | hs-CRP1 | Cor | 0.32802 | 4.15E-11 |
| B123 | B198 | hs-CRP1 | P | -0.26953 | 7.83E-08 |
| B124 | B139 | ACTH | ADA | -0.46337 | 0 |
| B124 | B181 | ACTH | APTT | -0.91884 | 0 |
| B124 | B182 | ACTH | TT | -0.91767 | 0 |
| B124 | B183 | ACTH | PTA | 0.918844 | 0 |
| B124 | B184 | ACTH | INR | 0.918843 | 0 |
| B124 | B185 | ACTH | PT | -0.91405 | 0 |
| B124 | B186 | ACTH | PFC | 0.918843 | 0 |
| B124 | B180 | ACTH | Tf | 0.367878 | 8.79E-14 |
| B124 | B195 | ACTH | Na | 0.33003 | 3.10E-11 |
| B124 | B200 | ACTH | OP | 0.318381 | 1.62E-10 |
| B124 | B198 | ACTH | P | 0.305813 | 8.88E-10 |
| B124 | B178 | ACTH | Ig-U | 0.301717 | 1.52E-09 |
| B124 | B158 | ACTH | IgM | 0.297082 | 2.77E-09 |
| B124 | B199 | ACTH | Mg | -0.26608 | 1.16E-07 |
| B124 | B179 | ACTH | α1-MG | 0.262369 | 1.76E-07 |
| B124 | B138 | ACTH | PAB | 0.260345 | 2.20E-07 |
| B126 | B127 | Alb | A-G ratio | 0.502764 | 0 |
| B126 | B188 | Alb | Ca | 0.261244 | 1.99E-07 |
| B126 | B129 | Alb | IBIL | 0.255727 | 3.65E-07 |
| B127 | B130 | A-G ratio | GLB | -0.9021 | 0 |
| B128 | B131 | ALT | AST | 0.727603 | 0 |
| B128 | B141 | ALT | AST/ALT | -0.49825 | 0 |
| B128 | B140 | ALT | r-GT | 0.381118 | 9.33E-15 |

| | | | | | |
|---|---|---|---|---|---|
| B128 | B216 | ALT | BMI | 0.257698 | 2.95E-07 |
| B129 | B132 | IBIL | Dbil | 0.746649 | 0 |
| B129 | B133 | IBIL | TBIL | 0.986583 | 0 |
| B131 | B140 | AST | r-GT | 0.428615 | 0 |
| B132 | B133 | Dbil | TBIL | 0.845167 | 0 |
| B134 | B136 | CHE | 5'NT | 0.515484 | 0 |
| B134 | B138 | CHE | PAB | 0.331354 | 2.56E-11 |
| B134 | B182 | CHE | TT | -0.26279 | 1.68E-07 |
| B134 | B197 | CHE | AG | -0.26154 | 1.93E-07 |
| B134 | B196 | CHE | TCO2 | 0.260018 | 2.28E-07 |
| B134 | B183 | CHE | PTA | 0.258861 | 2.59E-07 |
| B134 | B181 | CHE | APTT | -0.25873 | 2.63E-07 |
| B134 | B184 | CHE | INR | 0.258726 | 2.63E-07 |
| B134 | B186 | CHE | PFC | 0.258726 | 2.63E-07 |
| B134 | B185 | CHE | PT | -0.24894 | 7.54E-07 |
| B135 | B198 | TBA | P | -0.2877 | 8.99E-09 |
| B135 | B200 | TBA | OP | 0.284422 | 1.34E-08 |
| B135 | B137 | TBA | AFU | 0.255759 | 3.64E-07 |
| B136 | B139 | 5'NT | ADA | 0.355147 | 6.91E-13 |
| B136 | B168 | 5'NT | NMP22 | -0.25346 | 4.67E-07 |
| B137 | B178 | AFU | Ig-U | -0.37525 | 2.55E-14 |
| B138 | B193 | PAB | K | -0.49868 | 0 |
| B138 | B139 | PAB | ADA | -0.31427 | 2.85E-10 |
| B138 | B182 | PAB | TT | -0.30131 | 1.60E-09 |
| B138 | B197 | PAB | AG | -0.28927 | 7.40E-09 |
| B138 | B183 | PAB | PTA | 0.287621 | 9.08E-09 |
| B138 | B181 | PAB | APTT | -0.28718 | 9.59E-09 |
| B138 | B184 | PAB | INR | 0.287176 | 9.59E-09 |
| B138 | B186 | PAB | PFC | 0.287176 | 9.59E-09 |
| B138 | B196 | PAB | TCO2 | 0.286673 | 1.02E-08 |
| B138 | B158 | PAB | IgM | 0.28217 | 1.77E-08 |
| B138 | B152 | PAB | ACA-Ig(G-A-M) | -0.27703 | 3.27E-08 |
| B138 | B159 | PAB | β2-MG | -0.27703 | 3.27E-08 |
| B138 | B169 | PAB | U-CRY | -0.27702 | 3.27E-08 |
| B138 | B185 | PAB | PT | -0.25782 | 2.91E-07 |
| B139 | B158 | ADA | IgM | -0.43837 | 0 |
| B139 | B181 | ADA | APTT | 0.504109 | 0 |
| B139 | B182 | ADA | TT | 0.499469 | 0 |
| B139 | B183 | ADA | PTA | -0.50399 | 0 |
| B139 | B184 | ADA | INR | -0.50411 | 0 |
| B139 | B185 | ADA | PT | 0.509153 | 0 |
| B139 | B186 | ADA | PFC | -0.50411 | 0 |
| B139 | B157 | ADA | IgG | 0.394889 | 8.88E-16 |
| B139 | B199 | ADA | Mg | 0.315119 | 2.54E-10 |

| | | | | | |
|---|---|---|---|---|---|
| B14 | B73 | G-17 | ApoA | 0.454255 | 0 |
| B14 | B197 | G-17 | AG | -0.2689 | 8.42E-08 |
| B14 | B193 | G-17 | K | -0.24852 | 7.89E-07 |
| B140 | B151 | r-GT | Anti-CCP | 0.327353 | 4.56E-11 |
| B140 | B194 | r-GT | Cl | -0.24666 | 9.58E-07 |
| B141 | B222 | AST/ALT | NS | -0.25541 | 3.78E-07 |
| B142 | B144 | 25-(OH)D | Total-PINP | -0.92944 | 0 |
| B142 | B168 | 25-(OH)D | NMP22 | -0.27269 | 5.43E-08 |
| B143 | B145 | β-CTx | BGP | 0.835219 | 0 |
| B143 | B168 | β-CTx | NMP22 | -0.35316 | 9.45E-13 |
| B145 | B168 | BGP | NMP22 | -0.2813 | 1.96E-08 |
| B146 | B170 | CysC | U-GLU | 0.289078 | 7.58E-09 |
| B146 | B198 | CysC | P | 0.257329 | 3.07E-07 |
| B148 | B158 | Anti_TPO | IgM | 0.437196 | 0 |
| B15 | B156 | CD3-CD19+(B cell) | IgE | 0.901972 | 0 |
| B15 | B16 | CD3-CD19+(B cell) | CD3+T | -1 | 0 |
| B15 | B18 | CD3-CD19+(B cell) | CD3+CD4+helper(inducer)T | 1 | 0 |
| B15 | B22 | CD3-CD19+(B cell) | CD3+CD8+(T) | -1 | 0 |
| B15 | B23 | CD3-CD19+(B cell) | CD3-CD16+CD56+(NK) | -1 | 0 |
| B15 | B157 | CD3-CD19+(B cell) | IgG | 0.386637 | 3.55E-15 |
| B15 | B164 | CD3-CD19+(B cell) | COM | -0.35487 | 7.21E-13 |
| B15 | B78 | CD3-CD19+(B cell) | HBcAb | -0.26589 | 1.19E-07 |
| B151 | B155 | Anti-CCP | IgA | -0.44895 | 0 |
| B151 | B178 | Anti-CCP | Ig-U | -0.40715 | 0 |
| B151 | B180 | Anti-CCP | Tf | -0.42398 | 0 |
| B151 | B194 | Anti-CCP | Cl | -0.36575 | 1.25E-13 |
| B151 | B157 | Anti-CCP | IgG | -0.30832 | 6.36E-10 |
| B151 | B195 | Anti-CCP | Na | -0.27959 | 2.41E-08 |
| B153 | B170 | Glu | U-GLU | 0.575745 | 0 |
| B155 | B157 | IgA | IgG | 0.608636 | 0 |
| B155 | B180 | IgA | Tf | 0.279421 | 2.46E-08 |
| B155 | B178 | IgA | Ig-U | 0.258686 | 2.64E-07 |
| B156 | B164 | IgE | COM | -0.32027 | 1.24E-10 |
| B156 | B157 | IgE | IgG | 0.268345 | 8.96E-08 |
| B157 | B158 | IgG | IgM | -0.51312 | 0 |
| B157 | B200 | IgG | OP | -0.27124 | 6.43E-08 |
| B158 | B181 | IgM | APTT | -0.32227 | 9.40E-11 |
| B158 | B183 | IgM | PTA | 0.322262 | 9.40E-11 |
| B158 | B184 | IgM | INR | 0.322266 | 9.40E-11 |
| B158 | B186 | IgM | PFC | 0.322266 | 9.40E-11 |
| B158 | B182 | IgM | TT | -0.32171 | 1.02E-10 |
| B158 | B185 | IgM | PT | -0.32087 | 1.14E-10 |
| B158 | B200 | IgM | OP | 0.29321 | 4.52E-09 |
| B159 | B169 | β2-MG | U-CRY | 0.999997 | 0 |

| | | | | | |
|---|---|---|---|---|---|
| B16 | B156 | CD3+T | IgE | -0.90197 | 0 |
| B16 | B18 | CD3+T | CD3+CD4+helper(inducer)T | -1 | 0 |
| B16 | B19 | CD3+T | CD3+CD4+/CD3+CD8+ | -1 | 0 |
| B16 | B22 | CD3+T | CD3+CD8+(T) | 1 | 0 |
| B16 | B23 | CD3+T | CD3-CD16+CD56+(NK) | 1 | 0 |
| B16 | B157 | CD3+T | IgG | -0.38664 | 3.55E-15 |
| B16 | B164 | CD3+T | COM | 0.354875 | 7.21E-13 |
| B16 | B78 | CD3+T | HBcAb | 0.265888 | 1.19E-07 |
| B163 | B166 | SG | Pro | 0.372467 | 4.09E-14 |
| B163 | B176 | SG | Ucr | 0.365224 | 1.36E-13 |
| B163 | B221 | SG | Hei | 0.349637 | 1.64E-12 |
| B163 | B216 | SG | BMI | 0.344002 | 3.90E-12 |
| B163 | B222 | SG | NS | 0.306809 | 7.78E-10 |
| B167 | B171 | PC | EC | -0.42502 | 0 |
| B167 | B223 | PC | FH | 0.355529 | 6.50E-13 |
| B17 | B117 | CD3+CD16+CD56+NK | TT4 | -0.70278 | 0 |
| B17 | B21 | CD3+CD16+CD56+NK | CD3+CD4-CD8- | -0.51281 | 0 |
| B17 | B35 | CD3+CD16+CD56+NK | β2-MG | -0.46043 | 0 |
| B171 | B223 | EC | FH | 0.732695 | 0 |
| B174 | B190 | Urea/Cr | Cre | -0.47402 | 0 |
| B174 | B191 | Urea/Cr | Urea | 0.667803 | 0 |
| B174 | B221 | Urea/Cr | Hei | -0.41712 | 0 |
| B174 | B192 | Urea/Cr | UA | -0.26216 | 1.80E-07 |
| B175 | B176 | ALB-U | Ucr | 0.422013 | 0 |
| B175 | B177 | ALB-U | ACR | 0.744113 | 0 |
| B176 | B178 | Ucr | Ig-U | 0.524142 | 0 |
| B176 | B179 | Ucr | α1-MG | 0.342695 | 4.75E-12 |
| B177 | B180 | ACR | Tf | 0.330786 | 2.78E-11 |
| B178 | B179 | Ig-U | α1-MG | 0.335157 | 1.47E-11 |
| B178 | B181 | Ig-U | APTT | -0.32843 | 3.91E-11 |
| B178 | B183 | Ig-U | PTA | 0.328428 | 3.91E-11 |
| B178 | B184 | Ig-U | INR | 0.328428 | 3.91E-11 |
| B178 | B186 | Ig-U | PFC | 0.328428 | 3.91E-11 |
| B178 | B182 | Ig-U | TT | -0.32799 | 4.16E-11 |
| B178 | B185 | Ig-U | PT | -0.32675 | 4.97E-11 |
| B178 | B195 | Ig-U | Na | 0.277211 | 3.20E-08 |
| B179 | B181 | α1-MG | APTT | -0.28667 | 1.02E-08 |
| B179 | B183 | α1-MG | PTA | 0.286673 | 1.02E-08 |
| B179 | B184 | α1-MG | INR | 0.286671 | 1.02E-08 |
| B179 | B186 | α1-MG | PFC | 0.286671 | 1.02E-08 |
| B179 | B182 | α1-MG | TT | -0.28636 | 1.06E-08 |
| B179 | B185 | α1-MG | PT | -0.28506 | 1.24E-08 |
| B179 | B194 | α1-MG | Cl | -0.26048 | 2.17E-07 |
| B18 | B156 | CD3+CD4+helper(inducer)T | IgE | 0.901972 | 0 |

| | | | | | |
|---|---|---|---|---|---|
| B18 | B19 | CD3+CD4+helper(inducer)T | CD3+CD4+/CD3+CD8+ | 1 | 0 |
| B18 | B22 | CD3+CD4+helper(inducer)T | CD3+CD8+(T) | -1 | 0 |
| B18 | B23 | CD3+CD4+helper(inducer)T | CD3-CD16+CD56+(NK) | -1 | 0 |
| B18 | B157 | CD3+CD4+helper(inducer)T | IgG | 0.386637 | 3.55E-15 |
| B18 | B164 | CD3+CD4+helper(inducer)T | COM | -0.35487 | 7.21E-13 |
| B18 | B78 | CD3+CD4+helper(inducer)T | HBcAb | -0.26589 | 1.19E-07 |
| B180 | B181 | Tf | APTT | -0.39757 | 4.44E-16 |
| B180 | B182 | Tf | TT | -0.39656 | 4.44E-16 |
| B180 | B183 | Tf | PTA | 0.397559 | 4.44E-16 |
| B180 | B184 | Tf | INR | 0.397574 | 4.44E-16 |
| B180 | B185 | Tf | PT | -0.39647 | 4.44E-16 |
| B180 | B186 | Tf | PFC | 0.397574 | 4.44E-16 |
| B180 | B195 | Tf | Na | 0.280786 | 2.09E-08 |
| B180 | B194 | Tf | Cl | 0.255521 | 3.73E-07 |
| B181 | B182 | APTT | TT | 0.998634 | 0 |
| B181 | B183 | APTT | PTA | -1 | 0 |
| B181 | B185 | APTT | PT | 0.994954 | 0 |
| B181 | B195 | APTT | Na | -0.35797 | 4.41E-13 |
| B181 | B200 | APTT | OP | -0.34486 | 3.42E-12 |
| B181 | B198 | APTT | P | -0.33657 | 1.19E-11 |
| B181 | B199 | APTT | Mg | 0.291751 | 5.43E-09 |
| B182 | B183 | TT | PTA | -0.99872 | 0 |
| B182 | B184 | TT | INR | -0.99863 | 0 |
| B182 | B185 | TT | PT | 0.988352 | 0 |
| B182 | B186 | TT | PFC | -0.99863 | 0 |
| B182 | B195 | TT | Na | -0.34902 | 1.80E-12 |
| B182 | B200 | TT | OP | -0.33394 | 1.75E-11 |
| B182 | B198 | TT | P | -0.33332 | 1.92E-11 |
| B182 | B199 | TT | Mg | 0.285171 | 1.23E-08 |
| B183 | B184 | PTA | INR | 0.999999 | 0 |
| B183 | B185 | PTA | PT | -0.99479 | 0 |
| B183 | B186 | PTA | PFC | 0.999999 | 0 |
| B183 | B195 | PTA | Na | 0.357706 | 4.60E-13 |
| B183 | B200 | PTA | OP | 0.344543 | 3.59E-12 |
| B183 | B198 | PTA | P | 0.336481 | 1.21E-11 |
| B183 | B199 | PTA | Mg | -0.29156 | 5.56E-09 |
| B184 | B185 | INR | PT | -0.99495 | 0 |
| B184 | B195 | INR | Na | 0.357966 | 4.41E-13 |
| B184 | B200 | INR | OP | 0.344864 | 3.42E-12 |
| B184 | B198 | INR | P | 0.336567 | 1.19E-11 |
| B184 | B199 | INR | Mg | -0.29175 | 5.43E-09 |
| B185 | B186 | PT | PFC | -0.99495 | 0 |
| B185 | B195 | PT | Na | -0.3724 | 4.13E-14 |
| B185 | B200 | PT | OP | -0.3632 | 1.90E-13 |

| | | | | | |
|---|---|---|---|---|---|
| B185 | B198 | PT | P | -0.34021 | 6.91E-12 |
| B185 | B199 | PT | Mg | 0.302148 | 1.44E-09 |
| B186 | B195 | PFC | Na | 0.357966 | 4.41E-13 |
| B186 | B200 | PFC | OP | 0.344864 | 3.42E-12 |
| B186 | B198 | PFC | P | 0.336567 | 1.19E-11 |
| B186 | B199 | PFC | Mg | -0.29175 | 5.43E-09 |
| B189 | B190 | eGFB | Cre | -0.40213 | 2.22E-16 |
| B189 | B191 | eGFB | Urea | -0.29604 | 3.16E-09 |
| B189 | B220 | eGFB | SBP | -0.24943 | 7.17E-07 |
| B19 | B156 | CD3+CD4+/CD3+CD8+ | IgE | 0.901972 | 0 |
| B19 | B157 | CD3+CD4+/CD3+CD8+ | IgG | 0.386637 | 3.55E-15 |
| B19 | B164 | CD3+CD4+/CD3+CD8+ | COM | -0.35487 | 7.21E-13 |
| B19 | B78 | CD3+CD4+/CD3+CD8+ | HBcAb | -0.26589 | 1.19E-07 |
| B190 | B192 | Cre | UA | 0.538553 | 0 |
| B190 | B222 | Cre | NS | 0.415339 | 0 |
| B190 | B221 | Cre | Hei | 0.392829 | 1.33E-15 |
| B190 | B216 | Cre | BMI | 0.322789 | 8.73E-11 |
| B190 | B191 | Cre | Urea | 0.304792 | 1.02E-09 |
| B190 | B206 | Cre | Waist | 0.270998 | 6.61E-08 |
| B192 | B206 | UA | Waist | 0.405024 | 0 |
| B192 | B216 | UA | BMI | 0.428905 | 0 |
| B192 | B222 | UA | NS | 0.46681 | 0 |
| B192 | B221 | UA | Hei | 0.376164 | 2.18E-14 |
| B192 | B210 | UA | Hip | 0.353582 | 8.84E-13 |
| B192 | B207 | UA | WHR | 0.317359 | 1.86E-10 |
| B192 | B215 | UA | BMI | 0.269763 | 7.62E-08 |
| B193 | B196 | K | TCO2 | -0.34239 | 4.98E-12 |
| B193 | B197 | K | AG | 0.340975 | 6.16E-12 |
| B194 | B195 | Cl | Na | 0.346222 | 2.78E-12 |
| B194 | B196 | Cl | TCO2 | -0.28165 | 1.88E-08 |
| B195 | B200 | Na | OP | 0.872988 | 0 |
| B195 | B199 | Na | Mg | -0.32461 | 6.75E-11 |
| B195 | B197 | Na | AG | 0.323571 | 7.82E-11 |
| B196 | B197 | TCO2 | AG | -0.65366 | 0 |
| B196 | B213 | TCO2 | CVA-R | 0.334611 | 1.59E-11 |
| B196 | B200 | TCO2 | OP | 0.265689 | 1.21E-07 |
| B197 | B213 | AG | CVA-R | -0.32822 | 4.03E-11 |
| B197 | B200 | AG | OP | 0.320906 | 1.14E-10 |
| B197 | B199 | AG | Mg | -0.29565 | 3.32E-09 |
| B199 | B200 | Mg | OP | -0.26301 | 1.64E-07 |
| B2 | B189 | Age | eGFB | -0.52972 | 0 |
| B2 | B220 | Age | SBP | 0.426276 | 0 |
| B2 | B221 | Age | Hei | -0.41678 | 0 |
| B2 | B46 | Age | RBC Count | -0.37035 | 5.82E-14 |

| | | | | | |
|---|---|---|---|---|---|
| B2 | B51 | Age | Hb | -0.36849 | 7.95E-14 |
| B2 | B126 | Age | Alb | -0.34065 | 6.47E-12 |
| B2 | B72 | Age | HDL-C | 0.278308 | 2.81E-08 |
| B2 | B163 | Age | SG | -0.27148 | 6.25E-08 |
| B2 | B191 | Age | Urea | 0.267905 | 9.43E-08 |
| B2 | B219 | Age | OT | 0.25215 | 5.37E-07 |
| B20 | B38 | CD3+CD4+CD8+ | BAND% | 0.754732 | 0 |
| B20 | B171 | CD3+CD4+CD8+ | EC | 0.359139 | 3.65E-13 |
| B204 | B209 | Drinking | Smoking | 0.351972 | 1.14E-12 |
| B204 | B222 | Drinking | NS | 0.296331 | 3.04E-09 |
| B206 | B207 | Waist | WHR | 0.831564 | 0 |
| B206 | B210 | Waist | Hip | 0.796807 | 0 |
| B206 | B215 | Waist | BMI | 0.8172 | 0 |
| B206 | B216 | Waist | BMI | 0.804616 | 0 |
| B206 | B222 | Waist | NS | 0.714769 | 0 |
| B206 | B221 | Waist | Hei | 0.315823 | 2.30E-10 |
| B206 | B220 | Waist | SBP | 0.300297 | 1.83E-09 |
| B207 | B215 | WHR | BMI | 0.543829 | 0 |
| B207 | B216 | WHR | BMI | 0.516129 | 0 |
| B207 | B222 | WHR | NS | 0.535408 | 0 |
| B207 | B210 | WHR | Hip | 0.330518 | 2.89E-11 |
| B207 | B220 | WHR | SBP | 0.268831 | 8.48E-08 |
| B209 | B221 | Smoking | Hei | 0.28502 | 1.25E-08 |
| B209 | B222 | Smoking | NS | 0.256127 | 3.50E-07 |
| B21 | B116 | CD3+CD4-CD8- | TT3 | -0.84292 | 0 |
| B21 | B35 | CD3+CD4-CD8- | β2-MG | -0.4909 | 0 |
| B210 | B215 | Hip | BMI | 0.801318 | 0 |
| B210 | B216 | Hip | BMI | 0.806619 | 0 |
| B210 | B222 | Hip | NS | 0.637153 | 0 |
| B210 | B221 | Hip | Hei | 0.330854 | 2.75E-11 |
| B211 | B212 | IOP-R | IOP-L | 0.750434 | 0 |
| B213 | B214 | CVA-R | CVA-L | 0.832492 | 0 |
| B215 | B216 | BMI | BMI | 0.767777 | 0 |
| B215 | B222 | BMI | NS | 0.589897 | 0 |
| B215 | B220 | BMI | SBP | 0.316273 | 2.16E-10 |
| B216 | B221 | BMI | Hei | 0.669183 | 0 |
| B216 | B222 | BMI | NS | 0.81231 | 0 |
| B218 | B220 | DBP | SBP | 0.531154 | 0 |
| B22 | B156 | CD3+CD8+(T) | IgE | -0.90197 | 0 |
| B22 | B23 | CD3+CD8+(T) | CD3-CD16+CD56+(NK) | 1 | 0 |
| B22 | B157 | CD3+CD8+(T) | IgG | -0.38664 | 3.55E-15 |
| B22 | B164 | CD3+CD8+(T) | COM | 0.354875 | 7.21E-13 |
| B22 | B78 | CD3+CD8+(T) | HBcAb | 0.265888 | 1.19E-07 |
| B221 | B222 | Hei | NS | 0.594107 | 0 |

| | | | | | |
|---|---|---|---|---|---|
| B23 | B156 | CD3-CD16+CD56+(NK) | IgE | -0.90197 | 0 |
| B23 | B157 | CD3-CD16+CD56+(NK) | IgG | -0.38664 | 3.55E-15 |
| B23 | B164 | CD3-CD16+CD56+(NK) | COM | 0.354875 | 7.21E-13 |
| B23 | B78 | CD3-CD16+CD56+(NK) | HBcAb | 0.265888 | 1.19E-07 |
| B24 | B27 | αHBDH | LDH | 0.511115 | 0 |
| B24 | B25 | αHBDH | CK | 0.336451 | 1.21E-11 |
| B24 | B73 | αHBDH | ApoA | 0.286664 | 1.02E-08 |
| B24 | B60 | αHBDH | WBRVI | -0.28493 | 1.26E-08 |
| B24 | B59 | αHBDH | WBRV | -0.28447 | 1.34E-08 |
| B24 | B57 | αHBDH | RBCAI | -0.27198 | 5.90E-08 |
| B25 | B26 | CK | CKMB | 0.351474 | 1.23E-12 |
| B26 | B198 | CKMB | P | -0.35569 | 6.34E-13 |
| B26 | B59 | CKMB | WBRV | -0.30092 | 1.69E-09 |
| B26 | B57 | CKMB | RBCAI | -0.27944 | 2.45E-08 |
| B26 | B119 | CKMB | Hcv-rna | -0.269 | 8.31E-08 |
| B26 | B73 | CKMB | ApoA | 0.250973 | 6.08E-07 |
| B27 | B154 | LDH | RF | 0.318986 | 1.49E-10 |
| B27 | B134 | LDH | CHE | 0.308857 | 5.92E-10 |
| B27 | B139 | LDH | ADA | 0.249866 | 6.84E-07 |
| B29 | B32 | E2 | Testo | 1 | 0 |
| B3 | B89 | Hcy | CA 72-4 | 0.681022 | 0 |
| B3 | B81 | Hcy | TSGF | 0.348071 | 2.09E-12 |
| B30 | B31 | LH | FSH | 1 | 0 |
| B30 | B32 | LH | Testo | -1 | 0 |
| B31 | B32 | FSH | Testo | -1 | 0 |
| B32 | B33 | Testo | Prog | 1 | 0 |
| B35 | B116 | β2-MG | TT3 | -0.82857 | 0 |
| B36 | B227 | V-Lym% | OD | -0.32928 | 3.46E-11 |
| B37 | B167 | BAND | PC | -0.30305 | 1.28E-09 |
| B39 | B44 | Other cells | MCH | -0.27529 | 4.01E-08 |
| B39 | B43 | Other cells | MCV | -0.25315 | 4.82E-07 |
| B4 | B6 | WB-Ca | WB-Fe | -0.46894 | 0 |
| B4 | B7 | WB-Ca | WB-Cop | 0.622195 | 0 |
| B4 | B8 | WB-Ca | WB-Zinc | -0.94278 | 0 |
| B4 | B198 | WB-Ca | P | -0.32547 | 5.97E-11 |
| B4 | B5 | WB-Ca | WB-Mg | -0.26207 | 1.82E-07 |
| B40 | B42 | WBC | MON | 0.630434 | 0 |
| B40 | B207 | WBC | WHR | 0.268935 | 8.38E-08 |
| B40 | B51 | WBC | Hb | 0.251474 | 5.77E-07 |
| B40 | B53 | WBC | GR% | 0.249524 | 7.10E-07 |
| B41 | B42 | MON % | MON | 0.6341 | 0 |
| B41 | B53 | MON % | GR% | -0.27176 | 6.05E-08 |
| B42 | B207 | MON | WHR | 0.279538 | 2.42E-08 |
| B43 | B44 | MCV | MCH | 0.88824 | 0 |

| | | | | | |
|---|---|---|---|---|---|
| B43 | B46 | MCV | RBC Count | -0.48972 | 0 |
| B43 | B47 | MCV | RDW | 0.568311 | 0 |
| B44 | B45 | MCH | MCHC | 0.549158 | 0 |
| B44 | B47 | MCH | RDW | 0.365853 | 1.23E-13 |
| B44 | B51 | MCH | Hb | 0.35004 | 1.54E-12 |
| B44 | B46 | MCH | RBC Count | -0.33654 | 1.20E-11 |
| B45 | B51 | MCHC | Hb | 0.537521 | 0 |
| B45 | B222 | MCHC | NS | 0.304873 | 1.00E-09 |
| B45 | B216 | MCHC | BMI | 0.292092 | 5.21E-09 |
| B45 | B189 | MCHC | eGFB | 0.288862 | 7.79E-09 |
| B45 | B221 | MCHC | Hei | 0.276502 | 3.48E-08 |
| B46 | B216 | RBC Count | BMI | 0.427493 | 0 |
| B46 | B221 | RBC Count | Hei | 0.466479 | 0 |
| B46 | B222 | RBC Count | NS | 0.40791 | 0 |
| B46 | B51 | RBC Count | Hb | 0.760537 | 0 |
| B46 | B47 | RBC Count | RDW | -0.36723 | 9.79E-14 |
| B46 | B163 | RBC Count | SG | 0.352502 | 1.05E-12 |
| B46 | B192 | RBC Count | UA | 0.312014 | 3.87E-10 |
| B46 | B141 | RBC Count | AST/ALT | -0.29362 | 4.30E-09 |
| B46 | B72 | RBC Count | HDL-C | -0.29209 | 5.20E-09 |
| B46 | B206 | RBC Count | Waist | 0.266221 | 1.14E-07 |
| B46 | B210 | RBC Count | Hip | 0.26205 | 1.82E-07 |
| B46 | B174 | RBC Count | Urea/Cr | -0.26083 | 2.09E-07 |
| B47 | B72 | RDW | HDL-C | 0.24768 | 8.61E-07 |
| B48 | B53 | LYM% | GR% | -0.93652 | 0 |
| B49 | B50 | baso% | eo% | 0.349641 | 1.64E-12 |
| B49 | B154 | baso% | RF | 0.320454 | 1.21E-10 |
| B5 | B6 | WB-Mg | WB-Fe | 0.975255 | 0 |
| B5 | B7 | WB-Mg | WB-Cop | 0.592439 | 0 |
| B5 | B198 | WB-Mg | P | 0.291565 | 5.56E-09 |
| B50 | B154 | eo% | RF | 0.26528 | 1.27E-07 |
| B51 | B163 | Hb | SG | 0.405335 | 0 |
| B51 | B216 | Hb | BMI | 0.523612 | 0 |
| B51 | B221 | Hb | Hei | 0.578191 | 0 |
| B51 | B222 | Hb | NS | 0.533825 | 0 |
| B51 | B206 | Hb | Waist | 0.345662 | 3.03E-12 |
| B51 | B192 | Hb | UA | 0.325117 | 6.28E-11 |
| B51 | B210 | Hb | Hip | 0.302012 | 1.46E-09 |
| B51 | B209 | Hb | Smoking | 0.300853 | 1.70E-09 |
| B51 | B72 | Hb | HDL-C | -0.2934 | 4.41E-09 |
| B51 | B204 | Hb | Drinking | 0.286425 | 1.05E-08 |
| B51 | B189 | Hb | eGFB | 0.281739 | 1.86E-08 |
| B51 | B141 | Hb | AST/ALT | -0.27913 | 2.54E-08 |
| B51 | B207 | Hb | WHR | 0.269558 | 7.80E-08 |

| | | | | | |
|---|---|---|---|---|---|
| B51 | B174 | Hb | Urea/Cr | -0.26476 | 1.35E-07 |
| B51 | B190 | Hb | Cre | 0.259443 | 2.43E-07 |
| B54 | B63 | ESR | WBRSR(mPas_200) | 0.429704 | 0 |
| B54 | B69 | ESR | PV | 0.793306 | 0 |
| B54 | B66 | ESR | WBRSR(mPas_100) | 0.395587 | 6.66E-16 |
| B54 | B67 | ESR | WBRSR(mPas_50) | 0.363268 | 1.87E-13 |
| B54 | B61 | ESR | WBHSRV | 0.354665 | 7.45E-13 |
| B54 | B151 | ESR | Anti-CCP | -0.27899 | 2.59E-08 |
| B54 | B60 | ESR | WBRVI | -0.26195 | 1.84E-07 |
| B54 | B181 | ESR | APTT | -0.25535 | 3.80E-07 |
| B54 | B184 | ESR | INR | 0.255352 | 3.80E-07 |
| B54 | B186 | ESR | PFC | 0.255352 | 3.80E-07 |
| B54 | B183 | ESR | PTA | 0.255347 | 3.81E-07 |
| B54 | B182 | ESR | TT | -0.25486 | 4.01E-07 |
| B54 | B185 | ESR | PT | -0.25434 | 4.24E-07 |
| B55 | B56 | TK | IR | 0.830385 | 0 |
| B55 | B61 | TK | WBHSRV | 0.70655 | 0 |
| B55 | B62 | TK | WBHSRVI | 0.549846 | 0 |
| B55 | B98 | TK | anti-Jo-1 | -0.38606 | 4.00E-15 |
| B55 | B63 | TK | WBRSR(mPas_200) | 0.356512 | 5.56E-13 |
| B55 | B66 | TK | WBRSR(mPas_100) | 0.354207 | 8.01E-13 |
| B55 | B67 | TK | WBRSR(mPas_50) | 0.331689 | 2.44E-11 |
| B55 | B69 | TK | PV | -0.31399 | 2.96E-10 |
| B55 | B170 | TK | U-GLU | 0.28644 | 1.05E-08 |
| B55 | B139 | TK | ADA | -0.27718 | 3.21E-08 |
| B55 | B149 | TK | TGAb | 0.273988 | 4.67E-08 |
| B56 | B58 | IR | HCT | 0.452362 | 0 |
| B56 | B60 | IR | WBRVI | 0.567465 | 0 |
| B56 | B61 | IR | WBHSRV | 0.802405 | 0 |
| B56 | B62 | IR | WBHSRVI | 0.921193 | 0 |
| B56 | B63 | IR | WBRSR(mPas_200) | 0.663573 | 0 |
| B56 | B64 | IR | WBRSR(mPas_5) | 0.563472 | 0 |
| B56 | B65 | IR | WBRSR(mPas_1) | 0.534464 | 0 |
| B56 | B66 | IR | WBRSR(mPas_100) | 0.673093 | 0 |
| B56 | B67 | IR | WBRSR(mPas_50) | 0.670133 | 0 |
| B56 | B69 | IR | PV | -0.42896 | 0 |
| B56 | B98 | IR | anti-Jo-1 | -0.42279 | 0 |
| B56 | B59 | IR | WBRV | 0.359626 | 3.38E-13 |
| B56 | B100 | IR | anti-Ro52 | 0.332328 | 2.22E-11 |
| B56 | B101 | IR | anti-Sc70 | 0.332328 | 2.22E-11 |
| B56 | B102 | IR | anti-Sm | 0.332328 | 2.22E-11 |
| B56 | B103 | IR | anti-SSA/Ro | 0.332328 | 2.22E-11 |
| B56 | B104 | IR | anti-SSB | 0.332328 | 2.22E-11 |
| B56 | B105 | IR | anti-U1RNP | 0.332328 | 2.22E-11 |

| | | | | | |
|---|---|---|---|---|---|
| B56 | B106 | IR | anti-P | 0.332328 | 2.22E-11 |
| B56 | B107 | IR | AnuA | 0.332328 | 2.22E-11 |
| B56 | B108 | IR | Anti-Cyclin | 0.332328 | 2.22E-11 |
| B56 | B109 | IR | AMA-M2 | 0.332328 | 2.22E-11 |
| B56 | B110 | IR | AHA | 0.332328 | 2.22E-11 |
| B56 | B194 | IR | Cl | -0.33233 | 2.22E-11 |
| B56 | B95 | IR | anti-cenpb | 0.332328 | 2.22E-11 |
| B56 | B97 | IR | dsDNA | 0.332328 | 2.22E-11 |
| B56 | B99 | IR | anti-PM-Scl | 0.332328 | 2.22E-11 |
| B56 | B139 | IR | ADA | -0.30325 | 1.24E-09 |
| B56 | B138 | IR | PAB | 0.276472 | 3.49E-08 |
| B56 | B73 | IR | ApoA | -0.25666 | 3.30E-07 |
| B56 | B136 | IR | 5'NT | -0.24862 | 7.81E-07 |
| B57 | B59 | RBCAI | WBRV | 0.825507 | 0 |
| B57 | B60 | RBCAI | WBRVI | 0.815154 | 0 |
| B57 | B64 | RBCAI | WBRSR(mPas_5) | 0.595202 | 0 |
| B57 | B65 | RBCAI | WBRSR(mPas_1) | 0.720452 | 0 |
| B57 | B149 | RBCAI | TGAb | -0.39988 | 4.44E-16 |
| B57 | B58 | RBCAI | HCT | 0.337051 | 1.11E-11 |
| B57 | B198 | RBCAI | P | 0.316842 | 2.00E-10 |
| B57 | B69 | RBCAI | PV | -0.31035 | 4.85E-10 |
| B57 | B73 | RBCAI | ApoA | -0.25229 | 5.29E-07 |
| B57 | B139 | RBCAI | ADA | -0.25098 | 6.08E-07 |
| B58 | B60 | HCT | WBRVI | 0.692256 | 0 |
| B58 | B62 | HCT | WBHSRVI | 0.758109 | 0 |
| B58 | B63 | HCT | WBRSR(mPas_200) | 0.653981 | 0 |
| B58 | B64 | HCT | WBRSR(mPas_5) | 0.729575 | 0 |
| B58 | B65 | HCT | WBRSR(mPas_1) | 0.686145 | 0 |
| B58 | B66 | HCT | WBRSR(mPas_100) | 0.662844 | 0 |
| B58 | B67 | HCT | WBRSR(mPas_50) | 0.68846 | 0 |
| B58 | B78 | HCT | HBcAb | 0.377438 | 1.75E-14 |
| B58 | B59 | HCT | WBRV | 0.351729 | 1.18E-12 |
| B58 | B100 | HCT | anti-Ro52 | 0.351715 | 1.19E-12 |
| B58 | B101 | HCT | anti-Sc70 | 0.351715 | 1.19E-12 |
| B58 | B102 | HCT | anti-Sm | 0.351715 | 1.19E-12 |
| B58 | B103 | HCT | anti-SSA/Ro | 0.351715 | 1.19E-12 |
| B58 | B104 | HCT | anti-SSB | 0.351715 | 1.19E-12 |
| B58 | B105 | HCT | anti-U1RNP | 0.351715 | 1.19E-12 |
| B58 | B106 | HCT | anti-P | 0.351715 | 1.19E-12 |
| B58 | B107 | HCT | AnuA | 0.351715 | 1.19E-12 |
| B58 | B108 | HCT | Anti-Cyclin | 0.351715 | 1.19E-12 |
| B58 | B109 | HCT | AMA-M2 | 0.351715 | 1.19E-12 |
| B58 | B110 | HCT | AHA | 0.351715 | 1.19E-12 |
| B58 | B95 | HCT | anti-cenpb | 0.351715 | 1.19E-12 |

| | | | | | |
|---|---|---|---|---|---|
| B58 | B97 | HCT | dsDNA | 0.351715 | 1.19E-12 |
| B58 | B99 | HCT | anti-PM-Scl | 0.351715 | 1.19E-12 |
| B58 | B151 | HCT | Anti-CCP | -0.33291 | 2.04E-11 |
| B58 | B61 | HCT | WBHSRV | 0.331944 | 2.35E-11 |
| B59 | B60 | WBRV | WBRVI | 0.819096 | 0 |
| B59 | B62 | WBRV | WBHSRVI | 0.417432 | 0 |
| B59 | B63 | WBRV | WBRSR(mPas_200) | 0.449359 | 0 |
| B59 | B64 | WBRV | WBRSR(mPas_5) | 0.851442 | 0 |
| B59 | B65 | WBRV | WBRSR(mPas_1) | 0.891169 | 0 |
| B59 | B66 | WBRV | WBRSR(mPas_100) | 0.513759 | 0 |
| B59 | B67 | WBRV | WBRSR(mPas_50) | 0.59365 | 0 |
| B59 | B61 | WBRV | WBHSRV | 0.391819 | 1.33E-15 |
| B59 | B73 | WBRV | ApoA | -0.35405 | 8.21E-13 |
| B59 | B139 | WBRV | ADA | -0.34173 | 5.50E-12 |
| B59 | B198 | WBRV | P | 0.324701 | 6.66E-11 |
| B59 | B149 | WBRV | TGAb | -0.31474 | 2.67E-10 |
| B59 | B181 | WBRV | APTT | -0.28306 | 1.59E-08 |
| B59 | B183 | WBRV | PTA | 0.283058 | 1.59E-08 |
| B59 | B184 | WBRV | INR | 0.28306 | 1.59E-08 |
| B59 | B186 | WBRV | PFC | 0.28306 | 1.59E-08 |
| B59 | B182 | WBRV | TT | -0.28259 | 1.68E-08 |
| B59 | B185 | WBRV | PT | -0.28179 | 1.85E-08 |
| B59 | B124 | WBRV | ACTH | 0.260547 | 2.15E-07 |
| B6 | B7 | WB-Fe | WB-Cop | 0.399671 | 4.44E-16 |
| B6 | B198 | WB-Fe | P | 0.341406 | 5.78E-12 |
| B60 | B62 | WBRVI | WBHSRVI | 0.726996 | 0 |
| B60 | B63 | WBRVI | WBRSR(mPas_200) | 0.438732 | 0 |
| B60 | B64 | WBRVI | WBRSR(mPas_5) | 0.846142 | 0 |
| B60 | B65 | WBRVI | WBRSR(mPas_1) | 0.914495 | 0 |
| B60 | B66 | WBRVI | WBRSR(mPas_100) | 0.503631 | 0 |
| B60 | B67 | WBRVI | WBRSR(mPas_50) | 0.583139 | 0 |
| B60 | B69 | WBRVI | PV | -0.46456 | 0 |
| B60 | B139 | WBRVI | ADA | -0.32821 | 4.03E-11 |
| B60 | B61 | WBRVI | WBHSRV | 0.310398 | 4.81E-10 |
| B60 | B73 | WBRVI | ApoA | -0.30206 | 1.45E-09 |
| B60 | B194 | WBRVI | Cl | -0.28895 | 7.70E-09 |
| B60 | B149 | WBRVI | TGAb | -0.28748 | 9.24E-09 |
| B60 | B98 | WBRVI | anti-Jo-1 | -0.26413 | 1.45E-07 |
| B60 | B138 | WBRVI | PAB | 0.251327 | 5.86E-07 |
| B61 | B62 | WBHSRV | WBHSRVI | 0.709641 | 0 |
| B61 | B63 | WBHSRV | WBRSR(mPas_200) | 0.895519 | 0 |
| B61 | B64 | WBHSRV | WBRSR(mPas_5) | 0.630212 | 0 |
| B61 | B65 | WBHSRV | WBRSR(mPas_1) | 0.516503 | 0 |
| B61 | B66 | WBHSRV | WBRSR(mPas_100) | 0.889915 | 0 |

| | | | | | |
|---|---|---|---|---|---|
| B61 | B67 | WBHSRV | WBRSR(mPas_50) | 0.863181 | 0 |
| B61 | B100 | WBHSRV | anti-Ro52 | 0.385483 | 4.44E-15 |
| B61 | B101 | WBHSRV | anti-Sc70 | 0.385483 | 4.44E-15 |
| B61 | B102 | WBHSRV | anti-Sm | 0.385483 | 4.44E-15 |
| B61 | B103 | WBHSRV | anti-SSA/Ro | 0.385483 | 4.44E-15 |
| B61 | B104 | WBHSRV | anti-SSB | 0.385483 | 4.44E-15 |
| B61 | B105 | WBHSRV | anti-U1RNP | 0.385483 | 4.44E-15 |
| B61 | B106 | WBHSRV | anti-P | 0.385483 | 4.44E-15 |
| B61 | B107 | WBHSRV | AnuA | 0.385483 | 4.44E-15 |
| B61 | B108 | WBHSRV | Anti-Cyclin | 0.385483 | 4.44E-15 |
| B61 | B109 | WBHSRV | AMA-M2 | 0.385483 | 4.44E-15 |
| B61 | B110 | WBHSRV | AHA | 0.385483 | 4.44E-15 |
| B61 | B95 | WBHSRV | anti-cenpb | 0.385483 | 4.44E-15 |
| B61 | B97 | WBHSRV | dsDNA | 0.385483 | 4.44E-15 |
| B61 | B99 | WBHSRV | anti-PM-Scl | 0.385483 | 4.44E-15 |
| B61 | B151 | WBHSRV | Anti-CCP | -0.34405 | 3.87E-12 |
| B61 | B170 | WBHSRV | U-GLU | 0.319382 | 1.41E-10 |
| B61 | B98 | WBHSRV | anti-Jo-1 | -0.27801 | 2.91E-08 |
| B62 | B63 | WBHSRVI | WBRSR(mPas_200) | 0.74601 | 0 |
| B62 | B64 | WBHSRVI | WBRSR(mPas_5) | 0.72415 | 0 |
| B62 | B65 | WBHSRVI | WBRSR(mPas_1) | 0.689266 | 0 |
| B62 | B66 | WBHSRVI | WBRSR(mPas_100) | 0.762911 | 0 |
| B62 | B67 | WBHSRVI | WBRSR(mPas_50) | 0.774385 | 0 |
| B62 | B69 | WBHSRVI | PV | -0.42862 | 0 |
| B62 | B98 | WBHSRVI | anti-Jo-1 | -0.38918 | 2.22E-15 |
| B62 | B100 | WBHSRVI | anti-Ro52 | 0.365377 | 1.33E-13 |
| B62 | B101 | WBHSRVI | anti-Sc70 | 0.365377 | 1.33E-13 |
| B62 | B102 | WBHSRVI | anti-Sm | 0.365377 | 1.33E-13 |
| B62 | B103 | WBHSRVI | anti-SSA/Ro | 0.365377 | 1.33E-13 |
| B62 | B104 | WBHSRVI | anti-SSB | 0.365377 | 1.33E-13 |
| B62 | B105 | WBHSRVI | anti-U1RNP | 0.365377 | 1.33E-13 |
| B62 | B106 | WBHSRVI | anti-P | 0.365377 | 1.33E-13 |
| B62 | B107 | WBHSRVI | AnuA | 0.365377 | 1.33E-13 |
| B62 | B108 | WBHSRVI | Anti-Cyclin | 0.365377 | 1.33E-13 |
| B62 | B109 | WBHSRVI | AMA-M2 | 0.365377 | 1.33E-13 |
| B62 | B110 | WBHSRVI | AHA | 0.365377 | 1.33E-13 |
| B62 | B95 | WBHSRVI | anti-cenpb | 0.365377 | 1.33E-13 |
| B62 | B97 | WBHSRVI | dsDNA | 0.365377 | 1.33E-13 |
| B62 | B99 | WBHSRVI | anti-PM-Scl | 0.365377 | 1.33E-13 |
| B62 | B194 | WBHSRVI | Cl | -0.33765 | 1.01E-11 |
| B62 | B138 | WBHSRVI | PAB | 0.26879 | 8.52E-08 |
| B62 | B139 | WBHSRVI | ADA | -0.26697 | 1.05E-07 |
| B62 | B78 | WBHSRVI | HBcAb | 0.257917 | 2.88E-07 |
| B63 | B100 | WBRSR(mPas_200) | anti-Ro52 | 0.442489 | 0 |

| | | | | | |
|---|---|---|---|---|---|
| B63 | B101 | WBRSR(mPas_200) | anti-Sc70 | 0.442489 | 0 |
| B63 | B102 | WBRSR(mPas_200) | anti-Sm | 0.442489 | 0 |
| B63 | B103 | WBRSR(mPas_200) | anti-SSA/Ro | 0.442489 | 0 |
| B63 | B104 | WBRSR(mPas_200) | anti-SSB | 0.442489 | 0 |
| B63 | B105 | WBRSR(mPas_200) | anti-U1RNP | 0.442489 | 0 |
| B63 | B106 | WBRSR(mPas_200) | anti-P | 0.442489 | 0 |
| B63 | B107 | WBRSR(mPas_200) | AnuA | 0.442489 | 0 |
| B63 | B108 | WBRSR(mPas_200) | Anti-Cyclin | 0.442489 | 0 |
| B63 | B109 | WBRSR(mPas_200) | AMA-M2 | 0.442489 | 0 |
| B63 | B110 | WBRSR(mPas_200) | AHA | 0.442489 | 0 |
| B63 | B151 | WBRSR(mPas_200) | Anti-CCP | -0.45819 | 0 |
| B63 | B64 | WBRSR(mPas_200) | WBRSR(mPas_5) | 0.798504 | 0 |
| B63 | B65 | WBRSR(mPas_200) | WBRSR(mPas_1) | 0.665876 | 0 |
| B63 | B66 | WBRSR(mPas_200) | WBRSR(mPas_100) | 0.993152 | 0 |
| B63 | B67 | WBRSR(mPas_200) | WBRSR(mPas_50) | 0.979129 | 0 |
| B63 | B95 | WBRSR(mPas_200) | anti-cenpb | 0.442489 | 0 |
| B63 | B97 | WBRSR(mPas_200) | dsDNA | 0.442489 | 0 |
| B63 | B99 | WBRSR(mPas_200) | anti-PM-Scl | 0.442489 | 0 |
| B63 | B78 | WBRSR(mPas_200) | HBcAb | 0.361703 | 2.42E-13 |
| B63 | B69 | WBRSR(mPas_200) | PV | 0.268179 | 9.14E-08 |
| B63 | B178 | WBRSR(mPas_200) | Ig-U | 0.254561 | 4.14E-07 |
| B64 | B65 | WBRSR(mPas_5) | WBRSR(mPas_1) | 0.969594 | 0 |
| B64 | B66 | WBRSR(mPas_5) | WBRSR(mPas_100) | 0.843806 | 0 |
| B64 | B67 | WBRSR(mPas_5) | WBRSR(mPas_50) | 0.89515 | 0 |
| B64 | B78 | WBRSR(mPas_5) | HBcAb | 0.348893 | 1.84E-12 |
| B64 | B100 | WBRSR(mPas_5) | anti-Ro52 | 0.347261 | 2.37E-12 |
| B64 | B101 | WBRSR(mPas_5) | anti-Sc70 | 0.347261 | 2.37E-12 |
| B64 | B102 | WBRSR(mPas_5) | anti-Sm | 0.347261 | 2.37E-12 |
| B64 | B103 | WBRSR(mPas_5) | anti-SSA/Ro | 0.347261 | 2.37E-12 |
| B64 | B104 | WBRSR(mPas_5) | anti-SSB | 0.347261 | 2.37E-12 |
| B64 | B105 | WBRSR(mPas_5) | anti-U1RNP | 0.347261 | 2.37E-12 |
| B64 | B106 | WBRSR(mPas_5) | anti-P | 0.347261 | 2.37E-12 |
| B64 | B107 | WBRSR(mPas_5) | AnuA | 0.347261 | 2.37E-12 |
| B64 | B108 | WBRSR(mPas_5) | Anti-Cyclin | 0.347261 | 2.37E-12 |
| B64 | B109 | WBRSR(mPas_5) | AMA-M2 | 0.347261 | 2.37E-12 |
| B64 | B110 | WBRSR(mPas_5) | AHA | 0.347261 | 2.37E-12 |
| B64 | B95 | WBRSR(mPas_5) | anti-cenpb | 0.347261 | 2.37E-12 |
| B64 | B97 | WBRSR(mPas_5) | dsDNA | 0.347261 | 2.37E-12 |
| B64 | B99 | WBRSR(mPas_5) | anti-PM-Scl | 0.347261 | 2.37E-12 |
| B64 | B151 | WBRSR(mPas_5) | Anti-CCP | -0.34212 | 5.19E-12 |
| B64 | B73 | WBRSR(mPas_5) | ApoA | -0.29977 | 1.96E-09 |
| B64 | B139 | WBRSR(mPas_5) | ADA | -0.29645 | 3.00E-09 |
| B64 | B149 | WBRSR(mPas_5) | TGAb | -0.24889 | 7.58E-07 |
| B65 | B66 | WBRSR(mPas_1) | WBRSR(mPas_100) | 0.724735 | 0 |

| | | | | | |
|---|---|---|---|---|---|
| B65 | B67 | WBRSR(mPas_1) | WBRSR(mPas_50) | 0.791022 | 0 |
| B65 | B73 | WBRSR(mPas_1) | ApoA | -0.32895 | 3.63E-11 |
| B65 | B139 | WBRSR(mPas_1) | ADA | -0.31159 | 4.10E-10 |
| B65 | B100 | WBRSR(mPas_1) | anti-Ro52 | 0.303185 | 1.25E-09 |
| B65 | B101 | WBRSR(mPas_1) | anti-Sc70 | 0.303185 | 1.25E-09 |
| B65 | B102 | WBRSR(mPas_1) | anti-Sm | 0.303185 | 1.25E-09 |
| B65 | B103 | WBRSR(mPas_1) | anti-SSA/Ro | 0.303185 | 1.25E-09 |
| B65 | B104 | WBRSR(mPas_1) | anti-SSB | 0.303185 | 1.25E-09 |
| B65 | B105 | WBRSR(mPas_1) | anti-U1RNP | 0.303185 | 1.25E-09 |
| B65 | B106 | WBRSR(mPas_1) | anti-P | 0.303185 | 1.25E-09 |
| B65 | B107 | WBRSR(mPas_1) | AnuA | 0.303185 | 1.25E-09 |
| B65 | B108 | WBRSR(mPas_1) | Anti-Cyclin | 0.303185 | 1.25E-09 |
| B65 | B109 | WBRSR(mPas_1) | AMA-M2 | 0.303185 | 1.25E-09 |
| B65 | B110 | WBRSR(mPas_1) | AHA | 0.303185 | 1.25E-09 |
| B65 | B95 | WBRSR(mPas_1) | anti-cenpb | 0.303185 | 1.25E-09 |
| B65 | B97 | WBRSR(mPas_1) | dsDNA | 0.303185 | 1.25E-09 |
| B65 | B99 | WBRSR(mPas_1) | anti-PM-Scl | 0.303185 | 1.25E-09 |
| B65 | B151 | WBRSR(mPas_1) | Anti-CCP | -0.28772 | 8.97E-09 |
| B65 | B78 | WBRSR(mPas_1) | HBcAb | 0.280628 | 2.13E-08 |
| B66 | B100 | WBRSR(mPas_100) | anti-Ro52 | 0.420819 | 0 |
| B66 | B101 | WBRSR(mPas_100) | anti-Sc70 | 0.420819 | 0 |
| B66 | B102 | WBRSR(mPas_100) | anti-Sm | 0.420819 | 0 |
| B66 | B103 | WBRSR(mPas_100) | anti-SSA/Ro | 0.420819 | 0 |
| B66 | B104 | WBRSR(mPas_100) | anti-SSB | 0.420819 | 0 |
| B66 | B105 | WBRSR(mPas_100) | anti-U1RNP | 0.420819 | 0 |
| B66 | B106 | WBRSR(mPas_100) | anti-P | 0.420819 | 0 |
| B66 | B107 | WBRSR(mPas_100) | AnuA | 0.420819 | 0 |
| B66 | B108 | WBRSR(mPas_100) | Anti-Cyclin | 0.420819 | 0 |
| B66 | B109 | WBRSR(mPas_100) | AMA-M2 | 0.420819 | 0 |
| B66 | B110 | WBRSR(mPas_100) | AHA | 0.420819 | 0 |
| B66 | B151 | WBRSR(mPas_100) | Anti-CCP | -0.43307 | 0 |
| B66 | B67 | WBRSR(mPas_100) | WBRSR(mPas_50) | 0.994476 | 0 |
| B66 | B95 | WBRSR(mPas_100) | anti-cenpb | 0.420819 | 0 |
| B66 | B97 | WBRSR(mPas_100) | dsDNA | 0.420819 | 0 |
| B66 | B99 | WBRSR(mPas_100) | anti-PM-Scl | 0.420819 | 0 |
| B66 | B78 | WBRSR(mPas_100) | HBcAb | 0.352904 | 9.84E-13 |
| B66 | B69 | WBRSR(mPas_100) | PV | 0.247684 | 8.61E-07 |
| B67 | B100 | WBRSR(mPas_50) | anti-Ro52 | 0.414996 | 0 |
| B67 | B101 | WBRSR(mPas_50) | anti-Sc70 | 0.414996 | 0 |
| B67 | B102 | WBRSR(mPas_50) | anti-Sm | 0.414996 | 0 |
| B67 | B103 | WBRSR(mPas_50) | anti-SSA/Ro | 0.414996 | 0 |
| B67 | B104 | WBRSR(mPas_50) | anti-SSB | 0.414996 | 0 |
| B67 | B105 | WBRSR(mPas_50) | anti-U1RNP | 0.414996 | 0 |
| B67 | B106 | WBRSR(mPas_50) | anti-P | 0.414996 | 0 |

| | | | | | |
|---|---|---|---|---|---|
| B67 | B107 | WBRSR(mPas_50) | AnuA | 0.414996 | 0 |
| B67 | B108 | WBRSR(mPas_50) | Anti-Cyclin | 0.414996 | 0 |
| B67 | B109 | WBRSR(mPas_50) | AMA-M2 | 0.414996 | 0 |
| B67 | B110 | WBRSR(mPas_50) | AHA | 0.414996 | 0 |
| B67 | B151 | WBRSR(mPas_50) | Anti-CCP | -0.42499 | 0 |
| B67 | B95 | WBRSR(mPas_50) | anti-cenpb | 0.414996 | 0 |
| B67 | B97 | WBRSR(mPas_50) | dsDNA | 0.414996 | 0 |
| B67 | B99 | WBRSR(mPas_50) | anti-PM-Scl | 0.414996 | 0 |
| B67 | B78 | WBRSR(mPas_50) | HBcAb | 0.358574 | 4.00E-13 |
| B69 | B194 | PV | Cl | 0.347372 | 2.33E-12 |
| B69 | B98 | PV | anti-Jo-1 | 0.308586 | 6.14E-10 |
| B69 | B155 | PV | IgA | 0.29583 | 3.24E-09 |
| B69 | B151 | PV | Anti-CCP | -0.25076 | 6.22E-07 |
| B7 | B8 | WB-Cop | WB-Zinc | -0.84761 | 0 |
| B70 | B77 | LDL-C | TC | 0.850045 | 0 |
| B70 | B91 | LDL-C | TP | 0.252277 | 5.30E-07 |
| B70 | B76 | LDL-C | ApoB | 0.250531 | 6.38E-07 |
| B71 | B140 | TG | r-GT | 0.399827 | 4.44E-16 |
| B71 | B72 | TG | HDL-C | -0.37892 | 1.38E-14 |
| B71 | B77 | TG | TC | 0.308203 | 6.46E-10 |
| B71 | B128 | TG | ALT | 0.25518 | 3.88E-07 |
| B72 | B216 | HDL-C | BMI | -0.40299 | 2.22E-16 |
| B72 | B222 | HDL-C | NS | -0.34224 | 5.09E-12 |
| B72 | B192 | HDL-C | UA | -0.33622 | 1.25E-11 |
| B72 | B221 | HDL-C | Hei | -0.32829 | 3.99E-11 |
| B72 | B206 | HDL-C | Waist | -0.30959 | 5.37E-10 |
| B72 | B210 | HDL-C | Hip | -0.28612 | 1.09E-08 |
| B72 | B215 | HDL-C | BMI | -0.27332 | 5.05E-08 |
| B72 | B74 | HDL-C | ApoA1 | 0.2634 | 1.57E-07 |
| B72 | B77 | HDL-C | TC | 0.252611 | 5.11E-07 |
| B73 | B199 | ApoA | Mg | 0.331182 | 2.63E-11 |
| B73 | B76 | ApoA | ApoB | 0.278217 | 2.84E-08 |
| B73 | B75 | ApoA | ApoA1/B | -0.27554 | 3.89E-08 |
| B73 | B119 | ApoA | Hcv-rna | -0.26203 | 1.83E-07 |
| B73 | B79 | ApoA | hb V -dna | -0.2484 | 7.99E-07 |
| B73 | B197 | ApoA | AG | -0.24754 | 8.74E-07 |
| B74 | B196 | ApoA1 | TCO2 | -0.43407 | 0 |
| B74 | B75 | ApoA1 | ApoA1/B | 0.66891 | 0 |
| B74 | B197 | ApoA1 | AG | 0.361557 | 2.47E-13 |
| B74 | B213 | ApoA1 | CVA-R | -0.25473 | 4.07E-07 |
| B75 | B76 | ApoA1/B | ApoB | -0.77274 | 0 |
| B75 | B84 | ApoA1/B | C-PSA | 0.378405 | 1.49E-14 |
| B75 | B197 | ApoA1/B | AG | 0.330837 | 2.76E-11 |
| B75 | B196 | ApoA1/B | TCO2 | -0.32501 | 6.38E-11 |

| | | | | | |
|---|---|---|---|---|---|
| B75 | B213 | ApoA1/B | CVA-R | -0.31543 | 2.43E-10 |
| B75 | B180 | ApoA1/B | Tf | -0.31376 | 3.05E-10 |
| B75 | B134 | ApoA1/B | CHE | -0.29617 | 3.11E-09 |
| B76 | B180 | ApoB | Tf | 0.284615 | 1.31E-08 |
| B76 | B84 | ApoB | C-PSA | -0.26864 | 8.67E-08 |
| B76 | B136 | ApoB | 5'NT | 0.267065 | 1.04E-07 |
| B76 | B139 | ApoB | ADA | 0.246662 | 9.58E-07 |
| B77 | B91 | TC | TP | 0.271885 | 5.96E-08 |
| B78 | B175 | HBcAb | ALB-U | 0.269988 | 7.43E-08 |
| B78 | B119 | HBcAb | Hcv-rna | 0.255805 | 3.62E-07 |
| B79 | B119 | hb V -dna | Hcv-rna | 0.612221 | 0 |
| B79 | B138 | hb V -dna | PAB | 0.498508 | 0 |
| B79 | B181 | hb V -dna | APTT | -0.40869 | 0 |
| B79 | B182 | hb V -dna | TT | -0.42309 | 0 |
| B79 | B183 | hb V -dna | PTA | 0.409145 | 0 |
| B79 | B184 | hb V -dna | INR | 0.408688 | 0 |
| B79 | B186 | hb V -dna | PFC | 0.408688 | 0 |
| B79 | B185 | hb V -dna | PT | -0.3779 | 1.62E-14 |
| B79 | B124 | hb V -dna | ACTH | 0.372823 | 3.86E-14 |
| B79 | B90 | hb V -dna | CYFRA 21-1 | -0.34312 | 4.46E-12 |
| B79 | B198 | hb V -dna | P | 0.290124 | 6.66E-09 |
| B79 | B123 | hb V -dna | hs-CRP1 | -0.28622 | 1.08E-08 |
| B79 | B121 | hb V -dna | C3 | -0.28482 | 1.28E-08 |
| B79 | B122 | hb V -dna | C4 | -0.28482 | 1.28E-08 |
| B79 | B152 | hb V -dna | ACA-Ig(G-A-M) | -0.28482 | 1.28E-08 |
| B79 | B159 | hb V -dna | β2-MG | -0.28482 | 1.28E-08 |
| B79 | B169 | hb V -dna | U-CRY | -0.28482 | 1.28E-08 |
| B79 | B92 | hb V -dna | ALDH2-Glu504Glu | -0.28482 | 1.28E-08 |
| B79 | B93 | hb V -dna | ALDH2-Glu504Lys | 0.284817 | 1.28E-08 |
| B79 | B94 | hb V -dna | ALDH2-Lys504Lys | 0.284817 | 1.28E-08 |
| B79 | B180 | hb V -dna | Tf | 0.270703 | 6.84E-08 |
| B79 | B120 | hb V -dna | HCV-cAg | 0.248901 | 7.58E-07 |
| B81 | B89 | TSGF | CA 72-4 | 0.500038 | 0 |
| B81 | B182 | TSGF | TT | 0.355102 | 6.95E-13 |
| B81 | B183 | TSGF | PTA | -0.35511 | 6.95E-13 |
| B81 | B181 | TSGF | APTT | 0.355095 | 6.96E-13 |
| B81 | B184 | TSGF | INR | -0.35509 | 6.96E-13 |
| B81 | B186 | TSGF | PFC | -0.35509 | 6.96E-13 |
| B81 | B185 | TSGF | PT | 0.352356 | 1.07E-12 |
| B81 | B124 | TSGF | ACTH | -0.32357 | 7.82E-11 |
| B81 | B92 | TSGF | ALDH2-Glu504Glu | 0.25548 | 3.75E-07 |
| B81 | B93 | TSGF | ALDH2-Glu504Lys | -0.25548 | 3.75E-07 |
| B81 | B94 | TSGF | ALDH2-Lys504Lys | -0.25548 | 3.75E-07 |
| B82 | B98 | CEA | anti-Jo-1 | 0.60331 | 0 |

| | | | | | |
|---|---|---|---|---|---|
| B82 | B88 | CEA | CA 242 | 0.350005 | 1.55E-12 |
| B82 | B89 | CEA | CA 72-4 | 0.320357 | 1.23E-10 |
| B82 | B100 | CEA | anti-Ro52 | -0.3022 | 1.43E-09 |
| B82 | B101 | CEA | anti-Sc70 | -0.3022 | 1.43E-09 |
| B82 | B102 | CEA | anti-Sm | -0.3022 | 1.43E-09 |
| B82 | B103 | CEA | anti-SSA/Ro | -0.3022 | 1.43E-09 |
| B82 | B104 | CEA | anti-SSB | -0.3022 | 1.43E-09 |
| B82 | B105 | CEA | anti-U1RNP | -0.3022 | 1.43E-09 |
| B82 | B106 | CEA | anti-P | -0.3022 | 1.43E-09 |
| B82 | B107 | CEA | AnuA | -0.3022 | 1.43E-09 |
| B82 | B108 | CEA | Anti-Cyclin | -0.3022 | 1.43E-09 |
| B82 | B109 | CEA | AMA-M2 | -0.3022 | 1.43E-09 |
| B82 | B110 | CEA | AHA | -0.3022 | 1.43E-09 |
| B82 | B95 | CEA | anti-cenpb | -0.3022 | 1.43E-09 |
| B82 | B97 | CEA | dsDNA | -0.3022 | 1.43E-09 |
| B82 | B99 | CEA | anti-PM-Scl | -0.3022 | 1.43E-09 |
| B82 | B155 | CEA | IgA | 0.287459 | 9.26E-09 |
| B83 | B88 | AFP | CA 242 | 0.452814 | 0 |
| B83 | B87 | AFP | NSE | 0.327016 | 4.79E-11 |
| B84 | B139 | C-PSA | ADA | -0.25287 | 4.97E-07 |
| B84 | B136 | C-PSA | 5'NT | -0.25142 | 5.80E-07 |
| B85 | B98 | SCC | anti-Jo-1 | 0.30739 | 7.20E-10 |
| B85 | B89 | SCC | CA 72-4 | -0.29307 | 4.60E-09 |
| B86 | B87 | TBhCG | NSE | 0.422837 | 0 |
| B86 | B88 | TBhCG | CA 242 | 0.741413 | 0 |
| B86 | B90 | TBhCG | CYFRA 21-1 | 0.313652 | 3.10E-10 |
| B87 | B88 | NSE | CA 242 | 0.534397 | 0 |
| B87 | B98 | NSE | anti-Jo-1 | -0.37084 | 5.37E-14 |
| B87 | B89 | NSE | CA 72-4 | 0.298288 | 2.37E-09 |
| B87 | B90 | NSE | CYFRA 21-1 | 0.275444 | 3.94E-08 |
| B87 | B136 | NSE | 5'NT | 0.262222 | 1.79E-07 |
| B88 | B90 | CA 242 | CYFRA 21-1 | 0.254176 | 4.32E-07 |
| B90 | B124 | CYFRA 21-1 | ACTH | -0.63095 | 0 |
| B90 | B181 | CYFRA 21-1 | APTT | 0.69833 | 0 |
| B90 | B182 | CYFRA 21-1 | TT | 0.697361 | 0 |
| B90 | B183 | CYFRA 21-1 | PTA | -0.69833 | 0 |
| B90 | B184 | CYFRA 21-1 | INR | -0.69833 | 0 |
| B90 | B185 | CYFRA 21-1 | PT | 0.694833 | 0 |
| B90 | B186 | CYFRA 21-1 | PFC | -0.69833 | 0 |
| B90 | B92 | CYFRA 21-1 | ALDH2-Glu504Glu | 0.493013 | 0 |
| B90 | B93 | CYFRA 21-1 | ALDH2-Glu504Lys | -0.49301 | 0 |
| B90 | B94 | CYFRA 21-1 | ALDH2-Lys504Lys | -0.49301 | 0 |
| B90 | B139 | CYFRA 21-1 | ADA | 0.335153 | 1.47E-11 |
| B90 | B180 | CYFRA 21-1 | Tf | -0.27717 | 3.21E-08 |

| | | | | | |
|---|---|---|---|---|---|
| B91 | B126 | TP | Alb | 0.518706 | 0 |
| B91 | B127 | TP | A-G ratio | -0.4549 | 0 |
| B91 | B130 | TP | GLB | 0.768973 | 0 |
| B91 | B188 | TP | Ca | 0.275584 | 3.87E-08 |
| B92 | B121 | ALDH2-Glu504Glu | C3 | 0.497389 | 0 |
| B92 | B122 | ALDH2-Glu504Glu | C4 | 0.497389 | 0 |
| B92 | B124 | ALDH2-Glu504Glu | ACTH | -0.64962 | 0 |
| B92 | B152 | ALDH2-Glu504Glu | ACA-Ig(G-A-M) | 0.497398 | 0 |
| B92 | B159 | ALDH2-Glu504Glu | β2-MG | 0.49739 | 0 |
| B92 | B169 | ALDH2-Glu504Glu | U-CRY | 0.497388 | 0 |
| B92 | B181 | ALDH2-Glu504Glu | APTT | 0.706185 | 0 |
| B92 | B182 | ALDH2-Glu504Glu | TT | 0.731345 | 0 |
| B92 | B183 | ALDH2-Glu504Glu | PTA | -0.70698 | 0 |
| B92 | B184 | ALDH2-Glu504Glu | INR | -0.70619 | 0 |
| B92 | B185 | ALDH2-Glu504Glu | PT | 0.652454 | 0 |
| B92 | B186 | ALDH2-Glu504Glu | PFC | -0.70619 | 0 |
| B92 | B94 | ALDH2-Glu504Glu | ALDH2-Lys504Lys | -1 | 0 |
| B92 | B139 | ALDH2-Glu504Glu | ADA | 0.357376 | 4.85E-13 |
| B92 | B119 | ALDH2-Glu504Glu | Hcv-rna | -0.34892 | 1.83E-12 |
| B92 | B180 | ALDH2-Glu504Glu | Tf | -0.27628 | 3.57E-08 |
| B92 | B195 | ALDH2-Glu504Glu | Na | -0.27494 | 4.18E-08 |
| B92 | B200 | ALDH2-Glu504Glu | OP | -0.25845 | 2.71E-07 |
| B93 | B121 | ALDH2-Glu504Lys | C3 | -0.49739 | 0 |
| B93 | B122 | ALDH2-Glu504Lys | C4 | -0.49739 | 0 |
| B93 | B124 | ALDH2-Glu504Lys | ACTH | 0.649624 | 0 |
| B93 | B152 | ALDH2-Glu504Lys | ACA-Ig(G-A-M) | -0.4974 | 0 |
| B93 | B159 | ALDH2-Glu504Lys | β2-MG | -0.49739 | 0 |
| B93 | B169 | ALDH2-Glu504Lys | U-CRY | -0.49739 | 0 |
| B93 | B181 | ALDH2-Glu504Lys | APTT | -0.70619 | 0 |
| B93 | B182 | ALDH2-Glu504Lys | TT | -0.73134 | 0 |
| B93 | B183 | ALDH2-Glu504Lys | PTA | 0.706985 | 0 |
| B93 | B184 | ALDH2-Glu504Lys | INR | 0.706185 | 0 |
| B93 | B185 | ALDH2-Glu504Lys | PT | -0.65245 | 0 |
| B93 | B186 | ALDH2-Glu504Lys | PFC | 0.706185 | 0 |
| B93 | B94 | ALDH2-Glu504Lys | ALDH2-Lys504Lys | 1 | 0 |
| B93 | B139 | ALDH2-Glu504Lys | ADA | -0.35738 | 4.85E-13 |
| B93 | B119 | ALDH2-Glu504Lys | Hcv-rna | 0.348923 | 1.83E-12 |
| B93 | B180 | ALDH2-Glu504Lys | Tf | 0.276281 | 3.57E-08 |
| B93 | B195 | ALDH2-Glu504Lys | Na | 0.274943 | 4.18E-08 |
| B93 | B200 | ALDH2-Glu504Lys | OP | 0.258452 | 2.71E-07 |
| B94 | B121 | ALDH2-Lys504Lys | C3 | -0.49739 | 0 |
| B94 | B122 | ALDH2-Lys504Lys | C4 | -0.49739 | 0 |
| B94 | B124 | ALDH2-Lys504Lys | ACTH | 0.649624 | 0 |
| B94 | B152 | ALDH2-Lys504Lys | ACA-Ig(G-A-M) | -0.4974 | 0 |

| | | | | | |
|---|---|---|---|---|---|
| B94 | B159 | ALDH2-Lys504Lys | β2-MG | -0.49739 | 0 |
| B94 | B169 | ALDH2-Lys504Lys | U-CRY | -0.49739 | 0 |
| B94 | B181 | ALDH2-Lys504Lys | APTT | -0.70619 | 0 |
| B94 | B182 | ALDH2-Lys504Lys | TT | -0.73134 | 0 |
| B94 | B183 | ALDH2-Lys504Lys | PTA | 0.706985 | 0 |
| B94 | B184 | ALDH2-Lys504Lys | INR | 0.706185 | 0 |
| B94 | B185 | ALDH2-Lys504Lys | PT | -0.65245 | 0 |
| B94 | B186 | ALDH2-Lys504Lys | PFC | 0.706185 | 0 |
| B94 | B139 | ALDH2-Lys504Lys | ADA | -0.35738 | 4.85E-13 |
| B94 | B119 | ALDH2-Lys504Lys | Hcv-rna | 0.348923 | 1.83E-12 |
| B94 | B180 | ALDH2-Lys504Lys | Tf | 0.276281 | 3.57E-08 |
| B94 | B195 | ALDH2-Lys504Lys | Na | 0.274943 | 4.18E-08 |
| B94 | B200 | ALDH2-Lys504Lys | OP | 0.258452 | 2.71E-07 |
| B95 | B104 | anti-cenpb | anti-SSB | 1 | 0 |
| B95 | B106 | anti-cenpb | anti-P | 1 | 0 |
| B95 | B108 | anti-cenpb | Anti-Cyclin | 1 | 0 |
| B95 | B109 | anti-cenpb | AMA-M2 | 1 | 0 |
| B95 | B151 | anti-cenpb | Anti-CCP | -0.70526 | 0 |
| B95 | B178 | anti-cenpb | Ig-U | 0.453154 | 0 |
| B95 | B180 | anti-cenpb | Tf | 0.496456 | 0 |
| B95 | B98 | anti-cenpb | anti-Jo-1 | -0.47932 | 0 |
| B95 | B99 | anti-cenpb | anti-PM-Scl | 1 | 0 |
| B96 | B171 | dsDNA (IIF) | EC | 0.936586 | 0 |
| B97 | B100 | dsDNA | anti-Ro52 | 1 | 0 |
| B97 | B103 | dsDNA | anti-SSA/Ro | 1 | 0 |
| B97 | B105 | dsDNA | anti-U1RNP | 1 | 0 |
| B97 | B108 | dsDNA | Anti-Cyclin | 1 | 0 |
| B97 | B109 | dsDNA | AMA-M2 | 1 | 0 |
| B97 | B110 | dsDNA | AHA | 1 | 0 |
| B97 | B151 | dsDNA | Anti-CCP | -0.70526 | 0 |
| B97 | B178 | dsDNA | Ig-U | 0.453154 | 0 |
| B97 | B180 | dsDNA | Tf | 0.496456 | 0 |
| B97 | B98 | dsDNA | anti-Jo-1 | -0.47932 | 0 |
| B97 | B99 | dsDNA | anti-PM-Scl | 1 | 0 |
| B98 | B100 | anti-Jo-1 | anti-Ro52 | -0.47932 | 0 |
| B98 | B101 | anti-Jo-1 | anti-Sc70 | -0.47932 | 0 |
| B98 | B102 | anti-Jo-1 | anti-Sm | -0.47932 | 0 |
| B98 | B103 | anti-Jo-1 | anti-SSA/Ro | -0.47932 | 0 |
| B98 | B104 | anti-Jo-1 | anti-SSB | -0.47932 | 0 |
| B98 | B105 | anti-Jo-1 | anti-U1RNP | -0.47932 | 0 |
| B98 | B106 | anti-Jo-1 | anti-P | -0.47932 | 0 |
| B98 | B107 | anti-Jo-1 | AnuA | -0.47932 | 0 |
| B98 | B108 | anti-Jo-1 | Anti-Cyclin | -0.47932 | 0 |
| B98 | B109 | anti-Jo-1 | AMA-M2 | -0.47932 | 0 |

| | | | | | |
|---|---|---|---|---|---|
| B98 | B110 | anti-Jo-1 | AHA | -0.47932 | 0 |
| B98 | B139 | anti-Jo-1 | ADA | 0.513985 | 0 |
| B98 | B155 | anti-Jo-1 | IgA | 0.48677 | 0 |
| B98 | B99 | anti-Jo-1 | anti-PM-Scl | -0.47932 | 0 |
| B98 | B158 | anti-Jo-1 | IgM | -0.40005 | 4.44E-16 |
| B98 | B157 | anti-Jo-1 | IgG | 0.391939 | 1.33E-15 |
| B99 | B100 | anti-PM-Scl | anti-Ro52 | 1 | 0 |
| B99 | B103 | anti-PM-Scl | anti-SSA/Ro | 1 | 0 |
| B99 | B104 | anti-PM-Scl | anti-SSB | 1 | 0 |
| B99 | B105 | anti-PM-Scl | anti-U1RNP | 1 | 0 |
| B99 | B106 | anti-PM-Scl | anti-P | 1 | 0 |
| B99 | B109 | anti-PM-Scl | AMA-M2 | 1 | 0 |
| B99 | B110 | anti-PM-Scl | AHA | 1 | 0 |
| B99 | B151 | anti-PM-Scl | Anti-CCP | -0.70526 | 0 |
| B99 | B178 | anti-PM-Scl | Ig-U | 0.453154 | 0 |
| B99 | B180 | anti-PM-Scl | Tf | 0.496456 | 0 |

**Supplementary Table 23. PEI Correlations in Hepatitis B**

| Row | Column | Row | Column | Cor | *P* |
|---|---|---|---|---|---|
| B1 | B163 | Gender | SG | -0.38831 | 0 |
| B1 | B174 | Gender | Urea/Cr | 0.36025 | 0 |
| B1 | B190 | Gender | Cre | -0.63214 | 0 |
| B1 | B192 | Gender | UA | -0.50659 | 0 |
| B1 | B206 | Gender | Waist | -0.4416 | 0 |
| B1 | B207 | Gender | WHR | -0.45272 | 0 |
| B1 | B216 | Gender | BMI | -0.51769 | 0 |
| B1 | B221 | Gender | Hei | -0.60464 | 0 |
| B1 | B222 | Gender | NS | -0.61461 | 0 |
| B1 | B46 | Gender | RBC Count | -0.47012 | 0 |
| B1 | B51 | Gender | Hb | -0.62628 | 0 |
| B1 | B72 | Gender | HDL-C | 0.365087 | 0 |
| B1 | B45 | Gender | MCHC | -0.29938 | 4.44E-16 |
| B1 | B209 | Gender | Smoking | -0.27969 | 3.73E-14 |
| B1 | B210 | Gender | Hip | -0.25734 | 3.84E-12 |
| B1 | B215 | Gender | BMI | -0.25377 | 7.75E-12 |
| B1 | B161 | Gender | LEU | 0.247643 | 2.51E-11 |
| B1 | B218 | Gender | DBP | -0.24088 | 8.89E-11 |
| B1 | B204 | Gender | Drinking | -0.21915 | 4.00E-09 |
| B1 | B141 | Gender | AST/ALT | 0.196546 | 1.41E-07 |
| B1 | B140 | Gender | r-GT | -0.19515 | 1.73E-07 |
| B1 | B176 | Gender | Ucr | -0.19293 | 2.39E-07 |
| B1 | B128 | Gender | ALT | -0.1882 | 4.74E-07 |
| B1 | B44 | Gender | MCH | -0.18563 | 6.81E-07 |
| B1 | B42 | Gender | MON | -0.18489 | 7.55E-07 |
| B10 | B12 | PG I | PG II | 0.659891 | 0 |
| B10 | B14 | PG I | G-17 | 0.265023 | 8.20E-13 |
| B10 | B60 | PG I | WBRVI | 0.183049 | 9.75E-07 |
| B100 | B103 | anti-Ro52 | anti-SSA/Ro | 1 | 0 |
| B100 | B104 | anti-Ro52 | anti-SSB | 1 | 0 |
| B100 | B106 | anti-Ro52 | anti-P | 1 | 0 |
| B100 | B107 | anti-Ro52 | AnuA | 1 | 0 |
| B100 | B109 | anti-Ro52 | AMA-M2 | 1 | 0 |
| B100 | B151 | anti-Ro52 | Anti-CCP | 0.472831 | 0 |
| B100 | B186 | anti-Ro52 | PFC | 0.313552 | 0 |
| B100 | B142 | anti-Ro52 | 25-(OH)D | 0.249542 | 1.75E-11 |
| B100 | B184 | anti-Ro52 | INR | -0.22906 | 7.39E-10 |
| B100 | B185 | anti-Ro52 | PT | -0.19886 | 9.94E-08 |
| B100 | B175 | anti-Ro52 | ALB-U | 0.197112 | 1.29E-07 |
| B100 | B196 | anti-Ro52 | TCO2 | 0.193449 | 2.22E-07 |
| B100 | B137 | anti-Ro52 | AFU | -0.19228 | 2.63E-07 |
| B101 | B102 | anti-Sc70 | anti-Sm | 1 | 0 |

| | | | | | |
|---|---|---|---|---|---|
| B101 | B107 | anti-Sc70 | AnuA | 1 | 0 |
| B101 | B108 | anti-Sc70 | Anti-Cyclin | 1 | 0 |
| B101 | B151 | anti-Sc70 | Anti-CCP | 0.472831 | 0 |
| B101 | B186 | anti-Sc70 | PFC | 0.313552 | 0 |
| B101 | B142 | anti-Sc70 | 25-(OH)D | 0.249542 | 1.75E-11 |
| B101 | B184 | anti-Sc70 | INR | -0.22906 | 7.39E-10 |
| B101 | B185 | anti-Sc70 | PT | -0.19886 | 9.94E-08 |
| B101 | B175 | anti-Sc70 | ALB-U | 0.197112 | 1.29E-07 |
| B101 | B196 | anti-Sc70 | TCO2 | 0.193449 | 2.22E-07 |
| B101 | B137 | anti-Sc70 | AFU | -0.19228 | 2.63E-07 |
| B102 | B107 | anti-Sm | AnuA | 1 | 0 |
| B102 | B151 | anti-Sm | Anti-CCP | 0.472831 | 0 |
| B102 | B186 | anti-Sm | PFC | 0.313552 | 0 |
| B102 | B142 | anti-Sm | 25-(OH)D | 0.249542 | 1.75E-11 |
| B102 | B184 | anti-Sm | INR | -0.22906 | 7.39E-10 |
| B102 | B185 | anti-Sm | PT | -0.19886 | 9.94E-08 |
| B102 | B175 | anti-Sm | ALB-U | 0.197112 | 1.29E-07 |
| B102 | B196 | anti-Sm | TCO2 | 0.193449 | 2.22E-07 |
| B102 | B137 | anti-Sm | AFU | -0.19228 | 2.63E-07 |
| B103 | B104 | anti-SSA/Ro | anti-SSB | 1 | 0 |
| B103 | B105 | anti-SSA/Ro | anti-U1RNP | 1 | 0 |
| B103 | B106 | anti-SSA/Ro | anti-P | 1 | 0 |
| B103 | B107 | anti-SSA/Ro | AnuA | 1 | 0 |
| B103 | B109 | anti-SSA/Ro | AMA-M2 | 1 | 0 |
| B103 | B151 | anti-SSA/Ro | Anti-CCP | 0.472831 | 0 |
| B103 | B186 | anti-SSA/Ro | PFC | 0.313552 | 0 |
| B103 | B142 | anti-SSA/Ro | 25-(OH)D | 0.249542 | 1.75E-11 |
| B103 | B184 | anti-SSA/Ro | INR | -0.22906 | 7.39E-10 |
| B103 | B185 | anti-SSA/Ro | PT | -0.19886 | 9.94E-08 |
| B103 | B175 | anti-SSA/Ro | ALB-U | 0.197112 | 1.29E-07 |
| B103 | B196 | anti-SSA/Ro | TCO2 | 0.193449 | 2.22E-07 |
| B103 | B137 | anti-SSA/Ro | AFU | -0.19228 | 2.63E-07 |
| B104 | B105 | anti-SSB | anti-U1RNP | 1 | 0 |
| B104 | B106 | anti-SSB | anti-P | 1 | 0 |
| B104 | B107 | anti-SSB | AnuA | 1 | 0 |
| B104 | B108 | anti-SSB | Anti-Cyclin | 1 | 0 |
| B104 | B109 | anti-SSB | AMA-M2 | 1 | 0 |
| B104 | B151 | anti-SSB | Anti-CCP | 0.472831 | 0 |
| B104 | B186 | anti-SSB | PFC | 0.313552 | 0 |
| B104 | B142 | anti-SSB | 25-(OH)D | 0.249542 | 1.75E-11 |
| B104 | B184 | anti-SSB | INR | -0.22906 | 7.39E-10 |
| B104 | B185 | anti-SSB | PT | -0.19886 | 9.94E-08 |
| B104 | B175 | anti-SSB | ALB-U | 0.197112 | 1.29E-07 |
| B104 | B196 | anti-SSB | TCO2 | 0.193449 | 2.22E-07 |

| | | | | | |
|---|---|---|---|---|---|
| B104 | B137 | anti-SSB | AFU | -0.19228 | 2.63E-07 |
| B105 | B106 | anti-U1RNP | anti-P | 1 | 0 |
| B105 | B107 | anti-U1RNP | AnuA | 1 | 0 |
| B105 | B109 | anti-U1RNP | AMA-M2 | 1 | 0 |
| B105 | B151 | anti-U1RNP | Anti-CCP | 0.472831 | 0 |
| B105 | B186 | anti-U1RNP | PFC | 0.313552 | 0 |
| B105 | B142 | anti-U1RNP | 25-(OH)D | 0.249542 | 1.75E-11 |
| B105 | B184 | anti-U1RNP | INR | -0.22906 | 7.39E-10 |
| B105 | B185 | anti-U1RNP | PT | -0.19886 | 9.94E-08 |
| B105 | B175 | anti-U1RNP | ALB-U | 0.197112 | 1.29E-07 |
| B105 | B196 | anti-U1RNP | TCO2 | 0.193449 | 2.22E-07 |
| B105 | B137 | anti-U1RNP | AFU | -0.19228 | 2.63E-07 |
| B106 | B108 | anti-P | Anti-Cyclin | 1 | 0 |
| B106 | B109 | anti-P | AMA-M2 | 1 | 0 |
| B106 | B151 | anti-P | Anti-CCP | 0.472831 | 0 |
| B106 | B186 | anti-P | PFC | 0.313552 | 0 |
| B106 | B142 | anti-P | 25-(OH)D | 0.249542 | 1.75E-11 |
| B106 | B184 | anti-P | INR | -0.22906 | 7.39E-10 |
| B106 | B185 | anti-P | PT | -0.19886 | 9.94E-08 |
| B106 | B175 | anti-P | ALB-U | 0.197112 | 1.29E-07 |
| B106 | B196 | anti-P | TCO2 | 0.193449 | 2.22E-07 |
| B106 | B137 | anti-P | AFU | -0.19228 | 2.63E-07 |
| B107 | B108 | AnuA | Anti-Cyclin | 1 | 0 |
| B107 | B109 | AnuA | AMA-M2 | 1 | 0 |
| B107 | B151 | AnuA | Anti-CCP | 0.472831 | 0 |
| B107 | B186 | AnuA | PFC | 0.313552 | 0 |
| B107 | B142 | AnuA | 25-(OH)D | 0.249542 | 1.75E-11 |
| B107 | B184 | AnuA | INR | -0.22906 | 7.39E-10 |
| B107 | B185 | AnuA | PT | -0.19886 | 9.94E-08 |
| B107 | B175 | AnuA | ALB-U | 0.197112 | 1.29E-07 |
| B107 | B196 | AnuA | TCO2 | 0.193449 | 2.22E-07 |
| B107 | B137 | AnuA | AFU | -0.19228 | 2.63E-07 |
| B108 | B151 | Anti-Cyclin | Anti-CCP | 0.472831 | 0 |
| B108 | B186 | Anti-Cyclin | PFC | 0.313552 | 0 |
| B108 | B142 | Anti-Cyclin | 25-(OH)D | 0.249542 | 1.75E-11 |
| B108 | B184 | Anti-Cyclin | INR | -0.22906 | 7.39E-10 |
| B108 | B185 | Anti-Cyclin | PT | -0.19886 | 9.94E-08 |
| B108 | B175 | Anti-Cyclin | ALB-U | 0.197112 | 1.29E-07 |
| B108 | B196 | Anti-Cyclin | TCO2 | 0.193449 | 2.22E-07 |
| B108 | B137 | Anti-Cyclin | AFU | -0.19228 | 2.63E-07 |
| B109 | B151 | AMA-M2 | Anti-CCP | 0.472831 | 0 |
| B109 | B186 | AMA-M2 | PFC | 0.313552 | 0 |
| B109 | B142 | AMA-M2 | 25-(OH)D | 0.249542 | 1.75E-11 |
| B109 | B184 | AMA-M2 | INR | -0.22906 | 7.39E-10 |

| | | | | | |
|---|---|---|---|---|---|
| B109 | B185 | AMA-M2 | PT | -0.19886 | 9.94E-08 |
| B109 | B175 | AMA-M2 | ALB-U | 0.197112 | 1.29E-07 |
| B109 | B196 | AMA-M2 | TCO2 | 0.193449 | 2.22E-07 |
| B109 | B137 | AMA-M2 | AFU | -0.19228 | 2.63E-07 |
| B11 | B12 | PGI/PGII | PG II | -0.38802 | 0 |
| B110 | B157 | AHA | IgG | 0.351162 | 0 |
| B112 | B113 | FT4 | FT3 | 0.487005 | 0 |
| B112 | B114 | FT4 | TSH TT3 TT4(TT4) | 0.434034 | 0 |
| B113 | B115 | FT3 | TSH TT3 TT4(TT3) | 0.357752 | 0 |
| B114 | B115 | TSH TT3 TT4(TT4) | TSH TT3 TT4(TT3) | 0.499849 | 0 |
| B114 | B183 | TSH TT3 TT4(TT4) | PTA | -0.20889 | 2.11E-08 |
| B115 | B183 | TSH TT3 TT4(TT3) | PTA | -0.19825 | 1.09E-07 |
| B115 | B152 | TSH TT3 TT4(TT3) | ACA-Ig(G-A-M) | 0.185473 | 6.96E-07 |
| B119 | B120 | Hcv-rna | HCV-cAg | 0.439526 | 0 |
| B119 | B122 | Hcv-rna | C4 | -1 | 0 |
| B119 | B169 | Hcv-rna | U-CRY | -1 | 0 |
| B12 | B14 | PG II | G-17 | 0.290093 | 3.77E-15 |
| B120 | B121 | HCV-cAg | C3 | -0.43953 | 0 |
| B120 | B122 | HCV-cAg | C4 | -0.43953 | 0 |
| B120 | B159 | HCV-cAg | β2-MG | -0.43953 | 0 |
| B120 | B169 | HCV-cAg | U-CRY | -0.43953 | 0 |
| B121 | B122 | C3 | C4 | 1 | 0 |
| B121 | B169 | C3 | U-CRY | 0.999999 | 0 |
| B122 | B159 | C4 | β2-MG | 1 | 0 |
| B122 | B169 | C4 | U-CRY | 1 | 0 |
| B123 | B168 | hs-CRP1 | NMP22 | 0.191841 | 2.81E-07 |
| B124 | B125 | ACTH | Cor | 0.653675 | 0 |
| B125 | B181 | Cor | APTT | -0.28633 | 8.66E-15 |
| B125 | B182 | Cor | TT | -0.21863 | 4.36E-09 |
| B126 | B127 | Alb | A-G ratio | 0.499951 | 0 |
| B126 | B138 | Alb | PAB | 0.317646 | 0 |
| B126 | B188 | Alb | Ca | 0.478248 | 0 |
| B126 | B139 | Alb | ADA | -0.28083 | 2.91E-14 |
| B126 | B134 | Alb | CHE | 0.239514 | 1.14E-10 |
| B126 | B130 | Alb | GLB | -0.19608 | 1.51E-07 |
| B126 | B192 | Alb | UA | 0.183624 | 9.01E-07 |
| B127 | B130 | A-G ratio | GLB | -0.9159 | 0 |
| B127 | B139 | A-G ratio | ADA | -0.18456 | 7.90E-07 |
| B128 | B131 | ALT | AST | 0.819283 | 0 |
| B128 | B140 | ALT | r-GT | 0.33629 | 0 |
| B128 | B141 | ALT | AST/ALT | -0.43025 | 0 |
| B128 | B216 | ALT | BMI | 0.248875 | 1.99E-11 |
| B128 | B215 | ALT | BMI | 0.231303 | 4.99E-10 |
| B128 | B206 | ALT | Waist | 0.227576 | 9.56E-10 |

| | | | | | |
|---|---|---|---|---|---|
| B128 | B207 | ALT | WHR | 0.219876 | 3.54E-09 |
| B128 | B192 | ALT | UA | 0.195555 | 1.63E-07 |
| B128 | B222 | ALT | NS | 0.189606 | 3.87E-07 |
| B129 | B132 | IBIL | Dbil | 0.756319 | 0 |
| B129 | B133 | IBIL | TBIL | 0.985531 | 0 |
| B131 | B140 | AST | r-GT | 0.413774 | 0 |
| B132 | B133 | Dbil | TBIL | 0.856194 | 0 |
| B134 | B138 | CHE | PAB | 0.507609 | 0 |
| B134 | B139 | CHE | ADA | -0.39159 | 0 |
| B134 | B194 | CHE | Cl | -0.22148 | 2.71E-09 |
| B134 | B200 | CHE | OP | 0.206235 | 3.21E-08 |
| B134 | B188 | CHE | Ca | 0.186377 | 6.13E-07 |
| B135 | B139 | TBA | ADA | 0.283001 | 1.82E-14 |
| B135 | B199 | TBA | Mg | -0.23279 | 3.84E-10 |
| B135 | B138 | TBA | PAB | -0.19859 | 1.03E-07 |
| B136 | B138 | 5'NT | PAB | 0.424333 | 0 |
| B136 | B175 | 5'NT | ALB-U | 0.281031 | 2.80E-14 |
| B136 | B140 | 5'NT | r-GT | 0.2235 | 1.92E-09 |
| B136 | B184 | 5'NT | INR | -0.20589 | 3.39E-08 |
| B137 | B139 | AFU | ADA | 0.462806 | 0 |
| B137 | B151 | AFU | Anti-CCP | -0.28353 | 1.60E-14 |
| B137 | B199 | AFU | Mg | -0.26678 | 5.72E-13 |
| B137 | B150 | AFU | ALP | 0.264873 | 8.46E-13 |
| B137 | B186 | AFU | PFC | -0.26149 | 1.68E-12 |
| B137 | B138 | AFU | PAB | -0.21881 | 4.23E-09 |
| B137 | B198 | AFU | P | -0.19213 | 2.69E-07 |
| B138 | B139 | PAB | ADA | -0.596 | 0 |
| B138 | B199 | PAB | Mg | 0.316839 | 0 |
| B138 | B193 | PAB | K | 0.26314 | 1.20E-12 |
| B138 | B175 | PAB | ALB-U | 0.262646 | 1.33E-12 |
| B138 | B200 | PAB | OP | 0.250107 | 1.57E-11 |
| B138 | B194 | PAB | Cl | -0.23301 | 3.69E-10 |
| B138 | B188 | PAB | Ca | 0.18963 | 3.86E-07 |
| B138 | B184 | PAB | INR | -0.18651 | 6.01E-07 |
| B138 | B196 | PAB | TCO2 | 0.18625 | 6.24E-07 |
| B139 | B199 | ADA | Mg | -0.31379 | 0 |
| B139 | B150 | ADA | ALP | 0.278525 | 4.80E-14 |
| B139 | B198 | ADA | P | -0.24899 | 1.95E-11 |
| B139 | B196 | ADA | TCO2 | -0.23205 | 4.37E-10 |
| B139 | B193 | ADA | K | -0.20438 | 4.28E-08 |
| B139 | B181 | ADA | APTT | 0.197285 | 1.26E-07 |
| B139 | B186 | ADA | PFC | -0.18634 | 6.16E-07 |
| B14 | B143 | G-17 | β-CTx | 0.325853 | 0 |
| B14 | B144 | G-17 | Total-PINP | 0.472699 | 0 |

| | | | | | |
|---|---|---|---|---|---|
| B14 | B148 | G-17 | Anti_TPO | 0.441111 | 0 |
| B14 | B145 | G-17 | BGP | 0.250813 | 1.37E-11 |
| B14 | B24 | G-17 | αHBDH | 0.187512 | 5.22E-07 |
| B140 | B207 | r-GT | WHR | 0.29649 | 8.88E-16 |
| B140 | B192 | r-GT | UA | 0.287206 | 7.11E-15 |
| B140 | B206 | r-GT | Waist | 0.284161 | 1.40E-14 |
| B140 | B204 | r-GT | Drinking | 0.251931 | 1.11E-11 |
| B140 | B215 | r-GT | BMI | 0.246093 | 3.37E-11 |
| B140 | B216 | r-GT | BMI | 0.238586 | 1.35E-10 |
| B140 | B222 | r-GT | NS | 0.221946 | 2.50E-09 |
| B140 | B218 | r-GT | DBP | 0.203728 | 4.74E-08 |
| B140 | B209 | r-GT | Smoking | 0.191491 | 2.95E-07 |
| B141 | B216 | AST/ALT | BMI | -0.25009 | 1.58E-11 |
| B141 | B207 | AST/ALT | WHR | -0.24375 | 5.22E-11 |
| B141 | B206 | AST/ALT | Waist | -0.24167 | 7.68E-11 |
| B141 | B222 | AST/ALT | NS | -0.22444 | 1.64E-09 |
| B141 | B215 | AST/ALT | BMI | -0.21953 | 3.75E-09 |
| B141 | B163 | AST/ALT | SG | -0.20354 | 4.88E-08 |
| B141 | B192 | AST/ALT | UA | -0.20329 | 5.06E-08 |
| B142 | B143 | 25-(OH)D | β-CTx | 0.403402 | 0 |
| B142 | B183 | 25-(OH)D | PTA | -0.26607 | 6.63E-13 |
| B142 | B145 | 25-(OH)D | BGP | 0.198422 | 1.06E-07 |
| B143 | B144 | β-CTx | Total-PINP | 0.802848 | 0 |
| B143 | B145 | β-CTx | BGP | 0.720372 | 0 |
| B143 | B168 | β-CTx | NMP22 | -0.29461 | 1.33E-15 |
| B143 | B148 | β-CTx | Anti_TPO | 0.287249 | 7.11E-15 |
| B144 | B145 | Total-PINP | BGP | 0.795914 | 0 |
| B144 | B148 | Total-PINP | Anti_TPO | 0.48036 | 0 |
| B144 | B168 | Total-PINP | NMP22 | -0.25177 | 1.14E-11 |
| B145 | B148 | BGP | Anti_TPO | 0.217768 | 5.03E-09 |
| B148 | B149 | Anti_TPO | TGAb | 0.451927 | 0 |
| B149 | B175 | TGAb | ALB-U | 0.233265 | 3.52E-10 |
| B149 | B168 | TGAb | NMP22 | 0.202198 | 5.99E-08 |
| B149 | B224 | TGAb | pmh | 0.185527 | 6.91E-07 |
| B15 | B100 | CD3-CD19+(B cell) | anti-Ro52 | 0.389624 | 0 |
| B15 | B101 | CD3-CD19+(B cell) | anti-Sc70 | 0.389624 | 0 |
| B15 | B102 | CD3-CD19+(B cell) | anti-Sm | 0.389624 | 0 |
| B15 | B103 | CD3-CD19+(B cell) | anti-SSA/Ro | 0.389624 | 0 |
| B15 | B104 | CD3-CD19+(B cell) | anti-SSB | 0.389624 | 0 |
| B15 | B105 | CD3-CD19+(B cell) | anti-U1RNP | 0.389624 | 0 |
| B15 | B106 | CD3-CD19+(B cell) | anti-P | 0.389624 | 0 |
| B15 | B107 | CD3-CD19+(B cell) | AnuA | 0.389624 | 0 |
| B15 | B108 | CD3-CD19+(B cell) | Anti-Cyclin | 0.389624 | 0 |
| B15 | B109 | CD3-CD19+(B cell) | AMA-M2 | 0.389624 | 0 |

| | | | | | |
|---|---|---|---|---|---|
| B15 | B151 | CD3-CD19+(B cell) | Anti-CCP | 0.454231 | 0 |
| B15 | B16 | CD3-CD19+(B cell) | CD3+T | -0.56335 | 0 |
| B15 | B186 | CD3-CD19+(B cell) | PFC | 0.372108 | 0 |
| B15 | B19 | CD3-CD19+(B cell) | CD3+CD4+/CD3+CD8+ | 0.32526 | 0 |
| B15 | B22 | CD3-CD19+(B cell) | CD3+CD8+(T) | -0.50069 | 0 |
| B15 | B97 | CD3-CD19+(B cell) | dsDNA | 0.389624 | 0 |
| B15 | B98 | CD3-CD19+(B cell) | anti-Jo-1 | 0.389624 | 0 |
| B15 | B99 | CD3-CD19+(B cell) | anti-PM-Scl | -0.37021 | 0 |
| B15 | B95 | CD3-CD19+(B cell) | anti-cenpb | -0.23245 | 4.08E-10 |
| B15 | B17 | CD3-CD19+(B cell) | CD3+CD16+CD56+NK | -0.21777 | 5.03E-09 |
| B15 | B21 | CD3-CD19+(B cell) | CD3+CD4-CD8- | -0.20775 | 2.53E-08 |
| B15 | B193 | CD3-CD19+(B cell) | K | 0.203037 | 5.27E-08 |
| B15 | B18 | CD3-CD19+(B cell) | CD3+CD4+helper(inducer)T | 0.201672 | 6.49E-08 |
| B15 | B20 | CD3-CD19+(B cell) | CD3+CD4+CD8+ | -0.18777 | 5.03E-07 |
| B15 | B124 | CD3-CD19+(B cell) | ACTH | -0.18713 | 5.51E-07 |
| B151 | B186 | Anti-CCP | PFC | 0.37104 | 0 |
| B151 | B196 | Anti-CCP | TCO2 | 0.274661 | 1.10E-13 |
| B151 | B184 | Anti-CCP | INR | -0.26936 | 3.36E-13 |
| B151 | B177 | Anti-CCP | ACR | 0.261413 | 1.71E-12 |
| B151 | B175 | Anti-CCP | ALB-U | 0.245462 | 3.79E-11 |
| B151 | B185 | Anti-CCP | PT | -0.23598 | 2.17E-10 |
| B151 | B197 | Anti-CCP | AG | -0.23523 | 2.48E-10 |
| B151 | B157 | Anti-CCP | IgG | 0.207242 | 2.74E-08 |
| B153 | B170 | Glu | U-GLU | 0.445715 | 0 |
| B153 | B172 | Glu | KB | 0.236982 | 1.81E-10 |
| B153 | B220 | Glu | SBP | 0.23287 | 3.78E-10 |
| B153 | B218 | Glu | DBP | 0.193683 | 2.14E-07 |
| B155 | B157 | IgA | IgG | 0.354138 | 0 |
| B155 | B228 | IgA | OS | 0.654536 | 0 |
| B155 | B156 | IgA | IgE | 0.219843 | 3.56E-09 |
| B157 | B158 | IgG | IgM | 0.31565 | 0 |
| B157 | B183 | IgG | PTA | -0.27791 | 5.51E-14 |
| B157 | B170 | IgG | U-GLU | 0.220578 | 3.15E-09 |
| B157 | B182 | IgG | TT | 0.200381 | 7.90E-08 |
| B158 | B194 | IgM | Cl | -0.20334 | 5.03E-08 |
| B16 | B156 | CD3+T | IgE | -0.46831 | 0 |
| B16 | B18 | CD3+T | CD3+CD4+helper(inducer)T | 0.307531 | 0 |
| B16 | B183 | CD3+T | PTA | 0.340552 | 0 |
| B16 | B22 | CD3+T | CD3+CD8+(T) | 0.349024 | 0 |
| B16 | B23 | CD3+T | CD3-CD16+CD56+(NK) | -0.77511 | 0 |
| B16 | B151 | CD3+T | Anti-CCP | -0.27351 | 1.40E-13 |
| B16 | B100 | CD3+T | anti-Ro52 | -0.2312 | 5.07E-10 |
| B16 | B101 | CD3+T | anti-Sc70 | -0.2312 | 5.07E-10 |
| B16 | B102 | CD3+T | anti-Sm | -0.2312 | 5.07E-10 |

| | | | | | |
|---|---|---|---|---|---|
| B16 | B103 | CD3+T | anti-SSA/Ro | -0.2312 | 5.07E-10 |
| B16 | B104 | CD3+T | anti-SSB | -0.2312 | 5.07E-10 |
| B16 | B105 | CD3+T | anti-U1RNP | -0.2312 | 5.07E-10 |
| B16 | B106 | CD3+T | anti-P | -0.2312 | 5.07E-10 |
| B16 | B107 | CD3+T | AnuA | -0.2312 | 5.07E-10 |
| B16 | B108 | CD3+T | Anti-Cyclin | -0.2312 | 5.07E-10 |
| B16 | B109 | CD3+T | AMA-M2 | -0.2312 | 5.07E-10 |
| B16 | B97 | CD3+T | dsDNA | -0.2312 | 5.07E-10 |
| B16 | B98 | CD3+T | anti-Jo-1 | -0.2312 | 5.07E-10 |
| B16 | B21 | CD3+T | CD3+CD4-CD8- | 0.209181 | 2.02E-08 |
| B16 | B168 | CD3+T | NMP22 | -0.19726 | 1.26E-07 |
| B161 | B164 | LEU | COM | 0.186416 | 6.09E-07 |
| B163 | B166 | SG | Pro | 0.369216 | 0 |
| B163 | B176 | SG | Ucr | 0.381526 | 0 |
| B163 | B216 | SG | BMI | 0.338043 | 0 |
| B163 | B221 | SG | Hei | 0.315894 | 0 |
| B163 | B222 | SG | NS | 0.296943 | 8.88E-16 |
| B163 | B191 | SG | Urea | 0.270789 | 2.49E-13 |
| B163 | B206 | SG | Waist | 0.26415 | 9.80E-13 |
| B163 | B190 | SG | Cre | 0.261911 | 1.54E-12 |
| B163 | B210 | SG | Hip | 0.22724 | 1.01E-09 |
| B163 | B192 | SG | UA | 0.225325 | 1.41E-09 |
| B163 | B215 | SG | BMI | 0.210593 | 1.61E-08 |
| B163 | B207 | SG | WHR | 0.208762 | 2.16E-08 |
| B164 | B176 | COM | Ucr | 0.183774 | 8.82E-07 |
| B168 | B183 | NMP22 | PTA | -0.3647 | 0 |
| B168 | B175 | NMP22 | ALB-U | 0.265105 | 8.06E-13 |
| B168 | B197 | NMP22 | AG | 0.211514 | 1.39E-08 |
| B168 | B194 | NMP22 | Cl | -0.20593 | 3.36E-08 |
| B168 | B193 | NMP22 | K | 0.194216 | 1.98E-07 |
| B17 | B18 | CD3+CD16+CD56+NK | CD3+CD4+helper(inducer)T | -0.30543 | 0 |
| B17 | B19 | CD3+CD16+CD56+NK | CD3+CD4+/CD3+CD8+ | -0.38054 | 0 |
| B17 | B22 | CD3+CD16+CD56+NK | CD3+CD8+(T) | 0.407521 | 0 |
| B17 | B99 | CD3+CD16+CD56+NK | anti-PM-Scl | 0.443543 | 0 |
| B17 | B182 | CD3+CD16+CD56+NK | TT | 0.273201 | 1.50E-13 |
| B17 | B181 | CD3+CD16+CD56+NK | APTT | 0.247354 | 2.65E-11 |
| B17 | B175 | CD3+CD16+CD56+NK | ALB-U | 0.205944 | 3.36E-08 |
| B170 | B172 | U-GLU | KB | 0.213567 | 9.99E-09 |
| B172 | B179 | KB | α1-MG | 0.203368 | 5.01E-08 |
| B172 | B178 | KB | Ig-U | 0.198709 | 1.02E-07 |
| B174 | B190 | Urea/Cr | Cre | -0.4623 | 0 |
| B174 | B191 | Urea/Cr | Urea | 0.728631 | 0 |
| B174 | B221 | Urea/Cr | Hei | -0.27979 | 3.64E-14 |
| B174 | B192 | Urea/Cr | UA | -0.26182 | 1.57E-12 |

| | | | | | |
|---|---|---|---|---|---|
| B174 | B222 | Urea/Cr | NS | -0.2525 | 9.91E-12 |
| B174 | B216 | Urea/Cr | BMI | -0.22642 | 1.17E-09 |
| B174 | B189 | Urea/Cr | eGFB | 0.196676 | 1.38E-07 |
| B175 | B177 | ALB-U | ACR | 0.551605 | 0 |
| B176 | B177 | Ucr | ACR | -0.27869 | 4.62E-14 |
| B176 | B183 | Ucr | PTA | 0.240753 | 9.10E-11 |
| B176 | B216 | Ucr | BMI | 0.227184 | 1.02E-09 |
| B176 | B206 | Ucr | Waist | 0.204152 | 4.44E-08 |
| B176 | B198 | Ucr | P | -0.19081 | 3.26E-07 |
| B176 | B179 | Ucr | α1-MG | 0.187273 | 5.40E-07 |
| B177 | B178 | ACR | Ig-U | 0.357484 | 0 |
| B177 | B180 | ACR | Tf | 0.355035 | 0 |
| B178 | B179 | Ig-U | α1-MG | 0.411791 | 0 |
| B178 | B180 | Ig-U | Tf | 0.849848 | 0 |
| B18 | B156 | CD3+CD4+helper(inducer)T | IgE | -0.32839 | 0 |
| B18 | B181 | CD3+CD4+helper(inducer)T | APTT | -0.32592 | 0 |
| B18 | B19 | CD3+CD4+helper(inducer)T | CD3+CD4+/CD3+CD8+ | 0.896808 | 0 |
| B18 | B22 | CD3+CD4+helper(inducer)T | CD3+CD8+(T) | -0.67486 | 0 |
| B18 | B23 | CD3+CD4+helper(inducer)T | CD3-CD16+CD56+(NK) | -0.37734 | 0 |
| B18 | B20 | CD3+CD4+helper(inducer)T | CD3+CD4+CD8+ | 0.300745 | 4.44E-16 |
| B18 | B183 | CD3+CD4+helper(inducer)T | PTA | 0.231524 | 4.80E-10 |
| B181 | B182 | APTT | TT | 0.530822 | 0 |
| B181 | B184 | APTT | INR | 0.337321 | 0 |
| B181 | B185 | APTT | PT | 0.407327 | 0 |
| B181 | B186 | APTT | PFC | -0.35902 | 0 |
| B181 | B183 | APTT | PTA | -0.20759 | 2.59E-08 |
| B182 | B183 | TT | PTA | -0.29901 | 4.44E-16 |
| B182 | B185 | TT | PT | 0.191891 | 2.79E-07 |
| B183 | B194 | PTA | Cl | 0.231844 | 4.53E-10 |
| B184 | B185 | INR | PT | 0.915377 | 0 |
| B184 | B186 | INR | PFC | -0.40278 | 0 |
| B185 | B186 | PT | PFC | -0.40104 | 0 |
| B186 | B193 | PFC | K | 0.215984 | 6.74E-09 |
| B189 | B190 | eGFB | Cre | -0.51948 | 0 |
| B19 | B22 | CD3+CD4+/CD3+CD8+ | CD3+CD8+(T) | -0.87397 | 0 |
| B19 | B151 | CD3+CD4+/CD3+CD8+ | Anti-CCP | 0.284476 | 1.31E-14 |
| B19 | B100 | CD3+CD4+/CD3+CD8+ | anti-Ro52 | 0.242609 | 6.46E-11 |
| B19 | B101 | CD3+CD4+/CD3+CD8+ | anti-Sc70 | 0.242609 | 6.46E-11 |
| B19 | B102 | CD3+CD4+/CD3+CD8+ | anti-Sm | 0.242609 | 6.46E-11 |
| B19 | B103 | CD3+CD4+/CD3+CD8+ | anti-SSA/Ro | 0.242609 | 6.46E-11 |
| B19 | B104 | CD3+CD4+/CD3+CD8+ | anti-SSB | 0.242609 | 6.46E-11 |
| B19 | B105 | CD3+CD4+/CD3+CD8+ | anti-U1RNP | 0.242609 | 6.46E-11 |
| B19 | B106 | CD3+CD4+/CD3+CD8+ | anti-P | 0.242609 | 6.46E-11 |
| B19 | B107 | CD3+CD4+/CD3+CD8+ | AnuA | 0.242609 | 6.46E-11 |

| | | | | | |
|---|---|---|---|---|---|
| B19 | B108 | CD3+CD4+/CD3+CD8+ | Anti-Cyclin | 0.242609 | 6.46E-11 |
| B19 | B109 | CD3+CD4+/CD3+CD8+ | AMA-M2 | 0.242609 | 6.46E-11 |
| B19 | B97 | CD3+CD4+/CD3+CD8+ | dsDNA | 0.242609 | 6.46E-11 |
| B19 | B98 | CD3+CD4+/CD3+CD8+ | anti-Jo-1 | 0.242609 | 6.46E-11 |
| B19 | B186 | CD3+CD4+/CD3+CD8+ | PFC | 0.231806 | 4.56E-10 |
| B19 | B185 | CD3+CD4+/CD3+CD8+ | PT | -0.18707 | 5.56E-07 |
| B19 | B156 | CD3+CD4+/CD3+CD8+ | IgE | -0.18339 | 9.31E-07 |
| B190 | B192 | Cre | UA | 0.461507 | 0 |
| B190 | B216 | Cre | BMI | 0.379109 | 0 |
| B190 | B221 | Cre | Hei | 0.422194 | 0 |
| B190 | B222 | Cre | NS | 0.418573 | 0 |
| B190 | B206 | Cre | Waist | 0.290471 | 3.55E-15 |
| B190 | B207 | Cre | WHR | 0.25984 | 2.34E-12 |
| B190 | B191 | Cre | Urea | 0.229109 | 7.33E-10 |
| B190 | B210 | Cre | Hip | 0.216594 | 6.10E-09 |
| B190 | B209 | Cre | Smoking | 0.208017 | 2.43E-08 |
| B190 | B215 | Cre | BMI | 0.200168 | 8.16E-08 |
| B192 | B206 | UA | Waist | 0.41245 | 0 |
| B192 | B207 | UA | WHR | 0.369242 | 0 |
| B192 | B210 | UA | Hip | 0.304978 | 0 |
| B192 | B215 | UA | BMI | 0.340246 | 0 |
| B192 | B216 | UA | BMI | 0.460299 | 0 |
| B192 | B221 | UA | Hei | 0.363744 | 0 |
| B192 | B222 | UA | NS | 0.466429 | 0 |
| B192 | B218 | UA | DBP | 0.219409 | 3.83E-09 |
| B192 | B204 | UA | Drinking | 0.196403 | 1.44E-07 |
| B192 | B209 | UA | Smoking | 0.182943 | 9.90E-07 |
| B193 | B200 | K | OP | 0.218781 | 4.25E-09 |
| B194 | B197 | Cl | AG | -0.32174 | 0 |
| B194 | B196 | Cl | TCO2 | -0.26495 | 8.32E-13 |
| B194 | B200 | Cl | OP | 0.209555 | 1.90E-08 |
| B195 | B197 | Na | AG | 0.354654 | 0 |
| B195 | B200 | Na | OP | 0.899712 | 0 |
| B195 | B199 | Na | Mg | 0.214293 | 8.88E-09 |
| B195 | B212 | Na | IOP-L | 0.189039 | 4.20E-07 |
| B196 | B197 | TCO2 | AG | -0.64525 | 0 |
| B197 | B200 | AG | OP | 0.290078 | 3.77E-15 |
| B2 | B189 | Age | eGFB | -0.51682 | 0 |
| B2 | B220 | Age | SBP | 0.29375 | 1.78E-15 |
| B2 | B47 | Age | RDW | 0.265902 | 6.85E-13 |
| B2 | B46 | Age | RBC Count | -0.26112 | 1.81E-12 |
| B2 | B201 | Age | e-habits | 0.232557 | 4.00E-10 |
| B2 | B218 | Age | DBP | 0.218017 | 4.82E-09 |
| B2 | B52 | Age | PLT/BPC | -0.21525 | 7.60E-09 |

| | | | | | |
|---|---|---|---|---|---|
| B2 | B126 | Age | Alb | -0.20953 | 1.91E-08 |
| B2 | B209 | Age | Smoking | 0.206935 | 2.88E-08 |
| B2 | B221 | Age | Hei | -0.19872 | 1.02E-07 |
| B2 | B153 | Age | Glu | 0.196799 | 1.35E-07 |
| B2 | B207 | Age | WHR | 0.194181 | 1.99E-07 |
| B2 | B51 | Age | Hb | -0.18821 | 4.73E-07 |
| B20 | B181 | CD3+CD4+CD8+ | APTT | -0.33885 | 0 |
| B20 | B95 | CD3+CD4+CD8+ | anti-cenpb | 0.711817 | 0 |
| B20 | B185 | CD3+CD4+CD8+ | PT | -0.28118 | 2.71E-14 |
| B20 | B23 | CD3+CD4+CD8+ | CD3-CD16+CD56+(NK) | 0.25415 | 7.19E-12 |
| B20 | B183 | CD3+CD4+CD8+ | PTA | 0.250828 | 1.37E-11 |
| B20 | B184 | CD3+CD4+CD8+ | INR | -0.24159 | 7.80E-11 |
| B20 | B81 | CD3+CD4+CD8+ | TSGF | -0.22462 | 1.59E-09 |
| B20 | B125 | CD3+CD4+CD8+ | Cor | 0.21567 | 7.10E-09 |
| B20 | B205 | CD3+CD4+CD8+ | KDA | 0.199406 | 9.16E-08 |
| B20 | B99 | CD3+CD4+CD8+ | anti-PM-Scl | 0.195552 | 1.63E-07 |
| B20 | B136 | CD3+CD4+CD8+ | 5'NT | 0.192849 | 2.42E-07 |
| B204 | B209 | Drinking | Smoking | 0.3002 | 4.44E-16 |
| B204 | B222 | Drinking | NS | 0.191085 | 3.13E-07 |
| B206 | B207 | Waist | WHR | 0.864702 | 0 |
| B206 | B210 | Waist | Hip | 0.765646 | 0 |
| B206 | B215 | Waist | BMI | 0.815243 | 0 |
| B206 | B216 | Waist | BMI | 0.848832 | 0 |
| B206 | B221 | Waist | Hei | 0.405117 | 0 |
| B206 | B222 | Waist | NS | 0.660835 | 0 |
| B206 | B218 | Waist | DBP | 0.255906 | 5.10E-12 |
| B206 | B220 | Waist | SBP | 0.20978 | 1.84E-08 |
| B207 | B210 | WHR | Hip | 0.349359 | 0 |
| B207 | B215 | WHR | BMI | 0.615949 | 0 |
| B207 | B216 | WHR | BMI | 0.613874 | 0 |
| B207 | B222 | WHR | NS | 0.524663 | 0 |
| B207 | B221 | WHR | Hei | 0.262539 | 1.36E-12 |
| B207 | B218 | WHR | DBP | 0.239781 | 1.09E-10 |
| B207 | B209 | WHR | Smoking | 0.214124 | 9.13E-09 |
| B207 | B220 | WHR | SBP | 0.213372 | 1.03E-08 |
| B209 | B222 | Smoking | NS | 0.22912 | 7.31E-10 |
| B21 | B181 | CD3+CD4-CD8- | APTT | 0.452591 | 0 |
| B21 | B65 | CD3+CD4-CD8- | WBRSR(mPas_1) | -0.32584 | 0 |
| B21 | B59 | CD3+CD4-CD8- | WBRV | -0.29982 | 4.44E-16 |
| B21 | B60 | CD3+CD4-CD8- | WBRVI | -0.29569 | 8.88E-16 |
| B21 | B64 | CD3+CD4-CD8- | WBRSR(mPas_5) | -0.29443 | 1.33E-15 |
| B21 | B95 | CD3+CD4-CD8- | anti-cenpb | -0.26544 | 7.53E-13 |
| B21 | B57 | CD3+CD4-CD8- | RBCAI | -0.2353 | 2.45E-10 |
| B21 | B67 | CD3+CD4-CD8- | WBRSR(mPas_50) | -0.23156 | 4.77E-10 |

| | | | | | |
|---|---|---|---|---|---|
| B21 | B205 | CD3+CD4-CD8- | KDA | 0.22277 | 2.18E-09 |
| B21 | B195 | CD3+CD4-CD8- | Na | 0.2171 | 5.61E-09 |
| B21 | B66 | CD3+CD4-CD8- | WBRSR(mPas_100) | -0.21275 | 1.14E-08 |
| B21 | B62 | CD3+CD4-CD8- | WBHSRVI | -0.19656 | 1.40E-07 |
| B21 | B63 | CD3+CD4-CD8- | WBRSR(mPas_200) | -0.19293 | 2.39E-07 |
| B21 | B198 | CD3+CD4-CD8- | P | 0.191303 | 3.03E-07 |
| B210 | B215 | Hip | BMI | 0.743592 | 0 |
| B210 | B216 | Hip | BMI | 0.80648 | 0 |
| B210 | B221 | Hip | Hei | 0.42413 | 0 |
| B210 | B222 | Hip | NS | 0.573226 | 0 |
| B211 | B212 | IOP-R | IOP-L | 0.748024 | 0 |
| B213 | B214 | CVA-R | CVA-L | 0.762519 | 0 |
| B215 | B216 | BMI | BMI | 0.828294 | 0 |
| B215 | B222 | BMI | NS | 0.599239 | 0 |
| B215 | B218 | BMI | DBP | 0.234357 | 2.90E-10 |
| B215 | B220 | BMI | SBP | 0.187923 | 4.92E-07 |
| B216 | B221 | BMI | Hei | 0.646365 | 0 |
| B216 | B222 | BMI | NS | 0.714716 | 0 |
| B216 | B218 | BMI | DBP | 0.226987 | 1.06E-09 |
| B218 | B220 | DBP | SBP | 0.716888 | 0 |
| B218 | B222 | DBP | NS | 0.221915 | 2.52E-09 |
| B22 | B151 | CD3+CD8+(T) | Anti-CCP | -0.3398 | 0 |
| B22 | B100 | CD3+CD8+(T) | anti-Ro52 | -0.28931 | 4.44E-15 |
| B22 | B101 | CD3+CD8+(T) | anti-Sc70 | -0.28931 | 4.44E-15 |
| B22 | B102 | CD3+CD8+(T) | anti-Sm | -0.28931 | 4.44E-15 |
| B22 | B103 | CD3+CD8+(T) | anti-SSA/Ro | -0.28931 | 4.44E-15 |
| B22 | B104 | CD3+CD8+(T) | anti-SSB | -0.28931 | 4.44E-15 |
| B22 | B105 | CD3+CD8+(T) | anti-U1RNP | -0.28931 | 4.44E-15 |
| B22 | B106 | CD3+CD8+(T) | anti-P | -0.28931 | 4.44E-15 |
| B22 | B107 | CD3+CD8+(T) | AnuA | -0.28931 | 4.44E-15 |
| B22 | B108 | CD3+CD8+(T) | Anti-Cyclin | -0.28931 | 4.44E-15 |
| B22 | B109 | CD3+CD8+(T) | AMA-M2 | -0.28931 | 4.44E-15 |
| B22 | B97 | CD3+CD8+(T) | dsDNA | -0.28931 | 4.44E-15 |
| B22 | B98 | CD3+CD8+(T) | anti-Jo-1 | -0.28931 | 4.44E-15 |
| B22 | B66 | CD3+CD8+(T) | WBRSR(mPas_100) | 0.251338 | 1.24E-11 |
| B22 | B67 | CD3+CD8+(T) | WBRSR(mPas_50) | 0.243538 | 5.44E-11 |
| B22 | B63 | CD3+CD8+(T) | WBRSR(mPas_200) | 0.233817 | 3.19E-10 |
| B22 | B186 | CD3+CD8+(T) | PFC | -0.23194 | 4.46E-10 |
| B22 | B62 | CD3+CD8+(T) | WBHSRVI | 0.220324 | 3.29E-09 |
| B22 | B58 | CD3+CD8+(T) | HCT | 0.202811 | 5.45E-08 |
| B22 | B125 | CD3+CD8+(T) | Cor | -0.18414 | 8.38E-07 |
| B22 | B198 | CD3+CD8+(T) | P | -0.18352 | 9.13E-07 |
| B221 | B222 | Hei | NS | 0.467641 | 0 |
| B23 | B151 | CD3-CD16+CD56+(NK) | Anti-CCP | 0.304796 | 0 |

| | | | | | |
|---|---|---|---|---|---|
| B23 | B156 | CD3-CD16+CD56+(NK) | IgE | 0.399065 | 0 |
| B23 | B183 | CD3-CD16+CD56+(NK) | PTA | -0.29239 | 2.22E-15 |
| B23 | B142 | CD3-CD16+CD56+(NK) | 25-(OH)D | 0.265799 | 7.00E-13 |
| B23 | B100 | CD3-CD16+CD56+(NK) | anti-Ro52 | 0.258842 | 2.85E-12 |
| B23 | B101 | CD3-CD16+CD56+(NK) | anti-Sc70 | 0.258842 | 2.85E-12 |
| B23 | B102 | CD3-CD16+CD56+(NK) | anti-Sm | 0.258842 | 2.85E-12 |
| B23 | B103 | CD3-CD16+CD56+(NK) | anti-SSA/Ro | 0.258842 | 2.85E-12 |
| B23 | B104 | CD3-CD16+CD56+(NK) | anti-SSB | 0.258842 | 2.85E-12 |
| B23 | B105 | CD3-CD16+CD56+(NK) | anti-U1RNP | 0.258842 | 2.85E-12 |
| B23 | B106 | CD3-CD16+CD56+(NK) | anti-P | 0.258842 | 2.85E-12 |
| B23 | B107 | CD3-CD16+CD56+(NK) | AnuA | 0.258842 | 2.85E-12 |
| B23 | B108 | CD3-CD16+CD56+(NK) | Anti-Cyclin | 0.258842 | 2.85E-12 |
| B23 | B109 | CD3-CD16+CD56+(NK) | AMA-M2 | 0.258842 | 2.85E-12 |
| B23 | B97 | CD3-CD16+CD56+(NK) | dsDNA | 0.258842 | 2.85E-12 |
| B23 | B98 | CD3-CD16+CD56+(NK) | anti-Jo-1 | 0.258842 | 2.85E-12 |
| B23 | B58 | CD3-CD16+CD56+(NK) | HCT | -0.21916 | 3.99E-09 |
| B23 | B55 | CD3-CD16+CD56+(NK) | TK | 0.218963 | 4.12E-09 |
| B23 | B185 | CD3-CD16+CD56+(NK) | PT | -0.20451 | 4.20E-08 |
| B23 | B95 | CD3-CD16+CD56+(NK) | anti-cenpb | 0.203433 | 4.96E-08 |
| B23 | B136 | CD3-CD16+CD56+(NK) | 5'NT | 0.197143 | 1.29E-07 |
| B23 | B54 | CD3-CD16+CD56+(NK) | ESR | 0.19284 | 2.43E-07 |
| B23 | B200 | CD3-CD16+CD56+(NK) | OP | -0.1912 | 3.08E-07 |
| B23 | B184 | CD3-CD16+CD56+(NK) | INR | -0.18395 | 8.61E-07 |
| B24 | B27 | αHBDH | LDH | 0.66636 | 0 |
| B24 | B26 | αHBDH | CKMB | 0.243916 | 5.07E-11 |
| B24 | B114 | αHBDH | TSH TT3 TT4(TT4) | 0.226051 | 1.24E-09 |
| B24 | B112 | αHBDH | FT4 | 0.200228 | 8.09E-08 |
| B24 | B74 | αHBDH | ApoA1 | 0.194628 | 1.87E-07 |
| B25 | B69 | CK | PV | -0.19198 | 2.75E-07 |
| B25 | B26 | CK | CKMB | 0.183955 | 8.60E-07 |
| B26 | B57 | CKMB | RBCAI | -0.1973 | 1.26E-07 |
| B26 | B27 | CKMB | LDH | 0.194268 | 1.97E-07 |
| B26 | B170 | CKMB | U-GLU | 0.18891 | 4.28E-07 |
| B27 | B128 | LDH | ALT | 0.311646 | 0 |
| B27 | B131 | LDH | AST | 0.315013 | 0 |
| B27 | B150 | LDH | ALP | 0.192532 | 2.54E-07 |
| B27 | B139 | LDH | ADA | 0.192341 | 2.61E-07 |
| B27 | B114 | LDH | TSH TT3 TT4(TT4) | 0.191188 | 3.08E-07 |
| B28 | B135 | PRL | TBA | -0.50472 | 0 |
| B28 | B29 | PRL | E2 | 0.999997 | 0 |
| B28 | B30 | PRL | LH | -0.9126 | 0 |
| B28 | B31 | PRL | FSH | -0.99856 | 0 |
| B28 | B32 | PRL | Testo | 0.998827 | 0 |
| B28 | B33 | PRL | Prog | 0.999941 | 0 |

| | | | | | |
|---|---|---|---|---|---|
| B28 | B161 | PRL | LEU | -0.21378 | 9.65E-09 |
| B28 | B199 | PRL | Mg | 0.20084 | 7.37E-08 |
| B29 | B135 | E2 | TBA | -0.50604 | 0 |
| B29 | B30 | E2 | LH | -0.91156 | 0 |
| B29 | B31 | E2 | FSH | -0.99842 | 0 |
| B29 | B32 | E2 | Testo | 0.9987 | 0 |
| B29 | B33 | E2 | Prog | 0.999966 | 0 |
| B29 | B161 | E2 | LEU | -0.2134 | 1.03E-08 |
| B29 | B199 | E2 | Mg | 0.201375 | 6.79E-08 |
| B3 | B164 | Hcy | COM | 0.374255 | 0 |
| B3 | B176 | Hcy | Ucr | 0.316711 | 0 |
| B3 | B183 | Hcy | PTA | 0.277618 | 5.84E-14 |
| B3 | B61 | Hcy | WBHSRV | 0.239431 | 1.16E-10 |
| B3 | B56 | Hcy | IR | 0.235947 | 2.18E-10 |
| B3 | B18 | Hcy | CD3+CD4+helper(inducer)T | 0.231725 | 4.63E-10 |
| B3 | B62 | Hcy | WBHSRVI | 0.227623 | 9.48E-10 |
| B3 | B19 | Hcy | CD3+CD4+/CD3+CD8+ | 0.222464 | 2.29E-09 |
| B3 | B67 | Hcy | WBRSR(mPas_50) | 0.221882 | 2.53E-09 |
| B3 | B63 | Hcy | WBRSR(mPas_200) | 0.209934 | 1.79E-08 |
| B3 | B66 | Hcy | WBRSR(mPas_100) | 0.192094 | 2.70E-07 |
| B3 | B11 | Hcy | PGI/PGII | 0.184969 | 7.47E-07 |
| B30 | B31 | LH | FSH | 0.933204 | 0 |
| B30 | B32 | LH | Testo | -0.93133 | 0 |
| B30 | B33 | LH | Prog | -0.90812 | 0 |
| B30 | B161 | LH | LEU | 0.256204 | 4.81E-12 |
| B30 | B135 | LH | TBA | 0.248113 | 2.30E-11 |
| B30 | B129 | LH | IBIL | 0.192561 | 2.53E-07 |
| B30 | B133 | LH | TBIL | 0.190892 | 3.22E-07 |
| B30 | B219 | LH | OT | 0.190077 | 3.62E-07 |
| B31 | B135 | FSH | TBA | 0.47614 | 0 |
| B31 | B32 | FSH | Testo | -0.99999 | 0 |
| B31 | B33 | FSH | Prog | -0.99792 | 0 |
| B31 | B161 | FSH | LEU | 0.221487 | 2.70E-09 |
| B31 | B199 | FSH | Mg | -0.1893 | 4.04E-07 |
| B32 | B135 | Testo | TBA | -0.47896 | 0 |
| B32 | B33 | Testo | Prog | 0.998244 | 0 |
| B32 | B161 | Testo | LEU | -0.22077 | 3.05E-09 |
| B32 | B199 | Testo | Mg | 0.19044 | 3.43E-07 |
| B33 | B135 | Prog | TBA | -0.51032 | 0 |
| B33 | B161 | Prog | LEU | -0.21215 | 1.26E-08 |
| B33 | B199 | Prog | Mg | 0.203101 | 5.22E-08 |
| B34 | B125 | TK | Cor | -0.32696 | 0 |
| B34 | B79 | TK | hb V -dna | 0.417102 | 0 |
| B34 | B119 | TK | Hcv-rna | -0.28645 | 8.44E-15 |

| | | | | | |
|---|---|---|---|---|---|
| B34 | B121 | TK | C3 | 0.286445 | 8.44E-15 |
| B34 | B122 | TK | C4 | 0.286445 | 8.44E-15 |
| B34 | B159 | TK | β2-MG | 0.286445 | 8.44E-15 |
| B34 | B169 | TK | U-CRY | 0.286445 | 8.44E-15 |
| B34 | B124 | TK | ACTH | -0.26988 | 3.01E-13 |
| B34 | B194 | TK | Cl | -0.24312 | 5.87E-11 |
| B34 | B74 | TK | ApoA1 | 0.203582 | 4.84E-08 |
| B35 | B167 | β2-MG | PC | 0.496079 | 0 |
| B35 | B37 | β2-MG | BAND | 0.81127 | 0 |
| B35 | B38 | β2-MG | BAND% | -0.39364 | 0 |
| B35 | B96 | β2-MG | dsDNA (IIF) | 0.681874 | 0 |
| B37 | B38 | BAND | BAND% | 0.832193 | 0 |
| B38 | B96 | BAND% | dsDNA (IIF) | 0.897134 | 0 |
| B40 | B42 | WBC | MON | 0.69901 | 0 |
| B40 | B52 | WBC | PLT/BPC | 0.265248 | 7.83E-13 |
| B40 | B51 | WBC | Hb | 0.244609 | 4.45E-11 |
| B40 | B53 | WBC | GR% | 0.231064 | 5.20E-10 |
| B40 | B71 | WBC | TG | 0.224292 | 1.68E-09 |
| B40 | B46 | WBC | RBC Count | 0.205726 | 3.47E-08 |
| B40 | B48 | WBC | LYM% | -0.20468 | 4.09E-08 |
| B40 | B192 | WBC | UA | 0.199636 | 8.84E-08 |
| B40 | B70 | WBC | LDL-C | 0.194203 | 1.99E-07 |
| B40 | B215 | WBC | BMI | 0.193179 | 2.31E-07 |
| B40 | B41 | WBC | MON % | -0.18641 | 6.10E-07 |
| B41 | B42 | MON % | MON | 0.546946 | 0 |
| B41 | B131 | MON % | AST | 0.201647 | 6.52E-08 |
| B41 | B53 | MON % | GR% | -0.1907 | 3.31E-07 |
| B42 | B209 | MON | Smoking | 0.241711 | 7.63E-11 |
| B42 | B51 | MON | Hb | 0.235037 | 2.57E-10 |
| B42 | B192 | MON | UA | 0.187344 | 5.35E-07 |
| B42 | B206 | MON | Waist | 0.183406 | 9.28E-07 |
| B43 | B44 | MCV | MCH | 0.927114 | 0 |
| B43 | B45 | MCV | MCHC | 0.315807 | 0 |
| B43 | B46 | MCV | RBC Count | -0.53031 | 0 |
| B43 | B47 | MCV | RDW | 0.608609 | 0 |
| B43 | B51 | MCV | Hb | 0.231818 | 4.55E-10 |
| B43 | B52 | MCV | PLT/BPC | -0.18504 | 7.39E-07 |
| B44 | B45 | MCH | MCHC | 0.646606 | 0 |
| B44 | B46 | MCH | RBC Count | -0.43476 | 0 |
| B44 | B47 | MCH | RDW | 0.469058 | 0 |
| B44 | B51 | MCH | Hb | 0.390527 | 0 |
| B45 | B51 | MCHC | Hb | 0.522011 | 0 |
| B45 | B222 | MCHC | NS | 0.273446 | 1.42E-13 |
| B45 | B216 | MCHC | BMI | 0.223676 | 1.87E-09 |

| | | | | | |
|---|---|---|---|---|---|
| B45 | B192 | MCHC | UA | 0.215487 | 7.31E-09 |
| B45 | B72 | MCHC | HDL-C | -0.21269 | 1.15E-08 |
| B45 | B174 | MCHC | Urea/Cr | -0.21066 | 1.59E-08 |
| B45 | B221 | MCHC | Hei | 0.198484 | 1.05E-07 |
| B46 | B190 | RBC Count | Cre | 0.3546 | 0 |
| B46 | B192 | RBC Count | UA | 0.38243 | 0 |
| B46 | B216 | RBC Count | BMI | 0.350912 | 0 |
| B46 | B221 | RBC Count | Hei | 0.335202 | 0 |
| B46 | B222 | RBC Count | NS | 0.383816 | 0 |
| B46 | B47 | RBC Count | RDW | -0.45909 | 0 |
| B46 | B51 | RBC Count | Hb | 0.6529 | 0 |
| B46 | B126 | RBC Count | Alb | 0.286848 | 7.77E-15 |
| B46 | B206 | RBC Count | Waist | 0.26108 | 1.82E-12 |
| B46 | B72 | RBC Count | HDL-C | -0.24812 | 2.29E-11 |
| B46 | B163 | RBC Count | SG | 0.245786 | 3.57E-11 |
| B46 | B58 | RBC Count | HCT | 0.240308 | 9.87E-11 |
| B46 | B63 | RBC Count | WBRSR(mPas_200) | 0.237608 | 1.62E-10 |
| B46 | B64 | RBC Count | WBRSR(mPas_5) | 0.236394 | 2.01E-10 |
| B46 | B67 | RBC Count | WBRSR(mPas_50) | 0.228795 | 7.74E-10 |
| B46 | B66 | RBC Count | WBRSR(mPas_100) | 0.227854 | 9.11E-10 |
| B46 | B210 | RBC Count | Hip | 0.214512 | 8.57E-09 |
| B46 | B207 | RBC Count | WHR | 0.214385 | 8.75E-09 |
| B46 | B65 | RBC Count | WBRSR(mPas_1) | 0.212328 | 1.22E-08 |
| B46 | B215 | RBC Count | BMI | 0.210218 | 1.71E-08 |
| B46 | B141 | RBC Count | AST/ALT | -0.20882 | 2.14E-08 |
| B46 | B62 | RBC Count | WBHSRVI | 0.2044 | 4.27E-08 |
| B46 | B174 | RBC Count | Urea/Cr | -0.20224 | 5.95E-08 |
| B46 | B127 | RBC Count | A-G ratio | 0.196831 | 1.35E-07 |
| B46 | B134 | RBC Count | CHE | 0.190195 | 3.56E-07 |
| B47 | B52 | RDW | PLT/BPC | -0.22001 | 3.46E-09 |
| B48 | B53 | LYM% | GR% | -0.92929 | 0 |
| B49 | B50 | baso% | eo% | 0.409369 | 0 |
| B49 | B53 | baso% | GR% | -0.29755 | 6.66E-16 |
| B50 | B53 | eo% | GR% | -0.35405 | 0 |
| B51 | B190 | Hb | Cre | 0.460887 | 0 |
| B51 | B192 | Hb | UA | 0.439307 | 0 |
| B51 | B206 | Hb | Waist | 0.348537 | 0 |
| B51 | B207 | Hb | WHR | 0.327097 | 0 |
| B51 | B216 | Hb | BMI | 0.419642 | 0 |
| B51 | B221 | Hb | Hei | 0.409318 | 0 |
| B51 | B222 | Hb | NS | 0.498524 | 0 |
| B51 | B163 | Hb | SG | 0.294522 | 1.33E-15 |
| B51 | B174 | Hb | Urea/Cr | -0.29357 | 1.78E-15 |
| B51 | B58 | Hb | HCT | 0.292356 | 2.22E-15 |

| | | | | | |
|---|---|---|---|---|---|
| B51 | B126 | Hb | Alb | 0.274401 | 1.16E-13 |
| B51 | B72 | Hb | HDL-C | -0.27156 | 2.12E-13 |
| B51 | B215 | Hb | BMI | 0.256646 | 4.41E-12 |
| B51 | B63 | Hb | WBRSR(mPas_200) | 0.252407 | 1.01E-11 |
| B51 | B66 | Hb | WBRSR(mPas_100) | 0.2502 | 1.54E-11 |
| B51 | B67 | Hb | WBRSR(mPas_50) | 0.246365 | 3.20E-11 |
| B51 | B210 | Hb | Hip | 0.244451 | 4.58E-11 |
| B51 | B64 | Hb | WBRSR(mPas_5) | 0.243442 | 5.53E-11 |
| B51 | B141 | Hb | AST/ALT | -0.23673 | 1.89E-10 |
| B51 | B128 | Hb | ALT | 0.227799 | 9.20E-10 |
| B51 | B62 | Hb | WBHSRVI | 0.22763 | 9.47E-10 |
| B51 | B218 | Hb | DBP | 0.224002 | 1.77E-09 |
| B51 | B127 | Hb | A-G ratio | 0.222956 | 2.11E-09 |
| B51 | B65 | Hb | WBRSR(mPas_1) | 0.212835 | 1.13E-08 |
| B51 | B70 | Hb | LDL-C | 0.210675 | 1.59E-08 |
| B51 | B54 | Hb | ESR | -0.208 | 2.43E-08 |
| B51 | B209 | Hb | Smoking | 0.189636 | 3.86E-07 |
| B51 | B134 | Hb | CHE | 0.184954 | 7.48E-07 |
| B52 | B132 | PLT/BPC | Dbil | -0.23486 | 2.65E-10 |
| B52 | B133 | PLT/BPC | TBIL | -0.20582 | 3.42E-08 |
| B52 | B129 | PLT/BPC | IBIL | -0.1832 | 9.55E-07 |
| B54 | B151 | ESR | Anti-CCP | 0.451566 | 0 |
| B54 | B55 | ESR | TK | 0.389803 | 0 |
| B54 | B58 | ESR | HCT | -0.69687 | 0 |
| B54 | B60 | ESR | WBRVI | -0.40639 | 0 |
| B54 | B62 | ESR | WBHSRVI | -0.52136 | 0 |
| B54 | B63 | ESR | WBRSR(mPas_200) | -0.38371 | 0 |
| B54 | B64 | ESR | WBRSR(mPas_5) | -0.37688 | 0 |
| B54 | B65 | ESR | WBRSR(mPas_1) | -0.32503 | 0 |
| B54 | B66 | ESR | WBRSR(mPas_100) | -0.38625 | 0 |
| B54 | B67 | ESR | WBRSR(mPas_50) | -0.38559 | 0 |
| B54 | B69 | ESR | PV | 0.329632 | 0 |
| B54 | B59 | ESR | WBRV | 0.278445 | 4.88E-14 |
| B54 | B200 | ESR | OP | -0.22694 | 1.07E-09 |
| B54 | B56 | ESR | IR | -0.19682 | 1.35E-07 |
| B54 | B100 | ESR | anti-Ro52 | 0.183218 | 9.53E-07 |
| B54 | B101 | ESR | anti-Sc70 | 0.183218 | 9.53E-07 |
| B54 | B102 | ESR | anti-Sm | 0.183218 | 9.53E-07 |
| B54 | B103 | ESR | anti-SSA/Ro | 0.183218 | 9.53E-07 |
| B54 | B104 | ESR | anti-SSB | 0.183218 | 9.53E-07 |
| B54 | B105 | ESR | anti-U1RNP | 0.183218 | 9.53E-07 |
| B54 | B106 | ESR | anti-P | 0.183218 | 9.53E-07 |
| B54 | B107 | ESR | AnuA | 0.183218 | 9.53E-07 |
| B54 | B108 | ESR | Anti-Cyclin | 0.183218 | 9.53E-07 |

| | | | | | |
|---|---|---|---|---|---|
| B54 | B109 | ESR | AMA-M2 | 0.183218 | 9.53E-07 |
| B54 | B97 | ESR | dsDNA | 0.183218 | 9.53E-07 |
| B54 | B98 | ESR | anti-Jo-1 | 0.183218 | 9.53E-07 |
| B55 | B56 | TK | IR | 0.6373 | 0 |
| B55 | B58 | TK | HCT | -0.48019 | 0 |
| B55 | B59 | TK | WBRV | 0.32791 | 0 |
| B55 | B61 | TK | WBHSRV | 0.518848 | 0 |
| B55 | B69 | TK | PV | -0.38014 | 0 |
| B55 | B168 | TK | NMP22 | 0.219716 | 3.64E-09 |
| B55 | B139 | TK | ADA | 0.215361 | 7.47E-09 |
| B55 | B110 | TK | AHA | -0.20797 | 2.44E-08 |
| B55 | B151 | TK | Anti-CCP | 0.202095 | 6.09E-08 |
| B55 | B181 | TK | APTT | 0.193315 | 2.26E-07 |
| B55 | B164 | TK | COM | 0.187395 | 5.31E-07 |
| B55 | B183 | TK | PTA | -0.18675 | 5.82E-07 |
| B56 | B58 | IR | HCT | 0.347183 | 0 |
| B56 | B59 | IR | WBRV | 0.329968 | 0 |
| B56 | B60 | IR | WBRVI | 0.611407 | 0 |
| B56 | B61 | IR | WBHSRV | 0.874673 | 0 |
| B56 | B62 | IR | WBHSRVI | 0.85031 | 0 |
| B56 | B63 | IR | WBRSR(mPas_200) | 0.641808 | 0 |
| B56 | B64 | IR | WBRSR(mPas_5) | 0.586971 | 0 |
| B56 | B65 | IR | WBRSR(mPas_1) | 0.489306 | 0 |
| B56 | B66 | IR | WBRSR(mPas_100) | 0.579585 | 0 |
| B56 | B67 | IR | WBRSR(mPas_50) | 0.590816 | 0 |
| B56 | B69 | IR | PV | -0.47863 | 0 |
| B56 | B164 | IR | COM | 0.293538 | 1.78E-15 |
| B56 | B186 | IR | PFC | -0.25019 | 1.55E-11 |
| B56 | B110 | IR | AHA | -0.24584 | 3.53E-11 |
| B56 | B81 | IR | TSGF | 0.233022 | 3.68E-10 |
| B56 | B176 | IR | Ucr | 0.229514 | 6.83E-10 |
| B56 | B137 | IR | AFU | 0.220726 | 3.07E-09 |
| B56 | B182 | IR | TT | 0.192175 | 2.67E-07 |
| B57 | B110 | RBCAI | AHA | 0.336445 | 0 |
| B57 | B59 | RBCAI | WBRV | 0.743727 | 0 |
| B57 | B60 | RBCAI | WBRVI | 0.602261 | 0 |
| B57 | B65 | RBCAI | WBRSR(mPas_1) | 0.531799 | 0 |
| B57 | B64 | RBCAI | WBRSR(mPas_5) | 0.286284 | 8.88E-15 |
| B57 | B181 | RBCAI | APTT | -0.25904 | 2.74E-12 |
| B57 | B151 | RBCAI | Anti-CCP | 0.258743 | 2.91E-12 |
| B57 | B69 | RBCAI | PV | -0.24732 | 2.67E-11 |
| B57 | B61 | RBCAI | WBHSRV | -0.23832 | 1.42E-10 |
| B57 | B81 | RBCAI | TSGF | -0.21579 | 6.96E-09 |
| B57 | B180 | RBCAI | Tf | 0.213101 | 1.08E-08 |

| | | | | | |
|---|---|---|---|---|---|
| B58 | B151 | HCT | Anti-CCP | -0.35312 | 0 |
| B58 | B60 | HCT | WBRVI | 0.655335 | 0 |
| B58 | B61 | HCT | WBHSRV | 0.366996 | 0 |
| B58 | B62 | HCT | WBHSRVI | 0.778724 | 0 |
| B58 | B63 | HCT | WBRSR(mPas_200) | 0.796373 | 0 |
| B58 | B64 | HCT | WBRSR(mPas_5) | 0.804804 | 0 |
| B58 | B65 | HCT | WBRSR(mPas_1) | 0.72296 | 0 |
| B58 | B66 | HCT | WBRSR(mPas_100) | 0.804737 | 0 |
| B58 | B67 | HCT | WBRSR(mPas_50) | 0.806228 | 0 |
| B58 | B134 | HCT | CHE | 0.24527 | 3.93E-11 |
| B58 | B176 | HCT | Ucr | 0.225432 | 1.38E-09 |
| B58 | B200 | HCT | OP | 0.191816 | 2.82E-07 |
| B58 | B139 | HCT | ADA | -0.18806 | 4.83E-07 |
| B59 | B151 | WBRV | Anti-CCP | 0.363785 | 0 |
| B59 | B60 | WBRV | WBRVI | 0.617412 | 0 |
| B59 | B61 | WBRV | WBHSRV | 0.364442 | 0 |
| B59 | B64 | WBRV | WBRSR(mPas_5) | 0.516644 | 0 |
| B59 | B65 | WBRV | WBRSR(mPas_1) | 0.662313 | 0 |
| B59 | B67 | WBRV | WBRSR(mPas_50) | 0.336481 | 0 |
| B59 | B66 | WBRV | WBRSR(mPas_100) | 0.290333 | 3.55E-15 |
| B59 | B136 | WBRV | 5'NT | 0.256312 | 4.71E-12 |
| B59 | B175 | WBRV | ALB-U | 0.238723 | 1.32E-10 |
| B59 | B63 | WBRV | WBRSR(mPas_200) | 0.210769 | 1.57E-08 |
| B59 | B62 | WBRV | WBHSRVI | 0.200033 | 8.33E-08 |
| B59 | B177 | WBRV | ACR | 0.19414 | 2.01E-07 |
| B59 | B138 | WBRV | PAB | 0.190022 | 3.65E-07 |
| B59 | B181 | WBRV | APTT | -0.18995 | 3.68E-07 |
| B6 | B7 | WB-Fe | WB-Cop | 1 | 0 |
| B6 | B8 | WB-Fe | WB-Zinc | 1 | 0 |
| B60 | B61 | WBRVI | WBHSRV | 0.445784 | 0 |
| B60 | B62 | WBRVI | WBHSRVI | 0.775662 | 0 |
| B60 | B63 | WBRVI | WBRSR(mPas_200) | 0.571945 | 0 |
| B60 | B64 | WBRVI | WBRSR(mPas_5) | 0.837849 | 0 |
| B60 | B65 | WBRVI | WBRSR(mPas_1) | 0.908543 | 0 |
| B60 | B66 | WBRVI | WBRSR(mPas_100) | 0.631647 | 0 |
| B60 | B67 | WBRVI | WBRSR(mPas_50) | 0.673844 | 0 |
| B60 | B69 | WBRVI | PV | -0.45334 | 0 |
| B60 | B180 | WBRVI | Tf | 0.198812 | 1.00E-07 |
| B60 | B134 | WBRVI | CHE | 0.189163 | 4.13E-07 |
| B61 | B62 | WBHSRV | WBHSRVI | 0.764327 | 0 |
| B61 | B63 | WBHSRV | WBRSR(mPas_200) | 0.821475 | 0 |
| B61 | B64 | WBHSRV | WBRSR(mPas_5) | 0.674589 | 0 |
| B61 | B65 | WBHSRV | WBRSR(mPas_1) | 0.522956 | 0 |
| B61 | B66 | WBHSRV | WBRSR(mPas_100) | 0.745583 | 0 |

| | | | | | |
|---|---|---|---|---|---|
| B61 | B67 | WBHSRV | WBRSR(mPas_50) | 0.740074 | 0 |
| B61 | B176 | WBHSRV | Ucr | 0.281624 | 2.44E-14 |
| B61 | B110 | WBHSRV | AHA | -0.27961 | 3.82E-14 |
| B61 | B164 | WBHSRV | COM | 0.246398 | 3.18E-11 |
| B61 | B137 | WBHSRV | AFU | 0.238984 | 1.26E-10 |
| B61 | B81 | WBHSRV | TSGF | 0.218214 | 4.67E-09 |
| B61 | B182 | WBHSRV | TT | 0.212621 | 1.16E-08 |
| B61 | B186 | WBHSRV | PFC | -0.21214 | 1.26E-08 |
| B61 | B198 | WBHSRV | P | -0.19709 | 1.30E-07 |
| B62 | B151 | WBHSRVI | Anti-CCP | -0.30426 | 0 |
| B62 | B63 | WBHSRVI | WBRSR(mPas_200) | 0.853822 | 0 |
| B62 | B64 | WBHSRVI | WBRSR(mPas_5) | 0.828295 | 0 |
| B62 | B65 | WBHSRVI | WBRSR(mPas_1) | 0.720102 | 0 |
| B62 | B66 | WBHSRVI | WBRSR(mPas_100) | 0.828006 | 0 |
| B62 | B67 | WBHSRVI | WBRSR(mPas_50) | 0.835562 | 0 |
| B62 | B69 | WBHSRVI | PV | -0.36865 | 0 |
| B62 | B176 | WBHSRVI | Ucr | 0.285996 | 9.33E-15 |
| B62 | B186 | WBHSRVI | PFC | -0.26889 | 3.70E-13 |
| B62 | B164 | WBHSRVI | COM | 0.229519 | 6.82E-10 |
| B62 | B137 | WBHSRVI | AFU | 0.2278 | 9.20E-10 |
| B62 | B182 | WBHSRVI | TT | 0.211098 | 1.49E-08 |
| B62 | B94 | WBHSRVI | ALDH2-Lys504Lys | 0.20455 | 4.17E-08 |
| B62 | B110 | WBHSRVI | AHA | -0.18744 | 5.27E-07 |
| B63 | B176 | WBRSR(mPas_200) | Ucr | 0.320728 | 0 |
| B63 | B64 | WBRSR(mPas_200) | WBRSR(mPas_5) | 0.887487 | 0 |
| B63 | B65 | WBRSR(mPas_200) | WBRSR(mPas_1) | 0.733033 | 0 |
| B63 | B66 | WBRSR(mPas_200) | WBRSR(mPas_100) | 0.959666 | 0 |
| B63 | B67 | WBRSR(mPas_200) | WBRSR(mPas_50) | 0.949982 | 0 |
| B63 | B151 | WBRSR(mPas_200) | Anti-CCP | -0.22321 | 2.02E-09 |
| B63 | B182 | WBRSR(mPas_200) | TT | 0.216644 | 6.05E-09 |
| B63 | B137 | WBRSR(mPas_200) | AFU | 0.214513 | 8.57E-09 |
| B63 | B110 | WBRSR(mPas_200) | AHA | -0.21281 | 1.13E-08 |
| B63 | B94 | WBRSR(mPas_200) | ALDH2-Lys504Lys | 0.197729 | 1.18E-07 |
| B63 | B186 | WBRSR(mPas_200) | PFC | -0.19545 | 1.65E-07 |
| B63 | B134 | WBRSR(mPas_200) | CHE | 0.192873 | 2.41E-07 |
| B63 | B200 | WBRSR(mPas_200) | OP | 0.187471 | 5.25E-07 |
| B64 | B65 | WBRSR(mPas_5) | WBRSR(mPas_1) | 0.962389 | 0 |
| B64 | B66 | WBRSR(mPas_5) | WBRSR(mPas_100) | 0.916075 | 0 |
| B64 | B67 | WBRSR(mPas_5) | WBRSR(mPas_50) | 0.932378 | 0 |
| B64 | B176 | WBRSR(mPas_5) | Ucr | 0.281771 | 2.38E-14 |
| B64 | B134 | WBRSR(mPas_5) | CHE | 0.231013 | 5.25E-10 |
| B64 | B182 | WBRSR(mPas_5) | TT | 0.208247 | 2.34E-08 |
| B64 | B200 | WBRSR(mPas_5) | OP | 0.20371 | 4.75E-08 |
| B64 | B178 | WBRSR(mPas_5) | Ig-U | 0.196285 | 1.46E-07 |

| | | | | | |
|---|---|---|---|---|---|
| B64 | B94 | WBRSR(mPas_5) | ALDH2-Lys504Lys | 0.191616 | 2.90E-07 |
| B64 | B180 | WBRSR(mPas_5) | Tf | 0.189584 | 3.88E-07 |
| B64 | B138 | WBRSR(mPas_5) | PAB | 0.189573 | 3.89E-07 |
| B65 | B66 | WBRSR(mPas_1) | WBRSR(mPas_100) | 0.795956 | 0 |
| B65 | B67 | WBRSR(mPas_1) | WBRSR(mPas_50) | 0.827605 | 0 |
| B65 | B180 | WBRSR(mPas_1) | Tf | 0.240077 | 1.03E-10 |
| B65 | B134 | WBRSR(mPas_1) | CHE | 0.232517 | 4.02E-10 |
| B65 | B176 | WBRSR(mPas_1) | Ucr | 0.226023 | 1.25E-09 |
| B65 | B178 | WBRSR(mPas_1) | Ig-U | 0.225932 | 1.27E-09 |
| B65 | B138 | WBRSR(mPas_1) | PAB | 0.21499 | 7.93E-09 |
| B65 | B181 | WBRSR(mPas_1) | APTT | -0.19614 | 1.49E-07 |
| B65 | B200 | WBRSR(mPas_1) | OP | 0.192347 | 2.61E-07 |
| B66 | B67 | WBRSR(mPas_100) | WBRSR(mPas_50) | 0.991102 | 0 |
| B66 | B176 | WBRSR(mPas_100) | Ucr | 0.295909 | 8.88E-16 |
| B66 | B137 | WBRSR(mPas_100) | AFU | 0.239294 | 1.19E-10 |
| B66 | B182 | WBRSR(mPas_100) | TT | 0.226799 | 1.09E-09 |
| B66 | B186 | WBRSR(mPas_100) | PFC | -0.2241 | 1.74E-09 |
| B66 | B100 | WBRSR(mPas_100) | anti-Ro52 | -0.22355 | 1.91E-09 |
| B66 | B101 | WBRSR(mPas_100) | anti-Sc70 | -0.22355 | 1.91E-09 |
| B66 | B102 | WBRSR(mPas_100) | anti-Sm | -0.22355 | 1.91E-09 |
| B66 | B103 | WBRSR(mPas_100) | anti-SSA/Ro | -0.22355 | 1.91E-09 |
| B66 | B104 | WBRSR(mPas_100) | anti-SSB | -0.22355 | 1.91E-09 |
| B66 | B105 | WBRSR(mPas_100) | anti-U1RNP | -0.22355 | 1.91E-09 |
| B66 | B106 | WBRSR(mPas_100) | anti-P | -0.22355 | 1.91E-09 |
| B66 | B107 | WBRSR(mPas_100) | AnuA | -0.22355 | 1.91E-09 |
| B66 | B108 | WBRSR(mPas_100) | Anti-Cyclin | -0.22355 | 1.91E-09 |
| B66 | B109 | WBRSR(mPas_100) | AMA-M2 | -0.22355 | 1.91E-09 |
| B66 | B97 | WBRSR(mPas_100) | dsDNA | -0.22355 | 1.91E-09 |
| B66 | B98 | WBRSR(mPas_100) | anti-Jo-1 | -0.22355 | 1.91E-09 |
| B66 | B94 | WBRSR(mPas_100) | ALDH2-Lys504Lys | 0.215647 | 7.12E-09 |
| B66 | B134 | WBRSR(mPas_100) | CHE | 0.209172 | 2.02E-08 |
| B66 | B151 | WBRSR(mPas_100) | Anti-CCP | -0.20587 | 3.40E-08 |
| B66 | B200 | WBRSR(mPas_100) | OP | 0.198007 | 1.13E-07 |
| B67 | B176 | WBRSR(mPas_50) | Ucr | 0.295884 | 8.88E-16 |
| B67 | B100 | WBRSR(mPas_50) | anti-Ro52 | -0.22695 | 1.07E-09 |
| B67 | B101 | WBRSR(mPas_50) | anti-Sc70 | -0.22695 | 1.07E-09 |
| B67 | B102 | WBRSR(mPas_50) | anti-Sm | -0.22695 | 1.07E-09 |
| B67 | B103 | WBRSR(mPas_50) | anti-SSA/Ro | -0.22695 | 1.07E-09 |
| B67 | B104 | WBRSR(mPas_50) | anti-SSB | -0.22695 | 1.07E-09 |
| B67 | B105 | WBRSR(mPas_50) | anti-U1RNP | -0.22695 | 1.07E-09 |
| B67 | B106 | WBRSR(mPas_50) | anti-P | -0.22695 | 1.07E-09 |
| B67 | B107 | WBRSR(mPas_50) | AnuA | -0.22695 | 1.07E-09 |
| B67 | B108 | WBRSR(mPas_50) | Anti-Cyclin | -0.22695 | 1.07E-09 |
| B67 | B109 | WBRSR(mPas_50) | AMA-M2 | -0.22695 | 1.07E-09 |

| | | | | | |
|---|---|---|---|---|---|
| B67 | B97 | WBRSR(mPas_50) | dsDNA | -0.22695 | 1.07E-09 |
| B67 | B98 | WBRSR(mPas_50) | anti-Jo-1 | -0.22695 | 1.07E-09 |
| B67 | B137 | WBRSR(mPas_50) | AFU | 0.212646 | 1.16E-08 |
| B67 | B134 | WBRSR(mPas_50) | CHE | 0.211891 | 1.31E-08 |
| B67 | B182 | WBRSR(mPas_50) | TT | 0.205812 | 3.43E-08 |
| B67 | B200 | WBRSR(mPas_50) | OP | 0.200923 | 7.28E-08 |
| B67 | B186 | WBRSR(mPas_50) | PFC | -0.19515 | 1.73E-07 |
| B67 | B151 | WBRSR(mPas_50) | Anti-CCP | -0.19256 | 2.53E-07 |
| B69 | B78 | PV | HBcAb | 0.228343 | 8.37E-10 |
| B69 | B183 | PV | PTA | 0.213362 | 1.03E-08 |
| B69 | B91 | PV | TP | 0.194218 | 1.98E-07 |
| B7 | B8 | WB-Cop | WB-Zinc | 1 | 0 |
| B70 | B76 | LDL-C | ApoB | 0.351348 | 0 |
| B70 | B77 | LDL-C | TC | 0.862002 | 0 |
| B70 | B75 | LDL-C | ApoA1/B | -0.25369 | 7.86E-12 |
| B70 | B132 | LDL-C | Dbil | -0.23218 | 4.27E-10 |
| B70 | B215 | LDL-C | BMI | 0.226326 | 1.19E-09 |
| B70 | B206 | LDL-C | Waist | 0.201715 | 6.45E-08 |
| B70 | B207 | LDL-C | WHR | 0.184245 | 8.26E-07 |
| B71 | B140 | TG | r-GT | 0.364428 | 0 |
| B71 | B72 | TG | HDL-C | -0.41952 | 0 |
| B71 | B215 | TG | BMI | 0.300739 | 4.44E-16 |
| B71 | B206 | TG | Waist | 0.294922 | 1.33E-15 |
| B71 | B192 | TG | UA | 0.290864 | 3.11E-15 |
| B71 | B207 | TG | WHR | 0.284782 | 1.22E-14 |
| B71 | B216 | TG | BMI | 0.272506 | 1.74E-13 |
| B71 | B77 | TG | TC | 0.259763 | 2.37E-12 |
| B71 | B91 | TG | TP | 0.23905 | 1.24E-10 |
| B71 | B130 | TG | GLB | 0.236487 | 1.98E-10 |
| B71 | B222 | TG | NS | 0.209221 | 2.01E-08 |
| B71 | B210 | TG | Hip | 0.192885 | 2.41E-07 |
| B72 | B192 | HDL-C | UA | -0.32586 | 0 |
| B72 | B206 | HDL-C | Waist | -0.36433 | 0 |
| B72 | B207 | HDL-C | WHR | -0.35378 | 0 |
| B72 | B215 | HDL-C | BMI | -0.3484 | 0 |
| B72 | B216 | HDL-C | BMI | -0.39605 | 0 |
| B72 | B222 | HDL-C | NS | -0.34476 | 0 |
| B72 | B74 | HDL-C | ApoA1 | 0.372345 | 0 |
| B72 | B77 | HDL-C | TC | 0.291698 | 2.66E-15 |
| B72 | B75 | HDL-C | ApoA1/B | 0.28957 | 4.22E-15 |
| B72 | B190 | HDL-C | Cre | -0.25204 | 1.08E-11 |
| B72 | B141 | HDL-C | AST/ALT | 0.242181 | 6.99E-11 |
| B72 | B163 | HDL-C | SG | -0.24009 | 1.03E-10 |
| B72 | B210 | HDL-C | Hip | -0.23666 | 1.92E-10 |

| | | | | | |
|---|---|---|---|---|---|
| B72 | B221 | HDL-C | Hei | -0.23498 | 2.60E-10 |
| B72 | B174 | HDL-C | Urea/Cr | 0.18833 | 4.65E-07 |
| B73 | B76 | ApoA | ApoB | 0.239645 | 1.11E-10 |
| B74 | B75 | ApoA1 | ApoA1/B | 0.557938 | 0 |
| B74 | B194 | ApoA1 | Cl | -0.26807 | 4.39E-13 |
| B74 | B176 | ApoA1 | Ucr | -0.21066 | 1.60E-08 |
| B74 | B77 | ApoA1 | TC | 0.184683 | 7.77E-07 |
| B75 | B76 | ApoA1/B | ApoB | -0.68972 | 0 |
| B75 | B135 | ApoA1/B | TBA | 0.238412 | 1.40E-10 |
| B76 | B138 | ApoB | PAB | 0.307706 | 0 |
| B76 | B77 | ApoB | TC | 0.297591 | 6.66E-16 |
| B76 | B134 | ApoB | CHE | 0.288051 | 5.77E-15 |
| B76 | B194 | ApoB | Cl | -0.26099 | 1.86E-12 |
| B76 | B168 | ApoB | NMP22 | 0.256667 | 4.39E-12 |
| B76 | B92 | ApoB | ALDH2-Glu504Glu | -0.21086 | 1.55E-08 |
| B76 | B139 | ApoB | ADA | -0.20686 | 2.91E-08 |
| B76 | B175 | ApoB | ALB-U | 0.186384 | 6.12E-07 |
| B76 | B135 | ApoB | TBA | -0.18316 | 9.61E-07 |
| B77 | B132 | TC | Dbil | -0.21981 | 3.58E-09 |
| B78 | B178 | HBcAb | Ig-U | 0.21737 | 5.37E-09 |
| B78 | B180 | HBcAb | Tf | 0.183008 | 9.81E-07 |
| B81 | B125 | TSGF | Cor | -0.32104 | 0 |
| B81 | B93 | TSGF | ALDH2-Glu504Lys | -0.33812 | 0 |
| B81 | B139 | TSGF | ADA | 0.285753 | 9.77E-15 |
| B81 | B115 | TSGF | TSH TT3 TT4(TT3) | 0.269358 | 3.36E-13 |
| B81 | B137 | TSGF | AFU | 0.260214 | 2.17E-12 |
| B81 | B180 | TSGF | Tf | -0.25976 | 2.37E-12 |
| B81 | B181 | TSGF | APTT | 0.256995 | 4.11E-12 |
| B81 | B94 | TSGF | ALDH2-Lys504Lys | 0.241891 | 7.38E-11 |
| B81 | B123 | TSGF | hs-CRP1 | 0.240785 | 9.05E-11 |
| B81 | B178 | TSGF | Ig-U | -0.22471 | 1.57E-09 |
| B81 | B182 | TSGF | TT | 0.213823 | 9.59E-09 |
| B81 | B151 | TSGF | Anti-CCP | -0.21188 | 1.31E-08 |
| B81 | B198 | TSGF | P | -0.20546 | 3.62E-08 |
| B81 | B183 | TSGF | PTA | -0.19666 | 1.38E-07 |
| B81 | B154 | TSGF | RF | -0.1891 | 4.16E-07 |
| B82 | B85 | CEA | SCC | 0.471261 | 0 |
| B82 | B83 | CEA | AFP | 0.286187 | 8.88E-15 |
| B82 | B84 | CEA | C-PSA | 0.285195 | 1.11E-14 |
| B82 | B128 | CEA | ALT | 0.186649 | 5.90E-07 |
| B83 | B85 | AFP | SCC | 0.384305 | 0 |
| B83 | B119 | AFP | Hcv-rna | -0.27919 | 4.17E-14 |
| B83 | B121 | AFP | C3 | 0.279188 | 4.17E-14 |
| B83 | B122 | AFP | C4 | 0.279188 | 4.17E-14 |

| | | | | | |
|---|---|---|---|---|---|
| B83 | B159 | AFP | β2-MG | 0.279189 | 4.17E-14 |
| B83 | B169 | AFP | U-CRY | 0.279188 | 4.17E-14 |
| B83 | B153 | AFP | Glu | 0.267048 | 5.41E-13 |
| B83 | B170 | AFP | U-GLU | 0.202731 | 5.52E-08 |
| B84 | B87 | C-PSA | NSE | -0.19879 | 1.00E-07 |
| B86 | B90 | TBhCG | CYFRA 21-1 | 0.444342 | 0 |
| B90 | B139 | CYFRA 21-1 | ADA | 0.271296 | 2.24E-13 |
| B90 | B126 | CYFRA 21-1 | Alb | -0.21118 | 1.47E-08 |
| B90 | B138 | CYFRA 21-1 | PAB | -0.21001 | 1.77E-08 |
| B90 | B135 | CYFRA 21-1 | TBA | 0.201153 | 7.03E-08 |
| B90 | B186 | CYFRA 21-1 | PFC | -0.18451 | 7.96E-07 |
| B91 | B126 | TP | Alb | 0.463123 | 0 |
| B91 | B127 | TP | A-G ratio | -0.5077 | 0 |
| B91 | B130 | TP | GLB | 0.778264 | 0 |
| B91 | B188 | TP | Ca | 0.326421 | 0 |
| B92 | B93 | ALDH2-Glu504Glu | ALDH2-Glu504Lys | -0.70869 | 0 |
| B92 | B94 | ALDH2-Glu504Glu | ALDH2-Lys504Lys | -0.5045 | 0 |
| B93 | B94 | ALDH2-Glu504Lys | ALDH2-Lys504Lys | -0.25162 | 1.17E-11 |
| B93 | B139 | ALDH2-Glu504Lys | ADA | -0.21972 | 3.64E-09 |
| B93 | B137 | ALDH2-Glu504Lys | AFU | -0.2035 | 4.90E-08 |
| B93 | B115 | ALDH2-Glu504Lys | TSH TT3 TT4(TT3) | -0.18431 | 8.18E-07 |
| B94 | B184 | ALDH2-Lys504Lys | INR | -0.23797 | 1.51E-10 |
| B94 | B186 | ALDH2-Lys504Lys | PFC | -0.23414 | 3.02E-10 |
| B94 | B182 | ALDH2-Lys504Lys | TT | 0.215467 | 7.34E-09 |
| B94 | B115 | ALDH2-Lys504Lys | TSH TT3 TT4(TT3) | 0.215325 | 7.51E-09 |
| B94 | B197 | ALDH2-Lys504Lys | AG | 0.205393 | 3.66E-08 |
| B95 | B205 | anti-cenpb | KDA | 0.238326 | 1.42E-10 |
| B95 | B139 | anti-cenpb | ADA | 0.223034 | 2.08E-09 |
| B95 | B181 | anti-cenpb | APTT | -0.22164 | 2.64E-09 |
| B97 | B100 | dsDNA | anti-Ro52 | 1 | 0 |
| B97 | B102 | dsDNA | anti-Sm | 1 | 0 |
| B97 | B103 | dsDNA | anti-SSA/Ro | 1 | 0 |
| B97 | B104 | dsDNA | anti-SSB | 1 | 0 |
| B97 | B105 | dsDNA | anti-U1RNP | 1 | 0 |
| B97 | B106 | dsDNA | anti-P | 1 | 0 |
| B97 | B107 | dsDNA | AnuA | 1 | 0 |
| B97 | B108 | dsDNA | Anti-Cyclin | 1 | 0 |
| B97 | B109 | dsDNA | AMA-M2 | 1 | 0 |
| B97 | B151 | dsDNA | Anti-CCP | 0.472831 | 0 |
| B97 | B186 | dsDNA | PFC | 0.313552 | 0 |
| B97 | B142 | dsDNA | 25-(OH)D | 0.249542 | 1.75E-11 |
| B97 | B184 | dsDNA | INR | -0.22906 | 7.39E-10 |
| B97 | B99 | dsDNA | anti-PM-Scl | -0.20949 | 1.92E-08 |
| B97 | B185 | dsDNA | PT | -0.19886 | 9.94E-08 |

| | | | | | |
|---|---|---|---|---|---|
| B97 | B175 | dsDNA | ALB-U | 0.197112 | 1.29E-07 |
| B97 | B196 | dsDNA | TCO2 | 0.193449 | 2.22E-07 |
| B97 | B137 | dsDNA | AFU | -0.19228 | 2.63E-07 |
| B98 | B101 | anti-Jo-1 | anti-Sc70 | 1 | 0 |
| B98 | B102 | anti-Jo-1 | anti-Sm | 1 | 0 |
| B98 | B107 | anti-Jo-1 | AnuA | 1 | 0 |
| B98 | B108 | anti-Jo-1 | Anti-Cyclin | 1 | 0 |
| B98 | B151 | anti-Jo-1 | Anti-CCP | 0.472831 | 0 |
| B98 | B186 | anti-Jo-1 | PFC | 0.313552 | 0 |
| B98 | B142 | anti-Jo-1 | 25-(OH)D | 0.249542 | 1.75E-11 |
| B98 | B184 | anti-Jo-1 | INR | -0.22906 | 7.39E-10 |
| B98 | B99 | anti-Jo-1 | anti-PM-Scl | -0.20949 | 1.92E-08 |
| B98 | B185 | anti-Jo-1 | PT | -0.19886 | 9.94E-08 |
| B98 | B175 | anti-Jo-1 | ALB-U | 0.197112 | 1.29E-07 |
| B98 | B196 | anti-Jo-1 | TCO2 | 0.193449 | 2.22E-07 |
| B98 | B137 | anti-Jo-1 | AFU | -0.19228 | 2.63E-07 |
| B99 | B183 | anti-PM-Scl | PTA | 0.333792 | 0 |
| B99 | B186 | anti-PM-Scl | PFC | -0.31537 | 0 |
| B99 | B136 | anti-PM-Scl | 5'NT | 0.256415 | 4.61E-12 |
| B99 | B140 | anti-PM-Scl | r-GT | 0.240809 | 9.01E-11 |
| B99 | B100 | anti-PM-Scl | anti-Ro52 | -0.20949 | 1.92E-08 |
| B99 | B101 | anti-PM-Scl | anti-Sc70 | -0.20949 | 1.92E-08 |
| B99 | B102 | anti-PM-Scl | anti-Sm | -0.20949 | 1.92E-08 |
| B99 | B103 | anti-PM-Scl | anti-SSA/Ro | -0.20949 | 1.92E-08 |
| B99 | B104 | anti-PM-Scl | anti-SSB | -0.20949 | 1.92E-08 |
| B99 | B105 | anti-PM-Scl | anti-U1RNP | -0.20949 | 1.92E-08 |
| B99 | B106 | anti-PM-Scl | anti-P | -0.20949 | 1.92E-08 |
| B99 | B107 | anti-PM-Scl | AnuA | -0.20949 | 1.92E-08 |
| B99 | B108 | anti-PM-Scl | Anti-Cyclin | -0.20949 | 1.92E-08 |
| B99 | B109 | anti-PM-Scl | AMA-M2 | -0.20949 | 1.92E-08 |

**Supplementary Table 24. PEI Correlations in Hypertension+other diseases**

| Row | Column | Row | Column | Cor | *P* |
|---|---|---|---|---|---|
| B1 | B161 | Gender | LEU | 0.174573 | 0 |
| B1 | B163 | Gender | SG | -0.37282 | 0 |
| B1 | B174 | Gender | Urea/Cr | 0.408665 | 0 |
| B1 | B192 | Gender | UA | -0.34936 | 0 |
| B1 | B2 | Gender | Age | 0.180919 | 0 |
| B1 | B204 | Gender | Drinking | -0.29228 | 0 |
| B1 | B206 | Gender | Waist | -0.3332 | 0 |
| B1 | B209 | Gender | Smoking | -0.31641 | 0 |
| B1 | B210 | Gender | Hip | -0.25338 | 0 |
| B1 | B216 | Gender | BMI | -0.52976 | 0 |
| B1 | B218 | Gender | DBP | -0.16804 | 0 |
| B1 | B221 | Gender | Hei | -0.70011 | 0 |
| B1 | B222 | Gender | NS | -0.63696 | 0 |
| B1 | B41 | Gender | MON％ | -0.16696 | 0 |
| B1 | B42 | Gender | MON | -0.21134 | 0 |
| B1 | B45 | Gender | MCHC | -0.27825 | 0 |
| B1 | B46 | Gender | RBC Count | -0.35207 | 0 |
| B1 | B51 | Gender | Hb | -0.44605 | 0 |
| B1 | B72 | Gender | HDL-C | 0.29762 | 0 |
| B1 | B215 | Gender | BMI | -0.16496 | 4.44E-16 |
| B1 | B74 | Gender | ApoA1 | 0.163059 | 8.88E-16 |
| B1 | B48 | Gender | LYM% | 0.158 | 6.22E-15 |
| B1 | B141 | Gender | AST/ALT | 0.154172 | 2.78E-14 |
| B1 | B190 | Gender | Cre | -0.1499 | 1.41E-13 |
| B1 | B132 | Gender | Dbil | -0.12774 | 3.13E-10 |
| B1 | B44 | Gender | MCH | -0.12751 | 3.37E-10 |
| B1 | B127 | Gender | A-G ratio | -0.12743 | 3.45E-10 |
| B1 | B133 | Gender | TBIL | -0.12366 | 1.13E-09 |
| B1 | B3 | Gender | Hcy | -0.12242 | 1.66E-09 |
| B1 | B140 | Gender | r-GT | -0.12105 | 2.52E-09 |
| B1 | B130 | Gender | GLB | 0.117479 | 7.33E-09 |
| B1 | B219 | Gender | OT | 0.116188 | 1.07E-08 |
| B1 | B176 | Gender | Ucr | -0.11549 | 1.31E-08 |
| B1 | B166 | Gender | Pro | -0.11476 | 1.62E-08 |
| B1 | B128 | Gender | ALT | -0.11163 | 3.94E-08 |
| B1 | B129 | Gender | IBIL | -0.1106 | 5.26E-08 |
| B1 | B77 | Gender | TC | 0.106689 | 1.53E-07 |
| B1 | B113 | Gender | FT3 | -0.10594 | 1.87E-07 |
| B1 | B173 | Gender | NTT | 0.10482 | 2.52E-07 |
| B1 | B52 | Gender | PLT/BPC | 0.101932 | 5.35E-07 |
| B10 | B11 | PG Ⅰ | PGI/PGII | 0.168622 | 0 |
| B10 | B12 | PG Ⅰ | PG II | 0.587905 | 0 |

| | | | | | |
|---|---|---|---|---|---|
| B10 | B189 | PG I | eGFB | -0.19574 | 0 |
| B10 | B190 | PG I | Cre | 0.194114 | 0 |
| B10 | B191 | PG I | Urea | 0.184263 | 0 |
| B10 | B51 | PG I | Hb | -0.12359 | 1.16E-09 |
| B10 | B46 | PG I | RBC Count | -0.10566 | 2.02E-07 |
| B100 | B228 | anti-Ro52 | OS | 0.125436 | 6.50E-10 |
| B100 | B194 | anti-Ro52 | Cl | 0.111802 | 3.76E-08 |
| B100 | B198 | anti-Ro52 | P | 0.111318 | 4.31E-08 |
| B100 | B101 | anti-Ro52 | anti-Sc70 | 0.103467 | 3.59E-07 |
| B100 | B102 | anti-Ro52 | anti-Sm | 0.103467 | 3.59E-07 |
| B100 | B103 | anti-Ro52 | anti-SSA/Ro | 0.103467 | 3.59E-07 |
| B100 | B104 | anti-Ro52 | anti-SSB | 0.103467 | 3.59E-07 |
| B100 | B105 | anti-Ro52 | anti-U1RNP | 0.103467 | 3.59E-07 |
| B100 | B106 | anti-Ro52 | anti-P | 0.103467 | 3.59E-07 |
| B100 | B107 | anti-Ro52 | AnuA | 0.103466 | 3.59E-07 |
| B100 | B109 | anti-Ro52 | AMA-M2 | 0.103467 | 3.59E-07 |
| B100 | B110 | anti-Ro52 | AHA | 0.103467 | 3.59E-07 |
| B100 | B137 | anti-Ro52 | AFU | 0.103278 | 3.77E-07 |
| B101 | B103 | anti-Sc70 | anti-SSA/Ro | 1 | 0 |
| B101 | B104 | anti-Sc70 | anti-SSB | 1 | 0 |
| B101 | B107 | anti-Sc70 | AnuA | 1 | 0 |
| B101 | B109 | anti-Sc70 | AMA-M2 | 1 | 0 |
| B101 | B110 | anti-Sc70 | AHA | 1 | 0 |
| B101 | B120 | anti-Sc70 | HCV-cAg | 0.203435 | 0 |
| B101 | B156 | anti-Sc70 | IgE | 0.177741 | 0 |
| B101 | B168 | anti-Sc70 | NMP22 | -0.17131 | 0 |
| B101 | B155 | anti-Sc70 | IgA | -0.15872 | 4.66E-15 |
| B101 | B159 | anti-Sc70 | β2-MG | -0.15404 | 2.91E-14 |
| B101 | B134 | anti-Sc70 | CHE | -0.14987 | 1.43E-13 |
| B101 | B108 | anti-Sc70 | Anti-Cyclin | -0.14294 | 1.81E-12 |
| B101 | B136 | anti-Sc70 | 5'NT | -0.14092 | 3.71E-12 |
| B101 | B145 | anti-Sc70 | BGP | -0.14021 | 4.77E-12 |
| B101 | B193 | anti-Sc70 | K | 0.1384 | 8.96E-12 |
| B101 | B119 | anti-Sc70 | Hcv-rna | 0.128458 | 2.49E-10 |
| B101 | B135 | anti-Sc70 | TBA | 0.114445 | 1.77E-08 |
| B101 | B181 | anti-Sc70 | APTT | -0.11108 | 4.60E-08 |
| B101 | B198 | anti-Sc70 | P | 0.099814 | 9.17E-07 |
| B102 | B103 | anti-Sm | anti-SSA/Ro | 1 | 0 |
| B102 | B104 | anti-Sm | anti-SSB | 1 | 0 |
| B102 | B105 | anti-Sm | anti-U1RNP | 1 | 0 |
| B102 | B106 | anti-Sm | anti-P | 1 | 0 |
| B102 | B107 | anti-Sm | AnuA | 1 | 0 |
| B102 | B109 | anti-Sm | AMA-M2 | 1 | 0 |
| B102 | B110 | anti-Sm | AHA | 1 | 0 |

| | | | | | |
|---|---|---|---|---|---|
| B102 | B120 | anti-Sm | HCV-cAg | 0.203435 | 0 |
| B102 | B156 | anti-Sm | IgE | 0.177741 | 0 |
| B102 | B168 | anti-Sm | NMP22 | -0.17131 | 0 |
| B102 | B155 | anti-Sm | IgA | -0.15872 | 4.66E-15 |
| B102 | B159 | anti-Sm | β2-MG | -0.15404 | 2.91E-14 |
| B102 | B134 | anti-Sm | CHE | -0.14987 | 1.43E-13 |
| B102 | B108 | anti-Sm | Anti-Cyclin | -0.14294 | 1.81E-12 |
| B102 | B136 | anti-Sm | 5'NT | -0.14092 | 3.71E-12 |
| B102 | B145 | anti-Sm | BGP | -0.14021 | 4.77E-12 |
| B102 | B193 | anti-Sm | K | 0.1384 | 8.96E-12 |
| B102 | B119 | anti-Sm | Hcv-rna | 0.128458 | 2.49E-10 |
| B102 | B135 | anti-Sm | TBA | 0.114445 | 1.77E-08 |
| B102 | B181 | anti-Sm | APTT | -0.11108 | 4.60E-08 |
| B102 | B198 | anti-Sm | P | 0.099814 | 9.17E-07 |
| B103 | B104 | anti-SSA/Ro | anti-SSB | 1 | 0 |
| B103 | B105 | anti-SSA/Ro | anti-U1RNP | 1 | 0 |
| B103 | B106 | anti-SSA/Ro | anti-P | 1 | 0 |
| B103 | B107 | anti-SSA/Ro | AnuA | 0.999999 | 0 |
| B103 | B120 | anti-SSA/Ro | HCV-cAg | 0.203435 | 0 |
| B103 | B156 | anti-SSA/Ro | IgE | 0.177741 | 0 |
| B103 | B168 | anti-SSA/Ro | NMP22 | -0.17131 | 0 |
| B103 | B155 | anti-SSA/Ro | IgA | -0.15872 | 4.66E-15 |
| B103 | B159 | anti-SSA/Ro | β2-MG | -0.15404 | 2.91E-14 |
| B103 | B134 | anti-SSA/Ro | CHE | -0.14987 | 1.43E-13 |
| B103 | B108 | anti-SSA/Ro | Anti-Cyclin | -0.14294 | 1.81E-12 |
| B103 | B136 | anti-SSA/Ro | 5'NT | -0.14092 | 3.71E-12 |
| B103 | B145 | anti-SSA/Ro | BGP | -0.14021 | 4.77E-12 |
| B103 | B193 | anti-SSA/Ro | K | 0.1384 | 8.96E-12 |
| B103 | B119 | anti-SSA/Ro | Hcv-rna | 0.128458 | 2.49E-10 |
| B103 | B135 | anti-SSA/Ro | TBA | 0.114445 | 1.77E-08 |
| B103 | B181 | anti-SSA/Ro | APTT | -0.11108 | 4.60E-08 |
| B103 | B198 | anti-SSA/Ro | P | 0.099814 | 9.17E-07 |
| B104 | B106 | anti-SSB | anti-P | 1 | 0 |
| B104 | B107 | anti-SSB | AnuA | 0.999999 | 0 |
| B104 | B109 | anti-SSB | AMA-M2 | 1 | 0 |
| B104 | B110 | anti-SSB | AHA | 1 | 0 |
| B104 | B120 | anti-SSB | HCV-cAg | 0.203434 | 0 |
| B104 | B156 | anti-SSB | IgE | 0.177741 | 0 |
| B104 | B168 | anti-SSB | NMP22 | -0.17131 | 0 |
| B104 | B155 | anti-SSB | IgA | -0.15872 | 4.66E-15 |
| B104 | B159 | anti-SSB | β2-MG | -0.15404 | 2.91E-14 |
| B104 | B134 | anti-SSB | CHE | -0.14987 | 1.43E-13 |
| B104 | B108 | anti-SSB | Anti-Cyclin | -0.14294 | 1.81E-12 |
| B104 | B136 | anti-SSB | 5'NT | -0.14092 | 3.71E-12 |

| | | | | | |
|---|---|---|---|---|---|
| B104 | B145 | anti-SSB | BGP | -0.14021 | 4.77E-12 |
| B104 | B193 | anti-SSB | K | 0.1384 | 8.96E-12 |
| B104 | B119 | anti-SSB | Hcv-rna | 0.128458 | 2.49E-10 |
| B104 | B135 | anti-SSB | TBA | 0.114445 | 1.77E-08 |
| B104 | B181 | anti-SSB | APTT | -0.11108 | 4.60E-08 |
| B104 | B198 | anti-SSB | P | 0.099814 | 9.17E-07 |
| B105 | B107 | anti-U1RNP | AnuA | 1 | 0 |
| B105 | B120 | anti-U1RNP | HCV-cAg | 0.203435 | 0 |
| B105 | B156 | anti-U1RNP | IgE | 0.177741 | 0 |
| B105 | B168 | anti-U1RNP | NMP22 | -0.17131 | 0 |
| B105 | B155 | anti-U1RNP | IgA | -0.15872 | 4.66E-15 |
| B105 | B159 | anti-U1RNP | β2-MG | -0.15404 | 2.91E-14 |
| B105 | B134 | anti-U1RNP | CHE | -0.14987 | 1.43E-13 |
| B105 | B108 | anti-U1RNP | Anti-Cyclin | -0.14294 | 1.81E-12 |
| B105 | B136 | anti-U1RNP | 5'NT | -0.14092 | 3.71E-12 |
| B105 | B145 | anti-U1RNP | BGP | -0.14021 | 4.77E-12 |
| B105 | B193 | anti-U1RNP | K | 0.1384 | 8.96E-12 |
| B105 | B119 | anti-U1RNP | Hcv-rna | 0.128458 | 2.49E-10 |
| B105 | B135 | anti-U1RNP | TBA | 0.114445 | 1.77E-08 |
| B105 | B181 | anti-U1RNP | APTT | -0.11108 | 4.60E-08 |
| B105 | B198 | anti-U1RNP | P | 0.099814 | 9.17E-07 |
| B106 | B107 | anti-P | AnuA | 1 | 0 |
| B106 | B109 | anti-P | AMA-M2 | 1 | 0 |
| B106 | B120 | anti-P | HCV-cAg | 0.203434 | 0 |
| B106 | B156 | anti-P | IgE | 0.177741 | 0 |
| B106 | B168 | anti-P | NMP22 | -0.17131 | 0 |
| B106 | B155 | anti-P | IgA | -0.15872 | 4.66E-15 |
| B106 | B159 | anti-P | β2-MG | -0.15404 | 2.91E-14 |
| B106 | B134 | anti-P | CHE | -0.14987 | 1.43E-13 |
| B106 | B108 | anti-P | Anti-Cyclin | -0.14294 | 1.81E-12 |
| B106 | B136 | anti-P | 5'NT | -0.14092 | 3.71E-12 |
| B106 | B145 | anti-P | BGP | -0.14021 | 4.77E-12 |
| B106 | B193 | anti-P | K | 0.1384 | 8.96E-12 |
| B106 | B119 | anti-P | Hcv-rna | 0.128458 | 2.49E-10 |
| B106 | B135 | anti-P | TBA | 0.114445 | 1.77E-08 |
| B106 | B181 | anti-P | APTT | -0.11108 | 4.60E-08 |
| B106 | B198 | anti-P | P | 0.099814 | 9.17E-07 |
| B107 | B109 | AnuA | AMA-M2 | 0.999999 | 0 |
| B107 | B110 | AnuA | AHA | 0.999999 | 0 |
| B107 | B120 | AnuA | HCV-cAg | 0.203436 | 0 |
| B107 | B156 | AnuA | IgE | 0.177741 | 0 |
| B107 | B168 | AnuA | NMP22 | -0.17131 | 0 |
| B107 | B155 | AnuA | IgA | -0.15872 | 4.66E-15 |
| B107 | B159 | AnuA | β2-MG | -0.15404 | 2.93E-14 |

| | | | | | |
|---|---|---|---|---|---|
| B107 | B134 | AnuA | CHE | -0.14987 | 1.43E-13 |
| B107 | B108 | AnuA | Anti-Cyclin | -0.14294 | 1.81E-12 |
| B107 | B136 | AnuA | 5'NT | -0.14092 | 3.71E-12 |
| B107 | B145 | AnuA | BGP | -0.14021 | 4.77E-12 |
| B107 | B193 | AnuA | K | 0.1384 | 8.96E-12 |
| B107 | B119 | AnuA | Hcv-rna | 0.128458 | 2.49E-10 |
| B107 | B135 | AnuA | TBA | 0.114445 | 1.77E-08 |
| B107 | B181 | AnuA | APTT | -0.11108 | 4.60E-08 |
| B107 | B198 | AnuA | P | 0.099814 | 9.17E-07 |
| B108 | B181 | Anti-Cyclin | APTT | 0.391365 | 0 |
| B108 | B182 | Anti-Cyclin | TT | -0.42583 | 0 |
| B108 | B124 | Anti-Cyclin | ACTH | -0.16543 | 4.44E-16 |
| B108 | B120 | Anti-Cyclin | HCV-cAg | -0.14513 | 8.22E-13 |
| B108 | B109 | Anti-Cyclin | AMA-M2 | -0.14294 | 1.81E-12 |
| B108 | B110 | Anti-Cyclin | AHA | -0.14294 | 1.81E-12 |
| B108 | B137 | Anti-Cyclin | AFU | -0.11181 | 3.75E-08 |
| B108 | B134 | Anti-Cyclin | CHE | 0.100117 | 8.49E-07 |
| B109 | B120 | AMA-M2 | HCV-cAg | 0.203434 | 0 |
| B109 | B156 | AMA-M2 | IgE | 0.177741 | 0 |
| B109 | B168 | AMA-M2 | NMP22 | -0.17131 | 0 |
| B109 | B155 | AMA-M2 | IgA | -0.15872 | 4.66E-15 |
| B109 | B159 | AMA-M2 | β2-MG | -0.15404 | 2.91E-14 |
| B109 | B134 | AMA-M2 | CHE | -0.14987 | 1.43E-13 |
| B109 | B136 | AMA-M2 | 5'NT | -0.14092 | 3.71E-12 |
| B109 | B145 | AMA-M2 | BGP | -0.14021 | 4.77E-12 |
| B109 | B193 | AMA-M2 | K | 0.1384 | 8.96E-12 |
| B109 | B119 | AMA-M2 | Hcv-rna | 0.128458 | 2.49E-10 |
| B109 | B135 | AMA-M2 | TBA | 0.114445 | 1.77E-08 |
| B109 | B181 | AMA-M2 | APTT | -0.11108 | 4.60E-08 |
| B109 | B198 | AMA-M2 | P | 0.099814 | 9.17E-07 |
| B11 | B12 | PGI/PGII | PG II | -0.48342 | 0 |
| B11 | B14 | PGI/PGII | G-17 | -0.21957 | 0 |
| B11 | B169 | PGI/PGII | U-CRY | 0.175005 | 0 |
| B11 | B80 | PGI/PGII | anti-HP IgG | -0.15348 | 3.62E-14 |
| B110 | B120 | AHA | HCV-cAg | 0.203435 | 0 |
| B110 | B156 | AHA | IgE | 0.177741 | 0 |
| B110 | B168 | AHA | NMP22 | -0.17131 | 0 |
| B110 | B155 | AHA | IgA | -0.15872 | 4.66E-15 |
| B110 | B159 | AHA | β2-MG | -0.15404 | 2.91E-14 |
| B110 | B134 | AHA | CHE | -0.14987 | 1.43E-13 |
| B110 | B136 | AHA | 5'NT | -0.14092 | 3.71E-12 |
| B110 | B145 | AHA | BGP | -0.14021 | 4.77E-12 |
| B110 | B193 | AHA | K | 0.1384 | 8.96E-12 |
| B110 | B119 | AHA | Hcv-rna | 0.128458 | 2.49E-10 |

| | | | | | |
|---|---|---|---|---|---|
| B110 | B135 | AHA | TBA | 0.114445 | 1.77E-08 |
| B110 | B181 | AHA | APTT | -0.11108 | 4.60E-08 |
| B110 | B198 | AHA | P | 0.099814 | 9.17E-07 |
| B111 | B112 | TSH | FT4 | -0.21113 | 0 |
| B111 | B113 | TSH | FT3 | -0.18017 | 0 |
| B111 | B114 | TSH | TSH TT3 TT4(TT4) | -0.2798 | 0 |
| B111 | B115 | TSH | TSH TT3 TT4(TT3) | -0.17727 | 0 |
| B111 | B148 | TSH | Anti_TPO | 0.221981 | 0 |
| B112 | B113 | FT4 | FT3 | 0.228967 | 0 |
| B112 | B114 | FT4 | TSH TT3 TT4(TT4) | 0.538842 | 0 |
| B112 | B115 | FT4 | TSH TT3 TT4(TT3) | 0.209917 | 0 |
| B112 | B154 | FT4 | RF | 0.11282 | 2.82E-08 |
| B113 | B114 | FT3 | TSH TT3 TT4(TT4) | 0.199093 | 0 |
| B113 | B115 | FT3 | TSH TT3 TT4(TT3) | 0.528292 | 0 |
| B113 | B126 | FT3 | Alb | 0.130845 | 1.14E-10 |
| B113 | B176 | FT3 | Ucr | 0.107891 | 1.11E-07 |
| B114 | B115 | TSH TT3 TT4(TT4) | TSH TT3 TT4(TT3) | 0.412165 | 0 |
| B114 | B154 | TSH TT3 TT4(TT4) | RF | 0.184373 | 0 |
| B114 | B131 | TSH TT3 TT4(TT4) | AST | 0.158415 | 5.33E-15 |
| B114 | B128 | TSH TT3 TT4(TT4) | ALT | 0.129772 | 1.63E-10 |
| B114 | B148 | TSH TT3 TT4(TT4) | Anti_TPO | -0.12352 | 1.18E-09 |
| B114 | B165 | TSH TT3 TT4(TT4) | bilirubin | 0.110615 | 5.24E-08 |
| B115 | B128 | TSH TT3 TT4(TT3) | ALT | 0.145766 | 6.52E-13 |
| B115 | B131 | TSH TT3 TT4(TT3) | AST | 0.139366 | 6.41E-12 |
| B115 | B137 | TSH TT3 TT4(TT3) | AFU | 0.121979 | 1.90E-09 |
| B115 | B123 | TSH TT3 TT4(TT3) | hs-CRP1 | -0.11602 | 1.13E-08 |
| B115 | B134 | TSH TT3 TT4(TT3) | CHE | 0.103411 | 3.65E-07 |
| B116 | B117 | TT3 | TT4 | -0.8731 | 0 |
| B118 | B126 | CEA1 | Alb | -0.10181 | 5.52E-07 |
| B119 | B120 | Hcv-rna | HCV-cAg | 0.400438 | 0 |
| B119 | B124 | Hcv-rna | ACTH | -0.23254 | 0 |
| B119 | B125 | Hcv-rna | Cor | -0.20095 | 0 |
| B119 | B159 | Hcv-rna | β2-MG | -0.84477 | 0 |
| B119 | B186 | Hcv-rna | PFC | -0.1569 | 9.55E-15 |
| B119 | B139 | Hcv-rna | ADA | -0.1406 | 4.16E-12 |
| B119 | B185 | Hcv-rna | PT | -0.11307 | 2.63E-08 |
| B119 | B197 | Hcv-rna | AG | -0.11212 | 3.44E-08 |
| B119 | B156 | Hcv-rna | IgE | 0.110777 | 5.01E-08 |
| B119 | B194 | Hcv-rna | Cl | 0.10457 | 2.69E-07 |
| B119 | B137 | Hcv-rna | AFU | -0.10281 | 4.27E-07 |
| B12 | B14 | PG II | G-17 | 0.364621 | 0 |
| B12 | B191 | PG II | Urea | 0.17228 | 0 |
| B12 | B80 | PG II | anti-HP IgG | 0.15703 | 9.10E-15 |
| B12 | B189 | PG II | eGFB | -0.15484 | 2.13E-14 |

| | | | | | |
|---|---|---|---|---|---|
| B12 | B190 | PG II | Cre | 0.154302 | 2.62E-14 |
| B120 | B156 | HCV-cAg | IgE | 0.171095 | 0 |
| B120 | B159 | HCV-cAg | β2-MG | -0.33828 | 0 |
| B120 | B139 | HCV-cAg | ADA | -0.13741 | 1.26E-11 |
| B120 | B157 | HCV-cAg | IgG | -0.10083 | 7.08E-07 |
| B121 | B122 | C3 | C4 | 0.999989 | 0 |
| B121 | B155 | C3 | IgA | -0.70999 | 0 |
| B121 | B157 | C3 | IgG | -0.32037 | 0 |
| B121 | B130 | C3 | GLB | -0.11047 | 5.45E-08 |
| B122 | B155 | C4 | IgA | -0.71021 | 0 |
| B122 | B157 | C4 | IgG | -0.32038 | 0 |
| B122 | B130 | C4 | GLB | -0.11059 | 5.28E-08 |
| B123 | B138 | hs-CRP1 | PAB | -0.17598 | 0 |
| B123 | B179 | hs-CRP1 | α1-MG | 0.509714 | 0 |
| B123 | B186 | hs-CRP1 | PFC | 0.234861 | 0 |
| B123 | B139 | hs-CRP1 | ADA | 0.135328 | 2.57E-11 |
| B123 | B126 | hs-CRP1 | Alb | -0.11814 | 6.03E-09 |
| B123 | B134 | hs-CRP1 | CHE | -0.11224 | 3.32E-08 |
| B123 | B182 | hs-CRP1 | TT | -0.10565 | 2.02E-07 |
| B123 | B181 | hs-CRP1 | APTT | 0.100253 | 8.21E-07 |
| B124 | B182 | ACTH | TT | 0.173274 | 0 |
| B124 | B125 | ACTH | Cor | 0.166323 | 2.22E-16 |
| B124 | B195 | ACTH | Na | 0.140212 | 4.76E-12 |
| B124 | B200 | ACTH | OP | 0.136165 | 1.93E-11 |
| B124 | B197 | ACTH | AG | 0.129362 | 1.86E-10 |
| B124 | B199 | ACTH | Mg | 0.108216 | 1.01E-07 |
| B124 | B186 | ACTH | PFC | -0.1069 | 1.45E-07 |
| B125 | B139 | Cor | ADA | -0.1773 | 0 |
| B125 | B164 | Cor | COM | -0.20984 | 0 |
| B125 | B182 | Cor | TT | -0.1725 | 0 |
| B125 | B184 | Cor | INR | 0.197511 | 0 |
| B125 | B159 | Cor | β2-MG | 0.150013 | 1.35E-13 |
| B125 | B185 | Cor | PT | 0.148745 | 2.17E-13 |
| B125 | B136 | Cor | 5'NT | -0.11918 | 4.42E-09 |
| B125 | B197 | Cor | AG | -0.1168 | 8.94E-09 |
| B125 | B198 | Cor | P | -0.1045 | 2.74E-07 |
| B126 | B127 | Alb | A-G ratio | 0.520369 | 0 |
| B126 | B130 | Alb | GLB | -0.18882 | 0 |
| B126 | B188 | Alb | Ca | 0.385865 | 0 |
| B126 | B189 | Alb | eGFB | 0.263449 | 0 |
| B126 | B191 | Alb | Urea | -0.18033 | 0 |
| B126 | B138 | Alb | PAB | 0.157184 | 8.44E-15 |
| B126 | B141 | Alb | AST/ALT | -0.15453 | 2.40E-14 |
| B126 | B176 | Alb | Ucr | 0.122753 | 1.50E-09 |

| | | | | | |
|---|---|---|---|---|---|
| B126 | B134 | Alb | CHE | 0.121822 | 1.99E-09 |
| B126 | B146 | Alb | CysC | -0.11949 | 4.03E-09 |
| B126 | B218 | Alb | DBP | 0.118474 | 5.46E-09 |
| B126 | B190 | Alb | Cre | -0.11708 | 8.24E-09 |
| B126 | B221 | Alb | Hei | 0.115168 | 1.44E-08 |
| B126 | B129 | Alb | IBIL | 0.100955 | 6.86E-07 |
| B127 | B130 | A-G ratio | GLB | -0.8966 | 0 |
| B127 | B189 | A-G ratio | eGFB | 0.205409 | 0 |
| B127 | B221 | A-G ratio | Hei | 0.165241 | 4.44E-16 |
| B127 | B220 | A-G ratio | SBP | -0.13361 | 4.58E-11 |
| B127 | B216 | A-G ratio | BMI | 0.132048 | 7.71E-11 |
| B127 | B141 | A-G ratio | AST/ALT | -0.12738 | 3.51E-10 |
| B127 | B163 | A-G ratio | SG | 0.124732 | 8.11E-10 |
| B127 | B209 | A-G ratio | Smoking | 0.123791 | 1.09E-09 |
| B127 | B204 | A-G ratio | Drinking | 0.118347 | 5.67E-09 |
| B127 | B155 | A-G ratio | IgA | -0.11209 | 3.46E-08 |
| B127 | B138 | A-G ratio | PAB | 0.107906 | 1.10E-07 |
| B128 | B131 | ALT | AST | 0.850594 | 0 |
| B128 | B132 | ALT | Dbil | 0.293965 | 0 |
| B128 | B133 | ALT | TBIL | 0.174733 | 0 |
| B128 | B140 | ALT | r-GT | 0.346674 | 0 |
| B128 | B141 | ALT | AST/ALT | -0.40555 | 0 |
| B128 | B165 | ALT | bilirubin | 0.347602 | 0 |
| B128 | B215 | ALT | BMI | 0.176942 | 0 |
| B128 | B216 | ALT | BMI | 0.209281 | 0 |
| B128 | B222 | ALT | NS | 0.175872 | 0 |
| B128 | B206 | ALT | Waist | 0.160727 | 2.22E-15 |
| B128 | B210 | ALT | Hip | 0.140505 | 4.30E-12 |
| B128 | B189 | ALT | eGFB | 0.131235 | 1.01E-10 |
| B128 | B221 | ALT | Hei | 0.130101 | 1.46E-10 |
| B128 | B129 | ALT | IBIL | 0.114425 | 1.78E-08 |
| B128 | B163 | ALT | SG | 0.112579 | 3.02E-08 |
| B128 | B204 | ALT | Drinking | 0.108999 | 8.18E-08 |
| B128 | B192 | ALT | UA | 0.101839 | 5.47E-07 |
| B129 | B132 | IBIL | Dbil | 0.657246 | 0 |
| B129 | B133 | IBIL | TBIL | 0.97606 | 0 |
| B129 | B191 | IBIL | Urea | -0.1228 | 1.48E-09 |
| B129 | B131 | IBIL | AST | 0.10907 | 8.03E-08 |
| B129 | B221 | IBIL | Hei | 0.102163 | 5.04E-07 |
| B130 | B220 | GLB | SBP | 0.145585 | 6.96E-13 |
| B130 | B221 | GLB | Hei | -0.14039 | 4.48E-12 |
| B130 | B155 | GLB | IgA | 0.139691 | 5.72E-12 |
| B130 | B209 | GLB | Smoking | -0.1239 | 1.05E-09 |
| B130 | B189 | GLB | eGFB | -0.12067 | 2.83E-09 |

| | | | | | |
|---|---|---|---|---|---|
| B130 | B157 | GLB | IgG | 0.109516 | 7.10E-08 |
| B130 | B150 | GLB | ALP | 0.10447 | 2.76E-07 |
| B130 | B216 | GLB | BMI | -0.10064 | 7.43E-07 |
| B131 | B132 | AST | Dbil | 0.320309 | 0 |
| B131 | B133 | AST | TBIL | 0.178055 | 0 |
| B131 | B140 | AST | r-GT | 0.38223 | 0 |
| B131 | B165 | AST | bilirubin | 0.428065 | 0 |
| B132 | B133 | Dbil | TBIL | 0.805337 | 0 |
| B132 | B140 | Dbil | r-GT | 0.199823 | 0 |
| B132 | B165 | Dbil | bilirubin | 0.177781 | 0 |
| B132 | B185 | Dbil | PT | 0.125258 | 6.87E-10 |
| B133 | B140 | TBIL | r-GT | 0.118497 | 5.43E-09 |
| B133 | B191 | TBIL | Urea | -0.11232 | 3.25E-08 |
| B133 | B221 | TBIL | Hei | 0.106305 | 1.70E-07 |
| B134 | B136 | CHE | 5'NT | 0.180803 | 0 |
| B134 | B138 | CHE | PAB | 0.305622 | 0 |
| B134 | B168 | CHE | NMP22 | 0.271501 | 0 |
| B134 | B193 | CHE | K | -0.2418 | 0 |
| B134 | B146 | CHE | CysC | -0.15927 | 3.77E-15 |
| B134 | B156 | CHE | IgE | -0.14626 | 5.44E-13 |
| B134 | B197 | CHE | AG | 0.131314 | 9.81E-11 |
| B134 | B137 | CHE | AFU | 0.12038 | 3.09E-09 |
| B134 | B218 | CHE | DBP | 0.117156 | 8.07E-09 |
| B134 | B194 | CHE | Cl | -0.11335 | 2.42E-08 |
| B134 | B222 | CHE | NS | 0.101962 | 5.30E-07 |
| B135 | B137 | TBA | AFU | 0.159494 | 3.55E-15 |
| B135 | B193 | TBA | K | -0.11951 | 4.01E-09 |
| B135 | B187 | TBA | NSE | 0.117557 | 7.17E-09 |
| B135 | B138 | TBA | PAB | -0.10478 | 2.54E-07 |
| B135 | B139 | TBA | ADA | 0.101913 | 5.37E-07 |
| B136 | B150 | 5'NT | ALP | 0.278003 | 0 |
| B136 | B199 | 5'NT | Mg | 0.193018 | 0 |
| B136 | B200 | 5'NT | OP | 0.14374 | 1.36E-12 |
| B136 | B195 | 5'NT | Na | 0.142676 | 1.99E-12 |
| B136 | B140 | 5'NT | r-GT | 0.139049 | 7.15E-12 |
| B136 | B183 | 5'NT | PTA | -0.11401 | 2.01E-08 |
| B136 | B139 | 5'NT | ADA | 0.112693 | 2.92E-08 |
| B136 | B197 | 5'NT | AG | 0.11162 | 3.96E-08 |
| B136 | B152 | 5'NT | ACA-Ig(G-A-M) | -0.10783 | 1.12E-07 |
| B136 | B193 | 5'NT | K | -0.10026 | 8.19E-07 |
| B137 | B193 | AFU | K | -0.17283 | 0 |
| B137 | B139 | AFU | ADA | 0.139972 | 5.18E-12 |
| B137 | B198 | AFU | P | -0.13204 | 7.72E-11 |
| B138 | B139 | PAB | ADA | -0.48294 | 0 |

| | | | | | |
|---|---|---|---|---|---|
| B138 | B150 | PAB | ALP | -0.16526 | 4.44E-16 |
| B138 | B198 | PAB | P | -0.15059 | 1.09E-13 |
| B138 | B186 | PAB | PFC | -0.11231 | 3.26E-08 |
| B138 | B221 | PAB | Hei | 0.11077 | 5.02E-08 |
| B139 | B150 | ADA | ALP | 0.308368 | 0 |
| B139 | B200 | ADA | OP | 0.144351 | 1.09E-12 |
| B139 | B157 | ADA | IgG | 0.133998 | 4.02E-11 |
| B139 | B199 | ADA | Mg | -0.12798 | 2.90E-10 |
| B139 | B197 | ADA | AG | 0.1092 | 7.74E-08 |
| B139 | B146 | ADA | CysC | 0.105569 | 2.07E-07 |
| B139 | B195 | ADA | Na | 0.104114 | 3.03E-07 |
| B14 | B76 | G-17 | ApoB | 0.103641 | 3.43E-07 |
| B140 | B204 | r-GT | Drinking | 0.176265 | 0 |
| B140 | B165 | r-GT | bilirubin | 0.128363 | 2.56E-10 |
| B140 | B216 | r-GT | BMI | 0.115706 | 1.23E-08 |
| B140 | B222 | r-GT | NS | 0.115441 | 1.33E-08 |
| B140 | B192 | r-GT | UA | 0.112432 | 3.15E-08 |
| B140 | B141 | r-GT | AST/ALT | -0.11126 | 4.37E-08 |
| B140 | B163 | r-GT | SG | 0.105004 | 2.40E-07 |
| B140 | B221 | r-GT | Hei | 0.102279 | 4.89E-07 |
| B140 | B206 | r-GT | Waist | 0.102017 | 5.23E-07 |
| B140 | B153 | r-GT | Glu | 0.100066 | 8.60E-07 |
| B141 | B206 | AST/ALT | Waist | -0.21817 | 0 |
| B141 | B210 | AST/ALT | Hip | -0.23573 | 0 |
| B141 | B215 | AST/ALT | BMI | -0.25582 | 0 |
| B141 | B216 | AST/ALT | BMI | -0.31506 | 0 |
| B141 | B221 | AST/ALT | Hei | -0.2179 | 0 |
| B141 | B222 | AST/ALT | NS | -0.25574 | 0 |
| B141 | B218 | AST/ALT | DBP | -0.14335 | 1.56E-12 |
| B141 | B163 | AST/ALT | SG | -0.13493 | 2.94E-11 |
| B141 | B189 | AST/ALT | eGFB | -0.12272 | 1.51E-09 |
| B141 | B204 | AST/ALT | Drinking | -0.11088 | 4.87E-08 |
| B141 | B192 | AST/ALT | UA | -0.10785 | 1.12E-07 |
| B142 | B143 | 25-(OH)D | β-CTx | 0.310894 | 0 |
| B142 | B144 | 25-(OH)D | Total-PINP | 0.240978 | 0 |
| B142 | B151 | 25-(OH)D | Anti-CCP | 0.152644 | 4.97E-14 |
| B143 | B144 | β-CTx | Total-PINP | 0.748151 | 0 |
| B143 | B145 | β-CTx | BGP | 0.500277 | 0 |
| B143 | B151 | β-CTx | Anti-CCP | -0.20845 | 0 |
| B143 | B186 | β-CTx | PFC | 0.124629 | 8.37E-10 |
| B143 | B155 | β-CTx | IgA | 0.111183 | 4.47E-08 |
| B144 | B145 | Total-PINP | BGP | 0.36073 | 0 |
| B144 | B151 | Total-PINP | Anti-CCP | -0.20146 | 0 |
| B145 | B155 | BGP | IgA | 0.154673 | 2.29E-14 |

| | | | | | |
|---|---|---|---|---|---|
| B145 | B156 | BGP | IgE | -0.10982 | 6.54E-08 |
| B145 | B181 | BGP | APTT | 0.107503 | 1.23E-07 |
| B145 | B193 | BGP | K | -0.10184 | 5.47E-07 |
| B146 | B189 | CysC | eGFB | -0.26924 | 0 |
| B146 | B190 | CysC | Cre | 0.196558 | 0 |
| B146 | B191 | CysC | Urea | 0.309266 | 0 |
| B146 | B200 | CysC | OP | 0.233022 | 0 |
| B146 | B193 | CysC | K | 0.163637 | 6.66E-16 |
| B146 | B194 | CysC | Cl | 0.109768 | 6.62E-08 |
| B146 | B150 | CysC | ALP | 0.104011 | 3.12E-07 |
| B146 | B198 | CysC | P | 0.103953 | 3.16E-07 |
| B148 | B149 | Anti_TPO | TGAb | 0.477019 | 0 |
| B148 | B181 | Anti_TPO | APTT | -0.10418 | 2.98E-07 |
| B15 | B101 | CD3-CD19+(B cell) | anti-Sc70 | 0.358707 | 0 |
| B15 | B102 | CD3-CD19+(B cell) | anti-Sm | 0.358707 | 0 |
| B15 | B103 | CD3-CD19+(B cell) | anti-SSA/Ro | 0.358707 | 0 |
| B15 | B104 | CD3-CD19+(B cell) | anti-SSB | 0.358707 | 0 |
| B15 | B105 | CD3-CD19+(B cell) | anti-U1RNP | 0.358707 | 0 |
| B15 | B106 | CD3-CD19+(B cell) | anti-P | 0.358707 | 0 |
| B15 | B107 | CD3-CD19+(B cell) | AnuA | 0.358707 | 0 |
| B15 | B108 | CD3-CD19+(B cell) | Anti-Cyclin | -0.21515 | 0 |
| B15 | B109 | CD3-CD19+(B cell) | AMA-M2 | 0.358707 | 0 |
| B15 | B110 | CD3-CD19+(B cell) | AHA | 0.358707 | 0 |
| B15 | B156 | CD3-CD19+(B cell) | IgE | 0.198212 | 0 |
| B15 | B16 | CD3-CD19+(B cell) | CD3+T | 0.58108 | 0 |
| B15 | B168 | CD3-CD19+(B cell) | NMP22 | -0.18594 | 0 |
| B15 | B18 | CD3-CD19+(B cell) | CD3+CD4+helper(inducer) T | 0.507939 | 0 |
| B15 | B19 | CD3-CD19+(B cell) | CD3+CD4+/CD3+CD8+ | 0.377495 | 0 |
| B15 | B21 | CD3-CD19+(B cell) | CD3+CD4-CD8- | 0.408482 | 0 |
| B15 | B23 | CD3-CD19+(B cell) | CD3-CD16+CD56+(NK) | -0.67931 | 0 |
| B15 | B79 | CD3-CD19+(B cell) | hb V -dna | 0.215605 | 0 |
| B15 | B95 | CD3-CD19+(B cell) | anti-cenpb | 0.358707 | 0 |
| B15 | B98 | CD3-CD19+(B cell) | anti-Jo-1 | 0.358707 | 0 |
| B15 | B99 | CD3-CD19+(B cell) | anti-PM-Scl | 0.358707 | 0 |
| B15 | B134 | CD3-CD19+(B cell) | CHE | -0.16413 | 4.44E-16 |
| B15 | B120 | CD3-CD19+(B cell) | HCV-cAg | 0.159872 | 2.89E-15 |
| B15 | B181 | CD3-CD19+(B cell) | APTT | -0.14672 | 4.60E-13 |
| B15 | B94 | CD3-CD19+(B cell) | ALDH2-Lys504Lys | 0.131603 | 8.92E-11 |
| B15 | B145 | CD3-CD19+(B cell) | BGP | -0.12613 | 5.22E-10 |
| B15 | B64 | CD3-CD19+(B cell) | WBRSR(mPas_5) | -0.12438 | 9.05E-10 |
| B15 | B67 | CD3-CD19+(B cell) | WBRSR(mPas_50) | -0.12343 | 1.21E-09 |
| B15 | B65 | CD3-CD19+(B cell) | WBRSR(mPas_1) | -0.12264 | 1.55E-09 |
| B15 | B66 | CD3-CD19+(B cell) | WBRSR(mPas_100) | -0.12034 | 3.12E-09 |

| | | | | | |
|---|---|---|---|---|---|
| B15 | B22 | CD3-CD19+(B cell) | CD3+CD8+(T) | 0.117555 | 7.17E-09 |
| B15 | B92 | CD3-CD19+(B cell) | ALDH2-Glu504Glu | -0.11427 | 1.86E-08 |
| B15 | B61 | CD3-CD19+(B cell) | WBHSRV | -0.11393 | 2.05E-08 |
| B15 | B93 | CD3-CD19+(B cell) | ALDH2-Glu504Lys | 0.109737 | 6.68E-08 |
| B15 | B135 | CD3-CD19+(B cell) | TBA | 0.105101 | 2.34E-07 |
| B15 | B63 | CD3-CD19+(B cell) | WBRSR(mPas_200) | -0.1018 | 5.53E-07 |
| B15 | B60 | CD3-CD19+(B cell) | WBRVI | -0.1005 | 7.71E-07 |
| B150 | B200 | ALP | OP | 0.121454 | 2.23E-09 |
| B151 | B154 | Anti-CCP | RF | 0.178189 | 0 |
| B152 | B157 | ACA-Ig(G-A-M) | IgG | 0.126181 | 5.14E-10 |
| B152 | B156 | ACA-Ig(G-A-M) | IgE | 0.117753 | 6.76E-09 |
| B153 | B170 | Glu | U-GLU | 0.489316 | 0 |
| B153 | B206 | Glu | Waist | 0.174455 | 0 |
| B153 | B215 | Glu | BMI | 0.139998 | 5.14E-12 |
| B153 | B222 | Glu | NS | 0.129335 | 1.87E-10 |
| B153 | B175 | Glu | ALB-U | 0.124397 | 9.00E-10 |
| B153 | B216 | Glu | BMI | 0.111383 | 4.23E-08 |
| B153 | B210 | Glu | Hip | 0.105614 | 2.04E-07 |
| B153 | B160 | Glu | PH | -0.10159 | 5.84E-07 |
| B154 | B157 | RF | IgG | 0.131538 | 9.12E-11 |
| B154 | B156 | RF | IgE | -0.11747 | 7.36E-09 |
| B155 | B157 | IgA | IgG | 0.309072 | 0 |
| B155 | B158 | IgA | IgM | -0.13282 | 5.96E-11 |
| B156 | B186 | IgE | PFC | 0.148188 | 2.67E-13 |
| B156 | B182 | IgE | TT | -0.127 | 3.96E-10 |
| B156 | B195 | IgE | Na | -0.11948 | 4.05E-09 |
| B156 | B198 | IgE | P | 0.113479 | 2.34E-08 |
| B156 | B168 | IgE | NMP22 | -0.11077 | 5.02E-08 |
| B156 | B183 | IgE | PTA | 0.102646 | 4.45E-07 |
| B157 | B158 | IgG | IgM | -0.22973 | 0 |
| B157 | B185 | IgG | PT | 0.101416 | 6.10E-07 |
| B159 | B186 | β2-MG | PFC | 0.17118 | 0 |
| B159 | B198 | β2-MG | P | -0.12985 | 1.59E-10 |
| B159 | B193 | β2-MG | K | -0.1008 | 7.14E-07 |
| B16 | B138 | CD3+T | PAB | 0.199642 | 0 |
| B16 | B156 | CD3+T | IgE | 0.177942 | 0 |
| B16 | B17 | CD3+T | CD3+CD16+CD56+NK | 0.229013 | 0 |
| B16 | B18 | CD3+T | CD3+CD4+helper(inducer) T | 0.838252 | 0 |
| B16 | B19 | CD3+T | CD3+CD4+/CD3+CD8+ | 0.409564 | 0 |
| B16 | B20 | CD3+T | CD3+CD4+CD8+ | 0.359718 | 0 |
| B16 | B21 | CD3+T | CD3+CD4-CD8- | 0.373256 | 0 |
| B16 | B22 | CD3+T | CD3+CD8+(T) | 0.454745 | 0 |
| B16 | B23 | CD3+T | CD3-CD16+CD56+(NK) | -0.95751 | 0 |

| | | | | | |
|---|---|---|---|---|---|
| B16 | B34 | CD3+T | TK | -0.18568 | 0 |
| B16 | B96 | CD3+T | dsDNA (IIF) | -0.20128 | 0 |
| B16 | B97 | CD3+T | dsDNA | 0.202103 | 0 |
| B16 | B120 | CD3+T | HCV-cAg | 0.159641 | 3.11E-15 |
| B16 | B79 | CD3+T | hb V -dna | 0.153073 | 4.22E-14 |
| B16 | B157 | CD3+T | IgG | -0.14819 | 2.67E-13 |
| B16 | B168 | CD3+T | NMP22 | 0.114953 | 1.53E-08 |
| B16 | B65 | CD3+T | WBRSR(mPas_1) | -0.10591 | 1.89E-07 |
| B16 | B193 | CD3+T | K | -0.10334 | 3.71E-07 |
| B16 | B178 | CD3+T | Ig-U | -0.10225 | 4.93E-07 |
| B160 | B163 | PH | SG | -0.18538 | 0 |
| B160 | B192 | PH | UA | -0.22048 | 0 |
| B160 | B206 | PH | Waist | -0.10234 | 4.81E-07 |
| B161 | B173 | LEU | NTT | 0.252481 | 0 |
| B161 | B221 | LEU | Hei | -0.11339 | 2.40E-08 |
| B161 | B168 | LEU | NMP22 | 0.107946 | 1.09E-07 |
| B163 | B166 | SG | Pro | 0.255027 | 0 |
| B163 | B176 | SG | Ucr | 0.290616 | 0 |
| B163 | B204 | SG | Drinking | 0.182217 | 0 |
| B163 | B206 | SG | Waist | 0.228807 | 0 |
| B163 | B210 | SG | Hip | 0.196802 | 0 |
| B163 | B215 | SG | BMI | 0.183427 | 0 |
| B163 | B216 | SG | BMI | 0.325526 | 0 |
| B163 | B221 | SG | Hei | 0.325183 | 0 |
| B163 | B222 | SG | NS | 0.338965 | 0 |
| B163 | B209 | SG | Smoking | 0.14642 | 5.13E-13 |
| B163 | B189 | SG | eGFB | 0.12614 | 5.21E-10 |
| B163 | B170 | SG | U-GLU | 0.123355 | 1.24E-09 |
| B163 | B220 | SG | SBP | -0.11381 | 2.13E-08 |
| B163 | B188 | SG | Ca | -0.10968 | 6.79E-08 |
| B163 | B192 | SG | UA | 0.106523 | 1.60E-07 |
| B165 | B170 | bilirubin | U-GLU | 0.104192 | 2.97E-07 |
| B166 | B191 | Pro | Urea | 0.132235 | 7.24E-11 |
| B166 | B189 | Pro | eGFB | -0.12424 | 9.45E-10 |
| B166 | B176 | Pro | Ucr | 0.108723 | 8.82E-08 |
| B167 | B223 | PC | FH | 0.437891 | 0 |
| B168 | B183 | NMP22 | PTA | 0.208489 | 0 |
| B168 | B196 | NMP22 | TCO2 | 0.107238 | 1.32E-07 |
| B17 | B101 | CD3+CD16+CD56+NK | anti-Sc70 | -0.23281 | 0 |
| B17 | B102 | CD3+CD16+CD56+NK | anti-Sm | -0.23281 | 0 |
| B17 | B103 | CD3+CD16+CD56+NK | anti-SSA/Ro | -0.23281 | 0 |
| B17 | B104 | CD3+CD16+CD56+NK | anti-SSB | -0.23281 | 0 |
| B17 | B105 | CD3+CD16+CD56+NK | anti-U1RNP | -0.23281 | 0 |
| B17 | B106 | CD3+CD16+CD56+NK | anti-P | -0.23281 | 0 |

| | | | | | |
|---|---|---|---|---|---|
| B17 | B107 | CD3+CD16+CD56+NK | AnuA | -0.23281 | 0 |
| B17 | B109 | CD3+CD16+CD56+NK | AMA-M2 | -0.23281 | 0 |
| B17 | B110 | CD3+CD16+CD56+NK | AHA | -0.23281 | 0 |
| B17 | B134 | CD3+CD16+CD56+NK | CHE | 0.203947 | 0 |
| B17 | B168 | CD3+CD16+CD56+NK | NMP22 | 0.47432 | 0 |
| B17 | B182 | CD3+CD16+CD56+NK | TT | 0.173326 | 0 |
| B17 | B183 | CD3+CD16+CD56+NK | PTA | 0.195471 | 0 |
| B17 | B21 | CD3+CD16+CD56+NK | CD3+CD4-CD8- | 0.479454 | 0 |
| B17 | B22 | CD3+CD16+CD56+NK | CD3+CD8+(T) | 0.18064 | 0 |
| B17 | B23 | CD3+CD16+CD56+NK | CD3-CD16+CD56+(NK) | -0.23969 | 0 |
| B17 | B34 | CD3+CD16+CD56+NK | TK | 0.220005 | 0 |
| B17 | B95 | CD3+CD16+CD56+NK | anti-cenpb | -0.23281 | 0 |
| B17 | B98 | CD3+CD16+CD56+NK | anti-Jo-1 | -0.23281 | 0 |
| B17 | B99 | CD3+CD16+CD56+NK | anti-PM-Scl | -0.23281 | 0 |
| B17 | B20 | CD3+CD16+CD56+NK | CD3+CD4+CD8+ | 0.129124 | 2.00E-10 |
| B17 | B157 | CD3+CD16+CD56+NK | IgG | -0.12751 | 3.37E-10 |
| B17 | B193 | CD3+CD16+CD56+NK | K | -0.12368 | 1.12E-09 |
| B17 | B75 | CD3+CD16+CD56+NK | ApoA1/B | -0.10979 | 6.59E-08 |
| B17 | B74 | CD3+CD16+CD56+NK | ApoA1 | -0.10866 | 8.98E-08 |
| B17 | B61 | CD3+CD16+CD56+NK | WBHSRV | 0.104738 | 2.57E-07 |
| B17 | B55 | CD3+CD16+CD56+NK | TK | 0.104178 | 2.98E-07 |
| B170 | B225 | U-GLU | hPBG | 0.153466 | 3.64E-14 |
| B170 | B190 | U-GLU | Cre | 0.143208 | 1.65E-12 |
| B170 | B191 | U-GLU | Urea | 0.13844 | 8.84E-12 |
| B170 | B175 | U-GLU | ALB-U | 0.126737 | 4.31E-10 |
| B173 | B221 | NTT | Hei | -0.10847 | 9.47E-08 |
| B174 | B189 | Urea/Cr | eGFB | 0.223936 | 0 |
| B174 | B190 | Urea/Cr | Cre | -0.33203 | 0 |
| B174 | B192 | Urea/Cr | UA | -0.20618 | 0 |
| B174 | B216 | Urea/Cr | BMI | -0.22895 | 0 |
| B174 | B221 | Urea/Cr | Hei | -0.34644 | 0 |
| B174 | B222 | Urea/Cr | NS | -0.25343 | 0 |
| B174 | B191 | Urea/Cr | Urea | 0.164794 | 4.44E-16 |
| B174 | B209 | Urea/Cr | Smoking | -0.14114 | 3.43E-12 |
| B174 | B218 | Urea/Cr | DBP | -0.13684 | 1.53E-11 |
| B174 | B206 | Urea/Cr | Waist | -0.10857 | 9.21E-08 |
| B174 | B204 | Urea/Cr | Drinking | -0.10756 | 1.21E-07 |
| B174 | B210 | Urea/Cr | Hip | -0.10223 | 4.95E-07 |
| B175 | B177 | ALB-U | ACR | 0.602928 | 0 |
| B175 | B178 | ALB-U | Ig-U | 0.18126 | 0 |
| B175 | B180 | ALB-U | Tf | 0.204885 | 0 |
| B175 | B191 | ALB-U | Urea | 0.128183 | 2.72E-10 |
| B175 | B190 | ALB-U | Cre | 0.1128 | 2.84E-08 |
| B176 | B222 | Ucr | NS | 0.130923 | 1.12E-10 |

| | | | | | |
|---|---|---|---|---|---|
| B176 | B216 | Ucr | BMI | 0.123004 | 1.39E-09 |
| B176 | B177 | Ucr | ACR | -0.12277 | 1.49E-09 |
| B176 | B179 | Ucr | α1-MG | 0.1144 | 1.80E-08 |
| B176 | B221 | Ucr | Hei | 0.104523 | 2.72E-07 |
| B177 | B178 | ACR | Ig-U | 0.667516 | 0 |
| B177 | B180 | ACR | Tf | 0.815484 | 0 |
| B177 | B191 | ACR | Urea | 0.107331 | 1.29E-07 |
| B177 | B196 | ACR | TCO2 | 0.099489 | 9.95E-07 |
| B178 | B180 | Ig-U | Tf | 0.883872 | 0 |
| B178 | B198 | Ig-U | P | 0.12846 | 2.48E-10 |
| B178 | B196 | Ig-U | TCO2 | 0.105512 | 2.10E-07 |
| B18 | B108 | CD3+CD4+helper(inducer) T | Anti-Cyclin | -0.19775 | 0 |
| B18 | B136 | CD3+CD4+helper(inducer) T | 5'NT | 0.176886 | 0 |
| B18 | B138 | CD3+CD4+helper(inducer) T | PAB | 0.168502 | 0 |
| B18 | B19 | CD3+CD4+helper(inducer) T | CD3+CD4+/CD3+CD8+ | 0.760012 | 0 |
| B18 | B20 | CD3+CD4+helper(inducer) T | CD3+CD4+CD8+ | 0.378531 | 0 |
| B18 | B21 | CD3+CD4+helper(inducer) T | CD3+CD4-CD8- | 0.181505 | 0 |
| B18 | B23 | CD3+CD4+helper(inducer) T | CD3-CD16+CD56+(NK) | -0.8343 | 0 |
| B18 | B34 | CD3+CD4+helper(inducer) T | TK | -0.28874 | 0 |
| B18 | B156 | CD3+CD4+helper(inducer) T | IgE | 0.159844 | 3.11E-15 |
| B18 | B157 | CD3+CD4+helper(inducer) T | IgG | -0.15434 | 2.60E-14 |
| B18 | B79 | CD3+CD4+helper(inducer) T | hb V -dna | 0.140793 | 3.88E-12 |
| B18 | B193 | CD3+CD4+helper(inducer) T | K | -0.13812 | 9.89E-12 |
| B18 | B120 | CD3+CD4+helper(inducer) T | HCV-cAg | 0.134858 | 3.01E-11 |
| B18 | B199 | CD3+CD4+helper(inducer) T | Mg | 0.127985 | 2.89E-10 |
| B18 | B100 | CD3+CD4+helper(inducer) T | anti-Ro52 | 0.126288 | 4.97E-10 |
| B18 | B61 | CD3+CD4+helper(inducer) T | WBHSRV | -0.12564 | 6.10E-10 |
| B18 | B185 | CD3+CD4+helper(inducer) | PT | -0.10456 | 2.70E-07 |

| | | | | | |
|---|---|---|---|---|---|
| | | T | | | |
| B18 | B56 | CD3+CD4+helper(inducer) T | IR | -0.10429 | 2.90E-07 |
| B18 | B66 | CD3+CD4+helper(inducer) T | WBRSR(mPas_100) | -0.10345 | 3.61E-07 |
| B18 | B67 | CD3+CD4+helper(inducer) T | WBRSR(mPas_50) | -0.10255 | 4.56E-07 |
| B18 | B96 | CD3+CD4+helper(inducer) T | dsDNA (IIF) | -0.10027 | 8.18E-07 |
| B18 | B97 | CD3+CD4+helper(inducer) T | dsDNA | 0.099836 | 9.12E-07 |
| B180 | B196 | Tf | TCO2 | 0.156982 | 9.33E-15 |
| B180 | B198 | Tf | P | 0.109301 | 7.53E-08 |
| B180 | B197 | Tf | AG | -0.10813 | 1.04E-07 |
| B181 | B182 | APTT | TT | -0.23058 | 0 |
| B181 | B184 | APTT | INR | 0.287687 | 0 |
| B181 | B185 | APTT | PT | 0.202073 | 0 |
| B181 | B186 | APTT | PFC | 0.170237 | 0 |
| B181 | B199 | APTT | Mg | -0.12412 | 9.82E-10 |
| B181 | B187 | APTT | NSE | -0.12084 | 2.69E-09 |
| B182 | B183 | TT | PTA | -0.36502 | 0 |
| B182 | B184 | TT | INR | -0.20838 | 0 |
| B182 | B186 | TT | PFC | -0.31672 | 0 |
| B182 | B187 | TT | NSE | 0.298947 | 0 |
| B183 | B197 | PTA | AG | -0.13733 | 1.30E-11 |
| B183 | B185 | PTA | PT | -0.12957 | 1.73E-10 |
| B183 | B194 | PTA | Cl | 0.113532 | 2.30E-08 |
| B183 | B193 | PTA | K | -0.10616 | 1.77E-07 |
| B184 | B185 | INR | PT | 0.844178 | 0 |
| B184 | B187 | INR | NSE | -0.20854 | 0 |
| B186 | B198 | PFC | P | 0.115688 | 1.24E-08 |
| B187 | B194 | NSE | Cl | -0.15823 | 5.77E-15 |
| B187 | B204 | NSE | Drinking | 0.133501 | 4.75E-11 |
| B189 | B190 | eGFB | Cre | -0.54 | 0 |
| B189 | B191 | eGFB | Urea | -0.58098 | 0 |
| B189 | B220 | eGFB | SBP | -0.17508 | 0 |
| B189 | B192 | eGFB | UA | -0.16207 | 1.33E-15 |
| B189 | B204 | eGFB | Drinking | 0.132664 | 6.28E-11 |
| B189 | B216 | eGFB | BMI | 0.127807 | 3.06E-10 |
| B189 | B224 | eGFB | pmh | 0.123838 | 1.07E-09 |
| B189 | B215 | eGFB | BMI | 0.121924 | 1.93E-09 |
| B189 | B218 | eGFB | DBP | 0.11352 | 2.31E-08 |
| B189 | B209 | eGFB | Smoking | 0.112788 | 2.85E-08 |
| B189 | B210 | eGFB | Hip | 0.109667 | 6.81E-08 |

| | | | | | |
|---|---|---|---|---|---|
| B19 | B108 | CD3+CD4+/CD3+CD8+ | Anti-Cyclin | -0.23217 | 0 |
| B19 | B185 | CD3+CD4+/CD3+CD8+ | PT | -0.18689 | 0 |
| B19 | B21 | CD3+CD4+/CD3+CD8+ | CD3+CD4-CD8- | 0.296409 | 0 |
| B19 | B22 | CD3+CD4+/CD3+CD8+ | CD3+CD8+(T) | -0.60636 | 0 |
| B19 | B23 | CD3+CD4+/CD3+CD8+ | CD3-CD16+CD56+(NK) | -0.48142 | 0 |
| B19 | B34 | CD3+CD4+/CD3+CD8+ | TK | -0.24627 | 0 |
| B19 | B184 | CD3+CD4+/CD3+CD8+ | INR | -0.14749 | 3.46E-13 |
| B19 | B183 | CD3+CD4+/CD3+CD8+ | PTA | 0.145623 | 6.87E-13 |
| B19 | B55 | CD3+CD4+/CD3+CD8+ | TK | -0.14424 | 1.14E-12 |
| B19 | B157 | CD3+CD4+/CD3+CD8+ | IgG | -0.14307 | 1.73E-12 |
| B19 | B61 | CD3+CD4+/CD3+CD8+ | WBHSRV | -0.13519 | 2.69E-11 |
| B19 | B56 | CD3+CD4+/CD3+CD8+ | IR | -0.13471 | 3.17E-11 |
| B19 | B168 | CD3+CD4+/CD3+CD8+ | NMP22 | -0.13379 | 4.31E-11 |
| B19 | B136 | CD3+CD4+/CD3+CD8+ | 5'NT | 0.128645 | 2.34E-10 |
| B19 | B150 | CD3+CD4+/CD3+CD8+ | ALP | 0.125378 | 6.62E-10 |
| B19 | B135 | CD3+CD4+/CD3+CD8+ | TBA | 0.118286 | 5.78E-09 |
| B19 | B79 | CD3+CD4+/CD3+CD8+ | hb V -dna | 0.11093 | 4.80E-08 |
| B19 | B57 | CD3+CD4+/CD3+CD8+ | RBCAI | 0.099858 | 9.06E-07 |
| B190 | B191 | Cre | Urea | 0.713757 | 0 |
| B190 | B192 | Cre | UA | 0.107477 | 1.24E-07 |
| B190 | B221 | Cre | Hei | 0.103131 | 3.92E-07 |
| B191 | B192 | Urea | UA | 0.181896 | 0 |
| B192 | B206 | UA | Waist | 0.259668 | 0 |
| B192 | B210 | UA | Hip | 0.182143 | 0 |
| B192 | B215 | UA | BMI | 0.204021 | 0 |
| B192 | B216 | UA | BMI | 0.30417 | 0 |
| B192 | B221 | UA | Hei | 0.26881 | 0 |
| B192 | B222 | UA | NS | 0.322479 | 0 |
| B192 | B204 | UA | Drinking | 0.108415 | 9.60E-08 |
| B192 | B209 | UA | Smoking | 0.103173 | 3.88E-07 |
| B192 | B218 | UA | DBP | 0.103034 | 4.02E-07 |
| B193 | B198 | K | P | 0.162423 | 8.88E-16 |
| B193 | B194 | K | Cl | 0.139322 | 6.51E-12 |
| B193 | B216 | K | BMI | -0.10853 | 9.31E-08 |
| B194 | B195 | Cl | Na | 0.362749 | 0 |
| B194 | B196 | Cl | TCO2 | -0.20663 | 0 |
| B194 | B197 | Cl | AG | -0.3444 | 0 |
| B194 | B200 | Cl | OP | 0.348935 | 0 |
| B195 | B197 | Na | AG | 0.391611 | 0 |
| B195 | B200 | Na | OP | 0.869464 | 0 |
| B195 | B198 | Na | P | -0.10024 | 8.24E-07 |
| B196 | B197 | TCO2 | AG | -0.59453 | 0 |
| B197 | B200 | AG | OP | 0.382917 | 0 |
| B2 | B126 | Age | Alb | -0.25844 | 0 |

| | | | | | |
|---|---|---|---|---|---|
| B2 | B127 | Age | A-G ratio | -0.22291 | 0 |
| B2 | B128 | Age | ALT | -0.23584 | 0 |
| B2 | B141 | Age | AST/ALT | 0.341791 | 0 |
| B2 | B163 | Age | SG | -0.17555 | 0 |
| B2 | B189 | Age | eGFB | -0.34902 | 0 |
| B2 | B204 | Age | Drinking | -0.22245 | 0 |
| B2 | B209 | Age | Smoking | -0.1723 | 0 |
| B2 | B210 | Age | Hip | -0.20109 | 0 |
| B2 | B215 | Age | BMI | -0.23169 | 0 |
| B2 | B216 | Age | BMI | -0.40514 | 0 |
| B2 | B218 | Age | DBP | -0.39005 | 0 |
| B2 | B219 | Age | OT | 0.183354 | 0 |
| B2 | B220 | Age | SBP | 0.242934 | 0 |
| B2 | B221 | Age | Hei | -0.39067 | 0 |
| B2 | B222 | Age | NS | -0.27863 | 0 |
| B2 | B43 | Age | MCV | 0.173887 | 0 |
| B2 | B46 | Age | RBC Count | -0.30415 | 0 |
| B2 | B47 | Age | RDW | 0.25711 | 0 |
| B2 | B51 | Age | Hb | -0.27305 | 0 |
| B2 | B71 | Age | TG | -0.2184 | 0 |
| B2 | B72 | Age | HDL-C | 0.201298 | 0 |
| B2 | B45 | Age | MCHC | -0.16232 | 1.11E-15 |
| B2 | B130 | Age | GLB | 0.149323 | 1.75E-13 |
| B2 | B74 | Age | ApoA1 | 0.138099 | 9.95E-12 |
| B2 | B176 | Age | Ucr | -0.13404 | 3.97E-11 |
| B2 | B41 | Age | MON％ | 0.133798 | 4.30E-11 |
| B2 | B134 | Age | CHE | -0.13196 | 7.92E-11 |
| B2 | B174 | Age | Urea/Cr | 0.129419 | 1.82E-10 |
| B2 | B206 | Age | Waist | -0.12083 | 2.69E-09 |
| B2 | B191 | Age | Urea | 0.118245 | 5.85E-09 |
| B2 | B138 | Age | PAB | -0.11407 | 1.98E-08 |
| B2 | B146 | Age | CysC | 0.109823 | 6.52E-08 |
| B2 | B140 | Age | r-GT | -0.1067 | 1.53E-07 |
| B2 | B192 | Age | UA | -0.10465 | 2.63E-07 |
| B2 | B211 | Age | IOP-R | -0.10384 | 3.26E-07 |
| B2 | B193 | Age | K | 0.10201 | 5.24E-07 |
| B2 | B228 | Age | OS | 0.101711 | 5.66E-07 |
| B2 | B212 | Age | IOP-L | -0.10137 | 6.17E-07 |
| B20 | B100 | CD3+CD4+CD8+ | anti-Ro52 | 0.43309 | 0 |
| B20 | B145 | CD3+CD4+CD8+ | BGP | 0.177264 | 0 |
| B20 | B22 | CD3+CD4+CD8+ | CD3+CD8+(T) | 0.287506 | 0 |
| B20 | B23 | CD3+CD4+CD8+ | CD3-CD16+CD56+(NK) | -0.30956 | 0 |
| B20 | B34 | CD3+CD4+CD8+ | TK | -0.14955 | 1.61E-13 |
| B20 | B193 | CD3+CD4+CD8+ | K | -0.13301 | 5.61E-11 |

| | | | | | |
|---|---|---|---|---|---|
| B20 | B101 | CD3+CD4+CD8+ | anti-Sc70 | -0.12954 | 1.75E-10 |
| B20 | B102 | CD3+CD4+CD8+ | anti-Sm | -0.12954 | 1.75E-10 |
| B20 | B103 | CD3+CD4+CD8+ | anti-SSA/Ro | -0.12954 | 1.75E-10 |
| B20 | B104 | CD3+CD4+CD8+ | anti-SSB | -0.12954 | 1.75E-10 |
| B20 | B105 | CD3+CD4+CD8+ | anti-U1RNP | -0.12954 | 1.75E-10 |
| B20 | B106 | CD3+CD4+CD8+ | anti-P | -0.12954 | 1.75E-10 |
| B20 | B107 | CD3+CD4+CD8+ | AnuA | -0.12954 | 1.75E-10 |
| B20 | B109 | CD3+CD4+CD8+ | AMA-M2 | -0.12954 | 1.75E-10 |
| B20 | B110 | CD3+CD4+CD8+ | AHA | -0.12954 | 1.75E-10 |
| B20 | B95 | CD3+CD4+CD8+ | anti-cenpb | -0.12954 | 1.75E-10 |
| B20 | B98 | CD3+CD4+CD8+ | anti-Jo-1 | -0.12954 | 1.75E-10 |
| B20 | B99 | CD3+CD4+CD8+ | anti-PM-Scl | -0.12954 | 1.75E-10 |
| B20 | B181 | CD3+CD4+CD8+ | APTT | 0.107711 | 1.16E-07 |
| B20 | B138 | CD3+CD4+CD8+ | PAB | 0.100345 | 8.02E-07 |
| B201 | B204 | e-habits | Drinking | 0.110432 | 5.51E-08 |
| B202 | B223 | e-times | FH | 0.720422 | 0 |
| B203 | B216 | DH | BMI | 0.111089 | 4.59E-08 |
| B203 | B222 | DH | NS | 0.108377 | 9.70E-08 |
| B204 | B209 | Drinking | Smoking | 0.365163 | 0 |
| B204 | B216 | Drinking | BMI | 0.239461 | 0 |
| B204 | B221 | Drinking | Hei | 0.244496 | 0 |
| B204 | B222 | Drinking | NS | 0.250103 | 0 |
| B204 | B206 | Drinking | Waist | 0.165865 | 2.22E-16 |
| B204 | B215 | Drinking | BMI | 0.127978 | 2.90E-10 |
| B204 | B220 | Drinking | SBP | -0.12777 | 3.10E-10 |
| B204 | B210 | Drinking | Hip | 0.118124 | 6.06E-09 |
| B206 | B207 | Waist | WHR | 0.288734 | 0 |
| B206 | B210 | Waist | Hip | 0.766652 | 0 |
| B206 | B215 | Waist | BMI | 0.810302 | 0 |
| B206 | B216 | Waist | BMI | 0.787505 | 0 |
| B206 | B221 | Waist | Hei | 0.354862 | 0 |
| B206 | B222 | Waist | NS | 0.672992 | 0 |
| B206 | B209 | Waist | Smoking | 0.161339 | 1.78E-15 |
| B207 | B215 | WHR | BMI | 0.177509 | 0 |
| B207 | B222 | WHR | NS | 0.159976 | 2.66E-15 |
| B207 | B216 | WHR | BMI | 0.155481 | 1.67E-14 |
| B209 | B216 | Smoking | BMI | 0.21781 | 0 |
| B209 | B221 | Smoking | Hei | 0.249973 | 0 |
| B209 | B222 | Smoking | NS | 0.259497 | 0 |
| B209 | B210 | Smoking | Hip | 0.13017 | 1.43E-10 |
| B209 | B220 | Smoking | SBP | -0.12657 | 4.55E-10 |
| B21 | B168 | CD3+CD4-CD8- | NMP22 | 0.242911 | 0 |
| B21 | B183 | CD3+CD4-CD8- | PTA | 0.258563 | 0 |
| B21 | B23 | CD3+CD4-CD8- | CD3-CD16+CD56+(NK) | -0.40534 | 0 |

| | | | | | |
|---|---|---|---|---|---|
| B21 | B136 | CD3+CD4-CD8- | 5'NT | -0.16454 | 4.44E-16 |
| B21 | B79 | CD3+CD4-CD8- | hb V -dna | 0.144205 | 1.15E-12 |
| B21 | B178 | CD3+CD4-CD8- | Ig-U | -0.13421 | 3.75E-11 |
| B21 | B100 | CD3+CD4-CD8- | anti-Ro52 | -0.11824 | 5.86E-09 |
| B210 | B215 | Hip | BMI | 0.763198 | 0 |
| B210 | B216 | Hip | BMI | 0.78725 | 0 |
| B210 | B221 | Hip | Hei | 0.411415 | 0 |
| B210 | B222 | Hip | NS | 0.610502 | 0 |
| B210 | B218 | Hip | DBP | 0.122977 | 1.40E-09 |
| B211 | B212 | IOP-R | IOP-L | 0.80482 | 0 |
| B212 | B214 | IOP-L | CVA-L | 0.126753 | 4.29E-10 |
| B215 | B216 | BMI | BMI | 0.8041 | 0 |
| B215 | B222 | BMI | NS | 0.594225 | 0 |
| B215 | B221 | BMI | Hei | 0.162253 | 1.11E-15 |
| B215 | B218 | BMI | DBP | 0.137418 | 1.26E-11 |
| B216 | B218 | BMI | DBP | 0.228877 | 0 |
| B216 | B221 | BMI | Hei | 0.710876 | 0 |
| B216 | B222 | BMI | NS | 0.778671 | 0 |
| B216 | B219 | BMI | OT | -0.12661 | 4.48E-10 |
| B216 | B220 | BMI | SBP | -0.0995 | 9.93E-07 |
| B218 | B220 | DBP | SBP | 0.439644 | 0 |
| B218 | B221 | DBP | Hei | 0.216644 | 0 |
| B218 | B222 | DBP | NS | 0.19988 | 0 |
| B219 | B222 | OT | NS | -0.11787 | 6.54E-09 |
| B219 | B221 | OT | Hei | -0.10677 | 1.50E-07 |
| B22 | B108 | CD3+CD8+(T) | Anti-Cyclin | 0.187123 | 0 |
| B22 | B23 | CD3+CD8+(T) | CD3-CD16+CD56+(NK) | -0.34367 | 0 |
| B22 | B96 | CD3+CD8+(T) | dsDNA (IIF) | -0.24279 | 0 |
| B22 | B97 | CD3+CD8+(T) | dsDNA | 0.244195 | 0 |
| B22 | B124 | CD3+CD8+(T) | ACTH | 0.155672 | 1.55E-14 |
| B22 | B57 | CD3+CD8+(T) | RBCAI | -0.15489 | 2.09E-14 |
| B22 | B55 | CD3+CD8+(T) | TK | 0.143024 | 1.76E-12 |
| B22 | B168 | CD3+CD8+(T) | NMP22 | 0.133883 | 4.18E-11 |
| B22 | B137 | CD3+CD8+(T) | AFU | -0.1178 | 6.67E-09 |
| B22 | B185 | CD3+CD8+(T) | PT | 0.114734 | 1.63E-08 |
| B22 | B183 | CD3+CD8+(T) | PTA | -0.11379 | 2.14E-08 |
| B22 | B81 | CD3+CD8+(T) | TSGF | 0.110638 | 5.20E-08 |
| B22 | B148 | CD3+CD8+(T) | Anti_TPO | 0.106043 | 1.82E-07 |
| B22 | B156 | CD3+CD8+(T) | IgE | 0.104106 | 3.04E-07 |
| B22 | B56 | CD3+CD8+(T) | IR | 0.099776 | 9.25E-07 |
| B220 | B221 | SBP | Hei | -0.17178 | 0 |
| B221 | B222 | Hei | NS | 0.59343 | 0 |
| B227 | B228 | OD | OS | 0.158272 | 5.55E-15 |
| B23 | B138 | CD3-CD16+CD56+(NK) | PAB | -0.21055 | 0 |

| | | | | | |
|---|---|---|---|---|---|
| B23 | B120 | CD3-CD16+CD56+(NK) | HCV-cAg | -0.16642 | 2.22E-16 |
| B23 | B156 | CD3-CD16+CD56+(NK) | IgE | -0.16057 | 2.22E-15 |
| B23 | B79 | CD3-CD16+CD56+(NK) | hb V -dna | -0.15585 | 1.44E-14 |
| B23 | B157 | CD3-CD16+CD56+(NK) | IgG | 0.146358 | 5.25E-13 |
| B23 | B34 | CD3-CD16+CD56+(NK) | TK | 0.139482 | 6.15E-12 |
| B23 | B73 | CD3-CD16+CD56+(NK) | ApoA | 0.113106 | 2.60E-08 |
| B23 | B199 | CD3-CD16+CD56+(NK) | Mg | -0.11289 | 2.77E-08 |
| B23 | B100 | CD3-CD16+CD56+(NK) | anti-Ro52 | -0.10687 | 1.46E-07 |
| B23 | B65 | CD3-CD16+CD56+(NK) | WBRSR(mPas_1) | 0.106059 | 1.81E-07 |
| B23 | B181 | CD3-CD16+CD56+(NK) | APTT | 0.104774 | 2.55E-07 |
| B23 | B64 | CD3-CD16+CD56+(NK) | WBRSR(mPas_5) | 0.099688 | 9.46E-07 |
| B24 | B123 | αHBDH | hs-CRP1 | 0.307921 | 0 |
| B24 | B138 | αHBDH | PAB | -0.23447 | 0 |
| B24 | B139 | αHBDH | ADA | 0.230936 | 0 |
| B24 | B186 | αHBDH | PFC | 0.257983 | 0 |
| B24 | B26 | αHBDH | CKMB | 0.245089 | 0 |
| B24 | B27 | αHBDH | LDH | 0.796197 | 0 |
| B24 | B37 | αHBDH | BAND | -0.39063 | 0 |
| B24 | B38 | αHBDH | BAND% | -0.42824 | 0 |
| B24 | B39 | αHBDH | Other cells | 0.338369 | 0 |
| B24 | B73 | αHBDH | ApoA | 0.16955 | 0 |
| B24 | B187 | αHBDH | NSE | 0.134931 | 2.94E-11 |
| B24 | B134 | αHBDH | CHE | -0.12152 | 2.18E-09 |
| B24 | B146 | αHBDH | CysC | 0.110273 | 5.76E-08 |
| B24 | B157 | αHBDH | IgG | 0.102855 | 4.21E-07 |
| B24 | B193 | αHBDH | K | -0.10281 | 4.26E-07 |
| B24 | B199 | αHBDH | Mg | -0.10248 | 4.64E-07 |
| B24 | B150 | αHBDH | ALP | 0.102137 | 5.07E-07 |
| B24 | B184 | αHBDH | INR | -0.10183 | 5.49E-07 |
| B25 | B114 | CK | TSH TT3 TT4(TT4) | -0.13458 | 3.31E-11 |
| B25 | B168 | CK | NMP22 | 0.133446 | 4.84E-11 |
| B25 | B196 | CK | TCO2 | 0.115805 | 1.20E-08 |
| B25 | B179 | CK | α1-MG | -0.113 | 2.68E-08 |
| B25 | B187 | CK | NSE | 0.112041 | 3.51E-08 |
| B25 | B186 | CK | PFC | 0.110986 | 4.72E-08 |
| B26 | B27 | CKMB | LDH | 0.212069 | 0 |
| B26 | B186 | CKMB | PFC | 0.113461 | 2.35E-08 |
| B27 | B123 | LDH | hs-CRP1 | 0.275006 | 0 |
| B27 | B138 | LDH | PAB | -0.26504 | 0 |
| B27 | B139 | LDH | ADA | 0.28772 | 0 |
| B27 | B150 | LDH | ALP | 0.178723 | 0 |
| B27 | B186 | LDH | PFC | 0.172378 | 0 |
| B27 | B37 | LDH | BAND | -0.39784 | 0 |
| B27 | B38 | LDH | BAND% | -0.43284 | 0 |

| | | | | | |
|---|---|---|---|---|---|
| B27 | B39 | LDH | Other cells | 0.331287 | 0 |
| B27 | B73 | LDH | ApoA | 0.161386 | 1.55E-15 |
| B27 | B134 | LDH | CHE | -0.12519 | 7.02E-10 |
| B27 | B136 | LDH | 5'NT | 0.117395 | 7.52E-09 |
| B27 | B187 | LDH | NSE | 0.113425 | 2.38E-08 |
| B27 | B176 | LDH | Ucr | -0.11137 | 4.24E-08 |
| B27 | B194 | LDH | Cl | 0.103742 | 3.34E-07 |
| B28 | B125 | PRL | Cor | 0.449867 | 0 |
| B28 | B29 | PRL | E2 | 0.544185 | 0 |
| B28 | B30 | PRL | LH | 0.326351 | 0 |
| B28 | B31 | PRL | FSH | -0.90913 | 0 |
| B28 | B32 | PRL | Testo | 0.446104 | 0 |
| B28 | B33 | PRL | Prog | 0.413029 | 0 |
| B28 | B139 | PRL | ADA | -0.12896 | 2.11E-10 |
| B29 | B30 | E2 | LH | 0.962659 | 0 |
| B29 | B31 | E2 | FSH | -0.6359 | 0 |
| B29 | B32 | E2 | Testo | -0.40567 | 0 |
| B29 | B33 | E2 | Prog | -0.44299 | 0 |
| B29 | B125 | E2 | Cor | 0.134263 | 3.68E-11 |
| B3 | B26 | Hcy | CKMB | 0.247306 | 0 |
| B3 | B189 | Hcy | eGFB | -0.16014 | 2.66E-15 |
| B3 | B74 | Hcy | ApoA1 | -0.1528 | 4.71E-14 |
| B3 | B62 | Hcy | WBHSRVI | 0.137044 | 1.43E-11 |
| B3 | B192 | Hcy | UA | 0.135109 | 2.77E-11 |
| B3 | B175 | Hcy | ALB-U | 0.126992 | 3.97E-10 |
| B3 | B66 | Hcy | WBRSR(mPas_100) | 0.123735 | 1.11E-09 |
| B3 | B67 | Hcy | WBRSR(mPas_50) | 0.123174 | 1.31E-09 |
| B3 | B63 | Hcy | WBRSR(mPas_200) | 0.120469 | 3.00E-09 |
| B3 | B174 | Hcy | Urea/Cr | -0.11864 | 5.21E-09 |
| B3 | B64 | Hcy | WBRSR(mPas_5) | 0.114625 | 1.68E-08 |
| B3 | B60 | Hcy | WBRVI | 0.112531 | 3.06E-08 |
| B3 | B190 | Hcy | Cre | 0.112449 | 3.13E-08 |
| B3 | B191 | Hcy | Urea | 0.111991 | 3.56E-08 |
| B3 | B56 | Hcy | IR | 0.110185 | 5.90E-08 |
| B3 | B34 | Hcy | TK | 0.104453 | 2.78E-07 |
| B3 | B185 | Hcy | PT | 0.101542 | 5.91E-07 |
| B3 | B58 | Hcy | HCT | 0.101533 | 5.92E-07 |
| B3 | B65 | Hcy | WBRSR(mPas_1) | 0.099742 | 9.33E-07 |
| B30 | B31 | LH | FSH | -0.40323 | 0 |
| B30 | B32 | LH | Testo | -0.63796 | 0 |
| B30 | B33 | LH | Prog | -0.66916 | 0 |
| B31 | B125 | FSH | Cor | -0.484 | 0 |
| B31 | B32 | FSH | Testo | -0.44745 | 0 |
| B31 | B33 | FSH | Prog | -0.41021 | 0 |

| | | | | | |
|---|---|---|---|---|---|
| B31 | B139 | FSH | ADA | 0.1416 | 2.92E-12 |
| B32 | B125 | Testo | Cor | 0.417625 | 0 |
| B32 | B33 | Testo | Prog | 0.99915 | 0 |
| B32 | B139 | Testo | ADA | -0.12054 | 2.94E-09 |
| B33 | B125 | Prog | Cor | 0.403578 | 0 |
| B33 | B139 | Prog | ADA | -0.1164 | 1.01E-08 |
| B34 | B125 | TK | Cor | -0.17152 | 0 |
| B34 | B164 | TK | COM | 0.174645 | 0 |
| B34 | B186 | TK | PFC | 0.146863 | 4.36E-13 |
| B34 | B183 | TK | PTA | -0.14464 | 9.81E-13 |
| B34 | B138 | TK | PAB | -0.11949 | 4.04E-09 |
| B34 | B55 | TK | TK | 0.118639 | 5.20E-09 |
| B34 | B182 | TK | TT | 0.118636 | 5.21E-09 |
| B34 | B198 | TK | P | -0.11142 | 4.19E-08 |
| B34 | B139 | TK | ADA | 0.108485 | 9.42E-08 |
| B34 | B156 | TK | IgE | -0.10625 | 1.72E-07 |
| B34 | B57 | TK | RBCAI | -0.10258 | 4.53E-07 |
| B35 | B202 | β2-MG | e-times | 0.538982 | 0 |
| B35 | B223 | β2-MG | FH | 0.825439 | 0 |
| B37 | B123 | BAND | hs-CRP1 | -0.23548 | 0 |
| B37 | B138 | BAND | PAB | 0.186257 | 0 |
| B37 | B139 | BAND | ADA | -0.28484 | 0 |
| B37 | B157 | BAND | IgG | -0.17314 | 0 |
| B37 | B38 | BAND | BAND% | 0.993477 | 0 |
| B37 | B39 | BAND | Other cells | -0.50021 | 0 |
| B37 | B73 | BAND | ApoA | -0.1957 | 0 |
| B37 | B74 | BAND | ApoA1 | 0.162601 | 8.88E-16 |
| B37 | B199 | BAND | Mg | 0.123656 | 1.13E-09 |
| B37 | B134 | BAND | CHE | 0.117923 | 6.43E-09 |
| B37 | B120 | BAND | HCV-cAg | 0.113879 | 2.09E-08 |
| B38 | B123 | BAND% | hs-CRP1 | -0.25677 | 0 |
| B38 | B138 | BAND% | PAB | 0.202942 | 0 |
| B38 | B139 | BAND% | ADA | -0.31059 | 0 |
| B38 | B157 | BAND% | IgG | -0.18865 | 0 |
| B38 | B39 | BAND% | Other cells | -0.54505 | 0 |
| B38 | B73 | BAND% | ApoA | -0.21049 | 0 |
| B38 | B74 | BAND% | ApoA1 | 0.163908 | 4.44E-16 |
| B38 | B199 | BAND% | Mg | 0.134749 | 3.12E-11 |
| B38 | B134 | BAND% | CHE | 0.128344 | 2.58E-10 |
| B38 | B120 | BAND% | HCV-cAg | 0.124476 | 8.78E-10 |
| B39 | B123 | Other cells | hs-CRP1 | 0.19871 | 0 |
| B39 | B139 | Other cells | ADA | 0.241409 | 0 |
| B39 | B138 | Other cells | PAB | -0.15983 | 3.11E-15 |
| B39 | B73 | Other cells | ApoA | 0.157543 | 7.55E-15 |

| | | | | | |
|---|---|---|---|---|---|
| B39 | B157 | Other cells | IgG | 0.14846 | 2.42E-13 |
| B39 | B45 | Other cells | MCHC | -0.11906 | 4.59E-09 |
| B39 | B199 | Other cells | Mg | -0.10495 | 2.44E-07 |
| B39 | B134 | Other cells | CHE | -0.10408 | 3.06E-07 |
| B4 | B6 | WB-Ca | WB-Fe | 1 | 0 |
| B4 | B8 | WB-Ca | WB-Zinc | -0.9999 | 0 |
| B4 | B86 | WB-Ca | TBhCG | 0.116588 | 9.53E-09 |
| B40 | B41 | WBC | MON％ | -0.20705 | 0 |
| B40 | B42 | WBC | MON | 0.621687 | 0 |
| B40 | B46 | WBC | RBC Count | 0.170609 | 0 |
| B40 | B206 | WBC | Waist | 0.163778 | 4.44E-16 |
| B40 | B52 | WBC | PLT/BPC | 0.160443 | 2.22E-15 |
| B40 | B192 | WBC | UA | 0.145719 | 6.63E-13 |
| B40 | B215 | WBC | BMI | 0.140135 | 4.90E-12 |
| B40 | B51 | WBC | Hb | 0.132534 | 6.56E-11 |
| B40 | B222 | WBC | NS | 0.131583 | 8.98E-11 |
| B40 | B209 | WBC | Smoking | 0.131013 | 1.08E-10 |
| B40 | B53 | WBC | GR% | 0.125424 | 6.52E-10 |
| B40 | B49 | WBC | baso% | -0.12244 | 1.65E-09 |
| B40 | B160 | WBC | PH | -0.12032 | 3.15E-09 |
| B40 | B71 | WBC | TG | 0.118486 | 5.44E-09 |
| B40 | B153 | WBC | Glu | 0.117257 | 7.83E-09 |
| B40 | B216 | WBC | BMI | 0.113626 | 2.24E-08 |
| B40 | B123 | WBC | hs-CRP1 | 0.113299 | 2.46E-08 |
| B40 | B72 | WBC | HDL-C | -0.10524 | 2.26E-07 |
| B41 | B42 | MON％ | MON | 0.577218 | 0 |
| B41 | B53 | MON％ | GR% | -0.16256 | 8.88E-16 |
| B41 | B126 | MON％ | Alb | -0.16164 | 1.33E-15 |
| B41 | B47 | MON％ | RDW | 0.120588 | 2.90E-09 |
| B41 | B71 | MON％ | TG | -0.11312 | 2.59E-08 |
| B41 | B43 | MON％ | MCV | 0.109208 | 7.73E-08 |
| B41 | B49 | MON％ | baso% | 0.099836 | 9.12E-07 |
| B41 | B44 | MON％ | MCH | 0.099823 | 9.15E-07 |
| B42 | B48 | MON | LYM% | -0.1656 | 4.44E-16 |
| B42 | B123 | MON | hs-CRP1 | 0.158711 | 4.66E-15 |
| B42 | B209 | MON | Smoking | 0.157028 | 9.10E-15 |
| B42 | B192 | MON | UA | 0.155404 | 1.73E-14 |
| B42 | B222 | MON | NS | 0.154722 | 2.24E-14 |
| B42 | B206 | MON | Waist | 0.145337 | 7.62E-13 |
| B42 | B179 | MON | α1-MG | 0.144371 | 1.08E-12 |
| B42 | B51 | MON | Hb | 0.141194 | 3.37E-12 |
| B42 | B52 | MON | PLT/BPC | 0.138881 | 7.59E-12 |
| B42 | B46 | MON | RBC Count | 0.125786 | 5.82E-10 |
| B42 | B126 | MON | Alb | -0.1191 | 4.54E-09 |

| | | | | | |
|---|---|---|---|---|---|
| B42 | B127 | MON | A-G ratio | -0.11887 | 4.85E-09 |
| B42 | B216 | MON | BMI | 0.108748 | 8.76E-08 |
| B42 | B163 | MON | SG | 0.099991 | 8.77E-07 |
| B43 | B44 | MCV | MCH | 0.904668 | 0 |
| B43 | B46 | MCV | RBC Count | -0.52075 | 0 |
| B43 | B47 | MCV | RDW | 0.587671 | 0 |
| B43 | B45 | MCV | MCHC | 0.115941 | 1.15E-08 |
| B43 | B52 | MCV | PLT/BPC | -0.11502 | 1.50E-08 |
| B44 | B45 | MCH | MCHC | 0.525735 | 0 |
| B44 | B46 | MCH | RBC Count | -0.39641 | 0 |
| B44 | B47 | MCH | RDW | 0.402686 | 0 |
| B44 | B51 | MCH | Hb | 0.253385 | 0 |
| B44 | B204 | MCH | Drinking | 0.131498 | 9.24E-11 |
| B44 | B140 | MCH | r-GT | 0.123093 | 1.35E-09 |
| B44 | B132 | MCH | Dbil | 0.122753 | 1.50E-09 |
| B44 | B131 | MCH | AST | 0.105161 | 2.30E-07 |
| B44 | B133 | MCH | TBIL | 0.101598 | 5.82E-07 |
| B45 | B129 | MCHC | IBIL | 0.207315 | 0 |
| B45 | B133 | MCHC | TBIL | 0.203798 | 0 |
| B45 | B163 | MCHC | SG | 0.171746 | 0 |
| B45 | B216 | MCHC | BMI | 0.241256 | 0 |
| B45 | B221 | MCHC | Hei | 0.235166 | 0 |
| B45 | B222 | MCHC | NS | 0.255623 | 0 |
| B45 | B47 | MCHC | RDW | -0.23988 | 0 |
| B45 | B51 | MCHC | Hb | 0.47204 | 0 |
| B45 | B204 | MCHC | Drinking | 0.161536 | 1.55E-15 |
| B45 | B71 | MCHC | TG | 0.159123 | 4.00E-15 |
| B45 | B126 | MCHC | Alb | 0.150164 | 1.28E-13 |
| B45 | B189 | MCHC | eGFB | 0.148323 | 2.54E-13 |
| B45 | B193 | MCHC | K | -0.14802 | 2.84E-13 |
| B45 | B132 | MCHC | Dbil | 0.142521 | 2.10E-12 |
| B45 | B141 | MCHC | AST/ALT | -0.14161 | 2.91E-12 |
| B45 | B215 | MCHC | BMI | 0.140559 | 4.22E-12 |
| B45 | B210 | MCHC | Hip | 0.140522 | 4.27E-12 |
| B45 | B206 | MCHC | Waist | 0.136816 | 1.55E-11 |
| B45 | B218 | MCHC | DBP | 0.135901 | 2.11E-11 |
| B45 | B72 | MCHC | HDL-C | -0.12923 | 1.94E-10 |
| B45 | B153 | MCHC | Glu | 0.124956 | 7.56E-10 |
| B45 | B127 | MCHC | A-G ratio | 0.122515 | 1.61E-09 |
| B45 | B174 | MCHC | Urea/Cr | -0.11867 | 5.16E-09 |
| B45 | B46 | MCHC | RBC Count | 0.115343 | 1.37E-08 |
| B45 | B128 | MCHC | ALT | 0.110865 | 4.89E-08 |
| B45 | B140 | MCHC | r-GT | 0.110602 | 5.26E-08 |
| B45 | B192 | MCHC | UA | 0.108152 | 1.03E-07 |

| | | | | | |
|---|---|---|---|---|---|
| B45 | B209 | MCHC | Smoking | 0.102841 | 4.23E-07 |
| B45 | B134 | MCHC | CHE | 0.100999 | 6.79E-07 |
| B45 | B74 | MCHC | ApoA1 | -0.10005 | 8.65E-07 |
| B46 | B126 | RBC Count | Alb | 0.288487 | 0 |
| B46 | B129 | RBC Count | IBIL | 0.196415 | 0 |
| B46 | B133 | RBC Count | TBIL | 0.177596 | 0 |
| B46 | B141 | RBC Count | AST/ALT | -0.2484 | 0 |
| B46 | B163 | RBC Count | SG | 0.234795 | 0 |
| B46 | B189 | RBC Count | eGFB | 0.307872 | 0 |
| B46 | B190 | RBC Count | Cre | -0.22251 | 0 |
| B46 | B191 | RBC Count | Urea | -0.31416 | 0 |
| B46 | B206 | RBC Count | Waist | 0.23235 | 0 |
| B46 | B210 | RBC Count | Hip | 0.206514 | 0 |
| B46 | B215 | RBC Count | BMI | 0.208564 | 0 |
| B46 | B216 | RBC Count | BMI | 0.33451 | 0 |
| B46 | B218 | RBC Count | DBP | 0.265214 | 0 |
| B46 | B221 | RBC Count | Hei | 0.307261 | 0 |
| B46 | B222 | RBC Count | NS | 0.31818 | 0 |
| B46 | B47 | RBC Count | RDW | -0.4379 | 0 |
| B46 | B51 | RBC Count | Hb | 0.777017 | 0 |
| B46 | B58 | RBC Count | HCT | 0.196144 | 0 |
| B46 | B60 | RBC Count | WBRVI | 0.181835 | 0 |
| B46 | B61 | RBC Count | WBHSRV | 0.173638 | 0 |
| B46 | B62 | RBC Count | WBHSRVI | 0.198567 | 0 |
| B46 | B63 | RBC Count | WBRSR(mPas_200) | 0.206211 | 0 |
| B46 | B64 | RBC Count | WBRSR(mPas_5) | 0.206348 | 0 |
| B46 | B65 | RBC Count | WBRSR(mPas_1) | 0.193607 | 0 |
| B46 | B66 | RBC Count | WBRSR(mPas_100) | 0.206324 | 0 |
| B46 | B67 | RBC Count | WBRSR(mPas_50) | 0.207517 | 0 |
| B46 | B72 | RBC Count | HDL-C | -0.20677 | 0 |
| B46 | B209 | RBC Count | Smoking | 0.162533 | 8.88E-16 |
| B46 | B56 | RBC Count | IR | 0.15771 | 7.11E-15 |
| B46 | B128 | RBC Count | ALT | 0.154172 | 2.78E-14 |
| B46 | B174 | RBC Count | Urea/Cr | -0.14982 | 1.45E-13 |
| B46 | B192 | RBC Count | UA | 0.148221 | 2.64E-13 |
| B46 | B127 | RBC Count | A-G ratio | 0.146448 | 5.08E-13 |
| B46 | B91 | RBC Count | TP | 0.130043 | 1.49E-10 |
| B46 | B113 | RBC Count | FT3 | 0.127492 | 3.39E-10 |
| B46 | B70 | RBC Count | LDL-C | 0.12721 | 3.71E-10 |
| B46 | B204 | RBC Count | Drinking | 0.122982 | 1.40E-09 |
| B46 | B71 | RBC Count | TG | 0.122673 | 1.53E-09 |
| B46 | B74 | RBC Count | ApoA1 | -0.12261 | 1.57E-09 |
| B46 | B146 | RBC Count | CysC | -0.11471 | 1.65E-08 |
| B46 | B219 | RBC Count | OT | -0.11383 | 2.12E-08 |

| | | | | | |
|---|---|---|---|---|---|
| B47 | B126 | RDW | Alb | -0.21819 | 0 |
| B47 | B189 | RDW | eGFB | -0.21135 | 0 |
| B47 | B191 | RDW | Urea | 0.176988 | 0 |
| B47 | B51 | RDW | Hb | -0.20012 | 0 |
| B47 | B141 | RDW | AST/ALT | 0.15687 | 9.77E-15 |
| B47 | B52 | RDW | PLT/BPC | -0.15362 | 3.42E-14 |
| B47 | B127 | RDW | A-G ratio | -0.12938 | 1.84E-10 |
| B47 | B72 | RDW | HDL-C | 0.122047 | 1.86E-09 |
| B47 | B146 | RDW | CysC | 0.121146 | 2.45E-09 |
| B47 | B218 | RDW | DBP | -0.12062 | 2.87E-09 |
| B47 | B132 | RDW | Dbil | 0.115684 | 1.24E-08 |
| B47 | B216 | RDW | BMI | -0.11249 | 3.10E-08 |
| B47 | B190 | RDW | Cre | 0.105211 | 2.27E-07 |
| B47 | B71 | RDW | TG | -0.10436 | 2.85E-07 |
| B47 | B193 | RDW | K | 0.099832 | 9.13E-07 |
| B48 | B53 | LYM% | GR% | -0.94391 | 0 |
| B48 | B191 | LYM% | Urea | -0.15224 | 5.82E-14 |
| B48 | B190 | LYM% | Cre | -0.15152 | 7.64E-14 |
| B48 | B192 | LYM% | UA | -0.10223 | 4.95E-07 |
| B49 | B50 | baso% | eo% | 0.295193 | 0 |
| B49 | B53 | baso% | GR% | -0.15187 | 6.71E-14 |
| B49 | B52 | baso% | PLT/BPC | 0.101213 | 6.43E-07 |
| B5 | B8 | WB-Mg | WB-Zinc | -0.99988 | 0 |
| B5 | B86 | WB-Mg | TBhCG | 0.116588 | 9.53E-09 |
| B50 | B53 | eo% | GR% | -0.22844 | 0 |
| B50 | B190 | eo% | Cre | 0.141501 | 3.02E-12 |
| B51 | B113 | Hb | FT3 | 0.171537 | 0 |
| B51 | B126 | Hb | Alb | 0.319265 | 0 |
| B51 | B127 | Hb | A-G ratio | 0.169445 | 0 |
| B51 | B128 | Hb | ALT | 0.209081 | 0 |
| B51 | B129 | Hb | IBIL | 0.255918 | 0 |
| B51 | B133 | Hb | TBIL | 0.246451 | 0 |
| B51 | B141 | Hb | AST/ALT | -0.25757 | 0 |
| B51 | B163 | Hb | SG | 0.288489 | 0 |
| B51 | B174 | Hb | Urea/Cr | -0.17584 | 0 |
| B51 | B189 | Hb | eGFB | 0.327826 | 0 |
| B51 | B190 | Hb | Cre | -0.24165 | 0 |
| B51 | B191 | Hb | Urea | -0.32792 | 0 |
| B51 | B192 | Hb | UA | 0.169577 | 0 |
| B51 | B204 | Hb | Drinking | 0.215816 | 0 |
| B51 | B206 | Hb | Waist | 0.284972 | 0 |
| B51 | B209 | Hb | Smoking | 0.209257 | 0 |
| B51 | B210 | Hb | Hip | 0.257398 | 0 |
| B51 | B215 | Hb | BMI | 0.244856 | 0 |

| | | | | | |
|---|---|---|---|---|---|
| B51 | B216 | Hb | BMI | 0.395306 | 0 |
| B51 | B218 | Hb | DBP | 0.271206 | 0 |
| B51 | B221 | Hb | Hei | 0.367733 | 0 |
| B51 | B222 | Hb | NS | 0.389066 | 0 |
| B51 | B56 | Hb | IR | 0.173198 | 0 |
| B51 | B58 | Hb | HCT | 0.249977 | 0 |
| B51 | B60 | Hb | WBRVI | 0.200537 | 0 |
| B51 | B61 | Hb | WBHSRV | 0.177819 | 0 |
| B51 | B62 | Hb | WBHSRVI | 0.234971 | 0 |
| B51 | B63 | Hb | WBRSR(mPas_200) | 0.233136 | 0 |
| B51 | B64 | Hb | WBRSR(mPas_5) | 0.220944 | 0 |
| B51 | B65 | Hb | WBRSR(mPas_1) | 0.200984 | 0 |
| B51 | B66 | Hb | WBRSR(mPas_100) | 0.22857 | 0 |
| B51 | B67 | Hb | WBRSR(mPas_50) | 0.229256 | 0 |
| B51 | B72 | Hb | HDL-C | -0.20637 | 0 |
| B51 | B132 | Hb | Dbil | 0.157349 | 7.99E-15 |
| B51 | B71 | Hb | TG | 0.152589 | 5.11E-14 |
| B51 | B146 | Hb | CysC | -0.13864 | 8.25E-12 |
| B51 | B74 | Hb | ApoA1 | -0.13775 | 1.12E-11 |
| B51 | B91 | Hb | TP | 0.135808 | 2.18E-11 |
| B51 | B70 | Hb | LDL-C | 0.134604 | 3.28E-11 |
| B51 | B198 | Hb | P | -0.13423 | 3.73E-11 |
| B51 | B138 | Hb | PAB | 0.120743 | 2.77E-09 |
| B51 | B140 | Hb | r-GT | 0.116303 | 1.04E-08 |
| B51 | B134 | Hb | CHE | 0.115676 | 1.24E-08 |
| B51 | B188 | Hb | Ca | 0.114845 | 1.58E-08 |
| B51 | B219 | Hb | OT | -0.10761 | 1.20E-07 |
| B51 | B137 | Hb | AFU | 0.101837 | 5.48E-07 |
| B52 | B132 | PLT/BPC | Dbil | -0.15593 | 1.40E-14 |
| B52 | B70 | PLT/BPC | LDL-C | 0.141992 | 2.54E-12 |
| B52 | B77 | PLT/BPC | TC | 0.122526 | 1.61E-09 |
| B52 | B91 | PLT/BPC | TP | 0.103584 | 3.48E-07 |
| B53 | B153 | GR% | Glu | 0.117747 | 6.78E-09 |
| B53 | B191 | GR% | Urea | 0.113884 | 2.08E-08 |
| B53 | B190 | GR% | Cre | 0.106886 | 1.45E-07 |
| B54 | B123 | ESR | hs-CRP1 | 0.203915 | 0 |
| B54 | B56 | ESR | IR | -0.21395 | 0 |
| B54 | B58 | ESR | HCT | -0.31637 | 0 |
| B54 | B59 | ESR | WBRV | 0.21683 | 0 |
| B54 | B60 | ESR | WBRVI | -0.23027 | 0 |
| B54 | B62 | ESR | WBHSRVI | -0.28067 | 0 |
| B54 | B69 | ESR | PV | 0.529245 | 0 |
| B54 | B130 | ESR | GLB | 0.106664 | 1.54E-07 |
| B55 | B56 | TK | IR | 0.530737 | 0 |

| | | | | | |
|---|---|---|---|---|---|
| B55 | B58 | TK | HCT | -0.31162 | 0 |
| B55 | B59 | TK | WBRV | 0.184581 | 0 |
| B55 | B61 | TK | WBHSRV | 0.399602 | 0 |
| B55 | B69 | TK | PV | -0.30491 | 0 |
| B55 | B62 | TK | WBHSRVI | 0.143731 | 1.36E-12 |
| B55 | B57 | TK | RBCAI | -0.13343 | 4.87E-11 |
| B56 | B58 | IR | HCT | 0.608422 | 0 |
| B56 | B59 | IR | WBRV | 0.312782 | 0 |
| B56 | B60 | IR | WBRVI | 0.737687 | 0 |
| B56 | B61 | IR | WBHSRV | 0.869136 | 0 |
| B56 | B62 | IR | WBHSRVI | 0.912763 | 0 |
| B56 | B63 | IR | WBRSR(mPas_200) | 0.763198 | 0 |
| B56 | B64 | IR | WBRSR(mPas_5) | 0.671782 | 0 |
| B56 | B65 | IR | WBRSR(mPas_1) | 0.589148 | 0 |
| B56 | B66 | IR | WBRSR(mPas_100) | 0.722277 | 0 |
| B56 | B67 | IR | WBRSR(mPas_50) | 0.721271 | 0 |
| B56 | B69 | IR | PV | -0.3403 | 0 |
| B56 | B74 | IR | ApoA1 | -0.13734 | 1.29E-11 |
| B56 | B198 | IR | P | -0.11032 | 5.69E-08 |
| B57 | B59 | RBCAI | WBRV | 0.776236 | 0 |
| B57 | B60 | RBCAI | WBRVI | 0.520699 | 0 |
| B57 | B64 | RBCAI | WBRSR(mPas_5) | 0.33185 | 0 |
| B57 | B65 | RBCAI | WBRSR(mPas_1) | 0.522928 | 0 |
| B57 | B81 | RBCAI | TSGF | -0.14201 | 2.52E-12 |
| B57 | B61 | RBCAI | WBHSRV | -0.12267 | 1.54E-09 |
| B58 | B60 | HCT | WBRVI | 0.744063 | 0 |
| B58 | B61 | HCT | WBHSRV | 0.576455 | 0 |
| B58 | B62 | HCT | WBHSRVI | 0.859298 | 0 |
| B58 | B63 | HCT | WBRSR(mPas_200) | 0.830273 | 0 |
| B58 | B64 | HCT | WBRSR(mPas_5) | 0.785047 | 0 |
| B58 | B65 | HCT | WBRSR(mPas_1) | 0.716028 | 0 |
| B58 | B66 | HCT | WBRSR(mPas_100) | 0.775762 | 0 |
| B58 | B67 | HCT | WBRSR(mPas_50) | 0.780538 | 0 |
| B58 | B69 | HCT | PV | -0.16243 | 8.88E-16 |
| B58 | B74 | HCT | ApoA1 | -0.15365 | 3.40E-14 |
| B58 | B59 | HCT | WBRV | 0.130962 | 1.10E-10 |
| B58 | B187 | HCT | NSE | 0.114846 | 1.58E-08 |
| B58 | B136 | HCT | 5'NT | -0.11379 | 2.14E-08 |
| B58 | B133 | HCT | TBIL | 0.101158 | 6.52E-07 |
| B59 | B60 | WBRV | WBRVI | 0.65564 | 0 |
| B59 | B61 | WBRV | WBHSRV | 0.444501 | 0 |
| B59 | B62 | WBRV | WBHSRVI | 0.267066 | 0 |
| B59 | B63 | WBRV | WBRSR(mPas_200) | 0.369507 | 0 |
| B59 | B64 | WBRV | WBRSR(mPas_5) | 0.65545 | 0 |

| | | | | | |
|---|---|---|---|---|---|
| B59 | B65 | WBRV | WBRSR(mPas_1) | 0.769 | 0 |
| B59 | B66 | WBRV | WBRSR(mPas_100) | 0.456183 | 0 |
| B59 | B67 | WBRV | WBRSR(mPas_50) | 0.494438 | 0 |
| B59 | B69 | WBRV | PV | 0.234233 | 0 |
| B59 | B123 | WBRV | hs-CRP1 | 0.108435 | 9.55E-08 |
| B6 | B8 | WB-Fe | WB-Zinc | -0.9999 | 0 |
| B6 | B86 | WB-Fe | TBhCG | 0.116588 | 9.53E-09 |
| B60 | B61 | WBRVI | WBHSRV | 0.630579 | 0 |
| B60 | B62 | WBRVI | WBHSRVI | 0.830467 | 0 |
| B60 | B63 | WBRVI | WBRSR(mPas_200) | 0.720973 | 0 |
| B60 | B64 | WBRVI | WBRSR(mPas_5) | 0.876253 | 0 |
| B60 | B65 | WBRVI | WBRSR(mPas_1) | 0.906873 | 0 |
| B60 | B66 | WBRVI | WBRSR(mPas_100) | 0.733162 | 0 |
| B60 | B67 | WBRVI | WBRSR(mPas_50) | 0.763574 | 0 |
| B60 | B69 | WBRVI | PV | -0.25192 | 0 |
| B60 | B74 | WBRVI | ApoA1 | -0.15383 | 3.15E-14 |
| B61 | B62 | WBHSRV | WBHSRVI | 0.82252 | 0 |
| B61 | B63 | WBHSRV | WBRSR(mPas_200) | 0.901463 | 0 |
| B61 | B64 | WBHSRV | WBRSR(mPas_5) | 0.794197 | 0 |
| B61 | B65 | WBHSRV | WBRSR(mPas_1) | 0.69365 | 0 |
| B61 | B66 | WBHSRV | WBRSR(mPas_100) | 0.872947 | 0 |
| B61 | B67 | WBHSRV | WBRSR(mPas_50) | 0.866649 | 0 |
| B61 | B74 | WBHSRV | ApoA1 | -0.14461 | 9.93E-13 |
| B61 | B69 | WBHSRV | PV | 0.134281 | 3.66E-11 |
| B61 | B137 | WBHSRV | AFU | 0.124732 | 8.10E-10 |
| B61 | B157 | WBHSRV | IgG | 0.121091 | 2.49E-09 |
| B61 | B198 | WBHSRV | P | -0.11764 | 6.99E-09 |
| B61 | B134 | WBHSRV | CHE | 0.110108 | 6.03E-08 |
| B61 | B139 | WBHSRV | ADA | 0.107329 | 1.29E-07 |
| B61 | B186 | WBHSRV | PFC | 0.106786 | 1.49E-07 |
| B61 | B175 | WBHSRV | ALB-U | 0.106485 | 1.62E-07 |
| B61 | B94 | WBHSRV | ALDH2-Lys504Lys | -0.10431 | 2.88E-07 |
| B62 | B63 | WBHSRVI | WBRSR(mPas_200) | 0.894285 | 0 |
| B62 | B64 | WBHSRVI | WBRSR(mPas_5) | 0.814616 | 0 |
| B62 | B65 | WBHSRVI | WBRSR(mPas_1) | 0.726965 | 0 |
| B62 | B66 | WBHSRVI | WBRSR(mPas_100) | 0.860514 | 0 |
| B62 | B67 | WBHSRVI | WBRSR(mPas_50) | 0.861232 | 0 |
| B62 | B69 | WBHSRVI | PV | -0.25749 | 0 |
| B62 | B74 | WBHSRVI | ApoA1 | -0.17416 | 0 |
| B62 | B187 | WBHSRVI | NSE | 0.126062 | 5.34E-10 |
| B62 | B198 | WBHSRVI | P | -0.11039 | 5.58E-08 |
| B62 | B133 | WBHSRVI | TBIL | 0.104679 | 2.61E-07 |
| B62 | B132 | WBHSRVI | Dbil | 0.1019 | 5.39E-07 |
| B63 | B64 | WBRSR(mPas_200) | WBRSR(mPas_5) | 0.907341 | 0 |

| | | | | | |
|---|---|---|---|---|---|
| B63 | B65 | WBRSR(mPas_200) | WBRSR(mPas_1) | 0.804865 | 0 |
| B63 | B66 | WBRSR(mPas_200) | WBRSR(mPas_100) | 0.972232 | 0 |
| B63 | B67 | WBRSR(mPas_200) | WBRSR(mPas_50) | 0.968603 | 0 |
| B63 | B74 | WBRSR(mPas_200) | ApoA1 | -0.17352 | 0 |
| B63 | B69 | WBRSR(mPas_200) | PV | 0.159377 | 3.55E-15 |
| B63 | B137 | WBRSR(mPas_200) | AFU | 0.118333 | 5.70E-09 |
| B63 | B157 | WBRSR(mPas_200) | IgG | 0.111082 | 4.60E-08 |
| B63 | B198 | WBRSR(mPas_200) | P | -0.10734 | 1.29E-07 |
| B63 | B175 | WBRSR(mPas_200) | ALB-U | 0.102886 | 4.18E-07 |
| B64 | B65 | WBRSR(mPas_5) | WBRSR(mPas_1) | 0.972894 | 0 |
| B64 | B66 | WBRSR(mPas_5) | WBRSR(mPas_100) | 0.914794 | 0 |
| B64 | B67 | WBRSR(mPas_5) | WBRSR(mPas_50) | 0.932695 | 0 |
| B64 | B74 | WBRSR(mPas_5) | ApoA1 | -0.16183 | 1.33E-15 |
| B64 | B69 | WBRSR(mPas_5) | PV | 0.144645 | 9.80E-13 |
| B64 | B137 | WBRSR(mPas_5) | AFU | 0.113207 | 2.53E-08 |
| B64 | B101 | WBRSR(mPas_5) | anti-Sc70 | -0.11255 | 3.04E-08 |
| B64 | B102 | WBRSR(mPas_5) | anti-Sm | -0.11255 | 3.04E-08 |
| B64 | B103 | WBRSR(mPas_5) | anti-SSA/Ro | -0.11255 | 3.04E-08 |
| B64 | B104 | WBRSR(mPas_5) | anti-SSB | -0.11255 | 3.04E-08 |
| B64 | B105 | WBRSR(mPas_5) | anti-U1RNP | -0.11255 | 3.04E-08 |
| B64 | B106 | WBRSR(mPas_5) | anti-P | -0.11255 | 3.04E-08 |
| B64 | B107 | WBRSR(mPas_5) | AnuA | -0.11255 | 3.04E-08 |
| B64 | B109 | WBRSR(mPas_5) | AMA-M2 | -0.11255 | 3.04E-08 |
| B64 | B110 | WBRSR(mPas_5) | AHA | -0.11255 | 3.04E-08 |
| B64 | B95 | WBRSR(mPas_5) | anti-cenpb | -0.11255 | 3.04E-08 |
| B64 | B98 | WBRSR(mPas_5) | anti-Jo-1 | -0.11255 | 3.04E-08 |
| B64 | B99 | WBRSR(mPas_5) | anti-PM-Scl | -0.11255 | 3.04E-08 |
| B64 | B157 | WBRSR(mPas_5) | IgG | 0.112087 | 3.47E-08 |
| B64 | B134 | WBRSR(mPas_5) | CHE | 0.101338 | 6.23E-07 |
| B65 | B66 | WBRSR(mPas_1) | WBRSR(mPas_100) | 0.830239 | 0 |
| B65 | B67 | WBRSR(mPas_1) | WBRSR(mPas_50) | 0.858315 | 0 |
| B65 | B74 | WBRSR(mPas_1) | ApoA1 | -0.15023 | 1.24E-13 |
| B65 | B101 | WBRSR(mPas_1) | anti-Sc70 | -0.11573 | 1.22E-08 |
| B65 | B102 | WBRSR(mPas_1) | anti-Sm | -0.11573 | 1.22E-08 |
| B65 | B103 | WBRSR(mPas_1) | anti-SSA/Ro | -0.11573 | 1.22E-08 |
| B65 | B104 | WBRSR(mPas_1) | anti-SSB | -0.11573 | 1.22E-08 |
| B65 | B105 | WBRSR(mPas_1) | anti-U1RNP | -0.11573 | 1.22E-08 |
| B65 | B106 | WBRSR(mPas_1) | anti-P | -0.11573 | 1.22E-08 |
| B65 | B107 | WBRSR(mPas_1) | AnuA | -0.11573 | 1.22E-08 |
| B65 | B109 | WBRSR(mPas_1) | AMA-M2 | -0.11573 | 1.22E-08 |
| B65 | B110 | WBRSR(mPas_1) | AHA | -0.11573 | 1.22E-08 |
| B65 | B95 | WBRSR(mPas_1) | anti-cenpb | -0.11573 | 1.22E-08 |
| B65 | B98 | WBRSR(mPas_1) | anti-Jo-1 | -0.11573 | 1.22E-08 |
| B65 | B99 | WBRSR(mPas_1) | anti-PM-Scl | -0.11573 | 1.22E-08 |

| | | | | | |
|---|---|---|---|---|---|
| B65 | B69 | WBRSR(mPas_1) | PV | 0.11203 | 3.52E-08 |
| B65 | B157 | WBRSR(mPas_1) | IgG | 0.10531 | 2.21E-07 |
| B66 | B67 | WBRSR(mPas_100) | WBRSR(mPas_50) | 0.998347 | 0 |
| B66 | B69 | WBRSR(mPas_100) | PV | 0.204466 | 0 |
| B66 | B74 | WBRSR(mPas_100) | ApoA1 | -0.18097 | 0 |
| B66 | B137 | WBRSR(mPas_100) | AFU | 0.135181 | 2.70E-11 |
| B66 | B75 | WBRSR(mPas_100) | ApoA1/B | -0.11127 | 4.36E-08 |
| B66 | B175 | WBRSR(mPas_100) | ALB-U | 0.105561 | 2.07E-07 |
| B66 | B101 | WBRSR(mPas_100) | anti-Sc70 | -0.10482 | 2.52E-07 |
| B66 | B102 | WBRSR(mPas_100) | anti-Sm | -0.10482 | 2.52E-07 |
| B66 | B103 | WBRSR(mPas_100) | anti-SSA/Ro | -0.10482 | 2.52E-07 |
| B66 | B104 | WBRSR(mPas_100) | anti-SSB | -0.10482 | 2.52E-07 |
| B66 | B105 | WBRSR(mPas_100) | anti-U1RNP | -0.10482 | 2.52E-07 |
| B66 | B106 | WBRSR(mPas_100) | anti-P | -0.10482 | 2.52E-07 |
| B66 | B107 | WBRSR(mPas_100) | AnuA | -0.10482 | 2.52E-07 |
| B66 | B109 | WBRSR(mPas_100) | AMA-M2 | -0.10482 | 2.52E-07 |
| B66 | B110 | WBRSR(mPas_100) | AHA | -0.10482 | 2.52E-07 |
| B66 | B95 | WBRSR(mPas_100) | anti-cenpb | -0.10482 | 2.52E-07 |
| B66 | B98 | WBRSR(mPas_100) | anti-Jo-1 | -0.10482 | 2.52E-07 |
| B66 | B99 | WBRSR(mPas_100) | anti-PM-Scl | -0.10482 | 2.52E-07 |
| B66 | B157 | WBRSR(mPas_100) | IgG | 0.104757 | 2.56E-07 |
| B66 | B134 | WBRSR(mPas_100) | CHE | 0.104678 | 2.62E-07 |
| B66 | B187 | WBRSR(mPas_100) | NSE | 0.102768 | 4.31E-07 |
| B67 | B69 | WBRSR(mPas_50) | PV | 0.19479 | 0 |
| B67 | B74 | WBRSR(mPas_50) | ApoA1 | -0.18158 | 0 |
| B67 | B137 | WBRSR(mPas_50) | AFU | 0.133236 | 5.19E-11 |
| B67 | B75 | WBRSR(mPas_50) | ApoA1/B | -0.10997 | 6.26E-08 |
| B67 | B101 | WBRSR(mPas_50) | anti-Sc70 | -0.10602 | 1.83E-07 |
| B67 | B102 | WBRSR(mPas_50) | anti-Sm | -0.10602 | 1.83E-07 |
| B67 | B103 | WBRSR(mPas_50) | anti-SSA/Ro | -0.10602 | 1.83E-07 |
| B67 | B104 | WBRSR(mPas_50) | anti-SSB | -0.10602 | 1.83E-07 |
| B67 | B105 | WBRSR(mPas_50) | anti-U1RNP | -0.10602 | 1.83E-07 |
| B67 | B106 | WBRSR(mPas_50) | anti-P | -0.10602 | 1.83E-07 |
| B67 | B107 | WBRSR(mPas_50) | AnuA | -0.10602 | 1.83E-07 |
| B67 | B109 | WBRSR(mPas_50) | AMA-M2 | -0.10602 | 1.83E-07 |
| B67 | B110 | WBRSR(mPas_50) | AHA | -0.10602 | 1.83E-07 |
| B67 | B95 | WBRSR(mPas_50) | anti-cenpb | -0.10602 | 1.83E-07 |
| B67 | B98 | WBRSR(mPas_50) | anti-Jo-1 | -0.10602 | 1.83E-07 |
| B67 | B99 | WBRSR(mPas_50) | anti-PM-Scl | -0.10602 | 1.83E-07 |
| B67 | B134 | WBRSR(mPas_50) | CHE | 0.105319 | 2.21E-07 |
| B67 | B157 | WBRSR(mPas_50) | IgG | 0.104895 | 2.47E-07 |
| B67 | B175 | WBRSR(mPas_50) | ALB-U | 0.10422 | 2.95E-07 |
| B67 | B187 | WBRSR(mPas_50) | NSE | 0.099909 | 8.95E-07 |
| B69 | B123 | PV | hs-CRP1 | 0.192626 | 0 |

| | | | | | |
|---|---|---|---|---|---|
| B69 | B91 | PV | TP | 0.143951 | 1.26E-12 |
| B69 | B130 | PV | GLB | 0.135804 | 2.19E-11 |
| B69 | B157 | PV | IgG | 0.115922 | 1.16E-08 |
| B69 | B101 | PV | anti-Sc70 | -0.11178 | 3.78E-08 |
| B69 | B102 | PV | anti-Sm | -0.11178 | 3.78E-08 |
| B69 | B103 | PV | anti-SSA/Ro | -0.11178 | 3.78E-08 |
| B69 | B104 | PV | anti-SSB | -0.11178 | 3.78E-08 |
| B69 | B105 | PV | anti-U1RNP | -0.11178 | 3.78E-08 |
| B69 | B106 | PV | anti-P | -0.11178 | 3.78E-08 |
| B69 | B107 | PV | AnuA | -0.11178 | 3.78E-08 |
| B69 | B109 | PV | AMA-M2 | -0.11178 | 3.78E-08 |
| B69 | B110 | PV | AHA | -0.11178 | 3.78E-08 |
| B69 | B95 | PV | anti-cenpb | -0.11178 | 3.78E-08 |
| B69 | B98 | PV | anti-Jo-1 | -0.11178 | 3.78E-08 |
| B69 | B99 | PV | anti-PM-Scl | -0.11178 | 3.78E-08 |
| B69 | B186 | PV | PFC | 0.102484 | 4.64E-07 |
| B69 | B145 | PV | BGP | 0.102418 | 4.72E-07 |
| B69 | B137 | PV | AFU | 0.101562 | 5.88E-07 |
| B7 | B8 | WB-Cop | WB-Zinc | 0.999904 | 0 |
| B7 | B86 | WB-Cop | TBhCG | -0.11659 | 9.53E-09 |
| B70 | B132 | LDL-C | Dbil | -0.22049 | 0 |
| B70 | B75 | LDL-C | ApoA1/B | -0.20508 | 0 |
| B70 | B76 | LDL-C | ApoB | 0.32606 | 0 |
| B70 | B77 | LDL-C | TC | 0.816728 | 0 |
| B70 | B72 | LDL-C | HDL-C | 0.114166 | 1.92E-08 |
| B70 | B91 | LDL-C | TP | 0.109985 | 6.24E-08 |
| B71 | B140 | TG | r-GT | 0.184175 | 0 |
| B71 | B153 | TG | Glu | 0.200775 | 0 |
| B71 | B192 | TG | UA | 0.229258 | 0 |
| B71 | B206 | TG | Waist | 0.175396 | 0 |
| B71 | B215 | TG | BMI | 0.184636 | 0 |
| B71 | B216 | TG | BMI | 0.186317 | 0 |
| B71 | B222 | TG | NS | 0.188447 | 0 |
| B71 | B72 | TG | HDL-C | -0.40755 | 0 |
| B71 | B77 | TG | TC | 0.359255 | 0 |
| B71 | B91 | TG | TP | 0.2401 | 0 |
| B71 | B130 | TG | GLB | 0.165621 | 4.44E-16 |
| B71 | B209 | TG | Smoking | 0.154089 | 2.86E-14 |
| B71 | B74 | TG | ApoA1 | -0.14613 | 5.71E-13 |
| B71 | B126 | TG | Alb | 0.144137 | 1.18E-12 |
| B71 | B132 | TG | Dbil | -0.14409 | 1.20E-12 |
| B71 | B141 | TG | AST/ALT | -0.13735 | 1.29E-11 |
| B71 | B134 | TG | CHE | 0.136303 | 1.84E-11 |
| B71 | B204 | TG | Drinking | 0.130535 | 1.27E-10 |

| | | | | | |
|---|---|---|---|---|---|
| B71 | B170 | TG | U-GLU | 0.125482 | 6.41E-10 |
| B71 | B218 | TG | DBP | 0.122535 | 1.60E-09 |
| B71 | B210 | TG | Hip | 0.111467 | 4.13E-08 |
| B71 | B128 | TG | ALT | 0.111281 | 4.35E-08 |
| B71 | B188 | TG | Ca | 0.104701 | 2.60E-07 |
| B72 | B141 | HDL-C | AST/ALT | 0.207518 | 0 |
| B72 | B163 | HDL-C | SG | -0.17904 | 0 |
| B72 | B192 | HDL-C | UA | -0.29833 | 0 |
| B72 | B206 | HDL-C | Waist | -0.3469 | 0 |
| B72 | B210 | HDL-C | Hip | -0.25804 | 0 |
| B72 | B215 | HDL-C | BMI | -0.29481 | 0 |
| B72 | B216 | HDL-C | BMI | -0.38623 | 0 |
| B72 | B221 | HDL-C | Hei | -0.30365 | 0 |
| B72 | B222 | HDL-C | NS | -0.36357 | 0 |
| B72 | B74 | HDL-C | ApoA1 | 0.341241 | 0 |
| B72 | B77 | HDL-C | TC | 0.229735 | 0 |
| B72 | B75 | HDL-C | ApoA1/B | 0.152302 | 5.68E-14 |
| B72 | B209 | HDL-C | Smoking | -0.14486 | 9.06E-13 |
| B72 | B174 | HDL-C | Urea/Cr | 0.143694 | 1.38E-12 |
| B72 | B153 | HDL-C | Glu | -0.11737 | 7.57E-09 |
| B72 | B128 | HDL-C | ALT | -0.11519 | 1.43E-08 |
| B72 | B132 | HDL-C | Dbil | 0.106513 | 1.61E-07 |
| B72 | B207 | HDL-C | WHR | -0.10229 | 4.87E-07 |
| B72 | B73 | HDL-C | ApoA | 0.101925 | 5.36E-07 |
| B73 | B123 | ApoA | hs-CRP1 | 0.198462 | 0 |
| B73 | B74 | ApoA | ApoA1 | 0.17774 | 0 |
| B73 | B152 | ApoA | ACA-Ig(G-A-M) | -0.14836 | 2.51E-13 |
| B73 | B115 | ApoA | TSH TT3 TT4(TT3) | -0.12448 | 8.77E-10 |
| B73 | B138 | ApoA | PAB | -0.11948 | 4.05E-09 |
| B73 | B134 | ApoA | CHE | -0.10189 | 5.40E-07 |
| B73 | B199 | ApoA | Mg | -0.10177 | 5.57E-07 |
| B74 | B193 | ApoA1 | K | 0.219912 | 0 |
| B74 | B206 | ApoA1 | Waist | -0.18096 | 0 |
| B74 | B216 | ApoA1 | BMI | -0.21907 | 0 |
| B74 | B221 | ApoA1 | Hei | -0.18884 | 0 |
| B74 | B222 | ApoA1 | NS | -0.19591 | 0 |
| B74 | B75 | ApoA1 | ApoA1/B | 0.421613 | 0 |
| B74 | B123 | ApoA1 | hs-CRP1 | -0.15322 | 4.00E-14 |
| B74 | B198 | ApoA1 | P | 0.145934 | 6.13E-13 |
| B74 | B141 | ApoA1 | AST/ALT | 0.144602 | 9.96E-13 |
| B74 | B215 | ApoA1 | BMI | -0.14412 | 1.18E-12 |
| B74 | B101 | ApoA1 | anti-Sc70 | 0.139989 | 5.15E-12 |
| B74 | B102 | ApoA1 | anti-Sm | 0.139989 | 5.15E-12 |
| B74 | B103 | ApoA1 | anti-SSA/Ro | 0.139989 | 5.15E-12 |

| | | | | | |
|---|---|---|---|---|---|
| B74 | B104 | ApoA1 | anti-SSB | 0.139989 | 5.15E-12 |
| B74 | B105 | ApoA1 | anti-U1RNP | 0.139989 | 5.15E-12 |
| B74 | B106 | ApoA1 | anti-P | 0.139989 | 5.15E-12 |
| B74 | B107 | ApoA1 | AnuA | 0.139989 | 5.15E-12 |
| B74 | B109 | ApoA1 | AMA-M2 | 0.139989 | 5.15E-12 |
| B74 | B110 | ApoA1 | AHA | 0.139989 | 5.15E-12 |
| B74 | B95 | ApoA1 | anti-cenpb | 0.139989 | 5.15E-12 |
| B74 | B98 | ApoA1 | anti-Jo-1 | 0.139989 | 5.15E-12 |
| B74 | B99 | ApoA1 | anti-PM-Scl | 0.139989 | 5.15E-12 |
| B74 | B210 | ApoA1 | Hip | -0.13978 | 5.55E-12 |
| B74 | B199 | ApoA1 | Mg | 0.130921 | 1.12E-10 |
| B74 | B194 | ApoA1 | Cl | -0.12981 | 1.60E-10 |
| B74 | B192 | ApoA1 | UA | -0.11322 | 2.52E-08 |
| B74 | B200 | ApoA1 | OP | -0.10439 | 2.82E-07 |
| B74 | B150 | ApoA1 | ALP | -0.10373 | 3.36E-07 |
| B74 | B128 | ApoA1 | ALT | -0.10126 | 6.36E-07 |
| B75 | B172 | ApoA1/B | KB | 0.180422 | 0 |
| B75 | B76 | ApoA1/B | ApoB | -0.74181 | 0 |
| B75 | B92 | ApoA1/B | ALDH2-Glu504Glu | 0.134335 | 3.59E-11 |
| B75 | B93 | ApoA1/B | ALDH2-Glu504Lys | -0.13117 | 1.03E-10 |
| B75 | B198 | ApoA1/B | P | 0.129124 | 2.00E-10 |
| B75 | B194 | ApoA1/B | Cl | -0.1277 | 3.17E-10 |
| B75 | B101 | ApoA1/B | anti-Sc70 | 0.125208 | 6.98E-10 |
| B75 | B102 | ApoA1/B | anti-Sm | 0.125208 | 6.98E-10 |
| B75 | B103 | ApoA1/B | anti-SSA/Ro | 0.125208 | 6.98E-10 |
| B75 | B104 | ApoA1/B | anti-SSB | 0.125208 | 6.98E-10 |
| B75 | B105 | ApoA1/B | anti-U1RNP | 0.125208 | 6.98E-10 |
| B75 | B106 | ApoA1/B | anti-P | 0.125208 | 6.98E-10 |
| B75 | B107 | ApoA1/B | AnuA | 0.125208 | 6.98E-10 |
| B75 | B109 | ApoA1/B | AMA-M2 | 0.125208 | 6.98E-10 |
| B75 | B110 | ApoA1/B | AHA | 0.125208 | 6.98E-10 |
| B75 | B95 | ApoA1/B | anti-cenpb | 0.125208 | 6.98E-10 |
| B75 | B98 | ApoA1/B | anti-Jo-1 | 0.125208 | 6.98E-10 |
| B75 | B99 | ApoA1/B | anti-PM-Scl | 0.125208 | 6.98E-10 |
| B75 | B94 | ApoA1/B | ALDH2-Lys504Lys | -0.1229 | 1.43E-09 |
| B75 | B195 | ApoA1/B | Na | -0.11675 | 9.10E-09 |
| B75 | B222 | ApoA1/B | NS | -0.10604 | 1.82E-07 |
| B75 | B134 | ApoA1/B | CHE | -0.1053 | 2.22E-07 |
| B75 | B216 | ApoA1/B | BMI | -0.10484 | 2.51E-07 |
| B76 | B77 | ApoB | TC | 0.251173 | 0 |
| B76 | B134 | ApoB | CHE | 0.160038 | 2.66E-15 |
| B76 | B89 | ApoB | CA 72-4 | 0.124819 | 7.89E-10 |
| B76 | B168 | ApoB | NMP22 | 0.103225 | 3.83E-07 |
| B77 | B132 | TC | Dbil | -0.21504 | 0 |

| | | | | | |
|---|---|---|---|---|---|
| B77 | B91 | TC | TP | 0.189945 | 0 |
| B77 | B130 | TC | GLB | 0.152994 | 4.35E-14 |
| B77 | B188 | TC | Ca | 0.143302 | 1.59E-12 |
| B77 | B218 | TC | DBP | 0.114493 | 1.75E-08 |
| B77 | B127 | TC | A-G ratio | -0.11232 | 3.25E-08 |
| B77 | B220 | TC | SBP | 0.104218 | 2.95E-07 |
| B77 | B221 | TC | Hei | -0.10266 | 4.44E-07 |
| B77 | B133 | TC | TBIL | -0.1021 | 5.12E-07 |
| B78 | B80 | HBcAb | anti-HP IgG | -0.13003 | 1.49E-10 |
| B78 | B148 | HBcAb | Anti_TPO | 0.106519 | 1.60E-07 |
| B79 | B101 | hb V -dna | anti-Sc70 | 0.205607 | 0 |
| B79 | B102 | hb V -dna | anti-Sm | 0.205607 | 0 |
| B79 | B103 | hb V -dna | anti-SSA/Ro | 0.205607 | 0 |
| B79 | B104 | hb V -dna | anti-SSB | 0.205607 | 0 |
| B79 | B105 | hb V -dna | anti-U1RNP | 0.205607 | 0 |
| B79 | B106 | hb V -dna | anti-P | 0.205607 | 0 |
| B79 | B107 | hb V -dna | AnuA | 0.205606 | 0 |
| B79 | B109 | hb V -dna | AMA-M2 | 0.205607 | 0 |
| B79 | B110 | hb V -dna | AHA | 0.205607 | 0 |
| B79 | B119 | hb V -dna | Hcv-rna | 0.393279 | 0 |
| B79 | B120 | hb V -dna | HCV-cAg | 0.200223 | 0 |
| B79 | B159 | hb V -dna | β2-MG | -0.30977 | 0 |
| B79 | B81 | hb V -dna | TSGF | -0.21604 | 0 |
| B79 | B94 | hb V -dna | ALDH2-Lys504Lys | 0.214691 | 0 |
| B79 | B95 | hb V -dna | anti-cenpb | 0.205607 | 0 |
| B79 | B98 | hb V -dna | anti-Jo-1 | 0.205607 | 0 |
| B79 | B99 | hb V -dna | anti-PM-Scl | 0.205607 | 0 |
| B79 | B164 | hb V -dna | COM | -0.16569 | 2.22E-16 |
| B79 | B182 | hb V -dna | TT | -0.15482 | 2.15E-14 |
| B79 | B124 | hb V -dna | ACTH | -0.14796 | 2.91E-13 |
| B79 | B139 | hb V -dna | ADA | -0.14485 | 9.11E-13 |
| B79 | B193 | hb V -dna | K | 0.128139 | 2.75E-10 |
| B79 | B197 | hb V -dna | AG | -0.11603 | 1.12E-08 |
| B79 | B128 | hb V -dna | ALT | -0.11568 | 1.24E-08 |
| B79 | B183 | hb V -dna | PTA | 0.113183 | 2.54E-08 |
| B79 | B136 | hb V -dna | 5'NT | -0.10995 | 6.29E-08 |
| B79 | B156 | hb V -dna | IgE | 0.107757 | 1.15E-07 |
| B79 | B137 | hb V -dna | AFU | -0.10682 | 1.48E-07 |
| B79 | B131 | hb V -dna | AST | -0.10579 | 1.95E-07 |
| B79 | B125 | hb V -dna | Cor | 0.105489 | 2.11E-07 |
| B79 | B200 | hb V -dna | OP | -0.10324 | 3.81E-07 |
| B8 | B86 | WB-Zinc | TBhCG | -0.11658 | 9.56E-09 |
| B81 | B181 | TSGF | APTT | -0.2076 | 0 |
| B81 | B182 | TSGF | TT | 0.211125 | 0 |

| | | | | | |
|---|---|---|---|---|---|
| B81 | B184 | TSGF | INR | -0.18315 | 0 |
| B81 | B200 | TSGF | OP | 0.134398 | 3.52E-11 |
| B81 | B159 | TSGF | β2-MG | 0.134234 | 3.72E-11 |
| B81 | B108 | TSGF | Anti-Cyclin | -0.13397 | 4.07E-11 |
| B81 | B158 | TSGF | IgM | -0.12692 | 4.07E-10 |
| B81 | B199 | TSGF | Mg | 0.12437 | 9.08E-10 |
| B81 | B224 | TSGF | pmh | -0.1196 | 3.91E-09 |
| B81 | B197 | TSGF | AG | 0.106614 | 1.56E-07 |
| B81 | B164 | TSGF | COM | 0.100574 | 7.56E-07 |
| B82 | B84 | CEA | C-PSA | 0.310543 | 0 |
| B82 | B86 | CEA | TBhCG | -0.24912 | 0 |
| B82 | B87 | CEA | NSE | 0.185379 | 0 |
| B82 | B88 | CEA | CA 242 | 0.160261 | 2.66E-15 |
| B82 | B85 | CEA | SCC | 0.132707 | 6.19E-11 |
| B82 | B89 | CEA | CA 72-4 | 0.124449 | 8.85E-10 |
| B82 | B83 | CEA | AFP | 0.104902 | 2.47E-07 |
| B83 | B89 | AFP | CA 72-4 | 0.169774 | 0 |
| B83 | B140 | AFP | r-GT | 0.155248 | 1.82E-14 |
| B83 | B88 | AFP | CA 242 | 0.134806 | 3.07E-11 |
| B84 | B85 | C-PSA | SCC | 0.198643 | 0 |
| B84 | B86 | C-PSA | TBhCG | -0.25671 | 0 |
| B84 | B87 | C-PSA | NSE | 0.309448 | 0 |
| B84 | B135 | C-PSA | TBA | 0.100774 | 7.19E-07 |
| B85 | B87 | SCC | NSE | 0.178413 | 0 |
| B85 | B190 | SCC | Cre | 0.150249 | 1.24E-13 |
| B85 | B86 | SCC | TBhCG | -0.11474 | 1.63E-08 |
| B85 | B89 | SCC | CA 72-4 | 0.108682 | 8.92E-08 |
| B86 | B87 | TBhCG | NSE | -0.20518 | 0 |
| B86 | B90 | TBhCG | CYFRA 21-1 | 0.210006 | 0 |
| B86 | B137 | TBhCG | AFU | -0.12124 | 2.38E-09 |
| B87 | B90 | NSE | CYFRA 21-1 | 0.116761 | 9.06E-09 |
| B87 | B88 | NSE | CA 242 | 0.116732 | 9.14E-09 |
| B89 | B134 | CA 72-4 | CHE | 0.194081 | 0 |
| B9 | B25 | RH | CK | -0.22885 | 0 |
| B9 | B194 | RH | Cl | 0.118537 | 5.36E-09 |
| B91 | B126 | TP | Alb | 0.449048 | 0 |
| B91 | B127 | TP | A-G ratio | -0.4927 | 0 |
| B91 | B130 | TP | GLB | 0.792639 | 0 |
| B91 | B188 | TP | Ca | 0.322562 | 0 |
| B91 | B153 | TP | Glu | 0.109374 | 7.38E-08 |
| B91 | B192 | TP | UA | 0.107734 | 1.16E-07 |
| B91 | B157 | TP | IgG | 0.101524 | 5.94E-07 |
| B92 | B101 | ALDH2-Glu504Glu | anti-Sc70 | -0.18933 | 0 |
| B92 | B102 | ALDH2-Glu504Glu | anti-Sm | -0.18933 | 0 |

| | | | | | |
|---|---|---|---|---|---|
| B92 | B103 | ALDH2-Glu504Glu | anti-SSA/Ro | -0.18933 | 0 |
| B92 | B104 | ALDH2-Glu504Glu | anti-SSB | -0.18933 | 0 |
| B92 | B105 | ALDH2-Glu504Glu | anti-U1RNP | -0.18933 | 0 |
| B92 | B106 | ALDH2-Glu504Glu | anti-P | -0.18933 | 0 |
| B92 | B107 | ALDH2-Glu504Glu | AnuA | -0.18933 | 0 |
| B92 | B109 | ALDH2-Glu504Glu | AMA-M2 | -0.18933 | 0 |
| B92 | B110 | ALDH2-Glu504Glu | AHA | -0.18933 | 0 |
| B92 | B156 | ALDH2-Glu504Glu | IgE | -0.17928 | 0 |
| B92 | B93 | ALDH2-Glu504Glu | ALDH2-Glu504Lys | -0.99849 | 0 |
| B92 | B94 | ALDH2-Glu504Glu | ALDH2-Lys504Lys | -0.59047 | 0 |
| B92 | B95 | ALDH2-Glu504Glu | anti-cenpb | -0.18933 | 0 |
| B92 | B98 | ALDH2-Glu504Glu | anti-Jo-1 | -0.18933 | 0 |
| B92 | B99 | ALDH2-Glu504Glu | anti-PM-Scl | -0.18933 | 0 |
| B92 | B184 | ALDH2-Glu504Glu | INR | -0.13041 | 1.32E-10 |
| B92 | B125 | ALDH2-Glu504Glu | Cor | -0.11742 | 7.45E-09 |
| B92 | B124 | ALDH2-Glu504Glu | ACTH | 0.116055 | 1.11E-08 |
| B92 | B182 | ALDH2-Glu504Glu | TT | 0.114812 | 1.60E-08 |
| B92 | B197 | ALDH2-Glu504Glu | AG | 0.100569 | 7.57E-07 |
| B93 | B101 | ALDH2-Glu504Lys | anti-Sc70 | 0.181814 | 0 |
| B93 | B102 | ALDH2-Glu504Lys | anti-Sm | 0.181814 | 0 |
| B93 | B103 | ALDH2-Glu504Lys | anti-SSA/Ro | 0.181814 | 0 |
| B93 | B104 | ALDH2-Glu504Lys | anti-SSB | 0.181814 | 0 |
| B93 | B105 | ALDH2-Glu504Lys | anti-U1RNP | 0.181814 | 0 |
| B93 | B106 | ALDH2-Glu504Lys | anti-P | 0.181814 | 0 |
| B93 | B107 | ALDH2-Glu504Lys | AnuA | 0.181814 | 0 |
| B93 | B109 | ALDH2-Glu504Lys | AMA-M2 | 0.181814 | 0 |
| B93 | B110 | ALDH2-Glu504Lys | AHA | 0.181814 | 0 |
| B93 | B156 | ALDH2-Glu504Lys | IgE | 0.178095 | 0 |
| B93 | B94 | ALDH2-Glu504Lys | ALDH2-Lys504Lys | 0.545266 | 0 |
| B93 | B95 | ALDH2-Glu504Lys | anti-cenpb | 0.181814 | 0 |
| B93 | B98 | ALDH2-Glu504Lys | anti-Jo-1 | 0.181814 | 0 |
| B93 | B99 | ALDH2-Glu504Lys | anti-PM-Scl | 0.181814 | 0 |
| B93 | B184 | ALDH2-Glu504Lys | INR | 0.125182 | 7.04E-10 |
| B93 | B125 | ALDH2-Glu504Lys | Cor | 0.112701 | 2.92E-08 |
| B93 | B124 | ALDH2-Glu504Lys | ACTH | -0.11135 | 4.26E-08 |
| B93 | B182 | ALDH2-Glu504Lys | TT | -0.11017 | 5.93E-08 |
| B94 | B101 | ALDH2-Lys504Lys | anti-Sc70 | 0.218084 | 0 |
| B94 | B102 | ALDH2-Lys504Lys | anti-Sm | 0.218084 | 0 |
| B94 | B103 | ALDH2-Lys504Lys | anti-SSA/Ro | 0.218084 | 0 |
| B94 | B104 | ALDH2-Lys504Lys | anti-SSB | 0.218084 | 0 |
| B94 | B105 | ALDH2-Lys504Lys | anti-U1RNP | 0.218084 | 0 |
| B94 | B106 | ALDH2-Lys504Lys | anti-P | 0.218084 | 0 |
| B94 | B107 | ALDH2-Lys504Lys | AnuA | 0.218084 | 0 |
| B94 | B109 | ALDH2-Lys504Lys | AMA-M2 | 0.218084 | 0 |

| | | | | | |
|---|---|---|---|---|---|
| B94 | B110 | ALDH2-Lys504Lys | AHA | 0.218084 | 0 |
| B94 | B95 | ALDH2-Lys504Lys | anti-cenpb | 0.218084 | 0 |
| B94 | B98 | ALDH2-Lys504Lys | anti-Jo-1 | 0.218084 | 0 |
| B94 | B99 | ALDH2-Lys504Lys | anti-PM-Scl | 0.218084 | 0 |
| B94 | B184 | ALDH2-Lys504Lys | INR | 0.150933 | 9.55E-14 |
| B94 | B134 | ALDH2-Lys504Lys | CHE | -0.15032 | 1.20E-13 |
| B94 | B125 | ALDH2-Lys504Lys | Cor | 0.136158 | 1.94E-11 |
| B94 | B124 | ALDH2-Lys504Lys | ACTH | -0.13505 | 2.82E-11 |
| B94 | B182 | ALDH2-Lys504Lys | TT | -0.13351 | 4.73E-11 |
| B94 | B156 | ALDH2-Lys504Lys | IgE | 0.119377 | 4.17E-09 |
| B94 | B136 | ALDH2-Lys504Lys | 5'NT | -0.11146 | 4.14E-08 |
| B94 | B172 | ALDH2-Lys504Lys | KB | -0.10355 | 3.52E-07 |
| B94 | B193 | ALDH2-Lys504Lys | K | 0.101783 | 5.56E-07 |
| B95 | B101 | anti-cenpb | anti-Sc70 | 1 | 0 |
| B95 | B102 | anti-cenpb | anti-Sm | 1 | 0 |
| B95 | B103 | anti-cenpb | anti-SSA/Ro | 1 | 0 |
| B95 | B104 | anti-cenpb | anti-SSB | 1 | 0 |
| B95 | B105 | anti-cenpb | anti-U1RNP | 1 | 0 |
| B95 | B107 | anti-cenpb | AnuA | 1 | 0 |
| B95 | B109 | anti-cenpb | AMA-M2 | 1 | 0 |
| B95 | B110 | anti-cenpb | AHA | 1 | 0 |
| B95 | B120 | anti-cenpb | HCV-cAg | 0.203435 | 0 |
| B95 | B156 | anti-cenpb | IgE | 0.177741 | 0 |
| B95 | B168 | anti-cenpb | NMP22 | -0.17131 | 0 |
| B95 | B96 | anti-cenpb | dsDNA (IIF) | 0.175635 | 0 |
| B95 | B99 | anti-cenpb | anti-PM-Scl | 1 | 0 |
| B95 | B155 | anti-cenpb | IgA | -0.15872 | 4.66E-15 |
| B95 | B159 | anti-cenpb | β2-MG | -0.15404 | 2.91E-14 |
| B95 | B134 | anti-cenpb | CHE | -0.14987 | 1.43E-13 |
| B95 | B108 | anti-cenpb | Anti-Cyclin | -0.14294 | 1.81E-12 |
| B95 | B97 | anti-cenpb | dsDNA | -0.14114 | 3.44E-12 |
| B95 | B136 | anti-cenpb | 5'NT | -0.14092 | 3.71E-12 |
| B95 | B145 | anti-cenpb | BGP | -0.14021 | 4.77E-12 |
| B95 | B193 | anti-cenpb | K | 0.1384 | 8.96E-12 |
| B95 | B119 | anti-cenpb | Hcv-rna | 0.128458 | 2.49E-10 |
| B95 | B135 | anti-cenpb | TBA | 0.114445 | 1.77E-08 |
| B95 | B181 | anti-cenpb | APTT | -0.11108 | 4.60E-08 |
| B95 | B100 | anti-cenpb | anti-Ro52 | 0.103467 | 3.59E-07 |
| B95 | B198 | anti-cenpb | P | 0.099814 | 9.17E-07 |
| B96 | B101 | dsDNA (IIF) | anti-Sc70 | 0.175635 | 0 |
| B96 | B102 | dsDNA (IIF) | anti-Sm | 0.175635 | 0 |
| B96 | B103 | dsDNA (IIF) | anti-SSA/Ro | 0.175635 | 0 |
| B96 | B104 | dsDNA (IIF) | anti-SSB | 0.175635 | 0 |
| B96 | B105 | dsDNA (IIF) | anti-U1RNP | 0.175635 | 0 |

| | | | | | |
|---|---|---|---|---|---|
| B96 | B106 | dsDNA (IIF) | anti-P | 0.175635 | 0 |
| B96 | B107 | dsDNA (IIF) | AnuA | 0.175635 | 0 |
| B96 | B109 | dsDNA (IIF) | AMA-M2 | 0.175635 | 0 |
| B96 | B110 | dsDNA (IIF) | AHA | 0.175635 | 0 |
| B96 | B228 | dsDNA (IIF) | OS | -0.20433 | 0 |
| B96 | B97 | dsDNA (IIF) | dsDNA | -0.99939 | 0 |
| B96 | B98 | dsDNA (IIF) | anti-Jo-1 | 0.175635 | 0 |
| B96 | B99 | dsDNA (IIF) | anti-PM-Scl | 0.175635 | 0 |
| B97 | B228 | dsDNA | OS | 0.203306 | 0 |
| B97 | B101 | dsDNA | anti-Sc70 | -0.14114 | 3.44E-12 |
| B97 | B102 | dsDNA | anti-Sm | -0.14114 | 3.44E-12 |
| B97 | B103 | dsDNA | anti-SSA/Ro | -0.14114 | 3.44E-12 |
| B97 | B104 | dsDNA | anti-SSB | -0.14114 | 3.44E-12 |
| B97 | B105 | dsDNA | anti-U1RNP | -0.14114 | 3.44E-12 |
| B97 | B106 | dsDNA | anti-P | -0.14114 | 3.44E-12 |
| B97 | B107 | dsDNA | AnuA | -0.14114 | 3.44E-12 |
| B97 | B109 | dsDNA | AMA-M2 | -0.14114 | 3.44E-12 |
| B97 | B110 | dsDNA | AHA | -0.14114 | 3.44E-12 |
| B97 | B98 | dsDNA | anti-Jo-1 | -0.14114 | 3.44E-12 |
| B97 | B99 | dsDNA | anti-PM-Scl | -0.14114 | 3.44E-12 |
| B98 | B101 | anti-Jo-1 | anti-Sc70 | 1 | 0 |
| B98 | B104 | anti-Jo-1 | anti-SSB | 1 | 0 |
| B98 | B107 | anti-Jo-1 | AnuA | 0.999999 | 0 |
| B98 | B120 | anti-Jo-1 | HCV-cAg | 0.203435 | 0 |
| B98 | B156 | anti-Jo-1 | IgE | 0.177741 | 0 |
| B98 | B168 | anti-Jo-1 | NMP22 | -0.17131 | 0 |
| B98 | B155 | anti-Jo-1 | IgA | -0.15872 | 4.66E-15 |
| B98 | B159 | anti-Jo-1 | β2-MG | -0.15404 | 2.91E-14 |
| B98 | B134 | anti-Jo-1 | CHE | -0.14987 | 1.43E-13 |
| B98 | B108 | anti-Jo-1 | Anti-Cyclin | -0.14294 | 1.81E-12 |
| B98 | B136 | anti-Jo-1 | 5'NT | -0.14092 | 3.71E-12 |
| B98 | B145 | anti-Jo-1 | BGP | -0.14021 | 4.77E-12 |
| B98 | B193 | anti-Jo-1 | K | 0.1384 | 8.96E-12 |
| B98 | B119 | anti-Jo-1 | Hcv-rna | 0.128458 | 2.49E-10 |
| B98 | B135 | anti-Jo-1 | TBA | 0.114445 | 1.77E-08 |
| B98 | B181 | anti-Jo-1 | APTT | -0.11108 | 4.60E-08 |
| B98 | B100 | anti-Jo-1 | anti-Ro52 | 0.103467 | 3.59E-07 |
| B98 | B198 | anti-Jo-1 | P | 0.099814 | 9.17E-07 |
| B99 | B101 | anti-PM-Scl | anti-Sc70 | 1 | 0 |
| B99 | B102 | anti-PM-Scl | anti-Sm | 1 | 0 |
| B99 | B103 | anti-PM-Scl | anti-SSA/Ro | 1 | 0 |
| B99 | B104 | anti-PM-Scl | anti-SSB | 1 | 0 |
| B99 | B105 | anti-PM-Scl | anti-U1RNP | 1 | 0 |
| B99 | B107 | anti-PM-Scl | AnuA | 0.999999 | 0 |

| | | | | | |
|---|---|---|---|---|---|
| B99 | B110 | anti-PM-Scl | AHA | 1 | 0 |
| B99 | B120 | anti-PM-Scl | HCV-cAg | 0.203435 | 0 |
| B99 | B156 | anti-PM-Scl | IgE | 0.177741 | 0 |
| B99 | B168 | anti-PM-Scl | NMP22 | -0.17131 | 0 |
| B99 | B155 | anti-PM-Scl | IgA | -0.15872 | 4.66E-15 |
| B99 | B159 | anti-PM-Scl | β2-MG | -0.15404 | 2.91E-14 |
| B99 | B134 | anti-PM-Scl | CHE | -0.14987 | 1.43E-13 |
| B99 | B108 | anti-PM-Scl | Anti-Cyclin | -0.14294 | 1.81E-12 |
| B99 | B136 | anti-PM-Scl | 5'NT | -0.14092 | 3.71E-12 |
| B99 | B145 | anti-PM-Scl | BGP | -0.14021 | 4.77E-12 |
| B99 | B193 | anti-PM-Scl | K | 0.1384 | 8.96E-12 |
| B99 | B119 | anti-PM-Scl | Hcv-rna | 0.128458 | 2.49E-10 |
| B99 | B135 | anti-PM-Scl | TBA | 0.114445 | 1.77E-08 |
| B99 | B181 | anti-PM-Scl | APTT | -0.11108 | 4.60E-08 |
| B99 | B100 | anti-PM-Scl | anti-Ro52 | 0.103467 | 3.59E-07 |
| B99 | B198 | anti-PM-Scl | P | 0.099814 | 9.17E-07 |

**Supplementary Table 25. PEI Correlations in Coronary+others**

| Row | Column | Row | Column | Cor | *P* |
|---|---|---|---|---|---|
| B1 | B221 | Gender | Hei | -0.69258 | 8.88E-16 |
| B10 | B149 | PG I | TGAb | 0.626788 | 2.36E-12 |
| B10 | B137 | PG I | AFU | -0.62569 | 2.64E-12 |
| B10 | B134 | PG I | CHE | -0.61793 | 5.82E-12 |
| B10 | B138 | PG I | PAB | -0.60274 | 2.57E-11 |
| B10 | B14 | PG I | G-17 | 0.570817 | 4.59E-10 |
| B10 | B12 | PG I | PG II | 0.543064 | 4.43E-09 |
| B10 | B136 | PG I | 5'NT | 0.470415 | 6.91E-07 |
| B10 | B87 | PG I | NSE | -0.46489 | 9.70E-07 |
| B100 | B152 | anti-Ro52 | ACA-Ig(G-A-M) | 0.65263 | 1.42E-13 |
| B100 | B171 | anti-Ro52 | EC | 0.601929 | 2.78E-11 |
| B100 | B116 | anti-Ro52 | TT3 | 0.541736 | 4.91E-09 |
| B101 | B119 | anti-Sc70 | Hcv-rna | 0.749269 | 0 |
| B101 | B182 | anti-Sc70 | TT | 0.839146 | 0 |
| B101 | B159 | anti-Sc70 | β2-MG | 0.552052 | 2.17E-09 |
| B102 | B105 | anti-Sm | anti-U1RNP | 0.785674 | 0 |
| B103 | B104 | anti-SSA/Ro | anti-SSB | -0.75238 | 0 |
| B103 | B167 | anti-SSA/Ro | PC | -0.85716 | 0 |
| B103 | B183 | anti-SSA/Ro | PTA | -0.95482 | 0 |
| B103 | B184 | anti-SSA/Ro | INR | -0.87034 | 0 |
| B103 | B223 | anti-SSA/Ro | FH | -0.64258 | 4.37E-13 |
| B103 | B121 | anti-SSA/Ro | C3 | -0.55205 | 2.17E-09 |
| B103 | B125 | anti-SSA/Ro | Cor | -0.51854 | 2.78E-08 |
| B103 | B117 | anti-SSA/Ro | TT4 | -0.47336 | 5.76E-07 |
| B104 | B106 | anti-SSB | anti-P | -0.95443 | 0 |
| B105 | B159 | anti-U1RNP | β2-MG | 0.732467 | 0 |
| B105 | B169 | anti-U1RNP | U-CRY | 0.989949 | 0 |
| B105 | B109 | anti-U1RNP | AMA-M2 | 0.68853 | 1.78E-15 |
| B105 | B108 | anti-U1RNP | Anti-Cyclin | 0.666667 | 2.75E-14 |
| B105 | B182 | anti-U1RNP | TT | 0.527046 | 1.50E-08 |
| B106 | B121 | anti-P | C3 | -0.77033 | 0 |
| B106 | B183 | anti-P | PTA | -0.71058 | 0 |
| B106 | B223 | anti-P | FH | -0.84683 | 0 |
| B106 | B185 | anti-P | PT | -0.593 | 6.41E-11 |
| B106 | B110 | anti-P | AHA | -0.56614 | 6.83E-10 |
| B107 | B121 | AnuA | C3 | -0.96609 | 0 |
| B107 | B164 | AnuA | COM | -0.7303 | 0 |
| B107 | B185 | AnuA | PT | -0.95618 | 0 |
| B107 | B223 | AnuA | FH | -0.88354 | 0 |
| B107 | B225 | AnuA | hPBG | -0.85635 | 0 |
| B107 | B186 | AnuA | PFC | -0.67082 | 1.64E-14 |
| B107 | B183 | AnuA | PTA | -0.57289 | 3.84E-10 |

| | | | | | |
|---|---|---|---|---|---|
| B107 | B202 | AnuA | e-times | -0.47434 | 5.42E-07 |
| B109 | B116 | AMA-M2 | TT3 | 0.894427 | 0 |
| B109 | B152 | AMA-M2 | ACA-Ig(G-A-M) | 0.98382 | 0 |
| B11 | B12 | PGI/PGII | PG II | -0.708 | 0 |
| B110 | B117 | AHA | TT4 | -0.58926 | 9.03E-11 |
| B112 | B114 | FT4 | TSH TT3 TT4(TT4) | 0.513597 | 3.96E-08 |
| B113 | B197 | FT3 | AG | 0.518842 | 2.72E-08 |
| B113 | B196 | FT3 | TCO2 | -0.51528 | 3.52E-08 |
| B113 | B193 | FT3 | K | -0.51358 | 3.97E-08 |
| B115 | B135 | TSH TT3 TT4(TT3) | TBA | -0.57257 | 3.95E-10 |
| B115 | B136 | TSH TT3 TT4(TT3) | 5'NT | -0.52132 | 2.28E-08 |
| B115 | B194 | TSH TT3 TT4(TT3) | Cl | 0.475457 | 5.05E-07 |
| B116 | B171 | TT3 | EC | 0.833333 | 0 |
| B116 | B182 | TT3 | TT | 0.912871 | 0 |
| B117 | B121 | TT4 | C3 | -0.80178 | 0 |
| B117 | B125 | TT4 | Cor | -0.7303 | 0 |
| B117 | B183 | TT4 | PTA | -0.5547 | 1.76E-09 |
| B117 | B167 | TT4 | PC | -0.50518 | 7.14E-08 |
| B119 | B169 | Hcv-rna | U-CRY | 0.811107 | 0 |
| B119 | B159 | Hcv-rna | β2-MG | 0.69801 | 4.44E-16 |
| B119 | B122 | Hcv-rna | C4 | 0.629512 | 1.77E-12 |
| B12 | B134 | PG II | CHE | -0.57661 | 2.79E-10 |
| B12 | B137 | PG II | AFU | -0.57522 | 3.14E-10 |
| B12 | B149 | PG II | TGAb | 0.565404 | 7.26E-10 |
| B12 | B138 | PG II | PAB | -0.5485 | 2.89E-09 |
| B120 | B148 | HCV-cAg | Anti_TPO | -0.59998 | 3.34E-11 |
| B121 | B125 | C3 | Cor | -0.78072 | 0 |
| B121 | B183 | C3 | PTA | -0.74125 | 0 |
| B121 | B186 | C3 | PFC | -0.92582 | 0 |
| B121 | B223 | C3 | FH | -0.85912 | 0 |
| B122 | B159 | C4 | β2-MG | 0.985611 | 0 |
| B122 | B169 | C4 | U-CRY | 0.873128 | 0 |
| B122 | B152 | C4 | ACA-Ig(G-A-M) | 0.46676 | 8.65E-07 |
| B125 | B202 | Cor | e-times | -0.89443 | 0 |
| B125 | B185 | Cor | PT | -0.67612 | 8.66E-15 |
| B126 | B127 | Alb | A-G ratio | 0.608913 | 1.42E-11 |
| B127 | B130 | A-G ratio | GLB | -0.90521 | 0 |
| B128 | B131 | ALT | AST | 0.669669 | 1.91E-14 |
| B128 | B141 | ALT | AST/ALT | -0.61145 | 1.11E-11 |
| B129 | B133 | IBIL | TBIL | 0.957432 | 0 |
| B129 | B132 | IBIL | Dbil | 0.598676 | 3.78E-11 |
| B132 | B133 | Dbil | TBIL | 0.804273 | 0 |
| B132 | B140 | Dbil | r-GT | 0.546394 | 3.41E-09 |
| B134 | B137 | CHE | AFU | 0.811736 | 0 |

| | | | | | |
|---|---|---|---|---|---|
| B134 | B138 | CHE | PAB | 0.709096 | 0 |
| B134 | B149 | CHE | TGAb | -0.76297 | 0 |
| B134 | B136 | CHE | 5'NT | -0.62829 | 2.02E-12 |
| B134 | B198 | CHE | P | -0.601 | 3.04E-11 |
| B134 | B199 | CHE | Mg | -0.57182 | 4.22E-10 |
| B134 | B196 | CHE | TCO2 | -0.52187 | 2.19E-08 |
| B135 | B194 | TBA | Cl | -0.73944 | 0 |
| B135 | B136 | TBA | 5'NT | 0.638357 | 6.92E-13 |
| B135 | B199 | TBA | Mg | 0.625257 | 2.76E-12 |
| B135 | B200 | TBA | OP | -0.57671 | 2.76E-10 |
| B135 | B179 | TBA | α1-MG | 0.542175 | 4.75E-09 |
| B135 | B196 | TBA | TCO2 | 0.535306 | 8.05E-09 |
| B135 | B138 | TBA | PAB | 0.519166 | 2.66E-08 |
| B135 | B137 | TBA | AFU | 0.505899 | 6.79E-08 |
| B135 | B180 | TBA | Tf | 0.492421 | 1.69E-07 |
| B135 | B195 | TBA | Na | -0.47609 | 4.85E-07 |
| B136 | B196 | 5'NT | TCO2 | 0.829049 | 0 |
| B136 | B199 | 5'NT | Mg | 0.954757 | 0 |
| B136 | B197 | 5'NT | AG | -0.63143 | 1.45E-12 |
| B136 | B193 | 5'NT | K | 0.62101 | 4.26E-12 |
| B136 | B194 | 5'NT | Cl | -0.49958 | 1.05E-07 |
| B136 | B149 | 5'NT | TGAb | 0.498076 | 1.16E-07 |
| B137 | B138 | AFU | PAB | 0.986452 | 0 |
| B137 | B149 | AFU | TGAb | -0.65717 | 8.42E-14 |
| B138 | B149 | PAB | TGAb | -0.59069 | 7.93E-11 |
| B139 | B179 | ADA | α1-MG | -0.84609 | 0 |
| B139 | B180 | ADA | Tf | -0.7895 | 0 |
| B139 | B198 | ADA | P | 0.682261 | 4.00E-15 |
| B139 | B194 | ADA | Cl | 0.606996 | 1.71E-11 |
| B139 | B178 | ADA | Ig-U | -0.57305 | 3.79E-10 |
| B139 | B200 | ADA | OP | 0.482533 | 3.22E-07 |
| B139 | B161 | ADA | LEU | 0.478634 | 4.13E-07 |
| B139 | B196 | ADA | TCO2 | -0.46693 | 8.56E-07 |
| B141 | B170 | AST/ALT | U-GLU | 0.538024 | 6.54E-09 |
| B142 | B145 | 25-(OH)D | BGP | -0.98354 | 0 |
| B142 | B144 | 25-(OH)D | Total-PINP | -0.60776 | 1.59E-11 |
| B142 | B143 | 25-(OH)D | β-CTx | -0.4994 | 1.06E-07 |
| B143 | B144 | β-CTx | Total-PINP | 0.991518 | 0 |
| B143 | B151 | β-CTx | Anti-CCP | 0.755376 | 0 |
| B143 | B145 | β-CTx | BGP | 0.647705 | 2.47E-13 |
| B144 | B145 | Total-PINP | BGP | 0.741234 | 0 |
| B144 | B151 | Total-PINP | Anti-CCP | 0.6638 | 3.86E-14 |
| B148 | B149 | Anti_TPO | TGAb | 0.486432 | 2.51E-07 |
| B15 | B184 | CD3-CD19+(B cell) | INR | -0.80915 | 0 |

| | | | | | |
|---|---|---|---|---|---|
| B15 | B225 | CD3-CD19+(B cell) | hPBG | -0.80218 | 0 |
| B15 | B95 | CD3-CD19+(B cell) | anti-cenpb | -0.94324 | 0 |
| B15 | B185 | CD3-CD19+(B cell) | PT | -0.6482 | 2.34E-13 |
| B15 | B35 | CD3-CD19+(B cell) | β2-MG | -0.6482 | 2.34E-13 |
| B15 | B93 | CD3-CD19+(B cell) | ALDH2-Glu504Lys | -0.5145 | 3.72E-08 |
| B15 | B167 | CD3-CD19+(B cell) | PC | -0.48507 | 2.74E-07 |
| B151 | B168 | Anti-CCP | NMP22 | 0.571548 | 4.31E-10 |
| B152 | B182 | ACA-Ig(G-A-M) | TT | 0.860663 | 0 |
| B153 | B175 | Glu | ALB-U | 0.544447 | 3.97E-09 |
| B153 | B177 | Glu | ACR | 0.517687 | 2.96E-08 |
| B153 | B195 | Glu | Na | -0.48083 | 3.59E-07 |
| B155 | B156 | IgA | IgE | 0.845345 | 0 |
| B155 | B157 | IgA | IgG | -0.85163 | 0 |
| B155 | B158 | IgA | IgM | 0.564441 | 7.88E-10 |
| B156 | B157 | IgE | IgG | -0.99993 | 0 |
| B156 | B158 | IgE | IgM | 0.918132 | 0 |
| B157 | B158 | IgG | IgM | -0.91336 | 0 |
| B159 | B169 | β2-MG | U-CRY | 0.788851 | 0 |
| B159 | B182 | β2-MG | TT | 0.748331 | 0 |
| B16 | B102 | CD3+T | anti-Sm | 0.790569 | 0 |
| B16 | B18 | CD3+T | CD3+CD4+helper(inducer)T | 0.711512 | 0 |
| B16 | B30 | CD3+T | LH | 0.95 | 0 |
| B16 | B94 | CD3+T | ALDH2-Lys504Lys | 0.760639 | 0 |
| B16 | B152 | CD3+T | ACA-Ig(G-A-M) | 0.669439 | 1.95E-14 |
| B16 | B159 | CD3+T | β2-MG | 0.64254 | 4.39E-13 |
| B16 | B38 | CD3+T | BAND% | 0.581378 | 1.83E-10 |
| B16 | B33 | CD3+T | Prog | 0.53033 | 1.17E-08 |
| B16 | B169 | CD3+T | U-CRY | 0.505964 | 6.76E-08 |
| B161 | B170 | LEU | U-GLU | 0.479076 | 4.02E-07 |
| B161 | B200 | LEU | OP | 0.473303 | 5.78E-07 |
| B164 | B184 | COM | INR | 0.851257 | 0 |
| B164 | B225 | COM | hPBG | -0.92387 | 0 |
| B167 | B183 | PC | PTA | -0.8807 | 0 |
| B167 | B223 | PC | FH | -0.76332 | 0 |
| B169 | B182 | U-CRY | TT | 0.849706 | 0 |
| B17 | B152 | CD3+CD16+CD56+NK | ACA-Ig(G-A-M) | 0.746816 | 0 |
| B17 | B159 | CD3+CD16+CD56+NK | β2-MG | 0.927634 | 0 |
| B17 | B169 | CD3+CD16+CD56+NK | U-CRY | 0.727607 | 0 |
| B17 | B30 | CD3+CD16+CD56+NK | LH | 0.997054 | 0 |
| B17 | B29 | CD3+CD16+CD56+NK | E2 | 0.650791 | 1.75E-13 |
| B17 | B119 | CD3+CD16+CD56+NK | Hcv-rna | 0.629512 | 1.77E-12 |
| B17 | B102 | CD3+CD16+CD56+NK | anti-Sm | 0.485071 | 2.74E-07 |
| B171 | B182 | EC | TT | 0.63901 | 6.45E-13 |
| B175 | B177 | ALB-U | ACR | 0.982985 | 0 |

| | | | | | |
|---|---|---|---|---|---|
| B175 | B193 | ALB-U | K | 0.770979 | 0 |
| B175 | B195 | ALB-U | Na | -0.86974 | 0 |
| B175 | B197 | ALB-U | AG | -0.76693 | 0 |
| B175 | B200 | ALB-U | OP | 0.749569 | 0 |
| B175 | B194 | ALB-U | Cl | -0.61654 | 6.69E-12 |
| B175 | B196 | ALB-U | TCO2 | 0.527184 | 1.48E-08 |
| B177 | B193 | ACR | K | 0.763701 | 0 |
| B177 | B195 | ACR | Na | -0.84491 | 0 |
| B177 | B197 | ACR | AG | -0.75748 | 0 |
| B177 | B200 | ACR | OP | 0.761761 | 0 |
| B177 | B194 | ACR | Cl | -0.5839 | 1.46E-10 |
| B177 | B196 | ACR | TCO2 | 0.516384 | 3.25E-08 |
| B178 | B180 | Ig-U | Tf | 0.951058 | 0 |
| B179 | B194 | α1-MG | Cl | -0.61179 | 1.07E-11 |
| B179 | B198 | α1-MG | P | -0.56802 | 5.83E-10 |
| B179 | B200 | α1-MG | OP | -0.49038 | 1.94E-07 |
| B18 | B159 | CD3+CD4+helper(inducer)T | β2-MG | 0.896421 | 0 |
| B18 | B30 | CD3+CD4+helper(inducer)T | LH | 0.948683 | 0 |
| B18 | B33 | CD3+CD4+helper(inducer)T | Prog | 0.894427 | 0 |
| B18 | B98 | CD3+CD4+helper(inducer)T | anti-Jo-1 | 0.673575 | 1.18E-14 |
| B180 | B194 | Tf | Cl | -0.56936 | 5.20E-10 |
| B180 | B198 | Tf | P | -0.52933 | 1.26E-08 |
| B183 | B223 | PTA | FH | -0.83404 | 0 |
| B183 | B225 | PTA | hPBG | -0.66899 | 2.07E-14 |
| B183 | B184 | PTA | INR | -0.63439 | 1.06E-12 |
| B184 | B186 | INR | PFC | -0.83406 | 0 |
| B184 | B202 | INR | e-times | -0.73721 | 0 |
| B184 | B223 | INR | FH | -0.63666 | 8.31E-13 |
| B185 | B223 | PT | FH | -0.96003 | 0 |
| B185 | B225 | PT | hPBG | -0.83744 | 0 |
| B189 | B190 | eGFB | Cre | -0.5992 | 3.60E-11 |
| B19 | B169 | CD3+CD4+/CD3+CD8+ | U-CRY | 0.80829 | 0 |
| B19 | B159 | CD3+CD4+/CD3+CD8+ | β2-MG | 0.690066 | 1.33E-15 |
| B19 | B38 | CD3+CD4+/CD3+CD8+ | BAND% | 0.653197 | 1.33E-13 |
| B19 | B105 | CD3+CD4+/CD3+CD8+ | anti-U1RNP | -0.54433 | 4.01E-09 |
| B19 | B116 | CD3+CD4+/CD3+CD8+ | TT3 | -0.4714 | 6.50E-07 |
| B190 | B191 | Cre | Urea | 0.506992 | 6.30E-08 |
| B193 | B194 | K | Cl | -0.76885 | 0 |
| B193 | B195 | K | Na | -0.75303 | 0 |
| B193 | B196 | K | TCO2 | 0.918447 | 0 |
| B193 | B197 | K | AG | -0.99765 | 0 |
| B193 | B199 | K | Mg | 0.731296 | 0 |
| B194 | B195 | Cl | Na | 0.840388 | 0 |
| B194 | B196 | Cl | TCO2 | -0.78599 | 0 |

| | | | | | |
|---|---|---|---|---|---|
| B194 | B197 | Cl | AG | 0.797143 | 0 |
| B194 | B199 | Cl | Mg | -0.49299 | 1.63E-07 |
| B194 | B198 | Cl | P | 0.482485 | 3.23E-07 |
| B195 | B197 | Na | AG | 0.754191 | 0 |
| B195 | B196 | Na | TCO2 | -0.55019 | 2.52E-09 |
| B196 | B197 | TCO2 | AG | -0.93546 | 0 |
| B196 | B199 | TCO2 | Mg | 0.830432 | 0 |
| B197 | B199 | AG | Mg | -0.72346 | 0 |
| B20 | B93 | CD3+CD4+CD8+ | ALDH2-Glu504Lys | -0.75378 | 0 |
| B20 | B99 | CD3+CD4+CD8+ | anti-PM-Scl | -0.82916 | 0 |
| B20 | B223 | CD3+CD4+CD8+ | FH | -0.68926 | 1.55E-15 |
| B20 | B125 | CD3+CD4+CD8+ | Cor | -0.53936 | 5.90E-09 |
| B206 | B207 | Waist | WHR | 0.794081 | 0 |
| B206 | B210 | Waist | Hip | 0.737501 | 0 |
| B206 | B215 | Waist | BMI | 0.823895 | 0 |
| B206 | B216 | Waist | BMI | 0.765524 | 0 |
| B206 | B222 | Waist | NS | 0.631107 | 1.50E-12 |
| B207 | B215 | WHR | BMI | 0.537719 | 6.69E-09 |
| B21 | B107 | CD3+CD4-CD8- | AnuA | -0.84515 | 0 |
| B21 | B223 | CD3+CD4-CD8- | FH | -0.74673 | 0 |
| B21 | B35 | CD3+CD4-CD8- | β2-MG | -0.90914 | 0 |
| B21 | B92 | CD3+CD4-CD8- | ALDH2-Glu504Glu | -0.80178 | 0 |
| B21 | B97 | CD3+CD4-CD8- | dsDNA | -0.67445 | 1.07E-14 |
| B21 | B31 | CD3+CD4-CD8- | FSH | -0.59761 | 4.18E-11 |
| B21 | B96 | CD3+CD4-CD8- | dsDNA (IIF) | -0.56695 | 6.38E-10 |
| B21 | B23 | CD3+CD4-CD8- | CD3-CD16+CD56+(NK) | -0.56149 | 1.01E-09 |
| B21 | B95 | CD3+CD4-CD8- | anti-cenpb | -0.46771 | 8.16E-07 |
| B210 | B215 | Hip | BMI | 0.7484 | 0 |
| B210 | B216 | Hip | BMI | 0.774636 | 0 |
| B210 | B222 | Hip | NS | 0.583098 | 1.57E-10 |
| B211 | B212 | IOP-R | IOP-L | 0.82073 | 0 |
| B213 | B214 | CVA-R | CVA-L | 0.698652 | 4.44E-16 |
| B215 | B216 | BMI | BMI | 0.780053 | 0 |
| B215 | B222 | BMI | NS | 0.600002 | 3.34E-11 |
| B216 | B222 | BMI | NS | 0.804072 | 0 |
| B216 | B221 | BMI | Hei | 0.555081 | 1.70E-09 |
| B218 | B220 | DBP | SBP | 0.502075 | 8.83E-08 |
| B22 | B121 | CD3+CD8+(T) | C3 | -0.87482 | 0 |
| B22 | B223 | CD3+CD8+(T) | FH | -0.95038 | 0 |
| B22 | B35 | CD3+CD8+(T) | β2-MG | -0.90914 | 0 |
| B22 | B104 | CD3+CD8+(T) | anti-SSB | -0.69369 | 8.88E-16 |
| B22 | B183 | CD3+CD8+(T) | PTA | -0.5447 | 3.90E-09 |
| B221 | B222 | Hei | NS | 0.482624 | 3.20E-07 |
| B223 | B225 | FH | hPBG | -0.84416 | 0 |

| | | | | | |
|---|---|---|---|---|---|
| B23 | B107 | CD3-CD16+CD56+(NK) | AnuA | -0.9135 | 0 |
| B23 | B117 | CD3-CD16+CD56+(NK) | TT4 | -0.85769 | 0 |
| B23 | B183 | CD3-CD16+CD56+(NK) | PTA | -0.86232 | 0 |
| B23 | B184 | CD3-CD16+CD56+(NK) | INR | -0.89056 | 0 |
| B23 | B223 | CD3-CD16+CD56+(NK) | FH | -0.73374 | 0 |
| B23 | B35 | CD3-CD16+CD56+(NK) | β2-MG | -0.99258 | 0 |
| B23 | B36 | CD3-CD16+CD56+(NK) | V-Lym% | -0.87538 | 0 |
| B23 | B96 | CD3-CD16+CD56+(NK) | dsDNA (IIF) | -0.92848 | 0 |
| B23 | B225 | CD3-CD16+CD56+(NK) | hPBG | -0.64651 | 2.83E-13 |
| B23 | B92 | CD3-CD16+CD56+(NK) | ALDH2-Glu504Glu | -0.52523 | 1.71E-08 |
| B24 | B27 | αHBDH | LDH | 0.656053 | 9.57E-14 |
| B24 | B67 | αHBDH | WBRSR(mPas_50) | -0.53542 | 7.98E-09 |
| B24 | B66 | αHBDH | WBRSR(mPas_100) | -0.53354 | 9.20E-09 |
| B24 | B63 | αHBDH | WBRSR(mPas_200) | -0.51419 | 3.80E-08 |
| B24 | B57 | αHBDH | RBCAI | 0.501321 | 9.30E-08 |
| B24 | B61 | αHBDH | WBHSRV | -0.48289 | 3.15E-07 |
| B24 | B64 | αHBDH | WBRSR(mPas_5) | -0.46721 | 8.42E-07 |
| B25 | B134 | CK | CHE | 0.497166 | 1.23E-07 |
| B25 | B176 | CK | Ucr | 0.468903 | 7.59E-07 |
| B26 | B137 | CKMB | AFU | 0.698347 | 4.44E-16 |
| B26 | B138 | CKMB | PAB | 0.689824 | 1.33E-15 |
| B26 | B134 | CKMB | CHE | 0.543139 | 4.40E-09 |
| B26 | B87 | CKMB | NSE | 0.471651 | 6.40E-07 |
| B29 | B32 | E2 | Testo | 0.782624 | 0 |
| B29 | B79 | E2 | hb V -dna | 0.886593 | 0 |
| B29 | B100 | E2 | anti-Ro52 | 0.65938 | 6.51E-14 |
| B29 | B33 | E2 | Prog | 0.6 | 3.34E-11 |
| B30 | B101 | LH | anti-Sc70 | 0.912871 | 0 |
| B30 | B159 | LH | β2-MG | 0.907115 | 0 |
| B30 | B94 | LH | ALDH2-Lys504Lys | 0.760639 | 0 |
| B30 | B33 | LH | Prog | 0.565685 | 7.09E-10 |
| B30 | B119 | LH | Hcv-rna | 0.564288 | 7.98E-10 |
| B30 | B182 | LH | TT | 0.494975 | 1.43E-07 |
| B31 | B117 | FSH | TT4 | -0.91287 | 0 |
| B31 | B93 | FSH | ALDH2-Glu504Lys | -0.55902 | 1.23E-09 |
| B32 | B105 | Testo | anti-U1RNP | 0.824958 | 0 |
| B32 | B122 | Testo | C4 | 0.727607 | 0 |
| B32 | B169 | Testo | U-CRY | 1 | 0 |
| B32 | B38 | Testo | BAND% | -0.49497 | 1.43E-07 |
| B33 | B101 | Prog | anti-Sc70 | 0.903696 | 0 |
| B33 | B152 | Prog | ACA-Ig(G-A-M) | 0.989762 | 0 |
| B33 | B171 | Prog | EC | 0.730297 | 0 |
| B33 | B79 | Prog | hb V -dna | 0.784293 | 0 |
| B33 | B109 | Prog | AMA-M2 | 0.489898 | 2.00E-07 |

| | | | | | |
|---|---|---|---|---|---|
| B34 | B198 | TK | P | -0.66699 | 2.62E-14 |
| B34 | B187 | TK | NSE | -0.60477 | 2.12E-11 |
| B34 | B75 | TK | ApoA1/B | -0.59189 | 7.10E-11 |
| B34 | B135 | TK | TBA | 0.564426 | 7.88E-10 |
| B34 | B76 | TK | ApoB | 0.555771 | 1.61E-09 |
| B34 | B194 | TK | Cl | -0.52446 | 1.81E-08 |
| B34 | B139 | TK | ADA | -0.51557 | 3.44E-08 |
| B34 | B179 | TK | α1-MG | 0.511721 | 4.52E-08 |
| B34 | B134 | TK | CHE | 0.487389 | 2.35E-07 |
| B35 | B225 | β2-MG | hPBG | -0.74439 | 0 |
| B35 | B117 | β2-MG | TT4 | -0.61721 | 6.25E-12 |
| B35 | B183 | β2-MG | PTA | -0.59914 | 3.62E-11 |
| B35 | B125 | β2-MG | Cor | -0.59161 | 7.29E-11 |
| B35 | B95 | β2-MG | anti-cenpb | -0.5197 | 2.56E-08 |
| B36 | B103 | V-Lym% | anti-SSA/Ro | -0.98382 | 0 |
| B36 | B202 | V-Lym% | e-times | -0.75 | 0 |
| B36 | B95 | V-Lym% | anti-cenpb | -1 | 0 |
| B36 | B183 | V-Lym% | PTA | -0.64162 | 4.85E-13 |
| B36 | B164 | V-Lym% | COM | -0.57735 | 2.61E-10 |
| B36 | B99 | V-Lym% | anti-PM-Scl | 0.5 | 1.02E-07 |
| B36 | B167 | V-Lym% | PC | -0.4714 | 6.50E-07 |
| B37 | B98 | BAND | anti-Jo-1 | 0.907218 | 0 |
| B37 | B109 | BAND | AMA-M2 | 0.645497 | 3.16E-13 |
| B37 | B171 | BAND | EC | -0.48113 | 3.53E-07 |
| B38 | B100 | BAND% | anti-Ro52 | 0.917463 | 0 |
| B38 | B94 | BAND% | ALDH2-Lys504Lys | 0.831522 | 0 |
| B38 | B119 | BAND% | Hcv-rna | 0.596481 | 4.64E-11 |
| B38 | B79 | BAND% | hb V -dna | 0.56073 | 1.07E-09 |
| B38 | B98 | BAND% | anti-Jo-1 | 0.489898 | 2.00E-07 |
| B40 | B42 | WBC | MON | 0.654252 | 1.18E-13 |
| B41 | B42 | MON % | MON | 0.623488 | 3.31E-12 |
| B43 | B44 | MCV | MCH | 0.940905 | 0 |
| B43 | B47 | MCV | RDW | 0.598278 | 3.93E-11 |
| B43 | B89 | MCV | CA 72-4 | -0.49358 | 1.57E-07 |
| B44 | B45 | MCH | MCHC | 0.585214 | 1.30E-10 |
| B44 | B89 | MCH | CA 72-4 | -0.47542 | 5.06E-07 |
| B45 | B51 | MCHC | Hb | 0.483985 | 2.94E-07 |
| B46 | B51 | RBC Count | Hb | 0.662546 | 4.49E-14 |
| B46 | B47 | RBC Count | RDW | -0.5056 | 6.94E-08 |
| B48 | B53 | LYM% | GR% | -0.95701 | 0 |
| B5 | B122 | WB-Mg | C4 | 0.754594 | 0 |
| B5 | B16 | WB-Mg | CD3+T | 0.715542 | 0 |
| B5 | B169 | WB-Mg | U-CRY | 0.848528 | 0 |
| B5 | B182 | WB-Mg | TT | 0.822192 | 0 |

| | | | | | |
|---|---|---|---|---|---|
| B5 | B30 | WB-Mg | LH | 0.98387 | 0 |
| B5 | B94 | WB-Mg | ALDH2-Lys504Lys | 0.907115 | 0 |
| B5 | B7 | WB-Mg | WB-Cop | 0.682793 | 3.55E-15 |
| B5 | B98 | WB-Mg | anti-Jo-1 | 0.653197 | 1.33E-13 |
| B5 | B17 | WB-Mg | CD3+CD16+CD56+NK | 0.617395 | 6.14E-12 |
| B5 | B119 | WB-Mg | Hcv-rna | 0.596481 | 4.64E-11 |
| B5 | B171 | WB-Mg | EC | 0.57735 | 2.61E-10 |
| B5 | B38 | WB-Mg | BAND% | 0.56 | 1.14E-09 |
| B5 | B100 | WB-Mg | anti-Ro52 | 0.500435 | 9.88E-08 |
| B50 | B190 | eo% | Cre | 0.53295 | 9.62E-09 |
| B54 | B58 | ESR | HCT | -0.86704 | 0 |
| B54 | B62 | ESR | WBHSRVI | -0.80334 | 0 |
| B54 | B57 | ESR | RBCAI | 0.688692 | 1.78E-15 |
| B54 | B66 | ESR | WBRSR(mPas_100) | -0.68267 | 3.77E-15 |
| B54 | B63 | ESR | WBRSR(mPas_200) | -0.6814 | 4.44E-15 |
| B54 | B67 | ESR | WBRSR(mPas_50) | -0.67847 | 6.44E-15 |
| B54 | B59 | ESR | WBRV | 0.676647 | 7.99E-15 |
| B54 | B60 | ESR | WBRVI | -0.64899 | 2.14E-13 |
| B54 | B64 | ESR | WBRSR(mPas_5) | -0.55005 | 2.55E-09 |
| B54 | B56 | ESR | IR | -0.53641 | 7.40E-09 |
| B54 | B69 | ESR | PV | 0.467061 | 8.49E-07 |
| B55 | B56 | TK | IR | 0.754832 | 0 |
| B55 | B60 | TK | WBRVI | 0.726014 | 0 |
| B55 | B69 | TK | PV | -0.76325 | 0 |
| B55 | B61 | TK | WBHSRV | 0.552382 | 2.12E-09 |
| B56 | B60 | IR | WBRVI | 0.953202 | 0 |
| B56 | B61 | IR | WBHSRV | 0.81191 | 0 |
| B56 | B62 | IR | WBHSRVI | 0.897485 | 0 |
| B56 | B69 | IR | PV | -0.88302 | 0 |
| B56 | B57 | IR | RBCAI | -0.47233 | 6.14E-07 |
| B57 | B58 | RBCAI | HCT | -0.91068 | 0 |
| B57 | B59 | RBCAI | WBRV | 0.835044 | 0 |
| B57 | B62 | RBCAI | WBHSRVI | -0.77797 | 0 |
| B57 | B63 | RBCAI | WBRSR(mPas_200) | -0.79379 | 0 |
| B57 | B66 | RBCAI | WBRSR(mPas_100) | -0.83932 | 0 |
| B57 | B67 | RBCAI | WBRSR(mPas_50) | -0.81744 | 0 |
| B57 | B64 | RBCAI | WBRSR(mPas_5) | -0.55854 | 1.28E-09 |
| B57 | B61 | RBCAI | WBHSRV | -0.47163 | 6.41E-07 |
| B58 | B59 | HCT | WBRV | -0.81767 | 0 |
| B58 | B62 | HCT | WBHSRVI | 0.721986 | 0 |
| B58 | B63 | HCT | WBRSR(mPas_200) | 0.846678 | 0 |
| B58 | B66 | HCT | WBRSR(mPas_100) | 0.874946 | 0 |
| B58 | B67 | HCT | WBRSR(mPas_50) | 0.86328 | 0 |
| B58 | B64 | HCT | WBRSR(mPas_5) | 0.673035 | 1.27E-14 |

| | | | | | |
|---|---|---|---|---|---|
| B59 | B62 | WBRV | WBHSRVI | -0.56555 | 7.17E-10 |
| B59 | B66 | WBRV | WBRSR(mPas_100) | -0.48919 | 2.09E-07 |
| B6 | B103 | WB-Fe | anti-SSA/Ro | -0.83152 | 0 |
| B6 | B121 | WB-Fe | C3 | -0.7303 | 0 |
| B6 | B125 | WB-Fe | Cor | -0.80178 | 0 |
| B6 | B223 | WB-Fe | FH | -0.91077 | 0 |
| B6 | B36 | WB-Fe | V-Lym% | -0.95618 | 0 |
| B6 | B106 | WB-Fe | anti-P | -0.60945 | 1.35E-11 |
| B6 | B20 | WB-Fe | CD3+CD4+CD8+ | -0.5766 | 2.79E-10 |
| B6 | B183 | WB-Fe | PTA | -0.4872 | 2.38E-07 |
| B6 | B23 | WB-Fe | CD3-CD16+CD56+(NK) | -0.47082 | 6.74E-07 |
| B60 | B61 | WBRVI | WBHSRV | 0.75645 | 0 |
| B60 | B62 | WBRVI | WBHSRVI | 0.862277 | 0 |
| B60 | B69 | WBRVI | PV | -0.87094 | 0 |
| B61 | B62 | WBHSRV | WBHSRVI | 0.745909 | 0 |
| B61 | B63 | WBHSRV | WBRSR(mPas_200) | 0.729729 | 0 |
| B61 | B64 | WBHSRV | WBRSR(mPas_5) | 0.733236 | 0 |
| B61 | B67 | WBHSRV | WBRSR(mPas_50) | 0.697794 | 4.44E-16 |
| B61 | B66 | WBHSRV | WBRSR(mPas_100) | 0.692254 | 1.11E-15 |
| B61 | B65 | WBHSRV | WBRSR(mPas_1) | 0.613271 | 9.26E-12 |
| B62 | B69 | WBHSRVI | PV | -0.74732 | 0 |
| B62 | B63 | WBHSRVI | WBRSR(mPas_200) | 0.693325 | 8.88E-16 |
| B62 | B66 | WBHSRVI | WBRSR(mPas_100) | 0.694275 | 8.88E-16 |
| B62 | B67 | WBHSRVI | WBRSR(mPas_50) | 0.680272 | 5.11E-15 |
| B62 | B64 | WBHSRVI | WBRSR(mPas_5) | 0.512669 | 4.23E-08 |
| B63 | B64 | WBRSR(mPas_200) | WBRSR(mPas_5) | 0.945827 | 0 |
| B63 | B66 | WBRSR(mPas_200) | WBRSR(mPas_100) | 0.995582 | 0 |
| B63 | B67 | WBRSR(mPas_200) | WBRSR(mPas_50) | 0.997282 | 0 |
| B63 | B65 | WBRSR(mPas_200) | WBRSR(mPas_1) | 0.671654 | 1.51E-14 |
| B64 | B65 | WBRSR(mPas_5) | WBRSR(mPas_1) | 0.87471 | 0 |
| B64 | B66 | WBRSR(mPas_5) | WBRSR(mPas_100) | 0.919266 | 0 |
| B64 | B67 | WBRSR(mPas_5) | WBRSR(mPas_50) | 0.934064 | 0 |
| B65 | B67 | WBRSR(mPas_1) | WBRSR(mPas_50) | 0.644149 | 3.67E-13 |
| B65 | B66 | WBRSR(mPas_1) | WBRSR(mPas_100) | 0.613445 | 9.10E-12 |
| B66 | B67 | WBRSR(mPas_100) | WBRSR(mPas_50) | 0.999085 | 0 |
| B69 | B187 | PV | NSE | -0.51527 | 3.52E-08 |
| B7 | B102 | WB-Cop | anti-Sm | 0.80468 | 0 |
| B7 | B124 | WB-Cop | ACTH | 0.984798 | 0 |
| B7 | B17 | WB-Cop | CD3+CD16+CD56+NK | 0.810679 | 0 |
| B7 | B19 | WB-Cop | CD3+CD4+/CD3+CD8+ | 0.750479 | 0 |
| B7 | B30 | WB-Cop | LH | 0.763386 | 0 |
| B7 | B79 | WB-Cop | hb V -dna | 0.877866 | 0 |
| B7 | B182 | WB-Cop | TT | 0.539796 | 5.71E-09 |
| B7 | B101 | WB-Cop | anti-Sc70 | 0.536056 | 7.60E-09 |

| | | | | | |
|---|---|---|---|---|---|
| B7 | B169 | WB-Cop | U-CRY | 0.519947 | 2.51E-08 |
| B7 | B100 | WB-Cop | anti-Ro52 | 0.492827 | 1.65E-07 |
| B70 | B77 | LDL-C | TC | 0.931207 | 0 |
| B71 | B91 | TG | TP | 0.509598 | 5.25E-08 |
| B72 | B216 | HDL-C | BMI | -0.56054 | 1.09E-09 |
| B72 | B222 | HDL-C | NS | -0.47359 | 5.67E-07 |
| B73 | B78 | ApoA | HBcAb | 0.470971 | 6.68E-07 |
| B74 | B75 | ApoA1 | ApoA1/B | 0.548173 | 2.96E-09 |
| B74 | B198 | ApoA1 | P | 0.512396 | 4.31E-08 |
| B74 | B178 | ApoA1 | Ig-U | -0.49522 | 1.40E-07 |
| B74 | B180 | ApoA1 | Tf | -0.48929 | 2.08E-07 |
| B75 | B76 | ApoA1/B | ApoB | -0.77883 | 0 |
| B75 | B154 | ApoA1/B | RF | -0.59732 | 4.29E-11 |
| B75 | B134 | ApoA1/B | CHE | -0.58876 | 9.45E-11 |
| B75 | B198 | ApoA1/B | P | 0.542765 | 4.53E-09 |
| B75 | B149 | ApoA1/B | TGAb | 0.505024 | 7.22E-08 |
| B75 | B137 | ApoA1/B | AFU | -0.50081 | 9.63E-08 |
| B76 | B154 | ApoB | RF | 0.482859 | 3.16E-07 |
| B78 | B123 | HBcAb | hs-CRP1 | 0.643561 | 3.92E-13 |
| B79 | B109 | hb V -dna | AMA-M2 | 0.807427 | 0 |
| B79 | B159 | hb V -dna | β2-MG | 0.838445 | 0 |
| B79 | B124 | hb V -dna | ACTH | 0.646997 | 2.68E-13 |
| B79 | B98 | hb V -dna | anti-Jo-1 | 0.616316 | 6.84E-12 |
| B79 | B152 | hb V -dna | ACA-Ig(G-A-M) | 0.586967 | 1.11E-10 |
| B8 | B104 | WB-Zinc | anti-SSB | -0.96013 | 0 |
| B8 | B21 | WB-Zinc | CD3+CD4-CD8- | -0.79091 | 0 |
| B8 | B223 | WB-Zinc | FH | -0.8401 | 0 |
| B8 | B225 | WB-Zinc | hPBG | -0.72853 | 0 |
| B8 | B35 | WB-Zinc | β2-MG | -0.85534 | 0 |
| B8 | B97 | WB-Zinc | dsDNA | -0.95847 | 0 |
| B8 | B117 | WB-Zinc | TT4 | -0.67514 | 9.77E-15 |
| B8 | B22 | WB-Zinc | CD3+CD8+(T) | -0.65134 | 1.64E-13 |
| B8 | B125 | WB-Zinc | Cor | -0.6228 | 3.55E-12 |
| B8 | B95 | WB-Zinc | anti-cenpb | -0.60927 | 1.37E-11 |
| B8 | B121 | WB-Zinc | C3 | -0.5698 | 5.01E-10 |
| B8 | B103 | WB-Zinc | anti-SSA/Ro | -0.51378 | 3.91E-08 |
| B8 | B183 | WB-Zinc | PTA | -0.47305 | 5.87E-07 |
| B81 | B156 | TSGF | IgE | 0.672842 | 1.29E-14 |
| B81 | B157 | TSGF | IgG | -0.67271 | 1.31E-14 |
| B81 | B158 | TSGF | IgM | 0.620719 | 4.39E-12 |
| B81 | B155 | TSGF | IgA | 0.564792 | 7.65E-10 |
| B82 | B83 | CEA | AFP | 0.98185 | 0 |
| B82 | B86 | CEA | TBhCG | 0.686899 | 2.22E-15 |
| B82 | B134 | CEA | CHE | 0.563293 | 8.67E-10 |

| | | | | | |
|---|---|---|---|---|---|
| B82 | B89 | CEA | CA 72-4 | 0.555921 | 1.59E-09 |
| B82 | B137 | CEA | AFU | 0.525313 | 1.70E-08 |
| B82 | B138 | CEA | PAB | 0.479448 | 3.92E-07 |
| B82 | B84 | CEA | C-PSA | -0.46467 | 9.82E-07 |
| B83 | B86 | AFP | TBhCG | 0.704566 | 2.22E-16 |
| B83 | B134 | AFP | CHE | 0.685265 | 2.66E-15 |
| B83 | B137 | AFP | AFU | 0.640308 | 5.60E-13 |
| B83 | B138 | AFP | PAB | 0.584613 | 1.37E-10 |
| B83 | B89 | AFP | CA 72-4 | 0.563765 | 8.33E-10 |
| B83 | B149 | AFP | TGAb | -0.54364 | 4.23E-09 |
| B83 | B85 | AFP | SCC | 0.465779 | 9.18E-07 |
| B84 | B89 | C-PSA | CA 72-4 | -0.53709 | 7.03E-09 |
| B84 | B86 | C-PSA | TBhCG | -0.47401 | 5.53E-07 |
| B84 | B213 | C-PSA | CVA-R | 0.473101 | 5.85E-07 |
| B85 | B86 | SCC | TBhCG | 0.854844 | 0 |
| B85 | B87 | SCC | NSE | 0.881899 | 0 |
| B85 | B88 | SCC | CA 242 | 0.780969 | 0 |
| B85 | B134 | SCC | CHE | 0.602503 | 2.63E-11 |
| B85 | B90 | SCC | CYFRA 21-1 | 0.574308 | 3.40E-10 |
| B85 | B137 | SCC | AFU | 0.560736 | 1.07E-09 |
| B85 | B138 | SCC | PAB | 0.511586 | 4.57E-08 |
| B85 | B149 | SCC | TGAb | -0.48878 | 2.15E-07 |
| B86 | B88 | TBhCG | CA 242 | 0.893464 | 0 |
| B86 | B90 | TBhCG | CYFRA 21-1 | 0.59218 | 6.91E-11 |
| B86 | B87 | TBhCG | NSE | 0.556684 | 1.49E-09 |
| B86 | B134 | TBhCG | CHE | 0.483354 | 3.06E-07 |
| B87 | B134 | NSE | CHE | 0.73162 | 0 |
| B87 | B137 | NSE | AFU | 0.685483 | 2.66E-15 |
| B87 | B138 | NSE | PAB | 0.626171 | 2.51E-12 |
| B87 | B149 | NSE | TGAb | -0.57144 | 4.35E-10 |
| B87 | B89 | NSE | CA 72-4 | -0.48048 | 3.68E-07 |
| B88 | B90 | CA 242 | CYFRA 21-1 | 0.513665 | 3.94E-08 |
| B9 | B169 | RH | U-CRY | 0.816497 | 0 |
| B9 | B17 | RH | CD3+CD16+CD56+NK | 0.792118 | 0 |
| B9 | B79 | RH | hb V -dna | 0.996116 | 0 |
| B9 | B116 | RH | TT3 | 0.666667 | 2.75E-14 |
| B9 | B94 | RH | ALDH2-Lys504Lys | 0.654654 | 1.12E-13 |
| B9 | B32 | RH | Testo | -0.61237 | 1.01E-11 |
| B9 | B171 | RH | EC | -0.5 | 1.02E-07 |
| B90 | B187 | CYFRA 21-1 | NSE | 0.79911 | 0 |
| B91 | B130 | TP | GLB | 0.706414 | 0 |
| B91 | B126 | TP | Alb | 0.498333 | 1.14E-07 |
| B92 | B103 | ALDH2-Glu504Glu | anti-SSA/Ro | -0.94868 | 0 |
| B92 | B185 | ALDH2-Glu504Glu | PT | -0.94491 | 0 |

| | | | | | |
|---|---|---|---|---|---|
| B92 | B99 | ALDH2-Glu504Glu | anti-PM-Scl | 0.75 | 0 |
| B93 | B104 | ALDH2-Glu504Lys | anti-SSB | -0.89222 | 0 |
| B93 | B117 | ALDH2-Glu504Lys | TT4 | -0.71443 | 0 |
| B93 | B223 | ALDH2-Glu504Lys | FH | -0.9525 | 0 |
| B93 | B97 | ALDH2-Glu504Lys | dsDNA | -0.97502 | 0 |
| B93 | B99 | ALDH2-Glu504Lys | anti-PM-Scl | 0.75 | 0 |
| B93 | B103 | ALDH2-Glu504Lys | anti-SSA/Ro | -0.68516 | 2.66E-15 |
| B94 | B108 | ALDH2-Lys504Lys | Anti-Cyclin | -0.94491 | 0 |
| B94 | B105 | ALDH2-Lys504Lys | anti-U1RNP | 0.629941 | 1.70E-12 |
| B95 | B103 | anti-cenpb | anti-SSA/Ro | -0.94868 | 0 |
| B95 | B107 | anti-cenpb | AnuA | -0.94868 | 0 |
| B95 | B183 | anti-cenpb | PTA | -0.84921 | 0 |
| B95 | B225 | anti-cenpb | hPBG | -0.98473 | 0 |
| B95 | B97 | anti-cenpb | dsDNA | -0.7456 | 0 |
| B95 | B99 | anti-cenpb | anti-PM-Scl | -0.875 | 0 |
| B95 | B104 | anti-cenpb | anti-SSB | -0.68944 | 1.55E-15 |
| B95 | B202 | anti-cenpb | e-times | -0.625 | 2.83E-12 |
| B95 | B184 | anti-cenpb | INR | -0.58977 | 8.62E-11 |
| B95 | B110 | anti-cenpb | AHA | -0.57735 | 2.61E-10 |
| B95 | B125 | anti-cenpb | Cor | -0.55902 | 1.23E-09 |
| B95 | B167 | anti-cenpb | PC | -0.48614 | 2.55E-07 |
| B96 | B97 | dsDNA (IIF) | dsDNA | -0.64889 | 2.17E-13 |
| B96 | B186 | dsDNA (IIF) | PFC | -0.5 | 1.02E-07 |
| B97 | B103 | dsDNA | anti-SSA/Ro | -0.94312 | 0 |
| B97 | B110 | dsDNA | AHA | -0.72849 | 0 |
| B97 | B185 | dsDNA | PT | -0.86711 | 0 |
| B97 | B223 | dsDNA | FH | -0.87407 | 0 |
| B97 | B225 | dsDNA | hPBG | -0.81893 | 0 |
| B97 | B121 | dsDNA | C3 | -0.67585 | 8.88E-15 |
| B97 | B183 | dsDNA | PTA | -0.5455 | 3.66E-09 |
| B97 | B106 | dsDNA | anti-P | -0.49491 | 1.43E-07 |
| B98 | B108 | anti-Jo-1 | Anti-Cyclin | 0.952579 | 0 |
| B98 | B119 | anti-Jo-1 | Hcv-rna | 0.811708 | 0 |
| B98 | B122 | anti-Jo-1 | C4 | 0.840168 | 0 |
| B98 | B124 | anti-Jo-1 | ACTH | 0.816497 | 0 |
| B98 | B159 | anti-Jo-1 | β2-MG | 0.966092 | 0 |
| B98 | B100 | anti-Jo-1 | anti-Ro52 | 0.681005 | 4.66E-15 |
| B98 | B171 | anti-Jo-1 | EC | 0.628539 | 1.96E-12 |
| B98 | B182 | anti-Jo-1 | TT | 0.602464 | 2.64E-11 |
| B98 | B109 | anti-Jo-1 | AMA-M2 | 0.597319 | 4.29E-11 |
| B99 | B106 | anti-PM-Scl | anti-P | -0.58835 | 9.81E-11 |
| B99 | B223 | anti-PM-Scl | FH | -0.5715 | 4.33E-10 |
| B99 | B202 | anti-PM-Scl | e-times | -0.5 | 1.02E-07 |

**Supplementary Table 26. PEI Correlations in Diabetes+others**

| Row | Column | Row | Column | Cor | *P* |
|---|---|---|---|---|---|
| B1 | B216 | Gender | BMI | -0.443191974 | 0 |
| B1 | B221 | Gender | Hei | -0.655641509 | 0 |
| B1 | B222 | Gender | NS | -0.640859477 | 0 |
| B1 | B51 | Gender | Hb | -0.441414663 | 0 |
| B1 | B207 | Gender | WHR | -0.365030058 | 3.37E-13 |
| B1 | B72 | Gender | HDL-C | 0.338745074 | 1.82E-11 |
| B1 | B174 | Gender | Urea/Cr | 0.335696791 | 2.82E-11 |
| B1 | B206 | Gender | Waist | -0.335068543 | 3.08E-11 |
| B1 | B45 | Gender | MCHC | -0.310593529 | 8.75E-10 |
| B1 | B209 | Gender | Smoking | -0.30922387 | 1.05E-09 |
| B1 | B163 | Gender | SG | -0.303251609 | 2.25E-09 |
| B1 | B204 | Gender | Drinking | -0.264455116 | 2.18E-07 |
| B1 | B192 | Gender | UA | -0.262989113 | 2.56E-07 |
| B10 | B12 | PG I | PG II | 0.490712182 | 0 |
| B100 | B103 | anti-Ro52 | anti-SSA/Ro | 1 | 0 |
| B100 | B104 | anti-Ro52 | anti-SSB | 1 | 0 |
| B100 | B105 | anti-Ro52 | anti-U1RNP | 1 | 0 |
| B100 | B106 | anti-Ro52 | anti-P | 0.999999999 | 0 |
| B100 | B107 | anti-Ro52 | AnuA | 1 | 0 |
| B100 | B151 | anti-Ro52 | Anti-CCP | 0.589369501 | 0 |
| B100 | B193 | anti-Ro52 | K | 0.396577754 | 1.78E-15 |
| B100 | B157 | anti-Ro52 | IgG | 0.381317843 | 2.35E-14 |
| B100 | B198 | anti-Ro52 | P | -0.350329926 | 3.28E-12 |
| B100 | B185 | anti-Ro52 | PT | 0.30571844 | 1.64E-09 |
| B100 | B114 | anti-Ro52 | TSH TT3 TT4(TT4) | 0.304406136 | 1.94E-09 |
| B100 | B134 | anti-Ro52 | CHE | -0.262564376 | 2.68E-07 |
| B100 | B150 | anti-Ro52 | ALP | -0.252421184 | 7.86E-07 |
| B101 | B102 | anti-Sc70 | anti-Sm | 0.999999998 | 0 |
| B101 | B106 | anti-Sc70 | anti-P | 0.999999999 | 0 |
| B101 | B107 | anti-Sc70 | AnuA | 0.999999948 | 0 |
| B101 | B151 | anti-Sc70 | Anti-CCP | 0.589369504 | 0 |
| B101 | B193 | anti-Sc70 | K | 0.396577755 | 1.78E-15 |
| B101 | B157 | anti-Sc70 | IgG | 0.381317845 | 2.35E-14 |
| B101 | B198 | anti-Sc70 | P | -0.350329927 | 3.28E-12 |
| B101 | B185 | anti-Sc70 | PT | 0.305718442 | 1.64E-09 |
| B101 | B114 | anti-Sc70 | TSH TT3 TT4(TT4) | 0.304406137 | 1.94E-09 |
| B101 | B134 | anti-Sc70 | CHE | -0.262564377 | 2.68E-07 |
| B101 | B150 | anti-Sc70 | ALP | -0.252421186 | 7.86E-07 |
| B102 | B104 | anti-Sm | anti-SSB | 0.999999999 | 0 |
| B102 | B108 | anti-Sm | Anti-Cyclin | 1 | 0 |
| B102 | B151 | anti-Sm | Anti-CCP | 0.589369503 | 0 |
| B102 | B193 | anti-Sm | K | 0.396577755 | 1.78E-15 |

| | | | | | |
|---|---|---|---|---|---|
| B102 | B157 | anti-Sm | IgG | 0.381317844 | 2.35E-14 |
| B102 | B198 | anti-Sm | P | -0.350329926 | 3.28E-12 |
| B102 | B185 | anti-Sm | PT | 0.305718442 | 1.64E-09 |
| B102 | B114 | anti-Sm | TSH TT3 TT4(TT4) | 0.304406137 | 1.94E-09 |
| B102 | B134 | anti-Sm | CHE | -0.262564377 | 2.68E-07 |
| B102 | B150 | anti-Sm | ALP | -0.252421185 | 7.86E-07 |
| B103 | B104 | anti-SSA/Ro | anti-SSB | 1 | 0 |
| B103 | B105 | anti-SSA/Ro | anti-U1RNP | 1 | 0 |
| B103 | B106 | anti-SSA/Ro | anti-P | 1 | 0 |
| B103 | B151 | anti-SSA/Ro | Anti-CCP | 0.589369501 | 0 |
| B103 | B193 | anti-SSA/Ro | K | 0.396577754 | 1.78E-15 |
| B103 | B157 | anti-SSA/Ro | IgG | 0.381317843 | 2.35E-14 |
| B103 | B198 | anti-SSA/Ro | P | -0.350329926 | 3.28E-12 |
| B103 | B185 | anti-SSA/Ro | PT | 0.305718441 | 1.64E-09 |
| B103 | B114 | anti-SSA/Ro | TSH TT3 TT4(TT4) | 0.304406136 | 1.94E-09 |
| B103 | B134 | anti-SSA/Ro | CHE | -0.262564376 | 2.68E-07 |
| B103 | B150 | anti-SSA/Ro | ALP | -0.252421184 | 7.86E-07 |
| B104 | B105 | anti-SSB | anti-U1RNP | 0.999999999 | 0 |
| B104 | B107 | anti-SSB | AnuA | 0.999999997 | 0 |
| B104 | B109 | anti-SSB | AMA-M2 | 0.999999999 | 0 |
| B104 | B151 | anti-SSB | Anti-CCP | 0.589369501 | 0 |
| B104 | B193 | anti-SSB | K | 0.396577754 | 1.78E-15 |
| B104 | B157 | anti-SSB | IgG | 0.381317843 | 2.35E-14 |
| B104 | B198 | anti-SSB | P | -0.350329925 | 3.28E-12 |
| B104 | B185 | anti-SSB | PT | 0.305718441 | 1.64E-09 |
| B104 | B114 | anti-SSB | TSH TT3 TT4(TT4) | 0.304406136 | 1.94E-09 |
| B104 | B134 | anti-SSB | CHE | -0.262564376 | 2.68E-07 |
| B104 | B150 | anti-SSB | ALP | -0.252421184 | 7.86E-07 |
| B105 | B106 | anti-U1RNP | anti-P | 0.999999998 | 0 |
| B105 | B109 | anti-U1RNP | AMA-M2 | 0.999999999 | 0 |
| B105 | B110 | anti-U1RNP | AHA | 0.999999999 | 0 |
| B105 | B151 | anti-U1RNP | Anti-CCP | 0.589369501 | 0 |
| B105 | B193 | anti-U1RNP | K | 0.396577754 | 1.78E-15 |
| B105 | B157 | anti-U1RNP | IgG | 0.381317843 | 2.35E-14 |
| B105 | B198 | anti-U1RNP | P | -0.350329925 | 3.28E-12 |
| B105 | B185 | anti-U1RNP | PT | 0.30571844 | 1.64E-09 |
| B105 | B114 | anti-U1RNP | TSH TT3 TT4(TT4) | 0.304406136 | 1.94E-09 |
| B105 | B134 | anti-U1RNP | CHE | -0.262564376 | 2.68E-07 |
| B105 | B150 | anti-U1RNP | ALP | -0.252421184 | 7.86E-07 |
| B106 | B107 | anti-P | AnuA | 0.999999985 | 0 |
| B106 | B108 | anti-P | Anti-Cyclin | 1 | 0 |
| B106 | B109 | anti-P | AMA-M2 | 1 | 0 |
| B106 | B110 | anti-P | AHA | 0.999999998 | 0 |
| B106 | B151 | anti-P | Anti-CCP | 0.589369501 | 0 |

| | | | | | |
|---|---|---|---|---|---|
| B106 | B193 | anti-P | K | 0.396577754 | 1.78E-15 |
| B106 | B157 | anti-P | IgG | 0.381317843 | 2.35E-14 |
| B106 | B198 | anti-P | P | -0.350329925 | 3.28E-12 |
| B106 | B185 | anti-P | PT | 0.30571844 | 1.64E-09 |
| B106 | B114 | anti-P | TSH TT3 TT4(TT4) | 0.304406136 | 1.94E-09 |
| B106 | B134 | anti-P | CHE | -0.262564376 | 2.68E-07 |
| B106 | B150 | anti-P | ALP | -0.252421184 | 7.86E-07 |
| B107 | B108 | AnuA | Anti-Cyclin | 0.999999999 | 0 |
| B107 | B109 | AnuA | AMA-M2 | 0.999999998 | 0 |
| B107 | B151 | AnuA | Anti-CCP | 0.589369504 | 0 |
| B107 | B193 | AnuA | K | 0.396577755 | 1.78E-15 |
| B107 | B157 | AnuA | IgG | 0.381317845 | 2.35E-14 |
| B107 | B198 | AnuA | P | -0.350329926 | 3.28E-12 |
| B107 | B185 | AnuA | PT | 0.305718446 | 1.64E-09 |
| B107 | B114 | AnuA | TSH TT3 TT4(TT4) | 0.304406137 | 1.94E-09 |
| B107 | B134 | AnuA | CHE | -0.262564377 | 2.68E-07 |
| B107 | B150 | AnuA | ALP | -0.252421186 | 7.86E-07 |
| B108 | B110 | Anti-Cyclin | AHA | 1 | 0 |
| B108 | B151 | Anti-Cyclin | Anti-CCP | 0.589369501 | 0 |
| B108 | B193 | Anti-Cyclin | K | 0.396577754 | 1.78E-15 |
| B108 | B157 | Anti-Cyclin | IgG | 0.381317843 | 2.35E-14 |
| B108 | B198 | Anti-Cyclin | P | -0.350329926 | 3.28E-12 |
| B108 | B185 | Anti-Cyclin | PT | 0.305718441 | 1.64E-09 |
| B108 | B114 | Anti-Cyclin | TSH TT3 TT4(TT4) | 0.304406136 | 1.94E-09 |
| B108 | B134 | Anti-Cyclin | CHE | -0.262564376 | 2.68E-07 |
| B108 | B150 | Anti-Cyclin | ALP | -0.252421184 | 7.86E-07 |
| B109 | B151 | AMA-M2 | Anti-CCP | 0.589369501 | 0 |
| B109 | B193 | AMA-M2 | K | 0.396577754 | 1.78E-15 |
| B109 | B157 | AMA-M2 | IgG | 0.381317843 | 2.35E-14 |
| B109 | B198 | AMA-M2 | P | -0.350329926 | 3.28E-12 |
| B109 | B185 | AMA-M2 | PT | 0.30571844 | 1.64E-09 |
| B109 | B114 | AMA-M2 | TSH TT3 TT4(TT4) | 0.304406136 | 1.94E-09 |
| B109 | B134 | AMA-M2 | CHE | -0.262564376 | 2.68E-07 |
| B109 | B150 | AMA-M2 | ALP | -0.252421184 | 7.86E-07 |
| B11 | B12 | PGI/PGII | PG II | -0.565192792 | 0 |
| B11 | B21 | PGI/PGII | CD3+CD4-CD8- | 0.353818238 | 1.93E-12 |
| B11 | B19 | PGI/PGII | CD3+CD4+/CD3+CD8+ | -0.347975841 | 4.67E-12 |
| B11 | B14 | PGI/PGII | G-17 | -0.305573366 | 1.68E-09 |
| B11 | B185 | PGI/PGII | PT | -0.303568078 | 2.16E-09 |
| B11 | B22 | PGI/PGII | CD3+CD8+(T) | 0.301483661 | 2.82E-09 |
| B11 | B60 | PGI/PGII | WBRVI | 0.270283885 | 1.15E-07 |
| B110 | B151 | AHA | Anti-CCP | 0.589369501 | 0 |
| B110 | B193 | AHA | K | 0.396577754 | 1.78E-15 |
| B110 | B157 | AHA | IgG | 0.381317843 | 2.35E-14 |

| | | | | | |
|---|---|---|---|---|---|
| B110 | B198 | AHA | P | -0.350329926 | 3.28E-12 |
| B110 | B185 | AHA | PT | 0.305718441 | 1.64E-09 |
| B110 | B114 | AHA | TSH TT3 TT4(TT4) | 0.304406136 | 1.94E-09 |
| B110 | B134 | AHA | CHE | -0.262564376 | 2.68E-07 |
| B110 | B150 | AHA | ALP | -0.252421184 | 7.86E-07 |
| B111 | B120 | TSH | HCV-cAg | -0.43612265 | 0 |
| B111 | B112 | TSH | FT4 | -0.405041119 | 4.44E-16 |
| B112 | B114 | FT4 | TSH TT3 TT4(TT4) | 0.367015894 | 2.45E-13 |
| B112 | B113 | FT4 | FT3 | 0.356318543 | 1.32E-12 |
| B113 | B115 | FT3 | TSH TT3 TT4(TT3) | 0.352227049 | 2.46E-12 |
| B114 | B115 | TSH TT3 TT4(TT4) | TSH TT3 TT4(TT3) | 0.437866489 | 0 |
| B114 | B173 | TSH TT3 TT4(TT4) | NTT | 0.301485816 | 2.82E-09 |
| B116 | B117 | TT3 | TT4 | -0.692834742 | 0 |
| B119 | B120 | Hcv-rna | HCV-cAg | 0.466936012 | 0 |
| B119 | B121 | Hcv-rna | C3 | -0.706147397 | 0 |
| B119 | B122 | Hcv-rna | C4 | -0.706147821 | 0 |
| B119 | B124 | Hcv-rna | ACTH | 0.706149653 | 0 |
| B119 | B125 | Hcv-rna | Cor | -0.692337293 | 0 |
| B119 | B152 | Hcv-rna | ACA-Ig(G-A-M) | -0.706168268 | 0 |
| B119 | B158 | Hcv-rna | IgM | -0.505025144 | 0 |
| B119 | B169 | Hcv-rna | U-CRY | -0.70614412 | 0 |
| B119 | B180 | Hcv-rna | Tf | 0.471983653 | 0 |
| B119 | B184 | Hcv-rna | INR | 0.408050848 | 2.22E-16 |
| B119 | B194 | Hcv-rna | Cl | 0.332914152 | 4.19E-11 |
| B119 | B183 | Hcv-rna | PTA | 0.313179571 | 6.23E-10 |
| B119 | B178 | Hcv-rna | Ig-U | 0.311437849 | 7.83E-10 |
| B119 | B193 | Hcv-rna | K | -0.285808521 | 1.92E-08 |
| B119 | B186 | Hcv-rna | PFC | -0.274022467 | 7.54E-08 |
| B119 | B179 | Hcv-rna | α1-MG | 0.271301845 | 1.02E-07 |
| B119 | B182 | Hcv-rna | TT | -0.271267125 | 1.03E-07 |
| B119 | B195 | Hcv-rna | Na | 0.258797387 | 4.02E-07 |
| B120 | B124 | HCV-cAg | ACTH | 0.329752992 | 6.54E-11 |
| B120 | B121 | HCV-cAg | C3 | -0.329729576 | 6.56E-11 |
| B120 | B122 | HCV-cAg | C4 | -0.329730008 | 6.56E-11 |
| B120 | B152 | HCV-cAg | ACA-Ig(G-A-M) | -0.32973563 | 6.56E-11 |
| B120 | B169 | HCV-cAg | U-CRY | -0.329727389 | 6.57E-11 |
| B120 | B125 | HCV-cAg | Cor | -0.324552654 | 1.35E-10 |
| B120 | B180 | HCV-cAg | Tf | 0.322185777 | 1.86E-10 |
| B121 | B124 | C3 | ACTH | -0.999994007 | 0 |
| B121 | B125 | C3 | Cor | 0.979904206 | 0 |
| B121 | B158 | C3 | IgM | 0.709626721 | 0 |
| B121 | B169 | C3 | U-CRY | 0.999996144 | 0 |
| B121 | B182 | C3 | TT | 0.417386687 | 0 |
| B121 | B180 | C3 | Tf | -0.333294007 | 3.97E-11 |

| | | | | | |
|---|---|---|---|---|---|
| B121 | B194 | C3 | Cl | -0.270213457 | 1.16E-07 |
| B122 | B124 | C4 | ACTH | -0.999994639 | 0 |
| B122 | B125 | C4 | Cor | 0.979904731 | 0 |
| B122 | B158 | C4 | IgM | 0.709627145 | 0 |
| B122 | B169 | C4 | U-CRY | 0.999997501 | 0 |
| B122 | B182 | C4 | TT | 0.41738693 | 0 |
| B122 | B180 | C4 | Tf | -0.333294206 | 3.97E-11 |
| B122 | B194 | C4 | Cl | -0.270213632 | 1.16E-07 |
| B123 | B198 | hs-CRP1 | P | 0.341255756 | 1.26E-11 |
| B123 | B155 | hs-CRP1 | IgA | 0.299621336 | 3.56E-09 |
| B124 | B125 | ACTH | Cor | -0.980596928 | 0 |
| B124 | B158 | ACTH | IgM | -0.709602849 | 0 |
| B124 | B169 | ACTH | U-CRY | -0.999989342 | 0 |
| B124 | B182 | ACTH | TT | -0.417025457 | 0 |
| B124 | B180 | ACTH | Tf | 0.333314338 | 3.96E-11 |
| B124 | B194 | ACTH | Cl | 0.270192281 | 1.16E-07 |
| B125 | B152 | Cor | ACA-Ig(G-A-M) | 0.979933227 | 0 |
| B125 | B158 | Cor | IgM | 0.694260508 | 0 |
| B125 | B169 | Cor | U-CRY | 0.979899753 | 0 |
| B125 | B182 | Cor | TT | 0.388582947 | 6.66E-15 |
| B125 | B180 | Cor | Tf | -0.32787265 | 8.51E-11 |
| B125 | B194 | Cor | Cl | -0.26367413 | 2.38E-07 |
| B126 | B127 | Alb | A-G ratio | 0.555894025 | 0 |
| B126 | B188 | Alb | Ca | 0.377045006 | 4.80E-14 |
| B126 | B139 | Alb | ADA | -0.306846256 | 1.42E-09 |
| B126 | B138 | Alb | PAB | 0.304815921 | 1.85E-09 |
| B126 | B135 | Alb | TBA | -0.30309056 | 2.30E-09 |
| B126 | B134 | Alb | CHE | 0.289548807 | 1.23E-08 |
| B126 | B130 | Alb | GLB | -0.255659105 | 5.60E-07 |
| B127 | B130 | A-G ratio | GLB | -0.909483647 | 0 |
| B128 | B131 | ALT | AST | 0.804602391 | 0 |
| B128 | B141 | ALT | AST/ALT | -0.601919068 | 0 |
| B128 | B140 | ALT | r-GT | 0.377813613 | 4.22E-14 |
| B128 | B215 | ALT | BMI | 0.327383487 | 9.11E-11 |
| B128 | B216 | ALT | BMI | 0.288800009 | 1.34E-08 |
| B128 | B206 | ALT | Waist | 0.25111685 | 8.99E-07 |
| B129 | B132 | IBIL | Dbil | 0.740945069 | 0 |
| B129 | B133 | IBIL | TBIL | 0.974340431 | 0 |
| B129 | B139 | IBIL | ADA | 0.377436223 | 4.49E-14 |
| B129 | B135 | IBIL | TBA | 0.328571834 | 7.72E-11 |
| B129 | B131 | IBIL | AST | 0.279089967 | 4.22E-08 |
| B131 | B140 | AST | r-GT | 0.4557887 | 0 |
| B131 | B133 | AST | TBIL | 0.29735471 | 4.72E-09 |
| B131 | B135 | AST | TBA | 0.296687112 | 5.13E-09 |

| | | | | | |
|---|---|---|---|---|---|
| B131 | B132 | AST | Dbil | 0.286303569 | 1.81E-08 |
| B132 | B133 | Dbil | TBIL | 0.873000858 | 0 |
| B132 | B135 | Dbil | TBA | 0.591967395 | 0 |
| B132 | B136 | Dbil | 5'NT | 0.489480064 | 0 |
| B132 | B139 | Dbil | ADA | 0.67319391 | 0 |
| B132 | B134 | Dbil | CHE | -0.384589571 | 1.35E-14 |
| B132 | B138 | Dbil | PAB | -0.349508541 | 3.71E-12 |
| B132 | B137 | Dbil | AFU | -0.322466818 | 1.79E-10 |
| B133 | B135 | TBIL | TBA | 0.436363629 | 0 |
| B133 | B139 | TBIL | ADA | 0.499712364 | 0 |
| B133 | B136 | TBIL | 5'NT | 0.338958479 | 1.76E-11 |
| B133 | B134 | TBIL | CHE | -0.294640004 | 6.61E-09 |
| B133 | B137 | TBIL | AFU | -0.287740058 | 1.53E-08 |
| B133 | B138 | TBIL | PAB | -0.257546647 | 4.59E-07 |
| B134 | B135 | CHE | TBA | -0.596331361 | 0 |
| B134 | B138 | CHE | PAB | 0.565828705 | 0 |
| B134 | B139 | CHE | ADA | -0.513051543 | 0 |
| B134 | B150 | CHE | ALP | -0.421018989 | 0 |
| B134 | B185 | CHE | PT | -0.483522438 | 0 |
| B134 | B194 | CHE | Cl | -0.406735899 | 2.22E-16 |
| B134 | B181 | CHE | APTT | -0.383390751 | 1.64E-14 |
| B134 | B184 | CHE | INR | -0.353123678 | 2.15E-12 |
| B134 | B186 | CHE | PFC | -0.2982691 | 4.22E-09 |
| B134 | B136 | CHE | 5'NT | -0.273173035 | 8.30E-08 |
| B135 | B136 | TBA | 5'NT | 0.571427604 | 0 |
| B135 | B138 | TBA | PAB | -0.603451312 | 0 |
| B135 | B139 | TBA | ADA | 0.77959372 | 0 |
| B135 | B150 | TBA | ALP | 0.569463154 | 0 |
| B135 | B194 | TBA | Cl | 0.428843008 | 0 |
| B135 | B196 | TBA | TCO2 | -0.335390415 | 2.95E-11 |
| B135 | B193 | TBA | K | -0.319092343 | 2.83E-10 |
| B136 | B139 | 5'NT | ADA | 0.675842861 | 0 |
| B136 | B138 | 5'NT | PAB | -0.408982059 | 2.22E-16 |
| B136 | B150 | 5'NT | ALP | 0.272679424 | 8.78E-08 |
| B136 | B182 | 5'NT | TT | -0.26843809 | 1.41E-07 |
| B136 | B200 | 5'NT | OP | -0.263310823 | 2.47E-07 |
| B137 | B185 | AFU | PT | -0.419384796 | 0 |
| B137 | B184 | AFU | INR | -0.369112806 | 1.75E-13 |
| B137 | B181 | AFU | APTT | -0.366138263 | 2.82E-13 |
| B137 | B139 | AFU | ADA | -0.253156569 | 7.28E-07 |
| B138 | B139 | PAB | ADA | -0.595731174 | 0 |
| B138 | B150 | PAB | ALP | -0.477015461 | 0 |
| B138 | B155 | PAB | IgA | -0.31934846 | 2.74E-10 |
| B138 | B156 | PAB | IgE | -0.318484909 | 3.07E-10 |

| | | | | | |
|---|---|---|---|---|---|
| B138 | B193 | PAB | K | 0.303975281 | 2.05E-09 |
| B138 | B200 | PAB | OP | 0.255974158 | 5.42E-07 |
| B138 | B194 | PAB | CI | -0.25539691 | 5.76E-07 |
| B139 | B150 | ADA | ALP | 0.457923026 | 0 |
| B140 | B225 | r-GT | hPBG | 0.383305655 | 1.69E-14 |
| B140 | B206 | r-GT | Waist | 0.307621287 | 1.29E-09 |
| B140 | B215 | r-GT | BMI | 0.274845074 | 6.87E-08 |
| B140 | B216 | r-GT | BMI | 0.256201027 | 5.29E-07 |
| B141 | B215 | AST/ALT | BMI | -0.285042226 | 2.10E-08 |
| B142 | B151 | 25-(OH)D | Anti-CCP | -0.382528805 | 1.91E-14 |
| B142 | B143 | 25-(OH)D | β-CTx | -0.378619223 | 3.69E-14 |
| B143 | B144 | β-CTx | Total-PINP | 0.730010854 | 0 |
| B143 | B145 | β-CTx | BGP | 0.726070545 | 0 |
| B143 | B183 | β-CTx | PTA | -0.384171556 | 1.47E-14 |
| B143 | B151 | β-CTx | Anti-CCP | 0.368821286 | 1.83E-13 |
| B143 | B168 | β-CTx | NMP22 | 0.315492531 | 4.59E-10 |
| B144 | B145 | Total-PINP | BGP | 0.785828416 | 0 |
| B144 | B183 | Total-PINP | PTA | -0.363449592 | 4.32E-13 |
| B144 | B168 | Total-PINP | NMP22 | 0.296635183 | 5.17E-09 |
| B145 | B183 | BGP | PTA | -0.393525967 | 2.89E-15 |
| B148 | B149 | Anti_TPO | TGAb | 0.337812367 | 2.08E-11 |
| B149 | B177 | TGAb | ACR | 0.365920541 | 2.92E-13 |
| B149 | B175 | TGAb | ALB-U | 0.351268077 | 2.84E-12 |
| B149 | B168 | TGAb | NMP22 | -0.254164655 | 6.55E-07 |
| B15 | B100 | CD3-CD19+(B cell) | anti-Ro52 | 0.470036443 | 0 |
| B15 | B101 | CD3-CD19+(B cell) | anti-Sc70 | 0.470036445 | 0 |
| B15 | B102 | CD3-CD19+(B cell) | anti-Sm | 0.470036444 | 0 |
| B15 | B103 | CD3-CD19+(B cell) | anti-SSA/Ro | 0.470036443 | 0 |
| B15 | B104 | CD3-CD19+(B cell) | anti-SSB | 0.470036443 | 0 |
| B15 | B105 | CD3-CD19+(B cell) | anti-U1RNP | 0.470036443 | 0 |
| B15 | B106 | CD3-CD19+(B cell) | anti-P | 0.470036443 | 0 |
| B15 | B107 | CD3-CD19+(B cell) | AnuA | 0.470036445 | 0 |
| B15 | B108 | CD3-CD19+(B cell) | Anti-Cyclin | 0.470036443 | 0 |
| B15 | B109 | CD3-CD19+(B cell) | AMA-M2 | 0.470036443 | 0 |
| B15 | B110 | CD3-CD19+(B cell) | AHA | 0.470036443 | 0 |
| B15 | B143 | CD3-CD19+(B cell) | β-CTx | 0.42117639 | 0 |
| B15 | B145 | CD3-CD19+(B cell) | BGP | 0.432436807 | 0 |
| B15 | B16 | CD3-CD19+(B cell) | CD3+T | -0.865036921 | 0 |
| B15 | B18 | CD3-CD19+(B cell) | CD3+CD4+helper(inducer)T | -0.605956887 | 0 |
| B15 | B182 | CD3-CD19+(B cell) | TT | 0.461192457 | 0 |
| B15 | B183 | CD3-CD19+(B cell) | PTA | -0.651640256 | 0 |
| B15 | B23 | CD3-CD19+(B cell) | CD3-CD16+CD56+(NK) | 0.560329445 | 0 |
| B15 | B95 | CD3-CD19+(B cell) | anti-cenpb | 0.470036443 | 0 |
| B15 | B97 | CD3-CD19+(B cell) | dsDNA | 0.470036442 | 0 |

| | | | | | |
|---|---|---|---|---|---|
| B15 | B98 | CD3-CD19+(B cell) | anti-Jo-1 | 0.470036442 | 0 |
| B15 | B99 | CD3-CD19+(B cell) | anti-PM-Scl | 0.470036442 | 0 |
| B15 | B17 | CD3-CD19+(B cell) | CD3+CD16+CD56+NK | -0.404318843 | 4.44E-16 |
| B15 | B144 | CD3-CD19+(B cell) | Total-PINP | 0.398710564 | 1.11E-15 |
| B15 | B198 | CD3-CD19+(B cell) | P | -0.353294457 | 2.09E-12 |
| B15 | B151 | CD3-CD19+(B cell) | Anti-CCP | 0.346838487 | 5.53E-12 |
| B15 | B196 | CD3-CD19+(B cell) | TCO2 | 0.284605256 | 2.22E-08 |
| B15 | B22 | CD3-CD19+(B cell) | CD3+CD8+(T) | -0.282092806 | 2.98E-08 |
| B150 | B194 | ALP | Cl | 0.596188101 | 0 |
| B150 | B181 | ALP | APTT | 0.353613616 | 1.99E-12 |
| B150 | B182 | ALP | TT | 0.343227753 | 9.45E-12 |
| B150 | B193 | ALP | K | -0.330748282 | 5.69E-11 |
| B150 | B198 | ALP | P | 0.291932348 | 9.20E-09 |
| B150 | B196 | ALP | TCO2 | -0.278348058 | 4.60E-08 |
| B151 | B168 | Anti-CCP | NMP22 | 0.388530265 | 6.88E-15 |
| B151 | B157 | Anti-CCP | IgG | 0.283089815 | 2.65E-08 |
| B151 | B193 | Anti-CCP | K | 0.255737696 | 5.56E-07 |
| B152 | B158 | ACA-Ig(G-A-M) | IgM | 0.709647683 | 0 |
| B152 | B182 | ACA-Ig(G-A-M) | TT | 0.417398957 | 0 |
| B152 | B180 | ACA-Ig(G-A-M) | Tf | -0.333303864 | 3.97E-11 |
| B152 | B194 | ACA-Ig(G-A-M) | Cl | -0.270221487 | 1.16E-07 |
| B153 | B170 | Glu | U-GLU | 0.545040098 | 0 |
| B153 | B163 | Glu | SG | 0.278380552 | 4.58E-08 |
| B153 | B174 | Glu | Urea/Cr | 0.259211224 | 3.85E-07 |
| B154 | B187 | RF | NSE | -0.258438729 | 4.18E-07 |
| B155 | B156 | IgA | IgE | 0.577141426 | 0 |
| B155 | B178 | IgA | Ig-U | 0.46739412 | 0 |
| B155 | B180 | IgA | Tf | -0.34315795 | 9.54E-12 |
| B155 | B193 | IgA | K | -0.312702912 | 6.64E-10 |
| B155 | B157 | IgA | IgG | 0.290159776 | 1.14E-08 |
| B156 | B157 | IgE | IgG | 0.30305387 | 2.31E-09 |
| B156 | B178 | IgE | Ig-U | 0.273916042 | 7.63E-08 |
| B157 | B195 | IgG | Na | 0.279600299 | 3.98E-08 |
| B157 | B198 | IgG | P | -0.252807053 | 7.55E-07 |
| B158 | B169 | IgM | U-CRY | 0.709623431 | 0 |
| B158 | B194 | IgM | Cl | -0.399442643 | 8.88E-16 |
| B158 | B184 | IgM | INR | -0.300727062 | 3.10E-09 |
| B158 | B197 | IgM | AG | 0.274733288 | 6.96E-08 |
| B159 | B186 | β2-MG | PFC | 0.493874936 | 0 |
| B159 | B184 | β2-MG | INR | 0.28732765 | 1.60E-08 |
| B16 | B100 | CD3+T | anti-Ro52 | -0.75352045 | 0 |
| B16 | B101 | CD3+T | anti-Sc70 | -0.753520455 | 0 |
| B16 | B102 | CD3+T | anti-Sm | -0.753520452 | 0 |
| B16 | B103 | CD3+T | anti-SSA/Ro | -0.75352045 | 0 |

| | | | | | |
|---|---|---|---|---|---|
| B16 | B104 | CD3+T | anti-SSB | -0.753520449 | 0 |
| B16 | B105 | CD3+T | anti-U1RNP | -0.75352045 | 0 |
| B16 | B106 | CD3+T | anti-P | -0.75352045 | 0 |
| B16 | B107 | CD3+T | AnuA | -0.753520452 | 0 |
| B16 | B108 | CD3+T | Anti-Cyclin | -0.753520451 | 0 |
| B16 | B109 | CD3+T | AMA-M2 | -0.753520451 | 0 |
| B16 | B110 | CD3+T | AHA | -0.75352045 | 0 |
| B16 | B151 | CD3+T | Anti-CCP | -0.557153358 | 0 |
| B16 | B17 | CD3+T | CD3+CD16+CD56+NK | 0.410833619 | 0 |
| B16 | B18 | CD3+T | CD3+CD4+helper(inducer)T | 0.523698103 | 0 |
| B16 | B22 | CD3+T | CD3+CD8+(T) | 0.595960857 | 0 |
| B16 | B23 | CD3+T | CD3-CD16+CD56+(NK) | -0.897563668 | 0 |
| B16 | B95 | CD3+T | anti-cenpb | -0.753520451 | 0 |
| B16 | B97 | CD3+T | dsDNA | -0.753520448 | 0 |
| B16 | B98 | CD3+T | anti-Jo-1 | -0.753520448 | 0 |
| B16 | B99 | CD3+T | anti-PM-Scl | -0.753520448 | 0 |
| B16 | B198 | CD3+T | P | 0.406927469 | 2.22E-16 |
| B16 | B19 | CD3+T | CD3+CD4+/CD3+CD8+ | -0.36666811 | 2.59E-13 |
| B16 | B157 | CD3+T | IgG | -0.356830624 | 1.22E-12 |
| B16 | B145 | CD3+T | BGP | -0.346979023 | 5.42E-12 |
| B16 | B183 | CD3+T | PTA | 0.346167245 | 6.12E-12 |
| B16 | B143 | CD3+T | β-CTx | -0.337624499 | 2.14E-11 |
| B16 | B144 | CD3+T | Total-PINP | -0.319688669 | 2.61E-10 |
| B16 | B193 | CD3+T | K | -0.260934518 | 3.20E-07 |
| B16 | B182 | CD3+T | TT | -0.259400335 | 3.77E-07 |
| B163 | B176 | SG | Ucr | 0.441431812 | 0 |
| B163 | B170 | SG | U-GLU | 0.354125394 | 1.84E-12 |
| B163 | B166 | SG | Pro | 0.310753719 | 8.57E-10 |
| B167 | B223 | PC | FH | 0.451145631 | 0 |
| B168 | B175 | NMP22 | ALB-U | -0.268663846 | 1.38E-07 |
| B168 | B176 | NMP22 | Ucr | -0.261546198 | 2.99E-07 |
| B169 | B182 | U-CRY | TT | 0.417384763 | 0 |
| B169 | B180 | U-CRY | Tf | -0.33329246 | 3.97E-11 |
| B169 | B194 | U-CRY | Cl | -0.27021218 | 1.16E-07 |
| B17 | B182 | CD3+CD16+CD56+NK | TT | -0.563074336 | 0 |
| B17 | B20 | CD3+CD16+CD56+NK | CD3+CD4+CD8+ | 0.578669608 | 0 |
| B17 | B100 | CD3+CD16+CD56+NK | anti-Ro52 | -0.385859697 | 1.09E-14 |
| B17 | B101 | CD3+CD16+CD56+NK | anti-Sc70 | -0.385859699 | 1.09E-14 |
| B17 | B102 | CD3+CD16+CD56+NK | anti-Sm | -0.385859698 | 1.09E-14 |
| B17 | B103 | CD3+CD16+CD56+NK | anti-SSA/Ro | -0.385859697 | 1.09E-14 |
| B17 | B104 | CD3+CD16+CD56+NK | anti-SSB | -0.385859697 | 1.09E-14 |
| B17 | B105 | CD3+CD16+CD56+NK | anti-U1RNP | -0.385859697 | 1.09E-14 |
| B17 | B106 | CD3+CD16+CD56+NK | anti-P | -0.385859697 | 1.09E-14 |
| B17 | B107 | CD3+CD16+CD56+NK | AnuA | -0.385859699 | 1.09E-14 |

| | | | | | |
|---|---|---|---|---|---|
| B17 | B108 | CD3+CD16+CD56+NK | Anti-Cyclin | -0.385859697 | 1.09E-14 |
| B17 | B109 | CD3+CD16+CD56+NK | AMA-M2 | -0.385859697 | 1.09E-14 |
| B17 | B110 | CD3+CD16+CD56+NK | AHA | -0.385859697 | 1.09E-14 |
| B17 | B95 | CD3+CD16+CD56+NK | anti-cenpb | -0.385859697 | 1.09E-14 |
| B17 | B97 | CD3+CD16+CD56+NK | dsDNA | -0.385859696 | 1.09E-14 |
| B17 | B98 | CD3+CD16+CD56+NK | anti-Jo-1 | -0.385859696 | 1.09E-14 |
| B17 | B99 | CD3+CD16+CD56+NK | anti-PM-Scl | -0.385859696 | 1.09E-14 |
| B17 | B193 | CD3+CD16+CD56+NK | K | -0.311300027 | 7.98E-10 |
| B17 | B186 | CD3+CD16+CD56+NK | PFC | 0.287298662 | 1.61E-08 |
| B17 | B151 | CD3+CD16+CD56+NK | Anti-CCP | -0.287201857 | 1.63E-08 |
| B17 | B183 | CD3+CD16+CD56+NK | PTA | 0.282250287 | 2.92E-08 |
| B17 | B23 | CD3+CD16+CD56+NK | CD3-CD16+CD56+(NK) | -0.280772395 | 3.47E-08 |
| B173 | B228 | NTT | OS | 0.275649865 | 6.27E-08 |
| B174 | B191 | Urea/Cr | Urea | 0.325024124 | 1.26E-10 |
| B174 | B192 | Urea/Cr | UA | -0.3094913 | 1.01E-09 |
| B174 | B221 | Urea/Cr | Hei | -0.29059659 | 1.08E-08 |
| B174 | B190 | Urea/Cr | Cre | -0.288620527 | 1.37E-08 |
| B174 | B222 | Urea/Cr | NS | -0.274300948 | 7.31E-08 |
| B174 | B189 | Urea/Cr | eGFB | 0.269402371 | 1.27E-07 |
| B175 | B177 | ALB-U | ACR | 0.917329047 | 0 |
| B175 | B176 | ALB-U | Ucr | 0.255346817 | 5.79E-07 |
| B178 | B179 | Ig-U | α1-MG | 0.578667262 | 0 |
| B178 | B193 | Ig-U | K | -0.363549755 | 4.25E-13 |
| B178 | B180 | Ig-U | Tf | 0.323322602 | 1.59E-10 |
| B179 | B180 | α1-MG | Tf | 0.86451675 | 0 |
| B18 | B19 | CD3+CD4+helper(inducer)T | CD3+CD4+/CD3+CD8+ | 0.578693943 | 0 |
| B18 | B21 | CD3+CD4+helper(inducer)T | CD3+CD4-CD8- | -0.742526076 | 0 |
| B18 | B22 | CD3+CD4+helper(inducer)T | CD3+CD8+(T) | -0.346999309 | 5.40E-12 |
| B18 | B185 | CD3+CD4+helper(inducer)T | PT | 0.328858496 | 7.42E-11 |
| B18 | B23 | CD3+CD4+helper(inducer)T | CD3-CD16+CD56+(NK) | -0.325567272 | 1.17E-10 |
| B18 | B184 | CD3+CD4+helper(inducer)T | INR | 0.313978529 | 5.61E-10 |
| B18 | B183 | CD3+CD4+helper(inducer)T | PTA | 0.294987047 | 6.33E-09 |
| B18 | B181 | CD3+CD4+helper(inducer)T | APTT | 0.279277654 | 4.13E-08 |
| B181 | B184 | APTT | INR | 0.814544685 | 0 |
| B181 | B185 | APTT | PT | 0.837320044 | 0 |
| B181 | B194 | APTT | Cl | 0.514780263 | 0 |
| B181 | B182 | APTT | TT | 0.37593841 | 5.75E-14 |
| B181 | B186 | APTT | PFC | 0.3111712 | 8.11E-10 |
| B182 | B183 | TT | PTA | -0.473167224 | 0 |
| B182 | B186 | TT | PFC | -0.255380484 | 5.77E-07 |
| B183 | B196 | PTA | TCO2 | -0.423190322 | 0 |
| B183 | B197 | PTA | AG | 0.356385831 | 1.30E-12 |
| B183 | B184 | PTA | INR | 0.293362016 | 7.73E-09 |
| B184 | B185 | INR | PT | 0.899438001 | 0 |

| | | | | | |
|---|---|---|---|---|---|
| B184 | B186 | INR | PFC | 0.527966636 | 0 |
| B184 | B194 | INR | Cl | 0.469768985 | 0 |
| B184 | B195 | INR | Na | 0.349137561 | 3.92E-12 |
| B185 | B186 | PT | PFC | 0.583266934 | 0 |
| B185 | B194 | PT | Cl | 0.342323447 | 1.08E-11 |
| B185 | B195 | PT | Na | 0.330785427 | 5.66E-11 |
| B189 | B190 | eGFB | Cre | -0.494516064 | 0 |
| B189 | B191 | eGFB | Urea | -0.470856006 | 0 |
| B19 | B100 | CD3+CD4+/CD3+CD8+ | anti-Ro52 | 0.685480023 | 0 |
| B19 | B101 | CD3+CD4+/CD3+CD8+ | anti-Sc70 | 0.685480026 | 0 |
| B19 | B102 | CD3+CD4+/CD3+CD8+ | anti-Sm | 0.685480025 | 0 |
| B19 | B103 | CD3+CD4+/CD3+CD8+ | anti-SSA/Ro | 0.685480023 | 0 |
| B19 | B104 | CD3+CD4+/CD3+CD8+ | anti-SSB | 0.685480023 | 0 |
| B19 | B105 | CD3+CD4+/CD3+CD8+ | anti-U1RNP | 0.685480023 | 0 |
| B19 | B106 | CD3+CD4+/CD3+CD8+ | anti-P | 0.685480023 | 0 |
| B19 | B107 | CD3+CD4+/CD3+CD8+ | AnuA | 0.685480026 | 0 |
| B19 | B108 | CD3+CD4+/CD3+CD8+ | Anti-Cyclin | 0.685480023 | 0 |
| B19 | B109 | CD3+CD4+/CD3+CD8+ | AMA-M2 | 0.685480023 | 0 |
| B19 | B110 | CD3+CD4+/CD3+CD8+ | AHA | 0.685480023 | 0 |
| B19 | B151 | CD3+CD4+/CD3+CD8+ | Anti-CCP | 0.50947708 | 0 |
| B19 | B185 | CD3+CD4+/CD3+CD8+ | PT | 0.506572253 | 0 |
| B19 | B21 | CD3+CD4+/CD3+CD8+ | CD3+CD4-CD8- | -0.789285316 | 0 |
| B19 | B22 | CD3+CD4+/CD3+CD8+ | CD3+CD8+(T) | -0.95999777 | 0 |
| B19 | B23 | CD3+CD4+/CD3+CD8+ | CD3-CD16+CD56+(NK) | 0.569951807 | 0 |
| B19 | B95 | CD3+CD4+/CD3+CD8+ | anti-cenpb | 0.685480024 | 0 |
| B19 | B97 | CD3+CD4+/CD3+CD8+ | dsDNA | 0.685480022 | 0 |
| B19 | B98 | CD3+CD4+/CD3+CD8+ | anti-Jo-1 | 0.685480022 | 0 |
| B19 | B99 | CD3+CD4+/CD3+CD8+ | anti-PM-Scl | 0.685480022 | 0 |
| B19 | B134 | CD3+CD4+/CD3+CD8+ | CHE | -0.403325022 | 4.44E-16 |
| B19 | B186 | CD3+CD4+/CD3+CD8+ | PFC | 0.396274583 | 1.78E-15 |
| B19 | B195 | CD3+CD4+/CD3+CD8+ | Na | 0.384540892 | 1.38E-14 |
| B19 | B184 | CD3+CD4+/CD3+CD8+ | INR | 0.379971387 | 2.93E-14 |
| B19 | B157 | CD3+CD4+/CD3+CD8+ | IgG | 0.370279545 | 1.45E-13 |
| B19 | B34 | CD3+CD4+/CD3+CD8+ | TK | 0.358079579 | 1.00E-12 |
| B19 | B193 | CD3+CD4+/CD3+CD8+ | K | 0.317123862 | 3.69E-10 |
| B19 | B39 | CD3+CD4+/CD3+CD8+ | Other cells | 0.257379159 | 4.67E-07 |
| B19 | B60 | CD3+CD4+/CD3+CD8+ | WBRVI | -0.253323502 | 7.15E-07 |
| B190 | B191 | Cre | Urea | 0.689342277 | 0 |
| B192 | B221 | UA | Hei | 0.311083809 | 8.21E-10 |
| B192 | B216 | UA | BMI | 0.292977751 | 8.10E-09 |
| B192 | B222 | UA | NS | 0.286588969 | 1.75E-08 |
| B192 | B206 | UA | Waist | 0.258475478 | 4.16E-07 |
| B193 | B194 | K | Cl | -0.340416418 | 1.43E-11 |
| B193 | B198 | K | P | -0.329774934 | 6.52E-11 |

| | | | | | |
|---|---|---|---|---|---|
| B193 | B196 | K | TCO2 | 0.258865819 | 3.99E-07 |
| B194 | B197 | Cl | AG | -0.315407364 | 4.64E-10 |
| B195 | B200 | Na | OP | 0.531081869 | 0 |
| B195 | B197 | Na | AG | 0.378499728 | 3.77E-14 |
| B196 | B197 | TCO2 | AG | -0.738808975 | 0 |
| B196 | B198 | TCO2 | P | -0.380223459 | 2.82E-14 |
| B197 | B200 | AG | OP | 0.363983787 | 3.97E-13 |
| B2 | B189 | Age | eGFB | -0.461327898 | 0 |
| B2 | B216 | Age | BMI | -0.345319692 | 6.94E-12 |
| B2 | B141 | Age | AST/ALT | 0.316388155 | 4.07E-10 |
| B2 | B128 | Age | ALT | -0.293687758 | 7.43E-09 |
| B2 | B220 | Age | SBP | 0.281786085 | 3.09E-08 |
| B2 | B215 | Age | BMI | -0.278260386 | 4.65E-08 |
| B2 | B126 | Age | Alb | -0.271062361 | 1.05E-07 |
| B2 | B204 | Age | Drinking | -0.261809782 | 2.91E-07 |
| B20 | B181 | CD3+CD4+CD8+ | APTT | 0.537321297 | 0 |
| B20 | B184 | CD3+CD4+CD8+ | INR | 0.423402898 | 0 |
| B20 | B194 | CD3+CD4+CD8+ | Cl | 0.429509132 | 0 |
| B20 | B197 | CD3+CD4+CD8+ | AG | -0.460931839 | 0 |
| B20 | B23 | CD3+CD4+CD8+ | CD3-CD16+CD56+(NK) | -0.365375301 | 3.18E-13 |
| B20 | B185 | CD3+CD4+CD8+ | PT | 0.352398921 | 2.40E-12 |
| B20 | B100 | CD3+CD4+CD8+ | anti-Ro52 | -0.338122249 | 1.99E-11 |
| B20 | B101 | CD3+CD4+CD8+ | anti-Sc70 | -0.338122251 | 1.99E-11 |
| B20 | B102 | CD3+CD4+CD8+ | anti-Sm | -0.33812225 | 1.99E-11 |
| B20 | B103 | CD3+CD4+CD8+ | anti-SSA/Ro | -0.338122249 | 1.99E-11 |
| B20 | B104 | CD3+CD4+CD8+ | anti-SSB | -0.338122249 | 1.99E-11 |
| B20 | B105 | CD3+CD4+CD8+ | anti-U1RNP | -0.338122249 | 1.99E-11 |
| B20 | B106 | CD3+CD4+CD8+ | anti-P | -0.338122249 | 1.99E-11 |
| B20 | B107 | CD3+CD4+CD8+ | AnuA | -0.338122251 | 1.99E-11 |
| B20 | B108 | CD3+CD4+CD8+ | Anti-Cyclin | -0.338122249 | 1.99E-11 |
| B20 | B109 | CD3+CD4+CD8+ | AMA-M2 | -0.338122249 | 1.99E-11 |
| B20 | B110 | CD3+CD4+CD8+ | AHA | -0.338122249 | 1.99E-11 |
| B20 | B95 | CD3+CD4+CD8+ | anti-cenpb | -0.33812225 | 1.99E-11 |
| B20 | B97 | CD3+CD4+CD8+ | dsDNA | -0.338122248 | 1.99E-11 |
| B20 | B98 | CD3+CD4+CD8+ | anti-Jo-1 | -0.338122248 | 1.99E-11 |
| B20 | B99 | CD3+CD4+CD8+ | anti-PM-Scl | -0.338122248 | 1.99E-11 |
| B20 | B69 | CD3+CD4+CD8+ | PV | -0.312091894 | 7.19E-10 |
| B20 | B158 | CD3+CD4+CD8+ | IgM | -0.301471632 | 2.82E-09 |
| B20 | B196 | CD3+CD4+CD8+ | TCO2 | 0.273300569 | 8.18E-08 |
| B20 | B61 | CD3+CD4+CD8+ | WBHSRV | -0.256967791 | 4.88E-07 |
| B20 | B193 | CD3+CD4+CD8+ | K | -0.252297558 | 7.96E-07 |
| B20 | B151 | CD3+CD4+CD8+ | Anti-CCP | -0.251964334 | 8.24E-07 |
| B20 | B21 | CD3+CD4+CD8+ | CD3+CD4-CD8- | -0.251136802 | 8.97E-07 |
| B204 | B209 | Drinking | Smoking | 0.297109592 | 4.87E-09 |

| | | | | | |
|---|---|---|---|---|---|
| B204 | B221 | Drinking | Hei | 0.253132775 | 7.30E-07 |
| B206 | B207 | Waist | WHR | 0.764720694 | 0 |
| B206 | B210 | Waist | Hip | 0.759437879 | 0 |
| B206 | B215 | Waist | BMI | 0.791899549 | 0 |
| B206 | B216 | Waist | BMI | 0.826182575 | 0 |
| B206 | B222 | Waist | NS | 0.675089973 | 0 |
| B206 | B221 | Waist | Hei | 0.404201522 | 4.44E-16 |
| B207 | B215 | WHR | BMI | 0.44636984 | 0 |
| B207 | B216 | WHR | BMI | 0.509806205 | 0 |
| B207 | B222 | WHR | NS | 0.509617716 | 0 |
| B207 | B221 | WHR | Hei | 0.282855018 | 2.72E-08 |
| B21 | B181 | CD3+CD4-CD8- | APTT | -0.511472159 | 0 |
| B21 | B184 | CD3+CD4-CD8- | INR | -0.540148787 | 0 |
| B21 | B185 | CD3+CD4-CD8- | PT | -0.592211781 | 0 |
| B21 | B22 | CD3+CD4-CD8- | CD3+CD8+(T) | 0.69051836 | 0 |
| B21 | B134 | CD3+CD4-CD8- | CHE | 0.406001561 | 4.44E-16 |
| B21 | B195 | CD3+CD4-CD8- | Na | -0.403914624 | 4.44E-16 |
| B21 | B100 | CD3+CD4-CD8- | anti-Ro52 | -0.371097129 | 1.27E-13 |
| B21 | B101 | CD3+CD4-CD8- | anti-Sc70 | -0.371097131 | 1.27E-13 |
| B21 | B102 | CD3+CD4-CD8- | anti-Sm | -0.371097129 | 1.27E-13 |
| B21 | B103 | CD3+CD4-CD8- | anti-SSA/Ro | -0.371097129 | 1.27E-13 |
| B21 | B104 | CD3+CD4-CD8- | anti-SSB | -0.371097129 | 1.27E-13 |
| B21 | B105 | CD3+CD4-CD8- | anti-U1RNP | -0.371097129 | 1.27E-13 |
| B21 | B106 | CD3+CD4-CD8- | anti-P | -0.371097129 | 1.27E-13 |
| B21 | B107 | CD3+CD4-CD8- | AnuA | -0.37109713 | 1.27E-13 |
| B21 | B108 | CD3+CD4-CD8- | Anti-Cyclin | -0.371097129 | 1.27E-13 |
| B21 | B109 | CD3+CD4-CD8- | AMA-M2 | -0.371097129 | 1.27E-13 |
| B21 | B110 | CD3+CD4-CD8- | AHA | -0.371097129 | 1.27E-13 |
| B21 | B95 | CD3+CD4-CD8- | anti-cenpb | -0.371097129 | 1.27E-13 |
| B21 | B97 | CD3+CD4-CD8- | dsDNA | -0.371097128 | 1.27E-13 |
| B21 | B98 | CD3+CD4-CD8- | anti-Jo-1 | -0.371097128 | 1.27E-13 |
| B21 | B99 | CD3+CD4-CD8- | anti-PM-Scl | -0.371097128 | 1.27E-13 |
| B21 | B186 | CD3+CD4-CD8- | PFC | -0.317741842 | 3.40E-10 |
| B21 | B194 | CD3+CD4-CD8- | Cl | -0.282418522 | 2.87E-08 |
| B21 | B151 | CD3+CD4-CD8- | Anti-CCP | -0.277076185 | 5.32E-08 |
| B210 | B215 | Hip | BMI | 0.767388795 | 0 |
| B210 | B216 | Hip | BMI | 0.748138475 | 0 |
| B210 | B222 | Hip | NS | 0.516843999 | 0 |
| B210 | B221 | Hip | Hei | 0.326494513 | 1.03E-10 |
| B211 | B212 | IOP-R | IOP-L | 0.714908987 | 0 |
| B213 | B214 | CVA-R | CVA-L | 0.849387688 | 0 |
| B215 | B216 | BMI | BMI | 0.800267302 | 0 |
| B215 | B222 | BMI | NS | 0.531993971 | 0 |
| B216 | B221 | BMI | Hei | 0.685033616 | 0 |

| | | | | | |
|---|---|---|---|---|---|
| B216 | B222 | BMI | NS | 0.728984423 | 0 |
| B218 | B220 | DBP | SBP | 0.548116003 | 0 |
| B22 | B100 | CD3+CD8+(T) | anti-Ro52 | -0.80938203 | 0 |
| B22 | B101 | CD3+CD8+(T) | anti-Sc70 | -0.809382034 | 0 |
| B22 | B102 | CD3+CD8+(T) | anti-Sm | -0.809382032 | 0 |
| B22 | B103 | CD3+CD8+(T) | anti-SSA/Ro | -0.80938203 | 0 |
| B22 | B104 | CD3+CD8+(T) | anti-SSB | -0.809382029 | 0 |
| B22 | B105 | CD3+CD8+(T) | anti-U1RNP | -0.80938203 | 0 |
| B22 | B106 | CD3+CD8+(T) | anti-P | -0.80938203 | 0 |
| B22 | B107 | CD3+CD8+(T) | AnuA | -0.809382035 | 0 |
| B22 | B108 | CD3+CD8+(T) | Anti-Cyclin | -0.80938203 | 0 |
| B22 | B109 | CD3+CD8+(T) | AMA-M2 | -0.80938203 | 0 |
| B22 | B110 | CD3+CD8+(T) | AHA | -0.80938203 | 0 |
| B22 | B151 | CD3+CD8+(T) | Anti-CCP | -0.600969782 | 0 |
| B22 | B157 | CD3+CD8+(T) | IgG | -0.419960899 | 0 |
| B22 | B185 | CD3+CD8+(T) | PT | -0.454623318 | 0 |
| B22 | B23 | CD3+CD8+(T) | CD3-CD16+CD56+(NK) | -0.739398354 | 0 |
| B22 | B95 | CD3+CD8+(T) | anti-cenpb | -0.80938203 | 0 |
| B22 | B97 | CD3+CD8+(T) | dsDNA | -0.809382028 | 0 |
| B22 | B98 | CD3+CD8+(T) | anti-Jo-1 | -0.809382028 | 0 |
| B22 | B99 | CD3+CD8+(T) | anti-PM-Scl | -0.809382028 | 0 |
| B22 | B195 | CD3+CD8+(T) | Na | -0.388591887 | 6.66E-15 |
| B22 | B186 | CD3+CD8+(T) | PFC | -0.369574229 | 1.63E-13 |
| B22 | B134 | CD3+CD8+(T) | CHE | 0.365365082 | 3.19E-13 |
| B22 | B193 | CD3+CD8+(T) | K | -0.358212314 | 9.81E-13 |
| B22 | B34 | CD3+CD8+(T) | TK | -0.34458885 | 7.73E-12 |
| B22 | B184 | CD3+CD8+(T) | INR | -0.318782501 | 2.95E-10 |
| B22 | B198 | CD3+CD8+(T) | P | 0.272501079 | 8.96E-08 |
| B22 | B65 | CD3+CD8+(T) | WBRSR(mPas_1) | 0.255609668 | 5.63E-07 |
| B221 | B222 | Hei | NS | 0.567067707 | 0 |
| B227 | B228 | OD | OS | 0.306457291 | 1.50E-09 |
| B23 | B100 | CD3-CD16+CD56+(NK) | anti-Ro52 | 0.831639906 | 0 |
| B23 | B101 | CD3-CD16+CD56+(NK) | anti-Sc70 | 0.83163991 | 0 |
| B23 | B102 | CD3-CD16+CD56+(NK) | anti-Sm | 0.831639908 | 0 |
| B23 | B103 | CD3-CD16+CD56+(NK) | anti-SSA/Ro | 0.831639906 | 0 |
| B23 | B104 | CD3-CD16+CD56+(NK) | anti-SSB | 0.831639906 | 0 |
| B23 | B105 | CD3-CD16+CD56+(NK) | anti-U1RNP | 0.831639906 | 0 |
| B23 | B106 | CD3-CD16+CD56+(NK) | anti-P | 0.831639906 | 0 |
| B23 | B107 | CD3-CD16+CD56+(NK) | AnuA | 0.831639909 | 0 |
| B23 | B108 | CD3-CD16+CD56+(NK) | Anti-Cyclin | 0.831639906 | 0 |
| B23 | B109 | CD3-CD16+CD56+(NK) | AMA-M2 | 0.831639906 | 0 |
| B23 | B110 | CD3-CD16+CD56+(NK) | AHA | 0.831639906 | 0 |
| B23 | B151 | CD3-CD16+CD56+(NK) | Anti-CCP | 0.615322354 | 0 |
| B23 | B95 | CD3-CD16+CD56+(NK) | anti-cenpb | 0.831639907 | 0 |

| | | | | | |
|---|---|---|---|---|---|
| B23 | B97 | CD3-CD16+CD56+(NK) | dsDNA | 0.831639904 | 0 |
| B23 | B98 | CD3-CD16+CD56+(NK) | anti-Jo-1 | 0.831639904 | 0 |
| B23 | B99 | CD3-CD16+CD56+(NK) | anti-PM-Scl | 0.831639904 | 0 |
| B23 | B157 | CD3-CD16+CD56+(NK) | IgG | 0.393980959 | 2.66E-15 |
| B23 | B198 | CD3-CD16+CD56+(NK) | P | -0.379577089 | 3.15E-14 |
| B23 | B193 | CD3-CD16+CD56+(NK) | K | 0.313060282 | 6.33E-10 |
| B23 | B34 | CD3-CD16+CD56+(NK) | TK | 0.282714564 | 2.77E-08 |
| B23 | B114 | CD3-CD16+CD56+(NK) | TSH TT3 TT4(TT4) | 0.26072548 | 3.27E-07 |
| B23 | B158 | CD3-CD16+CD56+(NK) | IgM | 0.260160764 | 3.47E-07 |
| B24 | B26 | αHBDH | CKMB | 0.56628645 | 0 |
| B24 | B27 | αHBDH | LDH | 0.71248977 | 0 |
| B24 | B55 | αHBDH | TK | 0.325124484 | 1.24E-10 |
| B24 | B75 | αHBDH | ApoA1/B | 0.296146571 | 5.49E-09 |
| B24 | B25 | αHBDH | CK | 0.28732152 | 1.60E-08 |
| B24 | B134 | αHBDH | CHE | -0.279926812 | 3.83E-08 |
| B24 | B58 | αHBDH | HCT | -0.2768487 | 5.46E-08 |
| B25 | B26 | CK | CKMB | 0.407769074 | 2.22E-16 |
| B25 | B79 | CK | hb V -dna | -0.406701246 | 2.22E-16 |
| B25 | B73 | CK | ApoA | 0.327808604 | 8.58E-11 |
| B25 | B69 | CK | PV | -0.309992229 | 9.46E-10 |
| B25 | B135 | CK | TBA | 0.28343161 | 2.54E-08 |
| B25 | B55 | CK | TK | 0.258811774 | 4.01E-07 |
| B26 | B134 | CKMB | CHE | -0.464469972 | 0 |
| B26 | B135 | CKMB | TBA | 0.664681939 | 0 |
| B26 | B138 | CKMB | PAB | -0.44387 | 0 |
| B26 | B150 | CKMB | ALP | 0.502231593 | 0 |
| B26 | B194 | CKMB | Cl | 0.458358367 | 0 |
| B26 | B27 | CKMB | LDH | 0.428029421 | 0 |
| B26 | B55 | CKMB | TK | 0.431760876 | 0 |
| B26 | B196 | CKMB | TCO2 | -0.30299381 | 2.33E-09 |
| B26 | B58 | CKMB | HCT | -0.286283151 | 1.82E-08 |
| B26 | B193 | CKMB | K | -0.283937826 | 2.40E-08 |
| B26 | B139 | CKMB | ADA | 0.278315304 | 4.62E-08 |
| B26 | B56 | CKMB | IR | 0.270680464 | 1.10E-07 |
| B26 | B69 | CKMB | PV | -0.268689897 | 1.37E-07 |
| B27 | B132 | LDH | Dbil | 0.425838958 | 0 |
| B27 | B135 | LDH | TBA | 0.466776043 | 0 |
| B27 | B136 | LDH | 5'NT | 0.454547302 | 0 |
| B27 | B139 | LDH | ADA | 0.49542801 | 0 |
| B27 | B134 | LDH | CHE | -0.380495728 | 2.71E-14 |
| B27 | B133 | LDH | TBIL | 0.332394815 | 4.51E-11 |
| B27 | B150 | LDH | ALP | 0.296948868 | 4.97E-09 |
| B27 | B129 | LDH | IBIL | 0.261473908 | 3.02E-07 |
| B27 | B131 | LDH | AST | 0.261020656 | 3.17E-07 |

| | | | | | |
|---|---|---|---|---|---|
| B27 | B138 | LDH | PAB | -0.256701247 | 5.02E-07 |
| B28 | B116 | PRL | TT3 | -0.917911187 | 0 |
| B28 | B117 | PRL | TT4 | -0.624510605 | 0 |
| B28 | B29 | PRL | E2 | -0.452609736 | 0 |
| B28 | B33 | PRL | Prog | -0.668901276 | 0 |
| B29 | B116 | E2 | TT3 | -0.526423658 | 0 |
| B29 | B33 | E2 | Prog | -0.354107116 | 1.85E-12 |
| B3 | B124 | Hcy | ACTH | -0.31132333 | 7.95E-10 |
| B3 | B152 | Hcy | ACA-Ig(G-A-M) | 0.311236664 | 8.04E-10 |
| B3 | B121 | Hcy | C3 | 0.31122747 | 8.05E-10 |
| B3 | B122 | Hcy | C4 | 0.311227656 | 8.05E-10 |
| B3 | B169 | Hcy | U-CRY | 0.311226026 | 8.05E-10 |
| B3 | B125 | Hcy | Cor | 0.310541533 | 8.81E-10 |
| B3 | B119 | Hcy | Hcv-rna | -0.262502621 | 2.70E-07 |
| B30 | B116 | LH | TT3 | -0.55555781 | 0 |
| B31 | B223 | FSH | FH | -0.331294578 | 5.27E-11 |
| B33 | B116 | Prog | TT3 | -0.690865485 | 0 |
| B34 | B92 | TK | ALDH2-Glu504Glu | -0.382641055 | 1.87E-14 |
| B34 | B93 | TK | ALDH2-Glu504Lys | 0.382641055 | 1.87E-14 |
| B34 | B94 | TK | ALDH2-Lys504Lys | 0.382641053 | 1.87E-14 |
| B34 | B185 | TK | PT | 0.286776425 | 1.71E-08 |
| B34 | B100 | TK | anti-Ro52 | 0.275773255 | 6.18E-08 |
| B34 | B101 | TK | anti-Sc70 | 0.275773256 | 6.18E-08 |
| B34 | B102 | TK | anti-Sm | 0.275773255 | 6.18E-08 |
| B34 | B103 | TK | anti-SSA/Ro | 0.275773255 | 6.18E-08 |
| B34 | B104 | TK | anti-SSB | 0.275773255 | 6.18E-08 |
| B34 | B105 | TK | anti-U1RNP | 0.275773255 | 6.18E-08 |
| B34 | B106 | TK | anti-P | 0.275773255 | 6.18E-08 |
| B34 | B107 | TK | AnuA | 0.275773256 | 6.18E-08 |
| B34 | B108 | TK | Anti-Cyclin | 0.275773255 | 6.18E-08 |
| B34 | B109 | TK | AMA-M2 | 0.275773255 | 6.18E-08 |
| B34 | B110 | TK | AHA | 0.275773255 | 6.18E-08 |
| B34 | B95 | TK | anti-cenpb | 0.275773255 | 6.18E-08 |
| B34 | B97 | TK | dsDNA | 0.275773254 | 6.18E-08 |
| B34 | B98 | TK | anti-Jo-1 | 0.275773254 | 6.18E-08 |
| B34 | B99 | TK | anti-PM-Scl | 0.275773254 | 6.18E-08 |
| B34 | B157 | TK | IgG | 0.258897577 | 3.98E-07 |
| B35 | B159 | β2-MG | β2-MG | 0.999994519 | 0 |
| B35 | B186 | β2-MG | PFC | 0.493755911 | 0 |
| B35 | B184 | β2-MG | INR | 0.288146946 | 1.45E-08 |
| B36 | B167 | V-Lym% | PC | 0.626368234 | 0 |
| B36 | B223 | V-Lym% | FH | 0.698430296 | 0 |
| B36 | B37 | V-Lym% | BAND | 0.774596669 | 0 |
| B36 | B96 | V-Lym% | dsDNA (IIF) | 0.553636532 | 0 |

| | | | | | |
|---|---|---|---|---|---|
| B37 | B167 | BAND | PC | -0.379049022 | 3.42E-14 |
| B39 | B211 | Other cells | IOP-R | -0.389586429 | 5.77E-15 |
| B39 | B148 | Other cells | Anti_TPO | -0.349317572 | 3.82E-12 |
| B39 | B149 | Other cells | TGAb | -0.305275089 | 1.74E-09 |
| B39 | B195 | Other cells | Na | 0.286557285 | 1.76E-08 |
| B4 | B5 | WB-Ca | WB-Mg | 1 | 0 |
| B4 | B58 | WB-Ca | HCT | 0.484378161 | 0 |
| B4 | B6 | WB-Ca | WB-Fe | 1 | 0 |
| B4 | B8 | WB-Ca | WB-Zinc | 1 | 0 |
| B4 | B66 | WB-Ca | WBRSR(mPas_100) | 0.405797344 | 4.44E-16 |
| B4 | B67 | WB-Ca | WBRSR(mPas_50) | 0.404505679 | 4.44E-16 |
| B4 | B63 | WB-Ca | WBRSR(mPas_200) | 0.390698729 | 4.88E-15 |
| B4 | B62 | WB-Ca | WBHSRVI | 0.363067127 | 4.59E-13 |
| B4 | B64 | WB-Ca | WBRSR(mPas_5) | 0.312654121 | 6.68E-10 |
| B4 | B55 | WB-Ca | TK | -0.269580305 | 1.24E-07 |
| B40 | B42 | WBC | MON | 0.677889631 | 0 |
| B40 | B52 | WBC | PLT/BPC | 0.257137548 | 4.79E-07 |
| B40 | B53 | WBC | GR% | 0.255798016 | 5.52E-07 |
| B41 | B42 | MON % | MON | 0.601934531 | 0 |
| B43 | B44 | MCV | MCH | 0.914395091 | 0 |
| B43 | B46 | MCV | RBC Count | -0.615271089 | 0 |
| B43 | B47 | MCV | RDW | 0.620576617 | 0 |
| B44 | B45 | MCH | MCHC | 0.555825962 | 0 |
| B44 | B46 | MCH | RBC Count | -0.530474306 | 0 |
| B44 | B47 | MCH | RDW | 0.496486875 | 0 |
| B45 | B51 | MCHC | Hb | 0.42457375 | 0 |
| B45 | B72 | MCHC | HDL-C | -0.283594793 | 2.50E-08 |
| B46 | B47 | RBC Count | RDW | -0.476974378 | 0 |
| B46 | B51 | RBC Count | Hb | 0.695108729 | 0 |
| B46 | B126 | RBC Count | Alb | 0.287145405 | 1.64E-08 |
| B46 | B222 | RBC Count | NS | 0.259715762 | 3.64E-07 |
| B46 | B216 | RBC Count | BMI | 0.25295876 | 7.43E-07 |
| B48 | B53 | LYM% | GR% | -0.924629067 | 0 |
| B49 | B50 | baso% | eo% | 0.413105357 | 0 |
| B5 | B58 | WB-Mg | HCT | 0.484378161 | 0 |
| B5 | B6 | WB-Mg | WB-Fe | 1 | 0 |
| B5 | B8 | WB-Mg | WB-Zinc | 1 | 0 |
| B5 | B66 | WB-Mg | WBRSR(mPas_100) | 0.405797344 | 4.44E-16 |
| B5 | B67 | WB-Mg | WBRSR(mPas_50) | 0.404505679 | 4.44E-16 |
| B5 | B63 | WB-Mg | WBRSR(mPas_200) | 0.390698729 | 4.88E-15 |
| B5 | B62 | WB-Mg | WBHSRVI | 0.363067127 | 4.59E-13 |
| B5 | B64 | WB-Mg | WBRSR(mPas_5) | 0.312654121 | 6.68E-10 |
| B5 | B55 | WB-Mg | TK | -0.269580305 | 1.24E-07 |
| B50 | B225 | eo% | hPBG | 0.271296544 | 1.03E-07 |

| | | | | | |
|---|---|---|---|---|---|
| B51 | B222 | Hb | NS | 0.394333206 | 2.66E-15 |
| B51 | B216 | Hb | BMI | 0.34661984 | 5.72E-12 |
| B51 | B126 | Hb | Alb | 0.341896427 | 1.15E-11 |
| B51 | B221 | Hb | Hei | 0.334952608 | 3.14E-11 |
| B51 | B163 | Hb | SG | 0.288113844 | 1.46E-08 |
| B51 | B206 | Hb | Waist | 0.287803823 | 1.51E-08 |
| B51 | B71 | Hb | TG | 0.286637397 | 1.74E-08 |
| B51 | B207 | Hb | WHR | 0.285046687 | 2.10E-08 |
| B51 | B209 | Hb | Smoking | 0.25403149 | 6.64E-07 |
| B54 | B69 | ESR | PV | 0.433615147 | 0 |
| B54 | B114 | ESR | TSH TT3 TT4(TT4) | 0.30905663 | 1.07E-09 |
| B54 | B58 | ESR | HCT | -0.294961887 | 6.35E-09 |
| B54 | B62 | ESR | WBHSRVI | -0.275196895 | 6.60E-08 |
| B54 | B59 | ESR | WBRV | 0.274391201 | 7.23E-08 |
| B54 | B182 | ESR | TT | -0.26575661 | 1.89E-07 |
| B55 | B56 | TK | IR | 0.677254662 | 0 |
| B55 | B58 | TK | HCT | -0.585991115 | 0 |
| B55 | B61 | TK | WBHSRV | 0.55376848 | 0 |
| B55 | B135 | TK | TBA | 0.323286863 | 1.60E-10 |
| B55 | B59 | TK | WBRV | 0.27744856 | 5.10E-08 |
| B55 | B134 | TK | CHE | -0.274200548 | 7.39E-08 |
| B55 | B81 | TK | TSGF | -0.272576314 | 8.88E-08 |
| B56 | B60 | IR | WBRVI | 0.485006403 | 0 |
| B56 | B61 | IR | WBHSRV | 0.873021131 | 0 |
| B56 | B62 | IR | WBHSRVI | 0.764485802 | 0 |
| B56 | B63 | IR | WBRSR(mPas_200) | 0.605918505 | 0 |
| B56 | B64 | IR | WBRSR(mPas_5) | 0.472754365 | 0 |
| B56 | B66 | IR | WBRSR(mPas_100) | 0.598184502 | 0 |
| B56 | B67 | IR | WBRSR(mPas_50) | 0.587338233 | 0 |
| B56 | B65 | IR | WBRSR(mPas_1) | 0.384374427 | 1.40E-14 |
| B56 | B59 | IR | WBRV | 0.372612087 | 9.93E-14 |
| B56 | B81 | IR | TSGF | -0.317461939 | 3.53E-10 |
| B57 | B59 | RBCAI | WBRV | 0.739633774 | 0 |
| B57 | B60 | RBCAI | WBRVI | 0.649439613 | 0 |
| B57 | B64 | RBCAI | WBRSR(mPas_5) | 0.459645464 | 0 |
| B57 | B65 | RBCAI | WBRSR(mPas_1) | 0.613988382 | 0 |
| B57 | B119 | RBCAI | Hcv-rna | 0.363375254 | 4.37E-13 |
| B57 | B152 | RBCAI | ACA-Ig(G-A-M) | -0.284727084 | 2.18E-08 |
| B57 | B121 | RBCAI | C3 | -0.284718679 | 2.19E-08 |
| B57 | B122 | RBCAI | C4 | -0.284718846 | 2.19E-08 |
| B57 | B124 | RBCAI | ACTH | 0.284716229 | 2.19E-08 |
| B57 | B169 | RBCAI | U-CRY | -0.284717347 | 2.19E-08 |
| B57 | B125 | RBCAI | Cor | -0.278959341 | 4.29E-08 |
| B57 | B92 | RBCAI | ALDH2-Glu504Glu | -0.267555054 | 1.55E-07 |

| | | | | | |
|---|---|---|---|---|---|
| B57 | B93 | RBCAI | ALDH2-Glu504Lys | 0.267555054 | 1.55E-07 |
| B57 | B94 | RBCAI | ALDH2-Lys504Lys | 0.267555054 | 1.55E-07 |
| B57 | B79 | RBCAI | hb V -dna | 0.257061599 | 4.83E-07 |
| B58 | B60 | HCT | WBRVI | 0.674237158 | 0 |
| B58 | B62 | HCT | WBHSRVI | 0.769429636 | 0 |
| B58 | B63 | HCT | WBRSR(mPas_200) | 0.748361249 | 0 |
| B58 | B64 | HCT | WBRSR(mPas_5) | 0.749344295 | 0 |
| B58 | B65 | HCT | WBRSR(mPas_1) | 0.674367913 | 0 |
| B58 | B66 | HCT | WBRSR(mPas_100) | 0.784447457 | 0 |
| B58 | B67 | HCT | WBRSR(mPas_50) | 0.796502021 | 0 |
| B58 | B134 | HCT | CHE | 0.267040936 | 1.65E-07 |
| B59 | B60 | WBRV | WBRVI | 0.684611288 | 0 |
| B59 | B61 | WBRV | WBHSRV | 0.550036694 | 0 |
| B59 | B63 | WBRV | WBRSR(mPas_200) | 0.418896234 | 0 |
| B59 | B64 | WBRV | WBRSR(mPas_5) | 0.649761192 | 0 |
| B59 | B65 | WBRV | WBRSR(mPas_1) | 0.719146265 | 0 |
| B59 | B66 | WBRV | WBRSR(mPas_100) | 0.431772548 | 0 |
| B59 | B67 | WBRV | WBRSR(mPas_50) | 0.454691428 | 0 |
| B59 | B69 | WBRV | PV | 0.375205987 | 6.48E-14 |
| B59 | B119 | WBRV | Hcv-rna | 0.360068881 | 7.35E-13 |
| B59 | B92 | WBRV | ALDH2-Glu504Glu | -0.283825647 | 2.43E-08 |
| B59 | B93 | WBRV | ALDH2-Glu504Lys | 0.283825647 | 2.43E-08 |
| B59 | B94 | WBRV | ALDH2-Lys504Lys | 0.283825646 | 2.43E-08 |
| B59 | B197 | WBRV | AG | 0.253500508 | 7.02E-07 |
| B59 | B79 | WBRV | hb V -dna | 0.250558002 | 9.52E-07 |
| B6 | B58 | WB-Fe | HCT | 0.484378161 | 0 |
| B6 | B8 | WB-Fe | WB-Zinc | 1 | 0 |
| B6 | B66 | WB-Fe | WBRSR(mPas_100) | 0.405797344 | 4.44E-16 |
| B6 | B67 | WB-Fe | WBRSR(mPas_50) | 0.404505679 | 4.44E-16 |
| B6 | B63 | WB-Fe | WBRSR(mPas_200) | 0.390698729 | 4.88E-15 |
| B6 | B62 | WB-Fe | WBHSRVI | 0.363067127 | 4.59E-13 |
| B6 | B64 | WB-Fe | WBRSR(mPas_5) | 0.312654121 | 6.68E-10 |
| B6 | B55 | WB-Fe | TK | -0.269580305 | 1.24E-07 |
| B60 | B61 | WBRVI | WBHSRV | 0.470207077 | 0 |
| B60 | B62 | WBRVI | WBHSRVI | 0.73988826 | 0 |
| B60 | B63 | WBRVI | WBRSR(mPas_200) | 0.661893665 | 0 |
| B60 | B64 | WBRVI | WBRSR(mPas_5) | 0.891087364 | 0 |
| B60 | B65 | WBRVI | WBRSR(mPas_1) | 0.916037272 | 0 |
| B60 | B66 | WBRVI | WBRSR(mPas_100) | 0.713036301 | 0 |
| B60 | B67 | WBRVI | WBRSR(mPas_50) | 0.756780373 | 0 |
| B60 | B119 | WBRVI | Hcv-rna | 0.330837058 | 5.62E-11 |
| B60 | B186 | WBRVI | PFC | -0.298610085 | 4.04E-09 |
| B60 | B92 | WBRVI | ALDH2-Glu504Glu | -0.26833207 | 1.43E-07 |
| B60 | B93 | WBRVI | ALDH2-Glu504Lys | 0.26833207 | 1.43E-07 |

| | | | | | |
|---|---|---|---|---|---|
| B60 | B94 | WBRVI | ALDH2-Lys504Lys | 0.268332069 | 1.43E-07 |
| B61 | B62 | WBHSRV | WBHSRVI | 0.699310505 | 0 |
| B61 | B63 | WBHSRV | WBRSR(mPas_200) | 0.803517062 | 0 |
| B61 | B64 | WBHSRV | WBRSR(mPas_5) | 0.626090907 | 0 |
| B61 | B65 | WBHSRV | WBRSR(mPas_1) | 0.522605423 | 0 |
| B61 | B66 | WBHSRV | WBRSR(mPas_100) | 0.777441549 | 0 |
| B61 | B67 | WBHSRV | WBRSR(mPas_50) | 0.744736699 | 0 |
| B61 | B81 | WBHSRV | TSGF | -0.350446087 | 3.22E-12 |
| B61 | B69 | WBHSRV | PV | 0.289876704 | 1.18E-08 |
| B62 | B63 | WBHSRVI | WBRSR(mPas_200) | 0.858201983 | 0 |
| B62 | B64 | WBHSRVI | WBRSR(mPas_5) | 0.772039229 | 0 |
| B62 | B65 | WBHSRVI | WBRSR(mPas_1) | 0.667788919 | 0 |
| B62 | B66 | WBHSRVI | WBRSR(mPas_100) | 0.878736933 | 0 |
| B62 | B67 | WBHSRVI | WBRSR(mPas_50) | 0.880571016 | 0 |
| B63 | B64 | WBRSR(mPas_200) | WBRSR(mPas_5) | 0.862990848 | 0 |
| B63 | B65 | WBRSR(mPas_200) | WBRSR(mPas_1) | 0.74475926 | 0 |
| B63 | B66 | WBRSR(mPas_200) | WBRSR(mPas_100) | 0.99348703 | 0 |
| B63 | B67 | WBRSR(mPas_200) | WBRSR(mPas_50) | 0.971679882 | 0 |
| B63 | B69 | WBRSR(mPas_200) | PV | 0.427651803 | 0 |
| B63 | B81 | WBRSR(mPas_200) | TSGF | -0.250948627 | 9.15E-07 |
| B64 | B65 | WBRSR(mPas_5) | WBRSR(mPas_1) | 0.963608901 | 0 |
| B64 | B66 | WBRSR(mPas_5) | WBRSR(mPas_100) | 0.893577968 | 0 |
| B64 | B67 | WBRSR(mPas_5) | WBRSR(mPas_50) | 0.937612695 | 0 |
| B64 | B69 | WBRSR(mPas_5) | PV | 0.33888086 | 1.78E-11 |
| B64 | B92 | WBRSR(mPas_5) | ALDH2-Glu504Glu | -0.318258358 | 3.17E-10 |
| B64 | B93 | WBRSR(mPas_5) | ALDH2-Glu504Lys | 0.318258358 | 3.17E-10 |
| B64 | B94 | WBRSR(mPas_5) | ALDH2-Lys504Lys | 0.318258357 | 3.17E-10 |
| B64 | B119 | WBRSR(mPas_5) | Hcv-rna | 0.261924933 | 2.87E-07 |
| B65 | B66 | WBRSR(mPas_1) | WBRSR(mPas_100) | 0.787446501 | 0 |
| B65 | B67 | WBRSR(mPas_1) | WBRSR(mPas_50) | 0.834886076 | 0 |
| B65 | B92 | WBRSR(mPas_1) | ALDH2-Glu504Glu | -0.346995213 | 5.41E-12 |
| B65 | B93 | WBRSR(mPas_1) | ALDH2-Glu504Lys | 0.346995213 | 5.41E-12 |
| B65 | B94 | WBRSR(mPas_1) | ALDH2-Lys504Lys | 0.346995212 | 5.41E-12 |
| B65 | B69 | WBRSR(mPas_1) | PV | 0.306554729 | 1.48E-09 |
| B66 | B67 | WBRSR(mPas_100) | WBRSR(mPas_50) | 0.985021731 | 0 |
| B66 | B69 | WBRSR(mPas_100) | PV | 0.393664397 | 2.66E-15 |
| B67 | B69 | WBRSR(mPas_50) | PV | 0.349805196 | 3.55E-12 |
| B69 | B194 | PV | CI | -0.303618463 | 2.15E-09 |
| B69 | B197 | PV | AG | 0.273301559 | 8.18E-08 |
| B69 | B150 | PV | ALP | -0.265186002 | 2.02E-07 |
| B69 | B74 | PV | ApoA1 | 0.264457153 | 2.18E-07 |
| B7 | B58 | WB-Cop | HCT | 0.484378176 | 0 |
| B7 | B66 | WB-Cop | WBRSR(mPas_100) | 0.405797355 | 4.44E-16 |
| B7 | B67 | WB-Cop | WBRSR(mPas_50) | 0.40450569 | 4.44E-16 |

| | | | | | |
|---|---|---|---|---|---|
| B7 | B63 | WB-Cop | WBRSR(mPas_200) | 0.39069874 | 4.88E-15 |
| B7 | B62 | WB-Cop | WBHSRVI | 0.363067138 | 4.59E-13 |
| B7 | B64 | WB-Cop | WBRSR(mPas_5) | 0.312654131 | 6.68E-10 |
| B7 | B55 | WB-Cop | TK | -0.269580314 | 1.24E-07 |
| B70 | B76 | LDL-C | ApoB | 0.429527442 | 0 |
| B70 | B77 | LDL-C | TC | 0.762318444 | 0 |
| B70 | B74 | LDL-C | ApoA1 | 0.263898713 | 2.32E-07 |
| B70 | B72 | LDL-C | HDL-C | 0.253908611 | 6.73E-07 |
| B70 | B132 | LDL-C | Dbil | -0.25340365 | 7.09E-07 |
| B70 | B75 | LDL-C | ApoA1/B | -0.253154105 | 7.28E-07 |
| B71 | B72 | TG | HDL-C | -0.42793688 | 0 |
| B71 | B77 | TG | TC | 0.482233626 | 0 |
| B71 | B91 | TG | TP | 0.308431624 | 1.16E-09 |
| B71 | B153 | TG | Glu | 0.255990612 | 5.41E-07 |
| B71 | B130 | TG | GLB | 0.251784371 | 8.39E-07 |
| B72 | B216 | HDL-C | BMI | -0.43658543 | 0 |
| B72 | B222 | HDL-C | NS | -0.4194297 | 0 |
| B72 | B74 | HDL-C | ApoA1 | 0.402864399 | 4.44E-16 |
| B72 | B206 | HDL-C | Waist | -0.376565464 | 5.20E-14 |
| B72 | B215 | HDL-C | BMI | -0.34736446 | 5.12E-12 |
| B72 | B221 | HDL-C | Hei | -0.308888452 | 1.09E-09 |
| B72 | B207 | HDL-C | WHR | -0.304792821 | 1.85E-09 |
| B72 | B210 | HDL-C | Hip | -0.27123438 | 1.03E-07 |
| B72 | B192 | HDL-C | UA | -0.255307833 | 5.81E-07 |
| B73 | B79 | ApoA | hb V -dna | -0.291645753 | 9.53E-09 |
| B73 | B196 | ApoA | TCO2 | 0.275621693 | 6.29E-08 |
| B74 | B76 | ApoA1 | ApoB | 0.429954539 | 0 |
| B74 | B84 | ApoA1 | C-PSA | -0.335337317 | 2.97E-11 |
| B74 | B194 | ApoA1 | Cl | -0.311578884 | 7.69E-10 |
| B74 | B75 | ApoA1 | ApoA1/B | 0.283106306 | 2.64E-08 |
| B74 | B77 | ApoA1 | TC | 0.269070563 | 1.31E-07 |
| B75 | B76 | ApoA1/B | ApoB | -0.700656858 | 0 |
| B75 | B197 | ApoA1/B | AG | 0.272296137 | 9.17E-08 |
| B76 | B77 | ApoB | TC | 0.383745832 | 1.55E-14 |
| B76 | B84 | ApoB | C-PSA | -0.313745768 | 5.78E-10 |
| B76 | B196 | ApoB | TCO2 | 0.273248835 | 8.23E-08 |
| B76 | B199 | ApoB | Mg | 0.253580258 | 6.96E-07 |
| B77 | B91 | TC | TP | 0.293457383 | 7.64E-09 |
| B79 | B119 | hb V -dna | Hcv-rna | 0.705198175 | 0 |
| B79 | B120 | hb V -dna | HCV-cAg | 0.492507903 | 0 |
| B79 | B121 | hb V -dna | C3 | -0.497979684 | 0 |
| B79 | B122 | hb V -dna | C4 | -0.497979982 | 0 |
| B79 | B124 | hb V -dna | ACTH | 0.497990782 | 0 |
| B79 | B125 | hb V -dna | Cor | -0.488781991 | 0 |

| | | | | | |
|---|---|---|---|---|---|
| B79 | B152 | hb V -dna | ACA-Ig(G-A-M) | -0.497994395 | 0 |
| B79 | B169 | hb V -dna | U-CRY | -0.497977372 | 0 |
| B79 | B180 | hb V -dna | Tf | 0.498306465 | 0 |
| B79 | B155 | hb V -dna | IgA | -0.355222219 | 1.56E-12 |
| B79 | B158 | hb V -dna | IgM | -0.295572468 | 5.89E-09 |
| B79 | B184 | hb V -dna | INR | 0.283667385 | 2.47E-08 |
| B79 | B179 | hb V -dna | α1-MG | 0.262823873 | 2.61E-07 |
| B8 | B58 | WB-Zinc | HCT | 0.484378161 | 0 |
| B8 | B66 | WB-Zinc | WBRSR(mPas_100) | 0.405797344 | 4.44E-16 |
| B8 | B67 | WB-Zinc | WBRSR(mPas_50) | 0.404505679 | 4.44E-16 |
| B8 | B63 | WB-Zinc | WBRSR(mPas_200) | 0.390698729 | 4.88E-15 |
| B8 | B62 | WB-Zinc | WBHSRVI | 0.363067127 | 4.59E-13 |
| B8 | B64 | WB-Zinc | WBRSR(mPas_5) | 0.312654121 | 6.68E-10 |
| B8 | B55 | WB-Zinc | TK | -0.269580305 | 1.24E-07 |
| B80 | B156 | anti-HP IgG | IgE | 0.273911141 | 7.64E-08 |
| B81 | B121 | TSGF | C3 | 0.321649897 | 2.00E-10 |
| B81 | B122 | TSGF | C4 | 0.321650088 | 2.00E-10 |
| B81 | B124 | TSGF | ACTH | -0.321654233 | 2.00E-10 |
| B81 | B152 | TSGF | ACA-Ig(G-A-M) | 0.321659392 | 2.00E-10 |
| B81 | B169 | TSGF | U-CRY | 0.321648406 | 2.00E-10 |
| B81 | B125 | TSGF | Cor | 0.315547726 | 4.55E-10 |
| B82 | B85 | CEA | SCC | 0.560544337 | 0 |
| B82 | B89 | CEA | CA 72-4 | 0.654070848 | 0 |
| B82 | B88 | CEA | CA 242 | 0.294207373 | 6.97E-09 |
| B83 | B84 | AFP | C-PSA | 0.620364645 | 0 |
| B83 | B87 | AFP | NSE | 0.463401803 | 0 |
| B83 | B88 | AFP | CA 242 | 0.492584026 | 0 |
| B83 | B85 | AFP | SCC | 0.405847592 | 4.44E-16 |
| B83 | B90 | AFP | CYFRA 21-1 | 0.370789908 | 1.34E-13 |
| B84 | B85 | C-PSA | SCC | 0.417483967 | 0 |
| B84 | B87 | C-PSA | NSE | 0.649393489 | 0 |
| B84 | B88 | C-PSA | CA 242 | 0.359003153 | 8.68E-13 |
| B84 | B90 | C-PSA | CYFRA 21-1 | 0.275825908 | 6.14E-08 |
| B85 | B88 | SCC | CA 242 | 0.443821018 | 0 |
| B85 | B87 | SCC | NSE | 0.316357014 | 4.09E-10 |
| B85 | B89 | SCC | CA 72-4 | 0.274451612 | 7.18E-08 |
| B86 | B89 | TBhCG | CA 72-4 | 0.531607102 | 0 |
| B86 | B123 | TBhCG | hs-CRP1 | -0.270033649 | 1.18E-07 |
| B87 | B90 | NSE | CYFRA 21-1 | 0.364800776 | 3.49E-13 |
| B87 | B123 | NSE | hs-CRP1 | 0.281494172 | 3.19E-08 |
| B89 | B123 | CA 72-4 | hs-CRP1 | -0.263612484 | 2.39E-07 |
| B9 | B117 | RH | TT4 | -0.405095747 | 4.44E-16 |
| B9 | B29 | RH | E2 | -0.33028913 | 6.07E-11 |
| B91 | B127 | TP | A-G ratio | -0.500783305 | 0 |

| | | | | | |
|---|---|---|---|---|---|
| B91 | B130 | TP | GLB | 0.780472818 | 0 |
| B91 | B126 | TP | Alb | 0.404865761 | 4.44E-16 |
| B91 | B188 | TP | Ca | 0.308321676 | 1.18E-09 |
| B92 | B94 | ALDH2-Glu504Glu | ALDH2-Lys504Lys | -0.999999996 | 0 |
| B92 | B137 | ALDH2-Glu504Glu | AFU | 0.333474717 | 3.87E-11 |
| B92 | B157 | ALDH2-Glu504Glu | IgG | -0.294310245 | 6.88E-09 |
| B92 | B179 | ALDH2-Glu504Glu | α1-MG | -0.269814194 | 1.21E-07 |
| B92 | B158 | ALDH2-Glu504Glu | IgM | -0.264202788 | 2.25E-07 |
| B92 | B196 | ALDH2-Glu504Glu | TCO2 | -0.250773362 | 9.31E-07 |
| B93 | B94 | ALDH2-Glu504Lys | ALDH2-Lys504Lys | 0.999999996 | 0 |
| B93 | B137 | ALDH2-Glu504Lys | AFU | -0.333474717 | 3.87E-11 |
| B93 | B157 | ALDH2-Glu504Lys | IgG | 0.294310245 | 6.88E-09 |
| B93 | B179 | ALDH2-Glu504Lys | α1-MG | 0.269814194 | 1.21E-07 |
| B93 | B158 | ALDH2-Glu504Lys | IgM | 0.264202788 | 2.25E-07 |
| B93 | B196 | ALDH2-Glu504Lys | TCO2 | 0.250773362 | 9.31E-07 |
| B94 | B137 | ALDH2-Lys504Lys | AFU | -0.333474716 | 3.87E-11 |
| B94 | B157 | ALDH2-Lys504Lys | IgG | 0.294310244 | 6.88E-09 |
| B94 | B179 | ALDH2-Lys504Lys | α1-MG | 0.269814193 | 1.21E-07 |
| B94 | B158 | ALDH2-Lys504Lys | IgM | 0.264202787 | 2.25E-07 |
| B94 | B196 | ALDH2-Lys504Lys | TCO2 | 0.250773361 | 9.31E-07 |
| B95 | B104 | anti-cenpb | anti-SSB | 0.999999999 | 0 |
| B95 | B106 | anti-cenpb | anti-P | 0.999999999 | 0 |
| B95 | B108 | anti-cenpb | Anti-Cyclin | 0.999999999 | 0 |
| B95 | B109 | anti-cenpb | AMA-M2 | 0.999999999 | 0 |
| B95 | B151 | anti-cenpb | Anti-CCP | 0.589369502 | 0 |
| B95 | B98 | anti-cenpb | anti-Jo-1 | 0.999999998 | 0 |
| B95 | B193 | anti-cenpb | K | 0.396577754 | 1.78E-15 |
| B95 | B157 | anti-cenpb | IgG | 0.381317843 | 2.35E-14 |
| B95 | B198 | anti-cenpb | P | -0.350329926 | 3.28E-12 |
| B95 | B185 | anti-cenpb | PT | 0.305718441 | 1.64E-09 |
| B95 | B114 | anti-cenpb | TSH TT3 TT4(TT4) | 0.304406136 | 1.94E-09 |
| B95 | B134 | anti-cenpb | CHE | -0.262564376 | 2.68E-07 |
| B95 | B150 | anti-cenpb | ALP | -0.252421185 | 7.86E-07 |
| B97 | B100 | dsDNA | anti-Ro52 | 0.999999998 | 0 |
| B97 | B103 | dsDNA | anti-SSA/Ro | 0.999999999 | 0 |
| B97 | B105 | dsDNA | anti-U1RNP | 0.999999998 | 0 |
| B97 | B108 | dsDNA | Anti-Cyclin | 0.999999999 | 0 |
| B97 | B109 | dsDNA | AMA-M2 | 0.999999998 | 0 |
| B97 | B110 | dsDNA | AHA | 0.999999998 | 0 |
| B97 | B151 | dsDNA | Anti-CCP | 0.5893695 | 0 |
| B97 | B99 | dsDNA | anti-PM-Scl | 0.999999999 | 0 |
| B97 | B193 | dsDNA | K | 0.396577753 | 1.78E-15 |
| B97 | B157 | dsDNA | IgG | 0.381317842 | 2.35E-14 |
| B97 | B198 | dsDNA | P | -0.350329925 | 3.28E-12 |

| | | | | | |
|---|---|---|---|---|---|
| B97 | B185 | dsDNA | PT | 0.30571844 | 1.64E-09 |
| B97 | B114 | dsDNA | TSH TT3 TT4(TT4) | 0.304406135 | 1.94E-09 |
| B97 | B134 | dsDNA | CHE | -0.262564375 | 2.68E-07 |
| B97 | B150 | dsDNA | ALP | -0.252421184 | 7.86E-07 |
| B98 | B100 | anti-Jo-1 | anti-Ro52 | 0.999999998 | 0 |
| B98 | B101 | anti-Jo-1 | anti-Sc70 | 0.999999996 | 0 |
| B98 | B102 | anti-Jo-1 | anti-Sm | 0.999999998 | 0 |
| B98 | B103 | anti-Jo-1 | anti-SSA/Ro | 0.999999999 | 0 |
| B98 | B104 | anti-Jo-1 | anti-SSB | 0.999999998 | 0 |
| B98 | B105 | anti-Jo-1 | anti-U1RNP | 0.999999997 | 0 |
| B98 | B109 | anti-Jo-1 | AMA-M2 | 0.999999996 | 0 |
| B98 | B110 | anti-Jo-1 | AHA | 0.999999998 | 0 |
| B98 | B151 | anti-Jo-1 | Anti-CCP | 0.5893695 | 0 |
| B98 | B193 | anti-Jo-1 | K | 0.396577753 | 1.78E-15 |
| B98 | B157 | anti-Jo-1 | IgG | 0.381317842 | 2.35E-14 |
| B98 | B198 | anti-Jo-1 | P | -0.350329925 | 3.28E-12 |
| B98 | B185 | anti-Jo-1 | PT | 0.305718441 | 1.64E-09 |
| B98 | B114 | anti-Jo-1 | TSH TT3 TT4(TT4) | 0.304406136 | 1.94E-09 |
| B98 | B134 | anti-Jo-1 | CHE | -0.262564376 | 2.68E-07 |
| B98 | B150 | anti-Jo-1 | ALP | -0.252421184 | 7.86E-07 |
| B99 | B100 | anti-PM-Scl | anti-Ro52 | 0.999999997 | 0 |
| B99 | B103 | anti-PM-Scl | anti-SSA/Ro | 0.999999998 | 0 |
| B99 | B104 | anti-PM-Scl | anti-SSB | 1 | 0 |
| B99 | B105 | anti-PM-Scl | anti-U1RNP | 0.999999998 | 0 |
| B99 | B106 | anti-PM-Scl | anti-P | 1 | 0 |
| B99 | B109 | anti-PM-Scl | AMA-M2 | 0.999999998 | 0 |
| B99 | B110 | anti-PM-Scl | AHA | 0.999999999 | 0 |
| B99 | B151 | anti-PM-Scl | Anti-CCP | 0.5893695 | 0 |
| B99 | B193 | anti-PM-Scl | K | 0.396577753 | 1.78E-15 |
| B99 | B157 | anti-PM-Scl | IgG | 0.381317842 | 2.35E-14 |
| B99 | B198 | anti-PM-Scl | P | -0.350329925 | 3.28E-12 |
| B99 | B185 | anti-PM-Scl | PT | 0.30571844 | 1.64E-09 |
| B99 | B114 | anti-PM-Scl | TSH TT3 TT4(TT4) | 0.304406135 | 1.94E-09 |
| B99 | B134 | anti-PM-Scl | CHE | -0.262564375 | 2.68E-07 |
| B99 | B150 | anti-PM-Scl | ALP | -0.252421184 | 7.86E-07 |

**Supplementary Table 27. PEI Correlations in Bronchial disease**

| Row | Column | Row | Column | Cor | *P* |
|---|---|---|---|---|---|
| B1 | B163 | Gender | SG | -0.41257269 | 0 |
| B1 | B174 | Gender | Urea/Cr | 0.38883187 | 0 |
| B1 | B192 | Gender | UA | -0.42830993 | 0 |
| B1 | B216 | Gender | BMI | -0.39281319 | 0 |
| B1 | B221 | Gender | Hei | -0.64068402 | 0 |
| B1 | B222 | Gender | NS | -0.60274717 | 0 |
| B1 | B46 | Gender | RBC Count | -0.40887858 | 0 |
| B1 | B51 | Gender | Hb | -0.52611764 | 0 |
| B1 | B45 | Gender | MCHC | -0.29428682 | 6.24E-13 |
| B1 | B42 | Gender | MON | -0.28331701 | 4.67E-12 |
| B1 | B190 | Gender | Cre | -0.28264947 | 5.27E-12 |
| B1 | B209 | Gender | Smoking | -0.27493328 | 2.05E-11 |
| B1 | B41 | Gender | MON％ | -0.26476855 | 1.16E-10 |
| B1 | B204 | Gender | Drinking | -0.24497082 | 2.73E-09 |
| B1 | B206 | Gender | Waist | -0.24093287 | 5.03E-09 |
| B1 | B141 | Gender | AST/ALT | 0.235254029 | 1.17E-08 |
| B1 | B128 | Gender | ALT | -0.22648768 | 4.11E-08 |
| B1 | B52 | Gender | PLT/BPC | 0.219259903 | 1.12E-07 |
| B1 | B72 | Gender | HDL-C | 0.218326748 | 1.27E-07 |
| B1 | B44 | Gender | MCH | -0.2106098 | 3.54E-07 |
| B10 | B11 | PG I | PGI/PGII | 0.460740486 | 0 |
| B10 | B12 | PG I | PG II | 0.360651686 | 0 |
| B10 | B28 | PG I | PRL | -0.21422804 | 2.20E-07 |
| B10 | B29 | PG I | E2 | -0.21422804 | 2.20E-07 |
| B10 | B30 | PG I | LH | -0.21422804 | 2.20E-07 |
| B10 | B31 | PG I | FSH | -0.21422804 | 2.20E-07 |
| B10 | B32 | PG I | Testo | -0.21422804 | 2.20E-07 |
| B10 | B33 | PG I | Prog | -0.21422804 | 2.20E-07 |
| B100 | B103 | anti-Ro52 | anti-SSA/Ro | 1 | 0 |
| B100 | B104 | anti-Ro52 | anti-SSB | 0.999999997 | 0 |
| B100 | B105 | anti-Ro52 | anti-U1RNP | 1 | 0 |
| B100 | B106 | anti-Ro52 | anti-P | 0.999999998 | 0 |
| B100 | B107 | anti-Ro52 | AnuA | 0.999999976 | 0 |
| B100 | B151 | anti-Ro52 | Anti-CCP | 0.60849111 | 0 |

| | | | | | |
|---|---|---|---|---|---|
| | | | | 7 | |
| B100 | B179 | anti-Ro52 | α1-MG | -0.37132208 | 0 |
| B100 | B181 | anti-Ro52 | APTT | -0.49743253 | 0 |
| B100 | B182 | anti-Ro52 | TT | -0.36368895 | 0 |
| B100 | B186 | anti-Ro52 | PFC | 0.542615961 | 0 |
| B100 | B178 | anti-Ro52 | Ig-U | -0.28472773 | 3.62E-12 |
| B100 | B200 | anti-Ro52 | OP | -0.2737536 | 2.52E-11 |
| B100 | B195 | anti-Ro52 | Na | -0.27267879 | 3.03E-11 |
| B100 | B196 | anti-Ro52 | TCO2 | -0.25679436 | 4.27E-10 |
| B100 | B194 | anti-Ro52 | Cl | 0.240674745 | 5.23E-09 |
| B100 | B137 | anti-Ro52 | AFU | 0.236068598 | 1.04E-08 |
| B101 | B102 | anti-Sc70 | anti-Sm | 0.999999997 | 0 |
| B101 | B107 | anti-Sc70 | AnuA | 0.99999984 | 0 |
| B101 | B151 | anti-Sc70 | Anti-CCP | 0.608491124 | 0 |
| B101 | B179 | anti-Sc70 | α1-MG | -0.37132209 | 0 |
| B101 | B181 | anti-Sc70 | APTT | -0.49743254 | 0 |
| B101 | B182 | anti-Sc70 | TT | -0.36368896 | 0 |
| B101 | B186 | anti-Sc70 | PFC | 0.542615967 | 0 |
| B101 | B178 | anti-Sc70 | Ig-U | -0.28472774 | 3.62E-12 |
| B101 | B200 | anti-Sc70 | OP | -0.27375361 | 2.52E-11 |
| B101 | B195 | anti-Sc70 | Na | -0.2726788 | 3.03E-11 |
| B101 | B196 | anti-Sc70 | TCO2 | -0.25679437 | 4.27E-10 |
| B101 | B194 | anti-Sc70 | Cl | 0.240674749 | 5.23E-09 |
| B101 | B137 | anti-Sc70 | AFU | 0.236068601 | 1.04E-08 |
| B102 | B104 | anti-Sm | anti-SSB | 1 | 0 |
| B102 | B151 | anti-Sm | Anti-CCP | 0.60849112 | 0 |
| B102 | B179 | anti-Sm | α1-MG | -0.37132208 | 0 |
| B102 | B181 | anti-Sm | APTT | -0.49743253 | 0 |
| B102 | B182 | anti-Sm | TT | -0.36368895 | 0 |
| B102 | B186 | anti-Sm | PFC | 0.542615965 | 0 |
| B102 | B178 | anti-Sm | Ig-U | -0.28472773 | 3.62E-12 |
| B102 | B200 | anti-Sm | OP | -0.27375361 | 2.52E-11 |
| B102 | B195 | anti-Sm | Na | -0.27267879 | 3.03E-11 |
| B102 | B196 | anti-Sm | TCO2 | -0.25679436 | 4.27E-10 |
| B102 | B194 | anti-Sm | Cl | 0.24067474 | 5.23E-09 |

| | | | | | |
|---|---|---|---|---|---|
| | | | | 6 | |
| B102 | B137 | anti-Sm | AFU | 0.236068599 | 1.04E-08 |
| B103 | B104 | anti-SSA/Ro | anti-SSB | 0.999999997 | 0 |
| B103 | B105 | anti-SSA/Ro | anti-U1RNP | 1 | 0 |
| B103 | B106 | anti-SSA/Ro | anti-P | 0.999999998 | 0 |
| B103 | B107 | anti-SSA/Ro | AnuA | 0.999999987 | 0 |
| B103 | B108 | anti-SSA/Ro | Anti-Cyclin | 1 | 0 |
| B103 | B151 | anti-SSA/Ro | Anti-CCP | 0.608491116 | 0 |
| B103 | B179 | anti-SSA/Ro | α1-MG | -0.37132208 | 0 |
| B103 | B181 | anti-SSA/Ro | APTT | -0.49743253 | 0 |
| B103 | B182 | anti-SSA/Ro | TT | -0.36368895 | 0 |
| B103 | B186 | anti-SSA/Ro | PFC | 0.542615961 | 0 |
| B103 | B178 | anti-SSA/Ro | Ig-U | -0.28472773 | 3.62E-12 |
| B103 | B200 | anti-SSA/Ro | OP | -0.2737536 | 2.52E-11 |
| B103 | B195 | anti-SSA/Ro | Na | -0.27267879 | 3.03E-11 |
| B103 | B196 | anti-SSA/Ro | TCO2 | -0.25679436 | 4.27E-10 |
| B103 | B194 | anti-SSA/Ro | Cl | 0.240674745 | 5.23E-09 |
| B103 | B137 | anti-SSA/Ro | AFU | 0.236068598 | 1.04E-08 |
| B104 | B105 | anti-SSB | anti-U1RNP | 0.999999996 | 0 |
| B104 | B107 | anti-SSB | AnuA | 0.999999972 | 0 |
| B104 | B108 | anti-SSB | Anti-Cyclin | 0.999999997 | 0 |
| B104 | B109 | anti-SSB | AMA-M2 | 0.999999997 | 0 |
| B104 | B110 | anti-SSB | AHA | 0.999999999 | 0 |
| B104 | B151 | anti-SSB | Anti-CCP | 0.608491115 | 0 |
| B104 | B179 | anti-SSB | α1-MG | -0.37132208 | 0 |
| B104 | B181 | anti-SSB | APTT | -0.49743253 | 0 |
| B104 | B182 | anti-SSB | TT | -0.36368895 | 0 |
| B104 | B186 | anti-SSB | PFC | 0.542615959 | 0 |
| B104 | B178 | anti-SSB | Ig-U | -0.28472773 | 3.62E-12 |

| | | | | | |
|---|---|---|---|---|---|
| B104 | B200 | anti-SSB | OP | -0.2737536 | 2.52E-11 |
| B104 | B195 | anti-SSB | Na | -0.27267879 | 3.03E-11 |
| B104 | B196 | anti-SSB | TCO2 | -0.25679436 | 4.27E-10 |
| B104 | B194 | anti-SSB | Cl | 0.240674747 | 5.23E-09 |
| B104 | B137 | anti-SSB | AFU | 0.236068597 | 1.04E-08 |
| B105 | B106 | anti-U1RNP | anti-P | 0.999999995 | 0 |
| B105 | B107 | anti-U1RNP | AnuA | 0.999999973 | 0 |
| B105 | B108 | anti-U1RNP | Anti-Cyclin | 1 | 0 |
| B105 | B109 | anti-U1RNP | AMA-M2 | 0.999999998 | 0 |
| B105 | B151 | anti-U1RNP | Anti-CCP | 0.608491116 | 0 |
| B105 | B179 | anti-U1RNP | α1-MG | -0.37132208 | 0 |
| B105 | B181 | anti-U1RNP | APTT | -0.49743253 | 0 |
| B105 | B182 | anti-U1RNP | TT | -0.36368895 | 0 |
| B105 | B186 | anti-U1RNP | PFC | 0.54261596 | 0 |
| B105 | B178 | anti-U1RNP | Ig-U | -0.28472773 | 3.62E-12 |
| B105 | B200 | anti-U1RNP | OP | -0.2737536 | 2.52E-11 |
| B105 | B195 | anti-U1RNP | Na | -0.27267879 | 3.03E-11 |
| B105 | B196 | anti-U1RNP | TCO2 | -0.25679436 | 4.27E-10 |
| B105 | B194 | anti-U1RNP | Cl | 0.240674745 | 5.23E-09 |
| B105 | B137 | anti-U1RNP | AFU | 0.236068598 | 1.04E-08 |
| B106 | B108 | anti-P | Anti-Cyclin | 0.999999998 | 0 |
| B106 | B109 | anti-P | AMA-M2 | 0.999999998 | 0 |
| B106 | B110 | anti-P | AHA | 0.999999996 | 0 |
| B106 | B151 | anti-P | Anti-CCP | 0.608491115 | 0 |
| B106 | B179 | anti-P | α1-MG | -0.37132208 | 0 |
| B106 | B181 | anti-P | APTT | -0.49743253 | 0 |
| B106 | B182 | anti-P | TT | -0.36368895 | 0 |
| B106 | B186 | anti-P | PFC | 0.542615961 | 0 |
| B106 | B178 | anti-P | Ig-U | -0.28472773 | 3.62E-12 |
| B106 | B200 | anti-P | OP | -0.2737536 | 2.52E-11 |
| B106 | B195 | anti-P | Na | -0.27267879 | 3.03E-11 |

| | | | | | |
|---|---|---|---|---|---|
| B106 | B196 | anti-P | TCO2 | -0.25679436 | 4.27E-10 |
| B106 | B194 | anti-P | Cl | 0.240674748 | 5.23E-09 |
| B106 | B137 | anti-P | AFU | 0.236068598 | 1.04E-08 |
| B107 | B108 | AnuA | Anti-Cyclin | 0.999999959 | 0 |
| B107 | B109 | AnuA | AMA-M2 | 0.999999972 | 0 |
| B107 | B110 | AnuA | AHA | 0.999999989 | 0 |
| B107 | B151 | AnuA | Anti-CCP | 0.608491107 | 0 |
| B107 | B179 | AnuA | α1-MG | -0.37132207 | 0 |
| B107 | B181 | AnuA | APTT | -0.49743252 | 0 |
| B107 | B182 | AnuA | TT | -0.36368895 | 0 |
| B107 | B186 | AnuA | PFC | 0.542615952 | 0 |
| B107 | B178 | AnuA | Ig-U | -0.28472773 | 3.62E-12 |
| B107 | B200 | AnuA | OP | -0.2737536 | 2.52E-11 |
| B107 | B195 | AnuA | Na | -0.27267881 | 3.03E-11 |
| B107 | B196 | AnuA | TCO2 | -0.25679436 | 4.27E-10 |
| B107 | B194 | AnuA | Cl | 0.240674754 | 5.23E-09 |
| B107 | B137 | AnuA | AFU | 0.236068595 | 1.04E-08 |
| B108 | B110 | Anti-Cyclin | AHA | 0.999999999 | 0 |
| B108 | B151 | Anti-Cyclin | Anti-CCP | 0.608491117 | 0 |
| B108 | B179 | Anti-Cyclin | α1-MG | -0.37132208 | 0 |
| B108 | B181 | Anti-Cyclin | APTT | -0.49743253 | 0 |
| B108 | B182 | Anti-Cyclin | TT | -0.36368895 | 0 |
| B108 | B186 | Anti-Cyclin | PFC | 0.542615961 | 0 |
| B108 | B178 | Anti-Cyclin | Ig-U | -0.28472773 | 3.62E-12 |
| B108 | B200 | Anti-Cyclin | OP | -0.2737536 | 2.52E-11 |
| B108 | B195 | Anti-Cyclin | Na | -0.27267879 | 3.03E-11 |
| B108 | B196 | Anti-Cyclin | TCO2 | -0.25679436 | 4.27E-10 |
| B108 | B194 | Anti-Cyclin | Cl | 0.240674746 | 5.23E-09 |
| B108 | B137 | Anti-Cyclin | AFU | 0.236068598 | 1.04E-08 |
| B109 | B151 | AMA-M2 | Anti-CCP | 0.60849111 | 0 |

| | | | | | |
|---|---|---|---|---|---|
| | | | | 7 | |
| B109 | B179 | AMA-M2 | α1-MG | -0.37132208 | 0 |
| B109 | B181 | AMA-M2 | APTT | -0.49743253 | 0 |
| B109 | B182 | AMA-M2 | TT | -0.36368895 | 0 |
| B109 | B186 | AMA-M2 | PFC | 0.54261596 | 0 |
| B109 | B178 | AMA-M2 | Ig-U | -0.28472773 | 3.62E-12 |
| B109 | B200 | AMA-M2 | OP | -0.2737536 | 2.52E-11 |
| B109 | B195 | AMA-M2 | Na | -0.27267879 | 3.03E-11 |
| B109 | B196 | AMA-M2 | TCO2 | -0.25679436 | 4.27E-10 |
| B109 | B194 | AMA-M2 | Cl | 0.240674745 | 5.23E-09 |
| B109 | B137 | AMA-M2 | AFU | 0.236068598 | 1.04E-08 |
| B11 | B12 | PGI/PGII | PG II | -0.4521582 | 0 |
| B11 | B14 | PGI/PGII | G-17 | -0.23374642 | 1.46E-08 |
| B110 | B151 | AHA | Anti-CCP | 0.608491117 | 0 |
| B110 | B179 | AHA | α1-MG | -0.37132208 | 0 |
| B110 | B181 | AHA | APTT | -0.49743253 | 0 |
| B110 | B182 | AHA | TT | -0.36368895 | 0 |
| B110 | B186 | AHA | PFC | 0.542615961 | 0 |
| B110 | B178 | AHA | Ig-U | -0.28472773 | 3.62E-12 |
| B110 | B200 | AHA | OP | -0.2737536 | 2.52E-11 |
| B110 | B195 | AHA | Na | -0.27267879 | 3.03E-11 |
| B110 | B196 | AHA | TCO2 | -0.25679436 | 4.27E-10 |
| B110 | B194 | AHA | Cl | 0.240674746 | 5.23E-09 |
| B110 | B137 | AHA | AFU | 0.236068598 | 1.04E-08 |
| B111 | B148 | TSH | Anti_TPO | 0.283831159 | 4.26E-12 |
| B112 | B114 | FT4 | TSH TT3 TT4(TT4) | 0.367216607 | 0 |
| B113 | B115 | FT3 | TSH TT3 TT4(TT3) | 0.377672896 | 0 |
| B114 | B115 | TSH TT3 TT4(TT4) | TSH TT3 TT4(TT3) | 0.558506376 | 0 |
| B114 | B186 | TSH TT3 TT4(TT4) | PFC | 0.235509976 | 1.13E-08 |
| B115 | B200 | TSH TT3 TT4(TT3) | OP | 0.269919912 | 4.86E-11 |
| B115 | B195 | TSH TT3 TT4(TT3) | Na | 0.242671159 | 3.87E-09 |

| | | | | | |
|---|---|---|---|---|---|
| B115 | B179 | TSH TT3 TT4(TT3) | α1-MG | 0.223574072 | 6.18E-08 |
| B117 | B167 | TT4 | PC | -0.39911406 | 0 |
| B118 | B151 | CEA1 | Anti-CCP | -0.35233212 | 0 |
| B118 | B154 | CEA1 | RF | 0.332581363 | 2.22E-16 |
| B118 | B175 | CEA1 | ALB-U | 0.308196673 | 4.26E-14 |
| B118 | B132 | CEA1 | Dbil | 0.2943345 | 6.18E-13 |
| B118 | B140 | CEA1 | r-GT | 0.285987443 | 2.89E-12 |
| B119 | B169 | Hcv-rna | U-CRY | -0.99998704 | 0 |
| B119 | B120 | Hcv-rna | HCV-cAg | 0.239125419 | 6.60E-09 |
| B120 | B152 | HCV-cAg | ACA-Ig(G-A-M) | -0.23913935 | 6.58E-09 |
| B120 | B159 | HCV-cAg | β2-MG | -0.23912781 | 6.59E-09 |
| B120 | B121 | HCV-cAg | C3 | -0.23912578 | 6.60E-09 |
| B120 | B122 | HCV-cAg | C4 | -0.23912587 | 6.60E-09 |
| B120 | B169 | HCV-cAg | U-CRY | -0.23912226 | 6.60E-09 |
| B120 | B203 | HCV-cAg | DH | -0.21164297 | 3.09E-07 |
| B120 | B224 | HCV-cAg | pmh | -0.21110194 | 3.32E-07 |
| B121 | B169 | C3 | U-CRY | 0.999989917 | 0 |
| B122 | B169 | C4 | U-CRY | 0.999994043 | 0 |
| B123 | B155 | hs-CRP1 | IgA | 0.382710007 | 0 |
| B123 | B156 | hs-CRP1 | IgE | 0.467838674 | 0 |
| B123 | B139 | hs-CRP1 | ADA | 0.262149187 | 1.78E-10 |
| B123 | B227 | hs-CRP1 | OD | 0.242079546 | 4.23E-09 |
| B124 | B125 | ACTH | Cor | 0.74012183 | 0 |
| B124 | B139 | ACTH | ADA | -0.33145538 | 4.44E-16 |
| B124 | B199 | ACTH | Mg | -0.32589846 | 1.11E-15 |
| B124 | B193 | ACTH | K | -0.29388036 | 6.73E-13 |
| B125 | B187 | Cor | NSE | -0.39286204 | 0 |
| B125 | B199 | Cor | Mg | -0.35930848 | 0 |
| B125 | B139 | Cor | ADA | -0.31435759 | 1.24E-14 |
| B125 | B193 | Cor | K | -0.27317787 | 2.78E-11 |
| B125 | B137 | Cor | AFU | 0.217013778 | 1.52E-07 |
| B126 | B127 | Alb | A-G ratio | 0.51933405 | 0 |

| | | | | | |
|---|---|---|---|---|---|
| | | | | 9 | |
| B126 | B188 | Alb | Ca | 0.371679806 | 0 |
| B126 | B189 | Alb | eGFB | 0.263952703 | 1.32E-10 |
| B126 | B134 | Alb | CHE | 0.203786803 | 8.50E-07 |
| B127 | B130 | A-G ratio | GLB | -0.91083822 | 0 |
| B128 | B131 | ALT | AST | 0.617556625 | 0 |
| B128 | B140 | ALT | r-GT | 0.520621987 | 0 |
| B128 | B141 | ALT | AST/ALT | -0.68361227 | 0 |
| B128 | B216 | ALT | BMI | 0.350838683 | 0 |
| B128 | B222 | ALT | NS | 0.277744576 | 1.26E-11 |
| B128 | B192 | ALT | UA | 0.270588982 | 4.34E-11 |
| B128 | B215 | ALT | BMI | 0.266606735 | 8.51E-11 |
| B128 | B206 | ALT | Waist | 0.24672353 | 2.08E-09 |
| B128 | B221 | ALT | Hei | 0.236739768 | 9.39E-09 |
| B128 | B210 | ALT | Hip | 0.223804591 | 5.99E-08 |
| B129 | B132 | IBIL | Dbil | 0.698758042 | 0 |
| B129 | B133 | IBIL | TBIL | 0.984294021 | 0 |
| B131 | B140 | AST | r-GT | 0.386577485 | 0 |
| B131 | B192 | AST | UA | 0.241069939 | 4.93E-09 |
| B132 | B133 | Dbil | TBIL | 0.813988239 | 0 |
| B134 | B138 | CHE | PAB | 0.314316958 | 1.24E-14 |
| B134 | B136 | CHE | 5'NT | 0.278260654 | 1.15E-11 |
| B134 | B135 | CHE | TBA | 0.272263027 | 3.26E-11 |
| B134 | B199 | CHE | Mg | 0.238553357 | 7.18E-09 |

| | | | | | |
|---|---|---|---|---|---|
| B134 | B198 | CHE | P | 0.222160823 | 7.52E-08 |
| B135 | B136 | TBA | 5'NT | 0.866864464 | 0 |
| B135 | B150 | TBA | ALP | 0.356961864 | 0 |
| B135 | B197 | TBA | AG | 0.330840024 | 4.44E-16 |
| B135 | B196 | TBA | TCO2 | -0.27993233 | 8.55E-12 |
| B135 | B199 | TBA | Mg | 0.271261906 | 3.87E-11 |
| B135 | B198 | TBA | P | 0.264450048 | 1.22E-10 |
| B135 | B140 | TBA | r-GT | 0.245435414 | 2.54E-09 |
| B135 | B137 | TBA | AFU | -0.23759341 | 8.28E-09 |
| B135 | B188 | TBA | Ca | 0.221851781 | 7.84E-08 |
| B135 | B195 | TBA | Na | 0.217858032 | 1.35E-07 |
| B136 | B150 | 5'NT | ALP | 0.426842966 | 0 |
| B136 | B198 | 5'NT | P | 0.379514354 | 0 |
| B136 | B137 | 5'NT | AFU | -0.32305027 | 2.22E-15 |
| B136 | B140 | 5'NT | r-GT | 0.277651304 | 1.28E-11 |
| B136 | B195 | 5'NT | Na | 0.211493944 | 3.15E-07 |
| B136 | B199 | 5'NT | Mg | 0.209202111 | 4.25E-07 |
| B137 | B154 | AFU | RF | 0.30064116 | 1.87E-13 |
| B137 | B194 | AFU | Cl | -0.22716828 | 3.74E-08 |
| B137 | B196 | AFU | TCO2 | 0.226025521 | 4.39E-08 |
| B138 | B139 | PAB | ADA | -0.51928332 | 0 |
| B138 | B197 | PAB | AG | 0.268749222 | 5.93E-11 |
| B138 | B154 | PAB | RF | -0.24340082 | 3.47E-09 |
| B138 | B199 | PAB | Mg | 0.235040921 | 1.21E-08 |
| B138 | B198 | PAB | P | 0.205521385 | 6.82E-07 |
| B138 | B195 | PAB | Na | 0.20363783 | 8.66E-07 |

| | | | | | |
|---|---|---|---|---|---|
| | | | | 9 | |
| B139 | B154 | ADA | RF | 0.514590755 | 0 |
| B139 | B193 | ADA | K | 0.416280107 | 0 |
| B139 | B194 | ADA | Cl | -0.257287 | 3.94E-10 |
| B139 | B198 | ADA | P | -0.24510029 | 2.68E-09 |
| B139 | B200 | ADA | OP | 0.240728389 | 5.19E-09 |
| B139 | B195 | ADA | Na | 0.223601945 | 6.16E-08 |
| B139 | B197 | ADA | AG | 0.220607428 | 9.31E-08 |
| B139 | B150 | ADA | ALP | 0.204279365 | 7.99E-07 |
| B140 | B141 | r-GT | AST/ALT | -0.31543693 | 9.99E-15 |
| B141 | B216 | AST/ALT | BMI | -0.41549949 | 0 |
| B141 | B222 | AST/ALT | NS | -0.34119022 | 0 |
| B141 | B221 | AST/ALT | Hei | -0.32913331 | 4.44E-16 |
| B141 | B215 | AST/ALT | BMI | -0.29278234 | 8.26E-13 |
| B141 | B206 | AST/ALT | Waist | -0.2729892 | 2.87E-11 |
| B141 | B210 | AST/ALT | Hip | -0.27049322 | 4.41E-11 |
| B142 | B151 | 25-(OH)D | Anti-CCP | -0.49510376 | 0 |
| B142 | B168 | 25-(OH)D | NMP22 | -0.27766284 | 1.28E-11 |
| B142 | B175 | 25-(OH)D | ALB-U | -0.21763303 | 1.39E-07 |
| B143 | B144 | β-CTx | Total-PINP | 0.791075093 | 0 |
| B143 | B145 | β-CTx | BGP | 0.412089972 | 0 |
| B143 | B175 | β-CTx | ALB-U | 0.205267732 | 7.05E-07 |
| B144 | B145 | Total-PINP | BGP | 0.474318316 | 0 |
| B146 | B189 | CysC | eGFB | -0.24726922 | 1.92E-09 |
| B146 | B191 | CysC | Urea | 0.204885292 | 7.40E-07 |
| B15 | B100 | CD3-CD19+(B cell) | anti-Ro52 | 0.54435926 | 0 |
| B15 | B101 | CD3-CD19+(B cell) | anti-Sc70 | 0.544359266 | 0 |
| B15 | B102 | CD3-CD19+(B cell) | anti-Sm | 0.544359263 | 0 |
| B15 | B103 | CD3-CD19+(B cell) | anti-SSA/Ro | 0.54435926 | 0 |
| B15 | B104 | CD3-CD19+(B cell) | anti-SSB | 0.544359259 | 0 |

| | | | | | |
|---|---|---|---|---|---|
| B15 | B105 | CD3-CD19+(B cell) | anti-U1RNP | 0.54435926 | 0 |
| B15 | B106 | CD3-CD19+(B cell) | anti-P | 0.54435926 | 0 |
| B15 | B107 | CD3-CD19+(B cell) | AnuA | 0.544359251 | 0 |
| B15 | B108 | CD3-CD19+(B cell) | Anti-Cyclin | 0.544359261 | 0 |
| B15 | B109 | CD3-CD19+(B cell) | AMA-M2 | 0.54435926 | 0 |
| B15 | B110 | CD3-CD19+(B cell) | AHA | 0.544359261 | 0 |
| B15 | B151 | CD3-CD19+(B cell) | Anti-CCP | 0.452611948 | 0 |
| B15 | B16 | CD3-CD19+(B cell) | CD3+T | -0.46320258 | 0 |
| B15 | B17 | CD3-CD19+(B cell) | CD3+CD16+CD56+NK | 0.783021835 | 0 |
| B15 | B186 | CD3-CD19+(B cell) | PFC | 0.565105698 | 0 |
| B15 | B23 | CD3-CD19+(B cell) | CD3-CD16+CD56+(NK) | 0.553953165 | 0 |
| B15 | B34 | CD3-CD19+(B cell) | TK | 0.389855557 | 0 |
| B15 | B95 | CD3-CD19+(B cell) | anti-cenpb | 0.54435926 | 0 |
| B15 | B97 | CD3-CD19+(B cell) | dsDNA | 0.544359258 | 0 |
| B15 | B98 | CD3-CD19+(B cell) | anti-Jo-1 | 0.544359262 | 0 |
| B15 | B99 | CD3-CD19+(B cell) | anti-PM-Scl | 0.544359259 | 0 |
| B15 | B185 | CD3-CD19+(B cell) | PT | 0.301783381 | 1.50E-13 |
| B15 | B183 | CD3-CD19+(B cell) | PTA | 0.298143044 | 3.01E-13 |
| B15 | B182 | CD3-CD19+(B cell) | TT | -0.29144239 | 1.06E-12 |
| B15 | B157 | CD3-CD19+(B cell) | IgG | 0.278711981 | 1.06E-11 |
| B15 | B22 | CD3-CD19+(B cell) | CD3+CD8+(T) | -0.23867698 | 7.05E-09 |
| B15 | B59 | CD3-CD19+(B cell) | WBRV | 0.219184142 | 1.13E-07 |
| B15 | B18 | CD3-CD19+(B cell) | CD3+CD4+helper(inducer) T | -0.21844431 | 1.25E-07 |
| B15 | B55 | CD3-CD19+(B cell) | TK | 0.209688891 | 3.99E-07 |
| B150 | B200 | ALP | OP | 0.31311315 | 1.60E-14 |
| B150 | B195 | ALP | Na | 0.308980308 | 3.66E-14 |

| | | | | | |
|---|---|---|---|---|---|
| B151 | B181 | Anti-CCP | APTT | -0.36722914 | 0 |
| B151 | B186 | Anti-CCP | PFC | 0.461954386 | 0 |
| B151 | B168 | Anti-CCP | NMP22 | 0.293139773 | 7.73E-13 |
| B151 | B179 | Anti-CCP | α1-MG | -0.23261556 | 1.72E-08 |
| B151 | B182 | Anti-CCP | TT | -0.22955141 | 2.67E-08 |
| B151 | B178 | Anti-CCP | Ig-U | -0.2141462 | 2.22E-07 |
| B153 | B170 | Glu | U-GLU | 0.604781902 | 0 |
| B154 | B194 | RF | Cl | -0.32394426 | 1.78E-15 |
| B154 | B199 | RF | Mg | -0.22461719 | 5.35E-08 |
| B155 | B156 | IgA | IgE | 0.529745391 | 0 |
| B155 | B157 | IgA | IgG | 0.5585117 | 0 |
| B155 | B227 | IgA | OD | 0.232251789 | 1.81E-08 |
| B156 | B157 | IgE | IgG | 0.34099798 | 0 |
| B156 | B227 | IgE | OD | 0.270996005 | 4.05E-11 |
| B156 | B170 | IgE | U-GLU | 0.209777392 | 3.95E-07 |
| B157 | B185 | IgG | PT | 0.348250148 | 0 |
| B157 | B184 | IgG | INR | 0.222668637 | 7.01E-08 |
| B157 | B186 | IgG | PFC | 0.208299229 | 4.78E-07 |
| B158 | B179 | IgM | α1-MG | -0.21149354 | 3.16E-07 |
| B158 | B190 | IgM | Cre | -0.2055579 | 6.79E-07 |
| B159 | B169 | β2-MG | U-CRY | 0.999997539 | 0 |
| B16 | B157 | CD3+T | IgG | -0.40601811 | 0 |
| B16 | B178 | CD3+T | Ig-U | -0.52092874 | 0 |
| B16 | B179 | CD3+T | α1-MG | -0.57794078 | 0 |
| B16 | B18 | CD3+T | CD3+CD4+helper(inducer) T | 0.358781518 | 0 |
| B16 | B181 | CD3+T | APTT | -0.36284424 | 0 |
| B16 | B185 | CD3+T | PT | -0.64897252 | 0 |
| B16 | B186 | CD3+T | PFC | -0.51974058 | 0 |
| B16 | B20 | CD3+T | CD3+CD4+CD8+ | 0.783653986 | 0 |
| B16 | B21 | CD3+T | CD3+CD4-CD8- | 0.878161672 | 0 |

| | | | | | |
|---|---|---|---|---|---|
| B16 | B22 | CD3+T | CD3+CD8+(T) | 0.68524993 | 0 |
| B16 | B23 | CD3+T | CD3-CD16+CD56+(NK) | -0.87423379 | 0 |
| B16 | B79 | CD3+T | hb V -dna | 0.369853583 | 0 |
| B16 | B92 | CD3+T | ALDH2-Glu504Glu | 0.578852009 | 0 |
| B16 | B93 | CD3+T | ALDH2-Glu504Lys | -0.58991203 | 0 |
| B16 | B94 | CD3+T | ALDH2-Lys504Lys | 0.452627242 | 0 |
| B16 | B196 | CD3+T | TCO2 | -0.3236318 | 1.78E-15 |
| B16 | B200 | CD3+T | OP | -0.31778759 | 6.22E-15 |
| B16 | B57 | CD3+T | RBCAI | 0.266967 | 8.01E-11 |
| B16 | B139 | CD3+T | ADA | -0.26282572 | 1.60E-10 |
| B16 | B195 | CD3+T | Na | -0.25240269 | 8.59E-10 |
| B16 | B55 | CD3+T | TK | -0.25153642 | 9.85E-10 |
| B16 | B182 | CD3+T | TT | -0.24192557 | 4.33E-09 |
| B16 | B176 | CD3+T | Ucr | 0.239855387 | 5.91E-09 |
| B16 | B184 | CD3+T | INR | -0.23481458 | 1.25E-08 |
| B16 | B81 | CD3+T | TSGF | -0.2296538 | 2.63E-08 |
| B16 | B19 | CD3+T | CD3+CD4+/CD3+CD8+ | -0.22352928 | 6.22E-08 |
| B16 | B114 | CD3+T | TSH TT3 TT4(TT4) | -0.22167772 | 8.03E-08 |
| B163 | B222 | SG | NS | 0.32893084 | 4.44E-16 |
| B163 | B221 | SG | Hei | 0.327419625 | 8.88E-16 |
| B163 | B176 | SG | Ucr | 0.319563115 | 4.44E-15 |
| B163 | B166 | SG | Pro | 0.313019506 | 1.64E-14 |
| B163 | B216 | SG | BMI | 0.264850893 | 1.14E-10 |
| B163 | B191 | SG | Urea | 0.232993988 | 1.62E-08 |
| B165 | B172 | bilirubin | KB | 0.213389079 | 2.46E-07 |
| B166 | B175 | Pro | ALB-U | 0.214681725 | 2.07E-07 |
| B167 | B223 | PC | FH | -0.25205042 | 9.09E-10 |
| B17 | B100 | CD3+CD16+CD56+NK | anti-Ro52 | 0.471549329 | 0 |
| B17 | B101 | CD3+CD16+CD56+NK | anti-Sc70 | 0.471549334 | 0 |
| B17 | B102 | CD3+CD16+CD56+NK | anti-Sm | 0.471549331 | 0 |

| | | | | | |
|---|---|---|---|---|---|
| B17 | B103 | CD3+CD16+CD56+NK | anti-SSA/Ro | 0.471549329 | 0 |
| B17 | B104 | CD3+CD16+CD56+NK | anti-SSB | 0.471549327 | 0 |
| B17 | B105 | CD3+CD16+CD56+NK | anti-U1RNP | 0.471549329 | 0 |
| B17 | B106 | CD3+CD16+CD56+NK | anti-P | 0.471549328 | 0 |
| B17 | B107 | CD3+CD16+CD56+NK | AnuA | 0.471549323 | 0 |
| B17 | B108 | CD3+CD16+CD56+NK | Anti-Cyclin | 0.471549329 | 0 |
| B17 | B109 | CD3+CD16+CD56+NK | AMA-M2 | 0.471549329 | 0 |
| B17 | B110 | CD3+CD16+CD56+NK | AHA | 0.471549329 | 0 |
| B17 | B151 | CD3+CD16+CD56+NK | Anti-CCP | 0.390266852 | 0 |
| B17 | B184 | CD3+CD16+CD56+NK | INR | -0.3673096 | 0 |
| B17 | B198 | CD3+CD16+CD56+NK | P | -0.3423251 | 0 |
| B17 | B22 | CD3+CD16+CD56+NK | CD3+CD8+(T) | 0.356046826 | 0 |
| B17 | B34 | CD3+CD16+CD56+NK | TK | 0.336380335 | 0 |
| B17 | B95 | CD3+CD16+CD56+NK | anti-cenpb | 0.471549328 | 0 |
| B17 | B97 | CD3+CD16+CD56+NK | dsDNA | 0.471549327 | 0 |
| B17 | B98 | CD3+CD16+CD56+NK | anti-Jo-1 | 0.47154933 | 0 |
| B17 | B99 | CD3+CD16+CD56+NK | anti-PM-Scl | 0.471549328 | 0 |
| B17 | B18 | CD3+CD16+CD56+NK | CD3+CD4+helper(inducer) T | -0.31721123 | 7.11E-15 |
| B17 | B181 | CD3+CD16+CD56+NK | APTT | -0.29142829 | 1.06E-12 |
| B17 | B20 | CD3+CD16+CD56+NK | CD3+CD4+CD8+ | 0.279039575 | 1.00E-11 |
| B17 | B19 | CD3+CD16+CD56+NK | CD3+CD4+/CD3+CD8+ | -0.27217221 | 3.31E-11 |
| B17 | B21 | CD3+CD16+CD56+NK | CD3+CD4-CD8- | 0.255784013 | 5.02E-10 |
| B17 | B59 | CD3+CD16+CD56+NK | WBRV | 0.227764065 | 3.44E-08 |
| B17 | B182 | CD3+CD16+CD56+NK | TT | -0.22500181 | 5.07E-08 |
| B17 | B186 | CD3+CD16+CD56+NK | PFC | 0.219718067 | 1.05E-07 |

| | | | | | |
|---|---|---|---|---|---|
| B17 | B86 | CD3+CD16+CD56+NK | TBhCG | 0.213785258 | 2.33E-07 |
| B170 | B172 | U-GLU | KB | 0.233488143 | 1.51E-08 |
| B174 | B191 | Urea/Cr | Urea | 0.525054381 | 0 |
| B174 | B221 | Urea/Cr | Hei | -0.32512377 | 1.33E-15 |
| B174 | B190 | Urea/Cr | Cre | -0.29611054 | 4.43E-13 |
| B174 | B222 | Urea/Cr | NS | -0.25537844 | 5.36E-10 |
| B174 | B192 | Urea/Cr | UA | -0.2243485 | 5.55E-08 |
| B174 | B189 | Urea/Cr | eGFB | 0.208284249 | 4.79E-07 |
| B175 | B176 | ALB-U | Ucr | 0.293623441 | 7.06E-13 |
| B176 | B185 | Ucr | PT | -0.26804609 | 6.68E-11 |
| B176 | B184 | Ucr | INR | -0.2663029 | 8.95E-11 |
| B176 | B186 | Ucr | PFC | -0.20628112 | 6.19E-07 |
| B176 | B180 | Ucr | Tf | 0.206048359 | 6.38E-07 |
| B178 | B179 | Ig-U | α1-MG | 0.921111211 | 0 |
| B178 | B180 | Ig-U | Tf | 0.47044952 | 0 |
| B178 | B182 | Ig-U | TT | 0.508871602 | 0 |
| B178 | B195 | Ig-U | Na | 0.458340115 | 0 |
| B178 | B200 | Ig-U | OP | 0.502814832 | 0 |
| B178 | B181 | Ig-U | APTT | 0.31183043 | 2.09E-14 |
| B178 | B184 | Ig-U | INR | -0.27432682 | 2.28E-11 |
| B178 | B198 | Ig-U | P | -0.27296879 | 2.88E-11 |
| B178 | B197 | Ig-U | AG | 0.203541326 | 8.77E-07 |
| B179 | B181 | α1-MG | APTT | 0.343219917 | 0 |
| B179 | B182 | α1-MG | TT | 0.56345781 | 0 |
| B179 | B195 | α1-MG | Na | 0.504504503 | 0 |
| B179 | B200 | α1-MG | OP | 0.550202809 | 0 |
| B179 | B198 | α1-MG | P | -0.30407113 | 9.59E-14 |
| B179 | B184 | α1-MG | INR | -0.30304639 | 1.17E-13 |
| B179 | B197 | α1-MG | AG | 0.235529064 | 1.12E-08 |

| | | | | | |
|---|---|---|---|---|---|
| B179 | B196 | α1-MG | TCO2 | 0.221727495 | 7.98E-08 |
| B18 | B178 | CD3+CD4+helper(inducer) T | Ig-U | -0.57749666 | 0 |
| B18 | B179 | CD3+CD4+helper(inducer) T | α1-MG | -0.64176504 | 0 |
| B18 | B182 | CD3+CD4+helper(inducer) T | TT | -0.48023564 | 0 |
| B18 | B184 | CD3+CD4+helper(inducer) T | INR | 0.598520048 | 0 |
| B18 | B186 | CD3+CD4+helper(inducer) T | PFC | -0.46334742 | 0 |
| B18 | B19 | CD3+CD4+helper(inducer) T | CD3+CD4+/CD3+CD8+ | 0.82586328 | 0 |
| B18 | B195 | CD3+CD4+helper(inducer) T | Na | -0.41421783 | 0 |
| B18 | B197 | CD3+CD4+helper(inducer) T | AG | -0.34180893 | 0 |
| B18 | B198 | CD3+CD4+helper(inducer) T | P | 0.480064918 | 0 |
| B18 | B20 | CD3+CD4+helper(inducer) T | CD3+CD4+CD8+ | 0.543435031 | 0 |
| B18 | B200 | CD3+CD4+helper(inducer) T | OP | -0.45433248 | 0 |
| B18 | B22 | CD3+CD4+helper(inducer) T | CD3+CD8+(T) | -0.42017139 | 0 |
| B18 | B79 | CD3+CD4+helper(inducer) T | hb V -dna | 0.412287367 | 0 |
| B18 | B92 | CD3+CD4+helper(inducer) T | ALDH2-Glu504Glu | 0.642974144 | 0 |
| B18 | B93 | CD3+CD4+helper(inducer) T | ALDH2-Glu504Lys | -0.65530741 | 0 |
| B18 | B94 | CD3+CD4+helper(inducer) T | ALDH2-Lys504Lys | 0.503832848 | 0 |
| B18 | B55 | CD3+CD4+helper(inducer) T | TK | -0.32737373 | 8.88E-16 |
| B18 | B183 | CD3+CD4+helper(inducer) T | PTA | 0.31421232 | 1.29E-14 |
| B18 | B115 | CD3+CD4+helper(inducer) T | TSH TT3 TT4(TT3) | -0.29420026 | 6.34E-13 |
| B18 | B114 | CD3+CD4+helper(inducer) T | TSH TT3 TT4(TT4) | -0.26186297 | 1.87E-10 |
| B18 | B81 | CD3+CD4+helper(inducer) T | TSGF | -0.2543886 | 6.27E-10 |

| | | | | | |
|---|---|---|---|---|---|
| B18 | B59 | CD3+CD4+helper(inducer) T | WBRV | -0.21300629 | 2.59E-07 |
| B18 | B61 | CD3+CD4+helper(inducer) T | WBHSRV | -0.2084344 | 4.70E-07 |
| B18 | B185 | CD3+CD4+helper(inducer) T | PT | 0.205128323 | 7.17E-07 |
| B18 | B151 | CD3+CD4+helper(inducer) T | Anti-CCP | -0.20327653 | 9.07E-07 |
| B181 | B182 | APTT | TT | 0.678187749 | 0 |
| B181 | B185 | APTT | PT | 0.64982831 | 0 |
| B181 | B186 | APTT | PFC | -0.41615805 | 0 |
| B181 | B196 | APTT | TCO2 | 0.228770832 | 2.98E-08 |
| B181 | B200 | APTT | OP | 0.206054932 | 6.37E-07 |
| B182 | B183 | TT | PTA | -0.69198412 | 0 |
| B182 | B195 | TT | Na | 0.424603552 | 0 |
| B182 | B200 | TT | OP | 0.454512829 | 0 |
| B182 | B185 | TT | PT | 0.295167896 | 5.28E-13 |
| B182 | B197 | TT | AG | 0.218378212 | 1.26E-07 |
| B183 | B195 | PTA | Na | -0.22587717 | 4.48E-08 |
| B183 | B200 | PTA | OP | -0.22398375 | 5.84E-08 |
| B184 | B185 | INR | PT | 0.744583919 | 0 |
| B184 | B198 | INR | P | 0.437592791 | 0 |
| B184 | B197 | INR | AG | -0.31582366 | 9.33E-15 |
| B184 | B199 | INR | Mg | -0.23193144 | 1.89E-08 |
| B184 | B195 | INR | Na | -0.22491383 | 5.13E-08 |
| B184 | B200 | INR | OP | -0.21856646 | 1.23E-07 |
| B185 | B196 | PT | TCO2 | 0.28819414 | 1.93E-12 |
| B185 | B198 | PT | P | 0.24139616 | 4.69E-09 |
| B185 | B199 | PT | Mg | -0.22889198 | 2.93E-08 |
| B185 | B197 | PT | AG | -0.2161531 | 1.70E-07 |
| B186 | B199 | PFC | Mg | -0.24544165 | 2.54E-09 |
| B189 | B190 | eGFB | Cre | -0.44823645 | 0 |
| B189 | B191 | eGFB | Urea | -0.33947162 | 0 |
| B189 | B220 | eGFB | SBP | -0.29564754 | 4.83E-13 |
| B189 | B192 | eGFB | UA | -0.21246987 | 2.78E-07 |

| | | | | | |
|---|---|---|---|---|---|
| B19 | B179 | CD3+CD4+/CD3+CD8+ | α1-MG | -0.35125898 | 0 |
| B19 | B182 | CD3+CD4+/CD3+CD8+ | TT | -0.39818169 | 0 |
| B19 | B183 | CD3+CD4+/CD3+CD8+ | PTA | 0.42948332 | 0 |
| B19 | B184 | CD3+CD4+/CD3+CD8+ | INR | 0.76107524 3 | 0 |
| B19 | B185 | CD3+CD4+/CD3+CD8+ | PT | 0.60337781 4 | 0 |
| B19 | B197 | CD3+CD4+/CD3+CD8+ | AG | -0.38329476 | 0 |
| B19 | B198 | CD3+CD4+/CD3+CD8+ | P | 0.47990012 1 | 0 |
| B19 | B21 | CD3+CD4+/CD3+CD8+ | CD3+CD4-CD8- | -0.34790361 | 0 |
| B19 | B22 | CD3+CD4+/CD3+CD8+ | CD3+CD8+(T) | -0.84228591 | 0 |
| B19 | B92 | CD3+CD4+/CD3+CD8+ | ALDH2-Glu504Glu | 0.35197693 4 | 0 |
| B19 | B93 | CD3+CD4+/CD3+CD8+ | ALDH2-Glu504Lys | -0.3587421 | 0 |
| B19 | B23 | CD3+CD4+/CD3+CD8+ | CD3-CD16+CD56+(NK) | 0.33424279 1 | 2.22E-16 |
| B19 | B178 | CD3+CD4+/CD3+CD8+ | Ig-U | -0.31580936 | 9.33E-15 |
| B19 | B200 | CD3+CD4+/CD3+CD8+ | OP | -0.31181872 | 2.09E-14 |
| B19 | B195 | CD3+CD4+/CD3+CD8+ | Na | -0.30443782 | 8.95E-14 |
| B19 | B181 | CD3+CD4+/CD3+CD8+ | APTT | 0.28207327 7 | 5.84E-12 |
| B19 | B94 | CD3+CD4+/CD3+CD8+ | ALDH2-Lys504Lys | 0.27611165 1 | 1.67E-11 |
| B19 | B157 | CD3+CD4+/CD3+CD8+ | IgG | 0.26619285 4 | 9.12E-11 |
| B19 | B176 | CD3+CD4+/CD3+CD8+ | Ucr | -0.24598856 | 2.33E-09 |
| B19 | B79 | CD3+CD4+/CD3+CD8+ | hb V -dna | 0.22611094 8 | 4.34E-08 |
| B19 | B136 | CD3+CD4+/CD3+CD8+ | 5'NT | 0.22540839 3 | 4.79E-08 |
| B19 | B196 | CD3+CD4+/CD3+CD8+ | TCO2 | 0.21422506 | 2.20E-07 |
| B190 | B191 | Cre | Urea | 0.42512650 9 | 0 |
| B190 | B192 | Cre | UA | 0.22187607 8 | 7.82E-08 |
| B190 | B222 | Cre | NS | 0.22102336 8 | 8.79E-08 |
| B191 | B192 | Urea | UA | 0.22458241 9 | 5.37E-08 |
| B192 | B216 | UA | BMI | 0.34095907 | 0 |
| B192 | B222 | UA | NS | 0.39310168 4 | 0 |
| B192 | B221 | UA | Hei | 0.29809801 | 3.04E-13 |

| | | | | | |
|---|---|---|---|---|---|
| | | | | 3 | |
| B192 | B206 | UA | Waist | 0.271759505 | 3.55E-11 |
| B192 | B210 | UA | Hip | 0.231213553 | 2.10E-08 |
| B192 | B215 | UA | BMI | 0.226534856 | 4.09E-08 |
| B193 | B199 | K | Mg | 0.360422415 | 0 |
| B193 | B194 | K | Cl | -0.29194671 | 9.65E-13 |
| B193 | B195 | K | Na | -0.20339351 | 8.93E-07 |
| B194 | B197 | Cl | AG | -0.38455395 | 0 |
| B194 | B196 | Cl | TCO2 | -0.31902792 | 4.88E-15 |
| B195 | B197 | Na | AG | 0.519366667 | 0 |
| B195 | B200 | Na | OP | 0.965661844 | 0 |
| B196 | B197 | TCO2 | AG | -0.56681597 | 0 |
| B197 | B200 | AG | OP | 0.46220817 | 0 |
| B197 | B198 | AG | P | -0.32380485 | 1.78E-15 |
| B2 | B126 | Age | Alb | -0.33991482 | 0 |
| B2 | B189 | Age | eGFB | -0.62599871 | 0 |
| B2 | B220 | Age | SBP | 0.431546009 | 0 |
| B2 | B47 | Age | RDW | 0.412045714 | 0 |
| B2 | B41 | Age | MON％ | 0.312987636 | 1.64E-14 |
| B2 | B141 | Age | AST/ALT | 0.267694161 | 7.08E-11 |
| B2 | B221 | Age | Hei | -0.2664807 | 8.69E-11 |
| B2 | B128 | Age | ALT | -0.25084482 | 1.10E-09 |
| B2 | B216 | Age | BMI | -0.24724258 | 1.92E-09 |
| B2 | B191 | Age | Urea | 0.246833523 | 2.05E-09 |
| B2 | B207 | Age | WHR | 0.244914792 | 2.75E-09 |
| B2 | B46 | Age | RBC Count | -0.23568197 | 1.10E-08 |
| B2 | B43 | Age | MCV | 0.225329566 | 4.84E-08 |
| B2 | B146 | Age | CysC | 0.223075754 | 6.62E-08 |
| B2 | B45 | Age | MCHC | -0.20332611 | 9.01E-07 |
| B2 | B52 | Age | PLT/BPC | -0.20254649 | 9.94E-07 |

| | | | | | |
|---|---|---|---|---|---|
| B20 | B178 | CD3+CD4+CD8+ | Ig-U | -0.39802432 | 0 |
| B20 | B179 | CD3+CD4+CD8+ | α1-MG | -0.44239358 | 0 |
| B20 | B185 | CD3+CD4+CD8+ | PT | -0.39344512 | 0 |
| B20 | B186 | CD3+CD4+CD8+ | PFC | -0.69462698 | 0 |
| B20 | B21 | CD3+CD4+CD8+ | CD3+CD4-CD8- | 0.537364416 | 0 |
| B20 | B22 | CD3+CD4+CD8+ | CD3+CD8+(T) | 0.416990942 | 0 |
| B20 | B23 | CD3+CD4+CD8+ | CD3-CD16+CD56+(NK) | -0.83293325 | 0 |
| B20 | B92 | CD3+CD4+CD8+ | ALDH2-Glu504Glu | 0.443240741 | 0 |
| B20 | B93 | CD3+CD4+CD8+ | ALDH2-Glu504Lys | -0.45174613 | 0 |
| B20 | B94 | CD3+CD4+CD8+ | ALDH2-Lys504Lys | 0.34739616 | 0 |
| B20 | B182 | CD3+CD4+CD8+ | TT | -0.31791088 | 6.00E-15 |
| B20 | B200 | CD3+CD4+CD8+ | OP | -0.30620076 | 6.35E-14 |
| B20 | B114 | CD3+CD4+CD8+ | TSH TT3 TT4(TT4) | -0.29337079 | 7.41E-13 |
| B20 | B79 | CD3+CD4+CD8+ | hb V -dna | 0.284315899 | 3.90E-12 |
| B20 | B115 | CD3+CD4+CD8+ | TSH TT3 TT4(TT3) | -0.26255114 | 1.67E-10 |
| B20 | B157 | CD3+CD4+CD8+ | IgG | -0.25327983 | 7.48E-10 |
| B20 | B55 | CD3+CD4+CD8+ | TK | -0.23937814 | 6.35E-09 |
| B20 | B176 | CD3+CD4+CD8+ | Ucr | 0.223855411 | 5.95E-08 |
| B20 | B195 | CD3+CD4+CD8+ | Na | -0.21394468 | 2.28E-07 |
| B204 | B209 | Drinking | Smoking | 0.306199538 | 6.35E-14 |
| B204 | B221 | Drinking | Hei | 0.244027386 | 3.15E-09 |
| B204 | B222 | Drinking | NS | 0.224731056 | 5.26E-08 |
| B206 | B207 | Waist | WHR | 0.775269379 | 0 |
| B206 | B210 | Waist | Hip | 0.713741703 | 0 |
| B206 | B215 | Waist | BMI | 0.792769751 | 0 |
| B206 | B216 | Waist | BMI | 0.72149166 | 0 |
| B206 | B222 | Waist | NS | 0.663536122 | 0 |
| B206 | B220 | Waist | SBP | 0.293949494 | 6.64E-13 |
| B207 | B215 | WHR | BMI | 0.469871612 | 0 |
| B207 | B216 | WHR | BMI | 0.36132087 | 0 |

| | | | | | |
|---|---|---|---|---|---|
| B207 | B222 | WHR | NS | 0.416931783 | 0 |
| B207 | B220 | WHR | SBP | 0.24272175 | 3.84E-09 |
| B209 | B221 | Smoking | Hei | 0.203830617 | 8.45E-07 |
| B21 | B100 | CD3+CD4-CD8- | anti-Ro52 | 0.506334706 | 0 |
| B21 | B101 | CD3+CD4-CD8- | anti-Sc70 | 0.506334712 | 0 |
| B21 | B102 | CD3+CD4-CD8- | anti-Sm | 0.506334708 | 0 |
| B21 | B103 | CD3+CD4-CD8- | anti-SSA/Ro | 0.506334706 | 0 |
| B21 | B104 | CD3+CD4-CD8- | anti-SSB | 0.506334704 | 0 |
| B21 | B105 | CD3+CD4-CD8- | anti-U1RNP | 0.506334705 | 0 |
| B21 | B106 | CD3+CD4-CD8- | anti-P | 0.506334705 | 0 |
| B21 | B107 | CD3+CD4-CD8- | AnuA | 0.506334696 | 0 |
| B21 | B108 | CD3+CD4-CD8- | Anti-Cyclin | 0.506334706 | 0 |
| B21 | B109 | CD3+CD4-CD8- | AMA-M2 | 0.506334706 | 0 |
| B21 | B110 | CD3+CD4-CD8- | AHA | 0.506334706 | 0 |
| B21 | B151 | CD3+CD4-CD8- | Anti-CCP | 0.336887255 | 0 |
| B21 | B157 | CD3+CD4-CD8- | IgG | -0.34906475 | 0 |
| B21 | B178 | CD3+CD4-CD8- | Ig-U | -0.58575072 | 0 |
| B21 | B179 | CD3+CD4-CD8- | α1-MG | -0.64826392 | 0 |
| B21 | B181 | CD3+CD4-CD8- | APTT | -0.57372267 | 0 |
| B21 | B182 | CD3+CD4-CD8- | TT | -0.34025249 | 0 |
| B21 | B185 | CD3+CD4-CD8- | PT | -0.63227097 | 0 |
| B21 | B196 | CD3+CD4-CD8- | TCO2 | -0.39614098 | 0 |
| B21 | B200 | CD3+CD4-CD8- | OP | -0.38137179 | 0 |
| B21 | B22 | CD3+CD4-CD8- | CD3+CD8+(T) | 0.701704844 | 0 |
| B21 | B23 | CD3+CD4-CD8- | CD3-CD16+CD56+(NK) | -0.59765279 | 0 |
| B21 | B79 | CD3+CD4-CD8- | hb V -dna | 0.412480112 | 0 |
| B21 | B92 | CD3+CD4-CD8- | ALDH2-Glu504Glu | 0.648991053 | 0 |

| | | | | | |
|---|---|---|---|---|---|
| B21 | B93 | CD3+CD4-CD8- | ALDH2-Glu504Lys | -0.66131932 | 0 |
| B21 | B94 | CD3+CD4-CD8- | ALDH2-Lys504Lys | 0.505877664 | 0 |
| B21 | B95 | CD3+CD4-CD8- | anti-cenpb | 0.506334705 | 0 |
| B21 | B97 | CD3+CD4-CD8- | dsDNA | 0.506334703 | 0 |
| B21 | B98 | CD3+CD4-CD8- | anti-Jo-1 | 0.506334707 | 0 |
| B21 | B99 | CD3+CD4-CD8- | anti-PM-Scl | 0.506334705 | 0 |
| B21 | B195 | CD3+CD4-CD8- | Na | -0.32616615 | 1.11E-15 |
| B21 | B57 | CD3+CD4-CD8- | RBCAI | 0.308472659 | 4.04E-14 |
| B21 | B184 | CD3+CD4-CD8- | INR | -0.26597114 | 9.46E-11 |
| B21 | B81 | CD3+CD4-CD8- | TSGF | -0.25853506 | 3.22E-10 |
| B21 | B34 | CD3+CD4-CD8- | TK | 0.237738473 | 8.10E-09 |
| B21 | B139 | CD3+CD4-CD8- | ADA | -0.23245279 | 1.76E-08 |
| B21 | B76 | CD3+CD4-CD8- | ApoB | 0.214113209 | 2.23E-07 |
| B210 | B215 | Hip | BMI | 0.760110281 | 0 |
| B210 | B216 | Hip | BMI | 0.785392368 | 0 |
| B210 | B221 | Hip | Hei | 0.35662416 | 0 |
| B210 | B222 | Hip | NS | 0.590009874 | 0 |
| B211 | B212 | IOP-R | IOP-L | 0.799714215 | 0 |
| B213 | B214 | CVA-R | CVA-L | -0.25416037 | 6.50E-10 |
| B215 | B216 | BMI | BMI | 0.813268086 | 0 |
| B215 | B222 | BMI | NS | 0.572200902 | 0 |
| B215 | B220 | BMI | SBP | 0.203795455 | 8.49E-07 |
| B216 | B221 | BMI | Hei | 0.640046092 | 0 |
| B216 | B222 | BMI | NS | 0.740930595 | 0 |
| B218 | B220 | DBP | SBP | 0.570616869 | 0 |
| B22 | B157 | CD3+CD8+(T) | IgG | -0.39690239 | 0 |

| | | | | | |
|---|---|---|---|---|---|
| B22 | B181 | CD3+CD8+(T) | APTT | -0.38782031 | 0 |
| B22 | B183 | CD3+CD8+(T) | PTA | -0.33747462 | 0 |
| B22 | B184 | CD3+CD8+(T) | INR | -0.71944184 | 0 |
| B22 | B185 | CD3+CD8+(T) | PT | -0.78454545 | 0 |
| B22 | B198 | CD3+CD8+(T) | P | -0.38566949 | 0 |
| B22 | B23 | CD3+CD8+(T) | CD3-CD16+CD56+(NK) | -0.74157756 | 0 |
| B22 | B176 | CD3+CD8+(T) | Ucr | 0.32777325<br>9 | 8.88E-16 |
| B22 | B196 | CD3+CD8+(T) | TCO2 | -0.28278981 | 5.14E-12 |
| B22 | B197 | CD3+CD8+(T) | AG | 0.27769735<br>6 | 1.27E-11 |
| B22 | B57 | CD3+CD8+(T) | RBCAI | 0.21012226<br>4 | 3.77E-07 |
| B22 | B199 | CD3+CD8+(T) | Mg | 0.20607093<br>6 | 6.36E-07 |
| B22 | B186 | CD3+CD8+(T) | PFC | -0.20557538 | 6.78E-07 |
| B221 | B222 | Hei | NS | 0.52901351<br>1 | 0 |
| B227 | B228 | OD | OS | 0.20564864<br>1 | 6.71E-07 |
| B23 | B157 | CD3-CD16+CD56+(NK) | IgG | 0.40281625<br>1 | 0 |
| B23 | B184 | CD3-CD16+CD56+(NK) | INR | 0.40406069<br>3 | 0 |
| B23 | B185 | CD3-CD16+CD56+(NK) | PT | 0.65183803<br>6 | 0 |
| B23 | B186 | CD3-CD16+CD56+(NK) | PFC | 0.71666536<br>7 | 0 |
| B23 | B176 | CD3-CD16+CD56+(NK) | Ucr | -0.32752126 | 8.88E-16 |
| B23 | B100 | CD3-CD16+CD56+(NK) | anti-Ro52 | 0.24610583<br>8 | 2.29E-09 |
| B23 | B101 | CD3-CD16+CD56+(NK) | anti-Sc70 | 0.24610584<br>1 | 2.29E-09 |
| B23 | B102 | CD3-CD16+CD56+(NK) | anti-Sm | 0.24610584 | 2.29E-09 |
| B23 | B103 | CD3-CD16+CD56+(NK) | anti-SSA/Ro | 0.24610583<br>8 | 2.29E-09 |
| B23 | B104 | CD3-CD16+CD56+(NK) | anti-SSB | 0.24610583<br>8 | 2.29E-09 |
| B23 | B105 | CD3-CD16+CD56+(NK) | anti-U1RNP | 0.24610583<br>8 | 2.29E-09 |
| B23 | B106 | CD3-CD16+CD56+(NK) | anti-P | 0.24610583<br>8 | 2.29E-09 |
| B23 | B107 | CD3-CD16+CD56+(NK) | AnuA | 0.24610583<br>2 | 2.29E-09 |

| | | | | | |
|---|---|---|---|---|---|
| B23 | B108 | CD3-CD16+CD56+(NK) | Anti-Cyclin | 0.246105838 | 2.29E-09 |
| B23 | B109 | CD3-CD16+CD56+(NK) | AMA-M2 | 0.246105838 | 2.29E-09 |
| B23 | B110 | CD3-CD16+CD56+(NK) | AHA | 0.246105839 | 2.29E-09 |
| B23 | B95 | CD3-CD16+CD56+(NK) | anti-cenpb | 0.246105838 | 2.29E-09 |
| B23 | B97 | CD3-CD16+CD56+(NK) | dsDNA | 0.246105837 | 2.29E-09 |
| B23 | B98 | CD3-CD16+CD56+(NK) | anti-Jo-1 | 0.246105839 | 2.29E-09 |
| B23 | B99 | CD3-CD16+CD56+(NK) | anti-PM-Scl | 0.246105838 | 2.29E-09 |
| B23 | B151 | CD3-CD16+CD56+(NK) | Anti-CCP | 0.230654716 | 2.28E-08 |
| B23 | B199 | CD3-CD16+CD56+(NK) | Mg | -0.22061213 | 9.30E-08 |
| B23 | B93 | CD3-CD16+CD56+(NK) | ALDH2-Glu504Lys | 0.21793364 | 1.34E-07 |
| B23 | B114 | CD3-CD16+CD56+(NK) | TSH TT3 TT4(TT4) | 0.216033384 | 1.73E-07 |
| B23 | B92 | CD3-CD16+CD56+(NK) | ALDH2-Glu504Glu | -0.21379646 | 2.33E-07 |
| B23 | B179 | CD3-CD16+CD56+(NK) | α1-MG | 0.213246028 | 2.51E-07 |
| B24 | B27 | αHBDH | LDH | 0.664265241 | 0 |
| B24 | B154 | αHBDH | RF | 0.268296072 | 6.40E-11 |
| B24 | B139 | αHBDH | ADA | 0.246863799 | 2.04E-09 |
| B24 | B146 | αHBDH | CysC | 0.234912801 | 1.23E-08 |
| B25 | B60 | CK | WBRVI | 0.244834432 | 2.79E-09 |
| B25 | B56 | CK | IR | 0.241140495 | 4.88E-09 |
| B25 | B39 | CK | Other cells | 0.240250874 | 5.57E-09 |
| B25 | B123 | CK | hs-CRP1 | -0.23741819 | 8.50E-09 |
| B25 | B62 | CK | WBHSRVI | 0.234466171 | 1.31E-08 |
| B25 | B65 | CK | WBRSR(mPas_1) | 0.21381932 | 2.32E-07 |
| B25 | B154 | CK | RF | 0.203897216 | 8.38E-07 |
| B26 | B173 | CKMB | NTT | 0.26789464 | 6.85E-11 |

| | | | | | |
|---|---|---|---|---|---|
| | | | | 1 | |
| B26 | B154 | CKMB | RF | 0.241565131 | 4.58E-09 |
| B26 | B63 | CKMB | WBRSR(mPas_200) | 0.220999942 | 8.82E-08 |
| B26 | B67 | CKMB | WBRSR(mPas_50) | 0.213132473 | 2.54E-07 |
| B26 | B139 | CKMB | ADA | 0.211897333 | 2.99E-07 |
| B26 | B66 | CKMB | WBRSR(mPas_100) | 0.209272173 | 4.21E-07 |
| B27 | B124 | LDH | ACTH | -0.39971474 | 0 |
| B27 | B139 | LDH | ADA | 0.368749944 | 0 |
| B27 | B193 | LDH | K | 0.315555426 | 9.77E-15 |
| B27 | B125 | LDH | Cor | -0.28542465 | 3.20E-12 |
| B27 | B146 | LDH | CysC | 0.251372598 | 1.01E-09 |
| B27 | B131 | LDH | AST | 0.236356085 | 9.94E-09 |
| B27 | B199 | LDH | Mg | 0.204211619 | 8.06E-07 |
| B28 | B156 | PRL | IgE | -0.55181037 | 0 |
| B28 | B157 | PRL | IgG | -0.2124802 | 2.77E-07 |
| B29 | B156 | E2 | IgE | -0.55181037 | 0 |
| B29 | B30 | E2 | LH | 1 | 0 |
| B29 | B31 | E2 | FSH | 1 | 0 |
| B29 | B32 | E2 | Testo | 1 | 0 |
| B29 | B33 | E2 | Prog | 1 | 0 |
| B29 | B157 | E2 | IgG | -0.2124802 | 2.77E-07 |
| B30 | B156 | LH | IgE | -0.55181037 | 0 |
| B30 | B31 | LH | FSH | 1 | 0 |
| B30 | B157 | LH | IgG | -0.2124802 | 2.77E-07 |
| B31 | B156 | FSH | IgE | -0.55181037 | 0 |
| B31 | B32 | FSH | Testo | 1 | 0 |
| B31 | B33 | FSH | Prog | 1 | 0 |
| B31 | B157 | FSH | IgG | -0.2124802 | 2.77E-07 |
| B32 | B156 | Testo | IgE | -0.55181037 | 0 |
| B32 | B33 | Testo | Prog | 1 | 0 |
| B32 | B157 | Testo | IgG | -0.2124802 | 2.77E-07 |
| B33 | B156 | Prog | IgE | -0.55181037 | 0 |
| B33 | B157 | Prog | IgG | -0.2124802 | 2.77E-07 |
| B34 | B100 | TK | anti-Ro52 | 0.39833891 | 0 |

| | | | | | |
|---|---|---|---|---|---|
| | | | | 5 | |
| B34 | B101 | TK | anti-Sc70 | 0.39833892 | 0 |
| B34 | B102 | TK | anti-Sm | 0.398338918 | 0 |
| B34 | B103 | TK | anti-SSA/Ro | 0.398338915 | 0 |
| B34 | B104 | TK | anti-SSB | 0.398338914 | 0 |
| B34 | B105 | TK | anti-U1RNP | 0.398338915 | 0 |
| B34 | B106 | TK | anti-P | 0.398338915 | 0 |
| B34 | B107 | TK | AnuA | 0.398338909 | 0 |
| B34 | B108 | TK | Anti-Cyclin | 0.398338916 | 0 |
| B34 | B109 | TK | AMA-M2 | 0.398338915 | 0 |
| B34 | B110 | TK | AHA | 0.398338916 | 0 |
| B34 | B124 | TK | ACTH | -0.48054097 | 0 |
| B34 | B125 | TK | Cor | -0.35260478 | 0 |
| B34 | B95 | TK | anti-cenpb | 0.398338915 | 0 |
| B34 | B97 | TK | dsDNA | 0.398338913 | 0 |
| B34 | B98 | TK | anti-Jo-1 | 0.398338916 | 0 |
| B34 | B99 | TK | anti-PM-Scl | 0.398338914 | 0 |
| B34 | B93 | TK | ALDH2-Glu504Lys | -0.30918506 | 3.51E-14 |
| B34 | B92 | TK | ALDH2-Glu504Glu | 0.303416834 | 1.09E-13 |
| B34 | B179 | TK | α1-MG | -0.30305842 | 1.17E-13 |
| B34 | B151 | TK | Anti-CCP | 0.291968433 | 9.62E-13 |
| B34 | B178 | TK | Ig-U | -0.27374396 | 2.52E-11 |
| B34 | B182 | TK | TT | -0.26795225 | 6.78E-11 |
| B34 | B186 | TK | PFC | 0.253656307 | 7.05E-10 |
| B34 | B181 | TK | APTT | -0.23676042 | 9.37E-09 |
| B34 | B94 | TK | ALDH2-Lys504Lys | 0.23660831 | 9.58E-09 |
| B37 | B38 | BAND | BAND% | -1 | 0 |
| B37 | B40 | BAND | WBC | 0.29496132 | 5.50E-13 |

| | | | | | |
|---|---|---|---|---|---|
| | | | | 6 | |
| B38 | B40 | BAND% | WBC | -0.29496133 | 5.50E-13 |
| B39 | B76 | Other cells | ApoB | 0.250732327 | 1.12E-09 |
| B39 | B180 | Other cells | Tf | 0.203052435 | 9.32E-07 |
| B4 | B167 | WB-Ca | PC | -0.32659863 | 8.88E-16 |
| B40 | B42 | WBC | MON | 0.681168251 | 0 |
| B40 | B48 | WBC | LYM% | -0.35192973 | 0 |
| B40 | B53 | WBC | GR% | 0.391723272 | 0 |
| B40 | B52 | WBC | PLT/BPC | 0.234291254 | 1.35E-08 |
| B40 | B46 | WBC | RBC Count | 0.228606644 | 3.05E-08 |
| B40 | B51 | WBC | Hb | 0.216126621 | 1.71E-07 |
| B41 | B42 | MON % | MON | 0.544429176 | 0 |
| B41 | B189 | MON % | eGFB | -0.31379546 | 1.40E-14 |
| B41 | B47 | MON % | RDW | 0.234424497 | 1.32E-08 |
| B41 | B52 | MON % | PLT/BPC | -0.22106167 | 8.74E-08 |
| B42 | B48 | MON | LYM% | -0.32553138 | 1.33E-15 |
| B42 | B51 | MON | Hb | 0.249142293 | 1.43E-09 |
| B42 | B46 | MON | RBC Count | 0.219930534 | 1.02E-07 |
| B43 | B44 | MCV | MCH | 0.890855464 | 0 |
| B43 | B46 | MCV | RBC Count | -0.45406939 | 0 |
| B43 | B47 | MCV | RDW | 0.591636371 | 0 |
| B44 | B45 | MCH | MCHC | 0.578601013 | 0 |
| B44 | B46 | MCH | RBC Count | -0.33572273 | 0 |
| B44 | B47 | MCH | RDW | 0.371207373 | 0 |
| B44 | B51 | MCH | Hb | 0.289873765 | 1.42E-12 |
| B45 | B51 | MCHC | Hb | 0.443879216 | 0 |
| B45 | B221 | MCHC | Hei | 0.26764074 | 7.15E-11 |

| | | | | | |
|---|---|---|---|---|---|
| | | | | 2 | |
| B45 | B47 | MCHC | RDW | -0.2671926 | 7.71E-11 |
| B45 | B174 | MCHC | Urea/Cr | -0.20882541 | 4.47E-07 |
| B46 | B216 | RBC Count | BMI | 0.359902792 | 0 |
| B46 | B221 | RBC Count | Hei | 0.39425283 | 0 |
| B46 | B222 | RBC Count | NS | 0.375063921 | 0 |
| B46 | B47 | RBC Count | RDW | -0.3988947 | 0 |
| B46 | B51 | RBC Count | Hb | 0.754014302 | 0 |
| B46 | B141 | RBC Count | AST/ALT | -0.32495936 | 1.33E-15 |
| B46 | B163 | RBC Count | SG | 0.285713527 | 3.03E-12 |
| B46 | B128 | RBC Count | ALT | 0.275736933 | 1.79E-11 |
| B46 | B58 | RBC Count | HCT | 0.263842913 | 1.35E-10 |
| B46 | B65 | RBC Count | WBRSR(mPas_1) | 0.251378913 | 1.01E-09 |
| B46 | B64 | RBC Count | WBRSR(mPas_5) | 0.249649781 | 1.32E-09 |
| B46 | B67 | RBC Count | WBRSR(mPas_50) | 0.241709893 | 4.48E-09 |
| B46 | B66 | RBC Count | WBRSR(mPas_100) | 0.239732567 | 6.02E-09 |
| B46 | B60 | RBC Count | WBRVI | 0.237581811 | 8.29E-09 |
| B46 | B126 | RBC Count | Alb | 0.235025023 | 1.21E-08 |
| B46 | B63 | RBC Count | WBRSR(mPas_200) | 0.233862069 | 1.43E-08 |
| B46 | B62 | RBC Count | WBHSRVI | 0.231262481 | 2.09E-08 |
| B46 | B154 | RBC Count | RF | 0.219483256 | 1.08E-07 |
| B46 | B72 | RBC Count | HDL-C | -0.21572314 | 1.80E-07 |
| B46 | B192 | RBC Count | UA | 0.21063171 | 3.53E-07 |
| B46 | B206 | RBC Count | Waist | 0.204531689 | 7.74E-07 |
| B46 | B218 | RBC Count | DBP | 0.204200986 | 8.07E-07 |
| B47 | B189 | RDW | eGFB | -0.35053467 | 0 |
| B47 | B52 | RDW | PLT/BPC | -0.29720571 | 3.60E-13 |

| | | | | | |
|---|---|---|---|---|---|
| B47 | B126 | RDW | Alb | -0.25005555 | 1.24E-09 |
| B48 | B53 | LYM% | GR% | -0.92906706 | 0 |
| B49 | B50 | baso% | eo% | 0.419287606 | 0 |
| B49 | B53 | baso% | GR% | -0.27719052 | 1.39E-11 |
| B5 | B116 | WB-Mg | TT3 | -0.92754985 | 0 |
| B5 | B7 | WB-Mg | WB-Cop | -0.48986217 | 0 |
| B5 | B9 | WB-Mg | RH | 0.996927796 | 0 |
| B50 | B53 | eo% | GR% | -0.35748439 | 0 |
| B51 | B221 | Hb | Hei | 0.42131578 | 0 |
| B51 | B222 | Hb | NS | 0.382288975 | 0 |
| B51 | B216 | Hb | BMI | 0.308366049 | 4.13E-14 |
| B51 | B163 | Hb | SG | 0.292087646 | 9.41E-13 |
| B51 | B174 | Hb | Urea/Cr | -0.28119819 | 6.83E-12 |
| B51 | B141 | Hb | AST/ALT | -0.2777773 | 1.25E-11 |
| B51 | B128 | Hb | ALT | 0.247797839 | 1.77E-09 |
| B51 | B218 | Hb | DBP | 0.223900486 | 5.91E-08 |
| B51 | B126 | Hb | Alb | 0.220566579 | 9.36E-08 |
| B54 | B55 | ESR | TK | -0.39138371 | 0 |
| B54 | B56 | ESR | IR | -0.41554904 | 0 |
| B54 | B60 | ESR | WBRVI | -0.35387256 | 0 |
| B54 | B69 | ESR | PV | 0.628300835 | 0 |
| B54 | B59 | ESR | WBRV | -0.30497634 | 8.04E-14 |
| B54 | B57 | ESR | RBCAI | -0.30456562 | 8.73E-14 |
| B54 | B62 | ESR | WBHSRVI | -0.30369755 | 1.03E-13 |
| B54 | B123 | ESR | hs-CRP1 | 0.266555935 | 8.58E-11 |
| B54 | B61 | ESR | WBHSRV | -0.2509103 | 1.09E-09 |
| B54 | B65 | ESR | WBRSR(mPas_1) | -0.2318769 | 1.91E-08 |
| B55 | B56 | TK | IR | 0.373179313 | 0 |
| B55 | B58 | TK | HCT | -0.3853222 | 0 |
| B55 | B69 | TK | PV | -0.35458847 | 0 |
| B55 | B61 | TK | WBHSRV | 0.312578597 | 1.78E-14 |
| B55 | B57 | TK | RBCAI | -0.31103035 | 2.44E-14 |

| | | | | | |
|---|---|---|---|---|---|
| B55 | B123 | TK | hs-CRP1 | -0.27775656 | 1.25E-11 |
| B55 | B79 | TK | hb V -dna | -0.25879469 | 3.09E-10 |
| B55 | B93 | TK | ALDH2-Glu504Lys | 0.24771413<br>7 | 1.79E-09 |
| B55 | B92 | TK | ALDH2-Glu504Glu | -0.24293397 | 3.72E-09 |
| B55 | B179 | TK | α1-MG | 0.24198426<br>7 | 4.30E-09 |
| B55 | B186 | TK | PFC | 0.22626801<br>3 | 4.24E-08 |
| B55 | B178 | TK | Ig-U | 0.21534728 | 1.90E-07 |
| B55 | B154 | TK | RF | -0.21400042 | 2.27E-07 |
| B55 | B65 | TK | WBRSR(mPas_1) | -0.21199676 | 2.95E-07 |
| B55 | B196 | TK | TCO2 | 0.20782733<br>8 | 5.08E-07 |
| B56 | B154 | IR | RF | 0.53009269<br>3 | 0 |
| B56 | B58 | IR | HCT | 0.70001741<br>2 | 0 |
| B56 | B59 | IR | WBRV | 0.76850899<br>8 | 0 |
| B56 | B60 | IR | WBRVI | 0.86245081<br>3 | 0 |
| B56 | B61 | IR | WBHSRV | 0.93467525<br>8 | 0 |
| B56 | B62 | IR | WBHSRVI | 0.92946408<br>5 | 0 |
| B56 | B63 | IR | WBRSR(mPas_200) | 0.85658373<br>3 | 0 |
| B56 | B64 | IR | WBRSR(mPas_5) | 0.83336740<br>8 | 0 |
| B56 | B65 | IR | WBRSR(mPas_1) | 0.78820396<br>3 | 0 |
| B56 | B66 | IR | WBRSR(mPas_100) | 0.86394179<br>2 | 0 |
| B56 | B67 | IR | WBRSR(mPas_50) | 0.85362398<br>5 | 0 |
| B56 | B69 | IR | PV | -0.53317134 | 0 |
| B56 | B139 | IR | ADA | 0.32281794<br>1 | 2.22E-15 |
| B56 | B137 | IR | AFU | 0.31013738 | 2.91E-14 |
| B56 | B194 | IR | Cl | -0.2541539 | 6.51E-10 |
| B57 | B58 | RBCAI | HCT | 0.39808443<br>9 | 0 |
| B57 | B59 | RBCAI | WBRV | 0.47407024 | 0 |

| | | | | | |
|---|---|---|---|---|---|
| | | | | 9 | |
| B57 | B60 | RBCAI | WBRVI | 0.475243819 | 0 |
| B57 | B65 | RBCAI | WBRSR(mPas_1) | 0.446259788 | 0 |
| B57 | B69 | RBCAI | PV | -0.3241584 | 1.78E-15 |
| B57 | B64 | RBCAI | WBRSR(mPas_5) | 0.304375186 | 9.06E-14 |
| B57 | B196 | RBCAI | TCO2 | -0.27378165 | 2.51E-11 |
| B57 | B198 | RBCAI | P | -0.24560937 | 2.47E-09 |
| B57 | B137 | RBCAI | AFU | -0.23622244 | 1.01E-08 |
| B57 | B200 | RBCAI | OP | -0.21405963 | 2.25E-07 |
| B57 | B62 | RBCAI | WBHSRVI | 0.20949224 | 4.10E-07 |
| B57 | B93 | RBCAI | ALDH2-Glu504Lys | -0.20944791 | 4.12E-07 |
| B57 | B92 | RBCAI | ALDH2-Glu504Glu | 0.205422254 | 6.91E-07 |
| B57 | B179 | RBCAI | α1-MG | -0.2046864 | 7.59E-07 |
| B58 | B139 | HCT | ADA | 0.433030585 | 0 |
| B58 | B154 | HCT | RF | 0.644544611 | 0 |
| B58 | B59 | HCT | WBRV | 0.634856706 | 0 |
| B58 | B60 | HCT | WBRVI | 0.92713506 | 0 |
| B58 | B61 | HCT | WBHSRV | 0.67960685 | 0 |
| B58 | B62 | HCT | WBHSRVI | 0.904764482 | 0 |
| B58 | B63 | HCT | WBRSR(mPas_200) | 0.891104441 | 0 |
| B58 | B64 | HCT | WBRSR(mPas_5) | 0.945060612 | 0 |
| B58 | B65 | HCT | WBRSR(mPas_1) | 0.953701062 | 0 |
| B58 | B66 | HCT | WBRSR(mPas_100) | 0.907922828 | 0 |
| B58 | B67 | HCT | WBRSR(mPas_50) | 0.914185588 | 0 |
| B58 | B69 | HCT | PV | -0.2781223 | 1.18E-11 |
| B58 | B199 | HCT | Mg | -0.21754699 | 1.41E-07 |
| B58 | B75 | HCT | ApoA1/B | -0.20930504 | 4.20E-07 |
| B58 | B137 | HCT | AFU | 0.207184781 | 5.52E-07 |
| B59 | B154 | WBRV | RF | 0.366987662 | 0 |

| | | | | | |
|---|---|---|---|---|---|
| B59 | B60 | WBRV | WBRVI | 0.795497679 | 0 |
| B59 | B61 | WBRV | WBHSRV | 0.824752426 | 0 |
| B59 | B62 | WBRV | WBHSRVI | 0.739640819 | 0 |
| B59 | B63 | WBRV | WBRSR(mPas_200) | 0.778383728 | 0 |
| B59 | B64 | WBRV | WBRSR(mPas_5) | 0.790902506 | 0 |
| B59 | B65 | WBRV | WBRSR(mPas_1) | 0.81065079 | 0 |
| B59 | B66 | WBRV | WBRSR(mPas_100) | 0.7540783 | 0 |
| B59 | B67 | WBRV | WBRSR(mPas_50) | 0.786788164 | 0 |
| B59 | B139 | WBRV | ADA | 0.298781147 | 2.66E-13 |
| B59 | B100 | WBRV | anti-Ro52 | 0.297562635 | 3.36E-13 |
| B59 | B101 | WBRV | anti-Sc70 | 0.297562638 | 3.36E-13 |
| B59 | B102 | WBRV | anti-Sm | 0.297562637 | 3.36E-13 |
| B59 | B103 | WBRV | anti-SSA/Ro | 0.297562635 | 3.36E-13 |
| B59 | B104 | WBRV | anti-SSB | 0.297562634 | 3.36E-13 |
| B59 | B105 | WBRV | anti-U1RNP | 0.297562635 | 3.36E-13 |
| B59 | B106 | WBRV | anti-P | 0.297562635 | 3.36E-13 |
| B59 | B107 | WBRV | AnuA | 0.297562632 | 3.36E-13 |
| B59 | B108 | WBRV | Anti-Cyclin | 0.297562636 | 3.36E-13 |
| B59 | B109 | WBRV | AMA-M2 | 0.297562635 | 3.36E-13 |
| B59 | B110 | WBRV | AHA | 0.297562635 | 3.36E-13 |
| B59 | B95 | WBRV | anti-cenpb | 0.297562635 | 3.36E-13 |
| B59 | B97 | WBRV | dsDNA | 0.297562634 | 3.36E-13 |
| B59 | B98 | WBRV | anti-Jo-1 | 0.297562636 | 3.36E-13 |

| | | | | | |
|---|---|---|---|---|---|
| B59 | B99 | WBRV | anti-PM-Scl | 0.297562635 | 3.36E-13 |
| B59 | B186 | WBRV | PFC | 0.246912465 | 2.03E-09 |
| B59 | B151 | WBRV | Anti-CCP | 0.243047278 | 3.66E-09 |
| B6 | B167 | WB-Fe | PC | 0.517114501 | 0 |
| B6 | B223 | WB-Fe | FH | 0.990880714 | 0 |
| B6 | B8 | WB-Fe | WB-Zinc | 0.742610657 | 0 |
| B60 | B139 | WBRVI | ADA | 0.413489217 | 0 |
| B60 | B154 | WBRVI | RF | 0.626555594 | 0 |
| B60 | B61 | WBRVI | WBHSRV | 0.785291385 | 0 |
| B60 | B62 | WBRVI | WBHSRVI | 0.959401525 | 0 |
| B60 | B63 | WBRVI | WBRSR(mPas_200) | 0.888897511 | 0 |
| B60 | B64 | WBRVI | WBRSR(mPas_5) | 0.947842126 | 0 |
| B60 | B65 | WBRVI | WBRSR(mPas_1) | 0.96549689 | 0 |
| B60 | B66 | WBRVI | WBRSR(mPas_100) | 0.907273844 | 0 |
| B60 | B67 | WBRVI | WBRSR(mPas_50) | 0.916051307 | 0 |
| B60 | B69 | WBRVI | PV | -0.48411026 | 0 |
| B60 | B199 | WBRVI | Mg | -0.2130448 | 2.57E-07 |
| B60 | B138 | WBRVI | PAB | -0.20602488 | 6.40E-07 |
| B60 | B137 | WBRVI | AFU | 0.202856848 | 9.56E-07 |
| B61 | B137 | WBHSRV | AFU | 0.363154029 | 0 |
| B61 | B139 | WBHSRV | ADA | 0.352294713 | 0 |
| B61 | B154 | WBHSRV | RF | 0.511778062 | 0 |
| B61 | B62 | WBHSRV | WBHSRVI | 0.885086459 | 0 |
| B61 | B63 | WBHSRV | WBRSR(mPas_200) | 0.921926735 | 0 |

| | | | | | |
|---|---|---|---|---|---|
| B61 | B64 | WBHSRV | WBRSR(mPas_5) | 0.853323709 | 0 |
| B61 | B65 | WBHSRV | WBRSR(mPas_1) | 0.786271809 | 0 |
| B61 | B66 | WBHSRV | WBRSR(mPas_100) | 0.906348258 | 0 |
| B61 | B67 | WBHSRV | WBRSR(mPas_50) | 0.904618536 | 0 |
| B61 | B194 | WBHSRV | Cl | -0.2442821 | 3.03E-09 |
| B61 | B69 | WBHSRV | PV | -0.21900714 | 1.16E-07 |
| B61 | B186 | WBHSRV | PFC | 0.217339291 | 1.45E-07 |
| B62 | B139 | WBHSRVI | ADA | 0.43846854 | 0 |
| B62 | B154 | WBHSRVI | RF | 0.681216912 | 0 |
| B62 | B63 | WBHSRVI | WBRSR(mPas_200) | 0.952599303 | 0 |
| B62 | B64 | WBHSRVI | WBRSR(mPas_5) | 0.959500009 | 0 |
| B62 | B65 | WBHSRVI | WBRSR(mPas_1) | 0.932422513 | 0 |
| B62 | B66 | WBHSRVI | WBRSR(mPas_100) | 0.966869298 | 0 |
| B62 | B67 | WBHSRVI | WBRSR(mPas_50) | 0.961656249 | 0 |
| B62 | B69 | WBHSRVI | PV | -0.43196882 | 0 |
| B62 | B137 | WBHSRVI | AFU | 0.302673946 | 1.26E-13 |
| B62 | B194 | WBHSRVI | Cl | -0.241627 | 4.53E-09 |
| B62 | B199 | WBHSRVI | Mg | -0.21959811 | 1.07E-07 |
| B63 | B137 | WBRSR(mPas_200) | AFU | 0.342368323 | 0 |
| B63 | B139 | WBRSR(mPas_200) | ADA | 0.462373254 | 0 |
| B63 | B154 | WBRSR(mPas_200) | RF | 0.666921978 | 0 |
| B63 | B64 | WBRSR(mPas_200) | WBRSR(mPas_5) | 0.972061523 | 0 |
| B63 | B65 | WBRSR(mPas_200) | WBRSR(mPas_1) | 0.930281653 | 0 |
| B63 | B66 | WBRSR(mPas_200) | WBRSR(mPas_100) | 0.993422842 | 0 |
| B63 | B67 | WBRSR(mPas_200) | WBRSR(mPas_50) | 0.996017308 | 0 |

| | | | | | |
|---|---|---|---|---|---|
| B63 | B199 | WBRSR(mPas_200) | Mg | -0.23107501 | 2.14E-08 |
| B63 | B194 | WBRSR(mPas_200) | Cl | -0.22673991 | 3.97E-08 |
| B64 | B139 | WBRSR(mPas_5) | ADA | 0.470550485 | 0 |
| B64 | B154 | WBRSR(mPas_5) | RF | 0.660746417 | 0 |
| B64 | B65 | WBRSR(mPas_5) | WBRSR(mPas_1) | 0.984595223 | 0 |
| B64 | B66 | WBRSR(mPas_5) | WBRSR(mPas_100) | 0.977622186 | 0 |
| B64 | B67 | WBRSR(mPas_5) | WBRSR(mPas_50) | 0.983239778 | 0 |
| B64 | B137 | WBRSR(mPas_5) | AFU | 0.282729752 | 5.19E-12 |
| B64 | B69 | WBRSR(mPas_5) | PV | -0.25352945 | 7.19E-10 |
| B64 | B199 | WBRSR(mPas_5) | Mg | -0.23356648 | 1.49E-08 |
| B65 | B139 | WBRSR(mPas_1) | ADA | 0.474913644 | 0 |
| B65 | B154 | WBRSR(mPas_1) | RF | 0.656651372 | 0 |
| B65 | B66 | WBRSR(mPas_1) | WBRSR(mPas_100) | 0.93754514 | 0 |
| B65 | B67 | WBRSR(mPas_1) | WBRSR(mPas_50) | 0.950511141 | 0 |
| B65 | B69 | WBRSR(mPas_1) | PV | -0.28707438 | 2.37E-12 |
| B65 | B137 | WBRSR(mPas_1) | AFU | 0.241818935 | 4.40E-09 |
| B65 | B199 | WBRSR(mPas_1) | Mg | -0.23394415 | 1.41E-08 |
| B65 | B138 | WBRSR(mPas_1) | PAB | -0.21160053 | 3.11E-07 |
| B66 | B139 | WBRSR(mPas_100) | ADA | 0.471970026 | 0 |
| B66 | B154 | WBRSR(mPas_100) | RF | 0.673745686 | 0 |
| B66 | B67 | WBRSR(mPas_100) | WBRSR(mPas_50) | 0.995708663 | 0 |
| B66 | B137 | WBRSR(mPas_100) | AFU | 0.333587579 | 2.22E-16 |
| B66 | B199 | WBRSR(mPas_100) | Mg | -0.23227035 | 1.80E-08 |
| B66 | B194 | WBRSR(mPas_100) | Cl | -0.2239142 | 5.90E-08 |
| B66 | B69 | WBRSR(mPas_100) | PV | -0.21041652 | 3.63E-07 |
| B67 | B139 | WBRSR(mPas_50) | ADA | 0.468006976 | 0 |
| B67 | B154 | WBRSR(mPas_50) | RF | 0.665456157 | 0 |
| B67 | B137 | WBRSR(mPas_50) | AFU | 0.32216082 | 2.66E-15 |

| | | | | | |
|---|---|---|---|---|---|
| | | | | 3 | |
| B67 | B199 | WBRSR(mPas_50) | Mg | -0.22997399 | 2.51E-08 |
| B67 | B194 | WBRSR(mPas_50) | Cl | -0.21697059 | 1.52E-07 |
| B69 | B154 | PV | RF | -0.22952619 | 2.67E-08 |
| B69 | B79 | PV | hb V -dna | 0.2196535 | 1.06E-07 |
| B69 | B130 | PV | GLB | 0.213839922 | 2.32E-07 |
| B69 | B123 | PV | hs-CRP1 | 0.213538111 | 2.41E-07 |
| B69 | B91 | PV | TP | 0.211100252 | 3.32E-07 |
| B7 | B116 | WB-Cop | TT3 | -0.52784318 | 0 |
| B7 | B202 | WB-Cop | e-times | 0.338648106 | 0 |
| B7 | B35 | WB-Cop | β2-MG | 0.92966968 | 0 |
| B7 | B9 | WB-Cop | RH | 0.478091444 | 0 |
| B70 | B77 | LDL-C | TC | 0.868666204 | 0 |
| B70 | B76 | LDL-C | ApoB | 0.322195698 | 2.44E-15 |
| B70 | B132 | LDL-C | Dbil | -0.23156716 | 2.00E-08 |
| B70 | B75 | LDL-C | ApoA1/B | -0.20754922 | 5.26E-07 |
| B71 | B72 | TG | HDL-C | -0.40530791 | 0 |
| B71 | B91 | TG | TP | 0.327974724 | 8.88E-16 |
| B71 | B128 | TG | ALT | 0.29583311 | 4.66E-13 |
| B71 | B216 | TG | BMI | 0.267594223 | 7.20E-11 |
| B71 | B140 | TG | r-GT | 0.260231912 | 2.44E-10 |
| B71 | B130 | TG | GLB | 0.254083138 | 6.58E-10 |
| B71 | B215 | TG | BMI | 0.252231542 | 8.83E-10 |
| B71 | B192 | TG | UA | 0.216532646 | 1.62E-07 |
| B71 | B210 | TG | Hip | 0.208572653 | 4.61E-07 |
| B71 | B141 | TG | AST/ALT | -0.20774069 | 5.14E-07 |
| B72 | B206 | HDL-C | Waist | -0.3597257 | 0 |
| B72 | B215 | HDL-C | BMI | -0.38938974 | 0 |
| B72 | B216 | HDL-C | BMI | -0.47171487 | 0 |
| B72 | B222 | HDL-C | NS | -0.37675221 | 0 |

| | | | | | |
|---|---|---|---|---|---|
| B72 | B74 | HDL-C | ApoA1 | 0.33703332 | 0 |
| B72 | B210 | HDL-C | Hip | -0.31190684 | 2.04E-14 |
| B72 | B77 | HDL-C | TC | 0.30009496<br>3 | 2.07E-13 |
| B72 | B221 | HDL-C | Hei | -0.29718486 | 3.61E-13 |
| B72 | B192 | HDL-C | UA | -0.28665399 | 2.56E-12 |
| B72 | B141 | HDL-C | AST/ALT | 0.25151724<br>4 | 9.88E-10 |
| B72 | B207 | HDL-C | WHR | -0.24523645 | 2.62E-09 |
| B72 | B163 | HDL-C | SG | -0.22516198 | 4.96E-08 |
| B72 | B128 | HDL-C | ALT | -0.22314623 | 6.56E-08 |
| B73 | B196 | ApoA | TCO2 | -0.21509472 | 1.96E-07 |
| B74 | B135 | ApoA1 | TBA | 0.42596962<br>3 | 0 |
| B74 | B75 | ApoA1 | ApoA1/B | 0.52556259 | 0 |
| B74 | B136 | ApoA1 | 5'NT | 0.30891370<br>1 | 3.71E-14 |
| B74 | B198 | ApoA1 | P | 0.29298518<br>7 | 7.96E-13 |
| B74 | B134 | ApoA1 | CHE | 0.26279024<br>6 | 1.61E-10 |
| B74 | B199 | ApoA1 | Mg | 0.24560509<br>3 | 2.48E-09 |
| B75 | B76 | ApoA1/B | ApoB | -0.76356173 | 0 |
| B76 | B134 | ApoB | CHE | 0.27926537<br>8 | 9.62E-12 |
| B76 | B77 | ApoB | TC | 0.23347359 | 1.52E-08 |
| B76 | B136 | ApoB | 5'NT | 0.22840585<br>8 | 3.14E-08 |
| B77 | B91 | TC | TP | 0.26439891<br>6 | 1.23E-10 |
| B77 | B132 | TC | Dbil | -0.23333712 | 1.55E-08 |
| B79 | B134 | hb V -dna | CHE | -0.35237312 | 0 |
| B79 | B178 | hb V -dna | Ig-U | -0.34297203 | 0 |
| B79 | B179 | hb V -dna | α1-MG | -0.4181436 | 0 |
| B79 | B182 | hb V -dna | TT | -0.35891611 | 0 |
| B79 | B200 | hb V -dna | OP | -0.34913622 | 0 |
| B79 | B92 | hb V -dna | ALDH2-Glu504Glu | 0.52272128<br>2 | 0 |
| B79 | B93 | hb V -dna | ALDH2-Glu504Lys | -0.53258376 | 0 |
| B79 | B94 | hb V -dna | ALDH2-Lys504Lys | 0.40596325<br>9 | 0 |
| B79 | B195 | hb V -dna | Na | -0.32173908 | 2.66E-15 |
| B79 | B100 | hb V -dna | anti-Ro52 | 0.25335284 | 7.39E-10 |

| | | | | | |
|---|---|---|---|---|---|
| | | | | 6 | |
| B79 | B101 | hb V -dna | anti-Sc70 | 0.253352849 | 7.39E-10 |
| B79 | B102 | hb V -dna | anti-Sm | 0.253352848 | 7.39E-10 |
| B79 | B103 | hb V -dna | anti-SSA/Ro | 0.253352846 | 7.39E-10 |
| B79 | B104 | hb V -dna | anti-SSB | 0.253352846 | 7.39E-10 |
| B79 | B105 | hb V -dna | anti-U1RNP | 0.253352846 | 7.39E-10 |
| B79 | B106 | hb V -dna | anti-P | 0.253352846 | 7.39E-10 |
| B79 | B107 | hb V -dna | AnuA | 0.253352843 | 7.39E-10 |
| B79 | B108 | hb V -dna | Anti-Cyclin | 0.253352846 | 7.39E-10 |
| B79 | B109 | hb V -dna | AMA-M2 | 0.253352846 | 7.39E-10 |
| B79 | B110 | hb V -dna | AHA | 0.253352846 | 7.39E-10 |
| B79 | B95 | hb V -dna | anti-cenpb | 0.253352846 | 7.39E-10 |
| B79 | B97 | hb V -dna | dsDNA | 0.253352845 | 7.39E-10 |
| B79 | B98 | hb V -dna | anti-Jo-1 | 0.253352847 | 7.39E-10 |
| B79 | B99 | hb V -dna | anti-PM-Scl | 0.253352846 | 7.39E-10 |
| B79 | B181 | hb V -dna | APTT | -0.21555873 | 1.84E-07 |
| B79 | B81 | hb V -dna | TSGF | -0.20921862 | 4.24E-07 |
| B8 | B117 | WB-Zinc | TT4 | 0.419463975 | 0 |
| B8 | B223 | WB-Zinc | FH | 0.984891636 | 0 |
| B81 | B93 | TSGF | ALDH2-Glu504Lys | 0.331255212 | 4.44E-16 |
| B81 | B92 | TSGF | ALDH2-Glu504Glu | -0.32499417 | 1.33E-15 |
| B81 | B179 | TSGF | α1-MG | 0.324271841 | 1.55E-15 |
| B81 | B200 | TSGF | OP | 0.318663639 | 5.33E-15 |
| B81 | B195 | TSGF | Na | 0.308217588 | 4.26E-14 |

| | | | | | |
|---|---|---|---|---|---|
| B81 | B178 | TSGF | Ig-U | 0.291255973 | 1.10E-12 |
| B81 | B124 | TSGF | ACTH | -0.28333999 | 4.65E-12 |
| B81 | B94 | TSGF | ALDH2-Lys504Lys | -0.25526635 | 5.45E-10 |
| B81 | B125 | TSGF | Cor | -0.2446069 | 2.88E-09 |
| B81 | B182 | TSGF | TT | 0.224551259 | 5.40E-08 |
| B82 | B87 | CEA | NSE | 0.308508935 | 4.02E-14 |
| B82 | B88 | CEA | CA 242 | 0.284757062 | 3.61E-12 |
| B82 | B115 | CEA | TSH TT3 TT4(TT3) | 0.245905394 | 2.36E-09 |
| B83 | B84 | AFP | C-PSA | 0.268325547 | 6.37E-11 |
| B83 | B86 | AFP | TBhCG | 0.261409739 | 2.01E-10 |
| B83 | B88 | AFP | CA 242 | -0.25092226 | 1.09E-09 |
| B83 | B89 | AFP | CA 72-4 | 0.239723688 | 6.03E-09 |
| B84 | B87 | C-PSA | NSE | 0.352844412 | 0 |
| B84 | B85 | C-PSA | SCC | 0.303856246 | 1.00E-13 |
| B84 | B157 | C-PSA | IgG | 0.231013004 | 2.16E-08 |
| B84 | B86 | C-PSA | TBhCG | 0.218747151 | 1.20E-07 |
| B85 | B90 | SCC | CYFRA 21-1 | 0.332524399 | 2.22E-16 |
| B85 | B88 | SCC | CA 242 | 0.252026793 | 9.12E-10 |
| B85 | B89 | SCC | CA 72-4 | 0.239148514 | 6.57E-09 |
| B85 | B152 | SCC | ACA-Ig(G-A-M) | -0.2210497 | 8.76E-08 |
| B85 | B119 | SCC | Hcv-rna | 0.221034328 | 8.78E-08 |
| B85 | B121 | SCC | C3 | -0.22103442 | 8.78E-08 |
| B85 | B122 | SCC | C4 | -0.22103442 | 8.78E-08 |
| B85 | B159 | SCC | β2-MG | -0.22103563 | 8.78E-08 |
| B85 | B169 | SCC | U-CRY | -0.22103146 | 8.78E-08 |
| B85 | B87 | SCC | NSE | 0.207119806 | 5.56E-07 |
| B86 | B87 | TBhCG | NSE | 0.43282747 | 0 |

| | | | | | |
|---|---|---|---|---|---|
| | | | | 7 | |
| B86 | B90 | TBhCG | CYFRA 21-1 | 0.224004691 | 5.82E-08 |
| B86 | B119 | TBhCG | Hcv-rna | 0.214536875 | 2.11E-07 |
| B86 | B121 | TBhCG | C3 | -0.21453696 | 2.11E-07 |
| B86 | B122 | TBhCG | C4 | -0.21453696 | 2.11E-07 |
| B86 | B152 | TBhCG | ACA-Ig(G-A-M) | -0.2145518 | 2.11E-07 |
| B86 | B159 | TBhCG | β2-MG | -0.21453814 | 2.11E-07 |
| B86 | B169 | TBhCG | U-CRY | -0.2145341 | 2.11E-07 |
| B87 | B88 | NSE | CA 242 | 0.325873429 | 1.11E-15 |
| B87 | B90 | NSE | CYFRA 21-1 | 0.213964014 | 2.28E-07 |
| B87 | B119 | NSE | Hcv-rna | 0.212427353 | 2.79E-07 |
| B87 | B121 | NSE | C3 | -0.21242744 | 2.79E-07 |
| B87 | B122 | NSE | C4 | -0.21242744 | 2.79E-07 |
| B87 | B152 | NSE | ACA-Ig(G-A-M) | -0.21244215 | 2.79E-07 |
| B87 | B159 | NSE | β2-MG | -0.21242861 | 2.79E-07 |
| B87 | B169 | NSE | U-CRY | -0.2124246 | 2.79E-07 |
| B89 | B197 | CA 72-4 | AG | -0.2529303 | 7.91E-10 |
| B9 | B202 | RH | e-times | -0.875 | 0 |
| B90 | B187 | CYFRA 21-1 | NSE | 0.342086978 | 0 |
| B90 | B125 | CYFRA 21-1 | Cor | -0.23541258 | 1.14E-08 |
| B90 | B124 | CYFRA 21-1 | ACTH | -0.202889 | 9.52E-07 |
| B91 | B126 | TP | Alb | 0.41844389 | 0 |
| B91 | B127 | TP | A-G ratio | -0.53143882 | 0 |
| B91 | B130 | TP | GLB | 0.807592294 | 0 |
| B91 | B188 | TP | Ca | 0.226868745 | 3.90E-08 |
| B92 | B100 | ALDH2-Glu504Glu | anti-Ro52 | 0.404645902 | 0 |
| B92 | B101 | ALDH2-Glu504Glu | anti-Sc70 | 0.404645906 | 0 |
| B92 | B102 | ALDH2-Glu504Glu | anti-Sm | 0.404645903 | 0 |
| B92 | B103 | ALDH2-Glu504Glu | anti-SSA/Ro | 0.404645902 | 0 |
| B92 | B104 | ALDH2-Glu504Glu | anti-SSB | 0.4046459 | 0 |
| B92 | B105 | ALDH2-Glu504Glu | anti-U1RNP | 0.404645901 | 0 |

| | | | | | |
|---|---|---|---|---|---|
| B92 | B106 | ALDH2-Glu504Glu | anti-P | 0.404645901 | 0 |
| B92 | B107 | ALDH2-Glu504Glu | AnuA | 0.40464589 | 0 |
| B92 | B108 | ALDH2-Glu504Glu | Anti-Cyclin | 0.404645902 | 0 |
| B92 | B109 | ALDH2-Glu504Glu | AMA-M2 | 0.404645901 | 0 |
| B92 | B110 | ALDH2-Glu504Glu | AHA | 0.404645902 | 0 |
| B92 | B178 | ALDH2-Glu504Glu | Ig-U | -0.7370713 | 0 |
| B92 | B179 | ALDH2-Glu504Glu | α1-MG | -0.81746137 | 0 |
| B92 | B181 | ALDH2-Glu504Glu | APTT | -0.34325409 | 0 |
| B92 | B182 | ALDH2-Glu504Glu | TT | -0.56413099 | 0 |
| B92 | B195 | ALDH2-Glu504Glu | Na | -0.51666647 | 0 |
| B92 | B200 | ALDH2-Glu504Glu | OP | -0.54685089 | 0 |
| B92 | B93 | ALDH2-Glu504Glu | ALDH2-Glu504Lys | -0.99909906 | 0 |
| B92 | B94 | ALDH2-Glu504Glu | ALDH2-Lys504Lys | 0.338287376 | 0 |
| B92 | B95 | ALDH2-Glu504Glu | anti-cenpb | 0.404645901 | 0 |
| B92 | B97 | ALDH2-Glu504Glu | dsDNA | 0.4046459 | 0 |
| B92 | B98 | ALDH2-Glu504Glu | anti-Jo-1 | 0.404645902 | 0 |
| B92 | B99 | ALDH2-Glu504Glu | anti-PM-Scl | 0.404645901 | 0 |
| B92 | B194 | ALDH2-Glu504Glu | Cl | 0.318102219 | 5.77E-15 |
| B92 | B196 | ALDH2-Glu504Glu | TCO2 | -0.31160508 | 2.18E-14 |
| B92 | B198 | ALDH2-Glu504Glu | P | 0.305360491 | 7.46E-14 |
| B92 | B184 | ALDH2-Glu504Glu | INR | 0.303282832 | 1.12E-13 |
| B92 | B197 | ALDH2-Glu504Glu | AG | -0.25124799 | 1.03E-09 |
| B92 | B151 | ALDH2-Glu504Glu | Anti-CCP | 0.232062584 | 1.86E-08 |
| B92 | B115 | ALDH2-Glu504Glu | TSH TT3 TT4(TT3) | -0.23094741 | 2.18E-08 |
| B92 | B158 | ALDH2-Glu504Glu | IgM | 0.211493727 | 3.16E-07 |
| B92 | B139 | ALDH2-Glu504Glu | ADA | -0.20994747 | 3.86E-07 |
| B93 | B100 | ALDH2-Glu504Lys | anti-Ro52 | -0.41220681 | 0 |
| B93 | B101 | ALDH2-Glu504Lys | anti-Sc70 | -0.41220681 | 0 |
| B93 | B102 | ALDH2-Glu504Lys | anti-Sm | -0.41220681 | 0 |
| B93 | B103 | ALDH2-Glu504Lys | anti-SSA/Ro | -0.41220681 | 0 |
| B93 | B104 | ALDH2-Glu504Lys | anti-SSB | -0.41220681 | 0 |

| | | | | | |
|---|---|---|---|---|---|
| B93 | B105 | ALDH2-Glu504Lys | anti-U1RNP | -0.41220681 | 0 |
| B93 | B106 | ALDH2-Glu504Lys | anti-P | -0.41220681 | 0 |
| B93 | B107 | ALDH2-Glu504Lys | AnuA | -0.4122068 | 0 |
| B93 | B108 | ALDH2-Glu504Lys | Anti-Cyclin | -0.41220681 | 0 |
| B93 | B109 | ALDH2-Glu504Lys | AMA-M2 | -0.41220681 | 0 |
| B93 | B110 | ALDH2-Glu504Lys | AHA | -0.41220681 | 0 |
| B93 | B178 | ALDH2-Glu504Lys | Ig-U | 0.751234155 | 0 |
| B93 | B179 | ALDH2-Glu504Lys | α1-MG | 0.833083725 | 0 |
| B93 | B181 | ALDH2-Glu504Lys | APTT | 0.349689029 | 0 |
| B93 | B182 | ALDH2-Glu504Lys | TT | 0.574859426 | 0 |
| B93 | B195 | ALDH2-Glu504Lys | Na | 0.525040905 | 0 |
| B93 | B200 | ALDH2-Glu504Lys | OP | 0.557290589 | 0 |
| B93 | B94 | ALDH2-Glu504Lys | ALDH2-Lys504Lys | -0.37791954 | 0 |
| B93 | B95 | ALDH2-Glu504Lys | anti-cenpb | -0.41220681 | 0 |
| B93 | B97 | ALDH2-Glu504Lys | dsDNA | -0.4122068 | 0 |
| B93 | B98 | ALDH2-Glu504Lys | anti-Jo-1 | -0.41220681 | 0 |
| B93 | B99 | ALDH2-Glu504Lys | anti-PM-Scl | -0.41220681 | 0 |
| B93 | B194 | ALDH2-Glu504Lys | Cl | -0.31280165 | 1.69E-14 |
| B93 | B198 | ALDH2-Glu504Lys | P | -0.3111811 | 2.35E-14 |
| B93 | B184 | ALDH2-Glu504Lys | INR | -0.30901878 | 3.64E-14 |
| B93 | B196 | ALDH2-Glu504Lys | TCO2 | 0.308612851 | 3.93E-14 |
| B93 | B197 | ALDH2-Glu504Lys | AG | 0.254962671 | 5.72E-10 |
| B93 | B151 | ALDH2-Glu504Lys | Anti-CCP | -0.23627241 | 1.01E-08 |
| B93 | B115 | ALDH2-Glu504Lys | TSH TT3 TT4(TT3) | 0.23544689 | 1.14E-08 |
| B93 | B158 | ALDH2-Glu504Lys | IgM | -0.21545356 | 1.87E-07 |
| B93 | B139 | ALDH2-Glu504Lys | ADA | 0.213802497 | 2.33E-07 |
| B94 | B178 | ALDH2-Lys504Lys | Ig-U | -0.5781138 | 0 |
| B94 | B179 | ALDH2-Lys504Lys | α1-MG | -0.63927768 | 0 |
| B94 | B182 | ALDH2-Lys504Lys | TT | -0.44000064 | 0 |
| B94 | B195 | ALDH2-Lys504Lys | Na | -0.37079807 | 0 |
| B94 | B200 | ALDH2-Lys504Lys | OP | -0.42740729 | 0 |
| B94 | B100 | ALDH2-Lys504Lys | anti-Ro52 | 0.3126259 | 1.78E-14 |
| B94 | B101 | ALDH2-Lys504Lys | anti-Sc70 | 0.312625903 | 1.78E-14 |
| B94 | B102 | ALDH2-Lys504Lys | anti-Sm | 0.3126259 | 1.78E-14 |

| | | | | | |
|---|---|---|---|---|---|
| B94 | B103 | ALDH2-Lys504Lys | anti-SSA/Ro | 0.312625899 | 1.78E-14 |
| B94 | B104 | ALDH2-Lys504Lys | anti-SSB | 0.312625914 | 1.78E-14 |
| B94 | B105 | ALDH2-Lys504Lys | anti-U1RNP | 0.312625902 | 1.78E-14 |
| B94 | B106 | ALDH2-Lys504Lys | anti-P | 0.31262591 | 1.78E-14 |
| B94 | B107 | ALDH2-Lys504Lys | AnuA | 0.312625975 | 1.78E-14 |
| B94 | B108 | ALDH2-Lys504Lys | Anti-Cyclin | 0.312625904 | 1.78E-14 |
| B94 | B109 | ALDH2-Lys504Lys | AMA-M2 | 0.3126259 | 1.78E-14 |
| B94 | B110 | ALDH2-Lys504Lys | AHA | 0.312625898 | 1.78E-14 |
| B94 | B95 | ALDH2-Lys504Lys | anti-cenpb | 0.312625905 | 1.78E-14 |
| B94 | B97 | ALDH2-Lys504Lys | dsDNA | 0.312625906 | 1.78E-14 |
| B94 | B98 | ALDH2-Lys504Lys | anti-Jo-1 | 0.312625905 | 1.78E-14 |
| B94 | B99 | ALDH2-Lys504Lys | anti-PM-Scl | 0.312625907 | 1.78E-14 |
| B94 | B181 | ALDH2-Lys504Lys | APTT | -0.26566417 | 9.96E-11 |
| B94 | B198 | ALDH2-Lys504Lys | P | 0.238466088 | 7.28E-09 |
| B94 | B184 | ALDH2-Lys504Lys | INR | 0.235844471 | 1.07E-08 |
| B95 | B100 | anti-cenpb | anti-Ro52 | 0.999999999 | 0 |
| B95 | B103 | anti-cenpb | anti-SSA/Ro | 0.999999999 | 0 |
| B95 | B104 | anti-cenpb | anti-SSB | 0.999999997 | 0 |
| B95 | B105 | anti-cenpb | anti-U1RNP | 0.999999999 | 0 |
| B95 | B107 | anti-cenpb | AnuA | 0.999999984 | 0 |
| B95 | B108 | anti-cenpb | Anti-Cyclin | 0.999999998 | 0 |
| B95 | B109 | anti-cenpb | AMA-M2 | 0.999999997 | 0 |
| B95 | B110 | anti-cenpb | AHA | 1 | 0 |
| B95 | B151 | anti-cenpb | Anti-CCP | 0.608491116 | 0 |

| | | | | | |
|---|---|---|---|---|---|
| B95 | B179 | anti-cenpb | α1-MG | -0.37132208 | 0 |
| B95 | B181 | anti-cenpb | APTT | -0.49743253 | 0 |
| B95 | B182 | anti-cenpb | TT | -0.36368895 | 0 |
| B95 | B186 | anti-cenpb | PFC | 0.54261596<br>1 | 0 |
| B95 | B97 | anti-cenpb | dsDNA | 0.99999999<br>9 | 0 |
| B95 | B178 | anti-cenpb | Ig-U | -0.28472773 | 3.62E-12 |
| B95 | B200 | anti-cenpb | OP | -0.2737536 | 2.52E-11 |
| B95 | B195 | anti-cenpb | Na | -0.27267879 | 3.03E-11 |
| B95 | B196 | anti-cenpb | TCO2 | -0.25679436 | 4.27E-10 |
| B95 | B194 | anti-cenpb | Cl | 0.24067474<br>5 | 5.23E-09 |
| B95 | B137 | anti-cenpb | AFU | 0.23606859<br>8 | 1.04E-08 |
| B97 | B100 | dsDNA | anti-Ro52 | 0.99999999<br>6 | 0 |
| B97 | B103 | dsDNA | anti-SSA/Ro | 0.99999999<br>6 | 0 |
| B97 | B105 | dsDNA | anti-U1RNP | 0.99999999<br>5 | 0 |
| B97 | B108 | dsDNA | Anti-Cyclin | 0.99999999<br>8 | 0 |
| B97 | B109 | dsDNA | AMA-M2 | 0.99999999<br>6 | 0 |
| B97 | B110 | dsDNA | AHA | 0.99999999<br>6 | 0 |
| B97 | B151 | dsDNA | Anti-CCP | 0.60849111<br>3 | 0 |
| B97 | B179 | dsDNA | α1-MG | -0.37132208 | 0 |
| B97 | B181 | dsDNA | APTT | -0.49743252 | 0 |
| B97 | B182 | dsDNA | TT | -0.36368895 | 0 |
| B97 | B186 | dsDNA | PFC | 0.54261595<br>8 | 0 |
| B97 | B99 | dsDNA | anti-PM-Scl | 0.99999999<br>7 | 0 |
| B97 | B178 | dsDNA | Ig-U | -0.28472773 | 3.62E-12 |
| B97 | B200 | dsDNA | OP | -0.2737536 | 2.52E-11 |
| B97 | B195 | dsDNA | Na | -0.27267879 | 3.03E-11 |
| B97 | B196 | dsDNA | TCO2 | -0.25679436 | 4.27E-10 |
| B97 | B194 | dsDNA | Cl | 0.24067474<br>5 | 5.23E-09 |
| B97 | B137 | dsDNA | AFU | 0.23606859<br>7 | 1.04E-08 |

| | | | | | |
|---|---|---|---|---|---|
| B98 | B101 | anti-Jo-1 | anti-Sc70 | 0.99999998<br>7 | 0 |
| B98 | B102 | anti-Jo-1 | anti-Sm | 0.99999999<br>5 | 0 |
| B98 | B105 | anti-Jo-1 | anti-U1RNP | 0.99999999<br>9 | 0 |
| B98 | B107 | anti-Jo-1 | AnuA | 0.99999993<br>5 | 0 |
| B98 | B109 | anti-Jo-1 | AMA-M2 | 0.99999999<br>9 | 0 |
| B98 | B110 | anti-Jo-1 | AHA | 0.99999999<br>9 | 0 |
| B98 | B151 | anti-Jo-1 | Anti-CCP | 0.60849111<br>8 | 0 |
| B98 | B179 | anti-Jo-1 | α1-MG | -0.37132208 | 0 |
| B98 | B181 | anti-Jo-1 | APTT | -0.49743253 | 0 |
| B98 | B182 | anti-Jo-1 | TT | -0.36368895 | 0 |
| B98 | B186 | anti-Jo-1 | PFC | 0.54261596<br>1 | 0 |
| B98 | B178 | anti-Jo-1 | Ig-U | -0.28472773 | 3.62E-12 |
| B98 | B200 | anti-Jo-1 | OP | -0.2737536 | 2.52E-11 |
| B98 | B195 | anti-Jo-1 | Na | -0.27267879 | 3.03E-11 |
| B98 | B196 | anti-Jo-1 | TCO2 | -0.25679436 | 4.27E-10 |
| B98 | B194 | anti-Jo-1 | Cl | 0.24067474<br>7 | 5.23E-09 |
| B98 | B137 | anti-Jo-1 | AFU | 0.23606859<br>9 | 1.04E-08 |
| B99 | B100 | anti-PM-Scl | anti-Ro52 | 0.99999999<br>8 | 0 |
| B99 | B103 | anti-PM-Scl | anti-SSA/Ro | 0.99999999<br>8 | 0 |
| B99 | B104 | anti-PM-Scl | anti-SSB | 0.99999999<br>8 | 0 |
| B99 | B105 | anti-PM-Scl | anti-U1RNP | 0.99999999<br>8 | 0 |
| B99 | B106 | anti-PM-Scl | anti-P | 0.99999999<br>7 | 0 |
| B99 | B107 | anti-PM-Scl | AnuA | 0.99999996<br>8 | 0 |
| B99 | B109 | anti-PM-Scl | AMA-M2 | 0.99999999<br>7 | 0 |
| B99 | B151 | anti-PM-Scl | Anti-CCP | 0.60849111<br>5 | 0 |
| B99 | B179 | anti-PM-Scl | α1-MG | -0.37132208 | 0 |

| | | | | | |
|---|---|---|---|---|---|
| B99 | B181 | anti-PM-Scl | APTT | -0.49743253 | 0 |
| B99 | B182 | anti-PM-Scl | TT | -0.36368895 | 0 |
| B99 | B186 | anti-PM-Scl | PFC | 0.542615961 | 0 |
| B99 | B178 | anti-PM-Scl | Ig-U | -0.28472773 | 3.62E-12 |
| B99 | B200 | anti-PM-Scl | OP | -0.2737536 | 2.52E-11 |
| B99 | B195 | anti-PM-Scl | Na | -0.27267879 | 3.03E-11 |
| B99 | B196 | anti-PM-Scl | TCO2 | -0.25679436 | 4.27E-10 |
| B99 | B194 | anti-PM-Scl | Cl | 0.240674745 | 5.23E-09 |
| B99 | B137 | anti-PM-Scl | AFU | 0.236068597 | 1.04E-08 |

**Supplementary Table 28. PEI Correlations in Other disease conditions**

| Row | Column | Row | Column | Cor | *P* |
|---|---|---|---|---|---|
| B1 | B127 | Gender | A-G ratio | -0.17892 | 0 |
| B1 | B128 | Gender | ALT | -0.2238 | 0 |
| B1 | B140 | Gender | r-GT | -0.17911 | 0 |
| B1 | B141 | Gender | AST/ALT | 0.224985 | 0 |
| B1 | B153 | Gender | Glu | -0.17716 | 0 |
| B1 | B161 | Gender | LEU | 0.17101 | 0 |
| B1 | B163 | Gender | SG | -0.42484 | 0 |
| B1 | B166 | Gender | Pro | -0.17233 | 0 |
| B1 | B174 | Gender | Urea/Cr | 0.305315 | 0 |
| B1 | B190 | Gender | Cre | -0.24687 | 0 |
| B1 | B191 | Gender | Urea | -0.15699 | 0 |
| B1 | B192 | Gender | UA | -0.50875 | 0 |
| B1 | B204 | Gender | Drinking | -0.32405 | 0 |
| B1 | B206 | Gender | Waist | -0.50812 | 0 |
| B1 | B209 | Gender | Smoking | -0.35477 | 0 |
| B1 | B210 | Gender | Hip | -0.34623 | 0 |
| B1 | B215 | Gender | BMI | -0.30491 | 0 |
| B1 | B216 | Gender | BMI | -0.57912 | 0 |
| B1 | B218 | Gender | DBP | -0.25924 | 0 |
| B1 | B220 | Gender | SBP | -0.22569 | 0 |
| B1 | B221 | Gender | Hei | -0.63324 | 0 |
| B1 | B222 | Gender | NS | -0.73211 | 0 |
| B1 | B42 | Gender | MON | -0.24482 | 0 |
| B1 | B44 | Gender | MCH | -0.16839 | 0 |
| B1 | B45 | Gender | MCHC | -0.2903 | 0 |
| B1 | B46 | Gender | RBC Count | -0.46941 | 0 |
| B1 | B50 | Gender | eo% | -0.16196 | 0 |
| B1 | B51 | Gender | Hb | -0.58863 | 0 |
| B1 | B71 | Gender | TG | -0.17739 | 0 |
| B1 | B72 | Gender | HDL-C | 0.344249 | 0 |
| B1 | B41 | Gender | MON % | -0.14976 | 2.22E-15 |
| B1 | B132 | Gender | Dbil | -0.14575 | 1.18E-14 |
| B1 | B133 | Gender | TBIL | -0.14546 | 1.33E-14 |
| B1 | B130 | Gender | GLB | 0.144127 | 2.31E-14 |
| B1 | B170 | Gender | U-GLU | -0.13753 | 3.37E-13 |
| B1 | B129 | Gender | IBIL | -0.13283 | 2.10E-12 |
| B1 | B176 | Gender | Ucr | -0.13265 | 2.25E-12 |
| B1 | B2 | Gender | Age | -0.13044 | 5.18E-12 |
| B1 | B113 | Gender | FT3 | -0.11815 | 4.23E-10 |
| B1 | B3 | Gender | Hcy | -0.11617 | 8.25E-10 |
| B1 | B138 | Gender | PAB | -0.11292 | 2.42E-09 |
| B1 | B58 | Gender | HCT | -0.11176 | 3.52E-09 |

| | | | | | |
|---|---|---|---|---|---|
| B1 | B189 | Gender | eGFB | 0.1115 | 3.83E-09 |
| B1 | B62 | Gender | WBHSRVI | -0.11099 | 4.50E-09 |
| B1 | B52 | Gender | PLT/BPC | 0.110599 | 5.11E-09 |
| B1 | B148 | Gender | Anti_TPO | 0.107772 | 1.25E-08 |
| B1 | B126 | Gender | Alb | -0.10738 | 1.41E-08 |
| B1 | B198 | Gender | P | 0.102104 | 6.96E-08 |
| B1 | B131 | Gender | AST | -0.09669 | 3.31E-07 |
| B1 | B60 | Gender | WBRVI | -0.09617 | 3.83E-07 |
| B10 | B11 | PG Ⅰ | PGI/PGII | 0.17399 | 0 |
| B10 | B12 | PG Ⅰ | PG Ⅱ | 0.622188 | 0 |
| B10 | B189 | PG Ⅰ | eGFB | -0.10205 | 7.06E-08 |
| B100 | B101 | anti-Ro52 | anti-Sc70 | 0.246548 | 0 |
| B100 | B102 | anti-Ro52 | anti-Sm | 0.246548 | 0 |
| B100 | B103 | anti-Ro52 | anti-SSA/Ro | 0.391711 | 0 |
| B100 | B104 | anti-Ro52 | anti-SSB | 0.246548 | 0 |
| B100 | B105 | anti-Ro52 | anti-U1RNP | 0.666453 | 0 |
| B100 | B106 | anti-Ro52 | anti-P | 0.654074 | 0 |
| B100 | B107 | anti-Ro52 | AnuA | 0.639864 | 0 |
| B100 | B109 | anti-Ro52 | AMA-M2 | 0.582634 | 0 |
| B100 | B110 | anti-Ro52 | AHA | 0.462859 | 0 |
| B100 | B121 | anti-Ro52 | C3 | -0.47225 | 0 |
| B100 | B122 | anti-Ro52 | C4 | -0.3799 | 0 |
| B100 | B187 | anti-Ro52 | NSE | -0.09656 | 3.43E-07 |
| B101 | B104 | anti-Sc70 | anti-SSB | 1 | 0 |
| B101 | B121 | anti-Sc70 | C3 | 0.162823 | 0 |
| B101 | B180 | anti-Sc70 | Tf | -0.16234 | 0 |
| B101 | B186 | anti-Sc70 | PFC | 0.179834 | 0 |
| B101 | B122 | anti-Sc70 | C4 | 0.154227 | 4.44E-16 |
| B101 | B183 | anti-Sc70 | PTA | 0.146138 | 9.99E-15 |
| B101 | B120 | anti-Sc70 | HCV-cAg | -0.13654 | 4.98E-13 |
| B101 | B177 | anti-Sc70 | ACR | -0.13646 | 5.13E-13 |
| B101 | B158 | anti-Sc70 | IgM | 0.134079 | 1.30E-12 |
| B101 | B107 | anti-Sc70 | AnuA | -0.13254 | 2.35E-12 |
| B101 | B108 | anti-Sc70 | Anti-Cyclin | -0.12613 | 2.55E-11 |
| B101 | B178 | anti-Sc70 | Ig-U | -0.11829 | 4.03E-10 |
| B101 | B182 | anti-Sc70 | TT | 0.105557 | 2.47E-08 |
| B101 | B124 | anti-Sc70 | ACTH | -0.10554 | 2.48E-08 |
| B101 | B196 | anti-Sc70 | TCO2 | -0.09305 | 9.01E-07 |
| B102 | B104 | anti-Sm | anti-SSB | 1 | 0 |
| B102 | B121 | anti-Sm | C3 | 0.162823 | 0 |
| B102 | B180 | anti-Sm | Tf | -0.16234 | 0 |
| B102 | B186 | anti-Sm | PFC | 0.179834 | 0 |
| B102 | B122 | anti-Sm | C4 | 0.154227 | 4.44E-16 |
| B102 | B183 | anti-Sm | PTA | 0.146138 | 9.99E-15 |

| | | | | | |
|---|---|---|---|---|---|
| B102 | B120 | anti-Sm | HCV-cAg | -0.13654 | 4.98E-13 |
| B102 | B177 | anti-Sm | ACR | -0.13646 | 5.13E-13 |
| B102 | B158 | anti-Sm | IgM | 0.134079 | 1.30E-12 |
| B102 | B107 | anti-Sm | AnuA | -0.13254 | 2.35E-12 |
| B102 | B108 | anti-Sm | Anti-Cyclin | -0.12613 | 2.55E-11 |
| B102 | B178 | anti-Sm | Ig-U | -0.11829 | 4.03E-10 |
| B102 | B182 | anti-Sm | TT | 0.105557 | 2.47E-08 |
| B102 | B124 | anti-Sm | ACTH | -0.10554 | 2.48E-08 |
| B102 | B196 | anti-Sm | TCO2 | -0.09305 | 9.01E-07 |
| B103 | B105 | anti-SSA/Ro | anti-U1RNP | 0.63161 | 0 |
| B103 | B106 | anti-SSA/Ro | anti-P | 0.633219 | 0 |
| B103 | B107 | anti-SSA/Ro | AnuA | 0.633681 | 0 |
| B103 | B108 | anti-SSA/Ro | Anti-Cyclin | 0.396137 | 0 |
| B103 | B121 | anti-SSA/Ro | C3 | -0.48413 | 0 |
| B103 | B122 | anti-SSA/Ro | C4 | -0.39521 | 0 |
| B103 | B178 | anti-SSA/Ro | Ig-U | 0.284035 | 0 |
| B103 | B180 | anti-SSA/Ro | Tf | 0.333705 | 0 |
| B103 | B179 | anti-SSA/Ro | α1-MG | 0.138907 | 1.95E-13 |
| B103 | B157 | anti-SSA/Ro | IgG | 0.124852 | 4.05E-11 |
| B104 | B121 | anti-SSB | C3 | 0.162823 | 0 |
| B104 | B180 | anti-SSB | Tf | -0.16234 | 0 |
| B104 | B186 | anti-SSB | PFC | 0.179834 | 0 |
| B104 | B122 | anti-SSB | C4 | 0.154227 | 4.44E-16 |
| B104 | B183 | anti-SSB | PTA | 0.146138 | 9.99E-15 |
| B104 | B120 | anti-SSB | HCV-cAg | -0.13654 | 4.98E-13 |
| B104 | B177 | anti-SSB | ACR | -0.13646 | 5.13E-13 |
| B104 | B158 | anti-SSB | IgM | 0.134079 | 1.30E-12 |
| B104 | B107 | anti-SSB | AnuA | -0.13254 | 2.35E-12 |
| B104 | B108 | anti-SSB | Anti-Cyclin | -0.12613 | 2.55E-11 |
| B104 | B178 | anti-SSB | Ig-U | -0.11829 | 4.03E-10 |
| B104 | B182 | anti-SSB | TT | 0.105557 | 2.47E-08 |
| B104 | B124 | anti-SSB | ACTH | -0.10554 | 2.48E-08 |
| B104 | B196 | anti-SSB | TCO2 | -0.09305 | 9.01E-07 |
| B105 | B106 | anti-U1RNP | anti-P | 0.999099 | 0 |
| B105 | B107 | anti-U1RNP | AnuA | 0.99623 | 0 |
| B105 | B121 | anti-U1RNP | C3 | -0.77022 | 0 |
| B105 | B122 | anti-U1RNP | C4 | -0.63238 | 0 |
| B105 | B157 | anti-U1RNP | IgG | 0.161088 | 0 |
| B106 | B107 | anti-P | AnuA | 0.999014 | 0 |
| B106 | B121 | anti-P | C3 | -0.77494 | 0 |
| B106 | B122 | anti-P | C4 | -0.63712 | 0 |
| B106 | B157 | anti-P | IgG | 0.163233 | 0 |
| B107 | B121 | AnuA | C3 | -0.77837 | 0 |
| B107 | B122 | AnuA | C4 | -0.64086 | 0 |

| | | | | | |
|---|---|---|---|---|---|
| B107 | B157 | AnuA | IgG | 0.165161 | 0 |
| B108 | B177 | Anti-Cyclin | ACR | 0.166533 | 0 |
| B108 | B178 | Anti-Cyclin | Ig-U | 0.622547 | 0 |
| B108 | B180 | Anti-Cyclin | Tf | 0.809535 | 0 |
| B108 | B179 | Anti-Cyclin | α1-MG | 0.153717 | 4.44E-16 |
| B109 | B110 | AMA-M2 | AHA | 0.617891 | 0 |
| B109 | B155 | AMA-M2 | IgA | 0.10011 | 1.25E-07 |
| B109 | B136 | AMA-M2 | 5'NT | 0.097912 | 2.34E-07 |
| B109 | B187 | AMA-M2 | NSE | -0.09775 | 2.46E-07 |
| B109 | B198 | AMA-M2 | P | 0.094429 | 6.19E-07 |
| B11 | B12 | PGI/PGII | PG II | -0.4361 | 0 |
| B11 | B14 | PGI/PGII | G-17 | -0.18138 | 0 |
| B11 | B80 | PGI/PGII | anti-HP IgG | -0.14078 | 9.15E-14 |
| B110 | B125 | AHA | Cor | -0.13067 | 4.76E-12 |
| B110 | B115 | AHA | TSH TT3 TT4(TT3) | -0.11576 | 9.48E-10 |
| B110 | B187 | AHA | NSE | -0.10622 | 2.01E-08 |
| B110 | B124 | AHA | ACTH | -0.10311 | 5.15E-08 |
| B111 | B112 | TSH | FT4 | -0.23505 | 0 |
| B111 | B114 | TSH | TSH TT3 TT4(TT4) | -0.14508 | 1.55E-14 |
| B111 | B115 | TSH | TSH TT3 TT4(TT3) | -0.13494 | 9.31E-13 |
| B111 | B113 | TSH | FT3 | -0.1334 | 1.68E-12 |
| B111 | B173 | TSH | NTT | 0.09869 | 1.88E-07 |
| B112 | B113 | FT4 | FT3 | 0.396299 | 0 |
| B112 | B114 | FT4 | TSH TT3 TT4(TT4) | 0.578608 | 0 |
| B112 | B115 | FT4 | TSH TT3 TT4(TT3) | 0.222913 | 0 |
| B112 | B124 | FT4 | ACTH | 0.371197 | 0 |
| B112 | B125 | FT4 | Cor | 0.147833 | 4.88E-15 |
| B112 | B150 | FT4 | ALP | 0.138006 | 2.79E-13 |
| B113 | B114 | FT3 | TSH TT3 TT4(TT4) | 0.232665 | 0 |
| B113 | B115 | FT3 | TSH TT3 TT4(TT3) | 0.497472 | 0 |
| B113 | B124 | FT3 | ACTH | 0.411052 | 0 |
| B113 | B125 | FT3 | Cor | 0.176417 | 0 |
| B113 | B141 | FT3 | AST/ALT | -0.11076 | 4.85E-09 |
| B113 | B126 | FT3 | Alb | 0.108102 | 1.12E-08 |
| B113 | B150 | FT3 | ALP | 0.106164 | 2.05E-08 |
| B113 | B216 | FT3 | BMI | 0.101016 | 9.58E-08 |
| B113 | B210 | FT3 | Hip | 0.09576 | 4.29E-07 |
| B113 | B188 | FT3 | Ca | 0.095597 | 4.49E-07 |
| B113 | B222 | FT3 | NS | 0.093734 | 7.49E-07 |
| B114 | B115 | TSH TT3 TT4(TT4) | TSH TT3 TT4(TT3) | 0.390285 | 0 |
| B114 | B124 | TSH TT3 TT4(TT4) | ACTH | 0.369564 | 0 |
| B114 | B125 | TSH TT3 TT4(TT4) | Cor | 0.165796 | 0 |
| B114 | B150 | TSH TT3 TT4(TT4) | ALP | 0.11653 | 7.32E-10 |
| B114 | B138 | TSH TT3 TT4(TT4) | PAB | -0.11566 | 9.78E-10 |

| | | | | | |
|---|---|---|---|---|---|
| B115 | B124 | TSH TT3 TT4(TT3) | ACTH | 0.403439 | 0 |
| B115 | B125 | TSH TT3 TT4(TT3) | Cor | 0.187174 | 0 |
| B115 | B169 | TSH TT3 TT4(TT3) | U-CRY | 0.123689 | 6.14E-11 |
| B115 | B159 | TSH TT3 TT4(TT3) | β2-MG | 0.123687 | 6.15E-11 |
| B115 | B150 | TSH TT3 TT4(TT3) | ALP | 0.114014 | 1.69E-09 |
| B115 | B119 | TSH TT3 TT4(TT3) | Hcv-rna | -0.11072 | 4.92E-09 |
| B115 | B188 | TSH TT3 TT4(TT3) | Ca | 0.103975 | 3.98E-08 |
| B116 | B117 | TT3 | TT4 | 1 | 0 |
| B119 | B159 | Hcv-rna | β2-MG | -0.97783 | 0 |
| B119 | B169 | Hcv-rna | U-CRY | -0.97785 | 0 |
| B119 | B185 | Hcv-rna | PT | -0.19863 | 0 |
| B119 | B184 | Hcv-rna | INR | -0.14132 | 7.35E-14 |
| B119 | B181 | Hcv-rna | APTT | -0.13796 | 2.84E-13 |
| B119 | B194 | Hcv-rna | Cl | 0.118337 | 3.97E-10 |
| B119 | B197 | Hcv-rna | AG | -0.10907 | 8.30E-09 |
| B119 | B200 | Hcv-rna | OP | -0.10014 | 1.24E-07 |
| B119 | B158 | Hcv-rna | IgM | 0.094232 | 6.54E-07 |
| B12 | B14 | PG II | G-17 | 0.201397 | 0 |
| B12 | B80 | PG II | anti-HP IgG | 0.176859 | 0 |
| B12 | B156 | PG II | IgE | 0.151531 | 8.88E-16 |
| B12 | B186 | PG II | PFC | 0.144957 | 1.64E-14 |
| B12 | B93 | PG II | ALDH2-Glu504Lys | 0.127776 | 1.40E-11 |
| B12 | B169 | PG II | U-CRY | 0.110946 | 4.57E-09 |
| B12 | B159 | PG II | β2-MG | 0.110944 | 4.58E-09 |
| B12 | B92 | PG II | ALDH2-Glu504Glu | -0.10821 | 1.09E-08 |
| B12 | B119 | PG II | Hcv-rna | -0.10317 | 5.06E-08 |
| B12 | B37 | PG II | BAND | -0.09958 | 1.45E-07 |
| B12 | B164 | PG II | COM | 0.097804 | 2.42E-07 |
| B12 | B17 | PG II | CD3+CD16+CD56+NK | 0.093999 | 6.97E-07 |
| B120 | B134 | HCV-cAg | CHE | -0.1724 | 0 |
| B120 | B157 | HCV-cAg | IgG | -0.1932 | 0 |
| B120 | B175 | HCV-cAg | ALB-U | 0.336215 | 0 |
| B120 | B177 | HCV-cAg | ACR | 0.867363 | 0 |
| B120 | B178 | HCV-cAg | Ig-U | 0.439385 | 0 |
| B120 | B180 | HCV-cAg | Tf | 0.507883 | 0 |
| B120 | B195 | HCV-cAg | Na | 0.237934 | 0 |
| B120 | B199 | HCV-cAg | Mg | -0.18214 | 0 |
| B120 | B200 | HCV-cAg | OP | 0.226428 | 0 |
| B120 | B137 | HCV-cAg | AFU | 0.155303 | 2.22E-16 |
| B120 | B139 | HCV-cAg | ADA | 0.125962 | 2.71E-11 |
| B120 | B193 | HCV-cAg | K | -0.11619 | 8.20E-10 |
| B120 | B198 | HCV-cAg | P | 0.111555 | 3.76E-09 |
| B120 | B138 | HCV-cAg | PAB | -0.10827 | 1.07E-08 |
| B120 | B196 | HCV-cAg | TCO2 | 0.10744 | 1.38E-08 |

| | | | | | |
|---|---|---|---|---|---|
| B120 | B155 | HCV-cAg | IgA | -0.09658 | 3.42E-07 |
| B120 | B215 | HCV-cAg | BMI | 0.095951 | 4.07E-07 |
| B121 | B122 | C3 | C4 | 0.902188 | 0 |
| B121 | B157 | C3 | IgG | -0.13685 | 4.41E-13 |
| B122 | B157 | C4 | IgG | -0.14393 | 2.53E-14 |
| B123 | B156 | hs-CRP1 | IgE | 0.201231 | 0 |
| B123 | B186 | hs-CRP1 | PFC | 0.15833 | 0 |
| B123 | B125 | hs-CRP1 | Cor | -0.11924 | 2.92E-10 |
| B123 | B126 | hs-CRP1 | Alb | -0.10487 | 3.04E-08 |
| B123 | B164 | hs-CRP1 | COM | 0.096742 | 3.26E-07 |
| B123 | B139 | hs-CRP1 | ADA | 0.094798 | 5.60E-07 |
| B124 | B125 | ACTH | Cor | 0.601852 | 0 |
| B124 | B150 | ACTH | ALP | 0.187194 | 0 |
| B124 | B181 | ACTH | APTT | -0.16444 | 0 |
| B124 | B184 | ACTH | INR | -0.1255 | 3.21E-11 |
| B124 | B193 | ACTH | K | -0.11894 | 3.23E-10 |
| B124 | B185 | ACTH | PT | -0.10009 | 1.26E-07 |
| B124 | B195 | ACTH | Na | -0.09806 | 2.25E-07 |
| B124 | B200 | ACTH | OP | -0.09419 | 6.62E-07 |
| B125 | B156 | Cor | IgE | -0.24359 | 0 |
| B125 | B164 | Cor | COM | -0.17478 | 0 |
| B125 | B179 | Cor | α1-MG | -0.1584 | 0 |
| B125 | B186 | Cor | PFC | -0.19514 | 0 |
| B125 | B139 | Cor | ADA | -0.11249 | 2.78E-09 |
| B125 | B193 | Cor | K | -0.10147 | 8.38E-08 |
| B125 | B194 | Cor | Cl | -0.09356 | 7.86E-07 |
| B126 | B127 | Alb | A-G ratio | 0.51086 | 0 |
| B126 | B130 | Alb | GLB | -0.18659 | 0 |
| B126 | B188 | Alb | Ca | 0.403637 | 0 |
| B126 | B189 | Alb | eGFB | 0.191538 | 0 |
| B126 | B221 | Alb | Hei | 0.157291 | 0 |
| B126 | B128 | Alb | ALT | 0.153194 | 4.44E-16 |
| B126 | B138 | Alb | PAB | 0.145905 | 1.11E-14 |
| B126 | B192 | Alb | UA | 0.14572 | 1.20E-14 |
| B126 | B129 | Alb | IBIL | 0.145636 | 1.24E-14 |
| B126 | B141 | Alb | AST/ALT | -0.14266 | 4.24E-14 |
| B126 | B133 | Alb | TBIL | 0.139646 | 1.45E-13 |
| B126 | B218 | Alb | DBP | 0.129011 | 8.85E-12 |
| B126 | B139 | Alb | ADA | -0.12281 | 8.39E-11 |
| B126 | B134 | Alb | CHE | 0.114602 | 1.39E-09 |
| B126 | B216 | Alb | BMI | 0.101081 | 9.40E-08 |
| B126 | B199 | Alb | Mg | 0.099557 | 1.46E-07 |
| B126 | B191 | Alb | Urea | -0.09664 | 3.36E-07 |
| B126 | B222 | Alb | NS | 0.096194 | 3.80E-07 |

| | | | | | |
|---|---|---|---|---|---|
| B127 | B129 | A-G ratio | IBIL | 0.166519 | 0 |
| B127 | B130 | A-G ratio | GLB | -0.90713 | 0 |
| B127 | B133 | A-G ratio | TBIL | 0.169474 | 0 |
| B127 | B157 | A-G ratio | IgG | -0.18554 | 0 |
| B127 | B221 | A-G ratio | Hei | 0.205107 | 0 |
| B127 | B132 | A-G ratio | Dbil | 0.134535 | 1.09E-12 |
| B127 | B155 | A-G ratio | IgA | -0.12669 | 2.08E-11 |
| B127 | B139 | A-G ratio | ADA | -0.12136 | 1.40E-10 |
| B127 | B222 | A-G ratio | NS | 0.120647 | 1.79E-10 |
| B127 | B216 | A-G ratio | BMI | 0.107072 | 1.55E-08 |
| B127 | B209 | A-G ratio | Smoking | 0.100415 | 1.14E-07 |
| B127 | B163 | A-G ratio | SG | 0.093613 | 7.74E-07 |
| B128 | B131 | ALT | AST | 0.730974 | 0 |
| B128 | B140 | ALT | r-GT | 0.42337 | 0 |
| B128 | B141 | ALT | AST/ALT | -0.5135 | 0 |
| B128 | B163 | ALT | SG | 0.18528 | 0 |
| B128 | B192 | ALT | UA | 0.253042 | 0 |
| B128 | B206 | ALT | Waist | 0.28197 | 0 |
| B128 | B210 | ALT | Hip | 0.216015 | 0 |
| B128 | B215 | ALT | BMI | 0.268013 | 0 |
| B128 | B216 | ALT | BMI | 0.316044 | 0 |
| B128 | B218 | ALT | DBP | 0.182822 | 0 |
| B128 | B221 | ALT | Hei | 0.195109 | 0 |
| B128 | B222 | ALT | NS | 0.274878 | 0 |
| B128 | B137 | ALT | AFU | 0.121964 | 1.13E-10 |
| B128 | B204 | ALT | Drinking | 0.107446 | 1.38E-08 |
| B128 | B166 | ALT | Pro | 0.102453 | 6.27E-08 |
| B128 | B153 | ALT | Glu | 0.095108 | 5.14E-07 |
| B129 | B132 | IBIL | Dbil | 0.664958 | 0 |
| B129 | B133 | IBIL | TBIL | 0.979873 | 0 |
| B129 | B130 | IBIL | GLB | -0.12252 | 9.31E-11 |
| B129 | B168 | IBIL | NMP22 | -0.12055 | 1.85E-10 |
| B129 | B221 | IBIL | Hei | 0.108496 | 9.94E-09 |
| B130 | B157 | GLB | IgG | 0.208044 | 0 |
| B130 | B221 | GLB | Hei | -0.15915 | 0 |
| B130 | B155 | GLB | IgA | 0.139842 | 1.34E-13 |
| B130 | B139 | GLB | ADA | 0.128289 | 1.16E-11 |
| B130 | B133 | GLB | TBIL | -0.12526 | 3.49E-11 |
| B130 | B150 | GLB | ALP | 0.112554 | 2.72E-09 |
| B130 | B209 | GLB | Smoking | -0.10742 | 1.39E-08 |
| B130 | B132 | GLB | Dbil | -0.10117 | 9.16E-08 |
| B131 | B140 | AST | r-GT | 0.393193 | 0 |
| B131 | B192 | AST | UA | 0.142894 | 3.86E-14 |
| B131 | B206 | AST | Waist | 0.129482 | 7.43E-12 |

| | | | | | |
|---|---|---|---|---|---|
| B131 | B218 | AST | DBP | 0.127643 | 1.47E-11 |
| B131 | B216 | AST | BMI | 0.114138 | 1.62E-09 |
| B131 | B215 | AST | BMI | 0.111381 | 3.98E-09 |
| B131 | B132 | AST | Dbil | 0.105061 | 2.87E-08 |
| B131 | B222 | AST | NS | 0.104941 | 2.97E-08 |
| B131 | B204 | AST | Drinking | 0.101169 | 9.16E-08 |
| B131 | B135 | AST | TBA | 0.098405 | 2.04E-07 |
| B131 | B137 | AST | AFU | 0.097973 | 2.30E-07 |
| B131 | B220 | AST | SBP | 0.0964 | 3.59E-07 |
| B132 | B133 | Dbil | TBIL | 0.800566 | 0 |
| B132 | B140 | Dbil | r-GT | 0.152678 | 4.44E-16 |
| B132 | B135 | Dbil | TBA | 0.150129 | 1.78E-15 |
| B132 | B193 | Dbil | K | -0.12035 | 1.98E-10 |
| B132 | B139 | Dbil | ADA | 0.116591 | 7.17E-10 |
| B132 | B221 | Dbil | Hei | 0.113696 | 1.88E-09 |
| B132 | B134 | Dbil | CHE | -0.10681 | 1.68E-08 |
| B132 | B153 | Dbil | Glu | 0.101423 | 8.50E-08 |
| B132 | B204 | Dbil | Drinking | 0.098913 | 1.76E-07 |
| B132 | B155 | Dbil | IgA | 0.098053 | 2.25E-07 |
| B133 | B221 | TBIL | Hei | 0.117403 | 5.45E-10 |
| B133 | B168 | TBIL | NMP22 | -0.10404 | 3.91E-08 |
| B133 | B140 | TBIL | r-GT | 0.095841 | 4.19E-07 |
| B134 | B136 | CHE | 5'NT | 0.163868 | 0 |
| B134 | B138 | CHE | PAB | 0.397837 | 0 |
| B134 | B197 | CHE | AG | 0.155711 | 0 |
| B134 | B199 | CHE | Mg | 0.15247 | 6.66E-16 |
| B134 | B139 | CHE | ADA | -0.15114 | 1.33E-15 |
| B134 | B177 | CHE | ACR | -0.14856 | 3.55E-15 |
| B134 | B194 | CHE | Cl | -0.14404 | 2.40E-14 |
| B134 | B135 | CHE | TBA | -0.12165 | 1.26E-10 |
| B134 | B193 | CHE | K | 0.120511 | 1.88E-10 |
| B134 | B137 | CHE | AFU | 0.111894 | 3.37E-09 |
| B134 | B150 | CHE | ALP | 0.106005 | 2.15E-08 |
| B134 | B157 | CHE | IgG | -0.10011 | 1.25E-07 |
| B134 | B182 | CHE | TT | -0.09613 | 3.87E-07 |
| B135 | B137 | TBA | AFU | 0.21591 | 0 |
| B135 | B139 | TBA | ADA | 0.31002 | 0 |
| B135 | B140 | TBA | r-GT | 0.165622 | 0 |
| B135 | B150 | TBA | ALP | 0.254041 | 0 |
| B135 | B155 | TBA | IgA | 0.302625 | 0 |
| B135 | B157 | TBA | IgG | 0.198537 | 0 |
| B135 | B182 | TBA | TT | 0.280036 | 0 |
| B135 | B184 | TBA | INR | 0.237702 | 0 |
| B135 | B185 | TBA | PT | 0.206418 | 0 |

| | | | | | |
|---|---|---|---|---|---|
| B135 | B193 | TBA | K | -0.258 | 0 |
| B135 | B181 | TBA | APTT | 0.14955 | 2.22E-15 |
| B135 | B187 | TBA | NSE | -0.14602 | 1.07E-14 |
| B135 | B198 | TBA | P | -0.12201 | 1.11E-10 |
| B135 | B138 | TBA | PAB | -0.11012 | 5.94E-09 |
| B135 | B200 | TBA | OP | 0.105958 | 2.18E-08 |
| B135 | B158 | TBA | IgM | 0.105725 | 2.34E-08 |
| B135 | B156 | TBA | IgE | 0.093901 | 7.16E-07 |
| B136 | B139 | 5'NT | ADA | 0.215116 | 0 |
| B136 | B140 | 5'NT | r-GT | 0.242366 | 0 |
| B136 | B150 | 5'NT | ALP | 0.300025 | 0 |
| B136 | B194 | 5'NT | Cl | -0.14303 | 3.64E-14 |
| B136 | B173 | 5'NT | NTT | 0.114241 | 1.57E-09 |
| B136 | B197 | 5'NT | AG | 0.107921 | 1.19E-08 |
| B137 | B139 | AFU | ADA | 0.313989 | 0 |
| B137 | B150 | AFU | ALP | 0.202949 | 0 |
| B137 | B198 | AFU | P | -0.2072 | 0 |
| B137 | B200 | AFU | OP | 0.168542 | 0 |
| B137 | B177 | AFU | ACR | 0.132203 | 2.67E-12 |
| B137 | B140 | AFU | r-GT | 0.128003 | 1.29E-11 |
| B137 | B206 | AFU | Waist | 0.111146 | 4.29E-09 |
| B137 | B146 | AFU | CysC | 0.111003 | 4.49E-09 |
| B137 | B196 | AFU | TCO2 | -0.1053 | 2.67E-08 |
| B137 | B170 | AFU | U-GLU | 0.102426 | 6.32E-08 |
| B137 | B156 | AFU | IgE | 0.095835 | 4.20E-07 |
| B138 | B139 | PAB | ADA | -0.34856 | 0 |
| B138 | B193 | PAB | K | 0.19256 | 0 |
| B138 | B194 | PAB | Cl | -0.21793 | 0 |
| B138 | B197 | PAB | AG | 0.204542 | 0 |
| B138 | B199 | PAB | Mg | 0.292298 | 0 |
| B138 | B192 | PAB | UA | 0.124093 | 5.32E-11 |
| B138 | B222 | PAB | NS | 0.112666 | 2.62E-09 |
| B138 | B157 | PAB | IgG | -0.10816 | 1.10E-08 |
| B138 | B190 | PAB | Cre | 0.105076 | 2.86E-08 |
| B138 | B177 | PAB | ACR | -0.09982 | 1.36E-07 |
| B138 | B187 | PAB | NSE | 0.098517 | 1.97E-07 |
| B138 | B155 | PAB | IgA | -0.0958 | 4.24E-07 |
| B139 | B150 | ADA | ALP | 0.244693 | 0 |
| B139 | B155 | ADA | IgA | 0.208676 | 0 |
| B139 | B156 | ADA | IgE | 0.17116 | 0 |
| B139 | B157 | ADA | IgG | 0.326794 | 0 |
| B139 | B182 | ADA | TT | 0.17578 | 0 |
| B139 | B193 | ADA | K | -0.19419 | 0 |
| B139 | B200 | ADA | OP | 0.248529 | 0 |

| | | | | | |
|---|---|---|---|---|---|
| B139 | B199 | ADA | Mg | -0.14611 | 1.02E-14 |
| B139 | B177 | ADA | ACR | 0.136774 | 4.54E-13 |
| B139 | B153 | ADA | Glu | 0.132671 | 2.23E-12 |
| B139 | B140 | ADA | r-GT | 0.131982 | 2.90E-12 |
| B139 | B197 | ADA | AG | 0.129171 | 8.34E-12 |
| B139 | B170 | ADA | U-GLU | 0.122766 | 8.52E-11 |
| B139 | B187 | ADA | NSE | -0.10606 | 2.12E-08 |
| B14 | B112 | G-17 | FT4 | 0.135602 | 7.19E-13 |
| B14 | B156 | G-17 | IgE | 0.121768 | 1.21E-10 |
| B14 | B113 | G-17 | FT3 | 0.112556 | 2.72E-09 |
| B14 | B148 | G-17 | Anti_TPO | 0.104505 | 3.39E-08 |
| B140 | B141 | r-GT | AST/ALT | -0.19264 | 0 |
| B140 | B150 | r-GT | ALP | 0.169204 | 0 |
| B140 | B192 | r-GT | UA | 0.160968 | 0 |
| B140 | B204 | r-GT | Drinking | 0.2163 | 0 |
| B140 | B206 | r-GT | Waist | 0.199017 | 0 |
| B140 | B216 | r-GT | BMI | 0.19856 | 0 |
| B140 | B222 | r-GT | NS | 0.202377 | 0 |
| B140 | B215 | r-GT | BMI | 0.154669 | 2.22E-16 |
| B140 | B153 | r-GT | Glu | 0.152143 | 8.88E-16 |
| B140 | B218 | r-GT | DBP | 0.147425 | 5.77E-15 |
| B140 | B221 | r-GT | Hei | 0.144745 | 1.80E-14 |
| B140 | B170 | r-GT | U-GLU | 0.133727 | 1.49E-12 |
| B140 | B210 | r-GT | Hip | 0.129514 | 7.34E-12 |
| B140 | B209 | r-GT | Smoking | 0.121043 | 1.56E-10 |
| B140 | B163 | r-GT | SG | 0.115498 | 1.03E-09 |
| B140 | B166 | r-GT | Pro | 0.11434 | 1.52E-09 |
| B140 | B220 | r-GT | SBP | 0.10574 | 2.33E-08 |
| B140 | B155 | r-GT | IgA | 0.104659 | 3.24E-08 |
| B141 | B163 | AST/ALT | SG | -0.19186 | 0 |
| B141 | B192 | AST/ALT | UA | -0.20891 | 0 |
| B141 | B206 | AST/ALT | Waist | -0.25722 | 0 |
| B141 | B210 | AST/ALT | Hip | -0.2065 | 0 |
| B141 | B215 | AST/ALT | BMI | -0.23907 | 0 |
| B141 | B216 | AST/ALT | BMI | -0.30964 | 0 |
| B141 | B221 | AST/ALT | Hei | -0.23477 | 0 |
| B141 | B222 | AST/ALT | NS | -0.26736 | 0 |
| B141 | B218 | AST/ALT | DBP | -0.13808 | 2.70E-13 |
| B141 | B153 | AST/ALT | Glu | -0.12014 | 2.13E-10 |
| B141 | B224 | AST/ALT | pmh | 0.10199 | 7.20E-08 |
| B141 | B209 | AST/ALT | Smoking | -0.09325 | 8.53E-07 |
| B142 | B144 | 25-(OH)D | Total-PINP | -0.20949 | 0 |
| B142 | B145 | 25-(OH)D | BGP | 0.138667 | 2.14E-13 |
| B143 | B144 | β-CTx | Total-PINP | 0.755536 | 0 |

| | | | | | |
|---|---|---|---|---|---|
| B143 | B145 | β-CTx | BGP | 0.533108 | 0 |
| B143 | B151 | β-CTx | Anti-CCP | 0.200337 | 0 |
| B143 | B176 | β-CTx | Ucr | -0.0935 | 7.99E-07 |
| B144 | B145 | Total-PINP | BGP | 0.541227 | 0 |
| B144 | B151 | Total-PINP | Anti-CCP | 0.265107 | 0 |
| B144 | B154 | Total-PINP | RF | 0.09907 | 1.68E-07 |
| B145 | B183 | BGP | PTA | -0.14184 | 5.95E-14 |
| B145 | B176 | BGP | Ucr | -0.09293 | 9.31E-07 |
| B146 | B189 | CysC | eGFB | -0.28361 | 0 |
| B146 | B190 | CysC | Cre | 0.571458 | 0 |
| B146 | B191 | CysC | Urea | 0.272976 | 0 |
| B146 | B199 | CysC | Mg | 0.198519 | 0 |
| B146 | B200 | CysC | OP | 0.164142 | 0 |
| B146 | B193 | CysC | K | 0.140561 | 1.00E-13 |
| B146 | B197 | CysC | AG | 0.132843 | 2.09E-12 |
| B146 | B194 | CysC | Cl | -0.12598 | 2.70E-11 |
| B146 | B196 | CysC | TCO2 | -0.12191 | 1.15E-10 |
| B146 | B198 | CysC | P | 0.099729 | 1.39E-07 |
| B146 | B174 | CysC | Urea/Cr | -0.09902 | 1.71E-07 |
| B146 | B195 | CysC | Na | -0.09511 | 5.14E-07 |
| B148 | B149 | Anti_TPO | TGAb | 0.528489 | 0 |
| B15 | B101 | CD3-CD19+(B cell) | anti-Sc70 | 0.45898 | 0 |
| B15 | B102 | CD3-CD19+(B cell) | anti-Sm | 0.45898 | 0 |
| B15 | B104 | CD3-CD19+(B cell) | anti-SSB | 0.45898 | 0 |
| B15 | B109 | CD3-CD19+(B cell) | AMA-M2 | -0.20418 | 0 |
| B15 | B110 | CD3-CD19+(B cell) | AHA | -0.18835 | 0 |
| B15 | B16 | CD3-CD19+(B cell) | CD3+T | -0.46513 | 0 |
| B15 | B20 | CD3-CD19+(B cell) | CD3+CD4+CD8+ | -0.29004 | 0 |
| B15 | B22 | CD3-CD19+(B cell) | CD3+CD8+(T) | -0.33816 | 0 |
| B15 | B25 | CD3-CD19+(B cell) | CK | -0.19616 | 0 |
| B15 | B30 | CD3-CD19+(B cell) | LH | 0.171419 | 0 |
| B15 | B32 | CD3-CD19+(B cell) | Testo | -0.33841 | 0 |
| B15 | B33 | CD3-CD19+(B cell) | Prog | 0.182338 | 0 |
| B15 | B79 | CD3-CD19+(B cell) | hb V -dna | 0.164699 | 0 |
| B15 | B98 | CD3-CD19+(B cell) | anti-Jo-1 | 0.45898 | 0 |
| B15 | B99 | CD3-CD19+(B cell) | anti-PM-Scl | 0.45898 | 0 |
| B15 | B186 | CD3-CD19+(B cell) | PFC | 0.155157 | 2.22E-16 |
| B15 | B95 | CD3-CD19+(B cell) | anti-cenpb | -0.13116 | 3.96E-12 |
| B15 | B182 | CD3-CD19+(B cell) | TT | 0.126652 | 2.11E-11 |
| B15 | B62 | CD3-CD19+(B cell) | WBHSRVI | -0.12348 | 6.61E-11 |
| B15 | B57 | CD3-CD19+(B cell) | RBCAI | 0.112236 | 3.02E-09 |
| B15 | B31 | CD3-CD19+(B cell) | FSH | 0.109334 | 7.64E-09 |
| B15 | B18 | CD3-CD19+(B cell) | CD3+CD4+helper(inducer)T | -0.1079 | 1.20E-08 |
| B15 | B187 | CD3-CD19+(B cell) | NSE | -0.1027 | 5.83E-08 |

| | | | | | |
|---|---|---|---|---|---|
| B15 | B37 | CD3-CD19+(B cell) | BAND | -0.10027 | 1.19E-07 |
| B15 | B56 | CD3-CD19+(B cell) | IR | -0.09761 | 2.55E-07 |
| B15 | B23 | CD3-CD19+(B cell) | CD3-CD16+CD56+(NK) | 0.095688 | 4.38E-07 |
| B150 | B200 | ALP | OP | 0.151015 | 1.33E-15 |
| B150 | B155 | ALP | IgA | 0.110167 | 5.86E-09 |
| B150 | B157 | ALP | IgG | 0.101562 | 8.16E-08 |
| B150 | B187 | ALP | NSE | -0.09894 | 1.75E-07 |
| B150 | B193 | ALP | K | -0.09886 | 1.79E-07 |
| B151 | B154 | Anti-CCP | RF | 0.370182 | 0 |
| B153 | B163 | Glu | SG | 0.239155 | 0 |
| B153 | B170 | Glu | U-GLU | 0.653165 | 0 |
| B153 | B206 | Glu | Waist | 0.209923 | 0 |
| B153 | B222 | Glu | NS | 0.185968 | 0 |
| B153 | B215 | Glu | BMI | 0.147473 | 5.77E-15 |
| B153 | B220 | Glu | SBP | 0.147011 | 6.88E-15 |
| B153 | B216 | Glu | BMI | 0.138997 | 1.88E-13 |
| B153 | B160 | Glu | PH | -0.1268 | 2.00E-11 |
| B153 | B209 | Glu | Smoking | 0.12305 | 7.71E-11 |
| B153 | B172 | Glu | KB | 0.120242 | 2.06E-10 |
| B153 | B174 | Glu | Urea/Cr | 0.117092 | 6.06E-10 |
| B153 | B200 | Glu | OP | 0.113972 | 1.71E-09 |
| B153 | B214 | Glu | CVA-L | 0.111928 | 3.33E-09 |
| B153 | B218 | Glu | DBP | 0.105541 | 2.48E-08 |
| B153 | B166 | Glu | Pro | 0.099573 | 1.46E-07 |
| B153 | B191 | Glu | Urea | 0.093434 | 8.13E-07 |
| B154 | B228 | RF | OS | 0.289754 | 0 |
| B155 | B157 | IgA | IgG | 0.342222 | 0 |
| B155 | B182 | IgA | TT | 0.303975 | 0 |
| B155 | B193 | IgA | K | -0.16457 | 0 |
| B155 | B158 | IgA | IgM | 0.153924 | 4.44E-16 |
| B155 | B185 | IgA | PT | 0.127312 | 1.66E-11 |
| B155 | B184 | IgA | INR | 0.120419 | 1.94E-10 |
| B155 | B181 | IgA | APTT | 0.104384 | 3.52E-08 |
| B155 | B186 | IgA | PFC | -0.09686 | 3.15E-07 |
| B156 | B164 | IgE | COM | 0.271976 | 0 |
| B156 | B179 | IgE | α1-MG | 0.237238 | 0 |
| B156 | B182 | IgE | TT | 0.165587 | 0 |
| B156 | B186 | IgE | PFC | 0.456813 | 0 |
| B156 | B199 | IgE | Mg | -0.12934 | 7.84E-12 |
| B157 | B177 | IgG | ACR | -0.15259 | 6.66E-16 |
| B157 | B182 | IgG | TT | 0.145929 | 1.09E-14 |
| B157 | B195 | IgG | Na | -0.13092 | 4.34E-12 |
| B157 | B200 | IgG | OP | -0.10565 | 2.40E-08 |
| B157 | B186 | IgG | PFC | -0.09983 | 1.35E-07 |

| | | | | | |
|---|---|---|---|---|---|
| B158 | B199 | IgM | Mg | -0.10139 | 8.58E-08 |
| B159 | B169 | β2-MG | U-CRY | 0.999992 | 0 |
| B159 | B186 | β2-MG | PFC | 0.108465 | 1.00E-08 |
| B159 | B194 | β2-MG | Cl | -0.10168 | 7.88E-08 |
| B159 | B197 | β2-MG | AG | 0.100777 | 1.03E-07 |
| B16 | B109 | CD3+T | AMA-M2 | 0.232433 | 0 |
| B16 | B18 | CD3+T | CD3+CD4+helper(inducer)T | 0.433812 | 0 |
| B16 | B182 | CD3+T | TT | 0.170294 | 0 |
| B16 | B186 | CD3+T | PFC | -0.21541 | 0 |
| B16 | B187 | CD3+T | NSE | 0.172035 | 0 |
| B16 | B20 | CD3+T | CD3+CD4+CD8+ | 0.351775 | 0 |
| B16 | B22 | CD3+T | CD3+CD8+(T) | 0.60625 | 0 |
| B16 | B23 | CD3+T | CD3-CD16+CD56+(NK) | -0.85598 | 0 |
| B16 | B25 | CD3+T | CK | 0.230434 | 0 |
| B16 | B95 | CD3+T | anti-cenpb | 0.153911 | 4.44E-16 |
| B16 | B17 | CD3+T | CD3+CD16+CD56+NK | 0.14977 | 2.22E-15 |
| B16 | B110 | CD3+T | AHA | 0.130988 | 4.22E-12 |
| B16 | B101 | CD3+T | anti-Sc70 | -0.1238 | 5.91E-11 |
| B16 | B102 | CD3+T | anti-Sm | -0.1238 | 5.91E-11 |
| B16 | B104 | CD3+T | anti-SSB | -0.1238 | 5.91E-11 |
| B16 | B98 | CD3+T | anti-Jo-1 | -0.1238 | 5.91E-11 |
| B16 | B99 | CD3+T | anti-PM-Scl | -0.1238 | 5.91E-11 |
| B16 | B93 | CD3+T | ALDH2-Glu504Lys | -0.12074 | 1.74E-10 |
| B16 | B19 | CD3+T | CD3+CD4+/CD3+CD8+ | -0.10878 | 9.08E-09 |
| B16 | B155 | CD3+T | IgA | 0.100417 | 1.14E-07 |
| B16 | B157 | CD3+T | IgG | 0.097628 | 2.54E-07 |
| B16 | B138 | CD3+T | PAB | 0.093508 | 7.96E-07 |
| B160 | B163 | PH | SG | -0.15636 | 0 |
| B160 | B206 | PH | Waist | -0.14337 | 3.15E-14 |
| B160 | B192 | PH | UA | -0.13963 | 1.46E-13 |
| B160 | B215 | PH | BMI | -0.13648 | 5.09E-13 |
| B160 | B191 | PH | Urea | -0.12419 | 5.14E-11 |
| B160 | B216 | PH | BMI | -0.112 | 3.26E-09 |
| B160 | B222 | PH | NS | -0.11179 | 3.49E-09 |
| B160 | B210 | PH | Hip | -0.09745 | 2.67E-07 |
| B160 | B209 | PH | Smoking | -0.09707 | 2.97E-07 |
| B161 | B221 | LEU | Hei | -0.12865 | 1.01E-11 |
| B161 | B222 | LEU | NS | -0.10706 | 1.55E-08 |
| B161 | B173 | LEU | NTT | 0.101022 | 9.56E-08 |
| B163 | B166 | SG | Pro | 0.346249 | 0 |
| B163 | B170 | SG | U-GLU | 0.221686 | 0 |
| B163 | B176 | SG | Ucr | 0.302347 | 0 |
| B163 | B191 | SG | Urea | 0.239254 | 0 |
| B163 | B192 | SG | UA | 0.221765 | 0 |

| | | | | | |
|---|---|---|---|---|---|
| B163 | B204 | SG | Drinking | 0.176847 | 0 |
| B163 | B206 | SG | Waist | 0.271258 | 0 |
| B163 | B209 | SG | Smoking | 0.215343 | 0 |
| B163 | B210 | SG | Hip | 0.201094 | 0 |
| B163 | B215 | SG | BMI | 0.207139 | 0 |
| B163 | B216 | SG | BMI | 0.322372 | 0 |
| B163 | B221 | SG | Hei | 0.298883 | 0 |
| B163 | B222 | SG | NS | 0.370199 | 0 |
| B163 | B172 | SG | KB | 0.128587 | 1.04E-11 |
| B163 | B218 | SG | DBP | 0.126026 | 2.65E-11 |
| B163 | B190 | SG | Cre | 0.114334 | 1.52E-09 |
| B164 | B186 | COM | PFC | 0.208454 | 0 |
| B164 | B179 | COM | α1-MG | 0.111689 | 3.60E-09 |
| B166 | B176 | Pro | Ucr | 0.161678 | 0 |
| B166 | B222 | Pro | NS | 0.166094 | 0 |
| B166 | B216 | Pro | BMI | 0.152179 | 8.88E-16 |
| B166 | B221 | Pro | Hei | 0.122915 | 8.09E-11 |
| B166 | B206 | Pro | Waist | 0.120408 | 1.95E-10 |
| B166 | B215 | Pro | BMI | 0.111305 | 4.07E-09 |
| B166 | B192 | Pro | UA | 0.105128 | 2.81E-08 |
| B166 | B218 | Pro | DBP | 0.102381 | 6.41E-08 |
| B166 | B170 | Pro | U-GLU | 0.100579 | 1.09E-07 |
| B166 | B172 | Pro | KB | 0.096477 | 3.51E-07 |
| B168 | B183 | NMP22 | PTA | -0.09303 | 9.06E-07 |
| B169 | B186 | U-CRY | PFC | 0.108467 | 1.00E-08 |
| B169 | B194 | U-CRY | Cl | -0.10168 | 7.88E-08 |
| B169 | B197 | U-CRY | AG | 0.100779 | 1.03E-07 |
| B17 | B100 | CD3+CD16+CD56+NK | anti-Ro52 | -0.2728 | 0 |
| B17 | B110 | CD3+CD16+CD56+NK | AHA | -0.27545 | 0 |
| B17 | B19 | CD3+CD16+CD56+NK | CD3+CD4+/CD3+CD8+ | -0.29924 | 0 |
| B17 | B20 | CD3+CD16+CD56+NK | CD3+CD4+CD8+ | 0.38801 | 0 |
| B17 | B22 | CD3+CD16+CD56+NK | CD3+CD8+(T) | 0.273018 | 0 |
| B17 | B95 | CD3+CD16+CD56+NK | anti-cenpb | -0.37988 | 0 |
| B17 | B23 | CD3+CD16+CD56+NK | CD3-CD16+CD56+(NK) | -0.14203 | 5.51E-14 |
| B17 | B18 | CD3+CD16+CD56+NK | CD3+CD4+helper(inducer)T | -0.11676 | 6.77E-10 |
| B17 | B187 | CD3+CD16+CD56+NK | NSE | 0.099028 | 1.70E-07 |
| B17 | B73 | CD3+CD16+CD56+NK | ApoA | -0.0954 | 4.74E-07 |
| B170 | B172 | U-GLU | KB | 0.128443 | 1.09E-11 |
| B170 | B222 | U-GLU | NS | 0.128207 | 1.19E-11 |
| B170 | B200 | U-GLU | OP | 0.121422 | 1.37E-10 |
| B170 | B206 | U-GLU | Waist | 0.12106 | 1.55E-10 |
| B170 | B209 | U-GLU | Smoking | 0.119825 | 2.38E-10 |
| B170 | B216 | U-GLU | BMI | 0.110658 | 5.01E-09 |
| B170 | B214 | U-GLU | CVA-L | 0.104526 | 3.37E-08 |

| | | | | | |
|---|---|---|---|---|---|
| B170 | B194 | U-GLU | Cl | -0.09728 | 2.81E-07 |
| B170 | B174 | U-GLU | Urea/Cr | 0.09699 | 3.04E-07 |
| B173 | B213 | NTT | CVA-R | 0.133568 | 1.58E-12 |
| B174 | B189 | Urea/Cr | eGFB | 0.185543 | 0 |
| B174 | B190 | Urea/Cr | Cre | -0.29854 | 0 |
| B174 | B191 | Urea/Cr | Urea | 0.536679 | 0 |
| B174 | B192 | Urea/Cr | UA | -0.2192 | 0 |
| B174 | B216 | Urea/Cr | BMI | -0.21367 | 0 |
| B174 | B221 | Urea/Cr | Hei | -0.31133 | 0 |
| B174 | B222 | Urea/Cr | NS | -0.23527 | 0 |
| B174 | B210 | Urea/Cr | Hip | -0.13498 | 9.16E-13 |
| B174 | B206 | Urea/Cr | Waist | -0.10941 | 7.46E-09 |
| B174 | B200 | Urea/Cr | OP | 0.107825 | 1.23E-08 |
| B174 | B209 | Urea/Cr | Smoking | -0.10343 | 4.69E-08 |
| B175 | B177 | ALB-U | ACR | 0.699995 | 0 |
| B175 | B178 | ALB-U | Ig-U | 0.187014 | 0 |
| B175 | B180 | ALB-U | Tf | 0.182201 | 0 |
| B175 | B200 | ALB-U | OP | 0.095538 | 4.56E-07 |
| B175 | B199 | ALB-U | Mg | -0.09428 | 6.45E-07 |
| B176 | B221 | Ucr | Hei | 0.129754 | 6.71E-12 |
| B176 | B222 | Ucr | NS | 0.116837 | 6.60E-10 |
| B176 | B216 | Ucr | BMI | 0.116046 | 8.61E-10 |
| B177 | B178 | ACR | Ig-U | 0.531298 | 0 |
| B177 | B180 | ACR | Tf | 0.597639 | 0 |
| B177 | B195 | ACR | Na | 0.210991 | 0 |
| B177 | B199 | ACR | Mg | -0.17361 | 0 |
| B177 | B200 | ACR | OP | 0.201256 | 0 |
| B177 | B193 | ACR | K | -0.11918 | 2.98E-10 |
| B177 | B215 | ACR | BMI | 0.09979 | 1.37E-07 |
| B178 | B179 | Ig-U | α1-MG | 0.626289 | 0 |
| B178 | B180 | Ig-U | Tf | 0.867376 | 0 |
| B178 | B191 | Ig-U | Urea | 0.182838 | 0 |
| B178 | B200 | Ig-U | OP | 0.101079 | 9.40E-08 |
| B178 | B189 | Ig-U | eGFB | -0.09304 | 9.03E-07 |
| B179 | B180 | α1-MG | Tf | 0.241618 | 0 |
| B179 | B186 | α1-MG | PFC | 0.219459 | 0 |
| B179 | B191 | α1-MG | Urea | 0.228705 | 0 |
| B179 | B190 | α1-MG | Cre | 0.118491 | 3.77E-10 |
| B18 | B100 | CD3+CD4+helper(inducer)T | anti-Ro52 | 0.186491 | 0 |
| B18 | B109 | CD3+CD4+helper(inducer)T | AMA-M2 | 0.304354 | 0 |
| B18 | B110 | CD3+CD4+helper(inducer)T | AHA | 0.172064 | 0 |
| B18 | B181 | CD3+CD4+helper(inducer)T | APTT | -0.17459 | 0 |
| B18 | B182 | CD3+CD4+helper(inducer)T | TT | 0.17846 | 0 |
| B18 | B19 | CD3+CD4+helper(inducer)T | CD3+CD4+/CD3+CD8+ | 0.759247 | 0 |

| | | | | | |
|---|---|---|---|---|---|
| B18 | B20 | CD3+CD4+helper(inducer)T | CD3+CD4+CD8+ | 0.195508 | 0 |
| B18 | B21 | CD3+CD4+helper(inducer)T | CD3+CD4-CD8- | -0.35487 | 0 |
| B18 | B22 | CD3+CD4+helper(inducer)T | CD3+CD8+(T) | -0.39928 | 0 |
| B18 | B23 | CD3+CD4+helper(inducer)T | CD3-CD16+CD56+(NK) | -0.39998 | 0 |
| B18 | B25 | CD3+CD4+helper(inducer)T | CK | -0.41618 | 0 |
| B18 | B95 | CD3+CD4+helper(inducer)T | anti-cenpb | 0.266685 | 0 |
| B18 | B26 | CD3+CD4+helper(inducer)T | CKMB | -0.15074 | 1.33E-15 |
| B18 | B92 | CD3+CD4+helper(inducer)T | ALDH2-Glu504Glu | 0.15001 | 1.78E-15 |
| B18 | B37 | CD3+CD4+helper(inducer)T | BAND | -0.14236 | 4.80E-14 |
| B18 | B34 | CD3+CD4+helper(inducer)T | TK | 0.132423 | 2.45E-12 |
| B18 | B38 | CD3+CD4+helper(inducer)T | BAND% | -0.13087 | 4.41E-12 |
| B18 | B183 | CD3+CD4+helper(inducer)T | PTA | 0.127566 | 1.51E-11 |
| B18 | B186 | CD3+CD4+helper(inducer)T | PFC | -0.11967 | 2.51E-10 |
| B18 | B94 | CD3+CD4+helper(inducer)T | ALDH2-Lys504Lys | -0.11416 | 1.61E-09 |
| B18 | B157 | CD3+CD4+helper(inducer)T | IgG | -0.10177 | 7.69E-08 |
| B18 | B93 | CD3+CD4+helper(inducer)T | ALDH2-Glu504Lys | -0.09831 | 2.09E-07 |
| B18 | B138 | CD3+CD4+helper(inducer)T | PAB | 0.098001 | 2.28E-07 |
| B180 | B200 | Tf | OP | 0.117864 | 4.66E-10 |
| B180 | B195 | Tf | Na | 0.113059 | 2.31E-09 |
| B180 | B199 | Tf | Mg | -0.09818 | 2.17E-07 |
| B181 | B183 | APTT | PTA | -0.2939 | 0 |
| B181 | B184 | APTT | INR | 0.576674 | 0 |
| B181 | B185 | APTT | PT | 0.57069 | 0 |
| B181 | B186 | APTT | PFC | 0.093726 | 7.51E-07 |
| B182 | B183 | TT | PTA | -0.35109 | 0 |
| B182 | B193 | TT | K | -0.10405 | 3.89E-08 |
| B183 | B185 | PTA | PT | -0.14687 | 7.33E-15 |
| B183 | B184 | PTA | INR | -0.11877 | 3.43E-10 |
| B184 | B185 | INR | PT | 0.972359 | 0 |
| B184 | B186 | INR | PFC | -0.09474 | 5.69E-07 |
| B186 | B187 | PFC | NSE | -0.10969 | 6.82E-09 |
| B187 | B198 | NSE | P | 0.098453 | 2.01E-07 |
| B188 | B192 | Ca | UA | 0.11903 | 3.13E-10 |
| B188 | B193 | Ca | K | 0.11318 | 2.22E-09 |
| B189 | B190 | eGFB | Cre | -0.4983 | 0 |
| B189 | B191 | eGFB | Urea | -0.38868 | 0 |
| B189 | B220 | eGFB | SBP | -0.24669 | 0 |
| B189 | B206 | eGFB | Waist | -0.14165 | 6.44E-14 |
| B189 | B192 | eGFB | UA | -0.12677 | 2.02E-11 |
| B189 | B203 | eGFB | DH | 0.094343 | 6.34E-07 |
| B19 | B187 | CD3+CD4+/CD3+CD8+ | NSE | -0.15728 | 0 |
| B19 | B21 | CD3+CD4+/CD3+CD8+ | CD3+CD4-CD8- | -0.21537 | 0 |
| B19 | B22 | CD3+CD4+/CD3+CD8+ | CD3+CD8+(T) | -0.76931 | 0 |
| B19 | B25 | CD3+CD4+/CD3+CD8+ | CK | -0.31657 | 0 |

| | | | | | |
|---|---|---|---|---|---|
| B19 | B37 | CD3+CD4+/CD3+CD8+ | BAND | -0.18474 | 0 |
| B19 | B38 | CD3+CD4+/CD3+CD8+ | BAND% | -0.16983 | 0 |
| B19 | B92 | CD3+CD4+/CD3+CD8+ | ALDH2-Glu504Glu | 0.167766 | 0 |
| B19 | B94 | CD3+CD4+/CD3+CD8+ | ALDH2-Lys504Lys | -0.16784 | 0 |
| B19 | B95 | CD3+CD4+/CD3+CD8+ | anti-cenpb | 0.160278 | 0 |
| B19 | B183 | CD3+CD4+/CD3+CD8+ | PTA | 0.150984 | 1.33E-15 |
| B19 | B182 | CD3+CD4+/CD3+CD8+ | TT | 0.147491 | 5.77E-15 |
| B19 | B20 | CD3+CD4+/CD3+CD8+ | CD3+CD4+CD8+ | -0.14251 | 4.53E-14 |
| B19 | B34 | CD3+CD4+/CD3+CD8+ | TK | 0.132211 | 2.66E-12 |
| B19 | B26 | CD3+CD4+/CD3+CD8+ | CKMB | -0.12946 | 7.49E-12 |
| B19 | B181 | CD3+CD4+/CD3+CD8+ | APTT | -0.11727 | 5.70E-10 |
| B19 | B100 | CD3+CD4+/CD3+CD8+ | anti-Ro52 | 0.114599 | 1.39E-09 |
| B19 | B109 | CD3+CD4+/CD3+CD8+ | AMA-M2 | 0.108468 | 1.00E-08 |
| B19 | B124 | CD3+CD4+/CD3+CD8+ | ACTH | -0.10495 | 2.97E-08 |
| B19 | B157 | CD3+CD4+/CD3+CD8+ | IgG | -0.10392 | 4.05E-08 |
| B19 | B197 | CD3+CD4+/CD3+CD8+ | AG | 0.093405 | 8.19E-07 |
| B190 | B191 | Cre | Urea | 0.471002 | 0 |
| B190 | B192 | Cre | UA | 0.213767 | 0 |
| B190 | B193 | Cre | K | 0.158741 | 0 |
| B190 | B199 | Cre | Mg | 0.205102 | 0 |
| B190 | B222 | Cre | NS | 0.198671 | 0 |
| B190 | B221 | Cre | Hei | 0.149381 | 2.66E-15 |
| B190 | B206 | Cre | Waist | 0.14328 | 3.29E-14 |
| B190 | B216 | Cre | BMI | 0.133801 | 1.45E-12 |
| B190 | B194 | Cre | Cl | -0.12219 | 1.04E-10 |
| B190 | B200 | Cre | OP | 0.116187 | 8.21E-10 |
| B190 | B210 | Cre | Hip | 0.100241 | 1.20E-07 |
| B190 | B197 | Cre | AG | 0.099127 | 1.66E-07 |
| B190 | B195 | Cre | Na | -0.09887 | 1.78E-07 |
| B191 | B192 | Urea | UA | 0.166388 | 0 |
| B191 | B200 | Urea | OP | 0.214624 | 0 |
| B191 | B206 | Urea | Waist | 0.135772 | 6.73E-13 |
| B191 | B199 | Urea | Mg | 0.13487 | 9.56E-13 |
| B191 | B222 | Urea | NS | 0.121299 | 1.43E-10 |
| B191 | B197 | Urea | AG | 0.096824 | 3.19E-07 |
| B191 | B220 | Urea | SBP | 0.095939 | 4.08E-07 |
| B191 | B193 | Urea | K | 0.094501 | 6.07E-07 |
| B192 | B204 | UA | Drinking | 0.18817 | 0 |
| B192 | B206 | UA | Waist | 0.412521 | 0 |
| B192 | B210 | UA | Hip | 0.321961 | 0 |
| B192 | B215 | UA | BMI | 0.352994 | 0 |
| B192 | B216 | UA | BMI | 0.460991 | 0 |
| B192 | B218 | UA | DBP | 0.224235 | 0 |
| B192 | B220 | UA | SBP | 0.177214 | 0 |

| | | | | | |
|---|---|---|---|---|---|
| B192 | B221 | UA | Hei | 0.350204 | 0 |
| B192 | B222 | UA | NS | 0.479604 | 0 |
| B192 | B209 | UA | Smoking | 0.155245 | 2.22E-16 |
| B193 | B199 | K | Mg | 0.204594 | 0 |
| B193 | B198 | K | P | 0.122249 | 1.02E-10 |
| B194 | B195 | Cl | Na | 0.330048 | 0 |
| B194 | B197 | Cl | AG | -0.4321 | 0 |
| B194 | B199 | Cl | Mg | -0.15023 | 1.78E-15 |
| B195 | B197 | Na | AG | 0.238568 | 0 |
| B195 | B200 | Na | OP | 0.677972 | 0 |
| B195 | B198 | Na | P | 0.101117 | 9.30E-08 |
| B196 | B197 | TCO2 | AG | -0.72432 | 0 |
| B197 | B200 | AG | OP | 0.354244 | 0 |
| B197 | B199 | AG | Mg | 0.154496 | 2.22E-16 |
| B199 | B200 | Mg | OP | 0.152653 | 4.44E-16 |
| B2 | B126 | Age | Alb | -0.29389 | 0 |
| B2 | B153 | Age | Glu | 0.167532 | 0 |
| B2 | B189 | Age | eGFB | -0.5729 | 0 |
| B2 | B191 | Age | Urea | 0.259294 | 0 |
| B2 | B206 | Age | Waist | 0.216679 | 0 |
| B2 | B220 | Age | SBP | 0.401095 | 0 |
| B2 | B221 | Age | Hei | -0.22275 | 0 |
| B2 | B43 | Age | MCV | 0.201638 | 0 |
| B2 | B46 | Age | RBC Count | -0.17581 | 0 |
| B2 | B47 | Age | RDW | 0.251483 | 0 |
| B2 | B52 | Age | PLT/BPC | -0.18169 | 0 |
| B2 | B91 | Age | TP | -0.15545 | 2.22E-16 |
| B2 | B201 | Age | e-habits | 0.149409 | 2.44E-15 |
| B2 | B41 | Age | MON % | 0.147077 | 6.66E-15 |
| B2 | B44 | Age | MCH | 0.146948 | 7.11E-15 |
| B2 | B203 | Age | DH | -0.14377 | 2.71E-14 |
| B2 | B127 | Age | A-G ratio | -0.13156 | 3.40E-12 |
| B2 | B141 | Age | AST/ALT | 0.129363 | 7.77E-12 |
| B2 | B77 | Age | TC | 0.128382 | 1.12E-11 |
| B2 | B190 | Age | Cre | 0.113554 | 1.96E-09 |
| B2 | B219 | Age | OT | 0.109907 | 6.37E-09 |
| B2 | B160 | Age | PH | -0.10639 | 1.91E-08 |
| B2 | B176 | Age | Ucr | -0.09786 | 2.38E-07 |
| B2 | B128 | Age | ALT | -0.0975 | 2.63E-07 |
| B2 | B70 | Age | LDL-C | 0.095289 | 4.89E-07 |
| B2 | B27 | Age | LDH | 0.094996 | 5.30E-07 |
| B20 | B109 | CD3+CD4+CD8+ | AMA-M2 | 0.202697 | 0 |
| B20 | B21 | CD3+CD4+CD8+ | CD3+CD4-CD8- | -0.30634 | 0 |
| B20 | B22 | CD3+CD4+CD8+ | CD3+CD8+(T) | 0.335363 | 0 |

| | | | | | |
|---|---|---|---|---|---|
| B20 | B23 | CD3+CD4+CD8+ | CD3-CD16+CD56+(NK) | -0.26886 | 0 |
| B20 | B101 | CD3+CD4+CD8+ | anti-Sc70 | -0.14958 | 2.22E-15 |
| B20 | B102 | CD3+CD4+CD8+ | anti-Sm | -0.14958 | 2.22E-15 |
| B20 | B104 | CD3+CD4+CD8+ | anti-SSB | -0.14958 | 2.22E-15 |
| B20 | B98 | CD3+CD4+CD8+ | anti-Jo-1 | -0.14958 | 2.22E-15 |
| B20 | B99 | CD3+CD4+CD8+ | anti-PM-Scl | -0.14958 | 2.22E-15 |
| B20 | B187 | CD3+CD4+CD8+ | NSE | 0.149175 | 2.66E-15 |
| B20 | B186 | CD3+CD4+CD8+ | PFC | -0.13625 | 5.58E-13 |
| B20 | B25 | CD3+CD4+CD8+ | CK | -0.11006 | 6.06E-09 |
| B20 | B92 | CD3+CD4+CD8+ | ALDH2-Glu504Glu | 0.109172 | 8.04E-09 |
| B20 | B110 | CD3+CD4+CD8+ | AHA | -0.09779 | 2.43E-07 |
| B20 | B138 | CD3+CD4+CD8+ | PAB | 0.092935 | 9.30E-07 |
| B202 | B223 | e-times | FH | 0.638771 | 0 |
| B204 | B206 | Drinking | Waist | 0.221128 | 0 |
| B204 | B209 | Drinking | Smoking | 0.375612 | 0 |
| B204 | B210 | Drinking | Hip | 0.155642 | 0 |
| B204 | B215 | Drinking | BMI | 0.157058 | 0 |
| B204 | B216 | Drinking | BMI | 0.247284 | 0 |
| B204 | B218 | Drinking | DBP | 0.164406 | 0 |
| B204 | B221 | Drinking | Hei | 0.230209 | 0 |
| B204 | B222 | Drinking | NS | 0.278683 | 0 |
| B204 | B220 | Drinking | SBP | 0.099883 | 1.33E-07 |
| B206 | B209 | Waist | Smoking | 0.190562 | 0 |
| B206 | B210 | Waist | Hip | 0.761582 | 0 |
| B206 | B215 | Waist | BMI | 0.839688 | 0 |
| B206 | B216 | Waist | BMI | 0.831692 | 0 |
| B206 | B218 | Waist | DBP | 0.306469 | 0 |
| B206 | B220 | Waist | SBP | 0.310831 | 0 |
| B206 | B221 | Waist | Hei | 0.349561 | 0 |
| B206 | B222 | Waist | NS | 0.776635 | 0 |
| B209 | B216 | Smoking | BMI | 0.198068 | 0 |
| B209 | B221 | Smoking | Hei | 0.25083 | 0 |
| B209 | B222 | Smoking | NS | 0.265994 | 0 |
| B209 | B218 | Smoking | DBP | 0.122058 | 1.09E-10 |
| B209 | B210 | Smoking | Hip | 0.099016 | 1.71E-07 |
| B21 | B109 | CD3+CD4-CD8- | AMA-M2 | -0.21058 | 0 |
| B21 | B125 | CD3+CD4-CD8- | Cor | 0.219055 | 0 |
| B21 | B181 | CD3+CD4-CD8- | APTT | 0.263341 | 0 |
| B21 | B32 | CD3+CD4-CD8- | Testo | -0.18386 | 0 |
| B21 | B95 | CD3+CD4-CD8- | anti-cenpb | -0.16188 | 0 |
| B21 | B169 | CD3+CD4-CD8- | U-CRY | 0.136663 | 4.74E-13 |
| B21 | B159 | CD3+CD4-CD8- | β2-MG | 0.136661 | 4.75E-13 |
| B21 | B119 | CD3+CD4-CD8- | Hcv-rna | -0.13371 | 1.50E-12 |
| B21 | B79 | CD3+CD4-CD8- | hb V -dna | -0.12219 | 1.04E-10 |

| | | | | | |
|---|---|---|---|---|---|
| B21 | B33 | CD3+CD4-CD8- | Prog | 0.10003 | 1.28E-07 |
| B21 | B23 | CD3+CD4-CD8- | CD3-CD16+CD56+(NK) | 0.095064 | 5.20E-07 |
| B21 | B30 | CD3+CD4-CD8- | LH | 0.093557 | 7.86E-07 |
| B210 | B215 | Hip | BMI | 0.766541 | 0 |
| B210 | B216 | Hip | BMI | 0.818334 | 0 |
| B210 | B218 | Hip | DBP | 0.231876 | 0 |
| B210 | B220 | Hip | SBP | 0.176836 | 0 |
| B210 | B221 | Hip | Hei | 0.430077 | 0 |
| B210 | B222 | Hip | NS | 0.651327 | 0 |
| B211 | B212 | IOP-R | IOP-L | 0.732867 | 0 |
| B215 | B216 | BMI | BMI | 0.834991 | 0 |
| B215 | B218 | BMI | DBP | 0.297367 | 0 |
| B215 | B220 | BMI | SBP | 0.249239 | 0 |
| B215 | B222 | BMI | NS | 0.675331 | 0 |
| B215 | B221 | BMI | Hei | 0.137565 | 3.32E-13 |
| B216 | B218 | BMI | DBP | 0.304849 | 0 |
| B216 | B220 | BMI | SBP | 0.193801 | 0 |
| B216 | B221 | BMI | Hei | 0.65382 | 0 |
| B216 | B222 | BMI | NS | 0.813319 | 0 |
| B218 | B220 | DBP | SBP | 0.680661 | 0 |
| B218 | B222 | DBP | NS | 0.301672 | 0 |
| B218 | B221 | DBP | Hei | 0.136509 | 5.04E-13 |
| B22 | B157 | CD3+CD8+(T) | IgG | 0.180395 | 0 |
| B22 | B187 | CD3+CD8+(T) | NSE | 0.236624 | 0 |
| B22 | B23 | CD3+CD8+(T) | CD3-CD16+CD56+(NK) | -0.50924 | 0 |
| B22 | B25 | CD3+CD8+(T) | CK | 0.588828 | 0 |
| B22 | B26 | CD3+CD8+(T) | CKMB | 0.217506 | 0 |
| B22 | B101 | CD3+CD8+(T) | anti-Sc70 | -0.14636 | 9.10E-15 |
| B22 | B102 | CD3+CD8+(T) | anti-Sm | -0.14636 | 9.10E-15 |
| B22 | B104 | CD3+CD8+(T) | anti-SSB | -0.14636 | 9.10E-15 |
| B22 | B98 | CD3+CD8+(T) | anti-Jo-1 | -0.14636 | 9.10E-15 |
| B22 | B99 | CD3+CD8+(T) | anti-PM-Scl | -0.14636 | 9.10E-15 |
| B22 | B186 | CD3+CD8+(T) | PFC | -0.14225 | 5.04E-14 |
| B22 | B155 | CD3+CD8+(T) | IgA | 0.13924 | 1.70E-13 |
| B22 | B94 | CD3+CD8+(T) | ALDH2-Lys504Lys | 0.138531 | 2.26E-13 |
| B22 | B37 | CD3+CD8+(T) | BAND | 0.107402 | 1.40E-08 |
| B22 | B100 | CD3+CD8+(T) | anti-Ro52 | -0.1011 | 9.36E-08 |
| B22 | B38 | CD3+CD8+(T) | BAND% | 0.09873 | 1.86E-07 |
| B22 | B183 | CD3+CD8+(T) | PTA | -0.09784 | 2.39E-07 |
| B220 | B222 | SBP | NS | 0.260915 | 0 |
| B221 | B222 | Hei | NS | 0.541787 | 0 |
| B227 | B228 | OD | OS | 0.11759 | 5.12E-10 |
| B23 | B109 | CD3-CD16+CD56+(NK) | AMA-M2 | -0.17184 | 0 |
| B23 | B182 | CD3-CD16+CD56+(NK) | TT | -0.25993 | 0 |

| | | | | | |
|---|---|---|---|---|---|
| B23 | B186 | CD3-CD16+CD56+(NK) | PFC | 0.184915 | 0 |
| B23 | B187 | CD3-CD16+CD56+(NK) | NSE | -0.16707 | 0 |
| B23 | B25 | CD3-CD16+CD56+(NK) | CK | -0.21269 | 0 |
| B23 | B57 | CD3-CD16+CD56+(NK) | RBCAI | -0.1771 | 0 |
| B23 | B37 | CD3-CD16+CD56+(NK) | BAND | 0.137616 | 3.26E-13 |
| B23 | B94 | CD3-CD16+CD56+(NK) | ALDH2-Lys504Lys | -0.12794 | 1.32E-11 |
| B23 | B93 | CD3-CD16+CD56+(NK) | ALDH2-Glu504Lys | 0.127324 | 1.65E-11 |
| B23 | B38 | CD3-CD16+CD56+(NK) | BAND% | 0.126513 | 2.22E-11 |
| B23 | B59 | CD3-CD16+CD56+(NK) | WBRV | -0.12617 | 2.52E-11 |
| B23 | B95 | CD3-CD16+CD56+(NK) | anti-cenpb | -0.12385 | 5.81E-11 |
| B23 | B65 | CD3-CD16+CD56+(NK) | WBRSR(mPas_1) | -0.12362 | 6.29E-11 |
| B23 | B157 | CD3-CD16+CD56+(NK) | IgG | -0.12142 | 1.37E-10 |
| B23 | B32 | CD3-CD16+CD56+(NK) | Testo | 0.120555 | 1.85E-10 |
| B23 | B155 | CD3-CD16+CD56+(NK) | IgA | -0.11537 | 1.08E-09 |
| B23 | B124 | CD3-CD16+CD56+(NK) | ACTH | -0.10876 | 9.15E-09 |
| B23 | B60 | CD3-CD16+CD56+(NK) | WBRVI | -0.10439 | 3.51E-08 |
| B23 | B64 | CD3-CD16+CD56+(NK) | WBRSR(mPas_5) | -0.09582 | 4.21E-07 |
| B24 | B26 | αHBDH | CKMB | 0.499754 | 0 |
| B24 | B27 | αHBDH | LDH | 0.709971 | 0 |
| B24 | B54 | αHBDH | ESR | 0.156236 | 0 |
| B24 | B25 | αHBDH | CK | 0.126298 | 2.40E-11 |
| B24 | B131 | αHBDH | AST | 0.124228 | 5.07E-11 |
| B24 | B58 | αHBDH | HCT | -0.09873 | 1.86E-07 |
| B24 | B176 | αHBDH | Ucr | -0.09399 | 6.99E-07 |
| B25 | B155 | CK | IgA | 0.214381 | 0 |
| B25 | B157 | CK | IgG | 0.25367 | 0 |
| B25 | B193 | CK | K | -0.15579 | 0 |
| B25 | B26 | CK | CKMB | 0.353536 | 0 |
| B25 | B27 | CK | LDH | 0.154161 | 4.44E-16 |
| B25 | B54 | CK | ESR | 0.141933 | 5.73E-14 |
| B25 | B101 | CK | anti-Sc70 | -0.12084 | 1.67E-10 |
| B25 | B102 | CK | anti-Sm | -0.12084 | 1.67E-10 |
| B25 | B104 | CK | anti-SSB | -0.12084 | 1.67E-10 |
| B25 | B98 | CK | anti-Jo-1 | -0.12084 | 1.67E-10 |
| B25 | B99 | CK | anti-PM-Scl | -0.12084 | 1.67E-10 |
| B25 | B139 | CK | ADA | 0.107112 | 1.53E-08 |
| B25 | B134 | CK | CHE | -0.10204 | 7.10E-08 |
| B25 | B131 | CK | AST | 0.101587 | 8.10E-08 |
| B25 | B204 | CK | Drinking | 0.099741 | 1.39E-07 |
| B25 | B137 | CK | AFU | 0.099181 | 1.63E-07 |
| B25 | B187 | CK | NSE | 0.093913 | 7.13E-07 |
| B26 | B135 | CKMB | TBA | 0.198652 | 0 |
| B26 | B139 | CKMB | ADA | 0.211533 | 0 |
| B26 | B155 | CKMB | IgA | 0.18388 | 0 |

| | | | | | |
|---|---|---|---|---|---|
| B26 | B156 | CKMB | IgE | 0.232263 | 0 |
| B26 | B27 | CKMB | LDH | 0.380079 | 0 |
| B26 | B54 | CKMB | ESR | 0.203547 | 0 |
| B26 | B193 | CKMB | K | -0.1555 | 2.22E-16 |
| B26 | B137 | CKMB | AFU | 0.133307 | 1.75E-12 |
| B26 | B90 | CKMB | CYFRA 21-1 | 0.130991 | 4.22E-12 |
| B26 | B131 | CKMB | AST | 0.126952 | 1.89E-11 |
| B26 | B125 | CKMB | Cor | -0.12338 | 6.86E-11 |
| B26 | B157 | CKMB | IgG | 0.114864 | 1.28E-09 |
| B26 | B78 | CKMB | HBcAb | 0.113975 | 1.71E-09 |
| B26 | B186 | CKMB | PFC | 0.107759 | 1.25E-08 |
| B26 | B164 | CKMB | COM | 0.10607 | 2.11E-08 |
| B26 | B38 | CKMB | BAND% | 0.105319 | 2.65E-08 |
| B26 | B37 | CKMB | BAND | -0.10226 | 6.65E-08 |
| B26 | B81 | CKMB | TSGF | 0.099496 | 1.49E-07 |
| B26 | B179 | CKMB | α1-MG | 0.099308 | 1.57E-07 |
| B26 | B182 | CKMB | TT | 0.098017 | 2.27E-07 |
| B27 | B131 | LDH | AST | 0.17926 | 0 |
| B27 | B139 | LDH | ADA | 0.131218 | 3.87E-12 |
| B27 | B150 | LDH | ALP | 0.127191 | 1.73E-11 |
| B27 | B54 | LDH | ESR | 0.121463 | 1.35E-10 |
| B27 | B128 | LDH | ALT | 0.11594 | 8.92E-10 |
| B27 | B144 | LDH | Total-PINP | 0.111968 | 3.29E-09 |
| B27 | B197 | LDH | AG | 0.105719 | 2.35E-08 |
| B27 | B137 | LDH | AFU | 0.104556 | 3.34E-08 |
| B27 | B220 | LDH | SBP | 0.104 | 3.95E-08 |
| B27 | B177 | LDH | ACR | 0.099922 | 1.32E-07 |
| B27 | B176 | LDH | Ucr | -0.09389 | 7.18E-07 |
| B28 | B29 | PRL | E2 | 0.853113 | 0 |
| B28 | B30 | PRL | LH | -0.50838 | 0 |
| B28 | B31 | PRL | FSH | -0.59816 | 0 |
| B28 | B33 | PRL | Prog | 0.249792 | 0 |
| B29 | B30 | E2 | LH | -0.38219 | 0 |
| B29 | B31 | E2 | FSH | -0.41578 | 0 |
| B29 | B32 | E2 | Testo | -0.17331 | 0 |
| B3 | B192 | Hcy | UA | 0.130246 | 5.58E-12 |
| B3 | B27 | Hcy | LDH | 0.108071 | 1.13E-08 |
| B3 | B189 | Hcy | eGFB | -0.1069 | 1.63E-08 |
| B30 | B31 | LH | FSH | 0.958353 | 0 |
| B30 | B32 | LH | Testo | -0.49071 | 0 |
| B30 | B33 | LH | Prog | 0.290836 | 0 |
| B30 | B175 | LH | ALB-U | -0.13994 | 1.29E-13 |
| B31 | B175 | FSH | ALB-U | -0.19393 | 0 |
| B31 | B32 | FSH | Testo | -0.3112 | 0 |

| | | | | | |
|---|---|---|---|---|---|
| B31 | B177 | FSH | ACR | -0.09829 | 2.11E-07 |
| B32 | B33 | Testo | Prog | -0.51383 | 0 |
| B32 | B101 | Testo | anti-Sc70 | -0.13742 | 3.52E-13 |
| B32 | B102 | Testo | anti-Sm | -0.13742 | 3.52E-13 |
| B32 | B104 | Testo | anti-SSB | -0.13742 | 3.52E-13 |
| B32 | B98 | Testo | anti-Jo-1 | -0.13742 | 3.52E-13 |
| B32 | B99 | Testo | anti-PM-Scl | -0.13742 | 3.52E-13 |
| B32 | B75 | Testo | ApoA1/B | 0.128052 | 1.26E-11 |
| B32 | B76 | Testo | ApoB | -0.12497 | 3.88E-11 |
| B33 | B175 | Prog | ALB-U | 0.099449 | 1.51E-07 |
| B34 | B86 | TK | TBhCG | 0.4914 | 0 |
| B34 | B89 | TK | CA 72-4 | 0.155111 | 2.22E-16 |
| B34 | B101 | TK | anti-Sc70 | 0.147872 | 4.88E-15 |
| B34 | B102 | TK | anti-Sm | 0.147872 | 4.88E-15 |
| B34 | B104 | TK | anti-SSB | 0.147872 | 4.88E-15 |
| B34 | B98 | TK | anti-Jo-1 | 0.147872 | 4.88E-15 |
| B34 | B99 | TK | anti-PM-Scl | 0.147872 | 4.88E-15 |
| B34 | B181 | TK | APTT | -0.13037 | 5.33E-12 |
| B34 | B75 | TK | ApoA1/B | 0.128304 | 1.15E-11 |
| B34 | B182 | TK | TT | 0.118315 | 4.00E-10 |
| B34 | B94 | TK | ALDH2-Lys504Lys | -0.11384 | 1.79E-09 |
| B34 | B199 | TK | Mg | 0.105744 | 2.33E-08 |
| B34 | B79 | TK | hb V -dna | 0.098303 | 2.10E-07 |
| B35 | B202 | β2-MG | e-times | 0.508331 | 0 |
| B35 | B223 | β2-MG | FH | 0.2084 | 0 |
| B36 | B42 | V-Lym% | MON | -0.27711 | 0 |
| B36 | B37 | V-Lym% | BAND | -0.11166 | 3.63E-09 |
| B36 | B38 | V-Lym% | BAND% | 0.106819 | 1.67E-08 |
| B36 | B40 | V-Lym% | WBC | -0.10278 | 5.69E-08 |
| B37 | B139 | BAND | ADA | -0.20126 | 0 |
| B37 | B155 | BAND | IgA | -0.18335 | 0 |
| B37 | B156 | BAND | IgE | -0.27528 | 0 |
| B37 | B182 | BAND | TT | -0.23698 | 0 |
| B37 | B38 | BAND | BAND% | 0.610145 | 0 |
| B37 | B47 | BAND | RDW | -0.18949 | 0 |
| B37 | B57 | BAND | RBCAI | -0.17179 | 0 |
| B37 | B59 | BAND | WBRV | -0.18373 | 0 |
| B37 | B60 | BAND | WBRVI | -0.25613 | 0 |
| B37 | B62 | BAND | WBHSRVI | -0.17739 | 0 |
| B37 | B63 | BAND | WBRSR(mPas_200) | -0.18925 | 0 |
| B37 | B64 | BAND | WBRSR(mPas_5) | -0.25824 | 0 |
| B37 | B65 | BAND | WBRSR(mPas_1) | -0.27921 | 0 |
| B37 | B66 | BAND | WBRSR(mPas_100) | -0.23796 | 0 |
| B37 | B67 | BAND | WBRSR(mPas_50) | -0.21697 | 0 |

| | | | | | |
|---|---|---|---|---|---|
| B37 | B79 | BAND | hb V -dna | 0.165957 | 0 |
| B37 | B81 | BAND | TSGF | -0.21121 | 0 |
| B37 | B125 | BAND | Cor | 0.147191 | 6.44E-15 |
| B37 | B93 | BAND | ALDH2-Glu504Lys | -0.14441 | 2.04E-14 |
| B37 | B58 | BAND | HCT | -0.14236 | 4.80E-14 |
| B37 | B164 | BAND | COM | -0.14119 | 7.77E-14 |
| B37 | B92 | BAND | ALDH2-Glu504Glu | 0.132473 | 2.41E-12 |
| B37 | B61 | BAND | WBHSRV | -0.1272 | 1.73E-11 |
| B37 | B54 | BAND | ESR | -0.11731 | 5.63E-10 |
| B37 | B101 | BAND | anti-Sc70 | 0.111996 | 3.26E-09 |
| B37 | B102 | BAND | anti-Sm | 0.111996 | 3.26E-09 |
| B37 | B104 | BAND | anti-SSB | 0.111996 | 3.26E-09 |
| B37 | B98 | BAND | anti-Jo-1 | 0.111996 | 3.26E-09 |
| B37 | B99 | BAND | anti-PM-Scl | 0.111996 | 3.26E-09 |
| B37 | B56 | BAND | IR | -0.11005 | 6.09E-09 |
| B37 | B183 | BAND | PTA | 0.1095 | 7.24E-09 |
| B37 | B179 | BAND | α1-MG | -0.10332 | 4.85E-08 |
| B37 | B123 | BAND | hs-CRP1 | -0.093 | 9.14E-07 |
| B38 | B125 | BAND% | Cor | -0.19104 | 0 |
| B38 | B155 | BAND% | IgA | -0.18239 | 0 |
| B38 | B156 | BAND% | IgE | 0.250263 | 0 |
| B38 | B186 | BAND% | PFC | 0.309007 | 0 |
| B38 | B42 | BAND% | MON | -0.16621 | 0 |
| B38 | B47 | BAND% | RDW | -0.23644 | 0 |
| B38 | B57 | BAND% | RBCAI | -0.24435 | 0 |
| B38 | B58 | BAND% | HCT | -0.18508 | 0 |
| B38 | B59 | BAND% | WBRV | -0.23544 | 0 |
| B38 | B60 | BAND% | WBRVI | -0.31349 | 0 |
| B38 | B61 | BAND% | WBHSRV | -0.15876 | 0 |
| B38 | B62 | BAND% | WBHSRVI | -0.21941 | 0 |
| B38 | B63 | BAND% | WBRSR(mPas_200) | -0.22509 | 0 |
| B38 | B64 | BAND% | WBRSR(mPas_5) | -0.28544 | 0 |
| B38 | B65 | BAND% | WBRSR(mPas_1) | -0.31002 | 0 |
| B38 | B66 | BAND% | WBRSR(mPas_100) | -0.21293 | 0 |
| B38 | B67 | BAND% | WBRSR(mPas_50) | -0.25222 | 0 |
| B38 | B157 | BAND% | IgG | -0.1541 | 4.44E-16 |
| B38 | B56 | BAND% | IR | -0.15132 | 1.11E-15 |
| B38 | B182 | BAND% | TT | -0.14627 | 9.33E-15 |
| B38 | B179 | BAND% | α1-MG | 0.133358 | 1.71E-12 |
| B38 | B93 | BAND% | ALDH2-Glu504Lys | 0.1231 | 7.57E-11 |
| B38 | B164 | BAND% | COM | 0.118419 | 3.86E-10 |
| B38 | B124 | BAND% | ACTH | -0.11341 | 2.06E-09 |
| B38 | B92 | BAND% | ALDH2-Glu504Glu | -0.11317 | 2.23E-09 |
| B38 | B101 | BAND% | anti-Sc70 | 0.102996 | 5.34E-08 |

| | | | | | |
|---|---|---|---|---|---|
| B38 | B102 | BAND% | anti-Sm | 0.102996 | 5.34E-08 |
| B38 | B104 | BAND% | anti-SSB | 0.102996 | 5.34E-08 |
| B38 | B98 | BAND% | anti-Jo-1 | 0.102996 | 5.34E-08 |
| B38 | B99 | BAND% | anti-PM-Scl | 0.102996 | 5.34E-08 |
| B38 | B134 | BAND% | CHE | 0.099635 | 1.43E-07 |
| B39 | B171 | Other cells | EC | 0.207444 | 0 |
| B39 | B40 | Other cells | WBC | 0.548672 | 0 |
| B39 | B48 | Other cells | LYM% | 0.233286 | 0 |
| B39 | B53 | Other cells | GR% | -0.203 | 0 |
| B39 | B41 | Other cells | MON % | -0.13728 | 3.72E-13 |
| B39 | B44 | Other cells | MCH | -0.12967 | 6.93E-12 |
| B39 | B45 | Other cells | MCHC | -0.12521 | 3.56E-11 |
| B39 | B43 | Other cells | MCV | -0.10121 | 9.05E-08 |
| B4 | B10 | WB-Ca | PG I | -0.24725 | 0 |
| B4 | B12 | WB-Ca | PG II | -0.21229 | 0 |
| B4 | B136 | WB-Ca | 5'NT | -0.16644 | 0 |
| B4 | B143 | WB-Ca | β-CTx | -0.16642 | 0 |
| B4 | B5 | WB-Ca | WB-Mg | 0.266734 | 0 |
| B4 | B6 | WB-Ca | WB-Fe | 0.258763 | 0 |
| B4 | B7 | WB-Ca | WB-Cop | 0.933332 | 0 |
| B4 | B8 | WB-Ca | WB-Zinc | 0.302858 | 0 |
| B4 | B87 | WB-Ca | NSE | -0.15412 | 4.44E-16 |
| B4 | B140 | WB-Ca | r-GT | -0.11986 | 2.35E-10 |
| B4 | B58 | WB-Ca | HCT | 0.118092 | 4.31E-10 |
| B4 | B67 | WB-Ca | WBRSR(mPas_50) | 0.095072 | 5.19E-07 |
| B4 | B66 | WB-Ca | WBRSR(mPas_100) | 0.094782 | 5.62E-07 |
| B4 | B62 | WB-Ca | WBHSRVI | 0.094426 | 6.20E-07 |
| B40 | B171 | WBC | EC | 0.348303 | 0 |
| B40 | B41 | WBC | MON % | -0.16494 | 0 |
| B40 | B42 | WBC | MON | 0.263127 | 0 |
| B40 | B48 | WBC | LYM% | 0.158183 | 0 |
| B40 | B53 | WBC | GR% | -0.11778 | 4.80E-10 |
| B40 | B72 | WBC | HDL-C | -0.11159 | 3.72E-09 |
| B40 | B46 | WBC | RBC Count | 0.098963 | 1.74E-07 |
| B40 | B164 | WBC | COM | 0.094935 | 5.39E-07 |
| B41 | B126 | MON % | Alb | -0.15818 | 0 |
| B41 | B42 | MON % | MON | 0.562719 | 0 |
| B41 | B53 | MON % | GR% | -0.16401 | 0 |
| B41 | B47 | MON % | RDW | 0.11946 | 2.70E-10 |
| B41 | B43 | MON % | MCV | 0.117712 | 4.91E-10 |
| B41 | B189 | MON % | eGFB | -0.10546 | 2.54E-08 |
| B41 | B44 | MON % | MCH | 0.103813 | 4.18E-08 |
| B42 | B192 | MON | UA | 0.183515 | 0 |
| B42 | B206 | MON | Waist | 0.211648 | 0 |

| | | | | | |
|---|---|---|---|---|---|
| B42 | B209 | MON | Smoking | 0.204621 | 0 |
| B42 | B215 | MON | BMI | 0.170853 | 0 |
| B42 | B216 | MON | BMI | 0.191552 | 0 |
| B42 | B222 | MON | NS | 0.231287 | 0 |
| B42 | B46 | MON | RBC Count | 0.157089 | 0 |
| B42 | B51 | MON | Hb | 0.185604 | 0 |
| B42 | B52 | MON | PLT/BPC | 0.195939 | 0 |
| B42 | B210 | MON | Hip | 0.135714 | 6.88E-13 |
| B42 | B163 | MON | SG | 0.131793 | 3.12E-12 |
| B42 | B72 | MON | HDL-C | -0.13085 | 4.45E-12 |
| B42 | B220 | MON | SBP | 0.115264 | 1.12E-09 |
| B42 | B47 | MON | RDW | 0.112889 | 2.44E-09 |
| B42 | B48 | MON | LYM% | -0.11243 | 2.84E-09 |
| B42 | B221 | MON | Hei | 0.111704 | 3.58E-09 |
| B42 | B190 | MON | Cre | 0.109377 | 7.53E-09 |
| B42 | B191 | MON | Urea | 0.102677 | 5.87E-08 |
| B42 | B71 | MON | TG | 0.099682 | 1.41E-07 |
| B42 | B140 | MON | r-GT | 0.099651 | 1.42E-07 |
| B42 | B166 | MON | Pro | 0.096157 | 3.84E-07 |
| B42 | B218 | MON | DBP | 0.094488 | 6.10E-07 |
| B42 | B128 | MON | ALT | 0.094047 | 6.88E-07 |
| B43 | B189 | MCV | eGFB | -0.18861 | 0 |
| B43 | B44 | MCV | MCH | 0.921185 | 0 |
| B43 | B45 | MCV | MCHC | 0.235155 | 0 |
| B43 | B46 | MCV | RBC Count | -0.5183 | 0 |
| B43 | B47 | MCV | RDW | 0.570298 | 0 |
| B43 | B52 | MCV | PLT/BPC | -0.15282 | 4.44E-16 |
| B43 | B51 | MCV | Hb | 0.15074 | 1.33E-15 |
| B43 | B91 | MCV | TP | -0.11196 | 3.29E-09 |
| B43 | B59 | MCV | WBRV | -0.10561 | 2.43E-08 |
| B43 | B191 | MCV | Urea | 0.102207 | 6.75E-08 |
| B43 | B55 | MCV | TK | -0.10178 | 7.65E-08 |
| B44 | B45 | MCH | MCHC | 0.592677 | 0 |
| B44 | B46 | MCH | RBC Count | -0.38401 | 0 |
| B44 | B47 | MCH | RDW | 0.386888 | 0 |
| B44 | B51 | MCH | Hb | 0.345824 | 0 |
| B44 | B189 | MCH | eGFB | -0.14839 | 4.00E-15 |
| B44 | B52 | MCH | PLT/BPC | -0.14566 | 1.22E-14 |
| B44 | B209 | MCH | Smoking | 0.12577 | 2.91E-11 |
| B44 | B222 | MCH | NS | 0.118896 | 3.28E-10 |
| B44 | B204 | MCH | Drinking | 0.108286 | 1.06E-08 |
| B44 | B77 | MCH | TC | 0.104275 | 3.64E-08 |
| B44 | B130 | MCH | GLB | -0.10147 | 8.39E-08 |
| B44 | B127 | MCH | A-G ratio | 0.100075 | 1.26E-07 |

| | | | | | |
|---|---|---|---|---|---|
| B44 | B190 | MCH | Cre | 0.097728 | 2.47E-07 |
| B44 | B206 | MCH | Waist | 0.096661 | 3.34E-07 |
| B44 | B163 | MCH | SG | 0.095338 | 4.82E-07 |
| B44 | B192 | MCH | UA | 0.093177 | 8.71E-07 |
| B45 | B128 | MCHC | ALT | 0.158424 | 0 |
| B45 | B129 | MCHC | IBIL | 0.159583 | 0 |
| B45 | B133 | MCHC | TBIL | 0.157405 | 0 |
| B45 | B141 | MCHC | AST/ALT | -0.18012 | 0 |
| B45 | B153 | MCHC | Glu | 0.178336 | 0 |
| B45 | B163 | MCHC | SG | 0.186344 | 0 |
| B45 | B192 | MCHC | UA | 0.219436 | 0 |
| B45 | B206 | MCHC | Waist | 0.194676 | 0 |
| B45 | B210 | MCHC | Hip | 0.1774 | 0 |
| B45 | B215 | MCHC | BMI | 0.171837 | 0 |
| B45 | B216 | MCHC | BMI | 0.261805 | 0 |
| B45 | B221 | MCHC | Hei | 0.241986 | 0 |
| B45 | B222 | MCHC | NS | 0.275978 | 0 |
| B45 | B47 | MCHC | RDW | -0.22248 | 0 |
| B45 | B51 | MCHC | Hb | 0.562525 | 0 |
| B45 | B71 | MCHC | TG | 0.226994 | 0 |
| B45 | B72 | MCHC | HDL-C | -0.1554 | 2.22E-16 |
| B45 | B126 | MCHC | Alb | 0.149519 | 2.22E-15 |
| B45 | B170 | MCHC | U-GLU | 0.148989 | 3.11E-15 |
| B45 | B140 | MCHC | r-GT | 0.142265 | 5.00E-14 |
| B45 | B174 | MCHC | Urea/Cr | -0.13408 | 1.30E-12 |
| B45 | B218 | MCHC | DBP | 0.131856 | 3.04E-12 |
| B45 | B204 | MCHC | Drinking | 0.127558 | 1.51E-11 |
| B45 | B127 | MCHC | A-G ratio | 0.126368 | 2.34E-11 |
| B45 | B46 | MCHC | RBC Count | 0.12053 | 1.87E-10 |
| B45 | B209 | MCHC | Smoking | 0.118151 | 4.23E-10 |
| B45 | B188 | MCHC | Ca | 0.112118 | 3.14E-09 |
| B45 | B132 | MCHC | Dbil | 0.10963 | 6.95E-09 |
| B45 | B62 | MCHC | WBHSRVI | 0.093744 | 7.47E-07 |
| B46 | B113 | RBC Count | FT3 | 0.17528 | 0 |
| B46 | B126 | RBC Count | Alb | 0.289785 | 0 |
| B46 | B128 | RBC Count | ALT | 0.238034 | 0 |
| B46 | B141 | RBC Count | AST/ALT | -0.24506 | 0 |
| B46 | B163 | RBC Count | SG | 0.265054 | 0 |
| B46 | B174 | RBC Count | Urea/Cr | -0.16549 | 0 |
| B46 | B192 | RBC Count | UA | 0.307311 | 0 |
| B46 | B206 | RBC Count | Waist | 0.335214 | 0 |
| B46 | B209 | RBC Count | Smoking | 0.191282 | 0 |
| B46 | B210 | RBC Count | Hip | 0.292508 | 0 |
| B46 | B215 | RBC Count | BMI | 0.27306 | 0 |

| | | | | | |
|---|---|---|---|---|---|
| B46 | B216 | RBC Count | BMI | 0.423786 | 0 |
| B46 | B218 | RBC Count | DBP | 0.254189 | 0 |
| B46 | B221 | RBC Count | Hei | 0.392023 | 0 |
| B46 | B222 | RBC Count | NS | 0.438373 | 0 |
| B46 | B47 | RBC Count | RDW | -0.50942 | 0 |
| B46 | B51 | RBC Count | Hb | 0.721236 | 0 |
| B46 | B71 | RBC Count | TG | 0.162337 | 0 |
| B46 | B72 | RBC Count | HDL-C | -0.23262 | 0 |
| B46 | B60 | RBC Count | WBRVI | 0.154721 | 2.22E-16 |
| B46 | B188 | RBC Count | Ca | 0.153938 | 4.44E-16 |
| B46 | B204 | RBC Count | Drinking | 0.153681 | 4.44E-16 |
| B46 | B64 | RBC Count | WBRSR(mPas_5) | 0.154164 | 4.44E-16 |
| B46 | B65 | RBC Count | WBRSR(mPas_1) | 0.151814 | 8.88E-16 |
| B46 | B62 | RBC Count | WBHSRVI | 0.148877 | 3.11E-15 |
| B46 | B67 | RBC Count | WBRSR(mPas_50) | 0.147751 | 5.11E-15 |
| B46 | B66 | RBC Count | WBRSR(mPas_100) | 0.145883 | 1.11E-14 |
| B46 | B63 | RBC Count | WBRSR(mPas_200) | 0.145036 | 1.60E-14 |
| B46 | B140 | RBC Count | r-GT | 0.144176 | 2.26E-14 |
| B46 | B189 | RBC Count | eGFB | 0.143233 | 3.38E-14 |
| B46 | B127 | RBC Count | A-G ratio | 0.139267 | 1.69E-13 |
| B46 | B91 | RBC Count | TP | 0.138287 | 2.49E-13 |
| B46 | B153 | RBC Count | Glu | 0.134619 | 1.05E-12 |
| B46 | B170 | RBC Count | U-GLU | 0.131708 | 3.22E-12 |
| B46 | B58 | RBC Count | HCT | 0.127109 | 1.79E-11 |
| B46 | B115 | RBC Count | TSH TT3 TT4(TT3) | 0.125022 | 3.81E-11 |
| B46 | B61 | RBC Count | WBHSRV | 0.123922 | 5.65E-11 |
| B46 | B56 | RBC Count | IR | 0.123793 | 5.92E-11 |
| B46 | B220 | RBC Count | SBP | 0.117056 | 6.13E-10 |
| B46 | B75 | RBC Count | ApoA1/B | -0.11213 | 3.13E-09 |
| B46 | B59 | RBC Count | WBRV | 0.110361 | 5.51E-09 |
| B46 | B70 | RBC Count | LDL-C | 0.109915 | 6.35E-09 |
| B46 | B129 | RBC Count | IBIL | 0.10834 | 1.04E-08 |
| B46 | B176 | RBC Count | Ucr | 0.108046 | 1.14E-08 |
| B46 | B133 | RBC Count | TBIL | 0.104421 | 3.48E-08 |
| B46 | B50 | RBC Count | eo% | 0.097726 | 2.47E-07 |
| B46 | B166 | RBC Count | Pro | 0.093978 | 7.01E-07 |
| B47 | B126 | RDW | Alb | -0.15947 | 0 |
| B47 | B189 | RDW | eGFB | -0.20207 | 0 |
| B47 | B51 | RDW | Hb | -0.25916 | 0 |
| B47 | B91 | RDW | TP | -0.13675 | 4.58E-13 |
| B47 | B188 | RDW | Ca | -0.13131 | 3.74E-12 |
| B47 | B191 | RDW | Urea | 0.123292 | 7.07E-11 |
| B47 | B216 | RDW | BMI | -0.10378 | 4.23E-08 |
| B47 | B113 | RDW | FT3 | -0.1017 | 7.83E-08 |

| | | | | | |
|---|---|---|---|---|---|
| B47 | B134 | RDW | CHE | -0.09759 | 2.57E-07 |
| B47 | B221 | RDW | Hei | -0.09596 | 4.06E-07 |
| B48 | B53 | LYM% | GR% | -0.94773 | 0 |
| B49 | B50 | baso% | eo% | 0.299894 | 0 |
| B49 | B53 | baso% | GR% | -0.1406 | 9.86E-14 |
| B49 | B77 | baso% | TC | 0.10494 | 2.98E-08 |
| B49 | B52 | baso% | PLT/BPC | 0.100723 | 1.04E-07 |
| B5 | B6 | WB-Mg | WB-Fe | 0.966894 | 0 |
| B5 | B8 | WB-Mg | WB-Zinc | -0.14156 | 6.66E-14 |
| B5 | B65 | WB-Mg | WBRSR(mPas_1) | 0.129205 | 8.24E-12 |
| B5 | B64 | WB-Mg | WBRSR(mPas_5) | 0.127664 | 1.46E-11 |
| B5 | B60 | WB-Mg | WBRVI | 0.126832 | 1.98E-11 |
| B5 | B10 | WB-Mg | PG I | 0.122903 | 8.12E-11 |
| B5 | B67 | WB-Mg | WBRSR(mPas_50) | 0.120265 | 2.05E-10 |
| B5 | B66 | WB-Mg | WBRSR(mPas_100) | 0.11761 | 5.08E-10 |
| B5 | B63 | WB-Mg | WBRSR(mPas_200) | 0.117099 | 6.04E-10 |
| B5 | B62 | WB-Mg | WBHSRVI | 0.109233 | 7.88E-09 |
| B5 | B59 | WB-Mg | WBRV | 0.107784 | 1.24E-08 |
| B5 | B57 | WB-Mg | RBCAI | 0.107498 | 1.36E-08 |
| B5 | B12 | WB-Mg | PG II | 0.102125 | 6.91E-08 |
| B5 | B88 | WB-Mg | CA 242 | -0.10146 | 8.41E-08 |
| B5 | B58 | WB-Mg | HCT | 0.098817 | 1.81E-07 |
| B5 | B85 | WB-Mg | SCC | -0.09864 | 1.91E-07 |
| B5 | B61 | WB-Mg | WBHSRV | 0.095535 | 4.57E-07 |
| B50 | B53 | eo% | GR% | -0.21328 | 0 |
| B50 | B51 | eo% | Hb | 0.122946 | 8.00E-11 |
| B50 | B209 | eo% | Smoking | 0.122094 | 1.08E-10 |
| B50 | B190 | eo% | Cre | 0.111359 | 4.00E-09 |
| B50 | B222 | eo% | NS | 0.110531 | 5.22E-09 |
| B50 | B221 | eo% | Hei | 0.105102 | 2.83E-08 |
| B51 | B126 | Hb | Alb | 0.288153 | 0 |
| B51 | B127 | Hb | A-G ratio | 0.201324 | 0 |
| B51 | B128 | Hb | ALT | 0.277687 | 0 |
| B51 | B140 | Hb | r-GT | 0.206463 | 0 |
| B51 | B141 | Hb | AST/ALT | -0.29935 | 0 |
| B51 | B153 | Hb | Glu | 0.177602 | 0 |
| B51 | B163 | Hb | SG | 0.33615 | 0 |
| B51 | B170 | Hb | U-GLU | 0.163009 | 0 |
| B51 | B174 | Hb | Urea/Cr | -0.22713 | 0 |
| B51 | B188 | Hb | Ca | 0.180211 | 0 |
| B51 | B192 | Hb | UA | 0.36873 | 0 |
| B51 | B204 | Hb | Drinking | 0.234203 | 0 |
| B51 | B206 | Hb | Waist | 0.409147 | 0 |
| B51 | B209 | Hb | Smoking | 0.287986 | 0 |

| | | | | | |
|---|---|---|---|---|---|
| B51 | B210 | Hb | Hip | 0.334143 | 0 |
| B51 | B215 | Hb | BMI | 0.311828 | 0 |
| B51 | B216 | Hb | BMI | 0.488129 | 0 |
| B51 | B218 | Hb | DBP | 0.271466 | 0 |
| B51 | B221 | Hb | Hei | 0.458718 | 0 |
| B51 | B222 | Hb | NS | 0.527362 | 0 |
| B51 | B58 | Hb | HCT | 0.188283 | 0 |
| B51 | B62 | Hb | WBHSRVI | 0.168097 | 0 |
| B51 | B70 | Hb | LDL-C | 0.169806 | 0 |
| B51 | B71 | Hb | TG | 0.226453 | 0 |
| B51 | B72 | Hb | HDL-C | -0.22132 | 0 |
| B51 | B63 | Hb | WBRSR(mPas_200) | 0.155097 | 2.22E-16 |
| B51 | B66 | Hb | WBRSR(mPas_100) | 0.154419 | 2.22E-16 |
| B51 | B67 | Hb | WBRSR(mPas_50) | 0.15477 | 2.22E-16 |
| B51 | B113 | Hb | FT3 | 0.151174 | 1.11E-15 |
| B51 | B129 | Hb | IBIL | 0.149696 | 2.22E-15 |
| B51 | B60 | Hb | WBRVI | 0.148915 | 3.11E-15 |
| B51 | B64 | Hb | WBRSR(mPas_5) | 0.148044 | 4.44E-15 |
| B51 | B133 | Hb | TBIL | 0.147809 | 4.88E-15 |
| B51 | B65 | Hb | WBRSR(mPas_1) | 0.137537 | 3.36E-13 |
| B51 | B77 | Hb | TC | 0.135269 | 8.19E-13 |
| B51 | B166 | Hb | Pro | 0.13162 | 3.33E-12 |
| B51 | B220 | Hb | SBP | 0.129608 | 7.09E-12 |
| B51 | B176 | Hb | Ucr | 0.113379 | 2.08E-09 |
| B51 | B130 | Hb | GLB | -0.1128 | 2.51E-09 |
| B51 | B138 | Hb | PAB | 0.10571 | 2.35E-08 |
| B51 | B131 | Hb | AST | 0.105026 | 2.90E-08 |
| B51 | B132 | Hb | Dbil | 0.103242 | 4.96E-08 |
| B51 | B56 | Hb | IR | 0.102514 | 6.16E-08 |
| B51 | B219 | Hb | OT | -0.09965 | 1.42E-07 |
| B51 | B161 | Hb | LEU | -0.09672 | 3.28E-07 |
| B51 | B190 | Hb | Cre | 0.094817 | 5.57E-07 |
| B51 | B75 | Hb | ApoA1/B | -0.09463 | 5.86E-07 |
| B52 | B132 | PLT/BPC | Dbil | -0.13216 | 2.71E-12 |
| B52 | B91 | PLT/BPC | TP | 0.116064 | 8.56E-10 |
| B52 | B130 | PLT/BPC | GLB | 0.113178 | 2.22E-09 |
| B52 | B189 | PLT/BPC | eGFB | 0.11312 | 2.26E-09 |
| B52 | B191 | PLT/BPC | Urea | -0.10658 | 1.80E-08 |
| B52 | B133 | PLT/BPC | TBIL | -0.10578 | 2.30E-08 |
| B52 | B131 | PLT/BPC | AST | -0.09855 | 1.95E-07 |
| B54 | B108 | ESR | Anti-Cyclin | 0.16412 | 0 |
| B54 | B135 | ESR | TBA | 0.256734 | 0 |
| B54 | B138 | ESR | PAB | -0.18773 | 0 |
| B54 | B139 | ESR | ADA | 0.276795 | 0 |

| | | | | | |
|---|---|---|---|---|---|
| B54 | B155 | ESR | IgA | 0.182328 | 0 |
| B54 | B156 | ESR | IgE | 0.195034 | 0 |
| B54 | B157 | ESR | IgG | 0.180506 | 0 |
| B54 | B186 | ESR | PFC | 0.176022 | 0 |
| B54 | B56 | ESR | IR | -0.19826 | 0 |
| B54 | B58 | ESR | HCT | -0.45349 | 0 |
| B54 | B59 | ESR | WBRV | 0.171248 | 0 |
| B54 | B60 | ESR | WBRVI | -0.29367 | 0 |
| B54 | B62 | ESR | WBHSRVI | -0.35273 | 0 |
| B54 | B69 | ESR | PV | 0.575941 | 0 |
| B54 | B123 | ESR | hs-CRP1 | 0.152712 | 4.44E-16 |
| B54 | B175 | ESR | ALB-U | 0.152147 | 8.88E-16 |
| B54 | B193 | ESR | K | -0.14269 | 4.22E-14 |
| B54 | B180 | ESR | Tf | 0.140633 | 9.73E-14 |
| B54 | B213 | ESR | CVA-R | 0.13861 | 2.19E-13 |
| B54 | B55 | ESR | TK | 0.136609 | 4.85E-13 |
| B54 | B130 | ESR | GLB | 0.129047 | 8.74E-12 |
| B54 | B182 | ESR | TT | 0.128532 | 1.06E-11 |
| B54 | B125 | ESR | Cor | -0.11631 | 7.87E-10 |
| B54 | B63 | ESR | WBRSR(mPas_200) | -0.11536 | 1.08E-09 |
| B54 | B127 | ESR | A-G ratio | -0.11364 | 1.91E-09 |
| B54 | B177 | ESR | ACR | 0.111042 | 4.43E-09 |
| B54 | B81 | ESR | TSGF | 0.110061 | 6.07E-09 |
| B54 | B65 | ESR | WBRSR(mPas_1) | -0.10919 | 7.99E-09 |
| B54 | B199 | ESR | Mg | -0.10914 | 8.11E-09 |
| B54 | B67 | ESR | WBRSR(mPas_50) | -0.10854 | 9.79E-09 |
| B54 | B178 | ESR | Ig-U | 0.107945 | 1.18E-08 |
| B54 | B64 | ESR | WBRSR(mPas_5) | -0.09856 | 1.95E-07 |
| B54 | B179 | ESR | α1-MG | 0.093627 | 7.71E-07 |
| B54 | B150 | ESR | ALP | 0.09318 | 8.71E-07 |
| B55 | B107 | TK | AnuA | -0.15589 | 0 |
| B55 | B56 | TK | IR | 0.68765 | 0 |
| B55 | B58 | TK | HCT | -0.36251 | 0 |
| B55 | B59 | TK | WBRV | 0.4844 | 0 |
| B55 | B60 | TK | WBRVI | 0.199962 | 0 |
| B55 | B61 | TK | WBHSRV | 0.62213 | 0 |
| B55 | B62 | TK | WBHSRVI | 0.259768 | 0 |
| B55 | B63 | TK | WBRSR(mPas_200) | 0.16242 | 0 |
| B55 | B66 | TK | WBRSR(mPas_100) | 0.191466 | 0 |
| B55 | B67 | TK | WBRSR(mPas_50) | 0.192411 | 0 |
| B55 | B69 | TK | PV | -0.19592 | 0 |
| B55 | B105 | TK | anti-U1RNP | -0.15448 | 2.22E-16 |
| B55 | B106 | TK | anti-P | -0.15532 | 2.22E-16 |
| B55 | B64 | TK | WBRSR(mPas_5) | 0.147058 | 6.66E-15 |

| | | | | | |
|---|---|---|---|---|---|
| B55 | B65 | TK | WBRSR(mPas_1) | 0.131794 | 3.11E-12 |
| B55 | B121 | TK | C3 | 0.121977 | 1.13E-10 |
| B55 | B96 | TK | dsDNA (IIF) | 0.110864 | 4.69E-09 |
| B55 | B124 | TK | ACTH | -0.10765 | 1.29E-08 |
| B55 | B185 | TK | PT | 0.107558 | 1.33E-08 |
| B55 | B122 | TK | C4 | 0.101037 | 9.52E-08 |
| B55 | B184 | TK | INR | 0.09717 | 2.89E-07 |
| B55 | B100 | TK | anti-Ro52 | -0.09633 | 3.66E-07 |
| B56 | B100 | IR | anti-Ro52 | -0.15942 | 0 |
| B56 | B105 | IR | anti-U1RNP | -0.19061 | 0 |
| B56 | B106 | IR | anti-P | -0.18701 | 0 |
| B56 | B107 | IR | AnuA | -0.18288 | 0 |
| B56 | B57 | IR | RBCAI | 0.164719 | 0 |
| B56 | B58 | IR | HCT | 0.400067 | 0 |
| B56 | B59 | IR | WBRV | 0.607474 | 0 |
| B56 | B60 | IR | WBRVI | 0.764497 | 0 |
| B56 | B61 | IR | WBHSRV | 0.900204 | 0 |
| B56 | B62 | IR | WBHSRVI | 0.8772 | 0 |
| B56 | B63 | IR | WBRSR(mPas_200) | 0.751045 | 0 |
| B56 | B64 | IR | WBRSR(mPas_5) | 0.708865 | 0 |
| B56 | B65 | IR | WBRSR(mPas_1) | 0.6614 | 0 |
| B56 | B66 | IR | WBRSR(mPas_100) | 0.771235 | 0 |
| B56 | B67 | IR | WBRSR(mPas_50) | 0.774686 | 0 |
| B56 | B69 | IR | PV | -0.28477 | 0 |
| B56 | B121 | IR | C3 | 0.143402 | 3.13E-14 |
| B56 | B186 | IR | PFC | -0.14005 | 1.23E-13 |
| B56 | B96 | IR | dsDNA (IIF) | 0.129414 | 7.62E-12 |
| B56 | B137 | IR | AFU | 0.124814 | 4.11E-11 |
| B56 | B213 | IR | CVA-R | -0.12461 | 4.41E-11 |
| B56 | B103 | IR | anti-SSA/Ro | -0.12283 | 8.34E-11 |
| B56 | B122 | IR | C4 | 0.118048 | 4.38E-10 |
| B56 | B198 | IR | P | -0.10651 | 1.84E-08 |
| B56 | B74 | IR | ApoA1 | -0.10537 | 2.61E-08 |
| B56 | B75 | IR | ApoA1/B | -0.09953 | 1.48E-07 |
| B56 | B156 | IR | IgE | -0.09317 | 8.74E-07 |
| B57 | B110 | RBCAI | AHA | -0.19231 | 0 |
| B57 | B113 | RBCAI | FT3 | 0.20858 | 0 |
| B57 | B115 | RBCAI | TSH TT3 TT4(TT3) | 0.211808 | 0 |
| B57 | B124 | RBCAI | ACTH | 0.410973 | 0 |
| B57 | B125 | RBCAI | Cor | 0.273399 | 0 |
| B57 | B58 | RBCAI | HCT | 0.235927 | 0 |
| B57 | B59 | RBCAI | WBRV | 0.714845 | 0 |
| B57 | B60 | RBCAI | WBRVI | 0.649722 | 0 |
| B57 | B61 | RBCAI | WBHSRV | 0.193081 | 0 |

| | | | | | |
|---|---|---|---|---|---|
| B57 | B62 | RBCAI | WBHSRVI | 0.229207 | 0 |
| B57 | B63 | RBCAI | WBRSR(mPas_200) | 0.25668 | 0 |
| B57 | B64 | RBCAI | WBRSR(mPas_5) | 0.54793 | 0 |
| B57 | B65 | RBCAI | WBRSR(mPas_1) | 0.67118 | 0 |
| B57 | B66 | RBCAI | WBRSR(mPas_100) | 0.234554 | 0 |
| B57 | B67 | RBCAI | WBRSR(mPas_50) | 0.286129 | 0 |
| B57 | B112 | RBCAI | FT4 | 0.151803 | 8.88E-16 |
| B57 | B150 | RBCAI | ALP | 0.147073 | 6.66E-15 |
| B57 | B200 | RBCAI | OP | -0.11722 | 5.79E-10 |
| B57 | B114 | RBCAI | TSH TT3 TT4(TT4) | 0.116745 | 6.81E-10 |
| B57 | B92 | RBCAI | ALDH2-Glu504Glu | 0.113783 | 1.82E-09 |
| B57 | B185 | RBCAI | PT | -0.11344 | 2.04E-09 |
| B57 | B183 | RBCAI | PTA | -0.10732 | 1.44E-08 |
| B57 | B157 | RBCAI | IgG | 0.107159 | 1.51E-08 |
| B57 | B195 | RBCAI | Na | -0.10571 | 2.36E-08 |
| B57 | B199 | RBCAI | Mg | -0.09579 | 4.25E-07 |
| B57 | B76 | RBCAI | ApoB | -0.09507 | 5.19E-07 |
| B57 | B198 | RBCAI | P | -0.09284 | 9.55E-07 |
| B58 | B60 | HCT | WBRVI | 0.703559 | 0 |
| B58 | B61 | HCT | WBHSRV | 0.350111 | 0 |
| B58 | B62 | HCT | WBHSRVI | 0.776619 | 0 |
| B58 | B63 | HCT | WBRSR(mPas_200) | 0.73977 | 0 |
| B58 | B64 | HCT | WBRSR(mPas_5) | 0.696418 | 0 |
| B58 | B65 | HCT | WBRSR(mPas_1) | 0.653837 | 0 |
| B58 | B66 | HCT | WBRSR(mPas_100) | 0.721302 | 0 |
| B58 | B67 | HCT | WBRSR(mPas_50) | 0.725321 | 0 |
| B58 | B213 | HCT | CVA-R | -0.15204 | 8.88E-16 |
| B58 | B59 | HCT | WBRV | 0.142542 | 4.46E-14 |
| B58 | B69 | HCT | PV | -0.14188 | 5.86E-14 |
| B58 | B124 | HCT | ACTH | 0.122871 | 8.21E-11 |
| B58 | B110 | HCT | AHA | -0.11478 | 1.31E-09 |
| B58 | B186 | HCT | PFC | -0.11243 | 2.84E-09 |
| B58 | B138 | HCT | PAB | 0.108775 | 9.10E-09 |
| B58 | B100 | HCT | anti-Ro52 | -0.10733 | 1.43E-08 |
| B58 | B105 | HCT | anti-U1RNP | -0.10712 | 1.53E-08 |
| B58 | B221 | HCT | Hei | 0.105975 | 2.17E-08 |
| B58 | B198 | HCT | P | -0.10539 | 2.60E-08 |
| B58 | B106 | HCT | anti-P | -0.103 | 5.34E-08 |
| B58 | B125 | HCT | Cor | 0.099164 | 1.64E-07 |
| B58 | B107 | HCT | AnuA | -0.09849 | 1.99E-07 |
| B58 | B75 | HCT | ApoA1/B | -0.0983 | 2.10E-07 |
| B59 | B112 | WBRV | FT4 | 0.166323 | 0 |
| B59 | B113 | WBRV | FT3 | 0.156307 | 0 |
| B59 | B115 | WBRV | TSH TT3 TT4(TT3) | 0.185931 | 0 |

| | | | | | |
|---|---|---|---|---|---|
| B59 | B124 | WBRV | ACTH | 0.34699 | 0 |
| B59 | B125 | WBRV | Cor | 0.204477 | 0 |
| B59 | B150 | WBRV | ALP | 0.160662 | 0 |
| B59 | B60 | WBRV | WBRVI | 0.716101 | 0 |
| B59 | B61 | WBRV | WBHSRV | 0.765263 | 0 |
| B59 | B62 | WBRV | WBHSRVI | 0.478254 | 0 |
| B59 | B63 | WBRV | WBRSR(mPas_200) | 0.621065 | 0 |
| B59 | B64 | WBRV | WBRSR(mPas_5) | 0.782103 | 0 |
| B59 | B65 | WBRV | WBRSR(mPas_1) | 0.825781 | 0 |
| B59 | B66 | WBRV | WBRSR(mPas_100) | 0.605514 | 0 |
| B59 | B67 | WBRV | WBRSR(mPas_50) | 0.635937 | 0 |
| B59 | B69 | WBRV | PV | 0.315128 | 0 |
| B59 | B108 | WBRV | Anti-Cyclin | 0.145024 | 1.60E-14 |
| B59 | B110 | WBRV | AHA | -0.14081 | 9.06E-14 |
| B59 | B157 | WBRV | IgG | 0.133296 | 1.76E-12 |
| B59 | B114 | WBRV | TSH TT3 TT4(TT4) | 0.126551 | 2.19E-11 |
| B59 | B180 | WBRV | Tf | 0.12607 | 2.61E-11 |
| B59 | B195 | WBRV | Na | -0.12173 | 1.23E-10 |
| B59 | B198 | WBRV | P | -0.10095 | 9.76E-08 |
| B59 | B178 | WBRV | Ig-U | 0.100069 | 1.26E-07 |
| B59 | B100 | WBRV | anti-Ro52 | -0.09722 | 2.85E-07 |
| B59 | B155 | WBRV | IgA | 0.093773 | 7.41E-07 |
| B59 | B139 | WBRV | ADA | 0.092739 | 9.81E-07 |
| B6 | B10 | WB-Fe | PG I | 0.141562 | 6.66E-14 |
| B6 | B12 | WB-Fe | PG II | 0.115885 | 9.08E-10 |
| B6 | B64 | WB-Fe | WBRSR(mPas_5) | 0.109327 | 7.65E-09 |
| B6 | B65 | WB-Fe | WBRSR(mPas_1) | 0.109308 | 7.70E-09 |
| B6 | B60 | WB-Fe | WBRVI | 0.108416 | 1.02E-08 |
| B6 | B67 | WB-Fe | WBRSR(mPas_50) | 0.105381 | 2.60E-08 |
| B6 | B66 | WB-Fe | WBRSR(mPas_100) | 0.103484 | 4.62E-08 |
| B6 | B63 | WB-Fe | WBRSR(mPas_200) | 0.101884 | 7.42E-08 |
| B6 | B62 | WB-Fe | WBHSRVI | 0.097631 | 2.54E-07 |
| B6 | B58 | WB-Fe | HCT | 0.096168 | 3.83E-07 |
| B6 | B143 | WB-Fe | β-CTx | 0.095837 | 4.20E-07 |
| B60 | B124 | WBRVI | ACTH | 0.237131 | 0 |
| B60 | B125 | WBRVI | Cor | 0.1748 | 0 |
| B60 | B61 | WBRVI | WBHSRV | 0.704082 | 0 |
| B60 | B62 | WBRVI | WBHSRVI | 0.885661 | 0 |
| B60 | B63 | WBRVI | WBRSR(mPas_200) | 0.81157 | 0 |
| B60 | B64 | WBRVI | WBRSR(mPas_5) | 0.916571 | 0 |
| B60 | B65 | WBRVI | WBRSR(mPas_1) | 0.934416 | 0 |
| B60 | B66 | WBRVI | WBRSR(mPas_100) | 0.811778 | 0 |
| B60 | B67 | WBRVI | WBRSR(mPas_50) | 0.839005 | 0 |
| B60 | B69 | WBRVI | PV | -0.16986 | 0 |

| | | | | | |
|---|---|---|---|---|---|
| B60 | B110 | WBRVI | AHA | -0.14956 | 2.22E-15 |
| B60 | B198 | WBRVI | P | -0.14578 | 1.15E-14 |
| B60 | B213 | WBRVI | CVA-R | -0.13555 | 7.33E-13 |
| B60 | B100 | WBRVI | anti-Ro52 | -0.13222 | 2.65E-12 |
| B60 | B101 | WBRVI | anti-Sc70 | -0.1321 | 2.78E-12 |
| B60 | B102 | WBRVI | anti-Sm | -0.1321 | 2.78E-12 |
| B60 | B104 | WBRVI | anti-SSB | -0.1321 | 2.78E-12 |
| B60 | B98 | WBRVI | anti-Jo-1 | -0.1321 | 2.78E-12 |
| B60 | B99 | WBRVI | anti-PM-Scl | -0.1321 | 2.78E-12 |
| B60 | B115 | WBRVI | TSH TT3 TT4(TT3) | 0.121207 | 1.47E-10 |
| B60 | B105 | WBRVI | anti-U1RNP | -0.11908 | 3.07E-10 |
| B60 | B106 | WBRVI | anti-P | -0.11313 | 2.26E-09 |
| B60 | B186 | WBRVI | PFC | -0.10877 | 9.12E-09 |
| B60 | B107 | WBRVI | AnuA | -0.10669 | 1.74E-08 |
| B60 | B137 | WBRVI | AFU | 0.105119 | 2.82E-08 |
| B60 | B183 | WBRVI | PTA | -0.10396 | 4.00E-08 |
| B60 | B113 | WBRVI | FT3 | 0.100716 | 1.05E-07 |
| B61 | B100 | WBHSRV | anti-Ro52 | -0.16991 | 0 |
| B61 | B105 | WBHSRV | anti-U1RNP | -0.19451 | 0 |
| B61 | B106 | WBHSRV | anti-P | -0.19 | 0 |
| B61 | B107 | WBHSRV | AnuA | -0.18493 | 0 |
| B61 | B108 | WBHSRV | Anti-Cyclin | 0.15812 | 0 |
| B61 | B62 | WBHSRV | WBHSRVI | 0.783169 | 0 |
| B61 | B63 | WBHSRV | WBRSR(mPas_200) | 0.862998 | 0 |
| B61 | B64 | WBHSRV | WBRSR(mPas_5) | 0.817153 | 0 |
| B61 | B65 | WBHSRV | WBRSR(mPas_1) | 0.7582 | 0 |
| B61 | B66 | WBHSRV | WBRSR(mPas_100) | 0.865252 | 0 |
| B61 | B67 | WBHSRV | WBRSR(mPas_50) | 0.865441 | 0 |
| B61 | B69 | WBHSRV | PV | 0.151908 | 8.88E-16 |
| B61 | B121 | WBHSRV | C3 | 0.144729 | 1.82E-14 |
| B61 | B180 | WBHSRV | Tf | 0.134507 | 1.10E-12 |
| B61 | B96 | WBHSRV | dsDNA (IIF) | 0.130736 | 4.64E-12 |
| B61 | B75 | WBHSRV | ApoA1/B | -0.12134 | 1.41E-10 |
| B61 | B122 | WBHSRV | C4 | 0.119164 | 2.99E-10 |
| B61 | B74 | WBHSRV | ApoA1 | -0.11436 | 1.51E-09 |
| B61 | B137 | WBHSRV | AFU | 0.113629 | 1.92E-09 |
| B61 | B213 | WBHSRV | CVA-R | -0.1079 | 1.20E-08 |
| B61 | B71 | WBHSRV | TG | 0.107825 | 1.23E-08 |
| B61 | B124 | WBHSRV | ACTH | 0.107323 | 1.43E-08 |
| B61 | B178 | WBHSRV | Ig-U | 0.104704 | 3.20E-08 |
| B61 | B198 | WBHSRV | P | -0.10378 | 4.22E-08 |
| B61 | B101 | WBHSRV | anti-Sc70 | -0.09291 | 9.36E-07 |
| B61 | B102 | WBHSRV | anti-Sm | -0.09291 | 9.36E-07 |
| B61 | B104 | WBHSRV | anti-SSB | -0.09291 | 9.36E-07 |

| | | | | | |
|---|---|---|---|---|---|
| B61 | B98 | WBHSRV | anti-Jo-1 | -0.09291 | 9.36E-07 |
| B61 | B99 | WBHSRV | anti-PM-Scl | -0.09291 | 9.36E-07 |
| B62 | B100 | WBHSRVI | anti-Ro52 | -0.16391 | 0 |
| B62 | B105 | WBHSRVI | anti-U1RNP | -0.17035 | 0 |
| B62 | B106 | WBHSRVI | anti-P | -0.16468 | 0 |
| B62 | B107 | WBHSRVI | AnuA | -0.15844 | 0 |
| B62 | B186 | WBHSRVI | PFC | -0.16378 | 0 |
| B62 | B63 | WBHSRVI | WBRSR(mPas_200) | 0.890539 | 0 |
| B62 | B64 | WBHSRVI | WBRSR(mPas_5) | 0.843518 | 0 |
| B62 | B65 | WBHSRVI | WBRSR(mPas_1) | 0.789382 | 0 |
| B62 | B66 | WBHSRVI | WBRSR(mPas_100) | 0.900076 | 0 |
| B62 | B67 | WBHSRVI | WBRSR(mPas_50) | 0.903931 | 0 |
| B62 | B69 | WBHSRVI | PV | -0.26068 | 0 |
| B62 | B213 | WBHSRVI | CVA-R | -0.14004 | 1.23E-13 |
| B62 | B198 | WBHSRVI | P | -0.13138 | 3.65E-12 |
| B62 | B121 | WBHSRVI | C3 | 0.1244 | 4.76E-11 |
| B62 | B137 | WBHSRVI | AFU | 0.123891 | 5.71E-11 |
| B62 | B101 | WBHSRVI | anti-Sc70 | -0.12197 | 1.13E-10 |
| B62 | B102 | WBHSRVI | anti-Sm | -0.12197 | 1.13E-10 |
| B62 | B104 | WBHSRVI | anti-SSB | -0.12197 | 1.13E-10 |
| B62 | B98 | WBHSRVI | anti-Jo-1 | -0.12197 | 1.13E-10 |
| B62 | B99 | WBHSRVI | anti-PM-Scl | -0.12197 | 1.13E-10 |
| B62 | B74 | WBHSRVI | ApoA1 | -0.11656 | 7.24E-10 |
| B62 | B103 | WBHSRVI | anti-SSA/Ro | -0.1161 | 8.45E-10 |
| B62 | B75 | WBHSRVI | ApoA1/B | -0.11358 | 1.95E-09 |
| B62 | B96 | WBHSRVI | dsDNA (IIF) | 0.111762 | 3.52E-09 |
| B62 | B122 | WBHSRVI | C4 | 0.102009 | 7.15E-08 |
| B62 | B222 | WBHSRVI | NS | 0.100604 | 1.08E-07 |
| B62 | B156 | WBHSRVI | IgE | -0.09985 | 1.34E-07 |
| B62 | B79 | WBHSRVI | hb V -dna | -0.09684 | 3.17E-07 |
| B62 | B163 | WBHSRVI | SG | 0.096243 | 3.75E-07 |
| B62 | B221 | WBHSRVI | Hei | 0.096101 | 3.90E-07 |
| B62 | B110 | WBHSRVI | AHA | -0.0941 | 6.78E-07 |
| B63 | B100 | WBRSR(mPas_200) | anti-Ro52 | -0.16923 | 0 |
| B63 | B105 | WBRSR(mPas_200) | anti-U1RNP | -0.17681 | 0 |
| B63 | B106 | WBRSR(mPas_200) | anti-P | -0.17099 | 0 |
| B63 | B107 | WBRSR(mPas_200) | AnuA | -0.16459 | 0 |
| B63 | B124 | WBRSR(mPas_200) | ACTH | 0.183508 | 0 |
| B63 | B64 | WBRSR(mPas_200) | WBRSR(mPas_5) | 0.94472 | 0 |
| B63 | B65 | WBRSR(mPas_200) | WBRSR(mPas_1) | 0.879986 | 0 |
| B63 | B66 | WBRSR(mPas_200) | WBRSR(mPas_100) | 0.984381 | 0 |
| B63 | B67 | WBRSR(mPas_200) | WBRSR(mPas_50) | 0.984549 | 0 |
| B63 | B69 | WBRSR(mPas_200) | PV | 0.189701 | 0 |
| B63 | B108 | WBRSR(mPas_200) | Anti-Cyclin | 0.151764 | 8.88E-16 |

| | | | | | |
|---|---|---|---|---|---|
| B63 | B75 | WBRSR(mPas_200) | ApoA1/B | -0.13217 | 2.70E-12 |
| B63 | B180 | WBRSR(mPas_200) | Tf | 0.131909 | 2.98E-12 |
| B63 | B121 | WBRSR(mPas_200) | C3 | 0.128661 | 1.01E-11 |
| B63 | B213 | WBRSR(mPas_200) | CVA-R | -0.12646 | 2.26E-11 |
| B63 | B74 | WBRSR(mPas_200) | ApoA1 | -0.12571 | 2.98E-11 |
| B63 | B101 | WBRSR(mPas_200) | anti-Sc70 | -0.12489 | 3.99E-11 |
| B63 | B102 | WBRSR(mPas_200) | anti-Sm | -0.12489 | 3.99E-11 |
| B63 | B104 | WBRSR(mPas_200) | anti-SSB | -0.12489 | 3.99E-11 |
| B63 | B98 | WBRSR(mPas_200) | anti-Jo-1 | -0.12489 | 3.99E-11 |
| B63 | B99 | WBRSR(mPas_200) | anti-PM-Scl | -0.12489 | 3.99E-11 |
| B63 | B198 | WBRSR(mPas_200) | P | -0.12322 | 7.25E-11 |
| B63 | B96 | WBRSR(mPas_200) | dsDNA (IIF) | 0.11585 | 9.19E-10 |
| B63 | B110 | WBRSR(mPas_200) | AHA | -0.1096 | 7.03E-09 |
| B63 | B122 | WBRSR(mPas_200) | C4 | 0.105426 | 2.57E-08 |
| B63 | B125 | WBRSR(mPas_200) | Cor | 0.105194 | 2.75E-08 |
| B63 | B178 | WBRSR(mPas_200) | Ig-U | 0.104105 | 3.83E-08 |
| B63 | B137 | WBRSR(mPas_200) | AFU | 0.101801 | 7.61E-08 |
| B63 | B155 | WBRSR(mPas_200) | IgA | 0.098349 | 2.07E-07 |
| B63 | B163 | WBRSR(mPas_200) | SG | 0.098014 | 2.28E-07 |
| B63 | B71 | WBRSR(mPas_200) | TG | 0.097764 | 2.44E-07 |
| B64 | B124 | WBRSR(mPas_5) | ACTH | 0.301959 | 0 |
| B64 | B125 | WBRSR(mPas_5) | Cor | 0.166802 | 0 |
| B64 | B65 | WBRSR(mPas_5) | WBRSR(mPas_1) | 0.985334 | 0 |
| B64 | B66 | WBRSR(mPas_5) | WBRSR(mPas_100) | 0.929339 | 0 |
| B64 | B67 | WBRSR(mPas_5) | WBRSR(mPas_50) | 0.943965 | 0 |
| B64 | B69 | WBRSR(mPas_5) | PV | 0.195853 | 0 |
| B64 | B100 | WBRSR(mPas_5) | anti-Ro52 | -0.15385 | 4.44E-16 |
| B64 | B101 | WBRSR(mPas_5) | anti-Sc70 | -0.15119 | 1.11E-15 |
| B64 | B102 | WBRSR(mPas_5) | anti-Sm | -0.15119 | 1.11E-15 |
| B64 | B104 | WBRSR(mPas_5) | anti-SSB | -0.15119 | 1.11E-15 |
| B64 | B98 | WBRSR(mPas_5) | anti-Jo-1 | -0.15119 | 1.11E-15 |
| B64 | B99 | WBRSR(mPas_5) | anti-PM-Scl | -0.15119 | 1.11E-15 |
| B64 | B115 | WBRSR(mPas_5) | TSH TT3 TT4(TT3) | 0.149168 | 2.66E-15 |
| B64 | B110 | WBRSR(mPas_5) | AHA | -0.147 | 7.11E-15 |
| B64 | B198 | WBRSR(mPas_5) | P | -0.14318 | 3.42E-14 |
| B64 | B105 | WBRSR(mPas_5) | anti-U1RNP | -0.14008 | 1.21E-13 |
| B64 | B108 | WBRSR(mPas_5) | Anti-Cyclin | 0.134463 | 1.12E-12 |
| B64 | B113 | WBRSR(mPas_5) | FT3 | 0.133997 | 1.34E-12 |
| B64 | B106 | WBRSR(mPas_5) | anti-P | -0.13326 | 1.78E-12 |
| B64 | B112 | WBRSR(mPas_5) | FT4 | 0.127369 | 1.62E-11 |
| B64 | B213 | WBRSR(mPas_5) | CVA-R | -0.12616 | 2.53E-11 |
| B64 | B107 | WBRSR(mPas_5) | AnuA | -0.12587 | 2.81E-11 |
| B64 | B180 | WBRSR(mPas_5) | Tf | 0.117264 | 5.72E-10 |
| B64 | B74 | WBRSR(mPas_5) | ApoA1 | -0.10955 | 7.14E-09 |

| | | | | | |
|---|---|---|---|---|---|
| B64 | B150 | WBRSR(mPas_5) | ALP | 0.109539 | 7.15E-09 |
| B64 | B155 | WBRSR(mPas_5) | IgA | 0.105427 | 2.57E-08 |
| B64 | B137 | WBRSR(mPas_5) | AFU | 0.104441 | 3.46E-08 |
| B64 | B195 | WBRSR(mPas_5) | Na | -0.10108 | 9.41E-08 |
| B64 | B75 | WBRSR(mPas_5) | ApoA1/B | -0.10101 | 9.60E-08 |
| B64 | B121 | WBRSR(mPas_5) | C3 | 0.098828 | 1.80E-07 |
| B65 | B110 | WBRSR(mPas_1) | AHA | -0.15905 | 0 |
| B65 | B113 | WBRSR(mPas_1) | FT3 | 0.164487 | 0 |
| B65 | B115 | WBRSR(mPas_1) | TSH TT3 TT4(TT3) | 0.179001 | 0 |
| B65 | B124 | WBRSR(mPas_1) | ACTH | 0.357081 | 0 |
| B65 | B125 | WBRSR(mPas_1) | Cor | 0.19874 | 0 |
| B65 | B66 | WBRSR(mPas_1) | WBRSR(mPas_100) | 0.86154 | 0 |
| B65 | B67 | WBRSR(mPas_1) | WBRSR(mPas_50) | 0.88309 | 0 |
| B65 | B69 | WBRSR(mPas_1) | PV | 0.177068 | 0 |
| B65 | B101 | WBRSR(mPas_1) | anti-Sc70 | -0.15147 | 8.88E-16 |
| B65 | B102 | WBRSR(mPas_1) | anti-Sm | -0.15147 | 8.88E-16 |
| B65 | B104 | WBRSR(mPas_1) | anti-SSB | -0.15147 | 8.88E-16 |
| B65 | B98 | WBRSR(mPas_1) | anti-Jo-1 | -0.15147 | 8.88E-16 |
| B65 | B99 | WBRSR(mPas_1) | anti-PM-Scl | -0.15147 | 8.88E-16 |
| B65 | B112 | WBRSR(mPas_1) | FT4 | 0.145826 | 1.15E-14 |
| B65 | B198 | WBRSR(mPas_1) | P | -0.14376 | 2.71E-14 |
| B65 | B100 | WBRSR(mPas_1) | anti-Ro52 | -0.13922 | 1.72E-13 |
| B65 | B213 | WBRSR(mPas_1) | CVA-R | -0.12284 | 8.31E-11 |
| B65 | B108 | WBRSR(mPas_1) | Anti-Cyclin | 0.119045 | 3.12E-10 |
| B65 | B105 | WBRSR(mPas_1) | anti-U1RNP | -0.11878 | 3.41E-10 |
| B65 | B150 | WBRSR(mPas_1) | ALP | 0.117348 | 5.55E-10 |
| B65 | B106 | WBRSR(mPas_1) | anti-P | -0.11201 | 3.25E-09 |
| B65 | B195 | WBRSR(mPas_1) | Na | -0.10887 | 8.85E-09 |
| B65 | B107 | WBRSR(mPas_1) | AnuA | -0.1047 | 3.20E-08 |
| B65 | B180 | WBRSR(mPas_1) | Tf | 0.104701 | 3.20E-08 |
| B65 | B114 | WBRSR(mPas_1) | TSH TT3 TT4(TT4) | 0.102808 | 5.65E-08 |
| B65 | B74 | WBRSR(mPas_1) | ApoA1 | -0.09593 | 4.09E-07 |
| B66 | B100 | WBRSR(mPas_100) | anti-Ro52 | -0.16627 | 0 |
| B66 | B105 | WBRSR(mPas_100) | anti-U1RNP | -0.17001 | 0 |
| B66 | B106 | WBRSR(mPas_100) | anti-P | -0.16404 | 0 |
| B66 | B107 | WBRSR(mPas_100) | AnuA | -0.15748 | 0 |
| B66 | B108 | WBRSR(mPas_100) | Anti-Cyclin | 0.157802 | 0 |
| B66 | B67 | WBRSR(mPas_100) | WBRSR(mPas_50) | 0.995063 | 0 |
| B66 | B69 | WBRSR(mPas_100) | PV | 0.149327 | 2.66E-15 |
| B66 | B75 | WBRSR(mPas_100) | ApoA1/B | -0.13848 | 2.30E-13 |
| B66 | B180 | WBRSR(mPas_100) | Tf | 0.135601 | 7.19E-13 |
| B66 | B198 | WBRSR(mPas_100) | P | -0.13145 | 3.55E-12 |
| B66 | B101 | WBRSR(mPas_100) | anti-Sc70 | -0.12903 | 8.80E-12 |
| B66 | B102 | WBRSR(mPas_100) | anti-Sm | -0.12903 | 8.80E-12 |

| | | | | | |
|---|---|---|---|---|---|
| B66 | B104 | WBRSR(mPas_100) | anti-SSB | -0.12903 | 8.80E-12 |
| B66 | B98 | WBRSR(mPas_100) | anti-Jo-1 | -0.12903 | 8.80E-12 |
| B66 | B99 | WBRSR(mPas_100) | anti-PM-Scl | -0.12903 | 8.80E-12 |
| B66 | B213 | WBRSR(mPas_100) | CVA-R | -0.12527 | 3.49E-11 |
| B66 | B74 | WBRSR(mPas_100) | ApoA1 | -0.12505 | 3.77E-11 |
| B66 | B121 | WBRSR(mPas_100) | C3 | 0.123502 | 6.56E-11 |
| B66 | B137 | WBRSR(mPas_100) | AFU | 0.122265 | 1.02E-10 |
| B66 | B92 | WBRSR(mPas_100) | ALDH2-Glu504Glu | -0.11449 | 1.44E-09 |
| B66 | B110 | WBRSR(mPas_100) | AHA | -0.1122 | 3.06E-09 |
| B66 | B96 | WBRSR(mPas_100) | dsDNA (IIF) | 0.111026 | 4.46E-09 |
| B66 | B155 | WBRSR(mPas_100) | IgA | 0.109993 | 6.20E-09 |
| B66 | B178 | WBRSR(mPas_100) | Ig-U | 0.105992 | 2.16E-08 |
| B66 | B71 | WBRSR(mPas_100) | TG | 0.105531 | 2.49E-08 |
| B66 | B122 | WBRSR(mPas_100) | C4 | 0.101282 | 8.86E-08 |
| B66 | B163 | WBRSR(mPas_100) | SG | 0.097717 | 2.48E-07 |
| B67 | B100 | WBRSR(mPas_50) | anti-Ro52 | -0.1628 | 0 |
| B67 | B105 | WBRSR(mPas_50) | anti-U1RNP | -0.16417 | 0 |
| B67 | B106 | WBRSR(mPas_50) | anti-P | -0.15815 | 0 |
| B67 | B108 | WBRSR(mPas_50) | Anti-Cyclin | 0.155811 | 0 |
| B67 | B107 | WBRSR(mPas_50) | AnuA | -0.15155 | 8.88E-16 |
| B67 | B69 | WBRSR(mPas_50) | PV | 0.141693 | 6.31E-14 |
| B67 | B180 | WBRSR(mPas_50) | Tf | 0.13418 | 1.25E-12 |
| B67 | B101 | WBRSR(mPas_50) | anti-Sc70 | -0.13057 | 4.94E-12 |
| B67 | B102 | WBRSR(mPas_50) | anti-Sm | -0.13057 | 4.94E-12 |
| B67 | B104 | WBRSR(mPas_50) | anti-SSB | -0.13057 | 4.94E-12 |
| B67 | B98 | WBRSR(mPas_50) | anti-Jo-1 | -0.13057 | 4.94E-12 |
| B67 | B99 | WBRSR(mPas_50) | anti-PM-Scl | -0.13057 | 4.94E-12 |
| B67 | B213 | WBRSR(mPas_50) | CVA-R | -0.1263 | 2.39E-11 |
| B67 | B198 | WBRSR(mPas_50) | P | -0.12566 | 3.03E-11 |
| B67 | B75 | WBRSR(mPas_50) | ApoA1/B | -0.12536 | 3.38E-11 |
| B67 | B121 | WBRSR(mPas_50) | C3 | 0.118888 | 3.29E-10 |
| B67 | B110 | WBRSR(mPas_50) | AHA | -0.11887 | 3.30E-10 |
| B67 | B74 | WBRSR(mPas_50) | ApoA1 | -0.11685 | 6.57E-10 |
| B67 | B155 | WBRSR(mPas_50) | IgA | 0.114318 | 1.53E-09 |
| B67 | B137 | WBRSR(mPas_50) | AFU | 0.112296 | 2.96E-09 |
| B67 | B96 | WBRSR(mPas_50) | dsDNA (IIF) | 0.106823 | 1.67E-08 |
| B67 | B178 | WBRSR(mPas_50) | Ig-U | 0.104746 | 3.16E-08 |
| B67 | B71 | WBRSR(mPas_50) | TG | 0.102432 | 6.31E-08 |
| B67 | B156 | WBRSR(mPas_50) | IgE | -0.10067 | 1.06E-07 |
| B67 | B122 | WBRSR(mPas_50) | C4 | 0.097466 | 2.66E-07 |
| B67 | B186 | WBRSR(mPas_50) | PFC | -0.0969 | 3.12E-07 |
| B69 | B108 | PV | Anti-Cyclin | 0.312036 | 0 |
| B69 | B112 | PV | FT4 | 0.200942 | 0 |
| B69 | B124 | PV | ACTH | 0.265733 | 0 |

| | | | | | |
|---|---|---|---|---|---|
| B69 | B157 | PV | IgG | 0.177984 | 0 |
| B69 | B178 | PV | Ig-U | 0.216485 | 0 |
| B69 | B180 | PV | Tf | 0.273285 | 0 |
| B69 | B186 | PV | PFC | 0.182151 | 0 |
| B69 | B114 | PV | TSH TT3 TT4(TT4) | 0.146221 | 9.77E-15 |
| B69 | B113 | PV | FT3 | 0.14222 | 5.11E-14 |
| B69 | B130 | PV | GLB | 0.140902 | 8.70E-14 |
| B69 | B103 | PV | anti-SSA/Ro | 0.139624 | 1.46E-13 |
| B69 | B150 | PV | ALP | 0.125108 | 3.69E-11 |
| B69 | B139 | PV | ADA | 0.1198 | 2.40E-10 |
| B69 | B115 | PV | TSH TT3 TT4(TT3) | 0.118849 | 3.33E-10 |
| B69 | B127 | PV | A-G ratio | -0.11836 | 3.94E-10 |
| B69 | B135 | PV | TBA | 0.115901 | 9.04E-10 |
| B69 | B91 | PV | TP | 0.1108 | 4.79E-09 |
| B69 | B155 | PV | IgA | 0.105292 | 2.67E-08 |
| B69 | B195 | PV | Na | -0.10124 | 8.98E-08 |
| B69 | B182 | PV | TT | 0.097987 | 2.29E-07 |
| B69 | B177 | PV | ACR | 0.096872 | 3.14E-07 |
| B69 | B123 | PV | hs-CRP1 | 0.093705 | 7.55E-07 |
| B7 | B10 | WB-Cop | PG Ⅰ | -0.24312 | 0 |
| B7 | B12 | WB-Cop | PG Ⅱ | -0.21055 | 0 |
| B7 | B136 | WB-Cop | 5'NT | -0.16361 | 0 |
| B7 | B143 | WB-Cop | β-CTx | -0.16052 | 0 |
| B7 | B8 | WB-Cop | WB-Zinc | 0.392668 | 0 |
| B7 | B87 | WB-Cop | NSE | -0.14916 | 2.66E-15 |
| B7 | B58 | WB-Cop | HCT | 0.115724 | 9.59E-10 |
| B7 | B140 | WB-Cop | r-GT | -0.11115 | 4.29E-09 |
| B70 | B132 | LDL-C | Dbil | -0.16727 | 0 |
| B70 | B75 | LDL-C | ApoA1/B | -0.22456 | 0 |
| B70 | B76 | LDL-C | ApoB | 0.328064 | 0 |
| B70 | B77 | LDL-C | TC | 0.837962 | 0 |
| B70 | B218 | LDL-C | DBP | 0.120384 | 1.96E-10 |
| B70 | B206 | LDL-C | Waist | 0.119007 | 3.16E-10 |
| B70 | B215 | LDL-C | BMI | 0.110965 | 4.55E-09 |
| B70 | B188 | LDL-C | Ca | 0.108503 | 9.92E-09 |
| B70 | B91 | LDL-C | TP | 0.106063 | 2.11E-08 |
| B70 | B141 | LDL-C | AST/ALT | -0.09548 | 4.63E-07 |
| B71 | B128 | TG | ALT | 0.18488 | 0 |
| B71 | B130 | TG | GLB | 0.218675 | 0 |
| B71 | B140 | TG | r-GT | 0.241241 | 0 |
| B71 | B141 | TG | AST/ALT | -0.18708 | 0 |
| B71 | B153 | TG | Glu | 0.24666 | 0 |
| B71 | B170 | TG | U-GLU | 0.189889 | 0 |
| B71 | B192 | TG | UA | 0.272191 | 0 |

| | | | | | |
|---|---|---|---|---|---|
| B71 | B206 | TG | Waist | 0.276803 | 0 |
| B71 | B209 | TG | Smoking | 0.158343 | 0 |
| B71 | B210 | TG | Hip | 0.172206 | 0 |
| B71 | B215 | TG | BMI | 0.261185 | 0 |
| B71 | B216 | TG | BMI | 0.246309 | 0 |
| B71 | B218 | TG | DBP | 0.191631 | 0 |
| B71 | B222 | TG | NS | 0.259761 | 0 |
| B71 | B72 | TG | HDL-C | -0.41526 | 0 |
| B71 | B77 | TG | TC | 0.31956 | 0 |
| B71 | B91 | TG | TP | 0.294463 | 0 |
| B71 | B126 | TG | Alb | 0.154035 | 4.44E-16 |
| B71 | B220 | TG | SBP | 0.145327 | 1.42E-14 |
| B71 | B134 | TG | CHE | 0.139664 | 1.44E-13 |
| B71 | B188 | TG | Ca | 0.138083 | 2.70E-13 |
| B71 | B163 | TG | SG | 0.129536 | 7.28E-12 |
| B71 | B204 | TG | Drinking | 0.123472 | 6.64E-11 |
| B71 | B160 | TG | PH | -0.11945 | 2.71E-10 |
| B71 | B75 | TG | ApoA1/B | -0.11366 | 1.90E-09 |
| B71 | B138 | TG | PAB | 0.113041 | 2.32E-09 |
| B71 | B127 | TG | A-G ratio | -0.1114 | 3.95E-09 |
| B71 | B132 | TG | Dbil | -0.10245 | 6.29E-08 |
| B72 | B128 | HDL-C | ALT | -0.18443 | 0 |
| B72 | B141 | HDL-C | AST/ALT | 0.264635 | 0 |
| B72 | B153 | HDL-C | Glu | -0.16874 | 0 |
| B72 | B163 | HDL-C | SG | -0.20234 | 0 |
| B72 | B192 | HDL-C | UA | -0.32531 | 0 |
| B72 | B206 | HDL-C | Waist | -0.36744 | 0 |
| B72 | B209 | HDL-C | Smoking | -0.18013 | 0 |
| B72 | B210 | HDL-C | Hip | -0.26268 | 0 |
| B72 | B215 | HDL-C | BMI | -0.33804 | 0 |
| B72 | B216 | HDL-C | BMI | -0.39121 | 0 |
| B72 | B221 | HDL-C | Hei | -0.24951 | 0 |
| B72 | B222 | HDL-C | NS | -0.38959 | 0 |
| B72 | B74 | HDL-C | ApoA1 | 0.290673 | 0 |
| B72 | B75 | HDL-C | ApoA1/B | 0.211466 | 0 |
| B72 | B77 | HDL-C | TC | 0.23759 | 0 |
| B72 | B174 | HDL-C | Urea/Cr | 0.116736 | 6.83E-10 |
| B72 | B170 | HDL-C | U-GLU | -0.1163 | 7.91E-10 |
| B72 | B218 | HDL-C | DBP | -0.1161 | 8.46E-10 |
| B72 | B190 | HDL-C | Cre | -0.1152 | 1.14E-09 |
| B72 | B160 | HDL-C | PH | 0.107748 | 1.26E-08 |
| B72 | B176 | HDL-C | Ucr | -0.0999 | 1.33E-07 |
| B72 | B166 | HDL-C | Pro | -0.09626 | 3.73E-07 |
| B73 | B100 | ApoA | anti-Ro52 | 0.202486 | 0 |

| | | | | | |
|---|---|---|---|---|---|
| B73 | B109 | ApoA | AMA-M2 | 0.264609 | 0 |
| B73 | B110 | ApoA | AHA | 0.190559 | 0 |
| B73 | B76 | ApoA | ApoB | 0.215749 | 0 |
| B73 | B95 | ApoA | anti-cenpb | 0.297743 | 0 |
| B73 | B75 | ApoA | ApoA1/B | -0.10444 | 3.46E-08 |
| B73 | B156 | ApoA | IgE | 0.103436 | 4.68E-08 |
| B73 | B186 | ApoA | PFC | 0.098352 | 2.07E-07 |
| B74 | B75 | ApoA1 | ApoA1/B | 0.566001 | 0 |
| B74 | B194 | ApoA1 | Cl | -0.13318 | 1.84E-12 |
| B74 | B155 | ApoA1 | IgA | -0.12981 | 6.58E-12 |
| B74 | B169 | ApoA1 | U-CRY | 0.129414 | 7.62E-12 |
| B74 | B159 | ApoA1 | β2-MG | 0.129412 | 7.63E-12 |
| B74 | B139 | ApoA1 | ADA | -0.12661 | 2.15E-11 |
| B74 | B119 | ApoA1 | Hcv-rna | -0.12502 | 3.81E-11 |
| B74 | B197 | ApoA1 | AG | 0.124498 | 4.60E-11 |
| B74 | B199 | ApoA1 | Mg | 0.122932 | 8.04E-11 |
| B74 | B77 | ApoA1 | TC | 0.10785 | 1.22E-08 |
| B74 | B213 | ApoA1 | CVA-R | 0.106875 | 1.65E-08 |
| B74 | B216 | ApoA1 | BMI | -0.10479 | 3.11E-08 |
| B74 | B95 | ApoA1 | anti-cenpb | 0.103345 | 4.81E-08 |
| B74 | B125 | ApoA1 | Cor | 0.098361 | 2.06E-07 |
| B74 | B206 | ApoA1 | Waist | -0.09621 | 3.79E-07 |
| B74 | B89 | ApoA1 | CA 72-4 | 0.094231 | 6.54E-07 |
| B75 | B134 | ApoA1/B | CHE | -0.184 | 0 |
| B75 | B76 | ApoA1/B | ApoB | -0.74097 | 0 |
| B75 | B77 | ApoA1/B | TC | -0.16919 | 0 |
| B75 | B86 | ApoA1/B | TBhCG | 0.162557 | 0 |
| B75 | B110 | ApoA1/B | AHA | 0.140097 | 1.21E-13 |
| B75 | B139 | ApoA1/B | ADA | -0.13182 | 3.09E-12 |
| B75 | B89 | ApoA1/B | CA 72-4 | 0.125267 | 3.49E-11 |
| B75 | B206 | ApoA1/B | Waist | -0.11412 | 1.63E-09 |
| B75 | B200 | ApoA1/B | OP | -0.11333 | 2.11E-09 |
| B75 | B215 | ApoA1/B | BMI | -0.10995 | 6.28E-09 |
| B75 | B153 | ApoA1/B | Glu | -0.10601 | 2.14E-08 |
| B75 | B198 | ApoA1/B | P | 0.105523 | 2.49E-08 |
| B75 | B213 | ApoA1/B | CVA-R | 0.103923 | 4.05E-08 |
| B75 | B216 | ApoA1/B | BMI | -0.10359 | 4.46E-08 |
| B75 | B79 | ApoA1/B | hb V -dna | 0.102493 | 6.20E-08 |
| B75 | B150 | ApoA1/B | ALP | -0.09789 | 2.36E-07 |
| B75 | B170 | ApoA1/B | U-GLU | -0.09764 | 2.53E-07 |
| B75 | B222 | ApoA1/B | NS | -0.09624 | 3.76E-07 |
| B75 | B136 | ApoA1/B | 5'NT | -0.09579 | 4.25E-07 |
| B76 | B77 | ApoB | TC | 0.311362 | 0 |
| B76 | B134 | ApoB | CHE | 0.153232 | 4.44E-16 |

| | | | | | |
|---|---|---|---|---|---|
| B76 | B194 | ApoB | Cl | -0.15076 | 1.33E-15 |
| B76 | B138 | ApoB | PAB | 0.138585 | 2.21E-13 |
| B76 | B153 | ApoB | Glu | 0.133953 | 1.36E-12 |
| B76 | B170 | ApoB | U-GLU | 0.123824 | 5.85E-11 |
| B76 | B136 | ApoB | 5'NT | 0.118658 | 3.56E-10 |
| B76 | B150 | ApoB | ALP | 0.112226 | 3.03E-09 |
| B76 | B186 | ApoB | PFC | 0.111628 | 3.67E-09 |
| B76 | B200 | ApoB | OP | 0.106639 | 1.77E-08 |
| B76 | B86 | ApoB | TBhCG | -0.10605 | 2.12E-08 |
| B76 | B156 | ApoB | IgE | 0.10523 | 2.72E-08 |
| B76 | B179 | ApoB | α1-MG | 0.102857 | 5.56E-08 |
| B77 | B130 | TC | GLB | 0.158182 | 0 |
| B77 | B132 | TC | Dbil | -0.18271 | 0 |
| B77 | B188 | TC | Ca | 0.170981 | 0 |
| B77 | B91 | TC | TP | 0.196918 | 0 |
| B77 | B218 | TC | DBP | 0.133085 | 1.90E-12 |
| B77 | B153 | TC | Glu | 0.123557 | 6.44E-11 |
| B77 | B221 | TC | Hei | -0.12118 | 1.49E-10 |
| B77 | B220 | TC | SBP | 0.117062 | 6.12E-10 |
| B77 | B127 | TC | A-G ratio | -0.10785 | 1.22E-08 |
| B77 | B140 | TC | r-GT | 0.107807 | 1.23E-08 |
| B77 | B189 | TC | eGFB | -0.10211 | 6.95E-08 |
| B78 | B108 | HBcAb | Anti-Cyclin | 0.153783 | 4.44E-16 |
| B78 | B80 | HBcAb | anti-HP IgG | -0.12692 | 1.92E-11 |
| B78 | B180 | HBcAb | Tf | 0.123753 | 6.00E-11 |
| B78 | B185 | HBcAb | PT | -0.09939 | 1.54E-07 |
| B78 | B184 | HBcAb | INR | -0.09829 | 2.11E-07 |
| B78 | B145 | HBcAb | BGP | 0.09418 | 6.63E-07 |
| B79 | B101 | hb V -dna | anti-Sc70 | 0.30852 | 0 |
| B79 | B102 | hb V -dna | anti-Sm | 0.30852 | 0 |
| B79 | B104 | hb V -dna | anti-SSB | 0.30852 | 0 |
| B79 | B119 | hb V -dna | Hcv-rna | 0.309939 | 0 |
| B79 | B159 | hb V -dna | β2-MG | -0.28743 | 0 |
| B79 | B169 | hb V -dna | U-CRY | -0.28744 | 0 |
| B79 | B183 | hb V -dna | PTA | 0.200016 | 0 |
| B79 | B81 | hb V -dna | TSGF | -0.2599 | 0 |
| B79 | B98 | hb V -dna | anti-Jo-1 | 0.30852 | 0 |
| B79 | B99 | hb V -dna | anti-PM-Scl | 0.30852 | 0 |
| B79 | B92 | hb V -dna | ALDH2-Glu504Glu | 0.152433 | 6.66E-16 |
| B79 | B182 | hb V -dna | TT | -0.14702 | 6.88E-15 |
| B79 | B100 | hb V -dna | anti-Ro52 | 0.138942 | 1.92E-13 |
| B79 | B120 | hb V -dna | HCV-cAg | -0.1352 | 8.41E-13 |
| B79 | B181 | hb V -dna | APTT | -0.1297 | 6.85E-12 |
| B79 | B137 | hb V -dna | AFU | -0.12804 | 1.27E-11 |

| | | | | | |
|---|---|---|---|---|---|
| B79 | B158 | hb V -dna | IgM | 0.126611 | 2.14E-11 |
| B79 | B155 | hb V -dna | IgA | -0.11199 | 3.27E-09 |
| B79 | B177 | hb V -dna | ACR | -0.10932 | 7.67E-09 |
| B79 | B93 | hb V -dna | ALDH2-Glu504Lys | -0.10835 | 1.04E-08 |
| B79 | B139 | hb V -dna | ADA | -0.10328 | 4.90E-08 |
| B79 | B200 | hb V -dna | OP | -0.10031 | 1.18E-07 |
| B79 | B94 | hb V -dna | ALDH2-Lys504Lys | -0.0987 | 1.87E-07 |
| B79 | B185 | hb V -dna | PT | -0.09477 | 5.64E-07 |
| B8 | B88 | WB-Zinc | CA 242 | 0.103813 | 4.18E-08 |
| B8 | B59 | WB-Zinc | WBRV | -0.10083 | 1.01E-07 |
| B8 | B55 | WB-Zinc | TK | -0.10024 | 1.20E-07 |
| B8 | B85 | WB-Zinc | SCC | 0.093125 | 8.84E-07 |
| B81 | B125 | TSGF | Cor | -0.16307 | 0 |
| B81 | B156 | TSGF | IgE | 0.168617 | 0 |
| B81 | B182 | TSGF | TT | 0.162375 | 0 |
| B81 | B183 | TSGF | PTA | -0.17896 | 0 |
| B81 | B92 | TSGF | ALDH2-Glu504Glu | -0.16339 | 0 |
| B81 | B155 | TSGF | IgA | 0.154747 | 2.22E-16 |
| B81 | B94 | TSGF | ALDH2-Lys504Lys | 0.14974 | 2.22E-15 |
| B81 | B101 | TSGF | anti-Sc70 | -0.13617 | 5.76E-13 |
| B81 | B102 | TSGF | anti-Sm | -0.13617 | 5.76E-13 |
| B81 | B104 | TSGF | anti-SSB | -0.13617 | 5.76E-13 |
| B81 | B98 | TSGF | anti-Jo-1 | -0.13617 | 5.76E-13 |
| B81 | B99 | TSGF | anti-PM-Scl | -0.13617 | 5.76E-13 |
| B81 | B139 | TSGF | ADA | 0.126355 | 2.35E-11 |
| B81 | B138 | TSGF | PAB | -0.10795 | 1.18E-08 |
| B81 | B164 | TSGF | COM | 0.103346 | 4.81E-08 |
| B81 | B93 | TSGF | ALDH2-Glu504Lys | 0.093397 | 8.21E-07 |
| B82 | B83 | CEA | AFP | 0.293831 | 0 |
| B82 | B84 | CEA | C-PSA | 0.265755 | 0 |
| B82 | B88 | CEA | CA 242 | 0.312235 | 0 |
| B82 | B85 | CEA | SCC | 0.135692 | 6.94E-13 |
| B82 | B209 | CEA | Smoking | 0.103382 | 4.76E-08 |
| B82 | B159 | CEA | β2-MG | 0.095924 | 4.10E-07 |
| B82 | B169 | CEA | U-CRY | 0.095925 | 4.10E-07 |
| B82 | B119 | CEA | Hcv-rna | -0.09376 | 7.43E-07 |
| B83 | B84 | AFP | C-PSA | 0.1674 | 0 |
| B83 | B85 | AFP | SCC | 0.264576 | 0 |
| B83 | B88 | AFP | CA 242 | 0.269575 | 0 |
| B83 | B159 | AFP | β2-MG | 0.131111 | 4.03E-12 |
| B83 | B169 | AFP | U-CRY | 0.131113 | 4.03E-12 |
| B83 | B119 | AFP | Hcv-rna | -0.12846 | 1.08E-11 |
| B83 | B194 | AFP | CI | -0.11822 | 4.14E-10 |
| B83 | B140 | AFP | r-GT | 0.099287 | 1.58E-07 |

| | | | | | |
|---|---|---|---|---|---|
| B84 | B85 | C-PSA | SCC | 0.174683 | 0 |
| B84 | B87 | C-PSA | NSE | 0.161186 | 0 |
| B84 | B88 | C-PSA | CA 242 | 0.14414 | 2.31E-14 |
| B84 | B89 | C-PSA | CA 72-4 | 0.116752 | 6.79E-10 |
| B84 | B90 | C-PSA | CYFRA 21-1 | 0.114562 | 1.41E-09 |
| B85 | B87 | SCC | NSE | 0.171631 | 0 |
| B85 | B86 | SCC | TBhCG | -0.11926 | 2.90E-10 |
| B85 | B89 | SCC | CA 72-4 | -0.09456 | 5.98E-07 |
| B86 | B89 | TBhCG | CA 72-4 | 0.272662 | 0 |
| B86 | B149 | TBhCG | TGAb | 0.151656 | 8.88E-16 |
| B86 | B198 | TBhCG | P | -0.14972 | 2.22E-15 |
| B86 | B138 | TBhCG | PAB | -0.13093 | 4.31E-12 |
| B86 | B148 | TBhCG | Anti_TPO | 0.098998 | 1.72E-07 |
| B87 | B89 | NSE | CA 72-4 | 0.26511 | 0 |
| B87 | B90 | NSE | CYFRA 21-1 | 0.13943 | 1.58E-13 |
| B87 | B116 | NSE | TT3 | 0.10228 | 6.60E-08 |
| B87 | B117 | NSE | TT4 | 0.10228 | 6.60E-08 |
| B88 | B183 | CA 242 | PTA | 0.17819 | 0 |
| B89 | B90 | CA 72-4 | CYFRA 21-1 | 0.101291 | 8.84E-08 |
| B90 | B135 | CYFRA 21-1 | TBA | 0.169053 | 0 |
| B90 | B139 | CYFRA 21-1 | ADA | 0.152715 | 4.44E-16 |
| B90 | B155 | CYFRA 21-1 | IgA | 0.150764 | 1.33E-15 |
| B90 | B182 | CYFRA 21-1 | TT | 0.140032 | 1.24E-13 |
| B90 | B156 | CYFRA 21-1 | IgE | 0.123119 | 7.52E-11 |
| B90 | B150 | CYFRA 21-1 | ALP | 0.10528 | 2.68E-08 |
| B90 | B137 | CYFRA 21-1 | AFU | 0.094009 | 6.95E-07 |
| B91 | B126 | TP | Alb | 0.460772 | 0 |
| B91 | B127 | TP | A-G ratio | -0.49794 | 0 |
| B91 | B130 | TP | GLB | 0.785943 | 0 |
| B91 | B188 | TP | Ca | 0.306986 | 0 |
| B91 | B157 | TP | IgG | 0.139141 | 1.77E-13 |
| B91 | B189 | TP | eGFB | 0.136681 | 4.71E-13 |
| B91 | B218 | TP | DBP | 0.131645 | 3.30E-12 |
| B91 | B128 | TP | ALT | 0.115285 | 1.11E-09 |
| B91 | B131 | TP | AST | 0.106579 | 1.80E-08 |
| B91 | B209 | TP | Smoking | -0.10541 | 2.58E-08 |
| B91 | B192 | TP | UA | 0.104293 | 3.62E-08 |
| B91 | B140 | TP | r-GT | 0.102652 | 5.91E-08 |
| B91 | B150 | TP | ALP | 0.102536 | 6.12E-08 |
| B91 | B194 | TP | Cl | -0.09907 | 1.68E-07 |
| B92 | B125 | ALDH2-Glu504Glu | Cor | 0.189323 | 0 |
| B92 | B156 | ALDH2-Glu504Glu | IgE | -0.18372 | 0 |
| B92 | B186 | ALDH2-Glu504Glu | PFC | -0.1775 | 0 |
| B92 | B93 | ALDH2-Glu504Glu | ALDH2-Glu504Lys | -0.87173 | 0 |

| | | | | | |
|---|---|---|---|---|---|
| B92 | B94 | ALDH2-Glu504Glu | ALDH2-Lys504Lys | -0.34404 | 0 |
| B92 | B158 | ALDH2-Glu504Glu | IgM | 0.137499 | 3.41E-13 |
| B92 | B135 | ALDH2-Glu504Glu | TBA | 0.136611 | 4.84E-13 |
| B92 | B164 | ALDH2-Glu504Glu | COM | -0.12549 | 3.22E-11 |
| B92 | B184 | ALDH2-Glu504Glu | INR | 0.120685 | 1.77E-10 |
| B92 | B213 | ALDH2-Glu504Glu | CVA-R | 0.114556 | 1.41E-09 |
| B92 | B179 | ALDH2-Glu504Glu | α1-MG | -0.10094 | 9.80E-08 |
| B92 | B198 | ALDH2-Glu504Glu | P | 0.097988 | 2.29E-07 |
| B93 | B156 | ALDH2-Glu504Lys | IgE | 0.214856 | 0 |
| B93 | B181 | ALDH2-Glu504Lys | APTT | -0.1831 | 0 |
| B93 | B186 | ALDH2-Glu504Lys | PFC | 0.221013 | 0 |
| B93 | B94 | ALDH2-Glu504Lys | ALDH2-Lys504Lys | -0.16017 | 0 |
| B93 | B135 | ALDH2-Glu504Lys | TBA | -0.13855 | 2.25E-13 |
| B93 | B164 | ALDH2-Glu504Lys | COM | 0.136384 | 5.29E-13 |
| B93 | B158 | ALDH2-Glu504Lys | IgM | -0.13536 | 7.89E-13 |
| B93 | B179 | ALDH2-Glu504Lys | α1-MG | 0.124097 | 5.31E-11 |
| B93 | B213 | ALDH2-Glu504Lys | CVA-R | -0.11148 | 3.85E-09 |
| B93 | B125 | ALDH2-Glu504Lys | Cor | -0.11033 | 5.57E-09 |
| B93 | B193 | ALDH2-Glu504Lys | K | 0.104904 | 3.01E-08 |
| B93 | B198 | ALDH2-Glu504Lys | P | -0.0944 | 6.24E-07 |
| B94 | B125 | ALDH2-Lys504Lys | Cor | -0.16986 | 0 |
| B94 | B155 | ALDH2-Lys504Lys | IgA | 0.17988 | 0 |
| B94 | B181 | ALDH2-Lys504Lys | APTT | 0.184721 | 0 |
| B94 | B182 | ALDH2-Lys504Lys | TT | 0.165415 | 0 |
| B94 | B139 | ALDH2-Lys504Lys | ADA | -0.13628 | 5.52E-13 |
| B94 | B101 | ALDH2-Lys504Lys | anti-Sc70 | -0.11288 | 2.45E-09 |
| B94 | B102 | ALDH2-Lys504Lys | anti-Sm | -0.11288 | 2.45E-09 |
| B94 | B104 | ALDH2-Lys504Lys | anti-SSB | -0.11288 | 2.45E-09 |
| B94 | B98 | ALDH2-Lys504Lys | anti-Jo-1 | -0.11288 | 2.45E-09 |
| B94 | B99 | ALDH2-Lys504Lys | anti-PM-Scl | -0.11288 | 2.45E-09 |
| B95 | B100 | anti-cenpb | anti-Ro52 | 0.657405 | 0 |
| B95 | B109 | anti-cenpb | AMA-M2 | 0.893126 | 0 |
| B95 | B110 | anti-cenpb | AHA | 0.698732 | 0 |
| B95 | B136 | anti-cenpb | 5'NT | 0.105026 | 2.90E-08 |
| B95 | B187 | anti-cenpb | NSE | -0.10398 | 3.98E-08 |
| B95 | B155 | anti-cenpb | IgA | 0.098784 | 1.83E-07 |
| B96 | B100 | dsDNA (IIF) | anti-Ro52 | -0.4084 | 0 |
| B96 | B101 | dsDNA (IIF) | anti-Sc70 | 0.194341 | 0 |
| B96 | B102 | dsDNA (IIF) | anti-Sm | 0.194341 | 0 |
| B96 | B103 | dsDNA (IIF) | anti-SSA/Ro | -0.43215 | 0 |
| B96 | B104 | dsDNA (IIF) | anti-SSB | 0.194341 | 0 |
| B96 | B105 | dsDNA (IIF) | anti-U1RNP | -0.69501 | 0 |
| B96 | B106 | dsDNA (IIF) | anti-P | -0.70128 | 0 |
| B96 | B107 | dsDNA (IIF) | AnuA | -0.70649 | 0 |

| | | | | | |
|---|---|---|---|---|---|
| B96 | B121 | dsDNA (IIF) | C3 | 0.850944 | 0 |
| B96 | B122 | dsDNA (IIF) | C4 | 0.8061 | 0 |
| B96 | B97 | dsDNA (IIF) | dsDNA | -0.68735 | 0 |
| B96 | B98 | dsDNA (IIF) | anti-Jo-1 | 0.194341 | 0 |
| B96 | B99 | dsDNA (IIF) | anti-PM-Scl | 0.194341 | 0 |
| B96 | B157 | dsDNA (IIF) | IgG | -0.11669 | 6.94E-10 |
| B97 | B121 | dsDNA | C3 | -0.40648 | 0 |
| B97 | B122 | dsDNA | C4 | -0.48242 | 0 |
| B97 | B200 | dsDNA | OP | -0.09868 | 1.88E-07 |
| B97 | B193 | dsDNA | K | -0.09827 | 2.11E-07 |
| B98 | B100 | anti-Jo-1 | anti-Ro52 | 0.246548 | 0 |
| B98 | B101 | anti-Jo-1 | anti-Sc70 | 1 | 0 |
| B98 | B104 | anti-Jo-1 | anti-SSB | 1 | 0 |
| B98 | B121 | anti-Jo-1 | C3 | 0.162823 | 0 |
| B98 | B180 | anti-Jo-1 | Tf | -0.16234 | 0 |
| B98 | B186 | anti-Jo-1 | PFC | 0.179834 | 0 |
| B98 | B122 | anti-Jo-1 | C4 | 0.154227 | 4.44E-16 |
| B98 | B183 | anti-Jo-1 | PTA | 0.146138 | 9.99E-15 |
| B98 | B120 | anti-Jo-1 | HCV-cAg | -0.13654 | 4.98E-13 |
| B98 | B177 | anti-Jo-1 | ACR | -0.13646 | 5.13E-13 |
| B98 | B158 | anti-Jo-1 | IgM | 0.134079 | 1.30E-12 |
| B98 | B107 | anti-Jo-1 | AnuA | -0.13254 | 2.35E-12 |
| B98 | B108 | anti-Jo-1 | Anti-Cyclin | -0.12613 | 2.55E-11 |
| B98 | B178 | anti-Jo-1 | Ig-U | -0.11829 | 4.03E-10 |
| B98 | B182 | anti-Jo-1 | TT | 0.105557 | 2.47E-08 |
| B98 | B124 | anti-Jo-1 | ACTH | -0.10554 | 2.48E-08 |
| B98 | B196 | anti-Jo-1 | TCO2 | -0.09305 | 9.01E-07 |
| B99 | B100 | anti-PM-Scl | anti-Ro52 | 0.246548 | 0 |
| B99 | B101 | anti-PM-Scl | anti-Sc70 | 1 | 0 |
| B99 | B102 | anti-PM-Scl | anti-Sm | 1 | 0 |
| B99 | B121 | anti-PM-Scl | C3 | 0.162823 | 0 |
| B99 | B180 | anti-PM-Scl | Tf | -0.16234 | 0 |
| B99 | B186 | anti-PM-Scl | PFC | 0.179834 | 0 |
| B99 | B122 | anti-PM-Scl | C4 | 0.154227 | 4.44E-16 |
| B99 | B183 | anti-PM-Scl | PTA | 0.146138 | 9.99E-15 |
| B99 | B120 | anti-PM-Scl | HCV-cAg | -0.13654 | 4.98E-13 |
| B99 | B177 | anti-PM-Scl | ACR | -0.13646 | 5.13E-13 |
| B99 | B158 | anti-PM-Scl | IgM | 0.134079 | 1.30E-12 |
| B99 | B107 | anti-PM-Scl | AnuA | -0.13254 | 2.35E-12 |
| B99 | B108 | anti-PM-Scl | Anti-Cyclin | -0.12613 | 2.55E-11 |
| B99 | B178 | anti-PM-Scl | Ig-U | -0.11829 | 4.03E-10 |
| B99 | B182 | anti-PM-Scl | TT | 0.105557 | 2.47E-08 |
| B99 | B124 | anti-PM-Scl | ACTH | -0.10554 | 2.48E-08 |
| B99 | B196 | anti-PM-Scl | TCO2 | -0.09305 | 9.01E-07 |

**Supplementary Table 29. PEI Correlations in Brain diseases**

| Row | Column | Row | Column | Cor | *P* |
|---|---|---|---|---|---|
| B1 | B190 | Gender | Cre | -0.48631062 | 0 |
| B1 | B221 | Gender | Hei | -0.7259734 | 0 |
| B1 | B222 | Gender | NS | -0.58351537 | 0 |
| B1 | B51 | Gender | Hb | -0.42241064 | 1.52E-12 |
| B1 | B216 | Gender | BMI | -0.4113636 | 6.46E-12 |
| B1 | B163 | Gender | SG | -0.37511911 | 5.21E-10 |
| B1 | B42 | Gender | MON | -0.35904691 | 3.09E-09 |
| B1 | B192 | Gender | UA | -0.35423949 | 5.17E-09 |
| B1 | B46 | Gender | RBC Count | -0.32676714 | 8.28E-08 |
| B10 | B12 | PG I | PG II | 0.644868281 | 0 |
| B10 | B80 | PG I | anti-HP IgG | 0.517694815 | 0 |
| B10 | B14 | PG I | G-17 | 0.339413943 | 2.39E-08 |
| B100 | B101 | anti-Ro52 | anti-Sc70 | 0.999999993 | 0 |
| B100 | B103 | anti-Ro52 | anti-SSA/Ro | 1 | 0 |
| B100 | B105 | anti-Ro52 | anti-U1RNP | 0.999999999 | 0 |
| B100 | B106 | anti-Ro52 | anti-P | 0.999999998 | 0 |
| B100 | B107 | anti-Ro52 | AnuA | 0.999999835 | 0 |
| B100 | B109 | anti-Ro52 | AMA-M2 | 1 | 0 |
| B100 | B110 | anti-Ro52 | AHA | 1 | 0 |
| B100 | B198 | anti-Ro52 | P | 0.423843892 | 1.26E-12 |
| B100 | B136 | anti-Ro52 | 5'NT | 0.4004758 | 2.56E-11 |
| B100 | B199 | anti-Ro52 | Mg | 0.388322105 | 1.12E-10 |
| B101 | B102 | anti-Sc70 | anti-Sm | 0.999999983 | 0 |
| B101 | B103 | anti-Sc70 | anti-SSA/Ro | 0.999999993 | 0 |
| B101 | B104 | anti-Sc70 | anti-SSB | 0.999999994 | 0 |
| B101 | B105 | anti-Sc70 | anti-U1RNP | 0.999999996 | 0 |
| B101 | B106 | anti-Sc70 | anti-P | 0.999999988 | 0 |
| B101 | B107 | anti-Sc70 | AnuA | 0.99999989 | 0 |
| B101 | B108 | anti-Sc70 | Anti-Cyclin | 0.99999999 | 0 |
| B101 | B109 | anti-Sc70 | AMA-M2 | 0.999999994 | 0 |
| B101 | B110 | anti-Sc70 | AHA | 0.999999994 | 0 |
| B101 | B198 | anti-Sc70 | P | 0.423843888 | 1.26E-12 |
| B101 | B136 | anti-Sc70 | 5'NT | 0.400475796 | 2.56E-11 |
| B101 | B199 | anti-Sc70 | Mg | 0.388322101 | 1.12E-10 |
| B102 | B104 | anti-Sm | anti-SSB | 0.999999992 | 0 |
| B102 | B106 | anti-Sm | anti-P | 0.99999999 | 0 |
| B102 | B107 | anti-Sm | AnuA | 0.999999914 | 0 |
| B102 | B108 | anti-Sm | Anti-Cyclin | 0.999999995 | 0 |
| B102 | B198 | anti-Sm | P | 0.423843893 | 1.26E-12 |
| B102 | B136 | anti-Sm | 5'NT | 0.4004758 | 2.56E-11 |
| B102 | B199 | anti-Sm | Mg | 0.388322105 | 1.12E-10 |
| B103 | B105 | anti-SSA/Ro | anti-U1RNP | 0.999999999 | 0 |

| | | | | | |
|---|---|---|---|---|---|
| B103 | B107 | anti-SSA/Ro | AnuA | 0.999999846 | 0 |
| B103 | B110 | anti-SSA/Ro | AHA | 1 | 0 |
| B103 | B198 | anti-SSA/Ro | P | 0.423843892 | 1.26E-12 |
| B103 | B136 | anti-SSA/Ro | 5'NT | 0.4004758 | 2.56E-11 |
| B103 | B199 | anti-SSA/Ro | Mg | 0.388322105 | 1.12E-10 |
| B104 | B107 | anti-SSB | AnuA | 0.999999879 | 0 |
| B104 | B198 | anti-SSB | P | 0.423843895 | 1.26E-12 |
| B104 | B136 | anti-SSB | 5'NT | 0.400475803 | 2.56E-11 |
| B104 | B199 | anti-SSB | Mg | 0.388322108 | 1.12E-10 |
| B105 | B107 | anti-U1RNP | AnuA | 0.999999858 | 0 |
| B105 | B110 | anti-U1RNP | AHA | 0.999999999 | 0 |
| B105 | B198 | anti-U1RNP | P | 0.423843892 | 1.26E-12 |
| B105 | B136 | anti-U1RNP | 5'NT | 0.400475799 | 2.56E-11 |
| B105 | B199 | anti-U1RNP | Mg | 0.388322105 | 1.12E-10 |
| B106 | B107 | anti-P | AnuA | 0.999999839 | 0 |
| B106 | B108 | anti-P | Anti-Cyclin | 1 | 0 |
| B106 | B110 | anti-P | AHA | 0.999999997 | 0 |
| B106 | B198 | anti-P | P | 0.423843892 | 1.26E-12 |
| B106 | B136 | anti-P | 5'NT | 0.4004758 | 2.56E-11 |
| B106 | B199 | anti-P | Mg | 0.388322105 | 1.12E-10 |
| B107 | B108 | AnuA | Anti-Cyclin | 0.999999855 | 0 |
| B107 | B109 | AnuA | AMA-M2 | 0.999999843 | 0 |
| B107 | B110 | AnuA | AHA | 0.999999859 | 0 |
| B107 | B198 | AnuA | P | 0.423843823 | 1.26E-12 |
| B107 | B136 | AnuA | 5'NT | 0.400475735 | 2.56E-11 |
| B107 | B199 | AnuA | Mg | 0.388322045 | 1.12E-10 |
| B108 | B198 | Anti-Cyclin | P | 0.423843892 | 1.26E-12 |
| B108 | B136 | Anti-Cyclin | 5'NT | 0.4004758 | 2.56E-11 |
| B108 | B199 | Anti-Cyclin | Mg | 0.388322105 | 1.12E-10 |
| B109 | B110 | AMA-M2 | AHA | 1 | 0 |
| B109 | B198 | AMA-M2 | P | 0.423843892 | 1.26E-12 |
| B109 | B136 | AMA-M2 | 5'NT | 0.4004758 | 2.56E-11 |
| B109 | B199 | AMA-M2 | Mg | 0.388322105 | 1.12E-10 |
| B11 | B12 | PGI/PGII | PG II | -0.60141125 | 0 |
| B110 | B198 | AHA | P | 0.423843892 | 1.26E-12 |
| B110 | B136 | AHA | 5'NT | 0.4004758 | 2.56E-11 |
| B110 | B199 | AHA | Mg | 0.388322105 | 1.12E-10 |
| B111 | B112 | TSH | FT4 | -0.32471019 | 1.01E-07 |
| B112 | B114 | FT4 | TSH TT3 TT4(TT4) | 0.59921938 | 0 |
| B112 | B150 | FT4 | ALP | 0.320609345 | 1.49E-07 |
| B112 | B123 | FT4 | hs-CRP1 | 0.305673172 | 5.84E-07 |
| B113 | B115 | FT3 | TSH TT3 TT4(TT3) | 0.461737089 | 5.77E-15 |
| B114 | B123 | TSH TT3 TT4(TT4) | hs-CRP1 | 0.554529635 | 0 |
| B114 | B172 | TSH TT3 TT4(TT4) | KB | 0.501588492 | 0 |

| | | | | | |
|---|---|---|---|---|---|
| B115 | B156 | TSH TT3 TT4(TT3) | IgE | 0.362299056 | 2.17E-09 |
| B118 | B132 | CEA1 | Dbil | 0.651068689 | 0 |
| B118 | B140 | CEA1 | r-GT | 0.63169602 | 0 |
| B118 | B150 | CEA1 | ALP | 0.687552187 | 0 |
| B118 | B173 | CEA1 | NTT | 0.613446798 | 0 |
| B119 | B152 | Hcv-rna | ACA-Ig(G-A-M) | 0.703254243 | 0 |
| B119 | B167 | Hcv-rna | PC | 0.752578884 | 0 |
| B119 | B171 | Hcv-rna | EC | 0.558744237 | 0 |
| B12 | B197 | PG II | AG | -0.51003472 | 0 |
| B12 | B196 | PG II | TCO2 | 0.393403389 | 6.08E-11 |
| B12 | B156 | PG II | IgE | -0.39332119 | 6.14E-11 |
| B12 | B157 | PG II | IgG | 0.351553081 | 6.86E-09 |
| B12 | B14 | PG II | G-17 | 0.351523153 | 6.88E-09 |
| B12 | B66 | PG II | WBRSR(mPas_100) | -0.3477454 | 1.02E-08 |
| B12 | B63 | PG II | WBRSR(mPas_200) | -0.34559673 | 1.27E-08 |
| B12 | B67 | PG II | WBRSR(mPas_50) | -0.34230035 | 1.78E-08 |
| B12 | B86 | PG II | TBhCG | -0.33275283 | 4.63E-08 |
| B12 | B74 | PG II | ApoA1 | -0.3168792 | 2.11E-07 |
| B12 | B64 | PG II | WBRSR(mPas_5) | -0.31076766 | 3.69E-07 |
| B12 | B83 | PG II | AFP | -0.30528908 | 6.04E-07 |
| B120 | B149 | HCV-cAg | TGAb | -0.4352508 | 2.64E-13 |
| B120 | B148 | HCV-cAg | Anti_TPO | -0.34159426 | 1.92E-08 |
| B120 | B194 | HCV-cAg | Cl | 0.302518522 | 7.72E-07 |
| B121 | B152 | C3 | ACA-Ig(G-A-M) | 0.50914234 | 0 |
| B121 | B171 | C3 | EC | 0.404519917 | 1.54E-11 |
| B122 | B159 | C4 | β2-MG | -0.70521011 | 0 |
| B122 | B223 | C4 | FH | -0.67855114 | 0 |
| B123 | B172 | hs-CRP1 | KB | 0.83460675 | 0 |
| B124 | B180 | ACTH | Tf | -0.87287156 | 0 |
| B125 | B159 | Cor | β2-MG | -0.74069712 | 0 |
| B125 | B179 | Cor | α1-MG | -0.68494952 | 0 |
| B125 | B202 | Cor | e-times | 0.758175397 | 0 |
| B126 | B127 | Alb | A-G ratio | 0.555813345 | 0 |
| B126 | B188 | Alb | Ca | 0.432042865 | 4.12E-13 |
| B126 | B189 | Alb | eGFB | 0.360990499 | 2.51E-09 |
| B127 | B130 | A-G ratio | GLB | -0.90549754 | 0 |
| B127 | B189 | A-G ratio | eGFB | 0.362261502 | 2.18E-09 |
| B128 | B131 | ALT | AST | 0.775340815 | 0 |
| B128 | B141 | ALT | AST/ALT | -0.55106935 | 0 |
| B128 | B140 | ALT | r-GT | 0.385358257 | 1.59E-10 |
| B128 | B142 | ALT | 25-(OH)D | 0.351165152 | 7.14E-09 |
| B129 | B133 | IBIL | TBIL | 0.920330033 | 0 |
| B129 | B132 | IBIL | Dbil | 0.373121599 | 6.54E-10 |
| B131 | B140 | AST | r-GT | 0.564425842 | 0 |

| | | | | | |
|---|---|---|---|---|---|
| B131 | B150 | AST | ALP | 0.394853865 | 5.10E-11 |
| B131 | B132 | AST | Dbil | 0.366942846 | 1.30E-09 |
| B131 | B173 | AST | NTT | 0.308583717 | 4.50E-07 |
| B131 | B142 | AST | 25-(OH)D | 0.303079413 | 7.35E-07 |
| B132 | B133 | Dbil | TBIL | 0.706225666 | 0 |
| B132 | B140 | Dbil | r-GT | 0.776973209 | 0 |
| B132 | B150 | Dbil | ALP | 0.765618565 | 0 |
| B132 | B173 | Dbil | NTT | 0.694300371 | 0 |
| B133 | B140 | TBIL | r-GT | 0.341473444 | 1.94E-08 |
| B134 | B136 | CHE | 5'NT | 0.652471001 | 0 |
| B134 | B156 | CHE | IgE | 0.556664195 | 0 |
| B134 | B157 | CHE | IgG | 0.583581859 | 0 |
| B134 | B181 | CHE | APTT | 0.53405962 | 0 |
| B134 | B182 | CHE | TT | -0.53405962 | 0 |
| B134 | B183 | CHE | PTA | -0.53405962 | 0 |
| B134 | B184 | CHE | INR | 0.53405962 | 0 |
| B134 | B185 | CHE | PT | 0.53405962 | 0 |
| B134 | B186 | CHE | PFC | 0.53405962 | 0 |
| B134 | B158 | CHE | IgM | 0.437796191 | 1.85E-13 |
| B134 | B155 | CHE | IgA | 0.385378682 | 1.59E-10 |
| B134 | B197 | CHE | AG | -0.37920362 | 3.26E-10 |
| B134 | B135 | CHE | TBA | 0.366176723 | 1.42E-09 |
| B134 | B139 | CHE | ADA | 0.305834457 | 5.75E-07 |
| B135 | B198 | TBA | P | 0.548752322 | 0 |
| B135 | B136 | TBA | 5'NT | 0.391647538 | 7.52E-11 |
| B135 | B156 | TBA | IgE | 0.378531365 | 3.52E-10 |
| B136 | B198 | 5'NT | P | 0.598810107 | 0 |
| B136 | B199 | 5'NT | Mg | 0.489445587 | 0 |
| B136 | B193 | 5'NT | K | 0.470248073 | 1.55E-15 |
| B136 | B157 | 5'NT | IgG | 0.39804836 | 3.45E-11 |
| B136 | B139 | 5'NT | ADA | 0.362220395 | 2.19E-09 |
| B136 | B181 | 5'NT | APTT | 0.310314445 | 3.85E-07 |
| B136 | B182 | 5'NT | TT | -0.31031445 | 3.85E-07 |
| B136 | B183 | 5'NT | PTA | -0.31031445 | 3.85E-07 |
| B136 | B184 | 5'NT | INR | 0.310314445 | 3.85E-07 |
| B136 | B185 | 5'NT | PT | 0.310314445 | 3.85E-07 |
| B136 | B186 | 5'NT | PFC | 0.310314445 | 3.85E-07 |
| B137 | B138 | AFU | PAB | 0.648224945 | 0 |
| B137 | B139 | AFU | ADA | -0.37768419 | 3.89E-10 |
| B137 | B156 | AFU | IgE | 0.313366371 | 2.91E-07 |
| B138 | B139 | PAB | ADA | -0.58510429 | 0 |
| B138 | B156 | PAB | IgE | 0.473991677 | 8.88E-16 |
| B139 | B157 | ADA | IgG | 0.465343666 | 3.11E-15 |
| B139 | B200 | ADA | OP | -0.40768295 | 1.03E-11 |

| | | | | | |
|---|---|---|---|---|---|
| B139 | B197 | ADA | AG | -0.38982642 | 9.36E-11 |
| B139 | B198 | ADA | P | 0.359495175 | 2.94E-09 |
| B14 | B136 | G-17 | 5'NT | 0.344775399 | 1.39E-08 |
| B14 | B135 | G-17 | TBA | 0.337636559 | 2.85E-08 |
| B14 | B193 | G-17 | K | 0.319528676 | 1.65E-07 |
| B14 | B198 | G-17 | P | 0.305724628 | 5.81E-07 |
| B140 | B150 | r-GT | ALP | 0.877518086 | 0 |
| B140 | B173 | r-GT | NTT | 0.767884167 | 0 |
| B142 | B144 | 25-(OH)D | Total-PINP | -0.65792034 | 0 |
| B142 | B145 | 25-(OH)D | BGP | -0.46896449 | 1.78E-15 |
| B142 | B143 | 25-(OH)D | β-CTx | -0.44545157 | 6.26E-14 |
| B143 | B144 | β-CTx | Total-PINP | 0.648554193 | 0 |
| B143 | B145 | β-CTx | BGP | 0.589913754 | 0 |
| B143 | B151 | β-CTx | Anti-CCP | -0.4692113 | 1.78E-15 |
| B143 | B154 | β-CTx | RF | 0.373500423 | 6.26E-10 |
| B144 | B145 | Total-PINP | BGP | 0.378154133 | 3.68E-10 |
| B146 | B209 | CysC | Smoking | 0.378713419 | 3.45E-10 |
| B148 | B149 | Anti_TPO | TGAb | 0.867438084 | 0 |
| B15 | B123 | CD3-CD19+(B cell) | hs-CRP1 | -0.48685827 | 0 |
| B15 | B138 | CD3-CD19+(B cell) | PAB | 0.487477467 | 0 |
| B15 | B16 | CD3-CD19+(B cell) | CD3+T | -1 | 0 |
| B15 | B18 | CD3-CD19+(B cell) | CD3+CD4+helper(inducer)T | 1 | 0 |
| B15 | B19 | CD3-CD19+(B cell) | CD3+CD4+/CD3+CD8+ | 1 | 0 |
| B15 | B28 | CD3-CD19+(B cell) | PRL | 1 | 0 |
| B15 | B29 | CD3-CD19+(B cell) | E2 | -1 | 0 |
| B15 | B31 | CD3-CD19+(B cell) | FSH | 1 | 0 |
| B15 | B56 | CD3-CD19+(B cell) | IR | 0.714316492 | 0 |
| B15 | B60 | CD3-CD19+(B cell) | WBRVI | 0.51149382 | 0 |
| B15 | B61 | CD3-CD19+(B cell) | WBHSRV | 0.4932853 | 0 |
| B15 | B62 | CD3-CD19+(B cell) | WBHSRVI | 0.584027745 | 0 |
| B15 | B120 | CD3-CD19+(B cell) | HCV-cAg | 0.443705919 | 8.04E-14 |
| B15 | B156 | CD3-CD19+(B cell) | IgE | 0.43021394 | 5.30E-13 |
| B15 | B63 | CD3-CD19+(B cell) | WBRSR(mPas_200) | 0.417942309 | 2.75E-12 |
| B15 | B66 | CD3-CD19+(B cell) | WBRSR(mPas_100) | 0.415570122 | 3.75E-12 |
| B15 | B67 | CD3-CD19+(B cell) | WBRSR(mPas_50) | 0.40400005 | 1.65E-11 |
| B15 | B55 | CD3-CD19+(B cell) | TK | 0.40296789 | 1.87E-11 |
| B15 | B64 | CD3-CD19+(B cell) | WBRSR(mPas_5) | 0.342597209 | 1.73E-08 |
| B15 | B69 | CD3-CD19+(B cell) | PV | -0.32995546 | 6.09E-08 |
| B150 | B173 | ALP | NTT | 0.87085967 | 0 |
| B151 | B154 | Anti-CCP | RF | -0.82556948 | 0 |
| B153 | B170 | Glu | U-GLU | 0.481579155 | 2.22E-16 |
| B155 | B156 | IgA | IgE | 0.580394936 | 0 |
| B155 | B157 | IgA | IgG | 0.346570494 | 1.15E-08 |
| B155 | B181 | IgA | APTT | 0.33441334 | 3.93E-08 |

| | | | | | |
|---|---|---|---|---|---|
| B155 | B182 | IgA | TT | -0.33441334 | 3.93E-08 |
| B155 | B183 | IgA | PTA | -0.33441334 | 3.93E-08 |
| B155 | B184 | IgA | INR | 0.33441334 | 3.93E-08 |
| B155 | B185 | IgA | PT | 0.33441334 | 3.93E-08 |
| B155 | B186 | IgA | PFC | 0.33441334 | 3.93E-08 |
| B156 | B158 | IgE | IgM | 0.503407186 | 0 |
| B156 | B181 | IgE | APTT | 0.469036046 | 1.78E-15 |
| B156 | B182 | IgE | TT | -0.46903605 | 1.78E-15 |
| B156 | B183 | IgE | PTA | -0.46903605 | 1.78E-15 |
| B156 | B184 | IgE | INR | 0.469036046 | 1.78E-15 |
| B156 | B185 | IgE | PT | 0.469036046 | 1.78E-15 |
| B156 | B186 | IgE | PFC | 0.469036046 | 1.78E-15 |
| B157 | B196 | IgG | TCO2 | 0.493266374 | 0 |
| B157 | B197 | IgG | AG | -0.65389992 | 0 |
| B157 | B181 | IgG | APTT | 0.469698809 | 1.78E-15 |
| B157 | B182 | IgG | TT | -0.46969881 | 1.78E-15 |
| B157 | B183 | IgG | PTA | -0.46969881 | 1.78E-15 |
| B157 | B184 | IgG | INR | 0.469698809 | 1.78E-15 |
| B157 | B185 | IgG | PT | 0.469698809 | 1.78E-15 |
| B157 | B186 | IgG | PFC | 0.469698809 | 1.78E-15 |
| B157 | B200 | IgG | OP | -0.41310889 | 5.16E-12 |
| B157 | B195 | IgG | Na | -0.35180625 | 6.68E-09 |
| B158 | B181 | IgM | APTT | 0.378489003 | 3.54E-10 |
| B158 | B182 | IgM | TT | -0.378489 | 3.54E-10 |
| B158 | B183 | IgM | PTA | -0.378489 | 3.54E-10 |
| B158 | B184 | IgM | INR | 0.378489003 | 3.54E-10 |
| B158 | B185 | IgM | PT | 0.378489003 | 3.54E-10 |
| B158 | B186 | IgM | PFC | 0.378489003 | 3.54E-10 |
| B159 | B179 | β2-MG | α1-MG | -0.72566917 | 0 |
| B16 | B123 | CD3+T | hs-CRP1 | 0.486858266 | 0 |
| B16 | B138 | CD3+T | PAB | -0.48747747 | 0 |
| B16 | B18 | CD3+T | CD3+CD4+helper(inducer)T | -1 | 0 |
| B16 | B19 | CD3+T | CD3+CD4+/CD3+CD8+ | -1 | 0 |
| B16 | B22 | CD3+T | CD3+CD8+(T) | 1 | 0 |
| B16 | B23 | CD3+T | CD3-CD16+CD56+(NK) | -1 | 0 |
| B16 | B28 | CD3+T | PRL | -1 | 0 |
| B16 | B29 | CD3+T | E2 | 1 | 0 |
| B16 | B30 | CD3+T | LH | -1 | 0 |
| B16 | B31 | CD3+T | FSH | -1 | 0 |
| B16 | B32 | CD3+T | Testo | -1 | 0 |
| B16 | B33 | CD3+T | Prog | -1 | 0 |
| B16 | B56 | CD3+T | IR | -0.71431649 | 0 |
| B16 | B60 | CD3+T | WBRVI | -0.51149382 | 0 |
| B16 | B61 | CD3+T | WBHSRV | -0.4932853 | 0 |

| | | | | | |
|---|---|---|---|---|---|
| B16 | B62 | CD3+T | WBHSRVI | -0.58402775 | 0 |
| B16 | B120 | CD3+T | HCV-cAg | -0.44370591 | 8.04E-14 |
| B16 | B156 | CD3+T | IgE | -0.43021394 | 5.30E-13 |
| B16 | B63 | CD3+T | WBRSR(mPas_200) | -0.41794231 | 2.75E-12 |
| B16 | B66 | CD3+T | WBRSR(mPas_100) | -0.41557012 | 3.75E-12 |
| B16 | B67 | CD3+T | WBRSR(mPas_50) | -0.40400005 | 1.65E-11 |
| B16 | B55 | CD3+T | TK | -0.40296789 | 1.87E-11 |
| B16 | B64 | CD3+T | WBRSR(mPas_5) | -0.34259721 | 1.73E-08 |
| B16 | B69 | CD3+T | PV | 0.329955463 | 6.09E-08 |
| B163 | B166 | SG | Pro | 0.340766151 | 2.08E-08 |
| B163 | B191 | SG | Urea | 0.319371436 | 1.67E-07 |
| B163 | B176 | SG | Ucr | 0.311330398 | 3.51E-07 |
| B167 | B225 | PC | hPBG | 0.724437133 | 0 |
| B169 | B171 | U-CRY | EC | 0.702863496 | 0 |
| B17 | B122 | CD3+CD16+CD56+NK | C4 | -0.52633295 | 0 |
| B17 | B159 | CD3+CD16+CD56+NK | β2-MG | -0.60740116 | 0 |
| B17 | B179 | CD3+CD16+CD56+NK | α1-MG | -0.50467511 | 0 |
| B17 | B21 | CD3+CD16+CD56+NK | CD3+CD4-CD8- | -0.96394252 | 0 |
| B17 | B37 | CD3+CD16+CD56+NK | BAND | -0.55390031 | 0 |
| B171 | B225 | EC | hPBG | 0.85811633 | 0 |
| B172 | B191 | KB | Urea | 0.315766568 | 2.34E-07 |
| B174 | B191 | Urea/Cr | Urea | 0.634091022 | 0 |
| B175 | B177 | ALB-U | ACR | 0.732148934 | 0 |
| B178 | B223 | Ig-U | FH | 0.777817459 | 0 |
| B18 | B123 | CD3+CD4+helper(inducer)T | hs-CRP1 | -0.48685827 | 0 |
| B18 | B138 | CD3+CD4+helper(inducer)T | PAB | 0.487477467 | 0 |
| B18 | B19 | CD3+CD4+helper(inducer)T | CD3+CD4+/CD3+CD8+ | 1 | 0 |
| B18 | B28 | CD3+CD4+helper(inducer)T | PRL | 1 | 0 |
| B18 | B29 | CD3+CD4+helper(inducer)T | E2 | -1 | 0 |
| B18 | B30 | CD3+CD4+helper(inducer)T | LH | 1 | 0 |
| B18 | B31 | CD3+CD4+helper(inducer)T | FSH | 1 | 0 |
| B18 | B33 | CD3+CD4+helper(inducer)T | Prog | 1 | 0 |
| B18 | B56 | CD3+CD4+helper(inducer)T | IR | 0.714316492 | 0 |
| B18 | B60 | CD3+CD4+helper(inducer)T | WBRVI | 0.51149382 | 0 |
| B18 | B61 | CD3+CD4+helper(inducer)T | WBHSRV | 0.4932853 | 0 |
| B18 | B62 | CD3+CD4+helper(inducer)T | WBHSRVI | 0.584027745 | 0 |
| B18 | B120 | CD3+CD4+helper(inducer)T | HCV-cAg | 0.443705922 | 8.04E-14 |
| B18 | B156 | CD3+CD4+helper(inducer)T | IgE | 0.43021394 | 5.30E-13 |
| B18 | B63 | CD3+CD4+helper(inducer)T | WBRSR(mPas_200) | 0.417942309 | 2.75E-12 |
| B18 | B66 | CD3+CD4+helper(inducer)T | WBRSR(mPas_100) | 0.415570122 | 3.75E-12 |
| B18 | B67 | CD3+CD4+helper(inducer)T | WBRSR(mPas_50) | 0.40400005 | 1.65E-11 |
| B18 | B55 | CD3+CD4+helper(inducer)T | TK | 0.40296789 | 1.87E-11 |
| B18 | B64 | CD3+CD4+helper(inducer)T | WBRSR(mPas_5) | 0.342597209 | 1.73E-08 |
| B18 | B69 | CD3+CD4+helper(inducer)T | PV | -0.32995546 | 6.09E-08 |

| | | | | | |
|---|---|---|---|---|---|
| B181 | B197 | APTT | AG | -0.56842515 | 0 |
| B181 | B200 | APTT | OP | -0.47731041 | 4.44E-16 |
| B181 | B196 | APTT | TCO2 | 0.445267059 | 6.42E-14 |
| B181 | B195 | APTT | Na | -0.39781671 | 3.55E-11 |
| B182 | B184 | TT | INR | -1 | 0 |
| B182 | B185 | TT | PT | -1 | 0 |
| B182 | B197 | TT | AG | 0.56842515 | 0 |
| B182 | B200 | TT | OP | 0.477310415 | 4.44E-16 |
| B182 | B196 | TT | TCO2 | -0.44526706 | 6.42E-14 |
| B182 | B195 | TT | Na | 0.397816709 | 3.55E-11 |
| B183 | B184 | PTA | INR | -1 | 0 |
| B183 | B185 | PTA | PT | -1 | 0 |
| B183 | B197 | PTA | AG | 0.568425149 | 0 |
| B183 | B200 | PTA | OP | 0.477310412 | 4.44E-16 |
| B183 | B196 | PTA | TCO2 | -0.44526706 | 6.42E-14 |
| B183 | B195 | PTA | Na | 0.39781671 | 3.55E-11 |
| B184 | B186 | INR | PFC | 1 | 0 |
| B184 | B197 | INR | AG | -0.56842515 | 0 |
| B184 | B200 | INR | OP | -0.47731041 | 4.44E-16 |
| B184 | B196 | INR | TCO2 | 0.445267059 | 6.42E-14 |
| B184 | B195 | INR | Na | -0.39781671 | 3.55E-11 |
| B185 | B186 | PT | PFC | 1 | 0 |
| B185 | B197 | PT | AG | -0.56842515 | 0 |
| B185 | B200 | PT | OP | -0.47731041 | 4.44E-16 |
| B185 | B196 | PT | TCO2 | 0.445267059 | 6.42E-14 |
| B185 | B195 | PT | Na | -0.39781671 | 3.55E-11 |
| B186 | B197 | PFC | AG | -0.56842515 | 0 |
| B186 | B200 | PFC | OP | -0.47731041 | 4.44E-16 |
| B186 | B196 | PFC | TCO2 | 0.445267059 | 6.42E-14 |
| B186 | B195 | PFC | Na | -0.39781671 | 3.55E-11 |
| B189 | B190 | eGFB | Cre | -0.56574926 | 0 |
| B189 | B191 | eGFB | Urea | -0.44756616 | 4.62E-14 |
| B19 | B123 | CD3+CD4+/CD3+CD8+ | hs-CRP1 | -0.48685827 | 0 |
| B19 | B138 | CD3+CD4+/CD3+CD8+ | PAB | 0.487477467 | 0 |
| B19 | B23 | CD3+CD4+/CD3+CD8+ | CD3-CD16+CD56+(NK) | 1 | 0 |
| B19 | B28 | CD3+CD4+/CD3+CD8+ | PRL | 1 | 0 |
| B19 | B30 | CD3+CD4+/CD3+CD8+ | LH | 1 | 0 |
| B19 | B31 | CD3+CD4+/CD3+CD8+ | FSH | 1 | 0 |
| B19 | B32 | CD3+CD4+/CD3+CD8+ | Testo | 1 | 0 |
| B19 | B33 | CD3+CD4+/CD3+CD8+ | Prog | 1 | 0 |
| B19 | B56 | CD3+CD4+/CD3+CD8+ | IR | 0.714316492 | 0 |
| B19 | B60 | CD3+CD4+/CD3+CD8+ | WBRVI | 0.51149382 | 0 |
| B19 | B61 | CD3+CD4+/CD3+CD8+ | WBHSRV | 0.4932853 | 0 |
| B19 | B62 | CD3+CD4+/CD3+CD8+ | WBHSRVI | 0.584027745 | 0 |

| | | | | | |
|---|---|---|---|---|---|
| B19 | B120 | CD3+CD4+/CD3+CD8+ | HCV-cAg | 0.443705918 | 8.04E-14 |
| B19 | B156 | CD3+CD4+/CD3+CD8+ | IgE | 0.43021394 | 5.30E-13 |
| B19 | B63 | CD3+CD4+/CD3+CD8+ | WBRSR(mPas_200) | 0.417942309 | 2.75E-12 |
| B19 | B66 | CD3+CD4+/CD3+CD8+ | WBRSR(mPas_100) | 0.415570122 | 3.75E-12 |
| B19 | B67 | CD3+CD4+/CD3+CD8+ | WBRSR(mPas_50) | 0.40400005 | 1.65E-11 |
| B19 | B55 | CD3+CD4+/CD3+CD8+ | TK | 0.40296789 | 1.87E-11 |
| B19 | B64 | CD3+CD4+/CD3+CD8+ | WBRSR(mPas_5) | 0.342597209 | 1.73E-08 |
| B19 | B69 | CD3+CD4+/CD3+CD8+ | PV | -0.32995546 | 6.09E-08 |
| B190 | B191 | Cre | Urea | 0.57705182 | 0 |
| B190 | B192 | Cre | UA | 0.408900753 | 8.86E-12 |
| B190 | B221 | Cre | Hei | 0.330542457 | 5.75E-08 |
| B190 | B222 | Cre | NS | 0.318906561 | 1.75E-07 |
| B192 | B221 | UA | Hei | 0.305388181 | 5.99E-07 |
| B193 | B194 | K | Cl | 0.481614474 | 2.22E-16 |
| B194 | B195 | Cl | Na | 0.477668385 | 4.44E-16 |
| B194 | B200 | Cl | OP | 0.360196963 | 2.73E-09 |
| B194 | B199 | Cl | Mg | 0.345444583 | 1.29E-08 |
| B194 | B196 | Cl | TCO2 | -0.31785178 | 1.93E-07 |
| B195 | B199 | Na | Mg | 0.647422248 | 0 |
| B195 | B200 | Na | OP | 0.916593319 | 0 |
| B196 | B197 | TCO2 | AG | -0.69009177 | 0 |
| B196 | B198 | TCO2 | P | -0.44532702 | 6.35E-14 |
| B197 | B198 | AG | P | 0.398098607 | 3.43E-11 |
| B197 | B200 | AG | OP | 0.350231173 | 7.88E-09 |
| B199 | B200 | Mg | OP | 0.604268985 | 0 |
| B2 | B189 | Age | eGFB | -0.44290682 | 8.99E-14 |
| B2 | B47 | Age | RDW | 0.427567123 | 7.60E-13 |
| B2 | B46 | Age | RBC Count | -0.40092606 | 2.42E-11 |
| B2 | B141 | Age | AST/ALT | 0.351284156 | 7.05E-09 |
| B2 | B51 | Age | Hb | -0.34870634 | 9.23E-09 |
| B2 | B126 | Age | Alb | -0.32348752 | 1.13E-07 |
| B2 | B191 | Age | Urea | 0.316705046 | 2.14E-07 |
| B2 | B72 | Age | HDL-C | 0.306616906 | 5.37E-07 |
| B20 | B121 | CD3+CD4+CD8+ | C3 | 0.806059936 | 0 |
| B20 | B152 | CD3+CD4+CD8+ | ACA-Ig(G-A-M) | 0.794198533 | 0 |
| B20 | B169 | CD3+CD4+CD8+ | U-CRY | 0.878776058 | 0 |
| B20 | B96 | CD3+CD4+CD8+ | dsDNA (IIF) | 0.90623781 | 0 |
| B20 | B167 | CD3+CD4+CD8+ | PC | 0.477137852 | 4.44E-16 |
| B20 | B171 | CD3+CD4+CD8+ | EC | 0.38376645 | 1.92E-10 |
| B206 | B207 | Waist | WHR | 0.837641264 | 0 |
| B206 | B210 | Waist | Hip | 0.750375447 | 0 |
| B206 | B215 | Waist | BMI | 0.813300792 | 0 |
| B206 | B216 | Waist | BMI | 0.797057858 | 0 |
| B206 | B222 | Waist | NS | 0.702129993 | 0 |

| | | | | | |
|---|---|---|---|---|---|
| B207 | B215 | WHR | BMI | 0.539573771 | 0 |
| B207 | B216 | WHR | BMI | 0.489282036 | 0 |
| B207 | B222 | WHR | NS | 0.496224731 | 0 |
| B21 | B179 | CD3+CD4-CD8- | α1-MG | -0.54029547 | 0 |
| B21 | B122 | CD3+CD4-CD8- | C4 | -0.46491621 | 3.55E-15 |
| B210 | B215 | Hip | BMI | 0.783277312 | 0 |
| B210 | B216 | Hip | BMI | 0.81148641 | 0 |
| B210 | B222 | Hip | NS | 0.637503221 | 0 |
| B210 | B221 | Hip | Hei | 0.335352259 | 3.58E-08 |
| B211 | B212 | IOP-R | IOP-L | 0.865107492 | 0 |
| B213 | B214 | CVA-R | CVA-L | 0.883619514 | 0 |
| B215 | B216 | BMI | BMI | 0.797630221 | 0 |
| B215 | B222 | BMI | NS | 0.572561934 | 0 |
| B216 | B221 | BMI | Hei | 0.611630484 | 0 |
| B216 | B222 | BMI | NS | 0.774732218 | 0 |
| B218 | B220 | DBP | SBP | 0.539718511 | 0 |
| B22 | B123 | CD3+CD8+(T) | hs-CRP1 | 0.486858266 | 0 |
| B22 | B138 | CD3+CD8+(T) | PAB | -0.48747747 | 0 |
| B22 | B23 | CD3+CD8+(T) | CD3-CD16+CD56+(NK) | -1 | 0 |
| B22 | B32 | CD3+CD8+(T) | Testo | -1 | 0 |
| B22 | B56 | CD3+CD8+(T) | IR | -0.71431649 | 0 |
| B22 | B60 | CD3+CD8+(T) | WBRVI | -0.51149382 | 0 |
| B22 | B61 | CD3+CD8+(T) | WBHSRV | -0.4932853 | 0 |
| B22 | B62 | CD3+CD8+(T) | WBHSRVI | -0.58402775 | 0 |
| B22 | B120 | CD3+CD8+(T) | HCV-cAg | -0.44370592 | 8.04E-14 |
| B22 | B156 | CD3+CD8+(T) | IgE | -0.43021394 | 5.30E-13 |
| B22 | B63 | CD3+CD8+(T) | WBRSR(mPas_200) | -0.41794231 | 2.75E-12 |
| B22 | B66 | CD3+CD8+(T) | WBRSR(mPas_100) | -0.41557012 | 3.75E-12 |
| B22 | B67 | CD3+CD8+(T) | WBRSR(mPas_50) | -0.40400005 | 1.65E-11 |
| B22 | B55 | CD3+CD8+(T) | TK | -0.40296789 | 1.87E-11 |
| B22 | B64 | CD3+CD8+(T) | WBRSR(mPas_5) | -0.34259721 | 1.73E-08 |
| B22 | B69 | CD3+CD8+(T) | PV | 0.329955463 | 6.09E-08 |
| B221 | B222 | Hei | NS | 0.548290474 | 0 |
| B23 | B123 | CD3-CD16+CD56+(NK) | hs-CRP1 | -0.48685827 | 0 |
| B23 | B138 | CD3-CD16+CD56+(NK) | PAB | 0.487477467 | 0 |
| B23 | B28 | CD3-CD16+CD56+(NK) | PRL | 1 | 0 |
| B23 | B29 | CD3-CD16+CD56+(NK) | E2 | -1 | 0 |
| B23 | B30 | CD3-CD16+CD56+(NK) | LH | 1 | 0 |
| B23 | B31 | CD3-CD16+CD56+(NK) | FSH | 1 | 0 |
| B23 | B56 | CD3-CD16+CD56+(NK) | IR | 0.714316492 | 0 |
| B23 | B60 | CD3-CD16+CD56+(NK) | WBRVI | 0.51149382 | 0 |
| B23 | B61 | CD3-CD16+CD56+(NK) | WBHSRV | 0.4932853 | 0 |
| B23 | B62 | CD3-CD16+CD56+(NK) | WBHSRVI | 0.584027745 | 0 |
| B23 | B120 | CD3-CD16+CD56+(NK) | HCV-cAg | 0.443705922 | 8.04E-14 |

| | | | | | |
|---|---|---|---|---|---|
| B23 | B156 | CD3-CD16+CD56+(NK) | IgE | 0.43021394 | 5.30E-13 |
| B23 | B63 | CD3-CD16+CD56+(NK) | WBRSR(mPas_200) | 0.417942309 | 2.75E-12 |
| B23 | B66 | CD3-CD16+CD56+(NK) | WBRSR(mPas_100) | 0.415570122 | 3.75E-12 |
| B23 | B67 | CD3-CD16+CD56+(NK) | WBRSR(mPas_50) | 0.40400005 | 1.65E-11 |
| B23 | B55 | CD3-CD16+CD56+(NK) | TK | 0.40296789 | 1.87E-11 |
| B23 | B64 | CD3-CD16+CD56+(NK) | WBRSR(mPas_5) | 0.342597209 | 1.73E-08 |
| B23 | B69 | CD3-CD16+CD56+(NK) | PV | -0.32995546 | 6.09E-08 |
| B24 | B26 | αHBDH | CKMB | 0.562496621 | 0 |
| B24 | B27 | αHBDH | LDH | 0.440733418 | 1.23E-13 |
| B24 | B181 | αHBDH | APTT | -0.40470964 | 1.51E-11 |
| B24 | B182 | αHBDH | TT | 0.404709641 | 1.51E-11 |
| B24 | B183 | αHBDH | PTA | 0.404709641 | 1.51E-11 |
| B24 | B184 | αHBDH | INR | -0.40470964 | 1.51E-11 |
| B24 | B185 | αHBDH | PT | -0.40470964 | 1.51E-11 |
| B24 | B186 | αHBDH | PFC | -0.40470964 | 1.51E-11 |
| B24 | B25 | αHBDH | CK | 0.384228538 | 1.82E-10 |
| B24 | B87 | αHBDH | NSE | 0.363345141 | 1.94E-09 |
| B24 | B83 | αHBDH | AFP | -0.30357416 | 7.03E-07 |
| B25 | B26 | CK | CKMB | 0.45075432 | 2.89E-14 |
| B26 | B89 | CKMB | CA 72-4 | 0.330585213 | 5.72E-08 |
| B26 | B82 | CKMB | CEA | 0.30284968 | 7.50E-07 |
| B27 | B118 | LDH | CEA1 | 0.341744602 | 1.89E-08 |
| B27 | B131 | LDH | AST | 0.333540447 | 4.29E-08 |
| B28 | B123 | PRL | hs-CRP1 | -0.48685827 | 0 |
| B28 | B138 | PRL | PAB | 0.487477467 | 0 |
| B28 | B30 | PRL | LH | 1 | 0 |
| B28 | B32 | PRL | Testo | 1 | 0 |
| B28 | B56 | PRL | IR | 0.714316492 | 0 |
| B28 | B60 | PRL | WBRVI | 0.51149382 | 0 |
| B28 | B61 | PRL | WBHSRV | 0.4932853 | 0 |
| B28 | B62 | PRL | WBHSRVI | 0.584027745 | 0 |
| B28 | B120 | PRL | HCV-cAg | 0.443705918 | 8.04E-14 |
| B28 | B156 | PRL | IgE | 0.43021394 | 5.30E-13 |
| B28 | B63 | PRL | WBRSR(mPas_200) | 0.417942309 | 2.75E-12 |
| B28 | B66 | PRL | WBRSR(mPas_100) | 0.415570122 | 3.75E-12 |
| B28 | B67 | PRL | WBRSR(mPas_50) | 0.40400005 | 1.65E-11 |
| B28 | B55 | PRL | TK | 0.40296789 | 1.87E-11 |
| B28 | B64 | PRL | WBRSR(mPas_5) | 0.342597209 | 1.73E-08 |
| B28 | B69 | PRL | PV | -0.32995546 | 6.09E-08 |
| B29 | B123 | E2 | hs-CRP1 | 0.486858266 | 0 |
| B29 | B138 | E2 | PAB | -0.48747747 | 0 |
| B29 | B30 | E2 | LH | -1 | 0 |
| B29 | B32 | E2 | Testo | -1 | 0 |
| B29 | B56 | E2 | IR | -0.71431649 | 0 |

| | | | | | |
|---|---|---|---|---|---|
| B29 | B60 | E2 | WBRVI | -0.51149382 | 0 |
| B29 | B61 | E2 | WBHSRV | -0.4932853 | 0 |
| B29 | B62 | E2 | WBHSRVI | -0.58402775 | 0 |
| B29 | B120 | E2 | HCV-cAg | -0.44370592 | 8.04E-14 |
| B29 | B156 | E2 | IgE | -0.43021394 | 5.30E-13 |
| B29 | B63 | E2 | WBRSR(mPas_200) | -0.41794231 | 2.75E-12 |
| B29 | B66 | E2 | WBRSR(mPas_100) | -0.41557012 | 3.75E-12 |
| B29 | B67 | E2 | WBRSR(mPas_50) | -0.40400005 | 1.65E-11 |
| B29 | B55 | E2 | TK | -0.40296789 | 1.87E-11 |
| B29 | B64 | E2 | WBRSR(mPas_5) | -0.34259721 | 1.73E-08 |
| B29 | B69 | E2 | PV | 0.329955463 | 6.09E-08 |
| B3 | B26 | Hcy | CKMB | 0.350677147 | 7.52E-09 |
| B3 | B221 | Hcy | Hei | 0.305210044 | 6.08E-07 |
| B30 | B123 | LH | hs-CRP1 | -0.48685827 | 0 |
| B30 | B138 | LH | PAB | 0.487477467 | 0 |
| B30 | B31 | LH | FSH | 1 | 0 |
| B30 | B32 | LH | Testo | 1 | 0 |
| B30 | B33 | LH | Prog | 1 | 0 |
| B30 | B56 | LH | IR | 0.714316492 | 0 |
| B30 | B60 | LH | WBRVI | 0.51149382 | 0 |
| B30 | B61 | LH | WBHSRV | 0.4932853 | 0 |
| B30 | B62 | LH | WBHSRVI | 0.584027745 | 0 |
| B30 | B120 | LH | HCV-cAg | 0.443705919 | 8.04E-14 |
| B30 | B156 | LH | IgE | 0.43021394 | 5.30E-13 |
| B30 | B63 | LH | WBRSR(mPas_200) | 0.417942309 | 2.75E-12 |
| B30 | B66 | LH | WBRSR(mPas_100) | 0.415570122 | 3.75E-12 |
| B30 | B67 | LH | WBRSR(mPas_50) | 0.40400005 | 1.65E-11 |
| B30 | B55 | LH | TK | 0.40296789 | 1.87E-11 |
| B30 | B64 | LH | WBRSR(mPas_5) | 0.342597209 | 1.73E-08 |
| B30 | B69 | LH | PV | -0.32995546 | 6.09E-08 |
| B31 | B123 | FSH | hs-CRP1 | -0.48685827 | 0 |
| B31 | B138 | FSH | PAB | 0.487477467 | 0 |
| B31 | B32 | FSH | Testo | 1 | 0 |
| B31 | B56 | FSH | IR | 0.714316492 | 0 |
| B31 | B60 | FSH | WBRVI | 0.51149382 | 0 |
| B31 | B61 | FSH | WBHSRV | 0.4932853 | 0 |
| B31 | B62 | FSH | WBHSRVI | 0.584027745 | 0 |
| B31 | B120 | FSH | HCV-cAg | 0.443705918 | 8.04E-14 |
| B31 | B156 | FSH | IgE | 0.43021394 | 5.30E-13 |
| B31 | B63 | FSH | WBRSR(mPas_200) | 0.417942309 | 2.75E-12 |
| B31 | B66 | FSH | WBRSR(mPas_100) | 0.415570122 | 3.75E-12 |
| B31 | B67 | FSH | WBRSR(mPas_50) | 0.40400005 | 1.65E-11 |
| B31 | B55 | FSH | TK | 0.40296789 | 1.87E-11 |
| B31 | B64 | FSH | WBRSR(mPas_5) | 0.342597209 | 1.73E-08 |

| | | | | | |
|---|---|---|---|---|---|
| B31 | B69 | FSH | PV | -0.32995546 | 6.09E-08 |
| B32 | B123 | Testo | hs-CRP1 | -0.48685827 | 0 |
| B32 | B138 | Testo | PAB | 0.487477467 | 0 |
| B32 | B33 | Testo | Prog | 1 | 0 |
| B32 | B56 | Testo | IR | 0.714316492 | 0 |
| B32 | B60 | Testo | WBRVI | 0.51149382 | 0 |
| B32 | B61 | Testo | WBHSRV | 0.4932853 | 0 |
| B32 | B62 | Testo | WBHSRVI | 0.584027745 | 0 |
| B32 | B120 | Testo | HCV-cAg | 0.443705923 | 8.04E-14 |
| B32 | B156 | Testo | IgE | 0.43021394 | 5.30E-13 |
| B32 | B63 | Testo | WBRSR(mPas_200) | 0.417942309 | 2.75E-12 |
| B32 | B66 | Testo | WBRSR(mPas_100) | 0.415570122 | 3.75E-12 |
| B32 | B67 | Testo | WBRSR(mPas_50) | 0.40400005 | 1.65E-11 |
| B32 | B55 | Testo | TK | 0.40296789 | 1.87E-11 |
| B32 | B64 | Testo | WBRSR(mPas_5) | 0.342597209 | 1.73E-08 |
| B32 | B69 | Testo | PV | -0.32995546 | 6.09E-08 |
| B33 | B123 | Prog | hs-CRP1 | -0.48685827 | 0 |
| B33 | B138 | Prog | PAB | 0.487477467 | 0 |
| B33 | B56 | Prog | IR | 0.714316492 | 0 |
| B33 | B60 | Prog | WBRVI | 0.51149382 | 0 |
| B33 | B61 | Prog | WBHSRV | 0.4932853 | 0 |
| B33 | B62 | Prog | WBHSRVI | 0.584027745 | 0 |
| B33 | B120 | Prog | HCV-cAg | 0.443705918 | 8.04E-14 |
| B33 | B156 | Prog | IgE | 0.43021394 | 5.30E-13 |
| B33 | B63 | Prog | WBRSR(mPas_200) | 0.417942309 | 2.75E-12 |
| B33 | B66 | Prog | WBRSR(mPas_100) | 0.415570122 | 3.75E-12 |
| B33 | B67 | Prog | WBRSR(mPas_50) | 0.40400005 | 1.65E-11 |
| B33 | B55 | Prog | TK | 0.40296789 | 1.87E-11 |
| B33 | B64 | Prog | WBRSR(mPas_5) | 0.342597209 | 1.73E-08 |
| B33 | B69 | Prog | PV | -0.32995546 | 6.09E-08 |
| B35 | B119 | β2-MG | Hcv-rna | 0.897999131 | 0 |
| B35 | B121 | β2-MG | C3 | -0.67840053 | 0 |
| B35 | B169 | β2-MG | U-CRY | -0.90138782 | 0 |
| B35 | B171 | β2-MG | EC | 0.95032889 | 0 |
| B35 | B96 | β2-MG | dsDNA (IIF) | 0.624037721 | 0 |
| B35 | B167 | β2-MG | PC | 0.478796599 | 4.44E-16 |
| B35 | B225 | β2-MG | hPBG | 0.479701612 | 4.44E-16 |
| B35 | B38 | β2-MG | BAND% | 0.433012702 | 3.61E-13 |
| B35 | B94 | β2-MG | ALDH2-Lys504Lys | -0.4 | 2.71E-11 |
| B37 | B159 | BAND | β2-MG | -0.54739875 | 0 |
| B37 | B178 | BAND | Ig-U | 0.306186218 | 5.58E-07 |
| B38 | B180 | BAND% | Tf | 0.333333333 | 4.37E-08 |
| B39 | B79 | Other cells | hb V -dna | 0.404241831 | 1.60E-11 |
| B39 | B172 | Other cells | KB | -0.39399893 | 5.66E-11 |

| | | | | | |
|---|---|---|---|---|---|
| B39 | B168 | Other cells | NMP22 | 0.349230957 | 8.74E-09 |
| B39 | B57 | Other cells | RBCAI | -0.34730933 | 1.07E-08 |
| B39 | B54 | Other cells | ESR | -0.33508409 | 3.68E-08 |
| B39 | B123 | Other cells | hs-CRP1 | -0.31884897 | 1.75E-07 |
| B39 | B69 | Other cells | PV | -0.31569266 | 2.35E-07 |
| B4 | B17 | WB-Ca | CD3+CD16+CD56+NK | -0.74730284 | 0 |
| B4 | B21 | WB-Ca | CD3+CD4-CD8- | -0.70164642 | 0 |
| B40 | B42 | WBC | MON | 0.615627765 | 0 |
| B40 | B72 | WBC | HDL-C | -0.3528431 | 5.99E-09 |
| B40 | B222 | WBC | NS | 0.336823528 | 3.10E-08 |
| B40 | B46 | WBC | RBC Count | 0.316814106 | 2.12E-07 |
| B40 | B216 | WBC | BMI | 0.308100917 | 4.70E-07 |
| B41 | B42 | MON％ | MON | 0.64392944 | 0 |
| B42 | B222 | MON | NS | 0.361015836 | 2.50E-09 |
| B42 | B190 | MON | Cre | 0.341328948 | 1.97E-08 |
| B43 | B44 | MCV | MCH | 0.919617958 | 0 |
| B43 | B46 | MCV | RBC Count | -0.56066382 | 0 |
| B43 | B47 | MCV | RDW | 0.539250511 | 0 |
| B44 | B45 | MCH | MCHC | 0.589891544 | 0 |
| B44 | B46 | MCH | RBC Count | -0.39977997 | 2.79E-11 |
| B44 | B47 | MCH | RDW | 0.373376715 | 6.35E-10 |
| B44 | B51 | MCH | Hb | 0.367135563 | 1.28E-09 |
| B45 | B51 | MCHC | Hb | 0.631892085 | 0 |
| B46 | B47 | RBC Count | RDW | -0.59971213 | 0 |
| B46 | B51 | RBC Count | Hb | 0.694134525 | 0 |
| B46 | B126 | RBC Count | Alb | 0.340973129 | 2.04E-08 |
| B46 | B222 | RBC Count | NS | 0.32777037 | 7.52E-08 |
| B46 | B216 | RBC Count | BMI | 0.323998595 | 1.08E-07 |
| B46 | B72 | RBC Count | HDL-C | -0.31018671 | 3.89E-07 |
| B46 | B141 | RBC Count | AST/ALT | -0.30572545 | 5.81E-07 |
| B47 | B72 | RDW | HDL-C | 0.345611867 | 1.27E-08 |
| B47 | B51 | RDW | Hb | -0.31198452 | 3.31E-07 |
| B48 | B53 | LYM% | GR% | -0.936417 | 0 |
| B49 | B53 | baso% | GR% | -0.33169441 | 5.14E-08 |
| B49 | B50 | baso% | eo% | 0.31542664 | 2.41E-07 |
| B5 | B152 | WB-Mg | ACA-Ig(G-A-M) | 0.659212139 | 0 |
| B5 | B169 | WB-Mg | U-CRY | -0.78446454 | 0 |
| B5 | B38 | WB-Mg | BAND% | 0.816496581 | 0 |
| B5 | B92 | WB-Mg | ALDH2-Glu504Glu | 0.503115295 | 0 |
| B5 | B96 | WB-Mg | dsDNA (IIF) | 0.833493574 | 0 |
| B5 | B20 | WB-Mg | CD3+CD4+CD8+ | 0.455091027 | 1.51E-14 |
| B5 | B171 | WB-Mg | EC | 0.368932394 | 1.05E-09 |
| B5 | B167 | WB-Mg | PC | 0.360402923 | 2.67E-09 |
| B50 | B146 | eo% | CysC | 0.41181858 | 6.10E-12 |

| | | | | | |
|---|---|---|---|---|---|
| B51 | B216 | Hb | BMI | 0.427916713 | 7.25E-13 |
| B51 | B222 | Hb | NS | 0.420744244 | 1.90E-12 |
| B51 | B221 | Hb | Hei | 0.402303847 | 2.04E-11 |
| B51 | B126 | Hb | Alb | 0.363838285 | 1.84E-09 |
| B51 | B150 | Hb | ALP | -0.32469111 | 1.01E-07 |
| B51 | B173 | Hb | NTT | -0.31114561 | 3.57E-07 |
| B51 | B206 | Hb | Waist | 0.303225896 | 7.25E-07 |
| B51 | B174 | Hb | Urea/Cr | -0.3019339 | 8.12E-07 |
| B54 | B114 | ESR | TSH TT3 TT4(TT4) | 0.486962297 | 0 |
| B54 | B123 | ESR | hs-CRP1 | 0.849623165 | 0 |
| B54 | B172 | ESR | KB | 0.926783962 | 0 |
| B54 | B57 | ESR | RBCAI | 0.702605695 | 0 |
| B54 | B59 | ESR | WBRV | 0.71848184 | 0 |
| B54 | B64 | ESR | WBRSR(mPas_5) | 0.499403191 | 0 |
| B54 | B65 | ESR | WBRSR(mPas_1) | 0.578464712 | 0 |
| B54 | B69 | ESR | PV | 0.859164158 | 0 |
| B54 | B61 | ESR | WBHSRV | 0.419730519 | 2.17E-12 |
| B54 | B78 | ESR | HBcAb | -0.40055598 | 2.53E-11 |
| B54 | B67 | ESR | WBRSR(mPas_50) | 0.366385554 | 1.39E-09 |
| B54 | B191 | ESR | Urea | 0.329827347 | 6.16E-08 |
| B54 | B66 | ESR | WBRSR(mPas_100) | 0.326237911 | 8.72E-08 |
| B54 | B73 | ESR | ApoA | -0.3105133 | 3.78E-07 |
| B54 | B58 | ESR | HCT | -0.30474716 | 6.34E-07 |
| B55 | B56 | TK | IR | 0.694352174 | 0 |
| B55 | B58 | TK | HCT | -0.75134086 | 0 |
| B55 | B59 | TK | WBRV | 0.634312041 | 0 |
| B55 | B61 | TK | WBHSRV | 0.747792226 | 0 |
| B55 | B156 | TK | IgE | 0.37435174 | 5.69E-10 |
| B55 | B181 | TK | APTT | 0.307733088 | 4.86E-07 |
| B55 | B182 | TK | TT | -0.30773309 | 4.86E-07 |
| B55 | B183 | TK | PTA | -0.30773309 | 4.86E-07 |
| B55 | B184 | TK | INR | 0.307733088 | 4.86E-07 |
| B55 | B185 | TK | PT | 0.307733088 | 4.86E-07 |
| B55 | B186 | TK | PFC | 0.307733088 | 4.86E-07 |
| B56 | B156 | IR | IgE | 0.59512219 | 0 |
| B56 | B60 | IR | WBRVI | 0.601317719 | 0 |
| B56 | B61 | IR | WBHSRV | 0.80982541 | 0 |
| B56 | B62 | IR | WBHSRVI | 0.626976356 | 0 |
| B56 | B63 | IR | WBRSR(mPas_200) | 0.542406201 | 0 |
| B56 | B66 | IR | WBRSR(mPas_100) | 0.544675455 | 0 |
| B56 | B67 | IR | WBRSR(mPas_50) | 0.538112614 | 0 |
| B56 | B64 | IR | WBRSR(mPas_5) | 0.473986508 | 8.88E-16 |
| B56 | B123 | IR | hs-CRP1 | -0.44500354 | 6.66E-14 |
| B56 | B59 | IR | WBRV | 0.427107153 | 8.09E-13 |

| | | | | | |
|---|---|---|---|---|---|
| B56 | B65 | IR | WBRSR(mPas_1) | 0.418694243 | 2.49E-12 |
| B56 | B181 | IR | APTT | 0.396757119 | 4.04E-11 |
| B56 | B182 | IR | TT | -0.39675712 | 4.04E-11 |
| B56 | B183 | IR | PTA | -0.39675712 | 4.04E-11 |
| B56 | B184 | IR | INR | 0.396757119 | 4.04E-11 |
| B56 | B185 | IR | PT | 0.396757119 | 4.04E-11 |
| B56 | B186 | IR | PFC | 0.396757119 | 4.04E-11 |
| B56 | B138 | IR | PAB | 0.374257678 | 5.75E-10 |
| B56 | B155 | IR | IgA | 0.358033106 | 3.45E-09 |
| B56 | B120 | IR | HCV-cAg | 0.314111173 | 2.72E-07 |
| B56 | B115 | IR | TSH TT3 TT4(TT3) | 0.311304324 | 3.52E-07 |
| B57 | B114 | RBCAI | TSH TT3 TT4(TT4) | 0.551100917 | 0 |
| B57 | B123 | RBCAI | hs-CRP1 | 0.574979172 | 0 |
| B57 | B172 | RBCAI | KB | 0.679811755 | 0 |
| B57 | B59 | RBCAI | WBRV | 0.808801919 | 0 |
| B57 | B64 | RBCAI | WBRSR(mPas_5) | 0.656374849 | 0 |
| B57 | B65 | RBCAI | WBRSR(mPas_1) | 0.769984897 | 0 |
| B57 | B69 | RBCAI | PV | 0.716463643 | 0 |
| B57 | B60 | RBCAI | WBRVI | 0.475915947 | 6.66E-16 |
| B57 | B67 | RBCAI | WBRSR(mPas_50) | 0.465934962 | 3.11E-15 |
| B57 | B61 | RBCAI | WBHSRV | 0.44358623 | 8.17E-14 |
| B57 | B66 | RBCAI | WBRSR(mPas_100) | 0.411871089 | 6.06E-12 |
| B57 | B63 | RBCAI | WBRSR(mPas_200) | 0.307995466 | 4.74E-07 |
| B57 | B112 | RBCAI | FT4 | 0.303397865 | 7.14E-07 |
| B57 | B76 | RBCAI | ApoB | -0.30152053 | 8.42E-07 |
| B58 | B59 | HCT | WBRV | -0.52972716 | 0 |
| B58 | B60 | HCT | WBRVI | 0.52500069 | 0 |
| B58 | B62 | HCT | WBHSRVI | 0.739884969 | 0 |
| B58 | B63 | HCT | WBRSR(mPas_200) | 0.500072669 | 0 |
| B58 | B66 | HCT | WBRSR(mPas_100) | 0.446714196 | 5.20E-14 |
| B58 | B69 | HCT | PV | -0.4274432 | 7.73E-13 |
| B58 | B67 | HCT | WBRSR(mPas_50) | 0.415751649 | 3.66E-12 |
| B58 | B172 | HCT | KB | -0.3758227 | 4.81E-10 |
| B58 | B123 | HCT | hs-CRP1 | -0.35871693 | 3.20E-09 |
| B58 | B61 | HCT | WBHSRV | -0.31744565 | 2.00E-07 |
| B59 | B172 | WBRV | KB | 0.627046409 | 0 |
| B59 | B61 | WBRV | WBHSRV | 0.831718762 | 0 |
| B59 | B64 | WBRV | WBRSR(mPas_5) | 0.644006043 | 0 |
| B59 | B65 | WBRV | WBRSR(mPas_1) | 0.723734147 | 0 |
| B59 | B67 | WBRV | WBRSR(mPas_50) | 0.505970653 | 0 |
| B59 | B69 | WBRV | PV | 0.761481945 | 0 |
| B59 | B156 | WBRV | IgE | 0.468169416 | 2.22E-15 |
| B59 | B66 | WBRV | WBRSR(mPas_100) | 0.4626996 | 4.88E-15 |
| B59 | B123 | WBRV | hs-CRP1 | 0.454386695 | 1.69E-14 |

| | | | | | |
|---|---|---|---|---|---|
| B59 | B114 | WBRV | TSH TT3 TT4(TT4) | 0.408867594 | 8.90E-12 |
| B59 | B63 | WBRV | WBRSR(mPas_200) | 0.378632713 | 3.48E-10 |
| B59 | B60 | WBRV | WBRVI | 0.353718877 | 5.46E-09 |
| B59 | B134 | WBRV | CHE | 0.311225636 | 3.54E-07 |
| B59 | B73 | WBRV | ApoA | -0.30679885 | 5.28E-07 |
| B6 | B119 | WB-Fe | Hcv-rna | 0.702781928 | 0 |
| B6 | B171 | WB-Fe | EC | 0.521749195 | 0 |
| B6 | B94 | WB-Fe | ALDH2-Lys504Lys | -0.6 | 0 |
| B60 | B61 | WBRVI | WBHSRV | 0.513362026 | 0 |
| B60 | B62 | WBRVI | WBHSRVI | 0.82350199 | 0 |
| B60 | B63 | WBRVI | WBRSR(mPas_200) | 0.828391349 | 0 |
| B60 | B64 | WBRVI | WBRSR(mPas_5) | 0.854835162 | 0 |
| B60 | B65 | WBRVI | WBRSR(mPas_1) | 0.822399572 | 0 |
| B60 | B66 | WBRVI | WBRSR(mPas_100) | 0.856103761 | 0 |
| B60 | B67 | WBRVI | WBRSR(mPas_50) | 0.862877354 | 0 |
| B60 | B156 | WBRVI | IgE | 0.408145746 | 9.76E-12 |
| B61 | B156 | WBHSRV | IgE | 0.766335291 | 0 |
| B61 | B63 | WBHSRV | WBRSR(mPas_200) | 0.629523344 | 0 |
| B61 | B64 | WBHSRV | WBRSR(mPas_5) | 0.697217577 | 0 |
| B61 | B65 | WBHSRV | WBRSR(mPas_1) | 0.69095296 | 0 |
| B61 | B66 | WBHSRV | WBRSR(mPas_100) | 0.665767694 | 0 |
| B61 | B67 | WBHSRV | WBRSR(mPas_50) | 0.680165398 | 0 |
| B61 | B155 | WBHSRV | IgA | 0.452357581 | 2.29E-14 |
| B61 | B134 | WBHSRV | CHE | 0.43769059 | 1.88E-13 |
| B61 | B181 | WBHSRV | APTT | 0.431767907 | 4.28E-13 |
| B61 | B182 | WBHSRV | TT | -0.43176791 | 4.28E-13 |
| B61 | B183 | WBHSRV | PTA | -0.43176791 | 4.28E-13 |
| B61 | B184 | WBHSRV | INR | 0.431767907 | 4.28E-13 |
| B61 | B185 | WBHSRV | PT | 0.431767907 | 4.28E-13 |
| B61 | B186 | WBHSRV | PFC | 0.431767907 | 4.28E-13 |
| B61 | B69 | WBHSRV | PV | 0.419320818 | 2.29E-12 |
| B61 | B86 | WBHSRV | TBhCG | 0.403115374 | 1.84E-11 |
| B61 | B138 | WBHSRV | PAB | 0.388735718 | 1.07E-10 |
| B61 | B158 | WBHSRV | IgM | 0.376535226 | 4.43E-10 |
| B61 | B73 | WBHSRV | ApoA | -0.36605598 | 1.44E-09 |
| B61 | B115 | WBHSRV | TSH TT3 TT4(TT3) | 0.346060326 | 1.21E-08 |
| B61 | B83 | WBHSRV | AFP | 0.342723028 | 1.71E-08 |
| B62 | B123 | WBHSRVI | hs-CRP1 | -0.56821243 | 0 |
| B62 | B63 | WBHSRVI | WBRSR(mPas_200) | 0.737935397 | 0 |
| B62 | B64 | WBHSRVI | WBRSR(mPas_5) | 0.542651632 | 0 |
| B62 | B66 | WBHSRVI | WBRSR(mPas_100) | 0.701386435 | 0 |
| B62 | B67 | WBHSRVI | WBRSR(mPas_50) | 0.674252305 | 0 |
| B62 | B69 | WBHSRVI | PV | -0.46168793 | 5.77E-15 |
| B62 | B65 | WBHSRVI | WBRSR(mPas_1) | 0.432822694 | 3.70E-13 |

| | | | | | |
|---|---|---|---|---|---|
| B62 | B156 | WBHSRVI | IgE | 0.393156587 | 6.27E-11 |
| B62 | B172 | WBHSRVI | KB | -0.36310564 | 1.99E-09 |
| B62 | B114 | WBHSRVI | TSH TT3 TT4(TT4) | -0.34458703 | 1.41E-08 |
| B62 | B139 | WBHSRVI | ADA | -0.32602324 | 8.90E-08 |
| B63 | B156 | WBRSR(mPas_200) | IgE | 0.75334855 | 0 |
| B63 | B64 | WBRSR(mPas_200) | WBRSR(mPas_5) | 0.918656488 | 0 |
| B63 | B65 | WBRSR(mPas_200) | WBRSR(mPas_1) | 0.842220149 | 0 |
| B63 | B66 | WBRSR(mPas_200) | WBRSR(mPas_100) | 0.992598204 | 0 |
| B63 | B67 | WBRSR(mPas_200) | WBRSR(mPas_50) | 0.98397661 | 0 |
| B63 | B155 | WBRSR(mPas_200) | IgA | 0.432033154 | 4.13E-13 |
| B63 | B86 | WBRSR(mPas_200) | TBhCG | 0.411388824 | 6.44E-12 |
| B63 | B138 | WBRSR(mPas_200) | PAB | 0.391564808 | 7.60E-11 |
| B63 | B83 | WBRSR(mPas_200) | AFP | 0.33973145 | 2.31E-08 |
| B63 | B115 | WBRSR(mPas_200) | TSH TT3 TT4(TT3) | 0.333356829 | 4.36E-08 |
| B63 | B134 | WBRSR(mPas_200) | CHE | 0.324027538 | 1.08E-07 |
| B63 | B158 | WBRSR(mPas_200) | IgM | 0.323451692 | 1.14E-07 |
| B63 | B82 | WBRSR(mPas_200) | CEA | 0.31832349 | 1.84E-07 |
| B63 | B181 | WBRSR(mPas_200) | APTT | 0.314010716 | 2.75E-07 |
| B63 | B182 | WBRSR(mPas_200) | TT | -0.31401072 | 2.75E-07 |
| B63 | B183 | WBRSR(mPas_200) | PTA | -0.31401072 | 2.75E-07 |
| B63 | B184 | WBRSR(mPas_200) | INR | 0.314010716 | 2.75E-07 |
| B63 | B185 | WBRSR(mPas_200) | PT | 0.314010716 | 2.75E-07 |
| B63 | B186 | WBRSR(mPas_200) | PFC | 0.314010716 | 2.75E-07 |
| B64 | B156 | WBRSR(mPas_5) | IgE | 0.644809394 | 0 |
| B64 | B65 | WBRSR(mPas_5) | WBRSR(mPas_1) | 0.986612826 | 0 |
| B64 | B66 | WBRSR(mPas_5) | WBRSR(mPas_100) | 0.957678414 | 0 |
| B64 | B67 | WBRSR(mPas_5) | WBRSR(mPas_50) | 0.973283929 | 0 |
| B64 | B69 | WBRSR(mPas_5) | PV | 0.467047671 | 2.44E-15 |
| B64 | B155 | WBRSR(mPas_5) | IgA | 0.366103488 | 1.43E-09 |
| B64 | B86 | WBRSR(mPas_5) | TBhCG | 0.360752058 | 2.57E-09 |
| B64 | B138 | WBRSR(mPas_5) | PAB | 0.331903777 | 5.03E-08 |
| B64 | B115 | WBRSR(mPas_5) | TSH TT3 TT4(TT3) | 0.331780802 | 5.09E-08 |
| B64 | B172 | WBRSR(mPas_5) | KB | 0.319502081 | 1.65E-07 |
| B64 | B83 | WBRSR(mPas_5) | AFP | 0.305827224 | 5.76E-07 |
| B65 | B156 | WBRSR(mPas_1) | IgE | 0.569776923 | 0 |
| B65 | B66 | WBRSR(mPas_1) | WBRSR(mPas_100) | 0.89798915 | 0 |
| B65 | B67 | WBRSR(mPas_1) | WBRSR(mPas_50) | 0.922899343 | 0 |
| B65 | B69 | WBRSR(mPas_1) | PV | 0.552259574 | 0 |
| B65 | B172 | WBRSR(mPas_1) | KB | 0.42359369 | 1.30E-12 |
| B65 | B86 | WBRSR(mPas_1) | TBhCG | 0.32340655 | 1.14E-07 |
| B65 | B155 | WBRSR(mPas_1) | IgA | 0.322655197 | 1.23E-07 |
| B65 | B115 | WBRSR(mPas_1) | TSH TT3 TT4(TT3) | 0.311990915 | 3.30E-07 |
| B65 | B211 | WBRSR(mPas_1) | IOP-R | -0.30137497 | 8.53E-07 |
| B66 | B156 | WBRSR(mPas_100) | IgE | 0.737207261 | 0 |

| | | | | | |
|---|---|---|---|---|---|
| B66 | B67 | WBRSR(mPas_100) | WBRSR(mPas_50) | 0.998135817 | 0 |
| B66 | B155 | WBRSR(mPas_100) | IgA | 0.420503237 | 1.96E-12 |
| B66 | B86 | WBRSR(mPas_100) | TBhCG | 0.4020896 | 2.09E-11 |
| B66 | B138 | WBRSR(mPas_100) | PAB | 0.388661322 | 1.08E-10 |
| B66 | B115 | WBRSR(mPas_100) | TSH TT3 TT4(TT3) | 0.347298187 | 1.07E-08 |
| B66 | B83 | WBRSR(mPas_100) | AFP | 0.335763937 | 3.44E-08 |
| B66 | B158 | WBRSR(mPas_100) | IgM | 0.313547188 | 2.87E-07 |
| B66 | B82 | WBRSR(mPas_100) | CEA | 0.312355284 | 3.20E-07 |
| B66 | B134 | WBRSR(mPas_100) | CHE | 0.311272064 | 3.53E-07 |
| B66 | B181 | WBRSR(mPas_100) | APTT | 0.303740441 | 6.93E-07 |
| B66 | B182 | WBRSR(mPas_100) | TT | -0.30374044 | 6.93E-07 |
| B66 | B183 | WBRSR(mPas_100) | PTA | -0.30374044 | 6.93E-07 |
| B66 | B184 | WBRSR(mPas_100) | INR | 0.303740441 | 6.93E-07 |
| B66 | B185 | WBRSR(mPas_100) | PT | 0.303740441 | 6.93E-07 |
| B66 | B186 | WBRSR(mPas_100) | PFC | 0.303740441 | 6.93E-07 |
| B67 | B156 | WBRSR(mPas_50) | IgE | 0.722584925 | 0 |
| B67 | B155 | WBRSR(mPas_50) | IgA | 0.41188893 | 6.04E-12 |
| B67 | B86 | WBRSR(mPas_50) | TBhCG | 0.395664161 | 4.62E-11 |
| B67 | B138 | WBRSR(mPas_50) | PAB | 0.378671669 | 3.47E-10 |
| B67 | B115 | WBRSR(mPas_50) | TSH TT3 TT4(TT3) | 0.344924407 | 1.36E-08 |
| B67 | B83 | WBRSR(mPas_50) | AFP | 0.333225355 | 4.42E-08 |
| B67 | B69 | WBRSR(mPas_50) | PV | 0.323720785 | 1.11E-07 |
| B67 | B158 | WBRSR(mPas_50) | IgM | 0.308385371 | 4.58E-07 |
| B67 | B82 | WBRSR(mPas_50) | CEA | 0.307386664 | 5.01E-07 |
| B67 | B134 | WBRSR(mPas_50) | CHE | 0.305678938 | 5.83E-07 |
| B67 | B181 | WBRSR(mPas_50) | APTT | 0.304598876 | 6.42E-07 |
| B67 | B182 | WBRSR(mPas_50) | TT | -0.30459888 | 6.42E-07 |
| B67 | B183 | WBRSR(mPas_50) | PTA | -0.30459888 | 6.42E-07 |
| B67 | B184 | WBRSR(mPas_50) | INR | 0.304598876 | 6.42E-07 |
| B67 | B185 | WBRSR(mPas_50) | PT | 0.304598876 | 6.42E-07 |
| B67 | B186 | WBRSR(mPas_50) | PFC | 0.304598876 | 6.42E-07 |
| B69 | B114 | PV | TSH TT3 TT4(TT4) | 0.520271768 | 0 |
| B69 | B123 | PV | hs-CRP1 | 0.804950391 | 0 |
| B69 | B172 | PV | KB | 0.731077673 | 0 |
| B69 | B86 | PV | TBhCG | 0.340519907 | 2.14E-08 |
| B69 | B156 | PV | IgE | 0.308777814 | 4.42E-07 |
| B7 | B20 | WB-Cop | CD3+CD4+CD8+ | 0.552606552 | 0 |
| B7 | B225 | WB-Cop | hPBG | 0.738548946 | 0 |
| B7 | B180 | WB-Cop | Tf | 0.444444444 | 7.22E-14 |
| B70 | B77 | LDL-C | TC | 0.86138466 | 0 |
| B71 | B72 | TG | HDL-C | -0.39374784 | 5.83E-11 |
| B71 | B215 | TG | BMI | 0.361817939 | 2.29E-09 |
| B71 | B170 | TG | U-GLU | 0.301726069 | 8.27E-07 |
| B72 | B216 | HDL-C | BMI | -0.45416351 | 1.75E-14 |

| | | | | | |
|---|---|---|---|---|---|
| B72 | B215 | HDL-C | BMI | -0.44055849 | 1.26E-13 |
| B72 | B206 | HDL-C | Waist | -0.41835909 | 2.60E-12 |
| B72 | B77 | HDL-C | TC | 0.383456321 | 1.99E-10 |
| B72 | B222 | HDL-C | NS | -0.37302134 | 6.61E-10 |
| B72 | B141 | HDL-C | AST/ALT | 0.351240514 | 7.09E-09 |
| B72 | B210 | HDL-C | Hip | -0.3392516 | 2.43E-08 |
| B72 | B207 | HDL-C | WHR | -0.33787236 | 2.79E-08 |
| B73 | B100 | ApoA | anti-Ro52 | -0.57853689 | 0 |
| B73 | B101 | ApoA | anti-Sc70 | -0.57853689 | 0 |
| B73 | B102 | ApoA | anti-Sm | -0.5785369 | 0 |
| B73 | B103 | ApoA | anti-SSA/Ro | -0.57853689 | 0 |
| B73 | B104 | ApoA | anti-SSB | -0.5785369 | 0 |
| B73 | B105 | ApoA | anti-U1RNP | -0.57853689 | 0 |
| B73 | B106 | ApoA | anti-P | -0.57853689 | 0 |
| B73 | B107 | ApoA | AnuA | -0.5785368 | 0 |
| B73 | B108 | ApoA | Anti-Cyclin | -0.5785369 | 0 |
| B73 | B109 | ApoA | AMA-M2 | -0.57853689 | 0 |
| B73 | B110 | ApoA | AHA | -0.57853689 | 0 |
| B73 | B95 | ApoA | anti-cenpb | -0.5785369 | 0 |
| B73 | B97 | ApoA | dsDNA | -0.57853689 | 0 |
| B73 | B98 | ApoA | anti-Jo-1 | -0.57853689 | 0 |
| B73 | B99 | ApoA | anti-PM-Scl | -0.57853689 | 0 |
| B73 | B156 | ApoA | IgE | -0.34598803 | 1.22E-08 |
| B73 | B172 | ApoA | KB | -0.30373668 | 6.93E-07 |
| B74 | B135 | ApoA1 | TBA | 0.557501777 | 0 |
| B74 | B75 | ApoA1 | ApoA1/B | 0.535608195 | 0 |
| B74 | B156 | ApoA1 | IgE | 0.482487698 | 2.22E-16 |
| B74 | B86 | ApoA1 | TBhCG | 0.349522923 | 8.48E-09 |
| B74 | B197 | ApoA1 | AG | 0.347310644 | 1.07E-08 |
| B74 | B198 | ApoA1 | P | 0.343153404 | 1.64E-08 |
| B74 | B100 | ApoA1 | anti-Ro52 | -0.30095405 | 8.85E-07 |
| B74 | B101 | ApoA1 | anti-Sc70 | -0.30095405 | 8.85E-07 |
| B74 | B102 | ApoA1 | anti-Sm | -0.30095405 | 8.85E-07 |
| B74 | B103 | ApoA1 | anti-SSA/Ro | -0.30095405 | 8.85E-07 |
| B74 | B104 | ApoA1 | anti-SSB | -0.30095405 | 8.85E-07 |
| B74 | B105 | ApoA1 | anti-U1RNP | -0.30095405 | 8.85E-07 |
| B74 | B106 | ApoA1 | anti-P | -0.30095405 | 8.85E-07 |
| B74 | B107 | ApoA1 | AnuA | -0.300954 | 8.85E-07 |
| B74 | B108 | ApoA1 | Anti-Cyclin | -0.30095405 | 8.85E-07 |
| B74 | B109 | ApoA1 | AMA-M2 | -0.30095405 | 8.85E-07 |
| B74 | B110 | ApoA1 | AHA | -0.30095405 | 8.85E-07 |
| B74 | B95 | ApoA1 | anti-cenpb | -0.30095405 | 8.85E-07 |
| B74 | B97 | ApoA1 | dsDNA | -0.30095405 | 8.85E-07 |
| B74 | B98 | ApoA1 | anti-Jo-1 | -0.30095405 | 8.85E-07 |

| | | | | | |
|---|---|---|---|---|---|
| B74 | B99 | ApoA1 | anti-PM-Scl | -0.30095405 | 8.85E-07 |
| B75 | B76 | ApoA1/B | ApoB | -0.60751073 | 0 |
| B75 | B199 | ApoA1/B | Mg | -0.34343172 | 1.59E-08 |
| B78 | B176 | HBcAb | Ucr | 0.47792922 | 4.44E-16 |
| B78 | B79 | HBcAb | hb V -dna | 0.337212495 | 2.98E-08 |
| B78 | B172 | HBcAb | KB | -0.33019513 | 5.95E-08 |
| B78 | B123 | HBcAb | hs-CRP1 | -0.31985463 | 1.60E-07 |
| B8 | B93 | WB-Zinc | ALDH2-Glu504Lys | 0.938194187 | 0 |
| B81 | B155 | TSGF | IgA | -0.41604412 | 3.52E-12 |
| B82 | B195 | CEA | Na | -0.50176109 | 0 |
| B82 | B200 | CEA | OP | -0.50342988 | 0 |
| B82 | B89 | CEA | CA 72-4 | 0.531679382 | 0 |
| B82 | B196 | CEA | TCO2 | -0.4205472 | 1.95E-12 |
| B82 | B156 | CEA | IgE | 0.398098578 | 3.43E-11 |
| B82 | B199 | CEA | Mg | -0.3942545 | 5.49E-11 |
| B82 | B88 | CEA | CA 242 | 0.369018822 | 1.04E-09 |
| B82 | B85 | CEA | SCC | -0.34148303 | 1.94E-08 |
| B83 | B86 | AFP | TBhCG | 0.765717121 | 0 |
| B83 | B88 | AFP | CA 242 | -0.46166221 | 5.77E-15 |
| B83 | B181 | AFP | APTT | 0.456447525 | 1.24E-14 |
| B83 | B182 | AFP | TT | -0.45644753 | 1.24E-14 |
| B83 | B183 | AFP | PTA | -0.45644753 | 1.24E-14 |
| B83 | B184 | AFP | INR | 0.456447525 | 1.24E-14 |
| B83 | B185 | AFP | PT | 0.456447525 | 1.24E-14 |
| B83 | B186 | AFP | PFC | 0.456447525 | 1.24E-14 |
| B83 | B156 | AFP | IgE | 0.436918481 | 2.10E-13 |
| B84 | B116 | C-PSA | TT3 | 0.548023318 | 0 |
| B84 | B117 | C-PSA | TT4 | 0.548023318 | 0 |
| B84 | B134 | C-PSA | CHE | -0.6061178 | 0 |
| B84 | B157 | C-PSA | IgG | -0.59466375 | 0 |
| B84 | B86 | C-PSA | TBhCG | -0.52180613 | 0 |
| B84 | B85 | C-PSA | SCC | 0.458170618 | 9.77E-15 |
| B84 | B181 | C-PSA | APTT | -0.41665709 | 3.25E-12 |
| B84 | B182 | C-PSA | TT | 0.416657089 | 3.25E-12 |
| B84 | B183 | C-PSA | PTA | 0.416657088 | 3.25E-12 |
| B84 | B184 | C-PSA | INR | -0.41665709 | 3.25E-12 |
| B84 | B185 | C-PSA | PT | -0.41665709 | 3.25E-12 |
| B84 | B186 | C-PSA | PFC | -0.41665709 | 3.25E-12 |
| B84 | B200 | C-PSA | OP | 0.404253553 | 1.60E-11 |
| B84 | B195 | C-PSA | Na | 0.397990424 | 3.48E-11 |
| B84 | B156 | C-PSA | IgE | -0.38272786 | 2.17E-10 |
| B84 | B197 | C-PSA | AG | 0.37863969 | 3.48E-10 |
| B84 | B139 | C-PSA | ADA | -0.37318879 | 6.49E-10 |
| B84 | B136 | C-PSA | 5'NT | -0.36506401 | 1.61E-09 |

| | | | | | |
|---|---|---|---|---|---|
| B84 | B158 | C-PSA | IgM | -0.36125872 | 2.43E-09 |
| B84 | B196 | C-PSA | TCO2 | -0.3063683 | 5.49E-07 |
| B85 | B134 | SCC | CHE | -0.45629876 | 1.29E-14 |
| B85 | B157 | SCC | IgG | -0.43800922 | 1.80E-13 |
| B85 | B195 | SCC | Na | 0.421742409 | 1.66E-12 |
| B85 | B200 | SCC | OP | 0.403238273 | 1.81E-11 |
| B86 | B156 | TBhCG | IgE | 0.591282701 | 0 |
| B86 | B155 | TBhCG | IgA | 0.343174097 | 1.63E-08 |
| B86 | B116 | TBhCG | TT3 | -0.32872142 | 6.86E-08 |
| B86 | B117 | TBhCG | TT4 | -0.32872142 | 6.86E-08 |
| B86 | B181 | TBhCG | APTT | 0.328721423 | 6.86E-08 |
| B86 | B182 | TBhCG | TT | -0.32872142 | 6.86E-08 |
| B86 | B183 | TBhCG | PTA | -0.32872142 | 6.86E-08 |
| B86 | B184 | TBhCG | INR | 0.328721423 | 6.86E-08 |
| B86 | B185 | TBhCG | PT | 0.328721423 | 6.86E-08 |
| B86 | B186 | TBhCG | PFC | 0.328721423 | 6.86E-08 |
| B86 | B134 | TBhCG | CHE | 0.321702733 | 1.34E-07 |
| B86 | B158 | TBhCG | IgM | 0.314925614 | 2.53E-07 |
| B87 | B181 | NSE | APTT | -0.7031517 | 0 |
| B87 | B182 | NSE | TT | 0.703151697 | 0 |
| B87 | B183 | NSE | PTA | 0.703151696 | 0 |
| B87 | B184 | NSE | INR | -0.7031517 | 0 |
| B87 | B185 | NSE | PT | -0.7031517 | 0 |
| B87 | B186 | NSE | PFC | -0.7031517 | 0 |
| B87 | B197 | NSE | AG | 0.537699866 | 0 |
| B87 | B89 | NSE | CA 72-4 | 0.540383398 | 0 |
| B87 | B196 | NSE | TCO2 | -0.47518419 | 6.66E-16 |
| B87 | B157 | NSE | IgG | -0.32469469 | 1.01E-07 |
| B87 | B116 | NSE | TT3 | -0.31603124 | 2.28E-07 |
| B87 | B117 | NSE | TT4 | -0.31603124 | 2.28E-07 |
| B88 | B200 | CA 242 | OP | -0.38135081 | 2.54E-10 |
| B88 | B195 | CA 242 | Na | -0.35233967 | 6.31E-09 |
| B88 | B197 | CA 242 | AG | -0.32231339 | 1.27E-07 |
| B89 | B196 | CA 72-4 | TCO2 | -0.6314079 | 0 |
| B89 | B197 | CA 72-4 | AG | 0.41551159 | 3.78E-12 |
| B89 | B157 | CA 72-4 | IgG | -0.37778032 | 3.84E-10 |
| B89 | B199 | CA 72-4 | Mg | -0.36687664 | 1.31E-09 |
| B9 | B159 | RH | β2-MG | -0.71485073 | 0 |
| B9 | B178 | RH | Ig-U | -0.60878078 | 0 |
| B9 | B179 | RH | α1-MG | -0.77875499 | 0 |
| B9 | B21 | RH | CD3+CD4-CD8- | -0.72601344 | 0 |
| B91 | B130 | TP | GLB | 0.751948351 | 0 |
| B91 | B126 | TP | Alb | 0.475530994 | 6.66E-16 |
| B91 | B127 | TP | A-G ratio | -0.44120693 | 1.15E-13 |

| | | | | | |
|---|---|---|---|---|---|
| B91 | B118 | TP | CEA1 | -0.31381296 | 2.80E-07 |
| B92 | B167 | ALDH2-Glu504Glu | PC | 0.820537898 | 0 |
| B92 | B169 | ALDH2-Glu504Glu | U-CRY | 0.43852901 | 1.67E-13 |
| B92 | B171 | ALDH2-Glu504Glu | EC | -0.32998317 | 6.07E-08 |
| B93 | B152 | ALDH2-Glu504Lys | ACA-Ig(G-A-M) | 0.833003235 | 0 |
| B93 | B167 | ALDH2-Glu504Lys | PC | 0.882758227 | 0 |
| B93 | B180 | ALDH2-Glu504Lys | Tf | -0.70710678 | 0 |
| B94 | B119 | ALDH2-Lys504Lys | Hcv-rna | 0.999512076 | 0 |
| B94 | B167 | ALDH2-Lys504Lys | PC | 0.963771219 | 0 |
| B94 | B169 | ALDH2-Lys504Lys | U-CRY | 0.804315285 | 0 |
| B94 | B180 | ALDH2-Lys504Lys | Tf | 0.346410162 | 1.17E-08 |
| B95 | B101 | anti-cenpb | anti-Sc70 | 0.999999986 | 0 |
| B95 | B106 | anti-cenpb | anti-P | 0.999999996 | 0 |
| B95 | B107 | anti-cenpb | AnuA | 0.999999864 | 0 |
| B95 | B109 | anti-cenpb | AMA-M2 | 0.999999999 | 0 |
| B95 | B98 | anti-cenpb | anti-Jo-1 | 0.999999998 | 0 |
| B95 | B198 | anti-cenpb | P | 0.423843893 | 1.26E-12 |
| B95 | B136 | anti-cenpb | 5'NT | 0.400475801 | 2.56E-11 |
| B95 | B199 | anti-cenpb | Mg | 0.388322106 | 1.12E-10 |
| B96 | B152 | dsDNA (IIF) | ACA-Ig(G-A-M) | 0.897657101 | 0 |
| B96 | B167 | dsDNA (IIF) | PC | 0.788198329 | 0 |
| B96 | B169 | dsDNA (IIF) | U-CRY | -0.53846154 | 0 |
| B97 | B100 | dsDNA | anti-Ro52 | 0.999999997 | 0 |
| B97 | B101 | dsDNA | anti-Sc70 | 0.999999996 | 0 |
| B97 | B103 | dsDNA | anti-SSA/Ro | 0.999999999 | 0 |
| B97 | B106 | dsDNA | anti-P | 0.999999998 | 0 |
| B97 | B107 | dsDNA | AnuA | 0.999999926 | 0 |
| B97 | B108 | dsDNA | Anti-Cyclin | 0.999999999 | 0 |
| B97 | B109 | dsDNA | AMA-M2 | 0.999999998 | 0 |
| B97 | B110 | dsDNA | AHA | 1 | 0 |
| B97 | B98 | dsDNA | anti-Jo-1 | 0.999999999 | 0 |
| B97 | B198 | dsDNA | P | 0.423843891 | 1.26E-12 |
| B97 | B136 | dsDNA | 5'NT | 0.400475799 | 2.56E-11 |
| B97 | B199 | dsDNA | Mg | 0.388322104 | 1.12E-10 |
| B98 | B100 | anti-Jo-1 | anti-Ro52 | 0.999999993 | 0 |
| B98 | B101 | anti-Jo-1 | anti-Sc70 | 0.999999994 | 0 |
| B98 | B102 | anti-Jo-1 | anti-Sm | 0.999999999 | 0 |
| B98 | B103 | anti-Jo-1 | anti-SSA/Ro | 0.999999995 | 0 |
| B98 | B104 | anti-Jo-1 | anti-SSB | 0.999999993 | 0 |
| B98 | B105 | anti-Jo-1 | anti-U1RNP | 0.999999998 | 0 |
| B98 | B107 | anti-Jo-1 | AnuA | 0.999999914 | 0 |
| B98 | B108 | anti-Jo-1 | Anti-Cyclin | 0.999999997 | 0 |
| B98 | B109 | anti-Jo-1 | AMA-M2 | 0.999999991 | 0 |
| B98 | B110 | anti-Jo-1 | AHA | 0.999999993 | 0 |

| | | | | | |
|---|---|---|---|---|---|
| B98 | B99 | anti-Jo-1 | anti-PM-Scl | 0.999999995 | 0 |
| B98 | B198 | anti-Jo-1 | P | 0.423843889 | 1.26E-12 |
| B98 | B136 | anti-Jo-1 | 5'NT | 0.400475798 | 2.56E-11 |
| B98 | B199 | anti-Jo-1 | Mg | 0.388322103 | 1.12E-10 |
| B99 | B100 | anti-PM-Scl | anti-Ro52 | 0.999999995 | 0 |
| B99 | B103 | anti-PM-Scl | anti-SSA/Ro | 0.999999997 | 0 |
| B99 | B107 | anti-PM-Scl | AnuA | 0.99999987 | 0 |
| B99 | B108 | anti-PM-Scl | Anti-Cyclin | 0.999999999 | 0 |
| B99 | B109 | anti-PM-Scl | AMA-M2 | 0.999999999 | 0 |
| B99 | B110 | anti-PM-Scl | AHA | 0.999999999 | 0 |
| B99 | B198 | anti-PM-Scl | P | 0.42384389 | 1.26E-12 |
| B99 | B136 | anti-PM-Scl | 5'NT | 0.400475798 | 2.56E-11 |
| B99 | B199 | anti-PM-Scl | Mg | 0.388322104 | 1.12E-10 |

**Supplementary Table 30. PEI Correlations in Hepatic adipose infiltration**

| Row | Column | Row | Column | Cor | *P* |
|---|---|---|---|---|---|
| B1 | B190 | Gender | Cre | -0.48719 | 0 |
| B1 | B192 | Gender | UA | -0.47108 | 0 |
| B1 | B221 | Gender | Hei | -0.58144 | 0 |
| B1 | B46 | Gender | RBC Count | -0.50644 | 0 |
| B1 | B51 | Gender | Hb | -0.59397 | 0 |
| B1 | B222 | Gender | NS | -0.41933 | 4.29E-13 |
| B1 | B216 | Gender | BMI | -0.35189 | 2.09E-09 |
| B10 | B12 | PG Ⅰ | PG II | 0.619244 | 0 |
| B10 | B14 | PG Ⅰ | G-17 | 0.705459 | 0 |
| B10 | B74 | PG Ⅰ | ApoA1 | 0.319354 | 6.51E-08 |
| B100 | B101 | anti-Ro52 | anti-Sc70 | 1 | 0 |
| B100 | B103 | anti-Ro52 | anti-SSA/Ro | 1 | 0 |
| B100 | B105 | anti-Ro52 | anti-U1RNP | 1 | 0 |
| B100 | B106 | anti-Ro52 | anti-P | 1 | 0 |
| B100 | B107 | anti-Ro52 | AnuA | 1 | 0 |
| B100 | B110 | anti-Ro52 | AHA | 1 | 0 |
| B100 | B154 | anti-Ro52 | RF | 0.476063 | 0 |
| B100 | B183 | anti-Ro52 | PTA | 0.497714 | 0 |
| B100 | B198 | anti-Ro52 | P | -0.41547 | 7.35E-13 |
| B100 | B124 | anti-Ro52 | ACTH | 0.352095 | 2.04E-09 |
| B100 | B185 | anti-Ro52 | PT | 0.335754 | 1.21E-08 |
| B100 | B186 | anti-Ro52 | PFC | -0.33542 | 1.25E-08 |
| B100 | B134 | anti-Ro52 | CHE | -0.31897 | 6.76E-08 |
| B100 | B139 | anti-Ro52 | ADA | -0.3138 | 1.13E-07 |
| B100 | B182 | anti-Ro52 | TT | 0.29809 | 5.00E-07 |
| B101 | B102 | anti-Sc70 | anti-Sm | 1 | 0 |
| B101 | B103 | anti-Sc70 | anti-SSA/Ro | 1 | 0 |
| B101 | B104 | anti-Sc70 | anti-SSB | 1 | 0 |
| B101 | B105 | anti-Sc70 | anti-U1RNP | 1 | 0 |
| B101 | B106 | anti-Sc70 | anti-P | 1 | 0 |
| B101 | B107 | anti-Sc70 | AnuA | 1 | 0 |
| B101 | B108 | anti-Sc70 | Anti-Cyclin | 1 | 0 |
| B101 | B109 | anti-Sc70 | AMA-M2 | 1 | 0 |
| B101 | B110 | anti-Sc70 | AHA | 1 | 0 |
| B101 | B154 | anti-Sc70 | RF | 0.476063 | 0 |
| B101 | B183 | anti-Sc70 | PTA | 0.497714 | 0 |
| B101 | B198 | anti-Sc70 | P | -0.41547 | 7.35E-13 |
| B101 | B124 | anti-Sc70 | ACTH | 0.352095 | 2.04E-09 |
| B101 | B185 | anti-Sc70 | PT | 0.335754 | 1.21E-08 |
| B101 | B186 | anti-Sc70 | PFC | -0.33542 | 1.25E-08 |
| B101 | B134 | anti-Sc70 | CHE | -0.31897 | 6.76E-08 |
| B101 | B139 | anti-Sc70 | ADA | -0.3138 | 1.13E-07 |

| | | | | | |
|---|---|---|---|---|---|
| B101 | B182 | anti-Sc70 | TT | 0.29809 | 5.00E-07 |
| B102 | B104 | anti-Sm | anti-SSB | 1 | 0 |
| B102 | B106 | anti-Sm | anti-P | 1 | 0 |
| B102 | B107 | anti-Sm | AnuA | 1 | 0 |
| B102 | B108 | anti-Sm | Anti-Cyclin | 1 | 0 |
| B102 | B154 | anti-Sm | RF | 0.476063 | 0 |
| B102 | B183 | anti-Sm | PTA | 0.497714 | 0 |
| B102 | B198 | anti-Sm | P | -0.41547 | 7.35E-13 |
| B102 | B124 | anti-Sm | ACTH | 0.352095 | 2.04E-09 |
| B102 | B185 | anti-Sm | PT | 0.335754 | 1.21E-08 |
| B102 | B186 | anti-Sm | PFC | -0.33542 | 1.25E-08 |
| B102 | B134 | anti-Sm | CHE | -0.31897 | 6.76E-08 |
| B102 | B139 | anti-Sm | ADA | -0.3138 | 1.13E-07 |
| B102 | B182 | anti-Sm | TT | 0.29809 | 5.00E-07 |
| B103 | B105 | anti-SSA/Ro | anti-U1RNP | 1 | 0 |
| B103 | B107 | anti-SSA/Ro | AnuA | 1 | 0 |
| B103 | B110 | anti-SSA/Ro | AHA | 1 | 0 |
| B103 | B154 | anti-SSA/Ro | RF | 0.476063 | 0 |
| B103 | B183 | anti-SSA/Ro | PTA | 0.497714 | 0 |
| B103 | B198 | anti-SSA/Ro | P | -0.41547 | 7.35E-13 |
| B103 | B124 | anti-SSA/Ro | ACTH | 0.352095 | 2.04E-09 |
| B103 | B185 | anti-SSA/Ro | PT | 0.335754 | 1.21E-08 |
| B103 | B186 | anti-SSA/Ro | PFC | -0.33542 | 1.25E-08 |
| B103 | B134 | anti-SSA/Ro | CHE | -0.31897 | 6.76E-08 |
| B103 | B139 | anti-SSA/Ro | ADA | -0.3138 | 1.13E-07 |
| B103 | B182 | anti-SSA/Ro | TT | 0.29809 | 5.00E-07 |
| B104 | B107 | anti-SSB | AnuA | 1 | 0 |
| B104 | B154 | anti-SSB | RF | 0.476063 | 0 |
| B104 | B183 | anti-SSB | PTA | 0.497714 | 0 |
| B104 | B198 | anti-SSB | P | -0.41547 | 7.35E-13 |
| B104 | B124 | anti-SSB | ACTH | 0.352095 | 2.04E-09 |
| B104 | B185 | anti-SSB | PT | 0.335754 | 1.21E-08 |
| B104 | B186 | anti-SSB | PFC | -0.33542 | 1.25E-08 |
| B104 | B134 | anti-SSB | CHE | -0.31897 | 6.76E-08 |
| B104 | B139 | anti-SSB | ADA | -0.3138 | 1.13E-07 |
| B104 | B182 | anti-SSB | TT | 0.29809 | 5.00E-07 |
| B105 | B106 | anti-U1RNP | anti-P | 1 | 0 |
| B105 | B107 | anti-U1RNP | AnuA | 1 | 0 |
| B105 | B109 | anti-U1RNP | AMA-M2 | 1 | 0 |
| B105 | B110 | anti-U1RNP | AHA | 1 | 0 |
| B105 | B154 | anti-U1RNP | RF | 0.476063 | 0 |
| B105 | B183 | anti-U1RNP | PTA | 0.497714 | 0 |
| B105 | B198 | anti-U1RNP | P | -0.41547 | 7.35E-13 |
| B105 | B124 | anti-U1RNP | ACTH | 0.352095 | 2.04E-09 |

| | | | | | |
|---|---|---|---|---|---|
| B105 | B185 | anti-U1RNP | PT | 0.335754 | 1.21E-08 |
| B105 | B186 | anti-U1RNP | PFC | -0.33542 | 1.25E-08 |
| B105 | B134 | anti-U1RNP | CHE | -0.31897 | 6.76E-08 |
| B105 | B139 | anti-U1RNP | ADA | -0.3138 | 1.13E-07 |
| B105 | B182 | anti-U1RNP | TT | 0.29809 | 5.00E-07 |
| B106 | B107 | anti-P | AnuA | 1 | 0 |
| B106 | B108 | anti-P | Anti-Cyclin | 1 | 0 |
| B106 | B110 | anti-P | AHA | 1 | 0 |
| B106 | B154 | anti-P | RF | 0.476063 | 0 |
| B106 | B183 | anti-P | PTA | 0.497714 | 0 |
| B106 | B198 | anti-P | P | -0.41547 | 7.35E-13 |
| B106 | B124 | anti-P | ACTH | 0.352095 | 2.04E-09 |
| B106 | B185 | anti-P | PT | 0.335754 | 1.21E-08 |
| B106 | B186 | anti-P | PFC | -0.33542 | 1.25E-08 |
| B106 | B134 | anti-P | CHE | -0.31897 | 6.76E-08 |
| B106 | B139 | anti-P | ADA | -0.3138 | 1.13E-07 |
| B106 | B182 | anti-P | TT | 0.29809 | 5.00E-07 |
| B107 | B108 | AnuA | Anti-Cyclin | 1 | 0 |
| B107 | B109 | AnuA | AMA-M2 | 1 | 0 |
| B107 | B110 | AnuA | AHA | 1 | 0 |
| B107 | B154 | AnuA | RF | 0.476063 | 0 |
| B107 | B183 | AnuA | PTA | 0.497714 | 0 |
| B107 | B198 | AnuA | P | -0.41547 | 7.35E-13 |
| B107 | B124 | AnuA | ACTH | 0.352095 | 2.04E-09 |
| B107 | B185 | AnuA | PT | 0.335754 | 1.21E-08 |
| B107 | B186 | AnuA | PFC | -0.33542 | 1.25E-08 |
| B107 | B134 | AnuA | CHE | -0.31897 | 6.76E-08 |
| B107 | B139 | AnuA | ADA | -0.3138 | 1.13E-07 |
| B107 | B182 | AnuA | TT | 0.29809 | 5.00E-07 |
| B108 | B154 | Anti-Cyclin | RF | 0.476063 | 0 |
| B108 | B183 | Anti-Cyclin | PTA | 0.497714 | 0 |
| B108 | B198 | Anti-Cyclin | P | -0.41547 | 7.35E-13 |
| B108 | B124 | Anti-Cyclin | ACTH | 0.352095 | 2.04E-09 |
| B108 | B185 | Anti-Cyclin | PT | 0.335754 | 1.21E-08 |
| B108 | B186 | Anti-Cyclin | PFC | -0.33542 | 1.25E-08 |
| B108 | B134 | Anti-Cyclin | CHE | -0.31897 | 6.76E-08 |
| B108 | B139 | Anti-Cyclin | ADA | -0.3138 | 1.13E-07 |
| B108 | B182 | Anti-Cyclin | TT | 0.29809 | 5.00E-07 |
| B109 | B154 | AMA-M2 | RF | 0.476063 | 0 |
| B109 | B183 | AMA-M2 | PTA | 0.497714 | 0 |
| B109 | B198 | AMA-M2 | P | -0.41547 | 7.35E-13 |
| B109 | B124 | AMA-M2 | ACTH | 0.352095 | 2.04E-09 |
| B109 | B185 | AMA-M2 | PT | 0.335754 | 1.21E-08 |
| B109 | B186 | AMA-M2 | PFC | -0.33542 | 1.25E-08 |

| | | | | | |
|---|---|---|---|---|---|
| B109 | B134 | AMA-M2 | CHE | -0.31897 | 6.76E-08 |
| B109 | B139 | AMA-M2 | ADA | -0.3138 | 1.13E-07 |
| B109 | B182 | AMA-M2 | TT | 0.29809 | 5.00E-07 |
| B11 | B12 | PGI/PGII | PG II | -0.56967 | 0 |
| B11 | B123 | PGI/PGII | hs-CRP1 | -0.40536 | 2.92E-12 |
| B11 | B63 | PGI/PGII | WBRSR(mPas_200) | -0.35397 | 1.65E-09 |
| B11 | B61 | PGI/PGII | WBHSRV | -0.34915 | 2.83E-09 |
| B11 | B66 | PGI/PGII | WBRSR(mPas_100) | -0.34887 | 2.92E-09 |
| B11 | B67 | PGI/PGII | WBRSR(mPas_50) | -0.34661 | 3.75E-09 |
| B11 | B64 | PGI/PGII | WBRSR(mPas_5) | -0.32391 | 4.12E-08 |
| B11 | B25 | PGI/PGII | CK | 0.297714 | 5.17E-07 |
| B11 | B14 | PGI/PGII | G-17 | -0.29197 | 8.72E-07 |
| B110 | B154 | AHA | RF | 0.476063 | 0 |
| B110 | B183 | AHA | PTA | 0.497714 | 0 |
| B110 | B198 | AHA | P | -0.41547 | 7.35E-13 |
| B110 | B124 | AHA | ACTH | 0.352095 | 2.04E-09 |
| B110 | B185 | AHA | PT | 0.335754 | 1.21E-08 |
| B110 | B186 | AHA | PFC | -0.33542 | 1.25E-08 |
| B110 | B134 | AHA | CHE | -0.31897 | 6.76E-08 |
| B110 | B139 | AHA | ADA | -0.3138 | 1.13E-07 |
| B110 | B182 | AHA | TT | 0.29809 | 5.00E-07 |
| B111 | B114 | TSH | TSH TT3 TT4(TT4) | -0.50254 | 0 |
| B111 | B148 | TSH | Anti_TPO | 0.617706 | 0 |
| B111 | B149 | TSH | TGAb | 0.907258 | 0 |
| B111 | B115 | TSH | TSH TT3 TT4(TT3) | -0.3832 | 5.15E-11 |
| B111 | B112 | TSH | FT4 | -0.34509 | 4.43E-09 |
| B112 | B114 | FT4 | TSH TT3 TT4(TT4) | 0.483172 | 0 |
| B112 | B113 | FT4 | FT3 | 0.352952 | 1.85E-09 |
| B112 | B149 | FT4 | TGAb | -0.32213 | 4.93E-08 |
| B112 | B115 | FT4 | TSH TT3 TT4(TT3) | 0.304236 | 2.82E-07 |
| B112 | B148 | FT4 | Anti_TPO | -0.29196 | 8.73E-07 |
| B113 | B115 | FT3 | TSH TT3 TT4(TT3) | 0.489965 | 0 |
| B113 | B151 | FT3 | Anti-CCP | 0.319792 | 6.23E-08 |
| B113 | B125 | FT3 | Cor | 0.297578 | 5.24E-07 |
| B113 | B135 | FT3 | TBA | 0.29291 | 8.01E-07 |
| B114 | B115 | TSH TT3 TT4(TT4) | TSH TT3 TT4(TT3) | 0.543095 | 0 |
| B114 | B149 | TSH TT3 TT4(TT4) | TGAb | -0.4717 | 0 |
| B114 | B199 | TSH TT3 TT4(TT4) | Mg | 0.362437 | 6.27E-10 |
| B114 | B146 | TSH TT3 TT4(TT4) | CysC | 0.309544 | 1.70E-07 |
| B114 | B148 | TSH TT3 TT4(TT4) | Anti_TPO | -0.30557 | 2.49E-07 |
| B115 | B149 | TSH TT3 TT4(TT3) | TGAb | -0.43134 | 7.64E-14 |
| B115 | B124 | TSH TT3 TT4(TT3) | ACTH | 0.426761 | 1.49E-13 |
| B115 | B125 | TSH TT3 TT4(TT3) | Cor | 0.392318 | 1.62E-11 |
| B115 | B200 | TSH TT3 TT4(TT3) | OP | 0.38113 | 6.66E-11 |

| | | | | | |
|---|---|---|---|---|---|
| B115 | B156 | TSH TT3 TT4(TT3) | IgE | -0.34633 | 3.86E-09 |
| B115 | B183 | TSH TT3 TT4(TT3) | PTA | 0.3269 | 3.04E-08 |
| B115 | B151 | TSH TT3 TT4(TT3) | Anti-CCP | 0.30214 | 3.43E-07 |
| B115 | B137 | TSH TT3 TT4(TT3) | AFU | 0.292958 | 7.98E-07 |
| B116 | B142 | TT3 | 25-(OH)D | -0.77053 | 0 |
| B119 | B152 | Hcv-rna | ACA-Ig(G-A-M) | -1 | 0 |
| B119 | B169 | Hcv-rna | U-CRY | -1 | 0 |
| B119 | B120 | Hcv-rna | HCV-cAg | 0.461421 | 8.88E-16 |
| B119 | B172 | Hcv-rna | KB | -0.29888 | 4.64E-07 |
| B12 | B14 | PG II | G-17 | 0.806514 | 0 |
| B120 | B121 | HCV-cAg | C3 | -0.46142 | 8.88E-16 |
| B120 | B122 | HCV-cAg | C4 | -0.46142 | 8.88E-16 |
| B120 | B143 | HCV-cAg | β-CTx | -0.46142 | 8.88E-16 |
| B120 | B152 | HCV-cAg | ACA-Ig(G-A-M) | -0.46142 | 8.88E-16 |
| B120 | B159 | HCV-cAg | β2-MG | -0.46142 | 8.88E-16 |
| B120 | B169 | HCV-cAg | U-CRY | -0.46142 | 8.88E-16 |
| B120 | B136 | HCV-cAg | 5'NT | -0.3166 | 8.55E-08 |
| B120 | B199 | HCV-cAg | Mg | -0.30144 | 3.66E-07 |
| B121 | B152 | C3 | ACA-Ig(G-A-M) | 0.999997 | 0 |
| B121 | B169 | C3 | U-CRY | 1 | 0 |
| B121 | B172 | C3 | KB | 0.298883 | 4.64E-07 |
| B122 | B152 | C4 | ACA-Ig(G-A-M) | 0.999997 | 0 |
| B122 | B169 | C4 | U-CRY | 1 | 0 |
| B122 | B172 | C4 | KB | 0.298883 | 4.64E-07 |
| B123 | B136 | hs-CRP1 | 5'NT | 0.480732 | 0 |
| B123 | B139 | hs-CRP1 | ADA | 0.586482 | 0 |
| B123 | B150 | hs-CRP1 | ALP | 0.474845 | 0 |
| B123 | B138 | hs-CRP1 | PAB | -0.38153 | 6.34E-11 |
| B123 | B146 | hs-CRP1 | CysC | 0.349423 | 2.75E-09 |
| B123 | B194 | hs-CRP1 | Cl | 0.348979 | 2.89E-09 |
| B124 | B125 | ACTH | Cor | 0.917647 | 0 |
| B124 | B156 | ACTH | IgE | -0.79498 | 0 |
| B124 | B181 | ACTH | APTT | -0.6094 | 0 |
| B124 | B151 | ACTH | Anti-CCP | 0.429535 | 9.95E-14 |
| B124 | B184 | ACTH | INR | 0.390788 | 1.98E-11 |
| B124 | B200 | ACTH | OP | 0.381979 | 6.00E-11 |
| B124 | B183 | ACTH | PTA | 0.309502 | 1.71E-07 |
| B125 | B156 | Cor | IgE | -0.6385 | 0 |
| B125 | B181 | Cor | APTT | -0.43807 | 2.82E-14 |
| B125 | B151 | Cor | Anti-CCP | 0.330773 | 2.04E-08 |
| B125 | B200 | Cor | OP | 0.32383 | 4.15E-08 |
| B125 | B135 | Cor | TBA | 0.302618 | 3.28E-07 |
| B126 | B188 | Alb | Ca | 0.440412 | 2.00E-14 |
| B126 | B127 | Alb | A-G ratio | 0.330712 | 2.05E-08 |

| | | | | | |
|---|---|---|---|---|---|
| B127 | B130 | A-G ratio | GLB | -0.87291 | 0 |
| B128 | B131 | ALT | AST | 0.968904 | 0 |
| B129 | B132 | IBIL | Dbil | 0.680182 | 0 |
| B129 | B133 | IBIL | TBIL | 0.979295 | 0 |
| B132 | B133 | Dbil | TBIL | 0.814441 | 0 |
| B134 | B200 | CHE | OP | 0.434467 | 4.84E-14 |
| B134 | B137 | CHE | AFU | 0.430526 | 8.62E-14 |
| B134 | B138 | CHE | PAB | 0.393327 | 1.43E-11 |
| B134 | B195 | CHE | Na | 0.389541 | 2.32E-11 |
| B134 | B154 | CHE | RF | -0.31651 | 8.63E-08 |
| B135 | B157 | TBA | IgG | 0.448273 | 6.00E-15 |
| B135 | B185 | TBA | PT | -0.39211 | 1.67E-11 |
| B135 | B186 | TBA | PFC | -0.33852 | 9.01E-09 |
| B135 | B191 | TBA | Urea | 0.29295 | 7.99E-07 |
| B136 | B146 | 5'NT | CysC | 0.525504 | 0 |
| B136 | B194 | 5'NT | Cl | 0.508044 | 0 |
| B136 | B139 | 5'NT | ADA | 0.417077 | 5.88E-13 |
| B136 | B195 | 5'NT | Na | 0.406049 | 2.67E-12 |
| B136 | B193 | 5'NT | K | -0.38756 | 2.98E-11 |
| B136 | B150 | 5'NT | ALP | 0.373363 | 1.72E-10 |
| B136 | B140 | 5'NT | r-GT | 0.348624 | 3.00E-09 |
| B136 | B199 | 5'NT | Mg | 0.336736 | 1.09E-08 |
| B137 | B139 | AFU | ADA | 0.464108 | 4.44E-16 |
| B137 | B200 | AFU | OP | 0.329979 | 2.21E-08 |
| B138 | B196 | PAB | TCO2 | 0.519711 | 0 |
| B138 | B195 | PAB | Na | 0.393282 | 1.43E-11 |
| B138 | B139 | PAB | ADA | -0.34903 | 2.87E-09 |
| B138 | B150 | PAB | ALP | -0.33522 | 1.28E-08 |
| B138 | B194 | PAB | Cl | -0.3006 | 3.96E-07 |
| B139 | B146 | ADA | CysC | 0.547843 | 0 |
| B139 | B182 | ADA | TT | -0.47378 | 0 |
| B139 | B150 | ADA | ALP | 0.460163 | 8.88E-16 |
| B139 | B198 | ADA | P | 0.430488 | 8.66E-14 |
| B139 | B186 | ADA | PFC | 0.378079 | 9.70E-11 |
| B139 | B194 | ADA | Cl | 0.30233 | 3.37E-07 |
| B139 | B181 | ADA | APTT | -0.29152 | 9.09E-07 |
| B140 | B150 | r-GT | ALP | 0.314178 | 1.08E-07 |
| B142 | B223 | 25-(OH)D | FH | -0.50262 | 0 |
| B143 | B152 | β-CTx | ACA-Ig(G-A-M) | 0.999998 | 0 |
| B143 | B172 | β-CTx | KB | 0.298884 | 4.64E-07 |
| B145 | B167 | BGP | PC | 0.675441 | 0 |
| B145 | B225 | BGP | hPBG | 0.659912 | 0 |
| B146 | B199 | CysC | Mg | 0.489805 | 0 |
| B146 | B200 | CysC | OP | 0.334045 | 1.45E-08 |

| | | | | | |
|---|---|---|---|---|---|
| B146 | B156 | CysC | IgE | -0.3041 | 2.85E-07 |
| B148 | B149 | Anti_TPO | TGAb | 0.566041 | 0 |
| B15 | B100 | CD3-CD19+(B cell) | anti-Ro52 | 0.605529 | 0 |
| B15 | B101 | CD3-CD19+(B cell) | anti-Sc70 | 0.605529 | 0 |
| B15 | B102 | CD3-CD19+(B cell) | anti-Sm | 0.605529 | 0 |
| B15 | B103 | CD3-CD19+(B cell) | anti-SSA/Ro | 0.605529 | 0 |
| B15 | B104 | CD3-CD19+(B cell) | anti-SSB | 0.605529 | 0 |
| B15 | B105 | CD3-CD19+(B cell) | anti-U1RNP | 0.605529 | 0 |
| B15 | B106 | CD3-CD19+(B cell) | anti-P | 0.605529 | 0 |
| B15 | B107 | CD3-CD19+(B cell) | AnuA | 0.605529 | 0 |
| B15 | B108 | CD3-CD19+(B cell) | Anti-Cyclin | 0.605529 | 0 |
| B15 | B109 | CD3-CD19+(B cell) | AMA-M2 | 0.605529 | 0 |
| B15 | B110 | CD3-CD19+(B cell) | AHA | 0.605529 | 0 |
| B15 | B124 | CD3-CD19+(B cell) | ACTH | 0.609019 | 0 |
| B15 | B125 | CD3-CD19+(B cell) | Cor | 0.471845 | 0 |
| B15 | B151 | CD3-CD19+(B cell) | Anti-CCP | 0.648075 | 0 |
| B15 | B183 | CD3-CD19+(B cell) | PTA | 0.620002 | 0 |
| B15 | B19 | CD3-CD19+(B cell) | CD3+CD4+/CD3+CD8+ | -0.55073 | 0 |
| B15 | B22 | CD3-CD19+(B cell) | CD3+CD8+(T) | 0.656318 | 0 |
| B15 | B95 | CD3-CD19+(B cell) | anti-cenpb | 0.605529 | 0 |
| B15 | B97 | CD3-CD19+(B cell) | dsDNA | 0.605529 | 0 |
| B15 | B98 | CD3-CD19+(B cell) | anti-Jo-1 | 0.605529 | 0 |
| B15 | B156 | CD3-CD19+(B cell) | IgE | -0.46159 | 8.88E-16 |
| B15 | B23 | CD3-CD19+(B cell) | CD3-CD16+CD56+(NK) | -0.45184 | 3.55E-15 |
| B15 | B20 | CD3-CD19+(B cell) | CD3+CD4+CD8+ | -0.43181 | 7.15E-14 |
| B15 | B185 | CD3-CD19+(B cell) | PT | 0.420114 | 3.84E-13 |
| B15 | B181 | CD3-CD19+(B cell) | APTT | -0.40057 | 5.54E-12 |
| B15 | B198 | CD3-CD19+(B cell) | P | -0.37205 | 2.02E-10 |
| B15 | B200 | CD3-CD19+(B cell) | OP | 0.371359 | 2.19E-10 |
| B15 | B16 | CD3-CD19+(B cell) | CD3+T | 0.358699 | 9.65E-10 |
| B15 | B79 | CD3-CD19+(B cell) | hb V -dna | 0.354118 | 1.62E-09 |
| B15 | B154 | CD3-CD19+(B cell) | RF | 0.337262 | 1.03E-08 |
| B15 | B115 | CD3-CD19+(B cell) | TSH TT3 TT4(TT3) | 0.336019 | 1.17E-08 |
| B15 | B81 | CD3-CD19+(B cell) | TSGF | -0.33539 | 1.25E-08 |
| B15 | B17 | CD3-CD19+(B cell) | CD3+CD16+CD56+NK | -0.32472 | 3.79E-08 |
| B15 | B196 | CD3-CD19+(B cell) | TCO2 | 0.318838 | 6.85E-08 |
| B15 | B184 | CD3-CD19+(B cell) | INR | 0.316517 | 8.62E-08 |
| B150 | B196 | ALP | TCO2 | -0.33774 | 9.78E-09 |
| B150 | B194 | ALP | Cl | 0.299073 | 4.56E-07 |
| B151 | B156 | Anti-CCP | IgE | -0.50914 | 0 |
| B151 | B183 | Anti-CCP | PTA | 0.624794 | 0 |
| B151 | B198 | Anti-CCP | P | -0.47199 | 0 |
| B151 | B179 | Anti-CCP | α1-MG | 0.466454 | 4.44E-16 |
| B151 | B185 | Anti-CCP | PT | 0.424323 | 2.11E-13 |

| | | | | | |
|---|---|---|---|---|---|
| B151 | B186 | Anti-CCP | PFC | -0.4094 | 1.70E-12 |
| B151 | B195 | Anti-CCP | Na | 0.403074 | 3.97E-12 |
| B151 | B182 | Anti-CCP | TT | 0.359704 | 8.60E-10 |
| B151 | B196 | Anti-CCP | TCO2 | 0.345614 | 4.18E-09 |
| B151 | B200 | Anti-CCP | OP | 0.311319 | 1.43E-07 |
| B152 | B159 | ACA-Ig(G-A-M) | β2-MG | 0.999998 | 0 |
| B152 | B169 | ACA-Ig(G-A-M) | U-CRY | 0.999998 | 0 |
| B152 | B172 | ACA-Ig(G-A-M) | KB | 0.298882 | 4.64E-07 |
| B153 | B170 | Glu | U-GLU | 0.6347 | 0 |
| B154 | B183 | RF | PTA | 0.460307 | 8.88E-16 |
| B154 | B184 | RF | INR | 0.370735 | 2.36E-10 |
| B154 | B196 | RF | TCO2 | 0.320662 | 5.71E-08 |
| B154 | B185 | RF | PT | 0.320412 | 5.86E-08 |
| B154 | B156 | RF | IgE | 0.306788 | 2.21E-07 |
| B156 | B181 | IgE | APTT | 0.672397 | 0 |
| B156 | B200 | IgE | OP | -0.45267 | 3.11E-15 |
| B156 | B199 | IgE | Mg | -0.31534 | 9.68E-08 |
| B156 | B179 | IgE | α1-MG | -0.31264 | 1.26E-07 |
| B157 | B183 | IgG | PTA | -0.4816 | 0 |
| B157 | B184 | IgG | INR | -0.53046 | 0 |
| B157 | B185 | IgG | PT | -0.49012 | 0 |
| B157 | B197 | IgG | AG | 0.385492 | 3.87E-11 |
| B158 | B193 | IgM | K | -0.36242 | 6.28E-10 |
| B158 | B165 | IgM | bilirubin | -0.29389 | 7.33E-07 |
| B158 | B173 | IgM | NTT | -0.29389 | 7.33E-07 |
| B159 | B169 | β2-MG | U-CRY | 0.999999 | 0 |
| B159 | B172 | β2-MG | KB | 0.298883 | 4.64E-07 |
| B16 | B19 | CD3+T | CD3+CD4+/CD3+CD8+ | -0.51021 | 0 |
| B16 | B193 | CD3+T | K | -0.58794 | 0 |
| B16 | B199 | CD3+T | Mg | -0.54999 | 0 |
| B16 | B21 | CD3+T | CD3+CD4-CD8- | 0.499008 | 0 |
| B16 | B23 | CD3+T | CD3-CD16+CD56+(NK) | -0.94709 | 0 |
| B16 | B76 | CD3+T | ApoB | -0.54625 | 0 |
| B16 | B22 | CD3+T | CD3+CD8+(T) | 0.45741 | 1.33E-15 |
| B16 | B99 | CD3+T | anti-PM-Scl | 0.425906 | 1.68E-13 |
| B16 | B75 | CD3+T | ApoA1/B | 0.358737 | 9.61E-10 |
| B16 | B17 | CD3+T | CD3+CD16+CD56+NK | 0.350649 | 2.40E-09 |
| B16 | B134 | CD3+T | CHE | -0.32794 | 2.73E-08 |
| B16 | B196 | CD3+T | TCO2 | 0.326886 | 3.04E-08 |
| B16 | B154 | CD3+T | RF | 0.316664 | 8.50E-08 |
| B16 | B195 | CD3+T | Na | 0.311508 | 1.41E-07 |
| B16 | B18 | CD3+T | CD3+CD4+helper(inducer)T | -0.30731 | 2.11E-07 |
| B16 | B70 | CD3+T | LDL-C | -0.29928 | 4.48E-07 |
| B163 | B191 | SG | Urea | 0.443464 | 1.24E-14 |

| | | | | | |
|---|---|---|---|---|---|
| B163 | B166 | SG | Pro | 0.351096 | 2.28E-09 |
| B163 | B174 | SG | Urea/Cr | 0.309402 | 1.72E-07 |
| B167 | B225 | PC | hPBG | 0.732756 | 0 |
| B168 | B175 | NMP22 | ALB-U | -0.32283 | 4.59E-08 |
| B169 | B172 | U-CRY | KB | 0.298883 | 4.64E-07 |
| B17 | B137 | CD3+CD16+CD56+NK | AFU | 0.59136 | 0 |
| B17 | B146 | CD3+CD16+CD56+NK | CysC | 0.500563 | 0 |
| B17 | B18 | CD3+CD16+CD56+NK | CD3+CD4+helper(inducer)T | -0.73073 | 0 |
| B17 | B21 | CD3+CD16+CD56+NK | CD3+CD4-CD8- | 0.978039 | 0 |
| B17 | B94 | CD3+CD16+CD56+NK | ALDH2-Lys504Lys | -0.48991 | 0 |
| B17 | B99 | CD3+CD16+CD56+NK | anti-PM-Scl | 0.982422 | 0 |
| B17 | B22 | CD3+CD16+CD56+NK | CD3+CD8+(T) | -0.46856 | 2.22E-16 |
| B17 | B193 | CD3+CD16+CD56+NK | K | -0.43794 | 2.89E-14 |
| B17 | B100 | CD3+CD16+CD56+NK | anti-Ro52 | -0.4338 | 5.33E-14 |
| B17 | B101 | CD3+CD16+CD56+NK | anti-Sc70 | -0.4338 | 5.33E-14 |
| B17 | B102 | CD3+CD16+CD56+NK | anti-Sm | -0.4338 | 5.33E-14 |
| B17 | B103 | CD3+CD16+CD56+NK | anti-SSA/Ro | -0.4338 | 5.33E-14 |
| B17 | B104 | CD3+CD16+CD56+NK | anti-SSB | -0.4338 | 5.33E-14 |
| B17 | B105 | CD3+CD16+CD56+NK | anti-U1RNP | -0.4338 | 5.33E-14 |
| B17 | B106 | CD3+CD16+CD56+NK | anti-P | -0.4338 | 5.33E-14 |
| B17 | B107 | CD3+CD16+CD56+NK | AnuA | -0.4338 | 5.33E-14 |
| B17 | B108 | CD3+CD16+CD56+NK | Anti-Cyclin | -0.4338 | 5.33E-14 |
| B17 | B109 | CD3+CD16+CD56+NK | AMA-M2 | -0.4338 | 5.33E-14 |
| B17 | B110 | CD3+CD16+CD56+NK | AHA | -0.4338 | 5.33E-14 |
| B17 | B95 | CD3+CD16+CD56+NK | anti-cenpb | -0.4338 | 5.33E-14 |
| B17 | B97 | CD3+CD16+CD56+NK | dsDNA | -0.4338 | 5.33E-14 |
| B17 | B98 | CD3+CD16+CD56+NK | anti-Jo-1 | -0.4338 | 5.33E-14 |
| B17 | B23 | CD3+CD16+CD56+NK | CD3-CD16+CD56+(NK) | -0.41669 | 6.20E-13 |
| B17 | B200 | CD3+CD16+CD56+NK | OP | 0.383793 | 4.79E-11 |
| B17 | B92 | CD3+CD16+CD56+NK | ALDH2-Glu504Glu | 0.359638 | 8.66E-10 |
| B17 | B136 | CD3+CD16+CD56+NK | 5'NT | 0.347165 | 3.53E-09 |
| B17 | B158 | CD3+CD16+CD56+NK | IgM | 0.341695 | 6.40E-09 |
| B17 | B93 | CD3+CD16+CD56+NK | ALDH2-Glu504Lys | -0.33501 | 1.31E-08 |
| B17 | B157 | CD3+CD16+CD56+NK | IgG | -0.32351 | 4.29E-08 |
| B17 | B140 | CD3+CD16+CD56+NK | r-GT | 0.321469 | 5.27E-08 |
| B174 | B191 | Urea/Cr | Urea | 0.798031 | 0 |
| B174 | B190 | Urea/Cr | Cre | -0.4449 | 1.02E-14 |
| B175 | B177 | ALB-U | ACR | 0.759419 | 0 |
| B176 | B178 | Ucr | Ig-U | 0.386261 | 3.51E-11 |
| B178 | B179 | Ig-U | α1-MG | 0.302073 | 3.45E-07 |
| B178 | B180 | Ig-U | Tf | 0.294794 | 6.76E-07 |
| B179 | B184 | α1-MG | INR | 0.334012 | 1.45E-08 |
| B179 | B185 | α1-MG | PT | 0.30374 | 2.95E-07 |
| B18 | B124 | CD3+CD4+helper(inducer)T | ACTH | -0.59545 | 0 |

| | | | | | |
|---|---|---|---|---|---|
| B18 | B156 | CD3+CD4+helper(inducer)T | IgE | 0.600527 | 0 |
| B18 | B181 | CD3+CD4+helper(inducer)T | APTT | 0.509771 | 0 |
| B18 | B19 | CD3+CD4+helper(inducer)T | CD3+CD4+/CD3+CD8+ | 0.58056 | 0 |
| B18 | B21 | CD3+CD4+helper(inducer)T | CD3+CD4-CD8- | -0.75277 | 0 |
| B18 | B99 | CD3+CD4+helper(inducer)T | anti-PM-Scl | -0.69788 | 0 |
| B18 | B125 | CD3+CD4+helper(inducer)T | Cor | -0.45712 | 1.33E-15 |
| B18 | B200 | CD3+CD4+helper(inducer)T | OP | -0.43303 | 5.97E-14 |
| B18 | B184 | CD3+CD4+helper(inducer)T | INR | -0.41315 | 1.01E-12 |
| B18 | B146 | CD3+CD4+helper(inducer)T | CysC | -0.37808 | 9.71E-11 |
| B18 | B58 | CD3+CD4+helper(inducer)T | HCT | -0.36566 | 4.30E-10 |
| B18 | B115 | CD3+CD4+helper(inducer)T | TSH TT3 TT4(TT3) | -0.35959 | 8.71E-10 |
| B18 | B63 | CD3+CD4+helper(inducer)T | WBRSR(mPas_200) | -0.34828 | 3.12E-09 |
| B18 | B94 | CD3+CD4+helper(inducer)T | ALDH2-Lys504Lys | 0.347496 | 3.40E-09 |
| B18 | B64 | CD3+CD4+helper(inducer)T | WBRSR(mPas_5) | -0.34526 | 4.35E-09 |
| B18 | B67 | CD3+CD4+helper(inducer)T | WBRSR(mPas_50) | -0.33879 | 8.74E-09 |
| B18 | B62 | CD3+CD4+helper(inducer)T | WBHSRVI | -0.33783 | 9.70E-09 |
| B18 | B23 | CD3+CD4+helper(inducer)T | CD3-CD16+CD56+(NK) | 0.336158 | 1.16E-08 |
| B18 | B66 | CD3+CD4+helper(inducer)T | WBRSR(mPas_100) | -0.33451 | 1.38E-08 |
| B18 | B65 | CD3+CD4+helper(inducer)T | WBRSR(mPas_1) | -0.33212 | 1.77E-08 |
| B18 | B60 | CD3+CD4+helper(inducer)T | WBRVI | -0.31885 | 6.84E-08 |
| B18 | B137 | CD3+CD4+helper(inducer)T | AFU | -0.30779 | 2.01E-07 |
| B18 | B81 | CD3+CD4+helper(inducer)T | TSGF | 0.307099 | 2.15E-07 |
| B18 | B61 | CD3+CD4+helper(inducer)T | WBHSRV | -0.29404 | 7.23E-07 |
| B181 | B186 | APTT | PFC | -0.55711 | 0 |
| B181 | B184 | APTT | INR | -0.38112 | 6.67E-11 |
| B181 | B182 | APTT | TT | 0.350907 | 2.33E-09 |
| B181 | B185 | APTT | PT | -0.32739 | 2.89E-08 |
| B182 | B184 | TT | INR | -0.49325 | 0 |
| B182 | B186 | TT | PFC | -0.86375 | 0 |
| B182 | B198 | TT | P | -0.49147 | 0 |
| B183 | B184 | PTA | INR | 0.673239 | 0 |
| B183 | B185 | PTA | PT | 0.678248 | 0 |
| B183 | B196 | PTA | TCO2 | 0.386053 | 3.60E-11 |
| B183 | B198 | PTA | P | -0.36845 | 3.10E-10 |
| B183 | B197 | PTA | AG | -0.29329 | 7.74E-07 |
| B184 | B185 | INR | PT | 0.590427 | 0 |
| B184 | B186 | INR | PFC | 0.457796 | 1.33E-15 |
| B184 | B194 | INR | Cl | 0.300169 | 4.12E-07 |
| B184 | B197 | INR | AG | -0.29999 | 4.19E-07 |
| B185 | B197 | PT | AG | -0.37964 | 8.01E-11 |
| B185 | B196 | PT | TCO2 | 0.347362 | 3.45E-09 |
| B185 | B186 | PT | PFC | 0.330563 | 2.08E-08 |
| B185 | B187 | PT | NSE | 0.32227 | 4.86E-08 |
| B186 | B198 | PFC | P | 0.530807 | 0 |

| | | | | | |
|---|---|---|---|---|---|
| B186 | B194 | PFC | Cl | 0.305936 | 2.40E-07 |
| B189 | B190 | eGFB | Cre | -0.35409 | 1.63E-09 |
| B19 | B124 | CD3+CD4+/CD3+CD8+ | ACTH | -0.65565 | 0 |
| B19 | B125 | CD3+CD4+/CD3+CD8+ | Cor | -0.5064 | 0 |
| B19 | B181 | CD3+CD4+/CD3+CD8+ | APTT | 0.634809 | 0 |
| B19 | B22 | CD3+CD4+/CD3+CD8+ | CD3+CD8+(T) | -0.83474 | 0 |
| B19 | B79 | CD3+CD4+/CD3+CD8+ | hb V -dna | -0.5249 | 0 |
| B19 | B76 | CD3+CD4+/CD3+CD8+ | ApoB | 0.460461 | 8.88E-16 |
| B19 | B184 | CD3+CD4+/CD3+CD8+ | INR | -0.44174 | 1.62E-14 |
| B19 | B23 | CD3+CD4+/CD3+CD8+ | CD3-CD16+CD56+(NK) | 0.439797 | 2.18E-14 |
| B19 | B156 | CD3+CD4+/CD3+CD8+ | IgE | 0.417726 | 5.37E-13 |
| B19 | B75 | CD3+CD4+/CD3+CD8+ | ApoA1/B | -0.41373 | 9.36E-13 |
| B19 | B199 | CD3+CD4+/CD3+CD8+ | Mg | 0.395833 | 1.03E-11 |
| B19 | B81 | CD3+CD4+/CD3+CD8+ | TSGF | 0.353297 | 1.78E-09 |
| B19 | B134 | CD3+CD4+/CD3+CD8+ | CHE | 0.314689 | 1.03E-07 |
| B19 | B154 | CD3+CD4+/CD3+CD8+ | RF | -0.31173 | 1.38E-07 |
| B190 | B192 | Cre | UA | 0.359136 | 9.18E-10 |
| B190 | B221 | Cre | Hei | 0.310502 | 1.55E-07 |
| B192 | B221 | UA | Hei | 0.350057 | 2.56E-09 |
| B194 | B197 | Cl | AG | -0.52541 | 0 |
| B194 | B196 | Cl | TCO2 | -0.32298 | 4.53E-08 |
| B195 | B200 | Na | OP | 0.629106 | 0 |
| B195 | B196 | Na | TCO2 | 0.325483 | 3.51E-08 |
| B196 | B197 | TCO2 | AG | -0.48393 | 0 |
| B196 | B200 | TCO2 | OP | 0.528708 | 0 |
| B2 | B189 | Age | eGFB | -0.47845 | 0 |
| B2 | B141 | Age | AST/ALT | 0.365755 | 4.26E-10 |
| B2 | B46 | Age | RBC Count | -0.35223 | 2.01E-09 |
| B2 | B153 | Age | Glu | 0.331272 | 1.93E-08 |
| B2 | B126 | Age | Alb | -0.32759 | 2.83E-08 |
| B2 | B227 | Age | OD | -0.32142 | 5.29E-08 |
| B2 | B228 | Age | OS | -0.31303 | 1.21E-07 |
| B2 | B51 | Age | Hb | -0.29294 | 7.99E-07 |
| B20 | B100 | CD3+CD4+CD8+ | anti-Ro52 | -0.49147 | 0 |
| B20 | B101 | CD3+CD4+CD8+ | anti-Sc70 | -0.49147 | 0 |
| B20 | B102 | CD3+CD4+CD8+ | anti-Sm | -0.49147 | 0 |
| B20 | B103 | CD3+CD4+CD8+ | anti-SSA/Ro | -0.49147 | 0 |
| B20 | B104 | CD3+CD4+CD8+ | anti-SSB | -0.49147 | 0 |
| B20 | B105 | CD3+CD4+CD8+ | anti-U1RNP | -0.49147 | 0 |
| B20 | B106 | CD3+CD4+CD8+ | anti-P | -0.49147 | 0 |
| B20 | B107 | CD3+CD4+CD8+ | AnuA | -0.49147 | 0 |
| B20 | B108 | CD3+CD4+CD8+ | Anti-Cyclin | -0.49147 | 0 |
| B20 | B109 | CD3+CD4+CD8+ | AMA-M2 | -0.49147 | 0 |
| B20 | B110 | CD3+CD4+CD8+ | AHA | -0.49147 | 0 |

| | | | | | |
|---|---|---|---|---|---|
| B20 | B151 | CD3+CD4+CD8+ | Anti-CCP | -0.68064 | 0 |
| B20 | B182 | CD3+CD4+CD8+ | TT | -0.62768 | 0 |
| B20 | B183 | CD3+CD4+CD8+ | PTA | -0.6595 | 0 |
| B20 | B186 | CD3+CD4+CD8+ | PFC | 0.664016 | 0 |
| B20 | B198 | CD3+CD4+CD8+ | P | 0.595261 | 0 |
| B20 | B34 | CD3+CD4+CD8+ | TK | -0.66153 | 0 |
| B20 | B95 | CD3+CD4+CD8+ | anti-cenpb | -0.49147 | 0 |
| B20 | B97 | CD3+CD4+CD8+ | dsDNA | -0.49147 | 0 |
| B20 | B98 | CD3+CD4+CD8+ | anti-Jo-1 | -0.49147 | 0 |
| B20 | B185 | CD3+CD4+CD8+ | PT | -0.44605 | 8.44E-15 |
| B20 | B139 | CD3+CD4+CD8+ | ADA | 0.436603 | 3.51E-14 |
| B20 | B54 | CD3+CD4+CD8+ | ESR | 0.432061 | 6.88E-14 |
| B20 | B196 | CD3+CD4+CD8+ | TCO2 | -0.41398 | 9.04E-13 |
| B20 | B158 | CD3+CD4+CD8+ | IgM | -0.33639 | 1.13E-08 |
| B20 | B193 | CD3+CD4+CD8+ | K | 0.335555 | 1.23E-08 |
| B20 | B181 | CD3+CD4+CD8+ | APTT | -0.31218 | 1.32E-07 |
| B20 | B75 | CD3+CD4+CD8+ | ApoA1/B | 0.290496 | 9.96E-07 |
| B204 | B209 | Drinking | Smoking | 0.29349 | 7.60E-07 |
| B206 | B207 | Waist | WHR | 0.718831 | 0 |
| B206 | B210 | Waist | Hip | 0.803277 | 0 |
| B206 | B215 | Waist | BMI | 0.839032 | 0 |
| B206 | B216 | Waist | BMI | 0.834659 | 0 |
| B206 | B222 | Waist | NS | 0.667573 | 0 |
| B206 | B221 | Waist | Hei | 0.332844 | 1.64E-08 |
| B207 | B215 | WHR | BMI | 0.470108 | 2.22E-16 |
| B207 | B222 | WHR | NS | 0.467898 | 2.22E-16 |
| B207 | B216 | WHR | BMI | 0.449434 | 4.88E-15 |
| B21 | B137 | CD3+CD4-CD8- | AFU | 0.560152 | 0 |
| B21 | B193 | CD3+CD4-CD8- | K | -0.52735 | 0 |
| B21 | B23 | CD3+CD4-CD8- | CD3-CD16+CD56+(NK) | -0.54349 | 0 |
| B21 | B94 | CD3+CD4-CD8- | ALDH2-Lys504Lys | -0.48974 | 0 |
| B21 | B99 | CD3+CD4-CD8- | anti-PM-Scl | 0.982173 | 0 |
| B21 | B146 | CD3+CD4-CD8- | CysC | 0.441764 | 1.62E-14 |
| B21 | B200 | CD3+CD4-CD8- | OP | 0.425645 | 1.75E-13 |
| B21 | B158 | CD3+CD4-CD8- | IgM | 0.388662 | 2.59E-11 |
| B21 | B22 | CD3+CD4-CD8- | CD3+CD8+(T) | -0.35766 | 1.09E-09 |
| B21 | B92 | CD3+CD4-CD8- | ALDH2-Glu504Glu | 0.357541 | 1.10E-09 |
| B21 | B100 | CD3+CD4-CD8- | anti-Ro52 | -0.33401 | 1.45E-08 |
| B21 | B101 | CD3+CD4-CD8- | anti-Sc70 | -0.33401 | 1.45E-08 |
| B21 | B102 | CD3+CD4-CD8- | anti-Sm | -0.33401 | 1.45E-08 |
| B21 | B103 | CD3+CD4-CD8- | anti-SSA/Ro | -0.33401 | 1.45E-08 |
| B21 | B104 | CD3+CD4-CD8- | anti-SSB | -0.33401 | 1.45E-08 |
| B21 | B105 | CD3+CD4-CD8- | anti-U1RNP | -0.33401 | 1.45E-08 |
| B21 | B106 | CD3+CD4-CD8- | anti-P | -0.33401 | 1.45E-08 |

| | | | | | |
|---|---|---|---|---|---|
| B21 | B107 | CD3+CD4-CD8- | AnuA | -0.33401 | 1.45E-08 |
| B21 | B108 | CD3+CD4-CD8- | Anti-Cyclin | -0.33401 | 1.45E-08 |
| B21 | B109 | CD3+CD4-CD8- | AMA-M2 | -0.33401 | 1.45E-08 |
| B21 | B110 | CD3+CD4-CD8- | AHA | -0.33401 | 1.45E-08 |
| B21 | B95 | CD3+CD4-CD8- | anti-cenpb | -0.33401 | 1.45E-08 |
| B21 | B97 | CD3+CD4-CD8- | dsDNA | -0.33401 | 1.45E-08 |
| B21 | B98 | CD3+CD4-CD8- | anti-Jo-1 | -0.33401 | 1.45E-08 |
| B21 | B93 | CD3+CD4-CD8- | ALDH2-Glu504Lys | -0.33291 | 1.63E-08 |
| B21 | B136 | CD3+CD4-CD8- | 5'NT | 0.329948 | 2.22E-08 |
| B21 | B157 | CD3+CD4-CD8- | IgG | -0.3221 | 4.94E-08 |
| B21 | B140 | CD3+CD4-CD8- | r-GT | 0.321102 | 5.46E-08 |
| B210 | B215 | Hip | BMI | 0.785951 | 0 |
| B210 | B216 | Hip | BMI | 0.800216 | 0 |
| B210 | B222 | Hip | NS | 0.544947 | 0 |
| B210 | B221 | Hip | Hei | 0.340335 | 7.41E-09 |
| B211 | B212 | IOP-R | IOP-L | 0.776936 | 0 |
| B213 | B214 | CVA-R | CVA-L | 0.771791 | 0 |
| B215 | B216 | BMI | BMI | 0.831455 | 0 |
| B215 | B222 | BMI | NS | 0.670626 | 0 |
| B216 | B221 | BMI | Hei | 0.628858 | 0 |
| B216 | B222 | BMI | NS | 0.710175 | 0 |
| B218 | B220 | DBP | SBP | 0.593855 | 0 |
| B22 | B199 | CD3+CD8+(T) | Mg | -0.48742 | 0 |
| B22 | B75 | CD3+CD8+(T) | ApoA1/B | 0.555036 | 0 |
| B22 | B76 | CD3+CD8+(T) | ApoB | -0.54439 | 0 |
| B22 | B79 | CD3+CD8+(T) | hb V -dna | 0.558242 | 0 |
| B22 | B134 | CD3+CD8+(T) | CHE | -0.45888 | 1.11E-15 |
| B22 | B181 | CD3+CD8+(T) | APTT | -0.45133 | 3.55E-15 |
| B22 | B99 | CD3+CD8+(T) | anti-PM-Scl | -0.43792 | 2.89E-14 |
| B22 | B124 | CD3+CD8+(T) | ACTH | 0.406701 | 2.44E-12 |
| B22 | B154 | CD3+CD8+(T) | RF | 0.380244 | 7.44E-11 |
| B22 | B23 | CD3+CD8+(T) | CD3-CD16+CD56+(NK) | -0.38024 | 7.44E-11 |
| B22 | B137 | CD3+CD8+(T) | AFU | -0.32673 | 3.09E-08 |
| B22 | B125 | CD3+CD8+(T) | Cor | 0.31596 | 9.11E-08 |
| B221 | B222 | Hei | NS | 0.345572 | 4.20E-09 |
| B227 | B228 | OD | OS | 0.811111 | 0 |
| B23 | B193 | CD3-CD16+CD56+(NK) | K | 0.564134 | 0 |
| B23 | B99 | CD3-CD16+CD56+(NK) | anti-PM-Scl | -0.52362 | 0 |
| B23 | B76 | CD3-CD16+CD56+(NK) | ApoB | 0.437017 | 3.31E-14 |
| B23 | B200 | CD3-CD16+CD56+(NK) | OP | -0.4315 | 7.46E-14 |
| B23 | B63 | CD3-CD16+CD56+(NK) | WBRSR(mPas_200) | 0.396298 | 9.71E-12 |
| B23 | B199 | CD3-CD16+CD56+(NK) | Mg | 0.394431 | 1.24E-11 |
| B23 | B62 | CD3-CD16+CD56+(NK) | WBHSRVI | 0.393701 | 1.36E-11 |
| B23 | B66 | CD3-CD16+CD56+(NK) | WBRSR(mPas_100) | 0.392986 | 1.49E-11 |

| | | | | | |
|---|---|---|---|---|---|
| B23 | B67 | CD3-CD16+CD56+(NK) | WBRSR(mPas_50) | 0.391204 | 1.87E-11 |
| B23 | B64 | CD3-CD16+CD56+(NK) | WBRSR(mPas_5) | 0.36896 | 2.91E-10 |
| B23 | B196 | CD3-CD16+CD56+(NK) | TCO2 | -0.36124 | 7.20E-10 |
| B23 | B75 | CD3-CD16+CD56+(NK) | ApoA1/B | -0.35901 | 9.31E-10 |
| B23 | B58 | CD3-CD16+CD56+(NK) | HCT | 0.349633 | 2.68E-09 |
| B23 | B195 | CD3-CD16+CD56+(NK) | Na | -0.34546 | 4.25E-09 |
| B23 | B61 | CD3-CD16+CD56+(NK) | WBHSRV | 0.343788 | 5.10E-09 |
| B23 | B137 | CD3-CD16+CD56+(NK) | AFU | -0.33342 | 1.55E-08 |
| B23 | B65 | CD3-CD16+CD56+(NK) | WBRSR(mPas_1) | 0.332969 | 1.62E-08 |
| B23 | B56 | CD3-CD16+CD56+(NK) | IR | 0.322036 | 4.97E-08 |
| B23 | B60 | CD3-CD16+CD56+(NK) | WBRVI | 0.320106 | 6.04E-08 |
| B23 | B158 | CD3-CD16+CD56+(NK) | IgM | -0.31735 | 7.94E-08 |
| B24 | B27 | αHBDH | LDH | 0.655423 | 0 |
| B24 | B26 | αHBDH | CKMB | 0.394156 | 1.28E-11 |
| B24 | B89 | αHBDH | CA 72-4 | -0.33725 | 1.03E-08 |
| B24 | B168 | αHBDH | NMP22 | 0.303244 | 3.09E-07 |
| B25 | B55 | CK | TK | -0.44325 | 1.29E-14 |
| B25 | B198 | CK | P | -0.38934 | 2.38E-11 |
| B25 | B168 | CK | NMP22 | -0.38154 | 6.33E-11 |
| B25 | B59 | CK | WBRV | -0.33677 | 1.08E-08 |
| B26 | B148 | CKMB | Anti_TPO | -0.35282 | 1.88E-09 |
| B26 | B58 | CKMB | HCT | 0.297269 | 5.39E-07 |
| B26 | B27 | CKMB | LDH | 0.296965 | 5.54E-07 |
| B26 | B34 | CKMB | TK | -0.2952 | 6.51E-07 |
| B27 | B194 | LDH | Cl | 0.413822 | 9.24E-13 |
| B27 | B55 | LDH | TK | -0.33299 | 1.62E-08 |
| B27 | B150 | LDH | ALP | 0.325277 | 3.58E-08 |
| B28 | B202 | PRL | e-times | -0.76603 | 0 |
| B28 | B31 | PRL | FSH | -0.78446 | 0 |
| B29 | B116 | E2 | TT3 | -0.48146 | 0 |
| B29 | B223 | E2 | FH | -0.69529 | 0 |
| B29 | B202 | E2 | e-times | -0.37139 | 2.18E-10 |
| B29 | B37 | E2 | BAND | -0.29066 | 9.81E-07 |
| B3 | B62 | Hcy | WBHSRVI | 0.397911 | 7.86E-12 |
| B3 | B34 | Hcy | TK | -0.38921 | 2.42E-11 |
| B3 | B63 | Hcy | WBRSR(mPas_200) | 0.387259 | 3.10E-11 |
| B3 | B67 | Hcy | WBRSR(mPas_50) | 0.378898 | 8.78E-11 |
| B3 | B61 | Hcy | WBHSRV | 0.37883 | 8.85E-11 |
| B3 | B66 | Hcy | WBRSR(mPas_100) | 0.378297 | 9.45E-11 |
| B3 | B56 | Hcy | IR | 0.373579 | 1.68E-10 |
| B3 | B64 | Hcy | WBRSR(mPas_5) | 0.367449 | 3.49E-10 |
| B3 | B58 | Hcy | HCT | 0.360703 | 7.66E-10 |
| B3 | B65 | Hcy | WBRSR(mPas_1) | 0.33946 | 8.14E-09 |
| B3 | B60 | Hcy | WBRVI | 0.339362 | 8.23E-09 |

| | | | | | |
|---|---|---|---|---|---|
| B3 | B90 | Hcy | CYFRA 21-1 | 0.294764 | 6.77E-07 |
| B30 | B167 | LH | PC | 0.910823 | 0 |
| B30 | B225 | LH | hPBG | 0.641581 | 0 |
| B30 | B32 | LH | Testo | 0.429339 | 1.03E-13 |
| B31 | B142 | FSH | 25-(OH)D | -0.50262 | 0 |
| B31 | B33 | FSH | Prog | -0.96836 | 0 |
| B31 | B35 | FSH | β2-MG | -0.55816 | 0 |
| B32 | B144 | Testo | Total-PINP | -0.9759 | 0 |
| B32 | B167 | Testo | PC | 0.422248 | 2.84E-13 |
| B33 | B116 | Prog | TT3 | -0.72008 | 0 |
| B33 | B35 | Prog | β2-MG | -0.87831 | 0 |
| B34 | B186 | TK | PFC | -0.47067 | 0 |
| B34 | B151 | TK | Anti-CCP | 0.397899 | 7.87E-12 |
| B34 | B182 | TK | TT | 0.376428 | 1.19E-10 |
| B34 | B183 | TK | PTA | 0.373291 | 1.74E-10 |
| B34 | B181 | TK | APTT | 0.366046 | 4.11E-10 |
| B34 | B73 | TK | ApoA | 0.349489 | 2.73E-09 |
| B34 | B60 | TK | WBRVI | -0.32916 | 2.41E-08 |
| B34 | B198 | TK | P | -0.32744 | 2.87E-08 |
| B34 | B100 | TK | anti-Ro52 | 0.324409 | 3.91E-08 |
| B34 | B101 | TK | anti-Sc70 | 0.324409 | 3.91E-08 |
| B34 | B102 | TK | anti-Sm | 0.324409 | 3.91E-08 |
| B34 | B103 | TK | anti-SSA/Ro | 0.324409 | 3.91E-08 |
| B34 | B104 | TK | anti-SSB | 0.324409 | 3.91E-08 |
| B34 | B105 | TK | anti-U1RNP | 0.324409 | 3.91E-08 |
| B34 | B106 | TK | anti-P | 0.324409 | 3.91E-08 |
| B34 | B107 | TK | AnuA | 0.324409 | 3.91E-08 |
| B34 | B108 | TK | Anti-Cyclin | 0.324409 | 3.91E-08 |
| B34 | B109 | TK | AMA-M2 | 0.324409 | 3.91E-08 |
| B34 | B110 | TK | AHA | 0.324409 | 3.91E-08 |
| B34 | B95 | TK | anti-cenpb | 0.324409 | 3.91E-08 |
| B34 | B97 | TK | dsDNA | 0.324409 | 3.91E-08 |
| B34 | B98 | TK | anti-Jo-1 | 0.324409 | 3.91E-08 |
| B34 | B193 | TK | K | -0.29827 | 4.91E-07 |
| B35 | B37 | β2-MG | BAND | 0.3 | 4.19E-07 |
| B36 | B144 | V-Lym% | Total-PINP | 0.917663 | 0 |
| B36 | B145 | V-Lym% | BGP | 0.656042 | 0 |
| B36 | B167 | V-Lym% | PC | 0.779157 | 0 |
| B36 | B96 | V-Lym% | dsDNA (IIF) | 0.819513 | 0 |
| B37 | B116 | BAND | TT3 | -0.94868 | 0 |
| B37 | B142 | BAND | 25-(OH)D | -0.76277 | 0 |
| B4 | B6 | WB-Ca | WB-Fe | 0.685754 | 0 |
| B4 | B7 | WB-Ca | WB-Cop | -0.99502 | 0 |
| B4 | B8 | WB-Ca | WB-Zinc | -0.90132 | 0 |

| | | | | | |
|---|---|---|---|---|---|
| B4 | B75 | WB-Ca | ApoA1/B | 0.30166 | 3.59E-07 |
| B4 | B39 | WB-Ca | Other cells | -0.29614 | 5.97E-07 |
| B40 | B42 | WBC | MON | 0.628565 | 0 |
| B40 | B46 | WBC | RBC Count | 0.302677 | 3.26E-07 |
| B41 | B42 | MON％ | MON | 0.609449 | 0 |
| B41 | B49 | MON％ | baso% | 0.323963 | 4.10E-08 |
| B43 | B44 | MCV | MCH | 0.843341 | 0 |
| B43 | B46 | MCV | RBC Count | -0.54744 | 0 |
| B43 | B47 | MCV | RDW | 0.600698 | 0 |
| B43 | B72 | MCV | HDL-C | 0.29255 | 8.28E-07 |
| B44 | B45 | MCH | MCHC | 0.533862 | 0 |
| B44 | B46 | MCH | RBC Count | -0.42372 | 2.30E-13 |
| B44 | B47 | MCH | RDW | 0.326754 | 3.08E-08 |
| B45 | B51 | MCHC | Hb | 0.444025 | 1.15E-14 |
| B45 | B47 | MCHC | RDW | -0.34558 | 4.20E-09 |
| B46 | B51 | RBC Count | Hb | 0.797401 | 0 |
| B46 | B47 | RBC Count | RDW | -0.37566 | 1.30E-10 |
| B46 | B221 | RBC Count | Hei | 0.358662 | 9.69E-10 |
| B46 | B126 | RBC Count | Alb | 0.339084 | 8.48E-09 |
| B46 | B190 | RBC Count | Cre | 0.314347 | 1.07E-07 |
| B46 | B222 | RBC Count | NS | 0.300516 | 3.99E-07 |
| B46 | B216 | RBC Count | BMI | 0.296296 | 5.89E-07 |
| B48 | B53 | LYM% | GR% | -0.93812 | 0 |
| B49 | B50 | baso% | eo% | 0.359095 | 9.22E-10 |
| B49 | B53 | baso% | GR% | -0.2986 | 4.77E-07 |
| B5 | B6 | WB-Mg | WB-Fe | 0.742442 | 0 |
| B5 | B8 | WB-Mg | WB-Zinc | -0.45247 | 3.11E-15 |
| B51 | B190 | Hb | Cre | 0.378481 | 9.24E-11 |
| B51 | B192 | Hb | UA | 0.375159 | 1.39E-10 |
| B51 | B221 | Hb | Hei | 0.366946 | 3.70E-10 |
| B51 | B58 | Hb | HCT | 0.296099 | 6.00E-07 |
| B51 | B126 | Hb | Alb | 0.29207 | 8.65E-07 |
| B54 | B123 | ESR | hs-CRP1 | 0.580586 | 0 |
| B54 | B139 | ESR | ADA | 0.566014 | 0 |
| B54 | B60 | ESR | WBRVI | -0.48733 | 0 |
| B54 | B69 | ESR | PV | 0.626711 | 0 |
| B54 | B182 | ESR | TT | -0.46533 | 4.44E-16 |
| B54 | B58 | ESR | HCT | -0.45164 | 3.55E-15 |
| B54 | B56 | ESR | IR | -0.44007 | 2.09E-14 |
| B54 | B186 | ESR | PFC | 0.438007 | 2.84E-14 |
| B54 | B181 | ESR | APTT | -0.40264 | 4.21E-12 |
| B54 | B62 | ESR | WBHSRVI | -0.39711 | 8.73E-12 |
| B54 | B65 | ESR | WBRSR(mPas_1) | -0.373 | 1.80E-10 |
| B54 | B183 | ESR | PTA | -0.35702 | 1.17E-09 |

| | | | | | |
|---|---|---|---|---|---|
| B54 | B64 | ESR | WBRSR(mPas_5) | -0.3322 | 1.75E-08 |
| B54 | B146 | ESR | CysC | 0.328785 | 2.50E-08 |
| B54 | B198 | ESR | P | 0.319424 | 6.46E-08 |
| B54 | B59 | ESR | WBRV | -0.31657 | 8.58E-08 |
| B54 | B138 | ESR | PAB | -0.31461 | 1.04E-07 |
| B54 | B57 | ESR | RBCAI | -0.31073 | 1.52E-07 |
| B54 | B151 | ESR | Anti-CCP | -0.30991 | 1.64E-07 |
| B54 | B150 | ESR | ALP | 0.301949 | 3.49E-07 |
| B55 | B56 | TK | IR | 0.523329 | 0 |
| B55 | B59 | TK | WBRV | 0.621535 | 0 |
| B55 | B69 | TK | PV | -0.65588 | 0 |
| B55 | B57 | TK | RBCAI | 0.464298 | 4.44E-16 |
| B55 | B61 | TK | WBHSRV | 0.371582 | 2.13E-10 |
| B55 | B60 | TK | WBRVI | 0.336294 | 1.14E-08 |
| B56 | B58 | IR | HCT | 0.766533 | 0 |
| B56 | B59 | IR | WBRV | 0.643348 | 0 |
| B56 | B60 | IR | WBRVI | 0.884986 | 0 |
| B56 | B61 | IR | WBHSRV | 0.946335 | 0 |
| B56 | B62 | IR | WBHSRVI | 0.925801 | 0 |
| B56 | B63 | IR | WBRSR(mPas_200) | 0.822973 | 0 |
| B56 | B64 | IR | WBRSR(mPas_5) | 0.856112 | 0 |
| B56 | B65 | IR | WBRSR(mPas_1) | 0.832038 | 0 |
| B56 | B66 | IR | WBRSR(mPas_100) | 0.822885 | 0 |
| B56 | B67 | IR | WBRSR(mPas_50) | 0.833089 | 0 |
| B56 | B69 | IR | PV | -0.48288 | 0 |
| B56 | B185 | IR | PT | -0.40888 | 1.82E-12 |
| B56 | B183 | IR | PTA | -0.31808 | 7.38E-08 |
| B56 | B100 | IR | anti-Ro52 | -0.30306 | 3.15E-07 |
| B56 | B101 | IR | anti-Sc70 | -0.30306 | 3.15E-07 |
| B56 | B102 | IR | anti-Sm | -0.30306 | 3.15E-07 |
| B56 | B103 | IR | anti-SSA/Ro | -0.30306 | 3.15E-07 |
| B56 | B104 | IR | anti-SSB | -0.30306 | 3.15E-07 |
| B56 | B105 | IR | anti-U1RNP | -0.30306 | 3.15E-07 |
| B56 | B106 | IR | anti-P | -0.30306 | 3.15E-07 |
| B56 | B107 | IR | AnuA | -0.30306 | 3.15E-07 |
| B56 | B108 | IR | Anti-Cyclin | -0.30306 | 3.15E-07 |
| B56 | B109 | IR | AMA-M2 | -0.30306 | 3.15E-07 |
| B56 | B110 | IR | AHA | -0.30306 | 3.15E-07 |
| B56 | B95 | IR | anti-cenpb | -0.30306 | 3.15E-07 |
| B56 | B97 | IR | dsDNA | -0.30306 | 3.15E-07 |
| B56 | B98 | IR | anti-Jo-1 | -0.30306 | 3.15E-07 |
| B56 | B193 | IR | K | 0.292455 | 8.35E-07 |
| B57 | B59 | RBCAI | WBRV | 0.839864 | 0 |
| B57 | B60 | RBCAI | WBRVI | 0.517195 | 0 |

| | | | | | |
|---|---|---|---|---|---|
| B57 | B65 | RBCAI | WBRSR(mPas_1) | 0.458055 | 1.33E-15 |
| B57 | B69 | RBCAI | PV | -0.42387 | 2.25E-13 |
| B57 | B184 | RBCAI | INR | 0.399626 | 6.28E-12 |
| B57 | B182 | RBCAI | TT | -0.37646 | 1.18E-10 |
| B57 | B186 | RBCAI | PFC | 0.34817 | 3.16E-09 |
| B57 | B74 | RBCAI | ApoA1 | 0.32342 | 4.33E-08 |
| B57 | B157 | RBCAI | IgG | -0.31373 | 1.13E-07 |
| B57 | B123 | RBCAI | hs-CRP1 | -0.29519 | 6.52E-07 |
| B57 | B185 | RBCAI | PT | 0.294186 | 7.14E-07 |
| B58 | B60 | HCT | WBRVI | 0.779939 | 0 |
| B58 | B61 | HCT | WBHSRV | 0.812704 | 0 |
| B58 | B62 | HCT | WBHSRVI | 0.92397 | 0 |
| B58 | B63 | HCT | WBRSR(mPas_200) | 0.929012 | 0 |
| B58 | B64 | HCT | WBRSR(mPas_5) | 0.883152 | 0 |
| B58 | B65 | HCT | WBRSR(mPas_1) | 0.810612 | 0 |
| B58 | B66 | HCT | WBRSR(mPas_100) | 0.92182 | 0 |
| B58 | B67 | HCT | WBRSR(mPas_50) | 0.92106 | 0 |
| B58 | B124 | HCT | ACTH | 0.373647 | 1.66E-10 |
| B58 | B59 | HCT | WBRV | 0.327652 | 2.81E-08 |
| B58 | B182 | HCT | TT | 0.319256 | 6.57E-08 |
| B58 | B125 | HCT | Cor | 0.31021 | 1.59E-07 |
| B58 | B156 | HCT | IgE | -0.30726 | 2.12E-07 |
| B58 | B75 | HCT | ApoA1/B | -0.29314 | 7.85E-07 |
| B59 | B60 | WBRV | WBRVI | 0.799194 | 0 |
| B59 | B61 | WBRV | WBHSRV | 0.607835 | 0 |
| B59 | B64 | WBRV | WBRSR(mPas_5) | 0.658127 | 0 |
| B59 | B65 | WBRV | WBRSR(mPas_1) | 0.779505 | 0 |
| B59 | B62 | WBRV | WBHSRVI | 0.47015 | 2.22E-16 |
| B59 | B67 | WBRV | WBRSR(mPas_50) | 0.461819 | 6.66E-16 |
| B59 | B66 | WBRV | WBRSR(mPas_100) | 0.420001 | 3.90E-13 |
| B59 | B63 | WBRV | WBRSR(mPas_200) | 0.403142 | 3.94E-12 |
| B59 | B69 | WBRV | PV | -0.37079 | 2.34E-10 |
| B59 | B100 | WBRV | anti-Ro52 | -0.35392 | 1.66E-09 |
| B59 | B101 | WBRV | anti-Sc70 | -0.35392 | 1.66E-09 |
| B59 | B102 | WBRV | anti-Sm | -0.35392 | 1.66E-09 |
| B59 | B103 | WBRV | anti-SSA/Ro | -0.35392 | 1.66E-09 |
| B59 | B104 | WBRV | anti-SSB | -0.35392 | 1.66E-09 |
| B59 | B105 | WBRV | anti-U1RNP | -0.35392 | 1.66E-09 |
| B59 | B106 | WBRV | anti-P | -0.35392 | 1.66E-09 |
| B59 | B107 | WBRV | AnuA | -0.35392 | 1.66E-09 |
| B59 | B108 | WBRV | Anti-Cyclin | -0.35392 | 1.66E-09 |
| B59 | B109 | WBRV | AMA-M2 | -0.35392 | 1.66E-09 |
| B59 | B110 | WBRV | AHA | -0.35392 | 1.66E-09 |
| B59 | B95 | WBRV | anti-cenpb | -0.35392 | 1.66E-09 |

| | | | | | |
|---|---|---|---|---|---|
| B59 | B97 | WBRV | dsDNA | -0.35392 | 1.66E-09 |
| B59 | B98 | WBRV | anti-Jo-1 | -0.35392 | 1.66E-09 |
| B59 | B193 | WBRV | K | 0.312447 | 1.28E-07 |
| B59 | B186 | WBRV | PFC | 0.311893 | 1.36E-07 |
| B59 | B179 | WBRV | α1-MG | 0.29823 | 4.93E-07 |
| B6 | B7 | WB-Fe | WB-Cop | -0.60981 | 0 |
| B6 | B8 | WB-Fe | WB-Zinc | -0.93335 | 0 |
| B60 | B61 | WBRVI | WBHSRV | 0.840968 | 0 |
| B60 | B62 | WBRVI | WBHSRVI | 0.875765 | 0 |
| B60 | B63 | WBRVI | WBRSR(mPas_200) | 0.791517 | 0 |
| B60 | B64 | WBRVI | WBRSR(mPas_5) | 0.93303 | 0 |
| B60 | B65 | WBRVI | WBRSR(mPas_1) | 0.970271 | 0 |
| B60 | B66 | WBRVI | WBRSR(mPas_100) | 0.802081 | 0 |
| B60 | B67 | WBRVI | WBRSR(mPas_50) | 0.828358 | 0 |
| B60 | B69 | WBRVI | PV | -0.42048 | 3.65E-13 |
| B60 | B100 | WBRVI | anti-Ro52 | -0.35943 | 8.88E-10 |
| B60 | B101 | WBRVI | anti-Sc70 | -0.35943 | 8.88E-10 |
| B60 | B102 | WBRVI | anti-Sm | -0.35943 | 8.88E-10 |
| B60 | B103 | WBRVI | anti-SSA/Ro | -0.35943 | 8.88E-10 |
| B60 | B104 | WBRVI | anti-SSB | -0.35943 | 8.88E-10 |
| B60 | B105 | WBRVI | anti-U1RNP | -0.35943 | 8.88E-10 |
| B60 | B106 | WBRVI | anti-P | -0.35943 | 8.88E-10 |
| B60 | B107 | WBRVI | AnuA | -0.35943 | 8.88E-10 |
| B60 | B108 | WBRVI | Anti-Cyclin | -0.35943 | 8.88E-10 |
| B60 | B109 | WBRVI | AMA-M2 | -0.35943 | 8.88E-10 |
| B60 | B110 | WBRVI | AHA | -0.35943 | 8.88E-10 |
| B60 | B95 | WBRVI | anti-cenpb | -0.35943 | 8.88E-10 |
| B60 | B97 | WBRVI | dsDNA | -0.35943 | 8.88E-10 |
| B60 | B98 | WBRVI | anti-Jo-1 | -0.35943 | 8.88E-10 |
| B60 | B193 | WBRVI | K | 0.333717 | 1.50E-08 |
| B61 | B62 | WBHSRV | WBHSRVI | 0.932716 | 0 |
| B61 | B63 | WBHSRV | WBRSR(mPas_200) | 0.923887 | 0 |
| B61 | B64 | WBHSRV | WBRSR(mPas_5) | 0.919432 | 0 |
| B61 | B65 | WBHSRV | WBRSR(mPas_1) | 0.866761 | 0 |
| B61 | B66 | WBHSRV | WBRSR(mPas_100) | 0.924527 | 0 |
| B61 | B67 | WBHSRV | WBRSR(mPas_50) | 0.929664 | 0 |
| B61 | B100 | WBHSRV | anti-Ro52 | -0.36724 | 3.57E-10 |
| B61 | B101 | WBHSRV | anti-Sc70 | -0.36724 | 3.57E-10 |
| B61 | B102 | WBHSRV | anti-Sm | -0.36724 | 3.57E-10 |
| B61 | B103 | WBHSRV | anti-SSA/Ro | -0.36724 | 3.57E-10 |
| B61 | B104 | WBHSRV | anti-SSB | -0.36724 | 3.57E-10 |
| B61 | B105 | WBHSRV | anti-U1RNP | -0.36724 | 3.57E-10 |
| B61 | B106 | WBHSRV | anti-P | -0.36724 | 3.57E-10 |
| B61 | B107 | WBHSRV | AnuA | -0.36724 | 3.57E-10 |

| | | | | | |
|---|---|---|---|---|---|
| B61 | B108 | WBHSRV | Anti-Cyclin | -0.36724 | 3.57E-10 |
| B61 | B109 | WBHSRV | AMA-M2 | -0.36724 | 3.57E-10 |
| B61 | B110 | WBHSRV | AHA | -0.36724 | 3.57E-10 |
| B61 | B95 | WBHSRV | anti-cenpb | -0.36724 | 3.57E-10 |
| B61 | B97 | WBHSRV | dsDNA | -0.36724 | 3.57E-10 |
| B61 | B98 | WBHSRV | anti-Jo-1 | -0.36724 | 3.57E-10 |
| B61 | B183 | WBHSRV | PTA | -0.33875 | 8.78E-09 |
| B61 | B185 | WBHSRV | PT | -0.31692 | 8.28E-08 |
| B61 | B154 | WBHSRV | RF | -0.31353 | 1.16E-07 |
| B61 | B193 | WBHSRV | K | 0.297023 | 5.51E-07 |
| B62 | B63 | WBHSRVI | WBRSR(mPas_200) | 0.956036 | 0 |
| B62 | B64 | WBHSRVI | WBRSR(mPas_5) | 0.935097 | 0 |
| B62 | B65 | WBHSRVI | WBRSR(mPas_1) | 0.872101 | 0 |
| B62 | B66 | WBHSRVI | WBRSR(mPas_100) | 0.955945 | 0 |
| B62 | B67 | WBHSRVI | WBRSR(mPas_50) | 0.958346 | 0 |
| B62 | B185 | WBHSRVI | PT | -0.35495 | 1.48E-09 |
| B62 | B182 | WBHSRVI | TT | 0.320589 | 5.75E-08 |
| B62 | B100 | WBHSRVI | anti-Ro52 | -0.32037 | 5.88E-08 |
| B62 | B101 | WBHSRVI | anti-Sc70 | -0.32037 | 5.88E-08 |
| B62 | B102 | WBHSRVI | anti-Sm | -0.32037 | 5.88E-08 |
| B62 | B103 | WBHSRVI | anti-SSA/Ro | -0.32037 | 5.88E-08 |
| B62 | B104 | WBHSRVI | anti-SSB | -0.32037 | 5.88E-08 |
| B62 | B105 | WBHSRVI | anti-U1RNP | -0.32037 | 5.88E-08 |
| B62 | B106 | WBHSRVI | anti-P | -0.32037 | 5.88E-08 |
| B62 | B107 | WBHSRVI | AnuA | -0.32037 | 5.88E-08 |
| B62 | B108 | WBHSRVI | Anti-Cyclin | -0.32037 | 5.88E-08 |
| B62 | B109 | WBHSRVI | AMA-M2 | -0.32037 | 5.88E-08 |
| B62 | B110 | WBHSRVI | AHA | -0.32037 | 5.88E-08 |
| B62 | B95 | WBHSRVI | anti-cenpb | -0.32037 | 5.88E-08 |
| B62 | B97 | WBHSRVI | dsDNA | -0.32037 | 5.88E-08 |
| B62 | B98 | WBHSRVI | anti-Jo-1 | -0.32037 | 5.88E-08 |
| B62 | B124 | WBHSRVI | ACTH | 0.313109 | 1.20E-07 |
| B63 | B64 | WBRSR(mPas_200) | WBRSR(mPas_5) | 0.943873 | 0 |
| B63 | B65 | WBRSR(mPas_200) | WBRSR(mPas_1) | 0.858137 | 0 |
| B63 | B66 | WBRSR(mPas_200) | WBRSR(mPas_100) | 0.997992 | 0 |
| B63 | B67 | WBRSR(mPas_200) | WBRSR(mPas_50) | 0.995632 | 0 |
| B63 | B100 | WBRSR(mPas_200) | anti-Ro52 | -0.33304 | 1.61E-08 |
| B63 | B101 | WBRSR(mPas_200) | anti-Sc70 | -0.33304 | 1.61E-08 |
| B63 | B102 | WBRSR(mPas_200) | anti-Sm | -0.33304 | 1.61E-08 |
| B63 | B103 | WBRSR(mPas_200) | anti-SSA/Ro | -0.33304 | 1.61E-08 |
| B63 | B104 | WBRSR(mPas_200) | anti-SSB | -0.33304 | 1.61E-08 |
| B63 | B105 | WBRSR(mPas_200) | anti-U1RNP | -0.33304 | 1.61E-08 |
| B63 | B106 | WBRSR(mPas_200) | anti-P | -0.33304 | 1.61E-08 |
| B63 | B107 | WBRSR(mPas_200) | AnuA | -0.33304 | 1.61E-08 |

| | | | | | |
|---|---|---|---|---|---|
| B63 | B108 | WBRSR(mPas_200) | Anti-Cyclin | -0.33304 | 1.61E-08 |
| B63 | B109 | WBRSR(mPas_200) | AMA-M2 | -0.33304 | 1.61E-08 |
| B63 | B110 | WBRSR(mPas_200) | AHA | -0.33304 | 1.61E-08 |
| B63 | B95 | WBRSR(mPas_200) | anti-cenpb | -0.33304 | 1.61E-08 |
| B63 | B97 | WBRSR(mPas_200) | dsDNA | -0.33304 | 1.61E-08 |
| B63 | B98 | WBRSR(mPas_200) | anti-Jo-1 | -0.33304 | 1.61E-08 |
| B63 | B196 | WBRSR(mPas_200) | TCO2 | -0.32044 | 5.84E-08 |
| B63 | B124 | WBRSR(mPas_200) | ACTH | 0.30423 | 2.82E-07 |
| B63 | B156 | WBRSR(mPas_200) | IgE | -0.29125 | 9.30E-07 |
| B64 | B65 | WBRSR(mPas_5) | WBRSR(mPas_1) | 0.979301 | 0 |
| B64 | B66 | WBRSR(mPas_5) | WBRSR(mPas_100) | 0.951636 | 0 |
| B64 | B67 | WBRSR(mPas_5) | WBRSR(mPas_50) | 0.965843 | 0 |
| B64 | B100 | WBRSR(mPas_5) | anti-Ro52 | -0.40749 | 2.20E-12 |
| B64 | B101 | WBRSR(mPas_5) | anti-Sc70 | -0.40749 | 2.20E-12 |
| B64 | B102 | WBRSR(mPas_5) | anti-Sm | -0.40749 | 2.20E-12 |
| B64 | B103 | WBRSR(mPas_5) | anti-SSA/Ro | -0.40749 | 2.20E-12 |
| B64 | B104 | WBRSR(mPas_5) | anti-SSB | -0.40749 | 2.20E-12 |
| B64 | B105 | WBRSR(mPas_5) | anti-U1RNP | -0.40749 | 2.20E-12 |
| B64 | B106 | WBRSR(mPas_5) | anti-P | -0.40749 | 2.20E-12 |
| B64 | B107 | WBRSR(mPas_5) | AnuA | -0.40749 | 2.20E-12 |
| B64 | B108 | WBRSR(mPas_5) | Anti-Cyclin | -0.40749 | 2.20E-12 |
| B64 | B109 | WBRSR(mPas_5) | AMA-M2 | -0.40749 | 2.20E-12 |
| B64 | B110 | WBRSR(mPas_5) | AHA | -0.40749 | 2.20E-12 |
| B64 | B95 | WBRSR(mPas_5) | anti-cenpb | -0.40749 | 2.20E-12 |
| B64 | B97 | WBRSR(mPas_5) | dsDNA | -0.40749 | 2.20E-12 |
| B64 | B98 | WBRSR(mPas_5) | anti-Jo-1 | -0.40749 | 2.20E-12 |
| B64 | B193 | WBRSR(mPas_5) | K | 0.328496 | 2.58E-08 |
| B64 | B156 | WBRSR(mPas_5) | IgE | -0.29517 | 6.53E-07 |
| B65 | B66 | WBRSR(mPas_1) | WBRSR(mPas_100) | 0.869831 | 0 |
| B65 | B67 | WBRSR(mPas_1) | WBRSR(mPas_50) | 0.89349 | 0 |
| B65 | B100 | WBRSR(mPas_1) | anti-Ro52 | -0.41222 | 1.15E-12 |
| B65 | B101 | WBRSR(mPas_1) | anti-Sc70 | -0.41222 | 1.15E-12 |
| B65 | B102 | WBRSR(mPas_1) | anti-Sm | -0.41222 | 1.15E-12 |
| B65 | B103 | WBRSR(mPas_1) | anti-SSA/Ro | -0.41222 | 1.15E-12 |
| B65 | B104 | WBRSR(mPas_1) | anti-SSB | -0.41222 | 1.15E-12 |
| B65 | B105 | WBRSR(mPas_1) | anti-U1RNP | -0.41222 | 1.15E-12 |
| B65 | B106 | WBRSR(mPas_1) | anti-P | -0.41222 | 1.15E-12 |
| B65 | B107 | WBRSR(mPas_1) | AnuA | -0.41222 | 1.15E-12 |
| B65 | B108 | WBRSR(mPas_1) | Anti-Cyclin | -0.41222 | 1.15E-12 |
| B65 | B109 | WBRSR(mPas_1) | AMA-M2 | -0.41222 | 1.15E-12 |
| B65 | B110 | WBRSR(mPas_1) | AHA | -0.41222 | 1.15E-12 |
| B65 | B95 | WBRSR(mPas_1) | anti-cenpb | -0.41222 | 1.15E-12 |
| B65 | B97 | WBRSR(mPas_1) | dsDNA | -0.41222 | 1.15E-12 |
| B65 | B98 | WBRSR(mPas_1) | anti-Jo-1 | -0.41222 | 1.15E-12 |

| | | | | | |
|---|---|---|---|---|---|
| B65 | B193 | WBRSR(mPas_1) | K | 0.344781 | 4.58E-09 |
| B65 | B179 | WBRSR(mPas_1) | α1-MG | 0.310936 | 1.49E-07 |
| B66 | B67 | WBRSR(mPas_100) | WBRSR(mPas_50) | 0.998732 | 0 |
| B66 | B100 | WBRSR(mPas_100) | anti-Ro52 | -0.37503 | 1.41E-10 |
| B66 | B101 | WBRSR(mPas_100) | anti-Sc70 | -0.37503 | 1.41E-10 |
| B66 | B102 | WBRSR(mPas_100) | anti-Sm | -0.37503 | 1.41E-10 |
| B66 | B103 | WBRSR(mPas_100) | anti-SSA/Ro | -0.37503 | 1.41E-10 |
| B66 | B104 | WBRSR(mPas_100) | anti-SSB | -0.37503 | 1.41E-10 |
| B66 | B105 | WBRSR(mPas_100) | anti-U1RNP | -0.37503 | 1.41E-10 |
| B66 | B106 | WBRSR(mPas_100) | anti-P | -0.37503 | 1.41E-10 |
| B66 | B107 | WBRSR(mPas_100) | AnuA | -0.37503 | 1.41E-10 |
| B66 | B108 | WBRSR(mPas_100) | Anti-Cyclin | -0.37503 | 1.41E-10 |
| B66 | B109 | WBRSR(mPas_100) | AMA-M2 | -0.37503 | 1.41E-10 |
| B66 | B110 | WBRSR(mPas_100) | AHA | -0.37503 | 1.41E-10 |
| B66 | B95 | WBRSR(mPas_100) | anti-cenpb | -0.37503 | 1.41E-10 |
| B66 | B97 | WBRSR(mPas_100) | dsDNA | -0.37503 | 1.41E-10 |
| B66 | B98 | WBRSR(mPas_100) | anti-Jo-1 | -0.37503 | 1.41E-10 |
| B66 | B196 | WBRSR(mPas_100) | TCO2 | -0.3238 | 4.17E-08 |
| B66 | B154 | WBRSR(mPas_100) | RF | -0.30749 | 2.07E-07 |
| B67 | B100 | WBRSR(mPas_50) | anti-Ro52 | -0.38469 | 4.28E-11 |
| B67 | B101 | WBRSR(mPas_50) | anti-Sc70 | -0.38469 | 4.28E-11 |
| B67 | B102 | WBRSR(mPas_50) | anti-Sm | -0.38469 | 4.28E-11 |
| B67 | B103 | WBRSR(mPas_50) | anti-SSA/Ro | -0.38469 | 4.28E-11 |
| B67 | B104 | WBRSR(mPas_50) | anti-SSB | -0.38469 | 4.28E-11 |
| B67 | B105 | WBRSR(mPas_50) | anti-U1RNP | -0.38469 | 4.28E-11 |
| B67 | B106 | WBRSR(mPas_50) | anti-P | -0.38469 | 4.28E-11 |
| B67 | B107 | WBRSR(mPas_50) | AnuA | -0.38469 | 4.28E-11 |
| B67 | B108 | WBRSR(mPas_50) | Anti-Cyclin | -0.38469 | 4.28E-11 |
| B67 | B109 | WBRSR(mPas_50) | AMA-M2 | -0.38469 | 4.28E-11 |
| B67 | B110 | WBRSR(mPas_50) | AHA | -0.38469 | 4.28E-11 |
| B67 | B95 | WBRSR(mPas_50) | anti-cenpb | -0.38469 | 4.28E-11 |
| B67 | B97 | WBRSR(mPas_50) | dsDNA | -0.38469 | 4.28E-11 |
| B67 | B98 | WBRSR(mPas_50) | anti-Jo-1 | -0.38469 | 4.28E-11 |
| B67 | B196 | WBRSR(mPas_50) | TCO2 | -0.31507 | 9.94E-08 |
| B67 | B154 | WBRSR(mPas_50) | RF | -0.30141 | 3.67E-07 |
| B67 | B193 | WBRSR(mPas_50) | K | 0.290681 | 9.79E-07 |
| B67 | B156 | WBRSR(mPas_50) | IgE | -0.29068 | 9.80E-07 |
| B69 | B123 | PV | hs-CRP1 | 0.587248 | 0 |
| B69 | B136 | PV | 5'NT | 0.335673 | 1.22E-08 |
| B69 | B185 | PV | PT | 0.311388 | 1.42E-07 |
| B69 | B194 | PV | Cl | 0.303191 | 3.11E-07 |
| B69 | B186 | PV | PFC | 0.290678 | 9.80E-07 |
| B7 | B8 | WB-Cop | WB-Zinc | 0.853669 | 0 |
| B7 | B75 | WB-Cop | ApoA1/B | -0.3027 | 3.26E-07 |

| | | | | | |
|---|---|---|---|---|---|
| B7 | B39 | WB-Cop | Other cells | 0.294921 | 6.68E-07 |
| B70 | B77 | LDL-C | TC | 0.769078 | 0 |
| B70 | B76 | LDL-C | ApoB | 0.414942 | 7.92E-13 |
| B71 | B130 | TG | GLB | 0.551983 | 0 |
| B71 | B91 | TG | TP | 0.570989 | 0 |
| B71 | B77 | TG | TC | 0.444821 | 1.02E-14 |
| B71 | B72 | TG | HDL-C | -0.33357 | 1.52E-08 |
| B71 | B127 | TG | A-G ratio | -0.31404 | 1.10E-07 |
| B72 | B222 | HDL-C | NS | -0.30187 | 3.52E-07 |
| B73 | B123 | ApoA | hs-CRP1 | 0.446274 | 7.99E-15 |
| B73 | B134 | ApoA | CHE | -0.35913 | 9.19E-10 |
| B73 | B150 | ApoA | ALP | 0.355377 | 1.41E-09 |
| B73 | B138 | ApoA | PAB | -0.35493 | 1.48E-09 |
| B74 | B75 | ApoA1 | ApoA1/B | 0.463148 | 4.44E-16 |
| B74 | B193 | ApoA1 | K | 0.402032 | 4.56E-12 |
| B74 | B182 | ApoA1 | TT | -0.35419 | 1.61E-09 |
| B74 | B136 | ApoA1 | 5'NT | -0.30025 | 4.09E-07 |
| B75 | B134 | ApoA1/B | CHE | -0.48511 | 0 |
| B75 | B76 | ApoA1/B | ApoB | -0.77248 | 0 |
| B75 | B182 | ApoA1/B | TT | -0.37485 | 1.44E-10 |
| B75 | B138 | ApoA1/B | PAB | -0.33853 | 8.99E-09 |
| B76 | B134 | ApoB | CHE | 0.612775 | 0 |
| B76 | B199 | ApoB | Mg | 0.48968 | 0 |
| B76 | B193 | ApoB | K | 0.376292 | 1.21E-10 |
| B76 | B77 | ApoB | TC | 0.337624 | 9.91E-09 |
| B76 | B138 | ApoB | PAB | 0.31458 | 1.04E-07 |
| B76 | B154 | ApoB | RF | -0.30711 | 2.15E-07 |
| B77 | B91 | TC | TP | 0.369309 | 2.80E-10 |
| B77 | B130 | TC | GLB | 0.301351 | 3.69E-07 |
| B77 | B132 | TC | Dbil | -0.30129 | 3.71E-07 |
| B79 | B154 | hb V -dna | RF | 0.520042 | 0 |
| B79 | B199 | hb V -dna | Mg | -0.41628 | 6.58E-13 |
| B79 | B184 | hb V -dna | INR | 0.402943 | 4.04E-12 |
| B79 | B139 | hb V -dna | ADA | -0.37349 | 1.69E-10 |
| B79 | B185 | hb V -dna | PT | 0.368714 | 3.00E-10 |
| B79 | B137 | hb V -dna | AFU | -0.34555 | 4.21E-09 |
| B79 | B146 | hb V -dna | CysC | -0.33395 | 1.46E-08 |
| B79 | B81 | hb V -dna | TSGF | -0.3112 | 1.45E-07 |
| B79 | B134 | hb V -dna | CHE | -0.30752 | 2.06E-07 |
| B79 | B181 | hb V -dna | APTT | -0.29958 | 4.35E-07 |
| B79 | B120 | hb V -dna | HCV-cAg | 0.2953 | 6.45E-07 |
| B81 | B184 | TSGF | INR | -0.5519 | 0 |
| B81 | B185 | TSGF | PT | -0.46815 | 2.22E-16 |
| B81 | B181 | TSGF | APTT | 0.412366 | 1.13E-12 |

| | | | | | |
|---|---|---|---|---|---|
| B81 | B183 | TSGF | PTA | -0.40355 | 3.73E-12 |
| B81 | B157 | TSGF | IgG | 0.398824 | 6.97E-12 |
| B81 | B165 | TSGF | bilirubin | -0.39777 | 8.01E-12 |
| B81 | B173 | TSGF | NTT | -0.39777 | 8.01E-12 |
| B81 | B124 | TSGF | ACTH | -0.36611 | 4.08E-10 |
| B81 | B156 | TSGF | IgE | 0.347216 | 3.51E-09 |
| B81 | B151 | TSGF | Anti-CCP | -0.30573 | 2.45E-07 |
| B82 | B89 | CEA | CA 72-4 | 0.728018 | 0 |
| B82 | B84 | CEA | C-PSA | 0.439142 | 2.40E-14 |
| B82 | B88 | CEA | CA 242 | -0.39841 | 7.37E-12 |
| B82 | B168 | CEA | NMP22 | -0.32297 | 4.53E-08 |
| B82 | B87 | CEA | NSE | 0.292665 | 8.19E-07 |
| B83 | B87 | AFP | NSE | 0.652537 | 0 |
| B83 | B85 | AFP | SCC | 0.352006 | 2.06E-09 |
| B84 | B85 | C-PSA | SCC | 0.726849 | 0 |
| B84 | B88 | C-PSA | CA 242 | -0.55373 | 0 |
| B84 | B86 | C-PSA | TBhCG | -0.38641 | 3.44E-11 |
| B84 | B90 | C-PSA | CYFRA 21-1 | 0.357886 | 1.06E-09 |
| B85 | B87 | SCC | NSE | 0.554417 | 0 |
| B85 | B90 | SCC | CYFRA 21-1 | 0.452956 | 2.89E-15 |
| B85 | B88 | SCC | CA 242 | -0.31038 | 1.57E-07 |
| B85 | B86 | SCC | TBhCG | -0.30511 | 2.60E-07 |
| B86 | B88 | TBhCG | CA 242 | 0.871116 | 0 |
| B86 | B90 | TBhCG | CYFRA 21-1 | 0.451519 | 3.55E-15 |
| B88 | B89 | CA 242 | CA 72-4 | -0.5134 | 0 |
| B89 | B168 | CA 72-4 | NMP22 | -0.43854 | 2.62E-14 |
| B89 | B153 | CA 72-4 | Glu | 0.36207 | 6.54E-10 |
| B89 | B213 | CA 72-4 | CVA-R | -0.33984 | 7.82E-09 |
| B89 | B170 | CA 72-4 | U-GLU | 0.328452 | 2.59E-08 |
| B9 | B142 | RH | 25-(OH)D | -0.76784 | 0 |
| B9 | B29 | RH | E2 | -0.54139 | 0 |
| B9 | B31 | RH | FSH | -0.71348 | 0 |
| B9 | B33 | RH | Prog | -0.41166 | 1.24E-12 |
| B91 | B126 | TP | Alb | 0.529792 | 0 |
| B91 | B127 | TP | A-G ratio | -0.5826 | 0 |
| B91 | B130 | TP | GLB | 0.877304 | 0 |
| B91 | B188 | TP | Ca | 0.358229 | 1.02E-09 |
| B92 | B93 | ALDH2-Glu504Glu | ALDH2-Glu504Lys | -0.99849 | 0 |
| B92 | B99 | ALDH2-Glu504Glu | anti-PM-Scl | 0.363803 | 5.35E-10 |
| B92 | B149 | ALDH2-Glu504Glu | TGAb | 0.351731 | 2.12E-09 |
| B92 | B185 | ALDH2-Glu504Glu | PT | -0.29711 | 5.47E-07 |
| B92 | B158 | ALDH2-Glu504Glu | IgM | 0.297054 | 5.50E-07 |
| B93 | B99 | ALDH2-Glu504Lys | anti-PM-Scl | -0.33879 | 8.75E-09 |
| B93 | B149 | ALDH2-Glu504Lys | TGAb | -0.33037 | 2.12E-08 |

| | | | | | |
|---|---|---|---|---|---|
| B94 | B111 | ALDH2-Lys504Lys | TSH | -0.47883 | 0 |
| B94 | B99 | ALDH2-Lys504Lys | anti-PM-Scl | -0.49731 | 0 |
| B94 | B149 | ALDH2-Lys504Lys | TGAb | -0.43022 | 9.02E-14 |
| B94 | B185 | ALDH2-Lys504Lys | PT | 0.399939 | 6.02E-12 |
| B94 | B114 | ALDH2-Lys504Lys | TSH TT3 TT4(TT4) | 0.370053 | 2.56E-10 |
| B94 | B186 | ALDH2-Lys504Lys | PFC | 0.345908 | 4.05E-09 |
| B94 | B184 | ALDH2-Lys504Lys | INR | 0.33077 | 2.04E-08 |
| B94 | B137 | ALDH2-Lys504Lys | AFU | -0.32974 | 2.27E-08 |
| B94 | B200 | ALDH2-Lys504Lys | OP | -0.3253 | 3.57E-08 |
| B94 | B154 | ALDH2-Lys504Lys | RF | 0.304209 | 2.83E-07 |
| B94 | B112 | ALDH2-Lys504Lys | FT4 | 0.292319 | 8.45E-07 |
| B95 | B101 | anti-cenpb | anti-Sc70 | 1 | 0 |
| B95 | B106 | anti-cenpb | anti-P | 1 | 0 |
| B95 | B107 | anti-cenpb | AnuA | 1 | 0 |
| B95 | B109 | anti-cenpb | AMA-M2 | 1 | 0 |
| B95 | B154 | anti-cenpb | RF | 0.476063 | 0 |
| B95 | B183 | anti-cenpb | PTA | 0.497714 | 0 |
| B95 | B98 | anti-cenpb | anti-Jo-1 | 1 | 0 |
| B95 | B198 | anti-cenpb | P | -0.41547 | 7.35E-13 |
| B95 | B124 | anti-cenpb | ACTH | 0.352095 | 2.04E-09 |
| B95 | B185 | anti-cenpb | PT | 0.335754 | 1.21E-08 |
| B95 | B186 | anti-cenpb | PFC | -0.33542 | 1.25E-08 |
| B95 | B99 | anti-cenpb | anti-PM-Scl | -0.32678 | 3.07E-08 |
| B95 | B134 | anti-cenpb | CHE | -0.31897 | 6.76E-08 |
| B95 | B139 | anti-cenpb | ADA | -0.3138 | 1.13E-07 |
| B95 | B182 | anti-cenpb | TT | 0.29809 | 5.00E-07 |
| B96 | B167 | dsDNA (IIF) | PC | 0.754989 | 0 |
| B97 | B100 | dsDNA | anti-Ro52 | 1 | 0 |
| B97 | B103 | dsDNA | anti-SSA/Ro | 1 | 0 |
| B97 | B105 | dsDNA | anti-U1RNP | 1 | 0 |
| B97 | B106 | dsDNA | anti-P | 1 | 0 |
| B97 | B107 | dsDNA | AnuA | 1 | 0 |
| B97 | B108 | dsDNA | Anti-Cyclin | 1 | 0 |
| B97 | B109 | dsDNA | AMA-M2 | 1 | 0 |
| B97 | B110 | dsDNA | AHA | 1 | 0 |
| B97 | B154 | dsDNA | RF | 0.476063 | 0 |
| B97 | B183 | dsDNA | PTA | 0.497714 | 0 |
| B97 | B198 | dsDNA | P | -0.41547 | 7.35E-13 |
| B97 | B124 | dsDNA | ACTH | 0.352095 | 2.04E-09 |
| B97 | B185 | dsDNA | PT | 0.335754 | 1.21E-08 |
| B97 | B186 | dsDNA | PFC | -0.33542 | 1.25E-08 |
| B97 | B99 | dsDNA | anti-PM-Scl | -0.32678 | 3.07E-08 |
| B97 | B134 | dsDNA | CHE | -0.31897 | 6.76E-08 |
| B97 | B139 | dsDNA | ADA | -0.3138 | 1.13E-07 |

| | | | | | |
|---|---|---|---|---|---|
| B97 | B182 | dsDNA | TT | 0.29809 | 5.00E-07 |
| B98 | B100 | anti-Jo-1 | anti-Ro52 | 1 | 0 |
| B98 | B101 | anti-Jo-1 | anti-Sc70 | 1 | 0 |
| B98 | B102 | anti-Jo-1 | anti-Sm | 1 | 0 |
| B98 | B103 | anti-Jo-1 | anti-SSA/Ro | 1 | 0 |
| B98 | B104 | anti-Jo-1 | anti-SSB | 1 | 0 |
| B98 | B105 | anti-Jo-1 | anti-U1RNP | 1 | 0 |
| B98 | B107 | anti-Jo-1 | AnuA | 1 | 0 |
| B98 | B108 | anti-Jo-1 | Anti-Cyclin | 1 | 0 |
| B98 | B109 | anti-Jo-1 | AMA-M2 | 1 | 0 |
| B98 | B110 | anti-Jo-1 | AHA | 1 | 0 |
| B98 | B154 | anti-Jo-1 | RF | 0.476063 | 0 |
| B98 | B183 | anti-Jo-1 | PTA | 0.497714 | 0 |
| B98 | B198 | anti-Jo-1 | P | -0.41547 | 7.35E-13 |
| B98 | B124 | anti-Jo-1 | ACTH | 0.352095 | 2.04E-09 |
| B98 | B185 | anti-Jo-1 | PT | 0.335754 | 1.21E-08 |
| B98 | B186 | anti-Jo-1 | PFC | -0.33542 | 1.25E-08 |
| B98 | B99 | anti-Jo-1 | anti-PM-Scl | -0.32678 | 3.07E-08 |
| B98 | B134 | anti-Jo-1 | CHE | -0.31897 | 6.76E-08 |
| B98 | B139 | anti-Jo-1 | ADA | -0.3138 | 1.13E-07 |
| B98 | B182 | anti-Jo-1 | TT | 0.29809 | 5.00E-07 |
| B99 | B137 | anti-PM-Scl | AFU | 0.610793 | 0 |
| B99 | B146 | anti-PM-Scl | CysC | 0.519837 | 0 |
| B99 | B193 | anti-PM-Scl | K | -0.50236 | 0 |
| B99 | B200 | anti-PM-Scl | OP | 0.45934 | 8.88E-16 |
| B99 | B158 | anti-PM-Scl | IgM | 0.395352 | 1.10E-11 |
| B99 | B136 | anti-PM-Scl | 5'NT | 0.354348 | 1.58E-09 |
| B99 | B157 | anti-PM-Scl | IgG | -0.33669 | 1.09E-08 |
| B99 | B100 | anti-PM-Scl | anti-Ro52 | -0.32678 | 3.07E-08 |
| B99 | B101 | anti-PM-Scl | anti-Sc70 | -0.32678 | 3.07E-08 |
| B99 | B102 | anti-PM-Scl | anti-Sm | -0.32678 | 3.07E-08 |
| B99 | B103 | anti-PM-Scl | anti-SSA/Ro | -0.32678 | 3.07E-08 |
| B99 | B104 | anti-PM-Scl | anti-SSB | -0.32678 | 3.07E-08 |
| B99 | B105 | anti-PM-Scl | anti-U1RNP | -0.32678 | 3.07E-08 |
| B99 | B106 | anti-PM-Scl | anti-P | -0.32678 | 3.07E-08 |
| B99 | B107 | anti-PM-Scl | AnuA | -0.32678 | 3.07E-08 |
| B99 | B108 | anti-PM-Scl | Anti-Cyclin | -0.32678 | 3.07E-08 |
| B99 | B109 | anti-PM-Scl | AMA-M2 | -0.32678 | 3.07E-08 |
| B99 | B110 | anti-PM-Scl | AHA | -0.32678 | 3.07E-08 |
| B99 | B140 | anti-PM-Scl | r-GT | 0.315212 | 9.80E-08 |
| B99 | B115 | anti-PM-Scl | TSH TT3 TT4(TT3) | 0.294582 | 6.89E-07 |

**Supplementary Table 31. PEI Correlations in Asthma**

| Row | Column | Row | Column | Cor | *P* |
|---|---|---|---|---|---|
| B1 | B163 | Gender | SG | -0.41320024 | 0 |
| B1 | B174 | Gender | Urea/Cr | 0.329184431 | 0 |
| B1 | B190 | Gender | Cre | -0.65868278 | 0 |
| B1 | B192 | Gender | UA | -0.51110197 | 0 |
| B1 | B204 | Gender | Drinking | -0.38279287 | 0 |
| B1 | B206 | Gender | Waist | -0.47765615 | 0 |
| B1 | B207 | Gender | WHR | -0.46188452 | 0 |
| B1 | B210 | Gender | Hip | -0.29569796 | 0 |
| B1 | B216 | Gender | BMI | -0.57096145 | 0 |
| B1 | B221 | Gender | Hei | -0.63121386 | 0 |
| B1 | B222 | Gender | NS | -0.73201922 | 0 |
| B1 | B42 | Gender | MON | -0.28245501 | 0 |
| B1 | B46 | Gender | RBC Count | -0.47051242 | 0 |
| B1 | B51 | Gender | Hb | -0.58679322 | 0 |
| B1 | B72 | Gender | HDL-C | 0.327963236 | 0 |
| B1 | B128 | Gender | ALT | -0.26919178 | 4.44E-15 |
| B1 | B209 | Gender | Smoking | -0.26503563 | 1.20E-14 |
| B1 | B215 | Gender | BMI | -0.2580591 | 6.08E-14 |
| B1 | B166 | Gender | Pro | -0.22030431 | 1.80E-10 |
| B1 | B191 | Gender | Urea | -0.21959254 | 2.07E-10 |
| B1 | B141 | Gender | AST/ALT | 0.213953077 | 6.05E-10 |
| B1 | B41 | Gender | MON％ | -0.2103251 | 1.19E-09 |
| B1 | B218 | Gender | DBP | -0.20917269 | 1.47E-09 |
| B1 | B161 | Gender | LEU | 0.198982114 | 9.09E-09 |
| B1 | B189 | Gender | eGFB | 0.196677394 | 1.36E-08 |
| B1 | B127 | Gender | A-G ratio | -0.18933472 | 4.68E-08 |
| B1 | B133 | Gender | TBIL | -0.18201049 | 1.54E-07 |
| B1 | B132 | Gender | Dbil | -0.18029746 | 2.02E-07 |
| B1 | B140 | Gender | r-GT | -0.17978904 | 2.19E-07 |
| B1 | B71 | Gender | TG | -0.17808721 | 2.85E-07 |
| B10 | B12 | PG Ⅰ | PG Ⅱ | 0.656315674 | 0 |
| B10 | B197 | PG Ⅰ | AG | 0.241328393 | 2.49E-12 |
| B100 | B156 | anti-Ro52 | IgE | 0.433123851 | 0 |
| B100 | B158 | anti-Ro52 | IgM | 0.456094609 | 0 |
| B100 | B101 | anti-Ro52 | anti-Sc70 | -0.27098086 | 2.89E-15 |
| B100 | B102 | anti-Ro52 | anti-Sm | -0.27098086 | 2.89E-15 |
| B100 | B103 | anti-Ro52 | anti-SSA/Ro | -0.27098086 | 2.89E-15 |
| B100 | B104 | anti-Ro52 | anti-SSB | -0.27098086 | 2.89E-15 |
| B100 | B105 | anti-Ro52 | anti-U1RNP | -0.27098086 | 2.89E-15 |
| B100 | B106 | anti-Ro52 | anti-P | -0.27098086 | 2.89E-15 |
| B100 | B107 | anti-Ro52 | AnuA | -0.2709808 | 2.89E-15 |

| | | | | | |
|---|---|---|---|---|---|
| B100 | B108 | anti-Ro52 | Anti-Cyclin | -0.27098086 | 2.89E-15 |
| B100 | B109 | anti-Ro52 | AMA-M2 | -0.27098086 | 2.89E-15 |
| B100 | B110 | anti-Ro52 | AHA | -0.27098086 | 2.89E-15 |
| B100 | B151 | anti-Ro52 | Anti-CCP | -0.22834972 | 3.68E-11 |
| B101 | B102 | anti-Sc70 | anti-Sm | 0.999999992 | 0 |
| B101 | B105 | anti-Sc70 | anti-U1RNP | 0.999999999 | 0 |
| B101 | B107 | anti-Sc70 | AnuA | 0.999999682 | 0 |
| B101 | B108 | anti-Sc70 | Anti-Cyclin | 0.999999992 | 0 |
| B101 | B151 | anti-Sc70 | Anti-CCP | 0.452931832 | 0 |
| B101 | B156 | anti-Sc70 | IgE | -0.34171399 | 0 |
| B101 | B158 | anti-Sc70 | IgM | -0.28651441 | 0 |
| B101 | B186 | anti-Sc70 | PFC | 0.308748716 | 0 |
| B101 | B157 | anti-Sc70 | IgG | 0.273681126 | 1.33E-15 |
| B101 | B141 | anti-Sc70 | AST/ALT | -0.26503332 | 1.20E-14 |
| B101 | B181 | anti-Sc70 | APTT | -0.23523471 | 8.99E-12 |
| B101 | B154 | anti-Sc70 | RF | 0.177161152 | 3.29E-07 |
| B102 | B107 | anti-Sm | AnuA | 0.999999877 | 0 |
| B102 | B151 | anti-Sm | Anti-CCP | 0.452931831 | 0 |
| B102 | B156 | anti-Sm | IgE | -0.34171399 | 0 |
| B102 | B158 | anti-Sm | IgM | -0.28651441 | 0 |
| B102 | B186 | anti-Sm | PFC | 0.308748717 | 0 |
| B102 | B157 | anti-Sm | IgG | 0.273681126 | 1.33E-15 |
| B102 | B141 | anti-Sm | AST/ALT | -0.26503332 | 1.20E-14 |
| B102 | B181 | anti-Sm | APTT | -0.23523471 | 8.99E-12 |
| B102 | B154 | anti-Sm | RF | 0.177161151 | 3.29E-07 |
| B103 | B104 | anti-SSA/Ro | anti-SSB | 0.999999996 | 0 |
| B103 | B105 | anti-SSA/Ro | anti-U1RNP | 1 | 0 |
| B103 | B106 | anti-SSA/Ro | anti-P | 0.999999994 | 0 |
| B103 | B107 | anti-SSA/Ro | AnuA | 0.999999811 | 0 |
| B103 | B109 | anti-SSA/Ro | AMA-M2 | 1 | 0 |
| B103 | B151 | anti-SSA/Ro | Anti-CCP | 0.452931827 | 0 |
| B103 | B156 | anti-SSA/Ro | IgE | -0.34171399 | 0 |
| B103 | B158 | anti-SSA/Ro | IgM | -0.28651441 | 0 |
| B103 | B186 | anti-SSA/Ro | PFC | 0.308748713 | 0 |
| B103 | B157 | anti-SSA/Ro | IgG | 0.273681124 | 1.33E-15 |
| B103 | B141 | anti-SSA/Ro | AST/ALT | -0.26503332 | 1.20E-14 |
| B103 | B181 | anti-SSA/Ro | APTT | -0.23523471 | 8.99E-12 |
| B103 | B154 | anti-SSA/Ro | RF | 0.17716115 | 3.29E-07 |
| B104 | B105 | anti-SSB | anti-U1RNP | 0.999999995 | 0 |
| B104 | B106 | anti-SSB | anti-P | 0.999999998 | 0 |
| B104 | B107 | anti-SSB | AnuA | 0.999999839 | 0 |
| B104 | B108 | anti-SSB | Anti-Cyclin | 0.999999993 | 0 |
| B104 | B109 | anti-SSB | AMA-M2 | 0.999999998 | 0 |
| B104 | B110 | anti-SSB | AHA | 0.999999995 | 0 |

| | | | | | |
|---|---|---|---|---|---|
| B104 | B151 | anti-SSB | Anti-CCP | 0.452931825 | 0 |
| B104 | B156 | anti-SSB | IgE | -0.34171399 | 0 |
| B104 | B158 | anti-SSB | IgM | -0.28651441 | 0 |
| B104 | B186 | anti-SSB | PFC | 0.308748713 | 0 |
| B104 | B157 | anti-SSB | IgG | 0.273681124 | 1.33E-15 |
| B104 | B141 | anti-SSB | AST/ALT | -0.26503331 | 1.20E-14 |
| B104 | B181 | anti-SSB | APTT | -0.2352347 | 8.99E-12 |
| B104 | B154 | anti-SSB | RF | 0.177161149 | 3.29E-07 |
| B105 | B106 | anti-U1RNP | anti-P | 0.999999992 | 0 |
| B105 | B107 | anti-U1RNP | AnuA | 0.999999793 | 0 |
| B105 | B109 | anti-U1RNP | AMA-M2 | 0.999999998 | 0 |
| B105 | B151 | anti-U1RNP | Anti-CCP | 0.452931827 | 0 |
| B105 | B156 | anti-U1RNP | IgE | -0.34171399 | 0 |
| B105 | B158 | anti-U1RNP | IgM | -0.28651441 | 0 |
| B105 | B186 | anti-U1RNP | PFC | 0.308748713 | 0 |
| B105 | B157 | anti-U1RNP | IgG | 0.273681124 | 1.33E-15 |
| B105 | B141 | anti-U1RNP | AST/ALT | -0.26503332 | 1.20E-14 |
| B105 | B181 | anti-U1RNP | APTT | -0.23523471 | 8.99E-12 |
| B105 | B154 | anti-U1RNP | RF | 0.17716115 | 3.29E-07 |
| B106 | B107 | anti-P | AnuA | 1 | 0 |
| B106 | B108 | anti-P | Anti-Cyclin | 0.999999995 | 0 |
| B106 | B109 | anti-P | AMA-M2 | 0.999999995 | 0 |
| B106 | B110 | anti-P | AHA | 0.999999995 | 0 |
| B106 | B151 | anti-P | Anti-CCP | 0.452931824 | 0 |
| B106 | B156 | anti-P | IgE | -0.34171399 | 0 |
| B106 | B158 | anti-P | IgM | -0.2865144 | 0 |
| B106 | B186 | anti-P | PFC | 0.308748713 | 0 |
| B106 | B157 | anti-P | IgG | 0.273681123 | 1.33E-15 |
| B106 | B141 | anti-P | AST/ALT | -0.26503331 | 1.20E-14 |
| B106 | B181 | anti-P | APTT | -0.2352347 | 8.99E-12 |
| B106 | B154 | anti-P | RF | 0.177161149 | 3.29E-07 |
| B107 | B108 | AnuA | Anti-Cyclin | 0.999999778 | 0 |
| B107 | B109 | AnuA | AMA-M2 | 0.999999811 | 0 |
| B107 | B110 | AnuA | AHA | 0.999999833 | 0 |
| B107 | B151 | AnuA | Anti-CCP | 0.452931739 | 0 |
| B107 | B156 | AnuA | IgE | -0.34171392 | 0 |
| B107 | B158 | AnuA | IgM | -0.28651435 | 0 |
| B107 | B186 | AnuA | PFC | 0.308748659 | 0 |
| B107 | B157 | AnuA | IgG | 0.27368107 | 1.33E-15 |
| B107 | B141 | AnuA | AST/ALT | -0.26503326 | 1.20E-14 |
| B107 | B181 | AnuA | APTT | -0.23523466 | 8.99E-12 |
| B107 | B154 | AnuA | RF | 0.177161116 | 3.29E-07 |
| B108 | B110 | Anti-Cyclin | AHA | 0.999999998 | 0 |
| B108 | B151 | Anti-Cyclin | Anti-CCP | 0.452931828 | 0 |

| | | | | | |
|---|---|---|---|---|---|
| B108 | B156 | Anti-Cyclin | IgE | -0.34171399 | 0 |
| B108 | B158 | Anti-Cyclin | IgM | -0.28651441 | 0 |
| B108 | B186 | Anti-Cyclin | PFC | 0.308748714 | 0 |
| B108 | B157 | Anti-Cyclin | IgG | 0.273681124 | 1.33E-15 |
| B108 | B141 | Anti-Cyclin | AST/ALT | -0.26503332 | 1.20E-14 |
| B108 | B181 | Anti-Cyclin | APTT | -0.23523471 | 8.99E-12 |
| B108 | B154 | Anti-Cyclin | RF | 0.17716115 | 3.29E-07 |
| B109 | B110 | AMA-M2 | AHA | 1 | 0 |
| B109 | B151 | AMA-M2 | Anti-CCP | 0.452931827 | 0 |
| B109 | B156 | AMA-M2 | IgE | -0.34171399 | 0 |
| B109 | B158 | AMA-M2 | IgM | -0.28651441 | 0 |
| B109 | B186 | AMA-M2 | PFC | 0.308748713 | 0 |
| B109 | B157 | AMA-M2 | IgG | 0.273681124 | 1.33E-15 |
| B109 | B141 | AMA-M2 | AST/ALT | -0.26503332 | 1.20E-14 |
| B109 | B181 | AMA-M2 | APTT | -0.23523471 | 8.99E-12 |
| B109 | B154 | AMA-M2 | RF | 0.177161149 | 3.29E-07 |
| B11 | B12 | PGI/PGII | PG II | -0.45972514 | 0 |
| B11 | B14 | PGI/PGII | G-17 | -0.27340576 | 1.55E-15 |
| B11 | B183 | PGI/PGII | PTA | 0.213573715 | 6.50E-10 |
| B11 | B80 | PGI/PGII | anti-HP IgG | -0.17610905 | 3.87E-07 |
| B110 | B151 | AHA | Anti-CCP | 0.452931828 | 0 |
| B110 | B156 | AHA | IgE | -0.34171399 | 0 |
| B110 | B158 | AHA | IgM | -0.28651441 | 0 |
| B110 | B186 | AHA | PFC | 0.308748714 | 0 |
| B110 | B157 | AHA | IgG | 0.273681124 | 1.33E-15 |
| B110 | B141 | AHA | AST/ALT | -0.26503332 | 1.20E-14 |
| B110 | B181 | AHA | APTT | -0.23523471 | 8.99E-12 |
| B110 | B154 | AHA | RF | 0.17716115 | 3.29E-07 |
| B111 | B148 | TSH | Anti_TPO | 0.415764876 | 0 |
| B111 | B149 | TSH | TGAb | 0.216195142 | 3.96E-10 |
| B111 | B124 | TSH | ACTH | -0.19870047 | 9.55E-09 |
| B111 | B115 | TSH | TSH TT3 TT4(TT3) | -0.19361504 | 2.29E-08 |
| B112 | B113 | FT4 | FT3 | 0.572882472 | 0 |
| B112 | B114 | FT4 | TSH TT3 TT4(TT4) | 0.642482965 | 0 |
| B112 | B115 | FT4 | TSH TT3 TT4(TT3) | 0.493576043 | 0 |
| B113 | B114 | FT3 | TSH TT3 TT4(TT4) | 0.619849352 | 0 |
| B113 | B115 | FT3 | TSH TT3 TT4(TT3) | 0.811469227 | 0 |
| B114 | B115 | TSH TT3 TT4(TT4) | TSH TT3 TT4(TT3) | 0.689386864 | 0 |
| B116 | B202 | TT3 | e-times | 0.609749844 | 0 |
| B116 | B223 | TT3 | FH | -0.64051262 | 0 |
| B117 | B202 | TT4 | e-times | 0.970857462 | 0 |
| B117 | B223 | TT4 | FH | 0.941239114 | 0 |
| B118 | B189 | CEA1 | eGFB | -0.19176056 | 3.13E-08 |
| B119 | B120 | Hcv-rna | HCV-cAg | 0.297802586 | 0 |

| | | | | | |
|---|---|---|---|---|---|
| B119 | B122 | Hcv-rna | C4 | -0.99999928 | 0 |
| B119 | B125 | Hcv-rna | Cor | 0.533177151 | 0 |
| B119 | B169 | Hcv-rna | U-CRY | -0.99999281 | 0 |
| B119 | B179 | Hcv-rna | α1-MG | 0.312482516 | 0 |
| B119 | B180 | Hcv-rna | Tf | 0.195747761 | 1.59E-08 |
| B12 | B14 | PG II | G-17 | 0.250963991 | 3.04E-13 |
| B12 | B148 | PG II | Anti_TPO | 0.181187711 | 1.75E-07 |
| B120 | B121 | HCV-cAg | C3 | -0.29780338 | 0 |
| B120 | B122 | HCV-cAg | C4 | -0.29780272 | 0 |
| B120 | B159 | HCV-cAg | β2-MG | -0.29780574 | 0 |
| B120 | B169 | HCV-cAg | U-CRY | -0.29780192 | 0 |
| B121 | B122 | C3 | C4 | 0.999999385 | 0 |
| B121 | B125 | C3 | Cor | -0.53317735 | 0 |
| B121 | B159 | C3 | β2-MG | 0.99999993 | 0 |
| B121 | B169 | C3 | U-CRY | 0.99999567 | 0 |
| B121 | B179 | C3 | α1-MG | -0.31248263 | 0 |
| B121 | B180 | C3 | Tf | -0.19574784 | 1.59E-08 |
| B122 | B125 | C4 | Cor | -0.53317674 | 0 |
| B122 | B159 | C4 | β2-MG | 0.999998829 | 0 |
| B122 | B179 | C4 | α1-MG | -0.31248229 | 0 |
| B122 | B180 | C4 | Tf | -0.19574762 | 1.59E-08 |
| B123 | B148 | hs-CRP1 | Anti_TPO | 0.199231946 | 8.70E-09 |
| B124 | B155 | ACTH | IgA | -0.30816837 | 0 |
| B124 | B199 | ACTH | Mg | -0.5949463 | 0 |
| B124 | B178 | ACTH | Ig-U | 0.274666811 | 1.33E-15 |
| B124 | B179 | ACTH | α1-MG | 0.257492759 | 6.95E-14 |
| B124 | B180 | ACTH | Tf | 0.253850024 | 1.59E-13 |
| B124 | B200 | ACTH | OP | 0.230824838 | 2.23E-11 |
| B124 | B185 | ACTH | PT | -0.22507284 | 7.07E-11 |
| B124 | B148 | ACTH | Anti_TPO | -0.22235497 | 1.21E-10 |
| B124 | B186 | ACTH | PFC | 0.216986217 | 3.41E-10 |
| B124 | B195 | ACTH | Na | 0.216161439 | 3.99E-10 |
| B124 | B157 | ACTH | IgG | -0.18666879 | 7.26E-08 |
| B125 | B159 | Cor | β2-MG | -0.53317763 | 0 |
| B125 | B169 | Cor | U-CRY | -0.53317328 | 0 |
| B125 | B195 | Cor | Na | -0.2935092 | 0 |
| B125 | B199 | Cor | Mg | 0.387846119 | 0 |
| B125 | B200 | Cor | OP | -0.26830529 | 5.33E-15 |
| B125 | B157 | Cor | IgG | 0.260374535 | 3.57E-14 |
| B125 | B196 | Cor | TCO2 | -0.2331323 | 1.39E-11 |
| B125 | B139 | Cor | ADA | 0.198342664 | 1.02E-08 |
| B125 | B212 | Cor | IOP-L | -0.19491039 | 1.83E-08 |
| B126 | B127 | Alb | A-G ratio | 0.43658815 | 0 |
| B126 | B188 | Alb | Ca | 0.386149238 | 0 |

| | | | | | |
|---|---|---|---|---|---|
| B126 | B189 | Alb | eGFB | 0.193971174 | 2.15E-08 |
| B126 | B221 | Alb | Hei | 0.18382056 | 1.15E-07 |
| B127 | B130 | A-G ratio | GLB | -0.90133729 | 0 |
| B128 | B131 | ALT | AST | 0.728518447 | 0 |
| B128 | B140 | ALT | r-GT | 0.430464205 | 0 |
| B128 | B141 | ALT | AST/ALT | -0.39570698 | 0 |
| B128 | B192 | ALT | UA | 0.364067002 | 0 |
| B128 | B216 | ALT | BMI | 0.347765701 | 0 |
| B128 | B222 | ALT | NS | 0.329065963 | 0 |
| B128 | B215 | ALT | BMI | 0.272065605 | 2.22E-15 |
| B128 | B206 | ALT | Waist | 0.263817393 | 1.60E-14 |
| B128 | B221 | ALT | Hei | 0.224507276 | 7.91E-11 |
| B128 | B210 | ALT | Hip | 0.223192881 | 1.02E-10 |
| B128 | B207 | ALT | WHR | 0.202133522 | 5.23E-09 |
| B128 | B190 | ALT | Cre | 0.181366426 | 1.70E-07 |
| B129 | B132 | IBIL | Dbil | 0.685180016 | 0 |
| B129 | B133 | IBIL | TBIL | 0.981691474 | 0 |
| B131 | B140 | AST | r-GT | 0.347830673 | 0 |
| B131 | B192 | AST | UA | 0.272664383 | 1.78E-15 |
| B131 | B222 | AST | NS | 0.174944423 | 4.63E-07 |
| B132 | B133 | Dbil | TBIL | 0.811242368 | 0 |
| B132 | B184 | Dbil | INR | 0.211254533 | 1.00E-09 |
| B132 | B185 | Dbil | PT | 0.179713368 | 2.21E-07 |
| B134 | B137 | CHE | AFU | 0.40794927 | 0 |
| B134 | B138 | CHE | PAB | 0.651924016 | 0 |
| B134 | B199 | CHE | Mg | -0.26073333 | 3.29E-14 |
| B134 | B136 | CHE | 5'NT | 0.25387098 | 1.59E-13 |
| B134 | B198 | CHE | P | 0.253369987 | 1.77E-13 |
| B134 | B139 | CHE | ADA | -0.21978025 | 1.99E-10 |
| B134 | B152 | CHE | ACA-Ig(G-A-M) | -0.21337667 | 6.74E-10 |
| B134 | B183 | CHE | PTA | 0.195554462 | 1.64E-08 |
| B134 | B186 | CHE | PFC | 0.193803318 | 2.21E-08 |
| B135 | B138 | TBA | PAB | 0.267365589 | 6.88E-15 |
| B135 | B200 | TBA | OP | 0.252447376 | 2.18E-13 |
| B135 | B136 | TBA | 5'NT | 0.238376764 | 4.66E-12 |
| B135 | B198 | TBA | P | 0.223113542 | 1.04E-10 |
| B135 | B186 | TBA | PFC | 0.205236467 | 3.00E-09 |
| B136 | B138 | 5'NT | PAB | 0.372963892 | 0 |
| B136 | B193 | 5'NT | K | -0.24531006 | 1.06E-12 |
| B136 | B197 | 5'NT | AG | 0.21331316 | 6.82E-10 |
| B136 | B150 | 5'NT | ALP | 0.184568581 | 1.02E-07 |
| B136 | B199 | 5'NT | Mg | -0.16991719 | 9.89E-07 |
| B137 | B138 | AFU | PAB | 0.317972042 | 0 |
| B137 | B196 | AFU | TCO2 | 0.298735967 | 0 |

| | | | | | |
|---|---|---|---|---|---|
| B137 | B186 | AFU | PFC | 0.214344296 | 5.62E-10 |
| B137 | B197 | AFU | AG | -0.19028762 | 4.00E-08 |
| B138 | B139 | PAB | ADA | -0.24218979 | 2.07E-12 |
| B138 | B199 | PAB | Mg | -0.24103149 | 2.65E-12 |
| B138 | B186 | PAB | PFC | 0.217204349 | 3.27E-10 |
| B138 | B198 | PAB | P | 0.198403779 | 1.01E-08 |
| B138 | B181 | PAB | APTT | -0.17643596 | 3.68E-07 |
| B139 | B194 | ADA | Cl | -0.30180041 | 0 |
| B139 | B183 | ADA | PTA | -0.26215299 | 2.35E-14 |
| B139 | B157 | ADA | IgG | 0.212818676 | 7.48E-10 |
| B14 | B145 | G-17 | BGP | -0.20228219 | 5.09E-09 |
| B14 | B141 | G-17 | AST/ALT | 0.187680637 | 6.15E-08 |
| B140 | B192 | r-GT | UA | 0.320068349 | 0 |
| B140 | B173 | r-GT | NTT | 0.225241135 | 6.84E-11 |
| B140 | B222 | r-GT | NS | 0.190777165 | 3.68E-08 |
| B141 | B158 | AST/ALT | IgM | 0.460836531 | 0 |
| B141 | B222 | AST/ALT | NS | -0.2479031 | 5.99E-13 |
| B141 | B216 | AST/ALT | BMI | -0.24033865 | 3.08E-12 |
| B141 | B192 | AST/ALT | UA | -0.1975775 | 1.16E-08 |
| B141 | B215 | AST/ALT | BMI | -0.19558398 | 1.64E-08 |
| B141 | B151 | AST/ALT | Anti-CCP | -0.19153233 | 3.25E-08 |
| B141 | B206 | AST/ALT | Waist | -0.18624037 | 7.78E-08 |
| B142 | B151 | 25-(OH)D | Anti-CCP | -0.42847367 | 0 |
| B142 | B168 | 25-(OH)D | NMP22 | -0.21705518 | 3.36E-10 |
| B143 | B144 | β-CTx | Total-PINP | 0.340247327 | 0 |
| B143 | B145 | β-CTx | BGP | 0.223348833 | 9.94E-11 |
| B145 | B183 | BGP | PTA | 0.27612463 | 8.88E-16 |
| B145 | B186 | BGP | PFC | -0.23821667 | 4.82E-12 |
| B145 | B181 | BGP | APTT | 0.20741598 | 2.03E-09 |
| B146 | B189 | CysC | eGFB | -0.3059688 | 0 |
| B146 | B191 | CysC | Urea | 0.262741263 | 2.04E-14 |
| B146 | B190 | CysC | Cre | 0.250901269 | 3.08E-13 |
| B146 | B220 | CysC | SBP | 0.196191905 | 1.47E-08 |
| B148 | B149 | Anti_TPO | TGAb | 0.428318265 | 0 |
| B15 | B101 | CD3-CD19+(B cell) | anti-Sc70 | 0.692595234 | 0 |
| B15 | B102 | CD3-CD19+(B cell) | anti-Sm | 0.692595232 | 0 |
| B15 | B103 | CD3-CD19+(B cell) | anti-SSA/Ro | 0.692595227 | 0 |
| B15 | B104 | CD3-CD19+(B cell) | anti-SSB | 0.692595226 | 0 |
| B15 | B105 | CD3-CD19+(B cell) | anti-U1RNP | 0.692595228 | 0 |
| B15 | B106 | CD3-CD19+(B cell) | anti-P | 0.692595224 | 0 |
| B15 | B107 | CD3-CD19+(B cell) | AnuA | 0.692595097 | 0 |
| B15 | B108 | CD3-CD19+(B cell) | Anti-Cyclin | 0.692595229 | 0 |
| B15 | B109 | CD3-CD19+(B cell) | AMA-M2 | 0.692595227 | 0 |
| B15 | B110 | CD3-CD19+(B cell) | AHA | 0.692595229 | 0 |

| | | | | | |
|---|---|---|---|---|---|
| B15 | B141 | CD3-CD19+(B cell) | AST/ALT | -0.5366136 | 0 |
| B15 | B151 | CD3-CD19+(B cell) | Anti-CCP | 0.3867739 | 0 |
| B15 | B158 | CD3-CD19+(B cell) | IgM | -0.5234902 | 0 |
| B15 | B17 | CD3-CD19+(B cell) | CD3+CD16+CD56+NK | 0.656341752 | 0 |
| B15 | B181 | CD3-CD19+(B cell) | APTT | -0.42472342 | 0 |
| B15 | B183 | CD3-CD19+(B cell) | PTA | -0.37870934 | 0 |
| B15 | B186 | CD3-CD19+(B cell) | PFC | 0.320758848 | 0 |
| B15 | B19 | CD3-CD19+(B cell) | CD3+CD4+/CD3+CD8+ | -0.35527044 | 0 |
| B15 | B20 | CD3-CD19+(B cell) | CD3+CD4+CD8+ | 0.487537141 | 0 |
| B15 | B22 | CD3-CD19+(B cell) | CD3+CD8+(T) | 0.692102063 | 0 |
| B15 | B97 | CD3-CD19+(B cell) | dsDNA | 0.692595224 | 0 |
| B15 | B98 | CD3-CD19+(B cell) | anti-Jo-1 | 0.692595237 | 0 |
| B15 | B99 | CD3-CD19+(B cell) | anti-PM-Scl | 0.69259523 | 0 |
| B15 | B18 | CD3-CD19+(B cell) | CD3+CD4+helper(inducer)T | -0.27466122 | 1.33E-15 |
| B15 | B55 | CD3-CD19+(B cell) | TK | 0.25850083 | 5.51E-14 |
| B15 | B95 | CD3-CD19+(B cell) | anti-cenpb | -0.2251855 | 6.92E-11 |
| B15 | B193 | CD3-CD19+(B cell) | K | 0.216282486 | 3.90E-10 |
| B15 | B157 | CD3-CD19+(B cell) | IgG | 0.176708133 | 3.53E-07 |
| B15 | B156 | CD3-CD19+(B cell) | IgE | -0.17306855 | 6.16E-07 |
| B150 | B183 | ALP | PTA | -0.21506994 | 4.90E-10 |
| B151 | B168 | Anti-CCP | NMP22 | 0.291685227 | 0 |
| B151 | B158 | Anti-CCP | IgM | -0.2713119 | 2.66E-15 |
| B151 | B156 | Anti-CCP | IgE | -0.24802246 | 5.84E-13 |
| B151 | B154 | Anti-CCP | RF | 0.214706417 | 5.25E-10 |
| B151 | B176 | Anti-CCP | Ucr | -0.1838769 | 1.14E-07 |
| B152 | B181 | ACA-Ig(G-A-M) | APTT | 0.549162487 | 0 |
| B152 | B182 | ACA-Ig(G-A-M) | TT | 0.721954866 | 0 |
| B152 | B186 | ACA-Ig(G-A-M) | PFC | -0.30443682 | 0 |
| B152 | B183 | ACA-Ig(G-A-M) | PTA | -0.2467221 | 7.76E-13 |
| B153 | B170 | Glu | U-GLU | 0.58304124 | 0 |
| B153 | B220 | Glu | SBP | 0.237262556 | 5.89E-12 |
| B153 | B207 | Glu | WHR | 0.182465813 | 1.43E-07 |
| B153 | B206 | Glu | Waist | 0.170563343 | 8.98E-07 |
| B155 | B157 | IgA | IgG | 0.590326022 | 0 |
| B155 | B179 | IgA | α1-MG | -0.2958839 | 0 |
| B155 | B181 | IgA | APTT | 0.379718741 | 0 |
| B155 | B182 | IgA | TT | -0.35238061 | 0 |
| B155 | B184 | IgA | INR | 0.587771058 | 0 |
| B155 | B185 | IgA | PT | 0.659252564 | 0 |
| B155 | B186 | IgA | PFC | -0.50941347 | 0 |
| B155 | B195 | IgA | Na | -0.34247357 | 0 |
| B155 | B199 | IgA | Mg | 0.433344795 | 0 |
| B155 | B200 | IgA | OP | -0.36136877 | 0 |
| B155 | B178 | IgA | Ig-U | -0.26130702 | 2.89E-14 |

| | | | | | |
|---|---|---|---|---|---|
| B155 | B180 | IgA | Tf | -0.26000587 | 3.91E-14 |
| B155 | B183 | IgA | PTA | -0.23438403 | 1.07E-11 |
| B155 | B196 | IgA | TCO2 | -0.17974565 | 2.20E-07 |
| B156 | B184 | IgE | INR | 0.206480673 | 2.40E-09 |
| B156 | B198 | IgE | P | -0.19508017 | 1.78E-08 |
| B156 | B197 | IgE | AG | 0.172465976 | 6.75E-07 |
| B156 | B194 | IgE | Cl | -0.17004486 | 9.70E-07 |
| B157 | B182 | IgG | TT | -0.37170783 | 0 |
| B157 | B183 | IgG | PTA | -0.30281804 | 0 |
| B157 | B184 | IgG | INR | 0.553673001 | 0 |
| B157 | B185 | IgG | PT | 0.568782725 | 0 |
| B157 | B186 | IgG | PFC | -0.2599266 | 3.95E-14 |
| B157 | B199 | IgG | Mg | 0.239953747 | 3.34E-12 |
| B157 | B200 | IgG | OP | -0.1981217 | 1.06E-08 |
| B157 | B181 | IgG | APTT | 0.1823226 | 1.46E-07 |
| B157 | B195 | IgG | Na | -0.17756665 | 3.09E-07 |
| B158 | B184 | IgM | INR | -0.22582581 | 6.09E-11 |
| B158 | B185 | IgM | PT | -0.18849272 | 5.38E-08 |
| B158 | B193 | IgM | K | -0.17985203 | 2.16E-07 |
| B159 | B179 | β2-MG | α1-MG | -0.31248279 | 0 |
| B159 | B180 | β2-MG | Tf | -0.19574793 | 1.59E-08 |
| B16 | B100 | CD3+T | anti-Ro52 | 0.561504451 | 0 |
| B16 | B156 | CD3+T | IgE | 0.742877672 | 0 |
| B16 | B17 | CD3+T | CD3+CD16+CD56+NK | -0.59713082 | 0 |
| B16 | B18 | CD3+T | CD3+CD4+helper(inducer)T | 0.803005077 | 0 |
| B16 | B19 | CD3+T | CD3+CD4+/CD3+CD8+ | 0.790316618 | 0 |
| B16 | B21 | CD3+T | CD3+CD4-CD8- | -0.44642397 | 0 |
| B16 | B22 | CD3+T | CD3+CD8+(T) | -0.49572981 | 0 |
| B16 | B23 | CD3+T | CD3-CD16+CD56+(NK) | -0.94701323 | 0 |
| B16 | B92 | CD3+T | ALDH2-Glu504Glu | -0.7114502 | 0 |
| B16 | B93 | CD3+T | ALDH2-Glu504Lys | 0.711450199 | 0 |
| B16 | B94 | CD3+T | ALDH2-Lys504Lys | -0.71145019 | 0 |
| B16 | B95 | CD3+T | anti-cenpb | 0.564875211 | 0 |
| B16 | B20 | CD3+T | CD3+CD4+CD8+ | -0.25748003 | 6.97E-14 |
| B16 | B85 | CD3+T | SCC | 0.225537508 | 6.45E-11 |
| B16 | B101 | CD3+T | anti-Sc70 | -0.21249044 | 7.95E-10 |
| B16 | B102 | CD3+T | anti-Sm | -0.21249043 | 7.95E-10 |
| B16 | B103 | CD3+T | anti-SSA/Ro | -0.21249042 | 7.95E-10 |
| B16 | B104 | CD3+T | anti-SSB | -0.21249042 | 7.95E-10 |
| B16 | B105 | CD3+T | anti-U1RNP | -0.21249043 | 7.95E-10 |
| B16 | B106 | CD3+T | anti-P | -0.21249042 | 7.95E-10 |
| B16 | B107 | CD3+T | AnuA | -0.21249042 | 7.95E-10 |
| B16 | B108 | CD3+T | Anti-Cyclin | -0.21249043 | 7.95E-10 |
| B16 | B109 | CD3+T | AMA-M2 | -0.21249042 | 7.95E-10 |

| | | | | | |
|---|---|---|---|---|---|
| B16 | B110 | CD3+T | AHA | -0.21249043 | 7.95E-10 |
| B16 | B97 | CD3+T | dsDNA | -0.21249042 | 7.95E-10 |
| B16 | B98 | CD3+T | anti-Jo-1 | -0.21249043 | 7.95E-10 |
| B16 | B99 | CD3+T | anti-PM-Scl | -0.21249043 | 7.95E-10 |
| B16 | B141 | CD3+T | AST/ALT | -0.20291814 | 4.55E-09 |
| B16 | B69 | CD3+T | PV | -0.18817357 | 5.67E-08 |
| B16 | B186 | CD3+T | PFC | 0.183089434 | 1.29E-07 |
| B16 | B151 | CD3+T | Anti-CCP | -0.18014876 | 2.07E-07 |
| B160 | B163 | PH | SG | -0.18889114 | 5.04E-08 |
| B161 | B173 | LEU | NTT | 0.185362352 | 8.98E-08 |
| B163 | B166 | SG | Pro | 0.403971849 | 0 |
| B163 | B176 | SG | Ucr | 0.312828802 | 0 |
| B163 | B221 | SG | Hei | 0.33318889 | 0 |
| B163 | B222 | SG | NS | 0.315752831 | 0 |
| B163 | B190 | SG | Cre | 0.275958301 | 8.88E-16 |
| B163 | B216 | SG | BMI | 0.263300124 | 1.80E-14 |
| B163 | B191 | SG | Urea | 0.224828891 | 7.42E-11 |
| B163 | B192 | SG | UA | 0.217080838 | 3.35E-10 |
| B166 | B172 | Pro | KB | 0.188989862 | 4.96E-08 |
| B166 | B222 | Pro | NS | 0.178882525 | 2.52E-07 |
| B166 | B190 | Pro | Cre | 0.172847972 | 6.37E-07 |
| B168 | B183 | NMP22 | PTA | -0.19256574 | 2.73E-08 |
| B169 | B179 | U-CRY | α1-MG | -0.31248026 | 0 |
| B169 | B180 | U-CRY | Tf | -0.19574633 | 1.59E-08 |
| B17 | B100 | CD3+CD16+CD56+NK | anti-Ro52 | -0.63807499 | 0 |
| B17 | B101 | CD3+CD16+CD56+NK | anti-Sc70 | 0.742067595 | 0 |
| B17 | B102 | CD3+CD16+CD56+NK | anti-Sm | 0.742067592 | 0 |
| B17 | B103 | CD3+CD16+CD56+NK | anti-SSA/Ro | 0.742067587 | 0 |
| B17 | B104 | CD3+CD16+CD56+NK | anti-SSB | 0.742067584 | 0 |
| B17 | B105 | CD3+CD16+CD56+NK | anti-U1RNP | 0.742067587 | 0 |
| B17 | B106 | CD3+CD16+CD56+NK | anti-P | 0.742067583 | 0 |
| B17 | B107 | CD3+CD16+CD56+NK | AnuA | 0.742067445 | 0 |
| B17 | B108 | CD3+CD16+CD56+NK | Anti-Cyclin | 0.742067589 | 0 |
| B17 | B109 | CD3+CD16+CD56+NK | AMA-M2 | 0.742067587 | 0 |
| B17 | B110 | CD3+CD16+CD56+NK | AHA | 0.742067589 | 0 |
| B17 | B151 | CD3+CD16+CD56+NK | Anti-CCP | 0.458408632 | 0 |
| B17 | B156 | CD3+CD16+CD56+NK | IgE | -0.63429928 | 0 |
| B17 | B158 | CD3+CD16+CD56+NK | IgM | -0.33920453 | 0 |
| B17 | B18 | CD3+CD16+CD56+NK | CD3+CD4+helper(inducer)T | -0.86898465 | 0 |
| B17 | B181 | CD3+CD16+CD56+NK | APTT | -0.29246986 | 0 |
| B17 | B19 | CD3+CD16+CD56+NK | CD3+CD4+/CD3+CD8+ | -0.83450327 | 0 |
| B17 | B20 | CD3+CD16+CD56+NK | CD3+CD4+CD8+ | 0.402529282 | 0 |
| B17 | B21 | CD3+CD16+CD56+NK | CD3+CD4-CD8- | 0.550645247 | 0 |
| B17 | B22 | CD3+CD16+CD56+NK | CD3+CD8+(T) | 0.871764442 | 0 |

| | | | | | |
|---|---|---|---|---|---|
| B17 | B23 | CD3+CD16+CD56+NK | CD3-CD16+CD56+(NK) | 0.573011532 | 0 |
| B17 | B92 | CD3+CD16+CD56+NK | ALDH2-Glu504Glu | 0.510252497 | 0 |
| B17 | B93 | CD3+CD16+CD56+NK | ALDH2-Glu504Lys | -0.5102525 | 0 |
| B17 | B94 | CD3+CD16+CD56+NK | ALDH2-Lys504Lys | 0.51025249 | 0 |
| B17 | B95 | CD3+CD16+CD56+NK | anti-cenpb | -0.68908313 | 0 |
| B17 | B97 | CD3+CD16+CD56+NK | dsDNA | 0.742067583 | 0 |
| B17 | B98 | CD3+CD16+CD56+NK | anti-Jo-1 | 0.742067598 | 0 |
| B17 | B99 | CD3+CD16+CD56+NK | anti-PM-Scl | 0.74206759 | 0 |
| B17 | B55 | CD3+CD16+CD56+NK | TK | 0.26175026 | 2.60E-14 |
| B17 | B186 | CD3+CD16+CD56+NK | PFC | 0.248067312 | 5.78E-13 |
| B17 | B183 | CD3+CD16+CD56+NK | PTA | -0.24793828 | 5.94E-13 |
| B17 | B59 | CD3+CD16+CD56+NK | WBRV | 0.1853922 | 8.94E-08 |
| B17 | B157 | CD3+CD16+CD56+NK | IgG | 0.181790569 | 1.59E-07 |
| B17 | B193 | CD3+CD16+CD56+NK | K | 0.178584035 | 2.64E-07 |
| B174 | B190 | Urea/Cr | Cre | -0.48198546 | 0 |
| B174 | B191 | Urea/Cr | Urea | 0.603383404 | 0 |
| B174 | B221 | Urea/Cr | Hei | -0.36007544 | 0 |
| B174 | B216 | Urea/Cr | BMI | -0.26048037 | 3.49E-14 |
| B174 | B222 | Urea/Cr | NS | -0.25951171 | 4.35E-14 |
| B174 | B192 | Urea/Cr | UA | -0.21544819 | 4.56E-10 |
| B175 | B177 | ALB-U | ACR | 0.850583685 | 0 |
| B178 | B179 | Ig-U | α1-MG | 0.929890463 | 0 |
| B178 | B180 | Ig-U | Tf | 0.992488376 | 0 |
| B178 | B199 | Ig-U | Mg | -0.4302167 | 0 |
| B179 | B180 | α1-MG | Tf | 0.926041482 | 0 |
| B179 | B199 | α1-MG | Mg | -0.41886217 | 0 |
| B18 | B100 | CD3+CD4+helper(inducer)T | anti-Ro52 | 0.485675831 | 0 |
| B18 | B101 | CD3+CD4+helper(inducer)T | anti-Sc70 | -0.58699577 | 0 |
| B18 | B102 | CD3+CD4+helper(inducer)T | anti-Sm | -0.58699577 | 0 |
| B18 | B103 | CD3+CD4+helper(inducer)T | anti-SSA/Ro | -0.58699577 | 0 |
| B18 | B104 | CD3+CD4+helper(inducer)T | anti-SSB | -0.58699576 | 0 |
| B18 | B105 | CD3+CD4+helper(inducer)T | anti-U1RNP | -0.58699577 | 0 |
| B18 | B106 | CD3+CD4+helper(inducer)T | anti-P | -0.58699576 | 0 |
| B18 | B107 | CD3+CD4+helper(inducer)T | AnuA | -0.58699566 | 0 |
| B18 | B108 | CD3+CD4+helper(inducer)T | Anti-Cyclin | -0.58699577 | 0 |
| B18 | B109 | CD3+CD4+helper(inducer)T | AMA-M2 | -0.58699577 | 0 |
| B18 | B110 | CD3+CD4+helper(inducer)T | AHA | -0.58699577 | 0 |
| B18 | B151 | CD3+CD4+helper(inducer)T | Anti-CCP | -0.38776215 | 0 |
| B18 | B156 | CD3+CD4+helper(inducer)T | IgE | 0.719177507 | 0 |
| B18 | B19 | CD3+CD4+helper(inducer)T | CD3+CD4+/CD3+CD8+ | 0.959706032 | 0 |
| B18 | B20 | CD3+CD4+helper(inducer)T | CD3+CD4+CD8+ | -0.42334098 | 0 |
| B18 | B21 | CD3+CD4+helper(inducer)T | CD3+CD4-CD8- | -0.74818933 | 0 |
| B18 | B22 | CD3+CD4+helper(inducer)T | CD3+CD8+(T) | -0.81648698 | 0 |
| B18 | B23 | CD3+CD4+helper(inducer)T | CD3-CD16+CD56+(NK) | -0.82705391 | 0 |

| | | | | | |
|---|---|---|---|---|---|
| B18 | B92 | CD3+CD4+helper(inducer)T | ALDH2-Glu504Glu | -0.76606165 | 0 |
| B18 | B93 | CD3+CD4+helper(inducer)T | ALDH2-Glu504Lys | 0.766061646 | 0 |
| B18 | B94 | CD3+CD4+helper(inducer)T | ALDH2-Lys504Lys | -0.76606163 | 0 |
| B18 | B95 | CD3+CD4+helper(inducer)T | anti-cenpb | 0.526589976 | 0 |
| B18 | B97 | CD3+CD4+helper(inducer)T | dsDNA | -0.58699576 | 0 |
| B18 | B98 | CD3+CD4+helper(inducer)T | anti-Jo-1 | -0.58699578 | 0 |
| B18 | B99 | CD3+CD4+helper(inducer)T | anti-PM-Scl | -0.58699577 | 0 |
| B18 | B55 | CD3+CD4+helper(inducer)T | TK | -0.24892907 | 4.78E-13 |
| B18 | B85 | CD3+CD4+helper(inducer)T | SCC | 0.240637468 | 2.89E-12 |
| B18 | B59 | CD3+CD4+helper(inducer)T | WBRV | -0.17953009 | 2.28E-07 |
| B18 | B155 | CD3+CD4+helper(inducer)T | IgA | 0.177637164 | 3.06E-07 |
| B18 | B186 | CD3+CD4+helper(inducer)T | PFC | -0.17647233 | 3.66E-07 |
| B180 | B199 | Tf | Mg | -0.40362029 | 0 |
| B181 | B182 | APTT | TT | 0.301231001 | 0 |
| B181 | B183 | APTT | PTA | 0.326110245 | 0 |
| B181 | B184 | APTT | INR | 0.466797159 | 0 |
| B181 | B185 | APTT | PT | 0.666849473 | 0 |
| B181 | B186 | APTT | PFC | -0.79179173 | 0 |
| B181 | B193 | APTT | K | -0.20847811 | 1.67E-09 |
| B181 | B199 | APTT | Mg | 0.181838951 | 1.58E-07 |
| B182 | B184 | TT | INR | -0.62607168 | 0 |
| B182 | B185 | TT | PT | -0.36564761 | 0 |
| B183 | B186 | PTA | PFC | -0.2941229 | 0 |
| B183 | B199 | PTA | Mg | -0.26258913 | 2.13E-14 |
| B183 | B195 | PTA | Na | 0.220674947 | 1.68E-10 |
| B183 | B196 | PTA | TCO2 | 0.21338761 | 6.73E-10 |
| B183 | B200 | PTA | OP | 0.180910667 | 1.83E-07 |
| B184 | B185 | INR | PT | 0.922427885 | 0 |
| B184 | B186 | INR | PFC | -0.5255441 | 0 |
| B185 | B186 | PT | PFC | -0.67512296 | 0 |
| B185 | B199 | PT | Mg | 0.327045776 | 0 |
| B186 | B199 | PFC | Mg | -0.28531467 | 0 |
| B186 | B200 | PFC | OP | 0.26394803 | 1.55E-14 |
| B186 | B195 | PFC | Na | 0.186628813 | 7.31E-08 |
| B188 | B192 | Ca | UA | 0.171320614 | 8.02E-07 |
| B189 | B190 | eGFB | Cre | -0.59203061 | 0 |
| B189 | B191 | eGFB | Urea | -0.36676417 | 0 |
| B189 | B207 | eGFB | WHR | -0.29718052 | 0 |
| B189 | B220 | eGFB | SBP | -0.32385679 | 0 |
| B189 | B206 | eGFB | Waist | -0.22074121 | 1.65E-10 |
| B189 | B222 | eGFB | NS | -0.1733424 | 5.91E-07 |
| B19 | B100 | CD3+CD4+/CD3+CD8+ | anti-Ro52 | 0.339810957 | 0 |
| B19 | B101 | CD3+CD4+/CD3+CD8+ | anti-Sc70 | -0.60642117 | 0 |
| B19 | B102 | CD3+CD4+/CD3+CD8+ | anti-Sm | -0.60642116 | 0 |

| | | | | | |
|---|---|---|---|---|---|
| B19 | B103 | CD3+CD4+/CD3+CD8+ | anti-SSA/Ro | -0.60642116 | 0 |
| B19 | B104 | CD3+CD4+/CD3+CD8+ | anti-SSB | -0.60642116 | 0 |
| B19 | B105 | CD3+CD4+/CD3+CD8+ | anti-U1RNP | -0.60642116 | 0 |
| B19 | B106 | CD3+CD4+/CD3+CD8+ | anti-P | -0.60642116 | 0 |
| B19 | B107 | CD3+CD4+/CD3+CD8+ | AnuA | -0.60642104 | 0 |
| B19 | B108 | CD3+CD4+/CD3+CD8+ | Anti-Cyclin | -0.60642116 | 0 |
| B19 | B109 | CD3+CD4+/CD3+CD8+ | AMA-M2 | -0.60642116 | 0 |
| B19 | B110 | CD3+CD4+/CD3+CD8+ | AHA | -0.60642116 | 0 |
| B19 | B151 | CD3+CD4+/CD3+CD8+ | Anti-CCP | -0.40868358 | 0 |
| B19 | B156 | CD3+CD4+/CD3+CD8+ | IgE | 0.731968415 | 0 |
| B19 | B20 | CD3+CD4+/CD3+CD8+ | CD3+CD4+CD8+ | -0.63636062 | 0 |
| B19 | B21 | CD3+CD4+/CD3+CD8+ | CD3+CD4-CD8- | -0.56086303 | 0 |
| B19 | B22 | CD3+CD4+/CD3+CD8+ | CD3+CD8+(T) | -0.89765125 | 0 |
| B19 | B23 | CD3+CD4+/CD3+CD8+ | CD3-CD16+CD56+(NK) | -0.80489747 | 0 |
| B19 | B92 | CD3+CD4+/CD3+CD8+ | ALDH2-Glu504Glu | -0.89064576 | 0 |
| B19 | B93 | CD3+CD4+/CD3+CD8+ | ALDH2-Glu504Lys | 0.890645762 | 0 |
| B19 | B94 | CD3+CD4+/CD3+CD8+ | ALDH2-Lys504Lys | -0.89064575 | 0 |
| B19 | B95 | CD3+CD4+/CD3+CD8+ | anti-cenpb | 0.386882302 | 0 |
| B19 | B97 | CD3+CD4+/CD3+CD8+ | dsDNA | -0.60642116 | 0 |
| B19 | B98 | CD3+CD4+/CD3+CD8+ | anti-Jo-1 | -0.60642117 | 0 |
| B19 | B99 | CD3+CD4+/CD3+CD8+ | anti-PM-Scl | -0.60642116 | 0 |
| B19 | B85 | CD3+CD4+/CD3+CD8+ | SCC | 0.280241847 | 4.44E-16 |
| B19 | B55 | CD3+CD4+/CD3+CD8+ | TK | -0.22609291 | 5.78E-11 |
| B19 | B59 | CD3+CD4+/CD3+CD8+ | WBRV | -0.18253137 | 1.42E-07 |
| B19 | B196 | CD3+CD4+/CD3+CD8+ | TCO2 | -0.17917298 | 2.41E-07 |
| B190 | B191 | Cre | Urea | 0.35595249 | 0 |
| B190 | B192 | Cre | UA | 0.488643912 | 0 |
| B190 | B206 | Cre | Waist | 0.348360909 | 0 |
| B190 | B207 | Cre | WHR | 0.348995231 | 0 |
| B190 | B216 | Cre | BMI | 0.371038924 | 0 |
| B190 | B221 | Cre | Hei | 0.407376871 | 0 |
| B190 | B222 | Cre | NS | 0.509506629 | 0 |
| B190 | B204 | Cre | Drinking | 0.223717436 | 9.24E-11 |
| B190 | B210 | Cre | Hip | 0.203241863 | 4.29E-09 |
| B191 | B192 | Urea | UA | 0.178130326 | 2.83E-07 |
| B191 | B207 | Urea | WHR | 0.175836881 | 4.04E-07 |
| B192 | B206 | UA | Waist | 0.400448435 | 0 |
| B192 | B207 | UA | WHR | 0.331050202 | 0 |
| B192 | B210 | UA | Hip | 0.319864872 | 0 |
| B192 | B215 | UA | BMI | 0.323547084 | 0 |
| B192 | B216 | UA | BMI | 0.482827174 | 0 |
| B192 | B221 | UA | Hei | 0.396084032 | 0 |
| B192 | B222 | UA | NS | 0.498015442 | 0 |
| B192 | B218 | UA | DBP | 0.209394918 | 1.41E-09 |

| | | | | | |
|---|---|---|---|---|---|
| B192 | B204 | UA | Drinking | 0.191691524 | 3.16E-08 |
| B193 | B194 | K | Cl | -0.29940471 | 0 |
| B193 | B195 | K | Na | -0.28678888 | 0 |
| B193 | B200 | K | OP | -0.28858629 | 0 |
| B194 | B197 | Cl | AG | -0.30894644 | 0 |
| B194 | B200 | Cl | OP | 0.243584663 | 1.53E-12 |
| B194 | B195 | Cl | Na | 0.237745875 | 5.32E-12 |
| B194 | B199 | Cl | Mg | -0.18904309 | 4.91E-08 |
| B195 | B196 | Na | TCO2 | 0.466173366 | 0 |
| B195 | B200 | Na | OP | 0.956117947 | 0 |
| B195 | B197 | Na | AG | 0.223712998 | 9.25E-11 |
| B195 | B199 | Na | Mg | -0.20892979 | 1.54E-09 |
| B196 | B197 | TCO2 | AG | -0.59172863 | 0 |
| B196 | B200 | TCO2 | OP | 0.396353294 | 0 |
| B196 | B199 | TCO2 | Mg | -0.26352993 | 1.71E-14 |
| B197 | B200 | AG | OP | 0.255637036 | 1.06E-13 |
| B197 | B199 | AG | Mg | 0.212308583 | 8.23E-10 |
| B199 | B200 | Mg | OP | -0.2230835 | 1.05E-10 |
| B2 | B126 | Age | Alb | -0.33547656 | 0 |
| B2 | B189 | Age | eGFB | -0.64552045 | 0 |
| B2 | B191 | Age | Urea | 0.305901918 | 0 |
| B2 | B207 | Age | WHR | 0.31678284 | 0 |
| B2 | B220 | Age | SBP | 0.513363399 | 0 |
| B2 | B221 | Age | Hei | -0.29701143 | 0 |
| B2 | B47 | Age | RDW | 0.327865726 | 0 |
| B2 | B153 | Age | Glu | 0.268685375 | 4.88E-15 |
| B2 | B146 | Age | CysC | 0.225659154 | 6.30E-11 |
| B2 | B206 | Age | Waist | 0.201318132 | 6.04E-09 |
| B2 | B52 | Age | PLT/BPC | -0.19902365 | 9.03E-09 |
| B2 | B77 | Age | TC | 0.198650138 | 9.63E-09 |
| B2 | B46 | Age | RBC Count | -0.18463076 | 1.01E-07 |
| B2 | B118 | Age | CEA1 | 0.183934731 | 1.13E-07 |
| B2 | B190 | Age | Cre | 0.179367584 | 2.34E-07 |
| B2 | B201 | Age | e-habits | 0.178643946 | 2.62E-07 |
| B2 | B219 | Age | OT | 0.177434888 | 3.16E-07 |
| B2 | B41 | Age | MON％ | 0.174968364 | 4.62E-07 |
| B2 | B91 | Age | TP | -0.17359243 | 5.69E-07 |
| B20 | B101 | CD3+CD4+CD8+ | anti-Sc70 | 0.526944073 | 0 |
| B20 | B102 | CD3+CD4+CD8+ | anti-Sm | 0.526944072 | 0 |
| B20 | B103 | CD3+CD4+CD8+ | anti-SSA/Ro | 0.526944068 | 0 |
| B20 | B104 | CD3+CD4+CD8+ | anti-SSB | 0.526944067 | 0 |
| B20 | B105 | CD3+CD4+CD8+ | anti-U1RNP | 0.526944068 | 0 |
| B20 | B106 | CD3+CD4+CD8+ | anti-P | 0.526944066 | 0 |
| B20 | B107 | CD3+CD4+CD8+ | AnuA | 0.526943969 | 0 |

| | | | | | |
|---|---|---|---|---|---|
| B20 | B108 | CD3+CD4+CD8+ | Anti-Cyclin | 0.52694407 | 0 |
| B20 | B109 | CD3+CD4+CD8+ | AMA-M2 | 0.526944068 | 0 |
| B20 | B110 | CD3+CD4+CD8+ | AHA | 0.526944069 | 0 |
| B20 | B141 | CD3+CD4+CD8+ | AST/ALT | -0.48444422 | 0 |
| B20 | B151 | CD3+CD4+CD8+ | Anti-CCP | 0.402560588 | 0 |
| B20 | B156 | CD3+CD4+CD8+ | IgE | -0.35265467 | 0 |
| B20 | B22 | CD3+CD4+CD8+ | CD3+CD8+(T) | 0.785972976 | 0 |
| B20 | B23 | CD3+CD4+CD8+ | CD3-CD16+CD56+(NK) | 0.354776708 | 0 |
| B20 | B92 | CD3+CD4+CD8+ | ALDH2-Glu504Glu | 0.733264354 | 0 |
| B20 | B93 | CD3+CD4+CD8+ | ALDH2-Glu504Lys | -0.73326435 | 0 |
| B20 | B94 | CD3+CD4+CD8+ | ALDH2-Lys504Lys | 0.733264344 | 0 |
| B20 | B97 | CD3+CD4+CD8+ | dsDNA | 0.526944066 | 0 |
| B20 | B98 | CD3+CD4+CD8+ | anti-Jo-1 | 0.526944076 | 0 |
| B20 | B99 | CD3+CD4+CD8+ | anti-PM-Scl | 0.52694407 | 0 |
| B20 | B85 | CD3+CD4+CD8+ | SCC | -0.23055079 | 2.36E-11 |
| B20 | B158 | CD3+CD4+CD8+ | IgM | -0.20452012 | 3.42E-09 |
| B20 | B196 | CD3+CD4+CD8+ | TCO2 | 0.18961768 | 4.47E-08 |
| B202 | B225 | e-times | hPBG | 0.755318632 | 0 |
| B204 | B209 | Drinking | Smoking | 0.305750751 | 0 |
| B204 | B222 | Drinking | NS | 0.265736577 | 1.02E-14 |
| B204 | B221 | Drinking | Hei | 0.264994892 | 1.20E-14 |
| B204 | B216 | Drinking | BMI | 0.229468535 | 2.93E-11 |
| B204 | B207 | Drinking | WHR | 0.192284504 | 2.86E-08 |
| B204 | B206 | Drinking | Waist | 0.184632094 | 1.01E-07 |
| B206 | B207 | Waist | WHR | 0.850521347 | 0 |
| B206 | B210 | Waist | Hip | 0.762675767 | 0 |
| B206 | B215 | Waist | BMI | 0.822259367 | 0 |
| B206 | B216 | Waist | BMI | 0.810441512 | 0 |
| B206 | B220 | Waist | SBP | 0.297397685 | 0 |
| B206 | B221 | Waist | Hei | 0.322394261 | 0 |
| B206 | B222 | Waist | NS | 0.767763255 | 0 |
| B206 | B218 | Waist | DBP | 0.276008589 | 8.88E-16 |
| B207 | B210 | WHR | Hip | 0.316532466 | 0 |
| B207 | B215 | WHR | BMI | 0.577178879 | 0 |
| B207 | B216 | WHR | BMI | 0.538687645 | 0 |
| B207 | B218 | WHR | DBP | 0.284597956 | 0 |
| B207 | B220 | WHR | SBP | 0.309102823 | 0 |
| B207 | B222 | WHR | NS | 0.613819379 | 0 |
| B207 | B221 | WHR | Hei | 0.173700801 | 5.60E-07 |
| B209 | B222 | Smoking | NS | 0.173925605 | 5.41E-07 |
| B209 | B221 | Smoking | Hei | 0.171626227 | 7.66E-07 |
| B21 | B100 | CD3+CD4-CD8- | anti-Ro52 | -0.35150276 | 0 |
| B21 | B101 | CD3+CD4-CD8- | anti-Sc70 | 0.332420734 | 0 |
| B21 | B102 | CD3+CD4-CD8- | anti-Sm | 0.332420733 | 0 |

| | | | | | |
|---|---|---|---|---|---|
| B21 | B103 | CD3+CD4-CD8- | anti-SSA/Ro | 0.332420731 | 0 |
| B21 | B104 | CD3+CD4-CD8- | anti-SSB | 0.33242073 | 0 |
| B21 | B105 | CD3+CD4-CD8- | anti-U1RNP | 0.332420731 | 0 |
| B21 | B106 | CD3+CD4-CD8- | anti-P | 0.332420729 | 0 |
| B21 | B107 | CD3+CD4-CD8- | AnuA | 0.332420666 | 0 |
| B21 | B108 | CD3+CD4-CD8- | Anti-Cyclin | 0.332420732 | 0 |
| B21 | B109 | CD3+CD4-CD8- | AMA-M2 | 0.332420731 | 0 |
| B21 | B110 | CD3+CD4-CD8- | AHA | 0.332420732 | 0 |
| B21 | B141 | CD3+CD4-CD8- | AST/ALT | 0.517647567 | 0 |
| B21 | B156 | CD3+CD4-CD8- | IgE | -0.37433077 | 0 |
| B21 | B186 | CD3+CD4-CD8- | PFC | 0.330920899 | 0 |
| B21 | B22 | CD3+CD4-CD8- | CD3+CD8+(T) | 0.369027011 | 0 |
| B21 | B23 | CD3+CD4-CD8- | CD3-CD16+CD56+(NK) | 0.518649374 | 0 |
| B21 | B55 | CD3+CD4-CD8- | TK | 0.306949302 | 0 |
| B21 | B92 | CD3+CD4-CD8- | ALDH2-Glu504Glu | 0.338103263 | 0 |
| B21 | B93 | CD3+CD4-CD8- | ALDH2-Glu504Lys | -0.33810326 | 0 |
| B21 | B94 | CD3+CD4-CD8- | ALDH2-Lys504Lys | 0.338103258 | 0 |
| B21 | B95 | CD3+CD4-CD8- | anti-cenpb | -0.3724049 | 0 |
| B21 | B97 | CD3+CD4-CD8- | dsDNA | 0.332420729 | 0 |
| B21 | B98 | CD3+CD4-CD8- | anti-Jo-1 | 0.332420735 | 0 |
| B21 | B99 | CD3+CD4-CD8- | anti-PM-Scl | 0.332420732 | 0 |
| B21 | B155 | CD3+CD4-CD8- | IgA | -0.28026293 | 4.44E-16 |
| B21 | B158 | CD3+CD4-CD8- | IgM | 0.224347502 | 8.16E-11 |
| B21 | B181 | CD3+CD4-CD8- | APTT | -0.20902531 | 1.51E-09 |
| B21 | B56 | CD3+CD4-CD8- | IR | 0.193542455 | 2.31E-08 |
| B21 | B151 | CD3+CD4-CD8- | Anti-CCP | 0.18973338 | 4.38E-08 |
| B21 | B200 | CD3+CD4-CD8- | OP | 0.185047101 | 9.45E-08 |
| B21 | B61 | CD3+CD4-CD8- | WBHSRV | 0.176920553 | 3.42E-07 |
| B210 | B215 | Hip | BMI | 0.783265733 | 0 |
| B210 | B216 | Hip | BMI | 0.801876237 | 0 |
| B210 | B221 | Hip | Hei | 0.363581269 | 0 |
| B210 | B222 | Hip | NS | 0.636371287 | 0 |
| B211 | B212 | IOP-R | IOP-L | 0.803058931 | 0 |
| B215 | B216 | BMI | BMI | 0.801789017 | 0 |
| B215 | B222 | BMI | NS | 0.655226892 | 0 |
| B215 | B220 | BMI | SBP | 0.22964437 | 2.83E-11 |
| B215 | B218 | BMI | DBP | 0.197709131 | 1.13E-08 |
| B216 | B221 | BMI | Hei | 0.662789429 | 0 |
| B216 | B222 | BMI | NS | 0.808497203 | 0 |
| B216 | B218 | BMI | DBP | 0.196231625 | 1.46E-08 |
| B218 | B220 | DBP | SBP | 0.592801233 | 0 |
| B218 | B222 | DBP | NS | 0.222444548 | 1.19E-10 |
| B22 | B101 | CD3+CD8+(T) | anti-Sc70 | 0.782471377 | 0 |
| B22 | B102 | CD3+CD8+(T) | anti-Sm | 0.782471376 | 0 |

| | | | | | |
|---|---|---|---|---|---|
| B22 | B103 | CD3+CD8+(T) | anti-SSA/Ro | 0.78247137 | 0 |
| B22 | B104 | CD3+CD8+(T) | anti-SSB | 0.782471367 | 0 |
| B22 | B105 | CD3+CD8+(T) | anti-U1RNP | 0.78247137 | 0 |
| B22 | B106 | CD3+CD8+(T) | anti-P | 0.782471365 | 0 |
| B22 | B107 | CD3+CD8+(T) | AnuA | 0.782471221 | 0 |
| B22 | B108 | CD3+CD8+(T) | Anti-Cyclin | 0.782471371 | 0 |
| B22 | B109 | CD3+CD8+(T) | AMA-M2 | 0.782471369 | 0 |
| B22 | B110 | CD3+CD8+(T) | AHA | 0.782471371 | 0 |
| B22 | B141 | CD3+CD8+(T) | AST/ALT | -0.28957366 | 0 |
| B22 | B151 | CD3+CD8+(T) | Anti-CCP | 0.515287985 | 0 |
| B22 | B156 | CD3+CD8+(T) | IgE | -0.58707413 | 0 |
| B22 | B23 | CD3+CD8+(T) | CD3-CD16+CD56+(NK) | 0.539922124 | 0 |
| B22 | B55 | CD3+CD8+(T) | TK | 0.293448439 | 0 |
| B22 | B92 | CD3+CD8+(T) | ALDH2-Glu504Glu | 0.739655227 | 0 |
| B22 | B93 | CD3+CD8+(T) | ALDH2-Glu504Lys | -0.73965523 | 0 |
| B22 | B94 | CD3+CD8+(T) | ALDH2-Lys504Lys | 0.739655216 | 0 |
| B22 | B95 | CD3+CD8+(T) | anti-cenpb | -0.34271151 | 0 |
| B22 | B97 | CD3+CD8+(T) | dsDNA | 0.782471366 | 0 |
| B22 | B98 | CD3+CD8+(T) | anti-Jo-1 | 0.782471379 | 0 |
| B22 | B99 | CD3+CD8+(T) | anti-PM-Scl | 0.782471372 | 0 |
| B22 | B158 | CD3+CD8+(T) | IgM | -0.27914383 | 4.44E-16 |
| B22 | B100 | CD3+CD8+(T) | anti-Ro52 | -0.27719143 | 6.66E-16 |
| B22 | B186 | CD3+CD8+(T) | PFC | 0.23943537 | 3.73E-12 |
| B22 | B85 | CD3+CD8+(T) | SCC | -0.23101023 | 2.14E-11 |
| B22 | B59 | CD3+CD8+(T) | WBRV | 0.198183814 | 1.04E-08 |
| B22 | B181 | CD3+CD8+(T) | APTT | -0.19561774 | 1.63E-08 |
| B22 | B61 | CD3+CD8+(T) | WBHSRV | 0.172563541 | 6.65E-07 |
| B220 | B222 | SBP | NS | 0.2028551 | 4.60E-09 |
| B221 | B222 | Hei | NS | 0.523954494 | 0 |
| B223 | B225 | FH | hPBG | 0.825566644 | 0 |
| B23 | B100 | CD3-CD16+CD56+(NK) | anti-Ro52 | -0.54249207 | 0 |
| B23 | B156 | CD3-CD16+CD56+(NK) | IgE | -0.69986192 | 0 |
| B23 | B92 | CD3-CD16+CD56+(NK) | ALDH2-Glu504Glu | 0.721660028 | 0 |
| B23 | B93 | CD3-CD16+CD56+(NK) | ALDH2-Glu504Lys | -0.72166003 | 0 |
| B23 | B94 | CD3-CD16+CD56+(NK) | ALDH2-Lys504Lys | 0.721660016 | 0 |
| B23 | B95 | CD3-CD16+CD56+(NK) | anti-cenpb | -0.55214217 | 0 |
| B23 | B101 | CD3-CD16+CD56+(NK) | anti-Sc70 | 0.273136441 | 1.78E-15 |
| B23 | B102 | CD3-CD16+CD56+(NK) | anti-Sm | 0.273136441 | 1.78E-15 |
| B23 | B103 | CD3-CD16+CD56+(NK) | anti-SSA/Ro | 0.273136439 | 1.78E-15 |
| B23 | B104 | CD3-CD16+CD56+(NK) | anti-SSB | 0.273136438 | 1.78E-15 |
| B23 | B105 | CD3-CD16+CD56+(NK) | anti-U1RNP | 0.273136439 | 1.78E-15 |
| B23 | B106 | CD3-CD16+CD56+(NK) | anti-P | 0.273136438 | 1.78E-15 |
| B23 | B107 | CD3-CD16+CD56+(NK) | AnuA | 0.273136388 | 1.78E-15 |
| B23 | B108 | CD3-CD16+CD56+(NK) | Anti-Cyclin | 0.273136439 | 1.78E-15 |

| | | | | | |
|---|---|---|---|---|---|
| B23 | B109 | CD3-CD16+CD56+(NK) | AMA-M2 | 0.273136439 | 1.78E-15 |
| B23 | B110 | CD3-CD16+CD56+(NK) | AHA | 0.273136439 | 1.78E-15 |
| B23 | B97 | CD3-CD16+CD56+(NK) | dsDNA | 0.273136437 | 1.78E-15 |
| B23 | B98 | CD3-CD16+CD56+(NK) | anti-Jo-1 | 0.273136442 | 1.78E-15 |
| B23 | B99 | CD3-CD16+CD56+(NK) | anti-PM-Scl | 0.273136439 | 1.78E-15 |
| B23 | B151 | CD3-CD16+CD56+(NK) | Anti-CCP | 0.257302162 | 7.26E-14 |
| B23 | B85 | CD3-CD16+CD56+(NK) | SCC | -0.22841838 | 3.63E-11 |
| B23 | B181 | CD3-CD16+CD56+(NK) | APTT | 0.22316876 | 1.03E-10 |
| B23 | B183 | CD3-CD16+CD56+(NK) | PTA | 0.214306508 | 5.66E-10 |
| B23 | B196 | CD3-CD16+CD56+(NK) | TCO2 | 0.195237372 | 1.73E-08 |
| B23 | B141 | CD3-CD16+CD56+(NK) | AST/ALT | 0.19276157 | 2.64E-08 |
| B24 | B123 | αHBDH | hs-CRP1 | 0.309676228 | 0 |
| B24 | B27 | αHBDH | LDH | 0.391982091 | 0 |
| B24 | B199 | αHBDH | Mg | 0.230082107 | 2.59E-11 |
| B24 | B124 | αHBDH | ACTH | -0.19646857 | 1.40E-08 |
| B24 | B74 | αHBDH | ApoA1 | 0.178601212 | 2.63E-07 |
| B24 | B146 | αHBDH | CysC | 0.177514214 | 3.12E-07 |
| B25 | B26 | CK | CKMB | 0.320689289 | 0 |
| B25 | B146 | CK | CysC | 0.203078023 | 4.42E-09 |
| B25 | B118 | CK | CEA1 | 0.194651646 | 1.92E-08 |
| B25 | B57 | CK | RBCAI | 0.172065733 | 7.17E-07 |
| B27 | B131 | LDH | AST | 0.463331498 | 0 |
| B27 | B128 | LDH | ALT | 0.225801931 | 6.12E-11 |
| B27 | B75 | LDH | ApoA1/B | 0.197471685 | 1.18E-08 |
| B28 | B29 | PRL | E2 | -0.35950533 | 0 |
| B28 | B30 | PRL | LH | -0.49289568 | 0 |
| B28 | B31 | PRL | FSH | -0.77903587 | 0 |
| B28 | B32 | PRL | Testo | -0.82664639 | 0 |
| B28 | B33 | PRL | Prog | 0.743264174 | 0 |
| B29 | B30 | E2 | LH | 0.989115565 | 0 |
| B29 | B31 | E2 | FSH | -0.30499386 | 0 |
| B29 | B33 | E2 | Prog | -0.89147829 | 0 |
| B29 | B32 | E2 | Testo | -0.22791607 | 4.01E-11 |
| B3 | B25 | Hcy | CK | 0.173959937 | 5.38E-07 |
| B3 | B190 | Hcy | Cre | 0.172922118 | 6.30E-07 |
| B30 | B33 | LH | Prog | -0.94843914 | 0 |
| B31 | B32 | FSH | Testo | 0.996802098 | 0 |
| B32 | B33 | Testo | Prog | -0.23795692 | 5.09E-12 |
| B34 | B186 | TK | PFC | 0.276117531 | 8.88E-16 |
| B34 | B168 | TK | NMP22 | 0.187826817 | 6.01E-08 |
| B35 | B116 | β2-MG | TT3 | 0.995753696 | 0 |
| B35 | B225 | β2-MG | hPBG | 0.946923578 | 0 |
| B35 | B37 | β2-MG | BAND | 0.870819097 | 0 |
| B35 | B96 | β2-MG | dsDNA (IIF) | 0.834135464 | 0 |

| | | | | | |
|---|---|---|---|---|---|
| B37 | B202 | BAND | e-times | 0.269809071 | 3.77E-15 |
| B39 | B46 | Other cells | RBC Count | 0.243640874 | 1.52E-12 |
| B39 | B43 | Other cells | MCV | -0.23660347 | 6.76E-12 |
| B39 | B44 | Other cells | MCH | -0.23611801 | 7.48E-12 |
| B4 | B100 | WB-Ca | anti-Ro52 | -0.51901369 | 0 |
| B4 | B16 | WB-Ca | CD3+T | -0.29482445 | 0 |
| B4 | B17 | WB-Ca | CD3+CD16+CD56+NK | 0.363205652 | 0 |
| B4 | B23 | WB-Ca | CD3-CD16+CD56+(NK) | 0.288660284 | 0 |
| B4 | B5 | WB-Ca | WB-Mg | -0.48144802 | 0 |
| B4 | B8 | WB-Ca | WB-Zinc | 0.414410547 | 0 |
| B4 | B95 | WB-Ca | anti-cenpb | -0.52178872 | 0 |
| B4 | B18 | WB-Ca | CD3+CD4+helper(inducer)T | -0.27770445 | 4.44E-16 |
| B4 | B158 | WB-Ca | IgM | -0.24696267 | 7.36E-13 |
| B4 | B156 | WB-Ca | IgE | -0.23738202 | 5.74E-12 |
| B4 | B19 | WB-Ca | CD3+CD4+/CD3+CD8+ | -0.20531678 | 2.96E-09 |
| B4 | B198 | WB-Ca | P | -0.20009143 | 7.49E-09 |
| B4 | B21 | WB-Ca | CD3+CD4-CD8- | 0.195784181 | 1.58E-08 |
| B4 | B101 | WB-Ca | anti-Sc70 | 0.192795944 | 2.63E-08 |
| B4 | B102 | WB-Ca | anti-Sm | 0.192795946 | 2.63E-08 |
| B4 | B103 | WB-Ca | anti-SSA/Ro | 0.192795947 | 2.63E-08 |
| B4 | B104 | WB-Ca | anti-SSB | 0.192795949 | 2.63E-08 |
| B4 | B105 | WB-Ca | anti-U1RNP | 0.192795947 | 2.63E-08 |
| B4 | B106 | WB-Ca | anti-P | 0.192795949 | 2.63E-08 |
| B4 | B107 | WB-Ca | AnuA | 0.192795912 | 2.63E-08 |
| B4 | B108 | WB-Ca | Anti-Cyclin | 0.192795943 | 2.63E-08 |
| B4 | B109 | WB-Ca | AMA-M2 | 0.192795948 | 2.63E-08 |
| B4 | B110 | WB-Ca | AHA | 0.192795947 | 2.63E-08 |
| B4 | B97 | WB-Ca | dsDNA | 0.192795946 | 2.63E-08 |
| B4 | B98 | WB-Ca | anti-Jo-1 | 0.192795943 | 2.63E-08 |
| B4 | B99 | WB-Ca | anti-PM-Scl | 0.192795947 | 2.63E-08 |
| B4 | B138 | WB-Ca | PAB | 0.184401608 | 1.05E-07 |
| B4 | B22 | WB-Ca | CD3+CD8+(T) | 0.18368071 | 1.18E-07 |
| B40 | B42 | WBC | MON | 0.637954224 | 0 |
| B40 | B52 | WBC | PLT/BPC | 0.304505468 | 0 |
| B40 | B46 | WBC | RBC Count | 0.276688564 | 6.66E-16 |
| B40 | B48 | WBC | LYM% | -0.27652649 | 8.88E-16 |
| B40 | B53 | WBC | GR% | 0.269108711 | 4.44E-15 |
| B40 | B123 | WBC | hs-CRP1 | 0.242627396 | 1.89E-12 |
| B40 | B206 | WBC | Waist | 0.195159102 | 1.76E-08 |
| B40 | B51 | WBC | Hb | 0.185810166 | 8.35E-08 |
| B40 | B215 | WBC | BMI | 0.172044428 | 7.19E-07 |
| B40 | B222 | WBC | NS | 0.170990827 | 8.42E-07 |
| B41 | B42 | MON％ | MON | 0.610012605 | 0 |
| B41 | B190 | MON％ | Cre | 0.239397157 | 3.76E-12 |

| | | | | | |
|---|---|---|---|---|---|
| B41 | B189 | MON％ | eGFB | -0.18357413 | 1.20E-07 |
| B42 | B46 | MON | RBC Count | 0.273154364 | 1.78E-15 |
| B42 | B190 | MON | Cre | 0.272552591 | 2.00E-15 |
| B42 | B222 | MON | NS | 0.255970648 | 9.84E-14 |
| B42 | B48 | MON | LYM% | -0.25005169 | 3.73E-13 |
| B42 | B206 | MON | Waist | 0.238044327 | 5.00E-12 |
| B42 | B216 | MON | BMI | 0.205834296 | 2.70E-09 |
| B42 | B207 | MON | WHR | 0.204047548 | 3.72E-09 |
| B42 | B192 | MON | UA | 0.203295769 | 4.25E-09 |
| B42 | B51 | MON | Hb | 0.189910503 | 4.26E-08 |
| B42 | B210 | MON | Hip | 0.181019728 | 1.80E-07 |
| B42 | B215 | MON | BMI | 0.171769265 | 7.50E-07 |
| B43 | B44 | MCV | MCH | 0.88438728 | 0 |
| B43 | B46 | MCV | RBC Count | -0.51158784 | 0 |
| B43 | B47 | MCV | RDW | 0.479318945 | 0 |
| B43 | B51 | MCV | Hb | 0.215395214 | 4.61E-10 |
| B43 | B52 | MCV | PLT/BPC | -0.21435761 | 5.61E-10 |
| B43 | B45 | MCV | MCHC | 0.212020672 | 8.68E-10 |
| B44 | B45 | MCH | MCHC | 0.641730616 | 0 |
| B44 | B46 | MCH | RBC Count | -0.45840386 | 0 |
| B44 | B51 | MCH | Hb | 0.327395268 | 0 |
| B44 | B47 | MCH | RDW | 0.268261629 | 5.55E-15 |
| B44 | B52 | MCH | PLT/BPC | -0.18752693 | 6.31E-08 |
| B45 | B51 | MCHC | Hb | 0.351635465 | 0 |
| B45 | B47 | MCHC | RDW | -0.23838608 | 4.65E-12 |
| B46 | B192 | RBC Count | UA | 0.336212829 | 0 |
| B46 | B216 | RBC Count | BMI | 0.352119887 | 0 |
| B46 | B221 | RBC Count | Hei | 0.379906932 | 0 |
| B46 | B222 | RBC Count | NS | 0.412790532 | 0 |
| B46 | B47 | RBC Count | RDW | -0.38635634 | 0 |
| B46 | B51 | RBC Count | Hb | 0.675498703 | 0 |
| B46 | B126 | RBC Count | Alb | 0.279662755 | 4.44E-16 |
| B46 | B190 | RBC Count | Cre | 0.271772324 | 2.22E-15 |
| B46 | B206 | RBC Count | Waist | 0.26781796 | 6.22E-15 |
| B46 | B128 | RBC Count | ALT | 0.263273925 | 1.82E-14 |
| B46 | B174 | RBC Count | Urea/Cr | -0.25707886 | 7.64E-14 |
| B46 | B218 | RBC Count | DBP | 0.238458678 | 4.58E-12 |
| B46 | B207 | RBC Count | WHR | 0.222921502 | 1.08E-10 |
| B46 | B72 | RBC Count | HDL-C | -0.21729594 | 3.21E-10 |
| B46 | B204 | RBC Count | Drinking | 0.21133472 | 9.86E-10 |
| B46 | B71 | RBC Count | TG | 0.209575092 | 1.36E-09 |
| B46 | B141 | RBC Count | AST/ALT | -0.20926377 | 1.45E-09 |
| B46 | B58 | RBC Count | HCT | 0.208202079 | 1.76E-09 |
| B46 | B163 | RBC Count | SG | 0.207188321 | 2.11E-09 |

| | | | | | |
|---|---|---|---|---|---|
| B46 | B210 | RBC Count | Hip | 0.206119124 | 2.56E-09 |
| B46 | B66 | RBC Count | WBRSR(mPas_100) | 0.192139368 | 2.93E-08 |
| B46 | B67 | RBC Count | WBRSR(mPas_50) | 0.192126796 | 2.94E-08 |
| B46 | B63 | RBC Count | WBRSR(mPas_200) | 0.191274553 | 3.39E-08 |
| B46 | B91 | RBC Count | TP | 0.189768757 | 4.36E-08 |
| B46 | B62 | RBC Count | WBHSRVI | 0.188171766 | 5.67E-08 |
| B46 | B64 | RBC Count | WBRSR(mPas_5) | 0.185214601 | 9.20E-08 |
| B46 | B60 | RBC Count | WBRVI | 0.183276697 | 1.26E-07 |
| B46 | B65 | RBC Count | WBRSR(mPas_1) | 0.172215482 | 7.01E-07 |
| B47 | B189 | RDW | eGFB | -0.28097957 | 2.22E-16 |
| B47 | B52 | RDW | PLT/BPC | -0.25658269 | 8.57E-14 |
| B47 | B126 | RDW | Alb | -0.20388297 | 3.83E-09 |
| B47 | B191 | RDW | Urea | 0.173074318 | 6.16E-07 |
| B48 | B53 | LYM% | GR% | -0.91774906 | 0 |
| B49 | B50 | baso% | eo% | 0.378361194 | 0 |
| B49 | B53 | baso% | GR% | -0.27291013 | 1.78E-15 |
| B5 | B6 | WB-Mg | WB-Fe | 0.79765251 | 0 |
| B5 | B198 | WB-Mg | P | 0.253055572 | 1.91E-13 |
| B5 | B138 | WB-Mg | PAB | -0.23049216 | 2.38E-11 |
| B5 | B139 | WB-Mg | ADA | -0.21405913 | 5.93E-10 |
| B5 | B7 | WB-Mg | WB-Cop | 0.207943347 | 1.84E-09 |
| B5 | B193 | WB-Mg | K | 0.190579709 | 3.81E-08 |
| B5 | B73 | WB-Mg | ApoA | 0.183630522 | 1.19E-07 |
| B5 | B8 | WB-Mg | WB-Zinc | -0.17495149 | 4.63E-07 |
| B50 | B53 | eo% | GR% | -0.43665611 | 0 |
| B51 | B126 | Hb | Alb | 0.353330829 | 0 |
| B51 | B128 | Hb | ALT | 0.300248965 | 0 |
| B51 | B163 | Hb | SG | 0.295919676 | 0 |
| B51 | B190 | Hb | Cre | 0.386562727 | 0 |
| B51 | B192 | Hb | UA | 0.405906132 | 0 |
| B51 | B204 | Hb | Drinking | 0.289025785 | 0 |
| B51 | B206 | Hb | Waist | 0.303550862 | 0 |
| B51 | B216 | Hb | BMI | 0.409646917 | 0 |
| B51 | B221 | Hb | Hei | 0.445361603 | 0 |
| B51 | B222 | Hb | NS | 0.474132573 | 0 |
| B51 | B174 | Hb | Urea/Cr | -0.27118789 | 2.66E-15 |
| B51 | B207 | Hb | WHR | 0.254433021 | 1.39E-13 |
| B51 | B71 | Hb | TG | 0.245741033 | 9.61E-13 |
| B51 | B218 | Hb | DBP | 0.239007967 | 4.08E-12 |
| B51 | B210 | Hb | Hip | 0.236057296 | 7.58E-12 |
| B51 | B141 | Hb | AST/ALT | -0.23541934 | 8.65E-12 |
| B51 | B58 | Hb | HCT | 0.234753558 | 9.94E-12 |
| B51 | B62 | Hb | WBHSRVI | 0.218762134 | 2.42E-10 |
| B51 | B72 | Hb | HDL-C | -0.21779016 | 2.92E-10 |

| | | | | | |
|---|---|---|---|---|---|
| B51 | B66 | Hb | WBRSR(mPas_100) | 0.216679239 | 3.61E-10 |
| B51 | B67 | Hb | WBRSR(mPas_50) | 0.216292413 | 3.89E-10 |
| B51 | B63 | Hb | WBRSR(mPas_200) | 0.212488188 | 7.96E-10 |
| B51 | B60 | Hb | WBRVI | 0.208249681 | 1.74E-09 |
| B51 | B127 | Hb | A-G ratio | 0.207213023 | 2.10E-09 |
| B51 | B64 | Hb | WBRSR(mPas_5) | 0.205289641 | 2.98E-09 |
| B51 | B70 | Hb | LDL-C | 0.194117966 | 2.10E-08 |
| B51 | B215 | Hb | BMI | 0.192995213 | 2.54E-08 |
| B51 | B65 | Hb | WBRSR(mPas_1) | 0.185997559 | 8.10E-08 |
| B51 | B140 | Hb | r-GT | 0.180768901 | 1.87E-07 |
| B52 | B130 | PLT/BPC | GLB | 0.204298035 | 3.56E-09 |
| B52 | B190 | PLT/BPC | Cre | -0.18757301 | 6.26E-08 |
| B52 | B127 | PLT/BPC | A-G ratio | -0.18657752 | 7.37E-08 |
| B52 | B191 | PLT/BPC | Urea | -0.1798655 | 2.16E-07 |
| B52 | B189 | PLT/BPC | eGFB | 0.177523295 | 3.11E-07 |
| B52 | B91 | PLT/BPC | TP | 0.171444903 | 7.87E-07 |
| B54 | B123 | ESR | hs-CRP1 | 0.357362578 | 0 |
| B54 | B58 | ESR | HCT | -0.44101808 | 0 |
| B54 | B59 | ESR | WBRV | 0.285738779 | 0 |
| B54 | B60 | ESR | WBRVI | -0.31171209 | 0 |
| B54 | B62 | ESR | WBHSRVI | -0.33255881 | 0 |
| B54 | B69 | ESR | PV | 0.593140774 | 0 |
| B54 | B56 | ESR | IR | -0.22621396 | 5.64E-11 |
| B54 | B74 | ESR | ApoA1 | 0.211478518 | 9.60E-10 |
| B54 | B199 | ESR | Mg | 0.178257133 | 2.78E-07 |
| B55 | B101 | TK | anti-Sc70 | 0.294217593 | 0 |
| B55 | B102 | TK | anti-Sm | 0.294217593 | 0 |
| B55 | B103 | TK | anti-SSA/Ro | 0.294217592 | 0 |
| B55 | B104 | TK | anti-SSB | 0.294217591 | 0 |
| B55 | B105 | TK | anti-U1RNP | 0.294217591 | 0 |
| B55 | B106 | TK | anti-P | 0.29421759 | 0 |
| B55 | B107 | TK | AnuA | 0.294217537 | 0 |
| B55 | B108 | TK | Anti-Cyclin | 0.294217591 | 0 |
| B55 | B109 | TK | AMA-M2 | 0.294217591 | 0 |
| B55 | B110 | TK | AHA | 0.294217592 | 0 |
| B55 | B124 | TK | ACTH | -0.39073105 | 0 |
| B55 | B199 | TK | Mg | 0.404030982 | 0 |
| B55 | B56 | TK | IR | 0.512466909 | 0 |
| B55 | B57 | TK | RBCAI | -0.31180487 | 0 |
| B55 | B61 | TK | WBHSRV | 0.508602752 | 0 |
| B55 | B97 | TK | dsDNA | 0.29421759 | 0 |
| B55 | B98 | TK | anti-Jo-1 | 0.294217595 | 0 |
| B55 | B99 | TK | anti-PM-Scl | 0.294217593 | 0 |
| B55 | B59 | TK | WBRV | 0.255766675 | 1.03E-13 |

| | | | | | |
|---|---|---|---|---|---|
| B55 | B58 | TK | HCT | -0.24521222 | 1.08E-12 |
| B55 | B69 | TK | PV | -0.24517892 | 1.09E-12 |
| B55 | B183 | TK | PTA | -0.19995033 | 7.68E-09 |
| B55 | B179 | TK | α1-MG | -0.18301607 | 1.31E-07 |
| B55 | B178 | TK | Ig-U | -0.18040682 | 1.98E-07 |
| B55 | B180 | TK | Tf | -0.17223813 | 6.99E-07 |
| B56 | B57 | IR | RBCAI | -0.32517869 | 0 |
| B56 | B58 | IR | HCT | 0.679860665 | 0 |
| B56 | B59 | IR | WBRV | 0.299432322 | 0 |
| B56 | B60 | IR | WBRVI | 0.790670613 | 0 |
| B56 | B61 | IR | WBHSRV | 0.914725061 | 0 |
| B56 | B62 | IR | WBHSRVI | 0.928348727 | 0 |
| B56 | B63 | IR | WBRSR(mPas_200) | 0.860158007 | 0 |
| B56 | B64 | IR | WBRSR(mPas_5) | 0.741751347 | 0 |
| B56 | B65 | IR | WBRSR(mPas_1) | 0.653678455 | 0 |
| B56 | B66 | IR | WBRSR(mPas_100) | 0.844239372 | 0 |
| B56 | B67 | IR | WBRSR(mPas_50) | 0.830691299 | 0 |
| B56 | B69 | IR | PV | -0.46623269 | 0 |
| B56 | B137 | IR | AFU | 0.194365184 | 2.01E-08 |
| B56 | B101 | IR | anti-Sc70 | 0.178064644 | 2.86E-07 |
| B56 | B102 | IR | anti-Sm | 0.178064644 | 2.86E-07 |
| B56 | B103 | IR | anti-SSA/Ro | 0.178064642 | 2.86E-07 |
| B56 | B104 | IR | anti-SSB | 0.178064641 | 2.86E-07 |
| B56 | B105 | IR | anti-U1RNP | 0.178064642 | 2.86E-07 |
| B56 | B106 | IR | anti-P | 0.178064642 | 2.86E-07 |
| B56 | B107 | IR | AnuA | 0.178064606 | 2.86E-07 |
| B56 | B108 | IR | Anti-Cyclin | 0.178064642 | 2.86E-07 |
| B56 | B109 | IR | AMA-M2 | 0.178064642 | 2.86E-07 |
| B56 | B110 | IR | AHA | 0.178064642 | 2.86E-07 |
| B56 | B97 | IR | dsDNA | 0.178064641 | 2.86E-07 |
| B56 | B98 | IR | anti-Jo-1 | 0.178064644 | 2.86E-07 |
| B56 | B99 | IR | anti-PM-Scl | 0.178064642 | 2.86E-07 |
| B57 | B155 | RBCAI | IgA | -0.31313316 | 0 |
| B57 | B179 | RBCAI | α1-MG | 0.306946678 | 0 |
| B57 | B199 | RBCAI | Mg | -0.42782967 | 0 |
| B57 | B59 | RBCAI | WBRV | 0.649603472 | 0 |
| B57 | B65 | RBCAI | WBRSR(mPas_1) | 0.347543562 | 0 |
| B57 | B69 | RBCAI | PV | 0.314712914 | 0 |
| B57 | B81 | RBCAI | TSGF | -0.28889741 | 0 |
| B57 | B79 | RBCAI | hb V -dna | 0.280123489 | 4.44E-16 |
| B57 | B180 | RBCAI | Tf | 0.262522936 | 2.18E-14 |
| B57 | B185 | RBCAI | PT | -0.26205321 | 2.42E-14 |
| B57 | B178 | RBCAI | Ig-U | 0.259055035 | 4.84E-14 |
| B57 | B62 | RBCAI | WBHSRVI | -0.2520566 | 2.38E-13 |

| | | | | | |
|---|---|---|---|---|---|
| B57 | B181 | RBCAI | APTT | -0.21588286 | 4.20E-10 |
| B57 | B60 | RBCAI | WBRVI | 0.215053453 | 4.92E-10 |
| B57 | B61 | RBCAI | WBHSRV | -0.21352327 | 6.56E-10 |
| B57 | B186 | RBCAI | PFC | 0.207808964 | 1.89E-09 |
| B57 | B124 | RBCAI | ACTH | 0.192525976 | 2.75E-08 |
| B57 | B64 | RBCAI | WBRSR(mPas_5) | 0.177596354 | 3.08E-07 |
| B58 | B124 | HCT | ACTH | 0.345452881 | 0 |
| B58 | B199 | HCT | Mg | -0.31269089 | 0 |
| B58 | B60 | HCT | WBRVI | 0.873034754 | 0 |
| B58 | B61 | HCT | WBHSRV | 0.612882848 | 0 |
| B58 | B62 | HCT | WBHSRVI | 0.887578523 | 0 |
| B58 | B63 | HCT | WBRSR(mPas_200) | 0.871910545 | 0 |
| B58 | B64 | HCT | WBRSR(mPas_5) | 0.845069566 | 0 |
| B58 | B65 | HCT | WBRSR(mPas_1) | 0.782272488 | 0 |
| B58 | B66 | HCT | WBRSR(mPas_100) | 0.872818279 | 0 |
| B58 | B67 | HCT | WBRSR(mPas_50) | 0.872581967 | 0 |
| B58 | B69 | HCT | PV | -0.28662415 | 0 |
| B58 | B138 | HCT | PAB | 0.22138727 | 1.46E-10 |
| B58 | B134 | HCT | CHE | 0.186463677 | 7.51E-08 |
| B58 | B137 | HCT | AFU | 0.172787628 | 6.43E-07 |
| B58 | B59 | HCT | WBRV | 0.169900849 | 9.91E-07 |
| B59 | B155 | WBRV | IgA | -0.2863816 | 0 |
| B59 | B60 | WBRV | WBRVI | 0.540890709 | 0 |
| B59 | B61 | WBRV | WBHSRV | 0.535912677 | 0 |
| B59 | B63 | WBRV | WBRSR(mPas_200) | 0.442501085 | 0 |
| B59 | B64 | WBRV | WBRSR(mPas_5) | 0.636000329 | 0 |
| B59 | B65 | WBRV | WBRSR(mPas_1) | 0.720323513 | 0 |
| B59 | B66 | WBRV | WBRSR(mPas_100) | 0.446352645 | 0 |
| B59 | B67 | WBRV | WBRSR(mPas_50) | 0.479136228 | 0 |
| B59 | B69 | WBRV | PV | 0.400577601 | 0 |
| B59 | B186 | WBRV | PFC | 0.254503711 | 1.37E-13 |
| B59 | B181 | WBRV | APTT | -0.24707906 | 7.18E-13 |
| B59 | B185 | WBRV | PT | -0.2385323 | 4.51E-12 |
| B59 | B199 | WBRV | Mg | -0.2371071 | 6.08E-12 |
| B59 | B62 | WBRV | WBHSRVI | 0.221322467 | 1.48E-10 |
| B59 | B138 | WBRV | PAB | 0.193961953 | 2.16E-08 |
| B59 | B111 | WBRV | TSH | -0.17918277 | 2.40E-07 |
| B59 | B81 | WBRV | TSGF | -0.17888484 | 2.52E-07 |
| B6 | B100 | WB-Fe | anti-Ro52 | -0.47728728 | 0 |
| B6 | B17 | WB-Fe | CD3+CD16+CD56+NK | 0.335799418 | 0 |
| B6 | B95 | WB-Fe | anti-cenpb | -0.48011443 | 0 |
| B6 | B16 | WB-Fe | CD3+T | -0.27131202 | 2.66E-15 |
| B6 | B23 | WB-Fe | CD3-CD16+CD56+(NK) | 0.265850816 | 9.77E-15 |
| B6 | B18 | WB-Fe | CD3+CD4+helper(inducer)T | -0.25681354 | 8.13E-14 |

| | | | | | |
|---|---|---|---|---|---|
| B6 | B7 | WB-Fe | WB-Cop | 0.233376165 | 1.32E-11 |
| B6 | B158 | WB-Fe | IgM | -0.22681343 | 5.00E-11 |
| B6 | B156 | WB-Fe | IgE | -0.21903687 | 2.30E-10 |
| B6 | B19 | WB-Fe | CD3+CD4+/CD3+CD8+ | -0.19043132 | 3.90E-08 |
| B6 | B81 | WB-Fe | TSGF | 0.183435255 | 1.23E-07 |
| B6 | B21 | WB-Fe | CD3+CD4-CD8- | 0.180791516 | 1.87E-07 |
| B6 | B198 | WB-Fe | P | 0.180675205 | 1.90E-07 |
| B6 | B101 | WB-Fe | anti-Sc70 | 0.180216395 | 2.04E-07 |
| B6 | B102 | WB-Fe | anti-Sm | 0.180216393 | 2.04E-07 |
| B6 | B103 | WB-Fe | anti-SSA/Ro | 0.18021639 | 2.04E-07 |
| B6 | B104 | WB-Fe | anti-SSB | 0.180216394 | 2.04E-07 |
| B6 | B105 | WB-Fe | anti-U1RNP | 0.18021639 | 2.04E-07 |
| B6 | B106 | WB-Fe | anti-P | 0.180216395 | 2.04E-07 |
| B6 | B107 | WB-Fe | AnuA | 0.180216386 | 2.04E-07 |
| B6 | B108 | WB-Fe | Anti-Cyclin | 0.180216389 | 2.04E-07 |
| B6 | B109 | WB-Fe | AMA-M2 | 0.18021639 | 2.04E-07 |
| B6 | B110 | WB-Fe | AHA | 0.18021639 | 2.04E-07 |
| B6 | B97 | WB-Fe | dsDNA | 0.180216392 | 2.04E-07 |
| B6 | B98 | WB-Fe | anti-Jo-1 | 0.180216394 | 2.04E-07 |
| B6 | B99 | WB-Fe | anti-PM-Scl | 0.18021639 | 2.04E-07 |
| B6 | B22 | WB-Fe | CD3+CD8+(T) | 0.171143061 | 8.23E-07 |
| B60 | B199 | WBRVI | Mg | -0.31333886 | 0 |
| B60 | B61 | WBRVI | WBHSRV | 0.750962001 | 0 |
| B60 | B62 | WBRVI | WBHSRVI | 0.887923949 | 0 |
| B60 | B63 | WBRVI | WBRSR(mPas_200) | 0.887782308 | 0 |
| B60 | B64 | WBRVI | WBRSR(mPas_5) | 0.937854132 | 0 |
| B60 | B65 | WBRVI | WBRSR(mPas_1) | 0.925412947 | 0 |
| B60 | B66 | WBRVI | WBRSR(mPas_100) | 0.889470732 | 0 |
| B60 | B67 | WBRVI | WBRSR(mPas_50) | 0.901358914 | 0 |
| B60 | B69 | WBRVI | PV | -0.28509317 | 0 |
| B60 | B138 | WBRVI | PAB | 0.244562355 | 1.24E-12 |
| B60 | B124 | WBRVI | ACTH | 0.235082346 | 9.28E-12 |
| B60 | B155 | WBRVI | IgA | -0.21379189 | 6.24E-10 |
| B60 | B186 | WBRVI | PFC | 0.206668393 | 2.32E-09 |
| B60 | B134 | WBRVI | CHE | 0.197026482 | 1.28E-08 |
| B60 | B81 | WBRVI | TSGF | -0.17188701 | 7.36E-07 |
| B60 | B137 | WBRVI | AFU | 0.170650381 | 8.86E-07 |
| B61 | B62 | WBHSRV | WBHSRVI | 0.829479053 | 0 |
| B61 | B63 | WBHSRV | WBRSR(mPas_200) | 0.908444108 | 0 |
| B61 | B64 | WBHSRV | WBRSR(mPas_5) | 0.834642247 | 0 |
| B61 | B65 | WBHSRV | WBRSR(mPas_1) | 0.753287629 | 0 |
| B61 | B66 | WBHSRV | WBRSR(mPas_100) | 0.898474856 | 0 |
| B61 | B67 | WBHSRV | WBRSR(mPas_50) | 0.892199165 | 0 |
| B61 | B138 | WBHSRV | PAB | 0.194373574 | 2.01E-08 |

| | | | | | |
|---|---|---|---|---|---|
| B62 | B63 | WBHSRVI | WBRSR(mPas_200) | 0.933440081 | 0 |
| B62 | B64 | WBHSRVI | WBRSR(mPas_5) | 0.839857945 | 0 |
| B62 | B65 | WBHSRVI | WBRSR(mPas_1) | 0.746989403 | 0 |
| B62 | B66 | WBHSRVI | WBRSR(mPas_100) | 0.928167069 | 0 |
| B62 | B67 | WBHSRVI | WBRSR(mPas_50) | 0.918453496 | 0 |
| B62 | B69 | WBHSRVI | PV | -0.43492477 | 0 |
| B62 | B137 | WBHSRVI | AFU | 0.190356579 | 3.95E-08 |
| B62 | B138 | WBHSRVI | PAB | 0.186446609 | 7.53E-08 |
| B63 | B64 | WBRSR(mPas_200) | WBRSR(mPas_5) | 0.951609416 | 0 |
| B63 | B65 | WBRSR(mPas_200) | WBRSR(mPas_1) | 0.86924612 | 0 |
| B63 | B66 | WBRSR(mPas_200) | WBRSR(mPas_100) | 0.997346731 | 0 |
| B63 | B67 | WBRSR(mPas_200) | WBRSR(mPas_50) | 0.99522069 | 0 |
| B63 | B124 | WBRSR(mPas_200) | ACTH | 0.252038334 | 2.39E-13 |
| B63 | B138 | WBRSR(mPas_200) | PAB | 0.246308903 | 8.50E-13 |
| B63 | B134 | WBRSR(mPas_200) | CHE | 0.176955487 | 3.40E-07 |
| B64 | B124 | WBRSR(mPas_5) | ACTH | 0.297566999 | 0 |
| B64 | B138 | WBRSR(mPas_5) | PAB | 0.283889276 | 0 |
| B64 | B65 | WBRSR(mPas_5) | WBRSR(mPas_1) | 0.966822231 | 0 |
| B64 | B66 | WBRSR(mPas_5) | WBRSR(mPas_100) | 0.957965899 | 0 |
| B64 | B67 | WBRSR(mPas_5) | WBRSR(mPas_50) | 0.970083005 | 0 |
| B64 | B199 | WBRSR(mPas_5) | Mg | -0.28194177 | 2.22E-16 |
| B64 | B134 | WBRSR(mPas_5) | CHE | 0.200804082 | 6.61E-09 |
| B65 | B124 | WBRSR(mPas_1) | ACTH | 0.324416073 | 0 |
| B65 | B199 | WBRSR(mPas_1) | Mg | -0.34742107 | 0 |
| B65 | B66 | WBRSR(mPas_1) | WBRSR(mPas_100) | 0.87463193 | 0 |
| B65 | B67 | WBRSR(mPas_1) | WBRSR(mPas_50) | 0.893897099 | 0 |
| B65 | B138 | WBRSR(mPas_1) | PAB | 0.282258736 | 2.22E-16 |
| B65 | B155 | WBRSR(mPas_1) | IgA | -0.22126513 | 1.49E-10 |
| B65 | B134 | WBRSR(mPas_1) | CHE | 0.20385655 | 3.85E-09 |
| B65 | B186 | WBRSR(mPas_1) | PFC | 0.178937894 | 2.50E-07 |
| B65 | B179 | WBRSR(mPas_1) | α1-MG | 0.172593773 | 6.62E-07 |
| B65 | B111 | WBRSR(mPas_1) | TSH | -0.1718643 | 7.39E-07 |
| B65 | B81 | WBRSR(mPas_1) | TSGF | -0.17164878 | 7.63E-07 |
| B66 | B67 | WBRSR(mPas_100) | WBRSR(mPas_50) | 0.998904362 | 0 |
| B66 | B138 | WBRSR(mPas_100) | PAB | 0.253232365 | 1.83E-13 |
| B66 | B124 | WBRSR(mPas_100) | ACTH | 0.241106212 | 2.61E-12 |
| B66 | B134 | WBRSR(mPas_100) | CHE | 0.179731112 | 2.21E-07 |
| B66 | B199 | WBRSR(mPas_100) | Mg | -0.17349546 | 5.78E-07 |
| B67 | B138 | WBRSR(mPas_50) | PAB | 0.260223197 | 3.69E-14 |
| B67 | B124 | WBRSR(mPas_50) | ACTH | 0.252830957 | 2.00E-13 |
| B67 | B199 | WBRSR(mPas_50) | Mg | -0.19352701 | 2.32E-08 |
| B67 | B134 | WBRSR(mPas_50) | CHE | 0.18374052 | 1.17E-07 |
| B69 | B123 | PV | hs-CRP1 | 0.3604839 | 0 |
| B69 | B137 | PV | AFU | -0.2328099 | 1.48E-11 |

| | | | | | |
|---|---|---|---|---|---|
| B69 | B124 | PV | ACTH | 0.204088951 | 3.69E-09 |
| B69 | B89 | PV | CA 72-4 | -0.1954046 | 1.69E-08 |
| B7 | B8 | WB-Cop | WB-Zinc | 0.892367831 | 0 |
| B70 | B77 | LDL-C | TC | 0.893441316 | 0 |
| B70 | B132 | LDL-C | Dbil | -0.26011136 | 3.80E-14 |
| B70 | B76 | LDL-C | ApoB | 0.241520613 | 2.39E-12 |
| B70 | B71 | LDL-C | TG | 0.202720344 | 4.71E-09 |
| B71 | B192 | TG | UA | 0.326341638 | 0 |
| B71 | B72 | TG | HDL-C | -0.41369473 | 0 |
| B71 | B77 | TG | TC | 0.310146737 | 0 |
| B71 | B222 | TG | NS | 0.282388938 | 2.22E-16 |
| B71 | B206 | TG | Waist | 0.275575604 | 8.88E-16 |
| B71 | B128 | TG | ALT | 0.263804327 | 1.60E-14 |
| B71 | B91 | TG | TP | 0.260875115 | 3.18E-14 |
| B71 | B207 | TG | WHR | 0.256616203 | 8.48E-14 |
| B71 | B215 | TG | BMI | 0.244049067 | 1.39E-12 |
| B71 | B140 | TG | r-GT | 0.243582378 | 1.54E-12 |
| B71 | B216 | TG | BMI | 0.241457595 | 2.42E-12 |
| B71 | B119 | TG | Hcv-rna | -0.21644123 | 3.78E-10 |
| B71 | B121 | TG | C3 | 0.21644131 | 3.78E-10 |
| B71 | B122 | TG | C4 | 0.216441074 | 3.78E-10 |
| B71 | B159 | TG | β2-MG | 0.216441424 | 3.78E-10 |
| B71 | B169 | TG | U-CRY | 0.216439677 | 3.78E-10 |
| B71 | B126 | TG | Alb | 0.185589547 | 8.65E-08 |
| B71 | B210 | TG | Hip | 0.18335862 | 1.24E-07 |
| B71 | B130 | TG | GLB | 0.177526349 | 3.11E-07 |
| B71 | B132 | TG | Dbil | -0.17308826 | 6.14E-07 |
| B72 | B192 | HDL-C | UA | -0.34493972 | 0 |
| B72 | B206 | HDL-C | Waist | -0.36036171 | 0 |
| B72 | B207 | HDL-C | WHR | -0.28571207 | 0 |
| B72 | B210 | HDL-C | Hip | -0.29367155 | 0 |
| B72 | B215 | HDL-C | BMI | -0.34935235 | 0 |
| B72 | B216 | HDL-C | BMI | -0.43460148 | 0 |
| B72 | B221 | HDL-C | Hei | -0.29605607 | 0 |
| B72 | B222 | HDL-C | NS | -0.41221842 | 0 |
| B72 | B74 | HDL-C | ApoA1 | 0.27877564 | 4.44E-16 |
| B72 | B128 | HDL-C | ALT | -0.25397461 | 1.55E-13 |
| B72 | B77 | HDL-C | TC | 0.253217919 | 1.83E-13 |
| B72 | B163 | HDL-C | SG | -0.23144077 | 1.96E-11 |
| B72 | B141 | HDL-C | AST/ALT | 0.199606218 | 8.15E-09 |
| B73 | B139 | ApoA | ADA | -0.30692301 | 0 |
| B73 | B186 | ApoA | PFC | -0.26219481 | 2.35E-14 |
| B73 | B199 | ApoA | Mg | 0.239186931 | 3.93E-12 |
| B73 | B137 | ApoA | AFU | -0.2092403 | 1.45E-09 |

| | | | | | |
|---|---|---|---|---|---|
| B73 | B185 | ApoA | PT | 0.17942511 | 2.31E-07 |
| B74 | B212 | ApoA1 | IOP-L | -0.20227282 | 5.10E-09 |
| B74 | B138 | ApoA1 | PAB | -0.18032647 | 2.01E-07 |
| B74 | B216 | ApoA1 | BMI | -0.17488754 | 4.67E-07 |
| B74 | B199 | ApoA1 | Mg | 0.170300694 | 9.34E-07 |
| B75 | B194 | ApoA1/B | Cl | -0.33555557 | 0 |
| B75 | B200 | ApoA1/B | OP | -0.31827398 | 0 |
| B75 | B76 | ApoA1/B | ApoB | -0.59789844 | 0 |
| B75 | B197 | ApoA1/B | AG | 0.254499284 | 1.37E-13 |
| B75 | B196 | ApoA1/B | TCO2 | -0.24317471 | 1.68E-12 |
| B75 | B195 | ApoA1/B | Na | -0.2390697 | 4.03E-12 |
| B75 | B137 | ApoA1/B | AFU | -0.23581876 | 7.96E-12 |
| B75 | B135 | ApoA1/B | TBA | -0.22984899 | 2.72E-11 |
| B75 | B81 | ApoA1/B | TSGF | 0.228805057 | 3.35E-11 |
| B76 | B125 | ApoB | Cor | -0.30590618 | 0 |
| B76 | B200 | ApoB | OP | 0.322043475 | 0 |
| B76 | B195 | ApoB | Na | 0.255342819 | 1.13E-13 |
| B76 | B77 | ApoB | TC | 0.243749536 | 1.48E-12 |
| B76 | B194 | ApoB | Cl | 0.186499389 | 7.46E-08 |
| B76 | B196 | ApoB | TCO2 | 0.184174084 | 1.09E-07 |
| B76 | B137 | ApoB | AFU | 0.179005039 | 2.47E-07 |
| B77 | B132 | TC | Dbil | -0.27694617 | 6.66E-16 |
| B77 | B91 | TC | TP | 0.179214475 | 2.39E-07 |
| B78 | B180 | HBcAb | Tf | -0.24476042 | 1.19E-12 |
| B78 | B178 | HBcAb | Ig-U | -0.2423269 | 2.01E-12 |
| B78 | B179 | HBcAb | α1-MG | -0.23273298 | 1.51E-11 |
| B78 | B80 | HBcAb | anti-HP IgG | -0.18929615 | 4.71E-08 |
| B79 | B119 | hb V -dna | Hcv-rna | 0.285883774 | 0 |
| B79 | B121 | hb V -dna | C3 | -0.28588388 | 0 |
| B79 | B122 | hb V -dna | C4 | -0.28588357 | 0 |
| B79 | B159 | hb V -dna | β2-MG | -0.28588404 | 0 |
| B79 | B169 | hb V -dna | U-CRY | -0.28588172 | 0 |
| B79 | B178 | hb V -dna | Ig-U | 0.617492728 | 0 |
| B79 | B179 | hb V -dna | α1-MG | 0.728352077 | 0 |
| B79 | B180 | hb V -dna | Tf | 0.606877469 | 0 |
| B79 | B199 | hb V -dna | Mg | -0.38331483 | 0 |
| B79 | B81 | hb V -dna | TSGF | -0.36550661 | 0 |
| B79 | B155 | hb V -dna | IgA | -0.27035247 | 3.33E-15 |
| B79 | B124 | hb V -dna | ACTH | 0.235215112 | 9.03E-12 |
| B81 | B119 | TSGF | Hcv-rna | -0.37754949 | 0 |
| B81 | B121 | TSGF | C3 | 0.377549627 | 0 |
| B81 | B122 | TSGF | C4 | 0.377549217 | 0 |
| B81 | B155 | TSGF | IgA | 0.323546068 | 0 |
| B81 | B159 | TSGF | β2-MG | 0.377549827 | 0 |

| | | | | | |
|---|---|---|---|---|---|
| B81 | B169 | TSGF | U-CRY | 0.377546779 | 0 |
| B81 | B179 | TSGF | α1-MG | -0.40008948 | 0 |
| B81 | B180 | TSGF | Tf | -0.30482711 | 0 |
| B81 | B199 | TSGF | Mg | 0.332933976 | 0 |
| B81 | B178 | TSGF | Ig-U | -0.27336824 | 1.55E-15 |
| B81 | B125 | TSGF | Cor | -0.25800285 | 6.17E-14 |
| B81 | B124 | TSGF | ACTH | -0.20158643 | 5.76E-09 |
| B81 | B198 | TSGF | P | 0.171263 | 8.09E-07 |
| B82 | B84 | CEA | C-PSA | 0.327833572 | 0 |
| B82 | B85 | CEA | SCC | 0.290834528 | 0 |
| B82 | B88 | CEA | CA 242 | 0.582848484 | 0 |
| B82 | B89 | CEA | CA 72-4 | 0.289398861 | 0 |
| B82 | B83 | CEA | AFP | 0.261972223 | 2.46E-14 |
| B82 | B87 | CEA | NSE | 0.184029861 | 1.11E-07 |
| B83 | B84 | AFP | C-PSA | 0.526969974 | 0 |
| B83 | B85 | AFP | SCC | 0.269595958 | 4.00E-15 |
| B83 | B88 | AFP | CA 242 | 0.243855103 | 1.45E-12 |
| B84 | B85 | C-PSA | SCC | 0.32070822 | 0 |
| B84 | B86 | C-PSA | TBhCG | 0.31214265 | 0 |
| B84 | B89 | C-PSA | CA 72-4 | 0.379436729 | 0 |
| B84 | B90 | C-PSA | CYFRA 21-1 | 0.24047042 | 2.99E-12 |
| B84 | B87 | C-PSA | NSE | 0.230736854 | 2.27E-11 |
| B84 | B139 | C-PSA | ADA | -0.2180069 | 2.80E-10 |
| B84 | B198 | C-PSA | P | -0.1774788 | 3.14E-07 |
| B85 | B88 | SCC | CA 242 | 0.369075561 | 0 |
| B85 | B92 | SCC | ALDH2-Glu504Glu | -0.31652976 | 0 |
| B85 | B93 | SCC | ALDH2-Glu504Lys | 0.316529757 | 0 |
| B85 | B94 | SCC | ALDH2-Lys504Lys | -0.31652975 | 0 |
| B85 | B86 | SCC | TBhCG | -0.23550867 | 8.49E-12 |
| B85 | B196 | SCC | TCO2 | -0.18495644 | 9.59E-08 |
| B85 | B156 | SCC | IgE | 0.180076452 | 2.09E-07 |
| B85 | B139 | SCC | ADA | -0.17487224 | 4.68E-07 |
| B86 | B90 | TBhCG | CYFRA 21-1 | 0.387038187 | 0 |
| B86 | B88 | TBhCG | CA 242 | -0.24224757 | 2.05E-12 |
| B88 | B89 | CA 242 | CA 72-4 | 0.45440469 | 0 |
| B89 | B137 | CA 72-4 | AFU | 0.240727087 | 2.83E-12 |
| B9 | B38 | RH | BAND% | 0.396202908 | 0 |
| B91 | B126 | TP | Alb | 0.534225502 | 0 |
| B91 | B127 | TP | A-G ratio | -0.50395759 | 0 |
| B91 | B130 | TP | GLB | 0.804559562 | 0 |
| B91 | B188 | TP | Ca | 0.346188669 | 0 |
| B92 | B101 | ALDH2-Glu504Glu | anti-Sc70 | 0.376577446 | 0 |
| B92 | B102 | ALDH2-Glu504Glu | anti-Sm | 0.376577444 | 0 |
| B92 | B103 | ALDH2-Glu504Glu | anti-SSA/Ro | 0.376577443 | 0 |

| | | | | | |
|---|---|---|---|---|---|
| B92 | B104 | ALDH2-Glu504Glu | anti-SSB | 0.376577441 | 0 |
| B92 | B105 | ALDH2-Glu504Glu | anti-U1RNP | 0.376577443 | 0 |
| B92 | B106 | ALDH2-Glu504Glu | anti-P | 0.376577441 | 0 |
| B92 | B107 | ALDH2-Glu504Glu | AnuA | 0.376577378 | 0 |
| B92 | B108 | ALDH2-Glu504Glu | Anti-Cyclin | 0.376577443 | 0 |
| B92 | B109 | ALDH2-Glu504Glu | AMA-M2 | 0.376577442 | 0 |
| B92 | B110 | ALDH2-Glu504Glu | AHA | 0.376577443 | 0 |
| B92 | B156 | ALDH2-Glu504Glu | IgE | -0.62654051 | 0 |
| B92 | B93 | ALDH2-Glu504Glu | ALDH2-Glu504Lys | -1 | 0 |
| B92 | B94 | ALDH2-Glu504Glu | ALDH2-Lys504Lys | 0.999999985 | 0 |
| B92 | B97 | ALDH2-Glu504Glu | dsDNA | 0.376577442 | 0 |
| B92 | B98 | ALDH2-Glu504Glu | anti-Jo-1 | 0.376577448 | 0 |
| B92 | B99 | ALDH2-Glu504Glu | anti-PM-Scl | 0.376577444 | 0 |
| B92 | B151 | ALDH2-Glu504Glu | Anti-CCP | 0.272205993 | 2.22E-15 |
| B92 | B158 | ALDH2-Glu504Glu | IgM | 0.210741002 | 1.10E-09 |
| B92 | B196 | ALDH2-Glu504Glu | TCO2 | 0.21053142 | 1.14E-09 |
| B93 | B101 | ALDH2-Glu504Lys | anti-Sc70 | -0.37657745 | 0 |
| B93 | B102 | ALDH2-Glu504Lys | anti-Sm | -0.37657745 | 0 |
| B93 | B103 | ALDH2-Glu504Lys | anti-SSA/Ro | -0.37657744 | 0 |
| B93 | B104 | ALDH2-Glu504Lys | anti-SSB | -0.37657744 | 0 |
| B93 | B105 | ALDH2-Glu504Lys | anti-U1RNP | -0.37657744 | 0 |
| B93 | B106 | ALDH2-Glu504Lys | anti-P | -0.37657744 | 0 |
| B93 | B107 | ALDH2-Glu504Lys | AnuA | -0.37657737 | 0 |
| B93 | B108 | ALDH2-Glu504Lys | Anti-Cyclin | -0.37657744 | 0 |
| B93 | B109 | ALDH2-Glu504Lys | AMA-M2 | -0.37657744 | 0 |
| B93 | B110 | ALDH2-Glu504Lys | AHA | -0.37657744 | 0 |
| B93 | B156 | ALDH2-Glu504Lys | IgE | 0.626540505 | 0 |
| B93 | B94 | ALDH2-Glu504Lys | ALDH2-Lys504Lys | -0.99999998 | 0 |
| B93 | B97 | ALDH2-Glu504Lys | dsDNA | -0.37657744 | 0 |
| B93 | B98 | ALDH2-Glu504Lys | anti-Jo-1 | -0.37657745 | 0 |
| B93 | B99 | ALDH2-Glu504Lys | anti-PM-Scl | -0.37657744 | 0 |
| B93 | B151 | ALDH2-Glu504Lys | Anti-CCP | -0.27220599 | 2.22E-15 |
| B93 | B158 | ALDH2-Glu504Lys | IgM | -0.210741 | 1.10E-09 |
| B93 | B196 | ALDH2-Glu504Lys | TCO2 | -0.21053142 | 1.14E-09 |
| B94 | B101 | ALDH2-Lys504Lys | anti-Sc70 | 0.376577441 | 0 |
| B94 | B102 | ALDH2-Lys504Lys | anti-Sm | 0.376577438 | 0 |
| B94 | B103 | ALDH2-Lys504Lys | anti-SSA/Ro | 0.376577435 | 0 |
| B94 | B104 | ALDH2-Lys504Lys | anti-SSB | 0.37657746 | 0 |
| B94 | B105 | ALDH2-Lys504Lys | anti-U1RNP | 0.376577444 | 0 |
| B94 | B106 | ALDH2-Lys504Lys | anti-P | 0.376577471 | 0 |
| B94 | B107 | ALDH2-Lys504Lys | AnuA | 0.376577617 | 0 |
| B94 | B108 | ALDH2-Lys504Lys | Anti-Cyclin | 0.376577434 | 0 |
| B94 | B109 | ALDH2-Lys504Lys | AMA-M2 | 0.376577443 | 0 |
| B94 | B110 | ALDH2-Lys504Lys | AHA | 0.376577436 | 0 |

| | | | | | |
|---|---|---|---|---|---|
| B94 | B156 | ALDH2-Lys504Lys | IgE | -0.6265405 | 0 |
| B94 | B97 | ALDH2-Lys504Lys | dsDNA | 0.376577462 | 0 |
| B94 | B98 | ALDH2-Lys504Lys | anti-Jo-1 | 0.376577451 | 0 |
| B94 | B99 | ALDH2-Lys504Lys | anti-PM-Scl | 0.37657744 | 0 |
| B94 | B151 | ALDH2-Lys504Lys | Anti-CCP | 0.272205989 | 2.22E-15 |
| B94 | B158 | ALDH2-Lys504Lys | IgM | 0.210740999 | 1.10E-09 |
| B94 | B196 | ALDH2-Lys504Lys | TCO2 | 0.210531417 | 1.14E-09 |
| B95 | B100 | anti-cenpb | anti-Ro52 | 0.995876789 | 0 |
| B95 | B101 | anti-cenpb | anti-Sc70 | -0.35718554 | 0 |
| B95 | B102 | anti-cenpb | anti-Sm | -0.35718553 | 0 |
| B95 | B103 | anti-cenpb | anti-SSA/Ro | -0.35718553 | 0 |
| B95 | B104 | anti-cenpb | anti-SSB | -0.35718553 | 0 |
| B95 | B105 | anti-cenpb | anti-U1RNP | -0.35718553 | 0 |
| B95 | B106 | anti-cenpb | anti-P | -0.35718553 | 0 |
| B95 | B107 | anti-cenpb | AnuA | -0.35718546 | 0 |
| B95 | B108 | anti-cenpb | Anti-Cyclin | -0.35718553 | 0 |
| B95 | B109 | anti-cenpb | AMA-M2 | -0.35718553 | 0 |
| B95 | B110 | anti-cenpb | AHA | -0.35718553 | 0 |
| B95 | B156 | anti-cenpb | IgE | 0.452480836 | 0 |
| B95 | B158 | anti-cenpb | IgM | 0.469568124 | 0 |
| B95 | B97 | anti-cenpb | dsDNA | -0.35718553 | 0 |
| B95 | B98 | anti-cenpb | anti-Jo-1 | -0.35718554 | 0 |
| B95 | B99 | anti-cenpb | anti-PM-Scl | -0.35718553 | 0 |
| B95 | B151 | anti-cenpb | Anti-CCP | -0.26426195 | 1.44E-14 |
| B96 | B225 | dsDNA (IIF) | hPBG | 0.629389806 | 0 |
| B97 | B102 | dsDNA | anti-Sm | 0.999999997 | 0 |
| B97 | B103 | dsDNA | anti-SSA/Ro | 0.999999994 | 0 |
| B97 | B104 | dsDNA | anti-SSB | 0.999999997 | 0 |
| B97 | B105 | dsDNA | anti-U1RNP | 0.999999995 | 0 |
| B97 | B106 | dsDNA | anti-P | 0.999999996 | 0 |
| B97 | B107 | dsDNA | AnuA | 0.999999876 | 0 |
| B97 | B108 | dsDNA | Anti-Cyclin | 0.999999998 | 0 |
| B97 | B109 | dsDNA | AMA-M2 | 0.999999994 | 0 |
| B97 | B110 | dsDNA | AHA | 0.999999997 | 0 |
| B97 | B151 | dsDNA | Anti-CCP | 0.452931824 | 0 |
| B97 | B156 | dsDNA | IgE | -0.34171399 | 0 |
| B97 | B158 | dsDNA | IgM | -0.28651441 | 0 |
| B97 | B186 | dsDNA | PFC | 0.308748712 | 0 |
| B97 | B99 | dsDNA | anti-PM-Scl | 0.999999998 | 0 |
| B97 | B157 | dsDNA | IgG | 0.273681123 | 1.33E-15 |
| B97 | B100 | dsDNA | anti-Ro52 | -0.27098086 | 2.89E-15 |
| B97 | B141 | dsDNA | AST/ALT | -0.26503331 | 1.20E-14 |
| B97 | B181 | dsDNA | APTT | -0.2352347 | 8.99E-12 |
| B97 | B154 | dsDNA | RF | 0.177161149 | 3.29E-07 |

| | | | | | |
|---|---|---|---|---|---|
| B98 | B101 | anti-Jo-1 | anti-Sc70 | 0.999999977 | 0 |
| B98 | B102 | anti-Jo-1 | anti-Sm | 0.999999995 | 0 |
| B98 | B107 | anti-Jo-1 | AnuA | 0.999999687 | 0 |
| B98 | B108 | anti-Jo-1 | Anti-Cyclin | 0.999999998 | 0 |
| B98 | B151 | anti-Jo-1 | Anti-CCP | 0.452931833 | 0 |
| B98 | B156 | anti-Jo-1 | IgE | -0.34171399 | 0 |
| B98 | B158 | anti-Jo-1 | IgM | -0.28651441 | 0 |
| B98 | B186 | anti-Jo-1 | PFC | 0.308748717 | 0 |
| B98 | B99 | anti-Jo-1 | anti-PM-Scl | 0.999999999 | 0 |
| B98 | B157 | anti-Jo-1 | IgG | 0.273681127 | 1.33E-15 |
| B98 | B100 | anti-Jo-1 | anti-Ro52 | -0.27098086 | 2.89E-15 |
| B98 | B141 | anti-Jo-1 | AST/ALT | -0.26503332 | 1.20E-14 |
| B98 | B181 | anti-Jo-1 | APTT | -0.23523471 | 8.99E-12 |
| B98 | B154 | anti-Jo-1 | RF | 0.177161152 | 3.29E-07 |
| B99 | B101 | anti-PM-Scl | anti-Sc70 | 0.99999999 | 0 |
| B99 | B104 | anti-PM-Scl | anti-SSB | 0.999999998 | 0 |
| B99 | B106 | anti-PM-Scl | anti-P | 0.999999992 | 0 |
| B99 | B107 | anti-PM-Scl | AnuA | 0.999999791 | 0 |
| B99 | B151 | anti-PM-Scl | Anti-CCP | 0.452931829 | 0 |
| B99 | B156 | anti-PM-Scl | IgE | -0.34171399 | 0 |
| B99 | B158 | anti-PM-Scl | IgM | -0.28651441 | 0 |
| B99 | B186 | anti-PM-Scl | PFC | 0.308748715 | 0 |
| B99 | B157 | anti-PM-Scl | IgG | 0.273681125 | 1.33E-15 |
| B99 | B100 | anti-PM-Scl | anti-Ro52 | -0.27098086 | 2.89E-15 |
| B99 | B141 | anti-PM-Scl | AST/ALT | -0.26503332 | 1.20E-14 |
| B99 | B181 | anti-PM-Scl | APTT | -0.23523471 | 8.99E-12 |
| B99 | B154 | anti-PM-Scl | RF | 0.17716115 | 3.29E-07 |

**Supplementary Table 32. PEI Correlations in Other Cardiac diseases**

| Row | Column | Row | Column | Cor | *P* |
|---|---|---|---|---|---|
| B1 | B190 | Gender | Cre | -0.551863 | 0 |
| B1 | B216 | Gender | BMI | -0.486999 | 0 |
| B1 | B221 | Gender | Hei | -0.669205 | 0 |
| B1 | B222 | Gender | NS | -0.635298 | 0 |
| B1 | B46 | Gender | RBC Count | -0.468552 | 0 |
| B1 | B51 | Gender | Hb | -0.526159 | 0 |
| B1 | B174 | Gender | Urea/Cr | 0.3787769 | 3.53E-13 |
| B1 | B192 | Gender | UA | -0.362932 | 3.78E-12 |
| B1 | B204 | Gender | Drinking | -0.348494 | 2.94E-11 |
| B1 | B209 | Gender | Smoking | -0.34107 | 8.10E-11 |
| B1 | B206 | Gender | Waist | -0.29553 | 2.32E-08 |
| B10 | B12 | PG Ⅰ | PG II | 0.7065773 | 0 |
| B10 | B39 | PG Ⅰ | Other cells | 0.7236072 | 0 |
| B10 | B47 | PG Ⅰ | RDW | 0.3581102 | 7.59E-12 |
| B100 | B103 | anti-Ro52 | anti-SSA/Ro | 1 | 0 |
| B100 | B105 | anti-Ro52 | anti-U1RNP | 1 | 0 |
| B100 | B106 | anti-Ro52 | anti-P | 1 | 0 |
| B100 | B107 | anti-Ro52 | AnuA | 1 | 0 |
| B100 | B110 | anti-Ro52 | AHA | -0.677574 | 0 |
| B100 | B137 | anti-Ro52 | AFU | 0.4450762 | 0 |
| B100 | B156 | anti-Ro52 | IgE | -0.471088 | 0 |
| B100 | B154 | anti-Ro52 | RF | 0.3746809 | 6.60E-13 |
| B100 | B179 | anti-Ro52 | α1-MG | 0.3483944 | 2.98E-11 |
| B100 | B180 | anti-Ro52 | Tf | 0.3483708 | 2.99E-11 |
| B100 | B178 | anti-Ro52 | Ig-U | 0.3457582 | 4.28E-11 |
| B100 | B139 | anti-Ro52 | ADA | -0.327837 | 4.62E-10 |
| B100 | B151 | anti-Ro52 | Anti-CCP | -0.296133 | 2.16E-08 |
| B100 | B138 | anti-Ro52 | PAB | 0.2920171 | 3.45E-08 |
| B100 | B134 | anti-Ro52 | CHE | 0.2606755 | 9.48E-07 |
| B101 | B102 | anti-Sc70 | anti-Sm | 1 | 0 |
| B101 | B106 | anti-Sc70 | anti-P | 1 | 0 |
| B101 | B107 | anti-Sc70 | AnuA | 0.9999999 | 0 |
| B101 | B110 | anti-Sc70 | AHA | -0.677574 | 0 |
| B101 | B137 | anti-Sc70 | AFU | 0.4450762 | 0 |
| B101 | B156 | anti-Sc70 | IgE | -0.471088 | 0 |
| B101 | B154 | anti-Sc70 | RF | 0.3746809 | 6.60E-13 |
| B101 | B179 | anti-Sc70 | α1-MG | 0.3483944 | 2.98E-11 |
| B101 | B180 | anti-Sc70 | Tf | 0.3483708 | 2.99E-11 |
| B101 | B178 | anti-Sc70 | Ig-U | 0.3457582 | 4.28E-11 |
| B101 | B139 | anti-Sc70 | ADA | -0.327837 | 4.62E-10 |
| B101 | B151 | anti-Sc70 | Anti-CCP | -0.296133 | 2.16E-08 |
| B101 | B138 | anti-Sc70 | PAB | 0.2920171 | 3.45E-08 |

| | | | | | |
|---|---|---|---|---|---|
| B101 | B134 | anti-Sc70 | CHE | 0.2606755 | 9.48E-07 |
| B102 | B104 | anti-Sm | anti-SSB | 1 | 0 |
| B102 | B108 | anti-Sm | Anti-Cyclin | 1 | 0 |
| B102 | B110 | anti-Sm | AHA | -0.677574 | 0 |
| B102 | B137 | anti-Sm | AFU | 0.4450762 | 0 |
| B102 | B156 | anti-Sm | IgE | -0.471088 | 0 |
| B102 | B154 | anti-Sm | RF | 0.3746809 | 6.60E-13 |
| B102 | B179 | anti-Sm | α1-MG | 0.3483944 | 2.98E-11 |
| B102 | B180 | anti-Sm | Tf | 0.3483708 | 2.99E-11 |
| B102 | B178 | anti-Sm | Ig-U | 0.3457582 | 4.28E-11 |
| B102 | B139 | anti-Sm | ADA | -0.327837 | 4.62E-10 |
| B102 | B151 | anti-Sm | Anti-CCP | -0.296133 | 2.16E-08 |
| B102 | B138 | anti-Sm | PAB | 0.2920171 | 3.45E-08 |
| B102 | B134 | anti-Sm | CHE | 0.2606755 | 9.48E-07 |
| B103 | B105 | anti-SSA/Ro | anti-U1RNP | 1 | 0 |
| B103 | B106 | anti-SSA/Ro | anti-P | 1 | 0 |
| B103 | B107 | anti-SSA/Ro | AnuA | 1 | 0 |
| B103 | B110 | anti-SSA/Ro | AHA | -0.677574 | 0 |
| B103 | B137 | anti-SSA/Ro | AFU | 0.4450762 | 0 |
| B103 | B156 | anti-SSA/Ro | IgE | -0.471088 | 0 |
| B103 | B154 | anti-SSA/Ro | RF | 0.3746809 | 6.60E-13 |
| B103 | B179 | anti-SSA/Ro | α1-MG | 0.3483944 | 2.98E-11 |
| B103 | B180 | anti-SSA/Ro | Tf | 0.3483708 | 2.99E-11 |
| B103 | B178 | anti-SSA/Ro | Ig-U | 0.3457582 | 4.28E-11 |
| B103 | B139 | anti-SSA/Ro | ADA | -0.327837 | 4.62E-10 |
| B103 | B151 | anti-SSA/Ro | Anti-CCP | -0.296133 | 2.16E-08 |
| B103 | B138 | anti-SSA/Ro | PAB | 0.2920171 | 3.45E-08 |
| B103 | B134 | anti-SSA/Ro | CHE | 0.2606755 | 9.48E-07 |
| B104 | B105 | anti-SSB | anti-U1RNP | 1 | 0 |
| B104 | B107 | anti-SSB | AnuA | 1 | 0 |
| B104 | B109 | anti-SSB | AMA-M2 | 1 | 0 |
| B104 | B110 | anti-SSB | AHA | -0.677574 | 0 |
| B104 | B137 | anti-SSB | AFU | 0.4450762 | 0 |
| B104 | B156 | anti-SSB | IgE | -0.471088 | 0 |
| B104 | B154 | anti-SSB | RF | 0.3746809 | 6.60E-13 |
| B104 | B179 | anti-SSB | α1-MG | 0.3483944 | 2.98E-11 |
| B104 | B180 | anti-SSB | Tf | 0.3483708 | 2.99E-11 |
| B104 | B178 | anti-SSB | Ig-U | 0.3457582 | 4.28E-11 |
| B104 | B139 | anti-SSB | ADA | -0.327837 | 4.62E-10 |
| B104 | B151 | anti-SSB | Anti-CCP | -0.296133 | 2.16E-08 |
| B104 | B138 | anti-SSB | PAB | 0.2920171 | 3.45E-08 |
| B104 | B134 | anti-SSB | CHE | 0.2606755 | 9.48E-07 |
| B105 | B106 | anti-U1RNP | anti-P | 1 | 0 |
| B105 | B107 | anti-U1RNP | AnuA | 1 | 0 |

| | | | | | |
|---|---|---|---|---|---|
| B105 | B109 | anti-U1RNP | AMA-M2 | 1 | 0 |
| B105 | B110 | anti-U1RNP | AHA | -0.677574 | 0 |
| B105 | B137 | anti-U1RNP | AFU | 0.4450762 | 0 |
| B105 | B156 | anti-U1RNP | IgE | -0.471088 | 0 |
| B105 | B154 | anti-U1RNP | RF | 0.3746809 | 6.60E-13 |
| B105 | B179 | anti-U1RNP | α1-MG | 0.3483944 | 2.98E-11 |
| B105 | B180 | anti-U1RNP | Tf | 0.3483708 | 2.99E-11 |
| B105 | B178 | anti-U1RNP | Ig-U | 0.3457582 | 4.28E-11 |
| B105 | B139 | anti-U1RNP | ADA | -0.327837 | 4.62E-10 |
| B105 | B151 | anti-U1RNP | Anti-CCP | -0.296133 | 2.16E-08 |
| B105 | B138 | anti-U1RNP | PAB | 0.2920171 | 3.45E-08 |
| B105 | B134 | anti-U1RNP | CHE | 0.2606755 | 9.48E-07 |
| B106 | B107 | anti-P | AnuA | 1 | 0 |
| B106 | B108 | anti-P | Anti-Cyclin | 1 | 0 |
| B106 | B110 | anti-P | AHA | -0.677574 | 0 |
| B106 | B137 | anti-P | AFU | 0.4450762 | 0 |
| B106 | B156 | anti-P | IgE | -0.471088 | 0 |
| B106 | B154 | anti-P | RF | 0.3746809 | 6.60E-13 |
| B106 | B179 | anti-P | α1-MG | 0.3483944 | 2.98E-11 |
| B106 | B180 | anti-P | Tf | 0.3483708 | 2.99E-11 |
| B106 | B178 | anti-P | Ig-U | 0.3457582 | 4.28E-11 |
| B106 | B139 | anti-P | ADA | -0.327837 | 4.62E-10 |
| B106 | B151 | anti-P | Anti-CCP | -0.296133 | 2.16E-08 |
| B106 | B138 | anti-P | PAB | 0.2920171 | 3.45E-08 |
| B106 | B134 | anti-P | CHE | 0.2606755 | 9.48E-07 |
| B107 | B108 | AnuA | Anti-Cyclin | 1 | 0 |
| B107 | B109 | AnuA | AMA-M2 | 1 | 0 |
| B107 | B110 | AnuA | AHA | -0.677574 | 0 |
| B107 | B137 | AnuA | AFU | 0.4450762 | 0 |
| B107 | B156 | AnuA | IgE | -0.471088 | 0 |
| B107 | B154 | AnuA | RF | 0.3746809 | 6.60E-13 |
| B107 | B179 | AnuA | α1-MG | 0.3483944 | 2.98E-11 |
| B107 | B180 | AnuA | Tf | 0.3483708 | 2.99E-11 |
| B107 | B178 | AnuA | Ig-U | 0.3457582 | 4.28E-11 |
| B107 | B139 | AnuA | ADA | -0.327837 | 4.62E-10 |
| B107 | B151 | AnuA | Anti-CCP | -0.296133 | 2.16E-08 |
| B107 | B138 | AnuA | PAB | 0.2920171 | 3.45E-08 |
| B107 | B134 | AnuA | CHE | 0.2606755 | 9.48E-07 |
| B108 | B110 | Anti-Cyclin | AHA | -0.677574 | 0 |
| B108 | B137 | Anti-Cyclin | AFU | 0.4450762 | 0 |
| B108 | B156 | Anti-Cyclin | IgE | -0.471088 | 0 |
| B108 | B154 | Anti-Cyclin | RF | 0.3746809 | 6.60E-13 |
| B108 | B179 | Anti-Cyclin | α1-MG | 0.3483944 | 2.98E-11 |
| B108 | B180 | Anti-Cyclin | Tf | 0.3483708 | 2.99E-11 |

| | | | | | |
|---|---|---|---|---|---|
| B108 | B178 | Anti-Cyclin | Ig-U | 0.3457582 | 4.28E-11 |
| B108 | B139 | Anti-Cyclin | ADA | -0.327837 | 4.62E-10 |
| B108 | B151 | Anti-Cyclin | Anti-CCP | -0.296133 | 2.16E-08 |
| B108 | B138 | Anti-Cyclin | PAB | 0.2920171 | 3.45E-08 |
| B108 | B134 | Anti-Cyclin | CHE | 0.2606755 | 9.48E-07 |
| B109 | B110 | AMA-M2 | AHA | -0.677574 | 0 |
| B109 | B137 | AMA-M2 | AFU | 0.4450762 | 0 |
| B109 | B156 | AMA-M2 | IgE | -0.471088 | 0 |
| B109 | B154 | AMA-M2 | RF | 0.3746809 | 6.60E-13 |
| B109 | B179 | AMA-M2 | α1-MG | 0.3483944 | 2.98E-11 |
| B109 | B180 | AMA-M2 | Tf | 0.3483708 | 2.99E-11 |
| B109 | B178 | AMA-M2 | Ig-U | 0.3457582 | 4.28E-11 |
| B109 | B139 | AMA-M2 | ADA | -0.327837 | 4.62E-10 |
| B109 | B151 | AMA-M2 | Anti-CCP | -0.296133 | 2.16E-08 |
| B109 | B138 | AMA-M2 | PAB | 0.2920171 | 3.45E-08 |
| B109 | B134 | AMA-M2 | CHE | 0.2606755 | 9.48E-07 |
| B11 | B12 | PGI/PGII | PG II | -0.6235 | 0 |
| B11 | B14 | PGI/PGII | G-17 | -0.272673 | 2.80E-07 |
| B110 | B139 | AHA | ADA | 0.4730055 | 0 |
| B110 | B151 | AHA | Anti-CCP | 0.4117734 | 1.78E-15 |
| B110 | B157 | AHA | IgG | 0.3006494 | 1.28E-08 |
| B110 | B123 | AHA | hs-CRP1 | 0.2834881 | 8.85E-08 |
| B110 | B137 | AHA | AFU | -0.281236 | 1.13E-07 |
| B110 | B134 | AHA | CHE | -0.263735 | 6.98E-07 |
| B111 | B112 | TSH | FT4 | -0.27841 | 1.53E-07 |
| B112 | B114 | FT4 | TSH TT3 TT4(TT4) | 0.573777 | 0 |
| B112 | B198 | FT4 | P | 0.3429856 | 6.25E-11 |
| B113 | B115 | FT3 | TSH TT3 TT4(TT3) | 0.4355051 | 0 |
| B114 | B115 | TSH TT3 TT4(TT4) | TSH TT3 TT4(TT3) | 0.5715784 | 0 |
| B114 | B198 | TSH TT3 TT4(TT4) | P | 0.3647514 | 2.90E-12 |
| B114 | B138 | TSH TT3 TT4(TT4) | PAB | -0.311856 | 3.40E-09 |
| B115 | B172 | TSH TT3 TT4(TT3) | KB | 0.2825224 | 9.83E-08 |
| B119 | B124 | Hcv-rna | ACTH | 0.7060753 | 0 |
| B119 | B125 | Hcv-rna | Cor | 0.7060753 | 0 |
| B119 | B169 | Hcv-rna | U-CRY | -0.999998 | 0 |
| B119 | B120 | Hcv-rna | HCV-cAg | 0.4207534 | 4.44E-16 |
| B12 | B39 | PG II | Other cells | 0.5460093 | 0 |
| B12 | B47 | PG II | RDW | 0.3714917 | 1.07E-12 |
| B12 | B43 | PG II | MCV | 0.2742743 | 2.37E-07 |
| B12 | B46 | PG II | RBC Count | -0.265051 | 6.11E-07 |
| B120 | B121 | HCV-cAg | C3 | -0.420753 | 4.44E-16 |
| B120 | B122 | HCV-cAg | C4 | -0.420754 | 4.44E-16 |
| B120 | B152 | HCV-cAg | ACA-Ig(G-A-M) | -0.420755 | 4.44E-16 |
| B120 | B159 | HCV-cAg | β2-MG | -0.420754 | 4.44E-16 |

| | | | | | |
|---|---|---|---|---|---|
| B120 | B169 | HCV-cAg | U-CRY | -0.420752 | 4.44E-16 |
| B120 | B124 | HCV-cAg | ACTH | 0.2970836 | 1.94E-08 |
| B120 | B125 | HCV-cAg | Cor | 0.2970836 | 1.94E-08 |
| B120 | B138 | HCV-cAg | PAB | 0.2836953 | 8.65E-08 |
| B121 | B124 | C3 | ACTH | -0.706075 | 0 |
| B121 | B125 | C3 | Cor | -0.706075 | 0 |
| B121 | B169 | C3 | U-CRY | 0.9999985 | 0 |
| B122 | B124 | C4 | ACTH | -0.706076 | 0 |
| B122 | B125 | C4 | Cor | -0.706076 | 0 |
| B122 | B169 | C4 | U-CRY | 0.9999991 | 0 |
| B123 | B139 | hs-CRP1 | ADA | 0.3285782 | 4.20E-10 |
| B124 | B152 | ACTH | ACA-Ig(G-A-M) | -0.706081 | 0 |
| B124 | B159 | ACTH | β2-MG | -0.706076 | 0 |
| B124 | B169 | ACTH | U-CRY | -0.706074 | 0 |
| B124 | B138 | ACTH | PAB | 0.3303903 | 3.32E-10 |
| B124 | B134 | ACTH | CHE | 0.3080333 | 5.38E-09 |
| B124 | B150 | ACTH | ALP | -0.295635 | 2.29E-08 |
| B125 | B152 | Cor | ACA-Ig(G-A-M) | -0.706081 | 0 |
| B125 | B159 | Cor | β2-MG | -0.706076 | 0 |
| B125 | B169 | Cor | U-CRY | -0.706074 | 0 |
| B125 | B138 | Cor | PAB | 0.3303903 | 3.32E-10 |
| B125 | B134 | Cor | CHE | 0.3080333 | 5.38E-09 |
| B125 | B150 | Cor | ALP | -0.295635 | 2.29E-08 |
| B126 | B127 | Alb | A-G ratio | 0.420368 | 4.44E-16 |
| B126 | B188 | Alb | Ca | 0.383826 | 1.61E-13 |
| B127 | B130 | A-G ratio | GLB | -0.91246 | 0 |
| B128 | B131 | ALT | AST | 0.9136999 | 0 |
| B128 | B141 | ALT | AST/ALT | -0.374555 | 6.73E-13 |
| B129 | B132 | IBIL | Dbil | 0.4786094 | 0 |
| B129 | B133 | IBIL | TBIL | 0.9326676 | 0 |
| B132 | B133 | Dbil | TBIL | 0.763076 | 0 |
| B132 | B170 | Dbil | U-GLU | 0.3229079 | 8.66E-10 |
| B132 | B140 | Dbil | r-GT | 0.2706046 | 3.47E-07 |
| B134 | B138 | CHE | PAB | 0.6574514 | 0 |
| B134 | B139 | CHE | ADA | -0.57184 | 0 |
| B134 | B150 | CHE | ALP | -0.376045 | 5.36E-13 |
| B134 | B137 | CHE | AFU | 0.3526122 | 1.65E-11 |
| B134 | B135 | CHE | TBA | -0.284701 | 7.75E-08 |
| B135 | B138 | TBA | PAB | -0.293228 | 3.01E-08 |
| B136 | B195 | 5'NT | Na | 0.3431411 | 6.12E-11 |
| B136 | B140 | 5'NT | r-GT | 0.2983415 | 1.68E-08 |
| B136 | B196 | 5'NT | TCO2 | 0.2717436 | 3.08E-07 |
| B136 | B214 | 5'NT | CVA-L | -0.266576 | 5.24E-07 |
| B136 | B176 | 5'NT | Ucr | -0.264902 | 6.21E-07 |

| | | | | | |
|---|---|---|---|---|---|
| B137 | B180 | AFU | Tf | 0.3334046 | 2.24E-10 |
| B137 | B179 | AFU | α1-MG | 0.3328175 | 2.42E-10 |
| B137 | B178 | AFU | Ig-U | 0.3309255 | 3.10E-10 |
| B137 | B149 | AFU | TGAb | -0.299296 | 1.50E-08 |
| B137 | B148 | AFU | Anti_TPO | -0.270573 | 3.48E-07 |
| B137 | B194 | AFU | Cl | -0.268193 | 4.44E-07 |
| B138 | B139 | PAB | ADA | -0.558418 | 0 |
| B138 | B196 | PAB | TCO2 | 0.3830476 | 1.82E-13 |
| B138 | B197 | PAB | AG | -0.355803 | 1.05E-11 |
| B138 | B198 | PAB | P | -0.353231 | 1.52E-11 |
| B139 | B150 | ADA | ALP | 0.4562027 | 0 |
| B139 | B200 | ADA | OP | 0.446008 | 0 |
| B139 | B197 | ADA | AG | 0.345843 | 4.23E-11 |
| B139 | B198 | ADA | P | 0.3040158 | 8.67E-09 |
| B140 | B192 | r-GT | UA | 0.2614873 | 8.74E-07 |
| B142 | B143 | 25-(OH)D | β-CTx | -0.539114 | 0 |
| B142 | B151 | 25-(OH)D | Anti-CCP | -0.445942 | 0 |
| B142 | B144 | 25-(OH)D | Total-PINP | -0.361214 | 4.85E-12 |
| B142 | B168 | 25-(OH)D | NMP22 | -0.266039 | 5.53E-07 |
| B143 | B144 | β-CTx | Total-PINP | 0.2673843 | 4.83E-07 |
| B144 | B151 | Total-PINP | Anti-CCP | -0.292236 | 3.36E-08 |
| B145 | B167 | BGP | PC | 0.9123575 | 0 |
| B146 | B190 | CysC | Cre | 0.2917108 | 3.57E-08 |
| B146 | B189 | CysC | eGFB | -0.280694 | 1.20E-07 |
| B148 | B149 | Anti_TPO | TGAb | 0.8299072 | 0 |
| B149 | B199 | TGAb | Mg | -0.26972 | 3.80E-07 |
| B15 | B145 | CD3-CD19+(B cell) | BGP | 0.7619957 | 0 |
| B15 | B16 | CD3-CD19+(B cell) | CD3+T | 0.6552665 | 0 |
| B15 | B18 | CD3-CD19+(B cell) | CD3+CD4+helper(inducer)T | 0.5295491 | 0 |
| B15 | B20 | CD3-CD19+(B cell) | CD3+CD4+CD8+ | 0.6027691 | 0 |
| B15 | B36 | CD3-CD19+(B cell) | V-Lym% | 0.6611119 | 0 |
| B15 | B167 | CD3-CD19+(B cell) | PC | 0.2771327 | 1.75E-07 |
| B150 | B197 | ALP | AG | 0.4947299 | 0 |
| B150 | B200 | ALP | OP | 0.587617 | 0 |
| B150 | B195 | ALP | Na | 0.3824992 | 1.99E-13 |
| B150 | B194 | ALP | Cl | -0.267501 | 4.77E-07 |
| B151 | B168 | Anti-CCP | NMP22 | 0.406367 | 4.00E-15 |
| B153 | B206 | Glu | Waist | 0.2671246 | 4.95E-07 |
| B153 | B175 | Glu | ALB-U | 0.2663688 | 5.35E-07 |
| B153 | B215 | Glu | BMI | 0.2632708 | 7.31E-07 |
| B154 | B156 | RF | IgE | -0.275266 | 2.13E-07 |
| B157 | B158 | IgG | IgM | 0.6856963 | 0 |
| B159 | B169 | β2-MG | U-CRY | 0.9999977 | 0 |
| B16 | B167 | CD3+T | PC | 0.5321325 | 0 |

| | | | | | |
|---|---|---|---|---|---|
| B16 | B19 | CD3+T | CD3+CD4+/CD3+CD8+ | -0.707107 | 0 |
| B16 | B94 | CD3+T | ALDH2-Lys504Lys | 0.9274495 | 0 |
| B163 | B166 | SG | Pro | 0.350969 | 2.08E-11 |
| B163 | B191 | SG | Urea | 0.2618345 | 8.44E-07 |
| B167 | B223 | PC | FH | 0.8413753 | 0 |
| B17 | B116 | CD3+CD16+CD56+NK | TT3 | -0.971537 | 0 |
| B17 | B21 | CD3+CD16+CD56+NK | CD3+CD4-CD8- | -0.653275 | 0 |
| B17 | B23 | CD3+CD16+CD56+NK | CD3-CD16+CD56+(NK) | -0.773001 | 0 |
| B17 | B33 | CD3+CD16+CD56+NK | Prog | -0.785674 | 0 |
| B17 | B92 | CD3+CD16+CD56+NK | ALDH2-Glu504Glu | -0.848668 | 0 |
| B17 | B35 | CD3+CD16+CD56+NK | β2-MG | -0.322832 | 8.75E-10 |
| B170 | B173 | U-GLU | NTT | 0.3116847 | 3.47E-09 |
| B173 | B227 | NTT | OD | 0.4565176 | 0 |
| B174 | B191 | Urea/Cr | Urea | 0.6856339 | 0 |
| B174 | B221 | Urea/Cr | Hei | -0.340985 | 8.19E-11 |
| B174 | B190 | Urea/Cr | Cre | -0.333201 | 2.30E-10 |
| B175 | B177 | ALB-U | ACR | 0.9601111 | 0 |
| B175 | B225 | ALB-U | hPBG | 0.9403562 | 0 |
| B175 | B197 | ALB-U | AG | 0.3608084 | 5.15E-12 |
| B175 | B200 | ALB-U | OP | 0.3487657 | 2.83E-11 |
| B175 | B199 | ALB-U | Mg | -0.340682 | 8.53E-11 |
| B175 | B195 | ALB-U | Na | 0.2822361 | 1.01E-07 |
| B176 | B227 | Ucr | OD | 0.312856 | 3.01E-09 |
| B176 | B199 | Ucr | Mg | 0.2647621 | 6.30E-07 |
| B177 | B199 | ACR | Mg | -0.425904 | 0 |
| B177 | B225 | ACR | hPBG | 0.9934792 | 0 |
| B177 | B200 | ACR | OP | 0.3839004 | 1.59E-13 |
| B177 | B197 | ACR | AG | 0.3611841 | 4.87E-12 |
| B177 | B195 | ACR | Na | 0.3321689 | 2.64E-10 |
| B178 | B179 | Ig-U | α1-MG | 0.999964 | 0 |
| B178 | B180 | Ig-U | Tf | 0.9999732 | 0 |
| B178 | B193 | Ig-U | K | -0.264921 | 6.20E-07 |
| B179 | B180 | α1-MG | Tf | 0.9999908 | 0 |
| B179 | B193 | α1-MG | K | -0.266162 | 5.46E-07 |
| B18 | B145 | CD3+CD4+helper(inducer)T | BGP | 0.9730801 | 0 |
| B18 | B223 | CD3+CD4+helper(inducer)T | FH | 0.5039098 | 0 |
| B18 | B94 | CD3+CD4+helper(inducer)T | ALDH2-Lys504Lys | 0.8308996 | 0 |
| B18 | B167 | CD3+CD4+helper(inducer)T | PC | 0.4174236 | 6.66E-16 |
| B180 | B193 | Tf | K | -0.26684 | 5.10E-07 |
| B181 | B182 | APTT | TT | -0.987929 | 0 |
| B181 | B183 | APTT | PTA | -0.998964 | 0 |
| B181 | B185 | APTT | PT | 0.9999974 | 0 |
| B182 | B183 | TT | PTA | 0.9798542 | 0 |
| B182 | B184 | TT | INR | -0.987929 | 0 |

| | | | | | |
|---|---|---|---|---|---|
| B182 | B185 | TT | PT | -0.988279 | 0 |
| B182 | B186 | TT | PFC | 0.9879286 | 0 |
| B183 | B184 | PTA | INR | -0.998964 | 0 |
| B183 | B185 | PTA | PT | -0.998857 | 0 |
| B183 | B186 | PTA | PFC | 0.9989637 | 0 |
| B184 | B185 | INR | PT | 0.9999974 | 0 |
| B185 | B186 | PT | PFC | -0.999997 | 0 |
| B189 | B190 | eGFB | Cre | -0.632092 | 0 |
| B189 | B191 | eGFB | Urea | -0.453568 | 0 |
| B189 | B192 | eGFB | UA | -0.292128 | 3.40E-08 |
| B19 | B36 | CD3+CD4+/CD3+CD8+ | V-Lym% | 0.925 | 0 |
| B19 | B94 | CD3+CD4+/CD3+CD8+ | ALDH2-Lys504Lys | 0.4276995 | 0 |
| B19 | B30 | CD3+CD4+/CD3+CD8+ | LH | 0.41833 | 4.44E-16 |
| B19 | B223 | CD3+CD4+/CD3+CD8+ | FH | 0.4099458 | 2.22E-15 |
| B19 | B20 | CD3+CD4+/CD3+CD8+ | CD3+CD4+CD8+ | 0.3703986 | 1.26E-12 |
| B190 | B191 | Cre | Urea | 0.4266709 | 0 |
| B190 | B192 | Cre | UA | 0.4929016 | 0 |
| B190 | B222 | Cre | NS | 0.3707794 | 1.19E-12 |
| B190 | B221 | Cre | Hei | 0.3129576 | 2.97E-09 |
| B190 | B216 | Cre | BMI | 0.2743589 | 2.35E-07 |
| B192 | B222 | UA | NS | 0.3477821 | 3.24E-11 |
| B192 | B206 | UA | Waist | 0.3304513 | 3.30E-10 |
| B192 | B216 | UA | BMI | 0.3241101 | 7.44E-10 |
| B192 | B210 | UA | Hip | 0.2621237 | 8.20E-07 |
| B192 | B215 | UA | BMI | 0.2604655 | 9.68E-07 |
| B193 | B199 | K | Mg | 0.2938383 | 2.81E-08 |
| B194 | B196 | Cl | TCO2 | -0.444134 | 0 |
| B194 | B195 | Cl | Na | 0.3643735 | 3.06E-12 |
| B194 | B197 | Cl | AG | -0.358801 | 6.87E-12 |
| B195 | B200 | Na | OP | 0.7899762 | 0 |
| B195 | B197 | Na | AG | 0.3314028 | 2.91E-10 |
| B195 | B225 | Na | hPBG | 0.3219014 | 9.83E-10 |
| B195 | B198 | Na | P | 0.3129657 | 2.97E-09 |
| B196 | B197 | TCO2 | AG | -0.422465 | 2.22E-16 |
| B197 | B200 | AG | OP | 0.551586 | 0 |
| B197 | B225 | AG | hPBG | 0.360197 | 5.62E-12 |
| B198 | B200 | P | OP | 0.2742039 | 2.38E-07 |
| B199 | B225 | Mg | hPBG | -0.433257 | 0 |
| B2 | B189 | Age | eGFB | -0.562101 | 0 |
| B2 | B220 | Age | SBP | 0.4168992 | 6.66E-16 |
| B2 | B221 | Age | Hei | -0.352881 | 1.59E-11 |
| B2 | B207 | Age | WHR | 0.291169 | 3.79E-08 |
| B2 | B126 | Age | Alb | -0.263115 | 7.43E-07 |
| B20 | B38 | CD3+CD4+CD8+ | BAND% | 0.6625892 | 0 |

| | | | | | |
|---|---|---|---|---|---|
| B20 | B94 | CD3+CD4+CD8+ | ALDH2-Lys504Lys | 0.9575565 | 0 |
| B20 | B96 | CD3+CD4+CD8+ | dsDNA (IIF) | 0.9370426 | 0 |
| B20 | B145 | CD3+CD4+CD8+ | BGP | 0.4122831 | 1.55E-15 |
| B20 | B167 | CD3+CD4+CD8+ | PC | 0.2883549 | 5.19E-08 |
| B200 | B225 | OP | hPBG | 0.3785088 | 3.68E-13 |
| B204 | B209 | Drinking | Smoking | 0.5172652 | 0 |
| B204 | B222 | Drinking | NS | 0.2717253 | 3.09E-07 |
| B206 | B207 | Waist | WHR | 0.8199758 | 0 |
| B206 | B210 | Waist | Hip | 0.7311798 | 0 |
| B206 | B215 | Waist | BMI | 0.7845557 | 0 |
| B206 | B216 | Waist | BMI | 0.7479765 | 0 |
| B206 | B222 | Waist | NS | 0.6200392 | 0 |
| B206 | B218 | Waist | DBP | 0.3149052 | 2.34E-09 |
| B207 | B215 | WHR | BMI | 0.4929029 | 0 |
| B207 | B216 | WHR | BMI | 0.4254204 | 0 |
| B207 | B222 | WHR | NS | 0.4401768 | 0 |
| B207 | B220 | WHR | SBP | 0.3066292 | 6.36E-09 |
| B209 | B222 | Smoking | NS | 0.286818 | 6.15E-08 |
| B21 | B116 | CD3+CD4-CD8- | TT3 | -0.969697 | 0 |
| B21 | B29 | CD3+CD4-CD8- | E2 | -0.58191 | 0 |
| B21 | B31 | CD3+CD4-CD8- | FSH | -0.615457 | 0 |
| B21 | B33 | CD3+CD4-CD8- | Prog | -0.959403 | 0 |
| B21 | B35 | CD3+CD4-CD8- | β2-MG | -0.438019 | 0 |
| B21 | B92 | CD3+CD4-CD8- | ALDH2-Glu504Glu | -0.930484 | 0 |
| B210 | B215 | Hip | BMI | 0.7505311 | 0 |
| B210 | B216 | Hip | BMI | 0.771757 | 0 |
| B210 | B222 | Hip | NS | 0.5411742 | 0 |
| B210 | B221 | Hip | Hei | 0.3342811 | 2.00E-10 |
| B210 | B218 | Hip | DBP | 0.261549 | 8.69E-07 |
| B211 | B212 | IOP-R | IOP-L | 0.8828711 | 0 |
| B213 | B214 | CVA-R | CVA-L | 0.7410663 | 0 |
| B213 | B228 | CVA-R | OS | -0.271932 | 3.02E-07 |
| B215 | B216 | BMI | BMI | 0.7712224 | 0 |
| B215 | B222 | BMI | NS | 0.5145522 | 0 |
| B216 | B221 | BMI | Hei | 0.659667 | 0 |
| B216 | B222 | BMI | NS | 0.6965114 | 0 |
| B216 | B218 | BMI | DBP | 0.3289076 | 4.03E-10 |
| B218 | B220 | DBP | SBP | 0.5156754 | 0 |
| B218 | B222 | DBP | NS | 0.3512656 | 2.00E-11 |
| B221 | B222 | Hei | NS | 0.4834524 | 0 |
| B23 | B35 | CD3-CD16+CD56+(NK) | β2-MG | -0.807427 | 0 |
| B24 | B27 | αHBDH | LDH | 0.7833234 | 0 |
| B25 | B26 | CK | CKMB | 0.501067 | 0 |
| B25 | B58 | CK | HCT | 0.3458359 | 4.24E-11 |

| | | | | | |
|---|---|---|---|---|---|
| B25 | B62 | CK | WBHSRVI | 0.337243 | 1.35E-10 |
| B25 | B63 | CK | WBRSR(mPas_200) | 0.3212341 | 1.07E-09 |
| B25 | B61 | CK | WBHSRV | 0.3171331 | 1.78E-09 |
| B25 | B66 | CK | WBRSR(mPas_100) | 0.3137239 | 2.71E-09 |
| B25 | B67 | CK | WBRSR(mPas_50) | 0.3095037 | 4.51E-09 |
| B25 | B156 | CK | IgE | 0.3005047 | 1.31E-08 |
| B25 | B60 | CK | WBRVI | 0.2935884 | 2.89E-08 |
| B25 | B64 | CK | WBRSR(mPas_5) | 0.2795994 | 1.35E-07 |
| B25 | B56 | CK | IR | 0.2777663 | 1.64E-07 |
| B25 | B34 | CK | TK | -0.273042 | 2.69E-07 |
| B27 | B198 | LDH | P | 0.2957322 | 2.26E-08 |
| B29 | B116 | E2 | TT3 | -0.525596 | 0 |
| B29 | B202 | E2 | e-times | -0.67514 | 0 |
| B29 | B35 | E2 | β2-MG | -0.822217 | 0 |
| B3 | B227 | Hcy | OD | 0.6638989 | 0 |
| B3 | B78 | Hcy | HBcAb | 0.348971 | 2.75E-11 |
| B3 | B66 | Hcy | WBRSR(mPas_100) | 0.3039148 | 8.77E-09 |
| B3 | B173 | Hcy | NTT | 0.2996731 | 1.44E-08 |
| B3 | B67 | Hcy | WBRSR(mPas_50) | 0.2991224 | 1.53E-08 |
| B3 | B63 | Hcy | WBRSR(mPas_200) | 0.286014 | 6.71E-08 |
| B30 | B223 | LH | FH | -0.534522 | 0 |
| B33 | B116 | Prog | TT3 | -0.675136 | 0 |
| B33 | B35 | Prog | β2-MG | -0.944235 | 0 |
| B33 | B92 | Prog | ALDH2-Glu504Glu | -0.891056 | 0 |
| B34 | B86 | TK | TBhCG | 0.3351428 | 1.79E-10 |
| B34 | B85 | TK | SCC | -0.290857 | 3.92E-08 |
| B34 | B73 | TK | ApoA | -0.289882 | 4.38E-08 |
| B35 | B116 | β2-MG | TT3 | -0.849491 | 0 |
| B35 | B37 | β2-MG | BAND | -0.996116 | 0 |
| B35 | B92 | β2-MG | ALDH2-Glu504Glu | -0.285299 | 7.26E-08 |
| B36 | B145 | V-Lym% | BGP | 0.8428367 | 0 |
| B36 | B167 | V-Lym% | PC | 0.6714553 | 0 |
| B36 | B223 | V-Lym% | FH | 0.7267221 | 0 |
| B36 | B96 | V-Lym% | dsDNA (IIF) | 0.6166441 | 0 |
| B38 | B145 | BAND% | BGP | 0.394877 | 2.75E-14 |
| B39 | B47 | Other cells | RDW | 0.4036773 | 6.44E-15 |
| B39 | B43 | Other cells | MCV | 0.2958136 | 2.24E-08 |
| B39 | B46 | Other cells | RBC Count | -0.281627 | 1.08E-07 |
| B39 | B44 | Other cells | MCH | 0.2802694 | 1.25E-07 |
| B4 | B20 | WB-Ca | CD3+CD4+CD8+ | 0.5607673 | 0 |
| B4 | B223 | WB-Ca | FH | -0.534522 | 0 |
| B4 | B6 | WB-Ca | WB-Fe | 0.952381 | 0 |
| B4 | B94 | WB-Ca | ALDH2-Lys504Lys | 0.3067192 | 6.30E-09 |
| B40 | B42 | WBC | MON | 0.6462466 | 0 |

| | | | | | |
|---|---|---|---|---|---|
| B40 | B46 | WBC | RBC Count | 0.2748657 | 2.22E-07 |
| B41 | B42 | MON % | MON | 0.6510527 | 0 |
| B41 | B53 | MON % | GR% | -0.32748 | 4.84E-10 |
| B43 | B44 | MCV | MCH | 0.937983 | 0 |
| B43 | B46 | MCV | RBC Count | -0.605752 | 0 |
| B43 | B47 | MCV | RDW | 0.6618354 | 0 |
| B43 | B45 | MCV | MCHC | 0.2670495 | 4.99E-07 |
| B44 | B45 | MCH | MCHC | 0.582651 | 0 |
| B44 | B46 | MCH | RBC Count | -0.483135 | 0 |
| B44 | B47 | MCH | RDW | 0.5108088 | 0 |
| B45 | B51 | MCHC | Hb | 0.5561541 | 0 |
| B45 | B216 | MCHC | BMI | 0.266137 | 5.48E-07 |
| B46 | B47 | RBC Count | RDW | -0.596386 | 0 |
| B46 | B51 | RBC Count | Hb | 0.7222406 | 0 |
| B46 | B221 | RBC Count | Hei | 0.418991 | 4.44E-16 |
| B46 | B222 | RBC Count | NS | 0.4088116 | 2.66E-15 |
| B46 | B216 | RBC Count | BMI | 0.36046 | 5.41E-12 |
| B46 | B174 | RBC Count | Urea/Cr | -0.332059 | 2.68E-10 |
| B46 | B192 | RBC Count | UA | 0.3149997 | 2.32E-09 |
| B46 | B210 | RBC Count | Hip | 0.3017705 | 1.13E-08 |
| B47 | B51 | RDW | Hb | -0.307234 | 5.92E-09 |
| B47 | B132 | RDW | Dbil | 0.2783196 | 1.54E-07 |
| B48 | B53 | LYM% | GR% | -0.952141 | 0 |
| B49 | B50 | baso% | eo% | 0.2891129 | 4.77E-08 |
| B5 | B19 | WB-Mg | CD3+CD4+/CD3+CD8+ | 0.975 | 0 |
| B5 | B223 | WB-Mg | FH | -0.521749 | 0 |
| B5 | B6 | WB-Mg | WB-Fe | 0.876501 | 0 |
| B5 | B96 | WB-Mg | dsDNA (IIF) | 0.7589466 | 0 |
| B5 | B16 | WB-Mg | CD3+T | 0.3771236 | 4.55E-13 |
| B5 | B20 | WB-Mg | CD3+CD4+CD8+ | 0.3333587 | 2.26E-10 |
| B5 | B94 | WB-Mg | ALDH2-Lys504Lys | 0.3279029 | 4.58E-10 |
| B5 | B36 | WB-Mg | V-Lym% | 0.325 | 6.64E-10 |
| B50 | B135 | eo% | TBA | 0.2871127 | 5.95E-08 |
| B50 | B144 | eo% | Total-PINP | 0.2748426 | 2.23E-07 |
| B51 | B216 | Hb | BMI | 0.4634879 | 0 |
| B51 | B221 | Hb | Hei | 0.4849722 | 0 |
| B51 | B222 | Hb | NS | 0.4766995 | 0 |
| B51 | B174 | Hb | Urea/Cr | -0.371473 | 1.07E-12 |
| B51 | B192 | Hb | UA | 0.3505259 | 2.21E-11 |
| B51 | B210 | Hb | Hip | 0.3360695 | 1.58E-10 |
| B51 | B204 | Hb | Drinking | 0.322577 | 9.03E-10 |
| B51 | B206 | Hb | Waist | 0.3131364 | 2.91E-09 |
| B51 | B190 | Hb | Cre | 0.3042557 | 8.43E-09 |
| B51 | B126 | Hb | Alb | 0.2999789 | 1.39E-08 |

| | | | | | |
|---|---|---|---|---|---|
| B51 | B218 | Hb | DBP | 0.2809374 | 1.17E-07 |
| B54 | B59 | ESR | WBRV | 0.4369507 | 0 |
| B54 | B69 | ESR | PV | 0.4563678 | 0 |
| B54 | B57 | ESR | RBCAI | 0.3892538 | 6.84E-14 |
| B54 | B62 | ESR | WBHSRVI | -0.369402 | 1.46E-12 |
| B54 | B56 | ESR | IR | -0.349592 | 2.52E-11 |
| B54 | B55 | ESR | TK | -0.307885 | 5.48E-09 |
| B54 | B187 | ESR | NSE | 0.290904 | 3.90E-08 |
| B54 | B58 | ESR | HCT | -0.282473 | 9.88E-08 |
| B54 | B90 | ESR | CYFRA 21-1 | 0.271705 | 3.09E-07 |
| B55 | B148 | TK | Anti_TPO | -0.513873 | 0 |
| B55 | B149 | TK | TGAb | -0.617109 | 0 |
| B55 | B56 | TK | IR | 0.9261342 | 0 |
| B55 | B60 | TK | WBRVI | 0.7770793 | 0 |
| B55 | B61 | TK | WBHSRV | 0.793523 | 0 |
| B55 | B62 | TK | WBHSRVI | 0.7823046 | 0 |
| B55 | B69 | TK | PV | -0.931431 | 0 |
| B56 | B148 | IR | Anti_TPO | -0.462163 | 0 |
| B56 | B149 | IR | TGAb | -0.530381 | 0 |
| B56 | B58 | IR | HCT | 0.4565936 | 0 |
| B56 | B60 | IR | WBRVI | 0.919728 | 0 |
| B56 | B61 | IR | WBHSRV | 0.9476427 | 0 |
| B56 | B62 | IR | WBHSRVI | 0.9536138 | 0 |
| B56 | B63 | IR | WBRSR(mPas_200) | 0.5474995 | 0 |
| B56 | B64 | IR | WBRSR(mPas_5) | 0.4645448 | 0 |
| B56 | B66 | IR | WBRSR(mPas_100) | 0.527273 | 0 |
| B56 | B67 | IR | WBRSR(mPas_50) | 0.5195731 | 0 |
| B56 | B69 | IR | PV | -0.814583 | 0 |
| B56 | B65 | IR | WBRSR(mPas_1) | 0.3917342 | 4.62E-14 |
| B56 | B57 | IR | RBCAI | -0.314634 | 2.42E-09 |
| B56 | B59 | IR | WBRV | 0.2825487 | 9.80E-08 |
| B57 | B59 | RBCAI | WBRV | 0.5792043 | 0 |
| B57 | B62 | RBCAI | WBHSRVI | -0.374151 | 7.15E-13 |
| B57 | B58 | RBCAI | HCT | -0.369889 | 1.36E-12 |
| B57 | B69 | RBCAI | PV | 0.3235225 | 8.01E-10 |
| B57 | B61 | RBCAI | WBHSRV | -0.323517 | 8.02E-10 |
| B57 | B136 | RBCAI | 5'NT | 0.3194342 | 1.34E-09 |
| B57 | B63 | RBCAI | WBRSR(mPas_200) | -0.266836 | 5.10E-07 |
| B58 | B60 | HCT | WBRVI | 0.5878196 | 0 |
| B58 | B61 | HCT | WBHSRV | 0.591403 | 0 |
| B58 | B62 | HCT | WBHSRVI | 0.6953927 | 0 |
| B58 | B63 | HCT | WBRSR(mPas_200) | 0.8349531 | 0 |
| B58 | B64 | HCT | WBRSR(mPas_5) | 0.756765 | 0 |
| B58 | B65 | HCT | WBRSR(mPas_1) | 0.6641532 | 0 |

| | | | | | |
|---|---|---|---|---|---|
| B58 | B66 | HCT | WBRSR(mPas_100) | 0.830881 | 0 |
| B58 | B67 | HCT | WBRSR(mPas_50) | 0.8231658 | 0 |
| B58 | B78 | HCT | HBcAb | 0.3756964 | 5.66E-13 |
| B58 | B227 | HCT | OD | 0.3050067 | 7.71E-09 |
| B58 | B73 | HCT | ApoA | 0.2855761 | 7.04E-08 |
| B59 | B60 | WBRV | WBRVI | 0.4943886 | 0 |
| B59 | B61 | WBRV | WBHSRV | 0.4355207 | 0 |
| B59 | B63 | WBRV | WBRSR(mPas_200) | 0.5152659 | 0 |
| B59 | B64 | WBRV | WBRSR(mPas_5) | 0.7029655 | 0 |
| B59 | B65 | WBRV | WBRSR(mPas_1) | 0.8027181 | 0 |
| B59 | B66 | WBRV | WBRSR(mPas_100) | 0.530665 | 0 |
| B59 | B67 | WBRV | WBRSR(mPas_50) | 0.5598034 | 0 |
| B59 | B62 | WBRV | WBHSRVI | 0.2774 | 1.70E-07 |
| B59 | B75 | WBRV | ApoA1/B | -0.267076 | 4.98E-07 |
| B6 | B15 | WB-Fe | CD3-CD19+(B cell) | 0.9455144 | 0 |
| B6 | B16 | WB-Fe | CD3+T | 0.8639355 | 0 |
| B6 | B167 | WB-Fe | PC | 0.4807502 | 0 |
| B60 | B149 | WBRVI | TGAb | -0.44797 | 0 |
| B60 | B61 | WBRVI | WBHSRV | 0.9095782 | 0 |
| B60 | B62 | WBRVI | WBHSRVI | 0.934975 | 0 |
| B60 | B63 | WBRVI | WBRSR(mPas_200) | 0.6506764 | 0 |
| B60 | B64 | WBRVI | WBRSR(mPas_5) | 0.6614145 | 0 |
| B60 | B65 | WBRVI | WBRSR(mPas_1) | 0.6403458 | 0 |
| B60 | B66 | WBRVI | WBRSR(mPas_100) | 0.6459529 | 0 |
| B60 | B67 | WBRVI | WBRSR(mPas_50) | 0.6517527 | 0 |
| B60 | B69 | WBRVI | PV | -0.661233 | 0 |
| B60 | B148 | WBRVI | Anti_TPO | -0.385642 | 1.21E-13 |
| B60 | B78 | WBRVI | HBcAb | 0.2615164 | 8.72E-07 |
| B61 | B62 | WBHSRV | WBHSRVI | 0.9605797 | 0 |
| B61 | B63 | WBHSRV | WBRSR(mPas_200) | 0.7621804 | 0 |
| B61 | B64 | WBHSRV | WBRSR(mPas_5) | 0.6858802 | 0 |
| B61 | B65 | WBHSRV | WBRSR(mPas_1) | 0.606236 | 0 |
| B61 | B66 | WBHSRV | WBRSR(mPas_100) | 0.7445613 | 0 |
| B61 | B67 | WBHSRV | WBRSR(mPas_50) | 0.7382259 | 0 |
| B61 | B69 | WBHSRV | PV | -0.623792 | 0 |
| B61 | B149 | WBHSRV | TGAb | -0.403654 | 6.66E-15 |
| B61 | B148 | WBHSRV | Anti_TPO | -0.373123 | 8.35E-13 |
| B61 | B227 | WBHSRV | OD | 0.3496656 | 2.50E-11 |
| B61 | B78 | WBHSRV | HBcAb | 0.3357414 | 1.65E-10 |
| B61 | B75 | WBHSRV | ApoA1/B | -0.28421 | 8.18E-08 |
| B61 | B76 | WBHSRV | ApoB | 0.2768318 | 1.81E-07 |
| B62 | B149 | WBHSRVI | TGAb | -0.437738 | 0 |
| B62 | B63 | WBHSRVI | WBRSR(mPas_200) | 0.7304466 | 0 |
| B62 | B64 | WBHSRVI | WBRSR(mPas_5) | 0.6405917 | 0 |

| | | | | | |
|---|---|---|---|---|---|
| B62 | B65 | WBHSRVI | WBRSR(mPas_1) | 0.5505292 | 0 |
| B62 | B66 | WBHSRVI | WBRSR(mPas_100) | 0.71617 | 0 |
| B62 | B67 | WBHSRVI | WBRSR(mPas_50) | 0.7076264 | 0 |
| B62 | B69 | WBHSRVI | PV | -0.690807 | 0 |
| B62 | B148 | WBHSRVI | Anti_TPO | -0.414295 | 1.11E-15 |
| B62 | B227 | WBHSRVI | OD | 0.3471177 | 3.55E-11 |
| B62 | B78 | WBHSRVI | HBcAb | 0.3166295 | 1.90E-09 |
| B63 | B227 | WBRSR(mPas_200) | OD | 0.4753951 | 0 |
| B63 | B64 | WBRSR(mPas_200) | WBRSR(mPas_5) | 0.9559301 | 0 |
| B63 | B65 | WBRSR(mPas_200) | WBRSR(mPas_1) | 0.879555 | 0 |
| B63 | B66 | WBRSR(mPas_200) | WBRSR(mPas_100) | 0.9971702 | 0 |
| B63 | B67 | WBRSR(mPas_200) | WBRSR(mPas_50) | 0.9947287 | 0 |
| B63 | B78 | WBRSR(mPas_200) | HBcAb | 0.4855576 | 0 |
| B63 | B75 | WBRSR(mPas_200) | ApoA1/B | -0.321205 | 1.07E-09 |
| B63 | B74 | WBRSR(mPas_200) | ApoA1 | -0.277943 | 1.61E-07 |
| B64 | B227 | WBRSR(mPas_5) | OD | 0.4353726 | 0 |
| B64 | B65 | WBRSR(mPas_5) | WBRSR(mPas_1) | 0.9799979 | 0 |
| B64 | B66 | WBRSR(mPas_5) | WBRSR(mPas_100) | 0.9660892 | 0 |
| B64 | B67 | WBRSR(mPas_5) | WBRSR(mPas_50) | 0.9758884 | 0 |
| B64 | B78 | WBRSR(mPas_5) | HBcAb | 0.4618157 | 0 |
| B64 | B75 | WBRSR(mPas_5) | ApoA1/B | -0.321646 | 1.02E-09 |
| B64 | B74 | WBRSR(mPas_5) | ApoA1 | -0.277182 | 1.74E-07 |
| B65 | B66 | WBRSR(mPas_1) | WBRSR(mPas_100) | 0.8954464 | 0 |
| B65 | B67 | WBRSR(mPas_1) | WBRSR(mPas_50) | 0.9130382 | 0 |
| B65 | B78 | WBRSR(mPas_1) | HBcAb | 0.4180875 | 4.44E-16 |
| B65 | B227 | WBRSR(mPas_1) | OD | 0.3661495 | 2.36E-12 |
| B65 | B75 | WBRSR(mPas_1) | ApoA1/B | -0.31152 | 3.54E-09 |
| B65 | B74 | WBRSR(mPas_1) | ApoA1 | -0.26713 | 4.95E-07 |
| B66 | B227 | WBRSR(mPas_100) | OD | 0.5023004 | 0 |
| B66 | B67 | WBRSR(mPas_100) | WBRSR(mPas_50) | 0.9991207 | 0 |
| B66 | B78 | WBRSR(mPas_100) | HBcAb | 0.4928246 | 0 |
| B66 | B75 | WBRSR(mPas_100) | ApoA1/B | -0.312567 | 3.12E-09 |
| B66 | B74 | WBRSR(mPas_100) | ApoA1 | -0.271747 | 3.08E-07 |
| B67 | B227 | WBRSR(mPas_50) | OD | 0.4962085 | 0 |
| B67 | B78 | WBRSR(mPas_50) | HBcAb | 0.4900917 | 0 |
| B67 | B75 | WBRSR(mPas_50) | ApoA1/B | -0.315378 | 2.21E-09 |
| B67 | B74 | WBRSR(mPas_50) | ApoA1 | -0.272795 | 2.76E-07 |
| B69 | B148 | PV | Anti_TPO | 0.5624759 | 0 |
| B69 | B149 | PV | TGAb | 0.6535188 | 0 |
| B7 | B117 | WB-Cop | TT4 | 0.4870882 | 0 |
| B7 | B202 | WB-Cop | e-times | 0.5487955 | 0 |
| B7 | B22 | WB-Cop | CD3+CD8+(T) | 0.5423261 | 0 |
| B7 | B33 | WB-Cop | Prog | -0.960211 | 0 |
| B7 | B9 | WB-Cop | RH | -0.870927 | 0 |

| | | | | | |
|---|---|---|---|---|---|
| B7 | B93 | WB-Cop | ALDH2-Glu504Lys | 0.4348283 | 0 |
| B7 | B31 | WB-Cop | FSH | -0.271163 | 3.27E-07 |
| B70 | B77 | LDL-C | TC | 0.8464178 | 0 |
| B70 | B76 | LDL-C | ApoB | 0.2996321 | 1.45E-08 |
| B71 | B77 | TG | TC | 0.4312883 | 0 |
| B71 | B72 | TG | HDL-C | -0.371982 | 9.92E-13 |
| B71 | B91 | TG | TP | 0.3043009 | 8.38E-09 |
| B71 | B140 | TG | r-GT | 0.266725 | 5.16E-07 |
| B72 | B216 | HDL-C | BMI | -0.342662 | 6.53E-11 |
| B72 | B222 | HDL-C | NS | -0.304167 | 8.52E-09 |
| B72 | B192 | HDL-C | UA | -0.296183 | 2.15E-08 |
| B72 | B206 | HDL-C | Waist | -0.281317 | 1.12E-07 |
| B73 | B198 | ApoA | P | 0.2989067 | 1.57E-08 |
| B73 | B76 | ApoA | ApoB | 0.2831469 | 9.18E-08 |
| B74 | B75 | ApoA1 | ApoA1/B | 0.6160661 | 0 |
| B74 | B123 | ApoA1 | hs-CRP1 | -0.40708 | 3.55E-15 |
| B74 | B135 | ApoA1 | TBA | 0.2989816 | 1.56E-08 |
| B74 | B193 | ApoA1 | K | 0.2877388 | 5.55E-08 |
| B75 | B76 | ApoA1/B | ApoB | -0.74347 | 0 |
| B75 | B135 | ApoA1/B | TBA | 0.3325075 | 2.52E-10 |
| B75 | B134 | ApoA1/B | CHE | -0.260529 | 9.61E-07 |
| B76 | B137 | ApoB | AFU | 0.2951117 | 2.43E-08 |
| B76 | B194 | ApoB | Cl | -0.294741 | 2.53E-08 |
| B76 | B134 | ApoB | CHE | 0.2779818 | 1.60E-07 |
| B76 | B77 | ApoB | TC | 0.2602555 | 9.88E-07 |
| B77 | B188 | TC | Ca | 0.2719844 | 3.01E-07 |
| B78 | B227 | HBcAb | OD | 0.5071433 | 0 |
| B79 | B119 | hb V -dna | Hcv-rna | 0.4970761 | 0 |
| B79 | B121 | hb V -dna | C3 | -0.497076 | 0 |
| B79 | B122 | hb V -dna | C4 | -0.497076 | 0 |
| B79 | B124 | hb V -dna | ACTH | 0.7060754 | 0 |
| B79 | B125 | hb V -dna | Cor | 0.7060754 | 0 |
| B79 | B152 | hb V -dna | ACA-Ig(G-A-M) | -0.49708 | 0 |
| B79 | B159 | hb V -dna | β2-MG | -0.497077 | 0 |
| B79 | B169 | hb V -dna | U-CRY | -0.497075 | 0 |
| B79 | B114 | hb V -dna | TSH TT3 TT4(TT4) | -0.356655 | 9.34E-12 |
| B79 | B112 | hb V -dna | FT4 | -0.281076 | 1.15E-07 |
| B8 | B18 | WB-Zinc | CD3+CD4+helper(inducer)T | 0.9329904 | 0 |
| B8 | B145 | WB-Zinc | BGP | 0.4145133 | 8.88E-16 |
| B8 | B16 | WB-Zinc | CD3+T | -0.30429 | 8.39E-09 |
| B82 | B129 | CEA | IBIL | 0.3471683 | 3.53E-11 |
| B82 | B133 | CEA | TBIL | 0.2954756 | 2.33E-08 |
| B83 | B84 | AFP | C-PSA | 0.5570136 | 0 |
| B83 | B86 | AFP | TBhCG | 0.6485863 | 0 |

| | | | | | |
|---|---|---|---|---|---|
| B83 | B87 | AFP | NSE | 0.4157773 | 8.88E-16 |
| B84 | B86 | C-PSA | TBhCG | 0.6014747 | 0 |
| B84 | B85 | C-PSA | SCC | 0.393742 | 3.33E-14 |
| B84 | B87 | C-PSA | NSE | 0.3256834 | 6.09E-10 |
| B86 | B90 | TBhCG | CYFRA 21-1 | 0.3309367 | 3.10E-10 |
| B9 | B29 | RH | E2 | -0.489798 | 0 |
| B9 | B31 | RH | FSH | -0.860309 | 0 |
| B9 | B37 | RH | BAND | -0.794719 | 0 |
| B9 | B33 | RH | Prog | -0.353209 | 1.52E-11 |
| B90 | B187 | CYFRA 21-1 | NSE | 0.5976094 | 0 |
| B91 | B126 | TP | Alb | 0.4679088 | 0 |
| B91 | B127 | TP | A-G ratio | -0.576741 | 0 |
| B91 | B130 | TP | GLB | 0.8331217 | 0 |
| B91 | B188 | TP | Ca | 0.3846503 | 1.42E-13 |
| B92 | B116 | ALDH2-Glu504Glu | TT3 | -0.395456 | 2.53E-14 |
| B93 | B202 | ALDH2-Glu504Lys | e-times | 0.8451543 | 0 |
| B94 | B145 | ALDH2-Lys504Lys | BGP | 0.4669085 | 0 |
| B94 | B167 | ALDH2-Lys504Lys | PC | 0.5105442 | 0 |
| B94 | B223 | ALDH2-Lys504Lys | FH | 0.6163242 | 0 |
| B95 | B101 | anti-cenpb | anti-Sc70 | 1 | 0 |
| B95 | B104 | anti-cenpb | anti-SSB | 1 | 0 |
| B95 | B106 | anti-cenpb | anti-P | 1 | 0 |
| B95 | B107 | anti-cenpb | AnuA | 1 | 0 |
| B95 | B108 | anti-cenpb | Anti-Cyclin | 1 | 0 |
| B95 | B109 | anti-cenpb | AMA-M2 | 1 | 0 |
| B95 | B110 | anti-cenpb | AHA | -0.677574 | 0 |
| B95 | B137 | anti-cenpb | AFU | 0.4450762 | 0 |
| B95 | B156 | anti-cenpb | IgE | -0.471088 | 0 |
| B95 | B98 | anti-cenpb | anti-Jo-1 | 1 | 0 |
| B95 | B154 | anti-cenpb | RF | 0.3746809 | 6.60E-13 |
| B95 | B179 | anti-cenpb | α1-MG | 0.3483944 | 2.98E-11 |
| B95 | B180 | anti-cenpb | Tf | 0.3483708 | 2.99E-11 |
| B95 | B178 | anti-cenpb | Ig-U | 0.3457582 | 4.28E-11 |
| B95 | B139 | anti-cenpb | ADA | -0.327837 | 4.62E-10 |
| B95 | B151 | anti-cenpb | Anti-CCP | -0.296133 | 2.16E-08 |
| B95 | B138 | anti-cenpb | PAB | 0.2920171 | 3.45E-08 |
| B95 | B134 | anti-cenpb | CHE | 0.2606755 | 9.48E-07 |
| B97 | B100 | dsDNA | anti-Ro52 | 1 | 0 |
| B97 | B103 | dsDNA | anti-SSA/Ro | 1 | 0 |
| B97 | B105 | dsDNA | anti-U1RNP | 1 | 0 |
| B97 | B108 | dsDNA | Anti-Cyclin | 1 | 0 |
| B97 | B109 | dsDNA | AMA-M2 | 1 | 0 |
| B97 | B110 | dsDNA | AHA | -0.677574 | 0 |
| B97 | B137 | dsDNA | AFU | 0.4450762 | 0 |

| | | | | | |
|---|---|---|---|---|---|
| B97 | B156 | dsDNA | IgE | -0.471088 | 0 |
| B97 | B99 | dsDNA | anti-PM-Scl | 1 | 0 |
| B97 | B154 | dsDNA | RF | 0.3746809 | 6.60E-13 |
| B97 | B179 | dsDNA | α1-MG | 0.3483944 | 2.98E-11 |
| B97 | B180 | dsDNA | Tf | 0.3483708 | 2.99E-11 |
| B97 | B178 | dsDNA | Ig-U | 0.3457582 | 4.28E-11 |
| B97 | B139 | dsDNA | ADA | -0.327837 | 4.62E-10 |
| B97 | B151 | dsDNA | Anti-CCP | -0.296133 | 2.16E-08 |
| B97 | B138 | dsDNA | PAB | 0.2920171 | 3.45E-08 |
| B97 | B134 | dsDNA | CHE | 0.2606755 | 9.48E-07 |
| B98 | B100 | anti-Jo-1 | anti-Ro52 | 1 | 0 |
| B98 | B101 | anti-Jo-1 | anti-Sc70 | 1 | 0 |
| B98 | B102 | anti-Jo-1 | anti-Sm | 1 | 0 |
| B98 | B103 | anti-Jo-1 | anti-SSA/Ro | 1 | 0 |
| B98 | B104 | anti-Jo-1 | anti-SSB | 1 | 0 |
| B98 | B105 | anti-Jo-1 | anti-U1RNP | 1 | 0 |
| B98 | B107 | anti-Jo-1 | AnuA | 1 | 0 |
| B98 | B109 | anti-Jo-1 | AMA-M2 | 1 | 0 |
| B98 | B110 | anti-Jo-1 | AHA | -0.677574 | 0 |
| B98 | B137 | anti-Jo-1 | AFU | 0.4450762 | 0 |
| B98 | B156 | anti-Jo-1 | IgE | -0.471088 | 0 |
| B98 | B154 | anti-Jo-1 | RF | 0.3746809 | 6.60E-13 |
| B98 | B179 | anti-Jo-1 | α1-MG | 0.3483944 | 2.98E-11 |
| B98 | B180 | anti-Jo-1 | Tf | 0.3483708 | 2.99E-11 |
| B98 | B178 | anti-Jo-1 | Ig-U | 0.3457582 | 4.28E-11 |
| B98 | B139 | anti-Jo-1 | ADA | -0.327837 | 4.62E-10 |
| B98 | B151 | anti-Jo-1 | Anti-CCP | -0.296133 | 2.16E-08 |
| B98 | B138 | anti-Jo-1 | PAB | 0.2920171 | 3.45E-08 |
| B98 | B134 | anti-Jo-1 | CHE | 0.2606755 | 9.48E-07 |
| B99 | B100 | anti-PM-Scl | anti-Ro52 | 1 | 0 |
| B99 | B103 | anti-PM-Scl | anti-SSA/Ro | 1 | 0 |
| B99 | B104 | anti-PM-Scl | anti-SSB | 1 | 0 |
| B99 | B105 | anti-PM-Scl | anti-U1RNP | 1 | 0 |
| B99 | B106 | anti-PM-Scl | anti-P | 1 | 0 |
| B99 | B107 | anti-PM-Scl | AnuA | 1 | 0 |
| B99 | B109 | anti-PM-Scl | AMA-M2 | 1 | 0 |
| B99 | B110 | anti-PM-Scl | AHA | -0.677574 | 0 |
| B99 | B137 | anti-PM-Scl | AFU | 0.4450762 | 0 |
| B99 | B156 | anti-PM-Scl | IgE | -0.471088 | 0 |
| B99 | B154 | anti-PM-Scl | RF | 0.3746809 | 6.60E-13 |
| B99 | B179 | anti-PM-Scl | α1-MG | 0.3483944 | 2.98E-11 |
| B99 | B180 | anti-PM-Scl | Tf | 0.3483708 | 2.99E-11 |
| B99 | B178 | anti-PM-Scl | Ig-U | 0.3457582 | 4.28E-11 |
| B99 | B139 | anti-PM-Scl | ADA | -0.327837 | 4.62E-10 |

| | | | | | |
|---|---|---|---|---|---|
| B99 | B151 | anti-PM-Scl | Anti-CCP | -0.296133 | 2.16E-08 |
| B99 | B138 | anti-PM-Scl | PAB | 0.2920171 | 3.45E-08 |
| B99 | B134 | anti-PM-Scl | CHE | 0.2606755 | 9.48E-07 |

**Supplementary Table 33. PEI Correlations in Heart disease**

| Row | Column | Row | Column | Cor | *P* |
|---|---|---|---|---|---|
| B1 | B221 | Gender | Hei | -0.663065 | 0 |
| B1 | B222 | Gender | NS | -0.620303 | 0 |
| B1 | B216 | Gender | BMI | -0.492208 | 2.22E-15 |
| B1 | B51 | Gender | Hb | -0.47262 | 3.82E-14 |
| B1 | B174 | Gender | Urea/Cr | 0.4267699 | 1.49E-11 |
| B1 | B46 | Gender | RBC Count | -0.356285 | 2.94E-08 |
| B1 | B206 | Gender | Waist | -0.337545 | 1.65E-07 |
| B10 | B119 | PG Ⅰ | Hcv-rna | -0.717241 | 0 |
| B10 | B12 | PG Ⅰ | PG II | 0.8465091 | 0 |
| B10 | B120 | PG Ⅰ | HCV-cAg | -0.71724 | 0 |
| B10 | B121 | PG Ⅰ | C3 | 0.7172407 | 0 |
| B10 | B122 | PG Ⅰ | C4 | 0.7172411 | 0 |
| B10 | B137 | PG Ⅰ | AFU | 0.5854136 | 0 |
| B10 | B151 | PG Ⅰ | Anti-CCP | -0.717627 | 0 |
| B10 | B152 | PG Ⅰ | ACA-Ig(G-A-M) | 0.7172291 | 0 |
| B10 | B159 | PG Ⅰ | β2-MG | 0.7172416 | 0 |
| B10 | B169 | PG Ⅰ | U-CRY | 0.7172403 | 0 |
| B10 | B179 | PG Ⅰ | α1-MG | 0.7172409 | 0 |
| B10 | B198 | PG Ⅰ | P | -0.563346 | 0 |
| B10 | B79 | PG Ⅰ | hbＶ-dna | -0.717241 | 0 |
| B10 | B193 | PG Ⅰ | K | 0.5025705 | 4.44E-16 |
| B10 | B138 | PG Ⅰ | PAB | 0.4482896 | 1.01E-12 |
| B10 | B199 | PG Ⅰ | Mg | 0.3879271 | 1.22E-09 |
| B10 | B214 | PG Ⅰ | CVA-L | 0.3525425 | 4.19E-08 |
| B100 | B178 | anti-Ro52 | Ig-U | 0.4050957 | 1.87E-10 |
| B100 | B102 | anti-Ro52 | anti-Sm | -0.320256 | 7.39E-07 |
| B101 | B107 | anti-Sc70 | AnuA | 0.9709832 | 0 |
| B101 | B110 | anti-Sc70 | AHA | 0.8001713 | 0 |
| B102 | B125 | anti-Sm | Cor | -0.707107 | 0 |
| B103 | B105 | anti-SSA/Ro | anti-U1RNP | 0.9215122 | 0 |
| B103 | B107 | anti-SSA/Ro | AnuA | 0.9231862 | 0 |
| B103 | B156 | anti-SSA/Ro | IgE | 0.625 | 0 |
| B104 | B125 | anti-SSB | Cor | -1 | 0 |
| B104 | B180 | anti-SSB | Tf | -0.946729 | 0 |
| B104 | B108 | anti-SSB | Anti-Cyclin | 0.5070725 | 2.22E-16 |
| B104 | B202 | anti-SSB | e-times | -0.425926 | 1.66E-11 |
| B105 | B107 | anti-U1RNP | AnuA | 0.8892937 | 0 |
| B105 | B110 | anti-U1RNP | AHA | 0.5938631 | 0 |
| B106 | B223 | anti-P | FH | -0.442326 | 2.18E-12 |
| B107 | B110 | AnuA | AHA | 0.870153 | 0 |
| B108 | B117 | Anti-Cyclin | TT4 | -0.405096 | 1.87E-10 |
| B11 | B12 | PGI/PGII | PG II | -0.553566 | 0 |

| | | | | | |
|---|---|---|---|---|---|
| B11 | B144 | PGI/PGII | Total-PINP | 0.4528059 | 5.62E-13 |
| B11 | B143 | PGI/PGII | β-CTx | 0.4250319 | 1.84E-11 |
| B111 | B148 | TSH | Anti_TPO | 0.469054 | 6.26E-14 |
| B111 | B112 | TSH | FT4 | -0.451203 | 6.94E-13 |
| B112 | B114 | FT4 | TSH TT3 TT4(TT4) | 0.3406103 | 1.26E-07 |
| B112 | B148 | FT4 | Anti_TPO | -0.340268 | 1.30E-07 |
| B113 | B115 | FT3 | TSH TT3 TT4(TT3) | 0.6001771 | 0 |
| B114 | B115 | TSH TT3 TT4(TT4) | TSH TT3 TT4(TT3) | 0.4915197 | 2.44E-15 |
| B115 | B183 | TSH TT3 TT4(TT3) | PTA | -0.345643 | 7.95E-08 |
| B115 | B136 | TSH TT3 TT4(TT3) | 5'NT | 0.3314872 | 2.82E-07 |
| B116 | B117 | TT3 | TT4 | -0.532624 | 0 |
| B116 | B180 | TT3 | Tf | -0.521862 | 0 |
| B116 | B202 | TT3 | e-times | -0.926367 | 0 |
| B116 | B178 | TT3 | Ig-U | -0.414284 | 6.54E-11 |
| B118 | B148 | CEA1 | Anti_TPO | 0.5338012 | 0 |
| B118 | B149 | CEA1 | TGAb | 0.4352039 | 5.33E-12 |
| B118 | B187 | CEA1 | NSE | 0.3767751 | 3.89E-09 |
| B119 | B120 | Hcv-rna | HCV-cAg | 0.9999994 | 0 |
| B119 | B121 | Hcv-rna | C3 | -1 | 0 |
| B119 | B137 | Hcv-rna | AFU | -0.804069 | 0 |
| B119 | B138 | Hcv-rna | PAB | -0.61557 | 0 |
| B119 | B152 | Hcv-rna | ACA-Ig(G-A-M) | -0.999984 | 0 |
| B119 | B169 | Hcv-rna | U-CRY | -0.999999 | 0 |
| B119 | B198 | Hcv-rna | P | 0.579283 | 0 |
| B119 | B199 | Hcv-rna | Mg | -0.54099 | 0 |
| B119 | B193 | Hcv-rna | K | -0.500587 | 6.66E-16 |
| B119 | B200 | Hcv-rna | OP | -0.389346 | 1.05E-09 |
| B119 | B195 | Hcv-rna | Na | -0.340435 | 1.28E-07 |
| B12 | B119 | PG II | Hcv-rna | -0.838553 | 0 |
| B12 | B120 | PG II | HCV-cAg | -0.838552 | 0 |
| B12 | B121 | PG II | C3 | 0.8385527 | 0 |
| B12 | B122 | PG II | C4 | 0.8385531 | 0 |
| B12 | B137 | PG II | AFU | 0.6611052 | 0 |
| B12 | B138 | PG II | PAB | 0.5133206 | 0 |
| B12 | B151 | PG II | Anti-CCP | -0.839005 | 0 |
| B12 | B152 | PG II | ACA-Ig(G-A-M) | 0.8385391 | 0 |
| B12 | B159 | PG II | β2-MG | 0.8385537 | 0 |
| B12 | B169 | PG II | U-CRY | 0.8385522 | 0 |
| B12 | B179 | PG II | α1-MG | 0.8385529 | 0 |
| B12 | B198 | PG II | P | -0.51533 | 0 |
| B12 | B79 | PG II | hbＶ-dna | -0.838553 | 0 |
| B12 | B193 | PG II | K | 0.4929735 | 2.00E-15 |
| B12 | B199 | PG II | Mg | 0.4599735 | 2.17E-13 |
| B12 | B144 | PG II | Total-PINP | -0.34284 | 1.03E-07 |

| | | | | | |
|---|---|---|---|---|---|
| B120 | B121 | HCV-cAg | C3 | -0.999999 | 0 |
| B120 | B137 | HCV-cAg | AFU | -0.804068 | 0 |
| B120 | B138 | HCV-cAg | PAB | -0.61557 | 0 |
| B120 | B152 | HCV-cAg | ACA-Ig(G-A-M) | -0.999978 | 0 |
| B120 | B159 | HCV-cAg | β2-MG | -1 | 0 |
| B120 | B169 | HCV-cAg | U-CRY | -0.999998 | 0 |
| B120 | B179 | HCV-cAg | α1-MG | -0.999999 | 0 |
| B120 | B198 | HCV-cAg | P | 0.5792827 | 0 |
| B120 | B199 | HCV-cAg | Mg | -0.54099 | 0 |
| B120 | B193 | HCV-cAg | K | -0.500587 | 6.66E-16 |
| B120 | B200 | HCV-cAg | OP | -0.389345 | 1.05E-09 |
| B120 | B195 | HCV-cAg | Na | -0.340435 | 1.28E-07 |
| B121 | B137 | C3 | AFU | 0.8040689 | 0 |
| B121 | B138 | C3 | PAB | 0.6155703 | 0 |
| B121 | B152 | C3 | ACA-Ig(G-A-M) | 0.9999847 | 0 |
| B121 | B169 | C3 | U-CRY | 0.9999991 | 0 |
| B121 | B179 | C3 | α1-MG | 1 | 0 |
| B121 | B198 | C3 | P | -0.579283 | 0 |
| B121 | B199 | C3 | Mg | 0.5409901 | 0 |
| B121 | B193 | C3 | K | 0.5005868 | 6.66E-16 |
| B121 | B200 | C3 | OP | 0.3893457 | 1.05E-09 |
| B121 | B195 | C3 | Na | 0.3404348 | 1.28E-07 |
| B122 | B137 | C4 | AFU | 0.8040693 | 0 |
| B122 | B138 | C4 | PAB | 0.6155706 | 0 |
| B122 | B152 | C4 | ACA-Ig(G-A-M) | 0.9999861 | 0 |
| B122 | B169 | C4 | U-CRY | 0.9999998 | 0 |
| B122 | B198 | C4 | P | -0.579283 | 0 |
| B122 | B199 | C4 | Mg | 0.5409903 | 0 |
| B122 | B193 | C4 | K | 0.500587 | 6.66E-16 |
| B122 | B200 | C4 | OP | 0.3893459 | 1.05E-09 |
| B122 | B195 | C4 | Na | 0.3404349 | 1.28E-07 |
| B125 | B178 | Cor | Ig-U | -0.968963 | 0 |
| B125 | B202 | Cor | e-times | 0.5555556 | 0 |
| B126 | B127 | Alb | A-G ratio | 0.5665903 | 0 |
| B126 | B188 | Alb | Ca | 0.3375888 | 1.65E-07 |
| B127 | B130 | A-G ratio | GLB | -0.870423 | 0 |
| B128 | B131 | ALT | AST | 0.8480139 | 0 |
| B128 | B141 | ALT | AST/ALT | -0.469086 | 6.26E-14 |
| B129 | B132 | IBIL | Dbil | 0.65067 | 0 |
| B129 | B133 | IBIL | TBIL | 0.9759174 | 0 |
| B132 | B133 | Dbil | TBIL | 0.8005993 | 0 |
| B132 | B140 | Dbil | r-GT | 0.4040573 | 2.10E-10 |
| B134 | B138 | CHE | PAB | 0.6160091 | 0 |
| B134 | B195 | CHE | Na | 0.5806989 | 0 |

| | | | | | |
|---|---|---|---|---|---|
| B134 | B135 | CHE | TBA | 0.4750541 | 2.71E-14 |
| B134 | B139 | CHE | ADA | -0.402539 | 2.48E-10 |
| B134 | B200 | CHE | OP | 0.3990347 | 3.66E-10 |
| B134 | B197 | CHE | AG | 0.3753246 | 4.51E-09 |
| B135 | B136 | TBA | 5'NT | 0.4098303 | 1.09E-10 |
| B135 | B138 | TBA | PAB | 0.3678531 | 9.54E-09 |
| B135 | B137 | TBA | AFU | 0.3277762 | 3.90E-07 |
| B136 | B138 | 5'NT | PAB | 0.5094958 | 0 |
| B136 | B139 | 5'NT | ADA | -0.689059 | 0 |
| B136 | B198 | 5'NT | P | 0.5205801 | 0 |
| B137 | B138 | AFU | PAB | 0.6983825 | 0 |
| B137 | B151 | AFU | Anti-CCP | -0.804502 | 0 |
| B137 | B152 | AFU | ACA-Ig(G-A-M) | 0.8040558 | 0 |
| B137 | B159 | AFU | β2-MG | 0.8040698 | 0 |
| B137 | B169 | AFU | U-CRY | 0.8040684 | 0 |
| B137 | B179 | AFU | α1-MG | 0.804069 | 0 |
| B137 | B194 | AFU | Cl | -0.553965 | 0 |
| B137 | B195 | AFU | Na | 0.5961725 | 0 |
| B137 | B197 | AFU | AG | 0.5982739 | 0 |
| B137 | B198 | AFU | P | -0.608383 | 0 |
| B137 | B200 | AFU | OP | 0.6308868 | 0 |
| B137 | B199 | AFU | Mg | 0.5072971 | 2.22E-16 |
| B137 | B193 | AFU | K | 0.3283268 | 3.72E-07 |
| B138 | B139 | PAB | ADA | -0.602023 | 0 |
| B138 | B151 | PAB | Anti-CCP | -0.615902 | 0 |
| B138 | B152 | PAB | ACA-Ig(G-A-M) | 0.6155603 | 0 |
| B138 | B159 | PAB | β2-MG | 0.615571 | 0 |
| B138 | B169 | PAB | U-CRY | 0.6155699 | 0 |
| B138 | B179 | PAB | α1-MG | 0.6155704 | 0 |
| B138 | B195 | PAB | Na | 0.7477302 | 0 |
| B138 | B197 | PAB | AG | 0.5167095 | 0 |
| B138 | B200 | PAB | OP | 0.5514853 | 0 |
| B138 | B199 | PAB | Mg | 0.3331055 | 2.45E-07 |
| B139 | B199 | ADA | Mg | 0.5104448 | 0 |
| B139 | B228 | ADA | OS | 0.363958 | 1.40E-08 |
| B139 | B146 | ADA | CysC | 0.3361763 | 1.87E-07 |
| B14 | B196 | G-17 | TCO2 | -0.603695 | 0 |
| B14 | B200 | G-17 | OP | 0.5321141 | 0 |
| B14 | B25 | G-17 | CK | 0.8229728 | 0 |
| B14 | B198 | G-17 | P | 0.4567007 | 3.36E-13 |
| B14 | B195 | G-17 | Na | 0.4526963 | 5.71E-13 |
| B14 | B193 | G-17 | K | -0.447579 | 1.11E-12 |
| B14 | B194 | G-17 | Cl | 0.4420172 | 2.27E-12 |
| B14 | B114 | G-17 | TSH TT3 TT4(TT4) | -0.337114 | 1.72E-07 |

| | | | | | |
|---|---|---|---|---|---|
| B142 | B145 | 25-(OH)D | BGP | 0.6546522 | 0 |
| B142 | B143 | 25-(OH)D | β-CTx | 0.4600589 | 2.14E-13 |
| B142 | B144 | 25-(OH)D | Total-PINP | 0.3983107 | 3.97E-10 |
| B143 | B144 | β-CTx | Total-PINP | 0.9742981 | 0 |
| B143 | B145 | β-CTx | BGP | 0.8720692 | 0 |
| B143 | B214 | β-CTx | CVA-L | -0.493229 | 1.78E-15 |
| B144 | B145 | Total-PINP | BGP | 0.8128338 | 0 |
| B144 | B214 | Total-PINP | CVA-L | -0.511304 | 0 |
| B145 | B214 | BGP | CVA-L | -0.349947 | 5.34E-08 |
| B148 | B149 | Anti_TPO | TGAb | 0.6899245 | 0 |
| B149 | B164 | TGAb | COM | -0.328974 | 3.51E-07 |
| B15 | B16 | CD3-CD19+(B cell) | CD3+T | -1 | 0 |
| B15 | B18 | CD3-CD19+(B cell) | CD3+CD4+helper(inducer)T | -1 | 0 |
| B15 | B194 | CD3-CD19+(B cell) | CI | -0.708574 | 0 |
| B15 | B23 | CD3-CD19+(B cell) | CD3-CD16+CD56+(NK) | -1 | 0 |
| B15 | B137 | CD3-CD19+(B cell) | AFU | 0.4924971 | 2.22E-15 |
| B15 | B197 | CD3-CD19+(B cell) | AG | 0.3996106 | 3.44E-10 |
| B15 | B115 | CD3-CD19+(B cell) | TSH TT3 TT4(TT3) | -0.365236 | 1.24E-08 |
| B151 | B198 | Anti-CCP | P | 0.5795951 | 0 |
| B151 | B199 | Anti-CCP | Mg | -0.541282 | 0 |
| B151 | B193 | Anti-CCP | K | -0.500857 | 4.44E-16 |
| B151 | B200 | Anti-CCP | OP | -0.389555 | 1.02E-09 |
| B151 | B195 | Anti-CCP | Na | -0.340618 | 1.26E-07 |
| B152 | B159 | ACA-Ig(G-A-M) | β2-MG | 0.9999887 | 0 |
| B152 | B169 | ACA-Ig(G-A-M) | U-CRY | 0.9999891 | 0 |
| B152 | B179 | ACA-Ig(G-A-M) | α1-MG | 0.9999844 | 0 |
| B152 | B198 | ACA-Ig(G-A-M) | P | -0.579274 | 0 |
| B152 | B199 | ACA-Ig(G-A-M) | Mg | 0.5409813 | 0 |
| B152 | B193 | ACA-Ig(G-A-M) | K | 0.5005787 | 6.66E-16 |
| B152 | B200 | ACA-Ig(G-A-M) | OP | 0.3893395 | 1.05E-09 |
| B152 | B195 | ACA-Ig(G-A-M) | Na | 0.3404294 | 1.28E-07 |
| B153 | B170 | Glu | U-GLU | 0.4567001 | 3.36E-13 |
| B155 | B157 | IgA | IgG | -0.476156 | 2.31E-14 |
| B156 | B225 | IgE | hPBG | 0.6681531 | 0 |
| B157 | B158 | IgG | IgM | 0.6820303 | 0 |
| B159 | B169 | β2-MG | U-CRY | 0.9999978 | 0 |
| B159 | B198 | β2-MG | P | -0.579284 | 0 |
| B159 | B199 | β2-MG | Mg | 0.5409907 | 0 |
| B159 | B193 | β2-MG | K | 0.5005874 | 6.66E-16 |
| B159 | B200 | β2-MG | OP | 0.3893461 | 1.05E-09 |
| B159 | B195 | β2-MG | Na | 0.3404352 | 1.28E-07 |
| B16 | B18 | CD3+T | CD3+CD4+helper(inducer)T | 0.9999999 | 0 |
| B16 | B19 | CD3+T | CD3+CD4+/CD3+CD8+ | 0.9999998 | 0 |
| B16 | B194 | CD3+T | CI | 0.7085737 | 0 |

| | | | | | |
|---|---|---|---|---|---|
| B16 | B22 | CD3+T | CD3+CD8+(T) | -1 | 0 |
| B16 | B23 | CD3+T | CD3-CD16+CD56+(NK) | 0.9999998 | 0 |
| B16 | B137 | CD3+T | AFU | -0.492497 | 2.22E-15 |
| B16 | B197 | CD3+T | AG | -0.399611 | 3.44E-10 |
| B16 | B115 | CD3+T | TSH TT3 TT4(TT3) | 0.3652358 | 1.24E-08 |
| B163 | B176 | SG | Ucr | 0.4856148 | 6.00E-15 |
| B163 | B166 | SG | Pro | 0.3429436 | 1.02E-07 |
| B164 | B167 | COM | PC | 0.7044326 | 0 |
| B169 | B179 | U-CRY | α1-MG | 0.9999995 | 0 |
| B169 | B198 | U-CRY | P | -0.579283 | 0 |
| B169 | B199 | U-CRY | Mg | 0.5409897 | 0 |
| B169 | B193 | U-CRY | K | 0.5005865 | 6.66E-16 |
| B169 | B200 | U-CRY | OP | 0.3893454 | 1.05E-09 |
| B169 | B195 | U-CRY | Na | 0.3404346 | 1.28E-07 |
| B17 | B102 | CD3+CD16+CD56+NK | anti-Sm | -0.960119 | 0 |
| B17 | B104 | CD3+CD16+CD56+NK | anti-SSB | -0.959522 | 0 |
| B17 | B21 | CD3+CD16+CD56+NK | CD3+CD4-CD8- | -0.937338 | 0 |
| B17 | B37 | CD3+CD16+CD56+NK | BAND | -0.63408 | 0 |
| B17 | B95 | CD3+CD16+CD56+NK | anti-cenpb | -0.497059 | 1.11E-15 |
| B174 | B191 | Urea/Cr | Urea | 0.451382 | 6.78E-13 |
| B174 | B221 | Urea/Cr | Hei | -0.386676 | 1.39E-09 |
| B174 | B190 | Urea/Cr | Cre | -0.343134 | 1.00E-07 |
| B175 | B177 | ALB-U | ACR | 0.617387 | 0 |
| B175 | B227 | ALB-U | OD | 0.511082 | 0 |
| B176 | B177 | Ucr | ACR | -0.32632 | 4.42E-07 |
| B178 | B202 | Ig-U | e-times | -0.596285 | 0 |
| B178 | B180 | Ig-U | Tf | -0.46188 | 1.68E-13 |
| B179 | B198 | α1-MG | P | -0.579283 | 0 |
| B179 | B199 | α1-MG | Mg | 0.5409901 | 0 |
| B179 | B193 | α1-MG | K | 0.5005869 | 6.66E-16 |
| B179 | B200 | α1-MG | OP | 0.3893457 | 1.05E-09 |
| B179 | B195 | α1-MG | Na | 0.3404348 | 1.28E-07 |
| B18 | B19 | CD3+CD4+helper(inducer)T | CD3+CD4+/CD3+CD8+ | 1 | 0 |
| B18 | B194 | CD3+CD4+helper(inducer)T | Cl | 0.7085739 | 0 |
| B18 | B23 | CD3+CD4+helper(inducer)T | CD3-CD16+CD56+(NK) | 1 | 0 |
| B18 | B137 | CD3+CD4+helper(inducer)T | AFU | -0.492497 | 2.22E-15 |
| B18 | B197 | CD3+CD4+helper(inducer)T | AG | -0.399611 | 3.44E-10 |
| B18 | B115 | CD3+CD4+helper(inducer)T | TSH TT3 TT4(TT3) | 0.3652359 | 1.24E-08 |
| B180 | B202 | Tf | e-times | -0.946729 | 0 |
| B181 | B182 | APTT | TT | 0.7960301 | 0 |
| B181 | B184 | APTT | INR | 0.6794827 | 0 |
| B181 | B185 | APTT | PT | 0.6837592 | 0 |
| B181 | B186 | APTT | PFC | 0.8461272 | 0 |
| B181 | B183 | APTT | PTA | -0.467728 | 7.55E-14 |

| | | | | | |
|---|---|---|---|---|---|
| B182 | B183 | TT | PTA | -0.86806 | 0 |
| B182 | B184 | TT | INR | 0.566256 | 0 |
| B182 | B185 | TT | PT | 0.6638108 | 0 |
| B182 | B186 | TT | PFC | 0.9559545 | 0 |
| B183 | B184 | PTA | INR | -0.594726 | 0 |
| B183 | B185 | PTA | PT | -0.714416 | 0 |
| B183 | B186 | PTA | PFC | -0.693227 | 0 |
| B184 | B185 | INR | PT | 0.9873994 | 0 |
| B184 | B186 | INR | PFC | 0.4061828 | 1.65E-10 |
| B185 | B186 | PT | PFC | 0.4902613 | 3.11E-15 |
| B189 | B190 | eGFB | Cre | -0.623033 | 0 |
| B189 | B191 | eGFB | Urea | -0.589151 | 0 |
| B19 | B194 | CD3+CD4+/CD3+CD8+ | Cl | 0.7085739 | 0 |
| B19 | B23 | CD3+CD4+/CD3+CD8+ | CD3-CD16+CD56+(NK) | 1 | 0 |
| B19 | B137 | CD3+CD4+/CD3+CD8+ | AFU | -0.492497 | 2.22E-15 |
| B19 | B197 | CD3+CD4+/CD3+CD8+ | AG | -0.399611 | 3.44E-10 |
| B19 | B115 | CD3+CD4+/CD3+CD8+ | TSH TT3 TT4(TT3) | 0.3652359 | 1.24E-08 |
| B190 | B191 | Cre | Urea | 0.4937765 | 1.78E-15 |
| B191 | B192 | Urea | UA | 0.3569561 | 2.76E-08 |
| B193 | B198 | K | P | -0.764098 | 0 |
| B193 | B196 | K | TCO2 | 0.4841563 | 7.33E-15 |
| B193 | B199 | K | Mg | 0.4810079 | 1.15E-14 |
| B194 | B197 | Cl | AG | -0.570911 | 0 |
| B194 | B198 | Cl | P | 0.5852368 | 0 |
| B195 | B196 | Na | TCO2 | -0.638133 | 0 |
| B195 | B197 | Na | AG | 0.8632173 | 0 |
| B195 | B200 | Na | OP | 0.9281918 | 0 |
| B195 | B199 | Na | Mg | 0.4876943 | 4.44E-15 |
| B196 | B197 | TCO2 | AG | -0.772089 | 0 |
| B196 | B200 | TCO2 | OP | -0.700544 | 0 |
| B196 | B199 | TCO2 | Mg | -0.462539 | 1.53E-13 |
| B196 | B198 | TCO2 | P | -0.439894 | 2.96E-12 |
| B197 | B199 | AG | Mg | 0.5542742 | 0 |
| B197 | B200 | AG | OP | 0.8673368 | 0 |
| B198 | B199 | P | Mg | -0.382181 | 2.23E-09 |
| B199 | B200 | Mg | OP | 0.6513376 | 0 |
| B2 | B189 | Age | eGFB | -0.45582 | 3.78E-13 |
| B2 | B220 | Age | SBP | 0.3351505 | 2.05E-07 |
| B20 | B101 | CD3+CD4+CD8+ | anti-Sc70 | 0.6116357 | 0 |
| B20 | B103 | CD3+CD4+CD8+ | anti-SSA/Ro | 0.6073734 | 0 |
| B20 | B105 | CD3+CD4+CD8+ | anti-U1RNP | 0.5861606 | 0 |
| B20 | B107 | CD3+CD4+CD8+ | AnuA | 0.978709 | 0 |
| B20 | B110 | CD3+CD4+CD8+ | AHA | 0.7988126 | 0 |
| B20 | B225 | CD3+CD4+CD8+ | hPBG | 0.9149361 | 0 |

| | | | | | |
|---|---|---|---|---|---|
| B20 | B35 | CD3+CD4+CD8+ | β2-MG | 0.6535739 | 0 |
| B20 | B36 | CD3+CD4+CD8+ | V-Lym% | 0.8536585 | 0 |
| B20 | B38 | CD3+CD4+CD8+ | BAND% | 0.7682733 | 0 |
| B20 | B39 | CD3+CD4+CD8+ | Other cells | 0.7653197 | 0 |
| B20 | B92 | CD3+CD4+CD8+ | ALDH2-Glu504Glu | 0.9918366 | 0 |
| B20 | B96 | CD3+CD4+CD8+ | dsDNA (IIF) | 0.8598892 | 0 |
| B20 | B98 | CD3+CD4+CD8+ | anti-Jo-1 | 0.7363517 | 0 |
| B20 | B33 | CD3+CD4+CD8+ | Prog | 0.4734713 | 3.38E-14 |
| B204 | B209 | Drinking | Smoking | 0.4191471 | 3.71E-11 |
| B206 | B207 | Waist | WHR | 0.8011266 | 0 |
| B206 | B210 | Waist | Hip | 0.7595518 | 0 |
| B206 | B215 | Waist | BMI | 0.7929378 | 0 |
| B206 | B216 | Waist | BMI | 0.7687612 | 0 |
| B206 | B222 | Waist | NS | 0.7218839 | 0 |
| B206 | B221 | Waist | Hei | 0.3458199 | 7.82E-08 |
| B207 | B215 | WHR | BMI | 0.4851633 | 6.22E-15 |
| B207 | B222 | WHR | NS | 0.478715 | 1.60E-14 |
| B207 | B216 | WHR | BMI | 0.4378575 | 3.83E-12 |
| B21 | B100 | CD3+CD4-CD8- | anti-Ro52 | -0.84382 | 0 |
| B21 | B108 | CD3+CD4-CD8- | Anti-Cyclin | -0.690882 | 0 |
| B21 | B178 | CD3+CD4-CD8- | Ig-U | 0.7308817 | 0 |
| B21 | B223 | CD3+CD4-CD8- | FH | -0.826336 | 0 |
| B21 | B29 | CD3+CD4-CD8- | E2 | -0.792696 | 0 |
| B21 | B37 | CD3+CD4-CD8- | BAND | -0.521207 | 0 |
| B21 | B95 | CD3+CD4-CD8- | anti-cenpb | -0.634902 | 0 |
| B210 | B215 | Hip | BMI | 0.7755396 | 0 |
| B210 | B216 | Hip | BMI | 0.7809506 | 0 |
| B210 | B222 | Hip | NS | 0.6610933 | 0 |
| B210 | B221 | Hip | Hei | 0.389298 | 1.05E-09 |
| B211 | B212 | IOP-R | IOP-L | 0.3880219 | 1.20E-09 |
| B215 | B216 | BMI | BMI | 0.8187593 | 0 |
| B215 | B222 | BMI | NS | 0.6168676 | 0 |
| B216 | B221 | BMI | Hei | 0.695015 | 0 |
| B216 | B222 | BMI | NS | 0.7543289 | 0 |
| B22 | B194 | CD3+CD8+(T) | CI | -0.708574 | 0 |
| B22 | B137 | CD3+CD8+(T) | AFU | 0.4924971 | 2.22E-15 |
| B22 | B197 | CD3+CD8+(T) | AG | 0.3996106 | 3.44E-10 |
| B22 | B115 | CD3+CD8+(T) | TSH TT3 TT4(TT3) | -0.365236 | 1.24E-08 |
| B221 | B222 | Hei | NS | 0.5328143 | 0 |
| B23 | B194 | CD3-CD16+CD56+(NK) | CI | 0.7085739 | 0 |
| B23 | B137 | CD3-CD16+CD56+(NK) | AFU | -0.492497 | 2.22E-15 |
| B23 | B197 | CD3-CD16+CD56+(NK) | AG | -0.399611 | 3.44E-10 |
| B23 | B115 | CD3-CD16+CD56+(NK) | TSH TT3 TT4(TT3) | 0.3652359 | 1.24E-08 |
| B24 | B54 | αHBDH | ESR | 0.3820629 | 2.25E-09 |

| | | | | | |
|---|---|---|---|---|---|
| B24 | B26 | αHBDH | CKMB | 0.3335868 | 2.35E-07 |
| B25 | B193 | CK | K | -0.515078 | 0 |
| B25 | B196 | CK | TCO2 | -0.581952 | 0 |
| B25 | B198 | CK | P | 0.5010644 | 4.44E-16 |
| B25 | B200 | CK | OP | 0.4332124 | 6.81E-12 |
| B25 | B194 | CK | Cl | 0.386077 | 1.48E-09 |
| B25 | B195 | CK | Na | 0.3781376 | 3.38E-09 |
| B25 | B26 | CK | CKMB | 0.36402 | 1.39E-08 |
| B26 | B57 | CKMB | RBCAI | 0.6686722 | 0 |
| B26 | B74 | CKMB | ApoA1 | 0.4500251 | 8.09E-13 |
| B26 | B55 | CKMB | TK | -0.40645 | 1.60E-10 |
| B26 | B61 | CKMB | WBHSRV | -0.38526 | 1.61E-09 |
| B26 | B56 | CKMB | IR | -0.333775 | 2.31E-07 |
| B27 | B176 | LDH | Ucr | -0.431656 | 8.25E-12 |
| B28 | B202 | PRL | e-times | -0.596285 | 0 |
| B28 | B31 | PRL | FSH | -0.526648 | 0 |
| B28 | B95 | PRL | anti-cenpb | -0.501161 | 4.44E-16 |
| B28 | B178 | PRL | Ig-U | 0.4666667 | 8.73E-14 |
| B29 | B102 | E2 | anti-Sm | -0.779484 | 0 |
| B29 | B104 | E2 | anti-SSB | -0.888009 | 0 |
| B29 | B108 | E2 | Anti-Cyclin | -0.706073 | 0 |
| B29 | B117 | E2 | TT4 | -0.726241 | 0 |
| B29 | B178 | E2 | Ig-U | -0.903813 | 0 |
| B29 | B223 | E2 | FH | -0.630364 | 0 |
| B29 | B37 | E2 | BAND | -0.810891 | 0 |
| B29 | B116 | E2 | TT3 | -0.394549 | 5.98E-10 |
| B29 | B125 | E2 | Cor | -0.367452 | 9.93E-09 |
| B29 | B95 | E2 | anti-cenpb | -0.343149 | 9.99E-08 |
| B29 | B202 | E2 | e-times | -0.321521 | 6.64E-07 |
| B3 | B123 | Hcy | hs-CRP1 | 0.521957 | 0 |
| B30 | B103 | LH | anti-SSA/Ro | 0.8820529 | 0 |
| B30 | B124 | LH | ACTH | 0.5875697 | 0 |
| B30 | B225 | LH | hPBG | 0.59555 | 0 |
| B30 | B33 | LH | Prog | 0.652853 | 0 |
| B30 | B39 | LH | Other cells | 0.517906 | 0 |
| B30 | B96 | LH | dsDNA (IIF) | 0.9338366 | 0 |
| B30 | B97 | LH | dsDNA | 0.8816289 | 0 |
| B30 | B99 | LH | anti-PM-Scl | 0.6571028 | 0 |
| B30 | B156 | LH | IgE | 0.4951876 | 1.33E-15 |
| B30 | B32 | LH | Testo | 0.3551174 | 3.28E-08 |
| B30 | B101 | LH | anti-Sc70 | 0.3428305 | 1.03E-07 |
| B30 | B35 | LH | β2-MG | 0.3205453 | 7.21E-07 |
| B31 | B104 | FSH | anti-SSB | -0.878543 | 0 |
| B31 | B178 | FSH | Ig-U | -0.601884 | 0 |

| | | | | | |
|---|---|---|---|---|---|
| B31 | B180 | FSH | Tf | 0.4343722 | 5.90E-12 |
| B32 | B94 | Testo | ALDH2-Lys504Lys | 0.6377481 | 0 |
| B32 | B99 | Testo | anti-PM-Scl | 0.8458889 | 0 |
| B32 | B36 | Testo | V-Lym% | 0.5015655 | 4.44E-16 |
| B32 | B97 | Testo | dsDNA | 0.362209 | 1.66E-08 |
| B32 | B107 | Testo | AnuA | 0.3530939 | 3.97E-08 |
| B33 | B110 | Prog | AHA | 0.6202723 | 0 |
| B33 | B97 | Prog | dsDNA | 0.7969028 | 0 |
| B33 | B99 | Prog | anti-PM-Scl | 0.9355852 | 0 |
| B33 | B101 | Prog | anti-Sc70 | 0.4605389 | 2.01E-13 |
| B33 | B107 | Prog | AnuA | 0.4222003 | 2.58E-11 |
| B35 | B107 | β2-MG | AnuA | 0.7082641 | 0 |
| B35 | B124 | β2-MG | ACTH | 0.9327448 | 0 |
| B35 | B156 | β2-MG | IgE | 0.8219949 | 0 |
| B35 | B36 | β2-MG | V-Lym% | 0.9803609 | 0 |
| B35 | B38 | β2-MG | BAND% | 0.987763 | 0 |
| B35 | B96 | β2-MG | dsDNA (IIF) | 0.693395 | 0 |
| B35 | B99 | β2-MG | anti-PM-Scl | 0.8968544 | 0 |
| B35 | B103 | β2-MG | anti-SSA/Ro | 0.493197 | 1.78E-15 |
| B35 | B94 | β2-MG | ALDH2-Lys504Lys | -0.458535 | 2.63E-13 |
| B35 | B39 | β2-MG | Other cells | 0.444616 | 1.63E-12 |
| B35 | B225 | β2-MG | hPBG | 0.4393748 | 3.16E-12 |
| B35 | B93 | β2-MG | ALDH2-Glu504Lys | 0.4109975 | 9.56E-11 |
| B36 | B101 | V-Lym% | anti-Sc70 | 0.5745669 | 0 |
| B36 | B105 | V-Lym% | anti-U1RNP | 0.6350074 | 0 |
| B36 | B110 | V-Lym% | AHA | 0.9258964 | 0 |
| B36 | B124 | V-Lym% | ACTH | 0.8434347 | 0 |
| B36 | B39 | V-Lym% | Other cells | 0.6346554 | 0 |
| B36 | B94 | V-Lym% | ALDH2-Lys504Lys | 0.6695895 | 0 |
| B36 | B98 | V-Lym% | anti-Jo-1 | 0.9745832 | 0 |
| B36 | B92 | V-Lym% | ALDH2-Glu504Glu | -0.405751 | 1.73E-10 |
| B37 | B116 | BAND | TT3 | -0.983182 | 0 |
| B38 | B97 | BAND% | dsDNA | 0.4472136 | 1.17E-12 |
| B38 | B110 | BAND% | AHA | 0.3639127 | 1.41E-08 |
| B39 | B103 | Other cells | anti-SSA/Ro | 0.9296697 | 0 |
| B39 | B107 | Other cells | AnuA | 0.6033815 | 0 |
| B39 | B110 | Other cells | AHA | 0.55577 | 0 |
| B39 | B94 | Other cells | ALDH2-Lys504Lys | 0.655936 | 0 |
| B39 | B97 | Other cells | dsDNA | 0.4780914 | 1.78E-14 |
| B39 | B101 | Other cells | anti-Sc70 | 0.3688036 | 8.68E-09 |
| B4 | B135 | WB-Ca | TBA | 0.5615924 | 0 |
| B4 | B5 | WB-Ca | WB-Mg | 1 | 0 |
| B4 | B6 | WB-Ca | WB-Fe | 1 | 0 |
| B4 | B82 | WB-Ca | CEA | 0.56575 | 0 |

| | | | | | |
|---|---|---|---|---|---|
| B4 | B61 | WB-Ca | WBHSRV | -0.381188 | 2.47E-09 |
| B4 | B83 | WB-Ca | AFP | 0.3728675 | 5.78E-09 |
| B4 | B66 | WB-Ca | WBRSR(mPas_100) | -0.37132 | 6.75E-09 |
| B4 | B67 | WB-Ca | WBRSR(mPas_50) | -0.370193 | 7.56E-09 |
| B4 | B88 | WB-Ca | CA 242 | 0.3664185 | 1.10E-08 |
| B4 | B69 | WB-Ca | PV | -0.364999 | 1.26E-08 |
| B4 | B63 | WB-Ca | WBRSR(mPas_200) | -0.358225 | 2.44E-08 |
| B4 | B64 | WB-Ca | WBRSR(mPas_5) | -0.347112 | 6.94E-08 |
| B4 | B65 | WB-Ca | WBRSR(mPas_1) | -0.31899 | 8.22E-07 |
| B40 | B42 | WBC | MON | 0.3709695 | 7.00E-09 |
| B41 | B42 | MON％ | MON | 0.6253381 | 0 |
| B43 | B44 | MCV | MCH | 0.8745733 | 0 |
| B43 | B47 | MCV | RDW | 0.5089781 | 2.22E-16 |
| B43 | B46 | MCV | RBC Count | -0.491239 | 2.66E-15 |
| B43 | B52 | MCV | PLT/BPC | -0.316677 | 9.96E-07 |
| B44 | B45 | MCH | MCHC | 0.481672 | 1.07E-14 |
| B44 | B46 | MCH | RBC Count | -0.338956 | 1.46E-07 |
| B45 | B51 | MCHC | Hb | 0.4692928 | 6.08E-14 |
| B45 | B47 | MCHC | RDW | -0.405544 | 1.77E-10 |
| B46 | B51 | RBC Count | Hb | 0.8548587 | 0 |
| B46 | B47 | RBC Count | RDW | -0.406188 | 1.65E-10 |
| B46 | B216 | RBC Count | BMI | 0.4001133 | 3.25E-10 |
| B46 | B221 | RBC Count | Hei | 0.3820908 | 2.25E-09 |
| B46 | B222 | RBC Count | NS | 0.362275 | 1.65E-08 |
| B46 | B126 | RBC Count | Alb | 0.354186 | 3.59E-08 |
| B46 | B163 | RBC Count | SG | 0.3409021 | 1.22E-07 |
| B47 | B132 | RDW | Dbil | 0.3257328 | 4.65E-07 |
| B48 | B53 | LYM% | GR% | -0.951973 | 0 |
| B5 | B135 | WB-Mg | TBA | 0.5615924 | 0 |
| B5 | B7 | WB-Mg | WB-Cop | -1 | 0 |
| B5 | B8 | WB-Mg | WB-Zinc | -1 | 0 |
| B5 | B82 | WB-Mg | CEA | 0.56575 | 0 |
| B5 | B61 | WB-Mg | WBHSRV | -0.381188 | 2.47E-09 |
| B5 | B83 | WB-Mg | AFP | 0.3728675 | 5.78E-09 |
| B5 | B66 | WB-Mg | WBRSR(mPas_100) | -0.37132 | 6.75E-09 |
| B5 | B67 | WB-Mg | WBRSR(mPas_50) | -0.370193 | 7.56E-09 |
| B5 | B88 | WB-Mg | CA 242 | 0.3664185 | 1.10E-08 |
| B5 | B69 | WB-Mg | PV | -0.364999 | 1.26E-08 |
| B5 | B63 | WB-Mg | WBRSR(mPas_200) | -0.358225 | 2.44E-08 |
| B5 | B64 | WB-Mg | WBRSR(mPas_5) | -0.347112 | 6.94E-08 |
| B5 | B65 | WB-Mg | WBRSR(mPas_1) | -0.31899 | 8.22E-07 |
| B51 | B222 | Hb | NS | 0.4342891 | 5.97E-12 |
| B51 | B221 | Hb | Hei | 0.4139477 | 6.80E-11 |
| B51 | B216 | Hb | BMI | 0.3852086 | 1.62E-09 |

| | | | | | |
|---|---|---|---|---|---|
| B51 | B129 | Hb | IBIL | 0.3701757 | 7.57E-09 |
| B51 | B133 | Hb | TBIL | 0.3419741 | 1.11E-07 |
| B51 | B126 | Hb | Alb | 0.3317191 | 2.77E-07 |
| B54 | B56 | ESR | IR | -0.52792 | 0 |
| B54 | B69 | ESR | PV | 0.6307952 | 0 |
| B54 | B58 | ESR | HCT | -0.497827 | 8.88E-16 |
| B54 | B62 | ESR | WBHSRVI | -0.487008 | 4.88E-15 |
| B54 | B60 | ESR | WBRVI | -0.418211 | 4.14E-11 |
| B54 | B63 | ESR | WBRSR(mPas_200) | -0.373611 | 5.36E-09 |
| B54 | B66 | ESR | WBRSR(mPas_100) | -0.35274 | 4.11E-08 |
| B54 | B67 | ESR | WBRSR(mPas_50) | -0.350282 | 5.17E-08 |
| B54 | B64 | ESR | WBRSR(mPas_5) | -0.329054 | 3.49E-07 |
| B54 | B61 | ESR | WBHSRV | -0.323102 | 5.81E-07 |
| B55 | B57 | TK | RBCAI | -0.594907 | 0 |
| B55 | B58 | TK | HCT | -0.611678 | 0 |
| B55 | B59 | TK | WBRV | -0.716889 | 0 |
| B55 | B60 | TK | WBRVI | -0.784351 | 0 |
| B55 | B62 | TK | WBHSRVI | -0.532206 | 0 |
| B55 | B64 | TK | WBRSR(mPas_5) | -0.672293 | 0 |
| B55 | B65 | TK | WBRSR(mPas_1) | -0.761568 | 0 |
| B55 | B66 | TK | WBRSR(mPas_100) | -0.532868 | 0 |
| B55 | B67 | TK | WBRSR(mPas_50) | -0.557063 | 0 |
| B55 | B63 | TK | WBRSR(mPas_200) | -0.489733 | 3.11E-15 |
| B56 | B58 | IR | HCT | 0.9273673 | 0 |
| B56 | B60 | IR | WBRVI | 0.7788914 | 0 |
| B56 | B61 | IR | WBHSRV | 0.9569471 | 0 |
| B56 | B62 | IR | WBHSRVI | 0.9610589 | 0 |
| B56 | B63 | IR | WBRSR(mPas_200) | 0.950625 | 0 |
| B56 | B64 | IR | WBRSR(mPas_5) | 0.8660209 | 0 |
| B56 | B65 | IR | WBRSR(mPas_1) | 0.7888852 | 0 |
| B56 | B66 | IR | WBRSR(mPas_100) | 0.9327327 | 0 |
| B56 | B67 | IR | WBRSR(mPas_50) | 0.9241406 | 0 |
| B56 | B57 | IR | RBCAI | -0.460286 | 2.08E-13 |
| B57 | B59 | RBCAI | WBRV | 0.5851274 | 0 |
| B57 | B74 | RBCAI | ApoA1 | 0.5356244 | 0 |
| B57 | B61 | RBCAI | WBHSRV | -0.504284 | 4.44E-16 |
| B58 | B60 | HCT | WBRVI | 0.9255731 | 0 |
| B58 | B61 | HCT | WBHSRV | 0.8786996 | 0 |
| B58 | B62 | HCT | WBHSRVI | 0.9853699 | 0 |
| B58 | B63 | HCT | WBRSR(mPas_200) | 0.9574556 | 0 |
| B58 | B64 | HCT | WBRSR(mPas_5) | 0.9512071 | 0 |
| B58 | B65 | HCT | WBRSR(mPas_1) | 0.9197162 | 0 |
| B58 | B66 | HCT | WBRSR(mPas_100) | 0.9563708 | 0 |
| B58 | B67 | HCT | WBRSR(mPas_50) | 0.9576617 | 0 |

| | | | | | |
|---|---|---|---|---|---|
| B58 | B59 | HCT | WBRV | 0.4003887 | 3.15E-10 |
| B59 | B60 | WBRV | WBRVI | 0.6692073 | 0 |
| B59 | B64 | WBRV | WBRSR(mPas_5) | 0.6372358 | 0 |
| B59 | B65 | WBRV | WBRSR(mPas_1) | 0.7253219 | 0 |
| B59 | B67 | WBRV | WBRSR(mPas_50) | 0.5260117 | 0 |
| B59 | B66 | WBRV | WBRSR(mPas_100) | 0.5004054 | 6.66E-16 |
| B59 | B63 | WBRV | WBRSR(mPas_200) | 0.4452788 | 1.49E-12 |
| B59 | B62 | WBRV | WBHSRVI | 0.4237487 | 2.15E-11 |
| B6 | B135 | WB-Fe | TBA | 0.5615924 | 0 |
| B6 | B7 | WB-Fe | WB-Cop | -1 | 0 |
| B6 | B82 | WB-Fe | CEA | 0.56575 | 0 |
| B6 | B61 | WB-Fe | WBHSRV | -0.381188 | 2.47E-09 |
| B6 | B83 | WB-Fe | AFP | 0.3728675 | 5.78E-09 |
| B6 | B66 | WB-Fe | WBRSR(mPas_100) | -0.37132 | 6.75E-09 |
| B6 | B67 | WB-Fe | WBRSR(mPas_50) | -0.370193 | 7.56E-09 |
| B6 | B88 | WB-Fe | CA 242 | 0.3664185 | 1.10E-08 |
| B6 | B69 | WB-Fe | PV | -0.364999 | 1.26E-08 |
| B6 | B63 | WB-Fe | WBRSR(mPas_200) | -0.358225 | 2.44E-08 |
| B6 | B64 | WB-Fe | WBRSR(mPas_5) | -0.347112 | 6.94E-08 |
| B6 | B65 | WB-Fe | WBRSR(mPas_1) | -0.31899 | 8.22E-07 |
| B60 | B61 | WBRVI | WBHSRV | 0.7247258 | 0 |
| B60 | B62 | WBRVI | WBHSRVI | 0.9143487 | 0 |
| B60 | B63 | WBRVI | WBRSR(mPas_200) | 0.8661905 | 0 |
| B60 | B64 | WBRVI | WBRSR(mPas_5) | 0.9542674 | 0 |
| B60 | B65 | WBRVI | WBRSR(mPas_1) | 0.9781964 | 0 |
| B60 | B66 | WBRVI | WBRSR(mPas_100) | 0.8876169 | 0 |
| B60 | B67 | WBRVI | WBRSR(mPas_50) | 0.9011256 | 0 |
| B61 | B62 | WBHSRV | WBHSRVI | 0.925972 | 0 |
| B61 | B63 | WBHSRV | WBRSR(mPas_200) | 0.9681636 | 0 |
| B61 | B64 | WBHSRV | WBRSR(mPas_5) | 0.8787093 | 0 |
| B61 | B65 | WBHSRV | WBRSR(mPas_1) | 0.7940819 | 0 |
| B61 | B66 | WBHSRV | WBRSR(mPas_100) | 0.955353 | 0 |
| B61 | B67 | WBHSRV | WBRSR(mPas_50) | 0.945488 | 0 |
| B62 | B63 | WBHSRVI | WBRSR(mPas_200) | 0.9826381 | 0 |
| B62 | B64 | WBHSRVI | WBRSR(mPas_5) | 0.9632381 | 0 |
| B62 | B65 | WBHSRVI | WBRSR(mPas_1) | 0.921873 | 0 |
| B62 | B66 | WBHSRVI | WBRSR(mPas_100) | 0.9804478 | 0 |
| B62 | B67 | WBHSRVI | WBRSR(mPas_50) | 0.9801587 | 0 |
| B63 | B64 | WBRSR(mPas_200) | WBRSR(mPas_5) | 0.9650867 | 0 |
| B63 | B65 | WBRSR(mPas_200) | WBRSR(mPas_1) | 0.9120239 | 0 |
| B63 | B66 | WBRSR(mPas_200) | WBRSR(mPas_100) | 0.9977448 | 0 |
| B63 | B67 | WBRSR(mPas_200) | WBRSR(mPas_50) | 0.9951047 | 0 |
| B64 | B65 | WBRSR(mPas_5) | WBRSR(mPas_1) | 0.9874927 | 0 |
| B64 | B66 | WBRSR(mPas_5) | WBRSR(mPas_100) | 0.9791075 | 0 |

| | | | | | |
|---|---|---|---|---|---|
| B64 | B67 | WBRSR(mPas_5) | WBRSR(mPas_50) | 0.9854661 | 0 |
| B64 | B71 | WBRSR(mPas_5) | TG | 0.3232294 | 5.75E-07 |
| B65 | B66 | WBRSR(mPas_1) | WBRSR(mPas_100) | 0.9348065 | 0 |
| B65 | B67 | WBRSR(mPas_1) | WBRSR(mPas_50) | 0.946366 | 0 |
| B65 | B71 | WBRSR(mPas_1) | TG | 0.3274518 | 4.01E-07 |
| B66 | B67 | WBRSR(mPas_100) | WBRSR(mPas_50) | 0.9994109 | 0 |
| B7 | B135 | WB-Cop | TBA | -0.561592 | 0 |
| B7 | B8 | WB-Cop | WB-Zinc | 1 | 0 |
| B7 | B82 | WB-Cop | CEA | -0.56575 | 0 |
| B7 | B61 | WB-Cop | WBHSRV | 0.3811877 | 2.47E-09 |
| B7 | B83 | WB-Cop | AFP | -0.372868 | 5.78E-09 |
| B7 | B66 | WB-Cop | WBRSR(mPas_100) | 0.3713205 | 6.75E-09 |
| B7 | B67 | WB-Cop | WBRSR(mPas_50) | 0.3701925 | 7.56E-09 |
| B7 | B88 | WB-Cop | CA 242 | -0.366418 | 1.10E-08 |
| B7 | B69 | WB-Cop | PV | 0.3649991 | 1.26E-08 |
| B7 | B63 | WB-Cop | WBRSR(mPas_200) | 0.3582248 | 2.44E-08 |
| B7 | B64 | WB-Cop | WBRSR(mPas_5) | 0.3471122 | 6.94E-08 |
| B7 | B65 | WB-Cop | WBRSR(mPas_1) | 0.3189903 | 8.22E-07 |
| B70 | B77 | LDL-C | TC | 0.9101804 | 0 |
| B71 | B72 | TG | HDL-C | -0.462204 | 1.60E-13 |
| B72 | B192 | HDL-C | UA | -0.387809 | 1.23E-09 |
| B72 | B215 | HDL-C | BMI | -0.331369 | 2.85E-07 |
| B72 | B206 | HDL-C | Waist | -0.330657 | 3.04E-07 |
| B72 | B216 | HDL-C | BMI | -0.323456 | 5.64E-07 |
| B73 | B136 | ApoA | 5'NT | 0.3508996 | 4.88E-08 |
| B73 | B76 | ApoA | ApoB | 0.3188997 | 8.28E-07 |
| B74 | B76 | ApoA1 | ApoB | 0.5777042 | 0 |
| B74 | B139 | ApoA1 | ADA | -0.492535 | 2.22E-15 |
| B74 | B138 | ApoA1 | PAB | 0.4730073 | 3.62E-14 |
| B74 | B228 | ApoA1 | OS | -0.385651 | 1.55E-09 |
| B74 | B75 | ApoA1 | ApoA1/B | -0.371598 | 6.57E-09 |
| B74 | B136 | ApoA1 | 5'NT | 0.3531399 | 3.96E-08 |
| B74 | B134 | ApoA1 | CHE | 0.3370226 | 1.73E-07 |
| B74 | B130 | ApoA1 | GLB | -0.322602 | 6.06E-07 |
| B75 | B136 | ApoA1/B | 5'NT | -0.693777 | 0 |
| B75 | B139 | ApoA1/B | ADA | 0.78364 | 0 |
| B75 | B76 | ApoA1/B | ApoB | -0.912987 | 0 |
| B75 | B82 | ApoA1/B | CEA | 0.497883 | 8.88E-16 |
| B75 | B138 | ApoA1/B | PAB | -0.49014 | 3.11E-15 |
| B75 | B115 | ApoA1/B | TSH TT3 TT4(TT3) | -0.359479 | 2.16E-08 |
| B75 | B134 | ApoA1/B | CHE | -0.350045 | 5.29E-08 |
| B75 | B228 | ApoA1/B | OS | 0.3426937 | 1.04E-07 |
| B75 | B198 | ApoA1/B | P | -0.337695 | 1.63E-07 |
| B76 | B136 | ApoB | 5'NT | 0.6039959 | 0 |

| | | | | | |
|---|---|---|---|---|---|
| B76 | B139 | ApoB | ADA | -0.617354 | 0 |
| B76 | B138 | ApoB | PAB | 0.391273 | 8.52E-10 |
| B76 | B82 | ApoB | CEA | -0.37753 | 3.60E-09 |
| B76 | B228 | ApoB | OS | -0.375546 | 4.41E-09 |
| B76 | B115 | ApoB | TSH TT3 TT4(TT3) | 0.364594 | 1.32E-08 |
| B76 | B86 | ApoB | TBhCG | 0.3187456 | 8.38E-07 |
| B77 | B132 | TC | Dbil | -0.329872 | 3.25E-07 |
| B78 | B203 | HBcAb | DH | 0.3504142 | 5.11E-08 |
| B79 | B120 | hb V -dna | HCV-cAg | 0.9999994 | 0 |
| B79 | B121 | hb V -dna | C3 | -1 | 0 |
| B79 | B137 | hb V -dna | AFU | -0.804069 | 0 |
| B79 | B138 | hb V -dna | PAB | -0.61557 | 0 |
| B79 | B152 | hb V -dna | ACA-Ig(G-A-M) | -0.999984 | 0 |
| B79 | B169 | hb V -dna | U-CRY | -0.999999 | 0 |
| B79 | B198 | hb V -dna | P | 0.579283 | 0 |
| B79 | B199 | hb V -dna | Mg | -0.54099 | 0 |
| B79 | B193 | hb V -dna | K | -0.500587 | 6.66E-16 |
| B79 | B200 | hb V -dna | OP | -0.389346 | 1.05E-09 |
| B79 | B195 | hb V -dna | Na | -0.340435 | 1.28E-07 |
| B8 | B135 | WB-Zinc | TBA | -0.561592 | 0 |
| B8 | B82 | WB-Zinc | CEA | -0.56575 | 0 |
| B8 | B61 | WB-Zinc | WBHSRV | 0.3811877 | 2.47E-09 |
| B8 | B83 | WB-Zinc | AFP | -0.372868 | 5.78E-09 |
| B8 | B66 | WB-Zinc | WBRSR(mPas_100) | 0.3713205 | 6.75E-09 |
| B8 | B67 | WB-Zinc | WBRSR(mPas_50) | 0.3701925 | 7.56E-09 |
| B8 | B88 | WB-Zinc | CA 242 | -0.366418 | 1.10E-08 |
| B8 | B69 | WB-Zinc | PV | 0.3649991 | 1.26E-08 |
| B8 | B63 | WB-Zinc | WBRSR(mPas_200) | 0.3582248 | 2.44E-08 |
| B8 | B64 | WB-Zinc | WBRSR(mPas_5) | 0.3471122 | 6.94E-08 |
| B8 | B65 | WB-Zinc | WBRSR(mPas_1) | 0.3189903 | 8.22E-07 |
| B81 | B155 | TSGF | IgA | 0.7462566 | 0 |
| B81 | B158 | TSGF | IgM | 0.4084104 | 1.28E-10 |
| B82 | B88 | CEA | CA 242 | 0.5858643 | 0 |
| B82 | B139 | CEA | ADA | 0.4858355 | 5.77E-15 |
| B82 | B195 | CEA | Na | -0.378823 | 3.15E-09 |
| B82 | B136 | CEA | 5'NT | -0.375124 | 4.60E-09 |
| B82 | B138 | CEA | PAB | -0.357136 | 2.71E-08 |
| B82 | B83 | CEA | AFP | 0.3479453 | 6.43E-08 |
| B83 | B87 | AFP | NSE | 0.4937972 | 1.78E-15 |
| B83 | B88 | AFP | CA 242 | 0.4696708 | 5.77E-14 |
| B83 | B84 | AFP | C-PSA | 0.433101 | 6.91E-12 |
| B83 | B139 | AFP | ADA | -0.396438 | 4.87E-10 |
| B83 | B85 | AFP | SCC | 0.3916744 | 8.16E-10 |
| B85 | B87 | SCC | NSE | 0.5403509 | 0 |

| | | | | | |
|---|---|---|---|---|---|
| B85 | B89 | SCC | CA 72-4 | -0.490586 | 2.89E-15 |
| B85 | B86 | SCC | TBhCG | -0.466087 | 9.46E-14 |
| B85 | B88 | SCC | CA 242 | 0.3782342 | 3.35E-09 |
| B86 | B89 | TBhCG | CA 72-4 | 0.9406598 | 0 |
| B87 | B88 | NSE | CA 242 | 0.4294789 | 1.08E-11 |
| B88 | B177 | CA 242 | ACR | -0.320696 | 7.12E-07 |
| B89 | B193 | CA 72-4 | K | 0.3621366 | 1.67E-08 |
| B9 | B104 | RH | anti-SSB | -0.83205 | 0 |
| B9 | B117 | RH | TT4 | -0.935529 | 0 |
| B9 | B31 | RH | FSH | -0.591015 | 0 |
| B9 | B95 | RH | anti-cenpb | -0.828819 | 0 |
| B9 | B29 | RH | E2 | -0.484086 | 7.55E-15 |
| B9 | B178 | RH | Ig-U | -0.454794 | 4.33E-13 |
| B9 | B21 | RH | CD3+CD4-CD8- | -0.344488 | 8.84E-08 |
| B90 | B154 | CYFRA 21-1 | RF | -0.505116 | 4.44E-16 |
| B91 | B127 | TP | A-G ratio | -0.534914 | 0 |
| B91 | B130 | TP | GLB | 0.8206319 | 0 |
| B92 | B107 | ALDH2-Glu504Glu | AnuA | -0.904534 | 0 |
| B92 | B93 | ALDH2-Glu504Glu | ALDH2-Glu504Lys | -0.612372 | 0 |
| B92 | B98 | ALDH2-Glu504Glu | anti-Jo-1 | 0.5604485 | 0 |
| B92 | B97 | ALDH2-Glu504Glu | dsDNA | 0.412393 | 8.14E-11 |
| B92 | B103 | ALDH2-Glu504Glu | anti-SSA/Ro | -0.408248 | 1.31E-10 |
| B93 | B105 | ALDH2-Glu504Lys | anti-U1RNP | 0.8109308 | 0 |
| B93 | B156 | ALDH2-Glu504Lys | IgE | 0.5555556 | 0 |
| B93 | B94 | ALDH2-Glu504Lys | ALDH2-Lys504Lys | 0.4446486 | 1.62E-12 |
| B94 | B98 | ALDH2-Lys504Lys | anti-Jo-1 | 0.8323901 | 0 |
| B94 | B99 | ALDH2-Lys504Lys | anti-PM-Scl | 0.500639 | 6.66E-16 |
| B94 | B110 | ALDH2-Lys504Lys | AHA | -0.498408 | 8.88E-16 |
| B95 | B116 | anti-cenpb | TT3 | -0.817909 | 0 |
| B95 | B180 | anti-cenpb | Tf | 0.8198128 | 0 |
| B95 | B202 | anti-cenpb | e-times | -0.581068 | 0 |
| B95 | B104 | anti-cenpb | anti-SSB | -0.404672 | 1.96E-10 |
| B96 | B98 | dsDNA (IIF) | anti-Jo-1 | 0.8271702 | 0 |
| B97 | B101 | dsDNA | anti-Sc70 | 0.6781609 | 0 |
| B97 | B98 | dsDNA | anti-Jo-1 | 0.7924289 | 0 |
| B98 | B101 | anti-Jo-1 | anti-Sc70 | 0.7405965 | 0 |
| B98 | B99 | anti-Jo-1 | anti-PM-Scl | 0.3759047 | 4.25E-09 |
| B99 | B103 | anti-PM-Scl | anti-SSA/Ro | 0.9952332 | 0 |
| B99 | B107 | anti-PM-Scl | AnuA | 0.6533586 | 0 |
| B99 | B110 | anti-PM-Scl | AHA | 0.9574527 | 0 |

**Supplementary Table 34. PEI Correlations in Hepatopathy**

| Row | Column | Row | Column | Cor | *P* |
|---|---|---|---|---|---|
| B1 | B216 | Gender | BMI | -0.61231 | 0 |
| B1 | B221 | Gender | Hei | -0.68997 | 0 |
| B1 | B222 | Gender | NS | -0.74584 | 0 |
| B1 | B51 | Gender | Hb | -0.5987 | 0 |
| B1 | B46 | Gender | RBC Count | -0.55231 | 8.88E-16 |
| B1 | B206 | Gender | Waist | -0.52933 | 2.18E-14 |
| B1 | B207 | Gender | WHR | -0.4777 | 1.19E-11 |
| B1 | B192 | Gender | UA | -0.45438 | 1.48E-10 |
| B1 | B210 | Gender | Hip | -0.40361 | 1.93E-08 |
| B1 | B141 | Gender | AST/ALT | 0.390499 | 5.98E-08 |
| B1 | B215 | Gender | BMI | -0.39004 | 6.22E-08 |
| B1 | B72 | Gender | HDL-C | 0.369487 | 3.32E-07 |
| B1 | B209 | Gender | Smoking | -0.36358 | 5.25E-07 |
| B1 | B129 | Gender | IBIL | -0.35564 | 9.62E-07 |
| B10 | B12 | PG I | PG II | 0.454258 | 1.50E-10 |
| B100 | B103 | anti-Ro52 | anti-SSA/Ro | 0.944911 | 0 |
| B101 | B110 | anti-Sc70 | AHA | 0.884985 | 0 |
| B101 | B102 | anti-Sc70 | anti-Sm | -0.54661 | 2.22E-15 |
| B101 | B167 | anti-Sc70 | PC | 0.392695 | 4.97E-08 |
| B103 | B107 | anti-SSA/Ro | AnuA | 0.816497 | 0 |
| B104 | B106 | anti-SSB | anti-P | -0.86164 | 0 |
| B104 | B202 | anti-SSB | e-times | -0.69171 | 0 |
| B104 | B178 | anti-SSB | Ig-U | -0.44198 | 5.25E-10 |
| B104 | B180 | anti-SSB | Tf | -0.36927 | 3.37E-07 |
| B105 | B110 | anti-U1RNP | AHA | 0.584307 | 0 |
| B105 | B167 | anti-U1RNP | PC | 0.581231 | 0 |
| B106 | B180 | anti-P | Tf | -0.7 | 0 |
| B106 | B223 | anti-P | FH | -0.55216 | 8.88E-16 |
| B106 | B178 | anti-P | Ig-U | -0.47246 | 2.14E-11 |
| B107 | B110 | AnuA | AHA | 0.816497 | 0 |
| B107 | B167 | AnuA | PC | 0.806559 | 0 |
| B108 | B202 | Anti-Cyclin | e-times | -0.9934 | 0 |
| B108 | B117 | Anti-Cyclin | TT4 | 0.49029 | 2.83E-12 |
| B108 | B116 | Anti-Cyclin | TT3 | -0.45883 | 9.31E-11 |
| B109 | B110 | AMA-M2 | AHA | 0.388889 | 6.85E-08 |
| B11 | B12 | PGI/PGII | PG II | -0.66248 | 0 |
| B11 | B14 | PGI/PGII | G-17 | -0.53773 | 7.11E-15 |
| B11 | B158 | PGI/PGII | IgM | 0.361187 | 6.32E-07 |
| B112 | B114 | FT4 | TSH TT3 TT4(TT4) | 0.5305 | 1.87E-14 |
| B112 | B196 | FT4 | TCO2 | -0.41027 | 1.07E-08 |
| B112 | B194 | FT4 | Cl | -0.40419 | 1.83E-08 |
| B112 | B200 | FT4 | OP | -0.37505 | 2.13E-07 |

| | | | | | |
|---|---|---|---|---|---|
| B112 | B195 | FT4 | Na | -0.36431 | 4.97E-07 |
| B114 | B183 | TSH TT3 TT4(TT4) | PTA | -0.59306 | 0 |
| B114 | B186 | TSH TT3 TT4(TT4) | PFC | -0.6246 | 0 |
| B114 | B194 | TSH TT3 TT4(TT4) | Cl | -0.78964 | 0 |
| B114 | B195 | TSH TT3 TT4(TT4) | Na | -0.66102 | 0 |
| B114 | B196 | TSH TT3 TT4(TT4) | TCO2 | -0.75026 | 0 |
| B114 | B197 | TSH TT3 TT4(TT4) | AG | 0.625299 | 0 |
| B114 | B200 | TSH TT3 TT4(TT4) | OP | -0.64128 | 0 |
| B114 | B185 | TSH TT3 TT4(TT4) | PT | 0.476326 | 1.39E-11 |
| B114 | B193 | TSH TT3 TT4(TT4) | K | -0.41619 | 6.22E-09 |
| B114 | B184 | TSH TT3 TT4(TT4) | INR | 0.41356 | 7.91E-09 |
| B114 | B199 | TSH TT3 TT4(TT4) | Mg | 0.364325 | 4.96E-07 |
| B114 | B115 | TSH TT3 TT4(TT4) | TSH TT3 TT4(TT3) | 0.362396 | 5.76E-07 |
| B115 | B149 | TSH TT3 TT4(TT3) | TGAb | -0.4475 | 3.01E-10 |
| B115 | B156 | TSH TT3 TT4(TT3) | IgE | 0.389405 | 6.56E-08 |
| B116 | B223 | TT3 | FH | -0.5016 | 7.35E-13 |
| B116 | B178 | TT3 | Ig-U | 0.459855 | 8.35E-11 |
| B116 | B180 | TT3 | Tf | -0.35689 | 8.76E-07 |
| B117 | B202 | TT4 | e-times | -0.41562 | 6.56E-09 |
| B118 | B132 | CEA1 | Dbil | 0.529672 | 2.09E-14 |
| B118 | B153 | CEA1 | Glu | 0.497817 | 1.16E-12 |
| B118 | B139 | CEA1 | ADA | 0.479314 | 9.96E-12 |
| B118 | B135 | CEA1 | TBA | 0.471422 | 2.40E-11 |
| B118 | B137 | CEA1 | AFU | 0.458155 | 1.00E-10 |
| B118 | B170 | CEA1 | U-GLU | 0.448754 | 2.65E-10 |
| B118 | B133 | CEA1 | TBIL | 0.447585 | 2.99E-10 |
| B118 | B138 | CEA1 | PAB | -0.44215 | 5.17E-10 |
| B118 | B190 | CEA1 | Cre | 0.425842 | 2.53E-09 |
| B118 | B129 | CEA1 | IBIL | 0.368273 | 3.65E-07 |
| B118 | B165 | CEA1 | bilirubin | 0.35909 | 7.41E-07 |
| B119 | B121 | Hcv-rna | C3 | -1 | 0 |
| B119 | B152 | Hcv-rna | ACA-Ig(G-A-M) | -0.99999 | 0 |
| B119 | B169 | Hcv-rna | U-CRY | -1 | 0 |
| B119 | B120 | Hcv-rna | HCV-cAg | 0.402506 | 2.13E-08 |
| B12 | B14 | PG II | G-17 | 0.546319 | 2.22E-15 |
| B12 | B90 | PG II | CYFRA 21-1 | 0.41409 | 7.54E-09 |
| B12 | B79 | PG II | hb V -dna | -0.38308 | 1.11E-07 |
| B120 | B121 | HCV-cAg | C3 | -0.40251 | 2.13E-08 |
| B120 | B122 | HCV-cAg | C4 | -0.40251 | 2.13E-08 |
| B120 | B152 | HCV-cAg | ACA-Ig(G-A-M) | -0.4025 | 2.13E-08 |
| B120 | B159 | HCV-cAg | β2-MG | -0.40251 | 2.13E-08 |
| B120 | B169 | HCV-cAg | U-CRY | -0.40251 | 2.13E-08 |
| B120 | B179 | HCV-cAg | α1-MG | -0.40251 | 2.13E-08 |
| B121 | B152 | C3 | ACA-Ig(G-A-M) | 0.99999 | 0 |

| | | | | | |
|---|---|---|---|---|---|
| B121 | B169 | C3 | U-CRY | 0.999999 | 0 |
| B122 | B152 | C4 | ACA-Ig(G-A-M) | 0.999991 | 0 |
| B122 | B159 | C4 | β2-MG | 1 | 0 |
| B122 | B169 | C4 | U-CRY | 1 | 0 |
| B123 | B165 | hs-CRP1 | bilirubin | 0.67136 | 0 |
| B125 | B178 | Cor | Ig-U | -0.50395 | 5.52E-13 |
| B125 | B223 | Cor | FH | -0.36811 | 3.70E-07 |
| B126 | B127 | Alb | A-G ratio | 0.708017 | 0 |
| B126 | B188 | Alb | Ca | 0.577255 | 0 |
| B126 | B132 | Alb | Dbil | -0.38089 | 1.33E-07 |
| B127 | B130 | A-G ratio | GLB | -0.86444 | 0 |
| B127 | B188 | A-G ratio | Ca | 0.371294 | 2.88E-07 |
| B128 | B131 | ALT | AST | 0.686718 | 0 |
| B128 | B141 | ALT | AST/ALT | -0.55688 | 4.44E-16 |
| B128 | B140 | ALT | r-GT | 0.456957 | 1.13E-10 |
| B129 | B132 | IBIL | Dbil | 0.838371 | 0 |
| B129 | B133 | IBIL | TBIL | 0.975696 | 0 |
| B129 | B137 | IBIL | AFU | 0.419212 | 4.71E-09 |
| B129 | B139 | IBIL | ADA | 0.410469 | 1.05E-08 |
| B129 | B135 | IBIL | TBA | 0.404645 | 1.76E-08 |
| B131 | B140 | AST | r-GT | 0.572878 | 0 |
| B131 | B197 | AST | AG | 0.368378 | 3.62E-07 |
| B132 | B133 | Dbil | TBIL | 0.937424 | 0 |
| B132 | B135 | Dbil | TBA | 0.638676 | 0 |
| B132 | B137 | Dbil | AFU | 0.618722 | 0 |
| B132 | B139 | Dbil | ADA | 0.62065 | 0 |
| B132 | B165 | Dbil | bilirubin | 0.565628 | 0 |
| B132 | B138 | Dbil | PAB | -0.53895 | 6.00E-15 |
| B132 | B153 | Dbil | Glu | 0.38273 | 1.14E-07 |
| B132 | B170 | Dbil | U-GLU | 0.371648 | 2.80E-07 |
| B132 | B134 | Dbil | CHE | -0.36993 | 3.20E-07 |
| B133 | B137 | TBIL | AFU | 0.516824 | 1.11E-13 |
| B133 | B135 | TBIL | TBA | 0.514907 | 1.41E-13 |
| B133 | B139 | TBIL | ADA | 0.511718 | 2.11E-13 |
| B133 | B165 | TBIL | bilirubin | 0.439719 | 6.58E-10 |
| B133 | B138 | TBIL | PAB | -0.41983 | 4.45E-09 |
| B134 | B135 | CHE | TBA | -0.64327 | 0 |
| B134 | B139 | CHE | ADA | -0.60661 | 0 |
| B134 | B150 | CHE | ALP | -0.77589 | 0 |
| B134 | B158 | CHE | IgM | -0.51808 | 9.46E-14 |
| B134 | B198 | CHE | P | 0.50686 | 3.87E-13 |
| B134 | B138 | CHE | PAB | 0.501956 | 7.05E-13 |
| B134 | B200 | CHE | OP | -0.44993 | 2.35E-10 |
| B134 | B197 | CHE | AG | -0.44858 | 2.70E-10 |

| | | | | | |
|---|---|---|---|---|---|
| B134 | B136 | CHE | 5'NT | -0.43224 | 1.37E-09 |
| B134 | B195 | CHE | Na | -0.41897 | 4.81E-09 |
| B135 | B137 | TBA | AFU | 0.791212 | 0 |
| B135 | B138 | TBA | PAB | -0.85216 | 0 |
| B135 | B139 | TBA | ADA | 0.89689 | 0 |
| B135 | B150 | TBA | ALP | 0.599367 | 0 |
| B135 | B165 | TBA | bilirubin | 0.640952 | 0 |
| B135 | B136 | TBA | 5'NT | 0.446614 | 3.30E-10 |
| B135 | B170 | TBA | U-GLU | 0.376365 | 1.92E-07 |
| B136 | B150 | 5'NT | ALP | 0.778973 | 0 |
| B136 | B199 | 5'NT | Mg | -0.69899 | 0 |
| B136 | B140 | 5'NT | r-GT | 0.429875 | 1.72E-09 |
| B136 | B200 | 5'NT | OP | 0.362606 | 5.67E-07 |
| B136 | B194 | 5'NT | Cl | 0.357132 | 8.60E-07 |
| B137 | B138 | AFU | PAB | -0.76677 | 0 |
| B137 | B139 | AFU | ADA | 0.862487 | 0 |
| B137 | B165 | AFU | bilirubin | 0.640112 | 0 |
| B137 | B170 | AFU | U-GLU | 0.376622 | 1.88E-07 |
| B138 | B139 | PAB | ADA | -0.93493 | 0 |
| B138 | B165 | PAB | bilirubin | -0.56859 | 0 |
| B138 | B198 | PAB | P | 0.514741 | 1.44E-13 |
| B138 | B150 | PAB | ALP | -0.4211 | 3.95E-09 |
| B138 | B200 | PAB | OP | -0.37024 | 3.13E-07 |
| B139 | B165 | ADA | bilirubin | 0.631679 | 0 |
| B139 | B150 | ADA | ALP | 0.430213 | 1.67E-09 |
| B139 | B170 | ADA | U-GLU | 0.367591 | 3.85E-07 |
| B14 | B90 | G-17 | CYFRA 21-1 | 0.522237 | 5.53E-14 |
| B14 | B79 | G-17 | hb V -dna | -0.50149 | 7.46E-13 |
| B140 | B196 | r-GT | TCO2 | -0.43679 | 8.79E-10 |
| B140 | B150 | r-GT | ALP | 0.420934 | 4.01E-09 |
| B140 | B197 | r-GT | AG | 0.400034 | 2.64E-08 |
| B141 | B222 | AST/ALT | NS | -0.42148 | 3.81E-09 |
| B141 | B216 | AST/ALT | BMI | -0.39499 | 4.08E-08 |
| B141 | B221 | AST/ALT | Hei | -0.37171 | 2.78E-07 |
| B141 | B206 | AST/ALT | Waist | -0.36242 | 5.75E-07 |
| B142 | B143 | 25-(OH)D | β-CTx | -1 | 0 |
| B142 | B151 | 25-(OH)D | Anti-CCP | -0.70513 | 0 |
| B142 | B168 | 25-(OH)D | NMP22 | -0.70513 | 0 |
| B142 | B176 | 25-(OH)D | Ucr | -0.38458 | 9.81E-08 |
| B142 | B175 | 25-(OH)D | ALB-U | -0.36067 | 6.57E-07 |
| B143 | B144 | β-CTx | Total-PINP | 1 | 0 |
| B143 | B151 | β-CTx | Anti-CCP | 0.70513 | 0 |
| B143 | B168 | β-CTx | NMP22 | 0.70513 | 0 |
| B143 | B176 | β-CTx | Ucr | 0.384583 | 9.81E-08 |

| | | | | | |
|---|---|---|---|---|---|
| B143 | B175 | β-CTx | ALB-U | 0.360671 | 6.57E-07 |
| B144 | B151 | Total-PINP | Anti-CCP | 0.705129 | 0 |
| B144 | B168 | Total-PINP | NMP22 | 0.705129 | 0 |
| B144 | B176 | Total-PINP | Ucr | 0.38458 | 9.81E-08 |
| B144 | B175 | Total-PINP | ALB-U | 0.360668 | 6.57E-07 |
| B15 | B110 | CD3-CD19+(B cell) | AHA | 0.9245 | 0 |
| B15 | B167 | CD3-CD19+(B cell) | PC | 0.597763 | 0 |
| B15 | B35 | CD3-CD19+(B cell) | β2-MG | 0.57735 | 0 |
| B15 | B93 | CD3-CD19+(B cell) | ALDH2-Glu504Lys | 0.928191 | 0 |
| B15 | B94 | CD3-CD19+(B cell) | ALDH2-Lys504Lys | -0.73598 | 0 |
| B15 | B97 | CD3-CD19+(B cell) | dsDNA | 0.880705 | 0 |
| B15 | B99 | CD3-CD19+(B cell) | anti-PM-Scl | 0.537484 | 7.33E-15 |
| B15 | B19 | CD3-CD19+(B cell) | CD3+CD4+/CD3+CD8+ | 0.470871 | 2.55E-11 |
| B15 | B18 | CD3-CD19+(B cell) | CD3+CD4+helper(inducer) T | 0.413919 | 7.66E-09 |
| B15 | B124 | CD3-CD19+(B cell) | ACTH | 0.412021 | 9.10E-09 |
| B150 | B200 | ALP | OP | 0.53389 | 1.20E-14 |
| B150 | B195 | ALP | Na | 0.498762 | 1.04E-12 |
| B150 | B158 | ALP | IgM | 0.471962 | 2.26E-11 |
| B150 | B199 | ALP | Mg | -0.43803 | 7.77E-10 |
| B150 | B198 | ALP | P | -0.42342 | 3.18E-09 |
| B150 | B197 | ALP | AG | 0.409667 | 1.13E-08 |
| B151 | B168 | Anti-CCP | NMP22 | 0.494382 | 1.75E-12 |
| B151 | B154 | Anti-CCP | RF | 0.383092 | 1.11E-07 |
| B152 | B159 | ACA-Ig(G-A-M) | β2-MG | 0.999992 | 0 |
| B152 | B169 | ACA-Ig(G-A-M) | U-CRY | 0.999992 | 0 |
| B152 | B179 | ACA-Ig(G-A-M) | α1-MG | 0.999991 | 0 |
| B153 | B170 | Glu | U-GLU | 0.775793 | 0 |
| B153 | B190 | Glu | Cre | 0.429242 | 1.83E-09 |
| B154 | B156 | RF | IgE | -0.50006 | 8.86E-13 |
| B154 | B155 | RF | IgA | -0.45881 | 9.33E-11 |
| B155 | B157 | IgA | IgG | 0.772242 | 0 |
| B157 | B158 | IgG | IgM | 0.400624 | 2.51E-08 |
| B157 | B198 | IgG | P | -0.37421 | 2.28E-07 |
| B158 | B198 | IgM | P | -0.7157 | 0 |
| B158 | B200 | IgM | OP | 0.619256 | 0 |
| B158 | B195 | IgM | Na | 0.549842 | 1.33E-15 |
| B158 | B197 | IgM | AG | 0.407516 | 1.37E-08 |
| B159 | B169 | β2-MG | U-CRY | 0.999999 | 0 |
| B16 | B101 | CD3+T | anti-Sc70 | 0.729866 | 0 |
| B16 | B105 | CD3+T | anti-U1RNP | 0.874099 | 0 |
| B16 | B110 | CD3+T | AHA | 0.92966 | 0 |
| B16 | B18 | CD3+T | CD3+CD4+helper(inducer) T | 0.80353 | 0 |

| | | | | | |
|---|---|---|---|---|---|
| B16 | B19 | CD3+T | CD3+CD4+/CD3+CD8+ | 0.694709 | 0 |
| B16 | B20 | CD3+T | CD3+CD4+CD8+ | 0.738549 | 0 |
| B16 | B36 | CD3+T | V-Lym% | 0.88326 | 0 |
| B16 | B39 | CD3+T | Other cells | 0.767836 | 0 |
| B16 | B94 | CD3+T | ALDH2-Lys504Lys | 0.584685 | 0 |
| B16 | B97 | CD3+T | dsDNA | 0.522233 | 5.55E-14 |
| B16 | B99 | CD3+T | anti-PM-Scl | 0.449467 | 2.46E-10 |
| B16 | B109 | CD3+T | AMA-M2 | -0.40202 | 2.22E-08 |
| B16 | B32 | CD3+T | Testo | 0.371761 | 2.77E-07 |
| B163 | B166 | SG | Pro | 0.448324 | 2.77E-10 |
| B164 | B171 | COM | EC | 0.46171 | 6.86E-11 |
| B169 | B179 | U-CRY | α1-MG | 1 | 0 |
| B17 | B104 | CD3+CD16+CD56+NK | anti-SSB | -0.94164 | 0 |
| B17 | B117 | CD3+CD16+CD56+NK | TT4 | -0.96077 | 0 |
| B17 | B178 | CD3+CD16+CD56+NK | Ig-U | -0.6001 | 0 |
| B17 | B180 | CD3+CD16+CD56+NK | Tf | -0.75056 | 0 |
| B17 | B202 | CD3+CD16+CD56+NK | e-times | -0.86518 | 0 |
| B17 | B22 | CD3+CD16+CD56+NK | CD3+CD8+(T) | -0.59481 | 0 |
| B17 | B37 | CD3+CD16+CD56+NK | BAND | -0.95851 | 0 |
| B17 | B95 | CD3+CD16+CD56+NK | anti-cenpb | -0.60132 | 0 |
| B170 | B190 | U-GLU | Cre | 0.372618 | 2.59E-07 |
| B171 | B205 | EC | KDA | -0.43563 | 9.84E-10 |
| B174 | B191 | Urea/Cr | Urea | 0.487785 | 3.78E-12 |
| B175 | B176 | ALB-U | Ucr | 0.799283 | 0 |
| B175 | B177 | ALB-U | ACR | 0.694026 | 0 |
| B178 | B180 | Ig-U | Tf | -0.75593 | 0 |
| B18 | B101 | CD3+CD4+helper(inducer) T | anti-Sc70 | 0.998891 | 0 |
| B18 | B102 | CD3+CD4+helper(inducer) T | anti-Sm | 0.959403 | 0 |
| B18 | B110 | CD3+CD4+helper(inducer) T | AHA | 0.710669 | 0 |
| B18 | B35 | CD3+CD4+helper(inducer) T | β2-MG | 0.853486 | 0 |
| B18 | B36 | CD3+CD4+helper(inducer) T | V-Lym% | 0.818388 | 0 |
| B18 | B98 | CD3+CD4+helper(inducer) T | anti-Jo-1 | -0.8412 | 0 |
| B18 | B19 | CD3+CD4+helper(inducer) T | CD3+CD4+/CD3+CD8+ | 0.517088 | 1.07E-13 |
| B18 | B100 | CD3+CD4+helper(inducer) T | anti-Ro52 | 0.402911 | 2.05E-08 |
| B180 | B223 | Tf | FH | -0.50799 | 3.37E-13 |
| B181 | B182 | APTT | TT | 0.887069 | 0 |

| | | | | | |
|---|---|---|---|---|---|
| B181 | B183 | APTT | PTA | -0.94151 | 0 |
| B181 | B184 | APTT | INR | 0.996967 | 0 |
| B181 | B185 | APTT | PT | 0.986358 | 0 |
| B181 | B186 | APTT | PFC | -0.92208 | 0 |
| B181 | B212 | APTT | IOP-L | -0.44651 | 3.33E-10 |
| B181 | B211 | APTT | IOP-R | -0.38681 | 8.15E-08 |
| B181 | B194 | APTT | Cl | -0.38226 | 1.19E-07 |
| B181 | B196 | APTT | TCO2 | -0.36824 | 3.66E-07 |
| B182 | B183 | TT | PTA | -0.67969 | 0 |
| B182 | B184 | TT | INR | 0.848462 | 0 |
| B182 | B185 | TT | PT | 0.798986 | 0 |
| B182 | B186 | TT | PFC | -0.63936 | 0 |
| B182 | B212 | TT | IOP-L | -0.49654 | 1.35E-12 |
| B182 | B211 | TT | IOP-R | -0.42907 | 1.86E-09 |
| B183 | B184 | PTA | INR | -0.96488 | 0 |
| B183 | B185 | PTA | PT | -0.98408 | 0 |
| B183 | B186 | PTA | PFC | 0.99856 | 0 |
| B183 | B194 | PTA | Cl | 0.632419 | 0 |
| B183 | B196 | PTA | TCO2 | 0.612655 | 0 |
| B183 | B197 | PTA | AG | -0.54766 | 1.78E-15 |
| B183 | B195 | PTA | Na | 0.47263 | 2.10E-11 |
| B183 | B200 | PTA | OP | 0.43686 | 8.73E-10 |
| B184 | B185 | INR | PT | 0.996165 | 0 |
| B184 | B186 | INR | PFC | -0.9494 | 0 |
| B184 | B194 | INR | Cl | -0.44403 | 4.28E-10 |
| B184 | B196 | INR | TCO2 | -0.42855 | 1.95E-09 |
| B184 | B212 | INR | IOP-L | -0.42821 | 2.02E-09 |
| B184 | B197 | INR | AG | 0.382466 | 1.17E-07 |
| B184 | B211 | INR | IOP-R | -0.37114 | 2.91E-07 |
| B185 | B186 | PT | PFC | -0.97315 | 0 |
| B185 | B194 | PT | Cl | -0.50955 | 2.77E-13 |
| B185 | B196 | PT | TCO2 | -0.49299 | 2.06E-12 |
| B185 | B197 | PT | AG | 0.440322 | 6.20E-10 |
| B185 | B212 | PT | IOP-L | -0.40463 | 1.76E-08 |
| B185 | B195 | PT | Na | -0.3816 | 1.25E-07 |
| B186 | B194 | PFC | Cl | 0.665383 | 0 |
| B186 | B196 | PFC | TCO2 | 0.644971 | 0 |
| B186 | B197 | PFC | AG | -0.57668 | 0 |
| B186 | B195 | PFC | Na | 0.496994 | 1.28E-12 |
| B186 | B200 | PFC | OP | 0.459291 | 8.87E-11 |
| B189 | B190 | eGFB | Cre | -0.56633 | 0 |
| B189 | B191 | eGFB | Urea | -0.60992 | 0 |
| B19 | B101 | CD3+CD4+/CD3+CD8+ | anti-Sc70 | 0.909065 | 0 |
| B19 | B105 | CD3+CD4+/CD3+CD8+ | anti-U1RNP | 0.588663 | 0 |

| | | | | | |
|---|---|---|---|---|---|
| B19 | B30 | CD3+CD4+/CD3+CD8+ | LH | 0.634545 | 0 |
| B19 | B32 | CD3+CD4+/CD3+CD8+ | Testo | 0.717707 | 0 |
| B19 | B33 | CD3+CD4+/CD3+CD8+ | Prog | 0.685994 | 0 |
| B19 | B35 | CD3+CD4+/CD3+CD8+ | β2-MG | 0.854409 | 0 |
| B19 | B94 | CD3+CD4+/CD3+CD8+ | ALDH2-Lys504Lys | -0.6436 | 0 |
| B19 | B99 | CD3+CD4+/CD3+CD8+ | anti-PM-Scl | 0.795412 | 0 |
| B19 | B145 | CD3+CD4+/CD3+CD8+ | BGP | 0.485071 | 5.17E-12 |
| B19 | B38 | CD3+CD4+/CD3+CD8+ | BAND% | 0.438763 | 7.23E-10 |
| B19 | B167 | CD3+CD4+/CD3+CD8+ | PC | 0.392046 | 5.25E-08 |
| B19 | B103 | CD3+CD4+/CD3+CD8+ | anti-SSA/Ro | -0.3638 | 5.17E-07 |
| B190 | B191 | Cre | Urea | 0.583601 | 0 |
| B191 | B192 | Urea | UA | 0.374979 | 2.14E-07 |
| B192 | B222 | UA | NS | 0.441262 | 5.64E-10 |
| B192 | B216 | UA | BMI | 0.37197 | 2.73E-07 |
| B192 | B206 | UA | Waist | 0.357258 | 8.52E-07 |
| B193 | B194 | K | Cl | 0.567071 | 0 |
| B193 | B196 | K | TCO2 | 0.604598 | 0 |
| B193 | B197 | K | AG | -0.56565 | 0 |
| B193 | B198 | K | P | -0.44219 | 5.14E-10 |
| B193 | B200 | K | OP | 0.440635 | 6.01E-10 |
| B193 | B195 | K | Na | 0.37482 | 2.17E-07 |
| B194 | B195 | Cl | Na | 0.840105 | 0 |
| B194 | B196 | Cl | TCO2 | 0.703754 | 0 |
| B194 | B197 | Cl | AG | -0.60187 | 0 |
| B194 | B200 | Cl | OP | 0.801754 | 0 |
| B194 | B199 | Cl | Mg | -0.5221 | 5.64E-14 |
| B195 | B198 | Na | P | -0.70392 | 0 |
| B195 | B200 | Na | OP | 0.98714 | 0 |
| B195 | B196 | Na | TCO2 | 0.380158 | 1.41E-07 |
| B196 | B197 | TCO2 | AG | -0.94874 | 0 |
| B196 | B200 | TCO2 | OP | 0.370992 | 2.95E-07 |
| B198 | B200 | P | OP | -0.76191 | 0 |
| B2 | B189 | Age | eGFB | -0.62189 | 0 |
| B2 | B40 | Age | WBC | -0.42011 | 4.33E-09 |
| B2 | B220 | Age | SBP | 0.384781 | 9.65E-08 |
| B2 | B191 | Age | Urea | 0.364984 | 4.72E-07 |
| B20 | B100 | CD3+CD4+CD8+ | anti-Ro52 | 0.810093 | 0 |
| B20 | B103 | CD3+CD4+CD8+ | anti-SSA/Ro | 0.612372 | 0 |
| B20 | B110 | CD3+CD4+CD8+ | AHA | 0.765466 | 0 |
| B20 | B30 | CD3+CD4+CD8+ | LH | 0.866025 | 0 |
| B20 | B32 | CD3+CD4+CD8+ | Testo | 0.889282 | 0 |
| B20 | B36 | CD3+CD4+CD8+ | V-Lym% | 0.59797 | 0 |
| B20 | B38 | CD3+CD4+CD8+ | BAND% | 0.830868 | 0 |
| B20 | B93 | CD3+CD4+CD8+ | ALDH2-Glu504Lys | 0.829515 | 0 |

| | | | | | |
|---|---|---|---|---|---|
| B20 | B94 | CD3+CD4+CD8+ | ALDH2-Lys504Lys | 0.833333 | 0 |
| B20 | B96 | CD3+CD4+CD8+ | dsDNA (IIF) | 0.730297 | 0 |
| B20 | B167 | CD3+CD4+CD8+ | PC | 0.531596 | 1.60E-14 |
| B20 | B101 | CD3+CD4+CD8+ | anti-Sc70 | 0.526001 | 3.38E-14 |
| B20 | B105 | CD3+CD4+CD8+ | anti-U1RNP | 0.509196 | 2.90E-13 |
| B20 | B39 | CD3+CD4+CD8+ | Other cells | 0.47032 | 2.71E-11 |
| B20 | B98 | CD3+CD4+CD8+ | anti-Jo-1 | 0.469809 | 2.86E-11 |
| B20 | B33 | CD3+CD4+CD8+ | Prog | 0.396928 | 3.46E-08 |
| B206 | B207 | Waist | WHR | 0.807147 | 0 |
| B206 | B210 | Waist | Hip | 0.843022 | 0 |
| B206 | B215 | Waist | BMI | 0.825732 | 0 |
| B206 | B216 | Waist | BMI | 0.863636 | 0 |
| B206 | B222 | Waist | NS | 0.767709 | 0 |
| B206 | B221 | Waist | Hei | 0.563571 | 2.22E-16 |
| B207 | B216 | WHR | BMI | 0.568672 | 0 |
| B207 | B222 | WHR | NS | 0.591557 | 0 |
| B207 | B215 | WHR | BMI | 0.563659 | 2.22E-16 |
| B207 | B221 | WHR | Hei | 0.369099 | 3.42E-07 |
| B207 | B210 | WHR | Hip | 0.366707 | 4.12E-07 |
| B209 | B221 | Smoking | Hei | 0.375558 | 2.05E-07 |
| B21 | B108 | CD3+CD4-CD8- | Anti-Cyclin | -0.86603 | 0 |
| B21 | B202 | CD3+CD4-CD8- | e-times | -0.73923 | 0 |
| B21 | B223 | CD3+CD4-CD8- | FH | -0.8115 | 0 |
| B21 | B29 | CD3+CD4-CD8- | E2 | -0.52388 | 4.46E-14 |
| B21 | B125 | CD3+CD4-CD8- | Cor | -0.40825 | 1.28E-08 |
| B210 | B215 | Hip | BMI | 0.785418 | 0 |
| B210 | B216 | Hip | BMI | 0.841696 | 0 |
| B210 | B222 | Hip | NS | 0.675633 | 0 |
| B210 | B221 | Hip | Hei | 0.559125 | 4.44E-16 |
| B211 | B212 | IOP-R | IOP-L | 0.898572 | 0 |
| B213 | B214 | CVA-R | CVA-L | 0.620323 | 0 |
| B215 | B216 | BMI | BMI | 0.87358 | 0 |
| B215 | B222 | BMI | NS | 0.698743 | 0 |
| B216 | B221 | BMI | Hei | 0.732513 | 0 |
| B216 | B222 | BMI | NS | 0.805262 | 0 |
| B218 | B220 | DBP | SBP | 0.659372 | 0 |
| B22 | B117 | CD3+CD8+(T) | TT4 | -0.9871 | 0 |
| B22 | B178 | CD3+CD8+(T) | Ig-U | -0.83189 | 0 |
| B22 | B202 | CD3+CD8+(T) | e-times | -0.96491 | 0 |
| B22 | B29 | CD3+CD8+(T) | E2 | -0.84838 | 0 |
| B22 | B125 | CD3+CD8+(T) | Cor | -0.56195 | 2.22E-16 |
| B22 | B95 | CD3+CD8+(T) | anti-cenpb | -0.51212 | 2.01E-13 |
| B22 | B28 | CD3+CD8+(T) | PRL | -0.46732 | 3.75E-11 |
| B22 | B108 | CD3+CD8+(T) | Anti-Cyclin | -0.46359 | 5.61E-11 |

| | | | | | |
|---|---|---|---|---|---|
| B22 | B104 | CD3+CD8+(T) | anti-SSB | -0.36891 | 3.47E-07 |
| B221 | B222 | Hei | NS | 0.641371 | 0 |
| B227 | B228 | OD | OS | 0.77201 | 0 |
| B23 | B117 | CD3-CD16+CD56+(NK) | TT4 | 0.878275 | 0 |
| B23 | B178 | CD3-CD16+CD56+(NK) | Ig-U | -0.69293 | 0 |
| B23 | B29 | CD3-CD16+CD56+(NK) | E2 | -0.83034 | 0 |
| B23 | B31 | CD3-CD16+CD56+(NK) | FSH | 0.801784 | 0 |
| B23 | B95 | CD3-CD16+CD56+(NK) | anti-cenpb | -0.56415 | 0 |
| B24 | B25 | αHBDH | CK | 0.627691 | 0 |
| B24 | B27 | αHBDH | LDH | 0.574868 | 0 |
| B26 | B194 | CKMB | Cl | -0.59256 | 0 |
| B26 | B196 | CKMB | TCO2 | -0.58384 | 0 |
| B26 | B114 | CKMB | TSH TT3 TT4(TT4) | 0.561469 | 2.22E-16 |
| B26 | B123 | CKMB | hs-CRP1 | 0.538722 | 6.22E-15 |
| B26 | B197 | CKMB | AG | 0.524972 | 3.86E-14 |
| B26 | B140 | CKMB | r-GT | 0.496517 | 1.36E-12 |
| B26 | B186 | CKMB | PFC | -0.49093 | 2.62E-12 |
| B26 | B183 | CKMB | PTA | -0.46589 | 4.38E-11 |
| B26 | B195 | CKMB | Na | -0.43632 | 9.20E-10 |
| B26 | B200 | CKMB | OP | -0.40551 | 1.63E-08 |
| B26 | B165 | CKMB | bilirubin | 0.392229 | 5.17E-08 |
| B26 | B27 | CKMB | LDH | 0.388592 | 7.02E-08 |
| B26 | B185 | CKMB | PT | 0.373446 | 2.42E-07 |
| B27 | B165 | LDH | bilirubin | 0.621546 | 0 |
| B27 | B137 | LDH | AFU | 0.545219 | 2.66E-15 |
| B27 | B135 | LDH | TBA | 0.543277 | 3.33E-15 |
| B27 | B139 | LDH | ADA | 0.524808 | 3.95E-14 |
| B27 | B132 | LDH | Dbil | 0.500344 | 8.56E-13 |
| B27 | B138 | LDH | PAB | -0.47589 | 1.46E-11 |
| B27 | B133 | LDH | TBIL | 0.400301 | 2.58E-08 |
| B28 | B104 | PRL | anti-SSB | -0.54953 | 1.33E-15 |
| B28 | B202 | PRL | e-times | -0.51651 | 1.15E-13 |
| B28 | B95 | PRL | anti-cenpb | -0.44446 | 4.10E-10 |
| B28 | B31 | PRL | FSH | -0.38602 | 8.70E-08 |
| B29 | B178 | E2 | Ig-U | -0.83452 | 0 |
| B29 | B223 | E2 | FH | -0.57518 | 0 |
| B29 | B125 | E2 | Cor | -0.5284 | 2.46E-14 |
| B29 | B117 | E2 | TT4 | -0.50246 | 6.63E-13 |
| B29 | B202 | E2 | e-times | -0.40298 | 2.04E-08 |
| B29 | B95 | E2 | anti-cenpb | -0.39801 | 3.15E-08 |
| B3 | B88 | Hcy | CA 242 | 0.370607 | 3.04E-07 |
| B30 | B100 | LH | anti-Ro52 | 0.694879 | 0 |
| B30 | B105 | LH | anti-U1RNP | 0.877186 | 0 |
| B30 | B110 | LH | AHA | 0.848528 | 0 |

| | | | | | |
|---|---|---|---|---|---|
| B30 | B124 | LH | ACTH | 0.8141 | 0 |
| B30 | B145 | LH | BGP | 0.565685 | 0 |
| B30 | B167 | LH | PC | 0.838201 | 0 |
| B30 | B96 | LH | dsDNA (IIF) | 0.948683 | 0 |
| B30 | B97 | LH | dsDNA | 0.938971 | 0 |
| B30 | B98 | LH | anti-Jo-1 | 0.97648 | 0 |
| B30 | B99 | LH | anti-PM-Scl | 0.80111 | 0 |
| B30 | B33 | LH | Prog | 0.483333 | 6.31E-12 |
| B30 | B39 | LH | Other cells | 0.445896 | 3.54E-10 |
| B30 | B35 | LH | β2-MG | 0.430265 | 1.66E-09 |
| B31 | B117 | FSH | TT4 | -0.96362 | 0 |
| B31 | B178 | FSH | Ig-U | -0.88388 | 0 |
| B31 | B104 | FSH | anti-SSB | -0.52636 | 3.24E-14 |
| B32 | B102 | Testo | anti-Sm | -0.5754 | 0 |
| B32 | B103 | Testo | anti-SSA/Ro | -0.5754 | 0 |
| B32 | B105 | Testo | anti-U1RNP | 0.953205 | 0 |
| B32 | B107 | Testo | AnuA | 0.805387 | 0 |
| B32 | B110 | Testo | AHA | 0.931594 | 0 |
| B32 | B167 | Testo | PC | 0.649592 | 0 |
| B32 | B38 | Testo | BAND% | 0.842659 | 0 |
| B32 | B39 | Testo | Other cells | 0.91707 | 0 |
| B32 | B97 | Testo | dsDNA | 0.664411 | 0 |
| B32 | B99 | Testo | anti-PM-Scl | 0.85775 | 0 |
| B32 | B94 | Testo | ALDH2-Lys504Lys | 0.469809 | 2.86E-11 |
| B32 | B98 | Testo | anti-Jo-1 | 0.405405 | 1.65E-08 |
| B33 | B103 | Prog | anti-SSA/Ro | 0.942809 | 0 |
| B33 | B105 | Prog | anti-U1RNP | 0.826334 | 0 |
| B33 | B107 | Prog | AnuA | 0.57735 | 0 |
| B33 | B145 | Prog | BGP | -0.58926 | 0 |
| B33 | B167 | Prog | PC | 0.952501 | 0 |
| B33 | B36 | Prog | V-Lym% | 0.642857 | 0 |
| B33 | B94 | Prog | ALDH2-Lys504Lys | 0.866025 | 0 |
| B33 | B97 | Prog | dsDNA | 0.884538 | 0 |
| B33 | B99 | Prog | anti-PM-Scl | 0.702728 | 0 |
| B33 | B101 | Prog | anti-Sc70 | 0.460131 | 8.11E-11 |
| B33 | B93 | Prog | ALDH2-Glu504Lys | 0.422577 | 3.44E-09 |
| B33 | B100 | Prog | anti-Ro52 | 0.400892 | 2.45E-08 |
| B34 | B84 | TK | C-PSA | -0.88538 | 0 |
| B34 | B88 | TK | CA 242 | 0.372466 | 2.62E-07 |
| B35 | B102 | β2-MG | anti-Sm | 0.960769 | 0 |
| B35 | B103 | β2-MG | anti-SSA/Ro | -0.64051 | 0 |
| B35 | B109 | β2-MG | AMA-M2 | 0.907393 | 0 |
| B35 | B124 | β2-MG | ACTH | 0.921787 | 0 |
| B35 | B36 | β2-MG | V-Lym% | 0.905822 | 0 |

| | | | | | |
|---|---|---|---|---|---|
| B35 | B96 | β2-MG | dsDNA (IIF) | 0.644503 | 0 |
| B35 | B99 | β2-MG | anti-PM-Scl | 0.811597 | 0 |
| B35 | B94 | β2-MG | ALDH2-Lys504Lys | -0.55566 | 4.44E-16 |
| B35 | B93 | β2-MG | ALDH2-Glu504Lys | 0.516752 | 1.12E-13 |
| B35 | B107 | β2-MG | AnuA | 0.457604 | 1.06E-10 |
| B35 | B39 | β2-MG | Other cells | 0.388368 | 7.16E-08 |
| B36 | B103 | V-Lym% | anti-SSA/Ro | 0.909137 | 0 |
| B36 | B105 | V-Lym% | anti-U1RNP | 0.585698 | 0 |
| B36 | B107 | V-Lym% | AnuA | 0.701068 | 0 |
| B36 | B124 | V-Lym% | ACTH | 0.787839 | 0 |
| B36 | B167 | V-Lym% | PC | 0.925287 | 0 |
| B36 | B98 | V-Lym% | anti-Jo-1 | 0.996408 | 0 |
| B36 | B109 | V-Lym% | AMA-M2 | 0.404061 | 1.86E-08 |
| B36 | B110 | V-Lym% | AHA | 0.404061 | 1.86E-08 |
| B37 | B116 | BAND | TT3 | -0.71818 | 0 |
| B37 | B117 | BAND | TT4 | -0.96476 | 0 |
| B37 | B108 | BAND | Anti-Cyclin | -0.55902 | 4.44E-16 |
| B37 | B178 | BAND | Ig-U | -0.47809 | 1.14E-11 |
| B37 | B180 | BAND | Tf | -0.37947 | 1.49E-07 |
| B38 | B100 | BAND% | anti-Ro52 | -0.62678 | 0 |
| B38 | B167 | BAND% | PC | 0.649836 | 0 |
| B38 | B94 | BAND% | ALDH2-Lys504Lys | 0.892413 | 0 |
| B38 | B124 | BAND% | ACTH | 0.559893 | 4.44E-16 |
| B38 | B98 | BAND% | anti-Jo-1 | 0.520466 | 6.95E-14 |
| B39 | B103 | Other cells | anti-SSA/Ro | 0.970143 | 0 |
| B39 | B167 | Other cells | PC | 0.827653 | 0 |
| B39 | B94 | Other cells | ALDH2-Lys504Lys | 0.594089 | 0 |
| B39 | B97 | Other cells | dsDNA | 0.490098 | 2.89E-12 |
| B39 | B101 | Other cells | anti-Sc70 | 0.416655 | 5.96E-09 |
| B39 | B110 | Other cells | AHA | 0.363803 | 5.17E-07 |
| B4 | B101 | WB-Ca | anti-Sc70 | 0.585652 | 0 |
| B4 | B102 | WB-Ca | anti-Sm | 0.625 | 0 |
| B4 | B97 | WB-Ca | dsDNA | 0.866025 | 0 |
| B4 | B33 | WB-Ca | Prog | 0.53033 | 1.91E-14 |
| B4 | B105 | WB-Ca | anti-U1RNP | 0.47194 | 2.27E-11 |
| B4 | B99 | WB-Ca | anti-PM-Scl | 0.447214 | 3.10E-10 |
| B4 | B20 | WB-Ca | CD3+CD4+CD8+ | 0.357217 | 8.54E-07 |
| B40 | B42 | WBC | MON | 0.718622 | 0 |
| B40 | B52 | WBC | PLT/BPC | 0.464516 | 5.08E-11 |
| B41 | B42 | MON % | MON | 0.530778 | 1.80E-14 |
| B43 | B44 | MCV | MCH | 0.868732 | 0 |
| B43 | B46 | MCV | RBC Count | -0.59172 | 0 |
| B43 | B47 | MCV | RDW | 0.497735 | 1.17E-12 |
| B44 | B45 | MCH | MCHC | 0.52868 | 2.38E-14 |

| | | | | | |
|---|---|---|---|---|---|
| B44 | B46 | MCH | RBC Count | -0.41221 | 8.95E-09 |
| B45 | B51 | MCHC | Hb | 0.505687 | 4.47E-13 |
| B46 | B126 | RBC Count | Alb | 0.582513 | 0 |
| B46 | B47 | RBC Count | RDW | -0.57116 | 0 |
| B46 | B51 | RBC Count | Hb | 0.844473 | 0 |
| B46 | B222 | RBC Count | NS | 0.51219 | 1.99E-13 |
| B46 | B221 | RBC Count | Hei | 0.460815 | 7.55E-11 |
| B46 | B141 | RBC Count | AST/ALT | -0.44198 | 5.25E-10 |
| B46 | B216 | RBC Count | BMI | 0.416787 | 5.89E-09 |
| B46 | B127 | RBC Count | A-G ratio | 0.412223 | 8.94E-09 |
| B46 | B163 | RBC Count | SG | 0.370218 | 3.13E-07 |
| B46 | B91 | RBC Count | TP | 0.359596 | 7.13E-07 |
| B47 | B126 | RDW | Alb | -0.55138 | 8.88E-16 |
| B47 | B51 | RDW | Hb | -0.43559 | 9.88E-10 |
| B47 | B127 | RDW | A-G ratio | -0.40679 | 1.46E-08 |
| B48 | B53 | LYM% | GR% | -0.92319 | 0 |
| B5 | B100 | WB-Mg | anti-Ro52 | 0.777844 | 0 |
| B5 | B102 | WB-Mg | anti-Sm | 0.857493 | 0 |
| B5 | B107 | WB-Mg | AnuA | 0.840168 | 0 |
| B5 | B109 | WB-Mg | AMA-M2 | 0.571662 | 0 |
| B5 | B145 | WB-Mg | BGP | 0.771744 | 0 |
| B5 | B20 | WB-Mg | CD3+CD4+CD8+ | 0.665133 | 0 |
| B5 | B30 | WB-Mg | LH | 0.569959 | 0 |
| B5 | B7 | WB-Mg | WB-Cop | 0.703353 | 0 |
| B5 | B93 | WB-Mg | ALDH2-Glu504Lys | 0.737928 | 0 |
| B5 | B94 | WB-Mg | ALDH2-Lys504Lys | 0.735147 | 0 |
| B5 | B96 | WB-Mg | dsDNA (IIF) | 0.843661 | 0 |
| B5 | B15 | WB-Mg | CD3-CD19+(B cell) | 0.475651 | 1.50E-11 |
| B5 | B110 | WB-Mg | AHA | 0.45733 | 1.09E-10 |
| B5 | B35 | WB-Mg | β2-MG | 0.384465 | 9.91E-08 |
| B5 | B124 | WB-Mg | ACTH | 0.382158 | 1.20E-07 |
| B5 | B36 | WB-Mg | V-Lym% | 0.381127 | 1.30E-07 |
| B51 | B222 | Hb | NS | 0.54235 | 3.77E-15 |
| B51 | B221 | Hb | Hei | 0.534992 | 1.02E-14 |
| B51 | B126 | Hb | Alb | 0.508747 | 3.06E-13 |
| B51 | B216 | Hb | BMI | 0.487411 | 3.95E-12 |
| B51 | B141 | Hb | AST/ALT | -0.4343 | 1.12E-09 |
| B51 | B206 | Hb | Waist | 0.380175 | 1.41E-07 |
| B51 | B209 | Hb | Smoking | 0.378332 | 1.64E-07 |
| B51 | B91 | Hb | TP | 0.357907 | 8.11E-07 |
| B52 | B132 | PLT/BPC | Dbil | -0.36565 | 4.48E-07 |
| B54 | B57 | ESR | RBCAI | -1 | 0 |
| B54 | B63 | ESR | WBRSR(mPas_200) | -1 | 0 |
| B54 | B67 | ESR | WBRSR(mPas_50) | -1 | 0 |

| | | | | | |
|---|---|---|---|---|---|
| B56 | B57 | IR | RBCAI | 1 | 0 |
| B56 | B59 | IR | WBRV | 1 | 0 |
| B56 | B60 | IR | WBRVI | 1 | 0 |
| B56 | B61 | IR | WBHSRV | 1 | 0 |
| B56 | B62 | IR | WBHSRVI | 1 | 0 |
| B56 | B65 | IR | WBRSR(mPas_1) | 1 | 0 |
| B56 | B67 | IR | WBRSR(mPas_50) | 1 | 0 |
| B57 | B59 | RBCAI | WBRV | 1 | 0 |
| B57 | B60 | RBCAI | WBRVI | 1 | 0 |
| B57 | B61 | RBCAI | WBHSRV | 1 | 0 |
| B57 | B62 | RBCAI | WBHSRVI | 1 | 0 |
| B57 | B63 | RBCAI | WBRSR(mPas_200) | 1 | 0 |
| B57 | B64 | RBCAI | WBRSR(mPas_5) | 1 | 0 |
| B57 | B65 | RBCAI | WBRSR(mPas_1) | 1 | 0 |
| B57 | B66 | RBCAI | WBRSR(mPas_100) | 1 | 0 |
| B59 | B60 | WBRV | WBRVI | 1 | 0 |
| B59 | B61 | WBRV | WBHSRV | 1 | 0 |
| B59 | B62 | WBRV | WBHSRVI | 1 | 0 |
| B59 | B65 | WBRV | WBRSR(mPas_1) | 1 | 0 |
| B59 | B67 | WBRV | WBRSR(mPas_50) | 1 | 0 |
| B6 | B104 | WB-Fe | anti-SSB | -0.82061 | 0 |
| B6 | B202 | WB-Fe | e-times | -0.87415 | 0 |
| B6 | B223 | WB-Fe | FH | -0.95707 | 0 |
| B6 | B29 | WB-Fe | E2 | -0.98131 | 0 |
| B6 | B31 | WB-Fe | FSH | -0.89087 | 0 |
| B61 | B62 | WBHSRV | WBHSRVI | 1 | 0 |
| B61 | B64 | WBHSRV | WBRSR(mPas_5) | 1 | 0 |
| B61 | B65 | WBHSRV | WBRSR(mPas_1) | 1 | 0 |
| B61 | B67 | WBHSRV | WBRSR(mPas_50) | 1 | 0 |
| B62 | B63 | WBHSRVI | WBRSR(mPas_200) | 1 | 0 |
| B62 | B65 | WBHSRVI | WBRSR(mPas_1) | 1 | 0 |
| B62 | B67 | WBHSRVI | WBRSR(mPas_50) | 1 | 0 |
| B63 | B64 | WBRSR(mPas_200) | WBRSR(mPas_5) | 1 | 0 |
| B65 | B67 | WBRSR(mPas_1) | WBRSR(mPas_50) | 1 | 0 |
| B7 | B110 | WB-Cop | AHA | 0.565685 | 0 |
| B7 | B124 | WB-Cop | ACTH | 0.945406 | 0 |
| B7 | B18 | WB-Cop | CD3+CD4+helper(inducer) T | 0.663325 | 0 |
| B7 | B19 | WB-Cop | CD3+CD4+/CD3+CD8+ | 0.583095 | 0 |
| B7 | B33 | WB-Cop | Prog | 0.566667 | 0 |
| B7 | B36 | WB-Cop | V-Lym% | 0.614286 | 0 |
| B7 | B39 | WB-Cop | Other cells | 0.960392 | 0 |
| B7 | B93 | WB-Cop | ALDH2-Glu504Lys | 0.845154 | 0 |
| B7 | B98 | WB-Cop | anti-Jo-1 | 0.697486 | 0 |

| | | | | | |
|---|---|---|---|---|---|
| B7 | B105 | WB-Cop | anti-U1RNP | 0.533939 | 1.18E-14 |
| B7 | B101 | WB-Cop | anti-Sc70 | 0.485899 | 4.70E-12 |
| B7 | B30 | WB-Cop | LH | 0.48 | 9.22E-12 |
| B7 | B16 | WB-Cop | CD3+T | 0.447722 | 2.94E-10 |
| B7 | B20 | WB-Cop | CD3+CD4+CD8+ | 0.433013 | 1.27E-09 |
| B70 | B77 | LDL-C | TC | 0.761795 | 0 |
| B71 | B190 | TG | Cre | 0.553778 | 8.88E-16 |
| B71 | B77 | TG | TC | 0.493974 | 1.83E-12 |
| B71 | B153 | TG | Glu | 0.357102 | 8.62E-07 |
| B72 | B207 | HDL-C | WHR | -0.39782 | 3.20E-08 |
| B72 | B222 | HDL-C | NS | -0.37757 | 1.74E-07 |
| B72 | B206 | HDL-C | Waist | -0.37114 | 2.91E-07 |
| B72 | B74 | HDL-C | ApoA1 | 0.365097 | 4.67E-07 |
| B72 | B141 | HDL-C | AST/ALT | 0.359743 | 7.05E-07 |
| B73 | B76 | ApoA | ApoB | 0.557456 | 4.44E-16 |
| B73 | B187 | ApoA | NSE | 0.502352 | 6.71E-13 |
| B73 | B197 | ApoA | AG | -0.50085 | 8.06E-13 |
| B73 | B196 | ApoA | TCO2 | 0.480965 | 8.27E-12 |
| B73 | B150 | ApoA | ALP | -0.36232 | 5.79E-07 |
| B74 | B136 | ApoA1 | 5'NT | 0.843205 | 0 |
| B74 | B199 | ApoA1 | Mg | -0.79996 | 0 |
| B74 | B150 | ApoA1 | ALP | 0.528443 | 2.44E-14 |
| B74 | B76 | ApoA1 | ApoB | 0.473688 | 1.87E-11 |
| B75 | B134 | ApoA1/B | CHE | -0.60448 | 0 |
| B75 | B150 | ApoA1/B | ALP | 0.684879 | 0 |
| B75 | B158 | ApoA1/B | IgM | 0.801368 | 0 |
| B75 | B197 | ApoA1/B | AG | 0.602863 | 0 |
| B75 | B76 | ApoA1/B | ApoB | -0.65324 | 0 |
| B75 | B200 | ApoA1/B | OP | 0.53424 | 1.13E-14 |
| B75 | B198 | ApoA1/B | P | -0.52542 | 3.64E-14 |
| B75 | B195 | ApoA1/B | Na | 0.476309 | 1.40E-11 |
| B75 | B157 | ApoA1/B | IgG | 0.422667 | 3.41E-09 |
| B75 | B136 | ApoA1/B | 5'NT | 0.401665 | 2.29E-08 |
| B75 | B196 | ApoA1/B | TCO2 | -0.4014 | 2.34E-08 |
| B76 | B158 | ApoB | IgM | -0.698 | 0 |
| B76 | B197 | ApoB | AG | -0.59364 | 0 |
| B76 | B198 | ApoB | P | 0.609418 | 0 |
| B76 | B134 | ApoB | CHE | 0.417488 | 5.52E-09 |
| B76 | B199 | ApoB | Mg | -0.40213 | 2.20E-08 |
| B76 | B196 | ApoB | TCO2 | 0.371481 | 2.83E-07 |
| B76 | B157 | ApoB | IgG | -0.36618 | 4.30E-07 |
| B79 | B90 | hb V -dna | CYFRA 21-1 | -0.46071 | 7.63E-11 |
| B79 | B176 | hb V -dna | Ucr | -0.37343 | 2.43E-07 |
| B8 | B116 | WB-Zinc | TT3 | -0.93659 | 0 |

| | | | | | |
|---|---|---|---|---|---|
| B8 | B125 | WB-Zinc | Cor | 0.96225 | 0 |
| B8 | B180 | WB-Zinc | Tf | -0.80829 | 0 |
| B8 | B23 | WB-Zinc | CD3-CD16+CD56+(NK) | -0.57735 | 0 |
| B8 | B28 | WB-Zinc | PRL | -0.90972 | 0 |
| B8 | B29 | WB-Zinc | E2 | -0.98058 | 0 |
| B8 | B17 | WB-Zinc | CD3+CD16+CD56+NK | -0.5 | 8.93E-13 |
| B82 | B86 | CEA | TBhCG | 0.648946 | 0 |
| B82 | B88 | CEA | CA 242 | 0.768762 | 0 |
| B82 | B85 | CEA | SCC | 0.485184 | 5.11E-12 |
| B82 | B156 | CEA | IgE | -0.40863 | 1.24E-08 |
| B85 | B87 | SCC | NSE | 0.464764 | 4.95E-11 |
| B86 | B156 | TBhCG | IgE | -0.53178 | 1.58E-14 |
| B86 | B88 | TBhCG | CA 242 | 0.416037 | 6.31E-09 |
| B87 | B88 | NSE | CA 242 | 0.581305 | 0 |
| B88 | B156 | CA 242 | IgE | -0.38166 | 1.25E-07 |
| B88 | B89 | CA 242 | CA 72-4 | -0.37247 | 2.62E-07 |
| B89 | B115 | CA 72-4 | TSH TT3 TT4(TT3) | 0.546749 | 2.00E-15 |
| B89 | B156 | CA 72-4 | IgE | 0.522074 | 5.64E-14 |
| B9 | B123 | RH | hs-CRP1 | -0.95229 | 0 |
| B9 | B165 | RH | bilirubin | -0.70513 | 0 |
| B9 | B26 | RH | CKMB | -0.56325 | 2.22E-16 |
| B91 | B126 | TP | Alb | 0.613239 | 0 |
| B91 | B130 | TP | GLB | 0.548335 | 1.78E-15 |
| B91 | B188 | TP | Ca | 0.393728 | 4.55E-08 |
| B93 | B101 | ALDH2-Glu504Lys | anti-Sc70 | 0.597323 | 0 |
| B93 | B103 | ALDH2-Glu504Lys | anti-SSA/Ro | 0.597614 | 0 |
| B93 | B105 | ALDH2-Glu504Lys | anti-U1RNP | 0.902522 | 0 |
| B93 | B145 | ALDH2-Glu504Lys | BGP | 0.657376 | 0 |
| B93 | B167 | ALDH2-Glu504Lys | PC | 0.644008 | 0 |
| B93 | B110 | ALDH2-Glu504Lys | AHA | 0.557773 | 4.44E-16 |
| B94 | B109 | ALDH2-Lys504Lys | AMA-M2 | -0.95258 | 0 |
| B94 | B110 | ALDH2-Lys504Lys | AHA | -0.61237 | 0 |
| B94 | B98 | ALDH2-Lys504Lys | anti-Jo-1 | 0.570483 | 0 |
| B94 | B167 | ALDH2-Lys504Lys | PC | 0.513265 | 1.74E-13 |
| B94 | B97 | ALDH2-Lys504Lys | dsDNA | -0.41248 | 8.73E-09 |
| B95 | B117 | anti-cenpb | TT4 | -0.95686 | 0 |
| B95 | B223 | anti-cenpb | FH | -0.63259 | 0 |
| B95 | B202 | anti-cenpb | e-times | -0.46335 | 5.76E-11 |
| B95 | B104 | anti-cenpb | anti-SSB | -0.41665 | 5.96E-09 |
| B96 | B98 | dsDNA (IIF) | anti-Jo-1 | 0.661693 | 0 |
| B96 | B99 | dsDNA (IIF) | anti-PM-Scl | 0.866667 | 0 |
| B96 | B100 | dsDNA (IIF) | anti-Ro52 | 0.507093 | 3.76E-13 |
| B96 | B101 | dsDNA (IIF) | anti-Sc70 | 0.419058 | 4.78E-09 |
| B97 | B100 | dsDNA | anti-Ro52 | -0.87287 | 0 |

| | | | | | |
|---|---|---|---|---|---|
| B97 | B101 | dsDNA | anti-Sc70 | 0.811503 | 0 |
| B97 | B109 | dsDNA | AMA-M2 | -0.7698 | 0 |
| B97 | B98 | dsDNA | anti-Jo-1 | 0.854242 | 0 |
| B98 | B101 | anti-Jo-1 | anti-Sc70 | 0.872943 | 0 |
| B98 | B105 | anti-Jo-1 | anti-U1RNP | 0.886702 | 0 |
| B98 | B109 | anti-Jo-1 | AMA-M2 | -0.6028 | 0 |
| B98 | B145 | anti-Jo-1 | BGP | 0.904194 | 0 |
| B98 | B167 | anti-Jo-1 | PC | 0.531485 | 1.64E-14 |
| B98 | B103 | anti-Jo-1 | anti-SSA/Ro | -0.411 | 9.99E-09 |
| B99 | B110 | anti-PM-Scl | AHA | 0.919272 | 0 |
| B99 | B107 | anti-PM-Scl | AnuA | 0.426006 | 2.49E-09 |

**Supplementary Table 35. PEI Correlations in Pregnant**

| Row | Column | Row | Column | Cor | *P* |
|---|---|---|---|---|---|
| B1 | B211 | Gender | IOP-R | 0.861797 | 0 |
| B1 | B224 | Gender | pmh | -1 | 0 |
| B1 | B82 | Gender | CEA | 1 | 0 |
| B1 | B83 | Gender | AFP | -1 | 0 |
| B1 | B86 | Gender | TBhCG | 1 | 0 |
| B1 | B87 | Gender | NSE | 1 | 0 |
| B1 | B88 | Gender | CA 242 | -1 | 0 |
| B1 | B212 | Gender | IOP-L | 0.795336 | 2.45E-13 |
| B1 | B120 | Gender | HCV-cAg | 0.700649 | 1.84E-09 |
| B10 | B12 | PG I | PG II | 1 | 0 |
| B10 | B80 | PG I | anti-HP IgG | 0.700649 | 1.84E-09 |
| B100 | B152 | anti-Ro52 | ACA-Ig(G-A-M) | -0.86772 | 0 |
| B100 | B182 | anti-Ro52 | TT | -0.9 | 0 |
| B100 | B186 | anti-Ro52 | PFC | -0.86963 | 0 |
| B100 | B200 | anti-Ro52 | OP | -0.85812 | 0 |
| B100 | B137 | anti-Ro52 | AFU | -0.81989 | 1.09E-14 |
| B100 | B102 | anti-Ro52 | anti-Sm | -0.80498 | 7.59E-14 |
| B100 | B121 | anti-Ro52 | C3 | -0.79057 | 4.27E-13 |
| B100 | B125 | anti-Ro52 | Cor | -0.70711 | 1.12E-09 |
| B101 | B158 | anti-Sc70 | IgM | 1 | 0 |
| B101 | B175 | anti-Sc70 | ALB-U | 0.774597 | 2.48E-12 |
| B102 | B105 | anti-Sm | anti-U1RNP | -1 | 0 |
| B102 | B176 | anti-Sm | Ucr | -0.88752 | 0 |
| B102 | B177 | anti-Sm | ACR | -0.89815 | 0 |
| B102 | B185 | anti-Sm | PT | -0.84853 | 0 |
| B102 | B223 | anti-Sm | FH | -0.92164 | 0 |
| B102 | B198 | anti-Sm | P | -0.8165 | 1.73E-14 |
| B102 | B182 | anti-Sm | TT | -0.80498 | 7.59E-14 |
| B102 | B125 | anti-Sm | Cor | -0.6957 | 2.66E-09 |
| B102 | B136 | anti-Sm | 5'NT | -0.6532 | 4.85E-08 |
| B102 | B168 | anti-Sm | NMP22 | -0.6532 | 4.85E-08 |
| B102 | B186 | anti-Sm | PFC | -0.6364 | 1.35E-07 |
| B103 | B175 | anti-SSA/Ro | ALB-U | 0.894427 | 0 |
| B103 | B108 | anti-SSA/Ro | Anti-Cyclin | 0.771517 | 3.43E-12 |
| B103 | B110 | anti-SSA/Ro | AHA | 0.617213 | 4.06E-07 |
| B104 | B150 | anti-SSB | ALP | 0.979796 | 0 |
| B104 | B158 | anti-SSB | IgM | 0.948683 | 0 |
| B104 | B144 | anti-SSB | Total-PINP | 0.837708 | 8.88E-16 |
| B104 | B156 | anti-SSB | IgE | 0.816497 | 1.73E-14 |
| B104 | B159 | anti-SSB | β2-MG | 0.786991 | 6.41E-13 |
| B104 | B193 | anti-SSB | K | 0.759072 | 1.21E-11 |
| B104 | B197 | anti-SSB | AG | 0.700649 | 1.84E-09 |

| | | | | | |
|---|---|---|---|---|---|
| B105 | B121 | anti-U1RNP | C3 | -0.88388 | 0 |
| B105 | B106 | anti-U1RNP | anti-P | -0.68641 | 5.23E-09 |
| B105 | B151 | anti-U1RNP | Anti-CCP | -0.6455 | 7.82E-08 |
| B106 | B176 | anti-P | Ucr | -0.97907 | 0 |
| B106 | B185 | anti-P | PT | -0.9245 | 0 |
| B106 | B223 | anti-P | FH | -0.99887 | 0 |
| B106 | B167 | anti-P | PC | -0.83863 | 8.88E-16 |
| B106 | B135 | anti-P | TBA | -0.83205 | 2.00E-15 |
| B106 | B179 | anti-P | α1-MG | -0.78446 | 8.51E-13 |
| B106 | B182 | anti-P | TT | -0.70165 | 1.70E-09 |
| B106 | B183 | anti-P | PTA | -0.64194 | 9.71E-08 |
| B106 | B177 | anti-P | ACR | -0.64051 | 1.06E-07 |
| B106 | B186 | anti-P | PFC | -0.62404 | 2.77E-07 |
| B106 | B124 | anti-P | ACTH | -0.6191 | 3.65E-07 |
| B107 | B138 | AnuA | PAB | 0.896421 | 0 |
| B107 | B175 | AnuA | ALB-U | 0.919239 | 0 |
| B107 | B184 | AnuA | INR | 0.782624 | 1.04E-12 |
| B107 | B187 | AnuA | NSE | 0.744208 | 4.92E-11 |
| B107 | B194 | AnuA | Cl | 0.711512 | 7.92E-10 |
| B107 | B159 | AnuA | β2-MG | 0.681554 | 7.38E-09 |
| B107 | B134 | AnuA | CHE | 0.67082 | 1.54E-08 |
| B107 | B156 | AnuA | IgE | 0.636396 | 1.35E-07 |
| B108 | B134 | Anti-Cyclin | CHE | 0.872872 | 0 |
| B108 | B139 | Anti-Cyclin | ADA | 0.790912 | 4.10E-13 |
| B108 | B144 | Anti-Cyclin | Total-PINP | 0.778792 | 1.59E-12 |
| B108 | B110 | Anti-Cyclin | AHA | 0.761905 | 9.12E-12 |
| B108 | B138 | Anti-Cyclin | PAB | 0.758175 | 1.32E-11 |
| B108 | B197 | Anti-Cyclin | AG | 0.723747 | 2.93E-10 |
| B108 | B150 | Anti-Cyclin | ALP | 0.690066 | 4.02E-09 |
| B108 | B164 | Anti-Cyclin | COM | 0.637022 | 1.30E-07 |
| B108 | B199 | Anti-Cyclin | Mg | 0.635107 | 1.46E-07 |
| B108 | B158 | Anti-Cyclin | IgM | 0.62361 | 2.84E-07 |
| B109 | B159 | AMA-M2 | β2-MG | 0.931334 | 0 |
| B109 | B143 | AMA-M2 | β-CTx | 0.7698 | 4.10E-12 |
| B11 | B80 | PGI/PGII | anti-HP IgG | -0.70065 | 1.84E-09 |
| B110 | B159 | AHA | β2-MG | 0.720556 | 3.82E-10 |
| B110 | B138 | AHA | PAB | 0.699854 | 1.95E-09 |
| B110 | B195 | AHA | Na | 0.699854 | 1.95E-09 |
| B110 | B143 | AHA | β-CTx | 0.629941 | 1.97E-07 |
| B110 | B157 | AHA | IgG | 0.617213 | 4.06E-07 |
| B114 | B115 | TSH TT3 TT4(TT4) | TSH TT3 TT4(TT3) | 0.735587 | 1.07E-10 |
| B115 | B148 | TSH TT3 TT4(TT3) | Anti_TPO | -0.61901 | 3.67E-07 |
| B115 | B149 | TSH TT3 TT4(TT3) | TGAb | -0.61239 | 5.29E-07 |
| B116 | B197 | TT3 | AG | 0.94388 | 0 |

| | | | | | |
|---|---|---|---|---|---|
| B116 | B199 | TT3 | Mg | 0.976187 | 0 |
| B116 | B196 | TT3 | TCO2 | -0.7746 | 2.48E-12 |
| B117 | B199 | TT4 | Mg | 0.970143 | 0 |
| B12 | B80 | PG II | anti-HP IgG | 0.700649 | 1.84E-09 |
| B120 | B224 | HCV-cAg | pmh | -0.70065 | 1.84E-09 |
| B121 | B122 | C3 | C4 | -0.94868 | 0 |
| B121 | B182 | C3 | TT | -0.86963 | 0 |
| B121 | B186 | C3 | PFC | -0.875 | 0 |
| B121 | B202 | C3 | e-times | -0.86963 | 0 |
| B121 | B223 | C3 | FH | -0.85749 | 0 |
| B121 | B151 | C3 | Anti-CCP | -0.82158 | 8.66E-15 |
| B121 | B145 | C3 | BGP | -0.75378 | 2.01E-11 |
| B121 | B135 | C3 | TBA | -0.75 | 2.88E-11 |
| B121 | B225 | C3 | hPBG | -0.75 | 2.88E-11 |
| B121 | B125 | C3 | Cor | -0.67082 | 1.54E-08 |
| B122 | B145 | C4 | BGP | -0.95346 | 0 |
| B122 | B168 | C4 | NMP22 | -0.91287 | 0 |
| B122 | B176 | C4 | Ucr | -0.86824 | 0 |
| B122 | B185 | C4 | PT | -0.84327 | 4.44E-16 |
| B122 | B151 | C4 | Anti-CCP | -0.80829 | 5.02E-14 |
| B122 | B182 | C4 | TT | 0.8 | 1.40E-13 |
| B122 | B183 | C4 | PTA | -0.78072 | 1.29E-12 |
| B122 | B179 | C4 | α1-MG | -0.74536 | 4.43E-11 |
| B122 | B198 | C4 | P | 0.730297 | 1.69E-10 |
| B122 | B200 | C4 | OP | 0.667424 | 1.93E-08 |
| B124 | B152 | ACTH | ACA-Ig(G-A-M) | -0.94579 | 0 |
| B124 | B171 | ACTH | EC | -0.99038 | 0 |
| B124 | B179 | ACTH | α1-MG | -0.92848 | 0 |
| B124 | B198 | ACTH | P | -0.83391 | 1.33E-15 |
| B124 | B137 | ACTH | AFU | -0.80468 | 7.88E-14 |
| B124 | B186 | ACTH | PFC | -0.78784 | 5.83E-13 |
| B124 | B135 | ACTH | TBA | -0.7003 | 1.89E-09 |
| B125 | B225 | Cor | hPBG | -0.81989 | 1.09E-14 |
| B125 | B200 | Cor | OP | -0.80904 | 4.55E-14 |
| B125 | B152 | Cor | ACA-Ig(G-A-M) | -0.76696 | 5.49E-12 |
| B125 | B168 | Cor | NMP22 | -0.6455 | 7.82E-08 |
| B126 | B127 | Alb | A-G ratio | 0.784659 | 8.33E-13 |
| B127 | B130 | A-G ratio | GLB | -0.82337 | 6.66E-15 |
| B128 | B131 | ALT | AST | 0.865785 | 0 |
| B128 | B141 | ALT | AST/ALT | -0.66055 | 3.03E-08 |
| B129 | B133 | IBIL | TBIL | 0.987449 | 0 |
| B129 | B132 | IBIL | Dbil | 0.837792 | 8.88E-16 |
| B132 | B133 | Dbil | TBIL | 0.913369 | 0 |
| B134 | B159 | CHE | β2-MG | 0.889001 | 0 |

| | | | | | |
|---|---|---|---|---|---|
| B134 | B164 | CHE | COM | 0.834058 | 1.33E-15 |
| B134 | B197 | CHE | AG | 0.829156 | 3.11E-15 |
| B134 | B184 | CHE | INR | 0.8125 | 2.93E-14 |
| B134 | B144 | CHE | Total-PINP | 0.811107 | 3.49E-14 |
| B134 | B157 | CHE | IgG | 0.795495 | 2.40E-13 |
| B134 | B194 | CHE | Cl | 0.707107 | 1.12E-09 |
| B135 | B176 | TBA | Ucr | -0.91521 | 0 |
| B135 | B185 | TBA | PT | -1 | 0 |
| B136 | B168 | 5'NT | NMP22 | -1 | 0 |
| B136 | B198 | 5'NT | P | -1 | 0 |
| B136 | B151 | 5'NT | Anti-CCP | -0.84327 | 4.44E-16 |
| B136 | B152 | 5'NT | ACA-Ig(G-A-M) | -0.79212 | 3.57E-13 |
| B136 | B185 | 5'NT | PT | -0.7698 | 4.10E-12 |
| B137 | B151 | AFU | Anti-CCP | -0.86066 | 0 |
| B137 | B152 | AFU | ACA-Ig(G-A-M) | -0.8893 | 0 |
| B137 | B167 | AFU | PC | -0.7127 | 7.21E-10 |
| B138 | B199 | PAB | Mg | 0.907485 | 0 |
| B138 | B195 | PAB | Na | 0.714286 | 6.35E-10 |
| B139 | B143 | ADA | β-CTx | 0.923186 | 0 |
| B139 | B155 | ADA | IgA | 0.86164 | 0 |
| B139 | B159 | ADA | β2-MG | 0.974755 | 0 |
| B139 | B193 | ADA | K | 0.968665 | 0 |
| B139 | B199 | ADA | Mg | 0.982467 | 0 |
| B139 | B197 | ADA | AG | 0.771389 | 3.48E-12 |
| B139 | B157 | ADA | IgG | 0.603023 | 8.73E-07 |
| B14 | B80 | G-17 | anti-HP IgG | -0.70065 | 1.84E-09 |
| B142 | B197 | 25-(OH)D | AG | 0.9461 | 0 |
| B142 | B199 | 25-(OH)D | Mg | 0.951303 | 0 |
| B142 | B195 | 25-(OH)D | Na | 0.838628 | 8.88E-16 |
| B142 | B144 | 25-(OH)D | Total-PINP | 0.763542 | 7.75E-12 |
| B143 | B156 | β-CTx | IgE | 0.912871 | 0 |
| B143 | B158 | β-CTx | IgM | 0.942809 | 0 |
| B143 | B175 | β-CTx | ALB-U | 0.912871 | 0 |
| B143 | B178 | β-CTx | Ig-U | 1 | 0 |
| B143 | B195 | β-CTx | Na | 0.771517 | 3.43E-12 |
| B143 | B197 | β-CTx | AG | 0.73983 | 7.31E-11 |
| B143 | B150 | β-CTx | ALP | 0.730297 | 1.69E-10 |
| B143 | B164 | β-CTx | COM | 0.601929 | 9.24E-07 |
| B144 | B150 | Total-PINP | ALP | 0.923381 | 0 |
| B144 | B154 | Total-PINP | RF | -0.97333 | 0 |
| B144 | B159 | Total-PINP | β2-MG | 0.86529 | 0 |
| B144 | B193 | Total-PINP | K | 0.953821 | 0 |
| B144 | B175 | Total-PINP | ALB-U | 0.820783 | 9.77E-15 |
| B144 | B155 | Total-PINP | IgA | 0.749269 | 3.08E-11 |

| | | | | | |
|---|---|---|---|---|---|
| B144 | B164 | Total-PINP | COM | 0.744161 | 4.94E-11 |
| B144 | B197 | Total-PINP | AG | 0.684762 | 5.88E-09 |
| B145 | B167 | BGP | PC | -0.96866 | 0 |
| B145 | B176 | BGP | Ucr | -0.9461 | 0 |
| B145 | B186 | BGP | PFC | -0.97991 | 0 |
| B145 | B183 | BGP | PTA | -0.83744 | 8.88E-16 |
| B145 | B198 | BGP | P | -0.82687 | 4.00E-15 |
| B145 | B169 | BGP | U-CRY | 0.639602 | 1.12E-07 |
| B145 | B185 | BGP | PT | -0.60302 | 8.73E-07 |
| B148 | B149 | Anti_TPO | TGAb | 0.998881 | 0 |
| B15 | B124 | CD3-CD19+(B cell) | ACTH | -0.92705 | 0 |
| B15 | B136 | CD3-CD19+(B cell) | 5'NT | -0.90582 | 0 |
| B15 | B198 | CD3-CD19+(B cell) | P | -0.84921 | 0 |
| B15 | B223 | CD3-CD19+(B cell) | FH | -0.94174 | 0 |
| B15 | B28 | CD3-CD19+(B cell) | PRL | 0.915209 | 0 |
| B15 | B56 | CD3-CD19+(B cell) | IR | -0.87706 | 0 |
| B15 | B94 | CD3-CD19+(B cell) | ALDH2-Lys504Lys | -0.86824 | 0 |
| B15 | B185 | CD3-CD19+(B cell) | PT | -0.78446 | 8.51E-13 |
| B15 | B106 | CD3-CD19+(B cell) | anti-P | -0.7615 | 9.50E-12 |
| B15 | B93 | CD3-CD19+(B cell) | ALDH2-Glu504Lys | -0.74125 | 6.44E-11 |
| B15 | B137 | CD3-CD19+(B cell) | AFU | -0.7396 | 7.46E-11 |
| B15 | B145 | CD3-CD19+(B cell) | BGP | -0.70957 | 9.23E-10 |
| B15 | B97 | CD3-CD19+(B cell) | dsDNA | -0.66564 | 2.18E-08 |
| B15 | B135 | CD3-CD19+(B cell) | TBA | -0.65372 | 4.69E-08 |
| B15 | B176 | CD3-CD19+(B cell) | Ucr | -0.61538 | 4.49E-07 |
| B150 | B155 | ALP | IgA | 0.912871 | 0 |
| B150 | B159 | ALP | β2-MG | 0.883541 | 0 |
| B150 | B197 | ALP | AG | 0.858116 | 0 |
| B150 | B193 | ALP | K | 0.845154 | 2.22E-16 |
| B150 | B158 | ALP | IgM | 0.774597 | 2.48E-12 |
| B150 | B154 | ALP | RF | 0.632456 | 1.71E-07 |
| B150 | B199 | ALP | Mg | 0.613572 | 4.95E-07 |
| B151 | B185 | Anti-CCP | PT | -0.85201 | 0 |
| B151 | B225 | Anti-CCP | hPBG | -0.85201 | 0 |
| B151 | B167 | Anti-CCP | PC | -0.82808 | 3.55E-15 |
| B151 | B168 | Anti-CCP | NMP22 | -0.73786 | 8.71E-11 |
| B151 | B179 | Anti-CCP | α1-MG | -0.68853 | 4.49E-09 |
| B151 | B183 | Anti-CCP | PTA | -0.67612 | 1.08E-08 |
| B151 | B176 | Anti-CCP | Ucr | -0.6445 | 8.31E-08 |
| B152 | B168 | ACA-Ig(G-A-M) | NMP22 | -0.89113 | 0 |
| B152 | B182 | ACA-Ig(G-A-M) | TT | -0.86772 | 0 |
| B152 | B186 | ACA-Ig(G-A-M) | PFC | -0.68599 | 5.39E-09 |
| B152 | B176 | ACA-Ig(G-A-M) | Ucr | -0.60541 | 7.69E-07 |
| B154 | B175 | RF | ALB-U | -0.94868 | 0 |

| | | | | | |
|---|---|---|---|---|---|
| B154 | B195 | RF | Na | 0.801784 | 1.13E-13 |
| B154 | B199 | RF | Mg | -0.72761 | 2.12E-10 |
| B154 | B197 | RF | AG | -0.60302 | 8.73E-07 |
| B155 | B164 | IgA | COM | 0.842701 | 4.44E-16 |
| B156 | B194 | IgE | Cl | 0.782624 | 1.04E-12 |
| B156 | B199 | IgE | Mg | 0.766965 | 5.49E-12 |
| B157 | B197 | IgG | AG | 0.639602 | 1.12E-07 |
| B157 | B184 | IgG | INR | 0.618718 | 3.73E-07 |
| B158 | B164 | IgM | COM | 0.936382 | 0 |
| B158 | B187 | IgM | NSE | 0.905822 | 0 |
| B158 | B195 | IgM | Na | 0.872872 | 0 |
| B158 | B193 | IgM | K | 0.763763 | 7.58E-12 |
| B159 | B199 | β2-MG | Mg | 0.985666 | 0 |
| B159 | B164 | β2-MG | COM | 0.847403 | 2.22E-16 |
| B159 | B178 | β2-MG | Ig-U | 0.696574 | 2.49E-09 |
| B159 | B197 | β2-MG | AG | 0.650964 | 5.58E-08 |
| B159 | B175 | β2-MG | ALB-U | 0.602414 | 9.01E-07 |
| B16 | B110 | CD3+T | AHA | 0.948504 | 0 |
| B16 | B195 | CD3+T | Na | 0.909137 | 0 |
| B16 | B38 | CD3+T | BAND% | 0.92582 | 0 |
| B16 | B81 | CD3+T | TSGF | 0.981981 | 0 |
| B16 | B92 | CD3+T | ALDH2-Glu504Glu | 0.968665 | 0 |
| B16 | B95 | CD3+T | anti-cenpb | 0.910465 | 0 |
| B16 | B96 | CD3+T | dsDNA (IIF) | 0.845154 | 2.22E-16 |
| B16 | B142 | CD3+T | 25-(OH)D | 0.815374 | 2.00E-14 |
| B16 | B65 | CD3+T | WBRSR(mPas_1) | 0.801784 | 1.13E-13 |
| B16 | B175 | CD3+T | ALB-U | 0.776899 | 1.94E-12 |
| B16 | B57 | CD3+T | RBCAI | 0.771517 | 3.43E-12 |
| B16 | B164 | CD3+T | COM | 0.748705 | 3.25E-11 |
| B16 | B139 | CD3+T | ADA | 0.725241 | 2.59E-10 |
| B16 | B159 | CD3+T | β2-MG | 0.672022 | 1.42E-08 |
| B16 | B108 | CD3+T | Anti-Cyclin | 0.659829 | 3.18E-08 |
| B16 | B22 | CD3+T | CD3+CD8+(T) | 0.626783 | 2.37E-07 |
| B16 | B54 | CD3+T | ESR | 0.617213 | 4.06E-07 |
| B163 | B166 | SG | Pro | 0.612414 | 5.28E-07 |
| B164 | B199 | COM | Mg | 0.859727 | 0 |
| B164 | B184 | COM | INR | 0.834058 | 1.33E-15 |
| B164 | B194 | COM | Cl | 0.73721 | 9.23E-11 |
| B165 | B170 | bilirubin | U-GLU | 1 | 0 |
| B167 | B202 | PC | e-times | -0.95618 | 0 |
| B167 | B225 | PC | hPBG | -0.88192 | 0 |
| B167 | B223 | PC | FH | -0.84266 | 4.44E-16 |
| B167 | B171 | PC | EC | -0.7127 | 7.21E-10 |
| B167 | B177 | PC | ACR | -0.65465 | 4.42E-08 |

| | | | | | |
|---|---|---|---|---|---|
| B169 | B183 | U-CRY | PTA | -0.87287 | 0 |
| B169 | B202 | U-CRY | e-times | -0.89443 | 0 |
| B169 | B223 | U-CRY | FH | -0.97014 | 0 |
| B17 | B143 | CD3+CD16+CD56+NK | β-CTx | 0.96225 | 0 |
| B17 | B95 | CD3+CD16+CD56+NK | anti-cenpb | 0.917663 | 0 |
| B17 | B96 | CD3+CD16+CD56+NK | dsDNA (IIF) | 0.968963 | 0 |
| B17 | B134 | CD3+CD16+CD56+NK | CHE | 0.833333 | 1.78E-15 |
| B17 | B38 | CD3+CD16+CD56+NK | BAND% | 0.816497 | 1.73E-14 |
| B17 | B199 | CD3+CD16+CD56+NK | Mg | 0.808452 | 4.91E-14 |
| B17 | B164 | CD3+CD16+CD56+NK | COM | 0.764553 | 7.00E-12 |
| B17 | B184 | CD3+CD16+CD56+NK | INR | 0.75 | 2.88E-11 |
| B17 | B22 | CD3+CD16+CD56+NK | CD3+CD8+(T) | 0.703526 | 1.48E-09 |
| B17 | B139 | CD3+CD16+CD56+NK | ADA | 0.639602 | 1.12E-07 |
| B171 | B176 | EC | Ucr | -0.9245 | 0 |
| B171 | B179 | EC | α1-MG | -0.88889 | 0 |
| B171 | B185 | EC | PT | -0.94281 | 0 |
| B171 | B198 | EC | P | -0.88454 | 0 |
| B171 | B200 | EC | OP | -0.71067 | 8.47E-10 |
| B174 | B191 | Urea/Cr | Urea | 0.665553 | 2.19E-08 |
| B175 | B199 | ALB-U | Mg | 0.997054 | 0 |
| B175 | B193 | ALB-U | K | 0.760639 | 1.03E-11 |
| B175 | B180 | ALB-U | Tf | 0.67082 | 1.54E-08 |
| B175 | B187 | ALB-U | NSE | 0.613941 | 4.86E-07 |
| B176 | B183 | Ucr | PTA | -0.84732 | 2.22E-16 |
| B176 | B179 | Ucr | α1-MG | -0.64715 | 7.07E-08 |
| B176 | B223 | Ucr | FH | -0.60541 | 7.69E-07 |
| B177 | B198 | ACR | P | -0.66667 | 2.03E-08 |
| B178 | B199 | Ig-U | Mg | 0.910182 | 0 |
| B178 | B194 | Ig-U | Cl | 0.714435 | 6.28E-10 |
| B178 | B195 | Ig-U | Na | 0.694365 | 2.94E-09 |
| B178 | B193 | Ig-U | K | 0.617213 | 4.06E-07 |
| B179 | B225 | α1-MG | hPBG | -0.86424 | 0 |
| B179 | B183 | α1-MG | PTA | -0.80013 | 1.38E-13 |
| B18 | B109 | CD3+CD4+helper(inducer)T | AMA-M2 | -0.94281 | 0 |
| B18 | B150 | CD3+CD4+helper(inducer)T | ALP | 0.894427 | 0 |
| B18 | B54 | CD3+CD4+helper(inducer)T | ESR | 0.866025 | 0 |
| B18 | B57 | CD3+CD4+helper(inducer)T | RBCAI | -0.86603 | 0 |
| B18 | B142 | CD3+CD4+helper(inducer)T | 25-(OH)D | -0.83205 | 2.00E-15 |
| B18 | B195 | CD3+CD4+helper(inducer)T | Na | -0.75593 | 1.64E-11 |
| B18 | B134 | CD3+CD4+helper(inducer)T | CHE | -0.70711 | 1.12E-09 |
| B18 | B139 | CD3+CD4+helper(inducer)T | ADA | 0.603023 | 8.73E-07 |
| B182 | B183 | TT | PTA | -0.9759 | 0 |
| B183 | B223 | PTA | FH | -0.63511 | 1.46E-07 |
| B183 | B225 | PTA | hPBG | -0.61721 | 4.06E-07 |

| | | | | | |
|---|---|---|---|---|---|
| B183 | B200 | PTA | OP | -0.60481 | 7.94E-07 |
| B184 | B193 | INR | K | 0.868599 | 0 |
| B184 | B194 | INR | Cl | 0.883883 | 0 |
| B184 | B187 | INR | NSE | 0.83205 | 2.00E-15 |
| B184 | B195 | INR | Na | 0.801784 | 1.13E-13 |
| B185 | B225 | PT | hPBG | -0.88889 | 0 |
| B185 | B198 | PT | P | -0.7698 | 4.10E-12 |
| B186 | B200 | PFC | OP | -0.6784 | 9.19E-09 |
| B186 | B198 | PFC | P | -0.64952 | 6.10E-08 |
| B187 | B195 | NSE | Na | 0.889499 | 0 |
| B187 | B199 | NSE | Mg | 0.672673 | 1.36E-08 |
| B189 | B190 | eGFB | Cre | -0.80035 | 1.34E-13 |
| B189 | B192 | eGFB | UA | -0.68986 | 4.08E-09 |
| B19 | B168 | CD3+CD4+/CD3+CD8+ | NMP22 | -0.86164 | 0 |
| B19 | B169 | CD3+CD4+/CD3+CD8+ | U-CRY | -0.90453 | 0 |
| B19 | B176 | CD3+CD4+/CD3+CD8+ | Ucr | -0.91987 | 0 |
| B19 | B182 | CD3+CD4+/CD3+CD8+ | TT | -0.94388 | 0 |
| B19 | B28 | CD3+CD4+/CD3+CD8+ | PRL | -0.8528 | 0 |
| B19 | B56 | CD3+CD4+/CD3+CD8+ | IR | -0.85812 | 0 |
| B19 | B75 | CD3+CD4+/CD3+CD8+ | ApoA1/B | -0.89995 | 0 |
| B19 | B34 | CD3+CD4+/CD3+CD8+ | TK | -0.83984 | 4.44E-16 |
| B19 | B100 | CD3+CD4+/CD3+CD8+ | anti-Ro52 | -0.80904 | 4.55E-14 |
| B19 | B102 | CD3+CD4+/CD3+CD8+ | anti-Sm | -0.78393 | 9.04E-13 |
| B19 | B106 | CD3+CD4+/CD3+CD8+ | anti-P | -0.76871 | 4.59E-12 |
| B19 | B152 | CD3+CD4+/CD3+CD8+ | ACA-Ig(G-A-M) | -0.73127 | 1.55E-10 |
| B19 | B223 | CD3+CD4+/CD3+CD8+ | FH | -0.73127 | 1.55E-10 |
| B19 | B39 | CD3+CD4+/CD3+CD8+ | Other cells | -0.72392 | 2.89E-10 |
| B19 | B37 | CD3+CD4+/CD3+CD8+ | BAND | -0.71563 | 5.71E-10 |
| B19 | B225 | CD3+CD4+/CD3+CD8+ | hPBG | -0.71067 | 8.47E-10 |
| B190 | B191 | Cre | Urea | 0.63782 | 1.24E-07 |
| B195 | B196 | Na | TCO2 | 0.771517 | 3.43E-12 |
| B20 | B100 | CD3+CD4+CD8+ | anti-Ro52 | -0.9 | 0 |
| B20 | B106 | CD3+CD4+CD8+ | anti-P | -0.96476 | 0 |
| B20 | B145 | CD3+CD4+CD8+ | BGP | -0.85812 | 0 |
| B20 | B152 | CD3+CD4+CD8+ | ACA-Ig(G-A-M) | -0.86772 | 0 |
| B20 | B34 | CD3+CD4+CD8+ | TK | -0.99655 | 0 |
| B20 | B61 | CD3+CD4+CD8+ | WBHSRV | -0.99228 | 0 |
| B20 | B99 | CD3+CD4+CD8+ | anti-PM-Scl | -0.84515 | 2.22E-16 |
| B20 | B135 | CD3+CD4+CD8+ | TBA | -0.84327 | 4.44E-16 |
| B20 | B93 | CD3+CD4+CD8+ | ALDH2-Glu504Lys | -0.83666 | 8.88E-16 |
| B20 | B94 | CD3+CD4+CD8+ | ALDH2-Lys504Lys | -0.8 | 1.40E-13 |
| B20 | B63 | CD3+CD4+CD8+ | WBRSR(mPas_200) | -0.74536 | 4.43E-11 |
| B20 | B225 | CD3+CD4+CD8+ | hPBG | -0.73786 | 8.71E-11 |
| B20 | B97 | CD3+CD4+CD8+ | dsDNA | -0.71554 | 5.75E-10 |

| | | | | | |
|---|---|---|---|---|---|
| B20 | B223 | CD3+CD4+CD8+ | FH | -0.70502 | 1.32E-09 |
| B20 | B198 | CD3+CD4+CD8+ | P | -0.63901 | 1.16E-07 |
| B202 | B225 | e-times | hPBG | -0.94868 | 0 |
| B206 | B207 | Waist | WHR | 0.909226 | 0 |
| B206 | B210 | Waist | Hip | 0.74943 | 3.04E-11 |
| B206 | B222 | Waist | NS | 0.651473 | 5.40E-08 |
| B207 | B222 | WHR | NS | 0.712419 | 7.38E-10 |
| B21 | B121 | CD3+CD4-CD8- | C3 | -0.94491 | 0 |
| B21 | B122 | CD3+CD4-CD8- | C4 | -0.95618 | 0 |
| B21 | B225 | CD3+CD4-CD8- | hPBG | -0.88192 | 0 |
| B21 | B93 | CD3+CD4-CD8- | ALDH2-Glu504Lys | -1 | 0 |
| B21 | B97 | CD3+CD4-CD8- | dsDNA | -0.85524 | 0 |
| B21 | B100 | CD3+CD4-CD8- | anti-Ro52 | -0.83666 | 8.88E-16 |
| B21 | B171 | CD3+CD4-CD8- | EC | -0.80178 | 1.13E-13 |
| B21 | B179 | CD3+CD4-CD8- | α1-MG | -0.80178 | 1.13E-13 |
| B21 | B64 | CD3+CD4-CD8- | WBRSR(mPas_5) | -0.80178 | 1.13E-13 |
| B21 | B34 | CD3+CD4-CD8- | TK | -0.79407 | 2.84E-13 |
| B21 | B30 | CD3+CD4-CD8- | LH | -0.76064 | 1.03E-11 |
| B21 | B176 | CD3+CD4-CD8- | Ucr | -0.74125 | 6.44E-11 |
| B21 | B60 | CD3+CD4-CD8- | WBRVI | -0.73577 | 1.05E-10 |
| B21 | B223 | CD3+CD4-CD8- | FH | -0.71302 | 7.03E-10 |
| B21 | B124 | CD3+CD4-CD8- | ACTH | -0.69481 | 2.84E-09 |
| B21 | B102 | CD3+CD4-CD8- | anti-Sm | -0.64143 | 1.00E-07 |
| B211 | B212 | IOP-R | IOP-L | 0.992899 | 0 |
| B211 | B224 | IOP-R | pmh | -0.8618 | 0 |
| B212 | B224 | IOP-L | pmh | -0.79534 | 2.45E-13 |
| B213 | B214 | CVA-R | CVA-L | 0.933803 | 0 |
| B215 | B216 | BMI | BMI | 0.690658 | 3.85E-09 |
| B22 | B116 | CD3+CD8+(T) | TT3 | 0.94388 | 0 |
| B22 | B178 | CD3+CD8+(T) | Ig-U | 0.870388 | 0 |
| B22 | B54 | CD3+CD8+(T) | ESR | 0.923186 | 0 |
| B22 | B76 | CD3+CD8+(T) | ApoB | 0.934199 | 0 |
| B22 | B92 | CD3+CD8+(T) | ALDH2-Glu504Glu | 1 | 0 |
| B22 | B197 | CD3+CD8+(T) | AG | 0.727273 | 2.18E-10 |
| B22 | B184 | CD3+CD8+(T) | INR | 0.678401 | 9.19E-09 |
| B22 | B67 | CD3+CD8+(T) | WBRSR(mPas_50) | 0.639602 | 1.12E-07 |
| B223 | B225 | FH | hPBG | -0.62883 | 2.10E-07 |
| B227 | B228 | OD | OS | 0.859097 | 0 |
| B23 | B107 | CD3-CD16+CD56+(NK) | AnuA | 0.98387 | 0 |
| B23 | B117 | CD3-CD16+CD56+(NK) | TT4 | 1 | 0 |
| B23 | B143 | CD3-CD16+CD56+(NK) | β-CTx | 0.92376 | 0 |
| B23 | B156 | CD3-CD16+CD56+(NK) | IgE | 0.948683 | 0 |
| B23 | B157 | CD3-CD16+CD56+(NK) | IgG | 0.919239 | 0 |
| B23 | B193 | CD3-CD16+CD56+(NK) | K | 0.908688 | 0 |

| | | | | | |
|---|---|---|---|---|---|
| B23 | B194 | CD3-CD16+CD56+(NK) | Cl | 0.989949 | 0 |
| B23 | B54 | CD3-CD16+CD56+(NK) | ESR | 0.979796 | 0 |
| B23 | B65 | CD3-CD16+CD56+(NK) | WBRSR(mPas_1) | 0.989949 | 0 |
| B23 | B96 | CD3-CD16+CD56+(NK) | dsDNA (IIF) | 0.894427 | 0 |
| B23 | B187 | CD3-CD16+CD56+(NK) | NSE | 0.83205 | 2.00E-15 |
| B23 | B195 | CD3-CD16+CD56+(NK) | Na | 0.748331 | 3.36E-11 |
| B23 | B74 | CD3-CD16+CD56+(NK) | ApoA1 | 0.715542 | 5.75E-10 |
| B23 | B159 | CD3-CD16+CD56+(NK) | β2-MG | 0.711201 | 8.12E-10 |
| B23 | B79 | CD3-CD16+CD56+(NK) | hb V -dna | 0.7 | 1.93E-09 |
| B23 | B92 | CD3-CD16+CD56+(NK) | ALDH2-Glu504Glu | 0.663325 | 2.53E-08 |
| B23 | B67 | CD3-CD16+CD56+(NK) | WBRSR(mPas_50) | 0.659966 | 3.15E-08 |
| B23 | B55 | CD3-CD16+CD56+(NK) | TK | 0.604743 | 7.97E-07 |
| B23 | B98 | CD3-CD16+CD56+(NK) | anti-Jo-1 | 0.604743 | 7.97E-07 |
| B24 | B27 | αHBDH | LDH | 0.982027 | 0 |
| B24 | B26 | αHBDH | CKMB | 0.690276 | 3.96E-09 |
| B25 | B26 | CK | CKMB | 0.980308 | 0 |
| B25 | B27 | CK | LDH | 0.68381 | 6.29E-09 |
| B25 | B78 | CK | HBcAb | -0.61084 | 5.75E-07 |
| B26 | B27 | CKMB | LDH | 0.814433 | 2.26E-14 |
| B28 | B225 | PRL | hPBG | -1 | 0 |
| B28 | B34 | PRL | TK | -0.87538 | 0 |
| B28 | B33 | PRL | Prog | -0.75593 | 1.64E-11 |
| B28 | B97 | PRL | dsDNA | -0.75425 | 1.92E-11 |
| B28 | B29 | PRL | E2 | 0.673575 | 1.28E-08 |
| B28 | B121 | PRL | C3 | -0.66667 | 2.03E-08 |
| B28 | B102 | PRL | anti-Sm | -0.65997 | 3.15E-08 |
| B28 | B60 | PRL | WBRVI | -0.64889 | 6.35E-08 |
| B28 | B106 | PRL | anti-P | -0.64715 | 7.07E-08 |
| B28 | B151 | PRL | Anti-CCP | -0.60858 | 6.49E-07 |
| B29 | B135 | E2 | TBA | -0.86603 | 0 |
| B29 | B171 | E2 | EC | -0.95258 | 0 |
| B29 | B36 | E2 | V-Lym% | -0.92717 | 0 |
| B29 | B75 | E2 | ApoA1/B | -0.87039 | 0 |
| B29 | B64 | E2 | WBRSR(mPas_5) | 0.816497 | 1.73E-14 |
| B29 | B183 | E2 | PTA | -0.80178 | 1.13E-13 |
| B29 | B200 | E2 | OP | -0.78335 | 9.64E-13 |
| B29 | B125 | E2 | Cor | -0.7746 | 2.48E-12 |
| B29 | B56 | E2 | IR | -0.7746 | 2.48E-12 |
| B29 | B151 | E2 | Anti-CCP | -0.73786 | 8.71E-11 |
| B29 | B225 | E2 | hPBG | -0.67358 | 1.28E-08 |
| B29 | B168 | E2 | NMP22 | -0.66667 | 2.03E-08 |
| B29 | B59 | E2 | WBRV | -0.63901 | 1.16E-07 |
| B3 | B25 | Hcy | CK | 0.983797 | 0 |
| B3 | B26 | Hcy | CKMB | 0.929019 | 0 |

| | | | | | |
|---|---|---|---|---|---|
| B3 | B123 | Hcy | hs-CRP1 | -0.68996 | 4.05E-09 |
| B3 | B78 | Hcy | HBcAb | -0.61317 | 5.06E-07 |
| B30 | B179 | LH | α1-MG | -0.94868 | 0 |
| B30 | B202 | LH | e-times | -0.98995 | 0 |
| B30 | B37 | LH | BAND | -0.86603 | 0 |
| B30 | B59 | LH | WBRV | -0.98995 | 0 |
| B30 | B62 | LH | WBHSRVI | -0.9325 | 0 |
| B30 | B63 | LH | WBRSR(mPas_200) | -0.94868 | 0 |
| B30 | B64 | LH | WBRSR(mPas_5) | 0.948683 | 0 |
| B30 | B36 | LH | V-Lym% | -0.82078 | 9.77E-15 |
| B30 | B185 | LH | PT | -0.81989 | 1.09E-14 |
| B30 | B183 | LH | PTA | -0.75907 | 1.21E-11 |
| B30 | B97 | LH | dsDNA | -0.75895 | 1.22E-11 |
| B30 | B225 | LH | hPBG | -0.74536 | 4.43E-11 |
| B30 | B198 | LH | P | -0.6455 | 7.82E-08 |
| B30 | B102 | LH | anti-Sm | -0.63246 | 1.71E-07 |
| B31 | B124 | FSH | ACTH | -0.85769 | 0 |
| B31 | B125 | FSH | Cor | -0.91287 | 0 |
| B31 | B168 | FSH | NMP22 | -0.94281 | 0 |
| B31 | B185 | FSH | PT | -0.95258 | 0 |
| B31 | B62 | FSH | WBHSRVI | -0.85126 | 0 |
| B31 | B176 | FSH | Ucr | -0.80064 | 1.30E-13 |
| B31 | B60 | FSH | WBRVI | -0.79472 | 2.63E-13 |
| B31 | B33 | FSH | Prog | -0.77152 | 3.43E-12 |
| B31 | B152 | FSH | ACA-Ig(G-A-M) | -0.70014 | 1.91E-09 |
| B31 | B135 | FSH | TBA | -0.68041 | 7.99E-09 |
| B31 | B99 | FSH | anti-PM-Scl | -0.65465 | 4.42E-08 |
| B32 | B124 | Testo | ACTH | -0.99258 | 0 |
| B32 | B152 | Testo | ACA-Ig(G-A-M) | -0.90749 | 0 |
| B32 | B168 | Testo | NMP22 | -0.98198 | 0 |
| B32 | B177 | Testo | ACR | -0.98198 | 0 |
| B32 | B183 | Testo | PTA | -0.99146 | 0 |
| B32 | B61 | Testo | WBHSRV | -0.8895 | 0 |
| B32 | B167 | Testo | PC | -0.78571 | 7.41E-13 |
| B32 | B223 | Testo | FH | -0.77784 | 1.76E-12 |
| B32 | B135 | Testo | TBA | -0.75593 | 1.64E-11 |
| B32 | B58 | Testo | HCT | -0.74125 | 6.44E-11 |
| B32 | B59 | Testo | WBRV | -0.71714 | 5.05E-10 |
| B32 | B33 | Testo | Prog | -0.71429 | 6.35E-10 |
| B32 | B34 | Testo | TK | -0.69481 | 2.84E-09 |
| B32 | B60 | Testo | WBRVI | -0.67445 | 1.21E-08 |
| B32 | B121 | Testo | C3 | -0.66144 | 2.86E-08 |
| B32 | B62 | Testo | WBHSRVI | -0.63049 | 1.91E-07 |
| B32 | B75 | Testo | ApoA1/B | -0.62678 | 2.37E-07 |

| | | | | | |
|---|---|---|---|---|---|
| B32 | B171 | Testo | EC | -0.62361 | 2.84E-07 |
| B32 | B179 | Testo | α1-MG | -0.62361 | 2.84E-07 |
| B33 | B102 | Prog | anti-Sm | -0.85524 | 0 |
| B33 | B121 | Prog | C3 | -0.94491 | 0 |
| B33 | B125 | Prog | Cor | -0.92967 | 0 |
| B33 | B136 | Prog | 5'NT | -0.87287 | 0 |
| B33 | B182 | Prog | TT | -0.95618 | 0 |
| B33 | B75 | Prog | ApoA1/B | -0.91168 | 0 |
| B33 | B93 | Prog | ALDH2-Glu504Lys | -1 | 0 |
| B33 | B106 | Prog | anti-P | -0.83863 | 8.88E-16 |
| B33 | B59 | Prog | WBRV | -0.83666 | 8.88E-16 |
| B33 | B64 | Prog | WBRSR(mPas_5) | 0.801784 | 1.13E-13 |
| B33 | B176 | Prog | Ucr | -0.74125 | 6.44E-11 |
| B33 | B39 | Prog | Other cells | -0.73336 | 1.29E-10 |
| B33 | B100 | Prog | anti-Ro52 | -0.71714 | 5.05E-10 |
| B33 | B186 | Prog | PFC | -0.66144 | 2.86E-08 |
| B33 | B167 | Prog | PC | -0.64286 | 9.19E-08 |
| B33 | B60 | Prog | WBRVI | -0.61314 | 5.07E-07 |
| B34 | B106 | TK | anti-P | -0.98328 | 0 |
| B34 | B179 | TK | α1-MG | -0.92848 | 0 |
| B34 | B183 | TK | PTA | -0.89148 | 0 |
| B34 | B198 | TK | P | -0.94762 | 0 |
| B34 | B60 | TK | WBRVI | -0.93723 | 0 |
| B34 | B61 | TK | WBHSRV | -0.92705 | 0 |
| B34 | B93 | TK | ALDH2-Glu504Lys | -0.99258 | 0 |
| B34 | B97 | TK | dsDNA | -0.8542 | 0 |
| B34 | B200 | TK | OP | -0.8314 | 2.22E-15 |
| B34 | B75 | TK | ApoA1/B | -0.8314 | 2.22E-15 |
| B34 | B136 | TK | 5'NT | -0.7581 | 1.33E-11 |
| B34 | B167 | TK | PC | -0.74444 | 4.82E-11 |
| B34 | B177 | TK | ACR | -0.68229 | 7.01E-09 |
| B34 | B152 | TK | ACA-Ig(G-A-M) | -0.67557 | 1.12E-08 |
| B34 | B223 | TK | FH | -0.67557 | 1.12E-08 |
| B34 | B36 | TK | V-Lym% | -0.66272 | 2.63E-08 |
| B34 | B137 | TK | AFU | -0.61898 | 3.68E-07 |
| B34 | B135 | TK | TBA | -0.61276 | 5.18E-07 |
| B35 | B157 | β2-MG | IgG | 0.894427 | 0 |
| B35 | B159 | β2-MG | β2-MG | 0.883541 | 0 |
| B35 | B178 | β2-MG | Ig-U | 0.912871 | 0 |
| B35 | B67 | β2-MG | WBRSR(mPas_50) | 0.894427 | 0 |
| B35 | B98 | β2-MG | anti-Jo-1 | 0.956183 | 0 |
| B35 | B164 | β2-MG | COM | 0.791257 | 3.94E-13 |
| B35 | B134 | β2-MG | CHE | 0.790569 | 4.27E-13 |
| B35 | B158 | β2-MG | IgM | 0.774597 | 2.48E-12 |

| | | | | | |
|---|---|---|---|---|---|
| B35 | B199 | β2-MG | Mg | 0.766965 | 5.49E-12 |
| B35 | B142 | β2-MG | 25-(OH)D | 0.744208 | 4.92E-11 |
| B35 | B74 | β2-MG | ApoA1 | 0.707107 | 1.12E-09 |
| B35 | B175 | β2-MG | ALB-U | 0.7 | 1.93E-09 |
| B35 | B92 | β2-MG | ALDH2-Glu504Glu | 0.667424 | 1.93E-08 |
| B36 | B186 | V-Lym% | PFC | -0.86031 | 0 |
| B36 | B223 | V-Lym% | FH | -0.86558 | 0 |
| B36 | B39 | V-Lym% | Other cells | -0.94591 | 0 |
| B36 | B60 | V-Lym% | WBRVI | -0.89319 | 0 |
| B36 | B105 | V-Lym% | anti-U1RNP | -0.81111 | 3.49E-14 |
| B36 | B182 | V-Lym% | TT | -0.79802 | 1.78E-13 |
| B36 | B99 | V-Lym% | anti-PM-Scl | -0.73577 | 1.05E-10 |
| B36 | B59 | V-Lym% | WBRV | -0.72548 | 2.54E-10 |
| B36 | B102 | V-Lym% | anti-Sm | -0.71377 | 6.62E-10 |
| B36 | B152 | V-Lym% | ACA-Ig(G-A-M) | -0.7082 | 1.03E-09 |
| B36 | B200 | V-Lym% | OP | -0.69171 | 3.57E-09 |
| B36 | B151 | V-Lym% | Anti-CCP | -0.67017 | 1.61E-08 |
| B36 | B97 | V-Lym% | dsDNA | -0.64889 | 6.35E-08 |
| B36 | B145 | V-Lym% | BGP | -0.62254 | 3.01E-07 |
| B36 | B56 | V-Lym% | IR | -0.61559 | 4.44E-07 |
| B37 | B122 | BAND | C4 | -0.9798 | 0 |
| B37 | B124 | BAND | ACTH | -0.91539 | 0 |
| B37 | B145 | BAND | BGP | -0.97312 | 0 |
| B37 | B168 | BAND | NMP22 | -0.89443 | 0 |
| B37 | B182 | BAND | TT | -0.9798 | 0 |
| B37 | B58 | BAND | HCT | -0.86083 | 0 |
| B37 | B59 | BAND | WBRV | -0.89815 | 0 |
| B37 | B60 | BAND | WBRVI | -0.87959 | 0 |
| B37 | B93 | BAND | ALDH2-Glu504Lys | -0.87831 | 0 |
| B37 | B94 | BAND | ALDH2-Lys504Lys | -0.9798 | 0 |
| B37 | B202 | BAND | e-times | -0.8165 | 1.73E-14 |
| B37 | B75 | BAND | ApoA1/B | -0.7785 | 1.64E-12 |
| B37 | B66 | BAND | WBRSR(mPas_100) | -0.7746 | 2.48E-12 |
| B37 | B223 | BAND | FH | -0.75277 | 2.22E-11 |
| B37 | B102 | BAND | anti-Sm | -0.7303 | 1.69E-10 |
| B37 | B61 | BAND | WBHSRV | -0.70892 | 9.72E-10 |
| B37 | B125 | BAND | Cor | -0.69282 | 3.29E-09 |
| B37 | B39 | BAND | Other cells | -0.68885 | 4.39E-09 |
| B37 | B177 | BAND | ACR | -0.67082 | 1.54E-08 |
| B37 | B121 | BAND | C3 | -0.6455 | 7.82E-08 |
| B37 | B97 | BAND | dsDNA | -0.62075 | 3.33E-07 |
| B38 | B142 | BAND% | 25-(OH)D | 0.960769 | 0 |
| B38 | B81 | BAND% | TSGF | -0.94281 | 0 |
| B38 | B95 | BAND% | anti-cenpb | 0.936586 | 0 |

| | | | | | |
|---|---|---|---|---|---|
| B38 | B157 | BAND% | IgG | 0.721688 | 3.48E-10 |
| B38 | B155 | BAND% | IgA | -0.70711 | 1.12E-09 |
| B38 | B139 | BAND% | ADA | 0.696311 | 2.54E-09 |
| B38 | B158 | BAND% | IgM | 0.666667 | 2.03E-08 |
| B38 | B138 | BAND% | PAB | -0.65465 | 4.42E-08 |
| B38 | B156 | BAND% | IgE | -0.6455 | 7.82E-08 |
| B38 | B197 | BAND% | AG | 0.615457 | 4.47E-07 |
| B38 | B134 | BAND% | CHE | 0.612372 | 5.29E-07 |
| B38 | B184 | BAND% | INR | -0.61237 | 5.29E-07 |
| B39 | B125 | Other cells | Cor | -0.97619 | 0 |
| B39 | B168 | Other cells | NMP22 | -0.91018 | 0 |
| B39 | B56 | Other cells | IR | -0.97619 | 0 |
| B39 | B58 | Other cells | HCT | -0.99887 | 0 |
| B39 | B94 | Other cells | ALDH2-Lys504Lys | -0.99705 | 0 |
| B39 | B223 | Other cells | FH | -0.7487 | 3.25E-11 |
| B39 | B93 | Other cells | ALDH2-Glu504Lys | -0.73336 | 1.29E-10 |
| B39 | B145 | Other cells | BGP | -0.73127 | 1.55E-10 |
| B39 | B64 | Other cells | WBRSR(mPas_5) | -0.68599 | 5.39E-09 |
| B39 | B171 | Other cells | EC | -0.62883 | 2.10E-07 |
| B39 | B179 | Other cells | α1-MG | -0.62883 | 2.10E-07 |
| B39 | B75 | Other cells | ApoA1/B | -0.62158 | 3.18E-07 |
| B39 | B186 | Other cells | PFC | -0.60634 | 7.32E-07 |
| B4 | B121 | WB-Ca | C3 | -0.89767 | 0 |
| B4 | B13 | WB-Ca | PGI_PGII | -0.84867 | 0 |
| B4 | B145 | WB-Ca | BGP | -0.91168 | 0 |
| B4 | B29 | WB-Ca | E2 | -0.98198 | 0 |
| B4 | B31 | WB-Ca | FSH | -0.92582 | 0 |
| B4 | B33 | WB-Ca | Prog | -0.85714 | 0 |
| B4 | B125 | WB-Ca | Cor | -0.84515 | 2.22E-16 |
| B4 | B34 | WB-Ca | TK | -0.8437 | 4.44E-16 |
| B4 | B61 | WB-Ca | WBHSRV | -0.81537 | 2.00E-14 |
| B4 | B106 | WB-Ca | anti-P | -0.78621 | 7.00E-13 |
| B4 | B32 | WB-Ca | Testo | -0.78571 | 7.41E-13 |
| B4 | B63 | WB-Ca | WBRSR(mPas_200) | -0.7127 | 7.21E-10 |
| B4 | B62 | WB-Ca | WBHSRVI | -0.7093 | 9.43E-10 |
| B4 | B99 | WB-Ca | anti-PM-Scl | -0.70711 | 1.12E-09 |
| B4 | B102 | WB-Ca | anti-Sm | -0.69488 | 2.83E-09 |
| B4 | B39 | WB-Ca | Other cells | -0.68752 | 4.83E-09 |
| B4 | B200 | WB-Ca | OP | -0.68376 | 6.31E-09 |
| B4 | B75 | WB-Ca | ApoA1/B | -0.62678 | 2.37E-07 |
| B40 | B42 | WBC | MON | 0.61341 | 5.00E-07 |
| B40 | B53 | WBC | GR% | 0.601096 | 9.65E-07 |
| B41 | B42 | MON % | MON | 0.603126 | 8.68E-07 |
| B43 | B44 | MCV | MCH | 0.95261 | 0 |

| | | | | | |
|---|---|---|---|---|---|
| B43 | B47 | MCV | RDW | 0.622622 | 3.00E-07 |
| B44 | B45 | MCH | MCHC | 0.688901 | 4.37E-09 |
| B46 | B51 | RBC Count | Hb | 0.791866 | 3.67E-13 |
| B46 | B126 | RBC Count | Alb | 0.679233 | 8.67E-09 |
| B46 | B47 | RBC Count | RDW | -0.67739 | 9.85E-09 |
| B48 | B53 | LYM% | GR% | -0.97977 | 0 |
| B48 | B153 | LYM% | Glu | 0.652344 | 5.12E-08 |
| B5 | B144 | WB-Mg | Total-PINP | 0.917663 | 0 |
| B5 | B159 | WB-Mg | β2-MG | 0.987829 | 0 |
| B5 | B142 | WB-Mg | 25-(OH)D | 0.83205 | 2.00E-15 |
| B5 | B9 | WB-Mg | RH | 0.801784 | 1.13E-13 |
| B5 | B79 | WB-Mg | hb V -dna | 0.707107 | 1.12E-09 |
| B5 | B175 | WB-Mg | ALB-U | -0.67082 | 1.54E-08 |
| B51 | B126 | Hb | Alb | 0.600454 | 9.98E-07 |
| B53 | B153 | GR% | Glu | -0.68933 | 4.24E-09 |
| B54 | B104 | ESR | anti-SSB | 0.948683 | 0 |
| B54 | B107 | ESR | AnuA | 0.912871 | 0 |
| B54 | B143 | ESR | β-CTx | 0.942809 | 0 |
| B54 | B76 | ESR | ApoB | 0.948683 | 0 |
| B54 | B95 | ESR | anti-cenpb | 0.983415 | 0 |
| B54 | B96 | ESR | dsDNA (IIF) | 0.912871 | 0 |
| B54 | B199 | ESR | Mg | 0.841625 | 4.44E-16 |
| B54 | B196 | ESR | TCO2 | 0.824958 | 5.33E-15 |
| B54 | B193 | ESR | K | 0.818317 | 1.33E-14 |
| B54 | B175 | ESR | ALB-U | 0.774597 | 2.48E-12 |
| B54 | B67 | ESR | WBRSR(mPas_50) | 0.7698 | 4.10E-12 |
| B54 | B69 | ESR | PV | 0.7698 | 4.10E-12 |
| B54 | B138 | ESR | PAB | 0.763763 | 7.58E-12 |
| B54 | B142 | ESR | 25-(OH)D | 0.720577 | 3.81E-10 |
| B54 | B178 | ESR | Ig-U | 0.707107 | 1.12E-09 |
| B54 | B197 | ESR | AG | 0.677003 | 1.01E-08 |
| B54 | B108 | ESR | Anti-Cyclin | 0.668153 | 1.84E-08 |
| B54 | B195 | ESR | Na | 0.654654 | 4.42E-08 |
| B54 | B159 | ESR | β2-MG | 0.622171 | 3.08E-07 |
| B54 | B98 | ESR | anti-Jo-1 | 0.617213 | 4.06E-07 |
| B55 | B142 | TK | 25-(OH)D | 0.889499 | 0 |
| B55 | B144 | TK | Total-PINP | 0.981023 | 0 |
| B55 | B156 | TK | IgE | 0.956183 | 0 |
| B55 | B184 | TK | INR | 0.944911 | 0 |
| B55 | B194 | TK | Cl | 0.935414 | 0 |
| B55 | B98 | TK | anti-Jo-1 | 0.857143 | 0 |
| B55 | B110 | TK | AHA | 0.824786 | 5.55E-15 |
| B55 | B157 | TK | IgG | 0.801784 | 1.13E-13 |
| B55 | B154 | TK | RF | 0.755929 | 1.64E-11 |

| | | | | | |
|---|---|---|---|---|---|
| B55 | B175 | TK | ALB-U | 0.717137 | 5.05E-10 |
| B55 | B159 | TK | β2-MG | 0.672022 | 1.42E-08 |
| B55 | B139 | TK | ADA | 0.644658 | 8.23E-08 |
| B55 | B69 | TK | PV | 0.62361 | 2.84E-07 |
| B56 | B169 | IR | U-CRY | 0.948683 | 0 |
| B56 | B61 | IR | WBHSRV | -0.87706 | 0 |
| B56 | B168 | IR | NMP22 | -0.7746 | 2.48E-12 |
| B56 | B135 | IR | TBA | -0.74536 | 4.43E-11 |
| B56 | B225 | IR | hPBG | -0.74536 | 4.43E-11 |
| B56 | B223 | IR | FH | -0.69027 | 3.96E-09 |
| B56 | B62 | IR | WBHSRVI | -0.65275 | 4.99E-08 |
| B56 | B171 | IR | EC | -0.63246 | 1.71E-07 |
| B57 | B142 | RBCAI | 25-(OH)D | 0.960769 | 0 |
| B57 | B143 | RBCAI | β-CTx | 0.883883 | 0 |
| B57 | B150 | RBCAI | ALP | 0.903696 | 0 |
| B57 | B92 | RBCAI | ALDH2-Glu504Glu | 0.984732 | 0 |
| B57 | B156 | RBCAI | IgE | 0.839146 | 6.66E-16 |
| B57 | B158 | RBCAI | IgM | 0.833333 | 1.78E-15 |
| B57 | B159 | RBCAI | β2-MG | 0.829561 | 2.66E-15 |
| B57 | B81 | RBCAI | TSGF | 0.824958 | 5.33E-15 |
| B57 | B107 | RBCAI | AnuA | 0.821584 | 8.66E-15 |
| B57 | B144 | RBCAI | Total-PINP | 0.794719 | 2.63E-13 |
| B57 | B195 | RBCAI | Na | 0.763763 | 7.58E-12 |
| B57 | B157 | RBCAI | IgG | 0.721688 | 3.48E-10 |
| B57 | B79 | RBCAI | hb V -dna | 0.714435 | 6.28E-10 |
| B57 | B108 | RBCAI | Anti-Cyclin | 0.712697 | 7.21E-10 |
| B57 | B155 | RBCAI | IgA | 0.707107 | 1.12E-09 |
| B57 | B139 | RBCAI | ADA | 0.609272 | 6.26E-07 |
| B58 | B168 | HCT | NMP22 | -0.90582 | 0 |
| B58 | B177 | HCT | ACR | -0.90582 | 0 |
| B58 | B183 | HCT | PTA | -0.96836 | 0 |
| B58 | B198 | HCT | P | -0.90582 | 0 |
| B58 | B64 | HCT | WBRSR(mPas_5) | -0.97073 | 0 |
| B58 | B137 | HCT | AFU | -0.83205 | 2.00E-15 |
| B58 | B185 | HCT | PT | -0.78446 | 8.51E-13 |
| B58 | B145 | HCT | BGP | -0.76871 | 4.59E-12 |
| B58 | B93 | HCT | ALDH2-Glu504Lys | -0.74125 | 6.44E-11 |
| B58 | B135 | HCT | TBA | -0.71909 | 4.31E-10 |
| B58 | B100 | HCT | anti-Ro52 | -0.68219 | 7.05E-09 |
| B58 | B94 | HCT | ALDH2-Lys504Lys | 0.620174 | 3.44E-07 |
| B58 | B152 | HCT | ACA-Ig(G-A-M) | -0.60541 | 7.69E-07 |
| B59 | B136 | WBRV | 5'NT | -0.91287 | 0 |
| B59 | B145 | WBRV | BGP | -0.95346 | 0 |
| B59 | B198 | WBRV | P | -0.91287 | 0 |

| | | | | | |
|---|---|---|---|---|---|
| B59 | B75 | WBRV | ApoA1/B | -0.85812 | 0 |
| B59 | B167 | WBRV | PC | -0.83666 | 8.88E-16 |
| B59 | B93 | WBRV | ALDH2-Glu504Lys | -0.71714 | 5.05E-10 |
| B59 | B97 | WBRV | dsDNA | -0.71554 | 5.75E-10 |
| B59 | B61 | WBRV | WBHSRV | -0.68219 | 7.05E-09 |
| B59 | B62 | WBRV | WBHSRVI | -0.65938 | 3.27E-08 |
| B6 | B124 | WB-Fe | ACTH | -0.91915 | 0 |
| B6 | B62 | WB-Fe | WBHSRVI | -0.83406 | 1.33E-15 |
| B6 | B137 | WB-Fe | AFU | -0.82496 | 5.33E-15 |
| B6 | B151 | WB-Fe | Anti-CCP | -0.82158 | 8.66E-15 |
| B6 | B59 | WB-Fe | WBRV | -0.79057 | 4.27E-13 |
| B6 | B34 | WB-Fe | TK | -0.72219 | 3.34E-10 |
| B6 | B37 | WB-Fe | BAND | -0.71005 | 8.90E-10 |
| B6 | B106 | WB-Fe | anti-P | -0.62404 | 2.77E-07 |
| B6 | B13 | WB-Fe | PGI_PGII | -0.61237 | 5.29E-07 |
| B60 | B100 | WBRVI | anti-Ro52 | -0.87208 | 0 |
| B60 | B102 | WBRVI | anti-Sm | -0.91766 | 0 |
| B60 | B151 | WBRVI | Anti-CCP | -0.94776 | 0 |
| B60 | B171 | WBRVI | EC | -0.99413 | 0 |
| B60 | B177 | WBRVI | ACR | -0.93659 | 0 |
| B60 | B185 | WBRVI | PT | -0.97333 | 0 |
| B60 | B202 | WBRVI | e-times | -0.92338 | 0 |
| B60 | B93 | WBRVI | ALDH2-Glu504Lys | -0.8584 | 0 |
| B60 | B182 | WBRVI | TT | -0.82078 | 9.77E-15 |
| B60 | B183 | WBRVI | PTA | -0.801 | 1.24E-13 |
| B60 | B137 | WBRVI | AFU | -0.76472 | 6.89E-12 |
| B60 | B225 | WBRVI | hPBG | -0.75703 | 1.47E-11 |
| B60 | B62 | WBRVI | WBHSRVI | -0.74416 | 4.94E-11 |
| B60 | B179 | WBRVI | α1-MG | -0.68825 | 4.59E-09 |
| B60 | B200 | WBRVI | OP | -0.68476 | 5.88E-09 |
| B60 | B124 | WBRVI | ACTH | -0.68162 | 7.34E-09 |
| B60 | B152 | WBRVI | ACA-Ig(G-A-M) | -0.6677 | 1.90E-08 |
| B61 | B121 | WBHSRV | C3 | -0.98058 | 0 |
| B61 | B171 | WBHSRV | EC | -0.9245 | 0 |
| B61 | B176 | WBHSRV | Ucr | -0.92308 | 0 |
| B61 | B64 | WBHSRV | WBRSR(mPas_5) | -0.97073 | 0 |
| B61 | B66 | WBHSRV | WBRSR(mPas_100) | -0.78446 | 8.51E-13 |
| B61 | B200 | WBHSRV | OP | -0.76871 | 4.59E-12 |
| B61 | B106 | WBHSRV | anti-P | -0.7615 | 9.50E-12 |
| B61 | B179 | WBHSRV | α1-MG | -0.7396 | 7.46E-11 |
| B61 | B183 | WBHSRV | PTA | -0.72627 | 2.37E-10 |
| B61 | B105 | WBHSRV | anti-U1RNP | -0.69338 | 3.16E-09 |
| B61 | B223 | WBHSRV | FH | -0.67267 | 1.36E-08 |
| B61 | B75 | WBHSRV | ApoA1/B | -0.65044 | 5.76E-08 |

| | | | | | |
|---|---|---|---|---|---|
| B61 | B122 | WBHSRV | C4 | -0.62017 | 3.44E-07 |
| B61 | B62 | WBHSRV | WBHSRVI | -0.6134 | 5.00E-07 |
| B62 | B100 | WBHSRVI | anti-Ro52 | -0.98907 | 0 |
| B62 | B105 | WBHSRVI | anti-U1RNP | -0.88465 | 0 |
| B62 | B152 | WBHSRVI | ACA-Ig(G-A-M) | -0.92976 | 0 |
| B62 | B167 | WBHSRVI | PC | -0.86692 | 0 |
| B62 | B171 | WBHSRVI | EC | -0.88465 | 0 |
| B62 | B179 | WBHSRVI | α1-MG | -0.98295 | 0 |
| B62 | B168 | WBHSRVI | NMP22 | -0.8427 | 4.44E-16 |
| B62 | B202 | WBHSRVI | e-times | -0.79126 | 3.94E-13 |
| B62 | B106 | WBHSRVI | anti-P | -0.75181 | 2.43E-11 |
| B62 | B198 | WBHSRVI | P | -0.66212 | 2.74E-08 |
| B62 | B145 | WBHSRVI | BGP | -0.62869 | 2.12E-07 |
| B63 | B122 | WBRSR(mPas_200) | C4 | -0.89443 | 0 |
| B63 | B168 | WBRSR(mPas_200) | NMP22 | -0.95258 | 0 |
| B63 | B225 | WBRSR(mPas_200) | hPBG | -0.86424 | 0 |
| B63 | B75 | WBRSR(mPas_200) | ApoA1/B | -0.78174 | 1.15E-12 |
| B63 | B223 | WBRSR(mPas_200) | FH | -0.72761 | 2.12E-10 |
| B63 | B121 | WBRSR(mPas_200) | C3 | -0.70711 | 1.12E-09 |
| B63 | B66 | WBRSR(mPas_200) | WBRSR(mPas_100) | -0.70711 | 1.12E-09 |
| B63 | B100 | WBRSR(mPas_200) | anti-Ro52 | -0.67082 | 1.54E-08 |
| B64 | B135 | WBRSR(mPas_5) | TBA | -0.94281 | 0 |
| B64 | B145 | WBRSR(mPas_5) | BGP | -0.8528 | 0 |
| B64 | B97 | WBRSR(mPas_5) | dsDNA | 0.9 | 0 |
| B64 | B198 | WBRSR(mPas_5) | P | -0.8165 | 1.73E-14 |
| B64 | B66 | WBRSR(mPas_5) | WBRSR(mPas_100) | 0.707107 | 1.12E-09 |
| B64 | B177 | WBRSR(mPas_5) | ACR | 0.612372 | 5.29E-07 |
| B65 | B107 | WBRSR(mPas_1) | AnuA | 0.948683 | 0 |
| B65 | B108 | WBRSR(mPas_1) | Anti-Cyclin | 0.92582 | 0 |
| B65 | B184 | WBRSR(mPas_1) | INR | 0.883883 | 0 |
| B65 | B193 | WBRSR(mPas_1) | K | 0.944911 | 0 |
| B65 | B199 | WBRSR(mPas_1) | Mg | 0.857493 | 0 |
| B65 | B144 | WBRSR(mPas_1) | Total-PINP | 0.802955 | 9.77E-14 |
| B65 | B116 | WBRSR(mPas_1) | TT3 | -0.79057 | 4.27E-13 |
| B65 | B139 | WBRSR(mPas_1) | ADA | 0.753778 | 2.01E-11 |
| B65 | B157 | WBRSR(mPas_1) | IgG | -0.75 | 2.88E-11 |
| B65 | B164 | WBRSR(mPas_1) | COM | 0.73721 | 9.23E-11 |
| B65 | B95 | WBRSR(mPas_1) | anti-cenpb | 0.648886 | 6.35E-08 |
| B66 | B177 | WBRSR(mPas_100) | ACR | -0.86603 | 0 |
| B66 | B152 | WBRSR(mPas_100) | ACA-Ig(G-A-M) | -0.68599 | 5.39E-09 |
| B67 | B107 | WBRSR(mPas_50) | AnuA | 0.948683 | 0 |
| B67 | B142 | WBRSR(mPas_50) | 25-(OH)D | 0.9245 | 0 |
| B67 | B156 | WBRSR(mPas_50) | IgE | 0.968963 | 0 |
| B67 | B95 | WBRSR(mPas_50) | anti-cenpb | 0.865181 | 0 |

| | | | | | |
|---|---|---|---|---|---|
| B67 | B144 | WBRSR(mPas_50) | Total-PINP | 0.841191 | 4.44E-16 |
| B67 | B108 | WBRSR(mPas_50) | Anti-Cyclin | 0.822951 | 7.11E-15 |
| B67 | B110 | WBRSR(mPas_50) | AHA | 0.822951 | 7.11E-15 |
| B67 | B175 | WBRSR(mPas_50) | ALB-U | 0.819892 | 1.09E-14 |
| B67 | B139 | WBRSR(mPas_50) | ADA | 0.80403 | 8.55E-14 |
| B67 | B109 | WBRSR(mPas_50) | AMA-M2 | 0.785674 | 7.44E-13 |
| B67 | B184 | WBRSR(mPas_50) | INR | 0.707107 | 1.12E-09 |
| B69 | B108 | PV | Anti-Cyclin | 0.92582 | 0 |
| B69 | B143 | PV | β-CTx | 0.952579 | 0 |
| B69 | B159 | PV | β2-MG | 0.957895 | 0 |
| B69 | B76 | PV | ApoB | 0.852013 | 0 |
| B69 | B96 | PV | dsDNA (IIF) | 0.843274 | 4.44E-16 |
| B69 | B95 | PV | anti-cenpb | 0.757033 | 1.47E-11 |
| B69 | B142 | PV | 25-(OH)D | 0.7396 | 7.46E-11 |
| B69 | B110 | PV | AHA | 0.720082 | 3.97E-10 |
| B69 | B117 | PV | TT4 | -0.70711 | 1.12E-09 |
| B69 | B134 | PV | CHE | 0.707107 | 1.12E-09 |
| B69 | B184 | PV | INR | 0.707107 | 1.12E-09 |
| B69 | B164 | PV | COM | 0.688062 | 4.65E-09 |
| B69 | B196 | PV | TCO2 | 0.680414 | 7.99E-09 |
| B69 | B103 | PV | anti-SSA/Ro | 0.666667 | 2.03E-08 |
| B69 | B116 | PV | TT3 | 0.632456 | 1.71E-07 |
| B69 | B104 | PV | anti-SSB | 0.608581 | 6.49E-07 |
| B7 | B186 | WB-Cop | PFC | -0.875 | 0 |
| B7 | B19 | WB-Cop | CD3+CD4+/CD3+CD8+ | -0.8528 | 0 |
| B7 | B225 | WB-Cop | hPBG | -0.91667 | 0 |
| B7 | B62 | WB-Cop | WBHSRVI | -0.93831 | 0 |
| B7 | B223 | WB-Cop | FH | -0.77174 | 3.35E-12 |
| B7 | B93 | WB-Cop | ALDH2-Glu504Lys | -0.75593 | 1.64E-11 |
| B7 | B200 | WB-Cop | OP | -0.75378 | 2.01E-11 |
| B70 | B77 | LDL-C | TC | 0.837761 | 8.88E-16 |
| B71 | B77 | TG | TC | 0.730754 | 1.62E-10 |
| B71 | B127 | TG | A-G ratio | -0.65053 | 5.73E-08 |
| B73 | B110 | ApoA | AHA | 0.881917 | 0 |
| B73 | B138 | ApoA | PAB | 0.92582 | 0 |
| B73 | B175 | ApoA | ALB-U | 0.912871 | 0 |
| B73 | B164 | ApoA | COM | 0.842701 | 4.44E-16 |
| B73 | B143 | ApoA | β-CTx | 0.833333 | 1.78E-15 |
| B73 | B178 | ApoA | Ig-U | 0.833333 | 1.78E-15 |
| B73 | B194 | ApoA | Cl | 0.816497 | 1.73E-14 |
| B73 | B107 | ApoA | AnuA | 0.774597 | 2.48E-12 |
| B73 | B199 | ApoA | Mg | -0.70014 | 1.91E-09 |
| B73 | B96 | ApoA | dsDNA (IIF) | 0.645497 | 7.82E-08 |
| B73 | B139 | ApoA | ADA | 0.615457 | 4.47E-07 |

| | | | | | |
|---|---|---|---|---|---|
| B74 | B134 | ApoA1 | CHE | 0.894427 | 0 |
| B74 | B138 | ApoA1 | PAB | 0.956183 | 0 |
| B74 | B139 | ApoA1 | ADA | 0.858116 | 0 |
| B74 | B150 | ApoA1 | ALP | 0.989949 | 0 |
| B74 | B156 | ApoA1 | IgE | 0.989949 | 0 |
| B74 | B187 | ApoA1 | NSE | 0.992278 | 0 |
| B74 | B199 | ApoA1 | Mg | 0.867722 | 0 |
| B74 | B110 | ApoA1 | AHA | 0.78072 | 1.29E-12 |
| B74 | B178 | ApoA1 | Ig-U | 0.774597 | 2.48E-12 |
| B75 | B102 | ApoA1/B | anti-Sm | -0.89544 | 0 |
| B75 | B124 | ApoA1/B | ACTH | -0.98976 | 0 |
| B75 | B183 | ApoA1/B | PTA | -0.97701 | 0 |
| B75 | B198 | ApoA1/B | P | -0.82687 | 4.00E-15 |
| B75 | B202 | ApoA1/B | e-times | -0.76277 | 8.37E-12 |
| B75 | B125 | ApoA1/B | Cor | -0.74162 | 6.23E-11 |
| B75 | B100 | ApoA1/B | anti-Ro52 | -0.7151 | 5.95E-10 |
| B75 | B106 | ApoA1/B | anti-P | -0.71081 | 8.38E-10 |
| B76 | B199 | ApoB | Mg | 0.751469 | 2.51E-11 |
| B76 | B107 | ApoB | AnuA | 0.750555 | 2.73E-11 |
| B76 | B143 | ApoB | β-CTx | 0.745356 | 4.43E-11 |
| B76 | B156 | ApoB | IgE | 0.734847 | 1.14E-10 |
| B76 | B144 | ApoB | Total-PINP | 0.670166 | 1.61E-08 |
| B76 | B159 | ApoB | β2-MG | 0.655826 | 4.11E-08 |
| B76 | B187 | ApoB | NSE | 0.644503 | 8.31E-08 |
| B76 | B139 | ApoB | ADA | 0.60553 | 7.64E-07 |
| B77 | B127 | TC | A-G ratio | -0.62072 | 3.34E-07 |
| B79 | B104 | hbＶ-dna | anti-SSB | 0.968246 | 0 |
| B79 | B164 | hbＶ-dna | COM | 0.886186 | 0 |
| B79 | B178 | hbＶ-dna | Ig-U | 0.866025 | 0 |
| B79 | B184 | hbＶ-dna | INR | 0.875 | 0 |
| B79 | B187 | hbＶ-dna | NSE | 0.83205 | 2.00E-15 |
| B8 | B100 | WB-Zinc | anti-Ro52 | -1 | 0 |
| B8 | B125 | WB-Zinc | Cor | -0.98995 | 0 |
| B8 | B169 | WB-Zinc | U-CRY | -0.89443 | 0 |
| B8 | B183 | WB-Zinc | PTA | -0.87831 | 0 |
| B8 | B185 | WB-Zinc | PT | -0.94868 | 0 |
| B8 | B223 | WB-Zinc | FH | -0.97619 | 0 |
| B8 | B29 | WB-Zinc | E2 | -0.91287 | 0 |
| B8 | B34 | WB-Zinc | TK | -0.9135 | 0 |
| B8 | B61 | WB-Zinc | WBHSRV | -0.99228 | 0 |
| B8 | B64 | WB-Zinc | WBRSR(mPas_5) | 0.894427 | 0 |
| B8 | B225 | WB-Zinc | hPBG | -0.84327 | 4.44E-16 |
| B8 | B28 | WB-Zinc | PRL | 0.843274 | 4.44E-16 |
| B8 | B198 | WB-Zinc | P | -0.82158 | 8.66E-15 |

| | | | | | |
|---|---|---|---|---|---|
| B8 | B31 | WB-Zinc | FSH | -0.7746 | 2.48E-12 |
| B8 | B145 | WB-Zinc | BGP | -0.76277 | 8.37E-12 |
| B8 | B179 | WB-Zinc | α1-MG | -0.74536 | 4.43E-11 |
| B8 | B32 | WB-Zinc | Testo | -0.71714 | 5.05E-10 |
| B8 | B102 | WB-Zinc | anti-Sm | -0.71554 | 5.75E-10 |
| B8 | B99 | WB-Zinc | anti-PM-Scl | -0.67612 | 1.08E-08 |
| B8 | B124 | WB-Zinc | ACTH | -0.66436 | 2.37E-08 |
| B81 | B199 | TSGF | Mg | 0.840168 | 4.44E-16 |
| B81 | B150 | TSGF | ALP | 0.730297 | 1.69E-10 |
| B81 | B184 | TSGF | INR | 0.721688 | 3.48E-10 |
| B81 | B196 | TSGF | TCO2 | -0.66667 | 2.03E-08 |
| B81 | B187 | TSGF | NSE | 0.640513 | 1.06E-07 |
| B81 | B138 | TSGF | PAB | 0.617213 | 4.06E-07 |
| B81 | B139 | TSGF | ADA | 0.615457 | 4.47E-07 |
| B82 | B211 | CEA | IOP-R | 0.861797 | 0 |
| B82 | B224 | CEA | pmh | -1 | 0 |
| B82 | B83 | CEA | AFP | -1 | 0 |
| B82 | B84 | CEA | C-PSA | -1 | 0 |
| B82 | B85 | CEA | SCC | -1 | 0 |
| B82 | B86 | CEA | TBhCG | 1 | 0 |
| B82 | B87 | CEA | NSE | 1 | 0 |
| B82 | B88 | CEA | CA 242 | -1 | 0 |
| B82 | B89 | CEA | CA 72-4 | 1 | 0 |
| B82 | B90 | CEA | CYFRA 21-1 | 1 | 0 |
| B82 | B212 | CEA | IOP-L | 0.795336 | 2.45E-13 |
| B82 | B120 | CEA | HCV-cAg | 0.700649 | 1.84E-09 |
| B83 | B211 | AFP | IOP-R | -0.8618 | 0 |
| B83 | B224 | AFP | pmh | 1 | 0 |
| B83 | B86 | AFP | TBhCG | -1 | 0 |
| B83 | B87 | AFP | NSE | -1 | 0 |
| B83 | B90 | AFP | CYFRA 21-1 | -1 | 0 |
| B83 | B212 | AFP | IOP-L | -0.79534 | 2.45E-13 |
| B83 | B120 | AFP | HCV-cAg | -0.70065 | 1.84E-09 |
| B84 | B211 | C-PSA | IOP-R | -0.8618 | 0 |
| B84 | B85 | C-PSA | SCC | 1 | 0 |
| B84 | B86 | C-PSA | TBhCG | -1 | 0 |
| B84 | B87 | C-PSA | NSE | -1 | 0 |
| B84 | B212 | C-PSA | IOP-L | -0.79534 | 2.45E-13 |
| B84 | B120 | C-PSA | HCV-cAg | -0.70065 | 1.84E-09 |
| B85 | B211 | SCC | IOP-R | -0.8618 | 0 |
| B85 | B87 | SCC | NSE | -1 | 0 |
| B85 | B88 | SCC | CA 242 | 1 | 0 |
| B85 | B212 | SCC | IOP-L | -0.79534 | 2.45E-13 |
| B85 | B120 | SCC | HCV-cAg | -0.70065 | 1.84E-09 |

| | | | | | |
|---|---|---|---|---|---|
| B86 | B211 | TBhCG | IOP-R | 0.861797 | 0 |
| B86 | B224 | TBhCG | pmh | -1 | 0 |
| B86 | B87 | TBhCG | NSE | 1 | 0 |
| B86 | B88 | TBhCG | CA 242 | -1 | 0 |
| B86 | B212 | TBhCG | IOP-L | 0.795336 | 2.45E-13 |
| B86 | B120 | TBhCG | HCV-cAg | 0.700649 | 1.84E-09 |
| B87 | B211 | NSE | IOP-R | 0.861797 | 0 |
| B87 | B224 | NSE | pmh | -1 | 0 |
| B87 | B88 | NSE | CA 242 | -1 | 0 |
| B87 | B89 | NSE | CA 72-4 | 1 | 0 |
| B87 | B90 | NSE | CYFRA 21-1 | 1 | 0 |
| B87 | B212 | NSE | IOP-L | 0.795336 | 2.45E-13 |
| B87 | B120 | NSE | HCV-cAg | 0.700649 | 1.84E-09 |
| B88 | B211 | CA 242 | IOP-R | -0.8618 | 0 |
| B88 | B224 | CA 242 | pmh | 1 | 0 |
| B88 | B89 | CA 242 | CA 72-4 | -1 | 0 |
| B88 | B90 | CA 242 | CYFRA 21-1 | -1 | 0 |
| B88 | B212 | CA 242 | IOP-L | -0.79534 | 2.45E-13 |
| B88 | B120 | CA 242 | HCV-cAg | -0.70065 | 1.84E-09 |
| B89 | B211 | CA 72-4 | IOP-R | 0.861797 | 0 |
| B89 | B90 | CA 72-4 | CYFRA 21-1 | 1 | 0 |
| B89 | B212 | CA 72-4 | IOP-L | 0.795336 | 2.45E-13 |
| B89 | B120 | CA 72-4 | HCV-cAg | 0.700649 | 1.84E-09 |
| B9 | B156 | RH | IgE | 0.956183 | 0 |
| B9 | B22 | RH | CD3+CD8+(T) | 0.911685 | 0 |
| B9 | B98 | RH | anti-Jo-1 | 1 | 0 |
| B9 | B67 | RH | WBRSR(mPas_50) | 0.801784 | 1.13E-13 |
| B9 | B155 | RH | IgA | 0.763763 | 7.58E-12 |
| B9 | B154 | RH | RF | 0.755929 | 1.64E-11 |
| B9 | B196 | RH | TCO2 | 0.654654 | 4.42E-08 |
| B9 | B81 | RH | TSGF | -0.65465 | 4.42E-08 |
| B9 | B109 | RH | AMA-M2 | 0.629941 | 1.97E-07 |
| B9 | B38 | RH | BAND% | -0.61721 | 4.06E-07 |
| B90 | B211 | CYFRA 21-1 | IOP-R | 0.861797 | 0 |
| B90 | B212 | CYFRA 21-1 | IOP-L | 0.795336 | 2.45E-13 |
| B90 | B120 | CYFRA 21-1 | HCV-cAg | 0.700649 | 1.84E-09 |
| B91 | B126 | TP | Alb | 0.728761 | 1.92E-10 |
| B92 | B143 | ALDH2-Glu504Glu | β-CTx | 0.957427 | 0 |
| B92 | B157 | ALDH2-Glu504Glu | IgG | 0.959403 | 0 |
| B92 | B159 | ALDH2-Glu504Glu | β2-MG | 0.880716 | 0 |
| B92 | B150 | ALDH2-Glu504Glu | ALP | 0.76277 | 8.37E-12 |
| B92 | B197 | ALDH2-Glu504Glu | AG | 0.727273 | 2.18E-10 |
| B92 | B196 | ALDH2-Glu504Glu | TCO2 | 0.696311 | 2.54E-09 |
| B92 | B107 | ALDH2-Glu504Glu | AnuA | 0.6742 | 1.23E-08 |

| | | | | | |
|---|---|---|---|---|---|
| B92 | B175 | ALDH2-Glu504Glu | ALB-U | 0.667424 | 1.93E-08 |
| B92 | B199 | ALDH2-Glu504Glu | Mg | 0.658145 | 3.54E-08 |
| B92 | B184 | ALDH2-Glu504Glu | INR | 0.603023 | 8.73E-07 |
| B93 | B202 | ALDH2-Glu504Lys | e-times | -0.95618 | 0 |
| B93 | B125 | ALDH2-Glu504Lys | Cor | -0.84515 | 2.22E-16 |
| B93 | B183 | ALDH2-Glu504Lys | PTA | -0.8165 | 1.73E-14 |
| B93 | B200 | ALDH2-Glu504Lys | OP | -0.79772 | 1.84E-13 |
| B93 | B186 | ALDH2-Glu504Lys | PFC | -0.75593 | 1.64E-11 |
| B93 | B102 | ALDH2-Glu504Lys | anti-Sm | -0.74833 | 3.36E-11 |
| B93 | B99 | ALDH2-Glu504Lys | anti-PM-Scl | -0.60609 | 7.42E-07 |
| B94 | B186 | ALDH2-Lys504Lys | PFC | -0.94868 | 0 |
| B94 | B200 | ALDH2-Lys504Lys | OP | -0.95346 | 0 |
| B94 | B185 | ALDH2-Lys504Lys | PT | -0.84327 | 4.44E-16 |
| B94 | B106 | ALDH2-Lys504Lys | anti-P | -0.78935 | 4.91E-13 |
| B94 | B176 | ALDH2-Lys504Lys | Ucr | -0.74421 | 4.92E-11 |
| B95 | B103 | anti-cenpb | anti-SSA/Ro | 0.973329 | 0 |
| B95 | B107 | anti-cenpb | AnuA | 0.974679 | 0 |
| B95 | B144 | anti-cenpb | Total-PINP | 0.893188 | 0 |
| B95 | B150 | anti-cenpb | ALP | 0.870572 | 0 |
| B95 | B164 | anti-cenpb | COM | 0.95673 | 0 |
| B95 | B193 | anti-cenpb | K | 0.919709 | 0 |
| B95 | B116 | anti-cenpb | TT3 | 0.820783 | 9.77E-15 |
| B95 | B143 | anti-cenpb | β-CTx | 0.794719 | 2.63E-13 |
| B95 | B159 | anti-cenpb | β2-MG | 0.757532 | 1.40E-11 |
| B95 | B157 | anti-cenpb | IgG | 0.729996 | 1.73E-10 |
| B95 | B197 | anti-cenpb | AG | 0.691714 | 3.57E-09 |
| B95 | B155 | anti-cenpb | IgA | 0.662266 | 2.71E-08 |
| B95 | B187 | anti-cenpb | NSE | 0.636285 | 1.36E-07 |
| B96 | B104 | dsDNA (IIF) | anti-SSB | 0.866025 | 0 |
| B96 | B110 | dsDNA (IIF) | AHA | 0.87831 | 0 |
| B96 | B150 | dsDNA (IIF) | ALP | 0.989949 | 0 |
| B96 | B164 | dsDNA (IIF) | COM | 0.932505 | 0 |
| B96 | B116 | dsDNA (IIF) | TT3 | 0.8 | 1.40E-13 |
| B96 | B156 | dsDNA (IIF) | IgE | 0.777817 | 1.76E-12 |
| B96 | B199 | dsDNA (IIF) | Mg | 0.759257 | 1.18E-11 |
| B96 | B142 | dsDNA (IIF) | 25-(OH)D | 0.701646 | 1.70E-09 |
| B97 | B106 | dsDNA | anti-P | -0.98058 | 0 |
| B97 | B124 | dsDNA | ACTH | -0.96562 | 0 |
| B97 | B167 | dsDNA | PC | -0.96214 | 0 |
| B97 | B168 | dsDNA | NMP22 | -0.89815 | 0 |
| B97 | B202 | dsDNA | e-times | -0.89443 | 0 |
| B97 | B198 | dsDNA | P | -0.77567 | 2.22E-12 |
| B97 | B121 | dsDNA | C3 | -0.70711 | 1.12E-09 |
| B97 | B151 | dsDNA | Anti-CCP | -0.67132 | 1.49E-08 |

| | | | | | |
|---|---|---|---|---|---|
| B97 | B185 | dsDNA | PT | -0.65997 | 3.15E-08 |
| B97 | B152 | dsDNA | ACA-Ig(G-A-M) | -0.63059 | 1.90E-07 |
| B98 | B175 | anti-Jo-1 | ALB-U | 0.717137 | 5.05E-10 |
| B98 | B139 | anti-Jo-1 | ADA | 0.644658 | 8.23E-08 |
| B99 | B202 | anti-PM-Scl | e-times | -0.67612 | 1.08E-08 |

**Supplementary Table 35. P values of PEIs in healthy physical status vs 34 unhealthy physical status adjusted for age and sex.**

| PEI | PEI | Compare with normal | estimate | std.error | statistic | p.value |
|---|---|---|---|---|---|---|
| B3 | Hcy | (Intercept) | 14.90772996 | 0.00900448 | 1655.590351 | 0 |
| B3 | Hcy | Gender | -0.44012623 | 0.003978356 | -110.6301805 | 0 |
| B3 | Hcy | Age | 0.002704378 | 0.000152536 | 17.72944923 | 2.56E-70 |
| B3 | Hcy | Gout | 0.369748215 | 0.039468062 | 9.368289182 | 7.39E-21 |
| B3 | Hcy | Parkinson's syndrome | 0.677564136 | 0.129713144 | 5.223558039 | 1.76E-07 |
| B3 | Hcy | Hypertension+other diseases | 0.323902522 | 0.037293011 | 8.685341174 | 3.78E-18 |
| B3 | Hcy | Hypertension | 0.228543689 | 0.009191241 | 24.8653799 | 1.97E-136 |
| B3 | Hcy | Hypertension+Diabetes | 0.186173379 | 0.020136495 | 9.245570018 | 2.35E-20 |
| B3 | Hcy | Hypertension+Coronary heart disease | 0.409423438 | 0.040309846 | 10.15690906 | 3.10E-24 |
| B3 | Hcy | Hypertensive+Diabetes+Coronary Heart Disease | 0.235093742 | 0.059957225 | 3.921024409 | 8.82E-05 |
| B3 | Hcy | Coronary heart disease | 0.270798253 | 0.05020043 | 5.394341247 | 6.88E-08 |
| B4 | WB-Ca | (Intercept) | 1.65990543 | 2.54E-05 | 65446.43573 | 0 |
| B4 | WB-Ca | Gender | 8.57E-05 | 1.12E-05 | 7.647376102 | 2.05E-14 |
| B4 | WB-Ca | Diabetes | 0.000299086 | 4.82E-05 | 6.200761421 | 5.62E-10 |
| B4 | WB-Ca | Stomach trouble | 0.000477483 | 0.000142509 | 3.35055113 | 0.0008065 |
| B4 | WB-Ca | Hypertension | 0.000323794 | 2.59E-05 | 12.50708592 | 6.88E-36 |
| B4 | WB-Ca | Hepatic adipose infiltration | 0.001069587 | 0.000309656 | 3.454110402 | 0.0005521 |
| B4 | WB-Ca | Hypertension+Diabetes | 0.000299812 | 5.67E-05 | 5.285987438 | 1.25E-07 |
| B5 | WB-Mg | (Intercept) | 1.538188834 | 3.43E-05 | 44898.70994 | 0 |
| B5 | WB-Mg | Gender | -0.00014208 | 1.51E-05 | -9.386886831 | 6.19E-21 |
| B5 | WB-Mg | Cervical spondylopathy | 0.001486194 | 0.000388305 | 3.827392122 | 0.0001295 |
| B5 | WB-Mg | Diabetes | 0.000222551 | 6.52E-05 | 3.415861691 | 0.0006358 |
| B5 | WB-Mg | Gout | 0.000482709 | 0.000150163 | 3.214568462 | 0.0013065 |
| B5 | WB-Mg | Diabetes+other diseases | 0.001573524 | 0.000358712 | 4.386588577 | 1.15E-05 |
| B5 | WB-Mg | Hypertension | 0.000346223 | 3.50E-05 | 9.900680654 | 4.15E-23 |
| B5 | WB-Mg | Hypertension+Diabetes | 0.000319976 | 7.66E-05 | 4.176537322 | 2.96E-05 |
| B6 | WB-Fe | (Intercept) | 8.792245087 | 0.000201026 | 43736.94358 | 0 |
| B6 | WB-Fe | Gender | -0.0014585 | 8.88E-05 | -16.42138324 | 1.37E-60 |
| B6 | WB-Fe | Hyperthyroidism | 0.005372649 | 0.001467477 | 3.661147557 | 0.0002511 |
| B6 | WB-Fe | Cervical spondylopathy | 0.007811684 | 0.002278495 | 3.428440013 | 0.0006071 |
| B6 | WB-Fe | Diabetes | 0.00330766 | 0.0003823 | 8.651993567 | 5.07E-18 |
| B6 | WB-Fe | Gout | 0.00448833 | 0.000881127 | 5.093851135 | 3.51E-07 |
| B6 | WB-Fe | Cholecystolithiasis | 0.006322598 | 0.001296613 | 4.876241585 | 1.08E-06 |
| B6 | WB-Fe | Hypertension+other diseases | 0.002720534 | 0.000832569 | 3.267637924 | 0.0010845 |
| B6 | WB-Fe | Diabetes+other diseases | 0.010390655 | 0.002104854 | 4.936519621 | 7.95E-07 |
| B6 | WB-Fe | Other disease conditions | 0.003339518 | 0.000772897 | 4.320781994 | 1.55E-05 |
| B6 | WB-Fe | Hypertension | 0.003611496 | 0.000205195 | 17.60030635 | 2.52E-69 |
| B6 | WB-Fe | Hypertension+Diabetes | 0.00325419 | 0.000449549 | 7.238795121 | 4.53E-13 |
| B6 | WB-Fe | Hypertensive+Diabetes+Coronary Heart | 0.004331983 | 0.001338549 | 3.236327928 | 0.0012108 |

| | | | | | | |
|---|---|---|---|---|---|---|
| | | Disease | | | | |
| B7 | WB-Cop | (Intercept) | 16.4287723 | 0.000793879 | 20694.30553 | 0 |
| B8 | WB-Zinc | (Intercept) | 97.24811949 | 0.002601458 | 37382.16046 | 0 |
| B8 | WB-Zinc | Gender | -0.01333504 | 0.001149375 | -11.60199157 | 4.05E-31 |
| B8 | WB-Zinc | Hypothyroidism | 0.051647592 | 0.012920879 | 3.997219625 | 6.41E-05 |
| B8 | WB-Zinc | Heart disease | 0.153285152 | 0.034760812 | 4.409711477 | 1.04E-05 |
| B9 | RH | (Intercept) | 1.709743059 | 0.000194454 | 8792.531101 | 0 |
| B9 | RH | Gender | 0.000333034 | 8.59E-05 | 3.87638075 | 0.000106 |
| B10 | PG Ⅰ | (Intercept) | 95.18732833 | 0.10202058 | 933.0208549 | 0 |
| B10 | PG Ⅰ | Age | 0.088162392 | 0.001728229 | 51.0131462 | 0 |
| B10 | PG Ⅰ | Hypothyroidism | -1.71906152 | 0.506714159 | -3.392566586 | 0.0006924 |
| B10 | PG Ⅰ | Nephropathy | 7.224700752 | 0.859273536 | 8.407917211 | 4.18E-17 |
| B10 | PG Ⅰ | Diabetes | 0.628463836 | 0.194017578 | 3.239210805 | 0.0011987 |
| B10 | PG Ⅰ | Stomach trouble | 3.685718934 | 0.573234506 | 6.429687841 | 1.28E-10 |
| B10 | PG Ⅰ | Hypertension+other diseases | 3.305935398 | 0.422529078 | 7.824160682 | 5.12E-15 |
| B10 | PG Ⅰ | Hypertension | 0.974201165 | 0.104136575 | 9.35503367 | 8.38E-21 |
| B10 | PG Ⅰ | Hypertension+Diabetes | 1.797171873 | 0.228146097 | 7.877285181 | 3.35E-15 |
| B10 | PG Ⅰ | Hypertension+Coronary heart disease | 2.100241609 | 0.45670976 | 4.598635269 | 4.25E-06 |
| B10 | PG Ⅰ | Hypertensive+Diabetes+Coronary Heart Disease | 4.17952619 | 0.679314178 | 6.152567284 | 7.63E-10 |
| B10 | PG Ⅰ | Coronary heart disease+Diabetes | 8.103349214 | 1.233089902 | 6.571580224 | 4.98E-11 |
| B11 | PGI/PGII | (Intercept) | 17.64791607 | 0.566990893 | 31.12557235 | 1.43E-212 |
| B12 | PG Ⅱ | (Intercept) | 8.104031511 | 0.013259963 | 611.1654529 | 0 |
| B12 | PG Ⅱ | Gender | -0.0601347 | 0.005858512 | -10.2645006 | 1.02E-24 |
| B12 | PG Ⅱ | Age | 0.013282165 | 0.000224624 | 59.13070687 | 0 |
| B12 | PG Ⅱ | Rheumatoid arthritis | 0.67880678 | 0.134711741 | 5.038957829 | 4.68E-07 |
| B12 | PG Ⅱ | Stomach trouble | 0.245762621 | 0.074505248 | 3.298594782 | 0.0009717 |
| B12 | PG Ⅱ | Hypertension+other diseases | 0.24597313 | 0.054917548 | 4.478953234 | 7.50E-06 |
| B12 | PG Ⅱ | Hypertension | 0.064307243 | 0.013534987 | 4.751186314 | 2.02E-06 |
| B12 | PG Ⅱ | Coronary heart disease+Diabetes | 0.872432515 | 0.160268908 | 5.44355436 | 5.22E-08 |
| B14 | G-17 | (Intercept) | 4.360022564 | 0.026428596 | 164.9736751 | 0 |
| B14 | G-17 | Gender | 0.096497261 | 0.011676672 | 8.264106349 | 1.41E-16 |
| B14 | G-17 | Age | 0.009151495 | 0.0004477 | 20.44111166 | 7.59E-93 |
| B14 | G-17 | Rhinallergosis | 2.144013264 | 0.427567103 | 5.014448607 | 5.32E-07 |
| B14 | G-17 | Hypothyroidism | 0.879625378 | 0.13126512 | 6.701135649 | 2.07E-11 |
| B14 | G-17 | Hyperthyroidism | 1.169065209 | 0.192927417 | 6.059611576 | 1.37E-09 |
| B14 | G-17 | Diabetes | 0.264081 | 0.050260567 | 5.254238404 | 1.49E-07 |
| B14 | G-17 | Hypertension+other diseases | 0.372724208 | 0.109456839 | 3.405216248 | 0.0006611 |
| B14 | G-17 | Diabetes+other diseases | 0.92236901 | 0.276722701 | 3.333188814 | 0.0008586 |
| B14 | G-17 | Hypertension+Coronary heart disease | 0.385103322 | 0.1183114 | 3.254997585 | 0.001134 |
| B15 | CD3-CD19+(B cell) | (Intercept) | 16.73596953 | 0.002560678 | 6535.756015 | 0 |
| B15 | CD3-CD19+(B cell) | Gender | 0.009734903 | 0.001131358 | 8.604617981 | 7.67E-18 |
| B15 | CD3-CD19+(B cell) | Age | -0.00042658 | 4.34E-05 | -9.833944899 | 8.06E-23 |
| B15 | CD3-CD19+(B cell) | Cervical spondylopathy | -0.10501024 | 0.029023634 | -3.618094166 | 0.0002968 |

| | | | | | | |
|---|---|---|---|---|---|---|
| B15 | CD3-CD19+(B cell) | Diabetes | -0.01719166 | 0.004869769 | -3.530282256 | 0.0004151 |
| B15 | CD3-CD19+(B cell) | Stomach trouble | -0.0703432 | 0.014387972 | -4.889027997 | 1.01E-06 |
| B15 | CD3-CD19+(B cell) | Chronic pharyngitis | -0.12405915 | 0.01851127 | -6.701817429 | 2.06E-11 |
| B15 | CD3-CD19+(B cell) | Other disease conditions | -0.0350404 | 0.009845214 | -3.559130764 | 0.0003721 |
| B15 | CD3-CD19+(B cell) | Hypertension | -0.01028861 | 0.002613789 | -3.936282837 | 8.28E-05 |
| B16 | CD3+T | (Intercept) | 70.15141686 | 0.003951819 | 17751.67674 | 0 |
| B16 | CD3+T | Gender | 0.007785952 | 0.001745991 | 4.459330434 | 8.22E-06 |
| B16 | CD3+T | Age | -0.00028744 | 6.69E-05 | -4.293815874 | 1.76E-05 |
| B16 | CD3+T | Chronic pharyngitis | 0.122810184 | 0.028567894 | 4.298888181 | 1.72E-05 |
| B16 | CD3+T | Coronary heart disease | -0.0841138 | 0.022031592 | -3.817872221 | 0.0001346 |
| B17 | CD3+CD16+CD56+NK | (Intercept) | 7.759116553 | 0.000951329 | 8156.08113 | 0 |
| B17 | CD3+CD16+CD56+NK | Hepatic adipose infiltration | 0.108091793 | 0.011614839 | 9.306352927 | 1.33E-20 |
| B18 | CD3+CD4+helper(inducer)T | (Intercept) | 39.93211927 | 0.003415013 | 11693.10763 | 0 |
| B18 | CD3+CD4+helper(inducer)T | Gender | 0.010149746 | 0.00150882 | 6.726943516 | 1.73E-11 |
| B19 | CD3+CD4+/CD3+CD8+ | (Intercept) | 1.66830484 | 0.000297561 | 5606.594987 | 0 |
| B19 | CD3+CD4+/CD3+CD8+ | Age | 2.70E-05 | 5.04E-06 | 5.364443802 | 8.12E-08 |
| B19 | CD3+CD4+/CD3+CD8+ | Hypertension | 0.001423295 | 0.000303733 | 4.686010647 | 2.79E-06 |
| B19 | CD3+CD4+/CD3+CD8+ | Hypertension+Diabetes | 0.002249433 | 0.000665429 | 3.380426797 | 0.0007238 |
| B20 | CD3+CD4+CD8+ | (Intercept) | 1.209003986 | 0.000358142 | 3375.767892 | 0 |
| B20 | CD3+CD4+CD8+ | Gender | 0.000583758 | 0.000158234 | 3.689202869 | 0.000225 |
| B20 | CD3+CD4+CD8+ | Diabetes | 0.002530452 | 0.000681096 | 3.715263997 | 0.000203 |
| B20 | CD3+CD4+CD8+ | Hypertension | 0.001815471 | 0.00036557 | 4.966136984 | 6.83E-07 |
| B20 | CD3+CD4+CD8+ | Hepatic adipose infiltration | 0.01499519 | 0.004372579 | 3.429370147 | 0.000605 |
| B21 | CD3+CD4-CD8- | (Intercept) | 5.133434255 | 0.001135263 | 4521.801776 | 0 |
| B21 | CD3+CD4-CD8- | Age | -0.00011106 | 1.92E-05 | -5.77467555 | 7.71E-09 |
| B21 | CD3+CD4-CD8- | Stomach trouble | -0.02316956 | 0.00637883 | -3.632258636 | 0.000281 |
| B21 | CD3+CD4-CD8- | Hepatic adipose infiltration | 0.072084582 | 0.013860502 | 5.200719574 | 1.99E-07 |
| B22 | CD3+CD8+(T) | (Intercept) | 26.5222045 | 0.003259342 | 8137.287822 | 0 |
| B22 | CD3+CD8+(T) | Age | -0.00018371 | 5.52E-05 | -3.327223282 | 0.0008772 |
| B22 | CD3+CD8+(T) | Cervical spondylopathy | 0.129202146 | 0.036942534 | 3.497381777 | 0.0004699 |
| B22 | CD3+CD8+(T) | Chronic pharyngitis | 0.085580389 | 0.023561943 | 3.632144734 | 0.0002811 |
| B22 | CD3+CD8+(T) | Hepatitis B | 0.098043218 | 0.024798574 | 3.953582861 | 7.70E-05 |
| B23 | CD3-CD16+CD56+(NK) | (Intercept) | 15.8772081 | 0.003246558 | 4890.47416 | 0 |
| B23 | CD3-CD16+CD56+(NK) | Gender | -0.00759096 | 0.001434393 | -5.292107094 | 1.21E-07 |
| B23 | CD3-CD16+CD56+(NK) | Age | 0.000319542 | 5.50E-05 | 5.810195552 | 6.24E-09 |
| B23 | CD3-CD16+CD56+(NK) | Hypertension+other diseases | 0.042994084 | 0.013445965 | 3.197545355 | 0.0013861 |
| B24 | αHBDH | (Intercept) | 135.4643966 | 0.047482843 | 2852.912481 | 0 |
| B24 | αHBDH | Gender | 0.387140939 | 0.020978852 | 18.45386677 | 5.02E-76 |
| B24 | αHBDH | Age | 0.041103696 | 0.000804359 | 51.10115174 | 0 |
| B24 | αHBDH | Hypothyroidism | -1.25147987 | 0.235837016 | -5.306545559 | 1.12E-07 |
| B24 | αHBDH | Diabetes | -1.12002045 | 0.09030047 | -12.40326263 | 2.53E-35 |
| B24 | αHBDH | Cholecystolithiasis | -1.13475007 | 0.306263821 | -3.705139118 | 0.0002113 |
| B24 | αHBDH | Stomach trouble | -1.21068471 | 0.266797193 | -4.537846483 | 5.68E-06 |
| B24 | αHBDH | Other disease conditions | -0.64124889 | 0.182560508 | -3.512527945 | 0.0004439 |

| | | | | | | |
|---|---|---|---|---|---|---|
| B24 | αHBDH | Hypertension | -0.27081407 | 0.048467679 | -5.587518779 | 2.30E-08 |
| B24 | αHBDH | Hypertension+Diabetes | -1.05796784 | 0.106184708 | -9.963467086 | 2.21E-23 |
| B24 | αHBDH | Hyperlipidemia | -1.63351504 | 0.23223142 | -7.033996709 | 2.01E-12 |
| B25 | CK | (Intercept) | 128.2508858 | 0.521686196 | 245.8391399 | 0 |
| B25 | CK | Gender | -3.64266623 | 0.230491198 | -15.80392769 | 2.98E-56 |
| B25 | CK | Age | -0.07009037 | 0.008837366 | -7.931138425 | 2.17E-15 |
| B26 | CKMB | (Intercept) | 13.08522537 | 0.016527141 | 791.7416337 | 0 |
| B26 | CKMB | Gender | -0.0553398 | 0.007302015 | -7.578701839 | 3.49E-14 |
| B26 | CKMB | Age | 0.001800917 | 0.00027997 | 6.432540329 | 1.26E-10 |
| B26 | CKMB | Diabetes | 0.138997429 | 0.03143048 | 4.422376854 | 9.76E-06 |
| B26 | CKMB | Asthma | 0.713620115 | 0.116707906 | 6.114582477 | 9.69E-10 |
| B27 | LDH | (Intercept) | 165.0670552 | 0.073852284 | 2235.097493 | 0 |
| B27 | LDH | Gender | -0.15708842 | 0.032629388 | -4.814323197 | 1.48E-06 |
| B27 | LDH | Age | 0.078423412 | 0.001251058 | 62.68567834 | 0 |
| B27 | LDH | Rheumatoid arthritis | 3.368660698 | 0.750286368 | 4.489833268 | 7.13E-06 |
| B27 | LDH | Diabetes | -1.26808482 | 0.140448539 | -9.02882175 | 1.74E-19 |
| B27 | LDH | Hypertension+other diseases | 1.194335309 | 0.305867087 | 3.904752614 | 9.43E-05 |
| B27 | LDH | Brain diseases | 3.258159397 | 0.931617191 | 3.497315664 | 0.00047 |
| B27 | LDH | Hypertension | 1.003498149 | 0.075384045 | 13.31181092 | 1.99E-40 |
| B27 | LDH | Heart disease | 7.052807816 | 0.986817926 | 7.147020366 | 8.88E-13 |
| B27 | LDH | Hypertension+Coronary heart disease | 2.108277534 | 0.330610345 | 6.376925491 | 1.81E-10 |
| B27 | LDH | Coronary heart disease+Diabetes | -3.66369271 | 0.892628784 | -4.104385584 | 4.05E-05 |
| B28 | PRL | (Intercept) | 336.6778186 | 0.072970017 | 4613.919975 | 0 |
| B28 | PRL | Age | -0.00661037 | 0.001236112 | -5.347708466 | 8.91E-08 |
| B29 | E2 | (Intercept) | 342.1748281 | 0.094275687 | 3629.512977 | 0 |
| B29 | E2 | Age | -0.00776455 | 0.00159703 | -4.861864185 | 1.16E-06 |
| B30 | LH | (Intercept) | 13.12015031 | 0.003415026 | 3841.889263 | 0 |
| B30 | LH | Gender | 0.005209887 | 0.001508825 | 3.452942154 | 0.0005545 |
| B30 | LH | Age | 0.000772811 | 5.79E-05 | 13.3587542 | 1.06E-40 |
| B31 | FSH | (Intercept) | 22.86829194 | 0.007689627 | 2973.914385 | 0 |
| B31 | FSH | Age | 0.002288541 | 0.000130262 | 17.56871082 | 4.40E-69 |
| B31 | FSH | Hypothyroidism | 0.146608869 | 0.038192714 | 3.838660677 | 0.0001237 |
| B31 | FSH | Rheumatoid arthritis | 0.302966604 | 0.078121107 | 3.878165765 | 0.0001053 |
| B32 | Testo | (Intercept) | 1.72101631 | 0.001006374 | 1710.116099 | 0 |
| B32 | Testo | Gender | -0.00722969 | 0.000444636 | -16.25979866 | 1.94E-59 |
| B32 | Testo | Other disease conditions | 0.012548742 | 0.003869274 | 3.243177219 | 0.0011821 |
| B32 | Testo | Hypertension | 0.004631914 | 0.001027247 | 4.509056107 | 6.51E-06 |
| B33 | Prog | (Intercept) | 9.125998467 | 0.004794945 | 1903.25415 | 0 |
| B33 | Prog | Age | -0.00045073 | 8.12E-05 | -5.549052912 | 2.87E-08 |
| B34 | TK | (Intercept) | 2.072740635 | 0.019064695 | 108.7214159 | 0 |
| B34 | TK | Hepatitis B | 5.007969788 | 0.145052972 | 34.5251097 | 5.11E-261 |
| B35 | β2-MG | (Intercept) | 2.140036284 | 4.69E-05 | 45650.30865 | 0 |
| B35 | β2-MG | Rhinallergosis | 0.002499305 | 0.000758416 | 3.295426517 | 0.0009828 |
| B35 | β2-MG | Hypothyroidism | 0.002511445 | 0.000232837 | 10.78626267 | 4.01E-27 |

| | | | | | | |
|---|---|---|---|---|---|---|
| B35 | β2-MG | Hyperthyroidism | 0.002506347 | 0.000342214 | 7.323925996 | 2.41E-13 |
| B35 | β2-MG | Cervical spondylopathy | 0.002499769 | 0.000531342 | 4.704631871 | 2.54E-06 |
| B35 | β2-MG | Rheumatoid arthritis | 0.002507277 | 0.000476256 | 5.264555937 | 1.41E-07 |
| B35 | β2-MG | Chronic rhinitis | 0.002496658 | 0.000529604 | 4.714195028 | 2.43E-06 |
| B35 | β2-MG | Nephropathy | 0.002498545 | 0.00039484 | 6.327993785 | 2.48E-10 |
| B35 | β2-MG | Diabetes | 0.002502693 | 8.92E-05 | 28.07222215 | 2.57E-173 |
| B35 | β2-MG | Gout | 0.002486227 | 0.000205478 | 12.09974295 | 1.07E-33 |
| B35 | β2-MG | Parkinson's syndrome | 0.00249064 | 0.000675309 | 3.688146329 | 0.0002259 |
| B35 | β2-MG | Cholecystolithiasis | 0.002500054 | 0.000302368 | 8.268239108 | 1.36E-16 |
| B35 | β2-MG | Stomach trouble | 0.002497612 | 0.000263404 | 9.482066119 | 2.50E-21 |
| B35 | β2-MG | Chronic pharyngitis | 0.002497045 | 0.00033889 | 7.368307128 | 1.73E-13 |
| B35 | β2-MG | Lumbar disc protrusion | 0.002495409 | 0.000482995 | 5.166529456 | 2.39E-07 |
| B35 | β2-MG | Hepatitis B | 0.002493343 | 0.000356676 | 6.990494571 | 2.74E-12 |
| B35 | β2-MG | Hypertension+other diseases | 0.002495011 | 0.000194154 | 12.85068144 | 8.59E-38 |
| B35 | β2-MG | Diabetes+other diseases | 0.002109838 | 0.000490849 | 4.298342724 | 1.72E-05 |
| B35 | β2-MG | Bronchial disease | 0.002494141 | 0.000395785 | 6.301762054 | 2.94E-10 |
| B35 | β2-MG | Other disease conditions | 0.002498681 | 0.000180238 | 13.86319076 | 1.07E-43 |
| B35 | β2-MG | Brain diseases | 0.002491657 | 0.000591359 | 4.21344386 | 2.52E-05 |
| B35 | β2-MG | Hypertension | 0.002569441 | 4.79E-05 | 53.69646256 | 0 |
| B35 | β2-MG | Hepatic adipose infiltration | 0.002490433 | 0.000572348 | 4.351259414 | 1.35E-05 |
| B35 | β2-MG | Asthma | 0.002502899 | 0.00033104 | 7.560722069 | 4.01E-14 |
| B35 | β2-MG | Other Cardiac diseases | 0.002501036 | 0.000511005 | 4.894343624 | 9.87E-07 |
| B35 | β2-MG | Heart disease | 0.002493792 | 0.000626398 | 3.981161455 | 6.86E-05 |
| B35 | β2-MG | Hepatopathy | 0.002497168 | 0.000706101 | 3.536556801 | 0.0004054 |
| B35 | β2-MG | Hypertension+Diabetes | 0.002420735 | 0.000104834 | 23.09109315 | 6.18E-118 |
| B35 | β2-MG | Hypertension+Coronary heart disease | 0.00249424 | 0.00020986 | 11.88525155 | 1.42E-32 |
| B35 | β2-MG | Hypertensive+Diabetes+Coronary Heart Disease | 0.002492774 | 0.000312148 | 7.985876125 | 1.40E-15 |
| B35 | β2-MG | Hyperlipidemia | 0.00249342 | 0.000229278 | 10.87511307 | 1.52E-27 |
| B35 | β2-MG | Coronary heart disease | 0.002490363 | 0.000261352 | 9.528760058 | 1.60E-21 |
| B35 | β2-MG | Coronary heart disease+Diabetes | 0.002488912 | 0.00056661 | 4.392635736 | 1.12E-05 |
| B36 | V-Lym% | (Intercept) | 0.659495639 | 0.001271887 | 518.5176154 | 0 |
| B37 | BAND | (Intercept) | 0.509992491 | 0.00016194 | 3149.270519 | 0 |
| B38 | BAND% | (Intercept) | 0.03998483 | 3.31E-06 | 12088.33468 | 0 |
| B38 | BAND% | Age | 1.98E-07 | 5.60E-08 | 3.541635327 | 0.0003977 |
| B38 | BAND% | Rhinallergosis | -0.00029721 | 5.35E-05 | -5.554004519 | 2.79E-08 |
| B38 | BAND% | Hypothyroidism | -0.0003322 | 1.64E-05 | -20.22066284 | 6.77E-91 |
| B38 | BAND% | Hyperthyroidism | -0.00029933 | 2.41E-05 | -12.39666127 | 2.74E-35 |
| B38 | BAND% | Cervical spondylopathy | -0.00030043 | 3.75E-05 | -8.013443995 | 1.12E-15 |
| B38 | BAND% | Rheumatoid arthritis | -0.00030156 | 3.36E-05 | -8.97396826 | 2.87E-19 |
| B38 | BAND% | Chronic rhinitis | -0.00029683 | 3.74E-05 | -7.943473691 | 1.97E-15 |
| B38 | BAND% | Nephropathy | -0.0002813 | 2.79E-05 | -10.09726417 | 5.70E-24 |
| B38 | BAND% | Diabetes | -0.00029616 | 6.29E-06 | -47.08060128 | 0 |
| B38 | BAND% | Gout | -0.00029851 | 1.45E-05 | -20.5889937 | 3.63E-94 |

| | | | | | | |
|---|---|---|---|---|---|---|
| B38 | BAND% | Parkinson's syndrome | -0.00030301 | 4.76E-05 | -6.359137952 | 2.03E-10 |
| B38 | BAND% | Cholecystolithiasis | -0.00029701 | 2.13E-05 | -13.9214859 | 4.74E-44 |
| B38 | BAND% | Stomach trouble | -0.00029929 | 1.86E-05 | -16.1033416 | 2.47E-58 |
| B38 | BAND% | Chronic pharyngitis | -0.0002329 | 2.39E-05 | -9.739956547 | 2.04E-22 |
| B38 | BAND% | Lumbar disc protrusion | -0.0003006 | 3.41E-05 | -8.820588903 | 1.14E-18 |
| B38 | BAND% | Hepatitis B | -0.00029789 | 2.52E-05 | -11.83684247 | 2.53E-32 |
| B38 | BAND% | Hypertension+other diseases | -0.00032275 | 1.37E-05 | -23.55938601 | 1.10E-122 |
| B38 | BAND% | Coronary heart disease+other diseases | -0.00030303 | 6.65E-05 | -4.555289573 | 5.23E-06 |
| B38 | BAND% | Diabetes+other diseases | -0.00030155 | 3.46E-05 | -8.706685827 | 3.13E-18 |
| B38 | BAND% | Bronchial disease | -0.00032168 | 2.79E-05 | -11.51903346 | 1.06E-30 |
| B38 | BAND% | Other disease conditions | -0.00031704 | 1.27E-05 | -24.92929524 | 4.01E-137 |
| B38 | BAND% | Brain diseases | -0.00030301 | 4.17E-05 | -7.262034229 | 3.82E-13 |
| B38 | BAND% | Hypertension | -0.0003014 | 3.38E-06 | -89.26889317 | 0 |
| B38 | BAND% | Hepatic adipose infiltration | -0.00029732 | 4.04E-05 | -7.36231465 | 1.81E-13 |
| B38 | BAND% | Asthma | -0.0003002 | 2.34E-05 | -12.85211751 | 8.44E-38 |
| B38 | BAND% | Other Cardiac diseases | -0.00030189 | 3.61E-05 | -8.372773625 | 5.64E-17 |
| B38 | BAND% | Heart disease | -0.00030321 | 4.42E-05 | -6.860308207 | 6.88E-12 |
| B38 | BAND% | Hepatopathy | -0.00029968 | 4.98E-05 | -6.0150383 | 1.80E-09 |
| B38 | BAND% | Pregnant | -0.00029731 | 8.93E-05 | -3.328817225 | 0.0008722 |
| B38 | BAND% | Hypertension+Diabetes | -0.00031907 | 7.40E-06 | -43.13541113 | 0 |
| B38 | BAND% | Hypertension+Coronary heart disease | -0.00032077 | 1.48E-05 | -21.66271196 | 4.92E-104 |
| B38 | BAND% | Hypertensive+Diabetes+Coronary Heart Disease | -0.0003039 | 2.20E-05 | -13.79809007 | 2.65E-43 |
| B38 | BAND% | Hyperlipidemia | -0.00030181 | 1.62E-05 | -18.65584991 | 1.17E-77 |
| B38 | BAND% | Coronary heart disease | -0.00030261 | 1.84E-05 | -16.40990156 | 1.66E-60 |
| B38 | BAND% | Coronary heart disease+Diabetes | -0.00026594 | 4.00E-05 | -6.651869884 | 2.90E-11 |
| B39 | Other cells | (Intercept) | 1.089739481 | 0.001185528 | 919.2019865 | 0 |
| B39 | Other cells | Other disease conditions | 0.040029556 | 0.004558079 | 8.782110999 | 1.61E-18 |
| B39 | Other cells | Asthma | 0.031678919 | 0.008371712 | 3.784043078 | 0.0001543 |
| B40 | WBC | (Intercept) | 7.250137973 | 0.007565896 | 958.2655953 | 0 |
| B40 | WBC | Gender | -0.50286 | 0.003342761 | -150.4325152 | 0 |
| B40 | WBC | Age | -0.00957176 | 0.000128166 | -74.68232732 | 0 |
| B40 | WBC | Nephropathy | 0.373977572 | 0.063724146 | 5.868694896 | 4.39E-09 |
| B40 | WBC | Diabetes | 0.284943953 | 0.014388439 | 19.80367436 | 2.90E-87 |
| B40 | WBC | Gout | 0.453431515 | 0.033162521 | 13.67301089 | 1.49E-42 |
| B40 | WBC | Cholecystolithiasis | 0.32256083 | 0.048799948 | 6.609860182 | 3.85E-11 |
| B40 | WBC | Stomach trouble | -0.2504432 | 0.042511352 | -5.89120753 | 3.84E-09 |
| B40 | WBC | Hepatitis B | -0.43843851 | 0.057564819 | -7.616431635 | 2.61E-14 |
| B40 | WBC | Hypertension+other diseases | 0.474746438 | 0.031334963 | 15.1506939 | 7.61E-52 |
| B40 | WBC | Diabetes+other diseases | 0.257117934 | 0.079219314 | 3.245647082 | 0.0011719 |
| B40 | WBC | Other disease conditions | 0.155294168 | 0.029089114 | 5.338566453 | 9.37E-08 |
| B40 | WBC | Hypertension | 0.387951783 | 0.007722819 | 50.23447561 | 0 |
| B40 | WBC | Hepatic adipose infiltration | 0.617576678 | 0.09237253 | 6.685717898 | 2.30E-11 |
| B40 | WBC | Asthma | 0.384217984 | 0.053427263 | 7.191421789 | 6.42E-13 |

| | | | | | | |
|---|---|---|---|---|---|---|
| B40 | WBC | Pregnant | 2.349527975 | 0.204293652 | 11.50073901 | 1.31E-30 |
| B40 | WBC | Hypertension+Diabetes | 0.692281998 | 0.016919426 | 40.91639897 | 0 |
| B40 | WBC | Hypertension+Coronary heart disease | 0.491066262 | 0.033869819 | 14.49863831 | 1.25E-47 |
| B40 | WBC | Hypertensive+Diabetes+Coronary Heart Disease | 0.881566398 | 0.050378272 | 17.49894071 | 1.50E-68 |
| B40 | WBC | Hyperlipidemia | 0.119061237 | 0.037003656 | 3.217553344 | 0.0012929 |
| B40 | WBC | Coronary heart disease | 0.168786755 | 0.042180254 | 4.001558559 | 6.29E-05 |
| B40 | WBC | Coronary heart disease+Diabetes | 0.558964316 | 0.091446551 | 6.112470171 | 9.81E-10 |
| B41 | MON % | (Intercept) | 0.072566003 | 0.000608156 | 119.3212872 | 0 |
| B41 | MON % | Gender | -0.00575684 | 0.000268695 | -21.42514182 | 8.29E-102 |
| B41 | MON % | Age | 0.000121242 | 1.03E-05 | 11.76858004 | 5.70E-32 |
| B41 | MON % | Hypertension | 0.002384781 | 0.00062077 | 3.84164914 | 0.0001222 |
| B42 | MON | (Intercept) | 0.739755556 | 0.153093558 | 4.832048894 | 1.35E-06 |
| B43 | MCV | (Intercept) | 89.42501864 | 0.057958818 | 1542.906188 | 0 |
| B43 | MCV | Gender | -0.16744022 | 0.025607343 | -6.538757857 | 6.21E-11 |
| B43 | MCV | Age | 0.077135 | 0.000981823 | 78.56307916 | 0 |
| B43 | MCV | Diabetes | -0.78120792 | 0.110223148 | -7.087512331 | 1.37E-12 |
| B43 | MCV | Hypertension | -0.741358 | 0.059160934 | -12.5312085 | 5.08E-36 |
| B43 | MCV | Hypertension+Diabetes | -1.22239119 | 0.12961187 | -9.431167001 | 4.07E-21 |
| B44 | MCH | (Intercept) | 30.2863083 | 0.012145506 | 2493.622581 | 0 |
| B44 | MCH | Gender | -0.67018544 | 0.005366123 | -124.8919285 | 0 |
| B44 | MCH | Age | 0.018768724 | 0.000205745 | 91.22327501 | 0 |
| B44 | MCH | Rheumatoid arthritis | -0.50691129 | 0.123389652 | -4.108215569 | 3.99E-05 |
| B44 | MCH | Diabetes | -0.16698674 | 0.023097709 | -7.229580012 | 4.85E-13 |
| B44 | MCH | Hypertension | -0.13644839 | 0.012397414 | -11.00619732 | 3.58E-28 |
| B44 | MCH | Hepatopathy | 0.620585212 | 0.18293856 | 3.392314944 | 0.0006931 |
| B44 | MCH | Hypertension+Diabetes | -0.35114433 | 0.027160695 | -12.92839995 | 3.14E-38 |
| B44 | MCH | Hypertension+Coronary heart disease | -0.185829 | 0.0543711 | -3.417789969 | 0.0006313 |
| B45 | MCHC | (Intercept) | 338.1400052 | 0.048926795 | 6911.141485 | 0 |
| B45 | MCHC | Gender | -6.73968132 | 0.021616818 | -311.7795243 | 0 |
| B45 | MCHC | Age | -0.06798146 | 0.00082882 | -82.02197954 | 0 |
| B45 | MCHC | Rheumatoid arthritis | -3.63206674 | 0.497061236 | -7.307081068 | 2.73E-13 |
| B45 | MCHC | Diabetes | 1.001122595 | 0.093046505 | 10.75937882 | 5.37E-27 |
| B45 | MCHC | Parkinson's syndrome | -2.4777589 | 0.704810112 | -3.515498514 | 0.000439 |
| B45 | MCHC | Hypertension+other diseases | 0.725184719 | 0.202635525 | 3.578763979 | 0.0003452 |
| B45 | MCHC | Hypertension | 1.19038897 | 0.04994158 | 23.83562913 | 1.57E-125 |
| B45 | MCHC | Hepatic adipose infiltration | 3.227088879 | 0.597350505 | 5.402337244 | 6.58E-08 |
| B45 | MCHC | Hepatopathy | 2.782780193 | 0.736947265 | 3.776091351 | 0.0001593 |
| B45 | MCHC | Hypertension+Diabetes | 0.597362256 | 0.109413782 | 5.459661904 | 4.77E-08 |
| B45 | MCHC | Hyperlipidemia | 0.772210955 | 0.239293571 | 3.227044303 | 0.0012508 |
| B46 | RBC Count | (Intercept) | 6.056129881 | 0.002081744 | 2909.161454 | 0 |
| B46 | RBC Count | Gender | -0.61267124 | 0.000919755 | -666.1240898 | 0 |
| B46 | RBC Count | Age | -0.00742624 | 3.53E-05 | -210.5854047 | 0 |
| B46 | RBC Count | Hyperthyroidism | 0.066944839 | 0.015196627 | 4.405243135 | 1.06E-05 |

| | | | | | | |
|---|---|---|---|---|---|---|
| B46 | RBC Count | Rheumatoid arthritis | -0.07155716 | 0.021149031 | -3.383472369 | 0.0007158 |
| B46 | RBC Count | Nephropathy | -0.127817 | 0.017533596 | -7.289833652 | 3.11E-13 |
| B46 | RBC Count | Gout | -0.05704073 | 0.009124614 | -6.251302654 | 4.07E-10 |
| B46 | RBC Count | Parkinson's syndrome | -0.23813974 | 0.02998836 | -7.941072399 | 2.01E-15 |
| B46 | RBC Count | Cholecystolithiasis | 0.053301562 | 0.013427227 | 3.969662655 | 7.20E-05 |
| B46 | RBC Count | Other disease conditions | -0.03287311 | 0.008003823 | -4.107176192 | 4.01E-05 |
| B46 | RBC Count | Hypertension | 0.046953896 | 0.002124921 | 22.09676896 | 3.64E-108 |
| B46 | RBC Count | Hepatic adipose infiltration | 0.089047474 | 0.025416153 | 3.503577936 | 0.0004591 |
| B46 | RBC Count | Asthma | 0.075495696 | 0.014700426 | 5.135612938 | 2.81E-07 |
| B46 | RBC Count | Heart disease | -0.09926557 | 0.02781637 | -3.568602462 | 0.0003589 |
| B46 | RBC Count | Hepatopathy | -0.16061714 | 0.031355736 | -5.122416676 | 3.02E-07 |
| B46 | RBC Count | Pregnant | -0.45712845 | 0.05621107 | -8.132356356 | 4.22E-16 |
| B46 | RBC Count | Hypertension+Diabetes | -0.04574428 | 0.004655353 | -9.826167733 | 8.71E-23 |
| B46 | RBC Count | Hypertensive+Diabetes+Coronary Heart Disease | -0.07974894 | 0.013861501 | -5.75326919 | 8.76E-09 |
| B46 | RBC Count | Hyperlipidemia | -0.03850185 | 0.010181496 | -3.78155107 | 0.0001559 |
| B46 | RBC Count | Coronary heart disease | -0.0816631 | 0.011605829 | -7.036386963 | 1.97E-12 |
| B46 | RBC Count | Coronary heart disease+Diabetes | -0.1169195 | 0.025161372 | -4.646785565 | 3.37E-06 |
| B47 | RDW | (Intercept) | 40.48358908 | 0.017491398 | 2314.485671 | 0 |
| B47 | RDW | Gender | 0.545599138 | 0.007728043 | 70.59991266 | 0 |
| B47 | RDW | Age | 0.07881896 | 0.000296304 | 266.006787 | 0 |
| B47 | RDW | Hyperthyroidism | -0.49631341 | 0.127686321 | -3.886974003 | 0.0001015 |
| B47 | RDW | Rheumatoid arthritis | 1.595801116 | 0.177700085 | 8.980305865 | 2.71E-19 |
| B47 | RDW | Nephropathy | 0.732289804 | 0.147322189 | 4.970668784 | 6.67E-07 |
| B47 | RDW | Diabetes | -0.6677338 | 0.033264255 | -20.07361324 | 1.32E-89 |
| B47 | RDW | Gout | 0.254471261 | 0.076667566 | 3.319151399 | 0.000903 |
| B47 | RDW | Hypertension+other diseases | -0.31044979 | 0.072442483 | -4.285465824 | 1.82E-05 |
| B47 | RDW | Other disease conditions | 0.335476661 | 0.067250363 | 4.988473642 | 6.09E-07 |
| B47 | RDW | Hypertension | -0.50567098 | 0.017854184 | -28.32226695 | 2.22E-176 |
| B47 | RDW | Other Cardiac diseases | 0.655723588 | 0.190665719 | 3.43912682 | 0.0005836 |
| B47 | RDW | Hepatopathy | 1.218894445 | 0.263459675 | 4.62649339 | 3.72E-06 |
| B47 | RDW | Pregnant | 3.319899471 | 0.472301156 | 7.029200391 | 2.08E-12 |
| B47 | RDW | Hypertension+Diabetes | -0.85565405 | 0.03911558 | -21.87501899 | 4.81E-106 |
| B47 | RDW | Hypertension+Coronary heart disease | -0.43905208 | 0.078302751 | -5.607109332 | 2.06E-08 |
| B47 | RDW | Hypertensive+Diabetes+Coronary Heart Disease | -0.50306458 | 0.116468211 | -4.319329539 | 1.57E-05 |
| B47 | RDW | Hyperlipidemia | -0.67133037 | 0.085547785 | -7.847431355 | 4.25E-15 |
| B47 | RDW | Coronary heart disease | 0.331767219 | 0.097515426 | 3.402202422 | 0.0006685 |
| B47 | RDW | Coronary heart disease+Diabetes | -0.72477466 | 0.211412892 | -3.428242472 | 0.0006075 |
| B48 | LYM% | (Intercept) | 0.350378135 | 0.003072498 | 114.0369069 | 0 |
| B48 | LYM% | Age | -0.00036185 | 5.20E-05 | -6.952206808 | 3.60E-12 |
| B49 | baso% | (Intercept) | 0.005842706 | 8.78E-05 | 66.51031612 | 0 |
| B49 | baso% | Gender | -0.00018097 | 3.88E-05 | -4.662684849 | 3.12E-06 |
| B49 | baso% | Age | 5.83E-06 | 1.49E-06 | 3.919083254 | 8.89E-05 |

| | | | | | | |
|---|---|---|---|---|---|---|
| B50 | eo% | (Intercept) | 0.025594385 | 9.81E-05 | 261.0126106 | 0 |
| B50 | eo% | Gender | -0.00438072 | 4.33E-05 | -101.115311 | 0 |
| B50 | eo% | Age | 0.000107139 | 1.66E-06 | 64.49847011 | 0 |
| B50 | eo% | Rhinallergosis | 0.013766265 | 0.001586403 | 8.6776621 | 4.05E-18 |
| B50 | eo% | Chronic rhinitis | 0.00816929 | 0.00110779 | 7.374406086 | 1.65E-13 |
| B50 | eo% | Diabetes | -0.00109765 | 0.000186482 | -5.886088951 | 3.96E-09 |
| B50 | eo% | Bronchial disease | 0.003805074 | 0.000827875 | 4.596194456 | 4.30E-06 |
| B50 | eo% | Asthma | 0.016773775 | 0.000692446 | 24.22395526 | 1.38E-129 |
| B51 | Hb | (Intercept) | 182.502543 | 0.056992094 | 3202.243172 | 0 |
| B51 | Hb | Gender | -21.5323946 | 0.025180225 | -855.1311314 | 0 |
| B51 | Hb | Age | -0.13362261 | 0.000965446 | -138.4050297 | 0 |
| B51 | Hb | Rheumatoid arthritis | -4.54354845 | 0.578998896 | -7.847248894 | 4.26E-15 |
| B51 | Hb | Nephropathy | -4.29717977 | 0.480018817 | -8.952106909 | 3.49E-19 |
| B51 | Hb | Diabetes | -0.9052055 | 0.108384681 | -8.351784507 | 6.73E-17 |
| B51 | Hb | Gout | -2.04801474 | 0.249805375 | -8.198441433 | 2.44E-16 |
| B51 | Hb | Parkinson's syndrome | -8.42496189 | 0.82099397 | -10.26190472 | 1.05E-24 |
| B51 | Hb | Chronic pharyngitis | 1.620857345 | 0.411998636 | 3.934132794 | 8.35E-05 |
| B51 | Hb | Hepatitis B | 2.043539631 | 0.433622075 | 4.712720477 | 2.44E-06 |
| B51 | Hb | Hypertension+other diseases | -1.21068268 | 0.236038816 | -5.129167756 | 2.91E-07 |
| B51 | Hb | Other disease conditions | -1.32064102 | 0.219121368 | -6.026984169 | 1.67E-09 |
| B51 | Hb | Brain diseases | -4.15314028 | 0.718932595 | -5.776814555 | 7.62E-09 |
| B51 | Hb | Hypertension | 0.762736527 | 0.058174159 | 13.11125999 | 2.86E-39 |
| B51 | Hb | Hepatic adipose infiltration | 5.191115158 | 0.69582027 | 7.46042532 | 8.63E-14 |
| B51 | Hb | Asthma | 1.796974497 | 0.402454848 | 4.465033794 | 8.01E-06 |
| B51 | Hb | Other Cardiac diseases | -2.05639667 | 0.621244728 | -3.310123335 | 0.0009326 |
| B51 | Hb | Pregnant | -9.90813608 | 1.538895426 | -6.438472628 | 1.21E-10 |
| B51 | Hb | Hypertension+Diabetes | -3.04682231 | 0.127450009 | -23.90601882 | 2.92E-126 |
| B51 | Hb | Hypertension+Coronary heart disease | -1.18318645 | 0.255133283 | -4.637522912 | 3.53E-06 |
| B51 | Hb | Hypertensive+Diabetes+Coronary Heart Disease | -3.15013692 | 0.379487526 | -8.301028887 | 1.03E-16 |
| B51 | Hb | Coronary heart disease | -2.00203382 | 0.317733804 | -6.300978336 | 2.96E-10 |
| B51 | Hb | Coronary heart disease+Diabetes | -4.34538574 | 0.68884509 | -6.308219074 | 2.82E-10 |
| B52 | PLT/BPC | (Intercept) | 219.7440371 | 0.315817774 | 695.7937614 | 0 |
| B52 | PLT/BPC | Gender | 11.60720136 | 0.13953449 | 83.18517791 | 0 |
| B52 | PLT/BPC | Age | -0.93139883 | 0.005349954 | -174.0947387 | 0 |
| B52 | PLT/BPC | Diabetes | -6.81670731 | 0.60060627 | -11.34971052 | 7.48E-30 |
| B52 | PLT/BPC | Stomach trouble | -7.60641673 | 1.774520853 | -4.286462297 | 1.82E-05 |
| B52 | PLT/BPC | Hepatitis B | -27.3145574 | 2.402886954 | -11.36739177 | 6.11E-30 |
| B52 | PLT/BPC | Hypertension+other diseases | 7.96618779 | 1.307992892 | 6.090390732 | 1.13E-09 |
| B52 | PLT/BPC | Hypertension | 10.17041801 | 0.322368109 | 31.5490823 | 2.47E-218 |
| B52 | PLT/BPC | Asthma | 12.11415521 | 2.230175907 | 5.431928114 | 5.58E-08 |
| B52 | PLT/BPC | Other Cardiac diseases | -11.9735471 | 3.442585003 | -3.478068686 | 0.0005051 |
| B52 | PLT/BPC | Hepatopathy | -28.7298079 | 4.756923953 | -6.039576871 | 1.55E-09 |
| B52 | PLT/BPC | Hypertension+Diabetes | 6.202961738 | 0.706255474 | 8.782886604 | 1.60E-18 |

| | | | | | | |
|---|---|---|---|---|---|---|
| B52 | PLT/BPC | Hypertension+Coronary heart disease | 8.528356283 | 1.413803572 | 6.032207338 | 1.62E-09 |
| B52 | PLT/BPC | Hyperlipidemia | 7.918397415 | 1.544617064 | 5.126446936 | 2.95E-07 |
| B53 | GR% | (Intercept) | 0.549944073 | 0.003678714 | 149.4935767 | 0 |
| B53 | GR% | Gender | 0.014198079 | 0.001625328 | 8.735516662 | 2.43E-18 |
| B54 | ESR | (Intercept) | 8.137705712 | 0.00867674 | 937.8759955 | 0 |
| B54 | ESR | Gender | 0.200531934 | 0.003833554 | 52.30967013 | 0 |
| B54 | ESR | Age | 0.004530567 | 0.000146984 | 30.82353825 | 1.65E-208 |
| B54 | ESR | Rheumatoid arthritis | 0.338610907 | 0.088149466 | 3.84132683 | 0.0001224 |
| B54 | ESR | Hypertension | 0.085080515 | 0.008856703 | 9.606342071 | 7.54E-22 |
| B54 | ESR | Hypertension+Diabetes | 0.072317483 | 0.019403578 | 3.727017958 | 0.0001938 |
| B54 | ESR | Coronary heart disease | 0.192534921 | 0.048373262 | 3.980193076 | 6.89E-05 |
| B55 | TK | (Intercept) | 0.783130581 | 9.17E-05 | 8542.019413 | 0 |
| B55 | TK | Gender | 0.000594581 | 4.05E-05 | 14.67885682 | 8.93E-49 |
| B56 | IR | (Intercept) | 4.397176609 | 0.000720599 | 6102.110487 | 0 |
| B56 | IR | Gender | -0.01680636 | 0.000318375 | -52.78793614 | 0 |
| B56 | IR | Age | -0.00016656 | 1.22E-05 | -13.64502493 | 2.18E-42 |
| B57 | RBCAI | (Intercept) | 4.838005015 | 0.000552852 | 8750.993412 | 0 |
| B57 | RBCAI | Gender | -0.00508055 | 0.000244261 | -20.79967507 | 4.60E-96 |
| B57 | RBCAI | Age | -4.25E-05 | 9.37E-06 | -4.533203677 | 5.81E-06 |
| B58 | HCT | (Intercept) | 0.455797229 | 0.000402656 | 1131.978095 | 0 |
| B58 | HCT | Gender | -0.00248145 | 0.000177901 | -13.94846899 | 3.25E-44 |
| B59 | WBRV | (Intercept) | 40.88020179 | 0.006403175 | 6384.364423 | 0 |
| B59 | WBRV | Gender | -0.08034373 | 0.002829048 | -28.39956159 | 2.48E-177 |
| B59 | WBRV | Age | -0.00061015 | 0.00010847 | -5.625061467 | 1.85E-08 |
| B59 | WBRV | Hypertension | 0.023511374 | 0.006535982 | 3.597221133 | 0.0003217 |
| B60 | WBRVI | (Intercept) | 14.44305553 | 0.002407928 | 5998.127136 | 0 |
| B60 | WBRVI | Gender | -0.09786464 | 0.00106387 | -91.98931377 | 0 |
| B60 | WBRVI | Age | -0.00084145 | 4.08E-05 | -20.62878697 | 1.60E-94 |
| B60 | WBRVI | Gout | -0.03534264 | 0.010554328 | -3.348639449 | 0.0008121 |
| B61 | WBHSRV | (Intercept) | 6.018075629 | 0.000952376 | 6319.013004 | 0 |
| B61 | WBHSRV | Gender | -0.02133457 | 0.000420778 | -50.70262708 | 0 |
| B61 | WBHSRV | Age | -0.00017289 | 1.61E-05 | -10.71609014 | 8.59E-27 |
| B61 | WBHSRV | Hypertension | 0.005104175 | 0.000972129 | 5.250512123 | 1.52E-07 |
| B61 | WBHSRV | Hypertension+Diabetes | 0.00699641 | 0.002129775 | 3.285047126 | 0.0010197 |
| B62 | WBHSRVI | (Intercept) | 2.986842162 | 0.000409 | 7302.793705 | 0 |
| B62 | WBHSRVI | Gender | -0.01754075 | 0.000180704 | -97.06886968 | 0 |
| B62 | WBHSRVI | Age | -0.00015085 | 6.93E-06 | -21.77308193 | 4.46E-105 |
| B62 | WBHSRVI | Hypertension | 0.001599437 | 0.000417483 | 3.831144054 | 0.0001276 |
| B63 | WBRSR(mPas_200) | (Intercept) | 4.126641204 | 0.004962471 | 831.569898 | 0 |
| B63 | WBRSR(mPas_200) | Gender | -0.02467434 | 0.002192517 | -11.25388808 | 2.23E-29 |
| B64 | WBRSR(mPas_5) | (Intercept) | 9.026797304 | 0.001372648 | 6576.193232 | 0 |
| B64 | WBRSR(mPas_5) | Gender | -0.05492975 | 0.000606463 | -90.57398635 | 0 |
| B64 | WBRSR(mPas_5) | Age | -0.00040575 | 2.33E-05 | -17.44958438 | 3.56E-68 |
| B64 | WBRSR(mPas_5) | Diabetes | 0.012207972 | 0.002610432 | 4.67660951 | 2.92E-06 |

| | | | | | | |
|---|---|---|---|---|---|---|
| B64 | WBRSR(mPas_5) | Stomach trouble | -0.02886172 | 0.007712651 | -3.742127201 | 0.0001825 |
| B64 | WBRSR(mPas_5) | Hypertension | 0.009359945 | 0.001401118 | 6.680341305 | 2.39E-11 |
| B65 | WBRSR(mPas_1) | (Intercept) | 19.7792314 | 0.003233433 | 6117.099769 | 0 |
| B65 | WBRSR(mPas_1) | Gender | -0.12820217 | 0.001428594 | -89.74009657 | 0 |
| B65 | WBRSR(mPas_1) | Age | -0.00093845 | 5.48E-05 | -17.13307536 | 8.62E-66 |
| B65 | WBRSR(mPas_1) | Hypertension | 0.020869733 | 0.003300497 | 6.323209198 | 2.56E-10 |
| B66 | WBRSR(mPas_100) | (Intercept) | 4.396022859 | 0.000588521 | 7469.615762 | 0 |
| B66 | WBRSR(mPas_100) | Gender | -0.02415529 | 0.00026002 | -92.89782327 | 0 |
| B66 | WBRSR(mPas_100) | Age | -0.00016915 | 9.97E-06 | -16.96666779 | 1.49E-64 |
| B66 | WBRSR(mPas_100) | Stomach trouble | -0.01166958 | 0.003306787 | -3.528977895 | 0.0004172 |
| B66 | WBRSR(mPas_100) | Hypertension+other diseases | 0.00902277 | 0.002437421 | 3.701769821 | 0.0002141 |
| B66 | WBRSR(mPas_100) | Hypertension | 0.004529461 | 0.000600727 | 7.539965242 | 4.71E-14 |
| B67 | WBRSR(mPas_50) | (Intercept) | 4.85709514 | 0.000654031 | 7426.395631 | 0 |
| B67 | WBRSR(mPas_50) | Gender | -0.02707541 | 0.000288964 | -93.69826067 | 0 |
| B67 | WBRSR(mPas_50) | Age | -0.00018962 | 1.11E-05 | -17.11456533 | 1.19E-65 |
| B67 | WBRSR(mPas_50) | Stomach trouble | -0.01366874 | 0.003674879 | -3.719505813 | 0.0001996 |
| B67 | WBRSR(mPas_50) | Hypertension+other diseases | 0.009682048 | 0.00270874 | 3.574373271 | 0.0003511 |
| B67 | WBRSR(mPas_50) | Hypertension | 0.00508513 | 0.000667596 | 7.617070852 | 2.60E-14 |
| B69 | PV | (Intercept) | 1.368878115 | 8.20E-05 | 16693.09906 | 0 |
| B69 | PV | Gender | 0.000321452 | 3.62E-05 | 8.872451147 | 7.17E-19 |
| B69 | PV | Age | 1.73E-05 | 1.39E-06 | 12.41935361 | 2.07E-35 |
| B69 | PV | Diabetes | 0.000583573 | 0.000155948 | 3.742086932 | 0.0001825 |
| B69 | PV | Gout | 0.001471513 | 0.000359431 | 4.09401199 | 4.24E-05 |
| B69 | PV | Hepatitis B | -0.00226329 | 0.000623914 | -3.627573473 | 0.0002861 |
| B69 | PV | Hypertension | 0.000708751 | 8.37E-05 | 8.467404887 | 2.51E-17 |
| B69 | PV | Hypertension+Diabetes | 0.000757853 | 0.00018338 | 4.132681969 | 3.59E-05 |
| B70 | LDL-C | (Intercept) | 2.603604914 | 0.003637839 | 715.7009859 | 0 |
| B70 | LDL-C | Gender | -0.16712256 | 0.001607269 | -103.9792309 | 0 |
| B70 | LDL-C | Age | 0.010478597 | 6.16E-05 | 170.0380905 | 0 |
| B70 | LDL-C | Rheumatoid arthritis | -0.17129348 | 0.036957841 | -4.634834702 | 3.57E-06 |
| B70 | LDL-C | Diabetes | -0.21517333 | 0.006918258 | -31.10224023 | 2.95E-212 |
| B70 | LDL-C | Gout | -0.08831439 | 0.015945224 | -5.538610693 | 3.05E-08 |
| B70 | LDL-C | Parkinson's syndrome | -0.69022764 | 0.052404529 | -13.17114483 | 1.30E-39 |
| B70 | LDL-C | Chronic pharyngitis | 0.092270305 | 0.026298116 | 3.50862799 | 0.0004504 |
| B70 | LDL-C | Hepatitis B | -0.14724227 | 0.027678353 | -5.319762781 | 1.04E-07 |
| B70 | LDL-C | Hypertension+other diseases | -0.29963576 | 0.015066497 | -19.88755329 | 5.46E-88 |
| B70 | LDL-C | Coronary heart disease+other diseases | -0.53449345 | 0.073160652 | -7.305750185 | 2.76E-13 |
| B70 | LDL-C | Diabetes+other diseases | -0.31106874 | 0.03809028 | -8.166617145 | 3.18E-16 |
| B70 | LDL-C | Bronchial disease | -0.10422895 | 0.030713197 | -3.393620978 | 0.0006898 |
| B70 | LDL-C | Other disease conditions | -0.04792835 | 0.013986646 | -3.426722166 | 0.0006109 |
| B70 | LDL-C | Brain diseases | -0.55641218 | 0.045889891 | -12.12494004 | 7.84E-34 |
| B70 | LDL-C | Hypertension | -0.14715609 | 0.003713291 | -39.62956102 | 0 |
| B70 | LDL-C | Hepatic adipose infiltration | 0.204714918 | 0.044414618 | 4.609178863 | 4.04E-06 |
| B70 | LDL-C | Heart disease | -0.45421285 | 0.048608986 | -9.344215643 | 9.28E-21 |

| | | | | | | |
|---|---|---|---|---|---|---|
| B70 | LDL-C | Hypertension+Diabetes | -0.43848539 | 0.008135209 | -53.89970682 | 0 |
| B70 | LDL-C | Hypertension+Coronary heart disease | -0.71621303 | 0.016285308 | -43.97909075 | 0 |
| B70 | LDL-C | Hypertensive+Diabetes+Coronary Heart Disease | -0.87203125 | 0.024222912 | -36.00026413 | 1.35E-283 |
| B70 | LDL-C | Hyperlipidemia | -0.40606295 | 0.017792121 | -22.82262705 | 2.96E-115 |
| B70 | LDL-C | Coronary heart disease | -0.59616951 | 0.020281136 | -29.39527212 | 7.85E-190 |
| B70 | LDL-C | Coronary heart disease+Diabetes | -0.95105608 | 0.043969388 | -21.62995938 | 1.00E-103 |
| B71 | TG | (Intercept) | 2.305608458 | 0.006884744 | 334.8865988 | 0 |
| B71 | TG | Gender | -0.60550598 | 0.003041796 | -199.0620132 | 0 |
| B71 | TG | Age | 0.005177212 | 0.000116628 | 44.39074138 | 0 |
| B71 | TG | Diabetes | 0.295934926 | 0.013092696 | 22.60305433 | 4.37E-113 |
| B71 | TG | Gout | 0.674836952 | 0.030176067 | 22.36331729 | 9.64E-111 |
| B71 | TG | Parkinson's syndrome | -0.67983419 | 0.099174678 | -6.854917105 | 7.14E-12 |
| B71 | TG | Cholecystolithiasis | 0.194247386 | 0.04524012 | 4.293697428 | 1.76E-05 |
| B71 | TG | Hepatitis B | -0.29801439 | 0.052380809 | -5.689381187 | 1.28E-08 |
| B71 | TG | Hypertension+other diseases | 0.188247563 | 0.028513096 | 6.602143979 | 4.05E-11 |
| B71 | TG | Diabetes+other diseases | 0.38476933 | 0.072085206 | 5.337701769 | 9.42E-08 |
| B71 | TG | Other disease conditions | 0.106187298 | 0.02646949 | 4.011686591 | 6.03E-05 |
| B71 | TG | Brain diseases | -0.51271284 | 0.086845838 | -5.903712338 | 3.56E-09 |
| B71 | TG | Hypertension | 0.189938883 | 0.007027367 | 27.02845779 | 8.00E-161 |
| B71 | TG | Hepatic adipose infiltration | 0.996467758 | 0.084053904 | 11.85510378 | 2.04E-32 |
| B71 | TG | Pregnant | 0.640667623 | 0.185895948 | 3.446377559 | 0.0005682 |
| B71 | TG | Hypertension+Diabetes | 0.206593818 | 0.015395764 | 13.41887424 | 4.73E-41 |
| B71 | TG | Hypertension+Coronary heart disease | -0.20839783 | 0.030819682 | -6.761842413 | 1.36E-11 |
| B71 | TG | Hypertensive+Diabetes+Coronary Heart Disease | -0.15683263 | 0.045841457 | -3.421196345 | 0.0006235 |
| B71 | TG | Hyperlipidemia | 0.489588877 | 0.033671292 | 14.54024621 | 6.82E-48 |
| B71 | TG | Coronary heart disease | -0.40342444 | 0.038381711 | -10.51085075 | 7.73E-26 |
| B71 | TG | Coronary heart disease+Diabetes | -0.27597636 | 0.083211319 | -3.316572378 | 0.0009113 |
| B72 | HDL-C | (Intercept) | 0.943209142 | 0.001571916 | 600.0379009 | 0 |
| B72 | HDL-C | Gender | 0.249422755 | 0.000694503 | 359.1383261 | 0 |
| B72 | HDL-C | Age | 0.001476347 | 2.66E-05 | 55.44285147 | 0 |
| B72 | HDL-C | Hypothyroidism | -0.02687237 | 0.007807367 | -3.44192467 | 0.0005776 |
| B72 | HDL-C | Diabetes | -0.08492413 | 0.00298939 | -28.40851103 | 1.92E-177 |
| B72 | HDL-C | Gout | -0.11850163 | 0.006889957 | -17.19918352 | 2.76E-66 |
| B72 | HDL-C | Cholecystolithiasis | -0.06214821 | 0.010138841 | -6.129715189 | 8.81E-10 |
| B72 | HDL-C | Stomach trouble | 0.057596342 | 0.008832301 | 6.521102763 | 6.98E-11 |
| B72 | HDL-C | Hypertension+other diseases | -0.07240221 | 0.006510257 | -11.12125277 | 9.93E-29 |
| B72 | HDL-C | Diabetes+other diseases | -0.10612057 | 0.01645887 | -6.447621505 | 1.14E-10 |
| B72 | HDL-C | Other disease conditions | -0.02026072 | 0.006043652 | -3.35239735 | 0.0008012 |
| B72 | HDL-C | Brain diseases | 0.080579686 | 0.019829094 | 4.063709847 | 4.83E-05 |
| B72 | HDL-C | Hypertension | -0.06511105 | 0.001604519 | -40.57979866 | 0 |
| B72 | HDL-C | Hepatic adipose infiltration | -0.15995703 | 0.019191626 | -8.334730282 | 7.78E-17 |
| B72 | HDL-C | Pregnant | 0.156272998 | 0.042444734 | 3.68179946 | 0.0002316 |

| | | | | | | |
|---|---|---|---|---|---|---|
| B72 | HDL-C | Hypertension+Diabetes | -0.1212485 | 0.003515237 | -34.49227115 | 1.58E-260 |
| B72 | HDL-C | Hypertension+Coronary heart disease | -0.06740568 | 0.007036907 | -9.578878156 | 9.84E-22 |
| B72 | HDL-C | Hypertensive+Diabetes+Coronary Heart Disease | -0.13176836 | 0.010466759 | -12.58922259 | 2.44E-36 |
| B72 | HDL-C | Hyperlipidemia | -0.08649546 | 0.007688004 | -11.25070434 | 2.31E-29 |
| B72 | HDL-C | Coronary heart disease+Diabetes | -0.14708956 | 0.018999242 | -7.741864416 | 9.81E-15 |
| B73 | ApoA | (Intercept) | 183.6081447 | 0.37255827 | 492.8306773 | 0 |
| B73 | ApoA | Gender | 3.818599526 | 0.164603554 | 23.19876712 | 5.10E-119 |
| B73 | ApoA | Age | 0.10537012 | 0.006311138 | 16.69589842 | 1.44E-62 |
| B73 | ApoA | Diabetes | -3.72541791 | 0.708512476 | -5.258083707 | 1.46E-07 |
| B73 | ApoA | Hypertension+Diabetes | -3.73200564 | 0.833142841 | -4.479430722 | 7.49E-06 |
| B74 | ApoA1 | (Intercept) | 1.476689688 | 0.00043102 | 3426.03184 | 0 |
| B74 | ApoA1 | Gender | 0.018982959 | 0.000190433 | 99.68299089 | 0 |
| B74 | ApoA1 | Age | 0.000301394 | 7.30E-06 | 41.27845879 | 0 |
| B74 | ApoA1 | Hypothyroidism | -0.00786212 | 0.002140785 | -3.672541486 | 0.0002402 |
| B74 | ApoA1 | Diabetes | -0.01112641 | 0.000819693 | -13.57387443 | 5.78E-42 |
| B74 | ApoA1 | Gout | -0.01462813 | 0.001889231 | -7.742902772 | 9.73E-15 |
| B74 | ApoA1 | Cholecystolithiasis | -0.01378008 | 0.002780077 | -4.956725965 | 7.17E-07 |
| B74 | ApoA1 | Hypertension+other diseases | -0.01843698 | 0.001785117 | -10.32816421 | 5.27E-25 |
| B74 | ApoA1 | Hypertension | -0.00707583 | 0.00043996 | -16.08288559 | 3.43E-58 |
| B74 | ApoA1 | Hypertension+Diabetes | -0.01237836 | 0.00096388 | -12.84221321 | 9.59E-38 |
| B74 | ApoA1 | Hypertension+Coronary heart disease | -0.01256057 | 0.001929525 | -6.509672579 | 7.54E-11 |
| B74 | ApoA1 | Hypertensive+Diabetes+Coronary Heart Disease | -0.01528159 | 0.002869992 | -5.324609792 | 1.01E-07 |
| B74 | ApoA1 | Hyperlipidemia | -0.01156591 | 0.002108056 | -5.486528072 | 4.10E-08 |
| B74 | ApoA1 | Coronary heart disease+Diabetes | -0.01868052 | 0.005209605 | -3.58578484 | 0.0003361 |
| B75 | ApoA1/B | (Intercept) | 1.793048119 | 0.001765577 | 1015.559228 | 0 |
| B75 | ApoA1/B | Gender | 0.038736873 | 0.000780067 | 49.65841695 | 0 |
| B75 | ApoA1/B | Age | -0.00015256 | 2.99E-05 | -5.100829235 | 3.38E-07 |
| B75 | ApoA1/B | Hypertension | -0.00681144 | 0.001802197 | -3.779523202 | 0.0001571 |
| B76 | ApoB | (Intercept) | 0.893124116 | 0.000370686 | 2409.379147 | 0 |
| B76 | ApoB | Gender | -0.00762585 | 0.000163777 | -46.56254215 | 0 |
| B76 | ApoB | Age | 0.00022145 | 6.28E-06 | 35.26599557 | 3.07E-272 |
| B76 | ApoB | Diabetes | -0.0022783 | 0.000704953 | -3.231846769 | 0.00123 |
| B76 | ApoB | Hepatitis B | -0.01608235 | 0.002820353 | -5.702246938 | 1.18E-08 |
| B76 | ApoB | Bronchial disease | -0.01182712 | 0.003129596 | -3.779121233 | 0.0001574 |
| B76 | ApoB | Other disease conditions | -0.00463988 | 0.001425203 | -3.25559427 | 0.0011316 |
| B76 | ApoB | Heart disease | -0.01597174 | 0.00495313 | -3.224575107 | 0.0012616 |
| B76 | ApoB | Hypertension+Diabetes | -0.00763683 | 0.000828957 | -9.212576048 | 3.19E-20 |
| B76 | ApoB | Hypertension+Coronary heart disease | -0.01219954 | 0.001659431 | -7.351643184 | 1.96E-13 |
| B76 | ApoB | Hypertensive+Diabetes+Coronary Heart Disease | -0.01480743 | 0.002468252 | -5.999153636 | 1.98E-09 |
| B76 | ApoB | Hyperlipidemia | -0.00621932 | 0.001812971 | -3.430455591 | 0.0006026 |
| B76 | ApoB | Coronary heart disease | -0.01581443 | 0.002066596 | -7.652406298 | 1.97E-14 |

| | | | | | | |
|---|---|---|---|---|---|---|
| B76 | ApoB | Coronary heart disease+Diabetes | -0.02310132 | 0.004480367 | -5.156122784 | 2.52E-07 |
| B77 | TC | (Intercept) | 4.206762089 | 0.004952205 | 849.4724827 | 0 |
| B77 | TC | Gender | -0.05953377 | 0.002187981 | -27.20945082 | 5.89E-163 |
| B77 | TC | Age | 0.01570286 | 8.39E-05 | 187.183105 | 0 |
| B77 | TC | Rheumatoid arthritis | -0.16861341 | 0.050310862 | -3.351431584 | 0.000804 |
| B77 | TC | Diabetes | -0.20657234 | 0.009417853 | -21.93412072 | 1.31E-106 |
| B77 | TC | Parkinson's syndrome | -0.83086741 | 0.071338503 | -11.64682994 | 2.39E-31 |
| B77 | TC | Hepatitis B | -0.2570531 | 0.037678656 | -6.82224692 | 8.97E-12 |
| B77 | TC | Hypertension+other diseases | -0.31898001 | 0.020510085 | -15.55234998 | 1.56E-54 |
| B77 | TC | Coronary heart disease+other diseases | -0.6809831 | 0.0995939 | -6.83759848 | 8.06E-12 |
| B77 | TC | Diabetes+other diseases | -0.30278447 | 0.051852457 | -5.839346608 | 5.24E-09 |
| B77 | TC | Bronchial disease | -0.15508558 | 0.041810003 | -3.709293744 | 0.0002079 |
| B77 | TC | Brain diseases | -0.65347155 | 0.0624701 | -10.46054926 | 1.32E-25 |
| B77 | TC | Hypertension | -0.16668077 | 0.005054918 | -32.97397805 | 2.71E-238 |
| B77 | TC | Hepatic adipose infiltration | 0.198373611 | 0.060461804 | 3.280974067 | 0.0010345 |
| B77 | TC | Other Cardiac diseases | -0.20787698 | 0.053981723 | -3.850877108 | 0.0001177 |
| B77 | TC | Heart disease | -0.62608858 | 0.066171615 | -9.461588358 | 3.04E-21 |
| B77 | TC | Pregnant | 0.450894271 | 0.133719004 | 3.371953554 | 0.0007464 |
| B77 | TC | Hypertension+Diabetes | -0.50227743 | 0.011074494 | -45.35443582 | 0 |
| B77 | TC | Hypertension+Coronary heart disease | -0.8681306 | 0.022169257 | -39.15921001 | 0 |
| B77 | TC | Hypertensive+Diabetes+Coronary Heart Disease | -1.05983781 | 0.032974751 | -32.14088861 | 1.62E-226 |
| B77 | TC | Hyperlipidemia | -0.27398197 | 0.024220488 | -11.31199186 | 1.15E-29 |
| B77 | TC | Coronary heart disease | -0.73965091 | 0.027608795 | -26.79040938 | 4.86E-158 |
| B77 | TC | Coronary heart disease+Diabetes | -1.1935628 | 0.059855711 | -19.94066714 | 1.89E-88 |
| B78 | HBcAb | (Intercept) | 0.915864156 | 0.002872804 | 318.8049409 | 0 |
| B78 | HBcAb | Gender | -0.00541519 | 0.001269261 | -4.266413479 | 1.99E-05 |
| B78 | HBcAb | Age | 0.000439497 | 4.87E-05 | 9.031012799 | 1.70E-19 |
| B80 | anti-HP IgG | (Intercept) | 1.44240832 | 0.001393578 | 1035.039897 | 0 |
| B80 | anti-HP IgG | Gender | -0.00378232 | 0.00061571 | -6.143022744 | 8.10E-10 |
| B80 | anti-HP IgG | Age | 0.000523952 | 2.36E-05 | 22.19458462 | 4.16E-109 |
| B81 | TSGF | (Intercept) | 25.39937985 | 0.010648512 | 2385.251556 | 0 |
| B81 | TSGF | Other disease conditions | 0.150349562 | 0.040941056 | 3.672342044 | 0.0002404 |
| B81 | TSGF | Hepatic adipose infiltration | -0.48293849 | 0.130008393 | -3.714671615 | 0.0002035 |
| B82 | CEA | (Intercept) | 1.799743666 | 0.008169506 | 220.3001914 | 0 |
| B82 | CEA | Gender | -0.01936149 | 0.003609448 | -5.364114185 | 8.14E-08 |
| B82 | CEA | Age | 0.00100498 | 0.000138391 | 7.261865525 | 3.82E-13 |
| B83 | AFP | (Intercept) | 2.786623825 | 0.012609808 | 220.9886046 | 0 |
| B83 | AFP | Hepatopathy | 2.110849665 | 0.189931985 | 11.11371351 | 1.08E-28 |
| B84 | C-PSA | (Intercept) | 0.658884874 | 0.001484762 | 443.7647063 | 0 |
| B84 | C-PSA | Age | 0.000573596 | 2.52E-05 | 22.8052875 | 4.40E-115 |
| B84 | C-PSA | Hypertension | 0.00532953 | 0.001515557 | 3.516548832 | 0.0004372 |
| B84 | C-PSA | Hypertension+Coronary heart disease | 0.060541117 | 0.006646749 | 9.108380009 | 8.38E-20 |
| B85 | SCC | (Intercept) | 0.901116532 | 0.000806336 | 1117.545312 | 0 |

| | | | | | | |
|---|---|---|---|---|---|---|
| B85 | SCC | Gender | -0.00718829 | 0.000356255 | -20.17738182 | 1.62E-90 |
| B85 | SCC | Hypertension | 0.002813682 | 0.00082306 | 3.418564323 | 0.0006296 |
| B86 | TBhCG | (Intercept) | 22.68622878 | 0.616080814 | 36.82346256 | 1.33E-296 |
| B86 | TBhCG | Gender | 2.449312083 | 0.272196593 | 8.998320142 | 2.30E-19 |
| B86 | TBhCG | Age | -0.07066129 | 0.010436411 | -6.770650969 | 1.28E-11 |
| B87 | NSE | (Intercept) | 4.234024337 | 0.003198985 | 1323.552448 | 0 |
| B87 | NSE | Gender | -0.01427491 | 0.001413374 | -10.09988155 | 5.55E-24 |
| B87 | NSE | Age | 0.000413048 | 5.42E-05 | 7.622106441 | 2.50E-14 |
| B87 | NSE | Hypertension | 0.010561824 | 0.003265335 | 3.234530404 | 0.0012185 |
| B87 | NSE | Hypertension+Coronary heart disease | 0.05051683 | 0.014320715 | 3.527535495 | 0.0004195 |
| B88 | CA 242 | (Intercept) | 6.183770644 | 0.006280126 | 984.6570667 | 0 |
| B88 | CA 242 | Gender | 0.08321871 | 0.002774683 | 29.99215043 | 1.57E-197 |
| B88 | CA 242 | Diabetes | 0.059437939 | 0.011943226 | 4.976707045 | 6.47E-07 |
| B88 | CA 242 | Hypertension+Diabetes | 0.058610913 | 0.014044091 | 4.173350486 | 3.00E-05 |
| B88 | CA 242 | Coronary heart disease+Diabetes | 0.242922636 | 0.075905864 | 3.200314504 | 0.0013728 |
| B89 | CA 72-4 | (Intercept) | 4.752985955 | 0.012148029 | 391.2557261 | 0 |
| B89 | CA 72-4 | Gout | 0.271169885 | 0.053246736 | 5.092704356 | 3.53E-07 |
| B89 | CA 72-4 | Asthma | 0.611791471 | 0.085784411 | 7.131732454 | 9.92E-13 |
| B89 | CA 72-4 | Coronary heart disease+Diabetes | 0.602587746 | 0.146829317 | 4.104001549 | 4.06E-05 |
| B90 | CYFRA 21-1 | (Intercept) | 1.087818801 | 0.001324696 | 821.1835239 | 0 |
| B90 | CYFRA 21-1 | Gender | -0.01393057 | 0.000585277 | -23.80167505 | 3.53E-125 |
| B90 | CYFRA 21-1 | Age | 0.000987217 | 2.24E-05 | 43.99294279 | 0 |
| B90 | CYFRA 21-1 | Nephropathy | 0.035652485 | 0.011157322 | 3.195433741 | 0.0013963 |
| B90 | CYFRA 21-1 | Diabetes | 0.044583861 | 0.00251924 | 17.69734211 | 4.52E-70 |
| B90 | CYFRA 21-1 | Gout | 0.023036292 | 0.005806354 | 3.967428273 | 7.27E-05 |
| B90 | CYFRA 21-1 | Diabetes+other diseases | 0.07191712 | 0.013870338 | 5.184958165 | 2.16E-07 |
| B90 | CYFRA 21-1 | Brain diseases | 0.086200267 | 0.016710517 | 5.158443969 | 2.49E-07 |
| B90 | CYFRA 21-1 | Hypertension | 0.01067277 | 0.001352172 | 7.893058481 | 2.95E-15 |
| B90 | CYFRA 21-1 | Heart disease | 0.063785281 | 0.017700659 | 3.603554051 | 0.0003139 |
| B90 | CYFRA 21-1 | Hypertension+Diabetes | 0.061973678 | 0.002962386 | 20.92019277 | 3.71E-97 |
| B90 | CYFRA 21-1 | Hypertension+Coronary heart disease | 0.025485832 | 0.005930193 | 4.297639391 | 1.73E-05 |
| B90 | CYFRA 21-1 | Hypertensive+Diabetes+Coronary Heart Disease | 0.102641057 | 0.008820622 | 11.63648701 | 2.70E-31 |
| B91 | TP | (Intercept) | 75.98077023 | 0.019656559 | 3865.415706 | 0 |
| B91 | TP | Gender | -0.03184064 | 0.008684653 | -3.666311091 | 0.0002461 |
| B91 | TP | Age | -0.04421182 | 0.000332982 | -132.7753344 | 0 |
| B91 | TP | Nephropathy | -0.96543566 | 0.165558367 | -5.831391511 | 5.50E-09 |
| B91 | TP | Diabetes | 0.132785692 | 0.037381849 | 3.55214351 | 0.0003821 |
| B91 | TP | Gout | 0.837185804 | 0.08615781 | 9.716888148 | 2.56E-22 |
| B91 | TP | Cholecystolithiasis | 0.94259369 | 0.126784588 | 7.434607815 | 1.05E-13 |
| B91 | TP | Hypertension+other diseases | 0.904106011 | 0.081409727 | 11.10562632 | 1.18E-28 |
| B91 | TP | Hypertension | 0.929695631 | 0.020064253 | 46.33592121 | 0 |
| B91 | TP | Hepatic adipose infiltration | 0.929950648 | 0.239988233 | 3.874984347 | 0.0001066 |
| B91 | TP | Pregnant | -4.83744194 | 0.53076464 | -9.114099888 | 7.95E-20 |

| | | | | | | |
|---|---|---|---|---|---|---|
| B91 | TP | Hypertension+Diabetes | 0.982063195 | 0.043957476 | 22.34121008 | 1.58E-110 |
| B91 | TP | Hypertension+Coronary heart disease | 0.699140689 | 0.087995404 | 7.945195483 | 1.94E-15 |
| B91 | TP | Hypertensive+Diabetes+Coronary Heart Disease | 0.454717435 | 0.130885151 | 3.474171305 | 0.0005125 |
| B91 | TP | Coronary heart disease | -0.50728368 | 0.109586308 | -4.629078997 | 3.67E-06 |
| B92 | ALDH2-Glu504Glu | (Intercept) | 1.680196117 | 0.000149514 | 11237.72136 | 0 |
| B92 | ALDH2-Glu504Glu | Hypertension | 0.000763392 | 0.000152615 | 5.002075922 | 5.67E-07 |
| B92 | ALDH2-Glu504Glu | Hypertension+Diabetes | 0.001363495 | 0.000334354 | 4.07799489 | 4.54E-05 |
| B93 | ALDH2-Glu504Lys | (Intercept) | 1.295419889 | 0.000146276 | 8855.993008 | 0 |
| B93 | ALDH2-Glu504Lys | Hypertension | -0.00060574 | 0.00014931 | -4.056919311 | 4.97E-05 |
| B93 | ALDH2-Glu504Lys | Hypertension+Diabetes | -0.00120144 | 0.000327114 | -3.672865593 | 0.0002399 |
| B94 | ALDH2-Lys504Lys | (Intercept) | 1.024846638 | 4.99E-05 | 20557.77555 | 0 |
| B94 | ALDH2-Lys504Lys | Hypertension | -0.00016374 | 5.09E-05 | -3.217786229 | 0.0012919 |
| B95 | anti-cenpb | (Intercept) | 1.024765827 | 0.000130389 | 7859.268241 | 0 |
| B95 | anti-cenpb | Gender | 0.000241487 | 5.76E-05 | 4.191851794 | 2.77E-05 |
| B95 | anti-cenpb | Hepatitis B | 0.00331948 | 0.000992063 | 3.34603735 | 0.0008198 |
| B95 | anti-cenpb | Asthma | 0.003361228 | 0.000920757 | 3.650504702 | 0.0002617 |
| B96 | dsDNA (IIF) | (Intercept) | 1.012847772 | 1.06E-05 | 95260.18062 | 0 |
| B97 | dsDNA | (Intercept) | 1.018709873 | 0.000106906 | 9529.009416 | 0 |
| B98 | anti-Jo-1 | (Intercept) | 1.012218885 | 9.34E-05 | 10835.8271 | 0 |
| B98 | anti-Jo-1 | Coronary heart disease | 0.002870038 | 0.000520788 | 5.510950203 | 3.57E-08 |
| B99 | anti-PM-Scl | (Intercept) | 1.020082714 | 0.00011745 | 8685.229926 | 0 |
| B99 | anti-PM-Scl | Nephropathy | 0.006724248 | 0.000989231 | 6.797450189 | 1.07E-11 |
| B99 | anti-PM-Scl | Hepatitis B | 0.003486168 | 0.000893616 | 3.901192959 | 9.57E-05 |
| B100 | anti-Ro52 | (Intercept) | 1.086198543 | 0.000239068 | 4543.471201 | 0 |
| B100 | anti-Ro52 | Gender | 0.000771834 | 0.000105625 | 7.307310667 | 2.73E-13 |
| B100 | anti-Ro52 | Age | 1.34E-05 | 4.05E-06 | 3.300969494 | 0.0009636 |
| B100 | anti-Ro52 | Hypothyroidism | 0.003962597 | 0.001187399 | 3.337207653 | 0.0008463 |
| B100 | anti-Ro52 | Cholecystolithiasis | 0.005141605 | 0.001541986 | 3.334404288 | 0.0008549 |
| B101 | anti-Sc70 | (Intercept) | 1.010915605 | 8.94E-05 | 11311.26559 | 0 |
| B101 | anti-Sc70 | Gender | 0.00014871 | 3.95E-05 | 3.766086727 | 0.0001658 |
| B102 | anti-Sm | (Intercept) | 1.013746979 | 9.95E-05 | 10191.69159 | 0 |
| B103 | anti-SSA/Ro | (Intercept) | 1.100646591 | 0.00025753 | 4273.859065 | 0 |
| B103 | anti-SSA/Ro | Gender | 0.000947505 | 0.000113782 | 8.327392277 | 8.28E-17 |
| B103 | anti-SSA/Ro | Rheumatoid arthritis | 0.027576279 | 0.00261632 | 10.54010276 | 5.66E-26 |
| B104 | anti-SSB | (Intercept) | 1.015336311 | 0.000102658 | 9890.468887 | 0 |
| B104 | anti-SSB | Gender | 0.000165771 | 4.54E-05 | 3.654867014 | 0.0002573 |
| B104 | anti-SSB | Rheumatoid arthritis | 0.007195601 | 0.001042932 | 6.899393737 | 5.23E-12 |
| B104 | anti-SSB | Cholecystolithiasis | 0.004957549 | 0.000662143 | 7.48712423 | 7.05E-14 |
| B105 | anti-U1RNP | (Intercept) | 1.037956683 | 0.000162408 | 6391.048864 | 0 |
| B105 | anti-U1RNP | Gender | 0.000285424 | 7.18E-05 | 3.977754397 | 6.96E-05 |
| B106 | anti-P | (Intercept) | 1.013747792 | 9.26E-05 | 10947.41119 | 0 |
| B107 | AnuA | (Intercept) | 1.002031661 | 3.07E-05 | 32616.78848 | 0 |
| B107 | AnuA | Other disease conditions | 0.001046142 | 0.000118116 | 8.856869721 | 8.24E-19 |

| | | | | | | |
|---|---|---|---|---|---|---|
| B108 | Anti-Cyclin | (Intercept) | 1.023829062 | 0.00013499 | 7584.494489 | 0 |
| B109 | AMA-M2 | (Intercept) | 1.03773603 | 0.000152881 | 6787.873894 | 0 |
| B110 | AHA | (Intercept) | 1.03613541 | 0.000157665 | 6571.739209 | 0 |
| B110 | AHA | Chronic pharyngitis | 0.004067275 | 0.00113977 | 3.568503691 | 0.000359 |
| B112 | FT4 | (Intercept) | 12.9390186 | 0.004920447 | 2629.642954 | 0 |
| B112 | FT4 | Gender | -0.03872959 | 0.00217395 | -17.81530917 | 5.54E-71 |
| B112 | FT4 | Age | -0.00332459 | 8.34E-05 | -39.88600377 | 0 |
| B112 | FT4 | Hypothyroidism | 0.320659929 | 0.024438796 | 13.12093811 | 2.52E-39 |
| B112 | FT4 | Hyperthyroidism | 0.450330757 | 0.035919015 | 12.53739174 | 4.70E-36 |
| B112 | FT4 | Diabetes | 0.086886035 | 0.009357457 | 9.285218587 | 1.62E-20 |
| B112 | FT4 | Hypertension+other diseases | 0.117344337 | 0.020378554 | 5.758226837 | 8.50E-09 |
| B112 | FT4 | Hypertension | 0.077444854 | 0.005022501 | 15.41957869 | 1.23E-53 |
| B112 | FT4 | Hypertension+Diabetes | 0.158738058 | 0.011003474 | 14.42617685 | 3.59E-47 |
| B112 | FT4 | Hypertension+Coronary heart disease | 0.135655868 | 0.022027087 | 6.158593313 | 7.34E-10 |
| B112 | FT4 | Hypertensive+Diabetes+Coronary Heart Disease | 0.159984285 | 0.032763285 | 4.88303547 | 1.04E-06 |
| B112 | FT4 | Hyperlipidemia | -0.15649655 | 0.024065163 | -6.503033126 | 7.88E-11 |
| B113 | FT3 | (Intercept) | 4.76555051 | 0.002859229 | 1666.725701 | 0 |
| B113 | FT3 | Gender | -0.1060642 | 0.001263263 | -83.9604753 | 0 |
| B113 | FT3 | Age | -0.0016819 | 4.84E-05 | -34.72470197 | 5.11E-264 |
| B113 | FT3 | Hypothyroidism | -0.15274425 | 0.014201172 | -10.75574904 | 5.59E-27 |
| B113 | FT3 | Hyperthyroidism | 0.231037281 | 0.020872228 | 11.06912416 | 1.78E-28 |
| B113 | FT3 | Diabetes | -0.05841045 | 0.005437537 | -10.74207816 | 6.48E-27 |
| B113 | FT3 | Gout | -0.0569452 | 0.012532454 | -4.543818541 | 5.53E-06 |
| B113 | FT3 | Hypertension | 0.010152999 | 0.002918532 | 3.478803579 | 0.0005037 |
| B113 | FT3 | Hypertension+Diabetes | -0.04765205 | 0.006394023 | -7.452592535 | 9.16E-14 |
| B113 | FT3 | Hypertensive+Diabetes+Coronary Heart Disease | -0.06665798 | 0.019038461 | -3.501227531 | 0.0004631 |
| B113 | FT3 | Hyperlipidemia | -0.08420839 | 0.013984058 | -6.021742401 | 1.73E-09 |
| B114 | TSH TT3 TT4(TT4) | (Intercept) | 79.13080505 | 0.02554125 | 3098.157117 | 0 |
| B114 | TSH TT3 TT4(TT4) | Gender | 0.292628708 | 0.011284625 | 25.93162803 | 3.35E-148 |
| B114 | TSH TT3 TT4(TT4) | Hypothyroidism | 0.986528869 | 0.126857865 | 7.776647242 | 7.46E-15 |
| B114 | TSH TT3 TT4(TT4) | Hyperthyroidism | 1.335187525 | 0.186449837 | 7.16110857 | 8.01E-13 |
| B114 | TSH TT3 TT4(TT4) | Diabetes | 0.251380218 | 0.048573057 | 5.175301543 | 2.28E-07 |
| B114 | TSH TT3 TT4(TT4) | Hypertension | 0.154476027 | 0.026070998 | 5.925205822 | 3.12E-09 |
| B114 | TSH TT3 TT4(TT4) | Hypertension+Diabetes | 0.326181885 | 0.057117265 | 5.710740613 | 1.13E-08 |
| B115 | TSH TT3 TT4(TT3) | (Intercept) | 1.681810731 | 0.001040275 | 1616.697956 | 0 |
| B115 | TSH TT3 TT4(TT3) | Gender | -0.01205014 | 0.000459614 | -26.21796004 | 1.90E-151 |
| B115 | TSH TT3 TT4(TT3) | Age | -0.000123 | 1.76E-05 | -6.9798901 | 2.96E-12 |
| B115 | TSH TT3 TT4(TT3) | Hypothyroidism | -0.03451579 | 0.005166822 | -6.680274551 | 2.39E-11 |
| B115 | TSH TT3 TT4(TT3) | Diabetes | -0.00879741 | 0.001978343 | -4.446858973 | 8.71E-06 |
| B115 | TSH TT3 TT4(TT3) | Hypertension+Diabetes | -0.00867371 | 0.002326342 | -3.728476737 | 0.0001927 |
| B116 | TT3 | (Intercept) | 1.611263324 | 3.69E-05 | 43670.93362 | 0 |
| B116 | TT3 | Brain diseases | 0.001626318 | 0.000465423 | 3.494280483 | 0.0004754 |

| | | | | | | |
|---|---|---|---|---|---|---|
| B117 | TT4 | (Intercept) | 79.77576817 | 0.001443671 | 55258.9511 | 0 |
| B118 | CEA1 | (Intercept) | 1.51523311 | 0.025467861 | 59.49589251 | 0 |
| B118 | CEA1 | Gender | -0.16105695 | 0.011252201 | -14.31337321 | 1.83E-46 |
| B118 | CEA1 | Age | 0.011657752 | 0.000431426 | 27.021465 | 9.67E-161 |
| B118 | CEA1 | Diabetes | 0.424946621 | 0.04843349 | 8.773817942 | 1.73E-18 |
| B118 | CEA1 | Hypertension+Coronary heart disease | 1.26829653 | 0.114010532 | 11.12438038 | 9.59E-29 |
| B119 | Hcv-rna | (Intercept) | 42196.77632 | 307.819619 | 137.0828034 | 0 |
| B120 | HCV-cAg | (Intercept) | 1.001398698 | 3.45E-05 | 29021.6794 | 0 |
| B121 | C3 | (Intercept) | 1.156162266 | 2.17E-05 | 53209.41403 | 0 |
| B121 | C3 | Rheumatoid arthritis | -0.0009061 | 0.000220746 | -4.104701762 | 4.05E-05 |
| B121 | C3 | Other disease conditions | -0.00045478 | 8.35E-05 | -5.44379626 | 5.22E-08 |
| B122 | C4 | (Intercept) | 0.246553647 | 7.09E-06 | 34756.69765 | 0 |
| B122 | C4 | Rheumatoid arthritis | -0.00023354 | 7.21E-05 | -3.240526205 | 0.0011931 |
| B122 | C4 | Other disease conditions | -0.00018072 | 2.73E-05 | -6.626078996 | 3.45E-11 |
| B122 | C4 | Coronary heart disease | 0.000152498 | 3.95E-05 | 3.856047603 | 0.0001152 |
| B123 | hs-CRP1 | (Intercept) | 1.674520175 | 0.007707901 | 217.2472388 | 0 |
| B123 | hs-CRP1 | Gender | -0.05293574 | 0.003405502 | -15.54418117 | 1.77E-54 |
| B123 | hs-CRP1 | Age | 0.001741906 | 0.000130572 | 13.34059601 | 1.36E-40 |
| B123 | hs-CRP1 | Rheumatoid arthritis | 0.438508375 | 0.078306758 | 5.599879086 | 2.15E-08 |
| B123 | hs-CRP1 | Diabetes | 0.049100319 | 0.014658496 | 3.349615052 | 0.0008093 |
| B123 | hs-CRP1 | Gout | 0.353177548 | 0.03378495 | 10.45369442 | 1.41E-25 |
| B123 | hs-CRP1 | Hypertension+other diseases | 0.405693491 | 0.031923091 | 12.70846525 | 5.35E-37 |
| B123 | hs-CRP1 | Other disease conditions | 0.095701032 | 0.029635089 | 3.229314797 | 0.0012409 |
| B123 | hs-CRP1 | Hypertension | 0.064057018 | 0.007867769 | 8.141699972 | 3.90E-16 |
| B123 | hs-CRP1 | Hypertension+Diabetes | 0.07921283 | 0.017236988 | 4.595514619 | 4.32E-06 |
| B124 | ACTH | (Intercept) | 34.076131 | 0.009097736 | 3745.561704 | 0 |
| B124 | ACTH | Gender | -0.0600751 | 0.004019558 | -14.94569597 | 1.69E-50 |
| B125 | Cor | (Intercept) | 15.20055852 | 0.002134308 | 7122.006371 | 0 |
| B125 | Cor | Gender | -0.00839589 | 0.000942979 | -8.903575189 | 5.42E-19 |
| B126 | Alb | (Intercept) | 50.3349249 | 0.011419033 | 4407.985013 | 0 |
| B126 | Alb | Gender | -0.94662295 | 0.005045153 | -187.6301885 | 0 |
| B126 | Alb | Age | -0.06264905 | 0.000193438 | -323.8707215 | 0 |
| B126 | Alb | Rhinallergosis | 0.598741612 | 0.184739393 | 3.241006719 | 0.0011911 |
| B126 | Alb | Rheumatoid arthritis | -1.13568778 | 0.116009202 | -9.78963533 | 1.25E-22 |
| B126 | Alb | Nephropathy | -1.24989724 | 0.096177385 | -12.99574981 | 1.30E-38 |
| B126 | Alb | Diabetes | 0.188446422 | 0.021716139 | 8.677713101 | 4.04E-18 |
| B126 | Alb | Cholecystolithiasis | 0.4672629 | 0.073652636 | 6.344143599 | 2.24E-10 |
| B126 | Alb | Hypertension+other diseases | 0.280344054 | 0.047293138 | 5.927795568 | 3.07E-09 |
| B126 | Alb | Hypertension | 0.386459851 | 0.011655873 | 33.15580396 | 6.65E-241 |
| B126 | Alb | Hepatic adipose infiltration | 0.810033915 | 0.139415731 | 5.810204559 | 6.24E-09 |
| B126 | Alb | Hepatopathy | -1.31070279 | 0.171996241 | -7.620531616 | 2.53E-14 |
| B126 | Alb | Pregnant | -4.72672776 | 0.308335701 | -15.32981017 | 4.91E-53 |
| B126 | Alb | Hypertension+Diabetes | 0.421675336 | 0.0255361 | 16.51291044 | 3.03E-61 |
| B126 | Alb | Hypertension+Coronary heart disease | 0.624376497 | 0.051118938 | 12.2141915 | 2.63E-34 |

| | | | | | | |
|---|---|---|---|---|---|---|
| B126 | Alb | Hypertensive+Diabetes+Coronary Heart Disease | 0.399907631 | 0.076034765 | 5.259536613 | 1.44E-07 |
| B126 | Alb | Hyperlipidemia | 0.42959064 | 0.055848765 | 7.69203469 | 1.45E-14 |
| B127 | A-G ratio | (Intercept) | 1.970268436 | 0.001229434 | 1602.581814 | 0 |
| B127 | A-G ratio | Gender | -0.09290134 | 0.000543188 | -171.0297901 | 0 |
| B127 | A-G ratio | Age | -0.00323879 | 2.08E-05 | -155.5121247 | 0 |
| B127 | A-G ratio | Rheumatoid arthritis | -0.0954402 | 0.012490169 | -7.641225491 | 2.15E-14 |
| B127 | A-G ratio | Nephropathy | -0.05577048 | 0.01035497 | -5.385866234 | 7.21E-08 |
| B127 | A-G ratio | Diabetes | 0.014121739 | 0.002338075 | 6.039899179 | 1.54E-09 |
| B127 | A-G ratio | Gout | -0.05001381 | 0.005388804 | -9.281059238 | 1.68E-20 |
| B127 | A-G ratio | Stomach trouble | 0.029376796 | 0.006907959 | 4.252601652 | 2.11E-05 |
| B127 | A-G ratio | Lumbar disc protrusion | 0.041482406 | 0.012666907 | 3.274864645 | 0.0010572 |
| B127 | A-G ratio | Hypertension+other diseases | -0.02184353 | 0.005091831 | -4.289915555 | 1.79E-05 |
| B127 | A-G ratio | Hypertension | -0.01720265 | 0.001254933 | -13.70801661 | 9.19E-43 |
| B127 | A-G ratio | Hepatopathy | -0.08925381 | 0.018518032 | -4.819832272 | 1.44E-06 |
| B127 | A-G ratio | Pregnant | -0.16486472 | 0.033197065 | -4.966243919 | 6.83E-07 |
| B127 | A-G ratio | Hypertension+Diabetes | -0.01258114 | 0.002749353 | -4.576038808 | 4.74E-06 |
| B127 | A-G ratio | Hypertension+Coronary heart disease | 0.018479116 | 0.005503737 | 3.357557789 | 0.0007864 |
| B127 | A-G ratio | Hyperlipidemia | 0.027827756 | 0.006012976 | 4.627950858 | 3.69E-06 |
| B128 | ALT | (Intercept) | 52.19971764 | 0.121158392 | 430.8386454 | 0 |
| B128 | ALT | Gender | -13.5115349 | 0.053530155 | -252.4097837 | 0 |
| B128 | ALT | Age | -0.10815201 | 0.002052423 | -52.69478077 | 0 |
| B128 | ALT | Gout | 3.512777776 | 0.531056427 | 6.614697796 | 3.73E-11 |
| B128 | ALT | Parkinson's syndrome | -6.78177369 | 1.745335238 | -3.885656772 | 0.0001021 |
| B128 | ALT | Hepatitis B | 8.133587777 | 0.921828802 | 8.823317042 | 1.11E-18 |
| B128 | ALT | Bronchial disease | -3.70293739 | 1.022904432 | -3.620022824 | 0.0002946 |
| B128 | ALT | Hypertension | 1.761063311 | 0.123671323 | 14.239868 | 5.25E-46 |
| B128 | ALT | Hepatic adipose infiltration | 23.72327518 | 1.479230885 | 16.03757427 | 7.12E-58 |
| B128 | ALT | Hypertension+Diabetes | 1.418565832 | 0.270943515 | 5.235651547 | 1.64E-07 |
| B129 | IBIL | (Intercept) | 12.91676174 | 0.024007672 | 538.0264183 | 0 |
| B129 | IBIL | Gender | -1.45521293 | 0.010607061 | -137.192856 | 0 |
| B129 | IBIL | Age | 0.004392253 | 0.00040669 | 10.80000246 | 3.46E-27 |
| B129 | IBIL | Nephropathy | -0.91332592 | 0.202205842 | -4.516812717 | 6.28E-06 |
| B129 | IBIL | Diabetes | -0.40541865 | 0.045656576 | -8.879742648 | 6.71E-19 |
| B129 | IBIL | Cholecystolithiasis | -0.6891033 | 0.154849222 | -4.450156687 | 8.58E-06 |
| B129 | IBIL | Hypertension+other diseases | -0.46152029 | 0.099430326 | -4.641645169 | 3.46E-06 |
| B129 | IBIL | Diabetes+other diseases | -0.91912276 | 0.251374227 | -3.656392185 | 0.0002558 |
| B129 | IBIL | Other disease conditions | -0.60585668 | 0.092303924 | -6.563715271 | 5.25E-11 |
| B129 | IBIL | Brain diseases | -1.07084364 | 0.302847233 | -3.535920174 | 0.0004064 |
| B129 | IBIL | Hypertension | -0.07969948 | 0.024505612 | -3.252295155 | 0.0011448 |
| B129 | IBIL | Asthma | -0.60126354 | 0.169532357 | -3.546600514 | 0.0003903 |
| B129 | IBIL | Pregnant | -2.43142421 | 0.648253014 | -3.75073337 | 0.0001763 |
| B129 | IBIL | Hypertension+Diabetes | -0.90914366 | 0.053687763 | -16.9339085 | 2.59E-64 |
| B129 | IBIL | Hypertensive+Diabetes+Coronary Heart | -0.84401782 | 0.159857472 | -5.279814615 | 1.29E-07 |

| | | | | | | |
|---|---|---|---|---|---|---|
| | | Disease | | | | |
| B129 | IBIL | Hyperlipidemia | -0.77093531 | 0.1174179 | -6.565739204 | 5.18E-11 |
| B130 | GLB | (Intercept) | 25.64700733 | 0.016875402 | 1519.786484 | 0 |
| B130 | GLB | Gender | 0.914393511 | 0.007455884 | 122.6405295 | 0 |
| B130 | GLB | Age | 0.018423355 | 0.000285869 | 64.44676537 | 0 |
| B130 | GLB | Rheumatoid arthritis | 1.384135768 | 0.171442007 | 8.073492561 | 6.84E-16 |
| B130 | GLB | Gout | 0.864028223 | 0.073967559 | 11.68117804 | 1.60E-31 |
| B130 | GLB | Cholecystolithiasis | 0.473415386 | 0.108846156 | 4.349399211 | 1.37E-05 |
| B130 | GLB | Stomach trouble | -0.42224707 | 0.094819717 | -4.453156848 | 8.46E-06 |
| B130 | GLB | Lumbar disc protrusion | -0.60946298 | 0.173867943 | -3.505321163 | 0.0004561 |
| B130 | GLB | Hypertension+other diseases | 0.623966897 | 0.06989127 | 8.927680024 | 4.36E-19 |
| B130 | GLB | Hypertension | 0.543207506 | 0.017225412 | 31.53524002 | 3.82E-218 |
| B130 | GLB | Hepatopathy | 0.986317295 | 0.254181395 | 3.880367784 | 0.0001043 |
| B130 | GLB | Hypertension+Diabetes | 0.561072603 | 0.037738043 | 14.86755954 | 5.43E-50 |
| B130 | GLB | Coronary heart disease | -0.39242978 | 0.094081218 | -4.171180892 | 3.03E-05 |
| B131 | AST | (Intercept) | 30.64964398 | 0.069153702 | 443.2104603 | 0 |
| B131 | AST | Gender | -4.12559864 | 0.030553462 | -135.0288418 | 0 |
| B131 | AST | Age | 0.049228428 | 0.001171464 | 42.02300078 | 0 |
| B131 | AST | Diabetes | -1.63409955 | 0.131513012 | -12.42538299 | 1.92E-35 |
| B131 | AST | Gout | 1.268998744 | 0.30311163 | 4.186572274 | 2.83E-05 |
| B131 | AST | Hepatitis B | 4.733390301 | 0.526153185 | 8.996220944 | 2.34E-19 |
| B131 | AST | Hypertension+other diseases | 1.256754374 | 0.286407409 | 4.387995338 | 1.14E-05 |
| B131 | AST | Hypertension | 0.774683867 | 0.07058801 | 10.9747232 | 5.08E-28 |
| B131 | AST | Hepatic adipose infiltration | 7.676559708 | 0.844302152 | 9.092194884 | 9.73E-20 |
| B131 | AST | Other Cardiac diseases | 2.580745552 | 0.75381285 | 3.423589225 | 0.000618 |
| B131 | AST | Hypertension+Diabetes | -0.6868231 | 0.154646712 | -4.441239597 | 8.95E-06 |
| B131 | AST | Coronary heart disease+Diabetes | -2.79858655 | 0.835838531 | -3.34823826 | 0.0008133 |
| B132 | Dbil | (Intercept) | 4.483421958 | 0.007968022 | 562.6769166 | 0 |
| B132 | Dbil | Gender | -0.34417957 | 0.003520428 | -97.76638759 | 0 |
| B132 | Dbil | Hepatitis B | 0.291771062 | 0.060624377 | 4.812768022 | 1.49E-06 |
| B132 | Dbil | Hypertension | 0.052248317 | 0.008133286 | 6.424011148 | 1.33E-10 |
| B132 | Dbil | Asthma | -0.18386127 | 0.05626691 | -3.267662378 | 0.0010844 |
| B132 | Dbil | Other Cardiac diseases | 0.522490295 | 0.086855759 | 6.015609107 | 1.79E-09 |
| B132 | Dbil | Heart disease | 0.904024837 | 0.106469108 | 8.490959092 | 2.05E-17 |
| B132 | Dbil | Hepatopathy | 1.1949998 | 0.120016278 | 9.956980965 | 2.36E-23 |
| B132 | Dbil | Hypertension+Coronary heart disease | 0.409799467 | 0.035669993 | 11.48863328 | 1.51E-30 |
| B132 | Dbil | Hypertensive+Diabetes+Coronary Heart Disease | 0.439799593 | 0.053055866 | 8.289367897 | 1.14E-16 |
| B132 | Dbil | Coronary heart disease | 0.422864458 | 0.044422125 | 9.519230811 | 1.75E-21 |
| B132 | Dbil | Coronary heart disease+Diabetes | 0.350200742 | 0.096306915 | 3.636299038 | 0.0002766 |
| B133 | TBIL | (Intercept) | 17.3999144 | 0.029847518 | 582.9601865 | 0 |
| B133 | TBIL | Gender | -1.80001091 | 0.013187219 | -136.496627 | 0 |
| B133 | TBIL | Age | 0.004424835 | 0.000505617 | 8.751357239 | 2.11E-18 |
| B133 | TBIL | Nephropathy | -1.0380493 | 0.251392239 | -4.129201834 | 3.64E-05 |

| | | | | | | |
|---|---|---|---|---|---|---|
| B133 | TBIL | Diabetes | -0.3658958 | 0.056762499 | -6.446083279 | 1.15E-10 |
| B133 | TBIL | Cholecystolithiasis | -0.7841842 | 0.192516162 | -4.073342165 | 4.63E-05 |
| B133 | TBIL | Diabetes+other diseases | -1.02184614 | 0.312520792 | -3.269690114 | 0.0010767 |
| B133 | TBIL | Other disease conditions | -0.63580111 | 0.114756775 | -5.54042332 | 3.02E-08 |
| B133 | TBIL | Asthma | -0.78444493 | 0.210770957 | -3.721788522 | 0.0001978 |
| B133 | TBIL | Heart disease | 1.869723659 | 0.398824027 | 4.688091823 | 2.76E-06 |
| B133 | TBIL | Hepatopathy | 1.826543008 | 0.449570552 | 4.062861767 | 4.85E-05 |
| B133 | TBIL | Pregnant | -3.05127501 | 0.805940004 | -3.785982826 | 0.0001531 |
| B133 | TBIL | Hypertension+Diabetes | -0.90283979 | 0.066747265 | -13.52624396 | 1.11E-41 |
| B133 | TBIL | Hyperlipidemia | -0.68141416 | 0.145979703 | -4.667869169 | 3.04E-06 |
| B134 | CHE | (Intercept) | 9.415921376 | 0.002293683 | 4105.152882 | 0 |
| B134 | CHE | Gender | -0.06707219 | 0.001013394 | -66.1856698 | 0 |
| B134 | CHE | Age | -0.00030097 | 3.89E-05 | -7.745837659 | 9.51E-15 |
| B134 | CHE | Rhinallergosis | -0.13385292 | 0.03710767 | -3.607149677 | 0.0003096 |
| B134 | CHE | Diabetes | 0.023631173 | 0.004362011 | 5.417494724 | 6.05E-08 |
| B134 | CHE | Gout | 0.034897035 | 0.010053578 | 3.471105948 | 0.0005183 |
| B134 | CHE | Cholecystolithiasis | 0.076068247 | 0.014794233 | 5.141749837 | 2.72E-07 |
| B134 | CHE | Lumbar disc protrusion | -0.07562969 | 0.023631913 | -3.200320225 | 0.0013728 |
| B134 | CHE | Hypertension+other diseases | 0.03137905 | 0.009499534 | 3.303219938 | 0.0009559 |
| B134 | CHE | Hypertension | 0.021749604 | 0.002341256 | 9.289714559 | 1.55E-20 |
| B134 | CHE | Hepatic adipose infiltration | 0.100436442 | 0.028003735 | 3.586537426 | 0.0003351 |
| B134 | CHE | Hepatopathy | -0.11386376 | 0.034548017 | -3.29581171 | 0.0009814 |
| B135 | TBA | (Intercept) | 4.503741668 | 0.007159024 | 629.0999644 | 0 |
| B135 | TBA | Gender | -0.04850882 | 0.003162997 | -15.33634609 | 4.44E-53 |
| B135 | TBA | Age | 0.000665547 | 0.000121274 | 5.487968629 | 4.07E-08 |
| B135 | TBA | Cervical spondylopathy | 0.52367381 | 0.081142905 | 6.453722701 | 1.09E-10 |
| B135 | TBA | Hepatitis B | 0.354006818 | 0.054469148 | 6.499217098 | 8.08E-11 |
| B135 | TBA | Diabetes+other diseases | 0.387220795 | 0.074959125 | 5.165759241 | 2.40E-07 |
| B135 | TBA | Hepatopathy | 0.376949586 | 0.107830955 | 3.495745598 | 0.0004728 |
| B136 | 5'NT | (Intercept) | 2.457145102 | 0.008313868 | 295.5477782 | 0 |
| B137 | AFU | (Intercept) | 30.32314554 | 0.007896453 | 3840.097176 | 0 |
| B137 | AFU | Gender | -0.10674341 | 0.003488808 | -30.59595729 | 1.79E-205 |
| B137 | AFU | Age | 0.001239678 | 0.000133766 | 9.267514441 | 1.91E-20 |
| B137 | AFU | Diabetes | 0.131564731 | 0.015017074 | 8.761009556 | 1.94E-18 |
| B137 | AFU | Hepatitis B | 0.473639474 | 0.060079845 | 7.883500203 | 3.19E-15 |
| B137 | AFU | Hypertension+other diseases | 0.129977898 | 0.032703998 | 3.974373326 | 7.06E-05 |
| B137 | AFU | Other disease conditions | 0.172682117 | 0.030360027 | 5.687811707 | 1.29E-08 |
| B137 | AFU | Hypertension+Diabetes | 0.094333674 | 0.017658642 | 5.342068559 | 9.19E-08 |
| B137 | AFU | Coronary heart disease+Diabetes | 0.529598094 | 0.095441881 | 5.548906698 | 2.88E-08 |
| B138 | PAB | (Intercept) | 286.3073291 | 0.052875532 | 5414.741298 | 0 |
| B138 | PAB | Gender | -2.00888825 | 0.023361448 | -85.99159882 | 0 |
| B138 | PAB | Age | -0.02428702 | 0.000895712 | -27.11477942 | 7.72E-162 |
| B138 | PAB | Gout | 1.689438492 | 0.231761835 | 7.289545726 | 3.11E-13 |
| B138 | PAB | Hepatitis B | -5.60398342 | 0.402301382 | -13.92981399 | 4.22E-44 |

| | | | | | | |
|---|---|---|---|---|---|---|
| B138 | PAB | Hypertension | 0.590632304 | 0.053972217 | 10.94326559 | 7.19E-28 |
| B138 | PAB | Hypertension+Diabetes | 0.592706729 | 0.118244245 | 5.012563027 | 5.37E-07 |
| B138 | PAB | Hyperlipidemia | 1.548518595 | 0.258606248 | 5.987939606 | 2.13E-09 |
| B139 | ADA | (Intercept) | 8.46913963 | 0.002769004 | 3058.550824 | 0 |
| B139 | ADA | Gender | 0.029785707 | 0.0012234 | 24.34665606 | 6.97E-131 |
| B139 | ADA | Age | 0.001151535 | 4.69E-05 | 24.54935413 | 4.89E-133 |
| B139 | ADA | Diabetes | 0.083254681 | 0.005265952 | 15.80999665 | 2.70E-56 |
| B139 | ADA | Hepatitis B | 0.345250303 | 0.021067857 | 16.38753777 | 2.40E-60 |
| B139 | ADA | Hypertension+other diseases | 0.060532298 | 0.011468125 | 5.278308301 | 1.30E-07 |
| B139 | ADA | Diabetes+other diseases | 0.265901463 | 0.028993076 | 9.171205738 | 4.69E-20 |
| B139 | ADA | Other disease conditions | 0.037257688 | 0.010646178 | 3.499630484 | 0.0004659 |
| B139 | ADA | Hypertension | 0.010152715 | 0.002826436 | 3.592055916 | 0.0003281 |
| B139 | ADA | Hepatic adipose infiltration | 0.115845006 | 0.033806955 | 3.426661931 | 0.0006111 |
| B139 | ADA | Hepatopathy | 0.294970926 | 0.041707411 | 7.072386389 | 1.52E-12 |
| B139 | ADA | Hypertension+Diabetes | 0.079705036 | 0.006192255 | 12.87172979 | 6.55E-38 |
| B139 | ADA | Hypertensive+Diabetes+Coronary Heart Disease | 0.101902838 | 0.018437689 | 5.526876928 | 3.26E-08 |
| B139 | ADA | Coronary heart disease | 0.062445873 | 0.015437338 | 4.045119316 | 5.23E-05 |
| B139 | ADA | Coronary heart disease+Diabetes | 0.200684815 | 0.033468061 | 5.996308478 | 2.02E-09 |
| B140 | r-GT | (Intercept) | 55.99968187 | 0.192828294 | 290.4121629 | 0 |
| B140 | r-GT | Gender | -17.2755613 | 0.085195324 | -202.7759332 | 0 |
| B140 | r-GT | Age | 0.065313941 | 0.003266512 | 19.99501156 | 6.38E-89 |
| B140 | r-GT | Gout | 7.834159069 | 0.845196958 | 9.269033678 | 1.88E-20 |
| B140 | r-GT | Parkinson's syndrome | -13.5556738 | 2.777768915 | -4.880058127 | 1.06E-06 |
| B140 | r-GT | Hepatitis B | -5.32684285 | 1.467126392 | -3.630800238 | 0.0002826 |
| B140 | r-GT | Hypertension+other diseases | 4.268136615 | 0.798618882 | 5.344397324 | 9.07E-08 |
| B140 | r-GT | Hypertension | 3.755813802 | 0.19682772 | 19.08173198 | 3.72E-81 |
| B140 | r-GT | Hepatic adipose infiltration | 18.67790561 | 2.354253487 | 7.933685014 | 2.13E-15 |
| B140 | r-GT | Hepatopathy | 11.70612832 | 2.904426553 | 4.030443912 | 5.57E-05 |
| B140 | r-GT | Hypertension+Coronary heart disease | -3.85944599 | 0.863223519 | -4.470969459 | 7.79E-06 |
| B140 | r-GT | Coronary heart disease | -6.12881262 | 1.075027488 | -5.701075264 | 1.19E-08 |
| B140 | r-GT | Coronary heart disease+Diabetes | -8.7184216 | 2.330653511 | -3.740762647 | 0.0001835 |
| B141 | AST/ALT | (Intercept) | 0.552337317 | 0.00312441 | 176.7813434 | 0 |
| B141 | AST/ALT | Gender | 0.395233644 | 0.001380425 | 286.3129225 | 0 |
| B141 | AST/ALT | Age | 0.002590924 | 5.29E-05 | 48.95232064 | 0 |
| B141 | AST/ALT | Hypothyroidism | -0.05059616 | 0.015518267 | -3.260425972 | 0.0011125 |
| B141 | AST/ALT | Diabetes | -0.08457431 | 0.005941844 | -14.23368104 | 5.73E-46 |
| B141 | AST/ALT | Parkinson's syndrome | 0.356784608 | 0.045008373 | 7.927071935 | 2.25E-15 |
| B141 | AST/ALT | Stomach trouble | 0.064001038 | 0.017555471 | 3.645646233 | 0.0002667 |
| B141 | AST/ALT | Hepatitis B | -0.09064412 | 0.023771946 | -3.813071192 | 0.0001373 |
| B141 | AST/ALT | Bronchial disease | 0.105348861 | 0.026378465 | 3.993744984 | 6.50E-05 |
| B141 | AST/ALT | Brain diseases | 0.168048116 | 0.039413184 | 4.26375389 | 2.01E-05 |
| B141 | AST/ALT | Hypertension | -0.02686019 | 0.003189213 | -8.422200621 | 3.70E-17 |
| B141 | AST/ALT | Hepatic adipose infiltration | -0.2715988 | 0.038146124 | -7.11995788 | 1.08E-12 |

| | | | | | | |
|---|---|---|---|---|---|---|
| B141 | AST/ALT | Heart disease | 0.142463944 | 0.041748517 | 3.412431226 | 0.0006439 |
| B141 | AST/ALT | Hypertension+Diabetes | -0.06716382 | 0.00698704 | -9.612628563 | 7.09E-22 |
| B141 | AST/ALT | Hypertension+Coronary heart disease | 0.090508307 | 0.013986868 | 6.470949055 | 9.74E-11 |
| B141 | AST/ALT | Coronary heart disease | 0.106346893 | 0.017418741 | 6.105314497 | 1.03E-09 |
| B142 | 25-(OH)D | (Intercept) | 23.27813833 | 0.005842511 | 3984.269714 | 0 |
| B142 | 25-(OH)D | Gender | 0.025147551 | 0.002581336 | 9.74206878 | 2.00E-22 |
| B142 | 25-(OH)D | Age | 0.001504992 | 9.90E-05 | 15.20621813 | 3.27E-52 |
| B142 | 25-(OH)D | Diabetes+other diseases | 0.239460008 | 0.06117447 | 3.91437815 | 9.06E-05 |
| B142 | 25-(OH)D | Bronchial disease | 0.297258284 | 0.049326588 | 6.026329784 | 1.68E-09 |
| B142 | 25-(OH)D | Asthma | 0.177844932 | 0.04125742 | 4.310616897 | 1.63E-05 |
| B142 | 25-(OH)D | Hypertensive+Diabetes+Coronary Heart Disease | 0.137595325 | 0.038902939 | 3.536887691 | 0.0004049 |
| B142 | 25-(OH)D | Hyperlipidemia | 0.158075176 | 0.028574838 | 5.531971069 | 3.17E-08 |
| B142 | 25-(OH)D | Coronary heart disease+Diabetes | 0.231207246 | 0.070616546 | 3.274122847 | 0.0010599 |
| B143 | β-CTx | (Intercept) | 437.2280186 | 0.095761253 | 4565.813453 | 0 |
| B143 | β-CTx | Hypothyroidism | -2.28808335 | 0.475625439 | -4.810683289 | 1.50E-06 |
| B143 | β-CTx | Diabetes | -1.65485058 | 0.182113908 | -9.086898366 | 1.02E-19 |
| B143 | β-CTx | Gout | -1.59193261 | 0.419736743 | -3.792693015 | 0.000149 |
| B143 | β-CTx | Hepatitis B | 2.582673761 | 0.728595681 | 3.544728344 | 0.000393 |
| B143 | β-CTx | Diabetes+other diseases | -4.85428658 | 1.002675772 | -4.841332278 | 1.29E-06 |
| B143 | β-CTx | Heart disease | -4.22769574 | 1.279566671 | -3.304005828 | 0.0009532 |
| B143 | β-CTx | Hypertension+Diabetes | -2.23502042 | 0.214148521 | -10.43677727 | 1.69E-25 |
| B143 | β-CTx | Hypertensive+Diabetes+Coronary Heart Disease | -5.67477149 | 0.637635832 | -8.899706072 | 5.61E-19 |
| B143 | β-CTx | Hyperlipidemia | -4.45327676 | 0.468353836 | -9.508359753 | 1.94E-21 |
| B143 | β-CTx | Coronary heart disease+Diabetes | -4.3561843 | 1.157435441 | -3.763652073 | 0.0001675 |
| B144 | Total-PINP | (Intercept) | 47.69431054 | 0.008948135 | 5330.084123 | 0 |
| B144 | Total-PINP | Gender | 0.024634799 | 0.003953461 | 6.231197425 | 4.63E-10 |
| B144 | Total-PINP | Age | -0.00063182 | 0.000151581 | -4.168177976 | 3.07E-05 |
| B144 | Total-PINP | Hypothyroidism | -0.15936525 | 0.04444345 | -3.585798239 | 0.0003361 |
| B144 | Total-PINP | Diabetes | -0.17116499 | 0.01701711 | -10.05840548 | 8.46E-24 |
| B144 | Total-PINP | Gout | -0.14175682 | 0.039221092 | -3.614300597 | 0.0003012 |
| B144 | Total-PINP | Diabetes+other diseases | -0.3862025 | 0.093692151 | -4.122036844 | 3.76E-05 |
| B144 | Total-PINP | Hypertension | -0.03208602 | 0.009133727 | -3.51291695 | 0.0004432 |
| B144 | Total-PINP | Hypertension+Diabetes | -0.22262228 | 0.020010492 | -11.12527769 | 9.49E-29 |
| B144 | Total-PINP | Hypertensive+Diabetes+Coronary Heart Disease | -0.384601 | 0.059582045 | -6.454981481 | 1.08E-10 |
| B144 | Total-PINP | Hyperlipidemia | -0.42218549 | 0.043763976 | -9.646872404 | 5.08E-22 |
| B144 | Total-PINP | Coronary heart disease+Diabetes | -0.43598436 | 0.108153223 | -4.031173078 | 5.55E-05 |
| B145 | BGP | (Intercept) | 17.28278406 | 0.002336312 | 7397.463036 | 0 |
| B145 | BGP | Gender | 0.005297359 | 0.001032229 | 5.131963313 | 2.87E-07 |
| B145 | BGP | Diabetes | -0.04667089 | 0.004443081 | -10.50417409 | 8.29E-26 |
| B145 | BGP | Diabetes+other diseases | -0.11870908 | 0.024462543 | -4.85268762 | 1.22E-06 |
| B145 | BGP | Hypertension+Diabetes | -0.05463254 | 0.005224637 | -10.45671489 | 1.37E-25 |

| | | | | | | |
|---|---|---|---|---|---|---|
| B145 | BGP | Hypertension+Coronary heart disease | -0.04168505 | 0.010458837 | -3.98562901 | 6.73E-05 |
| B145 | BGP | Hypertensive+Diabetes+Coronary Heart Disease | -0.07843769 | 0.015556568 | -5.04209492 | 4.61E-07 |
| B145 | BGP | Hyperlipidemia | -0.09689352 | 0.011426551 | -8.479682051 | 2.26E-17 |
| B146 | CysC | (Intercept) | 0.79709918 | 0.00084029 | 948.5998748 | 0 |
| B146 | CysC | Gender | -0.01442626 | 0.000371257 | -38.85790708 | 0 |
| B146 | CysC | Age | 0.000696665 | 1.42E-05 | 48.94192011 | 0 |
| B146 | CysC | Rheumatoid arthritis | 0.040738667 | 0.008536747 | 4.772153297 | 1.82E-06 |
| B146 | CysC | Nephropathy | 0.040126065 | 0.007077387 | 5.669615636 | 1.43E-08 |
| B146 | CysC | Gout | 0.024238739 | 0.003683125 | 6.581025252 | 4.67E-11 |
| B146 | CysC | Hypertension+other diseases | 0.031618981 | 0.003480151 | 9.085519364 | 1.03E-19 |
| B146 | CysC | Other disease conditions | 0.011875915 | 0.003230721 | 3.675933579 | 0.000237 |
| B146 | CysC | Hypertension | 0.009379484 | 0.000857719 | 10.93538664 | 7.84E-28 |
| B146 | CysC | Hypertension+Diabetes | 0.022992926 | 0.00187912 | 12.2360061 | 2.01E-34 |
| B146 | CysC | Hypertension+Coronary heart disease | 0.019383721 | 0.00376168 | 5.152943194 | 2.56E-07 |
| B149 | TGAb | (Intercept) | 34.82676786 | 0.526643256 | 66.12971392 | 0 |
| B149 | TGAb | Gender | 11.81285796 | 0.232681325 | 50.7683974 | 0 |
| B149 | TGAb | Hyperthyroidism | 71.6439469 | 3.84446923 | 18.63558859 | 1.71E-77 |
| B149 | TGAb | Brain diseases | 24.18493186 | 6.643395214 | 3.640447554 | 0.0002722 |
| B149 | TGAb | Heart disease | 34.49634419 | 7.03703361 | 4.902114457 | 9.48E-07 |
| B150 | ALP | (Intercept) | 81.36751054 | 0.039245527 | 2073.293873 | 0 |
| B150 | ALP | Gender | -0.71406954 | 0.017339444 | -41.18180176 | 0 |
| B150 | ALP | Rhinallergosis | -2.06327766 | 0.634921977 | -3.24965544 | 0.0011555 |
| B150 | ALP | Hyperthyroidism | 0.98190452 | 0.286490369 | 3.427356124 | 0.0006095 |
| B150 | ALP | Stomach trouble | -0.76870726 | 0.220513257 | -3.485991163 | 0.0004903 |
| B150 | ALP | Hepatitis B | 1.257742268 | 0.298598028 | 4.212158651 | 2.53E-05 |
| B150 | ALP | Brain diseases | 4.92711373 | 0.495066713 | 9.952423783 | 2.47E-23 |
| B150 | ALP | Hypertension | 0.272599781 | 0.040059513 | 6.804870033 | 1.01E-11 |
| B150 | ALP | Hepatic adipose infiltration | 1.594032305 | 0.479151253 | 3.326783128 | 0.0008786 |
| B150 | ALP | Hepatopathy | 3.904862218 | 0.59112565 | 6.605807441 | 3.96E-11 |
| B150 | ALP | Hypertensive+Diabetes+Coronary Heart Disease | -0.91431949 | 0.261320245 | -3.498846733 | 0.0004673 |
| B150 | ALP | Hyperlipidemia | -1.12338882 | 0.191943948 | -5.852692019 | 4.84E-09 |
| B151 | Anti-CCP | (Intercept) | 8.106800143 | 0.020591675 | 393.6930893 | 0 |
| B151 | Anti-CCP | Age | 0.002552624 | 0.000348823 | 7.317819398 | 2.52E-13 |
| B151 | Anti-CCP | Rheumatoid arthritis | 8.869719042 | 0.209196685 | 42.39894637 | 0 |
| B152 | ACA-Ig(G-A-M) | (Intercept) | 3.909147996 | 0.000453948 | 8611.43654 | 0 |
| B152 | ACA-Ig(G-A-M) | Rhinallergosis | -0.02893774 | 0.007344068 | -3.940287269 | 8.14E-05 |
| B153 | Glu | (Intercept) | 4.684318986 | 0.024108769 | 194.2993817 | 0 |
| B153 | Glu | Gender | -0.15970858 | 0.010651727 | -14.99367878 | 8.20E-51 |
| B153 | Glu | Age | 0.012938292 | 0.000408403 | 31.68023873 | 3.90E-220 |
| B153 | Glu | Diabetes | 2.986924272 | 0.045848838 | 65.14721815 | 0 |
| B153 | Glu | Diabetes+other diseases | 2.321134582 | 0.252432775 | 9.195060259 | 3.76E-20 |
| B153 | Glu | Other disease conditions | 0.434427175 | 0.092692621 | 4.686750377 | 2.78E-06 |

| | | | | | | |
|---|---|---|---|---|---|---|
| B153 | Glu | Hypertension | 0.198933414 | 0.024608806 | 8.083830329 | 6.28E-16 |
| B153 | Glu | Hypertension+Diabetes | 2.20740417 | 0.053913844 | 40.94317878 | 0 |
| B153 | Glu | Hypertensive+Diabetes+Coronary Heart Disease | 1.846338014 | 0.160530639 | 11.50146801 | 1.30E-30 |
| B153 | Glu | Hyperlipidemia | 0.424959358 | 0.117912352 | 3.60402749 | 0.0003133 |
| B153 | Glu | Coronary heart disease+Diabetes | 2.282784333 | 0.291394934 | 7.833987704 | 4.73E-15 |
| B154 | RF | (Intercept) | 14.29784766 | 0.062017954 | 230.5436861 | 0 |
| B154 | RF | Age | 0.006645133 | 0.001050584 | 6.325178641 | 2.53E-10 |
| B154 | RF | Rheumatoid arthritis | 18.70910694 | 0.630058038 | 29.69425958 | 1.14E-193 |
| B154 | RF | Cholecystolithiasis | 1.53704282 | 0.400015124 | 3.842461764 | 0.0001218 |
| B154 | RF | Hypertension+other diseases | 0.828215647 | 0.25685395 | 3.224461403 | 0.0012621 |
| B155 | IgA | (Intercept) | 2.45418956 | 0.000844145 | 2907.307872 | 0 |
| B155 | IgA | Gender | -0.00140982 | 0.00037296 | -3.780072517 | 0.0001568 |
| B155 | IgA | Diabetes | 0.008562168 | 0.001605352 | 5.333512993 | 9.64E-08 |
| B155 | IgA | Gout | 0.013732692 | 0.003700021 | 3.711516706 | 0.000206 |
| B155 | IgA | Hypertension+other diseases | 0.021697615 | 0.003496116 | 6.206204667 | 5.43E-10 |
| B155 | IgA | Hypertension | 0.008734562 | 0.000861653 | 10.13697924 | 3.80E-24 |
| B155 | IgA | Hypertension+Diabetes | 0.007165075 | 0.001887741 | 3.795582166 | 0.0001473 |
| B156 | IgE | (Intercept) | 114.2591466 | 0.192301781 | 594.1658268 | 0 |
| B156 | IgE | Gender | -0.82099302 | 0.0849627 | -9.662981686 | 4.34E-22 |
| B156 | IgE | Diabetes+other diseases | 9.982538516 | 2.013510996 | 4.957777006 | 7.13E-07 |
| B156 | IgE | Other Cardiac diseases | 7.708021092 | 2.096193693 | 3.67715117 | 0.0002359 |
| B157 | IgG | (Intercept) | 12.20383121 | 0.002213425 | 5513.551048 | 0 |
| B157 | IgG | Gender | 0.029634696 | 0.000977935 | 30.30335272 | 1.33E-201 |
| B157 | IgG | Age | 0.000500695 | 3.75E-05 | 13.35350458 | 1.14E-40 |
| B157 | IgG | Diabetes | -0.03368908 | 0.00420938 | -8.003336482 | 1.21E-15 |
| B157 | IgG | Hypertension+Diabetes | -0.02230252 | 0.004949827 | -4.505717244 | 6.62E-06 |
| B158 | IgM | (Intercept) | 1.24978254 | 0.001073715 | 1163.979808 | 0 |
| B158 | IgM | Gender | 0.009885438 | 0.000474388 | 20.8382826 | 2.06E-96 |
| B159 | β2-MG | (Intercept) | 0.330669122 | 7.74E-06 | 42699.07095 | 0 |
| B159 | β2-MG | Diabetes+other diseases | -0.00027121 | 8.11E-05 | -3.344702824 | 0.0008237 |
| B160 | PH | (Intercept) | 6.08967121 | 0.002960394 | 2057.0474 | 0 |
| B160 | PH | Gender | 0.078278644 | 0.00130796 | 59.84787649 | 0 |
| B160 | PH | Age | -0.00402103 | 5.01E-05 | -80.18149114 | 0 |
| B160 | PH | Hypothyroidism | 0.057408557 | 0.014703638 | 3.904377787 | 9.45E-05 |
| B160 | PH | Diabetes | -0.18608855 | 0.005629928 | -33.05345039 | 1.97E-239 |
| B160 | PH | Gout | -0.10045526 | 0.012975877 | -7.741693607 | 9.82E-15 |
| B160 | PH | Hypertension+other diseases | 0.06524011 | 0.012260787 | 5.321037538 | 1.03E-07 |
| B160 | PH | Diabetes+other diseases | -0.12265493 | 0.030997042 | -3.956988257 | 7.59E-05 |
| B160 | PH | Hypertension | 0.083083147 | 0.003021795 | 27.49463049 | 2.41E-166 |
| B160 | PH | Hepatic adipose infiltration | -0.12004193 | 0.03614365 | -3.321245475 | 0.0008962 |
| B160 | PH | Pregnant | 0.529561226 | 0.0799363 | 6.624790299 | 3.48E-11 |
| B160 | PH | Hypertension+Diabetes | -0.07760186 | 0.006620256 | -11.72188155 | 9.90E-32 |
| B160 | PH | Hypertension+Coronary heart disease | 0.108296922 | 0.013252629 | 8.171731012 | 3.04E-16 |

| | | | | | | |
|---|---|---|---|---|---|---|
| B160 | PH | Coronary heart disease+Diabetes | -0.12881422 | 0.035781332 | -3.600039747 | 0.0003182 |
| B161 | LEU | (Intercept) | 0.811938439 | 0.003833442 | 211.8040005 | 0 |
| B161 | LEU | Gender | 0.277917087 | 0.00169369 | 164.0896936 | 0 |
| B161 | LEU | Age | 0.001116109 | 6.49E-05 | 17.18716109 | 3.40E-66 |
| B161 | LEU | Hypothyroidism | -0.0618775 | 0.01903988 | -3.249889021 | 0.0011545 |
| B161 | LEU | Hypertension | 0.026092498 | 0.003912951 | 6.668239914 | 2.59E-11 |
| B161 | LEU | Pregnant | 0.703922955 | 0.10351027 | 6.80051319 | 1.04E-11 |
| B161 | LEU | Hypertension+Diabetes | 0.055198436 | 0.008572632 | 6.438913237 | 1.20E-10 |
| B163 | SG | (Intercept) | 1.041649517 | 0.005485417 | 189.8943341 | 0 |
| B163 | SG | Gender | -0.00965895 | 0.002423565 | -3.985430995 | 6.74E-05 |
| B164 | COM | (Intercept) | 4.082390117 | 0.001807322 | 2258.806276 | 0 |
| B164 | COM | Gender | 0.009333452 | 0.00079851 | 11.68858073 | 1.47E-31 |
| B164 | COM | Age | -0.00019535 | 3.06E-05 | -6.380646548 | 1.76E-10 |
| B165 | bilirubin | (Intercept) | 1.001065586 | 0.000135437 | 7391.380589 | 0 |
| B165 | bilirubin | Gender | -0.00049606 | 5.98E-05 | -8.289868247 | 1.14E-16 |
| B165 | bilirubin | Hepatopathy | 0.010470886 | 0.002039983 | 5.132829391 | 2.85E-07 |
| B166 | Pro | (Intercept) | 2.547726319 | 0.007296244 | 349.1832709 | 0 |
| B166 | Pro | Gender | -0.52400173 | 0.003223624 | -162.5505243 | 0 |
| B166 | Pro | Age | -0.00388914 | 0.000123598 | -31.4659283 | 3.39E-217 |
| B166 | Pro | Chronic rhinitis | 0.276355323 | 0.082427742 | 3.352697949 | 0.0008003 |
| B166 | Pro | Nephropathy | 0.398642954 | 0.061452987 | 6.486958172 | 8.76E-11 |
| B166 | Pro | Diabetes | 0.156552436 | 0.013875628 | 11.28254802 | 1.61E-29 |
| B166 | Pro | Gout | -0.22437598 | 0.031980593 | -7.016004456 | 2.28E-12 |
| B166 | Pro | Parkinson's syndrome | 0.488574489 | 0.10510532 | 4.648427764 | 3.35E-06 |
| B166 | Pro | Stomach trouble | -0.1383097 | 0.040996226 | -3.373717925 | 0.0007416 |
| B166 | Pro | Hypertension | 0.172225199 | 0.007447574 | 23.12500564 | 2.82E-118 |
| B166 | Pro | Pregnant | 1.406715835 | 0.197012527 | 7.14023548 | 9.32E-13 |
| B166 | Pro | Hypertension+Diabetes | 0.319692364 | 0.01631641 | 19.59330323 | 1.84E-85 |
| B166 | Pro | Hypertension+Coronary heart disease | 0.176587207 | 0.032662683 | 5.406390185 | 6.43E-08 |
| B166 | Pro | Hypertensive+Diabetes+Coronary Heart Disease | 0.226677027 | 0.048582766 | 4.665790896 | 3.07E-06 |
| B166 | Pro | Hyperlipidemia | -0.14827153 | 0.035684828 | -4.155030047 | 3.25E-05 |
| B167 | PC | (Intercept) | 3.630720597 | 0.000104934 | 34600.20312 | 0 |
| B167 | PC | Asthma | 0.002881984 | 0.000740997 | 3.889330897 | 0.0001005 |
| B167 | PC | Heart disease | -0.00708993 | 0.001402127 | -5.056553791 | 4.27E-07 |
| B168 | NMP22 | (Intercept) | 1.007763484 | 8.38E-05 | 12031.25496 | 0 |
| B168 | NMP22 | Hypertension | 0.000417663 | 8.55E-05 | 4.884977136 | 1.03E-06 |
| B170 | U-GLU | (Intercept) | 1.068297081 | 0.002046855 | 521.9212417 | 0 |
| B170 | U-GLU | Gender | -0.04380891 | 0.000904341 | -48.4429313 | 0 |
| B170 | U-GLU | Age | 0.000779122 | 3.47E-05 | 22.47009778 | 8.77E-112 |
| B170 | U-GLU | Diabetes | 0.743761284 | 0.003892605 | 191.0703085 | 0 |
| B170 | U-GLU | Hypertension+other diseases | 0.027222064 | 0.008477267 | 3.211183865 | 0.0013219 |
| B170 | U-GLU | Diabetes+other diseases | 0.400863569 | 0.021431757 | 18.70418632 | 4.74E-78 |
| B170 | U-GLU | Other disease conditions | 0.107310678 | 0.007869682 | 13.63596099 | 2.47E-42 |

| | | | | | | |
|---|---|---|---|---|---|---|
| B170 | U-GLU | Hypertension | 0.016930095 | 0.002089309 | 8.103205166 | 5.36E-16 |
| B170 | U-GLU | Hypertension+Diabetes | 0.422061602 | 0.004577331 | 92.2069201 | 0 |
| B170 | U-GLU | Hypertensive+Diabetes+Coronary Heart Disease | 0.320887862 | 0.013629187 | 23.54416686 | 1.58E-122 |
| B170 | U-GLU | Hyperlipidemia | 0.053143784 | 0.010010859 | 5.308613932 | 1.10E-07 |
| B170 | U-GLU | Coronary heart disease+Diabetes | 0.338031606 | 0.024739677 | 13.66354164 | 1.69E-42 |
| B171 | EC | (Intercept) | 3.518721822 | 0.000891894 | 3945.225825 | 0 |
| B172 | KB | (Intercept) | 1.127634805 | 0.002308169 | 488.54076 | 0 |
| B172 | KB | Gender | 0.009799692 | 0.001019795 | 9.609477329 | 7.31E-22 |
| B172 | KB | Age | -0.00158775 | 3.91E-05 | -40.60708429 | 0 |
| B172 | KB | Diabetes | 0.135679241 | 0.00438956 | 30.90953293 | 1.16E-209 |
| B172 | KB | Parkinson's syndrome | 0.181583728 | 0.033250105 | 5.461147487 | 4.73E-08 |
| B172 | KB | Hypertension+Diabetes | 0.056066988 | 0.005161702 | 10.86211258 | 1.75E-27 |
| B173 | NTT | (Intercept) | 0.950710596 | 0.000727851 | 1306.189012 | 0 |
| B173 | NTT | Gender | 0.02288461 | 0.000321579 | 71.16332007 | 0 |
| B173 | NTT | Age | 0.000679516 | 1.23E-05 | 55.11174973 | 0 |
| B173 | NTT | Rheumatoid arthritis | 0.034142122 | 0.007394442 | 4.617267989 | 3.89E-06 |
| B173 | NTT | Diabetes | 0.007317941 | 0.00138419 | 5.286805016 | 1.25E-07 |
| B173 | NTT | Parkinson's syndrome | 0.040720258 | 0.010484981 | 3.883674783 | 0.0001029 |
| B173 | NTT | Hypertension | 0.002993377 | 0.000742947 | 4.029058863 | 5.60E-05 |
| B173 | NTT | Hypertension+Diabetes | 0.01862715 | 0.001627675 | 11.44402619 | 2.53E-30 |
| B173 | NTT | Hypertensive+Diabetes+Coronary Heart Disease | 0.049179022 | 0.004846466 | 10.14739776 | 3.41E-24 |
| B175 | ALB-U | (Intercept) | 28.38070352 | 0.23455716 | 120.9969607 | 0 |
| B175 | ALB-U | Gender | -1.11293383 | 0.103631956 | -10.73929196 | 6.68E-27 |
| B175 | ALB-U | Nephropathy | 21.73857614 | 1.975569649 | 11.0037002 | 3.68E-28 |
| B175 | ALB-U | Diabetes | 4.027267729 | 0.446068945 | 9.028352626 | 1.75E-19 |
| B175 | ALB-U | Hypertension+other diseases | 4.201411 | 0.97144342 | 4.324915806 | 1.53E-05 |
| B175 | ALB-U | Hypertension | 3.851683128 | 0.23942208 | 16.08741823 | 3.19E-58 |
| B175 | ALB-U | Hypertension+Diabetes | 15.06461191 | 0.524534374 | 28.71997083 | 2.62E-181 |
| B175 | ALB-U | Hypertensive+Diabetes+Coronary Heart Disease | 7.574404507 | 1.561822187 | 4.849722695 | 1.24E-06 |
| B177 | ACR | (Intercept) | 21.15084674 | 0.19681034 | 107.468168 | 0 |
| B177 | ACR | Gender | 0.42466969 | 0.08695467 | 4.883805419 | 1.04E-06 |
| B177 | ACR | Age | 0.011609233 | 0.003333968 | 3.482107363 | 0.0004975 |
| B177 | ACR | Nephropathy | 26.5770289 | 1.657645134 | 16.0330027 | 7.66E-58 |
| B177 | ACR | Diabetes | 2.886623991 | 0.374283952 | 7.712390492 | 1.24E-14 |
| B177 | ACR | Hypertension+other diseases | 6.021786419 | 0.815110952 | 7.387689254 | 1.50E-13 |
| B177 | ACR | Hypertension | 3.14422951 | 0.200892358 | 15.65131467 | 3.31E-55 |
| B177 | ACR | Hypertension+Diabetes | 14.91060682 | 0.440122094 | 33.87834198 | 2.04E-251 |
| B177 | ACR | Hypertension+Coronary heart disease | 3.877325653 | 0.881049723 | 4.400802308 | 1.08E-05 |
| B177 | ACR | Hypertensive+Diabetes+Coronary Heart Disease | 10.80017995 | 1.310481232 | 8.241384678 | 1.70E-16 |
| B177 | ACR | Hyperlipidemia | 5.694642939 | 0.962569669 | 5.916083921 | 3.30E-09 |

| | | | | | | |
|---|---|---|---|---|---|---|
| B178 | Ig-U | (Intercept) | 6.850460795 | 0.011709919 | 585.0134842 | 0 |
| B178 | Ig-U | Gender | -0.03174836 | 0.005173672 | -6.136523238 | 8.44E-10 |
| B178 | Ig-U | Nephropathy | 0.37611123 | 0.098627394 | 3.813456013 | 0.000137 |
| B178 | Ig-U | Hypertension | 0.059034074 | 0.011952793 | 4.938935368 | 7.86E-07 |
| B178 | Ig-U | Hypertension+Diabetes | 0.359437043 | 0.026186603 | 13.72598963 | 7.17E-43 |
| B179 | α1-MG | (Intercept) | 12.72305883 | 0.007727055 | 1646.559994 | 0 |
| B179 | α1-MG | Gender | -0.12825316 | 0.003413964 | -37.56722397 | 0.00E+00 |
| B179 | α1-MG | Age | 0.000621566 | 0.000130896 | 4.748534021 | 2.05E-06 |
| B179 | α1-MG | Diabetes | 0.115841022 | 0.014694921 | 7.883065091 | 3.20E-15 |
| B179 | α1-MG | Other disease conditions | 0.146233253 | 0.029708731 | 4.922231642 | 8.56E-07 |
| B179 | α1-MG | Hypertension+Diabetes | 0.158775839 | 0.017279821 | 9.188511909 | 3.99E-20 |
| B180 | Tf | (Intercept) | 3.177733045 | 0.006290352 | 505.1756971 | 0 |
| B180 | Tf | Gender | -0.00945526 | 0.002779201 | -3.402151171 | 0.0006686 |
| B180 | Tf | Nephropathy | 0.244085925 | 0.052980812 | 4.607062735 | 4.08E-06 |
| B180 | Tf | Diabetes | 0.045528446 | 0.011962674 | 3.805875455 | 0.0001413 |
| B180 | Tf | Hypertension | 0.025139805 | 0.00642082 | 3.91535768 | 9.03E-05 |
| B180 | Tf | Hypertension+Diabetes | 0.235212078 | 0.014066959 | 16.72089039 | 9.45E-63 |
| B181 | APTT | (Intercept) | 26.49553672 | 0.001614575 | 16410.22074 | 0 |
| B181 | APTT | Gender | 0.007519058 | 0.000713351 | 10.54047447 | 5.64E-26 |
| B181 | APTT | Age | -0.0001443 | 2.74E-05 | -5.275812997 | 1.32E-07 |
| B181 | APTT | Hypothyroidism | 0.025797605 | 0.008019246 | 3.216961287 | 0.0012956 |
| B181 | APTT | Hypertension | -0.00574602 | 0.001648063 | -3.486527473 | 0.0004894 |
| B181 | APTT | Hepatopathy | 0.091460318 | 0.024319125 | 3.760839244 | 0.0001694 |
| B182 | TT | (Intercept) | 18.85279165 | 0.00066805 | 28220.63533 | 0 |
| B182 | TT | Gender | -0.00234201 | 0.000295158 | -7.934782697 | 2.11E-15 |
| B182 | TT | Rhinallergosis | 0.042641865 | 0.010807844 | 3.945455156 | 7.97E-05 |
| B183 | PTA | (Intercept) | 27.65510108 | 0.020785182 | 1330.52001 | 0 |
| B183 | PTA | Hyperlipidemia | 0.478775646 | 0.10165719 | 4.709707671 | 2.48E-06 |
| B184 | INR | (Intercept) | 0.979965115 | 0.000765059 | 1280.901757 | 0 |
| B185 | PT | (Intercept) | 10.71616535 | 0.000497701 | 21531.32452 | 0 |
| B185 | PT | Gender | 0.001336364 | 0.000219894 | 6.077305747 | 1.22E-09 |
| B185 | PT | Age | 4.21E-05 | 8.43E-06 | 4.990636425 | 6.02E-07 |
| B185 | PT | Hepatitis B | 0.01278217 | 0.00378674 | 3.375508029 | 0.0007368 |
| B185 | PT | Other Cardiac diseases | 0.035104108 | 0.005425213 | 6.470549794 | 9.77E-11 |
| B185 | PT | Heart disease | 0.028419117 | 0.006650308 | 4.273353496 | 1.93E-05 |
| B185 | PT | Coronary heart disease | 0.013964669 | 0.002774709 | 5.032840469 | 4.83E-07 |
| B186 | PFC | (Intercept) | 2.617013713 | 0.000316059 | 8280.133517 | 0 |
| B186 | PFC | Gender | 0.000559353 | 0.000139641 | 4.005643355 | 6.19E-05 |
| B186 | PFC | Age | 4.79E-05 | 5.35E-06 | 8.948848262 | 3.60E-19 |
| B186 | PFC | Stomach trouble | -0.00597559 | 0.001775878 | -3.364866019 | 0.0007658 |
| B186 | PFC | Hepatitis B | -0.00906167 | 0.002404725 | -3.768278745 | 0.0001644 |
| B186 | PFC | Hypertension | 0.00156261 | 0.000322615 | 4.843578489 | 1.28E-06 |
| B186 | PFC | Hypertension+Diabetes | 0.003717615 | 0.000706796 | 5.259815975 | 1.44E-07 |
| B187 | NSE | (Intercept) | 13.43296953 | 0.010146827 | 1323.859165 | 0 |

| | | | | | | |
|---|---|---|---|---|---|---|
| B187 | NSE | Gender | -0.02817026 | 0.004483067 | -6.283703062 | 3.31E-10 |
| B187 | NSE | Age | 0.000635145 | 0.000171887 | 3.69512278 | 0.0002198 |
| B187 | NSE | Stomach trouble | 0.239769805 | 0.057013117 | 4.205520054 | 2.61E-05 |
| B188 | Ca | (Intercept) | 2.423776362 | 0.001619196 | 1496.901298 | 0 |
| B188 | Ca | Gender | -0.01800571 | 0.000715392 | -25.16899606 | 9.85E-140 |
| B188 | Ca | Age | -0.00095693 | 2.74E-05 | -34.88734524 | 1.78E-266 |
| B188 | Ca | Diabetes | 0.012818985 | 0.003079305 | 4.162947866 | 3.14E-05 |
| B188 | Ca | Hypertension | 0.009985747 | 0.001652779 | 6.04179044 | 1.52E-09 |
| B188 | Ca | Hypertension+Diabetes | 0.02227848 | 0.003620968 | 6.152631689 | 7.62E-10 |
| B189 | eGFB | (Intercept) | 125.504235 | 0.051642262 | 2430.262172 | 0 |
| B189 | eGFB | Gender | 4.674878243 | 0.022816565 | 204.8896638 | 0 |
| B189 | eGFB | Age | -0.61586271 | 0.00087482 | -703.987881 | 0 |
| B189 | eGFB | Hyperthyroidism | 1.226588439 | 0.376985908 | 3.253671859 | 0.0011393 |
| B189 | eGFB | Nephropathy | -13.4472105 | 0.434959584 | -30.91599999 | 9.51E-210 |
| B189 | eGFB | Diabetes | 2.746009186 | 0.098210641 | 27.96040374 | 5.89E-172 |
| B189 | eGFB | Gout | -5.68461806 | 0.226356214 | -25.11359404 | 3.97E-139 |
| B189 | eGFB | Stomach trouble | 1.309185028 | 0.290168185 | 4.511814517 | 6.43E-06 |
| B189 | eGFB | Lumbar disc protrusion | 2.976848649 | 0.532072302 | 5.5948198 | 2.21E-08 |
| B189 | eGFB | Hypertension+other diseases | -3.88502416 | 0.213881918 | -18.16434134 | 1.02E-73 |
| B189 | eGFB | Other disease conditions | 1.056754913 | 0.198552506 | 5.322294524 | 1.02E-07 |
| B189 | eGFB | Hypertension | -1.55088621 | 0.052713367 | -29.42111836 | 3.67E-190 |
| B189 | eGFB | Hepatic adipose infiltration | 2.678283493 | 0.630503814 | 4.24784662 | 2.16E-05 |
| B189 | eGFB | Other Cardiac diseases | -2.47472987 | 0.562928657 | -4.396169642 | 1.10E-05 |
| B189 | eGFB | Heart disease | -3.86799547 | 0.690046491 | -5.605412855 | 2.08E-08 |
| B189 | eGFB | Pregnant | 8.213947147 | 1.394439739 | 5.890499904 | 3.85E-09 |
| B189 | eGFB | Hypertension+Diabetes | -2.06235567 | 0.115486312 | -17.85800965 | 2.58E-71 |
| B189 | eGFB | Hypertension+Coronary heart disease | -4.02944209 | 0.231183993 | -17.42958951 | 5.06E-68 |
| B189 | eGFB | Hypertensive+Diabetes+Coronary Heart Disease | -4.46156951 | 0.343865137 | -12.97476548 | 1.72E-38 |
| B189 | eGFB | Hyperlipidemia | -3.45150455 | 0.252574507 | -13.66529264 | 1.65E-42 |
| B189 | eGFB | Coronary heart disease | -1.37213492 | 0.28790822 | -4.765876161 | 1.88E-06 |
| B190 | Cre | (Intercept) | 93.23874133 | 0.077687353 | 1200.179147 | 0 |
| B190 | Cre | Gender | -20.2944852 | 0.034323797 | -591.2657383 | 0 |
| B190 | Cre | Age | 0.107574312 | 0.001316024 | 81.74190993 | 0 |
| B190 | Cre | Hypothyroidism | 1.236185877 | 0.385856285 | 3.203746897 | 0.0013566 |
| B190 | Cre | Nephropathy | 53.78668523 | 0.654325696 | 82.20170102 | 0 |
| B190 | Cre | Diabetes | -4.11362658 | 0.147741879 | -27.8433346 | 1.55E-170 |
| B190 | Cre | Gout | 8.620716447 | 0.340515976 | 25.31662846 | 2.36E-141 |
| B190 | Cre | Hypertension+other diseases | 15.4815181 | 0.321750434 | 48.11654151 | 0 |
| B190 | Cre | Coronary heart disease+other diseases | 8.003861586 | 1.562371927 | 5.122891321 | 3.01E-07 |
| B190 | Cre | Diabetes+other diseases | 2.616756314 | 0.813431573 | 3.216934776 | 0.0012957 |
| B190 | Cre | Hypertension | 2.647701375 | 0.079298656 | 33.38898153 | 2.85E-244 |
| B190 | Cre | Other Cardiac diseases | 3.345807122 | 0.846834279 | 3.950958534 | 7.78E-05 |
| B190 | Cre | Heart disease | 10.24792792 | 1.038062311 | 9.872170301 | 5.51E-23 |

| | | | | | | |
|---|---|---|---|---|---|---|
| B190 | Cre | Pregnant | -8.69853099 | 2.097706976 | -4.146685448 | 3.37E-05 |
| B190 | Cre | Hypertension+Diabetes | 5.242407144 | 0.173730306 | 30.17554781 | 6.32E-200 |
| B190 | Cre | Hypertension+Coronary heart disease | 6.826699425 | 0.347778581 | 19.6294418 | 9.05E-86 |
| B190 | Cre | Hypertensive+Diabetes+Coronary Heart Disease | 9.353999631 | 0.51728897 | 18.08273551 | 4.50E-73 |
| B190 | Cre | Hyperlipidemia | 2.461919924 | 0.379957118 | 6.479467826 | 9.21E-11 |
| B190 | Cre | Coronary heart disease | 1.71304236 | 0.433110923 | 3.955204706 | 7.65E-05 |
| B191 | Urea | (Intercept) | 4.83489349 | 0.006389993 | 756.6351467 | 0 |
| B191 | Urea | Gender | -0.44549457 | 0.002823224 | -157.7963737 | 0 |
| B191 | Urea | Age | 0.019259045 | 0.000108247 | 177.9184051 | 0 |
| B191 | Urea | Nephropathy | 1.643571977 | 0.053820044 | 30.53828735 | 1.04E-204 |
| B191 | Urea | Diabetes | 0.219449684 | 0.012152166 | 18.05848307 | 6.98E-73 |
| B191 | Urea | Parkinson's syndrome | 0.363367512 | 0.092050416 | 3.947483651 | 7.90E-05 |
| B191 | Urea | Hypertension+other diseases | 0.417027528 | 0.026464837 | 15.75779722 | 6.18E-56 |
| B191 | Urea | Diabetes+other diseases | 0.397296671 | 0.06690693 | 5.938049667 | 2.89E-09 |
| B191 | Urea | Hypertension | 0.102843725 | 0.006522527 | 15.76746566 | 5.31E-56 |
| B191 | Urea | Heart disease | 0.631095029 | 0.085383411 | 7.391307337 | 1.46E-13 |
| B191 | Urea | Pregnant | -1.2528916 | 0.172542028 | -7.261370559 | 3.84E-13 |
| B191 | Urea | Hypertension+Diabetes | 0.513295158 | 0.014289784 | 35.92042853 | 2.39E-282 |
| B191 | Urea | Hypertension+Coronary heart disease | 0.213378323 | 0.028605722 | 7.459288277 | 8.71E-14 |
| B191 | Urea | Hypertensive+Diabetes+Coronary Heart Disease | 0.636088296 | 0.042548406 | 14.94975623 | 1.59E-50 |
| B191 | Urea | Coronary heart disease+Diabetes | 0.35069626 | 0.077233791 | 4.540710141 | 5.61E-06 |
| B192 | UA | (Intercept) | 496.6051344 | 0.347268188 | 1430.033476 | 0 |
| B192 | UA | Gender | -96.0645655 | 0.1534299 | -626.1137206 | 0 |
| B192 | UA | Age | -0.26422165 | 0.005882724 | -44.91484765 | 0 |
| B192 | UA | Rheumatoid arthritis | -20.3906787 | 3.527996335 | -5.779676841 | 7.49E-09 |
| B192 | UA | Nephropathy | 34.81904494 | 2.924884033 | 11.90441896 | 1.13E-32 |
| B192 | UA | Diabetes | -12.5528899 | 0.660417077 | -19.00751859 | 1.54E-80 |
| B192 | UA | Gout | 111.746858 | 1.522131481 | 73.414721 | 0 |
| B192 | UA | Parkinson's syndrome | -35.2126279 | 5.002537541 | -7.038953244 | 1.94E-12 |
| B192 | UA | Stomach trouble | -16.5492578 | 1.951234832 | -8.481428042 | 2.23E-17 |
| B192 | UA | Hepatitis B | -14.356383 | 2.642176177 | -5.433544925 | 5.53E-08 |
| B192 | UA | Hypertension+other diseases | 34.96351431 | 1.438248126 | 24.30979305 | 1.71E-130 |
| B192 | UA | Hypertension | 26.70132943 | 0.354470832 | 75.32729635 | 0 |
| B192 | UA | Hepatic adipose infiltration | 33.8177608 | 4.239820449 | 7.976224748 | 1.51E-15 |
| B192 | UA | Asthma | 9.561219309 | 2.452265864 | 3.898932595 | 9.66E-05 |
| B192 | UA | Other Cardiac diseases | 24.44142939 | 3.785411572 | 6.456742926 | 1.07E-10 |
| B192 | UA | Heart disease | 31.51351338 | 4.640214955 | 6.791390849 | 1.11E-11 |
| B192 | UA | Pregnant | -30.1775027 | 9.376904616 | -3.218279795 | 0.0012897 |
| B192 | UA | Hypertension+Diabetes | 22.52838578 | 0.776587256 | 29.00947137 | 6.15E-185 |
| B192 | UA | Hypertension+Coronary heart disease | 31.60664916 | 1.554595863 | 20.33110335 | 7.18E-92 |
| B192 | UA | Hypertensive+Diabetes+Coronary Heart Disease | 20.68683237 | 2.31231978 | 8.946354456 | 3.68E-19 |

| | | | | | | |
|---|---|---|---|---|---|---|
| B192 | UA | Hyperlipidemia | 13.55457302 | 1.698436292 | 7.980619046 | 1.46E-15 |
| B193 | K | (Intercept) | 4.066542121 | 0.000330953 | 12287.37427 | 0 |
| B193 | K | Gender | 0.000764295 | 0.000146221 | 5.226968328 | 1.72E-07 |
| B193 | K | Age | 6.56E-05 | 5.61E-06 | 11.6937547 | 1.38E-31 |
| B193 | K | Cervical spondylopathy | -0.01497294 | 0.003751137 | -3.991573778 | 6.56E-05 |
| B193 | K | Hepatitis B | -0.01584438 | 0.002518042 | -6.292339699 | 3.13E-10 |
| B193 | K | Hypertension+other diseases | -0.0088144 | 0.001370677 | -6.430688914 | 1.27E-10 |
| B193 | K | Hypertension | -0.00785067 | 0.000337817 | -23.23939167 | 1.98E-119 |
| B193 | K | Hypertension+Diabetes | -0.00326331 | 0.000740102 | -4.40927802 | 1.04E-05 |
| B194 | Cl | (Intercept) | 104.3894616 | 0.002417023 | 43189.26794 | 0 |
| B194 | Cl | Gender | 0.021386144 | 0.001067888 | 20.02657425 | 3.39E-89 |
| B194 | Cl | Age | 0.000719098 | 4.09E-05 | 17.56279346 | 4.88E-69 |
| B194 | Cl | Diabetes | -0.06007457 | 0.004596572 | -13.06942691 | 4.97E-39 |
| B194 | Cl | Hepatitis B | 0.06986079 | 0.018389825 | 3.798882915 | 0.0001454 |
| B194 | Cl | Hypertension | -0.01979776 | 0.002467154 | -8.024530597 | 1.02E-15 |
| B194 | Cl | Hypertension+Diabetes | -0.06359952 | 0.005405129 | -11.76651233 | 5.84E-32 |
| B194 | Cl | Hypertensive+Diabetes+Coronary Heart Disease | -0.06434211 | 0.016093989 | -3.997896589 | 6.39E-05 |
| B194 | Cl | Coronary heart disease+Diabetes | -0.11054075 | 0.029213781 | -3.783856404 | 0.0001544 |
| B195 | Na | (Intercept) | 141.3972769 | 0.002171545 | 65113.67638 | 0 |
| B195 | Na | Gender | -0.01473048 | 0.000959431 | -15.35334423 | 3.42E-53 |
| B195 | Na | Age | 0.000776008 | 3.68E-05 | 21.09520852 | 9.33E-99 |
| B195 | Na | Diabetes | -0.04334753 | 0.004129734 | -10.49644477 | 9.00E-26 |
| B195 | Na | Hypertension+other diseases | 0.035723511 | 0.008993684 | 3.972066338 | 7.13E-05 |
| B195 | Na | Hypertension | 0.011796541 | 0.002216585 | 5.321944484 | 1.03E-07 |
| B195 | Na | Hypertension+Diabetes | -0.02838236 | 0.004856172 | -5.844595339 | 5.08E-09 |
| B195 | Na | Hypertensive+Diabetes+Coronary Heart Disease | -0.06022781 | 0.014459448 | -4.16529099 | 3.11E-05 |
| B196 | TCO2 | (Intercept) | 26.43020406 | 0.002916879 | 9061.125131 | 0 |
| B196 | TCO2 | Gender | -0.00945646 | 0.001288734 | -7.337788959 | 2.17E-13 |
| B196 | TCO2 | Age | 0.000305599 | 4.94E-05 | 6.184711644 | 6.22E-10 |
| B197 | AG | (Intercept) | 10.57704226 | 0.003661054 | 2889.070493 | 0 |
| B197 | AG | Gender | -0.0264894 | 0.001617525 | -16.376497 | 2.88E-60 |
| B197 | AG | Age | -0.00024677 | 6.20E-05 | -3.978932699 | 6.92E-05 |
| B197 | AG | Stomach trouble | -0.0698323 | 0.020570774 | -3.394733421 | 0.000687 |
| B197 | AG | Chronic pharyngitis | 0.085456569 | 0.026465937 | 3.228926707 | 0.0012426 |
| B197 | AG | Hypertension+other diseases | 0.054765393 | 0.015162643 | 3.611863182 | 0.000304 |
| B197 | AG | Hypertension | 0.023454283 | 0.003736987 | 6.276254655 | 3.47E-10 |
| B197 | AG | Hypertension+Diabetes | 0.053036641 | 0.008187124 | 6.478055262 | 9.30E-11 |
| B198 | P | (Intercept) | 1.025998405 | 0.000155252 | 6608.597648 | 0 |
| B198 | P | Gender | 0.003638952 | 6.86E-05 | 53.0510471 | 0 |
| B198 | P | Age | -3.07E-05 | 2.63E-06 | -11.6663021 | 1.91E-31 |
| B198 | P | Diabetes | 0.001726999 | 0.000295251 | 5.849268221 | 4.94E-09 |
| B199 | Mg | (Intercept) | 0.86948253 | 7.30E-05 | 11918.61188 | 0 |

| | | | | | | |
|---|---|---|---|---|---|---|
| B199 | Mg | Gender | -0.00024568 | 3.22E-05 | -7.622381957 | 2.49E-14 |
| B199 | Mg | Age | 1.76E-05 | 1.24E-06 | 14.23647306 | 5.51E-46 |
| B199 | Mg | Cervical spondylopathy | -0.00273479 | 0.00082686 | -3.307441627 | 0.0009416 |
| B199 | Mg | Diabetes | -0.0021056 | 0.000138736 | -15.17702185 | 5.10E-52 |
| B199 | Mg | Gout | -0.00120015 | 0.000319759 | -3.753282467 | 0.0001745 |
| B199 | Mg | Hepatitis B | -0.00269661 | 0.00055505 | -4.858322294 | 1.18E-06 |
| B199 | Mg | Diabetes+other diseases | -0.00328182 | 0.000763846 | -4.296436171 | 1.74E-05 |
| B199 | Mg | Hypertension+Diabetes | -0.00262467 | 0.00016314 | -16.08845743 | 3.14E-58 |
| B199 | Mg | Hypertensive+Diabetes+Coronary Heart Disease | -0.00158518 | 0.000485756 | -3.263333418 | 0.0011011 |
| B200 | OP | (Intercept) | 282.4329022 | 0.00502425 | 56213.94083 | 0 |
| B200 | OP | Gender | -0.04801719 | 0.002219812 | -21.63119448 | 9.74E-104 |
| B200 | OP | Age | 0.00185557 | 8.51E-05 | 21.80181267 | 2.38E-105 |
| B200 | OP | Diabetes | 0.129203851 | 0.009554865 | 13.5223106 | 1.17E-41 |
| B200 | OP | Hypertension+other diseases | 0.115816514 | 0.020808466 | 5.565836148 | 2.61E-08 |
| B200 | OP | Hypertension | 0.035759167 | 0.005128457 | 6.972694442 | 3.11E-12 |
| B200 | OP | Hypertension+Diabetes | 0.123817355 | 0.011235606 | 11.02008661 | 3.07E-28 |
| B200 | OP | Coronary heart disease+Diabetes | 0.202493402 | 0.060726494 | 3.334514917 | 0.0008545 |
| B201 | e-habits | (Intercept) | 2.045813667 | 0.001768585 | 1156.75184 | 0 |
| B201 | e-habits | Gender | -0.03124466 | 0.000781395 | -39.98572417 | 0 |
| B201 | e-habits | Age | 0.002450691 | 3.00E-05 | 81.7992152 | 0 |
| B201 | e-habits | Rhinallergosis | 0.163665095 | 0.028612518 | 5.720052251 | 1.07E-08 |
| B201 | e-habits | Hypothyroidism | 0.037988189 | 0.008784178 | 4.324615092 | 1.53E-05 |
| B201 | e-habits | Cervical spondylopathy | 0.146452788 | 0.020045764 | 7.305921907 | 2.76E-13 |
| B201 | e-habits | Chronic rhinitis | 0.07604395 | 0.019980205 | 3.805964463 | 0.0001413 |
| B201 | e-habits | Diabetes | 0.037663017 | 0.003363405 | 11.1978834 | 4.20E-29 |
| B201 | e-habits | Gout | 0.056917114 | 0.007751987 | 7.342261195 | 2.10E-13 |
| B201 | e-habits | Cholecystolithiasis | 0.103977467 | 0.011407352 | 9.114951856 | 7.89E-20 |
| B201 | e-habits | Stomach trouble | 0.097129541 | 0.009937346 | 9.774193209 | 1.46E-22 |
| B201 | e-habits | Chronic pharyngitis | 0.112223031 | 0.012785186 | 8.777583164 | 1.67E-18 |
| B201 | e-habits | Lumbar disc protrusion | 0.147698857 | 0.0182218 | 8.105612894 | 5.25E-16 |
| B201 | e-habits | Hypertension+other diseases | 0.077162647 | 0.007324782 | 10.53446353 | 6.01E-26 |
| B201 | e-habits | Other disease conditions | 0.086235851 | 0.006799798 | 12.68212 | 7.49E-37 |
| B201 | e-habits | Brain diseases | 0.147524906 | 0.022309993 | 6.612503398 | 3.78E-11 |
| B201 | e-habits | Hypertension | 0.015444547 | 0.001805267 | 8.555271469 | 1.18E-17 |
| B201 | e-habits | Hepatopathy | 0.089110256 | 0.026638853 | 3.345123658 | 0.0008225 |
| B201 | e-habits | Hypertension+Diabetes | 0.027101286 | 0.003955042 | 6.852337705 | 7.27E-12 |
| B201 | e-habits | Hypertension+Coronary heart disease | 0.031195736 | 0.007917323 | 3.940187185 | 8.14E-05 |
| B201 | e-habits | Hyperlipidemia | 0.308070977 | 0.008649881 | 35.61563155 | 1.28E-277 |
| B201 | e-habits | Coronary heart disease | 0.057974977 | 0.009859949 | 5.879845281 | 4.11E-09 |
| B202 | e-times | (Intercept) | 3.24 | 1.51E-13 | 2.15E+13 | 0 |
| B203 | DH | (Intercept) | 2.374534659 | 0.003440044 | 690.2627824 | 0 |
| B203 | DH | Gender | -0.03528739 | 0.001519879 | -23.21723286 | 3.32E-119 |
| B203 | DH | Age | -0.00473002 | 5.83E-05 | -81.16802929 | 0 |

| | | | | | | |
|---|---|---|---|---|---|---|
| B203 | DH | Rhinallergosis | 0.415823498 | 0.055653727 | 7.471620017 | 7.93E-14 |
| B203 | DH | Hypothyroidism | -0.06090138 | 0.017085957 | -3.564411147 | 0.0003647 |
| B203 | DH | Cervical spondylopathy | 0.140498903 | 0.038990679 | 3.603397188 | 0.0003141 |
| B203 | DH | Chronic rhinitis | 0.198917638 | 0.03886316 | 5.118411251 | 3.08E-07 |
| B203 | DH | Cholecystolithiasis | 0.190394921 | 0.022188249 | 8.58088979 | 9.43E-18 |
| B203 | DH | Chronic pharyngitis | 0.223438812 | 0.02486825 | 8.984902969 | 2.59E-19 |
| B203 | DH | Other disease conditions | 0.058101279 | 0.013226172 | 4.392902031 | 1.12E-05 |
| B203 | DH | Hypertension | 0.017306262 | 0.003511394 | 4.928601735 | 8.28E-07 |
| B203 | DH | Hepatic adipose infiltration | 0.233057342 | 0.041999732 | 5.549019738 | 2.87E-08 |
| B203 | DH | Pregnant | 0.532303113 | 0.092887773 | 5.730604725 | 1.00E-08 |
| B204 | Drinking | (Intercept) | 2.735628848 | 0.00508733 | 537.7337587 | 0 |
| B204 | Drinking | Gender | -0.86632646 | 0.002247682 | -385.431058 | 0 |
| B204 | Drinking | Age | 0.001942683 | 8.62E-05 | 22.54231575 | 1.72E-112 |
| B204 | Drinking | Nephropathy | -0.23419485 | 0.042848293 | -5.465675222 | 4.61E-08 |
| B204 | Drinking | Diabetes | -0.09484561 | 0.009674826 | -9.803339841 | 1.09E-22 |
| B204 | Drinking | Gout | 0.093457635 | 0.022298571 | 4.191193858 | 2.78E-05 |
| B204 | Drinking | Parkinson's syndrome | -0.62535244 | 0.073285023 | -8.533154764 | 1.43E-17 |
| B204 | Drinking | Stomach trouble | 0.119579076 | 0.028584751 | 4.183317065 | 2.87E-05 |
| B204 | Drinking | Chronic pharyngitis | 0.150369702 | 0.036776555 | 4.088738169 | 4.34E-05 |
| B204 | Drinking | Hypertension+other diseases | -0.08075043 | 0.021069716 | -3.832535343 | 0.0001268 |
| B204 | Drinking | Coronary heart disease+other diseases | -0.57466611 | 0.102311387 | -5.616834335 | 1.95E-08 |
| B204 | Drinking | Diabetes+other diseases | -0.2967789 | 0.053267286 | -5.571504094 | 2.53E-08 |
| B204 | Drinking | Brain diseases | -0.30754073 | 0.064174639 | -4.792247126 | 1.65E-06 |
| B204 | Drinking | Hypertension | -0.02320905 | 0.005192845 | -4.469428126 | 7.84E-06 |
| B204 | Drinking | Hepatic adipose infiltration | 0.475857811 | 0.062111546 | 7.661342286 | 1.84E-14 |
| B204 | Drinking | Other Cardiac diseases | -0.21513886 | 0.055454651 | -3.879545807 | 0.0001047 |
| B204 | Drinking | Heart disease | -0.30672306 | 0.067977153 | -4.512149272 | 6.42E-06 |
| B204 | Drinking | Hypertension+Diabetes | -0.15689065 | 0.011376669 | -13.79056068 | 2.94E-43 |
| B204 | Drinking | Hypertension+Coronary heart disease | -0.25221377 | 0.022774161 | -11.07455815 | 1.67E-28 |
| B204 | Drinking | Hypertensive+Diabetes+Coronary Heart Disease | -0.27774016 | 0.03387449 | -8.199094908 | 2.43E-16 |
| B204 | Drinking | Hyperlipidemia | 1.165407603 | 0.024881361 | 46.83857938 | 0 |
| B204 | Drinking | Coronary heart disease | -0.28732078 | 0.02836212 | -10.13044111 | 4.06E-24 |
| B204 | Drinking | Coronary heart disease+Diabetes | -0.4189378 | 0.061488915 | -6.813224784 | 9.55E-12 |
| B205 | KDA | (Intercept) | 0.885653748 | 0.004665861 | 189.8157278 | 0 |
| B205 | KDA | Gender | 0.109493097 | 0.002061469 | 53.11411764 | 0 |
| B205 | KDA | Age | 0.004033484 | 7.90E-05 | 51.03111773 | 0 |
| B205 | KDA | Hypothyroidism | 0.166846929 | 0.023174321 | 7.19964684 | 6.04E-13 |
| B205 | KDA | Hyperthyroidism | 0.123211798 | 0.034060548 | 3.617434395 | 0.0002976 |
| B205 | KDA | Cervical spondylopathy | 0.209736172 | 0.052884513 | 3.965928 | 7.31E-05 |
| B205 | KDA | Nephropathy | 0.264214719 | 0.03929845 | 6.723286017 | 1.78E-11 |
| B205 | KDA | Gout | 0.159287333 | 0.020451207 | 7.788652064 | 6.78E-15 |
| B205 | KDA | Stomach trouble | 0.230896335 | 0.026216596 | 8.807258242 | 1.28E-18 |
| B205 | KDA | Chronic pharyngitis | 0.168051452 | 0.033729736 | 4.982293777 | 6.28E-07 |

| | | | | | | |
|---|---|---|---|---|---|---|
| B205 | KDA | Lumbar disc protrusion | 0.31535105 | 0.048072551 | 6.559898398 | 5.39E-11 |
| B205 | KDA | Hypertension+other diseases | 0.153576691 | 0.019324158 | 7.947393571 | 1.91E-15 |
| B205 | KDA | Coronary heart disease+other diseases | 0.361752969 | 0.093835219 | 3.855193951 | 0.0001156 |
| B205 | KDA | Bronchial disease | 0.224548179 | 0.03939248 | 5.700280273 | 1.20E-08 |
| B205 | KDA | Other disease conditions | 0.190933488 | 0.017939151 | 10.64339592 | 1.88E-26 |
| B205 | KDA | Hypertension | 0.052831898 | 0.004762635 | 11.09299816 | 1.36E-28 |
| B205 | KDA | Asthma | 0.524071512 | 0.032948399 | 15.90582614 | 5.88E-57 |
| B205 | KDA | Other Cardiac diseases | 0.180268665 | 0.050860412 | 3.544380734 | 0.0003936 |
| B205 | KDA | Heart disease | 0.252127869 | 0.062345465 | 4.044045054 | 5.25E-05 |
| B205 | KDA | Hypertension+Diabetes | 0.095322433 | 0.010434149 | 9.135621351 | 6.52E-20 |
| B205 | KDA | Hypertension+Coronary heart disease | 0.243387535 | 0.020887395 | 11.65236436 | 2.24E-31 |
| B205 | KDA | Hypertensive+Diabetes+Coronary Heart Disease | 0.132539206 | 0.031068098 | 4.266086949 | 1.99E-05 |
| B205 | KDA | Hyperlipidemia | 0.287250257 | 0.022820021 | 12.58764236 | 2.49E-36 |
| B205 | KDA | Coronary heart disease | 0.173112987 | 0.026012409 | 6.655015457 | 2.83E-11 |
| B205 | KDA | Coronary heart disease+Diabetes | 0.207348071 | 0.056394756 | 3.676726059 | 0.0002363 |
| B206 | Waist | (Intercept) | 87.79307359 | 0.040211749 | 2183.269239 | 0 |
| B206 | Waist | Gender | -10.2229022 | 0.01776634 | -575.4084641 | 0 |
| B206 | Waist | Age | 0.155595883 | 0.000681187 | 228.418709 | 0 |
| B206 | Waist | Hypothyroidism | 1.352609533 | 0.19972306 | 6.772425444 | 1.27E-11 |
| B206 | Waist | Hyperthyroidism | 0.951015558 | 0.293543738 | 3.239774643 | 0.0011963 |
| B206 | Waist | Diabetes | 1.428796447 | 0.07647267 | 18.6837527 | 6.96E-78 |
| B206 | Waist | Gout | 3.914511904 | 0.176254464 | 22.20943413 | 2.99E-109 |
| B206 | Waist | Parkinson's syndrome | -3.15185049 | 0.579266366 | -5.441107357 | 5.30E-08 |
| B206 | Waist | Cholecystolithiasis | 1.229904828 | 0.259365339 | 4.741978376 | 2.12E-06 |
| B206 | Waist | Stomach trouble | -1.76430795 | 0.225942275 | -7.808666853 | 5.79E-15 |
| B206 | Waist | Hypertension+other diseases | 3.108539106 | 0.166541232 | 18.66528226 | 9.83E-78 |
| B206 | Waist | Diabetes+other diseases | 1.799159771 | 0.421040291 | 4.273129692 | 1.93E-05 |
| B206 | Waist | Other disease conditions | 0.622284728 | 0.154604837 | 4.025001686 | 5.70E-05 |
| B206 | Waist | Brain diseases | -2.02356388 | 0.507255214 | -3.989242149 | 6.63E-05 |
| B206 | Waist | Hypertension | 3.340977112 | 0.041045775 | 81.39637032 | 0 |
| B206 | Waist | Hepatic adipose infiltration | 7.273957162 | 0.490947917 | 14.81614833 | 1.17E-49 |
| B206 | Waist | Asthma | 1.630686891 | 0.283958916 | 5.742686003 | 9.32E-09 |
| B206 | Waist | Hepatopathy | 2.113816125 | 0.605679114 | 3.489993423 | 0.0004831 |
| B206 | Waist | Pregnant | 6.765251161 | 1.085794043 | 6.230694672 | 4.65E-10 |
| B206 | Waist | Hypertension+Diabetes | 3.772186385 | 0.089924538 | 41.94835427 | 0 |
| B206 | Waist | Hypertension+Coronary heart disease | 1.872773349 | 0.180013661 | 10.40350682 | 2.40E-25 |
| B206 | Waist | Hypertensive+Diabetes+Coronary Heart Disease | 2.659801362 | 0.267753928 | 9.933752911 | 2.98E-23 |
| B206 | Waist | Hyperlipidemia | 1.399889792 | 0.196669592 | 7.117977798 | 1.10E-12 |
| B206 | Waist | Coronary heart disease | -0.90069507 | 0.224182531 | -4.017686229 | 5.88E-05 |
| B207 | WHR | (Intercept) | 0.882361812 | 0.002927709 | 301.3830149 | 0 |
| B207 | WHR | Gender | -0.07141942 | 0.001293519 | -55.21326245 | 0 |
| B207 | WHR | Age | 0.0016721 | 4.96E-05 | 33.71482286 | 5.12E-249 |

| | | | | | | |
|---|---|---|---|---|---|---|
| B207 | WHR | Diabetes | 0.018059824 | 0.005567769 | 3.243637392 | 0.0011802 |
| B207 | WHR | Hypertension | 0.019740422 | 0.002988432 | 6.605611201 | 3.96E-11 |
| B207 | WHR | Hypertension+Diabetes | 0.023631632 | 0.006547164 | 3.609445875 | 0.0003069 |
| B209 | Smoking | (Intercept) | 5.298185967 | 0.01939487 | 273.1746013 | 0 |
| B209 | Smoking | Gender | -3.08750539 | 0.008569034 | -360.3096088 | 0 |
| B209 | Smoking | Age | 0.04625438 | 0.000328549 | 140.7837434 | 0 |
| B209 | Smoking | Chronic rhinitis | -0.73802769 | 0.219109365 | -3.368307393 | 0.0007563 |
| B209 | Smoking | Diabetes | 0.255323625 | 0.036884183 | 6.922306552 | 4.45E-12 |
| B209 | Smoking | Parkinson's syndrome | -3.12365889 | 0.279390886 | -11.1802462 | 5.12E-29 |
| B209 | Smoking | Hypertension+other diseases | -0.67909026 | 0.080325918 | -8.454186081 | 2.81E-17 |
| B209 | Smoking | Coronary heart disease+other diseases | -1.73483127 | 0.390050628 | -4.44770792 | 8.68E-06 |
| B209 | Smoking | Bronchial disease | -0.78381248 | 0.163745147 | -4.78678298 | 1.70E-06 |
| B209 | Smoking | Other disease conditions | -0.27602189 | 0.074568773 | -3.701574769 | 0.0002143 |
| B209 | Smoking | Brain diseases | -1.10707956 | 0.244658575 | -4.524997974 | 6.04E-06 |
| B209 | Smoking | Hypertension | -0.44989046 | 0.019797137 | -22.72502645 | 2.74E-114 |
| B209 | Smoking | Asthma | -0.87420103 | 0.136958639 | -6.382956462 | 1.74E-10 |
| B209 | Smoking | Other Cardiac diseases | -1.006986 | 0.211414604 | -4.763086288 | 1.91E-06 |
| B209 | Smoking | Heart disease | -1.64701019 | 0.25915523 | -6.355303681 | 2.08E-10 |
| B209 | Smoking | Hypertension+Diabetes | -0.66172736 | 0.043372269 | -15.25692292 | 1.50E-52 |
| B209 | Smoking | Hypertension+Coronary heart disease | -1.84840613 | 0.086823919 | -21.28913484 | 1.52E-100 |
| B209 | Smoking | Hypertensive+Diabetes+Coronary Heart Disease | -1.40373478 | 0.129142674 | -10.86964319 | 1.61E-27 |
| B209 | Smoking | Hyperlipidemia | 1.580163793 | 0.094857383 | 16.65831102 | 2.69E-62 |
| B209 | Smoking | Coronary heart disease | -1.07480476 | 0.108127383 | -9.940171794 | 2.79E-23 |
| B209 | Smoking | Coronary heart disease+Diabetes | -1.15483875 | 0.234419554 | -4.926375503 | 8.38E-07 |
| B210 | Hip | (Intercept) | 100.7433562 | 0.029473942 | 3418.048247 | 0 |
| B210 | Hip | Gender | -4.58689571 | 0.013022166 | -352.2375402 | 0 |
| B210 | Hip | Hypothyroidism | 0.806401946 | 0.146390697 | 5.508560037 | 3.62E-08 |
| B210 | Hip | Diabetes | -0.45010907 | 0.056052052 | -8.030197833 | 9.74E-16 |
| B210 | Hip | Gout | 1.706808709 | 0.129188957 | 13.21172292 | 7.58E-40 |
| B210 | Hip | Parkinson's syndrome | -2.05949416 | 0.424583958 | -4.850616976 | 1.23E-06 |
| B210 | Hip | Cholecystolithiasis | 1.023849614 | 0.190106605 | 5.385660407 | 7.22E-08 |
| B210 | Hip | Stomach trouble | -1.47336268 | 0.165608554 | -8.896658087 | 5.77E-19 |
| B210 | Hip | Hypertension+other diseases | 1.541310811 | 0.122069465 | 12.6265058 | 1.52E-36 |
| B210 | Hip | Bronchial disease | -0.90935388 | 0.248839766 | -3.654375257 | 0.0002578 |
| B210 | Hip | Hypertension | 1.670662003 | 0.030085257 | 55.5309196 | 0 |
| B210 | Hip | Hepatic adipose infiltration | 3.931579558 | 0.359849323 | 10.92562722 | 8.73E-28 |
| B210 | Hip | Hypertension+Diabetes | 1.047760638 | 0.065911847 | 15.89639312 | 6.84E-57 |
| B210 | Hip | Hypertension+Coronary heart disease | 1.262376044 | 0.13194433 | 9.567489888 | 1.10E-21 |
| B210 | Hip | Hypertensive+Diabetes+Coronary Heart Disease | 0.7582231 | 0.196255176 | 3.863455307 | 0.0001118 |
| B211 | IOP-R | (Intercept) | 16.01192703 | 0.007249968 | 2208.551527 | 0 |
| B211 | IOP-R | Gender | -0.03741796 | 0.003203178 | -11.68151047 | 1.59E-31 |
| B211 | IOP-R | Age | -0.00418655 | 0.000122814 | -34.08841949 | 1.63E-254 |

| | | | | | | |
|---|---|---|---|---|---|---|
| B211 | IOP-R | Diabetes | 0.24992964 | 0.013787622 | 18.12710308 | 2.01E-73 |
| B211 | IOP-R | Hypertension+other diseases | 0.111094816 | 0.030026512 | 3.699890842 | 0.0002157 |
| B211 | IOP-R | Hypertension | 0.144741512 | 0.007400338 | 19.55876975 | 3.63E-85 |
| B211 | IOP-R | Hypertension+Diabetes | 0.240576638 | 0.016212923 | 14.83857264 | 8.37E-50 |
| B211 | IOP-R | Hyperlipidemia | 0.131662688 | 0.035458497 | 3.713149137 | 0.0002047 |
| B212 | IOP-L | (Intercept) | 16.19824456 | 0.007519186 | 2154.255148 | 0 |
| B212 | IOP-L | Gender | -0.0424369 | 0.003322124 | -12.77402791 | 2.31E-37 |
| B212 | IOP-L | Age | -0.00282993 | 0.000127375 | -22.21729876 | 2.51E-109 |
| B212 | IOP-L | Diabetes | 0.271085004 | 0.014299607 | 18.95751409 | 3.98E-80 |
| B212 | IOP-L | Hypertension+other diseases | 0.115481944 | 0.031141507 | 3.708296556 | 0.0002087 |
| B212 | IOP-L | Hypertension | 0.156860461 | 0.00767514 | 20.43747183 | 8.17E-93 |
| B212 | IOP-L | Hypertension+Diabetes | 0.246505464 | 0.016814969 | 14.65988219 | 1.18E-48 |
| B212 | IOP-L | Hyperlipidemia | 0.159102704 | 0.036775202 | 4.326358351 | 1.52E-05 |
| B213 | CVA-R | (Intercept) | 0.99341292 | 0.010389619 | 95.61591716 | 0 |
| B213 | CVA-R | Age | 0.003465525 | 0.000176 | 19.69046237 | 2.72E-86 |
| B213 | CVA-R | Coronary heart disease+other diseases | 3.993458211 | 0.208945831 | 19.11240914 | 2.07E-81 |
| B213 | CVA-R | Hyperlipidemia | 0.21636642 | 0.050814057 | 4.258003267 | 2.06E-05 |
| B214 | CVA-L | (Intercept) | 1.035688378 | 0.009966894 | 103.9128497 | 0 |
| B214 | CVA-L | Gender | -0.01540742 | 0.004403569 | -3.498847557 | 0.0004673 |
| B214 | CVA-L | Age | 0.002584785 | 0.000168839 | 15.30915387 | 6.75E-53 |
| B214 | CVA-L | Nephropathy | 0.294515019 | 0.083946675 | 3.508358371 | 0.0004509 |
| B214 | CVA-L | Coronary heart disease+other diseases | 1.947965956 | 0.20044441 | 9.718235376 | 2.53E-22 |
| B214 | CVA-L | Hypertension | 0.055394485 | 0.010173616 | 5.444915968 | 5.18E-08 |
| B214 | CVA-L | Hypertension+Diabetes | 0.116110547 | 0.022288719 | 5.2093863 | 1.90E-07 |
| B215 | BMI | (Intercept) | 24.83963749 | 0.014904329 | 1666.605498 | 0 |
| B215 | BMI | Gender | -2.13482836 | 0.006585025 | -324.1944083 | 0 |
| B215 | BMI | Age | 0.033853641 | 0.000252479 | 134.0847752 | 0 |
| B215 | BMI | Hypothyroidism | 0.352577628 | 0.074026582 | 4.762851651 | 1.91E-06 |
| B215 | BMI | Rheumatoid arthritis | -0.64062924 | 0.151417324 | -4.230884695 | 2.33E-05 |
| B215 | BMI | Diabetes | 0.139643488 | 0.0283443 | 4.926686772 | 8.37E-07 |
| B215 | BMI | Gout | 1.313672357 | 0.065328037 | 20.10886004 | 6.48E-90 |
| B215 | BMI | Parkinson's syndrome | -1.41069992 | 0.214702845 | -6.570476158 | 5.02E-11 |
| B215 | BMI | Cholecystolithiasis | 0.429034056 | 0.096132763 | 4.462932766 | 8.09E-06 |
| B215 | BMI | Stomach trouble | -0.8558995 | 0.083744633 | -10.22035048 | 1.61E-24 |
| B215 | BMI | Hypertension+other diseases | 0.949436342 | 0.061727865 | 15.38100073 | 2.23E-53 |
| B215 | BMI | Bronchial disease | -0.61093754 | 0.125832841 | -4.855151767 | 1.20E-06 |
| B215 | BMI | Brain diseases | -1.07328819 | 0.188012189 | -5.708609624 | 1.14E-08 |
| B215 | BMI | Hypertension | 1.233588294 | 0.015213458 | 81.08533075 | 0 |
| B215 | BMI | Hepatic adipose infiltration | 2.665063885 | 0.181967952 | 14.64578711 | 1.45E-48 |
| B215 | BMI | Hypertension+Diabetes | 0.995081704 | 0.033330183 | 29.85527252 | 9.46E-196 |
| B215 | BMI | Hypertension+Coronary heart disease | 0.477566151 | 0.066721369 | 7.157619182 | 8.22E-13 |
| B215 | BMI | Hypertensive+Diabetes+Coronary Heart Disease | 0.537858266 | 0.099241961 | 5.419665856 | 5.97E-08 |
| B215 | BMI | Coronary heart disease | -0.43996369 | 0.083092391 | -5.294873397 | 1.19E-07 |

| | | | | | | |
|---|---|---|---|---|---|---|
| B216 | BMI | (Intercept) | 85.41154394 | 0.046085949 | 1853.309885 | 0 |
| B216 | BMI | Gender | -14.45438 | 0.020361676 | -709.8816268 | 0 |
| B216 | BMI | Age | -0.03663223 | 0.000780696 | -46.92251668 | 0 |
| B216 | BMI | Hypothyroidism | 1.579311852 | 0.228898941 | 6.899603135 | 5.22E-12 |
| B216 | BMI | Diabetes | 0.442585574 | 0.087643926 | 5.049814563 | 4.42E-07 |
| B216 | BMI | Gout | 4.525308337 | 0.202002013 | 22.40229328 | 4.02E-111 |
| B216 | BMI | Parkinson's syndrome | -4.25509994 | 0.663886573 | -6.409377913 | 1.46E-10 |
| B216 | BMI | Cholecystolithiasis | 1.769177924 | 0.297253865 | 5.9517407 | 2.65E-09 |
| B216 | BMI | Stomach trouble | -2.61084163 | 0.258948303 | -10.08248212 | 6.62E-24 |
| B216 | BMI | Hypertension+other diseases | 2.969036878 | 0.190869856 | 15.55529479 | 1.49E-54 |
| B216 | BMI | Bronchial disease | -2.23492702 | 0.389090017 | -5.743984479 | 9.25E-09 |
| B216 | BMI | Brain diseases | -2.46168799 | 0.581355911 | -4.234390582 | 2.29E-05 |
| B216 | BMI | Hypertension | 3.473030667 | 0.047041811 | 73.82859169 | 0 |
| B216 | BMI | Hepatic adipose infiltration | 8.046568218 | 0.562666416 | 14.30077927 | 2.19E-46 |
| B216 | BMI | Pregnant | 4.213376871 | 1.244408667 | 3.385846612 | 0.0007096 |
| B216 | BMI | Hypertension+Diabetes | 2.712889107 | 0.103060866 | 26.32317391 | 1.20E-152 |
| B216 | BMI | Hypertension+Coronary heart disease | 1.562673224 | 0.20631036 | 7.574380778 | 3.61E-14 |
| B216 | BMI | Hypertensive+Diabetes+Coronary Heart Disease | 1.403726803 | 0.306867873 | 4.574368731 | 4.78E-06 |
| B216 | BMI | Hyperlipidemia | 1.335884617 | 0.225399418 | 5.926743865 | 3.09E-09 |
| B216 | BMI | Coronary heart disease | -1.03104699 | 0.256931493 | -4.012925682 | 6.00E-05 |
| B218 | DBP | (Intercept) | 74.99401517 | 0.050954069 | 1471.796402 | 0 |
| B218 | DBP | Gender | -5.63761754 | 0.022512507 | -250.4215733 | 0 |
| B218 | DBP | Age | 0.121667435 | 0.000863162 | 140.9554954 | 0 |
| B218 | DBP | Nephropathy | 1.426261999 | 0.429163244 | 3.323355431 | 0.0008895 |
| B218 | DBP | Gout | 2.476569777 | 0.223339755 | 11.08879954 | 1.43E-28 |
| B218 | DBP | Parkinson's syndrome | -5.58079468 | 0.734013799 | -7.603119567 | 2.89E-14 |
| B218 | DBP | Stomach trouble | -1.96807154 | 0.286301358 | -6.874125741 | 6.24E-12 |
| B218 | DBP | Lumbar disc protrusion | -2.80308992 | 0.524981823 | -5.339403748 | 9.33E-08 |
| B218 | DBP | Hepatitis B | -1.88594646 | 0.387682003 | -4.864673735 | 1.15E-06 |
| B218 | DBP | Hypertension+other diseases | 3.881880442 | 0.211031694 | 18.39477459 | 1.50E-75 |
| B218 | DBP | Coronary heart disease+other diseases | -5.39472816 | 1.024738306 | -5.264493512 | 1.41E-07 |
| B218 | DBP | Diabetes+other diseases | -2.39704903 | 0.533518605 | -4.492906165 | 7.03E-06 |
| B218 | DBP | Bronchial disease | -1.56642112 | 0.430190116 | -3.641229915 | 0.0002714 |
| B218 | DBP | Brain diseases | -3.27726616 | 0.642765312 | -5.098697922 | 3.42E-07 |
| B218 | DBP | Hypertension | 7.046683908 | 0.0520109 | 135.4847511 | 0 |
| B218 | DBP | Other Cardiac diseases | -2.97013493 | 0.555426982 | -5.34748045 | 8.92E-08 |
| B218 | DBP | Heart disease | -3.4793921 | 0.680850824 | -5.110358952 | 3.22E-07 |
| B218 | DBP | Hypertension+Diabetes | 1.470684502 | 0.113947323 | 12.90670513 | 4.16E-38 |
| B218 | DBP | Hypertension+Coronary heart disease | -1.43036361 | 0.228103199 | -6.270686347 | 3.60E-10 |
| B218 | DBP | Hypertensive+Diabetes+Coronary Heart Disease | -3.8554771 | 0.339282737 | -11.36361119 | 6.38E-30 |
| B218 | DBP | Hyperlipidemia | -1.99566012 | 0.24920866 | -8.007988668 | 1.17E-15 |
| B218 | DBP | Coronary heart disease | -3.06613349 | 0.28407151 | -10.7935269 | 3.71E-27 |

| | | | | | | |
|---|---|---|---|---|---|---|
| B218 | DBP | Coronary heart disease+Diabetes | -6.9294662 | 0.615865427 | -11.25159151 | 2.29E-29 |
| B219 | OT | (Intercept) | -0.10343083 | 0.011540596 | -8.962347223 | 3.18E-19 |
| B219 | OT | Gender | 0.33293848 | 0.005098862 | 65.29662793 | 0 |
| B219 | OT | Age | 0.031380817 | 0.000195498 | 160.5175557 | 0 |
| B219 | OT | Hypothyroidism | 0.381048995 | 0.057319645 | 6.64779055 | 2.98E-11 |
| B219 | OT | Hyperthyroidism | 0.347981487 | 0.084245769 | 4.130551474 | 3.62E-05 |
| B219 | OT | Cervical spondylopathy | 0.431893927 | 0.130805192 | 3.30181029 | 0.0009607 |
| B219 | OT | Nephropathy | 0.588188452 | 0.097201259 | 6.051243158 | 1.44E-09 |
| B219 | OT | Diabetes | 0.113844309 | 0.021947322 | 5.187161763 | 2.14E-07 |
| B219 | OT | Gout | 0.181223866 | 0.050584261 | 3.582613732 | 0.0003402 |
| B219 | OT | Stomach trouble | 0.381571424 | 0.064844445 | 5.884411899 | 4.00E-09 |
| B219 | OT | Hypertension+other diseases | 0.659750061 | 0.047796606 | 13.80328276 | 2.46E-43 |
| B219 | OT | Coronary heart disease+other diseases | 2.543222571 | 0.232093159 | 10.95776618 | 6.12E-28 |
| B219 | OT | Diabetes+other diseases | 0.391417909 | 0.120836723 | 3.239229763 | 0.0011986 |
| B219 | OT | Other disease conditions | 0.425806868 | 0.044370912 | 9.59653177 | 8.29E-22 |
| B219 | OT | Brain diseases | 0.874635432 | 0.145580029 | 6.007935559 | 1.88E-09 |
| B219 | OT | Hypertension | 0.253790174 | 0.011779958 | 21.54423418 | 6.39E-103 |
| B219 | OT | Asthma | 0.295384937 | 0.081494967 | 3.624578879 | 0.0002894 |
| B219 | OT | Other Cardiac diseases | 1.345794504 | 0.125798755 | 10.69799542 | 1.04E-26 |
| B219 | OT | Heart disease | 1.135227194 | 0.154206023 | 7.36175651 | 1.82E-13 |
| B219 | OT | Hepatopathy | 0.703986474 | 0.173827258 | 4.049919918 | 5.12E-05 |
| B219 | OT | Hypertension+Diabetes | 0.359527115 | 0.025807949 | 13.93086724 | 4.16E-44 |
| B219 | OT | Hypertension+Coronary heart disease | 2.322809984 | 0.051663134 | 44.96068705 | 0 |
| B219 | OT | Hypertensive+Diabetes+Coronary Heart Disease | 1.961474839 | 0.076844207 | 25.52534436 | 1.17E-143 |
| B219 | OT | Hyperlipidemia | 0.732816032 | 0.056443313 | 12.98322141 | 1.54E-38 |
| B219 | OT | Coronary heart disease | 3.112015072 | 0.064339406 | 48.36872528 | 0 |
| B219 | OT | Coronary heart disease+Diabetes | 3.73808917 | 0.139487469 | 26.79874542 | 3.89E-158 |
| B220 | SBP | (Intercept) | 111.235922 | 0.072664226 | 1530.820979 | 0 |
| B220 | SBP | Gender | -7.12179972 | 0.032104481 | -221.8319527 | 0 |
| B220 | SBP | Age | 0.389858158 | 0.001230932 | 316.7178217 | 0 |
| B220 | SBP | Rheumatoid arthritis | 3.218464305 | 0.73821655 | 4.359783458 | 1.30E-05 |
| B220 | SBP | Nephropathy | 2.154458035 | 0.612018153 | 3.520251851 | 0.0004312 |
| B220 | SBP | Diabetes | 2.356249838 | 0.138189151 | 17.05090326 | 3.53E-65 |
| B220 | SBP | Gout | 3.162468342 | 0.318498815 | 9.929293913 | 3.11E-23 |
| B220 | SBP | Stomach trouble | -3.63411504 | 0.408286662 | -8.900890925 | 5.55E-19 |
| B220 | SBP | Lumbar disc protrusion | -3.1771707 | 0.74866245 | -4.243795982 | 2.20E-05 |
| B220 | SBP | Hepatitis B | -4.83244825 | 0.552862871 | -8.740771912 | 2.32E-18 |
| B220 | SBP | Hypertension+other diseases | 11.75873082 | 0.300946619 | 39.07248018 | 0 |
| B220 | SBP | Other disease conditions | 0.92635607 | 0.279377079 | 3.315791241 | 0.0009139 |
| B220 | SBP | Hypertension | 15.09222416 | 0.074171345 | 203.4778264 | 0 |
| B220 | SBP | Hypertension+Diabetes | 13.6246466 | 0.162497211 | 83.84541803 | 0 |
| B220 | SBP | Hypertension+Coronary heart disease | 9.609783147 | 0.325291833 | 29.54203632 | 1.04E-191 |
| B220 | SBP | Hypertensive+Diabetes+Coronary Heart | 8.583157925 | 0.48384198 | 17.73958913 | 2.14E-70 |

| | | | | | | |
|---|---|---|---|---|---|---|
| | | Disease | | | | |
| B220 | SBP | Hyperlipidemia | 1.843783408 | 0.355389763 | 5.188059989 | 2.13E-07 |
| B221 | Hei | (Intercept) | 187.5874868 | 0.028618373 | 6554.792258 | 0 |
| B221 | Hei | Gender | -11.3503542 | 0.012644159 | -897.6757276 | 0 |
| B221 | Hei | Age | -0.16068176 | 0.000484795 | -331.4424896 | 0 |
| B221 | Hei | Hypothyroidism | 0.880552998 | 0.142141268 | 6.194914458 | 5.83E-10 |
| B221 | Hei | Gout | 0.875606126 | 0.12543886 | 6.980341844 | 2.95E-12 |
| B221 | Hei | Cholecystolithiasis | 0.772900429 | 0.18458819 | 4.187160769 | 2.82E-05 |
| B221 | Hei | Lumbar disc protrusion | 1.05829025 | 0.294856245 | 3.589173601 | 0.0003317 |
| B221 | Hei | Hypertension+other diseases | 0.497933123 | 0.118526033 | 4.20104437 | 2.66E-05 |
| B221 | Hei | Bronchial disease | -0.77494446 | 0.241616445 | -3.207333261 | 0.0013398 |
| B221 | Hei | Hypertension | 0.208000232 | 0.029211942 | 7.120383467 | 1.08E-12 |
| B221 | Hei | Hyperlipidemia | 1.366442922 | 0.13996814 | 9.76252823 | 1.63E-22 |
| B221 | Hei | Coronary heart disease+Diabetes | 1.160343971 | 0.345901056 | 3.354554578 | 0.000795 |
| B222 | NS | (Intercept) | 41.52027432 | 0.013203205 | 3144.711873 | 0 |
| B222 | NS | Gender | -5.2289509 | 0.005833435 | -896.375979 | 0 |
| B222 | NS | Age | 0.010525475 | 0.000223662 | 47.05966825 | 0 |
| B222 | NS | Hypothyroidism | 0.326357234 | 0.065577462 | 4.976667623 | 6.47E-07 |
| B222 | NS | Diabetes | 0.41417884 | 0.025109187 | 16.49511167 | 4.07E-61 |
| B222 | NS | Gout | 0.982784779 | 0.057871737 | 16.98211982 | 1.14E-64 |
| B222 | NS | Cholecystolithiasis | 0.536026602 | 0.085160526 | 6.294308287 | 3.09E-10 |
| B222 | NS | Stomach trouble | -0.47158821 | 0.074186331 | -6.356807288 | 2.06E-10 |
| B222 | NS | Hypertension+other diseases | 0.937805906 | 0.054682476 | 17.15002645 | 6.44E-66 |
| B222 | NS | Diabetes+other diseases | 0.467683525 | 0.138245198 | 3.383000159 | 0.000717 |
| B222 | NS | Other disease conditions | 0.211270541 | 0.050763256 | 4.161879224 | 3.16E-05 |
| B222 | NS | Hypertension | 0.923968801 | 0.013477051 | 68.55868038 | 0 |
| B222 | NS | Hepatic adipose infiltration | 2.103066797 | 0.161198805 | 13.04641681 | 6.72E-39 |
| B222 | NS | Pregnant | 1.960255784 | 0.356511752 | 5.49843244 | 3.83E-08 |
| B222 | NS | Hypertension+Diabetes | 1.03712607 | 0.029526 | 35.12585784 | 4.24E-270 |
| B222 | NS | Hypertension+Coronary heart disease | 0.523724027 | 0.05910604 | 8.860753166 | 7.96E-19 |
| B222 | NS | Hypertensive+Diabetes+Coronary Heart Disease | 0.826797692 | 0.087914851 | 9.404528116 | 5.24E-21 |
| B222 | NS | Hyperlipidemia | 0.377616538 | 0.064574881 | 5.847731087 | 4.99E-09 |
| B222 | NS | Coronary heart disease+Diabetes | 0.538106488 | 0.159582884 | 3.371956161 | 0.0007464 |
| B223 | FH | (Intercept) | 1.68 | 3.05E-14 | 5.52E+13 | 0 |
| B224 | pmh | (Intercept) | 0.981223972 | 0.011092521 | 88.45815653 | 0 |
| B224 | pmh | Gender | -0.04199674 | 0.004900894 | -8.569200902 | 1.04E-17 |
| B224 | pmh | Age | 0.005384402 | 0.000187907 | 28.65456001 | 1.71E-180 |
| B224 | pmh | Rhinallergosis | 1.492319237 | 0.179457024 | 8.315747169 | 9.13E-17 |
| B224 | pmh | Hypothyroidism | 0.895334226 | 0.055094154 | 16.25098418 | 2.24E-59 |
| B224 | pmh | Hyperthyroidism | 1.18903076 | 0.080974845 | 14.68395221 | 8.28E-49 |
| B224 | pmh | Cervical spondylopathy | 2.799280459 | 0.125726553 | 22.2648311 | 8.71E-110 |
| B224 | pmh | Rheumatoid arthritis | 1.284238951 | 0.112692079 | 11.39600021 | 4.40E-30 |
| B224 | pmh | Chronic rhinitis | 1.090012699 | 0.125315366 | 8.698156791 | 3.38E-18 |

| | | | | | | |
|---|---|---|---|---|---|---|
| B224 | pmh | Nephropathy | 1.968416907 | 0.093427325 | 21.06896351 | 1.62E-98 |
| B224 | pmh | Diabetes | 1.695349292 | 0.021095196 | 80.36660644 | 0 |
| B224 | pmh | Gout | 1.61722507 | 0.048620277 | 33.26235795 | 1.94E-242 |
| B224 | pmh | Parkinson's syndrome | 1.996031758 | 0.159792216 | 12.49142045 | 8.38E-36 |
| B224 | pmh | Cholecystolithiasis | 1.401335901 | 0.07154664 | 19.58632712 | 2.11E-85 |
| B224 | pmh | Stomach trouble | 3.484489907 | 0.062326796 | 55.90677065 | 0 |
| B224 | pmh | Chronic pharyngitis | 1.361144827 | 0.080188378 | 16.97434033 | 1.30E-64 |
| B224 | pmh | Lumbar disc protrusion | 3.314858044 | 0.114286692 | 29.00475972 | 7.05E-185 |
| B224 | pmh | Hepatitis B | 5.650828096 | 0.084397005 | 66.95531527 | 0 |
| B224 | pmh | Hypertension+other diseases | 2.069399792 | 0.045940856 | 45.04486816 | 0 |
| B224 | pmh | Coronary heart disease+other diseases | 3.919410979 | 0.223081916 | 17.56938008 | 4.35E-69 |
| B224 | pmh | Diabetes+other diseases | 2.780554357 | 0.116145119 | 23.94034609 | 1.28E-126 |
| B224 | pmh | Bronchial disease | 2.79666455 | 0.093650872 | 29.86266443 | 7.59E-196 |
| B224 | pmh | Other disease conditions | 2.162482264 | 0.042648168 | 50.70516155 | 0 |
| B224 | pmh | Brain diseases | 3.900537095 | 0.139927742 | 27.87536643 | 6.33E-171 |
| B224 | pmh | Hypertension | 0.230721058 | 0.01132259 | 20.37705721 | 2.81E-92 |
| B224 | pmh | Hepatic adipose infiltration | 9.947220293 | 0.13542933 | 73.44952758 | 0 |
| B224 | pmh | Asthma | 3.210217029 | 0.078330846 | 40.98279535 | 0 |
| B224 | pmh | Other Cardiac diseases | 2.884687546 | 0.120914496 | 23.85725157 | 9.36E-126 |
| B224 | pmh | Heart disease | 3.173478421 | 0.148218824 | 21.41076506 | 1.13E-101 |
| B224 | pmh | Hepatopathy | 4.475923527 | 0.167078245 | 26.78938553 | 5.00E-158 |
| B224 | pmh | Hypertension+Diabetes | 0.403471961 | 0.024805931 | 16.2651411 | 1.78E-59 |
| B224 | pmh | Hypertension+Coronary heart disease | 0.680248782 | 0.049657262 | 13.6988781 | 1.04E-42 |
| B224 | pmh | Hypertensive+Diabetes+Coronary Heart Disease | 0.900122176 | 0.073860656 | 12.18676125 | 3.68E-34 |
| B224 | pmh | Hyperlipidemia | 0.747460448 | 0.054251847 | 13.77760379 | 3.51E-43 |
| B224 | pmh | Coronary heart disease | 0.885269415 | 0.061841366 | 14.31516584 | 1.78E-46 |
| B224 | pmh | Coronary heart disease+Diabetes | 1.272823303 | 0.134071732 | 9.493599303 | 2.24E-21 |
| B225 | hPBG | (Intercept) | 7.604356604 | 0.000922572 | 8242.560168 | 0 |
| B225 | hPBG | Age | 5.61E-05 | 1.56E-05 | 3.591187086 | 0.0003292 |
| B225 | hPBG | Diabetes | 0.046685763 | 0.001754501 | 26.60913904 | 6.16E-156 |
| B225 | hPBG | Gout | 0.013089528 | 0.00404378 | 3.236953676 | 0.0012082 |
| B225 | hPBG | Hypertension+other diseases | 0.018622812 | 0.003820931 | 4.873894186 | 1.09E-06 |
| B225 | hPBG | Diabetes+other diseases | 0.052022358 | 0.009659865 | 5.385412664 | 7.23E-08 |
| B225 | hPBG | Hypertension | 0.005135315 | 0.000941707 | 5.453197544 | 4.95E-08 |
| B225 | hPBG | Hypertension+Diabetes | 0.065905797 | 0.002063125 | 31.94464097 | 8.70E-224 |
| B225 | hPBG | Hypertensive+Diabetes+Coronary Heart Disease | 0.068034061 | 0.006143039 | 11.07498516 | 1.67E-28 |
| B225 | hPBG | Coronary heart disease | 0.017953829 | 0.005143386 | 3.490662949 | 0.0004818 |
| B225 | hPBG | Coronary heart disease+Diabetes | 0.10829366 | 0.011150833 | 9.711710512 | 2.70E-22 |
| B227 | OD | (Intercept) | 0.750987957 | 0.0202091 | 37.16088136 | 5.11E-302 |
| B227 | OD | Gender | -0.03783715 | 0.008928777 | -4.237663169 | 2.26E-05 |
| B227 | OD | Age | 0.011154554 | 0.000342342 | 32.58305262 | 9.91E-233 |
| B227 | OD | Diabetes | 0.226246441 | 0.038432644 | 5.88683001 | 3.94E-09 |

| | | | | | | |
|---|---|---|---|---|---|---|
| B227 | OD | Parkinson's syndrome | 0.996343633 | 0.291120187 | 3.422447763 | 0.0006206 |
| B227 | OD | Diabetes+other diseases | 0.935826738 | 0.211600976 | 4.422601238 | 9.75E-06 |
| B227 | OD | Bronchial disease | 0.844404038 | 0.170619445 | 4.949049265 | 7.46E-07 |
| B227 | OD | Hypertension+Diabetes | 0.403449084 | 0.04519311 | 8.927225571 | 4.38E-19 |
| B227 | OD | Hypertension+Coronary heart disease | 0.634946461 | 0.090468934 | 7.01839221 | 2.25E-12 |
| B227 | OD | Hyperlipidemia | 0.408454047 | 0.098839656 | 4.132491577 | 3.59E-05 |
| B228 | OS | (Intercept) | 0.814346023 | 0.020010781 | 40.69536404 | 0 |
| B228 | OS | Gender | -0.02965087 | 0.008841156 | -3.353732595 | 0.0007973 |
| B228 | OS | Age | 0.010565329 | 0.000338983 | 31.16775372 | 3.84E-213 |
| B228 | OS | Diabetes | 0.187802016 | 0.038055492 | 4.934951785 | 8.02E-07 |
| B228 | OS | Parkinson's syndrome | 1.556822813 | 0.288263329 | 5.40069672 | 6.64E-08 |
| B228 | OS | Hypertension+other diseases | 0.547395376 | 0.082876778 | 6.60493072 | 3.98E-11 |
| B228 | OS | Brain diseases | 1.018865445 | 0.252428045 | 4.036260884 | 5.43E-05 |
| B228 | OS | Hypertension | 0.117119553 | 0.020425822 | 5.733896661 | 9.82E-09 |
| B228 | OS | Asthma | 0.620294757 | 0.141307949 | 4.389666405 | 1.14E-05 |
| B228 | OS | Heart disease | 2.629162941 | 0.267385062 | 9.832871462 | 8.15E-23 |
| B228 | OS | Hypertension+Diabetes | 0.22287428 | 0.044749615 | 4.980473719 | 6.34E-07 |
| B228 | OS | Hypertension+Coronary heart disease | 0.887877857 | 0.089581132 | 9.911438184 | 3.72E-23 |
| B228 | OS | Hypertensive+Diabetes+Coronary Heart Disease | 0.668201099 | 0.133243777 | 5.014876599 | 5.31E-07 |

**Supplementary Table 37. Optimal parameter combination of machine learning**

| | Automated feature selection | Parameters of Random forest | | |
|---|---|---|---|---|
| | SelectPercentile | n_estimators | max_depth | min_samples_leaf |
| Cholecystolithiasis | 15 | 100 | 10 | 4 |
| Hypertension | 16 | 100 | 20 | —— |
| Hypertension+Diabetes | 16 | 100 | 10 | —— |
| Hypertension+Coronary heart disease | 15 | 100 | 10 | —— |
| Hypertensive+Diabetes+Coronary Heart Disease | 15 | 100 | 10 | —— |
| Hyperlipidemia | 16 | 100 | 10 | —— |
| Coronary heart disease | 16 | 100 | 10 | —— |
| Coronary heart disease+Diabetes | 15 | 100 | 10 | —— |
| Rhinallergosis | 15 | 100 | 4 | —— |
| Hypothyroidism | 16 | 100 | 15 | —— |
| Hyperthyroidism | 16 | 100 | 6 | —— |
| Cervical spondylopathy | 16 | 100 | 6 | —— |
| Rheumatoid arthritis | 16 | 100 | 10 | —— |
| Chronic rhinitis | 16 | 100 | 10 | —— |
| Nephropathy | 16 | 100 | 10 | —— |
| Diabetes | 16 | 100 | 20 | —— |
| Gout | 16 | 100 | 10 | —— |
| Parkinson's syndrome | 16 | 100 | 10 | —— |
| Stomach trouble | 16 | 100 | 20 | —— |
| Chronic pharyngitis | 16 | 100 | 5 | —— |
| Lumbar disc protrusion | 16 | 100 | 5 | —— |
| Hepatitis B | 16 | 100 | 15 | —— |

| | | | | |
|---|---|---|---|---|
| Hypertension+other diseases | 16 | 100 | 15 | —— |
| Coronary heart disease+other diseases | 16 | 100 | 4 | —— |
| Diabetes+other diseases | 16 | 100 | 6 | —— |
| Bronchial disease | 16 | 100 | 8 | —— |
| Other disease conditions | 16 | 100 | 10 | 5 |
| Brain diseases | 16 | 100 | 20 | —— |
| Hepatic adipose infiltration | 16 | 100 | 10 | —— |
| Asthma | 16 | 100 | 10 | —— |
| Other Cardiac diseases | 16 | 100 | 6 | —— |
| Heart disease | 16 | 100 | _ | —— |
| Hepatopathy | 15 | 100 | 10 | —— |
| Pregnant | 16 | 70 | —— | —— |
| Unhealthy together | 16 | 100 | 20 | 10 |

**Supplementary codes for machine learning**

```
import numpy as np

from sklearn.ensemble import RandomForestClassifier

import pandas as pd

from imblearn.under_sampling import RandomUnderSampler

from sklearn.utils import shuffle

from sklearn.model_selection import train_test_split

from sklearn.feature_selection import SelectPercentile

from matplotlib import pyplot as plt

from sklearn.model_selection import GridSearchCV

from sklearn.metrics import f1_score

from sklearn.metrics import classification_report

from sklearn.metrics import roc_curve, auc

from sklearn.metrics import confusion_matrix


# Load data

dataset_original = np.loadtxt('C:\\Users\\dell\\Desktop\\Normal condition or Hypertension.csv',

                               encoding='utf-8', delimiter=',')

x_original = dataset_original[:, 1:209]

y_original = dataset_original[:, 0]


# Define a function that uses Random UnderSampler for undersampling
```

```
def RUS():

        # Establishing the model objects of RandomUnderSampler

        model_RandomUnderSampler = RandomUnderSampler()

        # Undersampling

        x_RandomUnderSampler_resampled, y_RandomUnderSampler_resampled =
model_RandomUnderSampler.fit_sample(x_original, y_original)

        # Converting under-sampled data into dataFrame

        x_resampled = pd.DataFrame(x_RandomUnderSampler_resampled)

        y_resampled = pd.DataFrame(y_RandomUnderSampler_resampled)

        RandomUnderSampler_resampled = pd.concat([y_resampled, x_resampled], axis=1)

        # shuffling

        res_data = shuffle(RandomUnderSampler_resampled)

        np.savetxt('res_data.csv', res_data, fmt='%10.4f', delimiter=',')

        return

RUS()


x_train = None

x_valid = None

y_train = None

y_valid = None

select = None

grid_search = None

y_score = None
```

```
# The purpose of defining the function is to divide x and y into x_trainval, y_trainval and
# test set by train_test_split method； Then x_trainval and y_trainval are randomly divided
# into training set and validation set, and relevant information is output.
def data_split():
    dataset = np.loadtxt('C:\\Users\\dell\\PycharmProjects\\Disease prediction\\res_data.csv', delimiter=',')
    x = dataset[:, 1:209]
    y = dataset[:, 0]
    print('the shape of x:', x.shape)
    print('the shape of y:', y.shape)
    x_trainval, x_test, y_trainval, y_test = train_test_split(x, y, test_size=0.3, random_state=1)
    np.savetxt('x_test_set.csv', x_test, fmt='%10.4f')
    np.savetxt('y_test_set.csv', y_test, fmt='%10.0f')
    global x_train, x_valid, y_train, y_valid
    x_train, x_valid, y_train, y_valid = train_test_split(x_trainval, y_trainval, test_size=0.3, random_state=2)
    print('the shape of x_train:', x_train.shape)
    print('the shape of y_train:', y_train.shape)
    y1 = pd.Series(y_train)
    print('y_train:\n', y1.value_counts())
    np.savetxt('y_valid_set.csv', y_valid, fmt='%10.0f')
    return
data_split()
```

```
# Define the function to automate feature selection

def select_feature():

    global select

    select = SelectPercentile(percentile=16)

    select.fit(x_train, y_train)

    # Transforming the training set

    global x_train_selected

    x_train_selected = select.transform(x_train)

    print('x_train_selected shape：{}'.format(x_train_selected.shape))

    mask = select.get_support()

    print(mask)

    return

select_feature()


# Define the function to select the best combination of parameters using grid search with cross
validation

def cross_validation_grid_search():

    forest = RandomForestClassifier(criterion='entropy', random_state=3)

    parameters = {'n_estimators': [80, 100],

                  'max_depth': [10, 20, 40]}

                  # 'min_samples_leaf': [20, 10]}

    global grid_search

    grid_search = GridSearchCV(estimator=forest, param_grid=parameters, cv=5)

    grid_search.fit(x_train_selected, y_train)
```

```
    x_valid_selected = select.transform(x_valid)
    print('valid score:{:.2f}'.format(grid_search.score(x_valid_selected, y_valid)))
    print('best parameters:{}'.format(grid_search.best_params_))
    # Best_score preserves the average accuracy of cross-validation, which is obtained by
    # cross-validation on the training set.
    print('best cross-validation score:{:.2f}'.format((grid_search.best_score_)))
    global y_valid_pred
    # Note that the validation set also needs to be transformed
    y_valid_pred = grid_search.predict(x_valid_selected)
    global y_score
    y_score1 = grid_search.predict_proba(x_valid_selected)
    y_score = y_score1[:, 1]
    np.savetxt('y_valid_pred.csv', y_valid_pred, fmt='%10.0f')
    print('the shape of y_validation_pred:', y_valid_pred.shape)
    return
cross_validation_grid_search()


print('f1_score: {:.2f}'.format(f1_score(y_valid, y_valid_pred)))
print(classification_report(y_valid, y_valid_pred.round()))


# Define the function to draw ROC curves
def ROC():
    # Calculate the true positive rate and false positive rate
    fpr, tpr, threshold = roc_curve(y_valid, y_score)
```

```
    # Calculate the value of Area Under Curve (AUC)
    roc_auc = auc(fpr, tpr)
    plt.figure()
    lw = 2
    plt.figure(figsize=(10, 10))
    plt.plot(fpr, tpr, color='darkorange',
             lw=lw, label='ROC curve (AUC = %0.2f)' % roc_auc)
    # Drawing curves in abscissa with false positivity and in ordinate with true positivity
    plt.plot([0, 1], [0, 1], color='navy', lw=lw, linestyle='--')
    plt.xticks([0, 0.2, 0.4, 0.6, 0.8, 1.0])
    plt.yticks([0, 0.2, 0.4, 0.6, 0.8, 1.0])
    plt.tick_params(labelsize=25)
    plt.xlabel('False Positive Rate', fontsize=25)
    plt.ylabel('True Positive Rate', fontsize=25)
    plt.title("Hypertension", fontsize=35)
    plt.legend(loc="lower right", fontsize=25)
    plt.show()
ROC()


# Compute and output Confusion Matrix
cm = confusion_matrix(y_valid, y_valid_pred)
print('confusion_matrix:\n', cm)
```
